# Deep-learning quantified cell-type-specific nuclear morphology predicts genomic instability and prognosis in multiple cancer types

**DOI:** 10.1101/2023.05.15.539600

**Authors:** John Abel, Suyog Jain, Deepta Rajan, Harshith Padigela, Kenneth Leidal, Aaditya Prakash, Jake Conway, Michael Nercessian, Christian Kirkup, Syed Ashar Javed, Raymond Biju, Natalia Harguindeguy, Daniel Shenker, Nicholas Indorf, Darpan Sanghavi, Robert Egger, Benjamin Trotter, Ylaine Gerardin, Jacqueline A. Brosnan-Cashman, Aditya Dhoot, Michael C. Montalto, Chintan Parmar, Ilan Wapinski, Archit Khosla, Michael G. Drage, Limin Yu, Amaro Taylor-Weiner

**Author notes:** Employee of PathAI at time of study (Suyog Jain, Deepta Rajan, Kenneth Leidal, Aaditya Prakash, Michael C. Montalto, Ilan Wapinski, Archit Khosla, Michael G. Drage, Amaro Taylor-Weiner). **Correspondence to:** John Abel, Ph.D. PathAI 1325 Boylston Street, Suite 10000 Boston, Massachusetts 02215. **Declaration of Competing interests:** The authors declare the following competing interests: J.A., S.J., D.R., H.P., K.L., A.P., J.C., M.N., C.K., S.A.J., R.B., N.H., D.Sh, N.I., D.Sa., R.E., B.T., Y.G., J.A.B., A.D., M.C.M., C.P., I.W., A.K., M.G.D., L.Y., and A.T.W. are currently, or were formerly, employed by and receive stock options from PathAI, Inc., a company that builds artificial intelligence tools for pathology. In addition, S.J. declares employment and stock from Meta, D.R. declares employment and stock from Microsoft, and A.P. declares employment and stock from Spring Discovery.

## Abstract

While alterations in nucleus size, shape, and color are ubiquitous in cancer, comprehensive quantification of nuclear morphology across a whole-slide histologic image remains a challenge. Here, we describe the development of a pan-tissue, deep learning-based digital pathology pipeline for exhaustive nucleus detection, segmentation, and classification and the utility of this pipeline for nuclear morphologic biomarker discovery. Manually-collected nucleus annotations were used to train an object detection and segmentation model for identifying nuclei, which was deployed to segment nuclei in H&E-stained slides from the BRCA, LUAD, and PRAD TCGA cohorts. Interpretable features describing the shape, size, color, and texture of each nucleus were extracted from segmented nuclei and compared to measurements of genomic instability, gene expression, and prognosis. The nuclear segmentation and classification model trained herein performed comparably to previously reported models. Features extracted from the model revealed differences sufficient to distinguish between BRCA, LUAD, and PRAD. Furthermore, cancer cell nuclear area was associated with increased aneuploidy score and homologous recombination deficiency. In BRCA, increased fibroblast nuclear area was indicative of poor progression-free and overall survival and was associated with gene expression signatures related to extracellular matrix remodeling and anti-tumor immunity. Thus, we developed a powerful pan-tissue approach for nucleus segmentation and featurization, enabling the construction of predictive models and the identification of features linking nuclear morphology with clinically-relevant prognostic biomarkers across multiple cancer types.

## INTRODUCTION

Histological assessment of tissue is central to the diagnosis and classification of malignancy, and critically informs patient management. Pathologists routinely report visible alterations in nuclear morphology. Altered nuclear features are ubiquitous in cancer, and changes in nuclear size, shape, coloration, texture, nucleoli, and nuclear-cytoplasmic ratio, as well as their intratumoral variance, are important features of histologic grade, which has prognostic relevance independent of disease stage^1,2^. The enumeration and morphologic features of mitoses also informs pathologist assessment of malignancy^3^. Furthermore, nuclear morphology can be important diagnostic features of certain cancers, such as nuclear clearing (“Ophan Annie Eyes”) and pseudoinclusions of papillary thyroid carcinoma^4^.

A complex interplay exists between nuclear morphology and the genetic, epigenetic, and transcriptomic milieu of cancer cells, reflecting the importance of the nucleus to the process of oncogenic transformation. Distorted nuclei can indicate dysregulated replication processes, aneuploidy, genomic instability, and genetic mutations that affect stability and function of the nuclear envelope^5^. Indeed, many cancers have altered expression of nuclear envelope components, resulting in nuclear rupture and micronuclei formation, further increasing genomic instability^5,6^. In addition, components of the nuclear envelope are known to bind to both chromatin and transcription factors, providing a spatial regulation to gene transcription and expression^5,7^. Therefore, the visual appearance of cancer cell nuclei has the potential to reveal key information about the biology of a tumor.

The quantitation of nuclear morphology has been a long sought-after goal^8^. Early studies used semi-quantitative approaches to enumerate features such as nuclear size and shape; these works revealed relationships between increased nuclear area and altered nuclear shape with poor prognosis and advanced disease in breast cancer and prostate cancer, respectively^9–11^. The use of computational approaches in pathology image analysis to identify and quantify nuclear changes has gained traction as modern computer vision methods have allowed for rapid, reproducible and cost-effective quantification of nuclear morphology. Using these methods, nuclear morphometric features have been shown to correlate with relevant clinical and pathological metrics, such as oligodendroglioma component in glioblastoma^12^, as well as stage^13^, disease aggressiveness^14^, recurrence^15–17^, and outcome^18^ in other cancer subtypes. In addition, increased nuclear size has been correlated with whole genome duplication^19,20^, and nuclear morphometric features have allowed for the prediction of relevant molecular information, such as ER status^21^ and Oncotype DX risk scores^22,23^ in breast cancer. Most recently, Nimgaonkar et al. described an AI-derived histologic signature, the main component of which was variance in nuclear morphology in cancer cells, that predicted response to gemcitabine in patients with pancreatic adenocarcinoma^24^.

Digitized whole slide images (WSIs) have enhanced the degree to which nuclear morphology can be studied in histological specimens^12,13^. However, the large size of WSIs - up to billions of pixels and containing thousands of nuclei - makes exhaustive manual annotation infeasible; thus studies have relied on manually-selected subregions of interest rather than entire slides^20,25^. Automated methods are, therefore, needed to fully quantify nuclear features in WSIs. We recently described a cell- and tissue-level computational pathology pipeline using WSIs for the automated computation of human interpretable features (HIFs), distinctive features with tangible methods for validation^26^. This pipeline allows the use of HIFs to predict treatment-relevant molecular phenotypes and allows for integration with current pathological methods. Given that morphological analysis of histology features is central to pathology workflows, we sought to extend this work to identify nuclear human interpretable features (nuHIFs) in multiple cancer types.

In this paper, we present a multi-tissue model for the exhaustive detection, segmentation, and classification of nuclei from entire hematoxylin and eosin (H&E)- stained WSIs, allowing for the exhaustive analysis of slide-level descriptors of nuclear size, shape, texture, and staining intensity. Furthermore, we demonstrate that these nuHIFs are predictive of clinically relevant information in multiple cancer types.

## MATERIALS AND METHODS

### Study Design

Manually collected annotations were used to train and validate an object detection and segmentation model to detect and segment nuclei from H&E stained tissue slides. Training data variation and number of annotations were selected to exceed previously used standards in the field^27^ and exhibit wide variation in tissue morphology as subjectively assessed by study pathologists (MGD and LY). This model was deployed on whole-slide H&E images from The Cancer Genome Atlas (TCGA) to extract features from each nucleus in each slide, and the resulting features were used to analyze the relationship between nuclear morphology and underlying molecular markers of cancer, and patient outcomes. Inclusion of TCGA slides was performed in accordance with literature norms (e.g. as by Saltz et al.^28^): TCGA slides were selected to be the DX1 (primary diagnostic) slide for each case in TCGA and no outlier exclusion was performed, to conservatively reflect real-world conditions where same-case replicates may not be available. Where multiple hypotheses were tested, all reported statistics were corrected to control false discovery rate as described below.

### Dataset Description and Annotation Collection

Over 29,000 manual annotations of cell nuclei were collected from H&E images from 21 tumor types at 40x and 20x magnification from TCGA^29^, as well as a proprietary set of H&E-stained tissue biopsies of skin, liver non-alcoholic steatohepatitis, colon inflammatory bowel disease, and kidney lupus. Board-certified pathologists (MGD and LY) selected 1000 x 1000 pixel patches that were exemplary of varied tissue and nuclear morphology from the training slides and trained collaborators to perform exhaustive manual annotation of nuclei in the patches. Annotations were checked for quality, adjusted, and confirmed by MGD and LY. This process resulted in 67 WSI patches exhaustively annotated for nuclei. These patches were split into training, validation, and held-out test data sets to ensure distribution of tissue types (Table 1). Following model training and initial testing, an additional two data sources were used to collect additional annotations for model testing. H&E-stained slides of ulcerative colitis were obtained from BioIVT (Westbury, NY), and H&E-stained breast cancer slides were generously provided by Cleveland Clinic Foundation (Cleveland, OH). An additional 14 512 x 512 pixel patches were identified from these data sources (seven patches from each source), and an additional 2,647 manual, exhaustive nucleus annotations were collected for model evaluation (Table 2).

**Table 1.** Samples used for training and evaluating the segmentation model.

| **Source** | **#**<br>**Frames** | **Magnificati**<br>**on** | **Microns**<br>**Per Pixel**<br>**(MPP)** | **Indication** | **Dataset** |
| --- | --- | --- | --- | --- | --- |
| TCGA-A2-A3XS-01Z-00-DX1 | 3 | 40X | 0.2456 | Breast, Infiltrating Ductal<br>Carcinoma | Train |
| TCGA-XF-AAMJ-01Z-00-DX1 | 2 | 40X | 0.2527 | Bladder, Urothelial<br>Carcinoma | Train |
| TCGA-UW-A72Q-01Z-00-DX1 | 3 | 40X | 0.2472 | Liver, Hepatocellular<br>Carcinoma | Train |
| TCGA-G3-A7M6-01Z-00-DX1 | 2 | 40X | 0.2527 | Liver, Hepatocellular<br>Carcinoma | Train |
| TCGA-<br>KM- | 1 | 40X | 0.2525 | Kidney, Renal Cell<br>Carcinoma | Train |
| A7QL-01Z-00-DX1 |  |  |  |  |  |
| TCGA-CV-A6JD-01Z-00-DX1 | 1 | 40X | 0.2527 | Head and Neck Squamous Cell Carcinoma | Train |
| TCGA-02-0009-01Z-00-DX1 | 1 | 20X | 0.5015 | Brain, Glioblastoma | Train |
| TCGA-LB-A8F3-01Z-00-DX1 | 3 | 40X | 0.2527 | Pancreas, Pancreatic Adenocarcinoma | Train |
| TCGA-36-2547-01A-01-TS1 | 1 | 20X | 0.5015 | Ovary, Ovarian Serous Cystadenocarcinoma | Train |
| TCGA-W5-AA30-01Z-00-DX1 | 3 | 40X | 0.2529 | Bile Duct, Cholangiocarcinoma | Train |
| TCGA-86-7701-01Z-00-DX1 | 2 | 40X | 0.252 | Lung, Lung Adenocarcinoma | Train |
| TCGA-B0-4823-01Z-00-DX1 | 3 | 40X | 0.252 | Kidney, Renal Clear Cell Carcinoma | Train |
| TCGA-02-0009-01Z-00-DX1 | 1 | 20X | 0.5015 | Brain, Glioblastoma multiforme | Train |
| TCGA-2G-AAKO-05Z-00-DX1 | 1 | 40X | 0.2277 | Testicle, Acute Myeloid Leukemia | Train |
| PathAI | 3 | 40X | 0.2511 | Liver, Hepatitis B | Train |
| PathAI | 3 | 40X | 0.2522 | Liver, Non-alcoholic steatohepatitis | Train |
| PathAI | 3 | 40X | 0.2522 | Kidney, Lupus | Train |
| PathAI | 1 | 40X | 0.2522 | Colon, Inflammatory Bowel Disease | Train |
| PathAI | 2 | 20X | 0.5023 | Small Intestine, Carcinoid | Train |
| PathAI | 1 | 40X | 0.2522 | Prostate, Prostate Adenocarcinoma | Train |
| TCGA-EJ-5495-01Z-00-DX1 | 2 | 40X | 0.252 | Prostate, Prostate Adenocarcinoma | Validation |
| TCGA-IG-A3YA-01Z-00-DX1 | 2 | 40X | 0.2465 | Esophagus, Esophageal Carcinoma | Validation |
| TCGA-GS-A9U3-01Z-00-DX1 | 2 | 40X | 0.2525 | Lymph, Diffuse Large B Cell Lymphoma | Validation |
| TCGA-LL-A5YM-01Z-00-DX1 | 2 | 40X | 0.2456 | Breast, Invasive Ductal Carcinoma | Validation |
| TCGA-P4-AAVO-01Z-00- | 3 | 40X | 0.2526 | Kidney, Renal Papillary Cell Carcinoma | Validation |
| DX1 |  |  |  |  |  |
| PathAI | 2 | 40X | 0.2522 | Liver, Non-alcoholic steatohepatitis | Validati<br>on |
| PathAI | 1 | 20X | 0.5023 | Colon, Human Papillomavirus | Validati<br>on |
| PathAI | 2 | 40X | 0.2511 | Brain, Glioma | Validati<br>on |
| PathAI | 2 | 40X | 0.2522 | Skin, Normal | Validati<br>on |
| TCGA-VQ-A8DV-01Z-00-DX1 | 1 | 40X | 0.2525 | Stomach, Stomach Adenocarcinoma | Test |
| TCGA-CV-6934-01Z-00-DX1 | 2 | 40X | 0.2525 | Head and Neck Squamous Cell Carcinoma | Test |
| TCGA-MQ-A6BR-01Z-00-DX1 | 1 | 40X | 0.2465 | Lung, Mesothelioma | Test |
| TCGA-<br>OR- | 1 | 40X | 0.2527 | Adrenal, Adrenal Cortical Carcinoma | Test |
| A5J8-01Z-00-DX1 |  |  |  |  |  |
| PathAI | 1 | 40X | 0.2522 | Colon, Inflammatory Bowel Disease | Test |
| PathAI | 1 | 20X | 0.5023 | Breast, Invasive Ductal Carcinoma | Test |
| PathAI | 2 | 40X | 0.2522 | Prostate, Prostate Adenocarcinoma | Test |

**Table 2:** Samples used for out-of-distribution (OOD) evaluation of model performance.

| Slide Identifier | # Frames | Magnification | Microns Per Pixel (MPP) | Indication | Dataset |
| --- | --- | --- | --- | --- | --- |
| 581805 | 1 | 40X | 0.2518 | Colon, Ulcerative Colitis | OOD-Test 1 |
| 581720 | 1 | 40X | 0.2518 | Colon, Ulcerative Colitis | OOD-Test-1 |
| 581810 | 1 | 40X | 0.2518 | Colon, Ulcerative Colitis | OOD-Test-1 |
| 581875 | 1 | 40X | 0.2518 | Colon, Ulcerative Colitis | OOD-Test-1 |
| 591979 | 1 | 40X | 0.2518 | Colon, Ulcerative Colitis | OOD-Test-1 |
| 581916 | 1 | 40X | 0.2518 | Colon, Ulcerative Colitis | OOD-Test-1 |
| 581956 | 1 | 40X | 0.2518 | Colon, Ulcerative Colitis | OOD-Test-1 |
| 591275 | 1 | 40X | 0.2521 | Breast, Breast Cancer | OOD-Test-2 |
| 591351 | 1 | 40X | 0.262385 | Breast, Breast Cancer | OOD-Test-2 |
| 591347 | 1 | 40X | 0.262385 | Breast, Breast Cancer | OOD-Test-2 |
| 592506 | 1 | 40X | 0.2521 | Breast, Breast Cancer | OOD-Test-2 |
| 592559 | 1 | 40X | 0.262125 | Breast, Breast Cancer | OOD-Test-2 |
| 597204 | 1 | 40X | 0.262125 | Breast, Breast Cancer | OOD-Test-2 |
| 597271 | 1 | 40X | 0.262125 | Breast, Breast Cancer | OOD-Test-2 |

### Nuclear Segmentation Model Architecture

A Mask-RCNN-style architecture was selected for nuclear segmentation. A ResNet50 backbone pretrained on the ImageNet dataset was used to produce the feature pyramid network. The first two of five modules that comprise ResNet50 were frozen during training to preserve the pretrained weights of early layers. Model development was performed using the PyTorch library^30^.

### Nuclear Segmentation Model Training

The manually-collected annotations were used to train the model for detecting and segmenting cellular nuclei (Figure 1A). During training, the annotated patches were augmented by crops, flips, rotations, and affine deformations.

**Figure 1.**
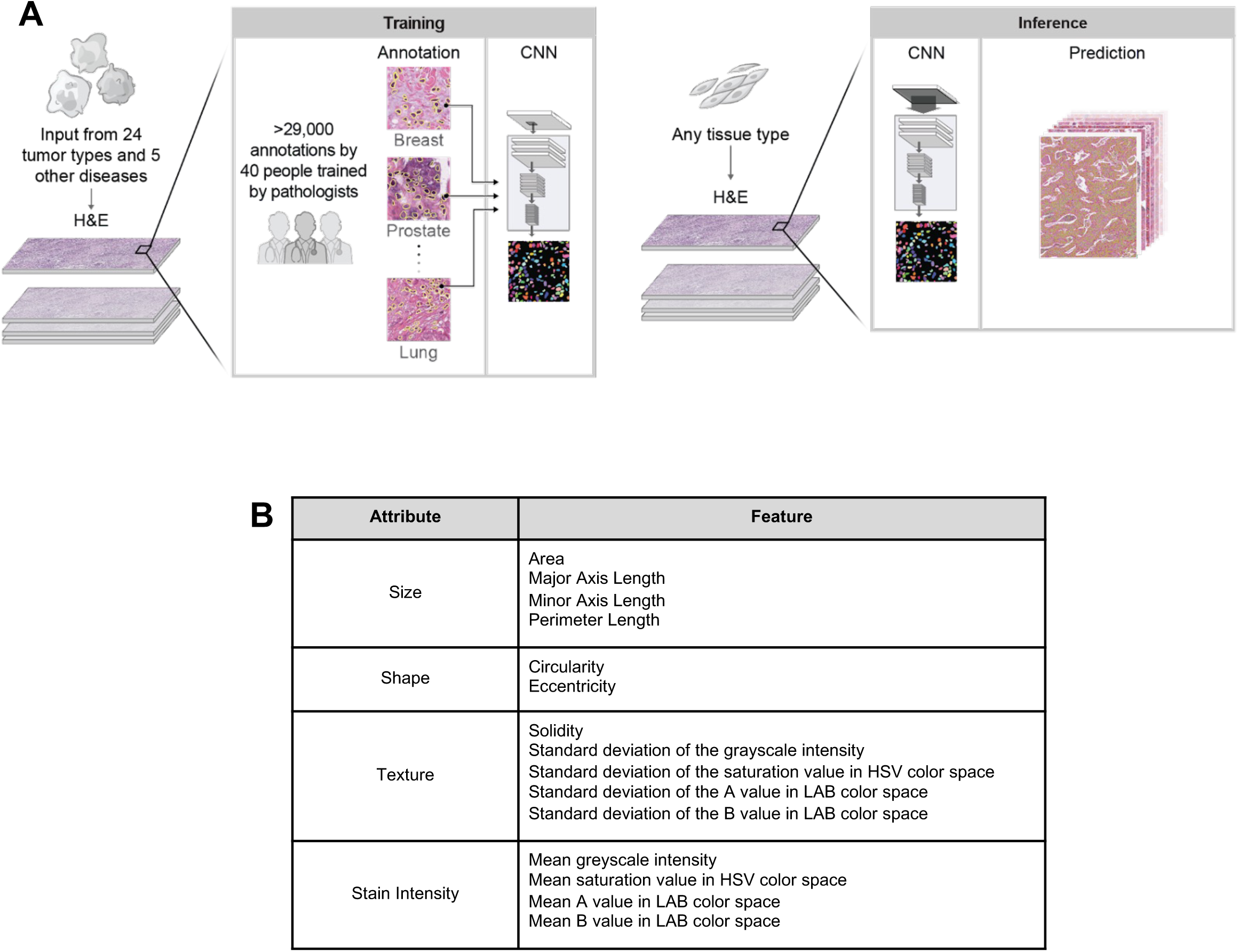
Machine learning model annotation collection, training, and application. **(A)** Model workflow. Briefly, pathologists trained expert annotators to perform exhaustive annotations of nuclei on H&E slide patches from diverse tissue sources. These were used to train a pan-H&E nucleus detection and segmentation model, which was subsequently evaluated on held-out patches and applied to exhaustively segment nuclei in three WSI datasets. **(B)** Features extracted from the model. Mean and standard deviation values were calculated for these features at the whole-slide level for cancer cells, lymphocytes, and fibroblasts.

### Cell Classification

Following nuclear segmentation, the cell class of each nucleus was assigned using PathExplore^TM^ (PathAI, Boston, MA)^31^ models specific to breast cancer (BRCA), lung adenocarcinoma (LUAD), and prostate adenocarcinoma (PRAD); PathExplore is for research use only and is not for use in diagnostic procedures. Cancer epithelial cells, fibroblasts, macrophages, lymphocytes and plasma cells were predicted for all three cancer types, while additional cell classes were predicted for LUAD (granulocytes and normal cells) and PRAD (smooth muscle cells, endothelial cells, and normal epithelial cells). Model performance for the prediction of cell types was assessed by comparing model predictions to pathologist annotations in nested pairwise fashion^32^. Model performance metrics for BRCA, LUAD, and PRAD are shown in Supplementary Figures S1-S3, respectively, and Supplementary Tables S1 and S2. Example prediction results are shown in Figure 3. The five pan-indication cell classes (cancer epithelial cells, fibroblasts, macrophages, lymphocytes, and plasma cells) were used for analyses assessing the biological implications of nuclear feature differences in BRCA, LUAD, and PRAD.

### Deployment Dataset and Feature Extraction

The nuclear segmentation model was deployed on publicly available images of H&E slides from the BRCA (N=886), PRAD (N=392), and LUAD (N=426) TCGA cohorts; a summary of clinicopathologic features of each cohort is shown in Table 3. Model performance was qualitatively assessed by board-certified pathologists and determined to be consistent with performance on the held-out test dataset. The features computed for each individual nucleus were: area, circularity, eccentricity, major and minor axis length, perimeter, solidity, and the mean and standard deviation of pixel grayscale intensity, pixel saturation, and pixel A and B channels in LAB colorspace. The mean and standard deviation of each feature from each nucleus class on the slides were used to summarize the nuclear morphology on each slide. This yielded 30 slide-level nuHIFs for each cell type, e.g. the mean area of cancer nuclei, the standard deviation of fibroblast nuclear eccentricity, or the mean pixel grayscale intensity of lymphocyte nuclei. Attributes and features described by nuHIFs are included in Figure 1B. Thus, the total number of features summarizing the morphology on each slide was 30 times the number of cell classes.

**Table 3.** Characteristics of patients in TCGA cohorts.

|  | TCGA BRCA<br>N=886 | TCGA LUAD<br>N=426 | TCGA PRAD<br>N=392 |
| --- | --- | --- | --- |
| Age at initial pathologic diagnosis (median, range) | 58 (26-90) | 66 (33-88) | 61 (41-77) |
| Sex (male), n (%) | 10 (1.1%) | 191 (44.8%) | 392 (100%) |
| American Joint Committee on Cancer (AJCC) Tumor Stage (n) | Stage I: 77<br>Stage IA: 76<br>Stage IB: 5<br>Stage II: 6<br>Stage IIA: 286<br>Stage IIB: 207<br>Stage III: 2<br>Stage IIIA: 128<br>Stage IIIB: 18<br>Stage IIIC: 51<br>Stage IV: 13<br>Unknown: 17 | Stage I: 5<br>Stage IA: 122<br>Stage IB: 108<br>Stage II: 1<br>Stage IIA: 47<br>Stage IIB: 58<br>Stage IIIA: 50<br>Stage IIIB: 5<br>Stage IV: 22<br>Unknown: 8 | Unknown: 392 (100%) |
| Whole genome doublings (WGD) | 0: 435, 56.8%<br>1: 298, 38.5%<br>2: 37, 4.8%<br>Unknown: 116 | 0: 168, 41.9%<br>1: 193, 48.1%<br>2: 40, 10.0%<br>Unknown: 25 | 0: 337, 92.1%<br>1: 29, 7.9%<br>Unknown: 26 |
| Progression-free survival (PFS, median) | 122.3 mo | 29.3 mo | 116.7 mo |
| Overall survival (OS, median) | 131.4 mo | 48.5 mo | Not Reached |

### Exploring Cancer Type and Nuclear Morphology

To compare the nuHIFs quantifying cancer cell, fibroblast, and lymphocyte morphology, uniform manifold approximation and projection (UMAP) analysis was performed. Nuclear HIFs were z-scored across all cancer types for standardization. UMAP was parameterized with 100 neighbors, an embedding dimension of 2, and the Euclidean distance metric. Features characteristic of each cancer type were evaluated by averaging each feature across the samples of each cancer type and z-scoring for visualization; hierarchical clustering (using Euclidean distance with average linkage) identified features that varied across cancer types.

### Classifying Cancer Type from Nuclear Morphology

Random forest (RF) binary classification models were trained and applied to each cell-type-specific nuHIF set to differentiate between pairs of cancer types. RF classification models were trained using 5-fold stratified cross-validation with balanced class weighting. The performance of each model was assessed using the area under the receiver operating characteristic curve (AUROC) on each held-out validation split. The mean AUROC on the held-out validation splits is reported. RF model training was performed in scikit-learn with default hyperparameters (100 trees)^33^.

### Classifying Breast Cancer Subtype from Nuclear Morphology

Characteristics of breast cancer molecular subtypes (luminal A, N=457; luminal B, N=159; HER-2, N=66; normal-like, N=31; basal-like, N=161) were obtained from a prior study by Berger and colleagues^34^. Random forest (RF) binary classification models were trained and applied to each cell-type-specific nuHIF set to differentiate between subtypes in a one-vs.-all manner. RF classification models and cross-validation schemes were identical to cancer type classification.

### Statistical Analysis

Spearman (rank-based) correlation was used to find the association between variation in cancer nuclear morphology and metrics of genomic instability. Variation in size was captured by the nuHIF “standard deviation of cancer cell nuclear area” for each slide. For metrics of genomic instability, previously published metrics were selected: aneuploidy score^35^ and homologous recombination deficiency (HRD) score^36^. RF binary classification models were trained in scikit-learn with default hyperparameters^33^ using 5-fold stratified cross-validation with balanced class weighting, and applied to the cancer nuHIF set from each cancer type to predict binarized whole-genome doubling (WGD; 1-2 doublings = 1; no doublings = 0). The performance of each model was evaluated using AUROC on each held-out validation split, and the mean AUROC is reported. The mean RF Gini importance (also called the mean decrease in impurity) of the top five features for each cancer type across the five splits are reported. Cox proportional hazard models were utilized to explore the relationship between BRCA fibroblast nuHIFs and overall and progression-free survival (OS and PFS, respectively). Ordinal tumor stage (1-4) and patient age were included as clinical covariates; 17 subjects missing tumor stage and the one missing survival data were excluded. Robust z-scoring (i.e. using the median and scaled interquartile range) of each nuHIF before modeling was performed for simple interpretation of the hazard ratios (HRs). The p-values associated with each nuHIF were corrected for false discovery rate (FDR) by the Benjamini-Hochberg procedure. Survival analyses were performed using the Lifelines library^37^. Gene expression data was acquired from the Genomic Data Commons (GDC)- processed TCGA BRCA cohort (release 18.0) from the UCSC Xena data portal^38^. Gene expression samples were paired to case-matched slides in our dataset, yielding 868 expression-nuHIF pairs. Spearman (rank-based) correlation was used to quantify the association between bulk RNAseq expression and the mean fibroblast nucleus area nuHIF for each gene and corrected for FDR via Benjamini-Hochberg procedure. Genes with corrected p < 0.05 and Spearman correlation greater than 0.15 or less than -0.15 were selected to comprise the significant positively and negatively associated gene sets, respectively, for gene set enrichment analysis (GSEA). GSEA^39^ was performed using the Molecular Signatures Database (MSigDB)^40^ and the REACTOME pathway database^41^, and the ten most significant pathway overlaps, with FDR-corrected p < 0.05, are reported.

## RESULTS

### Model development, performance, and nuclear feature extraction

We collected annotations and trained a machine learning (ML) model to detect and segment nuclei in H&E-stained WSIs as described in the Methods and shown in Figure 1. The model is not limited to sampling regions of interest from tissue samples, but rather can be utilized to exhaustively annotate WSIs. Application of the model to our held-out test data, including held-out tissue and disease types, demonstrated performance (mean Dice score=0.818, aggregated Jaccard index (AJI) =0.619) comparable to models reported previously in the literature^27,42^. Importantly, model speed was adequate to apply to multi-gigabyte WSIs at full resolution (approximately 0.25 μm/pixel; roughly 30 minutes per slide). Examples of model performance in mesothelioma, head and neck squamous cell carcinoma, and stomach adenocarcinoma are shown in Figure 2.

**Figure 2.**
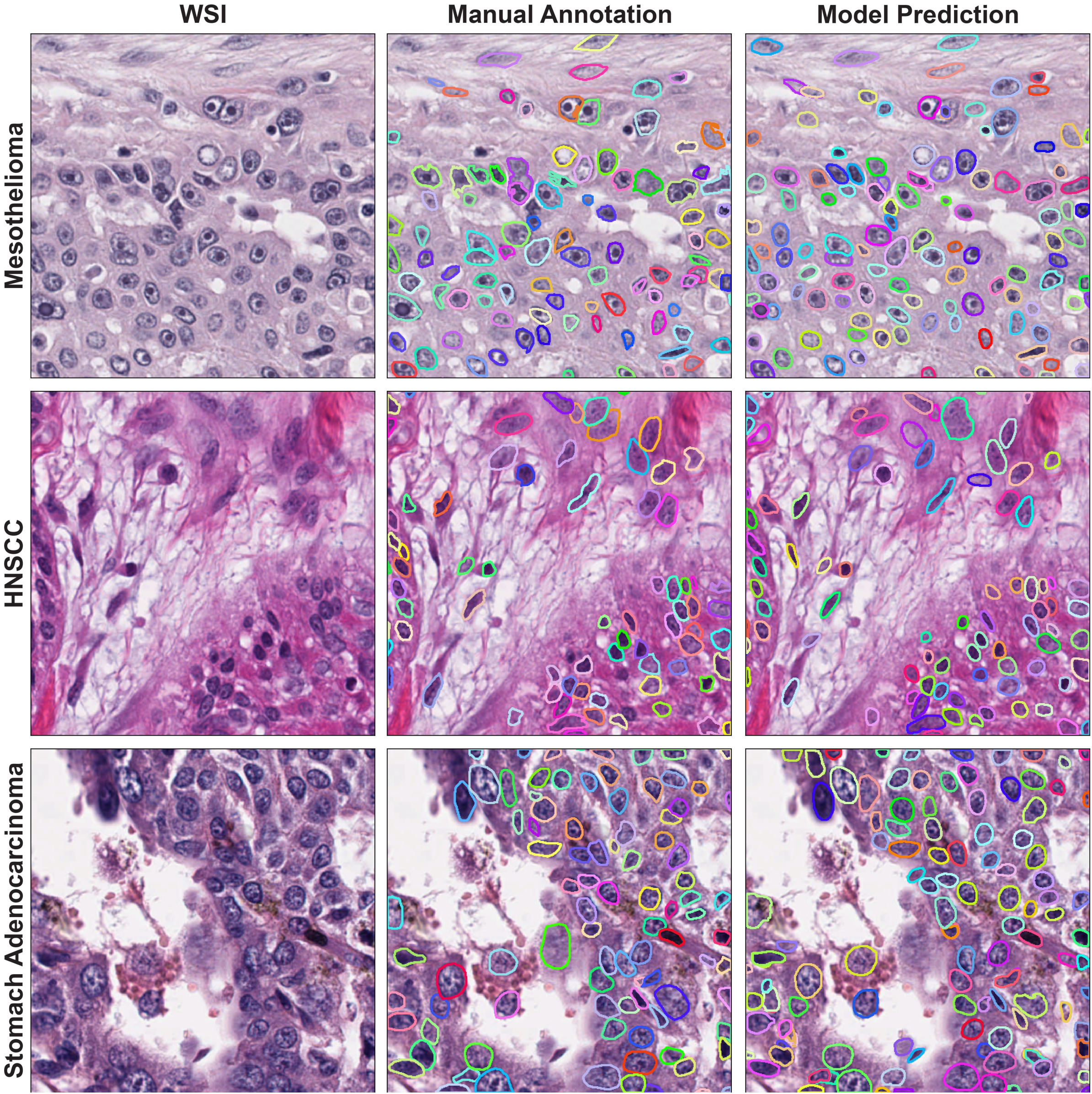
Example of model performance. Representative WSI patches from mesothelioma, head and neck squamous cell carcinoma (HNSCC), and stomach adenocarcinoma stained with H&E are shown in the left-most panel. Ground truth nuclei identified manually and nuclei predicted by the model are shown in the middle and right-most panels, respectively. Each color represents a nucleus instance.

We selected clinical samples from two additional datasets, designated OOD-Test-1 and OOD-Test-2, characterized in Table 2. We collected additional annotations on these datasets and characterized model performance. We found performance numerically comparable or superior to our initial held-out test data despite different sample origin, and one dataset containing non-cancer tissue samples (OOD-Test-1 mean Dice = 0.818, AJI = 0.628; OOD-Test-2 mean Dice = 0.826, AJI = 0.649).

Having evaluated our model’s performance, we deployed the resulting model on primary diagnostic (DX1) H&E slides from the breast cancer (BRCA; N=892), prostate adenocarcinoma (PRAD; N=392), and lung adenocarcinoma (LUAD; N=426) TCGA cohorts (Figure 3); model performance was visually assessed to be consistent with test data. The distribution of pixel sizes (microns per pixel; MPP) of these three cohorts are shown in Supplementary Figure 4. The median MPPs were 0.248, 0.252, and 0.252 for BRCA, LUAD, and PRAD datasets respectively. We extracted interpretable features describing the shape, size, staining intensity, and texture of every nucleus on each WSI (Supplementary Table S3). We performed further analysis on nuHIFs specific to cancer cells, fibroblasts, and lymphocytes, as these three cell classes are common across all cancer types and have been implicated in clinical outcomes.

**Figure 3.**
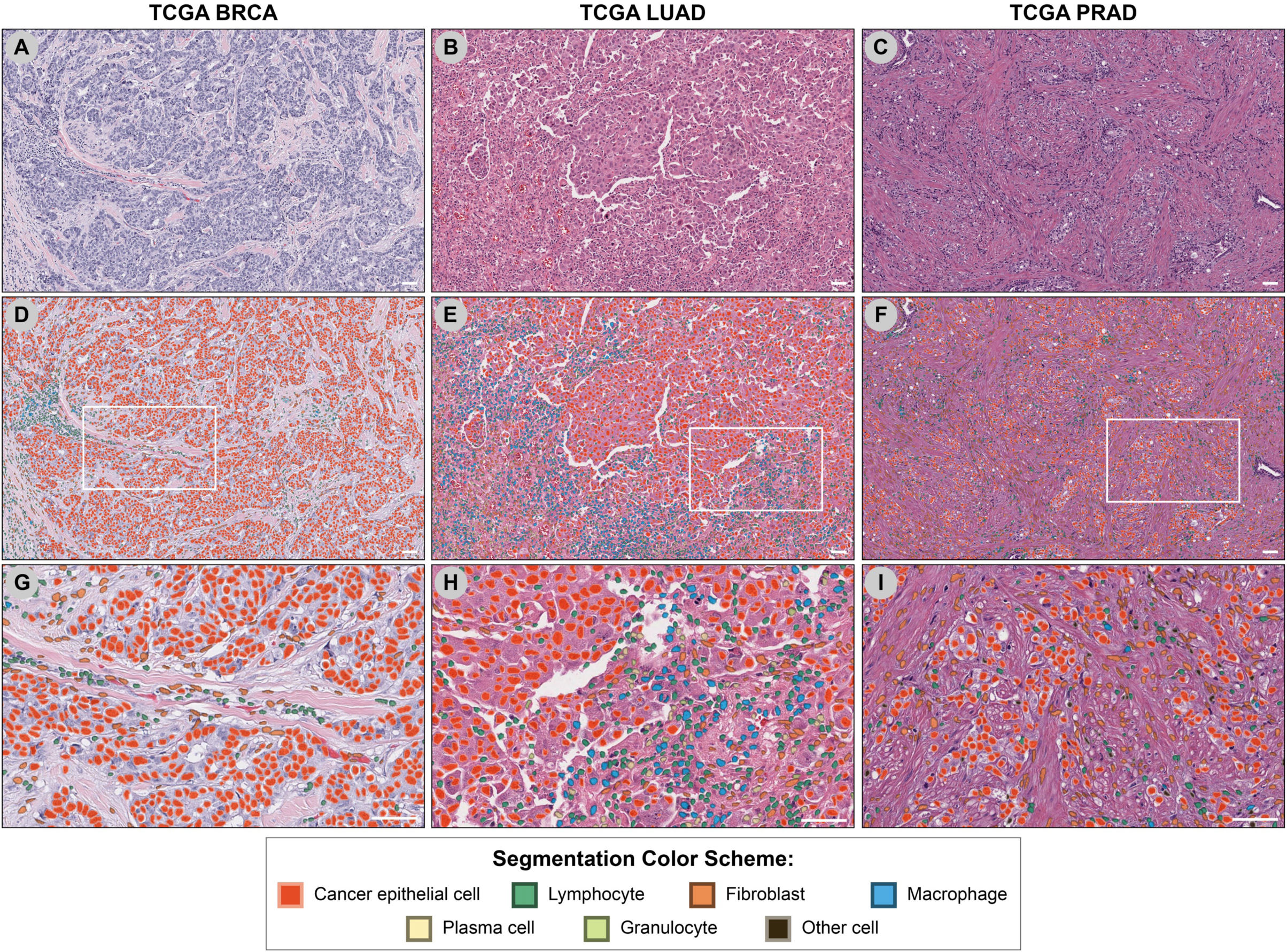
Nuclear segmentation and cell type identification in multiple cancer types. Representative H&E images of **(A)** breast cancer (TCGA BRCA), **(B)** lung adenocarcinoma (TCGA LUAD), and **(C)** prostate adenocarcinoma (TCGA PRAD) are shown at 40X magnification. **(D-F)** Nuclear segmentation and cell type identification masks are overlaid onto H&E images shown in panels A-C. **(H-I)** High-magnification images of BRCA **(G)**, LUAD **(H)**, and PRAD **(I)**. Magnified regions are indicated by dashed boxes in panels D-F. Scale bars indicate a distance of 50 μm.

### nuHIFs show within- and between-cancer type variation

To assess whether nuHIFs differ between cancer types, we performed UMAP to compare the nuHIFs from cancer cells (Figure 4A), fibroblasts (Figure 4B), and lymphocytes (Figure 4C) in BRCA, LUAD, and PRAD datasets. We observed notable inter- and intra-dataset variation in nuHIFs. For cancer cells, nuclear morphology was distinct between PRAD and LUAD datasets, while BRCA dataset cancer cells showed nuclear features similar to both PRAD and LUAD (Figure 4A). Unsupervised hierarchical clustering of z-scored features revealed specific nuHIFs differentially exhibited in these three cancer subtypes (Figure 4B). For example, features associated with nuclear size were higher in LUAD cancer nuclei relative to PRAD. Assessment of the distribution of three size-related features in BRCA, LUAD, and PRAD confirmed these observations – cancer nuclei in PRAD were smaller in area and major axis length than cancer nuclei in BRCA and LUAD (Supplementary Figure 5A), while fibroblast area and major axis length is larger in BRCA than in LUAD and PRAD (Supplementary Figure 5B). Minute variation in minor axis length was observed between the three cancer types for cancer cells and fibroblasts (Supplementary Figure 5C). In contrast, lymphocyte nucleus size parameters did not appear to differ between cancer types (Supplementary Figure 5). In addition to size features, features associated with nuclear staining were observed to differ between cancer types. In particular, notable differences in features relating to nucleus stain intensity, color, and shape between PRAD and LUAD were observed (Figure 4). The clearest distinction between cancer subtypes was discerned through nuHIFs of fibroblasts in BRCA, LUAD, and PRAD (Figure 4B,D). Unsupervised hierarchical clustering revealed specific features enriched in fibroblasts from these cancer subtypes. Interestingly, lymphocyte nuHIFs also differed between cancer types.

**Figure 4.**
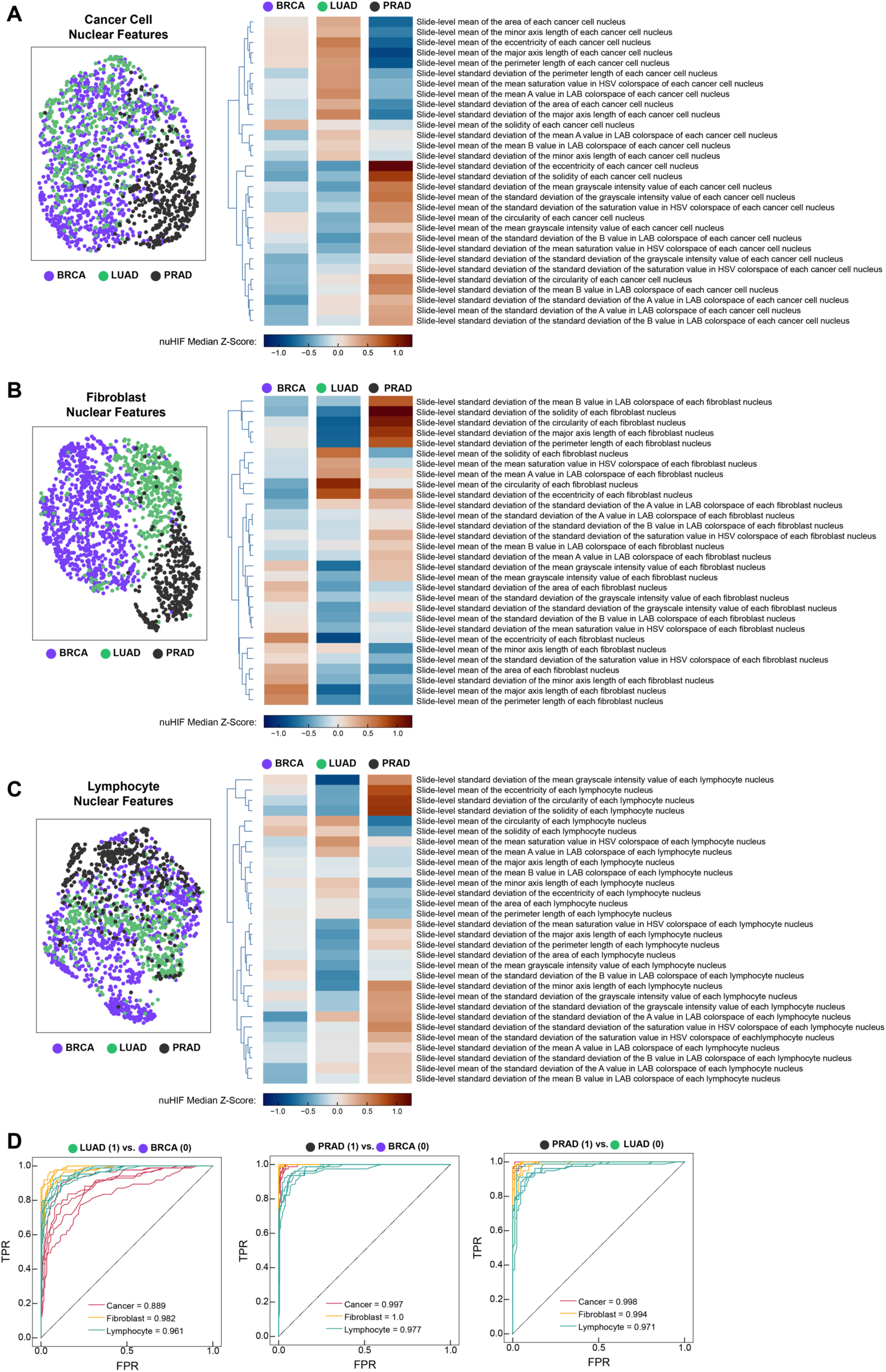
nuHIFs show variation within and between cancer types. Uniform manifold approximation and projection (UMAP) visualization of BRCA, LUAD, and PRAD defined by nuclear human interpretable feature (HIF) for **(A)** cancer cells, **(B)** fibroblasts, and **(C)** tumor-infiltrating lymphocytes. Clustered heatmaps of median Z-scores for all 30 nuHIFs are shown for each cell type. **(D)** Receiver operating characteristic (ROC) curves for binary classification between paired cancer types using nuHIFs from each of cancer, fibroblast, or lymphocyte nuclei. ROCs are shown for the five held-out validation splits and mean area under ROC (AUROC) is shown for each classification problem. In particular, fibroblast and lymphocyte nuclear features are highly able to differentiate between cancer types. Mean AUROC is shown for each class of nuclear HIF.

To ensure that the observed differences in nuclear features between cancer types were not biased by scanned image pixel size, we measured the Pearson correlation between nuclear size (using major axis length as a representative feature) and MPP for each cell type within BRCA, LUAD, and PRAD datasets individually, to remove the potential effect of possible between-cancer-type variation in nuclear size. The within-cancer-type variation in mean nuclear major axis length between slides at the same MPP is large for cancer cells (Supplementary Figure 6A), fibroblasts (Supplementary Figure 6B), and lymphocytes (Supplementary Figure 6C). In addition, the magnitude of the within-cancer-type Pearson correlations is low, although some rise to the level of significance, perhaps due to the high power of the large dataset. The within-cancer-type Pearson correlations also show an inconsistent sign, ranging from 0.206 to -0.151. Generally, these results suggest that there is an inconsistent directional effect of MPP on nuclear size, and other factors are likely driving the observed differences.

Because of the apparent association between nuclear morphology and cancer type, we hypothesized that nuHIF-quantified nuclear morphology could be a distinguishing feature of cancer types. To test this, we constructed a simple random forest binary classification model for differentiating between each pair of cancer types (BRCA, PRAD, LUAD) using cancer, fibroblast, or lymphocyte nuclear HIFs. We performed five-fold cross validation to estimate the extent to which cancer types may be differentiated by nuclear morphology. We found consistently strong performance for differentiating between cancer types using nuclear morphology (Figure 4D). Although lymphocyte nuclear morphology was less distinct between cancer types when visualized with UMAP, supervised analysis indicated that lymphocyte morphology differed between cancer types.

### Cancer nuclear morphology is associated with metrics of genomic instability in multiple cancer types

Cancer nuclear atypia is used clinically as a marker of malignancy. We therefore hypothesized that underlying levels of genomic instability may partially explain the observed heterogeneity in cancer nuclear morphology within cancer subtypes, as well as between cancer types with known differences in malignancy. We tested this hypothesis by assessing the relationship between cancer nuclear morphology and genomic instability in LUAD, BRCA, and PRAD cohorts using aneuploidy score and homologous recombination deficiency (HRD) score as metrics of genomic instability. Indeed, using the standard deviation of cancer nuclear area as a metric of nuclear atypia, we detected significant correlation between this nuHIF and both aneuploidy score (Figure 5A) and HRD score (Figure 5B). When assessed in a pan-cancer manner, the overall correlation increased, and the pattern observed in cancer nuHIF UMAP analysis persisted: PRAD displayed a lower level of genomic instability across both metrics compared to LUAD, while BRCA showed a wide range of genomic instability, with similarities to both PRAD and LUAD. These results confirm that cancer nuclear morphology, especially variability in nuclear size, is associated with the level of genomic instability.

**Figure 5.**
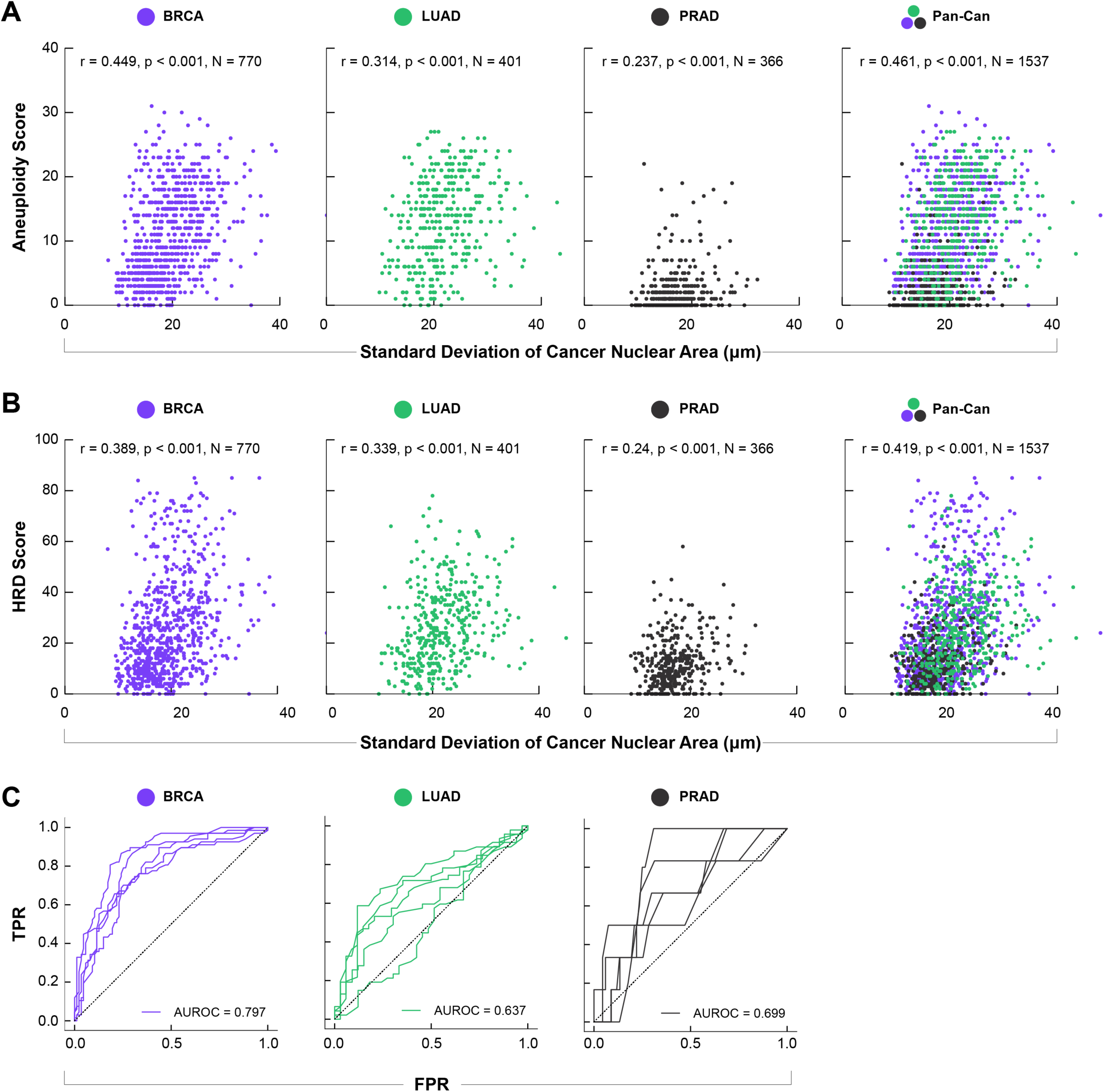
Variation in cancer nuclear size correlates with metrics of genomic instability. Standard deviation of cancer cell nuclear area was compared to **(A)** aneuploidy score and **(B)** homologous recombination deficiency (HRD) score for BRCA, LUAD, and PRAD. **(C)** Receiver operating characteristic (ROC) curves for prediction of whole-genome doublings in BRCA, LUAD, and PRAD. ROCs are shown for the five held-out validation splits; mean AUROC is shown for each cancer type.

Because aneuploidy score was correlated to variation in cancer nuclear area, we posited that cancer nuclear morphology was predictive of whole genome doubling. To address this hypothesis, we trained random forest models for predicting binarized whole-genome doubling using cancer nuHIFs from each of the BRCA, LUAD, and PRAD cancer types. We found that cancer nuclear morphology was predictive of WGD for each cancer type, with strongest predictive power in BRCA, and more variation in performance expected for PRAD, where WGD occurs less frequently (Figure 5C). The mean RF importance across the five splits is reported for the top five features for each cancer type in Supplementary Table S4. Briefly, variation in cancer nuclear dimensions were most important for predicting WGD in BRCA, mean cancer nuclear dimensions were most important for predicting WGD in LUAD, and a mix of color and shape features were found to be most important for PRAD.

### Nuclear morphology enables prediction of breast cancer molecular subtype

We hypothesized that nuclear morphology would differ in subtle but meaningful ways between molecular subtypes of breast cancer, and that these differences might enable classification of molecular subtypes of breast cancer from H&E images. To test this theory, we trained nuHIF-based classification models for predicting breast cancer subtype in a one-vs.-all manner (Figure 6). Briefly, we found that cell-type-specific nuclear morphology enabled classification of some but not all breast cancer molecular subtypes. Interestingly, the ability to predict subtype varied by subtype as well as by cell type being used to make the inference. Cancer nuclear morphology (Figure 6A) or lymphocyte nuclear morphology (Figure 6C) enabled moderate prediction (AUROC > 0.7) of luminal A and basal-like breast cancer subtypes. Cancer nuclear morphology but not lymphocyte or fibroblast nuclear morphology enabled moderate prediction of HER-2 breast cancer subtype. Interestingly, fibroblast nuclear morphology alone was a poor predictor of molecular subtype (Figure 6B). When aggregating cell types (Figure 6D), luminal A and basal-like prediction AUROC increased further to ≥ 0.80. These results suggest that altered nuclear morphology is a possible histological presentation of breast cancer molecular subtypes.

**Figure 6.**
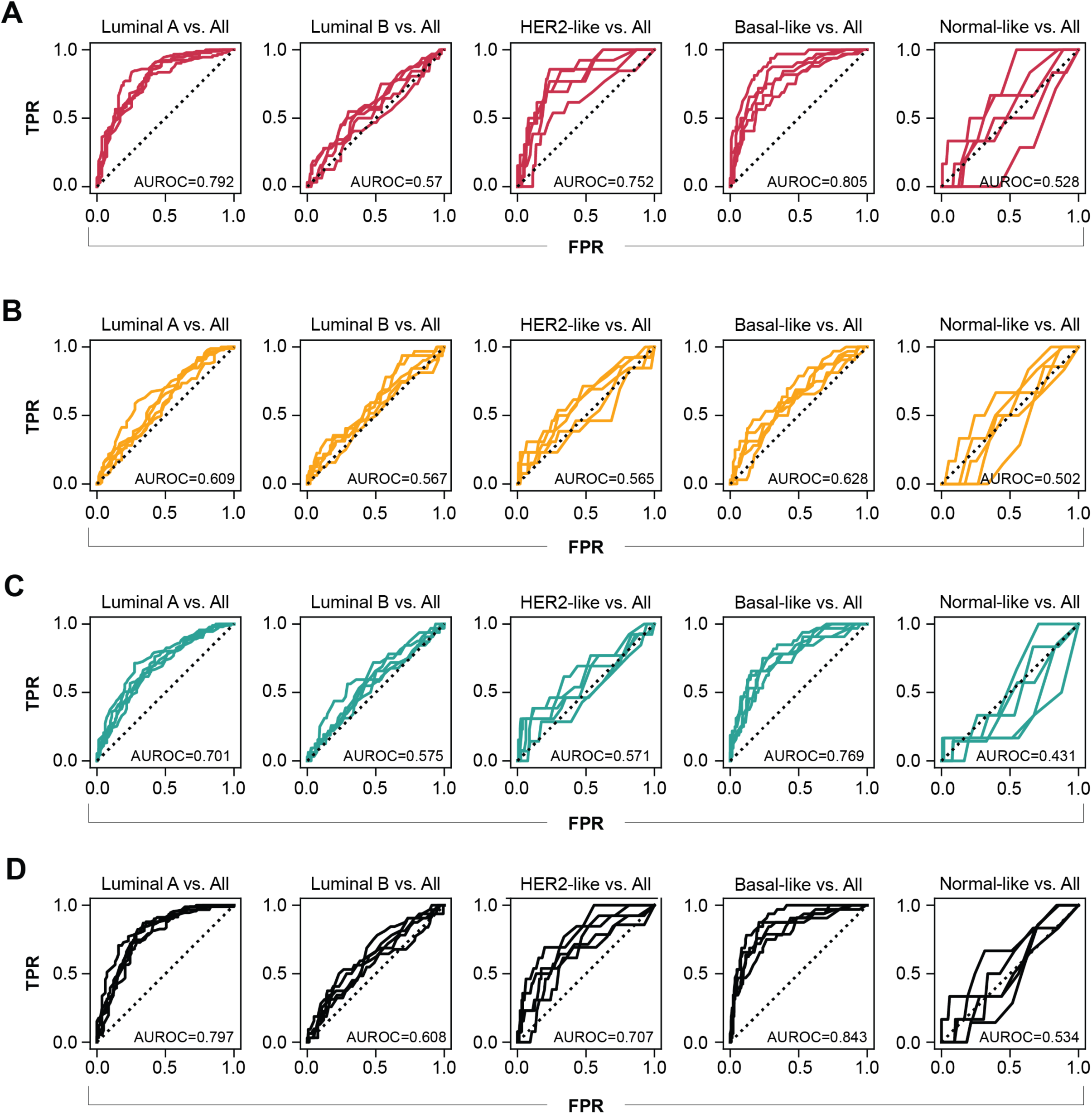
Cell-type-specific nuclear morphology enables classification of breast cancer molecular subtypes. One-vs-all binary classification of breast cancer molecular subtypes (luminal A, luminal B, HER2-like, basal-like, and normal-like)^34^ was performed using random forest classification on nuHIFs derived from A) cancer cells, B) fibroblasts, C) lymphocytes, and D) aggregated cell types. Five-fold stratified cross-validation was used, and mean AUROC for each of the iteratively held-out test sets is reported here.

### Fibroblast nuclear morphology is associated with survival and gene expression patterns in breast cancer

The interplay between fibroblasts and cancer cells is complex and prognostically relevant, as associations between cancer-associated fibroblasts (CAFs) and cancer progression have been recently described^43–46^. Notably, in breast cancer, CAFs have been shown to contribute to prognosis^47^, while CAF subset heterogeneity correlates with metastasis^48^. We therefore hypothesized that fibroblast nuHIFs in BRCA would be clinically prognostic, independent of further molecular testing. We sought to identify fibroblast nuHIFs that are associated with progression-free (PFS) and/or overall survival (OS). We performed regression between each fibroblast nuHIF and PFS and OS using Cox proportional hazards models with patient age and ordinal cancer stage as regression covariates. After FDR correction, multiple fibroblast nuHIFs were significantly prognostic of PFS (Supplementary Table S5) and OS (Supplementary Table S6). Features quantifying the same general attribute, *e.g.* nuclear area and nuclear axis length as measures of size, were indeed found to be correlated with one another (mean pairwise Pearson r = 0.90 for fibroblast nuclear area, major axis length, minor axis length, and perimeter). We selected the mean fibroblast nucleus area (“MEAN[FIBROBLAST_NUCLEUS_AREA]_H & E”) for further evaluation, and show Kaplan-Meier survival curves for PFS and OS for the population binarized by this feature median value. High nuclear area was prognostic of worse outcomes (Figure 7, PFS HR = 1.81, 95% CI [1.32-2.48], p = 0.0002; OS HR = 1.77, 95% CI [1.22, 2.56], p = 0.002).

**Figure 7.**
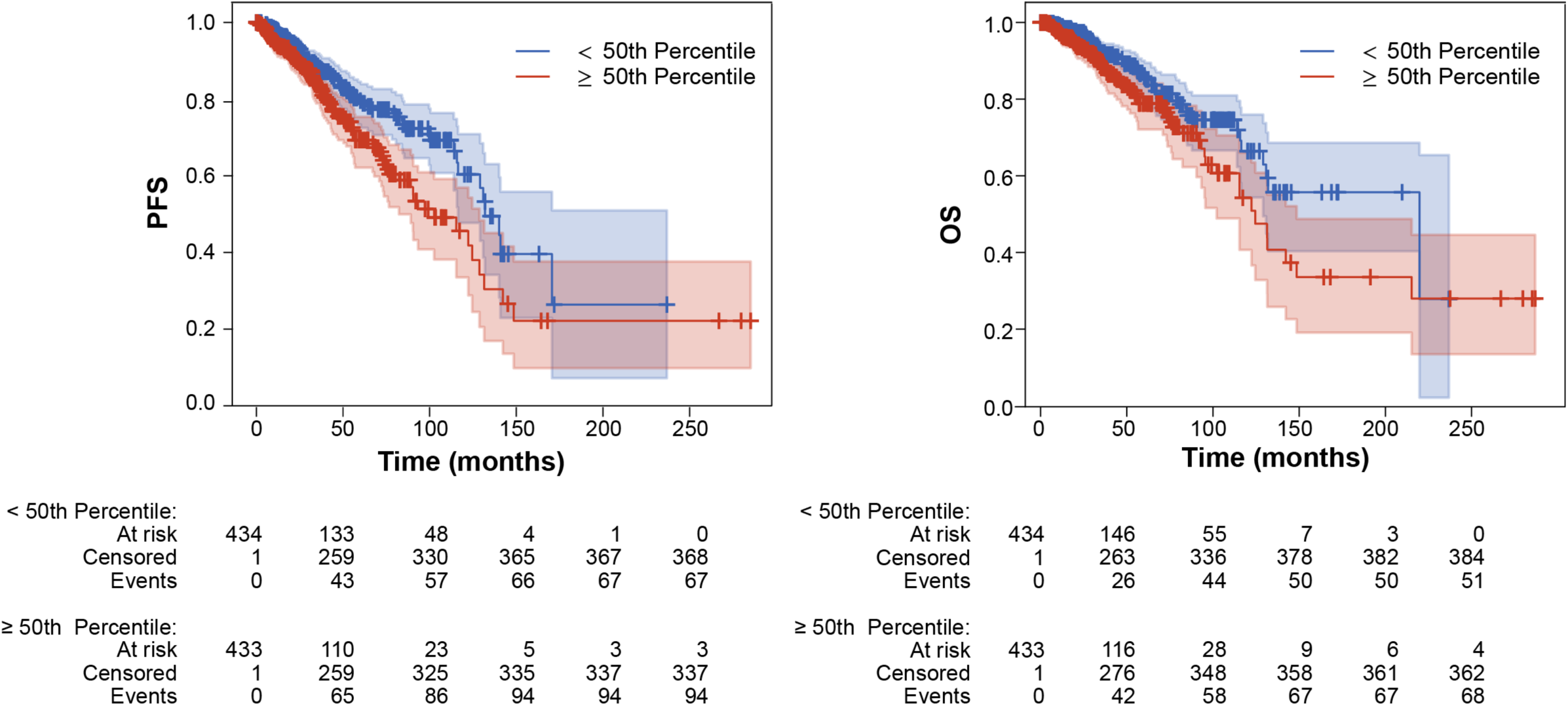
Association between fibroblast nuclear area and survival in breast cancer. Increased fibroblast nuclear area (≥ 50th percentile) corresponds to poor PFS (HR = 1.8163, 95% CI [1.3119-2.4823], p = 0.0002) and OS (HR = 1.7753, 95% CI [1.2206, 2.5620], p = 0.0022).

Having identified this relationship between fibroblast nuclear size and prognosis, we sought to assess whether mean fibroblast nuclear area was associated with differences in bulk gene expression in breast cancer. We computed the rank-based (Spearman) correlation between fibroblast mean nuclear area and each gene in TCGA bulk gene expression to identify genes associated with this nuHIF (see Methods for details). Fibroblast nuclear area was significantly, albeit weakly (absolute r > 0.15), associated with expression of numerous individual genes (Supplementary Table S7). In contrast to the weak associations observed at the individual gene level, gene set enrichment analysis performed on the genes associated with morphology revealed significant relationships between fibroblast nuclear size and levels of several previously identified expression pathways. Notably, larger fibroblast nuclear size showed positive association with gene expression in pathways associated with degradation and remodeling of the extracellular matrix (Supplementary Table S8) indicating higher fibroblast activity. Meanwhile, larger fibroblast nuclear size showed negative association with the expression of genes in pathways relating to immune response to the tumor, such as B cell receptor signaling and lymphoid cell interactions with non-lymphoid cells (Supplementary Table S9). Taken together, these results suggest that fibroblast nuclear morphology is indicative of underlying patterns of gene expression and is thus biologically grounded.

## DISCUSSION

In this study, we have presented a pan-tissue approach for nucleus segmentation, classification, and featurization on entire whole-slide pathology images. This method enabled the construction of predictive models and the identification of features linking nuclear morphology with quantitative biomarkers across BRCA, PRAD, and LUAD. These results highlight the potential of ML-enabled quantification of nuclear morphometry as a prognostic feature of many cancer types and a potential biomarker to be used by pathologists. Furthermore, this approach enables the quantitative testing of hypotheses and numerical quantification of histological relationships proposed by pathologists (*e.g.*, by establishing a numerical relationship between nuclear atypia and disease metrics). In addition, our approach enables the data-driven identification of sub-visual changes that may be clinically meaningful.

One particular strength of our approach is the ability to not only measure morphologic features associated with nuclei in a cancer specimen, but to assign a cell class to each nucleus, as well. To our knowledge, this work provides the first characterization of nuclear morphologies of specific cell types in different cancers at scale. As such, we were not only able to assess the associations of cancer cell nuclear morphology with clinically relevant metrics, but we were also able to examine these relationships using nuclear features of fibroblasts and lymphocytes. For example, fibroblast nuHIFs provided a clear separation of cancer types in both unsupervised and supervised analyses, indicating that the nuclear morphologies of fibroblasts differ in breast, lung, and prostate cancers. Given recent observations that CAFs can be classified into multiple functional subtypes based on gene expression^49^, the distinctive nuclear morphologies seen in fibroblasts of breast, lung, and prostate cancers suggests that fibroblasts may contribute to cancer progression differently in these three cancer types. Importantly, we cannot distinguish between the multiple known subtypes of intratumoral fibroblasts using the approach described herein. This caveat is particularly relevant to the associations of fibroblast nuclear morphology with gene expression in breast cancer. Increased nuclear size was positively associated with an extracellular matrix remodeling gene expression profile and negatively associated with the expression of genes relating to anti-tumor immune response (Supplementary Tables S4, S5). Interestingly, single-cell analysis of fibroblasts in breast cancer has revealed several disparate populations, including an immunosuppressive population characterized by the expression of genes involved in collagen production and extracellular matrix remodeling and a separate class with an inflammatory gene expression profile^50^. While our model cannot directly predict the presence of these fibroblast sub-populations, given the prognostic associations of nuclear morphology in our dataset and sc-RNAseq expression^50^, it will be of interest to test whether specific nuclear features of CAFs associate with functional subtypes.

Furthermore, nuclear features derived from our model were associated with PFS and OS in breast cancer. It is worth noting that this analysis, while incorporating patient age and clinical stage as regression covariates, was conducted on a large cohort of patients across study sites for whom relevant clinical information (e.g., treatment history) was not readily available. Therefore, while our result linking fibroblast nuclear morphology to prognosis in breast cancer is intriguing, further study in more controlled patient cohorts is needed to confirm this observation.

Herein, we observed that nuclear morphology differed between cancers as assessed using nucleus segmentation models. This result was observed not only for cancer epithelial cells and fibroblasts, but also, surprisingly, for lymphocytes. However, caution is warranted in interpretation – it is plausible that batch effects between slides from different tumor groups could drive variation in nuclear presentation, especially due to differences in pre-analytic variables such as slide preparation and staining. However, it is also plausible that this finding reflects the differences in genetic and epigenetic landscapes between tumor types, levels of genomic instability, and overall differences in cancer evolution between these cancer types that may manifest as disparate nuclear morphologies.

The observed relationship between greater variation in cancer nuclear area and genomic instability was consistent across cancer types, indicating a quantitative link between nuclear pleomorphism and genomic instability pertinent to numerous cancer histologies. Prior analyses have noted an association between increased variation in nuclear size and whole genome doubling, suggesting a direct link between variation in nuclear size and genomic instability^19,20^. Additional work has noted a correlation between nuclear morphology and HDR in luminal and triple-negative breast cancer^51^. Given that nuclear size reflects DNA content, variation in nuclear size features between cells may be linked to underlying genomic instability. Similarly, recent work identified a histologic signature based on variability in nuclear morphology in pancreatic cancer cells that was associated with improved response to gemcitabine but was not associated with a previously defined gene expression-based disease subtype^24^. Pancreatic cancer patients with BRCA1/2 mutations, associated with increased genomic instability, are known to respond more favorably to therapy regimens involving gemcitabine ^52^; thus, our result that nuclear variation is associated with genomic instability may explain this recent finding. To this end, our observation that variability in nuclear size (measured here by standard deviation of cancer cell nuclear area) is consistent with these prior hypotheses and allows for them to be tested on a larger scale for each case (all cells for each cell type in the WSI). While the biological result linking nuclear morphology with genomic instability is not novel, the observation of this expected result through the analyses of our novel model-derived nuclear features indicates that our approach supports the technical robustness and biological applicability of our approach.

One mitigation to potential batch effects is to analyze nuclear morphology within a single cancer type, and additionally to focus on size and shape features that are more likely to be robust to tissue preparation variabilities. For example, in breast cancer, we observed a clear relationship between fibroblast nuclear size, prognosis, and gene expression patterns. In breast cancer, increased fibroblast nuclear area was positively correlated with gene expression in extracellular matrix remodeling pathways and negatively correlated with genes in anti-tumor immune response pathways. The CAF subtypes present in a breast cancer sample may impact the tumor immune microenvironment^49^. While it would be interesting to posit that fibroblast nuclear morphology could reflect these subtypes, the ability to explore this is precluded by the use of bulk RNAseq data, since fibroblast nuclear features and the bulk expression profiling reflect a summarization of a whole slide. However, because nuclear morphology is quantified at single-cell resolution, this approach could be tied directly to single-cell expression analysis. Further work is necessary to delineate the functional relevance of nuclear morphology changes in fibroblasts in cancer.

As noted, batch effects have the potential to influence the interpretation of model outputs due to data that are aggregated across different sites, sources, and preparation laboratories. Pixel size variability, due to slide scanning with different MPP resolution, is one aspect of how these differences may manifest, but there are others to consider as well: differences in stain reagents, sample preparation, sample storage, or other pre-analytical variables. For the analyses described herein, the median MPP values were highly similar across the three indications, with the BRCA MPP slightly lower than that of LUAD and PRAD (Supplementary Figure S4). That said, to further ensure against differences in pixel dimension contributing to bias, the size-related features of the nuclei are reported here in units of microns or square microns, which is created by multiplying the size of the mask by the appropriate MPP conversion factor. Thus, differences in the MPP should not propagate into length-features, and the slide scan characteristics should not bias the features. Furthermore, we measured the Pearson correlation between nuclear size (using major axis length as a representative feature) and MPP for each cell type within BRCA, LUAD, and PRAD datasets individually to eliminate the potential effect of possible inter-cancer-type variation in nuclear size (Supplementary Figure S6). While the within-cancer-type variation in mean nuclear major axis length between slides at the same MPP is large, the magnitude of the within-cancer-type Pearson correlations is low, although some rise to the level of significance (likely due to the high power of the large datasets). Lastly, it is worth noting that cancer-type differences in nuclear size appear to be an outlier of relatively larger magnitude than expected if MPP bias was the primary driving factor (Supplementary Figure S5). Thus, we are confident that the observations noted in this study regarding nuclear size features are not biased by scan-specific metrics.

The approach that we undertook for nucleus segmentation and morphometry analysis in this paper has several key strengths. First, the ability to compute human-interpretable nuclear features at scale enables testing quantitative biological hypotheses, rather than relying on by-eye estimation of parameters such as variation in nuclear morphology.

The ability to perform these analyses on WSIs of H&E-stained cancer tissue additionally obviates the need to hand-select regions of interest, which may contribute to biased analyses. In addition, we were able to train and deploy our model on tissues from diverse cancer types, suggesting that the model can be readily deployed on samples from varied cancer indications^53^.

A particular strength of this approach is the interpretability of the predictions made. While HIF-based predictive clinical models are inherently less flexible than end-to-end black-box approaches (and, thus, can yield lower performance), they benefit from the lower dimensionality of features as a method of regularization, as the HIFs used herein directly map to low-dimensional representations of the tissue image. Furthermore, HIF-based models allow researchers and clinicians to learn from the features and generate novel hypotheses without discarding the wealth of known biology.

Although our results point to the potential of nuclear segmentation, classification, and feature analysis as a clinical screening tool, our study is limited in that our biomarker analysis was focused on academically curated datasets. These datasets were selected due to their size, completeness, and rich genomic and transcriptomic profiling data. Construction and validation of generalizable predictive machine-learning models requires the inclusion of a broad range of training and validation data, and future efforts should focus on validating these hypotheses in additional cohorts. The technical approaches we describe here have been validated by their application to other clinical data sets, showing their generalizability of this methodology and robustness of these models (data not shown).

In sum, this work highlights the power of ML-driven quantitative nuclear morphometry in multiple cancer types. The models and resulting features described herein have the potential not only to aid pathologists and research teams in discerning novel biomarkers but to provide meaningful prognostic information for cancer patients. The ability to measure these features robustly and consistently at scale may enable the development of improved clinical tools for advancing precision medicine.

## Supporting information

Supplementary figure S1 through S6

## ACKNOWLEDGEMENTS

The authors would like to thank the software engineering and machine learning operations teams at PathAI for developing the systems and pipelines used for model development and feature extraction. The authors also thank SciStories, LLC and GCI Health for assistance with figure design.

## FUNDING

This work was funded by PathAI, Inc.

## AUTHOR CONTRIBUTIONS

Conceptualization: J.A., S.J., D.J., A.P., M.G.D., L.Y., A.T.W.

Methodology: J.A., S.J., D.R., H.P., K.L., A.P., J.C., M.N., C.K., S.A.J., N.I., D.Sa., R.E., B.T., Y.G., A.D., C.P., I.W., A.K., M.G.D., L.Y., A.T.W.

Investigation: J.A., S.J., D.R., H.P., K.L., A.P., J.C., M.N., C.K., S.A.J., R.B., N.H., D.Sh., D.Sa., R.E., B.T., Y.G., A.D., C.P., M.G.D., L.Y., A.T.W.

Visualization: J.A., Y.G., J.A.B., I.W., A.T.W.

Funding acquisition: N/A

Project administration: J.A., D.Sa, C.P., I.W., A.T.W.

Supervision: M.C.M., I.W., A.K., L.Y., A.T.W.

Writing – original draft: J.A., Y.G., J.A.B.

Writing – review & editing: J.A., S.J., D.R., H.P., K.L., A.P., J.C., M.N., C.K., S.A.J., R.B., N.H., D.Sh., N.I., D.Sa., R.E., B.T., Y.G., J.A.B., A.D., M.C.M., C.P., I.W., A.K., M.G.D., L.Y., A.T.W.

## DATA AND MATERIALS AVAILABILITY

Histopathology images from the Cancer Genome Atlas dataset are available at https://www.cancer.gov/about-nci/organization/ccg/research/structural-genomics/tcga. Images and annotations of nuclei from the following datasets will be made available on GitHub prior to publication: training set, validation set, test set, OOD-2 dataset, and TCGA datasets. Images and annotations of nuclei from the OOD-1 dataset will be shared upon written request. Access to feature tables, cell-, tissue-, and nuclei-type heatmaps, as well as usage of cell- and tissue-type classification models, are available upon reasonable request to academic investigators without relevant conflicts of interest for non-commercial use who agree not to distribute the data. Access requests can be made to.

## CODE AVAILABILITY

Model parameters for nuclei, cell, and tissue models, and codes model training, inference, and feature extractions are not disclosed. Access requests for such a code will not be considered to safeguard PathAI’s intellectual property. All source code for reproducing correlational analyses and molecular predictions, will be deposited to GitHub prior to publication, and the link will be provided at that time.

## SUPPLEMENTARY FIGURE LEGENDS

**Supplementary Figure S1. Performance of cell classification model in breast cancer. A)** Comparison of model-predicted cell types to average pathologist annotations. The bar graph at the top left depicts the breakdown of average pathologist annotations for each class of model prediction (precision). The bar graph at the bottom right shows the breakdown of model predictions for each class of average pathologist annotation (recall). “Other Than Specified” refers to predictions of classes other than those listed, or background class. **B)** Agreement between model-derived cell counts and pathologist consensus counts. **C**) Precision, Recall, and F1 scores of model predictions compared to pathologists’ annotations in nested pairwise fashion^32^. **D**) Difference in the nested pairwise metric comparing mean difference between model and individual pathologist performance. Positive values indicate that the model out-performed pathologists when evaluated against held-out pathologists, while negative values indicate the model under-performed pathologists. Confidence intervals were obtained by bootstrapping.

**Supplementary Figure S2. Performance of cell classification model in NSCLC. A)** Comparison of model-predicted cell types to average pathologist annotations. The bar graph at the top left depicts the breakdown of average pathologist annotations for each class of model prediction (precision). The bar graph at the bottom right shows the breakdown of model predictions for each class of average pathologist annotation (recall). “Other Than Specified” refers to predictions of classes other than those listed, or background class. **B)** Agreement between model-derived cell counts and pathologist consensus counts. **C**) Precision, Recall, and F1 scores of model predictions compared to pathologists’ annotations in nested pairwise fashion^32^. **D**) Difference in the nested pairwise metric comparing mean difference between model and individual pathologist performance. Positive values indicate that the model out-performed pathologists when evaluated against held-out pathologists, while negative values indicate the model under-performed pathologists. Confidence intervals were obtained by bootstrapping.

**Supplementary Figure S3. Performance of cell classification model in prostate cancer. A)** Comparison of model-predicted cell types to average pathologist annotations. The bar graph at the top left depicts the breakdown of average pathologist annotations for each class of model prediction (precision). The bar graph at the bottom right shows the breakdown of model predictions for each class of average pathologist annotation (recall). “Other Than Specified” refers to predictions of classes other than those listed, or background class. **B)** Agreement between model-derived cell counts and pathologist consensus counts. **C**) Precision, Recall, and F1 scores of model predictions compared to pathologists’ annotations in nested pairwise fashion^32^. **D**) Difference in the nested pairwise metric comparing mean difference between model and individual pathologist performance. Positive values indicate that the model out-performed pathologists when evaluated against held-out pathologists, while negative values indicate the model under-performed pathologists. Confidence intervals were obtained by bootstrapping.

**Supplementary Figure S4. Distribution of pixel sizes (in MPP) across the three TCGA datasets (BRCA, LUAD, PRAD) used in this study.**

**Supplementary Figure S5. Distributions of nuclear size features in BRCA, LUAD, and PRAD datasets.** A) Distribution of area. B) Distribution of major axis length. C) Distribution of minor axis length.

**Supplementary Figure S6. Pearson correlation between nuclear size (using major axis length as a representative feature) and MPP for each cell type within BRCA, LUAD, and PRAD datasets.** A) Correlation between major axis length and MPP in cancer epithelial cells. B) Correlation between major axis length and MPP in fibroblasts. C) Correlation between major axis length and MPP in lymphocytes.

## Notes

### Summary of Updates

Updated cell segmentation and classification models (PathExplore, Boston, MA) were used to repeat all analyses. Supplemental figures (S1-S3) were added to provide model performance data for each of the cancer types of interest (BRCA, LUAD, PRAD). Table 2 has been added to show nuclear segmentation model performance in two out-of-distribution datasets. Figure 6 was added to show the utility of the model-derived nuclear features described herein for distinguishing BRCA molecular subtypes. An additional line of investigation was conducted to ensure that the original results presented were not due to batch effects and/or bias. Supplemental figures (S4-S6) were added to depict these new data. Additional text reflecting this work has been added to the results and discussion sections.

