## Supplementary Table S9 for "Deep-learning quantified cell-type-specific nuclear morphology predicts genomic instability and prognosis in multiple cancer types"

**Table S9**. Pathways enriched with diminished CAF nuclear size in fibroblasts.

| Gene Set Name | # Genes in Gene Set (K) | Description | # Genes in Overlap (k) | k/K | p-value | FDR q-value |
|---|---|---|---|---|---|---|
| REACTOME_IMMUNOREGULATORY_INTERACTIONS_BETWEEN_A_LYMPHOID_AND_A_NON_LYMPHOID_CELL | 191 | Immunoregulatory interactions between a Lymphoid and a non-Lymphoid cell | 59 | 0.3089 | 1.58E-75 | 2.67E-72 |
| REACTOME_FCGR_ACTIVATION | 69 | FCGR activation | 42 | 0.6087 | 6.63E-70 | 5.61E-67 |
| REACTOME_CD22_MEDIATED_BCR_REGULATION | 61 | CD22 mediated BCR regulation | 40 | 0.6557 | 1.34E-68 | 7.54E-66 |
| REACTOME_CREATION_OF_C4_AND_C2_ACTIVATORS | 71 | Creation of C4 and C2 activators | 41 | 0.5775 | 7.41E-67 | 3.13E-64 |
| REACTOME_ROLE_OF_PHOSPHOLIPIDS_IN_PHAGOCYTOSIS | 82 | Role of phospholipids in phagocytosis | 42 | 0.5122 | 2.23E-65 | 7.56E-63 |
| REACTOME_ANTIGEN_ACTIVATES_B_CELL_RECEPTOR_BCR_LEADING_TO_GENERATION_OF_SECOND_MESSENGERS | 86 | Antigen activates B Cell Receptor (BCR) leading to generation of second messengers | 42 | 0.4884 | 3.39E-64 | 9.57E-62 |
| REACTOME_INITIAL_TRIGGERING_OF_COMPLEMENT | 80 | Initial triggering of complement | 41 | 0.5125 | 7.59E-64 | 1.83E-61 |
| REACTOME_SCAVENGING_OF_HEME_FROM_PLASMA | 69 | Scavenging of heme from plasma | 39 | 0.5652 | 4.30E-63 | 9.10E-61 |
| REACTOME_FCGR3A_MEDIATED_IL10_SYNTHESIS | 95 | FCGR3A-mediated IL10 synthesis | 42 | 0.4421 | 8.42E-62 | 1.58E-59 |
| REACTOME_ROLE_OF_LAT2_NTAL_LAB_ON_CALCIUM_MOBILIZATION | 71 | Role of LAT2/NTAL/LAB on calcium mobilization | 38 | 0.5352 | 3.29E-60 | 5.57E-58 |
