## Supplementary Table S8 for "Deep-learning quantified cell-type-specific nuclear morphology predicts genomic instability and prognosis in multiple cancer types"

**Table S8**. Pathways enriched with increased CAF nuclear size in fibroblasts.

| Gene Set Name | # Genes in Gene Set (K) | Description | # Genes in Overlap (k) | k/K | p-value | FDR q-value |
|---|---|---|---|---|---|---|
| REACTOME_EXTRACELLULAR_MATRIX_ORGANIZATION | 300 | Extracellular matrix organization | 44 | 0.1467 | 1.51E-46 | 2.56E-43 |
| REACTOME_COLLAGEN_FORMATION | 90 | Collagen formation | 27 | 0.3 | 5.32E-38 | 4.50E-35 |
| REACTOME_COLLAGEN_BIOSYNTHESIS_AND_MODIFYING_ENZYMES | 67 | Collagen biosynthesis and modifying enzymes | 23 | 0.3433 | 4.16E-34 | 2.34E-31 |
| REACTOME_ASSEMBLY_OF_COLLAGEN_FIBRILS_AND_OTHER_MULTIMERIC_STRUCTURES | 61 | Assembly of collagen fibrils and other multimeric structures | 18 | 0.2951 | 1.50E-25 | 6.34E-23 |
| REACTOME_DEGRADATION_OF_THE_EXTRACELLULAR_MATRIX | 140 | Degradation of the extracellular matrix | 22 | 0.1571 | 2.28E-24 | 7.72E-22 |
| REACTOME_COLLAGEN_DEGRADATION | 64 | Collagen degradation | 17 | 0.2656 | 2.60E-23 | 7.34E-21 |
| REACTOME_INTEGRIN_CELL_SURFACE_INTERACTIONS | 85 | Integrin cell surface interactions | 16 | 0.1882 | 2.21E-19 | 5.33E-17 |
| REACTOME_COLLAGEN_CHAIN_TRIMERIZATION | 44 | Collagen chain trimerization | 13 | 0.2955 | 8.37E-19 | 1.77E-16 |
| REACTOME_SYNDECAN_INTERACTIONS | 27 | Syndecan interactions | 10 | 0.3704 | 6.60E-16 | 1.24E-13 |
| REACTOME_ECM_PROTEOGLYCANS | 76 | ECM proteoglycans | 13 | 0.1711 | 2.06E-15 | 3.48E-13 |
