## Supplementary Table S7 for "Deep-learning quantified cell-type-specific nuclear morphology predicts genomic instability and prognosis in multiple cancer types"

**Table S7.** Association between CAF nucleus area and gene expression levels

| | Spearman rho |
|---|---|
| ARSI | 0.254725677 |
| PTPRN | 0.253438842 |
| IGFL3 | 0.244728088 |
| TNFRSF12A | 0.24431962 |
| PPAPDC1A | 0.241117991 |
| RCN3 | 0.238874637 |
| MMP14 | 0.237794404 |
| MIR4435-1HG | 0.23526259 |
| BMP1 | 0.234741693 |
| GJB2 | 0.234232777 |
| RP11-145A3 | 0.228886516 |
| TPM2 | 0.227511794 |
| IL11 | 0.225049416 |
| NXN | 0.224221101 |
| NNMT | 0.222262875 |
| ADAMTS14 | 0.222177457 |
| SERPINH1 | 0.221100219 |
| LINC01561 | 0.218206811 |
| LOXL2 | 0.218072328 |
| TAX1BP3 | 0.217529514 |
| SUGCT | 0.217424337 |
| CERCAM | 0.217292089 |
| COL5A3 | 0.217107784 |
| MRC2 | 0.216412888 |
| UBTD1 | 0.213767339 |
| P3H1 | 0.213737589 |
| HHIPL1 | 0.212806553 |
| MFAP2 | 0.211912895 |
| COL7A1 | 0.211743976 |
| PPIC | 0.210350073 |
| SCARF2 | 0.210058525 |
| ANGPTL4 | 0.209509143 |
| LINC00632 | 0.208839398 |
| PTK7 | 0.208499015 |
| CLEC11A | 0.208255804 |
| COL1A1 | 0.207520758 |
| TLL2 | 0.20639643 |
| PDLIM7 | 0.206254079 |
| PLOD1 | 0.206037456 |
| SRPX2 | 0.20589557 |
| COL5A1 | 0.205450453 |
| CHST6 | 0.205252677 |

| | |
|---|---|
| FNDC4 | 0.204652446 |
| UCN2 | 0.203516026 |
| COL6A1 | 0.203513587 |
| APCDD1L | 0.203008225 |
| NPTX2 | 0.20299736 |
| COL6A2 | 0.202088613 |
| TMEM92 | 0.201018492 |
| RNF144A-AS1 | 0.198712214 |
| RP11-400N13 | 0.197881157 |
| C5orf46 | 0.197264396 |
| HOMER3 | 0.197235246 |
| EGLN3 | 0.196934805 |
| P4HA3 | 0.196808455 |
| SORCS2 | 0.196684729 |
| CD276 | 0.19630354 |
| RP11-417E7 | 0.196142789 |
| COL16A1 | 0.194469934 |
| CHPF | 0.194333928 |
| ADAMTS7 | 0.193130657 |
| FAM127A | 0.192752538 |
| WNT5A | 0.192042813 |
| LGALS1 | 0.191934064 |
| COL1A2 | 0.191841837 |
| CKAP4 | 0.190609616 |
| CILP2 | 0.190493257 |
| PCSK9 | 0.190207743 |
| SDC1 | 0.189211545 |
| CAMK2A | 0.188769488 |
| FHL3 | 0.18867714 |
| CTHRC1 | 0.188311524 |
| SPHK1 | 0.187914669 |
| EDN2 | 0.187774756 |
| CTD-2171N6 | 0.187067444 |
| RPSAP52 | 0.186881634 |
| NKAIN4 | 0.186726516 |
| P4HA2 | 0.186069007 |
| GPX8 | 0.185895734 |
| THBS2 | 0.185349013 |
| TMEM158 | 0.184851428 |
| SPON2 | 0.184344559 |
| FAM20C | 0.183041004 |
| PODNL1 | 0.182948684 |
| GDNF | 0.182665756 |
| MMP13 | 0.182357829 |
| S100A10 | 0.181578652 |

| | |
|---|---|
| WNT5A-AS1 | 0.181573873 |
| HTRA1 | 0.181291662 |
| MYL3 | 0.180855506 |
| HSD11B1L | 0.180664044 |
| PCOLCE | 0.180502493 |
| AF015262 | 0.180481486 |
| PITX3 | 0.179261255 |
| PRRX2 | 0.179106748 |
| RP11-863P13 | 0.179044419 |
| RP1-79C4 | 0.178954637 |
| FST | 0.178748686 |
| KIFC3 | 0.17858505 |
| ADM | 0.178205071 |
| THY1 | 0.178098462 |
| DACT3 | 0.17753747 |
| FKBP9P1 | 0.177449484 |
| ANXA2 | 0.177062063 |
| HMGA2 | 0.176847469 |
| GLIS1 | 0.17653008 |
| SMTN | 0.176009998 |
| FAM189A1 | 0.175968208 |
| CD99 | 0.175894197 |
| CA9 | 0.175425798 |
| LINC00511 | 0.175096302 |
| TMEM240 | 0.175093289 |
| SLC10A3 | 0.175086255 |
| HOXA11 | 0.174961282 |
| COPZ2 | 0.174833202 |
| VSNL1 | 0.174516035 |
| RPLP0P2 | 0.174288936 |
| COL18A1 | 0.17421044 |
| RP5-907D15 | 0.173984883 |
| GRP | 0.173594394 |
| CCNJL | 0.173262342 |
| FSCN1 | 0.173196014 |
| NKX3-2 | 0.173031204 |
| DDIT4 | 0.172944152 |
| MYL9 | 0.172902327 |
| SMCO2 | 0.172556454 |
| ZFHX4-AS1 | 0.172422583 |
| FLNC | 0.172349725 |
| FBLIM1 | 0.172240772 |
| APCDD1L-AS1 | 0.172207997 |
| EMILIN1 | 0.171927084 |
| PPAPDC3 | 0.171828158 |

| | |
|---|---|
| KCNMB4 | 0.171781347 |
| ACTN1-AS1 | 0.17176466 |
| SPSB1 | 0.171678105 |
| MMP2 | 0.171393163 |
| PAX1 | 0.171356346 |
| LMNA | 0.171212578 |
| PLAUR | 0.17082017 |
| MDFI | 0.170778996 |
| EFEMP2 | 0.170553554 |
| ACTN1 | 0.170501328 |
| PPP1R13L | 0.170487188 |
| MPZL2 | 0.170278323 |
| RP3-428L16 | 0.170229204 |
| COL13A1 | 0.170217111 |
| GPC1 | 0.169652305 |
| HOXA11-AS | 0.169500886 |
| ITGA5 | 0.169429969 |
| FLJ16779 | 0.168950167 |
| ZNF469 | 0.168894523 |
| IGF2-AS | 0.168776422 |
| COL5A2 | 0.16859771 |
| RARRES2 | 0.168521743 |
| S100A11 | 0.167944323 |
| PTH1R | 0.167821638 |
| BGN | 0.167627974 |
| LOX | 0.167482982 |
| WLS | 0.167388243 |
| S100A2 | 0.167188048 |
| PDGFRL | 0.167077903 |
| CDC42EP5 | 0.167000709 |
| MIR31HG | 0.1669921 |
| RP13-463N16 | 0.166850969 |
| MAP7D1 | 0.166788291 |
| COL3A1 | 0.166720641 |
| FN1 | 0.166675876 |
| FGF16 | 0.166602756 |
| SDCBP2 | 0.166388961 |
| AC027612 | 0.166146421 |
| C1QTNF3 | 0.165996814 |
| RFX8 | 0.165894261 |
| PPP1R3G | 0.165840454 |
| LINC00152 | 0.165797977 |
| ISLR | 0.165730495 |
| PXDN | 0.165647794 |
| BEAN1 | 0.165521202 |

| | |
|---|---|
| ARSE | 0.16525853 |
| AEBP1 | 0.165164722 |
| LINC01426 | 0.165156703 |
| VLDLR | 0.165101556 |
| CHSY3 | 0.164977793 |
| AP000695 | 0.164976472 |
| TMEM92-AS1 | 0.164885852 |
| ZYX | 0.164766955 |
| SPARC | 0.164379069 |
| KRT75 | 0.164272893 |
| LYPD1 | 0.164229909 |
| SYDE1 | 0.164222001 |
| CNIH3 | 0.164176827 |
| GXYLT2 | 0.164107931 |
| VWA1 | 0.16395678 |
| ACTB | 0.163940668 |
| RRBP1 | 0.163940556 |
| TNC | 0.163888945 |
| PLIN3 | 0.163884219 |
| SEPW1 | 0.163878228 |
| DCLK2 | 0.163768363 |
| ITGA11 | 0.163497095 |
| SYNDIG1 | 0.163453382 |
| PLS3 | 0.163207426 |
| P3H3 | 0.163158494 |
| LINC01503 | 0.163118027 |
| TP53I3 | 0.162783259 |
| DRD4 | 0.162748188 |
| IGLON5 | 0.162642177 |
| SPON1 | 0.162385065 |
| EFNB1 | 0.162361268 |
| TMEM45A | 0.162171139 |
| RP11-13P5 | 0.161793656 |
| PMEPA1 | 0.161732924 |
| GABRB2 | 0.161351956 |
| CORIN | 0.161108116 |
| POSTN | 0.160997469 |
| TGFBI | 0.160977729 |
| PTGFRN | 0.160840848 |
| RHBDL2 | 0.160785627 |
| PRKCDBP | 0.16075156 |
| OXT | 0.160619714 |
| VKORC1 | 0.160570771 |
| KLKP1 | 0.160493896 |
| CSMD2 | 0.160429649 |

| | |
|---|---|
| AC012487 | 0.160379468 |
| IFNWP19 | 0.160181023 |
| LINC01057 | 0.16017801 |
| IGFL2 | 0.15998493 |
| RP11-62F24 | 0.159606315 |
| GPX7 | 0.159449661 |
| TGFB1I1 | 0.158968226 |
| MXRA8 | 0.158954012 |
| COL11A1 | 0.158904763 |
| FAT3 | 0.158616386 |
| MGST3 | 0.158361223 |
| RPL39L | 0.158266502 |
| NREP | 0.157828009 |
| MMP11 | 0.157626846 |
| GALNT2 | 0.157252672 |
| FOXQ1 | 0.157216774 |
| RP1-198K11 | 0.157216001 |
| NKD2 | 0.157166045 |
| RP11-91J19 | 0.156517477 |
| TM6SF2 | 0.156371163 |
| PTRF | 0.156241241 |
| FAM101A | 0.156111896 |
| CCIN | 0.156100011 |
| SCT | 0.156005764 |
| GAS1 | 0.155773816 |
| RP11-22N19 | 0.155759836 |
| PQLC2L | 0.155752409 |
| EVA1A | 0.15565102 |
| DPYSL3 | 0.155426452 |
| SMYD3 | 0.155423513 |
| CLDN14 | 0.155004897 |
| KRT16 | 0.154834015 |
| LGALS7B | 0.154797159 |
| KIF26B | 0.154677657 |
| KLK4 | 0.154459615 |
| TGFB2 | 0.154423301 |
| CSDC2 | 0.154320181 |
| CD248 | 0.154305715 |
| FKBP10 | 0.153965402 |
| LINC01429 | 0.153681546 |
| NXPH4 | 0.153493158 |
| CSPG4 | 0.153489642 |
| RP11-815M8 | 0.153383257 |
| TNFRSF1A | 0.153370157 |
| AGPAT4 | 0.15330928 |

| | |
|---|---|
| CALB2 | 0.153157515 |
| ADAMTS16 | 0.152784829 |
| FIBIN | 0.152760921 |
| CYS1 | 0.152531776 |
| GLIS2 | 0.152303859 |
| C8orf58 | 0.151972049 |
| CLCA2 | 0.151875195 |
| ROS1 | 0.151825923 |
| RP11-380J14 | 0.151815151 |
| MFRP | 0.151582004 |
| CEBPD | 0.151525351 |
| GAP43 | 0.151396146 |
| JMJD1C-AS1 | 0.150891686 |
| PLAU | 0.150853414 |
| NMB | 0.150823832 |
| CLDN1 | 0.15081492 |
| CMTM3 | 0.15078383 |
| SMIM3 | 0.150713873 |
| AP2M1 | 0.15058704 |
| TUBB2A | 0.150525735 |
| KRT7 | 0.150380835 |
| CLTB | 0.150179933 |
| EFNA4 | 0.150117401 |
| ABCG4 | 0.14996755 |
| RP4-665J23 | 0.149897409 |
| FKBP9 | 0.149843975 |
| KANK4 | 0.149822095 |
| RP11-100E13 | 0.149400317 |
| MMP8 | 0.149336313 |
| H19 | 0.149196969 |
| PITX2 | 0.149081677 |
| TAGLN | 0.148976067 |
| SH3GL1 | 0.148924604 |
| GLIS3 | 0.1486861 |
| ELFN1 | 0.148682119 |
| FGF5 | 0.14863087 |
| COL8A1 | 0.148386701 |
| FKBP14 | 0.148378087 |
| CHPF2 | 0.148347983 |
| CNTN1 | 0.148258454 |
| DEPDC7 | 0.148231755 |
| RAB7B | 0.148163101 |
| NUDT11 | 0.147980711 |
| MAGEL2 | 0.147842441 |
| CREG2 | 0.147676894 |

| | |
|---|---|
| MYL6 | 0.147668381 |
| EMP1 | 0.147664772 |
| COL4A5 | 0.147654148 |
| PMP22 | 0.147457991 |
| SPP1 | 0.147340255 |
| RP11-531H8 | 0.147313513 |
| TNFAIP6 | 0.147178797 |
| C1QTNF6 | 0.147163764 |
| CST6 | 0.147138216 |
| FBLN1 | 0.147093919 |
| FAM109A | 0.147035219 |
| EN1 | 0.14700171 |
| OLFML2B | 0.146895938 |
| ASPN | 0.146788287 |
| TIMP2 | 0.146753848 |
| KDELC1 | 0.146652448 |
| SPOCD1 | 0.146396344 |
| AJ011932 | 0.146390222 |
| DMBT1 | 0.146363168 |
| TSPAN6 | 0.146142324 |
| CYP27C1 | 0.146111484 |
| PNMA6A | 0.145935915 |
| HEPH | 0.145907188 |
| RAET1L | 0.145778087 |
| AC093673 | 0.145744334 |
| RP11-107N15 | 0.145687504 |
| RP11-395A13 | 0.145632943 |
| CPXM1 | 0.145601984 |
| TMEM114 | 0.145576938 |
| ANTXR1 | 0.145495114 |
| GPR87 | 0.145470924 |
| PAEP | 0.14529635 |
| RPL17P50 | 0.145291012 |
| TUBB6 | 0.14523901 |
| RP3-495K2 | 0.144983428 |
| COL8A2 | 0.144898381 |
| RP11-401O9 | 0.144755138 |
| TGFB2-AS1 | 0.144606783 |
| NETO1 | 0.144581998 |
| FLNA | 0.144578032 |
| CTSK | 0.144474083 |
| HCFC1R1 | 0.144255952 |
| TPST1 | 0.144180824 |
| BCL7C | 0.144179931 |
| DGKI | 0.144145362 |

| | |
|---|---|
| MIR210HG | 0.143912886 |
| RP11-420L9 | 0.143794741 |
| HMGN1P13 | 0.143774915 |
| FOSL1 | 0.143750516 |
| WNT7B | 0.143600593 |
| TMEM215 | 0.143412007 |
| FAM225B | 0.143371132 |
| DSG4 | 0.143161665 |
| PLA2G5 | 0.142987841 |
| TRO | 0.142890162 |
| VASN | 0.142747142 |
| MSX1 | 0.14271475 |
| WDR13 | 0.142640495 |
| PTPRD | 0.142425081 |
| CDC42EP1 | 0.142398848 |
| RP4-598P13 | 0.142378664 |
| BARX1 | 0.1423785 |
| MAP3K6 | 0.142355534 |
| LAMB1 | 0.142349636 |
| MYH4 | 0.142301762 |
| TREM1 | 0.142210207 |
| PDPN | 0.14218161 |
| DACT1 | 0.142164679 |
| RAB3IL1 | 0.142149702 |
| SALL1 | 0.142112225 |
| HSPA8P4 | 0.142100643 |
| GIP | 0.141996246 |
| SPOCK1 | 0.141667113 |
| AKR1B15 | 0.141648872 |
| ACTA1 | 0.141581101 |
| EYA4 | 0.141525502 |
| C9orf3 | 0.141459699 |
| ANGPTL2 | 0.141236527 |
| NEFM | 0.141172099 |
| LRRC15 | 0.141131741 |
| SPINK1 | 0.141062937 |
| AGRN | 0.141006787 |
| RP11-703I16 | 0.140985372 |
| DPYSL4 | 0.140834984 |
| C11orf24 | 0.140807578 |
| OSMR | 0.140783335 |
| ADAM12 | 0.140610806 |
| LPPR5 | 0.140604945 |
| MYH13 | 0.140568914 |
| CCDC107 | 0.140529147 |

| | |
|---|---|
| LINC00857 | 0.140472097 |
| NR2F1-AS1 | 0.140299332 |
| ADAMTS12 | 0.140264521 |
| HSD17B6 | 0.140234511 |
| LRRC17 | 0.140010147 |
| NBL1 | 0.139946145 |
| PCDH7 | 0.13991498 |
| RP11-25L3 | 0.139860289 |
| ZNHIT1 | 0.13967804 |
| P4HA1 | 0.139621889 |
| LBP | 0.139468276 |
| HOXC13-AS | 0.139237786 |
| PSORS1C1 | 0.139230169 |
| DBN1 | 0.139164157 |
| CHST3 | 0.139099261 |
| WWTR1 | 0.139072283 |
| OLFML3 | 0.139065418 |
| RP11-191N8 | 0.138983702 |
| BCAR1 | 0.138983647 |
| WFDC1 | 0.138970585 |
| DIRC3 | 0.138943031 |
| RP11-91I8 | 0.138807583 |
| RAI14 | 0.13842232 |
| SHANK1 | 0.138390331 |
| ARHGEF38-IT1 | 0.138169367 |
| AC093850 | 0.137923526 |
| CTD-2269F5 | 0.13791635 |
| DES | 0.137792497 |
| SMO | 0.137656221 |
| GPR1 | 0.137566742 |
| SCG5 | 0.137527117 |
| DUOXA1 | 0.137293179 |
| FUT3 | 0.137287597 |
| SMOX | 0.137260576 |
| RP11-88G17 | 0.137172466 |
| LAMB3 | 0.136953734 |
| SHF | 0.136884038 |
| F2RL1 | 0.136804035 |
| LL21NC02-21A1 | 0.136639267 |
| ELN | 0.136601998 |
| CNN3 | 0.136427404 |
| RP11-390P2 | 0.136375997 |
| PRICKLE1 | 0.136332851 |
| FEZ1 | 0.135987701 |
| COL10A1 | 0.135873594 |

| | |
|---|---|
| AL589743 | 0.135873254 |
| MUM1L1 | 0.1357352 |
| LINC01119 | 0.135731336 |
| VSTM2L | 0.1356953 |
| METTL11B | 0.13565215 |
| RP5-1172A22 | 0.135518008 |
| ACTA2 | 0.135517876 |
| C14orf37 | 0.135457966 |
| SPSB4 | 0.135452944 |
| FSTL1 | 0.135419546 |
| TWIST2 | 0.135416402 |
| IGFL1 | 0.135275094 |
| INAFM1 | 0.135191889 |
| BBOX1-AS1 | 0.135125599 |
| TEAD2 | 0.13504485 |
| KRT1 | 0.13499322 |
| MGST1 | 0.134925924 |
| LRRC36 | 0.134846721 |
| SPANXB2 | 0.134843902 |
| AP001471 | 0.13483627 |
| KRT79 | 0.134710599 |
| KRT12 | 0.134646695 |
| B4GALT2 | 0.1345316 |
| SLC2A12 | 0.134410499 |
| FAM3A | 0.134374495 |
| ADAMTS2 | 0.134294808 |
| CD55 | 0.134174896 |
| RIN2 | 0.133799065 |
| COL6A3 | 0.133791642 |
| CTC-327F10 | 0.133757432 |
| RP11-395G23 | 0.13365731 |
| C9orf16 | 0.133489433 |
| CDH15 | 0.133290783 |
| MIR6815 | 0.132986274 |
| RP11-86H7 | 0.132945 |
| ACKR4 | 0.132921576 |
| NCKAP5L | 0.132577505 |
| HRH1 | 0.132515195 |
| FLG | 0.132441175 |
| JOSD2 | 0.132393646 |
| RP11-999E24 | 0.132367563 |
| AP000432 | 0.132333776 |
| ZNF593 | 0.132094786 |
| KLK12 | 0.131999832 |
| GFPT2 | 0.131918853 |

| | |
|---|---|
| VTCN1 | 0.131877232 |
| RP11-713C5 | 0.131758572 |
| KISS1 | 0.131719169 |
| PCDHB7 | 0.131691345 |
| SFRP2 | 0.13169021 |
| MRGPRF | 0.131505141 |
| SLN | 0.131257365 |
| UCHL1 | 0.130954513 |
| COL12A1 | 0.130948931 |
| LINC00941 | 0.13094381 |
| SLC29A4 | 0.130846973 |
| RP11-752L20 | 0.13064875 |
| FJX1 | 0.130525787 |
| CTD-3247H4 | 0.130463756 |
| GUK1 | 0.130315266 |
| LINC01048 | 0.13029749 |
| UBE2E3 | 0.130283804 |
| FAM101B | 0.130228062 |
| SOX11 | 0.13017485 |
| AL591893 | 0.13014513 |
| CNN2 | 0.130134216 |
| SBSN | 0.130074659 |
| LOXL1 | 0.130021225 |
| RP11-1060J15 | 0.130006862 |
| RP11-791G15 | 0.129972851 |
| EHD2 | 0.129955604 |
| PAM | 0.129898857 |
| KCNG2 | 0.129739045 |
| LINC01060 | 0.12959837 |
| RRAS | 0.12941504 |
| AC106786 | 0.129364819 |
| RP1-65J11 | 0.129300208 |
| C4orf26 | 0.129289987 |
| MYH1 | 0.129238591 |
| COLGALT1 | 0.129151103 |
| DIRC1 | 0.129137311 |
| LINC01096 | 0.128917937 |
| CENPB | 0.128891352 |
| CLTCL1 | 0.128882812 |
| KRT17 | 0.12888231 |
| SERPINB2 | 0.128558669 |
| C1QTNF1 | 0.128515689 |
| PRDM6 | 0.128396019 |
| CRYGN | 0.128290537 |
| DAPK3 | 0.128284051 |

| | |
|---|---|
| WISP1 | 0.12822896 |
| KRBOX1 | 0.128153407 |
| NTM | 0.128104136 |
| GLT8D2 | 0.127814505 |
| CCDC8 | 0.12779443 |
| SSC5D | 0.127667819 |
| CNN1 | 0.12758376 |
| MIR99AHG | 0.127549377 |
| NUPR1 | 0.12746982 |
| PTTG1IP | 0.127467346 |
| CNPY4 | 0.127276714 |
| TRIP6 | 0.127102455 |
| GLMP | 0.126872212 |
| COMP | 0.126826536 |
| CHRDL2 | 0.126725864 |
| FNDC1 | 0.126698084 |
| CORO6 | 0.126685655 |
| RP11-439M11 | 0.126598776 |
| RXFP3 | 0.126572142 |
| XXyac-YX65C7_A | 0.126533841 |
| PDLIM2 | 0.12649428 |
| SNAI2 | 0.12625853 |
| LIMK1 | 0.126254623 |
| INF2 | 0.126209579 |
| PGF | 0.126167475 |
| ZNF365 | 0.126159363 |
| DLX5 | 0.126095062 |
| LMX1A | 0.126063485 |
| AC112715 | 0.12601987 |
| UROD | 0.12591044 |
| NANOS3 | 0.125908247 |
| DFNA5 | 0.125818789 |
| ANXA2P1 | 0.125758584 |
| BCAT1 | 0.125708719 |
| NRP2 | 0.125508934 |
| MARVELD1 | 0.125495426 |
| MYPN | 0.125477931 |
| TM4SF19 | 0.125448307 |
| WBP5 | 0.125432912 |
| FBN2 | 0.125306618 |
| CTGF | 0.125189553 |
| PHLDA3 | 0.125139541 |
| UNC5B-AS1 | 0.125090927 |
| FAM57A | 0.125059444 |
| TMED1 | 0.124960054 |

| | |
|---|---|
| LZTS2 | 0.124951719 |
| RP11-184M15 | 0.124863227 |
| ST6GAL2-IT1 | 0.124856431 |
| FAM132A | 0.124826578 |
| GRAMD3 | 0.124815452 |
| GIPC1 | 0.1247393 |
| CCDC102A | 0.124588131 |
| KIF7 | 0.124585359 |
| EFNA3 | 0.124572316 |
| FOXF2 | 0.124557153 |
| AC011893 | 0.124429168 |
| CHMP4BP1 | 0.124388649 |
| EMX2 | 0.12435372 |
| NUCB1 | 0.124322631 |
| POR | 0.124200207 |
| LZTS3 | 0.124180057 |
| SH3PXD2B | 0.124095663 |
| PRRX1 | 0.124095086 |
| LCN1 | 0.124058945 |
| RDH8 | 0.124030365 |
| NKAIN3 | 0.124027914 |
| ARL4D | 0.123879039 |
| EGFL6 | 0.123876769 |
| TIMM8BP2 | 0.123814516 |
| KCNJ2-AS1 | 0.123810734 |
| SLC39A13 | 0.123756076 |
| CALD1 | 0.123729154 |
| RP11-732A21 | 0.123720346 |
| AC006262 | 0.123598178 |
| ST20-MTHFS | 0.123455858 |
| LINC00922 | 0.123442624 |
| CACNA1G | 0.123344369 |
| C17orf51 | 0.123306158 |
| CTD-2532N20 | 0.123287786 |
| HAGLROS | 0.123264928 |
| SLC44A3 | 0.123225094 |
| GDF6 | 0.123217889 |
| RP11-229O3 | 0.123073204 |
| MOSPD3 | 0.123063655 |
| FAP | 0.123013531 |
| ST6GAL2 | 0.123007522 |
| NIPAL4 | 0.122982201 |
| EDIL3 | 0.122927183 |
| NDN | 0.122841673 |
| KRT80 | 0.122835384 |

| | |
|---|---|
| AGPAT2 | 0.122780851 |
| KRT87P | 0.122770782 |
| APLN | 0.122767958 |
| SLC16A10 | 0.122765576 |
| IQCJ-SCHIP1 | 0.122633719 |
| TRIP10 | 0.122623003 |
| IGFBP6 | 0.122441878 |
| FBXL19 | 0.122426585 |
| RP11-323P17 | 0.122391218 |
| RFPL1 | 0.122378488 |
| AC007750 | 0.122367397 |
| MAGED1 | 0.122094086 |
| SH3PXD2A | 0.122088077 |
| RP11-473E2 | 0.121993632 |
| RP5-1011O1 | 0.121922466 |
| DUSP23 | 0.121868942 |
| GAL | 0.121794707 |
| AP2S1 | 0.121775263 |
| RP11-513O13 | 0.121706121 |
| LUM | 0.121681733 |
| GRIN2D | 0.121649137 |
| HS3ST3A1 | 0.121638404 |
| TTYH3 | 0.121613916 |
| TSPAN9 | 0.121385237 |
| FAM129B | 0.121365757 |
| GAREML | 0.121359208 |
| PGLS | 0.12135277 |
| DNM3OS | 0.121309847 |
| PCYT1B | 0.12111933 |
| CTC-537E7 | 0.121114085 |
| GRIA3 | 0.121103013 |
| CCDC67 | 0.121094885 |
| CCDC124 | 0.121023658 |
| MCAM | 0.120989163 |
| EMC10 | 0.120881308 |
| RP11-96B2 | 0.120667719 |
| AL589787 | 0.120597167 |
| C3orf80 | 0.120538948 |
| PGM2L1 | 0.120533888 |
| MYOZ1 | 0.120480043 |
| GDF5 | 0.120425785 |
| TENM3 | 0.120421986 |
| RP1-93H18 | 0.120401562 |
| PDIA5 | 0.120339386 |
| CTD-2263F21 | 0.120264461 |

| | |
|---|---|
| RP11-231D20 | 0.120250638 |
| LAMC2 | 0.120244442 |
| RP11-573D15 | 0.120243549 |
| LINC01559 | 0.120200609 |
| CD109 | 0.120020228 |
| LINC01212 | 0.119976431 |
| WISP2 | 0.11995576 |
| TMSB10 | 0.119924893 |
| RP11-783L4 | 0.119734892 |
| PMF1-BGLAP | 0.119681422 |
| FERMT2 | 0.119680455 |
| DOK5 | 0.119658035 |
| FTSJ1 | 0.119538942 |
| RP13-631K18 | 0.119518975 |
| KRT6A | 0.119479915 |
| SLC1A6 | 0.11947775 |
| ADAMTS6 | 0.119462659 |
| RP11-362K2 | 0.119417859 |
| RP1-293L8 | 0.119399923 |
| FKBP7 | 0.119394135 |
| SLC16A1 | 0.11931136 |
| PTPRD-AS1 | 0.119126767 |
| LRP1 | 0.11906039 |
| INSRR | 0.119003579 |
| C6orf15 | 0.118994495 |
| TSPO | 0.118988276 |
| KRT6B | 0.118962699 |
| GBX2 | 0.11889116 |
| AP000892 | 0.118831413 |
| FLJ26245 | 0.118773847 |
| LTBP1 | 0.118768805 |
| ANXA8 | 0.118756439 |
| PKM | 0.118636576 |
| ZNF358 | 0.118606715 |
| INHBA | 0.118456048 |
| RP11-167H9 | 0.118444685 |
| FHOD3 | 0.11843774 |
| BPIFB4 | 0.11839327 |
| AC004160 | 0.118359671 |
| RP5-952N6 | 0.118303716 |
| XXYLT1 | 0.118281863 |
| PCDHB17P | 0.118247743 |
| AKR1C2 | 0.118243136 |
| LAMA4 | 0.118196892 |
| SH3RF3 | 0.118185078 |

| | |
|---|---|
| ABCA13 | 0.118182723 |
| GABRA5 | 0.118108015 |
| RP11-372M18 | 0.118078209 |
| RP11-443C10 | 0.117911871 |
| AC007282 | 0.117874406 |
| RNASE1 | 0.117867371 |
| RPL7P9 | 0.117863947 |
| UPP1 | 0.117835983 |
| NGFRAP1 | 0.117834997 |
| SLED1 | 0.117709501 |
| CDR1 | 0.117687643 |
| GJA1 | 0.11765992 |
| DUOX2 | 0.117626942 |
| MLK7-AS1 | 0.117617681 |
| OSMR-AS1 | 0.117613292 |
| TPM1 | 0.117497159 |
| EPGN | 0.117475992 |
| ANXA2P2 | 0.11745005 |
| SHOX2 | 0.117385526 |
| AC108142 | 0.117310089 |
| CALU | 0.117223975 |
| RASAL2-AS1 | 0.117093588 |
| NOTUM | 0.117075175 |
| TNKS1BP1 | 0.117067949 |
| MSRB2 | 0.117066945 |
| NCS1 | 0.117054944 |
| ADAT3 | 0.116945581 |
| FAM180A | 0.11694279 |
| VIM | 0.116929674 |
| C10orf55 | 0.116924939 |
| GGN | 0.116843716 |
| SLC22A2 | 0.116825241 |
| RP4-580N22 | 0.116705084 |
| DCBLD1 | 0.116647075 |
| KCTD11 | 0.116630516 |
| HIC1 | 0.11660659 |
| PERM1 | 0.116513711 |
| TMEM53 | 0.11639538 |
| VAT1L | 0.116388449 |
| MAFB | 0.116370933 |
| CSMD1 | 0.116233399 |
| GLP2R | 0.116122307 |
| HOXA11-AS1_5 | 0.116113452 |
| RP11-14N7 | 0.116063776 |
| RP11-513N24 | 0.115981087 |

| | |
|---|---|
| CDK15 | 0.115928932 |
| ANKRD1 | 0.115927968 |
| VEPH1 | 0.115908587 |
| SH3RF3-AS1 | 0.115786238 |
| PXDC1 | 0.115750273 |
| RP11-626G11 | 0.115713639 |
| LINC00116 | 0.115706885 |
| SELM | 0.115695982 |
| KRTAP2-3 | 0.115694432 |
| PRDX5 | 0.115669265 |
| MIR503HG | 0.115651534 |
| AHNAK2 | 0.115639608 |
| PLIN2 | 0.115635738 |
| EDNRA | 0.115624537 |
| KRT81 | 0.115571772 |
| VCL | 0.115486466 |
| SEC16B | 0.115481082 |
| VSTM4 | 0.115446204 |
| GS1-257G1 | 0.115445373 |
| CDH11 | 0.115386108 |
| NACAD | 0.115353121 |
| SEMA3D | 0.115280932 |
| VLDLR-AS1 | 0.11527751 |
| RBP1 | 0.115211496 |
| TSPAN15 | 0.115120906 |
| CWH43 | 0.114980278 |
| RP11-200A1 | 0.114961741 |
| TMEM161A | 0.114961476 |
| BRI3 | 0.114837228 |
| SPANXD | 0.114815633 |
| RP11-465K1 | 0.114797987 |
| COMMD1 | 0.114705315 |
| NPC2 | 0.114625163 |
| RFPL4AP6 | 0.114621764 |
| LINC00704 | 0.114620424 |
| OBP2A | 0.114589834 |
| REP15 | 0.114588611 |
| MXRA5 | 0.114413898 |
| TRPM8 | 0.11433315 |
| DZIP1 | 0.114308163 |
| SFN | 0.114226373 |
| RAB20 | 0.114098517 |
| YIPF2 | 0.114093363 |
| BOK | 0.114012057 |
| CSTB | 0.113960464 |

| | |
|---|---|
| RP5-1198O20 | 0.113895903 |
| RPS20P14 | 0.113894043 |
| CRISPLD2 | 0.113879828 |
| FAM155A | 0.113763767 |
| RP11-20I20 | 0.113707095 |
| ROR2 | 0.113706034 |
| GPR161 | 0.113701941 |
| FSTL3 | 0.113686052 |
| EXTL3 | 0.113618254 |
| RP4-639F20 | 0.113551069 |
| LAMA1 | 0.113492667 |
| RP4-756H11 | 0.113392346 |
| MMP16 | 0.113238851 |
| GREM1 | 0.113230962 |
| VCAN | 0.113127125 |
| AC114803 | 0.113075225 |
| WWTR1-AS1 | 0.113070639 |
| WNT5B | 0.113059625 |
| NTMT1 | 0.113041856 |
| FLJ22447 | 0.112892082 |
| TRNP1 | 0.112847504 |
| TMED9 | 0.112847038 |
| TWF2 | 0.112795204 |
| THBS3 | 0.112765267 |
| SCX | 0.112754997 |
| DTX4 | 0.112682417 |
| PPL | 0.112600572 |
| HTRA3 | 0.112565166 |
| PLA2G4E | 0.112535479 |
| MT1X | 0.112518633 |
| CHCHD2 | 0.112484995 |
| HSD11B2 | 0.112363278 |
| SV2C | 0.1123489 |
| MIR210 | 0.112301517 |
| CRYAB | 0.112297452 |
| ADAMTS7P3 | 0.112252694 |
| RP4-668J24 | 0.112251207 |
| NOX4 | 0.112225169 |
| CTD-2297D10 | 0.112219681 |
| TGM5 | 0.112191717 |
| GAPDH | 0.112167176 |
| HHLA3 | 0.112113443 |
| SLC24A2 | 0.112086549 |
| LRP3 | 0.112002908 |
| PGRMC1 | 0.111920858 |

| | |
|---|---|
| HIF1A-AS1 | 0.111918698 |
| ADAMTS20 | 0.111912727 |
| EEF1G | 0.111849488 |
| APCDD1 | 0.11178182 |
| OXCT2P1 | 0.111709202 |
| CNFN | 0.111675527 |
| PDLIM4 | 0.111672326 |
| ZNF503 | 0.111633646 |
| SCGB3A2 | 0.11159563 |
| FKBP1A | 0.111518069 |
| ULBP2 | 0.111453472 |
| HAPLN1 | 0.111439147 |
| TIMM8AP1 | 0.111411806 |
| RP11-398G24 | 0.111383875 |
| FZD9 | 0.111364109 |
| PRADC1 | 0.111254913 |
| KLHDC8A | 0.111184789 |
| LTBR | 0.111137177 |
| LINC00887 | 0.111058478 |
| CTD-3035K23 | 0.111050662 |
| EPHB4 | 0.111049974 |
| KB-1980E6 | 0.111041189 |
| GGT6 | 0.111017825 |
| RHOV | 0.110996967 |
| C1QTNF2 | 0.110992018 |
| RP11-230G5 | 0.11094889 |
| RPS26P47 | 0.11090478 |
| SNHG23 | 0.11086211 |
| GJA3 | 0.110811396 |
| LIMCH1 | 0.110808661 |
| SLC12A4 | 0.110786929 |
| C1orf122 | 0.110714908 |
| TSR3 | 0.110615257 |
| HILS1 | 0.110517479 |
| FAM58A | 0.110511364 |
| RNU6-403P | 0.110457519 |
| COL22A1 | 0.110453724 |
| ST8SIA2 | 0.110419811 |
| NID1 | 0.110409592 |
| RP11-635L1 | 0.110408403 |
| RP11-19E11 | 0.11040819 |
| PEPD | 0.11039815 |
| RP1-240K6 | 0.110314792 |
| USE1 | 0.110267837 |
| RP11-308N19 | 0.110265201 |

| | |
|---|---|
| SGCD | 0.110162232 |
| RPLP1 | 0.110160539 |
| PLOD3 | 0.110158604 |
| HOXA-AS2 | 0.11014785 |
| ERCC1 | 0.110146287 |
| KIAA1644 | 0.110108575 |
| NALCN | 0.110018506 |
| ACTRT3 | 0.109982727 |
| RP11-89C3 | 0.109966592 |
| RP11-626H12 | 0.10981178 |
| KLK11 | 0.109739037 |
| FBLN2 | 0.109720892 |
| LINC00601 | 0.109712384 |
| RP1-191J18 | 0.10968022 |
| ADAMTSL2 | 0.109652721 |
| MYLK-AS1 | 0.109634153 |
| MYHAS | 0.109597254 |
| EVA1B | 0.109581779 |
| CTB-13F3 | 0.109484975 |
| RPL37 | 0.109478202 |
| PEX6 | 0.109441084 |
| FLG-AS1 | 0.109385528 |
| B3GNT7 | 0.1093667 |
| LAMB2 | 0.109327572 |
| FAM207A | 0.109277747 |
| VWC2 | 0.109256045 |
| PHC2 | 0.109194971 |
| SLPI | 0.109178208 |
| ODF3L1 | 0.10917484 |
| CLDN6 | 0.109115164 |
| IER5L | 0.109098148 |
| FNDC5 | 0.108988562 |
| NOTCH3 | 0.108968803 |
| HOXD11 | 0.108875093 |
| LINC00545 | 0.108860101 |
| HAS2-AS1 | 0.108821372 |
| TPTE2P4 | 0.108774998 |
| CCNA1 | 0.108767199 |
| SERPINF1 | 0.108750245 |
| LINC00460 | 0.108748929 |
| LINC01137 | 0.108747621 |
| MYO10 | 0.108739677 |
| GABRQ | 0.108649411 |
| CYB5R3 | 0.108637607 |
| IFNE | 0.108637279 |

| | |
|---|---|
| CLIP2 | 0.108583335 |
| EMX2OS | 0.108575521 |
| NINJ2 | 0.108555297 |
| NOG | 0.108493114 |
| MRPS6 | 0.108472521 |
| HHIP | 0.108468603 |
| LAMC1 | 0.108451813 |
| RP11-381O6 | 0.108443092 |
| PTP4A3 | 0.10843745 |
| ANKRD2 | 0.108383164 |
| CLDN2 | 0.108350949 |
| RNF217 | 0.108287155 |
| SGIP1 | 0.108235543 |
| QPRT | 0.108203021 |
| RP11-66N11 | 0.10820116 |
| RILP | 0.108153325 |
| RP11-438L7 | 0.10812707 |
| LINC01050 | 0.108103082 |
| BAIAP2L1 | 0.108092597 |
| FOXL1 | 0.10806627 |
| PLEKHA4 | 0.108054828 |
| KLK8 | 0.108022034 |
| ITFG3 | 0.107934618 |
| MVP | 0.107927567 |
| FAM46B | 0.107903603 |
| SRPX | 0.107894672 |
| FNDC1-IT1 | 0.107872751 |
| RP11-166A12 | 0.107858972 |
| RP11-323C15 | 0.107848623 |
| TAGLN2 | 0.107766722 |
| SLAMF9 | 0.10769732 |
| MBD3 | 0.107647443 |
| C1orf143 | 0.107644463 |
| C9orf170 | 0.107573559 |
| SLC30A10 | 0.107564809 |
| ARHGEF4 | 0.107544629 |
| NDUFA4L2 | 0.107534284 |
| CTD-2078B5 | 0.107494923 |
| OPCML | 0.107493905 |
| LZTS1 | 0.107475975 |
| FOXL2 | 0.107447555 |
| RAET1G | 0.107443253 |
| SMPD1 | 0.10736544 |
| KB-176G8 | 0.1073435 |
| PALM2 | 0.107328249 |

| | |
|---|---|
| CTD-2186M15 | 0.107321326 |
| PAPPA | 0.107319763 |
| ITPRIP | 0.107276989 |
| PRAF2 | 0.107251649 |
| GGCX | 0.107237881 |
| CLDN8 | 0.107184579 |
| P3H4 | 0.107166975 |
| ITGB6 | 0.107154733 |
| LYNX1 | 0.107150472 |
| RP11-1069G10 | 0.107143458 |
| SH2D5 | 0.107097081 |
| GALNT10 | 0.107031527 |
| PIANP | 0.107030998 |
| AVEN | 0.107025053 |
| MRPS21 | 0.10701748 |
| ERI3 | 0.106972604 |
| BAG2 | 0.106925178 |
| CASP14 | 0.106861896 |
| MEGF6 | 0.106796391 |
| FAM133A | 0.106720865 |
| TMEM2 | 0.106718676 |
| TMPRSS11E | 0.106687297 |
| PSG5 | 0.106624892 |
| CSRP2 | 0.106545515 |
| FAM19A5 | 0.106526333 |
| RCAN1 | 0.106517868 |
| SPRR2F | 0.106487836 |
| LIF | 0.106470517 |
| BEND6 | 0.106468191 |
| AHDC1 | 0.106448618 |
| GPC6 | 0.106413379 |
| CDH3 | 0.106331627 |
| CTC-518P12 | 0.106234886 |
| MIR6133 | 0.106193851 |
| NDNF | 0.106180393 |
| RP1-37C10 | 0.106123791 |
| SLC11A1 | 0.106123227 |
| AL603650 | 0.106122962 |
| PCBP4 | 0.106121311 |
| TMEM261 | 0.106117403 |
| RHOC | 0.106057047 |
| RP11-964E11 | 0.106049845 |
| TBC1D3 | 0.10603978 |
| SLC4A11 | 0.105978551 |
| RP11-1105O14 | 0.105938104 |

| | |
|---|---|
| STON1 | 0.105815362 |
| SNHG18 | 0.105804646 |
| CYR61 | 0.105803436 |
| PMF1 | 0.105775119 |
| KLK7 | 0.105760683 |
| TRPV4 | 0.105732196 |
| AKR1C1 | 0.10572414 |
| TPM4 | 0.105659244 |
| LINC01301 | 0.105655876 |
| RNU6-18P | 0.105522623 |
| SNTA1 | 0.105455774 |
| TBC1D2 | 0.105418954 |
| ZFP42 | 0.105382801 |
| PDE9A | 0.105331639 |
| MYH2 | 0.105329563 |
| DZIP1L | 0.105318913 |
| RP11-130L8 | 0.105306868 |
| RP5-864K19 | 0.105260491 |
| F2RL3 | 0.105257775 |
| MRPS26 | 0.105250854 |
| C1orf53 | 0.105233327 |
| TNNI2 | 0.105210666 |
| RP11-479G22 | 0.105111462 |
| TEX36 | 0.105094638 |
| ARF5 | 0.105044315 |
| FAM214B | 0.105038993 |
| ENO1 | 0.105008127 |
| C10orf10 | 0.105000945 |
| NLGN2 | 0.105000294 |
| SPTLC1P5 | 0.104976717 |
| LINC01142 | 0.104861793 |
| CAPN5 | 0.104842854 |
| SSR2 | 0.104826091 |
| CLDN10 | 0.104793563 |
| AC093627 | 0.104786833 |
| RP11-357P18 | 0.104747055 |
| SERPINB7 | 0.104731707 |
| SLC38A5 | 0.10470322 |
| KCNJ12 | 0.10469442 |
| KLK10 | 0.104631786 |
| PDGFRB | 0.104519529 |
| RP11-679B19 | 0.104424772 |
| ATP5EP2 | 0.104381075 |
| HSPB3 | 0.104352142 |
| LY6D | 0.104315462 |

| | |
|---|---|
| RP1-140K8 | 0.104233327 |
| AC004538 | 0.10415507 |
| RP11-9N12 | 0.104106293 |
| SEMA5B | 0.104082096 |
| PRKCSH | 0.104080849 |
| CPNE1 | 0.104075379 |
| TPRA1 | 0.104056755 |
| ANXA1 | 0.104028958 |
| ENC1 | 0.103995375 |
| FTH1P27 | 0.103970549 |
| ZFP36L2 | 0.103929252 |
| GCKR | 0.103907882 |
| MAD1L1 | 0.103823331 |
| MYH8 | 0.103820373 |
| LA16c-444G7 | 0.103723536 |
| SCEL | 0.103476786 |
| ALDH1B1 | 0.103458366 |
| MMP28 | 0.103378213 |
| DKKL1 | 0.103369913 |
| NAP1L3 | 0.103331272 |
| C7orf73 | 0.103311141 |
| AC007796 | 0.103216301 |
| KLF14 | 0.103157601 |
| GRHL3 | 0.103152771 |
| ASIP | 0.103114802 |
| LINC01105 | 0.103114027 |
| FAT1 | 0.103088991 |
| KCNJ15 | 0.103054776 |
| AKR1B1P2 | 0.102971517 |
| FAM8A3P | 0.102933808 |
| CH17-360D5 | 0.102928899 |
| HSPB7 | 0.102919384 |
| RP11-171N4 | 0.102865165 |
| KRT6C | 0.10280601 |
| RP11-66B24 | 0.102805965 |
| C20orf141 | 0.102803952 |
| RP11-126K15 | 0.102778654 |
| CNTN6 | 0.10272032 |
| APP | 0.102677755 |
| PLOD2 | 0.102628357 |
| RP11-1020M18 | 0.102577893 |
| TMUB1 | 0.102565675 |
| LMCD1 | 0.102542158 |
| CTC-546K23 | 0.102526989 |
| NDUFS6 | 0.102486007 |

| | |
|---|---|
| PARVA | 0.102467401 |
| RUSC2 | 0.102434712 |
| WTAPP1 | 0.102398036 |
| RPL29P19 | 0.102284396 |
| PDAP1 | 0.102263988 |
| FHL2 | 0.102245438 |
| FBN1 | 0.102244006 |
| YIF1B | 0.102208097 |
| RAD51AP2 | 0.102160987 |
| RP11-94A24 | 0.102145523 |
| MGAT1 | 0.10214015 |
| TCF7L2 | 0.102064928 |
| PPAN | 0.102033875 |
| LL22NC03-N14H11 | 0.101994583 |
| ANXA3 | 0.10198351 |
| MXRA5Y | 0.101951975 |
| RABGEF1 | 0.101911823 |
| ITPA | 0.101911805 |
| RP11-895K13 | 0.101894607 |
| MZT2B | 0.101877906 |
| CYP11A1 | 0.101794654 |
| MSANTD3 | 0.101791055 |
| FIS1 | 0.101736876 |
| RN7SKP231 | 0.101714294 |
| DNM1 | 0.101651142 |
| AKR1B10 | 0.101644098 |
| TBX5 | 0.101571883 |
| FHOD1 | 0.101554859 |
| GLYATL2 | 0.101545686 |
| TNFRSF19 | 0.101519062 |
| FETUB | 0.101506154 |
| RP11-767I20 | 0.101503531 |
| ZNF628 | 0.101474409 |
| RP11-411K7 | 0.101444474 |
| GAS6 | 0.101409253 |
| UNC5CL | 0.10136352 |
| VMO1 | 0.101319797 |
| PRKD1 | 0.10131403 |
| ATP13A5 | 0.101307123 |
| ARHGAP22 | 0.101303053 |
| NAT14 | 0.101276577 |
| RP11-14D22 | 0.101250175 |
| SPRR2C | 0.10123273 |
| RNF144A | 0.101212946 |
| FAM71E1 | 0.101199699 |

| | |
|---|---|
| C2orf54 | 0.101174232 |
| RP11-469N6 | 0.101128766 |
| HDGFRP2 | 0.101117798 |
| STOML1 | 0.101096755 |
| RTN4RL2 | 0.101090894 |
| NTNG2 | 0.101065386 |
| CCND2-AS1 | 0.101044558 |
| APOPT1 | 0.101006723 |
| INA | 0.100977407 |
| RP1-78O14 | 0.100942443 |
| RP11-279O9 | 0.10093296 |
| GLRX5 | 0.100923185 |
| CCL26 | 0.100887446 |
| MYO1B | 0.100876838 |
| CTD-2308L22 | 0.10085107 |
| CASC10 | 0.10083293 |
| RP11-98D18 | 0.100800593 |
| ST14 | 0.100784425 |
| ATAD3A | 0.100771587 |
| CPZ | 0.100742938 |
| RP11-86K22 | 0.100660308 |
| SCPEP1 | 0.100621962 |
| ALKBH4 | 0.100619934 |
| DIRAS3 | 0.100605161 |
| SERPINB11 | 0.100582484 |
| DKK3 | 0.100561253 |
| DPM3 | 0.100545643 |
| ANGPT2 | 0.100501157 |
| MIR9-1 | 0.10049176 |
| VWA5A | 0.100437954 |
| AP000240 | 0.10043605 |
| SLC35D3 | 0.100432251 |
| VASP | 0.100415497 |
| RP11-230B22 | 0.100369272 |
| CTD-2287O16 | 0.100358527 |
| SSTR2 | 0.100338954 |
| ARPC1B | 0.100331568 |
| SMAD5-AS1 | 0.100330235 |
| S100A16 | 0.1003082 |
| RP11-832A4 | 0.100302466 |
| EPS15P1 | 0.100248395 |
| S100A9 | 0.100230057 |
| MARCKSL1 | 0.100226987 |
| RPL39P40 | 0.100213677 |
| RPS4XP23 | 0.100185726 |

| | |
|---|---|
| FZD2 | 0.100114089 |
| ZFPM2 | 0.100091855 |
| NUDT10 | 0.100050772 |
| IGF2 | 0.099967292 |
| ANKRD35 | 0.099955812 |
| CXCL8 | 0.099937337 |
| RP11-429E11 | 0.099898361 |
| RASGRF2-AS1 | 0.099863599 |
| SEC61A1 | 0.099783656 |
| AL139099 | 0.099734502 |
| IL13RA2 | 0.099687169 |
| DSTN | 0.09966259 |
| ZSWIM4 | 0.099611909 |
| RBMS1 | 0.09956759 |
| RP11-685B14 | 0.09952381 |
| TMED3 | 0.099497392 |
| WT1 | 0.099495687 |
| JHDM1D-AS1 | 0.099483084 |
| MAN1B1 | 0.099448887 |
| TRIM29 | 0.099405872 |
| RP11-127L20 | 0.099389313 |
| SEMA7A | 0.099325998 |
| RHEBP2 | 0.099314743 |
| AURKAIP1 | 0.099303597 |
| MME | 0.099299151 |
| CTC-441N14 | 0.099289061 |
| LINC01265 | 0.099266565 |
| RP11-677M24 | 0.099166636 |
| FAM212A | 0.099141283 |
| KRT16P1 | 0.09913545 |
| MN1 | 0.099053261 |
| IL17D | 0.099007305 |
| ADRA1D | 0.09893438 |
| CDH2 | 0.098930241 |
| RP11-676J12 | 0.098921981 |
| CTB-178M22 | 0.098822253 |
| BFSP1 | 0.09881392 |
| MTHFD1L | 0.098738233 |
| EPN1 | 0.098651029 |
| LKAAEAR1 | 0.098645667 |
| LCAT | 0.098597594 |
| HHIP-AS1 | 0.098554668 |
| AP001626 | 0.098545362 |
| RP11-524D16__A | 0.098535224 |
| SHISA6 | 0.098496975 |

| | |
|---|---|
| HOXA3 | 0.098489385 |
| CCDC71L | 0.09848211 |
| CCK | 0.098476083 |
| STX4 | 0.098455969 |
| NLRX1 | 0.098428415 |
| CASC15 | 0.098389083 |
| HAND2 | 0.098387372 |
| DCN | 0.098373845 |
| LAMB4 | 0.098326884 |
| BCL2L12 | 0.098302418 |
| AGPAT4-IT1 | 0.098249444 |
| PPIB | 0.098239104 |
| LINC01235 | 0.098235457 |
| CEMIP | 0.098180255 |
| C7orf50 | 0.098033383 |
| CCDC85B | 0.098025662 |
| FAM127B | 0.09802276 |
| MFI2 | 0.098015578 |
| SULF2 | 0.098015262 |
| CD82 | 0.097995689 |
| FZD1 | 0.097969046 |
| RP11-9N20 | 0.097889835 |
| SNORD113-3 | 0.097810904 |
| PDLIM3 | 0.097805262 |
| TMEM256 | 0.097798248 |
| RP11-513G11 | 0.097778423 |
| RP11-313P22 | 0.097752646 |
| YBX3 | 0.097736682 |
| IGF2BP2 | 0.097733073 |
| RPL36 | 0.097723212 |
| SULF1 | 0.097671749 |
| NPAS2 | 0.097655283 |
| KRT3 | 0.097650794 |
| SAMD14 | 0.097629012 |
| RP11-849N15 | 0.09752791 |
| RP11-82L18 | 0.097507193 |
| NAA10 | 0.097493583 |
| EIF5A2 | 0.097483257 |
| MIF | 0.097475033 |
| RRAD | 0.097444074 |
| NXT1 | 0.097432966 |
| RPL3P4 | 0.097395625 |
| LINC01482 | 0.0973703 |
| JPH2 | 0.097348405 |
| RP11-499E18 | 0.097314412 |

| | |
|---|---|
| MMP19 | 0.097287211 |
| ATG101 | 0.097259191 |
| LINC01505 | 0.097252393 |
| ID3 | 0.097242502 |
| MSRB1 | 0.097181662 |
| DMKN | 0.097157419 |
| SLC6A8 | 0.097146051 |
| LRRC3 | 0.097143651 |
| KCNB2 | 0.097138636 |
| KIRREL | 0.097052466 |
| NENF | 0.097034567 |
| ADAM23 | 0.097007534 |
| SLC15A2 | 0.096980109 |
| SERPINB5 | 0.096933472 |
| BNC2 | 0.096904571 |
| RECK | 0.096891008 |
| RP11-445N18 | 0.096868486 |
| ITGB5 | 0.096826912 |
| RNF128 | 0.096812362 |
| SEMA4B | 0.096785905 |
| MSRB3 | 0.096775356 |
| KRT17P3 | 0.096727719 |
| DCBLD2 | 0.096724731 |
| GRID1 | 0.096708525 |
| SLC1A7 | 0.096678933 |
| C1orf86 | 0.096632801 |
| DMPK | 0.096626605 |
| AK5 | 0.09661014 |
| RP11-290H9 | 0.096609587 |
| TMEM59L | 0.096586715 |
| TRIOBP | 0.096512349 |
| RP11-215E13 | 0.09650777 |
| RP11-466H18 | 0.096487957 |
| GDA | 0.096461698 |
| SHOX | 0.096442102 |
| MTFP1 | 0.096431322 |
| IGFBP3 | 0.096413145 |
| RP11-290O12 | 0.096379023 |
| AB019441 | 0.096339802 |
| DEFB1 | 0.096338686 |
| MFGE8 | 0.096329625 |
| NDUFS4 | 0.096321494 |
| B4GALT7 | 0.096232058 |
| GNG5 | 0.0961888 |
| PHLDB1 | 0.096168892 |

| | |
|---|---|
| SMIM24 | 0.096148293 |
| CAV2 | 0.096143663 |
| RNF121 | 0.096143217 |
| ARHGAP28 | 0.096093094 |
| PIK3R2 | 0.096082786 |
| LCN2 | 0.096019965 |
| LINGO2 | 0.096005108 |
| CTB-102L5 | 0.095990875 |
| C1orf94 | 0.095979006 |
| SYT14 | 0.095978697 |
| FNDC3B | 0.095949887 |
| AP001434 | 0.095883076 |
| PNCK | 0.095857832 |
| NRK | 0.095845673 |
| MAOB | 0.095774438 |
| KLK2 | 0.095661105 |
| GADD45B | 0.095654563 |
| RP11-89M22 | 0.095628848 |
| POPDC3 | 0.095622361 |
| PBXIP1 | 0.095610244 |
| LAMC3 | 0.095603044 |
| RP4-673M15 | 0.095535023 |
| RP13-128O4 | 0.095523428 |
| DHDH | 0.095517665 |
| RP11-576I22 | 0.095491579 |
| OLFM2 | 0.095483932 |
| LHPP | 0.095477587 |
| RNF112 | 0.095382034 |
| RP11-14J7 | 0.095377957 |
| NDUFA11 | 0.095358457 |
| MYLPF | 0.095351858 |
| CASC18 | 0.095320006 |
| ZNF362 | 0.095310696 |
| RP11-91K9 | 0.095306522 |
| MEG9 | 0.095287042 |
| RPS2 | 0.095245856 |
| AKR1E2 | 0.095214767 |
| TSPAN18 | 0.095183789 |
| LINC00964 | 0.095108572 |
| NSFL1C | 0.095092752 |
| NFE2 | 0.095048863 |
| RN7SKP51 | 0.095036412 |
| ABC7-42391500H16 | 0.095034114 |
| DIRAS1 | 0.095001976 |
| KIAA1549L | 0.094967519 |

| | |
|---|---|
| PCSK1N | 0.094956282 |
| SH3RF2 | 0.094954699 |
| RP11-114M1 | 0.094945853 |
| CPN2 | 0.094937006 |
| NRP1 | 0.094936931 |
| TBX5-AS1 | 0.09491587 |
| ESM1 | 0.094914605 |
| GNAS-AS1 | 0.094893028 |
| MT1P3 | 0.094887527 |
| ADAMTS4 | 0.094877561 |
| GPR162 | 0.09487464 |
| TMEM217 | 0.094864779 |
| TWIST1 | 0.094857132 |
| UGT1A6 | 0.094837965 |
| SHBG | 0.094835578 |
| RP11-183I6 | 0.09482803 |
| MFAP5 | 0.094824759 |
| TSHZ3 | 0.094799065 |
| SLC52A2 | 0.094791008 |
| ADAD2 | 0.09475532 |
| DPP6 | 0.094752755 |
| PTGIR | 0.094744476 |
| FABP5 | 0.094732104 |
| TMEM35 | 0.094706711 |
| ART5 | 0.094686078 |
| FKBP8 | 0.094659096 |
| CTD-2154B17 | 0.094643499 |
| ISM1 | 0.094619745 |
| PRSS22 | 0.094594479 |
| DOHH | 0.094533788 |
| OSTC | 0.094530067 |
| XKR5 | 0.09452294 |
| SALL4 | 0.094501712 |
| GJC2 | 0.094460399 |
| KRT16P2 | 0.09445204 |
| MXRA7 | 0.094389689 |
| R3HDML | 0.094372686 |
| R3HCC1 | 0.094362394 |
| RBPMS | 0.094330914 |
| LINC00882 | 0.094328532 |
| TUBA1A | 0.094325537 |
| WNT11 | 0.094310094 |
| UNC93A | 0.094299352 |
| SLC15A1 | 0.094277355 |
| TXNDC17 | 0.094221811 |

| | |
|---|---|
| ZP1 | 0.094174295 |
| STEAP1 | 0.094173102 |
| FGL1 | 0.094172767 |
| DEXI | 0.094168004 |
| FAM26E | 0.09416019 |
| GPR3 | 0.09412603 |
| NKX2-8 | 0.09410755 |
| RP11-244F12 | 0.094088633 |
| AC010729 | 0.094070118 |
| IGFL1P1 | 0.094047769 |
| MYO1E | 0.094038343 |
| PDZK1IP1 | 0.094031292 |
| GPR153 | 0.094010193 |
| SYCE1L | 0.094002083 |
| IDH2 | 0.093969447 |
| MRPL28 | 0.09396247 |
| SLC43A2 | 0.093922208 |
| TCF7L1 | 0.093920068 |
| VWCE | 0.093907398 |
| RP11-557L19 | 0.09389871 |
| ZNF474 | 0.093875324 |
| KLC3 | 0.093868177 |
| SIX5 | 0.093839581 |
| KLK9 | 0.09382116 |
| NAA60 | 0.093777773 |
| CNTN3 | 0.093702579 |
| BX842568 | 0.093691984 |
| FGFRL1 | 0.093673173 |
| PLA2G15 | 0.093669173 |
| METRNL | 0.093663685 |
| RNU6-285P | 0.093658447 |
| B4GALT1-AS1 | 0.093519808 |
| MPZL1 | 0.093509873 |
| EIF4EBP1 | 0.093504961 |
| PTX3 | 0.093487493 |
| RP11-38H17 | 0.093483604 |
| COL23A1 | 0.09346215 |
| RUNX1 | 0.093455824 |
| RP11-42I10 | 0.09345575 |
| GABRE | 0.093405106 |
| KCNG1 | 0.093394972 |
| NDUFS5 | 0.093377235 |
| RAB40A | 0.09335422 |
| NRN1 | 0.093331837 |
| PNLIP | 0.093314048 |

| | |
|---|---|
| RP11-150C16 | 0.093312282 |
| EIF3K | 0.093281156 |
| PPEF1 | 0.093254048 |
| ALKBH7 | 0.093184594 |
| CTNNBIP1 | 0.093153504 |
| TAF7L | 0.093147966 |
| HOMER2P2 | 0.09314367 |
| DKFZp434J0226 | 0.093109453 |
| RP11-71E19 | 0.093062027 |
| MYH9 | 0.093028122 |
| ADORA2B | 0.093021387 |
| PQLC1 | 0.092998595 |
| MON1A | 0.092941699 |
| MCTS1 | 0.09293675 |
| PPP1R14B | 0.092919001 |
| RP11-309M7 | 0.092915176 |
| MMP12 | 0.092914145 |
| RP11-175B9 | 0.09288099 |
| REXO2 | 0.092874385 |
| RP11-1124B17 | 0.09286785 |
| KDELR3 | 0.092845044 |
| MASP1 | 0.092818622 |
| CADM4 | 0.092776632 |
| GAA | 0.092708089 |
| SIL1 | 0.092703922 |
| LINC00445 | 0.09267595 |
| ACOT9 | 0.092601852 |
| TNFRSF21 | 0.092583749 |
| FZD8 | 0.092578614 |
| NRG1 | 0.092569375 |
| ARHGAP23 | 0.092562204 |
| RFXANK | 0.092522928 |
| DUTP6 | 0.092505485 |
| NOS2 | 0.092489094 |
| GALNT15 | 0.092482665 |
| PPP1R14A | 0.092482126 |
| ITGB1 | 0.09245725 |
| RP11-115I9 | 0.09243098 |
| MIR4768 | 0.092421854 |
| PDGFC | 0.09241163 |
| RASGRF2 | 0.092387312 |
| EPHB2 | 0.092372595 |
| SYT8 | 0.092363529 |
| ZDHHC1 | 0.09232971 |
| ITGB1BP1 | 0.092326696 |

| | |
|---|---|
| TGM7 | 0.092257009 |
| UNC5B | 0.092242394 |
| ADAM33 | 0.092222747 |
| LYPLA2 | 0.092202039 |
| AC138647 | 0.092128147 |
| ATE1-AS1 | 0.092076296 |
| C1orf116 | 0.092045902 |
| CADPS | 0.092026058 |
| ERCC2 | 0.092009026 |
| RP11-809F4 | 0.09197088 |
| RGCC | 0.091958419 |
| MUC20 | 0.091951051 |
| PLXNA2 | 0.091922008 |
| TRAPPC3 | 0.091917785 |
| SHC1 | 0.091907961 |
| ISLR2 | 0.091907496 |
| AC007193 | 0.09190486 |
| ZNF428 | 0.09187382 |
| DPP4 | 0.091872406 |
| RP11-615I2 | 0.091863029 |
| LINC01449 | 0.091856795 |
| AC016999 | 0.091826833 |
| LSR | 0.091807994 |
| SERPINE1 | 0.091732437 |
| SRSF12 | 0.091724698 |
| HSPBP1 | 0.091722204 |
| HABP2 | 0.091694141 |
| MTSS1L | 0.091680695 |
| OAZ3 | 0.09165915 |
| ARSF | 0.091653251 |
| GPR110 | 0.091652519 |
| SNHG24 | 0.091638604 |
| KRT78 | 0.091614837 |
| BMP7 | 0.091595194 |
| BTG3 | 0.091582719 |
| B3GALT6 | 0.091527442 |
| FRRS1L | 0.091485738 |
| RN7SKP20 | 0.091469699 |
| HSPB2 | 0.091449522 |
| TMEM5-AS1 | 0.091425446 |
| LINC01133 | 0.091422234 |
| PUSL1 | 0.091421279 |
| CTA-293F17 | 0.091419028 |
| MLIP | 0.091402278 |
| SDC2 | 0.09138738 |

| | |
|---|---|
| MEG3 | 0.091385575 |
| AC007099 | 0.091382765 |
| NPR2 | 0.091366449 |
| SLC16A3 | 0.091355174 |
| GPR156 | 0.091343681 |
| ST3GAL3 | 0.091336941 |
| MYL6B | 0.091336624 |
| LAMTOR1 | 0.091329647 |
| CRHR1 | 0.091280633 |
| CTD-2024P10 | 0.091272173 |
| FAM229B | 0.091249588 |
| AC144836 | 0.091227798 |
| LIX1 | 0.091192228 |
| SPRR3 | 0.091158222 |
| NDUFA13 | 0.091108707 |
| UG0898H09 | 0.091096677 |
| KLHDC8B | 0.091083441 |
| IVL | 0.091080006 |
| C10orf25 | 0.09106184 |
| RP11-572P18 | 0.091052947 |
| F3 | 0.091032295 |
| XG | 0.091023531 |
| HOXA1 | 0.091015531 |
| SLC38A4 | 0.091015503 |
| KERA | 0.091014197 |
| DDRGK1 | 0.090987121 |
| RP11-680C21 | 0.090982669 |
| AC006115 | 0.090976828 |
| COL4A6 | 0.090936402 |
| LOXL4 | 0.090928625 |
| OST4 | 0.090914466 |
| EPYC | 0.090912604 |
| RP11-483A20 | 0.090900881 |
| LINC01405 | 0.090894041 |
| MYL7 | 0.090826275 |
| FBL | 0.090826258 |
| RPL29P14 | 0.090699967 |
| RP11-121L11 | 0.090684669 |
| CTC-451P13 | 0.090671963 |
| NCLN | 0.090671646 |
| RP11-221N13 | 0.090660832 |
| FAM195B | 0.090645394 |
| IL1RAP | 0.09063224 |
| ASIC4 | 0.090614656 |
| RP11-423E7 | 0.090604938 |

| | |
|---|---|
| S100A3 | 0.090585391 |
| LINC01186 | 0.090488864 |
| HRNR | 0.090486706 |
| RP3-473L9 | 0.090479655 |
| TMEM190 | 0.090463085 |
| KHDRBS3 | 0.090458037 |
| IL4R | 0.090440101 |
| SPINT2 | 0.090362349 |
| MIR4290 | 0.090360408 |
| ZNF524 | 0.090262363 |
| LIPH | 0.090239702 |
| RPS15A | 0.090225022 |
| LINC00162 | 0.090203959 |
| RP6-191P20 | 0.090195146 |
| RP11-536K7 | 0.090163326 |
| RUNX2 | 0.090150228 |
| SRM | 0.090144144 |
| S100A1 | 0.090139809 |
| RPL24 | 0.090135679 |
| RP11-177H13 | 0.090131233 |
| AC009166 | 0.090090998 |
| PDE4C | 0.090089183 |
| FAM20A | 0.090085295 |
| SNRPB | 0.090042837 |
| MRPS36P5 | 0.090027362 |
| MMP15 | 0.089922646 |
| KCP | 0.089871518 |
| MIR181A2HG | 0.089780094 |
| SPRR1A | 0.089778147 |
| ARL2 | 0.089766862 |
| ASB10 | 0.089747153 |
| CREB5 | 0.089631601 |
| STEAP3 | 0.089629275 |
| COX7A1 | 0.089623507 |
| RP11-479J7 | 0.089609165 |
| ACVR1 | 0.089609107 |
| C19orf81 | 0.089599249 |
| GLIPR2 | 0.089595766 |
| GREM2 | 0.089553242 |
| SNORA73B | 0.089552044 |
| NFIL3 | 0.089545494 |
| RP11-490G2 | 0.089524307 |
| PLAC9 | 0.089514461 |
| FAM89A | 0.089432503 |
| MUC12 | 0.089421047 |

| | |
|---|---|
| SLITRK3 | 0.089416029 |
| RP1-232P20 | 0.089373862 |
| AC004125 | 0.08936828 |
| TREX2 | 0.089356891 |
| DMWD | 0.089344425 |
| GAPDHP24 | 0.089339984 |
| HUNK | 0.089337671 |
| GPR78 | 0.089333977 |
| NOTCH1 | 0.089291976 |
| RP5-1059L7 | 0.08927938 |
| COL4A2 | 0.089278897 |
| STX10 | 0.089264552 |
| ETHE1 | 0.089245351 |
| OAF | 0.089227378 |
| LINC01091 | 0.089210136 |
| B3GNT5 | 0.089204289 |
| ECH1 | 0.089198056 |
| NPM3 | 0.089193479 |
| TSTA3 | 0.089185869 |
| BMP8B | 0.089160659 |
| GMPSP1 | 0.089096959 |
| CTBP2P8 | 0.08904705 |
| THOC6 | 0.089033863 |
| PTPN21 | 0.089031779 |
| SAPCD2P3 | 0.08902854 |
| CCM2 | 0.089020541 |
| OBP2B | 0.089015302 |
| RAB23 | 0.089000391 |
| FAM225A | 0.088997098 |
| RNH1 | 0.088986456 |
| RP11-95H3 | 0.088957183 |
| CYP26A1 | 0.088947657 |
| NSDHL | 0.08894024 |
| RBM42 | 0.08893144 |
| KRT42P | 0.088929758 |
| ALPK2 | 0.088896033 |
| GPAA1 | 0.088865576 |
| ZIC4 | 0.088851946 |
| LINC00520 | 0.088828129 |
| RPS16P9 | 0.088716808 |
| MID1 | 0.088685959 |
| PRADC1P1 | 0.088637287 |
| ANKRD65 | 0.088633213 |
| PROM1 | 0.0886163 |
| ZNF503-AS2 | 0.088596988 |

| | |
|---|---|
| ZFP36L1 | 0.088593527 |
| JUN | 0.088565154 |
| C21orf2 | 0.088563628 |
| HAS2 | 0.088550828 |
| CLMP | 0.088456405 |
| RNU6-725P | 0.088451728 |
| SLC22A16 | 0.088435361 |
| NDUFAB1 | 0.088400812 |
| LY6H | 0.088393818 |
| RP11-477I4 | 0.088392606 |
| CTB-40H15 | 0.088377903 |
| SDC3 | 0.088364215 |
| RND3 | 0.088327692 |
| NUAK1 | 0.088314687 |
| RAB25 | 0.088278816 |
| BOP1 | 0.088251968 |
| GYG2 | 0.088245921 |
| RP11-1081M5 | 0.088224185 |
| AC002454 | 0.088221359 |
| GPR143P | 0.088220966 |
| AC005256 | 0.088198194 |
| CYTH3 | 0.088188467 |
| NTAN1 | 0.088179425 |
| AC091801 | 0.088174868 |
| ANKFN1 | 0.08815913 |
| HAS1 | 0.088153771 |
| RP11-279O17 | 0.088132639 |
| UCHL3 | 0.088127758 |
| ZNF462 | 0.088112501 |
| TCEB2 | 0.088104947 |
| TPT1P5 | 0.088069174 |
| AGT | 0.088067327 |
| RP11-467L19 | 0.088049047 |
| PCDHB12 | 0.088046582 |
| CATSPERD | 0.088042081 |
| PCDHB16 | 0.088018041 |
| RPS19 | 0.08797443 |
| MUC1 | 0.087972886 |
| AC006145 | 0.087921738 |
| ADAMTS5 | 0.087854685 |
| YKT6 | 0.087848211 |
| AC009060 | 0.087830764 |
| GPX1 | 0.087808451 |
| MVB12A | 0.08780806 |
| AC007392 | 0.087780632 |

| | |
|---|---|
| RABAC1 | 0.087772896 |
| TMEM41A | 0.087766923 |
| ATP6V0D2 | 0.087733177 |
| AC010235 | 0.087731223 |
| PERP | 0.087693153 |
| PALLD | 0.087692781 |
| GDNF-AS1 | 0.087662957 |
| FZD7 | 0.087591046 |
| SLC25A1 | 0.087560477 |
| TEX29 | 0.087553767 |
| KRT86 | 0.087543955 |
| FABP6 | 0.087531013 |
| AC084357 | 0.087530477 |
| SCN2A | 0.087527997 |
| NEFL | 0.087512563 |
| KCNA1 | 0.087494536 |
| PDLIM1 | 0.087488958 |
| SPATA20P1 | 0.087467787 |
| CH17-13I23 | 0.087463556 |
| CDKN1A | 0.087462166 |
| KLK5 | 0.087425392 |
| LRRC8A | 0.087364878 |
| OGN | 0.087352319 |
| HSD3B7 | 0.087350775 |
| RP11-963H4 | 0.087331509 |
| AL158048 | 0.087293801 |
| RP11-157D23 | 0.087278161 |
| TMEM120A | 0.08727799 |
| PEMT | 0.087265841 |
| ST3GAL6-AS1 | 0.087254811 |
| S100A11P2 | 0.087231998 |
| HMCN1 | 0.087212666 |
| SNX7 | 0.087211066 |
| MYO1H | 0.087198415 |
| RP11-167P20 | 0.087190901 |
| SLC20A2 | 0.087167716 |
| ARMC5 | 0.087156869 |
| CGB8 | 0.08714726 |
| LEMD1 | 0.087070847 |
| RPS4Y1 | 0.087065254 |
| DTD1 | 0.087062762 |
| RP11-38M8 | 0.08705177 |
| UBE2QL1 | 0.087051469 |
| MAPK10 | 0.087042147 |
| RP11-252K23 | 0.087023855 |

| | |
|---|---|
| DENND2C | 0.087018183 |
| RGS20 | 0.086984888 |
| ZNF205 | 0.086944878 |
| MMP3 | 0.086934312 |
| ATP5J | 0.086889824 |
| MYL1 | 0.086860056 |
| MEG8 | 0.086827544 |
| SPTLC3 | 0.086815886 |
| TMEM242 | 0.086799941 |
| GLIS3-AS1 | 0.086788706 |
| FLJ20021 | 0.086783159 |
| ARHGEF40 | 0.08676874 |
| ATRAID | 0.086726171 |
| C12orf54 | 0.086700945 |
| ABCC6 | 0.086695714 |
| RP11-118A3 | 0.086687226 |
| CTC-260F20 | 0.086614891 |
| ACTBL2 | 0.086581988 |
| ADK | 0.086538051 |
| HSD17B2 | 0.086522856 |
| SERTAD1 | 0.086491332 |
| NUPR1L | 0.086477285 |
| NKX1-2 | 0.086471166 |
| AC073150 | 0.086465978 |
| RP3-416H24 | 0.086449651 |
| TENM4 | 0.086431776 |
| RP11-435J9 | 0.086421682 |
| ADPRHL2 | 0.086419255 |
| PRSS27 | 0.086395235 |
| NGEF | 0.086391942 |
| ZNF787 | 0.086368201 |
| SWI5 | 0.086277649 |
| RPL7P16 | 0.086274374 |
| CPA5 | 0.086263792 |
| B3GAT3 | 0.086229851 |
| IL36B | 0.086225742 |
| EDN1 | 0.086219693 |
| RP11-109J4 | 0.086218135 |
| RP11-166D19 | 0.086209776 |
| ZNF579 | 0.086207711 |
| GAL3ST1 | 0.08619131 |
| C19orf70 | 0.086108264 |
| SPRY3 | 0.086052988 |
| RP11-625H11 | 0.08604581 |
| BEX4 | 0.086022493 |

| | | |
|---|---|---|
| URGCP-MRPS24 | | 0.086014498 |
| RP11-541P9 | | 0.0859925 |
| RP11-185E8 | | 0.085976334 |
| MMP1 | | 0.085945076 |
| LINC01443 | | 0.085893664 |
| ARSJ | | 0.085881389 |
| PIGFP1 | | 0.08585854 |
| PCDHB14 | | 0.085853779 |
| FXYD5 | | 0.085849314 |
| RP11-329B9 | | 0.085846095 |
| C11orf87 | | 0.085830693 |
| PINX1 | | 0.085774761 |
| COMMD5 | | 0.085759393 |
| RP11-359D14 | | 0.085738003 |
| WDR83OS | | 0.085735225 |
| EXT2 | | 0.08570473 |
| SLC25A47 | | 0.085656042 |
| MPST | | 0.085649416 |
| LARP6 | | 0.085634867 |
| TACSTD2 | | 0.085567608 |
| ZFP92 | | 0.085552426 |
| LRRC43 | | 0.085474888 |
| AP000679 | | 0.085456887 |
| SNHG19 | | 0.085454096 |
| RN7SL228P | | 0.085424744 |
| OPN4 | | 0.085388865 |
| TNFSF9 | | 0.08537947 |
| FAM19A3 | | 0.085275998 |
| AC016735 | | 0.085268446 |
| RP11-173C1 | | 0.085266894 |
| AC006483 | | 0.085229896 |
| RP11-142C4 | | 0.085228561 |
| TRPC2 | | 0.085216735 |
| ATP10A | | 0.08520002 |
| ADAMTSL4 | | 0.085196169 |
| | 1-Dec | 0.08517006 |
| DAP | | 0.085138938 |
| IFNGR2 | | 0.085105839 |
| PDZD2 | | 0.085104499 |
| MBLAC1 | | 0.085078694 |
| AP000696 | | 0.085049676 |
| CIB2 | | 0.085035957 |
| FAM85B | | 0.08502595 |
| CDC42EP2 | | 0.085020347 |
| HOXA13 | | 0.084998858 |

| | |
|---|---|
| LRFN3 | 0.084995453 |
| RP11-290M5 | 0.084978613 |
| TRIML2 | 0.084972409 |
| LINC01082 | 0.084969069 |
| AK1 | 0.084936529 |
| KLHL4 | 0.084872634 |
| RSU1 | 0.084866033 |
| NFKBIB | 0.084855912 |
| SFRP4 | 0.084836488 |
| HSPA12B | 0.084780225 |
| APELA | 0.084772405 |
| RP4-630A11 | 0.084758972 |
| KLK3 | 0.084747927 |
| SCIN | 0.084732372 |
| MFSD2A | 0.084719255 |
| FAM162A | 0.084689225 |
| BACE2 | 0.084671066 |
| PORCN | 0.08466541 |
| ARHGEF38 | 0.084650507 |
| ZIC1 | 0.084634302 |
| COPE | 0.084600496 |
| KLF17 | 0.084570741 |
| ATP13A4 | 0.084570616 |
| C16orf91 | 0.084521739 |
| RP11-1152H15 | 0.084512952 |
| KDELR1 | 0.084474388 |
| LINC00163 | 0.084450936 |
| B4GALT1 | 0.084415185 |
| PLEKHM2 | 0.084411316 |
| PF4V1 | 0.084400981 |
| GATA6-AS1 | 0.084361652 |
| TMEM204 | 0.084350383 |
| SSPN | 0.084348857 |
| RP11-401P9 | 0.084328596 |
| KRT82 | 0.084324546 |
| IFNWP2 | 0.084316514 |
| ACTR1A | 0.084308985 |
| KLK13 | 0.084303139 |
| LGALS12 | 0.08426693 |
| ST5 | 0.084229503 |
| UNC93B6 | 0.084219575 |
| RP5-1121A15 | 0.084204336 |
| AC005330 | 0.084203415 |
| GJA5 | 0.084202934 |
| LINC01213 | 0.084186306 |

| | | |
|---|---|---|
| MSC | | 0.084184775 |
| TPTEP1 | | 0.084180794 |
| PPP1R14C | | 0.084163474 |
| PPP4C | | 0.084157593 |
| RP11-680A11 | | 0.084150932 |
| RP11-817I4 | | 0.084104102 |
| TMEM169 | | 0.084103953 |
| SYT3 | | 0.084083512 |
| GSDMC | | 0.084057309 |
| LRRC32 | | 0.084049086 |
| MACC1 | | 0.084032713 |
| GRAMD2 | | 0.08402782 |
| OXGR1 | | 0.08396761 |
| RNU6-44P | | 0.083956505 |
| HAUS7 | | 0.083947593 |
| NCK2 | | 0.083924801 |
| DMRT1 | | 0.083917883 |
| DMRTA1 | | 0.083909552 |
| JPH3 | | 0.083891702 |
| IRX6 | | 0.083883304 |
| RP11-547C5 | | 0.083877122 |
| RP11-59D5__B | | 0.083853173 |
| RPS26P4 | | 0.083852669 |
| RP11-376P6 | | 0.08383259 |
| SLC25A5 | | 0.083809447 |
| RP11-152P17 | | 0.08379871 |
| AC093901 | | 0.083796307 |
| RP11-134O21 | | 0.083795407 |
| INHBE | | 0.083779209 |
| TNNC2 | | 0.083774962 |
| | 11-Sep | 0.08376552 |
| RP11-247C2 | | 0.083759656 |
| CCDC9 | | 0.08372816 |
| P2RX6 | | 0.083720323 |
| NUTF2 | | 0.083693758 |
| PCAT6 | | 0.083691396 |
| C2CD4A | | 0.083679628 |
| ZNF812 | | 0.083647343 |
| AC037459 | | 0.083638603 |
| SPEG | | 0.083636416 |
| RP11-326A19 | | 0.0836301 |
| NRGN | | 0.08361182 |
| BAX | | 0.083598145 |
| LAYN | | 0.083548989 |
| LINC01354 | | 0.083542716 |

| | | |
|---|---|---|
| PLXDC2 | | 0.083540673 |
| HES4 | | 0.083443552 |
| RP1-138O17 | | 0.0834419 |
| PVRL4 | | 0.08344065 |
| ARHGEF25 | | 0.083435887 |
| ACTG2 | | 0.083419402 |
| CTD-2330J20 | | 0.083412353 |
| SDF4 | | 0.083355492 |
| RNU6-1234P | | 0.083318461 |
| SLC19A3 | | 0.083316653 |
| KDM5D | | 0.08329913 |
| NHSL1 | | 0.083284215 |
| MRVI1 | | 0.083258595 |
| LY6K | | 0.083256315 |
| RNU6-373P | | 0.083241783 |
| RP11-768F21 | | 0.083224995 |
| IFITM3 | | 0.083221291 |
| CTD-2532K18 | | 0.083210264 |
| ETV5 | | 0.083194369 |
| MRPS24 | | 0.083160302 |
| RPL35A | | 0.083160265 |
| FTL | | 0.083141883 |
| CMB9-22P13 | | 0.083119947 |
| ARL2BPP2 | | 0.08311638 |
| PADI1 | | 0.083095208 |
| CDH10 | | 0.083092659 |
| EGFEM1P | | 0.083088934 |
| MUCL1 | | 0.08308191 |
| OR11J5P | | 0.083070673 |
| AMDHD2 | | 0.083030269 |
| CORO2B | | 0.08303012 |
| ARPC1A | | 0.083021357 |
| BBC3 | | 0.08297622 |
| AP000473 | | 0.082973577 |
| KLK15 | | 0.082954132 |
| KRT17P6 | | 0.082944224 |
| GTF2E2 | | 0.082913259 |
| ERMN | | 0.082900998 |
| RPS3 | | 0.082884849 |
| MYL4 | | 0.082846577 |
| NCKAP5-IT1 | | 0.082825194 |
| | 4-Mar | 0.082822102 |
| TCEAL8 | | 0.082809962 |
| KCNH5 | | 0.082808934 |
| AP1M1 | | 0.082752415 |

| | |
|---|---|
| RIN1 | 0.082725735 |
| BTBD19 | 0.082720897 |
| VPS16 | 0.082699538 |
| LINC00326 | 0.082688048 |
| GALNT5 | 0.082687147 |
| FAM181B | 0.082672529 |
| RP11-297K8 | 0.082640366 |
| DNAH2 | 0.082620893 |
| AGTRAP | 0.082618139 |
| RNF122 | 0.082603366 |
| ZBTB18 | 0.08260119 |
| SOX21 | 0.082594585 |
| UGT1A7 | 0.082594366 |
| LINC00626 | 0.082590263 |
| FLRT3 | 0.082568258 |
| NAPRT | 0.082562155 |
| RNU6-1304P | 0.082543611 |
| XIRP2 | 0.082525551 |
| RP11-927P21 | 0.082519428 |
| RP11-44N22 | 0.082516626 |
| RNU6-860P | 0.082513973 |
| VAX1 | 0.082505387 |
| ACO1 | 0.082489817 |
| FAM171A2 | 0.082469277 |
| AC124861 | 0.082426157 |
| TEKT3 | 0.082415471 |
| SPECC1 | 0.082406093 |
| RPL7P6 | 0.08238991 |
| TYRO3 | 0.082384436 |
| TMEM160 | 0.082382501 |
| RP11-396O20 | 0.082369495 |
| RDH10 | 0.0823673 |
| PLEKHG2 | 0.082366835 |
| RAB32 | 0.08236503 |
| EEF1D | 0.082338276 |
| CTB-63M22 | 0.082331057 |
| SLC22A1 | 0.082306693 |
| BRK1 | 0.082295911 |
| SLC39A3 | 0.082283297 |
| DLX3 | 0.082275149 |
| FGF1 | 0.08224136 |
| RP13-614K11 | 0.082234251 |
| RP5-857K21 | 0.082230327 |
| NUDC | 0.082223461 |
| GRN | 0.082201488 |

| | |
|---|---|
| ZBTB47 | 0.082197339 |
| GSTA2 | 0.082176448 |
| NUTF2P5 | 0.082141912 |
| SLC38A8 | 0.082131006 |
| MMP27 | 0.082128409 |
| NAMPT | 0.082092702 |
| LINC01540 | 0.082074171 |
| RP11-93H12 | 0.082073006 |
| SLC43A3 | 0.082050803 |
| AL139815 | 0.082043758 |
| RP11-864N7 | 0.082042932 |
| AC113189 | 0.082041946 |
| TLCD2 | 0.08203897 |
| AMOTL2 | 0.082037983 |
| LBX1 | 0.082027325 |
| RP11-65J21 | 0.08197279 |
| CRISPLD1 | 0.081945886 |
| AC004471 | 0.081938447 |
| CDC34 | 0.081920322 |
| TUBB2B | 0.081905103 |
| RP11-427J23 | 0.081901478 |
| RP3-467K16 | 0.08189554 |
| RPL37P6 | 0.081861046 |
| IL12A-AS1 | 0.081860361 |
| SPRR4 | 0.081859811 |
| RILPL1 | 0.081798029 |
| S100A7 | 0.081749444 |
| TEAD4 | 0.081744371 |
| CLCA3P | 0.081733817 |
| RP6-91H8 | 0.081662674 |
| LINC00629 | 0.081651321 |
| GNG12-AS1 | 0.081604875 |
| CD24 | 0.081594783 |
| DPCD | 0.081584531 |
| IGSF3 | 0.081573703 |
| RNA5SP125 | 0.081564176 |
| TRIM6 | 0.081562353 |
| FBXO17 | 0.081532064 |
| LRG1 | 0.081508695 |
| APOOP2 | 0.081476728 |
| PRL | 0.081464543 |
| BICC1 | 0.081431352 |
| RP11-40C11 | 0.081390644 |
| HMGB3P7 | 0.081388767 |
| PLEKHS1 | 0.081366866 |

| | |
|---|---|
| CALCA | 0.081366698 |
| IL1RL2 | 0.081323106 |
| DDR2 | 0.081307384 |
| CTIF | 0.081262173 |
| GMDS | 0.081247772 |
| AC000403 | 0.081242879 |
| HOXA10 | 0.081241074 |
| MIR30C1 | 0.081230257 |
| HIST2H2AC | 0.081220273 |
| TXNRD3 | 0.081213668 |
| RP13-401N8 | 0.081201141 |
| JAM3 | 0.081200979 |
| RP1-20N18 | 0.081193367 |
| C1R | 0.081187751 |
| CD151 | 0.081164513 |
| RP11-661C8 | 0.081150531 |
| MIR5689 | 0.081139315 |
| RAET1E | 0.081124678 |
| AC002985 | 0.081107267 |
| TF | 0.081089719 |
| ETS2 | 0.081061215 |
| LINC00346 | 0.081057792 |
| C8B | 0.081057236 |
| MTMR11 | 0.081046312 |
| RP11-114F10 | 0.080970762 |
| MMP23B | 0.080926261 |
| RP11-2B6 | 0.080903348 |
| LAPTM4A | 0.080902585 |
| AC019117 | 0.080856063 |
| NID2 | 0.080833019 |
| AC107399 | 0.080807934 |
| MRAS | 0.08079752 |
| FRMPD4 | 0.080738249 |
| RNU6-952P | 0.080726846 |
| NMRK2 | 0.080716341 |
| GADD45GIP1 | 0.080677273 |
| SESN2 | 0.080675096 |
| AC245100 | 0.080656695 |
| TMEM11 | 0.080633866 |
| RP11-961A15 | 0.080631548 |
| STAC2 | 0.080587524 |
| CYB5R2 | 0.080586441 |
| HOXD13 | 0.080586356 |
| ASMTL | 0.0805587 |
| KRT83 | 0.080553161 |

| | |
|---|---|
| GSTP1 | 0.080547592 |
| PMVK | 0.08053989 |
| PYGL | 0.080517656 |
| RNA5SP104 | 0.080513299 |
| TIMP3 | 0.080502883 |
| RP11-803D5 | 0.080501561 |
| SLC2A14 | 0.080495077 |
| PCDHB11 | 0.080483162 |
| VCAN-AS1 | 0.080426343 |
| ACTG1 | 0.080424406 |
| IL20RB | 0.080420629 |
| CLEC18B | 0.080413179 |
| PLXDC1 | 0.08041047 |
| AP1S1 | 0.080368794 |
| TMEM39B | 0.080337704 |
| ALDH1L2 | 0.080269962 |
| PCDHB9 | 0.080258984 |
| MIR4731 | 0.080254623 |
| CLCN3P1 | 0.080253471 |
| CD63 | 0.080242202 |
| NECAB2 | 0.080235852 |
| PFDN2 | 0.080213792 |
| KRT5 | 0.080192507 |
| LINC00226 | 0.080139676 |
| RASD1 | 0.080110364 |
| KCNE2 | 0.08009881 |
| CTB-113I20 | 0.080078214 |
| AC005932 | 0.080069712 |
| C9orf69 | 0.080069153 |
| SNX12 | 0.080044184 |
| KLK6 | 0.080040194 |
| IL1R1 | 0.079980926 |
| UBE2E2 | 0.079966674 |
| ARFRP1 | 0.079948999 |
| RCN1 | 0.079932365 |
| AC106802 | 0.07991848 |
| RAB6B | 0.079899862 |
| GAS2L2 | 0.079895122 |
| AC092638 | 0.079893817 |
| SERPINB4 | 0.079889094 |
| GSG1L | 0.079885672 |
| LINC01545 | 0.079880565 |
| GOLM1 | 0.079878689 |
| LINC01420 | 0.079877126 |
| MUC21 | 0.079853707 |

| | |
|---|---|
| BAD | 0.079841627 |
| CXADR | 0.079821458 |
| CLCNKB | 0.07981229 |
| DENND2A | 0.079808286 |
| RP11-298E9 | 0.079803881 |
| MMP9 | 0.079802778 |
| GRB10 | 0.079770535 |
| RP11-141O11 | 0.079764455 |
| MIR507 | 0.079764267 |
| AC113208 | 0.079736115 |
| ESYT3 | 0.079733064 |
| FERMT1 | 0.07971805 |
| HMOX2 | 0.079707053 |
| APBA2 | 0.079700039 |
| PSCA | 0.079656472 |
| RP4-597N16 | 0.079652096 |
| RP11-483E23 | 0.079595782 |
| MYO3B | 0.079590199 |
| ANXA2P3 | 0.079586473 |
| LAMTOR4 | 0.079562191 |
| SLC2A1 | 0.079531064 |
| LINC00519 | 0.079526723 |
| MALSU1 | 0.079512217 |
| RP11-281O15 | 0.079484323 |
| RP11-411G2 | 0.07947808 |
| RP11-654E17 | 0.079478068 |
| JDP2 | 0.079472922 |
| SPACA6P | 0.079442521 |
| MTND1P36 | 0.079412997 |
| RP3-449M8 | 0.079402149 |
| TPPP3 | 0.079390537 |
| PRSS3 | 0.079366781 |
| IBSP | 0.07930703 |
| TMEM40 | 0.079303954 |
| MIR137HG | 0.079275994 |
| ARL4C | 0.079274011 |
| SF3B5 | 0.079267666 |
| WTIP | 0.079261694 |
| NGF | 0.07924241 |
| CTD-2600O9 | 0.079235803 |
| RP11-69I8 | 0.079230126 |
| LAMTOR2 | 0.079225711 |
| AKR1C3 | 0.079216315 |
| BSPRY | 0.079189114 |
| ZNF561-AS1 | 0.079173337 |

| | |
|---|---|
| SMAD6 | 0.079158955 |
| ABCG8 | 0.079150103 |
| EIF5A | 0.079143531 |
| PHLDA2 | 0.079135512 |
| GUCA1A | 0.079107023 |
| CTRB2 | 0.079071105 |
| RP11-90L1 | 0.079060755 |
| CDKN1C | 0.079044401 |
| NKX6-1 | 0.079043567 |
| LINC00972 | 0.079011816 |
| FAM41AY2 | 0.079011403 |
| EYA2 | 0.079010018 |
| AQP9 | 0.079001156 |
| RP11-815J4 | 0.079000473 |
| ANKH | 0.078977365 |
| RBAKDN | 0.078941888 |
| KIAA1024 | 0.07892473 |
| AC093875 | 0.078909532 |
| GYPA | 0.078903367 |
| B3GALTL | 0.078868244 |
| GTF3C6 | 0.078864895 |
| ILVBL | 0.078862886 |
| CYP26C1 | 0.0787944 |
| PLEKHN1 | 0.078782975 |
| EPPIN | 0.078778264 |
| RN7SKP197 | 0.078776738 |
| ZIC5 | 0.078749454 |
| PSMG3 | 0.078744425 |
| RHBDD2 | 0.078733931 |
| AC026877 | 0.078664262 |
| ASZ1 | 0.078663546 |
| SLURP1 | 0.078628146 |
| NRG4 | 0.078618857 |
| SOD2P1 | 0.078618269 |
| FAM50A | 0.078603135 |
| CTC-332L22 | 0.078578394 |
| SOWAHB | 0.078567971 |
| FDX1L | 0.078563636 |
| RNVU1-7 | 0.078563265 |
| PEBP4 | 0.078514951 |
| WNT2 | 0.078507187 |
| TMEM233 | 0.078476171 |
| AC100800 | 0.078474768 |
| LSM7 | 0.07842949 |
| HRASLS | 0.078425378 |

| | |
|---|---|
| GPR123 | 0.078400037 |
| SPACA3 | 0.078380478 |
| AMOTL1 | 0.078351068 |
| MRPL51 | 0.078337319 |
| RPS9 | 0.078326583 |
| ATP5O | 0.078306229 |
| ISY1 | 0.078284963 |
| LINC01014 | 0.078283717 |
| FADS3 | 0.078282786 |
| RP11-30P6 | 0.078263954 |
| PPP1R3A | 0.078260938 |
| MT1L | 0.078245315 |
| CHST14 | 0.078238728 |
| NLRP2 | 0.078232979 |
| RP11-179B15 | 0.078227055 |
| RNASE7 | 0.078225957 |
| RP5-1177I5 | 0.078203792 |
| DTNB | 0.078195489 |
| FUT6 | 0.078180668 |
| RP11-383H13 | 0.078180121 |
| SLC39A7 | 0.0781796 |
| FRMD6 | 0.078147766 |
| LPAR1 | 0.078126091 |
| AC112721 | 0.07812105 |
| PCDHB6 | 0.078116044 |
| RP11-651H2 | 0.078101523 |
| GHRHR | 0.078085367 |
| RP11-432I5 | 0.078085245 |
| EHHADH | 0.078066423 |
| CCDC92 | 0.078065921 |
| LINC01343 | 0.078049577 |
| DECR2 | 0.078000113 |
| C18orf63 | 0.077992518 |
| PTGER1 | 0.077972191 |
| ADCY2 | 0.077970642 |
| RP11-1008C21 | 0.077970586 |
| C1QL2 | 0.077951855 |
| LINC00543 | 0.077940981 |
| FRMD5 | 0.077918844 |
| RP11-553A21 | 0.077917562 |
| AC091057 | 0.077913815 |
| MIR199A2 | 0.077872726 |
| AADAC | 0.077833854 |
| SLC1A1 | 0.077822654 |
| SLC39A1 | 0.077812198 |

| | |
|---|---|
| RP11-575F12 | 0.077787789 |
| B3GNT9 | 0.077776047 |
| OPN3 | 0.077759526 |
| NXF3 | 0.077758265 |
| STX8 | 0.077755619 |
| RP11-753H16 | 0.077734304 |
| PDZRN3 | 0.077712063 |
| PADI3 | 0.077708872 |
| NHS-AS1 | 0.077703083 |
| RP11-432J24 | 0.077690941 |
| RP11-138H8 | 0.077679778 |
| EFS | 0.077634795 |
| SNRPGP20 | 0.077618962 |
| C19orf47 | 0.077614812 |
| PDZD3 | 0.077587659 |
| GSTM5 | 0.077586869 |
| AC008992 | 0.077580905 |
| MIB2 | 0.077565861 |
| KLK14 | 0.077565848 |
| DIO3 | 0.077557285 |
| ALG1L12P | 0.07749692 |
| ROR1-AS1 | 0.077466154 |
| COL11A2 | 0.07744634 |
| TMEM187 | 0.0774338 |
| RN7SKP224 | 0.077432279 |
| RNF217-AS1 | 0.077417413 |
| RP11-118K6 | 0.07739054 |
| C19orf68 | 0.077374783 |
| AC124789 | 0.077373044 |
| ITGB1BP2 | 0.077338391 |
| DEFA3 | 0.077318848 |
| RP11-57A1 | 0.077314419 |
| RP11-367F23 | 0.077300793 |
| AP000797 | 0.077258215 |
| TRIM15 | 0.077233134 |
| IDH3B | 0.077224414 |
| BNIP3P19 | 0.077204249 |
| UBXN6 | 0.077202776 |
| SNX21 | 0.077174868 |
| CCDC23 | 0.077152578 |
| TPI1 | 0.077115963 |
| AKT3 | 0.077107479 |
| SYNPO | 0.077083794 |
| DRAXIN | 0.077077747 |
| MIR3663HG | 0.077073712 |

| | |
|---|---|
| NUMBL | 0.077067681 |
| AC093620 | 0.077048445 |
| CLDN18 | 0.077023165 |
| MEIS1-AS2 | 0.07700077 |
| AGXT | 0.076989989 |
| NT5E | 0.07697298 |
| SHB | 0.076958188 |
| CH25H | 0.076908084 |
| ATP6V1G3 | 0.076871666 |
| C4B | 0.076865087 |
| F13A1 | 0.076865031 |
| OR7E90P | 0.076862257 |
| QSOX1 | 0.076833699 |
| SSC4D | 0.076819857 |
| WDR86 | 0.076810293 |
| SBSPON | 0.076798423 |
| EFNB3 | 0.076792693 |
| C15orf54 | 0.076777786 |
| TMEFF1 | 0.076720596 |
| MAP4K4 | 0.076703238 |
| AC004870 | 0.076690548 |
| RP5-858L17 | 0.076685591 |
| MCEMP1 | 0.076674412 |
| ADAMTSL4-AS1 | 0.076673785 |
| ZNF532 | 0.076666976 |
| RP11-396F22 | 0.076650771 |
| AC012462 | 0.076647666 |
| CYSTM1 | 0.076645728 |
| DGCR6 | 0.076643663 |
| RBMS3-AS2 | 0.076634989 |
| TRHDE | 0.07663466 |
| KCNE5 | 0.076633423 |
| WT1-AS | 0.076626784 |
| RP11-271M24 | 0.076626369 |
| RAP2B | 0.076623104 |
| RP11-346C4 | 0.076608584 |
| DSEL | 0.076590284 |
| MPG | 0.076568162 |
| RPL8 | 0.076476028 |
| ECHDC3 | 0.076446966 |
| CD9 | 0.076445738 |
| PRSS12 | 0.076440249 |
| FAM81B | 0.076416537 |
| LINC00242 | 0.076411169 |
| RP11-676B18 | 0.07639656 |

| | |
|---|---|
| C1orf123 | 0.076383931 |
| TMEM91 | 0.076380916 |
| PON2 | 0.076339129 |
| MIR511 | 0.076333541 |
| C5orf66-AS1 | 0.07628545 |
| TPRN | 0.076274586 |
| CLDN9 | 0.076223737 |
| CDC37 | 0.076214732 |
| FBXL15 | 0.076193504 |
| ENAH | 0.076188238 |
| PNLDC1 | 0.076152205 |
| PUF60 | 0.076121631 |
| RP11-302B13 | 0.076111258 |
| COPS6 | 0.076099825 |
| GAD1 | 0.076080776 |
| RP11-539E19 | 0.076025693 |
| PSG4 | 0.075989158 |
| RPL18 | 0.075970629 |
| WNT3A | 0.075967074 |
| RP11-6F2 | 0.075962886 |
| CDCP1 | 0.075958944 |
| ZC4H2 | 0.075918961 |
| HIF1A-AS2 | 0.075916792 |
| UFSP1 | 0.075912394 |
| DNAJB5 | 0.075907426 |
| TRIM16 | 0.075894923 |
| AC003092 | 0.075872923 |
| NDUFB7 | 0.075864224 |
| RP3-326L13 | 0.075835806 |
| RP11-4C20 | 0.075833793 |
| RPL28 | 0.07583066 |
| PHKG1 | 0.075827404 |
| PVRL1 | 0.075818194 |
| CITED4 | 0.075812705 |
| PITX1 | 0.075798455 |
| AC113617 | 0.075798143 |
| MIR3921 | 0.075731614 |
| GPRIN2 | 0.075728006 |
| DDX49 | 0.07572485 |
| DYNC1I1 | 0.075710227 |
| RP11-619A14 | 0.075699902 |
| LRRN4CL | 0.075672178 |
| ATP5D | 0.075668253 |
| JUNB | 0.075666318 |
| AC008427 | 0.075651416 |

| | |
|---|---|
| ZFPM2-AS1 | 0.075651059 |
| IRF6 | 0.075645573 |
| CA5BP1 | 0.075634019 |
| RN7SL675P | 0.07558726 |
| RPL15P3 | 0.075579467 |
| NT5DC2 | 0.075564825 |
| RP11-453D21 | 0.075553843 |
| LINC00160 | 0.075553069 |
| TIMM8B | 0.075552527 |
| AL513318 | 0.0755115 |
| RN7SL689P | 0.075508125 |
| NUDT1 | 0.075504729 |
| RP11-371I20 | 0.075497964 |
| CHCHD10 | 0.075479463 |
| RNU7-102P | 0.075464675 |
| SERPINB3 | 0.075446992 |
| AC021749 | 0.075414375 |
| PFKP | 0.075398473 |
| RP11-395L14 | 0.075334917 |
| AC144831 | 0.075312665 |
| RN7SL420P | 0.075311728 |
| GLB1L2 | 0.075303213 |
| AJUBA | 0.075292627 |
| C8orf82 | 0.075291641 |
| AFF2 | 0.075284422 |
| MESP2 | 0.075270003 |
| LUCAT1 | 0.075269013 |
| CPA4 | 0.075231006 |
| LTBP3 | 0.075218707 |
| KRTAP19-3 | 0.075205348 |
| RP4-794I6 | 0.075164063 |
| SPTB | 0.075136378 |
| GATA6 | 0.075117289 |
| SPRED3 | 0.075091055 |
| RNU6-733P | 0.075086889 |
| CTD-2566J3 | 0.075084386 |
| RP4-800M22 | 0.0750832 |
| PTPN14 | 0.07508218 |
| EIF1AY | 0.075045739 |
| CTA-941F9 | 0.075028764 |
| RP11-71N10 | 0.075028595 |
| ZNF668 | 0.07502281 |
| LA16c-335H7 | 0.075021181 |
| TMEM43 | 0.07501094 |
| FANCD2OS | 0.075008193 |

| | |
|---|---|
| RP11-315D16 | 0.075001428 |
| FASTK | 0.075000447 |
| SDSL | 0.074995739 |
| ANO7P1 | 0.074977208 |
| POLR2J | 0.074943514 |
| UPK1B | 0.074915701 |
| GAST | 0.074908702 |
| ECE2 | 0.074904945 |
| RPL7P15 | 0.074900844 |
| RP4-564O4 | 0.074899692 |
| AMTN | 0.074891547 |
| RP1-122P22 | 0.074881539 |
| PARL | 0.074875083 |
| ZFHX4 | 0.074834132 |
| EMD | 0.074787488 |
| ESRRA | 0.074759655 |
| MTMR2 | 0.074749161 |
| PLXNA1 | 0.074737886 |
| PCED1CP | 0.074725608 |
| FAM89B | 0.07472382 |
| BARHL2 | 0.074720313 |
| RP11-543P15 | 0.074715485 |
| FAM160B2 | 0.074714611 |
| SURF4 | 0.074708117 |
| LCE3D | 0.074701987 |
| PRPH2 | 0.074701792 |
| PANO1 | 0.074696843 |
| IKBIP | 0.074696024 |
| DEFB135 | 0.074680225 |
| RASL10B | 0.074626402 |
| CYP1B1-AS1 | 0.074619723 |
| TIMP1 | 0.074611927 |
| ABHD17AP1 | 0.074609182 |
| S100A12 | 0.074604526 |
| CDKL5 | 0.074598364 |
| PCDHGC5 | 0.074575265 |
| MIR3156-1 | 0.074552354 |
| LINC01370 | 0.074535611 |
| SGCE | 0.074484163 |
| NPY | 0.074480986 |
| SLC35B4 | 0.074472795 |
| PVRL3 | 0.074467344 |
| LINC01396 | 0.074462303 |
| CROCC | 0.074457278 |
| RP11-742B18 | 0.074439585 |

| | |
|---|---|
| CDC20P1 | 0.074438211 |
| PBDC1 | 0.074425091 |
| MRPL47 | 0.074416551 |
| OR2AT2P | 0.074412286 |
| RPL22L1 | 0.074390522 |
| IGFL4 | 0.074388145 |
| C12orf75 | 0.074387136 |
| EEF1GP8 | 0.074381 |
| SEC14L2 | 0.074376866 |
| ZNRF1 | 0.074364084 |
| TPGS1 | 0.074334724 |
| RP11-108K14 | 0.074333067 |
| ACTN2 | 0.074260237 |
| SORCS3 | 0.074258345 |
| RIT1 | 0.074246181 |
| CRNN | 0.07424285 |
| SEC13 | 0.074222329 |
| RP11-763K15 | 0.074221812 |
| ADAM15 | 0.074202998 |
| TOM1 | 0.07420296 |
| YIF1A | 0.074184653 |
| AC008045 | 0.074145048 |
| C11orf96 | 0.074072145 |
| MIER2 | 0.074065708 |
| IGSF9 | 0.074017892 |
| IQSEC3 | 0.074007566 |
| RPL32P33 | 0.07398716 |
| TSPY1 | 0.073977449 |
| AC016772 | 0.07396855 |
| SUMO3 | 0.073961499 |
| CTSLP4 | 0.073954617 |
| RP5-882O7 | 0.073954231 |
| FOXC2-AS1 | 0.073952651 |
| CD8BP | 0.073937769 |
| TMEM130 | 0.073937665 |
| AP000647 | 0.073932196 |
| RP11-745C15 | 0.073918879 |
| CNRIP1 | 0.073912957 |
| SMARCE1P4 | 0.073897819 |
| MIR3129 | 0.073887699 |
| RNU1-139P | 0.073864475 |
| ZPBP | 0.073858634 |
| RP11-259O2 | 0.073838143 |
| KRT71 | 0.073832152 |
| RN7SL310P | 0.073831569 |

| | |
|---|---|
| PCDHB5 | 0.073821827 |
| RP11-766F14 | 0.073819928 |
| RPL7P7 | 0.07380069 |
| AC087762 | 0.073798231 |
| C1QTNF4 | 0.073791761 |
| PXN | 0.07377042 |
| MIR99B | 0.073764387 |
| EPHA2 | 0.073758401 |
| RP11-313E19 | 0.073749726 |
| RPL18A | 0.073749024 |
| RP11-440G5 | 0.073717472 |
| LRFN1 | 0.073717376 |
| GLRX3 | 0.073684854 |
| EPHB3 | 0.073674937 |
| CTD-2538C1 | 0.07365516 |
| MESP1 | 0.073635289 |
| UGT1A3 | 0.073627573 |
| RPS26P3 | 0.073621862 |
| COMMD6 | 0.073616907 |
| LCN15 | 0.073573527 |
| CCDC80 | 0.073571137 |
| CAPG | 0.073533238 |
| RPL10AP6 | 0.073511414 |
| C1orf233 | 0.073491785 |
| RP11-57C13 | 0.073490451 |
| SRI | 0.073484324 |
| AHSG | 0.073468932 |
| C1RL | 0.073434015 |
| RHCE | 0.073427466 |
| RP11-545E8 | 0.073423147 |
| LRIG3 | 0.073408172 |
| TKT | 0.07340499 |
| OR14K1 | 0.073387942 |
| AC110810 | 0.073384197 |
| TSPAN10 | 0.073367519 |
| VTA1P1 | 0.073350657 |
| TBX18 | 0.073349509 |
| RP11-115C21 | 0.073347034 |
| KRT23 | 0.073336132 |
| NDST3 | 0.073315533 |
| AC241585 | 0.073312767 |
| SLC24A3 | 0.073306084 |
| GOLIM4 | 0.073302381 |
| RP11-689K5 | 0.073285143 |
| TLDC1 | 0.073283441 |

| | |
|---|---|
| RP11-114J13 | 0.073277989 |
| ALG3 | 0.073258789 |
| PWWP2B | 0.073252854 |
| EGFLAM | 0.073249318 |
| MIR1245A | 0.073238812 |
| CSPG5 | 0.073234099 |
| TIMM13 | 0.073217057 |
| AC079922 | 0.073213112 |
| TUBB3 | 0.073198656 |
| MRPL41 | 0.073196181 |
| RP11-320G24 | 0.073177377 |
| LINC00961 | 0.073149463 |
| SLC25A2 | 0.073148753 |
| AC090095 | 0.073148732 |
| CTB-1144G6 | 0.073130685 |
| CALN1 | 0.073100279 |
| SCN2B | 0.073091165 |
| KRT14 | 0.073081432 |
| C1orf106 | 0.073080567 |
| LINC00565 | 0.07307302 |
| RP11-54A9 | 0.073063159 |
| TPRXL | 0.073046986 |
| GRM6 | 0.073038711 |
| SIAE | 0.073036174 |
| COX7B | 0.073034351 |
| PCDHB8 | 0.073027582 |
| TBC1D8B | 0.073014294 |
| TIGD5 | 0.072989828 |
| REN | 0.072957042 |
| PDE10A | 0.072945119 |
| GLB1 | 0.072937119 |
| LINC00327 | 0.072931668 |
| RP11-358M14 | 0.072912811 |
| PHYHIP | 0.072911071 |
| NPR3 | 0.072904466 |
| RP13-188A5 | 0.072880949 |
| CTU2 | 0.072858083 |
| RPL13AP25 | 0.072839031 |
| AC090617 | 0.072838223 |
| INTS4P1 | 0.072797578 |
| PPFIBP1 | 0.072790563 |
| KRT4 | 0.072789604 |
| CTD-2288F12 | 0.07275102 |
| STUB1 | 0.072696234 |
| RN7SKP202 | 0.072690069 |

| | |
|---|---|
| HSPB1 | 0.072686931 |
| MRPL4 | 0.072682689 |
| MECR | 0.072676289 |
| ERICH5 | 0.07266637 |
| GOLT1A | 0.072661693 |
| MT1M | 0.07265666 |
| HECTD3 | 0.07265519 |
| CHSY1 | 0.072652957 |
| RP11-335I12 | 0.072648342 |
| MICAL2 | 0.072628901 |
| NFKBIL1 | 0.072614649 |
| NUBP2 | 0.072614128 |
| SLC6A2 | 0.072604229 |
| HLX | 0.072591764 |
| TMEM159 | 0.072588378 |
| SAMD1 | 0.072577215 |
| ANXA8L1 | 0.072567234 |
| LRRC4 | 0.072561828 |
| VGLL1 | 0.072555083 |
| FSCN1P1 | 0.072540846 |
| RRAGA | 0.072521398 |
| RP11-546K22 | 0.072521026 |
| LINC00853 | 0.072514607 |
| OR1C1 | 0.072500355 |
| SURF6 | 0.07249829 |
| RANBP3L | 0.072496442 |
| KDELR2 | 0.07248722 |
| ANKRD53 | 0.072473499 |
| ST3GAL2 | 0.072473247 |
| CDH17 | 0.072458449 |
| NEFH | 0.072424259 |
| ITIH5 | 0.072377559 |
| RP11-197K6 | 0.072369703 |
| SHFM1P1 | 0.072365111 |
| MEDAG | 0.072361856 |
| USP26 | 0.072357879 |
| LRPAP1 | 0.072352963 |
| TEDDM2P | 0.072342817 |
| RP11-373D23 | 0.07233058 |
| SYT16 | 0.072301772 |
| LINC01023 | 0.072300626 |
| AC092765 | 0.072295331 |
| TCEAL7 | 0.072285071 |
| GJB6 | 0.07227693 |
| SLC17A4 | 0.07226301 |

| | |
|---|---|
| AC073508 | 0.07225785 |
| TMEM9 | 0.07224226 |
| KRTAP20-4 | 0.072232905 |
| GRIN2A | 0.072223555 |
| RP11-213L8 | 0.072215006 |
| NFIA-AS2 | 0.072201018 |
| TIMM22 | 0.072190035 |
| CTNNAL1 | 0.072187653 |
| JRKL | 0.072178239 |
| BSG | 0.072168843 |
| GPC2 | 0.072166201 |
| SLC25A37 | 0.072153568 |
| RBMXP4 | 0.072133124 |
| RP5-1070A16 | 0.072131111 |
| RNA5SP133 | 0.072122311 |
| YTHDF3-AS1 | 0.072121064 |
| SNX3 | 0.072117492 |
| HOXC13 | 0.072114329 |
| GNAI1 | 0.072111222 |
| HYAL2 | 0.072109045 |
| FKBP1C | 0.07210391 |
| MTRNR2L6 | 0.072097621 |
| MRPS12 | 0.072091277 |
| YBX1 | 0.072088542 |
| CLIC4 | 0.072085714 |
| RP3-460G2 | 0.072077919 |
| FTH1P20 | 0.072055722 |
| EIF1P5 | 0.072043943 |
| RN7SKP42 | 0.072035096 |
| TSNAX-DISC1 | 0.071992224 |
| CD177P1 | 0.071989478 |
| PCBP3 | 0.071980184 |
| TMEM105 | 0.071978017 |
| STYXL1 | 0.071966658 |
| RGS4 | 0.071931475 |
| MEX3D | 0.071923065 |
| GPR124 | 0.071920814 |
| ZIM2-AS1 | 0.071911791 |
| NUTF2P3 | 0.07190881 |
| UBL4A | 0.071904925 |
| SLC25A6 | 0.071884552 |
| COX6B1P1 | 0.071882704 |
| RP11-369F10 | 0.071881053 |
| CDSN | 0.071875282 |
| RP11-339B21 | 0.071870579 |

| | |
|---|---|
| CEBPB | 0.071857146 |
| SLFN12 | 0.071849331 |
| ACTN4 | 0.071843099 |
| KRT76 | 0.071832473 |
| RP11-419J16 | 0.071830814 |
| CPPED1 | 0.071813665 |
| HYI-AS1 | 0.071807903 |
| AC009229 | 0.071807443 |
| EPS8L1 | 0.071806092 |
| FAM171A1 | 0.071794259 |
| VEGFC | 0.071777589 |
| ANXA13 | 0.071742481 |
| RP11-284H19 | 0.071741253 |
| LPCAT1 | 0.071724117 |
| TMC5 | 0.071720507 |
| PTGR1 | 0.071718963 |
| ALDH1A3 | 0.071711856 |
| RN7SL408P | 0.071696379 |
| COX6B1 | 0.071668226 |
| PROC | 0.071666812 |
| PRPF31 | 0.071665435 |
| OPN1SW | 0.071651816 |
| HNRNPA1P25 | 0.071624149 |
| KRBOX1-AS1 | 0.071624046 |
| RPS10 | 0.071607349 |
| RP11-574H6 | 0.071582727 |
| ITPRIPL2 | 0.071573822 |
| OR5B3 | 0.071572017 |
| RP1L1 | 0.071571314 |
| ENDOG | 0.071550044 |
| LYSMD4 | 0.071537076 |
| FIBCD1 | 0.071531468 |
| TRIM17 | 0.071527103 |
| CDKN2A | 0.071506619 |
| RP11-428J1 | 0.071505744 |
| OR13Z3P | 0.071500538 |
| RAB43 | 0.071489799 |
| VANGL2 | 0.071489297 |
| FNTB | 0.071455156 |
| OR7E161P | 0.0714535 |
| ALDH16A1 | 0.071450635 |
| NAA38 | 0.071443788 |
| SHH | 0.071434555 |
| RNU6-927P | 0.071406866 |
| SEL1L2 | 0.071376445 |

| | |
|---|---|
| PURG | 0.07137439 |
| SCAND1 | 0.071356249 |
| SRA1 | 0.071348602 |
| ROMO1 | 0.071336062 |
| ERGIC3 | 0.071315522 |
| S100A13 | 0.071291242 |
| AC093865 | 0.071266089 |
| ENPEP | 0.071261827 |
| TMEM37 | 0.071259352 |
| PSMB6 | 0.071245026 |
| RP11-499O7 | 0.071240266 |
| RP11-36B6 | 0.071225917 |
| FAM189A2 | 0.071219983 |
| HSD17B14 | 0.071217285 |
| S100A6 | 0.071195293 |
| EIF2B3 | 0.071187851 |
| AC098650 | 0.071174818 |
| C11orf74 | 0.071142696 |
| RP11-235G24 | 0.071140364 |
| RP11-649A16 | 0.071132846 |
| CTC-526N19 | 0.071125076 |
| RPL23AP82 | 0.071114434 |
| LSAMP | 0.071086712 |
| EIF2S2P2 | 0.071082983 |
| RPS3AP25 | 0.071076906 |
| RP11-26O3 | 0.071071129 |
| TMEM147 | 0.071062916 |
| PTCHD4 | 0.071057038 |
| AC090616 | 0.071055849 |
| FAM47C | 0.071017808 |
| TMEM79 | 0.0710175 |
| NKD1 | 0.07100429 |
| P4HB | 0.070980251 |
| TTTY10 | 0.070976382 |
| MVD | 0.070970204 |
| RP11-510P12 | 0.07096118 |
| MAP1S | 0.070923988 |
| RP11-442J21 | 0.070877832 |
| PHYHIPL | 0.070865588 |
| IGHD2OR15-2A | 0.070864205 |
| RP11-667F14 | 0.070863559 |
| SIX3 | 0.070837274 |
| LAGE3 | 0.070793062 |
| MIR8063 | 0.070772206 |
| RP11-174G17 | 0.070766886 |

| | |
|---|---|
| CNGB1 | 0.070739553 |
| ZNF865 | 0.070727998 |
| CRMP1 | 0.070713337 |
| MIR4740 | 0.070703467 |
| NUDT8 | 0.070697113 |
| AC060834 | 0.070689081 |
| CAPNS1 | 0.070674266 |
| RP1-301L19 | 0.070672349 |
| SPSB2 | 0.070650339 |
| KLF16 | 0.070629687 |
| OR2T6 | 0.07060557 |
| BCAS2P3 | 0.070586434 |
| MEIS2 | 0.070571992 |
| LINC01564 | 0.070553801 |
| RP6-43L17 | 0.070545026 |
| HNF1B | 0.070530773 |
| GBAP1 | 0.070514259 |
| CHI3L2 | 0.070509831 |
| RP11-416N13 | 0.070499428 |
| MIR8062 | 0.070496507 |
| SIRT2 | 0.070492546 |
| KCTD15 | 0.070479113 |
| RP11-514O12 | 0.070477017 |
| DEFA4 | 0.070438691 |
| ZNF697 | 0.07043046 |
| SNORD114-9 | 0.070408724 |
| LINC00284 | 0.07038243 |
| RNF175 | 0.070373453 |
| PINLYP | 0.070362494 |
| KANK2 | 0.070357508 |
| CTC-360J11 | 0.070298926 |
| ZNF598 | 0.070273709 |
| RNU1-3 | 0.070257703 |
| TMEM8C | 0.070238944 |
| LINC00452 | 0.070229877 |
| PRKG1 | 0.070225669 |
| RP11-139K1 | 0.07022061 |
| RP11-719J20 | 0.070217145 |
| CTD-2620I22 | 0.070187046 |
| WASF1 | 0.070183621 |
| RP11-634B7 | 0.070181526 |
| RN7SKP263 | 0.070178207 |
| AL354978 | 0.070160533 |
| GBGT1 | 0.070149201 |
| CAPN2 | 0.070143991 |

| | |
|---|---|
| TFAP2B | 0.070143923 |
| PRICKLE2 | 0.070141684 |
| ANKRD34C | 0.070139654 |
| SLC8B1 | 0.070108008 |
| PYGB | 0.070075133 |
| LINC01198 | 0.070073332 |
| WBP1LP2 | 0.070035856 |
| CCDC184 | 0.07001033 |
| PIN4 | 0.070003129 |
| DYTN | 0.069998208 |
| AC079305 | 0.069987222 |
| CTC-455F18 | 0.069976008 |
| FAM210CP | 0.069961198 |
| RP11-78O9 | 0.069959839 |
| AP3S1 | 0.069959165 |
| BACH1-IT2 | 0.069953644 |
| RNU6-977P | 0.069940191 |
| LINC00316 | 0.069919553 |
| PNMAL2 | 0.069916335 |
| MFSD12 | 0.069900148 |
| NME2 | 0.069885003 |
| RP5-881L22 | 0.069872957 |
| PES1 | 0.069865877 |
| PTPRS | 0.069829708 |
| RPS20 | 0.06978578 |
| CHRNA3 | 0.069779694 |
| PTPLA | 0.069744644 |
| NANOGNB | 0.06969026 |
| HS6ST2 | 0.069669115 |
| ERF | 0.069666687 |
| MT-TP | 0.069658184 |
| IDH3G | 0.06960367 |
| RN7SL143P | 0.06960183 |
| CTA-363E6 | 0.069587082 |
| MUC4 | 0.06957807 |
| MICALL1 | 0.069567352 |
| AC018464 | 0.069534711 |
| RN7SKP155 | 0.069534489 |
| ZNF837 | 0.069508894 |
| C4BPB | 0.069497206 |
| ABRACL | 0.069494568 |
| RPS8 | 0.069486288 |
| SAMD4A | 0.06947972 |
| PTP4A2P2 | 0.069479392 |
| AP001885 | 0.069470533 |

| | |
|---|---|
| LGMN | 0.069465041 |
| SPATA12 | 0.069459161 |
| NME4 | 0.069458436 |
| GNG11 | 0.069429523 |
| RN7SL318P | 0.069399643 |
| SHC3 | 0.069397094 |
| C9orf89 | 0.069394489 |
| C1orf105 | 0.069388982 |
| RP13-143G15 | 0.069373256 |
| TNFSF11 | 0.069370506 |
| RP1-170O19 | 0.069346549 |
| HOTAIRM1 | 0.069336347 |
| RP11-328C8 | 0.069334276 |
| UBA52 | 0.069332774 |
| CPQ | 0.069323416 |
| CAPNS2 | 0.069321581 |
| RP11-608O21 | 0.069274931 |
| RP11-424I19 | 0.069273977 |
| RP1-292L20 | 0.069270111 |
| YAP1 | 0.069268139 |
| SPANXN3 | 0.069235563 |
| DTX1 | 0.069208955 |
| SUN5 | 0.069203508 |
| PIWIL1 | 0.069200055 |
| TIMM23 | 0.069196694 |
| BHMT2 | 0.069195801 |
| ZNF454 | 0.069188582 |
| PLD3 | 0.069186089 |
| RAB13 | 0.069146999 |
| PCDHB10 | 0.069144841 |
| FUOM | 0.06914058 |
| NTN4 | 0.06913364 |
| CRTAP | 0.069118346 |
| LINC00547 | 0.069114259 |
| DKKL1P1 | 0.069096607 |
| FHAD1 | 0.069064874 |
| OPRK1 | 0.069064484 |
| KRT15 | 0.069053525 |
| DKK1 | 0.069038156 |
| MFSD10 | 0.069023198 |
| ATP1A1 | 0.069011588 |
| CTD-2541J13 | 0.068995843 |
| AGAP2-AS1 | 0.068988927 |
| RP4-718N17 | 0.068987786 |
| AZIN2 | 0.068957111 |

| | |
|---|---|
| LYPD4 | 0.068955749 |
| MIR22HG | 0.06895259 |
| C3orf30 | 0.06894235 |
| AC009228 | 0.068942261 |
| RP11-556E13 | 0.068941817 |
| HIST1H3A | 0.06894123 |
| MMP10 | 0.06893964 |
| LAMP2 | 0.068938152 |
| RP11-156K13 | 0.068932883 |
| RIPK4 | 0.068930453 |
| HOXB9 | 0.068913035 |
| MRPL55 | 0.068901816 |
| C4orf48 | 0.068900551 |
| PANX3 | 0.068877412 |
| WI2-1896O14 | 0.068867768 |
| DSE | 0.068866335 |
| RP11-8P13 | 0.068853829 |
| AC004699 | 0.068845298 |
| G0S2 | 0.068840511 |
| TBL3 | 0.068837757 |
| SPATS2L | 0.06882758 |
| NPPB | 0.068827355 |
| SLC9A5 | 0.06882652 |
| RP11-85O21 | 0.068805611 |
| RP11-378I6 | 0.068804019 |
| PCDHB2 | 0.068800807 |
| LMAN2 | 0.06879303 |
| RPL3L | 0.068767458 |
| TCHHL1 | 0.068765355 |
| SEC22A | 0.068689472 |
| BHMT | 0.068679006 |
| CALM2 | 0.068673099 |
| DTX2 | 0.068670587 |
| SP6 | 0.068662792 |
| RP1-127D3 | 0.068616081 |
| ERHP2 | 0.068604632 |
| POLR2H | 0.068561261 |
| KRT8P23 | 0.068533848 |
| RPS26P34 | 0.068531492 |
| RNA5SP93 | 0.068531026 |
| CSRP1 | 0.068500031 |
| FOLR3 | 0.068485539 |
| PFN2 | 0.068483621 |
| CTC-573N18 | 0.068470601 |
| SEMA6B | 0.068462522 |

| | |
|---|---|
| CLPSL1 | 0.068451491 |
| PSENEN | 0.068442373 |
| AL139188 | 0.068420786 |
| LINC01080 | 0.068418699 |
| RP5-1093G23 | 0.068417096 |
| RP5-1110E20 | 0.068414531 |
| TSPAN4 | 0.068411506 |
| UGT1A10 | 0.068387136 |
| NPM1P11 | 0.068357113 |
| ALDOA | 0.068354759 |
| CTB-32O4 | 0.06834495 |
| SLC38A10 | 0.068322237 |
| CTSB | 0.068318944 |
| AC092614 | 0.068318496 |
| CLCN1 | 0.068315222 |
| PABPC4L | 0.068312116 |
| FADS2P1 | 0.068296509 |
| RN7SL303P | 0.068295381 |
| RP11-10A14 | 0.068295306 |
| PEX14 | 0.068290682 |
| MYLK2 | 0.068272052 |
| CTD-2129N1 | 0.068269337 |
| PTCHD1 | 0.068252377 |
| MPP6 | 0.068213618 |
| AC109642 | 0.068210446 |
| AL035610 | 0.068202509 |
| ASL | 0.068192631 |
| NALCN-AS1 | 0.068190874 |
| ITGB8 | 0.068187831 |
| CDH23 | 0.068176389 |
| BCRP9 | 0.068167491 |
| MIR3689C | 0.068165598 |
| SMTNL2 | 0.068146446 |
| PRR5 | 0.068133112 |
| S100A5 | 0.068115403 |
| RP11-366L20 | 0.068110899 |
| C7orf57 | 0.068089095 |
| ORM1 | 0.068079296 |
| AC092699 | 0.068075852 |
| ERP27 | 0.068065947 |
| NEU2 | 0.06806278 |
| RN7SL448P | 0.068018677 |
| DCTN3 | 0.068010949 |
| DCHS1 | 0.068007898 |
| SNAI1 | 0.068006893 |

| | |
|---|---|
| RP11-87N24 | 0.06800454 |
| GBE1 | 0.067992157 |
| RPL35 | 0.067992046 |
| PRRG3 | 0.067989081 |
| AC079987 | 0.067955307 |
| RP11-400K9 | 0.067947746 |
| FAM224A | 0.067942503 |
| EGFR | 0.067910424 |
| RNU6-503P | 0.067852856 |
| GDPD5 | 0.06785204 |
| ZNF622 | 0.067834923 |
| DUSP10 | 0.067832169 |
| FOSL2 | 0.067829583 |
| SERINC2 | 0.067828355 |
| RP11-61J19 | 0.067827164 |
| TMEM132B | 0.067790964 |
| RP11-383C5 | 0.067777519 |
| ARFGAP1 | 0.067770529 |
| RP11-141M3 | 0.067764817 |
| TINAGL1 | 0.067745858 |
| APMAP | 0.067727681 |
| CPTP | 0.067713113 |
| SYNGR4 | 0.067709624 |
| MLF1 | 0.06770301 |
| DAW1 | 0.067701191 |
| CTD-3023L14 | 0.067677409 |
| RP11-456O19 | 0.067606476 |
| TMEM54 | 0.067599917 |
| PPP2R3A | 0.067579916 |
| RP11-406O23 | 0.06757687 |
| RPL21P133 | 0.06757369 |
| MYLK | 0.067547766 |
| SNRPGP9 | 0.067510123 |
| SHISA2 | 0.067500954 |
| CTD-2325P2 | 0.067488201 |
| AC068799 | 0.067481725 |
| C9orf92 | 0.067479451 |
| RRP1 | 0.067469734 |
| C10orf90 | 0.067449732 |
| CACNA1A | 0.067438552 |
| FOXC2 | 0.067436468 |
| MCU | 0.067419239 |
| MAD2L2 | 0.067417416 |
| TMEM132A | 0.067414346 |
| CAPSL | 0.067411587 |

| | |
|---|---|
| PPP1R14BP3 | 0.067410495 |
| C17orf64 | 0.067396332 |
| LINC00942 | 0.067385391 |
| MGMT | 0.067382568 |
| AC011738 | 0.067342383 |
| RP4-590F24 | 0.067334118 |
| SCN1A | 0.067320606 |
| RP3-523K23 | 0.06731578 |
| SEC11A | 0.067303625 |
| SYPL2 | 0.067302918 |
| RPL36A | 0.067300239 |
| RBPMS2 | 0.067293969 |
| TMEM220-AS1 | 0.067288406 |
| P2RX6P | 0.067268844 |
| LINC00844 | 0.067260091 |
| FGF18 | 0.067229501 |
| RPS28P7 | 0.067216942 |
| RNU6-302P | 0.067209141 |
| THEG | 0.06720523 |
| MRPL36 | 0.067203844 |
| ANPEP | 0.067177443 |
| RP4-541C22 | 0.067163012 |
| RP11-467J12 | 0.067160129 |
| NANOGP9 | 0.067158424 |
| SCAF1 | 0.067148958 |
| RPS19BP1 | 0.067148697 |
| KIAA1210 | 0.067136903 |
| CD58 | 0.067136008 |
| RP11-115J23 | 0.067125476 |
| GPSM1 | 0.06712397 |
| ZSWIM2 | 0.067116069 |
| U82695 | 0.067099193 |
| RP11-54I5 | 0.067099053 |
| UQCRC1 | 0.06707087 |
| RP11-249L21 | 0.067068898 |
| WNT9A | 0.067047893 |
| TLL1 | 0.067045484 |
| RP1-41C23 | 0.067037749 |
| RP4-761I2 | 0.067036897 |
| RXFP1 | 0.067035844 |
| DRAP1 | 0.066983778 |
| LAMP5 | 0.066938623 |
| CTD-2367A17 | 0.066929709 |
| CTB-14A14 | 0.066927699 |
| C4orf47 | 0.066924966 |

| | |
|---|---|
| MIR219A1 | 0.066905187 |
| AC000077 | 0.066872812 |
| DMRTA2 | 0.066865811 |
| EFNA1 | 0.066863419 |
| CDH16 | 0.06685753 |
| RP11-114G11 | 0.066854485 |
| SLC35C1 | 0.066834116 |
| ZMAT3 | 0.066821241 |
| HDGFL1 | 0.066820217 |
| LANCL3 | 0.066810506 |
| IL36RN | 0.066807932 |
| RP11-316O14 | 0.066804304 |
| RBPJP7 | 0.066796686 |
| SLC27A1 | 0.066770187 |
| MBNL1-AS1 | 0.06676695 |
| RNA5SP519 | 0.066757721 |
| NDUFS7 | 0.066719227 |
| AMPD2 | 0.066712362 |
| USP2 | 0.066710334 |
| RP11-181B18 | 0.066708797 |
| MAPK7 | 0.066708436 |
| NTRK1 | 0.066705329 |
| GAPDHP59 | 0.066702913 |
| C11orf39 | 0.066692343 |
| KRT127P | 0.066672389 |
| TUFM | 0.066663615 |
| RP11-201E8 | 0.066647332 |
| LINC00087 | 0.066639503 |
| ARPC4 | 0.0666262 |
| SERTAD4-AS1 | 0.066621902 |
| ENPP6 | 0.066604692 |
| RP11-429A20 | 0.066582768 |
| FAM25BP | 0.06658228 |
| RFPL3 | 0.066569052 |
| PLTP | 0.066541991 |
| KCNJ6 | 0.066538931 |
| RNU6-529P | 0.066528647 |
| C4orf17 | 0.066520929 |
| MTND1P17 | 0.066514089 |
| EGFL7 | 0.066506511 |
| NR1H2 | 0.066502548 |
| YBX1P6 | 0.066491844 |
| RP11-746M1 | 0.066478926 |
| RPS15AP17 | 0.0664524 |
| RP11-568K15 | 0.066442341 |

| | |
|---|---|
| PODN | 0.066441931 |
| AHRR | 0.06642647 |
| RHOD | 0.066417595 |
| HCAR2 | 0.066405669 |
| ETV1 | 0.066393222 |
| GOLGA6L2 | 0.066378082 |
| SLC35E4 | 0.066376645 |
| RP5-884M6 | 0.066370723 |
| TPST2 | 0.066341462 |
| TRPC4 | 0.066341145 |
| FURIN | 0.066339992 |
| NCCRP1 | 0.066305091 |
| RP11-571O6 | 0.066298123 |
| MYL6P4 | 0.066281221 |
| FAM131B | 0.066280938 |
| SIM2 | 0.066244285 |
| COMMD2 | 0.066224768 |
| TCTEX1D4 | 0.066210785 |
| TCP10L | 0.066184859 |
| LINC01423 | 0.066174651 |
| ITGAV | 0.066156486 |
| SNCAIP | 0.066136299 |
| SMCO4 | 0.066115684 |
| GLI2 | 0.066110438 |
| ETFB | 0.066099646 |
| IGF2BP1 | 0.06608998 |
| SNORD114-12 | 0.066074711 |
| KIF1C | 0.066058993 |
| PDF | 0.0660533 |
| AC011242 | 0.066028687 |
| KRT8P50 | 0.066013132 |
| HFE | 0.066000182 |
| LINC00636 | 0.065990444 |
| RP11-474O21 | 0.065949765 |
| ADAMTSL5 | 0.065919694 |
| STX1B | 0.065905256 |
| TDRP | 0.065894875 |
| TDRD1 | 0.065876726 |
| FAM86FP | 0.065844045 |
| AC012075 | 0.065835998 |
| RP11-710C12 | 0.065835106 |
| CGB7 | 0.065803266 |
| VPS37B | 0.06579314 |
| CHRD | 0.06579167 |
| PLCD3 | 0.065779149 |

| | |
|---|---|
| MIR485 | 0.065775118 |
| UQCC3 | 0.06577459 |
| TTTY9B | 0.065766103 |
| RP11-686D22 | 0.065737844 |
| KXD1 | 0.065719388 |
| PGM1 | 0.065710067 |
| RPL7P32 | 0.065708007 |
| AC008166 | 0.065690585 |
| RPL21P119 | 0.065689732 |
| RP11-29G8 | 0.065682977 |
| RP11-770E5 | 0.065680281 |
| REREP1Y | 0.065677326 |
| GLP1R | 0.065673251 |
| RN7SKP36 | 0.065669289 |
| RN7SKP113 | 0.065658488 |
| C16orf62 | 0.065649041 |
| APOBEC1 | 0.065638767 |
| RPL10 | 0.065635514 |
| C11orf88 | 0.065606434 |
| GNL1 | 0.06560342 |
| ATP1B3 | 0.06560089 |
| AC005523 | 0.065598962 |
| DBET | 0.065598871 |
| LINC01139 | 0.065597341 |
| RP11-517I3 | 0.065576516 |
| MT2P1 | 0.065568554 |
| FUNDC2 | 0.065565353 |
| P3H2 | 0.065557446 |
| MEGF8 | 0.065554469 |
| RP11-364M6 | 0.065552339 |
| UBE2Q2P6 | 0.065526877 |
| LINC00229 | 0.06552482 |
| SNORD113-4 | 0.065519605 |
| RASAL1 | 0.065500017 |
| P2RY1 | 0.065482336 |
| OR7E36P | 0.06548122 |
| TMEM200A | 0.065480308 |
| LIMK2 | 0.065460437 |
| AP000474 | 0.065447258 |
| RP11-123B3 | 0.0654261 |
| DCTPP1 | 0.065422612 |
| PSMA6P3 | 0.06540374 |
| MMP7 | 0.065354888 |
| P2RY11 | 0.065341325 |
| MIR4322 | 0.065339788 |

| | |
|---|---|
| ZNF90P3 | 0.065311089 |
| VCY1B | 0.065280036 |
| PPDPF | 0.065258493 |
| FAT2 | 0.065258196 |
| TUFT1 | 0.065250828 |
| GAPDHP2 | 0.065214486 |
| NAGLU | 0.065214454 |
| PLA2G2A | 0.065212013 |
| RSU1P2 | 0.065166785 |
| STK38L | 0.065152498 |
| ETV2 | 0.065152014 |
| LINC01272 | 0.065138004 |
| TFPT | 0.065132534 |
| AC003075 | 0.06512757 |
| PGK1 | 0.065122822 |
| EIF3KP1 | 0.065122517 |
| ZP4 | 0.065111829 |
| USP2-AS1 | 0.065107688 |
| ITGB3 | 0.065099379 |
| FOLH1 | 0.065084551 |
| RNU6-83P | 0.065082455 |
| RP11-792A8 | 0.065070131 |
| SYNDIG1L | 0.065066688 |
| LHX9 | 0.065053568 |
| RELB | 0.06503212 |
| LMO7-AS1 | 0.065028717 |
| FAM173A | 0.06501772 |
| LRP4 | 0.06495835 |
| SI | 0.064956208 |
| RP11-425D10 | 0.064951633 |
| VN1R10P | 0.064947063 |
| SNORA11B | 0.064942024 |
| RP11-567G11 | 0.064941572 |
| TMEM200C | 0.064928777 |
| ATP2B2 | 0.064914272 |
| RNU4-6P | 0.064909772 |
| ZNF556 | 0.064898642 |
| MT1H | 0.06489696 |
| AL391384 | 0.064892954 |
| EVPLL | 0.064875916 |
| LINC01444 | 0.064871014 |
| CALM1P1 | 0.064859132 |
| RPL13AP20 | 0.064855071 |
| DVL1 | 0.064834716 |
| PSG8 | 0.064829748 |

| | |
|---|---|
| RP11-414C23 | 0.064824395 |
| BCAR1P2 | 0.064813519 |
| RNU6-238P | 0.064806237 |
| PPIAP11 | 0.064802976 |
| G6PD | 0.064800352 |
| MGC32805 | 0.064787156 |
| CHCHD6 | 0.064782937 |
| QPCTL | 0.064769579 |
| DDX55P1 | 0.064769442 |
| MIR8053 | 0.064766511 |
| CLEC18A | 0.064764961 |
| CUEDC2 | 0.064742359 |
| MICA | 0.06473901 |
| RNF216P1 | 0.064720125 |
| RP1-231P7P | 0.064718602 |
| RP11-414H17 | 0.064716114 |
| ARHGEF17 | 0.064695641 |
| ADCY8 | 0.064691232 |
| AC007403 | 0.064670226 |
| GADD45A | 0.0646655 |
| TMEM132D | 0.064658332 |
| SCG2 | 0.06465616 |
| RPS5 | 0.06465229 |
| RN7SKP28 | 0.064650688 |
| RP11-729I10 | 0.06464543 |
| OSR1 | 0.064638819 |
| RIMBP2 | 0.064619016 |
| SOBP | 0.064595636 |
| EPPK1 | 0.064583859 |
| GALNS | 0.064582966 |
| CoTC_ribozyme | 0.064581723 |
| RAB11FIP5 | 0.064573068 |
| B9D2 | 0.064569179 |
| AC017060 | 0.064563148 |
| FABP5P10 | 0.064549333 |
| RP11-642D21 | 0.064536174 |
| ARMCX2 | 0.064532117 |
| RN7SL269P | 0.064526187 |
| KRT16P4 | 0.064520322 |
| RP11-315O6 | 0.064467488 |
| HOXA-AS3 | 0.064449891 |
| RP11-305M3 | 0.064447441 |
| RP5-933B4 | 0.064435725 |
| HOXA10-AS | 0.064434151 |
| CABP7 | 0.064429489 |

| | |
|---|---|
| LINC01193 | 0.06441715 |
| CFAP58 | 0.06441084 |
| CAHM | 0.064404044 |
| SNHG9 | 0.064381859 |
| FABP3 | 0.064377859 |
| ORM2 | 0.064363278 |
| LINC00456 | 0.064353766 |
| USH1G | 0.064348652 |
| HIST2H3D | 0.064334386 |
| CTD-2382E5 | 0.064324921 |
| HMGN2P26 | 0.064319735 |
| NUTM2F | 0.064309308 |
| FAM114A1 | 0.064303195 |
| CTC-806A22 | 0.064300416 |
| DGAT1 | 0.064289706 |
| MAP1B | 0.064281595 |
| LINC01563 | 0.064252374 |
| RNF7 | 0.064245649 |
| LINC00900 | 0.06423843 |
| LPAR3 | 0.0642379 |
| AC008132 | 0.064195489 |
| RP1-142L7 | 0.064182864 |
| RRAGC | 0.064156268 |
| LINC00615 | 0.064144488 |
| ALDH7A1 | 0.064142202 |
| JAG1 | 0.064138928 |
| NDUFB2 | 0.06412155 |
| GNG12 | 0.064111894 |
| GSN | 0.064107187 |
| COL26A1 | 0.064103912 |
| EREG | 0.064098124 |
| ZBED1 | 0.064092284 |
| CTD-2337A12 | 0.064087176 |
| TBC1D3D | 0.064077346 |
| RNU2-64P | 0.064066752 |
| BUD31 | 0.064036728 |
| RPS4XP7 | 0.064030692 |
| KCNK2 | 0.063995889 |
| PTHLH | 0.063994996 |
| RP11-215A19 | 0.063992483 |
| MUC15 | 0.063952183 |
| AL365502 | 0.063946836 |
| SGMS2 | 0.063939421 |
| MIR6072 | 0.063939202 |
| GCK | 0.063939129 |

| | |
|---|---|
| USP9Y | 0.063931525 |
| LINC00705 | 0.063930372 |
| ZNF536 | 0.063928635 |
| GNB1L | 0.063913243 |
| KCNS1 | 0.063900254 |
| ZNF502 | 0.063894359 |
| AC073109 | 0.063886816 |
| NME7 | 0.063881037 |
| UGT2B7 | 0.063876593 |
| IFNA1 | 0.063868033 |
| LRRTM1 | 0.06386728 |
| DLX1 | 0.063860687 |
| WBSCR22 | 0.063851734 |
| RNF24 | 0.06385019 |
| RARRES1 | 0.063843324 |
| PCDHGC3 | 0.063842152 |
| HSPG2 | 0.063837538 |
| RP11-564P9 | 0.063816796 |
| HOXC8 | 0.063811472 |
| BDH2 | 0.063811472 |
| CDH24 | 0.063803173 |
| TM4SF1 | 0.063794652 |
| TMEM38A | 0.063793852 |
| RP11-443P15 | 0.063785447 |
| EMC6 | 0.063783545 |
| GNB2 | 0.063779861 |
| AP001476 | 0.063777608 |
| RN7SKP97 | 0.063759647 |
| RP3-454B23 | 0.063753004 |
| PLD1 | 0.063714128 |
| RP11-451M19 | 0.063710724 |
| LINC01266 | 0.063708811 |
| TLN2 | 0.063650237 |
| OMD | 0.063649437 |
| ASPH | 0.063636562 |
| RP11-290L7 | 0.063636409 |
| RP11-1059L18 | 0.063631337 |
| ADCYAP1R1 | 0.063621231 |
| IL17RD | 0.063604262 |
| TSPAN2 | 0.063581899 |
| UQCRH | 0.063539162 |
| CD164L2 | 0.06353853 |
| CORO1C | 0.063531255 |
| RP5-1009N12 | 0.063503006 |
| SCHIP1 | 0.063486603 |

| | |
|---|---|
| SPP2 | 0.063477929 |
| SAMD11 | 0.063474731 |
| RPS26P43 | 0.063445429 |
| LINC00702 | 0.063444069 |
| OR3A2 | 0.063440058 |
| NWD1 | 0.063425795 |
| MIR4634 | 0.063421586 |
| SNORD114-24 | 0.063419213 |
| RP11-195F19 | 0.063408607 |
| PGAM1P5 | 0.063374307 |
| TPM4P1 | 0.06337202 |
| PNMAL1 | 0.06336708 |
| CHODL | 0.063360667 |
| CPNE2 | 0.063357424 |
| LUZP2 | 0.063356436 |
| MTHFD2P5 | 0.063356282 |
| SIRT4 | 0.063354224 |
| RP11-302L19 | 0.063349429 |
| AC007969 | 0.063346633 |
| SP8 | 0.063334739 |
| TMSB10P1 | 0.063327121 |
| RP11-865I6 | 0.063325986 |
| RP11-386B13 | 0.063314245 |
| RP11-295K3 | 0.063307819 |
| SCTR | 0.06330315 |
| CFI | 0.063300435 |
| RNU6-358P | 0.063280909 |
| RP11-793I11 | 0.063271381 |
| YWHAQP9 | 0.063265346 |
| PTS | 0.063259261 |
| DERA | 0.063254963 |
| CTD-2175A23 | 0.063253665 |
| JMJD4 | 0.063245437 |
| DGUOK | 0.063236097 |
| CTD-2201E9 | 0.06323147 |
| RP11-295G20 | 0.063214496 |
| CADM1 | 0.063209789 |
| RN7SL443P | 0.063195775 |
| LGALS3 | 0.063180914 |
| ASPG | 0.063172581 |
| RPL36AP40 | 0.063167337 |
| NEXN | 0.063155461 |
| MEA1 | 0.063145638 |
| KCNQ3 | 0.063136242 |
| BACH1-AS1 | 0.063134903 |

| | |
|---|---|
| UTF1 | 0.063128677 |
| FOXC1 | 0.063122883 |
| ZNF667-AS1 | 0.063110808 |
| SERPINA7 | 0.063096551 |
| RP11-61K9 | 0.063077449 |
| RNU6-961P | 0.063073755 |
| RP11-20J15 | 0.063068578 |
| SOX9 | 0.063063253 |
| HNRNPA1P73 | 0.063060627 |
| ARL4A | 0.063054024 |
| SLC27A5 | 0.063053913 |
| FGF11 | 0.063044478 |
| HEYL | 0.063025074 |
| AC016734 | 0.063011343 |
| DBI | 0.063004627 |
| GLG1 | 0.063002357 |
| RP11-563H6 | 0.062999148 |
| A4GALT | 0.062995659 |
| RNF187 | 0.062987026 |
| PPIE | 0.062984458 |
| SLC16A4 | 0.06297644 |
| RPS16 | 0.062965165 |
| EPHX4 | 0.062954076 |
| FAM47E-STBD1 | 0.062952122 |
| SNORD114-2 | 0.062946595 |
| PDE3A | 0.062942726 |
| RNU6-760P | 0.062937026 |
| RP5-1136G13 | 0.062915581 |
| UQCRQ | 0.062893775 |
| MAP1A | 0.062892194 |
| PI3 | 0.062878979 |
| TEAD3 | 0.062866463 |
| ZNF843 | 0.062858946 |
| HOXA5 | 0.062858109 |
| RP11-128A17 | 0.062853729 |
| CREB3L1 | 0.062829512 |
| KIAA1614 | 0.062829084 |
| SOX21-AS1 | 0.062825346 |
| GAS2L1 | 0.062824619 |
| AP001439 | 0.062811734 |
| INO80B | 0.06280099 |
| RNU6-34P | 0.062781045 |
| CLUH | 0.062778626 |
| RP11-62H20 | 0.062778579 |
| OR7E39P | 0.062770044 |

| | |
|---|---|
| SPRR2D | 0.062759602 |
| AC007229 | 0.062754504 |
| TAAR7P | 0.062747655 |
| MINOS1P3 | 0.062725708 |
| RP11-674P19 | 0.062705804 |
| RP11-351J23 | 0.062698861 |
| RPS15 | 0.062695739 |
| LYVE1 | 0.062692892 |
| TMEM8A | 0.062690529 |
| RNA5SP22 | 0.062689387 |
| AC015933 | 0.062682532 |
| C8orf48 | 0.062679025 |
| UBTFL3 | 0.062676356 |
| CGB5 | 0.062665471 |
| S100A8 | 0.062653132 |
| RP1-249H1 | 0.06264219 |
| PID1 | 0.062626917 |
| FTH1P3 | 0.062620275 |
| HDHD3 | 0.062619326 |
| FHL1 | 0.062615177 |
| OR6C6 | 0.062614022 |
| DEFB4A | 0.062595889 |
| ARID3C | 0.062577146 |
| WDR1 | 0.062575566 |
| RPL13P12 | 0.062567882 |
| LINC01549 | 0.062567842 |
| CLDN5 | 0.062559863 |
| RP13-81N3 | 0.062532352 |
| TEAD1 | 0.06252282 |
| SPRR2B | 0.062511026 |
| MAGIX | 0.0625068 |
| KNCN | 0.062502303 |
| SHMT2 | 0.062500344 |
| RP11-128A6 | 0.062486624 |
| BLACAT1 | 0.062480745 |
| RP11-387D10 | 0.062477138 |
| RP11-684N3 | 0.062468748 |
| PCSK5 | 0.062457849 |
| CLDN17 | 0.06242899 |
| RPL13 | 0.062422573 |
| NCBP2-AS2 | 0.062421085 |
| pRNA | 0.062396637 |
| CTA-298G8 | 0.06239474 |
| CUTA | 0.062378385 |
| GCNT2 | 0.062348096 |

| | |
|---|---|
| BCAP31 | 0.062342793 |
| ALS2CR11 | 0.06232337 |
| RPS23P4 | 0.062317433 |
| SUPT20HL1 | 0.062309603 |
| TCF15 | 0.062303895 |
| NANOGP1 | 0.062303583 |
| KB-1448A5 | 0.062302465 |
| RP11-296E3 | 0.062277369 |
| SFRP5 | 0.062272105 |
| SH2D7 | 0.062246774 |
| SPATA3 | 0.062242343 |
| SUPT20HL2 | 0.062236011 |
| FLJ43879 | 0.062219228 |
| IFNA2 | 0.062214499 |
| MT1JP | 0.062213418 |
| PENK | 0.062198356 |
| SNORD114-28 | 0.062197734 |
| TMEM182 | 0.06214111 |
| AP004782 | 0.062108652 |
| RPL7 | 0.062092754 |
| RNA5SP64 | 0.062082677 |
| HSD17B1 | 0.06207227 |
| MOGAT2 | 0.062064293 |
| SOX1 | 0.062062498 |
| NRBF2P4 | 0.062045866 |
| BAHCC1 | 0.062010202 |
| SLC35C2 | 0.062003243 |
| ZDHHC8 | 0.061998852 |
| PARD6G | 0.061993438 |
| GGCT | 0.061971744 |
| LINC00609 | 0.061971249 |
| SEC61B | 0.061966237 |
| UCN | 0.061957976 |
| RP11-715J22 | 0.061956023 |
| PCOLCE2 | 0.061928257 |
| RP11-154H12 | 0.061925002 |
| YPEL4 | 0.061883219 |
| RNU1-140P | 0.061871726 |
| RN7SL384P | 0.061856176 |
| ABHD11-AS1 | 0.061855572 |
| RPS2P28 | 0.061853146 |
| AC097461 | 0.061845631 |
| COX5B | 0.061841487 |
| GPC6-AS2 | 0.061835986 |
| NFIA | 0.061815179 |

| | |
|---|---|
| NWD2 | 0.061808364 |
| MIR3119-2 | 0.061804132 |
| RP11-783K16 | 0.061790899 |
| HABP4 | 0.061789671 |
| AC019100 | 0.06177288 |
| NRBP1 | 0.061768945 |
| RP11-427M20 | 0.061766886 |
| RNU6-883P | 0.061761683 |
| DAB2IP | 0.061759567 |
| EXTL1 | 0.061754581 |
| ANO1-AS2 | 0.061744278 |
| RP11-555G19 | 0.061743452 |
| OLIG2 | 0.061736817 |
| NOMO1 | 0.061731138 |
| COL27A1 | 0.061718003 |
| PLS1-AS1 | 0.061712189 |
| CIB1 | 0.06170803 |
| C1orf177 | 0.061705152 |
| TMEM31 | 0.061702318 |
| FCN2 | 0.061692391 |
| LINC00933 | 0.061688513 |
| CRYM | 0.061666903 |
| RP11-256I9 | 0.061666232 |
| RP11-353N4 | 0.061665942 |
| FAM127C | 0.061644511 |
| CCR10 | 0.061619431 |
| RNU6-1276P | 0.061616547 |
| CXADRP1 | 0.061610884 |
| RP11-466L17 | 0.061604909 |
| KRT39 | 0.061591625 |
| IFI27L2 | 0.061579969 |
| CLPSL2 | 0.061578491 |
| ACE2 | 0.061577899 |
| H1FX | 0.06157688 |
| RP1-309H15 | 0.061567751 |
| UGT1A2P | 0.061536597 |
| RP11-540D14 | 0.061525754 |
| CTD-2194D22 | 0.061496071 |
| WASH6P | 0.061495426 |
| MAP6D1 | 0.06149483 |
| AL096700 | 0.061491808 |
| CD99L2 | 0.061467704 |
| RP5-905G11 | 0.061467091 |
| RPN1 | 0.061437805 |
| RP1-205F14P | 0.061437383 |

| | |
|---|---|
| AP000351 | 0.061427731 |
| ROBO3 | 0.061416632 |
| BCHE | 0.06141486 |
| RPL21P42 | 0.061393774 |
| CDA | 0.061393765 |
| CTD-2119L1 | 0.061389338 |
| MYDGF | 0.061369113 |
| RP11-480C22 | 0.061367751 |
| RP1-107N3 | 0.061366227 |
| MIR346 | 0.061340622 |
| CCS | 0.061332591 |
| CBY1 | 0.061317762 |
| RFWD2P1 | 0.061316578 |
| PTRHD1 | 0.061297054 |
| GLYATL1 | 0.061296964 |
| PFKFB4 | 0.061290766 |
| ADA | 0.0612879 |
| SLC6A7 | 0.06127186 |
| AC022538 | 0.061261952 |
| LPHN2 | 0.06126122 |
| NLGN4X | 0.061260291 |
| DMGDH | 0.061250913 |
| CD207 | 0.06124828 |
| AP001596 | 0.06124617 |
| SOX6 | 0.061236772 |
| RP11-333I13 | 0.061235168 |
| SPTLC1P4 | 0.061223699 |
| RP11-394A14 | 0.061222663 |
| TNS4 | 0.061218334 |
| NEUROD1 | 0.061190006 |
| AC005082 | 0.061188469 |
| TOMM40 | 0.061181756 |
| COL9A3 | 0.06117677 |
| RP11-776B3 | 0.06116817 |
| HOXD9 | 0.061163318 |
| SLITRK4 | 0.061138461 |
| BOC | 0.061137587 |
| RNU6ATAC41P | 0.061130424 |
| ADCK5 | 0.061128526 |
| RPL32 | 0.061105716 |
| LRRTM4 | 0.061099054 |
| NES | 0.061092673 |
| C1orf35 | 0.061075742 |
| PMS2P10 | 0.061062682 |
| DGCR6L | 0.061055927 |

| | |
|---|---|
| AP000275 | 0.061042689 |
| MRGPRF-AS1 | 0.061042609 |
| DMRTC2 | 0.06103974 |
| SF3B6 | 0.061034624 |
| RN7SKP62 | 0.061010758 |
| KPTN | 0.061008353 |
| IL1RN | 0.060991366 |
| RN7SL496P | 0.060990961 |
| ARHGEF37 | 0.060987236 |
| GAPDHP51 | 0.060977569 |
| GYPB | 0.060959953 |
| CARHSP1 | 0.060958844 |
| KIAA1462 | 0.060949485 |
| NEDD9 | 0.060943681 |
| RP11-332M4 | 0.060938955 |
| MIR205HG | 0.060930085 |
| AC004009 | 0.06090883 |
| LDOC1 | 0.06087964 |
| RP11-402N8 | 0.060875242 |
| MAP7D2 | 0.06083983 |
| SPRR2E | 0.060833927 |
| CCKAR | 0.060825931 |
| AMPH | 0.060824029 |
| LDHA | 0.060821554 |
| PSMD4 | 0.060807879 |
| CLLU1OS | 0.060798664 |
| UBE2U | 0.06079486 |
| RP11-449O16 | 0.060789895 |
| ACTBP9 | 0.060787194 |
| RP11-108M9 | 0.060782352 |
| ACTR3BP6 | 0.060773481 |
| AL356693 | 0.060771971 |
| EDF1 | 0.060768584 |
| PTN | 0.060743207 |
| HMGN1P14 | 0.060742149 |
| VGLL2 | 0.060729604 |
| ASS1 | 0.060728192 |
| GLTSCR2 | 0.060726145 |
| MTAPP1 | 0.060713196 |
| RN7SL555P | 0.060711488 |
| RAD51B | 0.060702982 |
| CIB4 | 0.060697978 |
| INTS1 | 0.060695 |
| GAD2 | 0.06068876 |
| RP11-793H4 | 0.060687447 |

| | |
|---|---|
| LONP1 | 0.06068646 |
| RP11-395G17 | 0.06067905 |
| RP11-384C12 | 0.0606743 |
| GS1-174L6 | 0.060662959 |
| AL109659 | 0.060651615 |
| CER1 | 0.060634402 |
| C2orf76 | 0.060617341 |
| CDKN2B | 0.060615294 |
| TSPY14P | 0.060612915 |
| PRKAR1B | 0.060610494 |
| LINC01063 | 0.060586651 |
| NCAM1 | 0.060581432 |
| C8orf88 | 0.060577146 |
| ZNF667 | 0.060553803 |
| EVC2 | 0.060544686 |
| LCE1A | 0.060538506 |
| BAALC | 0.060531514 |
| RN7SKP238 | 0.06052971 |
| RPS3AP47 | 0.060525653 |
| CCDC127 | 0.060521578 |
| CLDN7 | 0.060487809 |
| UQCR11 | 0.060478507 |
| PCDHB3 | 0.06047659 |
| WWC2-AS1 | 0.060462474 |
| RP11-2L8 | 0.060457701 |
| ANKRD54P1 | 0.06045253 |
| C11orf73 | 0.060444887 |
| NAALADL2 | 0.0604431 |
| SNORA36B | 0.060378629 |
| HRCT1 | 0.060367824 |
| AC005083 | 0.060366967 |
| NTPCR | 0.060354762 |
| RPL7P10 | 0.060344264 |
| COL4A1 | 0.060341031 |
| INHBB | 0.060330426 |
| RNU7-56P | 0.060307546 |
| IRS4 | 0.060307094 |
| ACTBP7 | 0.060306689 |
| FLJ33581 | 0.060305459 |
| RPL39 | 0.060289642 |
| ZNF705D | 0.06028029 |
| NDUFB11 | 0.060278721 |
| PCNPP3 | 0.060257746 |
| LLGL1 | 0.060253846 |
| ZNF521 | 0.060253678 |

| | |
|---|---|
| ABHD17A | 0.060248431 |
| CACNA1S | 0.06024204 |
| HTR1A | 0.060229113 |
| RNA5SP432 | 0.060221049 |
| CDH22 | 0.060209684 |
| TRADD | 0.060208653 |
| C19orf60 | 0.060207016 |
| TMPRSS13 | 0.060177135 |
| EIF6 | 0.060158809 |
| PLA2G16 | 0.060155441 |
| EMP3 | 0.06015479 |
| KRT84 | 0.060150793 |
| JMJD8 | 0.06014385 |
| PGLYRP4 | 0.060129447 |
| LEPROT | 0.060126454 |
| PCDHB4 | 0.060119551 |
| SCRIB | 0.060099178 |
| IRAK1 | 0.060096685 |
| MIR181A1HG | 0.060081484 |
| IRX1 | 0.060078061 |
| HECW1 | 0.060064681 |
| RNU6ATAC39P | 0.060062284 |
| CTC-535M15 | 0.060048781 |
| EEF1B2P6 | 0.060035883 |
| PAPSS2 | 0.060035771 |
| SH3TC2 | 0.060011547 |
| LECT2 | 0.060010294 |
| AC016682 | 0.059995165 |
| BNIP3L | 0.05998922 |
| PKP2 | 0.059987825 |
| MOGS | 0.059973722 |
| ITPKC | 0.059973647 |
| RN7SKP45 | 0.059961686 |
| SEC14L3 | 0.059959873 |
| NMT2 | 0.059941981 |
| SRGAP2 | 0.059935339 |
| C20orf166-AS1 | 0.059917482 |
| C10orf107 | 0.059915294 |
| FLJ31356 | 0.059907152 |
| ARVCF | 0.05988276 |
| HN1 | 0.059873252 |
| C21orf33 | 0.059855298 |
| CSGALNACT1 | 0.059849102 |
| DRD2 | 0.059831453 |
| RP11-445L6 | 0.059805716 |

| | |
|---|---|
| SURF2 | 0.059800486 |
| PRSS56 | 0.059798314 |
| HECW1-IT1 | 0.059791007 |
| RPL31P28 | 0.05979044 |
| CTD-2353N24 | 0.059780543 |
| SVOPL | 0.059773583 |
| NCKAP5 | 0.059749265 |
| RP11-213H15 | 0.059749228 |
| HSPE1P1 | 0.05973926 |
| GPC4 | 0.059739218 |
| COX7BP2 | 0.059734975 |
| GCSHP2 | 0.05972366 |
| LINC00303 | 0.059718533 |
| RP11-490M8 | 0.059706324 |
| PTPLB | 0.059704928 |
| XXbac-BPG27H4 | 0.059696587 |
| RP11-344F13 | 0.059684371 |
| RP3-340H11 | 0.059681575 |
| RP11-575H3 | 0.059672589 |
| RBMX2 | 0.059656833 |
| DCUN1D5 | 0.05965181 |
| SMIM23 | 0.059642503 |
| RPL36A-HNRNPH2 | 0.059634532 |
| FKBP2 | 0.05962939 |
| SLC17A5 | 0.059618339 |
| RNA5SP45 | 0.059616721 |
| RNU6-433P | 0.05960756 |
| AC007016 | 0.059601682 |
| RP11-461F16 | 0.059592373 |
| PHGDH | 0.059592235 |
| POFUT2 | 0.059585853 |
| TTC23 | 0.059585798 |
| MIR663B | 0.059568848 |
| NINJ1 | 0.059568439 |
| TMEM110 | 0.059564532 |
| ZFYVE21 | 0.05956109 |
| RP11-325D8 | 0.059555989 |
| CP | 0.059549201 |
| EXT1 | 0.059533963 |
| FAM43A | 0.059525479 |
| C19orf48 | 0.059521497 |
| TSSC2 | 0.059500659 |
| SCARA3 | 0.059473309 |
| BTBD16 | 0.059465454 |
| EVC | 0.059464062 |

| | |
|---|---|
| AC007391 | 0.059456873 |
| RP11-348J12 | 0.059417198 |
| RP11-447E20 | 0.059416284 |
| AC005158 | 0.05941618 |
| AC002456 | 0.059408593 |
| PLEKHG5 | 0.059401827 |
| WDR46 | 0.059372579 |
| LINC01208 | 0.059359087 |
| RNA5SP239 | 0.059358416 |
| ACTG1P14 | 0.059346848 |
| BMP2 | 0.059341861 |
| CMSS1 | 0.059334996 |
| PLXNB3 | 0.059321451 |
| TULP2 | 0.059319853 |
| MAF | 0.059317004 |
| RP11-632L2 | 0.059315606 |
| TOMM7 | 0.059315293 |
| PRAME | 0.059314162 |
| CCDC89 | 0.059291146 |
| PTPRZ1 | 0.059279948 |
| SMIM20 | 0.059231084 |
| RP11-426C22 | 0.059228442 |
| PLB1 | 0.05922794 |
| RP11-227H15 | 0.059207694 |
| FAM132B | 0.059203526 |
| RP11-91A15 | 0.059180411 |
| LINC01254 | 0.059172871 |
| TGFB3 | 0.059160644 |
| SPRY1 | 0.059152848 |
| PNLIPRP2 | 0.059138068 |
| RPS2P1 | 0.059131057 |
| PRRG1 | 0.059095692 |
| SMIM12 | 0.059090483 |
| RPPH1 | 0.059085806 |
| BTBD2 | 0.059084622 |
| TMC7 | 0.059082873 |
| RNF144B | 0.059067524 |
| SLC22A17 | 0.059064212 |
| C19orf52 | 0.059053867 |
| RP11-643C9 | 0.059037135 |
| PROCR | 0.059033662 |
| EEPD1 | 0.059029643 |
| OR7G3 | 0.059027618 |
| RP11-182L21 | 0.05900927 |
| CTB-109A12 | 0.058992988 |

| | |
|---|---|
| LDLRAD2 | 0.058990181 |
| SMOC1 | 0.05897591 |
| NTF4 | 0.05897251 |
| SLC6A14 | 0.058969757 |
| ENTPD3 | 0.058952784 |
| CTC-277H1 | 0.058947946 |
| SVIL | 0.058945956 |
| RNA5SP289 | 0.058939355 |
| LINC00634 | 0.058928129 |
| RP11-799N11 | 0.058924419 |
| FMO4 | 0.058892037 |
| AC091939 | 0.058891494 |
| SHISA4 | 0.058861561 |
| ST3GAL4-AS1 | 0.058858305 |
| ALDH1L1 | 0.058856519 |
| RP11-440G9 | 0.058836943 |
| PPP2R1A | 0.058836072 |
| BHLHA9 | 0.058834776 |
| C1orf198 | 0.058789856 |
| ACADS | 0.05878046 |
| MPV17 | 0.058768199 |
| PRR5L | 0.058765594 |
| AC004946 | 0.058755013 |
| DOCK5 | 0.058747026 |
| KB-1410C5 | 0.05874431 |
| SERPINB10 | 0.058733407 |
| AC022537 | 0.058712357 |
| CETN4P | 0.058704875 |
| PRR32 | 0.058704503 |
| POMGNT2 | 0.058698428 |
| OR2T27 | 0.058697797 |
| AL138714 | 0.058696193 |
| RP1-32I10 | 0.058695771 |
| CHAC1 | 0.058687209 |
| EXOSC4 | 0.05866946 |
| GGH | 0.05866559 |
| FCGBP | 0.058661236 |
| CTD-2207P18 | 0.058657111 |
| CAMTA1 | 0.05864464 |
| RPL5P34 | 0.058642947 |
| TRIM54 | 0.058592929 |
| R3HDM4 | 0.058590293 |
| PGBD4P7 | 0.058569476 |
| TTTY15 | 0.058552336 |
| NDP | 0.058551616 |

| | |
|---|---|
| RP11-301O19 | 0.058548636 |
| DDX3P1 | 0.058528753 |
| SRD5A1P1 | 0.058522214 |
| AC023157 | 0.058518136 |
| CTD-2006M22 | 0.058494216 |
| RP11-436K8 | 0.05849201 |
| CENPP | 0.058476633 |
| C5orf38 | 0.058473767 |
| OR2W3 | 0.058472262 |
| TBC1D26 | 0.058470143 |
| GPI | 0.058460948 |
| RNU7-159P | 0.058441783 |
| IL1RAPL2 | 0.058437791 |
| RP11-315I14 | 0.058427017 |
| DGCR14 | 0.058427012 |
| TMEM246 | 0.058425319 |
| TYRP1 | 0.058422932 |
| CD59 | 0.05840703 |
| RBM17P3 | 0.058390125 |
| B3GNT8 | 0.058381428 |
| RP11-817J15 | 0.058374149 |
| UROC1 | 0.058368021 |
| SEPHS2P1 | 0.058359766 |
| C12orf57 | 0.058358097 |
| TRPC6P | 0.05834669 |
| AP5S1 | 0.058346153 |
| HEXB | 0.058346078 |
| AZGP1 | 0.058342673 |
| AP5Z1 | 0.058342338 |
| LINC00911 | 0.058324188 |
| NBEAP1 | 0.05831452 |
| RN7SL616P | 0.058297017 |
| MRPS15 | 0.058293406 |
| EVI5L | 0.058289629 |
| EPHX1 | 0.058283192 |
| TFDP3 | 0.058281266 |
| NKAIN2 | 0.058272105 |
| SSNA1 | 0.05823104 |
| ACAD9 | 0.058228082 |
| ALKBH3 | 0.058224249 |
| MAP3K13 | 0.058179299 |
| AP000697 | 0.058177623 |
| PYCRL | 0.058168842 |
| RP11-64K10 | 0.058156342 |
| C11orf53 | 0.058153757 |

| | |
|---|---|
| TEX26 | 0.058152043 |
| RNU6-1216P | 0.058151267 |
| MAMDC2 | 0.058148897 |
| RPL17P41 | 0.058135689 |
| SNORD114-6 | 0.058134166 |
| RP11-297J22 | 0.058132632 |
| DGKB | 0.05813213 |
| PLEKHF1 | 0.058116952 |
| NBAT1 | 0.058116587 |
| PWP2 | 0.058112353 |
| RP11-1152H14 | 0.058108063 |
| MYBPHL | 0.058106688 |
| RP11-360O19 | 0.058098841 |
| DGCR2 | 0.058089471 |
| PHF23 | 0.058087425 |
| RN7SKP181 | 0.058084984 |
| CTB-118P15 | 0.058051988 |
| PACSIN3 | 0.05805187 |
| MGP | 0.058050474 |
| LINC01218 | 0.058037073 |
| EVX1 | 0.058034632 |
| RP11-331K21 | 0.058030055 |
| ROM1 | 0.058018529 |
| LMO1 | 0.058010986 |
| PFN1 | 0.057994844 |
| AP000230 | 0.057993038 |
| AC004231 | 0.05798048 |
| APOBR | 0.057962768 |
| DPY19L1 | 0.057958656 |
| AC005703 | 0.057957668 |
| TSPY17P | 0.05793959 |
| SPANXC | 0.057936817 |
| ELSPBP1 | 0.057936804 |
| RP11-790J24 | 0.057909387 |
| CLEC2A | 0.057907344 |
| RGS2 | 0.057906803 |
| POMC | 0.057902492 |
| RP11-556O5 | 0.057894042 |
| RIN3 | 0.057880923 |
| POP7 | 0.057880104 |
| AC013275 | 0.057877009 |
| FAM43B | 0.057870467 |
| RPS27AP2 | 0.057854391 |
| TMEM174 | 0.057848694 |
| AP4M1 | 0.057848084 |

| | |
|---|---|
| CYYR1-AS1 | 0.057837227 |
| FOXRED1 | 0.057834186 |
| KRT18P53 | 0.057810355 |
| OR4F3 | 0.057805576 |
| S1PR2 | 0.057801924 |
| KCND2 | 0.057795989 |
| SSBP4 | 0.057786091 |
| MUC7 | 0.057784808 |
| CDK2AP1 | 0.057781458 |
| HM13 | 0.057769048 |
| RP11-707G18 | 0.057743098 |
| KB-1184D12 | 0.057741532 |
| RP11-381N20 | 0.057740072 |
| CTC-498J12 | 0.057734314 |
| LAD1 | 0.057728767 |
| CD81-AS1 | 0.057727903 |
| RP1-53O8 | 0.057727454 |
| MIR6726 | 0.057707617 |
| RP11-798L4 | 0.057704496 |
| UBALD1 | 0.057696245 |
| RNU1-14P | 0.05769392 |
| CATSPER1 | 0.05767457 |
| SHC2 | 0.057668504 |
| KCNJ2 | 0.057665509 |
| OFD1P1Y | 0.057663166 |
| CDY13P | 0.057663166 |
| RNU6-894P | 0.057663166 |
| MIA | 0.057614588 |
| C14orf166 | 0.057612632 |
| C1QL4 | 0.057611104 |
| RSPH6A | 0.057608921 |
| AC006077 | 0.057595769 |
| PNPLA4P1 | 0.057574579 |
| MIR1267 | 0.05757142 |
| RP11-556K13 | 0.057570621 |
| RP11-334G8 | 0.057554335 |
| AC079756 | 0.05752963 |
| RP11-10B2 | 0.057529002 |
| IFNA5 | 0.057520571 |
| RP11-255M6 | 0.057513909 |
| RN7SL251P | 0.057511224 |
| RP11-813I20 | 0.057510985 |
| ARHGEF19 | 0.057507399 |
| C16orf13 | 0.057503064 |
| WNT7A | 0.057501541 |

| | |
|---|---|
| TNFRSF10C | 0.057495101 |
| MFSD3 | 0.057478561 |
| RP1-144C9 | 0.057474763 |
| RP11-696F12 | 0.057473367 |
| HS6ST1 | 0.057468756 |
| PCDHA7 | 0.057467995 |
| RP11-14C10 | 0.057464406 |
| ATOX1 | 0.057461109 |
| PCDH17 | 0.057447992 |
| PDXK | 0.057417628 |
| FPGS | 0.0574116 |
| MBOAT7 | 0.057383766 |
| RP11-438P9 | 0.057376086 |
| TPT1P4 | 0.057372082 |
| PCDHGA9 | 0.057370141 |
| SRPK3 | 0.057370091 |
| TOP1MT | 0.057357309 |
| RNU1-119P | 0.057339973 |
| RP5-998H6 | 0.057335321 |
| RP11-332E4 | 0.057328912 |
| RP11-322E11 | 0.057312591 |
| DSG3 | 0.057304936 |
| RAB31 | 0.057302125 |
| LRAT | 0.057291672 |
| LCE2A | 0.057270506 |
| WWTR1-IT1 | 0.057258979 |
| AC069368 | 0.057255296 |
| GS1-44D20 | 0.057243073 |
| AC000089 | 0.057222103 |
| RNU7-93P | 0.057219781 |
| RP5-1116H23 | 0.057217899 |
| CTB-111H14 | 0.057217285 |
| NDUFB5 | 0.057215573 |
| RP11-416L21 | 0.057202525 |
| RNU6-308P | 0.057196355 |
| CTD-2014N11 | 0.057193775 |
| ARMCX6 | 0.0571928 |
| RP3-390M24 | 0.057190177 |
| RP1-16A9 | 0.057188647 |
| WDR41 | 0.057182232 |
| CYP4Z2P | 0.057177996 |
| CTD-2337I7 | 0.057175229 |
| OR10R3P | 0.05717207 |
| LINC01220 | 0.057168334 |
| RP3-481F12 | 0.057162151 |

| | |
|---|---|
| TBXA2R | 0.057115252 |
| KCNK1 | 0.057078599 |
| RN7SKP43 | 0.057060554 |
| SLC7A10 | 0.057051353 |
| HBB | 0.057050561 |
| SYCP1 | 0.057047648 |
| RP11-101E5 | 0.057029138 |
| SAMD5 | 0.057018169 |
| S100A4 | 0.05700561 |
| GNA11 | 0.057004047 |
| NDUFA1 | 0.056999098 |
| MIR548H4 | 0.056980762 |
| RNU7-57P | 0.056980762 |
| LINC00479 | 0.056967887 |
| RP11-292B8 | 0.056966943 |
| DHRS3 | 0.056952026 |
| RP11-431K24 | 0.056939962 |
| PRR7 | 0.056932379 |
| EGR1 | 0.05692462 |
| RP3-523C21 | 0.0569071 |
| PRR35 | 0.056904716 |
| AC016710 | 0.056904082 |
| ATP5F1P1 | 0.056903968 |
| RP11-4N23 | 0.0569016 |
| AC000111 | 0.056895642 |
| CD163L1 | 0.056893754 |
| KRT9 | 0.056892449 |
| KCNJ4 | 0.056881423 |
| TRIM51BP | 0.056880654 |
| ELOF1 | 0.05686752 |
| RP11-432I13 | 0.056859956 |
| AC004054 | 0.056858178 |
| COL25A1 | 0.05683838 |
| COX7A2L | 0.056795461 |
| RP11-170M17 | 0.056791201 |
| USB1 | 0.056787107 |
| RP11-763E3 | 0.05678147 |
| KCNS2 | 0.056781189 |
| MRGPRE | 0.056770102 |
| ROR1 | 0.05675758 |
| MLLT11 | 0.056751645 |
| NXF2B | 0.056735933 |
| CPAMD8 | 0.056730528 |
| SPRR2A | 0.056727438 |
| IFITM10 | 0.056724574 |

| | |
|---|---|
| NPM2 | 0.056718419 |
| PIGQ | 0.056710434 |
| STYK1 | 0.056681391 |
| BGLAP | 0.056663802 |
| TMEM133 | 0.056656367 |
| RN7SKP297 | 0.056648159 |
| FXYD1 | 0.056631233 |
| RP11-47P18 | 0.056624585 |
| C2orf71 | 0.056615432 |
| RP11-212D19 | 0.056615101 |
| UPK2 | 0.056606291 |
| MIR4644 | 0.05659432 |
| MRPL12 | 0.056585554 |
| RP11-20A20 | 0.056568026 |
| AC113171 | 0.056566302 |
| IFNA11P | 0.056562866 |
| NEXN-AS1 | 0.056562006 |
| RP11-457I16 | 0.056559696 |
| MYO3A | 0.056544932 |
| ALX3 | 0.05653363 |
| AC011294 | 0.05652264 |
| EIF3B | 0.056505476 |
| CTD-2015A6 | 0.056503488 |
| RNA5SP503 | 0.056502797 |
| FAS-AS1 | 0.056497585 |
| TMEM222 | 0.056495727 |
| HPGDS | 0.056495225 |
| KNDC1 | 0.056493704 |
| RP11-515C16 | 0.056489743 |
| RP11-114H7 | 0.056489701 |
| MID1IP1 | 0.056488787 |
| RP11-150D5 | 0.056485551 |
| PLEKHH3 | 0.05648488 |
| LSM4 | 0.056481996 |
| AC092043 | 0.05648107 |
| RP11-70D24 | 0.056466423 |
| MIR512-1 | 0.056434839 |
| PAQR7 | 0.056431278 |
| SPNS2 | 0.056428766 |
| RP11-361D15 | 0.056426064 |
| CYP1B1 | 0.056391667 |
| PRSS8 | 0.056388374 |
| RPL7P12 | 0.056387632 |
| RNU7-186P | 0.056385476 |
| CLDN23 | 0.056379759 |

| | |
|---|---|
| C16orf45 | 0.05637615 |
| SPAG7 | 0.056371554 |
| AC010127 | 0.056371097 |
| SCUBE3 | 0.056364614 |
| RN7SL679P | 0.05636173 |
| HNRNPA1P74 | 0.056360326 |
| RRAS2 | 0.056352335 |
| AC005682 | 0.056348037 |
| AL050316 | 0.056338791 |
| MT2A | 0.056333748 |
| TIMM50 | 0.056322045 |
| CACNA1C | 0.056315198 |
| RGS6 | 0.056306735 |
| CTC-786C10 | 0.056297246 |
| RN7SL208P | 0.056295396 |
| MRPS17P1 | 0.056290428 |
| RP11-51B23 | 0.056287606 |
| RP13-212L9 | 0.056281399 |
| RRP9 | 0.056260796 |
| EPHX3 | 0.05624993 |
| RN7SL705P | 0.056247248 |
| RP11-55L4 | 0.056245993 |
| RP11-701B6 | 0.056240176 |
| LINC00475 | 0.056232357 |
| AF013593 | 0.056232181 |
| RP11-395D3 | 0.056222069 |
| AKAP12 | 0.056218896 |
| SLC35B2 | 0.056217538 |
| LHCGR | 0.056216427 |
| IL13RA1 | 0.056210245 |
| RP11-557H15 | 0.056200136 |
| CTD-2282P23 | 0.056189656 |
| RP11-598F17 | 0.056184563 |
| RNA5SP410 | 0.056182829 |
| TRAPPC5 | 0.056179304 |
| RN7SL356P | 0.056165231 |
| EXOC3-AS1 | 0.056146744 |
| HEPHL1 | 0.056137341 |
| RP11-74C1 | 0.056137218 |
| KRTAP3-1 | 0.056137032 |
| ANAPC15 | 0.056134037 |
| RP11-626G3 | 0.056094952 |
| RNU6-923P | 0.056094821 |
| RABEP2 | 0.056094668 |
| SGK223 | 0.056087132 |

| | |
|---|---|
| ZBTB17 | 0.056084397 |
| RP4-798P15 | 0.056078369 |
| LINC01468 | 0.056073128 |
| C11orf98 | 0.056069253 |
| RNU6ATAC31P | 0.056068254 |
| AC107983 | 0.056060731 |
| AC004000 | 0.056058983 |
| RP11-742D12 | 0.056052921 |
| RP11-549L6 | 0.056049479 |
| THEGL | 0.05604462 |
| COMTD1 | 0.056035205 |
| RP11-298C3 | 0.056033995 |
| ZNF890P | 0.056029865 |
| MTND4P30 | 0.056027364 |
| NOP16 | 0.056024972 |
| ATP13A4-AS1 | 0.055990903 |
| AC098587 | 0.055990567 |
| DYSF | 0.05598097 |
| CXCL5 | 0.05597891 |
| AC016708 | 0.055972188 |
| RN7SL560P | 0.05597093 |
| SLC35F6 | 0.055955368 |
| OR51E2 | 0.05595524 |
| WASH4P | 0.05594828 |
| COX5BP1 | 0.055946132 |
| OXCT2 | 0.055942526 |
| KRT8P12 | 0.055936503 |
| SNRPD2 | 0.055933768 |
| AC005592 | 0.055932666 |
| ADAMTS3 | 0.055931646 |
| ZNF662 | 0.055930642 |
| DPM2 | 0.055904538 |
| RNU6-382P | 0.055888915 |
| ARSK | 0.055871328 |
| SNORA33 | 0.055852201 |
| LSM3P4 | 0.05585217 |
| ZNF185 | 0.055847252 |
| SDHA | 0.055840759 |
| RP5-968J1 | 0.055813202 |
| MIR539 | 0.055783252 |
| NKX2-2 | 0.055780345 |
| MEGF10 | 0.055733201 |
| ICAM5 | 0.055721276 |
| GOLGA6L6 | 0.055720383 |
| RNU2-57P | 0.055715348 |

| | |
|---|---|
| RPS17 | 0.055715023 |
| SNTB1 | 0.055701274 |
| GCNT6 | 0.05567497 |
| ENOX1-AS1 | 0.055672153 |
| CLTA | 0.055650611 |
| C19orf26 | 0.055645197 |
| NKX2-1 | 0.055640672 |
| MIR649 | 0.055612858 |
| RNF126P1 | 0.055610667 |
| LRRN3 | 0.055608618 |
| RPL41P1 | 0.055608135 |
| RP11-11N5 | 0.055604973 |
| RP11-1398P2 | 0.05559952 |
| ISPD | 0.055587017 |
| LINC00578 | 0.055585902 |
| TPD52L2 | 0.055582738 |
| RP11-109E12 | 0.055575157 |
| ZNF771 | 0.055560486 |
| SIK1 | 0.055546998 |
| CD99P1 | 0.055541136 |
| BEST3 | 0.05552595 |
| RN7SL181P | 0.055520529 |
| PEF1 | 0.055514884 |
| RN7SL401P | 0.055498521 |
| RP11-97N19 | 0.055494368 |
| SCARNA2 | 0.055486216 |
| MIR7855 | 0.055479473 |
| RP11-706C16 | 0.055469886 |
| SOSTDC1 | 0.055469678 |
| BAMBI | 0.055465486 |
| MAMLD1 | 0.055451067 |
| SCRN2 | 0.055437541 |
| AC093897 | 0.055436322 |
| RP11-72K17 | 0.055433786 |
| AC010980 | 0.055429919 |
| DHRS7B | 0.055429057 |
| AC079780 | 0.055426959 |
| RP11-280K24 | 0.055423354 |
| ADAMTS10 | 0.055411549 |
| B4GALNT1 | 0.055409968 |
| ACSF2 | 0.055397037 |
| BHLHE41 | 0.055394488 |
| RP11-34F13 | 0.055390357 |
| CCDC3 | 0.055388851 |
| OR7E89P | 0.055385768 |

| | |
|---|---|
| HOXD1 | 0.055378904 |
| RP11-556O9 | 0.055362769 |
| RP11-451J24 | 0.05536269 |
| CTB-78H18 | 0.055331246 |
| LRRTM3 | 0.055326034 |
| UBE2SP1 | 0.055311169 |
| CTC-552D5 | 0.055271615 |
| PTGER3 | 0.055262073 |
| RP11-344B2 | 0.055261212 |
| HMBS | 0.055254314 |
| TMEM255B | 0.055252454 |
| RP11-247A12 | 0.055245328 |
| LINC01093 | 0.055244515 |
| RP11-120I21 | 0.055244427 |
| AF064858 | 0.055242068 |
| RP11-385F7 | 0.05523636 |
| MEIS3 | 0.055227709 |
| RP11-93O17 | 0.055220273 |
| PARP4P3 | 0.055209224 |
| TRIM10 | 0.05519988 |
| PANX1 | 0.055182218 |
| PPP1R15A | 0.055171353 |
| RP11-439H9 | 0.055165153 |
| RP11-49G2 | 0.055162571 |
| CSNK1G2 | 0.055156115 |
| GRIP1 | 0.055146198 |
| CFL1 | 0.055136114 |
| PHLDA1 | 0.055125862 |
| SOD3 | 0.055125323 |
| RYK | 0.055116132 |
| DSCAM | 0.055114509 |
| F11R | 0.055101973 |
| SMARCA1 | 0.055098494 |
| ESRRG | 0.05509682 |
| WBSCR16 | 0.055092633 |
| FBN3 | 0.055080137 |
| RP11-697E14 | 0.055079287 |
| PLS3-AS1 | 0.055069991 |
| RP11-58C22 | 0.055054207 |
| NDUFV1 | 0.055053952 |
| AP2A1 | 0.055044054 |
| RPL31 | 0.055031365 |
| KRTAP19-5 | 0.05503075 |
| SNRPGP11 | 0.055026922 |
| CTC-293G12 | 0.055022993 |

| | |
|---|---|
| TBX15 | 0.055011978 |
| DMRT2 | 0.055001175 |
| RP11-193J6 | 0.054989601 |
| UBTD2 | 0.054988256 |
| U8 | 0.054981932 |
| RP1-300G12 | 0.054981458 |
| RP11-254I22 | 0.054980737 |
| GS1-57L11 | 0.054971845 |
| LRCH2 | 0.054968181 |
| LINC01377 | 0.054963473 |
| RP11-21L1 | 0.054953321 |
| VCY | 0.054942689 |
| ZNF219 | 0.054940087 |
| ADH1C | 0.054933333 |
| FAM126A | 0.054912904 |
| SPTSSA | 0.054898783 |
| RP11-403P13 | 0.054894898 |
| SPATA16 | 0.054880616 |
| FLJ40288 | 0.054875266 |
| TMEM239 | 0.054871647 |
| TMEM234 | 0.054860213 |
| PALM2-AKAP2 | 0.05483717 |
| GNAT3 | 0.05480262 |
| RNA5SP443 | 0.054800357 |
| RPS20P2 | 0.054790622 |
| DUSP27 | 0.054790602 |
| RP11-665C16 | 0.054773748 |
| TFG | 0.054773028 |
| MIR608 | 0.054764036 |
| SERPINA3 | 0.054758563 |
| RP11-565A3 | 0.054729286 |
| KRTAP1-5 | 0.054720074 |
| ELOVL1 | 0.0547187 |
| TPSAB1 | 0.054712951 |
| RNU7-119P | 0.054701501 |
| LRRC42 | 0.05469148 |
| MIR3913-2 | 0.054689828 |
| LL22NC03-102D1 | 0.054684547 |
| MIR873 | 0.054684407 |
| MIR6075 | 0.054683922 |
| ATOH1 | 0.054675071 |
| AC006288 | 0.054660587 |
| MSN | 0.05465667 |
| DNAJB2 | 0.054650474 |
| RP11-146G7 | 0.054649543 |

| | |
|---|---|
| TLX3 | 0.054646975 |
| RP11-185B14 | 0.054640951 |
| RP11-96H17 | 0.054632428 |
| FAM198A | 0.054630268 |
| ISOC2 | 0.054608109 |
| KRT8P38 | 0.054602354 |
| PCDHA5 | 0.054578107 |
| AL596330 | 0.054556372 |
| RP4-797C5 | 0.054554142 |
| RP11-168P13 | 0.05454885 |
| LINC00506 | 0.054537811 |
| PPP1R12C | 0.054537092 |
| CBLC | 0.054529204 |
| FOLR1 | 0.054528313 |
| PKIA | 0.054526841 |
| OR51B8P | 0.054525921 |
| RP11-717I24 | 0.054524106 |
| KDM4D | 0.054519752 |
| KIFAP3 | 0.054516831 |
| AP000320 | 0.05451674 |
| RNU6-1 | 0.054504696 |
| RP11-245P10 | 0.05449777 |
| HMGB1P36 | 0.05447772 |
| RP11-382D12 | 0.054477399 |
| HPD | 0.054472885 |
| PTGS2 | 0.054470578 |
| RP11-433C9 | 0.054446499 |
| FOXP2 | 0.054446018 |
| DDA1 | 0.054438018 |
| VMP1 | 0.054435209 |
| TFEB | 0.054433237 |
| CTD-3001H11 | 0.05443039 |
| SYNPR | 0.054427348 |
| TOMM40L | 0.054426818 |
| AC002465 | 0.05442249 |
| PSMD9 | 0.054418706 |
| CFDP1 | 0.054415208 |
| OR7E158P | 0.054399802 |
| RP11-589N15 | 0.054391672 |
| ZFY-AS1 | 0.0543903 |
| FAM118B | 0.054374406 |
| SLC25A39P2 | 0.054367392 |
| MAP2K2 | 0.054359857 |
| PGLYRP1 | 0.054358615 |
| C11orf68 | 0.054342553 |

| | |
|---|---|
| RPL7P35 | 0.054336168 |
| GSC | 0.054335558 |
| PAF1 | 0.054334181 |
| IFITM2 | 0.054330627 |
| NFYC | 0.054325939 |
| B3GNT4 | 0.054323036 |
| LINC01239 | 0.054320621 |
| AKR1B1 | 0.054280355 |
| PFKL | 0.054278737 |
| SRXN1 | 0.054276448 |
| GARS | 0.054274439 |
| SC5D | 0.054272857 |
| HMGB1P15 | 0.054254201 |
| RP11-119J18 | 0.054250023 |
| PSMC1P9 | 0.054248251 |
| GJB4 | 0.054244807 |
| RP11-46A10 | 0.054239963 |
| RP11-634H22 | 0.054226883 |
| ARMC10 | 0.054212966 |
| TBC1D25 | 0.054207459 |
| C1QTNF1-AS1 | 0.05420487 |
| TMEM177 | 0.054203589 |
| RP3-470B24 | 0.054203022 |
| RAB11B | 0.05419784 |
| AGRP | 0.054190239 |
| WT1-AS_2 | 0.054189125 |
| ATP6V0E1 | 0.054188389 |
| GNAS | 0.054181728 |
| RP11-483K5 | 0.054180567 |
| RPS20P35 | 0.054163187 |
| VAX2 | 0.054157094 |
| FBXW9 | 0.054156052 |
| OR7E102P | 0.054148575 |
| RP11-459D20 | 0.054140283 |
| ARSA | 0.054133372 |
| RP11-1007G5 | 0.054125095 |
| RP11-479I16 | 0.054120686 |
| TMOD1 | 0.054119325 |
| FRK | 0.054115567 |
| RN7SL815P | 0.05409898 |
| MED10 | 0.054087528 |
| CTA-331P3 | 0.054086878 |
| FABP5P1 | 0.054084396 |
| DIO3OS | 0.054080744 |
| SNORD114-3 | 0.054072854 |

| | |
|---|---|
| SLC39A4 | 0.054065872 |
| TARID | 0.054061628 |
| WDR91 | 0.054049666 |
| RPL7P44 | 0.054038176 |
| PITPNM3 | 0.054037628 |
| RALA | 0.054028679 |
| SGTA | 0.054027395 |
| C20orf85 | 0.054018409 |
| PYGO1 | 0.053991952 |
| NPHP3-AS1 | 0.053989684 |
| DAG1 | 0.053979114 |
| LINC00302 | 0.053968997 |
| RP11-352D3 | 0.053963206 |
| OAZ1 | 0.053960397 |
| HOXA4 | 0.053953348 |
| PLEKHG4B | 0.053951336 |
| RP11-3G21 | 0.053937894 |
| RN7SL418P | 0.053931667 |
| MROH9 | 0.053931295 |
| AC005754 | 0.053923133 |
| AC022819 | 0.053907218 |
| SCN5A | 0.053904092 |
| RPL17P1 | 0.053900463 |
| BBS9 | 0.053884747 |
| C19orf33 | 0.053881603 |
| RNY3P8 | 0.053876503 |
| RP11-995C19 | 0.053872928 |
| DDAH2 | 0.053867407 |
| MYH10 | 0.053848653 |
| AC100757 | 0.053841702 |
| RHBDF1 | 0.05383321 |
| ARPC5 | 0.053819275 |
| RP9P | 0.053813879 |
| NDUFB6 | 0.053801767 |
| RGMA | 0.05378638 |
| RP11-123J14 | 0.053779098 |
| TBX2 | 0.05377544 |
| LINC01583 | 0.053763468 |
| CCL7 | 0.053762467 |
| LGI3 | 0.053759084 |
| RPS6 | 0.053755737 |
| CCT3 | 0.05374662 |
| RP11-336A10 | 0.053730764 |
| SNORD115-1 | 0.053727819 |
| DDOST | 0.053707809 |

| | |
|---|---|
| AC024592 | 0.053705086 |
| SUGT1P | 0.053684687 |
| AC016730 | 0.053671727 |
| DARS-AS1 | 0.053667101 |
| AL022344 | 0.053630353 |
| LRP12 | 0.053621852 |
| CTD-2026G22 | 0.053618988 |
| AC079776 | 0.053606879 |
| RNA5SP305 | 0.053606109 |
| SRF | 0.053595823 |
| AC004012 | 0.053591747 |
| snoMBII-202 | 0.053583543 |
| SLC2A9 | 0.053579041 |
| PEA15 | 0.053571189 |
| AP003900 | 0.053563021 |
| CALML6 | 0.053562919 |
| RHOQ | 0.053561105 |
| LINC00982 | 0.053557435 |
| RP11-59H7 | 0.053550149 |
| FGFBP1 | 0.053518097 |
| RP11-561I11 | 0.053514482 |
| SLC38A11 | 0.05351344 |
| RNU6-806P | 0.05350769 |
| C2CD4B | 0.053504875 |
| AC010969 | 0.053500082 |
| MIR152 | 0.053461575 |
| RP11-93M12 | 0.053456668 |
| ARF4 | 0.053440393 |
| RP11-594C13 | 0.053440362 |
| TSPY11P | 0.05343226 |
| SLC25A18 | 0.053403926 |
| SLC45A2 | 0.053398864 |
| RNU6-229P | 0.053398817 |
| MTCL1P1 | 0.053398054 |
| RP11-538I12 | 0.053395534 |
| ABHD17AP9 | 0.053377261 |
| ACADVL | 0.053366752 |
| THEM7P | 0.053365538 |
| SLC39A14 | 0.053362176 |
| TRIM58 | 0.053358251 |
| ROPN1 | 0.053347027 |
| ALDH1A2 | 0.053334063 |
| SIVA1 | 0.053332537 |
| CEBPE | 0.053304502 |
| PRKRIRP6 | 0.053303491 |

| | |
|---|---|
| SYCE1 | 0.053303346 |
| GAPDHP28 | 0.053289773 |
| RP11-265D20 | 0.053285732 |
| AFAP1 | 0.053285633 |
| RAC1P2 | 0.053278563 |
| LINC00555 | 0.053265477 |
| SNORD28 | 0.053260523 |
| NCAM1-AS1 | 0.053240775 |
| PDGFRA | 0.053230337 |
| RPP40 | 0.053225202 |
| AC060835 | 0.053204203 |
| ABCB5 | 0.053189579 |
| AP001347 | 0.053188903 |
| BOLA3 | 0.05318014 |
| RP11-115A14 | 0.053178982 |
| XPR1 | 0.053172549 |
| CALML3 | 0.05317145 |
| BMS1P19 | 0.053169582 |
| RP11-354K1 | 0.053166569 |
| MAN1A2P1 | 0.053166506 |
| AC098591 | 0.053158793 |
| RP1-161N10 | 0.053141446 |
| LINC00222 | 0.053136424 |
| TMPRSS12 | 0.053134216 |
| FOXCUT | 0.05312539 |
| WIZ | 0.053112304 |
| MED8 | 0.05311089 |
| NTSR1 | 0.053105136 |
| CCHCR1 | 0.053099857 |
| RPS15AP1 | 0.05309541 |
| RP11-541F9 | 0.053067276 |
| GLYATL1P1 | 0.05306723 |
| ALK | 0.053063331 |
| RP11-659P15 | 0.053056303 |
| ANXA4 | 0.053041101 |
| TMEM102 | 0.053038775 |
| GABRR1 | 0.053020268 |
| SUSD5 | 0.05301176 |
| RP11-15J10 | 0.053009855 |
| H2BFM | 0.052993912 |
| PCDH18 | 0.052991387 |
| ZDHHC12 | 0.052976856 |
| SLCO6A1 | 0.052973947 |
| FAM84A | 0.052969135 |
| SLC9A4 | 0.052957164 |

| | |
|---|---|
| DEFA6 | 0.052949201 |
| RNA5SP293 | 0.052938841 |
| PFKFB2 | 0.052938343 |
| AC005281 | 0.052918352 |
| SRD5A1 | 0.052913356 |
| NAT2 | 0.052905076 |
| RPL18AP3 | 0.05289462 |
| AC016747 | 0.052890136 |
| MKRN4P | 0.05287663 |
| BACH1-IT1 | 0.052874057 |
| SLC25A3P1 | 0.052857903 |
| GGT2 | 0.052857615 |
| ITGBL1 | 0.052846525 |
| RP11-84A12 | 0.052841958 |
| PPP4R1-AS1 | 0.052824441 |
| SNORD114-11 | 0.052816772 |
| RP11-227F19 | 0.052811631 |
| KIAA2013 | 0.052800383 |
| PSMB4 | 0.052797927 |
| MIR3648-2 | 0.052786693 |
| GALE | 0.052782597 |
| HTR3C | 0.052780429 |
| GPR85 | 0.052769126 |
| FBLN7 | 0.052757461 |
| PGAM1 | 0.052752251 |
| RP11-205M3 | 0.052750544 |
| CTD-2544H17 | 0.05273129 |
| POLR3D | 0.052729273 |
| TANC1 | 0.052722892 |
| PDE11A | 0.052705365 |
| CPA1 | 0.052704456 |
| CEP170B | 0.052678964 |
| GJB5 | 0.05267602 |
| ZSWIM5P3 | 0.052669425 |
| AC133785 | 0.052662852 |
| CCL20 | 0.052658565 |
| AC114812 | 0.052656146 |
| RP11-277E18 | 0.052649879 |
| PPP2CB | 0.052647837 |
| GS1-184P14 | 0.052638107 |
| SNORA40 | 0.052626487 |
| LINC00622 | 0.052623595 |
| DDX43P3 | 0.052620912 |
| RP11-384O8 | 0.052616438 |
| RP11-619L19 | 0.052598786 |

| | |
|---|---|
| CTD-2336O2 | 0.052585527 |
| PRDM5 | 0.052576932 |
| C14orf1 | 0.052566587 |
| RP11-69H14 | 0.052558703 |
| GOLGA2P3Y | 0.052556328 |
| LINC01587 | 0.052552007 |
| CLCN4 | 0.052548689 |
| RP1-200K18 | 0.052543333 |
| PALM | 0.052531925 |
| LINC01554 | 0.052529217 |
| RP1-130H16 | 0.052527684 |
| TUT1 | 0.05252452 |
| SSBP1 | 0.05251959 |
| LRWD1 | 0.052517301 |
| RN7SKP145 | 0.052505963 |
| LRRC4C | 0.052502538 |
| LPP-AS2 | 0.052500947 |
| IQSEC3P1 | 0.052499826 |
| ARNTL2-AS1 | 0.05249306 |
| TMEM132C | 0.052492735 |
| CTC-573M9 | 0.05249023 |
| RPS4X | 0.052489895 |
| TECRP1 | 0.0524877 |
| MIR125B2 | 0.052485578 |
| RP11-318E3 | 0.052484351 |
| RNU2-1 | 0.052467159 |
| RP3-431A14 | 0.052454626 |
| HPN | 0.052440051 |
| SPDYE8P | 0.052434748 |
| AC145676 | 0.052419469 |
| AC000081 | 0.052406485 |
| LINC01241 | 0.052396213 |
| RPS2P5 | 0.052386914 |
| LINC01013 | 0.052380573 |
| AVPI1 | 0.052379602 |
| PSMB5 | 0.0523791 |
| RASL11B | 0.052370969 |
| NYNRIN | 0.052367434 |
| RPS23P8 | 0.052356606 |
| GRASP | 0.05235508 |
| CLRN3 | 0.052338866 |
| EBPL | 0.05233134 |
| CDK6 | 0.052328177 |
| UPK3B | 0.052316817 |
| ATP5L | 0.052295152 |

| | |
|---|---|
| ITGB1P1 | 0.052292589 |
| SERPINH1P1 | 0.05228871 |
| RPL36AP29 | 0.052280204 |
| CDHR4 | 0.052279449 |
| DOK4 | 0.052274779 |
| PCDHGA3 | 0.05226697 |
| HTR1F | 0.052259002 |
| FTLP2 | 0.052238061 |
| SPINK8 | 0.052227079 |
| ZBTB7C | 0.052225047 |
| RP11-475O6 | 0.052218721 |
| SMPX | 0.05221404 |
| RN7SKP172 | 0.052212826 |
| KLF8 | 0.052191147 |
| SHFM1 | 0.052186236 |
| ADCYAP1 | 0.052183567 |
| ARMC6 | 0.052169491 |
| DNPH1 | 0.052156969 |
| SOX9-AS1 | 0.052155667 |
| MED27 | 0.052148597 |
| RP11-851M3 | 0.052141222 |
| RP11-37O16 | 0.052117079 |
| C3orf22 | 0.052116104 |
| SERINC5 | 0.052114604 |
| TRMT61A | 0.052113563 |
| RN7SL782P | 0.052106549 |
| RP1-95L4 | 0.052105804 |
| MPRIPP1 | 0.052104435 |
| NEGR1-IT1 | 0.0520987 |
| RP11-15K3 | 0.052098156 |
| RAC3 | 0.052094473 |
| CTD-2201G3 | 0.052093177 |
| RPS17P13 | 0.05208548 |
| RAB40C | 0.052079868 |
| MRAP2 | 0.052075772 |
| TCEB1P28 | 0.052042675 |
| PPARG | 0.052039476 |
| MAP4 | 0.052036815 |
| XGY1 | 0.052028328 |
| MYO16-AS1 | 0.052027291 |
| TMEM117 | 0.052025912 |
| DYNLT1 | 0.052021726 |
| HAGLR | 0.052018358 |
| LINC01573 | 0.052010314 |
| RNU4-73P | 0.051993799 |

| | |
|---|---|
| C1orf50 | 0.051988627 |
| RPS10P16 | 0.051986778 |
| ELMOD1 | 0.05198485 |
| S100A7A | 0.051984142 |
| RPL31P1 | 0.051981457 |
| HOXC11 | 0.051981088 |
| RP11-231N3 | 0.051977695 |
| CTA-503F6 | 0.051972113 |
| RP5-1021I20 | 0.051963454 |
| KB-68A7 | 0.051949327 |
| LPAR2 | 0.051948755 |
| TNFSF4 | 0.05192535 |
| NHLH2 | 0.051909596 |
| PSMC4 | 0.051899823 |
| RNU6-614P | 0.051885924 |
| SAA1 | 0.051885897 |
| CCBE1 | 0.051883618 |
| CD81 | 0.051873645 |
| AC010091 | 0.051852777 |
| DEFB123 | 0.051852376 |
| RP11-299L17 | 0.051849683 |
| GPR111 | 0.051835437 |
| USH1C | 0.051824746 |
| RP11-810P12 | 0.051819485 |
| LINC00960 | 0.051817791 |
| RNU6-714P | 0.051812558 |
| TRAPPC1 | 0.051804675 |
| RNU6-1184P | 0.051794489 |
| RP4-715N11 | 0.05179119 |
| RP11-64D24 | 0.051790388 |
| UNC93B5 | 0.05178038 |
| SLC2A1-AS1 | 0.051778757 |
| RP11-800A3 | 0.051768524 |
| AC004461 | 0.051756968 |
| FKSG62 | 0.051748367 |
| ARGFXP1 | 0.051742725 |
| CTD-2315E11 | 0.05172674 |
| UBE2M | 0.051719871 |
| UBE2Q2L | 0.051719235 |
| RP11-115A15 | 0.05171869 |
| RPS11P6 | 0.051718283 |
| LINC01473 | 0.051716057 |
| AC002472 | 0.051699547 |
| TCEB3CL | 0.051688773 |
| ABHD8 | 0.051688204 |

| | |
|---|---|
| RP11-785G17 | 0.051680601 |
| RBP4 | 0.051672334 |
| TDRD10 | 0.051670439 |
| FGF14 | 0.051666678 |
| MIR3197 | 0.051662426 |
| COL17A1 | 0.051648483 |
| LINC01326 | 0.051637964 |
| AC009303 | 0.051629299 |
| OR8G5 | 0.051614527 |
| ATP2A1-AS1 | 0.051602824 |
| RNA5-8S5 | 0.051586465 |
| RP4-682E18 | 0.051582132 |
| ALPK3 | 0.051581632 |
| CH17-437K3 | 0.051580267 |
| TP53TG1 | 0.051565669 |
| PPFIA2 | 0.051554566 |
| ASGR1 | 0.05155125 |
| MIR4263 | 0.051549161 |
| MIR3713 | 0.051546285 |
| AC008914 | 0.051530341 |
| MSL3P1 | 0.051524869 |
| FOXN3-AS1 | 0.051518634 |
| RP11-223A3 | 0.051506579 |
| UBXN1 | 0.05150535 |
| RNU6-365P | 0.051492498 |
| RP11-442N24__B | 0.051489797 |
| METTL9 | 0.051464604 |
| SAT2 | 0.051439096 |
| TNKS2-AS1 | 0.051439069 |
| CLEC2L | 0.051437367 |
| ARTN | 0.051436677 |
| RN7SL58P | 0.051431946 |
| C20orf27 | 0.051431226 |
| CTD-2195M15 | 0.051427592 |
| PLK3 | 0.051426407 |
| DSTNP4 | 0.051404571 |
| TRGV8 | 0.051402817 |
| MIR1302-1 | 0.051385047 |
| AC104651 | 0.051376975 |
| AC003989 | 0.051360281 |
| NPM1P22 | 0.0513493 |
| SOHLH2 | 0.05134466 |
| CACNA2D3 | 0.05133528 |
| CTD-3007L5 | 0.051330083 |
| H1F0 | 0.051325398 |

| | |
|---|---|
| HEBP2 | 0.051309044 |
| ELOVL4 | 0.05130348 |
| CTD-2522B17 | 0.051299921 |
| DEFT1P | 0.051299469 |
| PGBD5 | 0.051295275 |
| AC004069 | 0.051294861 |
| TNNT2 | 0.051287963 |
| MIR933 | 0.051287881 |
| DSC3 | 0.051284485 |
| GS1-72M22 | 0.051280463 |
| TCF3 | 0.051278605 |
| TTTY8 | 0.051273987 |
| MB21D2 | 0.051267721 |
| EXOC3 | 0.051265953 |
| RP11-463H12 | 0.05126148 |
| GSTO1 | 0.051252502 |
| VENTXP1 | 0.05125159 |
| RNU6-1326P | 0.051248566 |
| DPP10-AS2 | 0.051248566 |
| UNC93B2 | 0.051244081 |
| ABCC6P2 | 0.05124338 |
| SMIM10 | 0.05122824 |
| PGAM4 | 0.051211664 |
| RP3-333A15 | 0.051205326 |
| LINC00552 | 0.051199564 |
| MTRNR2L1 | 0.051187465 |
| ADTRP | 0.051164349 |
| RP11-21B23 | 0.051162099 |
| IBA57 | 0.051159363 |
| RBP2 | 0.051159035 |
| RNU1-131P | 0.051154341 |
| LINC01350 | 0.05114669 |
| LINC00602 | 0.05114203 |
| PRSS48 | 0.051124371 |
| DLGAP4 | 0.051113016 |
| CCDC33 | 0.05111238 |
| SALRNA3 | 0.051111377 |
| ZNF350-AS1 | 0.051109017 |
| HDHD1P1 | 0.051083901 |
| CTU1 | 0.051083806 |
| TSC22D1 | 0.051081387 |
| FDXR | 0.051079973 |
| SNCB | 0.051067576 |
| RP11-138M12 | 0.051060297 |
| RP11-637O11 | 0.051056659 |

| | |
|---|---|
| DPPA2P4 | 0.051054594 |
| ZP3 | 0.051048325 |
| LINC01455 | 0.051037951 |
| AL121652 | 0.051034053 |
| NOP56P3 | 0.051033231 |
| RN7SKP33 | 0.051030245 |
| RPS2P7 | 0.051026185 |
| RP11-431M3 | 0.051018233 |
| NME1-NME2 | 0.051013179 |
| ARHGAP1 | 0.050999951 |
| AC023469 | 0.050999331 |
| SKI | 0.050957791 |
| POM121L8P | 0.050951288 |
| ZNF213 | 0.050948953 |
| AL451164 | 0.050945523 |
| RP11-183C12 | 0.050931507 |
| MRGPRX3 | 0.050922398 |
| HMX1 | 0.050917965 |
| RP11-315A17 | 0.050913194 |
| ZDHHC14 | 0.050911687 |
| RUVBL2 | 0.050906924 |
| RP11-377K22 | 0.050904582 |
| RP1-244F24 | 0.05090249 |
| KRT40 | 0.050885998 |
| HMGB3P18 | 0.05085716 |
| RP5-1125N11 | 0.050849346 |
| P2RY2 | 0.050845712 |
| RP11-642A1 | 0.050840132 |
| VN1R40P | 0.050839513 |
| NEK6 | 0.050839293 |
| PLN | 0.050833246 |
| MIR934 | 0.050832009 |
| RPL21P132 | 0.050817738 |
| NTN1 | 0.050817599 |
| STAU2-AS1 | 0.050816954 |
| OR8G3P | 0.050816495 |
| LRP4-AS1 | 0.050800189 |
| AC063932 | 0.050799178 |
| RP3-342P20 | 0.05079198 |
| HIST2H2BB | 0.050785368 |
| OR1E2 | 0.050778738 |
| ACTA2-AS1 | 0.050773876 |
| COX4I1 | 0.050773783 |
| RP11-115F18 | 0.050771748 |
| RP11-624M8 | 0.050769287 |

| | |
|---|---|
| APLP2 | 0.050690412 |
| RN7SKP111 | 0.050686982 |
| AC008163 | 0.050677531 |
| MIR1262 | 0.050668869 |
| SPATA2L | 0.050665927 |
| AC006355 | 0.050663003 |
| ATP5E | 0.050646354 |
| AC007559 | 0.050638664 |
| FTCD | 0.050629554 |
| MRPL2 | 0.05062626 |
| UBE2J2 | 0.050623079 |
| RTP1 | 0.050620468 |
| CINP | 0.050615971 |
| AC092309 | 0.050614098 |
| SH3BGRL3 | 0.050612287 |
| PLAT | 0.050609738 |
| RP3-429O6 | 0.050598144 |
| RP3-512B11 | 0.050596045 |
| ARHGEF5 | 0.050594705 |
| AC008598 | 0.050591202 |
| GCM2 | 0.050580456 |
| AC074212 | 0.050568453 |
| BRINP3 | 0.050562504 |
| SNORD72 | 0.05056041 |
| EPO | 0.050556685 |
| DESI2 | 0.050550164 |
| AC073465 | 0.050535998 |
| RNF215 | 0.05052339 |
| NDUFB10 | 0.050509325 |
| RP11-346M5 | 0.050501218 |
| PIN1 | 0.050500264 |
| LRRIQ4 | 0.05049347 |
| PRSS42 | 0.050474332 |
| APOA1BP | 0.050452411 |
| RPS17P11 | 0.05041774 |
| RP5-940F7 | 0.05041176 |
| CTD-2193P3 | 0.050409723 |
| RP11-679B17 | 0.050405451 |
| ZFAND2B | 0.050401841 |
| RP11-724O16 | 0.050396212 |
| IER2 | 0.050388929 |
| SFTPC | 0.050386393 |
| DNAJA1P4 | 0.050377673 |
| ZBTB11-AS1 | 0.050373021 |
| AC010086 | 0.050365686 |

| | |
|---|---|
| ALDH3A1 | 0.050359595 |
| FAM69C | 0.050353752 |
| RN7SL446P | 0.050350912 |
| OR4G11P | 0.050350635 |
| RP11-101E14 | 0.050337491 |
| UNC5D | 0.050333968 |
| HIST1H3C | 0.050315303 |
| HOTAIR | 0.050315203 |
| NFATC4 | 0.0503066 |
| CTSV | 0.050305074 |
| COL14A1 | 0.050296906 |
| LMF2 | 0.05029458 |
| RNU6-1235P | 0.0502932 |
| C15orf43 | 0.050288029 |
| ANP32BP2 | 0.050286527 |
| FAM86B1 | 0.05028645 |
| MIR8059 | 0.050280344 |
| PKDCC | 0.050276514 |
| ST6GALNAC6 | 0.05027043 |
| NEGR1 | 0.050270375 |
| RPL37P3 | 0.050259809 |
| DNAH11 | 0.050255371 |
| TMEM45B | 0.050239694 |
| STX1A | 0.050235862 |
| BTF3L4P1 | 0.050230073 |
| FAM197Y9 | 0.050213545 |
| PRNP | 0.050209665 |
| RPS6KA2-IT1 | 0.05020963 |
| RPS6P12 | 0.050208105 |
| DLL4 | 0.050201925 |
| RP11-437J19 | 0.0501951 |
| CHRNE | 0.050185217 |
| ASNSP1 | 0.050180109 |
| RP11-261B23 | 0.050173581 |
| APOBEC3AP1 | 0.050170412 |
| MDGA1 | 0.050148248 |
| GRM5 | 0.05014446 |
| AC010084 | 0.050141461 |
| SYT5 | 0.050134598 |
| MAB21L1 | 0.050129821 |
| ATP1B1 | 0.050129773 |
| RP11-193E15 | 0.05012348 |
| KCNAB1-AS2 | 0.050115659 |
| RNA5SP94 | 0.050106531 |
| CCDC61 | 0.050099837 |

| | |
|---|---|
| GALK1 | 0.05009113 |
| RN7SKP284 | 0.050075374 |
| HOXD-AS2 | 0.050075222 |
| DAPL1 | 0.050058871 |
| HPDL | 0.050051667 |
| RPL39P5 | 0.050050381 |
| MTATP6P3 | 0.05001694 |
| RFNG | 0.050012038 |
| RP11-368I23 | 0.050007695 |
| RP1-283K11 | 0.049987892 |
| GPRC5B | 0.049981599 |
| RN7SKP234 | 0.049977209 |
| RP11-505K9 | 0.049962808 |
| GPM6B | 0.049954175 |
| RPS15AP9 | 0.049928162 |
| EPHA5 | 0.049926425 |
| C1QTNF7 | 0.049925839 |
| CCKBR | 0.049914905 |
| AC009297 | 0.049909598 |
| RP11-276H7 | 0.04990803 |
| NRXN2 | 0.049905466 |
| RP11-745A24 | 0.049901122 |
| RETN | 0.049896288 |
| SEC23A | 0.049884795 |
| RPL17P11 | 0.049879141 |
| SERPINE2 | 0.049877539 |
| RP11-748L13 | 0.049862953 |
| MT3 | 0.049858544 |
| CCL24 | 0.049853438 |
| WFDC6 | 0.049843954 |
| AC093698 | 0.04983684 |
| AAMDC | 0.049833928 |
| PCDHA8 | 0.049828008 |
| RP1-276N6 | 0.049823518 |
| AC079250 | 0.049821313 |
| MICU3 | 0.049820327 |
| PQBP1 | 0.049816048 |
| AC073410 | 0.049809821 |
| LSM5 | 0.049805629 |
| RNU6-491P | 0.049803086 |
| TRPC1 | 0.049801703 |
| ANO10 | 0.049801219 |
| COX20 | 0.049793926 |
| C3P1 | 0.04978559 |
| FRMD8 | 0.049766911 |

| | |
|---|---|
| CLYBL | 0.04976505 |
| UTY | 0.049760037 |
| RP11-54D18 | 0.049750527 |
| LINC00868 | 0.049747241 |
| RPL35P5 | 0.049734872 |
| C14orf2 | 0.04972996 |
| IGFBP7 | 0.049725272 |
| CYGB | 0.049719262 |
| TMEM88B | 0.049697956 |
| PTPRK | 0.049688024 |
| RNU6-1159P | 0.049687971 |
| RPL22P11 | 0.049685798 |
| KCTD10 | 0.049675893 |
| AL136303 | 0.049674633 |
| C4orf3 | 0.049664022 |
| FDPS | 0.049659185 |
| AC104308 | 0.04965 |
| DHRS7C | 0.049648704 |
| RNA5SP346 | 0.049638878 |
| AC092171 | 0.049626998 |
| ZNF414 | 0.049623016 |
| RP11-51J9 | 0.049620039 |
| MIR3668 | 0.049610795 |
| TRIM60P8Y | 0.049610795 |
| CTSZ | 0.049598978 |
| RP11-24N18 | 0.049593769 |
| SLIRP | 0.049593303 |
| PAK4 | 0.049575423 |
| IGSF21 | 0.049570995 |
| SOX4 | 0.049559255 |
| VEGFA | 0.049545059 |
| PCDH12 | 0.049540445 |
| AP000568 | 0.049531714 |
| SLC6A3 | 0.049529612 |
| ARID5B | 0.049503029 |
| RNA5SP425 | 0.049493011 |
| MIR181A1 | 0.049484413 |
| RPA2P2 | 0.049482271 |
| OPCML-IT1 | 0.049481441 |
| RP11-600K15 | 0.04947778 |
| LEP | 0.049476674 |
| TTTY23 | 0.049474314 |
| AL133260 | 0.049468662 |
| AL356253 | 0.049467164 |
| LINC01043 | 0.049462723 |

| | |
|---|---|
| OR2V2 | 0.049452359 |
| RNU7-28P | 0.049445745 |
| A2ML1-AS2 | 0.049436552 |
| RP13-383K5 | 0.049430385 |
| RNU2-61P | 0.049426933 |
| MIEF2 | 0.049414225 |
| MIR4253 | 0.049408972 |
| KRTDAP | 0.049407604 |
| RP11-750B16 | 0.049390243 |
| CAPZB | 0.04938107 |
| ZNF90P2 | 0.049378658 |
| MIR6089 | 0.049377937 |
| AC016576 | 0.049376448 |
| PVR | 0.049368084 |
| BID | 0.049357851 |
| AC005498 | 0.049355635 |
| AC073135 | 0.049352706 |
| KRBA1 | 0.04934652 |
| RBMS3 | 0.049338743 |
| CTD-2292M14 | 0.049338233 |
| AL035467 | 0.049337834 |
| BABAM1 | 0.049332696 |
| KRT74 | 0.04932694 |
| PACERR | 0.04932249 |
| HS1BP3 | 0.049316212 |
| TMEM256-PLSCR3 | 0.049294332 |
| TMEM191A | 0.049287783 |
| AFAP1L1 | 0.049263596 |
| SAT1 | 0.049244004 |
| HEY2 | 0.049235966 |
| TSSC4 | 0.049229734 |
| GRIN2B | 0.049228694 |
| RP11-306G20 | 0.04922654 |
| RPL21P120 | 0.04921464 |
| TDGF1P7 | 0.0492068 |
| RP11-353M9 | 0.049201025 |
| CTD-2650P22 | 0.049200495 |
| RPL21P75 | 0.049195314 |
| DEFA8P | 0.049192642 |
| MIR502 | 0.049175845 |
| MIR181A2 | 0.0491668 |
| RP11-449J1 | 0.049166416 |
| RNF222 | 0.049155487 |
| PLIN4 | 0.049153116 |
| FAM131C | 0.049150193 |

| | |
|---|---|
| RP11-610J23 | 0.049137029 |
| STARD4-AS1 | 0.049126864 |
| MIR1272 | 0.049106506 |
| RP11-398J5 | 0.049100925 |
| AC007486 | 0.049094379 |
| PDRG1 | 0.049094211 |
| KPNA7 | 0.049093526 |
| AL590731 | 0.049091496 |
| DCAF12L1 | 0.049085319 |
| CTSD | 0.049082676 |
| TEX22 | 0.049078192 |
| HP | 0.049071683 |
| RP11-361D14 | 0.049071646 |
| AXL | 0.049063029 |
| GPX1P1 | 0.049049106 |
| VGLL4 | 0.049035251 |
| RPS24P1 | 0.04903371 |
| PPY2 | 0.049031531 |
| TES | 0.049031064 |
| RNU1-98P | 0.04902745 |
| CHL1-AS2 | 0.049017858 |
| FUNDC2P2 | 0.049008715 |
| NR2E1 | 0.049006717 |
| TCEAL5 | 0.049001738 |
| AC016700 | 0.04899577 |
| AC115099 | 0.048995118 |
| AC093388 | 0.048994539 |
| MIR4473 | 0.048994388 |
| RP11-91G21 | 0.048987429 |
| MAGOH3P | 0.048986976 |
| NF1P1 | 0.048986928 |
| ZNF688 | 0.048984551 |
| C19orf53 | 0.048984234 |
| PLEKHA8P1 | 0.048977648 |
| LINC01270 | 0.048969076 |
| RP11-585F1 | 0.048964791 |
| ARSH | 0.048948657 |
| AC011404 | 0.048945692 |
| DND1P1 | 0.0489382 |
| OR51A6P | 0.048932361 |
| RP11-345K20 | 0.048919525 |
| RN7SL189P | 0.048896141 |
| KRTCAP2 | 0.048888156 |
| SLC4A2 | 0.048885588 |
| RNU2-8P | 0.048867256 |

| | |
|---|---|
| LINC00205 | 0.048859001 |
| RASL11A | 0.048846349 |
| CRISP2 | 0.048843072 |
| BICD1 | 0.048840879 |
| TRPM1 | 0.048839775 |
| RP11-359P5 | 0.048833623 |
| CASC20 | 0.048832526 |
| RNF181 | 0.048789938 |
| NOP56P1 | 0.048789813 |
| TRIM43CP | 0.048788473 |
| RP11-535M15 | 0.04878711 |
| MTND1P34 | 0.048785202 |
| ASPRV1 | 0.048782328 |
| ANAPC10P1 | 0.048773668 |
| NDUFAF2 | 0.048772746 |
| RN7SL1 | 0.048764661 |
| RP11-757C15 | 0.04872373 |
| RNVU1-2 | 0.048720472 |
| RNU6ATAC17P | 0.04871244 |
| ALDH18A1 | 0.04871092 |
| RNU7-121P | 0.048710631 |
| POLR3GP1 | 0.048696335 |
| CDPF1 | 0.048695478 |
| TLR8-AS1 | 0.048674211 |
| PAPL | 0.048663122 |
| SART1 | 0.048651625 |
| TBC1D19 | 0.048649671 |
| CLDN16 | 0.048641796 |
| GRPEL2P1 | 0.04864113 |
| RP11-326I19 | 0.048640613 |
| GATAD2A | 0.048612851 |
| KRT20 | 0.048612554 |
| HNRNPCL1 | 0.048608328 |
| CPEB1 | 0.048599548 |
| DYNLL1 | 0.048595548 |
| CTD-3157E16 | 0.048578524 |
| SHARPIN | 0.048575212 |
| PRR18 | 0.048567468 |
| C1orf216 | 0.048561537 |
| LLNLR-304G9 | 0.048546349 |
| RP11-60L3 | 0.048542883 |
| FAM150A | 0.048536119 |
| AC079163 | 0.048524335 |
| AC140132 | 0.048518948 |
| CHST8 | 0.048510701 |

| | |
|---|---|
| TMEM121 | 0.048505888 |
| RNU2-52P | 0.048502113 |
| CLN3 | 0.048501832 |
| RP11-766H1 | 0.048499642 |
| UQCRHP3 | 0.048480625 |
| TSPY18P | 0.048475353 |
| RP11-395B7 | 0.048475174 |
| GAREM | 0.048461477 |
| ADRM1 | 0.048461309 |
| AC241952 | 0.048457836 |
| CDCA4P3 | 0.048454277 |
| AC073063 | 0.048449104 |
| DVL3 | 0.048444211 |
| APOD | 0.048438611 |
| C17orf49 | 0.048436583 |
| RP11-619J20 | 0.048434387 |
| CCDC12 | 0.048421568 |
| LINC00668 | 0.048419329 |
| FAM35CP | 0.048410568 |
| PPIH | 0.048408451 |
| MTND5P24 | 0.048395586 |
| ENOX1 | 0.048384897 |
| RN7SL216P | 0.048384568 |
| PIK3CD-AS2 | 0.048376357 |
| MIR548I4 | 0.048371517 |
| WNK3 | 0.048369714 |
| RNU6-974P | 0.048362361 |
| MAGEA4 | 0.048360873 |
| AP001048 | 0.048360379 |
| SH2D3A | 0.048355146 |
| GALNT18 | 0.04835297 |
| DAGLB | 0.048346253 |
| MSRA | 0.048343109 |
| PCDHGA2 | 0.048332373 |
| TULP1 | 0.048315618 |
| RPS10-NUDT3 | 0.048312572 |
| BACH1 | 0.048289767 |
| TXLNGY | 0.048289721 |
| AC046143 | 0.048285934 |
| RP11-229C3 | 0.048281964 |
| RP11-2K6 | 0.048277669 |
| WBP1L | 0.048267831 |
| TCF4 | 0.048257803 |
| SUB1P3 | 0.048254979 |
| HSPB6 | 0.048229001 |

| | |
|---|---|
| SLC5A1 | 0.048219642 |
| SDS | 0.048215364 |
| RPL6P27 | 0.048212535 |
| PHEX-AS1 | 0.048207409 |
| PTMS | 0.048204517 |
| OR7E16P | 0.048203186 |
| AC003666 | 0.048201458 |
| AC104655 | 0.048200405 |
| AC005076 | 0.04819419 |
| HSPE1P6 | 0.048191523 |
| AC027348 | 0.048188032 |
| RN7SKP118 | 0.048174882 |
| Z97356 | 0.048172818 |
| OR6K3 | 0.048170859 |
| RP11-345J18 | 0.04816593 |
| COPS8 | 0.048148849 |
| TSC22D2 | 0.048148663 |
| KCTD9P5 | 0.048145757 |
| HES7 | 0.048142903 |
| RN7SL70P | 0.048137637 |
| CTD-2256P15 | 0.04813549 |
| KLF9 | 0.048134244 |
| SLC39A2 | 0.048132692 |
| RP11-46B11 | 0.048124882 |
| RP11-63E5 | 0.048121238 |
| CHST13 | 0.048120893 |
| MYOZ3 | 0.048117443 |
| RNA5SP31 | 0.048109506 |
| LHB | 0.048108113 |
| RHEB | 0.048092828 |
| PRELID2 | 0.04809266 |
| CTD-2036A2 | 0.048092109 |
| GHR | 0.048084158 |
| TRNAU1AP | 0.048082279 |
| SPINT1 | 0.048075209 |
| CXCL2 | 0.048067804 |
| AL133232 | 0.048066228 |
| HSPA8P13 | 0.048063947 |
| COX4I2 | 0.04806265 |
| CXXC1P1 | 0.048050729 |
| CTD-2552K11 | 0.048036867 |
| LINC00473 | 0.048033784 |
| CTSL | 0.048025606 |
| SYF2P1 | 0.04802273 |
| AQP4 | 0.048015259 |

| | |
|---|---|
| LMOD1 | 0.048010927 |
| GNA12 | 0.048009475 |
| HIGD2A | 0.048008229 |
| ARHGAP20 | 0.048006647 |
| KRT8P9 | 0.048005568 |
| BNIP3P34 | 0.047993817 |
| MEIS1 | 0.047977307 |
| MTND1P6 | 0.047950626 |
| TUBB4BP5 | 0.047937494 |
| RP11-322L20 | 0.04792288 |
| MZT2A | 0.047917639 |
| KRTAP19-1 | 0.047896651 |
| RAB34 | 0.047889768 |
| CCNO | 0.0478896 |
| OOSP1P2 | 0.047873809 |
| OR10D4P | 0.04787173 |
| CD24P2 | 0.047859467 |
| LMOD3 | 0.047847131 |
| RP11-1053O12 | 0.047845003 |
| MIR4484 | 0.047833057 |
| RP11-266O8 | 0.047828163 |
| LHX4-AS1 | 0.047828128 |
| RP11-269F20 | 0.047809171 |
| ANKRD30BP3 | 0.047806499 |
| ADAM21 | 0.047800303 |
| BMS1P10 | 0.047795815 |
| BCL2L10 | 0.047778603 |
| HADHA | 0.047769149 |
| RPS4Y2 | 0.047763953 |
| AL158111 | 0.047763643 |
| RNY4P34 | 0.047760262 |
| AC006227 | 0.047750159 |
| EFCAB1 | 0.047736378 |
| FAR2P4 | 0.047724951 |
| SDHAF1 | 0.047720105 |
| STAC | 0.047716544 |
| ATP13A5-AS1 | 0.047709892 |
| RN7SL230P | 0.047701773 |
| RP11-383J24 | 0.047701376 |
| NPSR1 | 0.047698036 |
| RP11-1259L22 | 0.047669749 |
| LRRN1 | 0.047662744 |
| CMTM2 | 0.047658566 |
| MIR379 | 0.047650533 |
| TRIM28 | 0.047643078 |

| | |
|---|---|
| SAMD4B | 0.047642929 |
| MIR103A2 | 0.047635182 |
| TM2D2 | 0.047633105 |
| VENTXP5 | 0.04762963 |
| FGGY | 0.047608081 |
| RP11-124A7 | 0.047599141 |
| HGH1 | 0.04759582 |
| SSSCA1 | 0.047592694 |
| ACKR3 | 0.047583745 |
| SLC6A17 | 0.047582145 |
| RP13-444H2 | 0.047580335 |
| AC008625 | 0.047577233 |
| RP3-404F18 | 0.047562074 |
| RNU1-47P | 0.047539859 |
| IGFL1P2 | 0.047525901 |
| RP11-2F9 | 0.047523069 |
| POLR2I | 0.047514384 |
| RP11-162I7 | 0.047512171 |
| SNRPB2 | 0.047503109 |
| AC145123 | 0.047501333 |
| S100A14 | 0.047496504 |
| AL356213 | 0.047496198 |
| SPG21 | 0.047492504 |
| AL121761 | 0.047490025 |
| RP11-345N11 | 0.047476354 |
| VSIG2 | 0.047474197 |
| RBMS2 | 0.047472466 |
| HOMER1 | 0.047470233 |
| CTD-2169H9 | 0.047468448 |
| LINC00310 | 0.047466214 |
| OR10Q1 | 0.047466171 |
| RPS12P2 | 0.047459581 |
| RP11-363E6 | 0.047435241 |
| RP11-351M16 | 0.047422995 |
| SPNS1 | 0.047422752 |
| AC012668 | 0.047421055 |
| PROK1 | 0.04741531 |
| FTH1P19 | 0.047405672 |
| RGS17 | 0.047404035 |
| CLEC19A | 0.047399927 |
| SNRPD2P1 | 0.047393485 |
| EIF4A1P9 | 0.047393052 |
| MRPL9P1 | 0.047389659 |
| RP11-346D19 | 0.047388014 |
| RNU1-89P | 0.047385864 |

| | |
|---|---|
| FIGNL2 | 0.04738415 |
| MIR7641-1 | 0.047378886 |
| AC106878 | 0.047378184 |
| RP11-329E24 | 0.047355809 |
| SLC34A2 | 0.047347354 |
| UROS | 0.04733285 |
| AL158154 | 0.047319978 |
| CD36 | 0.047310896 |
| MS4A10 | 0.04730803 |
| RN7SL380P | 0.047304716 |
| RP11-312P21 | 0.047297123 |
| OFD1P11Y | 0.047294698 |
| CEACAMP4 | 0.047288419 |
| USP12-AS2 | 0.047271304 |
| RPL21P32 | 0.047269546 |
| ECHS1 | 0.047266652 |
| RP5-884C9 | 0.047254504 |
| FAM155A-IT1 | 0.047254504 |
| AC003088 | 0.047248 |
| C11orf91 | 0.047227907 |
| NANOGNBP3 | 0.047220676 |
| GPX3 | 0.047218241 |
| MIR208A | 0.047206055 |
| CES1P1 | 0.047204992 |
| LATS2 | 0.047198854 |
| ANKRD37 | 0.04719783 |
| GYS1 | 0.047195505 |
| ARSFP1 | 0.047187136 |
| RHBDD3 | 0.047185774 |
| ARHGAP42 | 0.047185495 |
| GS1-594A7 | 0.047181388 |
| AC004492 | 0.047179292 |
| NAV2-AS5 | 0.047177135 |
| IL17RC | 0.047168955 |
| HSPD1P14 | 0.047148476 |
| HIST1H2BH | 0.047145443 |
| RN7SKP162 | 0.047140556 |
| MGC45922 | 0.047139151 |
| C1orf110 | 0.047136235 |
| FAM188B | 0.04713273 |
| HOXD10 | 0.047118962 |
| UNC13D | 0.047118608 |
| RNU1-34P | 0.047105386 |
| FUCA2 | 0.047103724 |
| RP11-845M18 | 0.047093593 |

| | |
|---|---|
| BCKDK | 0.047091984 |
| RP11-404J24 | 0.047083285 |
| LAMP1 | 0.047079593 |
| RP11-390C19 | 0.047070739 |
| HOXC-AS3 | 0.047063234 |
| RP4-633O19__A | 0.047063174 |
| BMP8A | 0.047062159 |
| DKFZP434L187 | 0.047052543 |
| KIF26A | 0.047052261 |
| AC007461 | 0.047042752 |
| RP4-655L22 | 0.047034009 |
| CIC | 0.047023776 |
| PSMB2 | 0.0470232 |
| OSBPL3 | 0.047013153 |
| CIR1P3 | 0.047012928 |
| LMO7DN | 0.047011102 |
| FLYWCH2 | 0.047002678 |
| KRT77 | 0.046992082 |
| KLHDC3 | 0.04697263 |
| RN7SL749P | 0.046963928 |
| AC112497 | 0.046962899 |
| LINC00862 | 0.046960705 |
| CAV1 | 0.046950192 |
| RN7SL135P | 0.046948029 |
| PROS1 | 0.046941931 |
| MIR153-1 | 0.046927068 |
| SAMD13 | 0.046926842 |
| CTC-301O7 | 0.046921009 |
| AC109309 | 0.04691964 |
| RP11-78H24 | 0.046913688 |
| ID1 | 0.046913316 |
| MCAT | 0.046904794 |
| FAM195CP | 0.046904741 |
| C1orf159 | 0.046902599 |
| RP11-158I23 | 0.04690106 |
| RP11-59E19 | 0.046898596 |
| MIRLET7DHG | 0.046889772 |
| PPM1F | 0.046885668 |
| RP11-707M1 | 0.046885247 |
| DUX4L4 | 0.046881178 |
| TSPYL5 | 0.046879379 |
| RPS4XP6 | 0.046871695 |
| ARHGEF35 | 0.046871461 |
| LIMS2 | 0.046869946 |
| FLII | 0.046863118 |

| | |
|---|---|
| CFAP52 | 0.046856255 |
| HOXA9 | 0.046849573 |
| PLEC | 0.04684762 |
| RP11-3J1 | 0.04684068 |
| LINC01346 | 0.046834839 |
| SEC1P | 0.046831369 |
| MIR4450 | 0.046830285 |
| RIMS3 | 0.046827712 |
| MPV17L2 | 0.046806613 |
| METTL24 | 0.046783841 |
| RP11-805F19 | 0.046783229 |
| CLCA1 | 0.046776985 |
| STRA6 | 0.046774128 |
| GAPDHP27 | 0.046767255 |
| LINC00400 | 0.046766929 |
| RP11-358H9 | 0.046766584 |
| GAL3ST4 | 0.046757365 |
| RP11-725D20 | 0.046747101 |
| AL080286 | 0.046744697 |
| MRPS21P3 | 0.04674326 |
| METTL7AP1 | 0.046739162 |
| DPPA2P1 | 0.046731531 |
| HSFX1 | 0.046704336 |
| ZSCAN18 | 0.046702274 |
| ZNF608 | 0.046700878 |
| RP11-659E9 | 0.046699818 |
| FAM182A | 0.04668281 |
| KRT33B | 0.046654558 |
| ILK | 0.046648876 |
| RP11-276H19 | 0.046640485 |
| C4BPA | 0.04663598 |
| RARRES2P9 | 0.046617167 |
| RP11-102K13 | 0.046608929 |
| RP3-513G18 | 0.046599316 |
| PAGR1 | 0.046593432 |
| CES1 | 0.046587757 |
| RP13-349O20 | 0.046583005 |
| RN7SL483P | 0.046582706 |
| C1S | 0.046582101 |
| RNU6-770P | 0.046570842 |
| MIR1915 | 0.046566315 |
| IFT57 | 0.046563905 |
| GS1-24F4 | 0.046563331 |
| WI2-2221J1 | 0.046554833 |
| WWC3 | 0.046546155 |

| | |
|---|---|
| MAML2 | 0.046540276 |
| CGB | 0.046517153 |
| MIR412 | 0.046514429 |
| ABCG5 | 0.04651102 |
| C14orf180 | 0.046510332 |
| EBP | 0.046504033 |
| PSAT1 | 0.046502842 |
| TUBB8 | 0.04648588 |
| RMST_8 | 0.046484758 |
| NIFKP9 | 0.046471619 |
| AADACL2-AS1 | 0.046468995 |
| RP11-318G8 | 0.046466582 |
| BRE | 0.046463901 |
| RP11-15K2 | 0.046462512 |
| CARD18 | 0.046458539 |
| XKR8 | 0.046443658 |
| RP4-758J18 | 0.046441481 |
| DECR1 | 0.04643309 |
| OR10Z1 | 0.046426354 |
| LINC01309 | 0.046412949 |
| C8orf4 | 0.046412419 |
| SUPT5H | 0.046410559 |
| FBXO24 | 0.046410149 |
| MIR1972-1 | 0.046399038 |
| MURC | 0.046398595 |
| AC023672 | 0.046397589 |
| C19orf73 | 0.046397293 |
| RNU6-826P | 0.046396271 |
| RNU6-247P | 0.046392188 |
| AC118754 | 0.046370748 |
| RP11-536O18 | 0.046366545 |
| MEIKIN | 0.046365059 |
| RP11-397O8 | 0.046356651 |
| RP11-448K10 | 0.04635131 |
| LL22NC03-32F9 | 0.046351219 |
| RPS26 | 0.046348733 |
| RN7SKP266 | 0.046339665 |
| PRAC1 | 0.046337342 |
| MTND4P28 | 0.046331543 |
| SLC7A15P | 0.046315133 |
| RPL7P23 | 0.046313457 |
| RP11-217L21 | 0.046307418 |
| BOLA1 | 0.046302926 |
| EMILIN2 | 0.046291465 |
| ASB12 | 0.04629012 |

| | |
|---|---|
| B3GNTL1P1 | 0.046273234 |
| AF196972 | 0.046262296 |
| RP11-731N10 | 0.046259345 |
| STK19 | 0.046251389 |
| RP11-738B7 | 0.046251009 |
| RP11-405K6 | 0.046250297 |
| SLC22A14 | 0.046247105 |
| RN7SL542P | 0.046246906 |
| APOB | 0.046240814 |
| SERTM1 | 0.046238451 |
| RP5-859I17 | 0.046235202 |
| RP11-144A16 | 0.046234265 |
| EEFSEC | 0.046234253 |
| PDHA1 | 0.046222923 |
| RP11-982M15 | 0.046221457 |
| IRX4 | 0.046220866 |
| RP11-781P6 | 0.046219866 |
| RP4-794H19 | 0.046216057 |
| RNU6-140P | 0.046208254 |
| SLC6A11 | 0.046208047 |
| APOH | 0.046187619 |
| ABCD1 | 0.046168502 |
| APEX2 | 0.04616519 |
| ANKRD7 | 0.046164226 |
| ABCC1 | 0.046155775 |
| ST3GAL4 | 0.04615518 |
| TNFAIP8L3 | 0.046151868 |
| ACSL4 | 0.046150584 |
| LPL | 0.046149077 |
| EXOC3L1 | 0.046147905 |
| LRRC34P1 | 0.046137091 |
| RP11-102N11 | 0.046134399 |
| LINC00619 | 0.046133937 |
| DNAH9 | 0.046127726 |
| RP4-800F24 | 0.046127072 |
| SEC14L6 | 0.046101309 |
| SIX3-AS1 | 0.046099305 |
| EGLN3-AS1 | 0.04609908 |
| BCRP7 | 0.046090968 |
| POT1-AS1 | 0.046086228 |
| TUSC1 | 0.046083921 |
| BIN3 | 0.046079567 |
| TUBB | 0.046077893 |
| CTC-465D4 | 0.046073575 |
| C1orf137 | 0.046072933 |

| | |
|---|---|
| EEF1A1P27 | 0.046065588 |
| FKRP | 0.046063623 |
| RN7SKP242 | 0.046062303 |
| FBXO32 | 0.046061855 |
| AC025279 | 0.046054951 |
| DEFA5 | 0.046051598 |
| RN7SL178P | 0.046042456 |
| TRHDE-AS1 | 0.046034396 |
| RP11-402C9 | 0.046030308 |
| RNA5SP193 | 0.046026883 |
| TH | 0.046020171 |
| RP11-237D3 | 0.046013437 |
| LINC00927 | 0.046012351 |
| ANO3 | 0.046009953 |
| LHFPL5 | 0.046002629 |
| LCE3A | 0.045988439 |
| CRLF1 | 0.045977051 |
| CTD-2003C8 | 0.045956431 |
| KRT16P6 | 0.045941344 |
| PRAC2 | 0.045941056 |
| RNA5SP28 | 0.04593447 |
| TXNRD2 | 0.045930259 |
| USP17L17 | 0.045925812 |
| ERVV-1 | 0.045921224 |
| FGFBP3 | 0.045911634 |
| MATN3 | 0.045907839 |
| MAEL | 0.045904472 |
| C14orf80 | 0.045903039 |
| TST | 0.045881754 |
| AL159140 | 0.045876382 |
| RP13-644M16 | 0.045868815 |
| BNIP3P20 | 0.045857901 |
| RNA5SP270 | 0.045853089 |
| ID4 | 0.045848729 |
| TRAM2 | 0.045846311 |
| PDCL3P1 | 0.045840296 |
| THOC5 | 0.045833305 |
| AL161757 | 0.045832736 |
| KRTAP3-4P | 0.045830506 |
| LIPK | 0.045828961 |
| MIR205 | 0.045822245 |
| RHOBTB2 | 0.045818049 |
| CTD-2023N9 | 0.045817617 |
| AC090841 | 0.045809054 |
| RP11-470M17 | 0.045804215 |

| | |
|---|---|
| RP11-6C14 | 0.04580039 |
| CES3 | 0.045796113 |
| RNVU1-10 | 0.045795154 |
| AC090825 | 0.045789331 |
| RBMY1E | 0.045789331 |
| TREM2 | 0.045777024 |
| RP4-712E4 | 0.045771793 |
| RNU6-771P | 0.04577071 |
| SPANXN1 | 0.045758007 |
| PRRT3-AS1 | 0.045751367 |
| NKX2-5 | 0.045747548 |
| HSPB1P2 | 0.04574583 |
| RP11-945A11 | 0.045730031 |
| RP11-359E19 | 0.045728226 |
| ENKD1 | 0.045724854 |
| SPESP1 | 0.045718808 |
| RP11-82O18 | 0.04571837 |
| AC024198 | 0.045716506 |
| RP11-681N23 | 0.045714499 |
| STOML2 | 0.045711775 |
| REC114 | 0.045692691 |
| NAV2-AS2 | 0.045692278 |
| LXN | 0.045691383 |
| SEPN1 | 0.045678061 |
| ZC3HC1 | 0.045675326 |
| NTAN1P3 | 0.045673514 |
| CRB3 | 0.045673038 |
| MCM8-AS1 | 0.045668903 |
| TMEM214 | 0.045667289 |
| RP11-454P21 | 0.045664052 |
| RNU6-1133P | 0.045658249 |
| DYRK3 | 0.045657837 |
| CTB-11I22 | 0.045654208 |
| OR4Q1P | 0.045653667 |
| AC092156 | 0.04565285 |
| SHKBP1 | 0.045648665 |
| SULT1E1 | 0.045646029 |
| CACNG7 | 0.04563977 |
| C19orf43 | 0.045639436 |
| RP11-522M21 | 0.045637526 |
| ZNF653 | 0.045636255 |
| IFFO1 | 0.04562712 |
| AL358852 | 0.045624831 |
| RP4-760C5 | 0.045617954 |
| RNU6-55P | 0.045617509 |

| | |
|---|---|
| SLC22A31 | 0.045608953 |
| MIR4437 | 0.045604002 |
| DDN | 0.045596643 |
| TNFRSF10D | 0.045595621 |
| SRD5A2 | 0.04559381 |
| RP4-545L17 | 0.045587062 |
| OR13G1 | 0.045580991 |
| SHPK | 0.04558055 |
| EIF3KP3 | 0.045560017 |
| NPM1P47 | 0.04555307 |
| SLC9B2 | 0.045552065 |
| RP11-488L1 | 0.045541083 |
| IRGC | 0.04554072 |
| RP11-1L9 | 0.045537922 |
| OR13C9 | 0.045535935 |
| MIR4665 | 0.045535677 |
| CEACAM19 | 0.045526408 |
| ARNTL2 | 0.045520771 |
| MIR1252 | 0.045505496 |
| OR8B9P | 0.04550259 |
| RARRES2P4 | 0.045499713 |
| CYP11B2 | 0.045493409 |
| WNT6 | 0.045492407 |
| ANKS3 | 0.0454897 |
| AC055714 | 0.045472225 |
| CHCHD2P6 | 0.045468396 |
| GLYATL1P4 | 0.045459764 |
| LCE1C | 0.045450605 |
| FP671120 | 0.045448733 |
| MAGED4 | 0.045447583 |
| FGF19 | 0.045447234 |
| TNFRSF11B | 0.045436004 |
| RP11-144M17 | 0.045432026 |
| CARD14 | 0.045430981 |
| ERICH2 | 0.045427184 |
| DCD | 0.045423264 |
| TACC2 | 0.045416934 |
| CACNG2 | 0.045411888 |
| GSTA5 | 0.045405407 |
| OR6C4 | 0.045404901 |
| DLL1 | 0.045403798 |
| SLC39A11 | 0.0453971 |
| RNU4-13P | 0.045394837 |
| LIMA1 | 0.045394626 |
| HIST1H4C | 0.045394464 |

| | |
|---|---|
| RBM46 | 0.045393098 |
| IL1F10 | 0.045390012 |
| RP11-655G22 | 0.045388633 |
| RP11-141J10 | 0.045379888 |
| SNORD116-22 | 0.045377621 |
| HS6ST1P1 | 0.045364281 |
| DDX54 | 0.04536402 |
| RP11-25J3 | 0.045361888 |
| ALPL | 0.045344484 |
| RBMS1P1 | 0.04534172 |
| PTPRG | 0.045337098 |
| RP11-131L23 | 0.04533399 |
| RPL21P2 | 0.045325864 |
| FEZ2 | 0.04532359 |
| RP11-63B19 | 0.045305902 |
| LINC00244 | 0.045303387 |
| RP11-380D11 | 0.045301388 |
| MEPCE | 0.045300184 |
| AC021052 | 0.04529907 |
| RBMS3-AS3 | 0.045297678 |
| LINC01452 | 0.045290709 |
| CEND1 | 0.045277123 |
| RASAL2 | 0.045268369 |
| RP11-180I4 | 0.045267228 |
| JUND | 0.045266806 |
| PRKRIP1 | 0.045255587 |
| SERPINB13 | 0.045250026 |
| MTND4P17 | 0.045243408 |
| CTC-439O9 | 0.045239389 |
| RP11-51O6 | 0.045237856 |
| PJA1 | 0.045231921 |
| NR0B1 | 0.04522077 |
| TREML3P | 0.045217256 |
| RP11-547C13 | 0.045216464 |
| MICU1 | 0.045203845 |
| KRTAP2-2 | 0.045200269 |
| UPP2-IT1 | 0.045198756 |
| ARL6IP4 | 0.045185742 |
| RP11-434H6 | 0.045183342 |
| ANTXR2 | 0.045173128 |
| SVOP | 0.045166194 |
| WFDC2 | 0.045165054 |
| SCYL1 | 0.045164383 |
| TPTE2P1 | 0.045159875 |
| SPDYE9P | 0.045154605 |

| | |
|---|---|
| EHD1 | 0.045127767 |
| AL157702 | 0.045106927 |
| GALNT13 | 0.045104872 |
| RPL23AP53 | 0.045102315 |
| RNU6-570P | 0.045101756 |
| RP5-836J3 | 0.045101241 |
| RNU6-310P | 0.045098618 |
| CD300LD | 0.045095394 |
| LINC01007 | 0.045092713 |
| SNORA35 | 0.045083315 |
| GOT2P5 | 0.0450705 |
| QKI | 0.045069086 |
| RPS28 | 0.045065309 |
| TPT1P1 | 0.045063811 |
| TCEAL4 | 0.045049904 |
| RPS6P23 | 0.045042374 |
| SLC17A1 | 0.045036118 |
| AXIN1 | 0.045026275 |
| MARK2P6 | 0.045008037 |
| RP11-556I14 | 0.045002867 |
| RPS10L | 0.044998255 |
| MIR4799 | 0.044994671 |
| MAP2K3 | 0.044983203 |
| C11orf94 | 0.044977916 |
| RP11-1007I13 | 0.044962778 |
| RP11-145H9 | 0.044957714 |
| MACROD2-IT1 | 0.04494694 |
| RP11-121E16 | 0.044936292 |
| VAT1 | 0.044923275 |
| AL139147 | 0.044921716 |
| RNU6ATAC | 0.044898524 |
| OR52D1 | 0.04489794 |
| RP11-274H2 | 0.044892524 |
| RP11-413P11 | 0.044883618 |
| MIR1261 | 0.044880667 |
| RP11-475J5 | 0.044879533 |
| CITF22-92A6 | 0.044874901 |
| HNRNPA1P69 | 0.044870445 |
| AC093585 | 0.044867932 |
| LINC01123 | 0.044867886 |
| CEBPA-AS1 | 0.044866137 |
| EEF1B2P5 | 0.044858471 |
| RP11-380D23 | 0.044851787 |
| ZNF133 | 0.044850899 |
| RP1-137D17 | 0.044850779 |

| | |
|---|---|
| RNU4-2 | 0.044839675 |
| ANGPT1 | 0.044834638 |
| RNASE12 | 0.044827747 |
| TYR | 0.044817739 |
| ACAN | 0.044807921 |
| SERPINA15P | 0.044804611 |
| AC008063 | 0.044791665 |
| LINC01192 | 0.044789424 |
| RP11-66A2 | 0.044786954 |
| MORN5 | 0.044785033 |
| FOXO1B | 0.04477885 |
| RPS24 | 0.04477378 |
| IQCA1 | 0.044763807 |
| Six3os1_4 | 0.044751172 |
| RNA5SP325 | 0.044748329 |
| AC096587 | 0.04474657 |
| RP11-380L11 | 0.04474256 |
| VEGFB | 0.044737964 |
| ANGPTL6 | 0.044736346 |
| AC096669 | 0.044717657 |
| RP11-231E6 | 0.044711959 |
| RP1-39J2 | 0.04470084 |
| SNORD113-7 | 0.044697484 |
| MIR4758 | 0.044697484 |
| RP11-96G10 | 0.04469626 |
| GHc-1077H7 | 0.044693011 |
| NOTCH4 | 0.044690948 |
| LSAMP-AS1 | 0.044678956 |
| CHI3L1 | 0.044676845 |
| AC156455 | 0.044668161 |
| AC108463 | 0.044662519 |
| RNU6-223P | 0.044654539 |
| RNU7-158P | 0.044650247 |
| AADAT | 0.044644881 |
| RPLP2 | 0.044641476 |
| RN7SKP174 | 0.044600854 |
| RPS12 | 0.044590609 |
| RFKP1 | 0.044586167 |
| DMP1 | 0.044580777 |
| RPLP1P6 | 0.044579222 |
| RP11-756G12 | 0.044550438 |
| OR7E97P | 0.04454408 |
| VDAC2P1 | 0.044541912 |
| C9orf135 | 0.044530511 |
| CH17-478G19 | 0.04451416 |

| | |
|---|---|
| RN7SL299P | 0.044509837 |
| KLHL31 | 0.04449827 |
| PABPC1L2B-AS1 | 0.044498091 |
| CYCSP23 | 0.044495598 |
| RP11-785F11 | 0.044493369 |
| PPP1R3F | 0.044493247 |
| HSD17B10 | 0.044489321 |
| TCN1 | 0.044488584 |
| RP11-670E13 | 0.044481965 |
| OR7E23P | 0.044481422 |
| FKBPL | 0.044477785 |
| RPL22P19 | 0.044477383 |
| RP4-719C8 | 0.044477355 |
| RNU7-173P | 0.044477118 |
| ADAM11 | 0.04446372 |
| RP11-673F18 | 0.044457225 |
| RPL23AP68 | 0.044452349 |
| RP11-14I2 | 0.044441943 |
| PKP1 | 0.044435216 |
| RNU6-432P | 0.044428941 |
| RP11-214O14 | 0.044418321 |
| ILDR2 | 0.044406145 |
| RN7SL554P | 0.044402555 |
| AP003464 | 0.044399625 |
| ASB17 | 0.044395499 |
| RP11-248M19 | 0.044388688 |
| NT5C1A | 0.04436959 |
| RNU5A-1 | 0.04436624 |
| CLSTN1 | 0.04436418 |
| HSPB2-C11orf52 | 0.044360807 |
| RP11-507E23 | 0.044355256 |
| NUDT18 | 0.044351101 |
| DRP2 | 0.044349538 |
| UGP2 | 0.044331491 |
| CTD-2017D15 | 0.044329803 |
| ADAMTSL1 | 0.044329146 |
| QARS | 0.044325686 |
| GPR56 | 0.044324328 |
| TRPV6 | 0.044318711 |
| AWAT2 | 0.044316296 |
| C6orf226 | 0.044309071 |
| RP11-107E21 | 0.044302538 |
| FAM90A23P | 0.044288041 |
| RP11-231G15 | 0.044286413 |
| LINC01356 | 0.044285174 |

| | |
|---|---|
| LMNTD1 | 0.044282522 |
| RP11-547M24 | 0.044280079 |
| RP11-810P8 | 0.044262864 |
| EI24 | 0.044256492 |
| RP11-839D17 | 0.044254892 |
| OR9A4 | 0.044251335 |
| MIR4474 | 0.04425018 |
| RP11-317N12 | 0.044241588 |
| RNU6ATAC32P | 0.0442402 |
| SNX8 | 0.044237812 |
| RNU6-12P | 0.044220199 |
| GUSBP12 | 0.044219538 |
| SNRPEP5 | 0.04421522 |
| ZSCAN1 | 0.044197497 |
| RP4-604K5 | 0.044190871 |
| AC116562 | 0.04417659 |
| EIF4A3P1 | 0.044166927 |
| LINC01519 | 0.044166641 |
| C11orf70 | 0.044156171 |
| MYCNOS | 0.044152772 |
| MIR4528 | 0.044151561 |
| NR2F1 | 0.044138775 |
| MIR409 | 0.044133802 |
| MIR582 | 0.044133719 |
| CH507-210P18 | 0.044116449 |
| RPLP0 | 0.044115649 |
| OR8A1 | 0.044114042 |
| LINC00470 | 0.044106819 |
| RP5-1121H13 | 0.044095469 |
| SLC25A1P4 | 0.044095279 |
| PHBP7 | 0.044084971 |
| RP11-247L20 | 0.044080099 |
| CLDN4 | 0.044078568 |
| PEX16 | 0.044075907 |
| PRRC2CP1 | 0.044073474 |
| RP11-471L13 | 0.044069004 |
| KIAA1217 | 0.044068558 |
| CREM | 0.044062046 |
| SNCG | 0.044061321 |
| RP11-95M15 | 0.044046723 |
| AC023449 | 0.044035837 |
| HAO2-IT1 | 0.044029452 |
| LCE3E | 0.044024058 |
| SIRT6 | 0.044023161 |
| LA16c-60D12 | 0.044020916 |

| | |
|---|---|
| RP11-713C19 | 0.044017611 |
| USMG5 | 0.044013356 |
| AC079355 | 0.044006923 |
| ANKRD30BL | 0.043997579 |
| LYZL4 | 0.043990567 |
| RP11-1021N1 | 0.043975165 |
| SEMA6C | 0.043965949 |
| PTPRH | 0.043963437 |
| LINC00836 | 0.043934396 |
| MYPOP | 0.043931994 |
| PCDHGA12 | 0.043928954 |
| TMSB4X | 0.043928924 |
| TMPRSS5 | 0.043924228 |
| MIR370 | 0.043923374 |
| TMEM72-AS1 | 0.043920936 |
| RPS3AP44 | 0.043920278 |
| GPR115 | 0.043917113 |
| SF3A2 | 0.043913016 |
| PPP1R16A | 0.043903937 |
| RNU6-1339P | 0.043903888 |
| RP5-999L4 | 0.043901316 |
| XXbac-B33L19 | 0.043901201 |
| PROM2 | 0.043900737 |
| TMPRSS11D | 0.043899224 |
| RP11-329L6 | 0.043874112 |
| OR51S1 | 0.04387373 |
| THAP7-AS1 | 0.043872922 |
| LINC01394 | 0.043867933 |
| MIR4713 | 0.043859606 |
| MKL1 | 0.043856642 |
| DNM3 | 0.043852493 |
| AL390718 | 0.043846204 |
| RN7SL271P | 0.043845417 |
| ZNF511 | 0.04384399 |
| ZNF423 | 0.043840697 |
| IGFBP1 | 0.043838735 |
| KRTAP20-1 | 0.043837925 |
| REEP6 | 0.04383279 |
| PRDX6 | 0.04381597 |
| SLC44A2 | 0.043814501 |
| RP11-452F19 | 0.043806723 |
| MYL6P3 | 0.043798399 |
| IGFBP2 | 0.043796044 |
| RPL7AL2 | 0.043779637 |
| RP11-44L9 | 0.04377872 |

| | |
|---|---|
| FEZF1-AS1 | 0.043773706 |
| HBEGF | 0.043770126 |
| RNU4-35P | 0.04376049 |
| RPS27AP1 | 0.043749497 |
| LINC01151 | 0.043745324 |
| RP11-115C10 | 0.043744166 |
| MYO9B | 0.043743093 |
| LBX1-AS1 | 0.043727133 |
| CTD-2228K2 | 0.043725269 |
| KRT90P | 0.043725251 |
| RNU6-824P | 0.043722429 |
| SNORD114-1 | 0.043715513 |
| MRPL15P1 | 0.043714773 |
| RN7SL837P | 0.043708832 |
| RNA5SP164 | 0.0436904 |
| AC051649 | 0.043678735 |
| RNA5SP358 | 0.043675843 |
| AP000525 | 0.043674043 |
| LINC00359 | 0.043663875 |
| AC007274 | 0.043660269 |
| EOGT | 0.043658903 |
| C15orf39 | 0.0436552 |
| RNASEK | 0.043654568 |
| TMEM219 | 0.043649303 |
| URB1-AS1 | 0.043648986 |
| CHL1 | 0.043641016 |
| RP11-540K16 | 0.043627529 |
| NCALD | 0.043624222 |
| RPL7P45 | 0.043618374 |
| AC007318 | 0.043617766 |
| CIAPIN1 | 0.043606529 |
| RP3-388N13 | 0.04360594 |
| SND1 | 0.043595998 |
| TUSC3 | 0.043585449 |
| RP11-744A16 | 0.043584081 |
| ATL1 | 0.043577392 |
| MED16 | 0.043572927 |
| RNU6-132P | 0.043568994 |
| AC091705 | 0.043565429 |
| MEF2BNBP1 | 0.043558955 |
| C7orf65 | 0.043553207 |
| UQCRHP4 | 0.043546768 |
| LMNB2 | 0.043541689 |
| MROH5 | 0.043537219 |
| AC005094 | 0.043527018 |

| | |
|---|---|
| MIR323B | 0.043525501 |
| TIMM44 | 0.043525055 |
| ATP8B5P | 0.043516074 |
| RPL31P54 | 0.043513724 |
| RP11-22L13 | 0.043506466 |
| EMILIN3 | 0.043490969 |
| MIR8068 | 0.043479929 |
| MGARP | 0.043474586 |
| RPL12 | 0.043468885 |
| RNU6-7 | 0.043463323 |
| RPL21P131 | 0.043459746 |
| KCTD16 | 0.043459191 |
| MOB2 | 0.043455229 |
| AL050303 | 0.043453871 |
| RP11-129M6 | 0.043447004 |
| RP11-683L23 | 0.043432424 |
| ANKRD45 | 0.04342554 |
| RP13-126C7 | 0.04342432 |
| AL162711 | 0.043422478 |
| VPS25P1 | 0.043422302 |
| AC020931 | 0.043421877 |
| LINC01038 | 0.043421221 |
| LTC4S | 0.043416995 |
| RP11-264A11 | 0.043415921 |
| SPRY4 | 0.043415804 |
| SNRPBP1 | 0.043410368 |
| RP11-268J15 | 0.043393459 |
| TMEM115 | 0.043392398 |
| OR52A5 | 0.043360177 |
| RNU6-694P | 0.043357161 |
| AC092326 | 0.043346647 |
| RP11-259K5 | 0.043344936 |
| NPHS2 | 0.043336199 |
| DPYD-AS2 | 0.04333051 |
| SYT17 | 0.043315855 |
| AC006153 | 0.043315072 |
| RNU6-33P | 0.043304328 |
| UBL5 | 0.043285566 |
| GOLPH3 | 0.043277379 |
| RP11-66H6 | 0.043277028 |
| AP001331 | 0.043274344 |
| PKN3 | 0.043273267 |
| TMEM109 | 0.043272133 |
| MIR449A | 0.043270434 |
| USP54 | 0.043269677 |

| | |
|---|---|
| MXI1 | 0.043255109 |
| RP5-856G1 | 0.04325336 |
| PRND | 0.043246171 |
| LINC00998 | 0.043239685 |
| RPUSD1 | 0.043232261 |
| HSPD1P16 | 0.043226899 |
| RP11-977B10 | 0.043226233 |
| CAND2 | 0.043223907 |
| IFNWP15 | 0.043202665 |
| CRYBB2 | 0.043191072 |
| RP11-106M7 | 0.043182026 |
| ARRDC3 | 0.043178677 |
| CICP26 | 0.043164275 |
| RP11-130C6 | 0.043163848 |
| C11orf97 | 0.043160004 |
| PAX3 | 0.043159124 |
| RP11-14K3 | 0.043157516 |
| CC2D1A | 0.043156276 |
| CHAT | 0.04315368 |
| RP11-500C11 | 0.043141169 |
| RNU6-1032P | 0.043134174 |
| BAIAP2L2 | 0.043110172 |
| RN7SKP207 | 0.043106634 |
| RP11-523L1 | 0.043093514 |
| KCTD10P1 | 0.043090839 |
| PHEX | 0.04308338 |
| XXcos-LUCA11 | 0.043080204 |
| UBAC2 | 0.043076552 |
| RP11-977P2 | 0.043073509 |
| RP11-875O11 | 0.043063956 |
| OR4F7P | 0.043059714 |
| LRRC71 | 0.043034982 |
| RNA5SP105 | 0.043034698 |
| AC007003 | 0.043033932 |
| C1orf204 | 0.043030187 |
| RSL24D1P4 | 0.043028017 |
| ANKRD6 | 0.043026429 |
| CLDN24 | 0.043022521 |
| AC005178 | 0.043019546 |
| RP3-380B8 | 0.043014787 |
| ASTN1 | 0.042999047 |
| STK25 | 0.042996567 |
| AC016757 | 0.042995614 |
| TEX41 | 0.042988939 |
| RP11-51I5 | 0.042988245 |

| | |
|---|---|
| RPL13A | 0.042984827 |
| WNT4 | 0.042981496 |
| PDCD5 | 0.042981441 |
| WI2-85898F10 | 0.042978841 |
| MIR3162 | 0.042978191 |
| RN7SKP148 | 0.042973311 |
| RP5-915N17 | 0.042972518 |
| SHISA5 | 0.042957421 |
| DKFZP434A062 | 0.042947819 |
| FAM3C2 | 0.042944472 |
| OR7H2P | 0.042927794 |
| RP11-662I13 | 0.04292112 |
| WI2-89927D4 | 0.042917729 |
| ZNF488 | 0.042909938 |
| BCO1 | 0.042907391 |
| RP11-444P10 | 0.0429056 |
| CTD-2262B20 | 0.042905228 |
| AC079949 | 0.042899451 |
| AC007312 | 0.042899106 |
| DKFZp779M0652 | 0.042894911 |
| RNU6-319P | 0.042894874 |
| CCDC144NL-AS1 | 0.042882441 |
| CYP1D1P | 0.042874065 |
| NPAS4 | 0.042867961 |
| RP11-594N15 | 0.042867447 |
| AC093326 | 0.042862302 |
| PACRG | 0.042858954 |
| HIST1H4J | 0.042842419 |
| VSIG8 | 0.042834965 |
| KLHL30 | 0.04283026 |
| LINC00645 | 0.042821701 |
| AC011524 | 0.042810137 |
| RP1-290F12 | 0.042807469 |
| CISD1 | 0.042800819 |
| PGP | 0.042785916 |
| NOV | 0.042782027 |
| RORB-AS1 | 0.042776451 |
| CXCL1 | 0.042772105 |
| CTD-2007H13 | 0.042771767 |
| KCNK3 | 0.042769208 |
| RPL3 | 0.04275544 |
| LENG9 | 0.042750063 |
| RPL21P65 | 0.042739141 |
| LINC01065 | 0.042735562 |
| RNA5SP135 | 0.042733352 |

| | |
|---|---|
| GFRA2 | 0.042723383 |
| TRAPPC13P1 | 0.042719021 |
| RHOT2 | 0.042699028 |
| AP000997 | 0.042692634 |
| CHL1-AS1 | 0.042692301 |
| S100B | 0.042691437 |
| ARL5AP3 | 0.042682642 |
| RP11-325D15 | 0.042676319 |
| RASIP1 | 0.042675902 |
| TMEM47 | 0.04267525 |
| NINL | 0.042663771 |
| AC243772 | 0.042662769 |
| RNA5SP148 | 0.042650787 |
| SNORD113-2 | 0.042647236 |
| MEIS1-AS3 | 0.042636502 |
| RP11-838N2 | 0.042633987 |
| RN7SKP137 | 0.042631461 |
| RNU6-1049P | 0.042615872 |
| TTPA | 0.042607484 |
| CTD-2613O8 | 0.04260453 |
| MARK2P15 | 0.042604153 |
| RP11-448P19 | 0.042602966 |
| RXRG | 0.042595102 |
| RP11-114H20 | 0.042593505 |
| EMG1 | 0.042590893 |
| AC019048 | 0.042587498 |
| BX284613 | 0.042585289 |
| TM4SF19-AS1 | 0.042574662 |
| RP11-318I4 | 0.042569472 |
| CCDC153 | 0.042567841 |
| RN7SL824P | 0.042564967 |
| RNA5SP154 | 0.042561631 |
| NLGN1 | 0.042561126 |
| OR2S1P | 0.042550394 |
| SALL3 | 0.042542355 |
| RP11-392M9 | 0.042535866 |
| PDE1B | 0.042535821 |
| ECM2 | 0.042534761 |
| CYP21A2 | 0.042512389 |
| UQCR10 | 0.042505904 |
| AC006195 | 0.042499888 |
| MIR216B | 0.042499754 |
| ARG2 | 0.042491298 |
| RCAN2 | 0.042485977 |
| CTD-2021J15 | 0.042484362 |

| | |
|---|---|
| FGF13-AS1 | 0.042480978 |
| NAALADL1 | 0.04248047 |
| TRIM42 | 0.042457595 |
| AC006538 | 0.042456469 |
| AC104134 | 0.042454784 |
| B3GAT3P1 | 0.042452591 |
| MEMO1P5 | 0.042448069 |
| NAGK | 0.04244378 |
| RP11-572F4 | 0.042443071 |
| MUC19 | 0.04243898 |
| THBS1 | 0.042438347 |
| BMPER | 0.042435949 |
| RP11-169L17 | 0.042425588 |
| MRO | 0.042415797 |
| CCDC28B | 0.04241189 |
| TSPY15P | 0.042396028 |
| RP11-223J6 | 0.042380659 |
| MIAT_exon1 | 0.042377309 |
| IGHD5OR15-5B | 0.042377309 |
| IGHD1OR15-1A | 0.042377309 |
| RNU2-43P | 0.042371976 |
| SNAP91 | 0.04236889 |
| GSC2 | 0.042362131 |
| BRD7P6 | 0.042354104 |
| HSPD1P21 | 0.042329492 |
| NAALADL2-AS1 | 0.042329003 |
| MXD4 | 0.042326844 |
| FAM3C | 0.042323868 |
| PHB2 | 0.042322844 |
| RN7SL163P | 0.04231853 |
| SCNM1 | 0.042314658 |
| ADRA2A | 0.04231397 |
| AC009108 | 0.042313416 |
| SBNO2 | 0.042310751 |
| CPNE9 | 0.042306612 |
| PDZD11 | 0.042306378 |
| CAP1 | 0.042302416 |
| RNU6-809P | 0.042299291 |
| MIR711 | 0.042285825 |
| PON3 | 0.042281522 |
| KIRREL-IT1 | 0.042268958 |
| SC22CB-109E1 | 0.042259324 |
| PTGIS | 0.042238655 |
| RP11-90C1 | 0.042216741 |
| YBX1P2 | 0.042211277 |

| | |
|---|---|
| IFITM9P | 0.042209983 |
| MIR4310 | 0.04220625 |
| RNU1-18P | 0.042202392 |
| SLC35A2 | 0.042201351 |
| ULBP3 | 0.042193931 |
| NYAP1 | 0.042188263 |
| ERP29 | 0.042185182 |
| NFIA-AS1 | 0.042170759 |
| VPS72 | 0.042169535 |
| OR6F1 | 0.04216646 |
| RP11-552C15 | 0.042156928 |
| AC003681 | 0.042151458 |
| PGBD4P6 | 0.042144958 |
| RPS7P12 | 0.042144407 |
| RP11-399O19 | 0.042143078 |
| RNU4ATAC13P | 0.042142516 |
| CTD-2588E21 | 0.042134984 |
| AL354931 | 0.042134359 |
| LDLRAD4-AS1 | 0.042130725 |
| RP1-37E16 | 0.042127253 |
| TXN2 | 0.042126761 |
| SMURF1 | 0.04212064 |
| TENM2 | 0.042113068 |
| AC079135 | 0.042109124 |
| ACSL1 | 0.04208836 |
| C6orf118 | 0.042081014 |
| AC022616 | 0.042080676 |
| RP11-972L6 | 0.042077376 |
| MFAP4 | 0.042073903 |
| SYT12 | 0.042069512 |
| AL137852 | 0.04206925 |
| PLA2G2E | 0.042061907 |
| SEMG1 | 0.042056227 |
| RP11-73M7 | 0.042056192 |
| SNAPC1 | 0.04205554 |
| LINC01204 | 0.042055477 |
| RP11-416I2 | 0.042052335 |
| AC007967 | 0.042047647 |
| NACAP1 | 0.042046665 |
| TMEM208 | 0.042044953 |
| MYLK-AS2 | 0.042040696 |
| AL132671 | 0.042034221 |
| RP11-275O4 | 0.042031853 |
| SPPL2B | 0.042029492 |
| GAPDHP25 | 0.042014946 |

| | |
|---|---|
| RPS2P32 | 0.042012515 |
| DDX11L10 | 0.042005652 |
| ATP5G3 | 0.041981862 |
| ASB4 | 0.041976624 |
| HIGD1AP4 | 0.04197376 |
| WI2-89031B12 | 0.041952945 |
| CNIH4 | 0.041941953 |
| DBIP1 | 0.041939325 |
| RNU6-559P | 0.041935298 |
| RP11-380I20 | 0.041929696 |
| ALX4 | 0.041928636 |
| KRT8 | 0.041927385 |
| RGN | 0.041925115 |
| SLC45A3 | 0.041917357 |
| MEMO1P1 | 0.041901263 |
| KB-1582A10 | 0.041900439 |
| PAUPAR | 0.041894079 |
| HPGD | 0.041892267 |
| RHOG | 0.041882992 |
| ANKS6 | 0.041879643 |
| CFD | 0.041878527 |
| CCDC140 | 0.041878129 |
| CXCL16 | 0.041878099 |
| KCNA4 | 0.041877379 |
| AC010872 | 0.041877028 |
| TRBVB | 0.041875751 |
| CTD-2272G21 | 0.041872399 |
| AL359771 | 0.041860374 |
| LINC00906 | 0.041858282 |
| CPT1C | 0.041856796 |
| ECSIT | 0.041851326 |
| OR2T8 | 0.041846956 |
| RPL27 | 0.041840367 |
| COX8C | 0.041834795 |
| CTD-2311M21 | 0.041834641 |
| UTP11L | 0.041832497 |
| DAZ2 | 0.041831386 |
| LCE3B | 0.041831386 |
| RNU6-472P | 0.041827397 |
| OLFML1 | 0.041825278 |
| SFTPA2 | 0.041821344 |
| ST8SIA1 | 0.041820366 |
| SLC12A9 | 0.04181631 |
| MYEOV2 | 0.041815808 |
| HPSE2 | 0.041813817 |

| | |
|---|---|
| OR2M4 | 0.041812922 |
| RP4-718P11 | 0.04181075 |
| RP11-140C5 | 0.041791052 |
| WWC2-AS2 | 0.041786074 |
| RPL21P127 | 0.04177966 |
| AC073283 | 0.041776607 |
| AC226118 | 0.041765819 |
| CTD-2636A23 | 0.04175264 |
| RNY3P2 | 0.041750745 |
| RNU2-56P | 0.041745613 |
| RP11-497E19 | 0.041736455 |
| MIR3126 | 0.041730233 |
| RPL34P31 | 0.041729997 |
| AL135745 | 0.041724643 |
| ATRN | 0.04171666 |
| NDUFB9 | 0.041713255 |
| XAGE1A | 0.041709534 |
| STX19 | 0.041708051 |
| RPL23AP4 | 0.04170223 |
| RNF145 | 0.041686854 |
| TMEM185A | 0.041676825 |
| PSMD8 | 0.041670611 |
| PLCL1 | 0.041654034 |
| MIR376B | 0.041653018 |
| OR51D1 | 0.041650903 |
| RP3-510D11 | 0.041643094 |
| RP11-372K20 | 0.041638385 |
| DLX2-AS1 | 0.041629007 |
| RP11-278H7 | 0.041628851 |
| RP11-666A20 | 0.04162214 |
| RALY | 0.041621083 |
| SNORD116-10 | 0.041619338 |
| PRMT2 | 0.041614088 |
| ELL2P3 | 0.041613364 |
| GTPBP6 | 0.041612506 |
| ABL1 | 0.041611446 |
| OR10B1P | 0.041606421 |
| COL5A1-AS1 | 0.041597699 |
| POLR1E | 0.04158564 |
| NUTF2P2 | 0.041579182 |
| RPL39P6 | 0.041577839 |
| EBNA1BP2 | 0.041576505 |
| AL122003 | 0.041571886 |
| RP11-74B21 | 0.041565996 |
| AP002967 | 0.041560083 |

| | |
|---|---|
| RN7SKP200 | 0.041555608 |
| RPS23P1 | 0.041555259 |
| RP11-450H5 | 0.04154996 |
| CSRNP3 | 0.041535396 |
| TTTY14 | 0.041523367 |
| RP11-248E9 | 0.041522959 |
| MGAT4C | 0.04151319 |
| RP11-761B3 | 0.041510871 |
| RPL10A | 0.041509878 |
| PCDHA6 | 0.041496397 |
| Z69720 | 0.041490805 |
| RPSAP58 | 0.041485542 |
| GALR1 | 0.041479785 |
| RNA5SP461 | 0.041479511 |
| RP11-587P21 | 0.041474019 |
| ASS1P11 | 0.041468892 |
| RAC1 | 0.041465932 |
| PCDHGB2 | 0.041458008 |
| RP11-628M2 | 0.041452495 |
| APLF | 0.041446806 |
| THEM6 | 0.041442713 |
| TOR4A | 0.041442564 |
| KCNMB2 | 0.041434538 |
| MTATP6P9 | 0.041424601 |
| OR2D2 | 0.04142064 |
| FAM167B | 0.041406209 |
| FAM46D | 0.041403514 |
| EIF3G | 0.041400218 |
| PXDNL | 0.041392999 |
| GTF2H2 | 0.041384291 |
| RP11-227L6 | 0.041373946 |
| SUN2 | 0.041358746 |
| RN7SK | 0.04135843 |
| RP1-169K13 | 0.041350627 |
| EFHC2 | 0.041333424 |
| RP11-10D7 | 0.04132472 |
| REEP4 | 0.041312251 |
| IFNA13 | 0.041308955 |
| RP11-1182P23 | 0.041300334 |
| RP11-295M3 | 0.041300009 |
| MIR3675 | 0.041298542 |
| RNU6-1076P | 0.041285462 |
| RP11-172E9 | 0.041280691 |
| CRH | 0.04126158 |
| RN7SL98P | 0.041258786 |

| | |
|---|---|
| UGT1A8 | 0.041255793 |
| RNU1-148P | 0.041254884 |
| RP11-473M14 | 0.041252055 |
| CNGA1 | 0.041249657 |
| MMP20 | 0.041238829 |
| RP11-203I2 | 0.041225753 |
| MIR181B1 | 0.04122482 |
| CTD-2595P9 | 0.041216913 |
| PGAM1P2 | 0.041196639 |
| SUSD6 | 0.041196283 |
| LRRC14B | 0.041194607 |
| SRIP3 | 0.04118964 |
| AC016751 | 0.041185588 |
| PTDSS2 | 0.041184543 |
| MIR7158 | 0.041172272 |
| LINC01116 | 0.041172152 |
| PTCD1 | 0.041169733 |
| REG1A | 0.041169041 |
| AC114765 | 0.041163126 |
| AC131213 | 0.041153059 |
| SLC22A3 | 0.0411524 |
| HMGB1P16 | 0.041150971 |
| CNN2P3 | 0.04114613 |
| CTD-3094K11 | 0.041145783 |
| LGR5 | 0.041143085 |
| PKP3 | 0.041142662 |
| LA16c-4G1 | 0.041139723 |
| RPS28P1 | 0.041137263 |
| RP13-60M5 | 0.041132732 |
| RP11-335M9 | 0.041130448 |
| AC018359 | 0.041129842 |
| CTD-2031P19 | 0.04112789 |
| RP11-406O16 | 0.041114504 |
| MIR4735 | 0.041112076 |
| RN7SL180P | 0.041102807 |
| CCDC64B | 0.04110013 |
| CCDC82 | 0.041076985 |
| RP11-63N8 | 0.041070148 |
| RP11-140M13 | 0.041056599 |
| RNU1-46P | 0.041049967 |
| AC010150 | 0.041046956 |
| SERPING1 | 0.041041021 |
| AC121332 | 0.041040181 |
| TTC7A | 0.041029597 |
| RN7SKP286 | 0.041028579 |

| | |
|---|---|
| TLE1 | 0.04102128 |
| BCYRN1 | 0.041012387 |
| PHPT1 | 0.041011177 |
| RP11-702M1 | 0.041010945 |
| DPH3P2 | 0.041004641 |
| HMGB3P30 | 0.041003406 |
| KRT16P3 | 0.040985576 |
| MTND4P24 | 0.040984291 |
| OR10H1 | 0.040978722 |
| RP11-99H20 | 0.040957411 |
| AP001282 | 0.040951186 |
| RP11-843F11 | 0.04094983 |
| FLYWCH1 | 0.040949296 |
| MT-TA | 0.040948899 |
| C6 | 0.04094 |
| RPL35AP35 | 0.040932132 |
| TBCAP1 | 0.0409319 |
| AC234582 | 0.040929227 |
| PRSS45 | 0.040911867 |
| RNF26 | 0.040910875 |
| MIR7108 | 0.040907048 |
| TTC22 | 0.04090148 |
| NOC4L | 0.040893796 |
| PDSS1 | 0.040889181 |
| SMYD1 | 0.040885282 |
| CDH12 | 0.040876624 |
| RP11-176N18 | 0.040872278 |
| CLIC4P3 | 0.040866466 |
| CMTM7 | 0.040861664 |
| AC107079 | 0.040859713 |
| HILPDA | 0.040859673 |
| BCRP2 | 0.040854889 |
| FIZ1 | 0.040847896 |
| FCN3 | 0.040835095 |
| RP11-530C5 | 0.040834016 |
| MIR34C | 0.040833358 |
| SYN2 | 0.040833104 |
| RP11-388K12 | 0.040827546 |
| BANF1 | 0.040820788 |
| RNA5SP88 | 0.040812672 |
| AC007881 | 0.040809989 |
| AC096951 | 0.040798668 |
| PSMB1 | 0.040788303 |
| CHCHD2P2 | 0.040787748 |
| MIR4674 | 0.040782587 |

| | |
|---|---|
| RNU6-829P | 0.040778062 |
| TTLL11-IT1 | 0.040777036 |
| AL132988 | 0.040772957 |
| TGFB1 | 0.040770572 |
| SNORD114-20 | 0.040764489 |
| RP11-525J21 | 0.040763678 |
| XXbac-BCX196D17 | 0.040756823 |
| RP11-15I11 | 0.040749702 |
| LHX5 | 0.040739713 |
| AC023798 | 0.040739539 |
| ZMAT4 | 0.040737378 |
| SPRN | 0.040729025 |
| RP11-704M14 | 0.040708225 |
| PRKACA | 0.040706569 |
| SLC5A6 | 0.040697861 |
| RP11-174B4 | 0.040694789 |
| RN7SKP240 | 0.040680796 |
| AL023806 | 0.040675321 |
| PRTFDC1 | 0.040664837 |
| ZDHHC9 | 0.040644743 |
| FAM110A | 0.040638473 |
| LINC01582 | 0.040631339 |
| RP5-1097P24 | 0.040631185 |
| MIR8061 | 0.040629551 |
| RP11-699A5 | 0.040629371 |
| RNU6-37P | 0.040626505 |
| IMPA2 | 0.04062424 |
| FBXO10 | 0.040621542 |
| AP000859 | 0.040620387 |
| EXOSC7 | 0.040618751 |
| RPL7P20 | 0.040616778 |
| HNF4A | 0.040614858 |
| SNORD3B-2 | 0.040610853 |
| RP11-678D18 | 0.040607896 |
| TIMMDC1 | 0.040605541 |
| MIR5690 | 0.040601636 |
| ANKRD62P1 | 0.040596639 |
| RN7SL861P | 0.040594678 |
| C10orf120 | 0.040579543 |
| RP6-74O6 | 0.040574712 |
| TCEB1P2 | 0.040557731 |
| PPAN-P2RY11 | 0.040555179 |
| PRR14 | 0.040544627 |
| RP5-1185K9 | 0.040544518 |
| TMEM74B | 0.040543343 |

| | |
|---|---|
| SSR4P1 | 0.04054031 |
| RNU6-1161P | 0.040532442 |
| RP11-698N11 | 0.040532217 |
| CTD-2281E23 | 0.04052951 |
| PPP1R35 | 0.040527305 |
| NMRAL1 | 0.040516532 |
| MYH3 | 0.040483991 |
| USP9YP2 | 0.040481899 |
| ATP8A2 | 0.040479638 |
| GLDC | 0.040477908 |
| RARRES2P7 | 0.040477322 |
| OR2AT4 | 0.040472403 |
| PLA2R1 | 0.04047214 |
| TRIM11 | 0.040468568 |
| HMGN2P35 | 0.04046744 |
| RP11-1437A8 | 0.040466577 |
| SAA2-SAA4 | 0.040465204 |
| RPS15AP5 | 0.040462238 |
| AC006273 | 0.040454983 |
| RAB24 | 0.040433794 |
| RNU6-1290P | 0.040430655 |
| TBCB | 0.040428454 |
| FABP7 | 0.040422167 |
| RP11-75C9 | 0.040419647 |
| RNU4-80P | 0.04041701 |
| Z97055 | 0.040416355 |
| CETN2 | 0.040406332 |
| SNORD123 | 0.040398328 |
| SYPL1 | 0.040381866 |
| MIR3672 | 0.040360834 |
| RP11-5K16 | 0.04035986 |
| FOXE1 | 0.040345894 |
| GATSL3 | 0.040344599 |
| BVES | 0.040338999 |
| RNU6-901P | 0.040330096 |
| KRT8P36 | 0.040319155 |
| RP11-44F21 | 0.040306928 |
| JAG2 | 0.040306328 |
| CHODL-AS1 | 0.040304646 |
| RNU6-526P | 0.040299402 |
| CLIC1 | 0.040290178 |
| SLC44A1 | 0.040288541 |
| SNORD115-46 | 0.04028451 |
| RP11-789A21 | 0.040283805 |
| SDCBPP1 | 0.04028033 |

| | |
|---|---|
| RP11-397C12 | 0.040279931 |
| ATP5J2 | 0.040273992 |
| RN7SKP226 | 0.040273172 |
| HMGB3 | 0.040271517 |
| ATP12A | 0.040271473 |
| NPW | 0.040261882 |
| RP11-21M7 | 0.040253218 |
| UBOX5 | 0.040252893 |
| ARPIN | 0.040242865 |
| FRRS1 | 0.040242065 |
| ZC3H3 | 0.040237618 |
| OR9H1P | 0.040233291 |
| OR11K1P | 0.040229441 |
| C4A | 0.040200928 |
| PGC | 0.040184858 |
| FAM83H | 0.04018048 |
| MT-TK | 0.040176794 |
| RNU6-1247P | 0.040169439 |
| PSMD2 | 0.040142414 |
| LA16c-380F5 | 0.040135418 |
| RPL26 | 0.04013339 |
| TUBA4A | 0.040132702 |
| RNU6-160P | 0.040125036 |
| RP11-291H24 | 0.040123907 |
| PNKP | 0.040119008 |
| RAB40AL | 0.040118136 |
| RP11-87O11 | 0.04011799 |
| LINC00607 | 0.040106952 |
| AC002386 | 0.040105246 |
| RNU6-989P | 0.04008983 |
| RP11-293K19 | 0.040085175 |
| HNRNPA0 | 0.040078709 |
| MED15 | 0.040076532 |
| RNU11 | 0.040076325 |
| CTD-2046J7 | 0.040058038 |
| RP11-13K12 | 0.04005356 |
| OR1I1 | 0.040047685 |
| AC253572 | 0.040044738 |
| FGF8 | 0.040039196 |
| MIR362 | 0.040020437 |
| RP11-65M17 | 0.040017711 |
| UACA | 0.04001491 |
| XPNPEP1 | 0.040008212 |
| TMSB4Y | 0.040007321 |
| SNORD100 | 0.040001808 |

| | |
|---|---|
| RNU6-202P | 0.040000082 |
| MRPS11 | 0.039992379 |
| VN1R78P | 0.039988002 |
| HBZ | 0.039986332 |
| AC004967 | 0.039982556 |
| AC000067 | 0.039981633 |
| ANKRD9 | 0.039975411 |
| GPR21 | 0.039969885 |
| TP63 | 0.039955075 |
| C19orf12 | 0.039948526 |
| RP11-632K5 | 0.039945467 |
| BCRP4 | 0.039933291 |
| RP11-193H5 | 0.03993229 |
| RP3-455H14 | 0.039928034 |
| LPP | 0.03992473 |
| RNA5SP140 | 0.039911997 |
| ZNF282 | 0.039907427 |
| SACS-AS1 | 0.039906814 |
| RP11-545D22 | 0.039891761 |
| DNM1P28 | 0.039881183 |
| AC013701 | 0.039880513 |
| SGSM3 | 0.039880077 |
| PLA2G12B | 0.039877936 |
| MTND1P35 | 0.039877451 |
| RNU6-183P | 0.039873799 |
| ADIPOQ-AS1 | 0.039869251 |
| PKMP4 | 0.039867349 |
| CEACAM18 | 0.039863035 |
| OBSCN | 0.039855964 |
| AC093616 | 0.039852634 |
| MTND5P10 | 0.039841041 |
| AC010311 | 0.039839075 |
| RP11-483L5 | 0.039836281 |
| RP1-135L22 | 0.039831373 |
| GRXCR2 | 0.039813751 |
| GAPDHP14 | 0.039812161 |
| CTD-2162K18 | 0.039800569 |
| SCARNA17 | 0.039799327 |
| AP000479 | 0.039795885 |
| LINC01165 | 0.039793467 |
| CYCSP6 | 0.039793258 |
| COMMD4 | 0.039791812 |
| RP11-345F18 | 0.039790852 |
| PCDHGA5 | 0.039789542 |
| VGLL3 | 0.039788017 |

| | |
|---|---|
| TARBP2 | 0.039787514 |
| OR5M1 | 0.039786992 |
| RP11-127O4 | 0.039786848 |
| MT1E | 0.039786268 |
| PLEKHG3 | 0.039772444 |
| KANK1 | 0.039766676 |
| PRCC | 0.039756071 |
| RP11-671C19 | 0.039752519 |
| ENSAP3 | 0.039752381 |
| CACTIN | 0.039750824 |
| CTC-512J14 | 0.039749838 |
| SLC25A5P7 | 0.039739329 |
| SAA2 | 0.039737063 |
| AC078986 | 0.039729452 |
| C8G | 0.039694841 |
| RP4-806M20 | 0.039693699 |
| PHYH | 0.039689966 |
| MIR6810 | 0.039681042 |
| RNVU1-18 | 0.039675195 |
| TMEM44 | 0.039672886 |
| RPS5P7 | 0.039672684 |
| FZD10 | 0.039670951 |
| PSD | 0.039670058 |
| RP13-492C18 | 0.039655318 |
| RNU1-114P | 0.039654305 |
| RP11-709A23 | 0.039653347 |
| ADH4 | 0.039644137 |
| NXF2 | 0.039607565 |
| HPN-AS1 | 0.039605903 |
| SLC4A3 | 0.039594427 |
| CLEC16A | 0.039593441 |
| DGAT2 | 0.039592827 |
| LINC00869 | 0.039586929 |
| PABPC1P8 | 0.039586412 |
| TXNDC2 | 0.039569692 |
| RP11-503L19 | 0.039562708 |
| GABRA3 | 0.039561408 |
| RP11-873E20 | 0.039559207 |
| CTC-286N12 | 0.03955259 |
| LCE2B | 0.039548245 |
| CCDC181 | 0.039526331 |
| KEAP1 | 0.039525215 |
| GJB3 | 0.0395228 |
| RNA5S16 | 0.039511211 |
| GJB1 | 0.039509621 |

| | |
|---|---|
| RIPPLY1 | 0.039508354 |
| PCDHGA7 | 0.039495614 |
| S1PR5 | 0.039493957 |
| RP4-710M16 | 0.039492335 |
| RP11-513O17 | 0.039492194 |
| SLC35G2 | 0.039489027 |
| RPPH1-3P | 0.0394823 |
| UBASH3B | 0.039471835 |
| WFDC9 | 0.039461464 |
| CATSPERB | 0.039461044 |
| AC103881 | 0.03945169 |
| NAV1 | 0.039445099 |
| NLRP12 | 0.039441658 |
| HDAC8 | 0.039441025 |
| RP11-162A23 | 0.039435005 |
| MMD | 0.039434755 |
| OR13D3P | 0.039431974 |
| GABRD | 0.039429043 |
| MIR892A | 0.039427558 |
| PCDHB18P | 0.039424097 |
| PRR16 | 0.03942281 |
| RP11-309J13 | 0.039422251 |
| MIR921 | 0.039414375 |
| RP11-753N8 | 0.039395508 |
| RNU6-173P | 0.039393195 |
| RP13-43E11 | 0.039392089 |
| HS1BP3-IT1 | 0.039391674 |
| TMPRSS11A | 0.039385058 |
| FBXL12 | 0.039376966 |
| LINC01324 | 0.039370376 |
| SETP2 | 0.039365711 |
| MMP24-AS1 | 0.039364724 |
| NUDT22 | 0.03935343 |
| AC102948 | 0.039351662 |
| AP001172 | 0.039345192 |
| WDR63 | 0.039344965 |
| GRAMD4P1 | 0.039338009 |
| MIR4760 | 0.039335823 |
| AC009500 | 0.039323011 |
| MROH6 | 0.039315382 |
| LUZP4 | 0.039314522 |
| CELA2A | 0.039311902 |
| RPL12P38 | 0.039306864 |
| RNU6-1255P | 0.039303197 |
| CTD-2576N18 | 0.03930158 |

| | |
|---|---|
| RP1-155D22 | 0.039300524 |
| TARDBPP1 | 0.039291047 |
| RP11-554D15 | 0.039286846 |
| CTD-2541M15 | 0.039284478 |
| RP11-521D12 | 0.039281397 |
| RP11-106D4 | 0.039278261 |
| RP11-401N18 | 0.039274216 |
| HORMAD2 | 0.039274118 |
| ULK4P1 | 0.039273893 |
| MRPL11 | 0.039267157 |
| LDHB | 0.039265129 |
| OTOS | 0.039262358 |
| C16orf47 | 0.039260588 |
| UQCRFS1 | 0.039257538 |
| RNU6-1189P | 0.039257211 |
| RP11-350O14 | 0.039255656 |
| MLNR | 0.039251963 |
| RPS4XP20 | 0.039251761 |
| AC004448 | 0.039249671 |
| RP1-43E13 | 0.039239138 |
| MTND1P1 | 0.039237769 |
| KRTAP13-3 | 0.039232852 |
| RPL29 | 0.039230355 |
| RP11-120E11 | 0.039222913 |
| RPL7P40 | 0.03921646 |
| SLC25A15P5 | 0.039209523 |
| RP11-195L15 | 0.039205608 |
| DUOXA2 | 0.039201886 |
| DCTN1 | 0.039199637 |
| CTD-2325B11 | 0.039191598 |
| IP6K3 | 0.039186764 |
| WDR70 | 0.039184641 |
| RP11-313E4 | 0.039181999 |
| MIR605 | 0.039179881 |
| UGT8 | 0.039178191 |
| AC012360 | 0.039165627 |
| TBL1YP1 | 0.039159827 |
| NME2P1 | 0.039152488 |
| EEF1GP4 | 0.039122582 |
| PCAT7 | 0.039116749 |
| POLR2F | 0.039113189 |
| LY6E | 0.039104954 |
| RP1-309I22 | 0.039099262 |
| NEFMP1 | 0.039099065 |
| CYHR1 | 0.039096879 |

| | |
|---|---|
| RP1-45C12 | 0.039095979 |
| LINC01508 | 0.039095797 |
| RP11-418H16 | 0.039094023 |
| FRAT2 | 0.039093716 |
| BBOX1 | 0.039091762 |
| CDCA4 | 0.039085214 |
| BPIFB1 | 0.039083777 |
| MIR146B | 0.039079945 |
| RP11-849I19 | 0.039079135 |
| AC098870 | 0.039071597 |
| ZNF784 | 0.039064097 |
| RN7SKP108 | 0.039064031 |
| DLX2 | 0.039058391 |
| PLXNB2 | 0.039056413 |
| MIR7854 | 0.039055836 |
| RPL21P17 | 0.039052367 |
| RNU6-1196P | 0.039052057 |
| RN7SL670P | 0.039039511 |
| C1orf145 | 0.039038607 |
| LINC01315 | 0.039035016 |
| CTSE | 0.039033767 |
| UGT1A1 | 0.039029294 |
| MRAP | 0.039022844 |
| HSPA2 | 0.039011183 |
| HMGN1P19 | 0.039007628 |
| RP11-126L15 | 0.038990549 |
| C1orf21 | 0.038989154 |
| OC90 | 0.038988572 |
| HMGB4 | 0.038980224 |
| TRI-TAT2-2 | 0.038969251 |
| DEFB108P3 | 0.038965288 |
| MLF2 | 0.038960873 |
| RGS3 | 0.038960427 |
| RNU6-255P | 0.038952623 |
| CH17-335B8 | 0.038939226 |
| PTF1A | 0.038930118 |
| AQP8 | 0.038928302 |
| RNU6-938P | 0.038923806 |
| PDCL3P5 | 0.0389178 |
| MIR3666 | 0.038907232 |
| ATP11B | 0.038906973 |
| RP5-1155K23 | 0.038906487 |
| FAM138E | 0.038901805 |
| POMT2 | 0.03890115 |
| CRCT1 | 0.038885982 |

| | |
|---|---|
| SLC5A12 | 0.038878124 |
| AC099778 | 0.038870906 |
| CHST9 | 0.038861365 |
| RP11-162J8 | 0.038858148 |
| TGFBR1 | 0.038851678 |
| RP11-1090M7 | 0.038849098 |
| NRN1L | 0.038848106 |
| CTB-127C13 | 0.038841817 |
| SDC4P | 0.038837833 |
| ZFP64P1 | 0.038836617 |
| DPP10-AS1 | 0.038836192 |
| DYNLL1P7 | 0.038832888 |
| RP11-429D19 | 0.038827449 |
| ARPC2 | 0.038824012 |
| HOXA7 | 0.038822329 |
| RBM3 | 0.038820495 |
| SNORD114-4 | 0.038815882 |
| MDH2 | 0.038806895 |
| GAPDHP56 | 0.038803527 |
| RN7SKP280 | 0.038791683 |
| RP11-168A11 | 0.03878203 |
| LMO4 | 0.038776568 |
| RN7SKP156 | 0.038776298 |
| RPL10P15 | 0.038770522 |
| AC002310 | 0.038768474 |
| LINC00864 | 0.038767996 |
| RPL23AP12 | 0.038765118 |
| C20orf203 | 0.038748041 |
| DUSP14 | 0.038742203 |
| AC079398 | 0.038741556 |
| CTD-3065J16 | 0.03874118 |
| ZNF22 | 0.038734873 |
| HMGB1P5 | 0.038725924 |
| OLFM4 | 0.038724005 |
| CDY9P | 0.038721275 |
| MRPL20P1 | 0.038720872 |
| C8A | 0.038717223 |
| NAP1L1P2 | 0.038692764 |
| CALHM3 | 0.038691619 |
| RP11-87C7 | 0.038688164 |
| RPS2P46 | 0.038685754 |
| TMEM139 | 0.038680024 |
| MYCN | 0.038673029 |
| DAD1 | 0.038669754 |
| ADAM9 | 0.038669382 |

| | |
|---|---|
| RNU6-1124P | 0.038659974 |
| C17orf74 | 0.038643596 |
| NELFB | 0.038641138 |
| MIR3180-1 | 0.038640019 |
| RP1-182D15 | 0.038634449 |
| ANKRD34A | 0.038625008 |
| ZCCHC24 | 0.038623352 |
| CTD-2073O6 | 0.038620018 |
| AADACL2 | 0.038618677 |
| RP11-554F20 | 0.038616452 |
| MSLN | 0.038615891 |
| CTC-304I17 | 0.038603948 |
| RNA5SP330 | 0.038592463 |
| ANXA5 | 0.038592318 |
| HTR2A | 0.038590198 |
| NMUR2 | 0.038585349 |
| RPS3AP39 | 0.038582712 |
| RPL7P47 | 0.038579992 |
| RNU6-1010P | 0.038571839 |
| CDO1 | 0.038565954 |
| RN7SL632P | 0.038563875 |
| AC016724 | 0.038559209 |
| RNASE10 | 0.038556287 |
| AVP | 0.038554376 |
| MATN4 | 0.038539616 |
| RP1-37N7 | 0.03853477 |
| ARAP3 | 0.038532557 |
| MUC5AC | 0.038513481 |
| ZAN | 0.038513352 |
| MT-RNR1 | 0.038511942 |
| AOC4P | 0.03849845 |
| FTH1P7 | 0.038494397 |
| MRPS34 | 0.038490267 |
| RP11-397O4 | 0.038487844 |
| RNU6-129P | 0.038478346 |
| MIR659 | 0.038477176 |
| RP11-44N21 | 0.03846541 |
| CHRNA1 | 0.03844369 |
| HNRNPUL2-BSCL2 | 0.038443232 |
| RP11-636O21 | 0.038441906 |
| DPF1 | 0.038440869 |
| AC006050 | 0.038435135 |
| CUBN | 0.038428794 |
| ATP2B4 | 0.038423454 |
| RPL18AP14 | 0.038411217 |

| | |
|---|---|
| ANXA11 | 0.0384103 |
| GFOD2 | 0.038408849 |
| STARD13-AS | 0.038406676 |
| RP11-54A4 | 0.038402739 |
| MIR301A | 0.038396995 |
| NKAIN1P1 | 0.038395931 |
| HIST1H2AE | 0.038390231 |
| POLR2L | 0.038389946 |
| ATF3 | 0.038387695 |
| LCMT1 | 0.038387416 |
| DLX4 | 0.038385812 |
| LINC00885 | 0.038376388 |
| OR13J1 | 0.038376301 |
| OSBPL5 | 0.038376159 |
| SPRR1B | 0.038359901 |
| RN7SL786P | 0.038355373 |
| RNU4-59P | 0.038354519 |
| CDK16 | 0.038352717 |
| FAS | 0.038346093 |
| OR11H4 | 0.038338089 |
| OR9M1P | 0.038335197 |
| RP11-42O15 | 0.038334449 |
| XXbac-BPG308K3 | 0.038329733 |
| AC133106 | 0.038315486 |
| RP4-625H18 | 0.038305537 |
| SETD9 | 0.038305031 |
| LAMA5 | 0.038303486 |
| ZNF736P8Y | 0.038301141 |
| LRRC52 | 0.038299067 |
| TMEM189 | 0.03829396 |
| RP11-548K23 | 0.038287202 |
| RP11-449P1 | 0.038286631 |
| RPS4XP22 | 0.038279413 |
| UXS1 | 0.038278723 |
| AP003419 | 0.038276183 |
| NOL12 | 0.038275504 |
| AC005518 | 0.038262133 |
| FTH1 | 0.038259838 |
| RP11-45P15 | 0.03825861 |
| RP11-1028N23 | 0.038254872 |
| HIST1H3I | 0.038248394 |
| LINC00366 | 0.038241655 |
| RNU6-966P | 0.038240828 |
| ABHD17AP3 | 0.038237914 |
| PRAP1 | 0.038233373 |

| | |
|---|---|
| ADAMTS9 | 0.0382326 |
| PLEKHG6 | 0.038231614 |
| RPL11 | 0.038225232 |
| RP11-483F11 | 0.038224001 |
| CHMP6 | 0.038218757 |
| LINC01114 | 0.038213815 |
| PCDHB15 | 0.038206868 |
| FAM216B | 0.038193081 |
| SMAD1-AS1 | 0.038189989 |
| MIR6776 | 0.0381809 |
| TSSK1A | 0.03817676 |
| MIR4718 | 0.038174395 |
| IFNGR1 | 0.038172076 |
| C9orf84 | 0.038166282 |
| RP11-347D21 | 0.038164794 |
| RP11-714L20 | 0.03816154 |
| MFAP3L | 0.03815775 |
| SCGB3A1 | 0.038155303 |
| RALB | 0.038131777 |
| RNU7-195P | 0.038128153 |
| KRTAP21-3 | 0.038125256 |
| TMEM186 | 0.038125116 |
| RP11-401H2 | 0.038119505 |
| CEACAMP2 | 0.03811732 |
| RP11-119F19 | 0.038116582 |
| OR10V2P | 0.038105136 |
| RPL21P67 | 0.038098813 |
| GCNT3 | 0.03809507 |
| RN7SKP39 | 0.038091934 |
| RN7SKP101 | 0.038069629 |
| RP11-57C19 | 0.038046087 |
| DDIT4L | 0.03803914 |
| RP11-524H19 | 0.03803855 |
| MIR770 | 0.038034377 |
| AC035144 | 0.038034102 |
| OR2J2 | 0.038029476 |
| IFNA6 | 0.038025523 |
| RCC1 | 0.038021093 |
| SIGLEC31P | 0.038016612 |
| AL133269 | 0.038009922 |
| NRCAM | 0.038007734 |
| LL22NC03-22D1 | 0.038004399 |
| GAPDHP74 | 0.038000501 |
| USP9YP5 | 0.037997639 |
| RAX | 0.037997117 |

| | |
|---|---|
| NDUFC2 | 0.037995882 |
| FGF13 | 0.037994357 |
| TRBV25OR9-2 | 0.037986034 |
| RN7SKP244 | 0.037961069 |
| RN7SKP294 | 0.037948701 |
| ATP10B | 0.037945779 |
| RPL17P46 | 0.037945478 |
| RP11-547D24 | 0.037937361 |
| HOTTIP | 0.037930535 |
| MIR5096 | 0.037920313 |
| RNA5SP42 | 0.037911985 |
| RP11-83A16 | 0.037905202 |
| RP5-1136A10 | 0.037904733 |
| RP1-118J21 | 0.03790267 |
| SNORD126 | 0.037901915 |
| C4orf51 | 0.037896461 |
| DUSP5P1 | 0.037892813 |
| CYP4Z1 | 0.037887658 |
| ARHGAP40 | 0.03788316 |
| NSMCE1 | 0.03788278 |
| RP11-550I24 | 0.037876268 |
| IGHD4OR15-4A | 0.037873441 |
| USP9YP16 | 0.037873441 |
| OR7E38P | 0.037867691 |
| AC116311 | 0.037864422 |
| RP11-57P19 | 0.03786246 |
| KIAA0020 | 0.037856248 |
| KLC4 | 0.037854909 |
| KIF25 | 0.03785104 |
| RP11-312B18 | 0.037843694 |
| RNA5SP65 | 0.037841591 |
| GAPDHP64 | 0.037840831 |
| HIGD1AP18 | 0.03780825 |
| RP11-319E12 | 0.037796545 |
| MIIP | 0.037795557 |
| IGFBPL1 | 0.03779144 |
| ZDHHC15 | 0.037787761 |
| RNA5SP40 | 0.037787432 |
| RP11-699F16 | 0.037784077 |
| INPP5A | 0.037779836 |
| RNU6-1128P | 0.037778342 |
| AC022142 | 0.037768277 |
| LINC00635 | 0.037764488 |
| TNIP2 | 0.037760932 |
| ZFP36 | 0.037760691 |

| | |
|---|---|
| TBC1D10B | 0.037755853 |
| PKD1 | 0.037751109 |
| LINC01451 | 0.037750189 |
| RP11-272L14 | 0.037739761 |
| AC093270 | 0.03773238 |
| OR52E8 | 0.037731713 |
| TXNDC12 | 0.037729824 |
| RNASE4 | 0.037723573 |
| RNU6-1064P | 0.037722513 |
| SOX18 | 0.037720056 |
| ADSSL1 | 0.037703386 |
| TDRKH | 0.03769853 |
| PABPC5 | 0.037692967 |
| MIR520F | 0.037691137 |
| RP11-382M14 | 0.037690148 |
| RP11-245G13 | 0.037670607 |
| ARSB | 0.037664686 |
| AC010646 | 0.037661895 |
| FAM98A | 0.037660835 |
| HYPM | 0.037657452 |
| RP11-277J24 | 0.03765658 |
| RP11-248I9 | 0.037649193 |
| CTD-2227E11 | 0.037639647 |
| RP11-436D23 | 0.037618191 |
| HORMAD2-AS1 | 0.037613647 |
| DNAAF3 | 0.037613559 |
| TSKU | 0.037604498 |
| ADM5 | 0.037601614 |
| RP11-11A5 | 0.037600928 |
| AMOT | 0.037596795 |
| RP11-734J24 | 0.037595976 |
| LGI2 | 0.037591437 |
| RP1-269M15 | 0.03757792 |
| NIFKP3 | 0.037576037 |
| DNAH3 | 0.037574265 |
| CARM1 | 0.037572999 |
| CBX3P7 | 0.037570936 |
| PCDHGB4 | 0.037570645 |
| ASB11 | 0.037565487 |
| RP11-893F2 | 0.037564087 |
| RPL39P26 | 0.037555966 |
| LINC00482 | 0.03755432 |
| SCGB1A1 | 0.037546713 |
| IGSF10 | 0.037537388 |
| NBPF14 | 0.037533648 |

| | |
|---|---|
| SLC25A6P1 | 0.037528684 |
| DNMBP-AS1 | 0.037525804 |
| RP11-136I13 | 0.037519904 |
| HIST2H3C | 0.037512234 |
| RP11-114H23 | 0.037505651 |
| RN7SL124P | 0.037502699 |
| DMD-AS3 | 0.037498796 |
| SLC13A3 | 0.037495544 |
| RP11-505E24 | 0.03749391 |
| CTB-193M12 | 0.037480678 |
| RP11-311F12 | 0.037478579 |
| CPN1 | 0.037473492 |
| TMEM100 | 0.03746091 |
| RP11-356J5 | 0.037459542 |
| ZNF768 | 0.037459282 |
| RN7SL183P | 0.037456898 |
| RN7SKP72 | 0.037454311 |
| RNU6-791P | 0.037444439 |
| CYB5P4 | 0.037443872 |
| RP11-588P7 | 0.03744053 |
| TP53I13 | 0.037440286 |
| MT1XP1 | 0.037436687 |
| GALNT8 | 0.037434025 |
| LMTK3 | 0.03742862 |
| CYP24A1 | 0.037423526 |
| RP11-170L9 | 0.037423348 |
| CYCSP17 | 0.037422587 |
| PSMA7 | 0.037419485 |
| AC055876 | 0.037406239 |
| RP11-680N20 | 0.037404075 |
| AF241734 | 0.037393922 |
| RPS12P5 | 0.037389938 |
| FRMD6-AS2 | 0.037380783 |
| MIR3160-1 | 0.037373855 |
| PCDHA1 | 0.037371488 |
| SVIL-AS1 | 0.037369138 |
| LSM2 | 0.037356747 |
| MEX3A | 0.037349621 |
| CTD-2535L24 | 0.037346715 |
| RPL31P41 | 0.037346105 |
| AC009238 | 0.037334488 |
| RAB42P1 | 0.037313489 |
| HIST1H4L | 0.037313343 |
| TDH | 0.037308898 |
| CTD-2296D1 | 0.037301343 |

| | |
|---|---|
| CCZ1B | 0.03729347 |
| CBX3P6 | 0.037291442 |
| ADD3-AS1 | 0.037287524 |
| FAM83B | 0.037286381 |
| RP11-445O3 | 0.03728406 |
| RNU6-301P | 0.037274255 |
| OS9 | 0.037268204 |
| URM1 | 0.037265766 |
| EGF | 0.037263701 |
| CTC-501O10 | 0.037262595 |
| RNA5SP336 | 0.037255235 |
| GS1-393G12 | 0.037254414 |
| ADAM19 | 0.037252798 |
| RP11-482H16 | 0.037248807 |
| MIR7852 | 0.037247801 |
| AC112198 | 0.037245452 |
| RP11-495P10 | 0.037239198 |
| RP11-486P11 | 0.037225016 |
| KRTAP17-1 | 0.037224508 |
| RP13-514E23 | 0.037221063 |
| OLIG1 | 0.037204015 |
| HOXD12 | 0.037200617 |
| MIR411 | 0.03719939 |
| RBMY2TP | 0.037191036 |
| B4GALNT4 | 0.03718874 |
| YBX1P9 | 0.03718708 |
| KCNK12 | 0.037182175 |
| CDR2L | 0.037181893 |
| SNRNP70 | 0.037178842 |
| SNTG1 | 0.037176938 |
| AC135048 | 0.037176406 |
| RNU4-40P | 0.037175882 |
| RN7SKP143 | 0.03717154 |
| MIR8089 | 0.037170147 |
| AL512635 | 0.037165437 |
| GAPDHP44 | 0.037164411 |
| RNA5SP155 | 0.037159881 |
| RP11-791N19 | 0.037155291 |
| PNMA2 | 0.037155101 |
| MYH16 | 0.037140916 |
| RPL6P25 | 0.037139824 |
| RP11-356K23 | 0.037136893 |
| GGT5 | 0.037134058 |
| MIR5579 | 0.037131379 |
| RP11-63A1 | 0.037129293 |

| | |
|---|---|
| AC018742 | 0.037126009 |
| MAGEA10 | 0.037116553 |
| PRICKLE2-AS1 | 0.03711299 |
| RP11-158I9 | 0.037111453 |
| RP11-373J21 | 0.037102719 |
| BPIFA1 | 0.037101585 |
| ENTPD7 | 0.037101462 |
| CLIP3 | 0.037099582 |
| CKS1B | 0.037096829 |
| LPAR4 | 0.037094356 |
| DOCK6 | 0.0370812 |
| GSTA4 | 0.037069572 |
| ADAMTS18 | 0.037066411 |
| RP4-669K10 | 0.037057367 |
| RP11-154F14 | 0.037056968 |
| SCGB1C1 | 0.037055513 |
| PRR20A | 0.037054555 |
| ENKUR | 0.037041902 |
| CBLN2 | 0.037036951 |
| PCDHB13 | 0.037036826 |
| PRPF18 | 0.037032733 |
| AC109465 | 0.037028678 |
| EIF4G1 | 0.037026742 |
| LINC01085 | 0.037023003 |
| LPCAT2 | 0.037022351 |
| RP11-218E20 | 0.03701945 |
| RNU6-210P | 0.037019384 |
| RP11-532M24 | 0.037012844 |
| PLEKHG1 | 0.03700235 |
| RN7SL783P | 0.03700173 |
| CTD-2524L6 | 0.036997964 |
| NTF3 | 0.03699683 |
| AC011591 | 0.036989528 |
| RP11-157G21 | 0.036985705 |
| KB-7G2 | 0.036976964 |
| SNX19P1 | 0.03697312 |
| RP11-467L24 | 0.036967861 |
| LINC00973 | 0.03696217 |
| CTBP2P3 | 0.036950279 |
| FMN2 | 0.036948832 |
| UNC79 | 0.036945473 |
| RN7SL424P | 0.03694482 |
| RP11-578B16 | 0.036942617 |
| RCL1 | 0.036941789 |
| TMEM184B | 0.036937268 |

| | |
|---|---|
| RP11-287F9 | 0.036932514 |
| RPS14 | 0.036926998 |
| RNU6-447P | 0.036918075 |
| HOXC12 | 0.036915636 |
| OR6K6 | 0.036909186 |
| MIR1266 | 0.036907683 |
| ATP6V0CP2 | 0.036907156 |
| AC079150 | 0.036900488 |
| RNU6ATAC25P | 0.036886384 |
| RPL10AP2 | 0.036880819 |
| RNU6-1066P | 0.036877021 |
| AC079613 | 0.0368769 |
| FAM83C-AS1 | 0.036876678 |
| DUSP15 | 0.036876503 |
| RP3-436N22 | 0.036873745 |
| FGF14-AS1 | 0.036872682 |
| GAPDHP57 | 0.036869943 |
| NDUFA9 | 0.036863721 |
| CSDAP1 | 0.03686111 |
| KRTAP21-1 | 0.036856666 |
| USP17L8 | 0.036848235 |
| RPL31P35 | 0.036846524 |
| RNU5B-3P | 0.036844775 |
| ACADL | 0.036841308 |
| FBXW5 | 0.036826268 |
| WDR18 | 0.036802286 |
| RP4-612B15 | 0.0368021 |
| QTRT1 | 0.0368013 |
| AC009302 | 0.036796397 |
| RING1 | 0.036773856 |
| ATAD3C | 0.036767624 |
| HAPLN2 | 0.036765619 |
| MOB3B | 0.036763233 |
| RP11-478J18 | 0.036763205 |
| CHRM4 | 0.036756277 |
| AC006534 | 0.036743521 |
| QRFP | 0.036742599 |
| AP000354 | 0.036734916 |
| LCE3C | 0.036732918 |
| PCDH15 | 0.036732422 |
| BANF1P5 | 0.036731342 |
| AL160471 | 0.036729216 |
| PRDX1 | 0.036723212 |
| AC084882 | 0.036714421 |
| LINC00901 | 0.036711891 |

| | |
|---|---|
| RP11-6F6 | 0.03671162 |
| RPL10P3 | 0.0367063 |
| ECSCR | 0.036701667 |
| MYO1C | 0.036694113 |
| LSM3 | 0.03669296 |
| RNVU1-3 | 0.036683014 |
| CTD-2535I10 | 0.036681256 |
| RP3-398G3 | 0.036664784 |
| RNU4-28P | 0.036658132 |
| CCDC36 | 0.036651209 |
| AC009498 | 0.036648155 |
| CACNA1C-IT2 | 0.036647638 |
| AL359955 | 0.036645113 |
| FABP5P4 | 0.036643321 |
| WI2-87327B8 | 0.036643272 |
| RP11-404J23 | 0.036642368 |
| RP11-115H13 | 0.03663804 |
| RPL39P33 | 0.036637465 |
| RP11-663N22 | 0.036623819 |
| AC091320 | 0.036621347 |
| RPS6KA2 | 0.03659543 |
| RP11-5P18 | 0.036595054 |
| AC017083 | 0.036592649 |
| LL21NC02-1C16 | 0.036591411 |
| AC104012 | 0.036591188 |
| MAFK | 0.036581997 |
| CYCTP | 0.03657927 |
| MRPS2 | 0.036576471 |
| ARHGEF2 | 0.036576248 |
| BANF1P2 | 0.036573279 |
| RNA5SP274 | 0.036571103 |
| AC002523 | 0.036570564 |
| GLYAT | 0.036569102 |
| IL25 | 0.036568963 |
| COLEC10 | 0.036566987 |
| AC139099 | 0.036564062 |
| MRGPRG-AS1 | 0.036559422 |
| COL24A1 | 0.036547707 |
| AC010543 | 0.03654665 |
| UGT2B17 | 0.036544486 |
| XRCC4 | 0.036543539 |
| CH17-353B19 | 0.036540344 |
| RP11-553A10 | 0.036539936 |
| AC013439 | 0.0365337 |
| RP11-34N19 | 0.036524971 |

| | |
|---|---|
| CTC-444N24 | 0.036524915 |
| MIR4466 | 0.036524067 |
| RP11-3B12 | 0.03650751 |
| LDHAP7 | 0.036506179 |
| RP11-431D12 | 0.036488311 |
| RP11-680G10 | 0.036468988 |
| RP11-96L14 | 0.036454313 |
| RP11-210H10__A | 0.036453764 |
| RPL3P9 | 0.036447169 |
| RP3-431P23 | 0.03644573 |
| PRKAA1 | 0.036443293 |
| HOXA2 | 0.036442475 |
| RP4-593M8 | 0.03643727 |
| AC093702 | 0.036425457 |
| RAB9AP4 | 0.036425062 |
| GALNT1 | 0.036420557 |
| HNRNPA1P65 | 0.036416465 |
| AP001891 | 0.036406631 |
| PRKCA | 0.036406398 |
| AC002979 | 0.036404743 |
| AL008721 | 0.036401349 |
| RP11-65N13 | 0.036396331 |
| OCSTAMP | 0.036394539 |
| OSTCP4 | 0.036390998 |
| AC010983 | 0.036388612 |
| CTA-992D9 | 0.036381728 |
| RBM43P1 | 0.036380353 |
| RNU6-241P | 0.036369281 |
| MLST8 | 0.036368722 |
| RP11-70P17 | 0.036368631 |
| RGMB | 0.036364927 |
| TPSB2 | 0.036358584 |
| LINC01132 | 0.036358061 |
| CLPS | 0.036356747 |
| PRRG2 | 0.036348015 |
| AC002066 | 0.036347391 |
| SCN10A | 0.036345534 |
| GLRXP3 | 0.036335956 |
| AC093381 | 0.036324412 |
| SNORA23 | 0.036324309 |
| EEF2KMT | 0.036322451 |
| AC048346 | 0.036319006 |
| NRSN2-AS1 | 0.036307231 |
| PTPRU | 0.036306431 |
| NRBF2 | 0.036300273 |

| | |
|---|---|
| MIR4469 | 0.036294687 |
| KCNJ13 | 0.036292988 |
| LNP1 | 0.036288272 |
| AC010967 | 0.036274174 |
| CPA2 | 0.036267065 |
| SNORD114-5 | 0.036263155 |
| AC055733 | 0.036262814 |
| GS1-25M2 | 0.03626279 |
| RP11-736P16 | 0.036261397 |
| RP11-65F13 | 0.0362397 |
| LINC00415 | 0.036239265 |
| RNU6-937P | 0.036238366 |
| MIR4477B | 0.036237442 |
| NKAPL | 0.036236969 |
| RP11-219I21 | 0.036226677 |
| TRAPPC2B | 0.036224995 |
| RP5-935K16 | 0.036211525 |
| PRELID1P1 | 0.036207943 |
| TRPM4 | 0.036195878 |
| KRT19 | 0.036186668 |
| RNU5F-1 | 0.036174818 |
| TBC1D1 | 0.036170835 |
| ZNF93 | 0.036161774 |
| RNU1-43P | 0.036139735 |
| GPR4 | 0.036136861 |
| MAGI3 | 0.036133345 |
| RP11-430G17 | 0.036129435 |
| ZEB1 | 0.036124414 |
| RP11-382F24 | 0.036122841 |
| HOXC-AS1 | 0.036113009 |
| OR7E83P | 0.036106496 |
| RP11-333B11 | 0.036103079 |
| YIPF7 | 0.036100952 |
| MIR3141 | 0.036099189 |
| AC013477 | 0.036095223 |
| RN7SL582P | 0.036087375 |
| ANP32AP1 | 0.036084969 |
| AP000908 | 0.036083224 |
| CTD-3203P2 | 0.036073416 |
| AC091814 | 0.036072592 |
| TBKBP1 | 0.036071258 |
| RSPO2 | 0.036066812 |
| C17orf102 | 0.036065848 |
| RP11-111I12 | 0.036060877 |
| NRM | 0.0360587 |

| | |
|---|---|
| RPS27 | 0.036057546 |
| CTC-551A13 | 0.036046136 |
| RNA5SP377 | 0.036040581 |
| CRHR2 | 0.03603931 |
| SUMF1 | 0.036036652 |
| RP1-166H4 | 0.036022378 |
| C10orf71-AS1 | 0.036015926 |
| MIR4274 | 0.036008357 |
| RN7SL734P | 0.036007553 |
| SEC63P2 | 0.035999771 |
| CTD-2383M3 | 0.035995103 |
| CTSLP1 | 0.035979561 |
| MTND6P16 | 0.035978203 |
| RP13-884E18 | 0.035972707 |
| SUMO2P2 | 0.035968235 |
| MIR548AS | 0.035962709 |
| FAM78B | 0.0359565 |
| CYP8B1 | 0.035952566 |
| RP3-417G15 | 0.03594675 |
| FTH1P23 | 0.035939569 |
| RP11-7F17 | 0.035935903 |
| RN7SKP107 | 0.035935893 |
| KREMEN2 | 0.035935578 |
| CSMD2-AS1 | 0.035929017 |
| AC093117 | 0.035923508 |
| RP11-540O11 | 0.035918991 |
| C1orf140 | 0.035913369 |
| AL390065 | 0.035912712 |
| AC099754 | 0.035911322 |
| OTOR | 0.035905894 |
| RP11-252C24 | 0.03590503 |
| RP11-95L3 | 0.035904158 |
| PIAS3 | 0.035904125 |
| TMSB4XP1 | 0.035897455 |
| RNA5SP198 | 0.035892217 |
| CHCHD5 | 0.035884645 |
| HIST3H2A | 0.035868775 |
| GS1-519E5 | 0.035862849 |
| NOP10 | 0.035860682 |
| GLIPR1 | 0.03584602 |
| RBFOX2 | 0.035845406 |
| MAGEC1 | 0.035836465 |
| LEF1-AS1 | 0.035833574 |
| HELT | 0.035833407 |
| DHCR24 | 0.035828624 |

| | |
|---|---|
| RP1-261D10 | 0.035820159 |
| RAB42 | 0.035814689 |
| TXNP6 | 0.035812348 |
| RNA5SP401 | 0.035803617 |
| NDUFA3 | 0.035799209 |
| COX6B2 | 0.035791568 |
| MIR320A | 0.035787902 |
| RN7SL372P | 0.035779627 |
| NAPA-AS1 | 0.035769124 |
| PINK1 | 0.035768901 |
| FRMD7 | 0.035756508 |
| RNU6-219P | 0.035745335 |
| POU4F2 | 0.035743009 |
| RP3-413H6 | 0.035739085 |
| RPS27P26 | 0.035736571 |
| FAM159B | 0.035727999 |
| PLAGL1 | 0.035724303 |
| RNU6-994P | 0.035718671 |
| CABYRP1 | 0.035715965 |
| HSPD1P3 | 0.035711434 |
| RP11-168E17 | 0.035705057 |
| RPSAP34 | 0.035703062 |
| CASP1P1 | 0.03570168 |
| CCDC71 | 0.035700972 |
| RP11-775B15 | 0.035700841 |
| ATP6V1F | 0.035693344 |
| RPTN | 0.035683221 |
| SEZ6L2 | 0.035639686 |
| TIMM8A | 0.035638328 |
| NARFL | 0.035632448 |
| RP11-956J14 | 0.035630994 |
| RNU6-1308P | 0.035629811 |
| VPS13A-AS1 | 0.035629319 |
| RP11-449D8 | 0.035623334 |
| AC005062 | 0.035608074 |
| LINC01234 | 0.035607036 |
| RP11-443O13 | 0.035597563 |
| COMMD7 | 0.035597042 |
| UBA1 | 0.035589395 |
| CORT | 0.035583754 |
| NUP93 | 0.035580762 |
| HIST1H1A | 0.035569598 |
| YBX1P1 | 0.035556743 |
| RP11-736I10 | 0.035547323 |
| RP11-542C16 | 0.035547037 |

| | |
|---|---|
| RNY4P14 | 0.035544381 |
| HMCN2 | 0.035537449 |
| RPS27P22 | 0.035534308 |
| RNF183 | 0.03553323 |
| RP1-167F1 | 0.035531878 |
| PELP1 | 0.035527979 |
| ASS1P9 | 0.035526023 |
| CLCA4 | 0.035523744 |
| TLN1 | 0.035519029 |
| DNAJC5G | 0.035517972 |
| KLHL7 | 0.03551475 |
| RSPO4 | 0.035497448 |
| PYCARD | 0.035493577 |
| DEFB129 | 0.035488673 |
| RP11-307N16 | 0.035487205 |
| RNF151 | 0.035465029 |
| RN7SKP116 | 0.035464548 |
| TMEM51 | 0.035463045 |
| RNU6-428P | 0.03546284 |
| RPS14P8 | 0.035458538 |
| TSKS | 0.035456426 |
| DAB2 | 0.035451138 |
| CDH4 | 0.035449668 |
| ATP6V0C | 0.035448868 |
| RP11-136I14 | 0.035444291 |
| SRPRB | 0.035440496 |
| AC124997 | 0.035431191 |
| AP005118 | 0.035429165 |
| RP11-50D9 | 0.035425797 |
| UQCC2 | 0.035419918 |
| AC144835 | 0.035417921 |
| CCDC86 | 0.035417425 |
| AC108868 | 0.035417194 |
| RTN2 | 0.035408587 |
| SLC22A18AS | 0.035404959 |
| CHST12 | 0.035403359 |
| TMEM81 | 0.035391582 |
| PPP2R4 | 0.035391228 |
| VSX1 | 0.035390372 |
| CTD-2643K12 | 0.035390035 |
| GSTT2B | 0.035386578 |
| RNA5SP338 | 0.035374973 |
| AC005040 | 0.035372377 |
| NXPH2 | 0.035370472 |
| CLN6 | 0.035369627 |

| | |
|---|---|
| AL049791 | 0.035365883 |
| RNA5SP431 | 0.035361071 |
| NPFFR2 | 0.035353055 |
| C17orf82 | 0.035352849 |
| RP11-179B2 | 0.035351971 |
| C10orf35 | 0.035346091 |
| RNU7-163P | 0.035333104 |
| NLGN4Y | 0.035332387 |
| ZHX3 | 0.035328044 |
| PCMTD1P1 | 0.035314787 |
| IMPDH2 | 0.035309606 |
| MIR374A | 0.035307749 |
| PCDHA12 | 0.035295659 |
| TUBB4B | 0.03529234 |
| TMEM108 | 0.035291429 |
| RWDD1 | 0.035288303 |
| ZNF580 | 0.035283391 |
| FGF3 | 0.035270652 |
| RNU4-41P | 0.035270189 |
| AC012005 | 0.03526103 |
| PPP1R1B | 0.035253641 |
| LINC00871 | 0.035240688 |
| AC231645 | 0.035232739 |
| CTC-367J11 | 0.035227076 |
| ZMYM4-AS1 | 0.035226249 |
| RP11-496N12 | 0.035219524 |
| MIR8058 | 0.035218439 |
| ATP2B3 | 0.035202126 |
| CDC37L1 | 0.035201862 |
| RP1-43O17 | 0.03519927 |
| AL161626 | 0.035196335 |
| TMEM123 | 0.035194457 |
| TARS | 0.035192317 |
| GYPC | 0.035181414 |
| HIST1H3D | 0.03517901 |
| KB-1433G4 | 0.035175369 |
| FSHR | 0.035170865 |
| OR4F8P | 0.035167876 |
| RP11-359E8 | 0.035166935 |
| GET4 | 0.035164688 |
| RP1-102K2 | 0.035153151 |
| AC011530 | 0.035144962 |
| RP11-328L11 | 0.035139989 |
| DIRC3-AS1 | 0.035137674 |
| GADD45G | 0.035129654 |

| | |
|---|---|
| CHID1 | 0.035123924 |
| RP11-410K21 | 0.035123235 |
| UXT | 0.035115663 |
| EIF3KP2 | 0.035114122 |
| RP11-20D14 | 0.035112686 |
| RP11-35J1 | 0.035112299 |
| THRSP | 0.035093099 |
| OACYLP | 0.035091554 |
| PSAPL1 | 0.03508445 |
| GRXCR1 | 0.035080999 |
| ATG10-IT1 | 0.035078719 |
| MTND3P8 | 0.035070043 |
| LYPD3 | 0.035058321 |
| RNU2-17P | 0.035058056 |
| AC069213 | 0.035054265 |
| RP11-137F15 | 0.035052719 |
| PMM1 | 0.035048088 |
| AC139712 | 0.035034069 |
| HIST2H2AB | 0.035015589 |
| BRWD1-AS2 | 0.035006058 |
| RP11-152P23 | 0.034998392 |
| RNA5SP484 | 0.034996585 |
| RP11-52M17 | 0.034994667 |
| PRH1 | 0.034993961 |
| NAMA | 0.034990036 |
| RP11-347E10 | 0.034986726 |
| YEATS2 | 0.034981313 |
| RP11-628O18 | 0.034977105 |
| EMC8 | 0.034972196 |
| RPS3P7 | 0.034969362 |
| CYCSP55 | 0.034967894 |
| RP11-58O9 | 0.034959797 |
| CTD-2503I6 | 0.034944053 |
| KRTAP8-3P | 0.034942786 |
| HOXB5 | 0.034939618 |
| SLC19A1 | 0.034930185 |
| ODF3L2 | 0.034923116 |
| TBCAP2 | 0.03491884 |
| AC112504 | 0.034911056 |
| PTGES2 | 0.034905812 |
| LTBP4 | 0.03488996 |
| KLF7P1 | 0.034888704 |
| CTC-548K16 | 0.034887299 |
| WFDC5 | 0.034882683 |
| TMEM248 | 0.034867336 |

| | |
|---|---|
| PPBPP2 | 0.034864173 |
| RP11-151F5 | 0.0348639 |
| BAALC-AS2 | 0.034856435 |
| LINC01310 | 0.034856377 |
| STARD13-IT1 | 0.034855531 |
| ASPHD1 | 0.034853865 |
| ST20-AS1 | 0.034853214 |
| RPS10P28 | 0.034852269 |
| SPRY2 | 0.034849512 |
| AC092570 | 0.0348472 |
| CTC-575D19 | 0.03483859 |
| CCDC134 | 0.03483392 |
| AC004945 | 0.03483156 |
| AC134878 | 0.034828027 |
| RP11-91A18 | 0.034820678 |
| LINC00687 | 0.03481583 |
| GALNT12 | 0.034814589 |
| RP11-184J23 | 0.034807504 |
| AL591704 | 0.03480211 |
| RN7SKP227 | 0.034799452 |
| HAND2-AS1 | 0.034793704 |
| PEAK1 | 0.034791704 |
| AC023271 | 0.034779369 |
| RP11-646J21 | 0.03476917 |
| INTS10 | 0.034767778 |
| GUCY1A2 | 0.034765471 |
| MIR7845 | 0.034760101 |
| RAB38 | 0.034754456 |
| COX6A1P3 | 0.034749548 |
| MATN2 | 0.034746661 |
| MPDU1 | 0.034745581 |
| AC092569 | 0.034743021 |
| EPHA1 | 0.034742456 |
| RP11-223I10 | 0.034733423 |
| SMAGP | 0.034732818 |
| CCDC101 | 0.034730585 |
| IKBKG | 0.03472212 |
| TET1P1 | 0.034714911 |
| KB-1562D12 | 0.03471278 |
| CTD-3010D24 | 0.034708598 |
| CTB-54I1 | 0.034705754 |
| RP11-295G24 | 0.034703503 |
| DAAM2 | 0.034703496 |
| SUMO2P7 | 0.034699673 |
| RP11-532F12 | 0.034698696 |

| | |
|---|---|
| MIR583 | 0.034698445 |
| SNRPEP8 | 0.034693954 |
| AL359538 | 0.034686737 |
| AC020594 | 0.034685636 |
| RP11-769N22 | 0.034684588 |
| MIR450B | 0.03468391 |
| RNA5SP508 | 0.034675931 |
| CDH13 | 0.034669187 |
| VGF | 0.034654012 |
| MIR3678 | 0.034649084 |
| RP11-323I14__A | 0.034643056 |
| HSPE1P28 | 0.034640932 |
| LINC00331 | 0.034632463 |
| PRSS35 | 0.034631257 |
| CRP | 0.034624099 |
| LINC00278 | 0.034620125 |
| BCRP1 | 0.034609796 |
| SFTPD | 0.034602748 |
| C8orf34 | 0.034598937 |
| CBX1P3 | 0.034594851 |
| GLRA2 | 0.034586296 |
| RARS2 | 0.03458016 |
| RN7SL78P | 0.034574791 |
| ABCB8 | 0.034566522 |
| BRAT1 | 0.034557499 |
| ZMYND15 | 0.034551936 |
| SLC3A2 | 0.034550875 |
| NRSN2 | 0.034550652 |
| RP11-343H5 | 0.034548084 |
| MITF | 0.034537777 |
| RPF2 | 0.034523488 |
| IL22RA1 | 0.034521706 |
| DCAF4 | 0.034520865 |
| RP1-10C16 | 0.034515226 |
| LINC01479 | 0.034505096 |
| MIR6124 | 0.034503307 |
| PARK7 | 0.03450131 |
| VPS9D1 | 0.034495859 |
| FAM45B | 0.03449276 |
| RP11-408N14 | 0.03447409 |
| HYI | 0.034473439 |
| OPN1MW | 0.0344456 |
| RNU6-708P | 0.034440589 |
| PRYP2 | 0.034436937 |
| RNU6-436P | 0.034428077 |

| | |
|---|---|
| RN7SKP229 | 0.034427211 |
| CSN1S1 | 0.034427132 |
| EXOSC3P2 | 0.034423826 |
| CPE | 0.034423149 |
| OTP | 0.034422829 |
| FOLH1B | 0.03442151 |
| SLC12A8 | 0.034412916 |
| DLEU2_6 | 0.034404823 |
| MIR124-2 | 0.034398723 |
| RNU4-45P | 0.034383769 |
| AIM1L | 0.034379705 |
| BNIP3P16 | 0.034372446 |
| RP11-467N20 | 0.034370123 |
| RP11-9D8 | 0.034369996 |
| HES2 | 0.034331052 |
| HIST1H2AJ | 0.034330674 |
| TUSC8 | 0.034330185 |
| MED14P1 | 0.034324938 |
| RP11-413E6 | 0.034319814 |
| CYMP | 0.034313144 |
| ALDH1L1-AS1 | 0.034312863 |
| AEN | 0.034310976 |
| IL1R2 | 0.034310756 |
| AP000358 | 0.034310132 |
| CTA-520D8 | 0.034306225 |
| IRX3 | 0.034303739 |
| ADRA1B | 0.034298456 |
| RP11-807H17 | 0.034288978 |
| HSPB9 | 0.034288888 |
| DCAF12L2 | 0.034281509 |
| SSX4 | 0.034277754 |
| RP11-58O15 | 0.034275257 |
| AC007879 | 0.034265849 |
| NAMPTP3 | 0.034254487 |
| PLCB3 | 0.034249708 |
| TELO2 | 0.034249411 |
| RNY3P13 | 0.034249069 |
| AL589988 | 0.034233995 |
| PLVAP | 0.034233894 |
| TTC29 | 0.034226863 |
| GSTM4 | 0.034224814 |
| CRELD1 | 0.034221558 |
| RPL7P59 | 0.034221093 |
| DNAJC15 | 0.034218154 |
| EPB41L4A-AS2 | 0.034217 |

| | |
|---|---|
| AL049871 | 0.034211983 |
| RP5-933K21 | 0.034199697 |
| AC009313 | 0.034183522 |
| SNAP47 | 0.034179343 |
| RP11-90E5 | 0.034178938 |
| MIR548B | 0.034174928 |
| NOX5 | 0.034168962 |
| FBLN5 | 0.034168142 |
| RNU5E-5P | 0.034158394 |
| RP11-1038A11 | 0.034154009 |
| JAKMIP2-AS1 | 0.034145082 |
| RPL7P61 | 0.034134612 |
| FBXL22 | 0.034131099 |
| RP11-554A11 | 0.034126494 |
| ORAI3 | 0.03412507 |
| MIR126 | 0.034124929 |
| LPHN1 | 0.034121424 |
| CTD-2218K11 | 0.034119957 |
| RN7SL456P | 0.034117057 |
| RP11-104H15 | 0.034113568 |
| ATG9A | 0.034094427 |
| EPPIN-WFDC6 | 0.034089005 |
| RP1-90L14 | 0.034088518 |
| RP11-184I16 | 0.034073318 |
| RNU6-528P | 0.034072973 |
| IL17B | 0.034070937 |
| SNORD114-16 | 0.034070359 |
| MIR1290 | 0.03406796 |
| RNU6-916P | 0.034066245 |
| LOH12CR2 | 0.034063133 |
| SERPINF2 | 0.034054165 |
| GSTA7P | 0.034041239 |
| RP5-1068B5 | 0.034037015 |
| RP3-486I3 | 0.034034778 |
| STMN2 | 0.034024419 |
| KCNIP3 | 0.034022145 |
| CALML5 | 0.034020622 |
| ACPP | 0.034018666 |
| AC073370 | 0.034011766 |
| CTDNEP1 | 0.034004954 |
| PIK3R3 | 0.034004563 |
| RP11-492I21 | 0.034004273 |
| AL121819 | 0.034003056 |
| ARL2BPP5 | 0.034003006 |
| PDGFA | 0.033997418 |

| | |
|---|---|
| RN7SKP69 | 0.033997151 |
| RNU7-14P | 0.033994054 |
| ATOH7 | 0.033991987 |
| FOXO6 | 0.033989567 |
| AC013463 | 0.033986238 |
| FLRT2 | 0.033980562 |
| SLC46A2 | 0.033979896 |
| HYMAI | 0.033979863 |
| RPL35P9 | 0.033974244 |
| VWC2L-IT1 | 0.033965063 |
| RP13-942N8 | 0.033957212 |
| LINC00165 | 0.033953576 |
| CTD-2375G15 | 0.033953539 |
| KCND1 | 0.033951165 |
| ORAI1 | 0.033946458 |
| RP11-856B14 | 0.033934647 |
| NLRP10 | 0.033934271 |
| TTYH1 | 0.033931611 |
| GAR1 | 0.033914996 |
| LINC01138 | 0.033911628 |
| RP5-1030M6 | 0.033901092 |
| FAM83A | 0.033898653 |
| SLC17A7 | 0.033896977 |
| MT1P1 | 0.033891047 |
| DNAJB8 | 0.03388855 |
| OR8G1 | 0.033888058 |
| BATF3 | 0.033886772 |
| AC093159 | 0.033875773 |
| ZNF734P | 0.033875649 |
| MCL1 | 0.033871422 |
| RP11-116N8 | 0.033864397 |
| NR1D1 | 0.03384997 |
| LINC01020 | 0.033847643 |
| MIR548AG1 | 0.033839437 |
| SMG9 | 0.033838193 |
| SOX17 | 0.0338365 |
| AC025038 | 0.033825611 |
| CASKP1 | 0.033825491 |
| SNORD114-27 | 0.033823487 |
| RPL7P1 | 0.033822564 |
| ALDOAP1 | 0.033807594 |
| TRAF7 | 0.033805317 |
| PKMYT1 | 0.033796851 |
| LINC00113 | 0.033795081 |
| ADAM7 | 0.033793369 |

| | |
|---|---|
| FAM25E | 0.033792298 |
| TRIM60P3Y | 0.03379205 |
| KRTAP24-1 | 0.03378615 |
| DUXAP4 | 0.033782553 |
| RP11-501H19 | 0.033779421 |
| CTD-2381F24 | 0.033776086 |
| TBC1D10A | 0.033774153 |
| RNA5SP328 | 0.033772589 |
| RNU6-290P | 0.033767605 |
| RP11-575M22 | 0.033758493 |
| TUBA1C | 0.033755733 |
| TCEB1P7 | 0.033747151 |
| SULT4A1 | 0.033729864 |
| RP11-691G17 | 0.033729181 |
| RNF150 | 0.03371162 |
| TMEM125 | 0.033710503 |
| NDUFS8 | 0.03370868 |
| RPRM | 0.033708404 |
| SNORA57 | 0.033706136 |
| RPL24P2 | 0.033705368 |
| RN7SKP19 | 0.033698758 |
| CLIC3 | 0.033698577 |
| STEAP2 | 0.033697647 |
| ITGA2 | 0.033696791 |
| CDKL1 | 0.033695749 |
| RPL30P11 | 0.033694863 |
| RNF148 | 0.033692446 |
| RP11-89K22 | 0.033689791 |
| RNA5SP515 | 0.033689247 |
| ZCCHC16 | 0.033684977 |
| C2orf61 | 0.033679661 |
| SPRR2G | 0.033678267 |
| NDUFB11P1 | 0.033674841 |
| FGFBP2 | 0.03366755 |
| AL139377 | 0.033667116 |
| AC019084 | 0.033666772 |
| RP1-45I4 | 0.033664698 |
| MIR4771-1 | 0.033662915 |
| SELENBP1 | 0.033662464 |
| GNPTG | 0.033662334 |
| EIF4EBP2P1 | 0.033656989 |
| HCRTR2 | 0.033647616 |
| DLC1 | 0.033645868 |
| MIR1343 | 0.033641848 |
| RP5-1186P10 | 0.033639666 |

| | |
|---|---|
| RP1-27K12 | 0.033631925 |
| FAM9A | 0.033631583 |
| CYP4A27P | 0.033621879 |
| RP5-1051H14 | 0.033618289 |
| TWSG1 | 0.033609773 |
| EXD3 | 0.033607429 |
| AC128653 | 0.033605966 |
| TMEM141 | 0.03360421 |
| LINC01332 | 0.033600006 |
| TMEM178A | 0.033598219 |
| DUSP9 | 0.03358791 |
| RASL10A | 0.03355779 |
| ZNF114 | 0.033555836 |
| PEG3 | 0.033553976 |
| MTHFD2P6 | 0.033553775 |
| RP4-706L14 | 0.033551484 |
| TRMT112P3 | 0.03354542 |
| ATP6V1E1P1 | 0.033543873 |
| ATPIF1 | 0.03354017 |
| AC005901 | 0.033533933 |
| LYARP1 | 0.033533044 |
| FANK1 | 0.033528133 |
| PCOLCE-AS1 | 0.033526573 |
| RP4-777G9 | 0.033524349 |
| GPX6 | 0.033520483 |
| ADH6 | 0.033518844 |
| RPS7 | 0.033513658 |
| C4orf19 | 0.033509378 |
| AC105443 | 0.03350856 |
| PRG4 | 0.033508318 |
| H2AFB2 | 0.033503592 |
| RP11-403D15 | 0.03349025 |
| RNU1-138P | 0.033484722 |
| COL4A2-AS2 | 0.033475181 |
| CYP4F36P | 0.03346468 |
| LDHAP4 | 0.033459125 |
| USP5 | 0.03345639 |
| PIGPP4 | 0.0334542 |
| NGB | 0.033451561 |
| CTB-108O6 | 0.03344832 |
| RP11-672A2 | 0.033448151 |
| PRB2 | 0.033447837 |
| RN7SKP166 | 0.033438385 |
| CIART | 0.033434528 |
| RP11-744D14 | 0.033423545 |

| | |
|---|---|
| RP11-676J15 | 0.033421406 |
| SORBS1 | 0.033420686 |
| NKX2-4 | 0.033418189 |
| RP11-804L24 | 0.033414736 |
| RP11-816J8 | 0.033404309 |
| OR2W5 | 0.033398578 |
| NFIC | 0.033390675 |
| FAM183A | 0.033389231 |
| CLEC5A | 0.033383475 |
| RP11-12O16 | 0.033382516 |
| RP11-491H19 | 0.033380148 |
| EXOC3L2 | 0.033374768 |
| WIF1 | 0.033370763 |
| TVP23C | 0.033359455 |
| RP11-803A13 | 0.033357389 |
| SLC6A1-AS1 | 0.03335529 |
| RBMY2FP | 0.033354845 |
| RNU1-33P | 0.033347257 |
| NDUFA5P6 | 0.033340765 |
| RN7SKP257 | 0.033340448 |
| PER1 | 0.033338357 |
| RP11-832N8 | 0.033337841 |
| RPS4XP12 | 0.033337453 |
| CCDC103 | 0.033335431 |
| ZKSCAN7P1 | 0.03333162 |
| VPS28 | 0.033330133 |
| CTF1 | 0.033328421 |
| CTD-2126E3 | 0.033327084 |
| SRP68P1 | 0.033327003 |
| VIM-AS1 | 0.033305983 |
| AL392167 | 0.033288441 |
| OR2T33 | 0.033275809 |
| PPIAP6 | 0.033273234 |
| LINC00662 | 0.033267079 |
| RNU6-960P | 0.033243238 |
| AIMP2 | 0.03324146 |
| GOLGA2P7 | 0.033241015 |
| SCAP | 0.033235171 |
| GALNT3 | 0.033232529 |
| CTC-431G16 | 0.033228343 |
| FAM183DP | 0.03322231 |
| RP11-589M2 | 0.033222057 |
| RP11-94M14 | 0.033219257 |
| LL0XNC01-250H12 | 0.033217597 |
| KB-1732A1 | 0.033212379 |

| | |
|---|---|
| SMARCA4 | 0.033203728 |
| GNAT1 | 0.033201072 |
| AC005915 | 0.033193111 |
| WDYHV1 | 0.03318862 |
| RPL6P21 | 0.033180926 |
| RP11-324L17 | 0.033174241 |
| MRPS33P2 | 0.033173586 |
| PDIA6 | 0.033173512 |
| RPL29P22 | 0.033171285 |
| BX088702 | 0.033159226 |
| PLIN1 | 0.033149642 |
| FEZF1 | 0.03314428 |
| DLEU1-AS1 | 0.033138604 |
| RN7SL266P | 0.033136653 |
| RN7SL290P | 0.033133381 |
| LPPR2 | 0.033130441 |
| RP11-380M21 | 0.033130314 |
| MET | 0.033115017 |
| LURAP1L | 0.033112319 |
| CACNA2D1 | 0.033108821 |
| CTXN3 | 0.033105786 |
| RP11-432L12 | 0.033101616 |
| MED19 | 0.033099407 |
| SPA17P1 | 0.033096253 |
| LINC00313 | 0.033087118 |
| RP11-1223D19 | 0.033074383 |
| EPHA3 | 0.033064652 |
| GPR1-AS | 0.033062745 |
| PWRN1 | 0.033052593 |
| ARHGAP10 | 0.03305107 |
| AC073626 | 0.033039779 |
| PCDHGB6 | 0.033038623 |
| RFC2 | 0.033031236 |
| SUSD2 | 0.033020724 |
| BRD9 | 0.033012854 |
| RP5-1050K3 | 0.033007869 |
| PARD3-AS1 | 0.033007345 |
| SMYD2 | 0.033005858 |
| MARCO | 0.033003248 |
| RNU7-35P | 0.032997022 |
| RP4-736H5 | 0.032989775 |
| SH3D19 | 0.032955512 |
| FGF14-AS2 | 0.032955084 |
| FGF14-IT1 | 0.032936674 |
| RN7SKP189 | 0.032935509 |

| | |
|---|---|
| HNRNPMP1 | 0.032929437 |
| MTRNR2L11 | 0.032922933 |
| TMED11P | 0.032919625 |
| RP11-100E5 | 0.032918545 |
| MIR4449 | 0.032909043 |
| OR2T12 | 0.03289945 |
| RRP7A | 0.032894616 |
| RP4-640H8 | 0.032887851 |
| FOS | 0.032886151 |
| C11orf31 | 0.032882281 |
| MIR3173 | 0.032877963 |
| IGFBP4 | 0.032874002 |
| PLK2 | 0.032864717 |
| PIGCP2 | 0.032863781 |
| RNU6-1270P | 0.032862728 |
| C17orf97 | 0.032859768 |
| RN7SL414P | 0.032859249 |
| RN7SL519P | 0.032858616 |
| AL161891 | 0.032851444 |
| RP11-459F1 | 0.032845898 |
| RP1-17K7 | 0.032841702 |
| GPAA1P1 | 0.03283849 |
| RPS3AP33 | 0.032833859 |
| CTD-2514C3 | 0.032832783 |
| AP000477 | 0.03283144 |
| LCE1B | 0.032830042 |
| PELI2 | 0.032828437 |
| C2orf74 | 0.032825888 |
| KLHL13 | 0.032825832 |
| LINC01460 | 0.032815008 |
| RN7SKP191 | 0.032814951 |
| PPFIA4 | 0.032810278 |
| RNU6-1134P | 0.032805829 |
| OR5AM1P | 0.032804751 |
| CSPG4P12 | 0.032797443 |
| TMCC1P1 | 0.03278719 |
| BST1 | 0.032785253 |
| C4A-AS1 | 0.032779055 |
| PFKM | 0.032774648 |
| RP11-27K13 | 0.032774462 |
| CTD-2306M5 | 0.032774174 |
| STXBP6 | 0.032763513 |
| AC007204 | 0.032760934 |
| RP11-728C8 | 0.03275921 |
| MIR193A | 0.03275239 |

| | |
|---|---|
| SUCLG1 | 0.032745717 |
| RN7SL668P | 0.032745562 |
| RGPD4 | 0.032743788 |
| VENTX | 0.032734572 |
| UNC5C | 0.032731446 |
| AC092174 | 0.032722329 |
| MRPL24 | 0.032714776 |
| RNU6-659P | 0.032713312 |
| RUVBL1 | 0.032713027 |
| AKR1A1 | 0.032708227 |
| QRFPR | 0.032702587 |
| AC010930 | 0.032697288 |
| RNU6-1331P | 0.032697139 |
| C17orf59 | 0.032690124 |
| AC092066 | 0.032689007 |
| ACAP3 | 0.032686068 |
| RPL7P2 | 0.032685939 |
| CTB-66B24 | 0.032681161 |
| RP13-213K19 | 0.032666436 |
| RP11-527N22 | 0.032658643 |
| SPATA8 | 0.032650108 |
| MIRLET7A1 | 0.032649401 |
| HEY1 | 0.032648913 |
| RP11-1151B14 | 0.032648176 |
| CERS5 | 0.032643908 |
| QRSL1P2 | 0.032637398 |
| AP000261 | 0.032636502 |
| LINC00330 | 0.032628437 |
| GNRH2 | 0.032628216 |
| AC018495 | 0.032624937 |
| OR4K12P | 0.032620274 |
| CTBP2 | 0.032620204 |
| ADI1 | 0.032613376 |
| RP11-350J20 | 0.032610678 |
| RP11-67L3 | 0.032597543 |
| THAP3 | 0.03259518 |
| RNY4P30 | 0.032594966 |
| NANOS1 | 0.032588966 |
| ZBTB45P2 | 0.032582713 |
| RBMY2AP | 0.032577335 |
| RBPMS-AS1 | 0.032567923 |
| CCNE1 | 0.032564314 |
| RNY1P7 | 0.032563113 |
| PACS2 | 0.032555941 |
| ATAD3B | 0.032551885 |

| | |
|---|---|
| GSTM3 | 0.032548629 |
| DDX6P2 | 0.032546483 |
| RP1-153G14 | 0.032545756 |
| RP5-1051D14 | 0.032544912 |
| AC016575 | 0.032543555 |
| TCTEX1D1 | 0.032542843 |
| TMEFF2 | 0.032539287 |
| C11orf86 | 0.032536103 |
| MIR449C | 0.032535837 |
| DNAJC4 | 0.03253101 |
| RN7SL50P | 0.032528582 |
| RPL36AP6 | 0.032527555 |
| RP3-495O10 | 0.032522753 |
| ABCA4 | 0.032519679 |
| SLC16A8 | 0.032513967 |
| RPL41 | 0.032508032 |
| RP11-3G20 | 0.032507079 |
| RNA5SP60 | 0.032500738 |
| MTUS2-AS1 | 0.032491371 |
| RNU2-26P | 0.032486968 |
| HNRNPH3P1 | 0.032485852 |
| PDPR | 0.032483863 |
| GTDC1 | 0.032481315 |
| AL157407 | 0.03247465 |
| PAGE4 | 0.032473008 |
| RNU6-976P | 0.032469594 |
| RP11-18A15 | 0.032468172 |
| RP3-429G5 | 0.032462423 |
| AGBL1 | 0.032447498 |
| CLEC18C | 0.032446352 |
| PAK1IP1 | 0.032443024 |
| F8A1 | 0.032433498 |
| RNA5SP397 | 0.032433265 |
| RP11-419C19 | 0.032432592 |
| SPEM1 | 0.032426698 |
| RNA5SP26 | 0.032426454 |
| AC072052 | 0.032424453 |
| CITED1 | 0.032422633 |
| TADA3 | 0.032420475 |
| RP11-1072A3 | 0.03241653 |
| IRF5P1 | 0.032415003 |
| KANK3 | 0.032410725 |
| RP11-107D24 | 0.032404837 |
| ZNF212 | 0.03240332 |
| SNORD115-17 | 0.032383512 |

| | |
|---|---|
| AL137251 | 0.03238284 |
| AC080003 | 0.032378105 |
| LRRC75B | 0.03237677 |
| C19orf83 | 0.032374599 |
| RN7SL509P | 0.032367746 |
| IL17RE | 0.032360025 |
| RNU7-165P | 0.032357283 |
| RP11-435F13 | 0.032344772 |
| RN7SKP3 | 0.032344497 |
| TMPRSS4 | 0.032340154 |
| RNF6P1 | 0.032339093 |
| RNU2-28P | 0.032324819 |
| NRBF2P1 | 0.032324203 |
| PFDN5 | 0.032323056 |
| HAS3 | 0.032321866 |
| GNAI2 | 0.03231967 |
| GMPPA | 0.032312414 |
| ST20 | 0.032311335 |
| RNU6-1143P | 0.032307451 |
| WFDC8 | 0.032303885 |
| NDUFB4 | 0.032303223 |
| AC006386 | 0.032301592 |
| MIR557 | 0.032300152 |
| ABC7-481722F1 | 0.032290256 |
| RP11-420B22 | 0.032287886 |
| BACH2 | 0.032287576 |
| EGFLAM-AS1 | 0.032284807 |
| FIBP | 0.032278347 |
| AC106869 | 0.032276295 |
| DNAJC8P1 | 0.032275174 |
| AC019050 | 0.032266427 |
| RP4-730D4 | 0.032261278 |
| TSPAN9-IT1 | 0.032260419 |
| MLIP-AS1 | 0.032257546 |
| RTKN | 0.032255463 |
| RNU6-958P | 0.032254401 |
| RP11-372H2 | 0.032253979 |
| AMH | 0.032246584 |
| SYN3 | 0.032244147 |
| RMST_4 | 0.032241239 |
| HNRNPA1P32 | 0.032239663 |
| DSG1-AS1 | 0.032235807 |
| MTND5P3 | 0.032232683 |
| DIAPH2 | 0.032230922 |
| SNORD115-5 | 0.032228066 |

| | |
|---|---|
| RNU6-955P | 0.032226952 |
| RP11-470B22 | 0.032225593 |
| RANGAP1 | 0.032221471 |
| TTTY8B | 0.032218035 |
| RP1-317E23 | 0.032216149 |
| RNU2-71P | 0.032212721 |
| LINC00605 | 0.032212328 |
| CTD-2231H16 | 0.032195377 |
| HOXC6 | 0.032192148 |
| RPL37A | 0.032191572 |
| CHST7 | 0.032179348 |
| ST8SIA5 | 0.032178367 |
| DHRSX | 0.032172073 |
| RP1-99E18 | 0.03217004 |
| CTC-339O9 | 0.03216407 |
| RCVRN | 0.032156875 |
| RP11-387M24 | 0.03215373 |
| AGMO | 0.032149734 |
| RP11-655B23 | 0.032146559 |
| RP5-1065P14 | 0.03214139 |
| HTR5A-AS1 | 0.032133406 |
| TPT1-AS1 | 0.032127978 |
| GNAI2P1 | 0.032125021 |
| C20orf173 | 0.032121869 |
| RP11-26E5 | 0.03211406 |
| ARHGAP23P1 | 0.03211217 |
| FMOD | 0.032110545 |
| RNU6-644P | 0.03210893 |
| RNU5A-8P | 0.032105743 |
| PSORS1C2 | 0.032104132 |
| SLC22A12 | 0.032100884 |
| OR2E1P | 0.032097068 |
| BICD2 | 0.032096888 |
| RNY3 | 0.032096662 |
| RP11-113O24 | 0.032085786 |
| CHMP4B | 0.032084367 |
| LINC01483 | 0.032077897 |
| RP11-3B7 | 0.032075346 |
| OR1R1P | 0.032070616 |
| TBC1D24 | 0.032068292 |
| AC068570 | 0.032068168 |
| TMEM183AP1 | 0.032066378 |
| CTXN1 | 0.032058561 |
| RP11-299H22 | 0.032048587 |
| IFNWP4 | 0.03204625 |

| | |
|---|---|
| KCNK5 | 0.032034839 |
| AVPR1A | 0.032032923 |
| LINC01331 | 0.032025723 |
| RPL41P5 | 0.032025499 |
| GPR176 | 0.032024606 |
| RP11-113I22 | 0.032016992 |
| RPL32P14 | 0.032016135 |
| RNU6-1222P | 0.032014567 |
| RP11-25P11 | 0.032014076 |
| SNHG6 | 0.032009387 |
| REEP2 | 0.032007861 |
| RP11-192C21 | 0.032003326 |
| CTD-2272D18 | 0.031998429 |
| MIR183 | 0.031987925 |
| SEMA3E | 0.03198367 |
| SRC | 0.031980586 |
| ACKR2 | 0.03198053 |
| RP11-774D14 | 0.031973211 |
| SNORD96B | 0.031972524 |
| GGT1 | 0.031971171 |
| RP11-725G5 | 0.031966523 |
| PPP6R2P1 | 0.031955014 |
| CYCSP11 | 0.03194647 |
| RP4-569D19 | 0.031946375 |
| OVOL3 | 0.031942616 |
| MIR3943 | 0.031935896 |
| RP11-484O2 | 0.031935527 |
| GLB1L | 0.031935337 |
| RN7SL468P | 0.031934498 |
| RAB44 | 0.031928925 |
| CHCHD2P9 | 0.031927393 |
| CLDN11 | 0.031926109 |
| MIR6737 | 0.031925621 |
| RP1-180M12 | 0.031925426 |
| LINC00934 | 0.031916892 |
| AC140912 | 0.031916168 |
| DENND5B-AS1 | 0.031908009 |
| LONRF3 | 0.031904545 |
| PMPCA | 0.03189567 |
| MYRF | 0.03189541 |
| RP11-280G9 | 0.031883175 |
| RP11-45I20 | 0.031877127 |
| ITGA7 | 0.031873325 |
| MIR4803 | 0.031866963 |
| KIAA1161 | 0.031866274 |

| | |
|---|---|
| NR1H4 | 0.031863708 |
| ACPT | 0.031863196 |
| RNA5SP97 | 0.031858031 |
| RP11-282M16 | 0.031857248 |
| RFPL1S | 0.031845919 |
| SLC52A3 | 0.031842217 |
| AQP7P4 | 0.031835058 |
| HTR7 | 0.031824226 |
| NPY4R | 0.03180853 |
| SH3PXD2A-AS1 | 0.03180831 |
| RP11-204I15 | 0.031806974 |
| SCAMP3 | 0.03180201 |
| CTC-575I10 | 0.031800671 |
| TCOF1 | 0.031791535 |
| LINC00556 | 0.031790651 |
| RNU1-27P | 0.031786795 |
| RP11-1029M24 | 0.031783767 |
| EFEMP1 | 0.031776837 |
| RPL37P23 | 0.031766639 |
| EXOSC5 | 0.031762641 |
| FREM2 | 0.031755911 |
| ATF4P2 | 0.031753863 |
| AC067959 | 0.031752812 |
| CRIP2 | 0.031748482 |
| LINC00337 | 0.031747888 |
| AC007568 | 0.031744537 |
| RNU6-163P | 0.031743051 |
| GRM2 | 0.031742938 |
| NAB2 | 0.031721746 |
| OR5D18 | 0.031721587 |
| CTD-2378E21 | 0.031719896 |
| RP11-123O10 | 0.031711886 |
| RPL35P6 | 0.031709169 |
| TMEM261P1 | 0.031695801 |
| OSCAR | 0.03169408 |
| LRRC3-AS1 | 0.031690818 |
| PDP1 | 0.03168861 |
| ARHGEF7-AS1 | 0.031687131 |
| SERPINB8 | 0.031679251 |
| KCNK9 | 0.031672523 |
| RNU1-4 | 0.031668696 |
| HOXB7 | 0.031667325 |
| RP5-1099C19 | 0.031666615 |
| GOLGA8DP | 0.031666187 |
| WDR34 | 0.031665167 |

| | |
|---|---|
| AC097108 | 0.031663757 |
| F12 | 0.031661948 |
| MOGAT3 | 0.031661005 |
| RNU2-48P | 0.031660714 |
| CYP46A1 | 0.031648887 |
| AMMECR1 | 0.031644199 |
| AC007405 | 0.031642747 |
| LSM10 | 0.03164258 |
| CTD-2377O17 | 0.031636712 |
| ILF2 | 0.031636515 |
| GCG | 0.031634816 |
| RP11-822E23 | 0.031630473 |
| GFRA3 | 0.03162548 |
| HSFY3P | 0.03162209 |
| ASNSP6 | 0.031621275 |
| VPS51 | 0.031611174 |
| STAP2 | 0.031610672 |
| TEX26-AS1 | 0.031591719 |
| LINC01035 | 0.031591184 |
| RN7SL503P | 0.031587772 |
| MIR4308 | 0.031585314 |
| TMEM129 | 0.031585145 |
| AL645730 | 0.031584546 |
| KMT2E-AS1 | 0.031574968 |
| EDARADD | 0.031573684 |
| PDE6A | 0.031572679 |
| SIM1 | 0.031570109 |
| AC002553 | 0.031566994 |
| SGSH | 0.0315652 |
| ABO | 0.031563823 |
| CAMP | 0.031551906 |
| RPS3AP5 | 0.031523989 |
| MESDC2 | 0.031521626 |
| CH507-216K13 | 0.031521562 |
| AC022431 | 0.03151913 |
| RPL26L1 | 0.031514091 |
| RP11-115H18 | 0.031511289 |
| RBCK1 | 0.0315081 |
| XXbac-BPG55C20 | 0.031507125 |
| ZBTB8OSP2 | 0.03150315 |
| RP11-175G14 | 0.031500415 |
| RNF212 | 0.031497365 |
| RP11-161I6 | 0.031497084 |
| RP11-484K9 | 0.031493987 |
| RNU6-383P | 0.031482803 |

| | |
|---|---|
| GS1-600G8 | 0.031472494 |
| AC036178 | 0.03144923 |
| RP11-1079J22 | 0.03144779 |
| RP11-744J10 | 0.03144517 |
| RP11-7G12 | 0.031432942 |
| AC009158 | 0.031431905 |
| ITGA1 | 0.03142337 |
| VN1R31P | 0.031421436 |
| ADCK4 | 0.031416803 |
| SLC9A1 | 0.031407481 |
| RP11-195C7 | 0.031397315 |
| RP11-542B15 | 0.031396574 |
| AL359253 | 0.0313954 |
| RP11-459O1 | 0.03139459 |
| RNA5SP27 | 0.031383743 |
| PPIA | 0.031383313 |
| OR13C2 | 0.031380624 |
| MAGEH1 | 0.031380224 |
| RP3-412A9 | 0.031379313 |
| C19orf24 | 0.031373898 |
| KRT17P7 | 0.031370602 |
| RP11-447L10 | 0.031369374 |
| RP11-434D2 | 0.031361582 |
| OR9A3P | 0.031359633 |
| ERRFI1 | 0.031357284 |
| SEC61G | 0.031356856 |
| RNU6-753P | 0.031355701 |
| GOLGA2P9 | 0.031352406 |
| UHRF2P1 | 0.031347616 |
| PTENP1-AS | 0.031346785 |
| RNU6-376P | 0.031346148 |
| AC005701 | 0.031337515 |
| KCNQ4 | 0.031337246 |
| SARS2 | 0.031334678 |
| CTA-313A17 | 0.031329333 |
| AC005276 | 0.031325118 |
| CA5AP1 | 0.031324994 |
| AL136137 | 0.031322359 |
| CTC-340A15 | 0.031319434 |
| LYPD2 | 0.031318534 |
| RP11-78F17 | 0.031314286 |
| ILF2P1 | 0.031311392 |
| ZNF503-AS1 | 0.031304265 |
| CTD-2083E4 | 0.031301784 |
| AP000290 | 0.0312989 |

| | |
|---|---|
| PCDHB19P | 0.031295142 |
| FMO10P | 0.031293497 |
| RP11-414H23 | 0.031291176 |
| THPO | 0.031289794 |
| RP11-775J23 | 0.031280778 |
| RP11-177F15 | 0.03127792 |
| LHX8 | 0.031275033 |
| SMARCA5-AS1 | 0.031271414 |
| RP11-314C16 | 0.031266433 |
| SDAD1P1 | 0.031265094 |
| ROBO4 | 0.03125741 |
| LIX1L | 0.031255344 |
| RP4-734C18 | 0.03125318 |
| GSTZ1 | 0.031246805 |
| CAGE1 | 0.031245612 |
| SC22CB-1D7 | 0.031233707 |
| AHR | 0.031232069 |
| RP11-393K10 | 0.031228978 |
| MIR365B | 0.031227517 |
| CRYBB2P1 | 0.031216943 |
| C14orf178 | 0.031216588 |
| AC002429 | 0.031212872 |
| OR2F2 | 0.031211863 |
| RNA5SP376 | 0.03121083 |
| NREP-AS1 | 0.031206894 |
| LINC00315 | 0.031194107 |
| RP11-547I7 | 0.031190116 |
| SDR9C7 | 0.031186671 |
| TFAMP2 | 0.031173518 |
| CRISP3 | 0.031164638 |
| RP11-42F12 | 0.031163872 |
| MIR6771 | 0.031155937 |
| TRAPPC6A | 0.031152624 |
| LPCAT2BP | 0.031141483 |
| KRTAP1-3 | 0.031132729 |
| SNHG3 | 0.031120641 |
| MIR148A | 0.031112058 |
| MPP4 | 0.031110805 |
| TTTY7B | 0.031103054 |
| CDC20B | 0.031101525 |
| FAU | 0.031098947 |
| OR7G2 | 0.031093071 |
| AP000318 | 0.031085526 |
| RNA5SP192 | 0.031083845 |
| AC092597 | 0.03108208 |

| | |
|---|---|
| SF3B4 | 0.031078313 |
| TCEA2 | 0.031076323 |
| RP11-864I4 | 0.031061085 |
| RP11-472K22 | 0.031056139 |
| NOTCH2 | 0.03105154 |
| MIR5088 | 0.03104877 |
| RP11-1127D7 | 0.031045527 |
| PPP1R2P2 | 0.031044286 |
| ZBED3 | 0.031037716 |
| AP001059 | 0.031027647 |
| LINC01524 | 0.031024303 |
| MIR501 | 0.031022573 |
| RP11-254F7 | 0.031010515 |
| RP11-16N2 | 0.031010403 |
| LTBP2 | 0.03100685 |
| AC067969 | 0.031003325 |
| FILIP1L | 0.030998031 |
| RP11-187A9 | 0.030988536 |
| RP11-305O4 | 0.030986799 |
| CARM1P1 | 0.030986748 |
| OR52T1P | 0.03097949 |
| LINC01021 | 0.030978772 |
| RP11-69E9 | 0.030978636 |
| CTB-164L20 | 0.030977692 |
| SOCS5 | 0.030976914 |
| MAGI2-AS3 | 0.030970978 |
| AC006466 | 0.030970233 |
| RN7SKP211 | 0.030968291 |
| RP11-403A3 | 0.030957299 |
| FBXL6 | 0.030954438 |
| RP11-96K14 | 0.030951728 |
| AC005019 | 0.030946666 |
| LIPF | 0.030944754 |
| RNU4-71P | 0.030942781 |
| CTB-1H10 | 0.030938003 |
| LINC00577 | 0.030930817 |
| RP11-116D17 | 0.030929291 |
| RPL6P9 | 0.03092712 |
| RP11-321P16 | 0.030926583 |
| RP11-451G4 | 0.030924076 |
| AC068492 | 0.030919922 |
| RNU6-1172P | 0.0309189 |
| MIR4722 | 0.030915354 |
| OR9P1P | 0.030911304 |
| MMP24 | 0.030907348 |

| | |
|---|---|
| AC104088 | 0.03090734 |
| MPRIP | 0.030899292 |
| AC108022 | 0.030896053 |
| CTB-43I4 | 0.03089334 |
| GS1-115M3 | 0.030882073 |
| CENPBD1P1 | 0.030877709 |
| RNU6-71P | 0.030867549 |
| SENP3 | 0.030861802 |
| RP11-605F14 | 0.030860105 |
| RP11-348F1 | 0.030858486 |
| PSMC3 | 0.030858322 |
| CTD-2034I21 | 0.030849898 |
| RNU6-706P | 0.030843978 |
| AC087269 | 0.030831286 |
| RP11-589M4 | 0.030821117 |
| LHFPL2 | 0.030817148 |
| LDLR | 0.030812181 |
| AC003045 | 0.030798812 |
| RNU1-38P | 0.030791747 |
| AP000251 | 0.03078621 |
| AC012501 | 0.03078043 |
| RPL23AP67 | 0.030776536 |
| MIR1258 | 0.030776436 |
| PABPC4 | 0.030775044 |
| FXYD6 | 0.030774114 |
| RN7SL14P | 0.030772005 |
| C11orf1 | 0.03076803 |
| NUTM2E | 0.030766605 |
| INHBA-AS1 | 0.030759006 |
| FAM8A4P | 0.03075312 |
| ACOT1 | 0.030749462 |
| RP11-155N3 | 0.030746953 |
| LINC00551 | 0.030746137 |
| PCDHA11 | 0.030741814 |
| XYLT1 | 0.030733908 |
| ATP6V1G1P3 | 0.030727879 |
| OR2M3 | 0.030722328 |
| RP11-1082L8 | 0.030722106 |
| RP11-69E11 | 0.030713609 |
| CDK4 | 0.030710297 |
| C9orf50 | 0.0307034 |
| TMEM134 | 0.030700492 |
| SNAP25-AS1 | 0.030679721 |
| ITCH-IT1 | 0.030679077 |
| PSMG1 | 0.030671058 |

| | |
|---|---|
| AC026471 | 0.030669054 |
| CTD-2540M10 | 0.030668116 |
| RP11-272J7 | 0.030664942 |
| RNA5SP166 | 0.030663239 |
| RPL7AP34 | 0.030657 |
| PPP1R42 | 0.03065514 |
| RNA5SP179 | 0.030647173 |
| RN7SL649P | 0.030641068 |
| MIR4497 | 0.030640525 |
| AC074183 | 0.030640159 |
| CITF22-49E9 | 0.030638232 |
| RNU6-364P | 0.030637788 |
| KBTBD13 | 0.030636528 |
| BAI1 | 0.030619503 |
| RP11-115J16 | 0.030613237 |
| TPRX1P1 | 0.030611717 |
| SPANXA2-OT1 | 0.030607938 |
| RP11-165D7 | 0.030607891 |
| RN7SL346P | 0.030605578 |
| MIR378G | 0.030593169 |
| POF1B | 0.0305879 |
| RAB9BP1 | 0.030586101 |
| PEG10 | 0.030584822 |
| ANG | 0.030584468 |
| MGAT2 | 0.030581882 |
| GRIK2 | 0.030579783 |
| RP11-137D19 | 0.030579269 |
| UPK1A | 0.030576161 |
| AP000344 | 0.030560521 |
| RAB4A | 0.030560077 |
| HIGD1B | 0.030559649 |
| LAMA5-AS1 | 0.030557583 |
| RNA5SP382 | 0.030541074 |
| NPM1P2 | 0.030537033 |
| RP4-753F5 | 0.030533383 |
| MAPT-AS1 | 0.030529742 |
| RP11-212P7 | 0.030508279 |
| SH2D4A | 0.030505656 |
| EPHB6 | 0.03050307 |
| RP11-1057B6 | 0.030496102 |
| AC004895 | 0.030495774 |
| AC141930 | 0.030493632 |
| MTRF1LP2 | 0.030492508 |
| BRMS1 | 0.030491981 |
| RP13-49I15 | 0.030489527 |

| | |
|---|---|
| RP11-474D1 | 0.030488465 |
| RP11-506M13 | 0.030485172 |
| CLPB | 0.030482901 |
| PPM1G | 0.030482194 |
| MIR449B | 0.030481515 |
| RP11-21C17 | 0.030479663 |
| TEKT4P2 | 0.030473673 |
| RP11-351I24 | 0.030470861 |
| RNA5SP222 | 0.030469181 |
| DEFB136 | 0.03046739 |
| RTN4RL1 | 0.030465933 |
| NDP-AS1 | 0.030465524 |
| RP11-72J9 | 0.03046374 |
| AMIGO3 | 0.030449896 |
| RP11-529G21 | 0.030443354 |
| PRR23D2 | 0.030432186 |
| MIR3910-1 | 0.030431326 |
| PDIA4 | 0.030426229 |
| AC106827 | 0.030424488 |
| RP11-809C18 | 0.03041972 |
| PPP1CB | 0.030419419 |
| DDX26B-AS1 | 0.03040286 |
| C3orf14 | 0.030401856 |
| CACFD1 | 0.030401372 |
| HSF1 | 0.030390748 |
| AL365510 | 0.030380552 |
| MIR1976 | 0.030380092 |
| PLP1 | 0.030368346 |
| AC108683 | 0.030366993 |
| RP11-542G1 | 0.030358956 |
| RS1 | 0.030358313 |
| LAMA2 | 0.030357054 |
| CA13 | 0.030352756 |
| IMMP2L | 0.030350579 |
| IL34 | 0.03034643 |
| ISY1-RAB43 | 0.030344599 |
| RP11-680F8 | 0.030342802 |
| GALNT9 | 0.030330113 |
| FABP4 | 0.030313536 |
| AC006004 | 0.030308309 |
| HAUS1P3 | 0.030308192 |
| AC007228 | 0.030295674 |
| AC016994 | 0.030289613 |
| OLFM3 | 0.03028566 |
| TSPY24P | 0.030277467 |

| | |
|---|---|
| ESRG | 0.030272669 |
| AC019186 | 0.030269825 |
| AC005152 | 0.030267698 |
| SFRP1 | 0.030266799 |
| OR52J1P | 0.030259449 |
| RN7SKP104 | 0.030249225 |
| FAM168A | 0.030246909 |
| RN7SL656P | 0.030241644 |
| GJD4 | 0.030239257 |
| PLGRKT | 0.030238518 |
| OR11H1 | 0.030232043 |
| SSXP4 | 0.03020777 |
| OR7E110P | 0.030206238 |
| ARL5AP1 | 0.030206123 |
| RP11-468H14 | 0.03019896 |
| LINC00923 | 0.030195948 |
| SNRPGP2 | 0.030194461 |
| EXTL2 | 0.030194349 |
| ARHGEF7-AS2 | 0.030193298 |
| ISM2 | 0.030192666 |
| GRK4 | 0.03019113 |
| RP5-944M2 | 0.030190915 |
| B4GALT4 | 0.030190479 |
| ENDOU | 0.030187547 |
| HOXC10 | 0.030186795 |
| RBMY2JP | 0.030175251 |
| AP000619 | 0.030174953 |
| ASCL5 | 0.030166652 |
| ZBTB45 | 0.030161064 |
| TSTD1 | 0.030160413 |
| MIR4477A | 0.030156697 |
| C15orf52 | 0.030154738 |
| RNU6-281P | 0.030153257 |
| RP11-753A21 | 0.030152638 |
| FAM74A7 | 0.030143754 |
| STRA8 | 0.030143523 |
| SPIN2A | 0.030133204 |
| TBC1D20 | 0.030122476 |
| CAMKK1 | 0.030119295 |
| AC005477 | 0.030118267 |
| UBE2FP2 | 0.030117999 |
| RP11-339D20 | 0.030113875 |
| RP11-403P14 | 0.030106804 |
| RHEBP3 | 0.030101192 |
| RP11-77E14 | 0.030095957 |

| | |
|---|---|
| RP11-511B7 | 0.030095901 |
| AC005357 | 0.030095636 |
| RN7SKP57 | 0.030088786 |
| AC067960 | 0.030084889 |
| PKMP5 | 0.030082843 |
| TUB-AS1 | 0.030078598 |
| RP11-17M15 | 0.030075993 |
| CRTC1 | 0.030073506 |
| ALOX15B | 0.03007159 |
| TEKT1 | 0.03006652 |
| ARMC10P1 | 0.030050377 |
| KRT8P3 | 0.030044408 |
| RP11-85A1 | 0.030041587 |
| GFI1B | 0.030036856 |
| RP11-358H18 | 0.030036779 |
| RP11-435O11 | 0.030036104 |
| GULP1 | 0.030036016 |
| KRT34 | 0.030034478 |
| FAM86DP | 0.030033505 |
| GEM | 0.030031998 |
| MIRLET7F2 | 0.030027429 |
| SNORD114-10 | 0.030026122 |
| LA16c-380H5 | 0.030024611 |
| CGREF1 | 0.030024332 |
| KCNAB3 | 0.030019569 |
| OR7E29P | 0.03001306 |
| GSN-AS1 | 0.030010899 |
| MIR4535 | 0.03000781 |
| MIR320B1 | 0.030005462 |
| AL121657 | 0.029997426 |
| ZHX1-C8orf76 | 0.02999341 |
| RP11-357K6 | 0.029988258 |
| RP11-100N3 | 0.029987237 |
| RP11-430L3 | 0.029983979 |
| IMP4 | 0.029982228 |
| RNU6-488P | 0.029980586 |
| LINC00395 | 0.029980144 |
| CTC-246B18 | 0.029980051 |
| OR5BD1P | 0.029977237 |
| OR51L1 | 0.029968079 |
| AL121760 | 0.029964834 |
| CKLF | 0.029954729 |
| RP11-570G20 | 0.029950639 |
| PET117 | 0.029945892 |
| LINC00518 | 0.029945768 |

| | |
|---|---|
| ENO4 | 0.02994351 |
| HEG1 | 0.029937166 |
| RPL21P66 | 0.029918438 |
| TBC1D3P6 | 0.029917482 |
| COX8A | 0.029913258 |
| ILF2P2 | 0.029913164 |
| LINC01143 | 0.029912887 |
| AC231759 | 0.029907161 |
| FBXW4P1 | 0.029902559 |
| ADD3 | 0.029899936 |
| ST7-OT3_2 | 0.029899717 |
| EEF1GP2 | 0.029888508 |
| RP11-521C20 | 0.029888169 |
| CDK14 | 0.029880847 |
| MIR129-2 | 0.029880374 |
| AC006033 | 0.029872199 |
| RNU6-29P | 0.029863156 |
| TMEM238 | 0.029855934 |
| C2orf72 | 0.029854428 |
| CHTF18 | 0.029850204 |
| CTC-254B4 | 0.029846329 |
| NR2F6 | 0.029841775 |
| TMC1 | 0.029832248 |
| CBR3 | 0.029831152 |
| ZNF517 | 0.029830612 |
| RP11-747H12 | 0.029828645 |
| BSND | 0.029828602 |
| RP4-610C12 | 0.029822786 |
| CXCL6 | 0.029821829 |
| SLC25A43 | 0.029821384 |
| CTB-147C22 | 0.029821009 |
| RP11-355F22 | 0.029813374 |
| RBX1 | 0.029812993 |
| KB-1183D5 | 0.029807084 |
| RP11-326G21 | 0.029799383 |
| LTF | 0.02979835 |
| OR8T1P | 0.029767009 |
| AC062017 | 0.029766154 |
| KARSP1 | 0.029761859 |
| CTGLF10P | 0.029754988 |
| RP5-1158E12 | 0.029752861 |
| SLC7A14 | 0.029748715 |
| OSBPL10-AS1 | 0.029746135 |
| LINC00677 | 0.029745738 |
| PGAM1P12 | 0.029731829 |

| | |
|---|---|
| RP11-617I14 | 0.029731384 |
| CYP39A1 | 0.029731296 |
| HNRNPA1P62 | 0.029726332 |
| KCNMA1-AS1 | 0.029724017 |
| LINC01118 | 0.029722349 |
| AC009474 | 0.029721241 |
| MIR499A | 0.029718129 |
| NRTN | 0.029716461 |
| RN7SL831P | 0.029715534 |
| LL0XNC01-37G1 | 0.029709405 |
| CTNNB1 | 0.029704858 |
| SERBP1 | 0.029699965 |
| DEGS1 | 0.029695425 |
| ZNF71 | 0.029692039 |
| RP11-834C11 | 0.029690922 |
| AC016716 | 0.029690259 |
| RP11-30L3 | 0.029689493 |
| DUSP6 | 0.029680913 |
| DIO2 | 0.029671424 |
| OPLAH | 0.029668745 |
| OTOG | 0.029668324 |
| AC010641 | 0.029666043 |
| RP11-875H7 | 0.029661991 |
| MIR569 | 0.029656512 |
| OR8D4 | 0.029655889 |
| LINC00561 | 0.029655508 |
| GYG2P1 | 0.029653731 |
| ITGB4 | 0.029653321 |
| SH3D21 | 0.029650511 |
| RP11-287D1 | 0.029650176 |
| IER5 | 0.029650158 |
| NANOGNBP1 | 0.029647671 |
| RNASET2 | 0.029636818 |
| RPLP0P11 | 0.029632898 |
| PCDHA10 | 0.02962521 |
| GSDMD | 0.029615198 |
| AC134873 | 0.029613086 |
| AC068196 | 0.029610517 |
| MIR6786 | 0.029609589 |
| BAG3 | 0.029607812 |
| MIR6888 | 0.029600619 |
| EN2 | 0.029596956 |
| LINC01006 | 0.029596406 |
| HMGB1P42 | 0.029578451 |
| RP11-515O17 | 0.029567998 |

| | |
|---|---|
| ESAM | 0.029561372 |
| AC068580 | 0.029561279 |
| RNU6-985P | 0.029548108 |
| RNU6-284P | 0.029548108 |
| STK32A | 0.029543416 |
| ZNF513 | 0.029539585 |
| RP11-1C8 | 0.029537502 |
| CFL1P4 | 0.029534378 |
| RP11-124N14 | 0.029528444 |
| PCDHGA8 | 0.0295231 |
| CTD-2015G9 | 0.029521382 |
| AL034395 | 0.029515015 |
| PPP1R1A | 0.02950857 |
| RP11-48G14 | 0.029508226 |
| RP11-77O7 | 0.029492021 |
| RP11-26L20 | 0.029486696 |
| AC078917 | 0.029484564 |
| C8orf74 | 0.029482888 |
| SHROOM2 | 0.029482615 |
| LINC00535 | 0.0294815 |
| TNFRSF4 | 0.029477257 |
| OXER1 | 0.029471713 |
| CLEC4GP1 | 0.029471263 |
| LAGE3P1 | 0.029462122 |
| RP11-357D18 | 0.029452175 |
| RP11-109P6 | 0.029445105 |
| RRP12 | 0.029440102 |
| C20orf194 | 0.029432362 |
| RP11-392O17 | 0.029430297 |
| MIR548A1 | 0.029426633 |
| CHEK2P5 | 0.029424403 |
| PDE7B | 0.029424399 |
| RNU6-620P | 0.029411627 |
| HSD17B7 | 0.029405886 |
| SERAC1 | 0.029404528 |
| RP3-422G23 | 0.029400773 |
| AC068522 | 0.029399385 |
| FAM90A26 | 0.029393463 |
| IGF2R | 0.029392156 |
| AC073091 | 0.029392026 |
| AP000692 | 0.029390425 |
| FH | 0.0293873 |
| MIR6858 | 0.029383288 |
| ATP2C1 | 0.029377997 |
| RHOQP3 | 0.029357819 |

| | |
|---|---|
| LDHBP3 | 0.029355097 |
| AC068706 | 0.029354363 |
| NXF4 | 0.029345941 |
| AC092625 | 0.029338868 |
| GAS7 | 0.029333307 |
| OR4A44P | 0.029323618 |
| RP11-339D23 | 0.029322589 |
| PNP | 0.029315241 |
| C3orf33 | 0.029313827 |
| FMO9P | 0.029309518 |
| IL22RA2 | 0.029306436 |
| WDR24 | 0.029302496 |
| LRP5 | 0.029297714 |
| RN7SKP194 | 0.029292785 |
| RP11-419M24 | 0.029291024 |
| EIF4EBP3 | 0.029289416 |
| RP11-482E14 | 0.02928473 |
| RP11-138B9 | 0.029275596 |
| ACAA1 | 0.029270569 |
| RP11-395N6 | 0.029269269 |
| CACNA1C-AS3 | 0.029266781 |
| RP11-157P1 | 0.029253805 |
| RP4-726F1 | 0.029247426 |
| SNORA74A | 0.029246441 |
| FUT11 | 0.02924601 |
| ARF1 | 0.029244093 |
| STK25P1 | 0.029242376 |
| AC073427 | 0.029237718 |
| AC010096 | 0.029229116 |
| AC009965 | 0.029221917 |
| MROH2A | 0.029220776 |
| TPBGL | 0.029219385 |
| RARRES2P6 | 0.029216941 |
| SLFN13 | 0.029208352 |
| EPOR | 0.029207068 |
| MIR130B | 0.029206779 |
| LINC01179 | 0.029206421 |
| IFRD2 | 0.029206064 |
| MIR3922 | 0.029201105 |
| RNU6-800P | 0.029200534 |
| CTD-2014B16 | 0.029199073 |
| PIRT | 0.029197584 |
| MIR4304 | 0.029196492 |
| AC002486 | 0.029185349 |
| RN7SL708P | 0.029183035 |

| | |
|---|---|
| RP11-184D12 | 0.029169699 |
| LINC01338 | 0.029165373 |
| CALML4 | 0.02916396 |
| SV2A | 0.029163699 |
| SCNN1A | 0.029141707 |
| RP11-190J1 | 0.029138191 |
| FABP5P3 | 0.02913062 |
| C19orf67 | 0.02912478 |
| MIR132 | 0.029117805 |
| AC007319 | 0.029115017 |
| AMIGO2 | 0.029108218 |
| RP11-764K9 | 0.029107287 |
| RPS2P52 | 0.029082675 |
| OTX2-AS1 | 0.029077313 |
| ARMCX1 | 0.029072402 |
| RP11-12M9 | 0.029067881 |
| RP11-83C7 | 0.029067014 |
| PABPC5-AS1 | 0.029058247 |
| RP13-34L8 | 0.029052595 |
| AL591668 | 0.029048515 |
| RNU6-1320P | 0.029035566 |
| CPSF4 | 0.029032735 |
| DACT3-AS1 | 0.029032038 |
| RP11-2C24 | 0.029030893 |
| CTD-2325M2 | 0.029026502 |
| RP3-471C18 | 0.029022028 |
| PRKAB2 | 0.029019376 |
| NUDT14 | 0.029018018 |
| MTX1 | 0.02901532 |
| RP5-1039K5 | 0.029000641 |
| TMEM132E | 0.028993166 |
| CTD-2231E14 | 0.028982017 |
| RP11-182E7 | 0.028981467 |
| HCG16 | 0.028973061 |
| RP11-411H5 | 0.028971755 |
| MRPL34 | 0.028964193 |
| RNU6-1101P | 0.028962722 |
| MRPL57P7 | 0.028961848 |
| CCDC54 | 0.028959129 |
| RTN4R | 0.028957513 |
| RNU6-206P | 0.02895579 |
| SLC40A1 | 0.028946536 |
| RNASEH2CP1 | 0.028943718 |
| RNU6-1103P | 0.028937552 |
| SCN3B | 0.028936788 |

| | |
|---|---|
| FRG2DP | 0.028927071 |
| RP11-47J17 | 0.028922141 |
| AL035697 | 0.0289212 |
| PAM16 | 0.028915391 |
| OR4F29 | 0.028911568 |
| LL22NC03-80A10 | 0.028911167 |
| RNU6-15P | 0.02891099 |
| MIR6854 | 0.028907228 |
| RNU6-148P | 0.028903172 |
| RNA5S9 | 0.028902766 |
| CCT5 | 0.02889766 |
| CBLN1 | 0.028891373 |
| RN7SKP223 | 0.028890207 |
| LINC01282 | 0.028887411 |
| MKKS | 0.028884003 |
| SHC4 | 0.028881491 |
| RP11-508N22 | 0.028879705 |
| AL359851 | 0.028877735 |
| SLC39A12 | 0.028874578 |
| MAGEA2B | 0.028872674 |
| NEUROD2 | 0.02887227 |
| AC007271 | 0.028867941 |
| AL593844 | 0.028865703 |
| RP11-20O24 | 0.028860895 |
| UBXN8 | 0.028858569 |
| AACS | 0.028857044 |
| RP11-568A7 | 0.028852533 |
| XKR3 | 0.028849307 |
| RP11-282C5 | 0.028843255 |
| IQCJ | 0.028835987 |
| CALR | 0.028831368 |
| RP11-156J23 | 0.028815586 |
| RP11-125D12 | 0.028801872 |
| RNU6-1334P | 0.028801202 |
| LRRC29 | 0.028799832 |
| AC004108 | 0.028799463 |
| RNU6-1050P | 0.02879649 |
| AC073321 | 0.028794413 |
| RNU6-704P | 0.028792806 |
| NRG2 | 0.028789256 |
| RP11-321F6 | 0.028779716 |
| RP11-415H23 | 0.028774611 |
| RP11-128P17 | 0.028769361 |
| KCNS3 | 0.028767272 |
| RNU2-65P | 0.028766734 |

| | |
|---|---|
| RP11-292I17 | 0.028762378 |
| ZNF575 | 0.028760072 |
| AC113331 | 0.028754991 |
| RP11-889D3 | 0.028750843 |
| TNNT3 | 0.028748193 |
| RP11-92G12 | 0.028747294 |
| RP11-293A21 | 0.028745608 |
| SLC38A3 | 0.028733503 |
| CTC-350I8 | 0.028727248 |
| PWRN4 | 0.028724369 |
| CTC-369M3 | 0.028723173 |
| RP11-378I13 | 0.028722156 |
| PRSS16 | 0.028713919 |
| TMEM221 | 0.028713438 |
| RN7SKP93 | 0.028701586 |
| SNORA70C | 0.028698311 |
| CTAGE13P | 0.02868479 |
| ESYT2 | 0.028680683 |
| RP11-524L6 | 0.028678344 |
| DLG4 | 0.028678059 |
| LINC00592 | 0.028673281 |
| AC009081 | 0.028672564 |
| SNORD113-5 | 0.028662729 |
| HNRNPA1P66 | 0.02865853 |
| MROH7-TTC4 | 0.028644903 |
| RN7SKP47 | 0.028639492 |
| RPS23P6 | 0.028638679 |
| SPATA31E1 | 0.028636009 |
| RP11-48F14 | 0.028633942 |
| RP11-169F17 | 0.028627395 |
| APH1A | 0.028617387 |
| AC073264 | 0.028611315 |
| RP11-545H22 | 0.028610892 |
| AC098830 | 0.028602413 |
| LINC00839 | 0.028597163 |
| TBC1D7 | 0.028594576 |
| RNU7-8P | 0.028591956 |
| RSL24D1P11 | 0.028590553 |
| AL353751 | 0.028584638 |
| AC152010 | 0.028584259 |
| RP11-647K16 | 0.02857932 |
| AC008834 | 0.028578265 |
| UST | 0.028577106 |
| DVL2 | 0.028569757 |
| SULT1C3 | 0.028569732 |

| | |
|---|---|
| RP11-879F14 | 0.028566464 |
| MGC39584 | 0.028566423 |
| GNAL | 0.028564622 |
| RP11-10E18 | 0.028563659 |
| AGAP3 | 0.028549254 |
| RNU6-1299P | 0.028544928 |
| FRG2FP | 0.028544147 |
| RP11-111D3 | 0.028543836 |
| RN7SL643P | 0.0285422 |
| SERF2 | 0.028533476 |
| RN7SL865P | 0.028533179 |
| ANKRD33 | 0.028531563 |
| CTD-3066C23 | 0.028530971 |
| C16orf95 | 0.028518648 |
| HIST1H2BL | 0.028518606 |
| TATDN1P1 | 0.028517812 |
| AC069240 | 0.028516764 |
| AL022341 | 0.028516146 |
| RPL21P38 | 0.02851211 |
| TINCR | 0.028511522 |
| MIR542 | 0.028501936 |
| VMA21 | 0.028478832 |
| C6orf203 | 0.028476692 |
| KLF5 | 0.028475725 |
| HIST1H1B | 0.028468139 |
| FUT9 | 0.028464938 |
| FGF23 | 0.028460825 |
| FAM71E2 | 0.028460507 |
| MIR6077 | 0.028456921 |
| MIR655 | 0.028456312 |
| BTF3L4P3 | 0.028453697 |
| LRRC31 | 0.028453247 |
| SLC5A3 | 0.028451891 |
| RP11-158I13 | 0.028446152 |
| RNU6-984P | 0.028433246 |
| SNORD116-3 | 0.028432985 |
| SNORD114-18 | 0.028428151 |
| HMGN1P15 | 0.028425507 |
| FARP1 | 0.028423815 |
| SNRPGP12 | 0.028421725 |
| RNU2-3P | 0.028417669 |
| MIR3166 | 0.028407047 |
| RP11-116O11 | 0.028406725 |
| MIR375 | 0.028405731 |
| RNU6-1307P | 0.028405719 |

| | |
|---|---|
| ARMC2 | 0.02840294 |
| SLC37A3 | 0.028399256 |
| EDDM3DP | 0.028398896 |
| RPL23P11 | 0.028398756 |
| NT5C3B | 0.028384986 |
| AC027347 | 0.028384968 |
| RPP25L | 0.028379851 |
| ATP5F1P7 | 0.028378391 |
| RNU6-788P | 0.028375403 |
| ACOX3 | 0.028373506 |
| LEMD2 | 0.028370362 |
| SNORD6 | 0.028367522 |
| AC087897 | 0.02836516 |
| SIRPA | 0.028361227 |
| GP6 | 0.028351876 |
| MRPS21P9 | 0.028351773 |
| CENPW | 0.02835064 |
| ADAM30 | 0.028349684 |
| RASA3 | 0.028348947 |
| AC010295 | 0.028336434 |
| RP13-30A9 | 0.028333078 |
| RNU6-2 | 0.028331527 |
| IFNA8 | 0.028330961 |
| CTD-3076M17 | 0.028329818 |
| CTD-2007A10 | 0.028325911 |
| OTX2 | 0.028320085 |
| MT1B | 0.028309585 |
| AC072061 | 0.02830866 |
| RN7SL65P | 0.028303976 |
| CCDC169 | 0.02830105 |
| TMPRSS11F | 0.028300317 |
| WFDC3 | 0.028299345 |
| RNU1-95P | 0.028297038 |
| L1TD1 | 0.028296504 |
| RAG1 | 0.028295326 |
| RP11-516A11 | 0.028294132 |
| EFNA2 | 0.028293331 |
| FARP1-AS1 | 0.028283797 |
| NOXA1 | 0.028273465 |
| RP11-512N4 | 0.02826351 |
| GPX1P2 | 0.028256289 |
| HMGN5 | 0.028256069 |
| RP11-238F2 | 0.028254482 |
| POLR3H | 0.028252347 |
| CTAGE11P | 0.028250341 |

| | |
|---|---|
| LCMT1-AS1 | 0.028240009 |
| KSR1P1 | 0.028238844 |
| RP5-1056H1 | 0.028232109 |
| CTD-2060C23 | 0.02823162 |
| RP3-340B19 | 0.028231269 |
| RP6-1O2 | 0.028227683 |
| AC073900 | 0.028226808 |
| SUCLA2-AS1 | 0.028224663 |
| RNU6-929P | 0.028224043 |
| CCAR2 | 0.028223025 |
| PABPC1L2A | 0.028215137 |
| RNA5SP249 | 0.02821311 |
| RNU5E-9P | 0.028212688 |
| KRT16P5 | 0.028211314 |
| FERD3L | 0.028199708 |
| AC006007 | 0.028197646 |
| AC091565 | 0.028183299 |
| RNU6-349P | 0.028179941 |
| YDJC | 0.028177609 |
| CH17-189H20 | 0.028174614 |
| RPL5P20 | 0.028174504 |
| COMT | 0.028169497 |
| DPYD-AS1 | 0.028167616 |
| MTRNR2L10 | 0.028166262 |
| GLDCP1 | 0.028164183 |
| RN7SL267P | 0.02816243 |
| UBL5P2 | 0.028156393 |
| ICAM1 | 0.028152269 |
| OR1F1 | 0.028148924 |
| RPL30P15 | 0.028147646 |
| DGAT2L6 | 0.028147394 |
| AQPEP | 0.028145737 |
| STOML3 | 0.028139337 |
| RN7SL221P | 0.028131572 |
| FBXL8 | 0.028129384 |
| AL132709 | 0.028128947 |
| CTC-340D7 | 0.02812864 |
| RP11-345E19 | 0.028121295 |
| PTP4A1P3 | 0.028117472 |
| GALNT6 | 0.028114741 |
| MIR5190 | 0.028113688 |
| RP11-613D13 | 0.028112174 |
| RABL6 | 0.028109495 |
| FLAD1 | 0.028107578 |
| FMR1-AS1_2 | 0.028104973 |

| | |
|---|---|
| CTD-2542O7 | 0.028101434 |
| RP11-49E18 | 0.02809509 |
| RPL7P55 | 0.028093049 |
| RP11-81F13 | 0.028091995 |
| PRAMEF17 | 0.028091002 |
| GUCY1B3 | 0.028090554 |
| DUSP21 | 0.028082612 |
| TMEM203 | 0.028078107 |
| MIR6814 | 0.028078028 |
| RP11-114O8 | 0.028077314 |
| OR7E148P | 0.028064047 |
| IL11RA | 0.028058348 |
| SLCO4A1-AS1 | 0.028053559 |
| RP11-758N13 | 0.02804258 |
| LMO7 | 0.028035687 |
| OR7E41P | 0.028033996 |
| SNORA27 | 0.02803095 |
| SPSB3 | 0.02802391 |
| E2F6P3 | 0.028023439 |
| PTGES | 0.028017472 |
| RP1-124C6 | 0.028013438 |
| AC019129 | 0.028012399 |
| RP4-609E1 | 0.028012102 |
| RP11-33H15 | 0.02801015 |
| SPHKAP | 0.0280075 |
| SLC6A10P | 0.027992875 |
| RNU6-1252P | 0.027971857 |
| FES | 0.027970847 |
| GOLGA8CP | 0.027970776 |
| RP1-232L22__A | 0.027970554 |
| HIST1H3PS1 | 0.027954055 |
| ERO1L | 0.027953711 |
| RP11-126O22 | 0.027952842 |
| AL592549 | 0.027951025 |
| ITGA10 | 0.027944297 |
| RNU6-405P | 0.027943536 |
| DAZ1 | 0.027941081 |
| BVES-AS1 | 0.027934614 |
| RP11-536G4 | 0.027928703 |
| POMP | 0.027924631 |
| AC096664 | 0.027923428 |
| CNTNAP3 | 0.027914711 |
| ACEA_U3 | 0.027914222 |
| SERPINA11 | 0.027912136 |
| MFSD6L | 0.027910333 |

| | |
|---|---|
| IGHVIII-26-1 | 0.027906 |
| AL138498 | 0.027897179 |
| UBAP1 | 0.027896909 |
| EML1 | 0.027893522 |
| TMEM51-AS1 | 0.027884666 |
| DCLK3 | 0.027872238 |
| RPS26P13 | 0.02786503 |
| PPIAP16 | 0.027846075 |
| CD2BP2 | 0.027834004 |
| ACSM4 | 0.027833105 |
| FTLP14 | 0.027828284 |
| RP11-547L9 | 0.027827915 |
| RP11-217E22 | 0.027813544 |
| ADAMTS8 | 0.027813315 |
| RP11-434I12 | 0.027810804 |
| KIAA0930 | 0.027806653 |
| OR7E101P | 0.027806307 |
| BNIP3P1 | 0.027790481 |
| OR7E62P | 0.027787314 |
| RN7SL177P | 0.027783375 |
| RP11-651J20 | 0.02778189 |
| RN7SL275P | 0.027781561 |
| IFRD1 | 0.027780959 |
| RFPL4AP3 | 0.027776437 |
| RP11-293I14 | 0.027773856 |
| C8orf59P2 | 0.027766549 |
| TPSD1 | 0.027766108 |
| KRT10 | 0.027764028 |
| NPR1 | 0.027760289 |
| AC092289 | 0.027757833 |
| RP11-643N23 | 0.027754996 |
| MC3R | 0.027744906 |
| RN7SL292P | 0.027741487 |
| TCEB1P18 | 0.027732361 |
| SOX12 | 0.027729162 |
| FAM223A | 0.027726209 |
| PLA2G4A | 0.027722222 |
| INTS9 | 0.027716547 |
| SFTA3 | 0.027712632 |
| RPL17P25 | 0.02771012 |
| RP11-793K1 | 0.027703297 |
| RP4-796I8 | 0.027702222 |
| CTD-3064M3 | 0.027693048 |
| RP11-480G7 | 0.027686629 |
| RP11-68E19 | 0.027673107 |

| | |
|---|---|
| LSM1P2 | 0.027670729 |
| PIEZO1 | 0.02765714 |
| AP1B1 | 0.027651875 |
| LINC00970 | 0.027643815 |
| ALAS2 | 0.027643056 |
| MIR3140 | 0.027634876 |
| AL356140 | 0.027632494 |
| MIR3193 | 0.02763191 |
| RNU1-13P | 0.027631138 |
| C9orf66 | 0.02762659 |
| FAM50B | 0.027618627 |
| RP11-341D18 | 0.027611606 |
| RN7SL339P | 0.02760661 |
| NDUFS5P2 | 0.027604053 |
| PFDN1 | 0.027599128 |
| RP11-359N11 | 0.027598216 |
| RNU6-391P | 0.027598041 |
| AC007304 | 0.027594853 |
| PLD2 | 0.027590272 |
| ZNF101P1 | 0.027577396 |
| SNRPCP20 | 0.02757738 |
| OR51B5 | 0.027570193 |
| HNF1A | 0.027565704 |
| FAM90A14P | 0.027564075 |
| NMS | 0.027557123 |
| ABCF3 | 0.027556001 |
| SNORD115-35 | 0.027555373 |
| CHMP1A | 0.027550345 |
| CHCHD2P3 | 0.027546139 |
| NACAP7 | 0.02753841 |
| IGKV3D-34 | 0.027537665 |
| JUP | 0.027531925 |
| ANGPTL3 | 0.027525783 |
| TMEM237 | 0.027524799 |
| RP1-105O18 | 0.027519042 |
| IRAK2 | 0.027510045 |
| TMEM107 | 0.02750878 |
| ZNF714 | 0.027504966 |
| PPIAL4G | 0.027498228 |
| FIGN | 0.027497598 |
| RP4-745K6 | 0.027493168 |
| AC011357 | 0.027489753 |
| RP5-1103B4 | 0.027488948 |
| TGM1 | 0.027487644 |
| MTND2P15 | 0.027481144 |

| | |
|---|---|
| RP11-7D5 | 0.027473982 |
| GAPDHP17 | 0.027471949 |
| LINC01430 | 0.027471403 |
| ZNF705B | 0.027462637 |
| KRT8P22 | 0.027456179 |
| RP11-696L21 | 0.027450439 |
| AC009413 | 0.027448146 |
| PDCD6 | 0.0274466 |
| WFDC12 | 0.027445524 |
| KLHL40 | 0.0274449 |
| TNS1 | 0.027439921 |
| AK4 | 0.027438656 |
| PHLDB2 | 0.0274378 |
| RP4-612B18 | 0.027435074 |
| BANF1P3 | 0.027434578 |
| PARVB | 0.027424739 |
| OR7G1 | 0.027423485 |
| RP11-118M12 | 0.027418412 |
| NPY2R | 0.027417941 |
| LINC01571 | 0.027411568 |
| RP13-192B19 | 0.027411554 |
| SLC6A18 | 0.027401249 |
| ST13P11 | 0.027398626 |
| SSMEM1 | 0.02739775 |
| AMACR | 0.027397538 |
| RNY3P12 | 0.027390097 |
| CH17-408M7 | 0.027384368 |
| MTND6P8 | 0.027369945 |
| RP11-301L8 | 0.027352323 |
| CD1A | 0.027351109 |
| RNU6-13P | 0.027348731 |
| RP11-81B10 | 0.027344204 |
| RP11-275F13 | 0.027341302 |
| CCND2-AS2 | 0.027334832 |
| NOC2L | 0.027330651 |
| RARRES2P1 | 0.027329084 |
| GUCY1A3 | 0.027328586 |
| RN7SL516P | 0.027327939 |
| GOLGA6L1 | 0.02732494 |
| MIR1237 | 0.027320151 |
| GAPDHP40 | 0.027319625 |
| MIR8067 | 0.027318098 |
| H6PD | 0.027316623 |
| LINC01214 | 0.027315459 |
| LINC00642 | 0.027313974 |

| | |
|---|---|
| PKHD1 | 0.027312103 |
| AC084117 | 0.027308226 |
| RP11-96B5 | 0.027306527 |
| CPNE8 | 0.027303264 |
| FAM96AP2 | 0.027300726 |
| LRRC23 | 0.027294501 |
| SLC6A1 | 0.027281198 |
| AC068858 | 0.027279961 |
| LINC01073 | 0.027279219 |
| MTND1P22 | 0.027276987 |
| CTD-2314B22 | 0.027273353 |
| STOM | 0.027272044 |
| RBMX2P3 | 0.027271775 |
| ALS2CL | 0.027271728 |
| ARMC9 | 0.027270928 |
| RP9 | 0.027270649 |
| DSG2-AS1 | 0.027266267 |
| RP11-573N10 | 0.027262011 |
| CTC-436K13 | 0.027261604 |
| RP11-542F9 | 0.027253208 |
| RN7SKP77 | 0.027246708 |
| RP11-749H20 | 0.027243953 |
| CYP2J2 | 0.027234796 |
| CHIAP3 | 0.027234363 |
| KB-1958F4 | 0.027217589 |
| RP11-202D18 | 0.02721412 |
| CBR1 | 0.027212972 |
| EVX1-AS | 0.027212306 |
| RNF208 | 0.027208581 |
| ZNF711 | 0.027207539 |
| SKOR1 | 0.027204776 |
| FAH | 0.027202683 |
| BPESC1 | 0.027200601 |
| RP11-404G16 | 0.027200204 |
| LINC00681 | 0.027192852 |
| PRX | 0.027173007 |
| RP11-802H3 | 0.027171904 |
| TMSB10P2 | 0.027171528 |
| RP4-777L9 | 0.027170653 |
| EHBP1L1 | 0.027166067 |
| MIR30A | 0.02716328 |
| RN7SKP21 | 0.027154223 |
| KRT17P1 | 0.027145752 |
| RP11-92G19 | 0.027143556 |
| RN7SL546P | 0.027142466 |

| | |
|---|---|
| CTD-2339M3 | 0.027131686 |
| RP11-320A16 | 0.027130709 |
| ZNF560 | 0.027126996 |
| RP1-315G1 | 0.02711703 |
| NDEL1 | 0.027116316 |
| LINC00378 | 0.027115082 |
| RP11-711C17 | 0.027112989 |
| SNORA69 | 0.027112616 |
| FLG2 | 0.027105747 |
| DLG2-AS1_2 | 0.027097539 |
| ARF4P3 | 0.027096934 |
| U6atac | 0.027094677 |
| FEV | 0.027094585 |
| CRX | 0.027090844 |
| RP11-212D3 | 0.027085509 |
| MIR100HG | 0.027078956 |
| RP11-686G8 | 0.027075756 |
| MSTO1 | 0.027072705 |
| RP4-668E10 | 0.027066853 |
| PSPHP1 | 0.027055499 |
| MYO9A | 0.027044611 |
| BMS1P8 | 0.027037765 |
| HES1 | 0.027034899 |
| LINC01570 | 0.027018286 |
| FAM181A | 0.02700922 |
| SNORD114-21 | 0.027007931 |
| MFI2-AS1 | 0.026998302 |
| SLC25A26 | 0.026997371 |
| JADE3 | 0.026987343 |
| PPEF1-AS1 | 0.026979086 |
| RP11-298H24 | 0.026976753 |
| MIR4783 | 0.026970767 |
| APOL5 | 0.026967755 |
| PNMA6B | 0.026961985 |
| KRT8P14 | 0.026958069 |
| FGF9 | 0.026952376 |
| RPS3AP32 | 0.026949489 |
| CHRFAM7A | 0.026949208 |
| CTD-2318B16 | 0.026945889 |
| RP11-753G20 | 0.026943844 |
| PIGZ | 0.02694362 |
| RN7SKP296 | 0.02694298 |
| PAX9 | 0.026938933 |
| MTND3P2 | 0.02693602 |
| RN7SKP268 | 0.026931349 |

| | |
|---|---|
| JAZF1 | 0.026927936 |
| ADAM20P3 | 0.026926031 |
| CTD-2267G17 | 0.026914382 |
| GPR125 | 0.026909833 |
| AC135983 | 0.026906387 |
| AL591034 | 0.026904161 |
| RN7SL493P | 0.02689957 |
| RP11-145E5 | 0.026899349 |
| FGB | 0.026898402 |
| NIPA2P4 | 0.026895389 |
| AC066614 | 0.026880749 |
| TSEN34 | 0.026869552 |
| LCE5A | 0.026865771 |
| MPC2 | 0.026865719 |
| AL354720 | 0.026865553 |
| CICP27 | 0.026860863 |
| OR2H2 | 0.026860169 |
| KRT8P4 | 0.026859643 |
| PC | 0.026859003 |
| LIPE-AS1 | 0.026851746 |
| RNU6-1181P | 0.026848473 |
| PDE4A | 0.026845272 |
| GJC3 | 0.026844323 |
| FDX1P2 | 0.026842151 |
| SDHAF2 | 0.026840155 |
| RP11-481F24 | 0.026838036 |
| RP11-403I13 | 0.026837662 |
| MPP7 | 0.026833997 |
| MTND5P20 | 0.02683106 |
| NDRG1 | 0.026825438 |
| RN7SL190P | 0.026823337 |
| NAALADL2-AS2 | 0.02682139 |
| LINC01533 | 0.026820205 |
| STARD8 | 0.026818554 |
| MRPL9 | 0.026816768 |
| AL078471 | 0.026809691 |
| RN7SL283P | 0.02680561 |
| AL357973 | 0.0268001 |
| MIA-RAB4B | 0.026800043 |
| UNC93B8 | 0.026792797 |
| ATP5G1P6 | 0.02679098 |
| OR6K2 | 0.026789351 |
| RN7SL706P | 0.026789044 |
| TMEM216 | 0.02678739 |
| LDHAL6B | 0.026786752 |

| | | |
|---|---|---|
| AL138744 | | 0.026785334 |
| | 5-Sep | 0.02678339 |
| LINC00493 | | 0.026779036 |
| RP11-203F10 | | 0.026776428 |
| C9orf40 | | 0.026775948 |
| MAP3K11 | | 0.026773399 |
| HMGB3P5 | | 0.026767044 |
| IGDCC4 | | 0.026763798 |
| LINC01436 | | 0.026759725 |
| AC104135 | | 0.026753215 |
| ARRDC1 | | 0.02674856 |
| RANBP17 | | 0.026742551 |
| PLP2 | | 0.026737918 |
| TNIK | | 0.02672802 |
| CBWD5 | | 0.026724857 |
| RN7SL843P | | 0.026724316 |
| RP11-1217F2 | | 0.02672203 |
| RPS2P55 | | 0.02672121 |
| EEF1GP1 | | 0.02671394 |
| COQ9 | | 0.02671241 |
| SMPDL3B | | 0.02670627 |
| MAPK4 | | 0.026703972 |
| LINC00387 | | 0.026703446 |
| PRDX4 | | 0.026702958 |
| OR4B1 | | 0.026696016 |
| LLNLR-276E7 | | 0.026691876 |
| AC008781 | | 0.026691269 |
| KRT13 | | 0.02668554 |
| MAGEA11 | | 0.02668241 |
| PRRT4 | | 0.026680726 |
| AC090079 | | 0.026679734 |
| AC007106 | | 0.026673314 |
| CTC-228N24 | | 0.026673169 |
| RP11-761N21 | | 0.026669059 |
| COX20P1 | | 0.026665075 |
| RELT | | 0.026664054 |
| AC107993 | | 0.026654313 |
| FAM228A | | 0.026653632 |
| MME-AS1 | | 0.026650698 |
| OR52I2 | | 0.026641905 |
| MT-TC | | 0.026638349 |
| MARK2P13 | | 0.026637842 |
| GAGE12J | | 0.026635969 |
| CXorf22 | | 0.02663485 |
| RPSAP23 | | 0.026632667 |

| | |
|---|---|
| RP11-373N22 | 0.026631942 |
| RP11-626A5 | 0.026623694 |
| UBE2E1-AS1 | 0.026608431 |
| RP11-6J21 | 0.026599511 |
| RP1-52J10 | 0.026595221 |
| VDR | 0.026585409 |
| DNAJC22 | 0.026582414 |
| RP11-20G6 | 0.026570719 |
| CNN2P10 | 0.02656957 |
| AC005215 | 0.026565409 |
| PIK3C2G | 0.026565255 |
| AC011286 | 0.026563283 |
| SGCA | 0.026562664 |
| RP11-685F15 | 0.026548153 |
| NTHL1 | 0.026547324 |
| RP11-367H1 | 0.026544184 |
| AC096649 | 0.026527958 |
| RP11-260M19 | 0.026521738 |
| RPS6P16 | 0.026514593 |
| CCNY | 0.026512234 |
| CYCSP1 | 0.026510481 |
| SUMF2 | 0.026508978 |
| SNORD115 | 0.026497544 |
| RP4-601P9 | 0.026491315 |
| RPL17P26 | 0.026489056 |
| RN7SL220P | 0.026488885 |
| EEF1A1P6 | 0.026488363 |
| AL512655 | 0.026483521 |
| RP11-507F16 | 0.026470957 |
| MT-TT | 0.026468535 |
| MIR19B2 | 0.026464632 |
| GABARAPL1 | 0.02646319 |
| CES5A | 0.026463118 |
| SLC2A8 | 0.026456213 |
| RNU4-42P | 0.026453202 |
| LINC01271 | 0.026451312 |
| CX3CL1 | 0.026451282 |
| CTA-14H9 | 0.026448975 |
| NCOA7 | 0.026441905 |
| AC004791 | 0.026440247 |
| RP5-1185H19 | 0.026437949 |
| RP11-339A11 | 0.026433326 |
| PNLIPRP1 | 0.026432616 |
| NXNL1 | 0.026432198 |
| AAED1 | 0.026428788 |

| | |
|---|---|
| AC133104 | 0.026427495 |
| RPS7P9 | 0.026420042 |
| CRLS1 | 0.026415448 |
| RP11-463J17 | 0.02641544 |
| ACN9 | 0.026413085 |
| LARP7P4 | 0.026412361 |
| ARHGAP5-AS1 | 0.026401401 |
| EMC1 | 0.026394052 |
| SLC25A11 | 0.026389103 |
| FAM96B | 0.026384656 |
| MIR5706 | 0.026382111 |
| TPI1P4 | 0.026378282 |
| FEM1AP1 | 0.026371542 |
| LIMS1 | 0.026367093 |
| RP5-1112D6 | 0.026365586 |
| TMEM98 | 0.026365213 |
| AL445464 | 0.026364654 |
| RP11-120J4 | 0.026361847 |
| LEFTY2 | 0.026360197 |
| RP1-313L4 | 0.026360091 |
| PDCD5P1 | 0.026356656 |
| RP11-193H22 | 0.026351324 |
| DUSP7 | 0.026347557 |
| PCNXL2 | 0.026346441 |
| VN1R20P | 0.026343687 |
| RP11-5N19 | 0.026340542 |
| NBR2 | 0.026337938 |
| RPS29 | 0.026333286 |
| AC004988 | 0.026333244 |
| XX-DJ76P10__A | 0.026333033 |
| CDRT7 | 0.026325861 |
| AC009365 | 0.02632482 |
| SGK2 | 0.02632316 |
| RPP21 | 0.02632216 |
| RN7SL184P | 0.026318987 |
| CBX3P3 | 0.02631888 |
| TBC1D22A | 0.026317937 |
| RP1-292B18 | 0.026315807 |
| BEND3P3 | 0.026305768 |
| RHOBTB1 | 0.026301899 |
| LINC01230 | 0.026297059 |
| MIR520C | 0.026296922 |
| MIR6087 | 0.026293264 |
| SNX29 | 0.026292652 |
| RN7SKP291 | 0.026288973 |

| | |
|---|---|
| NANOS2 | 0.026287788 |
| FADS6 | 0.026284269 |
| IQCF5-AS1 | 0.026284217 |
| ACTBP8 | 0.0262771 |
| RPL19P11 | 0.02627636 |
| RP11-740C1 | 0.026260659 |
| MIR143HG | 0.026256539 |
| RP11-100F15 | 0.026252778 |
| IHH | 0.026248517 |
| RPS4XP19 | 0.026244909 |
| KB-1466C5 | 0.026237344 |
| MBL1P | 0.026236082 |
| MIR6853 | 0.026234921 |
| PSMA3P | 0.026222138 |
| CMAHP | 0.026221951 |
| B3GNT3 | 0.026204659 |
| RP11-295M18 | 0.026203956 |
| RP11-120E13 | 0.026193574 |
| FER1L6-AS2 | 0.026193195 |
| AL163953 | 0.026173671 |
| RPS24P7 | 0.026168302 |
| RP5-1195D24 | 0.02616813 |
| RP11-818F20 | 0.026162014 |
| HK3 | 0.026159883 |
| SNRPD2P2 | 0.026148126 |
| IGHD1OR15-1B | 0.026136086 |
| RP11-282K24 | 0.026106251 |
| RNU4-53P | 0.026095981 |
| RP11-116B13 | 0.026092705 |
| USP17L16P | 0.026090027 |
| AC004819 | 0.02608316 |
| AC084262 | 0.026073606 |
| RP11-232M24 | 0.026068553 |
| HIGD1A | 0.026067061 |
| RPL21P18 | 0.026058588 |
| RP11-395M19 | 0.026057116 |
| RNU4-1 | 0.026051996 |
| AC092155 | 0.026047922 |
| SCRG1 | 0.026047265 |
| KCNF1 | 0.026041367 |
| MPLKIP | 0.026039506 |
| SRRM3 | 0.026031134 |
| ACAT2 | 0.026028436 |
| RP11-305F5 | 0.026027823 |
| TNFSF15 | 0.026027338 |

| | |
|---|---|
| CTD-3093M3 | 0.026027121 |
| TGM4 | 0.026022587 |
| RP11-123G9 | 0.026017959 |
| ADAMTS17 | 0.026015003 |
| BCL6B | 0.026012937 |
| RPL27A | 0.02600957 |
| RP1-213J1P__B | 0.026003719 |
| RN7SL366P | 0.026003025 |
| TARS2P1 | 0.02600254 |
| RN7SL732P | 0.026001091 |
| AL445204 | 0.025999605 |
| LURAP1 | 0.025998035 |
| RP11-481A20 | 0.025995653 |
| LINC01427 | 0.025995377 |
| NDUFA3P3 | 0.025991404 |
| THUMPD3P1 | 0.025987533 |
| MIR8070 | 0.025985352 |
| RP11-407P15 | 0.025984721 |
| RP11-408E5 | 0.02597176 |
| LURAP1L-AS1 | 0.025970878 |
| OFD1P9Y | 0.025969039 |
| SLC9A7P1 | 0.025964267 |
| AADACL3 | 0.025962984 |
| ALDH7A1P1 | 0.025952746 |
| CCDC6 | 0.025952581 |
| RNU4-90P | 0.025952178 |
| OR2T4 | 0.025950847 |
| CYP4A22-AS1 | 0.025949653 |
| RN7SL591P | 0.025938408 |
| RP11-359H3 | 0.025934489 |
| LINC01428 | 0.025931864 |
| HHIPL2 | 0.025926405 |
| RP11-173M11 | 0.025925509 |
| SYN1 | 0.025918682 |
| NT5CP2 | 0.025914033 |
| RP11-84O12 | 0.025912518 |
| OBFC1 | 0.025910645 |
| AL928970 | 0.025906485 |
| TREH | 0.025902139 |
| RP11-815D16 | 0.025901263 |
| BZW2 | 0.025891686 |
| TCTE1 | 0.025890761 |
| RP11-659O3 | 0.025890573 |
| RP11-338O1 | 0.025886243 |
| RN7SL111P | 0.02588253 |

| | |
|---|---|
| MIR4435-2 | 0.025878184 |
| MRPL23 | 0.025877546 |
| RP6-114E22 | 0.025876638 |
| RNA5SP48 | 0.025872039 |
| OLFML2A | 0.025870959 |
| AC079340 | 0.025867788 |
| AP001065 | 0.025860452 |
| YAP1P2 | 0.025860169 |
| RP3-497J21 | 0.025859613 |
| OR2M2 | 0.025851726 |
| NUDT19P4 | 0.025849028 |
| RN7SL863P | 0.0258396 |
| AC021914 | 0.025836732 |
| CKM | 0.025834603 |
| AC068499 | 0.0258297 |
| PCP4L1 | 0.025825097 |
| SKINTL | 0.025820688 |
| ATP6V0E2 | 0.025812445 |
| RNU6-115P | 0.025810145 |
| MIR4761 | 0.025808866 |
| AP002414 | 0.025808366 |
| LINC00458 | 0.025793592 |
| EFNB2 | 0.025781913 |
| RARRES2P3 | 0.025769565 |
| MIR7152 | 0.025758281 |
| FTLP19 | 0.025755723 |
| RPL7AP69 | 0.025754602 |
| RP11-679I18 | 0.025751286 |
| MINOS1P4 | 0.025728637 |
| AC116176 | 0.025726643 |
| MIR1302-3 | 0.025723204 |
| KRTAP10-9 | 0.025719742 |
| KRTAP3-2 | 0.025717732 |
| MIR3179-3 | 0.025715995 |
| AC018738 | 0.025699082 |
| OSGIN1 | 0.025686281 |
| MSX2 | 0.025684235 |
| RPL5P8 | 0.025684088 |
| TMSB4XP4 | 0.02568323 |
| AC099339 | 0.025681385 |
| USP12-AS1 | 0.025673383 |
| RP11-401F2 | 0.025669344 |
| AL160269 | 0.025665653 |
| RN7SL72P | 0.025663573 |
| RP11-778D12 | 0.025663224 |

| | |
|---|---|
| DUTP1 | 0.025656775 |
| RP11-20J1 | 0.025641842 |
| CTD-2019O4 | 0.025632112 |
| BRAFP1 | 0.025627929 |
| AC008154 | 0.02562528 |
| RNF113B | 0.025620923 |
| RP11-429G19 | 0.02561818 |
| RWDD2B | 0.025615581 |
| RASGEF1C | 0.025614583 |
| RP11-560B16 | 0.025609911 |
| IGHVII-31-1 | 0.02560047 |
| GABRP | 0.025599115 |
| RASD2 | 0.025596026 |
| RP11-80P20 | 0.025595952 |
| OR51A9P | 0.025592652 |
| ZNF649 | 0.025592398 |
| RP11-368J21 | 0.025588365 |
| APOBEC3B-AS1 | 0.025585881 |
| MRPS5P3 | 0.025583153 |
| MAPT-IT1 | 0.025576213 |
| RN7SL464P | 0.025572173 |
| YARS | 0.025560304 |
| RP11-624J12 | 0.025559839 |
| UGT2B11 | 0.025558961 |
| AC026797 | 0.025551326 |
| RP11-208P4 | 0.025551056 |
| RNF223 | 0.025548062 |
| DPRXP7 | 0.025546653 |
| RN7SL598P | 0.025546626 |
| AC105398 | 0.025540731 |
| KRT8P24 | 0.025537959 |
| RNU4-66P | 0.025534458 |
| C1orf210 | 0.025526498 |
| TCEB3B | 0.025522914 |
| STXBP5-AS1 | 0.02552259 |
| RP5-827O9 | 0.025521029 |
| RP13-347D8 | 0.025520909 |
| SAC3D1 | 0.02552086 |
| CTD-2066L21 | 0.025519912 |
| GAPDHP32 | 0.025515518 |
| XRCC6P1 | 0.02550566 |
| RP13-565O16 | 0.025503611 |
| ATP5F1P3 | 0.025499307 |
| SLC26A10 | 0.025495765 |
| THRB-IT1 | 0.025494772 |

| | |
|---|---|
| GFER | 0.025486886 |
| FAM35DP | 0.02547733 |
| RN7SL371P | 0.025477135 |
| MEIS3P1 | 0.025473807 |
| MIR8074 | 0.025465103 |
| TMPRSS15 | 0.025464704 |
| AC127391 | 0.025463469 |
| RN7SKP6 | 0.025458744 |
| PPP1R11P2 | 0.025457299 |
| NTRK3-AS1 | 0.025456265 |
| RP11-25I15 | 0.025453573 |
| RP11-79P5 | 0.025450029 |
| STK40 | 0.02544255 |
| GNG10P1 | 0.025442324 |
| XKRYP1 | 0.025427326 |
| RHOA | 0.02542493 |
| RPS21 | 0.025413339 |
| SERPINA12 | 0.025407962 |
| OR4F28P | 0.025407279 |
| NRBP2 | 0.025398492 |
| GTF2IRD1 | 0.025382659 |
| RP5-1185I7 | 0.025373404 |
| ATP5C1 | 0.025369039 |
| PRPS1 | 0.025363458 |
| RP11-755O11 | 0.02536039 |
| RP11-1102P22 | 0.025359851 |
| RP11-63H19 | 0.025358007 |
| AC023158 | 0.025357787 |
| HENMT1 | 0.025355755 |
| AP000620 | 0.025353907 |
| RP11-445J9 | 0.025350286 |
| SCARNA18 | 0.025342269 |
| CRABP1 | 0.025341988 |
| LINC00319 | 0.025338366 |
| AC009009 | 0.025328754 |
| SSXP5 | 0.025317201 |
| RP11-464C19 | 0.025315386 |
| ULBP1 | 0.025312567 |
| MIR497HG | 0.025311028 |
| AP000662 | 0.025305987 |
| RNU6-567P | 0.025298819 |
| AC092620 | 0.025292518 |
| RPL13AP2 | 0.025292378 |
| RNU6ATAC5P | 0.025286161 |
| NME3 | 0.025283101 |

| | |
|---|---|
| AC011431 | 0.025279732 |
| MCC | 0.025279659 |
| RP11-521M14 | 0.025267202 |
| MIR381 | 0.025263888 |
| MIR652 | 0.02526222 |
| HPS6 | 0.025262039 |
| RPS17P2 | 0.025260049 |
| AL359218 | 0.025257561 |
| MTND2P19 | 0.025257042 |
| MRPL54 | 0.025256607 |
| VN1R36P | 0.025254206 |
| TSPY6P | 0.02524862 |
| MOSPD1 | 0.025247936 |
| FAM3D | 0.025242225 |
| RP11-453N3 | 0.025239925 |
| RNA5SP361 | 0.025237971 |
| BLID | 0.025232023 |
| RP11-111M22 | 0.025229331 |
| POLR2G | 0.025228512 |
| TSPY19P | 0.025228461 |
| KRTAP20-2 | 0.0252263 |
| AC068491 | 0.025216487 |
| PTPRG-AS1 | 0.025216009 |
| DSPP | 0.025215205 |
| HIGD1AP8 | 0.025211875 |
| ZNF777 | 0.025210856 |
| RPS20P32 | 0.025209261 |
| SYNM | 0.025204828 |
| ESPNL | 0.025201954 |
| MIR5000 | 0.025197828 |
| GSTA6P | 0.025196541 |
| FOXO3 | 0.025194948 |
| PIWIL2 | 0.02519465 |
| C7orf61 | 0.02518553 |
| HS3ST1 | 0.02518505 |
| CTD-3107M8 | 0.025180581 |
| NENFP3 | 0.025172988 |
| AC087073 | 0.025165273 |
| RP11-972K6 | 0.025151762 |
| FBXL21 | 0.025151553 |
| SMIM8 | 0.025145885 |
| CYP7B1 | 0.025145755 |
| C11orf52 | 0.025143713 |
| TMEM212-IT1 | 0.025143486 |
| SNORD88 | 0.02513717 |

| | |
|---|---|
| MIR3150A | 0.025134064 |
| EGR4 | 0.025127378 |
| RNVU1-17 | 0.025123278 |
| RP11-649G15 | 0.025120433 |
| HDGF | 0.02511928 |
| RN7SKP278 | 0.025117747 |
| HMGB1P1 | 0.025105977 |
| RP11-472G21 | 0.025101647 |
| MAP1LC3A | 0.025099781 |
| MIR30B | 0.025097041 |
| SHROOM2P1 | 0.025092888 |
| RP11-606P2 | 0.025090823 |
| RP11-268P4 | 0.025081046 |
| GUSBP1 | 0.02507872 |
| TFRC | 0.02507511 |
| RHOJ | 0.025071557 |
| RNU4-91P | 0.025069785 |
| AL121594 | 0.025067454 |
| RP11-707P20 | 0.025064996 |
| EIF3I | 0.025064394 |
| LINC01034 | 0.02506391 |
| ZNF90P1 | 0.025060095 |
| AC024896 | 0.025056385 |
| LRP10 | 0.025044783 |
| GUSB | 0.025043593 |
| RNU4-76P | 0.025042334 |
| PTPRF | 0.025039444 |
| RNY3P14 | 0.025035508 |
| MIR3926-2 | 0.02503529 |
| OXCT1-AS1 | 0.025035203 |
| PIP | 0.02502464 |
| HSD17B12 | 0.025016708 |
| MIR3917 | 0.025015246 |
| ABLIM1 | 0.025014047 |
| RP11-671M22 | 0.025007691 |
| RNU7-87P | 0.025006097 |
| RP11-105N14 | 0.025000302 |
| TIE1 | 0.024999516 |
| FTO-IT1 | 0.024996279 |
| RPSAP8 | 0.024995109 |
| AC078941 | 0.024990069 |
| RNU4-69P | 0.024986004 |
| TSLP | 0.02498284 |
| RP5-836N17 | 0.024981072 |
| RNU6-95P | 0.024980366 |

| | |
|---|---|
| RP11-263G22 | 0.024977445 |
| SLC12A2 | 0.024976855 |
| AC105914 | 0.024976821 |
| RP3-404K8 | 0.024973593 |
| RP4-644L1 | 0.024968288 |
| SNORA7 | 0.024958088 |
| AC034243 | 0.024944169 |
| RNY3P16 | 0.024943141 |
| RN7SKP250 | 0.024942739 |
| DDX43P2 | 0.024941669 |
| AC012499 | 0.024941116 |
| RP11-696D21 | 0.024941006 |
| RP11-278J20 | 0.024934686 |
| PXYLP1 | 0.024934379 |
| RNU2-36P | 0.024932242 |
| CTA-929C8 | 0.024931715 |
| CT45A11P | 0.024923688 |
| RP11-784B15 | 0.02491681 |
| SBK1 | 0.024901093 |
| RP11-618G20 | 0.024898775 |
| PRDX2P2 | 0.024897326 |
| GLI4 | 0.024894693 |
| C16orf90 | 0.024887184 |
| GPRC5C | 0.02488392 |
| AC007115 | 0.024882957 |
| EMID1 | 0.024881241 |
| RP4-718D20 | 0.024877291 |
| BCAM | 0.024875864 |
| AC079612 | 0.024869225 |
| KRTAP12-1 | 0.024860166 |
| snoU109 | 0.024856138 |
| KCNN4 | 0.024854561 |
| MIR589 | 0.024829215 |
| EIF1P7 | 0.024822503 |
| DHCR7 | 0.024816773 |
| TNN | 0.024813778 |
| AQP5 | 0.024810006 |
| STEAP3-AS1 | 0.024807973 |
| MIR548F3 | 0.02480794 |
| RP13-582O9 | 0.02480334 |
| CTA-331F8 | 0.024801663 |
| DNM1P35 | 0.02479846 |
| SOCS2P1 | 0.024793441 |
| MRPS21P6 | 0.024790841 |
| RP11-239A17 | 0.024786316 |

| | |
|---|---|
| BTD | 0.024785312 |
| C1orf234 | 0.024775585 |
| MIR6777 | 0.024770909 |
| ELOVL3 | 0.024766255 |
| HSPA8P6 | 0.024765143 |
| GDF11 | 0.024762241 |
| RP1-145M24 | 0.024761845 |
| RAP1BP3 | 0.024760511 |
| RP11-472G23 | 0.02475973 |
| RNF149 | 0.024752082 |
| CAPN6 | 0.024751952 |
| RNU7-96P | 0.024745192 |
| RN7SKP76 | 0.02474277 |
| RP11-442P12 | 0.024737065 |
| AC114737 | 0.02473074 |
| CFAP221 | 0.024726216 |
| RARSP1 | 0.024720846 |
| ZBTB7B | 0.024716825 |
| SLC25A6P5 | 0.024712536 |
| RP11-524P6 | 0.024711831 |
| RNU6-1243P | 0.024700573 |
| RN7SL634P | 0.024699635 |
| PPIAP14 | 0.024689558 |
| PON1 | 0.024685428 |
| HR | 0.02468166 |
| CIDEC | 0.024673252 |
| LL22NC03-13G6 | 0.024672747 |
| STEAP1B | 0.024670163 |
| EPHB1 | 0.024660525 |
| RP11-477N12 | 0.024657145 |
| RNU6-362P | 0.024651959 |
| MIR5704 | 0.02465114 |
| CTC-308K20 | 0.024637138 |
| BPIFA2 | 0.024636495 |
| SNORD115-39 | 0.024634797 |
| TMEM151B | 0.024632187 |
| AVPR1B | 0.024631755 |
| RNU6-674P | 0.024626992 |
| AC007367 | 0.024626916 |
| NDUFA6 | 0.024626086 |
| DMTN | 0.024620597 |
| RP11-93I21 | 0.024620333 |
| ALDH7A1P2 | 0.024615795 |
| RP11-520A21 | 0.024615155 |
| RP11-454H19 | 0.024606919 |

| | |
|---|---|
| LSM1 | 0.024606364 |
| CTD-2061E19 | 0.024600212 |
| RAP1BP1 | 0.024596886 |
| SKOR2 | 0.02458655 |
| SYNPO2L | 0.024584215 |
| RP11-149P14 | 0.024581373 |
| SH3BP4 | 0.024580986 |
| AC090666 | 0.024574804 |
| ERI1 | 0.024567404 |
| RP11-138I17 | 0.024565977 |
| FZD3 | 0.024553097 |
| PLBD1 | 0.024548129 |
| FAM83F | 0.024545487 |
| AC243945 | 0.02454416 |
| PRSS50 | 0.024538036 |
| CCER2 | 0.024529699 |
| UQCRFS1P1 | 0.024524278 |
| SPTLC1P3 | 0.024522027 |
| LCTL | 0.024518156 |
| SLC2A2 | 0.024514861 |
| RP1-169P22 | 0.024513867 |
| RAB11B-AS1 | 0.024513858 |
| ZNF391 | 0.024511253 |
| IPO13 | 0.024510676 |
| RPP25 | 0.024504239 |
| CTD-2651C21 | 0.024500848 |
| AC024067 | 0.024498316 |
| RP11-384C21 | 0.024492884 |
| AC004552 | 0.024492464 |
| RNU6-338P | 0.024479546 |
| AC009495 | 0.024474361 |
| AC004540 | 0.024468498 |
| MAMSTR | 0.024467995 |
| CTC-251D13 | 0.024463624 |
| RP11-46P12 | 0.024458893 |
| SNORD116-11 | 0.024457049 |
| MAGEA2 | 0.024452955 |
| DSCR4-IT1 | 0.024451236 |
| SPATA31A7 | 0.024448161 |
| KCNJ16 | 0.024445879 |
| YES1 | 0.024439213 |
| RPS6KA4 | 0.024435454 |
| AC092013 | 0.024434802 |
| CACNA2D3-AS1 | 0.024427831 |
| RNA5SP450 | 0.024427625 |

| | |
|---|---|
| SLC6A16 | 0.024424775 |
| AC097381 | 0.024423361 |
| ZSCAN5C | 0.024417624 |
| OBSL1 | 0.024413146 |
| RP11-180P8 | 0.024412026 |
| OR8R1P | 0.024402403 |
| GUSBP5 | 0.02439735 |
| TRPM3 | 0.024387541 |
| KRI1 | 0.024385889 |
| RP11-165A20 | 0.024384377 |
| RP11-378A12 | 0.024381105 |
| GPAA1P2 | 0.024374607 |
| PTPRM | 0.024370447 |
| RNA5SP308 | 0.02436907 |
| MIR885 | 0.024368238 |
| ZNHIT2 | 0.024367916 |
| GRID2IP | 0.024349832 |
| NDUFA5P2 | 0.024347464 |
| FRG2B | 0.024343999 |
| DEGS2 | 0.02434252 |
| RN7SKP221 | 0.024335486 |
| FLT4 | 0.024333292 |
| DMRT3 | 0.024332269 |
| AC008906 | 0.024325331 |
| RP1-121G13 | 0.024319586 |
| COPS7A | 0.024315114 |
| ATP6V0D1 | 0.024310946 |
| RP11-478H13 | 0.024292823 |
| AC013437 | 0.02429071 |
| AC010904 | 0.024289328 |
| GSS | 0.024289066 |
| RP11-10C8 | 0.024286838 |
| RP11-132G19 | 0.024280488 |
| AL138994 | 0.02427639 |
| GSTA9P | 0.024275135 |
| RP11-211C9 | 0.024274405 |
| RN7SL465P | 0.024271352 |
| CRYBA4 | 0.02426956 |
| RNF11 | 0.02426153 |
| RP11-34P13 | 0.024250814 |
| CTB-164N12 | 0.024250313 |
| KRTAP1-4 | 0.02424795 |
| WRAP53 | 0.024241567 |
| RP5-1007F24 | 0.024240238 |
| NIPAL2 | 0.02423686 |

| | |
|---|---|
| NOD1 | 0.024235222 |
| RNA5SP150 | 0.024233287 |
| UBE2A | 0.024230236 |
| MIR3935 | 0.024229923 |
| SNCA | 0.024228189 |
| RP11-179A10 | 0.024226781 |
| NUTF2P7 | 0.024225577 |
| RNU6-830P | 0.024225354 |
| RPL23AP62 | 0.024217948 |
| TC2N | 0.024212319 |
| RN7SKP152 | 0.02421013 |
| RP11-758M4 | 0.024204232 |
| CTD-2062F14 | 0.024203432 |
| OR4H6P | 0.02419641 |
| NAMPTP2 | 0.024193488 |
| RP4-743O11 | 0.024188039 |
| RP11-29B9 | 0.024185599 |
| SNRPEP3 | 0.024183102 |
| RN7SL191P | 0.024172175 |
| TCEAL3 | 0.0241592 |
| AC243654 | 0.024154358 |
| RP11-685A21 | 0.024148637 |
| MIR4721 | 0.024146869 |
| PPA2 | 0.024145972 |
| CHADL | 0.024145097 |
| MTND4P4 | 0.02413053 |
| USP17L3 | 0.024129317 |
| MRPS22 | 0.024121747 |
| C10orf99 | 0.024120426 |
| RP11-131K5 | 0.02411275 |
| MRPL45P1 | 0.024110457 |
| SNX18P27 | 0.024109556 |
| TSC22D3 | 0.024109468 |
| ZP2 | 0.024106731 |
| USP43 | 0.024103235 |
| AL139429 | 0.024090906 |
| SYNE4 | 0.024090155 |
| RN7SKP228 | 0.024082115 |
| PPP1R7 | 0.02407888 |
| BNIP3P18 | 0.024077675 |
| TMEM38B | 0.024077299 |
| RNU6-232P | 0.024074602 |
| SNORD15A | 0.024072668 |
| CTD-2332E11 | 0.024068061 |
| AC004453 | 0.024065838 |

| | |
|---|---|
| RP11-49K4 | 0.024065508 |
| PIGC | 0.02406314 |
| MIR4757 | 0.024058204 |
| COLEC11 | 0.024057149 |
| NDUFA3P4 | 0.024048777 |
| CTD-2542C24 | 0.024048662 |
| RNA5SP186 | 0.024047506 |
| GTF2H5 | 0.024045316 |
| CLEC14A | 0.024033446 |
| ABCA9-AS1 | 0.024029516 |
| RBFOX1 | 0.024026651 |
| EIF4A2P5 | 0.024023976 |
| GATA1 | 0.024022159 |
| FBXL13 | 0.02401845 |
| AC091185 | 0.024017053 |
| CASP3P1 | 0.024016865 |
| FER1L5 | 0.024014338 |
| FAM66E | 0.024010272 |
| PPIAP13 | 0.024010057 |
| CH17-140K24 | 0.02400769 |
| CYBRD1 | 0.023997314 |
| AC004386 | 0.023996364 |
| LDHAP3 | 0.0239953 |
| CTPS1 | 0.023990895 |
| RRS1 | 0.023979955 |
| BLVRB | 0.023976699 |
| TCHH | 0.023970699 |
| RNU6-547P | 0.023966608 |
| CTC-470E21 | 0.023960467 |
| GRM7 | 0.023958691 |
| C2CD4D | 0.023956029 |
| RP11-13J12 | 0.023955607 |
| CDY10P | 0.023952392 |
| PPAP2C | 0.023944642 |
| RP11-98L5 | 0.023941704 |
| C12orf77 | 0.023941675 |
| WDR38 | 0.023941417 |
| RFPL4B | 0.023930434 |
| RASSF7 | 0.023926353 |
| CTC-205M6 | 0.023921497 |
| RP5-1017F8 | 0.023920769 |
| RN7SKP49 | 0.023917795 |
| RP11-136C24 | 0.023915668 |
| MIR6081 | 0.023911425 |
| RN7SL577P | 0.023910193 |

| | |
|---|---|
| ASXL3 | 0.023901692 |
| LINC01586 | 0.023891977 |
| MAS1LP1 | 0.023886649 |
| AC104777 | 0.02388588 |
| KB-1517D11 | 0.02388372 |
| LINC01177 | 0.023876889 |
| RP11-1263C18 | 0.023876278 |
| FABP7P2 | 0.023875871 |
| RNU6-429P | 0.023874918 |
| RN7SKP295 | 0.023874646 |
| AC106765 | 0.023874503 |
| FAM108A8P | 0.023868804 |
| RPL23AP10 | 0.023865002 |
| LARGE-AS1 | 0.023864879 |
| ZNF32 | 0.023863373 |
| RP11-434E6 | 0.023857085 |
| RP11-647P12 | 0.023856475 |
| RP11-56I23 | 0.023856246 |
| LINC01442 | 0.023846298 |
| C17orf107 | 0.023845326 |
| RP11-263F14 | 0.023844098 |
| CYP4A22 | 0.023837723 |
| RNA5SP137 | 0.023836149 |
| RNA5-8SP4 | 0.023833934 |
| OR5BA1P | 0.023829774 |
| RP11-541G9 | 0.023829404 |
| ZNF571-AS1 | 0.023823334 |
| OR9K2 | 0.023822654 |
| EGLN3P1 | 0.023815258 |
| SLC6A20 | 0.023814553 |
| COX6CP4 | 0.023813047 |
| RN7SKP68 | 0.023809723 |
| OR13C7 | 0.023809429 |
| AC012066 | 0.023807418 |
| RP5-849H19 | 0.023795535 |
| RP11-467P22 | 0.02379537 |
| RN7SL645P | 0.023792903 |
| FBXW4 | 0.023792859 |
| FAM86JP | 0.023787724 |
| RP11-109D9 | 0.023784095 |
| CAMK2D | 0.023779109 |
| NTM-IT | 0.023774049 |
| RP4-724P12 | 0.023768471 |
| MEF2BNB | 0.023766737 |
| RP11-112L6 | 0.02376592 |

| | |
|---|---|
| PGAM1P1 | 0.023765743 |
| DLG5-AS1 | 0.02376495 |
| PRDM16 | 0.023762885 |
| DLK1 | 0.023762385 |
| PLXND1 | 0.02375989 |
| RP11-343P9 | 0.023750106 |
| TFDP2 | 0.023749247 |
| LINC01098 | 0.023746201 |
| RNU7-54P | 0.023743243 |
| MUL1 | 0.023742363 |
| RP11-96L7 | 0.023735559 |
| PICK1 | 0.023734028 |
| HOPX | 0.023721376 |
| C2CD4C | 0.023718251 |
| AP002490 | 0.023715453 |
| RP11-274J16 | 0.02371244 |
| IRF3 | 0.023696557 |
| STK11 | 0.023696501 |
| PALM3 | 0.023695665 |
| COA6 | 0.023694715 |
| RP4-647C14 | 0.023693747 |
| RP3-486D24 | 0.023692675 |
| PDLIM1P2 | 0.023690811 |
| OR52E3P | 0.023688165 |
| PTGES3L-AARSD1 | 0.023686695 |
| MIR4695 | 0.023685513 |
| HNRNPA1P33 | 0.023685187 |
| GPR88 | 0.023683175 |
| HSPA12A | 0.023682826 |
| AC018634 | 0.023681576 |
| RAET1M | 0.023679002 |
| MIR938 | 0.023677695 |
| RP11-789C17 | 0.023674448 |
| PPP1R1C | 0.023630423 |
| RNA5SP110 | 0.023621719 |
| ST8SIA6-AS1 | 0.023618116 |
| RNU6-707P | 0.023607225 |
| RP11-174M13 | 0.023604531 |
| KCNQ1 | 0.023604497 |
| B3GALT1 | 0.023602152 |
| RP11-673P17 | 0.023601468 |
| FGF7 | 0.02359631 |
| ELP5 | 0.023593873 |
| RP11-214N1 | 0.023589078 |
| RN7SL714P | 0.023585603 |

| | |
|---|---|
| RP11-298P3 | 0.023574554 |
| HOXB8 | 0.023560179 |
| GOLGA2P10 | 0.023554783 |
| C11orf95 | 0.023554281 |
| RP11-455B2 | 0.023553958 |
| GOLGA6L16P | 0.023550261 |
| CEL | 0.023541629 |
| ME3 | 0.023536866 |
| LRGUK | 0.023535564 |
| RP11-57A19 | 0.02352734 |
| GPNMB | 0.023524158 |
| PRAMEF10 | 0.023523335 |
| LGALS14 | 0.023520258 |
| ADARB1 | 0.02351973 |
| TK1 | 0.023516326 |
| PTPRA | 0.023511321 |
| RP11-1057N3 | 0.023509157 |
| RP11-456H18 | 0.023506481 |
| MAF1 | 0.023505386 |
| RPL5P22 | 0.023501744 |
| RNU6-1229P | 0.023500853 |
| SLC9A3 | 0.023498296 |
| SNRPGP19 | 0.023496953 |
| CTSA | 0.023487989 |
| RP11-54H7 | 0.023477263 |
| CTD-2298J14 | 0.023475196 |
| RNA5SP476 | 0.023474507 |
| RP11-240L7 | 0.023465687 |
| RP11-490G8 | 0.02345308 |
| RP11-7K24 | 0.023447225 |
| AC006272 | 0.02344467 |
| RP11-121P10 | 0.023444517 |
| RNU6-796P | 0.023443089 |
| RP11-318C24 | 0.023437373 |
| RPL4P5 | 0.023430275 |
| GPT2 | 0.023421754 |
| FTH1P15 | 0.023418585 |
| ANKRD62P1-PARP4P3 | 0.023414449 |
| AC024568 | 0.023407538 |
| LINC01170 | 0.023401408 |
| TSPEAR | 0.023394101 |
| TMPRSS11B | 0.02338235 |
| LINC00858 | 0.02337401 |
| FXYD4 | 0.023371881 |
| NR1H5P | 0.023363773 |

| | |
|---|---|
| AC131011 | 0.02336005 |
| RNU6-992P | 0.02335987 |
| EPDR1 | 0.023348504 |
| RP11-286O18 | 0.023335682 |
| NTN5 | 0.023312527 |
| AC020571 | 0.023310903 |
| SNORA14A | 0.023310844 |
| OR13Z1P | 0.023307054 |
| AC104169 | 0.023302117 |
| HYAL3 | 0.023296316 |
| AL513023 | 0.023291423 |
| RPL38 | 0.023280799 |
| AC107057 | 0.023277191 |
| GNAI3 | 0.023274324 |
| RP11-340L3 | 0.023272358 |
| MIR1286 | 0.023267534 |
| TSPAN17 | 0.023260445 |
| MARK4 | 0.023253951 |
| TFE3 | 0.023253077 |
| RPUSD2 | 0.023252165 |
| RNA5SP165 | 0.023240851 |
| LINC01107 | 0.023235046 |
| RP11-208K4 | 0.023228348 |
| RNA5SP161 | 0.023226293 |
| GS1-21A4 | 0.023219618 |
| PSMC1P8 | 0.023215346 |
| PYY3 | 0.023215235 |
| RSPO1 | 0.023210529 |
| RP11-806O11 | 0.023208734 |
| RP11-304M2 | 0.023208122 |
| MMP17 | 0.023207065 |
| RP13-497K6 | 0.023203354 |
| MIR28 | 0.02320315 |
| OR52N1 | 0.023202494 |
| RNA5SP373 | 0.02320112 |
| SNORA36C | 0.023194762 |
| RP11-941H19 | 0.023182241 |
| CTA-392E5 | 0.023175705 |
| OR7E163P | 0.023175175 |
| AC005159 | 0.023172147 |
| RP11-469H8 | 0.023161033 |
| NKX6-2 | 0.02315923 |
| TTTY25P | 0.023156723 |
| WNK2 | 0.023151733 |
| KCNH1-IT1 | 0.023150528 |

| | |
|---|---|
| RP11-21J18 | 0.023149315 |
| MIR548H1 | 0.023147843 |
| RP1-28O10 | 0.023138527 |
| BLCAP | 0.023133258 |
| BOLA2B | 0.023126256 |
| RP11-831A10 | 0.023123002 |
| BIN1 | 0.023116196 |
| RP11-586L23 | 0.02311581 |
| AC022909 | 0.023107018 |
| AL607076 | 0.02310665 |
| WDR81 | 0.023105182 |
| CCDC172 | 0.023101222 |
| OR4A3P | 0.023099927 |
| RP11-484L7 | 0.023097043 |
| LINC01365 | 0.023091988 |
| HSPE1P24 | 0.023091442 |
| LINC01323 | 0.02308479 |
| OR7E91P | 0.023079651 |
| RNU2-5P | 0.02307707 |
| C9orf142 | 0.023068976 |
| WWC1 | 0.023059245 |
| GSTM1 | 0.023056737 |
| PGLYRP3 | 0.023054904 |
| VN1R108P | 0.023052053 |
| RP13-258O15 | 0.023050838 |
| FOXJ1 | 0.023048182 |
| RP11-435O5 | 0.023045682 |
| RP4-753D10 | 0.023042593 |
| RP11-1K3 | 0.023037369 |
| KRTAP6-2 | 0.023034778 |
| ZFAT-AS1 | 0.023034446 |
| RPL32P35 | 0.023028841 |
| HIBADH | 0.023028509 |
| LINC01269 | 0.023026644 |
| EIF4A1P7 | 0.023024456 |
| NUTM2B | 0.023023562 |
| STK19B | 0.02302 |
| BEGAIN | 0.023013196 |
| RP1-276J11 | 0.02301 |
| RPL21P136 | 0.023001926 |
| CD93 | 0.022998889 |
| TTTY16 | 0.022997026 |
| FAM109B | 0.022995186 |
| AC005779 | 0.022993876 |
| SLC15A4 | 0.022993549 |

| | |
|---|---|
| AC006126 | 0.02298566 |
| FOCAD-AS1 | 0.022983104 |
| RPS26P6 | 0.022982416 |
| RN7SL587P | 0.02298171 |
| ACVR2A | 0.022977697 |
| AC073850 | 0.022973748 |
| NDRG4 | 0.022972748 |
| MIR4798 | 0.022968691 |
| RNU7-73P | 0.022967392 |
| ACVRL1 | 0.022965399 |
| AC018696 | 0.022961077 |
| RNU2-50P | 0.022958789 |
| NF1P8 | 0.022952317 |
| AL590226 | 0.022949707 |
| UPP2 | 0.022945311 |
| RP11-1033H12 | 0.022943754 |
| FAM195A | 0.022940635 |
| RP11-154D6 | 0.022935481 |
| RNU6-296P | 0.022932364 |
| RNF126 | 0.0229315 |
| RP1-163M9 | 0.02293084 |
| TRIM9 | 0.02292709 |
| FOXO1 | 0.022922755 |
| MIR762HG | 0.02292229 |
| GDF15 | 0.02291855 |
| MPTX1 | 0.022914734 |
| RP11-475I24 | 0.022912015 |
| AL121865 | 0.022899974 |
| SCARNA5 | 0.022888268 |
| CTD-2001E22 | 0.022884176 |
| AC106876 | 0.022884059 |
| RNU6-1026P | 0.022875293 |
| OR6P1 | 0.022873763 |
| CAPN15 | 0.022858287 |
| ARC | 0.022857543 |
| SLC22A11 | 0.022854008 |
| MIR7846 | 0.022850874 |
| FREM3 | 0.022850522 |
| RP11-555J4 | 0.022849339 |
| VPS18 | 0.022845543 |
| RP11-139F9 | 0.022844171 |
| MIR3189 | 0.022843509 |
| KCNMB2-AS1 | 0.022841386 |
| RP11-374A4 | 0.022838845 |
| ATP6V0CP3 | 0.0228376 |

| | |
|---|---|
| RP11-275I14 | 0.02283183 |
| SEPHS2 | 0.022830305 |
| RP11-37N22 | 0.022828705 |
| AURKC | 0.022819495 |
| RP11-424N24 | 0.022818453 |
| ZBTB45P1 | 0.022804132 |
| CTD-2550O8 | 0.022797799 |
| AC127383 | 0.022796955 |
| KRT18P48 | 0.022784277 |
| DUSP1 | 0.022783531 |
| RP11-563K23 | 0.022783115 |
| GALNT16 | 0.022780423 |
| ATOH8 | 0.022775828 |
| LINC01515 | 0.022758227 |
| SDK2 | 0.02275752 |
| AP000281 | 0.022753501 |
| RPL7AP38 | 0.022748889 |
| PLEKHJ1 | 0.022748608 |
| BLOC1S2 | 0.022748478 |
| ELF3 | 0.02274124 |
| COTL1 | 0.022740198 |
| AKAP6 | 0.022739808 |
| RBM17P1 | 0.022737076 |
| DBIL5P2 | 0.022736248 |
| AL118496 | 0.022734136 |
| KCNE4 | 0.022730821 |
| RP11-469A15 | 0.022728165 |
| MIR203A | 0.02272703 |
| RPS6P7 | 0.02272293 |
| RP11-692M12 | 0.022722722 |
| RP11-520H14 | 0.022718746 |
| SLITRK5 | 0.022718293 |
| ACSM1 | 0.022706988 |
| RP4-790G17 | 0.022702831 |
| MORF4L1P5 | 0.022698341 |
| RP13-324D24 | 0.022684991 |
| NDUFC1 | 0.02267854 |
| RPS20P1 | 0.022675242 |
| RP11-725M22 | 0.022668946 |
| RP11-463C14 | 0.022665602 |
| KRTAP19-4 | 0.022656299 |
| RNA5SP49 | 0.022654044 |
| RP11-983G14 | 0.022652414 |
| ERICD | 0.022648194 |
| LINC01257 | 0.022647147 |

| | |
|---|---|
| WDR4 | 0.022643785 |
| RP11-481H12 | 0.022640523 |
| AL732325 | 0.022632171 |
| RP11-778J15 | 0.022632003 |
| GABARAP | 0.022631877 |
| RNU6-666P | 0.022621525 |
| RP3-382I10 | 0.02261694 |
| RNA5SP428 | 0.022611668 |
| CTD-2114J12 | 0.022609161 |
| ARHGAP6 | 0.022599188 |
| STX6 | 0.022592285 |
| RP11-98N22 | 0.022591423 |
| RP11-855C21 | 0.022587584 |
| RP11-394B2 | 0.022580973 |
| RP5-1031J8 | 0.022579647 |
| RP11-577H5 | 0.022573253 |
| KCNK16 | 0.022567933 |
| CTD-2076M15 | 0.022567814 |
| GCNT1P2 | 0.022558726 |
| AP001092 | 0.022558545 |
| RP11-370B11 | 0.02254825 |
| RNU6-1123P | 0.022543151 |
| RP11-400D2 | 0.022539804 |
| GPER1 | 0.022533268 |
| SNORD42 | 0.022532665 |
| FLT1 | 0.022528729 |
| MRPS16P1 | 0.022527539 |
| RP13-487K5 | 0.022519903 |
| RN7SL25P | 0.022512463 |
| RP11-352E6 | 0.022508542 |
| RNU4-7P | 0.022508444 |
| SNORD8 | 0.022508279 |
| USP32P1 | 0.022502462 |
| RNU6-865P | 0.022495083 |
| RNU4-72P | 0.022492188 |
| FAM163B | 0.022486535 |
| RFX4 | 0.02247957 |
| RP11-446F17 | 0.022476198 |
| PDE8B | 0.022473396 |
| COA4 | 0.022460707 |
| CASC17 | 0.022459978 |
| CDIPT-AS1 | 0.022452287 |
| AP001634 | 0.022451468 |
| PHBP12 | 0.022448454 |
| RNU7-55P | 0.022437902 |

| | |
|---|---|
| RP11-143N13 | 0.022434535 |
| FAM170A | 0.022431382 |
| ESX1 | 0.022430708 |
| RN7SL127P | 0.022428626 |
| CTTNBP2NL | 0.022412091 |
| RP11-129M16 | 0.022408575 |
| GLYCAM1 | 0.022408112 |
| HGD | 0.0224045 |
| RP11-3P22 | 0.022396608 |
| LINGO1 | 0.022395402 |
| LRRC38 | 0.022393993 |
| RP11-236F9 | 0.02239349 |
| AC023513 | 0.022385139 |
| KLF6 | 0.022383997 |
| RNA5SP306 | 0.022372199 |
| ANAPC11 | 0.022363363 |
| RP11-33N16 | 0.022359976 |
| RP11-6I2 | 0.022358815 |
| TRIL | 0.022357465 |
| HSPE1P14 | 0.022356512 |
| MTHFD2P4 | 0.022355084 |
| EDA | 0.022354209 |
| MSMO1 | 0.022346283 |
| ETFA | 0.022344776 |
| RNU1-101P | 0.022338711 |
| RP11-425I13 | 0.022331841 |
| PRG3 | 0.0223238 |
| RNU1-42P | 0.022323695 |
| RP1-261G23 | 0.022322318 |
| CTD-2130O13 | 0.022322186 |
| RPS23 | 0.022313147 |
| AC002076 | 0.02231006 |
| SSBP3 | 0.022300607 |
| RP11-88E18 | 0.022299826 |
| RNU6ATAC19P | 0.022297101 |
| ITGA2B | 0.022287509 |
| LINC00703 | 0.022287231 |
| RP5-986I17 | 0.022283052 |
| RP11-627D16 | 0.022282888 |
| DIP2C | 0.022279583 |
| PELO | 0.02227776 |
| OR13C3 | 0.022277248 |
| RP4-669H2 | 0.02227611 |
| AC091487 | 0.0222758 |
| DNAH12 | 0.022272867 |

| | |
|---|---|
| LINC00323 | 0.022272463 |
| RP11-613E4 | 0.02226541 |
| CLU | 0.022262857 |
| MIR188 | 0.022262815 |
| LINC00301 | 0.022255857 |
| SNED1 | 0.022255638 |
| OOSP2 | 0.02225258 |
| HOXC-AS2 | 0.022242236 |
| LINC01183 | 0.022240939 |
| FAHD2P1 | 0.022238737 |
| PRSS38 | 0.022237434 |
| ZMAT5 | 0.0222372 |
| CEP112 | 0.022236753 |
| RNU6-680P | 0.022236204 |
| AP001464 | 0.02223483 |
| RP11-54O7 | 0.022232046 |
| H2AFX | 0.022222129 |
| MRPL37 | 0.022220455 |
| RP11-235D19 | 0.022218602 |
| RPS11 | 0.022214706 |
| SNORD115-20 | 0.022213024 |
| MARCKS | 0.022207766 |
| RNU6-1332P | 0.022189409 |
| RP11-260O18 | 0.022180207 |
| RP11-63E9 | 0.022179901 |
| RP11-323K10 | 0.022178141 |
| AC092684 | 0.02217134 |
| STARD13 | 0.022161699 |
| CYP1A1 | 0.022157461 |
| RP11-63E16 | 0.022145348 |
| DTYMK | 0.022142721 |
| CASC9 | 0.022132543 |
| RP11-711G10 | 0.022129593 |
| BTF3P14 | 0.022123125 |
| AC091114 | 0.022122079 |
| BPIFB3 | 0.022117437 |
| NACC1 | 0.02211738 |
| BOLA2P3 | 0.022114961 |
| HDAC2 | 0.022112896 |
| ZGPAT | 0.022108096 |
| VENTXP3 | 0.022106382 |
| RP11-219A15 | 0.022100129 |
| RNU6-574P | 0.022098375 |
| RP11-79C6 | 0.022096328 |
| ARL2BPP4 | 0.022095378 |

| | |
|---|---|
| RN7SKP32 | 0.022093443 |
| AC079907 | 0.022092932 |
| SNHG5 | 0.022092282 |
| GOLGA6C | 0.022086103 |
| RP11-161M6 | 0.022084672 |
| RN7SL128P | 0.022080486 |
| SNORA4 | 0.022079922 |
| DYNLL1P3 | 0.022079873 |
| AC011648 | 0.022071859 |
| RNU6-1014P | 0.022069306 |
| CDH9 | 0.022065919 |
| RP3-337H4 | 0.022059834 |
| LINC00311 | 0.022050682 |
| MAGEA5 | 0.022047117 |
| LINC01510 | 0.022044692 |
| RN7SL314P | 0.022039486 |
| RP11-308B16 | 0.022030234 |
| NSMF | 0.022029916 |
| VTRNA1-1 | 0.022029381 |
| LINC01497 | 0.022029365 |
| RP11-710E1 | 0.022014526 |
| ZBTB48 | 0.022010194 |
| TCF21 | 0.022009217 |
| AC008592 | 0.022001915 |
| APRT | 0.022000668 |
| PROKR1 | 0.021997948 |
| RP11-734E19 | 0.021993654 |
| CTC-830E23 | 0.021992521 |
| OR10G9 | 0.021989461 |
| SSR3 | 0.021988407 |
| CTD-3006G17 | 0.021986679 |
| ZNF660 | 0.021984426 |
| RP11-137L10 | 0.021983506 |
| MIR660 | 0.021982069 |
| KY | 0.021979559 |
| ARL3 | 0.021977188 |
| FSIP2 | 0.021962685 |
| FAHD2A | 0.021961876 |
| CICP2 | 0.021960428 |
| DHRS1 | 0.021959494 |
| RNU6-551P | 0.021954869 |
| GINM1 | 0.021949634 |
| RP11-157F20 | 0.021949564 |
| TBL1Y | 0.021946196 |
| MESTIT1 | 0.021945924 |

| | |
|---|---|
| AL356310 | 0.021942593 |
| RP11-165M1 | 0.02194029 |
| AP003068 | 0.021934321 |
| AC002044 | 0.021928079 |
| RMRP | 0.021924467 |
| RNU6-446P | 0.0219242 |
| MRGBP | 0.021922228 |
| RP11-383H17 | 0.021916337 |
| RP5-860P4 | 0.021914942 |
| RP11-236B18 | 0.021912841 |
| NFE2L3 | 0.021911567 |
| LINC01066 | 0.021908486 |
| RP11-231L11 | 0.02190602 |
| PTPRT | 0.021905782 |
| CXorf51B | 0.021905179 |
| TSC22D4 | 0.021886784 |
| PPIAP26 | 0.021885962 |
| RP11-356O9 | 0.021885302 |
| RP11-252M24 | 0.021885134 |
| C12orf45 | 0.021882356 |
| SLC35G5 | 0.021881301 |
| FRDAP | 0.021881109 |
| CHCHD2P7 | 0.021874407 |
| POLD1 | 0.021870672 |
| AC067717 | 0.021863527 |
| CXCL3 | 0.021862127 |
| SOWAHC | 0.021860792 |
| IL1RL1 | 0.02185431 |
| RP11-1018J11 | 0.021853525 |
| FUT4 | 0.02184829 |
| KLF7 | 0.02184481 |
| CYP26B1 | 0.021834615 |
| DEFA11P | 0.021824529 |
| LILRA2 | 0.021824177 |
| USP9YP35 | 0.021820095 |
| RNA5SP385 | 0.021806424 |
| TECTB | 0.021804733 |
| AC003658 | 0.021798668 |
| PPP1R8P1 | 0.021797546 |
| ENTHD1 | 0.021795527 |
| RNU7-136P | 0.021790248 |
| RP11-51E20 | 0.021788348 |
| RP11-486G15 | 0.02178716 |
| RPL30P13 | 0.021781869 |
| RP5-907C10 | 0.021779859 |

| | |
|---|---|
| FGG | 0.021767368 |
| RP11-528A4 | 0.02176537 |
| GAPDHP66 | 0.021765315 |
| GLIS2-AS1 | 0.021759158 |
| C8orf89 | 0.021757666 |
| RP11-52B19 | 0.021756174 |
| MTMR8 | 0.021755888 |
| RP11-368J22 | 0.021755404 |
| RP11-227D2 | 0.021752869 |
| RP11-265G8 | 0.021744229 |
| ECI2 | 0.021743987 |
| AC092662 | 0.02174199 |
| RNASE13 | 0.021737539 |
| SNORD56B | 0.021731135 |
| LCE1F | 0.021730221 |
| AP000237 | 0.021724556 |
| FTH1P10 | 0.021721912 |
| RN7SL188P | 0.02172163 |
| RP11-744H18 | 0.02171909 |
| AC079199 | 0.021710452 |
| RP11-389O22 | 0.021710323 |
| SQRDL | 0.021706702 |
| CDC42P2 | 0.021698561 |
| RN7SL552P | 0.021696149 |
| RPL7A | 0.021685845 |
| OR6C72P | 0.021682489 |
| RP11-368E13 | 0.02168226 |
| GAPDHP45 | 0.021680083 |
| CIDEA | 0.021677947 |
| SHROOM1 | 0.021676226 |
| ERCC8 | 0.021672263 |
| RN7SL791P | 0.02167013 |
| RN7SL309P | 0.021666795 |
| RP11-220C2 | 0.021665576 |
| RP11-139K4 | 0.021665498 |
| OR8X1P | 0.021663368 |
| ARHGEF9-IT1 | 0.021663039 |
| TCTN3 | 0.021654607 |
| CTD-2026J24 | 0.021654477 |
| LY6G6D | 0.021654182 |
| OR51B3P | 0.021648256 |
| AC099342 | 0.021644821 |
| ZNF300P1 | 0.021642494 |
| RBM20 | 0.021633173 |
| SNORD114-31 | 0.02163174 |

| | |
|---|---|
| TRGJ1 | 0.021623326 |
| AC005753 | 0.021618539 |
| RP3-483K16 | 0.021616795 |
| LINC01049 | 0.021612263 |
| RNU6-8 | 0.021601505 |
| RP3-406A7 | 0.021600688 |
| ZIM2 | 0.021596862 |
| MTRNR2L2 | 0.021596085 |
| RP11-595B24 | 0.021594238 |
| APOBEC4 | 0.021594006 |
| MRPL14 | 0.021588892 |
| RN7SL498P | 0.021588851 |
| RP11-49I11 | 0.021583794 |
| RN7SL293P | 0.02157968 |
| NR5A1 | 0.021578233 |
| MIRLET7C | 0.021576993 |
| RP11-383G10 | 0.021575949 |
| MAFF | 0.021575608 |
| KLF8P1 | 0.021573854 |
| RP11-157B13 | 0.021570746 |
| SCARF1 | 0.021569505 |
| TMSB4XP8 | 0.0215669 |
| RP11-84N19 | 0.021563816 |
| EIF2B2 | 0.021548239 |
| AC002395 | 0.021544687 |
| SEMA4F | 0.021540611 |
| AL159977 | 0.021539317 |
| SH2B1 | 0.021538639 |
| CLDN3 | 0.021526899 |
| THOP1 | 0.021524926 |
| RBPMSLP | 0.021521265 |
| RNU4-81P | 0.021515705 |
| FDX1P1 | 0.021514958 |
| C8orf31 | 0.02151341 |
| AC016866 | 0.02151009 |
| TNNI1 | 0.02150076 |
| RPS27P15 | 0.021498802 |
| ZFP37 | 0.021497428 |
| CUX2 | 0.021497267 |
| RBMY1F | 0.021495737 |
| SNORD105B | 0.021495649 |
| CAP2P1 | 0.02149484 |
| RP11-506H20 | 0.021488295 |
| RP4-651E10 | 0.021485542 |
| AC011322 | 0.021482172 |

| | |
|---|---|
| TNNT1 | 0.021479427 |
| UGT1A9 | 0.021473856 |
| RYR1 | 0.021471845 |
| RPL26P37 | 0.021466085 |
| CHRNB1 | 0.021464459 |
| AC068483 | 0.021459283 |
| RP11-428F18 | 0.021457879 |
| RPL7AP72 | 0.021456124 |
| VN1R2 | 0.021450389 |
| RNU6-1337P | 0.021450037 |
| CREB3L2 | 0.021447844 |
| CTSLP8 | 0.021446162 |
| RP11-501M7 | 0.021443644 |
| RP11-242G20 | 0.021439442 |
| LINC00652 | 0.021435737 |
| RNU6-1224P | 0.021433502 |
| FTLP3 | 0.021432588 |
| RP11-462D18 | 0.021431936 |
| OR4N1P | 0.021423612 |
| RPL7P5 | 0.021422369 |
| ZNF316 | 0.021416959 |
| TTLL2 | 0.021409089 |
| RP11-108B14 | 0.021407692 |
| GLI3 | 0.021406391 |
| POFUT1 | 0.021406186 |
| MIR365A | 0.02140036 |
| RP11-225N10 | 0.021400139 |
| SPATA31D5P | 0.021399139 |
| CYTL1 | 0.021398205 |
| CTD-2194F4 | 0.021397894 |
| PPP1R14BP2 | 0.021395186 |
| HTR6 | 0.021391417 |
| ISX | 0.021382512 |
| FAR2P2 | 0.021382198 |
| MIR549A | 0.021375695 |
| RP11-718O11 | 0.021374939 |
| PIH1D3 | 0.021366072 |
| RNU5B-6P | 0.021359451 |
| AC093160 | 0.021359256 |
| DEFA1 | 0.021359256 |
| ZNF717 | 0.021357477 |
| ATG10 | 0.021354612 |
| RP11-84A19 | 0.021354132 |
| AC092651 | 0.021341839 |
| RNU6ATAC10P | 0.021341483 |

| | |
|---|---|
| RP11-799D4 | 0.021338053 |
| NCSTNP1 | 0.02133593 |
| RPS7P13 | 0.021333932 |
| AC005618 | 0.021327393 |
| RP11-55C6 | 0.021325142 |
| ACYP2 | 0.021323969 |
| ZBTB22 | 0.021319894 |
| AL033403 | 0.021300909 |
| LHFP | 0.021298796 |
| RPSAP53 | 0.021298517 |
| OR2A14 | 0.021292903 |
| RP11-738O11 | 0.021286209 |
| SLC2A13 | 0.021284153 |
| AC079325 | 0.021283564 |
| MAP2 | 0.021273957 |
| EEF1B2P3 | 0.021268915 |
| TSPAN11 | 0.021266924 |
| ACP6 | 0.021259464 |
| MAP2K6 | 0.021259129 |
| RP11-445K13 | 0.02125878 |
| AC113618 | 0.021255581 |
| RNU6-227P | 0.021253674 |
| RPL21P39 | 0.021249525 |
| HSF2BP | 0.021249026 |
| RP11-483M24 | 0.021242273 |
| MIR5583-1 | 0.021240512 |
| RPL21P129 | 0.021229551 |
| PQLC2 | 0.021228672 |
| RP11-275N1 | 0.02122696 |
| RP11-489D6 | 0.021224541 |
| RP11-128D14 | 0.02122254 |
| RN7SKP186 | 0.021221674 |
| DPP10 | 0.021220884 |
| TPT1P11 | 0.021216653 |
| BX322559 | 0.021212297 |
| C6orf223 | 0.021205869 |
| RNU6-949P | 0.021203364 |
| MIR548AZ | 0.021199884 |
| RP11-142A22 | 0.021199412 |
| SEMA5A | 0.021197749 |
| RP11-91J3 | 0.021195563 |
| CTA-126B4 | 0.021185189 |
| CTA-31J9 | 0.021179117 |
| RP11-1H15 | 0.021177592 |
| RP11-451F20 | 0.021177569 |

| | |
|---|---|
| DNAJB11 | 0.021177321 |
| MIMT1 | 0.021177259 |
| RP11-629G13 | 0.021158984 |
| CXCL14 | 0.021158901 |
| HRASLS5 | 0.021153375 |
| CFB | 0.021152892 |
| RP11-678G15 | 0.021144519 |
| RP11-451B8 | 0.021140694 |
| CH507-236L23 | 0.021137184 |
| RNMTL1 | 0.021133281 |
| NAT16 | 0.021127841 |
| BMS1P3 | 0.021125935 |
| USP17L4 | 0.021119181 |
| RP11-283G6 | 0.021118881 |
| RNU6-1118P | 0.021118143 |
| WDR45 | 0.021116034 |
| AC006427 | 0.021115115 |
| ABCD1P2 | 0.021112778 |
| RP11-128M1 | 0.021108943 |
| RP6-149D17 | 0.021106132 |
| RP11-960B9 | 0.021102847 |
| RN7SKP112 | 0.021101189 |
| TAS2R2P | 0.021101071 |
| CACNA1C-AS2 | 0.021097952 |
| HOOK2 | 0.021095047 |
| AC092610 | 0.021090265 |
| DEFT1P2 | 0.021086294 |
| HMGN1P7 | 0.021084841 |
| RP11-476H24 | 0.021073717 |
| LINC00507 | 0.021064529 |
| TCEB2P4 | 0.021060231 |
| RNU6-641P | 0.021056497 |
| BCAR1P1 | 0.021055337 |
| ZNF883 | 0.021041519 |
| AC012512 | 0.021034437 |
| RP11-567C20 | 0.021030994 |
| RP11-745H7 | 0.021029582 |
| C2orf80 | 0.021024065 |
| STAT3 | 0.021021351 |
| RP11-330O11 | 0.021018281 |
| ABCC6P1 | 0.021017297 |
| AC007359 | 0.021016 |
| RAC1P4 | 0.021012733 |
| RN7SL369P | 0.021001704 |
| RP11-953B20 | 0.021001348 |

| | |
|---|---|
| ZBTB8B | 0.020998971 |
| ADCY10 | 0.020991788 |
| USP9YP15 | 0.020990708 |
| RP11-307C12 | 0.02098907 |
| BPGM | 0.020988531 |
| RP11-715H19 | 0.020988038 |
| RP13-415G19 | 0.020985563 |
| OR10AG1 | 0.020980412 |
| ARRDC3-AS1 | 0.020979284 |
| POU5F1 | 0.020978335 |
| HNMT | 0.020967265 |
| MIR5588 | 0.020955395 |
| MIR1249 | 0.020951634 |
| CYC1 | 0.020951115 |
| ATP5G1P5 | 0.020949709 |
| CRAT | 0.020949255 |
| AC092628 | 0.020945012 |
| MIR4643 | 0.020945012 |
| SLC2A5 | 0.020944845 |
| bP-21264C1 | 0.020944515 |
| ANO6 | 0.020944175 |
| FAM49A | 0.020943487 |
| C8orf59 | 0.020942836 |
| HSPB1P1 | 0.020940585 |
| AC116345 | 0.020934982 |
| RP11-292D4 | 0.020925461 |
| PTPRR | 0.020925319 |
| PTOV1 | 0.020924695 |
| BDH2P1 | 0.020920878 |
| AC011313 | 0.020918955 |
| RN7SKP261 | 0.020917692 |
| RSU1P3 | 0.020917186 |
| RNU4-83P | 0.020907276 |
| ZNF486 | 0.020897104 |
| SFTA2 | 0.020894375 |
| RNA5SP345 | 0.02089435 |
| C20orf62 | 0.020891978 |
| RN7SL445P | 0.020885232 |
| SYNC | 0.020885103 |
| OSBPL9 | 0.020882368 |
| LARGE | 0.020880861 |
| DSG2 | 0.020875019 |
| RPL30P3 | 0.020874668 |
| CHST11 | 0.020874107 |
| RNU6-981P | 0.020873619 |

| | |
|---|---|
| RNA5SP202 | 0.020871208 |
| RP11-864G5 | 0.02086935 |
| PIAS4 | 0.020858869 |
| RP11-83M16 | 0.020854555 |
| CNN2P1 | 0.020849012 |
| RP4-763G1 | 0.020846185 |
| RP11-835E18 | 0.020845857 |
| DAZAP1 | 0.020843464 |
| FABP5P7 | 0.020839547 |
| CTD-2644I21 | 0.020835444 |
| RNU6-1327P | 0.020830698 |
| GPR98 | 0.02082804 |
| ZNF32-AS3 | 0.02082768 |
| ZNF774 | 0.020824747 |
| XXbac-BPG13B8 | 0.020822378 |
| RNU6-46P | 0.020812124 |
| RNA5SP67 | 0.020808253 |
| LINC01117 | 0.020803535 |
| SOX7 | 0.020793434 |
| SCNN1D | 0.020783927 |
| RNU6-1260P | 0.020783547 |
| SLX1B | 0.02077868 |
| MIR941-4 | 0.020777732 |
| SLC35A4 | 0.020768893 |
| ADAMTS1 | 0.020767014 |
| RP11-19P22 | 0.020766491 |
| RPL23AP73 | 0.0207564 |
| RP11-32D16 | 0.020752236 |
| MYBBP1A | 0.020751088 |
| RNU5D-2P | 0.020747073 |
| PAMR1 | 0.020746102 |
| GSTK1 | 0.020744148 |
| CYB5A | 0.020738511 |
| BNIPL | 0.020726584 |
| CCDC167 | 0.020725319 |
| SOX30 | 0.020724685 |
| TP53INP2 | 0.020724594 |
| AC013268 | 0.020721007 |
| HSPA8P16 | 0.020718427 |
| RP11-435M3 | 0.020715748 |
| PADI2 | 0.020713858 |
| MRPS17 | 0.02070757 |
| TMEM229A | 0.02070379 |
| RP11-286B14 | 0.020701743 |
| RP11-307O1 | 0.020699976 |

| | |
|---|---|
| RNU7-161P | 0.020699013 |
| RP11-159M11 | 0.020697576 |
| RP11-1017G21 | 0.020692313 |
| RP11-396N11 | 0.020691895 |
| LYRM4 | 0.020690676 |
| ARL10 | 0.020690378 |
| RP11-552E10 | 0.020687725 |
| AC112719 | 0.020684051 |
| HTT-AS | 0.020677411 |
| WASH1 | 0.020676201 |
| MIR548AV | 0.020672663 |
| AC092168 | 0.020657646 |
| SEC63P1 | 0.020652349 |
| CTD-2306A12 | 0.020642337 |
| RP11-132M7 | 0.020640167 |
| RP11-18M17 | 0.02063832 |
| NFE4 | 0.020636242 |
| CASC8 | 0.020636182 |
| TSSC1-IT1 | 0.020630685 |
| CHMP2A | 0.020627696 |
| AC002069 | 0.020624672 |
| RP11-7C6 | 0.020622195 |
| CYP51A1-AS1 | 0.020621442 |
| GPD1 | 0.020619436 |
| RP11-63N3 | 0.020617571 |
| OR6M1 | 0.02061172 |
| KAZN | 0.020597518 |
| BCL7B | 0.020597369 |
| SPATA31B1P | 0.020596651 |
| RP5-1166A24 | 0.020587485 |
| RNU6-1092P | 0.020585148 |
| AC239798 | 0.020583534 |
| PRR22 | 0.020583118 |
| RN7SL504P | 0.020579204 |
| RNU4-4P | 0.020575668 |
| RP11-144C9 | 0.02057356 |
| CTD-2350C19 | 0.020569514 |
| MIR29A | 0.020568571 |
| RPE65 | 0.020567209 |
| RP4-534P7 | 0.020566851 |
| GYLTL1B | 0.020565945 |
| CTB-51J22 | 0.020561458 |
| DCTN2 | 0.020558912 |
| RP11-567C2 | 0.020556324 |
| ERMP1 | 0.020556047 |

| | |
|---|---|
| RNU7-181P | 0.020555271 |
| FAM184B | 0.020555098 |
| RP11-804N13 | 0.020554966 |
| MTUS2-AS2 | 0.020545773 |
| RNA5SP72 | 0.020539498 |
| RP11-174O3 | 0.020538997 |
| RP11-555M1 | 0.020538541 |
| TBX22 | 0.020538259 |
| RP11-23A22 | 0.020537477 |
| RNU6-194P | 0.020537097 |
| MLXIP | 0.020534502 |
| AC092470 | 0.020530273 |
| ALCAM | 0.020528529 |
| ZNF209P | 0.020521239 |
| FOXN3 | 0.020519264 |
| PART1 | 0.020518181 |
| ABCF2P2 | 0.020515838 |
| RN7SL553P | 0.020513333 |
| RP11-510N19 | 0.020508602 |
| RP11-166J5 | 0.02050784 |
| AC092431 | 0.020504572 |
| SLC36A1 | 0.020504565 |
| POLD2P1 | 0.020504041 |
| RNU6-78P | 0.020503729 |
| OR5I1 | 0.020500696 |
| LINC01228 | 0.020500049 |
| NAMPTP1 | 0.02049917 |
| RNU6-1082P | 0.020494815 |
| LYG1 | 0.020492118 |
| NPM1P28 | 0.020482262 |
| MIR2115 | 0.020480813 |
| C2orf81 | 0.020476974 |
| RP13-204A15 | 0.020474867 |
| RNU6-577P | 0.020470102 |
| AC092796 | 0.02046665 |
| HMG20B | 0.020461345 |
| KCMF1 | 0.020458331 |
| CLIP4 | 0.020450517 |
| RP11-452N17 | 0.020448423 |
| MIR619 | 0.020441933 |
| C19orf18 | 0.020441009 |
| EEF1DP5 | 0.020435178 |
| C17orf77 | 0.020434369 |
| RN7SL557P | 0.020434276 |
| DRD5P1 | 0.02043211 |

| | |
|---|---|
| RP11-218F4 | 0.0204254 |
| ACTBP2 | 0.020422367 |
| RNA5SP335 | 0.020420384 |
| RBMY1A3P | 0.02041991 |
| RNA5SP422 | 0.020406031 |
| GGTLC3 | 0.020400034 |
| RP11-263K19 | 0.020398924 |
| RN7SL702P | 0.020396974 |
| PKLR | 0.02039641 |
| C16orf59 | 0.020391891 |
| LRRC28 | 0.020389546 |
| RNU6-271P | 0.020389041 |
| RP11-739P6 | 0.020383804 |
| RP11-191L17 | 0.020380074 |
| PPIAP28 | 0.020379374 |
| AL645922 | 0.020365098 |
| ETNK2 | 0.02036402 |
| CABP5 | 0.020362778 |
| FAM27C | 0.020357174 |
| ZNF436 | 0.020356447 |
| THUMPD2 | 0.020355126 |
| POM121L2 | 0.02035401 |
| SNRPG | 0.020344149 |
| RP11-452C13 | 0.020344093 |
| CROT | 0.020341079 |
| RP11-314O13 | 0.020341059 |
| TRIM47 | 0.020328018 |
| HEPACAM | 0.020327969 |
| GKN1 | 0.020308506 |
| RP5-1029F21 | 0.020307072 |
| RP1-100J12 | 0.020297387 |
| MIR4297 | 0.020296408 |
| OR1A1 | 0.020292575 |
| SGK1 | 0.02028476 |
| RP11-1275H24 | 0.020282304 |
| GHET1 | 0.020281467 |
| RP11-178D12 | 0.020270022 |
| RN7SL67P | 0.020267552 |
| CXorf51A | 0.020267409 |
| LL0XNC01-220B11 | 0.020267409 |
| LIMS3L | 0.020267409 |
| OR4A14P | 0.02026635 |
| MIR128-1 | 0.020263821 |
| RP4-684O24 | 0.020263011 |
| RPF2P1 | 0.02026174 |

| | | |
|---|---|---|
| PCA3_1 | | 0.020261431 |
| KCNC3 | | 0.020259141 |
| RP11-347H15 | | 0.020257751 |
| RP11-480I12 | | 0.020255215 |
| RNA5SP369 | | 0.020252774 |
| AC005725 | | 0.020246042 |
| KPRP | | 0.020243916 |
| RPL5 | | 0.020233149 |
| OR7E85P | | 0.020230356 |
| RP11-121G22 | | 0.020230093 |
| PLCH1 | | 0.020228591 |
| SLC25A10 | | 0.020226432 |
| ORAI2 | | 0.020216999 |
| RP1-117P20 | | 0.020215383 |
| PABPC1P11 | | 0.020211163 |
| RP13-476E20 | | 0.020206506 |
| HMGN2P16 | | 0.020205094 |
| NDUFA5P4 | | 0.020201554 |
| RP11-381P6 | | 0.020197672 |
| CDRT1 | | 0.020187199 |
| AP005138 | | 0.020183786 |
| MAP7D3 | | 0.0201823 |
| RN7SL797P | | 0.02017046 |
| AC010677 | | 0.020169965 |
| AC073330 | | 0.020169356 |
| AL590126 | | 0.020163878 |
| CILP | | 0.020163323 |
| DUX4L26 | | 0.020160444 |
| | 2-Mar | 0.020154634 |
| RYR3 | | 0.020151508 |
| RNF130 | | 0.020136791 |
| RP11-629O19 | | 0.020134088 |
| RP11-79L21 | | 0.020132834 |
| RPS6KA2-AS1 | | 0.020131255 |
| C5AR1 | | 0.020128009 |
| DEPDC1-AS1 | | 0.020125125 |
| CNNM1 | | 0.020121224 |
| ADAP1 | | 0.020119655 |
| AC079610 | | 0.020119532 |
| TOR1AIP2 | | 0.02011506 |
| RPL36P1 | | 0.02011447 |
| MIR4486 | | 0.020110474 |
| LINC00640 | | 0.020109411 |
| RP11-321E2 | | 0.020107211 |
| RP11-452C8 | | 0.020105882 |

| | |
|---|---|
| SAR1A | 0.020105199 |
| RP11-90C4 | 0.020103872 |
| AL671883 | 0.020101379 |
| RNU6-861P | 0.020100662 |
| RP11-627G18 | 0.020100151 |
| DPPA5P2 | 0.020099404 |
| AC005863 | 0.020097908 |
| ZNF736P9Y | 0.020096781 |
| LRFN4 | 0.020092882 |
| RP11-587D21 | 0.020091479 |
| CTC-264K15 | 0.020087829 |
| AL391416 | 0.020079358 |
| NSUN5 | 0.020073626 |
| C10orf71 | 0.020071404 |
| CFTR | 0.020070953 |
| RPL24P4 | 0.020069681 |
| C20orf24 | 0.020063597 |
| MTND1P7 | 0.020063227 |
| LINC00390 | 0.020060873 |
| RP5-875O13 | 0.020058142 |
| ZNF471 | 0.020055467 |
| DNM1P49 | 0.020054276 |
| MIR3186 | 0.020052788 |
| TAF6L | 0.020049606 |
| RP11-215G15 | 0.02004672 |
| COX17P1 | 0.020040692 |
| FCER1A | 0.020035021 |
| CYP2C8 | 0.020028852 |
| RUNX1T1 | 0.020022107 |
| RPLP0P1 | 0.020019316 |
| PARK2 | 0.020008227 |
| RP11-353H3 | 0.020004172 |
| TMEM30A | 0.019996413 |
| RN7SL413P | 0.019990785 |
| RP11-139J15 | 0.019989385 |
| RN7SKP94 | 0.019969313 |
| FKSG63 | 0.019968786 |
| CH507-39O4 | 0.019967437 |
| SBK3 | 0.019953789 |
| CXCL17 | 0.019943709 |
| ZNF300 | 0.019940876 |
| PSG10P | 0.019932558 |
| MIR4500HG | 0.019931561 |
| CH17-118O6 | 0.019919982 |
| RP11-88I21 | 0.019903287 |

| | |
|---|---|
| INTS5 | 0.019902027 |
| RP11-346J10 | 0.019897196 |
| RP11-5P4 | 0.019885061 |
| AF121898 | 0.019878811 |
| OR4M2 | 0.019873855 |
| RP11-713D19 | 0.01987082 |
| DOK6 | 0.01986997 |
| RALYL | 0.019868653 |
| CTB-25B13 | 0.019867012 |
| CTD-2589H19 | 0.019865142 |
| LUZP4P1 | 0.019863251 |
| RP11-309P22 | 0.019856252 |
| AK2 | 0.019854602 |
| AL450226 | 0.0198518 |
| RP3-399L15 | 0.01984569 |
| RP11-374A22 | 0.019841725 |
| RNU6-777P | 0.019839558 |
| RP11-664D7 | 0.019839245 |
| OR9Q1 | 0.019835669 |
| RN7SKP279 | 0.01983275 |
| CTC-369A16 | 0.01983199 |
| RPL21P106 | 0.019830017 |
| SLC25A6P3 | 0.019827808 |
| GPR39 | 0.019813884 |
| MMD2 | 0.019811333 |
| AL031587 | 0.019808154 |
| RP11-25D10 | 0.019807384 |
| AC093763 | 0.019805121 |
| JTB | 0.019803623 |
| ZNF846 | 0.019799176 |
| AC004593 | 0.01978904 |
| TAF6 | 0.019787566 |
| CLCN5 | 0.019783287 |
| SH3BP5L | 0.019776589 |
| ZMIZ1-AS1 | 0.019776068 |
| TSSK1B | 0.019771361 |
| RPS29P20 | 0.019769886 |
| DDX11L9 | 0.019769738 |
| CFTRP3 | 0.019767199 |
| AC012307 | 0.019764287 |
| AL109947 | 0.019762537 |
| UGGT2 | 0.019755491 |
| FNDC8 | 0.019753689 |
| RN7SL168P | 0.019749183 |
| RP11-572B2 | 0.019745359 |

| | |
|---|---|
| MYO1D | 0.019738225 |
| AC015884 | 0.019737282 |
| SNORD53 | 0.01973695 |
| RP11-706P11 | 0.019735885 |
| SLC52A1 | 0.019730094 |
| AC004985 | 0.019729441 |
| PROX1-AS1 | 0.019725979 |
| AC006377 | 0.01972248 |
| POLR3GL | 0.019716531 |
| LINC00628 | 0.019713183 |
| RP11-706J10 | 0.019710015 |
| RP4-663N10 | 0.019696233 |
| MIR6879 | 0.019691834 |
| RPL7P37 | 0.019691413 |
| C18orf12 | 0.019680753 |
| RNU2-54P | 0.019670193 |
| AC002978 | 0.019668257 |
| BEST2 | 0.019667568 |
| OR6K1P | 0.019667549 |
| PIGR | 0.019660215 |
| RP11-237P21 | 0.019648714 |
| AC012354 | 0.019648229 |
| BECN1P1 | 0.019646348 |
| CTD-2050E21 | 0.019646326 |
| TBC1D3P7 | 0.019645198 |
| E2F6P2 | 0.019645073 |
| RP11-244H18 | 0.019643964 |
| GAPDHP65 | 0.019640928 |
| RP11-6K23 | 0.019639218 |
| RP11-6B4 | 0.019634641 |
| CFAP45 | 0.019614015 |
| NHS | 0.019612675 |
| AC013549 | 0.019609086 |
| RP4-788P17 | 0.01960094 |
| PMEL | 0.019596265 |
| API5P2 | 0.019589095 |
| RN7SL499P | 0.019586341 |
| AC013734 | 0.019585004 |
| RNU6ATAC23P | 0.019582696 |
| FAM205CP | 0.019578066 |
| HBA1 | 0.01957688 |
| RP11-43P8 | 0.019574709 |
| MALL | 0.019569979 |
| LINC00261 | 0.019565504 |
| RN7SL547P | 0.019564717 |

| | |
|---|---|
| RPL7AP71 | 0.019556935 |
| HNRNPA1P38 | 0.019542942 |
| RP5-984P4 | 0.019541784 |
| DPPA5P1 | 0.019541477 |
| RP11-376M2 | 0.0195388 |
| RP11-242P2 | 0.019536441 |
| RP11-993B23 | 0.019535792 |
| UGT2B28 | 0.01952824 |
| BAK1 | 0.019523034 |
| RP11-321M21 | 0.019521826 |
| FAM69B | 0.019520318 |
| MGAM | 0.019509975 |
| GNB2L1 | 0.01950774 |
| RNA5SP53 | 0.019507607 |
| MKRN5P | 0.019507316 |
| RP11-244G12 | 0.019506495 |
| AC007394 | 0.019505775 |
| LINC00407 | 0.019502224 |
| DYNAP | 0.019497831 |
| APOBEC3B | 0.019493023 |
| SORBS3 | 0.019481916 |
| COQ3 | 0.019479162 |
| HSPB11 | 0.019471627 |
| AC114273 | 0.01947139 |
| KDF1 | 0.01946411 |
| RN7SKP40 | 0.019464036 |
| KRT18P54 | 0.019457057 |
| RPL7AP14 | 0.019446406 |
| RPL35AP19 | 0.019443923 |
| MAGEB17 | 0.019435337 |
| MRPS17P9 | 0.019434557 |
| RP11-617F23 | 0.019431104 |
| RP11-266L9 | 0.019421932 |
| TRIM51GP | 0.019420451 |
| RP11-538P18 | 0.019419282 |
| ACOT2 | 0.019414248 |
| AC067968 | 0.019410723 |
| RNU6-406P | 0.019410532 |
| MTUS2 | 0.019404801 |
| KC6 | 0.019399673 |
| MAL2 | 0.019398787 |
| RP11-53L24 | 0.019391329 |
| CTC-370J7 | 0.019383966 |
| RNU6-207P | 0.019381613 |
| RP11-118H4 | 0.019380298 |

| | |
|---|---|
| HAUS5 | 0.019379381 |
| TRAV31 | 0.019378795 |
| CTC-207P7 | 0.019377284 |
| AP002802 | 0.019376228 |
| SLC4A9 | 0.019371666 |
| RN7SL618P | 0.019371269 |
| KRTAP1-1 | 0.019368242 |
| RCC2P3 | 0.019364761 |
| AP005273 | 0.019359506 |
| RN7SKP220 | 0.019354097 |
| GPATCH3 | 0.019354003 |
| MT1G | 0.019344626 |
| BCL3 | 0.019335695 |
| DNAJC17 | 0.019333574 |
| COLEC12 | 0.019332849 |
| NEDD4 | 0.019332216 |
| MROH3P | 0.019331846 |
| LINC01278 | 0.019331286 |
| PSMA2 | 0.0193231 |
| ADH1A | 0.019322433 |
| SLC16A2 | 0.019318783 |
| RP11-115N4 | 0.019317492 |
| AC020900 | 0.019316169 |
| SLC30A2 | 0.019314965 |
| ZNF280A | 0.019313785 |
| CTSF | 0.019311732 |
| RP1-59M18 | 0.019308821 |
| RP11-504E21 | 0.019306729 |
| GSTM3P2 | 0.019306377 |
| AC007126 | 0.019299748 |
| XXyac-YRM2039 | 0.019295063 |
| PPP1R26P1 | 0.019294992 |
| SNORD81 | 0.019293763 |
| OR7E46P | 0.019286087 |
| RP11-549A6 | 0.019284902 |
| MIR27A | 0.019280943 |
| PPIAP17 | 0.019278119 |
| LIMS3 | 0.019265474 |
| POM121L9P | 0.019263655 |
| MIRLET7A2 | 0.01926326 |
| FLOT1 | 0.019259041 |
| GSR | 0.019254483 |
| RP11-220H4 | 0.019248534 |
| RPL7AP65 | 0.019247406 |
| RNU4-65P | 0.019245535 |

| | |
|---|---|
| RP4-704D23 | 0.019244303 |
| SLC6A9 | 0.019243933 |
| MIR5694 | 0.019242227 |
| ACTBP11 | 0.01924151 |
| AL008707 | 0.019236606 |
| KCNAB1 | 0.019234445 |
| MRPL43 | 0.0192321 |
| MRPS36P4 | 0.01922773 |
| GBA | 0.019227114 |
| TCEAL2 | 0.01922199 |
| RP11-262K1 | 0.019205556 |
| TEX9 | 0.019203708 |
| PDLIM1P1 | 0.01919744 |
| ACSM6 | 0.019195089 |
| CUEDC1 | 0.019188508 |
| TGM6 | 0.019187125 |
| RNU1-70P | 0.019182692 |
| RNU6-1004P | 0.019182412 |
| AC107020 | 0.019180402 |
| SMIM2-IT1 | 0.019180373 |
| BANF1P4 | 0.019175334 |
| OR2G1P | 0.01917256 |
| RP11-217B1 | 0.019156999 |
| ATP11AUN | 0.019156804 |
| SNRPGP10 | 0.019155502 |
| LPO | 0.019154538 |
| HBA2 | 0.019150725 |
| SPAG16 | 0.019149566 |
| GSPT2 | 0.01914858 |
| RP11-485O14 | 0.019145367 |
| GS1-69O6 | 0.019141211 |
| HBE1 | 0.019138603 |
| UBE2Q2P12 | 0.019135732 |
| RP11-61K12 | 0.019133703 |
| LSP1 | 0.019132412 |
| RP11-343N15 | 0.019127864 |
| ZNF541 | 0.019124486 |
| PAICSP3 | 0.019124155 |
| SSX2 | 0.019122482 |
| RNU6-660P | 0.019119647 |
| SLC26A2 | 0.019112541 |
| RP11-38C18 | 0.019111998 |
| MIR876 | 0.019100759 |
| GAPDHP70 | 0.019100505 |
| MRPL23-AS1 | 0.019085515 |

| | |
|---|---|
| OR10V3P | 0.01908367 |
| LINC00570 | 0.019081004 |
| ZNF395 | 0.019076224 |
| OR9G2P | 0.019060686 |
| SNORD31 | 0.019059922 |
| RP11-399N22 | 0.019058875 |
| RNU6-552P | 0.019054939 |
| OR51P1P | 0.019053459 |
| GRIK1 | 0.019051768 |
| RP4-560B9 | 0.019049698 |
| ZSCAN16-AS1 | 0.019049525 |
| LINGO4 | 0.019046955 |
| HMBOX1-IT1 | 0.019044508 |
| RPS3AP50 | 0.019043 |
| RP1-223B1 | 0.01904284 |
| GRIN3B | 0.01904185 |
| RNU6-1288P | 0.019039081 |
| CCDC120 | 0.019037934 |
| KCNH1 | 0.019037001 |
| AC010987 | 0.019030454 |
| RBMY2QP | 0.019029788 |
| ALG9-IT1 | 0.019018744 |
| NEK4P3 | 0.019018328 |
| MRPL33 | 0.019013746 |
| MIR92B | 0.019012276 |
| FGA | 0.019006537 |
| MIR4458HG | 0.019000509 |
| AP000445 | 0.018999305 |
| C2orf82 | 0.018996406 |
| SNORD115-9 | 0.018996263 |
| RPS17P15 | 0.01899114 |
| AC090044 | 0.018989454 |
| DDT | 0.018985448 |
| MIR4254 | 0.018984528 |
| RBBP9 | 0.018984238 |
| PGCP1 | 0.018978669 |
| GRID1-AS1 | 0.018969458 |
| HOXB6 | 0.018968554 |
| RP11-1100L3 | 0.018967822 |
| MPND | 0.018965633 |
| LINC01312 | 0.018965084 |
| RP11-370K11 | 0.018960029 |
| UNC45A | 0.018958172 |
| STAB1 | 0.018956181 |
| TRGVB | 0.01895539 |

| | |
|---|---|
| RP11-104F15 | 0.018953877 |
| CTD-2331D11 | 0.018944731 |
| PPBP | 0.0189424 |
| ZFP57 | 0.018942164 |
| AC092635 | 0.018932881 |
| RP11-61G23 | 0.01893242 |
| DLEC1P1 | 0.018930327 |
| HTN3 | 0.018927784 |
| OFD1P3Y | 0.018926351 |
| WSCD1 | 0.018923864 |
| RP11-293B20 | 0.018911572 |
| AL354682 | 0.018904682 |
| RP11-1041F24 | 0.01889788 |
| RP5-1126H10 | 0.018895825 |
| METTL5 | 0.018887639 |
| OR9K1P | 0.018883416 |
| AC131097 | 0.018882261 |
| NDUFB1 | 0.018872122 |
| SNORD116-30 | 0.018868091 |
| TMEM200B | 0.018867135 |
| AHSA1 | 0.018866912 |
| SUMO2P18 | 0.018855886 |
| VWA5B2 | 0.018854967 |
| LRRC74B | 0.018852745 |
| FRMPD4-AS1 | 0.018849586 |
| JKAMPP1 | 0.018845775 |
| AC090559 | 0.01884405 |
| RP4-608O15 | 0.0188418 |
| IFT43 | 0.018839376 |
| CBWD1 | 0.018837553 |
| RNU4-11P | 0.018834594 |
| EEF1A1P21 | 0.018832305 |
| AC141002 | 0.01882984 |
| UBE2E2-AS1 | 0.018820594 |
| RP11-1037D22 | 0.018816201 |
| DNAJB5P1 | 0.018814324 |
| RIPPLY2 | 0.018812572 |
| RP11-560O20 | 0.018803266 |
| KCNQ1OT1 | 0.018800509 |
| AC004862 | 0.018796577 |
| GGTLC1 | 0.018796379 |
| VPS37D | 0.018794202 |
| CYP2C61P | 0.018788978 |
| RP1-146I3 | 0.018786245 |
| CTD-2044J15 | 0.018785857 |

| | |
|---|---|
| RP11-810K23 | 0.018782752 |
| RN7SL748P | 0.018779295 |
| RP11-416H1 | 0.018775817 |
| RN7SL199P | 0.018773464 |
| RNA5SP302 | 0.01877294 |
| RNU6-371P | 0.018762158 |
| AP006547 | 0.018758313 |
| HSBP1P2 | 0.018757006 |
| RNU6ATAC12P | 0.018755035 |
| AC087430 | 0.018749917 |
| RP11-130N24 | 0.018745204 |
| RNU6-928P | 0.018739087 |
| USF1 | 0.018735799 |
| RP11-246E12 | 0.018732869 |
| PCDHGA6 | 0.018729548 |
| RNU6-1233P | 0.01872792 |
| RP11-276E15 | 0.018725143 |
| AC068062 | 0.018720629 |
| RP11-567B20 | 0.018718888 |
| RP4-741O10 | 0.018713125 |
| RP5-866L20 | 0.018712899 |
| NPAS1 | 0.018710961 |
| AC142472 | 0.018709045 |
| RNU6-524P | 0.018707084 |
| RN7SKP282 | 0.018699371 |
| CNTNAP3B | 0.018698194 |
| L3MBTL3 | 0.018697454 |
| SOCS2-AS1 | 0.018696393 |
| OR6C74 | 0.018696348 |
| ANKRD30BP2 | 0.018688045 |
| AC093874 | 0.018686737 |
| RP11-588P8 | 0.018686692 |
| RP6-22P16 | 0.018686075 |
| NDST1 | 0.018684281 |
| OR5K1 | 0.018684015 |
| RP5-1100E15 | 0.01868051 |
| MIR4533 | 0.018678067 |
| RP11-34P1 | 0.018675575 |
| RNU4-54P | 0.018668551 |
| RNASE9 | 0.01866825 |
| SCMH1 | 0.01866575 |
| RN7SKP79 | 0.01866505 |
| LINC01207 | 0.01866049 |
| RPSAP14 | 0.018658813 |
| JOSD1 | 0.018653396 |

| | |
|---|---|
| DEFB4B | 0.018649606 |
| AC126177 | 0.01864761 |
| RSL24D1P6 | 0.018645865 |
| CTD-3030D20 | 0.018636465 |
| CTB-95B16 | 0.018632063 |
| C2orf40 | 0.018631746 |
| RP11-440G2 | 0.018631498 |
| SUGP1 | 0.018627478 |
| NKIRAS2 | 0.018627385 |
| RPIAP1 | 0.018620007 |
| CTD-2008L17 | 0.01861094 |
| METTL21C | 0.018603057 |
| RP11-272L13 | 0.018602775 |
| TRIM39-RPP21 | 0.018601721 |
| ECI1 | 0.018593951 |
| OR7E28P | 0.018588018 |
| SOX5P | 0.018586042 |
| GRM1 | 0.018575779 |
| SLC44A5 | 0.018573012 |
| OR10H5 | 0.01857182 |
| AP1S2P1 | 0.018568921 |
| NEU1 | 0.018568759 |
| OPA3 | 0.018567532 |
| SNORD115-33 | 0.018566393 |
| SNX17 | 0.018565429 |
| IRX2 | 0.018563662 |
| OR7E31P | 0.018560818 |
| AC022716 | 0.018559101 |
| RP11-747H7 | 0.018554564 |
| RP11-115M14 | 0.018550903 |
| RTCB | 0.018546675 |
| RP11-978I15 | 0.018546024 |
| RPL31P50 | 0.018542677 |
| CTB-77H17 | 0.018540713 |
| DBX2 | 0.018536402 |
| RNA5SP156 | 0.018529702 |
| AP000289 | 0.018524466 |
| SLC22A6 | 0.018516166 |
| SNORA50 | 0.018513825 |
| RP11-31F15 | 0.018512292 |
| MIR3689A | 0.01850959 |
| RIC1 | 0.01850485 |
| RNU6-435P | 0.018503083 |
| MIR606 | 0.018500117 |
| TUBB8P9 | 0.018485391 |

| | |
|---|---|
| RNA5SP122 | 0.018484126 |
| AC010518 | 0.018484116 |
| RP11-12P19 | 0.018478307 |
| RP11-113I24 | 0.018477337 |
| COX7C | 0.018474318 |
| BNIP1 | 0.018473388 |
| GAPDHP63 | 0.018472259 |
| ZNF518B | 0.018470355 |
| RP4-533D7 | 0.018460383 |
| C19orf25 | 0.018460066 |
| AC098477 | 0.018458087 |
| OR7E106P | 0.01845676 |
| DTNBP1 | 0.018456494 |
| CTC-255N20 | 0.018449894 |
| GAPDHP36 | 0.01844598 |
| NRON | 0.01844415 |
| RP11-362A1 | 0.018442959 |
| RNU6-1283P | 0.018438766 |
| KALRN | 0.018435284 |
| IGKV2-36 | 0.018430851 |
| LRRC37A7P | 0.018429442 |
| LINC01242 | 0.018427241 |
| AC012362 | 0.018425609 |
| TOPORS-AS1 | 0.018424679 |
| AC001226 | 0.018424381 |
| VN1R35P | 0.01841836 |
| MT-CO3 | 0.018417832 |
| LINC00559 | 0.018411716 |
| NPAP1 | 0.018406868 |
| GSTM2P1 | 0.018405351 |
| AC013271 | 0.018401382 |
| AC090957 | 0.018400604 |
| RP11-122M14 | 0.018391612 |
| RP11-169K16 | 0.018386668 |
| MTND3P1 | 0.018383644 |
| RPS26P31 | 0.018378462 |
| CTC-265F19 | 0.018376531 |
| RP11-109M19 | 0.018376423 |
| JRKL-AS1 | 0.018372527 |
| RP11-575B7 | 0.018365986 |
| ATP10D | 0.018361327 |
| PUS1 | 0.018360397 |
| RP3-425P12 | 0.01835845 |
| HDAC3 | 0.01835102 |
| FLJ42102 | 0.018340761 |

| | |
|---|---|
| RP11-474N24 | 0.018336116 |
| YWHAQ | 0.01833474 |
| RP11-391L3 | 0.018330397 |
| RNY4 | 0.018329641 |
| RPL4 | 0.018326163 |
| RPL13P | 0.018321692 |
| TRPV3 | 0.018320098 |
| RNU1-122P | 0.018318488 |
| OR3A1 | 0.018315876 |
| MIR4438 | 0.018314201 |
| CARD9 | 0.018311334 |
| MIR8069-1 | 0.018309654 |
| RNU6-386P | 0.01830536 |
| SLC25A39 | 0.018301734 |
| INGX | 0.018296599 |
| DEFB113 | 0.018295396 |
| LINC00028 | 0.018294091 |
| RNU6V | 0.018284893 |
| KRT18P47 | 0.018283518 |
| RNU6-98P | 0.018279496 |
| OR14A16 | 0.01827849 |
| PEX11G | 0.01827444 |
| ZNF649-AS1 | 0.018274146 |
| RNA5SP114 | 0.018273754 |
| SLC38A7 | 0.018273398 |
| AL365274 | 0.018271116 |
| TMA7 | 0.018268802 |
| CDY2A | 0.018263748 |
| HIST1H4A | 0.018257739 |
| RP11-797E24 | 0.018255948 |
| RP11-375I20 | 0.018253507 |
| CTNNA2 | 0.018253294 |
| RPS17P14 | 0.018253226 |
| AC234771 | 0.018244352 |
| XAGE3 | 0.018243757 |
| RN7SKP1 | 0.018238832 |
| NRSN1 | 0.01823848 |
| GNG7 | 0.018238401 |
| GCSAML | 0.018232681 |
| RP1-53C18 | 0.018228891 |
| PHBP11 | 0.0182219 |
| AC007511 | 0.018220759 |
| TSSK5P | 0.018215563 |
| ARL2-SNX15 | 0.018214172 |
| AC016993 | 0.018213823 |

| | |
|---|---|
| GLT8D1 | 0.018207999 |
| RAB5C | 0.018207367 |
| SRGAP3-AS2 | 0.01820354 |
| PCDHGA11 | 0.018201981 |
| CNTN4-AS2 | 0.018200688 |
| RBMS3-AS1 | 0.018199465 |
| DANCR | 0.018196501 |
| MIR2113 | 0.018195203 |
| METTL7B | 0.018194957 |
| GOT1 | 0.018190696 |
| MIR424 | 0.018189626 |
| PRKX | 0.018187775 |
| SNORD114-14 | 0.018185178 |
| NMI | 0.018185003 |
| AL133243 | 0.018181003 |
| TICAM1 | 0.01817957 |
| RGPD6 | 0.018178134 |
| BMS1P2 | 0.018178082 |
| SCARNA8 | 0.018177593 |
| ANKMY2 | 0.018175552 |
| PEX5 | 0.018169914 |
| ARSEP1 | 0.018166366 |
| DUSP12P1 | 0.018165083 |
| RFX2 | 0.018164742 |
| LINC01375 | 0.01816178 |
| RP11-421P23 | 0.018151312 |
| SPTSSB | 0.01815075 |
| RP11-367G18 | 0.018149225 |
| TMEM259 | 0.018149113 |
| CFAP74 | 0.018142341 |
| RP11-1026M7 | 0.018141184 |
| RP11-219G10 | 0.018137184 |
| TCAM1P | 0.018133447 |
| OLIG3 | 0.018132996 |
| AC106795 | 0.018127431 |
| RAB43P1 | 0.018125866 |
| COBLL1 | 0.018125745 |
| RNU6-267P | 0.018119007 |
| RP11-29O6 | 0.018117671 |
| RP11-1101K5 | 0.018117591 |
| NOS2P1 | 0.018113251 |
| SUMO2P5 | 0.018108121 |
| RP11-588D3 | 0.018105429 |
| RNU6-370P | 0.018100811 |
| RNU6-108P | 0.018099546 |

| | |
|---|---|
| LINC01159 | 0.018096828 |
| CLPP | 0.018090618 |
| HNRNPA1P71 | 0.018089529 |
| AC092275 | 0.018080735 |
| RP11-321E8 | 0.018080635 |
| MYBPH | 0.018073308 |
| LINC01185 | 0.018072223 |
| RP11-269C4 | 0.018071443 |
| RP11-281N10 | 0.018071197 |
| GLUD1P8 | 0.018070392 |
| RP1-309F20 | 0.018068685 |
| SPAG11A | 0.018065054 |
| MIR135A1 | 0.018055745 |
| AKT1S1 | 0.018054337 |
| AL121897 | 0.018052392 |
| GNL2 | 0.01804922 |
| RNA5SP498 | 0.01804632 |
| RNU6-855P | 0.018042248 |
| RPL29P12 | 0.018034244 |
| DHX32 | 0.018027415 |
| TUBB7P | 0.018027369 |
| AP001043 | 0.018026107 |
| TNNI3K | 0.018023244 |
| LCT | 0.018022226 |
| DISC2 | 0.018020687 |
| XRCC6P3 | 0.018016636 |
| RNA5SP162 | 0.018014255 |
| OR14I1 | 0.018013949 |
| MIR4327 | 0.018013808 |
| ZNF92P1Y | 0.018011519 |
| HOXB13-AS1_1 | 0.018010342 |
| PYY | 0.018008499 |
| RP1-171K16 | 0.018005392 |
| RP13-279N23 | 0.018004523 |
| AL157399 | 0.017993597 |
| LINC00032 | 0.01799227 |
| RNU6-1303P | 0.017990802 |
| HCG9 | 0.017985569 |
| RP11-361M4 | 0.017982384 |
| MIR4489 | 0.017979497 |
| RMI2 | 0.017978296 |
| AL359987 | 0.017972795 |
| LINC01500 | 0.017962365 |
| HIST1H2BG | 0.017960166 |
| CENPVP3 | 0.017955803 |

| | |
|---|---|
| CAP1P2 | 0.017952812 |
| RP11-156K23 | 0.017950351 |
| PTGER4P2 | 0.017947363 |
| CEP89 | 0.017944918 |
| TRIO | 0.017944304 |
| RP11-117L6 | 0.017938345 |
| ACAT1 | 0.017934462 |
| MTX1P1 | 0.017927169 |
| MAGT1 | 0.017920415 |
| CYCSP49 | 0.017908086 |
| BTBD11 | 0.017907242 |
| KLF2P4 | 0.017905088 |
| FBXL18 | 0.01790287 |
| KRT2 | 0.017895919 |
| RP11-365D23 | 0.017895791 |
| RP11-330L19 | 0.017892288 |
| MIR383 | 0.017885484 |
| CTB-83J4 | 0.017883978 |
| NLGN3 | 0.017876338 |
| RP11-673E1 | 0.017876204 |
| SMOC2 | 0.017873194 |
| RNU1-31P | 0.017868319 |
| GPR32P1 | 0.01786735 |
| CYCSP27 | 0.017865009 |
| SMARCB1 | 0.017863352 |
| AL591379 | 0.017859999 |
| BTBD3 | 0.017853379 |
| AC007292 | 0.017851519 |
| RN7SL278P | 0.017841951 |
| SHISA9 | 0.017839404 |
| SDHCP1 | 0.01783444 |
| RP11-1136G4 | 0.017830736 |
| RP13-685P2 | 0.017828782 |
| MIR6821 | 0.017816498 |
| CSPG4P4Y | 0.017810753 |
| CDC42BPG | 0.017802754 |
| RP11-444C12 | 0.017800732 |
| OR1M1 | 0.017799149 |
| RP11-379F4 | 0.017797005 |
| RP5-1048B16 | 0.017792679 |
| RNU7-95P | 0.017791723 |
| AC020934 | 0.017784621 |
| RN7SL700P | 0.017784011 |
| AL354833 | 0.017780483 |
| GPR150 | 0.017777369 |

| | |
|---|---|
| LINC00492 | 0.017771763 |
| ASB18 | 0.017771366 |
| RP11-296P7 | 0.017767709 |
| MIR519B | 0.017765611 |
| RP4-765H13 | 0.017759224 |
| RN7SKP247 | 0.017750222 |
| RP11-62I21 | 0.017750062 |
| CICP12 | 0.017748771 |
| BTBD6 | 0.017747961 |
| RNU6-121P | 0.017742099 |
| NDUFB8 | 0.017740667 |
| RP5-928E24 | 0.017739069 |
| MIR4787 | 0.01773722 |
| CCT4P2 | 0.017736481 |
| RP4-539M6 | 0.017724555 |
| KRT18P36 | 0.017724308 |
| VN1R66P | 0.017724207 |
| RP11-323D18 | 0.0177235 |
| AC005264 | 0.017720293 |
| GDI2 | 0.017709578 |
| RNU7-77P | 0.017708803 |
| LINC01548 | 0.017708063 |
| DUXAP6 | 0.017707525 |
| TMEM207 | 0.017704086 |
| RBPJP6 | 0.017701647 |
| RP4-800G7 | 0.017687642 |
| RP5-908M14 | 0.017683325 |
| IRF2BP1 | 0.017682469 |
| AC026241 | 0.017678366 |
| AC004911 | 0.017669696 |
| RN7SL653P | 0.017669311 |
| TMSB15A | 0.017666302 |
| WFDC11 | 0.017656937 |
| CCNB1IP1 | 0.017655101 |
| ZNF863P | 0.017652843 |
| RP11-61G24 | 0.01765012 |
| SNORD113-9 | 0.017648546 |
| PCDHA3 | 0.017635425 |
| AC010928 | 0.017632816 |
| GLRX2 | 0.017628495 |
| PRSS58 | 0.017627766 |
| SLC25A5P8 | 0.017625473 |
| CASC19 | 0.017622213 |
| MIR8082 | 0.017621037 |
| AC144838 | 0.017620324 |

| | |
|---|---|
| RP11-234N17 | 0.017615215 |
| LINC00347 | 0.017613521 |
| LINC01487 | 0.017609967 |
| RP11-366F6 | 0.017608293 |
| PGAM3P | 0.01759607 |
| GJC1 | 0.017591861 |
| RP11-127B16 | 0.017589625 |
| SNORD115-15 | 0.017587888 |
| RN7SKP86 | 0.017587267 |
| GPKOW | 0.017579972 |
| RNU6-188P | 0.01757986 |
| FUS | 0.017576139 |
| CTD-3080F16 | 0.01757553 |
| KCNJ9 | 0.017572923 |
| RP11-91P17 | 0.01756485 |
| LINC01456 | 0.017560567 |
| OSTCP2 | 0.01755731 |
| RP11-458D21 | 0.017555822 |
| AC009506 | 0.017550054 |
| CES1P2 | 0.017546369 |
| RNU6-1162P | 0.017541563 |
| STK32C | 0.017539058 |
| PMS2P5 | 0.017537719 |
| RNU6-841P | 0.017537336 |
| LINC00604 | 0.017536448 |
| LOXHD1 | 0.017534799 |
| RP11-713H12 | 0.017533833 |
| RPL7P58 | 0.017533631 |
| DEFB122 | 0.017529435 |
| BRI3P1 | 0.017524294 |
| SNX18P26 | 0.017516685 |
| RN7SL713P | 0.017511083 |
| AC013402 | 0.017509099 |
| KLRG2 | 0.017505371 |
| TAAR8 | 0.017493064 |
| RP11-404H14 | 0.01749237 |
| SIPA1L2 | 0.017492061 |
| PRAMEF5 | 0.017491953 |
| MED11 | 0.01748927 |
| PDLIM1P4 | 0.017485621 |
| HCG15 | 0.017485168 |
| PAK3 | 0.017481791 |
| LINC00693 | 0.017476837 |
| RP11-98L12 | 0.017470666 |
| OR11G1P | 0.017470048 |

| | |
|---|---|
| EHF | 0.01746739 |
| C5orf49 | 0.017466032 |
| RPL7P18 | 0.017450651 |
| RP11-509E10 | 0.017446137 |
| CTD-2036J7 | 0.017444245 |
| OR7E37P | 0.017442399 |
| FGD1 | 0.017432486 |
| RP13-526J3 | 0.017432108 |
| CYP2E1 | 0.017431798 |
| GNG5P3 | 0.017426981 |
| DCTN6 | 0.0174243 |
| RPL34P27 | 0.017421744 |
| CCDC85A | 0.017419972 |
| CCT8L1P | 0.017415859 |
| VN1R21P | 0.017415552 |
| GTF3C5 | 0.017413341 |
| MIR548X2 | 0.017401311 |
| RP11-148O7 | 0.017401311 |
| PRG2 | 0.017401045 |
| LGR6 | 0.017395927 |
| LINC01433 | 0.017395912 |
| RP11-444K7 | 0.017395361 |
| OR2A20P | 0.017393772 |
| CTC-326K19 | 0.017390049 |
| DCHS2 | 0.017386902 |
| AC008065 | 0.017384064 |
| RP1-257I9 | 0.017371865 |
| KIAA0226L | 0.017365358 |
| MAGEE2 | 0.017360783 |
| SMIM7 | 0.017357525 |
| CRYBG3 | 0.017350566 |
| ERVMER61-1 | 0.017350028 |
| DEDD | 0.017348315 |
| TIPRL | 0.017347887 |
| NELFCD | 0.017346808 |
| RNU6-1065P | 0.017342205 |
| RNU1-67P | 0.01733984 |
| COPS8P3 | 0.017335889 |
| MIR6825 | 0.01733207 |
| RP11-461C20 | 0.01733168 |
| YJEFN3 | 0.017327607 |
| DNAH10 | 0.017324835 |
| RP11-1129I3 | 0.017324198 |
| U91319 | 0.017311766 |
| AP003064 | 0.017305716 |

| | |
|---|---|
| RBMY2NP | 0.0172946 |
| RP11-149I23 | 0.017291712 |
| KANSL1-AS1 | 0.017291662 |
| RP1-76B20 | 0.017283702 |
| ATP6V0A4 | 0.017278261 |
| THRB-AS1 | 0.017277404 |
| RP11-383D22 | 0.017271022 |
| SLC25A51P2 | 0.0172685 |
| SMIM2-AS1 | 0.017266089 |
| AC008622 | 0.017265595 |
| RNA5SP235 | 0.017254574 |
| LINC00521 | 0.017252456 |
| IGFBP5 | 0.017252237 |
| MATK | 0.017246487 |
| BNIP3P33 | 0.017243648 |
| MIR3182 | 0.017241769 |
| RNU4-16P | 0.017236906 |
| STRC | 0.017231726 |
| CTD-2055G21 | 0.017228657 |
| MIR4650-1 | 0.017226819 |
| MIR6769B | 0.017226076 |
| DPRXP2 | 0.017224898 |
| AC068831 | 0.017224384 |
| RP5-1085F17 | 0.017220663 |
| MIR2278 | 0.017207818 |
| MIR147A | 0.017207661 |
| RN7SKP185 | 0.017206473 |
| CTD-3096P4 | 0.017202279 |
| NR2C2AP | 0.017191508 |
| AL160279 | 0.017189344 |
| SNRPE | 0.017185982 |
| TMEM136 | 0.017184252 |
| HLA-N | 0.017176325 |
| EPAS1 | 0.017167117 |
| LDHAL6CP | 0.017166641 |
| RP4-753D4 | 0.017164763 |
| COL18A1-AS1 | 0.017160285 |
| TNMD | 0.017157317 |
| RP11-624G19 | 0.017149342 |
| EIF4EP4 | 0.017149242 |
| RN7SKP102 | 0.017148682 |
| PPIAP23 | 0.017147677 |
| RPL31P53 | 0.017138965 |
| RNU6ATAC28P | 0.017138786 |
| RN7SL229P | 0.017137649 |

| | |
|---|---|
| MAGEF1 | 0.017134427 |
| PPOX | 0.017132138 |
| LINC00684 | 0.017128349 |
| RP5-1125M8 | 0.017117738 |
| AC003003 | 0.01711497 |
| AP1B1P2 | 0.017112253 |
| RP1-125I3 | 0.017111887 |
| TNFAIP8L1 | 0.017111449 |
| KRTAP13-1 | 0.017103424 |
| RP11-240B13 | 0.017095489 |
| MIR4470 | 0.017094221 |
| RP11-443G13 | 0.017094041 |
| SLC2A4RG | 0.017086927 |
| ZG16B | 0.017081624 |
| MTND5P21 | 0.017078377 |
| SMARCD3 | 0.01707742 |
| AC016745 | 0.017074702 |
| RP6-65G23 | 0.01707435 |
| TOP3BP1 | 0.017074139 |
| TRIM51 | 0.017074088 |
| MIR4460 | 0.017069439 |
| AL357514 | 0.017068306 |
| CENPT | 0.01706514 |
| CTD-2299E8 | 0.017061676 |
| AC108078 | 0.017061525 |
| MEF2BNB-MEF2B | 0.017059726 |
| HAPLN3 | 0.017052842 |
| RP11-569A11 | 0.017047895 |
| SPATA31C2 | 0.01703858 |
| RP11-1O7 | 0.017038278 |
| RNU6-1310P | 0.017035493 |
| GAPDHP19 | 0.017031727 |
| ALG1L2 | 0.017029659 |
| AL138925 | 0.017025357 |
| COA3 | 0.017020915 |
| SLC34A1 | 0.017019261 |
| AMHR2 | 0.017019148 |
| RP11-253M7 | 0.017018775 |
| RP11-259P15 | 0.017016288 |
| AC006987 | 0.01701237 |
| RNU6-139P | 0.017012361 |
| RNU7-196P | 0.017008249 |
| RP5-1024C24 | 0.017003012 |
| GFOD1 | 0.017002923 |
| HARS | 0.016999816 |

| | |
|---|---|
| H19_3 | 0.016995025 |
| RNA5SP391 | 0.016994659 |
| CTB-158D10 | 0.01698848 |
| AP000641 | 0.016981026 |
| RP5-1057B20 | 0.016979429 |
| AC068058 | 0.016975227 |
| DLGAP5P1 | 0.016973396 |
| MIR5192 | 0.016971319 |
| C10orf11 | 0.016968708 |
| MIR328 | 0.016966737 |
| CTC-507E12 | 0.016966001 |
| SLC41A3 | 0.016965843 |
| TSR2 | 0.016964354 |
| TMEM104 | 0.016963591 |
| HSPD1P10 | 0.016963369 |
| RP11-152C17 | 0.016958449 |
| ZNF295-AS1 | 0.01695727 |
| RP11-12G12 | 0.016953116 |
| MIR1225 | 0.016950896 |
| VARS | 0.01695027 |
| SLC41A2 | 0.01695014 |
| RN7SL558P | 0.016949988 |
| RP11-390G14 | 0.016944158 |
| VAC14 | 0.016943814 |
| METTL12 | 0.016935237 |
| RHBDF1P1 | 0.016934821 |
| RP11-25G10 | 0.016929292 |
| RN7SL819P | 0.01692791 |
| CCNG1P1 | 0.016927513 |
| CHCHD4P5 | 0.016926515 |
| RP11-349E4 | 0.016926453 |
| GNAI2P2 | 0.016915566 |
| ACTR3B | 0.016915422 |
| HIST1H1E | 0.016914997 |
| AC107083 | 0.016900642 |
| NMBR | 0.016897159 |
| RP4-799D16 | 0.016896638 |
| OR2B2 | 0.016895524 |
| UBXN2A | 0.016885579 |
| RP11-307C18 | 0.016883188 |
| SNORA41 | 0.01687974 |
| TMEM30C | 0.016876776 |
| RNU6-647P | 0.01687649 |
| ST3GAL6 | 0.016872164 |
| RP11-195E2 | 0.016871299 |

| | |
|---|---|
| LOC124685 | 0.016863494 |
| TRPC3 | 0.01686261 |
| RPL5P15 | 0.016862287 |
| RPL34P34 | 0.016862002 |
| RPS4XP15 | 0.016855133 |
| GPR149 | 0.016854068 |
| RP11-771F20 | 0.016850272 |
| KB-1742H10 | 0.016847422 |
| KCNMB3P1 | 0.016840826 |
| RP1-6P5 | 0.016825942 |
| TAS1R1 | 0.016821886 |
| ACR | 0.016818914 |
| TUSC7 | 0.016818837 |
| CSF2RBP1 | 0.016816599 |
| OR8Q1P | 0.016810275 |
| RP1-300I2 | 0.016795569 |
| DUSP22 | 0.016795044 |
| CCDC63 | 0.016792896 |
| LRRN2 | 0.0167927 |
| OR1F12 | 0.016785125 |
| RANP2 | 0.016780962 |
| RP11-136O12 | 0.016780286 |
| AC114813 | 0.016777463 |
| ARGFX | 0.016775017 |
| RIT2 | 0.016772983 |
| CTD-2587M23 | 0.016772685 |
| RP11-281J9 | 0.016772347 |
| OR5K4 | 0.016769288 |
| OR51V1 | 0.016763946 |
| AC000370 | 0.016763236 |
| RP11-531A24 | 0.01676228 |
| PKD2 | 0.016761741 |
| RP11-481G8 | 0.016760475 |
| ALG1L5P | 0.016754165 |
| OR6R2P | 0.016753287 |
| RPL34P26 | 0.016752576 |
| GTPBP8 | 0.016743042 |
| TUBB3P2 | 0.016742091 |
| KMO | 0.016738949 |
| SMUG1 | 0.01673787 |
| CCT4 | 0.016728828 |
| RP11-247I13 | 0.016728641 |
| RP11-826N14 | 0.016727652 |
| EEF1A1P20 | 0.016726354 |
| MIR500B | 0.016718906 |

| | |
|---|---|
| DCST2 | 0.016718334 |
| RP11-472F19 | 0.016717323 |
| ODF2L | 0.01671212 |
| CTD-2026G6 | 0.016702109 |
| Z84466 | 0.016697835 |
| IFNA17 | 0.016694757 |
| UGT1A5 | 0.016684148 |
| HIF3A | 0.016675819 |
| RP1-63P18 | 0.01667555 |
| RP11-72H11 | 0.016673087 |
| RP11-95P13 | 0.0166702 |
| RPL7P30 | 0.01666959 |
| CHIA | 0.016665892 |
| MTND1P16 | 0.016663368 |
| RNU6-1075P | 0.016647741 |
| AL591428 | 0.016646087 |
| MAPK6 | 0.016641307 |
| CNKSR2 | 0.016641078 |
| RP11-456I15 | 0.016639367 |
| RP11-882I15 | 0.016637944 |
| RNPS1 | 0.016629791 |
| AC010649 | 0.016626159 |
| RN7SL654P | 0.016624427 |
| PLD5 | 0.016622022 |
| MSGN1 | 0.016620898 |
| RPL9P32 | 0.016619186 |
| RAP1BP2 | 0.016618394 |
| MIR5100 | 0.016618292 |
| CTD-2007H18 | 0.016617032 |
| DSCAM-IT1 | 0.016608498 |
| PREP | 0.016605492 |
| RNU6-1107P | 0.016603841 |
| LINC01550 | 0.016600158 |
| MIR454 | 0.016598622 |
| AC104692 | 0.016593717 |
| MEST | 0.016592505 |
| PRKAG3 | 0.016590543 |
| LRTM1 | 0.016585114 |
| USP17L5 | 0.016582425 |
| MIR124-2HG | 0.016578808 |
| MIR1302-6 | 0.016577531 |
| CCDC105 | 0.016576137 |
| RN7SKP56 | 0.016575827 |
| TSACC | 0.016573342 |
| GPIHBP1 | 0.01657323 |

| | |
|---|---|
| TTC38 | 0.016572635 |
| COL20A1 | 0.016566619 |
| LYL1 | 0.016564188 |
| CCT5P1 | 0.016560703 |
| SNORD93 | 0.016558403 |
| RGS16 | 0.01655509 |
| FAM207BP | 0.01655411 |
| RP11-76K13 | 0.016552866 |
| RNU6-145P | 0.016552568 |
| NDUFAF3 | 0.016546978 |
| SBDS | 0.016545899 |
| RP11-449L23 | 0.01654386 |
| RPS3AP41 | 0.016541675 |
| XYLT2 | 0.016536 |
| CXADRP3 | 0.016532702 |
| CA3 | 0.016525865 |
| BANF2 | 0.016523154 |
| MIR4635 | 0.016521007 |
| RFWD2 | 0.016515702 |
| RP11-69C13 | 0.016515091 |
| CCDC94 | 0.016514213 |
| RP11-76G10 | 0.01651027 |
| SETP6 | 0.016507181 |
| FAM58CP | 0.016506551 |
| RNU6-692P | 0.01650342 |
| ALKBH2 | 0.016497264 |
| RP11-693L9 | 0.016496931 |
| RPL7P56 | 0.016492612 |
| PAFAH1B3 | 0.01648718 |
| HRAS | 0.016487087 |
| HRK | 0.016486538 |
| MIR5687 | 0.016472014 |
| AC012065 | 0.016469988 |
| P3H2-AS1 | 0.016469916 |
| BACE1 | 0.016465114 |
| AL117190 | 0.016457426 |
| CHML | 0.016447494 |
| AL732479 | 0.016445945 |
| RNA5SP171 | 0.016445945 |
| TBC1D3P1 | 0.016440207 |
| AC009502 | 0.016435833 |
| AC004832 | 0.016431642 |
| RP11-1055B8 | 0.016430228 |
| MIR6852 | 0.016422216 |
| RCBTB2P1 | 0.016413912 |

| | |
|---|---|
| C17orf50 | 0.016398269 |
| RP1-293L6 | 0.016389218 |
| KRT17P2 | 0.01638815 |
| PSMD10P2 | 0.016386705 |
| RP11-40H19 | 0.016380634 |
| CAPS | 0.016379826 |
| CNGA2 | 0.016378549 |
| MIR1273D | 0.016375495 |
| WI2-2373I1 | 0.016375038 |
| RNU7-20P | 0.016370822 |
| CETN1 | 0.016369166 |
| CAPN13 | 0.016367418 |
| BX571818 | 0.016365719 |
| ATP5F1P4 | 0.01636517 |
| LGALS7 | 0.016362448 |
| AC013480 | 0.016360942 |
| SMS | 0.016354616 |
| DIAPH2-AS1 | 0.016351783 |
| TMEM179 | 0.01635069 |
| MIR1285-2 | 0.01634825 |
| SLC30A3 | 0.016347017 |
| MIR708 | 0.01634524 |
| CCL11 | 0.01633832 |
| RP11-13N12 | 0.016329618 |
| TSGA10IP | 0.016326729 |
| PIN1P1 | 0.016324988 |
| WDR78 | 0.016322633 |
| RP11-192H6 | 0.016313558 |
| ABHD5 | 0.016311693 |
| MS4A15 | 0.016311216 |
| RP11-26C10 | 0.016309464 |
| OR8I4P | 0.016309464 |
| NOS2P3 | 0.016303713 |
| PCDHGB7 | 0.016301981 |
| RER1 | 0.016301292 |
| SLC39A5 | 0.016301067 |
| LINC00948 | 0.016298449 |
| CTD-2206G10 | 0.016294529 |
| BAGE2 | 0.016292193 |
| RP11-698E18 | 0.016289149 |
| RP11-789C1 | 0.016287085 |
| AC004969 | 0.016284342 |
| SHROOM4 | 0.016282408 |
| RP11-526K17 | 0.016273702 |
| RPS24P8 | 0.016272994 |

| | |
|---|---|
| GOLT1B | 0.016272714 |
| RP11-341G5 | 0.016270556 |
| AC012322 | 0.016247695 |
| LINC01351 | 0.016240456 |
| RPF2P2 | 0.016234046 |
| COMMD9 | 0.016231503 |
| ZNF732 | 0.016231333 |
| OR5AH1P | 0.016219307 |
| RNA5SP146 | 0.016210319 |
| LINC00574 | 0.016210118 |
| PCDHAC1 | 0.016207897 |
| AC013717 | 0.016204724 |
| DGKG | 0.016202014 |
| AC093732 | 0.01619653 |
| RP11-65L3 | 0.016193753 |
| KRTAP4-16P | 0.016188741 |
| AC079121 | 0.016183075 |
| TMLHE-AS1 | 0.016182394 |
| MAPKAPK2 | 0.016180543 |
| CTC-457E21 | 0.016179932 |
| CARD6 | 0.016172431 |
| RP4-796I17 | 0.01617196 |
| SCb-64M4 | 0.016170786 |
| RN7SL745P | 0.016167513 |
| C15orf59-AS1 | 0.016162569 |
| C10orf91 | 0.016157316 |
| UBE2L5P | 0.016152867 |
| RP11-174E22 | 0.016149625 |
| RN7SL479P | 0.016146275 |
| RASSF1 | 0.016145583 |
| RP11-449H15 | 0.016145496 |
| GAGE12H | 0.016144705 |
| AC108137 | 0.016136755 |
| LINC01252 | 0.016136485 |
| GRAMD4P6 | 0.016131347 |
| RNU6-170P | 0.016130711 |
| RP11-630C16 | 0.016121404 |
| MIR6068 | 0.01611936 |
| AC006373 | 0.01611396 |
| LINC01547 | 0.016081692 |
| CACNA1G-AS1 | 0.016078005 |
| CTD-2280E9 | 0.016077751 |
| GFRAL | 0.016073234 |
| RP11-609L23 | 0.016072826 |
| RNU6-1324P | 0.016071385 |

| | |
|---|---|
| RPL17P22 | 0.016069694 |
| RNU6-1217P | 0.01606942 |
| MAN1B1-AS1 | 0.016066994 |
| IGSF8 | 0.016062417 |
| RP11-409K15 | 0.016060232 |
| RGMB-AS1 | 0.016052091 |
| AC092646 | 0.0160493 |
| GAPDHP68 | 0.01603672 |
| AL021920 | 0.016036502 |
| RBMY2OP | 0.016036502 |
| MIR378C | 0.016036502 |
| ATP2B2-IT1 | 0.016036165 |
| MIR4266 | 0.01602461 |
| DGUOK-AS1 | 0.016020796 |
| RNU7-154P | 0.016017507 |
| KRT18P51 | 0.016015482 |
| RP11-141O15 | 0.016014816 |
| RNA5SP190 | 0.016009479 |
| PSD2 | 0.015977734 |
| TSPAN8 | 0.015965693 |
| RN7SKP165 | 0.015965144 |
| SLFN5 | 0.015958896 |
| AC004754 | 0.015958691 |
| SLC39A10 | 0.015958319 |
| RNA5SP471 | 0.015954709 |
| IGKV2-19 | 0.01595337 |
| LINC00382 | 0.015948637 |
| AC073621 | 0.015947264 |
| RP11-10O22 | 0.015946487 |
| RPL14P5 | 0.015945235 |
| BNIP3P24 | 0.015942796 |
| FDPSP7 | 0.015942289 |
| ARHGAP36 | 0.01594088 |
| AC134025 | 0.015936956 |
| SEC14L4 | 0.015936403 |
| AP000472 | 0.015936228 |
| HADHB | 0.015933928 |
| PSMB3 | 0.015927751 |
| ITLN2 | 0.015921409 |
| COX7B2 | 0.015918572 |
| TNFRSF10B | 0.015917127 |
| RP11-443F16 | 0.015917071 |
| RP11-5G9 | 0.01591614 |
| AC016821 | 0.015914645 |
| WAC-AS1 | 0.015911192 |

| | |
|---|---|
| RP11-486L19 | 0.015910922 |
| SNORD104 | 0.015907479 |
| SNX33 | 0.015902428 |
| C1orf74 | 0.015897424 |
| MIR4493 | 0.015896704 |
| SULT1D1P | 0.015893689 |
| TBX2-AS1 | 0.015892271 |
| CDC20 | 0.015890428 |
| MPPE1P1 | 0.015888704 |
| RP11-671P2 | 0.015873185 |
| RNA5SP444 | 0.015871941 |
| RP11-395E19 | 0.015865869 |
| TBC1D23 | 0.01586358 |
| RN7SKP264 | 0.015852032 |
| ABCC9 | 0.015847431 |
| PKIG | 0.01583837 |
| AQP7P1 | 0.015832305 |
| SRGAP2D | 0.015824455 |
| RP11-445O16 | 0.015824257 |
| MS4A2 | 0.015821436 |
| AP001505 | 0.015820341 |
| TMEM185AP1 | 0.015803848 |
| EIF4E2P2 | 0.015802071 |
| LINC01299 | 0.015801041 |
| BLOC1S1 | 0.015800433 |
| RHCG | 0.015797297 |
| RP11-20L24 | 0.015796115 |
| RP11-142A23 | 0.015792365 |
| RPS3AP30 | 0.015789768 |
| RP11-58A18 | 0.015788886 |
| RP11-613F7 | 0.015787191 |
| VN1R33P | 0.015784109 |
| PRKD3 | 0.015778386 |
| DUX4L16 | 0.015777595 |
| KLK1 | 0.015776866 |
| LINC01556 | 0.015772276 |
| LAP3P1 | 0.015771584 |
| FGF10-AS1 | 0.015771348 |
| RN7SL375P | 0.015770409 |
| LINC00936 | 0.015763408 |
| HCG4P8 | 0.015756255 |
| RN7SL580P | 0.015755397 |
| MIR30E | 0.015755133 |
| RP11-245J24 | 0.015749009 |
| CTD-3138B18 | 0.015737044 |

| | |
|---|---|
| RP11-252M18 | 0.015732517 |
| MIR15B | 0.015732156 |
| RNASEH2A | 0.015729602 |
| RP11-313C4 | 0.01572491 |
| NFIB | 0.015721416 |
| ALX1 | 0.015720628 |
| MPZL3 | 0.015720132 |
| DPT | 0.015713043 |
| RP11-1260E13 | 0.0157115 |
| RP1-89D4 | 0.015709493 |
| NPBWR1 | 0.015705238 |
| AC016577 | 0.015703655 |
| RP11-1058G23 | 0.015702586 |
| DLG1-AS1 | 0.015701293 |
| LINC00375 | 0.01569913 |
| RNU6-726P | 0.015694302 |
| RNA5SP256 | 0.015687288 |
| GDE1 | 0.015683517 |
| RP11-545G3 | 0.015682301 |
| SGCZ | 0.015678205 |
| RP11-638F5 | 0.015675389 |
| C7orf66 | 0.015672796 |
| KRTAP10-2 | 0.015671737 |
| AC103740 | 0.015664977 |
| RP11-10L7 | 0.015663659 |
| RN7SL870P | 0.01566129 |
| SERHL | 0.015661041 |
| RNU7-41P | 0.015661022 |
| RFTN1P1 | 0.015660009 |
| RP11-35J6 | 0.01565972 |
| RP11-711K1 | 0.015658958 |
| AC016819 | 0.015655405 |
| PRODH | 0.015650624 |
| MIR4771-2 | 0.01564884 |
| RN7SKP53 | 0.015637238 |
| KIRREL3 | 0.015636873 |
| SLC18B1 | 0.015636612 |
| CCDC51 | 0.015630584 |
| ZBBX | 0.015628599 |
| AC091038 | 0.015616901 |
| RP11-452G18 | 0.015616593 |
| LINC01070 | 0.0156129 |
| ZNF826P | 0.015611644 |
| VIM2P | 0.015610443 |
| CDC27P2 | 0.015609799 |

| | |
|---|---|
| MIR6069 | 0.015608412 |
| LINC01419 | 0.015607501 |
| RPS3AP22 | 0.015605701 |
| EEF1A1P25 | 0.015605395 |
| CTD-2176I21 | 0.015598469 |
| AC025594 | 0.015588752 |
| TMEM244 | 0.015585754 |
| WBSCR28 | 0.015584331 |
| RP11-719C20 | 0.015581461 |
| CTNNA1 | 0.015580238 |
| KLHL2P1 | 0.015575581 |
| STRN4 | 0.015572181 |
| PHBP4 | 0.015568821 |
| JPH1 | 0.015564907 |
| RPP30 | 0.015562469 |
| ZFP41 | 0.015559455 |
| XXbac-BPG34I8 | 0.015556706 |
| RP11-810B23 | 0.01554807 |
| CHST4 | 0.015545617 |
| AL136992 | 0.0155435 |
| GRHPR | 0.015537687 |
| RNA5SP32 | 0.015536905 |
| UCHL1-AS1 | 0.015535906 |
| ACE3P | 0.015535007 |
| FAM74A4 | 0.015529959 |
| RN7SL475P | 0.015525273 |
| FAM83C | 0.015522872 |
| AP001056 | 0.015514275 |
| CTD-2102P23 | 0.015513423 |
| GTF2F2P1 | 0.015510947 |
| P4HA2-AS1 | 0.015491268 |
| CEACAMP7 | 0.015469562 |
| GJA6P | 0.015468818 |
| HIP1 | 0.015464939 |
| MRPL52 | 0.015464381 |
| FIG4 | 0.01546159 |
| CTA-369K23 | 0.015452749 |
| AC138031 | 0.015444336 |
| ACTN3 | 0.015439816 |
| RP11-64C1 | 0.015437944 |
| DUS3L | 0.01543251 |
| AC021218 | 0.015428882 |
| MIR6078 | 0.015428443 |
| RPL23P4 | 0.015419734 |
| RNU6-1093P | 0.015418543 |

| | |
|---|---|
| PCNPP5 | 0.015417588 |
| RPL30P10 | 0.015412793 |
| RN7SKP235 | 0.015408987 |
| RP11-342M1 | 0.015403467 |
| CDH12P2 | 0.015402615 |
| MIR3689B | 0.015396282 |
| TOMM22P3 | 0.015395002 |
| MIR4299 | 0.015392545 |
| ATP8A2P3 | 0.01538673 |
| ACY1 | 0.015386369 |
| PFN1P11 | 0.015384801 |
| RP13-585F24 | 0.015377379 |
| TAF10 | 0.015374461 |
| RP11-5O23 | 0.01537436 |
| RP11-475C16 | 0.01537288 |
| ABHD14A | 0.015370777 |
| AP006261 | 0.015370394 |
| AC105235 | 0.015368701 |
| AIG1 | 0.01536553 |
| NASP | 0.015363577 |
| RP11-78A19 | 0.015357608 |
| TNRC18 | 0.015357605 |
| CTB-78O21 | 0.015343945 |
| RP11-349P19 | 0.015343185 |
| CTD-2152M20 | 0.015339431 |
| AC008967 | 0.015337987 |
| RNF103-CHMP3 | 0.015335852 |
| RP4-593H12 | 0.015334173 |
| CTA-276O3 | 0.01533115 |
| RNA5SP91 | 0.015321758 |
| ZSCAN5A | 0.015318477 |
| ARMS2 | 0.015306849 |
| MORC4 | 0.015304653 |
| RNU6-877P | 0.015302192 |
| ANO4 | 0.015301991 |
| AC003090 | 0.01529968 |
| ZNF676 | 0.015294229 |
| AC074363 | 0.015292688 |
| TRPM2-AS | 0.015289975 |
| NDUFS5P1 | 0.01528933 |
| ATP5G1P4 | 0.015283731 |
| RNA5SP151 | 0.015280709 |
| C15orf59 | 0.015278996 |
| NOL3 | 0.015278662 |
| AC092865 | 0.015266433 |

| | |
|---|---|
| RN7SL822P | 0.015265326 |
| AC012003 | 0.015260175 |
| SSTR1 | 0.015259447 |
| OR2A9P | 0.015259278 |
| MIR6829 | 0.015259236 |
| FAR2P1 | 0.015257752 |
| LINC00448 | 0.015255821 |
| SNORA17 | 0.015244092 |
| FGFR4 | 0.015242288 |
| AC110813 | 0.015236508 |
| CTD-3035D6 | 0.015236409 |
| LINC01201 | 0.015232611 |
| RP11-10G12 | 0.01523159 |
| RNU6-765P | 0.015225096 |
| TALDO1 | 0.015222417 |
| LEFTY1 | 0.015215084 |
| AC006988 | 0.015208949 |
| RN7SKP95 | 0.015201227 |
| RP11-545I10 | 0.015196026 |
| SPTLC1P2 | 0.01519443 |
| PMS2L2 | 0.015190991 |
| AC117444 | 0.015188915 |
| TUBBP9 | 0.015180287 |
| RP11-524K22 | 0.01517856 |
| NDFIP1P1 | 0.015177177 |
| RP11-526F3 | 0.015176261 |
| RP11-369C8 | 0.015175645 |
| RNU6-331P | 0.015175254 |
| AP001042 | 0.01516642 |
| RP11-872D17 | 0.015166359 |
| CBLN4 | 0.015161515 |
| MIR450A2 | 0.015160345 |
| XX-C2158C12 | 0.01515551 |
| SZT2-AS1 | 0.015154242 |
| DMBX1 | 0.015151063 |
| RNA5SP178 | 0.015150916 |
| RP11-284P20 | 0.015150451 |
| E2F6P1 | 0.01515011 |
| GTF2IRD1P1 | 0.015149913 |
| NODAL | 0.015149081 |
| VENTXP2 | 0.015147821 |
| DDC-AS1 | 0.015143751 |
| RNA5SP360 | 0.015143325 |
| RANBP20P | 0.015140661 |
| MIR3920 | 0.015140661 |

| | |
|---|---|
| MIR483 | 0.015140458 |
| CHRM3 | 0.015139865 |
| ARFGAP3 | 0.015136144 |
| RP11-567P19 | 0.015134352 |
| RETSAT | 0.015133855 |
| RNU2-21P | 0.01512711 |
| KRTAP5-9 | 0.015124359 |
| RCN1P1 | 0.015118983 |
| MAGEB1 | 0.015118133 |
| CSNK1G2P1 | 0.015115317 |
| ABCC3 | 0.015110152 |
| SMYD3-IT1 | 0.01510313 |
| RP11-301G7 | 0.01510297 |
| TAC1 | 0.015098367 |
| RP11-166B14 | 0.015093984 |
| RP3-324O17 | 0.015093234 |
| ADIPOQ | 0.015091805 |
| RP11-168L22 | 0.01509008 |
| ABCF2 | 0.015089128 |
| RP11-213G6 | 0.015087651 |
| IGFALS | 0.015086346 |
| CNPY2 | 0.015083099 |
| CTD-2270L9 | 0.015082485 |
| AF216667 | 0.015076118 |
| RP11-412H9 | 0.015070829 |
| AC011406 | 0.015070707 |
| RP5-1052M9 | 0.015069263 |
| RP5-988G17 | 0.01506554 |
| AC097359 | 0.015062593 |
| YBEY | 0.01506241 |
| RP11-962G15 | 0.015059287 |
| SNORD114-17 | 0.015057838 |
| TCEB1 | 0.015057052 |
| RP11-396M11 | 0.015056774 |
| AC093807 | 0.015048678 |
| RN7SL776P | 0.015036779 |
| AC084368 | 0.015035305 |
| AC104843 | 0.01503365 |
| RND2 | 0.015033051 |
| CFAP44-AS1 | 0.015032327 |
| ADAMTS13 | 0.015025962 |
| GRAMD4P7 | 0.015021096 |
| RP11-295D22 | 0.015016657 |
| MIR6511A2 | 0.015015237 |
| ERICH1-AS1 | 0.015008633 |

| | |
|---|---|
| RPS10P21 | 0.01500134 |
| RP11-330M2 | 0.014997735 |
| AC027811 | 0.014988044 |
| OFD1P5Y | 0.014985458 |
| CTC-471C19 | 0.014983099 |
| C1orf158 | 0.014978962 |
| CASP16 | 0.014975285 |
| AC114498 | 0.014972617 |
| SLC17A3 | 0.014970535 |
| GFAP | 0.014967561 |
| MIR648 | 0.014965554 |
| HNRNPDLP1 | 0.014961775 |
| HIST1H4B | 0.014961417 |
| REXO4 | 0.014958778 |
| RP11-541E12 | 0.014951314 |
| RPL12L3 | 0.014951169 |
| TBRG4 | 0.014949568 |
| RP11-301M17 | 0.014948228 |
| CTD-3224I3 | 0.01494818 |
| SRY | 0.01494818 |
| XKRYP3 | 0.014944655 |
| POLE4 | 0.014941419 |
| TUSC5 | 0.014937957 |
| RP5-1189B24 | 0.014937801 |
| TRABD2A | 0.014932004 |
| SELO | 0.014927855 |
| PGAM1P9 | 0.014925578 |
| RP11-486I11 | 0.014922981 |
| RP11-491H9 | 0.01492187 |
| RP11-328J2 | 0.014920216 |
| YIPF5 | 0.014918888 |
| RP11-154D17 | 0.014918233 |
| SMCP | 0.014917051 |
| ASCL4 | 0.014911407 |
| HNRNPA1P40 | 0.014904013 |
| DUXAP2 | 0.014899047 |
| SLC47A2 | 0.014895669 |
| RP11-211G3 | 0.014893976 |
| KRT125P | 0.014883617 |
| CPS1 | 0.014883314 |
| GAPDHP60 | 0.014880161 |
| SLC9C2 | 0.014879995 |
| SFTPA3P | 0.014871411 |
| CACYBPP1 | 0.014869432 |
| RP1-142F18 | 0.014867418 |

| | |
|---|---|
| GARSP1 | 0.014865692 |
| THBD | 0.014862494 |
| AP001992 | 0.01486234 |
| CEP131 | 0.014859164 |
| MIR489 | 0.014857982 |
| WDFY2 | 0.014857527 |
| SUFU | 0.014855276 |
| RP11-553L6 | 0.014854792 |
| TNFSF12-TNFSF13 | 0.01485392 |
| RP11-364L4 | 0.014852475 |
| CDC37L1-AS1 | 0.014851554 |
| DIRAS2 | 0.014851372 |
| RPS26P2 | 0.014848968 |
| MAGED4B | 0.014845937 |
| RNU6-485P | 0.014822194 |
| DDX3P2 | 0.014821787 |
| AC104037 | 0.014808174 |
| VSTM5 | 0.014805301 |
| MIR1263 | 0.014801289 |
| AC002467 | 0.014798622 |
| CNPY3 | 0.014797896 |
| TUBB1P2 | 0.014794605 |
| DUXAP8 | 0.014791887 |
| UQCRBP3 | 0.014790442 |
| AL590085 | 0.014786684 |
| RP11-661A12 | 0.014784631 |
| WI2-2118C23 | 0.01477165 |
| MIR320D2 | 0.0147692 |
| RP11-761E20 | 0.014766923 |
| PTCD2P1 | 0.014766251 |
| LINC00501 | 0.014760449 |
| ARL2BPP8 | 0.0147603 |
| VCX3A | 0.014754047 |
| TMPRSS9 | 0.01474515 |
| PPME1 | 0.014742378 |
| TAF13 | 0.014740852 |
| AC079586 | 0.014740339 |
| C19orf66 | 0.014738638 |
| MIR3165 | 0.014735584 |
| PPIAP22 | 0.014734433 |
| CTD-2243E23 | 0.014733269 |
| DEFA7P | 0.0147313 |
| KCNK7 | 0.014730956 |
| RP11-375A20 | 0.014722971 |
| GATM | 0.014718432 |

| | |
|---|---|
| ARID5A | 0.014717465 |
| RP5-1052I5 | 0.014710441 |
| RPL23AP60 | 0.014702271 |
| RP11-502H18 | 0.014699199 |
| RPL34P3 | 0.014695604 |
| PRICKLE2-AS3 | 0.014695461 |
| MARK3P3 | 0.014693221 |
| BX664724 | 0.014685768 |
| HMGB3P6 | 0.014685337 |
| C16orf86 | 0.014684496 |
| RP11-676F20 | 0.014683955 |
| DDX50P2 | 0.01467698 |
| RP1-281E18 | 0.014674788 |
| RP11-362B23 | 0.014668699 |
| SERPINC1 | 0.014667694 |
| RP11-455O6 | 0.014662858 |
| RNU1-62P | 0.014660538 |
| SCGB2A1 | 0.01465976 |
| CSNK2B | 0.014645983 |
| AC118550 | 0.014645131 |
| AL390036 | 0.01464251 |
| RNU6-797P | 0.014641959 |
| RP11-337L12 | 0.014637951 |
| CTDP1 | 0.014637219 |
| IFT22 | 0.014630112 |
| AC026799 | 0.014623469 |
| RNA5SP251 | 0.014616141 |
| NAT8B | 0.014612626 |
| AC022395 | 0.014600788 |
| METRN | 0.014597962 |
| TNFAIP1 | 0.014594836 |
| RPS3AP6 | 0.014589124 |
| RP11-211N11 | 0.014584112 |
| DMAP1 | 0.014578445 |
| PRDX2P4 | 0.014577288 |
| RP11-16L9 | 0.014576473 |
| LHX5-AS1 | 0.014575007 |
| LRRC4B | 0.014573552 |
| LINC00391 | 0.014572589 |
| CCDC81 | 0.014567747 |
| TLE1P1 | 0.014567464 |
| KRT17P5 | 0.014566216 |
| DPYSL2 | 0.014560193 |
| BARX1-AS1 | 0.014560148 |
| RAD1P1 | 0.014555596 |

| | |
|---|---|
| CHIT1 | 0.014554918 |
| NDUFA2 | 0.014554741 |
| GSTM2 | 0.014543429 |
| OR7A15P | 0.014542854 |
| FAT4 | 0.01453941 |
| IGSF11 | 0.014537326 |
| DLGAP1-AS2 | 0.014535354 |
| VWA3B | 0.014534992 |
| LINC00207 | 0.014523653 |
| RNU1-59P | 0.014520567 |
| ABHD1 | 0.014511056 |
| AL591925 | 0.014506768 |
| TRABD2B | 0.014506274 |
| MIR5002 | 0.014499185 |
| RNU6-266P | 0.01449797 |
| SMAD9 | 0.014497623 |
| RP11-728L1 | 0.014485043 |
| RNU4-29P | 0.014482311 |
| HIST3H2BB | 0.014480387 |
| RNA5SP475 | 0.014475045 |
| RNU7-21P | 0.014471983 |
| RP11-344A5 | 0.014469158 |
| KRTAP21-2 | 0.014466711 |
| STARD3NL | 0.014458439 |
| E2F3-IT1 | 0.014456749 |
| FOXR2 | 0.014451235 |
| LBR | 0.014449937 |
| AQP7P2 | 0.014443432 |
| PML | 0.014443164 |
| MIR519D | 0.014440602 |
| LRRC16A | 0.014434122 |
| RGAG4 | 0.014429545 |
| RPL22P5 | 0.014428768 |
| HMGN1P32 | 0.01442871 |
| AC019181 | 0.014428565 |
| RP11-385M4 | 0.014426027 |
| MAPK6PS4 | 0.014421473 |
| CTD-2335L22 | 0.014421001 |
| FGF7P1 | 0.014417699 |
| RP11-118F2 | 0.014411748 |
| RNA5SP442 | 0.014411329 |
| CC2D2A | 0.014406698 |
| RP11-193F5 | 0.014403404 |
| FAM163A | 0.014399427 |
| RPL35AP3 | 0.014394719 |

| | |
|---|---|
| FAM19A4 | 0.014394133 |
| RP11-366H4 | 0.014393296 |
| CA14 | 0.014392167 |
| BRDT | 0.014387586 |
| DYNLL1P4 | 0.014386791 |
| HIST2H3DP1 | 0.014382374 |
| Vault | 0.014381902 |
| TOMM22P2 | 0.014378604 |
| VDAC1P12 | 0.014372904 |
| RP3-325F22 | 0.014372724 |
| AC138649 | 0.014361534 |
| FAM83E | 0.014356872 |
| GEMIN2 | 0.014356705 |
| RN7SKP59 | 0.014354006 |
| RNU6-1027P | 0.014351046 |
| RP11-462G2 | 0.014349888 |
| AC243981 | 0.014343114 |
| RPL15P21 | 0.014336613 |
| HOXB-AS3 | 0.014334076 |
| RP11-735G4 | 0.01432603 |
| ATP11C | 0.014324462 |
| ELP3 | 0.01432381 |
| RN7SKP230 | 0.014321125 |
| N6AMT2 | 0.014320536 |
| RP11-521H3 | 0.014319644 |
| CYCSP5 | 0.014319445 |
| BDKRB2 | 0.014314936 |
| KLF2P2 | 0.014312413 |
| RN7SKP241 | 0.014304112 |
| ANP32E | 0.014300609 |
| THAP7 | 0.014289427 |
| FAM104B | 0.014288962 |
| PPARGC1A | 0.014288674 |
| RP11-397G5 | 0.014287739 |
| CCDC144NL | 0.014286371 |
| SLC2A4 | 0.014282153 |
| CTC-297N7 | 0.014281102 |
| HIGD1AP3 | 0.014276245 |
| RP13-297E16 | 0.014275515 |
| CCDC144A | 0.014275343 |
| RNU6-848P | 0.014274773 |
| RP11-697N18 | 0.014271938 |
| RP1-81P11 | 0.014264684 |
| ST13P13 | 0.01426269 |
| CTA-253N17 | 0.014262356 |

| | |
|---|---|
| NTF6G | 0.014258733 |
| FKBP1B | 0.014245928 |
| KATNA1 | 0.014232234 |
| LINC00951 | 0.014228362 |
| CHMP3 | 0.014227843 |
| LINC00202-1 | 0.014224039 |
| RP11-179G5 | 0.014216495 |
| RP5-839B4 | 0.014216137 |
| STK38 | 0.014212419 |
| RNU6-745P | 0.014211668 |
| KDM1A | 0.014209963 |
| AC239800 | 0.014206966 |
| IMPDH1P10 | 0.014202576 |
| SAFB | 0.01419 |
| RP4-575N6 | 0.014189851 |
| IGFN1 | 0.014187352 |
| LIPG | 0.014186074 |
| CDC42BPA | 0.014184176 |
| RP3-509I19 | 0.014177441 |
| RPL26P20 | 0.014172992 |
| RP11-744O11 | 0.014169966 |
| KB-1458E12 | 0.014162968 |
| RNU6-727P | 0.014159608 |
| ATP6V0B | 0.014158222 |
| THOC7 | 0.014148249 |
| ANKRD20A18P | 0.014147845 |
| CCDC169-SOHLH2 | 0.014144801 |
| GBP6 | 0.014144799 |
| ENOX1-AS2 | 0.014138711 |
| CTD-2523D13 | 0.014136565 |
| RP11-103J8 | 0.014135082 |
| OR4A17P | 0.014134743 |
| OR5D3P | 0.014125271 |
| MCF2L2 | 0.014121755 |
| CTC-258N23 | 0.014119925 |
| KRT18P12 | 0.014118964 |
| TMEM257 | 0.014113655 |
| DSC2 | 0.014111075 |
| CTD-3074O7 | 0.014105847 |
| RN7SL617P | 0.014104944 |
| AP000438 | 0.014103565 |
| FOXL2NB | 0.014102439 |
| AL356214 | 0.01410168 |
| AC000095 | 0.014099123 |
| GCNT1 | 0.014095317 |

| | |
|---|---|
| LA16c-83F12 | 0.014093988 |
| RP11-361I14 | 0.014093042 |
| RP11-492L8 | 0.014092202 |
| RNU4-34P | 0.014090294 |
| RP11-728B21 | 0.014088 |
| SH3BGRL | 0.014087688 |
| RP11-147G7 | 0.01408381 |
| MIR106A | 0.014081535 |
| CHCHD3 | 0.01407846 |
| OR13C8 | 0.014078232 |
| RPS3AP35 | 0.014078143 |
| AC092669 | 0.014076277 |
| MIR410 | 0.014074049 |
| CTD-2315M5 | 0.014071992 |
| RNA5SP52 | 0.014068835 |
| PPP1R37 | 0.014068822 |
| RP11-320M16 | 0.014060204 |
| CNKSR3 | 0.014059334 |
| PRR29 | 0.014055482 |
| RAB15 | 0.014052691 |
| OR5M4P | 0.014046395 |
| OR51B4 | 0.014044758 |
| ZFY | 0.014033455 |
| HIST1H3F | 0.014031879 |
| RNU5F-2P | 0.014019236 |
| RNF186 | 0.01401891 |
| LARP7P3 | 0.01401327 |
| TMEM201 | 0.014006606 |
| PNPLA3 | 0.014006048 |
| POTEB3 | 0.014001596 |
| MYH14 | 0.013996447 |
| RP3-492P14 | 0.013991225 |
| NIT2 | 0.013988558 |
| SLC29A2 | 0.01398841 |
| RP11-470P21 | 0.013987996 |
| WFDC21P | 0.013985842 |
| RP5-1097F14 | 0.013985285 |
| MTND5P16 | 0.0139851 |
| ASB9 | 0.013979684 |
| PHKG2 | 0.01397907 |
| KDELC2 | 0.013977172 |
| RP11-428O23 | 0.01397421 |
| RRAGD | 0.013964948 |
| AC016912 | 0.013962855 |
| GS1-115G20 | 0.01396148 |

| | |
|---|---|
| AC005752 | 0.013956664 |
| GRIK4 | 0.013952101 |
| FAM87A | 0.013951742 |
| MGC27382 | 0.013949372 |
| AC006041 | 0.013947664 |
| RP1-84O15 | 0.013946918 |
| H2AFJ | 0.013943515 |
| PSD3 | 0.013943198 |
| TECPR2 | 0.013940017 |
| LINC00086 | 0.013933635 |
| AC016894 | 0.013926384 |
| AC007098 | 0.013924579 |
| GTF2F1 | 0.013922714 |
| RNU6-261P | 0.013918739 |
| TEX36-AS1 | 0.013915649 |
| ATG4A | 0.013915216 |
| C1GALT1C1 | 0.013913839 |
| CTC-264O10 | 0.013910415 |
| RP11-563P16 | 0.013908831 |
| FOXP1-AS1 | 0.013904484 |
| AP000280 | 0.013904394 |
| AC108056 | 0.013903274 |
| CTB-138E5 | 0.013902581 |
| LINC01320 | 0.013897208 |
| CTD-2333M24 | 0.013886572 |
| CU463998 | 0.01388306 |
| RP11-454K7 | 0.013881145 |
| RP11-697G4 | 0.013880944 |
| HMGB3P8 | 0.01387356 |
| PHYHD1 | 0.013873279 |
| EIF3IP1 | 0.013868915 |
| AC114788 | 0.013866461 |
| OR4F21 | 0.013862665 |
| RNU1-49P | 0.013861964 |
| MTND5P26 | 0.013860917 |
| TOMM5 | 0.013853818 |
| PNLIPRP3 | 0.013853691 |
| NHLH1 | 0.013848864 |
| ADRA2C | 0.013848831 |
| AL137795 | 0.013842405 |
| AF241725 | 0.013839476 |
| RP11-786A20 | 0.013837168 |
| CHRNA5 | 0.013833835 |
| TLE2 | 0.013833445 |
| CEACAM1 | 0.013831138 |

| | |
|---|---|
| HIF1A | 0.013822467 |
| RN7SKP149 | 0.013822113 |
| RP11-175K6 | 0.013820272 |
| RNU6ATAC6P | 0.013818663 |
| MIR601 | 0.013816402 |
| PPP2R2A | 0.013811751 |
| AP000357 | 0.013807637 |
| RP11-481N16 | 0.01380684 |
| LRRC24 | 0.013806176 |
| CTD-2223O18 | 0.013796972 |
| PRSS36 | 0.01379294 |
| RNU6-1041P | 0.013789318 |
| CDKN2A-AS1 | 0.013788228 |
| LINC00364 | 0.013785384 |
| AL732374 | 0.013779764 |
| OR55B1P | 0.013778465 |
| RP11-1E6 | 0.0137778 |
| RNA5SP90 | 0.013770856 |
| HIST1H2BA | 0.013768611 |
| MRPL20 | 0.013766223 |
| OVAAL | 0.013760849 |
| RP11-1E22 | 0.013760216 |
| RP1-60O19 | 0.01375856 |
| AC114760 | 0.013755811 |
| KRT8P42 | 0.013753943 |
| MTNR1B | 0.013753144 |
| MIR2116 | 0.013752875 |
| AC108218 | 0.013747213 |
| HNRNPA1P54 | 0.013745296 |
| AC010975 | 0.013741258 |
| MEIG1 | 0.013735135 |
| AC004941 | 0.013732531 |
| RPS26P30 | 0.013730103 |
| MIR877 | 0.0137301 |
| RNU7-47P | 0.013728148 |
| AC005412 | 0.01372714 |
| RP11-702B10 | 0.013726091 |
| SPANXN5 | 0.013725423 |
| RPL34P29 | 0.013725042 |
| SPPL2C | 0.013723772 |
| CTD-2307P3 | 0.013723383 |
| RP11-543A18 | 0.013714907 |
| RNA5SP46 | 0.013713798 |
| RP11-346I3 | 0.013710602 |
| HOXB13 | 0.013709945 |

| | |
|---|---|
| MIR5688 | 0.013708477 |
| RP11-19N8 | 0.013701094 |
| MIR6085 | 0.013696605 |
| DIAPH3-AS2 | 0.013694186 |
| RNU6-1078P | 0.013688938 |
| RP11-319F12 | 0.013680666 |
| MAGEB4 | 0.013679444 |
| RNU6-1146P | 0.013671139 |
| RNU6-1335P | 0.013670553 |
| NPHP3-ACAD11 | 0.013670058 |
| HDLBP | 0.013667186 |
| RNA5SP237 | 0.013663654 |
| RP3-369A17 | 0.013659606 |
| PGRMC2 | 0.013657809 |
| MTND1P10 | 0.013649952 |
| RNU6-798P | 0.013649632 |
| AC128709 | 0.013648295 |
| FAM90A12P | 0.013636917 |
| POLD2 | 0.013634459 |
| AC020703 | 0.013629386 |
| IMPDH1P11 | 0.013628156 |
| OR8S21P | 0.01362706 |
| RP11-32B5 | 0.013624583 |
| LINC01248 | 0.013621973 |
| RPL5P32 | 0.013619526 |
| NCLP2 | 0.01361803 |
| RNFT1P3 | 0.013611139 |
| GAGE10 | 0.013609428 |
| RP11-255G12 | 0.013604693 |
| RNU6-1111P | 0.013604651 |
| MIR1293 | 0.013603735 |
| CCNJP2 | 0.013595197 |
| RP11-219B17 | 0.013593869 |
| RNA5SP340 | 0.013585555 |
| RP11-22P4 | 0.013571378 |
| MIR4734 | 0.013571354 |
| GPRASP2 | 0.013569898 |
| RN7SL488P | 0.013569052 |
| KLHL35 | 0.013567982 |
| TRIM6-TRIM34 | 0.013557163 |
| NDUFA4 | 0.013555553 |
| SPATA31D1 | 0.013554743 |
| MRPL39 | 0.0135528 |
| RP11-627G23 | 0.013550394 |
| MTND4P27 | 0.013548106 |

| | |
|---|---|
| HMGN1P34 | 0.013545272 |
| SEPHS1 | 0.013539832 |
| MORF4L2 | 0.013532464 |
| KRT8P49 | 0.013532138 |
| MAP1LC3P | 0.013531707 |
| RP11-112J1 | 0.013531552 |
| PTH2R | 0.013521358 |
| CDC14B | 0.013521319 |
| OR1D2 | 0.013517954 |
| AC079809 | 0.013516528 |
| RN7SL319P | 0.013513649 |
| RP11-82L20 | 0.013513195 |
| PEX11A | 0.013512296 |
| C4B-AS1 | 0.013507929 |
| MIR1270 | 0.013503944 |
| RNA5SP507 | 0.013491651 |
| RNU6-43P | 0.013484084 |
| RNA5SP96 | 0.013479653 |
| RP11-654C22 | 0.01346827 |
| RP11-209D20 | 0.013465049 |
| VSIG10L | 0.013464517 |
| CTSG | 0.013462104 |
| C12orf80 | 0.01346187 |
| RP5-1068H6 | 0.013459585 |
| RNU1-21P | 0.01345264 |
| CTD-2020P22 | 0.013448269 |
| MIR562 | 0.013443646 |
| RP4-683L5 | 0.01343894 |
| LINC01115 | 0.013435984 |
| CBX2 | 0.013433501 |
| RP11-661D19 | 0.013430057 |
| SPINK6 | 0.013419927 |
| RP11-27G22 | 0.013407359 |
| PTP4A1P2 | 0.013406632 |
| RP11-153F1 | 0.013402677 |
| RP11-722E23 | 0.013400309 |
| RP11-798K3 | 0.013399798 |
| RN7SL284P | 0.013398717 |
| AC078819 | 0.013389383 |
| B4GALT3 | 0.013387751 |
| RN7SL34P | 0.013387659 |
| RP11-692P14 | 0.013387527 |
| RNU2-60P | 0.013387418 |
| ZNF197-AS1 | 0.013387278 |
| TLE6 | 0.013386858 |

| | |
|---|---|
| RP11-114O18 | 0.013386418 |
| KRT18P66 | 0.013386002 |
| TSPY5P | 0.013385175 |
| KCNQ1-AS1 | 0.013383314 |
| AC080037 | 0.013381219 |
| MIR767 | 0.013381112 |
| TFPI2 | 0.013375546 |
| RPL32P31 | 0.013375073 |
| AC004019 | 0.013371865 |
| MTCH1 | 0.013371117 |
| NCRUPAR_1 | 0.013370528 |
| AC000041 | 0.013369493 |
| RP1-37J18 | 0.013364369 |
| RPS29P24 | 0.013362622 |
| PAPPA2 | 0.013362598 |
| RRM1-AS1 | 0.013351536 |
| MTHFD2P1 | 0.013351439 |
| RP1-144F13 | 0.01334955 |
| OR52U1P | 0.013344093 |
| RP11-1157N2__A | 0.013341478 |
| SUB1P1 | 0.01334112 |
| CTC-436P18 | 0.013340661 |
| RP11-387A1 | 0.013335357 |
| AC105009 | 0.013334502 |
| RP11-438O11 | 0.013333972 |
| BYSL | 0.013333534 |
| RN7SL792P | 0.013332812 |
| RP11-459A10 | 0.013328714 |
| MIR4717 | 0.013325361 |
| RP11-280O1 | 0.013324479 |
| RNA5SP219 | 0.013321999 |
| RPL9 | 0.01331811 |
| RP11-64D22 | 0.013311787 |
| MIR325HG | 0.013311291 |
| RP11-366N18 | 0.013310829 |
| SLC5A7 | 0.013306382 |
| LITAF | 0.013302835 |
| RP3-481A17 | 0.013302333 |
| ARHGAP44 | 0.013294333 |
| PELI1 | 0.013278462 |
| AC016292 | 0.013276825 |
| DNMBP | 0.013276527 |
| C16orf89 | 0.013274332 |
| RNU6-618P | 0.013271903 |
| AC010744 | 0.013270254 |

| | |
|---|---|
| FER | 0.01326875 |
| WIBG | 0.013268062 |
| TAAR6 | 0.013266342 |
| MIR4482 | 0.01326547 |
| RNU6-39P | 0.013262971 |
| RPS6P8 | 0.01326133 |
| OR52E7P | 0.013258066 |
| RP11-226M10 | 0.013252488 |
| RP5-1073F15 | 0.013251154 |
| URAD | 0.013241566 |
| RN7SL823P | 0.013240338 |
| PTRH1 | 0.01323763 |
| KRT8P19 | 0.013229908 |
| OR5P1P | 0.013226493 |
| UNC93B7 | 0.013224978 |
| AL009174 | 0.013222421 |
| RP11-762E8 | 0.013212328 |
| RP11-646E18 | 0.013208213 |
| CTD-2582M21 | 0.013207742 |
| DEFA9P | 0.013207637 |
| LUZP1 | 0.013203073 |
| TMEM173 | 0.013199966 |
| EML3 | 0.013194644 |
| FGF17 | 0.013194277 |
| ELAVL2 | 0.013186179 |
| RP11-794M8 | 0.013180417 |
| RP11-543C4 | 0.013180396 |
| ARMCX3-AS1 | 0.013178305 |
| RP11-411G9 | 0.013176459 |
| MIR4666B | 0.013170404 |
| CR383657 | 0.013170404 |
| RPS18 | 0.013162327 |
| RN7SL92P | 0.013151086 |
| FAM21EP | 0.013137042 |
| CD44 | 0.013132093 |
| RP11-325L12 | 0.013125484 |
| EIF3LP2 | 0.013117272 |
| RPL9P28 | 0.013112317 |
| CNN2P4 | 0.013112046 |
| BMS1P14 | 0.013110632 |
| LLNLR-285B5 | 0.013108738 |
| KCNA10 | 0.013108354 |
| FGFR1 | 0.013104482 |
| CYP7A1 | 0.013101409 |
| MIR4654 | 0.013097248 |

| | |
|---|---|
| RP11-15P13 | 0.013096687 |
| RP11-97I14 | 0.013085952 |
| RP1-184J9 | 0.013081415 |
| RP11-437L7 | 0.013072012 |
| ZNF696 | 0.013070899 |
| TXN | 0.013068704 |
| RNA5SP344 | 0.013067031 |
| RP1-66H9 | 0.013065258 |
| PACSIN2 | 0.013063476 |
| GFOD1-AS1 | 0.013063366 |
| RP11-235E17 | 0.013057373 |
| NDUFB8P1 | 0.01305313 |
| DEFB109P3 | 0.013047947 |
| AADACP1 | 0.013047779 |
| ENG | 0.013046657 |
| ZSCAN23 | 0.013043035 |
| RP11-110G21 | 0.013025911 |
| PSKH1 | 0.013025781 |
| FTH1P16 | 0.013024665 |
| RPL23AP40 | 0.013024657 |
| AC006461 | 0.013020678 |
| RNU6-120P | 0.013017926 |
| LINC01485 | 0.013017719 |
| RENBP | 0.01301406 |
| COX5A | 0.013012943 |
| INAFM2 | 0.013009204 |
| CTD-2351A16 | 0.013004239 |
| RP11-785D18 | 0.013000187 |
| LINC01209 | 0.012992457 |
| EHD4 | 0.012989128 |
| RP11-327L3 | 0.012985553 |
| OR8B6P | 0.012983968 |
| RP11-93N20 | 0.012979443 |
| RPL9P2 | 0.012979433 |
| TLR12P | 0.012974465 |
| RNA5SP366 | 0.012971908 |
| RP1-257C22 | 0.01296896 |
| SNORD113-1 | 0.012965455 |
| KRT18P50 | 0.012962541 |
| NECAB1 | 0.012962262 |
| TPT1P14 | 0.012959107 |
| PIM1 | 0.012952252 |
| AIM1 | 0.012949424 |
| CYCSP20 | 0.012949013 |
| C2orf27AP3 | 0.012946671 |

| | |
|---|---|
| CRIP1P1 | 0.012945973 |
| PKD1L2 | 0.012939433 |
| SNRPCP9 | 0.012937899 |
| LDHD | 0.012937219 |
| TTC1 | 0.012935489 |
| CRELD2 | 0.012928084 |
| C4orf27 | 0.012927321 |
| RP11-252I13 | 0.012925975 |
| PFKFB3 | 0.012925628 |
| ZNF385D-AS1 | 0.01292374 |
| TMBIM1 | 0.012917349 |
| SLC16A11 | 0.012917303 |
| RP11-883G14 | 0.012916687 |
| MED28P4 | 0.012915147 |
| RP11-217E13 | 0.012914944 |
| AL080272 | 0.01291449 |
| CTD-2128A3 | 0.012912551 |
| SERGEF | 0.012911227 |
| AC004221 | 0.012910395 |
| MIR4663 | 0.012909408 |
| RP13-444K19 | 0.01290904 |
| AL445384 | 0.012907917 |
| SIPA1L1 | 0.012904957 |
| FLJ31662 | 0.01290471 |
| MTND2P26 | 0.012904179 |
| AC010468 | 0.012903841 |
| EIF4A1P12 | 0.012903048 |
| RNA5SP483 | 0.012899023 |
| RPL31P12 | 0.012895848 |
| RP1-46F2 | 0.012890606 |
| MR1 | 0.012889943 |
| RNA5SP55 | 0.01288757 |
| RP11-176D17 | 0.012883378 |
| POLR2E | 0.012878835 |
| ITGA3 | 0.012872788 |
| TIMM17B | 0.012871821 |
| CPB2-AS1 | 0.012865837 |
| CTA-85E5 | 0.012865742 |
| RP13-52K8 | 0.012861712 |
| APBA1 | 0.012855746 |
| TSEN15 | 0.012854574 |
| CCDC130 | 0.01285409 |
| AC079298 | 0.012853152 |
| GPR37 | 0.012847913 |
| RN7SKP281 | 0.012838518 |

| | |
|---|---|
| VN2R11P | 0.012837165 |
| RNA5SP334 | 0.012835359 |
| ZNF135 | 0.012832117 |
| OTX1 | 0.01283063 |
| RNA5SP365 | 0.012829995 |
| OR2A1 | 0.012826468 |
| MIR4265 | 0.012823741 |
| UCP1 | 0.012815955 |
| PPIAP33 | 0.01280107 |
| RP11-758I14 | 0.01279913 |
| PTPRD-AS2 | 0.012799107 |
| FAM213A | 0.01279779 |
| ERAS | 0.01279775 |
| ZCCHC17 | 0.012795818 |
| SLITRK1 | 0.012790032 |
| LINC01553 | 0.012789654 |
| SPTAN1 | 0.012787054 |
| AL590683 | 0.012785377 |
| DBNDD1 | 0.012783129 |
| RNASEK-C17orf49 | 0.012783073 |
| AC090696 | 0.012781019 |
| SNORD66 | 0.012778783 |
| RNU6-295P | 0.01277703 |
| RP11-416K24 | 0.012775692 |
| KCNIP2 | 0.012773565 |
| MOXD1 | 0.012768635 |
| PRR9 | 0.012767367 |
| GPRIN1 | 0.01276376 |
| RNU7-43P | 0.012761785 |
| FAM224B | 0.012761628 |
| RASA2-IT1 | 0.012756473 |
| RNU6-178P | 0.012750263 |
| PANK2 | 0.012749695 |
| RP11-264E20 | 0.012744963 |
| RP11-420M1 | 0.012742133 |
| AC117507 | 0.012738033 |
| TRABD | 0.012736857 |
| CDIP1 | 0.012725675 |
| SALL4P5 | 0.012722554 |
| CTC-518B2 | 0.012721908 |
| RARA-AS1 | 0.012721842 |
| XX-FW80269A6 | 0.012719801 |
| RP11-123O22 | 0.012718477 |
| TFAP2E | 0.012717768 |
| ATF7 | 0.012714661 |

| | |
|---|---|
| RP11-108M12 | 0.012713384 |
| B3GNTL1P2 | 0.012711762 |
| DYNLL1P2 | 0.012706265 |
| RNA5SP76 | 0.012706176 |
| RP11-460I19 | 0.012701484 |
| PGAP1 | 0.012698511 |
| STX5 | 0.012697878 |
| RP4-781L3 | 0.012696857 |
| RP11-545A16 | 0.012694611 |
| DCAF15 | 0.012688018 |
| KLC1 | 0.01268638 |
| AC016738 | 0.012685772 |
| RPL32P9 | 0.012684008 |
| KLF2P1 | 0.012683583 |
| MTND6P4 | 0.012682276 |
| MEMO1 | 0.012680054 |
| HAUS4P1 | 0.012678111 |
| RN7SL274P | 0.012677265 |
| PTH | 0.012677083 |
| RP11-390F4 | 0.01267291 |
| RP11-413N13 | 0.012669393 |
| PAQR6 | 0.012667179 |
| MIR4471 | 0.012665107 |
| RP11-435I10 | 0.012663188 |
| AC017048 | 0.012655323 |
| SNAI3-AS1 | 0.012652332 |
| SLC18A3 | 0.012650246 |
| OR6N1 | 0.012647685 |
| RN7SL492P | 0.012645547 |
| RNU6-945P | 0.012643724 |
| OR7E33P | 0.012635569 |
| RP11-554K11 | 0.012632747 |
| FGD5P1 | 0.012632005 |
| RN7SL807P | 0.012629388 |
| UPK3BL | 0.012627525 |
| ALDH2 | 0.012626526 |
| C6orf52 | 0.01262502 |
| TBC1D3C | 0.012623723 |
| CNN2P6 | 0.012620565 |
| ASNSP4 | 0.012620549 |
| ZNF736P7Y | 0.012620166 |
| RP11-346K17 | 0.012620031 |
| KCTD17 | 0.012611865 |
| LINC00307 | 0.012611535 |
| RP11-12A2 | 0.012611256 |

| | |
|---|---|
| SERPINA2 | 0.01260969 |
| CYP51A1 | 0.012606209 |
| TLE4 | 0.01260446 |
| TM4SF1-AS1 | 0.012603347 |
| C2orf73 | 0.012590651 |
| YWHAEP2 | 0.012590223 |
| RN7SL419P | 0.01258728 |
| RNU2-42P | 0.012586293 |
| MIR651 | 0.012585577 |
| PCDHGB9P | 0.012583632 |
| MIR580 | 0.012580283 |
| RP3-468O1 | 0.012578495 |
| RP11-257I8 | 0.012570563 |
| DRAXINP1 | 0.012569985 |
| ENSA | 0.012569612 |
| RPS27P12 | 0.012567784 |
| COL18A1-AS2 | 0.012562752 |
| SNUPN | 0.012561389 |
| RND1 | 0.012558896 |
| RP11-424G14 | 0.012555384 |
| LSM3P2 | 0.012548779 |
| MPC1L | 0.012545415 |
| AC017002 | 0.012545064 |
| RN7SL150P | 0.012542529 |
| MRPL15 | 0.012539081 |
| RP11-20B7 | 0.012537697 |
| MYCBP2-AS1 | 0.012532553 |
| RNU6-1309P | 0.012529725 |
| AC012168 | 0.012526029 |
| CDY4P | 0.01251913 |
| OR2W4P | 0.012515777 |
| TMEM258 | 0.012512717 |
| EGFLAM-AS4 | 0.012512226 |
| AC013403 | 0.012512112 |
| DNER | 0.012511819 |
| RP11-122C5 | 0.012508463 |
| RPL36AP51 | 0.012507895 |
| ADSS | 0.012506633 |
| AL031721 | 0.012500109 |
| CDCA7 | 0.012493479 |
| GS1-164F24 | 0.012492891 |
| RP11-813N20 | 0.012491814 |
| SNORD109B | 0.012489559 |
| ADRB3 | 0.012487177 |
| TMEM86B | 0.012485702 |

| | |
|---|---|
| TECRL | 0.012484364 |
| BLOC1S4 | 0.012482892 |
| TIAM2 | 0.01247958 |
| RP11-764I5 | 0.012476965 |
| ARHGAP31-AS1 | 0.012474168 |
| RP11-5A19 | 0.012473608 |
| LRRC34 | 0.012471636 |
| CCDC122 | 0.01247065 |
| RP11-90P16 | 0.012468621 |
| RP11-181E10 | 0.012460962 |
| SNORD17 | 0.01245575 |
| POLR3KP1 | 0.012455179 |
| CTD-2170G1 | 0.012451923 |
| TMEM86A | 0.012449793 |
| TBX4 | 0.012449097 |
| RNA5SP229 | 0.012447436 |
| LINC00659 | 0.012443969 |
| NUDT4P2 | 0.012443591 |
| AL031681 | 0.012439199 |
| AQP1 | 0.012437904 |
| RP11-164A6 | 0.012435178 |
| CCDC178 | 0.012433739 |
| DDC | 0.012433431 |
| AC090051 | 0.012430193 |
| CCDC34P1 | 0.012429561 |
| PCDH10 | 0.01242559 |
| SNORA74B | 0.012418207 |
| PNPT1P2 | 0.012416342 |
| DEFB132 | 0.012415556 |
| AC006326 | 0.012414433 |
| RPL34 | 0.012412173 |
| RPL37P2 | 0.012411047 |
| PPP3CA | 0.012402889 |
| RP1-47A17 | 0.01240286 |
| PCDHGA10 | 0.012401642 |
| C9orf85 | 0.012399875 |
| GIF | 0.012398635 |
| RP11-12K22 | 0.012396744 |
| Z95114 | 0.012396526 |
| RP11-420N3 | 0.012396104 |
| SMPD2 | 0.012392767 |
| RP1-232N11 | 0.012389787 |
| TCEB3CL2 | 0.012382522 |
| ACTBP12 | 0.012378807 |
| IDH1-AS1 | 0.012368683 |

| | |
|---|---|
| FDPSP4 | 0.01236831 |
| MTND6P12 | 0.01236747 |
| AC133555 | 0.01236664 |
| CTC-458I2 | 0.012364312 |
| ARSD | 0.012360357 |
| PPP2R3B | 0.012351631 |
| VTI1BP2 | 0.012351465 |
| AC073578 | 0.012349716 |
| RP11-386I14 | 0.012348291 |
| RP11-350D17 | 0.012346831 |
| HIST1H2BE | 0.012343251 |
| GGT7 | 0.012341974 |
| RP11-43P5 | 0.012339976 |
| AC098973 | 0.012339907 |
| SDCBP | 0.012339835 |
| RP11-208G20 | 0.012338214 |
| RP5-1139I1 | 0.012336336 |
| MYL12A | 0.012326383 |
| SLC2A3 | 0.012317564 |
| AC003958 | 0.012316569 |
| ETV5-AS1 | 0.01230915 |
| ECE1 | 0.012308671 |
| YOD1 | 0.012302345 |
| CTD-2061E9 | 0.012299237 |
| LHX2 | 0.012296675 |
| ZNF576 | 0.012292316 |
| RP1-199J3 | 0.01228867 |
| BHMG1 | 0.012288087 |
| BMS1P17 | 0.012287221 |
| ABT1P1 | 0.012286726 |
| AC006499 | 0.012280092 |
| RP11-412H8 | 0.012275889 |
| PLA2G1B | 0.012273422 |
| EZR | 0.012271702 |
| RP11-11N6 | 0.012269834 |
| CBX1P1 | 0.012266943 |
| LINC00971 | 0.012266518 |
| SNORD99 | 0.012263273 |
| PUS3 | 0.012262622 |
| RPL23AP46 | 0.012256338 |
| OR52M1 | 0.012250196 |
| NPIPP1 | 0.012250082 |
| PTGDR2 | 0.012247696 |
| RP11-568A19 | 0.012246009 |
| LINC01415 | 0.012245369 |

| | |
|---|---|
| ALG1L10P | 0.012244318 |
| GEMIN8P3 | 0.012244298 |
| JAZF1-AS1 | 0.012243525 |
| RNU6-799P | 0.012243334 |
| RN7SL727P | 0.012241663 |
| RP11-663P9 | 0.012237024 |
| DPP7 | 0.012236965 |
| KALP | 0.012229463 |
| RBBP4P4 | 0.012229155 |
| LL0XNC01-240C2 | 0.0122269 |
| RN7SL438P | 0.012226787 |
| RP11-506B6 | 0.012225015 |
| RP4-683M8 | 0.012222294 |
| RP11-643M14 | 0.012222274 |
| RP11-73E6 | 0.012220688 |
| RN7SKP217 | 0.01221872 |
| USMG5P1 | 0.012213106 |
| RN7SL451P | 0.012204696 |
| RPS25 | 0.012204592 |
| MIR4431 | 0.012203607 |
| FGF7P2 | 0.012201054 |
| SNRPEP2 | 0.012201001 |
| RP11-122D10 | 0.012198721 |
| CTD-2373N4 | 0.012198191 |
| RP13-122B23 | 0.012191438 |
| PRMT1 | 0.012191363 |
| RP11-284F21 | 0.012188963 |
| AC104667 | 0.01218047 |
| TNNI3 | 0.012177914 |
| AL590007 | 0.012177771 |
| MIR4529 | 0.01217166 |
| RP11-244J10 | 0.012169608 |
| AC130709 | 0.012157973 |
| MIR4705 | 0.012157792 |
| MIR4293 | 0.012152312 |
| MIR7702 | 0.012149966 |
| KIAA1919 | 0.012141761 |
| RNPEPL1 | 0.012141128 |
| RNU6-823P | 0.012141021 |
| MIR6504 | 0.01214025 |
| AL109615 | 0.012137264 |
| RP11-361H10 | 0.012135299 |
| MIR4301 | 0.012131814 |
| MIR125B1 | 0.012131021 |
| ARHGAP24 | 0.012130709 |

| | |
|---|---|
| OR10AE3P | 0.012122911 |
| RNU6-344P | 0.01211598 |
| KLHL5 | 0.012114616 |
| AC010894 | 0.012113964 |
| BCAR3 | 0.012112364 |
| AL513123 | 0.012112224 |
| RP11-378J18 | 0.012109667 |
| PRELID1 | 0.012102838 |
| CTD-2060F24 | 0.012100647 |
| RP11-823E8 | 0.012093441 |
| CAST | 0.012092047 |
| RN7SKP60 | 0.012089547 |
| RP11-420O16 | 0.012086854 |
| TMEM196 | 0.012082227 |
| RNU6-119P | 0.012081929 |
| ZNF849P | 0.012078755 |
| AL589663 | 0.01207413 |
| HMGB3P11 | 0.012073742 |
| LINC00588 | 0.012072888 |
| SNORD115-42 | 0.012072664 |
| DEFB117 | 0.012071394 |
| APOE | 0.012058799 |
| AC012494 | 0.012055018 |
| TRIM43 | 0.012051274 |
| RPL24P8 | 0.01204572 |
| RP11-394O4 | 0.012044529 |
| AC007682 | 0.012043999 |
| RP11-200K12 | 0.012039094 |
| RASSF9 | 0.01203491 |
| GJB7 | 0.012034537 |
| LINC01108 | 0.012034323 |
| PLEKHO1 | 0.012028044 |
| CTD-2021H9 | 0.012026603 |
| FAM212B-AS1 | 0.012025789 |
| RNU6-127P | 0.0120257 |
| AD000091 | 0.012018188 |
| RASSF1-AS1 | 0.012017588 |
| RP11-501C14 | 0.012016339 |
| XXYLT1-AS1 | 0.012015346 |
| RNU6-243P | 0.01200835 |
| RP11-119B16 | 0.012006946 |
| MIR4259 | 0.012004876 |
| SNRPCP16 | 0.01199872 |
| BAIAP3 | 0.011996731 |
| HIST1H3B | 0.011986329 |

| | | |
|---|---|---|
| AKR1B10P1 | | 0.011985067 |
| ANKRD34C-AS1 | | 0.011984145 |
| RP11-15I20 | | 0.011983615 |
| AL008708 | | 0.011977917 |
| AL589947 | | 0.011975427 |
| MIR6765 | | 0.011973001 |
| CFC1 | | 0.011971851 |
| AC104304 | | 0.01197013 |
| SPIN1 | | 0.011968581 |
| GRIK1-AS1 | | 0.011968019 |
| RP11-364B6 | | 0.011966023 |
| CTD-2266L18 | | 0.011964593 |
| CAPZA3 | | 0.011963822 |
| COX6B1P7 | | 0.011957977 |
| RP11-532L16 | | 0.011956279 |
| MUC13 | | 0.011955131 |
| RNU6-665P | | 0.011947873 |
| CKLF-CMTM1 | | 0.011944376 |
| RAI2 | | 0.011943799 |
| AL355361 | | 0.011942076 |
| SMCO4P1 | | 0.011934049 |
| OR4A6P | | 0.01193303 |
| RP13-650J16 | | 0.011931474 |
| ZCCHC12 | | 0.011931326 |
| RNU6-781P | | 0.011929961 |
| AC055873 | | 0.011929343 |
| RNU6-240P | | 0.011925713 |
| FBXO39 | | 0.011918784 |
| RP11-102M11 | | 0.011918063 |
| | 8-Sep | 0.011917026 |
| WDR74 | | 0.011915128 |
| OR1A2 | | 0.011914677 |
| HIST1H2BPS2 | | 0.011914219 |
| LINC01448 | | 0.011913212 |
| CNTROB | | 0.011913118 |
| AC128677 | | 0.011912684 |
| RNU6-337P | | 0.011909145 |
| COX6CP5 | | 0.011906201 |
| BEST4 | | 0.01190617 |
| IFNA4 | | 0.011906105 |
| WFDC10B | | 0.011898531 |
| LRRC26 | | 0.011897707 |
| RP11-40F6 | | 0.011888974 |
| RNU6-608P | | 0.011884712 |
| POP5 | | 0.011880391 |

| | |
|---|---|
| MIR6804 | 0.011877168 |
| AL050335 | 0.011875634 |
| AC093106 | 0.011866215 |
| SLC27A6 | 0.01186303 |
| KRT126P | 0.011862979 |
| SPRED1 | 0.011860335 |
| AP000654 | 0.011859414 |
| RP11-305A4 | 0.011853702 |
| MRPL17 | 0.011851944 |
| CTNNBL1 | 0.011851683 |
| YEATS2-AS1 | 0.01184452 |
| RP3-354J5 | 0.011843223 |
| MLIP-IT1 | 0.011841047 |
| RP11-94H6 | 0.01184063 |
| SPAG17 | 0.011832567 |
| HOXD3 | 0.011830706 |
| POU6F2 | 0.011828722 |
| OR8A2P | 0.011827352 |
| LINC01144 | 0.01182584 |
| MIR6850 | 0.01182231 |
| WEE2 | 0.011821836 |
| RPSA | 0.01181944 |
| RPS12P24 | 0.011819403 |
| SP5 | 0.011816016 |
| NDUFAF4P1 | 0.011813361 |
| RP11-65D17 | 0.01180705 |
| FOXB2 | 0.011805654 |
| RP3-426I6 | 0.011804221 |
| D4S234E | 0.011803235 |
| RP11-739B23 | 0.011801099 |
| RNU6-750P | 0.011787502 |
| RP11-73M11 | 0.011787391 |
| POLR3KP2 | 0.011785486 |
| MBNL3 | 0.011784741 |
| RNU1-83P | 0.011782766 |
| NPM1P10 | 0.011780452 |
| CTB-32H22 | 0.011774941 |
| AC004052 | 0.011771299 |
| RP11-1081K18 | 0.011756458 |
| WWC2 | 0.01175447 |
| PSMD13 | 0.011746097 |
| IDUA | 0.011743567 |
| ATG5 | 0.01174312 |
| RP11-321C24 | 0.011735206 |
| AL583860 | 0.011734367 |

| | |
|---|---|
| AC108066 | 0.011727747 |
| TNXA | 0.011727018 |
| SULT1B1 | 0.011726462 |
| RP3-527G5 | 0.011718301 |
| HMGA1 | 0.011715621 |
| CHP2 | 0.01171469 |
| CRB3P1 | 0.011714068 |
| ROBO2P1 | 0.01170916 |
| TUBB8P3 | 0.011708625 |
| RP11-92J19 | 0.011701939 |
| RP11-815J21 | 0.011698235 |
| AC000367 | 0.01169253 |
| SLC25A13 | 0.01168522 |
| PAICSP2 | 0.011680065 |
| RP11-394D2 | 0.011676044 |
| ARHGEF7-IT1 | 0.011670076 |
| DNAJB3 | 0.011669212 |
| OR56A5 | 0.011669166 |
| OR51K1P | 0.011666208 |
| GABRA1 | 0.011658957 |
| RNU6-1045P | 0.011644339 |
| RNU6-1135P | 0.011641745 |
| RNA5SP41 | 0.011641431 |
| SRGAP2C | 0.011635618 |
| RP11-471J7 | 0.011627538 |
| CD300LB | 0.011627022 |
| HMGB3P23 | 0.011626777 |
| RP11-930O11 | 0.011622143 |
| GDF3 | 0.011619035 |
| A2ML1 | 0.01161504 |
| RP11-314N14 | 0.011614652 |
| CTC-278L1 | 0.011609769 |
| CSTF2 | 0.011609161 |
| TAS2R7 | 0.011607678 |
| BMP4 | 0.011605291 |
| MIR518A1 | 0.011604911 |
| DPY30 | 0.011600751 |
| SNORD115-36 | 0.011582927 |
| RP11-359E10 | 0.011582819 |
| RP11-692C23 | 0.011578599 |
| CBS | 0.011576781 |
| RP11-146E13 | 0.011576353 |
| AC004692 | 0.011571833 |
| D2HGDH | 0.011571355 |
| BNIP3 | 0.011570034 |

| | |
|---|---|
| OR52K1 | 0.011568214 |
| TMC2 | 0.01156569 |
| CATSPERG | 0.01156555 |
| SLC28A1 | 0.011547692 |
| LINC01280 | 0.011542662 |
| RP11-366I13 | 0.011541138 |
| AC097523 | 0.011536284 |
| AKIRIN1P1 | 0.011530104 |
| RP11-104D21 | 0.011526256 |
| RP11-559M23 | 0.011524803 |
| MIR6784 | 0.011519707 |
| IGSF23 | 0.011517867 |
| RP11-449M6 | 0.01151538 |
| C9orf135-AS1 | 0.011510448 |
| ISCUP1 | 0.011507985 |
| MIR103A1 | 0.011503219 |
| SNX18 | 0.011501249 |
| GYG1P2 | 0.011498262 |
| AC104389 | 0.011497392 |
| KRT8P48 | 0.01149692 |
| HDAC6 | 0.011489602 |
| GLRA4 | 0.011489252 |
| PPP1R2P6 | 0.01148858 |
| AF096876 | 0.011482671 |
| HIST1H2AD | 0.01147924 |
| B4GALT5 | 0.011467443 |
| RN7SL538P | 0.011465593 |
| AC002407 | 0.011464433 |
| OR2AT1P | 0.011464087 |
| HNRNPA1P64 | 0.011459412 |
| KLHL29 | 0.011459108 |
| RP11-497K21 | 0.01145754 |
| RP11-465B22 | 0.011452168 |
| RP11-326K13 | 0.011448112 |
| CYP51A1P3 | 0.011445354 |
| ALDH1L1-AS2 | 0.011442303 |
| TSPEAR-AS2 | 0.011440938 |
| RP4-798A17 | 0.011440691 |
| UBE2S | 0.01143693 |
| HMGN2P38 | 0.011434664 |
| ISYNA1 | 0.011433767 |
| RN7SL174P | 0.011432528 |
| AC005371 | 0.011432002 |
| BMS1P13 | 0.011429577 |
| RP11-447H19 | 0.011427792 |

| | |
|---|---|
| RP11-430K21 | 0.01142753 |
| CTD-2151A2 | 0.0114266 |
| RP11-118N24 | 0.011425635 |
| RP11-93L14 | 0.011424698 |
| RPL9P7 | 0.011423553 |
| RP11-340E6 | 0.011420187 |
| NR1I3 | 0.011416501 |
| RP11-736E3 | 0.011413032 |
| FLJ26850 | 0.011410575 |
| C3orf36 | 0.011409564 |
| NEK4P1 | 0.011409356 |
| RAB12 | 0.011406882 |
| RDH5 | 0.011402026 |
| AL117380 | 0.011394208 |
| CAPN1 | 0.01138623 |
| SNORA53 | 0.01138508 |
| RP4-614C15 | 0.011380564 |
| SOHLH1 | 0.011367449 |
| AC006390 | 0.011364713 |
| HIST1H2AL | 0.01136002 |
| RN7SL771P | 0.011358667 |
| RP1-202I21 | 0.011357625 |
| SOCS1 | 0.011353429 |
| MIR377 | 0.011351855 |
| AL356806 | 0.011350231 |
| RP11-64C12 | 0.01134967 |
| GOLGA7B | 0.011339028 |
| RNU6-695P | 0.01133844 |
| SLC35F3 | 0.011338345 |
| AC092287 | 0.01133544 |
| SND1-IT1 | 0.011335053 |
| RBM7 | 0.011327642 |
| TTC39DP | 0.011321092 |
| MIR124-3 | 0.011320803 |
| RP3-450M14 | 0.011320337 |
| OR10AA1P | 0.011319945 |
| MTND4P29 | 0.011317057 |
| RP3-522P13 | 0.01131624 |
| RP1-23E21 | 0.011313798 |
| RNU6-936P | 0.01131288 |
| TMSB15B | 0.011307699 |
| MIR5697 | 0.011307593 |
| AC011376 | 0.011302298 |
| TRIM60P13 | 0.011301156 |
| ATN1 | 0.011300626 |

| | |
|---|---|
| MIR7850 | 0.011300545 |
| AL161422 | 0.01129818 |
| AGBL4-IT1 | 0.011298029 |
| EMP2 | 0.011293073 |
| KNOP1P4 | 0.011289593 |
| AC104581 | 0.011288364 |
| RNF17 | 0.011285391 |
| PFN3 | 0.011272473 |
| RBM10 | 0.011270337 |
| TMEM164 | 0.011267285 |
| RNU7-182P | 0.011259672 |
| RNA5SP409 | 0.011259672 |
| BEX2 | 0.01125709 |
| MBL2 | 0.011256041 |
| F2RL2 | 0.011253796 |
| ALOX12P2 | 0.011252949 |
| RP11-595O22 | 0.011246208 |
| PIGU | 0.011236158 |
| BIRC7 | 0.011235966 |
| RPSAP15 | 0.01123389 |
| TMEM126A | 0.01123134 |
| AQP4-AS1 | 0.01123041 |
| KCNMB1 | 0.011226967 |
| AC021205 | 0.011225206 |
| ROPN1L | 0.011222465 |
| RPL21P111 | 0.011217325 |
| GPR75-ASB3 | 0.011216939 |
| RP11-95M5 | 0.011209994 |
| RRM2P3 | 0.011209373 |
| IGHVII-67-1 | 0.011208409 |
| LINC01541 | 0.011208156 |
| RNASEH1 | 0.011206985 |
| TAS2R13 | 0.011206938 |
| RP11-212E8 | 0.011203391 |
| AC008271 | 0.011203044 |
| RP1-40E16 | 0.011198669 |
| GCNT1P3 | 0.011193296 |
| RP11-572C15 | 0.01118696 |
| PGD | 0.01118624 |
| KRTAP19-9P | 0.011185379 |
| FECHP1 | 0.011181456 |
| RP3-419C19 | 0.011180131 |
| MRPS21P7 | 0.011177609 |
| AC107072 | 0.011173242 |
| CTC-305H11 | 0.011172231 |

| | | |
|---|---|---|
| SNRPCP5 | | 0.011170577 |
| RP11-626E13 | | 0.011167944 |
| ERC2 | | 0.011167882 |
| RPS10P14 | | 0.011165235 |
| AC123023 | | 0.01116398 |
| KLF4 | | 0.011161104 |
| NAV2-AS3 | | 0.011158647 |
| OPCML-IT2 | | 0.011150558 |
| RP1-19N1 | | 0.011143288 |
| KCTD21-AS1 | | 0.011143057 |
| RNA5SP315 | | 0.011138827 |
| RNU7-80P | | 0.011136574 |
| DPF3 | | 0.011119335 |
| FTH1P21 | | 0.011111138 |
| TMPRSS3 | | 0.01110966 |
| CD24P4 | | 0.01110749 |
| RPL15 | | 0.011107241 |
| | 11-Mar | 0.011105709 |
| DEFB108B | | 0.011104872 |
| MTND4P19 | | 0.011103377 |
| RP3-446N13 | | 0.011094927 |
| RP4-620E11 | | 0.011094441 |
| PPP2R2B-IT1 | | 0.011091614 |
| RNU6-956P | | 0.011089609 |
| LDHC | | 0.011088988 |
| SDHCP3 | | 0.011088035 |
| BCL2L1 | | 0.011087594 |
| RNU6-172P | | 0.011082222 |
| RP11-317J19 | | 0.011078618 |
| NDUFS3 | | 0.011074719 |
| MIR6738 | | 0.011070803 |
| LINC01327 | | 0.011068719 |
| HS2ST1 | | 0.011067798 |
| RP5-891H21 | | 0.011060369 |
| IPMKP1 | | 0.011058436 |
| MYL12B | | 0.011057974 |
| RPL35P2 | | 0.011049955 |
| ETV4 | | 0.011048708 |
| AP006748 | | 0.011048557 |
| TACR1 | | 0.011047379 |
| AC017081 | | 0.011041647 |
| GLA | | 0.011041434 |
| SUMO2P13 | | 0.01103587 |
| AC004158 | | 0.011023713 |
| TIMM9P3 | | 0.01102312 |

| | |
|---|---|
| CDIPT | 0.011020893 |
| RP11-353P15 | 0.011019709 |
| ACTL6B | 0.011017199 |
| NABP1 | 0.011000669 |
| AC140542 | 0.010995196 |
| RNA5SP157 | 0.010990349 |
| RP11-482L11 | 0.010984245 |
| MAP3K10 | 0.010982026 |
| RP1-149M18 | 0.010974023 |
| RNA5SP191 | 0.010972059 |
| CTC-575N7 | 0.010967211 |
| TIMM10 | 0.010966565 |
| VWDE | 0.010964965 |
| MIR1229 | 0.010964965 |
| RPL21P5 | 0.010960166 |
| ICMT | 0.010958118 |
| KB-1907C4 | 0.010952064 |
| GAS6-AS2 | 0.01095103 |
| UCA1 | 0.010950534 |
| PAPSS1 | 0.010950006 |
| RP11-686D16 | 0.010947916 |
| RPL7AP31 | 0.010946207 |
| PLCD1 | 0.01094342 |
| FAM230B | 0.010941447 |
| CCDC13 | 0.01094048 |
| TCF7L1-IT1 | 0.010939174 |
| CTD-2616J11 | 0.010938317 |
| RNA5SP204 | 0.010933 |
| SLC16A12-AS1 | 0.010930774 |
| RP11-395N21 | 0.010930549 |
| APBB1 | 0.010929857 |
| SNORA5 | 0.010929178 |
| DIO2-AS1 | 0.010926587 |
| AC073089 | 0.010918225 |
| TIGD7 | 0.010913409 |
| AP000962 | 0.01091166 |
| RP11-363G10 | 0.010910686 |
| LINC01410 | 0.010909707 |
| NT5M | 0.010907846 |
| EIF4A1P6 | 0.010907739 |
| RN7SL336P | 0.010906052 |
| RP11-574O7 | 0.01090031 |
| IFNWP5 | 0.010900154 |
| KLHL25 | 0.010898786 |
| ZNF408 | 0.010895064 |

| | |
|---|---|
| ITGB3BP | 0.010894134 |
| RP11-60A14 | 0.010889486 |
| AC011379 | 0.010881108 |
| RPL5P14 | 0.010875584 |
| LINC00189 | 0.010875539 |
| RNU7-97P | 0.01087506 |
| OR1E3 | 0.010872196 |
| NAIF1 | 0.010871975 |
| RP11-255M2 | 0.010868008 |
| RN7SL621P | 0.010867907 |
| DCLRE1B | 0.010866152 |
| SLC50A1 | 0.010864905 |
| RP11-316E14 | 0.010864774 |
| RN7SL518P | 0.0108605 |
| LGALS13 | 0.010857028 |
| GPAM | 0.010855453 |
| OR10A3 | 0.010854157 |
| CLUHP1 | 0.010851964 |
| KCNJ14 | 0.01084736 |
| AC008756 | 0.010841013 |
| AC015909 | 0.01083558 |
| Z99916 | 0.010831415 |
| RNU6-99P | 0.010830198 |
| PCDHGB5 | 0.010829128 |
| R3HDM2P1 | 0.010827814 |
| RP11-718M20 | 0.010822823 |
| RPS16P8 | 0.010821947 |
| RAB7A | 0.010821926 |
| MIR4658 | 0.010810737 |
| LINC01291 | 0.010804882 |
| RP11-379P1 | 0.010801296 |
| ACTC1 | 0.010795452 |
| RNU6-28P | 0.010795128 |
| DUOX1 | 0.01079493 |
| CTD-2308N23 | 0.010793943 |
| SYT14P1 | 0.010782102 |
| RN7SL435P | 0.010778836 |
| NPVF | 0.010777582 |
| C1QBPP | 0.010775091 |
| PFN1P12 | 0.010771128 |
| SCN8A | 0.010770631 |
| AL358815 | 0.010767961 |
| RP11-146F11 | 0.01076518 |
| MAN2A1 | 0.010764045 |
| PEX10 | 0.010762873 |

| | |
|---|---|
| RNU6-723P | 0.010760553 |
| BRIX1 | 0.010759952 |
| KB-1440D3 | 0.010757418 |
| MRPS9 | 0.010752974 |
| RP11-148E17 | 0.010750033 |
| RP11-556N21 | 0.010746375 |
| CDRT8 | 0.010744906 |
| ARL14 | 0.010738716 |
| TMEM202 | 0.010730586 |
| AC099665 | 0.010729598 |
| G6PC | 0.010727573 |
| KHDC1 | 0.010723615 |
| RNY4P18 | 0.010714046 |
| MIR8060 | 0.010713748 |
| AL356476 | 0.010713748 |
| RN7SL123P | 0.010713458 |
| MIR6500 | 0.01070422 |
| C19orf35 | 0.01070353 |
| MIR6515 | 0.010702561 |
| PROL1 | 0.010701072 |
| RP11-344H11 | 0.010694902 |
| GNAS-AS1_2 | 0.01068796 |
| PF4 | 0.010683978 |
| RP11-266J6 | 0.010667999 |
| RP11-384F7 | 0.010663631 |
| PABPC1P4 | 0.010662105 |
| CD300A | 0.010659352 |
| COL15A1 | 0.010658756 |
| H2AFY2 | 0.010656412 |
| AC234917 | 0.010655468 |
| SNORA54 | 0.010651184 |
| SMURF2P1 | 0.010649038 |
| RP11-621K7 | 0.010647418 |
| SAE1 | 0.010633825 |
| AC004485 | 0.010631489 |
| RNU7-127P | 0.010630373 |
| AC010900 | 0.010629429 |
| RP11-793A3 | 0.010625956 |
| TMC4 | 0.01062562 |
| THAP8 | 0.010622922 |
| RNU6-343P | 0.010617959 |
| RNU7-34P | 0.010613625 |
| LINC00533 | 0.010610465 |
| SNORD50 | 0.01061018 |
| MIR6806 | 0.010608983 |

| | | |
|---|---|---|
| SURF6P1 | | 0.01060668 |
| MIR3144 | | 0.010605602 |
| PSG11 | | 0.010600832 |
| NUDT15 | | 0.010596968 |
| FAM46A | | 0.010596837 |
| MIR6827 | | 0.010595942 |
| SPX | | 0.010584323 |
| VAMP3 | | 0.01057892 |
| OR5A2 | | 0.010574816 |
| ADH7 | | 0.010574597 |
| PIFO | | 0.010572687 |
| MRGPRG | | 0.010571998 |
| KRT8P6 | | 0.01056245 |
| TPBG | | 0.010552891 |
| RNU6-177P | | 0.010547524 |
| AC064853 | | 0.010538687 |
| TMPRSS11BNL | | 0.010537415 |
| HSD17B7P1 | | 0.010537041 |
| RNU6-1106P | | 0.010536632 |
| RP11-322A17 | | 0.010535214 |
| RPL7AP58 | | 0.010530952 |
| ADH5P2 | | 0.010530375 |
| RP11-561E1 | | 0.010529496 |
| CLCN7 | | 0.010528648 |
| RP4-673D20 | | 0.010524751 |
| RP5-968D22 | | 0.010524181 |
| SLCO1B7 | | 0.01051704 |
| TMEM150A | | 0.010514192 |
| TSPAN19 | | 0.010512467 |
| RN7SKP65 | | 0.010507832 |
| RP11-296C13 | | 0.010506747 |
| RNU4-58P | | 0.010506134 |
| RPS3AP15 | | 0.010505577 |
| RP11-622C24 | | 0.010503566 |
| RP11-25K19 | | 0.010502886 |
| VWA2 | | 0.010498731 |
| HSPE1P19 | | 0.010496695 |
| RPEP6 | | 0.010493052 |
| HOXC9 | | 0.010491921 |
| | 12-Sep | 0.010491527 |
| RP5-1177M21 | | 0.010490031 |
| RPSAP41 | | 0.010489516 |
| DHODH | | 0.010481781 |
| AL356756 | | 0.010480941 |
| CH17-12M21 | | 0.01047946 |

| | | |
|---|---|---|
| GCNT1P4 | | 0.010472269 |
| SOX2OT_exon4 | | 0.010468901 |
| PRSS23 | | 0.010468571 |
| OXTR | | 0.010466618 |
| NPS | | 0.010465745 |
| PRSS29P | | 0.010463362 |
| LCN8 | | 0.010459194 |
| PIGX | | 0.010458171 |
| MEX3B | | 0.010456757 |
| RP11-525G12 | | 0.010456413 |
| PCA3 | | 0.01045363 |
| AC090311 | | 0.010453628 |
| AC126917 | | 0.010451457 |
| ZEB1-AS1 | | 0.010451138 |
| RP1-40G4P | | 0.010450359 |
| AC068137 | | 0.010449127 |
| MED14OS | | 0.010443454 |
| ASCC2 | | 0.010442672 |
| RNU6-1178P | | 0.010439366 |
| RP11-488C13 | | 0.010432049 |
| CDH8 | | 0.01043005 |
| FAM53A | | 0.010426318 |
| DDO | | 0.010422206 |
| OR10H3 | | 0.010415538 |
| AC010890 | | 0.010414769 |
| STOX2 | | 0.010413462 |
| AL354932 | | 0.010413209 |
| LINC01140 | | 0.010407471 |
| RP11-405A12 | | 0.010398033 |
| KIRREL3-AS1 | | 0.01039727 |
| RP11-561B11 | | 0.010395056 |
| RP11-142A12 | | 0.01038936 |
| RP3-499B10 | | 0.010388799 |
| HDHD1P3 | | 0.010386823 |
| PLSCR4 | | 0.010386521 |
| RP11-156F23 | | 0.010383991 |
| LINC00334 | | 0.010382695 |
| RP1-117B12 | | 0.010378771 |
| RP11-359J6 | | 0.010374354 |
| NAGPA | | 0.010363785 |
| RNU6-1291P | | 0.010362133 |
| AC096553 | | 0.010361901 |
| AL138919 | | 0.010361463 |
| PCDH9-AS1 | | 0.010361154 |
| | 2-Mar | 0.010357385 |

| | |
|---|---|
| MIR3681 | 0.010356322 |
| RP11-881L2 | 0.010350319 |
| PCAT4 | 0.010347908 |
| RP11-26M5 | 0.010344488 |
| SOX2 | 0.010334622 |
| HAAO | 0.010332603 |
| MEGF9 | 0.01033037 |
| RP11-77G22 | 0.010314195 |
| LSM1P1 | 0.010313983 |
| RP11-390M11 | 0.010313613 |
| RORB | 0.010307692 |
| RP11-190G13 | 0.010305245 |
| AL355821 | 0.010304305 |
| ZBTB20-AS3 | 0.010304131 |
| RNU6-716P | 0.010303278 |
| CCER1 | 0.010301434 |
| RNF216 | 0.010300359 |
| PHKA2-AS1 | 0.010289753 |
| AC008641 | 0.010289596 |
| AC006989 | 0.010286375 |
| RP11-250B2 | 0.010282851 |
| LINC01152 | 0.010282722 |
| RN7SL169P | 0.010282619 |
| OR5H8 | 0.01028071 |
| RNU6-500P | 0.010278989 |
| RP11-448N11 | 0.010275765 |
| FZD5 | 0.010268507 |
| RTN4IP1 | 0.010267781 |
| TEX40 | 0.010260638 |
| RP11-14C22 | 0.010259006 |
| T | 0.010256032 |
| OR2M7 | 0.010252218 |
| RP11-475B2 | 0.010249166 |
| RPL5P27 | 0.010243166 |
| MORN1 | 0.010243129 |
| RNU6-1062P | 0.010241007 |
| EEF1GP5 | 0.010234702 |
| RP4-563E14 | 0.01023457 |
| RP11-471M2 | 0.01022908 |
| RECQL4 | 0.010228784 |
| KCNK15-AS1 | 0.010228252 |
| RP11-579D7 | 0.010219575 |
| CCDC147-AS1 | 0.01021788 |
| LINC00881 | 0.010209913 |
| UFL1-AS1 | 0.010209803 |

| | |
|---|---|
| AC019322 | 0.010208684 |
| SNORD3A | 0.010201701 |
| ANKRD54 | 0.010199722 |
| DPYD-IT1 | 0.010194223 |
| KATNBL1P3 | 0.010193127 |
| CSNK1E | 0.010192466 |
| U47924 | 0.010189564 |
| NOSIP | 0.010184094 |
| RP11-533K11 | 0.010181651 |
| RP11-524C21 | 0.010180775 |
| C1QL1 | 0.010177638 |
| RP11-672F9 | 0.010176979 |
| RNU1-104P | 0.010175043 |
| AC079741 | 0.010169511 |
| RBM22P7 | 0.010167825 |
| GPM6BP2 | 0.010167825 |
| CTD-2081C10 | 0.010160263 |
| RPS6KC1 | 0.01015959 |
| RP11-125O18 | 0.010157973 |
| RNU6-1081P | 0.010154231 |
| RRAGB | 0.010153841 |
| RP11-244O19 | 0.010152595 |
| HYDIN | 0.010143824 |
| COL6A4P2 | 0.010136567 |
| RP11-227F8 | 0.010135508 |
| OR5H15 | 0.010129868 |
| ZNF587P1 | 0.010110455 |
| RP11-288C17 | 0.010109672 |
| SLCO1B3 | 0.010108014 |
| RNU7-2P | 0.010105935 |
| TRIT1 | 0.010105913 |
| RP11-608B3 | 0.010104959 |
| MIR6511B1 | 0.010102039 |
| TRHR | 0.010099757 |
| MIR6880 | 0.010094415 |
| AC079584 | 0.010093317 |
| AC012146 | 0.01008675 |
| FABP5P11 | 0.01008502 |
| THAP4 | 0.010084238 |
| MIR204 | 0.010081294 |
| MIR190A | 0.01008045 |
| POP4 | 0.010079643 |
| RP11-21C4 | 0.010077025 |
| SRGAP3-AS4 | 0.010067182 |
| RP11-576C2 | 0.010064987 |

| | |
|---|---|
| SLC25A5P2 | 0.010063656 |
| SNORD114-30 | 0.010062035 |
| SSU72 | 0.010051362 |
| RP11-417D4 | 0.010048475 |
| SOCS3 | 0.010043753 |
| MPZ | 0.010043567 |
| FLJ37505 | 0.01003685 |
| MIR548AI | 0.010031344 |
| RNU6-1313P | 0.010030534 |
| OR4K5 | 0.010028962 |
| TET1 | 0.010021687 |
| DYNLL1P6 | 0.010021666 |
| RPL26P19 | 0.010017184 |
| AP000469 | 0.010017149 |
| RN7SKP115 | 0.01001392 |
| RNA5SP149 | 0.010013549 |
| HTN1 | 0.010009157 |
| CTBP2P1 | 0.010002587 |
| PABPC1P13 | 0.010002023 |
| MANBA | 0.010001127 |
| AC016995 | 0.009991713 |
| OR6U2P | 0.009990074 |
| RPS20P24 | 0.009988891 |
| RP13-786C16 | 0.009988709 |
| OSBPL6 | 0.009987304 |
| KCNJ8 | 0.009985108 |
| FAM181A-AS1 | 0.009978784 |
| MTRNR2L9 | 0.009977092 |
| HTR1D | 0.009976481 |
| MIR6074 | 0.009970203 |
| RP11-586K2 | 0.009965086 |
| CTD-2196P11 | 0.009954926 |
| RPS3A | 0.009948809 |
| TMPRSS2 | 0.009943395 |
| THG1L | 0.009943246 |
| RP11-598G2 | 0.009932337 |
| RAPSN | 0.009927985 |
| RP11-559N14 | 0.00992219 |
| OR7E15P | 0.009920132 |
| RNU6-410P | 0.009919673 |
| GPR112 | 0.009908782 |
| RP11-7M8 | 0.009908706 |
| AC004866 | 0.009908635 |
| SATB1-AS1 | 0.009903264 |
| RN7SL33P | 0.009902794 |

| | |
|---|---|
| RP11-509M23 | 0.00990015 |
| BMP10 | 0.009893974 |
| CTD-2534I21 | 0.009892168 |
| RN7SL223P | 0.009888953 |
| RP11-674E16 | 0.009882824 |
| RPSAP37 | 0.009878988 |
| MRPL35P2 | 0.009870563 |
| DNTTIP1 | 0.009870368 |
| ATP2B2-IT2 | 0.009869559 |
| RP11-134C1 | 0.00986936 |
| MRPL13 | 0.009863521 |
| AC008169 | 0.009862239 |
| OR4A16 | 0.009861545 |
| MTAP | 0.00985874 |
| RP11-402P6 | 0.009857815 |
| RP11-602N24 | 0.00985509 |
| CERS3 | 0.009854726 |
| AC134772 | 0.009850617 |
| A2ML1-AS1 | 0.009845008 |
| RP11-421K23 | 0.00983838 |
| RBPMS2P1 | 0.009838378 |
| ANAPC2 | 0.009837381 |
| RN7SL304P | 0.009829596 |
| RP11-461L18 | 0.009817351 |
| CELF5 | 0.009816995 |
| RNA5SP127 | 0.00981396 |
| ERMAP | 0.009809752 |
| LINC00437 | 0.009808628 |
| GEMIN4 | 0.009806849 |
| SLC16A9 | 0.009800988 |
| RP11-686O6 | 0.009799351 |
| BIK | 0.009795798 |
| RP11-59N23 | 0.0097927 |
| CTB-55B8 | 0.009791949 |
| RYKP1 | 0.009790324 |
| AKR1C6P | 0.009787641 |
| SCN1B | 0.009783425 |
| SUMO2P16 | 0.009781999 |
| PCBP2P3 | 0.009777797 |
| RNA5SP511 | 0.009777127 |
| RP11-85B7 | 0.009775772 |
| ELL2P4 | 0.00977558 |
| RP11-87P13 | 0.00977048 |
| RNU6-702P | 0.009767395 |
| RN7SKP140 | 0.00976722 |

| | |
|---|---|
| RP11-467C18 | 0.009767123 |
| FTH1P6 | 0.009765745 |
| RP11-105L10 | 0.009758382 |
| AC106883 | 0.009758382 |
| AC084125 | 0.009752261 |
| RP11-144G6 | 0.009748986 |
| CAMKMT | 0.009746809 |
| UBE2HP1 | 0.009739462 |
| RP11-37J13 | 0.009738257 |
| RNU6-1225P | 0.009738236 |
| RPL5P2 | 0.009737599 |
| CTA-339C12 | 0.009732457 |
| AC231657 | 0.009731453 |
| RP4-815D20 | 0.009730922 |
| GLTSCR1 | 0.009729171 |
| RP11-736G13 | 0.009726406 |
| TMSB4XP2 | 0.009723086 |
| CT47B1 | 0.009719324 |
| VDAC1P2 | 0.009718008 |
| RP11-100M12 | 0.009716791 |
| RNVU1-6 | 0.009711498 |
| TOMM22P6 | 0.009711261 |
| FOXA2 | 0.00970962 |
| OGFR | 0.009706938 |
| FTH1P2 | 0.00970558 |
| ATP6AP1L | 0.009704073 |
| RNU6-681P | 0.009696632 |
| TMEM220 | 0.00969263 |
| AC021451 | 0.00968194 |
| CTD-2290C23 | 0.009679139 |
| LINC00505 | 0.009677619 |
| LINC00560 | 0.009676713 |
| RP11-415D17 | 0.009675756 |
| AC037445 | 0.009675038 |
| RP11-398K22 | 0.009671792 |
| SLC17A8 | 0.009671525 |
| RP11-563N4 | 0.009664593 |
| AKR7A2P1 | 0.009664058 |
| RN7SKP180 | 0.009656086 |
| ARHGAP31 | 0.009655177 |
| PKN1 | 0.00965008 |
| RAB1A | 0.009645744 |
| RN7SL766P | 0.00964459 |
| ZNF783 | 0.009637837 |
| RP11-745L13 | 0.009633463 |

| | |
|---|---|
| FAHD2B | 0.009633297 |
| RP11-369K16 | 0.009625233 |
| RP11-220D10 | 0.009621909 |
| PSMC2 | 0.009620515 |
| RP11-156L14 | 0.009617965 |
| RN7SL432P | 0.009616546 |
| HSPA8P5 | 0.009615413 |
| LINC00616 | 0.00961117 |
| AC002398 | 0.009609054 |
| NAP1L4P2 | 0.009608586 |
| KCTD5 | 0.009605761 |
| AC005176 | 0.009605516 |
| RNA5SP363 | 0.009600583 |
| BAI2 | 0.009595231 |
| SNORD83 | 0.009593378 |
| AC004938 | 0.009593117 |
| UBE2Q2 | 0.009592905 |
| KCNRG | 0.009591584 |
| KLF10 | 0.009589388 |
| AC092107 | 0.009589296 |
| CLDN12 | 0.009583286 |
| RP11-587H10 | 0.009582884 |
| RP11-138H10 | 0.009581987 |
| TP53BP2 | 0.009581035 |
| ZNF192P2 | 0.009567472 |
| ECM1 | 0.009566894 |
| DNAJA1P2 | 0.009561147 |
| AL360074 | 0.009556421 |
| RN7SL2 | 0.009547545 |
| CD3EAP | 0.009547284 |
| MFHAS1 | 0.009542986 |
| STON1-GTF2A1L | 0.00953512 |
| RP11-424I3 | 0.009528471 |
| CDC26 | 0.009525162 |
| HMGCS2 | 0.00951774 |
| RP11-486M23 | 0.009514473 |
| AC073094 | 0.009513194 |
| RN7SKP203 | 0.009511591 |
| RPL36AP26 | 0.009505796 |
| CCDC58 | 0.009504659 |
| RP11-802F5 | 0.009502584 |
| RN7SL757P | 0.009500566 |
| MIR4427 | 0.009499583 |
| MIR581 | 0.009498488 |
| MYBPC2 | 0.009496953 |

| | |
|---|---|
| RPL7L1P1 | 0.009496593 |
| RP3-445O10 | 0.009494274 |
| FTLP15 | 0.009494055 |
| SNORA65 | 0.009493413 |
| RP11-710N8 | 0.009491694 |
| KRT38 | 0.009487798 |
| RRM2P4 | 0.009487683 |
| TRAF3IP2 | 0.009487617 |
| KLHDC4 | 0.009485291 |
| CYP2C60P | 0.009482884 |
| BCL2L12P1 | 0.009481987 |
| TONSL | 0.009480584 |
| CTRC | 0.009472945 |
| CHRNA2 | 0.009470959 |
| SGMS1 | 0.009466462 |
| SUMO4 | 0.009462497 |
| CHEK2P2 | 0.009461733 |
| HBM | 0.009459055 |
| CPSF1 | 0.009456359 |
| CES5AP1 | 0.00944665 |
| RP11-48L13 | 0.009443331 |
| RP5-1059H15 | 0.00944285 |
| C1orf185 | 0.009442371 |
| SLC6A6P1 | 0.009439111 |
| AC005363 | 0.009438285 |
| RP13-77O11 | 0.009438178 |
| ATXN1 | 0.009436768 |
| KIFC2 | 0.009436061 |
| CTC-470C15 | 0.009422423 |
| MSANTD3-TMEFF1 | 0.009421774 |
| RNA5SP147 | 0.009419855 |
| MPV17L | 0.009419669 |
| MIR892B | 0.009417185 |
| RNU7-194P | 0.009405916 |
| RAB30-AS1 | 0.009399669 |
| BCAP29 | 0.009395668 |
| SNORD116-17 | 0.009394935 |
| TMEM89 | 0.009390233 |
| BMP5 | 0.009389683 |
| SSX4B | 0.009388664 |
| RFC4 | 0.009387928 |
| AC025166 | 0.009387137 |
| ERC1 | 0.009383389 |
| BRSK1 | 0.00938164 |
| C20orf187 | 0.009381526 |

| | |
|---|---|
| RNU6-737P | 0.009381508 |
| RP4-725G10 | 0.009381191 |
| RN7SL460P | 0.00937972 |
| RP1-27C22 | 0.009375893 |
| AP001607 | 0.009375516 |
| U2 | 0.009374984 |
| CDR2 | 0.009373379 |
| CTD-2008N3 | 0.009373028 |
| DIAPH3-AS1 | 0.009370662 |
| CDT1 | 0.009368076 |
| RP11-561C5 | 0.009365219 |
| RP4-568F9 | 0.009356951 |
| CAMK2N2 | 0.009355499 |
| AC008173 | 0.009353965 |
| RP1-67A8 | 0.009351764 |
| RPL10AP3 | 0.009349203 |
| C11orf72 | 0.00934761 |
| RP11-259N19 | 0.009347331 |
| NDUFA8 | 0.009341787 |
| CNTNAP1 | 0.009339238 |
| SNORD39 | 0.009336521 |
| RNU7-4P | 0.009335185 |
| OR11H2 | 0.009334108 |
| LINC00993 | 0.009333482 |
| KCNK13 | 0.009330177 |
| MCIDAS | 0.00932579 |
| AC126763 | 0.009318655 |
| OR7E13P | 0.009318285 |
| RP11-346C16 | 0.009317438 |
| DNASE1L1 | 0.009316558 |
| SFPQP1 | 0.009301586 |
| RP11-363G2 | 0.009298674 |
| NDUFAF5 | 0.009298566 |
| MVK | 0.009286361 |
| RP11-92C4 | 0.009286207 |
| IGF1 | 0.00928437 |
| RP11-796G6 | 0.009276542 |
| MOCS1 | 0.009275719 |
| RP11-61L19 | 0.009273114 |
| RNU1-35P | 0.009271844 |
| CTD-2210P24 | 0.009271607 |
| AC093906 | 0.009268459 |
| AC041039 | 0.009268455 |
| RP11-301G21 | 0.009268284 |
| LINC00940 | 0.009264009 |

| | |
|---|---|
| RP11-113E21 | 0.009262901 |
| PRKG1-AS1 | 0.009259856 |
| ZNF582-AS1 | 0.009259662 |
| CNOT7P2 | 0.009252108 |
| OR6C70 | 0.009248657 |
| OR2T34 | 0.009248049 |
| MIR1193 | 0.009239506 |
| RP11-45L9 | 0.009232565 |
| RP5-1065J22 | 0.009227773 |
| RP11-255L13 | 0.009226278 |
| RNU6-1122P | 0.009224578 |
| CYP4F29P | 0.009223116 |
| DCSTAMP | 0.009222098 |
| ZKSCAN7 | 0.009220219 |
| RP6-102O10 | 0.009220091 |
| CXCL12 | 0.009219958 |
| PCCB | 0.009199046 |
| AC107081 | 0.009195846 |
| TMEM184A | 0.009194246 |
| SNORD113 | 0.009188233 |
| OR4K17 | 0.009187685 |
| RP11-310P5 | 0.009187397 |
| ACTBP14 | 0.009184149 |
| NFKB2 | 0.009183119 |
| TRMT2A | 0.009180887 |
| AC026951 | 0.009172703 |
| RP1-27O5 | 0.009169031 |
| MTND4P1 | 0.009168575 |
| MIR4498 | 0.009164761 |
| RP3-417O22 | 0.00916264 |
| SMAP1 | 0.009158132 |
| COX6CP1 | 0.009152549 |
| DUXAP10 | 0.00915226 |
| TMLHE | 0.00915082 |
| RP11-2P2 | 0.00915069 |
| CTD-3073N11 | 0.00914725 |
| CRK | 0.009146281 |
| RP11-98E6 | 0.009141999 |
| RN7SKP289 | 0.009140816 |
| KCTD14 | 0.009140178 |
| NCAPH2 | 0.00913882 |
| FLJ43681 | 0.009137134 |
| SNORA8 | 0.009133693 |
| PDC | 0.009133676 |
| RP11-214N9 | 0.009133238 |

| | |
|---|---|
| SNORD115-40 | 0.009130806 |
| MED13P1 | 0.009130316 |
| LINC01285 | 0.00913001 |
| RNU6-389P | 0.009129496 |
| RNU6ATAC30P | 0.009123534 |
| POM121L10P | 0.009123312 |
| DNAJC19P6 | 0.009122736 |
| AC092104 | 0.009122276 |
| ENPP7P14 | 0.009121815 |
| RP3-455E7 | 0.009120388 |
| RP11-461K13 | 0.009118587 |
| AC105399 | 0.009107357 |
| RP11-624L12 | 0.009104192 |
| POU5F1P5 | 0.009096211 |
| RNU6ATAC11P | 0.009089848 |
| LINC00051 | 0.009075478 |
| RP11-417O11 | 0.009073884 |
| RP3-520B18 | 0.009071314 |
| RP11-673C5 | 0.00907011 |
| RNU1-117P | 0.009065629 |
| RP11-123M21 | 0.009064902 |
| RP11-478C6 | 0.009064119 |
| AL442643 | 0.009062899 |
| RNU6-892P | 0.009049749 |
| AC064847 | 0.009043392 |
| AC013262 | 0.009040033 |
| CTSLP7 | 0.009039861 |
| RP11-446H18 | 0.009033202 |
| OR4K11P | 0.009024894 |
| RP11-142L1 | 0.009021558 |
| ETV6 | 0.009021326 |
| RP11-449G16 | 0.009020982 |
| CTD-2555C10 | 0.009020842 |
| RP11-690C23 | 0.00902003 |
| NARSP2 | 0.009016333 |
| BTF3P8 | 0.009013568 |
| CARSP1 | 0.009011658 |
| NELFE | 0.009009568 |
| RP11-425L10 | 0.009008563 |
| ACOT11 | 0.009002219 |
| MED15P1 | 0.009001411 |
| RP11-386J22 | 0.00899766 |
| SORT1 | 0.008995241 |
| RNU6ATAC2P | 0.008994188 |
| RN7SKP11 | 0.008978097 |

| | |
|---|---|
| SLC8A1-AS1 | 0.008974723 |
| AC007285 | 0.008969622 |
| CTD-2333K2 | 0.008968378 |
| RP11-160A9 | 0.008965605 |
| SPATA22 | 0.008964829 |
| FIRRE | 0.008964115 |
| IGLV3-7 | 0.008959472 |
| MIR4445 | 0.008956707 |
| CTD-3088G3 | 0.008948821 |
| CCDC17 | 0.008944709 |
| RP11-370F5 | 0.008938486 |
| AC124312 | 0.008935017 |
| RP1-287H17 | 0.008934625 |
| PDCD6IPP2 | 0.008934616 |
| RP11-146P2 | 0.008933976 |
| EEF1A1P19 | 0.00893269 |
| FABP5P5 | 0.00893253 |
| RP11-12K11 | 0.008921254 |
| RNU6-333P | 0.008909466 |
| OLA1 | 0.008901302 |
| OR5AP2 | 0.008899181 |
| RP11-69G7 | 0.00889704 |
| RP4-603I14 | 0.008889447 |
| HSPE1P11 | 0.008881606 |
| RP11-386I23 | 0.008874843 |
| RPS13 | 0.008866938 |
| HIST1H2AH | 0.008863702 |
| MIR541 | 0.008860695 |
| RP11-326L2 | 0.008855032 |
| FER1L6 | 0.008853976 |
| MMACHC | 0.008848054 |
| RP11-342F17 | 0.008840884 |
| LINC01397 | 0.008836977 |
| CBX1P4 | 0.008832263 |
| RP11-317P15 | 0.008827374 |
| RPL6P8 | 0.008826185 |
| RP5-875H18 | 0.008821642 |
| AC011994 | 0.008817934 |
| NCOR2 | 0.008816127 |
| RP11-680L20 | 0.008813443 |
| GPR128 | 0.008813306 |
| CTD-2579N5 | 0.008811038 |
| OR6R1P | 0.008803398 |
| TRIM50 | 0.008795223 |
| GS1-542M4 | 0.008787376 |

| | |
|---|---|
| RNU6-918P | 0.008785874 |
| VHLL | 0.008779775 |
| HMGB1P28 | 0.008776546 |
| RP11-322N21 | 0.008775423 |
| RNA5SP255 | 0.00877178 |
| RP11-326L9 | 0.008764016 |
| VAMP7 | 0.008762934 |
| RNU6-35P | 0.008759976 |
| KCNMA1-AS2 | 0.008757365 |
| RP11-284B18 | 0.008755768 |
| RP11-356B19 | 0.008753677 |
| AC097470 | 0.008750614 |
| TRIM48 | 0.00874542 |
| RN7SL259P | 0.008742896 |
| RNU7-169P | 0.008742667 |
| RP11-674I16 | 0.008740955 |
| RP11-354B3 | 0.008740727 |
| TEPP | 0.008739412 |
| LINC00392 | 0.008737229 |
| HMGB1P46 | 0.00873631 |
| AC005488 | 0.008729739 |
| LINC01077 | 0.008727595 |
| AC093267 | 0.008726289 |
| BNIP3P3 | 0.008725256 |
| RNU6-11P | 0.008724808 |
| RPL39P3 | 0.008724718 |
| BASP1 | 0.00872269 |
| RNA5SP479 | 0.008714724 |
| AC010731 | 0.008710973 |
| AL160165 | 0.00870616 |
| RP11-785H5 | 0.008705424 |
| CEACAMP9 | 0.008705413 |
| VSTM1 | 0.008704454 |
| RPL14 | 0.008702782 |
| RP11-565F4 | 0.008700195 |
| UBTFL2 | 0.008698485 |
| AQP7 | 0.008698175 |
| RP13-146A14 | 0.008694804 |
| NKPD1 | 0.008693591 |
| EEF1A1P37 | 0.008692651 |
| MTND3P5 | 0.008690853 |
| RP11-285C1 | 0.008689063 |
| RNU6-1148P | 0.008686951 |
| BEND5 | 0.008685795 |
| snoZ6 | 0.008680767 |

| | |
|---|---|
| SNORD88A | 0.00868036 |
| AC004066 | 0.008680195 |
| AC011239 | 0.008679078 |
| TMEM39A | 0.008669888 |
| AC011363 | 0.008666535 |
| RP4-594L9 | 0.008656842 |
| AL160037 | 0.008655148 |
| MIR4298 | 0.008653941 |
| LINC00211 | 0.008650214 |
| COX6CP11 | 0.008648593 |
| RP11-93K7 | 0.00864592 |
| RP11-269F19 | 0.00864557 |
| ARMC4P1 | 0.00864488 |
| RNLS | 0.008635951 |
| SLC5A8 | 0.008633069 |
| FRG1 | 0.008632751 |
| HDDC3 | 0.008629867 |
| SLC35F1 | 0.0086257 |
| ZNF626 | 0.008623207 |
| EI24P3 | 0.008621268 |
| RP11-546B8 | 0.008620755 |
| RP11-228O6 | 0.008620251 |
| MRPS21P1 | 0.008615008 |
| SAV1 | 0.008614778 |
| CMPK1 | 0.008613662 |
| AL445523 | 0.008610333 |
| PCED1A | 0.008599801 |
| CHMP1B2P | 0.008598919 |
| OR10P1 | 0.008597563 |
| CTC-232H10 | 0.008591496 |
| GAPDHP53 | 0.008589736 |
| OR2AL1P | 0.008584539 |
| CTD-2012K14 | 0.008582386 |
| NAT9 | 0.008580191 |
| KDM5B | 0.008578144 |
| RP11-81A22 | 0.008573046 |
| RP5-899B16 | 0.008572561 |
| CTA-29F11 | 0.008572451 |
| TTLL1 | 0.008568581 |
| NPIPB9 | 0.008560841 |
| AL355531 | 0.008560655 |
| MIR4739 | 0.008560339 |
| RP11-154P18 | 0.008558097 |
| TESK1 | 0.008550589 |
| AL354806 | 0.008546882 |

| | |
|---|---|
| TAZ | 0.008544859 |
| MIR3134 | 0.008544661 |
| GORASP2 | 0.008544487 |
| MGRN1 | 0.008544152 |
| RP11-188I24 | 0.00853795 |
| MIR4694 | 0.008536607 |
| PLCE1-AS2 | 0.008535659 |
| ANKRD66 | 0.008534972 |
| HIST1H2BM | 0.008533642 |
| BMPR2 | 0.008533286 |
| MIR4417 | 0.00853037 |
| AL080243 | 0.008530292 |
| AC069394 | 0.008528322 |
| GABRA6 | 0.008525993 |
| RNA5SP63 | 0.008524319 |
| RP5-991G20 | 0.008522718 |
| MIR3972 | 0.008519051 |
| ZNF707 | 0.00851669 |
| TMEM65 | 0.008515146 |
| RP11-797M17 | 0.008515063 |
| RP11-620J15 | 0.008513044 |
| OR8J3 | 0.008511312 |
| AGPAT1 | 0.008509713 |
| RN7SL325P | 0.008503363 |
| CCNYL1 | 0.008500913 |
| RP11-62J1 | 0.008495796 |
| ABCA17P | 0.008494754 |
| AC007567 | 0.008488864 |
| PRDX3P3 | 0.0084856 |
| RP4-565E6 | 0.008482856 |
| AC013251 | 0.008470964 |
| PRKY | 0.008465024 |
| AC104331 | 0.008456663 |
| CKMT2 | 0.008455999 |
| LINC00083 | 0.008454372 |
| CTD-2576F9 | 0.008449582 |
| RP11-363H12 | 0.008447812 |
| PROSP | 0.008447616 |
| RP11-177B4 | 0.008445463 |
| RPS3AP49 | 0.00844459 |
| AC118758 | 0.008433891 |
| AC004004 | 0.008431771 |
| RP11-28O3 | 0.008429201 |
| CTD-2027I19 | 0.008427589 |
| RP11-324C10 | 0.008425347 |

| | |
|---|---|
| NANOGP11 | 0.008421771 |
| RP11-447B18 | 0.008419247 |
| NFKBIE | 0.008418342 |
| RP11-351E7 | 0.008414059 |
| RNU6-730P | 0.00841225 |
| RPL11P5 | 0.008411788 |
| CTA-360L10 | 0.008410675 |
| RP3-354N19 | 0.008410307 |
| RP11-799M12 | 0.008408773 |
| RP11-745K9 | 0.008408014 |
| CA4 | 0.008404787 |
| PSTK | 0.008401727 |
| RP1-111B22 | 0.008399652 |
| RP11-885N19 | 0.00839846 |
| RP11-60H5 | 0.008398208 |
| RP11-173G21 | 0.008391427 |
| TRIM2 | 0.008388592 |
| HPS1 | 0.008387661 |
| KIAA1755 | 0.008386154 |
| AC067742 | 0.00838464 |
| LINC01097 | 0.008383724 |
| TSPYL6 | 0.008380093 |
| RP11-80H18 | 0.008376951 |
| RP5-978I12 | 0.008376573 |
| AC002368 | 0.008373333 |
| ADCK1 | 0.008370079 |
| CTB-105N12 | 0.00836816 |
| IQCA1-AS1 | 0.008367253 |
| RP11-394G3 | 0.008367197 |
| CACNB3 | 0.008361614 |
| LRRC19 | 0.008359576 |
| RP11-588K22 | 0.008354358 |
| RP11-723G8 | 0.008350914 |
| RP11-1394O16 | 0.008348128 |
| MIR5692B | 0.008342825 |
| RPEP1 | 0.008342353 |
| CYP4A43P | 0.008340109 |
| CHCHD1 | 0.008336868 |
| CTC-281M20 | 0.008334435 |
| BDKRB1 | 0.008334301 |
| MORF4L1 | 0.008334301 |
| CDX2 | 0.008332229 |
| RP5-1148A21 | 0.008330822 |
| AC073072 | 0.008330577 |
| AL008723 | 0.0083295 |

| | |
|---|---|
| RN7SKP195 | 0.008329329 |
| BRF1 | 0.008328347 |
| RPL7AP7 | 0.008323891 |
| SNRPF | 0.008323565 |
| PRIMA1 | 0.008317221 |
| LINC01184 | 0.008312849 |
| RNU6-274P | 0.008311635 |
| RP11-405P11 | 0.008301915 |
| RP11-23F23 | 0.008298365 |
| RP11-352B15 | 0.008283971 |
| CLASRP | 0.008281461 |
| RP11-510C10 | 0.008281304 |
| MYOM3 | 0.008277349 |
| NT5C1B-RDH14 | 0.008274462 |
| RP11-131M6 | 0.008272427 |
| AC073628 | 0.008271456 |
| LINC00491 | 0.008269638 |
| MIR5187 | 0.008262772 |
| MICG | 0.008258938 |
| FTH1P26 | 0.00825487 |
| RP11-101E13 | 0.008253851 |
| DCUN1D3 | 0.008253739 |
| DRC1 | 0.008250248 |
| FAM131A | 0.008247823 |
| RP11-406A20 | 0.008239823 |
| SORBS2 | 0.008239413 |
| RP11-274E6 | 0.008237305 |
| H2AFZP4 | 0.008234402 |
| UBE2L3 | 0.008231394 |
| AES | 0.008218612 |
| RP11-30L8 | 0.008213189 |
| EIF1P3 | 0.008212709 |
| GPR137B | 0.00821184 |
| AC108488 | 0.008211449 |
| ZNF723P | 0.008210494 |
| PLSCR5 | 0.008209724 |
| MIR381HG | 0.008206205 |
| AC002451 | 0.00820447 |
| RNU1-7P | 0.008204435 |
| RP11-260G13 | 0.008203059 |
| RN7SL646P | 0.008201051 |
| CTAG2 | 0.008196777 |
| RNA5SP469 | 0.008189931 |
| MRPS25 | 0.00818475 |
| RP11-119H12 | 0.008183373 |

| | |
|---|---|
| RP11-418I22 | 0.008181904 |
| RNY1P5 | 0.008177144 |
| RP4-694B14 | 0.008170703 |
| AC007742 | 0.008169169 |
| RP11-63A11 | 0.008168479 |
| RP11-92A5 | 0.008166518 |
| CCNB3P1 | 0.008163549 |
| ZNF835 | 0.008162182 |
| MAS1 | 0.008159321 |
| EHD4-AS1 | 0.008158837 |
| RP11-409C19 | 0.008156039 |
| RP11-675M1 | 0.00815524 |
| ULK4 | 0.008150888 |
| CTD-2572N17 | 0.008150546 |
| RP11-614F17 | 0.008142939 |
| RP11-11H9 | 0.008140605 |
| RP11-101P17 | 0.008140381 |
| RNU6-324P | 0.00813537 |
| RP11-294L13 | 0.008133377 |
| RN7SL400P | 0.008132829 |
| RP11-58O3 | 0.008130837 |
| LPP-AS1 | 0.008128544 |
| AL035106 | 0.008126908 |
| EPB41L4A-AS1 | 0.008125622 |
| TMCO5B | 0.00812154 |
| OR6C3 | 0.008121334 |
| UGT2A3 | 0.008117395 |
| ABC14-1080714F14 | 0.008115776 |
| SPINK13 | 0.008108442 |
| RP11-120J1 | 0.008108441 |
| PPIL6 | 0.008105733 |
| RP11-646D13 | 0.008094403 |
| RPL31P33 | 0.008092898 |
| RP11-426J5 | 0.008089294 |
| CTD-2086L14 | 0.008087199 |
| RP11-370P15 | 0.00808712 |
| RP13-1039J1 | 0.008087084 |
| CCDC68 | 0.008085304 |
| DAND5 | 0.008083522 |
| CTD-2314G24 | 0.008080073 |
| CTD-2016O11 | 0.008076898 |
| CTD-2299N12 | 0.008073845 |
| OR4F17 | 0.008067141 |
| MIR3156-2 | 0.008066082 |
| STARD4 | 0.008065694 |

| | |
|---|---|
| SC22CB-1E7 | 0.008064168 |
| MTND5P4 | 0.008052653 |
| AC092675 | 0.008051077 |
| RP5-1132H15 | 0.008038761 |
| LYPLA2P1 | 0.008036396 |
| RN7SL120P | 0.008034854 |
| RPS4XP10 | 0.008034069 |
| PCNXL3 | 0.008033171 |
| NOP56 | 0.008029115 |
| SEPT7P5 | 0.008028707 |
| LINC01010 | 0.008024888 |
| RP11-4H14 | 0.008024056 |
| HIGD1AP15 | 0.008014454 |
| NONO | 0.008013208 |
| MIR4478 | 0.008009568 |
| SCD5 | 0.008009133 |
| MIR524 | 0.008008015 |
| LA16c-60H5 | 0.008003925 |
| GABRA2 | 0.007998013 |
| PLG | 0.007997843 |
| CTD-2154H6 | 0.007988679 |
| MLLT1 | 0.007987923 |
| RAD21L1 | 0.007985851 |
| HOXC5 | 0.007982883 |
| RP11-775L16 | 0.007978791 |
| RP11-619L12 | 0.007977782 |
| MIR4804 | 0.007976197 |
| AC079917 | 0.007974508 |
| RP11-288H12 | 0.007969876 |
| FZD10-AS1 | 0.007967513 |
| CDK2AP2P2 | 0.007963104 |
| AL139333 | 0.007961297 |
| DUS2 | 0.007957819 |
| SALL4P6 | 0.007951515 |
| MIR525 | 0.007948938 |
| KATNBL1P2 | 0.007945495 |
| MT-RNR2 | 0.007938469 |
| LGI4 | 0.007937874 |
| CTC-527H23 | 0.007937083 |
| MAPRE1P3 | 0.007937046 |
| RP11-201K10 | 0.007934625 |
| MORN2 | 0.007934413 |
| LINC00266-4P | 0.007930638 |
| SLIT3 | 0.007926078 |
| CCNT2P1 | 0.007920482 |

| | |
|---|---|
| MIR656 | 0.007920228 |
| CDH19 | 0.007911387 |
| MIR3115 | 0.007908857 |
| TCFL5 | 0.007907361 |
| RP11-446H13 | 0.007902567 |
| AC005150 | 0.007900374 |
| RP11-24C14 | 0.007899464 |
| SNORD14 | 0.007892556 |
| BCAR4 | 0.007887988 |
| LINC01567 | 0.007887671 |
| RP11-103C3 | 0.007887293 |
| AQP10 | 0.007887085 |
| SLC25A5P6 | 0.007884807 |
| MIR4524B | 0.007883054 |
| AC002115 | 0.007879805 |
| SLITRK6 | 0.007877835 |
| ST6GALNAC2P1 | 0.007868834 |
| TEX38 | 0.007865531 |
| RP11-45H22 | 0.007864389 |
| AC007242 | 0.00785763 |
| OR51A10P | 0.007855653 |
| RP11-756H20 | 0.007850359 |
| RXRA | 0.007849293 |
| RP11-478B11 | 0.007848232 |
| AC104002 | 0.007848104 |
| AC091178 | 0.007841897 |
| AC096775 | 0.00784099 |
| MIR1305 | 0.007840513 |
| EHBP1 | 0.007840177 |
| RNU4-61P | 0.007839944 |
| LINC01471 | 0.007837255 |
| RP11-240E2 | 0.007833186 |
| AC011998 | 0.007831373 |
| TMEM191B | 0.007830339 |
| CUTC | 0.007823376 |
| AC004637 | 0.007820905 |
| ARHGAP42P5 | 0.007819463 |
| UQCRB | 0.007809794 |
| KCNE1 | 0.007809099 |
| RDH13 | 0.007808454 |
| LRRC1 | 0.007806966 |
| LYPD5 | 0.00780464 |
| S100A11P1 | 0.007796581 |
| RPS12P15 | 0.007791463 |
| SRGAP2B | 0.007782518 |

| | |
|---|---|
| VSIG4 | 0.007777123 |
| RP11-430H12 | 0.007775791 |
| THADA | 0.007771485 |
| ASS1P5 | 0.007770718 |
| RPSAP42 | 0.007770583 |
| RPS27A | 0.007766127 |
| RNU6-973P | 0.007758576 |
| ZDHHC4 | 0.007754182 |
| RP5-1101C3 | 0.007747005 |
| ZFAND3 | 0.007745531 |
| PLA2G3 | 0.007744556 |
| CTD-2156G10 | 0.007744169 |
| EFHD1 | 0.007740935 |
| AC016739 | 0.007737568 |
| AC009299 | 0.007730963 |
| RP11-573E11 | 0.00773056 |
| AC007679 | 0.007725494 |
| RSG1 | 0.007725381 |
| COX7A2 | 0.007723614 |
| RP11-19D2 | 0.007721889 |
| OR6C64P | 0.007720907 |
| RP11-487E1 | 0.007720795 |
| C4orf22 | 0.007718053 |
| AC100848 | 0.007713262 |
| RN7SL398P | 0.007712909 |
| AC024908 | 0.007711169 |
| TMEM101 | 0.007710962 |
| ZNF205-AS1 | 0.007710459 |
| STT3A | 0.007710385 |
| RP11-277P12 | 0.007707445 |
| AL611946 | 0.007702425 |
| RP11-289K10 | 0.007697638 |
| CTD-2525P14 | 0.00769606 |
| SLC26A3 | 0.007693753 |
| RASL12 | 0.007689286 |
| RGL1 | 0.007688338 |
| MIRLET7I | 0.007685009 |
| OR51F3P | 0.007675374 |
| AC104695 | 0.007670203 |
| AC006019 | 0.007664067 |
| RP11-1046B16 | 0.007657706 |
| RP11-509J21 | 0.007657635 |
| ARHGDIG | 0.007653651 |
| RNF2P1 | 0.007653312 |
| GALNTL5 | 0.007650011 |

| | |
|---|---|
| PRKACG | 0.007649266 |
| OLAH | 0.007647947 |
| CECR5 | 0.007647406 |
| MFSD2B | 0.007645731 |
| RP11-335E8 | 0.007641717 |
| RP11-384J4 | 0.007641483 |
| C22orf46 | 0.007639368 |
| TUBAP | 0.007637914 |
| MTRNR2L3 | 0.007635723 |
| MIR6859-2 | 0.007630931 |
| C6orf132 | 0.007629712 |
| RP11-420C9 | 0.007623663 |
| ATCAY | 0.007618343 |
| RP11-691H4 | 0.007617349 |
| RP11-163I18 | 0.007617112 |
| CTBP2P4 | 0.007616703 |
| RP1-209A6 | 0.007607241 |
| SPAG11B | 0.007604182 |
| MIR607 | 0.007599398 |
| TSPY25P | 0.007596357 |
| RP11-19C24 | 0.007594109 |
| CTD-2215L10 | 0.007591389 |
| MOS | 0.007586274 |
| LYPLAL1-AS1 | 0.007581718 |
| OR8B12 | 0.00757943 |
| GRB14 | 0.007579366 |
| PCDHGA1 | 0.007575272 |
| TGFA | 0.0075749 |
| AL354680 | 0.007567076 |
| RP11-478P10 | 0.007556305 |
| RP3-420J14 | 0.007555394 |
| LRRC40 | 0.007551959 |
| CCDC60 | 0.007549268 |
| AL590422 | 0.007548835 |
| RP11-102G14 | 0.007546341 |
| C15orf38-AP3S2 | 0.007546311 |
| RMDN2-AS1 | 0.007541208 |
| SPG20-AS1 | 0.007535592 |
| HMGB3P3 | 0.00753465 |
| AL512640 | 0.007534628 |
| RN7SKP139 | 0.007524072 |
| ING4 | 0.00752407 |
| CEACAM20 | 0.007522846 |
| UMODL1-AS1 | 0.007520669 |
| CTD-2382H12 | 0.007520282 |

| | |
|---|---|
| BX470187 | 0.007516935 |
| ITFG1 | 0.007515214 |
| SOCS2 | 0.007508441 |
| MOK | 0.00750673 |
| PLGLA | 0.007502242 |
| LINC01219 | 0.007501723 |
| AL121932 | 0.007500914 |
| RP11-797D24 | 0.007500409 |
| FADS1 | 0.007499994 |
| RPL35AP25 | 0.007493866 |
| RNU6-423P | 0.007492664 |
| MIR3928 | 0.007492196 |
| AL158839 | 0.007487424 |
| RBBP8NL | 0.007487417 |
| AC004673 | 0.007487214 |
| KRT8P8 | 0.007487053 |
| LZTR1 | 0.007483417 |
| NTF6B | 0.007483166 |
| LINC00454 | 0.007472824 |
| RP11-439K3 | 0.007471507 |
| ASNA1 | 0.007470747 |
| VAMP5 | 0.0074695 |
| LINC01004 | 0.007468756 |
| MIR7111 | 0.007466681 |
| RP11-252A24 | 0.007465984 |
| LINC01039 | 0.007464856 |
| AC069067 | 0.00745943 |
| IGLL4P | 0.007456293 |
| ERVMER34-1 | 0.007456141 |
| Z98941 | 0.00745426 |
| OR52E6 | 0.007453122 |
| RP11-34C15 | 0.007452076 |
| MTND4P7 | 0.007445645 |
| GPR116 | 0.007445406 |
| RP11-3K24 | 0.007441786 |
| RP11-403B2 | 0.007440385 |
| GVINP2 | 0.007437709 |
| ACTR6P1 | 0.007436388 |
| MAP4K5 | 0.007426652 |
| ZNF581 | 0.007426019 |
| AC084149 | 0.007425887 |
| CCT7P2 | 0.007423834 |
| RP5-898I4 | 0.00742248 |
| LINC00675 | 0.007422222 |
| HS3ST4 | 0.007418201 |

| | |
|---|---|
| AP006564 | 0.007415766 |
| RN7SL497P | 0.007415546 |
| MTHFSD | 0.007413758 |
| RP11-558A11 | 0.007412808 |
| HPX | 0.007405534 |
| LINC01311 | 0.007405367 |
| APOA4 | 0.007404644 |
| CTD-2195M18 | 0.007400791 |
| RP11-384P7 | 0.00739815 |
| MIR890 | 0.007391721 |
| PLBD1-AS1 | 0.00738972 |
| AC016909 | 0.007382887 |
| MPO | 0.007381981 |
| OR7E86P | 0.007381937 |
| UGDH | 0.007370742 |
| COX6CP17 | 0.007366866 |
| RP11-644C3 | 0.007364855 |
| PSG2 | 0.007363491 |
| AC026904 | 0.007361099 |
| FLJ13224 | 0.00735514 |
| AC069154 | 0.007353789 |
| RP11-93G23 | 0.007349321 |
| TMCO5A | 0.007344069 |
| AC005105 | 0.007343239 |
| RPS26P56 | 0.007342978 |
| RNU6-1198P | 0.007341422 |
| AL137797 | 0.007335116 |
| MORF4L2P1 | 0.007331389 |
| RN7SL83P | 0.007328451 |
| LINC01028 | 0.007325857 |
| RNU6-409P | 0.007325119 |
| RP11-32P22 | 0.007323831 |
| FAM122C | 0.007322926 |
| RP11-680F20 | 0.007321245 |
| APOLD1 | 0.007320787 |
| BDNF | 0.00731998 |
| AC013722 | 0.007318333 |
| NATD1 | 0.007317456 |
| TMEM198 | 0.007316935 |
| MED4-AS1 | 0.007316105 |
| AC096566 | 0.007312951 |
| RP11-20L19 | 0.007311711 |
| ITGA6 | 0.007310591 |
| RNA5SP95 | 0.007310304 |
| SLFNL1 | 0.007304455 |

| | |
|---|---|
| FRMPD1 | 0.00730166 |
| TMEM263 | 0.007300172 |
| RP11-45A16 | 0.007297977 |
| CTD-2144E22 | 0.0072948 |
| RNU6-1153P | 0.007294333 |
| KLHL9 | 0.007288804 |
| SNAP25 | 0.007288209 |
| AC015883 | 0.007285215 |
| MIR519C | 0.007280759 |
| RP11-261C10 | 0.007280348 |
| MIR6823 | 0.007279021 |
| ARL13B | 0.00727792 |
| RNU6-96P | 0.007276558 |
| C7orf62 | 0.007275786 |
| ZBED1P1 | 0.007273609 |
| ARF1P1 | 0.007268977 |
| CH17-38B12 | 0.007261006 |
| AL139151 | 0.007258846 |
| ZNF529-AS1 | 0.007256728 |
| RAD51L3-RFFL | 0.00725527 |
| VDAC1P7 | 0.007250773 |
| RP1-67K17 | 0.007247109 |
| CDCA4P2 | 0.007240781 |
| FZR1 | 0.007238495 |
| TUBB8P7 | 0.007236822 |
| LAMTOR3P1 | 0.007231498 |
| TEX261 | 0.007229973 |
| RNU6-1305P | 0.007223473 |
| SIGLEC15 | 0.00722294 |
| LRRC9 | 0.007222037 |
| ARMC4 | 0.007217949 |
| RP1-309K20 | 0.007217666 |
| OR7H1P | 0.007217611 |
| ATP6AP2 | 0.007213526 |
| PRR19 | 0.007211479 |
| RP11-477L16 | 0.007211005 |
| EFNA5 | 0.007207033 |
| RP11-285G1 | 0.007206952 |
| OR5B15P | 0.007202262 |
| MIR93 | 0.007201824 |
| RN7SKP132 | 0.00719781 |
| RNA5SP258 | 0.007196017 |
| GOLGA8J | 0.007195748 |
| RN7SKP288 | 0.007194645 |
| ABHD4 | 0.007190251 |

| | |
|---|---|
| UGT1A13P | 0.007186892 |
| AC010136 | 0.007185415 |
| DGCR5 | 0.007184157 |
| RP11-461F11 | 0.007180161 |
| COL21A1 | 0.007177431 |
| RP1-35C21 | 0.007176514 |
| Z73964 | 0.007176213 |
| NHP2L1 | 0.007175236 |
| CTD-2265D6 | 0.007171387 |
| RP11-541M12 | 0.007154336 |
| RP11-496I2 | 0.007153893 |
| LRP6 | 0.007152481 |
| FKBP1AP2 | 0.007148702 |
| LRRC3C | 0.007146749 |
| NAV2-IT1 | 0.007144844 |
| AC097463 | 0.007144337 |
| ZDHHC11 | 0.00714383 |
| REG3G | 0.007142586 |
| MIR6870 | 0.007142 |
| MCCD1P2 | 0.007141148 |
| DEF8 | 0.007126173 |
| MTRNR2L5 | 0.00712393 |
| HINT1 | 0.007121429 |
| RP11-307P5 | 0.007114513 |
| RP11-393J16 | 0.007109625 |
| RP11-428F8 | 0.007108437 |
| EDAR | 0.007107561 |
| RPS4XP13 | 0.007107531 |
| PSPN | 0.007106414 |
| RN7SL155P | 0.007105863 |
| SLC25A48 | 0.007103263 |
| RPL18AP16 | 0.007102747 |
| STX17-AS1 | 0.007102303 |
| RNU5E-3P | 0.007101166 |
| SDHAP1 | 0.007100647 |
| RP5-843L14 | 0.007097945 |
| RP11-492D6 | 0.007093894 |
| HNRNPCP10 | 0.00709309 |
| MYADM | 0.007088944 |
| FCF1 | 0.007083176 |
| SNORD115-8 | 0.00707879 |
| OR2L9P | 0.007077875 |
| PRPF19 | 0.007073594 |
| UCK1 | 0.007071008 |
| RP4-551E13 | 0.007068792 |

| | |
|---|---|
| RP11-70J12 | 0.007064841 |
| SLC16A1-AS1 | 0.007062505 |
| NAA50 | 0.007061519 |
| CELSR3-AS1 | 0.007055419 |
| RP11-560I19 | 0.007053583 |
| RP11-689C9 | 0.007052564 |
| LAPTM4B | 0.007050821 |
| EPS8L2 | 0.007050803 |
| TRMT1 | 0.007049221 |
| C12orf5 | 0.007048384 |
| DMD | 0.007047714 |
| RP11-47G4 | 0.007047146 |
| CTB-140J7 | 0.007046443 |
| LRRC6P1 | 0.007046291 |
| RP11-528A10 | 0.007045227 |
| EIF1P6 | 0.007044318 |
| RP11-106E7 | 0.007043536 |
| PRR15L | 0.007042486 |
| RP1-97J1 | 0.007034718 |
| RP11-843B15 | 0.007034199 |
| GCHFR | 0.007033797 |
| RP11-104O19 | 0.00702306 |
| RNU6-561P | 0.007019989 |
| AC007096 | 0.007014215 |
| RN7SL741P | 0.007012627 |
| SLC12A7 | 0.00701028 |
| XX-FYM637E10_5 | 0.007005609 |
| AC151960 | 0.007001263 |
| LINC00446 | 0.006999111 |
| CH17-340M24 | 0.006997126 |
| RNU6-1031P | 0.00699351 |
| AC093824 | 0.006991064 |
| MIR8066 | 0.006990637 |
| DIRC2 | 0.006985274 |
| IGHVII-62-1 | 0.006982823 |
| RN7SL810P | 0.006979717 |
| RN7SKP254 | 0.006976808 |
| MIR5692C1 | 0.006973907 |
| RP11-439I14 | 0.00696093 |
| SMIM9 | 0.006949523 |
| ARPC5L | 0.006946724 |
| PROSER2 | 0.00694371 |
| RP11-321G12 | 0.006943248 |
| RRP36 | 0.006942202 |
| CBWD3 | 0.006926555 |

| | |
|---|---|
| NAMA_1 | 0.006925993 |
| FANCC | 0.006925792 |
| RNU6-494P | 0.006922851 |
| PVRL2 | 0.006921662 |
| AC109351 | 0.006920965 |
| AC092380 | 0.006906587 |
| PROK2 | 0.006905313 |
| AC090060 | 0.00690102 |
| MIR6088 | 0.006894688 |
| THA1P | 0.006890988 |
| GEMIN7 | 0.006889903 |
| MED6P1 | 0.006886074 |
| TEX33 | 0.006885048 |
| RP11-268G13 | 0.006881634 |
| DCXR | 0.00687766 |
| RNF7P1 | 0.006877044 |
| XX-C2158C6 | 0.00687547 |
| CTD-2147F2 | 0.006874018 |
| SIX2 | 0.00687219 |
| AC010492 | 0.006869053 |
| GARNL3 | 0.006866832 |
| TAGLN2P1 | 0.006863725 |
| THOC7-AS1 | 0.006860416 |
| RN7SL117P | 0.006853807 |
| RPL23 | 0.006846459 |
| RP11-755B10 | 0.006845752 |
| CTC-458G6 | 0.006844751 |
| RPL30P12 | 0.006843539 |
| RNU6-820P | 0.006843499 |
| RP11-286O1 | 0.006834803 |
| VDAC1P3 | 0.006827467 |
| RPS3AP46 | 0.006811948 |
| HAP1 | 0.00680975 |
| RP3-405J10 | 0.006808336 |
| NUDT4P1 | 0.006807637 |
| AC093859 | 0.006807036 |
| RP11-69L16 | 0.006805322 |
| HOXA6 | 0.006804214 |
| RP11-223J15 | 0.006803778 |
| ERICH1 | 0.006800057 |
| BPIFB2 | 0.006798899 |
| MAGEA6 | 0.006795476 |
| SHANK3 | 0.006789433 |
| HNRNPA1P3 | 0.006789112 |
| MGLL | 0.00678694 |

| | |
|---|---|
| ACSM5P1 | 0.006784356 |
| TOLLIP | 0.006782754 |
| RNU6-1098P | 0.006782701 |
| RP3-332B22 | 0.006780657 |
| MTND1P3 | 0.006772834 |
| AC010196 | 0.006770659 |
| PRICKLE2-AS2 | 0.006768489 |
| AC128686 | 0.006765792 |
| AC012358 | 0.006764334 |
| AL390335 | 0.006763439 |
| RP11-363J17 | 0.006762685 |
| CALML3-AS1 | 0.006762502 |
| RP3-508D13 | 0.006760721 |
| RNU6-837P | 0.006758216 |
| RP11-1198D22 | 0.006756648 |
| AC091962 | 0.006753718 |
| ZNF710 | 0.006752185 |
| ATP5G1P3 | 0.006747113 |
| POU6F2-AS1 | 0.00674271 |
| CSN3 | 0.006729575 |
| RP11-335L23 | 0.006724334 |
| RP11-510J16 | 0.006722998 |
| RNU6-530P | 0.006715993 |
| PFN4 | 0.006715774 |
| CTD-2130F23 | 0.006705853 |
| CYP4F60P | 0.006695129 |
| CTC-448F2 | 0.006692926 |
| RP11-503P10 | 0.0066905 |
| MUC17 | 0.006690398 |
| WASH2P | 0.006681075 |
| RN7SKP110 | 0.006681062 |
| MIR3940 | 0.006679758 |
| KCNE3 | 0.006675456 |
| RN7SL52P | 0.006672868 |
| NPB | 0.006656731 |
| CDC14C | 0.006656057 |
| CYP4B1 | 0.006655288 |
| UBAC1 | 0.006654636 |
| MIR613 | 0.006651755 |
| CPA3 | 0.006651176 |
| SLC31A1 | 0.006649892 |
| OR2C3 | 0.006649053 |
| RPL19P18 | 0.006644118 |
| RN7SL241P | 0.00664358 |
| RP11-538C21 | 0.006641183 |

| | |
|---|---|
| RN7SKP44 | 0.006640987 |
| USP17L15 | 0.006633862 |
| OR4A10P | 0.006632293 |
| POU5F1P3 | 0.00663085 |
| RP11-289F5 | 0.006628764 |
| MIR5739 | 0.006628347 |
| WDR5 | 0.006627491 |
| GPR97 | 0.006626218 |
| CLDND2 | 0.006620235 |
| RNU6-1194P | 0.006619416 |
| RP11-91N2 | 0.006616959 |
| DPY19L2P1 | 0.006616892 |
| ZBTB20-AS2 | 0.006611406 |
| AC090018 | 0.006608828 |
| BET1 | 0.006602299 |
| TIMP4 | 0.00660122 |
| DDX11L8 | 0.006598629 |
| REG1P | 0.006597971 |
| CTD-2009A10 | 0.006596435 |
| CTA-228A9 | 0.006591266 |
| AP000745 | 0.006590734 |
| NAB1 | 0.006590299 |
| MOV10L1 | 0.006588848 |
| LMO3 | 0.006587061 |
| PGAM1P4 | 0.006584887 |
| AL133244 | 0.006583817 |
| OR4D12P | 0.006577281 |
| ATP5C1P1 | 0.006569256 |
| AC112219 | 0.006561967 |
| C15orf32 | 0.006559342 |
| TECR | 0.006553237 |
| RP11-191A15 | 0.006553098 |
| CTD-2203K17 | 0.006551934 |
| OR7G15P | 0.006548379 |
| RP11-157E16 | 0.006545885 |
| KNOP1 | 0.006544138 |
| AC087392 | 0.006543923 |
| FRMPD3-AS1 | 0.006542533 |
| OR5AR1 | 0.006542106 |
| KRTAP5-14P | 0.006541359 |
| CHCHD4P4 | 0.006538512 |
| CDH26 | 0.00653625 |
| ZC3H12B | 0.006533719 |
| RALGAPA2 | 0.006525421 |
| UBE2I | 0.006523859 |

| | |
|---|---|
| MORF4 | 0.00652357 |
| CUBNP2 | 0.00652349 |
| OTUD5 | 0.006523319 |
| AC092660 | 0.006519577 |
| SMIM4 | 0.006517347 |
| AP001601 | 0.006514024 |
| CPO | 0.006511213 |
| C9orf153 | 0.006504034 |
| RN7SKP89 | 0.006501182 |
| RP11-17E15 | 0.006497065 |
| AL356479 | 0.006497002 |
| OLA1P1 | 0.006490685 |
| RP11-308B5 | 0.006485995 |
| RPL5P6 | 0.006485539 |
| RPS27L | 0.00647995 |
| RNU6-164P | 0.006475933 |
| PIK3R1 | 0.006475001 |
| AL139340 | 0.006462965 |
| RP11-442G21 | 0.006461248 |
| AC116038 | 0.006451936 |
| UBTFL11 | 0.006448298 |
| SPANXA2 | 0.006447736 |
| BCL6 | 0.006439687 |
| HBG2 | 0.006436673 |
| RNU6-1242P | 0.00643641 |
| PCDHGA4 | 0.006435408 |
| LAMTOR5 | 0.006431613 |
| CTD-2571L23 | 0.006426816 |
| CTB-161C1 | 0.006420798 |
| RN7SL38P | 0.006420446 |
| AC087653 | 0.006407858 |
| RAI1 | 0.006404114 |
| IL4I1 | 0.00639909 |
| SDR16C5 | 0.006398555 |
| ATP5SL | 0.006397751 |
| RNU6-629P | 0.006383628 |
| UFC1 | 0.006380466 |
| ATP5LP2 | 0.00637631 |
| RN7SL87P | 0.006370732 |
| POLR3DP1 | 0.006367198 |
| KRASP1 | 0.006361407 |
| yR211F11 | 0.006360981 |
| BEND7 | 0.006357563 |
| VN1R5 | 0.006357112 |
| RPL6 | 0.00635654 |

| | |
|---|---|
| AC025300 | 0.006354725 |
| RP1-34H18 | 0.006354323 |
| RP11-307B6 | 0.006353682 |
| SETD1A | 0.006352614 |
| ZNF702P | 0.006349395 |
| RN7SKP71 | 0.006349043 |
| THEM4 | 0.006343572 |
| HAUS8P1 | 0.006334827 |
| FAM92A1P1 | 0.006325455 |
| OR4A8 | 0.006325147 |
| RNU7-116P | 0.006324671 |
| COLCA1 | 0.006322808 |
| PIH1D2 | 0.006320371 |
| STAT5A | 0.006319422 |
| TPTE | 0.006318905 |
| AL133391 | 0.006318744 |
| RP11-546O6 | 0.006316928 |
| TMEM167A | 0.006315905 |
| RN7SL549P | 0.006312254 |
| RNU6-673P | 0.006312249 |
| PPP4R4 | 0.006311648 |
| AC097711 | 0.006310411 |
| RP11-297A16 | 0.006309794 |
| PPP2R2D | 0.006309356 |
| AL360297 | 0.006308435 |
| C14orf119 | 0.006304612 |
| DNAH6 | 0.006302602 |
| RBMY2GP | 0.006300195 |
| FAM90A24P | 0.006297072 |
| RNU6-1190P | 0.006294331 |
| CTBP2P2 | 0.006288873 |
| AC008067 | 0.006287081 |
| SNORD115-11 | 0.006281476 |
| RNU4ATAC17P | 0.006277724 |
| SYTL4 | 0.006273782 |
| TLX2 | 0.006268869 |
| RP11-139E19 | 0.006268529 |
| RP11-510D21 | 0.006267345 |
| RN7SL306P | 0.006265453 |
| GPC5-AS2 | 0.006264268 |
| RP11-756P10 | 0.006256089 |
| FAM13A | 0.006248498 |
| SPATA33 | 0.006244851 |
| RP4-686C3 | 0.006244662 |
| OR1K1 | 0.006244277 |

| | |
|---|---|
| OR7A18P | 0.00624391 |
| ST13P16 | 0.006243714 |
| RP11-555K2 | 0.006242267 |
| CNGA3 | 0.006241937 |
| RP11-293A10 | 0.006241365 |
| IL6STP1 | 0.006237751 |
| RTN4 | 0.006234953 |
| RAB4B-EGLN2 | 0.006234749 |
| CRPP1 | 0.006232151 |
| SH3GL3 | 0.006229899 |
| C7orf60 | 0.006229706 |
| RP11-273G15 | 0.006215566 |
| TMEM161BP1 | 0.00621359 |
| RN7SKP91 | 0.006210897 |
| RP1-69B13 | 0.00621008 |
| AC092022 | 0.006209879 |
| LINC00865 | 0.006209538 |
| CCDC106 | 0.006207714 |
| RNU6-1086P | 0.006206265 |
| RP11-115D7 | 0.006200744 |
| HSPE1P27 | 0.006199471 |
| RP11-277J6 | 0.006198726 |
| RN7SL193P | 0.006190262 |
| C12orf50 | 0.006187439 |
| MIR101-1 | 0.006186389 |
| RP11-239L20 | 0.006183534 |
| DMBT1P1 | 0.0061831 |
| GHDC | 0.00617949 |
| KIF3C | 0.006171545 |
| ACTR3P3 | 0.006169316 |
| OR4K2 | 0.006168252 |
| SPTLC1P1 | 0.006166966 |
| SEMA3A | 0.006161033 |
| OR1L3 | 0.006159948 |
| AL596243 | 0.006155383 |
| RP11-136K14 | 0.006154089 |
| SMPD4P1 | 0.006151187 |
| RNU7-115P | 0.006150689 |
| TNP1 | 0.006143504 |
| IFI16 | 0.00614334 |
| BARHL1 | 0.006140505 |
| AC003986 | 0.006140254 |
| E4F1 | 0.006134502 |
| LHFPL3-AS1 | 0.006133592 |
| MIR7-3 | 0.006132971 |

| | |
|---|---|
| MIR4716 | 0.006130278 |
| RPL37P10 | 0.006120394 |
| COX6A1 | 0.006120045 |
| PCDH1 | 0.006115859 |
| DDX11L1 | 0.006115291 |
| OR14L1P | 0.006114236 |
| OGDHL | 0.006113446 |
| DHRS4L2 | 0.006109701 |
| HMGB3P20 | 0.006108501 |
| AP000577 | 0.006108465 |
| PPARGC1B | 0.006107561 |
| SNHG15 | 0.006107561 |
| RP11-479A21 | 0.00610314 |
| FDPSP1 | 0.006091626 |
| RP11-503L23 | 0.006089761 |
| AC007237 | 0.006086604 |
| RNU6-390P | 0.006085332 |
| AKR1B1P4 | 0.006084314 |
| CEACAMP10 | 0.006081879 |
| HMGB3P22 | 0.006077309 |
| GLTPD2 | 0.006077216 |
| RP11-168K9 | 0.006062693 |
| VASH2 | 0.006059522 |
| SIN3B | 0.006055391 |
| CYP4X1 | 0.006054424 |
| RPL36AP39 | 0.006051106 |
| PRM2 | 0.006050543 |
| LPHN3 | 0.006047168 |
| CTD-2015H3 | 0.006046898 |
| CTC-459M5 | 0.006045056 |
| MINOS1-NBL1 | 0.006043195 |
| SLC16A7 | 0.006040358 |
| GOLPH3L | 0.006039986 |
| RN7SL788P | 0.006037177 |
| RP11-145G20 | 0.006036473 |
| RN7SL214P | 0.00602392 |
| CRYBB1 | 0.006016822 |
| RP11-394I13 | 0.006016432 |
| RNU6-1047P | 0.006011884 |
| AL109805 | 0.006009363 |
| ARHGEF26-AS1 | 0.006009269 |
| OR4C1P | 0.006008277 |
| MIR602 | 0.006005703 |
| PITPNA-AS1 | 0.006004208 |
| CCDC154 | 0.006003464 |

| | |
|---|---|
| HSD3B1 | 0.006001623 |
| DEFB118 | 0.006001161 |
| CTBP1 | 0.00599738 |
| UBDP1 | 0.005997058 |
| FOXO3B | 0.00599671 |
| TPD52L3 | 0.005995163 |
| RPL18AP15 | 0.005994658 |
| RBM38 | 0.005992282 |
| RP11-224P11 | 0.005990429 |
| RP11-461M2 | 0.005982717 |
| RN7SKP136 | 0.005982277 |
| RBBP4P3 | 0.005969698 |
| MRPL18 | 0.005967462 |
| NAA20 | 0.00596376 |
| RP11-164H5 | 0.005958567 |
| AR | 0.005954996 |
| AC009411 | 0.005952123 |
| AOC3 | 0.005952113 |
| RP6-24A23 | 0.005950535 |
| AC005255 | 0.005949917 |
| RP11-384P14 | 0.005947217 |
| MIR3167 | 0.005946557 |
| DNAJC8 | 0.005945042 |
| MIR548X | 0.005936366 |
| RNU6-342P | 0.005933338 |
| OR4G1P | 0.005929115 |
| AC000003 | 0.005922474 |
| LACRT | 0.005910435 |
| RNU1-142P | 0.005907278 |
| SNRPGP13 | 0.005901314 |
| DUX4L17 | 0.005898654 |
| RP11-215A21 | 0.005892315 |
| SLC25A30-AS1 | 0.00589199 |
| SPATA17 | 0.005891849 |
| AL450327 | 0.005890961 |
| RN7SL100P | 0.005889857 |
| RP11-722P11 | 0.005884755 |
| RP11-923I11 | 0.005882993 |
| DLX6 | 0.005880467 |
| VN1R107P | 0.00588045 |
| D21S2088E | 0.005874868 |
| AC062031 | 0.005870007 |
| CDH18 | 0.005869844 |
| VKORC1L1 | 0.005869188 |
| RNU2-10P | 0.005868885 |

| | |
|---|---|
| CDRT15L2 | 0.005868333 |
| MUTYH | 0.005868072 |
| AC095056 | 0.005864686 |
| HCG17 | 0.005863713 |
| RN7SL836P | 0.00586266 |
| NPM1P12 | 0.005853535 |
| NANOG | 0.005852278 |
| ODF4 | 0.005851329 |
| ACA64 | 0.005850125 |
| CTB-167G5 | 0.005846553 |
| CRYBA1 | 0.005842382 |
| AC009800 | 0.005838121 |
| TBATA | 0.005837589 |
| NPRL3 | 0.005837503 |
| TMEM63C | 0.005836386 |
| HMGB1P12 | 0.005835172 |
| MARCKSL1P2 | 0.005826959 |
| TRIM16L | 0.005825223 |
| SLC7A11-AS1 | 0.005821721 |
| RN7SL712P | 0.005818843 |
| RNU6-690P | 0.005817425 |
| RN7SKP119 | 0.005817384 |
| CCT8 | 0.005817297 |
| MORN3 | 0.005816832 |
| SOCS5P3 | 0.005808082 |
| ANO7 | 0.005807734 |
| RP3-355L5 | 0.005806934 |
| TMEM249 | 0.005802974 |
| RNU4-70P | 0.005798105 |
| LINC01378 | 0.005797758 |
| RP4-587D13 | 0.00579676 |
| RNU6-755P | 0.005791141 |
| RNA5SP295 | 0.005789538 |
| ST6GAL1 | 0.005787826 |
| RN7SL681P | 0.005785412 |
| RPL21P112 | 0.00577573 |
| RP11-10G15 | 0.005774957 |
| PNMT | 0.005771979 |
| RP11-504A18 | 0.005771143 |
| BLOC1S5-TXNDC5 | 0.005757649 |
| RNU2-37P | 0.005755823 |
| DSP | 0.005754225 |
| OR1L6 | 0.005752675 |
| RN7SKP171 | 0.005750386 |
| RP11-277K23 | 0.005749014 |

| | |
|---|---|
| PCYT1B-AS1 | 0.005746233 |
| RP11-155D18 | 0.005742422 |
| RP4-761J14 | 0.005740345 |
| CTB-30L5 | 0.005734076 |
| C12orf56 | 0.005733303 |
| RN7SL263P | 0.005726691 |
| FTH1P5 | 0.005722484 |
| NPM1P45 | 0.005721473 |
| CYB561D2 | 0.005716232 |
| GSTA12P | 0.005714715 |
| CTD-2146O16 | 0.005712519 |
| RP11-390D11 | 0.005709073 |
| SLC25A1P5 | 0.005702907 |
| PPP1R32 | 0.005690836 |
| FTMT | 0.005687255 |
| CTD-2530H12 | 0.005682836 |
| DRD5 | 0.005682665 |
| RNU6-182P | 0.005682297 |
| HCG19P | 0.005679676 |
| RN7SL415P | 0.005673149 |
| AC010145 | 0.005664312 |
| RP11-539D10 | 0.005663956 |
| HPRT1 | 0.005658593 |
| NUTM2A | 0.005655637 |
| UBE2V2P1 | 0.005655534 |
| NSA2 | 0.005654499 |
| RP11-354E11 | 0.005652945 |
| COG4 | 0.005648992 |
| TRIML1 | 0.005629677 |
| HMGB3P27 | 0.005629175 |
| MRP63P3 | 0.005628577 |
| RP11-530I17 | 0.005627997 |
| RASSF8-AS1 | 0.005624545 |
| RP4-697K14 | 0.005623658 |
| RP3-523E19 | 0.005622428 |
| ULK4P2 | 0.005621064 |
| LRRC41 | 0.005619503 |
| OR7E99P | 0.005618278 |
| LPCAT3 | 0.005614851 |
| ARL5AP2 | 0.005614255 |
| BEX1 | 0.005612717 |
| NPIPA2 | 0.005603832 |
| KIF2B | 0.005595406 |
| MANF | 0.005595334 |
| LA16c-360A4 | 0.005590953 |

| | |
|---|---|
| GSTTP2 | 0.005587312 |
| RPSAP51 | 0.005584931 |
| FP325317 | 0.005582138 |
| MAPK6PS2 | 0.005579776 |
| RP11-758P17 | 0.005576787 |
| SDHDP1 | 0.005570594 |
| AC011754 | 0.005557735 |
| LINC00114 | 0.00555547 |
| ENPP1 | 0.005548672 |
| CTD-2377D24 | 0.005546764 |
| CCDC22 | 0.00554309 |
| RNU6-811P | 0.005538185 |
| AC104698 | 0.005535923 |
| RP11-513G19 | 0.005535111 |
| MIR4490 | 0.005527475 |
| RNU6-842P | 0.005527042 |
| LA16c-425C2 | 0.005526592 |
| NSUN5P1 | 0.005524373 |
| RP4-654H19 | 0.005524023 |
| RN7SL48P | 0.005521251 |
| Pinc | 0.005517126 |
| VN1R81P | 0.005516382 |
| REV3L-IT1 | 0.005514171 |
| CTNS | 0.005510214 |
| RP11-201M22 | 0.005501095 |
| MIR4446 | 0.005500387 |
| TCEB2P3 | 0.005499194 |
| RP4-605O3 | 0.005491292 |
| TUBA4B | 0.00549105 |
| A1BG | 0.005481915 |
| SKIL | 0.005480706 |
| LILRA6 | 0.005478901 |
| BST2 | 0.005474957 |
| MIR3919 | 0.005472959 |
| HTRA2 | 0.005468594 |
| PYROXD2 | 0.005467645 |
| PTPN3 | 0.005461114 |
| SULT2B1 | 0.005460686 |
| EEF1A1P2 | 0.00545984 |
| MIR3912 | 0.005458403 |
| FBXO46 | 0.005457505 |
| KCNA7 | 0.005457429 |
| AC010524 | 0.005456947 |
| RP11-126K1 | 0.005456165 |
| RP11-385D13 | 0.005454509 |

| | |
|---|---|
| RP11-293P20 | 0.005454149 |
| AL121756 | 0.00545345 |
| TSNAXIP1 | 0.005448742 |
| CDH6 | 0.005448239 |
| RPSAP35 | 0.005447564 |
| RP11-227H4 | 0.005442492 |
| RP11-447G11 | 0.005437959 |
| RNU6-1340P | 0.005437879 |
| MEP1AP4 | 0.005436346 |
| CT45A9 | 0.00543432 |
| RP13-16D10 | 0.005431919 |
| CHN1 | 0.00542969 |
| AC012640 | 0.005426273 |
| DNAL4 | 0.00541514 |
| RP11-525A16 | 0.005415021 |
| RP11-520D19 | 0.005413013 |
| RNF2 | 0.005411903 |
| PDZD7 | 0.005411307 |
| FAM71D | 0.005410733 |
| RP11-508N12 | 0.005406101 |
| RPL26P31 | 0.005391408 |
| AC009542 | 0.005391306 |
| MIR663A | 0.005390994 |
| RP11-492C2 | 0.005390994 |
| VWFP1 | 0.005380921 |
| ELL | 0.005379697 |
| RP11-680B3 | 0.005378657 |
| ABCA12 | 0.005377315 |
| MCFD2P1 | 0.005374471 |
| RP11-474J18 | 0.005368049 |
| KRTAP10-7 | 0.005366579 |
| MANBAL | 0.00536485 |
| PRKG2 | 0.005361073 |
| RN7SL41P | 0.005358736 |
| RP6-201G10 | 0.005358626 |
| MIR302E | 0.005353628 |
| RP11-552M6 | 0.005343931 |
| UGT2A1 | 0.005343378 |
| DUPD1 | 0.005332019 |
| RNU4-30P | 0.00533131 |
| RP11-552J9 | 0.005330002 |
| HSD3BP5 | 0.00532881 |
| PAQR9-AS1 | 0.005326082 |
| RP11-368N21 | 0.005321638 |
| GALP | 0.005319087 |

| | |
|---|---|
| C4orf45 | 0.005311246 |
| BRK1P2 | 0.005308727 |
| RP11-98G7 | 0.005307333 |
| CTC-329H14 | 0.005305797 |
| RP11-218E15 | 0.005304083 |
| NDUFA9P1 | 0.005302043 |
| AC112693 | 0.005297987 |
| GPR108 | 0.005296698 |
| UCP2 | 0.005293925 |
| ROPN1B | 0.005290503 |
| C12orf10 | 0.005287488 |
| RP3-366N23 | 0.005285971 |
| PES1P2 | 0.005284565 |
| RP11-147I11 | 0.005284305 |
| TRIM51JP | 0.005276822 |
| RP11-382N13 | 0.005276022 |
| AC105461 | 0.005273933 |
| SPATA18 | 0.005273273 |
| FAM160A1 | 0.005269683 |
| DDX19A | 0.005268045 |
| GJA4 | 0.005267915 |
| RPL7AP3 | 0.005264006 |
| YWHABP1 | 0.005263387 |
| FAM156A | 0.005262799 |
| MATN1 | 0.005260585 |
| RP5-1042I8 | 0.005259077 |
| RPS27P5 | 0.00525834 |
| BEAN1-AS1 | 0.005257604 |
| AP001482 | 0.005257483 |
| RBMX2P5 | 0.0052561 |
| TUBB4BP3 | 0.005252353 |
| HNRNPA1P41 | 0.005245612 |
| AC133749 | 0.00524006 |
| CACNA1B | 0.005232593 |
| SUCLA2P1 | 0.005228646 |
| HM13-AS1 | 0.005225862 |
| OR5W2 | 0.005224866 |
| RAB2A | 0.005222629 |
| AC012603 | 0.005221301 |
| SACS | 0.005221104 |
| CTD-3252C9 | 0.005218387 |
| RPL23AP35 | 0.005216771 |
| RP11-787B4 | 0.005200326 |
| ZMYND19 | 0.005197493 |
| RPS4XP2 | 0.005194014 |

| | |
|---|---|
| MAP1LC3C | 0.005191149 |
| MPDZ | 0.005186925 |
| LINC00115 | 0.005185679 |
| OR4A50P | 0.00518537 |
| ZNF99 | 0.00518229 |
| MIR4679-1 | 0.005181964 |
| AC078916 | 0.005174707 |
| R3HDM2P2 | 0.005174381 |
| RP11-1042B17 | 0.005174241 |
| WDPCP | 0.005172748 |
| CLDN19 | 0.005169624 |
| MUC22 | 0.00516691 |
| RP11-120B7 | 0.005164305 |
| TTLL11 | 0.005163222 |
| MIR4789 | 0.005163058 |
| MRPL22 | 0.005155743 |
| RP11-495O10 | 0.005154744 |
| RNA5SP459 | 0.005154455 |
| RNU6-9 | 0.005154337 |
| SNORA77 | 0.005152536 |
| AC104395 | 0.005143874 |
| MRPL42P4 | 0.005143516 |
| CCDC148-AS1 | 0.005137767 |
| LINC01173 | 0.005128981 |
| RP11-267L5 | 0.00511802 |
| RPL13P6 | 0.00511199 |
| RP11-31A23 | 0.005111823 |
| TM4SF4 | 0.005111548 |
| CTD-2309O5 | 0.005107613 |
| MIR568 | 0.005104364 |
| USP9YP4 | 0.005103962 |
| RP11-153M7 | 0.005103629 |
| EEF1DP6 | 0.005099359 |
| RNU1-115P | 0.00509384 |
| DARS | 0.005093749 |
| TMEM27 | 0.005090642 |
| MIR670HG | 0.005089804 |
| EEF1E1P1 | 0.005088088 |
| ARHGEF16 | 0.005086233 |
| ARL6IP5 | 0.005082567 |
| TRAPPC2P8 | 0.005081915 |
| RP5-1077I2 | 0.005077839 |
| PRDX3P4 | 0.005074855 |
| RP11-546M21 | 0.005069879 |
| FAM90A25P | 0.005067159 |

| | |
|---|---|
| VN2R19P | 0.005066432 |
| RP11-15B24 | 0.005063079 |
| RP11-463D19 | 0.005063013 |
| MIR3649 | 0.005056654 |
| SPAM1 | 0.005053898 |
| RP11-279E1 | 0.005051797 |
| PDE1C | 0.005051682 |
| LGALSL | 0.005046138 |
| RP1-7G5 | 0.005045193 |
| RP11-575A19 | 0.005043437 |
| RPSAP64 | 0.005042046 |
| RP11-402K9 | 0.005038278 |
| RNA5SP505 | 0.005038264 |
| RP11-36D19 | 0.005038208 |
| RNU6-1292P | 0.005038 |
| LENG1 | 0.005036035 |
| RP11-44N11 | 0.005031793 |
| ENTPD6 | 0.00503077 |
| RP11-506K19 | 0.005027926 |
| TVP23C-CDRT4 | 0.005026485 |
| AC004984 | 0.005021066 |
| RNU4-22P | 0.005020043 |
| TESC-AS1 | 0.005019854 |
| LSS | 0.005014099 |
| RNF40 | 0.005012127 |
| GPR113 | 0.005012062 |
| HULC | 0.005003435 |
| HMGB1P45 | 0.004999531 |
| C21orf140 | 0.004994344 |
| RP3-442L6 | 0.004993618 |
| AC091167 | 0.004989184 |
| MSNP1 | 0.004987311 |
| SLFN11 | 0.004977558 |
| ATP6V1D | 0.004977428 |
| TAF12 | 0.004976944 |
| SERPINB12 | 0.00497632 |
| CTD-2560K21 | 0.004975931 |
| RP11-314N13 | 0.004975809 |
| AC097499 | 0.004975195 |
| RP11-83J16 | 0.004970432 |
| OR8G7P | 0.004967847 |
| AC211469 | 0.004957933 |
| RP1-81D8 | 0.004952989 |
| RP1-149L1 | 0.00495004 |
| RP11-527H14 | 0.004949986 |

| | |
|---|---|
| SLC25A28 | 0.004947938 |
| ZNF629 | 0.004941761 |
| SPTLC1 | 0.00494111 |
| RN7SKP125 | 0.004939147 |
| RNA5SP145 | 0.004933667 |
| RP11-53B5 | 0.004933393 |
| RP1-190J20 | 0.004932346 |
| RP11-499A10 | 0.004930422 |
| TP53AIP1 | 0.004928944 |
| FRS3 | 0.004925072 |
| NUTM2A-AS1 | 0.004919323 |
| ASB1 | 0.004919081 |
| SLC22A18 | 0.004917016 |
| CFHR4 | 0.004911741 |
| RPS12P10 | 0.004911542 |
| RP3-389A20 | 0.00491001 |
| RNU6-699P | 0.004906735 |
| TSTD3 | 0.004905201 |
| MFAP1P1 | 0.004904035 |
| HFE2 | 0.004901453 |
| RP6-186E3 | 0.004899907 |
| RP11-735A19 | 0.004899297 |
| AC013409 | 0.004892015 |
| RP3-522D1 | 0.004890893 |
| SMAD1-AS2 | 0.00489083 |
| RPL19 | 0.004890205 |
| CTSL3P | 0.004889736 |
| RN7SKP117 | 0.004888503 |
| C1QTNF9 | 0.004881479 |
| RP11-601H16 | 0.004878316 |
| RP11-113C12 | 0.004868202 |
| MIR8076 | 0.004862238 |
| TEX37 | 0.004859711 |
| MAGEA3 | 0.00485375 |
| NCAPD2P1 | 0.004853093 |
| HIST1H4E | 0.004849705 |
| PEX3 | 0.00484691 |
| RP11-147K16 | 0.004844909 |
| ARHGEF10 | 0.004840622 |
| RNU6-1275P | 0.004832385 |
| PSMA6P1 | 0.004829347 |
| AC067956 | 0.004828401 |
| TMBIM7P | 0.004825929 |
| CCDC137 | 0.004824156 |
| HMHB1 | 0.004821996 |

| | |
|---|---|
| APOOP5 | 0.004815665 |
| RN7SKP232 | 0.004813932 |
| RPS27P20 | 0.00479418 |
| SLC35F5 | 0.00479368 |
| OR4F5 | 0.004793112 |
| LRRC37A5P | 0.004784937 |
| ESPNP | 0.004783265 |
| AC004549 | 0.004778832 |
| RNU6-864P | 0.00477781 |
| RNA5SP267 | 0.004773957 |
| pk | 0.004773847 |
| OR13C6P | 0.00476676 |
| LINC00499 | 0.004758887 |
| SNORA11C | 0.004757273 |
| RN7SL666P | 0.004756497 |
| TIGD2 | 0.00475459 |
| NEUROG2 | 0.00475208 |
| RBAK-RBAKDN | 0.004752078 |
| MTND1P30 | 0.004752051 |
| SLC26A9 | 0.004738004 |
| SNRPA | 0.004734403 |
| CLIC4P1 | 0.004731655 |
| MIR548H2 | 0.004728933 |
| SNORD12C | 0.004724162 |
| NFATC1 | 0.004721417 |
| ZNF652P1 | 0.004720995 |
| MIR1972-2 | 0.00471706 |
| HS3ST2 | 0.004715705 |
| SLC35G1 | 0.004712616 |
| TIMM17BP1 | 0.004709871 |
| TTTY20 | 0.00470859 |
| TAF9P1 | 0.00470859 |
| FAM197Y7 | 0.00470859 |
| CTD-2553L13 | 0.004702606 |
| NPHP1 | 0.004700765 |
| AC022201 | 0.004691838 |
| RPL7L1P2 | 0.004683937 |
| RP4-765C7 | 0.004683855 |
| ATP5I | 0.004681285 |
| MBD3L3 | 0.004678102 |
| SNORD113-8 | 0.004677013 |
| PDCL3 | 0.004673973 |
| ITSN1 | 0.004672205 |
| RPS24P17 | 0.004663984 |
| RN7SL461P | 0.004663656 |

| | |
|---|---|
| AC018450 | 0.004661756 |
| ARL9 | 0.004659786 |
| MICALL2 | 0.004658363 |
| MIR187 | 0.004652377 |
| RP11-155G14 | 0.004647972 |
| SHMT1 | 0.00464666 |
| CTD-2639E6 | 0.004646641 |
| RP11-356O24 | 0.004632219 |
| TOPORSLP | 0.004631878 |
| RP11-481J13 | 0.004631803 |
| PSMC1P2 | 0.004631192 |
| RP11-272B17 | 0.004626094 |
| PCSK7 | 0.004624594 |
| MIR8485 | 0.004623486 |
| OMP | 0.00461949 |
| CTA-989H11 | 0.004615775 |
| XAGE1B | 0.004612956 |
| RPS26P11 | 0.00461031 |
| RP5-991C6 | 0.004608114 |
| RP11-83N9 | 0.004607774 |
| SYT13 | 0.004606176 |
| RNU6-705P | 0.004600501 |
| CTD-2246P4 | 0.004600494 |
| EEF1A1P31 | 0.004598885 |
| PSORS1C3 | 0.004597521 |
| ARIH2OS | 0.004593262 |
| RP13-580F15 | 0.004591368 |
| C1orf229 | 0.004584732 |
| MIR141 | 0.004583355 |
| HNRNPA1P22 | 0.004582988 |
| UOX | 0.004579625 |
| CTTNBP2 | 0.004572461 |
| RP11-43A14 | 0.004550478 |
| RP11-3N2 | 0.004536402 |
| RNU6-245P | 0.004532008 |
| RPL26P30 | 0.004528744 |
| AC006366 | 0.004523676 |
| RP11-69C17 | 0.00452165 |
| CALM1P2 | 0.004518741 |
| RN7SL16P | 0.004518208 |
| AL928874 | 0.004517871 |
| LINC01525 | 0.004517833 |
| RP11-703G6 | 0.004513693 |
| CYCSP44 | 0.004511858 |
| RARS | 0.004509333 |

| | |
|---|---|
| LIPJ | 0.004505015 |
| MYLK3 | 0.004503398 |
| TPI1P3 | 0.004502734 |
| SIPA1 | 0.004500532 |
| AC107613 | 0.00449682 |
| ARMCX7P | 0.004495365 |
| AWAT1 | 0.004494001 |
| RPL7L1P8 | 0.004493683 |
| LINC00368 | 0.004493539 |
| PLEK2 | 0.004492179 |
| UBXN10-AS1 | 0.004487155 |
| SCARNA3 | 0.004485984 |
| FLJ41941 | 0.004474006 |
| AC007251 | 0.004472274 |
| RPL37AP8 | 0.004470186 |
| MIR1255B2 | 0.004466136 |
| GTF2F2 | 0.00446321 |
| AC007743 | 0.004462113 |
| ELOVL5 | 0.004456735 |
| AC073310 | 0.004455654 |
| ZSWIM7 | 0.004451879 |
| AC144525 | 0.004444545 |
| LINC01305 | 0.004443305 |
| PIGK | 0.004441646 |
| MYBPC1 | 0.004438287 |
| RP4-775D17 | 0.004437618 |
| RNU6-1236P | 0.004435628 |
| RP11-593F23 | 0.004434424 |
| OR4X2 | 0.004433143 |
| ENDOV | 0.004431395 |
| LRRC59 | 0.004429069 |
| RNU6-911P | 0.004427882 |
| RP11-545N8 | 0.004427269 |
| C11orf84 | 0.004418482 |
| RP5-1024N4 | 0.004415767 |
| GS1-383H3 | 0.004405776 |
| MIR551A | 0.004404494 |
| AC017074 | 0.004403375 |
| RP11-753B14 | 0.004394238 |
| ALG1L3P | 0.004391476 |
| RNU6-1210P | 0.004386374 |
| MLK4 | 0.00438622 |
| RP11-152O14 | 0.004378817 |
| RNU4-85P | 0.004377363 |
| RP11-1147P1 | 0.004377252 |

| | |
|---|---|
| AC079772 | 0.004375339 |
| RP11-58E21 | 0.004373978 |
| CASC23 | 0.004372792 |
| SMR3A | 0.004370576 |
| RNU6-1019P | 0.004370571 |
| AC012454 | 0.004366231 |
| AC019198 | 0.004364751 |
| RP11-397A16 | 0.00436422 |
| PI4K2A | 0.004360731 |
| OR2AF1P | 0.004358639 |
| RP11-494M8 | 0.004357126 |
| AL160291 | 0.004356326 |
| RLN3 | 0.004351429 |
| TRIM35 | 0.004350572 |
| RPL31P56 | 0.004350398 |
| IGKV3-34 | 0.004350349 |
| CTD-2001J20 | 0.004349524 |
| YIPF1 | 0.004342814 |
| EDA2R | 0.00434246 |
| DOCK4-AS1 | 0.004340025 |
| GM140 | 0.004335988 |
| FBXL7 | 0.004323167 |
| RP11-659F24 | 0.004319537 |
| RPL35P1 | 0.004318422 |
| AC107890 | 0.004317125 |
| AC104946 | 0.004315603 |
| CTB-113P19 | 0.004311482 |
| DEAF1 | 0.004310794 |
| RP5-934G17 | 0.00431004 |
| SSX3 | 0.004309319 |
| SQSTM1P1 | 0.004308825 |
| RP11-618M23 | 0.004307618 |
| TSPY10 | 0.00430582 |
| RP11-183A22 | 0.004300383 |
| RP1-278O22 | 0.004299517 |
| RNU6-579P | 0.004299481 |
| NKX3-1 | 0.004292728 |
| MIR26B | 0.004291097 |
| RP11-632C17__A | 0.004290272 |
| RP11-145M9 | 0.004283984 |
| RN7SL521P | 0.004279258 |
| RP11-283C24 | 0.004273695 |
| AP000860 | 0.004273368 |
| THSD1 | 0.004259964 |
| SRGAP3-AS1 | 0.004255096 |

| | |
|---|---|
| FAM185BP | 0.004254401 |
| DLGAP1-AS4 | 0.004251587 |
| FLJ36000 | 0.004250918 |
| RP11-300A12 | 0.004248131 |
| IQGAP1 | 0.004241898 |
| SNX33P1 | 0.004240239 |
| RP11-187E13 | 0.004239052 |
| RN7SKP144 | 0.004237833 |
| CLPTM1LP1 | 0.004226748 |
| RN7SKP133 | 0.004225709 |
| TRPM6 | 0.004224034 |
| KRTAP11-1 | 0.004222689 |
| RP3-380B4 | 0.004218025 |
| CYP4F62P | 0.004216258 |
| FAM196B | 0.004210879 |
| AC083829 | 0.004205195 |
| PSMC1P11 | 0.004200004 |
| MIR3116-1 | 0.00418846 |
| RP11-64J4 | 0.004186517 |
| SRP68P2 | 0.004185617 |
| MIR4456 | 0.004183594 |
| HDAC4 | 0.004176928 |
| CTD-2318H23 | 0.004176262 |
| SPATS1 | 0.004175366 |
| PTPLAD2 | 0.004169969 |
| RNU6-56P | 0.004167292 |
| COX6CP15 | 0.004161985 |
| GGT3P | 0.004158579 |
| SNRPCP6 | 0.004148138 |
| ST13P20 | 0.004140999 |
| RPS4XP5 | 0.004140672 |
| KBTBD12 | 0.004139467 |
| CDH20 | 0.004138601 |
| TFAP2D | 0.00413265 |
| AP000428 | 0.004132628 |
| RPLP0P7 | 0.004123206 |
| RP11-296I10 | 0.004121222 |
| CTD-2540B15 | 0.004118998 |
| AC096554 | 0.004117874 |
| AF124730 | 0.004117007 |
| RP11-219O3 | 0.004113031 |
| AC010723 | 0.004111558 |
| MIR1202 | 0.004109262 |
| CTD-2308B18 | 0.004109058 |
| CTC-432M15 | 0.00409729 |

| | |
|---|---|
| PCDH19 | 0.004097036 |
| CRYGB | 0.004085864 |
| OR1AC1P | 0.004085037 |
| CCDC110 | 0.004079528 |
| RPL31P7 | 0.004079309 |
| CDX1 | 0.00407845 |
| EEF1E1 | 0.004076272 |
| RP11-53M11 | 0.00407177 |
| MAFA | 0.00407102 |
| GALNT11 | 0.004057964 |
| NEIL2 | 0.004057406 |
| ABCC8 | 0.0040559 |
| ETF1P3 | 0.004050289 |
| RN7SL845P | 0.00404994 |
| GRM5-AS1 | 0.00404027 |
| AC023137 | 0.004036779 |
| RP11-318A15 | 0.004027825 |
| MIR487B | 0.004025606 |
| ALDOC | 0.004024902 |
| ADIPOR1P2 | 0.004024539 |
| CTB-46B19 | 0.004024075 |
| TPI1P1 | 0.004020493 |
| GTF2A1L | 0.004020054 |
| SCN4B | 0.004015451 |
| AC068286 | 0.004014205 |
| ACTR3 | 0.004007767 |
| AL080276 | 0.004005883 |
| AL645937 | 0.004003011 |
| CH17-76K2 | 0.004000889 |
| PCK1 | 0.003998431 |
| FAM195DP | 0.003997989 |
| RN7SKP270 | 0.003997901 |
| DNAH5 | 0.003994073 |
| ENTPD2 | 0.003989626 |
| MIR4508 | 0.003988789 |
| SEMA6D | 0.003988305 |
| TMEM61 | 0.003987152 |
| CELF2-AS2 | 0.00398284 |
| CTD-2384B9 | 0.003979645 |
| RP11-563N12 | 0.003976096 |
| ABCC4 | 0.003975337 |
| RP11-589P10 | 0.003973458 |
| RP11-472F14 | 0.003970511 |
| BCAS2 | 0.00396836 |
| E2F3 | 0.003965532 |

| | |
|---|---|
| HLFP1 | 0.003955081 |
| MIR4773-1 | 0.003950596 |
| MIR548U | 0.003945904 |
| CTD-2182N23 | 0.003940878 |
| AL354936 | 0.003940856 |
| RPL34P6 | 0.003936739 |
| AC093590 | 0.003935307 |
| DPRXP6 | 0.003931947 |
| CTB-92J24 | 0.003923949 |
| AARSP1 | 0.003920127 |
| RP11-302F12 | 0.003902251 |
| YWHAEP5 | 0.003900181 |
| MCHR1 | 0.003899409 |
| OR9N1P | 0.003890467 |
| AL161651 | 0.003889705 |
| RNA5SP510 | 0.003889705 |
| MT-CO1 | 0.003870812 |
| RNU6-198P | 0.003869188 |
| RP11-332H18 | 0.003864563 |
| RP11-106J23 | 0.003862807 |
| AC024569 | 0.003854995 |
| RP11-413M3 | 0.003854149 |
| DNAH14 | 0.003850737 |
| CLEC2B | 0.003849843 |
| RP4-811H24 | 0.003849642 |
| GRINA | 0.003840336 |
| PTGER2 | 0.003839927 |
| SZRD1 | 0.003836522 |
| CTD-2007F2 | 0.003829908 |
| CTA-326K9 | 0.003828279 |
| RP11-697K23 | 0.003828146 |
| ZNF788 | 0.003827926 |
| RNASE2 | 0.003824187 |
| RNA5SP89 | 0.003821034 |
| RN7SL363P | 0.003813048 |
| DHRS4 | 0.003807888 |
| APOC4-APOC2 | 0.003800471 |
| LINC01572 | 0.003798215 |
| MIR7154 | 0.003788911 |
| ECT2L | 0.003787869 |
| CYBA | 0.003785171 |
| RP11-459K23 | 0.003784472 |
| MIR3121 | 0.003780037 |
| RP11-78E6 | 0.003779774 |
| MFSD7 | 0.003777803 |

| | |
|---|---|
| MRPL48 | 0.003776389 |
| ZMYM3 | 0.003775217 |
| KRT8P7 | 0.003771857 |
| RP11-137P24 | 0.003767679 |
| EIF3EP2 | 0.00375378 |
| AC121157 | 0.003753224 |
| RP11-85D18 | 0.003744298 |
| RNU6-68P | 0.003742462 |
| PFN1P9 | 0.003742255 |
| LIN28B | 0.003741305 |
| MTND4P18 | 0.003741119 |
| A3GALT2 | 0.00373934 |
| RNU5A-7P | 0.003737889 |
| CUBNP3 | 0.003733701 |
| OR8I1P | 0.00373269 |
| RPS2P24 | 0.003732309 |
| MRPL30P1 | 0.003732278 |
| GNAQP1 | 0.003731759 |
| RNU6-162P | 0.003730788 |
| RP11-359P18 | 0.003725267 |
| AC008085 | 0.003717034 |
| SCGB1D4 | 0.003707488 |
| AL355490 | 0.003696491 |
| CTD-3093B17 | 0.003690446 |
| RP11-34H11 | 0.003686569 |
| ZGLP1 | 0.003680794 |
| POLR2LP | 0.003678871 |
| RSPRY1 | 0.003678431 |
| GM2AP2 | 0.003669193 |
| MIR548F4 | 0.003665126 |
| RN7SL236P | 0.003663957 |
| RP11-94H18 | 0.003663333 |
| RPS18P9 | 0.003657705 |
| FAM124B | 0.003653426 |
| KRT8P25 | 0.003651441 |
| RP11-755E23 | 0.003648472 |
| AF038458 | 0.003648066 |
| TRIM77 | 0.003643984 |
| VN1R25P | 0.003642359 |
| AC104183 | 0.003641007 |
| RNU6-1191P | 0.003640298 |
| WBP1LP6 | 0.003638239 |
| DHRS11 | 0.003635955 |
| RSPH14 | 0.003635101 |
| ACTR3BP1 | 0.003631869 |

| | |
|---|---|
| RP11-11P22 | 0.003628955 |
| RNA5SP169 | 0.003627462 |
| RNA5SP212 | 0.003626825 |
| IFNA10 | 0.00362669 |
| RHOQP2 | 0.003609467 |
| RNU6-1114P | 0.003606508 |
| ZFAS1 | 0.003598074 |
| RP11-332L8 | 0.003593608 |
| RP11-543H23 | 0.003592528 |
| BPIFB6 | 0.003590756 |
| RPL7L1P7 | 0.003589559 |
| RP3-518E13 | 0.003588057 |
| RP11-297M9 | 0.003587876 |
| RP11-887P2 | 0.003586967 |
| AL354749 | 0.003577066 |
| RP11-447M12 | 0.003566505 |
| BCAN | 0.003563319 |
| GAPDHP73 | 0.003559132 |
| AC005392 | 0.003558957 |
| SFXN3 | 0.003553105 |
| PRMT8 | 0.003549727 |
| RP1-203C2 | 0.003549486 |
| PCDHA4 | 0.00354882 |
| CTD-2588J1 | 0.003548732 |
| LINC00558 | 0.003547313 |
| ABCC11 | 0.003546184 |
| RPS11P1 | 0.003543951 |
| MIR6128 | 0.003539212 |
| ZNF677 | 0.00353569 |
| CNN2P2 | 0.003534463 |
| CTB-37A13 | 0.003532828 |
| TPRKBP1 | 0.003528517 |
| RP11-458K10 | 0.003528303 |
| MIR6788 | 0.003527784 |
| FAM229A | 0.003524471 |
| RP4-677H15 | 0.003522587 |
| CTD-2091N23 | 0.003520585 |
| CTD-2515A14 | 0.00351343 |
| AC008277 | 0.003510364 |
| RNA5-8SP2 | 0.003497224 |
| RPL7P38 | 0.003494903 |
| EPHA5-AS1 | 0.003494893 |
| RP11-565H13 | 0.003482167 |
| CTD-2320J21 | 0.003482097 |
| OR4A7P | 0.003480362 |

| | |
|---|---|
| CAMKV | 0.003480106 |
| MAPRE1P1 | 0.003479976 |
| RN7SKP206 | 0.003477671 |
| OCRL | 0.003471576 |
| RNU7-164P | 0.003468988 |
| VPS4A | 0.003468859 |
| CSDE1 | 0.003464357 |
| OR7E155P | 0.003463659 |
| RP11-343B18 | 0.003462682 |
| RP11-379P15 | 0.003462426 |
| RP11-11N9 | 0.003462329 |
| RNU6-787P | 0.003460176 |
| C22orf42 | 0.003455735 |
| ZNF354C | 0.003445305 |
| MAP3K4 | 0.003444561 |
| RP5-872K7 | 0.003441378 |
| KRT8P45 | 0.003440802 |
| RPL17P2 | 0.003431613 |
| CTRB1 | 0.003431364 |
| CYCSP32 | 0.003430614 |
| OR4L1 | 0.003424935 |
| DDX41 | 0.003421192 |
| RP11-184C23 | 0.00342024 |
| U52111 | 0.003420113 |
| RPUSD3 | 0.003410978 |
| RP11-331F4 | 0.003402903 |
| RNF10P1 | 0.003402711 |
| RPS14P4 | 0.003394162 |
| RP11-781E19 | 0.003392274 |
| FLJ37453 | 0.003387442 |
| FAM83A-AS1 | 0.0033848 |
| RNU6-345P | 0.003380729 |
| MECOM | 0.003379404 |
| SNX31 | 0.003375768 |
| SLC51B | 0.003374134 |
| MT-TD | 0.003370343 |
| RP11-351I21 | 0.003370285 |
| FTH1P11 | 0.003369525 |
| RP11-244B22 | 0.003363715 |
| RNA5SP139 | 0.003362476 |
| RP11-157I4 | 0.003357504 |
| CRIP1P3 | 0.003353908 |
| SYCE3 | 0.003351468 |
| RP11-98H4 | 0.003350666 |
| RP13-46H24 | 0.003349009 |

| | |
|---|---|
| RP11-45O22 | 0.003343781 |
| RSPH10B2 | 0.003342285 |
| NDUFV2 | 0.00334171 |
| LACTBL1 | 0.003341213 |
| AGBL3 | 0.003340872 |
| CTC-270D5 | 0.003331344 |
| PTPN20CP | 0.003328861 |
| EEF1A1P11 | 0.003326211 |
| RP11-297L6 | 0.003326024 |
| RP11-71J2 | 0.003325556 |
| CTD-2506P8 | 0.003324834 |
| PYDC1 | 0.003324508 |
| IGSF22 | 0.003322192 |
| EID2 | 0.003321299 |
| MIR3123 | 0.00332056 |
| RP5-1096D14 | 0.003318459 |
| CTC-313D10 | 0.003316032 |
| AC084809 | 0.003315178 |
| GMPPB | 0.003312145 |
| MYCBP | 0.003311364 |
| RP1-34M23 | 0.003308694 |
| CITED2 | 0.003308294 |
| AREG | 0.003304089 |
| DGCR9 | 0.003295283 |
| OR8K2P | 0.003293085 |
| DYRK4 | 0.003291698 |
| RP11-475O23 | 0.003286062 |
| NOTCH2NL | 0.003284191 |
| RPL36AP33 | 0.003283653 |
| OR51G2 | 0.003283358 |
| AP001412 | 0.003278252 |
| MIR3688-1 | 0.003276732 |
| RP11-45D17 | 0.00327552 |
| YBX1P4 | 0.003273935 |
| KCNK15 | 0.003273558 |
| AC000110 | 0.003273468 |
| FAM24A | 0.003268103 |
| LINC00251 | 0.003267746 |
| NLRP9 | 0.003261945 |
| OOEP | 0.003258109 |
| RNA5SP138 | 0.003257567 |
| GANAB | 0.003254934 |
| OR2A4 | 0.00324765 |
| ZNF839P1 | 0.003245192 |
| RP11-6N17 | 0.003238189 |

| | |
|---|---|
| RP3-430A16 | 0.003237159 |
| PIP5K1C | 0.003229909 |
| AC091947 | 0.003225301 |
| RPS4XP9 | 0.003223339 |
| ZNF648 | 0.00322248 |
| RP11-206P5 | 0.003219521 |
| TRPM5 | 0.003218182 |
| RP11-73B8 | 0.003211132 |
| LL22NC03-86D4 | 0.003210987 |
| MIR6770-1 | 0.0032073 |
| RNU11-2P | 0.003203691 |
| TRIM43B | 0.003197652 |
| PPP4R1 | 0.003196568 |
| AC026449 | 0.003193702 |
| C1orf87 | 0.003190569 |
| CTA-345G4 | 0.00319009 |
| RP11-706F1 | 0.003189303 |
| RP11-182J23 | 0.003184987 |
| RP11-798K23 | 0.003179135 |
| IFT52 | 0.003172883 |
| LINC01154 | 0.003165468 |
| ANKRD30BP1 | 0.003161124 |
| PCYOX1L | 0.003159711 |
| FAM231C | 0.003157917 |
| CTTN | 0.003156325 |
| SNX22 | 0.003153794 |
| RP11-160H22 | 0.003153593 |
| FAM171B | 0.003151952 |
| RP11-160O5 | 0.003148343 |
| CTB-5E10 | 0.003146347 |
| AMMECR1LP1 | 0.003146337 |
| RP11-63D14 | 0.003146233 |
| NPTN | 0.003145533 |
| DHDDS | 0.00314185 |
| PFDN6 | 0.003136156 |
| ACD | 0.003134072 |
| CRYBB3 | 0.003133822 |
| LCE1E | 0.003126793 |
| LINC01551 | 0.003124716 |
| RP11-604N13 | 0.003124262 |
| RNU4-9P | 0.003122182 |
| AC018361 | 0.003120665 |
| NIM1K | 0.003117216 |
| LINC01044 | 0.003105922 |
| RP11-168F9 | 0.003101708 |

| | |
|---|---|
| RN7SL808P | 0.003099826 |
| TDRD9 | 0.003092992 |
| ZIK1 | 0.00309234 |
| PSMD5 | 0.003092303 |
| RPL9P5 | 0.003089731 |
| PDZRN4 | 0.00308808 |
| AC099512 | 0.003086726 |
| RP11-356N1 | 0.003076582 |
| RNA5SP349 | 0.00307082 |
| RP11-62E14 | 0.003070332 |
| GCFC2 | 0.003067595 |
| RP11-177A2 | 0.003066233 |
| STEAP2-AS1 | 0.003065091 |
| UCN3 | 0.003062186 |
| RHPN1 | 0.003061009 |
| RN7SL340P | 0.003059764 |
| MIR3661 | 0.003048389 |
| FKSG56 | 0.003045971 |
| RNU6-235P | 0.00304517 |
| FARSBP1 | 0.003044236 |
| OR4F14P | 0.00304171 |
| RP11-388E23 | 0.003036004 |
| SERPIND1 | 0.003035765 |
| RN7SL240P | 0.003033482 |
| AP001442 | 0.003032658 |
| CTC-510F12 | 0.003031891 |
| RP11-729L2 | 0.003028955 |
| RP11-287J9 | 0.00302504 |
| MRPL21 | 0.00302257 |
| PDSS1P1 | 0.003021548 |
| OR2A3P | 0.003021419 |
| LINC01384 | 0.003014296 |
| PPP1R1AP2 | 0.003007603 |
| RP5-1033H22 | 0.003004262 |
| RP13-33H18 | 0.003002432 |
| LDHAL6FP | 0.002998787 |
| RP11-408E1 | 0.002993193 |
| BANF1P1 | 0.002992959 |
| AS3MT | 0.002991326 |
| OSTM1-AS1 | 0.002988365 |
| EGFR-AS1 | 0.002987158 |
| FTH1P8 | 0.00298681 |
| RPL36AP55 | 0.00298678 |
| AC104405 | 0.002978358 |
| RPS10P2 | 0.002977173 |

| | |
|---|---|
| RN7SL571P | 0.002974049 |
| MUC16 | 0.002971889 |
| PPP1R3C | 0.002970009 |
| SNORD115-23 | 0.002963253 |
| RN7SKP159 | 0.002963253 |
| RP11-201A3 | 0.00296005 |
| TCEB3C | 0.002954633 |
| CTD-2025J6 | 0.002952084 |
| LIPC | 0.002948706 |
| GACAT3 | 0.002948603 |
| PPP2R5B | 0.002946753 |
| DPH3 | 0.002945971 |
| AP001465 | 0.002944536 |
| TMEM256P1 | 0.002943296 |
| RNU6-469P | 0.002939413 |
| RP11-366M4 | 0.002938231 |
| MBD3L1 | 0.002936571 |
| UBE2R2 | 0.002936166 |
| CLSTN2-AS1 | 0.002934302 |
| ALG1L8P | 0.002930778 |
| AC008686 | 0.002926642 |
| AKT3-IT1 | 0.002923079 |
| CRYAA | 0.002921625 |
| FCF1P6 | 0.002912422 |
| HYDIN2 | 0.002911897 |
| RP11-861L17 | 0.002911456 |
| RPS20P23 | 0.002911393 |
| RP11-192P3 | 0.002909635 |
| PRPF19P1 | 0.002907634 |
| RP11-79N23 | 0.002906333 |
| GAPDHP52 | 0.002901598 |
| FDPSP6 | 0.002898335 |
| AL161740 | 0.002892635 |
| RP1-69E11 | 0.00288917 |
| CACNG8 | 0.002887913 |
| RP11-685N10 | 0.002884232 |
| AC024937 | 0.002883271 |
| ZNF691 | 0.002881857 |
| PCAT19 | 0.002880982 |
| KRTAP8-2P | 0.00287621 |
| CTD-3118D7 | 0.002875438 |
| RNU6-970P | 0.002873795 |
| AC093166 | 0.002868583 |
| GPR173 | 0.002867438 |
| KIAA1522 | 0.002866954 |

| | | |
|---|---|---|
| MSANTD2P1 | | 0.002865677 |
| UBE3AP2 | | 0.002863182 |
| GABRG1 | | 0.002857955 |
| MIR3660 | | 0.002856388 |
| AC105921 | | 0.00285355 |
| EIF5A2P1 | | 0.00285124 |
| MTND1P14 | | 0.002844068 |
| DERL2 | | 0.002838525 |
| SLC4A4 | | 0.002836683 |
| BAGE5 | | 0.002833611 |
| DUSP3 | | 0.002824403 |
| RPS3AP26 | | 0.00282431 |
| GOLGA6L17P | | 0.00281971 |
| RN7SKP269 | | 0.002819513 |
| | 2-Sep | 0.002818505 |
| SNORD38A | | 0.002817743 |
| PARD3 | | 0.002817482 |
| FBXO16 | | 0.002816905 |
| NUTF2P6 | | 0.002812367 |
| NME1 | | 0.002806746 |
| GPR53P | | 0.002804016 |
| TOMM22P1 | | 0.002803129 |
| RP11-100K18 | | 0.002802228 |
| RPL21P28 | | 0.002796755 |
| OR6C66P | | 0.002796663 |
| CDKN2B-AS | | 0.002796316 |
| RBM22P5 | | 0.002792886 |
| PCDHGC4 | | 0.002787338 |
| PKMP2 | | 0.002787102 |
| NDUFAF4 | | 0.002785834 |
| MROH7 | | 0.002781179 |
| GAPDHP67 | | 0.002777156 |
| OR10D1P | | 0.002771505 |
| RP11-516J2 | | 0.002765501 |
| RP11-196O2 | | 0.002761687 |
| LARP1BP3 | | 0.002761461 |
| RP11-90D11 | | 0.002759187 |
| WASF3 | | 0.002758298 |
| RP11-326E7 | | 0.002754014 |
| XXbac-B461K10 | | 0.002746714 |
| CUL7 | | 0.002743972 |
| RNU6-997P | | 0.00274194 |
| TRAPPC9 | | 0.002741627 |
| MAGI2 | | 0.002741348 |
| MIR6772 | | 0.002740444 |

| | |
|---|---|
| RP1-217P22 | 0.002734594 |
| MYLK4 | 0.002722743 |
| OR9I2P | 0.00271979 |
| SLC25A15P3 | 0.002714782 |
| SNORA31 | 0.00271413 |
| KRT8P28 | 0.002711457 |
| RP11-27M5 | 0.002710609 |
| AC104637 | 0.002708889 |
| RP11-307P22 | 0.002701837 |
| CNDP1 | 0.002700225 |
| AC018616 | 0.002699195 |
| RNU6-1042P | 0.002697067 |
| WDR25 | 0.002694072 |
| RHBDL1 | 0.002693309 |
| RNU6-36P | 0.002690893 |
| OR51E1 | 0.002688899 |
| RNU6-646P | 0.002687089 |
| RNA5SP259 | 0.002685849 |
| ARNT | 0.002683541 |
| RP11-701P16 | 0.002683055 |
| RRNAD1 | 0.002675448 |
| ESRRAP1 | 0.0026747 |
| C10orf67 | 0.002672639 |
| LEMD1-AS1 | 0.00266403 |
| MIR4436B2 | 0.002661377 |
| IFITM8P | 0.002656395 |
| TAB3P1 | 0.002645768 |
| TPT1 | 0.002645493 |
| BMP7-AS1 | 0.002644937 |
| RP11-483I24 | 0.002642837 |
| ATP8A2P1 | 0.002636996 |
| AC007677 | 0.002635875 |
| AC090945 | 0.002634247 |
| CYP4A26P | 0.002633568 |
| RN7SL738P | 0.002626941 |
| LINC01348 | 0.002619995 |
| SNORA49 | 0.002618447 |
| TMEM127 | 0.00261671 |
| RPL37P21 | 0.002612439 |
| C1orf112 | 0.002609808 |
| AC093724 | 0.002609305 |
| TMEM69 | 0.002608952 |
| RP11-486A14 | 0.002606199 |
| AC005510 | 0.002599142 |
| IGHVIII-76-1 | 0.002598323 |

| | |
|---|---|
| ERLIN1 | 0.002594607 |
| RP11-214N16 | 0.002593202 |
| MTUS1 | 0.002589211 |
| KRTAP6-1 | 0.002586466 |
| GOLGA6L3 | 0.002585816 |
| RNU6-887P | 0.002583456 |
| CTD-2207A17 | 0.002578956 |
| CD320 | 0.002573825 |
| AKAP14 | 0.002570832 |
| XAB2 | 0.002567796 |
| IAPP | 0.002564937 |
| RPL7P13 | 0.002563519 |
| EIF3FP3 | 0.002558047 |
| DGAT2L7P | 0.002555865 |
| PRSS37 | 0.002555571 |
| AL022400 | 0.002554977 |
| RP11-204L24 | 0.00255484 |
| CCDC155 | 0.002551111 |
| RPL21 | 0.002548484 |
| LINC00398 | 0.002547442 |
| MT-TR | 0.002543094 |
| RP1-303A1 | 0.002534686 |
| MTND1P15 | 0.002533972 |
| SRIP2 | 0.002530686 |
| MSLNL | 0.002530519 |
| NPM1P50 | 0.002529093 |
| RP11-48B14 | 0.002527311 |
| ASCC3 | 0.002525953 |
| MAST4-AS1 | 0.002519182 |
| NOMO3 | 0.002519143 |
| SNORD115-47 | 0.002518341 |
| APOM | 0.002516352 |
| MIR3651 | 0.002516014 |
| RP11-1057B8 | 0.002513132 |
| PGA5 | 0.002512791 |
| RP11-306K13 | 0.002512074 |
| ST8SIA6 | 0.002511329 |
| RP11-740D6 | 0.002510559 |
| CTC-448D22 | 0.002508269 |
| RP11-320M2 | 0.002505022 |
| LINC00265-2P | 0.002503776 |
| KAAG1 | 0.002500246 |
| ODF1 | 0.002495828 |
| CKS1BP7 | 0.002495054 |
| RPL21P50 | 0.002494511 |

| | |
|---|---|
| HCN1 | 0.002489914 |
| RP11-263E1 | 0.002488121 |
| RPL21P1 | 0.002488111 |
| MIR15A | 0.002477686 |
| IRAK3 | 0.002477634 |
| RN7SL311P | 0.002475871 |
| AC073133 | 0.002475473 |
| RNA5SP119 | 0.002473307 |
| RP11-544D21 | 0.002472821 |
| RP11-313I2 | 0.002471365 |
| RP11-319G6 | 0.002470415 |
| TCIRG1 | 0.002469671 |
| PSAT1P3 | 0.002465996 |
| RP5-926E3 | 0.002464869 |
| RN7SKP216 | 0.00245825 |
| SFR1 | 0.002455271 |
| ABCB10P3 | 0.002452533 |
| RP11-665E10 | 0.002450024 |
| MIR206 | 0.002449833 |
| BIN2P2 | 0.002449742 |
| RP4-612C19 | 0.002447704 |
| PLS1 | 0.002446303 |
| MIR874 | 0.0024449 |
| RNU7-48P | 0.002443568 |
| AL590708 | 0.002443278 |
| PLEKHA6 | 0.002441521 |
| CASKIN2 | 0.002441503 |
| SLC10A4 | 0.002434652 |
| RPL29P11 | 0.00243246 |
| RP11-36C20 | 0.002431809 |
| GSTO2 | 0.002430042 |
| RNU1-52P | 0.002429905 |
| PTGS1 | 0.002427288 |
| AC079248 | 0.002425884 |
| LYPLA2P2 | 0.002422998 |
| KRT18P60 | 0.002422487 |
| RP5-998N21 | 0.002407807 |
| RP11-15G16 | 0.00240534 |
| DAPK2 | 0.002399454 |
| TRDD1 | 0.002399085 |
| LINC00883 | 0.002392272 |
| AFP | 0.002391609 |
| GHc-602D8 | 0.002390248 |
| RP11-138J23 | 0.002388538 |
| USP17L7 | 0.002382623 |

| | |
|---|---|
| RN7SL597P | 0.002382582 |
| UBE2V1P10 | 0.002382107 |
| KRT18P9 | 0.002376296 |
| RN7SL51P | 0.002375239 |
| CTC-367F4 | 0.002367138 |
| TMCC1 | 0.002366281 |
| KCTD21 | 0.00235668 |
| TESK2 | 0.002352717 |
| MRPS10P2 | 0.002352106 |
| AL355098 | 0.002351979 |
| AC023669 | 0.002350754 |
| PAGE2 | 0.002349344 |
| GJD3 | 0.002344156 |
| SLAMF6P1 | 0.002343226 |
| RP11-179A9 | 0.00233845 |
| BCAP31P1 | 0.002336237 |
| RP11-351C8 | 0.002335425 |
| HERC2P3 | 0.002331247 |
| AC003991 | 0.002331054 |
| YWHAEP3 | 0.002329168 |
| MIR133B | 0.002329035 |
| MAP3K7CL | 0.002326688 |
| SNORD3B-1 | 0.002324209 |
| AL049798 | 0.002320274 |
| CTD-2247C11 | 0.002313898 |
| PPAP2A | 0.002310297 |
| GMEB2 | 0.002310167 |
| TSC22D1-AS1 | 0.002309218 |
| MRPS21P8 | 0.002309021 |
| RP13-824C8 | 0.002307703 |
| RNA5SP301 | 0.002304006 |
| SDHD | 0.002303041 |
| OR5M7P | 0.002300285 |
| RAD23BP2 | 0.002295874 |
| CICP4 | 0.002294185 |
| AL513327 | 0.002287086 |
| RN7SKP10 | 0.00228299 |
| RNU6-22P | 0.002282304 |
| RP5-836E8 | 0.002272688 |
| MIR6792 | 0.00227258 |
| AC010095 | 0.002270205 |
| BFAR | 0.002262797 |
| RP11-24J11 | 0.002257344 |
| RP11-33E15 | 0.002253784 |
| AP000350 | 0.002252409 |

| | | |
|---|---|---|
| AC004051 | | 0.002250621 |
| MTND1P33 | | 0.002248091 |
| RILPL2 | | 0.002245085 |
| LMF1 | | 0.002241233 |
| SEPT10P1 | | 0.00223889 |
| CEACAMP6 | | 0.002238865 |
| DOC2GP | | 0.002238024 |
| DLL3 | | 0.002235627 |
| AC084010 | | 0.00223538 |
| RP11-463J7 | | 0.002228846 |
| | 4-Sep | 0.002227223 |
| AL589675 | | 0.002226636 |
| HTR1B | | 0.002226414 |
| POLR2KP1 | | 0.002223451 |
| RP11-464F9 | | 0.002221437 |
| RNU6-1102P | | 0.002219662 |
| RABL3 | | 0.002217065 |
| REPS1 | | 0.00221312 |
| DHRS12 | | 0.002209809 |
| SNRPCP2 | | 0.002209117 |
| GSTA1 | | 0.002208691 |
| RNU6-212P | | 0.002206261 |
| C1orf146 | | 0.002205754 |
| HTT-AS1_1 | | 0.002204433 |
| KCNV2 | | 0.002202203 |
| PLSCR2 | | 0.002200681 |
| GHSR | | 0.002194844 |
| ZNF311 | | 0.002193045 |
| KRTAP19-6 | | 0.002190983 |
| RP11-453O22 | | 0.002188854 |
| KATNB1 | | 0.002187017 |
| RP11-666O2 | | 0.002185212 |
| FKSG68 | | 0.002183642 |
| VN1R80P | | 0.002177135 |
| TRIB2 | | 0.002175984 |
| AC140725 | | 0.002171685 |
| LONRF2P1 | | 0.002168166 |
| RNA5SP79 | | 0.002167951 |
| XRN2 | | 0.002165304 |
| PPIAP24 | | 0.002165211 |
| MNX1-AS2 | | 0.00215971 |
| AC016549 | | 0.002157869 |
| TSNARE1 | | 0.002157527 |
| ROBO1 | | 0.00215656 |
| GAL3ST3 | | 0.002153927 |

| | |
|---|---|
| CTC-400I9 | 0.002153084 |
| MS4A14 | 0.002149397 |
| NAP1L1P1 | 0.002147034 |
| CTC-209H22 | 0.002145577 |
| MTATP8P2 | 0.002144325 |
| RPGRIP1L | 0.002142048 |
| AC007136 | 0.002141007 |
| UGT3A2 | 0.002139783 |
| SNX9 | 0.002138773 |
| TBC1D3K | 0.002135743 |
| PRLHR | 0.002135447 |
| LAP3P2 | 0.002133904 |
| OR2S2 | 0.002132824 |
| ASCC1 | 0.002131517 |
| ZDHHC3 | 0.002130903 |
| AC239395 | 0.002130185 |
| AC063956 | 0.002130185 |
| AC017101 | 0.002129144 |
| CTC-265N9 | 0.002128888 |
| CTC-788C1 | 0.002126983 |
| PRDX4P1 | 0.002121964 |
| AL035045 | 0.002120085 |
| S100G | 0.002109348 |
| AL157882 | 0.002108886 |
| TTC28 | 0.002107609 |
| SARNP | 0.002104613 |
| RP11-506K6 | 0.002101417 |
| ATP5G1 | 0.002098734 |
| HES3 | 0.002098492 |
| ATP5G2 | 0.002097692 |
| RP11-399H11 | 0.002094999 |
| HMGCS1 | 0.00209291 |
| CCDC160 | 0.002092037 |
| RP11-696N14 | 0.002089245 |
| RP11-734C14 | 0.002086397 |
| RNU6-1300P | 0.002085707 |
| PPIAP15 | 0.002083445 |
| HYOU1 | 0.002081747 |
| AC125238 | 0.002080632 |
| AK3 | 0.002080426 |
| RN7SL233P | 0.00208004 |
| GPX4 | 0.002079347 |
| AC022384 | 0.00207386 |
| NIT1 | 0.002068202 |
| SEPW1P | 0.002063885 |

| | |
|---|---|
| RPA3 | 0.002052109 |
| EDDM3CP | 0.002049875 |
| RN7SL167P | 0.002048076 |
| SPDYC | 0.00204045 |
| RP11-178L8 | 0.002037596 |
| RP4-551N11 | 0.002034917 |
| AGGF1P3 | 0.002029764 |
| RNU6-1301P | 0.002026989 |
| RP11-480C16 | 0.002016956 |
| AL137190 | 0.002015638 |
| AC099537 | 0.002015504 |
| GAPDHP69 | 0.002013629 |
| RNU6-881P | 0.002010785 |
| RP11-789F5 | 0.00200764 |
| RN7SL752P | 0.002005761 |
| LRRN4 | 0.002002798 |
| HIVEP1 | 0.002002655 |
| RP11-529E15 | 0.002002052 |
| KRT18P62 | 0.002000253 |
| DCST1 | 0.001999102 |
| RP11-551G24 | 0.001995716 |
| RP4-553F4 | 0.001978124 |
| ADAM18 | 0.00197511 |
| MIR182 | 0.00197508 |
| ADAM3B | 0.001971222 |
| bP-2168N6 | 0.001967392 |
| RNU6-645P | 0.001967038 |
| RPS7P15 | 0.001964865 |
| RP11-479O17 | 0.001962389 |
| PGM2 | 0.00196124 |
| RP4-581F12 | 0.001952309 |
| HDHD1 | 0.001944532 |
| AC022486 | 0.001941045 |
| RNU6-387P | 0.001940141 |
| OR4X7P | 0.001937284 |
| HCFC2P1 | 0.001929407 |
| CORO7-PAM16 | 0.001921552 |
| AC025750 | 0.001921196 |
| MKX | 0.001916307 |
| LINC01297 | 0.001904955 |
| RNU6-822P | 0.001902821 |
| FADD | 0.001901051 |
| LINC01333 | 0.001899144 |
| RNA5SP310 | 0.001893232 |
| OAZ2 | 0.00189199 |

| | |
|---|---|
| INSIG2 | 0.001885013 |
| VDAC1 | 0.001873775 |
| OR56A3 | 0.001870396 |
| GMCL1P1 | 0.001868238 |
| RNF182 | 0.001867115 |
| C20orf96 | 0.001864491 |
| RNU7-107P | 0.001863134 |
| AL157883 | 0.001859963 |
| SUMO2P15 | 0.001857235 |
| DISP2 | 0.001852956 |
| RP11-874G11 | 0.001852512 |
| TPMTP4 | 0.001851892 |
| CDC123 | 0.001850891 |
| AC068057 | 0.00184838 |
| TTC23L | 0.001848379 |
| TMPRSS4-AS1 | 0.001846431 |
| RBMY2EP | 0.001842492 |
| FLJ22763 | 0.001837554 |
| RP11-421F16 | 0.001837364 |
| RP11-298D21 | 0.001836246 |
| TCP11 | 0.001835698 |
| KCTD8 | 0.001830944 |
| RP11-328D5 | 0.001830701 |
| LINC01560 | 0.001826834 |
| MACROD1 | 0.001823634 |
| NXNL2 | 0.001823392 |
| MIR6728 | 0.001823327 |
| EIF5AP4 | 0.001821159 |
| RP11-334N17 | 0.001820691 |
| TCP11X2 | 0.001820017 |
| WASH7P | 0.001819447 |
| SSX6 | 0.001819144 |
| CA10 | 0.001818563 |
| GGTLC2 | 0.001814805 |
| AC091654 | 0.001813578 |
| EIF3LP1 | 0.001807825 |
| C8orf76 | 0.001807447 |
| CYP21A1P | 0.001806119 |
| NIP7P3 | 0.001805803 |
| RN7SKP198 | 0.001804781 |
| CTD-2227C6 | 0.001803958 |
| PLA2G12AP2 | 0.001797874 |
| RP11-4O3 | 0.001792271 |
| RN7SL541P | 0.001790799 |
| RNU6-1028P | 0.001790516 |

| | |
|---|---|
| AC009264 | 0.001789041 |
| C17orf98 | 0.001786901 |
| KYNU | 0.001786441 |
| SMG1P3 | 0.001784636 |
| RP11-369J21 | 0.00178417 |
| RP11-87M18 | 0.0017839 |
| AXDND1 | 0.001779461 |
| RNA5SP326 | 0.001778761 |
| LDB2 | 0.001778069 |
| KCNIP2-AS1 | 0.001777891 |
| SOAT2 | 0.001769989 |
| GVQW1 | 0.00176791 |
| TNFSF12 | 0.001767073 |
| RP5-1025A1 | 0.001765247 |
| AGKP1 | 0.001757094 |
| OR7E7P | 0.001751891 |
| SNTN | 0.001748505 |
| HSBP1 | 0.001748467 |
| DPPA3P3 | 0.001745198 |
| OR7A11P | 0.001744265 |
| RPL12P23 | 0.001741868 |
| MTND4P16 | 0.001741399 |
| AC007358 | 0.001738914 |
| KIAA1549 | 0.001737286 |
| UQCRHP2 | 0.001736771 |
| ZNF72P | 0.001731575 |
| ELF5 | 0.001729667 |
| RPL21P104 | 0.001723556 |
| HSPE1P7 | 0.001723025 |
| SPG20 | 0.001717192 |
| TCEB1P20 | 0.001712483 |
| TOMM22 | 0.001707163 |
| RNU6-381P | 0.001706011 |
| RP11-399K21 | 0.001704078 |
| MRGPRX1 | 0.001703066 |
| CTB-56J15 | 0.001702386 |
| RP11-720L8 | 0.001698878 |
| AMELX | 0.001695082 |
| RPL39P25 | 0.001691236 |
| AC007562 | 0.001688614 |
| RP11-288A5 | 0.001681973 |
| GIPC3 | 0.001678064 |
| RP11-472K17 | 0.001677308 |
| SSTR4 | 0.001677141 |
| RN7SKP100 | 0.001676237 |

| | |
|---|---|
| RP1-91J24 | 0.001675639 |
| RP11-379J5 | 0.001674373 |
| RP4-656G21 | 0.001672312 |
| SOX14 | 0.001663427 |
| RAD1 | 0.001663366 |
| RPL5P7 | 0.001661707 |
| ACTBP1 | 0.001661474 |
| AC118063 | 0.001659703 |
| HIGD1AP1 | 0.001657458 |
| OR1D4 | 0.001656447 |
| OR2N1P | 0.001651953 |
| UFD1L | 0.00165144 |
| CHCHD2P1 | 0.001650702 |
| CTD-2244C20 | 0.00164998 |
| VTA1P2 | 0.001647924 |
| AC104441 | 0.00164585 |
| AC069286 | 0.001640832 |
| RP11-368P15 | 0.001639163 |
| FOXS1 | 0.001639142 |
| FBXO2 | 0.0016357 |
| AC130689 | 0.001634465 |
| FAM90A1 | 0.001634435 |
| SIGMAR1 | 0.0016341 |
| A2M | 0.00162569 |
| RNU6-972P | 0.00161555 |
| C4BPAP2 | 0.001609943 |
| AC002463 | 0.00160931 |
| OPHN1 | 0.001607457 |
| RNU6-1018P | 0.001604042 |
| MIR744 | 0.001603581 |
| KCNJ6-AS1 | 0.00160352 |
| RANBP1 | 0.001602749 |
| CTB-60E11 | 0.001597008 |
| CTD-2311A18 | 0.001593861 |
| CTD-2303H24 | 0.001591851 |
| BAALC-AS1 | 0.001584814 |
| HSPE1P21 | 0.001578334 |
| POTEF | 0.001576122 |
| RP11-135J2 | 0.001575744 |
| MARK2P17 | 0.001575405 |
| RP11-119K6 | 0.001573878 |
| MIR526A1 | 0.001572972 |
| AC091633 | 0.001570935 |
| C16orf82 | 0.00157074 |
| TGFBR3L | 0.001569786 |

| | |
|---|---|
| RP11-427G18 | 0.00156953 |
| RP11-28B23 | 0.00155438 |
| KRT19P6 | 0.001549464 |
| RN7SL841P | 0.001548621 |
| NF1P3 | 0.001546356 |
| RNU4-48P | 0.001542408 |
| CCDC70 | 0.001539564 |
| AC099759 | 0.001538049 |
| FO082796 | 0.001535249 |
| EDN3 | 0.001535219 |
| RP11-357J22 | 0.001535092 |
| DEFB119 | 0.001528444 |
| RP11-96P7 | 0.001520079 |
| ZNF705G | 0.001517492 |
| PLXNB1 | 0.001515639 |
| ABLIM2 | 0.001512364 |
| MIR99A | 0.001510513 |
| PABPC1P3 | 0.001509294 |
| RNU4ATAC5P | 0.001503838 |
| SNAPIN | 0.001499154 |
| CTD-2626G11 | 0.001497342 |
| ARHGAP39 | 0.001494726 |
| HIST1H3G | 0.001492399 |
| MIR134 | 0.001489899 |
| RP11-144G16 | 0.001484138 |
| ZNF285 | 0.001477795 |
| MIR4442 | 0.001470971 |
| RNMTL1P1 | 0.001465502 |
| CERS1 | 0.001465168 |
| AC097532 | 0.001463673 |
| RP5-892F13 | 0.00146315 |
| RNA5SP92 | 0.001455498 |
| RAPGEF4 | 0.001453701 |
| RP11-678L1 | 0.001450028 |
| DSG1 | 0.001448855 |
| GALNT14 | 0.001435449 |
| MIR520H | 0.001433049 |
| RNU1-40P | 0.001433049 |
| NDUFA5P7 | 0.001433049 |
| RNU1-41P | 0.001433049 |
| RNU1-97P | 0.001433049 |
| RNU6-256P | 0.001433049 |
| RNU6-546P | 0.001433049 |
| AC093680 | 0.001433049 |
| RNU1-86P | 0.001433049 |

| | |
|---|---|
| PRR20FP | 0.001433049 |
| SNORD115-18 | 0.001433049 |
| RNU1-45P | 0.001433049 |
| RNU1-128P | 0.001433049 |
| SNORD115-10 | 0.001433049 |
| SNORD115-27 | 0.001433049 |
| SNORD115-29 | 0.001433049 |
| SNORD115-19 | 0.001433049 |
| USP17L23 | 0.001433049 |
| SNORD115-34 | 0.001433049 |
| SNORD115-16 | 0.001433049 |
| FANK1-AS1 | 0.001424714 |
| IL36G | 0.001422157 |
| RP1-200G19 | 0.001421362 |
| MIR3915 | 0.001418983 |
| AC120104 | 0.001418817 |
| GAPDHP33 | 0.00141397 |
| HTR3A | 0.001412286 |
| C2orf70 | 0.001409332 |
| ERG | 0.001408918 |
| ZNF880 | 0.001408322 |
| RN7SL35P | 0.001397468 |
| CTD-2197M16 | 0.001394853 |
| DIP2A-IT1 | 0.001392825 |
| YWHAH | 0.001389345 |
| TAS2R40 | 0.001386253 |
| KRT18P11 | 0.001385921 |
| DEPTOR | 0.001385549 |
| RP11-65G9 | 0.001383295 |
| RN7SL494P | 0.00137891 |
| RP4-537K23 | 0.001377902 |
| PTCSC2 | 0.001376861 |
| C1orf68 | 0.001373904 |
| MEF2A | 0.001373251 |
| LA16c-313F4 | 0.001369623 |
| AC005631 | 0.001368354 |
| INO80E | 0.001367334 |
| PI4KAP1 | 0.001364674 |
| MTND5P18 | 0.001363402 |
| CTD-2033D24 | 0.001361439 |
| AC108032 | 0.001359009 |
| RP11-255E6 | 0.001357855 |
| RNU6-947P | 0.001353144 |
| AC005514 | 0.001350116 |
| SERPINA4 | 0.001349954 |

| | |
|---|---|
| C7orf49 | 0.001346701 |
| RP11-459C13 | 0.001344242 |
| CDK5RAP2 | 0.001338682 |
| CTD-2216M2 | 0.001338102 |
| MIR4754 | 0.001335308 |
| NMNAT1P1 | 0.001334294 |
| CSF3 | 0.001333755 |
| CTB-60B18 | 0.001328024 |
| RPL21P43 | 0.001327754 |
| RP11-140K17 | 0.001323537 |
| LINC00351 | 0.001323315 |
| RP11-368L12 | 0.001322798 |
| RP11-161D15 | 0.001322206 |
| REEP3 | 0.00132056 |
| RP1-229K20 | 0.001319861 |
| PLEKHM1 | 0.001319593 |
| AC127904 | 0.001317714 |
| RP11-272D12 | 0.001315437 |
| MIR191 | 0.001314366 |
| ZNF229 | 0.001313118 |
| FAM189B | 0.001311741 |
| TUBBP6 | 0.001309321 |
| RP11-720N19 | 0.001309204 |
| RP11-258F22 | 0.001308328 |
| CCT7 | 0.001306309 |
| DDX43 | 0.001306107 |
| RP11-68I18 | 0.00130417 |
| MRPL46 | 0.001301211 |
| MIR34B | 0.001298679 |
| MIR1231 | 0.001296001 |
| RP11-472I20 | 0.00129517 |
| RNU6-1186P | 0.001292969 |
| PTGES3P5 | 0.001289899 |
| DOCK4 | 0.001289005 |
| RP11-702C7 | 0.001286266 |
| FAM106CP | 0.001285357 |
| TEX30 | 0.001284149 |
| RP11-35J23 | 0.001282402 |
| RP11-111A24 | 0.001281907 |
| NPM1P48 | 0.001277693 |
| XIRP1 | 0.001274168 |
| RFX1 | 0.001271981 |
| MIR5091 | 0.001268755 |
| HMGB1P21 | 0.001267819 |
| PCCA | 0.001260297 |

| | |
|---|---|
| AC068292 | 0.001260229 |
| RN7SKP90 | 0.001258256 |
| OR10J2P | 0.001247324 |
| RN7SL156P | 0.001246199 |
| TRMT112P2 | 0.001245845 |
| ALG1L | 0.0012448 |
| KRTAP4-9 | 0.001244658 |
| AC074121 | 0.001242561 |
| CNOT10-AS1 | 0.001242242 |
| RP5-930J4 | 0.001225439 |
| RP11-262I2 | 0.001223687 |
| RP11-647O20 | 0.001222225 |
| GJD2 | 0.001220909 |
| RP11-89K21 | 0.001219177 |
| AC079233 | 0.00121345 |
| GFY | 0.001211057 |
| AP000705 | 0.001206995 |
| OR2T5 | 0.001206844 |
| PLXNA4 | 0.001204369 |
| MYO15A | 0.001203383 |
| DCTN5 | 0.001201262 |
| GH2 | 0.001199987 |
| LINC01398 | 0.001199689 |
| AL109750 | 0.001188689 |
| ZNF123P | 0.001188679 |
| RNU6-815P | 0.001184971 |
| RNU6-228P | 0.001179616 |
| ABCF1 | 0.001177521 |
| RP11-485B17 | 0.001175124 |
| NCSTN | 0.001174172 |
| AC005588 | 0.001167385 |
| RP11-384D10 | 0.001166059 |
| OR4A46P | 0.001165659 |
| C9orf9 | 0.0011626 |
| RP11-434C1 | 0.0011618 |
| OSBPL10 | 0.001160144 |
| LZTS1-AS1 | 0.001158063 |
| KLHL26 | 0.001157632 |
| RP11-98C8 | 0.00115657 |
| AL031655 | 0.001154517 |
| TCAP | 0.001154302 |
| TMEM246-AS1 | 0.001149283 |
| AC090707 | 0.001147367 |
| PDE4D | 0.001146804 |
| RP11-315F22 | 0.00114587 |

| | |
|---|---|
| TMEM171 | 0.001128216 |
| MIR130A | 0.001127013 |
| RP11-123C21 | 0.001125841 |
| MAGEB10 | 0.00112222 |
| RN7SL390P | 0.001120708 |
| CCDC37 | 0.001119921 |
| RPS12P26 | 0.001116571 |
| SNORD74 | 0.001116371 |
| PTP4A2P1 | 0.001114486 |
| AKR1D1 | 0.001107387 |
| OR5BC1P | 0.001104185 |
| AC105402 | 0.001099013 |
| RP11-248B24 | 0.001093132 |
| RP11-514D23 | 0.001092644 |
| OR2T1 | 0.001090506 |
| FAM99B | 0.001089365 |
| RP1-212G6 | 0.001083832 |
| GTF2IP2 | 0.001079682 |
| RP11-463I20 | 0.001079666 |
| CDY8P | 0.001078725 |
| HIST1H2AA | 0.00107628 |
| LINC00424 | 0.001075353 |
| TICAM2 | 0.001070414 |
| MIR1-1HG | 0.001066237 |
| RP11-85G20 | 0.001058562 |
| AC018463 | 0.001055736 |
| AGBL1-AS1 | 0.001055635 |
| RN7SL423P | 0.001050341 |
| PRDM9 | 0.001048885 |
| RP5-831C21 | 0.001045819 |
| RNU1-75P | 0.001044227 |
| STARP1 | 0.001043321 |
| PPIAP20 | 0.001043047 |
| RPL3P7 | 0.001039394 |
| GRIFIN | 0.00103884 |
| RP11-254F19 | 0.001037814 |
| RNU7-61P | 0.001029851 |
| RNA5SP19 | 0.001023137 |
| KCNK4 | 0.001022316 |
| RP11-490P13 | 0.001019935 |
| RP11-565P22 | 0.001011617 |
| RP11-294L11 | 0.001005818 |
| RP5-1106H14 | 0.001005122 |
| ZFAND5 | 0.001002853 |
| AL133475 | 0.000992224 |

| | |
|---|---|
| MIR16-2 | 0.00098669 |
| RNA5SP99 | 0.000985127 |
| SNORD78 | 0.00098177 |
| MIR625 | 0.000978046 |
| RP11-18B3 | 0.000975112 |
| USF2 | 0.000968173 |
| LINC00333 | 0.000966956 |
| RP11-507J18 | 0.000964587 |
| RMST_5 | 0.000963436 |
| MIR421 | 0.000962592 |
| RP11-90B22 | 0.000957312 |
| MIR4792 | 0.000957082 |
| TRAJ46 | 0.000956075 |
| CTC-543D15 | 0.000955595 |
| DGCR11 | 0.000954572 |
| RP11-848D3 | 0.000954012 |
| RP11-886D15 | 0.000953649 |
| RP1-86D1 | 0.000951528 |
| CHRAC1 | 0.000951316 |
| C17orf96 | 0.000950349 |
| AP006288 | 0.000938202 |
| CTD-2189E23 | 0.000936102 |
| AC062022 | 0.000930607 |
| RP1-49C23 | 0.000925662 |
| HRSP12 | 0.000924171 |
| SNRPEP4 | 0.000923408 |
| ANKRD50 | 0.000920152 |
| KIT | 0.000919185 |
| OR51I1 | 0.000915299 |
| HOTAIR_5 | 0.00091438 |
| RP11-326J18 | 0.000910126 |
| RNU6-1228P | 0.000905418 |
| AC104458 | 0.000901088 |
| RP11-662G23 | 0.000901011 |
| MIR3689F | 0.000896688 |
| SAP30L-AS1 | 0.000895611 |
| PTCHD3P2 | 0.000887343 |
| CUX1 | 0.000884337 |
| RNU4-84P | 0.00088388 |
| ASB6 | 0.000881918 |
| AC090625 | 0.000880735 |
| IL9RP4 | 0.000880383 |
| ANK3 | 0.000880206 |
| PLCL2-AS1 | 0.000878137 |
| AL158155 | 0.000876855 |

| | |
|---|---|
| OR10J1 | 0.000860505 |
| RP11-95J11 | 0.000858745 |
| RNU2-49P | 0.000856952 |
| ADH5P4 | 0.000846427 |
| EVA1C | 0.0008456 |
| GNA15 | 0.000844149 |
| FP236383 | 0.000838431 |
| RP11-650J17 | 0.000828889 |
| ALKBH6 | 0.000828408 |
| GNB4 | 0.000824669 |
| KRTAP22-1 | 0.000821391 |
| RNU6ATAC15P | 0.000819367 |
| CUL4AP1 | 0.000817055 |
| RP11-556I13 | 0.000814928 |
| CDC42EP3 | 0.000813599 |
| TTC3-AS1 | 0.0008107 |
| MIR5092 | 0.000807573 |
| CTD-2593A12 | 0.000803443 |
| HIST1H2APS1 | 0.000800979 |
| MIR6801 | 0.000792512 |
| RP11-723C11 | 0.000791997 |
| KRTAP4-2 | 0.000791044 |
| POLRMT | 0.00079036 |
| RN7SL383P | 0.00078299 |
| OR7E1P | 0.000778837 |
| MBL3P | 0.000774771 |
| RN7SL455P | 0.000771326 |
| RNU6-205P | 0.000765854 |
| RP11-958F21 | 0.000765418 |
| ACTRT1 | 0.000763178 |
| AC109496 | 0.000757782 |
| RP11-1B20 | 0.000756597 |
| RP13-820C6 | 0.000753669 |
| ADAM17 | 0.000751419 |
| RP11-127L21 | 0.000750702 |
| RNA5SP100 | 0.000750051 |
| GRK5 | 0.000749745 |
| OSTCP6 | 0.000748415 |
| BCL10 | 0.000747772 |
| WARS2 | 0.000743084 |
| LMAN2L | 0.000742284 |
| WBSCR17 | 0.000741893 |
| SFT2D1 | 0.000726804 |
| RNF39 | 0.00072539 |
| SNF8 | 0.000723995 |

| | |
|---|---|
| LMO7DN-IT1 | 0.00071793 |
| C2orf91 | 0.00071452 |
| AC008869 | 0.000712984 |
| FTSJ3 | 0.000711045 |
| NPC1L1 | 0.000710692 |
| MDK | 0.000709147 |
| NAP1L5 | 0.000706617 |
| RN7SL397P | 0.000705478 |
| BOLA3P4 | 0.000702331 |
| FAM182B | 0.000698766 |
| LINC01552 | 0.000698411 |
| RP13-714J12 | 0.00069565 |
| SNORA30 | 0.000693789 |
| AE000659 | 0.000693664 |
| CCNHP1 | 0.000689673 |
| ST7 | 0.000681853 |
| RP5-964N17 | 0.000680536 |
| SH2D4B | 0.000675773 |
| HDX | 0.000672104 |
| RSL24D1P5 | 0.000668495 |
| MIR4479 | 0.000667598 |
| MIR4676 | 0.000664815 |
| PRSS54 | 0.000663364 |
| RNA5SP474 | 0.000662808 |
| MAGED2 | 0.000661145 |
| CACNA1C-AS1 | 0.000657307 |
| AC009110 | 0.000656498 |
| OR7E125P | 0.000653473 |
| AC138942 | 0.000653169 |
| RP11-628E19 | 0.000650728 |
| DNAJA1P6 | 0.000650177 |
| RNA5SP47 | 0.000647552 |
| OR2A5 | 0.000647094 |
| RP11-554D13 | 0.000643091 |
| UPK1A-AS1 | 0.000641546 |
| ATP1B2 | 0.000641275 |
| PTMAP5 | 0.000641163 |
| GDF2 | 0.000640939 |
| AC007327 | 0.000636797 |
| AL355794 | 0.000636777 |
| EIF4H | 0.000634633 |
| RP11-448G4 | 0.000628979 |
| RNA5SP309 | 0.000627786 |
| RP11-601I15 | 0.000627173 |
| TAOK2 | 0.000625367 |

| | |
|---|---|
| RPL30 | 0.000624362 |
| RP11-203H2 | 0.00062436 |
| F10 | 0.000618967 |
| CDC25B | 0.000616678 |
| DEFB105B | 0.000614164 |
| RNA5SP106 | 0.000614164 |
| RNU6-906P | 0.000613975 |
| TUBB1P1 | 0.000612498 |
| ASRGL1 | 0.000608938 |
| OR2X1P | 0.000607814 |
| AP001062 | 0.000599971 |
| KDM4A | 0.000594724 |
| RNU6-128P | 0.000591237 |
| RPEL1 | 0.000588966 |
| RNU6-268P | 0.000587212 |
| OR6J1 | 0.000584399 |
| RPL3P3 | 0.000578209 |
| KIAA1211L | 0.000570425 |
| RP11-716O23 | 0.000567751 |
| RPL7P41 | 0.000566164 |
| MIR4719 | 0.000564748 |
| FKSG48 | 0.000561725 |
| MIR6822 | 0.000559832 |
| SPICE1 | 0.000559187 |
| RNU6-122P | 0.000559122 |
| LINC01352 | 0.000558634 |
| SNORD114-15 | 0.000551078 |
| SOST | 0.00055098 |
| RN7SKP109 | 0.000549707 |
| AC100767 | 0.000549226 |
| MIR550A1 | 0.000546268 |
| RSF1-IT2 | 0.000544334 |
| CTC-379B2 | 0.000543701 |
| THNSL2 | 0.000542033 |
| AC010240 | 0.000538957 |
| GK2 | 0.000536972 |
| RP11-88E10 | 0.000536433 |
| KIAA1045 | 0.00053368 |
| RN7SKP178 | 0.000531972 |
| RP11-1070A24 | 0.000531251 |
| ASAP3 | 0.000527149 |
| SYNCRIP | 0.000526367 |
| RP11-120C12 | 0.000526361 |
| MRPL16 | 0.000525884 |
| SNORD15B | 0.000520086 |

| | |
|---|---|
| CTB-162E12 | 0.000519734 |
| AC083826 | 0.000519407 |
| OR2AM1P | 0.000516296 |
| APCS | 0.000512865 |
| OR5AW1P | 0.000504196 |
| AC005775 | 0.000503203 |
| RNU6-843P | 0.000503112 |
| RNU6-808P | 0.000500395 |
| RNU6-656P | 0.000499278 |
| AC091729 | 0.000497696 |
| TRAIP | 0.000493361 |
| RP11-404K5 | 0.000493165 |
| RPL22 | 0.000492319 |
| ANP32B | 0.000488059 |
| OR6C75 | 0.000487586 |
| PRICKLE3 | 0.000485361 |
| TMEM213 | 0.000482014 |
| TP53 | 0.000479184 |
| RNU6-717P | 0.000477683 |
| MIR190B | 0.000475337 |
| PTCD2P2 | 0.000475015 |
| AKAP8L | 0.000474663 |
| CLPTM1L | 0.000473044 |
| RP11-349K16 | 0.00046871 |
| RP11-460B17 | 0.000468542 |
| TXNDC5 | 0.000467165 |
| MIR4287 | 0.000466962 |
| HNRNPA3P7 | 0.000461388 |
| HAUS8 | 0.000461304 |
| HSD17B8 | 0.00045976 |
| AC073071 | 0.000458627 |
| SPERT | 0.000455761 |
| RNA5SP280 | 0.000448507 |
| RN7SL623P | 0.000447993 |
| GS1-5L10 | 0.000446541 |
| SNAPC4 | 0.000440857 |
| RP11-168P6 | 0.000438941 |
| FCF1P5 | 0.000438476 |
| BRICD5 | 0.000432354 |
| RP11-75C23 | 0.00043068 |
| SYCN | 0.000421295 |
| AL021395 | 0.000417431 |
| HNRNPA1P29 | 0.000416806 |
| RASGRP4 | 0.000411051 |
| EXOGP1 | 0.000411017 |

| | |
|---|---|
| TDRD5 | 0.000410344 |
| OTUB1 | 0.000408241 |
| XXbac-BPG294E21 | 0.000406481 |
| RP11-589B3 | 0.000403681 |
| RP4-800O15 | 0.000396126 |
| PCDH9 | 0.000390064 |
| AC095030 | 0.000388965 |
| RP11-204E9 | 0.000376383 |
| ALG1L9P | 0.000375068 |
| ARL2BP | 0.000375049 |
| CYP4F27P | 0.000373053 |
| RP11-496H1 | 0.000369095 |
| PGBD4P5 | 0.000366137 |
| FAM58BP | 0.0003656 |
| GOLGA8A | 0.000365058 |
| GRWD1 | 0.000363179 |
| TMIE | 0.000362918 |
| TMCO1 | 0.00035222 |
| RP11-21L19 | 0.000346358 |
| RP4-813F11 | 0.000345936 |
| HSFY2 | 0.000342474 |
| RNA5SP117 | 0.000341202 |
| PPARD | 0.000338266 |
| PCNAP3 | 0.000337496 |
| RP11-475D12 | 0.000336244 |
| AC005355 | 0.000335971 |
| RIMS2 | 0.000334666 |
| RP11-202G11 | 0.000323298 |
| SDCCAG3P2 | 0.000322879 |
| PDGFB | 0.000322861 |
| MIR3118-3 | 0.000313381 |
| RP11-439C8 | 0.000308471 |
| GYG1 | 0.000305148 |
| AP3B2 | 0.000304953 |
| RP11-272P10 | 0.000303925 |
| DCTN1-AS1 | 0.000302424 |
| SNRPGP16 | 0.00030184 |
| PCSK1 | 0.000300013 |
| ZNF444 | 0.000298506 |
| TEX15 | 0.000297727 |
| RP1-34L19 | 0.00029261 |
| EBF4 | 0.00028498 |
| PDLIM5 | 0.000283064 |
| MIR1911 | 0.000274308 |
| AC073069 | 0.000273246 |

| | |
|---|---|
| SETP1 | 0.000272573 |
| GPR35 | 0.000265891 |
| SAMD12-AS1 | 0.000263454 |
| GPR12 | 0.000263419 |
| BSCL2 | 0.000262784 |
| KRT18P18 | 0.000262448 |
| RP11-216N21 | 0.000261605 |
| RP11-118E9 | 0.000261006 |
| RP4-559A3 | 0.000257861 |
| AL161935 | 0.000257375 |
| CTD-2110K23 | 0.000256635 |
| Z98052 | 0.000254832 |
| XXbac-BPGBPG34I8 | 0.000248215 |
| AC073218 | 0.000243637 |
| RP11-190A12 | 0.000242969 |
| LINC00371 | 0.000241354 |
| CXorf36 | 0.000241127 |
| GABRB3 | 0.000238504 |
| RN7SKP122 | 0.000238053 |
| SALL1P1 | 0.000234542 |
| RP5-1054A22 | 0.000233679 |
| RN7SL170P | 0.000232472 |
| RP11-166J22 | 0.000231859 |
| RP11-164O23 | 0.000226312 |
| RNU1-11P | 0.00022254 |
| RP1-131F15 | 0.000216292 |
| AC010132 | 0.000213042 |
| CORO7 | 0.000205535 |
| RNU6-654P | 0.000204721 |
| HES6 | 0.000203079 |
| CYCSP35 | 0.000203049 |
| RP11-852E15 | 0.00020185 |
| TBC1D28 | 0.000201731 |
| AC141928 | 0.000200573 |
| PCNAP1 | 0.000197075 |
| AC004816 | 0.000191811 |
| PSMF1 | 0.000189571 |
| PPM1AP1 | 0.000189019 |
| OR2K2 | 0.000186353 |
| RP11-163G10 | 0.00018439 |
| PDK4 | 0.000180361 |
| SLC9A2 | 0.000179229 |
| RNA5SP517 | 0.000176472 |
| AC116050 | 0.000167844 |
| RP11-582E3 | 0.000157923 |

| | |
|---|---|
| RN7SL687P | 0.000151972 |
| MED21 | 0.000151672 |
| YBX1P10 | 0.00015063 |
| LINC00443 | 0.000149399 |
| RNU4ATAC14P | 0.000149379 |
| RP11-769N19 | 0.000145622 |
| FOXH1 | 0.000138369 |
| RP11-60C6 | 0.000134242 |
| GRK1 | 0.000133506 |
| AC098795 | 0.000131479 |
| RNU7-46P | 0.000129442 |
| RN7SL101P | 0.000127533 |
| REG1B | 0.00012719 |
| GJA1P1 | 0.000126096 |
| SNHG12 | 0.000125829 |
| RPS29P22 | 0.000122067 |
| FKBP4P2 | 0.000116971 |
| COLGALT2 | 0.00011621 |
| RNA5SP481 | 0.000114512 |
| AC138744 | 0.00011414 |
| CCNC | 0.000107223 |
| CTC-529P8 | 0.00010304 |
| KATNBL1P5 | 0.000101968 |
| AC004878 | 9.96E-05 |
| RNU6-987P | 9.91E-05 |
| RP11-236L14 | 9.82E-05 |
| PNKD | 9.12E-05 |
| ZBTB8A | 8.76E-05 |
| RP11-527N12 | 7.34E-05 |
| PSG3 | 7.26E-05 |
| CKB | 7.06E-05 |
| RGS7 | 7.06E-05 |
| CTD-2138O14 | 7.03E-05 |
| C3orf56 | 6.94E-05 |
| GNGT1 | 6.90E-05 |
| RP11-249C24 | 6.33E-05 |
| RP11-154H23 | 5.53E-05 |
| HCCAT5 | 5.19E-05 |
| RP11-436H11 | 4.70E-05 |
| PMS2P6 | 4.45E-05 |
| YWHAE | 4.20E-05 |
| HNRNPFP1 | 3.86E-05 |
| RP11-46D6 | 3.60E-05 |
| MIR1208 | 3.38E-05 |
| RP11-406H21 | 3.11E-05 |

| | |
|---|---|
| RN7SL447P | 2.90E-05 |
| CASC6 | 2.82E-05 |
| MTCO3P2 | 2.68E-05 |
| RPL24P7 | 2.64E-05 |
| LINC01537 | 2.60E-05 |
| CCDC129 | 2.42E-05 |
| MOCS2 | 2.23E-05 |
| MIR101-2 | 1.97E-05 |
| RN7SKP153 | 1.38E-05 |
| CTC-353G13 | 1.07E-05 |
| GOPC | 9.75E-06 |
| OR5D16 | 9.47E-06 |
| UBTFL10 | 7.78E-06 |
| RPL10AP1 | 5.95E-06 |
| RP11-168O16 | 5.19E-06 |
| RP11-109E10 | 4.19E-06 |
| IFT46 | -2.79E-07 |
| RN7SL107P | -4.83E-07 |
| BANCR | -1.54E-06 |
| SNORD116-9 | -2.76E-06 |
| LINC01078 | -8.27E-06 |
| PGAM5P1 | -1.00E-05 |
| PRKRA | -1.24E-05 |
| RNU6-426P | -1.40E-05 |
| UBTFL8 | -1.67E-05 |
| AQP7P3 | -2.36E-05 |
| RIOK3P1 | -2.67E-05 |
| AF127577 | -2.71E-05 |
| RNU6-193P | -2.73E-05 |
| COMMD3-BMI1 | -3.46E-05 |
| EEF1B2 | -3.64E-05 |
| ZBTB9 | -4.03E-05 |
| IGKV2D-38 | -4.71E-05 |
| SNORD38 | -5.04E-05 |
| PHBP14 | -5.06E-05 |
| AC010376 | -5.55E-05 |
| RNU7-148P | -5.72E-05 |
| SLC9A3P1 | -6.38E-05 |
| AL391559 | -6.82E-05 |
| SNRPC | -7.04E-05 |
| LAS1L | -7.78E-05 |
| LINC00459 | -8.03E-05 |
| AC004074 | -8.28E-05 |
| RP13-436F16 | -8.33E-05 |
| RP11-61N20 | -8.37E-05 |

| | |
|---|---|
| SNORA76 | -8.68E-05 |
| TASP1 | -8.80E-05 |
| RNU6-1054P | -8.98E-05 |
| PDXP | -9.02E-05 |
| CTF2P | -9.10E-05 |
| AL121931 | -9.70E-05 |
| ACMSD | -9.87E-05 |
| AC083875 | -0.000100174 |
| AP001604 | -0.000107084 |
| RP11-90J7 | -0.000107479 |
| RN7SL97P | -0.000109975 |
| FTH1P4 | -0.000110405 |
| OR1E1 | -0.000113369 |
| CICP24 | -0.000114124 |
| ESPN | -0.00011941 |
| RP5-845O24 | -0.000120095 |
| MIR5682 | -0.000120499 |
| SOD1 | -0.000120563 |
| DCPS | -0.000120638 |
| RP11-62L18 | -0.000122846 |
| RFPL3S | -0.00012328 |
| RP11-123C5 | -0.000125061 |
| SNORA43 | -0.00013112 |
| CTD-3012A18 | -0.000134907 |
| SALL2 | -0.000135559 |
| RN7SKP258 | -0.00013663 |
| MIR6867 | -0.000138533 |
| DNLZ | -0.000139597 |
| MED31 | -0.00014436 |
| EIF5B | -0.000146276 |
| RP11-54J7 | -0.000153255 |
| AP005901 | -0.000157916 |
| RNU6-581P | -0.0001586 |
| RP11-351A20 | -0.000161041 |
| HSD3BP1 | -0.000161373 |
| OR2Q1P | -0.000162019 |
| RNU6-1187P | -0.000162691 |
| RN7SL202P | -0.000170879 |
| ELP6 | -0.000172342 |
| MOG | -0.000176386 |
| LINC00508 | -0.000176561 |
| RP11-108A14 | -0.000180205 |
| RP11-85L21 | -0.000185177 |
| SMAD1 | -0.000186259 |
| RP11-231I13 | -0.000186762 |

| | |
|---|---|
| MIRLET7F1 | -0.000191488 |
| ATP1B4 | -0.000194725 |
| MT-TQ | -0.000195695 |
| SNORD51 | -0.000196264 |
| RP11-197B12 | -0.000196372 |
| RP11-320J16 | -0.00019728 |
| MIR487A | -0.000200172 |
| RPL37P25 | -0.000201507 |
| PAX8-AS1 | -0.000214968 |
| GAPDHP23 | -0.000220138 |
| KB-1991G8 | -0.000220943 |
| LINC01251 | -0.000234036 |
| RNU7-62P | -0.000234193 |
| RP11-718B12 | -0.000234247 |
| ZNF877P | -0.000245142 |
| RNA5SP141 | -0.00024722 |
| VWA8-AS1 | -0.000248231 |
| RNU6-1323P | -0.000248277 |
| RP5-1180C18 | -0.000249662 |
| AC091042 | -0.000251668 |
| GDF10 | -0.0002517 |
| RP13-395E19 | -0.000254225 |
| DCT | -0.000257643 |
| RP11-61L23 | -0.000263137 |
| TUBGCP2 | -0.000263398 |
| OR2A12 | -0.000264942 |
| RNU5F-8P | -0.000266395 |
| RP11-221J22 | -0.000268409 |
| RP11-438B23 | -0.000268829 |
| RP5-1136G2 | -0.000272031 |
| NUP62CL | -0.00027231 |
| EVX2 | -0.000272653 |
| MTCO2P3 | -0.000274861 |
| AC004812 | -0.000277558 |
| STXBP1 | -0.000277612 |
| CTAG1B | -0.000280747 |
| MIR1273F | -0.000281527 |
| NPTX1 | -0.000284729 |
| TRIM8 | -0.000286468 |
| RPL13AP | -0.000287593 |
| RP1-297M16 | -0.000290334 |
| OR52A4 | -0.000296518 |
| ALG1 | -0.000297074 |
| RP5-1139B12 | -0.000299306 |
| ARMC3 | -0.0002994 |

| | |
|---|---|
| OR13Z2P | -0.000301491 |
| DEFB133 | -0.000304956 |
| RP11-760L24 | -0.000307068 |
| RNU1-19P | -0.000308048 |
| OGFR-AS1 | -0.000311984 |
| AC005324 | -0.000316451 |
| AF001548 | -0.000321502 |
| CNTN4 | -0.000327959 |
| TRBV24OR9-2 | -0.000333419 |
| CTD-2586B10 | -0.000336166 |
| AC107871 | -0.000337542 |
| CTD-2515C13 | -0.00034489 |
| RNA5SP438 | -0.00034497 |
| ASPSCR1 | -0.000358546 |
| ITIH3 | -0.000360165 |
| AC005783 | -0.00036103 |
| CCDC171 | -0.000364574 |
| DNMT3L | -0.000366331 |
| SERPINA7P1 | -0.000368301 |
| RP1-93L7 | -0.000368847 |
| CTD-2124B20 | -0.000373695 |
| IL9RP2 | -0.000376285 |
| RP11-20E24 | -0.000380339 |
| RN7SL273P | -0.000385735 |
| AC090578 | -0.000386609 |
| NAT6 | -0.000386826 |
| CABP1 | -0.000391217 |
| DDX18P6 | -0.000393064 |
| U51244 | -0.000393933 |
| MZF1-AS1 | -0.000395627 |
| RNA5SP427 | -0.000404035 |
| RP11-74C3 | -0.0004163 |
| HUNK-AS1 | -0.000417142 |
| FTLP5 | -0.000418338 |
| MMAB | -0.000421935 |
| RP11-665J16 | -0.000424553 |
| PROX1 | -0.000425172 |
| KRTAP9-11P | -0.000425925 |
| MIR4703 | -0.00042648 |
| SALL4P1 | -0.000426518 |
| PRAMEF2 | -0.000427084 |
| RP11-557F20 | -0.000430908 |
| MAPRE3 | -0.000432447 |
| MORC2 | -0.000434605 |
| WIPI2 | -0.000435833 |

| | |
|---|---|
| AOC1 | -0.000438959 |
| ME1 | -0.000444801 |
| RP11-88H10 | -0.000445452 |
| CTD-2514K5 | -0.000450739 |
| NOS3 | -0.00045122 |
| RP11-8L18 | -0.000455433 |
| RP11-713P14 | -0.000455499 |
| OR6C5P | -0.000461345 |
| RP11-281H11 | -0.000462685 |
| RP11-529H20 | -0.000463611 |
| TSC2 | -0.000464523 |
| AC011225 | -0.000467457 |
| SDHAP3 | -0.000469937 |
| RPL31P61 | -0.000470495 |
| RP11-45A17 | -0.000470588 |
| RP11-22D3 | -0.000472586 |
| DEFB104B | -0.000473983 |
| SCGB2A2 | -0.000475948 |
| C19orf57 | -0.000478737 |
| FRMD1 | -0.000479069 |
| CFL2 | -0.000479686 |
| RP11-416N4 | -0.000483232 |
| CCL22 | -0.000483668 |
| RP11-117L5 | -0.000484022 |
| FAM107A | -0.00048471 |
| RNA5SP225 | -0.000486841 |
| AMY2A | -0.000488569 |
| C16orf93 | -0.000493994 |
| PSRC1 | -0.000495426 |
| MAGEA13P | -0.000496102 |
| CTD-2057D4 | -0.000497831 |
| LINC00583 | -0.000499195 |
| RNA5SP390 | -0.000499711 |
| TGM2 | -0.000500394 |
| CTC-273B12 | -0.000503255 |
| KRTAP4-8 | -0.000503501 |
| TMCC3 | -0.000505994 |
| CRADD | -0.000512543 |
| WISP3 | -0.000513195 |
| AC008175 | -0.000518309 |
| SLC37A4 | -0.000532154 |
| AL162419 | -0.000534111 |
| RP11-317M11 | -0.000534607 |
| OVCH1-AS1 | -0.000536268 |
| RNU7-185P | -0.00054382 |

| | |
|---|---|
| OSTCP5 | -0.000544468 |
| SNORD115-32 | -0.000546917 |
| RP11-44N12 | -0.000551514 |
| FARS2 | -0.000554778 |
| CTC-347C20 | -0.0005574 |
| WBP2P1 | -0.000559039 |
| PPIEL | -0.000560053 |
| FKSG66 | -0.000561326 |
| FAM73B | -0.000564043 |
| RP11-23B5 | -0.000565489 |
| RP11-358M11 | -0.000567986 |
| FAM32A | -0.000570983 |
| TCERG1L | -0.000572544 |
| TPRKB | -0.000574574 |
| RP11-345M22 | -0.000583278 |
| LOC440461 | -0.000585239 |
| MIR378A | -0.000597258 |
| AC091895 | -0.000598421 |
| RP11-367G12 | -0.000598928 |
| AC005027 | -0.000603141 |
| AC105252 | -0.000603553 |
| ZMYND10-AS1 | -0.000606292 |
| CTD-2544L4 | -0.00061046 |
| PNPLA1 | -0.000613564 |
| RP11-925D8 | -0.000613924 |
| MDGA2 | -0.000619284 |
| GOLGA2P8 | -0.00062022 |
| RNU6-801P | -0.000621655 |
| PITRM1-AS1 | -0.000623637 |
| RPS26P15 | -0.000624724 |
| NEBL-AS1 | -0.000628251 |
| CHRM3-AS1 | -0.000634788 |
| DACT2 | -0.000636176 |
| RESP18 | -0.00063806 |
| RP11-138I1 | -0.000638707 |
| ATP4A | -0.000640075 |
| GS1-531I17 | -0.000640193 |
| SNORA71 | -0.000642916 |
| CECR7 | -0.000643842 |
| RP1-258N20 | -0.000644673 |
| RPL21P12 | -0.000645283 |
| RP11-430H10 | -0.000645704 |
| RMST_3 | -0.000646082 |
| SNORD115-3 | -0.000648455 |
| RPL3P6 | -0.000658166 |

| | |
|---|---|
| RNU4ATAC4P | -0.00065995 |
| NLRP5 | -0.000661176 |
| RIOK3 | -0.000662429 |
| MIR23C | -0.000664336 |
| RP11-470F18 | -0.000665304 |
| AC073973 | -0.000665622 |
| GYG1P1 | -0.000669011 |
| RP11-759A24 | -0.00067153 |
| RP11-63M22 | -0.000672278 |
| TSPAN3 | -0.000679081 |
| RP11-174I12 | -0.000688847 |
| TUBB4BP1 | -0.00069119 |
| OR4K14 | -0.000692559 |
| OR1S2 | -0.000693406 |
| MAFA-AS1 | -0.000694365 |
| BMP15 | -0.00069508 |
| RP11-492A10 | -0.000698753 |
| RP11-281P23 | -0.00069965 |
| HMP19 | -0.000701001 |
| DUX4L9 | -0.000705515 |
| RAB18 | -0.000709389 |
| OR5B10P | -0.000709699 |
| NANOGP7 | -0.000710033 |
| AF196779 | -0.000714251 |
| RPS4XP17 | -0.00071996 |
| RP11-59C5 | -0.000720218 |
| NIPSNAP3B | -0.000724088 |
| NUP210P2 | -0.000724184 |
| OR5G4P | -0.000724979 |
| RN7SL391P | -0.000725926 |
| MIR10B | -0.000727279 |
| CCDC157 | -0.000730246 |
| NCOR1P1 | -0.000731909 |
| POU3F2 | -0.000737312 |
| RNU6-696P | -0.000738969 |
| STAU2 | -0.000740907 |
| CPSF3 | -0.000742786 |
| CTAGE16P | -0.000743038 |
| AC092786 | -0.000744511 |
| RP11-753D20 | -0.00074542 |
| AC092316 | -0.000747072 |
| CTD-2526M8 | -0.000747971 |
| CCDC144B | -0.000751421 |
| ELAC2 | -0.000762285 |
| AC135584 | -0.000763335 |

| | |
|---|---|
| AL121875 | -0.000763335 |
| AC073255 | -0.000765235 |
| CDCA7L | -0.000770397 |
| RPL19P13 | -0.000772942 |
| RP11-981G7 | -0.000774415 |
| RPL41P2 | -0.000774583 |
| MIR1224 | -0.000776564 |
| CTD-3046C4 | -0.000776594 |
| NACA3P | -0.000781923 |
| RGS19 | -0.000782397 |
| SPATA45 | -0.000786101 |
| ZNF695 | -0.000786742 |
| CTC-774J1 | -0.000787258 |
| EEF2K | -0.000790807 |
| RNU5F-6P | -0.000795252 |
| NMNAT1P3 | -0.000797615 |
| AJ006995 | -0.000799982 |
| MAST1 | -0.00080331 |
| RP11-399F2 | -0.000804516 |
| ADAM21P1 | -0.000807215 |
| RNU1-63P | -0.000811021 |
| RP11-690G19 | -0.000811695 |
| RNU6-273P | -0.000812601 |
| AP001198 | -0.000821241 |
| FLJ12825 | -0.00082158 |
| RNF11B | -0.000821994 |
| RP1-4G17 | -0.000828126 |
| RNA5SP160 | -0.000830238 |
| FASN | -0.000833451 |
| RP11-583F24 | -0.00083562 |
| ZNF692 | -0.000840576 |
| RP11-530N7 | -0.000844166 |
| RNF133 | -0.000849167 |
| METTL1 | -0.000853526 |
| PPP4R1L | -0.000854289 |
| RNA5SP81 | -0.000865401 |
| FOXO4 | -0.000869787 |
| LLNLF-65H9 | -0.000870397 |
| SULT1A2 | -0.00087509 |
| RPL31P2 | -0.000881833 |
| SALRNA1 | -0.000883014 |
| AC005668 | -0.000883921 |
| HSP90B3P | -0.000884845 |
| AC007163 | -0.000885473 |
| AMER1 | -0.000886271 |

| | |
|---|---|
| AC024996 | -0.000887126 |
| NOMO2 | -0.000887146 |
| RNU6-231P | -0.000890418 |
| C17orf105 | -0.000898642 |
| RPS26P54 | -0.000899605 |
| YIPF3 | -0.000901193 |
| DIP2B | -0.000902068 |
| GUCY2F | -0.000904457 |
| AC136896 | -0.000905308 |
| SLC41A1 | -0.000906979 |
| RP11-592P9 | -0.000908295 |
| AC106753 | -0.00090859 |
| AC005042 | -0.000918087 |
| KLHL2 | -0.000920599 |
| TMEM99 | -0.000924468 |
| MIR8054 | -0.000926448 |
| OR7E104P | -0.000930914 |
| RNU6-1214P | -0.00093456 |
| RP11-640I15 | -0.000937486 |
| RP11-342F21 | -0.000937734 |
| RNA5SP485 | -0.000946741 |
| RP11-631M6 | -0.00094972 |
| RP11-403H13 | -0.000951776 |
| RPL21P33 | -0.000954186 |
| RPL30P5 | -0.000955622 |
| MAPK11 | -0.000955875 |
| FAM220A | -0.000956265 |
| OR5AK4P | -0.000958835 |
| RP11-139F4 | -0.000960114 |
| EXO5 | -0.000960768 |
| AC099799 | -0.000964672 |
| SCRT2 | -0.000967964 |
| CTD-2368P22 | -0.00096847 |
| RP5-1106E3 | -0.000970944 |
| RP11-105N13 | -0.000971641 |
| AC068718 | -0.00097587 |
| ATP5F1 | -0.000977122 |
| RP11-478L17 | -0.000978136 |
| AFAP1L2 | -0.000979708 |
| RP13-346H10 | -0.000981251 |
| RPL36AP41 | -0.000983308 |
| RP11-986E7 | -0.000986271 |
| RP11-689E3 | -0.000991142 |
| ASMTL-AS1 | -0.000992062 |
| AC004159 | -0.000992802 |

| | |
|---|---|
| TUBBP10 | -0.000995908 |
| PCDHB1 | -0.001004766 |
| MIR4684 | -0.00100583 |
| ESD | -0.001007449 |
| LGI1 | -0.001013569 |
| MIR519E | -0.00102238 |
| RP4-747G18 | -0.001023249 |
| RP11-770J1 | -0.001024212 |
| AC009312 | -0.001025631 |
| MIR1298 | -0.001031954 |
| RHPN2 | -0.001032083 |
| SRRT | -0.001036101 |
| RP11-49O14 | -0.001046923 |
| MRPL27 | -0.001047841 |
| CLLU1 | -0.001048932 |
| HMGN1P24 | -0.001049078 |
| HIST1H3H | -0.001053721 |
| SPIN2B | -0.001055228 |
| ANKRD30B | -0.001056198 |
| BPIFC | -0.001056596 |
| TUNAR | -0.001058301 |
| TGIF2LY | -0.001064448 |
| MIR4733 | -0.001065891 |
| RP11-440D17 | -0.001068773 |
| AC011196 | -0.001070311 |
| CYP1A2 | -0.001073312 |
| LCN1P1 | -0.001075918 |
| RLF | -0.001078503 |
| RP11-545P7 | -0.001079086 |
| NKX2-6 | -0.001083069 |
| CYP4F44P | -0.001102515 |
| RP11-334G22 | -0.001104321 |
| TOR1B | -0.001105146 |
| SNCA-AS1 | -0.001111397 |
| IL6 | -0.001115379 |
| CR848007 | -0.00111938 |
| RP11-378G13 | -0.00112152 |
| MLXIPL | -0.001122561 |
| LINC00920 | -0.001123789 |
| RPL34P18 | -0.001124068 |
| PSKH2 | -0.001129135 |
| RNU6-1001P | -0.001131472 |
| OR52B4 | -0.001133766 |
| HIGD1AP13 | -0.001135484 |
| RCOR2 | -0.001136738 |

| | |
|---|---|
| HSF4 | -0.00114018 |
| RP11-107F6 | -0.001140743 |
| RP11-430L16 | -0.001141875 |
| RNA5SP304 | -0.001144961 |
| OR4D2 | -0.001149421 |
| OR4P4 | -0.001154007 |
| RP11-318G21 | -0.001154329 |
| RN7SL173P | -0.001161409 |
| MT-TY | -0.001168907 |
| AC022872 | -0.001172611 |
| RNU2-16P | -0.001173551 |
| RNU6-457P | -0.001176167 |
| UBL5P1 | -0.001179381 |
| RP11-118H15 | -0.001183257 |
| EEF1B2P8 | -0.001192797 |
| CTC-391G2 | -0.001197149 |
| RNU6-718P | -0.001202449 |
| ABALON | -0.001203383 |
| RP11-165H23 | -0.001204533 |
| RN7SL37P | -0.001209557 |
| RP11-525E9 | -0.001214717 |
| FARSA | -0.001216239 |
| RN7SL794P | -0.001225294 |
| TOLLIP-AS1 | -0.001225691 |
| RNU2-53P | -0.001226527 |
| AL159997 | -0.001226559 |
| SDK1 | -0.00123745 |
| MIR154 | -0.00123899 |
| AC011752 | -0.001242072 |
| TMEM95 | -0.001243359 |
| RP11-434H14 | -0.00124406 |
| ELMO2P1 | -0.001244773 |
| PRM3 | -0.001253183 |
| RN7SL578P | -0.001253895 |
| AP000640 | -0.001257693 |
| AF131216 | -0.001259665 |
| CYP4F3 | -0.001261586 |
| E2F6P4 | -0.001265826 |
| OR7E59P | -0.001268364 |
| DUSP5P2 | -0.001269075 |
| PAWRP1 | -0.001270913 |
| RNA5-8SP6 | -0.001273221 |
| CTD-2192J16 | -0.00127667 |
| CETP | -0.001281191 |
| AL138885 | -0.001283009 |

| | |
|---|---|
| RP4-814D15 | -0.001289408 |
| AP001032 | -0.001291983 |
| DCAF13P3 | -0.001292031 |
| SIAH1 | -0.001294066 |
| RP11-75A9 | -0.001295853 |
| DRAM2 | -0.001296076 |
| AC140479 | -0.001298044 |
| LINC01205 | -0.001303614 |
| ANKK1 | -0.001305471 |
| MKRN2OS | -0.001312058 |
| SUPT3H | -0.001313695 |
| RP11-747D18 | -0.001316802 |
| RNA5SP298 | -0.00131765 |
| RPL36AP10 | -0.001323754 |
| RP11-90K18 | -0.001336071 |
| PGBD3 | -0.001344415 |
| OR52Z1 | -0.00134481 |
| AP001052 | -0.001346033 |
| CTD-2529P6 | -0.001347363 |
| CSNK2A3 | -0.001349849 |
| RNU1-15P | -0.001354998 |
| ZNF519P4 | -0.001362131 |
| TRIB3 | -0.001366274 |
| OR8I2 | -0.001368601 |
| PIH1D1 | -0.001369028 |
| ENPP3 | -0.001378219 |
| NLGN1-AS1 | -0.001381705 |
| RP11-5L12 | -0.001388455 |
| RP11-10L12 | -0.001395764 |
| RP11-162O12 | -0.001398609 |
| SCARNA10 | -0.001398849 |
| SLIT1-AS1 | -0.001400606 |
| LINC00968 | -0.001404684 |
| NANOGP2 | -0.001406288 |
| RPL37P15 | -0.001406548 |
| E2F4 | -0.001414946 |
| HPRTP4 | -0.001415467 |
| RP11-654D12 | -0.001417781 |
| MIR1276 | -0.001419434 |
| MTND5P11 | -0.001422425 |
| FCF1P9 | -0.001430399 |
| RNU6-53P | -0.001430569 |
| AMD1 | -0.001430872 |
| TBC1D3J | -0.001440062 |
| KRT18P6 | -0.001440129 |

| | |
|---|---|
| RP11-607P23 | -0.001445246 |
| TRIM46 | -0.001445254 |
| TBC1D3F | -0.001454905 |
| RP11-241N4 | -0.001458069 |
| RPS15AP3 | -0.001461381 |
| RP1-28O17 | -0.001461398 |
| CTC-463A16 | -0.001462018 |
| RN7SL376P | -0.001464864 |
| PPP1R14BP5 | -0.001467852 |
| MSL3 | -0.001468325 |
| GAPDHP48 | -0.001468987 |
| RNU6-259P | -0.001471857 |
| NUDT19 | -0.0014788 |
| RN7SL622P | -0.00148186 |
| RP11-669I1 | -0.00148844 |
| APBA3 | -0.001491731 |
| CYP2C58P | -0.001494137 |
| RP11-411B10 | -0.001496485 |
| ZNF273 | -0.001497442 |
| ADPRH | -0.001502782 |
| CADPS2 | -0.001502931 |
| OR4C7P | -0.00150369 |
| CTB-1048E9 | -0.00150545 |
| AC011523 | -0.001508322 |
| HMGA1P1 | -0.001508735 |
| RNASE11 | -0.001510677 |
| SPATA7 | -0.001519341 |
| RNA5SP177 | -0.001528974 |
| IFT172 | -0.001533221 |
| CTD-2568A17 | -0.001538322 |
| ALG1L7P | -0.001550407 |
| RP11-310I9 | -0.00155267 |
| LINC00994 | -0.00155763 |
| AL049868 | -0.001558193 |
| RP5-965G21 | -0.001560943 |
| SBF2 | -0.001561055 |
| SIGLEC27P | -0.001566779 |
| SOD1P2 | -0.001568188 |
| AL137164 | -0.001574541 |
| IKBKGP1 | -0.001574755 |
| RP11-438J1 | -0.001576645 |
| CFL1P2 | -0.001577463 |
| AC083760 | -0.001581152 |
| KRT18P34 | -0.001581539 |
| AC114783 | -0.001594617 |

| | |
|---|---|
| TCEB2P2 | -0.001597936 |
| RP11-692E14 | -0.001602374 |
| PRKCZ | -0.001606508 |
| RP5-1165K10 | -0.001609987 |
| MAGEA7P | -0.001610498 |
| HOXA10-HOXA9 | -0.001615091 |
| CTD-2158P22 | -0.001618033 |
| OR7E4P | -0.001620538 |
| AC079054 | -0.001631391 |
| ABCC13 | -0.001631784 |
| RNU4-20P | -0.001634016 |
| VIPR2 | -0.001645232 |
| IGFBP7-AS1 | -0.001648065 |
| OR5L2 | -0.001659033 |
| LINC01255 | -0.001659163 |
| RP11-629N8 | -0.001664058 |
| EEF1A1P32 | -0.001665303 |
| RP11-318C4 | -0.001673022 |
| BAI3 | -0.00167664 |
| MIR217HG | -0.001678472 |
| RP1-137F1 | -0.001678565 |
| GAS5 | -0.001678623 |
| RP11-847H18 | -0.00168526 |
| GPR123-AS1 | -0.001687424 |
| MRVI1-AS1 | -0.001690865 |
| TMED10 | -0.001693879 |
| SLC26A8 | -0.001695684 |
| RNA5SP350 | -0.001702248 |
| RP5-1193P9 | -0.001712029 |
| RP11-488I4 | -0.001712786 |
| ZSCAN4 | -0.001715721 |
| AC104654 | -0.00171663 |
| UBE2D3P3 | -0.001720715 |
| RNA5SP416 | -0.00172136 |
| AC144573 | -0.001724456 |
| RNU6-1035P | -0.001727519 |
| TTC7B | -0.001732318 |
| RP11-188P20 | -0.001732318 |
| MIR4764 | -0.001736569 |
| RP1-73A14 | -0.001744228 |
| LSM3P3 | -0.001748083 |
| LINC01029 | -0.001751397 |
| DLK2 | -0.001757491 |
| MTG1 | -0.001761194 |
| TRBV22-1 | -0.001764877 |

| | |
|---|---|
| CCL28 | -0.001765622 |
| RP11-1012E15 | -0.001766729 |
| RP11-625B21 | -0.001767371 |
| MED9 | -0.001781883 |
| IGKV2-38 | -0.001784672 |
| RP11-862P13 | -0.001786653 |
| C7orf55 | -0.001789604 |
| PAXBP1P1 | -0.001789922 |
| RP11-764E7 | -0.001798839 |
| RCC2P1 | -0.001802824 |
| AF246928 | -0.001803933 |
| RNF138P1 | -0.001810883 |
| RAB29 | -0.001813214 |
| RPL31P60 | -0.001818264 |
| SNX5 | -0.001819168 |
| RP5-1056L3 | -0.001819373 |
| RP11-493G17 | -0.001821054 |
| AL158035 | -0.001821961 |
| RP13-157F18 | -0.001825276 |
| RP11-490O24 | -0.00182915 |
| RP11-210L7 | -0.001830687 |
| RP11-101E19 | -0.00183225 |
| RP11-254A24 | -0.001832259 |
| MTRNR2L7 | -0.001833192 |
| AC064875 | -0.00183473 |
| AFAP1-AS1 | -0.001835496 |
| UBALD2 | -0.001835839 |
| PYGM | -0.001837476 |
| CTD-2545H1 | -0.001840105 |
| KRT8P37 | -0.001842225 |
| FNBP1L | -0.001849793 |
| AC006335 | -0.001849981 |
| MIR3648-1 | -0.001852786 |
| OR2AH1P | -0.001854899 |
| RP11-138C24 | -0.001859127 |
| KARS | -0.001860528 |
| RPL26P9 | -0.001864982 |
| EEF1A1P4 | -0.001866184 |
| RP1-290I10 | -0.001867571 |
| AC211476 | -0.001871778 |
| RN7SL868P | -0.00187401 |
| RPS4XP8 | -0.001874178 |
| RNA5SP435 | -0.001874741 |
| GUCY2C | -0.001876268 |
| AATBC | -0.001880659 |

| | |
|---|---|
| RNA5SP449 | -0.001883558 |
| RNU6-713P | -0.001884025 |
| SPTY2D1-AS1 | -0.001888195 |
| AC009478 | -0.001891744 |
| RPL35AP21 | -0.001896351 |
| CAD | -0.00189759 |
| FLT1P1 | -0.001899617 |
| RP4-562J12 | -0.00190289 |
| EEF1DP8 | -0.001906673 |
| RNU6-539P | -0.001913005 |
| RP11-367B6 | -0.001916602 |
| RP4-764D2 | -0.001917381 |
| ULK4P3 | -0.001917721 |
| LINC00664 | -0.001918605 |
| RNU6-325P | -0.001919699 |
| SCARB2 | -0.001919712 |
| FAM26D | -0.00192508 |
| RP11-909N17 | -0.001929326 |
| MIP | -0.001932678 |
| PDIA2 | -0.001936036 |
| RP11-534C12 | -0.001944532 |
| LDHBP2 | -0.001946242 |
| RP11-375A5 | -0.001947778 |
| ENDOD1 | -0.001950746 |
| RN7SKP192 | -0.001953867 |
| MIR521-2 | -0.001955435 |
| AL033528 | -0.001956237 |
| CNTNAP4 | -0.0019613 |
| IGLCOR22-2 | -0.001963693 |
| USHBP1 | -0.001964737 |
| HOXD8 | -0.001965314 |
| RP11-465O11 | -0.001966857 |
| RALY-AS1 | -0.001968124 |
| SAA4 | -0.001971803 |
| ALDH3B1 | -0.001971938 |
| HNRNPAB | -0.001973482 |
| MRTO4 | -0.001979138 |
| RP11-641A6 | -0.001980463 |
| MIR6499 | -0.001981097 |
| FAM205A | -0.001981457 |
| SNRNP25 | -0.00198645 |
| MAGOH | -0.001987864 |
| OR9A1P | -0.001988539 |
| KRTAP4-7 | -0.001988625 |
| PI15 | -0.001989018 |

| | |
|---|---|
| B3GNT6 | -0.001992979 |
| RP1-125N5 | -0.001993287 |
| RP11-314C9 | -0.002004799 |
| RP3-500L14 | -0.002005298 |
| TSPEAR-AS1 | -0.002006198 |
| RHOXF1 | -0.002011538 |
| AC005358 | -0.002011785 |
| AC005480 | -0.00201822 |
| ANKRD18CP | -0.002020443 |
| XRCC6P5 | -0.00202152 |
| KRTAP5-10 | -0.002022157 |
| AC000374 | -0.00203211 |
| RNA5SP118 | -0.002041335 |
| ANAPC13 | -0.002045336 |
| KCNQ5 | -0.002055365 |
| CLEC3B | -0.002056016 |
| RP11-309M23 | -0.002057987 |
| RP11-226P1 | -0.00205917 |
| CTD-2560E9 | -0.002061273 |
| H3F3AP1 | -0.002062431 |
| SEPT4-AS1 | -0.002064705 |
| SNORD23 | -0.002065954 |
| HIST2H3A | -0.002066995 |
| RP11-356I18 | -0.002068805 |
| ZNF343 | -0.002070621 |
| LSM11 | -0.002071291 |
| CYP4F11 | -0.002071604 |
| RNU4ATAC | -0.002073333 |
| TCP1 | -0.002075366 |
| DNASE2 | -0.00207771 |
| RP11-200A13 | -0.002079388 |
| RPS7P3 | -0.002080393 |
| HSPE1P12 | -0.002084382 |
| MIR4419B | -0.002086694 |
| RNA5SP144 | -0.002088964 |
| RNFT1P2 | -0.00209327 |
| AC092670 | -0.002100095 |
| CYP11B1 | -0.002106717 |
| MIR593 | -0.002109167 |
| RANGRF | -0.002110735 |
| RNU6-827P | -0.002115453 |
| AC007333 | -0.002115453 |
| RNU6-478P | -0.002115453 |
| MIR548AE1 | -0.002115453 |
| RNU6-1325P | -0.002115453 |

| | |
|---|---|
| WNT2B | -0.002115832 |
| TUBB4AP1 | -0.002119286 |
| RP11-80K6 | -0.002127073 |
| MTA1 | -0.002135517 |
| OR4A41P | -0.002137724 |
| TAB2 | -0.002138345 |
| TTLL4 | -0.002140559 |
| HHLA1 | -0.00214545 |
| CNKSR1 | -0.002145638 |
| RNU6-564P | -0.002146297 |
| FOXE3 | -0.00214775 |
| TRAPPC2P7 | -0.00214904 |
| OR7A5 | -0.002151665 |
| AC109829 | -0.002153371 |
| EMC4 | -0.002158867 |
| CTD-2006K23 | -0.00216025 |
| RCC2P6 | -0.002160442 |
| EIF2D | -0.002163611 |
| RP11-379B8 | -0.002164478 |
| RP5-1004I9 | -0.00216643 |
| RP11-164P12 | -0.002167463 |
| RP11-259K15 | -0.002171437 |
| MIR3201 | -0.002171628 |
| HIGD1AP6 | -0.002175313 |
| OR7E55P | -0.002178677 |
| AC005077 | -0.002179668 |
| RP11-244N9 | -0.002189609 |
| GS1-388B5 | -0.002193582 |
| CYCSP28 | -0.002199613 |
| RP11-155O18 | -0.002202516 |
| VAV2 | -0.002205194 |
| RP11-139G7 | -0.002213829 |
| CTC-235G5 | -0.002213897 |
| ART2BP | -0.002216915 |
| RP11-65E10 | -0.002227324 |
| PHKG1P4 | -0.002229398 |
| RPS8P3 | -0.002232691 |
| ATP5G1P7 | -0.002235893 |
| AP000282 | -0.002236827 |
| AC021192 | -0.00223851 |
| DPCR1 | -0.002242992 |
| PPFIA3 | -0.002245122 |
| PARP3 | -0.00224514 |
| RP11-536I6 | -0.002247576 |
| RPL27AP8 | -0.002251443 |

| | |
|---|---|
| HMGN1P10 | -0.002254437 |
| RAPGEF4-AS1 | -0.00225496 |
| RP11-298E10 | -0.002256998 |
| RNA5SP18 | -0.002262246 |
| FTX | -0.002266779 |
| PPA1 | -0.002267951 |
| RN7SL813P | -0.002268945 |
| CTD-2553C6 | -0.002270119 |
| AC006509 | -0.002272119 |
| KRTAP10-12 | -0.002281181 |
| CYCSP48 | -0.002286126 |
| MTND2P20 | -0.00229016 |
| RPL17P12 | -0.002290473 |
| ATP5HP2 | -0.00229243 |
| MIR4480 | -0.00229906 |
| RP11-449L13 | -0.002300188 |
| ECM1P2 | -0.00230278 |
| RP1-140A9 | -0.002306585 |
| TMEM151A | -0.00230666 |
| RP4-778K6 | -0.002309309 |
| RP3-469D22 | -0.002310964 |
| AC098614 | -0.002312604 |
| XXbac-BPG300A18 | -0.002313607 |
| RP11-514F8 | -0.002322358 |
| RNU6-950P | -0.002323878 |
| RP11-567N4 | -0.002331727 |
| RP5-827E24 | -0.00233422 |
| RNU6-993P | -0.002336205 |
| LINC00888 | -0.002338931 |
| RNU4-63P | -0.002343608 |
| MYOF | -0.002343787 |
| FAM25C | -0.002345026 |
| ARRDC1-AS1 | -0.002347117 |
| CTA-221G9 | -0.002352829 |
| TTC25 | -0.002360066 |
| RAB3B | -0.002364774 |
| DEFB126 | -0.002367628 |
| RNA5SP269 | -0.002370024 |
| RP11-572N21 | -0.002373814 |
| RP4-678D15 | -0.002374383 |
| B3GNT2 | -0.002374821 |
| ATP6V1C2 | -0.002375435 |
| TNFRSF11A | -0.00237763 |
| RP11-334A14 | -0.002379335 |
| FSIP2-AS1 | -0.002380148 |

| | |
|---|---|
| OR5B17 | -0.002381173 |
| SSFA2 | -0.002382505 |
| RAB9A | -0.002382542 |
| AC135724 | -0.002384821 |
| POLR3F | -0.002387528 |
| SNRPGP6 | -0.002390284 |
| PRR26 | -0.002395353 |
| GABRG3 | -0.002396019 |
| PDILT | -0.002396624 |
| RN7SL344P | -0.002404133 |
| SLC25A6P4 | -0.002406123 |
| RP1-20I3 | -0.002409248 |
| RP11-575I8 | -0.002409459 |
| MMRN2 | -0.002411715 |
| UQCRHL | -0.002413018 |
| RP3-449H6 | -0.002413846 |
| RPL13AP23 | -0.002415665 |
| AC010754 | -0.002415772 |
| SAPCD2P4 | -0.002418564 |
| STARD6 | -0.002422593 |
| AL121578 | -0.002423648 |
| PINK1-AS | -0.002427102 |
| RP11-704C2 | -0.002429164 |
| RP11-153K11 | -0.002432548 |
| RNU1-146P | -0.002433014 |
| LBH | -0.002436126 |
| RP11-1149M3 | -0.002443093 |
| FLJ20019 | -0.002446235 |
| AC078889 | -0.002450356 |
| RP11-640M9 | -0.002452216 |
| TMEM206 | -0.002453112 |
| MARK1 | -0.002454322 |
| RP11-453M23 | -0.00245796 |
| OR4C14P | -0.002458296 |
| NKX2-3 | -0.002460102 |
| NPSR1-AS1 | -0.002460271 |
| RP11-362J17 | -0.002461829 |
| OR4N2 | -0.002462223 |
| GAPDHP72 | -0.002469547 |
| IGLVVI-25-1 | -0.002470146 |
| AP001605 | -0.002471555 |
| MIR4749 | -0.002477983 |
| KRTAP22-2 | -0.002478234 |
| TMEM108-AS1 | -0.00248071 |
| ACVR1C | -0.002482565 |

| | |
|---|---|
| RP11-355N15 | -0.002485086 |
| RP5-1119A7 | -0.002485472 |
| AKR1B1P1 | -0.002487797 |
| NKX2-1-AS1 | -0.002488091 |
| RP11-259P6 | -0.002488476 |
| AC023115 | -0.002488597 |
| RP13-1016M1 | -0.002490066 |
| RGR | -0.00249118 |
| ASAP1-IT2 | -0.002495774 |
| RP11-390K5 | -0.002496575 |
| GUCA2B | -0.002497968 |
| AC009035 | -0.002499447 |
| TBX20 | -0.002500222 |
| TP73-AS1 | -0.002503087 |
| AL353782 | -0.002504049 |
| ZPLD1 | -0.002508171 |
| RPS15AP34 | -0.002510014 |
| AC024610 | -0.002510065 |
| RPL5P17 | -0.002511456 |
| MIR7975 | -0.002516198 |
| LINC01005 | -0.002517589 |
| ZNF806 | -0.002519788 |
| MGAT5 | -0.002522976 |
| RNU6-458P | -0.002524896 |
| RAB6C-AS1 | -0.002527689 |
| MIR4307 | -0.002528505 |
| INSC | -0.002535692 |
| RNU6-839P | -0.002542029 |
| RP3-393E18 | -0.002544415 |
| RP11-248J23 | -0.002545637 |
| AC091878 | -0.002545901 |
| RP11-715C4 | -0.002552074 |
| ZBTB16 | -0.002552888 |
| RP11-849F2 | -0.002553266 |
| RP11-498D10 | -0.002565127 |
| APOO | -0.002565955 |
| RP11-158N24 | -0.002567348 |
| RN7SL378P | -0.002568975 |
| ATG4B | -0.00257055 |
| RP11-52L5 | -0.002571526 |
| RN7SKP161 | -0.002571871 |
| TRIM7 | -0.002572355 |
| AC002530 | -0.002573825 |
| RP11-4L24 | -0.002575734 |
| SLC13A2 | -0.002577544 |

| | |
|---|---|
| GDI2P1 | -0.002577603 |
| AL133168 | -0.002578361 |
| AC018635 | -0.002579083 |
| RP11-1221G12 | -0.00258218 |
| MDH1B | -0.002584672 |
| MIR6754 | -0.002585448 |
| MIR3137 | -0.002592111 |
| TUBB8P4 | -0.002598936 |
| RP11-297H3 | -0.002599394 |
| RP11-543B16 | -0.002600756 |
| AC073115 | -0.002602934 |
| RNU7-24P | -0.00260515 |
| RNA5SP215 | -0.002634075 |
| FSTL5 | -0.002636042 |
| OR7E66P | -0.002636261 |
| AC027088 | -0.002641954 |
| CTD-2372A4 | -0.002646957 |
| MIR3196 | -0.002647625 |
| AC017053 | -0.002659245 |
| AC104000 | -0.002661377 |
| GOLGA2P6 | -0.00266236 |
| CCNT2-AS1 | -0.002662963 |
| PPM1L | -0.002663112 |
| AL353147 | -0.00266621 |
| KCNK6 | -0.002669699 |
| SUMO2P20 | -0.002671412 |
| ABC7-43041300I9 | -0.002678602 |
| RNU6-67P | -0.002681906 |
| PARP1P1 | -0.002683316 |
| EIF3M | -0.002683895 |
| RP11-177H2 | -0.002685841 |
| RP11-51M24 | -0.00268831 |
| RNU6-1044P | -0.002692349 |
| AL132772 | -0.002694574 |
| GRID2 | -0.002694874 |
| RP1-193H18 | -0.002695542 |
| SERPINA10 | -0.002696962 |
| RP11-382E9 | -0.0026973 |
| RP11-307I14 | -0.002698295 |
| RP11-195M16 | -0.002700421 |
| CTA-481E9 | -0.002702597 |
| FAM218A | -0.002704379 |
| FAM71A | -0.002704462 |
| GMNN | -0.0027049 |
| SLC35F4 | -0.002706818 |

| | |
|---|---|
| OR13I1P | -0.002707056 |
| RP11-243M5 | -0.002716495 |
| RP1-96H9 | -0.002725226 |
| FGF10 | -0.002726876 |
| S100P | -0.002726948 |
| NME8 | -0.002740195 |
| RP11-284G10 | -0.002744654 |
| RP11-748H22 | -0.002747878 |
| MIR646HG | -0.002755845 |
| MIR3147 | -0.002756891 |
| RP11-156G14 | -0.002767471 |
| RP11-162N7 | -0.002769047 |
| LINC01574 | -0.00277183 |
| RNU6-504P | -0.002777193 |
| AC010745 | -0.002779269 |
| OR5BN1P | -0.002786902 |
| LARP1BP1 | -0.002787289 |
| RN7SL615P | -0.002787972 |
| KDM8 | -0.002788122 |
| MIR520A | -0.002788127 |
| ACAA2 | -0.00278989 |
| BNIP3P42 | -0.00278997 |
| AL139318 | -0.002798799 |
| AC010878 | -0.002804565 |
| KB-1205A7 | -0.002804887 |
| RPL17P17 | -0.002806047 |
| RP11-613C6 | -0.002810294 |
| HIST1H2AM | -0.002810491 |
| NAV2 | -0.002810672 |
| NDUFA5P12 | -0.002810877 |
| AC007560 | -0.002812586 |
| RPL18AP7 | -0.002816842 |
| DCUN1D2-AS | -0.002819583 |
| PWRN3 | -0.002823587 |
| MIR9-3 | -0.002825457 |
| OR11J2P | -0.002833084 |
| PDSS2 | -0.00283378 |
| AC093822 | -0.002836478 |
| RP11-523J2 | -0.002838477 |
| ZNF48 | -0.002842804 |
| MIR6762 | -0.002846839 |
| AL162511 | -0.002847082 |
| ZPR1 | -0.002847139 |
| AP003176 | -0.002847312 |
| ORMDL2 | -0.002853242 |

| | |
|---|---|
| NDRG2 | -0.002862265 |
| RNU6-805P | -0.002863199 |
| OR5M12P | -0.002865141 |
| LONRF2P2 | -0.002866231 |
| CTC-320C6 | -0.00286963 |
| SIRT3 | -0.002876573 |
| TAF1D | -0.002876703 |
| CEBPA | -0.002877801 |
| DHX37 | -0.002883196 |
| EVPL | -0.002888685 |
| RNU4ATAC7P | -0.00288966 |
| RPL23AP54 | -0.002897189 |
| AARS2 | -0.002899495 |
| SEPHS1P1 | -0.00290035 |
| RP11-360P21 | -0.002901807 |
| MISP | -0.002902639 |
| NF2 | -0.00290917 |
| BOK-AS1 | -0.002909607 |
| BCAT2 | -0.002913133 |
| RP11-100N21 | -0.002913319 |
| CASK-AS1 | -0.002914357 |
| EGFLAM-AS2 | -0.002914559 |
| RP11-43D2 | -0.002918846 |
| PSG7 | -0.002919747 |
| SPHK2 | -0.002926677 |
| TCP11L1 | -0.002927905 |
| RNU6-113P | -0.002945722 |
| REXO1 | -0.002946381 |
| MIR31 | -0.002948141 |
| ALOX15 | -0.002949043 |
| PSMD12P | -0.002954767 |
| LINC01101 | -0.002956131 |
| KB-1299A7 | -0.002961715 |
| RP11-309L24 | -0.002968409 |
| RP11-99A1 | -0.002976697 |
| IPO9-AS1 | -0.002991387 |
| C17orf47 | -0.003000058 |
| PRR34 | -0.003003388 |
| MIR4255 | -0.003009037 |
| RP11-90K6 | -0.003013587 |
| C1orf228 | -0.003022291 |
| SLC25A5-AS1 | -0.003026161 |
| MIR3663 | -0.003027542 |
| MIR23A | -0.003028352 |
| U95743 | -0.003028863 |

| | |
|---|---|
| RASSF8 | -0.003031277 |
| KCNQ1DN | -0.003032505 |
| RP11-328K22 | -0.003033448 |
| RP11-175I6 | -0.003034139 |
| SPNS3 | -0.003037305 |
| RPL7L1P3 | -0.003039965 |
| AP005271 | -0.003044287 |
| RP11-242F24 | -0.003045783 |
| LINC00462 | -0.003050205 |
| UBXN7-AS1 | -0.003057527 |
| RP4-669B10 | -0.003064445 |
| RP11-550F7 | -0.003065791 |
| AC018866 | -0.003069004 |
| PLEKHH2 | -0.003070646 |
| AL353707 | -0.003071382 |
| PPP2CA | -0.003071484 |
| MTND4P21 | -0.003076417 |
| ACOT12 | -0.003082892 |
| GS1-259H13 | -0.003089596 |
| RP11-264M12 | -0.003090723 |
| RPS3AP52 | -0.003090781 |
| AC008938 | -0.003092382 |
| ELANE | -0.003093098 |
| PXMP2 | -0.003094238 |
| AC079140 | -0.003105102 |
| RP11-63A2 | -0.00310513 |
| KRAS | -0.003105401 |
| SLFNL1-AS1 | -0.003111058 |
| MTND2P6 | -0.003111985 |
| UNGP1 | -0.003114305 |
| RP11-468I15 | -0.003119606 |
| ZAR1 | -0.003122739 |
| RP11-134E15 | -0.003124251 |
| HPCA | -0.003125719 |
| RP11-195B3 | -0.003127884 |
| RP11-986G18 | -0.003129105 |
| PTPN13 | -0.003129309 |
| RNU6-175P | -0.00313026 |
| RPL4P4 | -0.003130872 |
| COX5BP8 | -0.003131783 |
| RSL24D1P8 | -0.003133452 |
| C1orf61 | -0.003136651 |
| SNHG11 | -0.003147989 |
| RP11-7O11 | -0.003149348 |
| CCDC109B | -0.003152194 |

| | |
|---|---|
| RP11-391A7 | -0.003153839 |
| RNU6-1052P | -0.003158204 |
| GAPDHP34 | -0.003161539 |
| AC008850 | -0.003164742 |
| RP11-42H13 | -0.003166557 |
| KRT8P21 | -0.003167382 |
| MCOLN3 | -0.003167655 |
| EFCAB8 | -0.003177145 |
| RP11-597M12 | -0.003178513 |
| ZNF286B | -0.003185814 |
| MIR7515HG | -0.003186407 |
| RP11-3M1 | -0.003196211 |
| OR52Y1P | -0.003196394 |
| RP11-14O19 | -0.003196946 |
| CISD1P1 | -0.00319739 |
| RP11-510M2 | -0.003203775 |
| MIR1183 | -0.003206693 |
| ZNF736P4Y | -0.0032073 |
| TTLL12 | -0.003208755 |
| BMPR1A | -0.003208885 |
| STK11IP | -0.003208941 |
| DUXAP1 | -0.003216296 |
| MS4A7 | -0.003221537 |
| MIR4504 | -0.003226412 |
| C11orf98P1 | -0.00322673 |
| AC013460 | -0.003229772 |
| ALOX12 | -0.003239621 |
| EEF1DP1 | -0.003240161 |
| NDUFAF4P4 | -0.003241489 |
| RP11-707O23 | -0.003246529 |
| ZNF565 | -0.003247864 |
| FOXK1 | -0.003259325 |
| ZFAND6 | -0.003260832 |
| AP000547 | -0.00326214 |
| RP11-173D14 | -0.003265588 |
| MIS18A | -0.003266488 |
| RNU6-1090P | -0.003271053 |
| RP11-496D24 | -0.003273002 |
| AL138807 | -0.003274968 |
| RP11-316N24 | -0.003277076 |
| CTSO | -0.003282284 |
| RP11-674N23 | -0.003284053 |
| RPL36AP43 | -0.00328569 |
| LRRC14 | -0.003286656 |
| DNAJC30 | -0.003286954 |

| | |
|---|---|
| HMGN2P30 | -0.003289123 |
| LINC00396 | -0.003290887 |
| CYP2A13 | -0.00329698 |
| RP11-46I1 | -0.003303321 |
| RP11-526I8 | -0.003304137 |
| C6orf1 | -0.003308852 |
| CTD-2383I20 | -0.003309563 |
| PRSS53 | -0.003310731 |
| ZNF470 | -0.003311048 |
| OR8K4P | -0.00331181 |
| RP11-640N11 | -0.003312384 |
| ABCC12 | -0.003314949 |
| RPL7AP61 | -0.003316926 |
| LA16c-312E8 | -0.003317141 |
| RN7SKP184 | -0.003326191 |
| SMC1B | -0.003326379 |
| LINC00349 | -0.003328117 |
| RP3-365O12 | -0.003332004 |
| CXorf49 | -0.003332781 |
| OR4A12P | -0.003334954 |
| RP1-146A15 | -0.003337509 |
| AL390966 | -0.003343781 |
| AC003968 | -0.003344059 |
| AC064837 | -0.003345383 |
| FTLP8 | -0.003348276 |
| RP11-196E1 | -0.003348782 |
| AP000936 | -0.003351963 |
| MYCL | -0.003357003 |
| ADCK2 | -0.003357338 |
| SLC2A6 | -0.003369692 |
| OR2T3 | -0.003370473 |
| SNRPGP1 | -0.0033708 |
| RN7SL329P | -0.003377013 |
| TMEM14A | -0.003377041 |
| NET1 | -0.003378362 |
| RP11-533F5 | -0.003378967 |
| SLC16A13 | -0.003382028 |
| ZNF256 | -0.003382362 |
| RP11-159D23 | -0.003384312 |
| RN7SKP15 | -0.003384404 |
| AC118562 | -0.003385122 |
| RP11-121A13 | -0.003388738 |
| RP3-418C23 | -0.003391198 |
| AL138810 | -0.003403235 |
| RP11-680H20 | -0.003404019 |

| | |
|---|---|
| RP11-61D1 | -0.003412841 |
| AC007050 | -0.003417413 |
| RP13-554M15 | -0.003420001 |
| CDHR3 | -0.003420187 |
| RPL6P29 | -0.003422436 |
| HOXD4 | -0.003425149 |
| RP11-6J24 | -0.003426138 |
| SUMO1P2 | -0.003428945 |
| MKI67P1 | -0.003436166 |
| RP11-481K9 | -0.003438522 |
| SESN1 | -0.003439016 |
| CNTNAP2 | -0.003440207 |
| CLYBL-AS2 | -0.003443907 |
| CACTIN-AS1 | -0.003447398 |
| RNU6-908P | -0.003449577 |
| NMU | -0.003451281 |
| MACC1-AS1 | -0.003453545 |
| RP11-379J13 | -0.003454239 |
| LINC00290 | -0.003456293 |
| SCNN1G | -0.003459287 |
| ACTL10 | -0.003459612 |
| MANSC1 | -0.003465901 |
| IGF2BP3 | -0.003467581 |
| SCARB1 | -0.003467855 |
| FOCAD | -0.003470106 |
| RP11-207F8 | -0.003473328 |
| RNU6-556P | -0.003473793 |
| RP11-262D11 | -0.003473985 |
| CCDC152 | -0.003475018 |
| LA16c-2F2 | -0.003475563 |
| CBX3 | -0.003476246 |
| LINC01018 | -0.003477706 |
| RPS2P48 | -0.003478757 |
| AL357140 | -0.003482877 |
| RP11-28P17 | -0.003487774 |
| RP11-618K16 | -0.003488225 |
| RNU6-609P | -0.00349222 |
| OR2T7 | -0.003492743 |
| RN7SKP87 | -0.003493332 |
| RN7SL526P | -0.003495371 |
| EEF1B2P7 | -0.003497702 |
| LY6G6C | -0.003499636 |
| ASPA | -0.003500731 |
| RP11-363I22 | -0.003508293 |
| MIR548AJ2 | -0.003509615 |

| | |
|---|---|
| NCRUPAR_2 | -0.003511888 |
| ATP13A2 | -0.003512768 |
| HSPA13 | -0.003516322 |
| RNU6-511P | -0.003516973 |
| LINCMD1 | -0.003520816 |
| HIST1H2BF | -0.003523967 |
| SLC6A21P | -0.003524197 |
| ANKRD20A4 | -0.003527842 |
| RGS21 | -0.003531739 |
| AC005597 | -0.003535548 |
| RNU6-1220P | -0.003537156 |
| MRPS5 | -0.003537774 |
| DLEU2 | -0.003537979 |
| AC012175 | -0.003538328 |
| RP11-280O24 | -0.00354214 |
| RP11-376N17 | -0.003543963 |
| AC137675 | -0.003546211 |
| CTC-394G3 | -0.00355043 |
| AC092484 | -0.003551105 |
| FRMD6-AS1 | -0.003553068 |
| MIR4780 | -0.00355504 |
| OR6L2P | -0.00355526 |
| TRIAP1 | -0.003558203 |
| LINC00376 | -0.003564846 |
| RNA5SP136 | -0.003565081 |
| RP11-76N22 | -0.003565345 |
| RNU6-959P | -0.003567057 |
| PANX2 | -0.003569013 |
| RNU7-30P | -0.003569839 |
| AC006925 | -0.003571488 |
| RP11-31E13 | -0.003578819 |
| RP1-90K10 | -0.003579511 |
| AC111155 | -0.003580584 |
| KB-1073A2 | -0.003584098 |
| MUPP | -0.00358438 |
| RP5-945I17 | -0.003592088 |
| CLNS1AP1 | -0.003597543 |
| DBNL | -0.0036004 |
| RP11-598C10 | -0.003604103 |
| RP11-806H10 | -0.003604735 |
| AC025263 | -0.003606407 |
| RN7SL517P | -0.003608537 |
| NAGS | -0.003612773 |
| AC083811 | -0.003615042 |
| RP11-256K9 | -0.003616743 |

| | |
|---|---|
| CNP | -0.003621889 |
| RP11-669M2 | -0.003622909 |
| AC093390 | -0.003626524 |
| RP11-810H18 | -0.003639824 |
| RNU2-18P | -0.003640382 |
| NOVA2 | -0.003640532 |
| TPSP2 | -0.003642089 |
| RP11-320P11 | -0.003643052 |
| SPARCL1 | -0.003646969 |
| AF067845 | -0.003649632 |
| AL360001 | -0.00365208 |
| RP11-1007J8 | -0.00366004 |
| AC011453 | -0.003660269 |
| OR4C3 | -0.0036605 |
| RP11-687D19 | -0.003663794 |
| RP11-116D2 | -0.003665429 |
| ITGA9 | -0.003665575 |
| RN7SKP204 | -0.003668634 |
| CYP51A1P1 | -0.003669095 |
| ZACN | -0.003672533 |
| CLEC4G | -0.003674915 |
| AC004744 | -0.003678591 |
| IQUB | -0.003679771 |
| TMEM165 | -0.003682524 |
| RN7SL73P | -0.003682616 |
| DNAJC25-GNG10 | -0.00368272 |
| CAMK1 | -0.00368325 |
| HMGA1P6 | -0.003685822 |
| ATP2A1 | -0.00368792 |
| KB-1683C8 | -0.003688091 |
| RNU1-36P | -0.003688884 |
| CTC-338M12 | -0.003689353 |
| PSMA6P2 | -0.003689572 |
| WI2-8325B5 | -0.003690453 |
| RP11-354E23 | -0.003695984 |
| AC019064 | -0.003697096 |
| HSPA8P14 | -0.003698006 |
| AL117187 | -0.003699996 |
| RNU6-627P | -0.003701954 |
| GTF2F2P2 | -0.003709623 |
| PIN4P1 | -0.003718295 |
| EPHA7 | -0.003721776 |
| AC013470 | -0.003722224 |
| TM4SF19-TCTEX1D2 | -0.003728429 |
| LINC00238 | -0.003728903 |

| | |
|---|---|
| AL033522 | -0.003734899 |
| CTD-2046I8 | -0.003735441 |
| SEC11B | -0.003738141 |
| GS1-54N10 | -0.003745131 |
| MIR586 | -0.003745787 |
| TAAR5 | -0.003747801 |
| DNAJC19P3 | -0.00375163 |
| RP11-296A18 | -0.00375289 |
| OR7E136P | -0.003754759 |
| RP11-114O13 | -0.003755628 |
| RPL13AP3 | -0.003757086 |
| RP3-503A6 | -0.003757618 |
| TMEM223 | -0.00376344 |
| AL445074 | -0.003763512 |
| TPT1P7 | -0.003764612 |
| KAT8 | -0.00376571 |
| IZUMO1 | -0.003772499 |
| STX18 | -0.003775105 |
| LA16c-306E5 | -0.003778336 |
| LINC00397 | -0.003778529 |
| RP11-789C2 | -0.003779831 |
| BOLA2P1 | -0.003780123 |
| NHP2 | -0.003781264 |
| GUSBP8 | -0.003781994 |
| SPINT3 | -0.003782813 |
| VPS25 | -0.003783534 |
| RP13-578N3 | -0.003783586 |
| COL6A4P1 | -0.003784391 |
| RP11-708L7 | -0.003789681 |
| RGPD3 | -0.003790078 |
| OR7E162P | -0.003790304 |
| NRG3-AS1 | -0.003792798 |
| AKR7A2 | -0.003796074 |
| AC010887 | -0.003804657 |
| KRTAP4-1 | -0.003804694 |
| MYCBP2-AS2 | -0.003805729 |
| MED25 | -0.003809302 |
| AC009878 | -0.003811469 |
| RNU6-464P | -0.003813858 |
| AC087393 | -0.00381489 |
| RP11-383I23 | -0.003815121 |
| HAT1 | -0.003817061 |
| WDR54 | -0.00382601 |
| BMP6P1 | -0.003831549 |
| HNRNPDLP2 | -0.003832395 |

| | |
|---|---|
| BEST1 | -0.003835908 |
| LINC01158 | -0.003838153 |
| RP11-515E23 | -0.003839005 |
| LRFN5 | -0.003839067 |
| POTEB | -0.003842843 |
| RP11-972P1 | -0.003845471 |
| RP4-646P19 | -0.003850106 |
| FAM60DP | -0.00385614 |
| KRTAP13-2 | -0.003856446 |
| SNN | -0.003856541 |
| CYP2C56P | -0.003856958 |
| RP11-459D22 | -0.003858884 |
| OR9R1P | -0.003864693 |
| KIF1A | -0.003889621 |
| AL163952 | -0.003889705 |
| RP11-545I5 | -0.003893064 |
| AP000593 | -0.003896673 |
| TAS2R62P | -0.003896738 |
| RP11-507K2 | -0.003898962 |
| RNA5SP250 | -0.003908784 |
| RP11-1J11 | -0.003912648 |
| COLQ | -0.003912786 |
| MTCL1 | -0.003915167 |
| CYP2D7 | -0.003916479 |
| PRKRIRP4 | -0.003917739 |
| RP11-699A7 | -0.003920243 |
| OR4F1P | -0.00392909 |
| RP11-570L14 | -0.003929879 |
| RP11-157H4 | -0.003931038 |
| CTD-2343P13 | -0.003934621 |
| ZNF286A | -0.00393528 |
| FABP1 | -0.00393779 |
| C16orf71 | -0.003942108 |
| CPS1-IT1 | -0.003945756 |
| LINC01203 | -0.003947178 |
| GPR64 | -0.003949773 |
| CTD-2026D23 | -0.00395682 |
| PRDX2P1 | -0.003957288 |
| ZNF92P3 | -0.003958248 |
| CTD-2334D19 | -0.003959571 |
| ABCD1P3 | -0.003961112 |
| RP11-704J17 | -0.003966095 |
| RP11-19F9 | -0.003968189 |
| AC139451 | -0.003977164 |
| RP11-217B7 | -0.0039773 |

| | |
|---|---|
| AC236972 | -0.003982436 |
| HDAC7 | -0.003985068 |
| NXPE2P1 | -0.003990335 |
| RP3-335N17 | -0.003991772 |
| MIR1238 | -0.003992227 |
| SPACA5B | -0.003997442 |
| RP11-1191J2 | -0.003998651 |
| RP11-438E8 | -0.004000223 |
| OR4C50P | -0.004001573 |
| TM9SF4 | -0.004008195 |
| OR5A1 | -0.004012589 |
| PGAM4P2 | -0.004016187 |
| UQCRC2 | -0.00401906 |
| RP11-127I20 | -0.00401973 |
| MYCLP1 | -0.004023594 |
| RP4-614N24 | -0.004025391 |
| MIR4444-2 | -0.004027967 |
| RN7SL39P | -0.004029074 |
| AC100802 | -0.004036113 |
| EIF4A1P4 | -0.004039485 |
| MAPK3 | -0.004040903 |
| RP11-577N1 | -0.00404125 |
| CTNND2 | -0.004047006 |
| CWC15 | -0.004047471 |
| DOT1L | -0.004048773 |
| EIF1P1 | -0.00404941 |
| AC115617 | -0.004049722 |
| RNU6-1279P | -0.004050367 |
| MIR3942 | -0.004050974 |
| AC021654 | -0.004054744 |
| AZGP1P1 | -0.004055231 |
| MLN | -0.004056471 |
| EBF1 | -0.004061109 |
| TUBBP8 | -0.00406324 |
| NT5C | -0.004065481 |
| BTF3P15 | -0.004066996 |
| AC072031 | -0.004071718 |
| KIAA1456 | -0.004071918 |
| RPL13AP17 | -0.004073929 |
| RP11-500G22 | -0.004074374 |
| MKX-AS1 | -0.004074908 |
| TMEM72 | -0.004077194 |
| TUBB1 | -0.004078515 |
| OR8V1P | -0.004079394 |
| NAALAD2 | -0.004083844 |

| | |
|---|---|
| RP11-342I1 | -0.004084051 |
| FOXD4L6 | -0.004090837 |
| RP11-535A19 | -0.004092329 |
| MIR5684 | -0.004092816 |
| CHORDC2P | -0.00409365 |
| CTB-2L9 | -0.004095054 |
| ECM1P1 | -0.004106315 |
| SLC30A1 | -0.004114543 |
| RP11-489M13 | -0.004122306 |
| AC006133 | -0.004122996 |
| RP11-756A22 | -0.004127808 |
| CTB-55O6 | -0.004129949 |
| RP11-203H19 | -0.004131066 |
| TXNL4A | -0.004137317 |
| SCN9A | -0.004137465 |
| VN1R96P | -0.004138195 |
| RNU6-715P | -0.004139109 |
| NGFR | -0.004141838 |
| MALRD1 | -0.004141952 |
| RP1-154J13 | -0.004144809 |
| AC069542 | -0.004150481 |
| MIR4662B | -0.004157518 |
| OR10A5 | -0.00416398 |
| RP11-329N22 | -0.004166393 |
| TRIM74 | -0.00416837 |
| MIR548AX | -0.004170382 |
| AC113607 | -0.00417064 |
| RP11-285A15 | -0.004171554 |
| RP11-416A17 | -0.004172792 |
| CYB5D2 | -0.004177151 |
| ZNF965P | -0.004178176 |
| RPS2P4 | -0.004179478 |
| RP3-332O11 | -0.004181311 |
| VPS9D1-AS1 | -0.004185077 |
| RNU6-1160P | -0.004185561 |
| MIR138-1 | -0.004186925 |
| WDR86-AS1 | -0.004187157 |
| RP11-335E14 | -0.004192106 |
| HIGD2B | -0.004192221 |
| SLIT2 | -0.004194473 |
| RPS2P44 | -0.004195225 |
| C10orf53 | -0.004196231 |
| LIPI | -0.00420067 |
| IQCG | -0.004200873 |
| RPL31P63 | -0.004205565 |

| | |
|---|---|
| SNRPCP17 | -0.004208833 |
| TRIM51HP | -0.004211603 |
| NTNG1 | -0.004213048 |
| IGHVII-78-1 | -0.004214504 |
| NXNP1 | -0.004215867 |
| LL22NC03-33B6 | -0.004215952 |
| AL034375 | -0.004223995 |
| RP11-142A5 | -0.004228181 |
| CMC2 | -0.004242921 |
| RP11-297C4 | -0.004247354 |
| RNU6-1000P | -0.004248305 |
| RP11-448A19 | -0.004252112 |
| RP11-208C17 | -0.004254281 |
| AC010103 | -0.004256012 |
| MIR181B2 | -0.004258223 |
| AC073834 | -0.004260022 |
| CIZ1 | -0.004266866 |
| RP11-15J23 | -0.004275218 |
| CYP2W1 | -0.00427661 |
| RP11-212J19 | -0.00428038 |
| RP11-125O5 | -0.004286269 |
| RN7SKP58 | -0.004289317 |
| COX7A2P2 | -0.00429017 |
| C11orf49 | -0.004294868 |
| FTX_2 | -0.004295004 |
| MRPL40 | -0.004295314 |
| RP11-780O17 | -0.0042959 |
| LINC01475 | -0.004296123 |
| IGF2BP2-AS1 | -0.004299232 |
| MAGEC3 | -0.004302473 |
| CTB-78F1 | -0.004308218 |
| TSHZ2 | -0.004309026 |
| AC093787 | -0.00430927 |
| LCA5 | -0.004324822 |
| DEFB103A | -0.004324858 |
| RNY3P5 | -0.004330994 |
| TRBV26OR9-2 | -0.004333804 |
| RPL31P47 | -0.004334653 |
| PLAC1 | -0.004335379 |
| AL031320 | -0.004343623 |
| RP11-268F1 | -0.004343843 |
| CTC-539A10 | -0.004349686 |
| UCK2 | -0.004353791 |
| OR4D7P | -0.004357274 |
| EQTN | -0.004359044 |

| | |
|---|---|
| F9 | -0.004362507 |
| FLT3LG | -0.004364136 |
| PMS2P1 | -0.004367894 |
| BTF3L4P4 | -0.004371491 |
| RN7SL817P | -0.004374124 |
| LINC01164 | -0.004374757 |
| HBG1 | -0.004375547 |
| ALOXE3 | -0.004379895 |
| ALKBH3-AS1 | -0.0043825 |
| RNU4-18P | -0.00438369 |
| RP3-406P24 | -0.00438676 |
| OR5AL2P | -0.004387531 |
| PCBD1 | -0.004389309 |
| FBP2 | -0.004395027 |
| FGF20 | -0.00439558 |
| SNORD115-38 | -0.004397296 |
| RNU6-20P | -0.004398594 |
| RP11-404I7 | -0.004400993 |
| GBAS | -0.004403133 |
| RP11-485F13 | -0.004417727 |
| UGT2B4 | -0.004422612 |
| RP11-364P22 | -0.004426924 |
| RNF25 | -0.004430464 |
| RNU6-321P | -0.004433784 |
| CTD-2517M22 | -0.004435227 |
| POLR2J3 | -0.004437813 |
| RNU6-667P | -0.004444886 |
| CYP2C18 | -0.004448507 |
| AKAP13 | -0.004449144 |
| RP11-527F13 | -0.004449443 |
| NDUFB1P1 | -0.004450321 |
| MIR300 | -0.004452275 |
| LRRC47 | -0.004453982 |
| CTD-2033C11 | -0.004455042 |
| MTG2 | -0.004455191 |
| RN7SKP88 | -0.004456513 |
| RASGRF1 | -0.004460735 |
| RP4-742J24 | -0.004465239 |
| MRPS35P3 | -0.004467793 |
| RNA5SP24 | -0.004473278 |
| DPRXP1 | -0.004474136 |
| RP4-529N6 | -0.004479523 |
| FLJ37035 | -0.004480011 |
| AL357936 | -0.004482043 |
| MIR548O | -0.004487575 |

| | | |
|---|---|---|
| MED28P2 | | -0.004487948 |
| DYRK1B | | -0.004489388 |
| RP11-687F6 | | -0.004495086 |
| AL049758 | | -0.004499396 |
| CFL1P1 | | -0.004500244 |
| COPS3 | | -0.004501928 |
| TRIM60 | | -0.004501977 |
| MED15P5 | | -0.004502499 |
| EXOSC1 | | -0.004504328 |
| COX6B1P4 | | -0.004506651 |
| RP11-442N1 | | -0.004507923 |
| SPINK14 | | -0.004517023 |
| RP11-187O7 | | -0.004521919 |
| CTC-506B8 | | -0.004523935 |
| RP11-778D9 | | -0.004524496 |
| RNU4-27P | | -0.004524979 |
| RP11-300M6 | | -0.004533488 |
| PLCE1 | | -0.004536385 |
| RP11-656G20 | | -0.004537879 |
| RP11-12M5 | | -0.004538897 |
| RNA5SP115 | | -0.004540463 |
| RNU4-33P | | -0.004541525 |
| PHLDB3 | | -0.004548907 |
| MIR6859-1 | | -0.004549656 |
| SEMA4G | | -0.004552758 |
| RNU6-66P | | -0.004555332 |
| CTD-3187F8 | | -0.004556269 |
| HTR3B | | -0.00456019 |
| AL117374 | | -0.004569363 |
| HAMP | | -0.004570787 |
| CYP4F25P | | -0.004571523 |
| | 7-Sep | -0.004572629 |
| RPLP0P6 | | -0.004573168 |
| RNASEH1-AS1 | | -0.004577894 |
| MIR4642 | | -0.004578003 |
| CTD-3214K23 | | -0.004579872 |
| ANGPTL1 | | -0.004582415 |
| RP11-215C7 | | -0.004583077 |
| RPL7P19 | | -0.004583134 |
| RP11-395C17 | | -0.004583419 |
| RP11-95P9 | | -0.004585823 |
| CTD-3233P19 | | -0.004588074 |
| RP11-540E16 | | -0.00459215 |
| RNU7-124P | | -0.004599819 |
| CTA-204B4 | | -0.004600351 |

| | |
|---|---|
| SELV | -0.004601716 |
| MIR4661 | -0.004601867 |
| RP11-707A18 | -0.004604569 |
| SMIM22 | -0.00460476 |
| STC2 | -0.004610323 |
| HMGN2P27 | -0.004610356 |
| RN7SL291P | -0.004612924 |
| JAGN1 | -0.004613058 |
| PCMTD1P3 | -0.004613722 |
| LLPH | -0.004615142 |
| RP11-288D15 | -0.00461563 |
| RP11-20F24 | -0.004619572 |
| PCDHA9 | -0.004623525 |
| RN7SL242P | -0.004627335 |
| GSTM5P1 | -0.004628849 |
| TTLL7-IT1 | -0.004629415 |
| TMEM60 | -0.004633115 |
| RP11-529O20 | -0.00463478 |
| RP11-24B13 | -0.004635445 |
| RP11-419L9 | -0.004636415 |
| FXYD3 | -0.004651404 |
| AC004595 | -0.004653357 |
| RN7SL665P | -0.004661548 |
| IFT20 | -0.00466601 |
| CFAP20 | -0.004667405 |
| ZFPL1 | -0.004671405 |
| RP11-467P9 | -0.004674391 |
| MIR450A1 | -0.004675716 |
| RP11-152L7 | -0.004676007 |
| RP5-1061H20 | -0.004677154 |
| MIR4748 | -0.004678308 |
| ANTXRL | -0.004684113 |
| TBC1D3G | -0.00468645 |
| EPB41L2 | -0.004689006 |
| DMRTC1 | -0.004689255 |
| TFPI | -0.004691332 |
| AC012627 | -0.00469182 |
| RP11-540H22 | -0.004698623 |
| RP11-182L7 | -0.004698894 |
| RP11-876F14 | -0.0046991 |
| ZNF876P | -0.004699295 |
| FAHD2CP | -0.004701825 |
| RPL17P49 | -0.00470523 |
| SLC22A23 | -0.004710253 |
| RP5-1016A21 | -0.004711722 |

| | |
|---|---|
| MIR329-1 | -0.004715613 |
| LINC01015 | -0.004719006 |
| POLL | -0.004731054 |
| EEF1A1P41 | -0.004733994 |
| CEACAMP1 | -0.004734985 |
| PGAP3 | -0.004736561 |
| RP11-393K19 | -0.004742615 |
| MIR6797 | -0.004742647 |
| RNA5SP74 | -0.004744306 |
| OR13K1P | -0.00474647 |
| RNU6-47P | -0.004747159 |
| AC104076 | -0.004747542 |
| MRPS28 | -0.004754572 |
| VPS45 | -0.004756097 |
| RP11-521O16 | -0.004760337 |
| RN7SL655P | -0.00476194 |
| PISRT1 | -0.004764394 |
| RNU6-412P | -0.004767829 |
| HOXA11-AS1_4 | -0.004769367 |
| OR7E140P | -0.00477248 |
| LRRC20 | -0.004772842 |
| RNY1P10 | -0.004772899 |
| AC019196 | -0.004778967 |
| OGFOD3 | -0.004781587 |
| AL139022 | -0.004782926 |
| NDUFAF4P2 | -0.00478713 |
| CTC-490E21 | -0.004791615 |
| OR6B1 | -0.004795319 |
| ATHL1 | -0.004796378 |
| HNRNPRP2 | -0.004798567 |
| RN7SL342P | -0.00479994 |
| CTB-23I7 | -0.004803819 |
| MIR4282 | -0.004807337 |
| KRT8P20 | -0.00481189 |
| HIRIP3 | -0.004815486 |
| ISL1 | -0.004818144 |
| MIR6730 | -0.004819875 |
| RP6-105D16 | -0.004827436 |
| PKN2-AS1 | -0.004828304 |
| RNU6-758P | -0.004829546 |
| CCDC102B | -0.004830277 |
| PTP4A1P1 | -0.00483053 |
| RAB11AP1 | -0.00483243 |
| RP1-137H15 | -0.004838659 |
| VIS1 | -0.004845218 |

| | |
|---|---|
| AC008836 | -0.00485037 |
| ARHGAP29 | -0.004859841 |
| AP000567 | -0.004865682 |
| RP11-16C1 | -0.004870339 |
| CBWD7 | -0.004876065 |
| RN7SL551P | -0.004876381 |
| CTD-2001C12 | -0.004877478 |
| CREB3 | -0.004878744 |
| SPATC1L | -0.004879107 |
| NIFKP8 | -0.004880211 |
| MEPE | -0.004880308 |
| TMEM212 | -0.004882266 |
| DNAH10OS | -0.004895601 |
| ATP5LP3 | -0.004898442 |
| MIR889 | -0.004910714 |
| DLX6-AS1 | -0.004912902 |
| RNU6-215P | -0.004913371 |
| RN7SL337P | -0.00491481 |
| RP11-16B13 | -0.004921294 |
| RP11-340F16 | -0.004921716 |
| MIR1185-2 | -0.00492693 |
| TMEM88 | -0.004927026 |
| SLC25A33 | -0.004928384 |
| GPT | -0.004933221 |
| PRAMEF9 | -0.004937848 |
| POMZP3 | -0.004937854 |
| RPL12P41 | -0.004943299 |
| OR6C7P | -0.004944508 |
| HMGB3P9 | -0.004945924 |
| SCO2 | -0.004954804 |
| LINC01467 | -0.004956516 |
| RN7SL237P | -0.004957122 |
| RPL21P11 | -0.004958625 |
| B3GALNT2P1 | -0.004960349 |
| RP1-267D11 | -0.004967604 |
| RP11-67L2 | -0.004979549 |
| RP11-790L1 | -0.004981552 |
| TRMT112P1 | -0.004982918 |
| PDK3 | -0.004983475 |
| LINC01341 | -0.004985558 |
| KRT8P40 | -0.004993351 |
| COQ2 | -0.004994377 |
| AGBL4 | -0.004996576 |
| RP11-216P16 | -0.004996748 |
| TBCCD1 | -0.004997968 |

| | |
|---|---|
| LCN1P2 | -0.005000262 |
| RNA5SP343 | -0.005004278 |
| AC087738 | -0.005007235 |
| SYT11 | -0.005013671 |
| CTA-963H5 | -0.005015364 |
| SQLE | -0.005018918 |
| MIR544B | -0.005019701 |
| AC003664 | -0.005021567 |
| DLG1 | -0.00502182 |
| KRT18P26 | -0.005022225 |
| RP11-16O13 | -0.005025494 |
| LINC01362 | -0.005027213 |
| CTD-2313J23 | -0.005029461 |
| RP11-17A4 | -0.005033764 |
| RNU6-60P | -0.005034884 |
| RP11-624D11 | -0.005035181 |
| AL592205 | -0.005041497 |
| CRTC3 | -0.005048798 |
| RP11-211F2 | -0.0050494 |
| LINC01090 | -0.005050532 |
| PDZD4 | -0.005053785 |
| NLE1 | -0.005054138 |
| USP22 | -0.005058901 |
| OR5F2P | -0.005062029 |
| RGS10 | -0.005063348 |
| CCL16 | -0.005063437 |
| SLC7A13 | -0.005063466 |
| CDKL3 | -0.005063757 |
| LSMEM1 | -0.005065115 |
| MIR500A | -0.005065863 |
| AF104455 | -0.005066049 |
| RP11-295K2 | -0.00506931 |
| CECR5-AS1 | -0.005071302 |
| SLC25A5P9 | -0.005071713 |
| AC005391 | -0.005081454 |
| SCCPDH | -0.005082623 |
| RNU6-463P | -0.005083814 |
| MIR4710 | -0.005083838 |
| AL139187 | -0.005086199 |
| RP11-332H21 | -0.005095306 |
| OR2B4P | -0.005095988 |
| AC117479 | -0.005100487 |
| AC007795 | -0.005103009 |
| AC011901 | -0.005103616 |
| RP11-553N16 | -0.00510417 |

| | |
|---|---|
| RP11-1236K1 | -0.005106069 |
| AC013399 | -0.005107188 |
| C12orf49 | -0.005117564 |
| RP11-575M4 | -0.005120554 |
| GLT1D1 | -0.005123253 |
| SNORD65 | -0.005124406 |
| CSNK2A1 | -0.005125323 |
| RNA5SP85 | -0.005130015 |
| RBMY3AP | -0.005133344 |
| PPIAL4F | -0.005135098 |
| CCDC148 | -0.005137305 |
| RP11-439M15 | -0.005145092 |
| FAM86EP | -0.005146793 |
| NANP | -0.005149473 |
| SKIDA1 | -0.005150459 |
| NHLRC4 | -0.005150905 |
| MRPS5P4 | -0.00515132 |
| SNORD116-5 | -0.005159662 |
| MAGI2-AS2 | -0.00516045 |
| RP11-385H1 | -0.005161067 |
| RP11-527D7 | -0.005168415 |
| AC021037 | -0.0051703 |
| RNU6-171P | -0.005178757 |
| RNF167 | -0.005179055 |
| BANP | -0.005183632 |
| NPAP1P2 | -0.005185016 |
| RP11-99A14 | -0.00518572 |
| FAM210B | -0.005188153 |
| CYP2G1P | -0.00519045 |
| SLC2A3P4 | -0.005199646 |
| C6orf47 | -0.005202926 |
| RAF1P1 | -0.005203765 |
| RP11-251P6 | -0.005208589 |
| OR6K5P | -0.00521057 |
| VAPA | -0.005210945 |
| AC019171 | -0.005211253 |
| RNA5SP419 | -0.005216102 |
| RP11-133F8 | -0.005220723 |
| TMEM167AP1 | -0.00522418 |
| MIR548Q | -0.005227601 |
| FARSB | -0.005229327 |
| CALCP | -0.005230301 |
| CCDC151 | -0.005233997 |
| OR6D1P | -0.005236603 |
| IQCF4 | -0.005251124 |

| | | |
|---|---|---|
| RP11-203P2 | | -0.005251713 |
| FAM106DP | | -0.005268261 |
| MIR6856 | | -0.005280892 |
| PLEKHA8 | | -0.005282055 |
| RP11-664H17 | | -0.005287371 |
| MGME1 | | -0.005291749 |
| RP11-124G5 | | -0.005292901 |
| RNU6-857P | | -0.00530155 |
| ZNF492 | | -0.005302024 |
| FUT8-AS1 | | -0.005302577 |
| OR7D2 | | -0.005304281 |
| RNA5SP62 | | -0.005304874 |
| AC007179 | | -0.00531931 |
| RP11-122F24 | | -0.005320817 |
| RP11-336F14 | | -0.005323425 |
| STRADB | | -0.005327713 |
| RP11-363N22 | | -0.005329071 |
| RNU6-722P | | -0.00533192 |
| KRT31 | | -0.005338954 |
| RP11-758H6 | | -0.005342145 |
| RP11-463M14 | | -0.005343159 |
| RP11-813P10 | | -0.005344451 |
| GFRA4 | | -0.005347453 |
| CYP27A1 | | -0.005352421 |
| AF238380 | | -0.005358653 |
| RP11-19O2 | | -0.005370138 |
| AIFM1P1 | | -0.005371734 |
| ISCA1P6 | | -0.005377465 |
| AF001550 | | -0.005381468 |
| RP11-144I2 | | -0.005384701 |
| RP4-802A10 | | -0.005386149 |
| PTTG1 | | -0.005389465 |
| ZNF98 | | -0.005389759 |
| RP11-57B24 | | -0.00539018 |
| | 10-Sep | -0.005391288 |
| RNU105C | | -0.005393413 |
| RP11-566K11 | | -0.005395329 |
| HTATSF1P | | -0.005396094 |
| SNORA74 | | -0.005398902 |
| UBE2V1P9 | | -0.005405133 |
| AC068724 | | -0.005409595 |
| RNA5SP80 | | -0.005411925 |
| RP4-655C5 | | -0.005415249 |
| MRPL42P3 | | -0.005415579 |
| RFKP2 | | -0.005418598 |

| | |
|---|---|
| AC115286 | -0.005420197 |
| ZNF738 | -0.005421522 |
| OCA2 | -0.005421935 |
| AC098848 | -0.005424317 |
| RNU2-12P | -0.005428083 |
| AC233287 | -0.005430323 |
| GAPDHP1 | -0.005438416 |
| H2BFWT | -0.005440584 |
| OR7E121P | -0.005440852 |
| TSSC1 | -0.00544156 |
| RP11-284L19 | -0.005448433 |
| HTR3E-AS1 | -0.005450403 |
| AC097724 | -0.005452723 |
| CCDC58P5 | -0.005453769 |
| AL158077 | -0.00545884 |
| AC067735 | -0.005461847 |
| HSPA8P17 | -0.00546516 |
| RP11-14N9 | -0.00546611 |
| AL590764 | -0.005471138 |
| LRRC75A-AS1 | -0.005475794 |
| CYTH2 | -0.005477822 |
| RP13-262C2 | -0.005481332 |
| RP11-343J18 | -0.005481751 |
| RP11-426M1 | -0.005489784 |
| ERN2 | -0.005491433 |
| SPATA31D3 | -0.005494306 |
| PRELP | -0.005495851 |
| RNU6-874P | -0.005499142 |
| RP11-327I22 | -0.005499832 |
| RP11-173A6 | -0.005507196 |
| RNU2-24P | -0.005507804 |
| RP4-706G24 | -0.005508692 |
| AC069063 | -0.005509416 |
| IARS2P1 | -0.005509706 |
| SUN1 | -0.005511907 |
| RP11-809H16 | -0.005516997 |
| ZNF885P | -0.005517738 |
| FAM197Y8 | -0.005517918 |
| RPS29P6 | -0.005519756 |
| RP11-467D6 | -0.005521321 |
| C17orf70 | -0.005529824 |
| ZBED3-AS1 | -0.005531071 |
| RP11-11L12 | -0.005532325 |
| CHEK2P3 | -0.005534526 |
| SNORD115-31 | -0.005536725 |

| | |
|---|---|
| GABRA4 | -0.005539604 |
| DUXAP7 | -0.005541458 |
| MIR1303 | -0.005544064 |
| RNU6-615P | -0.005545859 |
| HGFAC | -0.005546652 |
| BTG4 | -0.005547399 |
| C19orf44 | -0.005550607 |
| SLC20A1P1 | -0.005554851 |
| OR4C11 | -0.005557478 |
| RNU6-642P | -0.005558505 |
| LA16c-352F7 | -0.005564329 |
| RP11-256L11 | -0.005571441 |
| NPAS3 | -0.005571891 |
| AC007950 | -0.005572201 |
| RP11-269F21 | -0.005580084 |
| RNU6-849P | -0.005585869 |
| AP3B1 | -0.005590757 |
| LINC01492 | -0.005594766 |
| SUMO2P8 | -0.005595185 |
| BPI | -0.00559622 |
| WFIKKN2 | -0.005599575 |
| VDAC3P1 | -0.005602669 |
| LINC01287 | -0.005603408 |
| PCBP3-OT1 | -0.005608166 |
| CLSTN3 | -0.005608786 |
| TNNC1 | -0.005610601 |
| PSAT1P1 | -0.005612114 |
| AC007064 | -0.005631846 |
| RP11-456P18 | -0.005632824 |
| LINC01432 | -0.005633809 |
| RPL30P14 | -0.00564098 |
| RNU6-422P | -0.005643799 |
| TUBBP5 | -0.005644183 |
| RPL23AP77 | -0.005647723 |
| BRI3BPP1 | -0.005647806 |
| FDFT1 | -0.005648732 |
| MIR6729 | -0.005651038 |
| RP11-338L22 | -0.00565128 |
| RNU6-953P | -0.005651474 |
| LINC00890 | -0.005651658 |
| RP11-63A23 | -0.005652945 |
| RP11-94D20 | -0.005659753 |
| CTD-2538G9 | -0.005662759 |
| PMCHL1 | -0.005662864 |
| RNU6-995P | -0.005663956 |

| | |
|---|---|
| AL353763 | -0.005665183 |
| RNU6-473P | -0.005671602 |
| AC096750 | -0.005674113 |
| HMGB1P9 | -0.005681698 |
| RNU6-573P | -0.00568232 |
| RN7SKP124 | -0.005682355 |
| RP11-538D16 | -0.005682787 |
| AC009134 | -0.005684957 |
| BOLA3-AS1 | -0.005685961 |
| EXOC4 | -0.005685998 |
| SNORD91A | -0.005694061 |
| RNU7-103P | -0.005694252 |
| RNU7-81P | -0.005696606 |
| RPL18AP17 | -0.005698302 |
| PRSS30P | -0.005699943 |
| RP11-157J24 | -0.005700761 |
| GBX1 | -0.005702418 |
| MIDN | -0.005703209 |
| SLC25A12 | -0.005703395 |
| RP11-407P2 | -0.00570487 |
| AC007253 | -0.005707184 |
| AC117490 | -0.005708034 |
| CRISP1 | -0.005712018 |
| FAM150B | -0.005712352 |
| RP11-180K7 | -0.005713185 |
| PSMD7 | -0.005713442 |
| CTA-407F11 | -0.005715499 |
| RP11-260N14 | -0.005717353 |
| ASIC3 | -0.005718465 |
| MIR3138 | -0.005720755 |
| PHOX2A | -0.00572199 |
| MIR3180-2 | -0.005722325 |
| RP11-767C6 | -0.005723473 |
| LINC00374 | -0.005733762 |
| BRD7P2 | -0.005744606 |
| ANKRD60 | -0.005745179 |
| KRT18P8 | -0.005745766 |
| ZNF259P1 | -0.005747207 |
| AL359853 | -0.005748869 |
| MTND1P23 | -0.00574989 |
| IGKV2OR2-8 | -0.005756061 |
| AC063976 | -0.005760241 |
| SNRPCP8 | -0.005761292 |
| TTL | -0.00576937 |
| CSNK1A1L | -0.00578408 |

| | |
|---|---|
| AMZ2P2 | -0.005784168 |
| TMEM175 | -0.005785184 |
| LINC01128 | -0.005785687 |
| LINC00567 | -0.005791174 |
| LINC00682 | -0.00579184 |
| CTD-2228A4 | -0.00579422 |
| C2orf43 | -0.005798878 |
| MTCYBP3 | -0.005800516 |
| HEXA | -0.005801334 |
| RPL35AP | -0.005806177 |
| AL354923 | -0.005808668 |
| PPP1R18 | -0.005809223 |
| RP11-120A1 | -0.005812651 |
| AC097372 | -0.005814574 |
| RP11-524O1 | -0.005815275 |
| LINC01578 | -0.005815716 |
| RP11-301A5 | -0.005819769 |
| GAS2L1P2 | -0.005824311 |
| RN7SL754P | -0.005829898 |
| CYP4F34P | -0.005830807 |
| YPEL5P1 | -0.005832803 |
| ERGIC2 | -0.005833168 |
| WI2-2994D6 | -0.005834499 |
| RPL12P8 | -0.005835742 |
| MKNK1 | -0.005843215 |
| RP4-791M13 | -0.005848996 |
| RNU1-55P | -0.005850389 |
| RNA5SP205 | -0.005850827 |
| FOXD1 | -0.005851187 |
| RP11-231E4 | -0.005852859 |
| AC133141 | -0.005853878 |
| RNF216-IT1 | -0.00585455 |
| RAMP1 | -0.005855773 |
| RNU6-701P | -0.00585586 |
| RNU1-150P | -0.005856492 |
| ZNF969P | -0.005858066 |
| HDDC2 | -0.005858471 |
| RN7SL531P | -0.0058611 |
| AC098612 | -0.005863148 |
| EPB42 | -0.005864404 |
| RP11-490D19 | -0.005866162 |
| RBMY2DP | -0.005872092 |
| TERF1P2 | -0.005883351 |
| RP11-9G10 | -0.005884848 |
| AL139021 | -0.00588777 |

| | |
|---|---|
| RP11-349N19 | -0.005889561 |
| RP11-103J17 | -0.005891577 |
| RNA5SP321 | -0.005895851 |
| LMF1-AS1 | -0.005902111 |
| INHA | -0.005903087 |
| SGSM2 | -0.005905543 |
| SPATA6 | -0.005906622 |
| RNU6-505P | -0.005910034 |
| FAM110B | -0.005910101 |
| AC010423 | -0.005914707 |
| AL360157 | -0.005918286 |
| ENTPD4 | -0.005924446 |
| FMR1NB | -0.005928029 |
| ARL2BPP6 | -0.005931008 |
| PHB | -0.005931051 |
| TPRG1L | -0.005931833 |
| RP11-2N1 | -0.005933559 |
| ENO1-IT1 | -0.005938083 |
| LINC01393 | -0.005940979 |
| EEF2 | -0.005941935 |
| AC007364 | -0.005944196 |
| FTH1P14 | -0.005947322 |
| DYNC1H1 | -0.005953824 |
| SLC37A2 | -0.005957899 |
| HAUS1P2 | -0.005958901 |
| SOAT1 | -0.00595948 |
| LINC01527 | -0.005959849 |
| CTC-384G19 | -0.00596102 |
| NQO2 | -0.005965453 |
| TPM3 | -0.005966755 |
| VTRNA2-2P | -0.005967234 |
| OR5BQ1P | -0.005967933 |
| AL354873 | -0.005970181 |
| WDR77 | -0.005971909 |
| MTHFD2P3 | -0.00597339 |
| CFHR2 | -0.005979473 |
| RNA5SP387 | -0.005984979 |
| HNRNPA1P8 | -0.005985119 |
| MIR519A1 | -0.005988069 |
| DAB1-AS1 | -0.005988278 |
| SLC1A3 | -0.005991258 |
| RP11-548H3 | -0.005991984 |
| DNAH17 | -0.00599297 |
| RNASEH1P1 | -0.005993444 |
| SGCB | -0.005993528 |

| | |
|---|---|
| KRT72 | -0.005994408 |
| AGO3 | -0.005996524 |
| MIR548AO | -0.005996922 |
| CEP72 | -0.005997361 |
| KB-1836B5 | -0.005999576 |
| ARMC2-AS1 | -0.006004949 |
| C17orf112 | -0.00601714 |
| MIR3659 | -0.006018741 |
| AF212831 | -0.00602029 |
| AC008694 | -0.00602137 |
| AC010982 | -0.006022997 |
| AC097382 | -0.006024162 |
| NPAP1P3 | -0.006029687 |
| RP11-356C4 | -0.006031411 |
| TMEM254 | -0.006035372 |
| STEAP4 | -0.006036749 |
| CH17-333M13 | -0.006037093 |
| RP11-360D2 | -0.006037809 |
| HIST2H2BF | -0.006040005 |
| LINC00308 | -0.006041942 |
| CICP18 | -0.006042386 |
| FAM173B | -0.006044321 |
| MIR6718 | -0.006046723 |
| RP11-373E16 | -0.006057932 |
| RP11-385G16 | -0.006058647 |
| AC013448 | -0.00606074 |
| RP11-655M14 | -0.006066521 |
| RNA5SP287 | -0.006068123 |
| SRSF4 | -0.00607249 |
| MIR2355 | -0.006074789 |
| CCDC37-AS1 | -0.006075173 |
| MIR7-2 | -0.0060756 |
| PDYN | -0.006076367 |
| RP11-3L21 | -0.006079014 |
| AC108485 | -0.006079648 |
| RP11-477N3 | -0.006083225 |
| TMX4 | -0.006083449 |
| GPX5 | -0.006083723 |
| PDIA3P2 | -0.006085752 |
| WNT8A | -0.006086032 |
| HSCB | -0.006086909 |
| RP11-550A5 | -0.006090555 |
| MIR4637 | -0.00609345 |
| TUBB3P1 | -0.006096049 |
| RP11-106E15 | -0.006097022 |

| | |
|---|---|
| RNU6-679P | -0.006102679 |
| KCTD9P2 | -0.006106222 |
| AC106736 | -0.006108469 |
| SCARNA4 | -0.006112243 |
| AFMID | -0.006116045 |
| MIR585 | -0.006117083 |
| PPRC1 | -0.006120269 |
| SPG7 | -0.00612198 |
| RPS24P13 | -0.006122228 |
| CBX1P2 | -0.006127199 |
| LINC00937 | -0.006127341 |
| SPDYE4 | -0.006129519 |
| RNA5SP227 | -0.006131983 |
| GLOD5 | -0.006133902 |
| LYPD8 | -0.006134015 |
| CEACAM7 | -0.006134701 |
| SOX15 | -0.006136121 |
| DNM1P50 | -0.006139066 |
| OR13D2P | -0.00613922 |
| KCNMA1 | -0.006141405 |
| AL162853 | -0.006153304 |
| HAGHL | -0.00615534 |
| RP1-249F5 | -0.006159863 |
| NPC1 | -0.006161182 |
| LINC00408 | -0.006164405 |
| RP3-380C13 | -0.006164573 |
| H2AFY | -0.006166261 |
| FBXO25 | -0.006180178 |
| MC1R | -0.00618256 |
| AC004022 | -0.006183178 |
| LINC00112 | -0.006185463 |
| RP5-1099E6 | -0.006187439 |
| SP9 | -0.00618783 |
| EIF2B4 | -0.006189518 |
| MIR548L | -0.006190529 |
| MIR4525 | -0.006198 |
| RP11-131P10 | -0.006199869 |
| RP11-338K13 | -0.006201434 |
| NUCB1-AS1 | -0.006202148 |
| RP11-114H24 | -0.006202738 |
| RP11-552O4 | -0.006204725 |
| RP11-98G13 | -0.006209446 |
| IL10RB-AS1 | -0.006209798 |
| AC010209 | -0.006209832 |
| RP11-323H21 | -0.006211595 |

| | |
|---|---|
| RP11-5G18 | -0.006213436 |
| MGAT4B | -0.006214171 |
| RP11-401E5 | -0.006214608 |
| RP11-308D13 | -0.006215613 |
| SULT6B1 | -0.006226862 |
| RDX | -0.006239716 |
| RPL3P1 | -0.006242404 |
| AC026992 | -0.006244567 |
| AC010504 | -0.006248888 |
| KRT18P41 | -0.006256516 |
| MIR620 | -0.006265861 |
| RPL21P71 | -0.006266168 |
| PYCR1 | -0.006266322 |
| SULT1A4 | -0.006267489 |
| RP5-1022J11 | -0.006269241 |
| LMAN1L | -0.006273664 |
| CTD-2142D14 | -0.006275405 |
| TIGD6 | -0.006279811 |
| OR51B6 | -0.006282782 |
| CEP170 | -0.006283011 |
| CHRNA4 | -0.006286925 |
| SAR1P3 | -0.006288788 |
| LINC00410 | -0.006289928 |
| PLEKHG4 | -0.006291104 |
| RNA5SP264 | -0.0062915 |
| KCTD9 | -0.00629356 |
| WDR45BP1 | -0.00629985 |
| RP11-399E6 | -0.006307439 |
| EEF1A1P36 | -0.006308827 |
| RP11-108H9 | -0.006314818 |
| MIR508 | -0.006315367 |
| BNIP3P17 | -0.006316728 |
| RP11-618L22 | -0.006317093 |
| TMEM189-UBE2V1 | -0.006318405 |
| TRPC6 | -0.006322306 |
| TRAF2 | -0.006326064 |
| RP11-82L7 | -0.006326089 |
| MID1IP1-AS1 | -0.006326959 |
| GOLGA6D | -0.006328856 |
| ANKRD30A | -0.006329158 |
| LINC00838 | -0.006330299 |
| PTCHD1-AS | -0.00633146 |
| LINC01168 | -0.006333545 |
| ANKRD20A9P | -0.006334149 |
| KLF3P1 | -0.006335013 |

| | |
|---|---|
| MIR96 | -0.006336274 |
| NOS2P2 | -0.006336316 |
| PTEN | -0.006336501 |
| PTMAP8 | -0.00633786 |
| RNA5SP98 | -0.006345262 |
| MIR518E | -0.006353419 |
| FAM90A2P | -0.006354962 |
| LACTB | -0.006355516 |
| KIAA0125P2 | -0.006357001 |
| RTL1 | -0.006357708 |
| AC008264 | -0.006359258 |
| AP1B1P1 | -0.006364716 |
| RPL21P103 | -0.006366381 |
| MYEOV | -0.006366428 |
| RNU6-1338P | -0.006368798 |
| CALM2P4 | -0.006371483 |
| LRRK1 | -0.006377471 |
| RP11-882G5 | -0.006378698 |
| NDUFS2 | -0.006378959 |
| RP11-272D20 | -0.006382974 |
| IGKV2D-19 | -0.006384738 |
| F2 | -0.006384945 |
| RNU6-289P | -0.006386982 |
| PLBD2 | -0.006388299 |
| NUDT5 | -0.006389471 |
| TTTY1 | -0.006389662 |
| MEG3_1 | -0.006390986 |
| OTOP3 | -0.006393604 |
| RP11-541G7 | -0.006394724 |
| RNU4ATAC18P | -0.006396085 |
| MAGI2-IT1 | -0.006396478 |
| TBX3 | -0.006397379 |
| RP11-218D6 | -0.006404079 |
| AC091043 | -0.00640422 |
| DEFB110 | -0.006406814 |
| CTD-2328D6 | -0.00641058 |
| EXOC6B | -0.006411723 |
| MAP3K19 | -0.006414031 |
| FABP5P6 | -0.006414797 |
| METTL13 | -0.006416672 |
| RPS10P3 | -0.006417742 |
| RNU6-731P | -0.00642599 |
| RP4-789D17 | -0.006428572 |
| GNB5 | -0.006430999 |
| FBXW10 | -0.006431259 |

| | |
|---|---|
| SYAP1 | -0.006433771 |
| MRPL51P2 | -0.006434044 |
| AL359314 | -0.006435982 |
| RN7SL279P | -0.006437019 |
| UBE2D3P2 | -0.006438432 |
| AP000221 | -0.006440956 |
| RPS18P1 | -0.006443549 |
| MIR6851 | -0.006454279 |
| ABL2 | -0.006466628 |
| RPL7P22 | -0.006471196 |
| RP11-154D9 | -0.006472359 |
| RPS3AP14 | -0.006473811 |
| BMPR1B | -0.006475317 |
| RP1-161C16 | -0.006478646 |
| TTC39CP1 | -0.006481665 |
| REXO1L8P | -0.006488795 |
| RP11-548N1 | -0.006489088 |
| PRR31 | -0.006495008 |
| STMN1 | -0.006500267 |
| CHRND | -0.006504463 |
| CRACR2B | -0.006507207 |
| RNA5SP221 | -0.006509875 |
| ST6GALNAC1 | -0.006512048 |
| RN7SL638P | -0.006515762 |
| RN7SKP246 | -0.006515883 |
| RP11-190D6 | -0.006516447 |
| RNA5SP388 | -0.006523465 |
| RFT1 | -0.006526817 |
| AC092506 | -0.006528885 |
| PLA2G2F | -0.00653066 |
| RNU6-793P | -0.006531571 |
| RP11-472M19 | -0.006532189 |
| SLC7A8 | -0.00653225 |
| ZNF202 | -0.006533515 |
| mascRNA-menRNA | -0.006533804 |
| AMD1P1 | -0.006538126 |
| HK1 | -0.006544343 |
| PRCP | -0.006544566 |
| CTAGE6 | -0.006545493 |
| RP11-313A24 | -0.006547911 |
| AC091517 | -0.006550376 |
| RNU6-191P | -0.006553231 |
| LINC01017 | -0.006553948 |
| WDR11-AS1 | -0.006558048 |
| PIP5KL1 | -0.006558614 |

| | |
|---|---|
| CEACAMP8 | -0.006558999 |
| RP3-449O17 | -0.006571916 |
| RP5-988G15 | -0.006574675 |
| NACAP2 | -0.006579678 |
| ENY2 | -0.006580959 |
| TP53TG3D | -0.006582177 |
| MIR323A | -0.006603815 |
| ABCB9 | -0.006605313 |
| MYBL2 | -0.006614523 |
| FOXI1 | -0.006615636 |
| AC092265 | -0.006619322 |
| RP11-311B14 | -0.006627278 |
| RP11-1110J8 | -0.006635652 |
| RP11-955H22 | -0.006641112 |
| SLC26A6 | -0.006641501 |
| PCDH11Y | -0.006644542 |
| RNU4-62P | -0.006645046 |
| CASP12 | -0.006648361 |
| NLRP8 | -0.006650439 |
| AL133249 | -0.006650861 |
| GSTA3 | -0.006653043 |
| HMCES | -0.006655976 |
| AC117401 | -0.00666688 |
| RP1-149A16 | -0.006667642 |
| PCDHGB1 | -0.006668341 |
| NADK2 | -0.006674098 |
| YPEL5P3 | -0.006674654 |
| FAM212B | -0.006675716 |
| AC106706 | -0.006676462 |
| RP11-804H8 | -0.006679698 |
| RABIF | -0.006680591 |
| MROH1 | -0.006681428 |
| HMGN1P26 | -0.006683277 |
| BCL9L | -0.006685708 |
| RP11-408H20 | -0.006686826 |
| CTD-2293H3 | -0.006691812 |
| CD164 | -0.00670035 |
| AC138783 | -0.006705035 |
| DUXA | -0.006705766 |
| RP3-512E2 | -0.006715301 |
| CPSF3L | -0.006715402 |
| HPCAL4 | -0.006716318 |
| MIR4759 | -0.006716342 |
| CTD-2278B20 | -0.00671647 |
| RN7SL790P | -0.006718296 |

| | |
|---|---|
| RP11-382B18 | -0.006723243 |
| AC114271 | -0.006726565 |
| AC020956 | -0.006729485 |
| PHBP8 | -0.006733951 |
| OR7C1 | -0.006746286 |
| AC023380 | -0.006747706 |
| HINT1P1 | -0.006748118 |
| RP11-164C12 | -0.006748237 |
| OR7E156P | -0.006749066 |
| RP11-507B12 | -0.006755087 |
| AC103794 | -0.006755236 |
| USP17L13 | -0.006755803 |
| CITF22-62D4 | -0.00675609 |
| AL589863 | -0.006763653 |
| RNU1-93P | -0.006767756 |
| RPL29P25 | -0.006768789 |
| RP11-748C4 | -0.006772767 |
| AC100756 | -0.006776282 |
| RNU6-474P | -0.00677914 |
| CASP4 | -0.006779833 |
| C2-AS1 | -0.006781322 |
| RAB11FIP3 | -0.006783498 |
| KRT33A | -0.006784957 |
| RP4-633H17 | -0.006786478 |
| RP11-436J20 | -0.006787745 |
| CTB-26E19 | -0.006789255 |
| AC003005 | -0.006792095 |
| TIPARP-AS1 | -0.006793266 |
| LINC00237 | -0.006795808 |
| AC007131 | -0.00679648 |
| TMEM253 | -0.006796678 |
| WEE2-AS1 | -0.006797564 |
| PPIHP1 | -0.006798416 |
| DNAJC1 | -0.006801843 |
| RP1-140C12 | -0.00680231 |
| RN7SL795P | -0.006813188 |
| RP11-597K23 | -0.006815135 |
| RP11-240D10 | -0.006821511 |
| MIR222HG | -0.006821862 |
| AC092666 | -0.006831014 |
| OR7E117P | -0.006834164 |
| CRYGGP | -0.006837097 |
| RP5-906A24 | -0.006843441 |
| USP32P3 | -0.006844015 |
| AC005616 | -0.006845325 |

| | |
|---|---|
| HCRTR1 | -0.006847238 |
| GRHL1 | -0.006848375 |
| TREML4 | -0.006848446 |
| CNOT7P1 | -0.006849801 |
| FAIM | -0.006851873 |
| LMCD1-AS1 | -0.006852766 |
| ABCC2 | -0.006861101 |
| RP11-145E17 | -0.006863902 |
| DDX10 | -0.006868097 |
| LSM3P5 | -0.006869206 |
| CSTA | -0.006869232 |
| AC005375 | -0.006877676 |
| RNU6-413P | -0.006881502 |
| ELK3 | -0.006885195 |
| CSHL1 | -0.006886091 |
| RP11-360F5 | -0.006890526 |
| RNU6-684P | -0.006892284 |
| PMPCAP1 | -0.006892339 |
| RPL12P2 | -0.006895615 |
| PSMC1P10 | -0.006895995 |
| RP11-112N19 | -0.006899509 |
| RP11-167P23 | -0.006901082 |
| DRC7 | -0.006904468 |
| RP11-807E13 | -0.006904883 |
| PMM2 | -0.006910666 |
| NPEPL1 | -0.006911094 |
| EEF1A1P9 | -0.006912899 |
| RP5-875H3 | -0.006913572 |
| KLF4P1 | -0.006919774 |
| RP11-214J9 | -0.006926508 |
| HINT2P1 | -0.006926641 |
| RP11-465I4 | -0.006927198 |
| SSBP2 | -0.006929569 |
| CTD-2194M22 | -0.006931014 |
| RP3-406C18 | -0.006932783 |
| TTLL10 | -0.006934935 |
| TMEM17 | -0.006935411 |
| PNMA5 | -0.006937178 |
| PCDH8 | -0.006937891 |
| RSL24D1P2 | -0.006941828 |
| RP11-640I2 | -0.006947341 |
| MAEA | -0.006950352 |
| C11orf44 | -0.006950951 |
| RP11-291L19 | -0.006951937 |
| RPL5P26 | -0.0069578 |

| | |
|---|---|
| LINC01314 | -0.006960697 |
| SERF1AP1 | -0.006964014 |
| MTND2P5 | -0.006965013 |
| SNRPCP13 | -0.00697571 |
| MTND4P25 | -0.006977615 |
| RP11-338N12 | -0.006977932 |
| AC106900 | -0.006981172 |
| RP11-860B13 | -0.006984695 |
| RP4-675C20 | -0.006987912 |
| LINC00550 | -0.006990345 |
| CALM3 | -0.006992586 |
| RN7SL147P | -0.006993315 |
| RBM15B | -0.006994502 |
| RPL7AP33 | -0.006994742 |
| RNU6-990P | -0.007005626 |
| RP11-160N1 | -0.007011145 |
| RP4-635G19 | -0.007011707 |
| MYF5 | -0.007012025 |
| AL357115 | -0.007013052 |
| AIPL1 | -0.007014854 |
| ETF1 | -0.007016085 |
| RP11-458I4 | -0.00701692 |
| ANKRD13B | -0.007018597 |
| RNU2-63P | -0.007023485 |
| MLLT4-AS1 | -0.007024476 |
| RNU6-889P | -0.007025324 |
| LINC01372 | -0.007030476 |
| CHRNA7 | -0.007036239 |
| RNU6-543P | -0.007047478 |
| RN7SKP106 | -0.007049104 |
| ZNF90 | -0.007052179 |
| RAVER1 | -0.007052552 |
| RPL5P4 | -0.007053761 |
| AC097721 | -0.007053947 |
| RP11-315I20 | -0.007054449 |
| RNA5SP455 | -0.007057526 |
| RN7SL718P | -0.007062003 |
| GSTO3P | -0.007063699 |
| RP11-463H24 | -0.007066377 |
| KDELC1P1 | -0.007066573 |
| RP11-739N20 | -0.007070147 |
| UGT2A2 | -0.007073796 |
| RP11-369K17 | -0.007079559 |
| AL132640 | -0.007080794 |
| FRAS1 | -0.007081409 |

| | |
|---|---|
| DUSP4 | -0.007082004 |
| CTD-2249K22 | -0.007087813 |
| SPTLC2 | -0.007088144 |
| RNA5SP278 | -0.007092328 |
| MIR571 | -0.007093671 |
| SCAANT1 | -0.00709409 |
| MIR4636 | -0.007096216 |
| EID3 | -0.007097186 |
| ITM2C | -0.007101428 |
| RNU6-233P | -0.00710143 |
| FAM86KP | -0.007106536 |
| LINC01339 | -0.007108436 |
| TTTY11 | -0.007109075 |
| MTND4LP1 | -0.007112555 |
| HIST1H2AB | -0.007115683 |
| RP11-20G13 | -0.007122021 |
| RP11-113D19 | -0.007123092 |
| OR11I1P | -0.007124271 |
| AC004520 | -0.007126019 |
| TXNP7 | -0.00712649 |
| CTD-2376I4 | -0.007130081 |
| RNU4-60P | -0.007131698 |
| AC010480 | -0.007134163 |
| RP5-827C21 | -0.007137262 |
| UBE2CP3 | -0.007145218 |
| IFITM1 | -0.007146937 |
| RP11-316A16 | -0.007150657 |
| BTF3P12 | -0.007153151 |
| CCSER1 | -0.007156686 |
| FLJ44511 | -0.007156724 |
| NDUFS5P6 | -0.007162094 |
| PRDM14 | -0.007163525 |
| NPNT | -0.00716411 |
| BCO2 | -0.007166566 |
| KCNJ5 | -0.007171738 |
| KRTAP12-4 | -0.007171739 |
| RP11-222A5 | -0.00717394 |
| RP11-737F9 | -0.007175488 |
| ABHD17AP6 | -0.007175495 |
| RN7SKP55 | -0.007180099 |
| GAPDHP31 | -0.007184461 |
| RP11-157P23 | -0.007185101 |
| MIR4801 | -0.007191967 |
| RP11-18B16 | -0.007192521 |
| MIR34A | -0.0071985 |

| | |
|---|---|
| SPTBN5 | -0.007198642 |
| MKLN1-AS | -0.007202642 |
| RP11-137H2 | -0.00720272 |
| CD68 | -0.00720627 |
| MAN1A1 | -0.007207293 |
| MRPL42P6 | -0.007211234 |
| ZNF496 | -0.007211684 |
| RP11-177F11 | -0.007214944 |
| AL022393 | -0.007219013 |
| RP11-477J21 | -0.007221876 |
| RNU6-1261P | -0.007223441 |
| MIR6082 | -0.007225531 |
| KARSP3 | -0.007231194 |
| CCNG2P1 | -0.007231537 |
| AC132192 | -0.007232532 |
| ANKRD20A3 | -0.0072344 |
| CICP20 | -0.007238051 |
| RP11-70K10 | -0.007238416 |
| MIR4452 | -0.00723946 |
| MTHFS | -0.007245527 |
| RP11-322J23 | -0.007255839 |
| RN7SL864P | -0.00725615 |
| DAB1 | -0.007259029 |
| RNU7-12P | -0.007259144 |
| RP5-1027O11 | -0.007259919 |
| RP11-531F16 | -0.0072627 |
| AGL | -0.007267798 |
| RN7SL335P | -0.0072852 |
| RP11-307O10 | -0.007287428 |
| RNU6-480P | -0.007290628 |
| GNG5P1 | -0.007291237 |
| AC099050 | -0.00729558 |
| MIR494 | -0.00729884 |
| RNU6-1155P | -0.007298926 |
| RP11-677O4 | -0.007301234 |
| Z82205 | -0.007304158 |
| RN7SKP212 | -0.007309203 |
| NEO1 | -0.007310795 |
| CTC-260E6 | -0.007312712 |
| RP11-284A20 | -0.007315623 |
| EIF2S2P5 | -0.007316934 |
| FGFR3P5 | -0.007320063 |
| HMGB1P25 | -0.00732441 |
| LA16c-359F1 | -0.00732447 |
| MUM1 | -0.007325177 |

| | |
|---|---|
| PGAM1P10 | -0.007326232 |
| NGRN | -0.007333829 |
| HOXB-AS4 | -0.00733573 |
| RP11-140C18 | -0.007336552 |
| HIST1H2AI | -0.007337881 |
| RNU7-6P | -0.00733821 |
| DNAJA1P5 | -0.007343251 |
| RP11-2L13 | -0.007345554 |
| GRM7-AS3 | -0.007346053 |
| CTD-2015C24 | -0.007347213 |
| EIF4A2P1 | -0.007349688 |
| NSMCE2 | -0.007352695 |
| RP11-650L12 | -0.007354676 |
| LRRC30 | -0.007360124 |
| TNS2 | -0.007360639 |
| RP11-211G23 | -0.007360895 |
| RP11-384G23 | -0.007361233 |
| RNA5SP59 | -0.007366405 |
| GCSH | -0.007370054 |
| Z97632 | -0.007373927 |
| CTD-2358C21 | -0.007374118 |
| RPL35AP4 | -0.007378033 |
| COX17 | -0.00738185 |
| RP11-186F10 | -0.007383735 |
| DDR1-AS1 | -0.007385664 |
| AC117481 | -0.007387483 |
| RP11-315O8 | -0.007391976 |
| ATP6V1B1 | -0.007400251 |
| EIF4A1P10 | -0.007400399 |
| HMGCR | -0.007408232 |
| RP11-231G3 | -0.007412028 |
| MS4A18 | -0.007412993 |
| CYP2G2P | -0.007413422 |
| MAK | -0.007413721 |
| RP11-266N13 | -0.007415664 |
| AL590113 | -0.007419041 |
| RP1-278D1 | -0.007420974 |
| GAN | -0.007427731 |
| TGM3 | -0.007429413 |
| AC018730 | -0.007433327 |
| VN1R53P | -0.007438699 |
| OR14C36 | -0.007440425 |
| AP001046 | -0.007446299 |
| RNU6-143P | -0.007456353 |
| AC107622 | -0.007467233 |

| | |
|---|---|
| MRPS18CP6 | -0.00746793 |
| IGLVIV-59 | -0.007472793 |
| KRT18P21 | -0.007474682 |
| CMTM8 | -0.007481408 |
| SAMD11P1 | -0.007481573 |
| XX-FW83128A1 | -0.007482179 |
| RNU6-562P | -0.007487041 |
| RP11-23E19 | -0.007494463 |
| HMGB3P16 | -0.007495925 |
| MTND3P7 | -0.007500784 |
| IGBP1-AS2 | -0.007503698 |
| RPL21P47 | -0.00750434 |
| CTB-57H20 | -0.007510662 |
| RN7SL570P | -0.007519663 |
| DOC2A | -0.007520107 |
| RPL15P4 | -0.007525284 |
| AC013442 | -0.007526545 |
| WSCD2 | -0.007529006 |
| GIMD1 | -0.007530478 |
| CD70 | -0.007533225 |
| CH507-154B10 | -0.007539047 |
| FOSB | -0.007539178 |
| RP11-298A8 | -0.007539954 |
| AL118525 | -0.007543633 |
| NTF6A | -0.007544667 |
| PCTP | -0.007548145 |
| SGCG | -0.007549062 |
| IFT27 | -0.007549541 |
| CTD-2184D3 | -0.007550105 |
| RN7SL370P | -0.007551191 |
| SERBP1P3 | -0.007559135 |
| HMGN1P5 | -0.007561756 |
| GUSBP6 | -0.007563356 |
| KRT18P15 | -0.007565202 |
| RP11-502F1 | -0.007565275 |
| RP11-15B17 | -0.00756577 |
| GXYLT1P6 | -0.007570182 |
| HRASLS2 | -0.007575923 |
| AC097493 | -0.007578052 |
| BCR | -0.007578565 |
| AC018685 | -0.007582342 |
| HOOK3 | -0.007583626 |
| RP11-340C20 | -0.00758447 |
| RPL12P24 | -0.007593168 |
| LINC00896 | -0.007594307 |

| | |
|---|---|
| AC006942 | -0.007595981 |
| CUL4B | -0.007596538 |
| CCDC58P3 | -0.007598 |
| CTD-2501M5 | -0.007598478 |
| NXPE1 | -0.007602161 |
| AC008410 | -0.00760307 |
| AC018462 | -0.007603969 |
| BSPH1 | -0.007604212 |
| RP11-209A2 | -0.007605142 |
| MLLT10P2 | -0.007609214 |
| RP11-431J24 | -0.00760974 |
| OR1P1 | -0.007610424 |
| AK4P1 | -0.007612676 |
| TTTY21 | -0.007616922 |
| RP11-100G15 | -0.007617174 |
| MPHOSPH6 | -0.00762106 |
| CU013544 | -0.007622856 |
| AC006840 | -0.007623888 |
| MAPT | -0.007624074 |
| RP11-802D6 | -0.007624682 |
| CHRNB3 | -0.007630131 |
| ANKS4B | -0.007630588 |
| PSMB11 | -0.007632054 |
| MIR302A | -0.007639632 |
| KRTAP12-3 | -0.007644257 |
| EEF1A1P16 | -0.007644438 |
| ASS1P7 | -0.00764512 |
| CHRNA9 | -0.007646574 |
| RP11-787D11 | -0.007654923 |
| CTD-2589M5 | -0.007658364 |
| CH17-248H7 | -0.007671402 |
| SEPT7P4 | -0.007673315 |
| RP11-667K14 | -0.007674867 |
| KRT8P29 | -0.007675686 |
| AC111200 | -0.007676992 |
| GM2A | -0.007678867 |
| AC142293 | -0.007679578 |
| MIR941-5 | -0.007680496 |
| RNU6-111P | -0.007681208 |
| CTD-2534J5 | -0.007681959 |
| CTD-2014D20 | -0.007682194 |
| DNAJB5-AS1 | -0.007688519 |
| LINC00680 | -0.007688989 |
| SKP1 | -0.007690049 |
| AC005513 | -0.007694655 |

| | |
|---|---|
| TRIM68 | -0.007705976 |
| AL355795 | -0.007709458 |
| MTHFD1P1 | -0.007710284 |
| LINC00599 | -0.007711503 |
| CLCNKA | -0.007712342 |
| SOCS5P2 | -0.007714174 |
| MIR544A | -0.007717242 |
| VWA8P1 | -0.007717715 |
| RPL21P116 | -0.007717727 |
| VCX3B | -0.007718547 |
| CTAGE14P | -0.007724887 |
| RPS18P6 | -0.0077276 |
| SDCCAG3 | -0.007733177 |
| AP1M2 | -0.007738163 |
| AC005758 | -0.007739778 |
| VTRNA3-1P | -0.007740248 |
| RNU6-542P | -0.007741366 |
| AC013727 | -0.007741873 |
| RP11-94P11 | -0.007743719 |
| TEKT4 | -0.007744557 |
| SCGB1D2 | -0.007750258 |
| RP11-179G8 | -0.00775083 |
| TAGLN3 | -0.007752223 |
| LINC00639 | -0.007754723 |
| AC010642 | -0.00776475 |
| TTC36 | -0.007765849 |
| PNPLA6 | -0.007767504 |
| OR10J6P | -0.00776948 |
| OR5G3 | -0.007772835 |
| DHFRP1 | -0.007776729 |
| SNHG10 | -0.007781532 |
| RP11-466C23 | -0.007782653 |
| ACTR1B | -0.007789682 |
| NXPE2 | -0.007792943 |
| RP5-890O15 | -0.007794624 |
| RNU6-1336P | -0.007795135 |
| RP11-273P3 | -0.007795466 |
| OR8F1P | -0.007801771 |
| STK35 | -0.007803654 |
| PARP10 | -0.007805031 |
| RP11-788M5 | -0.007807482 |
| ALAD | -0.00781171 |
| MESTP4 | -0.007812157 |
| RP5-862P8 | -0.00781531 |
| RPS24P6 | -0.007823357 |

| | |
|---|---|
| RNA5SP331 | -0.00782749 |
| RP11-339A7 | -0.007828733 |
| MRPL50P2 | -0.007832143 |
| RP11-294O2 | -0.007841379 |
| TCEB1P14 | -0.00784765 |
| AC009127 | -0.00784765 |
| RP5-855F14 | -0.007852443 |
| AC096570 | -0.007864666 |
| AP000946 | -0.007865227 |
| RNU6ATAC33P | -0.007868106 |
| SAGE3P | -0.007868165 |
| USP35 | -0.007886932 |
| RP11-235P11 | -0.007890889 |
| RP11-54O15 | -0.007898581 |
| AGAP10 | -0.00789964 |
| RPS6KB2 | -0.007899696 |
| RNU7-9P | -0.007900686 |
| TRIM49 | -0.00790183 |
| MXD3 | -0.007902338 |
| RP5-892G5 | -0.007902385 |
| RP11-429J17 | -0.00790831 |
| COX14 | -0.007915734 |
| UBE2CP4 | -0.007915827 |
| KHSRP | -0.007916255 |
| TP53TG3 | -0.00791983 |
| PNRC1 | -0.007921725 |
| RNA5SP243 | -0.00792294 |
| ANKRD33B-AS1 | -0.007923002 |
| BNIP3P32 | -0.007924195 |
| RNU6-161P | -0.00792463 |
| AUTS2 | -0.00792632 |
| CEP57L1P1 | -0.007928864 |
| RPS7P2 | -0.007931022 |
| AC069276 | -0.007931153 |
| RP11-125N22 | -0.007937408 |
| FRG2C | -0.007943191 |
| CTC-425K20 | -0.007952415 |
| UBE2L2 | -0.00795724 |
| RNY1P1 | -0.007959583 |
| ACSM3 | -0.00796035 |
| COPRS | -0.007960908 |
| FXR2 | -0.007969355 |
| RNA5SP129 | -0.007972533 |
| AC011525 | -0.007973879 |
| PVRL3-AS1 | -0.007981524 |

| | |
|---|---|
| OTOP2 | -0.007981945 |
| TCTN2 | -0.007982062 |
| RP3-525L6 | -0.007982477 |
| BBX | -0.007990118 |
| TMX2-CTNND1 | -0.007990289 |
| MAPRE1 | -0.007991421 |
| CTD-3148I10 | -0.007996504 |
| FAM27B | -0.008001494 |
| SEC63 | -0.008004351 |
| RN7SL672P | -0.008006085 |
| CXCR2 | -0.008011432 |
| RP11-1D12 | -0.008012926 |
| RP11-675P14 | -0.008018864 |
| MIR1277 | -0.008021526 |
| AP002958 | -0.00802292 |
| RP11-300G22 | -0.008025686 |
| RYBP | -0.008025934 |
| RP11-575N15 | -0.008026672 |
| AC027238 | -0.008030985 |
| AC008626 | -0.008031201 |
| RP11-42O4 | -0.008037889 |
| SNRPN | -0.008040762 |
| MINOS1 | -0.008045563 |
| MYH11 | -0.008047386 |
| SULT1A1 | -0.008049953 |
| EEF1A1P28 | -0.008056375 |
| ECEL1P2 | -0.008056874 |
| RP11-686G23 | -0.008056961 |
| LINC01304 | -0.008057161 |
| RP5-1172N10 | -0.008058453 |
| LINC00536 | -0.008058584 |
| ZW10 | -0.008059721 |
| RP4-613B23 | -0.008059796 |
| C6orf25 | -0.008060093 |
| OR13F1 | -0.008063804 |
| OR4A5 | -0.008064708 |
| AMN | -0.008065948 |
| C20orf196 | -0.008066456 |
| RP11-774O3 | -0.008070196 |
| PCDHA2 | -0.00807458 |
| RNU6-555P | -0.008075593 |
| TBX6 | -0.008075982 |
| RNU6-728P | -0.008087354 |
| HMGXB3 | -0.008088634 |
| LA16c-380A1 | -0.008088654 |

| | |
|---|---|
| ZDHHC20P3 | -0.008091447 |
| GCSAML-AS1 | -0.008091621 |
| CADM2-AS2 | -0.008095897 |
| AL136446 | -0.008096489 |
| ISM1-AS1 | -0.008097763 |
| CSNK1A1 | -0.008097807 |
| BTBD7P1 | -0.008099235 |
| USP31 | -0.008101174 |
| EIF3LP3 | -0.008101385 |
| RNU6-831P | -0.008102235 |
| NRIP3 | -0.008103053 |
| TIMM9 | -0.00810456 |
| PRSS46 | -0.00811143 |
| C2orf66 | -0.008111439 |
| CPNE4 | -0.00811686 |
| PTP4A1P6 | -0.00811697 |
| KRTAP19-10P | -0.008120612 |
| RNA5S17 | -0.008120612 |
| RP11-167N24 | -0.008121687 |
| APLNR | -0.008121957 |
| B3GALT4 | -0.008130124 |
| LINC01072 | -0.00813069 |
| ATP5EP1 | -0.008131089 |
| FCF1P10 | -0.008131369 |
| COX6B1P3 | -0.008131784 |
| SAP30BP | -0.008133734 |
| MIR3179-4 | -0.008133933 |
| RNA5SP283 | -0.008134118 |
| KTN1-AS1 | -0.008134497 |
| NDUFB3 | -0.008134683 |
| SUB1P2 | -0.008135731 |
| ZSCAN10 | -0.008141185 |
| GPC3 | -0.008143409 |
| RP11-314A20 | -0.008145158 |
| ADH1B | -0.008147374 |
| AL109761 | -0.00814802 |
| RPL7P39 | -0.008148882 |
| RP4-597J3 | -0.008150762 |
| RNU6-557P | -0.008153077 |
| MTND2P24 | -0.008153286 |
| RP11-346D14 | -0.008155193 |
| DHX34 | -0.008156712 |
| PCCA-AS1 | -0.008165291 |
| ZIC2 | -0.008177728 |
| GPX2 | -0.008181355 |

| | |
|---|---|
| RP11-389G6 | -0.008182232 |
| RP11-351O2 | -0.008184434 |
| HMGN3-AS1 | -0.008186443 |
| ANP32D | -0.008186782 |
| RP11-466A19 | -0.008189755 |
| RP11-958J22 | -0.008192669 |
| NPM1P42 | -0.008193514 |
| ONECUT3 | -0.008196878 |
| RPS27AP14 | -0.008197336 |
| SCARNA21 | -0.008198229 |
| RARG | -0.008199262 |
| BOLA3P1 | -0.008203537 |
| CDHR2 | -0.008207616 |
| PKD1P5 | -0.008209021 |
| TTTY7 | -0.008212266 |
| PABPC1 | -0.008213309 |
| RN7SL5P | -0.008214208 |
| HNRNPA1P57 | -0.0082147 |
| RN7SL789P | -0.008217627 |
| CTB-22K14 | -0.008221743 |
| RP11-34D15 | -0.008224026 |
| LBX2 | -0.008227264 |
| BNIP3P13 | -0.00823343 |
| KRTAP19-8 | -0.008233431 |
| RP11-35O7 | -0.008234388 |
| SLC7A3 | -0.008236178 |
| AC124853 | -0.008246597 |
| AC096546 | -0.008247395 |
| AC108025 | -0.008248198 |
| LINC00269 | -0.008248991 |
| CNTFR | -0.008253202 |
| CIDECP | -0.008257851 |
| HNRNPA1P18 | -0.008259797 |
| AC108448 | -0.008264255 |
| KRT18P10 | -0.00826734 |
| AC005540 | -0.008272132 |
| OR3D1P | -0.008276854 |
| AC068138 | -0.008277092 |
| AL161643 | -0.008285912 |
| HMGA1P7 | -0.008287352 |
| AC018832 | -0.008287797 |
| MPL | -0.008293857 |
| RP11-340A13 | -0.008298364 |
| PLCXD1 | -0.008298541 |
| AC074008 | -0.008300594 |

| | |
|---|---|
| RP11-425P5 | -0.008305071 |
| OPRD1 | -0.008305816 |
| AC079834 | -0.008308973 |
| RP11-468D11 | -0.008310804 |
| RP5-1069C8 | -0.008312426 |
| PRR13P7 | -0.00831397 |
| ACSL5 | -0.008318151 |
| RNU6-700P | -0.008323926 |
| AL161652 | -0.00833078 |
| RP11-102D18 | -0.008331839 |
| CYCSP19 | -0.008338745 |
| MIR644A | -0.00834062 |
| RN7SL690P | -0.00834607 |
| RP11-299M14 | -0.008346965 |
| RNU7-69P | -0.008348351 |
| AC087491 | -0.00836283 |
| TIPARP | -0.008369837 |
| PSAP | -0.008372033 |
| RP11-379I19 | -0.008373859 |
| SLC10A6 | -0.008374211 |
| RP11-550A9 | -0.008375021 |
| CTD-2330K9 | -0.00838156 |
| RPS3AP42 | -0.008386436 |
| NTN3 | -0.008390303 |
| PLA2G12AP1 | -0.008391249 |
| SLC25A14 | -0.00839462 |
| AC008265 | -0.008401078 |
| RNA5SP120 | -0.008401526 |
| BX470102 | -0.008403397 |
| AP004372 | -0.008403434 |
| BMS1P11 | -0.008403459 |
| RPL5P19 | -0.008407171 |
| CACNB4 | -0.008409876 |
| DKK4 | -0.008412004 |
| ZNF790-AS1 | -0.008414323 |
| CTD-2071N1 | -0.008414488 |
| RPS3AP12 | -0.008419008 |
| USP32P2 | -0.008422746 |
| SREBF1 | -0.008425058 |
| PLEKHA1 | -0.008425914 |
| SHROOM3 | -0.008431421 |
| RNU6ATAC20P | -0.008433759 |
| ARF4P1 | -0.008436527 |
| RP11-841C19 | -0.008438888 |
| AUP1 | -0.008440073 |

| | |
|---|---|
| TBC1D3P5 | -0.008442197 |
| LINC01088 | -0.008455442 |
| RP11-397D12 | -0.008457153 |
| ESYT1 | -0.008468204 |
| SPEF1 | -0.008469953 |
| TTTY2B | -0.008470002 |
| LIPE | -0.008473693 |
| LINC00694 | -0.008474766 |
| F2R | -0.00847747 |
| RP11-3J11 | -0.008482439 |
| PDHA2 | -0.008482907 |
| MIR6803 | -0.008483886 |
| MAST4-IT1 | -0.008484075 |
| NUBP1 | -0.008486419 |
| RP4-810F7 | -0.008487417 |
| WI2-1528O10 | -0.008488413 |
| KRT8P27 | -0.008488951 |
| MTND2P18 | -0.008495859 |
| SCOCP1 | -0.008496698 |
| SIGLEC20P | -0.008497262 |
| AL391650 | -0.008497836 |
| YBX1P3 | -0.008498222 |
| CDC42P1 | -0.008500824 |
| RP5-826L7 | -0.008502445 |
| F11-AS1 | -0.008512093 |
| SPIDR | -0.008513602 |
| ACOT7 | -0.008514625 |
| RP11-351O1 | -0.008518374 |
| HTR3E | -0.00852008 |
| TMEM161B-AS1 | -0.008523053 |
| SMIM2 | -0.008526913 |
| SERPINE4P | -0.008528533 |
| BCORP1 | -0.008531634 |
| DIXDC1 | -0.008533454 |
| RN7SL735P | -0.008536054 |
| RP11-468N14 | -0.008538258 |
| ELL2P2 | -0.008541148 |
| C22orf15 | -0.008550576 |
| RP11-238I10 | -0.008551555 |
| LINC01450 | -0.008552226 |
| RP5-1181K21 | -0.008556501 |
| EEF1A1 | -0.008560115 |
| RP11-208N14 | -0.008560321 |
| TRIM32 | -0.008563148 |
| PTCSC3 | -0.008564094 |

| | |
|---|---|
| MIR4260 | -0.008564383 |
| CTD-2165H16 | -0.008570581 |
| KCNIP4-IT1 | -0.008572397 |
| AC007322 | -0.008574139 |
| LINC00904 | -0.00858026 |
| SNHG21 | -0.008580377 |
| ASB9P1 | -0.008582563 |
| RN7SL491P | -0.00858592 |
| RNU6-317P | -0.008586217 |
| KCNAB1-AS1 | -0.008596818 |
| LINC01016 | -0.008599257 |
| IMPDH1 | -0.008599615 |
| SYF2P2 | -0.008600033 |
| SAGE4P | -0.008602645 |
| RP11-616K6 | -0.008604758 |
| RP11-1G11 | -0.008605438 |
| HNRNPCL2 | -0.008605898 |
| LINC01273 | -0.008606879 |
| KRTAP9-8 | -0.008607786 |
| DNAJB13 | -0.008610247 |
| ULK1 | -0.008615876 |
| XKR4 | -0.008619905 |
| OR51I2 | -0.008621472 |
| MIR563 | -0.008624178 |
| RNU6ATAC27P | -0.008626231 |
| PMS2P11 | -0.008627699 |
| RP11-629E24 | -0.00863684 |
| CDK5 | -0.008637663 |
| RP11-541H12 | -0.008638434 |
| CD34 | -0.008640156 |
| CTB-161J7 | -0.00864246 |
| RNU6-1263P | -0.008643011 |
| FOXP4-AS1 | -0.008645291 |
| RP11-467L13 | -0.008645831 |
| RP5-1066H13 | -0.008655924 |
| RP11-312B8 | -0.008658678 |
| RNU6-1104P | -0.008659722 |
| OR4B2P | -0.00866448 |
| DSTNP2 | -0.008666874 |
| AC078852 | -0.008667199 |
| IFNWP18 | -0.008671995 |
| AC098828 | -0.008672563 |
| AC079807 | -0.008675283 |
| PNPLA4 | -0.008677348 |
| LINC00239 | -0.008680573 |

| | |
|---|---|
| CSPG4P1Y | -0.008681791 |
| UQCRBP2 | -0.008690332 |
| LINC01296 | -0.008690468 |
| ZYG11A | -0.008691396 |
| RP11-171A24 | -0.008691424 |
| FLJ21408 | -0.008691991 |
| RP11-402F9 | -0.00869284 |
| RP11-484N16 | -0.008699834 |
| RN7SL720P | -0.008703023 |
| PSMG3-AS1 | -0.008704903 |
| RP11-112L18 | -0.008712023 |
| RP11-302I18 | -0.008712833 |
| APEH | -0.008716197 |
| RP11-744N12 | -0.008721405 |
| RP11-50C13 | -0.008722392 |
| TAB1 | -0.008723081 |
| AP000704 | -0.008724327 |
| RPS3AP53 | -0.008724996 |
| RHOT1P2 | -0.008729215 |
| FAM71F1 | -0.008742856 |
| ZNF299P | -0.008742949 |
| RP11-521C22 | -0.008748552 |
| BSN-AS1 | -0.008749565 |
| RPL7AP8 | -0.008751656 |
| RP11-332M2 | -0.008757985 |
| CH17-431G21 | -0.008758041 |
| RP11-797H7 | -0.008758264 |
| BOLA3P3 | -0.008758834 |
| PDCD10 | -0.008760906 |
| LL09NC01-254D11 | -0.008763489 |
| RP11-143E21 | -0.008763635 |
| RNU6-1126P | -0.008769524 |
| RNY4P16 | -0.008770354 |
| MIR6842 | -0.008772808 |
| RP11-415J8 | -0.008774246 |
| C6orf120 | -0.008781241 |
| RP11-470L19 | -0.008784152 |
| SCGB1D1 | -0.008789025 |
| SIK2 | -0.008791884 |
| NAALADL2-AS3 | -0.008792102 |
| ILDR1 | -0.008792144 |
| AC217773 | -0.008794226 |
| AC018737 | -0.008796941 |
| SRIP1 | -0.008804797 |
| SNRPCP15 | -0.008807776 |

| | |
|---|---|
| RP11-180N14 | -0.008808261 |
| CLCN2 | -0.008812238 |
| GABRG3-AS1 | -0.00881443 |
| MICAL3 | -0.008819662 |
| AGBL5-AS1 | -0.008820252 |
| OR1L1 | -0.008821638 |
| RP11-420A23 | -0.008826304 |
| KHDC1L | -0.008827221 |
| RP1-48J14 | -0.008836618 |
| AL357075 | -0.008837274 |
| RP11-432M24 | -0.008840398 |
| AC005229 | -0.008841951 |
| DNAJC18 | -0.008846007 |
| MIR4675 | -0.008851546 |
| AL669914 | -0.008853309 |
| CTC-435M10 | -0.008855235 |
| ANKRD20A11P | -0.00885836 |
| RP11-77A13 | -0.008860894 |
| PSPC1P2 | -0.008862493 |
| MIR4295 | -0.008866742 |
| RN7SL459P | -0.008867525 |
| EEF1DP4 | -0.008876197 |
| RP11-5K23 | -0.008882219 |
| OSTN | -0.008882234 |
| RP11-358N2 | -0.008885925 |
| CCT7P1 | -0.008887958 |
| TM7SF3 | -0.008892056 |
| AF196970 | -0.008892296 |
| U1 | -0.008893153 |
| DPP3P1 | -0.00889332 |
| IGLVI-42 | -0.008894953 |
| RP11-225H22 | -0.008896279 |
| CFL1P3 | -0.008898347 |
| CCND3P1 | -0.008898423 |
| AP003096 | -0.008912831 |
| MRPL3 | -0.008914717 |
| ABHD12B | -0.008915658 |
| ZNF385D | -0.008916205 |
| RTCA-AS1 | -0.008917564 |
| PPIAP2 | -0.008921049 |
| NQO1 | -0.00892361 |
| RP11-473M10 | -0.008927112 |
| HSPE1P4 | -0.008929511 |
| RP11-780O24 | -0.008929773 |
| RPS15AP38 | -0.008933361 |

| | |
|---|---|
| KITLG | -0.008936244 |
| RAP1GAP2 | -0.008938941 |
| PSMC1P12 | -0.008939877 |
| PUS7 | -0.00894002 |
| RP11-1325J9 | -0.008941789 |
| RPS6P22 | -0.008942248 |
| RP11-1086F11 | -0.008943995 |
| RNU6-222P | -0.008946791 |
| RP11-58K22 | -0.008950668 |
| CSPG4P3Y | -0.008952814 |
| DPP3 | -0.008954049 |
| RP11-1114I9 | -0.008967954 |
| MCFD2 | -0.00897111 |
| MIR6752 | -0.008971902 |
| RNA5SP395 | -0.008979462 |
| CALCB | -0.0089806 |
| AL049872 | -0.008983375 |
| IGKV2D-23 | -0.008986677 |
| RP11-316M20 | -0.008989891 |
| KRT8P11 | -0.00899236 |
| RP11-15E18 | -0.008993883 |
| RNU6-270P | -0.0089959 |
| RP11-502M1 | -0.008996792 |
| ITGA9-AS1 | -0.009001158 |
| TSPAN12 | -0.00900713 |
| Evf-2_5p | -0.009007404 |
| OR13C1P | -0.009011327 |
| INPP5F | -0.009012191 |
| ALB | -0.009012294 |
| TUBBP1 | -0.009014405 |
| PDE4B | -0.009018145 |
| ATF1P1 | -0.009018221 |
| PRMT5P1 | -0.009019202 |
| RNU6-225P | -0.009021039 |
| STMN1P2 | -0.009021887 |
| ELK1 | -0.00902514 |
| RN7SL162P | -0.009025489 |
| FAR1P1 | -0.00902636 |
| MIR3132 | -0.009026769 |
| RP13-465B17 | -0.009029641 |
| RNU6-1009P | -0.009031658 |
| RP11-756D7 | -0.009033566 |
| RP13-612N21 | -0.00903357 |
| GS1-304P7 | -0.009039214 |
| LANCL2 | -0.00904263 |

| | |
|---|---|
| BNIP3P30 | -0.00904534 |
| ACOT4 | -0.009046388 |
| AC025442 | -0.009047716 |
| PAK1 | -0.009048583 |
| RP11-888D10 | -0.009057477 |
| AC025811 | -0.009058833 |
| HMGN2P34 | -0.009059253 |
| MTND4P10 | -0.009061455 |
| CLVS2 | -0.009063383 |
| RP11-57J16 | -0.009065038 |
| AKR1B1P8 | -0.009065661 |
| RP11-760D2 | -0.009066768 |
| GSTA10P | -0.00906865 |
| TRAJ53 | -0.009068728 |
| MIR6794 | -0.009069605 |
| PRR36 | -0.009070984 |
| AC015818 | -0.009071579 |
| PITRM1 | -0.009073459 |
| RP11-266K4 | -0.00907891 |
| RN7SKP273 | -0.009080029 |
| LINC00476 | -0.009086985 |
| MAGEA9B | -0.009088554 |
| RP11-317M22 | -0.009092066 |
| MSTN | -0.009096071 |
| RP11-159J3 | -0.009096213 |
| LINC01277 | -0.009096706 |
| RHOB | -0.009096995 |
| SNRPD1 | -0.009098706 |
| BCL11A | -0.009099637 |
| AC137695 | -0.009100156 |
| POTEA | -0.00910524 |
| LINC01361 | -0.009110495 |
| TPM3P2 | -0.009111057 |
| NOBOX | -0.009113344 |
| MED22 | -0.009115396 |
| C1orf52 | -0.009117163 |
| RP11-445F12 | -0.009124304 |
| NBEAL1 | -0.009124494 |
| AE000661 | -0.00912659 |
| MRPS35P2 | -0.00912663 |
| RPS7P7 | -0.009127557 |
| RP11-457D20 | -0.009129061 |
| MIR5590 | -0.009134447 |
| LINC00572 | -0.009137285 |
| RP11-95H11 | -0.009137651 |

| | |
|---|---|
| RP11-493K23 | -0.009143695 |
| RP11-778J16 | -0.009144666 |
| RP11-393N4 | -0.009147357 |
| RP11-213N20 | -0.009148822 |
| RP11-903H12 | -0.009148906 |
| SORD2P | -0.00914922 |
| COLCA2 | -0.009151865 |
| CPXCR1 | -0.009155166 |
| CAPZA2 | -0.009156402 |
| RP11-89N17 | -0.009162212 |
| C2orf48 | -0.009165003 |
| CKS1BP5 | -0.009165756 |
| KIF5A | -0.009166124 |
| TEX264 | -0.0091695 |
| GATM-AS1 | -0.00917019 |
| EEF1DP3 | -0.009171273 |
| STARD9 | -0.009179603 |
| RN7SKP277 | -0.009192654 |
| LDHAP5 | -0.009196039 |
| PFN1P8 | -0.009200085 |
| OR5G1P | -0.009202967 |
| RP11-514P8 | -0.009207174 |
| AC003688 | -0.009207235 |
| CTD-2588C8 | -0.009211063 |
| RP3-340N1 | -0.0092147 |
| PSMC1P7 | -0.009215317 |
| DRG2 | -0.009215549 |
| PIGT | -0.0092178 |
| FLJ46066 | -0.009218445 |
| KIRREL3-AS3 | -0.009230673 |
| OR2AP1 | -0.009232771 |
| CTD-2043I16 | -0.009239588 |
| PRDM12 | -0.009245453 |
| ZNF204P | -0.009245969 |
| chr22-38_28785274-29006793 | -0.009249597 |
| DKK2 | -0.009252834 |
| AC111194 | -0.009253803 |
| CTD-2134A5 | -0.009254983 |
| CGB2 | -0.009256716 |
| RNU6-838P | -0.009257124 |
| SPATA31D2P | -0.00926061 |
| HEBP1 | -0.009264332 |
| SRSF10P1 | -0.00926638 |
| FGF12 | -0.009268109 |
| RP4-755D9 | -0.009271153 |

| | |
|---|---|
| IER3 | -0.00927224 |
| RP3-438O4 | -0.009273301 |
| HNRNPA1P60 | -0.009273983 |
| RP11-630A13 | -0.009274303 |
| CLK3P2 | -0.009274879 |
| LRCOL1 | -0.009278843 |
| ALYREF | -0.009291254 |
| CYP2F2P | -0.00929346 |
| AC136932 | -0.009297313 |
| NDUFV2P1 | -0.009312018 |
| RP11-449H3 | -0.009313239 |
| MIR3180-4 | -0.009314381 |
| RPL7P25 | -0.009315693 |
| SNORD115-30 | -0.009316542 |
| MRPS18BP2 | -0.009320135 |
| MIR6071 | -0.009322254 |
| RP11-123O1 | -0.009322923 |
| RP11-430E17 | -0.009323125 |
| RP5-981O7 | -0.00932576 |
| AC020689 | -0.009330219 |
| MEIS3P2 | -0.009331152 |
| NDUFB4P2 | -0.009332174 |
| RP11-1J7 | -0.009335917 |
| AC004112 | -0.00934214 |
| RP11-807H7 | -0.009344795 |
| CHURC1 | -0.009348448 |
| GRM8 | -0.009349694 |
| SERPINA13P | -0.009352727 |
| FAM172A | -0.009354383 |
| PAK7 | -0.0093583 |
| GNMT | -0.009359276 |
| RP11-555H23 | -0.009360211 |
| CTD-3118D11 | -0.009360489 |
| PSEN2 | -0.009361006 |
| USP17L14P | -0.009364981 |
| MAGOHB | -0.009365974 |
| H3F3A | -0.009369435 |
| SNORA72 | -0.009370403 |
| RP11-12A20 | -0.009373455 |
| OR2M5 | -0.009378073 |
| TCF25 | -0.0093795 |
| RP5-1100H13 | -0.009382835 |
| MTCO1P3 | -0.009383652 |
| MORF4L1P4 | -0.00938679 |
| ICE2P2 | -0.009391014 |

| | |
|---|---|
| FAM53C | -0.009392394 |
| KRT18 | -0.009393398 |
| RN7SKP34 | -0.009394036 |
| ERVV-2 | -0.009402737 |
| AC005034 | -0.009402888 |
| RNU6-586P | -0.009405946 |
| EMR1 | -0.009411799 |
| ARHGAP35 | -0.00941299 |
| RBMX2P4 | -0.009415161 |
| MBD3L5 | -0.009419385 |
| AC069157 | -0.009419521 |
| PRAMEF4 | -0.009422404 |
| OR4E2 | -0.009426442 |
| CCDC173 | -0.009426851 |
| TAS2R38 | -0.009426883 |
| RBP3 | -0.009428245 |
| C16orf58 | -0.009429549 |
| PARM1 | -0.009429772 |
| PSMC5 | -0.00943314 |
| RP11-392P7 | -0.009434591 |
| AC005237 | -0.009435871 |
| RP11-617J18 | -0.00943682 |
| RP5-858B6 | -0.009438256 |
| GNB3 | -0.00944354 |
| AC110283 | -0.009444451 |
| RP11-1112C15 | -0.009446608 |
| SNHG7 | -0.009447838 |
| HNRNPUL2 | -0.009448433 |
| CTC-493L21 | -0.009451105 |
| CEP164 | -0.009451838 |
| RP11-212E4 | -0.009453337 |
| SS18L2 | -0.009454424 |
| AC090505 | -0.009457072 |
| COX5AP2 | -0.009457438 |
| RP11-96C21 | -0.009464044 |
| C1QBP | -0.009465997 |
| RP11-344N17 | -0.009466724 |
| NDUFA5P8 | -0.009468618 |
| RN7SL215P | -0.009469651 |
| RP11-74M13 | -0.009470163 |
| CYP4F22 | -0.009477588 |
| snoU2-30 | -0.009478766 |
| AC011233 | -0.009484309 |
| RP11-25O3 | -0.009485415 |
| MIR1323 | -0.00948542 |

| | |
|---|---|
| LINC00598 | -0.009488387 |
| NUDT4 | -0.009489049 |
| TCEB1P24 | -0.009491276 |
| RP11-192I3 | -0.009491366 |
| CTD-2651B20 | -0.009494463 |
| RNU6-38P | -0.009500246 |
| AL360178 | -0.00950169 |
| RP13-512J5 | -0.00950224 |
| BET1L | -0.009509962 |
| CH17-472G23 | -0.009517255 |
| LINC01523 | -0.009517377 |
| AC010148 | -0.009523721 |
| RP11-300M24 | -0.009524935 |
| AC087348 | -0.009528571 |
| CTC-529L17 | -0.009529043 |
| LINC01149 | -0.009530555 |
| HSPE1P22 | -0.009535897 |
| C22orf23 | -0.0095372 |
| GAPDHP21 | -0.009539137 |
| RP11-310J24 | -0.00953981 |
| MIR514B | -0.009540347 |
| AL450063 | -0.009541 |
| RN7SKP85 | -0.009541654 |
| RP11-441M10 | -0.009543442 |
| ZSCAN31 | -0.009549498 |
| CTD-2270N23 | -0.009550131 |
| YBX2 | -0.009551284 |
| RP11-774G5 | -0.009553919 |
| RNA5SP87 | -0.009560308 |
| RP1-154K9 | -0.009561273 |
| SNORD92 | -0.009562865 |
| RNA5SP291 | -0.009564739 |
| TAS2R39 | -0.009567139 |
| RNU6ATAC4P | -0.009569563 |
| NT5C3AP1 | -0.009569687 |
| ATF4P1 | -0.00957127 |
| RP11-716D16 | -0.009572967 |
| RP11-30J20 | -0.00957521 |
| NACA | -0.009580011 |
| RP1-34B20 | -0.009580821 |
| SNORD111B | -0.009581763 |
| RPL27P12 | -0.009583092 |
| AC006000 | -0.009583699 |
| SNAPC2 | -0.009584272 |
| WBP1 | -0.009586114 |

| | |
|---|---|
| RN7SL426P | -0.009586818 |
| GINS3 | -0.009586933 |
| CTD-2527I21 | -0.009589761 |
| RNU6-1151P | -0.009594363 |
| LINC01195 | -0.009601052 |
| ATP5A1P8 | -0.009603344 |
| FOXP1-IT1 | -0.009606183 |
| LINC01232 | -0.009608645 |
| RP11-715L17 | -0.009614985 |
| SCD | -0.009615566 |
| ENPP7P12 | -0.009617046 |
| RP11-793B23 | -0.009617469 |
| KRT8P2 | -0.009618151 |
| ADAMTS9-AS1 | -0.009618621 |
| AC133552 | -0.009619855 |
| RNU6-394P | -0.00962039 |
| ATF6 | -0.009624516 |
| PCGF7P | -0.009627529 |
| UGT1A12P | -0.009629649 |
| LINC01141 | -0.009634431 |
| OR10H2 | -0.009634633 |
| RNU1-64P | -0.009636456 |
| RP11-777J24 | -0.009637228 |
| SLC25A23 | -0.009637409 |
| RP13-131K19 | -0.00964647 |
| MIR5004 | -0.009646535 |
| MIR6817 | -0.009647398 |
| GAPDHS | -0.009653836 |
| ZFPM1 | -0.009654266 |
| MIR4281 | -0.009660981 |
| AC011243 | -0.009669358 |
| RP11-92B11 | -0.009674996 |
| GEMIN2P2 | -0.00968056 |
| ENOX2 | -0.009681579 |
| AC073284 | -0.009685541 |
| FHP1 | -0.009686298 |
| SLC43A1 | -0.00968757 |
| SASH1 | -0.009699328 |
| RNU6-854P | -0.009709752 |
| LINC00929 | -0.009713993 |
| LLPHP1 | -0.009716075 |
| EI24P4 | -0.009716321 |
| RNU6-352P | -0.009718231 |
| NOL4 | -0.009721443 |
| MIR4325 | -0.009722084 |

| | |
|---|---|
| TTLL7 | -0.009726399 |
| RP11-437J2 | -0.009735524 |
| LINC00548 | -0.009737873 |
| SNORD19B | -0.009739286 |
| XRCC1 | -0.0097432 |
| OR5M9 | -0.009761644 |
| RP11-141A19 | -0.009765919 |
| RN7SKP292 | -0.009770109 |
| FAM41C | -0.009770336 |
| RP5-827L5 | -0.009771277 |
| RPS13P2 | -0.009774476 |
| AL079334 | -0.009775377 |
| OR9I3P | -0.009776333 |
| PPP1R3B | -0.009777155 |
| LCE6A | -0.009778167 |
| ACTL6A | -0.00978056 |
| STK31 | -0.009781333 |
| CTC-507E2 | -0.00978463 |
| RD3 | -0.009786593 |
| RP11-567E21 | -0.009788515 |
| PGAM1P13 | -0.009789843 |
| AL136234 | -0.009792046 |
| RDH16 | -0.009794849 |
| RP11-40C6 | -0.009802105 |
| RP11-916L7 | -0.009802937 |
| PIR | -0.009803947 |
| RP11-703H8 | -0.009806935 |
| RP11-173L6 | -0.00980912 |
| RP11-637O19 | -0.009811039 |
| AKIRIN1P2 | -0.009817146 |
| HSD17B13 | -0.009825915 |
| BCKDHB | -0.009828115 |
| AC005187 | -0.009828569 |
| ATP5J2P3 | -0.009829671 |
| AC018767 | -0.009829699 |
| CT867977 | -0.009836428 |
| ZNF319 | -0.009836878 |
| MBIP | -0.00983993 |
| CTNNAP1 | -0.009845245 |
| CRIPT | -0.009846758 |
| RP3-375P9 | -0.009853587 |
| COL9A2 | -0.009858107 |
| RNA5SP400 | -0.009859737 |
| RNU6-1108P | -0.009860343 |
| RP11-787I22 | -0.009865665 |

| | |
|---|---|
| CLN5 | -0.009873364 |
| NMD3 | -0.009878201 |
| RP1-257A15 | -0.009878471 |
| GS1-256O22 | -0.009882415 |
| PDE2A | -0.009882555 |
| SPATA17-AS1 | -0.009890601 |
| AL096864 | -0.009894901 |
| snoZ196 | -0.009896264 |
| MIR6090 | -0.009899592 |
| ARRDC2 | -0.009905179 |
| KIAA0355 | -0.009908528 |
| RP1-273P12 | -0.009911987 |
| OR8J2 | -0.009920933 |
| RP11-753C18 | -0.009924477 |
| RP11-354O24 | -0.009927629 |
| OR56A7P | -0.009927832 |
| HIRAP1 | -0.009928365 |
| CTD-2265O21 | -0.009929258 |
| VN2R3P | -0.009935383 |
| AC108925 | -0.009936161 |
| FAM170B-AS1 | -0.009937154 |
| ADORA3 | -0.009937776 |
| RP4-775C13 | -0.009949689 |
| AC004470 | -0.009957169 |
| MIR526B | -0.009957948 |
| RNU6-399P | -0.009962765 |
| RP11-413G15 | -0.009970287 |
| TK2 | -0.009974912 |
| PRLR | -0.009978875 |
| POU4F3 | -0.009978942 |
| RP11-386D6 | -0.009979285 |
| SAFB2 | -0.009981666 |
| DSCR4 | -0.009982014 |
| TRIM41 | -0.009987378 |
| EEF1A1P22 | -0.009993816 |
| RNU6-416P | -0.009998321 |
| RP11-453D16 | -0.009999898 |
| RP11-24I21 | -0.010000615 |
| SLC8A2 | -0.010003141 |
| RP11-254G11 | -0.010007324 |
| RP11-68L18 | -0.010007705 |
| VN2R17P | -0.010007707 |
| SPACA4 | -0.01000974 |
| AC073387 | -0.010010939 |
| RN7SKP2 | -0.010011938 |

| | |
|---|---|
| RBM22P2 | -0.010016145 |
| SNORD116-8 | -0.010017805 |
| RP4-630C24 | -0.01001803 |
| AC018735 | -0.010020411 |
| LINC00685 | -0.010020514 |
| RP11-507A3 | -0.010021706 |
| TMEM14C | -0.010022431 |
| SLC25A34 | -0.010022766 |
| RNU6-501P | -0.01002411 |
| HORMAD1 | -0.010024712 |
| PDCD5P2 | -0.010027804 |
| RP11-467D10 | -0.010028913 |
| OR10Y1P | -0.010029374 |
| RP11-89M20 | -0.010030551 |
| RP11-557C9 | -0.010032422 |
| VN1R1 | -0.010035343 |
| RNA5SP523 | -0.010038359 |
| HSPD1P8 | -0.010039339 |
| KCTD12 | -0.010042599 |
| RNU4-25P | -0.010048443 |
| CDK2AP2P3 | -0.010048725 |
| RP11-266E8 | -0.010052224 |
| SMC2-AS1 | -0.010057195 |
| RPL30P1 | -0.010061392 |
| RP11-313F23 | -0.010062397 |
| MIR133A2 | -0.010062611 |
| RNU6-1226P | -0.010063442 |
| TAAR1 | -0.010063695 |
| RP11-402L1 | -0.010070698 |
| MIR4648 | -0.010071438 |
| AP000478 | -0.010076965 |
| AL772307 | -0.010078785 |
| IQCK | -0.010080926 |
| RNASE3 | -0.010081136 |
| HIST1H2BI | -0.010085331 |
| OR8B4 | -0.010086037 |
| RP5-956O18 | -0.010095598 |
| SOX13 | -0.010095774 |
| MIR3939 | -0.010104049 |
| RP11-460E7 | -0.010104333 |
| RP11-543N12 | -0.010105282 |
| LRIT1 | -0.010105345 |
| RN7SKP75 | -0.010107744 |
| RP11-966I7 | -0.010109596 |
| RN7SL204P | -0.010109667 |

| | |
|---|---|
| RN7SKP255 | -0.010110181 |
| KRT18P13 | -0.01011381 |
| AC024082 | -0.010117016 |
| RNU6-721P | -0.010119051 |
| RP11-344N10 | -0.010126156 |
| MIR1273A | -0.010129813 |
| RP4-568B10 | -0.010129855 |
| RP11-90D4 | -0.010130005 |
| FOXI2 | -0.010132558 |
| HMGA1P3 | -0.010132842 |
| RP11-348M3 | -0.010135112 |
| SLC16A14P1 | -0.01013527 |
| ZNF385A | -0.010135906 |
| AC025458 | -0.010138958 |
| OR8S1 | -0.010139561 |
| RNU6-710P | -0.010141391 |
| RAET1E-AS1 | -0.010148798 |
| ITGA8 | -0.010155013 |
| CCDC186 | -0.010155106 |
| CATIP | -0.010159088 |
| EGLN1 | -0.010161302 |
| PELI3 | -0.010161544 |
| RP11-657O9 | -0.01016736 |
| INSL5 | -0.010167959 |
| MIR3177 | -0.010171618 |
| CCNB1IP1P1 | -0.010172199 |
| ANGPT4 | -0.01017513 |
| RPL13P2 | -0.010175737 |
| HCG14 | -0.010176418 |
| SST | -0.010177834 |
| RP11-874J12 | -0.010180754 |
| VWF | -0.010181414 |
| LL0XNC01-138C3 | -0.010182285 |
| XX-FW83563B9 | -0.010187443 |
| TMEM42 | -0.010187964 |
| RNU7-75P | -0.010188098 |
| CTB-31C7 | -0.010189751 |
| CTD-2160D9 | -0.010190237 |
| RP11-564H18 | -0.010194341 |
| RNA5SP488 | -0.010194362 |
| RP5-1086K13 | -0.01019442 |
| AC010455 | -0.010197061 |
| TYROBP | -0.010198215 |
| CLCF1 | -0.010200057 |
| EIF4E2 | -0.010200857 |

| | |
|---|---|
| LDHAP2 | -0.010201084 |
| GNA14-AS1 | -0.010201355 |
| FAM69A | -0.010203481 |
| AC092048 | -0.010205052 |
| MAP3K1 | -0.010215983 |
| TMEM97P1 | -0.010216054 |
| RNU4-12P | -0.010221425 |
| PTPN2P2 | -0.010221555 |
| MIR4261 | -0.010222336 |
| SNORD5 | -0.010232322 |
| RN7SL471P | -0.010237553 |
| ADAMTS9-AS2 | -0.010237642 |
| RP11-257P3 | -0.010240853 |
| DRGX | -0.010245584 |
| RPS26P8 | -0.010252761 |
| BCKDHA | -0.010258943 |
| PPT1 | -0.010260134 |
| OR7E18P | -0.010261633 |
| FAM83G | -0.010261697 |
| MIR506 | -0.010265012 |
| MOBP | -0.010266209 |
| RP11-179A18 | -0.010269041 |
| GSPT1 | -0.010269325 |
| RP3-393B19 | -0.010273298 |
| CXorf49B | -0.010274228 |
| SPAG8 | -0.010275502 |
| RNA5SP66 | -0.010277638 |
| RP11-472F21 | -0.010280581 |
| KCNN2 | -0.0102806 |
| ADSL | -0.010284024 |
| TMEM44-AS1 | -0.010284061 |
| TRMU | -0.010294331 |
| RPL5P23 | -0.010296843 |
| RP11-1315F8 | -0.010298003 |
| RP11-545E17 | -0.010298685 |
| MIR6871 | -0.01029998 |
| AC092896 | -0.010301198 |
| GAFA3 | -0.010301422 |
| RP11-523L20 | -0.01030615 |
| RP4-705O1 | -0.01030677 |
| AC118345 | -0.010306977 |
| TBPL1 | -0.010307076 |
| CTB-33G10 | -0.010309662 |
| RP11-580P21 | -0.010312056 |
| BTF3P2 | -0.010318635 |

| | |
|---|---|
| CFHR1 | -0.010319313 |
| RNU6-147P | -0.010321779 |
| RP11-78O22 | -0.010325297 |
| GPAT2 | -0.010331002 |
| RNU6-636P | -0.010331595 |
| JTBP1 | -0.010331952 |
| RPL30P9 | -0.01033283 |
| RP1-172B20 | -0.010333671 |
| RP11-33A14 | -0.01033882 |
| HTR1DP1 | -0.010339064 |
| OR4H12P | -0.010342257 |
| RP11-46F15 | -0.010342848 |
| GPR42 | -0.010344013 |
| RP11-196I18 | -0.010345822 |
| C12orf74 | -0.010350413 |
| RP11-325I22 | -0.010351028 |
| DNMT3B | -0.010356938 |
| RNU6-1137P | -0.01035936 |
| RP13-313G19 | -0.010363789 |
| MIR769 | -0.010364449 |
| MIR1273E | -0.01036952 |
| RP11-555H7 | -0.010372111 |
| ATP6V1E1 | -0.010372623 |
| GOLGA6L7P | -0.010377043 |
| ZBTB20 | -0.010378521 |
| RNU6-1265P | -0.010381753 |
| LINC01289 | -0.010384629 |
| TRDMT1 | -0.010390354 |
| OR2L1P | -0.010390769 |
| TMEM56 | -0.010392326 |
| RP11-54F2 | -0.01039716 |
| RN7SL764P | -0.010398056 |
| MIR566 | -0.010399529 |
| CH17-125A10 | -0.010401147 |
| SOS1-IT1 | -0.010402801 |
| LINC00352 | -0.01040652 |
| RP11-443B20 | -0.010409797 |
| CRLF2 | -0.010412887 |
| ACOT6 | -0.010415967 |
| RP11-775H9 | -0.010416322 |
| RN7SL160P | -0.01041793 |
| AC079112 | -0.010420971 |
| ELAC1 | -0.010422411 |
| RN7SKP245 | -0.010425974 |
| CASK | -0.010427695 |

| | |
|---|---|
| GPN1 | -0.010435788 |
| RP11-602M11 | -0.010443673 |
| AK3P4 | -0.010444415 |
| AC073127 | -0.010447798 |
| OR8B7P | -0.010451343 |
| DYRK2 | -0.01045244 |
| RN7SL798P | -0.010452548 |
| MXD1 | -0.010452682 |
| HNRNPM | -0.010456775 |
| BOD1P2 | -0.010459807 |
| AC008865 | -0.010461698 |
| AC026700 | -0.010462727 |
| SPC24 | -0.010464739 |
| AC023824 | -0.010465424 |
| MIR6509 | -0.010465737 |
| PCGF1 | -0.01046846 |
| PDE6H | -0.010470493 |
| FAM185A | -0.010474097 |
| KIAA2012 | -0.010476617 |
| HCG4P5 | -0.010476665 |
| KB-1043D8 | -0.010478861 |
| LINGO1-AS2 | -0.010480382 |
| RP4-547N15 | -0.010484166 |
| LINC00379 | -0.010485071 |
| AC068135 | -0.010488947 |
| ELMO3 | -0.010496628 |
| CLDN10-AS1 | -0.010498277 |
| RP11-651K21 | -0.01049875 |
| MRPL35P3 | -0.01049935 |
| VN2R1P | -0.010502095 |
| CTD-2015H6 | -0.010503438 |
| SERTAD2 | -0.010507792 |
| USP6NL | -0.010512145 |
| RNU6-1077P | -0.010514513 |
| RP11-535E8 | -0.010519952 |
| SFXN2 | -0.010523178 |
| ATF7IP2 | -0.010523513 |
| RP11-310I24 | -0.010525963 |
| RP11-44M6 | -0.010526211 |
| ACTR3C | -0.010526434 |
| PRPF40B | -0.010533765 |
| RNU5A-6P | -0.010533805 |
| AC006552 | -0.010533916 |
| LMO2 | -0.010533932 |
| LINC00698 | -0.010533996 |

| | |
|---|---|
| RP11-7N14 | -0.010534838 |
| RNU6-1056P | -0.01054101 |
| DYNLT3 | -0.010543793 |
| SNORD114-7 | -0.010549721 |
| AC104066 | -0.010552817 |
| MIR3175 | -0.01055713 |
| MYL6P2 | -0.010558765 |
| CSF2 | -0.010558966 |
| LINC01247 | -0.010559085 |
| RP11-270F18 | -0.010568852 |
| AC064850 | -0.010572822 |
| TMEM231 | -0.010577264 |
| DAOA-AS1_2 | -0.010577267 |
| RP11-558O12 | -0.010586993 |
| RP11-347N5 | -0.010588501 |
| NPM1P26 | -0.010590656 |
| RPL17P19 | -0.01059715 |
| PCDHA13 | -0.010600244 |
| RNA5SP113 | -0.010600282 |
| ADAM20 | -0.010603236 |
| RNU7-187P | -0.010604446 |
| RNU6-185P | -0.010605648 |
| RN7SKP29 | -0.010607214 |
| NCMAP | -0.010609973 |
| MIR7843 | -0.010612849 |
| RP11-713M15 | -0.01062175 |
| LINC01258 | -0.010629919 |
| RNA5SP23 | -0.010635456 |
| DNAJC2 | -0.010636039 |
| RP11-276M12 | -0.010636128 |
| Z99756 | -0.010642778 |
| FAM25G | -0.010653923 |
| BHLHE22 | -0.010656542 |
| RNU2-69P | -0.010664391 |
| LDHAL6EP | -0.010665762 |
| DGKH | -0.010672376 |
| RP11-78C3 | -0.010672685 |
| RP5-1189K21 | -0.010673882 |
| RP11-763F8 | -0.010676474 |
| TCERG1L-AS1 | -0.010686233 |
| LINC01288 | -0.010687446 |
| OFCC1 | -0.010689801 |
| MIR5699 | -0.010690423 |
| RN7SKP25 | -0.010691827 |
| CLDN25 | -0.010697067 |

| | |
|---|---|
| ARMC12 | -0.010698946 |
| SNX18P5 | -0.010701556 |
| UBE2R2-AS1 | -0.010706488 |
| AC012370 | -0.010709012 |
| AC022210 | -0.010710675 |
| AC126323 | -0.010713748 |
| HECW2 | -0.01071407 |
| CHST2 | -0.010716861 |
| HMGB3P13 | -0.010717384 |
| GABBR2 | -0.010719614 |
| TRIM51FP | -0.010721335 |
| FLJ30679 | -0.010723707 |
| GPR101 | -0.010728995 |
| AARD | -0.010730071 |
| RN7SL69P | -0.010730582 |
| RP11-267A15 | -0.010740206 |
| AC005102 | -0.010740224 |
| KRT18P55 | -0.010740726 |
| OR7E2P | -0.010741685 |
| RN7SKP167 | -0.010745941 |
| RP11-146E23 | -0.010754009 |
| TRIM60P14 | -0.010756965 |
| CTH | -0.010757123 |
| FAM157B | -0.010757828 |
| ZNF728 | -0.010760849 |
| RP5-965F6 | -0.010762547 |
| FAM90A11P | -0.010763627 |
| OR4A15 | -0.010764823 |
| AC130360 | -0.010765554 |
| NUDT17 | -0.01076637 |
| TAS2R60 | -0.010769293 |
| CCDC144CP | -0.010776399 |
| RNU6-1281P | -0.010776788 |
| RP11-796A5 | -0.010777947 |
| MESDC1 | -0.010779599 |
| AC079354 | -0.010779724 |
| LINC00343 | -0.010779847 |
| RP11-44D19 | -0.010781192 |
| RNA5SP466 | -0.010783544 |
| POLR1C | -0.010785757 |
| CTB-70D19 | -0.010788079 |
| SNORA76A | -0.010788106 |
| SMG5 | -0.010791804 |
| PLAC8L1 | -0.010792696 |
| MIR3181 | -0.010793519 |

| | |
|---|---|
| MIR517B | -0.010794206 |
| IGHVIII-67-3 | -0.010797211 |
| EIF4A2P2 | -0.01079793 |
| G3BP1 | -0.010802986 |
| RP11-358B23 | -0.010803544 |
| SERPINA6 | -0.010803836 |
| RP11-589F4 | -0.010807043 |
| RN7SL332P | -0.010814005 |
| SECTM1 | -0.010814819 |
| SSU72P2 | -0.010815319 |
| HSD52 | -0.010816156 |
| OGFOD1P1 | -0.010824994 |
| RP3-526F5 | -0.010834593 |
| SSX2B | -0.010834796 |
| RP11-357H3 | -0.010835563 |
| RP11-488I20 | -0.010837531 |
| LY96 | -0.01083815 |
| BLACE | -0.010840742 |
| ACTL8 | -0.01084509 |
| RP11-107M16 | -0.010848629 |
| SNORD46 | -0.010850434 |
| CCDC182 | -0.010850616 |
| RP11-36I17 | -0.010854213 |
| KLHDC7A | -0.010859878 |
| RP11-32A1 | -0.01086042 |
| RP11-739G5 | -0.01086193 |
| RP11-754I20 | -0.010865871 |
| RP5-979D14 | -0.010867046 |
| RN7SL91P | -0.010867091 |
| RP11-259F16 | -0.010869488 |
| RNU4-88P | -0.010876349 |
| RP13-735L24 | -0.010877712 |
| AC092332 | -0.010878689 |
| MIR4269 | -0.01087953 |
| ATP6V1E2 | -0.010881129 |
| RP1-30E17 | -0.010884143 |
| AC122714 | -0.010885605 |
| CD14 | -0.010886822 |
| RP11-263I1 | -0.010887656 |
| PPP1R2P5 | -0.010888077 |
| UMODL1 | -0.010888611 |
| AL442203 | -0.010889958 |
| PEX13 | -0.01089099 |
| LRRIQ1 | -0.010891346 |
| RP11-296E23 | -0.01089235 |

| | |
|---|---|
| ATP13A1 | -0.010900255 |
| RP1-140J1 | -0.010902864 |
| TM9SF1 | -0.010904516 |
| RNA5SP472 | -0.010904704 |
| KIR3DP1 | -0.010907808 |
| MIR8075 | -0.010911714 |
| MIR5586 | -0.01092035 |
| OR4S1 | -0.010922793 |
| HNRNPDP2 | -0.010923696 |
| FZD4 | -0.010927345 |
| RP11-90E9 | -0.010927911 |
| FAM167A-AS1 | -0.010936467 |
| BNIP3P29 | -0.010937297 |
| GRPEL2P2 | -0.010940918 |
| RP1-50A13 | -0.010942259 |
| LINC00867 | -0.010943516 |
| P2RX2 | -0.010944537 |
| RP11-13A1 | -0.010953507 |
| STRAP | -0.010954193 |
| AGAP1-IT1 | -0.010955236 |
| RP11-286M16 | -0.010961339 |
| RNU7-174P | -0.010962966 |
| RNU6-445P | -0.010965943 |
| RP11-15M15 | -0.010967812 |
| CDCP2 | -0.010970188 |
| TJP1 | -0.010972259 |
| TMF1P1 | -0.010976207 |
| AC005262 | -0.010981064 |
| EXPH5 | -0.010984389 |
| ZFYVE1 | -0.010986641 |
| AL591509 | -0.01098671 |
| RPS29P28 | -0.01098671 |
| RP11-809N15 | -0.010987214 |
| MTRNR2L12 | -0.010987422 |
| RP11-277A4 | -0.010989215 |
| RP11-419L4 | -0.010990276 |
| C9orf114 | -0.010990622 |
| RP11-324D17 | -0.010990824 |
| RNU6-1003P | -0.010993109 |
| RP11-548L20 | -0.010995886 |
| OR52E2 | -0.011007512 |
| RPS15AP16 | -0.011012743 |
| RP11-867O8 | -0.011013042 |
| DKC1 | -0.011013228 |
| CTD-2554C21 | -0.011014698 |

| | | |
|---|---|---|
| CDC42P5 | | -0.01102418 |
| LINC00708 | | -0.011024236 |
| RP5-1102E8 | | -0.011025723 |
| MRPL50P4 | | -0.011027142 |
| PRAM1 | | -0.01103228 |
| EIF4A3 | | -0.011033657 |
| AC132812 | | -0.011034904 |
| RP11-296L20 | | -0.01103846 |
| DCAF13 | | -0.011038606 |
| KLHL38 | | -0.011045754 |
| RP11-141E13 | | -0.011046649 |
| RP11-476J6 | | -0.011047313 |
| ELAVL4 | | -0.011047814 |
| TCF24 | | -0.011049915 |
| FGFR3P6 | | -0.011050101 |
| CLN8 | | -0.011051462 |
| RNU6-1029P | | -0.011051941 |
| AKT1 | | -0.011053323 |
| IFT140 | | -0.011054457 |
| RP11-429K17 | | -0.011056289 |
| CRABP2 | | -0.011059127 |
| RP11-326L17 | | -0.011059406 |
| NSUN2 | | -0.011061118 |
| HAUS6P1 | | -0.011066697 |
| RP11-789C7 | | -0.011066965 |
| RP11-158A14 | | -0.011067689 |
| AL021394 | | -0.011074875 |
| MIR618 | | -0.011075325 |
| RP11-570J4 | | -0.011075376 |
| RP1-68D18 | | -0.011076035 |
| SOX8 | | -0.011076356 |
| DUXAP3 | | -0.011082072 |
| PSMD10 | | -0.011085417 |
| ADHFE1 | | -0.011085957 |
| LRRC3B | | -0.011086079 |
| CTC-461H2 | | -0.011086802 |
| RP11-130C19 | | -0.011088892 |
| UBE2V1P12 | | -0.011090902 |
| PPIAP30 | | -0.011092651 |
| TYRL | | -0.011093767 |
| CTC-454I21 | | -0.011095874 |
| RP11-390N6 | | -0.011101417 |
| RP11-47A17 | | -0.011102192 |
| OR2G3 | | -0.011104194 |
| | 15-Sep | -0.011111651 |

| | |
|---|---|
| RP11-738G5 | -0.011115495 |
| RNU5F-7P | -0.011115792 |
| KRT8P18 | -0.011118924 |
| RP11-899L11 | -0.011119157 |
| RP1-92O14 | -0.011119186 |
| PCP4 | -0.011119382 |
| KIF1B | -0.0111198 |
| POT1 | -0.011131782 |
| WRN | -0.011131949 |
| KIAA1614-AS1 | -0.011137786 |
| LINC00903 | -0.011138194 |
| FRMD4A | -0.011142461 |
| EFCAB6-AS1 | -0.011147904 |
| RP11-998D10 | -0.011150686 |
| RP11-181G12 | -0.011158778 |
| LINC00856 | -0.011159061 |
| RNA5SP493 | -0.011171406 |
| RNU6-42P | -0.011179106 |
| RP11-289I10 | -0.01117984 |
| RP1-18D14 | -0.011180083 |
| RNU6-1144P | -0.011181344 |
| ERICH6 | -0.011187224 |
| RNU6-217P | -0.011188506 |
| RP11-13E5 | -0.011190369 |
| AC010888 | -0.011192319 |
| AC138304 | -0.011192736 |
| RPS3AP54 | -0.011194154 |
| IFNL4 | -0.011201587 |
| RP11-570H19 | -0.011204752 |
| RP11-78A18 | -0.011206201 |
| AC008740 | -0.01120851 |
| TECTA | -0.011208957 |
| MIR5189 | -0.011209886 |
| LL22NC01-81G9 | -0.011211725 |
| SLC10A5P1 | -0.011211814 |
| MTND4P22 | -0.011211817 |
| MIR3691 | -0.011212562 |
| RP11-128P10 | -0.011212812 |
| TTC39B | -0.011214446 |
| U85056 | -0.011218343 |
| RNU6-62P | -0.011220155 |
| RP11-172F4 | -0.011223027 |
| MIR4769 | -0.011223392 |
| SNAI1P1 | -0.011225874 |
| MIR128-2 | -0.011226213 |

| | |
|---|---|
| RP11-359P14 | -0.011231946 |
| MIR4447 | -0.011236146 |
| RP11-457D2 | -0.011239479 |
| AC007395 | -0.011243631 |
| PPP1R12BP2 | -0.011245076 |
| FADS2 | -0.011250596 |
| RAB4B | -0.011252159 |
| RN7SL130P | -0.011257156 |
| RP11-697E2 | -0.011261722 |
| SQSTM1 | -0.011265536 |
| FABP5P15 | -0.011267286 |
| BMP3 | -0.011268475 |
| SLC31A2 | -0.011271658 |
| AC026423 | -0.011276459 |
| ATL2 | -0.0112767 |
| ANO8 | -0.01127856 |
| RP11-21J7 | -0.011281944 |
| JMJD6 | -0.011282988 |
| RPS2P35 | -0.011284113 |
| AC092798 | -0.011287077 |
| BNIP3P7 | -0.011290829 |
| WI2-80269A6 | -0.011292081 |
| TNRC18P2 | -0.011296081 |
| VSTM2A | -0.011304722 |
| AQP12A | -0.011304737 |
| RN7SL63P | -0.011305766 |
| PRKRIRP3 | -0.011306791 |
| NOX1 | -0.01130805 |
| RNU6-792P | -0.011310632 |
| LINC01056 | -0.011320221 |
| EIF4EP3 | -0.011321386 |
| LL0XNC01-105G4 | -0.011322264 |
| RP3-326I13 | -0.011328674 |
| RP11-254A17 | -0.011331143 |
| AC138811 | -0.011334006 |
| AP002530 | -0.011334419 |
| NFE2L1 | -0.011340349 |
| B4GALNT3 | -0.0113434 |
| AC011518 | -0.011345542 |
| NRG1-IT3 | -0.011349267 |
| OFD1P6Y | -0.0113514 |
| TMEM30B | -0.01135261 |
| AP001350 | -0.011354229 |
| RP11-399J13 | -0.011356807 |
| RP11-463J10 | -0.01135946 |

| | |
|---|---|
| FOXA3 | -0.01135976 |
| LDLRAD4 | -0.011363197 |
| CRY1 | -0.011369709 |
| LINC01344 | -0.011371259 |
| NDUFA5P1 | -0.011374278 |
| AL583784 | -0.011375566 |
| FAM72D | -0.01137704 |
| RP3-521E19 | -0.011378199 |
| AQP12B | -0.011387464 |
| IMP3P1 | -0.011395126 |
| LPIN3 | -0.011395533 |
| RP11-204E4 | -0.011397349 |
| RP11-218I7 | -0.011398895 |
| ILF3-AS1 | -0.011399645 |
| AOC2 | -0.01140264 |
| RP11-451O18 | -0.011403971 |
| RP11-442O1 | -0.011406553 |
| DDX11 | -0.011409282 |
| SYT4 | -0.011411871 |
| EPB41L4A | -0.011412966 |
| AP000673 | -0.011414732 |
| NOP2 | -0.011418715 |
| OR2A41P | -0.01142038 |
| NCOA7-AS1 | -0.011423305 |
| RPL7AP28 | -0.01142446 |
| HEXIM1 | -0.011428241 |
| RP11-753E22 | -0.011429397 |
| RP11-474N8 | -0.011432276 |
| RNY1P15 | -0.011432334 |
| CTD-2325A15 | -0.01143386 |
| AKAP17BP | -0.01143449 |
| CTD-2316B1 | -0.011442139 |
| TBX19 | -0.011442363 |
| RP11-2N5 | -0.011444491 |
| RPL36AP45 | -0.011445772 |
| H1FX-AS1 | -0.011449786 |
| RN7SL254P | -0.01145416 |
| LINC01580 | -0.011456588 |
| RNU6-819P | -0.011458677 |
| RP5-828K20 | -0.011460587 |
| AP002884 | -0.011461193 |
| OR6V1 | -0.011466159 |
| HARS2 | -0.01146709 |
| PCAT18 | -0.011470131 |
| RNU7-52P | -0.011472804 |

| | |
|---|---|
| RP11-67H24 | -0.011478282 |
| POU3F4 | -0.011481637 |
| GALNTL6 | -0.011482767 |
| TAS2R12 | -0.011484547 |
| DOLPP1 | -0.011490644 |
| MED15P6 | -0.011493057 |
| COX6A1P2 | -0.011493416 |
| AC011306 | -0.011497593 |
| AFF2-IT1 | -0.011500696 |
| RP11-209E8 | -0.011513609 |
| AC010883 | -0.011515074 |
| RPS20P15 | -0.011524979 |
| OR1J2 | -0.011526124 |
| KRTAP10-13P | -0.011529294 |
| LINC01307 | -0.011530572 |
| SMIM13 | -0.011530906 |
| FYN | -0.011531688 |
| RNU1-17P | -0.011531977 |
| SNX3P1Y | -0.011532633 |
| RN7SKP38 | -0.011533296 |
| AC004510 | -0.011534965 |
| SCML2P1 | -0.011547015 |
| GJE1 | -0.011551069 |
| ZNF251 | -0.011551428 |
| RFC5P1 | -0.011557655 |
| OR52A1 | -0.011558596 |
| RP11-77P16 | -0.011560303 |
| RP11-167N4 | -0.011560458 |
| AC002075 | -0.011562045 |
| RP13-379L11 | -0.011565183 |
| STK3 | -0.011566926 |
| MARCKSL1P1 | -0.011567538 |
| RP11-529H2 | -0.011577686 |
| RP11-597M17 | -0.011579399 |
| P2RY4 | -0.011590585 |
| RPL31P20 | -0.01159252 |
| TTC9 | -0.011592528 |
| ZNF618 | -0.011598053 |
| VN1R13P | -0.01159952 |
| AC007339 | -0.011607284 |
| RP11-586K12 | -0.011607684 |
| KCNK18 | -0.01161124 |
| PCSK4 | -0.011612882 |
| AL353805 | -0.011618769 |
| RP11-465K4 | -0.011620224 |

| | |
|---|---|
| RPL14P3 | -0.011620231 |
| EEF1A1P26 | -0.011620793 |
| RP3-395C13 | -0.011631539 |
| RIMKLB | -0.011634111 |
| RP11-534I8 | -0.011639437 |
| MCTP2 | -0.011640065 |
| RNU6-27P | -0.011647616 |
| RPS21P8 | -0.011647933 |
| AC011551 | -0.011648069 |
| MIR633 | -0.011652269 |
| RP11-324F21 | -0.011659525 |
| RP11-135D11 | -0.011659597 |
| TRPM2 | -0.011661163 |
| OR6L1P | -0.011661203 |
| LINC01425 | -0.011664084 |
| MIR3670-2 | -0.011669114 |
| MT-TI | -0.011669406 |
| MIR200B | -0.011671858 |
| LRRC37A12P | -0.011675233 |
| RN7SL164P | -0.011679357 |
| MTND2P21 | -0.011679364 |
| TPRG1-AS1 | -0.011681606 |
| MIR4433B | -0.011681652 |
| DISP1 | -0.01168496 |
| RP11-354P17 | -0.011686873 |
| MIR3664 | -0.011688249 |
| MIR3667 | -0.011688972 |
| ACSS2 | -0.011689071 |
| RP11-157D18 | -0.01170323 |
| CTD-2116N24 | -0.011704436 |
| THYN1 | -0.011708142 |
| PRR33 | -0.011713235 |
| CPEB1-AS1 | -0.011713893 |
| RN7SL854P | -0.011717978 |
| CTD-2258A20 | -0.011722781 |
| AD001527 | -0.011726106 |
| MIR4277 | -0.011729762 |
| C7orf26 | -0.011729873 |
| RP4-667H12 | -0.011739686 |
| PPP1R36 | -0.011740013 |
| OR6N2 | -0.011740192 |
| EMR2 | -0.011747158 |
| DPH2 | -0.011747362 |
| RP11-692D12 | -0.011747511 |
| NANS | -0.011747641 |

| | |
|---|---|
| RP11-213G21 | -0.011748489 |
| RP11-313H3 | -0.011749918 |
| SMAD9-IT1 | -0.011757101 |
| RP4-552O12 | -0.011759695 |
| CASC22 | -0.011766375 |
| AC117945 | -0.011772294 |
| LINC01003 | -0.01177328 |
| AC092846 | -0.011773411 |
| RP11-635N19 | -0.011775996 |
| CTD-3131K8 | -0.011778415 |
| OR11L1 | -0.011780104 |
| RNU6-396P | -0.011782007 |
| GCSHP1 | -0.011785499 |
| LINC00686 | -0.011786163 |
| LARS2-AS1 | -0.011789973 |
| RP11-59O2 | -0.011794614 |
| ACOT8 | -0.01179542 |
| RP11-309H21 | -0.011795452 |
| FAM47B | -0.011796 |
| XPO6 | -0.011796871 |
| SLITRK2 | -0.011797433 |
| MTND6P11 | -0.011803192 |
| RPL7AP73 | -0.011804389 |
| DEFB107B | -0.011805595 |
| AC093756 | -0.011805595 |
| RN7SL148P | -0.011809069 |
| SLC31A1P1 | -0.011811045 |
| RP1-52D1 | -0.011816194 |
| KLF7-IT1 | -0.011816539 |
| OR10AK1P | -0.011817171 |
| RP11-78L16 | -0.011823077 |
| RP13-104F24 | -0.011823198 |
| LRSAM1 | -0.011829022 |
| SNORD115-43 | -0.011830014 |
| TERC | -0.011830342 |
| FBRSL1 | -0.011832631 |
| FABP5P14 | -0.01183458 |
| NPM1P38 | -0.011835767 |
| RPL12P11 | -0.011838468 |
| CH17-302M23 | -0.011842184 |
| CNBD1 | -0.011846445 |
| ERBB2 | -0.011855907 |
| C1orf168 | -0.011857885 |
| AC108057 | -0.011858442 |
| AL160010 | -0.011861435 |

| | |
|---|---|
| RNU6-45P | -0.011862162 |
| SRMP1 | -0.011862373 |
| TECRP2 | -0.01186614 |
| RNU6-554P | -0.011866516 |
| RP11-216M21 | -0.011870969 |
| LINC01511 | -0.01187255 |
| AC003988 | -0.011872888 |
| RNA5SP168 | -0.011874308 |
| CHST15 | -0.011887629 |
| EIF3D | -0.011887964 |
| MIR1228 | -0.011890591 |
| RFTN2 | -0.011898718 |
| OR56B2P | -0.011899052 |
| RP1-128O3 | -0.011900014 |
| ZMYND12 | -0.011903499 |
| WDR6 | -0.011908914 |
| PSMB7 | -0.0119091 |
| AC016910 | -0.011911535 |
| RN7SL308P | -0.011914145 |
| RP4-753D5 | -0.011920235 |
| RP1-134O19 | -0.011920425 |
| RNU6-766P | -0.011921186 |
| RN7SL8P | -0.011925279 |
| RP11-362F19 | -0.011928171 |
| MIR4522 | -0.01193296 |
| AC135068 | -0.011933053 |
| LINC00671 | -0.011934212 |
| PRPF6 | -0.011934738 |
| IFT80 | -0.011935575 |
| ARL6IP1P1 | -0.011939285 |
| NOL6 | -0.011940636 |
| RP11-307L14 | -0.011940673 |
| SERPINA5 | -0.011943017 |
| LINC00370 | -0.011949986 |
| AE000662 | -0.011952244 |
| ALMS1-IT1 | -0.011952841 |
| RAP2CP1 | -0.011955135 |
| CERS4 | -0.011955409 |
| RNU4-67P | -0.011961436 |
| AC139085 | -0.011962099 |
| MTND4P31 | -0.011962432 |
| FTH1P12 | -0.011965214 |
| RP11-461H3 | -0.011965993 |
| RNA5SP128 | -0.011967057 |
| GPC5-IT1 | -0.011969546 |

| | |
|---|---|
| KNOP1P5 | -0.011977084 |
| RP11-767C4 | -0.011977866 |
| ANKRD16 | -0.01198875 |
| ADRA1A | -0.011992277 |
| MIR8073 | -0.011993639 |
| CYCS | -0.011997104 |
| RP11-359D24 | -0.011998716 |
| OR8D2 | -0.012002271 |
| AL592310 | -0.012008377 |
| RP11-48D4 | -0.012011439 |
| NXT1P1 | -0.01201219 |
| RP11-2C15 | -0.012013612 |
| SLC15A3 | -0.012015318 |
| RP3-322G13 | -0.012023574 |
| RP3-433F14 | -0.012025444 |
| SPATA42 | -0.012025739 |
| RNU1-69P | -0.012028273 |
| RP11-171L9 | -0.012028659 |
| RP11-473I1 | -0.012030075 |
| RP11-339F13 | -0.012031797 |
| KRT18P3 | -0.012033473 |
| RPL5P13 | -0.012034976 |
| DEFA10P | -0.012037528 |
| MIR3680-2 | -0.012041061 |
| AC009234 | -0.01204175 |
| FABP9 | -0.012043283 |
| VPS26AP1 | -0.012051424 |
| MADCAM1 | -0.012052093 |
| MRPS36P2 | -0.01205239 |
| RNU6-1212P | -0.012053356 |
| LL22NC03-22A12 | -0.012053724 |
| GAPDHP38 | -0.012057868 |
| PANK1 | -0.012064046 |
| ITPKB | -0.012064325 |
| VN1R37P | -0.012066318 |
| FAM21FP | -0.012070483 |
| VN1R6P | -0.012071782 |
| SLC9B1P2 | -0.012078461 |
| HMGB1P27 | -0.01208333 |
| RP1-120G22 | -0.012085164 |
| ENPP5 | -0.012085814 |
| MIR24-2 | -0.012088073 |
| ZNF277 | -0.012088401 |
| AC018717 | -0.012089918 |
| MIR548A3 | -0.012093201 |

| | |
|---|---|
| RP11-545P6 | -0.012093923 |
| OR51F2 | -0.012094398 |
| NOTCH2P1 | -0.012095169 |
| SPATA31D4 | -0.012100424 |
| RP11-476H16 | -0.012106586 |
| SALL4P2 | -0.012111118 |
| RP11-379L18 | -0.012113157 |
| LINC01445 | -0.012116248 |
| SDR16C6P | -0.01212697 |
| IFFO2 | -0.012127379 |
| ALDH7A1P3 | -0.012128065 |
| BEND3 | -0.012129649 |
| RP11-439L18 | -0.012132213 |
| HSD3BP4 | -0.012143718 |
| TMEM230 | -0.012149631 |
| RP11-736K20 | -0.012151417 |
| VN1R12P | -0.012161195 |
| RP11-189G24 | -0.012162759 |
| RP11-631N16 | -0.01216593 |
| AC009227 | -0.012168948 |
| RP11-136H13 | -0.012171229 |
| RP11-744K17 | -0.012172716 |
| ALPPL2 | -0.012174649 |
| AKIRIN2P1 | -0.012182023 |
| CHMP2B | -0.012182116 |
| CNTN4-AS1 | -0.012182507 |
| AL512359 | -0.012183357 |
| LINC00358 | -0.012191946 |
| NUP210P1 | -0.012194429 |
| SUCLG2 | -0.012198136 |
| OR7A1P | -0.012202803 |
| TERF1P1 | -0.012203315 |
| RP11-39M21 | -0.012204368 |
| GUSBP3 | -0.012209348 |
| RP5-834N19 | -0.012212537 |
| FAM90A8P | -0.012215038 |
| DAP3P1 | -0.012215187 |
| ABC12-47964100C23 | -0.012215351 |
| AC004980 | -0.012217355 |
| NEK10 | -0.012219662 |
| SNORD14E | -0.012225986 |
| PPIAP7 | -0.012226035 |
| ANKRD26P3 | -0.012226316 |
| AC005086 | -0.012233134 |
| TLR3 | -0.012233784 |

| | |
|---|---|
| OR4U1P | -0.012234377 |
| RP5-1103G7 | -0.012237037 |
| RNY4P9 | -0.012237634 |
| REPS2 | -0.012240723 |
| RP11-794P6 | -0.012242608 |
| AC093602 | -0.012243778 |
| TRAJ43 | -0.012244031 |
| SNRPD3 | -0.012249189 |
| ARL5AP4 | -0.012253961 |
| RP11-1049H7 | -0.012256233 |
| RP11-222N13 | -0.012265483 |
| RNA5SP252 | -0.012266978 |
| OCIAD2 | -0.012267776 |
| AGTR2 | -0.012270594 |
| RP11-23E10 | -0.012271403 |
| AC007557 | -0.012272612 |
| RP1-158P9 | -0.012272646 |
| ARHGAP19-SLIT1 | -0.012276317 |
| FAM86B2 | -0.012276459 |
| RP11-759A9 | -0.012277029 |
| RPL9P18 | -0.012286876 |
| TBCA | -0.012296763 |
| AC004057 | -0.012297655 |
| IFNA16 | -0.012298067 |
| RNU6-123P | -0.012298307 |
| SUB1 | -0.012303684 |
| SIGLEC30P | -0.012310207 |
| GOLGA2P11 | -0.012310551 |
| MIR521-1 | -0.01231339 |
| RNU6-287P | -0.012314956 |
| DICER1-AS1 | -0.012317006 |
| ZNF117 | -0.012324076 |
| MYT1L | -0.012324979 |
| AP002495 | -0.012335241 |
| TAAR4P | -0.012338804 |
| LINC00393 | -0.012339472 |
| RP11-804M7 | -0.012339938 |
| PLAA | -0.012344821 |
| VDAC2 | -0.012346254 |
| YY1P1 | -0.012349228 |
| AC073043 | -0.012353024 |
| RP11-153I24 | -0.012353641 |
| RP11-393I23 | -0.012353739 |
| FAM86C2P | -0.012360059 |
| MIR3617 | -0.012361181 |

| | |
|---|---|
| RP11-756J15 | -0.012361556 |
| TMEM25 | -0.012362124 |
| FCF1P4 | -0.012362267 |
| CU104787 | -0.012362915 |
| MTATP6P7 | -0.01236398 |
| NMRK1 | -0.01236778 |
| PRR30 | -0.012369024 |
| PICALM | -0.012373362 |
| FAUP2 | -0.012373455 |
| TSPY2 | -0.012373963 |
| AC128715 | -0.012381566 |
| CDK10 | -0.012381641 |
| AC012074 | -0.012385809 |
| FTLP17 | -0.012390209 |
| SIRPAP1 | -0.012392674 |
| RP11-49C9 | -0.012394824 |
| SUMO1 | -0.012396954 |
| PRPS2 | -0.012398907 |
| COX19 | -0.01239941 |
| KIF4B | -0.012399741 |
| TCEB1P5 | -0.012406709 |
| TYK2 | -0.012411689 |
| RP11-707M3 | -0.012412005 |
| OFD1P4Y | -0.012413128 |
| PRELID1P2 | -0.012413864 |
| AL390071 | -0.012419599 |
| MIR4747 | -0.012426817 |
| RP11-956E11 | -0.012427858 |
| RP11-519G16 | -0.012432304 |
| AC010323 | -0.012448079 |
| RP6-204F4 | -0.012450101 |
| C17orf104 | -0.012453663 |
| RP11-359B12 | -0.012456156 |
| SENP8 | -0.012456249 |
| RNU6-26P | -0.012462305 |
| UBL5P4 | -0.01246388 |
| CALHM2 | -0.012465031 |
| AC112229 | -0.012465742 |
| MIR20B | -0.012468882 |
| GPR6 | -0.012471133 |
| PTPN23 | -0.012472659 |
| SDHCP4 | -0.012476656 |
| SNORA70E | -0.01247769 |
| NFIX | -0.012479618 |
| TAF1A | -0.012480101 |

| | |
|---|---|
| KYNUP2 | -0.012481299 |
| MBOAT2 | -0.012482334 |
| RP4-650F12 | -0.012482679 |
| LL22NC03-63E9 | -0.012483252 |
| RPL12P4 | -0.01248425 |
| SEMA3F | -0.012484306 |
| ZNF438 | -0.012484492 |
| AC007056 | -0.012487977 |
| CYCSP45 | -0.012491626 |
| RP11-708H21 | -0.012496221 |
| MTATP6P1 | -0.012496753 |
| C14orf105 | -0.012498342 |
| RP11-312P12 | -0.012507209 |
| RP11-428P16 | -0.012508872 |
| CTD-2517O10 | -0.012509721 |
| RP11-392C20 | -0.012513647 |
| CRYGA | -0.012514128 |
| RNU6-675P | -0.012522269 |
| RP11-249M12 | -0.012526306 |
| OR11H13P | -0.012528365 |
| LINC00891 | -0.012529523 |
| CASP6 | -0.012532234 |
| PHTF2 | -0.012532681 |
| HMGCL | -0.012533555 |
| LINC00440 | -0.012539211 |
| C14orf132 | -0.012543881 |
| AC007611 | -0.012544445 |
| NF1P2 | -0.012545083 |
| KLB | -0.012546449 |
| RNU6-1251P | -0.012553546 |
| POLD4 | -0.012554319 |
| KCTD1 | -0.012560272 |
| RN7SKP67 | -0.012561639 |
| GPR179 | -0.012562964 |
| AC019172 | -0.012565866 |
| RAC1P6 | -0.012566103 |
| DLD | -0.012566691 |
| APOOP4 | -0.012566804 |
| MIR5585 | -0.012567572 |
| MIR4284 | -0.012568561 |
| TG | -0.012573892 |
| AC025540 | -0.012576141 |
| CUBNP1 | -0.012578383 |
| UBE2V1P1 | -0.012580322 |
| PHBP1 | -0.012581484 |

| | |
|---|---|
| RPS3AP17 | -0.012585134 |
| RN7SL563P | -0.012585517 |
| RNU7-90P | -0.012590629 |
| RPS3AP1 | -0.012594439 |
| RP1-69D17 | -0.012599748 |
| SMIM1 | -0.012601614 |
| GOT2P2 | -0.012604317 |
| BTBD9-AS1 | -0.012605415 |
| TUBG2 | -0.012608628 |
| UBE2H | -0.012611326 |
| RP11-285F16 | -0.012612989 |
| RNU6-107P | -0.012617603 |
| CTB-88F18 | -0.012619596 |
| KIF13B | -0.012622917 |
| ISCA1 | -0.01262314 |
| FAM99A | -0.012623629 |
| MC5R | -0.012624277 |
| RP11-431N15 | -0.012625184 |
| ECEL1 | -0.012625415 |
| CCM2L | -0.012625671 |
| AP5B1 | -0.012627047 |
| MIR551B | -0.012627291 |
| RNU6-1317P | -0.012627386 |
| ATP6AP1 | -0.012628759 |
| HOXC4 | -0.012628871 |
| RP11-94B19 | -0.012629672 |
| ZMAT2 | -0.012631941 |
| RP11-314N2 | -0.012637706 |
| AC068754 | -0.012638427 |
| RP11-362M19 | -0.012638941 |
| RP11-567M21 | -0.012642595 |
| C15orf26 | -0.012642726 |
| VNN3 | -0.012647518 |
| CTC-280C13 | -0.012648723 |
| LINC00959 | -0.012649579 |
| TXNDC9 | -0.012652463 |
| RP6-60B16 | -0.012670949 |
| MIR609 | -0.012672802 |
| MIR4422 | -0.012673058 |
| LINC01477 | -0.012681686 |
| FP325318 | -0.012683532 |
| AC137527 | -0.012685444 |
| AC104794 | -0.012687829 |
| RP5-967N21 | -0.01268932 |
| RP11-162K11 | -0.012692591 |

| | |
|---|---|
| PNKDP1 | -0.012694108 |
| HMGN1P17 | -0.01269802 |
| PODXL2 | -0.012699088 |
| OR7E122P | -0.012700381 |
| HYAL1 | -0.012702139 |
| OR6C71P | -0.0127024 |
| KIAA0922 | -0.01270266 |
| C5orf52 | -0.012705474 |
| ZNF479 | -0.012706272 |
| MYOM1 | -0.012709209 |
| MRI1 | -0.012723275 |
| RNU1-108P | -0.012723626 |
| RN7SL140P | -0.012726294 |
| RP5-844F9 | -0.012727224 |
| RPL5P21 | -0.01273269 |
| CTDSP1 | -0.012733992 |
| POM121L3P | -0.012734671 |
| AC108938 | -0.012735616 |
| CCDC24 | -0.012736001 |
| ST7-AS1 | -0.012739462 |
| RPS17P1 | -0.012744846 |
| RP11-491F9 | -0.012747588 |
| NUDT16L1 | -0.012754011 |
| AC006478 | -0.012760961 |
| BX323860 | -0.012760961 |
| AC116165 | -0.012760961 |
| EPS8 | -0.0127675 |
| PITPNM2 | -0.012767798 |
| CTD-2277K2 | -0.012769423 |
| MRPS6P4 | -0.01277208 |
| VENTXP7 | -0.012772536 |
| RN7SL146P | -0.012775831 |
| MIR2392 | -0.012775934 |
| RP11-157L10 | -0.012777739 |
| LRRC45 | -0.012778403 |
| TPTE2P3 | -0.012779981 |
| KRTAP3-3 | -0.012780241 |
| PRPH | -0.012782568 |
| RNU1-78P | -0.012783438 |
| CTD-2281M20 | -0.012789607 |
| XIRP2-AS1 | -0.012789776 |
| RP11-79C23 | -0.012794935 |
| RP11-134K1 | -0.012798748 |
| LINC00240 | -0.012799315 |
| RNU6-74P | -0.012800126 |

| | |
|---|---|
| RP11-324P9 | -0.012803366 |
| KRT18P14 | -0.012804678 |
| APOA2 | -0.012808372 |
| LILRB5 | -0.012810218 |
| TGFA-IT1 | -0.012812503 |
| RP11-706O15 | -0.012812897 |
| RP11-788H18 | -0.012818077 |
| OR5M8 | -0.012829836 |
| CTC-463N11 | -0.012831277 |
| PTGFR | -0.012835395 |
| VTRNA1-2 | -0.01283568 |
| RPL29P28 | -0.012836445 |
| RP11-615J4 | -0.012842554 |
| AL391827 | -0.012844178 |
| SUSD1 | -0.01284648 |
| RP4-751H13 | -0.012849346 |
| RNU6-1094P | -0.012851761 |
| AC145124 | -0.012859664 |
| AC011366 | -0.012867196 |
| CH17-270A2 | -0.012867439 |
| RP11-580I1 | -0.012872097 |
| RP11-219D15 | -0.01287847 |
| Z83001 | -0.012890425 |
| RNU6-873P | -0.012891811 |
| AATK | -0.012892845 |
| RP11-131O15 | -0.012893618 |
| ACSM2B | -0.012894557 |
| RP11-214F16 | -0.012896458 |
| AC122136 | -0.012897363 |
| AC079790 | -0.012901114 |
| AC004775 | -0.012901628 |
| GIGYF1 | -0.01290933 |
| UAP1 | -0.01291026 |
| PPIAP27 | -0.01291044 |
| RP11-422P22 | -0.012910765 |
| MTND1P4 | -0.012917448 |
| ZNF177 | -0.01292344 |
| INS-IGF2 | -0.012928298 |
| KRTAP2-4 | -0.012928324 |
| RP13-582L3 | -0.012928334 |
| AC022173 | -0.012929966 |
| SH3TC1 | -0.012933014 |
| NAP1L4P1 | -0.012934789 |
| RP11-455B3 | -0.012934905 |
| SNORD114-23 | -0.012938095 |

| | |
|---|---|
| NDUFS5P3 | -0.012940445 |
| AC012671 | -0.012946782 |
| AL353997 | -0.012952973 |
| HMGB1P26 | -0.01295307 |
| IL23A | -0.012955899 |
| CHEK2 | -0.012957071 |
| ZNF705A | -0.012957254 |
| MIR4463 | -0.012963922 |
| RP11-465L10 | -0.012965968 |
| MAPK15 | -0.012968346 |
| SFTA1P | -0.012970303 |
| AC093826 | -0.012973927 |
| LAMP5-AS1 | -0.012976761 |
| MIR4666A | -0.012977286 |
| CTD-2374C24 | -0.0129803 |
| FBRS | -0.012982877 |
| RPL30P4 | -0.012983798 |
| PDE4DIP | -0.012986766 |
| RN7SKP22 | -0.012999287 |
| RP11-351D16 | -0.012999771 |
| AL024509 | -0.013000572 |
| AURKB | -0.013005018 |
| RP11-546M4 | -0.013005605 |
| SKP1P3 | -0.013008228 |
| RP11-293E1 | -0.013009564 |
| AL031601 | -0.013010795 |
| NOX3 | -0.013011326 |
| MIR548AB | -0.013012455 |
| SSX7 | -0.013013507 |
| PPAP2B | -0.013016032 |
| MT-TE | -0.013018272 |
| SNORD36C | -0.013021828 |
| RP11-782E2 | -0.013022826 |
| RP5-1028L10 | -0.013025964 |
| TRAPPC2L | -0.013027716 |
| TXNDC15 | -0.013035828 |
| AARS | -0.013037689 |
| RP11-212F11 | -0.013038034 |
| RP11-476M19 | -0.013040018 |
| RN7SL691P | -0.013043149 |
| LINC01290 | -0.013043729 |
| RP4-666O22 | -0.013044867 |
| MIR373 | -0.013050267 |
| AC233702 | -0.013054852 |
| RPS3P1 | -0.013056357 |

| | |
|---|---|
| NPM1P46 | -0.013056861 |
| MIR646 | -0.013058339 |
| FNTAL1 | -0.013060889 |
| RP4-813D12 | -0.013065916 |
| RNU6-213P | -0.013066869 |
| RP11-12A16 | -0.013069678 |
| ACTG1P12 | -0.013070833 |
| RP11-4K16 | -0.013072748 |
| KAZALD1 | -0.013074155 |
| RN7SKP205 | -0.013076159 |
| RP11-24J19 | -0.013076377 |
| GTPBP4 | -0.013076686 |
| TRAF6P1 | -0.013079547 |
| OVCH2 | -0.013079833 |
| VN1R70P | -0.013079965 |
| FAM71BP1 | -0.013080645 |
| MIR4777 | -0.013091098 |
| NRXN1 | -0.013094691 |
| DCAF6 | -0.013096445 |
| RNU6-668P | -0.013098749 |
| MIR6745 | -0.013099765 |
| AC005487 | -0.013105385 |
| RP5-1119O21 | -0.013105762 |
| SLC29A1 | -0.013107087 |
| ARAFP1 | -0.013107884 |
| HNRNPH1P3 | -0.013109261 |
| LINC01069 | -0.013115597 |
| ANKRD26P1 | -0.013118678 |
| RP11-338L18 | -0.013119376 |
| CDRT15P1 | -0.01312734 |
| RNF19BPY | -0.013128357 |
| MC4R | -0.013133343 |
| SPATA41 | -0.013135274 |
| IL27RA | -0.013135274 |
| CTD-2516K3 | -0.013139628 |
| RP11-10H3 | -0.013149274 |
| F8 | -0.013153229 |
| RP11-129G17 | -0.01315346 |
| RNA5SP500 | -0.013155069 |
| BNC1 | -0.013160006 |
| MRPS16P3 | -0.013172043 |
| PLXNA3 | -0.013173974 |
| SLC36A2 | -0.013175054 |
| PRB1 | -0.013179439 |
| AC005926 | -0.013179861 |

| | |
|---|---|
| AC006452 | -0.013184015 |
| RP11-300I6 | -0.013185297 |
| MAGEB5 | -0.013188637 |
| RP11-617O8 | -0.013193514 |
| PSMC1P5 | -0.013194402 |
| NIFKP1 | -0.013203617 |
| AC011380 | -0.013206249 |
| RPSAP9 | -0.013207259 |
| RP11-301G19 | -0.013207571 |
| RPL36AP53 | -0.013209365 |
| HDAC1P1 | -0.013209406 |
| RP11-588H23 | -0.013211581 |
| CACNA1I | -0.013213473 |
| MIR1254-2 | -0.013213686 |
| CA2 | -0.013214441 |
| RARRES3 | -0.013226013 |
| FAM133CP | -0.013227069 |
| MT-ND6 | -0.013228562 |
| RNA5SP185 | -0.013228772 |
| TMEM97P2 | -0.01322964 |
| RP11-528M18 | -0.01323084 |
| DYNLL1P5 | -0.013231656 |
| OXLD1 | -0.013232767 |
| MIR617 | -0.013233239 |
| AL035419 | -0.013242037 |
| CTD-2564P9 | -0.013243715 |
| PGM3 | -0.0132494 |
| ATP6V0E2-AS1 | -0.013249419 |
| RP11-615I16 | -0.013249745 |
| OR9Q2 | -0.013253025 |
| SLC3A1 | -0.01326017 |
| RP11-8J23 | -0.013260836 |
| CDKL4 | -0.013263747 |
| RNU6-932P | -0.013265664 |
| ZNF232 | -0.013267615 |
| CENPUP1 | -0.013271367 |
| SNORD116-26 | -0.013275397 |
| SERBP1P5 | -0.013277404 |
| RP1-92C8 | -0.013278345 |
| PABPC1P6 | -0.013278858 |
| MIR491 | -0.013280534 |
| A4GNT | -0.013281932 |
| AP000435 | -0.013289009 |
| RP11-169I9 | -0.013293016 |
| ORC5 | -0.0132977 |

| | |
|---|---|
| TUSC2 | -0.013302035 |
| SDR42E1 | -0.013304556 |
| RNU6-425P | -0.013312829 |
| CNDP2 | -0.013313292 |
| HHAT | -0.013313664 |
| EFHB | -0.01331385 |
| RNU6-565P | -0.013318748 |
| GSTA11P | -0.013323273 |
| RPL3P13 | -0.013324194 |
| AL022336 | -0.013325754 |
| RP11-573C10 | -0.01332579 |
| AC004129 | -0.013327991 |
| RNU6-514P | -0.013330097 |
| RP11-44F14 | -0.013341851 |
| MIR3611 | -0.013343034 |
| AC007317 | -0.013344581 |
| RP4-781B1 | -0.013347836 |
| EFCAB10 | -0.013348902 |
| CTD-2320B12 | -0.013349204 |
| THOC3 | -0.013349386 |
| RNU6-234P | -0.013354504 |
| RN7SL392P | -0.013354914 |
| CTB-167B5 | -0.013357496 |
| AC111186 | -0.013360365 |
| RP11-108F13 | -0.013360683 |
| RP11-618P17 | -0.013362491 |
| RP11-289H16 | -0.013364172 |
| RP11-158L12 | -0.013364289 |
| CATIP-AS1 | -0.013365506 |
| MIR95 | -0.013365848 |
| MIR212 | -0.013368291 |
| TM2D3 | -0.013374969 |
| C1DP3 | -0.013390399 |
| RP11-433B3 | -0.013392981 |
| AC099326 | -0.013393183 |
| LRP11 | -0.013395639 |
| RP11-279L11 | -0.013401743 |
| RP11-141J13 | -0.013404237 |
| RP11-1072N2 | -0.01340466 |
| MIR361 | -0.013405696 |
| LINC00612 | -0.013409778 |
| CCT6P2 | -0.013418425 |
| RP3-347M6 | -0.013422147 |
| SIGLEC28P | -0.013423625 |
| RP11-844G16 | -0.013432338 |

| | |
|---|---|
| GH1 | -0.013435684 |
| PSMG4 | -0.013444665 |
| CARS2 | -0.013445986 |
| RNU6-187P | -0.013447801 |
| AC084872 | -0.013448051 |
| LINC00846 | -0.013455623 |
| ALG14 | -0.013459251 |
| MIR339 | -0.013459555 |
| RP11-484L8 | -0.013460233 |
| RN7SL399P | -0.013460499 |
| RNU6-795P | -0.013461451 |
| OR9G1 | -0.013471084 |
| KRT17P4 | -0.013471723 |
| ACAD8 | -0.01347328 |
| EEF1A1P12 | -0.013475066 |
| NF1P7 | -0.013475658 |
| CRTAC1 | -0.013476346 |
| AL590822 | -0.013486007 |
| SLC45A1 | -0.013491141 |
| SCNN1B | -0.013492946 |
| DDX6P1 | -0.013495449 |
| AL359737 | -0.013500079 |
| AOX1 | -0.013505337 |
| RNU6-1163P | -0.01350867 |
| RNU6-424P | -0.013509254 |
| RP11-734K21 | -0.013509864 |
| AL137792 | -0.013528181 |
| SYNPR-AS1 | -0.0135294 |
| RNU6-291P | -0.013532678 |
| SLC25A5P4 | -0.01353517 |
| RNA5SP378 | -0.013535488 |
| RN7SL300P | -0.013536644 |
| ITIH4-AS1 | -0.013536692 |
| RNU6-601P | -0.01353763 |
| UFSP2 | -0.013539813 |
| FGD5 | -0.013542399 |
| CTD-3199J23 | -0.013543627 |
| RNA5SP282 | -0.013545576 |
| RP11-16L21 | -0.013547986 |
| SYNJ2BP-COX16 | -0.013548539 |
| SLC9A3R2 | -0.013562084 |
| PPIAP10 | -0.013578202 |
| AC009406 | -0.013579846 |
| SLC25A3 | -0.01358121 |
| MIR503 | -0.013583194 |

| | |
|---|---|
| RPS10P27 | -0.013584707 |
| C19orf40 | -0.013584913 |
| RP11-699L21 | -0.013586772 |
| RP11-375D13 | -0.013587345 |
| PITHD1 | -0.013589955 |
| DEFB106B | -0.013591276 |
| CIRBP-AS1 | -0.013592541 |
| EEF1A1P29 | -0.013595793 |
| RNU6-550P | -0.013596716 |
| OR4A4P | -0.013600321 |
| AC012451 | -0.013603179 |
| NPM1P43 | -0.013603637 |
| KB-1247B1 | -0.01360522 |
| KRTAP10-4 | -0.01360543 |
| RP11-85G21 | -0.013610079 |
| RP3-439F8 | -0.013611407 |
| LINC01566 | -0.013613465 |
| SEMA4C | -0.013615444 |
| HEATR4 | -0.013625975 |
| RN7SL457P | -0.013630022 |
| CDK7 | -0.013633231 |
| POU5F2 | -0.013633811 |
| SYT6 | -0.013634664 |
| RNU5E-8P | -0.013641156 |
| IGBP1P5 | -0.013642887 |
| LCE1D | -0.013643684 |
| SLC7A5 | -0.013648357 |
| CNPY1 | -0.013651089 |
| RP4-594A5 | -0.013660582 |
| TEKT5 | -0.01366214 |
| RP11-131M11 | -0.013663521 |
| RPS26P52 | -0.013666141 |
| RPL5P3 | -0.013671777 |
| RNU7-23P | -0.013673493 |
| MIR1287 | -0.013676594 |
| PP7080 | -0.013676954 |
| AC125421 | -0.013677533 |
| AC078899 | -0.013678155 |
| RNU6-380P | -0.013681177 |
| RP11-243A14 | -0.013682862 |
| ARMCX4 | -0.013683224 |
| HNRNPMP2 | -0.013686639 |
| WDR7-OT1 | -0.013687269 |
| RP11-359B20 | -0.013689756 |
| RGS14 | -0.013691355 |

| | |
|---|---|
| AL355512 | -0.013693505 |
| RN7SL307P | -0.013702641 |
| RP11-460I13 | -0.01370981 |
| TMED7-TICAM2 | -0.013709849 |
| IPO4 | -0.013713216 |
| FTO | -0.01371437 |
| AC004460 | -0.013716327 |
| TCEB1P29 | -0.013717079 |
| RNF113A | -0.013720193 |
| CTB-95D12 | -0.013720696 |
| AK7 | -0.0137221 |
| BMF | -0.013726128 |
| GOLGA6B | -0.013731787 |
| RP11-166P13 | -0.013731896 |
| RNU2-23P | -0.013732293 |
| ASH1L-AS1 | -0.013734612 |
| GPR19 | -0.013734873 |
| AL844165 | -0.013735561 |
| FAM231D | -0.013738991 |
| SEC14L5 | -0.013742791 |
| FGFR2 | -0.013742985 |
| RP11-192N10 | -0.013744068 |
| ATP6V0E1P1 | -0.013747897 |
| KL | -0.013749199 |
| SOCS5P5 | -0.013749606 |
| FAM154B | -0.013752474 |
| RP11-408P14 | -0.013752883 |
| PBX2P1 | -0.01375696 |
| HIP1R | -0.01375893 |
| RAB28P1 | -0.013764011 |
| RP11-226F19 | -0.013764394 |
| CTA-397C4 | -0.013767317 |
| RN7SKP74 | -0.01376737 |
| NLRP13 | -0.013768489 |
| HNRNPCP3 | -0.013770717 |
| RNU6-1183P | -0.013773561 |
| RP11-639F1 | -0.013775546 |
| USP9YP32 | -0.013776018 |
| RAB41 | -0.013776605 |
| COTL1P1 | -0.013778068 |
| RPL7P24 | -0.013779182 |
| AP000887 | -0.013779409 |
| MIR4455 | -0.013787893 |
| CYP46A4P | -0.013800629 |
| MAPKAPK5P1 | -0.013810252 |

| | |
|---|---|
| CTD-2555I5 | -0.013812573 |
| RP11-136H19 | -0.013813568 |
| RP11-648L3 | -0.013816754 |
| RP11-566H8 | -0.013822942 |
| KCTD19 | -0.013823683 |
| AC144652 | -0.013825574 |
| PCDHAC2 | -0.013829034 |
| ZNF487 | -0.013834375 |
| RP11-359E3 | -0.013840254 |
| OLA1P2 | -0.013840397 |
| RP5-1027G4 | -0.01384829 |
| CCDC25 | -0.013848515 |
| MIR1283-2 | -0.013849111 |
| SYT9 | -0.013849393 |
| AKR1B1P7 | -0.013850551 |
| LINC00467 | -0.013850897 |
| KB-1000E4 | -0.013851393 |
| PI4KB | -0.013851585 |
| MST1R | -0.013856143 |
| PGBD4P3 | -0.013856676 |
| RNU6-430P | -0.013858966 |
| SLC20A1 | -0.013859027 |
| RP13-488H8 | -0.013862376 |
| RP11-686N4 | -0.013869044 |
| ZNF341 | -0.013869855 |
| PLK1 | -0.013870767 |
| AC092117 | -0.013876524 |
| RNU6-61P | -0.013877099 |
| AC139453 | -0.013878844 |
| RP11-36N20 | -0.013881222 |
| RNU6-167P | -0.013883677 |
| AC079466 | -0.013884179 |
| TMEM185B | -0.01388727 |
| C11orf16 | -0.013888175 |
| FAM66D | -0.013890639 |
| ATP5J2-PTCD1 | -0.013898677 |
| ELFN2 | -0.013899923 |
| SNORD116-7 | -0.013905288 |
| RNU6-541P | -0.013906415 |
| RP11-305P22 | -0.013906534 |
| AP000688 | -0.013906659 |
| SPINK9 | -0.013907722 |
| DESI1 | -0.013911141 |
| RPS20P10 | -0.013912474 |
| BNIP3P11 | -0.013916165 |

| | |
|---|---|
| RNA5SP421 | -0.013917364 |
| RP11-524K14 | -0.013918521 |
| HGSNAT | -0.013919365 |
| CEP70 | -0.01392143 |
| AC010141 | -0.013923106 |
| SLC7A9 | -0.013927293 |
| HN1L | -0.013927309 |
| GYG1P3 | -0.013927747 |
| KRTCAP3 | -0.013928723 |
| MIR1269B | -0.013932438 |
| OR51F5P | -0.013932777 |
| RP11-545J16 | -0.013936895 |
| C5orf34 | -0.013938882 |
| PET100 | -0.013944315 |
| RN7SKP158 | -0.013946554 |
| RP11-261N11 | -0.013947805 |
| FAM124A | -0.013951422 |
| AC063944 | -0.013966307 |
| AP000265 | -0.013970948 |
| RP11-557J10 | -0.013973095 |
| SLC35F2 | -0.0139768 |
| PLEKHA5 | -0.013980409 |
| RP11-548H18 | -0.013982325 |
| AC017079 | -0.013982358 |
| RPS19P3 | -0.01398587 |
| U91328 | -0.013989507 |
| ATP6V1B2 | -0.013989972 |
| RNA5SP226 | -0.013995798 |
| RNU6-971P | -0.013999096 |
| RP11-730N24 | -0.01400111 |
| RP11-338I24 | -0.014002474 |
| EEF1B2P2 | -0.014002581 |
| FER1L6-AS1 | -0.014003163 |
| ABCD4 | -0.014003815 |
| CADM2-AS1 | -0.014005632 |
| ZNF69 | -0.014007443 |
| RP11-534L20 | -0.01401388 |
| MIR3151 | -0.014016414 |
| RP11-11K13 | -0.014017599 |
| OFD1P2Y | -0.014020073 |
| RNU6-214P | -0.014022581 |
| VN1R48P | -0.014029766 |
| H3F3AP6 | -0.014029974 |
| SYMPK | -0.01403016 |
| YARS2 | -0.014031946 |

| | |
|---|---|
| RPS12P31 | -0.014034662 |
| MIR378D1 | -0.014037395 |
| PPIAL4D | -0.014037407 |
| SNORA58 | -0.014038306 |
| AC016732 | -0.014048527 |
| RN7SKP121 | -0.014049901 |
| CTD-2315A10 | -0.014050276 |
| TINAG | -0.014051128 |
| UBTFL9 | -0.014054567 |
| FZD6 | -0.014055333 |
| DST | -0.014067539 |
| RP11-375H17 | -0.014070159 |
| DNAJB8-AS1 | -0.014074106 |
| OR10J3 | -0.014075134 |
| EFHD2 | -0.014075241 |
| VPS35 | -0.014075706 |
| TXNP4 | -0.014076443 |
| TMEM8B | -0.014078218 |
| MIR378E | -0.014078728 |
| RN7SL806P | -0.014086653 |
| AC009052 | -0.014087298 |
| NRAP | -0.014087864 |
| KCNQ5-AS1 | -0.014088193 |
| OR7E130P | -0.014089113 |
| AC008937 | -0.014090379 |
| RP11-97E7 | -0.014091211 |
| PPM1H | -0.014091688 |
| PAQR4 | -0.014092247 |
| TMEM168 | -0.014092358 |
| RP11-688D15 | -0.014094038 |
| AC243980 | -0.01409594 |
| RP11-379B18 | -0.014097363 |
| ART3 | -0.014100712 |
| HMGN1P9 | -0.014114126 |
| HNRNPLP1 | -0.014115265 |
| GAPDHP46 | -0.014118006 |
| RPL15P2 | -0.014119269 |
| CCNJP1 | -0.014123949 |
| OR2T29 | -0.014124061 |
| MIR520B | -0.01412577 |
| IAH1 | -0.014126723 |
| CA1 | -0.014133018 |
| KRTAP27-1 | -0.014141931 |
| RP1-197J16 | -0.014144284 |
| ACTRT2 | -0.01414452 |

| | |
|---|---|
| GS1-204I12 | -0.014146336 |
| CRIM1 | -0.014149738 |
| CTC-339D2 | -0.01415437 |
| RRM2P2 | -0.014156383 |
| AC020922 | -0.014157995 |
| AL139295 | -0.014159749 |
| PMM2P1 | -0.014160012 |
| AC002389 | -0.01416184 |
| NDFIP1 | -0.014162594 |
| IQGAP2 | -0.014166836 |
| AP000255 | -0.014167833 |
| RN7SL849P | -0.01416835 |
| RP11-698F20 | -0.014168441 |
| RNU6-522P | -0.014169304 |
| RP11-781M16 | -0.014173192 |
| RP11-759F5 | -0.014179716 |
| RAC1P1 | -0.014184293 |
| RP11-354G16 | -0.01418572 |
| NHSL2 | -0.014185916 |
| RNU6-443P | -0.014192487 |
| SNORD58 | -0.014193281 |
| B9D1 | -0.014198763 |
| AC104809 | -0.014204944 |
| WDR92 | -0.014206317 |
| RP11-149F8 | -0.014211547 |
| ATP5LP5 | -0.014216838 |
| CACNG1 | -0.014217085 |
| RNU2-39P | -0.014217151 |
| FAM231A | -0.014220467 |
| RNU7-175P | -0.014221952 |
| RNU6ATAC8P | -0.014224874 |
| RP11-638L3 | -0.014228289 |
| ABHD17B | -0.014229164 |
| TMEM119 | -0.014233741 |
| RP11-708J19 | -0.014235341 |
| RP11-126F18 | -0.014236431 |
| RP11-989F5 | -0.014236483 |
| KLF2P3 | -0.014240537 |
| RP11-332O19 | -0.014241382 |
| RP11-521A24 | -0.014243853 |
| OR7E96P | -0.014246937 |
| AC017028 | -0.014248278 |
| ZNF670 | -0.01425551 |
| PCDHGB3 | -0.014258432 |
| LINC00280 | -0.014262251 |

| | |
|---|---|
| RP11-375B1 | -0.014266 |
| RPL5P33 | -0.014276664 |
| RNU6-1195P | -0.014280726 |
| CYP19A1 | -0.014281311 |
| BCRP8 | -0.014282138 |
| RP11-134L4 | -0.014296839 |
| TMEM64 | -0.014298488 |
| AP001627 | -0.014303109 |
| AC006328 | -0.014304015 |
| PRUNE2 | -0.014309149 |
| TEX13B | -0.01431372 |
| DSTNP5 | -0.014314395 |
| RP11-589C21 | -0.014325079 |
| AL138721 | -0.014329338 |
| RN7SKP99 | -0.014333896 |
| ZNF138 | -0.014334229 |
| RP11-77B22 | -0.014334321 |
| SNHG8 | -0.014337764 |
| RNA5SP480 | -0.014346739 |
| AL133247 | -0.014349616 |
| MIR6790 | -0.014350878 |
| RP11-275A14 | -0.014356055 |
| RNU6-408P | -0.014362881 |
| RPS14P7 | -0.014363084 |
| HIST2H3PS2 | -0.014363181 |
| RP11-473A10 | -0.014365448 |
| RP11-291L22 | -0.014366528 |
| RNU1-77P | -0.01436656 |
| AC083855 | -0.014372007 |
| HMGN2P39 | -0.014374678 |
| RN7SL36P | -0.014378282 |
| RP11-81H3 | -0.014381411 |
| AP004242 | -0.014382715 |
| RP11-508O18 | -0.014383077 |
| AC005235 | -0.014386905 |
| TXNP1 | -0.014387559 |
| LINC01074 | -0.014389508 |
| CHRNB4 | -0.014396287 |
| AC114973 | -0.014399129 |
| GLRXP | -0.014403952 |
| CDX4 | -0.014410155 |
| MIR576 | -0.01441049 |
| EIF4EBP2P3 | -0.014411898 |
| RNU6-1073P | -0.014412324 |
| RNU6-70P | -0.014413846 |

| | |
|---|---|
| CTD-2270F17 | -0.014419311 |
| RP11-400F19 | -0.014421284 |
| ZFAND2A | -0.014421526 |
| IGSF1 | -0.01442188 |
| OPN5 | -0.014423133 |
| RP11-264F23 | -0.014428166 |
| RP4-535B20 | -0.014434175 |
| RP11-410L14 | -0.014434792 |
| RP6-99M1 | -0.014435148 |
| RP4-737E23 | -0.014437345 |
| L3HYPDH | -0.014439964 |
| RP4-585I14 | -0.01444564 |
| RPL17P37 | -0.014448727 |
| RP11-473P24 | -0.014449379 |
| C17orf62 | -0.014449639 |
| AC073065 | -0.014452284 |
| AP000322 | -0.014454066 |
| CCDC11P1 | -0.014454357 |
| HMOX1 | -0.014458272 |
| OR10J8P | -0.014459519 |
| CTD-2010I16 | -0.014459947 |
| RP11-554D14 | -0.01446412 |
| VN2R9P | -0.014467858 |
| SEPT7P9 | -0.014470678 |
| LINC01431 | -0.01447297 |
| CHP1 | -0.014480952 |
| RP5-912I13 | -0.014482741 |
| RNU6-493P | -0.014484388 |
| LINC01534 | -0.014485548 |
| RP11-1281K21 | -0.014485665 |
| BPIFA4P | -0.014486589 |
| CFC1B | -0.01448699 |
| RP5-874C20 | -0.014487984 |
| MIR4664 | -0.014488172 |
| HIPK1-AS1 | -0.014491712 |
| MIR5702 | -0.014498269 |
| HSP90AB6P | -0.014502178 |
| LINC01474 | -0.014502251 |
| AC112249 | -0.014507239 |
| SEC31A | -0.014509028 |
| CLC | -0.014512038 |
| ZNF727 | -0.014516327 |
| RP11-396J6 | -0.01451883 |
| MTND5P31 | -0.014526698 |
| LILRA5 | -0.014531485 |

| | |
|---|---|
| RP3-465N24 | -0.01453395 |
| XPNPEP2 | -0.014534445 |
| USP9YP20 | -0.014535212 |
| NNAT | -0.014535876 |
| RP11-503G7 | -0.014538606 |
| LYN | -0.014542946 |
| TAS2R9 | -0.014545958 |
| RN7SKP128 | -0.01454716 |
| AL138963 | -0.014548039 |
| BPNT1 | -0.014551746 |
| NR2F2-AS1 | -0.01455489 |
| SYCP2L | -0.014556956 |
| MT-CYB | -0.014558928 |
| DSC1 | -0.014569589 |
| RPL21P23 | -0.014570707 |
| FGF21 | -0.014572525 |
| CKS2 | -0.014572807 |
| RP4-616B8 | -0.014573101 |
| MTNR1A | -0.014573192 |
| SLC38A2 | -0.014574017 |
| RPL8P2 | -0.014575648 |
| AC087793 | -0.014577223 |
| EZH2P1 | -0.01457837 |
| AL121970 | -0.014579719 |
| OPALIN | -0.014582512 |
| OR5BT1P | -0.014582779 |
| RNA5SP477 | -0.014583345 |
| RN7SL601P | -0.014583959 |
| AL023876 | -0.014587085 |
| RP11-163F15 | -0.014587288 |
| SOX3 | -0.014588547 |
| KRTAP9-3 | -0.014591054 |
| BCCIP | -0.014591524 |
| EIF3EP1 | -0.014591562 |
| AK4P5 | -0.014593748 |
| PDCL3P3 | -0.01459659 |
| LINC01222 | -0.014599141 |
| RNU6-1060P | -0.014600182 |
| AL118509 | -0.01460161 |
| GS1-309P15 | -0.014604352 |
| RPL21P54 | -0.014606849 |
| HMGB1P30 | -0.01460708 |
| CYCSP8 | -0.014610128 |
| TSPAN1 | -0.014616791 |
| OR10D5P | -0.014618713 |

| | |
|---|---|
| AC013406 | -0.014621257 |
| GUCY2GP | -0.014625299 |
| RNU6-258P | -0.014625682 |
| AKT2 | -0.014628847 |
| PDE5A | -0.014631321 |
| GPR137C | -0.014633033 |
| ENO1P2 | -0.014633467 |
| ZFYVE9P2 | -0.014633724 |
| RNU6-757P | -0.014638108 |
| NDOR1 | -0.014653127 |
| OR10C1 | -0.014653822 |
| OSR2 | -0.014655806 |
| RP11-445N20 | -0.014658133 |
| WARS2-IT1 | -0.014659007 |
| TUBB8P8 | -0.014662136 |
| RNF32 | -0.014667435 |
| RP11-401I19 | -0.01467588 |
| AP000457 | -0.014680543 |
| AC006460 | -0.014691517 |
| RP11-425D17 | -0.014693037 |
| MIR3125 | -0.014695757 |
| RNU2-70P | -0.014696249 |
| HNRNPCL3 | -0.014698073 |
| SNORD115-25 | -0.014702118 |
| RP4-564F22 | -0.014702869 |
| PCDH9-AS3 | -0.014703854 |
| FAAHP1 | -0.014704125 |
| RN7SL767P | -0.014709362 |
| RNU6-821P | -0.01471032 |
| MKNK2 | -0.014716516 |
| AC010525 | -0.014720748 |
| RP11-430C1 | -0.014724559 |
| TGIF1P1 | -0.014724933 |
| BTC | -0.014725409 |
| NPM1P17 | -0.014728392 |
| KRTAP10-11 | -0.014729281 |
| RP5-832C2 | -0.014730619 |
| RP11-666F17 | -0.014733501 |
| AC005599 | -0.014733906 |
| STAR | -0.014737404 |
| OR10AC1 | -0.014737502 |
| RP11-43A1 | -0.014738828 |
| CTA-212D2 | -0.014740709 |
| ARHGAP12 | -0.014751997 |
| MRPS36P1 | -0.014753003 |

| | |
|---|---|
| NTRK3 | -0.014759495 |
| RP11-103C16 | -0.014763331 |
| RP11-563J2 | -0.014763913 |
| KRCC1 | -0.01476796 |
| RP11-6N13 | -0.014773287 |
| RP11-252P19 | -0.014774968 |
| ZNF460 | -0.014775458 |
| RP11-867G2 | -0.014775661 |
| RNU7-99P | -0.014778524 |
| RPL6P30 | -0.0147809 |
| RP11-324I22 | -0.014791775 |
| RP11-486B10 | -0.014792705 |
| MFSD5 | -0.014798213 |
| MTND5P6 | -0.014798811 |
| AC090958 | -0.014800681 |
| DNAJC19P4 | -0.014805009 |
| C6orf222 | -0.014806242 |
| MIR591 | -0.014806941 |
| MIR3118-4 | -0.014808174 |
| LINC01579 | -0.014809834 |
| GALR2 | -0.014810649 |
| HGS | -0.014811441 |
| LHFPL1 | -0.014811551 |
| RP11-92F20 | -0.014812212 |
| LETM1P2 | -0.014813057 |
| BMS1P15 | -0.014816353 |
| RP11-282M24 | -0.014817898 |
| BRDTP1 | -0.014818803 |
| LINC00453 | -0.01481962 |
| ALOX12B | -0.014819838 |
| TCEAL3-AS1 | -0.014821288 |
| AF224669 | -0.014823606 |
| OR5M2P | -0.014825392 |
| MIR222 | -0.014825405 |
| AF003626 | -0.01483375 |
| RAD51AP1P1 | -0.014839092 |
| MIR4488 | -0.014841382 |
| OR1AA1P | -0.014842135 |
| OR10AF1P | -0.014845135 |
| AC005606 | -0.014846276 |
| MIR5006 | -0.014847412 |
| AL050327 | -0.014854399 |
| THRB | -0.014854643 |
| MNT | -0.014855424 |
| CTD-2342N23 | -0.014856439 |

| | |
|---|---|
| RP5-879K22 | -0.014857579 |
| RAC1P7 | -0.014859066 |
| C5orf15 | -0.014860132 |
| RNU6-79P | -0.014863636 |
| RP11-15G8 | -0.014864959 |
| RNU6-482P | -0.014865131 |
| AL359546 | -0.014866369 |
| RP11-1E4 | -0.01487209 |
| CCT8L2 | -0.014872863 |
| AC010146 | -0.014872897 |
| RP11-147K6 | -0.014876797 |
| CDK9 | -0.014877583 |
| ZNF189 | -0.0148787 |
| SMIM17 | -0.014885558 |
| RP11-424M22 | -0.014889229 |
| RP11-810O3 | -0.014891238 |
| DEFB103B | -0.014892075 |
| GNPAT | -0.014892152 |
| RP11-115D19 | -0.014893404 |
| AC093382 | -0.014894116 |
| RP1-197B17 | -0.014894459 |
| GSDMA | -0.014896472 |
| RP1-60N8 | -0.014897167 |
| RNU6-1170P | -0.014898255 |
| RIIAD1 | -0.014901078 |
| RDH12 | -0.014902218 |
| MYL5 | -0.014902571 |
| RP11-796E10 | -0.014909576 |
| POU1F1 | -0.01491069 |
| AC008072 | -0.014911692 |
| RN7SL814P | -0.014912776 |
| HMGB1P39 | -0.014913112 |
| RP11-651P23 | -0.014915148 |
| TSSK6 | -0.014916581 |
| OR3B1P | -0.014924788 |
| OR2T35 | -0.014931469 |
| SCARNA24 | -0.014938362 |
| ADD2 | -0.014939484 |
| PDCD11 | -0.014946275 |
| AC068282 | -0.014947577 |
| RP5-837J1 | -0.014952751 |
| RNU6-269P | -0.014952874 |
| RPL9P3 | -0.01495457 |
| AC083810 | -0.014956327 |
| SNORA64 | -0.01495755 |

| | |
|---|---|
| FOXD3-AS1 | -0.014960213 |
| RP11-473M20 | -0.014962015 |
| PRR25 | -0.014964298 |
| RAB5CP2 | -0.014964928 |
| FBXL3 | -0.014968787 |
| RNA5SP342 | -0.014969618 |
| FNDC3CP | -0.014973708 |
| AC104653 | -0.014975485 |
| AC108667 | -0.014976674 |
| VN1R65P | -0.014977114 |
| AC007040 | -0.014981509 |
| CTD-2037L6 | -0.014984552 |
| CRYGFP | -0.014985738 |
| ZNF446 | -0.014988826 |
| AP002364 | -0.014989419 |
| RABEPK | -0.01498996 |
| FKBP4P1 | -0.014991728 |
| RNU12 | -0.014993737 |
| RP11-77H9 | -0.014999356 |
| RP11-332G1 | -0.014999575 |
| DISC1-IT1 | -0.015002271 |
| CEACAM8 | -0.015004152 |
| MGC16275 | -0.015004957 |
| RP11-40H20 | -0.015005544 |
| RP11-7I15 | -0.015014269 |
| RP11-521F1 | -0.01501878 |
| PATE2 | -0.015020464 |
| RP11-54G14 | -0.015020973 |
| CTD-2049O17 | -0.015023355 |
| MIR3680-1 | -0.015025633 |
| HTR5BP | -0.015027967 |
| AL162381 | -0.015029167 |
| RP11-115L11 | -0.015030458 |
| RP11-44I10 | -0.015031458 |
| AC133965 | -0.015032101 |
| MIR4440 | -0.015032278 |
| RIMS4 | -0.015034787 |
| PDIA3 | -0.015039581 |
| RP11-90P13 | -0.015040499 |
| RP11-14I17 | -0.015046119 |
| COASY | -0.015047656 |
| RP11-424D14 | -0.015048773 |
| MIR6876 | -0.015057012 |
| PABPC1P12 | -0.01505792 |
| VN1R46P | -0.015059228 |

| | |
|---|---|
| PARS2 | -0.015062522 |
| RP1-31B8 | -0.015062763 |
| TEX11 | -0.015064621 |
| RP13-577H12 | -0.015070767 |
| MIR4530 | -0.015073989 |
| TBC1D21 | -0.015074099 |
| RP5-916O11 | -0.015074137 |
| CASC1 | -0.01507495 |
| RNASEH1P2 | -0.015075669 |
| FAM66B | -0.015077289 |
| SF3A3P2 | -0.015081259 |
| RP11-710M11 | -0.015085686 |
| RP11-5O17 | -0.015085797 |
| RP11-397E7 | -0.01508669 |
| AP003558 | -0.015087985 |
| FUT8 | -0.015088625 |
| RPL13AP5 | -0.015088644 |
| INO80C | -0.015095212 |
| LINC01210 | -0.015095655 |
| DUTP5 | -0.01510685 |
| ATP5A1P6 | -0.015107021 |
| HMGB1P48 | -0.015107202 |
| ZDHHC18 | -0.015107826 |
| RP13-329D4 | -0.015108754 |
| CST13P | -0.015110109 |
| RSRC1 | -0.015111975 |
| AC020626 | -0.015114104 |
| CTD-2125J1 | -0.015117014 |
| GAS2L1P1 | -0.015118542 |
| HNRNPA1P7 | -0.015121592 |
| RP11-1148L6 | -0.015125092 |
| RP5-1096J16 | -0.015127555 |
| LRP1B | -0.015130763 |
| AC074051 | -0.015132563 |
| APITD1-CORT | -0.015136906 |
| CTD-2036A18 | -0.015137054 |
| OR4R3P | -0.015140225 |
| RP11-331K15 | -0.015142062 |
| SKIV2L | -0.015143977 |
| GAPDHP54 | -0.015145231 |
| RNU1-20P | -0.015146371 |
| TRMT10C | -0.015148386 |
| KCNJ1 | -0.015149178 |
| CTD-3064H18 | -0.015157081 |
| RN7SL716P | -0.015158644 |

| | |
|---|---|
| ANKRD20A19P | -0.015159161 |
| MIR4765 | -0.015160164 |
| RP11-333B6 | -0.015160748 |
| OVGP1 | -0.015161838 |
| RP11-716H6 | -0.015167381 |
| PDDC1 | -0.015170396 |
| MIR6848 | -0.015170691 |
| RPL7AP30 | -0.015170843 |
| RP11-317F20 | -0.015174447 |
| RNU1-96P | -0.015178726 |
| TTC3P1 | -0.015178918 |
| ZNF404 | -0.015192146 |
| CIRH1A | -0.01519516 |
| RP11-466I1 | -0.015197026 |
| HAUS1P1 | -0.015198918 |
| OFD1P7Y | -0.015200538 |
| GLUL | -0.015201616 |
| RP11-102F4 | -0.015204342 |
| SLC16A1P1 | -0.015206594 |
| C15orf61 | -0.015210714 |
| GRAMD4P2 | -0.015211527 |
| RP11-478B9 | -0.015214993 |
| SCARNA13 | -0.015222065 |
| MAB21L3 | -0.015223254 |
| TULP3 | -0.015226603 |
| ELTD1 | -0.01522798 |
| AC097466 | -0.015230453 |
| AL138999 | -0.015231565 |
| RP11-1105G2 | -0.015233283 |
| RP11-334E6 | -0.015236716 |
| SMCO1 | -0.015237255 |
| CD302 | -0.015237767 |
| CTD-2647E9 | -0.015240692 |
| RNA5SP486 | -0.015243962 |
| LDLRAP1 | -0.015251423 |
| RP13-7D7 | -0.015257168 |
| DOCK9-AS2 | -0.015257507 |
| LGMNP1 | -0.0152583 |
| OR10R1P | -0.015262009 |
| PGLYRP2 | -0.015262323 |
| EPHA8 | -0.015267056 |
| CLCP1 | -0.015267292 |
| AC073409 | -0.015267906 |
| RPS15P5 | -0.015271263 |
| KB-208E9 | -0.015274034 |

| | |
|---|---|
| SLC14A2-AS1 | -0.015275579 |
| MIR218-2 | -0.015284013 |
| DAAM1 | -0.015284652 |
| FAF1 | -0.015287276 |
| CALM1 | -0.015289639 |
| GGT8P | -0.015292207 |
| NARF | -0.015298141 |
| TRBV20OR9-2 | -0.015298803 |
| RP11-363J20 | -0.015301286 |
| EEF1GP3 | -0.015301362 |
| UTP14A | -0.015301435 |
| RNU6-31P | -0.015306192 |
| PSPH | -0.015307426 |
| MIR30C2 | -0.0153153 |
| MT-TS1 | -0.015316132 |
| DDX39B-AS1 | -0.015320075 |
| RNU6-1120P | -0.015324309 |
| KCTD4 | -0.015324369 |
| UBE2DNL | -0.015325048 |
| CTD-2234B20 | -0.015326427 |
| FAM218BP | -0.015326977 |
| RNU6-509P | -0.015328867 |
| AC139103 | -0.015334831 |
| RP5-1120P11 | -0.015336487 |
| AC006014 | -0.015340085 |
| TRBV29OR9-2 | -0.015342704 |
| NOL10 | -0.015342869 |
| RAB14 | -0.015346218 |
| SYTL3 | -0.015347558 |
| RP11-82H13 | -0.015350814 |
| AC079467 | -0.015352168 |
| IGHD3OR15-3B | -0.015354098 |
| AC007009 | -0.015356126 |
| RNU6-1150P | -0.015357193 |
| CCL17 | -0.015363546 |
| GNG10 | -0.015367596 |
| NEFLP1 | -0.015369214 |
| MIR5591 | -0.015369762 |
| OR7E21P | -0.01537052 |
| RP3-336K20__B | -0.015372768 |
| RPS20P20 | -0.015374572 |
| RP11-566F5 | -0.015379422 |
| SIAH1P1 | -0.015379486 |
| BGLT3 | -0.015381278 |
| XXyac-YR21CG7 | -0.015381312 |

| | |
|---|---|
| MIR4786 | -0.01538778 |
| RP11-370I10 | -0.015388155 |
| SSXP10 | -0.015390657 |
| RP11-309N24 | -0.01539677 |
| IGLVIV-53 | -0.015399427 |
| OR10G7 | -0.015403267 |
| PAFAH1B2 | -0.01540369 |
| ASGR2 | -0.015404369 |
| RPSAP1 | -0.015407812 |
| TRDD2 | -0.015411753 |
| COX4I1P1 | -0.015418437 |
| MIR761 | -0.015425162 |
| RP11-177J6 | -0.015428042 |
| AC012493 | -0.015429334 |
| PPP3R1 | -0.015429887 |
| PXT1 | -0.015432194 |
| CYP2F1 | -0.015436057 |
| MIR920 | -0.015438575 |
| SNRPFP3 | -0.015440353 |
| RP11-189E14 | -0.015441012 |
| AL118520 | -0.015442074 |
| LL0XNC01-16G2 | -0.015442719 |
| RNA5SP356 | -0.01544372 |
| KRT18P65 | -0.015448347 |
| MS4A6E | -0.01544863 |
| TBCE | -0.015448902 |
| MED28P7 | -0.015451239 |
| RP5-878I13 | -0.015456028 |
| LINC01075 | -0.015456837 |
| DHRS4L1 | -0.015457944 |
| RP11-731D1 | -0.015458206 |
| LINC01476 | -0.015460382 |
| OSBPL8 | -0.015462391 |
| HS3ST5 | -0.015463479 |
| TUBA3E | -0.015464496 |
| MMP21 | -0.015467124 |
| AC007535 | -0.01547882 |
| UBE2Q2P9 | -0.015491844 |
| RP11-116P24 | -0.015497872 |
| RP11-457K10 | -0.015498246 |
| ASS1P10 | -0.015498256 |
| AC005625 | -0.015503718 |
| UBL4B | -0.015504302 |
| TYMP | -0.015504811 |
| AC078842 | -0.015512231 |

| | |
|---|---|
| AC026367 | -0.015516477 |
| VDAC1P10 | -0.015516636 |
| LINC00332 | -0.015520893 |
| TMEM212-AS1 | -0.015525585 |
| MZT1P1 | -0.015526393 |
| ZNF213-AS1 | -0.015531324 |
| SCGB1B2P | -0.015533612 |
| EIF2B5-IT1 | -0.015533774 |
| UGT1A11P | -0.01553553 |
| WDR82P1 | -0.015537513 |
| AC012368 | -0.015539498 |
| CTD-2275D24 | -0.015539776 |
| RP4-607I7 | -0.015543229 |
| RP11-390E23 | -0.015543343 |
| MTCYBP1 | -0.015545688 |
| OR7E108P | -0.015546041 |
| YY2 | -0.015547659 |
| MIR7162 | -0.015547809 |
| ASCL2 | -0.015549594 |
| RP11-886P16 | -0.01555021 |
| MT1F | -0.015556069 |
| ARL2BPP3 | -0.015560692 |
| CAMSAP2 | -0.015566246 |
| AL137855 | -0.015567397 |
| PBX3 | -0.015569391 |
| RNU6-40P | -0.015571715 |
| RP11-16D22 | -0.01557218 |
| AL390776 | -0.015574313 |
| AC002064 | -0.015577207 |
| RNU2-45P | -0.015579548 |
| ABLIM3 | -0.015579698 |
| MEIOB | -0.015581103 |
| RP11-31F19 | -0.015583373 |
| RP11-375N15 | -0.015584969 |
| RP11-781A6 | -0.015585731 |
| AC002519 | -0.015586057 |
| FAM135A | -0.01558768 |
| AC016597 | -0.015590107 |
| EEF1DP7 | -0.015594871 |
| TRIM64EP | -0.015596489 |
| HNRNPA1P58 | -0.015597237 |
| NFE2L2 | -0.015598024 |
| NFE2L3P1 | -0.015598203 |
| RP11-1H8 | -0.015598351 |
| RNU2-15P | -0.01560238 |

| | |
|---|---|
| RP3-329E20 | -0.015605251 |
| RP11-143H17 | -0.015610911 |
| ALDH4A1 | -0.015613002 |
| MTCO1P5 | -0.015613169 |
| ZNF449 | -0.015615197 |
| RCC2 | -0.015616128 |
| CICP7 | -0.01561709 |
| AC009014 | -0.015619975 |
| RP1-76C18 | -0.015622892 |
| AL360078 | -0.015627059 |
| CDY2B | -0.015627059 |
| PRIM2 | -0.015632072 |
| AC024571 | -0.015635475 |
| CCDC136 | -0.015638566 |
| CTAGE8 | -0.015638867 |
| ARHGEF1 | -0.015639198 |
| HIGD1AP10 | -0.015639225 |
| UBE2Q2P11 | -0.015639767 |
| RP11-5P15 | -0.015644262 |
| HMGB3P31 | -0.015646981 |
| RP11-252O18 | -0.015649315 |
| CTD-2545M3 | -0.015649743 |
| RMST | -0.015655275 |
| AC114296 | -0.015659408 |
| EPS8L3 | -0.015661626 |
| CTD-2653B5 | -0.015663995 |
| RP11-629F19 | -0.015664789 |
| LINC01440 | -0.015667488 |
| ZNF467 | -0.015668948 |
| OR5BB1P | -0.015671051 |
| OR7E159P | -0.015675154 |
| GTPBP3 | -0.015681116 |
| RP11-332K15 | -0.015681446 |
| AL360217 | -0.015682819 |
| RNU6-1312P | -0.015682942 |
| AC013401 | -0.015683847 |
| LINC00538 | -0.015684008 |
| AP006222 | -0.015690587 |
| RP11-257K9 | -0.015692702 |
| AC226119 | -0.015693771 |
| RP11-254B13 | -0.015694512 |
| LINC01509 | -0.015700307 |
| AC005394 | -0.015707609 |
| NUPL1P1 | -0.015713889 |
| CTD-2623H2 | -0.015714399 |

| | |
|---|---|
| RN7SKP14 | -0.015714672 |
| GATA5 | -0.015715144 |
| CD27-AS1 | -0.01571749 |
| RP11-80F22 | -0.015719731 |
| RP11-678K21 | -0.015722289 |
| DNAJC21 | -0.015725137 |
| CTB-51A17 | -0.015725558 |
| MRPS33P4 | -0.01572646 |
| RN7SKP78 | -0.015728463 |
| MIR6855 | -0.015729194 |
| RNU6-320P | -0.015730869 |
| MTND4P13 | -0.015731616 |
| RN7SKP96 | -0.015731894 |
| FAM135B | -0.015731935 |
| RP11-1096D5 | -0.015734869 |
| SMIM19 | -0.015735091 |
| RP11-660M18 | -0.015735276 |
| C16orf74 | -0.015736486 |
| FTLP10 | -0.015741596 |
| PILRA | -0.015744319 |
| CCT6B | -0.015746868 |
| POLR3C | -0.01574858 |
| PDE7A | -0.015749119 |
| C20orf181 | -0.015752336 |
| EXTL3-AS1 | -0.015753613 |
| POTEM | -0.015754922 |
| TBL1XR1-AS1 | -0.015756073 |
| AC004951 | -0.015760673 |
| MAPKAP1 | -0.015762162 |
| MIR6886 | -0.01576354 |
| MIR6893 | -0.015763687 |
| GK-IT1 | -0.01576802 |
| FAR2P3 | -0.015768561 |
| LINC00540 | -0.015772066 |
| RP1-288M22 | -0.015772449 |
| RP11-710F7 | -0.015774967 |
| GPR62 | -0.01578025 |
| AC004850 | -0.015783379 |
| OVOL2 | -0.015786554 |
| RNA5SP181 | -0.015791264 |
| RP11-697H10 | -0.01579552 |
| RP1-266L20 | -0.015798573 |
| ODAM | -0.015801281 |
| SCAMP4 | -0.015802908 |
| RP11-231C18 | -0.015803684 |

| | |
|---|---|
| MYH7B | -0.015807243 |
| CTC-329D1 | -0.015813197 |
| RP11-79E3 | -0.015813352 |
| RP11-435F17 | -0.015814239 |
| TMEM218 | -0.015817011 |
| AC007161 | -0.015822161 |
| HERC2P8 | -0.015824386 |
| RPA4 | -0.015826011 |
| RP11-67P15 | -0.01582804 |
| AC003006 | -0.015828443 |
| OR10W1 | -0.015828679 |
| ZNF793-AS1 | -0.015830351 |
| VASH1 | -0.015830556 |
| CTD-2236F14 | -0.015838625 |
| RN7SL740P | -0.015842224 |
| RP11-571E6 | -0.015842571 |
| PGA3 | -0.015843394 |
| RP11-945C19 | -0.015850748 |
| AL138710 | -0.015853199 |
| AL359091 | -0.015854484 |
| RNU7-45P | -0.015857025 |
| ZNF672 | -0.015858575 |
| RP11-556H2 | -0.015861743 |
| OR13D1 | -0.015862794 |
| AP000949 | -0.015867227 |
| CPB1 | -0.01586875 |
| ZFP28 | -0.015870036 |
| RP11-232C2 | -0.015870971 |
| AC119403 | -0.01587217 |
| ATP6V0E1P2 | -0.01587534 |
| RN7SL362P | -0.01587619 |
| SNORD105 | -0.015879568 |
| RP11-167B3 | -0.015883012 |
| SWAP70 | -0.015883618 |
| KLHL24 | -0.015885125 |
| AC005559 | -0.015887299 |
| RP11-737F10 | -0.015890364 |
| RNU6-506P | -0.01589704 |
| INPP1 | -0.015898745 |
| FAM98C | -0.015901294 |
| GPR68 | -0.015911359 |
| RP11-395F4 | -0.015913943 |
| SNORD114-29 | -0.01591672 |
| TMA16P1 | -0.015917626 |
| AL596092 | -0.015921462 |

| | |
|---|---|
| AC243562 | -0.015923886 |
| RP11-1220K2 | -0.015926207 |
| RP11-1113N2 | -0.015928953 |
| SNRPCP1 | -0.015929035 |
| KRT8P35 | -0.015930786 |
| PRKAA2 | -0.015936421 |
| TAS2R1 | -0.015938138 |
| CORO1B | -0.01594016 |
| EIF4BP4 | -0.015945762 |
| ZDHHC16 | -0.015946077 |
| EIF3E | -0.015946579 |
| GS1-39E22 | -0.015947862 |
| LINC00700 | -0.015948844 |
| RP11-7M10 | -0.015949659 |
| RN7SL567P | -0.015950629 |
| RP11-498J9 | -0.015955497 |
| SLC2A7 | -0.01595734 |
| SRSF1P1 | -0.015957419 |
| AC006277 | -0.015959045 |
| RP11-76P2 | -0.015959543 |
| LINC01217 | -0.015961693 |
| VNN1 | -0.015963622 |
| LATS2-AS1 | -0.015967638 |
| RP5-981L23 | -0.015969239 |
| RPS12P27 | -0.01597625 |
| TMEM247 | -0.01598408 |
| EIF1AXP1 | -0.015986172 |
| KRT8P31 | -0.015988485 |
| RN7SL20P | -0.015999366 |
| HPCAL1 | -0.015999512 |
| HSP90B2P | -0.015999689 |
| GPHB5 | -0.016000756 |
| AL355593 | -0.016001796 |
| RP1-263J7 | -0.016009339 |
| FEM1A | -0.016014526 |
| AC069155 | -0.016021895 |
| CTNND1 | -0.016027606 |
| AC026185 | -0.016027753 |
| AC073657 | -0.016027787 |
| C7orf43 | -0.01602796 |
| RP11-839G9 | -0.016035262 |
| POLRMTP1 | -0.016038957 |
| CMP21-97G8 | -0.016040871 |
| RP11-30K9 | -0.016045049 |
| LINC01498 | -0.016046534 |

| | |
|---|---|
| ST6GALNAC4 | -0.016057096 |
| PTCHD3P3 | -0.0160687 |
| TANC2 | -0.016072724 |
| AC008060 | -0.016075969 |
| PCGEM1 | -0.016079219 |
| RP11-381O7 | -0.016079801 |
| CTD-2547L16 | -0.016081357 |
| RNASEH2C | -0.01608346 |
| RN7SL357P | -0.016089168 |
| POMGNT1 | -0.016089209 |
| UGGT1 | -0.01609293 |
| OPRL1 | -0.016094362 |
| G6PC3 | -0.016094976 |
| MIR3689D2 | -0.01609715 |
| GPATCH2 | -0.016104949 |
| MIR4781 | -0.016107065 |
| LLNLR-470E3 | -0.016109815 |
| NCKAP1 | -0.016111777 |
| RP4-735C1 | -0.016114344 |
| MAK16 | -0.016115294 |
| AKAP4 | -0.016116547 |
| RP11-697E23 | -0.016123782 |
| MIR4270 | -0.016129138 |
| RP11-427E4 | -0.016129982 |
| BAP1 | -0.016136225 |
| RNU6-190P | -0.01613848 |
| AC100821 | -0.01614642 |
| CYP51A1P2 | -0.01614914 |
| AC099850 | -0.016156281 |
| TVP23CP1 | -0.016156391 |
| AP005227 | -0.016158245 |
| AC004915 | -0.016165062 |
| RP11-218L14 | -0.016165254 |
| MEAF6P1 | -0.016165353 |
| RP11-527J8 | -0.016170124 |
| RPL39P29 | -0.016172758 |
| MCCD1 | -0.01617968 |
| CD47 | -0.016180115 |
| LINC00837 | -0.016181842 |
| RP11-388M20 | -0.01618223 |
| BNIP3P23 | -0.016184319 |
| RP11-795F19 | -0.016184729 |
| AC011498 | -0.016186558 |
| FAM86HP | -0.016187111 |
| RN7SL648P | -0.016191758 |

| | | |
|---|---|---|
| LINC00265-3P | | -0.016195574 |
| RP11-731J8 | | -0.016196198 |
| RNA5SP351 | | -0.016198904 |
| UBA52P5 | | -0.016199047 |
| AC010082 | | -0.016201599 |
| KLHL3 | | -0.016203632 |
| RP11-3P17 | | -0.016206032 |
| EPHA6 | | -0.016206127 |
| CBX3P1 | | -0.016208922 |
| RNA5SP415 | | -0.016210475 |
| RP11-928F19 | | -0.016213438 |
| CTD-2336H13 | | -0.016214456 |
| CTD-2023J5 | | -0.016219458 |
| RN7SL122P | | -0.016225728 |
| RP11-262M14 | | -0.016225816 |
| CTD-2004A9 | | -0.016226492 |
| INSL4 | | -0.016227827 |
| RP4-555L14 | | -0.016229265 |
| RN7SKP177 | | -0.01623348 |
| STARD3 | | -0.016233959 |
| CTC-430J12 | | -0.016235181 |
| RP11-45F15 | | -0.016235863 |
| MGAT3 | | -0.016237048 |
| RNU6-1322P | | -0.016237483 |
| RN7SL677P | | -0.016239961 |
| RP11-26L16 | | -0.01624155 |
| TMCC2 | | -0.01624276 |
| RP11-720L2 | | -0.016247932 |
| RP11-296A16 | | -0.016248084 |
| FOXG1-AS1 | | -0.01625356 |
| | 6-Mar | -0.016261179 |
| CTD-3018O17 | | -0.016263117 |
| PRKAG2-AS1 | | -0.01626464 |
| RP11-242F11 | | -0.016265004 |
| DNAJB12 | | -0.01626611 |
| AC093802 | | -0.016269282 |
| RP11-281B1 | | -0.016272081 |
| AC004016 | | -0.016275032 |
| AC018865 | | -0.016276558 |
| CTAG1A | | -0.016278953 |
| RP11-709P2 | | -0.016279053 |
| FABP7P1 | | -0.016279319 |
| AL353135 | | -0.016281783 |
| SMIM6 | | -0.016282832 |
| RP11-451O13 | | -0.016290764 |

| | |
|---|---|
| GLI1 | -0.01629372 |
| RP11-24C3 | -0.016299276 |
| RNA5SP261 | -0.016299807 |
| TNRC18P1 | -0.016301872 |
| RP11-630H24 | -0.016302259 |
| RP11-536P16 | -0.016306395 |
| RP11-603J24 | -0.016309795 |
| RP11-358N4 | -0.016312038 |
| GS1-279B7 | -0.016312251 |
| PIGFP3 | -0.01631375 |
| TAF9P2 | -0.016314948 |
| PCNA | -0.016316177 |
| NUCKS1 | -0.016317144 |
| RNU6-719P | -0.016317296 |
| RP11-85E16 | -0.016318333 |
| MIR6799 | -0.016319679 |
| RP11-571L19 | -0.016321219 |
| AC027124 | -0.016325284 |
| RP11-173D3 | -0.01632858 |
| VN1R88P | -0.016331321 |
| AC136611 | -0.016335546 |
| NEAT1 | -0.016336401 |
| CST9LP1 | -0.01633666 |
| TRAJ57 | -0.016337211 |
| RNU6-720P | -0.016339058 |
| HOXB13-AS1_2 | -0.016341346 |
| TXLNA | -0.016343062 |
| TMEM179B | -0.016345797 |
| AC068533 | -0.016348904 |
| ADAD1P2 | -0.01634939 |
| KRT18P68 | -0.016352613 |
| RP4-735N21 | -0.016353156 |
| LINC01568 | -0.016354975 |
| RP11-384L8 | -0.016358802 |
| CYP4F23P | -0.016359069 |
| CHTOP | -0.016365202 |
| RN7SL692P | -0.016367119 |
| ZSCAN2 | -0.016368514 |
| RP11-17E13 | -0.016369175 |
| EARS2 | -0.016370896 |
| RP11-397H6 | -0.01637202 |
| ASUN | -0.016372756 |
| PFN1P10 | -0.016372773 |
| FOXD4L1 | -0.016376467 |
| AC068490 | -0.016378517 |

| | |
|---|---|
| FAIM2 | -0.016380254 |
| RNU6-48P | -0.01638222 |
| ECHDC2 | -0.016383287 |
| RP11-343H19 | -0.016384805 |
| TACC1P1 | -0.016388562 |
| AIFM1 | -0.016395529 |
| CYP2A7P1 | -0.016398659 |
| GLYATL1P3 | -0.016405712 |
| FAM108A9P | -0.016407117 |
| SPA17 | -0.016407158 |
| CCND2 | -0.016410562 |
| KRTAP4-4 | -0.016412178 |
| CDKN2AIPNLP2 | -0.01641606 |
| RN7SL449P | -0.016418919 |
| ATP7A | -0.016421744 |
| RP11-193I22 | -0.016422114 |
| SHE | -0.016423661 |
| RN7SL234P | -0.016424625 |
| XKR9 | -0.01642656 |
| MIR4715 | -0.016428813 |
| RP11-196H14 | -0.016429913 |
| RP13-608F4 | -0.016439291 |
| MAPKBP1 | -0.016444182 |
| CTB-33O18 | -0.016444797 |
| HMGB1P18 | -0.016447413 |
| RPS4XP18 | -0.016449532 |
| OR2B11 | -0.016449593 |
| SH3GLB1 | -0.016451904 |
| CRYGC | -0.016456157 |
| AC034228 | -0.016456446 |
| LINC00840 | -0.016458088 |
| TAF9BP2 | -0.016468799 |
| FAM110D | -0.016473914 |
| PAWR | -0.016477151 |
| RN7SL480P | -0.016478172 |
| AC006026 | -0.016478525 |
| NEK11 | -0.016487273 |
| METTL22 | -0.016489226 |
| HNRNPA1P55 | -0.01648948 |
| RNA5SP130 | -0.016490218 |
| RP11-344B23 | -0.016492825 |
| RNU2-55P | -0.016496481 |
| RN7SL326P | -0.016496622 |
| MIR548D2 | -0.01649784 |
| ERCC6-PGBD3 | -0.016500396 |

| | |
|---|---|
| RP11-156E8 | -0.016501177 |
| RNU1-65P | -0.016505308 |
| CTB-99A3 | -0.016509421 |
| ZNF101P2 | -0.016513042 |
| EEF1A1P7 | -0.016514362 |
| RP11-718I15 | -0.016514698 |
| PRKCA-AS1 | -0.016528549 |
| RN7SL566P | -0.016529711 |
| CTD-2340E1 | -0.016537847 |
| AC007272 | -0.016543709 |
| OR51A5P | -0.016543872 |
| ZNF174 | -0.016544205 |
| PPP1R1AP1 | -0.016548493 |
| RP11-159A18 | -0.016557718 |
| OR2A42 | -0.016560428 |
| SNX18P23 | -0.016562314 |
| AL049539 | -0.01657534 |
| RP11-813B8 | -0.016586244 |
| AL133153 | -0.016590265 |
| CDK8 | -0.016594087 |
| NANOGP3 | -0.016595543 |
| RN7SL11P | -0.016595979 |
| RNA5SP77 | -0.016602865 |
| MIR8057 | -0.016606473 |
| VAC14-AS1 | -0.016609586 |
| RNU6-220P | -0.016609807 |
| REG4 | -0.016616059 |
| LRCH1 | -0.016625828 |
| MIR3690 | -0.016632113 |
| RP11-465N4 | -0.016636358 |
| KRTAP29-1 | -0.016642611 |
| GPR107 | -0.01664408 |
| AC007365 | -0.016645568 |
| STX12 | -0.016646963 |
| MYT1L-AS1 | -0.016648797 |
| ABHD11 | -0.016654052 |
| AC109335 | -0.016654245 |
| RP11-563M4 | -0.016657401 |
| ZNF322P1 | -0.016657461 |
| CACNA1H | -0.016659318 |
| RNMTL1P2 | -0.01666353 |
| RP11-190P13 | -0.016663929 |
| MT4 | -0.016664475 |
| FAM169B | -0.01667322 |
| RP11-25I9 | -0.016679716 |

| | |
|---|---|
| RP11-83B20 | -0.016681161 |
| ARHGEF28 | -0.016681179 |
| HNRNPCP2 | -0.016682965 |
| RP11-181C3 | -0.016689849 |
| ATP5G1P1 | -0.016690536 |
| EID2B | -0.016695784 |
| MIR3941 | -0.016698213 |
| METTL21B | -0.016701161 |
| KIF22 | -0.016701291 |
| RP11-419K12 | -0.016701574 |
| RP11-262H14 | -0.016702854 |
| CTC-554D6 | -0.016704328 |
| MIR3146 | -0.01671048 |
| GUCA1C | -0.016713125 |
| RAP1B | -0.016717329 |
| RNU6-769P | -0.016721173 |
| RP11-266E6 | -0.016722152 |
| NCF2 | -0.016728102 |
| CXCL1P | -0.016729155 |
| TRAJ51 | -0.016730292 |
| AC009531 | -0.016730748 |
| OR10A4 | -0.016733894 |
| RP11-95O2 | -0.016738988 |
| SNORD116-15 | -0.016739884 |
| MED4 | -0.01674094 |
| AL157904 | -0.016741514 |
| RNU6-318P | -0.016742392 |
| AC034236 | -0.016746493 |
| NDUFB4P1 | -0.016749386 |
| PNO1 | -0.01674987 |
| RNU6-939P | -0.01675094 |
| AL356217 | -0.016753593 |
| MED26 | -0.016754522 |
| CTB-7E3 | -0.016764292 |
| KRTAP5-3 | -0.016765607 |
| ISCA2P1 | -0.016766379 |
| CMB9-14B22 | -0.01677194 |
| MIR6839 | -0.016772322 |
| AC007326 | -0.016774198 |
| LINC01153 | -0.016775359 |
| OSCP1 | -0.016776811 |
| NFYAP1 | -0.016780102 |
| LPAL2 | -0.016781669 |
| AC104073 | -0.016782387 |
| GRIK5 | -0.016784495 |

| | |
|---|---|
| FCF1P7 | -0.016789375 |
| RP3-477M7 | -0.016790507 |
| RP4-666F24 | -0.016791317 |
| GAPDHP58 | -0.016791806 |
| C17orf61-PLSCR3 | -0.016796184 |
| LINC01047 | -0.016809615 |
| RP1-78B3 | -0.016810871 |
| PRSS21 | -0.016813901 |
| RP1-178F10 | -0.016821892 |
| RPL31P3 | -0.016825934 |
| SNORA7B | -0.016828253 |
| KRT19P4 | -0.016832841 |
| SUV39H2 | -0.016833595 |
| RP11-24P14 | -0.01683425 |
| COMMD3 | -0.016840832 |
| AC133633 | -0.016841795 |
| LIN37 | -0.016844144 |
| ALG10 | -0.016850135 |
| AC018867 | -0.016852404 |
| RP11-380G5 | -0.016859169 |
| DPRX | -0.016860773 |
| TCEAL1 | -0.016861596 |
| TVP23BP1 | -0.016863737 |
| CTD-2012B7 | -0.016864015 |
| CTB-55P19 | -0.016865573 |
| SNORD115-37 | -0.016865723 |
| AC093304 | -0.016866759 |
| TRIM51DP | -0.01686898 |
| XX-2136C77 | -0.016869558 |
| RPS15AP11 | -0.01687513 |
| CENPN | -0.016884834 |
| NIPA2P5 | -0.016886497 |
| SLC7A4 | -0.016888328 |
| UBTFL5 | -0.016893344 |
| LINC00488 | -0.016893836 |
| SRP54-AS1 | -0.016893951 |
| RP11-243E13 | -0.016894595 |
| RNU6-709P | -0.016897041 |
| DNM2 | -0.016902305 |
| MIR4515 | -0.016907752 |
| AP001137 | -0.016910711 |
| OR5J1P | -0.016911248 |
| SGPP2 | -0.016914864 |
| GOLGA7 | -0.016914994 |
| LCE2C | -0.016916651 |

| | |
|---|---|
| RP11-104G3 | -0.01692084 |
| RP11-638I8 | -0.016921301 |
| RNU6-946P | -0.016929534 |
| MIR3692 | -0.016929961 |
| MTND4P3 | -0.016932184 |
| SH3RF1 | -0.016932948 |
| GID8 | -0.016933376 |
| PRAMEF14 | -0.016933783 |
| RP11-410C4 | -0.016939611 |
| AC009196 | -0.016941099 |
| AC099332 | -0.016941526 |
| C19orf71 | -0.016946772 |
| AC009065 | -0.01694929 |
| RPS12P23 | -0.016950641 |
| CEP83-AS1 | -0.016952763 |
| RPA1 | -0.016955275 |
| POTEJ | -0.016957786 |
| RP11-63G10 | -0.016959507 |
| TTR | -0.016959584 |
| RP11-80I15 | -0.016960266 |
| RP11-774I5 | -0.016962076 |
| RP11-99E15 | -0.016966467 |
| RP11-157E14 | -0.016969092 |
| RP11-251G23 | -0.016971108 |
| GPRC5A | -0.01697548 |
| SSXP3 | -0.016976837 |
| IFNA12P | -0.016976849 |
| MARK2P9 | -0.016986315 |
| GPR146 | -0.016987574 |
| CCDC159 | -0.01699096 |
| OR4C4P | -0.016993425 |
| SLCO1A2 | -0.016993939 |
| ERC2-IT1 | -0.016996052 |
| LACE1 | -0.016996486 |
| RP11-403E24 | -0.016997593 |
| RP11-98J9 | -0.016998589 |
| RPL3P10 | -0.017000041 |
| AC011410 | -0.017001326 |
| MIR6129 | -0.017004476 |
| MTHFD2P2 | -0.017009422 |
| PIEZO1P1 | -0.017010906 |
| DUX4L27 | -0.017012124 |
| AC092642 | -0.017013911 |
| XXyac-YR29IB3 | -0.017014863 |
| RP11-159K7 | -0.017017103 |

| | |
|---|---|
| RP11-428G2 | -0.017019437 |
| CLK2 | -0.017019501 |
| AC008940 | -0.017020688 |
| RNU5B-4P | -0.017023983 |
| UBE2L4 | -0.01702861 |
| TPGS2 | -0.017028748 |
| ALG6 | -0.017030683 |
| RP11-14J16 | -0.017033844 |
| SLC5A2 | -0.017037467 |
| MIR1265 | -0.017040343 |
| KRT18P19 | -0.017041141 |
| CTD-2552B11 | -0.017044597 |
| AIP | -0.017045586 |
| RNU4-31P | -0.017049377 |
| RBM17 | -0.01705007 |
| PRELID1P3 | -0.017054478 |
| AC004543 | -0.017060259 |
| CICP16 | -0.017069091 |
| AL109618 | -0.017074449 |
| RP11-327L9 | -0.017076435 |
| RNU6-986P | -0.0170772 |
| APOOP1 | -0.017084772 |
| CCDC42B | -0.017085578 |
| RP11-351M8 | -0.017086695 |
| Six3os1_5 | -0.017093005 |
| RP11-175D17 | -0.017099697 |
| RP11-47F1 | -0.017100095 |
| AMY1B | -0.017101581 |
| PRAMEF12 | -0.017102399 |
| KLHL7-AS1 | -0.017103794 |
| SLC25A44 | -0.017105346 |
| TRDN | -0.017106929 |
| HSD17B1P1 | -0.017108575 |
| APOC3 | -0.017110401 |
| AL132796 | -0.017119564 |
| RP11-133M8 | -0.017120255 |
| RP11-304C12 | -0.01712347 |
| AC090150 | -0.017126646 |
| RP5-1046G13 | -0.017127612 |
| LCE4A | -0.017133396 |
| ZNF584 | -0.017133924 |
| C14orf79 | -0.017138446 |
| MTND5P19 | -0.01713883 |
| MIR548J | -0.017139746 |
| TNK1 | -0.017143897 |

| | |
|---|---|
| CMTM4 | -0.017147134 |
| RNU6-1237P | -0.01714846 |
| RP11-143M1 | -0.01715472 |
| EFCC1 | -0.017156511 |
| GIMAP1-GIMAP5 | -0.017156788 |
| ZNF793 | -0.017159712 |
| SMCR2 | -0.017160688 |
| AC097455 | -0.017165558 |
| RP13-152O15 | -0.017165965 |
| RNA5SP233 | -0.017166135 |
| RP5-936J12 | -0.017166511 |
| LRRC72 | -0.017167614 |
| RNU6-251P | -0.017171421 |
| AL590723 | -0.017172241 |
| RP3-424M6 | -0.017180847 |
| RECQL | -0.017187508 |
| YWHAG | -0.017188308 |
| SUMO2P4 | -0.01718946 |
| BRWD1-IT1 | -0.017191117 |
| SPRED2 | -0.017192327 |
| TAF9BP1 | -0.017193069 |
| CTD-2528A14 | -0.017194629 |
| RP11-736N17 | -0.01719667 |
| RP4-630J13 | -0.017200879 |
| RP13-672B3 | -0.017201542 |
| RP11-767L7 | -0.017206588 |
| RN7SL636P | -0.017212385 |
| RP11-18D7 | -0.017213583 |
| RP11-79O8 | -0.01721663 |
| NDUFB1P2 | -0.017221259 |
| RP11-513D4 | -0.017221656 |
| ZNRF3 | -0.017223692 |
| LINC00955 | -0.017225954 |
| OFD1P17 | -0.01722946 |
| MIR548AL | -0.017230666 |
| AL158141 | -0.017232379 |
| NATP | -0.017239883 |
| LINC01054 | -0.017241203 |
| TNK2 | -0.017241613 |
| MIR6859-4 | -0.017247175 |
| LUC7L2 | -0.017247307 |
| RAB11FIP1 | -0.017247883 |
| SSR4 | -0.017250097 |
| MTND5P29 | -0.017256086 |
| RNA5SP228 | -0.017257044 |

| | |
|---|---|
| MIR548AG2 | -0.017257676 |
| AP000436 | -0.017264273 |
| ANKDD1B | -0.017264584 |
| MIR657 | -0.017269082 |
| PIWIL3 | -0.017272888 |
| VN1R28P | -0.0172754 |
| RP11-119D18 | -0.01727645 |
| TXNP3 | -0.017279653 |
| RPL31P40 | -0.017280642 |
| RN7SKP265 | -0.017283696 |
| ZNF879 | -0.017283977 |
| RP11-359G22 | -0.017284258 |
| BIVM | -0.017285559 |
| IFI27L1 | -0.017294397 |
| SMAD5-AS1_2 | -0.01729594 |
| AP000361 | -0.01729648 |
| AEBP2 | -0.017297001 |
| ZNF324 | -0.017301448 |
| RP11-806J6 | -0.017301669 |
| ERICH6-AS1 | -0.017304462 |
| DBX1 | -0.017305012 |
| WDR93 | -0.017305727 |
| LINC00383 | -0.017310776 |
| RP11-74D3 | -0.017316849 |
| S1PR3 | -0.017317337 |
| HMGB3P26 | -0.017319517 |
| WFS1 | -0.017326231 |
| PARP4P2 | -0.017329401 |
| CTB-51E13 | -0.017338386 |
| RPS7P10 | -0.017339422 |
| RN7SL846P | -0.017341227 |
| ZG16 | -0.017341967 |
| PSMC1P4 | -0.017342406 |
| CEACAM6 | -0.01734505 |
| WFDC10A | -0.017348212 |
| RNU7-143P | -0.017351238 |
| RP11-148I19 | -0.017351659 |
| RN7SL81P | -0.01735308 |
| PABPC1P2 | -0.017358555 |
| RP1-85F18 | -0.017362286 |
| CPEB2-AS1 | -0.01736322 |
| RTN1 | -0.017373247 |
| ZNF70P1 | -0.017377461 |
| MIR6802 | -0.017380219 |
| SDF2 | -0.017381396 |

| | |
|---|---|
| RNA5SP199 | -0.017381923 |
| ZNF699 | -0.017386308 |
| RP11-727F15 | -0.017386549 |
| MAN2B2 | -0.01738895 |
| TRPT1 | -0.017398401 |
| BCL2L15 | -0.01740482 |
| RP11-670N15 | -0.017406865 |
| HIST1H2BPS3 | -0.01740869 |
| RP1-136J15 | -0.017411276 |
| AC008103 | -0.017412747 |
| RPL21P41 | -0.017417362 |
| SMARCE1P7 | -0.01742 |
| OR4A47 | -0.017421915 |
| DPH6-AS1 | -0.017429551 |
| AL158069 | -0.017431079 |
| POLR2J2 | -0.017431205 |
| RPL35P4 | -0.017432213 |
| INS | -0.017435969 |
| PABPC1L2B | -0.017437355 |
| RP11-973D8 | -0.017440329 |
| GMPS | -0.017440636 |
| RP1-102H19 | -0.017441132 |
| RP11-1008M1 | -0.01744995 |
| PSG9 | -0.017451949 |
| HIGD1AP9 | -0.017454625 |
| SNRPFP1 | -0.017456758 |
| RP5-898J17 | -0.017457823 |
| IGDCC3 | -0.017459504 |
| AGPAT5 | -0.017464302 |
| RNU4ATAC9P | -0.017464745 |
| CTA-398F10 | -0.017465995 |
| MIR1288 | -0.01746688 |
| MLLT3 | -0.017468934 |
| RP11-125H8 | -0.017472461 |
| AL161668 | -0.017472482 |
| RP5-897D18 | -0.017475908 |
| DUSP11 | -0.017476321 |
| SDCBPP3 | -0.017478765 |
| EGR2 | -0.017482386 |
| FUT2 | -0.017486814 |
| RP5-1000K24 | -0.017487521 |
| HSPA6 | -0.017490182 |
| CHCHD3P2 | -0.017492436 |
| RP11-52L11 | -0.017495227 |
| RP11-109M17 | -0.017496501 |

| | |
|---|---|
| SLC28A2 | -0.017500136 |
| KB-1471A8 | -0.017500381 |
| RP11-795A2 | -0.017501364 |
| OR7E111P | -0.017504184 |
| RP11-807H22 | -0.017504941 |
| VN1R68P | -0.017509088 |
| NUTM2G | -0.017510221 |
| MAGI1-AS1 | -0.017517458 |
| KRTAP9-12P | -0.017519022 |
| CYP4A44P | -0.017524309 |
| RP11-267N12 | -0.017527054 |
| RPS4XP11 | -0.017527393 |
| AMYP1 | -0.017529606 |
| PRYP6 | -0.017531528 |
| RP11-22C8 | -0.017536671 |
| RPL12P34 | -0.017536708 |
| AC091096 | -0.017543689 |
| RP11-327P2 | -0.017546575 |
| RP11-111F16 | -0.017547238 |
| LINC00648 | -0.017548504 |
| RP11-158O16 | -0.017550328 |
| SAR1AP4 | -0.017551384 |
| RNF152P1 | -0.017552663 |
| RNU6-237P | -0.017558349 |
| RP11-554J4 | -0.017561292 |
| RNA5SP514 | -0.01756381 |
| RPL7P3 | -0.017570121 |
| OR4A1P | -0.017574159 |
| AC138472 | -0.017574662 |
| RP11-349G13 | -0.01757665 |
| RP11-406D1 | -0.017579721 |
| RP11-588G21 | -0.017582757 |
| TOM1L2 | -0.017592158 |
| GAPDHP50 | -0.01759307 |
| CTC-218H9 | -0.017594949 |
| MIR4465 | -0.017609011 |
| RP11-465L8 | -0.017609114 |
| RP11-26P13 | -0.017611829 |
| MGAT2P1 | -0.017623316 |
| RP3-384D21 | -0.017624597 |
| ANKEF1 | -0.017628923 |
| OR7E109P | -0.017632786 |
| CRB1 | -0.017634435 |
| OR52E5 | -0.017639889 |
| SPCS2P3 | -0.017640481 |

| | |
|---|---|
| RNU6-625P | -0.017642249 |
| C12orf4 | -0.017642319 |
| RN7SKP248 | -0.017646266 |
| RP1-269O5 | -0.017648412 |
| TNFRSF18 | -0.017652608 |
| RNU6-954P | -0.017656395 |
| RNU1-51P | -0.017657774 |
| RP11-421L21 | -0.017659585 |
| RP11-461A8 | -0.01766032 |
| AP000998 | -0.017662135 |
| RNU6-339P | -0.017663868 |
| LINC00845 | -0.017664284 |
| RP11-360A18 | -0.017668775 |
| AC007383 | -0.01766926 |
| DYNC1I2 | -0.017677985 |
| OR4C10P | -0.017687015 |
| CCDC58P1 | -0.017687968 |
| MIR6808 | -0.017688218 |
| CDH12P3 | -0.017694998 |
| ZBTB8OS | -0.017695065 |
| MIR27B | -0.017695494 |
| XXbac-BPG157A10 | -0.017699583 |
| AL591708 | -0.017700741 |
| RPL7L1P5 | -0.017701232 |
| RN7SL671P | -0.017706303 |
| RP3-399J4 | -0.017706497 |
| CXCR1 | -0.017706511 |
| MARK2P4 | -0.017715919 |
| RP3-437I16 | -0.017721126 |
| OR14J1 | -0.017726232 |
| PQLC3 | -0.017728034 |
| RPL7L1P9 | -0.01773162 |
| RANP6 | -0.017732638 |
| RP1-21P18 | -0.017733259 |
| AC006150 | -0.017734708 |
| IGKV3D-31 | -0.017736035 |
| TEN1 | -0.017738323 |
| VDAC3 | -0.017739681 |
| PRUNEP1 | -0.017740568 |
| RP11-624A21 | -0.017748837 |
| RP11-406A9 | -0.017750516 |
| RP11-317J10 | -0.017754232 |
| RNA5SP354 | -0.017757453 |
| TOPAZ1 | -0.01775841 |
| MIOXP1 | -0.017762458 |

| | |
|---|---|
| ARHGEF18 | -0.017762957 |
| GRPR | -0.017765424 |
| URB1 | -0.01776731 |
| TM4SF18 | -0.017772762 |
| RN7SL4P | -0.017773891 |
| AP000356 | -0.017775073 |
| RN7SL524P | -0.017777277 |
| GTF3C1 | -0.017777971 |
| RP4-614O4 | -0.017780334 |
| RPS29P12 | -0.017781374 |
| FREM2-AS1 | -0.017782476 |
| RARRES2P8 | -0.017788362 |
| RN7SL743P | -0.017790137 |
| RP11-56H16 | -0.017792088 |
| RP11-341B24 | -0.017797707 |
| ZNF497 | -0.017798735 |
| EHD3 | -0.017803647 |
| C6orf48 | -0.017804745 |
| MIR32 | -0.017806235 |
| CTB-179I1 | -0.017806308 |
| ARX | -0.017809676 |
| COX6CP3 | -0.017809925 |
| RNU6-772P | -0.017810753 |
| USP9YP11 | -0.017810753 |
| CALM2P3 | -0.017813263 |
| KATNBL1P6 | -0.017814821 |
| RP11-771K4 | -0.017815617 |
| EBF3 | -0.017820336 |
| RP4-672N11 | -0.017824212 |
| CICP9 | -0.01783071 |
| COX5BP3 | -0.017831927 |
| AP1S3 | -0.017832616 |
| TSHB | -0.017832783 |
| FTLP1 | -0.017834854 |
| SPDYE2B | -0.017837572 |
| SNORD53_SNORD92 | -0.017843231 |
| RP11-320H14 | -0.017843445 |
| CD200R1L | -0.017843571 |
| CTC-321K16 | -0.017846561 |
| GS1-122H1 | -0.01784809 |
| SEMA3B | -0.017850533 |
| RSU1P1 | -0.01785251 |
| MAP10 | -0.017854012 |
| HSD3BP2 | -0.017854186 |
| HMGN1P37 | -0.017856077 |

| | |
|---|---|
| RP11-215H22 | -0.017860112 |
| ACTG1P2 | -0.017861538 |
| C7orf31 | -0.017862384 |
| RP11-143P4 | -0.017868421 |
| LINC01514 | -0.017869599 |
| FO538757 | -0.017874124 |
| RGAG1 | -0.017880302 |
| MTCO2P4 | -0.017881152 |
| RP11-351N6 | -0.017885086 |
| AC113188 | -0.017887521 |
| EFTUD2 | -0.017889548 |
| MIR4267 | -0.017890548 |
| RNF212B | -0.017893791 |
| GNRH1 | -0.017899744 |
| PRTG | -0.017902944 |
| RNY4P28 | -0.017905401 |
| MIR548H5 | -0.017906614 |
| RP11-305D15 | -0.017906939 |
| TIMM17A | -0.017907726 |
| VSIG1 | -0.017919559 |
| AL158147 | -0.017928252 |
| VN1R87P | -0.017931625 |
| ADH5 | -0.017932229 |
| DUTP3 | -0.017934294 |
| PRSS44 | -0.017936735 |
| RP11-654C20 | -0.017939361 |
| PARN | -0.017941011 |
| HSPE1P5 | -0.017942367 |
| RP11-331N16 | -0.017946717 |
| RP11-165D6 | -0.017950721 |
| NSUN7 | -0.017952621 |
| CTD-2045M21 | -0.017953686 |
| ARHGEF9 | -0.017958202 |
| GRIK3 | -0.017961385 |
| AP005242 | -0.017963706 |
| NPM1P51 | -0.017964093 |
| DUS1L | -0.017968808 |
| LA16c-329F2 | -0.017974092 |
| PAIP1 | -0.017976957 |
| AC012047 | -0.017982766 |
| PDZD8 | -0.017983822 |
| NFU1P2 | -0.017987858 |
| RP11-332J15 | -0.017994275 |
| GULOP | -0.01800124 |
| RP11-1081L13 | -0.018005144 |

| | |
|---|---|
| RP11-272G22 | -0.018005181 |
| SERBP1P1 | -0.018010505 |
| RP11-932A10 | -0.018011234 |
| ADARB2-AS1 | -0.018017816 |
| RP11-627H22 | -0.018022223 |
| RP11-285F7 | -0.018025703 |
| VN1R85P | -0.018028203 |
| OR1B1 | -0.018032577 |
| RPL7P34 | -0.018033822 |
| RP11-463M16 | -0.018034192 |
| RP11-350A1 | -0.018037542 |
| GMNC | -0.018049231 |
| AC002056 | -0.018056766 |
| NXT2 | -0.018061091 |
| OR6C2 | -0.018062589 |
| AC005780 | -0.018062779 |
| RP11-162D16 | -0.018062818 |
| GXYLT1P2 | -0.018066227 |
| OSBPL9P1 | -0.018066717 |
| RP11-33E24 | -0.018067577 |
| SH3BGRL2 | -0.018071919 |
| CTD-2501O3 | -0.01807359 |
| MIR6889 | -0.018077318 |
| OR51A7 | -0.018080158 |
| CASC2 | -0.018080254 |
| COG5 | -0.018082208 |
| TSPY13P | -0.018083715 |
| RP11-96J15 | -0.018085675 |
| RP11-35J10 | -0.018087166 |
| HYKK | -0.018088422 |
| GPR17 | -0.018089897 |
| LIN28A | -0.018091426 |
| LINC00433 | -0.018099009 |
| NEDD8-MDP1 | -0.018109509 |
| RNA5SP276 | -0.018110659 |
| RP11-95I19 | -0.018112464 |
| RNU6-794P | -0.018112559 |
| PDE1A | -0.018113037 |
| RP11-8L2 | -0.018114032 |
| HDHD1P2 | -0.018114769 |
| DNAJB1 | -0.01811633 |
| RP11-78I14 | -0.018118134 |
| DDX11-AS1 | -0.018120256 |
| AC134882 | -0.018120333 |
| LRIT2 | -0.018120334 |

| | |
|---|---|
| RP11-706D8 | -0.018121295 |
| HTR2B | -0.018122228 |
| FOXG1 | -0.018129573 |
| TMEM106B | -0.018131215 |
| UBE2B | -0.018132554 |
| AP000807 | -0.018132692 |
| PCYT1A | -0.018133596 |
| HNRNPA1P49 | -0.01813554 |
| HMGN1P35 | -0.018140391 |
| RP11-605F22 | -0.018144759 |
| MRPS18A | -0.018151048 |
| AC026495 | -0.01815278 |
| NOSTRIN | -0.018153262 |
| BNIP3P15 | -0.018157116 |
| GREB1L | -0.018157579 |
| AC023385 | -0.01816493 |
| RP11-494G9 | -0.018168624 |
| AC068538 | -0.018168939 |
| PRRT3 | -0.018173989 |
| RAC1P9 | -0.018174768 |
| RP11-2A4 | -0.01817649 |
| MIR941-1 | -0.018177218 |
| RNU6-1164P | -0.018182472 |
| RHOBTB3 | -0.01818331 |
| IGHVIII-67-4 | -0.01818702 |
| RNA5SP294 | -0.018189841 |
| SLC17A6 | -0.018191973 |
| MIR4669 | -0.018194388 |
| LINC00411 | -0.018206177 |
| AC108059 | -0.018208025 |
| PTGES2-AS1 | -0.018208274 |
| RP11-349I1 | -0.018209908 |
| SERPINI2 | -0.018212685 |
| RNU2-20P | -0.018215966 |
| CTD-2540F13 | -0.018217116 |
| COG1 | -0.018218325 |
| STH | -0.018219283 |
| MNX1 | -0.018221105 |
| RP11-119N19 | -0.018222643 |
| RNU6-999P | -0.018223468 |
| RP5-1044H5 | -0.018227137 |
| EBF2 | -0.018228614 |
| RNU6-448P | -0.018231778 |
| RP1-97D16 | -0.018235334 |
| AC022073 | -0.018236717 |

| | |
|---|---|
| AC044907 | -0.018238612 |
| NDUFB2-AS1 | -0.018242382 |
| RNU6-1046P | -0.018248233 |
| OR51B2 | -0.018249697 |
| TMPRSS11CP | -0.018253171 |
| HRC | -0.018254401 |
| KCNJ18 | -0.018256453 |
| SLC25A25 | -0.018257769 |
| GRAMD4P5 | -0.018259262 |
| PGPEP1L | -0.018262454 |
| RP11-228G3 | -0.018263961 |
| AC011073 | -0.01826627 |
| IL31RA | -0.018268497 |
| AC144833 | -0.018270073 |
| RNA5SP230 | -0.018270376 |
| RP11-747I9 | -0.01827209 |
| AL139296 | -0.018272427 |
| RNU4-57P | -0.018273249 |
| RP11-507D14 | -0.018275789 |
| AC103816 | -0.018279584 |
| ZFR2 | -0.018281882 |
| ATP2B1 | -0.018282496 |
| GPR45 | -0.018283293 |
| AC008753 | -0.01829405 |
| FDPSP5 | -0.018297325 |
| AP000857 | -0.01829921 |
| AGAP7P | -0.018299849 |
| SLC28A3 | -0.018301846 |
| LINC00575 | -0.018303819 |
| RP11-96D24 | -0.018310397 |
| MIR6508 | -0.018312554 |
| MIR944 | -0.018313194 |
| OR4D8P | -0.018313496 |
| RP11-117O7 | -0.018314021 |
| NEUROD4 | -0.018314601 |
| LINC01349 | -0.018316474 |
| AL158801 | -0.018319372 |
| RP11-636N17 | -0.018320302 |
| TACO1 | -0.018325474 |
| PRR34-AS1 | -0.0183318 |
| RPL29P2 | -0.018338565 |
| RP11-13J10 | -0.018338707 |
| OSBPL9P4 | -0.018339947 |
| SERBP1P4 | -0.018342277 |
| CTD-3057O21 | -0.018345487 |

| | |
|---|---|
| AP001171 | -0.018353063 |
| RNA5SP232 | -0.018356677 |
| RP11-21G20 | -0.018357703 |
| WDR27 | -0.018358741 |
| HNRNPCP6 | -0.018369373 |
| CFL1P8 | -0.018374066 |
| HMGCLL1 | -0.018377445 |
| RP11-328P23 | -0.018381351 |
| LINC00637 | -0.018381495 |
| RP5-902P8 | -0.018382389 |
| RP11-185H22 | -0.018385197 |
| CDRT4 | -0.018390742 |
| AC009501 | -0.018390761 |
| GOLGA6L4 | -0.018400561 |
| RNU6-677P | -0.018406173 |
| KLHL30-AS1 | -0.018406627 |
| AL109763 | -0.018412569 |
| GLRB | -0.018413069 |
| NETO2 | -0.018413441 |
| CLIC6 | -0.018416585 |
| TGFBR2 | -0.018423116 |
| RNU6-1020P | -0.018424358 |
| AC018755 | -0.018430819 |
| USP9YP10 | -0.018431398 |
| HNRNPA1P28 | -0.018433716 |
| PPP1R26P3 | -0.01843564 |
| SKP2 | -0.018439377 |
| FAR2 | -0.018443768 |
| RP1-56J10 | -0.01844429 |
| RP11-662M24 | -0.018452639 |
| TRIM72 | -0.018457592 |
| OR5AL1 | -0.018458286 |
| RFPL3-AS1_1 | -0.018463639 |
| INPPL1 | -0.018472253 |
| AUNIP | -0.018474206 |
| RP11-47A8 | -0.018493184 |
| RP11-715I4 | -0.018501032 |
| RP11-205A8 | -0.01850131 |
| RP11-687M24 | -0.018501811 |
| MT-TS2 | -0.018511042 |
| TSPAN14 | -0.018513725 |
| SURF1 | -0.018514115 |
| RRN3P3 | -0.018521036 |
| RN7SL717P | -0.018523023 |
| RN7SL454P | -0.018524482 |

| | |
|---|---|
| RCE1 | -0.0185322 |
| RITA1 | -0.018533335 |
| AC108051 | -0.018534106 |
| AC009245 | -0.018534153 |
| HMGN1P28 | -0.018538574 |
| FTCD-AS1 | -0.018539129 |
| RP11-320N7 | -0.018539242 |
| RP11-593F5 | -0.018540099 |
| OR8C1P | -0.018543402 |
| UBE2Q2P1 | -0.018547903 |
| PYURF | -0.018550228 |
| COPS8P2 | -0.018555404 |
| PTP4A1P5 | -0.018556107 |
| EPN2 | -0.018558322 |
| RP3-410C9 | -0.018561643 |
| RP11-446P9 | -0.018564746 |
| MIR3914-1 | -0.018566824 |
| RP5-896L10 | -0.018567037 |
| ZNRF4 | -0.01856882 |
| ERV3-1 | -0.018573392 |
| RP11-43N22 | -0.018574274 |
| RP11-253E3 | -0.018577541 |
| CATSPER3 | -0.018579513 |
| ARID1B | -0.018579793 |
| RP11-126D17 | -0.018580771 |
| CH507-42P11 | -0.018581041 |
| CTC-498M16 | -0.018581806 |
| TERF1P3 | -0.018586264 |
| RP11-788A1 | -0.018588698 |
| AIFM3 | -0.018594044 |
| BX322557 | -0.018595607 |
| RP11-286E11 | -0.018599455 |
| RNU6-1084P | -0.018600359 |
| AL080285 | -0.018601879 |
| AC104592 | -0.018605629 |
| VN1R110P | -0.018606607 |
| RPS26P49 | -0.018612333 |
| FUZ | -0.01861291 |
| GDPD3 | -0.018614343 |
| RP11-419N10 | -0.018617134 |
| OR4K16P | -0.018618276 |
| RP1-203P18 | -0.018622836 |
| COX6A2 | -0.018622923 |
| PGR | -0.018625339 |
| TSPY4 | -0.018629638 |

| | |
|---|---|
| CRIP3 | -0.01863528 |
| PARD3B | -0.018638456 |
| CTC-458A3 | -0.018639587 |
| C1GALT1 | -0.018650344 |
| RPL18P11 | -0.018650979 |
| OR8D1 | -0.018656983 |
| RP11-66D17 | -0.018664001 |
| CABP4 | -0.01866655 |
| RPL31P9 | -0.018669705 |
| AC027288 | -0.018676155 |
| SEMA3C | -0.018677174 |
| RNU6-604P | -0.018677177 |
| CTD-2410N18 | -0.018677378 |
| ACTR3P2 | -0.018678055 |
| HNRNPCL4 | -0.018685912 |
| RP11-219G17 | -0.018686044 |
| CTC-203F4 | -0.018687239 |
| RNU6-563P | -0.018692537 |
| RP11-107N7 | -0.01869528 |
| AC005786 | -0.018698384 |
| RP11-867G23 | -0.018700207 |
| ARV1 | -0.01870231 |
| RNA5SP234 | -0.018703162 |
| CTD-3051D23 | -0.018704879 |
| HMBOX1 | -0.018705082 |
| PSG1 | -0.018706656 |
| RN7SL211P | -0.018708613 |
| GOLGA8G | -0.018709594 |
| RP5-998G20 | -0.018712947 |
| IVNS1ABP | -0.018714087 |
| SNHG17 | -0.018714143 |
| AC018643 | -0.018717735 |
| RPS17P8 | -0.018719504 |
| RP3-453P22 | -0.018720386 |
| FMO1 | -0.018720394 |
| CCT6A | -0.018729957 |
| RN7SL66P | -0.018733956 |
| RP11-446E24 | -0.01873553 |
| AC087237 | -0.018737758 |
| RP11-976B16 | -0.018740469 |
| SLC6A19 | -0.018749262 |
| SLC25A4 | -0.018751372 |
| KRTAP10-5 | -0.018758154 |
| CDY12P | -0.018766119 |
| AC079603 | -0.018766119 |

| | |
|---|---|
| USP9YP8 | -0.018766119 |
| RP11-729M20 | -0.018766411 |
| CSAG4 | -0.018768096 |
| RNU6-150P | -0.018769102 |
| RPS7P4 | -0.018770583 |
| MIR185 | -0.018776219 |
| ATXN8OS | -0.01877801 |
| UBXN11 | -0.018782778 |
| ATP5L2 | -0.01878523 |
| RNU6-1087P | -0.018786261 |
| RP11-122L9 | -0.018786896 |
| RNU6-153P | -0.018786989 |
| AC009517 | -0.018787264 |
| FIGLA | -0.018789982 |
| RNY3P1 | -0.018792699 |
| AL365181 | -0.018794909 |
| MIR5571 | -0.018796229 |
| RP4-740C4 | -0.018805328 |
| RP11-6D1 | -0.018818703 |
| U2AF2 | -0.018821087 |
| MIR466 | -0.018830415 |
| FAM230C | -0.018831034 |
| RNU6-764P | -0.018831778 |
| GPANK1 | -0.018835729 |
| RP11-11A9 | -0.018839384 |
| RNU6-1277P | -0.018841055 |
| CBX3P9 | -0.018842709 |
| OR8J1 | -0.018844561 |
| AP002856 | -0.01885235 |
| LONRF1 | -0.018858558 |
| AC011155 | -0.018863553 |
| RP11-174F8 | -0.018863937 |
| FAM178B | -0.018865293 |
| CAMK1D | -0.018876122 |
| RAN | -0.01887815 |
| RP11-137O10 | -0.018882035 |
| CTB-129O4 | -0.018882727 |
| HMX2 | -0.018891715 |
| AP000889 | -0.018894007 |
| RP1-241P17 | -0.018899404 |
| RPL7P8 | -0.01890247 |
| AC190387 | -0.01890288 |
| RNU4-26P | -0.018904854 |
| RNU6-606P | -0.018906396 |
| SCOC-AS1 | -0.018909444 |

| | |
|---|---|
| CTD-2354A18 | -0.018911264 |
| PKDREJ | -0.018911613 |
| ACTR3BP2 | -0.018911784 |
| MIR3118-1 | -0.018916613 |
| RN7SL217P | -0.018919619 |
| SETMAR | -0.018919621 |
| FKSG51 | -0.018922461 |
| RP11-3J10 | -0.01892251 |
| GABARAPL3 | -0.018928266 |
| RP11-404E12 | -0.01892934 |
| AC092116 | -0.018932386 |
| CHRM1 | -0.018932463 |
| KRT19P2 | -0.018932764 |
| HLX-AS1 | -0.018934866 |
| RP11-48B3 | -0.018936124 |
| TRPC7-AS2 | -0.01893906 |
| MIR4472-2 | -0.01893912 |
| RNA5SP54 | -0.018942205 |
| LINC01493 | -0.018943167 |
| PTRH2 | -0.01896314 |
| AC006547 | -0.018964498 |
| RP11-27G13 | -0.018968105 |
| RP11-227D13 | -0.018972764 |
| RHO | -0.018973789 |
| PRPF38AP1 | -0.018976073 |
| RNU6-154P | -0.018977762 |
| SETP22 | -0.018977907 |
| SPATA6L | -0.018984257 |
| EIF4BP8 | -0.018989414 |
| RN7SL470P | -0.01899008 |
| FUNDC1 | -0.018991959 |
| AC093004 | -0.018992514 |
| DHH | -0.0189942 |
| KB-1125A3 | -0.018994211 |
| ATP5LP4 | -0.01899456 |
| LINC00472 | -0.018995135 |
| GRIA2 | -0.019001789 |
| RP5-1086L22 | -0.019010639 |
| MROH2B | -0.019016239 |
| TLR5 | -0.019016761 |
| GCSHP5 | -0.019018876 |
| RP4-583K8 | -0.019021245 |
| RP11-159L20 | -0.019024112 |
| AL354828 | -0.019033342 |
| RNU6-1088P | -0.019033662 |

| | |
|---|---|
| AL162614 | -0.019036002 |
| SLC25A1P1 | -0.019036186 |
| FAM200A | -0.019038715 |
| AC007422 | -0.01904041 |
| ARL8A | -0.019044315 |
| CCDC13-AS1 | -0.019045063 |
| RYR2 | -0.019045115 |
| RN7SL317P | -0.019045314 |
| RNA5SP362 | -0.019049693 |
| DNM1P51 | -0.019052025 |
| IGBP1 | -0.019053711 |
| LINC00401 | -0.019060903 |
| CYP2A7 | -0.019062535 |
| HSPBAP1 | -0.019062958 |
| CTD-2006H14 | -0.019068949 |
| LRRC37A6P | -0.019070158 |
| RP11-432J3 | -0.019071455 |
| MIR202HG | -0.019075522 |
| KRT18P58 | -0.019079664 |
| FAM162B | -0.01908016 |
| RP11-325O24 | -0.019081562 |
| AL354775 | -0.019085889 |
| KIAA0895L | -0.01908614 |
| BCAP31P2 | -0.01908625 |
| SF3A3 | -0.019088633 |
| KRTAP9-7 | -0.019100067 |
| GRIA4 | -0.01910085 |
| FMO6P | -0.019100966 |
| SLMO2P1 | -0.019102101 |
| GOT2P1 | -0.019103332 |
| RP11-93P10 | -0.01910361 |
| STK32B | -0.019106178 |
| CYP3A43 | -0.019108387 |
| RP11-375H19 | -0.019116787 |
| PCDH11X | -0.01911791 |
| CD300C | -0.019118644 |
| RP11-438D14 | -0.019119201 |
| MIR4305 | -0.019119644 |
| MTMR9 | -0.019119667 |
| CTB-85P21 | -0.019120588 |
| RP11-221H10 | -0.019125461 |
| MYC | -0.019127407 |
| CMA1 | -0.01912753 |
| AC005013 | -0.019130506 |
| RP11-142L4 | -0.019131463 |

| | |
|---|---|
| NDUFA10 | -0.019134366 |
| RP11-81M19 | -0.01913838 |
| MIR5186 | -0.019144045 |
| RP11-439E19 | -0.019149306 |
| RNU1-57P | -0.019154405 |
| MIR3616 | -0.019156495 |
| P2RY6 | -0.019159055 |
| MRPL37P1 | -0.019167539 |
| RNF180 | -0.019173642 |
| CTCFL | -0.019175383 |
| RP11-628J14 | -0.019176084 |
| GLS | -0.019179112 |
| RNF138P2 | -0.019179135 |
| HSD17B3 | -0.019183451 |
| CA6 | -0.019185098 |
| CLDN15 | -0.019186294 |
| OCEL1 | -0.019193512 |
| AC073928 | -0.019197798 |
| AC004070 | -0.019199901 |
| LAMTOR3P2 | -0.019200037 |
| RP11-303E16 | -0.019202015 |
| TSFM | -0.019210499 |
| RP11-213O5 | -0.019212966 |
| UBE2E1 | -0.019218183 |
| RNU1-44P | -0.01922437 |
| RP1-278E11 | -0.019225756 |
| KRTAP19-2 | -0.019232966 |
| RP5-955M13 | -0.019233908 |
| ARHGEF15 | -0.019250873 |
| ANGPTL7 | -0.019251706 |
| PLCH1-AS1 | -0.019252277 |
| RNU6-896P | -0.019256394 |
| HSP90AA5P | -0.019262783 |
| KHK | -0.019263023 |
| RP11-360L9 | -0.01926466 |
| MIR543 | -0.019266092 |
| SNRPEP7 | -0.019266612 |
| AC009005 | -0.019266669 |
| MTCO1P4 | -0.019267389 |
| FMO2 | -0.019275656 |
| PSMB3P2 | -0.019280052 |
| RP11-122L4 | -0.01928234 |
| ARF1P2 | -0.019286253 |
| ZNF646 | -0.019288884 |
| AC079154 | -0.019291373 |

| | |
|---|---|
| CTC-276P9 | -0.01929436 |
| IDI2 | -0.01929701 |
| RNU6ATAC29P | -0.019298122 |
| AC025280 | -0.019301642 |
| AL357060 | -0.019301709 |
| CCDC73 | -0.019307602 |
| EGOT | -0.019312626 |
| PATE4 | -0.01931441 |
| ICT1 | -0.019315397 |
| RP11-552E20 | -0.019325132 |
| RP4-718J7 | -0.019325924 |
| RP11-572H4 | -0.01932882 |
| OR5AZ1P | -0.019332493 |
| DACH2 | -0.019338465 |
| RPS20P5 | -0.019339342 |
| LINC00879 | -0.019344763 |
| LINC00202-2 | -0.019348951 |
| GUCY2D | -0.019352723 |
| RP4-595K12 | -0.019353821 |
| STARD7-AS1 | -0.019354413 |
| MIR3158-1 | -0.019355142 |
| RBMS2P1 | -0.019359349 |
| RPLP1P13 | -0.019360455 |
| RP11-289E15 | -0.019361106 |
| TAAR9 | -0.019361186 |
| HIST1H1C | -0.019371734 |
| RNU6-328P | -0.019372525 |
| RP11-514A9 | -0.019374727 |
| RPL34P21 | -0.019375105 |
| RNU6-628P | -0.01937715 |
| EIF1P4 | -0.019377841 |
| HMGB1P10 | -0.019377911 |
| RP11-164C1 | -0.019378419 |
| RP11-128L5 | -0.01938074 |
| RNU6-1197P | -0.019383556 |
| RP5-877J2 | -0.01938424 |
| AC123768 | -0.019385045 |
| RP11-793L10 | -0.019385746 |
| RN7SL761P | -0.019385775 |
| RN7SL379P | -0.019385775 |
| LINC00958 | -0.019388618 |
| CNOT6LP1 | -0.019392 |
| FAM187B | -0.019393643 |
| RNY1 | -0.019398219 |
| RP11-790I12 | -0.019398272 |

| | |
|---|---|
| RP11-18O11 | -0.01940151 |
| EFCAB11 | -0.019405224 |
| DLGAP3 | -0.01940821 |
| RCBTB2 | -0.01941929 |
| AP3S2 | -0.019421281 |
| ABCB10P1 | -0.019422757 |
| RP11-56O18 | -0.019427714 |
| RP4-676L2 | -0.019430384 |
| CTA-440B3 | -0.019436757 |
| GOLGA2 | -0.019437319 |
| RNA5SP487 | -0.01943855 |
| AC073333 | -0.019441966 |
| AC119674 | -0.019443163 |
| RP11-138E2 | -0.019443408 |
| RN7SL152P | -0.019446442 |
| ADAMTSL3 | -0.019447905 |
| NPM1P35 | -0.019450753 |
| RP1-137K24 | -0.019452517 |
| USH2A | -0.019453329 |
| RP11-72M10 | -0.019454434 |
| RN7SL26P | -0.01945492 |
| RP11-65D24 | -0.01945567 |
| LINC00235 | -0.019461655 |
| Z83826 | -0.019466976 |
| CU459202 | -0.019468821 |
| PADI4 | -0.019470252 |
| AL139239 | -0.019470745 |
| CNTN2 | -0.019470972 |
| OR5M3 | -0.019471986 |
| MIR5584 | -0.019474767 |
| CDC42 | -0.019478213 |
| RP11-80I3 | -0.019480527 |
| NDUFA3P6 | -0.019480748 |
| GYG2-AS1 | -0.019482036 |
| FKBP4P7 | -0.019486337 |
| FAM19A2 | -0.019489209 |
| YBX1P8 | -0.01949474 |
| RP11-181B11 | -0.01949494 |
| RP1-218B13 | -0.019495015 |
| RPL7P43 | -0.019498622 |
| RNU4-68P | -0.019499809 |
| TUBA3C | -0.019502781 |
| AL121967 | -0.019508327 |
| HMGA1P5 | -0.019510785 |
| RNA5SP405 | -0.019510785 |

| | |
|---|---|
| LINC00603 | -0.019514059 |
| MIR548V | -0.019515686 |
| PTBP1 | -0.019515964 |
| XXbac-BPG308J9 | -0.019521365 |
| RP11-537E18 | -0.019528538 |
| MIR3135A | -0.019529384 |
| CDCA2 | -0.019536076 |
| RNY3P9 | -0.01953698 |
| ISX-AS1 | -0.019540859 |
| PAFAH1B1 | -0.019543426 |
| RP5-859M6 | -0.019543846 |
| OIT3 | -0.019548784 |
| MMS19 | -0.019552375 |
| RP11-963H1 | -0.019554431 |
| AC012306 | -0.019562645 |
| MIR4797 | -0.019563655 |
| RP11-129O7 | -0.019563974 |
| MRPL49P1 | -0.019564655 |
| CEP57L1 | -0.019566459 |
| NUTM2B-AS1 | -0.019566961 |
| CACNA1C-AS4 | -0.019569231 |
| MIR532 | -0.019572337 |
| AC005160 | -0.019573183 |
| ZNF512B | -0.019573994 |
| DPP3P2 | -0.019575816 |
| RP11-367N14 | -0.019578702 |
| RNU6-738P | -0.019581766 |
| RGS1 | -0.019582571 |
| AP000235 | -0.01958451 |
| CDY3P | -0.019585004 |
| RP11-159D8 | -0.019587629 |
| RP5-1077B9 | -0.019596247 |
| PDGFD | -0.019597735 |
| SHC1P2 | -0.019599915 |
| SUMO2P12 | -0.019601164 |
| RNU7-70P | -0.019606258 |
| RP11-526P5 | -0.019610183 |
| RNU6-242P | -0.019612741 |
| RP11-641C17 | -0.019613127 |
| UPK3A | -0.019613545 |
| CGB1 | -0.019614023 |
| RP11-485C11 | -0.019617111 |
| RPL21P109 | -0.019619642 |
| RP11-386B1 | -0.019619796 |
| LINC01262 | -0.019621255 |

| | |
|---|---|
| C3orf49 | -0.019627005 |
| RPL10L | -0.019630627 |
| FAM72C | -0.019631036 |
| LARP7P1 | -0.019632027 |
| RPS19P1 | -0.019632616 |
| CTD-2301A4 | -0.019634864 |
| ADRA2B | -0.019634964 |
| ZNF747 | -0.019644602 |
| RN7SKP146 | -0.019648267 |
| C10orf131 | -0.019648814 |
| RNU4-50P | -0.019648838 |
| MED20 | -0.019650184 |
| AC009403 | -0.019657886 |
| MIR1299 | -0.01965806 |
| CHST10 | -0.019659431 |
| ARL2BPP1 | -0.019660911 |
| AC017078 | -0.019663575 |
| KB-1568E2 | -0.019666847 |
| KDR | -0.019670092 |
| RP11-439A17 | -0.019670212 |
| AC098823 | -0.019671813 |
| ZYG11AP1 | -0.019674007 |
| RP11-44B19 | -0.019675249 |
| RP11-664H7 | -0.019677378 |
| MFN1 | -0.019678278 |
| RP11-484P15 | -0.019679915 |
| FAM47DP | -0.019682484 |
| RP1-256G22 | -0.019684763 |
| RP11-399L7 | -0.019686059 |
| AC009362 | -0.019687733 |
| RUSC1 | -0.019689423 |
| GDI2P2 | -0.019689666 |
| RP11-6L6 | -0.019697959 |
| MRGPRX4 | -0.019698054 |
| TMTC3 | -0.019698967 |
| ARMCX3 | -0.019710521 |
| AF277315 | -0.019711507 |
| BIRC8 | -0.019711966 |
| KRT18P44 | -0.019712565 |
| RPL36AP21 | -0.019714126 |
| PSMC1P6 | -0.019714525 |
| AC009260 | -0.019717018 |
| RPL5P28 | -0.019718363 |
| SNRPCP11 | -0.019720412 |
| AL033378 | -0.019721312 |

| | |
|---|---|
| L3MBTL4 | -0.019726485 |
| CTD-3025N20 | -0.019726913 |
| KLF17P2 | -0.019727153 |
| LINC01194 | -0.019728313 |
| RBM8A | -0.019731694 |
| GTF3AP1 | -0.019735123 |
| MIR3679 | -0.019736715 |
| ANKRD20A17P | -0.019738037 |
| MIR194-1 | -0.019742498 |
| BBS5 | -0.019742541 |
| AC005284 | -0.019743767 |
| RNA5SP184 | -0.01974535 |
| TPCN1 | -0.0197455 |
| MIR6510 | -0.019749935 |
| CYFIP1 | -0.019749946 |
| AL354764 | -0.01975098 |
| RNA5SP327 | -0.019752766 |
| RP11-245D16 | -0.019755574 |
| MAP3K15 | -0.019756446 |
| HIST1H2BPS1 | -0.019762678 |
| SLC36A4 | -0.019763751 |
| ATP5F1P6 | -0.019768472 |
| RP3-437C15 | -0.019768581 |
| OR4Q2 | -0.019768692 |
| RP11-51P8 | -0.019770179 |
| RTFDC1 | -0.019771287 |
| CTC-447K7 | -0.019774208 |
| LL22NC03-121E8 | -0.019775434 |
| NEURL2 | -0.019776924 |
| SNORD90 | -0.019778449 |
| RPS27P27 | -0.019778869 |
| RPL29P7 | -0.019781325 |
| ARL6IP1P2 | -0.019788358 |
| RP11-12N13 | -0.019789857 |
| CTD-2376I20 | -0.019792091 |
| RNU6-767P | -0.019792346 |
| RP11-307E17 | -0.019794874 |
| ANKRD26P4 | -0.019797124 |
| ZNF280C | -0.019798488 |
| TPT1P12 | -0.019799139 |
| CCDC138 | -0.019800721 |
| ACSM2A | -0.019802136 |
| MCRS1 | -0.019803214 |
| C5orf17 | -0.019803931 |
| MIR6131 | -0.019809182 |

| | |
|---|---|
| RP11-327J17 | -0.019811301 |
| RN7SKP208 | -0.019812629 |
| RNA5SP111 | -0.019814381 |
| Clostridiales-1 | -0.019814588 |
| TRI-TAT2-3 | -0.019817664 |
| GAGE1 | -0.019818708 |
| POLR2C | -0.019821614 |
| UQCRFS1P2 | -0.019821859 |
| AC000032 | -0.019823507 |
| RPSAP2 | -0.019825865 |
| AC010880 | -0.019833302 |
| RP11-700F16 | -0.019835228 |
| ZSCAN25 | -0.019835941 |
| OR5AK2 | -0.019838733 |
| ZCCHC9 | -0.019845355 |
| RP11-33O4 | -0.019845913 |
| RN7SL698P | -0.019848542 |
| ZSCAN21 | -0.019849951 |
| GORAB | -0.019851067 |
| RNU6-57P | -0.019851257 |
| AC005593 | -0.019852776 |
| RNU7-129P | -0.019854182 |
| DOCK3 | -0.019861133 |
| HNRNPA1P30 | -0.019861613 |
| NCAM2 | -0.019861858 |
| RP11-182I10 | -0.019862739 |
| RP11-574O16 | -0.019864603 |
| HSPA5 | -0.019865282 |
| RP11-612J15 | -0.019867819 |
| AASS | -0.019868277 |
| OR5BS1P | -0.019871538 |
| MOB1AP2 | -0.019872474 |
| FOXJ2 | -0.019878622 |
| RP11-1267H10 | -0.019882466 |
| SCARNA1 | -0.019882497 |
| CTD-2357A8 | -0.019884827 |
| KRT8P47 | -0.019886651 |
| AC142119 | -0.019887458 |
| RP3-414A15 | -0.019889283 |
| RP11-446J8 | -0.019892395 |
| CHEK2P4 | -0.019899479 |
| HOMER2 | -0.019902604 |
| CHIAP2 | -0.019905502 |
| TRIAP1P1 | -0.019905738 |
| C2CD2 | -0.019908595 |

| | |
|---|---|
| RNU1-24P | -0.019910682 |
| RPS15AP6 | -0.0199114 |
| LINC01347 | -0.019913358 |
| RNU7-134P | -0.019914916 |
| RP3-395P12 | -0.019915749 |
| NACAP3 | -0.019916128 |
| HMGB3P21 | -0.019916166 |
| KLHL10 | -0.019922469 |
| RP11-486O13 | -0.019923104 |
| UHRF2 | -0.019923126 |
| TAS2R16 | -0.019924689 |
| RPL4P6 | -0.019926326 |
| FTHL17 | -0.019928527 |
| SARS | -0.019929787 |
| MIR598 | -0.019938512 |
| RP1-228H13 | -0.01993868 |
| RP11-324F11 | -0.019939597 |
| GATA2-AS1 | -0.019940559 |
| TPPP | -0.019941564 |
| CENPL | -0.01994175 |
| TPM3P6 | -0.019942084 |
| FTSJ2 | -0.019942513 |
| RP11-1084I9 | -0.019947323 |
| SNORA71D | -0.019951057 |
| RPL12P33 | -0.019951253 |
| RPS20P33 | -0.019953659 |
| OR8H1 | -0.019954962 |
| TMEM71 | -0.019965956 |
| RN7SKP293 | -0.01997163 |
| ARHGAP42P2 | -0.019974576 |
| RN7SL288P | -0.019979746 |
| AC114877 | -0.019980385 |
| C1orf115 | -0.019980952 |
| CTD-2116N17 | -0.019981538 |
| RP11-108E14 | -0.019987899 |
| UBE2D2 | -0.019988282 |
| RNU6-1105P | -0.019989882 |
| RP11-1082I11 | -0.019990003 |
| RP11-481E4 | -0.019991264 |
| AC008440 | -0.019991384 |
| MIR4779 | -0.019997225 |
| AC124914 | -0.019997439 |
| RP11-38F22 | -0.019999761 |
| RP11-489C13 | -0.020009786 |
| ACSL3 | -0.020019781 |

| | |
|---|---|
| ZDHHC24 | -0.020027112 |
| RNU6-276P | -0.020030178 |
| TMTC2 | -0.020034312 |
| CTD-2074D8 | -0.020037248 |
| RP11-270C12 | -0.020039274 |
| OR51H1 | -0.020040479 |
| RP11-697E22 | -0.02004142 |
| CHCHD3P3 | -0.020043468 |
| HDAC5 | -0.020043931 |
| OGDH | -0.02004769 |
| DEFB125 | -0.020051268 |
| ASS1P6 | -0.020060453 |
| PRSS33 | -0.020066975 |
| TLR2 | -0.020073328 |
| OR10A7 | -0.020076537 |
| CTD-2105E13 | -0.020077577 |
| LMOD2 | -0.020086603 |
| AC091770 | -0.02009245 |
| TAL1 | -0.020092547 |
| ZNF679 | -0.020093165 |
| PEX12P1 | -0.020100909 |
| NMNAT1 | -0.020106967 |
| FRMD3 | -0.020107115 |
| GOLGA2P2Y | -0.020107482 |
| HNRNPCP7 | -0.020111525 |
| RP11-553N19 | -0.020112912 |
| RP11-258O13 | -0.020117174 |
| PDE8A | -0.020118037 |
| CTB-191D16 | -0.020124159 |
| RN7SKP253 | -0.020133237 |
| NISCH | -0.020135749 |
| RPL7P52 | -0.020139839 |
| RP11-344A16 | -0.020140233 |
| RNU6ATAC18P | -0.020143591 |
| AL451006 | -0.020143692 |
| CCBL2 | -0.020153462 |
| RBBP7 | -0.020153797 |
| SNORD62A | -0.020156331 |
| RNU2-59P | -0.020161588 |
| RPL5P16 | -0.020163144 |
| C20orf202 | -0.020163176 |
| KRTAP7-1 | -0.020163195 |
| RP11-393N21 | -0.020164301 |
| MCM7 | -0.020168904 |
| LLfos-48D6 | -0.020169918 |

| | |
|---|---|
| RP11-275O18 | -0.020173367 |
| NEUROG3 | -0.020176206 |
| AL606923 | -0.020177218 |
| ZNF329 | -0.020179584 |
| TYW1B | -0.020182802 |
| SNX30 | -0.020182933 |
| RNA5SP436 | -0.020186528 |
| RNA5SP247 | -0.020189835 |
| CA7 | -0.020190826 |
| AC007450 | -0.020193506 |
| MIR4526 | -0.02019526 |
| TUB | -0.020198543 |
| SYTL5 | -0.020202335 |
| CXorf40B | -0.020204627 |
| CMTM1 | -0.020207064 |
| AC006480 | -0.020208933 |
| RP3-455J7 | -0.020219668 |
| SLC18A2 | -0.020223464 |
| AC004745 | -0.020224244 |
| FAM210A | -0.020224311 |
| KRTAP9-6 | -0.02022642 |
| REG3A | -0.02022808 |
| COPB2 | -0.020229874 |
| NLGN4Y-AS1 | -0.02023016 |
| ICE1 | -0.020235772 |
| SMIM21 | -0.020244255 |
| RNY4P6 | -0.020250444 |
| RP13-238F13 | -0.020256442 |
| RNU1-23P | -0.020257636 |
| RAB9AP1 | -0.020265708 |
| FAM90A5P | -0.020267409 |
| FAM83D | -0.020274955 |
| CTD-3137H5 | -0.020281025 |
| TOE1 | -0.020283235 |
| TBCEL | -0.020285095 |
| TAF11 | -0.020287347 |
| ZNF284 | -0.0202917 |
| KLF3P2 | -0.02029187 |
| GLUD1P9 | -0.020294291 |
| FAM160B1 | -0.020297449 |
| RP11-537P22 | -0.020297886 |
| ADPRHL1 | -0.020300873 |
| EEF1A2 | -0.02030305 |
| RPL30P7 | -0.020305503 |
| BOD1P1 | -0.020308318 |

| | |
|---|---|
| RBFA | -0.020311962 |
| TOR1A | -0.020317673 |
| AC099796 | -0.020321371 |
| PIF1 | -0.020322716 |
| GEMIN8 | -0.020325302 |
| GAGE2E | -0.020327043 |
| AC091643 | -0.020328986 |
| CTB-50E14 | -0.020331015 |
| TTC13 | -0.020331125 |
| AL121933 | -0.020331951 |
| RP11-467I17 | -0.020332366 |
| AL356289 | -0.020332735 |
| CTD-2122P11 | -0.020333675 |
| RNU6-687P | -0.020336091 |
| RP11-98O2 | -0.020336495 |
| RP11-355I22 | -0.020338297 |
| RNU6-754P | -0.020342823 |
| PRKRIRP9 | -0.020343081 |
| MIR4289 | -0.020343148 |
| FXYD7 | -0.02034527 |
| ZEB2 | -0.020345451 |
| RP11-88H12 | -0.020346632 |
| PEAR1 | -0.020347089 |
| AC144449 | -0.020349374 |
| RP11-169K19 | -0.020352109 |
| LINC01243 | -0.020359183 |
| SNORD121A | -0.020359729 |
| OR4V1P | -0.020361592 |
| PPARA | -0.020366792 |
| MIR4326 | -0.020373469 |
| HBQ1 | -0.020374019 |
| SERPINA1 | -0.020376969 |
| RP11-685B24 | -0.020377463 |
| IDNK | -0.020383611 |
| MAFIP | -0.020386695 |
| RP11-229M1 | -0.020388305 |
| CAMSAP3 | -0.020391723 |
| SAYSD1 | -0.020392579 |
| RNU6-1080P | -0.020392914 |
| NUS1P1 | -0.020393168 |
| SIX1 | -0.020397137 |
| GRAMD4 | -0.0203987 |
| AL079306 | -0.020398909 |
| RP11-348P10 | -0.020400952 |
| RNA5SP332 | -0.020405119 |

| | |
|---|---|
| RNU4ATAC11P | -0.020406198 |
| POLB | -0.020409491 |
| PRELID2P1 | -0.020410059 |
| TUBB8P6 | -0.0204126 |
| OR4A21P | -0.020417259 |
| RP11-147I3 | -0.020421492 |
| DUSP12 | -0.02042326 |
| GABRG2 | -0.020429692 |
| RP11-571F15 | -0.020433327 |
| SPPL3 | -0.020436451 |
| OTUD7A | -0.020437921 |
| MIR181C | -0.020442497 |
| RP11-3K16 | -0.02044328 |
| USP11 | -0.020443428 |
| AC002116 | -0.020444191 |
| RNU6-106P | -0.020445721 |
| VN1R17P | -0.020445738 |
| CTC-575C13 | -0.02044575 |
| AP003354 | -0.020446207 |
| ATG4AP1 | -0.020452295 |
| PRR23B | -0.020455489 |
| CTD-2544M6 | -0.020459075 |
| NAT8L | -0.020463838 |
| KRT19P3 | -0.020464372 |
| AC004932 | -0.020464802 |
| TCAF2P1 | -0.020468993 |
| RNU6-1248P | -0.020470892 |
| RP11-236P24 | -0.020475878 |
| HMGN1P12 | -0.020477819 |
| RP3-468B3 | -0.020478724 |
| LINC00309 | -0.020488889 |
| LINC01395 | -0.020489238 |
| MIR3907 | -0.020496422 |
| RP11-537A6 | -0.020497779 |
| KCNC4 | -0.020499635 |
| CBWD2 | -0.020505031 |
| AC015688 | -0.020508555 |
| ZNF334 | -0.020512119 |
| RN7SKP17 | -0.020513449 |
| RNU6-871P | -0.020517118 |
| RP3-323K23 | -0.020520816 |
| RP11-739A13 | -0.020524912 |
| FANCG | -0.020527152 |
| EXOSC8 | -0.020530669 |
| PRH1-PRR4 | -0.020531294 |

| | |
|---|---|
| MRPS6P2 | -0.020531935 |
| RNU6-369P | -0.020535725 |
| KRTAP9-2 | -0.020542523 |
| AC096643 | -0.020542708 |
| NDUFV3 | -0.020543525 |
| ELOVL6 | -0.020545795 |
| OXSM | -0.020548214 |
| SNORD115-45 | -0.020548258 |
| RP11-346D6 | -0.020550636 |
| TNS3 | -0.020553144 |
| LINC01381 | -0.020554586 |
| ABCG2 | -0.020555935 |
| AC009336 | -0.020557731 |
| RP11-360I2 | -0.020558148 |
| RN7SKP123 | -0.020558828 |
| RPL23AP39 | -0.020562348 |
| RNU2-29P | -0.020563824 |
| RP11-469J4 | -0.020564666 |
| RP11-304L19 | -0.02056587 |
| RP11-138E9 | -0.020568078 |
| HES5 | -0.020570977 |
| RP11-229A12 | -0.020572012 |
| RP11-358M3 | -0.020577782 |
| ZNF749 | -0.020581592 |
| RP13-362E11 | -0.020582069 |
| RP11-846F4 | -0.020584702 |
| VTRNA2-1 | -0.020585313 |
| CSF1 | -0.020585555 |
| AC006548 | -0.020595818 |
| MIR513A1 | -0.020601329 |
| CDH5 | -0.020602058 |
| ACTR10 | -0.020606598 |
| RP11-97F8 | -0.020608681 |
| SUGT1P2 | -0.020611356 |
| RP11-316K19 | -0.02061498 |
| AC096732 | -0.020622962 |
| CTD-2653D5 | -0.020623091 |
| AC138430 | -0.020625373 |
| CCL13 | -0.020628531 |
| AC007966 | -0.020633137 |
| AL353731 | -0.020639894 |
| YBX1P5 | -0.020644759 |
| ARL15 | -0.020644888 |
| RP11-404O13 | -0.020649315 |
| RP11-122A21 | -0.020661524 |

| | |
|---|---|
| RP11-69H7 | -0.020664674 |
| RN7SL693P | -0.020665462 |
| SPOPL | -0.020665745 |
| RN7SL775P | -0.020668001 |
| RNU6-484P | -0.020669256 |
| LA16c-325D7 | -0.020676937 |
| RP11-627K11 | -0.02067715 |
| RPS26P55 | -0.020678274 |
| RNU5B-1 | -0.020681467 |
| CSPG4P5 | -0.020683507 |
| RP11-319G9 | -0.02068545 |
| LINC01031 | -0.020687926 |
| AC104981 | -0.020691321 |
| AL627171 | -0.020693919 |
| GCLM | -0.020699737 |
| CHERP | -0.020701262 |
| LINC01182 | -0.020705343 |
| FNTAP2 | -0.020707564 |
| RP11-529K1 | -0.0207077 |
| RP5-996D20 | -0.020712105 |
| ASS1P1 | -0.020712288 |
| RNU6-1241P | -0.020714372 |
| ZCCHC13 | -0.020717822 |
| RNU2-68P | -0.020719305 |
| AC002543 | -0.020720892 |
| H1FOO | -0.020721261 |
| LRRC37A3 | -0.020725356 |
| RP11-552M14 | -0.020727403 |
| MDFIC | -0.020727552 |
| RP11-716D19 | -0.020728766 |
| LYZL2 | -0.02073391 |
| FAM205BP | -0.020738837 |
| PHF19 | -0.020739162 |
| PTPRVP | -0.020739614 |
| RP13-467H17 | -0.020741975 |
| CTD-2369P2 | -0.02074225 |
| MRPS14 | -0.020747404 |
| AC104802 | -0.020754985 |
| RPL7P48 | -0.02076114 |
| RP5-1163L11 | -0.020761235 |
| RP11-606D9 | -0.020763284 |
| C1orf101 | -0.020764056 |
| CAAP1 | -0.020765879 |
| IL10RB | -0.020766065 |
| BCL2L2-PABPN1 | -0.020766549 |

| | |
|---|---|
| FAM174A | -0.020766791 |
| TFF2 | -0.020774634 |
| MED28P8 | -0.020776853 |
| RP11-215I16 | -0.020780203 |
| MTND2P11 | -0.020783699 |
| RPS20P25 | -0.020787504 |
| ATP5F1P2 | -0.02078916 |
| RP11-1167A19 | -0.020791292 |
| THUMPD1P1 | -0.020793084 |
| AC104387 | -0.020797868 |
| G6PC2 | -0.020806218 |
| FAM45A | -0.020809565 |
| RNU6-1139P | -0.020809605 |
| POU3F3 | -0.020810209 |
| GTF2IRD2P1 | -0.020810805 |
| OLR1 | -0.020812839 |
| RP11-332E19 | -0.020813525 |
| MT1HL1 | -0.020817378 |
| LINC00665 | -0.020831426 |
| CSPG4P8 | -0.020834961 |
| TRIM71 | -0.020835022 |
| PDE6B | -0.020838031 |
| SH3KBP1 | -0.020845008 |
| TAS2R42 | -0.020847322 |
| GAPDHP76 | -0.02084787 |
| CDKN2AIPNLP3 | -0.020851282 |
| RP11-373A6 | -0.020852379 |
| ATG12P2 | -0.020853062 |
| RP11-124N2 | -0.020854731 |
| RNU6-1131P | -0.020855178 |
| RP11-296K13 | -0.020858586 |
| AL513487 | -0.020859027 |
| OR1D3P | -0.020862487 |
| RP1-249C1 | -0.020864206 |
| GOLGA8T | -0.02086601 |
| VIPAS39 | -0.020872582 |
| CACNG4 | -0.020875707 |
| AC018628 | -0.020876356 |
| OR5BM1P | -0.020881189 |
| MTM1 | -0.020882219 |
| RPL32P26 | -0.020884723 |
| MAVS | -0.020886015 |
| IPO8P1 | -0.020889326 |
| HSFY4P | -0.02088934 |
| RNA5SP281 | -0.020894336 |

| | |
|---|---|
| AC090602 | -0.020895144 |
| ATP5HP1 | -0.020898829 |
| AC004053 | -0.020899217 |
| RP11-457D13 | -0.020905551 |
| MTND1P31 | -0.020907625 |
| GDAP1L1 | -0.020907688 |
| NPPC | -0.020908087 |
| RNA5SP448 | -0.02091043 |
| FMO8P | -0.020912187 |
| TEX2 | -0.020912676 |
| KB-1083B1 | -0.020916744 |
| ZEB2P1 | -0.020918903 |
| RP11-500G10 | -0.020920715 |
| RNU6-703P | -0.02092294 |
| SNRPFP2 | -0.020923579 |
| RP11-662J14 | -0.02092821 |
| GLUD1P7 | -0.020928603 |
| RNA5SP134 | -0.020930243 |
| LDLRAD1 | -0.020930867 |
| HAPLN4 | -0.020935112 |
| LINC00353 | -0.020935607 |
| NMNAT1P2 | -0.020937356 |
| NAPSA | -0.020942668 |
| SLC25A20 | -0.020948566 |
| TTTY5 | -0.020949813 |
| TRIP13 | -0.020950948 |
| H3F3BP2 | -0.020951061 |
| RP11-215P8 | -0.020953218 |
| ARHGAP27 | -0.02095826 |
| MYL2 | -0.020958481 |
| OR9G3P | -0.020958579 |
| STMN4 | -0.020967779 |
| JAK1 | -0.020973479 |
| SNORA19 | -0.020974895 |
| PRKRIR | -0.020976661 |
| RNA5SP518 | -0.020978008 |
| CENPVP1 | -0.020978656 |
| RP11-90M2 | -0.020983446 |
| OR8G2P | -0.020983835 |
| SH2B2 | -0.020985889 |
| RP3-473B4 | -0.020994195 |
| HMGN3 | -0.02099601 |
| RNA5SP413 | -0.020996785 |
| SNORA80E | -0.020999319 |
| UBQLN3 | -0.021000624 |

| | |
|---|---|
| KRTAP5-8 | -0.021002748 |
| RP4-568C11 | -0.021008922 |
| MPC1 | -0.021012178 |
| RPL6P12 | -0.021013127 |
| CBX3P8 | -0.021015058 |
| AGPAT9 | -0.021015658 |
| RNU6-1089P | -0.021017035 |
| UNC80 | -0.021018531 |
| RP11-644L4 | -0.02102156 |
| RPL35AP26 | -0.021022859 |
| RAPH1 | -0.021026802 |
| RPL7P53 | -0.021027447 |
| RNU6-752P | -0.02102995 |
| RP11-524O24 | -0.021032924 |
| MIR363 | -0.021033596 |
| RP11-51C14 | -0.021034311 |
| MIR6740 | -0.021038021 |
| MT-TL2 | -0.021044495 |
| ITLN1 | -0.021046776 |
| RP11-14N4 | -0.021046992 |
| OR52N2 | -0.021047139 |
| AC108205 | -0.021047366 |
| FOXI3 | -0.021047991 |
| OR14A2 | -0.021048415 |
| GNAT2 | -0.021049756 |
| RP11-162D9 | -0.021049944 |
| Z83851 | -0.021052459 |
| BMI1P1 | -0.021053329 |
| RP1-209B9 | -0.021061113 |
| FAM166A | -0.021064529 |
| RNU6-926P | -0.021065324 |
| OR1G1 | -0.021066876 |
| RP11-344E13 | -0.021068189 |
| SWT1 | -0.021069874 |
| MND1 | -0.021070041 |
| RNU6-1266P | -0.021072956 |
| CLUL1 | -0.021075456 |
| BNIP3P22 | -0.021075828 |
| CLP1 | -0.02107633 |
| POLR3G | -0.021077558 |
| MIR7150 | -0.021082901 |
| TMEM26 | -0.021084516 |
| LINC00689 | -0.02108564 |
| RNA5SP522 | -0.021086294 |
| RN7SKP52 | -0.021091152 |

| | |
|---|---|
| RNU6-1316P | -0.021093616 |
| AC002550 | -0.021095698 |
| RP11-129I19 | -0.021096206 |
| RP11-336N8 | -0.021097845 |
| FFAR4 | -0.021100201 |
| AL138726 | -0.021101691 |
| KIAA0087 | -0.021103138 |
| FAM92A1 | -0.021107587 |
| RP11-977G19 | -0.021115476 |
| CLNS1A | -0.021117299 |
| SPOCK3 | -0.021129904 |
| RP11-1016B18 | -0.021131133 |
| RN7SKP114 | -0.021135895 |
| SNX18P8 | -0.021139447 |
| HIGD1AP17 | -0.021141691 |
| CYP2A6 | -0.021144573 |
| RP3-358H7 | -0.021147148 |
| CCT5P2 | -0.021149046 |
| FAM209B | -0.021150628 |
| RP11-10C24 | -0.021151143 |
| TRIM59 | -0.021152519 |
| PAX8 | -0.021153599 |
| AKAP8P1 | -0.021153923 |
| AC061992 | -0.021159219 |
| NDUFB8P2 | -0.021160429 |
| CHCHD2P5 | -0.021160643 |
| IMMP1LP1 | -0.021161593 |
| MTND6P17 | -0.02116209 |
| RP11-203P23 | -0.021166096 |
| RP11-259G18 | -0.021166606 |
| RP11-25L19 | -0.021169919 |
| GLTP | -0.021170623 |
| AL139319 | -0.021171889 |
| AC119751 | -0.021177095 |
| DET1 | -0.02118067 |
| RN7SL564P | -0.021181094 |
| COX7A2P1 | -0.021184034 |
| TATDN2P1 | -0.02118772 |
| PHBP13 | -0.021188896 |
| CISH | -0.021190735 |
| RP4-569M23 | -0.02119161 |
| CTD-2292P10 | -0.021201768 |
| EIF2S2 | -0.021204596 |
| MIR196A1 | -0.021206906 |
| AK2P2 | -0.021208111 |

| | |
|---|---|
| AC099757 | -0.021208226 |
| RBM8B | -0.021209283 |
| RP11-241G9 | -0.021210329 |
| ZNF66 | -0.021215071 |
| RPL23AP83 | -0.021221556 |
| OR12D2 | -0.021221747 |
| RP5-1118D24 | -0.021225723 |
| ELK1P1 | -0.021226224 |
| ZNF613 | -0.021226532 |
| RP11-653B10 | -0.021233049 |
| RNA5SP408 | -0.021237231 |
| OR7E26P | -0.021242244 |
| BACH1-IT3 | -0.021247268 |
| SNORD116-27 | -0.021247293 |
| MIR942 | -0.021250951 |
| RP11-223P11 | -0.021254853 |
| CR381670 | -0.021267328 |
| SNORD30 | -0.021273354 |
| RNU6-49P | -0.021280531 |
| DNAJC25 | -0.021280916 |
| SRR | -0.021282311 |
| LELP1 | -0.021289074 |
| NEK4P2 | -0.021292183 |
| AHCY | -0.021296228 |
| P2RX4 | -0.02129807 |
| DBIL5P | -0.021302405 |
| CTB-31N19 | -0.021306815 |
| PTGER4P3 | -0.021308008 |
| RP1-39G22 | -0.021309326 |
| MSI1 | -0.021311745 |
| RP11-145B3 | -0.021312669 |
| IMP3 | -0.02131288 |
| RP11-801I18 | -0.021316138 |
| ADAM32 | -0.021317829 |
| TMEM59 | -0.021322797 |
| EPB41L4B | -0.02132609 |
| RP11-799O21 | -0.021328873 |
| RP11-174C7 | -0.021333418 |
| ZNF274 | -0.021337458 |
| RCC2P5 | -0.021339192 |
| TDGF1P1 | -0.021343398 |
| RP11-1391J7 | -0.021345067 |
| NCOR1P3 | -0.021348372 |
| LL22NC03-N64E9 | -0.021349405 |
| RP11-1180F24 | -0.021350133 |

| | |
|---|---|
| RP3-344J20 | -0.02135272 |
| MIR7159 | -0.021360603 |
| CROCCP2 | -0.02136291 |
| PRR27 | -0.021366006 |
| CTAGE4 | -0.021366528 |
| RP11-766N7 | -0.021367316 |
| RN7SL250P | -0.021368344 |
| AC113610 | -0.021370212 |
| RP11-739P1 | -0.021371555 |
| RP11-129H15 | -0.021379467 |
| AC090573 | -0.021380903 |
| C5orf28 | -0.021380939 |
| AC243836 | -0.021385302 |
| RP11-728E14 | -0.021386023 |
| POU6F1 | -0.021389869 |
| AC019055 | -0.021393683 |
| RP11-435D7 | -0.021396375 |
| MIR6811 | -0.021399617 |
| RNA5SP512 | -0.021402799 |
| RP11-776H12 | -0.021403274 |
| Z70272 | -0.021403781 |
| RP11-702L6 | -0.021406682 |
| CFTRP1 | -0.021407567 |
| DAPK1 | -0.021408214 |
| RP11-241F15 | -0.021410236 |
| RP11-181K3 | -0.021412105 |
| OR56B3P | -0.021412321 |
| RP11-208N20 | -0.021414215 |
| MGAT4EP | -0.021416719 |
| SUB1P4 | -0.021417632 |
| L3MBTL4-AS1 | -0.021424699 |
| RNU6-141P | -0.021432345 |
| TJP2 | -0.021439304 |
| RP11-32D4 | -0.021441952 |
| PKNOX2-AS1 | -0.021442164 |
| RNY1P4 | -0.021446015 |
| AC104058 | -0.021447981 |
| RP11-701H16 | -0.021453763 |
| EIF3F | -0.021457035 |
| RP11-561O23 | -0.021457112 |
| C3orf67-AS1 | -0.021462033 |
| RNU7-176P | -0.021465579 |
| RP11-812I20 | -0.021466786 |
| RP11-684B2 | -0.021469599 |
| RP11-1415C14 | -0.021471571 |

| | |
|---|---|
| RNU6-421P | -0.021474105 |
| OR4K3 | -0.021476575 |
| ACVR1B | -0.021477706 |
| GPN3P1 | -0.0214852 |
| RN7SKP147 | -0.021487462 |
| C9orf172 | -0.021490488 |
| AC002127 | -0.021490753 |
| RP11-323I1 | -0.021491993 |
| GEMIN2P1 | -0.02149221 |
| ZNF804B | -0.021495728 |
| AC079145 | -0.021495978 |
| SKAP2 | -0.021497372 |
| PHOX2B | -0.021507062 |
| OR2Y1 | -0.021511425 |
| CKMT2-AS1 | -0.021511772 |
| OR2AG1 | -0.021514158 |
| SERPINB8P1 | -0.021515253 |
| ZSCAN20 | -0.021515866 |
| AC063979 | -0.02152281 |
| TAF9B | -0.021523364 |
| PALD1 | -0.021525875 |
| RNA5SP272 | -0.021529654 |
| SRPR | -0.021531848 |
| MIR522 | -0.021535213 |
| RP11-561G1 | -0.021538498 |
| AL136985 | -0.021538687 |
| BCAS3 | -0.021541243 |
| FGF2 | -0.021550007 |
| KRT18P17 | -0.021555037 |
| STON2 | -0.021563644 |
| LINC00976 | -0.021569534 |
| RP11-445P19 | -0.021570586 |
| RP11-410D17 | -0.021576453 |
| MIR1268A | -0.021576845 |
| RNU6-905P | -0.021577795 |
| MIR597 | -0.021579471 |
| RTCA | -0.021579515 |
| MIR4699 | -0.021579526 |
| IL20RA | -0.021580585 |
| MIR4641 | -0.021587865 |
| RNA5SP30 | -0.021590081 |
| TMEM205 | -0.021591515 |
| RP11-1E1 | -0.02159799 |
| RN7SL510P | -0.021602461 |
| RP11-99J16__A | -0.021611238 |

| | |
|---|---|
| RN7SL608P | -0.021613171 |
| DNAJC5 | -0.02161907 |
| MARVELD3 | -0.021620205 |
| GM2AP1 | -0.021623631 |
| RNA5SP262 | -0.021623691 |
| RNU6-951P | -0.021628192 |
| RNU6-1008P | -0.021628697 |
| QRSL1P1 | -0.021629321 |
| CNIH2 | -0.02163308 |
| LACC1 | -0.021635424 |
| NCKIPSD | -0.021640727 |
| SIKE1 | -0.021645546 |
| GPS1 | -0.021647388 |
| RP11-165D20 | -0.021652096 |
| CXorf58 | -0.021654663 |
| RPL36AP15 | -0.021655994 |
| HHATL | -0.021659027 |
| RNU6-306P | -0.021671189 |
| PAQR9 | -0.021672717 |
| MAP6 | -0.021676077 |
| CYP2D6 | -0.021676282 |
| WRBP1 | -0.02167646 |
| KIF2C | -0.021679705 |
| RP11-318C2 | -0.021683115 |
| AP001063 | -0.021684992 |
| CTD-2049O4 | -0.021686498 |
| LINC00477 | -0.021686625 |
| MLKL | -0.02168845 |
| DNAJC19 | -0.021689194 |
| RP11-694I15 | -0.021693157 |
| ENGASE | -0.021699334 |
| RP11-219J21 | -0.021707264 |
| RP11-372E1 | -0.021707614 |
| AC003080 | -0.021710365 |
| RP11-570L15 | -0.021717797 |
| RNU4-14P | -0.021721863 |
| PNPLA8 | -0.021721996 |
| ALKBH5 | -0.021723428 |
| RNU1-56P | -0.021730947 |
| LA16c-306A4 | -0.021731503 |
| RNU6ATAC21P | -0.021737631 |
| REM1 | -0.021738685 |
| DHX38 | -0.021740062 |
| CACNB1 | -0.021741103 |
| RP11-315D13 | -0.021747976 |

| | |
|---|---|
| GIPC2 | -0.021753513 |
| RP11-570P14 | -0.021762783 |
| RN7SL525P | -0.021764479 |
| PIDD1 | -0.021764863 |
| RP11-433A10 | -0.0217684 |
| MIR6507 | -0.021768698 |
| OR2A2 | -0.021771496 |
| SCARNA14 | -0.021771656 |
| C9 | -0.021772944 |
| RP11-114M5 | -0.021775727 |
| MIR6736 | -0.021780094 |
| NME1P1 | -0.021780864 |
| POC1A | -0.02178291 |
| RP11-1113C11 | -0.021785943 |
| CTD-2139B15 | -0.021786431 |
| FEM1AP2 | -0.021786487 |
| SPATA31E2P | -0.021786949 |
| RNU6-1319P | -0.021794244 |
| RP11-28A22 | -0.021799561 |
| XKRX | -0.021799897 |
| RNU6-1286P | -0.021800225 |
| KCNJ11 | -0.021801943 |
| RN7SKP48 | -0.021804898 |
| CH17-451C3 | -0.021805191 |
| RP4-660H19 | -0.021813367 |
| RNU6-919P | -0.021822028 |
| AC110056 | -0.021822846 |
| ASNS | -0.021823247 |
| EIF4A1P8 | -0.021825111 |
| RP11-138B4 | -0.021828841 |
| KPNA4 | -0.02183428 |
| AC063980 | -0.021839735 |
| RP11-497K15 | -0.021846526 |
| DNAJB6 | -0.021846764 |
| RFX3 | -0.021849238 |
| GHRH | -0.021852348 |
| AC005251 | -0.021852541 |
| LINC01465 | -0.021854653 |
| TANGO2 | -0.021861927 |
| INSM2 | -0.021870668 |
| UBTFL7 | -0.021874952 |
| NUFIP1 | -0.0218751 |
| RCCD1 | -0.021878207 |
| RP5-968P14 | -0.021880421 |
| CTD-2154I11 | -0.021882208 |

| | |
|---|---|
| HS3ST3B1 | -0.021882356 |
| OR4K8P | -0.021886324 |
| MIR3180-5 | -0.021888716 |
| CTB-25J19 | -0.021893529 |
| RN7SL744P | -0.021894361 |
| SETP12 | -0.021896974 |
| RP11-10N16 | -0.021896985 |
| RNU7-113P | -0.021902096 |
| SNORA18 | -0.021905206 |
| CTD-2210A23 | -0.021906186 |
| ERVFRD-3 | -0.021908975 |
| PRPF38AP2 | -0.021911624 |
| MARK2P14 | -0.021912061 |
| RHEBP1 | -0.02191332 |
| SPRYD7 | -0.021917316 |
| PRDM13 | -0.021920567 |
| PKD1P1 | -0.021920655 |
| BCL9 | -0.021922097 |
| AC018690 | -0.021923278 |
| RPL17P40 | -0.021923294 |
| SCAND3P1 | -0.021925322 |
| MST1P2 | -0.021927121 |
| AC068039 | -0.021933279 |
| MYOM2 | -0.021934117 |
| LINC01529 | -0.021938556 |
| LINC00917 | -0.021938677 |
| RN7SL482P | -0.021948769 |
| RNU6-492P | -0.021949325 |
| EIF4A1P13 | -0.021949408 |
| HSPA1A | -0.021952275 |
| RNA5SP420 | -0.021952603 |
| TKTL2 | -0.021954112 |
| PHACTR1 | -0.021954341 |
| GTF2H2B | -0.021956315 |
| MIR4316 | -0.021956741 |
| RP11-603K19 | -0.02195852 |
| TCEAL4P1 | -0.021959112 |
| MIR5188 | -0.021959966 |
| RNU1-8P | -0.021960911 |
| RP11-111A21 | -0.02196195 |
| RP11-526H11 | -0.021964394 |
| LSM6 | -0.021965634 |
| OR52N5 | -0.021973139 |
| EIF2AP4 | -0.021973713 |
| RGS12 | -0.021974174 |

| | |
|---|---|
| SLC44A4 | -0.021975718 |
| GNAQ | -0.021981114 |
| RN7SL635P | -0.021986304 |
| RP13-895J2 | -0.021987647 |
| ARMC8P1 | -0.021989144 |
| RGS8 | -0.021989635 |
| RP11-274J15 | -0.02199725 |
| BOD1 | -0.021998808 |
| RP11-252P18 | -0.02200309 |
| MIR548I3 | -0.022003093 |
| MIR3131 | -0.022004629 |
| PYCARD-AS1 | -0.022006382 |
| RNA5SP121 | -0.022006812 |
| AL109612 | -0.022008951 |
| ANK2 | -0.022009487 |
| RP11-104E19 | -0.022010989 |
| SLC6A13 | -0.022014645 |
| RN7SL466P | -0.022014842 |
| CTD-2127O16 | -0.02201614 |
| RPL12P3 | -0.022017932 |
| ZNF519P1 | -0.022017947 |
| RNA5-8SP7 | -0.022018047 |
| RP11-511H9 | -0.022022634 |
| PITPNA | -0.022024632 |
| RP11-613M10 | -0.022029991 |
| SUCLG2-AS1 | -0.022030884 |
| RP11-131J4 | -0.022034668 |
| RPS26P24 | -0.022035719 |
| ATL3 | -0.022044131 |
| NANOGP8 | -0.022047284 |
| FOXP1 | -0.02204975 |
| GCM1 | -0.022053677 |
| CTC-412M14 | -0.022057017 |
| RP11-61I13 | -0.02206298 |
| AL137224 | -0.022070767 |
| RP11-436P7 | -0.022075259 |
| RNU6-1176P | -0.022079721 |
| GCOM2 | -0.02208511 |
| DUSP13 | -0.022085925 |
| RNU6-643P | -0.022090996 |
| AP002498 | -0.022095635 |
| GGACT | -0.02210311 |
| RP11-818C3 | -0.022105613 |
| RP3-403L10 | -0.022108364 |
| RNA5SP460 | -0.022108662 |

| | |
|---|---|
| ATP5A1P10 | -0.022109385 |
| RP11-1036F1 | -0.022115757 |
| HSPE1 | -0.022116952 |
| RP11-383B24 | -0.022117664 |
| SENCR | -0.022118153 |
| USP24P1 | -0.022118857 |
| AL133228 | -0.022119483 |
| RNU6ATAC3P | -0.02212226 |
| RNU7-25P | -0.022127864 |
| CTA-415G2 | -0.022129268 |
| RP4-675G8 | -0.022129987 |
| AC009494 | -0.022131187 |
| AC244230 | -0.022131475 |
| CTB-187M2 | -0.022134333 |
| EEF1A1P24 | -0.022135614 |
| RP11-319E16 | -0.022139184 |
| RP11-676M6 | -0.022140042 |
| CEP19 | -0.022149828 |
| IMMP1LP2 | -0.022152957 |
| GLYATL3 | -0.022158767 |
| CTD-2366F13 | -0.022162554 |
| RP11-406H23 | -0.022166884 |
| CTD-2108O9 | -0.022168309 |
| OCLN | -0.022171429 |
| RP11-180O5 | -0.022174169 |
| EIF2A | -0.022176955 |
| ARHGAP22-IT1 | -0.022181192 |
| BAAT | -0.022181392 |
| RNU6-603P | -0.022181804 |
| NEDD8 | -0.022182518 |
| OR7E157P | -0.022183183 |
| MIR3156-3 | -0.022184494 |
| DPYSL5 | -0.022186812 |
| NIPSNAP3A | -0.022191244 |
| RBMXL3 | -0.022194802 |
| MTCH2 | -0.022203319 |
| RP11-15N24 | -0.022205747 |
| RP11-74K19 | -0.022214025 |
| SLC27A3 | -0.022214985 |
| ATP9A | -0.022215543 |
| OR2P1P | -0.022217081 |
| RP11-785H20 | -0.022218647 |
| MIR599 | -0.022225482 |
| ISPD-AS1 | -0.022225702 |
| RN7SL151P | -0.0222303 |

| | |
|---|---|
| GLRA3 | -0.022232626 |
| CHD3 | -0.022238521 |
| TMPRSS7 | -0.0222395 |
| AC009987 | -0.022239678 |
| MTND4P5 | -0.022243751 |
| ACER2 | -0.022245516 |
| TFAP2C | -0.02224613 |
| ANKS1B | -0.022246856 |
| AC060226 | -0.022249124 |
| C1QTNF8 | -0.022251019 |
| RN7SKP142 | -0.022251686 |
| RP11-881M11 | -0.022259359 |
| ISCA1P3 | -0.022259928 |
| RP11-341G2 | -0.022262981 |
| AP003305 | -0.022265856 |
| WBP1LP10 | -0.022272172 |
| RP11-291I6 | -0.022274541 |
| RPLP2P1 | -0.0222749 |
| SREK1IP1 | -0.022278671 |
| SLC25A15P2 | -0.022278969 |
| AC073055 | -0.02227968 |
| SNORA68 | -0.022280162 |
| RNU6-263P | -0.02228062 |
| RPL39P31 | -0.022283538 |
| PPIAP32 | -0.022284429 |
| MTND5P7 | -0.022286167 |
| RP11-101O6 | -0.022287848 |
| snoMe28S-Am2634 | -0.022288325 |
| AKIRIN2 | -0.022289797 |
| RP11-973F15 | -0.022293808 |
| RP11-4B16 | -0.022294411 |
| AL121753 | -0.022299333 |
| ATP8B3 | -0.022300068 |
| DEFB114 | -0.022305931 |
| RP11-726G1 | -0.022309687 |
| CTD-2568P8 | -0.022314836 |
| AC083867 | -0.022315935 |
| FAM41AY1 | -0.02232023 |
| TBR1 | -0.022322673 |
| RGL3 | -0.022329557 |
| RNU4ATAC16P | -0.02233 |
| F8A2 | -0.02233041 |
| KNOP1P3 | -0.022337779 |
| OR5D15P | -0.02233967 |
| ARAFP3 | -0.022341487 |

| | |
|---|---|
| RP11-66C24 | -0.022344471 |
| RNU6-575P | -0.022344486 |
| RP11-717H13 | -0.022352438 |
| TAF13P2 | -0.022352932 |
| HIST2H2BC | -0.02235594 |
| FAM60A | -0.022356275 |
| TDGF1P6 | -0.022358289 |
| AL157359 | -0.022358791 |
| RNU6-747P | -0.022361749 |
| LINC00581 | -0.022363261 |
| PKMP1 | -0.022365235 |
| ZNF519P3 | -0.02236674 |
| WIPI1 | -0.022371587 |
| RPS20P4 | -0.022374855 |
| RP11-572M11 | -0.02237777 |
| RPS27AP3 | -0.022381164 |
| C22orf34 | -0.022385076 |
| RNY4P20 | -0.022387688 |
| TMEM191C | -0.022388481 |
| TMEM225 | -0.022391001 |
| KCTD7 | -0.022392797 |
| RP11-84C13 | -0.022396312 |
| TOR3A | -0.022396611 |
| ATP6V1A | -0.022398249 |
| MIR3152 | -0.022401859 |
| FGD3 | -0.022405207 |
| RP11-7F18 | -0.022408928 |
| MIR6826 | -0.022416583 |
| WDR83 | -0.022423087 |
| AL158051 | -0.022423692 |
| RP11-428I12 | -0.022428311 |
| ZNF736P10Y | -0.022429563 |
| AC008691 | -0.022434431 |
| RP11-522I20 | -0.022435962 |
| U40455 | -0.02244036 |
| PHF5A | -0.022443385 |
| AC083862 | -0.02244887 |
| RP11-305K5 | -0.022449004 |
| BMP6 | -0.022450251 |
| CXorf57 | -0.022450493 |
| RP11-148M9 | -0.022451379 |
| RNU6-1287P | -0.022457833 |
| CCDC116 | -0.022460499 |
| RREB1 | -0.022461079 |
| RPP40P1 | -0.022464738 |

| | |
|---|---|
| SHQ1 | -0.022465358 |
| RP11-395N3 | -0.022466834 |
| ENPP7P1 | -0.022471703 |
| OSM | -0.022479089 |
| RN7SL596P | -0.02248182 |
| NDUFA12 | -0.022482178 |
| AC092835 | -0.022491713 |
| RP1-302D9 | -0.02249254 |
| RP11-717D12 | -0.022493046 |
| RP11-251M1 | -0.022493311 |
| PRB3 | -0.022496892 |
| RPS3AP43 | -0.02249824 |
| RP4-613A2 | -0.022500417 |
| RP11-95P2 | -0.022504566 |
| AC009404 | -0.022505323 |
| CTD-2533K21 | -0.02250554 |
| SSPO | -0.022506514 |
| RP11-613M5 | -0.022509537 |
| RP1-265C24 | -0.022512448 |
| CKS1BP6 | -0.022513368 |
| RP11-111G20 | -0.022514677 |
| ABTB2 | -0.022522552 |
| TRAJ55 | -0.022526189 |
| LINC01504 | -0.022527465 |
| PTPMT1 | -0.022529585 |
| RP11-433J22 | -0.022538243 |
| AL354820 | -0.022541993 |
| ARL4P | -0.022542349 |
| MIR545 | -0.022552915 |
| LINC01127 | -0.022553299 |
| HIST3H3 | -0.022553812 |
| GNS | -0.022554051 |
| RP3-393K13 | -0.022554951 |
| FAM86GP | -0.022555446 |
| LL22NC01-116C6 | -0.022558728 |
| RP11-669C19 | -0.02255917 |
| RN7SKP252 | -0.022561585 |
| MTND4P6 | -0.022562471 |
| MIR4487 | -0.022565424 |
| PPP5D1 | -0.02257167 |
| MIR8083 | -0.022574304 |
| CYB5R1 | -0.022575391 |
| MFSD11 | -0.022577512 |
| C15orf62 | -0.022584099 |
| JAM2 | -0.02258488 |

| | |
|---|---|
| EIF4BP5 | -0.022588266 |
| MKNK2P1 | -0.022589247 |
| RNU6-133P | -0.022593853 |
| C21orf91 | -0.022600862 |
| TDO2 | -0.022602323 |
| RP11-562L8 | -0.02260303 |
| RP11-218I21 | -0.02260639 |
| CLMN | -0.022607374 |
| TOMM20 | -0.022607821 |
| TTTY2 | -0.022615094 |
| MRPS33 | -0.022618723 |
| RP11-1029J19 | -0.022618804 |
| RNU6-297P | -0.022624111 |
| RBM22P4 | -0.022629416 |
| HGF | -0.022633012 |
| RP13-26D14 | -0.022648976 |
| AC106901 | -0.02265131 |
| RNU6-818P | -0.022651757 |
| RP11-503E24 | -0.022651798 |
| VANGL1 | -0.022653106 |
| RP13-13A3 | -0.022653322 |
| RP11-408H1 | -0.022656951 |
| AC093376 | -0.022663786 |
| ERP44 | -0.022665795 |
| C1orf56 | -0.022674949 |
| WDR61 | -0.022680661 |
| MIR518C | -0.022682011 |
| ZFYVE16 | -0.022684047 |
| MIR572 | -0.022684753 |
| HAUS6P2 | -0.022685105 |
| AC008671 | -0.022685911 |
| MIR3178 | -0.02268662 |
| RIC8A | -0.022688922 |
| METTL18 | -0.022690913 |
| NCK1 | -0.022701406 |
| PHOSPHO1 | -0.02270456 |
| LINC00939 | -0.022710468 |
| RP11-533N14 | -0.022713556 |
| C3orf70 | -0.022714095 |
| PRSS57 | -0.022715491 |
| SNORA70 | -0.02271925 |
| RAB6A | -0.022719584 |
| OR8K1 | -0.022721106 |
| DNAJB4 | -0.022727175 |
| MT-TH | -0.022727193 |

| | |
|---|---|
| FAM120AOS | -0.022735175 |
| RP3-348I23 | -0.022735208 |
| IGKV2D-14 | -0.022740415 |
| MAP2K7 | -0.022744757 |
| CHEK1 | -0.022744961 |
| GOLGA6L11P | -0.022749836 |
| LHFPL3-AS2 | -0.022749903 |
| CCDC96 | -0.022751325 |
| AL163760 | -0.022754091 |
| BTF3P5 | -0.022755492 |
| ZNF684 | -0.022755864 |
| RP11-381I15 | -0.02275869 |
| KRT8P13 | -0.022759055 |
| RPL18AP2 | -0.022761656 |
| USP28 | -0.022768218 |
| RP13-1032I1 | -0.022768739 |
| SNX18P13 | -0.02277249 |
| RP11-313C15 | -0.022777601 |
| KCNMB3 | -0.022777633 |
| AC133041 | -0.022781482 |
| OR51J1 | -0.022782308 |
| MIR520E | -0.022785943 |
| LARGE-IT1 | -0.022786614 |
| NME5 | -0.022789561 |
| SNORD116-18 | -0.022789913 |
| MRPL38 | -0.022792331 |
| RP11-451F14 | -0.022793949 |
| RNU6-176P | -0.022800938 |
| MIR301B | -0.022802106 |
| DIAPH1 | -0.022805894 |
| GTF3AP2 | -0.02280809 |
| BAATP1 | -0.022809421 |
| EP300-AS1 | -0.022813776 |
| SUCLG2P4 | -0.022817 |
| LYSMD1 | -0.022817262 |
| HPVC1 | -0.02282108 |
| RP11-885B4 | -0.022828003 |
| RP11-465K16 | -0.022828142 |
| RP11-72I2 | -0.022828987 |
| PCBD2 | -0.022829858 |
| LINC01059 | -0.022831974 |
| RP11-189B4 | -0.022832438 |
| TMEM256P2 | -0.022833752 |
| RP11-183E9 | -0.02283438 |
| RP11-398B16 | -0.022837579 |

| | |
|---|---|
| VIPR1 | -0.022838621 |
| CLVS1 | -0.022841137 |
| RNU6-775P | -0.022848879 |
| CFH | -0.022852334 |
| RNU6-553P | -0.02285238 |
| RNU1-72P | -0.022853292 |
| RP4-736I12 | -0.022854359 |
| UNC13B | -0.022860464 |
| RNU7-66P | -0.022860545 |
| ASTN2-AS1 | -0.022861158 |
| RP11-522B15 | -0.022862139 |
| RP11-4G2 | -0.022863146 |
| RP11-152K4 | -0.022864232 |
| NRAS | -0.022866195 |
| SPINK2 | -0.022873498 |
| AC092910 | -0.022873826 |
| RP11-386G11 | -0.022879665 |
| PBX4 | -0.022883479 |
| RP5-925F19 | -0.022884694 |
| RP5-1057I20 | -0.022885321 |
| MED28P1 | -0.02288585 |
| RP11-124O11 | -0.022888762 |
| SLC9A7 | -0.022891052 |
| HNRNPA1P43 | -0.022902628 |
| AC121336 | -0.022902745 |
| CTB-162J7 | -0.022905469 |
| AC108519 | -0.022907187 |
| PDCD4-AS1 | -0.022908466 |
| OR52H2P | -0.022913015 |
| ARRDC4 | -0.022913546 |
| RNU6-711P | -0.022915484 |
| RNA5SP245 | -0.022920978 |
| SLC16A5 | -0.022925862 |
| RP11-332P22 | -0.022928182 |
| CLUHP5 | -0.022933675 |
| MIR1278 | -0.022936932 |
| BRD9P1 | -0.022937986 |
| TNIP1 | -0.022947128 |
| KRT18P7 | -0.02294906 |
| AL589843 | -0.022949307 |
| MIR4464 | -0.022950592 |
| BRINP1 | -0.022951736 |
| AC005626 | -0.022952352 |
| PRRC2B | -0.022956431 |
| STK4-AS1 | -0.022958441 |

| | |
|---|---|
| C11orf85 | -0.022960831 |
| IGHVIII-22-2 | -0.022962653 |
| AB019440 | -0.022968072 |
| CCDC185 | -0.022974111 |
| RNU6-1328P | -0.022975414 |
| RN7SKP46 | -0.02297641 |
| MAN2B1 | -0.022977493 |
| CPB2 | -0.022977528 |
| DNAAF5 | -0.022979093 |
| KRT8P32 | -0.022980909 |
| RPL35AP30 | -0.022983062 |
| CTC-250I14 | -0.022983532 |
| HIST2H2BE | -0.022983707 |
| FAM204BP | -0.022984036 |
| COQ7 | -0.022985921 |
| AC026403 | -0.022986345 |
| MIR550A2 | -0.022987755 |
| DYNC1I2P1 | -0.022988182 |
| C5orf66-AS2 | -0.022988298 |
| LINC00365 | -0.022988415 |
| CTD-2024D23 | -0.022996669 |
| AL357095 | -0.022997026 |
| TEX28 | -0.022999189 |
| CLPTM1 | -0.023000154 |
| ST18 | -0.023002556 |
| RP11-452D2 | -0.023006188 |
| MCMDC2 | -0.023014555 |
| AL133167 | -0.023020805 |
| FAM177A1P1 | -0.023023301 |
| RN7SL561P | -0.023023983 |
| DCDC2 | -0.023024875 |
| CH17-228N4 | -0.023027887 |
| RSPH3 | -0.023028546 |
| AP001372 | -0.023029588 |
| AC020636 | -0.023030017 |
| TIFA | -0.02303009 |
| AC005776 | -0.023035043 |
| LINC01585 | -0.023038549 |
| AC096633 | -0.023041882 |
| BMPR1B-AS1 | -0.023042624 |
| ARPP21 | -0.023048169 |
| TMEM62 | -0.023049812 |
| BTF3P9 | -0.023055351 |
| RP11-707E21 | -0.023057111 |
| RP4-555N2 | -0.023058862 |

| | |
|---|---|
| MIR4279 | -0.023059252 |
| RN7SKP83 | -0.023065038 |
| RPL7P29 | -0.023067229 |
| SNX18P24 | -0.023070042 |
| HNRNPA1P2 | -0.023072676 |
| ZNF415 | -0.023073255 |
| RP11-394J1 | -0.023083671 |
| FOXD4L4 | -0.023086109 |
| VN1R3 | -0.023087627 |
| AC118463 | -0.023092858 |
| DNAJC28 | -0.02309454 |
| TBC1D13 | -0.023100493 |
| IDH1 | -0.023103117 |
| CTD-2245E15 | -0.023103226 |
| TTC28-AS1_3 | -0.023114233 |
| OR4A9P | -0.023121954 |
| DNAJC27-AS1 | -0.023122076 |
| RP11-178C3 | -0.023124066 |
| UNC93B1 | -0.023126243 |
| RP11-114H21 | -0.023126325 |
| AC145141 | -0.023126459 |
| RP13-766D20 | -0.023127583 |
| LINC01317 | -0.023133761 |
| RP11-384A12 | -0.023144285 |
| IQCF3 | -0.023144432 |
| RP11-562F9 | -0.023145781 |
| FXYD6P3 | -0.023147072 |
| C6orf99 | -0.023147733 |
| PSME4 | -0.023151398 |
| C8orf33 | -0.023153928 |
| AL121988 | -0.023154647 |
| SLC26A4-AS1 | -0.023158773 |
| RP11-46E17 | -0.023160353 |
| ALG8 | -0.023162412 |
| ZNF154 | -0.023165966 |
| RP11-17E3 | -0.02316636 |
| TAS2R31 | -0.023167225 |
| VCP | -0.023169464 |
| ZNF530 | -0.02317339 |
| KRTAP16-1 | -0.023179667 |
| AC021078 | -0.023181806 |
| RP11-93O14 | -0.023183471 |
| RPSAP31 | -0.023186104 |
| CTC-311G1 | -0.023188599 |
| RP11-776A13 | -0.023191808 |

| | |
|---|---|
| NAP1L6 | -0.023192449 |
| SSU72P1 | -0.023196345 |
| RP11-16F15 | -0.023206675 |
| RELN | -0.023207072 |
| AP001055 | -0.023215317 |
| OR52K3P | -0.023216705 |
| AC002059 | -0.023217468 |
| MTURN | -0.023218359 |
| RP11-16O9 | -0.023218754 |
| RP11-946L16 | -0.023218768 |
| OR4C12 | -0.023221921 |
| PRSS55 | -0.02322471 |
| RNU7-167P | -0.023225542 |
| MORC1-AS1 | -0.023226019 |
| LINC01231 | -0.023227326 |
| ATRNL1 | -0.023231668 |
| RPL7AP66 | -0.023237876 |
| LINC01124 | -0.023240092 |
| FOXN4 | -0.023252464 |
| RNU6-3P | -0.023257423 |
| TCAF2 | -0.023262268 |
| RP11-501J20 | -0.023262416 |
| ADAM10 | -0.023263254 |
| HSPA8P7 | -0.023265571 |
| COX6CP13 | -0.023265984 |
| ENPP7P4 | -0.023267339 |
| TMEM52 | -0.023268501 |
| SLC27A4 | -0.023271682 |
| CTD-2275D10 | -0.023277068 |
| CALR3 | -0.023280668 |
| RP5-1128N12 | -0.023285494 |
| RP11-478K15 | -0.023290097 |
| NIFK | -0.023291813 |
| INHBC | -0.023298007 |
| AC018902 | -0.023299415 |
| RP11-432B6 | -0.023299648 |
| AL031768 | -0.023299817 |
| PLCH1-AS2 | -0.023301786 |
| RN7SL40P | -0.023302304 |
| AC104057 | -0.023306375 |
| CTD-2396E7 | -0.02330893 |
| RP11-39C10 | -0.023311884 |
| AC022007 | -0.023313693 |
| CHMP7 | -0.023314661 |
| GRM7-AS2 | -0.023315136 |

| | |
|---|---|
| RP11-408O19 | -0.02331557 |
| AP006216 | -0.023316614 |
| HAND1 | -0.023317585 |
| NLRP1 | -0.023318419 |
| RN7SL416P | -0.02331979 |
| SLC38A6 | -0.023321005 |
| SESN3 | -0.023323015 |
| MIR4774 | -0.023323769 |
| RPS23P2 | -0.023331225 |
| AC016753 | -0.023331728 |
| DEPDC4 | -0.023333099 |
| UBE2D1 | -0.023333694 |
| RP11-399B17 | -0.02333656 |
| CTD-3116E22 | -0.023336674 |
| AC097465 | -0.023338593 |
| RP11-812D23 | -0.023343264 |
| DHX40P1 | -0.023346238 |
| AF241726 | -0.023346323 |
| RPS2P2 | -0.023347277 |
| OR1Q1 | -0.023349895 |
| MIR548AD | -0.023352301 |
| AL645728 | -0.023354506 |
| PLEKHO2 | -0.023357453 |
| RP11-553K8 | -0.023357815 |
| FO393414 | -0.023358339 |
| ORC3 | -0.023361602 |
| PSMD14 | -0.023368933 |
| RNA5SP389 | -0.023373061 |
| MIR6070 | -0.023373105 |
| RP11-422J15 | -0.023378402 |
| TFAP2A-AS1 | -0.023381845 |
| DDAH1 | -0.02338231 |
| ENTPD1 | -0.023382813 |
| XRCC6 | -0.023384673 |
| IGHVII-65-1 | -0.023387026 |
| RNU7-105P | -0.023390177 |
| NKIRAS1 | -0.023394962 |
| OR4A2P | -0.023397132 |
| PLAC4 | -0.023404944 |
| C5orf42 | -0.023406311 |
| RP4-713A8 | -0.023409511 |
| RP11-27N21 | -0.02340992 |
| RP11-292B1 | -0.023411387 |
| RP11-626P14 | -0.023412717 |
| AC073107 | -0.023415007 |

| | |
|---|---|
| STRA13 | -0.023419131 |
| RP11-696F10 | -0.023419255 |
| TUBA1B | -0.023421438 |
| MIR2681 | -0.023424871 |
| POU5F1P4 | -0.023424885 |
| CTD-2033A16 | -0.023427373 |
| AL359983 | -0.02342797 |
| RNA5SP353 | -0.023428355 |
| CYP2AB1P | -0.023431323 |
| NUTF2P8 | -0.023432708 |
| C2orf83 | -0.02343344 |
| RN7SL201P | -0.023437793 |
| ATP1B1P1 | -0.023441185 |
| XXbac-BPG116M5 | -0.023442107 |
| RP11-408B11 | -0.023442749 |
| C6orf165 | -0.023443374 |
| RP11-258B16 | -0.023443909 |
| AC147651 | -0.023444006 |
| RP11-273B20 | -0.023444657 |
| C5orf30 | -0.023445848 |
| RPS27P16 | -0.023448914 |
| PIGA | -0.023456211 |
| RP11-111N20 | -0.023457024 |
| MYOCD | -0.02345711 |
| RP3-514P16 | -0.023457895 |
| RP11-328K15 | -0.02345882 |
| RP11-1292F20 | -0.023459285 |
| CTD-2345H1 | -0.023460637 |
| TRAJ50 | -0.023460776 |
| GSTT2 | -0.023461551 |
| KIAA0101 | -0.023467226 |
| PIGPP2 | -0.02347421 |
| TXNL1P1 | -0.023476295 |
| RHOU | -0.023476975 |
| VSTM2A-OT1 | -0.023483408 |
| RNA5SP371 | -0.023486784 |
| RP5-1006K12 | -0.023487469 |
| AC009973 | -0.023487777 |
| AC104451 | -0.02348931 |
| CTD-2537L20 | -0.023489747 |
| RPL7P11 | -0.023490068 |
| ATP6V1B1-AS1 | -0.023492724 |
| RP11-363E7 | -0.023492864 |
| TRPC5 | -0.023493967 |
| C17orf89 | -0.023497869 |

| | |
|---|---|
| RNU6-693P | -0.023504572 |
| RP11-452J6 | -0.023505399 |
| TRAPPC12 | -0.023510093 |
| ZCCHC14 | -0.023513646 |
| SNORD116-6 | -0.023514737 |
| PYDC2 | -0.023516654 |
| CASP7 | -0.02351746 |
| RP11-80A15 | -0.023523662 |
| ZNF716 | -0.02352497 |
| RP11-944L7 | -0.023525848 |
| DRD5P2 | -0.023530233 |
| MIR4802 | -0.023538626 |
| APTX | -0.023540289 |
| TMPRSS6 | -0.02354122 |
| HMGN2P47 | -0.023541552 |
| RPL9P31 | -0.023544717 |
| RP11-91K11 | -0.023547075 |
| CD177 | -0.023548774 |
| RPL9P30 | -0.023551995 |
| UBE2SP2 | -0.023552649 |
| AC003009 | -0.023555481 |
| ABCF2P1 | -0.023558761 |
| RP11-680E19 | -0.023558965 |
| RP11-796L2 | -0.023560296 |
| C15orf27 | -0.023561202 |
| RPL4P3 | -0.023562528 |
| RN7SL47P | -0.023563467 |
| FAM86LP | -0.023564267 |
| FBXO44 | -0.023566337 |
| MIR4778 | -0.023568948 |
| MYO5BP3 | -0.023572579 |
| FOXP4 | -0.023575807 |
| CDK5RAP3 | -0.023585147 |
| CTB-120L21 | -0.02358899 |
| AC008278 | -0.023589591 |
| GGA1 | -0.023589985 |
| ZNF570 | -0.023590003 |
| HIBCH | -0.023592385 |
| RP11-69A21 | -0.023592448 |
| ATP6V0A1 | -0.023593799 |
| RP11-38L15 | -0.023596264 |
| FAM25A | -0.023600887 |
| SMAD5 | -0.023604478 |
| SPATA31A6 | -0.023605753 |
| AC079804 | -0.02360701 |

| | |
|---|---|
| OR5H1 | -0.023608506 |
| CTB-187L3 | -0.023618385 |
| HSPA8P9 | -0.02361867 |
| RP11-540N6 | -0.023630069 |
| NEURL3 | -0.023631735 |
| WDR75 | -0.023634898 |
| DDTL | -0.023638861 |
| RP4-646N3 | -0.02363888 |
| RP1-272E8 | -0.023639705 |
| HLA-DPB2 | -0.023641212 |
| RP11-234K24 | -0.023641774 |
| MTND2P25 | -0.023645996 |
| PPIAP19 | -0.023648571 |
| RNU6ATAC22P | -0.023650076 |
| FKSG61 | -0.023650337 |
| LIPA | -0.023651885 |
| CTA-249B10 | -0.023655371 |
| TRIM64FP | -0.023656554 |
| VSX2 | -0.023660884 |
| SLC25A27 | -0.023670844 |
| ABHD17AP4 | -0.023673471 |
| NOP14-AS1 | -0.023677598 |
| AC104417 | -0.023680533 |
| RP11-77I22 | -0.023683154 |
| CREB3L3 | -0.023690867 |
| HOXB-AS1 | -0.023691943 |
| RP4-808O4 | -0.023693592 |
| AC097635 | -0.023698321 |
| LA16c-17H1 | -0.023709615 |
| AC105342 | -0.023709909 |
| CTC-241F20 | -0.023715817 |
| NELFA | -0.023718995 |
| HBP1 | -0.023720223 |
| ZMYND10 | -0.023722009 |
| PTENP1 | -0.023722083 |
| TBCAP3 | -0.023723016 |
| TPMTP3 | -0.02372319 |
| DYX1C1 | -0.023723423 |
| RP11-184A2 | -0.023728383 |
| TBC1D3H | -0.02372889 |
| EPCAM | -0.023729079 |
| OR7E105P | -0.023733256 |
| SMIM5 | -0.02373721 |
| AC097452 | -0.023741378 |
| RN7SKP134 | -0.023745581 |

| | |
|---|---|
| RNU1-88P | -0.023748999 |
| RP11-25O10 | -0.02375475 |
| ARHGAP26 | -0.023767369 |
| MS4A13 | -0.023769757 |
| CTD-2325K12 | -0.023772408 |
| PSMD10P3 | -0.023774535 |
| OR7K1P | -0.023775049 |
| RNU6-158P | -0.023775125 |
| FAM168B | -0.023775146 |
| AC073257 | -0.023776059 |
| REEP5 | -0.023777286 |
| SDAD1P2 | -0.02378341 |
| PRDX2 | -0.023786719 |
| ST13P14 | -0.023789584 |
| RP11-297D21 | -0.023791705 |
| AL161908 | -0.023797986 |
| GRPEL2P3 | -0.023802897 |
| ASAP1 | -0.023811092 |
| RP11-408J6 | -0.023811533 |
| STK33 | -0.023812705 |
| CTD-2526A2 | -0.023813998 |
| CTD-2050N2 | -0.023815567 |
| RP11-69M1 | -0.02382486 |
| MRPL19 | -0.023826888 |
| RNU6-1306P | -0.023828715 |
| RP1-191L6 | -0.023829087 |
| LRRC10 | -0.023839909 |
| LYPLAL1 | -0.023841847 |
| DSCR8 | -0.023843296 |
| QRICH2 | -0.02384754 |
| RP11-672L10 | -0.023852731 |
| LRRC8E | -0.023855764 |
| MIR6885 | -0.023856072 |
| IMPAD1 | -0.023856471 |
| KRTAP15-1 | -0.023857925 |
| RP5-1111A8 | -0.023858366 |
| RP11-300E4 | -0.023859068 |
| LUC7L | -0.02386088 |
| CRYZP1 | -0.023863899 |
| LINC01581 | -0.023864078 |
| PIGUP1 | -0.023864541 |
| TRIM53AP | -0.023867776 |
| SUSD4 | -0.023881142 |
| FAM64A | -0.023886388 |
| AL157823 | -0.023886798 |

| | |
|---|---|
| NPM1P37 | -0.02388729 |
| MIR604 | -0.023887297 |
| MT-ND5 | -0.023895821 |
| TYRO3P | -0.023897155 |
| OR12D1 | -0.023901575 |
| MAP2K1P1 | -0.023901794 |
| MTND1P11 | -0.02390425 |
| RNA5SP159 | -0.023905229 |
| HADHAP1 | -0.023905572 |
| GOLGA8Q | -0.023906644 |
| CYP4F10P | -0.0239089 |
| AK4P3 | -0.023910202 |
| FAM103A2P | -0.023911946 |
| RP11-25C15 | -0.023916907 |
| PGDP1 | -0.02391715 |
| TDGF1P5 | -0.023919264 |
| GAMT | -0.023920381 |
| LNX1-AS2 | -0.023921578 |
| RP11-33M22 | -0.023930117 |
| RP11-58B2 | -0.023930246 |
| AC023283 | -0.023930346 |
| OR8B10P | -0.023930856 |
| MIR208B | -0.023939974 |
| AL592528 | -0.023940495 |
| PM20D2 | -0.023941014 |
| RP11-2A1 | -0.02394647 |
| RPPH1-2P | -0.023946739 |
| SCGN | -0.023947739 |
| UBBP2 | -0.023948153 |
| DSCAS | -0.023949118 |
| SSX5 | -0.023949708 |
| RP11-744I24 | -0.023951866 |
| TRIM49D2 | -0.023952392 |
| MIR105-1 | -0.023952392 |
| PTP4A1P4 | -0.023954314 |
| RP11-185O21 | -0.023958523 |
| OR7E100P | -0.023968617 |
| RMDN2 | -0.023968848 |
| RPS6P20 | -0.023971788 |
| RNY3P10 | -0.023976554 |
| RP11-850F7 | -0.023985229 |
| CSRNP1 | -0.023986169 |
| PDCD2L | -0.023989742 |
| RP5-842K16 | -0.02399005 |
| AC005532 | -0.023996786 |

| | |
|---|---|
| KLF2 | -0.0240031 |
| AC095040 | -0.024007246 |
| AC092933 | -0.024011335 |
| MAN1A2 | -0.024014096 |
| RNU6-549P | -0.024019525 |
| RP11-545M17 | -0.024023256 |
| MIR4306 | -0.02402477 |
| PAK6 | -0.024025682 |
| MRPS17P3 | -0.024026253 |
| GK5 | -0.024030916 |
| RP11-739N10 | -0.02403258 |
| RP11-347L18 | -0.024034394 |
| NABP2 | -0.024036516 |
| RNU6-686P | -0.024044034 |
| IGKV2-10 | -0.024044149 |
| PTGR2 | -0.024050712 |
| RP13-507P19 | -0.024052724 |
| RP11-656D10 | -0.024054693 |
| NDUFA5P10 | -0.024056568 |
| RP11-509A17 | -0.024057694 |
| AC024590 | -0.024058756 |
| LINC01495 | -0.024059766 |
| POU5F1P7 | -0.024064124 |
| CDNF | -0.02406474 |
| EGLN2 | -0.02406688 |
| FCGR1C | -0.02406734 |
| SERF1A | -0.024069089 |
| TSPAN7 | -0.024073913 |
| RP11-616M22 | -0.024076254 |
| RP11-8H2 | -0.024079552 |
| MRPL48P1 | -0.024080518 |
| SDR42E2 | -0.024081821 |
| RP3-423B22 | -0.024085941 |
| RNU6-545P | -0.024086931 |
| TTC12 | -0.024090732 |
| LINC01412 | -0.024091563 |
| FAM154A | -0.02409517 |
| RNGTTP1 | -0.024095459 |
| FAM228B | -0.024105524 |
| HIVEP2 | -0.024106845 |
| DPY19L2P4 | -0.024108993 |
| AC008759 | -0.024110539 |
| AL022574 | -0.024112019 |
| MIR4685 | -0.02411211 |
| AC006037 | -0.024117898 |

| | |
|---|---|
| RP11-308K2 | -0.024119505 |
| LRRC49 | -0.024121152 |
| RP1-21O18 | -0.024122993 |
| RP11-159N11 | -0.024130604 |
| PGAM4P1 | -0.024133439 |
| RP11-334L9 | -0.024136734 |
| RP11-401E14 | -0.024137606 |
| CACHD1 | -0.024139981 |
| ZNF425 | -0.024149618 |
| RP11-622A1 | -0.02415042 |
| MIR221 | -0.024150506 |
| LARP7P2 | -0.024151097 |
| MRFAP1 | -0.024158735 |
| AC008074 | -0.024163015 |
| OR2M1P | -0.024163512 |
| WBP1LP3 | -0.024164772 |
| AC116614 | -0.024168548 |
| CTD-2195B23 | -0.024169308 |
| RP11-190C22 | -0.024173228 |
| AC008565 | -0.02418497 |
| MRPL22P1 | -0.02418618 |
| MIATNB | -0.024186606 |
| RNU6-803P | -0.024189556 |
| RNU2-6P | -0.02419534 |
| WASF2 | -0.024197509 |
| RNU4-56P | -0.024199114 |
| TCEB1P3 | -0.024204926 |
| DRG1 | -0.024208635 |
| RPL38P1 | -0.024209144 |
| SUV420H2 | -0.024212524 |
| RP11-399C16 | -0.02421701 |
| SMPD4 | -0.024217491 |
| CTD-2194A8 | -0.024223394 |
| DMRTB1 | -0.024228634 |
| C11orf45 | -0.024230496 |
| A1CF | -0.02423153 |
| RNU4ATAC15P | -0.02423473 |
| AL354928 | -0.024235428 |
| ZMPSTE24 | -0.024237473 |
| AC084854 | -0.024239512 |
| POMT1 | -0.02424259 |
| IL3RA | -0.024243502 |
| RP11-599J14 | -0.024244721 |
| TMEM70 | -0.024249139 |
| RP11-133L14 | -0.024252589 |

| | |
|---|---|
| RP11-6O2 | -0.024255986 |
| ATG10-AS1 | -0.024259694 |
| RNA5SP109 | -0.024262488 |
| AC092652 | -0.024264338 |
| RP11-74E22 | -0.024264377 |
| AC005740 | -0.024268842 |
| A2M-AS1 | -0.024273605 |
| RP11-442J17 | -0.024282101 |
| UPF1 | -0.024300732 |
| AC003984 | -0.024302699 |
| CTD-2224J9 | -0.024306762 |
| AC025016 | -0.024308869 |
| TPTE2P6 | -0.024309039 |
| KATNAL1 | -0.024310426 |
| OR4C2P | -0.024313901 |
| RP11-385N17 | -0.024314604 |
| RP11-72M17 | -0.02431649 |
| RNY4P24 | -0.024316614 |
| CCDC149 | -0.024322259 |
| RP11-147G16 | -0.024323101 |
| MIR4682 | -0.024325525 |
| AC106870 | -0.024326001 |
| MIR3183 | -0.024329349 |
| CELA2B | -0.024333813 |
| PHKA1P1 | -0.0243351 |
| AP001255 | -0.024337231 |
| RP11-65E22 | -0.024344695 |
| CH507-254M2 | -0.02434901 |
| POLR1A | -0.024350185 |
| NUMB | -0.024352251 |
| ANKHD1-EIF4EBP3 | -0.024355153 |
| CH17-78J1 | -0.024355301 |
| PTPN5 | -0.024361012 |
| AC007952 | -0.024364613 |
| XXyac-YR12DB5 | -0.024371768 |
| RP11-265D19 | -0.02437236 |
| RP11-144C15 | -0.024373799 |
| HIST1H2BO | -0.024375711 |
| RNA5SP68 | -0.024376489 |
| RNU6-682P | -0.024380005 |
| RP11-544P1 | -0.024381753 |
| RPL10P1 | -0.024383832 |
| CACNA1C-IT1 | -0.024395447 |
| RNU6-935P | -0.024397163 |
| ARAP1 | -0.024400513 |

| | |
|---|---|
| AP001205 | -0.024403356 |
| RIPK2 | -0.024406262 |
| COX6CP2 | -0.024406466 |
| RNU4-8P | -0.024406548 |
| RP11-244C20 | -0.024408856 |
| RP1-106C24 | -0.024414895 |
| MIR1269A | -0.0244192 |
| MIR320D1 | -0.02442524 |
| MRPS21P2 | -0.024426633 |
| RP11-358D14 | -0.024426698 |
| RNA5SP285 | -0.024428232 |
| RNA5SP396 | -0.024428339 |
| DDX28 | -0.024431901 |
| RP11-543D10 | -0.024449453 |
| C15orf56 | -0.024451097 |
| RP1-90L6 | -0.024453157 |
| RPA2 | -0.024456199 |
| CT45A5 | -0.024459395 |
| AC000362 | -0.024461397 |
| CYP4F8 | -0.024462361 |
| DYNC2H1 | -0.024463567 |
| MIR6793 | -0.024466728 |
| RP11-14K2 | -0.024467708 |
| SFXN5 | -0.024468516 |
| RP11-35P15 | -0.024469302 |
| SNORD12B | -0.024471017 |
| AC007390 | -0.024472608 |
| LARP1BP2 | -0.024475256 |
| RPL30P6 | -0.024477807 |
| LDB1 | -0.024478489 |
| PNMA3 | -0.024482272 |
| BSX | -0.024485349 |
| GS1-18A18 | -0.024488363 |
| C1orf54 | -0.024488871 |
| RPL12P15 | -0.024489626 |
| ABCA9 | -0.024490294 |
| ARL6 | -0.024491773 |
| HFM1 | -0.024492241 |
| KHDRBS2 | -0.024494405 |
| CTD-2350J17 | -0.024496026 |
| KLHL17 | -0.024498899 |
| RP13-977E11 | -0.024499715 |
| AC079779 | -0.02450011 |
| FOXD3 | -0.024500339 |
| PRSS51 | -0.024504981 |

| | |
|---|---|
| LINC00670 | -0.024508604 |
| AC144634 | -0.024509028 |
| RN7SL108P | -0.024512912 |
| AC015923 | -0.024513765 |
| OTUD7B | -0.024514323 |
| PRAMEF26 | -0.024514845 |
| RP11-667F9 | -0.024516811 |
| IBA57-AS1 | -0.024522026 |
| RNU4-87P | -0.024522033 |
| RP5-1092L12 | -0.024525025 |
| AC013267 | -0.024527172 |
| RNU6-828P | -0.024540617 |
| MTCYBP4 | -0.024545257 |
| RN7SL507P | -0.024548695 |
| RP11-304C16 | -0.024549036 |
| MIR4513 | -0.024549069 |
| SCP2 | -0.024550138 |
| RP11-71J4 | -0.024553999 |
| C3orf58 | -0.024555832 |
| RP11-13E1 | -0.024559837 |
| RP13-57D9 | -0.024560504 |
| LINC01111 | -0.024560907 |
| BNIP3P28 | -0.024561648 |
| TAS2R8 | -0.024563298 |
| NPM1 | -0.024565283 |
| OR52B6 | -0.024566233 |
| FILIP1 | -0.024568353 |
| DUX4L19 | -0.024573368 |
| RP11-123K3 | -0.024577988 |
| RNU6-246P | -0.02458533 |
| RP11-793H13 | -0.024587349 |
| AC017084 | -0.024587618 |
| MIR4708 | -0.024589865 |
| TAF1A-AS1 | -0.024591536 |
| TPT1P9 | -0.024591722 |
| STX3 | -0.024594903 |
| AC002551 | -0.024595272 |
| LRRC63 | -0.024595617 |
| IFNA22P | -0.024597396 |
| DOCK1 | -0.024601545 |
| LINC00449 | -0.024603193 |
| OR4C15 | -0.024604292 |
| DEAR | -0.024613769 |
| RNU1-143P | -0.024615249 |
| MAPK8IP3 | -0.024615332 |

| | |
|---|---|
| SRRM4 | -0.024615828 |
| RAB26 | -0.024616876 |
| TMEM235 | -0.024621062 |
| SCGB1C2 | -0.024627188 |
| LINC01032 | -0.02463154 |
| ROGDI | -0.024641584 |
| RP6-109B7 | -0.024642459 |
| AL139398 | -0.024650056 |
| AL162759 | -0.024656112 |
| AP000781 | -0.02466439 |
| AC024162 | -0.024668382 |
| FASTKD5 | -0.024671223 |
| SLC35E1 | -0.024678051 |
| PDE6D | -0.024679186 |
| RNU6-102P | -0.024683104 |
| RP3-417L20 | -0.024683613 |
| RP11-1080G15 | -0.02469167 |
| RPS6P21 | -0.024692143 |
| AC000099 | -0.024692687 |
| RNU1-82P | -0.024693465 |
| RP11-277B15 | -0.024699963 |
| LLNLR-268E12 | -0.0247015 |
| RP11-54C4 | -0.024704564 |
| ST6GALNAC3 | -0.024708359 |
| RN7SL277P | -0.024710637 |
| RPL19P14 | -0.024713733 |
| LINC00314 | -0.024714699 |
| SCARNA23 | -0.024717595 |
| RP11-121M22 | -0.0247185 |
| MTND4P20 | -0.024721103 |
| LLNLF-18A12 | -0.024722476 |
| RP11-310E22 | -0.024726178 |
| RP13-238N7 | -0.024733656 |
| AC090324 | -0.024735906 |
| PRSS3P4 | -0.024736048 |
| RP11-354A14 | -0.024736342 |
| OR5P2 | -0.024736807 |
| RP11-149P24 | -0.024742679 |
| RP11-154B12 | -0.024746811 |
| RNU2-44P | -0.024747681 |
| GLIDR | -0.024747728 |
| GOLGA8H | -0.024748949 |
| CTD-2507G9 | -0.024750083 |
| AC011933 | -0.024750652 |
| COX7CP1 | -0.024761875 |

| | |
|---|---|
| AC007036 | -0.024763209 |
| MT-TW | -0.02476662 |
| SPINK5 | -0.024767079 |
| TAPBPL | -0.024772995 |
| RNU6-341P | -0.024775786 |
| XXbac-BPG118E17 | -0.024777108 |
| DKFZp313A047 | -0.024779165 |
| RORA-AS2 | -0.024781633 |
| SLC17A2 | -0.024783225 |
| AAAS | -0.024784084 |
| CTD-2561J22 | -0.024792102 |
| RIOK1 | -0.024792828 |
| SDF2L1 | -0.024793275 |
| LRRC61 | -0.024793293 |
| RPS28P8 | -0.024799002 |
| CAMK2N1 | -0.024799545 |
| TRIM49B | -0.024801406 |
| AL513547 | -0.024802876 |
| KCNK17 | -0.02480658 |
| RP11-286H14 | -0.024808196 |
| RP11-84C10 | -0.024808234 |
| SPATA9 | -0.024808922 |
| MEMO1P3 | -0.024811572 |
| RP11-433P17 | -0.024814318 |
| RP4-586O15 | -0.02481663 |
| MIR1471 | -0.024819957 |
| AC073326 | -0.024820936 |
| FAM8A6P | -0.024821864 |
| RP3-331H24 | -0.024822802 |
| RP11-544I20 | -0.024827425 |
| HLA-DQB3 | -0.024829706 |
| OR5M10 | -0.024830162 |
| HMGN2P40 | -0.024836348 |
| PLCB4 | -0.024845054 |
| AL603766 | -0.024847618 |
| RP11-462L8 | -0.024847659 |
| AC010633 | -0.024849706 |
| AGFG1 | -0.024852645 |
| CAPN9 | -0.024852942 |
| AC129778 | -0.024857583 |
| EIF5P2 | -0.024859151 |
| RP11-408A13 | -0.024859715 |
| RP3-365I19 | -0.024866685 |
| RP11-324H9 | -0.024866711 |
| RP11-24D15 | -0.024872177 |

| | |
|---|---|
| TRAV28 | -0.024872933 |
| RP3-400B16 | -0.02487498 |
| AC100828 | -0.024881132 |
| RPL7P54 | -0.024889181 |
| RP11-339B9 | -0.024890959 |
| CMC4 | -0.024893639 |
| KRT8P30 | -0.024894321 |
| RDH11 | -0.024895363 |
| RP11-203L2 | -0.024895474 |
| FOXN1 | -0.024896291 |
| RP11-286N3 | -0.024899139 |
| RNA5SP337 | -0.024903816 |
| RN7SL835P | -0.024910894 |
| RP11-894J14 | -0.024912685 |
| NOXO1 | -0.024913362 |
| C18orf15 | -0.02491906 |
| ABCA3 | -0.024919252 |
| EEF1A1P17 | -0.024923184 |
| TCEB1P32 | -0.024923279 |
| MTND2P2 | -0.024923628 |
| RN7SKP103 | -0.024925018 |
| DDI1 | -0.024927696 |
| DLG5 | -0.024929113 |
| ARL5C | -0.024931313 |
| SKA2P1 | -0.02493185 |
| RPL36P16 | -0.024935117 |
| SLC25A41 | -0.024937212 |
| C14orf39 | -0.024939817 |
| PLA2G4D | -0.024943181 |
| RP11-128I7 | -0.024944313 |
| AC137723 | -0.024944451 |
| SLC7A2 | -0.024946509 |
| RGPD1 | -0.024946575 |
| PRRC1 | -0.024948593 |
| GRAPL | -0.024949498 |
| XRCC3 | -0.02495237 |
| FKBP5 | -0.024952482 |
| SYS1-DBNDD2 | -0.024956761 |
| PFN1P1 | -0.024957693 |
| RBM34 | -0.02496158 |
| FABP5P2 | -0.024963751 |
| GLRX3P2 | -0.024971106 |
| RP11-800A18 | -0.024973695 |
| RNU6ATAC38P | -0.024974107 |
| DDX19B | -0.024974157 |

| | |
|---|---|
| RP11-1094H24 | -0.024977065 |
| RP11-777B9 | -0.024977112 |
| ALOX5AP | -0.024979887 |
| RP11-199F6 | -0.024983253 |
| ADAD1 | -0.024984623 |
| RN7SL79P | -0.024987386 |
| RHPN1-AS1 | -0.024988427 |
| MIR5680 | -0.024988999 |
| SRCAP | -0.024991367 |
| OR13C4 | -0.024993243 |
| RNA5SP238 | -0.02499341 |
| RP11-456A14 | -0.025003058 |
| RN7SKP73 | -0.025004731 |
| RN7SL586P | -0.025005606 |
| CTD-2026A21 | -0.025005817 |
| MRPS35 | -0.025006177 |
| CTC-250P20 | -0.025007032 |
| RNU6-527P | -0.025008053 |
| MIR340 | -0.02502118 |
| CPXM2 | -0.025023889 |
| RNU7-50P | -0.025025858 |
| EEF1A1P42 | -0.025025891 |
| PHF20 | -0.025032187 |
| DPPA2P2 | -0.025037384 |
| FTH1P18 | -0.025045296 |
| ZNF559-ZNF177 | -0.025048314 |
| CCDC90B | -0.025048616 |
| NUTM2D | -0.025049202 |
| RP11-439H13 | -0.02504945 |
| RP11-574E24 | -0.025049723 |
| AJ009632 | -0.025051534 |
| UBQLN4P2 | -0.025054302 |
| SLC6A4 | -0.025062366 |
| NEPNP | -0.025067036 |
| NR4A1 | -0.025069808 |
| MIR548W | -0.025072036 |
| C2orf16 | -0.025072133 |
| DBIP2 | -0.025075736 |
| HMGB3P24 | -0.025077755 |
| RABL2A | -0.025084822 |
| RP11-128B16 | -0.025085152 |
| OR7A19P | -0.025085568 |
| CCNI2 | -0.025085939 |
| AC005307 | -0.025087685 |
| RP11-452L6 | -0.025088618 |

| | |
|---|---|
| ZNF962P | -0.025090968 |
| RP11-476C8 | -0.025093205 |
| BNIP3P35 | -0.025093648 |
| CABS1 | -0.025096094 |
| ADCY7 | -0.025098144 |
| ERICH3-AS1 | -0.025098531 |
| RP11-490H24 | -0.025099343 |
| MIPEPP2 | -0.025100669 |
| TRBV12-1 | -0.025102688 |
| AC106875 | -0.025104403 |
| ATP5LP6 | -0.025107609 |
| TRAPPC2P3 | -0.025109477 |
| RP11-624A4 | -0.025111713 |
| TRMT11 | -0.025111949 |
| RAB17 | -0.025114628 |
| RORC | -0.025114982 |
| SNORA80B | -0.02511732 |
| OR5B1P | -0.025119105 |
| RNU1-105P | -0.025119237 |
| SYNE1-AS1 | -0.025120784 |
| PSMD7P1 | -0.025125616 |
| TRIM64DP | -0.025128848 |
| LINC00526 | -0.02512968 |
| RP3-352A20 | -0.025134314 |
| SAR1P1 | -0.025136642 |
| RNU2-9P | -0.02514078 |
| AL031229 | -0.025141144 |
| RP11-61F5 | -0.025143242 |
| AC004062 | -0.025147781 |
| SERHL2 | -0.0251505 |
| AC091069 | -0.025150972 |
| EIF3S5P1 | -0.025151796 |
| CT83 | -0.025158538 |
| RN7SL12P | -0.025165303 |
| CTD-2312P21 | -0.025165344 |
| RP11-344A7 | -0.025170545 |
| RBBP8P1 | -0.025173649 |
| MIR6816 | -0.025177535 |
| IL36A | -0.025179036 |
| RP11-231N9 | -0.025183593 |
| RP11-11C20 | -0.02518549 |
| RPS18P13 | -0.025187488 |
| AC009955 | -0.025188556 |
| FITM1 | -0.025189648 |
| DPAGT1 | -0.025190948 |

| | |
|---|---|
| RAB1B | -0.025196995 |
| SNPH | -0.025198092 |
| RPL6P5 | -0.025198727 |
| NECAP2 | -0.025208828 |
| ROCK2 | -0.025209609 |
| AC097483 | -0.02520985 |
| RP11-1006G14 | -0.025212326 |
| CRYZ | -0.025213461 |
| C2orf27AP2 | -0.025214704 |
| AC116407 | -0.02521627 |
| RP11-778H2 | -0.025219443 |
| CMYA5 | -0.02522027 |
| MIR3139 | -0.025223404 |
| RPL21P134 | -0.025230304 |
| FBXO18 | -0.025230317 |
| AL512791 | -0.025232706 |
| AL139328 | -0.025234911 |
| IFNNP1 | -0.025238658 |
| AC009236 | -0.025241313 |
| RPS15AP7 | -0.025243308 |
| AVPR2 | -0.025248848 |
| CYP4F2 | -0.025250781 |
| CSPG4P13 | -0.025258441 |
| HNRNPA1P10 | -0.025258988 |
| RNU6-944P | -0.025263834 |
| CPLX3 | -0.025264156 |
| SDCCAG8 | -0.025264533 |
| NBPF10 | -0.025265947 |
| HIGD1AP14 | -0.025266526 |
| DROSHA | -0.025267919 |
| VRK1 | -0.025270003 |
| RN7SKP188 | -0.025277645 |
| RP11-473J6 | -0.025278661 |
| CTD-2013N24 | -0.025281091 |
| PACS1 | -0.025283938 |
| HSPD1P18 | -0.025287014 |
| KDM6B | -0.025287622 |
| RPL32P34 | -0.02528858 |
| RP11-683O4 | -0.025290065 |
| SNX2P2 | -0.02529333 |
| CDK2AP2 | -0.025296757 |
| HSPE1P3 | -0.025296882 |
| RNU6-401P | -0.025298424 |
| GPN2 | -0.025300553 |
| RP11-814M22 | -0.025303678 |

| | |
|---|---|
| PHBP3 | -0.025305093 |
| UBA52P9 | -0.025306813 |
| RP5-1018K9 | -0.025307453 |
| LINC01447 | -0.025310534 |
| PABPC1P5 | -0.02531545 |
| IGLVVII-41-1 | -0.025318656 |
| VENTXP4 | -0.025321215 |
| RN7SL42P | -0.02532301 |
| AL391380 | -0.025326224 |
| TUBA3FP | -0.0253268 |
| SYNJ2-IT1 | -0.025326974 |
| SUSD3 | -0.025327735 |
| EDEM2 | -0.025328256 |
| HNF4A-AS1 | -0.025328803 |
| RP11-118F19 | -0.025330012 |
| CERS3-AS1 | -0.025332614 |
| CCDC58P2 | -0.025333961 |
| CICP23 | -0.025334176 |
| MIR4532 | -0.025339045 |
| RP11-779P15 | -0.0253392 |
| CDK20 | -0.025341578 |
| CTC-480C2 | -0.025343622 |
| TPT1P3 | -0.025344551 |
| KCNC4-AS1 | -0.025347207 |
| RP11-443I9 | -0.025347328 |
| RP11-694D5 | -0.025348129 |
| MRGPRD | -0.025350289 |
| MTND4P12 | -0.025352369 |
| HIST1H1PS1 | -0.025353127 |
| WDFY3-AS2 | -0.025353653 |
| DRD3 | -0.025354036 |
| REM2 | -0.025356518 |
| RP11-1024P17 | -0.025357523 |
| RNU5E-7P | -0.025358941 |
| RNU6-780P | -0.025362806 |
| CACYBP | -0.025365709 |
| GPBAR1 | -0.025374249 |
| SENP3-EIF4A1 | -0.025374422 |
| RNU6-1097P | -0.02537785 |
| RP11-756K15 | -0.025379765 |
| SNX4 | -0.025385394 |
| RNU6-470P | -0.025391691 |
| RP11-549B18 | -0.02539215 |
| PCBP1 | -0.025392222 |
| AC016723 | -0.025397011 |

| | |
|---|---|
| RP11-19G24 | -0.025400088 |
| RP11-160C18 | -0.025401585 |
| MYO5B | -0.025402883 |
| AC096574 | -0.025404636 |
| TM9SF3 | -0.025406753 |
| EEF1A1P13 | -0.025408985 |
| PPP1R17 | -0.02540899 |
| PRR7-AS1 | -0.02540903 |
| AC010608 | -0.02541102 |
| RN7SL44P | -0.025411638 |
| CKBP1 | -0.025412004 |
| OR11G2 | -0.025412498 |
| KIAA0825 | -0.025414418 |
| PTX4 | -0.025415368 |
| PLRG1 | -0.025417097 |
| ZNF92P2 | -0.025423269 |
| DOCK7 | -0.025424688 |
| POTEE | -0.025426187 |
| GOT1L1 | -0.025426524 |
| UBQLNL | -0.025438777 |
| SERP2 | -0.025442494 |
| TMEM26-AS1 | -0.025446469 |
| ZFYVE9 | -0.025450941 |
| RP11-740N7 | -0.025451626 |
| FERP1 | -0.025453978 |
| RNU6-1059P | -0.02545471 |
| KDM4E | -0.025456398 |
| UBE3D | -0.02546082 |
| RP11-313P13 | -0.025464867 |
| RP13-317D12 | -0.025465402 |
| SPDYE5 | -0.025469025 |
| RP11-618P13 | -0.025469115 |
| CTC-398G3 | -0.025474254 |
| RNA5SP279 | -0.025477061 |
| FAM27L | -0.025477574 |
| GALC | -0.025480356 |
| AC007773 | -0.025482942 |
| BOLA3P2 | -0.025485456 |
| ZNF296 | -0.025496133 |
| PHKBP1 | -0.02549782 |
| HUS1B | -0.025502015 |
| RN7SKP24 | -0.025503807 |
| ATP4B | -0.025506985 |
| NCEH1 | -0.025511222 |
| MTFR1 | -0.025518032 |

| | |
|---|---|
| TCF12 | -0.025520209 |
| ZNF436-AS1 | -0.025520637 |
| GOLGA6L22 | -0.025521901 |
| LRP2 | -0.025522982 |
| CTA-268H5 | -0.025530322 |
| OR5M6P | -0.025532633 |
| RP11-687E1 | -0.025534594 |
| RP11-305L7 | -0.025535093 |
| BMS1P16 | -0.025535288 |
| RN7SL612P | -0.025535559 |
| AC093084 | -0.025542501 |
| RP11-337C18 | -0.025544712 |
| ANKRD22 | -0.025550927 |
| TP53TG3C | -0.025555296 |
| RPS23P10 | -0.025556398 |
| SNORD116-23 | -0.025556547 |
| FATE1 | -0.02555787 |
| CTB-52I2 | -0.025559262 |
| AC069285 | -0.025561817 |
| ECEL1P1 | -0.025564137 |
| U2AF1L4 | -0.025565383 |
| MNX1-AS1 | -0.025566109 |
| SPRTN | -0.025567392 |
| HHATL-AS1 | -0.025567973 |
| RP11-717A5 | -0.025568474 |
| BX682235 | -0.025570065 |
| NEURL1-AS1 | -0.025570282 |
| MIR548AR | -0.025572091 |
| RP11-155G15 | -0.025573293 |
| FAM169A | -0.025575895 |
| SAMD7 | -0.02557683 |
| RP11-608O8 | -0.025580494 |
| RP11-59J16 | -0.025581504 |
| RP3-377O6 | -0.025585144 |
| HMX3 | -0.025586297 |
| TPI1P2 | -0.025587505 |
| RP11-139H14 | -0.025587781 |
| RN7SL523P | -0.025588945 |
| AC091856 | -0.025590163 |
| LINC00841 | -0.025594939 |
| GSTP1P1 | -0.025595247 |
| COL6A6 | -0.025595785 |
| RP11-814P5 | -0.025598018 |
| AC092535 | -0.025608045 |
| ADIPOR2 | -0.025610948 |

| | |
|---|---|
| CYSRT1 | -0.025611971 |
| ATXN7L3 | -0.02561426 |
| HSPE1P2 | -0.025616554 |
| MIR548N | -0.025624292 |
| ACACA | -0.025630818 |
| CTD-2050I18 | -0.025630887 |
| CTD-2335O3 | -0.025636878 |
| RP1-90J20 | -0.025639174 |
| AL022578 | -0.025644291 |
| POLR1D | -0.025648512 |
| OR6C69P | -0.025649196 |
| SPDYA | -0.025651843 |
| RRP15 | -0.025652177 |
| RP5-994D16 | -0.025656513 |
| HSD17B11 | -0.02566215 |
| TRAF3IP2-AS1 | -0.025662708 |
| AC011477 | -0.025663111 |
| LINC01478 | -0.025666238 |
| RP11-385E5 | -0.025672274 |
| MIR5010 | -0.02567411 |
| AC019109 | -0.025675265 |
| AL357874 | -0.025678258 |
| SORCS1 | -0.025678688 |
| RP13-130D24 | -0.025679047 |
| OSTM1 | -0.025681965 |
| ATP5H | -0.025683063 |
| RN7SL57P | -0.025685512 |
| AC073934 | -0.02568599 |
| NIPSNAP1 | -0.025686095 |
| TEX12 | -0.025687487 |
| CASC11 | -0.025687998 |
| RNU7-157P | -0.02569235 |
| MERTK | -0.025694003 |
| MTND6P3 | -0.025694894 |
| CBX5P1 | -0.025695852 |
| RP11-16P20 | -0.025696437 |
| RNU6-672P | -0.025705271 |
| RP11-600L4 | -0.025713215 |
| ZNF355P | -0.025715801 |
| WDR26 | -0.025717483 |
| RP11-661G16 | -0.025721272 |
| KRT19P1 | -0.025721704 |
| CALCOCO1 | -0.025722674 |
| PCGF2 | -0.025723343 |
| LINC01125 | -0.025724999 |

| | |
|---|---|
| RBMY1J | -0.025726643 |
| AC069082 | -0.025726643 |
| ZSCAN22 | -0.025726934 |
| NIPA2P2 | -0.02573607 |
| FLJ42969 | -0.025743821 |
| RP11-70F11 | -0.025749625 |
| EIF4E2P1 | -0.025751744 |
| DAP3 | -0.02575607 |
| ANKRD20A7P | -0.025756606 |
| NOB1 | -0.025758731 |
| CTD-2582D11 | -0.025764038 |
| RP11-27P7 | -0.025764721 |
| HCG11 | -0.025773746 |
| LINC00176 | -0.025777113 |
| DPY19L3 | -0.025779848 |
| LOXL1-AS1 | -0.025781802 |
| GAPDHP55 | -0.025783549 |
| FLJ35934 | -0.025784029 |
| USP9YP1 | -0.025785734 |
| AC116035 | -0.025795013 |
| SNRPEP10 | -0.025795535 |
| AC010139 | -0.025802481 |
| RP11-193P11 | -0.025807096 |
| FOXD4L5 | -0.0258129 |
| RP3-323A16 | -0.025813294 |
| OR2T32P | -0.02582397 |
| KF511026 | -0.025828272 |
| MTND6P1 | -0.025828889 |
| CLUAP1 | -0.025832855 |
| RP11-31K23 | -0.025833198 |
| RP11-206F17 | -0.025833358 |
| SLC29A4P2 | -0.025834546 |
| MIR5700 | -0.0258382 |
| PROKR2 | -0.025838969 |
| RP11-654F3 | -0.025839106 |
| OR51A4 | -0.025839116 |
| TTLL8 | -0.025844935 |
| RP11-108N13 | -0.025848769 |
| FMNL3 | -0.025852354 |
| RNF115 | -0.025854214 |
| RN7SL832P | -0.025858624 |
| RP11-22A3 | -0.025858969 |
| AC009276 | -0.025860073 |
| NANOGP4 | -0.025862019 |
| OR5AU1 | -0.025862121 |

| | |
|---|---|
| RN7SL683P | -0.025863327 |
| NCOR1P2 | -0.02586455 |
| STAC3 | -0.025864819 |
| CAPN10 | -0.02586575 |
| AC022681 | -0.025867098 |
| EFCAB3 | -0.025869881 |
| OR51F4P | -0.025874715 |
| BTF3L4P2 | -0.025878866 |
| AC005757 | -0.02588524 |
| RNF8 | -0.025886271 |
| RP11-701I24 | -0.025887713 |
| TRMT61B | -0.025905919 |
| RP11-655H13 | -0.025906957 |
| OR52P1P | -0.025909319 |
| ZBTB38 | -0.0259099 |
| RP11-191H23 | -0.025910271 |
| CRY2 | -0.025912021 |
| OR6W1P | -0.025916567 |
| RP11-248J18 | -0.02591697 |
| APC2 | -0.025918161 |
| RNU6-272P | -0.02591819 |
| RP11-245C17 | -0.025921065 |
| AIF1L | -0.025921529 |
| MAPK12 | -0.02592218 |
| RP11-266K22 | -0.025923288 |
| RP11-473C19 | -0.025929723 |
| ZCCHC5 | -0.025933102 |
| RP11-492O8 | -0.025933598 |
| KRTAP5-7 | -0.025935269 |
| RNU7-152P | -0.025936963 |
| LIN7B | -0.025940823 |
| NUTF2P4 | -0.025941918 |
| RP11-187C18 | -0.025948089 |
| RP11-1145L24 | -0.025951216 |
| WTAP | -0.025959726 |
| TMEM167B | -0.025962889 |
| RPL5P29 | -0.025971333 |
| FLVCR2 | -0.025972489 |
| ERVFRD-1 | -0.02597351 |
| MRPL1 | -0.025979987 |
| AL389915 | -0.025982081 |
| RP4-704D21 | -0.025986156 |
| PPP1R26 | -0.025996267 |
| AC068312 | -0.025996992 |
| RP11-787P24 | -0.026000999 |

| | |
|---|---|
| TUFMP1 | -0.026002814 |
| WHAMMP2 | -0.026010258 |
| AL137229 | -0.026011298 |
| OR5V1 | -0.026018707 |
| EIF4E1B | -0.026023863 |
| RPL21P4 | -0.026023926 |
| SMARCD2 | -0.026023952 |
| ZNF574 | -0.026024975 |
| RP11-439H8 | -0.02602696 |
| RP11-626K17 | -0.026031926 |
| USP9YP7 | -0.026032184 |
| RNU4-38P | -0.026032543 |
| GPR27 | -0.026033348 |
| AL590762 | -0.026040213 |
| CTD-3145H4 | -0.026044655 |
| RELA | -0.026046744 |
| SSR1P2 | -0.026049674 |
| RP11-367E12 | -0.026057359 |
| TOR2A | -0.026057832 |
| RP11-233G1 | -0.026059005 |
| RP11-578C11 | -0.026059602 |
| snoU83B | -0.026064924 |
| C8orf87 | -0.026065821 |
| AC016907 | -0.026069399 |
| TFR2 | -0.026072829 |
| MIR320C1 | -0.026075769 |
| IL1RAPL1 | -0.026077982 |
| RP11-791O21 | -0.026079583 |
| NUDT19P3 | -0.026082524 |
| RNU6-1127P | -0.026082624 |
| CITF22-1A6 | -0.026085945 |
| RPL36AL | -0.026093388 |
| LL0XNC01-73E8 | -0.026095695 |
| MIR6769A | -0.02609622 |
| SAMD15 | -0.026098225 |
| MYOC | -0.026101357 |
| TBPL2 | -0.026101898 |
| RP11-269M20 | -0.026108367 |
| GML | -0.026108879 |
| AC004603 | -0.026110874 |
| CST4 | -0.026111031 |
| DUSP16 | -0.026114151 |
| RP11-46O21 | -0.026116227 |
| RP11-321A17 | -0.026118951 |
| RP11-380N8 | -0.026120439 |

| | |
|---|---|
| RP11-479O16 | -0.026122885 |
| OR5H2 | -0.026128359 |
| EIF2B5 | -0.026129296 |
| COX16 | -0.026130375 |
| IPO11 | -0.026131361 |
| CHMP1AP1 | -0.026131579 |
| RP4-671G15 | -0.026135648 |
| MIR3174 | -0.026140399 |
| OPN1LW | -0.026143313 |
| SLC8A3 | -0.02614578 |
| RP5-1164C1 | -0.026146541 |
| AL512505 | -0.026146896 |
| AL358354 | -0.026149503 |
| ZNF773 | -0.026153186 |
| CTC-505O3 | -0.02615471 |
| XXbac-BPGBPG24O18 | -0.02615712 |
| RP11-801G16 | -0.026157667 |
| GLULP3 | -0.026166686 |
| RNU6-836P | -0.026169447 |
| RP11-588F10 | -0.026172511 |
| MS4A8 | -0.026179427 |
| TMEM183A | -0.026184443 |
| RNU6-613P | -0.026186636 |
| CSRP3 | -0.026195115 |
| SLC9B1P3 | -0.02619631 |
| RP11-335O4 | -0.026197563 |
| RP11-424M24 | -0.026200254 |
| RNU7-133P | -0.026200629 |
| TRMT2B-AS1 | -0.026202611 |
| OR4A18P | -0.026203202 |
| AP000580 | -0.026203441 |
| RCN2 | -0.026205132 |
| TWISTNB | -0.026208834 |
| RP11-569A2 | -0.026210955 |
| OR2Z1 | -0.026212222 |
| RP11-30H9 | -0.026213803 |
| FAM138D | -0.026215113 |
| RP11-312J18 | -0.026217077 |
| PRB4 | -0.026217605 |
| MIR6763 | -0.026220856 |
| RP11-167P22 | -0.026221937 |
| FAM206BP | -0.026223121 |
| ZNRD1 | -0.026223905 |
| MTND5P22 | -0.026226315 |
| OR5BN2P | -0.026226315 |

| | |
|---|---|
| AL596268 | -0.026226315 |
| CCNH | -0.026231831 |
| RP5-991O23 | -0.026232629 |
| AC007387 | -0.026235107 |
| AC234031 | -0.026235628 |
| RN7SKP30 | -0.026239779 |
| NANOGNBP2 | -0.026244356 |
| RN7SL774P | -0.026249935 |
| RP11-114B7 | -0.026250929 |
| AC109589 | -0.026253194 |
| ZNF385D-AS2 | -0.026255481 |
| LINC01387 | -0.026264255 |
| TPD52L1 | -0.026264595 |
| PBOV1 | -0.0262664 |
| HIST1H1D | -0.026266641 |
| WBP1LP12 | -0.026272567 |
| GRK7 | -0.026277841 |
| RP5-1053E7 | -0.026278186 |
| GRAP | -0.026283517 |
| RP11-843A23 | -0.026288433 |
| RP11-576D8 | -0.026289114 |
| POM121L1P | -0.026291316 |
| RN7SKP151 | -0.026291642 |
| USP29 | -0.02629219 |
| HIST1H4K | -0.026294519 |
| CSNK2A2 | -0.026296001 |
| SLC23A2 | -0.026302811 |
| MKRN3 | -0.026304315 |
| PSEN1 | -0.026304504 |
| LINC00587 | -0.02630702 |
| U5 | -0.026311181 |
| DYDC1 | -0.026313397 |
| RP11-310A13 | -0.026320767 |
| IMPG1 | -0.026320941 |
| NDFIP2-AS1 | -0.02633042 |
| RP11-474D14 | -0.026330958 |
| HIPK4 | -0.026332073 |
| CLEC4M | -0.026339544 |
| ATP5JP1 | -0.02633966 |
| KCNT1 | -0.026340043 |
| TMEM78 | -0.026346791 |
| RP11-543F8 | -0.026348377 |
| RP11-700N1 | -0.026349026 |
| MTND3P6 | -0.026352332 |
| RTP3 | -0.02635428 |

| | |
|---|---|
| KB-1592A4 | -0.026354923 |
| HNRNPDLP3 | -0.026359786 |
| OR4A48P | -0.026360655 |
| FAM27E3 | -0.026360826 |
| RP11-434J24 | -0.026361358 |
| OR7E154P | -0.02636278 |
| COPG1 | -0.026363725 |
| LINC00630 | -0.026364562 |
| RP11-264J4 | -0.026367763 |
| RP11-104D3 | -0.026368786 |
| AC013467 | -0.02636944 |
| DDTP1 | -0.026375171 |
| OR7E129P | -0.026378989 |
| ADAM22 | -0.026383707 |
| RNU6-388P | -0.026383779 |
| CISTR | -0.026384394 |
| ARHGAP26-IT1 | -0.026387208 |
| AC095061 | -0.02638732 |
| MED28P6 | -0.026390902 |
| TPMTP2 | -0.026393709 |
| RP11-554I8 | -0.026395698 |
| GMPR | -0.026398257 |
| OR10R2 | -0.026400552 |
| AC090955 | -0.026400611 |
| RP11-193M21 | -0.026405468 |
| RP11-38J22 | -0.026408676 |
| TCEB1P12 | -0.026409048 |
| OR7E10P | -0.026421405 |
| NDUFA5P9 | -0.026429002 |
| KRT18P61 | -0.026432683 |
| TYMSOS | -0.02643463 |
| KRTAP19-11P | -0.026447891 |
| STK24P1 | -0.026447979 |
| CTD-2530N21 | -0.026449907 |
| NOS1AP | -0.026450333 |
| TLR1 | -0.026453013 |
| PRMT7 | -0.026455059 |
| AC064848 | -0.026455528 |
| HNRNPA1P15 | -0.026456181 |
| RNU6-1249P | -0.026456757 |
| HSFX2 | -0.026458725 |
| RP11-728K20 | -0.026460377 |
| AF186192 | -0.026462874 |
| RNU6-593P | -0.026467596 |
| CTD-2540L5 | -0.026479356 |

| | |
|---|---|
| RN7SL652P | -0.026481059 |
| DHRS2 | -0.026481684 |
| PDS5B | -0.026485461 |
| HMGN2P17 | -0.026485498 |
| ADRB1 | -0.026486854 |
| IFIT1B | -0.026494695 |
| RP11-497D6 | -0.026498292 |
| RNA5SP492 | -0.026501832 |
| MAMDC2-AS1 | -0.026502671 |
| SNRPEP9 | -0.026505406 |
| RP11-642M2 | -0.026509879 |
| RN7SL703P | -0.026510074 |
| TJAP1 | -0.02651082 |
| RN7SL756P | -0.02651438 |
| RP11-421N8 | -0.026519598 |
| CCZ1 | -0.026521741 |
| LINC00866 | -0.026527789 |
| NFU1P1 | -0.026528861 |
| RP5-837I24 | -0.02652907 |
| DLST | -0.02653177 |
| LTV1P1 | -0.026533884 |
| HMGB1P37 | -0.026535957 |
| TRIM62 | -0.026538226 |
| AL158089 | -0.026540927 |
| RP11-62N21 | -0.026545066 |
| MIR3133 | -0.026548601 |
| RP4-581O6 | -0.02655134 |
| LINC00974 | -0.026551862 |
| RNU6-1121P | -0.026552422 |
| CTD-3179P9 | -0.026552851 |
| RP11-811I15 | -0.026557879 |
| RP11-341N2 | -0.026558178 |
| NCAN | -0.026558607 |
| RSL24D1P1 | -0.026564653 |
| LL0YNC03-29C1 | -0.026574809 |
| SNORA42 | -0.026577575 |
| PTH2 | -0.026577679 |
| CDCA4P4 | -0.02658012 |
| HSP90AB7P | -0.026580368 |
| MIR504 | -0.026580606 |
| PLCB1-IT1 | -0.026581017 |
| RP5-903E17 | -0.026581663 |
| CXorf67 | -0.026582134 |
| RP11-397P13 | -0.026582868 |
| AC002314 | -0.02658771 |

| | |
|---|---|
| PRAMEF25 | -0.02658816 |
| RNA5SP197 | -0.026593108 |
| RP11-38M15 | -0.026593141 |
| HIGD1AP5 | -0.026593286 |
| SIGLEC17P | -0.026596164 |
| NDUFA5P3 | -0.026597121 |
| EFCAB14P1 | -0.026599856 |
| TJP3 | -0.026602991 |
| AL158151 | -0.026605017 |
| ZNRF2P1 | -0.026611447 |
| AC104434 | -0.026612118 |
| RNU6-309P | -0.026612118 |
| RNU6-978P | -0.026612118 |
| TVP23BP2 | -0.026612146 |
| AC068643 | -0.026613058 |
| CHCHD2P11 | -0.026614161 |
| RN7SL231P | -0.02661723 |
| CTC-366B18 | -0.026619569 |
| PXMP4 | -0.026627271 |
| CTB-47B11 | -0.026629922 |
| RP11-885L14 | -0.026630945 |
| INTS4P2 | -0.026632854 |
| TSEN15P2 | -0.026632927 |
| RP11-109G23 | -0.026634164 |
| ABCB4 | -0.026634342 |
| RN7SL520P | -0.026637105 |
| ZNF687 | -0.02663797 |
| RP13-221M14 | -0.026638146 |
| PAPD7 | -0.026641133 |
| DNAJB7 | -0.026642127 |
| HNRNPA1P45 | -0.026649305 |
| EFCAB2 | -0.026654361 |
| PRUNE | -0.026660426 |
| STT3A-AS1 | -0.026662501 |
| IPO9 | -0.026667329 |
| CTC-508F8 | -0.026669095 |
| RP11-523H20 | -0.026669655 |
| TMIGD1 | -0.026674314 |
| USP53 | -0.026683237 |
| AC135999 | -0.026683475 |
| AP000719 | -0.026686365 |
| RAD17P2 | -0.026687228 |
| COPA | -0.026689265 |
| RNU6-917P | -0.026692428 |
| IFNB1 | -0.0266976 |

| | |
|---|---|
| USP12PY | -0.026699027 |
| FEM1C | -0.026700037 |
| MIR33B | -0.026704181 |
| EIF4EP5 | -0.026709939 |
| MIR6780A | -0.026714784 |
| MAGEC2 | -0.026718214 |
| VIMP | -0.026721415 |
| ZBTB4 | -0.026725266 |
| RP1-169I5 | -0.026725467 |
| CUL1P1 | -0.026726925 |
| CPSF1P1 | -0.026727661 |
| TGOLN2 | -0.026728597 |
| RP11-55K13 | -0.026730576 |
| RP11-31H5 | -0.026730978 |
| ZRANB3 | -0.026731257 |
| SRGNP1 | -0.026732303 |
| HEATR1 | -0.02673682 |
| NDE1P2 | -0.02673704 |
| CYP3A7 | -0.026740227 |
| NPIPA1 | -0.026745007 |
| SNORD32B | -0.026754765 |
| RNU6-1254P | -0.026755307 |
| DCAF13P1 | -0.026756926 |
| YRDCP3 | -0.026757375 |
| RP11-24M17 | -0.026765378 |
| CTD-3037G24 | -0.026765782 |
| ODF2 | -0.026765994 |
| NPM1P33 | -0.026772712 |
| MEF2AP1 | -0.026775715 |
| CYP4A11 | -0.026775966 |
| RSL24D1P3 | -0.026777898 |
| AC008171 | -0.026778046 |
| TRPC7 | -0.026780966 |
| AC118282 | -0.026786428 |
| AP003550 | -0.02678787 |
| RP11-371M22 | -0.026789792 |
| HIST1H2BJ | -0.026790516 |
| LINC01584 | -0.026792164 |
| TBC1D8 | -0.0267958 |
| TGFBRAP1 | -0.026796209 |
| KIAA0040 | -0.026799037 |
| STX2 | -0.026801158 |
| RNU2-47P | -0.02680292 |
| LA16c-429E7 | -0.026803742 |
| AC018512 | -0.026804201 |

| | |
|---|---|
| AC005037 | -0.026804579 |
| KRTAP10-10 | -0.026805128 |
| AC004447 | -0.026806982 |
| TMTC1 | -0.026807596 |
| MRPS27 | -0.026807633 |
| HMGN2P36 | -0.026808763 |
| RP11-33I11 | -0.026813426 |
| RPN2 | -0.026816694 |
| ADPRM | -0.026816954 |
| HMGN2P25 | -0.026817086 |
| AL353783 | -0.026819564 |
| ASXL1 | -0.02682395 |
| AC005522 | -0.026828388 |
| RP11-315A16 | -0.026830678 |
| GTF2H4 | -0.026834667 |
| RP3-364H10 | -0.026838832 |
| FXN | -0.026840434 |
| IL1A | -0.026841599 |
| MIOX | -0.026841905 |
| RN7SKP135 | -0.026843299 |
| XXbac-B135H6 | -0.026844267 |
| LINC01399 | -0.02684802 |
| AC093843 | -0.026853201 |
| DPPA3P2 | -0.026854827 |
| ZNF215 | -0.026861375 |
| RP11-539E17 | -0.026861947 |
| RP11-624D20 | -0.026863901 |
| RP11-361M10 | -0.026864819 |
| RP11-121A14 | -0.026865026 |
| H2BFS | -0.026865373 |
| RP11-454C18 | -0.026865854 |
| AC010319 | -0.026867405 |
| RP11-135A24 | -0.026868889 |
| RP11-123N4 | -0.026872675 |
| RP11-313J2 | -0.026875457 |
| LAMTOR5P1 | -0.026880361 |
| RP11-397G17 | -0.026886355 |
| RNU6-1017P | -0.026887216 |
| OGFRL1 | -0.026888083 |
| EWSAT1 | -0.026890725 |
| C22orf24 | -0.026897881 |
| CRCP | -0.02690613 |
| RP3-333B15 | -0.026907627 |
| AC020915 | -0.026911875 |
| SMU1 | -0.026914242 |

| | |
|---|---|
| RP4-811M8 | -0.026919081 |
| RNU6-1091P | -0.026921521 |
| OR1F2P | -0.02692463 |
| AC018799 | -0.026927879 |
| OR7A3P | -0.026928411 |
| RP11-415G4 | -0.026932919 |
| NSL1 | -0.026933555 |
| TUBA3D | -0.026936662 |
| ATPAF2 | -0.026941276 |
| ITGAM | -0.026942616 |
| PYCR2 | -0.026949183 |
| OR2L5 | -0.026949243 |
| OR7L1P | -0.026951299 |
| OR4A40P | -0.02695184 |
| CTB-139P11 | -0.026953111 |
| SERBP1P2 | -0.026956547 |
| RP11-755J8 | -0.026957201 |
| RP11-398M15 | -0.026959533 |
| CBR3-AS1 | -0.026962747 |
| RP11-241J12 | -0.02696391 |
| AC000354 | -0.026964193 |
| MT1CP | -0.026964587 |
| RNU6-169P | -0.026967051 |
| OR4T1P | -0.026968184 |
| AP006621 | -0.026979901 |
| ABHD12 | -0.026984738 |
| RP11-15A1 | -0.026984962 |
| CYCSP4 | -0.026988719 |
| RN7SL619P | -0.026989306 |
| RP11-756H6 | -0.026991005 |
| BIRC5 | -0.026991548 |
| TXNL1 | -0.026991715 |
| RNA5SP347 | -0.026997921 |
| CELA3B | -0.026998926 |
| DDX39AP1 | -0.027000412 |
| NFRKB | -0.027000813 |
| ZNF790 | -0.027003586 |
| TTC4 | -0.02700513 |
| RN7SL785P | -0.027007071 |
| OR10T1P | -0.02700723 |
| GABRR3 | -0.027010256 |
| KRT73 | -0.027016815 |
| C8orf37-AS1 | -0.027027006 |
| NCBP2 | -0.027028666 |
| PNPLA2 | -0.027029019 |

| | |
|---|---|
| RP11-92K15 | -0.027029559 |
| AC143336 | -0.027032432 |
| PLGLB2 | -0.027033159 |
| ARHGAP8 | -0.027035531 |
| POLN | -0.027037001 |
| CLUHP4 | -0.027037902 |
| RPL38P3 | -0.027039651 |
| PAIP2B | -0.027041131 |
| P2RX3 | -0.027048057 |
| CDKN2AIPNL | -0.027050025 |
| CDKN2B-AS_3 | -0.027053083 |
| ARHGAP21 | -0.027062211 |
| RP11-297L17 | -0.027062388 |
| MAP3K7 | -0.027062881 |
| RP11-444D3 | -0.027064265 |
| RNU7-155P | -0.027065313 |
| TMEM135 | -0.027065988 |
| RN7SL302P | -0.027079142 |
| RP11-292F9 | -0.027079713 |
| GS1-124K5 | -0.027080743 |
| HCG4B | -0.027085991 |
| FLJ38576 | -0.027087768 |
| PRRC2A | -0.027090529 |
| RP11-65B23 | -0.027090624 |
| CST2P1 | -0.027091452 |
| CDK2 | -0.027094157 |
| RP11-464I4 | -0.027096597 |
| MFSD9 | -0.027098269 |
| AC023274 | -0.02710097 |
| ABCB7 | -0.027108372 |
| RPL5P35 | -0.027114049 |
| RP11-688H10 | -0.027121811 |
| RP1-167A14 | -0.027122921 |
| AC076972 | -0.027127824 |
| GOLGA8UP | -0.027129754 |
| RNU6-315P | -0.027129912 |
| CLRN1 | -0.027132324 |
| RNU6-934P | -0.027133088 |
| AC106873 | -0.027134613 |
| LY6G6F | -0.027134627 |
| RP3-453I5 | -0.02713576 |
| RPL7AP25 | -0.027136123 |
| CLIC4P2 | -0.027136764 |
| STS | -0.027137564 |
| AL121952 | -0.027138994 |

| | |
|---|---|
| HMGB1P38 | -0.027142134 |
| PCDH20 | -0.027148396 |
| IQCF6 | -0.027149079 |
| FIGF | -0.027149592 |
| MTND5P17 | -0.027149845 |
| STSP1 | -0.027153401 |
| AC093515 | -0.027154036 |
| MTND2P32 | -0.027155335 |
| MIR3689E | -0.027159243 |
| WI2-88277B6 | -0.02716251 |
| OR5AQ1P | -0.027171119 |
| RNU6-124P | -0.027171639 |
| PSMA2P2 | -0.027172857 |
| IRF2BPL | -0.027173937 |
| ABCA8 | -0.027174254 |
| RP11-506O24 | -0.027185383 |
| SGOL1-AS1 | -0.027186463 |
| RP4-687K1 | -0.027187328 |
| RP11-307A17 | -0.027189327 |
| ZNF29P | -0.027190164 |
| RP13-259N13 | -0.027208415 |
| SLC13A5 | -0.027209935 |
| FKBP3 | -0.027212376 |
| KCNB1 | -0.027213074 |
| TUBB2BP1 | -0.027213716 |
| PPIF | -0.027216544 |
| ZNF324B | -0.027220116 |
| LINC00884 | -0.027220339 |
| FUK | -0.027227577 |
| MEMO1P4 | -0.027229262 |
| RP11-306O13 | -0.027235858 |
| AAK1 | -0.027236731 |
| RP11-367J11 | -0.027236842 |
| OR52E4 | -0.027245945 |
| IGLVI-68 | -0.027247394 |
| SNORD115-24 | -0.027251077 |
| RP1-102E24 | -0.02725128 |
| TRAPPC2P2 | -0.027268597 |
| GPR37L1 | -0.027268602 |
| RP11-317J9 | -0.027271333 |
| ZDBF2 | -0.027276342 |
| CAPN8 | -0.027278667 |
| AC117422 | -0.027279571 |
| TAX1BP1 | -0.027281402 |
| FLJ40194 | -0.027291769 |

| | |
|---|---|
| OR52S1P | -0.027296931 |
| AJAP1 | -0.027299822 |
| CAMTA1-IT1 | -0.027300254 |
| NAMA_2 | -0.027303625 |
| MIR8081 | -0.027305273 |
| CRYM-AS1 | -0.027317246 |
| EFCAB5 | -0.027326555 |
| GBF1 | -0.027329293 |
| RP11-152L20 | -0.027339948 |
| DTX3 | -0.027345089 |
| RNA5SP468 | -0.027347605 |
| SEPP1 | -0.027349499 |
| RNA5SP286 | -0.027350418 |
| RIMS1 | -0.027351605 |
| RBKS | -0.027352792 |
| RP11-552E4 | -0.027353083 |
| RPS7P8 | -0.027353866 |
| TRAJ52 | -0.027354776 |
| MAMDC4 | -0.027360736 |
| LINC01052 | -0.027363078 |
| AC012445 | -0.027363346 |
| UTRN | -0.027364476 |
| AGPAT6 | -0.027368997 |
| TTTY12 | -0.027369418 |
| RP4-736L20 | -0.027369699 |
| RP11-338C15 | -0.027369946 |
| RP11-421N10 | -0.027371868 |
| NMNAT2 | -0.027377574 |
| LINC00184 | -0.027386362 |
| RNU6-419P | -0.027386775 |
| OR4A42P | -0.027388356 |
| RNU6-965P | -0.027393811 |
| POTEH-AS1 | -0.027398527 |
| TACR2 | -0.027403026 |
| RPL7AP4 | -0.027408048 |
| RNA5SP502 | -0.027408173 |
| DYDC2 | -0.027408252 |
| RNU6-749P | -0.027408646 |
| RP11-88H1 | -0.027409666 |
| CTD-3229J4 | -0.027417013 |
| RPL23AP2 | -0.027417762 |
| AC005017 | -0.02741919 |
| CDC45 | -0.02742219 |
| FKSG59 | -0.027423122 |
| AP003307 | -0.027424816 |

| | |
|---|---|
| AP000870 | -0.027424916 |
| UBL7 | -0.027426562 |
| ATG7 | -0.027429446 |
| RPL17P51 | -0.027430566 |
| SFT2D2 | -0.02743834 |
| MIR4491 | -0.027439023 |
| SLC25A1P3 | -0.02744071 |
| KRTAP9-1 | -0.027441096 |
| FAM222B | -0.027443028 |
| RP3-463P15 | -0.027447053 |
| CTD-2193G5 | -0.027448362 |
| ETV3L | -0.027452071 |
| LMNTD2 | -0.027454843 |
| OR2T10 | -0.027454947 |
| RGPD4-AS1 | -0.027461167 |
| RP11-320N21 | -0.027466352 |
| TTYH2 | -0.027468164 |
| IZUMO4 | -0.027468387 |
| MYL6P1 | -0.027469193 |
| UBBP3 | -0.027470804 |
| RNA5SP208 | -0.027474471 |
| ZFHX2 | -0.0274781 |
| RP11-275E15 | -0.027478166 |
| RP11-8J9 | -0.027478309 |
| CTD-2323K18 | -0.027481207 |
| CTD-2384A14 | -0.027484325 |
| C20orf78 | -0.027485301 |
| EFTUD1P2 | -0.027488447 |
| RP4-799P18 | -0.027488686 |
| MTND2P17 | -0.02749001 |
| RP11-12D5 | -0.027494675 |
| CDHR1 | -0.027495376 |
| RN7SL592P | -0.027497206 |
| KCND3-IT1 | -0.027503586 |
| C8orf17 | -0.027503672 |
| C20orf195 | -0.02750586 |
| MIR5681A | -0.027507596 |
| ZNF720 | -0.027513264 |
| RNA5SP71 | -0.027514043 |
| AC012442 | -0.027516129 |
| RP11-271K21 | -0.027522311 |
| RAB28 | -0.027524818 |
| AC231656 | -0.027527005 |
| PLXNC1 | -0.027531051 |
| RP5-1049G16 | -0.027536167 |

| | |
|---|---|
| RP11-457P14 | -0.027540451 |
| KIAA1324 | -0.027544242 |
| CTD-2248H3 | -0.027552568 |
| VAMP8 | -0.027552633 |
| HIST1H2BC | -0.027556968 |
| FMNL2 | -0.027558122 |
| ZFHX3 | -0.027558382 |
| CTD-2300H10 | -0.027560601 |
| LINC01019 | -0.027560885 |
| RP11-722M1 | -0.027561146 |
| IL20 | -0.027570122 |
| TBL2 | -0.027570438 |
| CTB-105L4 | -0.027572791 |
| RP11-1103G16 | -0.027574464 |
| RP11-347C12 | -0.027579499 |
| CTD-2533G20 | -0.027582408 |
| RP11-118G23 | -0.027583303 |
| CCNB1 | -0.027587146 |
| FAM136A | -0.027594812 |
| SNORA32 | -0.027597762 |
| RNU6-698P | -0.027602526 |
| CDCA3 | -0.027608822 |
| LINC01037 | -0.02761179 |
| LINC00899 | -0.02761952 |
| MS4A5 | -0.027619891 |
| AC019185 | -0.027622651 |
| OGFOD2 | -0.027623966 |
| RP11-390B4 | -0.027625371 |
| XXyac-YM21GA2 | -0.027629995 |
| RP11-542A14 | -0.027631465 |
| HINFP | -0.027631911 |
| LINC01167 | -0.027634828 |
| AC025156 | -0.027636505 |
| AC009237 | -0.027640916 |
| RP11-817G13 | -0.027642071 |
| PLIN5 | -0.027642368 |
| CCAT1 | -0.027645393 |
| TMOD4 | -0.027646835 |
| MIR30D | -0.02764828 |
| CTA-392C11 | -0.027648582 |
| RP11-2I17 | -0.027654176 |
| PLA2G10 | -0.027656033 |
| RP11-471B18 | -0.027658256 |
| USP25 | -0.027665754 |
| RP11-180D15 | -0.027680821 |

| | |
|---|---|
| RP11-183G22 | -0.027681201 |
| RP11-522N14 | -0.02768305 |
| AC010884 | -0.027684481 |
| NPTXR | -0.027685774 |
| AC008798 | -0.02768579 |
| TAF9P3 | -0.027686941 |
| BRWD1-AS1 | -0.027689752 |
| COPS5 | -0.027690239 |
| RP11-150O12 | -0.027693383 |
| C10orf111 | -0.027694893 |
| RP11-762J3 | -0.027695329 |
| MYADML2 | -0.027697007 |
| ZBTB46 | -0.027702147 |
| CTB-161M19 | -0.027704301 |
| RP11-272M24 | -0.027706788 |
| MIR5580 | -0.027708368 |
| RP11-111J6 | -0.027709815 |
| AC023992 | -0.027709982 |
| AMT | -0.027711431 |
| CTD-3032H12 | -0.027712212 |
| RNA5SP497 | -0.027720073 |
| OR10AE1P | -0.027720134 |
| TUBB4BP4 | -0.027721855 |
| GP9 | -0.02772539 |
| RP11-534P6 | -0.027726399 |
| RP11-460N11 | -0.027729254 |
| KRT32 | -0.027731883 |
| RP11-306B9 | -0.027741399 |
| AC083939 | -0.027742482 |
| RPS26P21 | -0.027744067 |
| EXOSC3 | -0.027749088 |
| C1RL-AS1 | -0.027754205 |
| TMEM63A | -0.027759637 |
| AC115522 | -0.027761109 |
| ANP32C | -0.027764891 |
| AC106053 | -0.027764951 |
| RP4-730K3 | -0.027767545 |
| RP11-404L6 | -0.027767624 |
| C16orf72 | -0.027769647 |
| FAM207CP | -0.027772515 |
| AC078855 | -0.027782694 |
| AC092664 | -0.027785409 |
| MIR4711 | -0.027786901 |
| RP11-34F20 | -0.027787434 |
| PGBD4P8 | -0.02778764 |

| | |
|---|---|
| TPSG1 | -0.027790808 |
| TEF | -0.02779309 |
| CEACAMP3 | -0.027795775 |
| MIR6768 | -0.027798517 |
| CDS2 | -0.027805872 |
| MIR3179-1 | -0.027810277 |
| KYNUP1 | -0.027810443 |
| TEKT2 | -0.027812719 |
| RP11-356M17 | -0.027814653 |
| RP11-826F13 | -0.027814896 |
| FNIP1 | -0.027816217 |
| MTND1P19 | -0.027816672 |
| ABCC5 | -0.027822077 |
| EHMT1-IT1 | -0.02783832 |
| RPL21P108 | -0.027839998 |
| MRPS36 | -0.027843455 |
| RP11-322I2 | -0.027846487 |
| RAB3GAP2 | -0.027847288 |
| AP002806 | -0.027853137 |
| LINC01513 | -0.02785713 |
| ZNF726 | -0.027857484 |
| MIR5095 | -0.02785884 |
| IFNA7 | -0.027860198 |
| RP1-41P2 | -0.027864195 |
| GAPDHP26 | -0.027864614 |
| RNU4ATAC10P | -0.027865977 |
| RPL21P135 | -0.027866554 |
| PABPC1P9 | -0.027868367 |
| NIPAL1 | -0.027869261 |
| CFAP57 | -0.027869782 |
| IGHVII-53-1 | -0.027870644 |
| DUSP8P4 | -0.027873551 |
| RNY1P3 | -0.027875343 |
| RPH3AL | -0.027875345 |
| THSD7A | -0.027876219 |
| RBM48P1 | -0.027877759 |
| KIAA0408 | -0.027878816 |
| HIST1H1PS2 | -0.027878946 |
| RNU6-159P | -0.027885992 |
| AC104133 | -0.027888703 |
| MIR6868 | -0.027894817 |
| SNORD91B | -0.027898308 |
| ZNF888 | -0.027902397 |
| CREBBP | -0.027904518 |
| PXN-AS1 | -0.027906565 |

| | |
|---|---|
| SNORD116-29 | -0.027908179 |
| OR7E19P | -0.027908468 |
| Hammerhead_HH10 | -0.027912388 |
| OTULIN | -0.027913933 |
| KDM4B | -0.027915626 |
| LINC00524 | -0.027919405 |
| OR52B3P | -0.02792358 |
| ZC3H11B | -0.027923731 |
| RN7SKP8 | -0.02792462 |
| RN7SL630P | -0.027926082 |
| LHX1 | -0.027926922 |
| RP11-195B21 | -0.027928287 |
| AL133230 | -0.027930789 |
| KCTD20 | -0.027930845 |
| PIWIL4 | -0.027931291 |
| RP11-124B13 | -0.027933545 |
| RNU6-669P | -0.027935555 |
| CAMSAP1 | -0.027943571 |
| CTB-54O9 | -0.027946101 |
| RP11-179D22 | -0.027951266 |
| DGKK | -0.027952848 |
| AF178030 | -0.027958569 |
| POU4F1 | -0.027961435 |
| DCAF5 | -0.02796253 |
| RP11-129B9 | -0.027965191 |
| YWHAZP8 | -0.027967856 |
| RP11-216L13 | -0.027968968 |
| B3GALNT2 | -0.027969619 |
| NDUFA7 | -0.027974791 |
| AC012593 | -0.027977948 |
| RNU6-10P | -0.027984592 |
| AC104781 | -0.027993566 |
| RNA5SP463 | -0.027998628 |
| HCAR3 | -0.028012843 |
| ELFN1-AS1 | -0.028014902 |
| PGAP2 | -0.028015277 |
| CTC-339F2 | -0.028015759 |
| RPL6P2 | -0.028019533 |
| CYP17A1 | -0.028022041 |
| RP11-561P12 | -0.028022198 |
| MTPN | -0.028022291 |
| HEXDC | -0.028037454 |
| SNORA13 | -0.028038305 |
| AC073648 | -0.028039695 |
| KRTAP5-11 | -0.028041556 |

| | |
|---|---|
| RP11-526P6 | -0.028046599 |
| MAP3K8 | -0.028046627 |
| MIR5693 | -0.028051189 |
| ANKS1A | -0.028051911 |
| FBXO34 | -0.028054832 |
| CTD-2313J17 | -0.028057027 |
| RP11-1378G18 | -0.028061496 |
| GS1-293C5 | -0.028063088 |
| CNN2P11 | -0.028066407 |
| RP11-195O1 | -0.028067213 |
| RP11-641D5 | -0.028069232 |
| TRMT112 | -0.028073512 |
| KRTAP19-7 | -0.028074842 |
| RN7SL294P | -0.028079129 |
| CDYL | -0.028081773 |
| NARS2 | -0.028082052 |
| MAGEB18 | -0.028082964 |
| CCDC14 | -0.028088619 |
| EXOSC10 | -0.028089159 |
| DNAI2 | -0.028091456 |
| AL356019 | -0.028099224 |
| XAGE2B | -0.02810317 |
| EFCAB9 | -0.028103441 |
| TMCO3 | -0.028103615 |
| AC139143 | -0.028104789 |
| RMST_9 | -0.028107692 |
| MTND1P27 | -0.02810944 |
| CTD-2287N17 | -0.02811475 |
| RN7SL751P | -0.028118099 |
| RP1-65P5 | -0.028122536 |
| RP11-384B12 | -0.028127184 |
| FMO11P | -0.028136402 |
| CHAC2 | -0.028140231 |
| CCDC113 | -0.028143096 |
| RP11-169D4 | -0.028147553 |
| STIM1 | -0.028159934 |
| PRKCI | -0.028161943 |
| RP1-272L16 | -0.028162698 |
| GXYLT1P5 | -0.028162797 |
| RP11-528N21 | -0.028167403 |
| RP11-422P24 | -0.028169888 |
| FAM86C1 | -0.028176139 |
| KB-173C10 | -0.028178195 |
| RP11-556G22 | -0.028179699 |
| AC144522 | -0.028182775 |

| | |
|---|---|
| PKNOX2 | -0.028187073 |
| AC103863 | -0.028188778 |
| KCNN1 | -0.028190933 |
| KLF12 | -0.02819988 |
| RBM22P3 | -0.028202377 |
| RNU6-748P | -0.028214046 |
| RPS12P18 | -0.028216516 |
| OR8L1P | -0.028221195 |
| RPL36P18 | -0.028221218 |
| PCNPP1 | -0.028221895 |
| RP11-361F19 | -0.02822398 |
| AC098617 | -0.028228766 |
| ZNF321P | -0.028233386 |
| AP002815 | -0.02823519 |
| CD209 | -0.028241352 |
| RP11-690P14 | -0.028249007 |
| RP11-507M3 | -0.028263832 |
| RP11-401L13 | -0.028269093 |
| RNU6-1165P | -0.028271395 |
| FNIP2 | -0.028273911 |
| RP11-190J23 | -0.02827428 |
| AL442639 | -0.028274844 |
| RAB40B | -0.028276404 |
| RPL31P57 | -0.028276494 |
| DEFB127 | -0.028276945 |
| FTCDNL1 | -0.028279753 |
| AL449212 | -0.028281434 |
| LRRC18 | -0.02828182 |
| RP11-112N23 | -0.028284669 |
| RP11-775G23 | -0.028287191 |
| CH17-53B9 | -0.028289589 |
| AC008391 | -0.028290118 |
| ZNF254 | -0.028293354 |
| AC005048 | -0.028293385 |
| BTF3P11 | -0.028297637 |
| OR10A6 | -0.028298113 |
| RN7SL513P | -0.028300432 |
| AF213884 | -0.028301661 |
| NXPH1 | -0.028307322 |
| CBX6 | -0.028311141 |
| NACC2 | -0.028314304 |
| TAS2R18 | -0.028314869 |
| CAP2 | -0.028318267 |
| RP11-542E6 | -0.028319957 |
| AC006322 | -0.028320608 |

| | |
|---|---|
| RP11-690J15 | -0.028322812 |
| INADL | -0.028335979 |
| KB-1208A12 | -0.028336705 |
| RNA5SP434 | -0.02833701 |
| CTC-325H20 | -0.028337988 |
| RNU7-172P | -0.028341097 |
| RP11-794A8 | -0.028342202 |
| RP11-13L2 | -0.028342984 |
| CD46P1 | -0.028345307 |
| SNORA14B | -0.02835193 |
| SNORA34 | -0.028354291 |
| LINC00328-2P | -0.028355426 |
| MIR548A2 | -0.028362472 |
| RP11-723O4 | -0.028362901 |
| MIR5685 | -0.028362904 |
| MIR4511 | -0.028367549 |
| POGK | -0.028378567 |
| RNA5SP430 | -0.02838043 |
| FOPNL | -0.028385525 |
| AC104763 | -0.028387531 |
| ABCC5-AS1 | -0.028387973 |
| RP11-580J4 | -0.028388532 |
| AC090774 | -0.028389773 |
| RP11-114L10 | -0.028391924 |
| RP11-58G13 | -0.028393307 |
| CCDC27 | -0.028394208 |
| RNU6-379P | -0.028394635 |
| SUCLA2P3 | -0.02839636 |
| QPCT | -0.0283968 |
| UBQLN4 | -0.028400112 |
| RNU6-1231P | -0.028405273 |
| RNU6-336P | -0.028405719 |
| HLA-DQB2 | -0.028405805 |
| MMGT1 | -0.02840588 |
| EIF4A1P3 | -0.028406564 |
| TCTEX1D2 | -0.028407145 |
| TENM1 | -0.028407678 |
| EYA1 | -0.028408092 |
| IGSF5 | -0.028409712 |
| NAP1L1P3 | -0.02841484 |
| AC093165 | -0.028416875 |
| AC110781 | -0.02841754 |
| OR5T3 | -0.028417816 |
| RPS27P29 | -0.028418826 |
| CTD-2027G2 | -0.028419012 |

| | |
|---|---|
| RNU6-903P | -0.02841953 |
| MRPL49P2 | -0.028427956 |
| CEP170P1 | -0.028428578 |
| CFAP69 | -0.028429006 |
| RP11-125A15 | -0.028432488 |
| CTD-2571E19 | -0.028432818 |
| MTX2 | -0.02843282 |
| MBOAT4 | -0.028435006 |
| RP11-124D9 | -0.028443844 |
| BNIP3P38 | -0.028446863 |
| LINC01237 | -0.028450375 |
| RNA5SP210 | -0.028453891 |
| XKRYP2 | -0.028456261 |
| AL589736 | -0.02846892 |
| RP11-424E7 | -0.028470753 |
| BRMS1L | -0.02847245 |
| DCDC2B | -0.028474736 |
| PPCDC | -0.028479148 |
| RPL34P1 | -0.028482025 |
| SRSF8 | -0.028483986 |
| SNORA47 | -0.028485881 |
| RP11-96A1 | -0.028488601 |
| THUMPD1 | -0.02848966 |
| SPATA2P1 | -0.028490434 |
| HIATL1 | -0.028490944 |
| RP11-168J5 | -0.02849259 |
| BPHL | -0.028493177 |
| PARD6G-AS1 | -0.028497251 |
| HSP90AA6P | -0.028497825 |
| AC016727 | -0.028502039 |
| XXbac-BPG283O16 | -0.028505103 |
| CTD-2363C16 | -0.02850565 |
| RP11-705O24 | -0.028508242 |
| LINC01120 | -0.028511591 |
| ABCD1P5 | -0.028513794 |
| PUM1 | -0.028520582 |
| RP11-1O2 | -0.028522934 |
| RNU6-264P | -0.02852338 |
| RP11-29P20 | -0.028523539 |
| RN7SL282P | -0.028523914 |
| RNA5SP188 | -0.028525296 |
| FAM186B | -0.028526778 |
| FAM177A1 | -0.028528062 |
| MIR4671 | -0.028531452 |
| MIR217 | -0.028541131 |

| | |
|---|---|
| RP11-643G5 | -0.02854424 |
| AC003002 | -0.028545838 |
| HMGB3P17 | -0.028546735 |
| TMCC1-AS1 | -0.028548435 |
| FAM201B | -0.028551457 |
| RNU2-19P | -0.028553342 |
| KRT18P64 | -0.028554886 |
| TMEM211 | -0.028555782 |
| PSAT1P2 | -0.028556145 |
| GSX1 | -0.028560704 |
| RNA5SP333 | -0.028562854 |
| PCGF6 | -0.028564603 |
| AMBP | -0.028568409 |
| CENPM | -0.028568603 |
| TRAV8-5 | -0.028569235 |
| AC006003 | -0.02857272 |
| RP11-293K20 | -0.028572853 |
| RP11-769N21 | -0.028574862 |
| CTB-61M7 | -0.02857696 |
| RP11-30G8 | -0.028578307 |
| PIGF | -0.028579562 |
| NUS1 | -0.028582539 |
| RNA5SP183 | -0.028583237 |
| KIF20A | -0.028587562 |
| RPL17P35 | -0.028588235 |
| SPACA6P-AS | -0.028591607 |
| FAM90A21P | -0.028592742 |
| FAM197Y3 | -0.028592742 |
| RNU6-1158P | -0.028598717 |
| STPG2-AS1 | -0.028598836 |
| GS1-120K12 | -0.028599659 |
| RP11-273B19 | -0.028607219 |
| RP11-158G18 | -0.028609405 |
| UBLCP1 | -0.028609554 |
| HMGN2P22 | -0.028610524 |
| GINS1 | -0.028611731 |
| MIR4693 | -0.028612474 |
| AC018880 | -0.028613147 |
| RP11-9G1 | -0.028617294 |
| TCAF1P1 | -0.028618364 |
| GAGE2A | -0.028619676 |
| RNU6-773P | -0.02862319 |
| OR10Q2P | -0.028623602 |
| ARL11 | -0.028628513 |
| RNU6-441P | -0.028631786 |

| | |
|---|---|
| MKLN1 | -0.028632532 |
| RP5-849L7 | -0.028634851 |
| RP11-529P9 | -0.028636757 |
| RP1-159G19 | -0.028638523 |
| SLC25A5P3 | -0.028641575 |
| CNN3P1 | -0.028647746 |
| RPL21P122 | -0.028648642 |
| LYST | -0.02864963 |
| RP11-324J13 | -0.028651391 |
| CABIN1 | -0.028651677 |
| SNX18P2 | -0.028653646 |
| AL512290 | -0.028654185 |
| AC005288 | -0.028654186 |
| RP11-665G4 | -0.028659111 |
| RP11-30O15 | -0.028665494 |
| GPR61 | -0.028666082 |
| TRAV37 | -0.028666162 |
| AC010907 | -0.028667166 |
| MIR153-2 | -0.028667311 |
| TUBB8P5 | -0.028668824 |
| SFTPA1 | -0.028671424 |
| FAM149B1P1 | -0.028674472 |
| VAPB | -0.028676961 |
| LINC00678 | -0.02867698 |
| KRTAP21-4P | -0.028678336 |
| LINC01491 | -0.028678448 |
| VN1R91P | -0.02868383 |
| AC084219 | -0.028684022 |
| AL136380 | -0.028684667 |
| CALCR | -0.02869285 |
| TRIM73 | -0.028696344 |
| RNU7-197P | -0.028698838 |
| RP3-402G11 | -0.028701576 |
| CTD-2135D7 | -0.028705956 |
| TNFRSF10A | -0.028707195 |
| TFP1 | -0.028709742 |
| RP11-752P2 | -0.028710721 |
| MIR26A2 | -0.028711254 |
| LINC01518 | -0.028711789 |
| RNA5SP489 | -0.028714586 |
| RPL21P40 | -0.028717999 |
| LSM12P1 | -0.028721298 |
| SPZ1 | -0.028721823 |
| C6orf89 | -0.02872448 |
| RNA5SP143 | -0.028732115 |

| | |
|---|---|
| C19orf80 | -0.028735951 |
| RMDN3 | -0.028736425 |
| FTLP16 | -0.02873672 |
| ZNF775 | -0.028738155 |
| MIR5094 | -0.028742894 |
| AK4P2 | -0.028744214 |
| AC016908 | -0.028750023 |
| AC114494 | -0.028753095 |
| RP5-1037N22 | -0.028759897 |
| BLZF2P | -0.028761352 |
| ELMOD3 | -0.028769207 |
| RP11-192H23 | -0.02876971 |
| RP11-418B12 | -0.028770752 |
| SNORA70F | -0.028770955 |
| RP11-63P12 | -0.028778557 |
| AC018731 | -0.028780686 |
| AC245470 | -0.028785328 |
| RPL35AP18 | -0.028789158 |
| RP11-745O10 | -0.028789841 |
| RPS15AP12 | -0.028795236 |
| RNU6-580P | -0.0287988 |
| MBNL2 | -0.028798883 |
| SMG6 | -0.028801097 |
| RPL7P36 | -0.028801859 |
| AC006946 | -0.028806679 |
| OR8B5P | -0.02880727 |
| RNA5SP384 | -0.028812531 |
| RP11-20F18 | -0.028817896 |
| RP11-480O10 | -0.028819214 |
| RP3-467N11 | -0.028823386 |
| TMEM138 | -0.028825638 |
| NAV2-AS4 | -0.028831784 |
| HMGB1P31 | -0.028832131 |
| SUMO2P14 | -0.028843233 |
| RP1-149C7 | -0.028844246 |
| RP11-134J21 | -0.028845356 |
| EPB41L5 | -0.028848355 |
| RNY4P3 | -0.028848595 |
| MARK2P11 | -0.028849723 |
| ADCY9 | -0.028850085 |
| AC005609 | -0.028871402 |
| C3orf35 | -0.028871984 |
| OR2AG2 | -0.028874357 |
| KTI12 | -0.028877584 |
| AC113608 | -0.028880939 |

| | |
|---|---|
| CTC-503K11 | -0.028881693 |
| COX6CP18 | -0.028884082 |
| CTD-2024I7 | -0.028887945 |
| RP11-4B14 | -0.028888803 |
| RN7SKP262 | -0.028889011 |
| AL136456 | -0.028890156 |
| OR10G3 | -0.028890486 |
| ZNF680P1 | -0.028890781 |
| RNU6-536P | -0.028890916 |
| RN7SL252P | -0.028893046 |
| RP11-44N17 | -0.028901662 |
| AFM | -0.028903542 |
| RP11-165E7 | -0.028906687 |
| CXXC5 | -0.028907092 |
| MIF-AS1 | -0.028909995 |
| AC091492 | -0.028910433 |
| OR7E93P | -0.028911554 |
| RP11-476K15 | -0.028913558 |
| MAT1A | -0.028915833 |
| UBE2V1P6 | -0.028918783 |
| RNU6-1169P | -0.028918853 |
| PTPRE | -0.028922442 |
| RP11-52J3 | -0.02892259 |
| RNU6ATAC34P | -0.028923391 |
| RN7SL182P | -0.028925266 |
| HIGD1AP2 | -0.028928703 |
| MIR1301 | -0.02893219 |
| CTA-109P11 | -0.028932216 |
| OR51N1P | -0.02893501 |
| HOTAIR_4 | -0.02893634 |
| RP5-855F16 | -0.02894517 |
| RN7SL285P | -0.028946349 |
| RNU6-279P | -0.028947182 |
| GS1-166A23 | -0.028947908 |
| NUDCD3 | -0.028948397 |
| RP11-436F23 | -0.028955281 |
| APOA1 | -0.028960149 |
| GPR50 | -0.028962087 |
| BPIFA3 | -0.028962246 |
| PLCXD3 | -0.028963186 |
| LCNL1 | -0.028965709 |
| TFB2M | -0.028966183 |
| MIR3684 | -0.028970296 |
| RP11-24P24 | -0.028971133 |
| GRIP2 | -0.028971598 |

| | |
|---|---|
| LTB4R | -0.028973942 |
| RP11-342A23 | -0.028979284 |
| RNU4-64P | -0.028986135 |
| EEF1A1P38 | -0.028989386 |
| CDHR5 | -0.028989614 |
| UBL5P3 | -0.02899 |
| RP11-1102P16 | -0.028999955 |
| UBE2Q2P10 | -0.029003633 |
| CYCSP25 | -0.029005916 |
| NOL8P1 | -0.029006244 |
| RNA5SP207 | -0.029007724 |
| RNU6-516P | -0.029010515 |
| NPIPA7 | -0.029015769 |
| RNU6-983P | -0.029020227 |
| AAR2 | -0.029021144 |
| FAM83H-AS1 | -0.02902267 |
| LONP2 | -0.029023916 |
| RP11-380O24 | -0.029027207 |
| CA5A | -0.029029785 |
| AP000711 | -0.029031534 |
| ACRV1 | -0.029034449 |
| HSPD1P2 | -0.029039085 |
| RPS6KA5 | -0.029039898 |
| RP11-469L4 | -0.02904091 |
| AC090952 | -0.029043472 |
| SNORA80A | -0.029050944 |
| RNU6-314P | -0.029054825 |
| KDM5C | -0.029058039 |
| RP11-441O15 | -0.029058243 |
| CTD-2203A3 | -0.029061785 |
| RP11-180C16 | -0.029066107 |
| TDGF1 | -0.029067599 |
| ZDHHC11B | -0.029071977 |
| ANKRD34B | -0.029073591 |
| ASS1P4 | -0.029074271 |
| RFC3P1 | -0.029080854 |
| RP11-17J14 | -0.029081169 |
| SLC35A3 | -0.029081482 |
| TSPYL4 | -0.029082505 |
| RP11-677N16 | -0.029084722 |
| SIGLEC9 | -0.029085296 |
| AC007631 | -0.029088341 |
| RP11-552I14 | -0.029089619 |
| C6orf195 | -0.029089828 |
| RNU6-1055P | -0.029091051 |

| | |
|---|---|
| RP11-63I8 | -0.029091194 |
| USP21 | -0.029099845 |
| RAB6C | -0.029101894 |
| FTH1P25 | -0.02910554 |
| AC002539 | -0.029110447 |
| RP4-705F19 | -0.02911778 |
| RPIA | -0.029119065 |
| HSD3BP3 | -0.02912222 |
| RP11-651L5 | -0.029122432 |
| ANKRD19P | -0.029131493 |
| ZNF720P1 | -0.029131978 |
| ANP32BP3 | -0.029133587 |
| RP11-331O9 | -0.029133936 |
| AC004691 | -0.029134082 |
| BATF | -0.029136758 |
| CYP4F30P | -0.029136918 |
| UMPS | -0.029140256 |
| FLVCR1 | -0.029141745 |
| RP11-298E2 | -0.029146726 |
| POU2F3 | -0.029149708 |
| TMEM241 | -0.02915501 |
| EEF1A1P5 | -0.029155755 |
| RP11-805J14 | -0.029156418 |
| RP11-466J24 | -0.029156648 |
| DDX10P1 | -0.029158681 |
| RP4-710M3 | -0.029168718 |
| SNRNP40 | -0.029171011 |
| AC015726 | -0.029173653 |
| DXO | -0.029174137 |
| LINC01249 | -0.02917471 |
| CTB-119C2 | -0.029178881 |
| RP11-407P18 | -0.029184401 |
| KRT18P35 | -0.029185077 |
| DMRTC1B | -0.029187287 |
| AC138123 | -0.029189854 |
| RP11-140H17 | -0.02919486 |
| RNU6-657P | -0.029195003 |
| RP11-1148O4 | -0.029195651 |
| RP5-1033K19 | -0.02919659 |
| MARK2P10 | -0.029201371 |
| HSPE1P9 | -0.029203179 |
| URI1 | -0.029205375 |
| AC024158 | -0.029205564 |
| AC010184 | -0.029205689 |
| KCNH7 | -0.029206844 |

| | |
|---|---|
| C8orf44-SGK3 | -0.029209939 |
| TRIP4 | -0.029215646 |
| CTD-2034I4 | -0.029218011 |
| RP11-391M7 | -0.029218173 |
| MPP2 | -0.029218995 |
| AC090420 | -0.029220029 |
| TDRD12 | -0.029229358 |
| UBE2V1P11 | -0.029233189 |
| FAM219A | -0.029236484 |
| RP11-508M8 | -0.029239941 |
| SLC35A1 | -0.029242028 |
| SPANXN2 | -0.029242698 |
| AL096855 | -0.029244163 |
| ZSCAN5B | -0.029245257 |
| RP1-59B16 | -0.029247769 |
| OR52X1P | -0.029247789 |
| AMY1C | -0.029252519 |
| RP3-334F4 | -0.029253435 |
| MIR4421 | -0.029255421 |
| LA16c-60G3 | -0.029260585 |
| RNU11-5P | -0.029260748 |
| CCDC183 | -0.029265731 |
| CICP13 | -0.029270492 |
| RNU7-170P | -0.029271993 |
| CADM2 | -0.029272413 |
| LINC01531 | -0.029276965 |
| AC007250 | -0.029279591 |
| AC087650 | -0.029285113 |
| MIR1256 | -0.029295389 |
| ZDHHC21 | -0.029298607 |
| CSN1S2AP | -0.029300375 |
| LINC00638 | -0.029301119 |
| AC105179 | -0.02930218 |
| RP11-158J3 | -0.029306414 |
| AC092812 | -0.029306534 |
| SCGB1D5P | -0.029310934 |
| LLPHP2 | -0.029311811 |
| LINC00935 | -0.0293149 |
| KLHL22 | -0.029316506 |
| B3GALT5-AS1 | -0.029318179 |
| SAPCD2P1 | -0.029318822 |
| RP11-371A19 | -0.029320168 |
| MIR653 | -0.029320209 |
| RNU6-466P | -0.029322067 |
| AC062032 | -0.029327093 |

| | |
|---|---|
| RPS11P7 | -0.029331105 |
| RNU6-420P | -0.029337121 |
| BMP2KL | -0.029340428 |
| PGK2 | -0.029341156 |
| SNORD75 | -0.029341215 |
| IPMK | -0.02934354 |
| MINK1 | -0.029347503 |
| RP5-855D21 | -0.029349642 |
| MIR4767 | -0.029359816 |
| IMMP1L | -0.029361233 |
| RP11-789L4 | -0.029361791 |
| MIR8052 | -0.029364706 |
| AL732579 | -0.029367243 |
| TPT1P8 | -0.029367526 |
| HCG20 | -0.02936797 |
| RP11-567J20 | -0.029371478 |
| ESRRB | -0.029378929 |
| PAPPA-AS2 | -0.029381082 |
| PPP1R10P1 | -0.029382492 |
| AC005772 | -0.029383097 |
| OR8B1P | -0.029383344 |
| RNU6-50P | -0.029392367 |
| MIR3142 | -0.029392939 |
| MIR4689 | -0.029393607 |
| C15orf41 | -0.029403542 |
| RP11-379K17 | -0.029410181 |
| MRPL10 | -0.029412677 |
| RP11-269G24 | -0.029419897 |
| RNU6-532P | -0.02942172 |
| AL021917 | -0.029425727 |
| RNY4P27 | -0.029426622 |
| AC013436 | -0.029427141 |
| PHACTR2 | -0.029442893 |
| MT1A | -0.029443228 |
| ZNF12 | -0.029444586 |
| CYCSP42 | -0.029445006 |
| TTC39A-AS1 | -0.029447877 |
| CTD-2054N24 | -0.029456561 |
| ZNF70 | -0.029458205 |
| MIR4439 | -0.029465269 |
| AMELY | -0.029470932 |
| RP11-160E2 | -0.029471571 |
| MRRFP1 | -0.029474662 |
| CPOX | -0.029475806 |
| RNU6-756P | -0.029476383 |

| | | |
|---|---|---|
| GNPATP | | -0.029480238 |
| YWHAQP7 | | -0.029480759 |
| PHKG1P2 | | -0.029486479 |
| FABP2 | | -0.029487782 |
| AP000569 | | -0.029488824 |
| KLHL36 | | -0.029493332 |
| RNA5SP174 | | -0.029500731 |
| AL050310 | | -0.029500809 |
| PPP6R2 | | -0.029501165 |
| INO80B-WBP1 | | -0.029502267 |
| AC079988 | | -0.029502363 |
| PDZK1P2 | | -0.029503728 |
| AC026366 | | -0.029508588 |
| TTK | | -0.029518208 |
| SAP25 | | -0.029521779 |
| KCND3-AS1 | | -0.029522112 |
| PSAT1P4 | | -0.029529079 |
| ARPC3P2 | | -0.029532609 |
| FPGT-TNNI3K | | -0.029536396 |
| SPIN4-AS1 | | -0.029536869 |
| NT5C1B | | -0.029537057 |
| AC007790 | | -0.029540469 |
| CCDC42 | | -0.02954343 |
| NLN | | -0.02954619 |
| C10orf105 | | -0.029547871 |
| CTD-3247F14 | | -0.029548967 |
| RP11-107G24 | | -0.029553194 |
| TAS2R41 | | -0.029554839 |
| HADH | | -0.029556684 |
| ASCL3 | | -0.029557095 |
| PIGG | | -0.029557651 |
| RP11-712C7 | | -0.029564737 |
| GNG5P2 | | -0.029564797 |
| RP11-665I14 | | -0.029566072 |
| SYNE3 | | -0.029567866 |
| HAX1 | | -0.029570154 |
| GSX2 | | -0.029572829 |
| | 5-Mar | -0.029579345 |
| RP11-574K11 | | -0.029580127 |
| CTA-722E9 | | -0.029581257 |
| AC010971 | | -0.029583488 |
| AC096922 | | -0.029584731 |
| UGT2B15 | | -0.029585369 |
| NLRP9P | | -0.029591309 |
| RNU4-51P | | -0.029600917 |

| | |
|---|---|
| RP11-23I7 | -0.029603785 |
| EMC3 | -0.029604463 |
| BMS1P20 | -0.029604481 |
| NAA35 | -0.029610714 |
| RP3-327A19 | -0.029617317 |
| RPA2P3 | -0.029620132 |
| DALRD3 | -0.029624166 |
| ARHGEF34P | -0.029628073 |
| TCAF1 | -0.029628799 |
| H2AFZP1 | -0.029631657 |
| MIR518B | -0.029634655 |
| AC005741 | -0.029639207 |
| ZCWPW1 | -0.02964145 |
| LINC01363 | -0.029643232 |
| RP11-1O10 | -0.0296467 |
| AC023128 | -0.029658443 |
| RNU6-1200P | -0.029659035 |
| AC107977 | -0.02965988 |
| RP11-190I17 | -0.029662177 |
| OR1N2 | -0.029669297 |
| LINC01202 | -0.029670928 |
| C11orf65 | -0.029671498 |
| ACP1 | -0.029674996 |
| CFAP43 | -0.029675703 |
| HNRNPA1P56 | -0.029678305 |
| RP11-535N6 | -0.029679589 |
| FOLR2 | -0.029683461 |
| POLR3GP2 | -0.029683965 |
| RNU7-51P | -0.029684589 |
| CECR9 | -0.029684617 |
| EHHADH-AS1 | -0.029691032 |
| RP11-767N15 | -0.029692583 |
| NDC1 | -0.029698178 |
| RP11-168P8 | -0.029698577 |
| TMEM18 | -0.029709323 |
| LMBR1 | -0.029710142 |
| ZNF840P | -0.02971583 |
| RP11-93H10 | -0.029716171 |
| SPATA1 | -0.029720096 |
| RP11-299G20 | -0.029720747 |
| ZNF772 | -0.029722031 |
| MINA | -0.029723296 |
| CTA-299D3 | -0.029735885 |
| SLC25A21 | -0.02973619 |
| YAE1D1 | -0.029737157 |

| | |
|---|---|
| RP11-452H21 | -0.029745338 |
| RP11-234A1 | -0.029750683 |
| MTMR9LP | -0.02975193 |
| RP11-84A14 | -0.029752249 |
| RN7SL474P | -0.029753421 |
| DIO1 | -0.029754526 |
| MORF4L1P6 | -0.029758285 |
| RP11-93B21 | -0.029761662 |
| RP11-98C1 | -0.029764105 |
| RP11-632F7 | -0.029765438 |
| ACTG1P1 | -0.029765456 |
| RP11-603B13 | -0.029765493 |
| RPL7AP10 | -0.029769593 |
| GPR50-AS1 | -0.02977068 |
| AL137006 | -0.029773104 |
| EIF4BP7 | -0.029773772 |
| RNU6-65P | -0.029778068 |
| OR2L13 | -0.029779641 |
| HIST2H2BA | -0.029781358 |
| TBC1D22B | -0.029781512 |
| RP11-64P14 | -0.029782469 |
| H3F3BP1 | -0.029783059 |
| HLF | -0.029783484 |
| RP11-363G15 | -0.029785175 |
| RP11-303G3 | -0.02979586 |
| FSBP | -0.029802511 |
| RP11-295B17 | -0.029808105 |
| PMS2P4 | -0.029808658 |
| RN7SKP120 | -0.029809288 |
| MIR2909 | -0.029812218 |
| WBP1LP5 | -0.029815214 |
| AC016831 | -0.029815988 |
| RP11-109N23 | -0.02981757 |
| AP000721 | -0.029817644 |
| LA16c-360H6 | -0.029818295 |
| RNU2-41P | -0.029825208 |
| RN7SL505P | -0.029834433 |
| H2AFB3 | -0.029836074 |
| KB-1254G8 | -0.029836927 |
| RN7SKP163 | -0.029838983 |
| CELF4 | -0.029843804 |
| CTXN2 | -0.029847712 |
| HAO1 | -0.029851135 |
| CYLC1 | -0.029852931 |
| C7orf33 | -0.02985512 |

| | |
|---|---|
| ARF4P2 | -0.02986267 |
| MAPKAPK3 | -0.029870186 |
| RPS18P12 | -0.029876121 |
| KRT8P44 | -0.029877423 |
| TNPO1P2 | -0.029880065 |
| RP11-267C16 | -0.029884182 |
| FUT10 | -0.029887805 |
| CMB9-52H4 | -0.029890279 |
| FAM198B | -0.029890745 |
| NUS1P2 | -0.029896836 |
| MIR184 | -0.029900541 |
| RNU6-1289P | -0.029902089 |
| AC133680 | -0.029902775 |
| RP11-171I2 | -0.029904848 |
| RNU6-774P | -0.029914107 |
| AC138965 | -0.029914682 |
| TCEA1P4 | -0.029916191 |
| RP11-455J15 | -0.029921163 |
| THEMIS2 | -0.02993257 |
| IL6RP1 | -0.029933013 |
| CBL | -0.029941538 |
| SNORA20 | -0.029947513 |
| BX088563 | -0.029949788 |
| RP11-517A5 | -0.029955071 |
| RFPL4AP1 | -0.029961654 |
| MIR670 | -0.029966359 |
| XXyac-YX60D10 | -0.029967209 |
| ATP2C2 | -0.029973986 |
| AIG1P1 | -0.029975383 |
| RPL35AP22 | -0.029977645 |
| BEND2 | -0.029979342 |
| RNA5SP418 | -0.029979346 |
| HIGD1AP12 | -0.029981862 |
| AC010747 | -0.029981876 |
| IRF7 | -0.029982991 |
| OR6A2 | -0.029990797 |
| RNU6-1211P | -0.029993649 |
| MIR1296 | -0.029998161 |
| EPHX2 | -0.02999888 |
| CTB-91J4 | -0.029999383 |
| RN7SKP154 | -0.030001072 |
| AC134300 | -0.030002261 |
| RP11-644K8 | -0.03000292 |
| MIR4725 | -0.030005771 |
| SNRPCP14 | -0.030009614 |

| | |
|---|---|
| AK4P4 | -0.030010976 |
| MIR548AW | -0.030015045 |
| ZDHHC8P1 | -0.030015681 |
| FGF12-AS1 | -0.030018099 |
| AC092375 | -0.030023611 |
| WDR90 | -0.030026546 |
| RNA5SP307 | -0.030026694 |
| RANP3 | -0.030027948 |
| AC005052 | -0.030034755 |
| QDPR | -0.030041598 |
| INPP4B | -0.030043831 |
| RP11-3L23 | -0.030043922 |
| PLCB1 | -0.030044742 |
| RNU6ATAC7P | -0.03004666 |
| COQ10BP2 | -0.030047338 |
| CTB-158E9 | -0.030048003 |
| MYRFL | -0.030053413 |
| RP11-32K4 | -0.030053423 |
| RP11-577G20 | -0.030053581 |
| COX6CP7 | -0.030063336 |
| CHRNB2 | -0.030064148 |
| RP5-1100I6 | -0.030066684 |
| TAF1C | -0.030070269 |
| LLNLR-276H7 | -0.030073523 |
| RPL5P5 | -0.030074998 |
| RP11-65B7 | -0.030079698 |
| PAPLN | -0.03008 |
| SHCBP1L | -0.0300886 |
| CEP164P1 | -0.030093469 |
| CTC-251I16 | -0.030098401 |
| NEU4 | -0.030098721 |
| AC074375 | -0.030100843 |
| MEI4 | -0.030101983 |
| AC092573 | -0.030102898 |
| MRPL45 | -0.030103982 |
| ZCCHC18 | -0.03010614 |
| VN2R10P | -0.030109554 |
| OPTN | -0.030109731 |
| AC018688 | -0.030110063 |
| RNU6-540P | -0.030110426 |
| MTAPP2 | -0.030120195 |
| SRCIN1 | -0.03012082 |
| AC087499 | -0.030122815 |
| HMGN1P20 | -0.030126474 |
| RAB22A | -0.030129955 |

| | |
|---|---|
| PRDM7 | -0.030133073 |
| EI24P1 | -0.030133082 |
| CDRT15 | -0.030133298 |
| RP11-642D6 | -0.030134164 |
| MTND5P8 | -0.030136463 |
| TM9SF2 | -0.030138867 |
| AC073873 | -0.030140006 |
| RP11-488P3 | -0.030143345 |
| WDR66 | -0.030143817 |
| CHD1L | -0.030148133 |
| RNA5SP38 | -0.030148149 |
| NAA11 | -0.030148784 |
| RP11-536N17 | -0.030153906 |
| RN7SL118P | -0.030154266 |
| ALDH6A1 | -0.030158422 |
| RN7SKP35 | -0.030166104 |
| AC012497 | -0.030167949 |
| RN7SKP225 | -0.03017086 |
| AC093074 | -0.030174417 |
| RNA5SP290 | -0.030175686 |
| TAF5L | -0.030177772 |
| RPL3P8 | -0.030180667 |
| IQCE | -0.030181381 |
| ATP5A1P2 | -0.030181598 |
| DENND1A | -0.030182274 |
| SNORD94 | -0.030186723 |
| RPL17 | -0.030192172 |
| GGTA2P | -0.030192844 |
| MIR548F2 | -0.030197677 |
| RN7SL862P | -0.030197678 |
| ST13P21 | -0.03019828 |
| FAAH2 | -0.030201121 |
| SFXN4 | -0.03020127 |
| MIR196A2 | -0.030204455 |
| EEF1A1P8 | -0.03020501 |
| RP4-799G3 | -0.030212295 |
| TBL1XR1 | -0.030215634 |
| AC009967 | -0.03022259 |
| RP11-331H2 | -0.030224388 |
| CHRNG | -0.030225523 |
| AC133644 | -0.030231746 |
| RNA5SP404 | -0.030235148 |
| OSBPL9P3 | -0.030237315 |
| KIF6 | -0.030238574 |
| QRSL1 | -0.03023917 |

| | |
|---|---|
| RPS27P1 | -0.030241155 |
| PCDH8P1 | -0.030242413 |
| HNRNPRP1 | -0.030246788 |
| SWSAP1 | -0.030248342 |
| NAPEPLD | -0.030252584 |
| RP5-1173P7 | -0.030254711 |
| AC073476 | -0.030255649 |
| ENO3 | -0.030258836 |
| Hammerhead_HH9 | -0.030266787 |
| RP1-57E3 | -0.030268016 |
| CTA-21C21 | -0.030269486 |
| TOMM34 | -0.03027359 |
| RP11-552D4 | -0.030277951 |
| AL355885 | -0.030278333 |
| GC | -0.030278737 |
| CPLX4 | -0.030279321 |
| OR4S2 | -0.030281717 |
| UPF3AP3 | -0.030282957 |
| PPP2R5D | -0.030283023 |
| CD300E | -0.030287321 |
| AC093375 | -0.030290674 |
| AL022237 | -0.030295639 |
| PCYOX1 | -0.030304828 |
| RP11-80N2 | -0.030313562 |
| RUFY1 | -0.030315489 |
| AC104837 | -0.030315965 |
| CBY3 | -0.030316981 |
| CT978678 | -0.030317688 |
| RP11-268I9 | -0.03031842 |
| UQCRHP1 | -0.030323339 |
| ARPC3 | -0.030324066 |
| RP11-380F14 | -0.030328006 |
| RPL36AP35 | -0.030333567 |
| RP11-331H13 | -0.030336355 |
| PPIL4 | -0.030339658 |
| RN7SKP18 | -0.030345672 |
| ACTBP13 | -0.030350943 |
| AC117947 | -0.030351174 |
| EDDM3B | -0.030355052 |
| LINC01555 | -0.030355805 |
| ABTB1 | -0.030367603 |
| AJ003147 | -0.030371436 |
| RP5-919F19 | -0.030372526 |
| LINC00208 | -0.03038408 |
| RP11-459O16 | -0.030385995 |

| | |
|---|---|
| RP11-253D19 | -0.03038608 |
| TTC39A | -0.030388869 |
| ARCN1 | -0.030397502 |
| RP11-342D11 | -0.030398028 |
| HLA-W | -0.030398237 |
| RP11-712B9 | -0.03040167 |
| HSD17B4 | -0.03040433 |
| ODF3 | -0.030405017 |
| MIR4730 | -0.03040724 |
| PFAS | -0.030412461 |
| LINC01532 | -0.030414907 |
| MIR3122 | -0.030415727 |
| HNRNPKP3 | -0.030419341 |
| MZT1P2 | -0.030420216 |
| HSPB8 | -0.030422024 |
| UBE2V1P3 | -0.030422193 |
| UCMA | -0.030423679 |
| NAT1 | -0.030432592 |
| KHSRPP1 | -0.030435214 |
| TXNDC8 | -0.030443026 |
| GGPS1 | -0.030448276 |
| FBXO40 | -0.030454075 |
| CTD-2525I3 | -0.030457096 |
| APOC1P1 | -0.030460456 |
| RPL10P14 | -0.030463187 |
| RNU6-208P | -0.030463589 |
| SLC35G4 | -0.030464761 |
| AC019178 | -0.030470345 |
| RP11-64I5 | -0.030473601 |
| LINC00554 | -0.030474674 |
| SREBF2 | -0.03047998 |
| HNRNPH2 | -0.03048212 |
| RPSAP4 | -0.030485769 |
| RP11-35L17 | -0.030488843 |
| MAPK6PS3 | -0.030498468 |
| RP5-1098D14 | -0.030500516 |
| AC005351 | -0.030502754 |
| RP11-461L13 | -0.030503053 |
| SRPK2 | -0.030506363 |
| FSCN3 | -0.03050938 |
| RP11-92K2 | -0.030512763 |
| RP11-240A16 | -0.030517193 |
| OR11H7 | -0.030518874 |
| LINC01546 | -0.030520392 |
| RP4-597A16 | -0.030521237 |

| | |
|---|---|
| AC004702 | -0.030522214 |
| RP11-214D15 | -0.030527729 |
| RP11-635O16 | -0.030531057 |
| RP11-290K4 | -0.030532121 |
| VN1R67P | -0.03053348 |
| CSE1L-AS1 | -0.030534085 |
| KRTAP9-10P | -0.030534997 |
| AL356270 | -0.030545659 |
| LINC00428 | -0.030551086 |
| LINC01267 | -0.030552947 |
| MTND2P29 | -0.030556474 |
| HIRA | -0.030558588 |
| AP003025 | -0.03056103 |
| GPR89A | -0.0305611 |
| RP11-496D1 | -0.030561813 |
| DOLK | -0.030562551 |
| RNU2-27P | -0.030563559 |
| MGC15885 | -0.030569337 |
| RP11-536P6 | -0.030569867 |
| RP1-273G13 | -0.030573919 |
| AC093928 | -0.030575904 |
| AL353626 | -0.030576063 |
| RP11-119F7 | -0.030578477 |
| OR7E145P | -0.030580837 |
| YWHAQP6 | -0.030587117 |
| TGIF2LX | -0.030587555 |
| RNU6-876P | -0.030589533 |
| AC000068 | -0.030591408 |
| AC078927 | -0.030602975 |
| LINC00052 | -0.030606563 |
| RPS10P18 | -0.030608804 |
| TDGF1P2 | -0.030612221 |
| SLC4A8 | -0.030612339 |
| ATG3 | -0.030612916 |
| RP1-116K23 | -0.030615444 |
| GLDN | -0.030616619 |
| KATNAL2 | -0.030621438 |
| RP11-72L22 | -0.03062399 |
| RP11-107C16 | -0.030625665 |
| RNA5SP424 | -0.030626627 |
| GDPD2 | -0.030627731 |
| OR2A13P | -0.030630147 |
| RP11-180D21 | -0.030633412 |
| CLYBL-AS1 | -0.03063505 |
| AC078785 | -0.030635364 |

| | |
|---|---|
| MTND2P8 | -0.030639342 |
| MTND6P2 | -0.030639634 |
| FLNB | -0.030644173 |
| LSMEM2 | -0.030645514 |
| FYCO1 | -0.030645587 |
| LINC01012 | -0.030646304 |
| FMR1-AS1 | -0.030646543 |
| AC092198 | -0.030650005 |
| RP4-697P8 | -0.030653395 |
| RHOXF2 | -0.03065374 |
| DGKA | -0.030654723 |
| VPS26A | -0.03065582 |
| RP1-319M7 | -0.030657735 |
| AC092017 | -0.030657836 |
| RAB10 | -0.030660155 |
| C21orf62 | -0.030662599 |
| RN7SL610P | -0.030663423 |
| ARAF | -0.030669216 |
| ZNF606 | -0.030670109 |
| RPL5P9 | -0.030680997 |
| RNU6-847P | -0.030682381 |
| AL118506 | -0.030687728 |
| SORCS3-AS1 | -0.030688517 |
| HOXB1 | -0.030689629 |
| AC016644 | -0.030690548 |
| SYNGR1 | -0.030690929 |
| MIR4500 | -0.030695376 |
| AL161645 | -0.03069788 |
| RP3-363L9 | -0.030698704 |
| RNU6-260P | -0.030698937 |
| RPL29P3 | -0.030701379 |
| SCGB2B2 | -0.03070185 |
| RN7SL530P | -0.030702101 |
| RP11-732A19 | -0.030705581 |
| RNA5SP84 | -0.030709044 |
| SLK | -0.030711246 |
| TPM3P3 | -0.030711593 |
| EFCAB12 | -0.030713404 |
| RNU7-193P | -0.030713542 |
| SRP9 | -0.030714986 |
| ZDHHC20P1 | -0.030714991 |
| SNORA25 | -0.030716885 |
| AC090043 | -0.030718369 |
| FP236240 | -0.030724484 |
| AC007402 | -0.030725521 |

| | |
|---|---|
| INTS7 | -0.030725591 |
| AC005537 | -0.030726905 |
| RP11-172H24 | -0.030730773 |
| AL731561 | -0.030734045 |
| AP000343 | -0.030734898 |
| ABHD14B | -0.030735415 |
| LINC00571 | -0.030735716 |
| HMGB3P1 | -0.030737687 |
| RP11-518K17 | -0.030739635 |
| FAM71F2 | -0.03074122 |
| HMGN2P10 | -0.030743047 |
| CSGALNACT2 | -0.030743731 |
| ALDOAP2 | -0.030745714 |
| ALDH9A1 | -0.030747657 |
| TAT-AS1 | -0.030748684 |
| RNU6-996P | -0.030748768 |
| STK39 | -0.030750541 |
| RP11-356O22 | -0.030751498 |
| MRPL50P1 | -0.030753898 |
| BNIP3P31 | -0.030754618 |
| RPL21P10 | -0.030754982 |
| RN7SKP193 | -0.030757733 |
| SRGAP2-AS1 | -0.030763299 |
| KCTD13 | -0.030763527 |
| CDC27P1 | -0.030764104 |
| RNU4ATAC3P | -0.030776436 |
| RNU6-776P | -0.030776436 |
| RNU7-183P | -0.030776436 |
| AATF | -0.030776644 |
| SETD7 | -0.030788794 |
| AL136090 | -0.030789114 |
| SEPT7P7 | -0.030795111 |
| AC009022 | -0.030795644 |
| HNRNPA1P72 | -0.03079618 |
| AL772400 | -0.030796196 |
| IFITM5 | -0.030801645 |
| CTB-186G2 | -0.030803281 |
| UBE2V2P4 | -0.030809369 |
| RP11-477G18 | -0.030810185 |
| RN7SL584P | -0.030815288 |
| CTC-215O4 | -0.030819383 |
| OR51M1 | -0.030829164 |
| RP11-134G8 | -0.030830489 |
| CNGA4 | -0.030832079 |
| RPS2P41 | -0.030840879 |

| | |
|---|---|
| RNU6-1264P | -0.030845124 |
| AC002365 | -0.030845927 |
| SKP1P1 | -0.030846694 |
| RNA5SP182 | -0.030847566 |
| KLLN | -0.03084848 |
| KB-1674E1 | -0.030855295 |
| AC007006 | -0.030855948 |
| RNU6-1258P | -0.030855948 |
| AC096763 | -0.030855948 |
| RNA5SP406 | -0.030855948 |
| LINC00898 | -0.030858016 |
| EBAG9P1 | -0.030858102 |
| MIR3621 | -0.030860635 |
| RPS7P6 | -0.03087181 |
| RP5-1023B21 | -0.030872707 |
| GEMIN6 | -0.030876109 |
| PPIL1 | -0.030876742 |
| NPIPB7 | -0.030881743 |
| UBAP2L | -0.030886231 |
| LIPN | -0.030888519 |
| CTD-2310F14 | -0.03089198 |
| PRDM1 | -0.030893487 |
| OR2L3 | -0.030895766 |
| RNA5SP394 | -0.030898474 |
| AC017104 | -0.030898864 |
| AC013286 | -0.030900073 |
| SLC35D1 | -0.03090171 |
| ABCC10 | -0.030903515 |
| SLC25A42 | -0.030904687 |
| MRPS10 | -0.030905078 |
| AKIRIN1 | -0.03090571 |
| RP11-5N11 | -0.030909331 |
| ZNF528-AS1 | -0.030913097 |
| AC105760 | -0.030919702 |
| RP3-368B9 | -0.030921541 |
| RP11-423F24 | -0.030921633 |
| SETP21 | -0.030923489 |
| RNA5SP263 | -0.030925593 |
| AC253578 | -0.030926161 |
| PPP1R9A | -0.030931814 |
| LA16c-321D4 | -0.030932058 |
| OR5B21 | -0.030933403 |
| APOL4 | -0.030933581 |
| RNU6-236P | -0.030933682 |
| CTD-2380F24 | -0.030941574 |

| | |
|---|---|
| HSPE1P8 | -0.030941819 |
| GALNT4 | -0.030942019 |
| RP11-307L3 | -0.030955862 |
| ELOVL2-AS1 | -0.030963773 |
| OR9L1P | -0.030965656 |
| RP1-225L15 | -0.030966006 |
| SPI1 | -0.030968095 |
| FBXO27 | -0.030969416 |
| EEF1A1P1 | -0.030973069 |
| RP1-111J24 | -0.030973563 |
| RP11-492M19 | -0.030977275 |
| RPL5P12 | -0.030977943 |
| CH17-351M24 | -0.030984809 |
| RNU4-36P | -0.030986423 |
| DEFB124 | -0.030988808 |
| RP11-138H11 | -0.030989347 |
| RP11-219E7 | -0.030989661 |
| AC123900 | -0.030993535 |
| HLA-F-AS1 | -0.030997417 |
| AP002022 | -0.030999198 |
| MPP1 | -0.031011315 |
| EEF1A1P10 | -0.03101485 |
| IL32 | -0.031015073 |
| RPL7AP57 | -0.031021645 |
| AC074391 | -0.031022389 |
| SIGLEC11 | -0.031022986 |
| RP11-466P24 | -0.031026066 |
| AL353602 | -0.031030562 |
| RP11-402J6 | -0.031033009 |
| TSPAN5 | -0.031040526 |
| bP-21201H5 | -0.031042869 |
| RP11-76E17 | -0.031043107 |
| RP11-173A16 | -0.031045248 |
| RP11-1078H9 | -0.031046665 |
| ATP5HP3 | -0.031054856 |
| RP11-398J16 | -0.031055098 |
| LINC00336 | -0.031055552 |
| LINC00919 | -0.031057988 |
| TUBA8P2 | -0.031059926 |
| RPS6KL1 | -0.031061196 |
| RP5-850E9 | -0.031062792 |
| PPIL1P1 | -0.031066701 |
| MCCC1-AS1 | -0.031070297 |
| RP11-179A16 | -0.031072312 |
| MIR548I1 | -0.031073368 |

| | |
|---|---|
| LINC01293 | -0.031074684 |
| PDE6C | -0.031079318 |
| MIR1200 | -0.031080306 |
| SLC30A7 | -0.03108503 |
| ARPP21-AS1 | -0.031088941 |
| CTB-36O1 | -0.031095077 |
| EMC7 | -0.031097849 |
| RP11-345K9 | -0.031102526 |
| ARHGAP26-AS1 | -0.031102765 |
| OR7E43P | -0.031103548 |
| NPIPB6 | -0.031106979 |
| LPPR4 | -0.031107989 |
| CACNG5 | -0.031108216 |
| RP11-294J22 | -0.031109138 |
| RP11-466F5 | -0.031112882 |
| THBS4 | -0.031117348 |
| CTA-732E4 | -0.031117446 |
| CAB39L | -0.031117924 |
| OR7E14P | -0.031119562 |
| LINC00589 | -0.031119821 |
| MECP2 | -0.031123636 |
| RNU6-1285P | -0.031125973 |
| MIR555 | -0.031129468 |
| IFNWP9 | -0.031131262 |
| SRRM2-AS1 | -0.031134651 |
| GS1-433O24 | -0.031141468 |
| KTN1 | -0.031152549 |
| RP4-803J11 | -0.031156474 |
| FAM186A | -0.031162224 |
| OR11K2P | -0.031164937 |
| IFNA20P | -0.031168014 |
| PCNP | -0.031170727 |
| ZYG11B | -0.031180699 |
| SLC22A9 | -0.031183417 |
| AC004840 | -0.031185851 |
| RPL3P2 | -0.03118736 |
| RP11-680G24 | -0.031195658 |
| AC118465 | -0.031196034 |
| ZNF16 | -0.031196291 |
| ALG1L6P | -0.031199716 |
| RNU6-374P | -0.031201069 |
| RP11-265N7 | -0.031202642 |
| RPS3AP4 | -0.031205219 |
| RP1-305G21 | -0.031207988 |
| CACUL1 | -0.031211659 |

| | |
|---|---|
| AL390119 | -0.031223505 |
| RP1-62D2 | -0.03122543 |
| DHRSX-IT1 | -0.031225595 |
| RP11-474B16 | -0.03122618 |
| MIR5194 | -0.031238711 |
| MIR367 | -0.031243565 |
| HLTF-AS1 | -0.031243841 |
| AC226150 | -0.031245703 |
| AC002366 | -0.03124721 |
| CTA-215D11 | -0.031248277 |
| RPS4XP4 | -0.031249974 |
| MIR148B | -0.031250775 |
| MIR4653 | -0.031251118 |
| RNU6-483P | -0.03125363 |
| AMER2 | -0.031254996 |
| DHX33 | -0.031255586 |
| RNA5SP355 | -0.031255827 |
| RP11-314A15 | -0.031257815 |
| CNOT7 | -0.031265336 |
| ZNF484 | -0.031266619 |
| S100PBP | -0.03126675 |
| VN1R64P | -0.031268197 |
| AC067957 | -0.031269562 |
| AC016737 | -0.031269649 |
| XDH | -0.031270768 |
| EDNRB-AS1 | -0.031271568 |
| MIR5705 | -0.031276404 |
| PLCG1 | -0.031279587 |
| MIR4311 | -0.031280048 |
| STK17B | -0.03128275 |
| RP11-124N19 | -0.031283779 |
| AL589986 | -0.031287017 |
| RNU6-4P | -0.03128938 |
| RP11-733O18 | -0.031293876 |
| C9orf129 | -0.031305351 |
| NANOGP10 | -0.031306422 |
| RP11-373N24 | -0.031307245 |
| MIR661 | -0.031310538 |
| TARS2 | -0.031317059 |
| AC107218 | -0.031322619 |
| VIT | -0.031325162 |
| RP5-997D24 | -0.031327236 |
| AC083843 | -0.031328817 |
| SSXP9 | -0.031328904 |
| MAS1L | -0.031329066 |

| | |
|---|---|
| CHFR | -0.031332501 |
| MTOR-AS1 | -0.031333621 |
| LINC01522 | -0.031335508 |
| HSP90AB2P | -0.031344334 |
| LYZL1 | -0.03134493 |
| RP11-288G3 | -0.031350489 |
| HCFC1 | -0.031350902 |
| AC016722 | -0.031354241 |
| SNRNP35 | -0.031354791 |
| SNORA79 | -0.031355626 |
| MTND4P15 | -0.03135681 |
| RP11-440K22 | -0.031363171 |
| AC005730 | -0.03136477 |
| SPACA1 | -0.031366213 |
| OR7E160P | -0.031367245 |
| MIR6715A | -0.031369559 |
| TRIM63 | -0.031381074 |
| RP11-543H12 | -0.031382617 |
| KRTAP4-3 | -0.031385113 |
| TPM3P7 | -0.031387506 |
| RP11-395P13 | -0.031395718 |
| AC007375 | -0.031396405 |
| RP11-51G5 | -0.031398098 |
| RP11-351K23 | -0.031398113 |
| PA2G4P2 | -0.031406562 |
| KIAA2026 | -0.031409937 |
| OR4C9P | -0.031410938 |
| RP11-472N13 | -0.031418973 |
| RNU6-1206P | -0.03141961 |
| MDH1 | -0.031427371 |
| HPR | -0.03142927 |
| B4GALNT2 | -0.031430013 |
| LGALS4 | -0.031430975 |
| SUMO2P10 | -0.031431035 |
| SMC5-AS1 | -0.031431687 |
| RP11-1246C19 | -0.031432636 |
| CGN | -0.031433157 |
| PSMC1P1 | -0.03143418 |
| MTND1P32 | -0.03143485 |
| CHCHD3P1 | -0.031435951 |
| AC131160 | -0.031438483 |
| LIN28AP1 | -0.031438861 |
| IFNW1 | -0.031451398 |
| OR6Q1 | -0.031451801 |
| PRAMEF11 | -0.031453287 |

| | |
|---|---|
| SESTD1 | -0.031457214 |
| NIP7 | -0.031459949 |
| PKD1P6 | -0.031460005 |
| FOXD4 | -0.031461587 |
| TMEM245 | -0.031461605 |
| PODXL | -0.0314654 |
| OR11N1P | -0.031465555 |
| OR2L6P | -0.031466845 |
| RP11-707F2 | -0.031467409 |
| SDCBPP2 | -0.031471307 |
| MICD | -0.031472002 |
| CCL18 | -0.03147268 |
| MIR6760 | -0.031474688 |
| RP11-172C16 | -0.03147586 |
| RP11-335E6 | -0.031476242 |
| RPL5P1 | -0.031481702 |
| ZEB2-AS1 | -0.03148238 |
| ZMIZ2 | -0.031490127 |
| SYT1 | -0.03149115 |
| YAP1P1 | -0.031491947 |
| SLC39A12-AS1 | -0.031492598 |
| RN7SKP16 | -0.031493447 |
| ST7-OT4 | -0.031497225 |
| RP11-453A12 | -0.031498233 |
| CTD-2201G16 | -0.031500395 |
| RNU1-112P | -0.031500744 |
| RP11-619I22 | -0.031505643 |
| NPHP4 | -0.031508211 |
| RP11-342C20 | -0.031511524 |
| IMMP1LP3 | -0.031513015 |
| AC009970 | -0.031517504 |
| FAM13A-AS1_2 | -0.031520615 |
| ZNF41 | -0.031535989 |
| IZUMO3 | -0.031540049 |
| ZNF398 | -0.031545906 |
| ERCC6L | -0.031546613 |
| CSK | -0.031550669 |
| ABCA7 | -0.031551525 |
| BTF3 | -0.031552548 |
| GAB1 | -0.031555562 |
| TPCN2 | -0.031558167 |
| RP11-348M17 | -0.031563169 |
| AZU1P1 | -0.031571026 |
| AC104162 | -0.031571739 |
| EIF4EBP1P1 | -0.031576523 |

| | |
|---|---|
| LINC01438 | -0.031578661 |
| RP11-1141N12 | -0.031579408 |
| C17orf99 | -0.031581792 |
| COL2A1 | -0.031584494 |
| AC069303 | -0.031589949 |
| BECN1P2 | -0.031589949 |
| AC069070 | -0.031595321 |
| PITPNM1 | -0.031601518 |
| RP4-681N20 | -0.031602931 |
| MIR320C2 | -0.031604026 |
| TMED7 | -0.031604327 |
| IKBKE | -0.03160535 |
| RP11-659G9 | -0.031608654 |
| GORASP1 | -0.0316119 |
| MUSK | -0.031627079 |
| RP4-714D9 | -0.031628421 |
| OR8K5 | -0.031629548 |
| MTRF1L | -0.031631696 |
| FTX_1 | -0.031636821 |
| RP11-758B24 | -0.031642025 |
| RP11-780M14 | -0.031643749 |
| SLC25A16 | -0.03164751 |
| FGFR1OP2P1 | -0.03165297 |
| ZNF275 | -0.031652999 |
| RP11-407N8 | -0.031658568 |
| SNORD114-19 | -0.03166196 |
| SETD8 | -0.031662153 |
| RN7SL323P | -0.031666905 |
| RPL7P33 | -0.031667775 |
| RP3-461P17 | -0.031669781 |
| RP11-444J21 | -0.031677055 |
| RP11-204K16 | -0.031680306 |
| WASIR1 | -0.031682461 |
| AL356137 | -0.031683783 |
| BAG1 | -0.031684219 |
| RN7SL472P | -0.031685725 |
| RN7SL405P | -0.031686163 |
| ERVK3-1 | -0.031687456 |
| N4BP3 | -0.031689094 |
| RBM11P1 | -0.031694331 |
| RP11-632B21 | -0.031694557 |
| H1FNT | -0.031695273 |
| PAICSP5 | -0.031695657 |
| RP11-326C3 | -0.031697392 |
| WBP1LP8 | -0.031698299 |

| | |
|---|---|
| KRTAP10-3 | -0.031698734 |
| RP11-669B18 | -0.031699935 |
| SNORD116-25 | -0.031702282 |
| SAMSN1-AS1 | -0.03170967 |
| ATP6V1G2 | -0.03171676 |
| PEBP1P1 | -0.031718406 |
| AGTR1 | -0.031720221 |
| ZNF157 | -0.03172126 |
| FUT5 | -0.031728826 |
| RNU6-592P | -0.031731802 |
| AC112179 | -0.031731802 |
| FREM1 | -0.031734696 |
| TUBAL3 | -0.03173759 |
| RP4-580O19 | -0.031738001 |
| ARMC8 | -0.031739068 |
| RP11-803B1 | -0.031743158 |
| CCDC39-AS1 | -0.031746585 |
| RNU6-346P | -0.031749053 |
| RP11-96O20 | -0.03174931 |
| AK8 | -0.031750399 |
| CICP5 | -0.031750687 |
| RP1-159A19 | -0.031752985 |
| PHKA1 | -0.031757692 |
| RP11-616M17 | -0.031762364 |
| RP11-613H2 | -0.031763643 |
| RPL36P2 | -0.031764417 |
| IGBP1P3 | -0.031774368 |
| LRRC66 | -0.031780595 |
| SGOL1P1 | -0.03178118 |
| CACNA1F | -0.031782084 |
| AC007041 | -0.031784784 |
| AC114969 | -0.031785737 |
| RP11-349H17 | -0.031786378 |
| HMGN2P32 | -0.031787831 |
| PRRX2-AS1 | -0.03178851 |
| RP11-727M10 | -0.031795149 |
| RP11-934B9 | -0.031797253 |
| AC084121 | -0.031801243 |
| YWHAEP1 | -0.031802043 |
| CTAGE12P | -0.031802708 |
| PSMA4 | -0.03180281 |
| RP11-612A1 | -0.031806965 |
| ANKRD24 | -0.031808485 |
| AA06 | -0.031810274 |
| SLC9B1P4 | -0.031815494 |

| | |
|---|---|
| RNU6-1294P | -0.031824894 |
| CTC-503J8 | -0.031825501 |
| LINC00487 | -0.031829147 |
| RPL12P43 | -0.031829855 |
| RN7SL825P | -0.031832443 |
| AP006294 | -0.031833156 |
| RP11-505O17 | -0.031837447 |
| RP1-101K10 | -0.031838248 |
| SNORD114-13 | -0.03183864 |
| RP11-170J3 | -0.03184131 |
| RP4-621B10 | -0.031842359 |
| MAP7 | -0.031843166 |
| MRPS16P2 | -0.031843521 |
| C10orf95 | -0.031847984 |
| GNL2P1 | -0.031851066 |
| LINC00850 | -0.031851466 |
| RN7SL851P | -0.031851598 |
| AC215219 | -0.031860627 |
| RP11-175E9 | -0.031863468 |
| RNY3P11 | -0.031865324 |
| RN7SL393P | -0.031867616 |
| ZNF281 | -0.031872209 |
| PRR23D1 | -0.031872451 |
| MIR4728 | -0.031882027 |
| RP11-26H16 | -0.031882788 |
| AC008697 | -0.031884286 |
| PRSS3P2 | -0.031885545 |
| TMEM262 | -0.031885828 |
| GACAT1 | -0.031886686 |
| POM121L12 | -0.031888801 |
| CTD-2234N14 | -0.031890212 |
| RANBP6 | -0.031896098 |
| AC004535 | -0.031896553 |
| GNG8 | -0.0318989 |
| SOWAHA | -0.031899168 |
| CTD-2286N8 | -0.031904005 |
| MYL10 | -0.031904292 |
| SMIM15 | -0.031908285 |
| RNU6-890P | -0.031908971 |
| RP11-421B23 | -0.031909295 |
| TTC30A | -0.031918183 |
| GAPDHP71 | -0.031920768 |
| RP11-304F15 | -0.031921253 |
| RAX2 | -0.031923903 |
| CTB-73N10 | -0.031930295 |

| | |
|---|---|
| TRBVA | -0.031935047 |
| CD300LG | -0.031935327 |
| OOSP1P1 | -0.031936222 |
| SLC22A4 | -0.031936509 |
| RP11-168O22 | -0.031941068 |
| CPNE6 | -0.03194526 |
| RN7SL28P | -0.031946443 |
| RPL23AP76 | -0.031946539 |
| CTC-471F3 | -0.031947598 |
| ZNF337-AS1 | -0.031955133 |
| ZNF221 | -0.03195651 |
| RNU6-461P | -0.031958415 |
| MIR202 | -0.031961247 |
| MIR6766 | -0.031964482 |
| CDK11B | -0.031965422 |
| RP11-492M23 | -0.031965813 |
| AC108479 | -0.03196589 |
| LINC00403 | -0.031965916 |
| RP1-22N22 | -0.031978817 |
| RARA | -0.031980883 |
| CYP2T3P | -0.031985703 |
| RNU6-332P | -0.031996452 |
| ANKRD20A8P | -0.031999982 |
| HSP90AA1 | -0.032000679 |
| OR5AC4P | -0.03200184 |
| RP11-107K17 | -0.032003617 |
| AC114752 | -0.032007951 |
| KRT18P2 | -0.032010466 |
| RP13-75G22 | -0.032011271 |
| EIF3H | -0.032016494 |
| OR2U2P | -0.032027354 |
| MS4A3 | -0.032031852 |
| GAPDHP20 | -0.03203489 |
| MLLT4 | -0.032034951 |
| RP11-398H6 | -0.032035762 |
| RNU6-135P | -0.032038435 |
| PLCXD2-AS1 | -0.03204001 |
| TP53I11 | -0.032043733 |
| C8orf22 | -0.032047293 |
| NDUFA4P2 | -0.032049427 |
| AP000470 | -0.032052606 |
| MIR548AK | -0.032054298 |
| RP11-751K21 | -0.032055261 |
| TRAV11 | -0.032060511 |
| GRK5-IT1 | -0.032065967 |

| | |
|---|---|
| SDAD1P3 | -0.032068997 |
| SLC5A9 | -0.032069148 |
| ACTG1P10 | -0.032074916 |
| AC007078 | -0.032078834 |
| RNF5P1 | -0.032080553 |
| Six3os1_7 | -0.032081732 |
| RP11-89B16 | -0.032087902 |
| IMP3P2 | -0.032089742 |
| STXBP2 | -0.032094395 |
| ATP6V0E1P3 | -0.032096228 |
| SRP68P3 | -0.032106397 |
| PPCS | -0.032107419 |
| MOGAT1 | -0.03210821 |
| RN7SL602P | -0.032111669 |
| AC008536 | -0.03211362 |
| RP11-76E16 | -0.032113942 |
| ATP1A2 | -0.032119325 |
| CFAP53 | -0.032121578 |
| IP6K2 | -0.032122489 |
| RP11-450I1 | -0.032124109 |
| ZNF208 | -0.032128681 |
| AC124276 | -0.032134659 |
| RP11-564C4 | -0.032136509 |
| HNRNPA3P13 | -0.032137036 |
| MIR4425 | -0.03214096 |
| MIR302B | -0.032142634 |
| LRRC37A11P | -0.032150404 |
| HNRNPA1P42 | -0.03215059 |
| AL591610 | -0.032154633 |
| RP4-725K1 | -0.032156048 |
| RP11-330M19 | -0.032161603 |
| TAL2 | -0.032162903 |
| RP11-697M17 | -0.032164226 |
| IQCJ-SCHIP1-AS1 | -0.03216775 |
| KRTAP13-6P | -0.032168998 |
| RP11-168J19 | -0.03217003 |
| ASS1P12 | -0.032176541 |
| ABCB10P4 | -0.032177121 |
| MIR193B | -0.03217915 |
| RP11-686F15 | -0.03218071 |
| FABP12 | -0.032181877 |
| PTMA | -0.032185674 |
| HNRNPA1 | -0.032190809 |
| MIR2276 | -0.032192852 |
| ZNF542P | -0.032195181 |

| | |
|---|---|
| TDRD3 | -0.03219987 |
| RP11-267M23 | -0.032200986 |
| RP11-284H18 | -0.032203175 |
| PPT2 | -0.032207033 |
| COPS5P1 | -0.032208579 |
| MEF2B | -0.032208615 |
| SETP4 | -0.032210078 |
| OR2T2 | -0.032215907 |
| RPL15P20 | -0.032216115 |
| TNPO1P3 | -0.032217511 |
| PIGS | -0.032222178 |
| CTD-2058B24 | -0.032223203 |
| RP5-1114G22 | -0.032226458 |
| Z83819 | -0.03222913 |
| DDX39A | -0.032230717 |
| RNU6-840P | -0.032235038 |
| CFP | -0.032235629 |
| H2AFZP5 | -0.032236406 |
| C1orf226 | -0.032238234 |
| RNA5SP103 | -0.032242868 |
| CLEC3A | -0.032246837 |
| SUCLA2P2 | -0.032247051 |
| DIEXF | -0.032247704 |
| PRSS41 | -0.032249788 |
| RP11-810F22 | -0.03225255 |
| SLC45A4 | -0.03225403 |
| RNA5SP277 | -0.032257941 |
| CA12 | -0.032259054 |
| LGALS9 | -0.032262068 |
| KIZ | -0.032264226 |
| RP1-139G21 | -0.032264478 |
| CDC42P6 | -0.032265124 |
| OR52Q1P | -0.032266609 |
| RP11-669N7 | -0.032267335 |
| MIR574 | -0.032268005 |
| PMPCB | -0.032270626 |
| RP11-474L11 | -0.032272355 |
| MIR548AQ | -0.03227294 |
| AC010891 | -0.032277725 |
| SCARNA11 | -0.032279158 |
| AC012485 | -0.032281519 |
| RP13-379O24 | -0.032283017 |
| FTH1P1 | -0.03228696 |
| GLULP4 | -0.032291079 |
| PRPF4 | -0.032297641 |

| | |
|---|---|
| RPSAP63 | -0.032307704 |
| C9orf116 | -0.032308414 |
| OR6M2P | -0.032312755 |
| DEFB131 | -0.032313179 |
| RNU6-590P | -0.032314963 |
| RP11-462G22 | -0.032316212 |
| HRG | -0.032322222 |
| RP11-63L7 | -0.032322671 |
| AC067718 | -0.0323237 |
| IFIT1P1 | -0.032324158 |
| CTD-2012I17 | -0.032325276 |
| WNK4 | -0.032325419 |
| RP11-506N2 | -0.032337439 |
| STK33P1 | -0.032344364 |
| OR51R1P | -0.032344402 |
| FDX1 | -0.032344899 |
| RP11-342H21 | -0.032346436 |
| KCTD3 | -0.032351783 |
| AL353141 | -0.032354566 |
| RNU6-304P | -0.032355184 |
| IRGQ | -0.032357923 |
| RP11-30M20 | -0.032359072 |
| RP11-996F15 | -0.032364863 |
| CTD-2251F13 | -0.032365291 |
| ARHGDIA | -0.032371579 |
| RP11-757G1 | -0.032371896 |
| POLE2 | -0.03237638 |
| RN7SL7P | -0.032380413 |
| SAPCD1 | -0.032380968 |
| PTK6 | -0.032381682 |
| WFDC13 | -0.032385948 |
| RP11-324J3 | -0.032388639 |
| RP11-317B7 | -0.03239283 |
| GOLGA3 | -0.032394018 |
| AC018816 | -0.032394967 |
| SPINK7 | -0.032396134 |
| RP11-750I4 | -0.032400189 |
| AC073342 | -0.032400341 |
| RNU6-1257P | -0.032403505 |
| NFAM1 | -0.032403804 |
| SPATA24 | -0.032405627 |
| RNU6-334P | -0.032412262 |
| WBP2NL | -0.032412418 |
| RPRML | -0.03241308 |
| CTB-184G21 | -0.032415826 |

| | |
|---|---|
| MIR4495 | -0.032416271 |
| SNORD127 | -0.032417716 |
| RN7SL685P | -0.03241922 |
| RNF170 | -0.03243134 |
| C10orf2 | -0.03243508 |
| ZNF146 | -0.032436233 |
| PLA2G7 | -0.032436494 |
| GDF5OS | -0.032438534 |
| DNM1P31 | -0.032441507 |
| CTC-137K3 | -0.032448491 |
| MAPRE2 | -0.032448736 |
| AC092299 | -0.032449099 |
| RP11-61D3 | -0.032449646 |
| GAPDHP37 | -0.032449881 |
| RSL1D1 | -0.032452569 |
| RPL7P21 | -0.032453406 |
| RP11-317O24 | -0.03245575 |
| RN7SL515P | -0.032455922 |
| PPP2R5E | -0.032456848 |
| RP11-168L7 | -0.032458636 |
| RP11-222G7 | -0.032458837 |
| RP11-775D22 | -0.032466039 |
| LRRC6 | -0.032466691 |
| AC010609 | -0.032468307 |
| PAX2 | -0.032471371 |
| EIF4A1P2 | -0.032472238 |
| CTC-529G1 | -0.03247389 |
| RP11-475F14 | -0.032476888 |
| RP11-2G1 | -0.03248232 |
| RP11-654G14 | -0.032483385 |
| RP11-361F15 | -0.032484029 |
| AL356135 | -0.032486074 |
| HCRT | -0.032486958 |
| OR2T11 | -0.03248712 |
| RP11-648O15 | -0.032487896 |
| RP11-402G3 | -0.032491407 |
| MIR627 | -0.032492589 |
| RP11-270L13 | -0.032493173 |
| RNU1-120P | -0.032495816 |
| SCAMP1 | -0.03249752 |
| GTF2A2 | -0.032499548 |
| AC068506 | -0.03250101 |
| KRTAP5-4 | -0.032503027 |
| RP11-366L18 | -0.032505134 |
| MFSD1 | -0.032507753 |

| | |
|---|---|
| RNA5SP288 | -0.032510435 |
| RN7SL138P | -0.032510674 |
| MYO16-AS2 | -0.032513562 |
| PATE3 | -0.032516812 |
| ANXA7 | -0.032527196 |
| SCHLAP1 | -0.032530697 |
| KRT27 | -0.032533765 |
| RP11-196D18 | -0.032535756 |
| RP11-719K4 | -0.032539109 |
| PRDM8 | -0.032540861 |
| CTC-233O10 | -0.032541975 |
| TRAJ49 | -0.032542636 |
| RNA5SP170 | -0.032544659 |
| RNA5SP200 | -0.032546777 |
| MBTPS1 | -0.032558695 |
| LRRC2-AS1 | -0.032560589 |
| MTMR12 | -0.032563532 |
| DAOA-AS1 | -0.032563633 |
| RP11-310N16 | -0.032564741 |
| NF1P4 | -0.032565437 |
| RP11-177G23 | -0.032571786 |
| OSTF1 | -0.032572332 |
| FEZF2 | -0.032580942 |
| AKAP7 | -0.032583403 |
| VN1R11P | -0.032586717 |
| KLHDC9 | -0.032590175 |
| RP11-451H23 | -0.03259695 |
| GOLGA8F | -0.032606273 |
| RP11-411G7 | -0.032611095 |
| SPRYD3 | -0.032614902 |
| CTA-256D12 | -0.032617204 |
| ENTPD8 | -0.032618846 |
| AC015922 | -0.032619943 |
| AC021134 | -0.032626881 |
| RNU6-537P | -0.032627921 |
| RP11-762L8 | -0.032629146 |
| CTD-3105H18 | -0.032629302 |
| RNA5SP299 | -0.032631062 |
| AC004079 | -0.032633529 |
| AL157392 | -0.03263423 |
| FOXR1 | -0.032642672 |
| RP5-963E22 | -0.032651369 |
| ZBTB34 | -0.032653508 |
| EVI5 | -0.032653713 |
| CTD-3193O13 | -0.032654676 |

| | |
|---|---|
| LINC00412 | -0.032657993 |
| AC007128 | -0.032658205 |
| GPR141 | -0.032662535 |
| SETP17 | -0.032664412 |
| OR52L2P | -0.032671525 |
| OR56A1 | -0.032676876 |
| FRA10AC1 | -0.032679574 |
| IDI1 | -0.032684765 |
| VMAC | -0.032690719 |
| C2orf27A | -0.032695166 |
| NPM1P14 | -0.032697693 |
| TMED10P2 | -0.032699482 |
| B4GALT6 | -0.032699966 |
| RP11-629P16 | -0.032702056 |
| PPFIBP2 | -0.032703762 |
| LINC00895 | -0.03270415 |
| RP11-856F16 | -0.032707693 |
| MCEE | -0.032708655 |
| LINC00930 | -0.03271027 |
| LINC01046 | -0.032710881 |
| MIR7160 | -0.032714168 |
| FSD2 | -0.032714987 |
| RP11-10K17 | -0.032718793 |
| RP11-410B16 | -0.032719071 |
| AC024619 | -0.032720717 |
| RP1-178F15 | -0.032721 |
| RAC1P5 | -0.032721234 |
| RP11-556N4 | -0.032723097 |
| MIR936 | -0.032725102 |
| RP11-364C11 | -0.032730809 |
| DPH5 | -0.032739224 |
| UBQLN2 | -0.032741903 |
| RP11-452K12 | -0.032747336 |
| C22orf29 | -0.032749308 |
| KRTAP4-12 | -0.032752486 |
| MRPS15P1 | -0.032754043 |
| SNORD101 | -0.03275567 |
| ITGAE | -0.032755708 |
| ARHGAP5 | -0.032765383 |
| TRIM45 | -0.032768639 |
| OR2G6 | -0.032781514 |
| NPM1P6 | -0.032784398 |
| ILKAP | -0.032786072 |
| RN7SL442P | -0.032789677 |
| PIPSL | -0.032790444 |

| | |
|---|---|
| RNU6-1282P | -0.032792301 |
| B4GAT1 | -0.032792305 |
| RP11-502I4 | -0.0327929 |
| RP11-757O6 | -0.032792923 |
| ATP6V1G2-DDX39B | -0.032793365 |
| AP000362 | -0.032795675 |
| RNU7-19P | -0.032796183 |
| KRTAP13-5P | -0.032796397 |
| SNX14 | -0.032796789 |
| CTB-41I6 | -0.032802683 |
| ANO2 | -0.032811264 |
| NPBWR2 | -0.032818287 |
| TLR6 | -0.032821739 |
| GLB1L3 | -0.032822661 |
| ATXN3 | -0.032824771 |
| ZNF512 | -0.032824809 |
| ANKDD1A | -0.032827711 |
| RN7SL68P | -0.032827777 |
| C9orf173-AS1 | -0.032829441 |
| PDCL2P1 | -0.032830131 |
| CTSLP2 | -0.032847748 |
| RNA5SP175 | -0.032851764 |
| PDCD2 | -0.032852159 |
| ZNF596 | -0.032856475 |
| ANXA6 | -0.032859247 |
| PIGFP2 | -0.032859644 |
| RNA5SP116 | -0.032860505 |
| LRRC46 | -0.032864792 |
| CDC37P1 | -0.032864839 |
| CHCHD4P2 | -0.032868218 |
| OR6C68 | -0.03287093 |
| CCDC175 | -0.032872635 |
| MIR496 | -0.032872684 |
| PAQR8 | -0.032872922 |
| HMGB1P7 | -0.032873025 |
| RP11-5A1 | -0.032875258 |
| RP11-697B24 | -0.032875758 |
| RNU6-1067P | -0.032888252 |
| ASH2L | -0.03289523 |
| VTI1BP4 | -0.03290185 |
| RNF219-AS1 | -0.03290421 |
| AC107068 | -0.032906246 |
| RP11-211N8 | -0.032908633 |
| RN7SL60P | -0.032910695 |
| CTD-2206N4 | -0.032912106 |

| | |
|---|---|
| RP11-271F18 | -0.032916219 |
| AC093609 | -0.032917756 |
| RPP38 | -0.032920255 |
| KRTAP10-8 | -0.032922228 |
| AMBRA1 | -0.032923027 |
| IQCD | -0.032925092 |
| AC104301 | -0.032926609 |
| LINGO3 | -0.032927901 |
| LINC00200 | -0.032932115 |
| FAM3B | -0.032935418 |
| CXXC4 | -0.032942916 |
| RP11-897M7 | -0.032942938 |
| AC005300 | -0.032943144 |
| AIMP1P1 | -0.032947001 |
| LL22NC03-123E1 | -0.032948523 |
| RP1-60G11 | -0.032949256 |
| RP11-13J8 | -0.032950902 |
| FARSA-AS1 | -0.03295223 |
| RP4-771M4 | -0.032953675 |
| AP000770 | -0.03296124 |
| SLC15A5 | -0.032961316 |
| RP11-402D21 | -0.032962948 |
| GHc-857G6 | -0.032971479 |
| AC005777 | -0.03297201 |
| MYCT1 | -0.032974434 |
| CTD-3162L10 | -0.032974862 |
| OR1L4 | -0.032988618 |
| OR5AN1 | -0.032989376 |
| FRMPD3 | -0.032990035 |
| RP11-181K12 | -0.032991174 |
| RNU6-1154P | -0.032995883 |
| RP11-451L19 | -0.032997525 |
| RP11-684B21 | -0.032999522 |
| CTD-2151L9 | -0.033005193 |
| FSHB | -0.033005401 |
| OR7E149P | -0.033006942 |
| AC084859 | -0.033009693 |
| RP5-1024G6 | -0.033010268 |
| MIR1197 | -0.033011919 |
| C6orf106 | -0.033014584 |
| BROX | -0.033014771 |
| FAM120B | -0.033017264 |
| CHTF8 | -0.033019385 |
| RAB8B | -0.033023236 |
| SIAH2-AS1 | -0.033023456 |

| | |
|---|---|
| PDXDC1 | -0.033026678 |
| BNIP3P36 | -0.033032934 |
| HCN4 | -0.033033684 |
| CTD-3194G12 | -0.033035875 |
| SNRPA1 | -0.033037916 |
| RNU6-519P | -0.033042942 |
| AL158210 | -0.033045314 |
| HCG4P3 | -0.033047107 |
| PTTG3P | -0.033058606 |
| MIR1468 | -0.033059311 |
| RNA5SP453 | -0.033061825 |
| RP11-252C15 | -0.033062952 |
| PPP2R2DP1 | -0.033064087 |
| RP11-118M9 | -0.033077572 |
| AC007618 | -0.033083967 |
| HDAC1P2 | -0.03308436 |
| RP11-53B2 | -0.033085062 |
| OR52P2P | -0.033090125 |
| BLMH | -0.03309256 |
| RNU6-920P | -0.033096995 |
| STXBP5 | -0.033099072 |
| MIR4540 | -0.033104061 |
| AL954742 | -0.033110351 |
| RN7SL417P | -0.033114661 |
| AP001437 | -0.033122094 |
| RTP2 | -0.0331224 |
| FAM120C | -0.033125213 |
| ZNF736P6Y | -0.033130943 |
| FAM220CP | -0.033134519 |
| TMEM120B | -0.03313619 |
| RP11-367H5 | -0.033138952 |
| IGIP | -0.03313965 |
| RNU6-902P | -0.033144025 |
| RP11-93H24 | -0.033145679 |
| RP11-889L3 | -0.033146553 |
| AC114801 | -0.033148444 |
| ADAT2 | -0.033161475 |
| BPIFB5P | -0.03316292 |
| THEM5 | -0.033165196 |
| RNA5SP513 | -0.033165537 |
| RPS21P1 | -0.033166889 |
| LIME1 | -0.033168414 |
| MRPL35P1 | -0.033173798 |
| DUX4L18 | -0.033174118 |
| CISD3 | -0.033179968 |

| | |
|---|---|
| RPS6KA3 | -0.03318062 |
| LINC00500 | -0.0331827 |
| SRD5A3P1 | -0.033182911 |
| GXYLT1P3 | -0.033183931 |
| MIR548AN | -0.033194455 |
| RP11-400G3 | -0.033194963 |
| TFCP2L1 | -0.033203318 |
| RP11-22N12 | -0.033203608 |
| SMN1 | -0.033207858 |
| PCNPP4 | -0.03320877 |
| Y10196 | -0.033210858 |
| AL355149 | -0.033221198 |
| CTD-2329C7 | -0.03322542 |
| RP11-75N6 | -0.033228166 |
| GAMTP2 | -0.033228647 |
| TBX10 | -0.03322927 |
| SDCBP2P1 | -0.033231361 |
| MIR1324 | -0.033231957 |
| CDC42BPB | -0.03323558 |
| RP11-126M14 | -0.033236203 |
| RNU6-1100P | -0.033238921 |
| CH17-262O2 | -0.03323941 |
| OR4C49P | -0.033246366 |
| RP11-48O20 | -0.033251715 |
| TRAV8-7 | -0.033262376 |
| RNU6-347P | -0.033264881 |
| RP11-216F5 | -0.0332656 |
| PAX6 | -0.033268382 |
| OR2AK2 | -0.03327114 |
| RN7SKP168 | -0.033278413 |
| RP11-10F11 | -0.033285434 |
| OR7M1P | -0.033289628 |
| MST1 | -0.033293276 |
| OR5G5P | -0.033294114 |
| PYGO2 | -0.033294262 |
| RP11-328N19 | -0.03329544 |
| RP11-1112J20 | -0.033297804 |
| RFESD | -0.033299174 |
| AL096803 | -0.033299347 |
| FAHD1 | -0.033300216 |
| DHX57 | -0.033302355 |
| HSPA8P18 | -0.033303915 |
| TRBV23OR9-2 | -0.033307038 |
| BMX | -0.033308312 |
| SLC25A38P1 | -0.033310377 |

| | |
|---|---|
| RP11-88L20 | -0.033312144 |
| CENPI | -0.033312532 |
| DDX11L2 | -0.033315522 |
| RP11-159F24 | -0.033319993 |
| AP000897 | -0.033323965 |
| TSPY22P | -0.033331702 |
| FAM90A22P | -0.033332438 |
| RP4-782L23 | -0.033333674 |
| CHRM2 | -0.033334371 |
| OLA1P3 | -0.033338658 |
| TAS2R20 | -0.033339752 |
| DGKQ | -0.033340143 |
| HCST | -0.033346301 |
| DOK1 | -0.033349092 |
| MIR6747 | -0.033351743 |
| CEND1P1 | -0.033352421 |
| RNY3P7 | -0.033352477 |
| RNU6-23P | -0.03335955 |
| AC073143 | -0.033360472 |
| TSEN15P3 | -0.033363666 |
| TNP2 | -0.03336472 |
| AC087174 | -0.033369572 |
| RP11-57H14 | -0.033370805 |
| PTDSS1 | -0.033375828 |
| AC084082 | -0.033380893 |
| AC004899 | -0.033381548 |
| TRBV22OR9-2 | -0.033382355 |
| LINC00355 | -0.033383521 |
| TDRG1 | -0.03338506 |
| RPS15AP25 | -0.033389233 |
| UQCRFS1P3 | -0.033393624 |
| AC009262 | -0.033394344 |
| RP11-320G10 | -0.033396929 |
| AC073635 | -0.033398124 |
| RP1-92C4 | -0.033398138 |
| RP11-320P7 | -0.033401286 |
| LINC01121 | -0.033401795 |
| RP13-653N12 | -0.03340719 |
| RP11-598D14 | -0.033408352 |
| RP11-705O1 | -0.033420261 |
| AC013269 | -0.033423008 |
| RP11-907D1 | -0.033424989 |
| FOXD2-AS1 | -0.033426007 |
| XXyac-YX155B6 | -0.033429849 |
| HNRNPLP2 | -0.03343051 |

| | |
|---|---|
| RP11-35N6 | -0.033432129 |
| RP11-728F11 | -0.033433839 |
| DRD1 | -0.033437559 |
| C15orf65 | -0.033439719 |
| AK3P3 | -0.03343989 |
| RANP1 | -0.033443505 |
| RP11-186B7 | -0.033444241 |
| TSPAN16 | -0.033448531 |
| MB | -0.033452836 |
| IFT74 | -0.033455813 |
| NUTM2HP | -0.033456984 |
| LRRC55 | -0.03346212 |
| RP11-30B1 | -0.033463834 |
| SERPINA9 | -0.033464234 |
| AC019080 | -0.033476309 |
| OR5M13P | -0.033478654 |
| CTC-558O2 | -0.033479347 |
| AC000036 | -0.033481228 |
| RP11-132J14 | -0.033482148 |
| AC005024 | -0.033488578 |
| CHMP4C | -0.033490698 |
| RP11-598O8 | -0.033495631 |
| RP11-1029F8 | -0.033497203 |
| AC092069 | -0.033501811 |
| ZFYVE9P1 | -0.033510311 |
| RP11-215H18 | -0.033515193 |
| SNORD95 | -0.03351828 |
| AC078794 | -0.03352413 |
| RP11-101E3 | -0.033525912 |
| LCE2D | -0.033527635 |
| RNU6-316P | -0.033528291 |
| RP1-298M8 | -0.03354036 |
| TNFSF10 | -0.033546906 |
| MFAP3 | -0.033550254 |
| RP1-296G17 | -0.033550714 |
| AC108044 | -0.033551215 |
| ACOX2 | -0.033552487 |
| TRIB1 | -0.033563725 |
| RP4-769N13 | -0.033564525 |
| RN7SKP157 | -0.033564538 |
| TFAMP1 | -0.033569105 |
| TISP43 | -0.033569483 |
| UBBP4 | -0.03357113 |
| AC074011 | -0.033571564 |
| RP11-417J1 | -0.033580192 |

| | |
|---|---|
| RN7SL838P | -0.033589524 |
| ZNF713 | -0.033591317 |
| MIR588 | -0.033591887 |
| MIR1251 | -0.033594885 |
| NR2E3 | -0.033595998 |
| ACTG1P3 | -0.033596288 |
| RP11-72I18 | -0.033597548 |
| C14orf142 | -0.033600992 |
| RPL7P26 | -0.033604504 |
| U91324 | -0.033604706 |
| MOB3C | -0.033606741 |
| AC005539 | -0.033611515 |
| BRINP2 | -0.033612394 |
| LINC00444 | -0.033619394 |
| LOH12CR1 | -0.033619913 |
| KSR1 | -0.033620788 |
| RP11-470E16 | -0.033624648 |
| CTD-3138F19 | -0.033632259 |
| TREX1 | -0.033635449 |
| C16orf52 | -0.033636547 |
| CRYGS | -0.033637216 |
| RNA5SP319 | -0.03363992 |
| RP4-659I19 | -0.033641888 |
| CICP10 | -0.033643908 |
| LINC00863 | -0.033648045 |
| SLCO2A1 | -0.033649459 |
| MIR4726 | -0.033649882 |
| RNU6-1284P | -0.033661759 |
| TBC1D3E | -0.033674023 |
| AC112518 | -0.033676248 |
| SEBOX | -0.033677531 |
| CDC42EP4 | -0.033678632 |
| ARHGEF12 | -0.033679376 |
| RPL12P37 | -0.033683724 |
| SPATA8-AS1 | -0.033690355 |
| RP11-329A14 | -0.033692655 |
| CSMD3 | -0.033695387 |
| BRD9P2 | -0.033703438 |
| RP11-357H14 | -0.033705575 |
| RNA5SP82 | -0.033715356 |
| COX6CP8 | -0.033717578 |
| VN1R51P | -0.03371871 |
| CTD-2123J17 | -0.033720422 |
| HIC2 | -0.033724569 |
| CIR1P1 | -0.033725148 |

| | | |
|---|---|---|
| CTD-2339L15 | | -0.033726181 |
| MIR4476 | | -0.033726762 |
| CENPA | | -0.033734318 |
| GNPTAB | | -0.033735918 |
| MIR518F | | -0.033740766 |
| RP11-382J24 | | -0.033741308 |
| BAG6 | | -0.033746989 |
| AADACL4 | | -0.033750251 |
| RNU6-689P | | -0.033752022 |
| AC011467 | | -0.033752486 |
| MIR493 | | -0.03375329 |
| PAGE2B | | -0.033758864 |
| CHD5 | | -0.033759862 |
| LINC01104 | | -0.033765722 |
| OTOL1 | | -0.033769717 |
| RNU6-1138P | | -0.033771042 |
| RNU7-130P | | -0.033776009 |
| RP11-79M19 | | -0.033776265 |
| PTPN20A | | -0.033779658 |
| DLGAP2 | | -0.033785492 |
| TIMM8BP1 | | -0.033785607 |
| TCN2 | | -0.033785632 |
| RP13-487C10 | | -0.033790269 |
| UBA3 | | -0.033792442 |
| CR769776 | | -0.033796429 |
| C18orf21 | | -0.033797149 |
| RP1-15D7 | | -0.033799294 |
| NTRK2 | | -0.033801986 |
| MIR520D | | -0.033803508 |
| A1BG-AS1 | | -0.033803587 |
| CTD-2017F17 | | -0.033803921 |
| ENHO | | -0.033809671 |
| ZDHHC22 | | -0.033811687 |
| TMEM243 | | -0.033811922 |
| ALDH8A1 | | -0.033812092 |
| FAM106A | | -0.033818663 |
| RP11-840I19 | | -0.033819077 |
| | 10-Mar | -0.033821052 |
| YBX1P7 | | -0.033824745 |
| RP11-40A13 | | -0.033825586 |
| RNU6-116P | | -0.033827642 |
| LMLN-AS1 | | -0.033830911 |
| RP11-650P15 | | -0.033835115 |
| KRT85 | | -0.033850836 |
| RP11-584P21 | | -0.0338589 |

| | |
|---|---|
| NPM1P4 | -0.033859258 |
| RP11-510I6 | -0.03386393 |
| MIR1250 | -0.03386943 |
| TSPO2 | -0.033873129 |
| CROCCP5 | -0.033876327 |
| RPS6KA1 | -0.033876669 |
| RNU6-1013P | -0.03387868 |
| RNU6-335P | -0.033881636 |
| RBM22P12 | -0.033882451 |
| RP11-648M2 | -0.033883186 |
| RN7SL728P | -0.033883613 |
| MIR548AJ1 | -0.033884098 |
| ZNF214 | -0.033889469 |
| OR8B8 | -0.03389603 |
| LOXL3 | -0.033896632 |
| RNU11-3P | -0.033898958 |
| FKBP1AP3 | -0.033902815 |
| RP1-127L4 | -0.033906517 |
| RNU6-982P | -0.033911523 |
| RNU7-63P | -0.033916271 |
| SNRPGP18 | -0.033919829 |
| BRD4 | -0.033925173 |
| AC066692 | -0.033925372 |
| C6orf10 | -0.033925375 |
| SNRPCP10 | -0.033925379 |
| APTR | -0.033926234 |
| RP11-805I24 | -0.033926723 |
| ISCA1P4 | -0.033930468 |
| RNU6-498P | -0.033942481 |
| MIR151B | -0.033942749 |
| RP4-809F18 | -0.033943136 |
| ASAH2B | -0.033949639 |
| HSPD1P15 | -0.033956044 |
| RANBP3 | -0.033956523 |
| CR1L | -0.033960221 |
| GCA | -0.033960691 |
| RNU5E-10P | -0.033961032 |
| SLC34A3 | -0.03396119 |
| RPL17P45 | -0.033966361 |
| AC009541 | -0.033968514 |
| CHTF8P1 | -0.033969769 |
| RNU6-434P | -0.033975204 |
| MIR554 | -0.033975334 |
| TFB1M | -0.033976134 |
| TUBB4BP6 | -0.033977219 |

| | |
|---|---|
| FAM58DP | -0.033978555 |
| AP000295 | -0.033978809 |
| USP9X | -0.03397885 |
| RNU6-878P | -0.033979677 |
| LINC00320 | -0.033986875 |
| ZNF500 | -0.033993065 |
| RNY1P14 | -0.033994277 |
| RP11-122K13 | -0.033995539 |
| TPT1P6 | -0.033996988 |
| ATP5A1P7 | -0.033997073 |
| LL22NC03-48A11 | -0.03400039 |
| RP11-209K10 | -0.034001888 |
| NUDCD2 | -0.034005921 |
| OR5T1 | -0.034013071 |
| CTA-351J1 | -0.03401517 |
| SRRM1P2 | -0.034016699 |
| AC026703 | -0.034032253 |
| OR8B2 | -0.034038115 |
| CCDC177 | -0.034039749 |
| MED15P7 | -0.034048071 |
| USP6 | -0.03404996 |
| RP11-548K12 | -0.034056738 |
| IL20RB-AS1 | -0.034065978 |
| MIR4323 | -0.034072774 |
| RP11-369E15 | -0.034078819 |
| CBLB | -0.034087004 |
| AC016142 | -0.034091576 |
| HOXB4 | -0.034098706 |
| GADL1 | -0.03410938 |
| RLIM | -0.034113051 |
| HKDC1 | -0.034125108 |
| LDLRAD3 | -0.034127489 |
| AC104996 | -0.034127722 |
| RNU6-497P | -0.034128004 |
| RP11-185J20 | -0.034139554 |
| RP11-730B22 | -0.034143185 |
| MTND5P1 | -0.034143607 |
| RP11-959I15 | -0.034143772 |
| MYLKP1 | -0.034143935 |
| RNY1P2 | -0.034145803 |
| RP11-23D5 | -0.034169435 |
| TTI2 | -0.034173891 |
| RP5-1033E15 | -0.034175019 |
| ATP1B3P1 | -0.034176678 |
| CBX3P5 | -0.034176704 |

| | |
|---|---|
| PPIAP1 | -0.034181502 |
| RP11-367O10 | -0.034188263 |
| ZNF615 | -0.034188627 |
| RP11-501E14 | -0.034191963 |
| CROCCP4 | -0.034195544 |
| CTC-450M9 | -0.034200889 |
| RNU6-969P | -0.034202927 |
| AP001189 | -0.034206137 |
| RP4-545K15 | -0.034207842 |
| RP11-396C23 | -0.034208962 |
| EEA1 | -0.03420939 |
| NUP133 | -0.034211437 |
| ZNF304 | -0.034218656 |
| LA16c-385E7 | -0.034224835 |
| RP11-338H14 | -0.034228948 |
| GBA3 | -0.034233804 |
| LINC00967 | -0.03423935 |
| MAGEB2 | -0.034242591 |
| RNU6-444P | -0.034247132 |
| PA2G4P1 | -0.034248566 |
| ADAM6 | -0.034257439 |
| ARPC3P5 | -0.034261243 |
| RP11-535C7 | -0.034266822 |
| NEB | -0.034269914 |
| RP11-530A18 | -0.034270903 |
| OR9S24P | -0.034272365 |
| RN7SL624P | -0.034272941 |
| RP11-2G17 | -0.034276735 |
| RP1-156L9 | -0.034276983 |
| CHD4 | -0.034280166 |
| PTPN12 | -0.034291459 |
| RP11-527L4 | -0.034297806 |
| RP11-136B18 | -0.034303239 |
| AC090810 | -0.034305744 |
| RP11-567P17 | -0.034307597 |
| RP11-94I2 | -0.034308208 |
| DHRS7 | -0.034311311 |
| PIP5K1P2 | -0.034314898 |
| RP11-335O13 | -0.034316647 |
| ZAR1L | -0.034323919 |
| RP11-115H15 | -0.034324586 |
| RNU5D-1 | -0.034326234 |
| SLC12A1 | -0.034328601 |
| CALCRL | -0.034336168 |
| FAM9B | -0.034336347 |

| | |
|---|---|
| RP11-95I16 | -0.034338897 |
| RUVBL1-AS1 | -0.034341542 |
| LINC00159 | -0.034344931 |
| GCDH | -0.034348745 |
| TTC33 | -0.034350215 |
| CHM | -0.034357322 |
| AL161752 | -0.034357599 |
| CTB-189B5 | -0.034366915 |
| RP11-568G11 | -0.034373535 |
| RP1-164F3 | -0.034373573 |
| EEF1A1P3 | -0.034375143 |
| CAMK2G | -0.034377212 |
| AC011288 | -0.034380477 |
| RNU6-17P | -0.034382655 |
| TDP1 | -0.034385026 |
| Vax2os1_3 | -0.034389683 |
| BEND7P1 | -0.034390905 |
| MSH5-SAPCD1 | -0.034397771 |
| ZC3H18 | -0.034405176 |
| HSBP1L1 | -0.034408208 |
| RP11-144O23 | -0.034411588 |
| AL117334 | -0.034412796 |
| FAM90A10P | -0.034418547 |
| LINC00029 | -0.034419604 |
| AMZ1 | -0.03442233 |
| GAPDHP15 | -0.034424168 |
| DEFB108P1 | -0.034431329 |
| NT5DC4 | -0.034433446 |
| RNU6-197P | -0.034434375 |
| RP11-429P7 | -0.034434508 |
| AP005135 | -0.03443632 |
| RNF19B | -0.034437847 |
| RNU7-156P | -0.034440158 |
| RNU6-893P | -0.03444227 |
| RP11-274M17 | -0.034444526 |
| RP11-844P9 | -0.034446047 |
| MGAT5B | -0.034453196 |
| AC002981 | -0.034453231 |
| KLHDC10 | -0.034455485 |
| MIR4318 | -0.034461419 |
| SUMO1P3 | -0.034461792 |
| PRR4 | -0.034463318 |
| ZNF733P | -0.034466337 |
| RP11-205K6 | -0.034470398 |
| RP11-70C1 | -0.034471364 |

| | |
|---|---|
| RP11-396B14 | -0.034472825 |
| SUMO2P6 | -0.034480359 |
| RNU6-1208P | -0.034482291 |
| SCRT1 | -0.034482679 |
| GTPBP2 | -0.034483933 |
| MIR3168 | -0.034486316 |
| RNU6-226P | -0.034488324 |
| LIPT2 | -0.034488677 |
| RP11-415K20 | -0.034491857 |
| DCUN1D1 | -0.034495543 |
| RP11-91I20 | -0.034496907 |
| RNU6-761P | -0.03449774 |
| RP11-317B3 | -0.034500621 |
| CTB-131B5 | -0.034502278 |
| OSBPL7 | -0.034504027 |
| CTC-236F12 | -0.034508174 |
| TRIM31 | -0.034517819 |
| RP1-181J22 | -0.034522371 |
| DAPK1-IT1 | -0.034524687 |
| RNU6-21P | -0.034526317 |
| KPNA6 | -0.034526539 |
| OR7E11P | -0.034528829 |
| KIF19 | -0.03452905 |
| LIPM | -0.034529292 |
| ADIRF-AS1 | -0.034530446 |
| RNA5SP322 | -0.034534611 |
| PRR3 | -0.034535414 |
| LBX2-AS1 | -0.034536698 |
| LLPH-AS1 | -0.034543408 |
| RP11-444B24 | -0.034546127 |
| RP11-567M16 | -0.034553945 |
| AL121989 | -0.034555118 |
| OR7E116P | -0.034555777 |
| ATIC | -0.034557127 |
| CMIP | -0.034560085 |
| RP11-149B9 | -0.034560085 |
| HAVCR2 | -0.034562541 |
| MED15P8 | -0.034566263 |
| RP11-312A15 | -0.034567293 |
| AL157773 | -0.034567394 |
| UGT2A3P7 | -0.034569346 |
| DNM3-IT1 | -0.034570689 |
| APBB2 | -0.034580049 |
| RP11-381K7 | -0.034581753 |
| OR52N3P | -0.034584169 |

| | |
|---|---|
| RP5-867C24 | -0.034585479 |
| ZNF148 | -0.034586077 |
| FDPSP3 | -0.034586947 |
| ELDR | -0.03458705 |
| MIR518A2 | -0.034590031 |
| CTC-204F22 | -0.034594719 |
| NBAS | -0.034595324 |
| MKRNP2 | -0.034598748 |
| RP11-440E5 | -0.034601649 |
| RP11-731K22 | -0.034614654 |
| CDC73 | -0.034615659 |
| IGHV3-33-2 | -0.034615687 |
| SPATA31A3 | -0.034620639 |
| MRPL2P1 | -0.034621003 |
| SP2-AS1 | -0.034622227 |
| AC004837 | -0.034626118 |
| RP11-467I20 | -0.034630216 |
| LRRC39 | -0.034633316 |
| KRT128P | -0.034635373 |
| GUCA2A | -0.034638079 |
| SLC16A12 | -0.034639715 |
| ADRB2 | -0.03464033 |
| MIR548I2 | -0.034643963 |
| MUC5B | -0.034647869 |
| RP11-412B14 | -0.034648086 |
| SPTBN2 | -0.034658136 |
| SAR1B | -0.034659512 |
| MRPS18CP4 | -0.034660451 |
| CYB5B | -0.034665559 |
| RP11-74M11 | -0.034667775 |
| RP11-397K18 | -0.034668183 |
| DNM1P34 | -0.03467379 |
| TBC1D16 | -0.034675383 |
| ZFYVE19 | -0.034675829 |
| FRG1B | -0.034678471 |
| RP11-291C6 | -0.034678818 |
| C16orf97 | -0.034679231 |
| LINC00690 | -0.034690082 |
| RP11-143J24 | -0.034692945 |
| MTND3P9 | -0.034697168 |
| RP3-510L9 | -0.034697845 |
| RP11-278L15 | -0.034701457 |
| SERTAD3 | -0.034703477 |
| OR1AB1P | -0.034704046 |
| RP11-377H23 | -0.034704983 |

| | |
|---|---|
| RP1-142O9 | -0.034708629 |
| RP4-742N3 | -0.034714104 |
| HEIH | -0.034715534 |
| AC074290 | -0.034718939 |
| AC005204 | -0.034719198 |
| PRKAR2A | -0.034721059 |
| AL354984 | -0.034723144 |
| RNU6-979P | -0.034726821 |
| LINC00710 | -0.034729928 |
| RNU6-14P | -0.034733836 |
| RN7SKP61 | -0.034734381 |
| MIR515-2 | -0.034734381 |
| AKR1C8P | -0.034736341 |
| CCDC162P | -0.034736855 |
| PHRF1 | -0.034749935 |
| RP4-734G22 | -0.034752037 |
| RNU6-634P | -0.034752727 |
| RN7SL74P | -0.034758009 |
| RP11-231I16 | -0.03476002 |
| UGT3A1 | -0.034766652 |
| KB-1930G5 | -0.034767357 |
| RP11-850A17 | -0.03476798 |
| AP000266 | -0.034771958 |
| CTD-2026K11 | -0.034777415 |
| OR2D3 | -0.034779778 |
| AC003682 | -0.034784334 |
| RP4-555D20 | -0.034792513 |
| LYZL6 | -0.034794517 |
| RPL34P33 | -0.034795109 |
| RP11-561N12 | -0.034796186 |
| RP5-884M20 | -0.034799097 |
| CHRDL1 | -0.034799426 |
| MTND4P14 | -0.034800541 |
| CERS2 | -0.034800933 |
| RP11-29O13 | -0.034803218 |
| RNASEL | -0.034803221 |
| RNU6-664P | -0.034805061 |
| RP11-67M9 | -0.034814929 |
| IFT122 | -0.034815259 |
| RNA5SP329 | -0.03481681 |
| SLC9A9-AS2 | -0.034818859 |
| TRIM60P17 | -0.0348217 |
| PDCD4 | -0.034822515 |
| ZNF704 | -0.034826348 |
| CENPH | -0.034837102 |

| | |
|---|---|
| HERC2P11 | -0.034839953 |
| ZNF415P1 | -0.034844432 |
| AC068042 | -0.034846794 |
| RNF224 | -0.034850572 |
| MIR3202-1 | -0.034855206 |
| RNA5SP25 | -0.034856504 |
| RNA5-8SP5 | -0.034858085 |
| AC021021 | -0.034858608 |
| KIAA0513 | -0.03486021 |
| PNMA1 | -0.034861773 |
| PTTG2 | -0.034862843 |
| RNA5SP180 | -0.034864038 |
| FBXO30 | -0.03486955 |
| SNURF | -0.034874183 |
| AL121584 | -0.034875971 |
| ARAFP2 | -0.034876701 |
| CTC-344H19 | -0.034880795 |
| RP11-806N19 | -0.034880968 |
| OR7E87P | -0.034882016 |
| KLF15 | -0.034884639 |
| HINT2 | -0.034886332 |
| AC007563 | -0.034886759 |
| MIR516A2 | -0.034890122 |
| DDX10P2 | -0.034890984 |
| PCYT2 | -0.034895262 |
| IRS1 | -0.034901867 |
| RP1-111C20 | -0.03490263 |
| RP11-163M18 | -0.03490319 |
| AC013404 | -0.03490699 |
| RP11-815N9 | -0.034907431 |
| CTC-360G5 | -0.034910872 |
| MIR6877 | -0.034911312 |
| NCOA1 | -0.034915487 |
| LINC00498 | -0.03491568 |
| STOX1 | -0.034916305 |
| RPS3P6 | -0.034917724 |
| RNU7-110P | -0.034921271 |
| AC016582 | -0.034925079 |
| VN1R93P | -0.034925097 |
| RAD23A | -0.034925236 |
| AC110620 | -0.034926033 |
| LINC00092 | -0.034926442 |
| TMED2 | -0.034926706 |
| KRTAP20-3 | -0.034930775 |
| TAPBP | -0.034931041 |

| | |
|---|---|
| RP11-25D3 | -0.034934343 |
| RN7SKP169 | -0.03493682 |
| IGLVVI-22-1 | -0.034938557 |
| MIR4280 | -0.034940641 |
| SNORD79 | -0.034943874 |
| RP11-423G4 | -0.034945182 |
| GABBR1 | -0.034945627 |
| RNA5SP284 | -0.034950529 |
| EML2 | -0.034950949 |
| RP3-425C14 | -0.034953237 |
| TUBG1 | -0.034953777 |
| LINC01446 | -0.03495678 |
| POM121L7 | -0.034957423 |
| CH507-338C24 | -0.034957763 |
| RPL18AP8 | -0.034962985 |
| MAP2K5 | -0.034964326 |
| RP11-512M8 | -0.034964545 |
| TRIM49D1 | -0.034965552 |
| RP11-316M21 | -0.034966298 |
| PDZRN3-AS1 | -0.034966693 |
| QTRT1P1 | -0.034966882 |
| ENPP7P10 | -0.034969063 |
| RERGL | -0.034970321 |
| PLA2G2C | -0.034970735 |
| AC126755 | -0.034973287 |
| CYP4F12 | -0.034987501 |
| TPX2 | -0.034988048 |
| CTD-2194L12 | -0.03499208 |
| MIR5692C2 | -0.034996411 |
| RPL5P25 | -0.034999667 |
| RP11-520B13 | -0.0350015 |
| OR2B3 | -0.035007017 |
| RP11-342A1 | -0.035007161 |
| TPRX2P | -0.035009996 |
| RP3-341O5 | -0.035013164 |
| SSU72P8 | -0.035017836 |
| DISC1FP1 | -0.035021093 |
| CNN2P8 | -0.035021095 |
| BAK1P2 | -0.035021148 |
| STK24-AS1 | -0.035027082 |
| RP11-148K1 | -0.035027994 |
| SOGA3 | -0.035032075 |
| RPL17P20 | -0.035037438 |
| EXO1 | -0.035041185 |
| EIF4EBP2 | -0.035044553 |

| | | |
|---|---|---|
| MAP1LC3B | | -0.035046618 |
| AP000855 | | -0.035047559 |
| RP3-337O18 | | -0.03504858 |
| AL672296 | | -0.035051781 |
| MIR3929 | | -0.035052184 |
| MARK2P12 | | -0.035053969 |
| AC137590 | | -0.035055239 |
| UBB | | -0.035057428 |
| PEX11B | | -0.035057502 |
| RP11-643A5 | | -0.035060639 |
| AC008991 | | -0.03507019 |
| OR2L2 | | -0.035071347 |
| PKMP3 | | -0.035071865 |
| RP11-237N19 | | -0.035076051 |
| TIGD1 | | -0.035085429 |
| RP11-407A16 | | -0.035085779 |
| SNORD11 | | -0.03508662 |
| AC123886 | | -0.0350874 |
| MIR4673 | | -0.035089078 |
| CTD-3154N5 | | -0.035089772 |
| RAB36 | | -0.035091866 |
| RP11-219C24 | | -0.035097121 |
| IFNA14 | | -0.035097341 |
| ONECUT1 | | -0.03510449 |
| | 14-Sep | -0.03510838 |
| AC073464 | | -0.035112809 |
| SEC14L1P1 | | -0.035117365 |
| TMEM248P1 | | -0.03512126 |
| PGGT1BP1 | | -0.035124423 |
| BIRC2 | | -0.035127254 |
| CCDC74B | | -0.035128761 |
| RBPJP5 | | -0.035131593 |
| TAAR2 | | -0.035131953 |
| LINC01169 | | -0.035132563 |
| RP11-191G24 | | -0.03513438 |
| RP11-388B24 | | -0.035147323 |
| RP11-1250I15 | | -0.035147898 |
| CT45A10 | | -0.035163865 |
| RP11-325B23 | | -0.035164 |
| RP11-474I16 | | -0.035164046 |
| FAM174B | | -0.035164093 |
| VTA1 | | -0.035164335 |
| CMTM5 | | -0.0351648 |
| RP11-51L5 | | -0.035164801 |
| AP001630 | | -0.03516519 |

| | |
|---|---|
| RP11-372B4 | -0.035166158 |
| RNU6-744P | -0.035172619 |
| SH3BGR | -0.035177638 |
| NPHP3 | -0.035183294 |
| SLC25A20P1 | -0.035184324 |
| CXorf23 | -0.035187517 |
| NBPF17P | -0.035188099 |
| GUSBP2 | -0.035191145 |
| MIR4286 | -0.035195102 |
| MIR548F5 | -0.035196228 |
| NMD3P2 | -0.03519756 |
| CHIAP1 | -0.035200715 |
| AC099791 | -0.03520304 |
| RNU6ATAC9P | -0.035204589 |
| RPL14P1 | -0.035206178 |
| RBM22 | -0.03520709 |
| RP11-28H5 | -0.035208911 |
| RP11-278I4 | -0.035209729 |
| RP11-347J14 | -0.035214648 |
| SPTA1 | -0.035220987 |
| PAK2 | -0.0352299 |
| RNU6ATAC35P | -0.035236963 |
| KIAA1715 | -0.035237305 |
| CTDSPL | -0.035240412 |
| PRRT1 | -0.035241064 |
| RP11-101C11 | -0.03524143 |
| RP11-165F24 | -0.035241817 |
| RN7SL659P | -0.035245412 |
| TMCO2 | -0.035246488 |
| CFAP61 | -0.035249417 |
| AC079781 | -0.035249434 |
| RP11-173A8 | -0.035255566 |
| RFPL4AL1 | -0.035255848 |
| RP11-239C9 | -0.035256088 |
| MIR4309 | -0.03525649 |
| CTA-38K21 | -0.035259701 |
| RNA5SP465 | -0.035262531 |
| MAP3K9 | -0.035263855 |
| RP11-591L14 | -0.035268362 |
| RN7SL62P | -0.035268412 |
| RPL35AP33 | -0.035269931 |
| RP13-348B13 | -0.035270776 |
| CH17-219N22 | -0.035273574 |
| AC010740 | -0.035280719 |
| MIRLET7BHG | -0.03528354 |

| | |
|---|---|
| HMGN1P16 | -0.035292456 |
| AC023347 | -0.035294375 |
| RPL6P7 | -0.03529914 |
| OR5BK1P | -0.035299411 |
| SNORA45B | -0.035301759 |
| AC007948 | -0.035310445 |
| SRBD1 | -0.035311597 |
| AC007680 | -0.035311673 |
| RP11-846C15 | -0.035313444 |
| AC098872 | -0.035313681 |
| OR12D3 | -0.035314526 |
| RNU6-572P | -0.035325898 |
| SLC9A9-AS1 | -0.035328597 |
| RP4-745E8 | -0.035330362 |
| MROH4P | -0.035335214 |
| LINC00606 | -0.035344243 |
| CCSER2 | -0.035347617 |
| OR10T2 | -0.035348137 |
| RN7SL213P | -0.035349491 |
| RP11-29B2 | -0.035356329 |
| SMG1P1 | -0.035360256 |
| FABP5P12 | -0.035361963 |
| FOXRED2 | -0.035363134 |
| CECR2 | -0.035363823 |
| EEF1GP6 | -0.035364668 |
| RHBDL3 | -0.03536892 |
| HSPD1P7 | -0.035372177 |
| RP11-753B7 | -0.035373807 |
| RNFT2 | -0.0353756 |
| RP11-871F6 | -0.035379192 |
| SEPHS1P4 | -0.035381336 |
| MIR6861 | -0.035384135 |
| ACSF3 | -0.035384177 |
| DUSP8 | -0.035385144 |
| ANP32A | -0.035385982 |
| RP11-232L2 | -0.035388353 |
| OR5AS1 | -0.035390563 |
| KLHL21 | -0.035390819 |
| RP11-483G21 | -0.035391731 |
| SDHDP6 | -0.035392911 |
| CTC-565M22 | -0.035395797 |
| DCTD | -0.03539988 |
| HMGB1P13 | -0.035401822 |
| GPCPD1 | -0.035402857 |
| AC024995 | -0.035403309 |

| | |
|---|---|
| AC098649 | -0.035410001 |
| C4orf33 | -0.035410894 |
| AL020997 | -0.035416631 |
| CTC-248O19 | -0.035422949 |
| CCT8P1 | -0.035425667 |
| RP11-257I14 | -0.035427581 |
| OTOA | -0.035430523 |
| ETS1 | -0.035430616 |
| RP11-57G22 | -0.035433752 |
| HSPE1P10 | -0.035436903 |
| AC002306 | -0.035437375 |
| RP11-447M4 | -0.03543771 |
| Z69890 | -0.035439808 |
| FAM24B | -0.035441426 |
| RP11-39H3 | -0.035441872 |
| HUS1 | -0.035444663 |
| MIR3148 | -0.035444838 |
| MTND5P27 | -0.035446413 |
| OR7E22P | -0.035454945 |
| ERVWE2 | -0.035455114 |
| CT867976 | -0.035460109 |
| RP11-45J1 | -0.035462044 |
| SVILP1 | -0.035463474 |
| MIR3923 | -0.035464948 |
| KLF17P1 | -0.035465743 |
| RP11-436D10 | -0.035467309 |
| RP11-269C23 | -0.035471179 |
| MEF2D | -0.03547166 |
| UBC | -0.035474283 |
| AL590558 | -0.035475212 |
| VIL1 | -0.035478726 |
| DDX39BP2 | -0.035482294 |
| POLHP1 | -0.035484814 |
| XXbac-BPG181B23 | -0.035486339 |
| TRAF4 | -0.035488628 |
| HNRNPH1P2 | -0.035492111 |
| SIGLEC19P | -0.035495623 |
| RNU5A-4P | -0.03549709 |
| RP5-1014O16 | -0.035499913 |
| AC016561 | -0.035508295 |
| TMEM14B | -0.03550941 |
| RPS17P5 | -0.035512835 |
| MTA2 | -0.035516778 |
| MIR1294 | -0.035516877 |
| RP11-113A11 | -0.035518023 |

| | | |
|---|---|---|
| INPP5K | | -0.035523216 |
| RP4-738P15 | | -0.035526171 |
| AC004947 | | -0.03552648 |
| UCHL5 | | -0.035527309 |
| | 9-Mar | -0.03552956 |
| RNU6-863P | | -0.035533691 |
| IQGAP3 | | -0.035535067 |
| AIFM2 | | -0.035535849 |
| RP11-144L1 | | -0.035536761 |
| AC025284 | | -0.035538006 |
| UBE2N | | -0.035539402 |
| AC093639 | | -0.035543544 |
| RP11-576N17 | | -0.035551015 |
| RFESDP1 | | -0.035551958 |
| ZNF736P2Y | | -0.035553266 |
| RNF220 | | -0.035561357 |
| GALR3 | | -0.035562251 |
| OR52M2P | | -0.035562335 |
| RNU6-456P | | -0.035563134 |
| KDSR | | -0.035563552 |
| RNU7-140P | | -0.035564008 |
| PHF6 | | -0.035564464 |
| RPL7AP11 | | -0.035569376 |
| MIR1273G | | -0.035572043 |
| KCNQ5-IT1 | | -0.035577496 |
| WRAP73 | | -0.035578902 |
| MIR6076 | | -0.035579481 |
| OR8H3 | | -0.035582031 |
| IGHVIII-5-2 | | -0.035583231 |
| RNA5SP303 | | -0.035586203 |
| RNA5SP73 | | -0.035588407 |
| RPL36P19 | | -0.035589777 |
| EIF3A | | -0.035591405 |
| AL139383 | | -0.035597205 |
| RP11-420H19 | | -0.035597229 |
| AC019011 | | -0.035607843 |
| ZNF729 | | -0.035612334 |
| CTA-972D3 | | -0.035617767 |
| C6orf57 | | -0.035619052 |
| RPL5P11 | | -0.035621945 |
| LINC00517 | | -0.035624001 |
| AC104984 | | -0.03562598 |
| AC023818 | | -0.035631487 |
| PPP1R12BP1 | | -0.035633674 |
| RP11-2F20 | | -0.035636302 |

| | |
|---|---|
| KIAA1024L | -0.035645099 |
| SCARNA16 | -0.035646119 |
| ADAM29 | -0.035646517 |
| AC140076 | -0.035646993 |
| CYP3A54P | -0.035647964 |
| GON4L | -0.035650682 |
| AMBN | -0.03565479 |
| RP11-1406H17 | -0.035657701 |
| AL020993 | -0.035661357 |
| TACC1 | -0.035665659 |
| MIR3976HG | -0.035665919 |
| KRT89P | -0.035666216 |
| SCUBE1 | -0.035668189 |
| OR10H4 | -0.035669068 |
| USP17L2 | -0.035670096 |
| AC006196 | -0.035670443 |
| AC002984 | -0.035674233 |
| SNORA2B | -0.035680103 |
| MIR23B | -0.035681118 |
| CFL1P5 | -0.035682367 |
| RP11-425A23 | -0.035686336 |
| TUBB4BP2 | -0.035686428 |
| REREP2Y | -0.035689747 |
| AC239859 | -0.035689747 |
| TAS2R15P | -0.035692413 |
| AC104623 | -0.035694748 |
| CASC4 | -0.035704023 |
| RP11-872J21 | -0.035704954 |
| RP11-518I13 | -0.035710121 |
| SNORD27 | -0.035713571 |
| CTD-2131I18 | -0.035715801 |
| AL356791 | -0.035718148 |
| SNORD1B | -0.035723931 |
| SNORA62 | -0.035726717 |
| OR2U1P | -0.035727016 |
| DDX4 | -0.035727695 |
| SHANK2-AS3 | -0.035733249 |
| SLC23A3 | -0.035734314 |
| SIGLEC16 | -0.03573761 |
| RP11-454P7 | -0.035743776 |
| RNA5SP482 | -0.03574498 |
| RP11-760H22 | -0.035745923 |
| SBF1 | -0.035750091 |
| SEZ6L | -0.035750524 |
| RP11-199F11 | -0.035756584 |

| | |
|---|---|
| F13B | -0.035758054 |
| RP11-452K20 | -0.035761924 |
| AIDA | -0.035767431 |
| HSPH1 | -0.035768789 |
| MICAL1 | -0.035790632 |
| ENO2 | -0.035791525 |
| RP11-617B3 | -0.035795473 |
| MTND5P30 | -0.035797308 |
| RP11-767C1 | -0.035798986 |
| GAPDHP29 | -0.035799039 |
| MUT | -0.035799842 |
| L1CAM | -0.035801386 |
| UBE2V1P5 | -0.035802288 |
| LGALS8-AS1 | -0.035804868 |
| TREML5P | -0.035809203 |
| RPL12P9 | -0.035813417 |
| CYP2U1 | -0.03581841 |
| HMGB3P19 | -0.035819153 |
| RP11-148L24 | -0.035820661 |
| ZNF7 | -0.035822057 |
| RNU6-786P | -0.035822582 |
| HSFY5P | -0.035826228 |
| RP11-796E2 | -0.035829223 |
| MPPED2 | -0.035832904 |
| SLC30A5 | -0.035836922 |
| CH507-152C13 | -0.035836949 |
| HSPE1P20 | -0.035837426 |
| CTB-20D2 | -0.035839508 |
| RP11-577B7 | -0.035840854 |
| SLC7A11 | -0.035843062 |
| MIR577 | -0.035844407 |
| NMTRQ-TTG12-1 | -0.03584446 |
| RP11-141C7 | -0.035850197 |
| RP3-431C21 | -0.035854065 |
| ZNF317P1 | -0.035854316 |
| RP11-282I1 | -0.035857781 |
| AP000402 | -0.035860285 |
| HNRNPA1P63 | -0.035863704 |
| RP4-681L3 | -0.035866413 |
| AGK | -0.035867156 |
| SPATA32 | -0.035870077 |
| RP11-967K21 | -0.035873036 |
| RD3L | -0.035884868 |
| RP11-261P13 | -0.035898978 |
| OR52B2 | -0.035900663 |

| | |
|---|---|
| INSIG1 | -0.035903735 |
| INMT-FAM188B | -0.035909519 |
| FSD1 | -0.03591124 |
| TNR-IT1 | -0.035911661 |
| RP5-1166F10 | -0.03591208 |
| PAICSP6 | -0.035913088 |
| ABCA6 | -0.035913595 |
| MIR6775 | -0.035915856 |
| MIR3619 | -0.035919352 |
| MIR664A | -0.035922549 |
| DAK | -0.035923531 |
| CTB-147N14 | -0.035928219 |
| RP11-586E1 | -0.035929388 |
| RP11-317B17 | -0.035929669 |
| IGSF11-AS1 | -0.035929772 |
| EPB41L1 | -0.035934527 |
| TRIM60P15 | -0.035937949 |
| CTD-2353F22 | -0.035939282 |
| CHCHD7 | -0.035939755 |
| CTD-2117L12 | -0.035940177 |
| MIR516B1 | -0.035942579 |
| AC004076 | -0.035945653 |
| RNU4-10P | -0.035947405 |
| ETFDH | -0.035950471 |
| SMCO3 | -0.035955219 |
| PPM1J | -0.035961858 |
| TSPY8 | -0.035962709 |
| OFD1P16Y | -0.035962709 |
| NPY5R | -0.035967122 |
| RP11-343B5 | -0.035969176 |
| SDHC | -0.035975254 |
| LINC01435 | -0.035977026 |
| MIR4475 | -0.035977311 |
| MIR4696 | -0.035977834 |
| RP1-251I12 | -0.035980811 |
| RPSAP54 | -0.03598238 |
| AC079630 | -0.035986837 |
| ZNF419 | -0.035992166 |
| FAM149A | -0.035996427 |
| GPR143 | -0.035997432 |
| USP20 | -0.035998399 |
| CCNG2 | -0.036000688 |
| RP5-933E2 | -0.03600204 |
| SDCCAG3P1 | -0.036005694 |
| GS1-421I3 | -0.036012149 |

| | |
|---|---|
| ERVH48-1 | -0.036016585 |
| RNU1-87P | -0.036018082 |
| KRTAP10-1 | -0.036019106 |
| SHD | -0.03601922 |
| RAMP2 | -0.03602281 |
| LRRC53 | -0.036028917 |
| CRNDE | -0.036037043 |
| AC105339 | -0.036039167 |
| RP11-253A20 | -0.03603972 |
| RP11-470C13 | -0.036040149 |
| LRRC7 | -0.036042606 |
| FKSG52 | -0.036042796 |
| CCDC15 | -0.036043536 |
| GPR20 | -0.03604671 |
| RP3-514A23 | -0.036048257 |
| DYNLRB1 | -0.036048522 |
| AQP3 | -0.036051015 |
| PDE4DIP1 | -0.03605328 |
| CHIC1 | -0.03605416 |
| RPL18P10 | -0.036055322 |
| SRD5A3 | -0.036058272 |
| RNU6-299P | -0.036061181 |
| ZNF610 | -0.036067332 |
| KLF3-AS1 | -0.036067518 |
| SNORA59A | -0.036067784 |
| RNY1P8 | -0.036067983 |
| RP11-1157N2__B | -0.03606979 |
| CDK3 | -0.036072375 |
| MIR632 | -0.036072747 |
| MYO5BP2 | -0.036074674 |
| PRPS1L1 | -0.036081905 |
| GS1-345D13 | -0.036084211 |
| NDUFS5P5 | -0.036087939 |
| FHL5 | -0.036089185 |
| TGIF1 | -0.036091054 |
| BCRP3 | -0.036091684 |
| HSPA9P1 | -0.036092329 |
| ALOX15P2 | -0.036093399 |
| LPGAT1 | -0.036093659 |
| PTMAP1 | -0.036094253 |
| RP3-467L1 | -0.036095055 |
| C3 | -0.036100022 |
| WDR44 | -0.036101622 |
| ZNF639 | -0.036103799 |
| EEF1GP7 | -0.036107519 |

| | |
|---|---|
| ABCA1 | -0.036108674 |
| RP1-196A12 | -0.036109007 |
| AC004490 | -0.036109776 |
| AL596220 | -0.036116878 |
| ACTR3BP3 | -0.036122756 |
| UBE2C | -0.036123688 |
| RNF43 | -0.036132582 |
| ELOVL2 | -0.0361326 |
| AL590684 | -0.036132862 |
| KRT18P37 | -0.036135355 |
| LINC01330 | -0.036135564 |
| DFNB59 | -0.036135931 |
| RP4-657D16 | -0.036137456 |
| ANKRD28 | -0.036147857 |
| RN7SL154P | -0.036148461 |
| PRH2 | -0.036152089 |
| RP5-1182A14 | -0.036152312 |
| SCARA5 | -0.036156386 |
| SLC13A1 | -0.036157179 |
| PIK3R4 | -0.036161811 |
| RP11-536C12 | -0.036161837 |
| HLA-V | -0.036162301 |
| ACTG1P13 | -0.036163694 |
| PRKRIRP7 | -0.036163956 |
| LGR4 | -0.036166686 |
| SNHG22 | -0.036168193 |
| SEMG2 | -0.036169167 |
| RP11-452N4 | -0.036171762 |
| RNU6-439P | -0.036173706 |
| AF228730 | -0.036185382 |
| AC005621 | -0.036189076 |
| GDI1 | -0.036194575 |
| NUTM1 | -0.036199979 |
| LINC01562 | -0.036199988 |
| FCHSD1 | -0.036204938 |
| CTD-2233K9 | -0.036206092 |
| CCDC85C | -0.036210836 |
| PTCH2 | -0.036211339 |
| Z85986 | -0.036212025 |
| AP000244 | -0.036213995 |
| UBA52P6 | -0.036226824 |
| MICB | -0.036228363 |
| MARK3P1 | -0.036232003 |
| TRAV9-1 | -0.036233653 |
| RP11-597G23 | -0.036233726 |

| | |
|---|---|
| XXbac-BPG254F23 | -0.03623485 |
| GAPDHP49 | -0.036235786 |
| COG7 | -0.036236121 |
| SIGLEC26P | -0.036241674 |
| PMCH | -0.036242684 |
| AP000563 | -0.036243422 |
| RP11-558F24 | -0.036244258 |
| CWC27 | -0.036244438 |
| ARMC7 | -0.036246131 |
| CDK19 | -0.03624654 |
| SRP19 | -0.036246559 |
| RP11-553E24 | -0.036251733 |
| AL445199 | -0.036252864 |
| TMEM82 | -0.036256188 |
| KLF11 | -0.036259769 |
| RP11-443B9 | -0.036261019 |
| MIR200C | -0.036268174 |
| DEFB121 | -0.036271074 |
| RP11-617D20 | -0.0362714 |
| C10orf12 | -0.036274858 |
| ASIC1 | -0.03627843 |
| RP13-494C23 | -0.036280062 |
| HNRNPA3P16 | -0.036286402 |
| NSUN5P2 | -0.036290784 |
| ZNF551 | -0.036292049 |
| YWHAEP4 | -0.036308992 |
| RP11-1348G14 | -0.036309836 |
| ZFP64 | -0.036316292 |
| OR2R1P | -0.036317182 |
| RANP5 | -0.036318334 |
| ATAT1 | -0.036320758 |
| SNORD82 | -0.03632316 |
| MT-ND3 | -0.036323455 |
| AC016903 | -0.036326884 |
| RP5-824H1 | -0.036328375 |
| HBZP1 | -0.036330824 |
| GABPA | -0.036331158 |
| RP11-863K10 | -0.036334674 |
| RAP2C | -0.036337056 |
| MIR4742 | -0.036338725 |
| IGKV1D-37 | -0.036339244 |
| AC096753 | -0.036351546 |
| LRIG1 | -0.036351847 |
| CHAF1B | -0.036352722 |
| RPL36AP48 | -0.036358993 |

| | |
|---|---|
| OTOP1 | -0.036359259 |
| GLRX | -0.036360183 |
| AL360091 | -0.03636261 |
| RP1-163G9 | -0.036365624 |
| RP11-796I2 | -0.036371046 |
| U4atac | -0.036372103 |
| CTC-451A6 | -0.036372942 |
| CCDC114 | -0.036378137 |
| RP5-1115A15 | -0.036386586 |
| RNA5SP242 | -0.036388509 |
| CLRN2 | -0.036396524 |
| PTPRQ | -0.036400436 |
| RP3-418A9 | -0.036404709 |
| RP11-498E11 | -0.036409146 |
| CAV3 | -0.036414514 |
| TTLL9 | -0.036420204 |
| PGA4 | -0.036422546 |
| CH17-258A22 | -0.036422995 |
| AKR1C5P | -0.036427085 |
| IGLVI-38 | -0.036428647 |
| DTWD2 | -0.036435572 |
| ZNF85 | -0.036436521 |
| RP11-22H5 | -0.036438013 |
| B4GALT4-AS1 | -0.036445999 |
| RNU6-1024P | -0.036452349 |
| PGM5P3-AS1 | -0.036453242 |
| RNU6-662P | -0.036456217 |
| NRG3 | -0.036456801 |
| RP11-153K16 | -0.036457241 |
| RBMX2P1 | -0.03645918 |
| OR2B6 | -0.036460392 |
| RP11-101K23 | -0.036464696 |
| RPSAP55 | -0.036469075 |
| DCANP1 | -0.036472423 |
| CTD-3022L24 | -0.036472813 |
| CTC-513N18 | -0.036473644 |
| RP11-42D20 | -0.036474455 |
| OR2AJ1 | -0.036476423 |
| ZNF805 | -0.036479239 |
| RPL7L1P10 | -0.036482868 |
| LINC00111 | -0.036486582 |
| ZIM3 | -0.036488037 |
| AMMECR1-IT1 | -0.036488105 |
| RPS3AP29 | -0.036489165 |
| HIAT1 | -0.036489881 |

| | | |
|---|---|---|
| RASEF | | -0.036490421 |
| | 1-Mar | -0.036490737 |
| IPO5 | | -0.036494923 |
| RN7SKP267 | | -0.036497282 |
| CYP17A1-AS1 | | -0.036501702 |
| TCF3P1 | | -0.036502551 |
| KRT18P43 | | -0.036507973 |
| RNU6-582P | | -0.036513547 |
| GREB1 | | -0.036515799 |
| RP11-449I17 | | -0.036518022 |
| CTD-2201I18 | | -0.036519641 |
| AC073137 | | -0.036520486 |
| EIF2S2P4 | | -0.036521154 |
| EEF1A1P30 | | -0.036521947 |
| FXR1 | | -0.036523222 |
| MSH6 | | -0.036529009 |
| CCNYL2 | | -0.036530384 |
| PIGHP1 | | -0.036534081 |
| HDAC10 | | -0.036535372 |
| AC062028 | | -0.036537544 |
| RNA5SP501 | | -0.036539096 |
| RP11-159H22 | | -0.036539235 |
| SNX25 | | -0.036539446 |
| RP11-324E6 | | -0.036539446 |
| NF1P6 | | -0.036539484 |
| APPL2 | | -0.036542777 |
| RP11-968A15 | | -0.036544405 |
| ZNF785 | | -0.03654447 |
| RNU1-109P | | -0.036544531 |
| RP11-185E12 | | -0.036545033 |
| RN7SKP82 | | -0.036547736 |
| CREB3L4 | | -0.036553698 |
| MUS81 | | -0.036560322 |
| AC021016 | | -0.036568675 |
| ADGB | | -0.036569915 |
| RBMXP1 | | -0.03657165 |
| RP11-407B7 | | -0.036573029 |
| PGM5 | | -0.036574238 |
| FEN1P1 | | -0.036582312 |
| RP11-416A14 | | -0.036582838 |
| AC097506 | | -0.03658431 |
| RARB | | -0.036587188 |
| RAB35 | | -0.036591393 |
| NUP205 | | -0.036596453 |
| RP11-723D22 | | -0.03660787 |

| | |
|---|---|
| ZNF333 | -0.036609328 |
| RNA5SP246 | -0.036610996 |
| RP11-317L10 | -0.036614083 |
| RNU6-602P | -0.036614117 |
| MIR766 | -0.036617095 |
| MLEC | -0.036617143 |
| RP11-45P22 | -0.036617865 |
| LINC01099 | -0.036617926 |
| RN7SL801P | -0.036620615 |
| EPRS | -0.036623524 |
| RP11-478H16 | -0.03662593 |
| RPL21P126 | -0.036627591 |
| PLA2G4E-AS1 | -0.036627933 |
| RP11-397C18 | -0.036629018 |
| AP000866 | -0.036631695 |
| PPP1R27 | -0.036636216 |
| NMNAT3 | -0.036636269 |
| CH17-132F21 | -0.036637223 |
| RNU6-417P | -0.036638819 |
| RPL12P44 | -0.036639741 |
| ITPKA | -0.036640213 |
| AC092601 | -0.036646725 |
| MIR548AY | -0.036652972 |
| TRBV21OR9-2 | -0.036659045 |
| NDFIP2 | -0.036659954 |
| PALMD | -0.036667563 |
| RFPL2 | -0.036675064 |
| RP11-71H17 | -0.036677196 |
| TCTE3 | -0.036681704 |
| SNX24 | -0.036688755 |
| RP11-182J1 | -0.036691862 |
| LINC00950 | -0.03669322 |
| SPIN4 | -0.03669469 |
| OR10J7P | -0.036699124 |
| RNU6-535P | -0.03670191 |
| RNU6-869P | -0.036702012 |
| RP5-905H7 | -0.036706792 |
| SNX6P1 | -0.036713123 |
| RP2 | -0.03671644 |
| NDUFA5P5 | -0.036716933 |
| SNORD115-48 | -0.036717585 |
| RAD23BLP | -0.036719656 |
| RNU6-661P | -0.03672793 |
| AC093523 | -0.036729578 |
| DEFB128 | -0.036729875 |

| | |
|---|---|
| POU6F2-AS2 | -0.036730188 |
| RP11-349F21 | -0.036730241 |
| RP11-316M1 | -0.036734056 |
| ZFRP1 | -0.036734542 |
| SLCO4A1 | -0.036734562 |
| RP11-55L3 | -0.036744742 |
| LINC01366 | -0.03674596 |
| ABAT | -0.036746339 |
| ISG15 | -0.036753465 |
| RNA5SP37 | -0.036761841 |
| NSRP1P1 | -0.036763288 |
| OR7E53P | -0.03676406 |
| TRMT2B | -0.0367682 |
| UBE2G1 | -0.036768219 |
| RP11-316I3 | -0.036769002 |
| MIR4462 | -0.036775146 |
| RNF166 | -0.036775475 |
| RN7SL402P | -0.036775519 |
| RP11-815I9 | -0.036779382 |
| RP11-1149O23 | -0.036785578 |
| SLC6A6 | -0.036788034 |
| EIF4EBP2P2 | -0.036788961 |
| AC005041 | -0.03678977 |
| MTND6P9 | -0.036790093 |
| LINC00613 | -0.036790842 |
| AC083906 | -0.036792179 |
| RP11-567O16 | -0.036797195 |
| TRAP1 | -0.03679916 |
| ASMT | -0.0367995 |
| LL09NC01-139C3 | -0.036800705 |
| MTND1P21 | -0.03680161 |
| OR5H7P | -0.036804257 |
| RP11-95G6 | -0.03680714 |
| RP1-172I22 | -0.036808166 |
| URGCP | -0.036816091 |
| SUV39H1 | -0.036817114 |
| MIR135B | -0.036822062 |
| HNRNPA1P24 | -0.036828854 |
| LCN12 | -0.036829747 |
| RPL3P12 | -0.036834076 |
| KIF9-AS1 | -0.036840576 |
| BX255923 | -0.036840973 |
| RNU1-85P | -0.036843163 |
| WDR72 | -0.036847179 |
| ADCY4 | -0.036847925 |

| | |
|---|---|
| UBE2T | -0.036848018 |
| TMEM52B | -0.036849041 |
| ADAMTS15 | -0.036849451 |
| RP13-516M14 | -0.036850623 |
| RP11-174J11 | -0.036850859 |
| CPEB3_ribozyme | -0.036854919 |
| ADAM2 | -0.0368561 |
| METTL8 | -0.036865005 |
| TRGV11 | -0.036865042 |
| AFF4 | -0.036868056 |
| LEPR | -0.036868521 |
| RP11-632K21 | -0.036869868 |
| RP11-263C24 | -0.036880028 |
| RNU6-957P | -0.036880278 |
| CSRNP2 | -0.036883312 |
| ICE2P1 | -0.036885784 |
| C21orf59 | -0.036893341 |
| ABC7-42418200C9 | -0.036893504 |
| MIR6865 | -0.036894995 |
| TROVE2 | -0.036896467 |
| AC009941 | -0.036899523 |
| RP11-417O18 | -0.036899605 |
| TTC4P1 | -0.036901677 |
| DPH7 | -0.036906067 |
| OR5J7P | -0.036911682 |
| ZNF341-AS1 | -0.036915835 |
| RP4-705D16 | -0.036924304 |
| MIR4423 | -0.036924853 |
| AC099684 | -0.036931115 |
| THAP2 | -0.036934515 |
| RP11-647F2 | -0.036936244 |
| ARHGAP42P3 | -0.03693977 |
| RNA5SP172 | -0.036941584 |
| RGS9BP | -0.036952826 |
| ZNF33BP1 | -0.036953846 |
| IARS2 | -0.036956134 |
| RP11-542C10 | -0.036975237 |
| KRTAP10-6 | -0.036978891 |
| RP11-244H3 | -0.036982107 |
| DENND1B | -0.036985587 |
| CDY5P | -0.036988378 |
| TRAM1L1 | -0.036992229 |
| SEMA3G | -0.03699342 |
| CAT | -0.036996117 |
| CDY22P | -0.037002192 |

| | |
|---|---|
| DCAF8L2 | -0.037004968 |
| HMGB1P22 | -0.037010897 |
| MIR6781 | -0.037019353 |
| OR5H14 | -0.037028791 |
| KB-1090H4 | -0.037035896 |
| NRIP1 | -0.037038073 |
| SVEP1 | -0.03703865 |
| ATP7B | -0.037040901 |
| UBOX5-AS1 | -0.037044743 |
| RNU1-100P | -0.037046292 |
| OR5M5P | -0.037050016 |
| HTR2C | -0.037053696 |
| MPI | -0.037056864 |
| ZNF669 | -0.037057199 |
| MIR510 | -0.037057367 |
| RP11-822P1 | -0.037058496 |
| ALKBH1 | -0.037062316 |
| AP001342 | -0.037063434 |
| PLEKHM1P | -0.037064679 |
| RP11-73E17 | -0.037066669 |
| RP11-597A11 | -0.037067803 |
| AC104306 | -0.037069581 |
| HSPE1P16 | -0.03707089 |
| NECAP1P1 | -0.0370711 |
| TET3 | -0.037073516 |
| ALOX5 | -0.037077181 |
| RP5-951N9 | -0.037086984 |
| RP11-458I7 | -0.037087506 |
| SMARCE1P3 | -0.037089882 |
| RP11-560L11 | -0.037093951 |
| RP11-321L2 | -0.037094596 |
| THAP11 | -0.037094764 |
| GPR158 | -0.037097409 |
| SMCR5 | -0.03710143 |
| RP11-39K24 | -0.037105124 |
| TMEM181 | -0.037107936 |
| RP1-122K4 | -0.03710843 |
| MIR3136 | -0.03711004 |
| RNU6-1227P | -0.037112928 |
| MIR5708 | -0.03711787 |
| RP11-256I23 | -0.037118738 |
| TUBAP2 | -0.037121742 |
| MRPS18B | -0.037123565 |
| RN7SKP183 | -0.037125242 |
| RP11-61K20 | -0.037125242 |

| | |
|---|---|
| RNU6-533P | -0.037125242 |
| OR4R1P | -0.037125242 |
| RNA5SP142 | -0.037125242 |
| RNA5SP126 | -0.037125242 |
| RNU6-832P | -0.037125242 |
| RNU6-655P | -0.037125242 |
| RNU6-812P | -0.037125242 |
| SNORD115-44 | -0.037125242 |
| RNU2-66P | -0.037125242 |
| AC012119 | -0.037125242 |
| RNU6-1012P | -0.037125242 |
| RNU6-1130P | -0.037125242 |
| SNORD115-2 | -0.037125242 |
| NUDT7 | -0.037129016 |
| RP11-354I10 | -0.037131279 |
| NPDC1 | -0.037132291 |
| AKIP1 | -0.037133407 |
| FBXO4 | -0.037138952 |
| RASGEF1A | -0.037145259 |
| RPS7P1 | -0.037149222 |
| RP5-916L7 | -0.037149868 |
| AC016696 | -0.037151541 |
| RN7SL750P | -0.03715489 |
| BBS12 | -0.03715925 |
| PROB1 | -0.03716206 |
| OR51Q1 | -0.037170824 |
| SEC62 | -0.037173986 |
| ACBD4 | -0.037176758 |
| USP9YP14 | -0.037183579 |
| COX6CP12 | -0.037186918 |
| MIR4291 | -0.037191036 |
| AL117341 | -0.037192214 |
| RP11-116O18 | -0.03719534 |
| CTD-2666L21 | -0.037195512 |
| ITPR2 | -0.037199791 |
| RP11-334J6 | -0.037205176 |
| MTND6P15 | -0.037205463 |
| RP11-1035H13 | -0.03720597 |
| AC068715 | -0.037216726 |
| RP4-604A21 | -0.037218577 |
| C3orf18 | -0.037222806 |
| RP11-983C2 | -0.03722856 |
| ST6GALNAC5 | -0.037229914 |
| AKR1C4 | -0.03723194 |
| KIAA1671 | -0.037241151 |

| | |
|---|---|
| RNU6-1174P | -0.037243658 |
| C1orf141 | -0.037247437 |
| HDAC9 | -0.037248873 |
| AL359815 | -0.03725228 |
| OCM | -0.037257001 |
| EIF2S1 | -0.037258361 |
| AF165138 | -0.037264773 |
| DDX18 | -0.037265152 |
| AF146191 | -0.037265379 |
| ZNF74 | -0.037265822 |
| RP11-83J21 | -0.037269933 |
| GEMIN8P2 | -0.037270179 |
| SPATA19 | -0.037272556 |
| AC092841 | -0.037272766 |
| ABCB11 | -0.037277163 |
| AC024704 | -0.037279561 |
| ZKSCAN5 | -0.037283907 |
| RPL23AP6 | -0.037285165 |
| MACF1 | -0.037285358 |
| PLCE1-AS1 | -0.037288524 |
| FAM65A | -0.037294996 |
| FCF1P3 | -0.037295043 |
| PGM5-AS1 | -0.03729528 |
| RXRB | -0.037299126 |
| MARK2 | -0.037300484 |
| RP11-357N13 | -0.03730649 |
| RP11-384E22 | -0.037307951 |
| MIR7856 | -0.037312677 |
| TPH2 | -0.037313514 |
| COX11P1 | -0.037313704 |
| AC013737 | -0.03731526 |
| RNU6-254P | -0.037315644 |
| RP11-423J7 | -0.037316424 |
| AL591806 | -0.037322037 |
| RP11-1J4 | -0.037327517 |
| DCLK1 | -0.037328783 |
| AC083900 | -0.037330902 |
| AP2A2 | -0.037331444 |
| GRK6P1 | -0.037343116 |
| GART | -0.03734644 |
| RP11-566J3 | -0.037349668 |
| RPL21P3 | -0.037349691 |
| XYLB | -0.037351928 |
| RP11-10E6 | -0.037352232 |
| AC027320 | -0.037353329 |

| | |
|---|---|
| CX3CR1 | -0.037353603 |
| AL157884 | -0.037356552 |
| GAK | -0.037357566 |
| PMFBP1 | -0.037360096 |
| PSMG2 | -0.037363836 |
| RN7SL811P | -0.037363864 |
| AC092944 | -0.037367771 |
| DPPA4 | -0.037368847 |
| RP11-375O18 | -0.037373487 |
| RP11-121J20 | -0.037374824 |
| CARD17 | -0.037384924 |
| RPS29P2 | -0.037393335 |
| AC093430 | -0.03740004 |
| Z93241 | -0.037401122 |
| TICRR | -0.037402386 |
| MRPL35P4 | -0.037402418 |
| RP11-489E7 | -0.037402798 |
| RP11-262A16 | -0.037405643 |
| DPM1 | -0.03740767 |
| DHX36 | -0.037412824 |
| FFAR3 | -0.037413292 |
| MIR4736 | -0.037413432 |
| DNAL1 | -0.037413903 |
| RNU7-126P | -0.037414594 |
| MIR6782 | -0.037414956 |
| RP11-35G22 | -0.037417382 |
| AKR1B10P2 | -0.037417707 |
| PABPC1P10 | -0.037418229 |
| CCDC75P1 | -0.037421891 |
| RP1-202O8 | -0.037423847 |
| AC008449 | -0.037424986 |
| SEPT2P1 | -0.037425261 |
| EIF4A1P1 | -0.037430099 |
| RP4-543J13 | -0.037432743 |
| RNU4-79P | -0.037433491 |
| AC011343 | -0.037435569 |
| MIR4257 | -0.037435746 |
| ARL5B | -0.03743889 |
| ASNSP3 | -0.037439522 |
| RNU6-600P | -0.03744128 |
| MYO7B | -0.037448361 |
| XPO7 | -0.037450928 |
| RNPEP | -0.03745303 |
| SNORA67 | -0.037456156 |
| RP11-365O16 | -0.037457347 |

| | |
|---|---|
| LLNLR-246C6 | -0.037457366 |
| HNRNPA3P1 | -0.037459682 |
| MIR548G | -0.03746172 |
| RP3-445N2 | -0.037464731 |
| ZNF705E | -0.037465759 |
| RP11-38O23 | -0.037467745 |
| FGF6 | -0.037467867 |
| PRKCE | -0.037470501 |
| RP11-430B1 | -0.03747372 |
| ZHX1 | -0.037474966 |
| OR5M14P | -0.037475168 |
| RNU6-1218P | -0.037476399 |
| TSPAN33 | -0.037477478 |
| MIR4753 | -0.037479352 |
| RP11-398J10 | -0.037482157 |
| PTCH1 | -0.037494446 |
| RHAG | -0.03749534 |
| ENO1P3 | -0.037495384 |
| SDPR | -0.037499228 |
| AP000708 | -0.037500431 |
| PSMD1 | -0.037504605 |
| SLC5A11 | -0.037505685 |
| MTND4P2 | -0.0375107 |
| TACC3 | -0.037512121 |
| RNU7-137P | -0.037512519 |
| ROPN1L-AS1 | -0.037512809 |
| ACTL9 | -0.037513699 |
| AC135050 | -0.037519394 |
| DYNLL1P1 | -0.037521648 |
| RP11-479F13 | -0.037521904 |
| LINC00945 | -0.037521933 |
| CFAP46 | -0.037522157 |
| CSRP2BP | -0.037522429 |
| RP5-850O15 | -0.037522446 |
| KIAA0556 | -0.037522652 |
| GPR151 | -0.0375233 |
| LSM6P2 | -0.037523903 |
| PPIAL4B | -0.037526933 |
| PHF13 | -0.03753216 |
| RNU11-6P | -0.037533887 |
| CTD-2006O16 | -0.037535603 |
| SNORA37 | -0.037542793 |
| OTUB2 | -0.0375447 |
| PCSK6 | -0.037544867 |
| AC067903 | -0.037549806 |

| | |
|---|---|
| MIR8065 | -0.03755474 |
| NAV3 | -0.037561165 |
| MIR8080 | -0.037564482 |
| RNU6-114P | -0.0375691 |
| AL022476 | -0.037578152 |
| PRKX-AS1 | -0.037581466 |
| NBPF1 | -0.037591883 |
| SNORA51 | -0.037593266 |
| SUPV3L1 | -0.037598321 |
| SLC25A36 | -0.037604107 |
| AL163193 | -0.037605753 |
| GMEB1 | -0.03761047 |
| RETNLB | -0.037610819 |
| FCF1P2 | -0.037611177 |
| RNU6-189P | -0.037614992 |
| TMX1 | -0.037619419 |
| RNU7-11P | -0.037626816 |
| RP11-532N4 | -0.037629379 |
| MID2 | -0.037632517 |
| CCNB1IP1P2 | -0.03763533 |
| CNTNAP5 | -0.037636706 |
| POC1B-GALNT4 | -0.037637336 |
| SF3B3 | -0.037645058 |
| MIR758 | -0.037645112 |
| RNU7-26P | -0.037653479 |
| SLMAP | -0.037655532 |
| AC005686 | -0.037656276 |
| SNORD29 | -0.037657797 |
| OR4F4 | -0.03766423 |
| LRRC37A13P | -0.037665278 |
| SNORD62 | -0.037667671 |
| AC005336 | -0.037670237 |
| AC131012 | -0.037672554 |
| RNU6-637P | -0.03767446 |
| MIR4794 | -0.037675687 |
| DDX56 | -0.037676761 |
| RP11-9B6 | -0.037678427 |
| RP11-420K10 | -0.037681332 |
| HSPA4 | -0.037690287 |
| LINC00210 | -0.037694137 |
| RN7SL382P | -0.037698636 |
| SLC1A5 | -0.037699274 |
| FAM221B | -0.037701068 |
| LINC00563 | -0.037701496 |
| XIAP-AS1 | -0.037702629 |

| | |
|---|---|
| AF131217 | -0.037713494 |
| AC079235 | -0.03771422 |
| AC006486 | -0.037716558 |
| TEN1-CDK3 | -0.037722307 |
| AC022153 | -0.037722611 |
| RPUSD4 | -0.037733806 |
| BMS1P7 | -0.037735264 |
| TSN | -0.037736801 |
| RNU6-1145P | -0.037736966 |
| FAM161B | -0.037738122 |
| RP11-362I1 | -0.037739121 |
| MIR4681 | -0.037740589 |
| RP5-1077H22 | -0.037745237 |
| HNRNPA1P35 | -0.037750163 |
| PCDHGB8P | -0.037754365 |
| RN7SKP81 | -0.037755129 |
| RN7SKP182 | -0.037759195 |
| TSGA13 | -0.037768005 |
| CTD-2345M20 | -0.037768803 |
| MIR548T | -0.037773928 |
| OR11M1P | -0.037777901 |
| RN7SL869P | -0.03777819 |
| KCNU1 | -0.037778421 |
| AC024560 | -0.037782496 |
| MTND6P7 | -0.037784579 |
| RP11-568G20 | -0.037787766 |
| AL121748 | -0.037791801 |
| RPS8P6 | -0.037792858 |
| PIEZO1P2 | -0.037793364 |
| RP11-452I5 | -0.037793492 |
| MCTS2P | -0.037794107 |
| TTPAL | -0.037794125 |
| PLK5 | -0.037809638 |
| EEF1A1P14 | -0.037811745 |
| AC019046 | -0.037818165 |
| RPL32P28 | -0.037819539 |
| RNU6-130P | -0.037820736 |
| ZNF124 | -0.037829345 |
| BAG4 | -0.037838908 |
| FTH1P24 | -0.037839786 |
| FEM1AP4 | -0.037840798 |
| HEMK1 | -0.037843615 |
| LRRC58 | -0.037851169 |
| MESTP3 | -0.037853257 |
| DDX39BP1 | -0.037867076 |

| | |
|---|---|
| ACAD11 | -0.037877589 |
| RNA5SP21 | -0.037880969 |
| KRT18P49 | -0.037896807 |
| SNORD114-22 | -0.037898251 |
| MRPS16 | -0.03790319 |
| HSPA8P8 | -0.037905298 |
| DDX3Y | -0.037907685 |
| AQP2 | -0.0379084 |
| ZBED9 | -0.037916238 |
| GAL3ST2 | -0.037921139 |
| HMGN2P41 | -0.037923808 |
| RP11-613G13 | -0.037924518 |
| GOLGA2P4 | -0.037924652 |
| RP11-487E13 | -0.037938266 |
| DNASE1L2 | -0.037943043 |
| CTC-336P14 | -0.037944909 |
| EME2 | -0.037946857 |
| TRIM53CP | -0.03795175 |
| RP11-719N22 | -0.037954664 |
| ACCS | -0.037959211 |
| UBE2NP1 | -0.0379602 |
| STKLD1 | -0.037961388 |
| VTI1B | -0.037964997 |
| FBXL2 | -0.037965834 |
| RNU6-790P | -0.037966719 |
| RP11-365N19 | -0.037987842 |
| NPM1P13 | -0.037991349 |
| ERO1LB | -0.037992608 |
| SCG3 | -0.037993578 |
| ISCA1P1 | -0.037993632 |
| RP5-1173A5 | -0.037994739 |
| RP11-140L24 | -0.037995426 |
| RNU2-58P | -0.037999339 |
| LL09NC01-251B2 | -0.038000005 |
| RP11-6E9 | -0.038000121 |
| HNRNPCP9 | -0.038004951 |
| KPNA3 | -0.038005334 |
| ZNF350 | -0.03800846 |
| THRA | -0.038008869 |
| OR10G4 | -0.038009295 |
| RP11-160K17 | -0.038009922 |
| MPPED1 | -0.038014537 |
| DEK | -0.038018432 |
| ABHD15-AS1 | -0.038019277 |
| MIA3 | -0.038022544 |

| | |
|---|---|
| RP11-434D9 | -0.038026151 |
| WDR49 | -0.038031933 |
| CYP3A52P | -0.038034429 |
| AC233299 | -0.03803617 |
| C17orf53 | -0.038040573 |
| RP5-1063M23 | -0.038042458 |
| RNU11-4P | -0.038046437 |
| PIGM | -0.038049094 |
| CTC-428G20 | -0.038053243 |
| N4BP1 | -0.038054341 |
| BTBD7 | -0.03806329 |
| HMGB1 | -0.038071644 |
| RP11-765K14 | -0.03807486 |
| SSR1P1 | -0.038075946 |
| RSPH9 | -0.038076444 |
| NBPF21P | -0.038081805 |
| CYYR1 | -0.038082695 |
| KRTAP23-1 | -0.038083649 |
| AC074117 | -0.038085635 |
| RPL35AP15 | -0.038088408 |
| CBX3P4 | -0.038089006 |
| AC009161 | -0.038096432 |
| MIR199A1 | -0.038096632 |
| RP5-1049N15 | -0.038097824 |
| VBP1 | -0.038102659 |
| RP11-110L15 | -0.038103206 |
| SUCLG2P2 | -0.038106777 |
| LINC01506 | -0.038109834 |
| RP5-964H19 | -0.03811839 |
| SNRPGP7 | -0.038135345 |
| DDX11L5 | -0.038139373 |
| SPAG4 | -0.038145079 |
| RP11-307F22 | -0.038145967 |
| XXbac-BPG249D20 | -0.038148124 |
| RPL3P11 | -0.038149924 |
| RN7SL218P | -0.038150018 |
| RP11-1252I4 | -0.038150261 |
| GCSHP4 | -0.038158557 |
| RPL23P10 | -0.038160713 |
| MIR548AP | -0.038160894 |
| KCTD9P4 | -0.038168398 |
| C1QTNF9B | -0.038170233 |
| TMEM143 | -0.038171443 |
| ENO1-AS1 | -0.038177927 |
| MIR4429 | -0.038181622 |

| | |
|---|---|
| PAX7 | -0.038184587 |
| RP11-814H16 | -0.038185409 |
| RP3-359N14 | -0.038185804 |
| RP1-66C13 | -0.038186291 |
| NPIPB15 | -0.038195621 |
| RP11-529H22 | -0.038195726 |
| COPG2 | -0.038196207 |
| NEURL4 | -0.038202905 |
| RPL36P20 | -0.03820351 |
| AC005387 | -0.038205307 |
| GLYATL1P2 | -0.038207137 |
| KCNN3 | -0.038218125 |
| RPS2P39 | -0.03821836 |
| HSPA8P1 | -0.038223131 |
| LINC01364 | -0.03822557 |
| SIGLEC29P | -0.038225859 |
| ZNF663P | -0.038227959 |
| ADAM5 | -0.038231439 |
| CXorf56 | -0.038242126 |
| LINC00654 | -0.038247651 |
| LINC00297 | -0.038247726 |
| GLYCTK | -0.038248061 |
| RP11-121C1 | -0.038251244 |
| RP11-728G15 | -0.038253935 |
| RN7SKP141 | -0.038258548 |
| SPDEF | -0.038258815 |
| RP11-444A22 | -0.038262756 |
| UBE2F | -0.03826395 |
| RP11-618I10 | -0.038269234 |
| RPL15P1 | -0.038275032 |
| MIR139 | -0.038277153 |
| RP11-42A4 | -0.038286027 |
| CTD-2200P10 | -0.038290535 |
| RPS29P25 | -0.038293471 |
| HOXB3 | -0.038294854 |
| AC090514 | -0.038296294 |
| RNA5SP402 | -0.038296646 |
| RP11-434O22 | -0.038297833 |
| ZDHHC19 | -0.038302295 |
| RP11-132A1 | -0.038305682 |
| RP5-1108M17 | -0.038309723 |
| LINC00534 | -0.038317652 |
| RP11-379H8 | -0.038317745 |
| UBFD1 | -0.038320231 |
| RP4-564M11 | -0.038321765 |

| | |
|---|---|
| OR10G5P | -0.038325258 |
| CTD-3193K9 | -0.038329132 |
| RNA5SP300 | -0.038331889 |
| LINC00908 | -0.038332736 |
| PANK4 | -0.038339916 |
| RP11-204M4 | -0.038342587 |
| RP11-288E14 | -0.038342753 |
| RP11-105C20 | -0.038349788 |
| CTC-348L5 | -0.038350675 |
| RN7SKP210 | -0.038351129 |
| RP11-603B24 | -0.038354339 |
| DAGLA | -0.038357759 |
| MIR575 | -0.038360786 |
| MS4A12 | -0.038367519 |
| MRPS18BP1 | -0.038368378 |
| RP11-662B19 | -0.038370596 |
| RNU6-239P | -0.038370733 |
| KRT18P24 | -0.038378505 |
| ERLIN2 | -0.038379694 |
| CTD-2547G23 | -0.038380029 |
| PRRG4 | -0.038380215 |
| DEFB115 | -0.038381819 |
| RN7SL265P | -0.038382743 |
| DLAT | -0.038383248 |
| RGS9 | -0.038384104 |
| ZNF264 | -0.038390002 |
| OR9G4 | -0.038394805 |
| FAM76A | -0.038396328 |
| RP1-177I10 | -0.03839732 |
| FBF1 | -0.038400309 |
| RP11-161I10 | -0.038403714 |
| AC013472 | -0.038404416 |
| POLM | -0.03840911 |
| CTD-2355J17 | -0.038410191 |
| STAU1 | -0.038412608 |
| MIR638 | -0.038413889 |
| RAI1-AS1 | -0.038414062 |
| GRIN2C | -0.038415994 |
| DUSP8P3 | -0.038419055 |
| LRTOMT | -0.038424106 |
| KRTAP5-5 | -0.038425106 |
| MIR92A2 | -0.038426181 |
| FAM166B | -0.038428199 |
| RP5-1002M8 | -0.03843062 |
| MIR643 | -0.038431836 |

| | |
|---|---|
| OR5S1P | -0.038437837 |
| RN7SL149P | -0.03844708 |
| SNORA44 | -0.038451197 |
| AC132068 | -0.038454278 |
| VILL | -0.038456237 |
| RP11-134N8 | -0.038459835 |
| LL0XNC01-39B3 | -0.038459943 |
| ORC6 | -0.038463791 |
| AKNAD1 | -0.038464892 |
| CTB-175E5 | -0.038468733 |
| KRTAP4-11 | -0.038469832 |
| IKZF2 | -0.038470173 |
| RP11-124N3 | -0.038470691 |
| MTRNR2L8 | -0.038475308 |
| RN7SKP199 | -0.03847722 |
| SMG1P6 | -0.038477243 |
| RN7SL724P | -0.038479061 |
| LINC01489 | -0.038483728 |
| AC090133 | -0.038484828 |
| RP11-285J16 | -0.038487322 |
| ARHGEF26 | -0.038490043 |
| MIR4258 | -0.038491528 |
| CDY7P | -0.038491989 |
| RNU4-86P | -0.038492118 |
| PRR11 | -0.038492611 |
| RP11-164D18 | -0.038499905 |
| SLC25A15P1 | -0.038502413 |
| MIR150 | -0.038503096 |
| RP11-475D10 | -0.038504539 |
| AL157871 | -0.038509348 |
| CTD-3076O17 | -0.038510607 |
| SEC22B | -0.038511607 |
| RP11-162K6 | -0.038512739 |
| HMGN1P33 | -0.038514069 |
| RP4-607J2 | -0.038514648 |
| SSX9 | -0.038521822 |
| GPR126 | -0.038525859 |
| HSFY1 | -0.038525874 |
| PRMT6 | -0.038527199 |
| RP13-635I23 | -0.038531319 |
| FAM153B | -0.038531636 |
| SHISA3 | -0.03853259 |
| GLO1 | -0.038535645 |
| LINC00614 | -0.038537639 |
| PDIA3P1 | -0.038543125 |

| | |
|---|---|
| PES1P1 | -0.038546424 |
| RP11-1E11 | -0.038547814 |
| CTD-3126B10 | -0.038551857 |
| RN7SKP13 | -0.038555845 |
| RNU6-804P | -0.038555845 |
| MIR6130 | -0.038555845 |
| CAPN12 | -0.038559777 |
| IDS | -0.038560819 |
| RNU6-427P | -0.038564362 |
| RP11-148B18 | -0.038566328 |
| RP11-567G24 | -0.038571604 |
| FNTA | -0.038574624 |
| OR5BJ1P | -0.038577252 |
| AC107982 | -0.038582778 |
| RNU7-60P | -0.038583185 |
| RPL27AP5 | -0.038583567 |
| RP11-265F19 | -0.038603553 |
| RP11-293M10 | -0.038606421 |
| KRTAP6-3 | -0.038608066 |
| B3GALNT1P1 | -0.038613206 |
| RP11-529A4 | -0.03861684 |
| RP11-365H8 | -0.038617463 |
| OSBP2 | -0.038618421 |
| FAM60CP | -0.038620046 |
| RNU6-1110P | -0.038620342 |
| UBTFL6 | -0.038620516 |
| SNORD7 | -0.038628032 |
| RN7SL353P | -0.038628915 |
| CTD-2555O16 | -0.038631892 |
| LINC00466 | -0.03863945 |
| MIR4514 | -0.038643578 |
| RP11-446E9 | -0.038643613 |
| RP5-1033H2 | -0.038648015 |
| MIR2114 | -0.03864925 |
| AC116154 | -0.038654869 |
| RNU6-1085P | -0.038655108 |
| RNU6-1177P | -0.038657902 |
| HMGN2P23 | -0.038665105 |
| AC108456 | -0.038669143 |
| RNU6-597P | -0.038674662 |
| FLCN | -0.038677996 |
| MIR1302-4 | -0.03867876 |
| ARIH2P1 | -0.038679831 |
| UBE2Q1 | -0.038684843 |
| SRPK2P | -0.038688536 |

| | |
|---|---|
| RN7SL658P | -0.038689381 |
| SARAF | -0.038689941 |
| RBBP4P5 | -0.038690443 |
| HNRNPUP1 | -0.03869122 |
| RP5-837M10 | -0.038693215 |
| COL4A2-AS1 | -0.038705212 |
| GNB1 | -0.038705476 |
| GINS4 | -0.038706462 |
| RNA5SP260 | -0.038707969 |
| PSMD6 | -0.038709272 |
| HIST2H4B | -0.038710665 |
| RP11-440J4 | -0.038711754 |
| TPTE2P2 | -0.038713745 |
| MSMB | -0.038720655 |
| RP5-914M6 | -0.038726155 |
| RP11-661P17 | -0.038727902 |
| AC055764 | -0.038728952 |
| PSMC1 | -0.03873318 |
| HIST1H2AC | -0.038747562 |
| VN1R4 | -0.03874777 |
| USP27X | -0.038750836 |
| RP11-341A11 | -0.038752582 |
| LALBA | -0.038755005 |
| AC007528 | -0.038765487 |
| KRT18P67 | -0.038765562 |
| MTFR2 | -0.038775377 |
| AL135784 | -0.038778533 |
| RP4-639J15 | -0.038786481 |
| SUPT4H1P | -0.038793998 |
| RP11-100N20 | -0.038798679 |
| FYTTD1P1 | -0.038803429 |
| RP11-142G1 | -0.038803751 |
| RP11-318K15 | -0.038807062 |
| GAGE13 | -0.038807205 |
| AK3P2 | -0.038808843 |
| RP11-958N24 | -0.038809871 |
| RLN2 | -0.038816221 |
| Z95704 | -0.038817641 |
| DYNLL1-AS1 | -0.03881843 |
| RN7SL159P | -0.038820596 |
| AC006482 | -0.038822433 |
| RP11-485O10 | -0.038824696 |
| CSTL1 | -0.038834996 |
| CASQ2 | -0.03884293 |
| HNRNPA3P5 | -0.038843204 |

| | |
|---|---|
| LINC00502 | -0.038844496 |
| AP000868 | -0.038844625 |
| PQLC1P1 | -0.038844762 |
| RP11-115P21 | -0.038853854 |
| KLHL20 | -0.038857836 |
| NDNL2 | -0.038866507 |
| SNX1 | -0.038867046 |
| RP11-215D10 | -0.038867861 |
| RP11-112L7 | -0.038871561 |
| RLBP1 | -0.038876295 |
| CTD-2555K7 | -0.038879087 |
| SALRNA2 | -0.038883839 |
| YY1AP1 | -0.038887307 |
| RP11-382A18 | -0.03888895 |
| AC006458 | -0.03889199 |
| RP1-97G4 | -0.038894906 |
| RP11-517H2 | -0.038896917 |
| NKRF | -0.038904704 |
| SLC39A8 | -0.038909336 |
| RP4-604G5 | -0.038912576 |
| MIR6873 | -0.038916109 |
| DNAJC13 | -0.038916425 |
| EIF2AK1 | -0.038919104 |
| RP11-151G12 | -0.038920013 |
| FTLP4 | -0.038922195 |
| LLNLR-284B4 | -0.038932268 |
| RP11-700P18 | -0.038933616 |
| CTD-2062A1 | -0.038934823 |
| MLC1 | -0.0389437 |
| PFN1P3 | -0.03894465 |
| MYF6 | -0.038949297 |
| VSTM2B | -0.038951332 |
| RFX3-AS1 | -0.038957926 |
| LINC00322 | -0.038962297 |
| CTD-2631K10 | -0.038966694 |
| MGAT4D | -0.038970332 |
| RNY4P37 | -0.03897177 |
| AC010243 | -0.038975396 |
| RP11-39E3 | -0.03897983 |
| MIR4776-1 | -0.03898299 |
| RP11-697H9 | -0.038986502 |
| AC092415 | -0.038995914 |
| ATP8B2 | -0.03899734 |
| DONSON | -0.039001322 |
| DDX43P1 | -0.03900267 |

| | |
|---|---|
| AC012353 | -0.039004358 |
| OR4A43P | -0.039007307 |
| KAL1 | -0.039009117 |
| PHKBP2 | -0.03901474 |
| AC011284 | -0.039014931 |
| PRR13P1 | -0.03901877 |
| ABHD6 | -0.039019146 |
| AC012613 | -0.039025555 |
| RP11-553D4 | -0.039032251 |
| RNA5SP320 | -0.039032539 |
| RNU6-525P | -0.039034124 |
| SLC35G3 | -0.039040154 |
| PA2G4P4 | -0.039043432 |
| GSTCD | -0.039045268 |
| TRMT44 | -0.039052766 |
| RP11-622O11 | -0.039053447 |
| SLC7A1 | -0.039059929 |
| RP11-643G16 | -0.039067802 |
| APBB3 | -0.03906899 |
| PPT2-EGFL8 | -0.039069697 |
| LRRD1 | -0.039076229 |
| RN7SL359P | -0.039076427 |
| RPL9P15 | -0.039077781 |
| bP-2171C21 | -0.039079322 |
| PLCZ1 | -0.039081087 |
| MBD5 | -0.039084246 |
| AC242988 | -0.039085708 |
| GNG2 | -0.039087595 |
| OR52H1 | -0.03909709 |
| RP11-468O2 | -0.039099662 |
| RP1-278C19 | -0.039100492 |
| RNA5SP458 | -0.039108302 |
| H2AFV | -0.039111354 |
| TOMM20P1 | -0.039112878 |
| SAMM50 | -0.039114834 |
| MIR802 | -0.039116081 |
| CTD-2537O9 | -0.039116457 |
| MYNN | -0.039118183 |
| C7orf71 | -0.039118535 |
| OXCT1 | -0.039128565 |
| SIDT1 | -0.039130295 |
| PAICSP7 | -0.039132665 |
| PIGV | -0.039138295 |
| MCHR2-AS1 | -0.039138454 |
| RN7SL674P | -0.039139598 |

| | |
|---|---|
| AP001340 | -0.039140603 |
| RP5-945F2 | -0.03914558 |
| CMAS | -0.039145886 |
| AC080032 | -0.039150706 |
| RP11-346E8 | -0.03915138 |
| CST1 | -0.039154046 |
| LINC00661 | -0.039155432 |
| C12orf60 | -0.039160082 |
| RP11-25E2 | -0.039162292 |
| RP11-432J9 | -0.039168855 |
| RP11-714M23 | -0.039174625 |
| RP1-86C11 | -0.03917627 |
| RP11-299J5 | -0.039176294 |
| MPP3 | -0.039177348 |
| SLC51A | -0.039179469 |
| RP11-15F12 | -0.039180443 |
| RPEP3 | -0.039190825 |
| FAM104A | -0.039192139 |
| RP11-79M21 | -0.039195428 |
| RP11-883G10 | -0.039200454 |
| LINC01360 | -0.03920641 |
| OR4F2P | -0.039206568 |
| GPR137 | -0.03921735 |
| UBE4A | -0.03922068 |
| ICA1 | -0.039220829 |
| ZBTB5 | -0.039222317 |
| FAM180B | -0.039231732 |
| UBE2NL | -0.039235093 |
| MIR3157 | -0.039235246 |
| AC004166 | -0.03923603 |
| HIST1H2APS6 | -0.039238249 |
| FAM197Y6 | -0.039238249 |
| MIR6134 | -0.039238249 |
| OR51F1 | -0.039238249 |
| TTTY13 | -0.039238249 |
| RNA5SP417 | -0.039244532 |
| MOXD2P | -0.039246992 |
| ZNRF2 | -0.039252068 |
| KRT18P31 | -0.039253328 |
| RNU6-440P | -0.039257258 |
| RP11-24P4 | -0.039258622 |
| LINC00442 | -0.039261113 |
| RBM22P13 | -0.039262233 |
| LINC00596 | -0.039263459 |
| SEMA6A | -0.039269017 |

| | |
|---|---|
| RP11-393I2 | -0.039271027 |
| AC105081 | -0.039276649 |
| SLC11A2 | -0.039277632 |
| AC012078 | -0.039289042 |
| DHX40 | -0.039289502 |
| HSBP1P1 | -0.039289871 |
| ST13P12 | -0.039292725 |
| CALR4P | -0.039296531 |
| AL021546 | -0.039299866 |
| AC048380 | -0.039300033 |
| RP11-499P1 | -0.039303011 |
| POTEC | -0.039303424 |
| RNU6-277P | -0.039318843 |
| C14orf93 | -0.039323289 |
| RAB3GAP1 | -0.039325113 |
| RN7SL452P | -0.03932977 |
| RP11-650K20 | -0.039339395 |
| NR6A1 | -0.039341709 |
| PVALB | -0.039344554 |
| AL358340 | -0.039349666 |
| ADAMTS19-AS1 | -0.039350283 |
| RP11-412D9 | -0.039350844 |
| RP11-421M1 | -0.039354379 |
| NIP7P1 | -0.03935962 |
| RNU6-152P | -0.039361372 |
| RP11-550P17 | -0.039362097 |
| OR4K13 | -0.039362251 |
| RN7SL644P | -0.039364093 |
| TMEM126B | -0.039369133 |
| RBMX2P2 | -0.039371977 |
| RP11-192M23 | -0.039372636 |
| CELF3 | -0.039373306 |
| MTCYBP2 | -0.03937473 |
| KIF3B | -0.039383943 |
| RN7SL125P | -0.03938477 |
| MRPS11P1 | -0.039384886 |
| RP11-299D14 | -0.039385497 |
| RNU2-35P | -0.039385868 |
| CTB-118N6 | -0.039387764 |
| DCX | -0.03938845 |
| RP11-460G11 | -0.03939827 |
| RP1-138B7 | -0.039399703 |
| ZPBP2 | -0.039401404 |
| LINC01517 | -0.039402423 |
| EIF4A1P11 | -0.039403742 |

| | |
|---|---|
| RP11-191F9 | -0.039403927 |
| STX18-AS1 | -0.039405656 |
| FAM156B | -0.039407655 |
| LTN1 | -0.039409582 |
| RP11-573G6 | -0.03941881 |
| RP11-57G10 | -0.039420365 |
| SLC5A4P1 | -0.039420623 |
| KRT8P17 | -0.039422129 |
| RP11-815I22 | -0.039429112 |
| DEPDC1 | -0.039431685 |
| CYCSP12 | -0.039437073 |
| RP11-637N19 | -0.039438372 |
| MIR342 | -0.039439012 |
| ATXN10 | -0.039441434 |
| RPS19P6 | -0.039443894 |
| WFIKKN1 | -0.039444485 |
| ELP4 | -0.039446662 |
| HIST1H2BB | -0.039447696 |
| FAM91A3P | -0.039449927 |
| LINC00471 | -0.039450327 |
| RP3-507I15 | -0.039451027 |
| KCNT2 | -0.039452855 |
| ZC3HAV1L | -0.039455463 |
| MIR378F | -0.039459918 |
| ARHGEF10L | -0.039463947 |
| RNU7-125P | -0.039464302 |
| STC1 | -0.039465007 |
| IPO8 | -0.039465603 |
| PPP1R14D | -0.039469523 |
| AC106874 | -0.039471792 |
| LZIC | -0.039473882 |
| AC007182 | -0.039477215 |
| ZFP82 | -0.039477436 |
| AC009977 | -0.039480356 |
| CTD-2090I13 | -0.039481177 |
| EDEM3 | -0.039482478 |
| RP11-236J17 | -0.039482594 |
| PYY2 | -0.039483139 |
| NMNAT1P4 | -0.039484868 |
| MT-TF | -0.039491421 |
| MREGP1 | -0.039493931 |
| AGO1 | -0.039497771 |
| RP11-691F15 | -0.03949866 |
| RPS29P13 | -0.039509474 |
| MIR4436B1 | -0.039511211 |

| | |
|---|---|
| STK16 | -0.039511633 |
| AAMP | -0.039512191 |
| MSH3 | -0.039516619 |
| CDC42P3 | -0.039519698 |
| AC011513 | -0.039522519 |
| NPIPA8 | -0.039527064 |
| HINT1P2 | -0.039528059 |
| GPRC6A | -0.03952813 |
| MEF2C-AS1 | -0.039528768 |
| CASP1 | -0.039531671 |
| RP11-794C22 | -0.039532168 |
| AL596329 | -0.039536509 |
| SLC9A6 | -0.039540043 |
| CTBP2P7 | -0.039541253 |
| CTD-2218G20 | -0.039543684 |
| RP11-216P6 | -0.039543878 |
| C3orf67 | -0.039546425 |
| CTC-550M4 | -0.039559177 |
| LINC-ROR | -0.039559678 |
| TBX1 | -0.039560565 |
| RP11-512G4 | -0.039561093 |
| C7 | -0.039566186 |
| AC104772 | -0.039567427 |
| IMMT | -0.039568156 |
| GPM6BP1 | -0.039572854 |
| LINC01374 | -0.039577091 |
| BLZF1 | -0.039578798 |
| MIR100 | -0.03958352 |
| RP11-15D14 | -0.039592719 |
| RP11-167N5 | -0.039598227 |
| LINC01400 | -0.039610365 |
| KPNA1 | -0.039613814 |
| TERF1P4 | -0.039618893 |
| GS1-114I9 | -0.039620189 |
| MIR6761 | -0.039628325 |
| PPP4R2 | -0.039628326 |
| SMAD3 | -0.039629498 |
| MIR488 | -0.03963296 |
| RNU6-591P | -0.03963296 |
| AC010739 | -0.03963296 |
| AC098583 | -0.03963296 |
| CYP2AC1P | -0.039635009 |
| SLC26A1 | -0.03963668 |
| RAP2C-AS1 | -0.039638726 |
| RP11-319C21 | -0.039640267 |

| | |
|---|---|
| DYNC2LI1 | -0.039643117 |
| STATH | -0.039643658 |
| LHX6 | -0.039648829 |
| SHCBP1 | -0.039649704 |
| RP11-280H21 | -0.039659236 |
| SRMP2 | -0.039667958 |
| NFKBIA | -0.039669128 |
| ARF1P3 | -0.03966927 |
| GP2 | -0.039669352 |
| GS1-590J6 | -0.039670663 |
| KLHL34 | -0.039676537 |
| IL18R1 | -0.039677686 |
| RNU7-200P | -0.039677689 |
| CST2 | -0.039679705 |
| PALB2 | -0.039683603 |
| RP11-318K12 | -0.039685061 |
| FBXO28 | -0.039691101 |
| PGBD1 | -0.039701632 |
| OR5BE1P | -0.039702746 |
| NLRP6 | -0.03971543 |
| RP11-718L23 | -0.039726838 |
| MALT1 | -0.039729782 |
| WWOX | -0.039730712 |
| RP11-17E4 | -0.039735886 |
| MED7 | -0.039736424 |
| SIX6 | -0.039736831 |
| AC091697 | -0.039743269 |
| ZNF582 | -0.039743717 |
| RNU6-925P | -0.039745571 |
| AHNAK | -0.039746675 |
| UQCRC2P1 | -0.039747198 |
| USP17L1 | -0.039748563 |
| MIR4999 | -0.039753328 |
| YTHDF1P1 | -0.039754533 |
| KB-1589B1 | -0.03975977 |
| CMBL | -0.039760183 |
| RP11-505C13 | -0.03976407 |
| CCR3 | -0.039768101 |
| APLP1 | -0.039776965 |
| SLC25A3P2 | -0.039777258 |
| RP4-792G4 | -0.039778122 |
| LINC00989 | -0.039779835 |
| SNORD116-24 | -0.03978765 |
| RP11-395B20 | -0.039788907 |
| ADAM20P2 | -0.039791894 |

| | |
|---|---|
| RC3H1-IT1 | -0.039792459 |
| EIF5AP3 | -0.039799419 |
| CRYGD | -0.039801067 |
| RP11-893G17 | -0.039803211 |
| RNU6-1298P | -0.03981123 |
| FBXW8 | -0.039818772 |
| RP5-1186N24 | -0.039819224 |
| SNHG4 | -0.039826718 |
| AC008088 | -0.039827668 |
| TPT1P13 | -0.039829726 |
| CYCSP24 | -0.039830778 |
| CTBP2P5 | -0.039853357 |
| RNU1-28P | -0.039856393 |
| APOBEC3A | -0.039858264 |
| RP11-139I14 | -0.039861844 |
| RP11-252I14 | -0.039866107 |
| FLJ39080 | -0.039868788 |
| PLD4 | -0.039870141 |
| RPL19P1 | -0.039873516 |
| MORF4L1P1 | -0.039876281 |
| RNU6-980P | -0.039879453 |
| RP11-230L22 | -0.039880841 |
| NUP210L | -0.039883376 |
| PPP2R5C | -0.039884523 |
| BIRC6-AS1 | -0.039889118 |
| GSK3A | -0.039897398 |
| SNORD56 | -0.039897575 |
| MAST2 | -0.039900171 |
| RP11-159H3 | -0.039906547 |
| PPAPDC2 | -0.039907669 |
| RP3-509L4 | -0.039911924 |
| TRIM49C | -0.039912985 |
| PPP1R26P4 | -0.039913297 |
| RP11-56P9 | -0.039920607 |
| CDKN2C | -0.039922385 |
| RNU6-1057P | -0.039926207 |
| RP11-265B8 | -0.039926556 |
| KDM4C | -0.03993193 |
| RSPH1 | -0.039932265 |
| AC092839 | -0.039939278 |
| SNORA2 | -0.03994821 |
| RP11-3D4 | -0.039950126 |
| LINC01094 | -0.039953345 |
| RNA5SP478 | -0.039953802 |
| SNRPCP19 | -0.039957523 |

| | |
|---|---|
| RP11-98J23 | -0.039958191 |
| ATF4 | -0.039959354 |
| RP11-513I15 | -0.039964192 |
| AL021807 | -0.039966251 |
| EEF1B2P4 | -0.039969216 |
| MICALCL | -0.039969904 |
| RP11-1C1 | -0.039970243 |
| AL136231 | -0.039970312 |
| IGHV1OR16-4 | -0.039970359 |
| CWF19L1 | -0.039977886 |
| RP11-1360M22 | -0.039980489 |
| CEBPG | -0.03998557 |
| TCEB1P31 | -0.039987769 |
| OR6C1 | -0.039990496 |
| RN7SKP26 | -0.040001727 |
| ONECUT2 | -0.040009299 |
| Z97352 | -0.040015485 |
| RP11-496H15 | -0.040025132 |
| LINC00404 | -0.04002928 |
| RP11-224O19 | -0.040030235 |
| RP11-300D11 | -0.040033282 |
| FRG2 | -0.040036284 |
| RP13-216E22 | -0.040036924 |
| IGBP1P4 | -0.040038134 |
| UBAC2-AS1 | -0.040041758 |
| AC016987 | -0.040043759 |
| AC008562 | -0.040047291 |
| RNU4-75P | -0.040047794 |
| FXYD6P2 | -0.040050576 |
| RPSAP56 | -0.040054006 |
| AL590964 | -0.040057135 |
| MTRNR2L13 | -0.040063395 |
| ZNF3 | -0.040063582 |
| ANLN | -0.040063936 |
| AC011465 | -0.040064262 |
| GPR26 | -0.040064471 |
| CXorf40A | -0.040071285 |
| RP11-364B14 | -0.040076177 |
| FAM213B | -0.04007789 |
| AC104297 | -0.040078662 |
| AGO2 | -0.040080271 |
| RP11-291J9 | -0.040081035 |
| FTH1P9 | -0.040082012 |
| SLMO2P11 | -0.040084855 |
| AC009729 | -0.040088186 |

| | |
|---|---|
| RP11-233E12 | -0.040088672 |
| HS6ST3 | -0.040089113 |
| SLC20A1P3 | -0.040092338 |
| PABPN1 | -0.040096309 |
| OSBPL9P5 | -0.040097883 |
| RP11-180C1 | -0.040102242 |
| SLC25A5P1 | -0.040102465 |
| TCAIM | -0.040104626 |
| CLEC4F | -0.040109581 |
| GAS6-AS1 | -0.040110356 |
| RNA5SP176 | -0.040115213 |
| RNA5SP462 | -0.04011788 |
| RNU1-129P | -0.040118255 |
| CTD-2335A18 | -0.040123676 |
| RP11-11M20 | -0.040124461 |
| ERI3-IT1 | -0.04013403 |
| IGLVIVOR22-2 | -0.040136452 |
| ZNF140 | -0.040140516 |
| MYO6 | -0.04014353 |
| TMBIM4 | -0.040144367 |
| BTF3P4 | -0.040144978 |
| RNA5SP29 | -0.040164645 |
| AC006070 | -0.04017179 |
| CCDC26 | -0.040172732 |
| IQCF2 | -0.040178298 |
| RP11-67A1 | -0.040179065 |
| CCND3 | -0.040180871 |
| IFI30 | -0.040184016 |
| RP11-194G10 | -0.040194752 |
| MIR584 | -0.0402048 |
| RP11-263C16 | -0.040206783 |
| C1QTNF9B-AS1 | -0.040208428 |
| DDX1 | -0.040208612 |
| RAB9B | -0.040210733 |
| PNPT1P1 | -0.040212252 |
| N6AMT1 | -0.040220222 |
| MED30 | -0.040224668 |
| ABT1 | -0.040224948 |
| RP13-39P12 | -0.040225878 |
| MYOZ2 | -0.040225881 |
| RNMT | -0.040227515 |
| L34079 | -0.040231819 |
| CSN2 | -0.040232417 |
| RN7SL404P | -0.040232838 |
| RP13-455A7 | -0.040238002 |

| | |
|---|---|
| RCN1P2 | -0.040240353 |
| MIR4324 | -0.040240753 |
| RP11-148B3 | -0.040242532 |
| RP11-151H2 | -0.040244246 |
| WDR3 | -0.040256298 |
| TMPRSS11GP | -0.040258836 |
| RN7SL697P | -0.040268262 |
| RP11-323F24 | -0.040273573 |
| ZDHHC20P2 | -0.040276015 |
| YPEL3 | -0.04027868 |
| OR5AP1P | -0.04027975 |
| AARSD1 | -0.040284708 |
| RP11-420K8 | -0.040288006 |
| CTAGE9 | -0.040288755 |
| RNA5SP398 | -0.040297303 |
| AC021087 | -0.04029907 |
| AC092832 | -0.040304217 |
| RP11-642C5 | -0.040308101 |
| CNTFR-AS1 | -0.040310238 |
| LAMA3 | -0.040314365 |
| MIR626 | -0.040314789 |
| RP11-386M24 | -0.040320896 |
| RAB11FIP1P1 | -0.040326525 |
| ZNF736P3Y | -0.040329014 |
| GATA4 | -0.04032957 |
| FA2H | -0.040339539 |
| MIR3677 | -0.040340128 |
| RNU6-942P | -0.040343433 |
| RP11-148B6 | -0.040343562 |
| AL713999 | -0.040351432 |
| MIR3145 | -0.040353643 |
| HIST1H2APS4 | -0.040362008 |
| IGHVIII-38-1 | -0.040363602 |
| HDGFP1 | -0.040365623 |
| RP11-719L21 | -0.040369104 |
| RP11-72I8 | -0.040372521 |
| XK | -0.040374514 |
| RP11-552F3 | -0.04037874 |
| ODF3B | -0.040379503 |
| AC109815 | -0.040387725 |
| LINC00504 | -0.040389115 |
| RP11-15H20 | -0.040398183 |
| IMPDH1P4 | -0.040404231 |
| RP11-329J18 | -0.040404372 |
| DENND6B | -0.040408342 |

| | |
|---|---|
| CENPV | -0.040417421 |
| USP17L6P | -0.04042382 |
| TARM1 | -0.040427844 |
| PRR20E | -0.040431612 |
| LINC01558 | -0.040436123 |
| RNU7-128P | -0.040438779 |
| RP13-147D17 | -0.040440761 |
| PRR12 | -0.04044706 |
| OR4N3P | -0.040452701 |
| AC016543 | -0.040458372 |
| PRKAR2B | -0.040460809 |
| HERC2P4 | -0.040463675 |
| RP11-123I22 | -0.040466577 |
| OR5T2 | -0.040471576 |
| KIF4A | -0.04047173 |
| RP11-713M6 | -0.040471962 |
| ATP5G1P8 | -0.040475217 |
| MAGEA9 | -0.040475486 |
| ARL13A | -0.04047658 |
| CTD-2215E18 | -0.040476679 |
| ITGB5-AS1 | -0.040477466 |
| KAT2A | -0.040480847 |
| SLX4 | -0.040487824 |
| BCL2A1 | -0.040490038 |
| RNU6-192P | -0.040492742 |
| MSANTD4 | -0.040494448 |
| ACBD3 | -0.040494541 |
| DFFBP1 | -0.040495628 |
| FAM87B | -0.040498372 |
| KDM4A-AS1 | -0.040504941 |
| RNU7-92P | -0.040505736 |
| RP11-809N8 | -0.040507233 |
| OXSR1 | -0.040508495 |
| BCL2L11 | -0.040511528 |
| KDM5C-IT1 | -0.040516316 |
| ENPP7P13 | -0.040517508 |
| NFS1 | -0.040517984 |
| U51561 | -0.040518472 |
| RP11-245A18 | -0.040522246 |
| AL133335 | -0.040528007 |
| CYB5D1 | -0.040528459 |
| RP11-472E5 | -0.040530172 |
| SPCS1 | -0.040534487 |
| RP11-646I6 | -0.040542152 |
| NUP155 | -0.040545706 |

| | |
|---|---|
| GFM1 | -0.040555064 |
| MIR4516 | -0.040564032 |
| NPM1P18 | -0.040566845 |
| AL136001 | -0.040568038 |
| AC008780 | -0.04057422 |
| C6orf58 | -0.040578491 |
| BRD7P4 | -0.040579201 |
| MIR4428 | -0.040586822 |
| OR2F1 | -0.040590092 |
| ZNF134 | -0.040591847 |
| AC011450 | -0.040593069 |
| ENTPD1-AS1 | -0.04059328 |
| RN7SL22P | -0.040597085 |
| C1orf189 | -0.040599059 |
| PCP2 | -0.04059968 |
| SRSF9P1 | -0.040599736 |
| LINC00363 | -0.040601205 |
| AC008581 | -0.040605963 |
| RP11-305F18 | -0.040608533 |
| ZNF8 | -0.040609299 |
| RP11-409O11 | -0.040615012 |
| MIR429 | -0.040618369 |
| S1PR1 | -0.0406189 |
| RP11-372K14 | -0.040619439 |
| AC118344 | -0.040620043 |
| HERC3 | -0.040621132 |
| AL031905 | -0.040626021 |
| AL365232 | -0.040626271 |
| RP11-26J3 | -0.040629933 |
| VAMP4 | -0.040630119 |
| RP11-437B10 | -0.040631515 |
| SMC5 | -0.040642622 |
| ZNF346 | -0.040645357 |
| RP11-543E8 | -0.040648726 |
| CDC16 | -0.040649469 |
| RN7SKP256 | -0.040667662 |
| CTA-444M12 | -0.04066773 |
| C4orf36 | -0.040668 |
| RP11-483H20 | -0.040669832 |
| RNU6-166P | -0.040673333 |
| CASP9 | -0.040673711 |
| NPM1P30 | -0.040673913 |
| RNU6-1141P | -0.04067884 |
| RP11-26N15 | -0.040681116 |
| LYPLA1P2 | -0.040684875 |

| | |
|---|---|
| UAP1L1 | -0.040685712 |
| LINC00362 | -0.040695243 |
| RP11-820K3 | -0.040696805 |
| FFAR1 | -0.040699245 |
| RP11-241I20 | -0.04070068 |
| DNAH7 | -0.040700913 |
| RPS24P14 | -0.040701678 |
| VDAC1P4 | -0.040703131 |
| PHF1 | -0.040703424 |
| PCNAP4 | -0.040707789 |
| CH507-528H12 | -0.040708563 |
| CTD-2561F5 | -0.040715309 |
| RP11-74J13 | -0.040720653 |
| AC090338 | -0.040723981 |
| SNORA66 | -0.04072773 |
| AC124864 | -0.040729052 |
| LINC01109 | -0.040734409 |
| MIR4790 | -0.040738159 |
| RP1-207H1 | -0.04073848 |
| HIST1H4D | -0.040743717 |
| MIR548F1 | -0.040752896 |
| CRTC2 | -0.040752952 |
| NTS | -0.040755023 |
| CIRBP | -0.040755464 |
| AC108105 | -0.040759008 |
| TRMT112P4 | -0.04075936 |
| TRBV6-7 | -0.040766805 |
| RNU2-72P | -0.040771077 |
| AC098680 | -0.040773166 |
| CCDC180 | -0.040774386 |
| RN7SL219P | -0.040783086 |
| RP11-295P9 | -0.040783763 |
| RP11-166N6 | -0.040786308 |
| KCNQ2 | -0.040786427 |
| KRT28 | -0.040788948 |
| MTND5P12 | -0.040799372 |
| ANKRD27 | -0.040804768 |
| ZNF18 | -0.040805066 |
| RNU7-27P | -0.040805671 |
| Z84721 | -0.040807704 |
| AL606500 | -0.040809053 |
| RP11-290C10 | -0.040810221 |
| RPL21P9 | -0.040812253 |
| PTCD2 | -0.040813885 |
| AKAP9 | -0.040813941 |

| | |
|---|---|
| HECA | -0.040814573 |
| RP13-539F13 | -0.040821305 |
| AC073415 | -0.040828626 |
| CWF19L2 | -0.040829086 |
| BRD3 | -0.040832509 |
| MRC1 | -0.040835933 |
| SEMA4A | -0.040838258 |
| RP11-244M2 | -0.04084363 |
| CSAG2 | -0.040845476 |
| LL22NC03-27C5 | -0.040851049 |
| AKR1C7P | -0.040853582 |
| RNU7-141P | -0.040856605 |
| MAP3K5 | -0.040856696 |
| PPIAL4E | -0.040856911 |
| RNU6-402P | -0.04086384 |
| AC093766 | -0.040864825 |
| AP4B1 | -0.040871022 |
| MIB1 | -0.040872102 |
| RNA5SP153 | -0.04087602 |
| KDM3A | -0.040878706 |
| LINC01166 | -0.040885109 |
| AP000787 | -0.040885261 |
| CRB2 | -0.040885445 |
| CTD-2184C24 | -0.040886729 |
| AL034449 | -0.040886819 |
| MIR517A | -0.040896242 |
| HNRNPA1P46 | -0.040898565 |
| TFAP4 | -0.040901442 |
| RIBC1 | -0.040906522 |
| ADM2 | -0.040907471 |
| CYB561A3 | -0.040907526 |
| ACSM5 | -0.04090855 |
| RPS27P25 | -0.040908703 |
| TUBBP11 | -0.040911811 |
| RPL21P121 | -0.040927965 |
| RP11-285A18 | -0.040936891 |
| ZNF192P1 | -0.040942766 |
| EAF2 | -0.040944123 |
| AP001631 | -0.040945493 |
| XXbac-BPG154L12 | -0.040946467 |
| FAM199X | -0.04094857 |
| ARSDP1 | -0.040952865 |
| FGD4 | -0.040953296 |
| RP1-134E15 | -0.040955885 |
| NPRL2 | -0.040957743 |

| | |
|---|---|
| AL160056 | -0.040958196 |
| FOXF1 | -0.040960496 |
| AC105052 | -0.040964557 |
| CTSLP6 | -0.040965424 |
| AGXT2 | -0.040971431 |
| KIAA1841 | -0.040973725 |
| ZDHHC7 | -0.040974618 |
| RP11-388K2 | -0.040979011 |
| AHCYP5 | -0.040989837 |
| CHAF1A | -0.040997168 |
| RGP1 | -0.040998526 |
| AC021231 | -0.04099887 |
| SLC35G6 | -0.041000527 |
| NBEAP3 | -0.041001189 |
| RANBP2 | -0.041003605 |
| RNA5SP189 | -0.041005822 |
| RP11-285E9 | -0.041009704 |
| RNU6-1244P | -0.041012501 |
| RP11-21M24 | -0.041014498 |
| MIR6894 | -0.041026761 |
| RP11-467H10 | -0.041029411 |
| DLGAP5 | -0.041029839 |
| COL4A3BP | -0.041036574 |
| FGF12-AS3 | -0.041038688 |
| RP3-337D23 | -0.041048533 |
| CLIP1 | -0.041055105 |
| IL5 | -0.041056585 |
| SNORD60 | -0.041058001 |
| RNU1-94P | -0.041067243 |
| KB-226F1 | -0.04107438 |
| AP000462 | -0.041078024 |
| MIR3135B | -0.041088174 |
| HCG4P7 | -0.04109053 |
| RNU6-740P | -0.041090892 |
| XX-CR54 | -0.041091575 |
| LINC01588 | -0.04109319 |
| CNIH1 | -0.041094809 |
| ZNF432 | -0.041097991 |
| INPP5J | -0.04110147 |
| RNU1-16P | -0.041102686 |
| RP11-328K4 | -0.04110693 |
| MUC6 | -0.041114205 |
| RP11-182M20 | -0.04112148 |
| H2AFZ | -0.041123406 |
| RNU6-293P | -0.041123739 |

| | |
|---|---|
| CBX5 | -0.041124745 |
| MSANTD2 | -0.041130494 |
| TDGF1P3 | -0.041130936 |
| SRPK1 | -0.041134439 |
| HNRNPA1P47 | -0.041135179 |
| CAPN11 | -0.041136727 |
| RP11-745I13 | -0.041139699 |
| DLG2-AS1_1 | -0.041146503 |
| AC017035 | -0.041150559 |
| ZNF705C | -0.041157705 |
| WDR64 | -0.041160315 |
| DTX2P1-UPK3BP1-PMS2P11 | -0.041161658 |
| MPP5 | -0.041161882 |
| SLC47A1 | -0.041165082 |
| AATK-AS1 | -0.041165548 |
| AC068946 | -0.041168332 |
| RP3-368A4 | -0.041168654 |
| AP001187 | -0.041170386 |
| MCCC1 | -0.041173771 |
| ATP5G2P3 | -0.041175841 |
| MIR3976 | -0.041176171 |
| CKMT1B | -0.041181942 |
| CTB-147C13 | -0.041189231 |
| RASA1 | -0.041190311 |
| RP11-234B24 | -0.041199379 |
| RN7SL350P | -0.041204455 |
| IGKV3-31 | -0.04121629 |
| RP11-631F7 | -0.041218689 |
| ST6GALNAC4P1 | -0.041220987 |
| HNRNPA1P37 | -0.041222256 |
| CTA-390C10 | -0.041223138 |
| MYOD1 | -0.041237025 |
| RP11-759L5 | -0.041237025 |
| RPL22P22 | -0.041242545 |
| AC025335 | -0.041246946 |
| PRPSAP1 | -0.041248323 |
| OR11Q1P | -0.041248694 |
| CCPG1 | -0.041250127 |
| GPRC5D | -0.041250691 |
| BAG5 | -0.041251355 |
| RP11-164N20 | -0.041256788 |
| BHLHB9 | -0.041256937 |
| RPS15AP27 | -0.041259607 |
| LRRC25 | -0.041261328 |
| MIR5692A2 | -0.041272621 |

| | |
|---|---|
| RN7SKP164 | -0.041272621 |
| SERPINE3 | -0.041274578 |
| MIR513B | -0.041276421 |
| LINC00608 | -0.041282681 |
| LINGO1-AS1 | -0.041287314 |
| RN7SL583P | -0.041288321 |
| KRT18P57 | -0.041288585 |
| USP19 | -0.041294725 |
| RNU6-1201P | -0.041296318 |
| RP11-171G2 | -0.041296542 |
| PTGDS | -0.041298967 |
| RP11-489G11 | -0.041305854 |
| CAMK2B | -0.04130639 |
| PPP1R2P3 | -0.041308538 |
| CHD7 | -0.041317535 |
| YPEL1 | -0.04132001 |
| RNU6-1280P | -0.041321901 |
| RP11-886H22 | -0.04132415 |
| STMN3 | -0.041325628 |
| RP11-354I13 | -0.041329331 |
| AC116533 | -0.041330137 |
| CTC-534B23 | -0.041334801 |
| RNU6-910P | -0.041338413 |
| RP11-86A5 | -0.041338697 |
| RP11-62C3 | -0.041340301 |
| OR4M1 | -0.041348296 |
| RP11-188D8 | -0.041349643 |
| ZSWIM5P2 | -0.041349873 |
| FAM183B | -0.041350327 |
| MIR5698 | -0.041353083 |
| RP11-395P16 | -0.041354298 |
| AC139100 | -0.041354913 |
| CASC16 | -0.041356596 |
| OTX2P1 | -0.041357145 |
| AL901608 | -0.041358257 |
| LINC00399 | -0.041359517 |
| TMA16P2 | -0.041360228 |
| RP11-509P12 | -0.041363678 |
| SLC6A15 | -0.041366392 |
| RN7SL272P | -0.041370893 |
| RNA5SP312 | -0.041372835 |
| ADH5P3 | -0.041376448 |
| RP11-352G9 | -0.041377929 |
| SOD2 | -0.041380589 |
| MIR4499 | -0.041395013 |

| | |
|---|---|
| RP11-264B14 | -0.041396883 |
| LIN28AP2 | -0.041398436 |
| RP11-476B13 | -0.041400184 |
| SNORD41 | -0.04140531 |
| HIST1H2APS3 | -0.041407268 |
| USP46-AS1 | -0.041414414 |
| AF254982 | -0.041417989 |
| KATNBL1P4 | -0.041420465 |
| RP11-664I21 | -0.041421423 |
| EGFLAM-AS3 | -0.041422815 |
| CTD-2128F4 | -0.04143509 |
| SNORD11B | -0.041438535 |
| VN1R76P | -0.041438764 |
| DDR1 | -0.041439029 |
| RP11-209D14 | -0.041440754 |
| MYZAP | -0.041442545 |
| RNU6-392P | -0.041450116 |
| TM2D1 | -0.04145075 |
| TCEB3-AS1 | -0.041451587 |
| CTD-2588J6 | -0.041455084 |
| PSMC1P13 | -0.041457996 |
| AL132768 | -0.041459868 |
| MORN4 | -0.041460127 |
| RNU6-104P | -0.041468136 |
| OPA1-AS1 | -0.041474967 |
| LINC01544 | -0.041475834 |
| C9orf147 | -0.041480212 |
| RP11-543G18 | -0.041480241 |
| AL158086 | -0.041487089 |
| LINC01507 | -0.041491375 |
| H2AFZP6 | -0.041491832 |
| MIR6895 | -0.041494119 |
| RP11-140I24 | -0.041500247 |
| MED29 | -0.041502473 |
| HAO2 | -0.04150513 |
| AKAP17A | -0.041511199 |
| RP11-70O5 | -0.041511302 |
| EHMT2 | -0.04151172 |
| RP11-112N13 | -0.041520707 |
| LINC00624 | -0.041524996 |
| GMIP | -0.041525172 |
| NAP1L4P3 | -0.041530064 |
| ST7-OT4_2 | -0.041531868 |
| CTD-2385L22 | -0.041532567 |
| MTMR1 | -0.041533972 |

| | |
|---|---|
| TBC1D3P3 | -0.041534288 |
| ZC3H12A | -0.041535238 |
| RNU7-120P | -0.04153612 |
| FOSL1P1 | -0.041536121 |
| LA16c-3G11 | -0.041538091 |
| RNU5E-1 | -0.041540792 |
| RP11-34A14 | -0.041543852 |
| DDX50 | -0.041544875 |
| CT55 | -0.041544899 |
| RP11-259P20 | -0.041545132 |
| REV1 | -0.04154588 |
| AC098826 | -0.041548273 |
| RP11-556O15 | -0.041549297 |
| PI16 | -0.041550726 |
| RP4-803A2 | -0.041557294 |
| HAUS6P3 | -0.04155813 |
| SLC26A4 | -0.041560225 |
| RP11-160H12 | -0.041560914 |
| MAN2A2 | -0.041562495 |
| RN7SL209P | -0.041565901 |
| SMDT1 | -0.041566309 |
| LINC01337 | -0.041566873 |
| RP11-593P24 | -0.041567551 |
| PIGPP3 | -0.041571538 |
| TRRAP | -0.041575946 |
| TAF7 | -0.041580561 |
| C9orf171 | -0.041581474 |
| RN7SL444P | -0.041584043 |
| CTD-2291D10 | -0.041590314 |
| KRT18P40 | -0.041595662 |
| DGKZ | -0.041602031 |
| SERP1 | -0.041602366 |
| HCG9P5 | -0.041604226 |
| AC005014 | -0.041608166 |
| AC009505 | -0.041608618 |
| MIR219A2 | -0.041613832 |
| WDR31 | -0.041614311 |
| RP11-25H12 | -0.041624389 |
| RP11-256K7 | -0.041624413 |
| RP11-440L16 | -0.041625759 |
| MIR5787 | -0.041626199 |
| CNOT4P1 | -0.041627011 |
| RP3-388M5 | -0.041627052 |
| CTA-833B7 | -0.041627778 |
| STAG3L4 | -0.041631484 |

| | |
|---|---|
| AC016743 | -0.041634964 |
| PIK3CA | -0.04163593 |
| PIEZO2 | -0.041638275 |
| SNX18P25 | -0.041638536 |
| LINC01024 | -0.041653661 |
| PITPNB | -0.041654759 |
| RNU6-851P | -0.041658298 |
| RP11-301J16 | -0.041659661 |
| RNU6-534P | -0.041661232 |
| RPS23P3 | -0.041661782 |
| XX-FW81066F1 | -0.041668997 |
| PSMA3 | -0.041670816 |
| RP11-140A10 | -0.041677163 |
| PPP1R12B | -0.041679095 |
| LINS | -0.041694426 |
| snoZ40 | -0.041694905 |
| HNRNPA1P26 | -0.041696057 |
| SAMD12 | -0.041696714 |
| RNA5SP20 | -0.041704673 |
| RP1-12G14 | -0.041715599 |
| TRIM26BP | -0.041720083 |
| RP11-575G13 | -0.041720775 |
| AL049691 | -0.041724334 |
| C3orf84 | -0.041725543 |
| RP11-326A13 | -0.041725622 |
| CCDC115 | -0.04172626 |
| SETDB1 | -0.041733423 |
| RNA5SP374 | -0.041736014 |
| RP11-165I9 | -0.041740383 |
| NCBP2L | -0.041742272 |
| ALPI | -0.041743979 |
| HIST1H4PS1 | -0.041744439 |
| RADIL | -0.041754019 |
| RP11-820I16 | -0.041755726 |
| MASTL | -0.041758708 |
| RPL23P2 | -0.041768215 |
| ZBTB21 | -0.041776737 |
| C3orf17 | -0.041783993 |
| FP325331 | -0.041791533 |
| AC004824 | -0.041795279 |
| RN7SL847P | -0.041797193 |
| AC004067 | -0.04179724 |
| AC015691 | -0.041797569 |
| HCG18 | -0.041797779 |
| RPL7P27 | -0.041797832 |

| | |
|---|---|
| RNA5SP70 | -0.041799448 |
| RP13-580B18 | -0.041799721 |
| RP13-483F6 | -0.041802735 |
| Y_RNA | -0.041809464 |
| RP11-134N1 | -0.041811906 |
| RP11-38C17 | -0.041812804 |
| PAICSP4 | -0.041814393 |
| PPP1R2 | -0.041819957 |
| EHMT1 | -0.041827102 |
| MYO1A | -0.041828216 |
| CDK5PS | -0.041828624 |
| CTB-10G5 | -0.041837919 |
| SPATA31A1 | -0.041838074 |
| TDGF1P4 | -0.041838586 |
| CTC-499B15 | -0.041843778 |
| MED24 | -0.041845391 |
| SNORA36A | -0.041855307 |
| MIR4319 | -0.0418559 |
| MTND2P14 | -0.041856882 |
| ZNF850 | -0.041856908 |
| AC090587 | -0.041858824 |
| AC010207 | -0.04185898 |
| CA11 | -0.041859717 |
| IL19 | -0.041861391 |
| AC016656 | -0.041870856 |
| LINC00350 | -0.041872009 |
| AC007277 | -0.041872963 |
| SNORD116-19 | -0.04187368 |
| SLCO3A1 | -0.041876443 |
| AC093151 | -0.041878411 |
| LINC01176 | -0.04188035 |
| LY6G6E | -0.041884209 |
| RP11-3L10 | -0.041887538 |
| ALG5 | -0.041890156 |
| ITM2B | -0.041905133 |
| ASTE1 | -0.041909226 |
| RN7SKP12 | -0.041919039 |
| RP11-18F14 | -0.041924652 |
| GPHA2 | -0.041932514 |
| HCFC2 | -0.041933078 |
| AP000403 | -0.041934932 |
| KRTAP5-1 | -0.041936542 |
| CERS6-AS1 | -0.041937795 |
| RNU6-1136P | -0.041938588 |
| RP11-735B13 | -0.041946207 |

| | |
|---|---|
| MIR7155 | -0.041947197 |
| RN7SL569P | -0.041948363 |
| RN7SKP98 | -0.041948656 |
| GOLGA8VP | -0.041952884 |
| CADM3 | -0.041960389 |
| CCDC126 | -0.041964317 |
| C8orf44 | -0.041967554 |
| AC000029 | -0.041968876 |
| AL138751 | -0.041971074 |
| HMGN2P11 | -0.041975696 |
| AC083873 | -0.041978183 |
| ZNF17 | -0.041980243 |
| SARDH | -0.041983332 |
| C2orf47 | -0.041984634 |
| RP11-1023P17 | -0.041987811 |
| RNA5SP372 | -0.041987883 |
| SMR3B | -0.041991996 |
| RP11-94C24 | -0.041996486 |
| SLC2A11 | -0.041997732 |
| KLF13 | -0.041998179 |
| NCDN | -0.041999109 |
| RASSF6 | -0.042004542 |
| MIR3908 | -0.042007657 |
| RP11-339N8 | -0.042008065 |
| RBMY2KP | -0.042008501 |
| SLC25A6P6 | -0.042009635 |
| REPIN1 | -0.04201351 |
| RP11-388G22 | -0.042014356 |
| CTD-2269E23 | -0.042017432 |
| CNN2P12 | -0.042018486 |
| PRPF3 | -0.042021957 |
| LA16c-395F10 | -0.042024627 |
| CHURC1-FNTB | -0.042024641 |
| LYPLA2P3 | -0.042027427 |
| NPM1P52 | -0.042027906 |
| TRPC7-AS1 | -0.042031946 |
| CTBP1-AS2 | -0.042031966 |
| PANDAR | -0.042034679 |
| CD72 | -0.042037548 |
| CTD-2118P12 | -0.042037866 |
| LL0XNC01-177E8 | -0.042039009 |
| RP11-293F5 | -0.042043895 |
| XRCC5 | -0.042044265 |
| CTD-2008A1 | -0.042045259 |
| RP11-124I4 | -0.042045371 |

| | |
|---|---|
| RP11-331F9 | -0.042048495 |
| RNU7-85P | -0.042050043 |
| SSR1 | -0.042050423 |
| RP11-161H23 | -0.042051795 |
| ATP5G2P1 | -0.04205389 |
| CH507-396I9 | -0.042056316 |
| RP11-456J20 | -0.042057076 |
| HMGN2P9 | -0.042060833 |
| CCDC150P1 | -0.042064709 |
| OR10AB1P | -0.042066342 |
| RP11-772C9 | -0.042070497 |
| MBD4 | -0.042070815 |
| LINC01187 | -0.042074565 |
| RNU6-1016P | -0.042074796 |
| MIR5703 | -0.042075685 |
| BCRP6 | -0.042079946 |
| AL358133 | -0.042080198 |
| DBR1 | -0.042083429 |
| INTS12 | -0.042096434 |
| FHIT | -0.042099616 |
| RP11-292K15 | -0.04210707 |
| IRF2BP2 | -0.042107635 |
| RNU6-783P | -0.042114268 |
| MMADHC | -0.042119152 |
| AC142528 | -0.042119374 |
| PPM1N | -0.042123654 |
| HS3ST6 | -0.042130589 |
| UNC5A | -0.042137925 |
| AC007740 | -0.042145663 |
| AL355075 | -0.042145797 |
| MIR1179 | -0.042149176 |
| HSPE1-MOB4 | -0.042155025 |
| ZBTB33 | -0.042157981 |
| RP11-2J18 | -0.042159912 |
| FBP2P1 | -0.042162948 |
| RP11-622J9 | -0.042165927 |
| OR7A10 | -0.042167327 |
| ZNF35 | -0.042170466 |
| RNF5 | -0.042170596 |
| MIR6846 | -0.042175447 |
| RP1-55C23 | -0.042179239 |
| RP11-15L13 | -0.042179975 |
| IGHD | -0.042183032 |
| OR10D3 | -0.042185211 |
| RP11-43N5 | -0.042187826 |

| | |
|---|---|
| RNU6-496P | -0.042188619 |
| RP11-239E10 | -0.042188857 |
| NFASC | -0.042190578 |
| LINC01221 | -0.042193483 |
| CTD-2134P3 | -0.042196434 |
| MTND6P5 | -0.042197872 |
| C1orf95 | -0.04219923 |
| OR4F6 | -0.042199262 |
| AC011385 | -0.042204379 |
| RNU6-517P | -0.04220566 |
| CYP4F24P | -0.042206219 |
| SNORD70 | -0.042210244 |
| RP11-67H2 | -0.04221853 |
| RBFOX3 | -0.042218804 |
| AK3P5 | -0.042219045 |
| AC103724 | -0.042224907 |
| RBM17P4 | -0.042232107 |
| ARHGEF33 | -0.042235454 |
| SLC39A9 | -0.042235659 |
| RP11-776K3 | -0.042236875 |
| CTD-2132H18 | -0.042243033 |
| RP11-420K14 | -0.042243453 |
| RP11-544M22 | -0.042251957 |
| RP11-118D22 | -0.042254112 |
| MIR4459 | -0.042266676 |
| IGHVII-49-1 | -0.042270391 |
| DOCK11 | -0.042274321 |
| DHX29 | -0.042275624 |
| GPATCH4 | -0.042275735 |
| MIR1180 | -0.04227741 |
| GNL3LP1 | -0.042277698 |
| RP11-517M22 | -0.042285204 |
| AC016959 | -0.042285327 |
| SATL1 | -0.04228781 |
| PCNXL4 | -0.042287847 |
| FAM200B | -0.04229449 |
| ADCY3 | -0.042297113 |
| AC064852 | -0.042309889 |
| SATB1 | -0.04231423 |
| RP4-635A23 | -0.04231706 |
| LHFPL4 | -0.042318837 |
| MTFMT | -0.042322807 |
| BNIP3P26 | -0.042325546 |
| PRY2 | -0.042335897 |
| SNORD115-41 | -0.042335986 |

| | |
|---|---|
| RP11-16C18 | -0.042339365 |
| UPRT | -0.042341989 |
| TOB2P1 | -0.042342049 |
| LINC00969 | -0.042343534 |
| RP11-762N20 | -0.042343581 |
| RP11-385N23 | -0.042345765 |
| LINC00523 | -0.042346754 |
| IQCB1 | -0.042347348 |
| ZFP69B | -0.04234958 |
| MIR7847 | -0.042350754 |
| ERBB4 | -0.042350813 |
| RP11-256L6 | -0.04235129 |
| DYNC1LI2 | -0.042354474 |
| PA2G4 | -0.042355274 |
| RP1-8B22 | -0.042355542 |
| RP11-3N13 | -0.042355686 |
| MIR4303 | -0.042361529 |
| AC132825 | -0.04237068 |
| UBE2Q2P8 | -0.042377309 |
| RNU7-147P | -0.042377309 |
| ARHGEF3 | -0.042378605 |
| RP11-219B4 | -0.042386666 |
| MRPL57 | -0.042387963 |
| CTD-3060P21 | -0.04238815 |
| RNU6-510P | -0.042390132 |
| SNORD59 | -0.042390605 |
| AC072062 | -0.042398324 |
| SSXP1 | -0.042399779 |
| HIPK2 | -0.042401397 |
| JARID2-AS1 | -0.042405657 |
| SGSM1 | -0.042406253 |
| THAP9-AS1 | -0.042408225 |
| KIDINS220 | -0.042408802 |
| ZIK1P1 | -0.042412642 |
| CTC-482H14 | -0.042418744 |
| MAGEE1 | -0.042426681 |
| RN7SL239P | -0.042427848 |
| CYCSP39 | -0.042430079 |
| RP11-60I3 | -0.042431173 |
| SMG7-AS1 | -0.042431519 |
| WASH3P | -0.042434533 |
| AP004290 | -0.042438696 |
| RNU4ATAC6P | -0.042443887 |
| SNX19 | -0.042446831 |
| TAF1L | -0.042455991 |

| | |
|---|---|
| RNU6-126P | -0.042459171 |
| TMEM147-AS1 | -0.042459743 |
| EFTUD1P1 | -0.042464153 |
| RP11-809O17 | -0.042469939 |
| RP11-356M6 | -0.042477716 |
| INSL3 | -0.042478474 |
| FLRT1 | -0.042481828 |
| RP11-474G23 | -0.042485045 |
| LINC01135 | -0.042486162 |
| GPR142 | -0.042487079 |
| TATDN1 | -0.042487968 |
| RP11-420J11 | -0.04249035 |
| RP11-90P5 | -0.042495035 |
| ZNF260 | -0.042497661 |
| RP11-12D24 | -0.042498665 |
| RP11-490N5 | -0.042503435 |
| TMEM9B | -0.042504583 |
| AC005624 | -0.042509334 |
| MIR129-1 | -0.04251379 |
| LSP1P3 | -0.042524537 |
| DNASE2B | -0.042529257 |
| RNU6-249P | -0.042534503 |
| AC068207 | -0.042535672 |
| NUP43 | -0.042538668 |
| GUCA1B | -0.042539151 |
| LINC00457 | -0.042539202 |
| AC079896 | -0.042546104 |
| ZNF815P | -0.04254797 |
| PPP1R11 | -0.042551692 |
| CAMLG | -0.042553571 |
| MIR3130-1 | -0.042556885 |
| MIR3978 | -0.042559511 |
| FANCE | -0.042560343 |
| GTF2I | -0.042561999 |
| RP4-749H3 | -0.042564254 |
| PKP4P1 | -0.042566726 |
| RP4-739H11 | -0.042573288 |
| NACAP5 | -0.042582262 |
| RPL7P4 | -0.042588549 |
| AC110769 | -0.042595263 |
| RP11-73B2 | -0.042598634 |
| GOSR2 | -0.042598782 |
| RASSF4 | -0.042600196 |
| RP3-525N14 | -0.042600908 |
| C1orf27 | -0.042614485 |

| | |
|---|---|
| LINC00469 | -0.042617545 |
| MIR7-3HG | -0.042619537 |
| RP11-701B16 | -0.042625309 |
| CCDC150 | -0.042633611 |
| DNAJA1P3 | -0.042640074 |
| AC068279 | -0.042642584 |
| EIF1 | -0.042643472 |
| MIR6750 | -0.042644139 |
| AC096509 | -0.042645009 |
| SETD3 | -0.042647212 |
| TMEM55A | -0.042651715 |
| RP4-752I6 | -0.042655048 |
| E2F1 | -0.042659436 |
| B3GALT5 | -0.042666152 |
| RP11-547D23 | -0.042670647 |
| CSNK2B-LY6G5B-562 | -0.042676607 |
| RNU1-141P | -0.042677501 |
| RP11-282O18 | -0.042685911 |
| HNRNPCP4 | -0.042694954 |
| RP11-554D20 | -0.042697636 |
| MIR345 | -0.042706225 |
| FTX_4 | -0.042706761 |
| C6orf201 | -0.042712226 |
| WI2-3308P17 | -0.042712664 |
| RP11-502N13 | -0.042713168 |
| SSH3 | -0.042715829 |
| KRT18P42 | -0.042716811 |
| MTND5P28 | -0.042717736 |
| CLIC1P1 | -0.042718177 |
| PTP4A1 | -0.042719457 |
| RP11-248G5 | -0.042725368 |
| RNU6-899P | -0.042732735 |
| CPHL1P | -0.042732834 |
| RP11-629B11 | -0.042733393 |
| RP11-728D14 | -0.042737371 |
| AC092854 | -0.042738156 |
| MT-TN | -0.042738434 |
| UGT2B29P | -0.042738779 |
| SMN2 | -0.042739997 |
| RP3-403A15 | -0.04274251 |
| RN7SL132P | -0.042743425 |
| TTC30B | -0.042745598 |
| KIAA1586 | -0.042747384 |
| RPS15AP40 | -0.042752691 |
| UBAP2 | -0.042764147 |

| | |
|---|---|
| LINC00656 | -0.042764671 |
| RP11-381K20 | -0.042778012 |
| CC2D2B | -0.042781201 |
| PCNT | -0.042781357 |
| ANO1 | -0.042786437 |
| TMEM57 | -0.042790065 |
| C1orf174 | -0.042793246 |
| RP11-349A8 | -0.042794229 |
| RP3-525N10 | -0.042801038 |
| BHLHE23 | -0.04280478 |
| KIFC1 | -0.042806214 |
| RP4-669P10 | -0.042809228 |
| LINC00851 | -0.042812088 |
| CDCA4P1 | -0.042816821 |
| B3GAT1 | -0.042818312 |
| MAZ | -0.04281855 |
| RNA5SP50 | -0.042826596 |
| RP11-1299A16 | -0.042830116 |
| AC009414 | -0.042834946 |
| AGAP1 | -0.042837806 |
| HIST1H2AG | -0.042838309 |
| KCNH6 | -0.042839237 |
| SLC25A39P1 | -0.042843417 |
| RP11-20P5 | -0.042852711 |
| GYPE | -0.042853645 |
| SLC4A1APP1 | -0.042853796 |
| AC144450 | -0.04285557 |
| SPANXA1 | -0.042859987 |
| PAGE5 | -0.042860653 |
| BAHD1 | -0.042861063 |
| GSK3B | -0.042862905 |
| MIAT | -0.042868766 |
| LINC01103 | -0.042872571 |
| HNRNPA1P67 | -0.042880061 |
| PRKD2 | -0.042880134 |
| AQP6 | -0.042881902 |
| CDRT15P2 | -0.042891174 |
| SSU72P5 | -0.042891339 |
| C9orf173 | -0.04289446 |
| RNU6-361P | -0.042896766 |
| RN7SL105P | -0.042900325 |
| MTCO3P1 | -0.042900925 |
| RN7SL411P | -0.042906221 |
| CC2D1B | -0.042907447 |
| EML4 | -0.042907912 |

| | |
|---|---|
| RNU6-298P | -0.042908269 |
| IL12A | -0.042910838 |
| OR2H1 | -0.04291341 |
| RNU6-624P | -0.042917212 |
| SULT1A3 | -0.042917765 |
| CTB-191K22 | -0.04291969 |
| AC007256 | -0.042922502 |
| AC093642 | -0.042923857 |
| RP11-797A18 | -0.042936419 |
| AC080002 | -0.042937578 |
| LINC00692 | -0.04293819 |
| CTB-43E15 | -0.042938318 |
| COL6A5 | -0.042939129 |
| AHSP | -0.042943232 |
| IQSEC2 | -0.042944751 |
| ASF1B | -0.042945104 |
| RP11-520P18 | -0.042946558 |
| RP11-374I15 | -0.042952461 |
| SNORD71 | -0.042958454 |
| VCX | -0.042958875 |
| RP11-179K3 | -0.042967751 |
| BMS1 | -0.042968435 |
| RPL36AP13 | -0.04297023 |
| RP11-359H18 | -0.042972704 |
| MTCP1 | -0.042976715 |
| RP11-574F11 | -0.042977012 |
| RN7SL667P | -0.042978235 |
| RP11-374N7 | -0.042985629 |
| CHKB-AS1 | -0.042989702 |
| CTD-2161E19 | -0.042991514 |
| SLC35B3 | -0.042992809 |
| RPL34-AS1 | -0.042998317 |
| LLGL2 | -0.042998334 |
| AMER3 | -0.042999809 |
| OR2C1 | -0.043005009 |
| MSR1 | -0.043010019 |
| RP1-253P7 | -0.043010652 |
| FUNDC2P3 | -0.043016735 |
| MTND1P20 | -0.043018465 |
| CTD-2089O24 | -0.043022709 |
| IGLVV-66 | -0.043024488 |
| RN7SKP272 | -0.043026508 |
| RINT1 | -0.043027805 |
| LINC00667 | -0.043031452 |
| RP11-153F5 | -0.04303356 |

| | | |
|---|---|---|
| CTB-1I21 | | -0.043040162 |
| RPL37AP1 | | -0.043057416 |
| RP11-432J22 | | -0.043058914 |
| RNU6ATAC16P | | -0.043059212 |
| CES2 | | -0.043062319 |
| DPPA2 | | -0.043064726 |
| TRIM5 | | -0.043071733 |
| RP11-861E23 | | -0.043074001 |
| ATXN2L | | -0.04307685 |
| NT5DC3 | | -0.043095641 |
| LPHN3-AS1 | | -0.043100118 |
| PTPRB | | -0.043104795 |
| RNU7-151P | | -0.043108965 |
| MIR16-1 | | -0.043115017 |
| RP11-351C21 | | -0.043115141 |
| AP000783 | | -0.043116674 |
| RN7SL574P | | -0.043117131 |
| OSGEP | | -0.043118116 |
| RP1-28H20 | | -0.043118265 |
| MKNK1-AS1 | | -0.043118993 |
| IGHVIII-25-1 | | -0.043119942 |
| CNN2P7 | | -0.043127939 |
| SLCO4C1 | | -0.043131222 |
| KRTAP2-5P | | -0.04313248 |
| WIPF3 | | -0.043132536 |
| RP11-316J7 | | -0.043132745 |
| SMAD7 | | -0.043136071 |
| COA7 | | -0.043138173 |
| KRT18P29 | | -0.043142212 |
| RNU6-313P | | -0.043156377 |
| | 8-Mar | -0.043156574 |
| RP11-526K21 | | -0.04316065 |
| RP11-480A16 | | -0.043161653 |
| TMPPE | | -0.04316223 |
| RNU7-106P | | -0.043165572 |
| RP11-226L15 | | -0.043166323 |
| RP11-63C8 | | -0.043168388 |
| GMFBP1 | | -0.04317201 |
| SNORD69 | | -0.043177869 |
| SLC30A6 | | -0.043185022 |
| SRMS | | -0.04318519 |
| ATF4P3 | | -0.043187875 |
| RP11-802O23 | | -0.043190369 |
| DHRS9 | | -0.043190622 |
| MIR25 | | -0.043193866 |

| | | |
|---|---|---|
| RPS27P14 | | -0.043200552 |
| RAB11A | | -0.043202976 |
| RP11-576E20 | | -0.043203673 |
| HNRNPA1P39 | | -0.043208266 |
| SCRN1 | | -0.043211721 |
| SNORD116-4 | | -0.043213025 |
| | 7-Mar | -0.043218809 |
| SDHDP2 | | -0.043219164 |
| HBBP1 | | -0.043219642 |
| CTC-471J1 | | -0.04322026 |
| RP11-519C12 | | -0.043224737 |
| AL050321 | | -0.043225673 |
| RNU6-103P | | -0.043240096 |
| RNU6-415P | | -0.043243438 |
| CHST1 | | -0.043250811 |
| BNIP3P39 | | -0.043261786 |
| CTD-2587H19 | | -0.043262255 |
| KHDC3L | | -0.04326503 |
| CTD-2057J6 | | -0.043265393 |
| RNU6-76P | | -0.043265461 |
| AC012531 | | -0.043265758 |
| RP11-182B22 | | -0.043266897 |
| RP11-235C23 | | -0.043269433 |
| NANOGP6 | | -0.043270733 |
| AL391001 | | -0.043271682 |
| RP11-64K7 | | -0.043279649 |
| CEP41 | | -0.043286049 |
| RP11-120D5 | | -0.043286535 |
| RP11-108K3 | | -0.043288321 |
| MIR6838 | | -0.043288357 |
| RP11-519M16 | | -0.043301725 |
| C19orf54 | | -0.043309455 |
| CTC-756D1 | | -0.043310692 |
| RP11-598P20 | | -0.043315004 |
| RP5-886K2 | | -0.043318337 |
| LINC01033 | | -0.043325352 |
| AC007792 | | -0.043328628 |
| ALG1L15P | | -0.04333151 |
| RNA5SP163 | | -0.043335949 |
| LLNLR-249E10 | | -0.04334286 |
| OR56B1 | | -0.043346069 |
| AC074019 | | -0.04335119 |
| RP11-398C13 | | -0.043367225 |
| SALL4P7 | | -0.043369346 |
| SNX19P2 | | -0.043378905 |

| | |
|---|---|
| GOLGA8K | -0.043379891 |
| AC004869 | -0.043380273 |
| RP11-461G12 | -0.04338601 |
| TCF7 | -0.043387952 |
| PCMT1 | -0.043388249 |
| RP4-738P11 | -0.043390967 |
| NRD1 | -0.043391189 |
| MCM5 | -0.043397087 |
| ALS2CR12 | -0.043400325 |
| ZNF619P1 | -0.043400561 |
| RNU2-11P | -0.043402442 |
| RP11-118B18 | -0.043406987 |
| SCUBE2 | -0.043407878 |
| ANKRD33B | -0.043409608 |
| URAHP | -0.043409608 |
| AL021398 | -0.043413982 |
| MIR937 | -0.04341474 |
| FGF22 | -0.043421371 |
| RP11-717F1 | -0.043421775 |
| SP7 | -0.043422279 |
| CTD-2536I1 | -0.04342568 |
| RP11-429B14 | -0.04342782 |
| RNU6-518P | -0.04343428 |
| RP1-223H12 | -0.043434378 |
| RP11-863N1 | -0.043437182 |
| MARS | -0.043437758 |
| CTC-287O8 | -0.043440837 |
| C1orf131 | -0.043440903 |
| ZNF789 | -0.043442782 |
| C2orf50 | -0.043443526 |
| RP11-499F3 | -0.04344577 |
| IGFLR1 | -0.043448029 |
| EXOSC3P1 | -0.043449382 |
| OSTCP8 | -0.043450973 |
| RHD | -0.04345098 |
| AF230666 | -0.043457089 |
| MIR4646 | -0.043462259 |
| RP11-70L8 | -0.043464204 |
| SULT1C4 | -0.043464366 |
| SNORA71A | -0.043464381 |
| RNU6-77P | -0.043466299 |
| MGC50722 | -0.043467711 |
| RN7SKP5 | -0.043471235 |
| GLT6D1 | -0.04347806 |
| RP11-379H18 | -0.043478281 |

| | |
|---|---|
| AC012181 | -0.043484518 |
| ZNF548 | -0.043486226 |
| GAB2 | -0.043491807 |
| RP11-129J12 | -0.043493726 |
| AC005884 | -0.043494343 |
| ZER1 | -0.043496887 |
| CTD-2185K10 | -0.043497868 |
| GPR133 | -0.043501966 |
| CTD-2168K21 | -0.043513319 |
| RPL12P29 | -0.043517536 |
| NHP2P2 | -0.043518743 |
| ZNF287 | -0.043520032 |
| C6orf229 | -0.043520064 |
| GRAMD1A | -0.043520478 |
| OR52W1 | -0.043522656 |
| DENND3 | -0.043523939 |
| RP11-529E10 | -0.043524274 |
| RNU4-47P | -0.043526043 |
| RP11-660L16 | -0.043533037 |
| ING2 | -0.043534786 |
| CCDC168 | -0.043536692 |
| OGFOD1 | -0.043537614 |
| ASAH1 | -0.043538563 |
| SCFD1 | -0.043541354 |
| CSH1 | -0.043542543 |
| C2CD2L | -0.04354407 |
| AJ239318 | -0.043552753 |
| RP11-640A1 | -0.043554035 |
| TMEM198B | -0.043554787 |
| MIR1285-1 | -0.043557708 |
| C5orf66 | -0.043559364 |
| RP11-266O15 | -0.043559739 |
| ZNF577 | -0.043578751 |
| CTC-563A5 | -0.043580242 |
| STAT5B | -0.043585467 |
| AC092755 | -0.043591892 |
| RHBDF2 | -0.043592184 |
| RP11-498M15 | -0.043595023 |
| PTER | -0.043595626 |
| SLC38A9 | -0.04359797 |
| RNU6-742P | -0.043598978 |
| VTRNA1-3 | -0.043601357 |
| C10orf113 | -0.043601436 |
| ZWILCH | -0.043606808 |
| IQCF5 | -0.043610309 |

| | |
|---|---|
| MIR4296 | -0.043611413 |
| PRAMEF7 | -0.043612302 |
| MIR640 | -0.043623909 |
| RNU1-92P | -0.043625718 |
| CENPO | -0.043629804 |
| MC2R | -0.043635316 |
| OR2J3 | -0.043636064 |
| ATP5J2P6 | -0.043641052 |
| C6orf3 | -0.04364413 |
| CTA-414D7 | -0.043647342 |
| DTD2 | -0.043647628 |
| RP11-84H6 | -0.043649919 |
| GSKIP | -0.043652949 |
| SRSF9 | -0.043653917 |
| SNORA70B | -0.043656501 |
| SH2D6 | -0.043673399 |
| RP13-24G8 | -0.043677439 |
| KRT25 | -0.043678886 |
| KCTD2 | -0.043680243 |
| AC026188 | -0.043680858 |
| RNA5SP56 | -0.043683888 |
| CCDC93 | -0.043684132 |
| RP11-395I14 | -0.043689479 |
| PAOX | -0.043690421 |
| TEX101 | -0.043694569 |
| FLVCR1-AS1 | -0.043695184 |
| KRT35 | -0.043695401 |
| MIR505 | -0.043696894 |
| RNU6-400P | -0.043698728 |
| RP11-89F3 | -0.043705047 |
| MIR4762 | -0.04371676 |
| LSM14B | -0.043720264 |
| CTA-254O6 | -0.043721444 |
| ASB15 | -0.043722263 |
| RP11-348B17 | -0.043728446 |
| PRKRIRP8 | -0.043729393 |
| AC004257 | -0.043730719 |
| ETNPPL | -0.043739673 |
| PGBD2 | -0.043741511 |
| Z97634 | -0.043742981 |
| KRT8P15 | -0.04374833 |
| DKFZP434K028 | -0.043754511 |
| AC137488 | -0.043755299 |
| TROAP | -0.043760024 |
| LINC01225 | -0.043760332 |

| | |
|---|---|
| OR4D6 | -0.043763413 |
| PRINS | -0.043765189 |
| TPP1 | -0.043775634 |
| TMEM252 | -0.043776026 |
| TRIM60P16 | -0.043777565 |
| HNRNPA1P76 | -0.043778047 |
| LRRC73 | -0.043785011 |
| UBA5 | -0.043792695 |
| RBM44 | -0.0437945 |
| AC016764 | -0.043796631 |
| LRP2BP | -0.043798165 |
| MIR1537 | -0.043798583 |
| LPA | -0.043804536 |
| TBC1D2B | -0.043807933 |
| OR52K2 | -0.043813738 |
| AC004006 | -0.04381882 |
| RP4-631H13 | -0.043827551 |
| PRR13P3 | -0.043830538 |
| POLDIP2 | -0.043831394 |
| DCAKD | -0.043831711 |
| CIDEB | -0.043836269 |
| RP11-428C6 | -0.043842723 |
| FAM13C | -0.043849182 |
| DNAJB1P1 | -0.043849217 |
| GRTP1-AS1 | -0.043855046 |
| RPS2P45 | -0.043857568 |
| KMT2C | -0.043860214 |
| AC006926 | -0.043863633 |
| AC068056 | -0.043869887 |
| TLK2P1 | -0.043876196 |
| DNAJC19P5 | -0.04389338 |
| MED12 | -0.043895937 |
| CYP2D8P | -0.043901789 |
| RNA5SP470 | -0.043903224 |
| RNU6-431P | -0.043903581 |
| RP11-386I8 | -0.043906204 |
| AC114808 | -0.043909043 |
| ZNF735 | -0.043910658 |
| ZNF526 | -0.043910747 |
| VAMP1 | -0.043916477 |
| RP11-480G3 | -0.043917957 |
| SNORA70G | -0.043918004 |
| RP11-485M7 | -0.0439191 |
| ATXN1L | -0.043921742 |
| FAM8A2P | -0.043924734 |

| | |
|---|---|
| RNU6-211P | -0.04393168 |
| MT-TV | -0.043936867 |
| HIST1H2APS2 | -0.043938125 |
| C1orf64 | -0.04393976 |
| RNU6-612P | -0.043947892 |
| SULT1C2P1 | -0.043947914 |
| SPEF2 | -0.043951046 |
| AL450992 | -0.043952609 |
| MRPS31P1 | -0.043952999 |
| DLGAP2-AS1 | -0.043958738 |
| PSMC1P3 | -0.043960398 |
| ZNF418 | -0.043960479 |
| RNA5SP211 | -0.043962029 |
| MLYCD | -0.043964479 |
| LINC01521 | -0.0439714 |
| RP4-591N18 | -0.043975039 |
| PARP1 | -0.043976349 |
| AC008674 | -0.043977032 |
| EPHA1-AS1 | -0.043981714 |
| RP11-1079H9 | -0.043982149 |
| ZNHIT1P1 | -0.043986476 |
| HMGN1 | -0.043987048 |
| RP5-921G16 | -0.04398791 |
| CTB-180A7 | -0.043995114 |
| AC099506 | -0.043996694 |
| RNU6-743P | -0.044004339 |
| CTD-2183H9 | -0.044005318 |
| KRTAP26-1 | -0.044006061 |
| MIR1289-1 | -0.044006893 |
| PPIAP31 | -0.044008857 |
| WASL | -0.044013337 |
| PABPC3 | -0.044015347 |
| ZC2HC1B | -0.044015784 |
| MIR875 | -0.044016955 |
| GPM6A | -0.044019467 |
| MAB21L2 | -0.044027676 |
| RP11-145M4 | -0.044033961 |
| SDHB | -0.044034175 |
| ZNF444P1 | -0.044036091 |
| MIR587 | -0.044036195 |
| RP11-293D9 | -0.044037858 |
| RP11-633G11 | -0.044038879 |
| RP11-99C10 | -0.044041978 |
| PKD1L3 | -0.044049536 |
| HNRNPA1P52 | -0.044049649 |

| | |
|---|---|
| RNU6-867P | -0.044060778 |
| RNU7-38P | -0.044061447 |
| CUL1 | -0.044062623 |
| TM4SF5 | -0.044067473 |
| MTX3 | -0.044080391 |
| GTF2IP1 | -0.044083647 |
| RP11-206B11 | -0.044089408 |
| RN7SKP190 | -0.044093413 |
| RPL35AP12 | -0.04409356 |
| APEX1 | -0.044095536 |
| LL0XNC01-131B10 | -0.044096131 |
| SNURFL | -0.04409815 |
| ADAMTS7P1 | -0.044098842 |
| PARP16 | -0.044109546 |
| YY1P2 | -0.044114466 |
| HSPD1P6 | -0.044125585 |
| RANP4 | -0.044128775 |
| TTBK2 | -0.044131724 |
| SERTAD4 | -0.044133119 |
| HERC4 | -0.044138403 |
| RPS2P25 | -0.044140145 |
| ARHGEF39 | -0.044140301 |
| MEN1 | -0.044140878 |
| DDX27 | -0.044143427 |
| ST13P2 | -0.044144085 |
| ACADM | -0.044148748 |
| RP11-25C19 | -0.044150106 |
| MIR3687-2 | -0.044151561 |
| DUT | -0.044155446 |
| RP11-72B4 | -0.04415651 |
| C10orf126 | -0.044162663 |
| EEF1A1P15 | -0.044167166 |
| LYRM9 | -0.044174777 |
| AC098820 | -0.044175372 |
| AC011193 | -0.044175581 |
| CCNYL3 | -0.044181086 |
| RP11-552D8 | -0.04418603 |
| RP11-255B23 | -0.044187198 |
| XAGE5 | -0.044188437 |
| HIST2H4A | -0.04419272 |
| EIF5AL1 | -0.044197755 |
| RNU6-64P | -0.044197907 |
| OR6X1 | -0.04420177 |
| CTD-2313P7 | -0.044210646 |
| CELF6 | -0.04421488 |

| | |
|---|---|
| AC114763 | -0.044214932 |
| RP11-566E18 | -0.044215225 |
| LRRIQ3 | -0.044220193 |
| RP11-803P9 | -0.044220683 |
| BTBD9 | -0.044220956 |
| DCAF8L1 | -0.044221498 |
| ERVW-1 | -0.044222098 |
| ARHGAP42P4 | -0.044226465 |
| PRTN3 | -0.044229662 |
| RP11-366L5 | -0.044230391 |
| AC078881 | -0.044235081 |
| RP11-78B10 | -0.044236494 |
| ZIC3 | -0.044236499 |
| ZFYVE27 | -0.04423757 |
| AC005033 | -0.044239544 |
| RP4-798A10 | -0.044239728 |
| OR1M4P | -0.044242153 |
| SNORA11 | -0.044242315 |
| ZNF654 | -0.044242612 |
| RNU6-155P | -0.044251887 |
| AC145343 | -0.044254017 |
| LINC00706 | -0.044256213 |
| MIR4645 | -0.044256413 |
| LINC01391 | -0.044257741 |
| ELL2 | -0.044262111 |
| NUS1P4 | -0.044263533 |
| CRYGEP | -0.044277313 |
| DERL1 | -0.044281795 |
| AC105393 | -0.044283685 |
| TXNP2 | -0.044283925 |
| AGER | -0.044286224 |
| MIR3937 | -0.044286264 |
| OR8U1 | -0.044286346 |
| RP11-453N18 | -0.044287417 |
| GAGE12C | -0.044288041 |
| MTND1P24 | -0.044291721 |
| SUCO | -0.044292103 |
| CDKN3 | -0.044295563 |
| SLMO1 | -0.04429614 |
| RP11-204C16 | -0.044296343 |
| RP11-798M19 | -0.044298801 |
| RNU6-30P | -0.044301904 |
| RNU5B-2P | -0.044316718 |
| RP11-297P16 | -0.04432224 |
| RNU2-13P | -0.044323745 |

| | |
|---|---|
| RP11-169K17 | -0.044325453 |
| RP11-164N3 | -0.044326614 |
| SNORA5A | -0.044333825 |
| CSF3R | -0.044336179 |
| CYCSP30 | -0.044345306 |
| RP3-466P17 | -0.044354309 |
| AC104457 | -0.044355851 |
| LINC01413 | -0.044357479 |
| RP11-231E19 | -0.044365571 |
| HMGB3P10 | -0.044365599 |
| TTN-AS1 | -0.044366767 |
| ERP29P1 | -0.044368531 |
| AC024132 | -0.044370416 |
| RP4-536B24 | -0.044373252 |
| TULP3P1 | -0.044375108 |
| RN7SL262P | -0.044377173 |
| ABHD14A-ACY1 | -0.044377626 |
| RP13-1056D16 | -0.044379815 |
| RP11-1324A7 | -0.044382292 |
| AZU1 | -0.044382872 |
| RP1-161P9 | -0.044395368 |
| RN7SL387P | -0.044396699 |
| HDAC11 | -0.044401689 |
| RP11-240G22 | -0.044402189 |
| CEP63 | -0.044403196 |
| ZNF836 | -0.044412313 |
| FOXN3-AS2 | -0.044412364 |
| SNX29P2 | -0.044418701 |
| PRPF38A | -0.044418769 |
| CTC-229L21 | -0.044421042 |
| RNU6-329P | -0.044422636 |
| RP11-474P12 | -0.044424053 |
| RP11-31L22 | -0.044446473 |
| GRIA1 | -0.044447205 |
| LINC00529 | -0.044447452 |
| RP11-85I21 | -0.044449106 |
| TDPX2 | -0.044452945 |
| RP11-242C24 | -0.044454728 |
| HNRNPA1P17 | -0.04445621 |
| NDUFA5P11 | -0.044459073 |
| FBXO42 | -0.044462548 |
| OR7A17 | -0.044464555 |
| LINC00276 | -0.044467848 |
| IFI35 | -0.044468873 |
| CTC-558O19 | -0.044476208 |

| | |
|---|---|
| OR2W2P | -0.044477278 |
| AGA | -0.044480241 |
| RNU5F-3P | -0.044484351 |
| AUH | -0.044486177 |
| E2F3P2 | -0.044499012 |
| CEP55 | -0.044503089 |
| RP11-102N12 | -0.044503908 |
| LRRFIP2 | -0.04450467 |
| RNU6-1215P | -0.044511145 |
| TOB2 | -0.044516038 |
| ACTR3BP5 | -0.04451859 |
| IPPK | -0.044518624 |
| COG2 | -0.044520187 |
| RP11-359K18 | -0.04452709 |
| MIR548AM | -0.044531007 |
| MIR4435-1 | -0.044534682 |
| CTC-224D3 | -0.044534692 |
| TAB3 | -0.044540039 |
| MAPK14 | -0.044540914 |
| CORO2A | -0.044544765 |
| ZFP91-CNTF | -0.044545845 |
| PIGN | -0.044553659 |
| CCNF | -0.044558254 |
| ATG4D | -0.044559259 |
| HNRNPU | -0.044559538 |
| RP5-1174J21 | -0.044560088 |
| AC110086 | -0.044561003 |
| USP9YP9 | -0.044561003 |
| CTD-2254N19 | -0.044561564 |
| RPL29P33 | -0.044564081 |
| PLCG1-AS1 | -0.044568801 |
| TOMM20P3 | -0.044570438 |
| TMBIM6 | -0.044571873 |
| MIR4714 | -0.044585149 |
| KRT26 | -0.044587693 |
| RSPO3 | -0.044588302 |
| AC090453 | -0.044588862 |
| RP1-101D8 | -0.044589862 |
| RP11-621H8 | -0.044590516 |
| AP001136 | -0.044599741 |
| MIR6749 | -0.044605573 |
| KIAA1257 | -0.044605791 |
| RNU6-360P | -0.044614289 |
| MIA2 | -0.044614594 |
| RNU6-921P | -0.044614917 |

| | |
|---|---|
| EEF1A1P34 | -0.044619953 |
| AC093700 | -0.044623567 |
| RP11-483P21 | -0.044625438 |
| BTF3P13 | -0.044627869 |
| RP11-1101H11 | -0.044633652 |
| RP11-773H22 | -0.044635722 |
| IDI2-AS1 | -0.044638102 |
| MIR107 | -0.044640643 |
| RP11-302K17 | -0.044640899 |
| OR6K4P | -0.044643274 |
| AC106801 | -0.044644962 |
| PDX1 | -0.044645643 |
| ZNF816-ZNF321P | -0.044650146 |
| RNU6-110P | -0.044650345 |
| GTF2IP3 | -0.044652146 |
| DCUN1D4 | -0.04465264 |
| RP11-354M20 | -0.044656419 |
| ACTN4P1 | -0.044660199 |
| NOS1 | -0.044660307 |
| RP11-419L20 | -0.044660402 |
| ATF1 | -0.044661384 |
| UBXN10 | -0.044664547 |
| AL607148 | -0.044665452 |
| RP11-334E15 | -0.044667772 |
| ASNSD1 | -0.044669347 |
| RP11-12K6 | -0.044671495 |
| AP001468 | -0.044671683 |
| RP11-188C12 | -0.044674222 |
| AC114798 | -0.044676797 |
| RP11-537I16 | -0.04468227 |
| AC006128 | -0.044686818 |
| KIAA1598 | -0.044686911 |
| AP001178 | -0.044689768 |
| RP11-1110F20 | -0.044699023 |
| LINC01122 | -0.044702744 |
| CTD-2245F17 | -0.044702754 |
| NME6 | -0.044705163 |
| LA16c-13E4 | -0.044705213 |
| ST7-AS2 | -0.044707195 |
| STPG1 | -0.044707396 |
| RP11-460N20 | -0.044708884 |
| SCEL-AS1 | -0.044709472 |
| AP3D1 | -0.044709554 |
| LL0XNC01-221F2 | -0.044710503 |
| SEC24A | -0.044710838 |

| | |
|---|---|
| RP11-138E16 | -0.044712091 |
| CCBL1 | -0.04472535 |
| RP11-131K17 | -0.044726574 |
| AKAP1 | -0.044727229 |
| IFNL4P1 | -0.044730092 |
| MIR888 | -0.044733848 |
| NAAA | -0.044736197 |
| RPSAP32 | -0.044738037 |
| C17orf100 | -0.044739378 |
| MIR603 | -0.044741161 |
| ADD1 | -0.044742392 |
| SNORD115-21 | -0.044745062 |
| RN7SL93P | -0.044746562 |
| RP11-252E2 | -0.044747223 |
| RP11-19J3 | -0.044748115 |
| ARHGAP33 | -0.044750318 |
| SLC29A4P1 | -0.044755504 |
| RP11-68L1 | -0.044757629 |
| RNU6-607P | -0.044769039 |
| POM121C | -0.044790004 |
| RGPD2 | -0.044795066 |
| C9orf47 | -0.044795818 |
| RP11-345I18 | -0.044806979 |
| LYRM1 | -0.044814675 |
| EPHA4 | -0.044814842 |
| RN7SKP64 | -0.044817813 |
| RNF185 | -0.044818321 |
| C16orf87 | -0.044824777 |
| RN7SL614P | -0.044825283 |
| OR51A1P | -0.044825766 |
| TSPY16P | -0.044836892 |
| OR2H5P | -0.0448382 |
| SOCS5P4 | -0.04483834 |
| ARHGEF3-AS1 | -0.044839344 |
| ZNF252P-AS1 | -0.044839773 |
| RNU6-630P | -0.044849259 |
| KDM7A | -0.044849355 |
| AC011718 | -0.044852546 |
| PSMB7P1 | -0.044852622 |
| SMAD2 | -0.044853486 |
| GS1-410F4 | -0.044853786 |
| C10orf54 | -0.044857225 |
| GPR152 | -0.044858677 |
| OR52B1P | -0.044859536 |
| AL021918 | -0.044864733 |

| | |
|---|---|
| RNU6-1074P | -0.044866527 |
| IL31 | -0.044866748 |
| WDR95P | -0.044868826 |
| CTD-2653M23 | -0.044871791 |
| LINC01538 | -0.044874283 |
| NRDE2 | -0.044875477 |
| RP11-685B13 | -0.04488225 |
| RP11-29B11 | -0.044882641 |
| RP11-141P6 | -0.044885874 |
| ADAM3A | -0.044888308 |
| RP11-741G21 | -0.044889963 |
| ACTL7A | -0.044890132 |
| RP11-68D16 | -0.044894314 |
| CTC-479C5 | -0.044899032 |
| MINOS1P2 | -0.044907718 |
| LINC01076 | -0.044913498 |
| POTED | -0.044913527 |
| CCDC132 | -0.044923052 |
| RNU6-1204P | -0.044925591 |
| TKTL1 | -0.044934875 |
| AC021660 | -0.044935269 |
| RPL7P14 | -0.044941262 |
| CTD-2140B24 | -0.044945446 |
| RP11-142O6 | -0.044946312 |
| AC007362 | -0.044947145 |
| ZNF568 | -0.04495429 |
| PLEKHB1 | -0.044955034 |
| IGHVIV-44-1 | -0.044956543 |
| CLEC1A | -0.044960114 |
| AK9 | -0.044962644 |
| RP11-432M8 | -0.044965918 |
| AANAT | -0.044966839 |
| GPR135 | -0.044974179 |
| RP11-693I21 | -0.044977963 |
| RP11-333A23 | -0.044981846 |
| RNU6-520P | -0.044982035 |
| RP11-194N12 | -0.044983788 |
| RP11-456K23 | -0.044994069 |
| RP11-255I10 | -0.044994979 |
| RP11-576C12 | -0.044995619 |
| MIR1226 | -0.045000151 |
| SNORD116-2 | -0.045003718 |
| AC064865 | -0.045004928 |
| MINPP1 | -0.045007092 |
| AC093844 | -0.045008233 |

| | |
|---|---|
| LIFR-AS1 | -0.045008823 |
| RNU6-544P | -0.045010654 |
| RP4-561L24 | -0.045016767 |
| CD200 | -0.045018088 |
| AP000925 | -0.045018093 |
| TMEM176A | -0.045021828 |
| FRZB | -0.045022684 |
| RP11-674C21 | -0.045026215 |
| Z83844 | -0.045046489 |
| MOV10 | -0.045046815 |
| RP11-454E5 | -0.045056732 |
| CTD-3244O18 | -0.045059216 |
| AC018693 | -0.045065661 |
| MCM4 | -0.045066295 |
| RNU6-437P | -0.045066589 |
| MIPEP | -0.045075467 |
| CDV3P1 | -0.045076262 |
| NMUR1 | -0.045077879 |
| PTCRA | -0.045079342 |
| RP11-756G20 | -0.045083018 |
| DSCC1 | -0.045088361 |
| VPS53 | -0.045092622 |
| TMEM154 | -0.045096696 |
| RP11-22M7 | -0.045098971 |
| TMEM131 | -0.045105404 |
| RP1-138A5 | -0.045108654 |
| AC004449 | -0.045112026 |
| RP11-379K22 | -0.045114457 |
| GTF2E1 | -0.045114986 |
| RAB27B | -0.045115488 |
| RP13-487P22 | -0.045115585 |
| ZNF737 | -0.045126372 |
| MOB1AP1 | -0.045129002 |
| ZNF263 | -0.045129293 |
| RNU5F-4P | -0.045134446 |
| SNRPCP3 | -0.045134921 |
| RP11-1114A5 | -0.045136233 |
| KCNG4 | -0.045137498 |
| AL357932 | -0.045144338 |
| NIP7P2 | -0.045144633 |
| LGALS2 | -0.045145312 |
| FP565210 | -0.045145599 |
| AF121897 | -0.045151607 |
| LINC00924 | -0.045154594 |
| XRCC6P2 | -0.045157946 |

| | |
|---|---|
| TAOK3 | -0.045161481 |
| OR10J9P | -0.045161814 |
| AB015752 | -0.04516376 |
| LGALS3BP | -0.045167025 |
| NUP85 | -0.04516736 |
| RP1-72E17 | -0.045176244 |
| H2AFZP3 | -0.045181401 |
| RNU6-1157P | -0.045181618 |
| RN7SL535P | -0.045182881 |
| RP11-84D1 | -0.045183123 |
| RP11-153N17 | -0.045184149 |
| HNRNPDLP4 | -0.045187927 |
| AC068014 | -0.045188285 |
| RP11-527D15 | -0.045194709 |
| CACNA1D | -0.045195231 |
| RP11-854K16 | -0.045196264 |
| RP11-346C20 | -0.045197836 |
| AL589822 | -0.045199649 |
| AC018442 | -0.045200215 |
| RP11-64B16 | -0.045200631 |
| RP5-1050E16 | -0.04520137 |
| MIR3188 | -0.045205686 |
| RNU6-1274P | -0.045206153 |
| MIR4312 | -0.045217711 |
| DIABLO | -0.045221074 |
| RP11-526A4 | -0.045222949 |
| RP11-338N10 | -0.045223022 |
| MTND1P26 | -0.045225224 |
| HMGB1P47 | -0.045226493 |
| PKNOX1 | -0.045228442 |
| RP11-234G16 | -0.045229948 |
| RP11-685G9 | -0.045230086 |
| RGPD8 | -0.045232851 |
| LLNLF-187D8 | -0.045234296 |
| NPM1P41 | -0.045240685 |
| IPO7P1 | -0.045243727 |
| ABCA11P | -0.045247959 |
| H3F3AP5 | -0.045248964 |
| MIR4655 | -0.045252367 |
| EIF1AXP2 | -0.045252911 |
| RP11-610P16 | -0.045253491 |
| Six3os1_2 | -0.045255432 |
| DNMT1 | -0.045255792 |
| EIF3L | -0.045258601 |
| METTL21EP | -0.04525963 |

| | |
|---|---|
| RP11-1136G11 | -0.0452607 |
| RNU6-438P | -0.045267303 |
| RNA5SP495 | -0.045268219 |
| LRP5L | -0.045268276 |
| OR5L1 | -0.045271245 |
| GABRB1 | -0.045273203 |
| NIFKP2 | -0.045277176 |
| RNU6-1156P | -0.045281351 |
| KLHL18 | -0.045283793 |
| MIR6753 | -0.045290469 |
| CCNB2 | -0.045292333 |
| MIR181D | -0.045293678 |
| RP11-438C19 | -0.0452966 |
| RP6-127F18 | -0.0452983 |
| NDUFAF4P3 | -0.045303298 |
| AC009310 | -0.045303631 |
| RP5-865N13 | -0.045304453 |
| MIR4502 | -0.045312656 |
| RNU6-817P | -0.045316643 |
| ANKRD11 | -0.045318994 |
| SAG | -0.04532662 |
| SSSCA1-AS1 | -0.045330381 |
| AC012513 | -0.045335608 |
| YES1P1 | -0.045336017 |
| RNU6-998P | -0.045337831 |
| RBM22P1 | -0.045341311 |
| AC027119 | -0.045342553 |
| TAS2R19 | -0.045345453 |
| RP11-220I1 | -0.045352221 |
| AC013738 | -0.045352798 |
| ENPP7P7 | -0.045356319 |
| RBP7 | -0.045365508 |
| GNG5P5 | -0.045371973 |
| FAM157C | -0.045373329 |
| RPL21P107 | -0.045375682 |
| APOOP3 | -0.045377867 |
| RP11-110A12 | -0.045385474 |
| HSPD1P11 | -0.045390595 |
| AL121588 | -0.045395453 |
| RP11-40B20 | -0.045397273 |
| RP11-429P3 | -0.045401393 |
| RP11-689P11 | -0.045404189 |
| ZNF827 | -0.045407054 |
| MFN2 | -0.045412747 |
| CAPN14 | -0.045415477 |

| | |
|---|---|
| RP11-463O9 | -0.045417045 |
| TRBV7-5 | -0.045419343 |
| ALDH3A2 | -0.045420748 |
| SNTB2 | -0.045423371 |
| RP5-1027O15 | -0.045424221 |
| RP11-989E6 | -0.045431298 |
| RPS23P7 | -0.045432638 |
| RNU1-76P | -0.04544191 |
| AC138035 | -0.045442503 |
| LINC01191 | -0.045442857 |
| LINC00992 | -0.045443005 |
| VTI1BP3 | -0.045443836 |
| TMOD3 | -0.045445177 |
| NHLRC1 | -0.045445623 |
| RP11-248N22 | -0.045448261 |
| RP11-350D23 | -0.045460463 |
| AF064860 | -0.045460732 |
| DOCK9-AS1 | -0.045461829 |
| ZNRF2P3 | -0.045462123 |
| GINS2 | -0.045466834 |
| AC004987 | -0.045467243 |
| AC008810 | -0.04546786 |
| DDHD2 | -0.04547115 |
| ECHDC1 | -0.045471224 |
| ALDH3B2 | -0.045476025 |
| RP11-326I11 | -0.045477401 |
| OR5H3P | -0.045480133 |
| RNU6-621P | -0.045485464 |
| RP11-104J23 | -0.045486522 |
| AKAP11 | -0.045492249 |
| CTD-3098H1 | -0.045492422 |
| KRT18P28 | -0.045494735 |
| RNA5SP494 | -0.045502153 |
| RP11-24H2 | -0.045502761 |
| CCNI | -0.045504565 |
| SYT2 | -0.045508008 |
| PPP2R2C | -0.045511301 |
| AL121893 | -0.045513775 |
| RNASE8 | -0.045514787 |
| AL136529 | -0.045516369 |
| ZNF366 | -0.045518855 |
| DPP9 | -0.045519133 |
| STRIP2 | -0.045519636 |
| RP11-435I3 | -0.045525129 |
| RP11-218M22 | -0.045525757 |

| | |
|---|---|
| RNU6-218P | -0.045527199 |
| STAG1 | -0.045530799 |
| ARPC3P1 | -0.045534521 |
| MZF1 | -0.045538874 |
| MARVELD2 | -0.045540493 |
| URB2 | -0.045541702 |
| MED6 | -0.045546986 |
| RP11-490E15 | -0.045549899 |
| HSPA4L | -0.045558652 |
| AC007283 | -0.045560563 |
| HNRNPH1P1 | -0.045561815 |
| CTDSP2 | -0.045561889 |
| CTGLF8P | -0.045567713 |
| RP11-23B7 | -0.045570457 |
| SIAH2 | -0.045575322 |
| CTPS2 | -0.045576569 |
| TERF1P5 | -0.045579085 |
| USP40 | -0.045581611 |
| XKR6 | -0.045581612 |
| AC106782 | -0.045582645 |
| RP11-117N2 | -0.045583535 |
| RP11-383C6 | -0.045584812 |
| RP1-182O16 | -0.045586644 |
| CTD-2124B8 | -0.04558669 |
| TMEM232 | -0.045589434 |
| AC005550 | -0.04559295 |
| RP11-60E8 | -0.045602459 |
| RPL30P16 | -0.045602963 |
| RP11-111E14 | -0.045603437 |
| C10orf62 | -0.0456068 |
| PSMB10 | -0.045607249 |
| ZNF678 | -0.045610635 |
| ACBD7 | -0.045612877 |
| EIF2B1 | -0.045613705 |
| CAB39 | -0.045617705 |
| CASD1 | -0.045617724 |
| NOTO | -0.045623585 |
| CCNG1 | -0.045637836 |
| SLCO1C1 | -0.045638839 |
| NOLC1 | -0.045641799 |
| BLVRA | -0.045647381 |
| VWC2L | -0.045653699 |
| RP11-317G6 | -0.045655614 |
| NRBF2P3 | -0.045656066 |
| CCDC77 | -0.045657279 |

| | |
|---|---|
| CNTLN | -0.045658507 |
| SLC26A5 | -0.045661636 |
| OR52J3 | -0.045663299 |
| ZNF285B | -0.045668249 |
| AC087380 | -0.04567378 |
| CCDC88B | -0.045675103 |
| RP11-56N19 | -0.045676418 |
| CNTN5 | -0.045684716 |
| EIF4EP1 | -0.04568939 |
| NPAP1P7 | -0.045693198 |
| RP4-734P14 | -0.045695859 |
| SLC25A51 | -0.045697151 |
| DDX51 | -0.045698267 |
| RP11-361A23 | -0.045698313 |
| ADIRF | -0.045699993 |
| RP11-554L12 | -0.04571086 |
| MIR561 | -0.045711385 |
| CTA-125H2 | -0.045711452 |
| KIF9 | -0.045713542 |
| KRT37 | -0.045720289 |
| KIAA1467 | -0.045721449 |
| JMY | -0.04572411 |
| RP11-1113L8 | -0.04572478 |
| RP11-936I5 | -0.045730635 |
| RSAD2 | -0.045736278 |
| RP11-181C21 | -0.04573705 |
| VTI1BP1 | -0.045738395 |
| OR2G2 | -0.045738862 |
| CTGLF12P | -0.045743981 |
| CTD-2342J14 | -0.045751609 |
| RP11-96A15 | -0.045751613 |
| RP5-888M10 | -0.045756158 |
| RP11-450H6 | -0.045759422 |
| ZSWIM8 | -0.045770047 |
| ZBTB8OSP1 | -0.045771533 |
| METTL6 | -0.04577347 |
| RP11-109I13 | -0.045778151 |
| RPL5P10 | -0.045781991 |
| RP11-585K6 | -0.04578319 |
| METTL8P1 | -0.045783398 |
| RBMY2WP | -0.045789331 |
| AL031653 | -0.045789331 |
| RP11-434D11 | -0.045791905 |
| AC009961 | -0.045800802 |
| RP11-429O1 | -0.045801629 |

| | |
|---|---|
| RAPGEFL1 | -0.045802067 |
| FAM151B | -0.045802383 |
| MIR578 | -0.045803154 |
| ATP11A-AS1 | -0.045803967 |
| ZFP2 | -0.045805007 |
| LINC01470 | -0.045806716 |
| RP11-462B18 | -0.045808665 |
| AC004655 | -0.045811161 |
| TTC28-AS1 | -0.045813677 |
| ZNF534 | -0.045815138 |
| YTHDF3 | -0.045815872 |
| ANXA9 | -0.045817863 |
| PTPRJ-AS1 | -0.04582277 |
| LMBRD2 | -0.045825435 |
| DSCR9 | -0.045826962 |
| KIF23 | -0.045827221 |
| CTC-239J10 | -0.045828908 |
| TSTD2 | -0.045829342 |
| TAF15 | -0.04583044 |
| CXXC1 | -0.045832133 |
| TNR | -0.04583624 |
| RPL34P23 | -0.045841264 |
| COQ4 | -0.045842124 |
| KRT8P1 | -0.045847146 |
| RP11-539G18 | -0.045847632 |
| IFNAR2 | -0.045847706 |
| RP11-498P14 | -0.045848785 |
| RPS24P12 | -0.045858698 |
| RP11-211A18 | -0.045862664 |
| ACP2 | -0.045863018 |
| CDKN2D | -0.045864916 |
| RNA5SP152 | -0.045866605 |
| HMGN3P1 | -0.045869636 |
| MIR4503 | -0.04587364 |
| RPL34P22 | -0.045874062 |
| RNU6-450P | -0.045877368 |
| CCDC163P | -0.045878145 |
| RP11-19H6 | -0.045878412 |
| FMO5 | -0.045884843 |
| PGM5P4 | -0.04588542 |
| AL139821 | -0.04588558 |
| RBMXP2 | -0.045886538 |
| MIR4441 | -0.045895972 |
| RP11-135M8__A | -0.045898916 |
| AL845472 | -0.045899095 |

| | |
|---|---|
| CKS1BP3 | -0.045901259 |
| AC083884 | -0.045903728 |
| MAOA | -0.045906871 |
| DLGAP1 | -0.04590796 |
| RNU6-1329P | -0.045918456 |
| RP11-22P6 | -0.045920124 |
| RP5-1012F16 | -0.045924444 |
| RP11-24H1 | -0.045929455 |
| RP11-56H7 | -0.045941841 |
| RN7SL396P | -0.0459433 |
| KLRF2 | -0.045943409 |
| CTC-499J9 | -0.045944422 |
| RNA5SP173 | -0.04594749 |
| FAM65C | -0.045950148 |
| AL592063 | -0.04596577 |
| CDK5RAP1 | -0.045968381 |
| CTD-2113L7 | -0.045968565 |
| RP11-905K4 | -0.0459705 |
| RAD18 | -0.045975767 |
| RNU6-25P | -0.045986207 |
| DEFB105A | -0.045987665 |
| LIG1 | -0.045992233 |
| TUBB8P1 | -0.045993231 |
| PPP1R11P1 | -0.045995549 |
| RNU6-499P | -0.045998807 |
| AZGP1P2 | -0.045999432 |
| CST5 | -0.046006578 |
| RNU6-1272P | -0.046007673 |
| PTCHD3 | -0.046008482 |
| RNU2-14P | -0.046011962 |
| RP11-323A7 | -0.04601302 |
| RP3-452M16 | -0.046020101 |
| RP11-368I7 | -0.046020997 |
| STAT6 | -0.046024849 |
| THOC2 | -0.046028179 |
| AL117209 | -0.046029249 |
| TRGJ2 | -0.046034216 |
| ZBTB12 | -0.04604297 |
| RP11-517C16 | -0.046043436 |
| RCOR1 | -0.046045668 |
| YWHAB | -0.046045743 |
| RP11-36B15 | -0.046047276 |
| IK | -0.046047585 |
| TMEM251 | -0.04604952 |
| TRMT5 | -0.046053185 |

| | |
|---|---|
| ZNF485 | -0.046054524 |
| OR4K1 | -0.046055395 |
| AC016629 | -0.046058877 |
| ALPP | -0.046059157 |
| SPATA3-AS1 | -0.046061899 |
| MIR6857 | -0.046062293 |
| YWHAZP2 | -0.046062923 |
| AC092421 | -0.046067559 |
| RP11-217C7 | -0.046068566 |
| TTC3 | -0.046073167 |
| RP11-137J7 | -0.04607761 |
| SPACA7 | -0.04607802 |
| AC011748 | -0.046079336 |
| RP11-463O12 | -0.04608033 |
| HEXIM2 | -0.046086954 |
| SLC46A3 | -0.046087605 |
| GLIPR1L2 | -0.046096591 |
| AC002070 | -0.046098585 |
| RNY4P17 | -0.04610136 |
| EPM2A | -0.046105131 |
| RP11-270B14 | -0.046105924 |
| RP11-900F13 | -0.046107981 |
| RP11-177N22 | -0.046110544 |
| HEATR5A | -0.046110862 |
| TMEM210 | -0.046113524 |
| RP11-87C12 | -0.046113627 |
| HSF2 | -0.046115792 |
| AC026954 | -0.046118423 |
| MFF | -0.04611849 |
| MBTPS2 | -0.046119606 |
| PGAM1P6 | -0.046119722 |
| RN7SL330P | -0.046122376 |
| RP11-751L19 | -0.046122876 |
| SEC24B-AS1 | -0.046122937 |
| EIF4A2P4 | -0.046127702 |
| LCN6 | -0.04613431 |
| RP11-550C4 | -0.046135001 |
| ATP2A2 | -0.046136295 |
| RP11-219H23 | -0.046140906 |
| PRDM2 | -0.046141393 |
| ZNF114P1 | -0.046142289 |
| PRR13P4 | -0.046143205 |
| LA16c-23H5 | -0.046143911 |
| RP11-65I12 | -0.046145128 |
| RNU6-922P | -0.046149322 |

| | |
|---|---|
| LIMD2 | -0.046150677 |
| RP11-553P9 | -0.04615075 |
| RP11-8P11 | -0.046153271 |
| RP11-1023L17 | -0.046153524 |
| EEF1E1-BLOC1S5 | -0.046156873 |
| C4BPAP1 | -0.046169546 |
| LINC01040 | -0.046173533 |
| DCP1B | -0.046174976 |
| MAFG-AS1 | -0.046175981 |
| RNA5SP375 | -0.046179727 |
| RPL21P13 | -0.046181057 |
| RP1-234P15 | -0.046183572 |
| IGHVIII-13-1 | -0.04618585 |
| WDR82 | -0.046190828 |
| MTRR | -0.046191256 |
| RP11-22K10 | -0.046191537 |
| RP11-349J5 | -0.046194566 |
| AC003101 | -0.046194587 |
| DPYD | -0.046195107 |
| SNORD36 | -0.046198774 |
| HK2P1 | -0.046202452 |
| SMIM18 | -0.046206904 |
| OR52J2P | -0.046210275 |
| RP1-90G24 | -0.046213186 |
| C1QTNF9-AS1 | -0.046217069 |
| RORA | -0.046224839 |
| AC093817 | -0.046229216 |
| CTAGE1 | -0.046230225 |
| RP11-565F19 | -0.046230309 |
| SNX18P14 | -0.046234159 |
| RP11-415C15 | -0.046235103 |
| DFFA | -0.046242105 |
| ZMYM6NB | -0.046242198 |
| ADCY6 | -0.046243928 |
| PUS10 | -0.046244077 |
| RP11-63B13 | -0.04624699 |
| RP11-297N6 | -0.046251571 |
| VDAC1P6 | -0.046252921 |
| RP1-240B8 | -0.046255815 |
| RP5-894A10 | -0.046262217 |
| RP11-159G9 | -0.04626538 |
| ATG12 | -0.046270385 |
| RANP9 | -0.046278885 |
| PRR23C | -0.046281705 |
| FYTTD1 | -0.046285604 |

| | |
|---|---|
| RNU6-1333P | -0.046289269 |
| RNA5SP223 | -0.04629432 |
| SLC25A36P1 | -0.046296642 |
| AC131263 | -0.046304051 |
| PKI55 | -0.046309531 |
| EIF4BP3 | -0.046310238 |
| NEUROD6 | -0.046313275 |
| PMP2 | -0.046316825 |
| FCER1G | -0.046320359 |
| RELL2 | -0.046321978 |
| BRD7P1 | -0.046326385 |
| BIRC6-AS2 | -0.046331247 |
| AP3M2 | -0.046339411 |
| CMTM6 | -0.046341086 |
| AC131056 | -0.046353887 |
| MRPL35 | -0.046354779 |
| SRD5A3-AS1 | -0.046355021 |
| RNA5SP499 | -0.046355635 |
| XIAP | -0.046361161 |
| AC004541 | -0.046361473 |
| ANGPTL5 | -0.046363822 |
| MRPL53P1 | -0.046365923 |
| HZGJ | -0.04636881 |
| AC090954 | -0.046370565 |
| HCCS | -0.046370929 |
| DPRXP5 | -0.046377739 |
| LL22NC03-75A1 | -0.046378534 |
| PRKAB1 | -0.04638066 |
| MCM2 | -0.046382837 |
| OR10AH1P | -0.046389748 |
| ZNF410 | -0.046390297 |
| TBC1D3I | -0.046392542 |
| CTD-2545G14 | -0.046393815 |
| TRUB2 | -0.046411042 |
| C9orf24 | -0.046413862 |
| PARP4P1 | -0.046427275 |
| RP11-436M15 | -0.04644319 |
| RP11-454H13 | -0.046444497 |
| RN7SL200P | -0.046455104 |
| FAM102A | -0.046456123 |
| MIR548S | -0.046458484 |
| DDB1 | -0.046468831 |
| RPL23AP3 | -0.046476541 |
| CCL23 | -0.046480169 |
| DLEU1 | -0.046489092 |

| | |
|---|---|
| ERH | -0.046490116 |
| CMB9-94B1 | -0.046493153 |
| OR2J1 | -0.046494816 |
| IGLVV-58 | -0.046497763 |
| DKFZp434P228 | -0.04650116 |
| RP11-2E11 | -0.046505112 |
| RPL18P13 | -0.046506111 |
| RHOA-IT1 | -0.046510824 |
| CPED1 | -0.046534248 |
| SH3BP5 | -0.046534583 |
| FBXO31 | -0.046536146 |
| LINC01206 | -0.046539543 |
| RPS4XP21 | -0.046546706 |
| OR9I1 | -0.046547543 |
| AC016712 | -0.046554772 |
| BRF2 | -0.046555142 |
| RP11-202K23 | -0.046556799 |
| AC021224 | -0.046558588 |
| CEACAMP5 | -0.046560165 |
| RP11-101O21 | -0.046566611 |
| ARMCX5 | -0.046566826 |
| RP11-157L3 | -0.046568775 |
| CTD-3075F15 | -0.046570244 |
| RP1-28C20 | -0.046570615 |
| RP11-500M10 | -0.046577542 |
| RP11-73M14 | -0.046578254 |
| UHRF1 | -0.046579441 |
| OR5B2 | -0.046579803 |
| MTRF1LP1 | -0.046579857 |
| HNRNPA1P48 | -0.04658359 |
| MTND2P33 | -0.046593958 |
| ZSCAN12 | -0.046594083 |
| RP11-290L1 | -0.046600828 |
| YTHDF2 | -0.046603665 |
| RP11-340I6 | -0.046605277 |
| FBXO15 | -0.046613563 |
| STAM2 | -0.046613675 |
| UBE2V2P2 | -0.046615856 |
| RP11-336K24 | -0.046618828 |
| C5orf47 | -0.046626863 |
| OR8A3P | -0.046629507 |
| EMCN | -0.04663148 |
| MIR7849 | -0.046633388 |
| RP11-23N2 | -0.046634164 |
| RNU6-1315P | -0.046644378 |

| | |
|---|---|
| RP11-2L4 | -0.046645034 |
| CTD-3253I12 | -0.046646539 |
| RPS7P5 | -0.046648387 |
| FAM72A | -0.046652132 |
| MIR6859-3 | -0.046652903 |
| RP11-136F16 | -0.046665347 |
| AC097713 | -0.046667626 |
| RP11-365F18 | -0.046678282 |
| PRKRIRP1 | -0.046680363 |
| SLIT2-IT1 | -0.046680597 |
| AC027806 | -0.04668137 |
| RN7SL477P | -0.046690799 |
| AC078816 | -0.04669901 |
| CHGA | -0.046702182 |
| GTF2IRD2B | -0.046703427 |
| CHCHD2P8 | -0.046707116 |
| ISCA2 | -0.046713642 |
| IGLVIV-64 | -0.046718029 |
| PFDN4 | -0.046719521 |
| RNU6-697P | -0.046720766 |
| SUMO2P1 | -0.046722729 |
| SPACA5 | -0.04672503 |
| SPATA31C1 | -0.04672534 |
| FAM110C | -0.04672901 |
| EIF2B5-AS1 | -0.046730408 |
| SCAF8 | -0.046734089 |
| RNA5SP433 | -0.046743095 |
| RNU6-294P | -0.046744697 |
| DBP | -0.046745643 |
| BMI1 | -0.046746276 |
| SYNJ2 | -0.046753737 |
| SLC25A21-AS1 | -0.046756342 |
| RP11-459E5 | -0.046759464 |
| GRPEL2 | -0.04676317 |
| RP11-203I16 | -0.046764974 |
| OR51A2 | -0.04677031 |
| LHFPL3 | -0.046773496 |
| RP11-103H7 | -0.046774257 |
| AC092159 | -0.046774788 |
| RP11-46H11 | -0.046776044 |
| RP11-386P4 | -0.046778775 |
| ATP6V1G1P4 | -0.046781099 |
| KCNH3 | -0.04678248 |
| CCT6P4 | -0.046783444 |
| RNU6-1199P | -0.046787124 |

| | |
|---|---|
| RP11-423H2 | -0.046787319 |
| RN7SKP209 | -0.046787395 |
| RP11-863H1 | -0.046796196 |
| RP11-579O24 | -0.046803274 |
| C16orf78 | -0.046808757 |
| RP11-101B14 | -0.046818126 |
| ZC3H7B | -0.046818167 |
| FLJ27354 | -0.046819954 |
| AC004901 | -0.046820682 |
| RASSF10 | -0.046824927 |
| RP11-455I9 | -0.046827675 |
| RP11-514F3 | -0.046827799 |
| PNPLA5 | -0.046838895 |
| COPZ1 | -0.046842503 |
| FCN1 | -0.046842933 |
| FCAR | -0.04684561 |
| TBC1D32 | -0.046849778 |
| IL15RA | -0.046855415 |
| TMEM106A | -0.046858057 |
| RNU6-640P | -0.046863411 |
| WDR45B | -0.0468655 |
| LINC00266-3 | -0.046865622 |
| RNU6ATAC14P | -0.046865908 |
| ABHD2 | -0.046869407 |
| MIR374B | -0.046871279 |
| PRKAR1AP | -0.04687452 |
| AC005088 | -0.046884911 |
| KRT18P4 | -0.046891864 |
| C11orf71 | -0.04689231 |
| SHANK2 | -0.046894617 |
| RNU6ATAC13P | -0.046895484 |
| CCL2 | -0.046898282 |
| METAP1 | -0.046900348 |
| CTA-342B11 | -0.046904414 |
| MAP3K3 | -0.046906655 |
| LINC01569 | -0.046909576 |
| DAZAP2 | -0.046910767 |
| MIR3973 | -0.046914925 |
| LSM8 | -0.046915213 |
| RP3-391O22 | -0.046917383 |
| RP11-19E18 | -0.046918186 |
| RP11-467M13 | -0.046920351 |
| RRP7B | -0.04692446 |
| RNU6-862P | -0.046924862 |
| RP11-459I19 | -0.046925171 |

| | |
|---|---|
| PRKRIRP5 | -0.046927309 |
| LINC01150 | -0.046932195 |
| RP11-726G23 | -0.046934069 |
| RERE | -0.046934972 |
| LINC01160 | -0.046953843 |
| RNU4-15P | -0.046957038 |
| ISCA1P2 | -0.046962402 |
| ACO2 | -0.046964313 |
| BUD13 | -0.046965653 |
| RP11-143K11 | -0.046969134 |
| STIP1P3 | -0.046972477 |
| WSB2 | -0.0469725 |
| LINC01376 | -0.046972983 |
| BCL2L14 | -0.046979608 |
| SRGAP3-AS3 | -0.046980226 |
| RNA5SP393 | -0.046981592 |
| FKBP6 | -0.0469878 |
| RN7SKP27 | -0.046988591 |
| SOWAHD | -0.046989933 |
| TGFBR3 | -0.046999198 |
| RP11-724N1 | -0.047001563 |
| PRAMEF19 | -0.04700358 |
| FAM86MP | -0.047003888 |
| NDUFS5P4 | -0.047004881 |
| RNU6-452P | -0.047011121 |
| HIF1AP1 | -0.047011418 |
| CEACAM5 | -0.047011819 |
| BC040587 | -0.047017659 |
| RAB30 | -0.047018213 |
| AC109357 | -0.047020995 |
| OR6E1P | -0.047022864 |
| RP11-571M6 | -0.047038549 |
| PLEKHA2 | -0.047039312 |
| HTRA4 | -0.047042812 |
| RP11-286H15 | -0.047049039 |
| RP11-16B9 | -0.047055923 |
| WDR62 | -0.047056001 |
| RP11-802E16 | -0.047057675 |
| AC091866 | -0.047059662 |
| ADAM8 | -0.047061769 |
| SLMO2P2 | -0.047064214 |
| RP1-215K18 | -0.047064936 |
| LINC00963 | -0.047066327 |
| ODC1 | -0.047073751 |
| IGKV2-23 | -0.047073878 |

| | |
|---|---|
| AC079117 | -0.047076625 |
| FPR1 | -0.047077211 |
| CASP2 | -0.047083072 |
| ORC1 | -0.047088207 |
| SNRK-AS1 | -0.047090369 |
| RRH | -0.047095263 |
| AC080008 | -0.047099486 |
| ZNF709 | -0.047100282 |
| KRT8P33 | -0.047104133 |
| RCC2P4 | -0.047108808 |
| MUC3A | -0.047108822 |
| HELB | -0.047113101 |
| TSEN15P1 | -0.047115602 |
| TRAK2 | -0.04712259 |
| AC130462 | -0.047130249 |
| RNU4-23P | -0.047135258 |
| NEK2 | -0.047136935 |
| LYRM5 | -0.047142461 |
| IGLV3-2 | -0.047145275 |
| RNU6-479P | -0.04714575 |
| RPL23A | -0.047147019 |
| RP11-344L21 | -0.047148195 |
| RSF1-IT1 | -0.047150095 |
| TRIM55 | -0.047150937 |
| ATP6V1G1P7 | -0.047152605 |
| APH1B | -0.047154089 |
| HPRTP3 | -0.047155951 |
| AC068657 | -0.047166063 |
| MAPKAPK5-AS1 | -0.04716622 |
| AC097467 | -0.047171595 |
| OR5AO1P | -0.04719015 |
| AC002511 | -0.047191084 |
| RP11-116B19 | -0.047192025 |
| RP11-234O6 | -0.047196549 |
| RAD51 | -0.047199393 |
| RNU6-257P | -0.047199869 |
| AC015600 | -0.047200493 |
| RP11-715F3 | -0.047201943 |
| GTF3C4 | -0.047202798 |
| MX2 | -0.047204194 |
| RP11-435P24 | -0.047214443 |
| RP3-434P1 | -0.047228353 |
| RNU6-41P | -0.047231207 |
| CTC-472C24 | -0.047232236 |
| SCO1 | -0.047232288 |

| | |
|---|---|
| TBC1D3P1-DHX40P1 | -0.047235824 |
| IL6ST | -0.0472404 |
| RP11-443B7 | -0.047251898 |
| CTAGE5 | -0.047252624 |
| CTNNA3 | -0.047252795 |
| CRYZL1 | -0.047254652 |
| SNX5P1 | -0.047255364 |
| RP11-572O6 | -0.047261269 |
| RP11-432N13 | -0.0472618 |
| RASSF2 | -0.047263266 |
| EAF1-AS1 | -0.047263612 |
| RNF141 | -0.047278094 |
| HTATSF1 | -0.047278299 |
| RP11-716A19 | -0.047278404 |
| ACTL7B | -0.047280921 |
| KIAA1191 | -0.047280997 |
| RNU7-153P | -0.047282181 |
| CD5L | -0.047283977 |
| PRDX3 | -0.047284644 |
| RNA5SP370 | -0.04728479 |
| RP11-506E9 | -0.04729001 |
| AL136221 | -0.047290418 |
| RNU6-1115P | -0.047292815 |
| OR8H2 | -0.047296329 |
| RP11-380P13 | -0.047298825 |
| CTD-2140G10 | -0.047304225 |
| RABGAP1L-IT1 | -0.047308987 |
| USP51 | -0.047314784 |
| LINC01486 | -0.047314827 |
| EDDM3A | -0.047315716 |
| GRB7 | -0.047322134 |
| KCNC2 | -0.04732423 |
| ZMYND11 | -0.047328757 |
| RP11-500G9 | -0.047331858 |
| PFKFB1 | -0.047337502 |
| NUDT16 | -0.047337725 |
| KIF27 | -0.047338785 |
| CABYR | -0.047340572 |
| OTUD6A | -0.047346208 |
| MCPH1 | -0.047351288 |
| FGF12-AS2 | -0.047351419 |
| OR2AD1P | -0.047352382 |
| ELL2P1 | -0.047352666 |
| MIR628 | -0.047354196 |
| AL441988 | -0.047357782 |

| | |
|---|---|
| RP11-446N19 | -0.047362575 |
| NHP2P1 | -0.047364969 |
| RN7SL604P | -0.047365022 |
| RNU7-179P | -0.047366462 |
| RP11-298J20 | -0.047367766 |
| RNU6-578P | -0.047374648 |
| USP17L10 | -0.047374907 |
| RP11-788A4 | -0.047375017 |
| LZTFL1 | -0.047375401 |
| RP1-102G20 | -0.047376426 |
| RP11-262A12 | -0.047377209 |
| RP11-546B15 | -0.047378161 |
| WBSCR27 | -0.047382397 |
| FAM118A | -0.047385653 |
| HSDL1 | -0.047386899 |
| CCNB2P1 | -0.047390486 |
| MIR892C | -0.047392266 |
| AC092657 | -0.047397783 |
| ZCWPW2 | -0.047399346 |
| AP001962 | -0.047401611 |
| C20orf144 | -0.047402417 |
| OR51C4P | -0.047405916 |
| DNAJA1P1 | -0.047410641 |
| RP13-140E4 | -0.047413315 |
| AP000345 | -0.047414137 |
| ST13P10 | -0.047414529 |
| RP3-435K13 | -0.047416274 |
| GOLGA6L10 | -0.047434462 |
| PHKA2 | -0.047435999 |
| CA5B | -0.047436204 |
| SSU72P7 | -0.047442321 |
| FUT7 | -0.047443077 |
| AC074191 | -0.047446453 |
| AC092634 | -0.04745057 |
| RP11-373F14 | -0.047451956 |
| NPIPA3 | -0.047454063 |
| C5orf22 | -0.047456298 |
| CTC-295J13 | -0.047460188 |
| TPD52 | -0.04746147 |
| CTA-397H3 | -0.047463831 |
| OR51C1P | -0.047465315 |
| RP11-392B6 | -0.047478089 |
| DDIAS | -0.047478364 |
| TRAJ54 | -0.047480962 |
| HNRNPA1P11 | -0.047481147 |

| | |
|---|---|
| IFNL2 | -0.047495525 |
| BTNL10 | -0.04749696 |
| CDK5R2 | -0.047511302 |
| RPL7P28 | -0.047512357 |
| ADAM20P1 | -0.047514042 |
| U82670 | -0.047514182 |
| MSANTD1 | -0.047523873 |
| CCDC53 | -0.047524077 |
| RP5-1014D13 | -0.047524133 |
| RP11-355E10 | -0.047525511 |
| SH2D2A | -0.047526068 |
| PPIAP29 | -0.047527259 |
| RP11-5A11 | -0.047528862 |
| SERPINB9P1 | -0.047530068 |
| CDH12P1 | -0.047530267 |
| ATPAF1 | -0.047534348 |
| RP11-355K23B | -0.047539442 |
| AL355977 | -0.047541916 |
| RNA5SP399 | -0.047543241 |
| RPS3AP9 | -0.047544797 |
| REV3L | -0.047545641 |
| DPRXP4 | -0.047563062 |
| RNU6-599P | -0.047563582 |
| RP11-385F5 | -0.047566516 |
| C1orf111 | -0.047582953 |
| SNORD83A | -0.047583758 |
| RP11-2H3 | -0.047585465 |
| MIR5589 | -0.047587715 |
| RNA5SP33 | -0.047591234 |
| RP11-438D8 | -0.047591758 |
| PHKB | -0.04759342 |
| DZANK1 | -0.04759476 |
| NIFKP4 | -0.047598548 |
| VDAC1P11 | -0.047598902 |
| RNA5SP206 | -0.04760809 |
| C16orf96 | -0.047609445 |
| AL603910 | -0.047615012 |
| ZNF592 | -0.047616267 |
| bP-2189O9 | -0.047616927 |
| OAT | -0.047619933 |
| LYPLA1P3 | -0.047626086 |
| UCKL1 | -0.047628789 |
| RP11-184B22 | -0.047632421 |
| RFPL4AP5 | -0.047633746 |
| SAP30 | -0.04763958 |

| | |
|---|---|
| SLC22A13 | -0.047640931 |
| TREML1 | -0.047647667 |
| LRRC34P2 | -0.047650923 |
| KRT8P10 | -0.047657685 |
| CELP | -0.047662902 |
| RNA5SP464 | -0.047668786 |
| ENSAP2 | -0.047674924 |
| RP11-544H14 | -0.047678016 |
| ZNF630-AS1 | -0.047678136 |
| RP11-1030E3 | -0.047678396 |
| ITIH6 | -0.047682464 |
| DICER1 | -0.047684252 |
| LA16c-361A3 | -0.047684575 |
| NDUFV2-AS1 | -0.047686559 |
| RNU6-898P | -0.047694138 |
| RNU1-22P | -0.047698934 |
| RPS3AP2 | -0.047705901 |
| RPL35AP31 | -0.047707857 |
| SAPCD2 | -0.047712439 |
| RP11-327O17 | -0.047716758 |
| RP11-453O5 | -0.047719811 |
| RP11-387H17 | -0.047719907 |
| F11 | -0.047723569 |
| ZNF718 | -0.047729407 |
| SCARNA6 | -0.047732888 |
| LDB3 | -0.047734821 |
| TMX2 | -0.047735063 |
| LINC00272 | -0.047741083 |
| HSPA14 | -0.047741538 |
| HMGN2P20 | -0.047743079 |
| RP3-508I15 | -0.047746413 |
| AC005549 | -0.047747769 |
| XXbac-B476C20 | -0.04775899 |
| USP9YP3 | -0.047759438 |
| RP11-551L14 | -0.047760144 |
| OR4G4P | -0.047765989 |
| LL0XNC01-237H1 | -0.04776724 |
| GCLC | -0.047775121 |
| MT-CO2 | -0.047776033 |
| UTP15 | -0.047781465 |
| TRMT112P5 | -0.047782578 |
| TOR1AIP1 | -0.047784368 |
| ZNF30-AS1 | -0.04778743 |
| OR1N1 | -0.047797726 |
| MTF1 | -0.047801522 |

| | |
|---|---|
| PRDM11 | -0.047802713 |
| RBBP4P6 | -0.047805733 |
| AC034220 | -0.047805932 |
| RP11-25B7 | -0.047810642 |
| RP11-592N21 | -0.047824574 |
| CTD-2270P14 | -0.04782677 |
| RP11-241K18 | -0.047829812 |
| ACTR5 | -0.047833003 |
| COG6 | -0.047834361 |
| ANKHD1 | -0.0478381 |
| RNU6-904P | -0.04783828 |
| RNU6-651P | -0.047847335 |
| PGBD4P4 | -0.04785384 |
| RP11-348J24 | -0.047855181 |
| AC127030 | -0.047858913 |
| CTB-50L17 | -0.047863925 |
| GDF7 | -0.0478695 |
| CTC-820M8 | -0.047871558 |
| ASB13 | -0.047873004 |
| HNRNPCP8 | -0.047874585 |
| OR5C1 | -0.047882971 |
| RP11-632P5 | -0.047883942 |
| YTHDF1 | -0.047884186 |
| RN7SL487P | -0.047886508 |
| KRT24 | -0.047886903 |
| HMGB3P14 | -0.047891271 |
| PPP1R26P5 | -0.047892108 |
| RP1-66N13 | -0.0478936 |
| ZRANB2-AS1 | -0.047894939 |
| RNU6-560P | -0.047897194 |
| ACY3 | -0.047898469 |
| RP11-326E22 | -0.047899921 |
| C10orf32 | -0.047919016 |
| RNU6-359P | -0.047919054 |
| SNRPEP6 | -0.047919536 |
| SNIP1 | -0.047923183 |
| MAGEB16 | -0.04792943 |
| RNU6-196P | -0.047932953 |
| PRKDC | -0.047933937 |
| LIM2 | -0.0479387 |
| OR4A45P | -0.047939792 |
| RP11-354M1 | -0.047941424 |
| RP11-436F21 | -0.047946316 |
| C9orf62 | -0.04795145 |
| AC006378 | -0.047954059 |

| | |
|---|---|
| MRRF | -0.047955389 |
| STXBP5L | -0.047955656 |
| LRRC56 | -0.047959166 |
| OR2A25 | -0.047961381 |
| LINC00564 | -0.047962309 |
| RP11-122C21 | -0.04796407 |
| C1DP1 | -0.047965441 |
| TRIM31-AS1 | -0.047968137 |
| AC104978 | -0.047972363 |
| OR2AS2P | -0.047973025 |
| RP5-865E18 | -0.047973025 |
| ACAD10 | -0.047974125 |
| RDH14 | -0.047979055 |
| TUBBP2 | -0.047982945 |
| ZNF394 | -0.047983372 |
| VTI1A | -0.047984228 |
| RP5-1092A3 | -0.047992098 |
| RNA5SP224 | -0.047992295 |
| PACRG-AS1 | -0.047994747 |
| CD46 | -0.047995056 |
| RP11-365H22 | -0.048009777 |
| ATP6V1G1P2 | -0.048015104 |
| HSP90B1 | -0.048015597 |
| ACOT13 | -0.04801649 |
| RP11-344G13 | -0.04802732 |
| SAAL1 | -0.048027876 |
| RP11-134L10 | -0.048027969 |
| RAG2 | -0.048031743 |
| SH2D3C | -0.04803249 |
| TFCP2 | -0.048035728 |
| VPS37A | -0.048036993 |
| IGLV3-26 | -0.048037907 |
| RNU1-54P | -0.048038026 |
| IGHV2OR16-5 | -0.048040177 |
| HSPE1P26 | -0.048043199 |
| RP11-16L14 | -0.048044232 |
| RNU6-739P | -0.048045071 |
| CTD-2230M5 | -0.048046679 |
| POTEI | -0.048047171 |
| MTND5P2 | -0.048052489 |
| ACLY | -0.04805331 |
| RP1-71H19 | -0.048058167 |
| RN7SL500P | -0.048060004 |
| RP11-145O15 | -0.048060643 |
| RP5-1042K10 | -0.048063208 |

| | |
|---|---|
| RAD54B | -0.048063394 |
| ART1 | -0.048065137 |
| RP11-714G18 | -0.048065555 |
| SERBP1P6 | -0.04807121 |
| F8A3 | -0.048071855 |
| HLA-DRB5 | -0.04807452 |
| AC073987 | -0.048081227 |
| MT-ATP6 | -0.048084958 |
| FARP2 | -0.048085497 |
| RP11-252M21 | -0.048085886 |
| ALPK1 | -0.04808613 |
| HSPD1P9 | -0.048089773 |
| AP000459 | -0.048101728 |
| RN7SL205P | -0.048104361 |
| LINC01233 | -0.048104993 |
| RPL30P2 | -0.048107218 |
| MIR4652 | -0.048108055 |
| RXFP2 | -0.048109752 |
| CTD-3051L14 | -0.048110201 |
| TET2 | -0.048111452 |
| PRR23A | -0.048111727 |
| CCDC88A | -0.04811454 |
| HMGB1P4 | -0.048114656 |
| RP11-3D23 | -0.048117827 |
| NRBF2P2 | -0.048118347 |
| LINC00905 | -0.048119778 |
| SMIM14 | -0.048127322 |
| CDC23 | -0.04813002 |
| RP11-261P24 | -0.048133341 |
| SSRP1 | -0.048133369 |
| TRPS1 | -0.048133406 |
| C12orf71 | -0.048148061 |
| OSBPL2 | -0.048153705 |
| SUN3 | -0.048154804 |
| CNBP | -0.048155156 |
| RNU6-1015P | -0.048156553 |
| PDZD9 | -0.048161304 |
| MCM3 | -0.048162319 |
| SNORD116-14 | -0.04816253 |
| LSINCT5 | -0.048163202 |
| HMGN1P8 | -0.048165425 |
| CDC42P4 | -0.048169597 |
| SCML1 | -0.048171008 |
| MYOT | -0.048173329 |
| DSCAM-AS1 | -0.048174655 |

| | |
|---|---|
| U82671 | -0.04817607 |
| RNA5SP324 | -0.048176429 |
| AC009969 | -0.048179301 |
| FAM114A2 | -0.048187065 |
| EWSR1 | -0.048192442 |
| RNF165 | -0.048195046 |
| FAM60BP | -0.048197269 |
| USP41 | -0.048206287 |
| TCF23 | -0.048209191 |
| CTD-2319I12 | -0.048209745 |
| RP11-644F5 | -0.048210582 |
| RP11-692N5 | -0.048213727 |
| HMGN2P46 | -0.048218154 |
| FTLP13 | -0.048220004 |
| RP11-423P10 | -0.048224517 |
| HLA-S | -0.048224644 |
| C1orf100 | -0.048229184 |
| AC109471 | -0.048235259 |
| AL356309 | -0.048236367 |
| MRPL32P1 | -0.048241944 |
| ZNF599 | -0.048242397 |
| RPS24P18 | -0.048242733 |
| TESC | -0.048243755 |
| FAM74A1 | -0.048245987 |
| AC011372 | -0.048245987 |
| AC135178 | -0.048247741 |
| AC091153 | -0.048253197 |
| RP11-146N23 | -0.048258849 |
| CROCCP1 | -0.048263496 |
| TYW1 | -0.048263701 |
| SH3BP1 | -0.048268947 |
| MOB3A | -0.048272073 |
| MED15P4 | -0.048279329 |
| FPGT | -0.048282511 |
| ALDH1A1 | -0.048283497 |
| SNORD3C | -0.048292068 |
| RECQL5 | -0.048293506 |
| LINC00557 | -0.048298124 |
| OR51A8P | -0.048300568 |
| ATP6V0E1P4 | -0.048300644 |
| HDAC11-AS1 | -0.048301815 |
| RP11-548M13 | -0.048304468 |
| C14orf183 | -0.048305486 |
| MIR4766 | -0.04830906 |
| ANKUB1 | -0.048311223 |

| | |
|---|---|
| LRCH3 | -0.048312763 |
| NELL1 | -0.04831646 |
| FAM90A20P | -0.048323224 |
| WDR55 | -0.04832761 |
| LRRC8B | -0.048328113 |
| GIT1 | -0.048331071 |
| CTA-544A11 | -0.04833534 |
| DNPEP | -0.048343946 |
| LA16c-313D11 | -0.048352504 |
| GKN2 | -0.048353574 |
| OR4A13P | -0.048362236 |
| ZNF43 | -0.048368152 |
| RP11-183K14 | -0.048370188 |
| MYH15 | -0.048371755 |
| RP11-866E20 | -0.048371897 |
| CTB-134H23 | -0.048380607 |
| RP11-861A13 | -0.048388136 |
| HMGB3P12 | -0.048389274 |
| MIR298 | -0.048394321 |
| AL035461 | -0.048398994 |
| KIAA0141 | -0.048403707 |
| RP11-167J8 | -0.048405518 |
| GOT2P3 | -0.048406868 |
| ZNF385B | -0.048407138 |
| FAM209A | -0.048408459 |
| TTC37 | -0.04841007 |
| DYX1C1-CCPG1 | -0.048410745 |
| RNU6-1203P | -0.048419252 |
| LINC00489 | -0.048419649 |
| RNU105B | -0.04841972 |
| AC083864 | -0.0484204 |
| MIR548H3 | -0.048432142 |
| RP11-573J24 | -0.048443661 |
| NAE1 | -0.048443857 |
| CCDC158 | -0.04844665 |
| RPS26P5 | -0.048446764 |
| ICAM2 | -0.048451244 |
| LINC00167 | -0.048456474 |
| RNA5SP194 | -0.04845914 |
| SIPA1L3 | -0.048464305 |
| TOMM70A | -0.048471058 |
| MIR2467 | -0.048472405 |
| RNA5SP473 | -0.048475196 |
| DNAJC6 | -0.048475524 |
| MIR138-2 | -0.048485424 |

| | |
|---|---|
| ASB5 | -0.048489658 |
| PPP3R2 | -0.048494141 |
| RNU7-7P | -0.048501748 |
| EBLN1 | -0.048504631 |
| RNU6-1037P | -0.048506423 |
| CTD-2349B8 | -0.048508445 |
| RNU7-190P | -0.048508688 |
| AC003084 | -0.048512057 |
| TNFRSF25 | -0.048515619 |
| LPAR6 | -0.048522763 |
| HDC | -0.04852308 |
| INTS6P1 | -0.0485258 |
| AC103564 | -0.048529018 |
| RP11-816B4 | -0.048529793 |
| AY269186 | -0.0485327 |
| AC008521 | -0.048534552 |
| MIR369 | -0.04853504 |
| ENPP2 | -0.048543527 |
| OR10AD1 | -0.048550876 |
| PRAMEF6 | -0.048553143 |
| RP11-169E6 | -0.048556147 |
| RP11-550H2 | -0.048556403 |
| SP3 | -0.048564328 |
| HSD17B7P2 | -0.048573705 |
| AC064829 | -0.048574474 |
| AC080094 | -0.048582588 |
| SNHG16 | -0.048584998 |
| CICP1 | -0.048585459 |
| GNE | -0.048590301 |
| DENND6A | -0.048613316 |
| GAPDHP16 | -0.048614115 |
| WDR35 | -0.048615549 |
| BLOC1S3 | -0.048616256 |
| OR4A11P | -0.048616686 |
| AKAP3 | -0.048617186 |
| TRAJ31 | -0.04861832 |
| VPS13A | -0.048618656 |
| AC003956 | -0.048621991 |
| OR7E8P | -0.048622949 |
| RP11-96H19 | -0.048628219 |
| DEFB134 | -0.048634927 |
| MICF | -0.048635542 |
| CT45A1 | -0.048635952 |
| RP11-93G5 | -0.048638157 |
| ADORA1 | -0.048643885 |

| | |
|---|---|
| PGK1P2 | -0.048649572 |
| KRTAP13-4 | -0.048650457 |
| SLC5A5 | -0.048650687 |
| AC003973 | -0.048662816 |
| RP1-212P9 | -0.04866338 |
| ABI2 | -0.048664946 |
| AC073215 | -0.048666709 |
| RP11-572C21 | -0.048669589 |
| RN7SL804P | -0.048669726 |
| AC002480 | -0.048683042 |
| RP5-1050D4 | -0.048685591 |
| APPL1 | -0.048689598 |
| AC005162 | -0.048690595 |
| MTND1P2 | -0.048690696 |
| PREB | -0.048694268 |
| RNU6-1297P | -0.048698992 |
| AC055758 | -0.048699324 |
| NUP210P3 | -0.048712169 |
| PARD6A | -0.048716595 |
| RN7SL134P | -0.048721933 |
| RPEP2 | -0.048729043 |
| C2orf44 | -0.048731126 |
| EGR3 | -0.048732502 |
| CUL2 | -0.048734903 |
| AC108676 | -0.048737181 |
| RP11-452J21 | -0.048737881 |
| MMAA | -0.048747368 |
| RPL35P3 | -0.048756764 |
| SPDL1 | -0.048764039 |
| RNU6ATAC26P | -0.048766453 |
| LINC00483 | -0.048768085 |
| AC005702 | -0.048770848 |
| NUBPL | -0.04877336 |
| ZNF655 | -0.048774421 |
| DLG2 | -0.048775835 |
| CABLES2 | -0.048780821 |
| RP11-46J23 | -0.048782384 |
| RP11-596D21 | -0.048785784 |
| MIR3687-1 | -0.04879191 |
| SCN3A | -0.048796952 |
| TUBB4A | -0.04879702 |
| FGFR3P1 | -0.048798185 |
| POM121L4P | -0.048809596 |
| CLTC | -0.048809994 |
| CLGN | -0.048816078 |

| | |
|---|---|
| PBLD | -0.048821865 |
| CTD-2187J20 | -0.048822312 |
| TRMT6 | -0.048823018 |
| RP11-99H8 | -0.048824003 |
| TTC26 | -0.048824916 |
| OR10N1P | -0.048829374 |
| RP11-413H22 | -0.04883478 |
| KIN | -0.048835447 |
| ZNF595 | -0.048840135 |
| CATSPER4 | -0.048842191 |
| RP11-369G6 | -0.048845289 |
| TNKS | -0.048846219 |
| FAT1P1 | -0.048852337 |
| GAPDHP39 | -0.048856625 |
| FENDRR | -0.048871212 |
| IGHVIII-2-1 | -0.048884315 |
| RBP5 | -0.048896975 |
| RN7SKP92 | -0.048898579 |
| SPCS2 | -0.04889891 |
| MCOLN1 | -0.048901515 |
| UNC93B3 | -0.048901991 |
| RP5-906C1 | -0.048903719 |
| SLC22A20 | -0.048904826 |
| IP6K1 | -0.048911859 |
| ERVH-1 | -0.048912667 |
| MIR4659B | -0.048920185 |
| ZNF426 | -0.048924716 |
| NEK7 | -0.048933646 |
| HIST1H2BD | -0.048936121 |
| CH507-513H4 | -0.048938446 |
| AC009695 | -0.048944154 |
| ARHGDIB | -0.048948382 |
| LINC01284 | -0.048949239 |
| RP11-141B14 | -0.048960717 |
| OPRM1 | -0.048961603 |
| RNU6-16P | -0.048965801 |
| SLC25A6P2 | -0.048967546 |
| CEP162 | -0.048967583 |
| RNA5SP447 | -0.048968684 |
| AP000959 | -0.048973748 |
| LDOC1L | -0.04897415 |
| TM7SF2 | -0.048975285 |
| COX10 | -0.048976309 |
| AC104534 | -0.048977154 |
| RP11-361A21 | -0.048978398 |

| | |
|---|---|
| AC007899 | -0.048978636 |
| RP11-323I15 | -0.04898161 |
| CSF2RA | -0.048983007 |
| AC022098 | -0.048983361 |
| LNX1-AS1 | -0.048993848 |
| RP11-639B1 | -0.048997467 |
| CALM2P1 | -0.048997576 |
| DPH1 | -0.048999565 |
| SIGLECL1 | -0.049004386 |
| RNA5SP86 | -0.049009482 |
| CSF1R | -0.049010375 |
| RPSAP47 | -0.049012151 |
| RP11-274E7 | -0.049015641 |
| FBXW2 | -0.049017501 |
| BRSK2 | -0.04901939 |
| RBM11 | -0.049020208 |
| NR3C1 | -0.049027306 |
| ILF3 | -0.04902846 |
| MRM1 | -0.049039083 |
| MAGEB3 | -0.049040141 |
| CHRM5 | -0.04905221 |
| RAB27A | -0.049056889 |
| FAM160A2 | -0.049058749 |
| SERINC1 | -0.049061503 |
| MIR610 | -0.04906303 |
| MTERF3 | -0.049064052 |
| DUX4L8 | -0.049064872 |
| RP11-38O14 | -0.049067556 |
| HKR1 | -0.049068406 |
| PCNPP2 | -0.049069295 |
| CCAT2 | -0.049073253 |
| AC009480 | -0.049075741 |
| RP11-479O9 | -0.049076887 |
| SNRPCP18 | -0.049079416 |
| PCSK2 | -0.049082114 |
| CYCSP2 | -0.049092867 |
| NUP88 | -0.049103812 |
| RP11-434P11 | -0.049105717 |
| RP11-775A3 | -0.049115645 |
| DUS4L | -0.049117729 |
| RNU6-179P | -0.049119057 |
| RP13-392I16 | -0.049125013 |
| LINC00339 | -0.049128706 |
| SHOC2 | -0.049133413 |
| AL161731 | -0.049134861 |

| | |
|---|---|
| CTD-2318O12 | -0.049135669 |
| ZNF81 | -0.049136055 |
| AC020983 | -0.049138628 |
| C1QL3 | -0.049139851 |
| PCAT2 | -0.049140111 |
| XKRY | -0.049140225 |
| RNU2-34P | -0.049141133 |
| PEX2 | -0.049145562 |
| LIPT1P1 | -0.049153012 |
| UBQLN4P1 | -0.049156835 |
| TMEM67 | -0.049169526 |
| CCDC28A | -0.049170047 |
| MIR378J | -0.049171106 |
| MIR4706 | -0.049184738 |
| RNU6-460P | -0.049185101 |
| AC078882 | -0.049193534 |
| RNU1-80P | -0.049196952 |
| CTC-467M3 | -0.049197117 |
| CCNL2 | -0.049200151 |
| CTB-89H12 | -0.049200467 |
| TUBA3GP | -0.049210936 |
| MTERF1 | -0.049216338 |
| CCL14 | -0.04921801 |
| RPH3A | -0.049224882 |
| NDUFAF7 | -0.049225975 |
| RP13-270P17 | -0.049226682 |
| CPA6 | -0.049228326 |
| POM121L14P | -0.049228678 |
| FAM227B | -0.049232469 |
| AF254983 | -0.049234766 |
| RP11-143A12 | -0.049244458 |
| HIST2H2AA4 | -0.049248565 |
| RP11-345J4 | -0.049257139 |
| AMFR | -0.049258907 |
| FAM204A | -0.049259484 |
| HLA-Z | -0.049260292 |
| CTD-2012J19 | -0.049262629 |
| RP11-823P9 | -0.049265553 |
| RNU6-311P | -0.049268282 |
| AC093668 | -0.049269398 |
| ARPC4-TTLL3 | -0.049285272 |
| RP11-182E14 | -0.049289394 |
| LINC00997 | -0.049293048 |
| RP5-965K10 | -0.049293172 |
| TBC1D9B | -0.049297104 |

| | |
|---|---|
| AD000671 | -0.04929776 |
| DEPDC1B | -0.04929889 |
| SPINT5P | -0.049299763 |
| RP11-536C5 | -0.049303988 |
| IRX1P1 | -0.049312711 |
| MELK | -0.049312844 |
| RP11-517P14 | -0.0493177 |
| RP11-781P14 | -0.049324996 |
| C4orf46 | -0.049325254 |
| RP11-583F2 | -0.04932932 |
| DUTP4 | -0.049332725 |
| VAV3 | -0.049336082 |
| ZBED6 | -0.049336161 |
| PIGO | -0.049337459 |
| OFD1P8Y | -0.049337834 |
| ALG1L14P | -0.049338258 |
| ZRSR1 | -0.049340239 |
| RP11-588L15 | -0.049350722 |
| AC005224 | -0.049352176 |
| AC005227 | -0.049353224 |
| SEC14L1 | -0.049356604 |
| FDPSP2 | -0.049361687 |
| MYCBP2 | -0.049369237 |
| RP11-709D24 | -0.049371237 |
| THSD7B | -0.049372234 |
| RP11-754N21 | -0.049372622 |
| CAND1 | -0.049379973 |
| SMYD5 | -0.049381182 |
| HTR7P1 | -0.049385889 |
| STAG3L5P | -0.049391992 |
| RP1-127B20 | -0.04939393 |
| SNORD121B | -0.049394999 |
| RPL37P4 | -0.049395034 |
| ZNF211 | -0.049397201 |
| CTA-250D10 | -0.049397936 |
| LINC00633 | -0.049397942 |
| LTV1 | -0.049398504 |
| RN7SL222P | -0.049402916 |
| RP11-317G22 | -0.049404971 |
| DYM | -0.049405313 |
| AC026271 | -0.049407527 |
| RNU6-63P | -0.049420897 |
| WDR53 | -0.049427324 |
| RNU6-455P | -0.049428348 |
| RIOK2 | -0.049434152 |

| | |
|---|---|
| AC073046 | -0.049443858 |
| RNU6-1079P | -0.049451245 |
| YME1L1 | -0.049457502 |
| RN7SKP275 | -0.049458549 |
| RPS5P3 | -0.04946121 |
| RNU6-975P | -0.049467114 |
| CTB-181H17 | -0.04946942 |
| DISC1 | -0.04946967 |
| RP11-244K5 | -0.049472593 |
| RAE1 | -0.049473447 |
| RN7SL606P | -0.049474053 |
| RP11-510I5 | -0.049474231 |
| ZNF616 | -0.049475661 |
| MIR3945 | -0.049484274 |
| CST11 | -0.049485013 |
| RP11-27M24 | -0.049489943 |
| TMEM74 | -0.049492048 |
| PPATP1 | -0.049492292 |
| AC003104 | -0.049503246 |
| DHX8 | -0.049510546 |
| SLC14A2 | -0.049511674 |
| AC091199 | -0.049511923 |
| RNU4-44P | -0.049512524 |
| DCAF13P2 | -0.049519799 |
| SNORA84 | -0.049522553 |
| AC005785 | -0.049524314 |
| MIRLET7D | -0.049532118 |
| DNM1L | -0.049535477 |
| CBX1 | -0.04953637 |
| AC010974 | -0.049538141 |
| CALY | -0.049539365 |
| RN7SKP260 | -0.049542003 |
| CXorf66 | -0.049542753 |
| HIST1H4F | -0.049546476 |
| RNU6-1068P | -0.049547265 |
| LOR | -0.04954928 |
| RP11-960H2 | -0.049553267 |
| DNAJC12 | -0.049559609 |
| GLUDP5 | -0.049560085 |
| DDX47 | -0.04956266 |
| KIRREL2 | -0.049566719 |
| RABEPKP1 | -0.049580747 |
| RNA5SP187 | -0.049588916 |
| KIAA1875 | -0.049598466 |
| GFM2 | -0.049598531 |

| | |
|---|---|
| MAP2K4P1 | -0.049601183 |
| ITCH-AS1 | -0.04960258 |
| CFL1P6 | -0.049602839 |
| ATP5LP7 | -0.049609986 |
| RP11-557B9 | -0.049610795 |
| RP11-443K8 | -0.049616983 |
| RP11-605F20 | -0.049621971 |
| MIR548AC | -0.049622349 |
| HMGN1P30 | -0.049624732 |
| HMGN1P36 | -0.049625622 |
| AC064846 | -0.049626983 |
| RPS15P9 | -0.049637427 |
| FXYD2 | -0.049639975 |
| WDFY3-AS1 | -0.049641651 |
| CTA-292E10 | -0.049642868 |
| RNU2-62P | -0.049642948 |
| ASS1P2 | -0.049643047 |
| AC005808 | -0.049646067 |
| NBPF13P | -0.049647408 |
| RNU6-471P | -0.049653428 |
| H3F3B | -0.049660246 |
| RNU7-138P | -0.049661565 |
| IYD | -0.049664882 |
| AC018878 | -0.049665122 |
| MITD1 | -0.049667688 |
| XPO5 | -0.049667744 |
| CREG1 | -0.049668023 |
| POLG | -0.049668358 |
| RP11-544L8__B | -0.049671184 |
| RNU6-1278P | -0.04967371 |
| POU5F1P2 | -0.049679842 |
| KCTD9P6 | -0.049681627 |
| OR7E128P | -0.049683226 |
| NOM1 | -0.049692638 |
| RP11-328J6 | -0.049695949 |
| PTPDC1 | -0.049701736 |
| NPAP1P6 | -0.049706621 |
| CTD-2008E3 | -0.049707559 |
| PIGB | -0.049708787 |
| AC021876 | -0.049716395 |
| RP11-168C9 | -0.049718562 |
| HINT3 | -0.049719653 |
| THAP5P1 | -0.049719703 |
| AC068446 | -0.049721897 |
| EIF5P1 | -0.049722278 |

| | |
|---|---|
| ZBTB10 | -0.049722872 |
| CTB-39G8 | -0.04972371 |
| RN7SL187P | -0.049731381 |
| TMCO4 | -0.049736081 |
| SLC25A51P3 | -0.049740065 |
| PP12613 | -0.049746692 |
| ANKRD18A | -0.049752681 |
| ATG9B | -0.049755859 |
| SCN4A | -0.049756938 |
| USP39 | -0.049757534 |
| HNRNPA1P31 | -0.049757994 |
| TIMM21 | -0.049759878 |
| SMARCC1 | -0.049764752 |
| AC093162 | -0.049766762 |
| AC064871 | -0.049767777 |
| MEF2C | -0.049768436 |
| RP3-357I16 | -0.049771328 |
| OR7E94P | -0.049775008 |
| RP11-428O18 | -0.049776457 |
| HIPK1 | -0.049781162 |
| RP11-360A10 | -0.049781749 |
| SRRM5 | -0.049783953 |
| CTD-2501B8 | -0.049796123 |
| RN7SL351P | -0.049816324 |
| ATP5HP4 | -0.049821435 |
| AC011999 | -0.049831561 |
| RP11-114G22 | -0.049831701 |
| GNPDA1 | -0.049834635 |
| ZNF823 | -0.049837035 |
| RP11-210G22 | -0.049839869 |
| MTND2P22 | -0.049840303 |
| ZSWIM3 | -0.049840923 |
| CTD-2600H12 | -0.049845437 |
| TCTA | -0.049849035 |
| NUDT6 | -0.049854896 |
| MIR4660 | -0.049856963 |
| MAP9 | -0.049857054 |
| NPM1P21 | -0.049865548 |
| ENTPD5 | -0.049871455 |
| RP11-97C18 | -0.04987355 |
| GS1-306C12 | -0.049875149 |
| BNIP3P41 | -0.049879488 |
| TLCD1 | -0.049880627 |
| INPP5B | -0.049882395 |
| RP11-787D18 | -0.049883364 |

| | |
|---|---|
| LINC00279 | -0.049883757 |
| AP001506 | -0.049883757 |
| RP11-61E11 | -0.049885418 |
| RP11-206M11 | -0.049892727 |
| RP11-2O17 | -0.049897806 |
| MAFG | -0.049905373 |
| AC007349 | -0.049906287 |
| LSP1P2 | -0.049908897 |
| CDCA5 | -0.049912164 |
| FAM196A | -0.049913987 |
| ANAPC16 | -0.04992275 |
| LINC01335 | -0.049924331 |
| RP11-433A19 | -0.049926867 |
| RP11-609N14 | -0.049929218 |
| OR51H2P | -0.049934647 |
| MIR6514 | -0.049935976 |
| RN7SL594P | -0.049941368 |
| RP11-521P1 | -0.049942634 |
| RP11-166B2 | -0.049943756 |
| CTD-2013M15 | -0.049945322 |
| LINC00293 | -0.049947859 |
| ZNF501 | -0.049949002 |
| IGHD1-20 | -0.049959414 |
| DRICH1 | -0.049966066 |
| GPR119 | -0.049966634 |
| FAM138B | -0.049976101 |
| RP11-214N15 | -0.049979382 |
| NRG1-IT1 | -0.049982128 |
| MSRB1P1 | -0.049986147 |
| HNRNPABP1 | -0.049986938 |
| FCGR3B | -0.049991819 |
| DLGAP1-AS1 | -0.049993097 |
| DHX30 | -0.049998809 |
| VWA3A | -0.049999335 |
| KANSL1L | -0.049999888 |
| MAPK1 | -0.050016261 |
| DACH1 | -0.050019294 |
| RNU6-1072P | -0.050020238 |
| KRTAP2-1 | -0.050028143 |
| GLULP6 | -0.050029136 |
| RP1-274L7 | -0.050032598 |
| TXNIP | -0.050041583 |
| RP11-654A16 | -0.050042565 |
| LRRFIP1P1 | -0.050044328 |
| AC090644 | -0.050045494 |

| | |
|---|---|
| SCRN3 | -0.050045565 |
| EIF4A1P5 | -0.050060434 |
| RP5-943J3 | -0.050060934 |
| PMS2P9 | -0.05006431 |
| RNU6-1318P | -0.050070445 |
| RP11-311H10 | -0.05007225 |
| CTC-484P3 | -0.050072629 |
| IGKV2OR2-7 | -0.050072844 |
| CCSAP | -0.050076952 |
| FAM105A | -0.05008218 |
| LA16c-OS12 | -0.050082962 |
| RP11-162G10 | -0.050083949 |
| AC018618 | -0.050085288 |
| NEK2P4 | -0.050085599 |
| ATG3P1 | -0.050088552 |
| FAM206A | -0.050103446 |
| GPALPP1 | -0.050106944 |
| RNU6-59P | -0.050107195 |
| NEBL | -0.050110293 |
| SLC25A29 | -0.050114907 |
| FCGR1B | -0.050121977 |
| C5orf51 | -0.050124843 |
| INSL6 | -0.050127968 |
| RP11-298O21 | -0.050128898 |
| BAIAP2-AS1 | -0.050129978 |
| RP11-392L5 | -0.050130404 |
| TRAM1 | -0.050134257 |
| BMS1P12 | -0.050136676 |
| ATF4P4 | -0.050140619 |
| TSR1 | -0.05014302 |
| RNU6-617P | -0.050150405 |
| RP1-177A13 | -0.050155503 |
| RP4-781K5 | -0.050157228 |
| SSX8 | -0.050157732 |
| RBM28 | -0.050158128 |
| RP11-732M18 | -0.050163998 |
| MIR1284 | -0.050165124 |
| NSUN4 | -0.050170649 |
| RP11-649E7 | -0.050172922 |
| ACTR3BP4 | -0.050178446 |
| BCORL1 | -0.050178594 |
| MUSTN1 | -0.050178771 |
| RP4-592A1 | -0.050179283 |
| NUPL2 | -0.050180361 |
| EIF4BP9 | -0.050189432 |

| | |
|---|---|
| CDK11A | -0.050192436 |
| RP11-110J1 | -0.05019389 |
| MIR548K | -0.050194362 |
| RPL32P3 | -0.050197385 |
| CTC-490G23 | -0.050200154 |
| RNA5SP445 | -0.050200771 |
| MIR509-3 | -0.050204759 |
| PDX1-AS1 | -0.050216068 |
| KIAA2018 | -0.050217703 |
| RP11-256G5 | -0.050218219 |
| RP11-746P2 | -0.050219099 |
| RP11-550A18 | -0.050219603 |
| AC004458 | -0.050222088 |
| MIR4795 | -0.050222967 |
| MICE | -0.050226336 |
| BTF3L4 | -0.050232736 |
| RP11-30P21 | -0.05023443 |
| RNU6-1022P | -0.050242535 |
| RCC2P7 | -0.050248149 |
| RP11-622J8 | -0.050248355 |
| RP5-882C2 | -0.050253723 |
| CD101 | -0.050258839 |
| DENND5A | -0.0502623 |
| CLEC1B | -0.050265888 |
| TIPIN | -0.05026617 |
| LRP8 | -0.050272161 |
| AC027506 | -0.050274127 |
| RPS27P18 | -0.050274892 |
| AL022345 | -0.050277375 |
| RNU6-1180P | -0.050280243 |
| RNU6-348P | -0.050282881 |
| RAP1A | -0.050284719 |
| ALG11 | -0.050288534 |
| NRXN3 | -0.050288906 |
| HSP90AA2P | -0.050295734 |
| RNU6-418P | -0.050295788 |
| CCDC50 | -0.050300553 |
| LRRC48 | -0.050301985 |
| RP11-183E24 | -0.050303091 |
| MIR425 | -0.050310414 |
| AC083863 | -0.050311145 |
| RN7SL165P | -0.050311783 |
| RP11-166O4 | -0.050311939 |
| SRFBP1 | -0.050315251 |
| RN7SL502P | -0.050317639 |

| | |
|---|---|
| MTND2P4 | -0.050320823 |
| RNU5A-5P | -0.05032089 |
| CTC-492K19 | -0.050323332 |
| LINC01180 | -0.050325238 |
| MOCOS | -0.050327921 |
| BNIP3P4 | -0.050331504 |
| JRK | -0.050333801 |
| IFT74-AS1 | -0.050334213 |
| KPNA2 | -0.050336517 |
| HOXB2 | -0.050336685 |
| SLC38A1 | -0.050338173 |
| PAFAH2 | -0.05033901 |
| RNA5SP201 | -0.050339041 |
| ZNF767P | -0.050339289 |
| AP000487 | -0.050339885 |
| SEC24D | -0.05034035 |
| PTPLAD1 | -0.05034381 |
| RP11-421E17 | -0.050351669 |
| OR5K2 | -0.05035205 |
| TACR3 | -0.0503533 |
| RNU2-38P | -0.05035667 |
| AJ239322 | -0.050357694 |
| FALEC | -0.05035957 |
| OR5AC2 | -0.050359632 |
| RP1-32B1 | -0.050360325 |
| BHLHE40 | -0.050365263 |
| MED15P3 | -0.050368735 |
| RP11-10A7 | -0.050370152 |
| MIR6779 | -0.050371937 |
| NPAP1P4 | -0.050381025 |
| RP11-960D24 | -0.050383041 |
| TXNDC16 | -0.050383328 |
| NDST4 | -0.050384637 |
| SNORD37 | -0.050386993 |
| RP11-433O3 | -0.050391572 |
| NMD3P1 | -0.050393063 |
| TIMM9P2 | -0.050394428 |
| RP11-109D24 | -0.0504083 |
| TNPO3 | -0.050410009 |
| RNF169 | -0.050413693 |
| OR5K3 | -0.050430043 |
| DNAI1 | -0.05043074 |
| SLX1A-SULT1A3 | -0.050431265 |
| GAPDHP47 | -0.050433655 |
| PRKCD | -0.050435126 |

| | |
|---|---|
| RP11-133L19 | -0.050436182 |
| NPM1P9 | -0.050437387 |
| TUBB4BP7 | -0.050438624 |
| RN7SL678P | -0.050441691 |
| RP4-566L20 | -0.05044635 |
| C8orf46 | -0.050447136 |
| CTD-2547H18 | -0.050460322 |
| OR5AK1P | -0.050463995 |
| EFTUD1 | -0.050465099 |
| CTC-296K1 | -0.050467083 |
| LILRB3 | -0.050467183 |
| RP11-359I18 | -0.050471496 |
| AC068489 | -0.050474399 |
| AL136123 | -0.050476966 |
| RP11-108M11 | -0.050482635 |
| RP11-945F5 | -0.050486799 |
| TMEM87A | -0.050487296 |
| BNIP3P10 | -0.050488358 |
| OR7E126P | -0.05049158 |
| ZNF250 | -0.050496226 |
| SEC23IP | -0.05049885 |
| AC004920 | -0.050499589 |
| IGKV1D-32 | -0.050504607 |
| RP11-182I21 | -0.05050546 |
| AC096559 | -0.050511107 |
| CTC-422A18 | -0.050511234 |
| ZNF558 | -0.050511743 |
| ANKRD36P1 | -0.050513167 |
| OR1X1P | -0.050514523 |
| AC024475 | -0.050527861 |
| RNU6-454P | -0.050531887 |
| ABCB6 | -0.050535577 |
| RP11-203F8 | -0.050536858 |
| CTA-373H7 | -0.050537862 |
| AP006477 | -0.050538425 |
| LINC01136 | -0.050539365 |
| CH17-80A12 | -0.050544052 |
| RN7SL280P | -0.050547729 |
| FAM214A | -0.050548173 |
| ARRDC5 | -0.050551331 |
| ZNF33AP1 | -0.050552647 |
| PDCL | -0.050563411 |
| STT3B | -0.050564192 |
| CHKA | -0.050565327 |
| MIR17HG | -0.050566276 |

| | |
|---|---|
| DAP3P2 | -0.050567584 |
| RP11-2E17 | -0.050570035 |
| RN7SKP50 | -0.050570903 |
| FRG2GP | -0.050572984 |
| AC009831 | -0.050586714 |
| PTPN9 | -0.050591356 |
| CIR1P2 | -0.050597044 |
| C2orf88 | -0.050604529 |
| OR7E35P | -0.050605277 |
| PIBF1 | -0.050606668 |
| ZNF175 | -0.050609683 |
| PAQR5 | -0.050609906 |
| AC093433 | -0.050614163 |
| IGBP1P1 | -0.050623589 |
| SNORA63 | -0.050624159 |
| RP11-255H23 | -0.050625441 |
| CDKAL1 | -0.050629274 |
| AC110615 | -0.050629317 |
| ST8SIA3 | -0.050638134 |
| RPA2P1 | -0.050638679 |
| RP11-265N6 | -0.050641218 |
| RP11-518L10 | -0.050641516 |
| AC114945 | -0.050641544 |
| RTP4 | -0.050644679 |
| RP11-274J7 | -0.050649723 |
| AL162151 | -0.05065322 |
| RP11-44K6 | -0.050653821 |
| LINC01106 | -0.050668254 |
| RP11-145P16 | -0.050669239 |
| TRAPPC4 | -0.050672216 |
| ZDHHC5 | -0.050674392 |
| TDGP1 | -0.050675704 |
| RP11-131N11 | -0.050678961 |
| RP11-464D20 | -0.050679955 |
| TMC6 | -0.050684458 |
| RP11-120K19 | -0.050687091 |
| OR51A3P | -0.050687897 |
| ZFP1 | -0.05068803 |
| UBR4 | -0.050690895 |
| RNASEH2B-AS1 | -0.050691454 |
| FSTL4 | -0.050691484 |
| FAM201A | -0.050695232 |
| CDC42EP3P1 | -0.050698672 |
| MAML3 | -0.050709854 |
| RP11-749H17 | -0.050713698 |

| | |
|---|---|
| COMMD10 | -0.050716348 |
| DDIT3 | -0.050728255 |
| ABCD1P4 | -0.050733127 |
| AHCYP3 | -0.05073443 |
| RP11-660M5 | -0.050739484 |
| EIF4B | -0.050739902 |
| PHF2P2 | -0.050740225 |
| SMIM11P1 | -0.050749167 |
| AC091736 | -0.050749457 |
| AC069181 | -0.050755306 |
| RP11-775C24 | -0.050760778 |
| TCP11L2 | -0.050761857 |
| RN7SKP243 | -0.05076236 |
| LINC01526 | -0.050762767 |
| RP11-493P1 | -0.050765935 |
| SMKR1 | -0.050767029 |
| OR7E5P | -0.050769382 |
| HLA-C | -0.050772406 |
| CTD-2547E10 | -0.050776002 |
| SMUG1P1 | -0.050776496 |
| HECTD2 | -0.0507877 |
| RNU6-468P | -0.05079274 |
| SNRPGP5 | -0.050795308 |
| EIF4EP2 | -0.050798651 |
| RP11-204C23 | -0.050801182 |
| AL109658 | -0.05080164 |
| AKR1D1P1 | -0.05080415 |
| RNA5SP292 | -0.050821239 |
| TRY2P | -0.050821978 |
| WNT9B | -0.05082668 |
| LOC401913 | -0.050829151 |
| AC114730 | -0.050834102 |
| RNA5SP504 | -0.050839123 |
| AC016970 | -0.050839123 |
| RP11-433J8 | -0.050840319 |
| RP11-71C5 | -0.050840321 |
| AC090627 | -0.050842077 |
| AC011499 | -0.050847531 |
| UMAD1 | -0.050847721 |
| COX6B1P2 | -0.050848728 |
| RSF1 | -0.050850084 |
| CTC-1337H24 | -0.050851833 |
| IRX5 | -0.05085334 |
| AL353664 | -0.0508552 |
| RNU4-39P | -0.050860182 |

| | |
|---|---|
| PPIAL4C | -0.050860359 |
| PHKG1P1 | -0.050861255 |
| ATAD1 | -0.050863759 |
| METTL21AP1 | -0.050864293 |
| CCDC57 | -0.050865787 |
| EPN3 | -0.050870234 |
| GPR89B | -0.050871201 |
| CTD-2591A1 | -0.050879779 |
| AC053503 | -0.050880429 |
| MIR8086 | -0.050885172 |
| RNU7-171P | -0.050885737 |
| DPYS | -0.05088735 |
| MIR548Y | -0.050888767 |
| RNU6-1246P | -0.050893251 |
| CCDC87 | -0.050893993 |
| RP13-150K15 | -0.050895141 |
| ABCD3 | -0.0509027 |
| CYCSP46 | -0.050910436 |
| CTB-111F10 | -0.05091097 |
| CTD-2384B11 | -0.050916691 |
| RP11-500K19 | -0.050923333 |
| FKBP4 | -0.050927204 |
| NXPH3 | -0.050933287 |
| CXADRP2 | -0.050939215 |
| AC019070 | -0.050947874 |
| UBE2V2 | -0.050948042 |
| RNU6-411P | -0.050949427 |
| Six3os1_1 | -0.050949834 |
| RBM24 | -0.050951735 |
| RPS4XP14 | -0.050952062 |
| RP11-120K18 | -0.050952131 |
| RP11-1085N6 | -0.050956835 |
| KIAA1958 | -0.050958423 |
| CTC-340I23 | -0.050961099 |
| CHCHD2P4 | -0.050961433 |
| AL354896 | -0.050966176 |
| RP11-265E18 | -0.050977091 |
| CTAGE15 | -0.050982585 |
| RP11-243J18 | -0.050985035 |
| AF111168 | -0.050997647 |
| RN7SKP179 | -0.051002423 |
| CYP3A4 | -0.051002821 |
| INTS6-AS1 | -0.051010128 |
| RP11-308D16 | -0.051012472 |
| MIR1539 | -0.051012897 |

| | |
|---|---|
| C1DP2 | -0.051013819 |
| RP11-325K19 | -0.051020164 |
| LRRC10B | -0.051026947 |
| FAF2 | -0.051028008 |
| LSM14A | -0.051030743 |
| RP11-426A6 | -0.05103377 |
| RP11-345P4 | -0.051036008 |
| AL157372 | -0.051049465 |
| CCDC30 | -0.051055246 |
| MIR8055 | -0.051057123 |
| FBXO22 | -0.051059005 |
| SDCBP2-AS1 | -0.051061833 |
| AC078974 | -0.051062172 |
| ASB16-AS1 | -0.051065014 |
| YIPF6 | -0.051066465 |
| RNU6ATAC42P | -0.051068733 |
| AE000660 | -0.051070428 |
| CTD-2331H12 | -0.051074038 |
| RP5-1009E24 | -0.05107429 |
| SH3YL1 | -0.051087564 |
| AC024084 | -0.051106551 |
| NR0B2 | -0.051110305 |
| AC091304 | -0.051112085 |
| HMGN2P24 | -0.051136573 |
| ENPP7P9 | -0.051138886 |
| HSPE1P25 | -0.051147232 |
| MIR5003 | -0.051154342 |
| BPIFB9P | -0.051154578 |
| RP11-430A19 | -0.05115835 |
| SNORA60 | -0.051173328 |
| RP11-648K4 | -0.051174503 |
| RP11-147C23 | -0.051181332 |
| ATF6B | -0.05118688 |
| OR10G8 | -0.051193068 |
| IGHVII-1-1 | -0.051195621 |
| C12orf40 | -0.051197971 |
| ERBB2IP | -0.051198881 |
| BTG1 | -0.051200071 |
| LAPTM5 | -0.051206044 |
| RN7SL15P | -0.051214389 |
| ABCA10 | -0.051216556 |
| TMCO6 | -0.051222082 |
| GRIPAP1 | -0.051223868 |
| PEBP1P3 | -0.051230548 |
| RP11-202D1 | -0.051230685 |

| | |
|---|---|
| RNU6-605P | -0.051237906 |
| TMEM63B | -0.051238361 |
| RP11-629O1 | -0.051238938 |
| FBP1 | -0.051241822 |
| MIR8056 | -0.051245823 |
| RP11-476H20 | -0.051258597 |
| MRPL32 | -0.051265563 |
| CTD-2647L4 | -0.051266772 |
| RP11-152N13 | -0.051269302 |
| JMJD1C | -0.05127174 |
| RP11-255C15 | -0.051279452 |
| RP11-447D11 | -0.051279975 |
| SNX27 | -0.051284261 |
| AF015720 | -0.051291114 |
| AC010297 | -0.051293603 |
| RP11-498M14 | -0.051293856 |
| RNU1-32P | -0.051295789 |
| SCFD2 | -0.051296522 |
| UTP14C | -0.051298811 |
| OR10S1 | -0.051311877 |
| VN1R79P | -0.051316453 |
| RNA5SP248 | -0.051321611 |
| MIR6864 | -0.0513244 |
| AL035457 | -0.051329409 |
| SGOL1P2 | -0.051332009 |
| RP11-106A1 | -0.051332598 |
| RNA5SP437 | -0.051338995 |
| PSMD8P1 | -0.051342699 |
| CCL1 | -0.051345677 |
| RP11-343D24 | -0.051349932 |
| PLAGL2 | -0.05135085 |
| RP11-128N14 | -0.051354233 |
| CENPIP1 | -0.051356296 |
| RP11-214O1 | -0.051356641 |
| TTC21A | -0.051359706 |
| RP11-325L7 | -0.051365118 |
| BRWD3 | -0.051367353 |
| GRIN1 | -0.051381633 |
| RNU6-746P | -0.051385047 |
| SNORD48 | -0.051386549 |
| RCOR3 | -0.051387503 |
| RP1-151F17 | -0.051393233 |
| TSIX | -0.051394821 |
| NICN1 | -0.051398927 |
| CH17-52D20 | -0.051402122 |

| | |
|---|---|
| ANKRD20A2 | -0.051403432 |
| SEPHS1P7 | -0.051406787 |
| ST3GAL5-AS1 | -0.051408805 |
| KB-1507C5 | -0.051409736 |
| RP11-569G13 | -0.051410692 |
| FAM216A | -0.051412025 |
| C18orf42 | -0.05141576 |
| RP11-616K22 | -0.051419101 |
| RNU4-77P | -0.051421116 |
| RP11-480D4 | -0.051421914 |
| RPL7P60 | -0.05142344 |
| ENPP7P6 | -0.05142685 |
| NFU1 | -0.051433179 |
| RNU6-203P | -0.051441436 |
| RNA5SP132 | -0.051441445 |
| C2orf57 | -0.051442569 |
| RAD1P2 | -0.051449553 |
| RP5-1168A5 | -0.051459497 |
| SNORD115-22 | -0.051463061 |
| SERPINB6 | -0.051472307 |
| HSD3B2 | -0.051475336 |
| RP13-228J13 | -0.051480868 |
| RP5-973N23 | -0.051484309 |
| STAG3L5P-PVRIG2P-PILRB | -0.051486502 |
| MCM6 | -0.051487061 |
| IMPA1P | -0.051487225 |
| ACTBP6 | -0.051488555 |
| HAR1B | -0.051497414 |
| DOC2B | -0.051498094 |
| SSH1 | -0.051499043 |
| RP11-296O14 | -0.051500475 |
| MNAT1 | -0.05150241 |
| PWP1 | -0.051503006 |
| MRPL57P6 | -0.051506396 |
| RNU6-751P | -0.051521527 |
| RPL7AP2 | -0.051531487 |
| INTS4 | -0.051532663 |
| SAGE2P | -0.051533021 |
| AC005822 | -0.051543786 |
| KRT43P | -0.051548339 |
| NXF5 | -0.051548376 |
| RMDN1 | -0.051562338 |
| MIR1273C | -0.051564953 |
| RP11-392A14 | -0.051569193 |
| NR1I2 | -0.051572719 |

| | |
|---|---|
| TUBGCP5 | -0.051577427 |
| RNU2-33P | -0.051577545 |
| CYP20A1 | -0.051579009 |
| PRDX3P2 | -0.051579728 |
| HNRNPA1P19 | -0.051586933 |
| RP11-1277A3 | -0.051591475 |
| RNU1-84P | -0.05159257 |
| RNU6-80P | -0.051592865 |
| RN7SKP37 | -0.051594605 |
| LNX2 | -0.051602098 |
| SEH1L | -0.051602415 |
| OR4N5 | -0.051603544 |
| AL512506 | -0.051606451 |
| RNU6-924P | -0.051606744 |
| RP11-471G13 | -0.051610071 |
| LINC00377 | -0.051619335 |
| RNY1P16 | -0.051620357 |
| TRIM53BP | -0.051626214 |
| TMUB2 | -0.05162662 |
| RTEL1P1 | -0.051626626 |
| ETF1P2 | -0.051630056 |
| PGM5P4-AS1 | -0.051630639 |
| SORD | -0.051633393 |
| AC092162 | -0.051633925 |
| RPS23P5 | -0.051635984 |
| MAP4K2 | -0.051640835 |
| SAA3P | -0.051641096 |
| AC012506 | -0.051644129 |
| AC144530 | -0.051644854 |
| LL22NC03-104C7 | -0.05164511 |
| ADCY1 | -0.051648035 |
| TNXB | -0.051651086 |
| IQCH | -0.051657338 |
| ZNF674 | -0.051659012 |
| RP11-2C7 | -0.051664959 |
| TRAJ27 | -0.051668482 |
| RP11-718G2 | -0.051671127 |
| CTD-2302E22 | -0.051673432 |
| ZNF658 | -0.05167358 |
| AP001271 | -0.051677668 |
| TNPO2 | -0.051678455 |
| RP11-688I9 | -0.051682576 |
| EIF2S3 | -0.051683665 |
| RPL23AP69 | -0.051683741 |
| SETP5 | -0.051684774 |

| | |
|---|---|
| RP11-276H1 | -0.051688303 |
| CTD-2005H7 | -0.051694879 |
| SS18L2P1 | -0.05169579 |
| RP11-421H10 | -0.051700692 |
| CSNK1D | -0.05170361 |
| TCEB1P33 | -0.051706228 |
| RP11-328K2 | -0.051707993 |
| CPD | -0.051711796 |
| MIR4670 | -0.051717813 |
| RP11-274A11 | -0.051717938 |
| LINC00427 | -0.051718947 |
| FUBP3 | -0.05172508 |
| CYCSP29 | -0.051728194 |
| THRAP3 | -0.051729248 |
| U73166 | -0.051730234 |
| RPSAP6 | -0.051730662 |
| ANKRD26P2 | -0.051731839 |
| TAS2R14 | -0.051737676 |
| RPS15AP10 | -0.051738482 |
| SLC30A4 | -0.051747965 |
| RP11-231P20 | -0.051749512 |
| GPR132 | -0.051760021 |
| RP11-503D12 | -0.051765012 |
| KB-1042C11 | -0.05177931 |
| RP3-441A12 | -0.051784447 |
| OR4K7P | -0.051788712 |
| RP13-619I2 | -0.051789049 |
| RBM19 | -0.051789809 |
| RABGAP1 | -0.05179033 |
| RP11-272K23 | -0.051792006 |
| ZC3H4 | -0.051792321 |
| RN7SKP285 | -0.051795195 |
| RP1-288L1 | -0.051797762 |
| RNU6-1038P | -0.051799978 |
| ADORA2A-AS1 | -0.051802892 |
| TTC39C-AS1 | -0.05180331 |
| AL391095 | -0.051807161 |
| RN7SL49P | -0.05180969 |
| MT-TG | -0.051838188 |
| AC005578 | -0.051843724 |
| CHAD | -0.051856212 |
| USP32 | -0.051863673 |
| RP4-635E18 | -0.051872212 |
| TRIM65 | -0.051874613 |
| ZNF781 | -0.051876771 |

| | | |
|---|---|---|
| AP003471 | | -0.051876901 |
| CYCSP26 | | -0.051879012 |
| RP11-196B3 | | -0.051883196 |
| RP11-84G21 | | -0.051886632 |
| RP11-350G13 | | -0.051886908 |
| ELMOD2 | | -0.051893144 |
| SLC25A31 | | -0.051893475 |
| RP11-46D1 | | -0.05189504 |
| MAN2C1 | | -0.051895265 |
| KIAA1524 | | -0.051896493 |
| MRPL40P1 | | -0.051901865 |
| SUOX | | -0.051904139 |
| RNU6-1036P | | -0.051906887 |
| RPL7AP56 | | -0.051908081 |
| RP1-223D17 | | -0.051913449 |
| CYB561 | | -0.051914168 |
| C1orf194 | | -0.051924815 |
| RP11-549J18 | | -0.051925926 |
| CARTPT | | -0.051928147 |
| RN7SL581P | | -0.051930446 |
| MIR1295A | | -0.051931568 |
| FANCD2P2 | | -0.051936677 |
| RP11-350N15 | | -0.051937462 |
| ARHGAP4 | | -0.051939601 |
| COPS7B | | -0.051940941 |
| LINC00621 | | -0.051942163 |
| CNOT11 | | -0.051960681 |
| ELK4 | | -0.051965351 |
| RNA5SP516 | | -0.051966622 |
| RPS17P17 | | -0.05196734 |
| RP11-192B18 | | -0.051969465 |
| SLC26A11 | | -0.051969593 |
| CDC25C | | -0.051976143 |
| | 9-Sep | -0.051976366 |
| SPDYE6 | | -0.051978645 |
| HNRNPA1P53 | | -0.051985182 |
| NPPA | | -0.051986132 |
| SLC9B1P1 | | -0.051986835 |
| RP11-197P3 | | -0.05198848 |
| RP11-21L23 | | -0.051990562 |
| OPA1 | | -0.051991641 |
| LINC00441 | | -0.051993014 |
| RP5-1107A17 | | -0.051994474 |
| RN7SL543P | | -0.051996798 |
| RP11-185N2 | | -0.051997284 |

| | |
|---|---|
| TCTN1 | -0.05200673 |
| MIR6165 | -0.05200755 |
| TEX10 | -0.052008367 |
| WARSP1 | -0.052012285 |
| PAAF1 | -0.052015047 |
| RNA5SP339 | -0.052018239 |
| POU5F1B | -0.052027382 |
| OR5D2P | -0.052028731 |
| ISL2 | -0.052031265 |
| RNA5SP209 | -0.052034864 |
| RNA5SP467 | -0.052037393 |
| PRKAR2A-AS1 | -0.052037634 |
| AC110926 | -0.052040797 |
| MKRN2 | -0.052043104 |
| OSBPL11 | -0.05204755 |
| DUSP5 | -0.052049225 |
| PGPEP1 | -0.052057802 |
| RP11-142J21 | -0.052058947 |
| RP11-202H2 | -0.052060535 |
| RP11-384K6 | -0.052060537 |
| FAM19A1 | -0.052062359 |
| MYO18A | -0.052068519 |
| RP3-433G13 | -0.052071297 |
| C1QA | -0.052073486 |
| C19orf45 | -0.052083078 |
| PCMTD1P2 | -0.052088479 |
| AL158040 | -0.052090881 |
| ANKIB1 | -0.052091608 |
| RP4-620F22 | -0.052093454 |
| MTND5P23 | -0.052095764 |
| EIF4G2 | -0.052100409 |
| SZT2 | -0.052105041 |
| AC096558 | -0.052106635 |
| RP11-5P22 | -0.052109172 |
| NPM1P34 | -0.052109273 |
| RNU6-5P | -0.052109867 |
| MTND6P20 | -0.052112791 |
| TTLL10-AS1 | -0.052127266 |
| CTD-2033D15 | -0.052130084 |
| PHACTR3 | -0.052136373 |
| CTD-2373H9 | -0.052145785 |
| PPP5C | -0.052146606 |
| CICP8 | -0.052146862 |
| AC009974 | -0.052148144 |
| RNF10 | -0.052151387 |

| | |
|---|---|
| MIR595 | -0.052153935 |
| REXO1L1P | -0.052155005 |
| CTB-70G10 | -0.052157304 |
| TSPY26P | -0.052158848 |
| OOSP1 | -0.05216022 |
| FBXO36 | -0.052165137 |
| HMGN2P28 | -0.052167275 |
| RNU6-101P | -0.05217275 |
| MMEL1 | -0.052185101 |
| SLC10A2 | -0.052187363 |
| AC007389 | -0.052209635 |
| RP11-524F11 | -0.052209771 |
| RP11-50E11 | -0.052216045 |
| PPP1R2P10 | -0.05221929 |
| RP11-142M10 | -0.052221597 |
| CTD-2002J20 | -0.052221725 |
| AC004237 | -0.052232892 |
| RP11-585P4 | -0.052234033 |
| RP11-946P6 | -0.052234981 |
| AC018647 | -0.052235312 |
| RP11-429O2 | -0.052242606 |
| RP11-775E10 | -0.052252783 |
| RP11-158I3 | -0.052256548 |
| ATP5B | -0.052259895 |
| RPS4XP3 | -0.052260472 |
| CTD-2116N20 | -0.052262848 |
| ZNF573 | -0.052266611 |
| RP11-62G11 | -0.052274357 |
| HDGFRP3 | -0.052279114 |
| KRBA2 | -0.052286381 |
| BEX5 | -0.052289412 |
| CENPJ | -0.052291952 |
| RP11-104C4 | -0.052296511 |
| HNF4G | -0.052299462 |
| ARL6IP1 | -0.052299636 |
| IFNAR1 | -0.052302706 |
| ISG20 | -0.05231986 |
| MESTP1 | -0.052327429 |
| VN1R7P | -0.052328032 |
| RP11-1212A22 | -0.052338805 |
| SLC2A3P2 | -0.052344826 |
| RP11-503C24 | -0.05235007 |
| C14orf159 | -0.052352792 |
| LY6G5C | -0.052354094 |
| SUCLA2 | -0.052355471 |

| | |
|---|---|
| OR7E25P | -0.052358363 |
| MIR448 | -0.052363404 |
| SLC22A7 | -0.05237018 |
| ANKRD20A5P | -0.052371655 |
| OR5H4P | -0.052374898 |
| AL110292 | -0.052376451 |
| EIF4A2P3 | -0.052377542 |
| FAUP1 | -0.052383779 |
| RPRD2 | -0.052386375 |
| RP11-486O12 | -0.052387528 |
| CDH1 | -0.052389686 |
| MRPS17P5 | -0.052392783 |
| POLR3B | -0.05239391 |
| MTND1P5 | -0.052396929 |
| AC011995 | -0.052397662 |
| MAGI1-IT1 | -0.05240848 |
| RNU6-595P | -0.052411714 |
| MTND6P6 | -0.05241672 |
| AL109800 | -0.052418114 |
| PEBP1 | -0.052418469 |
| ACKR1 | -0.052433285 |
| GOLGA8EP | -0.052436563 |
| GRM7-AS1 | -0.052443657 |
| AL161898 | -0.052447826 |
| DHFRP2 | -0.052452165 |
| PHF2 | -0.052455271 |
| MIR4677 | -0.052456752 |
| PJA2 | -0.052459475 |
| CEACAM3 | -0.052461633 |
| AC022021 | -0.052466323 |
| TRIM33 | -0.052468406 |
| CCNB3 | -0.052472276 |
| RP5-1121E10 | -0.052473849 |
| MIR8077 | -0.052476893 |
| MIR337 | -0.052478695 |
| RP11-445J14 | -0.052479552 |
| AL928742 | -0.05248487 |
| OR2V1 | -0.052493567 |
| EMC9 | -0.05249611 |
| GJA10 | -0.052498238 |
| MTND1P29 | -0.052502401 |
| MIR516B2 | -0.052506216 |
| TRIM24 | -0.052508259 |
| AL138479 | -0.052509613 |
| RN7SL664P | -0.052514605 |

| | |
|---|---|
| TMEM155 | -0.052515197 |
| RP11-42L13 | -0.052518524 |
| ZNF627 | -0.052518715 |
| C5orf60 | -0.052519829 |
| AL137024 | -0.052522181 |
| RNU6-589P | -0.052522747 |
| RP11-198M15 | -0.0525282 |
| LINC00380 | -0.05252851 |
| RP4-591L5 | -0.0525288 |
| KCNV1 | -0.052538135 |
| BCDIN3D-AS1 | -0.052538586 |
| LRIG2 | -0.052542363 |
| CTC-525D6 | -0.052546496 |
| MED28 | -0.052546661 |
| AC091813 | -0.052548702 |
| NDE1 | -0.052567908 |
| RP11-231C14 | -0.05257261 |
| PAN3-AS1 | -0.052573285 |
| RP11-445F6 | -0.052573811 |
| RPL7AL3 | -0.052580035 |
| OSTCP1 | -0.052580191 |
| KRTAP8-1 | -0.052586204 |
| XXbac-B444P24 | -0.052588356 |
| SSBP3-AS1 | -0.052596951 |
| EIF4BP1 | -0.052599776 |
| ZBTB42 | -0.052600803 |
| USP27X-AS1 | -0.05260244 |
| LIFR | -0.052605621 |
| GCN1L1 | -0.052606273 |
| MRPL3P1 | -0.052609146 |
| ADAMTS7P4 | -0.052610775 |
| MIR596 | -0.052614363 |
| ALDH7A1P4 | -0.052615547 |
| PREX2 | -0.052619036 |
| AC002542 | -0.052620206 |
| GSTA8P | -0.052624017 |
| RP11-329A6 | -0.052626082 |
| CHRNA6 | -0.052629832 |
| HLA-A | -0.052635669 |
| EFR3A | -0.05263699 |
| MAGOH2P | -0.052640323 |
| AC073869 | -0.052647431 |
| PIM3 | -0.052648619 |
| SCYL3 | -0.052658945 |
| RP5-823G15 | -0.052662339 |

| | |
|---|---|
| RP11-467K18 | -0.052664419 |
| ANKRD63 | -0.052668541 |
| PGBD4P2 | -0.052669453 |
| VN1R104P | -0.052673805 |
| AP001469 | -0.052673829 |
| NEDD4L | -0.052673866 |
| LRRC69 | -0.052675243 |
| RP11-330C7 | -0.05267612 |
| RPL7AP15 | -0.05267758 |
| ITFG1-AS1 | -0.052681805 |
| RP11-1228E12 | -0.052684326 |
| RNA5SP491 | -0.052684833 |
| RPL7AP23 | -0.052685119 |
| RP11-65J3 | -0.052686607 |
| MIR149 | -0.052689291 |
| B3GNTL1 | -0.052690332 |
| RP11-425A6 | -0.052695433 |
| HSPA8 | -0.052696193 |
| CTD-2014E2 | -0.052699473 |
| AC104938 | -0.052703086 |
| SETP11 | -0.052703347 |
| AC104841 | -0.052703785 |
| ZC2HC1A | -0.052705123 |
| COL28A1 | -0.052716287 |
| IGHMBP2 | -0.052716547 |
| RP1-276E15 | -0.052721866 |
| LINC00618 | -0.052723064 |
| ANKRD13A | -0.052728231 |
| MIR3671 | -0.052732883 |
| RP11-76H14 | -0.052734596 |
| RP11-80H5 | -0.052736609 |
| ZNF491 | -0.05274399 |
| CTD-2161F6 | -0.052745302 |
| CRAMP1L | -0.052751693 |
| RNU6-853P | -0.052754968 |
| PSD4 | -0.052776606 |
| KIAA1731NL | -0.052777995 |
| RAD23B | -0.052794504 |
| GS1-358P8 | -0.052797946 |
| RPL19P21 | -0.052799374 |
| MIR4755 | -0.052803791 |
| HLCS | -0.052806728 |
| TBC1D17 | -0.052814449 |
| AC062016 | -0.052825507 |
| RNU6-991P | -0.052826193 |

| | |
|---|---|
| POM121L13P | -0.052828193 |
| STK26 | -0.05282911 |
| RP11-334C17 | -0.052843269 |
| CTD-2026D20 | -0.052848981 |
| NCAPG2 | -0.052851344 |
| MIR629 | -0.05285652 |
| RP11-138A23 | -0.052874246 |
| FERMT3 | -0.052875624 |
| KRBOX4 | -0.052878247 |
| RP13-210D15 | -0.052885641 |
| REXO1L6P | -0.052886336 |
| AP001001 | -0.052886336 |
| C7orf72 | -0.05288656 |
| WDR36 | -0.052889094 |
| RNU6-1096P | -0.05289302 |
| RP11-253I19 | -0.052897711 |
| RP11-678B3 | -0.052902892 |
| AC105389 | -0.05290739 |
| AGBL5 | -0.052908574 |
| RP11-350G24 | -0.052908985 |
| EPS15L1 | -0.052915477 |
| AL022326 | -0.052923137 |
| RP11-506H21 | -0.052929504 |
| CTD-2356P16 | -0.052932127 |
| DHX35 | -0.052941729 |
| RNA5SP451 | -0.052948745 |
| AC073636 | -0.052950435 |
| SHANK2-AS1 | -0.052954957 |
| LGSN | -0.052955335 |
| FKSG54 | -0.052964678 |
| RNU6-477P | -0.052968757 |
| AJ239321 | -0.052972463 |
| C3orf52 | -0.052979126 |
| RNU6-195P | -0.052981057 |
| CHST5 | -0.052985158 |
| EIF1P2 | -0.052987161 |
| RP11-426K3 | -0.052987899 |
| CHMP1B | -0.052990103 |
| RNA5SP454 | -0.052991369 |
| SNX32 | -0.052995052 |
| C14orf28 | -0.053003295 |
| PSTPIP2 | -0.053003481 |
| RPL9P29 | -0.053003834 |
| TCEA1P2 | -0.053004338 |
| ZNF483 | -0.053009993 |

| | |
|---|---|
| NFAT5 | -0.053011779 |
| AC096971 | -0.053013321 |
| RPL23AP50 | -0.053015475 |
| RPS15AP18 | -0.053015931 |
| RP11-75D3 | -0.053022817 |
| RP11-1096G20 | -0.053024174 |
| RP11-61O11 | -0.053034484 |
| CYCSP43 | -0.053034887 |
| MIR1283-1 | -0.053044891 |
| UQCRBP1 | -0.053054061 |
| LINC01229 | -0.053060575 |
| MTATP6P4 | -0.053065577 |
| RP11-170K4 | -0.053066843 |
| RP11-49G10 | -0.053068758 |
| RP11-557B23 | -0.05307109 |
| TMEM116 | -0.053073047 |
| RP11-335F8 | -0.053077239 |
| AC002383 | -0.053084027 |
| RPL7P46 | -0.053085992 |
| RPL17P33 | -0.053092948 |
| OR4D11 | -0.053095443 |
| FKTN | -0.053097922 |
| RN7SL629P | -0.053099486 |
| RP11-494O16 | -0.053100284 |
| PLEKHG7 | -0.053107782 |
| GPR84 | -0.053109644 |
| DPEP3 | -0.053113186 |
| ECD | -0.05311809 |
| TAS2R46 | -0.053118966 |
| RPL23AP22 | -0.053130401 |
| MIR143 | -0.053132871 |
| TCEB1P30 | -0.053133693 |
| PPIL2 | -0.053137515 |
| SSX2IP | -0.053142408 |
| RNU6-393P | -0.053160352 |
| RP11-75A5 | -0.053162907 |
| RP11-737G21 | -0.053163506 |
| ZNF32-AS1 | -0.05316949 |
| RNU6-32P | -0.053175557 |
| MORC3 | -0.053179823 |
| C22orf39 | -0.053184661 |
| GOT2 | -0.053190242 |
| GCNT1P1 | -0.053193005 |
| ZNF572 | -0.053193666 |
| AC068287 | -0.053195277 |

| | |
|---|---|
| KB-1639H6 | -0.053195659 |
| RP5-890O3 | -0.053199638 |
| TMEM144 | -0.053201034 |
| BTBD17 | -0.053205646 |
| RP11-73K9 | -0.053209257 |
| FAM220BP | -0.0532111 |
| RNU6-879P | -0.053211356 |
| NKX6-3 | -0.053212025 |
| RP13-278H16 | -0.053216069 |
| KIAA0319 | -0.053218345 |
| BCL9P1 | -0.053220797 |
| AC007248 | -0.053223603 |
| RP11-653J6 | -0.053223863 |
| PTP4A1P7 | -0.053225382 |
| OR5AC1 | -0.053225766 |
| MIR654 | -0.053231503 |
| KRT18P39 | -0.05324146 |
| RP11-1144P22 | -0.053256476 |
| MIR6813 | -0.053260441 |
| MIR1227 | -0.053262298 |
| BMPR1APS1 | -0.05326438 |
| SP3P | -0.053277251 |
| PDCL3P7 | -0.053280557 |
| CCDC78 | -0.053283753 |
| RP11-423C15 | -0.053285568 |
| snosnR66 | -0.053295779 |
| EFR3B | -0.05329648 |
| CTB-47B8 | -0.053305248 |
| LRRC27 | -0.053307513 |
| ADIG | -0.053307866 |
| ACTG1P11 | -0.053308335 |
| ATRX | -0.053314657 |
| ADAP2 | -0.05331862 |
| HTR3D | -0.053321351 |
| AL135878 | -0.053323914 |
| RNU6-363P | -0.053328957 |
| AL161452 | -0.053330701 |
| DTWD1 | -0.053331439 |
| CYTH1 | -0.053332128 |
| KCNC1 | -0.053336558 |
| CR392039 | -0.053336968 |
| ZNF20 | -0.053340277 |
| AP001462 | -0.053355136 |
| RNA5SP112 | -0.053355322 |
| CTA-276F8 | -0.053355422 |

| | |
|---|---|
| RNA5SP311 | -0.053356099 |
| RP11-275H4 | -0.053360261 |
| RP11-94P14 | -0.05336997 |
| ME2P1 | -0.053371266 |
| RP11-76E12 | -0.05337462 |
| NVL | -0.053375944 |
| OR52I1 | -0.053376017 |
| RP1-34P24 | -0.05337753 |
| HID1-AS1 | -0.053378714 |
| KRT18P23 | -0.053381163 |
| RP11-811J10 | -0.053382375 |
| LINC01216 | -0.053386765 |
| RP11-946L20 | -0.05339238 |
| CYP3A5 | -0.053395405 |
| FAM66C | -0.053397563 |
| SNAPC3 | -0.053400131 |
| HLA-DRB9 | -0.053402079 |
| RP11-270M14 | -0.053405103 |
| ZNF112 | -0.05340921 |
| OR2A7 | -0.053410307 |
| ASH2LP1 | -0.053412599 |
| FEN1 | -0.053415796 |
| RPL5P18 | -0.053417427 |
| PRDX3P1 | -0.053419754 |
| MAGEB6P1 | -0.05342058 |
| RP11-427L15 | -0.053424244 |
| EXOC7P1 | -0.053428423 |
| AC096507 | -0.053430963 |
| RPL10AP5 | -0.053432622 |
| RNU6-490P | -0.053437877 |
| RP5-828H9 | -0.053445419 |
| AF131215 | -0.053452338 |
| KBTBD3 | -0.053454961 |
| RP11-466M21 | -0.053458514 |
| RNA5SP403 | -0.053462176 |
| POMK | -0.053465343 |
| AMMECR1L | -0.053466887 |
| OR51G1 | -0.053471446 |
| NPIPA5 | -0.053473018 |
| DSTYK | -0.053477027 |
| DBT | -0.053477232 |
| NIPA2P1 | -0.053488977 |
| RP11-1109M24 | -0.053489188 |
| RP5-905N1 | -0.053504468 |
| AP000223 | -0.053507266 |

| | |
|---|---|
| AC137674 | -0.05351048 |
| FMO3 | -0.053513829 |
| C9orf41-AS1 | -0.053518824 |
| ZFP69 | -0.053520192 |
| RN7SL434P | -0.053522938 |
| AC096921 | -0.053526644 |
| HNRNPKP5 | -0.053527764 |
| CTA-223H9 | -0.053532226 |
| PIK3R6 | -0.053536118 |
| MIR567 | -0.053538942 |
| KLHL1 | -0.053541627 |
| AC021613 | -0.05354529 |
| HLA-H | -0.053547207 |
| RP5-821D11 | -0.053549421 |
| GPSM2 | -0.053557886 |
| RP11-149I9 | -0.053567312 |
| RP11-41O4 | -0.05356838 |
| USP9YP34 | -0.05356874 |
| UBE2V1P7 | -0.05356874 |
| CT45A6 | -0.05356874 |
| AC009508 | -0.05357039 |
| RP11-1079K10 | -0.053577099 |
| CHD9 | -0.053584474 |
| AL034553 | -0.053591556 |
| LINC00880 | -0.053596031 |
| EYA3 | -0.0535964 |
| RP4-785G19 | -0.053597274 |
| VWA7 | -0.053599042 |
| AMDP1 | -0.053602403 |
| CTA-796E4 | -0.053607597 |
| ZNF429 | -0.053610279 |
| RPS3AP23 | -0.05361512 |
| EBLN2 | -0.053615526 |
| AC078828 | -0.053618309 |
| ALG9 | -0.053619452 |
| MIR4752 | -0.05362 |
| MNS1 | -0.053620773 |
| SETP20 | -0.053621145 |
| ST13 | -0.05362174 |
| AL136040 | -0.053627931 |
| MMP23A | -0.053628213 |
| CCDC166 | -0.053633678 |
| LINC00317 | -0.05363414 |
| RNU6-652P | -0.053649929 |
| MT-ND4 | -0.053653407 |

| | |
|---|---|
| RP11-67L14 | -0.053659477 |
| RP11-111K18 | -0.053662129 |
| RNU7-67P | -0.053662743 |
| AP003498 | -0.053667497 |
| RP11-105C19 | -0.053678929 |
| NAP1L2 | -0.053680771 |
| TLX1NB | -0.053682161 |
| ZNRD1-AS1_2 | -0.053682511 |
| DLG3-AS1 | -0.053684079 |
| MIR8078 | -0.053695555 |
| C10orf82 | -0.053697314 |
| RPL29P32 | -0.053700869 |
| TXNP5 | -0.053704411 |
| USP46 | -0.053709502 |
| CTRL | -0.053709774 |
| MRPL42 | -0.053709837 |
| SLC25A17 | -0.05371114 |
| RNU6-19P | -0.053724437 |
| UGT2B10 | -0.053727751 |
| RP11-285E23 | -0.053729777 |
| LA16c-349E10 | -0.053730807 |
| GTPBP10 | -0.053738229 |
| TMPOP1 | -0.053748352 |
| ART2P | -0.0537534 |
| OR7E24 | -0.053757555 |
| RP11-333O1 | -0.053759588 |
| TECPR1 | -0.053761523 |
| RNU6-100P | -0.053764262 |
| OR5B19P | -0.053764465 |
| ATF5 | -0.053766417 |
| SNX2 | -0.053773877 |
| RP11-15J22 | -0.053777283 |
| LINC01089 | -0.053779738 |
| CICP6 | -0.053789094 |
| LINC01499 | -0.053789499 |
| RNA5SP296 | -0.053793882 |
| SNRPGP15 | -0.053795405 |
| UBE4B | -0.053797376 |
| RNU6-1256P | -0.053800808 |
| TUG1 | -0.053802437 |
| UBE2D4 | -0.053805358 |
| PSMA2P1 | -0.053806908 |
| GUCY1B2 | -0.053811555 |
| AC110754 | -0.05381173 |
| GPR158-AS1 | -0.053814328 |

| | |
|---|---|
| ANKLE2 | -0.053814828 |
| AC141586 | -0.05381734 |
| RPL27AP | -0.053818796 |
| RP13-359K18 | -0.053828314 |
| ASH1L | -0.05384069 |
| SNORD115-12 | -0.053841702 |
| AC093611 | -0.053841702 |
| AL391867 | -0.053841702 |
| RNU6-1023P | -0.053841702 |
| MIR5011 | -0.053841702 |
| AC016194 | -0.053841702 |
| RMRPP5 | -0.053841702 |
| RNU6-566P | -0.053841702 |
| AC105245 | -0.053841702 |
| FABP12P1 | -0.053841702 |
| MIR3169 | -0.053841702 |
| AC091321 | -0.053841702 |
| AC134729 | -0.053841702 |
| AC092925 | -0.053841702 |
| RNU6-180P | -0.053841702 |
| RNA5SP407 | -0.053841702 |
| MIR526A2 | -0.053841702 |
| RNA5SP380 | -0.053841702 |
| MIR4770 | -0.053841702 |
| KF459411 | -0.053841702 |
| AL590726 | -0.053841702 |
| MIR297 | -0.053841702 |
| AC016113 | -0.053841702 |
| RNU6-1007P | -0.053841702 |
| AL356739 | -0.053841702 |
| AL354741 | -0.053841702 |
| RNU6ATAC37P | -0.053841702 |
| MIR513A2 | -0.053841702 |
| RNU6-224P | -0.053841702 |
| RNA5SP446 | -0.053841702 |
| AC104687 | -0.053841702 |
| RNU6-1230P | -0.053841702 |
| MIR516A1 | -0.053841702 |
| RNU4-55P | -0.053841702 |
| AC069243 | -0.053841702 |
| MIR514A3 | -0.053841702 |
| AC078988 | -0.053841702 |
| RNU6-330P | -0.053841702 |
| AC019051 | -0.053841702 |
| RNU4ATAC8P | -0.053841702 |

| | |
|---|---|
| AP005638 | -0.053841702 |
| MIR4330 | -0.053841702 |
| AF186996 | -0.053841702 |
| AC098847 | -0.053841702 |
| MIR4268 | -0.053841702 |
| AC026305 | -0.053841702 |
| AL109922 | -0.053841702 |
| RP11-656A10 | -0.053841702 |
| RNU6-741P | -0.053841702 |
| MIR7151 | -0.053841702 |
| RP11-569H17 | -0.053841702 |
| PPATP2 | -0.053841702 |
| DEFB108P4 | -0.053841702 |
| RP11-13M3 | -0.053841702 |
| AP000960 | -0.053841702 |
| BNIP3P6 | -0.053841702 |
| OR6C73P | -0.053841702 |
| MIR105-2 | -0.053841702 |
| RNU6-283P | -0.053841702 |
| AC068797 | -0.053841702 |
| SNORD2 | -0.053841702 |
| RN7SKP233 | -0.053841702 |
| RP4-657M3 | -0.053841702 |
| RP11-736K12 | -0.053841702 |
| AC093816 | -0.053841702 |
| AP003972 | -0.053841702 |
| AC097361 | -0.053841702 |
| MIR382 | -0.053841702 |
| AC034205 | -0.053841702 |
| AC004769 | -0.053841702 |
| RNU6-639P | -0.053841702 |
| AL591415 | -0.053841702 |
| RNU6-72P | -0.053841702 |
| RPEP5 | -0.053841702 |
| AC108739 | -0.053841702 |
| AC108211 | -0.053841702 |
| MIR3924 | -0.053841702 |
| MIR1912 | -0.053841702 |
| AC106860 | -0.053841702 |
| RP11-534P19 | -0.053841702 |
| C10orf32-ASMT | -0.053842518 |
| LINC01496 | -0.053849799 |
| RNU6-622P | -0.053857591 |
| NUDT19P5 | -0.053861107 |
| HMGN2P15 | -0.053872459 |

| | |
|---|---|
| C11orf80 | -0.053878291 |
| RPL7P31 | -0.053878891 |
| IGHVII-74-1 | -0.053879417 |
| RNU1-132P | -0.053884361 |
| MIR1271 | -0.053890169 |
| RNU4-24P | -0.053892527 |
| RP13-16H11 | -0.0538944 |
| RP11-751A18 | -0.053896388 |
| RPL15P18 | -0.053898347 |
| CLUHP2 | -0.053906047 |
| FAM120A | -0.053908358 |
| AC005538 | -0.053909357 |
| RN7SL198P | -0.053917318 |
| FBXO11 | -0.053921475 |
| ATXN7L2 | -0.053931708 |
| AC073343 | -0.05393533 |
| HTR4 | -0.053936903 |
| OR10J5 | -0.053940352 |
| RP11-412P11 | -0.053946961 |
| FDPSP8 | -0.053950112 |
| RP11-338E21 | -0.053954026 |
| RP11-398E10 | -0.053958098 |
| MIR3128 | -0.053967171 |
| DDX18P1 | -0.053980402 |
| DLGAP1-AS3 | -0.053983495 |
| LINC01189 | -0.053995239 |
| LHX3 | -0.053995571 |
| RNU7-74P | -0.054002722 |
| RPL9P16 | -0.054003561 |
| AC021887 | -0.054004111 |
| RP11-428K3 | -0.054004121 |
| ZNF322 | -0.054004455 |
| MAP3K2 | -0.054008623 |
| RP13-890H12 | -0.054013852 |
| EVL | -0.054016623 |
| MACROD2 | -0.054016828 |
| SGPP1 | -0.054023451 |
| PDK2 | -0.054026056 |
| RP5-1159O4 | -0.054028065 |
| RPL37P1 | -0.054030183 |
| CLRN1-AS1 | -0.054033226 |
| AC013444 | -0.054037312 |
| KATNBL1 | -0.05404215 |
| TMEM194B | -0.054051527 |
| RP11-528I4 | -0.054052073 |

| | |
|---|---|
| STARD5 | -0.054052308 |
| AC092881 | -0.05405289 |
| CTD-2086O20 | -0.054057899 |
| OR4A49P | -0.054061443 |
| YRDCP2 | -0.05406588 |
| AC062029 | -0.054068514 |
| EIF4BP6 | -0.054068607 |
| ROBO2 | -0.054080676 |
| RP4-728D4 | -0.054085605 |
| RP11-344J7 | -0.054089336 |
| UPF3AP2 | -0.054092824 |
| MIR3657 | -0.054114974 |
| TNPO1 | -0.054115325 |
| RPL17P44 | -0.054117811 |
| AC118653 | -0.054123034 |
| SYT15 | -0.054130191 |
| RP11-151A6 | -0.054130194 |
| Mico1 | -0.054132081 |
| MIRLET7G | -0.054148416 |
| EFCAB6 | -0.054150806 |
| RP11-440I14 | -0.054152075 |
| AP1S2 | -0.054160238 |
| RP11-363L24 | -0.054173803 |
| PHLPP1 | -0.054185858 |
| RP11-596C23 | -0.054185941 |
| AL356865 | -0.054186468 |
| RP11-671J11 | -0.05418848 |
| C21orf62-AS1 | -0.054190435 |
| RNU6-144P | -0.054193151 |
| AL121923 | -0.054197013 |
| RPL17-C18orf32 | -0.054201422 |
| RP11-417F21 | -0.054205737 |
| LRTM2 | -0.054211441 |
| TMEM50A | -0.054213208 |
| KLHL28 | -0.054217041 |
| RP11-202A13 | -0.054226698 |
| MCCC2 | -0.054230846 |
| AC068121 | -0.054230975 |
| AC132807 | -0.054231936 |
| CT45A3 | -0.054232485 |
| COQ6 | -0.054243777 |
| LYSMD3 | -0.054250382 |
| DEFB106A | -0.054253801 |
| RP11-585K14 | -0.054254458 |
| RP11-227P3 | -0.054261397 |

| | |
|---|---|
| RP11-174N3 | -0.054263341 |
| CD163 | -0.054273397 |
| CTC-359D24 | -0.054274755 |
| RP11-50B3 | -0.054280553 |
| GHRL | -0.054298979 |
| PREX1 | -0.054301584 |
| RMND5B | -0.054306403 |
| RP11-322L17 | -0.054314828 |
| CENPF | -0.054316115 |
| APC | -0.054317455 |
| MIR3612 | -0.054317455 |
| RP13-88F20 | -0.054323217 |
| TANGO6 | -0.05433539 |
| RP11-476F14 | -0.054335694 |
| RN7SL76P | -0.054337637 |
| GS1-214D18 | -0.054339798 |
| KLHL6-AS1 | -0.054341397 |
| GLUD2 | -0.054344563 |
| KRT8P51 | -0.054345983 |
| AC007246 | -0.054348154 |
| CTC-261N6 | -0.054352543 |
| DTNA | -0.054355056 |
| PRM1 | -0.054358595 |
| KRT36 | -0.054359744 |
| AC010329 | -0.054364334 |
| RN7SL368P | -0.054364487 |
| CTB-3M24 | -0.054364713 |
| RP11-578O24 | -0.054365592 |
| VN1R32P | -0.054368761 |
| GAS2 | -0.054376322 |
| AC005828 | -0.054376776 |
| RP11-197O8 | -0.054379327 |
| HLA-K | -0.054379755 |
| RNU7-94P | -0.054387626 |
| CTD-2515H24 | -0.054394715 |
| ZNF420 | -0.054397161 |
| AC016697 | -0.054406645 |
| TXNDC12-AS1 | -0.054407081 |
| RP11-197M22 | -0.054409361 |
| RP11-203B9 | -0.054413924 |
| AC241377 | -0.054420151 |
| RP11-161N3 | -0.054421142 |
| AP000560 | -0.054423753 |
| MIR1275 | -0.054425193 |
| MIR7848 | -0.054432585 |

| | |
|---|---|
| ARHGAP17 | -0.05443279 |
| RP11-264K23 | -0.054433953 |
| SCARNA12 | -0.054436417 |
| USP49 | -0.054438 |
| HSPE1P23 | -0.05443933 |
| CCNB1IP1P3 | -0.054441386 |
| KDM6A | -0.054444214 |
| ANKRD42 | -0.054460177 |
| AC093911 | -0.054463457 |
| AC067940 | -0.054476026 |
| AC090607 | -0.054480114 |
| PNPO | -0.054481797 |
| ZBTB43 | -0.054483602 |
| RPL13AP6 | -0.054488314 |
| ALDOB | -0.054489922 |
| AC010368 | -0.054491475 |
| RP11-338I21 | -0.054502269 |
| SPRY4-IT1 | -0.054503492 |
| ZNF57 | -0.054509649 |
| RABGGTB | -0.054517333 |
| ZNF862 | -0.054531306 |
| MIR3927 | -0.054535108 |
| OR6M3P | -0.054539541 |
| RUSC1-AS1 | -0.054541911 |
| NPM1P32 | -0.054546057 |
| SV2B | -0.054547455 |
| LINC00271 | -0.054563772 |
| COX6B1P6 | -0.054564767 |
| HMGN2P13 | -0.054569337 |
| AXIN2 | -0.054570898 |
| RNA5SP35 | -0.054577383 |
| SLCO1B1 | -0.054580609 |
| PMS2CL | -0.054581708 |
| SP140L | -0.054582099 |
| AC074035 | -0.054582134 |
| RNU6-1025P | -0.054589918 |
| PAQR3 | -0.05459369 |
| FAM27E4 | -0.054603038 |
| RP11-109D20 | -0.054603665 |
| RP3-341D10 | -0.054615041 |
| RP11-55K22 | -0.054617178 |
| RP11-123K19 | -0.054619359 |
| SEL1L | -0.054622156 |
| TMEM260 | -0.05462545 |
| NF1P5 | -0.054627416 |

| | |
|---|---|
| RNU4-49P | -0.054630644 |
| CTD-2320G14 | -0.054634233 |
| RP11-188P17 | -0.054635932 |
| MIR3683 | -0.054637691 |
| RP11-668G10 | -0.054646638 |
| AC012476 | -0.054646878 |
| RN7SL647P | -0.054647424 |
| RP11-299H21 | -0.054653842 |
| STMN1P1 | -0.054656126 |
| MGAT2P2 | -0.054656914 |
| RP4-785J1 | -0.054659966 |
| MIR218-1 | -0.054668628 |
| AC015977 | -0.054685104 |
| GS1-541M1 | -0.054691176 |
| AL121899 | -0.054700702 |
| FAM92A1P2 | -0.054701051 |
| TDRD6 | -0.054702263 |
| RP11-486E2 | -0.054711345 |
| KIAA0895 | -0.054717844 |
| TPMTP1 | -0.054722484 |
| AC083822 | -0.054724693 |
| CH507-9B2 | -0.054729212 |
| TIMM9P1 | -0.054732991 |
| RP1-15D23 | -0.054734737 |
| HIST3H2BA | -0.054735825 |
| TLR9 | -0.054736999 |
| RNU6-326P | -0.054737619 |
| MIR641 | -0.054739559 |
| KIF13A | -0.054742273 |
| MESTP2 | -0.054750148 |
| LINC00102 | -0.054750484 |
| AL035250 | -0.054751386 |
| PDCL2P2 | -0.054784299 |
| ITIH2 | -0.054785168 |
| CCDC40 | -0.054790424 |
| SNORA3 | -0.054791602 |
| MIR558 | -0.054794952 |
| MIR592 | -0.054796642 |
| RN7SL316P | -0.054796673 |
| ZSCAN5D | -0.054800035 |
| TRAV7 | -0.054802873 |
| RP5-893G23 | -0.054817279 |
| STAB2 | -0.054827551 |
| GID4 | -0.0548289 |
| AC005301 | -0.054833415 |

| | |
|---|---|
| FAM63B | -0.054836808 |
| AC092687 | -0.054842502 |
| RP4-754E20__A | -0.054843801 |
| RP11-522L3 | -0.054851506 |
| CTD-2141G3 | -0.054852572 |
| ECT2 | -0.054855432 |
| ZSCAN12P1 | -0.054864865 |
| SNX13 | -0.054867656 |
| PTMAP4 | -0.054869386 |
| LARP4P | -0.054871035 |
| MIR4672 | -0.054872635 |
| SNORA26 | -0.054872717 |
| RP11-288C18 | -0.054874428 |
| ENPP7P2 | -0.054880318 |
| RP11-11N7 | -0.054887284 |
| SHMT1P1 | -0.054889684 |
| AC073130 | -0.054894596 |
| ETAA1 | -0.054899266 |
| SCARNA15 | -0.054900495 |
| ARFIP1 | -0.054903806 |
| CTD-2587H24 | -0.054905481 |
| ISG20L2 | -0.054907899 |
| RP11-392O1 | -0.054908287 |
| RP11-166N17 | -0.054909146 |
| MTCO2P1 | -0.054910679 |
| CTD-2024F21 | -0.054914017 |
| MIR6812 | -0.054914383 |
| RP11-325P15 | -0.054935354 |
| AP001258 | -0.054935379 |
| RP11-305P14 | -0.054938934 |
| OR10U1P | -0.054941892 |
| OR4G2P | -0.05494572 |
| RP11-466G12 | -0.054955788 |
| RP11-343C2 | -0.054961353 |
| KIZ-AS1 | -0.054962909 |
| AKAP8 | -0.054966041 |
| RP11-481J2 | -0.054972776 |
| RNU6-73P | -0.054976623 |
| SCARNA22 | -0.054981243 |
| OR10J4 | -0.054983069 |
| RP1-197O17 | -0.054995311 |
| PPIAP25 | -0.054995695 |
| RP11-125M16 | -0.054996494 |
| USP34 | -0.054998043 |
| MMS22L | -0.05499821 |

| | |
|---|---|
| ATP9B | -0.054998768 |
| MBNL1 | -0.054999903 |
| RPL12P18 | -0.055001584 |
| RNU6ATAC36P | -0.055003619 |
| TRIM51CP | -0.055008114 |
| RCC2P2 | -0.055009315 |
| ITGB2 | -0.055017095 |
| SIDT1-AS1 | -0.055019153 |
| BTBD7P2 | -0.055024426 |
| TMEM106C | -0.05503691 |
| FRMPD2 | -0.05504397 |
| STRCP1 | -0.055044907 |
| AC245036 | -0.055045876 |
| CHCHD4 | -0.055047868 |
| ATP8B4 | -0.055051682 |
| POTEB2 | -0.055055887 |
| AC084193 | -0.055058436 |
| C5orf24 | -0.055060222 |
| CCDC183-AS1 | -0.055060706 |
| AL645859 | -0.055061764 |
| OR4D10 | -0.055063451 |
| GS1-466O4 | -0.055065129 |
| AC114302 | -0.055072151 |
| PDCD6IP | -0.05507386 |
| RP11-757H14 | -0.055077139 |
| CYB561D1 | -0.055079014 |
| FAM157A | -0.055084665 |
| CYP2B7P | -0.055087664 |
| AC004156 | -0.05508888 |
| LYPLA1P1 | -0.055106014 |
| KCTD9P3 | -0.055107268 |
| PTK2 | -0.055109266 |
| C1orf167 | -0.055114609 |
| AL713997 | -0.055115366 |
| C17orf85 | -0.055116522 |
| KANTR | -0.055127369 |
| UPK3BP1 | -0.055135172 |
| CTB-114C7 | -0.0551356 |
| RNVU1-20 | -0.055139111 |
| VPS26B | -0.055140095 |
| ZNF750 | -0.055141919 |
| RP11-333E1 | -0.05516925 |
| ZNF402P | -0.055170873 |
| ANKRD18B | -0.055172661 |
| LYAR | -0.055177046 |

| | |
|---|---|
| MEAF6 | -0.055179148 |
| MKS1 | -0.055180916 |
| RP11-293B7 | -0.055184081 |
| RP11-382H24 | -0.055185429 |
| RP11-139B1 | -0.055186938 |
| PIP5K1P1 | -0.055196043 |
| RP11-151M7 | -0.055199522 |
| BRAF | -0.055199559 |
| ZCCHC3 | -0.055202349 |
| IGHVII-26-2 | -0.055209003 |
| RN7SL30P | -0.055210394 |
| RP11-769O8 | -0.055217614 |
| UPF3B | -0.055218276 |
| MOB4 | -0.055219094 |
| MIR4729 | -0.055223657 |
| RNU4-17P | -0.055227025 |
| LINC01441 | -0.055228441 |
| AC018804 | -0.055232211 |
| RP11-267D19 | -0.055234154 |
| BTNL8 | -0.055236338 |
| AHCYL1 | -0.055242816 |
| UMOD | -0.055247894 |
| HMGB1P44 | -0.055250826 |
| SH3GLB2 | -0.055258891 |
| RP11-286B5 | -0.05526311 |
| CPVL | -0.055264026 |
| HMGN1P11 | -0.055271872 |
| SBDSP1 | -0.055272343 |
| AC027319 | -0.055273104 |
| IL3 | -0.055274256 |
| UBL3 | -0.055275711 |
| RP11-152C15 | -0.055280005 |
| NUP50-AS1 | -0.055280064 |
| IPPKP1 | -0.055280399 |
| MAP2K4 | -0.055280492 |
| MIR325 | -0.055282259 |
| CDC40 | -0.055283469 |
| TMC3 | -0.055285684 |
| RP11-486M3 | -0.055287079 |
| RNU6-633P | -0.055288582 |
| ZNF34 | -0.055290204 |
| RFC3 | -0.055298837 |
| RNU6-136P | -0.055303047 |
| MTERF2 | -0.055308214 |
| RP1-150O5 | -0.05531517 |

| | |
|---|---|
| SNORA73 | -0.055315339 |
| RP11-476I15 | -0.055316838 |
| MIR3199-2 | -0.055317424 |
| RP1-159M24 | -0.05531784 |
| ARL5A | -0.055323322 |
| RP11-137N23 | -0.055326378 |
| MIR3170 | -0.055326765 |
| PPP1R8 | -0.055326857 |
| CSE1L | -0.055330895 |
| NBEA | -0.055334244 |
| RP11-1152B5 | -0.055340941 |
| AC016683 | -0.05534701 |
| ZBED4 | -0.055347993 |
| HNRNPA3P3 | -0.055352505 |
| HYALP1 | -0.055361531 |
| STIP1 | -0.055362319 |
| ZDHHC20-IT1 | -0.055363612 |
| RP11-342D14 | -0.055369837 |
| RP4-591C20 | -0.055370226 |
| PRAMEF20 | -0.055371636 |
| RP11-54P19 | -0.055376199 |
| MARK2P5 | -0.055384146 |
| RN7SL110P | -0.055384279 |
| OTOGL | -0.055389365 |
| C6orf163 | -0.055391381 |
| FANCB | -0.055396609 |
| VDAC1P13 | -0.055404748 |
| RP11-10O17 | -0.055410098 |
| RNU6-782P | -0.055416489 |
| GABARAPL2 | -0.055417633 |
| ATG4C | -0.05541888 |
| RP11-348N5 | -0.055419289 |
| RIPPLY3 | -0.055426508 |
| CPT2 | -0.055430062 |
| ZNF561 | -0.055432387 |
| AMDHD1 | -0.055436574 |
| ABI1 | -0.055439476 |
| RP11-1430O6 | -0.055440208 |
| RP11-463C8 | -0.05544705 |
| ALOX12-AS1 | -0.055447532 |
| AC010092 | -0.055448995 |
| GOLGA8O | -0.055456503 |
| RP11-6C10 | -0.055465481 |
| NR1H3 | -0.055468463 |
| RP11-43D4 | -0.055468908 |

| | |
|---|---|
| FAM133DP | -0.05547075 |
| RP11-21A7A | -0.05547247 |
| MTND6P14 | -0.055473461 |
| OR5AN2P | -0.055482757 |
| RNA5SP341 | -0.055486814 |
| PMS2 | -0.055489153 |
| RP11-1193F23 | -0.055492232 |
| RP11-557C18 | -0.055493303 |
| AC006994 | -0.055496515 |
| RP11-471N19 | -0.055502813 |
| RP11-814E24 | -0.055508479 |
| MED17 | -0.055515647 |
| ZNF292 | -0.055520261 |
| RP11-734B5 | -0.055520418 |
| POLD3 | -0.055529638 |
| RP11-202I11 | -0.055532021 |
| PIGL | -0.055542755 |
| CDC42SE1 | -0.055545285 |
| RP1-186E20 | -0.055546579 |
| SNX3P1X | -0.055548062 |
| ST13P4 | -0.055555044 |
| RP5-894D12 | -0.055557154 |
| SLX4IP | -0.055558384 |
| DLG3 | -0.055558681 |
| MAP3K12 | -0.05555976 |
| AC069513 | -0.055560749 |
| AC124074 | -0.05556229 |
| ZFX-AS1 | -0.055564097 |
| AC090206 | -0.055581343 |
| RP11-181D18 | -0.055581841 |
| PIGW | -0.05558819 |
| RGSL1 | -0.05559321 |
| EXOC5P1 | -0.055598003 |
| TSPAN31 | -0.055599595 |
| IL1B | -0.055609307 |
| SIRT5 | -0.055609995 |
| MIR1302-11 | -0.055612803 |
| LANCL1 | -0.055613139 |
| NPIPB1P | -0.055614187 |
| RP4-641G12 | -0.05561541 |
| RP11-420A21 | -0.055618627 |
| POLK | -0.055626294 |
| AC093690 | -0.055627642 |
| ENTPD3-AS1 | -0.055627708 |
| FGR | -0.055630387 |

| | |
|---|---|
| RNU6-616P | -0.055634812 |
| MTND2P23 | -0.055635098 |
| RP11-314D7 | -0.05564132 |
| PCSK6-AS1 | -0.055649146 |
| RNA5SP34 | -0.055650969 |
| AC235953 | -0.055659093 |
| BRD2-IT1 | -0.055665487 |
| RNU6-495P | -0.055665901 |
| RP11-60A24 | -0.055672356 |
| RP11-487I5 | -0.055673876 |
| SMARCD1 | -0.055675784 |
| AC097662 | -0.05568345 |
| MUC2 | -0.055687513 |
| DSCR10 | -0.055695148 |
| RNU6-157P | -0.055697958 |
| ZNF385C | -0.055704064 |
| RP11-90L20 | -0.055710621 |
| TSPY20P | -0.055716045 |
| PRKACB | -0.055716288 |
| ACVR2B-AS1 | -0.055726056 |
| RP11-9M16 | -0.055729615 |
| CTC-360P9 | -0.055739142 |
| TCEB1P4 | -0.055739171 |
| IL23R | -0.055748895 |
| MILR1 | -0.055754113 |
| BTBD10 | -0.055759937 |
| RP11-355F16 | -0.055762839 |
| RP11-246A10 | -0.055770447 |
| ADCK3 | -0.055770486 |
| AC103686 | -0.055771888 |
| KIF28P | -0.055774484 |
| KISS1R | -0.055774577 |
| SH3BP2 | -0.055780458 |
| AP006285 | -0.055782804 |
| CTB-181F24 | -0.055783138 |
| BMS1P9 | -0.055783346 |
| ANKFY1 | -0.055786096 |
| CACNA1C-IT3 | -0.055794845 |
| RNU6-288P | -0.055798385 |
| AC062020 | -0.055798915 |
| LANCL1-AS1 | -0.055799865 |
| OR7D1P | -0.055803551 |
| CCDC65 | -0.055806432 |
| HSPE1P13 | -0.05581167 |
| DLEU2L | -0.055812134 |

| | |
|---|---|
| RAF1 | -0.055813483 |
| RNU6-286P | -0.055814515 |
| AL450423 | -0.055814773 |
| CTC-244M17 | -0.05582048 |
| C2orf68 | -0.05582247 |
| CBFB | -0.055844778 |
| SERF1B | -0.055858229 |
| TPRG1-AS2 | -0.055859549 |
| RNU6-588P | -0.055860013 |
| LINC00388 | -0.055864311 |
| EML5 | -0.055865764 |
| CDC27 | -0.055869132 |
| PRRC2C | -0.055869634 |
| GS1-278J22 | -0.055875199 |
| DGCR8 | -0.055877486 |
| RP11-442O18 | -0.055878434 |
| PBX2 | -0.055881468 |
| PCAT1 | -0.055885189 |
| LINC00474 | -0.055887015 |
| RNU2-7P | -0.055887285 |
| PPIAP9 | -0.055887917 |
| AL845259 | -0.055893206 |
| EIF4BP2 | -0.055902318 |
| OR2A1-AS1 | -0.055904464 |
| GLCE | -0.055905859 |
| RP11-300J18 | -0.055911399 |
| ZNF569 | -0.055913581 |
| IGKV3-25 | -0.055919107 |
| SNORD73 | -0.055919248 |
| RP11-64K12 | -0.055921804 |
| RPS4XP1 | -0.055926379 |
| GOLGA8M | -0.05592834 |
| NCAPD2 | -0.05593133 |
| RP11-313L6 | -0.055932739 |
| AFG3L2P1 | -0.05593431 |
| FAM153C | -0.055951729 |
| RP11-141M1 | -0.055952181 |
| AC074389 | -0.055957236 |
| AC138623 | -0.05595844 |
| RP1-245M18 | -0.055959442 |
| GTF2IRD2 | -0.055964206 |
| RP11-332H14 | -0.055966978 |
| SLC25A52 | -0.055969697 |
| RNU6-632P | -0.055975995 |
| OR4C13 | -0.055981088 |

| | |
|---|---|
| RNF152 | -0.055981937 |
| RP11-296L22 | -0.055982271 |
| CCDC47 | -0.05598363 |
| EID1 | -0.055989026 |
| DNALI1 | -0.055993472 |
| SCYL2 | -0.055996617 |
| GAS8 | -0.055997287 |
| CSPP1 | -0.056003352 |
| RN7SKP222 | -0.056007459 |
| SPIN3 | -0.056010757 |
| MIR320B2 | -0.056021547 |
| HPYR1 | -0.056036535 |
| U7 | -0.056038152 |
| AC093107 | -0.056040823 |
| VEZF1P1 | -0.056042023 |
| CTC-548H10 | -0.056044694 |
| RNA5SP57 | -0.056046914 |
| TRUB1 | -0.05605301 |
| GOLGA6L5P | -0.056054079 |
| METAP1D | -0.056057271 |
| RP11-373D7 | -0.056063493 |
| RP11-708B6 | -0.056071017 |
| ENSAP1 | -0.056071334 |
| VENTXP6 | -0.056073162 |
| ADORA2A | -0.056075643 |
| MTHFR | -0.056084881 |
| RP11-497H16 | -0.056085678 |
| DDX31 | -0.056091542 |
| AC087651 | -0.056107743 |
| MIR5587 | -0.056113055 |
| BLOC1S2P1 | -0.056114841 |
| AC073551 | -0.056115084 |
| PHBP10 | -0.056126816 |
| MEGF11 | -0.056141153 |
| C18orf54 | -0.056144716 |
| CHUK | -0.0561452 |
| MIR6807 | -0.056146758 |
| RP11-81K2 | -0.056152348 |
| UXT-AS1 | -0.05615642 |
| RP11-1134I14 | -0.05615734 |
| LMBRD1 | -0.05616001 |
| RNU6-891P | -0.056161915 |
| RPL5P24 | -0.056165253 |
| SMTNL1 | -0.056175998 |
| SETBP1 | -0.056180997 |

| | |
|---|---|
| RP11-17M16 | -0.056181717 |
| TRIM77BP | -0.056183468 |
| PLEKHM3 | -0.056194598 |
| HMGB3P4 | -0.056200575 |
| BCS1L | -0.056203472 |
| RP1-286D6 | -0.056203714 |
| PRAMEF8 | -0.056206485 |
| CRHR1-IT1 | -0.056210263 |
| AL357515 | -0.056211166 |
| EEF1A1P40 | -0.056216328 |
| RNU1-91P | -0.056217205 |
| RP1-29C18 | -0.056219575 |
| AL136160 | -0.056222477 |
| NPM1P5 | -0.056228254 |
| MGST2 | -0.056230115 |
| SNORA36 | -0.056238796 |
| RNU6-366P | -0.056242311 |
| SAPCD2P2 | -0.056244103 |
| RP11-711D18 | -0.056257212 |
| TRBV8-1 | -0.05626269 |
| HMGN2P3 | -0.056266061 |
| RP11-175M2 | -0.056269582 |
| DIMT1 | -0.056272573 |
| RAD54L | -0.056281243 |
| CARD10 | -0.056281448 |
| RP11-424C20 | -0.05629521 |
| RP11-218M11 | -0.056295961 |
| MIR3689D1 | -0.056298358 |
| RP11-449J10 | -0.056300242 |
| NKAIN1 | -0.056306175 |
| AC066612 | -0.056315231 |
| RP11-75N20 | -0.056318928 |
| CSH2 | -0.056322124 |
| CDH23-AS1 | -0.056326499 |
| ZKSCAN1 | -0.056328948 |
| RP11-956A19 | -0.056334339 |
| AIRE | -0.056335868 |
| MIR3682 | -0.056342312 |
| GACAT2 | -0.056346936 |
| RNU6-397P | -0.056348074 |
| MRPS23 | -0.05634919 |
| MIR3609 | -0.056358159 |
| NUP98 | -0.056359944 |
| CSNK1G3 | -0.056369694 |
| AC007787 | -0.056374355 |

| | |
|---|---|
| PMAIP1 | -0.056374996 |
| MIR760 | -0.056377967 |
| POGZ | -0.056378159 |
| AJ011931 | -0.05638153 |
| RNU6-117P | -0.05638156 |
| RP11-333J10 | -0.056382031 |
| RP11-460H9 | -0.056382332 |
| RP11-307O13 | -0.05638395 |
| EXOC5 | -0.056390904 |
| CTD-2622I13 | -0.056399759 |
| MRPL44 | -0.056400076 |
| UNC119 | -0.056401118 |
| HIVEP3 | -0.0564027 |
| BARX2 | -0.056418793 |
| IGKV1-22 | -0.056427468 |
| RP11-1072C15 | -0.056444786 |
| CRYBA2 | -0.056446322 |
| KIF18B | -0.056448078 |
| RP11-162A12 | -0.056457177 |
| MIR4300 | -0.056469137 |
| RPL7P57 | -0.056469758 |
| SEPT7P8 | -0.056472946 |
| DNMT3A | -0.056476582 |
| ADAMTS19 | -0.056479896 |
| METTL7A | -0.056486127 |
| DSCR3 | -0.056493699 |
| LRRTM2 | -0.056493935 |
| CTB-113D17 | -0.056496508 |
| SIGLEC1 | -0.056498611 |
| NYX | -0.056503205 |
| RASA3-IT1 | -0.05650482 |
| RP11-28G8 | -0.056505936 |
| SYNGR3 | -0.05651504 |
| OR10V1 | -0.056521462 |
| OTUD1 | -0.056522389 |
| RNU7-160P | -0.056524382 |
| HEATR3 | -0.056529254 |
| HCP5B | -0.056537349 |
| C14orf177 | -0.056547502 |
| MIR6746 | -0.056548748 |
| RP4-583P15 | -0.056553646 |
| ALMS1 | -0.056560567 |
| CDKN2B-AS1 | -0.056561795 |
| CTD-2285G11 | -0.056570562 |
| OR4F16 | -0.056571319 |

| | |
|---|---|
| XXbac-BPG181M17 | -0.056575508 |
| ABHD17C | -0.056578354 |
| RNU1-136P | -0.056587276 |
| RP11-116K4 | -0.056591757 |
| CHPT1 | -0.056594838 |
| RP11-368M16 | -0.056594961 |
| LINC00977 | -0.056607213 |
| CTD-2265M8 | -0.056608393 |
| LY86 | -0.056609202 |
| WDR59 | -0.056613407 |
| PRKAG1 | -0.0566199 |
| RP13-254B10 | -0.05662209 |
| RP11-307C19 | -0.056623031 |
| SGTB | -0.056629966 |
| RP11-398A8 | -0.056642604 |
| CSAG3 | -0.056642854 |
| RP11-335K5 | -0.056648318 |
| RP11-763B22 | -0.056648954 |
| WDR37 | -0.056649204 |
| LINC00294 | -0.056653018 |
| GBP1 | -0.05665499 |
| KNOP1P1 | -0.056658467 |
| SLC7A6OS | -0.056661037 |
| RBM47 | -0.056661781 |
| RNU6-1273P | -0.056671431 |
| PASD1 | -0.056674689 |
| KRT18P38 | -0.056679992 |
| SS18L2P2 | -0.056682285 |
| COL9A1 | -0.056696519 |
| MIEN1 | -0.056698062 |
| RAPGEF2 | -0.056698899 |
| CARD16 | -0.05670009 |
| RP11-513M1 | -0.056702178 |
| RP11-346L1 | -0.056708289 |
| JKAMP | -0.056711699 |
| RP11-366O20 | -0.056714873 |
| OSBPL9P2 | -0.056716723 |
| PHACTR2P1 | -0.056733034 |
| RP11-359N5 | -0.056735293 |
| MIR6833 | -0.056737093 |
| AC008280 | -0.05674283 |
| RP5-1170K4 | -0.05674332 |
| DOCK11P1 | -0.056743381 |
| UBTF | -0.056744352 |
| SLC4A1 | -0.056745001 |

| | |
|---|---|
| SP2 | -0.056745766 |
| SYCE2 | -0.056746715 |
| CTBS | -0.056750938 |
| TTC6 | -0.056752969 |
| MMRN1 | -0.056753116 |
| RP11-715G18 | -0.05675495 |
| SELT | -0.056762623 |
| AL356261 | -0.056769475 |
| THNSL1 | -0.056769507 |
| PSME3 | -0.056770288 |
| VOPP1 | -0.056775404 |
| HNRNPA3P4 | -0.056779107 |
| CTB-151G24 | -0.05678577 |
| DHFRL1 | -0.056800913 |
| HCG23 | -0.056801811 |
| IGHVIII-82 | -0.056801861 |
| AC109826 | -0.056809341 |
| HLA-DPA2 | -0.056813428 |
| TGS1 | -0.056819406 |
| TCP11X3P | -0.056826806 |
| RP4-726N1 | -0.056829552 |
| ZRANB2-AS2 | -0.056837491 |
| AC013248 | -0.056858252 |
| AL392111 | -0.056860715 |
| PI4KAP2 | -0.056863966 |
| HNRNPA3P8 | -0.056874506 |
| MIR4292 | -0.056877772 |
| RN7SKP126 | -0.056882286 |
| OR2AS1P | -0.056885785 |
| DTX2P1 | -0.056887949 |
| RNU6-138P | -0.056894709 |
| CCDC176 | -0.05689967 |
| CDK1 | -0.056903652 |
| RP11-2H8 | -0.056903976 |
| RP5-983L19 | -0.056906019 |
| SEMA3B-AS1 | -0.056928959 |
| MCM10 | -0.05693236 |
| MCMBP | -0.056934667 |
| IGLJ7 | -0.056945183 |
| DNTTIP2 | -0.056946445 |
| RP11-795H16 | -0.05695057 |
| ERCC4 | -0.056952138 |
| AC073610 | -0.056954966 |
| RAB5CP1 | -0.056960425 |
| RP11-539L10 | -0.056969832 |

| | |
|---|---|
| AC009491 | -0.056970166 |
| SNX6 | -0.056971599 |
| CABP2 | -0.056982945 |
| SLC23A1 | -0.056984418 |
| MIR5008 | -0.056995914 |
| SYNGAP1 | -0.057000661 |
| AC069277 | -0.057003491 |
| RNU7-149P | -0.057005856 |
| CICP11 | -0.057007725 |
| UNC50 | -0.057008159 |
| RP11-342M3 | -0.057009617 |
| GPHN | -0.057011824 |
| CTA-363E19 | -0.057015285 |
| NPY1R | -0.057020494 |
| OR4R2P | -0.057027229 |
| RP11-944C7 | -0.05702984 |
| MIR3646 | -0.057031052 |
| RAC1P8 | -0.05703434 |
| RP11-206L10 | -0.057035379 |
| AC104590 | -0.057037097 |
| DKFZP434H168 | -0.057039653 |
| CTA-305I2 | -0.057040202 |
| MIR8069-2 | -0.057044498 |
| AC112778 | -0.05704751 |
| AREL1 | -0.057049575 |
| MIR3153 | -0.05705732 |
| CTD-2313N18 | -0.057058248 |
| ZNF730 | -0.057063502 |
| PDPK2P | -0.057063654 |
| RP11-17G11 | -0.057065607 |
| PANK3 | -0.057066189 |
| ZNF886P | -0.057070193 |
| ZNF529 | -0.057072571 |
| RP11-641J8 | -0.05707348 |
| OR4Q3 | -0.057081803 |
| NEFHP1 | -0.057084698 |
| OR5AK3P | -0.057085271 |
| AC079080 | -0.057086996 |
| RP11-405F3 | -0.057091696 |
| RP11-492E3 | -0.057096703 |
| ZNF407 | -0.057100442 |
| FAM183CP | -0.057106581 |
| MIR3171 | -0.05710908 |
| AC009305 | -0.057118166 |
| PLAG1 | -0.057122751 |

| | |
|---|---|
| RHOQP1 | -0.057126778 |
| RP11-103B5 | -0.057128489 |
| AC108462 | -0.057146766 |
| RNU7-40P | -0.057148895 |
| DDX3P3 | -0.05714973 |
| RP11-133K1 | -0.05714983 |
| MTND3P3 | -0.05715122 |
| AC002128 | -0.057160778 |
| CCP110 | -0.057166994 |
| BIVM-ERCC5 | -0.057168165 |
| PPY | -0.057168527 |
| MTFR1L | -0.057180315 |
| MIR302C | -0.05718365 |
| OR5P4P | -0.057184919 |
| PPM1M | -0.057188799 |
| PPIAP3 | -0.057190748 |
| RNU7-18P | -0.057218154 |
| RP11-824M15 | -0.057218443 |
| GNA14 | -0.057219498 |
| AC091177 | -0.057219517 |
| CLEC7A | -0.057227536 |
| TFF3 | -0.057230736 |
| RP11-33N14 | -0.057241137 |
| CARD8-AS1 | -0.057242365 |
| RNU6-856P | -0.057244168 |
| MTPAP | -0.057257602 |
| APOBEC3F | -0.057258756 |
| MIR4288 | -0.057264278 |
| LEUTX | -0.057266793 |
| PIGBOS1 | -0.057267482 |
| AC020951 | -0.057268334 |
| GRAMD1B | -0.057268803 |
| KYNUP3 | -0.057270541 |
| CTD-2630F21 | -0.057279538 |
| OGFRP1 | -0.057285362 |
| KRT223P | -0.057290835 |
| TMEM255A | -0.057295074 |
| AP2B1 | -0.057296078 |
| RN7SL709P | -0.057296271 |
| HIST1H2BK | -0.057297623 |
| RP1-172N19 | -0.057301721 |
| MIR3180-3 | -0.057313936 |
| AC087501 | -0.057314877 |
| KRT18P46 | -0.057317905 |
| RN7SL395P | -0.0573214 |

| | |
|---|---|
| ATP13A3 | -0.057332266 |
| IMPACT | -0.057334908 |
| AC099539 | -0.05734025 |
| ZNF682 | -0.057352751 |
| ORAOV1P1 | -0.057353915 |
| RTN3P1 | -0.057359363 |
| AC011742 | -0.057362016 |
| LINC01502 | -0.05737157 |
| RNU6-1119P | -0.05737339 |
| GLMN | -0.057376045 |
| MTMR6 | -0.057377254 |
| PROSER3 | -0.057382668 |
| RNF41 | -0.057382854 |
| DPY19L1P2 | -0.057391434 |
| PIK3CB | -0.057393366 |
| CPNE7 | -0.057395543 |
| SRRD | -0.057399023 |
| PPAPDC1B | -0.057408958 |
| GOLGA5P1 | -0.057414925 |
| SLC35D2 | -0.057420921 |
| C4orf32 | -0.057423656 |
| MIR4632 | -0.057424253 |
| TSEN2P1 | -0.057425811 |
| RP11-16E18 | -0.057426726 |
| ZNF223 | -0.057431191 |
| CTC-661I16 | -0.057434486 |
| MIR548C | -0.057435407 |
| RNU6-351P | -0.057435656 |
| BTNL3 | -0.057444773 |
| RN7SL388P | -0.057446517 |
| SLC25A35 | -0.057446894 |
| HMGN2P21 | -0.057455687 |
| RNA5SP124 | -0.057457354 |
| IARS | -0.057460179 |
| TAMM41 | -0.057475733 |
| POGLUT1 | -0.057479417 |
| FEM1B | -0.05748511 |
| SIGLEC22P | -0.057485852 |
| POLR3E | -0.057487194 |
| RN7SKP129 | -0.057488935 |
| WDTC1 | -0.057493017 |
| ESRP2 | -0.057496292 |
| RPL21P24 | -0.05750081 |
| DNAJC19P8 | -0.05750335 |
| RP11-775A1 | -0.057504098 |

| | |
|---|---|
| DARS2 | -0.057507883 |
| ATG16L1 | -0.057515437 |
| MRPL53 | -0.057515642 |
| MIR6881 | -0.057517405 |
| UTS2R | -0.057518646 |
| NBPF7 | -0.057518829 |
| L3MBTL2 | -0.057522228 |
| RNU6-1099P | -0.057525876 |
| AC009947 | -0.057526686 |
| CDKN2AIPNLP1 | -0.057529645 |
| RNU7-180P | -0.057530136 |
| TBXAS1 | -0.057533466 |
| KIF16B | -0.05754221 |
| ABHD13 | -0.057551215 |
| RP11-352D13 | -0.057553578 |
| FKBP4P6 | -0.0575642 |
| SYBU | -0.057566211 |
| LMBR1L | -0.05756729 |
| DUTP7 | -0.057569792 |
| RP11-370B6 | -0.057570472 |
| CTD-2013N17 | -0.057574119 |
| CELSR1 | -0.057574733 |
| SUMO2P17 | -0.057575942 |
| RP11-10I6 | -0.057584151 |
| RNU6-378P | -0.057584799 |
| AL139184 | -0.057595117 |
| RP11-507K13 | -0.057597463 |
| BTBD8 | -0.057598539 |
| SRP14-AS1 | -0.05759998 |
| DCAF12 | -0.057608185 |
| AL353803 | -0.057611445 |
| ZNF859P | -0.057612417 |
| AL035422 | -0.057624826 |
| ZFR | -0.057634233 |
| ZNF746 | -0.057634568 |
| AC011298 | -0.057648138 |
| TAF4B | -0.05765068 |
| RP11-127B20 | -0.057657806 |
| KRT8P41 | -0.057659691 |
| ZNF257 | -0.057660225 |
| SLC6A12 | -0.057661397 |
| MIR324 | -0.057677918 |
| JPH4 | -0.057692524 |
| ADAM1B | -0.057693231 |
| KBTBD11 | -0.057697119 |

| | |
|---|---|
| RP4-701O16 | -0.057712678 |
| LINC00620 | -0.057712989 |
| RP11-119D9 | -0.057718711 |
| MIR6809 | -0.057718748 |
| OR5D14 | -0.057721975 |
| RP1-63G5 | -0.057724985 |
| RRN3P2 | -0.057725307 |
| RP11-624L4 | -0.057739042 |
| MIR5695 | -0.057739644 |
| FSCN2 | -0.057741456 |
| POU5F1P6 | -0.05774217 |
| PITPNC1 | -0.057750647 |
| GPC5-AS1 | -0.057757673 |
| RP11-285E18 | -0.057761233 |
| ZNF847P | -0.057764159 |
| CTC-497E21 | -0.057768462 |
| RP11-699C17 | -0.05777226 |
| POTENP | -0.057773715 |
| MIR4285 | -0.057778594 |
| TAS2R6P | -0.057780731 |
| KIAA1429 | -0.057781495 |
| APOF | -0.057786724 |
| PCDH9-AS2 | -0.057789767 |
| ADPGK-AS1 | -0.057796496 |
| ACTR2 | -0.057800026 |
| SMARCAL1 | -0.057804436 |
| CTD-2104P17 | -0.057805273 |
| RP11-768G7 | -0.057810895 |
| H3F3AP4 | -0.057816622 |
| MED28P5 | -0.05782723 |
| RP11-1365D11 | -0.057829957 |
| MIR1185-1 | -0.0578322 |
| CCNK | -0.057833088 |
| AC069294 | -0.057834707 |
| CYP2B6 | -0.057839 |
| RN7SL246P | -0.057840508 |
| CTD-2008P7 | -0.057851303 |
| RP11-18H7 | -0.057853108 |
| PRYP5 | -0.057856104 |
| RALGPS1 | -0.057863713 |
| RNU6-648P | -0.057864598 |
| CYP4F32P | -0.057866591 |
| SRP14 | -0.057871936 |
| ACA59 | -0.057874096 |
| MIR1207 | -0.05788085 |

| | |
|---|---|
| RP5-1173I20 | -0.057881385 |
| SIGLEC7 | -0.057889742 |
| TARSL2 | -0.057892867 |
| UNC119B | -0.057901928 |
| PGGT1BP2 | -0.057904626 |
| RP11-498E2 | -0.057911369 |
| OR52V1P | -0.05791751 |
| SLC2A10 | -0.057921576 |
| AC012507 | -0.057922776 |
| RP11-341A22 | -0.05792301 |
| RP11-533K9 | -0.05792944 |
| AC009487 | -0.057934004 |
| GALK2 | -0.057936888 |
| KMT2B | -0.0579506 |
| PHB2P1 | -0.057954278 |
| FASTKD3 | -0.057973615 |
| RN7SL826P | -0.057984358 |
| RP11-142I2 | -0.057985745 |
| HMGN2P7 | -0.057988626 |
| RNU6-355P | -0.057991674 |
| GNG4 | -0.057994379 |
| SDC4 | -0.057995179 |
| RN7SL297P | -0.058002498 |
| PLEKHB2 | -0.058002733 |
| APOOL | -0.058003123 |
| MCM8 | -0.058008389 |
| OR2AI1P | -0.058009041 |
| TLK1 | -0.058018845 |
| MLX | -0.058020371 |
| TPPP2 | -0.058022206 |
| ZNF75A | -0.05803092 |
| RP11-305E6 | -0.058037916 |
| RP11-572O17 | -0.058042251 |
| AC091132 | -0.058045058 |
| KRTAP5-AS1 | -0.058048843 |
| LINC00566 | -0.058049156 |
| RP11-295G12 | -0.058051439 |
| RN7SKP9 | -0.058053647 |
| TWF1 | -0.058055386 |
| VDAC1P5 | -0.058056394 |
| RP11-214K3 | -0.058057284 |
| RNU6-1113P | -0.058064404 |
| RP11-305O6 | -0.058066045 |
| MIR548E | -0.058070564 |
| RP11-626I20 | -0.058076037 |

| | | |
|---|---|---|
| MAPK13 | | -0.058078476 |
| IGLVIV-66-1 | | -0.058080552 |
| RNU4-78P | | -0.05808103 |
| RP11-43F13 | | -0.058085974 |
| RHEBL1 | | -0.05809029 |
| LINC00434 | | -0.058091094 |
| MICC | | -0.058091424 |
| PEX7 | | -0.058093174 |
| ARL16 | | -0.058094718 |
| LINC00418 | | -0.058098958 |
| RP11-58H15 | | -0.058104494 |
| VWA8 | | -0.058121157 |
| ARG1 | | -0.058123176 |
| MIPOL1 | | -0.058123222 |
| | 3-Sep | -0.058129343 |
| HSP90AA4P | | -0.058130175 |
| RPS20P22 | | -0.058134242 |
| CTD-3184A7 | | -0.058135204 |
| SYTL1 | | -0.058143427 |
| COG8 | | -0.0581462 |
| SLBP | | -0.058147334 |
| EZR-AS1 | | -0.058147932 |
| ANKRD23 | | -0.058148432 |
| MARK2P8 | | -0.058148972 |
| RP11-552M11 | | -0.058150721 |
| THEMIS3P | | -0.058151296 |
| RP11-397A15 | | -0.058153757 |
| VN1R42P | | -0.058160805 |
| FBXO5 | | -0.058161791 |
| AL139139 | | -0.058168337 |
| MCHR2 | | -0.058168378 |
| ARL8B | | -0.058177364 |
| OR10K2 | | -0.05818483 |
| RP11-673D15 | | -0.0581874 |
| VCX2 | | -0.058193911 |
| AGGF1P4 | | -0.058197664 |
| RP1-104O17 | | -0.058198282 |
| ZDHHC23 | | -0.058198462 |
| RPL23AP78 | | -0.058199422 |
| ZSCAN30 | | -0.058199634 |
| RPS15AP36 | | -0.058206371 |
| RNU1-48P | | -0.05820909 |
| NBPF8 | | -0.058211226 |
| GCNT1P5 | | -0.058211521 |
| DPP9-AS1 | | -0.058214328 |

| | |
|---|---|
| RP11-315L6 | -0.058217339 |
| NPM1P24 | -0.058218377 |
| UBE2WP1 | -0.058219719 |
| RP11-154H17 | -0.058221761 |
| AC104383 | -0.058245595 |
| MIR133A1HG | -0.058246369 |
| DSN1 | -0.058248604 |
| CHRNA10 | -0.058250893 |
| SCN7A | -0.058254878 |
| RN7SL96P | -0.058256755 |
| NIFKP6 | -0.058258135 |
| RPS6P2 | -0.058263028 |
| CDCA8 | -0.058271303 |
| AC010931 | -0.058281405 |
| RP11-175I17 | -0.058283923 |
| PTTG4P | -0.058287294 |
| TMED6 | -0.058291583 |
| ZNF165 | -0.058298578 |
| C5 | -0.058305369 |
| STRN | -0.058309593 |
| INTS3 | -0.058312588 |
| RP11-574F21 | -0.058319789 |
| HMGB1P23 | -0.058325085 |
| AC005329 | -0.058330231 |
| RP11-151E14 | -0.058331391 |
| POLR2CP | -0.058331411 |
| CNEP1R1 | -0.058332291 |
| AC020550 | -0.058336588 |
| RP11-611L7 | -0.058343278 |
| RP11-29A2 | -0.058344392 |
| RP11-110F24 | -0.058345571 |
| NSMCE4A | -0.058346078 |
| SETP15 | -0.058350119 |
| LINC01181 | -0.058353523 |
| RP11-671E7 | -0.058362097 |
| CFAP99 | -0.058367139 |
| AC092377 | -0.058369695 |
| TRAPPC2 | -0.058369912 |
| CAPZA1 | -0.058375214 |
| TTLL5 | -0.058379921 |
| LINC01359 | -0.058382413 |
| NOL7 | -0.058384666 |
| BTNL9 | -0.058386005 |
| AC008073 | -0.05839195 |
| PWAR1 | -0.058393907 |

| | |
|---|---|
| SATB2 | -0.058394843 |
| POLR2A | -0.058395922 |
| AC016691 | -0.058396002 |
| DDX25 | -0.05839719 |
| PLSCR5-AS1 | -0.058403122 |
| CTD-2010I22 | -0.05840553 |
| RAVER2 | -0.058407848 |
| AC132479 | -0.058422761 |
| AC004846 | -0.058423828 |
| VN1R69P | -0.058425064 |
| RP5-837O21 | -0.058428646 |
| LPCAT4 | -0.05843291 |
| FAM149B1 | -0.05843371 |
| IL17A | -0.058434216 |
| RP11-354H21 | -0.058436794 |
| SLC25A18P1 | -0.058439701 |
| TAF1 | -0.05844091 |
| ZNF25 | -0.058443366 |
| TPRG1 | -0.058444389 |
| MICU2 | -0.058444482 |
| RP11-89A16 | -0.058444644 |
| RP11-505P4 | -0.058445398 |
| GPN3 | -0.058447943 |
| ZNF821 | -0.058449524 |
| ZZZ3 | -0.058451032 |
| RP11-380D15 | -0.058455931 |
| RNA5SP217 | -0.058459826 |
| CELF2 | -0.058462511 |
| BCL2L2 | -0.058465581 |
| RP4-622L5 | -0.05847686 |
| RP4-655J12 | -0.058478516 |
| KRTAP9-9 | -0.058478918 |
| PAPOLB | -0.058480097 |
| AL603632 | -0.058482052 |
| RP11-733C7 | -0.058488583 |
| PPP3CC | -0.058490177 |
| RHBG | -0.058491717 |
| RP11-184E9 | -0.058499612 |
| TTC8 | -0.058500969 |
| ITGAX | -0.058510383 |
| CTD-3020H12 | -0.058530174 |
| RIPK1 | -0.058543054 |
| PSMA1 | -0.058544115 |
| AC132803 | -0.058547858 |
| MIPEPP3 | -0.058555185 |

| | |
|---|---|
| RAP1GAP | -0.058561846 |
| AC067961 | -0.058567634 |
| UBE2MP1 | -0.058573046 |
| AP001595 | -0.058575514 |
| RP11-204P2 | -0.0585773 |
| CERS6 | -0.058586926 |
| TRAJ59 | -0.058596087 |
| TRBV5-3 | -0.058596467 |
| RNU6-658P | -0.058604342 |
| OR56A4 | -0.058605524 |
| YY1 | -0.058606592 |
| AL160011 | -0.05860844 |
| RP11-313D6 | -0.058609116 |
| HAR1A | -0.058611033 |
| MAGI1 | -0.058613271 |
| RNU6-778P | -0.058618532 |
| AC116606 | -0.058618532 |
| NSMAF | -0.058618723 |
| FAM107B | -0.058620341 |
| RP11-452D21 | -0.058622916 |
| MIR635 | -0.058624323 |
| DPP10-AS3 | -0.058624393 |
| RP1-128M12 | -0.058629226 |
| DOPEY1 | -0.058629904 |
| CH507-24F1 | -0.05863242 |
| AC104071 | -0.058633744 |
| AL513011 | -0.058633744 |
| AC078991 | -0.058633744 |
| NIPA2 | -0.058636286 |
| CSPG4P11 | -0.058642971 |
| RP11-76I14 | -0.058661373 |
| OCIAD1-AS1 | -0.058662186 |
| PRDM4 | -0.058666725 |
| RP1-101G11 | -0.058667223 |
| DENR | -0.058676772 |
| RP11-656A15 | -0.058676934 |
| AL162430 | -0.058686578 |
| GS1-111G14 | -0.058687332 |
| AC074264 | -0.058689313 |
| TRDJ4 | -0.05869308 |
| ITPR1-AS1 | -0.058694583 |
| SPATA4 | -0.05869636 |
| DEFB109P1 | -0.058699168 |
| ATP5BP1 | -0.058702979 |
| RP11-21G15 | -0.058704255 |

| | |
|---|---|
| AC073587 | -0.058704643 |
| RP11-536K17 | -0.058706292 |
| AGGF1P2 | -0.058715034 |
| RP11-342M21 | -0.058719566 |
| RNA5SP271 | -0.058725411 |
| YWHAQP5 | -0.058726914 |
| RN7SL354P | -0.058726997 |
| AC004656 | -0.058727899 |
| AC022120 | -0.058730195 |
| DUX4 | -0.058740397 |
| ATP5A1 | -0.058750003 |
| RP5-1074L1 | -0.058776497 |
| IMPDH1P6 | -0.058776864 |
| FEM1AP3 | -0.058779694 |
| HLA-DRB6 | -0.058786621 |
| RPL5P31 | -0.058792513 |
| OR2AQ1P | -0.058793333 |
| SLC25A24 | -0.058795437 |
| RP11-455G16 | -0.058796757 |
| GPR160 | -0.058801372 |
| SEC23B | -0.058808015 |
| HLA-J | -0.058809931 |
| AC011816 | -0.058814408 |
| PLCH2 | -0.058820964 |
| RP11-306I1 | -0.058823932 |
| CENPBD1 | -0.058824741 |
| MIR891A | -0.058826646 |
| ARL14EPL | -0.05883543 |
| IGLJ5 | -0.05883668 |
| MIR8064 | -0.058836873 |
| RP11-400N9 | -0.058841956 |
| RNU6-915P | -0.05884236 |
| DYNLRB2 | -0.058843919 |
| MB21D1 | -0.058849821 |
| RP11-417N10 | -0.05886132 |
| TTLL13P | -0.05886273 |
| LINC01300 | -0.058865855 |
| PHKA1-AS1 | -0.058867748 |
| KCND3 | -0.058872724 |
| RNA5SP131 | -0.058874313 |
| RN7SL89P | -0.058876693 |
| AP000302 | -0.058876911 |
| FLYWCH1P1 | -0.05887833 |
| RP11-65L19 | -0.058884393 |
| RNU6-913P | -0.058889054 |

| | |
|---|---|
| DR1 | -0.058897674 |
| RP11-394B5 | -0.058901095 |
| RN7SL833P | -0.05891438 |
| AL354981 | -0.0589161 |
| SLC26A7 | -0.058917773 |
| AC093419 | -0.058923096 |
| BACE1-AS | -0.058932746 |
| UBQLN1 | -0.058937081 |
| TPM3P4 | -0.058937307 |
| USP10 | -0.058947407 |
| RAC1P3 | -0.058951778 |
| DRAM1 | -0.058954849 |
| RP11-9H20 | -0.058959472 |
| C9orf152 | -0.058965231 |
| ZNF623 | -0.058970775 |
| RP11-600F24 | -0.058977957 |
| PEX19 | -0.058978106 |
| AHCYP7 | -0.058986547 |
| PROSER2-AS1 | -0.058988116 |
| RP5-901A4 | -0.058988618 |
| SETP16 | -0.058992579 |
| AC064874 | -0.059001109 |
| RP13-560N11 | -0.059004896 |
| TRIM56 | -0.059005177 |
| MIR6783 | -0.059010242 |
| RP11-513M16 | -0.059013586 |
| RP11-349A22 | -0.059014281 |
| DIS3 | -0.059019094 |
| HERC2P10 | -0.059023213 |
| ZNF563 | -0.059024192 |
| RLN1 | -0.059045243 |
| RIBC2 | -0.059048007 |
| SPINK4 | -0.059052077 |
| AKTIP | -0.059055467 |
| KRT18P27 | -0.059060618 |
| RP11-883A18 | -0.059068682 |
| RABGAP1L | -0.059072566 |
| AF003625 | -0.059073377 |
| RPS2P6 | -0.059074694 |
| HIST1H3E | -0.059085046 |
| ZNF131 | -0.059086836 |
| HNRNPA1P5 | -0.059093222 |
| ZNF384 | -0.059096697 |
| CTD-2227I18 | -0.059098469 |
| RNU6-475P | -0.059098875 |

| | |
|---|---|
| FOXJ3 | -0.059099507 |
| AL590084 | -0.059101021 |
| RPL17P43 | -0.059101924 |
| GATB | -0.059105386 |
| UVRAG | -0.059109274 |
| RP11-432F4 | -0.059112761 |
| RPL23AP81 | -0.059112871 |
| AC015969 | -0.059113625 |
| PVT1 | -0.059113702 |
| KCNK10 | -0.059121455 |
| AC068134 | -0.059141021 |
| RP5-940J5 | -0.059144941 |
| AC109638 | -0.059145559 |
| TRAM2-AS1 | -0.059149834 |
| PTAR1 | -0.059156867 |
| SYT10 | -0.059158496 |
| KLF1 | -0.059167718 |
| UGDH-AS1 | -0.05916937 |
| LRRC70 | -0.059176255 |
| CTD-3220F14 | -0.059182208 |
| VPS8 | -0.059190711 |
| KRT8P5 | -0.059206037 |
| AC087163 | -0.059208566 |
| EDRF1 | -0.059208981 |
| AC122718 | -0.059212552 |
| RP11-350F16 | -0.0592153 |
| RP11-433J20 | -0.059217668 |
| SF3A3P1 | -0.059221978 |
| RP11-172E10 | -0.059234484 |
| BAZ2B | -0.059237559 |
| WDR60 | -0.059248387 |
| AC005534 | -0.059261082 |
| ZNF439 | -0.059261635 |
| AKAP2 | -0.059261644 |
| RNU2-31P | -0.059264304 |
| RPL39P36 | -0.059267199 |
| CUZD1 | -0.059275849 |
| CUL5 | -0.059282473 |
| RN7SKP249 | -0.059282798 |
| GPS2P2 | -0.059289543 |
| RP11-23P13 | -0.059289752 |
| RP11-16K12 | -0.059290226 |
| RP5-1087E8 | -0.059291466 |
| LPPR3 | -0.059296816 |
| OR5E1P | -0.059318248 |

| | |
|---|---|
| RP11-553K23 | -0.059319777 |
| HIST1H2BN | -0.059330103 |
| RNA5SP102 | -0.059337144 |
| RAB1C | -0.059341475 |
| RNU6-54P | -0.059343657 |
| IFNK | -0.059346148 |
| RNU6-968P | -0.059346746 |
| CTD-2309H9 | -0.059347428 |
| BUB1P1 | -0.05935454 |
| LINC01322 | -0.059355822 |
| SKIV2L2 | -0.059359295 |
| FOXK2 | -0.059363555 |
| NPFFR1 | -0.05936446 |
| LINC01256 | -0.059366113 |
| RNU6-1125P | -0.05936622 |
| RP11-172F10 | -0.059375316 |
| CTBP2P6 | -0.059379468 |
| RP11-179H18 | -0.059380029 |
| AC097262 | -0.059382401 |
| ZSCAN9 | -0.05938257 |
| AC015987 | -0.059384605 |
| RP11-50O21 | -0.059389794 |
| TRIM52-AS1 | -0.059390236 |
| RP11-512F24 | -0.059392567 |
| TOX4 | -0.059392915 |
| GOLGA5 | -0.059394329 |
| CTD-2015B23 | -0.059399521 |
| RP3-416J7 | -0.059401017 |
| RPL21P7 | -0.05940621 |
| RPS7P14 | -0.059411944 |
| RN7SL809P | -0.059418007 |
| RP11-315E17 | -0.059422492 |
| OR5J2 | -0.059424348 |
| CTD-2326C4 | -0.0594255 |
| PPP1CA | -0.059436117 |
| RAET1K | -0.059438145 |
| LINC01516 | -0.0594384 |
| PRPSAP2 | -0.059441977 |
| SAP130 | -0.059443447 |
| TIAF1 | -0.059450889 |
| FCGRT | -0.059452043 |
| DDX46 | -0.059455448 |
| SATB2-AS1 | -0.059460205 |
| CTD-3113P16 | -0.059471658 |
| RP11-52A20 | -0.059475198 |

| | |
|---|---|
| OR4P1P | -0.059477203 |
| ZNF689 | -0.059478184 |
| AC078950 | -0.059480476 |
| SETD6 | -0.059481421 |
| LECT1 | -0.059486099 |
| NAGA | -0.059486779 |
| TRMT12 | -0.059487244 |
| RP3-378P9 | -0.059489551 |
| CYP2C59P | -0.059498431 |
| RP11-111F5 | -0.059498784 |
| RNU6-81P | -0.0595018 |
| TET2-AS1 | -0.059501928 |
| CANT1 | -0.05950918 |
| ZBTB41 | -0.059510799 |
| RP11-96J19 | -0.059515659 |
| RP11-1258F18 | -0.059520125 |
| RP11-20I23 | -0.059521758 |
| TYMS | -0.059523413 |
| NR4A3 | -0.059523674 |
| RP11-152H18 | -0.059527208 |
| RP5-937E21 | -0.059530577 |
| TRIM38 | -0.059534651 |
| OR9A2 | -0.05953481 |
| MIR615 | -0.059538275 |
| RP3-472M2 | -0.059539313 |
| TMEM50B | -0.059552271 |
| MIR548AT | -0.059553795 |
| SLC9A3P2 | -0.059558794 |
| CTD-2210P15 | -0.059572981 |
| GRPEL1 | -0.059575769 |
| RP11-388P9 | -0.059582303 |
| RNY4P19 | -0.059582332 |
| RN7SL704P | -0.059582708 |
| RNU6-650P | -0.059584682 |
| ALAS1 | -0.059585853 |
| AZIN1 | -0.059587007 |
| CTB-107G13 | -0.059587422 |
| FCHO1 | -0.059600868 |
| RP11-17E2 | -0.05960226 |
| TATDN2P3 | -0.059604581 |
| TCEAL6 | -0.059609788 |
| AL109843 | -0.059613666 |
| GXYLT1P1 | -0.05961565 |
| CNGB3 | -0.059618896 |
| TUBB8P10 | -0.059622229 |

| | |
|---|---|
| CLASP1 | -0.059628553 |
| RNU5E-4P | -0.059641061 |
| UBE2CP1 | -0.059645592 |
| TIAL1 | -0.059646135 |
| CH17-60O17 | -0.059652942 |
| NOXRED1 | -0.059653968 |
| DUSP8P5 | -0.059658899 |
| AC078864 | -0.05966087 |
| RNU6-663P | -0.0596616 |
| KRT8P26 | -0.059662299 |
| RARRES2P5 | -0.059664289 |
| RP11-135N5 | -0.059665551 |
| RP11-173B14 | -0.05966804 |
| LINC00430 | -0.059668944 |
| AC079768 | -0.059668999 |
| ELMO1-AS1 | -0.059669763 |
| CASZ1 | -0.059673057 |
| ABI3 | -0.059673634 |
| C1orf43 | -0.059676555 |
| KLHL33 | -0.059684236 |
| SPECC1L | -0.059685542 |
| SNORD116-16 | -0.059685692 |
| RNU1-1 | -0.059700904 |
| RNU6-75P | -0.059702149 |
| IGLV5-37 | -0.059714009 |
| RP11-74K11 | -0.059718209 |
| RPP40P2 | -0.059719392 |
| LIN9 | -0.059730548 |
| RP11-78J21 | -0.059734736 |
| TRH | -0.059744513 |
| AC092578 | -0.05974477 |
| RP11-136G2 | -0.059746543 |
| AC005517 | -0.059748471 |
| NXF1 | -0.0597504 |
| MED28P3 | -0.059759314 |
| RP11-715G15 | -0.059766734 |
| RP11-677I18 | -0.059769597 |
| TPO | -0.059774146 |
| RP11-165N19 | -0.059779747 |
| MAGI2-AS1 | -0.059780216 |
| MIR299 | -0.059780823 |
| HCG4P11 | -0.059785497 |
| NUP107 | -0.059796188 |
| ASTN2 | -0.059797491 |
| TRIM36 | -0.05980177 |

| | |
|---|---|
| DNASE1 | -0.059817343 |
| RNU4-74P | -0.059818899 |
| RP11-503I22 | -0.059819998 |
| PPM1A | -0.059822813 |
| SEPT7P6 | -0.059825238 |
| RP11-285M22 | -0.059826085 |
| NOP58 | -0.059826106 |
| RP11-388G3 | -0.059826597 |
| AC195454 | -0.059831464 |
| ACSS3 | -0.059834441 |
| MT1DP | -0.059841419 |
| AC002072 | -0.059849827 |
| MRPS33P3 | -0.059854565 |
| RP11-191L9 | -0.059858209 |
| LINC00461 | -0.059864318 |
| OR4C45 | -0.059866543 |
| YBX2P1 | -0.05987119 |
| RSL24D1P9 | -0.059872643 |
| RP4-584D14 | -0.059878331 |
| CUL9 | -0.059878555 |
| RP11-454L9 | -0.059880952 |
| AP003356 | -0.059884757 |
| RP11-768G3 | -0.059887603 |
| MAPK8IP1 | -0.059891783 |
| RP11-573M3 | -0.05989354 |
| PZP | -0.059900201 |
| INTS6 | -0.059903839 |
| RNA5SP36 | -0.059904599 |
| RUNDC3A | -0.059910398 |
| ZNF607 | -0.059914147 |
| AL391994 | -0.059914392 |
| EDNRB | -0.059916417 |
| CELA3A | -0.059918657 |
| UBA2 | -0.05992464 |
| MTND2P30 | -0.059926947 |
| HMGB1P11 | -0.059927853 |
| LINC01067 | -0.059930798 |
| RHOF | -0.059932045 |
| ACACB | -0.059936771 |
| RN7SKP160 | -0.059949281 |
| C3orf38 | -0.059951749 |
| MYOG | -0.059955995 |
| CCND3P2 | -0.059956655 |
| RNA5SP423 | -0.059961158 |
| FSCB | -0.059962984 |

| | |
|---|---|
| FAM151A | -0.059965555 |
| RPS15AP32 | -0.059966017 |
| HLA-DRB1 | -0.059976661 |
| PTMAP6 | -0.059977919 |
| RPS11P5 | -0.059978261 |
| NMT1 | -0.059984736 |
| GOLGA8S | -0.059992417 |
| ZMYM4 | -0.060000923 |
| GZF1 | -0.060002542 |
| CFHR3 | -0.060008152 |
| RP11-797J4 | -0.060010337 |
| SPCS2P1 | -0.060013861 |
| OMG | -0.060018734 |
| IL17RA | -0.060031733 |
| ANAPC10 | -0.060036273 |
| RP11-700A24 | -0.060039425 |
| RN7SKP105 | -0.060043432 |
| CBLL1 | -0.060046134 |
| SNORD64 | -0.060047333 |
| CTD-2017C7 | -0.060047651 |
| CD97 | -0.060053279 |
| GPR63 | -0.060057186 |
| RNU6-165P | -0.060059029 |
| TMX2P1 | -0.060062972 |
| CDON | -0.060064107 |
| SYS1 | -0.060067958 |
| OVCH1 | -0.060068152 |
| RPL31P59 | -0.060068455 |
| ERAL1 | -0.060081708 |
| C17orf78 | -0.060082956 |
| NR2F2 | -0.060085001 |
| TRMT112P6 | -0.060088338 |
| AC091493 | -0.06010815 |
| MAP4K3 | -0.060108221 |
| RP13-977J11 | -0.060114528 |
| NENFP1 | -0.060116338 |
| TRBV26 | -0.060116359 |
| MIR550A3 | -0.060136603 |
| AC010547 | -0.060136917 |
| RP11-425M5 | -0.060142716 |
| SNORD115-14 | -0.060143845 |
| SNORD111 | -0.060147722 |
| RAMP2-AS1 | -0.060148148 |
| LY86-AS1 | -0.060148705 |
| FBXW11P1 | -0.060157852 |

| | |
|---|---|
| TRIP11 | -0.060162567 |
| DHTKD1 | -0.060169135 |
| AL137800 | -0.060176841 |
| AC093157 | -0.060176955 |
| CTD-3028N15 | -0.060181218 |
| ERAP1 | -0.060192652 |
| AC007842 | -0.060193015 |
| AOX2P | -0.060196484 |
| EFCAB14 | -0.060205434 |
| CTD-2531D15 | -0.060209363 |
| AC114755 | -0.060220389 |
| TNFRSF14 | -0.060228579 |
| RNF123 | -0.060230477 |
| RP11-565J7 | -0.060242651 |
| FFAR2 | -0.060244078 |
| C9orf78 | -0.060245808 |
| KRTAP25-1 | -0.060252621 |
| SRRM2 | -0.060257548 |
| RP5-942I16 | -0.060261032 |
| RN7SKP276 | -0.06026141 |
| FAM73A | -0.060263874 |
| NEK5 | -0.06026406 |
| RSPH4A | -0.060268302 |
| MIR140 | -0.060274529 |
| DLGAP1-AS5 | -0.060286971 |
| FBXW11 | -0.060288545 |
| KIAA1683 | -0.060312062 |
| CARS | -0.060314481 |
| BRD7 | -0.060316899 |
| RP11-451N19 | -0.060329509 |
| GHRLOS | -0.060341459 |
| SNORD89 | -0.060345146 |
| HS6ST2-AS1 | -0.060345301 |
| TCP10L2 | -0.060347264 |
| CTC-484M2 | -0.060348861 |
| SLC6A5 | -0.060353426 |
| RP5-827A12 | -0.060361971 |
| ASPDH | -0.060367641 |
| AL122019 | -0.060369856 |
| RPS4XP16 | -0.060370725 |
| CTD-3080P12 | -0.060371268 |
| AL160231 | -0.060384084 |
| HSP90AB1 | -0.060387247 |
| LL22NC03-86G7 | -0.060388531 |
| RIMKLA | -0.060389498 |

| | |
|---|---|
| NUDT3 | -0.060393089 |
| ZNF764 | -0.060395582 |
| PURA | -0.060402522 |
| RCBTB1 | -0.060405759 |
| RP1-59D14 | -0.060410299 |
| AC007556 | -0.060413783 |
| RP11-496B10 | -0.060426285 |
| ARHGAP18 | -0.060428216 |
| POP1 | -0.060430728 |
| CICP17 | -0.060431249 |
| CTD-2213F21 | -0.060432737 |
| UBQLN1P1 | -0.060437007 |
| DGKD | -0.060443045 |
| POLE3 | -0.060443231 |
| PKIB | -0.060444273 |
| QTRTD1 | -0.060444756 |
| MIR4505 | -0.060449313 |
| MTTP | -0.060459678 |
| KLC2 | -0.06047326 |
| MCCD1P1 | -0.060483546 |
| C1orf195 | -0.06048427 |
| MRPS7 | -0.06048526 |
| SNORD116 | -0.060489834 |
| RP11-52I18 | -0.060510665 |
| AC018761 | -0.060512464 |
| WASF3-AS1 | -0.06051772 |
| JADRR | -0.060519491 |
| C10orf76 | -0.060522136 |
| DIAPH3 | -0.060532835 |
| AC007279 | -0.060535403 |
| MRE11A | -0.060543012 |
| KLHL15 | -0.060547477 |
| RP11-163O19 | -0.06054873 |
| AP002381 | -0.060550095 |
| RP11-413E1 | -0.06055013 |
| GDAP2 | -0.060550696 |
| CTC-209L16 | -0.060556766 |
| KCNA5 | -0.060559046 |
| MTND4P11 | -0.060562385 |
| RNU6-1179P | -0.060562758 |
| CTB-32P11 | -0.060562758 |
| CBX3P2 | -0.060563962 |
| RPS3AP24 | -0.06057073 |
| MIR3936 | -0.060572771 |
| MTHFD2P7 | -0.060582339 |

| | |
|---|---|
| AC007238 | -0.060583832 |
| RP11-474P2 | -0.060589711 |
| HIST1H3J | -0.06059233 |
| KRT8P39 | -0.060595523 |
| RPL7P42 | -0.060598126 |
| LINC01318 | -0.060613438 |
| RP3-323P24 | -0.060618253 |
| RP11-290D2 | -0.060620526 |
| FAM8A1 | -0.06062229 |
| RN7SKP259 | -0.060622424 |
| SLC10A5 | -0.060623108 |
| CDV3 | -0.06063029 |
| FAM134C | -0.06063362 |
| LINC01036 | -0.060635263 |
| HNRNPA3P14 | -0.060638358 |
| CELF2-AS1 | -0.060643506 |
| C10orf88 | -0.060643872 |
| CTD-2235C13 | -0.06064538 |
| MTATP8P1 | -0.060656076 |
| YPEL5 | -0.060658477 |
| RP11-472N19 | -0.060659109 |
| MIR4264 | -0.060659209 |
| RP5-1125A11 | -0.060661845 |
| EEF1A1P39 | -0.0606648 |
| PTOV1-AS2 | -0.060665399 |
| ZNF614 | -0.060667482 |
| RP11-307I2 | -0.060677091 |
| RN7SL430P | -0.060681644 |
| UEVLD | -0.060682906 |
| PIGH | -0.060685399 |
| RP11-21K12 | -0.060686686 |
| SLC48A1 | -0.060687111 |
| AC084290 | -0.060688081 |
| LINC00354 | -0.060691022 |
| IGHD4-17 | -0.060693949 |
| OR6B2 | -0.060695069 |
| TRBV6-8 | -0.060698586 |
| IFNA21 | -0.060711767 |
| NBPF3 | -0.060718368 |
| RNF225 | -0.060732934 |
| RP11-56D16 | -0.060734453 |
| NBPF25P | -0.060734462 |
| ZCCHC11 | -0.060734592 |
| AC108039 | -0.060735513 |
| FITM2 | -0.06073569 |

| | |
|---|---|
| POM121L6P | -0.060736038 |
| ATP5G2P4 | -0.060738802 |
| RP4-717M23 | -0.060740333 |
| EIF2AK3 | -0.06074222 |
| MTND5P9 | -0.060743153 |
| TUBB8P2 | -0.0607432 |
| BNIP3P8 | -0.060763278 |
| OR4D5 | -0.060765464 |
| GEMIN8P4 | -0.060766482 |
| CLHC1 | -0.060766854 |
| RNU6-453P | -0.060772669 |
| LINC01411 | -0.060773078 |
| RP11-406H4 | -0.060774489 |
| GUSBP10 | -0.060777071 |
| GPRASP1 | -0.060777236 |
| SCAF4 | -0.060784213 |
| RP11-701H24 | -0.060786604 |
| GPR34 | -0.060796139 |
| RN7SL361P | -0.060799168 |
| SSU72P4 | -0.060803198 |
| THRAP3P1 | -0.060815314 |
| RP1-20B11 | -0.06081692 |
| SLC24A1 | -0.060817294 |
| TRPV2 | -0.060819191 |
| EMC2 | -0.060819601 |
| RNU6-933P | -0.060820727 |
| SAPCD1-AS1 | -0.06082493 |
| RP1-224A6 | -0.060829598 |
| PAXIP1-AS1 | -0.060830857 |
| RP11-402L5 | -0.060831413 |
| RP11-1293J14 | -0.060839515 |
| RP11-443N24 | -0.060842784 |
| LINC01490 | -0.060843254 |
| CETN3 | -0.060853463 |
| TIMM23B | -0.060860384 |
| ALS2 | -0.060873426 |
| ZNF527 | -0.060875064 |
| HAUS6 | -0.060879287 |
| ARID1A | -0.060879306 |
| AC006978 | -0.060879417 |
| RAB11FIP2 | -0.060880422 |
| ABR | -0.060886004 |
| AF238378 | -0.060904667 |
| MCOLN2 | -0.060909856 |
| LINC00987 | -0.060912014 |

| | |
|---|---|
| GATA2 | -0.060916758 |
| CTD-2373J6 | -0.060918392 |
| RP13-235E23 | -0.060925012 |
| OR5M11 | -0.060926572 |
| SUGT1 | -0.060927289 |
| RP11-526N18 | -0.060929923 |
| GDAP1 | -0.060933652 |
| CTA-714B7 | -0.060934632 |
| AC090181 | -0.060938038 |
| LY75 | -0.060941411 |
| RP11-685M7 | -0.060942471 |
| RP11-27I1 | -0.060960946 |
| XX-2136C48 | -0.060961979 |
| RAMP3 | -0.060967291 |
| B3GALNT1 | -0.060969486 |
| VN1R71P | -0.060971466 |
| CDADC1 | -0.060972724 |
| RNY4P23 | -0.060972759 |
| RPL17P3 | -0.060974012 |
| RP11-439L8 | -0.060978042 |
| LRRC74A | -0.060988484 |
| AC016706 | -0.060990738 |
| PPP2R5A | -0.060993655 |
| RPL7AP6 | -0.060994715 |
| ASIC5 | -0.06099577 |
| RP11-725P16 | -0.060999955 |
| STPG2 | -0.061001295 |
| RP11-354P11 | -0.061005885 |
| RP11-107C10 | -0.061006409 |
| C11orf63 | -0.061011367 |
| AL355537 | -0.061011831 |
| RP11-452D12 | -0.061015821 |
| SPRYD4 | -0.061017023 |
| DPEP2 | -0.061021675 |
| MIR6891 | -0.061022403 |
| CES4A | -0.061031992 |
| BNIP3P27 | -0.061034324 |
| SRP9P1 | -0.061038062 |
| GOLGB1 | -0.061040596 |
| MANSC4 | -0.061043131 |
| DEFB104A | -0.061045011 |
| TCEB1P21 | -0.061047948 |
| INMT | -0.061049137 |
| AP001619 | -0.061058193 |
| C11orf54 | -0.061058309 |

| | |
|---|---|
| ANKRD52 | -0.061062086 |
| RP11-554E23 | -0.061068189 |
| AC099668 | -0.061069519 |
| RP11-484D2 | -0.061071097 |
| LST1 | -0.061071835 |
| ANAPC1 | -0.061075854 |
| LOC100421166 | -0.061078371 |
| HNRNPA1P1 | -0.061082699 |
| FAM47E | -0.061083482 |
| SIRT7 | -0.061084431 |
| RP4-700A9 | -0.061095566 |
| GTF2H2C | -0.061098608 |
| CU459211 | -0.061098973 |
| OSER1-AS1 | -0.061099315 |
| EPHA10 | -0.061101827 |
| HSPD1 | -0.061112693 |
| RP11-31I22 | -0.061116289 |
| OR1J4 | -0.061120401 |
| ZNF431 | -0.061130033 |
| KIAA0232 | -0.061138591 |
| PURB | -0.06113995 |
| WDFY1 | -0.061143875 |
| MIR331 | -0.061144915 |
| TSPYL2 | -0.061152025 |
| RN7SL333P | -0.061153838 |
| FUCA1P1 | -0.061155505 |
| ATP6V0A2 | -0.061158927 |
| RP11-91I11 | -0.061175006 |
| ANKRD31 | -0.061175496 |
| CPNE3 | -0.061176714 |
| AMZ2P1 | -0.061177272 |
| CTB-174D11 | -0.061185123 |
| RP11-486F17 | -0.061186033 |
| RP4-566D2 | -0.061190136 |
| CFAP44 | -0.061197701 |
| AL592546 | -0.061199073 |
| OR52B5P | -0.061201839 |
| CTC-550B14 | -0.061203301 |
| RP11-143G3 | -0.061204835 |
| AC096670 | -0.061213747 |
| RBBP5 | -0.061215599 |
| ATG2B | -0.061220921 |
| MARK3 | -0.061227749 |
| RNU6-356P | -0.061234927 |
| Z98949 | -0.061241441 |

| | |
|---|---|
| RP11-325N19 | -0.061242932 |
| FBXL19-AS1 | -0.061242949 |
| AC106775 | -0.061248522 |
| CTSC | -0.061249033 |
| AGFG2 | -0.061249443 |
| SLC33A1 | -0.061253313 |
| TBC1D3B | -0.061255339 |
| CCDC74A | -0.061256885 |
| LINC01355 | -0.06125709 |
| RN7SL255P | -0.061261362 |
| RP11-589F5 | -0.061265405 |
| AC007091 | -0.061270243 |
| RAB33A | -0.061274355 |
| SPATA21 | -0.061289796 |
| RNU6-1311P | -0.061290763 |
| NPTN-IT1 | -0.061292507 |
| TNFSF13 | -0.061293966 |
| NUS1P3 | -0.061294108 |
| CDKN1B | -0.061306915 |
| COX6CP16 | -0.061326361 |
| LINC00707 | -0.061328894 |
| AL590096 | -0.061331865 |
| AC109333 | -0.061336136 |
| RP11-414J4 | -0.061337233 |
| RP11-365P13 | -0.061339984 |
| SRSF3 | -0.06134141 |
| RP11-274B18 | -0.061344303 |
| FAM215B | -0.061346294 |
| AC012123 | -0.061350343 |
| RN7SL121P | -0.061351013 |
| RP11-521C10 | -0.061356618 |
| MIR3143 | -0.061358526 |
| AC099344 | -0.061358652 |
| RP11-279N21 | -0.061361532 |
| RP1-91N13 | -0.061366192 |
| RP11-158C21 | -0.061367732 |
| AC138517 | -0.061369636 |
| SPATA25 | -0.061372369 |
| SNORD113-6 | -0.061372647 |
| LINC00691 | -0.06137595 |
| SHISA7 | -0.061376012 |
| PRICKLE4 | -0.061378453 |
| RP5-1068E13 | -0.061393077 |
| MIR200A | -0.061394526 |
| SYT7 | -0.06139704 |

| | |
|---|---|
| HNRNPA1P51 | -0.061399166 |
| RP11-144G7 | -0.061409968 |
| POLR2K | -0.061413115 |
| ENPP7P8 | -0.061413813 |
| AHCTF1 | -0.061430976 |
| RP11-661C3 | -0.061443369 |
| MAGEA8 | -0.061462374 |
| RP11-196G18 | -0.061466903 |
| NPM1P19 | -0.061472152 |
| FHDC1 | -0.061485379 |
| RNU6-131P | -0.061492418 |
| DENND4C | -0.06149457 |
| CTC-512J12 | -0.061498908 |
| HNRNPA1P6 | -0.061500827 |
| EXOC6 | -0.061502235 |
| KRT18P1 | -0.061509985 |
| HNRNPC | -0.061511798 |
| CYCSP40 | -0.061514183 |
| RP11-219F10 | -0.06151463 |
| CTD-2619J13 | -0.061519464 |
| AP005530 | -0.061519498 |
| SLC9C1 | -0.061521021 |
| NPIPB3 | -0.06152685 |
| SPATS2 | -0.061527967 |
| RP11-9E13 | -0.061537476 |
| ACTN4P2 | -0.061537979 |
| AC005253 | -0.061544656 |
| AC010528 | -0.061545883 |
| RP11-32D17 | -0.06155028 |
| PSMA5 | -0.061557698 |
| PILRB | -0.061572266 |
| IRGM | -0.06158244 |
| AP004371 | -0.061582861 |
| RP11-121C6 | -0.061588042 |
| MIR6819 | -0.061597883 |
| SNORD124 | -0.061600687 |
| HMGB1P19 | -0.061607556 |
| RP11-939C17 | -0.061617294 |
| HMGB1P8 | -0.061625375 |
| RP11-259A24 | -0.061627537 |
| RP11-490B18 | -0.06163625 |
| LINC00854 | -0.061637088 |
| RP11-33E12 | -0.061642111 |
| RP11-234K19 | -0.061656631 |
| DNAJA3 | -0.061661312 |

| | |
|---|---|
| OSBP | -0.061662652 |
| ACER3 | -0.061663098 |
| SNAP47-AS1 | -0.061664486 |
| LINC01276 | -0.061671494 |
| RNU6-886P | -0.061673317 |
| RNU6-1147P | -0.061674762 |
| UBP1 | -0.061682392 |
| AC007249 | -0.061683052 |
| RP13-643D4 | -0.061685446 |
| BBS7 | -0.06168602 |
| RNU6-895P | -0.061686932 |
| RP11-380B4 | -0.061691658 |
| RN7SL784P | -0.061702245 |
| RP11-299J3 | -0.061709277 |
| RP11-323J4 | -0.061720659 |
| SSTR5-AS1 | -0.061721361 |
| CANX | -0.061724459 |
| SSTR5 | -0.061724562 |
| RP11-314M24 | -0.061727949 |
| AC244021 | -0.06172819 |
| PIP5K1A | -0.061728199 |
| NAP1L1 | -0.061732645 |
| RP1-167G20 | -0.06173365 |
| RP11-414C16 | -0.061740321 |
| ALG2 | -0.061741762 |
| RPS12P20 | -0.06174529 |
| RTN3 | -0.061752088 |
| NIFKP7 | -0.061756332 |
| OR5P3 | -0.061759777 |
| UBE2F-SCLY | -0.061767001 |
| MTND1P28 | -0.061768221 |
| LEKR1 | -0.061769093 |
| RNA5SP218 | -0.061772207 |
| RNU6-1267P | -0.061772428 |
| S100A7L2 | -0.061775566 |
| SIX4 | -0.061776257 |
| SLC9A3R1 | -0.061783066 |
| AC087294 | -0.06178543 |
| RGS11 | -0.061799886 |
| RNU6-512P | -0.061802902 |
| RNA5SP123 | -0.061804294 |
| SPINT4 | -0.061818322 |
| ACE | -0.061825189 |
| RP3-394A18 | -0.06182757 |
| CTD-2302A16 | -0.06183356 |

| | |
|---|---|
| RP11-560J1 | -0.061833952 |
| GABRR2 | -0.06183821 |
| PARP8 | -0.061841227 |
| RNY4P36 | -0.061845902 |
| OR10G6 | -0.061851258 |
| SOCS2P2 | -0.061853004 |
| RP11-498M5 | -0.061863461 |
| KIAA2022 | -0.061875808 |
| PRELID1P4 | -0.061887328 |
| CCDC179 | -0.061899963 |
| AC010999 | -0.061904465 |
| HMGB1P41 | -0.061911173 |
| RP11-529F4 | -0.06192484 |
| GPC5 | -0.061925001 |
| NRADDP | -0.061928869 |
| AC007435 | -0.061929203 |
| MAP3K14 | -0.061929621 |
| PHTF1 | -0.061932784 |
| RP11-102J14 | -0.061937534 |
| RP11-417P24 | -0.061945071 |
| AC064834 | -0.061949866 |
| RP13-20L14 | -0.061950367 |
| YPEL5P2 | -0.061955336 |
| AC004381 | -0.061957083 |
| HMGN2P31 | -0.061957711 |
| Z82214 | -0.061957768 |
| RP11-702H23 | -0.061971931 |
| RP4-753M9 | -0.061972668 |
| CTD-2510F5 | -0.061977177 |
| IZUMO2 | -0.061979829 |
| EIF1B-AS1 | -0.061982182 |
| IZUMO1R | -0.061988714 |
| STYX | -0.061999206 |
| NFKB1 | -0.062002425 |
| RP11-62L10 | -0.062006756 |
| PGAM5 | -0.062009104 |
| CSAD | -0.062024026 |
| SLC1A2 | -0.062031598 |
| RP11-20E23 | -0.06203196 |
| EZH2 | -0.062037701 |
| AC103996 | -0.062040142 |
| EDC3 | -0.062041775 |
| GAS5-AS1 | -0.062043133 |
| RP3-453C12 | -0.062043321 |
| ZNF280B | -0.062043745 |

| | |
|---|---|
| XXbac-BPG32J3 | -0.062044987 |
| ELAVL3 | -0.062050333 |
| GRM4 | -0.062053422 |
| SLC9A3P3 | -0.062055355 |
| TXNL4B | -0.062056418 |
| APOC2 | -0.062062967 |
| PKP4 | -0.062063116 |
| KRT17P8 | -0.062063477 |
| DCDC1 | -0.062076487 |
| UNK | -0.0620852 |
| RXFP4 | -0.062086196 |
| RP11-377D9 | -0.062091393 |
| RP11-77M5 | -0.06209723 |
| OR6Y1 | -0.062110277 |
| TRAF5 | -0.06212003 |
| AC092384 | -0.062135943 |
| RP5-892K4 | -0.062148811 |
| SETP8 | -0.062149308 |
| SLC10A7 | -0.062151417 |
| AGBL5-IT1 | -0.062155309 |
| DAOA | -0.062156111 |
| CNN2P9 | -0.062159 |
| UBR7 | -0.062164311 |
| NADSYN1 | -0.062170451 |
| PDCD1LG2 | -0.062171251 |
| RNF219 | -0.062172255 |
| ZBTB20-AS4 | -0.06217883 |
| SLMO2P3 | -0.062182916 |
| SLIT1 | -0.062183028 |
| KRT18P56 | -0.062186779 |
| RAB5A | -0.06218753 |
| IQCB2P | -0.062189356 |
| CGNL1 | -0.062192573 |
| RP3-380E11 | -0.062194906 |
| CTD-2008O4 | -0.062220956 |
| ENPP7P11 | -0.06222107 |
| PHF10 | -0.062221244 |
| BTF3P10 | -0.062224525 |
| MIER1 | -0.062226416 |
| NTSR2 | -0.062228899 |
| KIF5B | -0.062231514 |
| RP11-864J10 | -0.06223824 |
| RNU6-930P | -0.062238894 |
| AC026254 | -0.062241704 |
| CTD-2609K8 | -0.062242994 |

| | |
|---|---|
| ZNF853 | -0.062247161 |
| RP11-413B19 | -0.062249927 |
| RP6-159A1 | -0.062252259 |
| TAPT1-AS1 | -0.062257915 |
| RNU6-275P | -0.06226309 |
| NRARP | -0.062267887 |
| HSPA9 | -0.062272688 |
| ELAVL1 | -0.062278102 |
| RNU6-845P | -0.062280765 |
| AC010970 | -0.062299263 |
| EIF4E3 | -0.062301563 |
| RNU6-253P | -0.06230482 |
| KRT18P25 | -0.06230876 |
| MIR7974 | -0.062311291 |
| OGG1 | -0.062314531 |
| RP5-972B16 | -0.062316462 |
| TP53INP1 | -0.062321415 |
| AC058791 | -0.062321471 |
| OCM2 | -0.062322953 |
| ZNF585A | -0.062323816 |
| RP11-37B2 | -0.06232562 |
| RP5-1154L15 | -0.062334106 |
| RP1-85D24 | -0.062334318 |
| AC025048 | -0.062338008 |
| RP11-496I9 | -0.062345249 |
| EPM2AIP1 | -0.062346663 |
| GPR180 | -0.062350626 |
| RP11-260A9 | -0.062351503 |
| RP4-809F4 | -0.062353273 |
| RP11-513D5 | -0.062357379 |
| CCDC97 | -0.062357789 |
| KB-1269D1 | -0.062361944 |
| AC024060 | -0.062362465 |
| TEK | -0.062363166 |
| RP11-291O7 | -0.06237181 |
| RNY4P29 | -0.062375126 |
| RP11-365H23 | -0.062387807 |
| RP11-691N7 | -0.062388629 |
| CEACAMP11 | -0.062395925 |
| AC009410 | -0.062400164 |
| FAM193B | -0.06240114 |
| ZNF671 | -0.062406517 |
| RP5-899E9 | -0.062409188 |
| DNHD1 | -0.062414722 |
| PRMT9 | -0.062415094 |

| | |
|---|---|
| RP11-163E9 | -0.062415299 |
| MIR6731 | -0.062415823 |
| LPIN1 | -0.062417568 |
| FNDC3A | -0.062422499 |
| ACYP1 | -0.062425885 |
| CCDC108 | -0.062427614 |
| UBR1 | -0.062428676 |
| RP11-391J13 | -0.062431094 |
| CTC-459F4 | -0.062431206 |
| HRH2 | -0.062434642 |
| RP11-362L22 | -0.062440552 |
| TPT1P10 | -0.062447431 |
| SENP5 | -0.062452975 |
| AC005020 | -0.062457947 |
| GP5 | -0.062460754 |
| RP11-343B22 | -0.062470573 |
| EGFL8 | -0.062475338 |
| RPS26P35 | -0.062495235 |
| GABPB1-AS1 | -0.062497498 |
| HIF1AN | -0.062500381 |
| ST13P19 | -0.062507202 |
| RNASE6 | -0.062519192 |
| NBPF5P | -0.062521053 |
| AC004383 | -0.062531034 |
| RP11-399D15 | -0.06253487 |
| OLMALINC | -0.062540197 |
| PDK1 | -0.062544253 |
| RNU2-30P | -0.062545691 |
| CST3 | -0.062549798 |
| UNCX | -0.062554869 |
| SULT2A1 | -0.062568318 |
| HCN3 | -0.062571529 |
| SYK | -0.062571864 |
| AC115283 | -0.062577648 |
| NUDT12 | -0.062581036 |
| NXPE4 | -0.062581794 |
| C2orf42 | -0.062582078 |
| RP4-773N10 | -0.062586227 |
| AC010287 | -0.062587394 |
| GPR139 | -0.062587682 |
| ATP1B3-AS1 | -0.062601534 |
| TANK | -0.062602618 |
| RNU1-130P | -0.062604873 |
| TNRC18P3 | -0.062605343 |
| CD1C | -0.062613208 |

| | |
|---|---|
| ZBTB37 | -0.062614898 |
| TMEM75 | -0.062622723 |
| ELOVL7 | -0.062649114 |
| AC004813 | -0.062649137 |
| PSMA2P3 | -0.062654872 |
| CTC-421K24 | -0.062657865 |
| AC116618 | -0.062662245 |
| NOL4L | -0.062663142 |
| CTD-2591A6 | -0.062664819 |
| ACAP2 | -0.062667756 |
| RP5-1022P6 | -0.062668984 |
| TOMM22P5 | -0.062669973 |
| NACA2 | -0.062674805 |
| EIF3C | -0.062675589 |
| DAZL | -0.062676444 |
| ATF7IP | -0.062677152 |
| RP11-214L13 | -0.062681302 |
| RP11-747E23 | -0.062683578 |
| PCM1 | -0.062688241 |
| EXD1 | -0.062689932 |
| RBM5-AS1 | -0.062690776 |
| BLNK | -0.062691757 |
| RP5-997D16 | -0.062692762 |
| LINC00431 | -0.062694607 |
| AC011753 | -0.062703028 |
| MIR5007 | -0.062703487 |
| RPL34P17 | -0.062706747 |
| APOA5 | -0.062722696 |
| MPHOSPH10 | -0.062725694 |
| ZNF550 | -0.062727108 |
| RP11-241M13 | -0.062728286 |
| PKD2L1 | -0.062729341 |
| RP11-675F6 | -0.062730706 |
| MIR6719 | -0.062731235 |
| LNX1 | -0.062738103 |
| COTL1P2 | -0.062738355 |
| YWHAZP7 | -0.062738878 |
| MIR106B | -0.062741022 |
| TNFAIP8 | -0.062741452 |
| RNU6-763P | -0.062743232 |
| AC009473 | -0.062757407 |
| NAPA | -0.062762235 |
| KIAA0319L | -0.062773752 |
| GPR52 | -0.062784902 |
| SUMO2P3 | -0.062792117 |

| | |
|---|---|
| TMEM178B | -0.062793725 |
| AC136289 | -0.062794086 |
| RP11-301N24 | -0.062801165 |
| RP11-739L10 | -0.062805967 |
| EXOG | -0.06280659 |
| KIAA1107 | -0.06281219 |
| MTND2P16 | -0.062814911 |
| MIR5696 | -0.062829653 |
| TAS1R2 | -0.062846958 |
| DNM1P38 | -0.062851563 |
| AC138761 | -0.062851635 |
| RP6-29D12 | -0.062852429 |
| RNU6-653P | -0.062857352 |
| AC016768 | -0.062862367 |
| TXLNB | -0.062864211 |
| AL590410 | -0.062864839 |
| RNU6-462P | -0.062877574 |
| AC091180 | -0.062878635 |
| XXbac-BPG248L24 | -0.062878686 |
| BSDC1 | -0.062880845 |
| FAXDC2 | -0.062882166 |
| ATXN3L | -0.062883455 |
| NCRNA00250 | -0.062883724 |
| ZNF440 | -0.062890724 |
| RNU6-611P | -0.062890783 |
| RP11-80B9 | -0.062894561 |
| CTD-3032J10 | -0.062895944 |
| AC139452 | -0.062898308 |
| RP11-806K15 | -0.062909734 |
| TTC19 | -0.062922242 |
| MKL2 | -0.06293454 |
| RP11-481J8 | -0.062935011 |
| TBRG1 | -0.062940978 |
| MIR1827 | -0.062970081 |
| PCIF1 | -0.062971472 |
| RP11-401N16 | -0.062973625 |
| RP11-161P17 | -0.0629775 |
| RP11-209M4 | -0.062979614 |
| STX7 | -0.062983528 |
| TFIP11 | -0.063001743 |
| RP11-757F18 | -0.063006804 |
| MIR6796 | -0.063009965 |
| RP5-884G6 | -0.063010078 |
| C19orf38 | -0.063010357 |
| SMARCA2 | -0.063011995 |

| | |
|---|---|
| RP11-571I18 | -0.063017063 |
| RP11-712C19 | -0.063025057 |
| AC074289 | -0.06302513 |
| RP11-37L2 | -0.063031442 |
| UNGP3 | -0.063033937 |
| NUP54 | -0.063042935 |
| RP11-621L6 | -0.063045133 |
| RP11-250H24 | -0.063049835 |
| FXYD6P1 | -0.063049956 |
| CTB-35F21 | -0.063050011 |
| ENPP7P3 | -0.063053985 |
| DCUN1D2 | -0.063058992 |
| SNORA55 | -0.063060073 |
| TMEM176B | -0.063061057 |
| UBR5 | -0.063070862 |
| UBN1 | -0.063084258 |
| KIAA1328 | -0.063090993 |
| GNL3 | -0.063092798 |
| TRAPPC3L | -0.063097179 |
| RP11-490O6 | -0.063097716 |
| MIR765 | -0.063097789 |
| AC098817 | -0.063098648 |
| RNA5SP43 | -0.063100232 |
| ARR3 | -0.063101732 |
| RN7SL352P | -0.063107189 |
| RP11-279O22 | -0.06310862 |
| AC068535 | -0.063109813 |
| RP11-473C18 | -0.063111593 |
| NUP214 | -0.063118343 |
| GATA3-AS1 | -0.063119696 |
| SSX1 | -0.063122332 |
| RPL3P5 | -0.063125752 |
| POTEG | -0.063143321 |
| Z69666 | -0.063153159 |
| KB-1552D7 | -0.063154618 |
| RP11-109F19 | -0.063160891 |
| QSER1 | -0.063168169 |
| AC137932 | -0.063169248 |
| AURKA | -0.063177937 |
| LINC01494 | -0.06318115 |
| RP11-778O17 | -0.063181424 |
| RP11-131L12 | -0.063183835 |
| RERG-AS1 | -0.063187797 |
| GOLGA6A | -0.063194976 |
| RP6-14C6 | -0.063212205 |

| | |
|---|---|
| CDS1 | -0.063212245 |
| SMARCE1P2 | -0.063219026 |
| IFITM4P | -0.063219233 |
| TTC21B | -0.063225957 |
| TRIQK | -0.063227706 |
| KLHL42 | -0.063230907 |
| QRSL1P3 | -0.06323172 |
| SDAD1P4 | -0.063231745 |
| LAMTOR3 | -0.063232302 |
| OR11H3P | -0.063233425 |
| ALG1L11P | -0.063235379 |
| SORL1 | -0.063235744 |
| PRC1-AS1 | -0.063247942 |
| FAM192A | -0.063248005 |
| TCEB1P22 | -0.063251559 |
| AC005007 | -0.063256673 |
| RNA5SP490 | -0.06325913 |
| HLA-DMA | -0.063263541 |
| CRNKL1 | -0.063271206 |
| RP11-69J7 | -0.063275912 |
| NCOA4 | -0.063276248 |
| RP11-379F12 | -0.063276892 |
| RNU2-40P | -0.063281023 |
| SF3A1 | -0.063281085 |
| MMP25-AS1 | -0.063281848 |
| AL133405 | -0.063290186 |
| SUPT7L | -0.063296323 |
| DDX21 | -0.063300212 |
| TMEM163 | -0.063306389 |
| RP11-681H10 | -0.063308768 |
| ZNF337 | -0.063313571 |
| PRR21 | -0.063326623 |
| RP11-498B4 | -0.063330172 |
| C19orf84 | -0.063331225 |
| KIAA1377 | -0.063341237 |
| CTD-2037K23 | -0.063345279 |
| TMEM56-RWDD3 | -0.063349758 |
| MIR6503 | -0.063354943 |
| RP11-332H17 | -0.063359819 |
| AIRN | -0.063360813 |
| RP3-496C20 | -0.06336518 |
| RSPH10B | -0.063365371 |
| GCAT | -0.063367043 |
| DPY19L4 | -0.063375415 |
| BMS1P4 | -0.063383118 |

| | |
|---|---|
| LINC01264 | -0.063397746 |
| SNORA5C | -0.063398448 |
| ZFAT | -0.063402672 |
| VN1R38P | -0.063402994 |
| OR13H1 | -0.063403617 |
| RP1-302G2 | -0.063419173 |
| MIR4727 | -0.063425219 |
| LINC00345 | -0.063433738 |
| HTR1E | -0.063449262 |
| AC062021 | -0.063451327 |
| RP11-115E19 | -0.063455385 |
| RP11-521B24 | -0.063457614 |
| RP11-29H23 | -0.063461893 |
| FCHO2 | -0.063462117 |
| FAM72B | -0.06346554 |
| TRPC5OS | -0.063472701 |
| LINC00910 | -0.063481299 |
| PLA2G4F | -0.06349224 |
| PPBPP1 | -0.063497228 |
| GTF2H3 | -0.063499681 |
| SLC22A5 | -0.063504128 |
| NDST2 | -0.063509021 |
| IL27 | -0.06351176 |
| ZNF674-AS1 | -0.063519105 |
| KCTD18 | -0.063528594 |
| MTND6P18 | -0.063528625 |
| MED12L | -0.063529431 |
| RP11-384C4 | -0.063533897 |
| ZSWIM8-AS1 | -0.063534459 |
| SRP72P1 | -0.063543748 |
| NSUN6 | -0.063544651 |
| MTRNR2L4 | -0.063547837 |
| RP11-87N3 | -0.063553964 |
| ZNF77 | -0.063558084 |
| CD1B | -0.06356232 |
| MFSD4 | -0.063564168 |
| TMSB4XP3 | -0.063567096 |
| AP001628 | -0.063571642 |
| RN7SKP170 | -0.063576576 |
| CECR3 | -0.063576886 |
| AE000658 | -0.063581281 |
| ATP1A4 | -0.063587099 |
| BUB1 | -0.063608784 |
| TEX43 | -0.063609442 |
| RP11-666G4 | -0.063610399 |

| | |
|---|---|
| FBXO47 | -0.063612788 |
| NSUN3 | -0.063615668 |
| EIF4HP2 | -0.063624729 |
| IFI27 | -0.063631761 |
| LACTB2 | -0.063639799 |
| RP11-61A14 | -0.063641639 |
| RNU7-88P | -0.06364918 |
| SNHG1 | -0.06365139 |
| NEK2P2 | -0.063652115 |
| E2F4P1 | -0.063652379 |
| AL109620 | -0.063656779 |
| PRR13 | -0.063659 |
| POLR2KP2 | -0.063660706 |
| RP11-267J23 | -0.063663223 |
| SNAP29 | -0.063671968 |
| SSUH2 | -0.063677103 |
| OVOL1-AS1 | -0.063677367 |
| RWDD2A | -0.06368594 |
| TRIM4 | -0.06368821 |
| LCN10 | -0.06368898 |
| AL157931 | -0.063706228 |
| SPATA13-AS1 | -0.063710116 |
| CTD-2655K5 | -0.063714835 |
| TRIM60P19 | -0.063716803 |
| RP11-49L2 | -0.063719099 |
| NPM1P36 | -0.063719298 |
| AL391261 | -0.063728377 |
| TRIM67 | -0.063733316 |
| MIR6512 | -0.063735671 |
| SNORD115-6 | -0.063740266 |
| RP3-471M13 | -0.06374878 |
| JAKMIP3 | -0.063749069 |
| FDCSP | -0.063750979 |
| PGAM1P3 | -0.063757087 |
| BEND3P2 | -0.063765447 |
| DUTP8 | -0.063773784 |
| CCNA2 | -0.063777833 |
| TNPO1P1 | -0.06378236 |
| OR7C2 | -0.063784651 |
| MIR4793 | -0.063784818 |
| RRN3P1 | -0.063786484 |
| PHF14 | -0.063791526 |
| AC017006 | -0.063797684 |
| RP11-690D19 | -0.06379988 |
| IFIT6P | -0.063804155 |

| | |
|---|---|
| SCARNA20 | -0.063818242 |
| AC006539 | -0.063819915 |
| TPR | -0.063823416 |
| AC090286 | -0.063838986 |
| RP11-433M22 | -0.063845436 |
| ZNF724P | -0.06385218 |
| CCR9 | -0.063856517 |
| NUP37 | -0.063857018 |
| UBE2Q2P2 | -0.063859399 |
| RP11-6B19 | -0.063859518 |
| VWA9 | -0.063863232 |
| KCTD9P1 | -0.063866258 |
| PIP4K2C | -0.063867511 |
| CELSR3 | -0.063882991 |
| RP11-81B23 | -0.063884371 |
| GOLGA8R | -0.063889076 |
| FAXC | -0.063895792 |
| APOL2 | -0.063896536 |
| IQCC | -0.063898954 |
| RN7SKP283 | -0.063904215 |
| PYROXD1 | -0.063904983 |
| TWF1P1 | -0.063909393 |
| RP11-570K4 | -0.063915488 |
| WNT3 | -0.063918304 |
| FPR2 | -0.063922315 |
| KIAA1407 | -0.063922714 |
| C1orf127 | -0.06392334 |
| IWS1 | -0.063931123 |
| RUNDC3B | -0.063938305 |
| ZMIZ1 | -0.063940035 |
| RP1-74B13 | -0.063947745 |
| MIR3159 | -0.063950002 |
| RN7SL18P | -0.063953274 |
| AC009263 | -0.063954718 |
| C12orf73 | -0.063962399 |
| MTOR | -0.063962474 |
| MTR | -0.063963087 |
| TFDP1 | -0.063966548 |
| RP4-562D20 | -0.06397178 |
| RNU7-79P | -0.063973686 |
| ZNF248 | -0.063974921 |
| CCT2 | -0.063984037 |
| PIGQP1 | -0.063991167 |
| CTD-2179L22 | -0.063994571 |
| SERPINI1 | -0.063998252 |

| | |
|---|---|
| LRRC37A17P | -0.064007927 |
| KCNAB2 | -0.064010215 |
| CTD-2011G17 | -0.06401316 |
| BTRC | -0.064015536 |
| SYDE2 | -0.064020337 |
| ARNTL | -0.064020951 |
| MTND4P8 | -0.064020987 |
| KRT222 | -0.064028502 |
| ANAPC5 | -0.064034235 |
| CSTF3-AS1 | -0.064035309 |
| AC073842 | -0.064036133 |
| Metazoa_SRP | -0.064046142 |
| TPT1P2 | -0.064053057 |
| PLLP | -0.064053361 |
| APOBEC2 | -0.06406123 |
| ARHGEF11 | -0.064062087 |
| RP11-47L3 | -0.064064078 |
| XXbac-BPG16N22 | -0.064074621 |
| TP53TG5 | -0.064078348 |
| PDSS1P2 | -0.06408369 |
| CST8 | -0.064087451 |
| CTD-2529O21 | -0.064093694 |
| GPC6-AS1 | -0.064098308 |
| M6PR | -0.064098796 |
| RP3-462C17 | -0.064099247 |
| LINC00643 | -0.064108071 |
| BNIP2 | -0.064112489 |
| RP11-109A6 | -0.064115026 |
| CTD-3224K15 | -0.064116581 |
| MT-ND4L | -0.064117792 |
| RP11-462G12 | -0.064138332 |
| TLDC2 | -0.064145254 |
| RP11-1399P15 | -0.064152058 |
| LINC01126 | -0.064154777 |
| RNA5SP317 | -0.064157658 |
| RP11-91P24 | -0.064172176 |
| WNT16 | -0.064178077 |
| AC092071 | -0.064179857 |
| SYP | -0.064180492 |
| RP11-20I6 | -0.064182818 |
| MTND2P7 | -0.064186751 |
| ACCSL | -0.064189732 |
| PRPF8 | -0.064193367 |
| CEP68 | -0.064193684 |
| STAM-AS1 | -0.064196031 |

| | |
|---|---|
| NEUROG1 | -0.064196689 |
| RNGTT | -0.06419893 |
| AL450244 | -0.064200167 |
| HAGH | -0.064202075 |
| FDXACB1 | -0.064210987 |
| NFYA | -0.064215675 |
| MRPS18C | -0.064216829 |
| FAM35A | -0.064218094 |
| ZBTB46-AS1 | -0.064225003 |
| SLC25A5P5 | -0.064225802 |
| LINC00174 | -0.064226429 |
| XRRA1 | -0.064230411 |
| NR3C2 | -0.064231378 |
| RNU6-262P | -0.064238719 |
| AL031594 | -0.064249841 |
| CPEB4 | -0.064254356 |
| PTGES3L | -0.064254431 |
| AP5M1 | -0.06426258 |
| RP1-137K2 | -0.064263087 |
| ZDHHC6 | -0.06426524 |
| MEOX2 | -0.064267343 |
| PCBP2 | -0.064273873 |
| LTB | -0.064274617 |
| PPP2R1B | -0.064277483 |
| PNLIPP1 | -0.064280182 |
| PI4K2B | -0.064292274 |
| RP3-428A13 | -0.064294535 |
| CTC-359M8 | -0.06430303 |
| CNOT3 | -0.06430554 |
| RP11-207C16 | -0.064307584 |
| VN1R105P | -0.064308657 |
| CACYBPP2 | -0.064309296 |
| LRPPRC | -0.064311624 |
| CTD-2085J24 | -0.064311656 |
| RP11-497J7 | -0.064312109 |
| IQSEC1 | -0.064318359 |
| IL17C | -0.064319742 |
| AF106564 | -0.064324334 |
| RP11-410E4 | -0.06432738 |
| HIST1H4I | -0.064329652 |
| TCEA1 | -0.064330992 |
| RN7SL225P | -0.064334057 |
| RNU1-58P | -0.064338793 |
| RP11-1003J3 | -0.064348242 |
| GUCD1 | -0.06434863 |

| | |
|---|---|
| CLDND1 | -0.064365747 |
| BCL2L13 | -0.064381878 |
| C16orf70 | -0.064382362 |
| AC104131 | -0.064385478 |
| RP11-216B9 | -0.064390809 |
| RNU1-103P | -0.064391006 |
| HERC2P7 | -0.064392345 |
| PECAM1 | -0.064401172 |
| ZNF416 | -0.064402084 |
| FBXO38 | -0.064402307 |
| RP11-285A1 | -0.064402767 |
| RP11-156P1 | -0.064403832 |
| DCAF4L2 | -0.064415728 |
| EIF2S3L | -0.064417899 |
| LINC01197 | -0.064425285 |
| SNORA81 | -0.064432504 |
| RP11-419C23 | -0.064436449 |
| AL593848 | -0.064452265 |
| RP4-591B8 | -0.064464072 |
| NKX1-1 | -0.064464198 |
| HLCS-IT1 | -0.06446627 |
| EED | -0.064468542 |
| WNK1 | -0.064489492 |
| FN3KRP | -0.064496376 |
| HEXA-AS1 | -0.064499094 |
| RP11-1336O20 | -0.064501552 |
| RNU6-204P | -0.064504924 |
| RP11-713P17 | -0.064508803 |
| FAM63A | -0.064508842 |
| LEAP2 | -0.064510907 |
| RNU7-59P | -0.064511161 |
| ZNF706 | -0.064517679 |
| FIGNL1 | -0.06451874 |
| MAST4 | -0.064522759 |
| SERINC3 | -0.06452408 |
| NOL9 | -0.064537494 |
| NUDCD1 | -0.064538908 |
| RIPK3 | -0.064539262 |
| OR2AE1 | -0.064541653 |
| ADAL | -0.064542685 |
| Nkx2-2as | -0.064542952 |
| RP11-317N8 | -0.064553811 |
| ZNF645 | -0.064557108 |
| RP11-740P5 | -0.064565422 |
| MIR3198-1 | -0.064567699 |

| | |
|---|---|
| AP000944 | -0.064570362 |
| RP11-16P6 | -0.064572993 |
| RP11-44H4 | -0.064575959 |
| AGGF1 | -0.064577849 |
| LGALS9B | -0.064583487 |
| IGBP1-AS1 | -0.064589583 |
| CTC-218B8 | -0.064590607 |
| HSDL2 | -0.064595785 |
| RPF1 | -0.064602223 |
| RP11-230F18 | -0.064605478 |
| COX15 | -0.064606092 |
| SCP2D1 | -0.064614638 |
| RP5-1055C14 | -0.064629033 |
| PIP5K1B | -0.064632587 |
| TRERF1 | -0.064638913 |
| AC006159 | -0.064642145 |
| TUBB8P11 | -0.064647321 |
| IL4 | -0.064652407 |
| XPOT | -0.064660886 |
| RNA5SP297 | -0.064677265 |
| RP11-186O14 | -0.064683008 |
| GNL3L | -0.064683026 |
| LINC01535 | -0.064688888 |
| CTD-2339F6 | -0.064699283 |
| CRYL1 | -0.064699418 |
| ZNF32-AS2 | -0.064699725 |
| RDM1 | -0.064704621 |
| MIR4432 | -0.064710137 |
| SPTY2D1 | -0.064716218 |
| CH17-373J23 | -0.064716753 |
| AL358134 | -0.064717534 |
| LINC00305 | -0.064723126 |
| RHOT1P1 | -0.064723386 |
| IGHD2-8 | -0.06472897 |
| FRY | -0.064731568 |
| CTD-2207O23 | -0.064731783 |
| RP11-27G24 | -0.064734138 |
| RP11-612B6 | -0.064734191 |
| AL133493 | -0.064737207 |
| AC016717 | -0.064737552 |
| RP11-667M19 | -0.064741988 |
| SYTL2 | -0.064742173 |
| RP11-393B14 | -0.064745285 |
| AL445183 | -0.064747127 |
| RP3-510H16 | -0.064749211 |

| | |
|---|---|
| DCTN4 | -0.064755848 |
| NCF4 | -0.064757708 |
| MFAP1 | -0.064766286 |
| RP11-266A24 | -0.064767957 |
| PRKCQ-AS1 | -0.064769805 |
| GLOD4 | -0.064769969 |
| OSGEPL1-AS1 | -0.064771368 |
| HMGB1P32 | -0.064773013 |
| DDB2 | -0.064774323 |
| APITD1 | -0.06477436 |
| AHI1 | -0.06478597 |
| UBE2W | -0.064786956 |
| NBPF26 | -0.064788091 |
| HOGA1 | -0.064791625 |
| CTB-152G17 | -0.064792947 |
| TRIP12 | -0.064793263 |
| AC012314 | -0.064793784 |
| RP11-428L21 | -0.064798829 |
| NLRC4 | -0.064808259 |
| RP11-1379J22 | -0.064811776 |
| COPS2 | -0.064812222 |
| RP11-97C16 | -0.064819162 |
| RP11-383M4 | -0.064833656 |
| WDR43 | -0.064841712 |
| HMGN2P18 | -0.064848264 |
| KLKB1 | -0.064861012 |
| SPAG9 | -0.064863406 |
| FAM215A | -0.064864362 |
| RP13-616I3 | -0.064864991 |
| RAP1AP | -0.064865674 |
| WASF5P | -0.064867713 |
| MIR4688 | -0.064868449 |
| CCR1 | -0.064869527 |
| RP5-1170D6 | -0.064881269 |
| OR2A15P | -0.064882544 |
| BRI3BP | -0.064883779 |
| AC079127 | -0.064888241 |
| RP11-430L17 | -0.064888424 |
| RP11-475A13 | -0.064891034 |
| RP11-314B1 | -0.064895033 |
| ADCY5 | -0.064895221 |
| TRIM27 | -0.064896245 |
| FAM20B | -0.064898794 |
| RP11-124D2 | -0.06490231 |
| MIR548M | -0.064903039 |

| | |
|---|---|
| RP11-355B11 | -0.064907427 |
| ASTL | -0.064911957 |
| IGLVIV-65 | -0.064914653 |
| AC099676 | -0.06492659 |
| MIR4252 | -0.064931859 |
| TRGJP | -0.064936087 |
| MDC1 | -0.064941568 |
| SLC24A5 | -0.064946705 |
| RP11-99L13 | -0.064948652 |
| SUPT4H1 | -0.064950498 |
| RNU6-964P | -0.064960691 |
| PP2672 | -0.064966 |
| SOCS5P1 | -0.064971551 |
| AC023481 | -0.064979466 |
| AC092905 | -0.064993119 |
| SNORD116-12 | -0.064993744 |
| PATL1 | -0.064997682 |
| AL592466 | -0.064998252 |
| MIR8084 | -0.064999489 |
| FBXO7 | -0.065001998 |
| TRAJ61 | -0.065008042 |
| NHLRC3 | -0.06501798 |
| LDHAP1 | -0.065034601 |
| IGHD6-6 | -0.065035324 |
| SNORA28 | -0.065038847 |
| ABRA | -0.065040237 |
| KLF3 | -0.065048568 |
| MIR199B | -0.065061736 |
| SMPD4P2 | -0.065064618 |
| NTAN1P2 | -0.065075936 |
| CPEB2 | -0.065076811 |
| LEPROTL1 | -0.065083751 |
| RPL7L1P12 | -0.06509651 |
| RP11-1000B6 | -0.065105352 |
| AC012365 | -0.065105596 |
| RNU5A-2P | -0.065108427 |
| GBA2 | -0.065108942 |
| SUGT1P3 | -0.065109408 |
| OR1H1P | -0.065113744 |
| MYO18B | -0.065116129 |
| WASF4P | -0.065116153 |
| MAPK9 | -0.065117222 |
| RP5-1142A6 | -0.065120441 |
| RNU6-768P | -0.065121293 |
| PGS1 | -0.065127976 |

| | |
|---|---|
| RP11-455P21 | -0.065131185 |
| RP11-360K13 | -0.065131501 |
| MIR2053 | -0.065138986 |
| DNAJA2 | -0.065139883 |
| EIF4A1 | -0.065144888 |
| FBXL17 | -0.065146135 |
| AD000092 | -0.065147716 |
| ZNF555 | -0.065148144 |
| RP11-511B23 | -0.065154978 |
| RBAK | -0.065156405 |
| ZNF225 | -0.065160331 |
| C1QTNF3-AMACR | -0.065163684 |
| RP11-845C23 | -0.065172822 |
| SNHG14 | -0.065177429 |
| MLPH | -0.06518195 |
| IGKV2-14 | -0.06518765 |
| RP11-703C10 | -0.065189034 |
| RTEL1-TNFRSF6B | -0.065201058 |
| AC011516 | -0.065205323 |
| AICDA | -0.065205765 |
| KIF2A | -0.065207124 |
| RP11-302M6 | -0.06521128 |
| RNU1-134P | -0.065225962 |
| EDC4 | -0.065230659 |
| RP11-436I24 | -0.06523212 |
| WWC3-AS1 | -0.065234798 |
| RNU6-880P | -0.065244183 |
| STK10 | -0.065256521 |
| SLC25A32 | -0.065264075 |
| AIF1 | -0.065265042 |
| INCENP | -0.065268168 |
| RUFY4 | -0.065275202 |
| AC110611 | -0.065278285 |
| AC083899 | -0.065284169 |
| TBP | -0.065284336 |
| RNA5SP203 | -0.065293271 |
| RP11-524N5 | -0.065294344 |
| TBL1X | -0.065294439 |
| RP11-151N17 | -0.065295613 |
| TBC1D30 | -0.065300616 |
| HNRNPA1P23 | -0.06530376 |
| CTD-2036P10 | -0.065305052 |
| ERHP1 | -0.065305846 |
| AC093230 | -0.065311328 |
| AFF1 | -0.065322087 |

| | |
|---|---|
| RP11-108L7 | -0.065322105 |
| PHAX | -0.065328375 |
| RP11-950K24 | -0.065330061 |
| SOCS7 | -0.065339408 |
| MTSS1 | -0.065346869 |
| ZNF107 | -0.065350572 |
| RNU6-859P | -0.065350787 |
| MIR552 | -0.065350979 |
| AJ006998 | -0.065371617 |
| BBS2 | -0.065373382 |
| RP11-127H13 | -0.065374737 |
| NT5C2 | -0.065375149 |
| RN7SL762P | -0.065379031 |
| RP11-401E9 | -0.065379941 |
| LINC00381 | -0.065385501 |
| HIPK3 | -0.065388843 |
| CDC7 | -0.065393457 |
| TCF19 | -0.065393829 |
| AK4P6 | -0.065404509 |
| WDR47 | -0.065404751 |
| RP5-903G2 | -0.065407585 |
| ESF1 | -0.065426128 |
| RP11-17P16 | -0.065426203 |
| ICOSLG | -0.065435896 |
| FN3K | -0.065445106 |
| CCDC43 | -0.06544626 |
| CARSP2 | -0.065449855 |
| FAM170B | -0.065456048 |
| TADA1 | -0.065460325 |
| SLC7A5P1 | -0.06546161 |
| ZNF76 | -0.065464344 |
| SMYD4 | -0.065468623 |
| RP11-375N9 | -0.065468867 |
| RAB3IP | -0.065469758 |
| C1D | -0.065471284 |
| ZDHHC4P1 | -0.065474154 |
| KIAA1244 | -0.065476828 |
| AP001432 | -0.065483445 |
| RP11-600G3 | -0.06548464 |
| RP4-635E8 | -0.065487926 |
| ZDHHC20 | -0.065490857 |
| SLC16A14 | -0.065501388 |
| FGD6 | -0.065502932 |
| C1orf109 | -0.065506151 |
| MAP1LC3B2 | -0.065508086 |

| | |
|---|---|
| CDC5L | -0.065512476 |
| MIR6882 | -0.065512761 |
| SNORA71B | -0.065514018 |
| RNU4ATAC12P | -0.065514909 |
| EIF3CL | -0.065521259 |
| MIR3176 | -0.065531411 |
| RN7SL55P | -0.065532671 |
| RP11-298J23 | -0.065533193 |
| ODCP | -0.065534606 |
| RNU6-340P | -0.065539881 |
| ZNF252P | -0.065540143 |
| EPSTI1 | -0.065541408 |
| AP002954 | -0.065541606 |
| SNX25P1 | -0.065547371 |
| ATP6V1H | -0.065551157 |
| HNRNPA1P14 | -0.065552532 |
| CTB-180C19 | -0.065563059 |
| RNU6-1039P | -0.065564973 |
| RN7SL141P | -0.065567469 |
| SPATA31A5 | -0.065568582 |
| RP11-422N16 | -0.065575628 |
| GUSBP4 | -0.065582079 |
| LINC00847 | -0.065582656 |
| RP11-264L1 | -0.065585788 |
| PXK | -0.065596703 |
| C9orf64 | -0.065598713 |
| OSBPL1A | -0.065600573 |
| AL445232 | -0.065607811 |
| RP11-277I20 | -0.065608598 |
| RP11-712P20 | -0.065611805 |
| AP001786 | -0.065615081 |
| FTOP1 | -0.065618656 |
| MRGPRX2 | -0.06562495 |
| IGHV1OR16-2 | -0.06562645 |
| CGA | -0.065628795 |
| RPS3AP34 | -0.065633721 |
| RP5-1047A19 | -0.065646101 |
| GAPDHP61 | -0.065648867 |
| RP11-861E21 | -0.065653673 |
| TRIM64B | -0.065667413 |
| RPS20P31 | -0.065674402 |
| RP11-1021O19 | -0.06567841 |
| TBC1D14 | -0.065692336 |
| RP11-344B5 | -0.065692754 |
| RP11-477H21 | -0.065695517 |

| | |
|---|---|
| AC079799 | -0.065697626 |
| DENND5B | -0.065697954 |
| AC025165 | -0.065698493 |
| CXCR4 | -0.065700317 |
| RP11-296E7 | -0.065701894 |
| RP11-599B13 | -0.065710083 |
| STAM | -0.065712057 |
| RALGPS2 | -0.065715518 |
| HSFY1P1 | -0.065727549 |
| LINC00264 | -0.065734172 |
| AC005796 | -0.065734421 |
| RP11-388N2 | -0.065737602 |
| FER1L4 | -0.0657379 |
| TIAM1 | -0.065738161 |
| ITPR3 | -0.065740059 |
| RP11-638I2 | -0.065740877 |
| MIR4276 | -0.06574126 |
| LINC00676 | -0.065742679 |
| FCF1P1 | -0.065742839 |
| MIR3161 | -0.065746768 |
| RP11-703P11 | -0.065754329 |
| PDCL3P4 | -0.065754924 |
| AC078883 | -0.065758287 |
| AC009477 | -0.065778629 |
| TMTC4 | -0.065780395 |
| JAK2 | -0.065798108 |
| AL161790 | -0.065800866 |
| RNU1-74P | -0.065807076 |
| AC093838 | -0.065810052 |
| MREG | -0.065810294 |
| MIR33A | -0.065818772 |
| RNVU1-15 | -0.065822245 |
| CCL21 | -0.065831414 |
| RP11-548P2 | -0.065831857 |
| SNORA59B | -0.065835215 |
| AC136365 | -0.065841354 |
| RP11-983P16 | -0.065845049 |
| C1orf186 | -0.065853104 |
| XXbac-BPG258E24 | -0.065866064 |
| RP1-77H15 | -0.065872579 |
| ASIC2 | -0.065873918 |
| AP001257 | -0.065876295 |
| METTL15P1 | -0.065884382 |
| CEBPZOS | -0.06588626 |
| SYNGR2 | -0.065889944 |

| | |
|---|---|
| CAMKK2 | -0.065893051 |
| RNVU1-8 | -0.065895892 |
| ZNF442 | -0.065901833 |
| NEIL1 | -0.065903898 |
| NEK1 | -0.065910782 |
| LINC01162 | -0.065911737 |
| IL33 | -0.065913889 |
| RP3-511B24 | -0.065920619 |
| RNU6-858P | -0.065927687 |
| DNAJC10 | -0.065929425 |
| ZFP90 | -0.065933314 |
| AC006487 | -0.065935174 |
| FAM161A | -0.065942709 |
| AP004289 | -0.065949987 |
| RP11-1M18 | -0.065956033 |
| ZBED8 | -0.065956496 |
| RP1-253B10 | -0.065963003 |
| LINC00094 | -0.065963492 |
| RP11-459F6 | -0.065967138 |
| RP11-511I11 | -0.065972728 |
| ZBTB12P1 | -0.065976353 |
| AP003385 | -0.06599247 |
| ZNF347 | -0.066005782 |
| RP11-477E3 | -0.066010823 |
| RBM43 | -0.066013671 |
| TATDN2P2 | -0.066020722 |
| RP5-837G17 | -0.066025563 |
| TRAJ34 | -0.066029072 |
| RAP2A | -0.066035941 |
| CYCSP53 | -0.066043453 |
| RP11-417L19 | -0.066046044 |
| RP11-563D10 | -0.06604667 |
| MIR3938 | -0.066060447 |
| CTD-2562J17 | -0.066062287 |
| RPL22P18 | -0.066066875 |
| RP11-60A8 | -0.066068515 |
| AC096772 | -0.066078771 |
| DNAAF1 | -0.066081953 |
| GCOM1 | -0.066082978 |
| CD33 | -0.06608519 |
| RP4-724E16 | -0.066085283 |
| MIR495 | -0.066086518 |
| OR5B12 | -0.066098469 |
| AP000279 | -0.066110498 |
| MGA | -0.066111219 |

| | |
|---|---|
| FGD5-AS1 | -0.066111517 |
| RP11-611E13 | -0.066112187 |
| UPF3AP1 | -0.066118705 |
| RNU6-724P | -0.066125603 |
| C6orf183 | -0.066133353 |
| ZBTB24 | -0.066133936 |
| RPL13AP7 | -0.066137453 |
| DNAJB9 | -0.066141602 |
| PDZK1 | -0.066145918 |
| MIR5009 | -0.066145934 |
| MDC1-AS1 | -0.066146712 |
| RNU2-51P | -0.066146857 |
| DNAJC19P9 | -0.066150641 |
| TUBA8 | -0.066154481 |
| GPR33 | -0.066163653 |
| PAXIP1-AS2 | -0.066164059 |
| RNU6-1250P | -0.066167109 |
| AC097533 | -0.066168038 |
| RNU6-649P | -0.066177848 |
| RP11-521L9 | -0.066178726 |
| RNU6-897P | -0.066183059 |
| GPD1L | -0.066211744 |
| LINC00921 | -0.066222703 |
| RFPL4AP7 | -0.066230325 |
| BAK1P1 | -0.066246792 |
| AL121656 | -0.066253215 |
| C6orf136 | -0.066253551 |
| SETP7 | -0.066253668 |
| SLC25A51P1 | -0.066260329 |
| RP5-1142J19 | -0.066263522 |
| CNTF | -0.06626561 |
| TRIM40 | -0.066273269 |
| AL590763 | -0.066286384 |
| LIG4 | -0.066287227 |
| DDX53 | -0.066293411 |
| ESCO2 | -0.066298148 |
| RN7SL258P | -0.06630112 |
| AC010976 | -0.066303395 |
| RP11-232D9 | -0.066305178 |
| KRT18P59 | -0.066307352 |
| RNU7-123P | -0.066312282 |
| CTD-3239E11 | -0.06632335 |
| RP11-166D18 | -0.066323497 |
| PHGR1 | -0.066332062 |
| RP11-932O9 | -0.06633255 |

| | |
|---|---|
| RP1-188K17 | -0.066335583 |
| RP11-178F10 | -0.066339755 |
| GTF3AP5 | -0.06634746 |
| AC021523 | -0.066349799 |
| ZC3H14 | -0.066353518 |
| ITPK1-AS1 | -0.066353947 |
| MIR186 | -0.066356153 |
| NUF2 | -0.066357388 |
| RP11-74C13 | -0.066358515 |
| AC004953 | -0.066359573 |
| RP5-1154E9 | -0.066359924 |
| RNU6-596P | -0.066374704 |
| MSH4 | -0.066374867 |
| CFL1P7 | -0.066375546 |
| KBTBD4 | -0.066384868 |
| ZNF703 | -0.066388329 |
| RN7SL850P | -0.066389099 |
| TRAV32 | -0.06639322 |
| GLUD1P4 | -0.066394095 |
| NRROS | -0.066401688 |
| VIPR1-AS1 | -0.066409582 |
| RP11-23J9 | -0.0664098 |
| MX1 | -0.066411325 |
| ZC3H11A | -0.066413911 |
| CTA-315H11 | -0.066417483 |
| C5AR2 | -0.066421726 |
| RGS17P1 | -0.066433578 |
| GTF3C2 | -0.066439364 |
| ZNF547 | -0.066446397 |
| AC102953 | -0.066448912 |
| SLC9A9 | -0.066455271 |
| TRIM64C | -0.066456538 |
| RP11-340M11 | -0.066459242 |
| RP11-176H8 | -0.066459513 |
| MAP1LC3BP1 | -0.066462154 |
| OR4C5 | -0.066474596 |
| FAAH | -0.066499217 |
| RP1-288H2 | -0.066502401 |
| CTD-2623N2 | -0.066504185 |
| FAR1-IT1 | -0.066505783 |
| EPC2 | -0.066507181 |
| ARFGEF1 | -0.066508148 |
| ZNF564 | -0.066508688 |
| VN1R8P | -0.066516894 |
| RNA5SP51 | -0.066517691 |

| | |
|---|---|
| TERF1 | -0.066519497 |
| MIR7853 | -0.066527373 |
| HCG4 | -0.066530495 |
| RNU6-1051P | -0.066540191 |
| HLA-B | -0.066547536 |
| CTD-2135J3 | -0.066557123 |
| TIMD4 | -0.066559722 |
| ARRB1 | -0.066564783 |
| RPL12P39 | -0.066565622 |
| MAML1 | -0.066567648 |
| CTD-3222D19 | -0.06656936 |
| KIF14 | -0.066569416 |
| METTL15 | -0.066569583 |
| RAD9B | -0.066575835 |
| IL9 | -0.066576212 |
| SEPT7P3 | -0.066577282 |
| RP1-56K13 | -0.066582048 |
| RP11-344L13 | -0.066584578 |
| AC123788 | -0.066595729 |
| RP11-108O10 | -0.066597032 |
| RP11-950C14 | -0.066597194 |
| RP11-159H20 | -0.06660229 |
| MKI67 | -0.066604357 |
| VRTN | -0.066607764 |
| OR8B3 | -0.066612581 |
| RN7SKP213 | -0.066620053 |
| RP3-407E4 | -0.06662246 |
| FKBP15 | -0.066625865 |
| ZNF160 | -0.06665288 |
| CCL15-CCL14 | -0.066654014 |
| CATSPER2P1 | -0.066661922 |
| AC005481 | -0.066664059 |
| RP11-634B22 | -0.066669582 |
| NKAP | -0.066671244 |
| NFYC-AS1 | -0.066676342 |
| MRPS10P1 | -0.066692193 |
| LY75-CD302 | -0.066692683 |
| RP1-50J22 | -0.066692865 |
| RP11-991C1 | -0.066693921 |
| AL590431 | -0.0666986 |
| CLEC4A | -0.06670425 |
| UBE2G2 | -0.066708789 |
| OR7E12P | -0.066718483 |
| CYP2T1P | -0.066729125 |
| RP11-325E14 | -0.066731175 |

| | |
|---|---|
| ZNF680 | -0.066734316 |
| RP11-511H23 | -0.066734763 |
| AC109925 | -0.066741335 |
| TINF2 | -0.066742558 |
| SLC24A4 | -0.066745647 |
| RP11-25I11 | -0.06675024 |
| EYS | -0.06675374 |
| XRCC6BP1 | -0.066757889 |
| RPSAP57 | -0.066764267 |
| TP53BP1 | -0.066765871 |
| MYO15B | -0.066773574 |
| AC093899 | -0.066775788 |
| RTKN2 | -0.066779602 |
| RP4-801G22 | -0.066784918 |
| AC092038 | -0.06678614 |
| CAPNS1P1 | -0.066790275 |
| BDH1 | -0.066791454 |
| OR2W6P | -0.066803144 |
| APOC1 | -0.066804868 |
| AC006372 | -0.066808156 |
| NBPF6 | -0.066829056 |
| SGPL1 | -0.066829371 |
| TMPOP2 | -0.066832475 |
| SSB | -0.066835418 |
| POLH | -0.066836572 |
| LMAN1 | -0.066838674 |
| RP11-703M24 | -0.066841681 |
| AC023050 | -0.066843303 |
| RNF135 | -0.066845651 |
| RP11-1022B3 | -0.066854127 |
| AC005304 | -0.066854974 |
| MIR7976 | -0.066857914 |
| MIR4668 | -0.066859097 |
| AC093110 | -0.066863431 |
| ATP11A | -0.066865801 |
| FOXN3P1 | -0.066875217 |
| IGHVIII-44 | -0.066876309 |
| RP11-122G18 | -0.066879366 |
| LRFN2 | -0.066883093 |
| RP11-1074O12 | -0.066885862 |
| MIR4443 | -0.066887571 |
| RRM2 | -0.066890397 |
| RP11-265P11 | -0.066899487 |
| RN7SL172P | -0.066901325 |
| LYST-AS1 | -0.066901978 |

| | |
|---|---|
| RP11-203J24 | -0.066904649 |
| OR10K1 | -0.066905329 |
| HRH4 | -0.066910315 |
| RNA5SP318 | -0.066943283 |
| RUFY2 | -0.0669448 |
| RP11-138I18 | -0.066956801 |
| USP44 | -0.066958867 |
| FBXL4 | -0.066961917 |
| APOC4 | -0.066962313 |
| AC005794 | -0.066965396 |
| ABBA01017803 | -0.066971879 |
| RP11-155L15 | -0.066973181 |
| POTEKP | -0.06697617 |
| AC106788 | -0.066976928 |
| IL12RB2 | -0.066983819 |
| LAIR2 | -0.066986573 |
| ZNF761 | -0.066988541 |
| SPO11 | -0.066988569 |
| FBXO8 | -0.066989118 |
| KB-318B8 | -0.066992572 |
| COQ10B | -0.066993639 |
| LINC01058 | -0.067000692 |
| SF3B2 | -0.067000765 |
| PAXIP1 | -0.06700283 |
| RP11-500M8 | -0.067006088 |
| LETM1 | -0.067007351 |
| RN7SL32P | -0.067016239 |
| AC034187 | -0.067024636 |
| CPSF4L | -0.067026602 |
| RNA5SP452 | -0.067031436 |
| AC009120 | -0.067040357 |
| R3HDM1 | -0.067042367 |
| ZNF630 | -0.06704927 |
| SHISA8 | -0.067050524 |
| RNU6-729P | -0.067050686 |
| LGALS9C | -0.067051041 |
| FAM122A | -0.067055 |
| OR4K6P | -0.067057214 |
| RP11-521E5 | -0.067062172 |
| RP5-1067M6 | -0.067064484 |
| AC090804 | -0.067072012 |
| RTEL1 | -0.067073494 |
| ARF3 | -0.067075131 |
| RP11-779O18 | -0.067077533 |
| RP1-3E10 | -0.067078324 |

| | |
|---|---|
| RNU6-1116P | -0.067079877 |
| AC079753 | -0.067095729 |
| CTD-2560C21 | -0.067099943 |
| CYP4F35P | -0.067099992 |
| RBM41 | -0.067102351 |
| LARS | -0.067110221 |
| RP1-37M3 | -0.067111774 |
| LGALS8 | -0.067111877 |
| ANKRD62 | -0.067113701 |
| RP11-246K15 | -0.067131228 |
| CTD-3234P18 | -0.067131469 |
| AC010886 | -0.067132678 |
| MCF2L-AS1 | -0.067138957 |
| MIR5197 | -0.067141677 |
| NLRP3 | -0.067150837 |
| POM121 | -0.06716213 |
| PSTPIP1 | -0.067188159 |
| IKBKB | -0.067192904 |
| RP5-862K6 | -0.067195647 |
| MIR4791 | -0.067204739 |
| SNORD116-13 | -0.067205732 |
| FGFR1OP | -0.067212272 |
| RNU7-71P | -0.067218033 |
| RP11-88H9 | -0.067219523 |
| SNORA1 | -0.06722488 |
| NCAPH | -0.067231957 |
| FCF1P8 | -0.067236944 |
| RPL7L1P13 | -0.067242755 |
| RP13-461N9 | -0.067251395 |
| SFMBT1 | -0.067253837 |
| RP11-90K17 | -0.067256668 |
| RP11-816J6 | -0.067258868 |
| EMR3 | -0.067261199 |
| TRIM39 | -0.067265633 |
| HSPA8P11 | -0.067269474 |
| MTND5P13 | -0.06727265 |
| FBXO3-AS1 | -0.067283963 |
| ZRANB1 | -0.067286899 |
| GNRHR2P1 | -0.067298074 |
| RP11-56L13 | -0.067299065 |
| AC139887 | -0.067300332 |
| MIR5691 | -0.067302759 |
| ZNF507 | -0.067304183 |
| AC009677 | -0.06730891 |
| RP11-890B15 | -0.067339868 |

| | |
|---|---|
| ST13P3 | -0.067340746 |
| UCP3 | -0.067342343 |
| RCC2P8 | -0.067343836 |
| ANKRD13C | -0.067345524 |
| ZNF268 | -0.067354343 |
| RN7SL104P | -0.067361145 |
| TPM3P1 | -0.06736128 |
| RP11-444D13 | -0.067361303 |
| DOPEY2 | -0.067370809 |
| RNU1-2 | -0.067376611 |
| XKRY2 | -0.06737918 |
| RP11-155I9 | -0.067382113 |
| RP11-371F15 | -0.067383958 |
| CTC-232P5 | -0.067385044 |
| RP11-31H15 | -0.067385337 |
| RP11-158M2 | -0.067388131 |
| DEFB112 | -0.067396775 |
| FBXW12 | -0.067401343 |
| TLK1P1 | -0.067409193 |
| OR3A5P | -0.067413863 |
| MFSD1P1 | -0.067422137 |
| BTBD1 | -0.06742257 |
| SKA1 | -0.067438459 |
| LINC01227 | -0.067441348 |
| NT5DC1 | -0.067447185 |
| TAF9 | -0.067447892 |
| RP11-762B21 | -0.067448338 |
| PGAM1P11 | -0.067450982 |
| RNU6-354P | -0.067457277 |
| NOP14 | -0.067459501 |
| INE1 | -0.067466516 |
| AC006022 | -0.067471137 |
| MIR211 | -0.067472388 |
| RP11-684B20 | -0.067473496 |
| RP11-230C9 | -0.067486145 |
| CTC-351M12 | -0.067493791 |
| RP11-322D14 | -0.067497334 |
| AC096967 | -0.067502775 |
| PPP1R13B | -0.067503057 |
| RP11-494H4 | -0.06750936 |
| CCDC146 | -0.067510108 |
| NOP56P2 | -0.067510564 |
| DHX9P1 | -0.067515117 |
| LINC00658 | -0.067529947 |
| DUSP18 | -0.067544696 |

| | |
|---|---|
| RN7SKP130 | -0.067549811 |
| RAPGEF3 | -0.067555971 |
| TOP3A | -0.067562036 |
| DLEC1 | -0.067564325 |
| AP4S1 | -0.067576269 |
| RP11-383G6 | -0.067579045 |
| TOPORS | -0.067579172 |
| RP11-1036E20 | -0.067580747 |
| YLPM1 | -0.067581907 |
| RP11-45M22 | -0.067587377 |
| WDR87 | -0.067591826 |
| RP11-445L13__B | -0.067596257 |
| IQCH-AS1 | -0.067600289 |
| HNRNPA3 | -0.067608401 |
| PIGCP1 | -0.067608578 |
| RP11-495K9 | -0.067610639 |
| IMPA1 | -0.067613611 |
| YWHAZP5 | -0.067617129 |
| CYLC2 | -0.067625597 |
| ST13P6 | -0.067629331 |
| SLC4A7 | -0.067636719 |
| RP11-389J22 | -0.06763967 |
| WNT8B | -0.067660827 |
| ACSBG1 | -0.067660923 |
| ZNF345 | -0.067661687 |
| CTA-280A3__B | -0.067662668 |
| RP1-225E12 | -0.067669888 |
| RP11-629O4 | -0.067669965 |
| VPS41 | -0.067683642 |
| AP000867 | -0.067693233 |
| RP11-393M18 | -0.067694589 |
| AC011247 | -0.067701602 |
| MIR6757 | -0.067701934 |
| MORC2-AS1 | -0.067703697 |
| GRTP1 | -0.067707568 |
| RP11-218F10 | -0.067712647 |
| RP11-321F8 | -0.067712948 |
| CNR1 | -0.067718176 |
| FAM103A1 | -0.067721076 |
| CFAP54 | -0.067732658 |
| RP11-448G15 | -0.067745077 |
| ZNHIT3 | -0.06774863 |
| NEURL1B | -0.067755142 |
| VN1R90P | -0.067769729 |
| CTD-2522E6 | -0.06777027 |

| | |
|---|---|
| CTD-2062O1 | -0.067779338 |
| DDX3X | -0.067785897 |
| AC004875 | -0.067796559 |
| RNU6-449P | -0.067796559 |
| IFI44 | -0.067796781 |
| PWAR6 | -0.067802065 |
| ZNF624 | -0.067803033 |
| NT5CP1 | -0.067816028 |
| LA16c-358B7 | -0.06781801 |
| ANGEL1 | -0.067823871 |
| HMGB1P29 | -0.067826051 |
| ITIH4 | -0.067831538 |
| RP11-217O12 | -0.067834868 |
| AL359709 | -0.067838512 |
| CST12P | -0.067840704 |
| MTND2P9 | -0.067841787 |
| PARVG | -0.067843835 |
| SNRNP200 | -0.067844449 |
| RNU6-1296P | -0.067866187 |
| SYNE2 | -0.067875948 |
| AC013733 | -0.067876149 |
| PAPD4 | -0.067878199 |
| IFT81 | -0.067880878 |
| FAM188A | -0.067887576 |
| ATMIN | -0.067893511 |
| DNAJC9-AS1 | -0.067895177 |
| OR2B7P | -0.067896461 |
| RPL7P50 | -0.067898114 |
| ATP5J2LP | -0.067903178 |
| LINC00421 | -0.067903525 |
| RPS23P9 | -0.067914152 |
| TOP3B | -0.067923826 |
| RP11-1007O24 | -0.06794583 |
| SLC25A22 | -0.067947672 |
| DHPS | -0.067948546 |
| DIS3L2 | -0.067951114 |
| RNY1P13 | -0.067956505 |
| OMA1 | -0.067964547 |
| KRT18P5 | -0.067986124 |
| EIF1B | -0.067992511 |
| DPPA3P1 | -0.068000503 |
| RNU6-82P | -0.068002697 |
| FLNB-AS1 | -0.068020103 |
| MBD6 | -0.068023322 |
| RP11-681L4 | -0.068029445 |

| | |
|---|---|
| RPS7P11 | -0.068036736 |
| AC068669 | -0.06804041 |
| GMCL1 | -0.068042578 |
| C3AR1 | -0.068044941 |
| RIMKLBP2 | -0.06804769 |
| SNORD115-4 | -0.068049791 |
| RPSAP43 | -0.068054282 |
| F7 | -0.068060081 |
| GBP2 | -0.068064737 |
| RNA5SP456 | -0.068072851 |
| STX17 | -0.068078264 |
| EPB41L3 | -0.068090059 |
| GS1-165B14 | -0.068090845 |
| SKA3 | -0.068108963 |
| RASA2 | -0.06811514 |
| INSR | -0.068131122 |
| RP11-560F18 | -0.068131192 |
| TNFRSF8 | -0.068133522 |
| CASP3 | -0.068143829 |
| MRPS36P3 | -0.068144058 |
| CTA-280A3 | -0.068149573 |
| RP11-806L2 | -0.068168767 |
| ANKLE1 | -0.068188538 |
| LA16c-349E11 | -0.068189476 |
| LL0XNC01-36H8 | -0.068199834 |
| ZNF239 | -0.068214679 |
| DCP1A | -0.068218716 |
| LINC00657 | -0.06823174 |
| RP11-415I12 | -0.068238801 |
| RP11-598O12 | -0.068241475 |
| RNU6-943P | -0.06824836 |
| CTD-2283N19 | -0.068251999 |
| UBXN7 | -0.068257955 |
| DNAJA1 | -0.068274347 |
| AL449083 | -0.068278812 |
| AC097639 | -0.068279146 |
| RP11-819M15 | -0.068283199 |
| OTOF | -0.06828737 |
| MBD2 | -0.068302125 |
| RP11-504P24 | -0.068303576 |
| MIR623 | -0.068305164 |
| AASDHPPT | -0.068321381 |
| CDC42-IT1 | -0.068321551 |
| FRMD8P1 | -0.06833596 |
| ATG13 | -0.068340191 |

| | |
|---|---|
| DNM1P32 | -0.068340639 |
| AURKAPS1 | -0.068340742 |
| DNASE1L3 | -0.068348025 |
| CTD-2173L22 | -0.06835301 |
| TBCK | -0.068366797 |
| GLTPP1 | -0.068377753 |
| USP4 | -0.06838077 |
| IL17F | -0.068384523 |
| LARS2 | -0.068400492 |
| ATF2 | -0.068403636 |
| TMEM150B | -0.068408903 |
| POLR2M | -0.068410427 |
| ZBED6CL | -0.068416902 |
| ZNF514 | -0.068421944 |
| SNORA46 | -0.068426941 |
| SLC12A5 | -0.068429516 |
| FNDC7 | -0.068431994 |
| TRAJ14 | -0.068439823 |
| FAM74A6 | -0.068447784 |
| TPST2P1 | -0.068452164 |
| SNRPGP8 | -0.068453433 |
| NFX1 | -0.06845402 |
| MANEA-AS1 | -0.068469096 |
| FAM84B | -0.068477462 |
| LETM2 | -0.068481742 |
| KHDRBS1 | -0.068482039 |
| GPR83 | -0.068487228 |
| ZNF765 | -0.068490598 |
| NFKBIZ | -0.068490933 |
| PDCL2 | -0.068491864 |
| EFHC1 | -0.068503305 |
| ATP6V1G1P6 | -0.06850634 |
| DUXAP5 | -0.068514602 |
| IGHVII-15-1 | -0.068521004 |
| RP11-359M6 | -0.068524322 |
| RP11-444E17 | -0.068530328 |
| VN1R83P | -0.068536476 |
| CRYZP2 | -0.068543963 |
| ZNF652 | -0.068545447 |
| INTU | -0.068546024 |
| RP11-389C8 | -0.068552145 |
| SLC37A1 | -0.068558173 |
| RNU7-144P | -0.068558567 |
| RP11-313M3 | -0.068560859 |
| HLA-E | -0.068562024 |

| | |
|---|---|
| COMMD8 | -0.068562787 |
| AC005306 | -0.068564841 |
| SCN11A | -0.068566663 |
| ATRIP | -0.068574649 |
| HNRNPUL1 | -0.068581262 |
| RP11-483C6 | -0.068586238 |
| RNA5SP367 | -0.068586728 |
| ITPK1 | -0.068595123 |
| RP11-45A12 | -0.068612679 |
| LINC00298 | -0.068623009 |
| CARKD | -0.068624818 |
| PPP1R3E | -0.068625506 |
| MIR7851 | -0.068627684 |
| XPNPEP3 | -0.068632595 |
| CHDH | -0.068633525 |
| MIEF1 | -0.068637656 |
| NDUFA3P1 | -0.068639912 |
| AGR3 | -0.0686407 |
| AC008703 | -0.068655056 |
| AC093484 | -0.068655864 |
| CTB-4E7 | -0.068657115 |
| RP11-30P16 | -0.068657678 |
| UBXN2B | -0.068664317 |
| AC004906 | -0.068668442 |
| APPBP2 | -0.068668708 |
| ARFIP2 | -0.068681471 |
| EHMT2-AS1 | -0.068700337 |
| RP11-343J3 | -0.068700486 |
| EFCAB13 | -0.068702589 |
| HUWE1 | -0.068707296 |
| HIST1H4H | -0.068709026 |
| ZNF121 | -0.068710105 |
| ZKSCAN3 | -0.06871178 |
| CTD-2290P7 | -0.068719003 |
| ZNF559 | -0.068729529 |
| KIF5C | -0.068730143 |
| RP11-829H16 | -0.068731073 |
| RP11-567F11 | -0.068734689 |
| ETV3 | -0.068736097 |
| RP11-55J15 | -0.068736243 |
| RP11-295D4 | -0.068736897 |
| RNU6-1167P | -0.068737982 |
| RP5-974N19 | -0.068755504 |
| RP3-415N12 | -0.068759784 |
| TMF1 | -0.068770406 |

| | |
|---|---|
| RP11-526D8 | -0.068775095 |
| AL365202 | -0.068784609 |
| RP11-418J17 | -0.068786927 |
| CTB-75G16 | -0.068789048 |
| RP11-289A15 | -0.068790396 |
| ZC3H15 | -0.06879143 |
| MIOS | -0.06879463 |
| RGS22 | -0.068796427 |
| ZBED5-AS1 | -0.068797439 |
| RP11-142L16 | -0.068809132 |
| RP11-746B8 | -0.068812091 |
| CEP78 | -0.068817924 |
| RP11-488L18 | -0.068827729 |
| RP11-341G23 | -0.068829342 |
| MTMR7 | -0.068830538 |
| RNU1-106P | -0.068832735 |
| NUDT21 | -0.06883279 |
| RGS13 | -0.068834327 |
| CEP57 | -0.068834985 |
| FGFR3 | -0.068848567 |
| UBE2J1 | -0.068850874 |
| CPM | -0.068860735 |
| EPT1 | -0.06886561 |
| RP11-404E16 | -0.068866482 |
| RAPGEF1 | -0.068876178 |
| AC067805 | -0.068881646 |
| RN7SL328P | -0.068888713 |
| IL7 | -0.06889389 |
| AC005008 | -0.068895198 |
| RNF207 | -0.068896699 |
| RP11-46C24 | -0.0689047 |
| CACNA2D4 | -0.068905946 |
| HNRNPA3P10 | -0.068918647 |
| MYADML | -0.06892391 |
| RP11-799B12 | -0.068929166 |
| RNU6-888P | -0.068929388 |
| PRAMEF1 | -0.068931687 |
| ZSWIM6 | -0.06893205 |
| ITPR1 | -0.068952795 |
| WAC | -0.068953148 |
| HMGN2P19 | -0.068963691 |
| MIR7973-1 | -0.068968787 |
| EMBP1 | -0.06897183 |
| RSBN1 | -0.06897594 |
| MEX3C | -0.06897821 |

| | |
|---|---|
| METTL16 | -0.068981168 |
| RN7SL481P | -0.068982031 |
| RNU6-312P | -0.068989375 |
| CSTF1 | -0.06899942 |
| RP11-531A21 | -0.069002445 |
| TAS2R10 | -0.069020348 |
| RP11-87E22 | -0.069022451 |
| STXBP4 | -0.069024798 |
| MIR527 | -0.069028977 |
| RP11-340F14 | -0.069033424 |
| RNU6-558P | -0.069034217 |
| LINC00486 | -0.069039574 |
| COPS4 | -0.069040836 |
| MIR1307 | -0.069049978 |
| CYB5AP5 | -0.06905513 |
| AC008268 | -0.069057523 |
| AC079172 | -0.069059838 |
| CTHRC1P1 | -0.0690609 |
| ZNF417 | -0.069065228 |
| RP11-403A21 | -0.069068545 |
| PPP1R21 | -0.069074549 |
| DNAJC14 | -0.069088615 |
| KB-67B5 | -0.06910226 |
| CYTH4 | -0.06910335 |
| ORC4 | -0.069116858 |
| RNU6-1043P | -0.069126643 |
| PTPN4 | -0.0691275 |
| MIR5707 | -0.069129987 |
| PCAT14 | -0.069130288 |
| GRHL2 | -0.069133268 |
| LL22NC03-84E4 | -0.069134587 |
| PIK3CD | -0.069138905 |
| DPPA3 | -0.069139909 |
| TMEM170B | -0.069140692 |
| FAM217A | -0.069148871 |
| HOMER2P1 | -0.069153169 |
| RP11-138A9 | -0.069154821 |
| CTB-160O22 | -0.069155052 |
| UBE2K | -0.069161306 |
| CTB-176F20 | -0.069162063 |
| FBXL5 | -0.069177735 |
| HJURP | -0.069188638 |
| NUP35 | -0.069189866 |
| SYNJ2BP | -0.069200248 |
| DAPP1 | -0.069217904 |

| | |
|---|---|
| RBSN | -0.069233124 |
| NADK2-AS1 | -0.069237076 |
| THAP1 | -0.069239189 |
| ZNF184 | -0.069270055 |
| GSG1 | -0.069285185 |
| CCDC88C | -0.069287693 |
| ANAPC4 | -0.069289368 |
| SIRPB2 | -0.069291061 |
| IKBKAP | -0.069302243 |
| PROSC | -0.069307843 |
| ITM2BP1 | -0.069310641 |
| MIR1255A | -0.069311255 |
| C7orf69 | -0.069313628 |
| NDUFA6-AS1 | -0.069324514 |
| RP11-161I2 | -0.069325472 |
| GYS2 | -0.069329618 |
| ZNF280D | -0.069333165 |
| LNPEP | -0.069337817 |
| CIPC | -0.069345221 |
| RP4-791C19 | -0.069352869 |
| GATAD1 | -0.069357538 |
| FAM95B1 | -0.069359951 |
| RP11-196G11 | -0.069373744 |
| RN7SL334P | -0.069377818 |
| RRM1 | -0.069379456 |
| PPP1CC | -0.06938433 |
| PCBP2-OT1 | -0.069389408 |
| CFAP97 | -0.069393745 |
| RP11-299A16 | -0.069402566 |
| AC007880 | -0.069407965 |
| NDC80 | -0.069419532 |
| RP11-258F1 | -0.069430389 |
| AC007919 | -0.069434429 |
| AC008752 | -0.069435529 |
| DYNLL2 | -0.06943557 |
| PRR13P5 | -0.069444575 |
| LLNLF-158E9 | -0.069444818 |
| RP11-441A11 | -0.069449144 |
| DDX26B | -0.069450975 |
| ENPP7P5 | -0.069455024 |
| RP11-359J14 | -0.069456385 |
| RNU6-909P | -0.069461475 |
| CTD-2538A21 | -0.069467485 |
| RP11-777F6 | -0.069469718 |
| RP11-255P5 | -0.069472107 |

| | |
|---|---|
| MRPS31P4 | -0.069481259 |
| MYD88 | -0.069481265 |
| RNU6-307P | -0.06948777 |
| RP11-351A11 | -0.069498322 |
| SOS2 | -0.069500131 |
| MPPE1 | -0.069500298 |
| RP11-414B7 | -0.069502302 |
| RPE | -0.069510885 |
| RP11-264E18 | -0.069511124 |
| TEDDM1 | -0.06951436 |
| HBS1L | -0.069520541 |
| RP11-100L22 | -0.069535241 |
| DNAJC11 | -0.069535425 |
| HMGN4 | -0.069538253 |
| SLC30A9 | -0.069542309 |
| PHF5CP | -0.069548592 |
| DHRS4-AS1 | -0.069554607 |
| CKAP2P1 | -0.069556551 |
| RNU6-486P | -0.069559387 |
| FAM227A | -0.069561417 |
| PTPN2P1 | -0.069563887 |
| CYCSP10 | -0.069567326 |
| CELA1 | -0.069573829 |
| TRBJ2-4 | -0.069575176 |
| ZNF141 | -0.069589288 |
| RP11-15H7 | -0.069595314 |
| PEX26 | -0.069598814 |
| SH3GL2 | -0.069599606 |
| PRCD | -0.069601791 |
| TSPYL1 | -0.069602405 |
| RP11-51F16 | -0.069614964 |
| RP6-218J18 | -0.069616203 |
| HOXB-AS2 | -0.069627883 |
| RP11-720L3 | -0.069633175 |
| RP11-149I2 | -0.069637702 |
| AQP11 | -0.069637979 |
| PECR | -0.069638518 |
| U3 | -0.069642816 |
| OOEP-AS1 | -0.069642976 |
| RAB5B | -0.069646016 |
| RP11-5C23 | -0.069650965 |
| RP11-12J10 | -0.069654519 |
| UBBP1 | -0.06965589 |
| VCAM1 | -0.069659821 |
| CTB-12O2 | -0.069665079 |

| | |
|---|---|
| PDCD7 | -0.069670296 |
| SNX18P4 | -0.069673667 |
| SET | -0.069675692 |
| SPDYE2 | -0.069688756 |
| ADCY10P1 | -0.069696902 |
| HYAL4 | -0.069697079 |
| PCED1B | -0.069699432 |
| NCOA2 | -0.06970133 |
| SCART1 | -0.069712847 |
| ZNF736P11Y | -0.06971529 |
| SSH2 | -0.069718782 |
| MIR216A | -0.069725573 |
| KRT18P32 | -0.069726005 |
| STK17A | -0.069741611 |
| PBX1 | -0.069746281 |
| EEF1A1P33 | -0.069748721 |
| TRMT1L | -0.069750021 |
| RP11-106M3 | -0.069757165 |
| TOMM22P4 | -0.069770808 |
| PLA2G12A | -0.069778468 |
| PLET1 | -0.069782671 |
| RP13-60P5 | -0.069783098 |
| RNU6-142P | -0.069785946 |
| YWHAZ | -0.069789315 |
| GTF2A1 | -0.069790227 |
| GSG2 | -0.069790562 |
| RP11-734K2 | -0.069792292 |
| UBE2V1 | -0.069792795 |
| C14orf169 | -0.069794432 |
| GHITM | -0.069808553 |
| DEDD2 | -0.069816405 |
| SIGLEC14 | -0.069829618 |
| WDR12 | -0.069833801 |
| TRIM51EP | -0.069835785 |
| DNM1P17 | -0.069844729 |
| ANXA10 | -0.06984602 |
| PRC1 | -0.069851737 |
| AC013429 | -0.069860752 |
| RP1-20C7 | -0.06987544 |
| NOVA1-AS1 | -0.06989104 |
| HACL1 | -0.069893041 |
| ZFYVE26 | -0.069897413 |
| AC018641 | -0.069913844 |
| AL353733 | -0.069914227 |
| AL136115 | -0.069922544 |

| | |
|---|---|
| RNA5SP158 | -0.069929083 |
| NCBP2-AS1 | -0.069935268 |
| KIF25-AS1 | -0.069939889 |
| RNU6-1061P | -0.069952296 |
| UBE3A | -0.06995429 |
| AC007652 | -0.069956172 |
| MYO1F | -0.069961062 |
| BMP2K | -0.069969919 |
| IGHD1-7 | -0.06997165 |
| NCAPG | -0.069974477 |
| KCNH8 | -0.069974599 |
| LCMT1-AS2 | -0.069978379 |
| RP11-383B4 | -0.069981946 |
| CTD-2621I17 | -0.069982663 |
| NOL8 | -0.06998417 |
| FAM133B | -0.069991892 |
| RP11-98F14 | -0.069992257 |
| MIR3164 | -0.069993155 |
| MIR4283-1 | -0.069999278 |
| TBC1D9 | -0.070009232 |
| NBPF19 | -0.070012451 |
| RNF19A | -0.070013028 |
| NUP188 | -0.070015539 |
| LYPLA1 | -0.070022479 |
| RP11-382A20 | -0.070025346 |
| RPL13P4 | -0.070029682 |
| HTATIP2 | -0.070034349 |
| POM121B | -0.070038982 |
| MIR6721 | -0.070042879 |
| ELF1 | -0.070051039 |
| HIGD1AP16 | -0.070056416 |
| RP11-261P9 | -0.070057846 |
| GLYR1 | -0.07006129 |
| SOX2-OT | -0.070062027 |
| PPIAP21 | -0.070067235 |
| MRPL42P2 | -0.070074821 |
| RIF1 | -0.07007584 |
| UTP20 | -0.070082966 |
| RP11-1012A1 | -0.070083915 |
| PRDX2P3 | -0.070098506 |
| UBE2FP1 | -0.07010835 |
| STAG2 | -0.070109441 |
| RN7SKP80 | -0.070115422 |
| TMEM184C | -0.070126726 |
| ZNF519 | -0.070127823 |

| | |
|---|---|
| CD40 | -0.070130019 |
| ZNF84 | -0.07014734 |
| RNA5SP359 | -0.070149451 |
| MAP2K1 | -0.070151955 |
| OR5BH1P | -0.070157568 |
| AC005795 | -0.070165188 |
| MTND5P5 | -0.070169703 |
| GMFB | -0.070170746 |
| TRPA1 | -0.070175639 |
| TTN | -0.070176049 |
| RP11-810D13 | -0.070181479 |
| AGTPBP1 | -0.070181705 |
| RP13-452N2 | -0.07019041 |
| GMPR2 | -0.070192012 |
| ZNF594 | -0.070194375 |
| RP5-1157M23 | -0.070206003 |
| RNU6-199P | -0.070207218 |
| AC006557 | -0.070215589 |
| ERAP2 | -0.070227195 |
| AP001058 | -0.070228965 |
| FECH | -0.070230749 |
| DNM1P41 | -0.070235695 |
| AL138696 | -0.070241363 |
| ATP5F1P5 | -0.070241649 |
| CTD-3064C13 | -0.070243923 |
| SOS1 | -0.070246117 |
| SMC6 | -0.07024634 |
| IGHEP2 | -0.070249391 |
| RP11-297B17 | -0.070249916 |
| TEX35 | -0.070250109 |
| HPSE | -0.070251847 |
| RP4-654C18 | -0.070252881 |
| PARPBP | -0.070264648 |
| ANKRD10 | -0.070273132 |
| AMN1 | -0.070280332 |
| DDX6 | -0.070282825 |
| SCARNA7 | -0.070285425 |
| CSNK1G2-AS1 | -0.070291611 |
| SLC25A46 | -0.070291663 |
| PBK | -0.070296798 |
| RNU6-282P | -0.070296941 |
| SMG6-IT1 | -0.070298728 |
| RNU6-97P | -0.070299217 |
| ABC12-47043100G14 | -0.070304277 |
| RP11-169N13 | -0.070305207 |

| | |
|---|---|
| RNF214 | -0.070307217 |
| CCDC38 | -0.070309272 |
| MIR6744 | -0.07031416 |
| PSMA6P4 | -0.07031668 |
| FAM129A | -0.070318306 |
| EML6 | -0.070319795 |
| CTC-242N15 | -0.070326785 |
| RP5-1099D15 | -0.070334597 |
| EPC1 | -0.070335163 |
| NUMA1 | -0.070352559 |
| RPL21P123 | -0.070353738 |
| C12orf43 | -0.070363369 |
| AC093323 | -0.070364987 |
| PAH | -0.0703672 |
| AMIGO1 | -0.070367629 |
| DNAJC9 | -0.070368485 |
| CNOT6 | -0.070375443 |
| DCAF17 | -0.070384895 |
| TRAPPC13 | -0.070390123 |
| RP11-1396O13 | -0.070391135 |
| MTL5 | -0.07040538 |
| TTC17 | -0.070406105 |
| MIR624 | -0.070410643 |
| CFAP70 | -0.070412059 |
| RNU6-1238P | -0.070420427 |
| RPL39P38 | -0.07042252 |
| OR5BP1P | -0.070431723 |
| RP3-323N1 | -0.070432397 |
| RP11-419I17 | -0.070432438 |
| RP11-390F10 | -0.070440049 |
| SP1 | -0.070445512 |
| AL009050 | -0.070455945 |
| OR4F15 | -0.070457514 |
| FBXL16 | -0.070460229 |
| ASF1A | -0.07046261 |
| FABP3P2 | -0.070471936 |
| TEX19 | -0.070474909 |
| RWDD4P1 | -0.070484022 |
| NADK | -0.070489551 |
| MTND4P32 | -0.070489606 |
| AC004041 | -0.070490923 |
| CASKIN1 | -0.070491467 |
| RP11-535A5 | -0.070496302 |
| RP11-498C9 | -0.070497998 |
| UNC93B4 | -0.07050627 |

| | |
|---|---|
| ZXDB | -0.070507561 |
| AC114814 | -0.070509349 |
| UBL7-AS1 | -0.070510054 |
| AC004063 | -0.070516867 |
| TRMT112P7 | -0.070526088 |
| POPDC2 | -0.070532027 |
| RP5-881P19 | -0.070534491 |
| AL391730 | -0.070536895 |
| CTD-2532D12 | -0.070541088 |
| CHIC2 | -0.070543581 |
| RP11-757G14 | -0.070544176 |
| CTD-2201E18 | -0.070550781 |
| RP5-1065O2 | -0.070550815 |
| TCEB1P19 | -0.070557277 |
| AC027307 | -0.070580245 |
| MIR3944 | -0.070580288 |
| HMG20A | -0.070589146 |
| SNTG2 | -0.070592851 |
| CTD-3096M3 | -0.070596646 |
| KIRREL3-AS2 | -0.070600254 |
| ATP6V1C1 | -0.070606765 |
| C21orf91-OT1 | -0.070609024 |
| MCTP1 | -0.070610579 |
| ANGEL2 | -0.070610784 |
| CTB-22K21 | -0.070612331 |
| RP11-90H3 | -0.070633074 |
| AC099545 | -0.070638888 |
| RP11-545D19 | -0.070639121 |
| NYAP2 | -0.070639273 |
| GTSE1 | -0.070642711 |
| CNOT10 | -0.070650507 |
| PCLO | -0.070651158 |
| NEK9 | -0.0706586 |
| RP11-895M11 | -0.070661925 |
| KIF21B | -0.070668591 |
| ASLP1 | -0.070685018 |
| RNU4-21P | -0.070685897 |
| C3orf20 | -0.070687747 |
| ORMDL3 | -0.070694378 |
| MAN1C1 | -0.070695811 |
| LAMP3 | -0.070697765 |
| RP11-541N10 | -0.070707439 |
| RNF146 | -0.070718677 |
| KIAA0430 | -0.070736576 |
| METTL17 | -0.070741413 |

| | |
|---|---|
| XX-C00717C00720L | -0.070742409 |
| RP11-63K6 | -0.070743068 |
| TRIM64 | -0.070744256 |
| CDH12P4 | -0.070744632 |
| IDO1 | -0.070751088 |
| AC006960 | -0.070753944 |
| NUDT9 | -0.070759367 |
| RP11-578F21 | -0.070774427 |
| SPANXN4 | -0.070784562 |
| AGAP5 | -0.070787482 |
| LSM12 | -0.070789173 |
| RP11-201O14 | -0.070789952 |
| LINC00709 | -0.070792657 |
| AC019330 | -0.070801475 |
| TCP1P1 | -0.070802401 |
| AC007566 | -0.070812951 |
| RP1-14D6 | -0.070825291 |
| ERI2 | -0.070828896 |
| MLLT10P1 | -0.070829526 |
| CMTR2 | -0.070838813 |
| RP11-88I18 | -0.070840052 |
| RNA5SP195 | -0.07084055 |
| PDCL3P6 | -0.070845528 |
| SIGLEC21P | -0.070847919 |
| LINC01011 | -0.070857251 |
| ASAP2 | -0.070866256 |
| B4GALNT2P1 | -0.070866767 |
| OR6B3 | -0.070868979 |
| GATS | -0.07087167 |
| POLR3K | -0.070884117 |
| KBTBD2 | -0.070885364 |
| AC092301 | -0.070886114 |
| TBC1D12 | -0.070888917 |
| RP11-472B18 | -0.070890444 |
| ZNF766 | -0.070895243 |
| SPCS2P4 | -0.07089569 |
| IL2RA | -0.070900415 |
| HLA-T | -0.070905404 |
| AHCYP2 | -0.07090742 |
| CTD-2299I21 | -0.070913062 |
| LINC00273 | -0.070920961 |
| PTGES3P4 | -0.07092332 |
| RP11-89K11 | -0.070926279 |
| C20orf197 | -0.070926891 |
| SLC25A14P1 | -0.070941385 |

| | |
|---|---|
| RP11-438N5 | -0.070943407 |
| KRT18P16 | -0.070948528 |
| RP11-542P2 | -0.07095227 |
| AC022816 | -0.070959727 |
| RABEP1 | -0.070964269 |
| RP11-360N9 | -0.070976384 |
| ARL2BPP7 | -0.070978164 |
| DHX16 | -0.070979228 |
| MGEA5 | -0.070982652 |
| ING1 | -0.070982856 |
| RP4-657E11 | -0.070986243 |
| SNORD14A | -0.070986875 |
| RP11-499E14 | -0.07098968 |
| ZNF549 | -0.071000476 |
| IMPG2 | -0.071006309 |
| RP11-132N15 | -0.071011671 |
| WBP1LP1 | -0.071027902 |
| RP3-461F17 | -0.071028905 |
| RPL7L1P11 | -0.071033938 |
| RP11-348H3 | -0.071041624 |
| ATR | -0.071048013 |
| ADNP2 | -0.071054766 |
| AC253576 | -0.071055601 |
| GMDS-AS1 | -0.071056552 |
| CTD-2292M16 | -0.071062469 |
| MIR6835 | -0.071063174 |
| IL10 | -0.071063446 |
| RNF103 | -0.07106485 |
| RNPS1P1 | -0.071071494 |
| CTD-2095E4 | -0.071072925 |
| RP11-96F8 | -0.071075882 |
| ABCE1 | -0.071083419 |
| RP11-353J17 | -0.07108468 |
| SPTBN4 | -0.071102694 |
| OR7D4 | -0.07110645 |
| OR11H5P | -0.071112132 |
| ZNF776 | -0.071112778 |
| WWP2 | -0.071116462 |
| CTB-58E17 | -0.07111942 |
| SLC22A25 | -0.071123541 |
| AGGF1P1 | -0.071125025 |
| AL157788 | -0.071125727 |
| AF011889 | -0.071127091 |
| METTL15P3 | -0.071128549 |
| DOK7 | -0.071135694 |

| | |
|---|---|
| RNA5-8SP3 | -0.071136618 |
| RP11-21B21 | -0.071140035 |
| PGGT1B | -0.071140724 |
| RP11-1280N14 | -0.071150343 |
| MS4A4E | -0.071155219 |
| MED23 | -0.071175553 |
| RP11-493E12 | -0.071185973 |
| USP12PX | -0.071187939 |
| RP11-310H4 | -0.071190364 |
| KRTAP5-2 | -0.071201493 |
| UBTFL1 | -0.071204935 |
| AC098824 | -0.071206585 |
| RP11-234P3 | -0.071207725 |
| KIF18A | -0.07121509 |
| RP11-126H7 | -0.071217074 |
| CEBPZ | -0.071217918 |
| ZNF571 | -0.071220429 |
| SKA2 | -0.071225118 |
| RN7SL45P | -0.071227371 |
| DBNDD2 | -0.071247631 |
| LINC00644 | -0.071254893 |
| RP11-568J23 | -0.071266069 |
| CTD-2370N5 | -0.071270577 |
| CTD-2197I11 | -0.071277727 |
| KIAA1143 | -0.071281809 |
| NHEG1 | -0.071283608 |
| RP4-694A7 | -0.071294493 |
| BHLHB9P1 | -0.071301257 |
| CAMK4 | -0.071305996 |
| UTP6 | -0.07130607 |
| OR51T1 | -0.071308275 |
| ATP6V0CP1 | -0.071319431 |
| RP1-247F3 | -0.07132128 |
| UBR3 | -0.071328099 |
| IGHD5-18 | -0.071329379 |
| RFTN1 | -0.071330946 |
| PCGF3 | -0.071332862 |
| AC024028 | -0.071336993 |
| RPAP2 | -0.071339504 |
| C15orf40 | -0.071341625 |
| RP11-282A11 | -0.071349058 |
| MTMR3 | -0.071349309 |
| RP11-10N23 | -0.07134957 |
| MIR509-1 | -0.071355623 |
| RPL19P20 | -0.071360759 |

| | |
|---|---|
| SNORA16 | -0.071360889 |
| GNRHR2 | -0.071367171 |
| RP11-700E23 | -0.071372896 |
| PPP1R15B | -0.071382688 |
| AHCYP4 | -0.071388158 |
| RP13-225O21 | -0.071393275 |
| CTD-3216D2 | -0.071395165 |
| ZNF578 | -0.071404012 |
| SRP54 | -0.071408977 |
| ODF2-AS1 | -0.071414631 |
| LMX1B | -0.071420494 |
| LINC01481 | -0.071433143 |
| RP11-500B12 | -0.071435183 |
| HARBI1 | -0.071438504 |
| RN7SL385P | -0.071443962 |
| LINC01226 | -0.071449526 |
| CBX1P5 | -0.071460829 |
| RP11-711M9 | -0.071467466 |
| RP11-154D3 | -0.071476426 |
| CTD-2526L21 | -0.071480826 |
| RP11-630D6 | -0.071491706 |
| RARRES2P2 | -0.071498492 |
| HNRNPA1P4 | -0.071506935 |
| C2orf27B | -0.071507778 |
| RNU6-912P | -0.071509102 |
| AC121251 | -0.071513442 |
| UBE3C | -0.071523066 |
| MIR6505 | -0.071533694 |
| ZNF813 | -0.071542081 |
| KRTAP4-5 | -0.071544855 |
| CTD-2351A8 | -0.071547406 |
| RP5-1031D4 | -0.071549022 |
| ROCK1 | -0.071550435 |
| TNF | -0.071553337 |
| CCL25 | -0.071556475 |
| LINC00894 | -0.071569896 |
| AC159540 | -0.071581869 |
| TSNAX | -0.071586845 |
| RIMKLBP1 | -0.071591926 |
| RP11-44D5 | -0.0715952 |
| ETV7 | -0.071609358 |
| AC010522 | -0.071612573 |
| SCAMP1-AS1 | -0.071613712 |
| AC098869 | -0.071614308 |
| SLAIN2 | -0.071616633 |

| | |
|---|---|
| HCK | -0.0716168 |
| MTND3P4 | -0.071617338 |
| PHOSPHO2 | -0.07161814 |
| LINC01087 | -0.071625858 |
| C15orf48 | -0.071631406 |
| KRIT1 | -0.071632354 |
| MIR455 | -0.071635914 |
| AC138781 | -0.071638292 |
| RP11-112J3 | -0.071655708 |
| HLA-DPB1 | -0.071657025 |
| AL355598 | -0.07166782 |
| ALG12 | -0.071669677 |
| PRAMEF18 | -0.071673035 |
| OR2W1 | -0.071679601 |
| LINC00674 | -0.071689976 |
| LINC01163 | -0.071692548 |
| AC112220 | -0.071698569 |
| HNRNPF | -0.071699632 |
| GFRA1 | -0.071702042 |
| TTC27 | -0.071703167 |
| RP13-926M18 | -0.071706458 |
| FAM187B2P | -0.071710735 |
| CTD-2347I17 | -0.071717037 |
| MOCS1P1 | -0.071730512 |
| CGRRF1 | -0.071731577 |
| NPM1P39 | -0.071741318 |
| ZMYM1 | -0.071763597 |
| CBX8 | -0.071764249 |
| WHAMML1 | -0.071784045 |
| PLCL2 | -0.071790985 |
| RP11-59O6 | -0.07179311 |
| CD79B | -0.071795078 |
| FAM9C | -0.071806684 |
| IGF1R | -0.07180773 |
| RNU7-13P | -0.071809358 |
| NME9 | -0.071815042 |
| FAM219B | -0.07181733 |
| RANP8 | -0.071827379 |
| MAGEA12 | -0.071829126 |
| CASC4P1 | -0.071829527 |
| ZXDA | -0.071838819 |
| LLNLF-136C3 | -0.071839873 |
| PRPS1P1 | -0.071845309 |
| RPL29P29 | -0.071849758 |
| CHORDC1 | -0.07186016 |

| | |
|---|---|
| LINC01342 | -0.071867372 |
| AL162426 | -0.07187255 |
| FUNDC2P4 | -0.071883813 |
| RNU7-53P | -0.071886502 |
| HNRNPU-AS1 | -0.071887863 |
| MIR215 | -0.071891604 |
| RP11-121L10 | -0.071899487 |
| RBBP8 | -0.071910432 |
| AC130464 | -0.071926756 |
| FOXD2 | -0.071928181 |
| MIS18A-AS1 | -0.071929205 |
| ZNF19 | -0.071931549 |
| GTF3C3 | -0.071935214 |
| CTC-529I10 | -0.071936126 |
| RGS5 | -0.071953076 |
| CGGBP1 | -0.071954006 |
| IPO7 | -0.071955569 |
| BLOC1S6 | -0.071959439 |
| RP11-61G19 | -0.071973713 |
| GPLD1 | -0.07198184 |
| RNU6-300P | -0.071982474 |
| RNF14 | -0.071998882 |
| LINC01095 | -0.072000005 |
| RP5-854E16 | -0.072020802 |
| VDAC1P1 | -0.072029752 |
| HAUS4 | -0.072030939 |
| METTL23 | -0.072039163 |
| CH17-409J4 | -0.072050701 |
| CCL3 | -0.07205855 |
| SLC16A6P1 | -0.072059074 |
| KIAA0753 | -0.072068541 |
| VN1R92P | -0.072070188 |
| MIR1273H | -0.072071652 |
| SUDS3 | -0.072072188 |
| RP11-76C10 | -0.072076785 |
| MYCBPAP | -0.072077937 |
| RP11-724M22 | -0.072097726 |
| AC009784 | -0.072098174 |
| HEATR6 | -0.072102645 |
| MTND5P15 | -0.072102675 |
| RP11-374P20 | -0.072104398 |
| AC009499 | -0.072113296 |
| LRRC75A | -0.072114162 |
| CTD-2311B13 | -0.072117244 |
| AC093142 | -0.072125378 |

| | |
|---|---|
| VPS13D | -0.072125902 |
| DCAF10 | -0.072126348 |
| MIR1302-8 | -0.072136625 |
| RNU6ATAC24P | -0.072146318 |
| TARDBP | -0.072151987 |
| LINC-PINT | -0.072153754 |
| WDR11 | -0.072159299 |
| AC130469 | -0.07216274 |
| RP11-510I21 | -0.072164678 |
| OR4K4P | -0.072166399 |
| RP11-1334A24 | -0.072177923 |
| PHF11 | -0.072191598 |
| RP11-545L5 | -0.072196099 |
| HMGN2P6 | -0.072199167 |
| SLC36A3 | -0.072201367 |
| PRR5-ARHGAP8 | -0.07221292 |
| RP11-259O18 | -0.072222707 |
| TTC34 | -0.07222284 |
| RP11-513H8 | -0.072224553 |
| RAPGEF5 | -0.072226669 |
| FOXM1 | -0.072246688 |
| SLC25A15 | -0.072256475 |
| RRM2B | -0.072258894 |
| OR51AB1P | -0.072261862 |
| RP1-305B16 | -0.072266331 |
| RP11-713N11 | -0.072266998 |
| METTL20 | -0.072281183 |
| TRIM36-IT1 | -0.072289395 |
| EIF4G3 | -0.072292086 |
| IFI6 | -0.072294207 |
| FAM86B3P | -0.072297295 |
| RP11-458J1 | -0.072297742 |
| AC063957 | -0.072300725 |
| RPL29P30 | -0.072304434 |
| RP11-109L13 | -0.072308782 |
| TUBE1 | -0.072318506 |
| RP11-462P6 | -0.072319638 |
| RN7SKP219 | -0.072325783 |
| WHSC1L1 | -0.072329576 |
| SNAP23P | -0.072334488 |
| AC092641 | -0.072335241 |
| UBE2L6 | -0.07233899 |
| RP11-475E11 | -0.072340699 |
| AC137934 | -0.072342358 |
| CTD-2313D3 | -0.072363182 |

| | |
|---|---|
| NOA1 | -0.072371326 |
| RN7SL116P | -0.072374493 |
| AP000347 | -0.072376257 |
| EXOC8 | -0.07237968 |
| WNT10A | -0.07238024 |
| RPL4P2 | -0.072384941 |
| SLC22A8 | -0.072387331 |
| PAICSP1 | -0.072387924 |
| DPPA5 | -0.072391663 |
| ZNF562 | -0.0724005 |
| KNSTRN | -0.072401988 |
| RPP14 | -0.072408816 |
| HMGA1P4 | -0.072414833 |
| RN7SL394P | -0.072419495 |
| SNX16 | -0.072423812 |
| GPBP1 | -0.072427775 |
| VAV3-AS1 | -0.072428736 |
| HMGN2P4 | -0.072430893 |
| RRP1B | -0.072439609 |
| RP1-310O13 | -0.072441318 |
| LINC00243 | -0.072441573 |
| TCEA1P1 | -0.072447459 |
| OR4A19P | -0.072452798 |
| RPL7L1 | -0.072458661 |
| TTLL6 | -0.072462459 |
| SRL | -0.072483322 |
| RP11-132E11 | -0.072487858 |
| RPS16P5 | -0.072491463 |
| RWDD3 | -0.072495927 |
| ZNF778 | -0.072499295 |
| SDE2 | -0.072507872 |
| ZFAND4 | -0.072508802 |
| HNRNPKP2 | -0.072522291 |
| TMEM68 | -0.072523891 |
| LYG2 | -0.072527376 |
| RP11-420A6 | -0.072540558 |
| CTC-428H11 | -0.072546627 |
| RP11-555K12 | -0.072549524 |
| IER3IP1 | -0.072568935 |
| TAS2R3 | -0.07257494 |
| AC090154 | -0.072579034 |
| CNOT2 | -0.072579298 |
| NPLOC4 | -0.072581494 |
| THAP5 | -0.072582238 |
| PPP6C | -0.072582964 |

| | |
|---|---|
| ADIPOR1 | -0.072587894 |
| GCC1 | -0.072588787 |
| STK24 | -0.072591411 |
| LRRC37A4P | -0.072596899 |
| RPL21P8 | -0.072611305 |
| HNRNPA1P9 | -0.072615574 |
| RP11-25L5 | -0.072619093 |
| FBXO36P1 | -0.07262521 |
| ZHX2 | -0.072625589 |
| TULP4 | -0.072635505 |
| BCAS2P1 | -0.072636314 |
| KLHL8 | -0.072637254 |
| IGKV3D-25 | -0.072638873 |
| EME1 | -0.072639617 |
| PACRGL | -0.072656195 |
| RP11-116A1 | -0.072658555 |
| RABL2B | -0.072660939 |
| RP11-1109F11 | -0.072662205 |
| RPL23AP80 | -0.072675403 |
| SYNE1 | -0.07269002 |
| C2 | -0.072690336 |
| HMGB2P1 | -0.072696025 |
| MTA3 | -0.072696364 |
| ZNF554 | -0.072699322 |
| NCBP1 | -0.072706392 |
| RP1-122O8 | -0.072719634 |
| OR11H6 | -0.072721005 |
| RNA5SP429 | -0.072727254 |
| YAF2 | -0.072728254 |
| RP11-228B15 | -0.072738896 |
| RUNX3 | -0.072744906 |
| AC092636 | -0.072748871 |
| SLC5A10 | -0.072750636 |
| RP11-616L12 | -0.072751394 |
| XKR7 | -0.072774314 |
| RPS24P16 | -0.072776217 |
| RP11-102C16 | -0.072777661 |
| KRT18P33 | -0.072789251 |
| IGLVIVOR22-1 | -0.072794454 |
| PTPRO | -0.072794489 |
| AL592294 | -0.072804395 |
| GAPDHP22 | -0.072807508 |
| ARFGAP2 | -0.072820053 |
| CS | -0.072822714 |
| EIF2AK4 | -0.072824221 |

| | |
|---|---|
| ZNF878 | -0.072826568 |
| LINC00683 | -0.072829863 |
| GNAZ | -0.072830844 |
| OR6C76 | -0.072834968 |
| RP11-342K6 | -0.07283771 |
| TPP2 | -0.072838882 |
| LRRC8D | -0.072839384 |
| AC019097 | -0.072841896 |
| PDE6G | -0.072847645 |
| RP13-539J13 | -0.072853438 |
| C9orf117 | -0.07286225 |
| RN7SKP218 | -0.072876142 |
| AP001537 | -0.072876602 |
| U52112 | -0.072883155 |
| AC234775 | -0.07288478 |
| COX5BP6 | -0.072884893 |
| DPPA5P4 | -0.072892532 |
| IGLV3-4 | -0.072893658 |
| RP1-296L11 | -0.07289512 |
| TPTE2 | -0.072895307 |
| POLR2J4 | -0.072898084 |
| ACOX1 | -0.072909341 |
| AC078942 | -0.072914809 |
| RNU6-882P | -0.072920562 |
| ZC2HC1C | -0.072930309 |
| RP11-216N14 | -0.072931222 |
| RP11-78H18 | -0.072936227 |
| CST9 | -0.072938492 |
| VN1R84P | -0.072940568 |
| UPB1 | -0.072943836 |
| RGL2 | -0.072947426 |
| DLEU2_4 | -0.072958588 |
| RP4-744I24 | -0.072963709 |
| SNORA15 | -0.072983451 |
| PRPS1P2 | -0.072990052 |
| AC008069 | -0.072997663 |
| HNRNPKP1 | -0.07300367 |
| USP16 | -0.073004471 |
| ZDHHC2 | -0.073007187 |
| H2AFB1 | -0.073011453 |
| AC094019 | -0.073011606 |
| RP1-209B5 | -0.073020167 |
| WDSUB1 | -0.073020862 |
| RP11-82O2 | -0.073027483 |
| AL137019 | -0.073027642 |

| | |
|---|---|
| PTPN11 | -0.073030555 |
| RP11-805L22 | -0.07304017 |
| RP11-424M21 | -0.073041196 |
| MEOX2-AS1 | -0.073042 |
| RP11-222O23 | -0.073056825 |
| METTL10 | -0.073058947 |
| TMEM180 | -0.073061496 |
| ZNF433 | -0.073064436 |
| RNA5SP509 | -0.073069495 |
| HCG2040054 | -0.073071784 |
| RP11-392E22 | -0.07308134 |
| RP11-540A21 | -0.073082093 |
| FLJ31104 | -0.073082205 |
| RP1-187B23 | -0.073082606 |
| LYRM2 | -0.073083134 |
| AC026357 | -0.073101346 |
| RP11-173E2 | -0.073102177 |
| GRAMD4P8 | -0.073103954 |
| FAM159A | -0.073104568 |
| NIPBL | -0.073107508 |
| TMEM97 | -0.073111917 |
| RP11-630I5 | -0.073122783 |
| FAM98B | -0.07312442 |
| RPL31P58 | -0.073126616 |
| IL6R | -0.073128067 |
| CTD-2196E14 | -0.073129443 |
| RP11-592H6 | -0.073130923 |
| MIR6722 | -0.073133131 |
| SNX19P3 | -0.073134522 |
| RNU1-133P | -0.073142866 |
| MOCS3 | -0.073148291 |
| RP11-342K2 | -0.073152849 |
| FAM95C | -0.073155529 |
| BCDIN3D | -0.073162971 |
| SENP1 | -0.073167659 |
| OPTC | -0.073175819 |
| FLJ45079 | -0.073180956 |
| RP11-85G18 | -0.07318392 |
| IGKV2OR2-2 | -0.073188281 |
| RP11-96K19 | -0.073193037 |
| CTD-3185P2 | -0.073195288 |
| RP11-6B6 | -0.073196587 |
| DOK2 | -0.073196665 |
| UQCC1 | -0.073198265 |
| MIR4633 | -0.073199779 |

| | |
|---|---|
| SCOC | -0.073208014 |
| ZCRB1 | -0.073213559 |
| FBXL14 | -0.073224796 |
| SPC25 | -0.073225187 |
| OASL | -0.073234695 |
| CTD-3099C6 | -0.073236853 |
| UBXN4 | -0.07323836 |
| AMY2B | -0.073248072 |
| BOLA2P2 | -0.073248685 |
| RP11-791M20 | -0.073252302 |
| GAPDHP75 | -0.073254207 |
| AP002852 | -0.07326115 |
| RN7SL364P | -0.073269182 |
| SNORD77 | -0.073273991 |
| AC002331 | -0.073276459 |
| SLMO2 | -0.073277896 |
| AKR1B1P6 | -0.073278585 |
| GEMIN5 | -0.073279962 |
| RP5-966M1 | -0.073284576 |
| AL139351 | -0.073294248 |
| LINC01030 | -0.073304304 |
| RSAD1 | -0.073308223 |
| MTRF1 | -0.073313749 |
| METTL2A | -0.07331481 |
| RP11-484D4 | -0.073316117 |
| CTB-79E8 | -0.073327015 |
| RN7SKP196 | -0.073332047 |
| CTD-2520I13 | -0.07334816 |
| SPDYE3 | -0.073353025 |
| FASTKD2 | -0.073359556 |
| ARF6 | -0.073366124 |
| ANHX | -0.073369178 |
| RP11-164J13 | -0.073370367 |
| RNU6-807P | -0.073379043 |
| RNU6-610P | -0.07338471 |
| RP11-782K4 | -0.073390699 |
| ASPM | -0.073397585 |
| GNPNAT1 | -0.07340393 |
| CYSLTR1 | -0.073407446 |
| RNU6-548P | -0.073413523 |
| MSTO2P | -0.073419205 |
| CCL15 | -0.07342029 |
| MIR422A | -0.073421396 |
| YIPF4 | -0.073424563 |
| CITF22-24E5 | -0.073429035 |

| | |
|---|---|
| ZNF583 | -0.073433085 |
| CPSF2 | -0.073434759 |
| VPS52 | -0.073446053 |
| BAIAP2 | -0.073459523 |
| RNF19BPX | -0.073461907 |
| RP5-1178H5 | -0.073463656 |
| PRMT5 | -0.073463672 |
| NR1D2 | -0.073467914 |
| ACTG1P9 | -0.0734691 |
| WBP11 | -0.073477905 |
| CLSTN2 | -0.07350053 |
| RP11-445P17 | -0.073513783 |
| NPM1P25 | -0.073517274 |
| CCDC83 | -0.073529932 |
| KATNBL1P1 | -0.073529944 |
| MIR3653 | -0.073534054 |
| MLH3 | -0.073534354 |
| RP11-1112G13 | -0.073536385 |
| RP11-157E21 | -0.073542046 |
| HPS3 | -0.073543992 |
| RNU6-735P | -0.073546426 |
| BZRAP1 | -0.073547211 |
| SRRM1P3 | -0.073558914 |
| RNU7-3P | -0.073563134 |
| TFAP2A | -0.073564439 |
| RP11-642E20 | -0.073566416 |
| THUMPD3-AS1 | -0.073582394 |
| RP11-539I5 | -0.073592571 |
| RP11-47G11 | -0.073632304 |
| SENP2 | -0.073634656 |
| RRS1-AS1 | -0.073638006 |
| RP11-560G2 | -0.073639632 |
| SLC25A47P1 | -0.073648945 |
| AP3M1 | -0.073654713 |
| CTD-2063L20 | -0.073665113 |
| RP11-780K2 | -0.073667651 |
| RP11-197K3 | -0.073668482 |
| RP11-525G13 | -0.073668935 |
| RP3-474G15 | -0.073674568 |
| U2AF1 | -0.073686193 |
| RP4-706A16 | -0.073690566 |
| GGTA1P | -0.073698473 |
| TPRX1 | -0.07370593 |
| UBE2V1P4 | -0.073711441 |
| RAB21 | -0.07371533 |

| | |
|---|---|
| RP11-652G5 | -0.073716258 |
| XPC | -0.073723944 |
| ACTR8 | -0.0737248 |
| MIR4639 | -0.073732082 |
| RNA5SP75 | -0.0737333 |
| IGHA2 | -0.073737806 |
| TRGV6 | -0.073740189 |
| MAPK8 | -0.073754643 |
| RP11-798G7 | -0.073771686 |
| RP11-279F6 | -0.073773643 |
| AC092338 | -0.073776291 |
| ZNF132 | -0.073784728 |
| DCLRE1A | -0.073789621 |
| USP50 | -0.07379038 |
| GGTLC5P | -0.073793348 |
| UBE2V2P3 | -0.073797445 |
| PTOV1-AS1 | -0.073808097 |
| PIAS2 | -0.073811576 |
| RP11-297K7 | -0.073828505 |
| ZNF829 | -0.073836935 |
| YPEL2 | -0.073847503 |
| RNU6-181P | -0.073853897 |
| FLJ33360 | -0.07385587 |
| LINC00641 | -0.073856601 |
| AC008984 | -0.073864951 |
| AMZ2 | -0.073872899 |
| MIR8088 | -0.073872952 |
| AC007000 | -0.073878538 |
| RP11-229D13 | -0.073878761 |
| RP11-1086I4 | -0.073881446 |
| RN7SL508P | -0.073882244 |
| IREB2 | -0.073883132 |
| RNU6-1188P | -0.073888029 |
| HNF4GP1 | -0.073892466 |
| RNF125 | -0.073896863 |
| RN7SL799P | -0.073901731 |
| RP11-14P20 | -0.073903127 |
| SNW1 | -0.073912045 |
| SAMD8 | -0.073923804 |
| TTC9B | -0.073928469 |
| CHD6 | -0.07395707 |
| RP11-856M7 | -0.073965118 |
| C11orf40 | -0.073969126 |
| AP001525 | -0.073969403 |
| RP11-749I16 | -0.073971296 |

| | |
|---|---|
| STRN3 | -0.073973332 |
| AC109631 | -0.073983835 |
| RNU1-124P | -0.073988889 |
| RP11-461O14 | -0.073990619 |
| SASS6 | -0.074001352 |
| EIF4HP1 | -0.074004218 |
| RP11-89H19 | -0.074008142 |
| ANKRD36C | -0.074009817 |
| ST6GALNAC2 | -0.074010543 |
| HACE1 | -0.074015138 |
| RP3-408N23 | -0.07401641 |
| RP11-284M14 | -0.074022959 |
| AC026318 | -0.074035349 |
| MPRIP-AS1 | -0.074038542 |
| RP1-72A23 | -0.074052375 |
| SHC1P1 | -0.074057328 |
| KMT2E | -0.074061205 |
| RAD21-AS1 | -0.074061447 |
| RP11-143A22 | -0.074078102 |
| RNA5SP323 | -0.074083187 |
| ERICH3 | -0.074105903 |
| RP11-82O19 | -0.07410874 |
| C1QB | -0.074130176 |
| CSTF2T | -0.074136092 |
| RNU6-1207P | -0.07413666 |
| RP11-485G7 | -0.074137089 |
| C3orf79 | -0.074145745 |
| SEPHS1P6 | -0.074147473 |
| RNU6-1095P | -0.074158279 |
| MTIF3 | -0.074165135 |
| ZNF644 | -0.074170215 |
| MTBP | -0.074170568 |
| ECEL1P3 | -0.074176446 |
| HMGB1P35 | -0.074185658 |
| SMG1P5 | -0.074195481 |
| SCDP1 | -0.074201184 |
| MIR4494 | -0.074214011 |
| RP1-251M9 | -0.07421752 |
| MST1L | -0.074217528 |
| PHC3 | -0.074221212 |
| SLX1B-SULT1A4 | -0.074231557 |
| RP11-139H15 | -0.074237641 |
| RP11-150D20 | -0.074238919 |
| RP11-25H11 | -0.074243202 |
| RNU6-1048P | -0.074245132 |

| | |
|---|---|
| RP11-455J20 | -0.074251014 |
| TOP1 | -0.074262107 |
| LIN7A | -0.074262405 |
| ZNF271P | -0.074266182 |
| DAO | -0.074267647 |
| AC115115 | -0.074275726 |
| RNA5SP231 | -0.074280555 |
| VEZT | -0.074295597 |
| TMOD2 | -0.074297625 |
| SMURF2 | -0.074304323 |
| OR1L8 | -0.074319008 |
| DPY19L2 | -0.074331524 |
| RP11-352M15 | -0.074337887 |
| TOMM20P4 | -0.074341686 |
| CTD-2089N3 | -0.074360771 |
| RP1-192P9 | -0.07436231 |
| RP1-206D15 | -0.074366493 |
| AC013410 | -0.074368313 |
| ERGIC1 | -0.074370428 |
| CASP5 | -0.074378896 |
| RP11-35O15 | -0.07438191 |
| CYSLTR2 | -0.07438588 |
| HEPACAM2 | -0.074386342 |
| CA15P1 | -0.074389748 |
| TRA2B | -0.074395992 |
| REEP1 | -0.074403695 |
| MT-ND1 | -0.074409165 |
| METTL25 | -0.074410876 |
| FAM35BP | -0.074427611 |
| LCLAT1 | -0.074436961 |
| MIR26A1 | -0.074438041 |
| SNORA71C | -0.074443408 |
| AC104113 | -0.074450209 |
| STIL | -0.074454413 |
| AC009960 | -0.07445623 |
| OTUD3 | -0.074458804 |
| RP11-41L14 | -0.07446071 |
| MIR600HG | -0.074466656 |
| RN7SL753P | -0.074471549 |
| RP11-343L5 | -0.074472888 |
| CTA-445C9 | -0.074474135 |
| LDHBP1 | -0.074495906 |
| CTD-2233C11 | -0.074509704 |
| FGF4 | -0.074512546 |
| CAPS2 | -0.074519923 |

| | |
|---|---|
| MIR759 | -0.074520432 |
| ZNF430 | -0.074524556 |
| ZNF780A | -0.074531347 |
| MIR5582 | -0.074538309 |
| PGAM1P7 | -0.074543709 |
| RNU2-22P | -0.074550791 |
| RP11-175O19 | -0.074550976 |
| FAM47A | -0.074560459 |
| EXD2 | -0.074563925 |
| CT62 | -0.074565817 |
| CWC22 | -0.074569767 |
| RP11-1338A24 | -0.074576812 |
| FRMD4B | -0.074583889 |
| LINC01392 | -0.074585828 |
| IGHD4-4 | -0.074588072 |
| TSEN54 | -0.074588447 |
| TMEM254-AS1 | -0.074612188 |
| MIR4262 | -0.074614063 |
| KRT41P | -0.074617672 |
| HIST4H4 | -0.074624858 |
| ASB7 | -0.074658087 |
| RP3-370M22 | -0.074658092 |
| BBS10 | -0.07465859 |
| HSP90AB3P | -0.074660748 |
| PATZ1 | -0.074668823 |
| RN7SL144P | -0.074681749 |
| WBP2 | -0.074695354 |
| RP11-1174L13 | -0.074698266 |
| LIG3 | -0.074711448 |
| TNRC6A | -0.074713792 |
| AC006028 | -0.07471778 |
| AC244258 | -0.074720825 |
| RP1-199H16 | -0.074724022 |
| RP11-407N17 | -0.074729476 |
| GJA8 | -0.074733206 |
| AC010105 | -0.074734878 |
| GALT | -0.074735188 |
| LILRA4 | -0.074743971 |
| BX649553 | -0.074745762 |
| TPTE2P5 | -0.074764548 |
| RNU6-250P | -0.074764763 |
| BRPF1 | -0.074764939 |
| ZNF567 | -0.074770855 |
| RP11-436A7 | -0.074771774 |
| NIFK-AS1 | -0.074776065 |

| | |
|---|---|
| RP11-582J16 | -0.07477733 |
| CKMT1A | -0.074778767 |
| ZNF516 | -0.07478347 |
| RP11-279N8 | -0.074794382 |
| RGS7BP | -0.074802287 |
| IPO5P1 | -0.07480241 |
| HHLA2 | -0.074813528 |
| FPR3 | -0.074813536 |
| YRDC | -0.074813815 |
| RP4-530I15 | -0.074819007 |
| CARS-AS1 | -0.074822526 |
| ARRB2 | -0.074823453 |
| HMGN2P8 | -0.074823665 |
| DDX23 | -0.074838802 |
| AL121655 | -0.074841044 |
| BNIP3P37 | -0.074846532 |
| OR10A2 | -0.074852671 |
| ARL6IP1P3 | -0.074890327 |
| JAKMIP2 | -0.07489127 |
| AC234063 | -0.074891426 |
| RP1-111D6 | -0.074901682 |
| GOLGA8N | -0.074902693 |
| TAT | -0.074906658 |
| LARP4B | -0.074912108 |
| HBD | -0.07491367 |
| C2orf78 | -0.074915817 |
| ANKRD39 | -0.074929839 |
| PPEF2 | -0.074930599 |
| FSIP1 | -0.074938955 |
| STAMBP | -0.074947905 |
| ADAM24P | -0.074960384 |
| EXOSC6 | -0.074979013 |
| DBF4P1 | -0.074982711 |
| MKRN9P | -0.074984745 |
| LINC01250 | -0.074986818 |
| AC009945 | -0.074992689 |
| GOLGA8B | -0.075004409 |
| PREPL | -0.075007386 |
| RNU6-24P | -0.075009738 |
| MAU2 | -0.075010084 |
| RP11-210M15 | -0.07501411 |
| MT-ND2 | -0.075022568 |
| RP11-10J18 | -0.075022723 |
| RP11-391J2 | -0.075026889 |
| AKR7A3 | -0.075030122 |

| | |
|---|---|
| TMEM110-MUSTN1 | -0.075034682 |
| IGKV3OR22-2 | -0.07503681 |
| RP11-574M7 | -0.075040297 |
| SNORA24 | -0.075043382 |
| DNAJC19P7 | -0.075046917 |
| RPS15AP24 | -0.07505291 |
| SMNDC1 | -0.075053714 |
| CYP27B1 | -0.075057677 |
| RP4-550H1 | -0.07506266 |
| CTC-453G23 | -0.075083595 |
| BARD1 | -0.075090944 |
| HIGD1C | -0.075092128 |
| RP13-991F5 | -0.075093939 |
| LINC00221 | -0.07509501 |
| ELF4 | -0.07509593 |
| RP11-232A1 | -0.075099625 |
| RNU6-523P | -0.075102071 |
| RP3-393P12 | -0.075109351 |
| RP11-97O12 | -0.075115521 |
| SNAPC5 | -0.075118089 |
| RAB28P3 | -0.075121098 |
| HIST1H1T | -0.07513442 |
| PSMC6 | -0.075140508 |
| TCL1B | -0.075144693 |
| RIC3 | -0.075149322 |
| THTPA | -0.075153942 |
| RPL26P34 | -0.075156247 |
| DGKZP1 | -0.075157049 |
| SEC24B | -0.075157328 |
| MIR887 | -0.075176458 |
| HNRNPR | -0.07517824 |
| RP11-407H12 | -0.075179513 |
| RP11-374M1 | -0.0751808 |
| AF003529 | -0.07518273 |
| RASA4DP | -0.075183402 |
| TAS2R50 | -0.075183644 |
| IGHD1-26 | -0.075192247 |
| RP11-528G1 | -0.075198421 |
| LGALS16 | -0.075199541 |
| NPEPPS | -0.075203823 |
| LINC00348 | -0.075204838 |
| RP11-1140I5 | -0.075205281 |
| LLNLR-260G6 | -0.075207579 |
| RP11-218C23 | -0.075217629 |
| RP11-1149M10 | -0.075239127 |

| | |
|---|---|
| GNRHR | -0.075252891 |
| MIR621 | -0.075259621 |
| CD2AP | -0.075283157 |
| CTC-325J23 | -0.075291105 |
| PTMAP2 | -0.0753098 |
| TRAJ26 | -0.075310206 |
| TMC3-AS1 | -0.075318079 |
| ZNHIT6 | -0.075333987 |
| CCNL2P1 | -0.075341356 |
| ADO | -0.075348387 |
| RP1-249I4 | -0.07534865 |
| HLA-L | -0.075351513 |
| RP11-90B9 | -0.075364019 |
| PTPN18 | -0.075366323 |
| YWHAZP3 | -0.075375261 |
| CENPCP1 | -0.075378568 |
| TBC1D4 | -0.075383087 |
| HLA-F | -0.075386008 |
| RP11-438L19 | -0.075388808 |
| RP11-405L18 | -0.075394337 |
| BCAS4 | -0.075395236 |
| U4 | -0.075395767 |
| OR6T1 | -0.075396627 |
| PNPLA7 | -0.075409544 |
| FAM21C | -0.075413265 |
| SCAPER | -0.075414251 |
| ATG2A | -0.075417414 |
| ATP5S | -0.075428856 |
| GPATCH2L | -0.075432596 |
| RP11-504G3 | -0.075434134 |
| WDR33 | -0.075436168 |
| AL049831 | -0.075439364 |
| UNC45B | -0.07544489 |
| TTBK1 | -0.075447241 |
| PLA2G4C | -0.075462141 |
| ISCU | -0.07546629 |
| RP11-489P6 | -0.075466937 |
| SCLY | -0.075470476 |
| RN7SKP271 | -0.075473526 |
| GPR22 | -0.075480115 |
| RP11-96C23 | -0.075480338 |
| KHDC1P1 | -0.075484109 |
| CNOT4 | -0.075490031 |
| NLRP14 | -0.075493593 |
| ZNF860 | -0.075494124 |

| | |
|---|---|
| AC068587 | -0.07549825 |
| LINC00877 | -0.075511007 |
| SNORA38 | -0.075513 |
| EIF5AP2 | -0.075514878 |
| MYCNUT | -0.075517575 |
| RP11-546J1 | -0.075548967 |
| PRNT | -0.075559502 |
| CTD-2240E14 | -0.075569175 |
| RP11-225B17 | -0.075575969 |
| XXcos-LUCA16 | -0.075582844 |
| RP11-818O24 | -0.075590825 |
| SNRPGP17 | -0.075593134 |
| MIER3 | -0.07559917 |
| ZFAND1 | -0.075600212 |
| RP11-67K19 | -0.07560156 |
| HID1 | -0.075607524 |
| BNIP3P2 | -0.075609064 |
| KRT73-AS1 | -0.075611054 |
| RP11-143I21 | -0.075621369 |
| PIPOX | -0.075621553 |
| HNRNPCP1 | -0.075623302 |
| MAD2L1 | -0.07562546 |
| RP11-13N13 | -0.075632656 |
| MTND6P10 | -0.075645517 |
| PAIP2 | -0.07564721 |
| SYF2 | -0.075649275 |
| AGPAT3 | -0.075652419 |
| C8orf37 | -0.075657368 |
| C2orf27AP1 | -0.075659929 |
| HMMR-AS1 | -0.075661483 |
| TOP2B | -0.075664773 |
| RP11-649A18 | -0.075679583 |
| AC006116 | -0.075687565 |
| CEACAM4 | -0.07568804 |
| CTA-242H14 | -0.075700866 |
| PPP2R3C | -0.075703491 |
| SLC22A24 | -0.07571864 |
| CPSF7 | -0.075742191 |
| AC133528 | -0.075757801 |
| AGPS | -0.075759512 |
| MED14 | -0.075766862 |
| SNORD44 | -0.075771184 |
| ALMS1P | -0.075775489 |
| STMND1 | -0.075782167 |
| MIR1302-5 | -0.075786223 |

| | |
|---|---|
| CIB3 | -0.075792195 |
| ZNF45 | -0.075803514 |
| CADM3-AS1 | -0.075814824 |
| AC087884 | -0.075816831 |
| BASP1P1 | -0.07582364 |
| C5orf63 | -0.075824892 |
| TNKS2 | -0.075827646 |
| HNRNPDP1 | -0.075831544 |
| RP11-374F3 | -0.07583946 |
| OSGEPL1 | -0.075848465 |
| RP11-848P1 | -0.075849265 |
| PRODH2 | -0.075849909 |
| DDX24 | -0.075867815 |
| RP11-69I13 | -0.075869145 |
| RP11-483I13 | -0.075872206 |
| RP11-1197K16 | -0.075873603 |
| MIR3655 | -0.075874115 |
| MAGEA1 | -0.075874914 |
| RAB33B | -0.075887202 |
| PM20D1 | -0.075889738 |
| RP11-268G12 | -0.075898464 |
| ZC3H12C | -0.075900616 |
| AC006995 | -0.075902449 |
| TNFAIP3 | -0.075910998 |
| DDX50P1 | -0.075927298 |
| RP11-525K10 | -0.075930317 |
| RNF4 | -0.075946423 |
| CMB9-55F22 | -0.075950888 |
| ZNF197 | -0.075955037 |
| SNRK | -0.075959614 |
| STAMBPL1 | -0.075960805 |
| RP11-564D11 | -0.075972069 |
| RP11-394O2 | -0.075972281 |
| ZNF461 | -0.075981327 |
| ENPP4 | -0.075983299 |
| HP1BP3 | -0.075993272 |
| KLHDC1 | -0.075995188 |
| PPP1R9B | -0.076009123 |
| LINC01565 | -0.076014704 |
| RN7SL23P | -0.076019215 |
| RP4-640E24 | -0.07602667 |
| HNRNPA1P12 | -0.0760331 |
| RP11-449J3 | -0.076040348 |
| RAB3D | -0.076043897 |
| IGHVII-51-2 | -0.07604509 |

| | |
|---|---|
| RP11-804F13 | -0.076053149 |
| AC090283 | -0.076056305 |
| RPS5P2 | -0.07607171 |
| TRAV15 | -0.076078259 |
| EXOSC2 | -0.076083173 |
| RNU6-670P | -0.076087664 |
| TAS2R43 | -0.076092787 |
| BDNF-AS | -0.076093853 |
| ARL1 | -0.076094969 |
| ERBB3 | -0.076099806 |
| RP11-305B6 | -0.076099903 |
| RNY1P11 | -0.076109928 |
| ALOX12P1 | -0.076123539 |
| RP11-74E24 | -0.076123999 |
| MIR559 | -0.076141175 |
| DNM1P47 | -0.076150842 |
| OR1X5P | -0.076151769 |
| CERK | -0.076155083 |
| SLC7A7 | -0.076155734 |
| GTF2H1 | -0.076157967 |
| RNU7-84P | -0.076158582 |
| UNKL | -0.076159977 |
| KLHDC7B | -0.076167028 |
| FTLP6 | -0.076167661 |
| MTND6P19 | -0.076168945 |
| RP11-417B4 | -0.076187045 |
| RRP8 | -0.076189448 |
| EP400 | -0.076190713 |
| UBE2FP3 | -0.076191209 |
| LENEP | -0.076192409 |
| CTC-493P15 | -0.076196612 |
| SLC7A6 | -0.076200313 |
| RP11-386G21 | -0.076200336 |
| CNST | -0.076206025 |
| RP11-403P17 | -0.076206025 |
| AMY1A | -0.076206307 |
| AC078851 | -0.076210024 |
| MACROD2-AS1 | -0.076218137 |
| OCLM | -0.076223353 |
| KB-1027C11 | -0.076225384 |
| ANKRD20A10P | -0.076230124 |
| STAG3L3 | -0.076232129 |
| INIP | -0.076233059 |
| CTD-3022G6 | -0.076248598 |
| EIF4A2 | -0.076259404 |

| | |
|---|---|
| PTGES3P3 | -0.076261581 |
| RP11-645N11 | -0.076263327 |
| RP11-434B12 | -0.07626947 |
| RP11-707G14 | -0.076287691 |
| ZNF200 | -0.076289712 |
| RP11-337A23 | -0.076290338 |
| ZWINT | -0.076294401 |
| CAPRIN1 | -0.076295834 |
| MTMR14 | -0.07630082 |
| WSB1 | -0.07630415 |
| RP11-804A23 | -0.07630539 |
| TRIM34 | -0.076307881 |
| RP11-37C7 | -0.076308548 |
| TVP23B | -0.076311258 |
| GPR55 | -0.0763234 |
| RP11-73O6 | -0.076333163 |
| CD1E | -0.076334988 |
| RP11-446F3 | -0.076339121 |
| BRPF3 | -0.076345064 |
| RP11-38G5 | -0.076350931 |
| IGHD7-27 | -0.076353288 |
| ATP5A1P3 | -0.076363977 |
| TMEM5 | -0.076364376 |
| RP11-534L6 | -0.076371344 |
| RNU6-907P | -0.076373439 |
| ESRRAP2 | -0.076380399 |
| RP1-45N11 | -0.076384028 |
| CCL4L1 | -0.07638692 |
| MIR6739 | -0.076394487 |
| MEP1B | -0.076410543 |
| PABPC1L | -0.076415095 |
| C7orf25 | -0.076420509 |
| AC104650 | -0.076436846 |
| GALM | -0.076448603 |
| FAM177B | -0.076456994 |
| ERICH4 | -0.076457584 |
| DNAJC19P1 | -0.076459167 |
| LLNLR-307A6 | -0.07646423 |
| RP11-278J6 | -0.076467807 |
| RNU6-367P | -0.076469095 |
| RP3-333H23 | -0.076473061 |
| PIAS1 | -0.076477814 |
| TCEA3 | -0.076479544 |
| SIGIRR | -0.07649774 |
| FLJ42393 | -0.076498234 |

| | |
|---|---|
| TAS2R4 | -0.076511648 |
| RP11-543D5 | -0.076518684 |
| MIR579 | -0.076524805 |
| ZXDC | -0.076527788 |
| FRAT1 | -0.0765279 |
| NDRG3 | -0.076531304 |
| LSG1 | -0.076538747 |
| TATDN3 | -0.076541649 |
| NOC3L | -0.076541872 |
| UNG | -0.076554375 |
| ZDHHC20P4 | -0.076564299 |
| AL356458 | -0.076570218 |
| LRBA | -0.076570376 |
| BOD1L2 | -0.076576294 |
| RP11-259P1 | -0.076587285 |
| RP11-503N18 | -0.076589875 |
| ZNF770 | -0.076595698 |
| RNU6-1040P | -0.076604451 |
| AC012594 | -0.076606532 |
| SERINC4 | -0.076618075 |
| RERG-IT1 | -0.076619014 |
| MIR4420 | -0.076626596 |
| SNORA48 | -0.076635275 |
| RNU6-252P | -0.076641911 |
| AC105053 | -0.076642077 |
| VWA5B1 | -0.076659438 |
| DQX1 | -0.076659998 |
| TNFRSF9 | -0.076669097 |
| RNU6-759P | -0.076675296 |
| RP11-144H23 | -0.076675386 |
| USPL1 | -0.076676408 |
| HTT-AS1_2 | -0.076685635 |
| KRTAP4-6 | -0.076685778 |
| SRGAP1 | -0.076690791 |
| ARF4P4 | -0.076711012 |
| PWAR5 | -0.076711794 |
| DHX9 | -0.076722792 |
| CCND2P1 | -0.076723288 |
| MON1B | -0.076727164 |
| SLC22A15 | -0.076727332 |
| FLJ46284 | -0.076728061 |
| HNF1A-AS1 | -0.076740992 |
| RELL1 | -0.076741621 |
| UBE2D3P4 | -0.076742215 |
| DIDO1 | -0.076742272 |

| | |
|---|---|
| ZNF780B | -0.076743463 |
| PRORY | -0.076743728 |
| RP11-57H12 | -0.076746663 |
| RNU6-875P | -0.076751981 |
| KCNG3 | -0.076753396 |
| RP11-517B11 | -0.076761398 |
| RP11-132F7 | -0.076773118 |
| SLCO2B1 | -0.076775204 |
| PER3 | -0.076775929 |
| RP11-618N24 | -0.076776992 |
| OR52L1 | -0.076777185 |
| CAPN3 | -0.076781474 |
| METAP2 | -0.076788376 |
| SAMD10 | -0.07679059 |
| VPS33B | -0.076796804 |
| AC009892 | -0.076805785 |
| RP5-1041C10 | -0.076809365 |
| RAD51C | -0.076817326 |
| RN7SL431P | -0.076819245 |
| RN7SKP287 | -0.076820859 |
| PCNX | -0.076822722 |
| COCH | -0.076829167 |
| RP5-859D4 | -0.076834866 |
| DEFB109P1B | -0.076851981 |
| AC100787 | -0.076863734 |
| RAD9A | -0.076865942 |
| UBFD1P1 | -0.076867392 |
| ZNF665 | -0.076875747 |
| NCOR1 | -0.076878092 |
| GPM6BP3 | -0.076880193 |
| HTR2A-AS1 | -0.076886993 |
| EPS15 | -0.076888529 |
| SLC13A4 | -0.076889218 |
| TUBGCP6 | -0.076909758 |
| CLCP2 | -0.076909768 |
| PLSCR1 | -0.076917126 |
| RP11-497G19 | -0.076923053 |
| USP1 | -0.076928029 |
| RP11-89K18 | -0.076928356 |
| EEF1B2P1 | -0.076928426 |
| RP1-5O6 | -0.076948788 |
| CABLES1 | -0.076956179 |
| RNA5SP496 | -0.076965598 |
| CTD-2324A24 | -0.076966062 |
| SLCO5A1 | -0.076967324 |

| | |
|---|---|
| SNORA70D | -0.076974745 |
| NAA15 | -0.076976254 |
| RP11-930P14 | -0.076995697 |
| SMC2 | -0.077000758 |
| TDRD15 | -0.077005549 |
| TMEM150C | -0.07701927 |
| ST13P18 | -0.0770355 |
| ACSBG2 | -0.07703732 |
| HNRNPA3P6 | -0.077062509 |
| C7orf34 | -0.077067679 |
| RP11-405O10 | -0.077070645 |
| RP11-90O23 | -0.077075108 |
| NCK1-AS1 | -0.077078491 |
| MAATS1 | -0.077091645 |
| SLC14A1 | -0.077094008 |
| RP11-849H4 | -0.077095404 |
| AC006296 | -0.07711879 |
| ZNF267 | -0.077124521 |
| GPBP1L1 | -0.077126865 |
| RNA5SP441 | -0.077132915 |
| AC079395 | -0.077133318 |
| RSL24D1 | -0.077145527 |
| RP5-1171I10 | -0.077148991 |
| ZNF641 | -0.07715109 |
| AL807772 | -0.077152275 |
| AC079796 | -0.077152275 |
| DCAF8 | -0.077156783 |
| CHGB | -0.07715808 |
| PRR13P2 | -0.077159466 |
| RP11-22C11 | -0.077170458 |
| MLH1 | -0.07718084 |
| SGOL1 | -0.077188524 |
| REL | -0.077190478 |
| MTATP6P2 | -0.077191291 |
| FAF2P1 | -0.077193992 |
| COX20P2 | -0.077196479 |
| SNHG20 | -0.077202366 |
| REST | -0.077207762 |
| GS1-251I9 | -0.077221971 |
| RP11-1195F20 | -0.077225837 |
| ZRSR2 | -0.077226219 |
| GNG13 | -0.077234119 |
| RP11-144F15 | -0.077242393 |
| AC011290 | -0.077253178 |
| snoU13 | -0.077261095 |

| | |
|---|---|
| PSMC3IP | -0.077264118 |
| SEPT7-AS1 | -0.077264844 |
| RP11-506M12 | -0.077265646 |
| PAGE3 | -0.077279836 |
| ZNF605 | -0.077280807 |
| LCA5L | -0.077284137 |
| PTGER4 | -0.077300454 |
| OSGIN2 | -0.077314316 |
| CCDC142 | -0.077330149 |
| TUBD1 | -0.077333088 |
| SMARCE1P6 | -0.077340636 |
| FBXO45 | -0.077352568 |
| AP004550 | -0.077353658 |
| SPTBN1 | -0.077357052 |
| VRK2 | -0.077370225 |
| RLIMP1 | -0.077373415 |
| PROX2 | -0.077377774 |
| LAT2 | -0.077404273 |
| ZNF597 | -0.077422841 |
| WDR73 | -0.077428665 |
| PHC1P1 | -0.077429972 |
| VPS13C | -0.077440163 |
| AC245028 | -0.077441224 |
| OR5W1P | -0.077461304 |
| NUP160 | -0.077464052 |
| LRRK2 | -0.07746876 |
| C18orf32 | -0.07746902 |
| SCLT1 | -0.07747289 |
| PTPN2 | -0.077476592 |
| MIR6798 | -0.077479777 |
| WBP11P1 | -0.07748162 |
| C16orf92 | -0.077485259 |
| EPX | -0.077486105 |
| RAD51AP1 | -0.077487012 |
| XIST | -0.077489359 |
| SPR | -0.077491235 |
| B3GNT2P1 | -0.077491616 |
| C8orf59P1 | -0.077498327 |
| P4HTM | -0.077499552 |
| RP11-15E1 | -0.077501273 |
| MIR4756 | -0.077513258 |
| ASH1L-IT1 | -0.077515996 |
| CCL8 | -0.077518529 |
| ARAP1-AS1 | -0.077529687 |
| RP11-734I18 | -0.07753106 |

| | |
|---|---|
| RP4-733B9 | -0.077531988 |
| RP11-295I5 | -0.077538159 |
| BOLL | -0.077540096 |
| RPL26P35 | -0.077546426 |
| LYPD6 | -0.07754947 |
| RP11-428C19 | -0.077549859 |
| AC078993 | -0.077555573 |
| TNFAIP2 | -0.077563759 |
| COQ5 | -0.077566866 |
| SHPRH | -0.077572187 |
| MIR3194 | -0.077574391 |
| UFL1 | -0.07757762 |
| PDE12 | -0.077585007 |
| HMGA1P8 | -0.07758528 |
| MIR5581 | -0.07759218 |
| NSD1 | -0.077594998 |
| TDG | -0.077596579 |
| RP11-210N13 | -0.077599252 |
| ZNF543 | -0.0776043 |
| CLUHP3 | -0.077604393 |
| RN7SL425P | -0.077609216 |
| RAB11AP2 | -0.077613023 |
| OTC | -0.077615766 |
| RP11-26F2 | -0.077619224 |
| RNU6-712P | -0.077619798 |
| TSEN2 | -0.077620059 |
| AC018766 | -0.07764086 |
| ZNF33A | -0.077680378 |
| C8orf49 | -0.077685329 |
| AP000253 | -0.077686257 |
| AC003985 | -0.07769322 |
| ABCA2 | -0.077693365 |
| RNU6-1005P | -0.077696012 |
| PPP6R1 | -0.07769943 |
| RP11-104N10 | -0.077702165 |
| RP11-129K12 | -0.077716896 |
| RNU6-1205P | -0.077719111 |
| TAS2R5 | -0.077720721 |
| ZNF786 | -0.07772371 |
| RN7SL75P | -0.077723968 |
| RP13-15M17 | -0.077744049 |
| OR7E115P | -0.077744755 |
| AC110299 | -0.077751618 |
| APOL1 | -0.07775627 |
| LINC01424 | -0.077759897 |

| | |
|---|---|
| RP11-542K23 | -0.077767199 |
| RP11-1137G4 | -0.077774914 |
| SLC4A1APP2 | -0.077780482 |
| RP11-185P18 | -0.077796847 |
| TRAF3 | -0.077803658 |
| LINC01336 | -0.077817573 |
| CTD-2143L24 | -0.077846307 |
| KIF17 | -0.07784952 |
| TP53RK | -0.077861205 |
| RPL26P3 | -0.077879863 |
| RP11-173M1 | -0.077884294 |
| OR4F13P | -0.077885551 |
| AC006380 | -0.077896503 |
| HNRNPA1P44 | -0.07790645 |
| FAM66A | -0.07791421 |
| AC080125 | -0.077921515 |
| NOL11 | -0.077923459 |
| FCRLB | -0.077937041 |
| GABPB2 | -0.077940706 |
| CCT4P1 | -0.07794403 |
| RP11-143J12 | -0.077944894 |
| MIR4419A | -0.077946221 |
| RNU6-200P | -0.077949776 |
| FMR1-IT1 | -0.077951016 |
| CMB9-55A18 | -0.077956515 |
| LINC00282 | -0.077958409 |
| GCGR | -0.077958479 |
| PDHX | -0.077959479 |
| AP1G1 | -0.077964763 |
| TMEM33 | -0.077969116 |
| LATS1 | -0.077971144 |
| CTD-2199O4 | -0.077977762 |
| RN7SL305P | -0.077984253 |
| CREB1 | -0.077991406 |
| TNFRSF1B | -0.077991778 |
| MSL2 | -0.07800162 |
| HELLS | -0.078002495 |
| HLA-G | -0.078003406 |
| RBBP4P1 | -0.078013429 |
| MTHFD1 | -0.078046571 |
| CLK3 | -0.078060488 |
| ST3GAL1 | -0.078062386 |
| TMX3 | -0.078072842 |
| RP11-133N21 | -0.078073048 |
| AC010226 | -0.078094368 |

| | | |
|---|---|---|
| TCEB1P23 | | -0.078095394 |
| CLASP2 | | -0.078104378 |
| VPS26BP1 | | -0.078105627 |
| PTBP2 | | -0.078116248 |
| PIK3C3 | | -0.078118835 |
| RABGGTA | | -0.078123821 |
| ADORA2BP1 | | -0.078138778 |
| LRR1 | | -0.078141868 |
| NCOA6 | | -0.078149087 |
| ST3GAL5 | | -0.078153757 |
| HCG24 | | -0.078160477 |
| RP11-737O24 | | -0.078166709 |
| IGKV2OR2-10 | | -0.078168925 |
| DUSP19 | | -0.078174372 |
| MAL | | -0.078186356 |
| TIA1 | | -0.078187563 |
| AC011899 | | -0.078194354 |
| XPOTP1 | | -0.078198179 |
| TPM3P8 | | -0.078200502 |
| TXNRD1 | | -0.078211341 |
| RP11-333E13 | | -0.078228216 |
| RP11-857B24 | | -0.078233146 |
| RP11-393K12 | | -0.07824091 |
| MIR556 | | -0.078243263 |
| RP11-146N18 | | -0.078249509 |
| PAGE1 | | -0.078253238 |
| | 6-Sep | -0.078270227 |
| RP11-481C4 | | -0.078275474 |
| MRPL50 | | -0.078277707 |
| RP11-258C19 | | -0.078287605 |
| INPP5E | | -0.078291289 |
| SNORD116-1 | | -0.078291459 |
| RP11-473L15 | | -0.078293647 |
| MOSPD2 | | -0.078294712 |
| LINC00254 | | -0.078295117 |
| RNU6-1223P | | -0.078301364 |
| RP11-378N18 | | -0.078309683 |
| VRK3 | | -0.078316053 |
| RP1-257A7 | | -0.078345654 |
| KIAA0907 | | -0.07835704 |
| AC018890 | | -0.078374882 |
| AGR2 | | -0.078384707 |
| FCGR2A | | -0.078393582 |
| AC097374 | | -0.0784018 |
| RACGAP1 | | -0.078406922 |

| | |
|---|---|
| AC008079 | -0.078419611 |
| RP11-1099M24 | -0.078422346 |
| MEOX1 | -0.078426151 |
| RP11-492I2 | -0.078430704 |
| CCL3L3 | -0.078437118 |
| KMT2A | -0.07845604 |
| PHACTR4 | -0.078464431 |
| ATP8A2P2 | -0.078464936 |
| RPARP-AS1 | -0.078476004 |
| ALLC | -0.078480641 |
| BCOR | -0.078486497 |
| SNRPFP4 | -0.078490175 |
| DNAJC3 | -0.078496544 |
| RP13-941N14 | -0.078515108 |
| SLC19A2 | -0.078529476 |
| AC083949 | -0.078529698 |
| RP11-569G9 | -0.078541812 |
| CCDC84 | -0.078549942 |
| SOX5 | -0.07856131 |
| RP11-107P7 | -0.078561408 |
| H3F3AP3 | -0.078564669 |
| ZBTB44 | -0.078565961 |
| HM13-IT1 | -0.078567319 |
| ETNK1 | -0.078575915 |
| ACBD5 | -0.07859132 |
| F5 | -0.078592939 |
| RUNDC3A-AS1 | -0.078597592 |
| CNNM4 | -0.078601981 |
| CH17-164J11 | -0.078603284 |
| AC000120 | -0.07860663 |
| ANAPC7 | -0.078607712 |
| POU2F2 | -0.078611917 |
| ZNF354A | -0.078613424 |
| RP11-81K13 | -0.078616257 |
| CRIP1P4 | -0.07862006 |
| HNRNPA1P36 | -0.078628518 |
| AC010364 | -0.078628914 |
| RP11-702F3 | -0.07863602 |
| LA16c-431H6 | -0.078639434 |
| RP1-136B1 | -0.078639665 |
| PRKRIRP2 | -0.078640147 |
| RP11-347P5 | -0.078644061 |
| SFI1 | -0.078649258 |
| WIPF1 | -0.078659993 |
| CSTP1 | -0.078663299 |

| | |
|---|---|
| SIGLEC6 | -0.078664147 |
| SPDYE1 | -0.078672219 |
| AC015971 | -0.078672244 |
| OR1D5 | -0.078673719 |
| RP11-337N6 | -0.078676943 |
| PARPG1 | -0.07868219 |
| CTD-2023M8 | -0.078688637 |
| GAS2L3 | -0.078692757 |
| SETD4 | -0.078693911 |
| RPL5P30 | -0.078703197 |
| RP11-43N16 | -0.078710098 |
| RALBP1 | -0.078717763 |
| IGHVII-44-2 | -0.078724537 |
| KB-288A10 | -0.078727364 |
| BRCA1 | -0.078730136 |
| AC006445 | -0.07875047 |
| SNORD112 | -0.078756135 |
| BTG2 | -0.078766565 |
| CSNK1A1P3 | -0.078773086 |
| WDFY3 | -0.078773747 |
| LYPD6B | -0.078776631 |
| CYCSP34 | -0.07877789 |
| MORF4L2-AS1 | -0.078788148 |
| RP11-331D5 | -0.078797303 |
| RP11-566K19 | -0.078803496 |
| RNA5SP268 | -0.078804265 |
| KIAA0196-AS1 | -0.078807652 |
| BTF3P6 | -0.078810677 |
| RP5-971N18 | -0.078830053 |
| GATAD2B | -0.078831573 |
| PHF20L1 | -0.078850923 |
| AC005071 | -0.078854054 |
| SMIM11 | -0.078857623 |
| RP11-281A20 | -0.078866948 |
| TRGJP1 | -0.07886727 |
| RP11-165J3 | -0.078873751 |
| HCN2 | -0.078887259 |
| DPH6 | -0.078898236 |
| CTC-312O10 | -0.078905934 |
| VEZF1 | -0.07891033 |
| HTR5A | -0.078912399 |
| AC000124 | -0.078914903 |
| HCG22 | -0.078927671 |
| PHLPP2 | -0.078928935 |
| MDN1 | -0.078939708 |

| | |
|---|---|
| RP11-613F22 | -0.078947855 |
| LAMTOR5-AS1 | -0.078951876 |
| NPM1P27 | -0.078962648 |
| IVD | -0.078969458 |
| LINC00886 | -0.078970146 |
| RP11-855O10 | -0.078974049 |
| CTB-174O21 | -0.07898664 |
| GPR157 | -0.078988715 |
| AL132875 | -0.079004271 |
| PSMD12 | -0.079020511 |
| MIR4424 | -0.079021465 |
| RNY1P9 | -0.079026014 |
| SMARCC2 | -0.079026707 |
| FAM138A | -0.079035085 |
| RAD52 | -0.0790356 |
| RMI1 | -0.07903614 |
| AFF3 | -0.079036773 |
| RP11-197N18 | -0.07904174 |
| KPNA5 | -0.079046968 |
| RBM15 | -0.079050652 |
| RNU2-25P | -0.079062834 |
| FAM32B | -0.079063831 |
| AP001057 | -0.079087609 |
| HERC6 | -0.079116292 |
| GSDMB | -0.079121 |
| KCNIP1 | -0.079126311 |
| GIMAP1 | -0.079126972 |
| HMGB3P2 | -0.079131597 |
| C11orf57 | -0.079134042 |
| BTN2A3P | -0.079138731 |
| GATSL2 | -0.079142582 |
| RP11-371E8 | -0.079143565 |
| AP1G2 | -0.079148684 |
| ANKZF1 | -0.079184333 |
| ZNF382 | -0.079192835 |
| PKD1L1 | -0.07919557 |
| AC073254 | -0.079199552 |
| SENP6 | -0.079204706 |
| TMIGD2 | -0.07920716 |
| MIR4698 | -0.079207926 |
| SMG1P2 | -0.079213413 |
| CYP2C9 | -0.079221433 |
| PARG | -0.079223665 |
| RP11-403N16 | -0.079224008 |
| RANP7 | -0.079224488 |

| | |
|---|---|
| RP11-77P6 | -0.079238735 |
| SS18 | -0.079247535 |
| M1AP | -0.07926015 |
| RNY3P4 | -0.079268604 |
| PRSS3P1 | -0.07928092 |
| IGHV3-30-2 | -0.079281291 |
| BSN | -0.0792843 |
| RN7SL568P | -0.079287536 |
| CLEC4C | -0.079291511 |
| MIR4294 | -0.079292637 |
| ZFP3 | -0.079300319 |
| RNF168 | -0.079306384 |
| ESRP1 | -0.079307594 |
| MIR5572 | -0.079309272 |
| TADA2B | -0.079310738 |
| RASGRP2 | -0.079314013 |
| POLDIP3 | -0.079332376 |
| RP11-669M16 | -0.079332845 |
| RP11-567O18 | -0.079336839 |
| RP11-767N6 | -0.079338293 |
| C1orf132 | -0.07934793 |
| RAP1GDS1 | -0.079352098 |
| ACHE | -0.079356619 |
| POU3F1 | -0.079361973 |
| GPR144 | -0.079370336 |
| ATE1 | -0.079371318 |
| GMFG | -0.079374927 |
| IL13 | -0.079379676 |
| HIST2H2BD | -0.079383083 |
| THSD4 | -0.079396621 |
| RN7SL403P | -0.079407136 |
| AQR | -0.079411412 |
| RNU6-377P | -0.079416582 |
| C21orf58 | -0.079428232 |
| LINC01340 | -0.079442285 |
| RP11-681L8 | -0.079444161 |
| SLC27A2 | -0.079444418 |
| CDK18 | -0.079446111 |
| EAF1 | -0.079452586 |
| PAX4 | -0.079455082 |
| C12orf65 | -0.079456289 |
| MIR642A | -0.079462461 |
| SEC62-AS1 | -0.079465671 |
| SCAND2P | -0.079467508 |
| PBRM1 | -0.079471527 |

| | |
|---|---|
| TRIM13 | -0.07947242 |
| RP11-1020A11 | -0.079474187 |
| AC020743 | -0.079474861 |
| LTB4R2 | -0.079496197 |
| NECAP1P2 | -0.079498417 |
| RNU6-784P | -0.079504721 |
| AC012456 | -0.079516614 |
| PDXDC2P | -0.079520478 |
| NF1 | -0.079523566 |
| CPT1B | -0.07952418 |
| AC068641 | -0.07953302 |
| ZMYND8 | -0.07953512 |
| RP11-536L3 | -0.079538092 |
| TYW3 | -0.079541148 |
| RNU1-61P | -0.0795596 |
| ZNF546 | -0.079565503 |
| HECTD1 | -0.079566935 |
| MTND2P31 | -0.079568413 |
| MED18 | -0.079569261 |
| AC019118 | -0.07957172 |
| RP11-560A7 | -0.079576384 |
| TEX21P | -0.079579871 |
| XRCC2 | -0.079592202 |
| DNAJC27 | -0.079596016 |
| PRKAG2 | -0.079598639 |
| CTD-2021A8 | -0.079600037 |
| GDPGP1 | -0.079602174 |
| EFCAB14-AS1 | -0.07960223 |
| ZNF585B | -0.079609244 |
| RP11-473O4 | -0.079614402 |
| RP11-757A13 | -0.079620425 |
| RP11-67C2 | -0.079625519 |
| LINC01100 | -0.07963239 |
| MIR498 | -0.079635657 |
| XPA | -0.079637506 |
| RNU6-1321P | -0.079639461 |
| MIR320E | -0.079642282 |
| THAP10 | -0.079651739 |
| RN7SKP66 | -0.07965177 |
| OR5D17P | -0.079658562 |
| HNRNPKP4 | -0.079667053 |
| OR4C46 | -0.079672243 |
| AP001425 | -0.079675322 |
| QRICH1 | -0.079676056 |
| RNF20 | -0.079680931 |

| | |
|---|---|
| WDHD1 | -0.079685601 |
| TLX1 | -0.079703264 |
| TERT | -0.07970725 |
| HMGN2P5 | -0.079708188 |
| HERPUD2 | -0.079716133 |
| RP4-621N11 | -0.079722166 |
| FAM230A | -0.079726794 |
| AC113133 | -0.07973542 |
| TEX13A | -0.079737988 |
| NONOP2 | -0.079753967 |
| SYP-AS1 | -0.079756023 |
| CICP3 | -0.07976652 |
| RP11-97N10 | -0.079768966 |
| GTSCR1 | -0.079770983 |
| RBPJ | -0.079772098 |
| CCNDBP1 | -0.079801122 |
| NIN | -0.079804657 |
| NT5C3A | -0.079815765 |
| KRTAP9-4 | -0.07981621 |
| CAPRIN2 | -0.079823561 |
| SNRPCP4 | -0.079825725 |
| RNA5SP213 | -0.07982573 |
| RP11-30L15 | -0.0798331 |
| RP11-353N14 | -0.079841924 |
| SUGP2 | -0.079845422 |
| RP11-113K21 | -0.079849292 |
| OXR1 | -0.079861739 |
| CHMP4A | -0.079869014 |
| RP1-166H1 | -0.079874479 |
| IST1 | -0.079891192 |
| CASP8AP2 | -0.079905834 |
| LMLN | -0.079908439 |
| MIR5089 | -0.079912316 |
| OR5BR1P | -0.079915061 |
| MTND4P26 | -0.079925577 |
| TMA16 | -0.079930133 |
| COL19A1 | -0.079930467 |
| TTC9C | -0.079932775 |
| RP11-512N21 | -0.079941432 |
| XRN1 | -0.079947473 |
| BCRP5 | -0.079947691 |
| AC006947 | -0.079957939 |
| RP11-712L6 | -0.079976646 |
| AC093373 | -0.07998356 |
| CYP2C19 | -0.079986847 |

| | |
|---|---|
| FAM90A28P | -0.079993794 |
| RP4-576H24 | -0.079998584 |
| RP11-168J18 | -0.080005526 |
| ZFP91 | -0.080015904 |
| ZZEF1 | -0.0800365 |
| AC092958 | -0.080042883 |
| SYNPO2 | -0.080050585 |
| RNA5SP44 | -0.080051656 |
| DZIP3 | -0.08005412 |
| AC092106 | -0.080054974 |
| BCAS2P2 | -0.080056784 |
| COX6CP6 | -0.08005894 |
| AC093716 | -0.080073807 |
| HMGB1P14 | -0.080078164 |
| RP11-80H8 | -0.080081066 |
| RPL23AP72 | -0.080084851 |
| CD83 | -0.080103778 |
| DHFR | -0.080108764 |
| STX18-IT1 | -0.080114586 |
| RNY4P13 | -0.080128837 |
| LRRC37A15P | -0.080129652 |
| RP11-10A2 | -0.080142821 |
| LINC01385 | -0.080150571 |
| ARNT2 | -0.080164376 |
| AP005718 | -0.080171925 |
| HELZ2 | -0.080176618 |
| CCDC39 | -0.0801806 |
| RP4-758J24 | -0.080190665 |
| MIR4518 | -0.080200079 |
| SOGA1 | -0.080211615 |
| C9orf106 | -0.080215915 |
| SLC25A19 | -0.08022795 |
| ZCCHC6 | -0.080228378 |
| COL11A2P1 | -0.080237484 |
| SCAMP5 | -0.080250891 |
| SIRPB1 | -0.080252342 |
| RNU6-584P | -0.08025264 |
| LINC00893 | -0.08030053 |
| CR1 | -0.080302776 |
| RB1 | -0.080308884 |
| WBP4 | -0.080315564 |
| TARBP1 | -0.080323601 |
| AC010655 | -0.080330929 |
| PRPF4B | -0.080332085 |
| RP11-598D12 | -0.080335501 |

| | |
|---|---|
| AC026150 | -0.080343787 |
| IGHVIII-5-1 | -0.080349926 |
| SRGAP3 | -0.080354021 |
| RP11-751H17 | -0.080358243 |
| FAM96AP1 | -0.080363885 |
| RP11-564C24 | -0.080370886 |
| OARD1 | -0.080371045 |
| POLA1 | -0.080374617 |
| RP11-22B10 | -0.080374862 |
| SRSF2 | -0.08038619 |
| RNU6-1021P | -0.08038982 |
| CH17-264B6 | -0.08039053 |
| PWWP2A | -0.080403958 |
| VPREB1 | -0.080425401 |
| TYSND1 | -0.080430006 |
| AC008985 | -0.080432497 |
| CLIP1-AS1 | -0.080433254 |
| LILRA1 | -0.080438323 |
| STRIP1 | -0.080447904 |
| ATG14 | -0.080460277 |
| SGK494 | -0.080462268 |
| DMXL1 | -0.08046277 |
| CUX2P1 | -0.08050798 |
| CACNG6 | -0.080509742 |
| KDM5A | -0.080512577 |
| HDAC1 | -0.080516335 |
| AL671972 | -0.080516677 |
| SLC30A8 | -0.080528405 |
| USP7 | -0.080529843 |
| IDE | -0.080532206 |
| MTHFD2 | -0.080533303 |
| RP11-158D2 | -0.080534931 |
| GPATCH11 | -0.080537062 |
| RNU6-834P | -0.080540752 |
| COG3 | -0.080547369 |
| UBE2O | -0.080552988 |
| RP11-417L14 | -0.080556777 |
| E2F2 | -0.080560226 |
| LAIR1 | -0.080562328 |
| RP11-87F15 | -0.080564446 |
| RNU6-1112P | -0.080569613 |
| SUMO1P1 | -0.080569893 |
| RP11-535C21 | -0.080572805 |
| TLR4 | -0.08057284 |
| STK36 | -0.08057431 |

| | |
|---|---|
| RP1-232L24 | -0.080575153 |
| FTLP18 | -0.080581713 |
| CTD-2308G16 | -0.080586761 |
| AC140481 | -0.080587476 |
| TSPAN13 | -0.080588041 |
| HNRNPA1P20 | -0.080591539 |
| PWWP2AP1 | -0.080600538 |
| ARSG | -0.080611539 |
| IGHD5-5 | -0.080622824 |
| ZC3H10 | -0.080628936 |
| PMS1 | -0.080636024 |
| DFNB31 | -0.080644229 |
| RP13-650G11 | -0.080658867 |
| RP11-356M20 | -0.080672648 |
| SPDYE7P | -0.08067279 |
| ACADSB | -0.080678538 |
| RP11-361L15 | -0.080684535 |
| HNRNPDL | -0.080694911 |
| CCT6P3 | -0.08069798 |
| RP11-1082A3 | -0.080719359 |
| ZBTB3 | -0.080720493 |
| SLC35A5 | -0.080735303 |
| FAM26F | -0.080739136 |
| TRERNA1 | -0.08074341 |
| RFK | -0.080752048 |
| PROP1 | -0.080761632 |
| WDR20 | -0.080762281 |
| GK-AS1 | -0.080767697 |
| TOB1-AS1 | -0.080771379 |
| PRORSD1P | -0.080774654 |
| RP4-633I8 | -0.080780998 |
| UBA6-AS1 | -0.08078338 |
| SNORD18 | -0.080784424 |
| RP11-331G2 | -0.080787881 |
| RP11-140I16 | -0.080802882 |
| INCA1 | -0.080805557 |
| NUDT2 | -0.080813539 |
| E2F7 | -0.08082193 |
| RP11-733D4 | -0.08082405 |
| B3GAT2 | -0.080825938 |
| CELSR2 | -0.080829856 |
| RP11-598F7 | -0.080844443 |
| RP11-4O1 | -0.08085155 |
| DENND4B | -0.080860388 |
| PRKAR1A | -0.080861225 |

| | |
|---|---|
| CH507-145C22 | -0.080868509 |
| MYB | -0.080872742 |
| BCLAF1P1 | -0.080875405 |
| MYO19 | -0.080880128 |
| MTND5P25 | -0.080880934 |
| KRR1P1 | -0.080891898 |
| KRT8P46 | -0.08089347 |
| MIR553 | -0.080893578 |
| IGHV3-16 | -0.080894595 |
| C1DP4 | -0.080898218 |
| LRRC37A2 | -0.080903162 |
| RP11-505K1 | -0.080910899 |
| CTC-524C5 | -0.080922344 |
| TM6SF1 | -0.080924018 |
| LENG8 | -0.080942307 |
| OR2H4P | -0.080943832 |
| USP3 | -0.080960857 |
| RP11-77G23 | -0.080962518 |
| CBX4 | -0.080965732 |
| ELMO1 | -0.080965825 |
| FLOT2 | -0.080973937 |
| AZI2 | -0.080975407 |
| HOMEZ | -0.080977323 |
| CTD-2267D19 | -0.080977937 |
| DNAJC16 | -0.08098283 |
| ZSCAN32 | -0.080996059 |
| CLEC4E | -0.080997549 |
| MTDH | -0.080997938 |
| RP11-720D4 | -0.081012796 |
| GPR25 | -0.081022246 |
| PRIMPOL | -0.081027948 |
| RP11-437F6 | -0.081045196 |
| AL360176 | -0.081049898 |
| RP11-666A8 | -0.081062834 |
| RBM4 | -0.081064285 |
| KIAA1324L | -0.081068452 |
| KAT5 | -0.081073067 |
| PACSIN1 | -0.081076304 |
| ZUFSP | -0.08108023 |
| SEZ6 | -0.081083032 |
| TMEM161B | -0.08109037 |
| HMMR | -0.081097496 |
| FAM81A | -0.081108622 |
| RP11-136K7 | -0.0811205 |
| FAM134A | -0.081123971 |

| | |
|---|---|
| LINC01102 | -0.081197472 |
| RFC1 | -0.081208552 |
| CITF22-49D8 | -0.081208761 |
| TRIM75P | -0.081210056 |
| ZKSCAN2 | -0.081236832 |
| RNU12-2P | -0.081239902 |
| CEPT1 | -0.081243902 |
| NRL | -0.081244758 |
| AC006130 | -0.081265323 |
| IFIT2 | -0.081265577 |
| WDR48 | -0.081274787 |
| FAM175A | -0.081279401 |
| AC034110 | -0.081282405 |
| FCGR1A | -0.081290806 |
| RNU6-813P | -0.081292192 |
| HERC2 | -0.081305561 |
| LRRC37A | -0.081310677 |
| RP11-638O6 | -0.081311492 |
| ZNF155 | -0.081313896 |
| AC000123 | -0.081314528 |
| TMED4 | -0.081315626 |
| WWP1P1 | -0.081320769 |
| KBTBD7 | -0.081331441 |
| RP11-819C21 | -0.08133265 |
| OR7E47P | -0.081335407 |
| CPT1A | -0.081342065 |
| LYRM7 | -0.081349897 |
| TAPT1 | -0.081359461 |
| RP11-274B21 | -0.081359647 |
| FBXO36-IT1 | -0.081371056 |
| RNU6-635P | -0.081374868 |
| UHMK1 | -0.081388132 |
| ARIH1 | -0.081388578 |
| AC006042 | -0.081390848 |
| MIR1289-2 | -0.08139365 |
| RP11-677M14 | -0.081403425 |
| RP11-146I2 | -0.081409536 |
| ANKRD44-IT1 | -0.081412397 |
| ID2-AS1 | -0.081412542 |
| MIR590 | -0.08141299 |
| RP3-510O8 | -0.081418464 |
| ENOSF1 | -0.081424319 |
| LCMT2 | -0.081425808 |
| OXA1L | -0.081433752 |
| CTAGE10P | -0.081442356 |

| | |
|---|---|
| SVIP | -0.08144406 |
| CDAN1 | -0.0814542 |
| AC016405 | -0.081458455 |
| MIR4436A | -0.081460806 |
| MIR3163 | -0.081460826 |
| RP1-308E4 | -0.081463328 |
| KRT8P43 | -0.081478644 |
| ANAPC1P1 | -0.081481223 |
| AFG3L2 | -0.081484061 |
| TRAJ21 | -0.081489337 |
| HLA-DQB1-AS1 | -0.081496838 |
| HERC2P9 | -0.081498946 |
| CTD-3065B20 | -0.08149972 |
| CDC25A | -0.081499802 |
| HNRNPK | -0.081499988 |
| SLC35E1P1 | -0.08150129 |
| RP11-10J21 | -0.081513138 |
| LA16c-316G12 | -0.081531193 |
| CTD-2561B21 | -0.08153705 |
| RP11-428G5 | -0.081547043 |
| RUNDC1 | -0.081563265 |
| RP11-419C5 | -0.081572459 |
| PSMD3 | -0.081580642 |
| RN7SL320P | -0.0815907 |
| AF127936 | -0.081597518 |
| RP11-351N4 | -0.081598013 |
| HIATL2 | -0.08160096 |
| ATP6V1G1 | -0.081610207 |
| RP11-408I18 | -0.08161533 |
| SFSWAP | -0.081618932 |
| SBF1P1 | -0.081629633 |
| HK2 | -0.081631045 |
| RP11-429G17 | -0.081631794 |
| PPIP5K2 | -0.081632459 |
| GLRA1 | -0.081635492 |
| STAT2 | -0.081638152 |
| YWHABP2 | -0.081641446 |
| AAGAB | -0.081642413 |
| ZNF283 | -0.081645929 |
| NOVA1 | -0.081651139 |
| MLECP1 | -0.081654262 |
| GLYCTK-AS1 | -0.081658078 |
| KIF11 | -0.081660032 |
| TRIM22 | -0.08166312 |
| CLOCK | -0.081678675 |

| | |
|---|---|
| NCR2 | -0.081679061 |
| COIL | -0.081689819 |
| IRS2 | -0.081691791 |
| RP11-401G5 | -0.081694606 |
| CFHR5 | -0.081709606 |
| IL5RA | -0.081725169 |
| SFPQ | -0.081741933 |
| SNORA9 | -0.081748791 |
| TRDJ3 | -0.081749456 |
| CTD-2503O16 | -0.081752577 |
| CHD2 | -0.081757841 |
| DCP2 | -0.081757971 |
| IFNL3 | -0.081769057 |
| RPL7AP60 | -0.081797257 |
| RP11-415A20 | -0.081799846 |
| ZCCHC10 | -0.081802308 |
| PATE1 | -0.081836334 |
| RN7SL441P | -0.081838517 |
| KLRF1 | -0.081869124 |
| MFNG | -0.081883335 |
| OR1J1 | -0.081887759 |
| FMR1 | -0.081891019 |
| RP11-563E2 | -0.081898224 |
| HMGN2 | -0.081904936 |
| NNT | -0.081907782 |
| AC073957 | -0.081910455 |
| C18orf65 | -0.081912933 |
| ITFG2 | -0.081919206 |
| LINC00106 | -0.081944491 |
| RP11-56F10 | -0.081961392 |
| FAM204CP | -0.081963136 |
| EAPP | -0.081964064 |
| OIP5 | -0.08198293 |
| ASB14 | -0.081984232 |
| BRD7P5 | -0.081996399 |
| TRAJ28 | -0.082003835 |
| PHF21B | -0.082008212 |
| PIP4K2A | -0.082008847 |
| KRTAP5-6 | -0.082013427 |
| NPIPB8 | -0.082015822 |
| RNU6-1109P | -0.082042299 |
| DBF4 | -0.082047137 |
| DDX42 | -0.082051714 |
| IFIT1 | -0.082056012 |
| CTC-268N12 | -0.082061033 |

| | |
|---|---|
| RFX7 | -0.082064254 |
| MVB12B | -0.082068552 |
| TVP23CP2 | -0.082072944 |
| BCL7A | -0.082087083 |
| CICP14 | -0.08209834 |
| RN7SKP239 | -0.082099737 |
| RN7SL19P | -0.082101863 |
| PCMTD1 | -0.082102786 |
| CD180 | -0.08210487 |
| NPM1P29 | -0.082108993 |
| RPGRIP1 | -0.082111512 |
| TEC | -0.082116294 |
| CTD-2145A24 | -0.082118396 |
| MBD3L2 | -0.082119258 |
| LILRP2 | -0.082122298 |
| TOM1L1 | -0.082124741 |
| RNU6-1011P | -0.082128697 |
| RP1-29G21 | -0.082136688 |
| RP11-242J7 | -0.082142721 |
| DPPA2P3 | -0.082148425 |
| LINC00870 | -0.082149677 |
| PRDM15 | -0.082156351 |
| SBK2 | -0.082166488 |
| ESPL1 | -0.082167626 |
| JAK3 | -0.082172185 |
| ULK3 | -0.082177599 |
| ARL17B | -0.082190269 |
| AP001625 | -0.082222769 |
| 5_8S_rRNA | -0.08223232 |
| ZNF320 | -0.082233546 |
| ARL6IP6 | -0.082234755 |
| RNU6-1034P | -0.082248703 |
| RPTOR | -0.082251835 |
| TIGD4 | -0.082253528 |
| KCNIP4 | -0.082253937 |
| PLEKHA7 | -0.082257342 |
| C16orf46 | -0.082257751 |
| RP11-130F10 | -0.082265307 |
| RP11-455F5 | -0.082265342 |
| PLCXD2 | -0.082270123 |
| NPM1P8 | -0.082272506 |
| PDPK1 | -0.08227511 |
| PKD2L2 | -0.082278159 |
| MAD2L1BP | -0.082285976 |
| RNU6-626P | -0.082295109 |

| | |
|---|---|
| UBAP1L | -0.082296879 |
| APOBEC3C | -0.082301772 |
| RNU6-619P | -0.082303043 |
| HLA-P | -0.082307279 |
| SAGE1 | -0.082312193 |
| RFWD3 | -0.08233182 |
| RP11-665C14 | -0.082343547 |
| TMEM55B | -0.082345271 |
| GFPT1 | -0.082346834 |
| RNU6-278P | -0.082352001 |
| RP11-40E6 | -0.082364881 |
| OR2AO1P | -0.082365046 |
| SMARCE1P1 | -0.082368341 |
| GEN1 | -0.082371598 |
| AC027601 | -0.082408362 |
| H3F3C | -0.082431303 |
| SH2B3 | -0.082436159 |
| KB-431C1 | -0.082439527 |
| SELE | -0.082441387 |
| RP11-1094M14 | -0.082443545 |
| SNORD42B | -0.082458481 |
| DAXX | -0.082479789 |
| MIR4472-1 | -0.082489942 |
| SPAG1 | -0.08249032 |
| TRAJ10 | -0.082522223 |
| ARGFXP2 | -0.082527346 |
| NLRP11 | -0.082530003 |
| RP11-524G24 | -0.082540216 |
| MIPEPP1 | -0.082544806 |
| CHKB | -0.082549522 |
| MIR7156 | -0.082561172 |
| RP11-242D8 | -0.082563011 |
| RPL35P8 | -0.082581197 |
| AC015820 | -0.082586857 |
| NFYB | -0.082593301 |
| CTB-17P3 | -0.082603425 |
| RBM5 | -0.082614474 |
| C2CD3 | -0.082630754 |
| MIR1205 | -0.082634137 |
| RP11-452B18 | -0.082636158 |
| POLR1B | -0.082638549 |
| RP11-504I13 | -0.08263907 |
| FAM49B | -0.082645601 |
| RNU7-192P | -0.082648469 |
| RNU6-137P | -0.082656365 |

| | |
|---|---|
| RP11-214C8 | -0.082661537 |
| ZKSCAN8 | -0.082664727 |
| ERCC6L2 | -0.0826779 |
| AL672043 | -0.08268505 |
| RFX6 | -0.082691396 |
| RNU6-353P | -0.082696796 |
| GPR79 | -0.082708022 |
| KBTBD8 | -0.082713269 |
| OR56B4 | -0.082714601 |
| RNU6-1213P | -0.082736244 |
| AP000355 | -0.082738253 |
| PPM1E | -0.082741177 |
| RP11-79J24 | -0.082741315 |
| IGLV3-31 | -0.082759819 |
| UBE2V1P8 | -0.082764683 |
| AC092574 | -0.082765411 |
| KDM3B | -0.082772825 |
| AC069282 | -0.082774853 |
| AC074338 | -0.082776286 |
| PTBP3 | -0.082785737 |
| MIR5191 | -0.082798915 |
| RP11-162P23 | -0.082801408 |
| PIP4K2B | -0.082807189 |
| ZNF451 | -0.082808213 |
| MOB1A | -0.082808789 |
| TMEM145 | -0.082827283 |
| RP11-637C24 | -0.082843618 |
| RP11-98P2 | -0.082847998 |
| RP11-22B23 | -0.082879788 |
| TRBV17 | -0.08288074 |
| RNA5SP241 | -0.082887202 |
| PCK2 | -0.08290524 |
| PSG6 | -0.082915509 |
| TAS2R64P | -0.082926843 |
| RP11-631M21 | -0.082939846 |
| IFIT3 | -0.082943344 |
| ZNF814 | -0.082961745 |
| CTD-2342I9 | -0.082967242 |
| RP11-494I9 | -0.082975799 |
| CEACAM22P | -0.082985265 |
| TRIM23 | -0.082987997 |
| HLTF | -0.082998565 |
| SNORA14 | -0.083017358 |
| RP11-758M2 | -0.083029988 |
| CTA-984G1 | -0.083034333 |

| | |
|---|---|
| SS18L1 | -0.083037265 |
| AC018892 | -0.08303981 |
| ZFP14 | -0.08304413 |
| MYB-AS1 | -0.083061621 |
| DYNC1LI1 | -0.083061898 |
| MATR3 | -0.083062921 |
| MTND3P10 | -0.083070189 |
| TMEM14E | -0.083079867 |
| RP11-693J15 | -0.083100019 |
| KIAA0196 | -0.083117454 |
| RPL7P17 | -0.083121401 |
| AC092568 | -0.08313579 |
| CD74 | -0.083163428 |
| AF240627 | -0.0831677 |
| RNU7-29P | -0.083184183 |
| SLC2A13P1 | -0.083185103 |
| RTBDN | -0.083185802 |
| RP11-382J12 | -0.083188888 |
| ABI3BP | -0.083190778 |
| RP11-180M15 | -0.08320116 |
| TNRC6C | -0.083207002 |
| RP11-523O18 | -0.083216718 |
| ZNF28 | -0.083217552 |
| TMEM140 | -0.083222333 |
| GPR32 | -0.08322511 |
| SH3GL1P2 | -0.083245513 |
| ANKAR | -0.083246539 |
| EXOC7 | -0.083250539 |
| NAT8 | -0.083251376 |
| RP11-217F16 | -0.083251573 |
| TMEM128 | -0.083251618 |
| AL365361 | -0.083252965 |
| SNORD114-25 | -0.083254384 |
| RP11-449P15 | -0.083264474 |
| NECAB3 | -0.083268121 |
| MYBPC3 | -0.083279176 |
| IGHVII-60-1 | -0.083285229 |
| MORF4L1P7 | -0.083285586 |
| RNU6-571P | -0.083289541 |
| RNU2-32P | -0.083298974 |
| INTS8 | -0.08330269 |
| SLC16A6 | -0.08332857 |
| RP1-230L10 | -0.083341783 |
| NBPF11 | -0.083344404 |
| API5 | -0.083346301 |

| | |
|---|---|
| FBLL1 | -0.083350998 |
| SNORD19 | -0.083357576 |
| ZNF83 | -0.083365279 |
| DGCR12 | -0.083369742 |
| RP11-758H9 | -0.083377503 |
| WRB | -0.083383177 |
| AC022154 | -0.083397656 |
| LDHAL6DP | -0.083401991 |
| ISOC1 | -0.083403811 |
| MON2 | -0.083403978 |
| RP11-373I8 | -0.083411653 |
| AC091849 | -0.083413502 |
| RGPD5 | -0.083429016 |
| RP11-933H2 | -0.083429421 |
| TCP10 | -0.083433661 |
| OR6C65 | -0.083434672 |
| SIGLEC18P | -0.083436503 |
| EIF4E | -0.083436594 |
| MFSD6 | -0.083451087 |
| RPL7L1P4 | -0.083461524 |
| BIRC6 | -0.083465358 |
| KLHDC2 | -0.083481377 |
| TMEM87B | -0.083487647 |
| MSI2 | -0.08349708 |
| SMG1 | -0.083505843 |
| CYCSP52 | -0.083518403 |
| USP3-AS1 | -0.083542645 |
| VIP | -0.083550525 |
| DDX52 | -0.083560971 |
| RNU2-46P | -0.083561691 |
| AC110602 | -0.083573869 |
| CTD-2232E5 | -0.083574186 |
| RP11-21I4 | -0.08357572 |
| RP1-63M2 | -0.083576675 |
| C1QC | -0.083580693 |
| RP11-159D12 | -0.083589363 |
| GIGYF2 | -0.083597382 |
| WDR19 | -0.083604713 |
| ANKRD13D | -0.083612676 |
| RP11-678G14 | -0.083623061 |
| RP11-482D24 | -0.083627247 |
| AJ271736 | -0.083639584 |
| ARAP2 | -0.083643003 |
| C2orf49 | -0.083653012 |
| YTHDC2 | -0.083667785 |

| | |
|---|---|
| RP11-9L18 | -0.083672809 |
| SCML2P2 | -0.08367991 |
| RP11-120M18 | -0.083690512 |
| IGKV2D-10 | -0.083695765 |
| AC002117 | -0.083702718 |
| SLAMF8 | -0.083712587 |
| RP4-713B5 | -0.083716015 |
| TOX3 | -0.08371827 |
| RP11-74D7 | -0.083724523 |
| TRBJ2-2P | -0.08372888 |
| RP11-173P15 | -0.083737173 |
| ZMYM6 | -0.083766487 |
| POLR2D | -0.083784107 |
| PAPOLA | -0.083791809 |
| RP11-438F14 | -0.083803038 |
| LINC00701 | -0.083803356 |
| ZNF79 | -0.083805447 |
| RP1-30M3 | -0.08380567 |
| NDUFAF6 | -0.083808964 |
| ENTHD2 | -0.083813168 |
| LINC01268 | -0.083813299 |
| ARMC1 | -0.083824536 |
| GOLGA2P5 | -0.083824611 |
| RP11-575C1 | -0.083836103 |
| RNA5SP101 | -0.083873571 |
| RP11-523H24 | -0.083883479 |
| RN7SL253P | -0.083884589 |
| SNORD117 | -0.08388676 |
| SMLR1 | -0.083888316 |
| HNRNPA3P15 | -0.083891526 |
| RP11-609L3 | -0.083893719 |
| SMARCE1P5 | -0.083899863 |
| ARGLU1 | -0.083902047 |
| AC007551 | -0.083902598 |
| RNU6-476P | -0.083907173 |
| ANXA2R | -0.083907684 |
| RP3-388E23 | -0.083908208 |
| LINC01134 | -0.083918615 |
| PSIP1 | -0.083925917 |
| MS4A4A | -0.083949193 |
| U66059 | -0.083950354 |
| TOX2 | -0.08395483 |
| PP14571 | -0.083976346 |
| RP11-120E5 | -0.083979335 |
| PSMB8 | -0.083987185 |

| | |
|---|---|
| TCL6 | -0.0839889 |
| RNA5SP214 | -0.084013126 |
| BRCC3 | -0.084016414 |
| RP11-362K14 | -0.084020567 |
| SEC22C | -0.084020731 |
| LCOR | -0.084026871 |
| VARS2 | -0.084032211 |
| CASP8 | -0.084033978 |
| DDX20 | -0.084053458 |
| C10orf128 | -0.084054928 |
| AL079339 | -0.084057501 |
| ASB16 | -0.084060751 |
| NHEJ1 | -0.0840609 |
| TFAM | -0.084063021 |
| RN7SL131P | -0.084067607 |
| TAF5 | -0.084075654 |
| FAM208B | -0.084076975 |
| MCF2 | -0.084083052 |
| AC005562 | -0.084094167 |
| AC006076 | -0.084099039 |
| TXLNG | -0.084108549 |
| ZNF302 | -0.084114912 |
| ZNF816 | -0.08411694 |
| TAF8 | -0.084117405 |
| SAP30L | -0.084119154 |
| AC097468 | -0.084121662 |
| CD80 | -0.084126075 |
| AC073316 | -0.084127934 |
| RP11-68I3 | -0.084134429 |
| RP11-342L8 | -0.084139362 |
| MRPL49 | -0.084144625 |
| ARL17A | -0.084158077 |
| FLJ38122 | -0.084164152 |
| BRD1 | -0.084165668 |
| RP11-1376P16 | -0.084165811 |
| TRPC4AP | -0.084168812 |
| AC011897 | -0.084172748 |
| AC025627 | -0.08417845 |
| MIR4418 | -0.084178463 |
| SELP | -0.084181896 |
| GUCY2EP | -0.084185576 |
| CD1D | -0.084189445 |
| RP11-354K4 | -0.084193012 |
| KIAA0586 | -0.084199939 |
| MIR663AHG | -0.084222429 |

| | |
|---|---|
| MBD1 | -0.084233689 |
| AC004623 | -0.084234597 |
| SOD1P1 | -0.084248073 |
| WARS2P1 | -0.084253035 |
| RP11-575L7 | -0.084261481 |
| RP11-666A1 | -0.084262532 |
| UBE2Q1-AS1 | -0.084264091 |
| C9orf41 | -0.084264891 |
| ARHGAP19 | -0.084269728 |
| L29074 | -0.084270462 |
| RP1-58B11 | -0.084284663 |
| RP11-203E8 | -0.084288555 |
| ERCC6 | -0.084292817 |
| CTC-487M23 | -0.084297487 |
| CELF1 | -0.084305841 |
| ERCC5 | -0.084309079 |
| HSPA7 | -0.084309209 |
| POC5 | -0.084309562 |
| SPAG5 | -0.084309637 |
| MRS2 | -0.084316353 |
| ZNF861P | -0.084318088 |
| OVOL1 | -0.084323237 |
| CECR6 | -0.084328354 |
| RP11-186N15 | -0.084330661 |
| RP11-263K4 | -0.084337923 |
| RP3-395M20 | -0.084339015 |
| TIMELESS | -0.084341638 |
| IGKV1-35 | -0.084352145 |
| POTEH | -0.084358541 |
| KCNA2 | -0.084374991 |
| RP11-59H1 | -0.084387975 |
| SNORD63 | -0.084394326 |
| TP73 | -0.084395296 |
| CCDC7 | -0.084407911 |
| LINC00595 | -0.084418982 |
| IFNL1 | -0.084420147 |
| MED13 | -0.084425493 |
| MAPK1IP1L | -0.084426665 |
| PARP9 | -0.084430702 |
| ING5 | -0.084432265 |
| RP11-876N24 | -0.084445978 |
| NCL | -0.084458518 |
| FAM91A1 | -0.084459764 |
| RP11-271C24 | -0.084460374 |
| CTD-2020K17 | -0.084460918 |

| | |
|---|---|
| LINC00944 | -0.084465106 |
| FCGR3A | -0.084466388 |
| PTP4A2 | -0.084512697 |
| PTPRJ | -0.084519878 |
| RNU6-125P | -0.084524628 |
| GPR183 | -0.084530353 |
| RP11-694O4 | -0.084537196 |
| RP11-416O18 | -0.08454189 |
| BRCA2 | -0.084544884 |
| HMHA1 | -0.084561778 |
| RAD17P1 | -0.084562712 |
| SLC1A4 | -0.084572755 |
| PABPC1P7 | -0.084580616 |
| EVI2A | -0.084582839 |
| ZNF701 | -0.084585946 |
| KAT6A | -0.0845895 |
| RP11-245J9 | -0.084590989 |
| RP11-896J10 | -0.084591301 |
| SETP10 | -0.084592581 |
| RP11-229P13 | -0.084602077 |
| CALHM1 | -0.08462005 |
| AL591377 | -0.084639878 |
| MDS2 | -0.084641686 |
| NPM1P40 | -0.084648894 |
| CCDC18 | -0.084651754 |
| AP000439 | -0.084656162 |
| RP11-358L22 | -0.084661485 |
| WDR7 | -0.084664555 |
| CACNB2 | -0.084668815 |
| KBTBD11-OT1 | -0.084678963 |
| ARIH2 | -0.084680816 |
| CTD-3128G10 | -0.084681783 |
| IGHV1OR15-4 | -0.084684093 |
| MRPS31 | -0.084684351 |
| MDP1 | -0.084700333 |
| RP1-152L7 | -0.084728427 |
| LILRB4 | -0.084730604 |
| GPR155 | -0.084739311 |
| PRR15 | -0.084740018 |
| ZNF445 | -0.084756931 |
| OR4D9 | -0.084767506 |
| UBIAD1 | -0.084779425 |
| AC013476 | -0.084779807 |
| BTAF1 | -0.084781397 |
| RP11-344P13 | -0.084784677 |

| | |
|---|---|
| GUSBP11 | -0.084786625 |
| ZNF544 | -0.084805714 |
| NPM1P49 | -0.084806641 |
| NDUFA5 | -0.084811817 |
| RP11-467D18 | -0.084821469 |
| CSNK1A1P1 | -0.084824831 |
| DPP8 | -0.084828134 |
| RP11-547D13 | -0.084832921 |
| CLCC1 | -0.084834013 |
| SRSF5 | -0.084850442 |
| RP11-350G8 | -0.084876731 |
| SLC35B1 | -0.084880806 |
| ADIPOR1P1 | -0.084883323 |
| DIS3L2P1 | -0.084886748 |
| ZNF609 | -0.084889197 |
| SEC16A | -0.084890276 |
| COX6CP14 | -0.084912317 |
| AC090132 | -0.084915234 |
| RBM48 | -0.084921961 |
| RBBP6 | -0.084927208 |
| AC092667 | -0.084927834 |
| SLC25A24P2 | -0.084938637 |
| PIK3IP1 | -0.084941255 |
| RP11-727A23 | -0.084952604 |
| SEC24C | -0.084976103 |
| 7SK | -0.084993179 |
| GPATCH1 | -0.084998672 |
| AC074367 | -0.085002902 |
| LINC00943 | -0.085012611 |
| CFLAR-AS1 | -0.085014282 |
| CTD-2200A16 | -0.085055234 |
| CASQ1 | -0.085056836 |
| GAPVD1 | -0.08506167 |
| RP11-358D17 | -0.085061972 |
| RHNO1 | -0.085069372 |
| ACVR2B | -0.085073987 |
| DKFZP434I0714 | -0.085074638 |
| RP11-382D8 | -0.085079789 |
| INSM1 | -0.085088149 |
| RP11-195B17 | -0.085088261 |
| GNPDA2 | -0.085097299 |
| RP11-902B17 | -0.085118975 |
| AC009274 | -0.085121665 |
| ZNF586 | -0.085123775 |
| FOXA1 | -0.085140855 |

| | |
|---|---|
| AC007463 | -0.08514808 |
| RP11-378E13 | -0.085156064 |
| METTL2B | -0.085159404 |
| SF3B1 | -0.085159907 |
| RNASEH2B | -0.085160055 |
| MYCLP2 | -0.085170866 |
| AC005189 | -0.085172307 |
| AC046176 | -0.085196699 |
| TVP23A | -0.085204336 |
| CACNA1E | -0.08521875 |
| CXCL11 | -0.08523284 |
| VPS39 | -0.085233435 |
| IDSP1 | -0.08523895 |
| TPMT | -0.085240747 |
| NPIPB4 | -0.085242199 |
| PRPF40A | -0.085242422 |
| L2HGDH | -0.085248078 |
| RPL7P49 | -0.085267801 |
| LINC00324 | -0.085276693 |
| TOPBP1 | -0.08527954 |
| IGJP1 | -0.085284989 |
| NR2C2 | -0.08528888 |
| KBTBD6 | -0.085303392 |
| NAP1L4 | -0.085315895 |
| RP11-264I13 | -0.085328677 |
| ZNF839 | -0.085329923 |
| PLCB2 | -0.085337477 |
| HSPD1P1 | -0.085354986 |
| RP11-884K10 | -0.08537893 |
| DUTP2 | -0.085380349 |
| SUMO2 | -0.085382595 |
| RBM45 | -0.085383526 |
| DDX59 | -0.085393201 |
| MIR5692A1 | -0.085396587 |
| HERC5 | -0.085403396 |
| RP11-113D6 | -0.085406737 |
| MBD3L4 | -0.085413698 |
| SMARCA5 | -0.085413964 |
| SLC25A24P1 | -0.085414633 |
| RP11-638D14 | -0.085426181 |
| SEPSECS-AS1 | -0.085438914 |
| LPIN2 | -0.085447417 |
| CTBP1-AS | -0.085458431 |
| FAM13A-AS1 | -0.085473762 |
| HAL | -0.085475875 |

| | |
|---|---|
| RP11-107I14 | -0.085480876 |
| COBL | -0.085482749 |
| NRF1 | -0.08548567 |
| AC073641 | -0.085487171 |
| TFF1 | -0.085496249 |
| HLA-DPA3 | -0.085514124 |
| RP11-754B17 | -0.085516371 |
| NOD2 | -0.085527178 |
| TTLL3 | -0.085533951 |
| SCAMP2 | -0.085537672 |
| AC091053 | -0.085538342 |
| RP11-120K24 | -0.085541133 |
| IMMTP1 | -0.085551399 |
| FUBP1 | -0.085557226 |
| VHL | -0.085560315 |
| RPAIN | -0.085560799 |
| RP11-397D21 | -0.085567021 |
| TAP2 | -0.085574939 |
| GTF3A | -0.085575571 |
| RPL39P28 | -0.085587034 |
| NPAT | -0.085602624 |
| EMC3-AS1 | -0.085608447 |
| RP13-314C10 | -0.085613893 |
| AC132008 | -0.085619975 |
| CEACAM16 | -0.085646451 |
| RP1-232L22__B | -0.0856563 |
| CTD-2028E8 | -0.085656961 |
| USP13 | -0.085665305 |
| P2RX5-TAX1BP3 | -0.085698033 |
| VPS36 | -0.085698758 |
| LA16c-390E6 | -0.085701413 |
| LL0XNC01-116E7 | -0.085701651 |
| HCG21 | -0.085705354 |
| FAM221A | -0.085709921 |
| NSFP1 | -0.085712726 |
| RP11-862G15 | -0.085716663 |
| ZNF830 | -0.085717978 |
| GPR148 | -0.085722236 |
| RP11-357F12 | -0.085727835 |
| AC018712 | -0.085733128 |
| CCNJ | -0.085738053 |
| CTA-384D8 | -0.085738165 |
| AC139530 | -0.085739974 |
| RP11-146D12 | -0.085741931 |
| RP11-380I10 | -0.085760405 |

| | |
|---|---|
| RP11-474B12 | -0.085764082 |
| CTD-2132N18 | -0.085784176 |
| MIR645 | -0.085801427 |
| FBXO3 | -0.085801702 |
| GTF3C2-AS1 | -0.085802442 |
| TAF2 | -0.085804158 |
| RP11-692C24 | -0.085809533 |
| RP11-16E23 | -0.085823712 |
| RP11-271K11 | -0.085836447 |
| TAOK1 | -0.085853146 |
| SMPDL3A | -0.0858575 |
| PRMT5-AS1 | -0.0858859 |
| ASCL1 | -0.085893311 |
| AGMAT | -0.085896032 |
| RP4-682C21 | -0.085912016 |
| CTC-241N9 | -0.085914079 |
| HELLPAR | -0.0859218 |
| RP11-689B22 | -0.085946737 |
| USP12 | -0.085955421 |
| RP11-78O7 | -0.085962137 |
| MAG | -0.085966643 |
| ARHGAP32 | -0.08597157 |
| TRAJ29 | -0.085971923 |
| TMEM80 | -0.08598145 |
| BZW1P1 | -0.085985899 |
| TTC32 | -0.085987068 |
| AC245407 | -0.085987673 |
| NUP153 | -0.086036522 |
| ZNF230 | -0.086039331 |
| SMEK1 | -0.08604562 |
| RNU6-230P | -0.086046881 |
| GAPDHP42 | -0.086051122 |
| CLCN6 | -0.086056765 |
| ADPGK | -0.086063704 |
| AC068044 | -0.086078942 |
| OAS2 | -0.08607924 |
| TCEB2P1 | -0.086086379 |
| KIAA0391 | -0.086096692 |
| CTB-179K24 | -0.086098348 |
| RP11-56A10 | -0.086104861 |
| BLOC1S5 | -0.086114032 |
| TRAV33 | -0.086131829 |
| RP3-470L22 | -0.086132734 |
| KIAA0368 | -0.086135354 |
| ARPP19 | -0.08613861 |

| | |
|---|---|
| ADNP | -0.086146462 |
| BUB1B | -0.086147801 |
| CH17-224D4 | -0.086149958 |
| NAT10 | -0.086157011 |
| PRLH | -0.086165258 |
| RP11-536C10 | -0.086166306 |
| RP11-395C3 | -0.086170405 |
| SEMA3F-AS1 | -0.086176454 |
| PDE3B | -0.086177142 |
| CDKL2 | -0.08618196 |
| CEBPB-AS1 | -0.086183282 |
| PPP3CB | -0.086187524 |
| PLEKHD1 | -0.086193393 |
| CD274 | -0.086194557 |
| C7orf55-LUC7L2 | -0.086196231 |
| ZNF106 | -0.086207822 |
| SETP3 | -0.086221201 |
| MTF2 | -0.086227879 |
| CIAPIN1P | -0.086228102 |
| UBE2V1P2 | -0.086230049 |
| C17orf67 | -0.086238466 |
| ATP8B1 | -0.086239582 |
| RPS6KB1 | -0.086240642 |
| AC096582 | -0.086241119 |
| RP11-702L15 | -0.086245269 |
| RAD54L2 | -0.086250336 |
| HCLS1 | -0.086252829 |
| PIGP | -0.086259639 |
| RP11-8L21 | -0.086269259 |
| RP11-33B1 | -0.086269351 |
| ZNF26 | -0.086276086 |
| IGLV10-67 | -0.086288901 |
| RP11-563N6 | -0.086310816 |
| TRIM44 | -0.086315008 |
| HNRNPA1L2 | -0.086320088 |
| UPF2 | -0.08632606 |
| SPPL2A | -0.086328851 |
| TGIF2-C20orf24 | -0.086340838 |
| AC009963 | -0.086346499 |
| SPATA5L1 | -0.086347884 |
| PRSS1 | -0.086352472 |
| ROCK1P1 | -0.086373616 |
| PSMA6 | -0.086378918 |
| EIF1AX | -0.086380462 |
| TAF4 | -0.086389263 |

| | |
|---|---|
| GDF9 | -0.086392482 |
| RP11-118E18 | -0.086403654 |
| LINC00907 | -0.086405067 |
| DBH-AS1 | -0.086406737 |
| KIAA1033 | -0.086408706 |
| CTD-2574D22 | -0.086424409 |
| RPL31P49 | -0.086442644 |
| MRPS31P5 | -0.086451932 |
| RP11-238K6 | -0.086472092 |
| KANSL1 | -0.086485118 |
| AL662800 | -0.086488212 |
| AL691477 | -0.086496113 |
| RP1-168L15 | -0.086500386 |
| KIF12 | -0.086508208 |
| CICP22 | -0.086510378 |
| TTC5 | -0.086511891 |
| ASXL2 | -0.086516841 |
| AC008464 | -0.08654409 |
| RP3-323B6 | -0.086545562 |
| SELPLG | -0.08655407 |
| SPEN | -0.086556433 |
| SNAP23 | -0.086557382 |
| CTD-2284J15 | -0.086560005 |
| LILRP1 | -0.086586936 |
| STARD10 | -0.086595225 |
| ACTR6 | -0.086599933 |
| RNU6-940P | -0.086603994 |
| CEP85L | -0.086619245 |
| RNY1P12 | -0.08662031 |
| SMC1A | -0.086623562 |
| LL22NC03-2H8 | -0.08662678 |
| STX11 | -0.086648716 |
| ZNF137P | -0.086652214 |
| RP11-831H9 | -0.086658127 |
| TNFSF18 | -0.086662077 |
| RP11-1070N10 | -0.086666537 |
| ZMYND19P1 | -0.086672501 |
| GOLGA1 | -0.086673889 |
| RP4-816N1 | -0.086676141 |
| TBC1D3P4 | -0.086683743 |
| RNU6-487P | -0.086690599 |
| AC125634 | -0.086699016 |
| TRBJ2-1 | -0.086701767 |
| RIC8B | -0.08671551 |
| UBE2CP2 | -0.08671612 |

| | |
|---|---|
| C12orf29 | -0.08671711 |
| ARHGAP30 | -0.086720812 |
| PSMD11 | -0.08673032 |
| RP11-533E19 | -0.086742655 |
| RP11-299P2 | -0.086757832 |
| RP11-73M18 | -0.08676647 |
| RP11-212I21 | -0.086766731 |
| ANKRA2 | -0.08676794 |
| VAMP2 | -0.086769875 |
| RN7SL589P | -0.086776248 |
| OR52N4 | -0.086779211 |
| NBPF12 | -0.086803141 |
| RP11-645C24 | -0.086817226 |
| RN7SL440P | -0.086822082 |
| PLA2G6 | -0.086822361 |
| RNU6-1209P | -0.08682278 |
| CTD-3149D2 | -0.086822905 |
| ACIN1 | -0.086830101 |
| RWDD4P2 | -0.086839571 |
| SLC12A6 | -0.086849916 |
| MIR4690 | -0.086851547 |
| RFC5 | -0.086852409 |
| SLC5A4 | -0.086854004 |
| RP11-313P18 | -0.086855295 |
| CKAP2 | -0.086856372 |
| PAPOLG | -0.08686413 |
| MIR4314 | -0.086867071 |
| MIR4539 | -0.086871316 |
| LCN9 | -0.086878588 |
| AC025918 | -0.086896216 |
| ZNF818P | -0.086899668 |
| CYCSP22 | -0.086918252 |
| CTD-3214H19 | -0.086924505 |
| LIN52 | -0.086924858 |
| CASP1P2 | -0.086952184 |
| CPLX1 | -0.086953623 |
| NPM1P31 | -0.086954978 |
| CTD-2050B12 | -0.086955415 |
| LINC01520 | -0.086966526 |
| NOP9 | -0.086967167 |
| CTD-2575K13 | -0.086974945 |
| PHF3 | -0.086976954 |
| LINC00485 | -0.086977117 |
| GATC | -0.086981735 |
| FAM184A | -0.086987187 |

| | |
|---|---|
| FAM155B | -0.086992826 |
| RALGAPB | -0.087000564 |
| RP11-266I3 | -0.087017124 |
| DNAJC24 | -0.087021793 |
| C2orf15 | -0.087050204 |
| PRMT3 | -0.087052362 |
| SETSIP | -0.087054531 |
| ZNF317 | -0.087061013 |
| SLA | -0.087066893 |
| VPS37C | -0.087071897 |
| LRRFIP1 | -0.087088549 |
| AC074286 | -0.087089703 |
| CCDC125 | -0.087090391 |
| TRAJ1 | -0.087092625 |
| TRDV3 | -0.087098278 |
| RERG | -0.087124309 |
| FCGR2B | -0.087124607 |
| MSH2 | -0.08712937 |
| CLDN20 | -0.087132742 |
| CST7 | -0.087136663 |
| MIR155HG | -0.087147268 |
| HIST1H4G | -0.087147321 |
| RNU7-49P | -0.087148486 |
| RP4-676J13 | -0.08716196 |
| SON | -0.087194787 |
| NDUFA3P2 | -0.087205686 |
| B3GALT2 | -0.087228677 |
| AC113167 | -0.087229679 |
| SRP68 | -0.087256594 |
| AC011625 | -0.087258784 |
| ZNF807 | -0.087271297 |
| HLA-DQB1 | -0.087273878 |
| RP11-47I22 | -0.087281843 |
| RP11-135A1 | -0.087283155 |
| RP11-311P8 | -0.087284313 |
| SH3GL1P1 | -0.087288679 |
| ZNF792 | -0.087289879 |
| ZBTB40 | -0.087302289 |
| RP11-222K16 | -0.087302427 |
| GAPDHP62 | -0.087308469 |
| RP11-83A24 | -0.087317583 |
| TRAPPC6B | -0.087323015 |
| RN7SL688P | -0.08733019 |
| ALG3P1 | -0.087345913 |
| RNU6-948P | -0.087348041 |

| | |
|---|---|
| POLR3A | -0.087356543 |
| RNU6-151P | -0.087362242 |
| STAG3L1 | -0.087366461 |
| CTD-2371O3 | -0.087381474 |
| C1orf162 | -0.08738473 |
| IRF8 | -0.08739273 |
| COX10-AS1 | -0.087395874 |
| S100Z | -0.087396747 |
| AC013474 | -0.087397814 |
| FAM138C | -0.087408979 |
| UBA6 | -0.087414368 |
| UGT1A4 | -0.087430701 |
| RP5-1070G24 | -0.087434399 |
| XXbac-BPG252P9 | -0.087439393 |
| AC124944 | -0.087449947 |
| MAP3K14-AS1 | -0.087482799 |
| CTD-2649C14 | -0.087482929 |
| PLK4 | -0.087494781 |
| PPP1R2P1 | -0.087506649 |
| ZNF638-IT1 | -0.087521064 |
| CUL3 | -0.08753642 |
| TCEA1P3 | -0.087536899 |
| CCDC91 | -0.087539378 |
| FKSG70 | -0.087545947 |
| FBXO48 | -0.08754816 |
| PIK3C2B | -0.087549146 |
| AP000476 | -0.087549906 |
| LINC01389 | -0.087561579 |
| RP11-95D17 | -0.087564719 |
| CASC3 | -0.087592739 |
| NMNAT1P5 | -0.08759361 |
| CKAP2L | -0.087598042 |
| RP11-1123I8 | -0.087602948 |
| MIR4506 | -0.087627169 |
| PPTC7 | -0.087630508 |
| LINC00463 | -0.087630699 |
| USP37 | -0.087633671 |
| HEXDC-IT1 | -0.087642776 |
| AC079448 | -0.087646526 |
| EBAG9 | -0.087648276 |
| ZNF620 | -0.087649988 |
| DLGAP4-AS1 | -0.087651532 |
| WDR88 | -0.087667384 |
| AC019206 | -0.087675407 |
| MED15P9 | -0.087678827 |

| | |
|---|---|
| DDX12P | -0.08768011 |
| AC096589 | -0.087689605 |
| SPECC1L-ADORA2A | -0.087705155 |
| CTD-2003D5 | -0.087710792 |
| TIRAP | -0.087715647 |
| TLK2 | -0.087722735 |
| LILRB2 | -0.087725805 |
| ADNP-AS1 | -0.087728243 |
| MKRN3-AS1 | -0.087739442 |
| ERLEC1 | -0.087752076 |
| NEK3 | -0.087755462 |
| GBP3 | -0.087763295 |
| CH17-262A2 | -0.087786523 |
| LFNG | -0.087787036 |
| FAM134B | -0.087787334 |
| AL672294 | -0.087788292 |
| AGAP6 | -0.087791445 |
| MIR3685 | -0.087793729 |
| HERC1 | -0.087799315 |
| SNX18P3 | -0.087804924 |
| MANEA | -0.087857867 |
| RP11-862L9 | -0.087859115 |
| POLA2 | -0.087861607 |
| SNORD67 | -0.087868167 |
| RNU6-835P | -0.087868409 |
| UBA7 | -0.087900045 |
| LINC00928 | -0.087900879 |
| RBM12 | -0.087940289 |
| AC025674 | -0.087941835 |
| RP11-62H7 | -0.087944146 |
| HNRNPA1P13 | -0.08795272 |
| FRY-AS1 | -0.087970812 |
| MTND1P9 | -0.087973772 |
| NDUFA12P1 | -0.087975055 |
| ZNF625 | -0.087976874 |
| RP11-457M11 | -0.087978833 |
| FNDC9 | -0.087981482 |
| DIP2A | -0.087990896 |
| SUZ12P1 | -0.088011027 |
| HNRNPLL | -0.088018432 |
| RP11-294N21 | -0.088030975 |
| RNA5SP61 | -0.088044711 |
| ZBTB1 | -0.088050006 |
| CENPU | -0.08806969 |
| SMAD4 | -0.088090975 |

| | |
|---|---|
| QSOX2 | -0.088091105 |
| GBP1P1 | -0.088092054 |
| RNF139 | -0.088092966 |
| FAM179B | -0.088096333 |
| CTB-31O20 | -0.088102808 |
| PHC1 | -0.088116241 |
| TRBV12-2 | -0.088130256 |
| CNOT8 | -0.088135442 |
| RPAP3 | -0.08816015 |
| ALG13 | -0.088164783 |
| LMNB1 | -0.088169062 |
| HLA-DOB | -0.088180895 |
| RP11-738E22 | -0.088188072 |
| MIR634 | -0.08820624 |
| MTND6P21 | -0.088225392 |
| HSPD1P5 | -0.088226571 |
| RPS29P26 | -0.088230868 |
| OR2BH1P | -0.088233965 |
| CYP2S1 | -0.088234516 |
| uc_338 | -0.088239781 |
| RNU6-638P | -0.088267849 |
| TERF2 | -0.088271783 |
| OFD1 | -0.088273476 |
| MCM9 | -0.088274518 |
| UHRF1BP1L | -0.088280639 |
| AC069499 | -0.088290256 |
| LINC00384 | -0.088299099 |
| RP11-178H8 | -0.088301198 |
| RP11-666E17 | -0.088305687 |
| RPL31P13 | -0.088308028 |
| RP11-480N24 | -0.088336935 |
| KLHL23 | -0.088340418 |
| BRCC3P1 | -0.088340708 |
| BLM | -0.088342763 |
| C9orf163 | -0.088349163 |
| DPY19L2P2 | -0.088357573 |
| EIF5 | -0.088378597 |
| ZNF490 | -0.088383229 |
| PIK3AP1 | -0.088385109 |
| TMEM19 | -0.088390951 |
| CDC6 | -0.088397202 |
| ALG10B | -0.088416552 |
| XXbac-BPGBPG55C20 | -0.088422636 |
| SUV420H1 | -0.088425594 |
| WIPF2 | -0.088425817 |

| | |
|---|---|
| MIR548P | -0.088432264 |
| BBS1 | -0.088433818 |
| AP000304 | -0.088440742 |
| AC134915 | -0.088443829 |
| FAM53B | -0.088447567 |
| XBP1P1 | -0.088448498 |
| RP11-173D9 | -0.088468385 |
| SPATC1 | -0.088481139 |
| HMGA1P2 | -0.088482816 |
| RP11-242C19 | -0.088484665 |
| ELK2BP | -0.088487071 |
| OTUD4 | -0.088488164 |
| RP11-314P15 | -0.088489277 |
| AL161448 | -0.088490014 |
| FABP5P13 | -0.088494179 |
| LINC01260 | -0.088500969 |
| C11orf30 | -0.0885069 |
| CTD-2528L19 | -0.088518752 |
| MIR8087 | -0.088522096 |
| AC069257 | -0.088526887 |
| AC008269 | -0.088527352 |
| ARID3A | -0.088540613 |
| ZNF91 | -0.088545339 |
| NHLRC2 | -0.088563796 |
| RNF38 | -0.088569266 |
| AC093495 | -0.088573266 |
| CLINT1 | -0.088594904 |
| USP33 | -0.088596299 |
| PTGES3 | -0.088596634 |
| RP11-9E17 | -0.088597192 |
| ST7L | -0.088605435 |
| RP11-108P20 | -0.088615536 |
| AC007191 | -0.088639576 |
| HNRNPA1P50 | -0.088643758 |
| PHBP9 | -0.088656782 |
| SOCS4 | -0.088661586 |
| AC079951 | -0.088668121 |
| R3HCC1L | -0.088701197 |
| RNU6-58P | -0.088708505 |
| RP11-454F8 | -0.088713282 |
| RP11-59K5 | -0.088716495 |
| RNU6-280P | -0.08872428 |
| AC092170 | -0.088728723 |
| MIR4650-2 | -0.088730524 |
| RP11-8L8 | -0.088731209 |

| | |
|---|---|
| ATG16L2 | -0.088783731 |
| KMT2D | -0.088791917 |
| RP5-890E16 | -0.08879588 |
| RP11-474C8 | -0.088796699 |
| UBE3B | -0.088808941 |
| BZW1 | -0.088853911 |
| AC007875 | -0.088857745 |
| RP11-165H4 | -0.08886109 |
| NLRP4 | -0.088875598 |
| GPR75 | -0.0888766 |
| MORC1 | -0.088881346 |
| CREBZF | -0.088884368 |
| NEU3 | -0.088893689 |
| GLUD1P3 | -0.088895289 |
| NCF1C | -0.08889715 |
| ZNRF2P2 | -0.088907841 |
| DPF2 | -0.088914025 |
| RBM27 | -0.088928928 |
| ZNF891 | -0.088940035 |
| RP1-80N2 | -0.088966232 |
| KIF3A | -0.088968037 |
| TMEM236 | -0.088969569 |
| GNGT2 | -0.08898266 |
| LBHD1 | -0.088991814 |
| RP3-339A18 | -0.088996379 |
| FAM71C | -0.089000038 |
| ZNF845 | -0.089002326 |
| RP11-355O1 | -0.089010587 |
| CDK5R1 | -0.089020243 |
| PSME2P2 | -0.089020746 |
| ALG1L13P | -0.089020827 |
| RP13-923O23 | -0.089022774 |
| ITGB2-AS1 | -0.089023165 |
| LMTK2 | -0.089024672 |
| RP3-468K18 | -0.089029962 |
| COA1 | -0.089035165 |
| CAMTA2 | -0.089035853 |
| RNU6-322P | -0.089056751 |
| IGHVII-33-1 | -0.089058223 |
| LINC01259 | -0.089061164 |
| RP1-102D24 | -0.089113573 |
| ACER1 | -0.089114658 |
| AP000437 | -0.089136358 |
| RNA5SP275 | -0.089168298 |
| IGSF9B | -0.089177941 |

| | |
|---|---|
| TPH1 | -0.089183023 |
| RP11-404P21 | -0.089184495 |
| WNT10B | -0.089185859 |
| HTT | -0.08918626 |
| ERMARD | -0.08920801 |
| MTND6P13 | -0.08921311 |
| PAICS | -0.089229369 |
| AC008281 | -0.089235674 |
| RNU6-678P | -0.089248665 |
| RP11-427P5 | -0.089258441 |
| KRTAP12-2 | -0.08925845 |
| AC093788 | -0.089259211 |
| TMEM41B | -0.089263194 |
| RP13-266I11 | -0.089264254 |
| RP11-114N19 | -0.089273866 |
| MYO16 | -0.089280586 |
| IPP | -0.089281762 |
| RP11-693M3 | -0.089345613 |
| AC013264 | -0.089347454 |
| CCDC34 | -0.089353542 |
| CACNA2D2 | -0.089366008 |
| KB-1572G7 | -0.089368174 |
| PTGDR | -0.089374998 |
| NCF1B | -0.089391274 |
| RP11-27M9 | -0.089402154 |
| OR2J4P | -0.089403403 |
| SMCR8 | -0.089404409 |
| CEP350 | -0.089409321 |
| AC113192 | -0.089413153 |
| LINC00562 | -0.089413908 |
| MGAT3-AS1 | -0.089414424 |
| EI24P2 | -0.089435872 |
| RP5-851M4 | -0.089452625 |
| IL37 | -0.089462607 |
| AD000684 | -0.089468524 |
| YWHAEP7 | -0.089473632 |
| RNU6-502P | -0.089488867 |
| PARP11 | -0.08949219 |
| UBE2Z | -0.089492525 |
| RBM23 | -0.08949446 |
| FKBP11 | -0.08952194 |
| STARD7 | -0.089529959 |
| ANKRD36B | -0.089543113 |
| ZNF2 | -0.089551188 |
| GKAP1 | -0.089557588 |

| | |
|---|---|
| CASP10 | -0.089565998 |
| UBN2 | -0.089572417 |
| GOLGA6L9 | -0.089575338 |
| CTAGE3P | -0.089579395 |
| SNRPGP14 | -0.089588632 |
| RPRD1B | -0.089594576 |
| FAM208A | -0.089597441 |
| RALGAPA1 | -0.089598111 |
| FBXO21 | -0.089620977 |
| AC025171 | -0.089621833 |
| ANKMY1 | -0.089621889 |
| TEX14 | -0.089624977 |
| LRIT3 | -0.089635565 |
| RICTOR | -0.089655657 |
| PPP1R12A | -0.08966429 |
| MIR4448 | -0.089673541 |
| RPL38P4 | -0.089674316 |
| RFPL4A | -0.089679016 |
| LINC01224 | -0.089689273 |
| AC026167 | -0.089713034 |
| RBBP4 | -0.089720516 |
| CTD-3092A11 | -0.089720944 |
| RB1CC1 | -0.089724944 |
| RNU6-168P | -0.089734127 |
| TTC39C | -0.089737391 |
| AC244157 | -0.089758642 |
| RALGAPA1P | -0.08977306 |
| RRN3 | -0.08979291 |
| SGK3 | -0.089792929 |
| RNVU1-19 | -0.089799568 |
| ZNF621 | -0.089805543 |
| C9orf156 | -0.089837693 |
| G2E3 | -0.089848801 |
| FANCF | -0.08985641 |
| TRAJ45 | -0.089860284 |
| ZNF92 | -0.089862922 |
| ZNF195 | -0.089869155 |
| MYT1 | -0.089880288 |
| SNORD62B | -0.08988681 |
| CAPN7 | -0.089886905 |
| TNRC6B | -0.089888579 |
| RP11-669E14 | -0.089897237 |
| KLHL12 | -0.089899687 |
| C3orf62 | -0.089936935 |
| TMEM192 | -0.089958796 |

| | |
|---|---|
| CALB1 | -0.089980548 |
| MIR3925 | -0.08998721 |
| AGAP11 | -0.089993508 |
| SLC25A30 | -0.090012585 |
| RNU6-789P | -0.090018047 |
| RP11-537H15 | -0.090023567 |
| C17orf58 | -0.090026167 |
| RBM12B-AS1 | -0.090031278 |
| RP11-434D12 | -0.090039729 |
| XPO1 | -0.090042595 |
| DDX18P5 | -0.090050072 |
| BDP1 | -0.090051005 |
| CTD-2340D6 | -0.090052492 |
| GIPR | -0.090057313 |
| USP45 | -0.090057536 |
| RP11-93K22 | -0.090059115 |
| CTD-2072I24 | -0.090059849 |
| RN7SL559P | -0.090061681 |
| IBTK | -0.090091081 |
| PIK3CD-AS1 | -0.090099147 |
| USP42 | -0.090105017 |
| SLC10A1 | -0.090105521 |
| GTPBP1 | -0.09013045 |
| LENG8-AS1 | -0.090134264 |
| DNAJC7 | -0.090145167 |
| BMPR1APS2 | -0.090149254 |
| CCR12P | -0.09015838 |
| OR5F1 | -0.090166397 |
| RP3-377H17 | -0.090189703 |
| SPATA20 | -0.090207421 |
| PLAC8 | -0.090210472 |
| DUSP2 | -0.090210603 |
| TRAJ56 | -0.090221186 |
| TRIM37 | -0.090231776 |
| RP5-1071N3 | -0.090232999 |
| RP11-386O9 | -0.09023451 |
| RP11-1252D15 | -0.090258003 |
| RBMXL2 | -0.090268547 |
| CSNK1G1 | -0.090269303 |
| RP4-570O12 | -0.09027511 |
| ZBTB40-IT1 | -0.09028101 |
| IGKV1OR2-1 | -0.090282493 |
| HERC2P5 | -0.090309156 |
| IPO7P2 | -0.090323585 |
| FBXW7 | -0.090331855 |

| | |
|---|---|
| RP11-109P11 | -0.090336373 |
| FANCD2 | -0.090337287 |
| SUMO1P4 | -0.090358688 |
| AL138898 | -0.09036096 |
| BMS1P1 | -0.090361791 |
| RP11-175P13 | -0.090370154 |
| TRMT13 | -0.090380787 |
| RCAN3 | -0.090382982 |
| RP11-305E17 | -0.090405071 |
| MLANA | -0.090405732 |
| RP11-338K17 | -0.090449739 |
| TRPV1 | -0.090452543 |
| AP000146 | -0.090459263 |
| RP3-476K8 | -0.090460363 |
| SENP7 | -0.09046107 |
| ZNF75D | -0.09046373 |
| SLC9B1 | -0.090463823 |
| NAA40 | -0.090469219 |
| RP11-292E2 | -0.09047254 |
| RP11-609D21 | -0.090476423 |
| ATXN7L1 | -0.090481796 |
| CRTC3-AS1 | -0.090486189 |
| DTL | -0.090491824 |
| KRT18P20 | -0.090491835 |
| SNRNP27 | -0.090506895 |
| BECN1 | -0.090532924 |
| TCEB3 | -0.090542059 |
| RAB2B | -0.090557483 |
| IGLJ6 | -0.090559126 |
| CEP295 | -0.090561669 |
| PMCHL2 | -0.090565551 |
| SREK1 | -0.090575363 |
| NUDT13 | -0.090584573 |
| RP11-91K8 | -0.090616948 |
| AC005104 | -0.090621323 |
| LEF1 | -0.090627235 |
| RP11-490K7 | -0.09062737 |
| ANP32BP1 | -0.090628814 |
| TATDN2 | -0.090654343 |
| ZNF441 | -0.090661599 |
| RP11-367J7 | -0.090672635 |
| SLFN14 | -0.090679415 |
| TSC1 | -0.090690121 |
| CROCCP3 | -0.090693657 |
| JPX | -0.090698047 |

| | |
|---|---|
| RP11-453E17 | -0.090704299 |
| Z98750 | -0.090708922 |
| TRAJ8 | -0.090711722 |
| ZC3H7A | -0.090713453 |
| RP11-17A1 | -0.090721049 |
| COX11 | -0.090721453 |
| ZNF675 | -0.09072696 |
| RP1-281H8 | -0.090727816 |
| MIR3909 | -0.090750971 |
| WDR89 | -0.090753752 |
| ZBTB6 | -0.090757641 |
| RP11-755F10 | -0.090757678 |
| RP11-762I7 | -0.090774311 |
| RP11-136L23 | -0.090779811 |
| TRAJ20 | -0.090788901 |
| ZNF841 | -0.090789996 |
| CTD-2017D11 | -0.090791372 |
| RP11-218C14 | -0.090798152 |
| STAG3L2 | -0.090808787 |
| LGALS9DP | -0.090812309 |
| PFN1P2 | -0.090820578 |
| FAM90A27P | -0.090824711 |
| EXOC1 | -0.090830035 |
| XXbac-BPG170G13 | -0.090838336 |
| RUFY3 | -0.090854798 |
| MANEAL | -0.090855189 |
| GXYLT1 | -0.090857347 |
| TPRKBP2 | -0.090872531 |
| RBM25 | -0.090877292 |
| KIF4CP | -0.09091026 |
| ASB2 | -0.090914838 |
| KDM1B | -0.090938318 |
| NBPF9 | -0.090938635 |
| ELF2 | -0.090961575 |
| RPAP1 | -0.090962747 |
| ATP1A1-AS1 | -0.090972162 |
| LINC01402 | -0.090977168 |
| PA2G4P3 | -0.090993942 |
| XRCC6P4 | -0.090994073 |
| EP300 | -0.091007177 |
| SNORD9 | -0.091027204 |
| RP11-730A19 | -0.091037002 |
| RP13-726E6 | -0.09104404 |
| FAM167A | -0.091045858 |
| RP11-168K11 | -0.091050878 |

| | |
|---|---|
| UNC13C | -0.091056683 |
| BSN-AS2 | -0.09106388 |
| AC114776 | -0.091078116 |
| CMPK2 | -0.091078957 |
| KNG1 | -0.09107999 |
| DNAJB14 | -0.091088055 |
| ICK | -0.091095907 |
| KIAA1109 | -0.091097656 |
| AC008547 | -0.091104015 |
| PIK3C2A | -0.091118568 |
| HSD11B1 | -0.091129918 |
| FAM13B | -0.091132336 |
| MYBL1 | -0.091141118 |
| RP11-66N24 | -0.091148132 |
| ZFX | -0.091156021 |
| TRAJ48 | -0.091168379 |
| ABHD10 | -0.091174533 |
| AC064836 | -0.091179378 |
| NR5A2 | -0.091180766 |
| GHc-210E9 | -0.091205849 |
| CXCL10 | -0.091213735 |
| TRAF6 | -0.091221661 |
| VPRBP | -0.091226871 |
| TRIM61 | -0.091232301 |
| DYNLT3P2 | -0.091235356 |
| SPAST | -0.091240694 |
| RP1-267L14 | -0.091241723 |
| RP1-69M21 | -0.091242829 |
| CCDC79 | -0.091248123 |
| FBXO43 | -0.091255635 |
| GNA13 | -0.09127331 |
| LGALS17A | -0.091274006 |
| SLC25A45 | -0.091279952 |
| LINC01053 | -0.091281844 |
| AC010746 | -0.091286671 |
| ZNF468 | -0.091290557 |
| SIDT2 | -0.091294595 |
| PER2 | -0.091307395 |
| AL096869 | -0.091308187 |
| ZNF525 | -0.091308195 |
| ZNF589 | -0.091318149 |
| SPIB | -0.091326545 |
| COX6C | -0.091337034 |
| SMG8 | -0.091345518 |
| AC005722 | -0.091346868 |

| | |
|---|---|
| RAB19 | -0.091350314 |
| OGT | -0.091379491 |
| PGM5P2 | -0.09138946 |
| AC138969 | -0.091416088 |
| DCC | -0.091421666 |
| CDYL2 | -0.091458565 |
| MIR3155A | -0.091461667 |
| RP11-79H23 | -0.091479677 |
| SNORA22 | -0.091481612 |
| MIR7107 | -0.091482186 |
| RP1-130L23 | -0.091494341 |
| GIMAP3P | -0.091495141 |
| PAG1 | -0.091498045 |
| VCPKMT | -0.091499255 |
| HNRNPA1P27 | -0.091518741 |
| CARNS1 | -0.091523795 |
| TTI1 | -0.091526009 |
| OR5H5P | -0.091527286 |
| CTC-429L19 | -0.091532975 |
| ENPP7 | -0.091535678 |
| DLEU7 | -0.091547666 |
| RP11-494K3 | -0.091555395 |
| TMEM14D | -0.091562697 |
| NNT-AS1 | -0.091570997 |
| USF1P1 | -0.09159517 |
| TLE3 | -0.091605287 |
| GOLGA2P1 | -0.091629692 |
| RP11-409K20 | -0.091635544 |
| ADARB2 | -0.091644894 |
| AC011515 | -0.091646884 |
| CENPK | -0.091654592 |
| LARP4 | -0.091663504 |
| TAS2R30 | -0.091678756 |
| AC097495 | -0.091688048 |
| CYCSP51 | -0.091708756 |
| ITCH | -0.09172386 |
| AKR7L | -0.091725441 |
| ZBTB26 | -0.091730297 |
| RP11-730G20 | -0.09174894 |
| CHN2 | -0.091774095 |
| GSE1 | -0.091777016 |
| AC087350 | -0.091785074 |
| ARHGEF7 | -0.091795342 |
| HSPA1B | -0.091800607 |
| CNOT1 | -0.091820385 |

| | |
|---|---|
| LRRC37A16P | -0.091822301 |
| DYNLT3P1 | -0.09183244 |
| RNU4ATAC2P | -0.091838085 |
| BTF3P7 | -0.091842158 |
| ZSWIM5 | -0.0918442 |
| MYO5C | -0.09188223 |
| RC3H2 | -0.091885002 |
| GRAMD1C | -0.09188582 |
| VCPIP1 | -0.09188943 |
| RP11-138P22 | -0.09189424 |
| JARID2 | -0.091915626 |
| GTF2B | -0.09193025 |
| RPL7AP26 | -0.091939103 |
| SETX | -0.091947721 |
| MCUR1 | -0.091948874 |
| SNORD45 | -0.091958532 |
| CEP250 | -0.091962642 |
| SDHCP2 | -0.091971837 |
| RP11-611O2 | -0.091971933 |
| AIMP1 | -0.091984281 |
| RP1-313I6 | -0.09199723 |
| TRAV39 | -0.092029339 |
| GPS2P1 | -0.092031923 |
| COA5 | -0.092042516 |
| IGLV4-60 | -0.092048698 |
| RN7SL381P | -0.092082298 |
| TMED8 | -0.09209169 |
| LRCH4 | -0.092094369 |
| VPS54 | -0.092101569 |
| RP11-111H3 | -0.092103753 |
| RP4-549F15 | -0.09211127 |
| ABHD16A | -0.092141813 |
| TRAJ23 | -0.092142054 |
| IL21R | -0.092142743 |
| TRAFD1 | -0.092150037 |
| AC000078 | -0.0921527 |
| RNU4-52P | -0.092170153 |
| UIMC1 | -0.09218094 |
| RP11-625I7 | -0.092182171 |
| RP11-477D19 | -0.092193369 |
| RP11-968O1 | -0.09219404 |
| ZSCAN26 | -0.092205946 |
| C6orf141 | -0.092209277 |
| RP5-947P14 | -0.09221239 |
| RP11-158H5 | -0.092221519 |

| | |
|---|---|
| ZNF800 | -0.092221798 |
| GTSF1 | -0.092222836 |
| CTB-134F13 | -0.092237291 |
| ASPHD2 | -0.092242599 |
| RNU6-414P | -0.092242963 |
| AL157902 | -0.092252002 |
| OR13A1 | -0.09225395 |
| RNU6-442P | -0.092254318 |
| TRAJ25 | -0.092255127 |
| RP11-153M3 | -0.092273177 |
| AL589931 | -0.092275172 |
| FOXP3 | -0.092297411 |
| MIR4772 | -0.092298299 |
| DOCK8 | -0.092303178 |
| RAB3C | -0.092311185 |
| FTLP12 | -0.09231864 |
| RPL31P11 | -0.092343658 |
| ZNF10 | -0.092343813 |
| ACOXL | -0.092349291 |
| RP11-177C12 | -0.092353196 |
| RP11-288I21 | -0.0923788 |
| SHANK2-AS2 | -0.092404254 |
| RN7SL153P | -0.092414239 |
| C12orf79 | -0.092423431 |
| IGLVI-56 | -0.09242467 |
| CREBL2 | -0.092428356 |
| CTSH | -0.092431221 |
| CCNT2 | -0.092439817 |
| MTND1P18 | -0.092443268 |
| RP11-471B22 | -0.092446831 |
| COPB1 | -0.092472786 |
| RP11-174G6 | -0.092477232 |
| AP000431 | -0.092480984 |
| RAD23BP1 | -0.092487593 |
| CTD-2659N19 | -0.092493039 |
| CCDC112 | -0.092495038 |
| MIR302D | -0.092506622 |
| SEPT7P2 | -0.092510034 |
| LINC01543 | -0.092511792 |
| MYH7 | -0.092515882 |
| NUCB2 | -0.092523058 |
| IGKV1D-35 | -0.092542086 |
| GRM3 | -0.092547346 |
| ALKBH8 | -0.092554613 |
| RP11-326N17 | -0.092557814 |

| | |
|---|---|
| RANBP10 | -0.092571692 |
| SLC25A25-AS1 | -0.092573776 |
| SLC8A1 | -0.092578874 |
| CNBD2 | -0.092594224 |
| TMPO-AS1 | -0.092597889 |
| NR2C1 | -0.09260027 |
| HNRNPA1P68 | -0.092606491 |
| ABHD15 | -0.092630021 |
| USP30 | -0.092635156 |
| AC104986 | -0.09264448 |
| NDUFS1 | -0.09264524 |
| RASA4B | -0.092687527 |
| RP1-90J4 | -0.092688791 |
| CYB5R4 | -0.092698321 |
| AOX3P | -0.092707329 |
| ZNF136 | -0.092713913 |
| TDP2 | -0.092717894 |
| RP11-411B6 | -0.092740811 |
| ST8SIA4 | -0.092742639 |
| RNF13 | -0.092744556 |
| WDR5B | -0.092746286 |
| ZC3H8 | -0.092746789 |
| MIR6083 | -0.09275498 |
| ZNF506 | -0.09278106 |
| MTND2P28 | -0.092785302 |
| ARID2 | -0.092821266 |
| MIR1254-1 | -0.092830545 |
| CRIPAK | -0.092841732 |
| RALGDS | -0.092865938 |
| RP11-12L8 | -0.092866608 |
| IGHJ2P | -0.092883013 |
| RPS6KA6 | -0.092893195 |
| RAD17 | -0.092927615 |
| ATM | -0.092928955 |
| AC004014 | -0.092947069 |
| SCAF11 | -0.092951486 |
| LINC01353 | -0.092960884 |
| OR11H12 | -0.092976016 |
| MIR6795 | -0.09298091 |
| BUB3 | -0.092982576 |
| MIR3613 | -0.092983892 |
| CORO1A | -0.09298745 |
| HNRNPA1P16 | -0.093005963 |
| GCNT4 | -0.093012456 |
| PP2D1 | -0.093027584 |

| | | |
|---|---|---|
| WRNIP1 | | -0.093030466 |
| | 3-Mar | -0.093040439 |
| ADAT1 | | -0.093072849 |
| MIR647 | | -0.093074055 |
| AC004893 | | -0.093077463 |
| SIK3-IT1 | | -0.093077742 |
| RP11-257A22 | | -0.093090612 |
| GATA3 | | -0.093103883 |
| PRR14L | | -0.093119214 |
| LRRC8C | | -0.093121819 |
| SPATA5 | | -0.09312247 |
| RP11-973N13 | | -0.093122714 |
| ZNF782 | | -0.093125149 |
| CTC-425O23 | | -0.093132461 |
| HLA-DQA2 | | -0.093143643 |
| RP5-1184F4 | | -0.093164785 |
| RASGRP3 | | -0.093166826 |
| TBC1D15 | | -0.093180073 |
| SIN3A | | -0.09319825 |
| UCKL1-AS1 | | -0.093202604 |
| EDRF1-AS1 | | -0.093202949 |
| C9orf131 | | -0.093206341 |
| N4BP2L2 | | -0.093207683 |
| COX4I1P2 | | -0.09321786 |
| RPL21P110 | | -0.093236674 |
| CENPQ | | -0.093239387 |
| CTD-2555A7 | | -0.093241444 |
| TTF2 | | -0.093242029 |
| LPAR5 | | -0.093255704 |
| RP11-397J20 | | -0.093274591 |
| RP11-56M3 | | -0.093288177 |
| MIR6780B | | -0.09329074 |
| RN7SL842P | | -0.093311074 |
| BTNL2 | | -0.09331882 |
| SNX11 | | -0.093327056 |
| RP13-638C3 | | -0.093333551 |
| KRT18P63 | | -0.093336136 |
| TRAF3IP1 | | -0.093336154 |
| ZNF23 | | -0.093366594 |
| MKRN1 | | -0.09336756 |
| PUM2 | | -0.09339865 |
| RP11-533O20 | | -0.093416317 |
| SNAI3 | | -0.093426 |
| RP1-239B22 | | -0.093436679 |
| CCL27 | | -0.093445667 |

| | |
|---|---|
| NAPB | -0.093469723 |
| RP11-10K16 | -0.093484998 |
| U2SURP | -0.093488049 |
| RP11-117D22 | -0.093495796 |
| RNF111 | -0.093498896 |
| RP11-350E12 | -0.093517925 |
| JADE2 | -0.093536628 |
| IRF5 | -0.093549112 |
| ZNF619 | -0.093550266 |
| RPS3AP40 | -0.093551971 |
| FAM193A | -0.093555698 |
| SAMD9L | -0.093557596 |
| PPIL3 | -0.093559792 |
| FANCA | -0.093561075 |
| RNU6-844P | -0.093567966 |
| RP11-77K12 | -0.093570192 |
| RAD50 | -0.093593077 |
| TRAPPC10 | -0.093619069 |
| PPHLN1 | -0.093622008 |
| RNU6ATAC40P | -0.093623814 |
| IGHVIII-11-1 | -0.093628961 |
| IL15 | -0.093632669 |
| CFAP36 | -0.093653359 |
| TAS2R63P | -0.093676996 |
| RNU6-209P | -0.093691887 |
| RP11-115N12 | -0.09369795 |
| MAGEB6 | -0.09370877 |
| STX8P1 | -0.093711213 |
| RPRD1A | -0.093720561 |
| API5P1 | -0.093727204 |
| ELMO2 | -0.093730236 |
| PRKXP1 | -0.09376673 |
| MRFAP1L1 | -0.093788323 |
| MIR4744 | -0.093814815 |
| EMR4P | -0.093828415 |
| MRPS18AP1 | -0.093852908 |
| TRPV5 | -0.093861158 |
| HNRNPA3P12 | -0.093899608 |
| CTD-2341M24 | -0.093906979 |
| RP1-283E3 | -0.093907751 |
| KIF24 | -0.093920161 |
| PFN1P4 | -0.093957246 |
| MMP26 | -0.09396455 |
| FANCM | -0.093965819 |
| ABCB10 | -0.093974005 |

| | |
|---|---|
| TUBGCP4 | -0.093981075 |
| PEX5L-AS1 | -0.093992878 |
| NUP62 | -0.094005281 |
| DOCK9 | -0.094012277 |
| RNU6-1006P | -0.09401761 |
| LL22NC03-30E12 | -0.094018841 |
| GPATCH8 | -0.094019719 |
| GZMB | -0.094023719 |
| CBR4 | -0.094034808 |
| SYCP3 | -0.094034888 |
| RP11-383F6 | -0.094055861 |
| DPEP1 | -0.094065256 |
| CCRL2 | -0.094078047 |
| EPG5 | -0.094082754 |
| RP11-110I1 | -0.094101025 |
| MTHFD2L | -0.094107146 |
| RBPJL | -0.094107177 |
| ZC3H13 | -0.094114811 |
| ITPKB-IT1 | -0.094119955 |
| RNU6-1245P | -0.094124536 |
| RMND5A | -0.094141901 |
| ZNF566 | -0.094149604 |
| RP1-117O3 | -0.09415502 |
| RP11-46I8 | -0.094180462 |
| ZBED5 | -0.094180768 |
| AC007381 | -0.094184723 |
| SUPT6H | -0.094185512 |
| RNU6-1053P | -0.094189936 |
| AP001429 | -0.094191569 |
| RP11-529J17 | -0.094196864 |
| POLQ | -0.094200694 |
| ZNF611 | -0.094227133 |
| CBLN3 | -0.094237998 |
| HPS4 | -0.09424464 |
| KIAA1279 | -0.094254129 |
| CEP104 | -0.094256287 |
| RNU6-508P | -0.094262679 |
| RP11-550I15 | -0.094286607 |
| LINC01001 | -0.094290319 |
| GNG3 | -0.094304547 |
| RAB39A | -0.0943053 |
| ZNF493 | -0.094311415 |
| FAM96A | -0.094314932 |
| THSD4-AS1 | -0.094315243 |
| FANCI | -0.094320625 |

| | |
|---|---|
| CENPUP2 | -0.094324853 |
| SLX1A | -0.094330321 |
| HYLS1 | -0.094331472 |
| SBF2-AS1 | -0.094340812 |
| CHMP5 | -0.094347101 |
| CASC5 | -0.094355101 |
| ZNF234 | -0.094360199 |
| MTIF2 | -0.094366078 |
| RAB11FIP4 | -0.094382358 |
| CBFA2T2 | -0.094387568 |
| GGNBP1 | -0.09438886 |
| TGIF2 | -0.094394452 |
| SNX29P1 | -0.094394725 |
| NRIP2 | -0.094407122 |
| CTA-797E19 | -0.094426807 |
| RP11-661O13 | -0.094486594 |
| RBM14 | -0.094494977 |
| DCLRE1C | -0.094495014 |
| APBB1IP | -0.094523387 |
| CTAGE7P | -0.094549032 |
| FLI1 | -0.094551351 |
| RP4-534N18 | -0.094553762 |
| RP11-295H24 | -0.094554235 |
| RPL31P52 | -0.094562706 |
| RNU6-762P | -0.094570865 |
| LIAS | -0.094581492 |
| CLEC12B | -0.094582375 |
| RP11-391H12 | -0.09459047 |
| EMX1 | -0.094613766 |
| SYCP2 | -0.094626332 |
| CYCSP38 | -0.094629196 |
| RP4-724E13 | -0.094647328 |
| TRIM3 | -0.094661682 |
| BBIP1 | -0.094665887 |
| GRK6 | -0.094680939 |
| AP000442 | -0.094689181 |
| CTB-129P6 | -0.094691525 |
| EIF3J | -0.094704623 |
| ZNF335 | -0.094711917 |
| FCRL4 | -0.094713009 |
| ANKRD10-IT1 | -0.094727638 |
| AC005009 | -0.094732551 |
| RP11-49K24 | -0.094745565 |
| SLC25A40 | -0.094753016 |
| CTD-2005D20 | -0.094760358 |

| | |
|---|---|
| RIMBP3C | -0.094768487 |
| ZNF510 | -0.094780292 |
| UVSSA | -0.094783455 |
| SLC4A1AP | -0.094790841 |
| CD86 | -0.094800516 |
| LEMD3 | -0.094805837 |
| RBM33 | -0.094841132 |
| ZNF736 | -0.094856221 |
| ZNF227 | -0.094878119 |
| NENFP2 | -0.094878285 |
| TSSK3 | -0.094884684 |
| UTS2B | -0.094925455 |
| BRAP | -0.094925824 |
| RP11-817O13 | -0.094927145 |
| BCL11B | -0.094935759 |
| RP11-66N5 | -0.094938012 |
| LIMD1 | -0.094966551 |
| DNA2 | -0.094971258 |
| PLEKHH1 | -0.094991929 |
| GAPT | -0.094994162 |
| RP11-495O11 | -0.094994402 |
| OLFM1 | -0.095004171 |
| CIR1 | -0.09502527 |
| ARFGEF2 | -0.095037401 |
| VN1R34P | -0.095041601 |
| DBH | -0.095043927 |
| EIF4ENIF1 | -0.095050741 |
| GJA9 | -0.09506147 |
| MTND2P12 | -0.095072835 |
| RP11-981P6 | -0.095074807 |
| IGLV3-17 | -0.095080201 |
| MYO5A | -0.09508291 |
| FAM57B | -0.095098371 |
| KSR2 | -0.095099991 |
| ZNF480 | -0.095127544 |
| RP11-62C7 | -0.09514537 |
| OR11A1 | -0.095155845 |
| TBC1D31 | -0.09515655 |
| CLPX | -0.095176198 |
| SLC39A6 | -0.095176811 |
| FAM175B | -0.095199417 |
| RWDD4 | -0.095200738 |
| DNM1P33 | -0.095213249 |
| DCAF4L1 | -0.095228516 |
| AL355480 | -0.095228674 |

| | |
|---|---|
| GIT2 | -0.095250731 |
| SART3 | -0.095257857 |
| RP11-894P9 | -0.095260945 |
| RPL21P44 | -0.095275136 |
| CTC-343N3 | -0.095277429 |
| SLU7 | -0.095322716 |
| MTO1 | -0.095322976 |
| CTC-559E9 | -0.095323441 |
| TSHZ1 | -0.095340056 |
| PEX1 | -0.095341972 |
| DCAF7 | -0.095379034 |
| CTC-378H22 | -0.095382538 |
| RP11-227G15 | -0.095393995 |
| HYPK | -0.095396486 |
| KCNH2 | -0.095403668 |
| NARS | -0.095419352 |
| BRIP1 | -0.095426422 |
| AC098592 | -0.09543484 |
| RP11-293N14 | -0.095440051 |
| RP11-734K23 | -0.095471136 |
| IGKV1-32 | -0.095472962 |
| ZC3H6 | -0.095486406 |
| PDHB | -0.095530892 |
| NAA30 | -0.095540046 |
| CD7 | -0.095577536 |
| PCID2 | -0.095605072 |
| HCAR1 | -0.095609221 |
| MIR4701 | -0.09561995 |
| AHSA2 | -0.095621222 |
| PRDM10 | -0.095632311 |
| USP8P1 | -0.095640971 |
| ADAR | -0.095644534 |
| CTC-461F20 | -0.095702574 |
| RP11-705C15 | -0.095706453 |
| MIS12 | -0.095709281 |
| RHOT1 | -0.095736724 |
| STRADBP1 | -0.095737321 |
| SRSF11 | -0.095745674 |
| RP11-428L9 | -0.09574689 |
| PPM1B | -0.095757711 |
| TRMT10A | -0.09576493 |
| KIF20B | -0.095780838 |
| METTL14 | -0.095788038 |
| ATXN2 | -0.095788522 |
| ITPRIPL1 | -0.095791871 |

| | |
|---|---|
| PLCG2 | -0.095802495 |
| AC005614 | -0.095806855 |
| GOLGA4 | -0.095821714 |
| RP11-107E5 | -0.095837031 |
| RP11-517O13 | -0.095858731 |
| ZNF833P | -0.09587154 |
| CTD-2562G15 | -0.095879158 |
| NRBF2P5 | -0.095888258 |
| E2F5 | -0.095896266 |
| CTB-49A3 | -0.095903943 |
| PFN1P7 | -0.095908262 |
| THUMPD3 | -0.095917923 |
| RP11-843P14 | -0.095921769 |
| SFMBT2 | -0.095924863 |
| NPIPB11 | -0.095925272 |
| FAM76B | -0.095940603 |
| TSSK4 | -0.095942985 |
| EIF2AK2 | -0.095944417 |
| CA8 | -0.095944566 |
| ZCCHC4 | -0.095944882 |
| C8orf86 | -0.095945096 |
| RP4-594I10 | -0.09594954 |
| IGHD4-11 | -0.095956194 |
| PPM1D | -0.09595813 |
| CLEC4D | -0.095959182 |
| RNU6-451P | -0.095976661 |
| AL138935 | -0.096007852 |
| MIR570 | -0.096014774 |
| EXOSC9 | -0.096015416 |
| RP11-1084A12 | -0.096017525 |
| MEFV | -0.096025481 |
| RN7SKP70 | -0.096028552 |
| RPS3P2 | -0.096034411 |
| LCP1 | -0.096050003 |
| RP11-274H24 | -0.096058795 |
| RNU6-583P | -0.096071296 |
| BOD1L1 | -0.096102824 |
| KIAA1147 | -0.096105485 |
| AC127526 | -0.09612425 |
| TRMT10B | -0.096133114 |
| DDX55 | -0.096135681 |
| AIMP1P2 | -0.096153146 |
| ZNF638 | -0.096176948 |
| ZNF253 | -0.0961856 |
| FSD1L | -0.096193377 |

| | |
|---|---|
| MYLIP | -0.096198233 |
| APIP | -0.096212038 |
| TRAJ44 | -0.096230225 |
| TNIP3 | -0.096236571 |
| ZNF181 | -0.096252989 |
| SPATA13 | -0.096260747 |
| AP001610 | -0.096261202 |
| GPD2 | -0.09627859 |
| SGMS1-AS1 | -0.096306163 |
| RASGEF1B | -0.096339914 |
| ULK2 | -0.096345216 |
| NFKBID | -0.096369366 |
| RP4-669L17 | -0.096372631 |
| LINC00161 | -0.096378328 |
| FAM92B | -0.096395972 |
| C9orf43 | -0.096396121 |
| RP11-40G16 | -0.096399748 |
| RP11-549K20 | -0.096405927 |
| RP11-364P2 | -0.096413761 |
| ZNF528 | -0.096415433 |
| IGKV1D-22 | -0.096425528 |
| FANCL | -0.096428904 |
| RPL23AP23 | -0.096439476 |
| RP11-730K11 | -0.096440715 |
| MASP2 | -0.096441353 |
| RP11-101E7 | -0.096456736 |
| NSF | -0.096467566 |
| RP11-544A12 | -0.096525334 |
| C18orf25 | -0.09653222 |
| MYEF2 | -0.096535811 |
| IL17RB | -0.09653622 |
| RP11-493E3 | -0.096558755 |
| PHF8 | -0.096572966 |
| VDAC1P8 | -0.096584873 |
| RP11-321N4 | -0.096585518 |
| RP11-75C10 | -0.096591315 |
| SNORD115-28 | -0.096600177 |
| RHOXF2B | -0.096603279 |
| ZNF518A | -0.096608428 |
| CTC-571O20 | -0.096612091 |
| AL356585 | -0.096667611 |
| TRGVA | -0.096676933 |
| RP11-1143G9 | -0.096681116 |
| DSCAML1 | -0.096682869 |
| RP11-23D24 | -0.096692225 |

| | |
|---|---|
| RP11-501O2 | -0.096705563 |
| TMEM229B | -0.09672914 |
| LINC00266-1 | -0.09674339 |
| GLUD1 | -0.096755839 |
| PCDH9-AS4 | -0.096783059 |
| L3MBTL1 | -0.096786817 |
| PNISR | -0.096789347 |
| BICD1P1 | -0.096799248 |
| TCF20 | -0.096803711 |
| PUS7L | -0.096805999 |
| RP11-831F12 | -0.096809156 |
| LINC01022 | -0.09683576 |
| RP11-489O18 | -0.096843714 |
| GLS2 | -0.096844347 |
| TOX | -0.096874635 |
| CTC-534A2 | -0.096875063 |
| TRNT1 | -0.0968775 |
| AVL9 | -0.096878561 |
| MBLAC2 | -0.096880198 |
| ZNF330 | -0.096902208 |
| RP11-1058N17 | -0.096905297 |
| RP11-71B7 | -0.096907722 |
| MSS51 | -0.096908274 |
| ZMYM2 | -0.096914078 |
| KNOP1P2 | -0.096918547 |
| TSG101 | -0.09694234 |
| FUT1 | -0.096944647 |
| PRRT2 | -0.096951996 |
| RP13-753N3 | -0.096953886 |
| AL513548 | -0.096983689 |
| TBC1D29 | -0.096991287 |
| SEL1L3 | -0.096998529 |
| FGFR1OP2 | -0.09700264 |
| ACP5 | -0.097028428 |
| ABHD3 | -0.097057973 |
| AC093063 | -0.097059471 |
| RP11-328J14 | -0.097060032 |
| VDAC1P9 | -0.097064081 |
| CCT6P1 | -0.097072913 |
| MRPS30 | -0.097086998 |
| CNTD1 | -0.097089677 |
| PCBP1-AS1 | -0.097096803 |
| SARM1 | -0.097097138 |
| ID2 | -0.097100468 |
| TLR7 | -0.097104636 |

| | |
|---|---|
| MYSM1 | -0.097114664 |
| RP11-324H7 | -0.097120097 |
| HNRNPA2B1 | -0.097152247 |
| CTD-2542L18 | -0.097156222 |
| TBC1D5 | -0.097159057 |
| IGLV8OR8-1 | -0.097169246 |
| MRPL30 | -0.09717729 |
| RNU6-375P | -0.097196856 |
| RP11-167P11 | -0.097211146 |
| RNU6-1175P | -0.097238434 |
| FMNL1 | -0.097251377 |
| AVIL | -0.097256549 |
| CXCL13 | -0.097264942 |
| RFXAP | -0.09728323 |
| GPR82 | -0.097284495 |
| TBCC | -0.097286765 |
| LINC00484 | -0.097295335 |
| RNA5SP273 | -0.097303494 |
| RP11-580I16 | -0.097304273 |
| CHD8 | -0.097305872 |
| KLRAP1 | -0.097314319 |
| CKAP5 | -0.097338841 |
| RHBDD1 | -0.097351847 |
| RP11-562A8 | -0.097358098 |
| FUCA1 | -0.097366191 |
| RNU6-531P | -0.097384498 |
| RP11-700H6 | -0.097417516 |
| CTD-2329K10 | -0.097432548 |
| ZNF169 | -0.097436204 |
| RP1-127H14 | -0.09744463 |
| RP1-101A2 | -0.097478754 |
| RP11-376O6 | -0.097492801 |
| CYP4F26P | -0.097512012 |
| SETD2 | -0.097512337 |
| RP11-320L11 | -0.097515649 |
| RP11-134K13 | -0.097526403 |
| RP11-45K10 | -0.097565626 |
| RNPC3 | -0.097575075 |
| LINC00672 | -0.097599802 |
| IGHD6-13 | -0.097600571 |
| OR4D1 | -0.097639273 |
| RP11-134D3 | -0.097640189 |
| PNPT1 | -0.097660865 |
| TTF1 | -0.097666632 |
| CTB-36H16 | -0.097703397 |

| | |
|---|---|
| SLC35E3 | -0.097711565 |
| KIAA0100 | -0.097732179 |
| HMGB3P15 | -0.097755928 |
| LTK | -0.097772638 |
| RP11-288L9 | -0.097773525 |
| RP11-16N11 | -0.097847645 |
| AC131254 | -0.097851009 |
| RP11-959F10 | -0.097855946 |
| EXTL2P1 | -0.097863661 |
| TRAJ17 | -0.097880593 |
| VPS13B | -0.097893266 |
| AC099805 | -0.097933328 |
| RP5-1153D9 | -0.097943769 |
| RP1 | -0.097944867 |
| IRG1 | -0.097944943 |
| CENPE | -0.09795498 |
| RP11-276E17 | -0.097979094 |
| RP11-370A5 | -0.097985156 |
| RNU6-481P | -0.097999687 |
| AC005519 | -0.098006041 |
| KRT18P52 | -0.098013681 |
| AGO4 | -0.098030127 |
| POLR2B | -0.098035318 |
| IGHV3-65 | -0.098039861 |
| NEK8 | -0.098043468 |
| PPP1R26-AS1 | -0.098057589 |
| RP11-19J5 | -0.09806396 |
| IL7R | -0.098072715 |
| FAM111A | -0.098075971 |
| EPN2-AS1 | -0.098097743 |
| PRMT1P1 | -0.098168946 |
| MTND4P9 | -0.098174059 |
| HLA-DPA1 | -0.098183399 |
| BDP1P | -0.098188579 |
| SRP72P2 | -0.098197484 |
| OR5BL1P | -0.098199075 |
| AFTPH | -0.098214377 |
| RP11-579E24 | -0.098216024 |
| MIR616 | -0.098216504 |
| CTD-2535P7 | -0.098244085 |
| FAM179A | -0.098266883 |
| AASDH | -0.09827505 |
| IGHVII-46-1 | -0.098278037 |
| PARP4 | -0.098300056 |
| RP11-348A7 | -0.098302285 |

| | |
|---|---|
| MATN1-AS1 | -0.098304484 |
| DYRK1A | -0.098314642 |
| RSRC2 | -0.098324019 |
| AC007038 | -0.098333239 |
| IKZF5 | -0.098343462 |
| AC007308 | -0.098355871 |
| SOD1P3 | -0.09836248 |
| SEC31B | -0.098404339 |
| MAPK8IP2 | -0.098415633 |
| APAF1 | -0.098422238 |
| LHX4 | -0.09842408 |
| NCAPD3 | -0.098435764 |
| CTSS | -0.098436024 |
| IGLVI-63 | -0.098444042 |
| HNRNPD | -0.098444657 |
| MIR4686 | -0.098449565 |
| COQ10A | -0.098456732 |
| KIF15 | -0.098458128 |
| JMJD7-PLA2G4B | -0.098476696 |
| MTND5P14 | -0.098488827 |
| TIMM10B | -0.098489329 |
| XPO4 | -0.09849052 |
| ZBTB11 | -0.098506446 |
| CTC-429P9 | -0.098509069 |
| ARAP1-AS2 | -0.098511378 |
| MIR1203 | -0.098539884 |
| IFNL3P1 | -0.098553722 |
| TRBJ2-3 | -0.0985544 |
| HSPA8P3 | -0.098571875 |
| MAPKAPK5 | -0.098580738 |
| CLSPN | -0.098591008 |
| AC061961 | -0.09861727 |
| RP11-390P24 | -0.09861953 |
| RNA5SP107 | -0.098634005 |
| RP11-135F9 | -0.098635289 |
| RP11-85I17 | -0.098668238 |
| MIR6832 | -0.098670309 |
| CLCN3 | -0.098670621 |
| ZNF799 | -0.098673281 |
| IGKV2D-36 | -0.098673517 |
| MAGEA8-AS1 | -0.098681005 |
| FOXD4L3 | -0.098687136 |
| RP11-324O2 | -0.098689115 |
| RP11-516C1 | -0.09871906 |
| OTUD6B | -0.098755741 |

| | |
|---|---|
| TRAJ38 | -0.098771883 |
| RP11-288K12 | -0.098781715 |
| RP11-431J17 | -0.0987825 |
| 5S_rRNA | -0.098787261 |
| EBLN3 | -0.098794608 |
| EXOC2 | -0.098811427 |
| XXbac-BPG246D15 | -0.098813297 |
| AC092295 | -0.098819296 |
| CSAG1 | -0.098823966 |
| ITM2A | -0.098855373 |
| COX6B1P5 | -0.098885982 |
| MIR326 | -0.098893114 |
| PMS2P3 | -0.098896584 |
| CUL4A | -0.098901477 |
| PTMAP3 | -0.098909962 |
| AC100830 | -0.098922289 |
| SELK | -0.0989332 |
| EP400NL | -0.098943265 |
| IGHD3-16 | -0.098944984 |
| RP11-750H9 | -0.098950926 |
| GRPEL2-AS1 | -0.098958917 |
| FAM53B-AS1 | -0.098967802 |
| NLK | -0.098973964 |
| PKN2 | -0.098982281 |
| CTC-303L1 | -0.09899053 |
| NBPF4 | -0.098995836 |
| USP38 | -0.098997296 |
| RP3-462D8 | -0.099006332 |
| UBA52P8 | -0.099015924 |
| RNA5SP457 | -0.099019839 |
| PROSER1 | -0.099034804 |
| FRS2 | -0.099039102 |
| LCORL | -0.099050526 |
| NUDT16P1 | -0.099050619 |
| SNORA16B | -0.099077455 |
| RP11-526I2 | -0.099079104 |
| CTSW | -0.099086081 |
| MT-TM | -0.099095872 |
| TNFAIP8L2 | -0.099118733 |
| HNRNPA3P9 | -0.099121293 |
| NBPF15 | -0.099151646 |
| NFATC3 | -0.099152335 |
| ZNF346-IT1 | -0.099159172 |
| C6orf62 | -0.099160614 |
| PLEKHA3 | -0.099170308 |

| | |
|---|---|
| DTX3L | -0.099177266 |
| TRAJ58 | -0.09918172 |
| IGHV1OR15-6 | -0.099198594 |
| IGLV1-41 | -0.099208459 |
| TCERG1 | -0.099214979 |
| CEP83 | -0.099220189 |
| TRBV1 | -0.0992525 |
| SEC61A2 | -0.09928168 |
| STRADA | -0.099293662 |
| RP11-409I10 | -0.099302166 |
| CTA-113A6 | -0.099320794 |
| THOC1 | -0.099335115 |
| GIN1 | -0.099345999 |
| AC004854 | -0.099350888 |
| LDHAL6A | -0.099358868 |
| TRIM21 | -0.099361423 |
| AC098784 | -0.099400327 |
| SETD5 | -0.099400606 |
| CSF2RB | -0.09941257 |
| NPL | -0.099427007 |
| TTC31 | -0.099477726 |
| RNU6-216P | -0.099492295 |
| CD4 | -0.099524072 |
| SRGN | -0.099524984 |
| CNTD2 | -0.099550306 |
| C1orf220 | -0.099552557 |
| SEMA6A-AS1 | -0.099557378 |
| AL136419 | -0.099559609 |
| CTD-2521M24 | -0.099560892 |
| PHYKPL | -0.099567646 |
| GALNT7 | -0.0995704 |
| PABPC1P1 | -0.099583665 |
| CNNM2 | -0.099584763 |
| IRAK1BP1 | -0.099620169 |
| SP4 | -0.099632542 |
| SP110 | -0.099649361 |
| ZNF587B | -0.09968713 |
| KLHL11 | -0.099711039 |
| PLGLB1 | -0.099715451 |
| DNTT | -0.09971568 |
| ZNF318 | -0.099735784 |
| AF129075 | -0.099743087 |
| SNRPGP3 | -0.099747005 |
| OR2B8P | -0.09974786 |
| ZNF540 | -0.099751989 |

| | |
|---|---|
| AC067945 | -0.099754906 |
| VPS29 | -0.099773571 |
| YWHAZP4 | -0.099783709 |
| GLIPR1L1 | -0.099792516 |
| RP11-260E18 | -0.099801842 |
| RP11-182N22 | -0.099816429 |
| CISD2 | -0.099826783 |
| LY6G5B | -0.099842207 |
| ZNF207 | -0.099842356 |
| NKAPP1 | -0.099850431 |
| RP11-95G17 | -0.099861664 |
| PDLIM1P3 | -0.099888632 |
| ZSWIM5P1 | -0.099891801 |
| RP11-91H12 | -0.099892466 |
| RMND1 | -0.099912908 |
| TRAJ42 | -0.099923097 |
| CTD-2378E12 | -0.09992535 |
| NDUFAF1 | -0.099927122 |
| ETF1P1 | -0.099929544 |
| RP1-8B1 | -0.099930459 |
| IGHV3OR16-16 | -0.099931086 |
| S1PR4 | -0.099931234 |
| MED1 | -0.099933709 |
| RP11-794G24 | -0.099952521 |
| TRAJ12 | -0.099962343 |
| LL22NC03-23C6 | -0.09997771 |
| AC012361 | -0.099990002 |
| RP3-443C4 | -0.099994852 |
| RNA5SP426 | -0.100005441 |
| AC021188 | -0.100007395 |
| VAV1 | -0.100020801 |
| IGHV7-34-1 | -0.100034427 |
| ZNF664 | -0.100041732 |
| FAM136BP | -0.10004437 |
| IGHV1-24 | -0.100100879 |
| HELZ | -0.100110591 |
| CEP85 | -0.100120377 |
| DBF4B | -0.100129308 |
| BBS4 | -0.100154909 |
| UFM1 | -0.100157532 |
| AL672183 | -0.100186073 |
| UTS2 | -0.100194206 |
| AL138706 | -0.100221175 |
| RP11-762H8 | -0.100225261 |
| OR4K15 | -0.100234804 |

| | |
|---|---|
| RP4-665N4 | -0.100244384 |
| PSMA8 | -0.100254564 |
| GPSM3 | -0.100259974 |
| COX6CP10 | -0.100277526 |
| LL22NC03-75H12 | -0.100288695 |
| IKZF4 | -0.100290022 |
| INTS2 | -0.100290171 |
| TTC14 | -0.100324721 |
| RNF44 | -0.10036329 |
| RSRP1 | -0.100381989 |
| RP11-761I4 | -0.100389037 |
| KRT18P22 | -0.100410843 |
| IGHD5-24 | -0.100418543 |
| RP11-439C15 | -0.100448895 |
| PI4KA | -0.100456597 |
| KIR3DL3 | -0.10046956 |
| CTB-171A8 | -0.100487724 |
| RBM26-AS1 | -0.10049057 |
| BNIP3P40 | -0.100498973 |
| CFLAR | -0.100506943 |
| HMGXB4 | -0.100510627 |
| RP11-242O24 | -0.100556297 |
| SPIRE1 | -0.100578518 |
| ACSL6 | -0.100580528 |
| CTD-2583P5 | -0.100589108 |
| ZNF182 | -0.100592231 |
| SMC4 | -0.100595673 |
| HMGN1P2 | -0.100633617 |
| CTD-2587M2 | -0.10064545 |
| LONRF2 | -0.100652457 |
| CYP4F9P | -0.100660095 |
| NAPSB | -0.100678876 |
| TRBV5-7 | -0.100698256 |
| FAM222A | -0.100706673 |
| SNORD59A | -0.100722072 |
| DDX5 | -0.100729502 |
| RNU6-1262P | -0.10074473 |
| LINC00265 | -0.100755215 |
| MSH5 | -0.100764313 |
| ZNF808 | -0.100767866 |
| UTP18 | -0.100782788 |
| FICD | -0.1007853 |
| FAM78A | -0.100792798 |
| RP3-343K2 | -0.100794093 |
| TEFM | -0.100795905 |

| | |
|---|---|
| XXbac-B562F10 | -0.100805453 |
| RNF34 | -0.10082238 |
| SNRPGP4 | -0.100838314 |
| CCRN4L | -0.100839572 |
| OR10G2 | -0.100849891 |
| LINC00593 | -0.100851928 |
| CIITA | -0.100882978 |
| EIF2S2P3 | -0.10088964 |
| RP11-85K15 | -0.100912505 |
| CYCSP33 | -0.100916777 |
| TAF3 | -0.100955502 |
| HNRNPA1P21 | -0.10096115 |
| OSER1 | -0.100965605 |
| WWP1 | -0.10096648 |
| RP11-278C7 | -0.100971243 |
| RP11-343D2 | -0.100992046 |
| RN7SL377P | -0.100998284 |
| AFG3L1P | -0.101001123 |
| UBA52P7 | -0.101026935 |
| FOXN2 | -0.101034334 |
| RP5-887A10 | -0.101036127 |
| RNY4P25 | -0.10103738 |
| TEP1 | -0.10105266 |
| SBNO1 | -0.101074429 |
| AKAP10 | -0.101090094 |
| RP11-640B6 | -0.101116134 |
| RP11-347I19 | -0.10111711 |
| RP11-301G23 | -0.101119889 |
| RP11-410N8 | -0.101129009 |
| CTC-420A11 | -0.101129059 |
| RNFT1 | -0.101166972 |
| C12orf76 | -0.101182266 |
| SF1 | -0.101199848 |
| TXNDC11 | -0.101226138 |
| AC009133 | -0.101227477 |
| TMEM9B-AS1 | -0.101238119 |
| IGLV3-15 | -0.10125453 |
| CTD-3236F5 | -0.101263213 |
| IGKV2OR2-7D | -0.101279813 |
| RP11-170N16 | -0.101291487 |
| CCR6 | -0.101296771 |
| CLIC2 | -0.101307536 |
| DDX11L16 | -0.101319336 |
| RP11-848G14 | -0.10132925 |
| ZNF14 | -0.101358348 |

| | |
|---|---|
| RP11-239H6 | -0.101364625 |
| MIR6892 | -0.101369973 |
| RP4-545C24 | -0.101386115 |
| AP000648 | -0.101423076 |
| ME2 | -0.101424397 |
| RP4-764O22 | -0.101433262 |
| CIAO1 | -0.101439207 |
| MBOAT1 | -0.101442277 |
| SUDS3P1 | -0.101454552 |
| FAM111B | -0.101455227 |
| PPP3CB-AS1 | -0.10148907 |
| BTBD18 | -0.10149314 |
| PEX12 | -0.101502838 |
| KRT8P34 | -0.101513545 |
| ZMAT1 | -0.101521146 |
| AC009159 | -0.101528193 |
| CYP4V2 | -0.101552273 |
| VPS4B | -0.101574488 |
| HAUS2 | -0.10158113 |
| CLEC6A | -0.101586218 |
| ARHGEF6 | -0.101595624 |
| ANKRD17 | -0.101611606 |
| CLEC10A | -0.101615964 |
| VPREB3 | -0.101619712 |
| SAMM50P1 | -0.101623584 |
| UBR2 | -0.101646956 |
| KIAA1468 | -0.10166638 |
| OR13C5 | -0.101667184 |
| ATAD2 | -0.101681637 |
| PADI6 | -0.101698638 |
| GTSE1-AS1 | -0.10171215 |
| MIF4GD | -0.101716615 |
| MCM3AP | -0.101718401 |
| KCNJ3 | -0.101729694 |
| RP11-752D24 | -0.101739242 |
| HLA-U | -0.101750545 |
| MIR4692 | -0.101754341 |
| RNF139-AS1 | -0.101755091 |
| AMICA1 | -0.101764896 |
| CTA-150C2 | -0.101772686 |
| GUSBP9 | -0.101782239 |
| RN7SL338P | -0.101782574 |
| GBP7 | -0.10178805 |
| IGKV2D-28 | -0.10179827 |
| MIR4457 | -0.101814219 |

| | |
|---|---|
| AC008155 | -0.101846449 |
| UTP23 | -0.101880231 |
| ANKRD36 | -0.101883729 |
| CASC21 | -0.101897317 |
| CD84 | -0.10190907 |
| ATXN7 | -0.101924791 |
| RP11-473N11 | -0.101934675 |
| PIK3IP1-AS1 | -0.101939588 |
| SETP14 | -0.101946653 |
| SGOL2 | -0.101953035 |
| AP000254 | -0.101962561 |
| LRIF1 | -0.101976161 |
| RNU6-833P | -0.101978457 |
| XX-C283C717 | -0.101988282 |
| WAS | -0.102000497 |
| BTN2A1 | -0.102006376 |
| REC8 | -0.102039457 |
| RP11-567L7 | -0.102056658 |
| LINC01422 | -0.102056692 |
| ZRANB2 | -0.102071551 |
| RNU4-82P | -0.102072741 |
| TMEM209 | -0.102084873 |
| CTA-243E7 | -0.102085794 |
| E2F8 | -0.102087775 |
| BAZ1B | -0.102128205 |
| RP11-4K3__A | -0.102133369 |
| SPOCK2 | -0.102136131 |
| TRAF1 | -0.102173547 |
| RP11-378A13 | -0.102215111 |
| ERN1 | -0.102275653 |
| RP4-756G23 | -0.10229465 |
| TRIM66 | -0.102296454 |
| IGHV3-50 | -0.102308772 |
| MSL1 | -0.102311376 |
| RP4-753P9 | -0.102347523 |
| NPY6R | -0.102387328 |
| RP11-690I21 | -0.102395471 |
| RP11-61F12 | -0.102407498 |
| CST9L | -0.102443623 |
| SCML2 | -0.102494268 |
| ANKRD20A1 | -0.102519588 |
| TRIM14 | -0.102521711 |
| RP11-50I19 | -0.102529599 |
| SMARCE1 | -0.102530474 |
| RP11-131J3 | -0.102535651 |

| | |
|---|---|
| INPP5D | -0.102549414 |
| DCDC2C | -0.102553669 |
| RP11-286N22 | -0.102557173 |
| DFFB | -0.102575425 |
| ZKSCAN4 | -0.102591314 |
| MLLT10 | -0.10260871 |
| MALAT1 | -0.102613454 |
| MAT2B | -0.102628134 |
| NBR1 | -0.102629827 |
| AC092580 | -0.102654683 |
| TRAJ40 | -0.102662714 |
| RP11-640N20 | -0.102683169 |
| NACAP6 | -0.102688612 |
| RP11-431M7 | -0.102703293 |
| RP11-597D13 | -0.102707096 |
| HCG25 | -0.102729404 |
| BRWD1 | -0.102735581 |
| RP11-345J13 | -0.1027396 |
| NCOA3 | -0.102743265 |
| RP11-156E6 | -0.102763935 |
| RP11-284J1 | -0.102764716 |
| CCDC170 | -0.102765833 |
| AC005154 | -0.102797593 |
| LINC00216 | -0.102801872 |
| AL356154 | -0.102804253 |
| RNF138 | -0.102811826 |
| ASB8 | -0.102824719 |
| OIP5-AS1 | -0.102837297 |
| RP11-478C1 | -0.102837614 |
| RAB3A | -0.102860888 |
| SUPT20H | -0.102878247 |
| ICA1L | -0.102881354 |
| BRS3 | -0.102883668 |
| RP11-356I2 | -0.102883922 |
| RPL7P51 | -0.102887868 |
| SETD1B | -0.102934008 |
| R3HDM2 | -0.102940352 |
| RP11-17G12 | -0.102955463 |
| SUMO2P19 | -0.102965308 |
| CEP44 | -0.102977935 |
| CYP2R1 | -0.102985117 |
| RP11-343K8 | -0.10300206 |
| ZFP62 | -0.103010495 |
| MT-ATP8 | -0.103015332 |
| RP11-499P20 | -0.103034012 |

| | |
|---|---|
| CEP76 | -0.103044859 |
| CARF | -0.103053101 |
| PISD | -0.103054999 |
| AHCYL2 | -0.103066125 |
| RP11-658F2 | -0.103076154 |
| IGLV1-36 | -0.103077195 |
| AC010733 | -0.103083856 |
| Z84812 | -0.103090873 |
| C11orf42 | -0.103100471 |
| AL109767 | -0.103102349 |
| WARS | -0.103125458 |
| KLHL41 | -0.103139524 |
| NAA25 | -0.103162595 |
| ZBTB32 | -0.103163525 |
| KB-1615E4 | -0.103167548 |
| AOAH-IT1 | -0.103180573 |
| FMO7P | -0.103188284 |
| RP11-73G16 | -0.103193545 |
| YWHAZP6 | -0.103201986 |
| RP11-407G23 | -0.103208048 |
| CTCF | -0.103212253 |
| FASTKD1 | -0.103217239 |
| HOOK1 | -0.103220439 |
| POLI | -0.103261129 |
| RP11-298I3 | -0.103261241 |
| SRSF7 | -0.103287195 |
| AC005954 | -0.103299483 |
| IGHV7-40 | -0.103304084 |
| ZGRF1 | -0.103324592 |
| RP11-264B17 | -0.103330379 |
| ZNF37A | -0.10334197 |
| CCR4 | -0.103350868 |
| SOCS6 | -0.103354938 |
| NSRP1 | -0.103369246 |
| SLC46A1 | -0.103370846 |
| RCSD1 | -0.103383442 |
| CD52 | -0.103386511 |
| CTA-217C2 | -0.103398817 |
| RP11-673E11 | -0.103420284 |
| RP11-28F1 | -0.103420996 |
| RP4-777O23 | -0.103422651 |
| RP11-243J16 | -0.103441141 |
| CTC1 | -0.103447612 |
| AGAP2 | -0.103447668 |
| RP11-770G2 | -0.103457505 |

| | |
|---|---|
| BCLAF1 | -0.103459984 |
| ZADH2 | -0.103481716 |
| IGHV1-2 | -0.103494644 |
| RP11-579A4 | -0.103504229 |
| ELMSAN1 | -0.103505028 |
| CEACAM21 | -0.103511987 |
| KB-1460A1 | -0.103516303 |
| EMB | -0.103519596 |
| FAM21A | -0.103538872 |
| WDR17 | -0.103547779 |
| RP11-363D14 | -0.103548365 |
| AC129492 | -0.103550877 |
| TRAJ36 | -0.103562856 |
| CEP192 | -0.103569626 |
| ZNF180 | -0.103576287 |
| AC100832 | -0.103613457 |
| RN7SKP237 | -0.103619477 |
| IGLJ4 | -0.103626655 |
| DLEU7-AS1 | -0.103664405 |
| CALM2P2 | -0.103706 |
| RP11-240M16 | -0.103732342 |
| AB019438 | -0.103734771 |
| CTD-2047H16 | -0.103747234 |
| WHSC1 | -0.103750844 |
| HNRNPL | -0.103751272 |
| OR2I1P | -0.103758485 |
| RP11-379C10 | -0.103773395 |
| OSTF1P1 | -0.103791652 |
| RP11-400L8 | -0.103793115 |
| GGNBP2 | -0.10379866 |
| MIR6774 | -0.103852844 |
| RASAL3 | -0.103866347 |
| RP11-53O19 | -0.103871016 |
| MMP25 | -0.103908674 |
| PLD6 | -0.103944396 |
| FCER2 | -0.103954252 |
| PPP1R10 | -0.103956788 |
| LRRC37BP1 | -0.103960267 |
| CDK13 | -0.103962258 |
| RBM18 | -0.103975784 |
| CLEC12A | -0.103980859 |
| PPID | -0.104027842 |
| RFX5 | -0.104032121 |
| STIM2 | -0.104046633 |
| PPFIA1 | -0.104083993 |

| | |
|---|---|
| RBM12B | -0.104104776 |
| CATIP-AS2 | -0.104113687 |
| ZNF473 | -0.104118432 |
| OTUD4P1 | -0.104124511 |
| CTDSPL2 | -0.104144405 |
| METTL21A | -0.104148461 |
| NAA16 | -0.104150545 |
| USP8 | -0.104156369 |
| GRAP2 | -0.104168909 |
| LINC00925 | -0.104212204 |
| NAF1 | -0.104231516 |
| AC007255 | -0.104246582 |
| RPEP4 | -0.104261597 |
| LRRC37A14P | -0.104288486 |
| SMC3 | -0.104295723 |
| RP5-1091N2 | -0.104311503 |
| IGKV1-9 | -0.104312362 |
| CTD-2260A17 | -0.104317864 |
| RNU6-884P | -0.104369167 |
| IGKV1OR2-118 | -0.104419511 |
| HNRNPH1 | -0.104432604 |
| DDX58 | -0.104436009 |
| RBMX | -0.104443637 |
| ANKRD18EP | -0.104445777 |
| PLA2G4B | -0.104462614 |
| CCDC69 | -0.104470875 |
| MIR4520-1 | -0.104491633 |
| ORAOV1 | -0.104493853 |
| DHX15 | -0.104514747 |
| TRAK1 | -0.104517203 |
| TTC21B-AS1 | -0.104521636 |
| POC1B | -0.104523845 |
| TSPAN32 | -0.104530832 |
| SERPINB1 | -0.10454926 |
| PROZ | -0.104551808 |
| LINC01539 | -0.104558239 |
| RP11-689J19 | -0.104558793 |
| PRKCG | -0.104560627 |
| KDM2A | -0.104581857 |
| ZNF552 | -0.104587104 |
| TGDS | -0.104593039 |
| FAM71B | -0.104625132 |
| EIF1AD | -0.104653488 |
| AC011747 | -0.104658625 |
| OCIAD1 | -0.104683759 |

| | |
|---|---|
| GPA33 | -0.104739569 |
| RP5-1147A1 | -0.104751592 |
| DEFB116 | -0.104759373 |
| ZBTB7A | -0.104776433 |
| KIR2DP1 | -0.104779374 |
| NBPF8P | -0.104780117 |
| TRIM60P18 | -0.104802425 |
| RN7SKP173 | -0.104815289 |
| NUFIP2 | -0.104817142 |
| AL390877 | -0.10485941 |
| NARF-IT1 | -0.104877554 |
| TMED5 | -0.104887973 |
| RNU6-1140P | -0.104892901 |
| TRBV5-2 | -0.104897603 |
| CDC42SE2 | -0.10490602 |
| RP11-274J2 | -0.104926748 |
| HNRNPH3 | -0.104938449 |
| AC017076 | -0.104969693 |
| AC005220 | -0.104977611 |
| MOAP1 | -0.104993782 |
| RP11-478C19 | -0.104995743 |
| PARP2 | -0.105003382 |
| GRIN3A | -0.105046808 |
| IGHV3-32 | -0.105065516 |
| RPS27P23 | -0.105100872 |
| DDX17 | -0.105105861 |
| GLE1 | -0.105129732 |
| CRBN | -0.105133918 |
| ZNF681 | -0.105163092 |
| CTD-2288O8 | -0.105166979 |
| RP11-298C2 | -0.105176158 |
| TCL1A | -0.105192215 |
| HLA-DRA | -0.10521221 |
| LINC00342 | -0.10522702 |
| DNAAF2 | -0.105288883 |
| RP11-329N15 | -0.105328207 |
| RP11-456D7 | -0.105334008 |
| RP1-164L12 | -0.10540465 |
| MIR573 | -0.105422796 |
| TRAJ7 | -0.105422833 |
| SIRT1 | -0.10544863 |
| C15orf53 | -0.105450603 |
| NKTR | -0.105452798 |
| AC099552 | -0.105454014 |
| RP13-36G14 | -0.105474138 |

| | |
|---|---|
| IGHV1OR16-3 | -0.105486667 |
| CDK12 | -0.10549546 |
| ZNF224 | -0.105502418 |
| IFI44L | -0.105503181 |
| IGHV1OR15-1 | -0.105503844 |
| KAT6B | -0.105511591 |
| BORA | -0.105527164 |
| TGIF2P1 | -0.105554385 |
| MIR5196 | -0.10556714 |
| AC002400 | -0.105573033 |
| HLA-DOA | -0.105584636 |
| RP11-632K20 | -0.105612228 |
| RPL23AP64 | -0.105623635 |
| RP4-800J21 | -0.105670643 |
| TRAJ9 | -0.105703946 |
| SCAI | -0.105720512 |
| AC007906 | -0.105754153 |
| CCND1 | -0.105784589 |
| PTCHD2 | -0.105793075 |
| VNN2 | -0.105797743 |
| ZNF30 | -0.105806748 |
| RP11-625L16 | -0.105812631 |
| CCDC62 | -0.105826372 |
| FAM117B | -0.105829298 |
| PLCB2-AS1 | -0.105838013 |
| DOK3 | -0.105864425 |
| DNAH8 | -0.105909875 |
| RP11-285B24 | -0.105911138 |
| OR3A3 | -0.105925646 |
| RP11-347C18 | -0.105926065 |
| POLE | -0.105929432 |
| GRB2 | -0.105968876 |
| ACAP2-IT1 | -0.105978153 |
| AACSP1 | -0.105985529 |
| TRAJ18 | -0.10603786 |
| RNF114 | -0.10607355 |
| BEND3P1 | -0.106081301 |
| LLNLR-304A6 | -0.106087297 |
| ZNF763 | -0.106117385 |
| NIPA1 | -0.106120827 |
| CRKL | -0.106130855 |
| PARP6 | -0.106150689 |
| CRIP1P2 | -0.106160431 |
| CTC-523E23 | -0.106160587 |
| MCF2L | -0.106253354 |

| | |
|---|---|
| ZDHHC17 | -0.106285206 |
| CERKL | -0.106309449 |
| ZNFX1 | -0.106322157 |
| CSTF3 | -0.106327515 |
| HECTD4 | -0.106341618 |
| LINC00926 | -0.106364652 |
| BAZ1A | -0.106374903 |
| AKAP5 | -0.106401416 |
| GPS2 | -0.106437566 |
| PRKCQ | -0.106456619 |
| LINC00423 | -0.106469422 |
| PSMB9 | -0.10646994 |
| TAS1R3 | -0.106540566 |
| TAC3 | -0.106546715 |
| IGKV3OR2-5 | -0.106552727 |
| RP11-837J7 | -0.106555079 |
| DERL3 | -0.106575954 |
| ZNF740 | -0.10659082 |
| GGA2 | -0.106604048 |
| AC009518 | -0.106646073 |
| PABPN1L | -0.106688011 |
| INVS | -0.106690284 |
| LEO1 | -0.106706229 |
| BNIP3P25 | -0.106762005 |
| PHF7 | -0.106767013 |
| IL18 | -0.106767869 |
| RP11-159H10 | -0.106784345 |
| RP11-293G6__B | -0.106788421 |
| SLC4A5 | -0.106799294 |
| LINC01298 | -0.106817335 |
| CEP97 | -0.106835072 |
| CNPPD1 | -0.106837026 |
| MYO1G | -0.106860134 |
| EIF3J-AS1 | -0.106865604 |
| PTPN22 | -0.10688047 |
| AP1AR | -0.106915243 |
| ATAD2B | -0.106920062 |
| RHOT1P3 | -0.106954684 |
| USP30-AS1 | -0.106974493 |
| SIMC1 | -0.106987991 |
| RAC2 | -0.107034337 |
| COL4A4 | -0.107043881 |
| COX18 | -0.107054877 |
| FLT3 | -0.107057697 |
| RP11-203B7 | -0.107083897 |

| | |
|---|---|
| RP11-535I24 | -0.107084062 |
| GCC2 | -0.107143086 |
| CYB5RL | -0.107165636 |
| GTF3AP6 | -0.107178823 |
| USP18 | -0.107180687 |
| ANKRD26 | -0.107256337 |
| TOMM20L | -0.107258094 |
| EIF1AX-AS1 | -0.107287256 |
| HRH3 | -0.107296445 |
| NUSAP1 | -0.107329308 |
| RP11-93B14 | -0.107331206 |
| ITGA4 | -0.107349011 |
| ZNRD1-AS1 | -0.107352211 |
| RP13-870H17 | -0.107354572 |
| GANC | -0.10735956 |
| RCHY1 | -0.107370928 |
| STRBP | -0.107381664 |
| GPR182 | -0.107395131 |
| ZC3HAV1 | -0.107414112 |
| ZNF143 | -0.107431805 |
| RP11-110H1 | -0.107494716 |
| RP5-961K14 | -0.107503243 |
| LARP1P1 | -0.107520963 |
| DSTNP1 | -0.107536108 |
| USP14 | -0.10758757 |
| RP1-20N2 | -0.107588239 |
| SUPT16H | -0.107643815 |
| ING3 | -0.107658792 |
| ADRBK1 | -0.107664988 |
| LASP1 | -0.107668002 |
| RP11-179A7 | -0.107701719 |
| SFTPB | -0.107733328 |
| TOP2A | -0.107743875 |
| MOB1B | -0.107787486 |
| MRPS31P2 | -0.10780788 |
| CTD-2240J17 | -0.107808673 |
| PIK3R5 | -0.107823767 |
| KCNH4 | -0.107826208 |
| RPS3AP3 | -0.107844727 |
| CREBRF | -0.107873295 |
| DNAJA4 | -0.107881127 |
| SRP72 | -0.107895937 |
| PPAT | -0.10789951 |
| AL928761 | -0.107908865 |
| NCF1 | -0.107908924 |

| | |
|---|---|
| CTD-2349P21 | -0.107917669 |
| PTAFR | -0.107924664 |
| SLC22A10 | -0.107943329 |
| RP11-104L21 | -0.107993019 |
| AC099535 | -0.107993667 |
| RP11-365D9 | -0.10803308 |
| DMTF1 | -0.108038344 |
| LA16c-381G6 | -0.108045155 |
| RP11-568N6 | -0.108055665 |
| ACAP1 | -0.108063312 |
| ADRBK2 | -0.108111928 |
| RP11-210K20 | -0.108112295 |
| ZCCHC7 | -0.108123185 |
| TRDJ2 | -0.108127867 |
| RNY4P10 | -0.108128708 |
| FNBP1 | -0.108131818 |
| SPOP | -0.1081462 |
| ANKRD61 | -0.10815061 |
| IGHD3-3 | -0.108162175 |
| DAZAP2P1 | -0.108162664 |
| PTCD3 | -0.108167614 |
| RNA5SP78 | -0.108168018 |
| RP11-25K21 | -0.108181815 |
| UBD | -0.108194891 |
| AC023590 | -0.108242011 |
| P2RY12 | -0.10824973 |
| RP11-357G3 | -0.108250674 |
| AC024361 | -0.108265771 |
| AC015720 | -0.108274903 |
| TNRC6C-AS1 | -0.108281685 |
| RP11-482M8 | -0.108287143 |
| LRRCC1 | -0.108287248 |
| RNU6-407P | -0.108325804 |
| RP11-830F9 | -0.108335175 |
| HCG27 | -0.10833832 |
| SLC29A3 | -0.108357465 |
| TRBV11-1 | -0.108389565 |
| ANKRD18DP | -0.108393098 |
| RP11-34E5 | -0.108414455 |
| ZNF266 | -0.10843691 |
| IFT88 | -0.10845092 |
| AC074366 | -0.108462569 |
| RN7SL268P | -0.10847452 |
| CHAMP1 | -0.108502643 |
| PCBP2P1 | -0.108521805 |

| | |
|---|---|
| GZMM | -0.108540842 |
| AC005197 | -0.108574148 |
| RP11-565N2 | -0.108579465 |
| P2RX5 | -0.108583331 |
| AL049794 | -0.108589296 |
| HLA-DQA1 | -0.108596377 |
| AC017080 | -0.108604114 |
| NGDN | -0.108608211 |
| TRBV7-1 | -0.108608905 |
| RP11-111A22 | -0.108617928 |
| PHIP | -0.108621774 |
| RP11-644A7 | -0.108638016 |
| PSPC1P1 | -0.108642971 |
| RBM6 | -0.108668064 |
| RP11-79L9 | -0.108679227 |
| CTD-2022H16 | -0.108720829 |
| CH17-264L24 | -0.108735267 |
| CTB-131K11 | -0.108762115 |
| METTL3 | -0.108786916 |
| KANSL3 | -0.108824034 |
| RP11-126O1 | -0.108826709 |
| C9orf91 | -0.108843067 |
| FAM129C | -0.10885805 |
| GAB3 | -0.108865152 |
| FAM178A | -0.108872185 |
| C2orf69 | -0.10888733 |
| TRBV11-3 | -0.108907695 |
| PPM1K | -0.108918029 |
| PDP2 | -0.10891855 |
| LINC00173 | -0.108919144 |
| PLCD4 | -0.108927127 |
| NECAP1 | -0.108940337 |
| TRIM25 | -0.108948542 |
| CTD-2516F10 | -0.108980208 |
| BPTF | -0.108990423 |
| RP1-168P16 | -0.109000816 |
| SPATA2 | -0.109002107 |
| RPL29P24 | -0.109039391 |
| CTD-2007L18 | -0.109051541 |
| RP11-426D19 | -0.109051835 |
| TRAPPC8 | -0.109053235 |
| HEATR5B | -0.109063021 |
| MSX2P1 | -0.109072864 |
| RP11-461O7 | -0.109093913 |
| USP36 | -0.109104344 |

| | |
|---|---|
| INO80D | -0.10911601 |
| C12orf66 | -0.10913402 |
| RP11-114F3 | -0.109156165 |
| INPP4A | -0.109170933 |
| C5orf64 | -0.109176662 |
| CCNT1 | -0.109199678 |
| UBE2D3 | -0.109202711 |
| LL0XNC01-7P3 | -0.109210432 |
| RP1-130G2 | -0.109252402 |
| UBE2D3P1 | -0.10926325 |
| RP13-93L13 | -0.109279939 |
| IGLCOR22-1 | -0.109291412 |
| AC145110 | -0.109297279 |
| AGBL2 | -0.109343964 |
| ACRC | -0.109353601 |
| MIR4512 | -0.109368922 |
| ARHGAP42P1 | -0.109394016 |
| RET | -0.10940499 |
| F10-AS1 | -0.109431393 |
| FAM222A-AS1 | -0.109442376 |
| GDPD1 | -0.109462927 |
| RN7SL605P | -0.109468252 |
| ZNF33B | -0.109518092 |
| C5orf58 | -0.109531711 |
| USP47 | -0.109536382 |
| PPWD1 | -0.109557406 |
| SAMD9 | -0.109581165 |
| GOLGA8IP | -0.109597489 |
| E2F6 | -0.109599919 |
| PSME2 | -0.109619641 |
| CPSF6 | -0.109627492 |
| IGHV1-45 | -0.109634518 |
| DPY19L1P1 | -0.109640051 |
| PARP14 | -0.109686676 |
| USP48 | -0.10970037 |
| ZNF226 | -0.109743107 |
| RP11-1084E5 | -0.109824521 |
| RNA5SP379 | -0.109829 |
| RP11-142E9 | -0.10984406 |
| NAPG | -0.109871913 |
| RP11-58A17 | -0.109877576 |
| RP11-540B6 | -0.109883987 |
| SDR39U1 | -0.109911635 |
| CCDC59 | -0.109914426 |
| ESR2 | -0.109932492 |

| | |
|---|---|
| SSTR3 | -0.109952732 |
| AC069363 | -0.10996623 |
| UGCG | -0.10997761 |
| HMGB1P3 | -0.110014564 |
| SNX18P9 | -0.110042293 |
| PCGF5 | -0.11006087 |
| RP11-147L13 | -0.110083345 |
| TRIM52 | -0.110084759 |
| RNU1-73P | -0.110101931 |
| DNAJC3-AS1 | -0.11013077 |
| RP11-71G12 | -0.110137183 |
| FAN1 | -0.110156297 |
| TREML2 | -0.110156929 |
| HNRNPA1P59 | -0.110159312 |
| WEE1 | -0.110168558 |
| LARP7 | -0.110223147 |
| AC004924 | -0.110239247 |
| AC090960 | -0.110264543 |
| BCAS1 | -0.110280786 |
| IGLVI-20 | -0.110303462 |
| LRRC37B | -0.110312788 |
| PEX5L-AS2 | -0.110326834 |
| ZNF383 | -0.110342817 |
| GCNT7 | -0.110342881 |
| RP11-330H6 | -0.110369607 |
| SMG1P7 | -0.110420662 |
| RP5-991B18 | -0.110436356 |
| RNA5SP39 | -0.110451796 |
| RP11-404F10 | -0.110471756 |
| AC012363 | -0.110476995 |
| CHKB-CPT1B | -0.110480088 |
| DCAF16 | -0.110522081 |
| ESR1 | -0.11054117 |
| RP11-561O4 | -0.110558927 |
| GPR31 | -0.110563746 |
| ALG13-AS1 | -0.110570452 |
| TARDBPP2 | -0.110578204 |
| RP11-700H13 | -0.110579733 |
| C5orf45 | -0.110590214 |
| HHEX | -0.110593153 |
| NEIL3 | -0.110601824 |
| RP11-158K1 | -0.110603145 |
| APOA1-AS | -0.110604945 |
| RP11-1319K7 | -0.110649444 |
| SNX15 | -0.110669734 |

| | |
|---|---|
| CCNL1 | -0.110673417 |
| FAM153A | -0.11071204 |
| GLTSCR1L | -0.11072034 |
| IL9R | -0.110724979 |
| LETMD1 | -0.110729085 |
| AC108004 | -0.110742909 |
| GUF1 | -0.110770687 |
| MAST3 | -0.110801795 |
| NXPE3 | -0.110814968 |
| FAM126B | -0.11082375 |
| RNA5SP240 | -0.110876984 |
| NFATC2IP | -0.110891362 |
| KIAA1211 | -0.110923401 |
| HDHD2 | -0.110939252 |
| RP11-476D10 | -0.110945642 |
| MAT2A | -0.1109589 |
| EFCAB7 | -0.110961821 |
| SMARCAD1 | -0.110963309 |
| AL022314 | -0.11097692 |
| CCDC117 | -0.111001023 |
| TRPM7 | -0.111003553 |
| CDC14A | -0.111008372 |
| ORC2 | -0.111025917 |
| LINC00824 | -0.11102613 |
| RP11-521I2 | -0.111034733 |
| TAAR3 | -0.111060401 |
| RNA5SP69 | -0.111077932 |
| ZSWIM1 | -0.111085491 |
| DHRS13 | -0.111140191 |
| AC073052 | -0.111147615 |
| FAM102B | -0.111206185 |
| RP11-213G2 | -0.111212623 |
| RP11-679C8 | -0.111245169 |
| CHD1 | -0.111246317 |
| RP4-717I23 | -0.111254429 |
| PEX5L | -0.111255994 |
| PROCA1 | -0.111259173 |
| TRAJ16 | -0.111268096 |
| RBBP4P2 | -0.111270143 |
| NEK4 | -0.111272681 |
| KRT18P45 | -0.111275657 |
| TRAJ47 | -0.11128804 |
| LINC00843 | -0.111324991 |
| MARS2 | -0.111325223 |
| RP4-784A16 | -0.111380679 |

| | |
|---|---|
| MIR98 | -0.111392364 |
| SAMHD1 | -0.111430753 |
| UHRF1BP1 | -0.111452187 |
| MDM1 | -0.111509026 |
| RN7SKP214 | -0.111527841 |
| RP11-12D16 | -0.1115333 |
| RPL12P1 | -0.111555206 |
| PTGES3P2 | -0.111587404 |
| NPPA-AS1 | -0.111592203 |
| TRAJ19 | -0.111595918 |
| RP11-131H24 | -0.111608045 |
| P2RY14 | -0.111611096 |
| KIAA0226 | -0.111630408 |
| SAMSN1 | -0.111633367 |
| AIM2 | -0.111633739 |
| AC009948 | -0.111684569 |
| SFXN1 | -0.111706616 |
| PSMD5-AS1 | -0.111715045 |
| SSU72P3 | -0.111783277 |
| CRHBP | -0.111821365 |
| GPR15 | -0.111860146 |
| RP4-680D5 | -0.111882932 |
| NIPAL3 | -0.111912895 |
| PHF12 | -0.111919072 |
| RP11-752G15 | -0.111945771 |
| ZFP30 | -0.111947799 |
| SNORA12 | -0.111948109 |
| MFSD8 | -0.111971558 |
| ST13P15 | -0.112012817 |
| RBM39 | -0.112029384 |
| RP11-260M2 | -0.112038873 |
| TRIM26 | -0.112074223 |
| SPAG6 | -0.112090417 |
| CRIP1 | -0.112096103 |
| BET1P1 | -0.112138066 |
| ACRBP | -0.112183754 |
| MCM3AP-AS1 | -0.112187028 |
| METTL4 | -0.112200722 |
| GOSR1 | -0.11220208 |
| SLC25A38 | -0.112216555 |
| SMG7 | -0.112226825 |
| TERF2IP | -0.112248203 |
| RP11-50D16 | -0.112250034 |
| NELL2 | -0.112252724 |
| CD300LF | -0.112275925 |

| | |
|---|---|
| SH3BP5-AS1 | -0.112318494 |
| ZNF142 | -0.112344635 |
| RTF1 | -0.112350049 |
| DPY19L2P3 | -0.112435002 |
| ZSCAN16 | -0.112443523 |
| LTA4H | -0.112495079 |
| C15orf57 | -0.112550616 |
| KHNYN | -0.112556682 |
| SPG11 | -0.112575027 |
| HNRNPA1P61 | -0.112609816 |
| BNIP3P5 | -0.112612485 |
| FCAMR | -0.112630049 |
| RP11-56B16 | -0.112656093 |
| MED13L | -0.112664761 |
| NUFIP1P | -0.112675394 |
| RBMXL1 | -0.112678399 |
| RN7SL684P | -0.112697281 |
| WASIR2 | -0.112745254 |
| ZNF600 | -0.112768523 |
| FBXL20 | -0.112770012 |
| AC007278 | -0.112797603 |
| RP3-329A5 | -0.112801325 |
| RBL1 | -0.112843559 |
| WAPAL | -0.112848415 |
| CBX7 | -0.112850815 |
| TOB1 | -0.112873086 |
| RPL35AP32 | -0.112908852 |
| NEDD1 | -0.112942131 |
| RBM4B | -0.112960811 |
| SP100 | -0.113027716 |
| ZNF331 | -0.113028014 |
| POU2F1 | -0.113050061 |
| SNRNP48 | -0.113069262 |
| CTB-12A17 | -0.113081961 |
| FAR1 | -0.113101152 |
| AC015849 | -0.113111962 |
| C4orf29 | -0.113124018 |
| DDX60L | -0.113154196 |
| WDR82P2 | -0.113156951 |
| HMGB1P20 | -0.113164296 |
| SMEK2 | -0.113173881 |
| IQCF1 | -0.113174347 |
| FMN1 | -0.113189621 |
| PTBP1P | -0.11320481 |
| CD37 | -0.113230572 |

| | |
|---|---|
| IDO2 | -0.113252676 |
| G3BP2 | -0.113267578 |
| TRBJ2-7 | -0.113274113 |
| RASA4 | -0.113295337 |
| MIR1913 | -0.11330315 |
| MGAT4A | -0.113321273 |
| IGHEP1 | -0.113339595 |
| TRGV1 | -0.113356145 |
| NCR3LG1 | -0.113361405 |
| AC002401 | -0.11336715 |
| AC023593 | -0.113380316 |
| TRAJ35 | -0.113414938 |
| HP09025 | -0.113456909 |
| RC3H1 | -0.113465782 |
| AC126365 | -0.113475677 |
| C2CD5 | -0.113502174 |
| PPIP5K1 | -0.113506193 |
| SRSF1 | -0.113509207 |
| RP11-511P7 | -0.113513538 |
| TRAJ22 | -0.113556213 |
| ICE2 | -0.113579964 |
| IL18BP | -0.113602681 |
| RAD21 | -0.113603053 |
| RNU6-118P | -0.113612908 |
| CTD-3195I5 | -0.113661854 |
| RP11-81A1 | -0.113668154 |
| N4BP2L2-IT2 | -0.113670944 |
| KDM2B | -0.113700397 |
| FBXO6 | -0.113723021 |
| KCNMA1-AS3 | -0.113742565 |
| MADD | -0.113744306 |
| RP4-695O20 | -0.113747049 |
| LINC00954 | -0.113749878 |
| IL24 | -0.113768381 |
| DNAJC5B | -0.113776303 |
| NRIR | -0.113785596 |
| RP1-228P16 | -0.113816774 |
| RNU6-1152P | -0.113846369 |
| OR6S1 | -0.113862527 |
| SRRM1 | -0.113869911 |
| RP11-813F20 | -0.113902317 |
| ANKRD29 | -0.113925597 |
| AC116609 | -0.113948776 |
| MIR4697HG | -0.113978913 |
| LINC00158 | -0.113987982 |

| | |
|---|---|
| NUPL1 | -0.113998568 |
| GGA3 | -0.114059929 |
| CNNM3 | -0.114072283 |
| RAD51-AS1 | -0.114087409 |
| RP3-448I9 | -0.114111027 |
| CH17-260O16 | -0.114138195 |
| ZNF844 | -0.114138686 |
| CLK4 | -0.114164194 |
| RBM14-RBM4 | -0.114175822 |
| CIT | -0.114181367 |
| DENND4A | -0.114198633 |
| ZNF235 | -0.114214522 |
| C9orf57 | -0.114236139 |
| IGHE | -0.114258067 |
| TMEM170A | -0.114267045 |
| AC104532 | -0.114285223 |
| AP003774 | -0.114292944 |
| CCDC174 | -0.114361133 |
| IGHV3-79 | -0.114375291 |
| MNDA | -0.114388353 |
| RP11-532F6 | -0.114451835 |
| RNF213 | -0.11451887 |
| SNORA2A | -0.114546911 |
| ANKRD40 | -0.114556769 |
| CTC-231O11 | -0.114562575 |
| LINC01480 | -0.114627638 |
| TRAV26-2 | -0.114633156 |
| KNTC1 | -0.114653778 |
| RNU6-598P | -0.114668868 |
| RP11-20B24 | -0.114675685 |
| SLC35E2 | -0.1146759 |
| ARHGAP11A | -0.11472407 |
| GNLY | -0.114732461 |
| RN7SL145P | -0.114745722 |
| RP11-458F8 | -0.114761411 |
| ABCB1 | -0.1147757 |
| HCFC1-AS1 | -0.114776298 |
| RP11-98I9 | -0.114779272 |
| RP11-75L1 | -0.114794053 |
| TAF1B | -0.114825823 |
| KRR1 | -0.11483027 |
| TAP1 | -0.114863294 |
| UNQ6494 | -0.114891272 |
| LINC01501 | -0.11489724 |
| PDS5A | -0.114898012 |

| | |
|---|---|
| RN7SKP127 | -0.114901489 |
| RP11-157K17 | -0.114912466 |
| SIRPD | -0.114948158 |
| AC008982 | -0.114980691 |
| CAPN10-AS1 | -0.115036158 |
| CCR8 | -0.115036753 |
| RP11-257O5 | -0.115105612 |
| RTTN | -0.115131827 |
| ARID4B | -0.115133464 |
| CTD-2509G16 | -0.11516423 |
| CYFIP2 | -0.115217096 |
| IGLV3-30 | -0.115250505 |
| ZNF222 | -0.115251423 |
| SECISBP2L | -0.115255963 |
| RP3-432I18 | -0.11526237 |
| RP11-118B22 | -0.115285644 |
| TRAJ5 | -0.115287173 |
| CTD-2562J15 | -0.115318965 |
| ZNF100 | -0.115356321 |
| DOCK10 | -0.115388397 |
| TM4SF20 | -0.115413307 |
| UBA52P3 | -0.115421068 |
| COL4A3 | -0.115439617 |
| SLC25A53 | -0.115445776 |
| PTK2B | -0.115538617 |
| C17orf80 | -0.115545464 |
| TNFSF8 | -0.115554916 |
| SEC13P1 | -0.115558249 |
| MDM2 | -0.115577763 |
| IGKV2-4 | -0.115587103 |
| IFNLR1 | -0.115598824 |
| LINC00304 | -0.115649391 |
| PRSS3P3 | -0.115651598 |
| RASGRP1 | -0.11570402 |
| PIKFYVE | -0.115723425 |
| RSBN1L | -0.115740803 |
| KAT7 | -0.115755185 |
| TNFSF14 | -0.115767604 |
| TRAJ3 | -0.115770622 |
| PLEKHA3P1 | -0.115789703 |
| ZNF217 | -0.115791112 |
| SYNJ1 | -0.115857757 |
| IRAK4 | -0.115860343 |
| TCHP | -0.115863506 |
| HERPUD1 | -0.115880734 |

| | |
|---|---|
| HAUS1 | -0.115903266 |
| RP11-384M15 | -0.115945515 |
| RP11-497H17 | -0.115959374 |
| UPF3A | -0.115975734 |
| HLA-DMB | -0.116014154 |
| CTLA4 | -0.116125606 |
| TBCD | -0.116226982 |
| ACSS1 | -0.116242778 |
| RP11-290F24 | -0.116243225 |
| ZNF276 | -0.116288529 |
| ST13P5 | -0.116301535 |
| RP11-281P11 | -0.116312584 |
| RP11-384M20 | -0.116323901 |
| PAXBP1 | -0.116331173 |
| C6orf211 | -0.11633962 |
| CCAR1 | -0.116344718 |
| STXBP3 | -0.116355174 |
| LINC01358 | -0.116358419 |
| FAM65B | -0.116393817 |
| BRD8 | -0.1164106 |
| TRAV23DV6 | -0.116433052 |
| TPM3P9 | -0.116433242 |
| CMC1 | -0.116434898 |
| MT-TL1 | -0.116495606 |
| TAC4 | -0.116507579 |
| NLRP7 | -0.116516205 |
| SLC32A1 | -0.116539263 |
| A2MP1 | -0.116577156 |
| TNFSF13B | -0.116580263 |
| RPL7AP64 | -0.116657505 |
| ZNF725P | -0.116663313 |
| RANBP9 | -0.116669569 |
| RP11-483E17 | -0.116686526 |
| IDH3A | -0.116695431 |
| CRLF3 | -0.116707859 |
| RP5-1057J7 | -0.116721143 |
| NCKAP1L | -0.116741498 |
| SIK3 | -0.116743117 |
| RP11-381E24 | -0.116760501 |
| RBM26 | -0.116770708 |
| ZNF326 | -0.116821166 |
| TRAV18 | -0.116830133 |
| JSRP1 | -0.116838684 |
| IGKV6-21 | -0.116838898 |
| FBXO9 | -0.116850731 |

| | |
|---|---|
| LARP1B | -0.116855419 |
| AC079767 | -0.116866384 |
| IGHD2-15 | -0.116899406 |
| DEPDC5 | -0.116914733 |
| CAMK1G | -0.116953057 |
| RP1-179N16 | -0.116973843 |
| TLR8 | -0.117024301 |
| ITPKB-AS1 | -0.117043577 |
| RN7SKP150 | -0.117064214 |
| C11orf58 | -0.117092899 |
| PSMA3-AS1 | -0.11710784 |
| IGHD6-19 | -0.117141775 |
| SECISBP2 | -0.117171061 |
| TPK1 | -0.117184866 |
| IL16 | -0.117196215 |
| CYLD | -0.117279866 |
| IGHV3-6 | -0.117299913 |
| RP11-109P14 | -0.117308657 |
| IGKV2OR2-1 | -0.117395685 |
| RP11-367G6 | -0.117404297 |
| ZBTB49 | -0.117416858 |
| UBBP5 | -0.117439399 |
| LINC00528 | -0.117457842 |
| MYRIP | -0.117461888 |
| ZNF24 | -0.117479614 |
| RP11-855A2 | -0.117507987 |
| AC092566 | -0.117551323 |
| ZNF443 | -0.117556343 |
| RP11-795J1 | -0.117587756 |
| SMPD3 | -0.117603322 |
| RP3-475N16 | -0.117628402 |
| HMGN1P4 | -0.117677282 |
| PGBD4P1 | -0.117737679 |
| IL9RP3 | -0.117751513 |
| TRAPPC11 | -0.117771255 |
| CTC-542B22 | -0.117774531 |
| TRBV12-5 | -0.117776082 |
| LINC01386 | -0.117801572 |
| hsa-mir-4538 | -0.117842281 |
| BFSP2-AS1 | -0.117850513 |
| NBN | -0.117901995 |
| LINC00525 | -0.117902031 |
| CTD-2576D5 | -0.117917063 |
| ART4 | -0.117942947 |
| RTP5 | -0.117964802 |

| | |
|---|---|
| SETP9 | -0.117967873 |
| TRAJ33 | -0.117972737 |
| RP11-791G16 | -0.117984906 |
| LRRC57 | -0.117996158 |
| LINC01079 | -0.118008911 |
| MTND4P23 | -0.118024031 |
| RP11-265O12 | -0.118048195 |
| GTSF1L | -0.118093475 |
| RP11-18C24 | -0.118123372 |
| NUP50 | -0.118125838 |
| PCMTD2 | -0.118146285 |
| GLULP5 | -0.11816291 |
| SIRPB3P | -0.11820644 |
| BZW1P2 | -0.118228484 |
| IGLVI-70 | -0.118232728 |
| ARHGAP9 | -0.118238085 |
| RP11-1084J3 | -0.11830814 |
| TRBV10-1 | -0.118379682 |
| RGS18 | -0.11843054 |
| CPLX2 | -0.118444138 |
| RP11-474L23 | -0.118485418 |
| LINC01146 | -0.11849209 |
| KIR2DS4 | -0.118499656 |
| GIMAP8 | -0.118510673 |
| SAP18 | -0.118546824 |
| P2RX7 | -0.118591942 |
| ZNF791 | -0.118594137 |
| LINC00957 | -0.118700561 |
| ZNF721 | -0.118716431 |
| AC144568 | -0.1187241 |
| RP11-1060G2 | -0.118730264 |
| RP3-522J7 | -0.118778553 |
| KCTD6 | -0.118816547 |
| KANSL2 | -0.118842781 |
| CARD11 | -0.118853181 |
| RP11-175P19 | -0.118875323 |
| SMG1P4 | -0.11892593 |
| ZNF625-ZNF20 | -0.118943427 |
| PSMD6-AS2 | -0.118945 |
| SLC35E2B | -0.118953446 |
| HVCN1 | -0.118957018 |
| FGD2 | -0.118974545 |
| RP11-586D19 | -0.118990005 |
| SLC18A1 | -0.11907189 |
| CATSPER2 | -0.119079256 |

| | |
|---|---|
| CASR | -0.119090339 |
| DDHD1 | -0.119139092 |
| CEP128 | -0.119140394 |
| IGLV3-21 | -0.119158749 |
| AP4E1 | -0.119187763 |
| N4BP2 | -0.119217514 |
| CCDC121 | -0.119233458 |
| USP24 | -0.119250148 |
| AC011997 | -0.119288271 |
| LTA | -0.119353222 |
| TCEANC2 | -0.119365334 |
| DHX58 | -0.119427067 |
| AKNA | -0.119438714 |
| HSPE1P18 | -0.119547605 |
| AP000233 | -0.119558081 |
| PCF11 | -0.119602889 |
| RP11-23J18 | -0.119624463 |
| NCOA5 | -0.11969506 |
| DNM1P46 | -0.119795217 |
| OAS1 | -0.119822694 |
| RP11-396K3 | -0.119827047 |
| RP11-421E14 | -0.119869048 |
| MLLT6 | -0.119915442 |
| CASS4 | -0.119959816 |
| ZNF354B | -0.119964672 |
| ANKRD46 | -0.120029959 |
| AL133245 | -0.120046218 |
| RP11-474I11 | -0.120082142 |
| SRSF6 | -0.120082184 |
| EDEM1 | -0.120093812 |
| CALCOCO2 | -0.120135079 |
| PVRIG | -0.120145402 |
| APOL6 | -0.120153722 |
| KPNB1 | -0.120180179 |
| USP15 | -0.120199566 |
| RP11-973H7 | -0.120266341 |
| BFSP2 | -0.120294616 |
| B2M | -0.120339293 |
| TOX4P1 | -0.120378418 |
| RP11-440L14 | -0.120397268 |
| BTK | -0.120402403 |
| MZT1 | -0.12044914 |
| TCEANC | -0.12045221 |
| TRDC | -0.120452619 |
| MAX | -0.120461643 |

| | |
|---|---|
| TRAJ24 | -0.120504551 |
| CTA-246H3 | -0.120524432 |
| PRKCH | -0.120526948 |
| HMGN1P38 | -0.120526959 |
| LRRC2 | -0.120545974 |
| IGKV2-18 | -0.120561805 |
| CEP120 | -0.120564773 |
| PPP2R2B | -0.12056654 |
| TYW5 | -0.120596997 |
| SIAH3 | -0.120602683 |
| RP11-58A11 | -0.120610984 |
| AC011330 | -0.120614984 |
| AC055866 | -0.120620236 |
| ANO5 | -0.120644526 |
| TRAV38-1 | -0.120714406 |
| GIMAP6 | -0.120814886 |
| MIR5093 | -0.120834315 |
| PLEK | -0.120854106 |
| NGLY1 | -0.120880414 |
| RP11-449J21 | -0.120922984 |
| AC006160 | -0.12093452 |
| CXorf21 | -0.120952604 |
| PEBP1P2 | -0.120960071 |
| STX16 | -0.120977442 |
| RP11-330A16 | -0.120991602 |
| IGHV1-68 | -0.121014177 |
| RP11-655C2 | -0.121019414 |
| RP11-1018N14 | -0.12105501 |
| TMEM199 | -0.121119327 |
| SNORA75 | -0.121123187 |
| RLTPR | -0.121123848 |
| DDX39B | -0.121157841 |
| DOCK2 | -0.121158883 |
| PAXBP1-AS1 | -0.121224039 |
| RP11-388C12 | -0.121236523 |
| MDM4 | -0.121300061 |
| LINC01068 | -0.121303619 |
| AC073325 | -0.121315619 |
| LARP1 | -0.121324825 |
| KIAA0195 | -0.121330853 |
| U62631 | -0.121361854 |
| EBI3 | -0.121382502 |
| EXOC3L4 | -0.121412373 |
| WI2-1959D15 | -0.121412461 |
| RNY4P7 | -0.121447596 |

| | |
|---|---|
| RP1-257I20 | -0.121456265 |
| RP11-35G9 | -0.121499344 |
| LILRB1 | -0.121517745 |
| TTC16 | -0.121535532 |
| CEP290 | -0.121536499 |
| DMC1 | -0.12154957 |
| ZDHHC13 | -0.121570398 |
| C9orf72 | -0.121584204 |
| CTD-2537I9 | -0.121636969 |
| NUB1 | -0.12164495 |
| N4BP2L1 | -0.12164536 |
| RP3-466I7 | -0.121646434 |
| C4orf50 | -0.1216524 |
| CTD-2278I10 | -0.121653595 |
| RP11-353K11 | -0.121669231 |
| VPS11 | -0.121671184 |
| IGLV3-9 | -0.121706588 |
| NEMF | -0.121728266 |
| FNBP1P1 | -0.12174316 |
| NLRC5 | -0.121748732 |
| KIR3DL1 | -0.121782372 |
| LPXN | -0.121791282 |
| IGHV3-25 | -0.121793774 |
| ESCO1 | -0.121845052 |
| RN7SL812P | -0.121863539 |
| NIF3L1 | -0.121878784 |
| RP11-415F23 | -0.121961225 |
| RP11-102L12 | -0.122003316 |
| IGKV1-37 | -0.122024731 |
| ERCC3 | -0.122025246 |
| TUBGCP3 | -0.122038958 |
| WHAMM | -0.122087277 |
| CMKLR1 | -0.122113603 |
| JMJD7 | -0.12213353 |
| APOBEC3D | -0.122142814 |
| NBEAL2 | -0.122174202 |
| FBXO41 | -0.122185216 |
| XAF1 | -0.122196212 |
| MROH8 | -0.122213013 |
| IGHD4-23 | -0.122221514 |
| CTD-2252P21 | -0.12229882 |
| HAVCR1 | -0.122319359 |
| DNAH1 | -0.122356851 |
| AC009950 | -0.122384034 |
| TRAJ2 | -0.122395712 |

| | |
|---|---|
| NFXL1 | -0.12241918 |
| MYL8P | -0.122474203 |
| PTPN1 | -0.122498625 |
| CCDC66 | -0.122509528 |
| EZH1 | -0.122514998 |
| RP5-908D6 | -0.122516431 |
| CD6 | -0.122535501 |
| IGHD2-21 | -0.122588041 |
| IGLV9-49 | -0.122632596 |
| AC008746 | -0.122633459 |
| IFIH1 | -0.122639468 |
| HSPA8P19 | -0.122652582 |
| RP11-392O18 | -0.122671397 |
| IGHA1 | -0.122707081 |
| MIR378H | -0.122707747 |
| NBPF22P | -0.122717477 |
| ANO9 | -0.122792052 |
| TRBV21-1 | -0.122797651 |
| IGLL3P | -0.12283122 |
| ZNF708 | -0.122841152 |
| TRAJ13 | -0.122842088 |
| FAM122B | -0.122863553 |
| TRBJ2-2 | -0.122969255 |
| IFIT5 | -0.122981381 |
| RP11-542M13 | -0.122994979 |
| IL22 | -0.123082155 |
| RP11-564A8 | -0.123094335 |
| TNK2-AS1 | -0.12316128 |
| ELP2 | -0.123250881 |
| WDFY4 | -0.123326866 |
| AOAH | -0.123333843 |
| C1orf147 | -0.123357909 |
| PSPC1 | -0.123418889 |
| RP11-642P15 | -0.123490613 |
| PPIG | -0.123546262 |
| TSGA10 | -0.123604404 |
| RP11-24F11 | -0.123657091 |
| CTC-457L16 | -0.123684794 |
| ZNF396 | -0.12368744 |
| CLEC9A | -0.123727462 |
| HSPA1L | -0.123803631 |
| AC105265 | -0.12381768 |
| IGKV1OR9-1 | -0.123826727 |
| LINC01281 | -0.123864836 |
| TRANK1 | -0.123889402 |

| | |
|---|---|
| DDI2 | -0.123956996 |
| NEURL1 | -0.123978764 |
| STAT4 | -0.124008012 |
| RAD51D | -0.124025948 |
| MTMR4 | -0.124050768 |
| AL365331 | -0.124055237 |
| PRPF39 | -0.124065559 |
| RP11-283I3 | -0.124108519 |
| ANKRD32 | -0.124109598 |
| RP11-723J4 | -0.12411461 |
| SEPSECS | -0.124125636 |
| ATAD5 | -0.124209658 |
| CTD-2002H8 | -0.124211966 |
| IGLV3-12 | -0.124238014 |
| MIR548AA1 | -0.124240622 |
| RP11-430C7 | -0.124254832 |
| RP11-325E5 | -0.124298999 |
| POLG2 | -0.124299486 |
| IGHV3-54 | -0.12430785 |
| FRMPD2L2 | -0.12431109 |
| LYZ | -0.124329571 |
| MBTD1 | -0.124392141 |
| RP11-377G16 | -0.124405954 |
| ZFC3H1 | -0.12441763 |
| IL12B | -0.124569946 |
| ARID3B | -0.124593601 |
| MBP | -0.124641547 |
| CCNE2 | -0.124665325 |
| AC093818 | -0.124668265 |
| OTUD6B-AS1 | -0.124670758 |
| CYBB | -0.124703261 |
| BANK1 | -0.124709534 |
| PHF21A | -0.124735226 |
| KLRC4 | -0.124789736 |
| RP11-335G20 | -0.12480398 |
| STAG3 | -0.124824402 |
| ZBTB14 | -0.124845575 |
| LIMD1-AS1 | -0.124872909 |
| SIT1 | -0.124875455 |
| IGHV3-57 | -0.124917294 |
| RP13-126P21 | -0.124951361 |
| TRGV5P | -0.124999 |
| SDAD1 | -0.125035927 |
| IGKV1OR-3 | -0.12504 |
| IGKV6D-41 | -0.125068971 |

| | |
|---|---|
| LINC00996 | -0.125098047 |
| TDRD7 | -0.125177701 |
| RP11-693N9 | -0.125180752 |
| TRGJP2 | -0.125199144 |
| PNOC | -0.125203346 |
| GAS8-AS1 | -0.125219932 |
| TRBV8-2 | -0.125222671 |
| TBK1 | -0.125265426 |
| RQCD1 | -0.125272552 |
| TRBV10-2 | -0.125296503 |
| RNU6-1149P | -0.12533217 |
| AL353662 | -0.125363534 |
| VPS33A | -0.125403925 |
| RP11-57K17 | -0.125417775 |
| CTD-2026C7 | -0.125521672 |
| HMGB1P17 | -0.125547168 |
| MYH6 | -0.12559351 |
| ELK2AP | -0.125667148 |
| CNTRL | -0.125705315 |
| AC244205 | -0.125718146 |
| RP11-278A23 | -0.125737856 |
| RP11-468E2 | -0.125786379 |
| ZMYM5 | -0.125787979 |
| AC020595 | -0.12581656 |
| SIGLEC5 | -0.125876443 |
| TBC1D3L | -0.125883515 |
| ZNF44 | -0.125889397 |
| TNFRSF13C | -0.125913948 |
| IGHV1OR15-3 | -0.125916193 |
| UNC13A | -0.125941309 |
| ZFYVE28 | -0.125985885 |
| PPP6R3 | -0.126046018 |
| TIFAB | -0.12604685 |
| TRAV25 | -0.126067552 |
| PSMD10P1 | -0.126085105 |
| CD53 | -0.126086131 |
| NCR3 | -0.126094998 |
| TRGC1 | -0.12610806 |
| SNX20 | -0.126165912 |
| RP11-164H13 | -0.126176772 |
| SERPINB9 | -0.126186508 |
| ZNF804A | -0.126188297 |
| RP11-493L12 | -0.126202672 |
| PDCD1 | -0.126229294 |
| CDK17 | -0.126247943 |

| | |
|---|---|
| GBP5 | -0.126255925 |
| IGHD3-10 | -0.126258005 |
| ZBED2 | -0.126307275 |
| ZNF852 | -0.126309397 |
| STX16-NPEPL1 | -0.126310384 |
| IGHV3-47 | -0.126312483 |
| RBL2 | -0.126359129 |
| ORMDL1 | -0.12641316 |
| IL26 | -0.126430833 |
| GDPD4 | -0.126436258 |
| TRDV2 | -0.126453626 |
| TRAV35 | -0.12645383 |
| ZNF557 | -0.126515118 |
| RP11-154J22 | -0.126565932 |
| PAN2 | -0.126605745 |
| MIR4737 | -0.126628478 |
| ANKRD12 | -0.126648947 |
| ANKRD49 | -0.126660278 |
| IGHV1-14 | -0.126686181 |
| IGKV1OR2-11 | -0.126798705 |
| XCL1 | -0.127025997 |
| AC022182 | -0.127059979 |
| RP11-10J5 | -0.127072871 |
| RASSF5 | -0.12709239 |
| IGHD1-1 | -0.127110542 |
| RP5-1029K10 | -0.127202727 |
| RP11-82K18 | -0.127204148 |
| AC104634 | -0.127267754 |
| TMEM194A | -0.127289924 |
| CPEB3 | -0.127293924 |
| KIF21A | -0.127294482 |
| GLCCI1 | -0.127372867 |
| HSPA8P15 | -0.127437174 |
| KLRB1 | -0.12745862 |
| C9orf139 | -0.127459308 |
| CMTR1 | -0.127498193 |
| RNU6-1069P | -0.127516023 |
| ARHGAP15 | -0.127569936 |
| TRAV30 | -0.127575965 |
| RPL9P33 | -0.127577453 |
| PAX5 | -0.127579253 |
| PHBP5 | -0.127593142 |
| IGLV8-61 | -0.127606901 |
| CEP95 | -0.127661252 |
| LINC01388 | -0.12767676 |

| | |
|---|---|
| DENND1C | -0.127765052 |
| IGKV1OR9-2 | -0.127770833 |
| LINC00649 | -0.127775713 |
| BAZ2A | -0.127806877 |
| XCR1 | -0.127809363 |
| snoU2_19 | -0.127826816 |
| LAT | -0.127840348 |
| GRSF1 | -0.127852218 |
| PRF1 | -0.127906137 |
| PGBD4 | -0.127922752 |
| SASH3 | -0.127944613 |
| CPNE5 | -0.127948669 |
| IGHVII-30-1 | -0.127961365 |
| TSHR | -0.127961568 |
| MPHOSPH9 | -0.127965172 |
| DIS3L | -0.127973824 |
| RP11-4M23 | -0.12801336 |
| TRAJ30 | -0.128029636 |
| BATF2 | -0.128046273 |
| CXorf38 | -0.128106685 |
| RP4-671O14 | -0.128118537 |
| ABCA5 | -0.128180568 |
| RP11-1166P10 | -0.128250557 |
| KARSP2 | -0.128306772 |
| SLC9A8 | -0.128307289 |
| RNA5SP383 | -0.128342401 |
| RP11-426L16 | -0.12834826 |
| ARHGAP25 | -0.128423592 |
| CWC25 | -0.128438923 |
| RP11-203M5 | -0.128440927 |
| FNBP4 | -0.128459464 |
| RP11-624C23 | -0.128497158 |
| NPIPB5 | -0.128559431 |
| RP5-1028K7 | -0.128588802 |
| INO80 | -0.128613982 |
| ELL3 | -0.128641406 |
| RNF157 | -0.128696907 |
| ZBTB39 | -0.128697576 |
| IRF2 | -0.128721726 |
| CTB-133G6 | -0.128733822 |
| NPHS1 | -0.128776323 |
| TRAJ39 | -0.128841217 |
| FBXO33 | -0.128845415 |
| HPS5 | -0.128849918 |
| ANK1 | -0.128879064 |

| | | |
|---|---|---|
| DUSP26 | | -0.128888154 |
| LAG3 | | -0.12889351 |
| P2RY8 | | -0.128912339 |
| AC005546 | | -0.128950669 |
| CRACR2A | | -0.12896222 |
| TRAV8-4 | | -0.12896312 |
| HERC2P2 | | -0.129011209 |
| IGLV3-13 | | -0.129059041 |
| DEF6 | | -0.129071881 |
| CD160 | | -0.129091231 |
| CACNG3 | | -0.129169894 |
| TRAJ4 | | -0.129190328 |
| IGHV3OR15-7 | | -0.129229601 |
| IGLV7-35 | | -0.129245936 |
| PNN | | -0.129267369 |
| CLNK | | -0.129280772 |
| RP11-53I6 | | -0.129367394 |
| ZBTB25 | | -0.12946561 |
| MEP1A | | -0.12946885 |
| XXbac-BPG299F13 | | -0.129491341 |
| MTMR10 | | -0.129528087 |
| NCLP1 | | -0.129533566 |
| RAB8A | | -0.129599904 |
| IGHVIII-47-1 | | -0.129636288 |
| IGKV1OR22-1 | | -0.129724179 |
| IGLV1-50 | | -0.1297706 |
| IGHVII-22-1 | | -0.12984348 |
| FIP1L1 | | -0.129871153 |
| CDH7 | | -0.129874189 |
| TRAJ37 | | -0.129933695 |
| TRBV16 | | -0.129959192 |
| APOBEC3G | | -0.129972311 |
| IGLJCOR18 | | -0.130019764 |
| | 1-Sep | -0.130042007 |
| GIMAP2 | | -0.130046324 |
| CCL5 | | -0.130084874 |
| FAM117A | | -0.130089637 |
| TRBV4-1 | | -0.130092253 |
| WNT1 | | -0.130099818 |
| YEATS4 | | -0.130118811 |
| SACM1L | | -0.130123871 |
| PLEKHF2 | | -0.130124337 |
| ITSN2 | | -0.130146105 |
| IGKV2D-26 | | -0.130160835 |
| IGLV1-62 | | -0.130177734 |

| | |
|---|---|
| ERICH6B | -0.13024145 |
| TRAJ11 | -0.130363513 |
| ARL14EP | -0.130374487 |
| HAUS3 | -0.130389204 |
| RP11-87H9 | -0.130417149 |
| NAIP | -0.13042743 |
| BNIP3P9 | -0.130440033 |
| ARID4A | -0.130584189 |
| CLK1 | -0.130587222 |
| RP5-1073O3 | -0.130676918 |
| IGKV2D-18 | -0.130799254 |
| YTHDC1 | -0.130808385 |
| P2RX1 | -0.130818435 |
| AL590222 | -0.130886688 |
| RP1-151B14 | -0.130966373 |
| CD69 | -0.130984449 |
| RP11-152F13 | -0.131001366 |
| SMCHD1 | -0.131020971 |
| ZNF587 | -0.131052005 |
| PPP1R16B | -0.131071169 |
| RP11-223C24 | -0.131085499 |
| AC010761 | -0.131113887 |
| BCL2 | -0.131123953 |
| LINC00544 | -0.131130663 |
| MIR8071-1 | -0.131154691 |
| JADE1 | -0.131202542 |
| OAS3 | -0.131257558 |
| GABPB1 | -0.131339795 |
| FRYL | -0.131352688 |
| NKG7 | -0.131387518 |
| RP11-731C17 | -0.131416691 |
| IGKV2-28 | -0.131444859 |
| PIK3CG | -0.131468303 |
| IGKV1D-12 | -0.131477978 |
| THAP9 | -0.13150209 |
| AC009095 | -0.131524456 |
| ITIH1 | -0.131539526 |
| IGKV2D-24 | -0.13157574 |
| AL359753 | -0.131586089 |
| SUCNR1 | -0.131600011 |
| U6 | -0.131620995 |
| AL928768 | -0.13162711 |
| IGHV1OR15-9 | -0.131732626 |
| PTPN6 | -0.131736928 |
| IGHV1-17 | -0.131761012 |

| | |
|---|---|
| TXK | -0.131776006 |
| HESX1 | -0.131907094 |
| CTD-2583A14 | -0.131959896 |
| IGKV2-24 | -0.131961664 |
| MAP4K1 | -0.131988753 |
| C17orf75 | -0.1319972 |
| AC246787 | -0.132013412 |
| BHLHA15 | -0.132102953 |
| PRIM1 | -0.13213583 |
| IGHV7-56 | -0.132161762 |
| NR4A2 | -0.132175031 |
| SYNRG | -0.132210958 |
| SPCS3 | -0.132248374 |
| TRBV23-1 | -0.13225152 |
| IGHV3OR16-11 | -0.132347278 |
| ZAP70 | -0.132422317 |
| RP11-327E2 | -0.132426391 |
| C18orf8 | -0.132433573 |
| AGAP4 | -0.132449108 |
| TRBV28 | -0.132476328 |
| XXYLT1-AS2 | -0.132539291 |
| CD28 | -0.132541503 |
| FCHSD2 | -0.132596017 |
| MS4A6A | -0.13259892 |
| SLAIN1 | -0.132616839 |
| E2F3P1 | -0.132638463 |
| PDIK1L | -0.132668448 |
| RP11-121C2 | -0.132668969 |
| KIR2DL1 | -0.132758923 |
| IGHV3-36 | -0.132902782 |
| ITGAD | -0.132941536 |
| IGHJ1 | -0.133033792 |
| CTA-212A2 | -0.133065117 |
| SRSF10 | -0.133087741 |
| IGHV3OR16-15 | -0.133160353 |
| IGLV3-29 | -0.133172517 |
| RNF31 | -0.133244567 |
| RNF6 | -0.133299676 |
| IGLV6-57 | -0.133303877 |
| AP4B1-AS1 | -0.13331478 |
| BIRC3 | -0.133328812 |
| AP001107 | -0.13338451 |
| WDR76 | -0.133395271 |
| CACYBPP3 | -0.13348193 |
| LINC01215 | -0.133601004 |

| | |
|---|---|
| ZNF101 | -0.133701201 |
| CHRM3-AS2 | -0.133701802 |
| CTR9 | -0.133721853 |
| FAIM3 | -0.1337986 |
| LCK | -0.133848016 |
| RP11-61O1 | -0.133933965 |
| LINC00892 | -0.133967129 |
| IGKV1D-42 | -0.133989551 |
| CH17-385C13 | -0.134059383 |
| SLC17A9 | -0.134069998 |
| IGKV1OR2-2 | -0.134147145 |
| CD22 | -0.134150783 |
| MTERF4 | -0.134152551 |
| CTD-3014M21 | -0.134181196 |
| ATP2A3 | -0.13424608 |
| ZSCAN29 | -0.134303236 |
| NANOGP5 | -0.134316479 |
| CD200R1 | -0.134332075 |
| IGHJ2 | -0.13437387 |
| ATXN7L3B | -0.134377528 |
| TRAV13-2 | -0.134398467 |
| SNX10 | -0.134425865 |
| ZBTB20-AS1 | -0.134431183 |
| P2RY13 | -0.13446462 |
| TRAF3IP3 | -0.134596868 |
| SELL | -0.134658285 |
| RPGR | -0.134666657 |
| NUP210 | -0.134716371 |
| SPIRE2 | -0.134747535 |
| AC125232 | -0.13483005 |
| RAB39B | -0.134878499 |
| PRAMENP | -0.134884586 |
| RP11-96D1 | -0.134908361 |
| TRA2A | -0.134924678 |
| RP11-512C24 | -0.134927039 |
| TRAV8-1 | -0.134935784 |
| RP11-731F5 | -0.134940248 |
| XCL2 | -0.135105994 |
| LUC7L3 | -0.135109207 |
| RASSF3 | -0.13524512 |
| KLHL32 | -0.135435387 |
| SIGLEC10 | -0.135511438 |
| AC026202 | -0.135529246 |
| RNU6-850P | -0.135562848 |
| ZBTB2 | -0.135633174 |

| | |
|---|---|
| TRGV9 | -0.135652412 |
| RP1-71H24 | -0.135663987 |
| CD244 | -0.135741961 |
| IL12RB1 | -0.13578706 |
| DUSP28 | -0.135919475 |
| RFFL | -0.135940666 |
| RAPGEF6 | -0.135957393 |
| SIGLEC8 | -0.136003348 |
| RP5-1180D12 | -0.136101883 |
| IGHD2-2 | -0.136112163 |
| MIS18BP1 | -0.136265946 |
| HNRNPA3P11 | -0.136293656 |
| ZNF236 | -0.136420353 |
| GK | -0.136462364 |
| PSMB8-AS1 | -0.136550572 |
| SNX18P7 | -0.136563863 |
| TRBV6-6 | -0.136569881 |
| RP11-352G18 | -0.136622448 |
| MPEG1 | -0.136662819 |
| IGHV7-27 | -0.136702691 |
| IGLV7-46 | -0.136703596 |
| MIXL1 | -0.136810733 |
| KEL | -0.13683986 |
| TRAV34 | -0.13686946 |
| TRAPPC12-AS1 | -0.13695598 |
| IGKV5-2 | -0.13705114 |
| HEMGN | -0.137194453 |
| IRF1 | -0.137210564 |
| AC104699 | -0.137244951 |
| IGHD5-12 | -0.137268817 |
| APOBEC3H | -0.137276706 |
| HSPA8P20 | -0.137278276 |
| USO1 | -0.137285153 |
| IGKV1D-27 | -0.137303571 |
| LINC00852 | -0.137341454 |
| ZNF700 | -0.137392172 |
| PARP12 | -0.137414406 |
| RP1-167O22 | -0.137473645 |
| IGLV2-33 | -0.137495395 |
| SIGLEC12 | -0.137552906 |
| BTN3A3 | -0.137608535 |
| IGHV4-61 | -0.137629941 |
| GIMAP5 | -0.137649802 |
| ITGB7 | -0.13765116 |
| RP5-1007H16 | -0.137720883 |

| | |
|---|---|
| RP11-327F22 | -0.137819184 |
| RP11-361C13 | -0.137827262 |
| IGHV4-55 | -0.137830243 |
| HNRNPA3P2 | -0.137880083 |
| RP11-71L14 | -0.137922914 |
| IGKV1OR2-6 | -0.138060843 |
| ARHGAP11B | -0.138064406 |
| PRPF38B | -0.138067085 |
| GPR65 | -0.138099105 |
| IGHV4-31 | -0.138145227 |
| SUZ12 | -0.138198812 |
| C5orf56 | -0.138266778 |
| AC244250 | -0.138399692 |
| BIN2 | -0.138469354 |
| LRMP | -0.13854586 |
| CD3D | -0.138577675 |
| ATP1A3 | -0.138593918 |
| HNRNPA1P34 | -0.138679071 |
| ADAMDEC1 | -0.138679075 |
| HMSD | -0.138692625 |
| TRAV20 | -0.138712686 |
| CCR7 | -0.13871623 |
| TRIM69 | -0.138790671 |
| RP11-707P17 | -0.138954048 |
| SEC11C | -0.138975981 |
| IGLV5-48 | -0.139004896 |
| LAP3 | -0.139121104 |
| HCP5 | -0.139143207 |
| KLHL14 | -0.139157796 |
| KLRC1 | -0.139168409 |
| IGKV2-29 | -0.139180331 |
| RP1-148H17 | -0.139252909 |
| TRAJ41 | -0.139320664 |
| SPN | -0.13936584 |
| RP11-416N2 | -0.139388752 |
| SYVN1 | -0.139487557 |
| AC116366 | -0.139508265 |
| IGLV3-32 | -0.13958231 |
| IGHV1-69 | -0.139588873 |
| PTPRN2 | -0.139618614 |
| GFI1 | -0.139653034 |
| TMC8 | -0.139728126 |
| ABCD2 | -0.139747209 |
| BTN3A2 | -0.139788649 |
| TFEC | -0.139866848 |

| | |
|---|---|
| LYSMD2 | -0.139882979 |
| AMPD3 | -0.139925716 |
| RP11-325K4 | -0.139935558 |
| NFATC2 | -0.139974481 |
| MIR650 | -0.140040028 |
| CCL19 | -0.140118717 |
| CTD-2006C1 | -0.140173206 |
| TRBV15 | -0.140209042 |
| HSPC047 | -0.140282457 |
| FAM172BP | -0.140295692 |
| KLHL6 | -0.140339036 |
| ALDH5A1 | -0.140431133 |
| IGHV4-4 | -0.14064907 |
| OXNAD1 | -0.140769362 |
| PAIP1P1 | -0.140831052 |
| TESPA1 | -0.140852194 |
| RINL | -0.140856268 |
| CDKN2AIP | -0.14087666 |
| SMAP2 | -0.140969817 |
| CTC-281F24 | -0.140990905 |
| IGHVIII-51-1 | -0.141009545 |
| RP11-275I4 | -0.141013171 |
| PAPD5 | -0.141077208 |
| TRAV40 | -0.141113337 |
| CD247 | -0.141120447 |
| IGKV2D-30 | -0.141146207 |
| HMGB2 | -0.141154328 |
| KLRG1 | -0.141200469 |
| CEP152 | -0.141224173 |
| RP11-812E19 | -0.141262872 |
| IL2 | -0.141304196 |
| SKAP1 | -0.141371156 |
| SLFN12L | -0.141383114 |
| RP11-395I6 | -0.141435475 |
| RP1-38C16 | -0.141463114 |
| RP11-121A8 | -0.141499552 |
| PSME1 | -0.14150746 |
| CNR2 | -0.141522236 |
| IGKV3D-7 | -0.141567315 |
| TRAV8-6 | -0.141694078 |
| IGKV3D-20 | -0.14174461 |
| RPL32P1 | -0.141812267 |
| ADAM28 | -0.141934306 |
| TUBG1P | -0.141968383 |
| IGLV3-6 | -0.1420382 |

| | |
|---|---|
| CXorf65 | -0.142069367 |
| HMGB3P32 | -0.142214432 |
| ZNF233 | -0.14227194 |
| IGHV3-22 | -0.142315902 |
| IGLV2-28 | -0.14240702 |
| CCL4 | -0.142450255 |
| BEND4 | -0.142499625 |
| TRAV26-1 | -0.142645603 |
| TRAJ6 | -0.142673481 |
| RP11-560A15 | -0.142763846 |
| BTN3A1 | -0.142819201 |
| CD8B | -0.142839034 |
| CR2 | -0.142847674 |
| RP11-476B1 | -0.142850012 |
| ABCG1 | -0.142860449 |
| IGHVII-40-1 | -0.142873864 |
| KIR2DL4 | -0.142904784 |
| SLC4A10 | -0.142930306 |
| IGKV3-15 | -0.142997739 |
| SCARNA9 | -0.143016214 |
| IGHVIII-67-2 | -0.143136915 |
| ZNF367 | -0.143262365 |
| TRAV2 | -0.143272472 |
| IGKV2D-40 | -0.143278307 |
| PTPN7 | -0.14329738 |
| CNOT6L | -0.143400938 |
| GPRIN3 | -0.143452047 |
| PAN3 | -0.143526172 |
| SETDB2 | -0.143575848 |
| LIN7C | -0.143610882 |
| PARD6B | -0.143874782 |
| CTD-2035E11 | -0.143929482 |
| FCGR2C | -0.143953911 |
| SCML4 | -0.143993491 |
| IL21R-AS1 | -0.144019409 |
| RNA5SP364 | -0.144024748 |
| GZMAP1 | -0.14404575 |
| MPHOSPH8 | -0.144059497 |
| IGHV1OR16-1 | -0.144099591 |
| STAP1 | -0.144108129 |
| KLRD1 | -0.144128003 |
| IL2RG | -0.144178089 |
| CD48 | -0.144204564 |
| AC006129 | -0.144221644 |
| PRKCB | -0.144256027 |

| | | |
|---|---|---|
| DCLRE1CP1 | | -0.144259849 |
| IGKV7-3 | | -0.144280113 |
| IKZF1 | | -0.14429828 |
| IGKV1OR1-1 | | -0.144305097 |
| TMPO | | -0.144365371 |
| IGHV2-70 | | -0.144489838 |
| RP11-640L9 | | -0.144527741 |
| IGHV3OR16-7 | | -0.144595599 |
| TADA2A | | -0.144675264 |
| LCP2 | | -0.144693683 |
| IGLC7 | | -0.144774318 |
| TRBV5-1 | | -0.144967831 |
| IL18RAP | | -0.145028607 |
| KAT2B | | -0.145072714 |
| TRAJ60 | | -0.145104394 |
| IGLC3 | | -0.145145211 |
| TRAV29DV5 | | -0.145207271 |
| ZNF37BP | | -0.145293412 |
| CLECL1 | | -0.145300985 |
| RP11-204N11 | | -0.145398979 |
| LL22NC03-24A12 | | -0.145401957 |
| IGLV3-27 | | -0.145406461 |
| GCH1 | | -0.145424339 |
| KIAA1551 | | -0.145491411 |
| TRGV10 | | -0.145578396 |
| DDX60 | | -0.145707998 |
| ZBP1 | | -0.145710323 |
| PTGES3P1 | | -0.145716686 |
| IGHV4-28 | | -0.145762641 |
| MIR5195 | | -0.145778663 |
| IGLV4-3 | | -0.145885049 |
| RP5-1092A11 | | -0.14590673 |
| RP11-17M24 | | -0.145933062 |
| CCDC141 | | -0.145996124 |
| CXCR6 | | -0.146008848 |
| IGLV5-45 | | -0.146010093 |
| LINC01055 | | -0.146011722 |
| FOXB1 | | -0.146036913 |
| EVI2B | | -0.146076777 |
| IGHV3OR16-6 | | -0.146119188 |
| UTP3 | | -0.146368268 |
| | 1-Mar | -0.146379153 |
| IL10RA | | -0.146382464 |
| IGHD1-14 | | -0.146420628 |
| RP11-405M12 | | -0.146489353 |

| | |
|---|---|
| ZC3H12D | -0.146497837 |
| DENND2D | -0.14653596 |
| GPR171 | -0.146557791 |
| IGLJ2 | -0.146574251 |
| CTB-96E2 | -0.146634797 |
| CBFA2T3 | -0.146655295 |
| IPCEF1 | -0.146660356 |
| SLC12A3 | -0.146664987 |
| IGLV5-52 | -0.146670796 |
| SCIMP | -0.146749364 |
| RIMBP3 | -0.146800368 |
| IGKV2D-29 | -0.146835773 |
| LIPT1 | -0.14686736 |
| TRAV14DV4 | -0.146972516 |
| TRAV6 | -0.147026122 |
| RNU6-623P | -0.147046387 |
| TRAV1-2 | -0.147085328 |
| RHOH | -0.147086402 |
| OR4C6 | -0.14713866 |
| RP11-652L8 | -0.147139865 |
| IGLV3-25 | -0.147156407 |
| DCAF11 | -0.147343158 |
| TAGAP | -0.147436762 |
| TRAC | -0.14744253 |
| CTD-2011F17 | -0.147467649 |
| SNORA38B | -0.147490898 |
| HELQ | -0.147569158 |
| TRBV19 | -0.147959358 |
| CD5 | -0.148053328 |
| HSF5 | -0.148070107 |
| TRBV3-1 | -0.148180754 |
| RP11-291B21 | -0.148181349 |
| CEP135 | -0.1481978 |
| IGHV3-76 | -0.148236724 |
| AD000864 | -0.148277413 |
| TRBV2 | -0.14829849 |
| ZCCHC2 | -0.148360114 |
| IGHV2-26 | -0.148368504 |
| IGHJ1P | -0.148532641 |
| IGHV1-3 | -0.148572474 |
| TRAV41 | -0.14857879 |
| IGHV3-60 | -0.148610104 |
| CENPC | -0.148693208 |
| IGKV1OR2-108 | -0.148701778 |
| TRAV21 | -0.148805913 |

| | |
|---|---|
| IGLV2-18 | -0.148871591 |
| IGHV3-49 | -0.149000476 |
| TRBV29-1 | -0.149025368 |
| IGHV3-72 | -0.149098873 |
| VTN | -0.149110222 |
| IGHV6-1 | -0.149138907 |
| ENOPH1 | -0.149205714 |
| TRBV24-1 | -0.149219447 |
| IGHV1-12 | -0.149298549 |
| CTD-2501E16 | -0.149310529 |
| CD96 | -0.149310649 |
| CTB-186H2 | -0.149396186 |
| RIMBP3B | -0.149405169 |
| TRBC2 | -0.149433891 |
| IGHJ3 | -0.149613568 |
| PNRC2 | -0.149791879 |
| IGLV11-55 | -0.149791954 |
| RP3-477O4 | -0.149874947 |
| CTD-2506J14 | -0.150143599 |
| AC010458 | -0.150265191 |
| RN7SL834P | -0.15030514 |
| IGHV3OR16-12 | -0.150326751 |
| ANTXRLP1 | -0.15033194 |
| IGHV1-58 | -0.150459138 |
| IGKV1OR-2 | -0.150461774 |
| IGHV3-20 | -0.150700141 |
| FCRLA | -0.150731274 |
| TRBV6-4 | -0.1511391 |
| FGL2 | -0.151141129 |
| TBX21 | -0.151168147 |
| LINC00402 | -0.15139224 |
| AC096579 | -0.15141711 |
| LINC00909 | -0.151482112 |
| RP11-362A9 | -0.15157705 |
| AC103563 | -0.151671487 |
| KLRC2 | -0.151698461 |
| MYO7A | -0.151702531 |
| IGLV10-54 | -0.151745575 |
| IGHV3-53 | -0.151814291 |
| ICAM3 | -0.151879896 |
| JAKMIP1 | -0.151988655 |
| ST3GAL1P1 | -0.152095888 |
| IGKV3-7 | -0.152128507 |
| RP4-647J21 | -0.152140206 |
| NBPF2P | -0.152217852 |

| | |
|---|---|
| CXCR3 | -0.152317441 |
| ASAH2 | -0.152399322 |
| IL17REL | -0.152484457 |
| RP11-325F22 | -0.152504826 |
| TBC1D10C | -0.152546363 |
| RP11-436I9 | -0.152707517 |
| ZNF683 | -0.152749042 |
| HEATR9 | -0.152757849 |
| UBASH3A | -0.152779213 |
| ICOS | -0.152917847 |
| RP11-546D6 | -0.15306419 |
| IGHG4 | -0.153190745 |
| NUAK2 | -0.153310062 |
| IGHV3-13 | -0.153333139 |
| MIR8071-2 | -0.153407972 |
| TBC1D27 | -0.153445259 |
| RP11-700J17 | -0.153586877 |
| RP11-87G24 | -0.153710779 |
| LIN54 | -0.153737485 |
| TRBV6-5 | -0.153793693 |
| IGKV2OR22-3 | -0.153838893 |
| STAT1 | -0.153956062 |
| KIR3DL2 | -0.15396674 |
| PATL2 | -0.154061834 |
| SLA2 | -0.154111232 |
| IGKV3D-11 | -0.154234727 |
| IGHV1-67 | -0.154268105 |
| IGHV3OR16-8 | -0.154290758 |
| IGHV1-69-2 | -0.154355596 |
| CD19 | -0.154385988 |
| TRAV3 | -0.154399132 |
| MIR4537 | -0.154425073 |
| IGHV3-62 | -0.154428117 |
| IGKV3D-15 | -0.154478504 |
| IGLV3-24 | -0.154596675 |
| IGLL1 | -0.15462538 |
| C11orf21 | -0.154630098 |
| IGLV1-40 | -0.154746388 |
| IGHV4-34 | -0.154886199 |
| ITGAL | -0.154967679 |
| CYTIP | -0.154982954 |
| IGKV2OR22-4 | -0.155088763 |
| RP11-294C11 | -0.155146778 |
| TTC24 | -0.155195079 |
| ANKRD44 | -0.155259598 |

| | |
|---|---|
| TRBV13 | -0.15527873 |
| IGKV1-6 | -0.155309289 |
| BIN2P1 | -0.155309989 |
| IGHV3-33 | -0.155340081 |
| IGLV1-47 | -0.1554292 |
| ZNF397 | -0.15550867 |
| IGHV3-11 | -0.155606679 |
| CCDC64 | -0.155610684 |
| RP11-18H21 | -0.155628514 |
| SLAMF1 | -0.155686455 |
| IGHV5-51 | -0.155763349 |
| FTH1P22 | -0.15577996 |
| GNAO1 | -0.155791101 |
| FAM217B | -0.155850973 |
| IGKV2-40 | -0.155939702 |
| SEMA4D | -0.155996059 |
| GIMAP4 | -0.156060396 |
| IGKV1OR2-9 | -0.156153055 |
| RP11-148O21 | -0.156217404 |
| TRBV6-1 | -0.156254893 |
| APOL3 | -0.156258228 |
| TRBV20-1 | -0.15633491 |
| KLRK1 | -0.156364331 |
| TRBV5-6 | -0.156407702 |
| SIRPG | -0.156509744 |
| IGKV1D-33 | -0.156565765 |
| TRBV4-2 | -0.156637025 |
| FCRL6 | -0.156645504 |
| TLR10 | -0.156698825 |
| TRAV27 | -0.156738184 |
| TRBV25-1 | -0.156753712 |
| SULT1C2 | -0.156779548 |
| IGHV3-52 | -0.156858481 |
| CD3E | -0.156889568 |
| GZMH | -0.157093725 |
| ZCCHC8 | -0.157125448 |
| PCED1B-AS1 | -0.157131811 |
| LINC00299 | -0.157165029 |
| TRAV24 | -0.157202625 |
| IGKV1D-16 | -0.157275731 |
| IGLV3-1 | -0.157303097 |
| CD40LG | -0.157379221 |
| CRTAM | -0.157402651 |
| IGKV1-17 | -0.157499168 |
| PTPRC | -0.157529056 |

| | |
|---|---|
| BHLHE40-AS1 | -0.157607906 |
| RP11-441F2 | -0.157634643 |
| IGHG3 | -0.157661453 |
| IGLV3-19 | -0.157734219 |
| TRGV7 | -0.157872996 |
| CD2 | -0.15800303 |
| RP11-89K10 | -0.158018397 |
| TRBV11-2 | -0.158031254 |
| LINC00494 | -0.158067152 |
| IGKV2-26 | -0.158157334 |
| TRBV7-6 | -0.158214439 |
| CXCR5 | -0.158270483 |
| PLA1A | -0.158326096 |
| BLK | -0.158343521 |
| LRRC16B | -0.158409095 |
| PPP1R2P9 | -0.15846525 |
| BTN2A2 | -0.158494754 |
| IGHV3OR16-9 | -0.158593359 |
| IGKV1D-13 | -0.158704141 |
| IGLV3-10 | -0.158810743 |
| IGHV3-7 | -0.158820265 |
| IGHV3-73 | -0.158823351 |
| BTLA | -0.158929077 |
| IGHV5-78 | -0.158939125 |
| TRG-AS1 | -0.159049495 |
| CCR5 | -0.159261914 |
| CTD-2313F11 | -0.159391109 |
| TRBV5-5 | -0.15956374 |
| CARD8 | -0.159617259 |
| IGKV6D-21 | -0.159789009 |
| IGKV1OR10-1 | -0.159913107 |
| FYB | -0.159966447 |
| IGLV2-8 | -0.160068772 |
| IGLV3-22 | -0.160158327 |
| TRAV17 | -0.160164818 |
| IGKV3-11 | -0.160178231 |
| TRAV12-2 | -0.160328484 |
| TRBV18 | -0.16039139 |
| AMPD1 | -0.160439643 |
| SLTM | -0.160440086 |
| KLRC3 | -0.160492806 |
| IGSF6 | -0.16053276 |
| IGHV4OR15-8 | -0.160554253 |
| IGLV7-43 | -0.160700081 |
| TRGV2 | -0.160711096 |

| | |
|---|---|
| IGKV1OR22-5 | -0.161055037 |
| TIGIT | -0.161068784 |
| RP11-202G18 | -0.161105458 |
| KLRC4-KLRK1 | -0.161307051 |
| ENAM | -0.161407366 |
| IGKV1-13 | -0.161450856 |
| FCRL3 | -0.161452958 |
| IGHV2-5 | -0.161453928 |
| IGKV1-5 | -0.161458743 |
| IGKV1OR2-3 | -0.161472313 |
| TRAV12-3 | -0.161519025 |
| GZMA | -0.161529362 |
| RP11-960L18 | -0.16153988 |
| TIGD3 | -0.161541065 |
| PARP15 | -0.161573364 |
| EPB41 | -0.161736329 |
| TRAV22 | -0.161775724 |
| TRAV13-1 | -0.161804882 |
| IGKV1-8 | -0.161860068 |
| CXCL9 | -0.161957995 |
| CD3G | -0.162197931 |
| ANKRD55 | -0.162205059 |
| CLEC17A | -0.162231146 |
| RP11-170L3 | -0.162298875 |
| GSAP | -0.162347072 |
| CLIC5 | -0.162353361 |
| IGHG1 | -0.162491079 |
| CTD-2547L24 | -0.162658788 |
| SPAG5-AS1 | -0.162660035 |
| EOMES | -0.162761413 |
| IGLJ3 | -0.162867636 |
| TRAV4 | -0.1628785 |
| IGHV1OR21-1 | -0.162923816 |
| IGLC2 | -0.163027596 |
| KIR2DL3 | -0.163187698 |
| IL2RB | -0.163213771 |
| RP1-13D10 | -0.163432919 |
| IGHV1OR15-2 | -0.16350586 |
| TRAV16 | -0.163578076 |
| IGKV1D-43 | -0.163847478 |
| TNFRSF13B | -0.163917389 |
| THAP6 | -0.164025044 |
| RP11-637A17 | -0.164246384 |
| SIRPG-AS1 | -0.164254861 |
| LINC00426 | -0.164271304 |

| | |
|---|---|
| TRBV12-4 | -0.164309075 |
| TRBV10-3 | -0.164359875 |
| IGKV1-33 | -0.164380217 |
| IGHV3OR16-13 | -0.164728981 |
| PPP1R3D | -0.164887649 |
| TRBV9 | -0.164961457 |
| IGKV1-27 | -0.164988705 |
| IGKV1D-8 | -0.165045407 |
| STK4 | -0.165245004 |
| RP11-255A11 | -0.16525271 |
| IGLV4-69 | -0.16538017 |
| IGHV7-81 | -0.16543126 |
| CECR1 | -0.165710792 |
| IGLV2-11 | -0.165756382 |
| TRBV30 | -0.165921305 |
| IGLV3-16 | -0.16609821 |
| IGJ | -0.166185436 |
| IGHV3-48 | -0.166195327 |
| KIAA0125 | -0.166338072 |
| LL22NC03-88E1 | -0.16667935 |
| IFNG-AS1 | -0.166743134 |
| PCAT29 | -0.166811837 |
| IGHV3OR16-10 | -0.166886228 |
| IGHD3-22 | -0.166899344 |
| TRAV8-3 | -0.167012622 |
| MS4A1 | -0.167076754 |
| IGKV3OR2-268 | -0.167104832 |
| RP1-47M23 | -0.167131235 |
| DTHD1 | -0.167306977 |
| IGKV1-16 | -0.167328159 |
| HSH2D | -0.167440561 |
| SPIC | -0.167491447 |
| SH2D1B | -0.167560772 |
| SP140 | -0.167602838 |
| TRBV7-9 | -0.167646323 |
| PYHIN1 | -0.167959207 |
| TRBV5-4 | -0.167998061 |
| VSIG10 | -0.16835755 |
| GIMAP7 | -0.168360992 |
| TRAV9-2 | -0.168416846 |
| FCRL1 | -0.168482843 |
| CCR2 | -0.168634586 |
| IGHV4-39 | -0.168703358 |
| RP11-109E24 | -0.168744724 |
| TRAV36DV7 | -0.168839254 |

| | |
|---|---|
| IGLL5 | -0.169004926 |
| IGHV3-19 | -0.169177483 |
| P2RY10 | -0.169693058 |
| RP11-402J7 | -0.169856565 |
| IFNG | -0.169915507 |
| IL21-AS1 | -0.169928943 |
| IGHG2 | -0.17002073 |
| CD27 | -0.170030349 |
| GCSAM | -0.170045438 |
| DCK | -0.170101943 |
| GBP4 | -0.170163192 |
| IGLC6 | -0.170398113 |
| IL21 | -0.170495053 |
| ITK | -0.170513068 |
| TRAV8-2 | -0.170765116 |
| POU2AF1 | -0.171058342 |
| SAMD3 | -0.171157546 |
| IGHV3-64 | -0.171249364 |
| RGL4 | -0.171289995 |
| IGHM | -0.171395042 |
| IGHV3-21 | -0.171712573 |
| IGHV3-37 | -0.171770013 |
| GPR114 | -0.171794464 |
| AL122127 | -0.171823912 |
| TRGC2 | -0.171971971 |
| CXCR2P1 | -0.17200736 |
| IRF4 | -0.172358358 |
| PLA2G2D | -0.172463593 |
| IGHV3-71 | -0.172466579 |
| IGHD6-25 | -0.172513831 |
| CD226 | -0.172619598 |
| LY9 | -0.172653962 |
| BRD2 | -0.172852036 |
| CD79A | -0.172861004 |
| GPR174 | -0.173116384 |
| IGHV3-43 | -0.173148423 |
| NCR1 | -0.173251073 |
| TRBV7-4 | -0.173360385 |
| IGKJ5 | -0.173440129 |
| AC010149 | -0.173456381 |
| PKHD1L1 | -0.17350346 |
| IGLV2-14 | -0.173804515 |
| IGHV3-41 | -0.173959369 |
| OSTN-AS1 | -0.174081285 |
| TRAV1-1 | -0.174358341 |

| | |
|---|---|
| IGHV3-35 | -0.174511214 |
| DGKE | -0.17460229 |
| IGHV1-46 | -0.174721129 |
| TRGV5 | -0.17472413 |
| IGKV1-39 | -0.174819461 |
| DMXL2 | -0.174906452 |
| AC104820 | -0.174930207 |
| TRBV27 | -0.174951944 |
| TRAV12-1 | -0.175146539 |
| CD38 | -0.175153588 |
| IGKV3-20 | -0.175377282 |
| TNFRSF17 | -0.175420751 |
| IGLC5 | -0.175455211 |
| IGHV3-63 | -0.175471841 |
| GZMK | -0.175736692 |
| BTN1A1 | -0.17578025 |
| ZNF831 | -0.175886879 |
| IGKC | -0.175943725 |
| RP11-81H14 | -0.176077527 |
| TRAV10 | -0.176118273 |
| RP11-664D1 | -0.17630372 |
| LINC00582 | -0.176435255 |
| IGLV2-23 | -0.17659817 |
| TMEM156 | -0.177077933 |
| IGLV2-5 | -0.177241133 |
| IGHV3-75 | -0.177252424 |
| RPL4P1 | -0.177285192 |
| IGHV3-38 | -0.17742443 |
| IRF9 | -0.177428293 |
| RORA-AS1 | -0.177491313 |
| FCRL2 | -0.17762094 |
| IGLV2-34 | -0.177802296 |
| IGKV4-1 | -0.178054478 |
| RP11-295P22 | -0.178259256 |
| SH2D1A | -0.178468226 |
| KCNA3 | -0.178481415 |
| C12orf42 | -0.178607362 |
| IGHV1-18 | -0.178626415 |
| LINC00861 | -0.178794353 |
| LA16c-390H2 | -0.179116561 |
| TRAT1 | -0.179265696 |
| IGHV3-30 | -0.179617289 |
| IGKV1-12 | -0.179763299 |
| ZNF80 | -0.179913015 |
| TOMM20P2 | -0.179978589 |

| | |
|---|---|
| RASA4CP | -0.180049357 |
| IGLV1-44 | -0.180060955 |
| IGKV1D-39 | -0.180146959 |
| TRGV4 | -0.180671361 |
| IGHV3-66 | -0.180919142 |
| GVINP1 | -0.180975276 |
| TRDV1 | -0.181241331 |
| LINC01484 | -0.181310512 |
| IGHV3-23 | -0.182432311 |
| IKZF3 | -0.182917445 |
| IGHJ3P | -0.18331594 |
| KIR3DX1 | -0.183555833 |
| FCRL5 | -0.184263624 |
| RP4-742C19 | -0.184454462 |
| MEI1 | -0.184497198 |
| BZRAP1-AS1 | -0.184598226 |
| CD8A | -0.184678397 |
| AC006369 | -0.184853408 |
| TRBV7-7 | -0.184905378 |
| SLAMF6 | -0.185022859 |
| TRAV19 | -0.185159056 |
| IGLC4 | -0.185392102 |
| GP1BA | -0.186144973 |
| RP11-768B22 | -0.186418725 |
| RP1-153P14 | -0.186741604 |
| GPR18 | -0.186770753 |
| C16orf54 | -0.186840259 |
| IGHGP | -0.187459542 |
| PASK | -0.187814888 |
| SLAMF7 | -0.188075532 |
| TRAV5 | -0.188180833 |
| IGKV2-30 | -0.188453468 |
| IGHV3-74 | -0.188507342 |
| TRBV14 | -0.188510435 |
| TRBV7-3 | -0.18927101 |
| IGKV1D-17 | -0.189629101 |
| IGHV3-42 | -0.191122086 |
| NUGGC | -0.191151774 |
| LINC00539 | -0.191394208 |
| RP11-290F5 | -0.191714687 |
| TRGV3 | -0.192015523 |
| AC104024 | -0.192872434 |
| IGLJ1 | -0.192875308 |
| HNRNPA1P70 | -0.193779956 |
| RP11-284N8 | -0.194206816 |

| | |
|---|---|
| CLEC2D | -0.194703488 |
| IGLV1-51 | -0.195256629 |
| THEMIS | -0.195973892 |
| LAX1 | -0.195985769 |
| IGHD3-9 | -0.196244453 |
| NLRC3 | -0.196355766 |
| IGHV3-15 | -0.196375683 |
| RP3-492J12 | -0.197501669 |
| ANKRD36BP2 | -0.198317949 |
| FAM46C | -0.19923299 |
| IGHV4-59 | -0.200028015 |
| MZB1 | -0.20414221 |
| TRBV12-3 | -0.205693664 |
| FASLG | -0.205912356 |
| ATP8A1 | -0.206015297 |
| RP11-16E12 | -0.206990056 |
| RAB37 | -0.20833832 |
| KCNJ10 | -0.216417708 |
| AC007386 | -0.224490962 |
| PIM2 | -0.230296195 |

| Spearman p | Spearman adj p |
|---|---|
| 2.91E-14 | 9.41E-10 |
| 3.95E-14 | 9.41E-10 |
| 3.00E-13 | 3.92E-09 |
| 3.30E-13 | 3.92E-09 |
| 6.80E-13 | 6.48E-09 |
| 1.12E-12 | 8.91E-09 |
| 1.43E-12 | 9.71E-09 |
| 2.49E-12 | 1.48E-08 |
| 2.79E-12 | 1.48E-08 |
| 3.12E-12 | 1.49E-08 |
| 9.85E-12 | 3.91E-08 |
| 1.32E-11 | 4.83E-08 |
| 2.21E-11 | 7.51E-08 |
| 2.62E-11 | 7.81E-08 |
| 3.93E-11 | 1.06E-07 |
| 4.00E-11 | 1.06E-07 |
| 5.00E-11 | 1.25E-07 |
| 9.00E-11 | 2.10E-07 |
| 9.24E-11 | 2.10E-07 |
| 1.03E-10 | 2.14E-07 |
| 1.05E-10 | 2.14E-07 |
| 1.08E-10 | 2.14E-07 |
| 1.12E-10 | 2.14E-07 |
| 1.29E-10 | 2.28E-07 |
| 2.18E-10 | 3.61E-07 |
| 2.20E-10 | 3.61E-07 |
| 2.64E-10 | 4.19E-07 |
| 3.14E-10 | 4.83E-07 |
| 3.25E-10 | 4.84E-07 |
| 4.26E-10 | 6.15E-07 |
| 4.51E-10 | 6.32E-07 |
| 5.02E-10 | 6.83E-07 |
| 5.71E-10 | 7.56E-07 |
| 6.10E-10 | 7.80E-07 |
| 6.39E-10 | 7.80E-07 |
| 7.36E-10 | 8.76E-07 |
| 9.12E-10 | 1.02E-06 |
| 9.37E-10 | 1.02E-06 |
| 9.77E-10 | 1.02E-06 |
| 1.00E-09 | 1.02E-06 |
| 1.09E-09 | 1.06E-06 |
| 1.13E-09 | 1.08E-06 |

| | |
|---|---|
| 1.27E-09 | 1.19E-06 |
| 1.57E-09 | 1.39E-06 |
| 1.57E-09 | 1.39E-06 |
| 1.73E-09 | 1.47E-06 |
| 1.73E-09 | 1.47E-06 |
| 2.05E-09 | 1.72E-06 |
| 2.51E-09 | 2.06E-06 |
| 3.83E-09 | 2.99E-06 |
| 4.46E-09 | 3.37E-06 |
| 4.99E-09 | 3.62E-06 |
| 5.02E-09 | 3.62E-06 |
| 5.30E-09 | 3.77E-06 |
| 5.42E-09 | 3.80E-06 |
| 5.54E-09 | 3.83E-06 |
| 5.94E-09 | 3.92E-06 |
| 6.11E-09 | 3.93E-06 |
| 8.26E-09 | 4.98E-06 |
| 8.46E-09 | 5.04E-06 |
| 1.05E-08 | 6.02E-06 |
| 1.12E-08 | 6.21E-06 |
| 1.27E-08 | 6.92E-06 |
| 1.30E-08 | 6.94E-06 |
| 1.32E-08 | 6.98E-06 |
| 1.64E-08 | 8.21E-06 |
| 1.67E-08 | 8.29E-06 |
| 1.76E-08 | 8.63E-06 |
| 2.09E-08 | 9.96E-06 |
| 2.26E-08 | 1.06E-05 |
| 2.29E-08 | 1.07E-05 |
| 2.44E-08 | 1.10E-05 |
| 2.62E-08 | 1.14E-05 |
| 2.68E-08 | 1.15E-05 |
| 3.03E-08 | 1.28E-05 |
| 3.13E-08 | 1.30E-05 |
| 3.21E-08 | 1.30E-05 |
| 3.60E-08 | 1.42E-05 |
| 3.71E-08 | 1.45E-05 |
| 4.07E-08 | 1.56E-05 |
| 4.43E-08 | 1.63E-05 |
| 4.83E-08 | 1.72E-05 |
| 6.02E-08 | 2.08E-05 |
| 6.11E-08 | 2.09E-05 |
| 6.41E-08 | 2.16E-05 |
| 6.75E-08 | 2.25E-05 |
| 7.69E-08 | 2.53E-05 |

| | |
|---|---|
| 7.69E-08 | 2.53E-05 |
| 8.06E-08 | 2.61E-05 |
| 8.67E-08 | 2.74E-05 |
| 8.95E-08 | 2.79E-05 |
| 9.20E-08 | 2.84E-05 |
| 9.23E-08 | 2.84E-05 |
| 1.13E-07 | 3.28E-05 |
| 1.16E-07 | 3.32E-05 |
| 1.17E-07 | 3.34E-05 |
| 1.19E-07 | 3.37E-05 |
| 1.23E-07 | 3.44E-05 |
| 1.26E-07 | 3.48E-05 |
| 1.34E-07 | 3.61E-05 |
| 1.37E-07 | 3.66E-05 |
| 1.50E-07 | 3.91E-05 |
| 1.52E-07 | 3.91E-05 |
| 1.62E-07 | 4.04E-05 |
| 1.68E-07 | 4.16E-05 |
| 1.77E-07 | 4.33E-05 |
| 1.92E-07 | 4.60E-05 |
| 1.93E-07 | 4.60E-05 |
| 1.96E-07 | 4.60E-05 |
| 2.11E-07 | 4.81E-05 |
| 2.23E-07 | 4.94E-05 |
| 2.23E-07 | 4.94E-05 |
| 2.23E-07 | 4.94E-05 |
| 2.28E-07 | 4.99E-05 |
| 2.32E-07 | 5.02E-05 |
| 2.44E-07 | 5.15E-05 |
| 2.53E-07 | 5.29E-05 |
| 2.57E-07 | 5.34E-05 |
| 2.66E-07 | 5.49E-05 |
| 2.83E-07 | 5.77E-05 |
| 2.99E-07 | 5.94E-05 |
| 3.02E-07 | 5.97E-05 |
| 3.10E-07 | 6.05E-05 |
| 3.14E-07 | 6.11E-05 |
| 3.16E-07 | 6.12E-05 |
| 3.34E-07 | 6.34E-05 |
| 3.41E-07 | 6.38E-05 |
| 3.45E-07 | 6.40E-05 |
| 3.51E-07 | 6.49E-05 |
| 3.53E-07 | 6.49E-05 |
| 3.69E-07 | 6.71E-05 |
| 3.75E-07 | 6.73E-05 |

| | |
|---|---|
| 3.78E-07 | 6.73E-05 |
| 3.79E-07 | 6.73E-05 |
| 3.84E-07 | 6.77E-05 |
| 4.02E-07 | 7.03E-05 |
| 4.04E-07 | 7.05E-05 |
| 4.13E-07 | 7.13E-05 |
| 4.40E-07 | 7.50E-05 |
| 4.42E-07 | 7.52E-05 |
| 4.58E-07 | 7.71E-05 |
| 4.62E-07 | 7.71E-05 |
| 4.63E-07 | 7.71E-05 |
| 4.79E-07 | 7.91E-05 |
| 4.82E-07 | 7.94E-05 |
| 4.83E-07 | 7.94E-05 |
| 5.28E-07 | 8.38E-05 |
| 5.40E-07 | 8.55E-05 |
| 5.46E-07 | 8.62E-05 |
| 5.89E-07 | 9.19E-05 |
| 5.94E-07 | 9.24E-05 |
| 6.05E-07 | 9.35E-05 |
| 6.22E-07 | 9.49E-05 |
| 6.29E-07 | 9.58E-05 |
| 6.88E-07 | 0.000102413 |
| 7.01E-07 | 0.000104047 |
| 7.23E-07 | 0.00010654 |
| 7.39E-07 | 0.000107616 |
| 7.50E-07 | 0.000108534 |
| 7.73E-07 | 0.000110919 |
| 7.86E-07 | 0.00011149 |
| 7.96E-07 | 0.00011195 |
| 7.97E-07 | 0.00011195 |
| 8.14E-07 | 0.0001134 |
| 8.22E-07 | 0.00011383 |
| 8.31E-07 | 0.000114353 |
| 8.36E-07 | 0.000114479 |
| 8.46E-07 | 0.000115438 |
| 8.74E-07 | 0.000118941 |
| 9.07E-07 | 0.000122043 |
| 9.28E-07 | 0.000124163 |
| 9.43E-07 | 0.000125418 |
| 9.51E-07 | 0.0001261 |
| 9.57E-07 | 0.000126568 |
| 9.67E-07 | 0.000127171 |
| 9.79E-07 | 0.000128077 |
| 9.98E-07 | 0.000130213 |

| | |
|---|---|
| 1.04E-06 | 0.000133968 |
| 1.05E-06 | 0.000135049 |
| 1.05E-06 | 0.000135049 |
| 1.06E-06 | 0.00013582 |
| 1.08E-06 | 0.000136906 |
| 1.08E-06 | 0.000136906 |
| 1.10E-06 | 0.000137757 |
| 1.12E-06 | 0.000139895 |
| 1.19E-06 | 0.000147201 |
| 1.21E-06 | 0.000147662 |
| 1.21E-06 | 0.000147662 |
| 1.22E-06 | 0.000147662 |
| 1.22E-06 | 0.000148295 |
| 1.24E-06 | 0.000149467 |
| 1.27E-06 | 0.00015166 |
| 1.27E-06 | 0.00015166 |
| 1.27E-06 | 0.00015166 |
| 1.28E-06 | 0.00015166 |
| 1.28E-06 | 0.00015166 |
| 1.28E-06 | 0.00015166 |
| 1.30E-06 | 0.00015343 |
| 1.36E-06 | 0.000158649 |
| 1.36E-06 | 0.000159305 |
| 1.42E-06 | 0.000164099 |
| 1.43E-06 | 0.000164509 |
| 1.44E-06 | 0.000165111 |
| 1.51E-06 | 0.000171542 |
| 1.52E-06 | 0.000171626 |
| 1.54E-06 | 0.000173139 |
| 1.60E-06 | 0.000178843 |
| 1.61E-06 | 0.000178843 |
| 1.65E-06 | 0.00018149 |
| 1.75E-06 | 0.000190245 |
| 1.77E-06 | 0.000190667 |
| 1.87E-06 | 0.000196436 |
| 1.94E-06 | 0.000202416 |
| 1.97E-06 | 0.000204338 |
| 1.98E-06 | 0.000204492 |
| 2.02E-06 | 0.000208228 |
| 2.03E-06 | 0.000209483 |
| 2.04E-06 | 0.000210087 |
| 2.08E-06 | 0.000212846 |
| 2.10E-06 | 0.000213933 |
| 2.12E-06 | 0.000214574 |
| 2.14E-06 | 0.000215215 |

| | |
|---|---|
| 2.16E-06 | 0.000215905 |
| 2.22E-06 | 0.00022029 |
| 2.22E-06 | 0.00022029 |
| 2.29E-06 | 0.000225059 |
| 2.42E-06 | 0.000235519 |
| 2.47E-06 | 0.00024002 |
| 2.66E-06 | 0.000255413 |
| 2.66E-06 | 0.000255413 |
| 2.68E-06 | 0.00025581 |
| 2.80E-06 | 0.000264179 |
| 2.90E-06 | 0.000271511 |
| 2.94E-06 | 0.00027314 |
| 3.14E-06 | 0.000286673 |
| 3.23E-06 | 0.000292925 |
| 3.41E-06 | 0.000304618 |
| 3.43E-06 | 0.000305096 |
| 3.43E-06 | 0.000305096 |
| 3.45E-06 | 0.000305642 |
| 3.79E-06 | 0.000327472 |
| 3.87E-06 | 0.0003326 |
| 3.95E-06 | 0.000336125 |
| 4.02E-06 | 0.000340589 |
| 4.03E-06 | 0.000340589 |
| 4.08E-06 | 0.000343885 |
| 4.22E-06 | 0.000351168 |
| 4.23E-06 | 0.000351168 |
| 4.23E-06 | 0.000351168 |
| 4.29E-06 | 0.000355076 |
| 4.43E-06 | 0.00036244 |
| 4.44E-06 | 0.00036244 |
| 4.71E-06 | 0.000378954 |
| 4.83E-06 | 0.000385706 |
| 4.85E-06 | 0.00038709 |
| 4.94E-06 | 0.000392429 |
| 5.09E-06 | 0.000401458 |
| 5.12E-06 | 0.000401544 |
| 5.19E-06 | 0.000404812 |
| 5.20E-06 | 0.000404982 |
| 5.46E-06 | 0.000420115 |
| 5.69E-06 | 0.000433295 |
| 5.84E-06 | 0.000443102 |
| 5.84E-06 | 0.000443102 |
| 5.93E-06 | 0.000447681 |
| 5.94E-06 | 0.000447801 |
| 5.99E-06 | 0.000449538 |

| | |
|---|---|
| 6.12E-06 | 0.000457837 |
| 6.45E-06 | 0.000479711 |
| 6.48E-06 | 0.000479711 |
| 6.69E-06 | 0.000492312 |
| 6.91E-06 | 0.000504438 |
| 7.23E-06 | 0.000523655 |
| 7.33E-06 | 0.0005292 |
| 7.38E-06 | 0.000531318 |
| 7.40E-06 | 0.000531318 |
| 7.64E-06 | 0.000544321 |
| 7.70E-06 | 0.00054745 |
| 7.84E-06 | 0.000554414 |
| 8.41E-06 | 0.000590974 |
| 8.46E-06 | 0.000593255 |
| 8.49E-06 | 0.000594689 |
| 8.50E-06 | 0.000594689 |
| 8.54E-06 | 0.00059639 |
| 8.62E-06 | 0.000600459 |
| 8.78E-06 | 0.000609354 |
| 8.85E-06 | 0.00061367 |
| 9.03E-06 | 0.000623414 |
| 9.29E-06 | 0.000636409 |
| 9.37E-06 | 0.000640104 |
| 9.56E-06 | 0.000652584 |
| 9.66E-06 | 0.000658003 |
| 9.73E-06 | 0.000660991 |
| 9.76E-06 | 0.000662049 |
| 1.03E-05 | 0.000695207 |
| 1.04E-05 | 0.00069998 |
| 1.06E-05 | 0.000707565 |
| 1.08E-05 | 0.000714887 |
| 1.10E-05 | 0.000722336 |
| 1.10E-05 | 0.000726459 |
| 1.14E-05 | 0.000744884 |
| 1.14E-05 | 0.000744884 |
| 1.15E-05 | 0.000749113 |
| 1.19E-05 | 0.000769017 |
| 1.19E-05 | 0.000769017 |
| 1.20E-05 | 0.000769254 |
| 1.21E-05 | 0.000775595 |
| 1.21E-05 | 0.000776344 |
| 1.23E-05 | 0.000779484 |
| 1.26E-05 | 0.000795971 |
| 1.28E-05 | 0.000808989 |
| 1.31E-05 | 0.000825548 |

| | |
|---|---|
| 1.31E-05 | 0.000825548 |
| 1.31E-05 | 0.000825548 |
| 1.31E-05 | 0.000825653 |
| 1.35E-05 | 0.000843561 |
| 1.37E-05 | 0.000852709 |
| 1.38E-05 | 0.0008547 |
| 1.40E-05 | 0.000868516 |
| 1.40E-05 | 0.000868516 |
| 1.41E-05 | 0.000868516 |
| 1.42E-05 | 0.000871396 |
| 1.43E-05 | 0.000875046 |
| 1.44E-05 | 0.000876825 |
| 1.46E-05 | 0.000887224 |
| 1.48E-05 | 0.000895688 |
| 1.49E-05 | 0.000898139 |
| 1.51E-05 | 0.000904271 |
| 1.56E-05 | 0.000925196 |
| 1.56E-05 | 0.000925196 |
| 1.57E-05 | 0.000925196 |
| 1.61E-05 | 0.000952028 |
| 1.62E-05 | 0.000953635 |
| 1.66E-05 | 0.000967295 |
| 1.67E-05 | 0.000968363 |
| 1.69E-05 | 0.000981694 |
| 1.70E-05 | 0.000983771 |
| 1.72E-05 | 0.000986536 |
| 1.73E-05 | 0.000992606 |
| 1.73E-05 | 0.000995547 |
| 1.74E-05 | 0.000996505 |
| 1.76E-05 | 0.001005611 |
| 1.77E-05 | 0.001007175 |
| 1.81E-05 | 0.001022015 |
| 1.81E-05 | 0.001022015 |
| 1.82E-05 | 0.001027953 |
| 1.89E-05 | 0.001056275 |
| 1.91E-05 | 0.001065865 |
| 1.94E-05 | 0.001083947 |
| 1.98E-05 | 0.001101788 |
| 1.99E-05 | 0.00110217 |
| 1.99E-05 | 0.00110217 |
| 2.02E-05 | 0.001113692 |
| 2.08E-05 | 0.00113735 |
| 2.10E-05 | 0.001142647 |
| 2.10E-05 | 0.001142647 |
| 2.11E-05 | 0.001146329 |

| | |
|---|---|
| 2.17E-05 | 0.001167693 |
| 2.21E-05 | 0.00118351 |
| 2.21E-05 | 0.001185299 |
| 2.22E-05 | 0.001187814 |
| 2.27E-05 | 0.001208985 |
| 2.32E-05 | 0.001234081 |
| 2.34E-05 | 0.001238025 |
| 2.40E-05 | 0.001265809 |
| 2.46E-05 | 0.001289542 |
| 2.49E-05 | 0.001301994 |
| 2.54E-05 | 0.001315265 |
| 2.55E-05 | 0.001319463 |
| 2.57E-05 | 0.00132812 |
| 2.65E-05 | 0.001361938 |
| 2.66E-05 | 0.001363706 |
| 2.66E-05 | 0.001364424 |
| 2.66E-05 | 0.001364424 |
| 2.67E-05 | 0.001366679 |
| 2.67E-05 | 0.001366679 |
| 2.69E-05 | 0.001372372 |
| 2.72E-05 | 0.001384553 |
| 2.73E-05 | 0.001388287 |
| 2.74E-05 | 0.001389899 |
| 2.75E-05 | 0.001391157 |
| 2.76E-05 | 0.001396541 |
| 2.76E-05 | 0.001397183 |
| 2.80E-05 | 0.001411979 |
| 2.93E-05 | 0.001464973 |
| 2.93E-05 | 0.001466936 |
| 2.96E-05 | 0.001478462 |
| 2.98E-05 | 0.001485226 |
| 3.01E-05 | 0.001491192 |
| 3.09E-05 | 0.001525809 |
| 3.12E-05 | 0.00153394 |
| 3.14E-05 | 0.001537286 |
| 3.17E-05 | 0.001544797 |
| 3.19E-05 | 0.001551387 |
| 3.20E-05 | 0.001552555 |
| 3.26E-05 | 0.001574475 |
| 3.27E-05 | 0.001577699 |
| 3.28E-05 | 0.001581089 |
| 3.36E-05 | 0.001611716 |
| 3.36E-05 | 0.001611716 |
| 3.38E-05 | 0.001617659 |
| 3.39E-05 | 0.001624412 |

| | |
|---|---|
| 3.42E-05 | 0.001634862 |
| 3.50E-05 | 0.001666067 |
| 3.51E-05 | 0.001669483 |
| 3.53E-05 | 0.001674327 |
| 3.63E-05 | 0.001715205 |
| 3.66E-05 | 0.001726061 |
| 3.68E-05 | 0.001727909 |
| 3.70E-05 | 0.001735049 |
| 3.79E-05 | 0.001769475 |
| 3.82E-05 | 0.001777891 |
| 3.89E-05 | 0.001803935 |
| 4.01E-05 | 0.001849383 |
| 4.02E-05 | 0.001849409 |
| 4.05E-05 | 0.001859544 |
| 4.08E-05 | 0.001868217 |
| 4.10E-05 | 0.001872778 |
| 4.10E-05 | 0.001872778 |
| 4.14E-05 | 0.001886732 |
| 4.14E-05 | 0.001886732 |
| 4.15E-05 | 0.001886732 |
| 4.17E-05 | 0.001889826 |
| 4.24E-05 | 0.001921198 |
| 4.45E-05 | 0.001997682 |
| 4.47E-05 | 0.002002119 |
| 4.60E-05 | 0.002053792 |
| 4.75E-05 | 0.00210553 |
| 4.75E-05 | 0.00210553 |
| 4.83E-05 | 0.00212919 |
| 4.91E-05 | 0.002160304 |
| 4.97E-05 | 0.002177274 |
| 4.99E-05 | 0.002184276 |
| 5.15E-05 | 0.002232473 |
| 5.15E-05 | 0.002232473 |
| 5.17E-05 | 0.002232998 |
| 5.22E-05 | 0.002249985 |
| 5.37E-05 | 0.002307105 |
| 5.42E-05 | 0.0023255 |
| 5.48E-05 | 0.002340777 |
| 5.59E-05 | 0.002381606 |
| 5.62E-05 | 0.002388598 |
| 5.74E-05 | 0.002431096 |
| 5.78E-05 | 0.002442604 |
| 5.81E-05 | 0.002453783 |
| 6.07E-05 | 0.002549477 |
| 6.16E-05 | 0.002575157 |

| | |
|---|---|
| 6.16E-05 | 0.002575157 |
| 6.27E-05 | 0.002612407 |
| 6.27E-05 | 0.002612407 |
| 6.30E-05 | 0.002621986 |
| 6.33E-05 | 0.002629361 |
| 6.44E-05 | 0.002662358 |
| 6.44E-05 | 0.002662358 |
| 6.49E-05 | 0.002677349 |
| 6.49E-05 | 0.002677349 |
| 6.52E-05 | 0.002682696 |
| 6.52E-05 | 0.002682696 |
| 6.64E-05 | 0.00272834 |
| 6.71E-05 | 0.002752209 |
| 6.77E-05 | 0.002772783 |
| 6.83E-05 | 0.002793762 |
| 6.88E-05 | 0.002809471 |
| 6.94E-05 | 0.002821578 |
| 7.01E-05 | 0.002838117 |
| 7.01E-05 | 0.002838117 |
| 7.01E-05 | 0.002838211 |
| 7.13E-05 | 0.00287359 |
| 7.18E-05 | 0.002889283 |
| 7.29E-05 | 0.002926156 |
| 7.40E-05 | 0.00296071 |
| 7.43E-05 | 0.002964298 |
| 7.51E-05 | 0.002983654 |
| 7.62E-05 | 0.003020979 |
| 7.98E-05 | 0.003136933 |
| 7.99E-05 | 0.003137056 |
| 8.02E-05 | 0.003147811 |
| 8.12E-05 | 0.003176693 |
| 8.29E-05 | 0.003238178 |
| 8.50E-05 | 0.003297183 |
| 8.83E-05 | 0.003404224 |
| 8.87E-05 | 0.003417541 |
| 8.90E-05 | 0.003423202 |
| 9.28E-05 | 0.003545712 |
| 9.36E-05 | 0.003564432 |
| 9.44E-05 | 0.003588164 |
| 9.50E-05 | 0.00359797 |
| 9.53E-05 | 0.003606656 |
| 9.57E-05 | 0.003615915 |
| 9.85E-05 | 0.0037001 |
| 9.97E-05 | 0.00373581 |
| 0.000100658 | 0.003760022 |

| | |
|---|---|
| 0.000101172 | 0.003773324 |
| 0.000102653 | 0.003819573 |
| 0.000103149 | 0.003829061 |
| 0.000103501 | 0.00383667 |
| 0.000103515 | 0.00383667 |
| 0.000105884 | 0.00389862 |
| 0.000109135 | 0.00398909 |
| 0.000113237 | 0.004097243 |
| 0.000113314 | 0.004097243 |
| 0.000113385 | 0.004097243 |
| 0.000114728 | 0.00413951 |
| 0.000117525 | 0.004224427 |
| 0.000119292 | 0.004278262 |
| 0.000120193 | 0.00430733 |
| 0.000122376 | 0.004362576 |
| 0.00012264 | 0.00436871 |
| 0.000122843 | 0.004372687 |
| 0.000123675 | 0.004395731 |
| 0.000124474 | 0.004417541 |
| 0.000124923 | 0.004423568 |
| 0.000125088 | 0.004424518 |
| 0.000125992 | 0.004431783 |
| 0.000126809 | 0.004448155 |
| 0.000127029 | 0.00445182 |
| 0.000127553 | 0.004463879 |
| 0.000127819 | 0.004466337 |
| 0.000128698 | 0.004490471 |
| 0.000131205 | 0.004561231 |
| 0.000133449 | 0.004625757 |
| 0.000136429 | 0.004708472 |
| 0.000137256 | 0.004730159 |
| 0.000138327 | 0.004763619 |
| 0.000138497 | 0.004766032 |
| 0.000139355 | 0.004781755 |
| 0.000140828 | 0.004818408 |
| 0.000141062 | 0.004822934 |
| 0.000144825 | 0.004921752 |
| 0.000145288 | 0.004921752 |
| 0.000145437 | 0.004921752 |
| 0.000145445 | 0.004921752 |
| 0.000151194 | 0.005067489 |
| 0.000151973 | 0.005090024 |
| 0.000154163 | 0.005141665 |
| 0.000156117 | 0.005188686 |
| 0.000156238 | 0.005189084 |

| | |
|---|---|
| 0.000157269 | 0.005216044 |
| 0.000158693 | 0.005255954 |
| 0.000159628 | 0.005275921 |
| 0.000165231 | 0.005402329 |
| 0.000165626 | 0.005406626 |
| 0.000168138 | 0.005469896 |
| 0.000169826 | 0.005503606 |
| 0.000170521 | 0.005513516 |
| 0.000172139 | 0.005550633 |
| 0.00017219 | 0.005550633 |
| 0.000176127 | 0.005648766 |
| 0.0001798 | 0.005747218 |
| 0.000184764 | 0.00588612 |
| 0.000185764 | 0.005914017 |
| 0.000187985 | 0.005976754 |
| 0.000188603 | 0.005992384 |
| 0.00018888 | 0.005993183 |
| 0.000190825 | 0.006034824 |
| 0.000191426 | 0.006049792 |
| 0.000192292 | 0.006069117 |
| 0.000193191 | 0.006089412 |
| 0.000198629 | 0.00620619 |
| 0.00019872 | 0.00620619 |
| 0.000199776 | 0.006226909 |
| 0.000200767 | 0.006234482 |
| 0.000200959 | 0.00623522 |
| 0.000202483 | 0.006269453 |
| 0.000203236 | 0.006277246 |
| 0.00020428 | 0.006297235 |
| 0.000206921 | 0.006355613 |
| 0.000206974 | 0.006355613 |
| 0.000209158 | 0.006403695 |
| 0.000210639 | 0.006434963 |
| 0.000211874 | 0.00645869 |
| 0.000216888 | 0.006587843 |
| 0.000217231 | 0.006594056 |
| 0.000217676 | 0.006603357 |
| 0.000218432 | 0.006622059 |
| 0.000218825 | 0.006629772 |
| 0.00022208 | 0.00670703 |
| 0.000225136 | 0.00676503 |
| 0.000226454 | 0.006791776 |
| 0.000227742 | 0.006821813 |
| 0.00022858 | 0.006838305 |
| 0.000231244 | 0.006899007 |

| | |
|---|---|
| 0.000231469 | 0.006899007 |
| 0.000233867 | 0.006952795 |
| 0.000234052 | 0.006953958 |
| 0.000234867 | 0.0069669 |
| 0.000235171 | 0.006969822 |
| 0.000237265 | 0.00701773 |
| 0.000241472 | 0.007101417 |
| 0.00024155 | 0.007101417 |
| 0.000241916 | 0.007105375 |
| 0.000242343 | 0.00711156 |
| 0.000245972 | 0.007213603 |
| 0.000247131 | 0.007229798 |
| 0.000248135 | 0.007254701 |
| 0.000249031 | 0.007271983 |
| 0.000252591 | 0.007330981 |
| 0.000253181 | 0.007343641 |
| 0.000255668 | 0.007389272 |
| 0.000255685 | 0.007389272 |
| 0.000256757 | 0.007409298 |
| 0.000257608 | 0.007417144 |
| 0.000257681 | 0.007417144 |
| 0.000262157 | 0.007518977 |
| 0.000262226 | 0.007518977 |
| 0.000264119 | 0.007553842 |
| 0.000264235 | 0.007553842 |
| 0.000265908 | 0.007592575 |
| 0.000266736 | 0.007608583 |
| 0.000267008 | 0.007610281 |
| 0.000270799 | 0.007695321 |
| 0.00027528 | 0.007808664 |
| 0.0002757 | 0.007815926 |
| 0.000278838 | 0.007890798 |
| 0.000280067 | 0.007911485 |
| 0.00028066 | 0.007923534 |
| 0.000281399 | 0.007939699 |
| 0.000282692 | 0.007966733 |
| 0.000282927 | 0.007968625 |
| 0.000287673 | 0.008083173 |
| 0.000287988 | 0.008087277 |
| 0.000289651 | 0.008129188 |
| 0.000289852 | 0.00813001 |
| 0.000290696 | 0.008140073 |
| 0.000292538 | 0.008181264 |
| 0.000295422 | 0.00824281 |
| 0.000295636 | 0.00824281 |

| | |
|---|---|
| 0.00029749 | 0.008276035 |
| 0.000297834 | 0.00827604 |
| 0.00029793 | 0.00827604 |
| 0.000298012 | 0.00827604 |
| 0.000302549 | 0.008359254 |
| 0.000302921 | 0.008363593 |
| 0.000309266 | 0.008499363 |
| 0.000309807 | 0.008509332 |
| 0.000311063 | 0.008529058 |
| 0.000311516 | 0.008531657 |
| 0.000311911 | 0.008537571 |
| 0.000321799 | 0.008744719 |
| 0.00032202 | 0.008744719 |
| 0.000325507 | 0.008813476 |
| 0.000328157 | 0.008875154 |
| 0.000330164 | 0.0089193 |
| 0.000332966 | 0.008973063 |
| 0.000333704 | 0.00897928 |
| 0.00033634 | 0.009024702 |
| 0.000337274 | 0.009039927 |
| 0.000338526 | 0.009039927 |
| 0.00033894 | 0.009039927 |
| 0.000339884 | 0.009058576 |
| 0.000348827 | 0.009232639 |
| 0.000349599 | 0.00924167 |
| 0.000349859 | 0.00924167 |
| 0.000350115 | 0.009243295 |
| 0.000351823 | 0.009267857 |
| 0.000359502 | 0.009418044 |
| 0.000359716 | 0.009418457 |
| 0.000360167 | 0.009425091 |
| 0.000360498 | 0.009428589 |
| 0.000363417 | 0.009493839 |
| 0.000364838 | 0.009510747 |
| 0.000369315 | 0.009591684 |
| 0.000378333 | 0.009798194 |
| 0.000381357 | 0.00983924 |
| 0.000383869 | 0.009871956 |
| 0.000384088 | 0.009871956 |
| 0.000386426 | 0.009910788 |
| 0.000388796 | 0.009943706 |
| 0.000388963 | 0.009943706 |
| 0.000389859 | 0.009955399 |
| 0.000392598 | 0.010004518 |
| 0.000395922 | 0.010072966 |

| | |
|---|---|
| 0.000396539 | 0.010080538 |
| 0.000396815 | 0.010080538 |
| 0.000396855 | 0.010080538 |
| 0.000398776 | 0.010119738 |
| 0.000406943 | 0.010271049 |
| 0.000408949 | 0.01031622 |
| 0.000409899 | 0.010323782 |
| 0.000411322 | 0.010354135 |
| 0.00042018 | 0.010543692 |
| 0.000422705 | 0.010591456 |
| 0.000422751 | 0.010591456 |
| 0.000423814 | 0.010612505 |
| 0.000429502 | 0.010726744 |
| 0.000430463 | 0.010745101 |
| 0.000432348 | 0.010783464 |
| 0.000432453 | 0.010783464 |
| 0.000433183 | 0.010796027 |
| 0.000435358 | 0.010833237 |
| 0.000436232 | 0.010849311 |
| 0.000436515 | 0.010850666 |
| 0.000440571 | 0.010934374 |
| 0.000449743 | 0.011109913 |
| 0.000453085 | 0.011169285 |
| 0.000455963 | 0.011234426 |
| 0.000456425 | 0.011236163 |
| 0.000456741 | 0.011236163 |
| 0.000458045 | 0.011256592 |
| 0.000461709 | 0.011311629 |
| 0.00046479 | 0.011363733 |
| 0.000467777 | 0.011419172 |
| 0.000468039 | 0.011419724 |
| 0.000468683 | 0.011429589 |
| 0.000474971 | 0.011553348 |
| 0.00047655 | 0.011585831 |
| 0.00048459 | 0.011733381 |
| 0.000485201 | 0.011736964 |
| 0.000485575 | 0.011739338 |
| 0.000487977 | 0.011785435 |
| 0.000489799 | 0.011817449 |
| 0.000492847 | 0.011878956 |
| 0.000494042 | 0.01190174 |
| 0.000495914 | 0.011940787 |
| 0.000496167 | 0.011940846 |
| 0.000498715 | 0.011971926 |
| 0.000499368 | 0.011978653 |

| | |
|---|---|
| 0.000499498 | 0.011978653 |
| 0.000503647 | 0.012047809 |
| 0.000505311 | 0.012075487 |
| 0.000514692 | 0.012232155 |
| 0.000516827 | 0.012264539 |
| 0.000517229 | 0.012266495 |
| 0.000517425 | 0.012266495 |
| 0.000519026 | 0.012281333 |
| 0.000519082 | 0.012281333 |
| 0.000526324 | 0.012409515 |
| 0.000527595 | 0.012433324 |
| 0.000529211 | 0.012459077 |
| 0.000531139 | 0.012492124 |
| 0.000531682 | 0.012498596 |
| 0.000531939 | 0.012498596 |
| 0.000538791 | 0.012622236 |
| 0.000540049 | 0.012639267 |
| 0.000541594 | 0.012656772 |
| 0.000545455 | 0.012715807 |
| 0.000550002 | 0.012811277 |
| 0.000555235 | 0.012912195 |
| 0.000563247 | 0.013054883 |
| 0.000564387 | 0.013070628 |
| 0.000564835 | 0.013070628 |
| 0.000564897 | 0.013070628 |
| 0.000565642 | 0.013077586 |
| 0.000572472 | 0.013201088 |
| 0.000572647 | 0.013201088 |
| 0.000573471 | 0.0132137 |
| 0.000573769 | 0.01321417 |
| 0.000578902 | 0.013288326 |
| 0.000580075 | 0.013294651 |
| 0.000587761 | 0.013419599 |
| 0.000591505 | 0.013466353 |
| 0.000592578 | 0.013484332 |
| 0.000594131 | 0.013513224 |
| 0.000600197 | 0.013618674 |
| 0.000608009 | 0.013745888 |
| 0.000608469 | 0.013747426 |
| 0.000609635 | 0.01376722 |
| 0.000618855 | 0.013896382 |
| 0.000626398 | 0.014045882 |
| 0.000627003 | 0.014052828 |
| 0.000630406 | 0.01412246 |
| 0.000636109 | 0.014236828 |

| | |
|---|---|
| 0.000639731 | 0.014285823 |
| 0.000639798 | 0.014285823 |
| 0.000641149 | 0.014309276 |
| 0.000649738 | 0.01444681 |
| 0.000652283 | 0.014476364 |
| 0.000654885 | 0.01451382 |
| 0.000655366 | 0.014515487 |
| 0.000656142 | 0.014517117 |
| 0.000656253 | 0.014517117 |
| 0.000658049 | 0.0145501 |
| 0.000659318 | 0.014561259 |
| 0.000660172 | 0.014569355 |
| 0.00066045 | 0.014569355 |
| 0.000661253 | 0.014580331 |
| 0.000665052 | 0.01463698 |
| 0.000671236 | 0.014739025 |
| 0.000671628 | 0.014740838 |
| 0.000674173 | 0.014777601 |
| 0.000674234 | 0.014777601 |
| 0.000678579 | 0.014838699 |
| 0.000681009 | 0.014878174 |
| 0.000686355 | 0.014966197 |
| 0.000686609 | 0.014966197 |
| 0.000691534 | 0.015032257 |
| 0.000698345 | 0.015152647 |
| 0.000709041 | 0.015349773 |
| 0.000710462 | 0.015367012 |
| 0.000710483 | 0.015367012 |
| 0.000720077 | 0.015518159 |
| 0.000721757 | 0.015533275 |
| 0.000723133 | 0.015555423 |
| 0.000730396 | 0.015648473 |
| 0.000736732 | 0.015728607 |
| 0.000737002 | 0.015728607 |
| 0.000737108 | 0.015728607 |
| 0.000739541 | 0.015768468 |
| 0.000739638 | 0.015768468 |
| 0.000753678 | 0.016024744 |
| 0.00076025 | 0.016135662 |
| 0.000762294 | 0.016171846 |
| 0.000769022 | 0.016271083 |
| 0.000779649 | 0.01643024 |
| 0.00078008 | 0.016432056 |
| 0.000786912 | 0.016546697 |
| 0.000791276 | 0.016609128 |

| | |
|---|---|
| 0.000796769 | 0.01669097 |
| 0.000796927 | 0.01669097 |
| 0.000798142 | 0.016709053 |
| 0.00080812 | 0.016858672 |
| 0.000813035 | 0.0169067 |
| 0.000813127 | 0.0169067 |
| 0.000813483 | 0.0169067 |
| 0.000814867 | 0.016925272 |
| 0.000820794 | 0.01701871 |
| 0.000826708 | 0.017096667 |
| 0.000831881 | 0.01716637 |
| 0.000840836 | 0.017313661 |
| 0.000854711 | 0.017508525 |
| 0.00085543 | 0.017508918 |
| 0.000864943 | 0.017640532 |
| 0.000869735 | 0.01769819 |
| 0.00087016 | 0.01769819 |
| 0.00087118 | 0.017708789 |
| 0.000872829 | 0.017727169 |
| 0.000886841 | 0.017965783 |
| 0.000891051 | 0.018012675 |
| 0.000891095 | 0.018012675 |
| 0.000896014 | 0.018082407 |
| 0.000898867 | 0.018116943 |
| 0.000899848 | 0.018129034 |
| 0.000906805 | 0.018244242 |
| 0.000914711 | 0.018343134 |
| 0.000918151 | 0.01839662 |
| 0.000921044 | 0.01843133 |
| 0.00092269 | 0.018456504 |
| 0.000925984 | 0.01850685 |
| 0.000937994 | 0.018723318 |
| 0.000939422 | 0.01873612 |
| 0.000944143 | 0.018806649 |
| 0.000944549 | 0.018806873 |
| 0.00094903 | 0.018857772 |
| 0.00094918 | 0.018857772 |
| 0.000951796 | 0.018887995 |
| 0.000952348 | 0.01888946 |
| 0.000955167 | 0.018915489 |
| 0.000957647 | 0.018940977 |
| 0.000963099 | 0.019032987 |
| 0.000965838 | 0.019063389 |
| 0.000974403 | 0.019192672 |
| 0.000982873 | 0.019304265 |

| | |
|---|---|
| 0.000983097 | 0.019304265 |
| 0.000983716 | 0.019304265 |
| 0.000990296 | 0.019409395 |
| 0.000997382 | 0.019524171 |
| 0.001005038 | 0.019655164 |
| 0.001008607 | 0.01970197 |
| 0.001008947 | 0.01970197 |
| 0.001013062 | 0.019756663 |
| 0.001017121 | 0.019795707 |
| 0.00102545 | 0.019909949 |
| 0.001032435 | 0.02002378 |
| 0.001033989 | 0.020043067 |
| 0.001036963 | 0.02008436 |
| 0.001040009 | 0.020125564 |
| 0.00104217 | 0.020152421 |
| 0.00105418 | 0.020295067 |
| 0.001061961 | 0.020402583 |
| 0.001067273 | 0.020479869 |
| 0.001076109 | 0.020616188 |
| 0.00107699 | 0.020617971 |
| 0.001077068 | 0.020617971 |
| 0.001078059 | 0.020628654 |
| 0.0010807 | 0.020645999 |
| 0.001083062 | 0.020657985 |
| 0.001083623 | 0.020660415 |
| 0.001088525 | 0.020711184 |
| 0.00109356 | 0.020775019 |
| 0.001098451 | 0.020842991 |
| 0.00110429 | 0.020926429 |
| 0.001104605 | 0.020926429 |
| 0.001107117 | 0.020957336 |
| 0.001115478 | 0.021063336 |
| 0.001127143 | 0.021243554 |
| 0.001138699 | 0.021351497 |
| 0.001139425 | 0.021356707 |
| 0.001145839 | 0.02143911 |
| 0.001146292 | 0.02143911 |
| 0.00115035 | 0.021485378 |
| 0.001151575 | 0.021486129 |
| 0.001151718 | 0.021486129 |
| 0.001151744 | 0.021486129 |
| 0.001152949 | 0.021500194 |
| 0.001163001 | 0.021641326 |
| 0.001168699 | 0.02170787 |
| 0.001169019 | 0.02170787 |

| | |
|---|---|
| 0.001181607 | 0.021876903 |
| 0.001181815 | 0.021876903 |
| 0.001182053 | 0.021876903 |
| 0.001183375 | 0.021882678 |
| 0.001183568 | 0.021882678 |
| 0.001188216 | 0.021943073 |
| 0.001199387 | 0.022097957 |
| 0.001203852 | 0.022163061 |
| 0.00120587 | 0.022175335 |
| 0.001225394 | 0.022438226 |
| 0.001234668 | 0.022590673 |
| 0.001236991 | 0.022624493 |
| 0.001238082 | 0.02263575 |
| 0.001242214 | 0.022685165 |
| 0.001245756 | 0.022741139 |
| 0.001248154 | 0.022758733 |
| 0.00125293 | 0.02280218 |
| 0.001254938 | 0.022822739 |
| 0.001267569 | 0.023007048 |
| 0.001268457 | 0.023014395 |
| 0.001273334 | 0.02306773 |
| 0.001280666 | 0.02316531 |
| 0.001283159 | 0.023192799 |
| 0.001288356 | 0.023260239 |
| 0.001295001 | 0.023344816 |
| 0.001297905 | 0.023379423 |
| 0.00130611 | 0.02348287 |
| 0.001308371 | 0.023505066 |
| 0.001308826 | 0.023505066 |
| 0.001316905 | 0.023605654 |
| 0.001319217 | 0.023629312 |
| 0.001334197 | 0.023799176 |
| 0.001336915 | 0.023829802 |
| 0.001349873 | 0.023978239 |
| 0.001351957 | 0.023986306 |
| 0.001357354 | 0.02404116 |
| 0.001363842 | 0.024138123 |
| 0.001364936 | 0.024148512 |
| 0.001367317 | 0.024161259 |
| 0.001367502 | 0.024161259 |
| 0.001367685 | 0.024161259 |
| 0.001368803 | 0.024172031 |
| 0.001381556 | 0.024346425 |
| 0.001383232 | 0.024346425 |
| 0.001383279 | 0.024346425 |

| | |
|---|---|
| 0.001390962 | 0.024396553 |
| 0.001392078 | 0.024396553 |
| 0.00139497 | 0.024434812 |
| 0.001403899 | 0.024555082 |
| 0.001406867 | 0.024588941 |
| 0.001407433 | 0.024589798 |
| 0.001409858 | 0.024607348 |
| 0.001411119 | 0.024615662 |
| 0.001411936 | 0.024615662 |
| 0.001419815 | 0.024715466 |
| 0.001424509 | 0.024779073 |
| 0.001433848 | 0.024887014 |
| 0.001441445 | 0.024955176 |
| 0.001446251 | 0.025024953 |
| 0.001446526 | 0.025024953 |
| 0.001453623 | 0.025081827 |
| 0.001457532 | 0.025096839 |
| 0.001461111 | 0.025122146 |
| 0.001462678 | 0.025140016 |
| 0.00146662 | 0.025197206 |
| 0.001468337 | 0.02520089 |
| 0.001473266 | 0.025249094 |
| 0.00148648 | 0.025438939 |
| 0.00148755 | 0.025446519 |
| 0.001491194 | 0.025460557 |
| 0.001492554 | 0.025460557 |
| 0.001495897 | 0.025490197 |
| 0.001498002 | 0.025516935 |
| 0.001499585 | 0.025534764 |
| 0.001512165 | 0.025684695 |
| 0.001522901 | 0.025830205 |
| 0.00153066 | 0.025916028 |
| 0.001531125 | 0.025916028 |
| 0.001542221 | 0.026074312 |
| 0.001543595 | 0.026088287 |
| 0.001546769 | 0.026132659 |
| 0.001548398 | 0.02614164 |
| 0.001554613 | 0.02620283 |
| 0.001554774 | 0.02620283 |
| 0.001557612 | 0.026241385 |
| 0.001562121 | 0.026308044 |
| 0.001562805 | 0.026310254 |
| 0.001575217 | 0.026463091 |
| 0.001578733 | 0.026475465 |
| 0.001581181 | 0.026488541 |

| | |
|---|---|
| 0.001582294 | 0.026492751 |
| 0.001582545 | 0.026492751 |
| 0.001589436 | 0.02658008 |
| 0.001593531 | 0.026620525 |
| 0.00159576 | 0.026648416 |
| 0.001604417 | 0.026780467 |
| 0.001607286 | 0.026797555 |
| 0.001609284 | 0.026817829 |
| 0.001609979 | 0.026820039 |
| 0.001611126 | 0.026821382 |
| 0.001618722 | 0.026937411 |
| 0.001629516 | 0.027042895 |
| 0.001629603 | 0.027042895 |
| 0.001630585 | 0.027049769 |
| 0.001631837 | 0.02706111 |
| 0.001639274 | 0.027156071 |
| 0.001647167 | 0.027267858 |
| 0.001657754 | 0.027390885 |
| 0.001668778 | 0.02752039 |
| 0.001681572 | 0.027670454 |
| 0.001681944 | 0.027670454 |
| 0.001687288 | 0.027720059 |
| 0.00169796 | 0.027876166 |
| 0.001711625 | 0.028032863 |
| 0.001714942 | 0.02806789 |
| 0.001716408 | 0.028082232 |
| 0.001721618 | 0.028157791 |
| 0.001724629 | 0.028174955 |
| 0.001725033 | 0.028174955 |
| 0.001728443 | 0.02821129 |
| 0.001734596 | 0.028292337 |
| 0.00174895 | 0.028477692 |
| 0.001766076 | 0.028697686 |
| 0.001773387 | 0.028806654 |
| 0.001775791 | 0.028835866 |
| 0.001785933 | 0.028910828 |
| 0.001786035 | 0.028910828 |
| 0.001786082 | 0.028910828 |
| 0.001786379 | 0.028910828 |
| 0.001787081 | 0.028910828 |
| 0.001797961 | 0.029047322 |
| 0.001799264 | 0.029058506 |
| 0.001801085 | 0.02907382 |
| 0.001812202 | 0.029219098 |
| 0.001819581 | 0.029317043 |

| | |
|---|---|
| 0.00184214 | 0.029575983 |
| 0.001844122 | 0.029575983 |
| 0.001844346 | 0.029575983 |
| 0.001849593 | 0.029620228 |
| 0.001852273 | 0.029653175 |
| 0.001857572 | 0.029718028 |
| 0.001859073 | 0.029732056 |
| 0.001871205 | 0.029885948 |
| 0.001871837 | 0.029886013 |
| 0.001896985 | 0.030206515 |
| 0.001909717 | 0.030309394 |
| 0.001916763 | 0.030369153 |
| 0.001923705 | 0.030458874 |
| 0.001933568 | 0.030584871 |
| 0.001933969 | 0.030584871 |
| 0.001936028 | 0.030603129 |
| 0.00193836 | 0.030629815 |
| 0.001947361 | 0.03074979 |
| 0.001947889 | 0.03074979 |
| 0.001949236 | 0.030760857 |
| 0.001952651 | 0.030804535 |
| 0.001957074 | 0.030843662 |
| 0.001976545 | 0.031078543 |
| 0.001979865 | 0.031099937 |
| 0.001989825 | 0.031225497 |
| 0.001990881 | 0.031231775 |
| 0.001997015 | 0.031310955 |
| 0.001998445 | 0.031310955 |
| 0.001998575 | 0.031310955 |
| 0.002003276 | 0.031343636 |
| 0.002026336 | 0.031608127 |
| 0.00203016 | 0.031649893 |
| 0.00203355 | 0.031692366 |
| 0.002040142 | 0.031753545 |
| 0.002041508 | 0.03176443 |
| 0.002049602 | 0.03182797 |
| 0.002052732 | 0.03185581 |
| 0.002058554 | 0.031935757 |
| 0.002060356 | 0.0319533 |
| 0.002073221 | 0.032110988 |
| 0.002096462 | 0.032328675 |
| 0.002116266 | 0.032523844 |
| 0.002125456 | 0.032644009 |
| 0.002131561 | 0.032716674 |
| 0.002139323 | 0.032806415 |

| | |
|---|---|
| 0.0021568 | 0.033003638 |
| 0.002173573 | 0.033211659 |
| 0.002184087 | 0.033291883 |
| 0.002189321 | 0.033338755 |
| 0.00218959 | 0.033338755 |
| 0.002190775 | 0.033346134 |
| 0.002194813 | 0.033396927 |
| 0.002200853 | 0.033449779 |
| 0.002208171 | 0.033546584 |
| 0.002213594 | 0.033596831 |
| 0.002222643 | 0.033669795 |
| 0.002227339 | 0.033719481 |
| 0.002246008 | 0.033958054 |
| 0.002246663 | 0.033958054 |
| 0.002268226 | 0.034164702 |
| 0.002324028 | 0.034785186 |
| 0.002328243 | 0.034818409 |
| 0.002346668 | 0.034979158 |
| 0.002348584 | 0.034979158 |
| 0.002357521 | 0.035058226 |
| 0.002362189 | 0.035102644 |
| 0.002384295 | 0.035342927 |
| 0.002398072 | 0.035436851 |
| 0.002399209 | 0.035442655 |
| 0.002408163 | 0.035533544 |
| 0.002408346 | 0.035533544 |
| 0.002414267 | 0.035576821 |
| 0.00242238 | 0.035625087 |
| 0.002442228 | 0.035822606 |
| 0.002451266 | 0.035899894 |
| 0.002452445 | 0.035906116 |
| 0.002454731 | 0.035928546 |
| 0.002467797 | 0.036053302 |
| 0.002482124 | 0.036166361 |
| 0.002482135 | 0.036166361 |
| 0.002482624 | 0.036166361 |
| 0.002488776 | 0.036226442 |
| 0.002503013 | 0.036344822 |
| 0.002513449 | 0.036440805 |
| 0.00252561 | 0.036562264 |
| 0.002538089 | 0.036695202 |
| 0.002541118 | 0.036719006 |
| 0.002546959 | 0.036758797 |
| 0.002550733 | 0.036768699 |
| 0.002560955 | 0.036871405 |

| | |
|---|---|
| 0.002565608 | 0.036927232 |
| 0.002573802 | 0.03702279 |
| 0.002583023 | 0.037131082 |
| 0.002611787 | 0.037456066 |
| 0.002616984 | 0.037508002 |
| 0.002621715 | 0.037558453 |
| 0.002622081 | 0.037558453 |
| 0.002631264 | 0.037667326 |
| 0.002643355 | 0.037817686 |
| 0.002647335 | 0.037863255 |
| 0.00264872 | 0.037871679 |
| 0.002668167 | 0.03804695 |
| 0.002676234 | 0.038116323 |
| 0.002686472 | 0.038227851 |
| 0.002689364 | 0.038246145 |
| 0.002697615 | 0.038306296 |
| 0.002708154 | 0.03838612 |
| 0.002708159 | 0.03838612 |
| 0.002712685 | 0.038417197 |
| 0.002717086 | 0.038445237 |
| 0.002739124 | 0.03871106 |
| 0.00274008 | 0.038711753 |
| 0.002754514 | 0.03880191 |
| 0.002760551 | 0.038837595 |
| 0.002777497 | 0.039010273 |
| 0.002779393 | 0.039013893 |
| 0.002798899 | 0.03918376 |
| 0.002803515 | 0.03923515 |
| 0.002806005 | 0.039248641 |
| 0.002813245 | 0.039315266 |
| 0.002816761 | 0.039330872 |
| 0.002817476 | 0.039330872 |
| 0.002825425 | 0.039404549 |
| 0.002833618 | 0.039480955 |
| 0.002843285 | 0.039549392 |
| 0.002855881 | 0.039678253 |
| 0.002867972 | 0.039801567 |
| 0.00286957 | 0.039810375 |
| 0.002871485 | 0.039813751 |
| 0.002872615 | 0.039817815 |
| 0.002879969 | 0.039873342 |
| 0.002887321 | 0.03994029 |
| 0.002892187 | 0.039984385 |
| 0.002897716 | 0.040025958 |
| 0.002901423 | 0.04005393 |

| | |
|---|---|
| 0.002908562 | 0.040129218 |
| 0.002921346 | 0.04025896 |
| 0.002924438 | 0.040261436 |
| 0.002930378 | 0.040325086 |
| 0.002932035 | 0.040327688 |
| 0.002939254 | 0.040400529 |
| 0.00294516 | 0.040458366 |
| 0.002955917 | 0.040547661 |
| 0.002964277 | 0.040615543 |
| 0.002974276 | 0.040682315 |
| 0.002976993 | 0.040707788 |
| 0.002979796 | 0.040734425 |
| 0.002990065 | 0.04082793 |
| 0.002993119 | 0.040846211 |
| 0.00300055 | 0.040917093 |
| 0.003005791 | 0.04094875 |
| 0.003015155 | 0.041036693 |
| 0.003019847 | 0.041041056 |
| 0.003023577 | 0.041073562 |
| 0.003031917 | 0.041128195 |
| 0.003056087 | 0.041374421 |
| 0.003067362 | 0.041475369 |
| 0.00306796 | 0.041475369 |
| 0.003072315 | 0.041522456 |
| 0.003079011 | 0.041558042 |
| 0.003085292 | 0.041615182 |
| 0.003089918 | 0.04165398 |
| 0.003103134 | 0.041793596 |
| 0.003105933 | 0.041803261 |
| 0.003122 | 0.041951669 |
| 0.003122571 | 0.041951669 |
| 0.003123708 | 0.041955116 |
| 0.003128729 | 0.041998865 |
| 0.00314262 | 0.042137831 |
| 0.003145857 | 0.042157498 |
| 0.003151762 | 0.042224742 |
| 0.003153993 | 0.042236261 |
| 0.003154395 | 0.042236261 |
| 0.00316106 | 0.042301712 |
| 0.003162796 | 0.042313056 |
| 0.003179212 | 0.042449205 |
| 0.003184797 | 0.042483746 |
| 0.003185733 | 0.042483746 |
| 0.003189793 | 0.042507065 |
| 0.003198335 | 0.042585152 |

| | |
|---|---|
| 0.003220322 | 0.042818062 |
| 0.003227175 | 0.042873261 |
| 0.003239871 | 0.043029929 |
| 0.003265809 | 0.04325381 |
| 0.003269391 | 0.043277177 |
| 0.003275163 | 0.043292226 |
| 0.003287369 | 0.043370507 |
| 0.003298291 | 0.043462435 |
| 0.003323534 | 0.043678238 |
| 0.003339141 | 0.043786694 |
| 0.003354234 | 0.043924297 |
| 0.003362095 | 0.044014983 |
| 0.003378359 | 0.04417932 |
| 0.003392639 | 0.044305236 |
| 0.003406799 | 0.044392784 |
| 0.00341537 | 0.04448734 |
| 0.003415924 | 0.04448734 |
| 0.00342002 | 0.04451634 |
| 0.003431157 | 0.044636908 |
| 0.003445213 | 0.044782229 |
| 0.003450638 | 0.044816895 |
| 0.003471452 | 0.045013576 |
| 0.003475164 | 0.045037185 |
| 0.003478844 | 0.04506647 |
| 0.003480313 | 0.045067117 |
| 0.003483648 | 0.045085787 |
| 0.003491095 | 0.04515762 |
| 0.003524347 | 0.045476591 |
| 0.00353283 | 0.045524371 |
| 0.003534784 | 0.045537232 |
| 0.003562423 | 0.04579371 |
| 0.003577963 | 0.045911461 |
| 0.003602749 | 0.046081438 |
| 0.00360416 | 0.046081965 |
| 0.003606978 | 0.046093254 |
| 0.003641163 | 0.046428249 |
| 0.003644032 | 0.046428249 |
| 0.00367019 | 0.046663149 |
| 0.00370054 | 0.046923757 |
| 0.003702414 | 0.04693502 |
| 0.00371925 | 0.047085782 |
| 0.003734342 | 0.047205779 |
| 0.003737621 | 0.047230479 |
| 0.003741196 | 0.047250587 |
| 0.003754713 | 0.047357279 |

| | |
|---|---|
| 0.003757401 | 0.047357279 |
| 0.003759979 | 0.047374736 |
| 0.003762115 | 0.047376589 |
| 0.003769254 | 0.047428865 |
| 0.003779053 | 0.047476911 |
| 0.003793081 | 0.047585514 |
| 0.003793693 | 0.047585514 |
| 0.003798529 | 0.047633627 |
| 0.003815361 | 0.047781805 |
| 0.003824157 | 0.047827157 |
| 0.003843266 | 0.048005046 |
| 0.003847006 | 0.04802657 |
| 0.003848326 | 0.048030456 |
| 0.003868355 | 0.048179427 |
| 0.003922104 | 0.048708752 |
| 0.003924949 | 0.048718668 |
| 0.003926018 | 0.048719249 |
| 0.003928666 | 0.048722705 |
| 0.003928783 | 0.048722705 |
| 0.003936008 | 0.048766975 |
| 0.003945862 | 0.048850942 |
| 0.003975291 | 0.049125894 |
| 0.004004814 | 0.049413802 |
| 0.004006932 | 0.049427131 |
| 0.004009567 | 0.049446821 |
| 0.004017021 | 0.049525923 |
| 0.004026732 | 0.049594301 |
| 0.004032757 | 0.049650933 |
| 0.00403412 | 0.049650933 |
| 0.004037847 | 0.049679808 |
| 0.004057347 | 0.049881073 |
| 0.004063604 | 0.049919342 |
| 0.004065312 | 0.049927439 |
| 0.004073606 | 0.049990628 |
| 0.004112304 | 0.050394724 |
| 0.004120275 | 0.050433391 |
| 0.004125519 | 0.050475328 |
| 0.004129502 | 0.050507388 |
| 0.004132676 | 0.050520265 |
| 0.004144646 | 0.050640597 |
| 0.004148948 | 0.050656159 |
| 0.004163441 | 0.050805073 |
| 0.004173296 | 0.05089925 |
| 0.004181834 | 0.050951189 |
| 0.00419512 | 0.051074309 |

| | |
|---|---|
| 0.004205779 | 0.051118119 |
| 0.004216784 | 0.051204887 |
| 0.004219458 | 0.051204887 |
| 0.004223352 | 0.051234225 |
| 0.004247372 | 0.051473173 |
| 0.004256977 | 0.051529779 |
| 0.004261488 | 0.051550863 |
| 0.004262441 | 0.051550863 |
| 0.004264433 | 0.051561861 |
| 0.00429879 | 0.051793189 |
| 0.004305958 | 0.051866425 |
| 0.004316804 | 0.051970784 |
| 0.004327833 | 0.052037789 |
| 0.004346646 | 0.052198109 |
| 0.004358341 | 0.052257822 |
| 0.00436384 | 0.052299096 |
| 0.004372984 | 0.052329681 |
| 0.00438991 | 0.052479475 |
| 0.004395847 | 0.052537265 |
| 0.004406662 | 0.052574171 |
| 0.004410981 | 0.052599347 |
| 0.004430532 | 0.052779627 |
| 0.004431761 | 0.052781068 |
| 0.004438432 | 0.052847297 |
| 0.004450636 | 0.052923057 |
| 0.004469722 | 0.053113635 |
| 0.004472291 | 0.053130906 |
| 0.004479125 | 0.053167489 |
| 0.004479354 | 0.053167489 |
| 0.004488863 | 0.053204175 |
| 0.004519906 | 0.053469833 |
| 0.004521824 | 0.053479238 |
| 0.00453013 | 0.053564174 |
| 0.00454116 | 0.053667949 |
| 0.004549407 | 0.053711849 |
| 0.00455395 | 0.05373781 |
| 0.004561619 | 0.053762987 |
| 0.004576047 | 0.053856535 |
| 0.004592682 | 0.053974398 |
| 0.004593156 | 0.053974398 |
| 0.004597007 | 0.054006328 |
| 0.004600465 | 0.054033627 |
| 0.004638661 | 0.054384182 |
| 0.004657238 | 0.054539038 |
| 0.00466581 | 0.054599145 |

| | |
|---|---|
| 0.004674694 | 0.054675026 |
| 0.004676693 | 0.054675026 |
| 0.004676886 | 0.054675026 |
| 0.004698578 | 0.054847834 |
| 0.00470305 | 0.054886584 |
| 0.004730388 | 0.055097066 |
| 0.004736875 | 0.055119263 |
| 0.004743096 | 0.055178176 |
| 0.00474829 | 0.055210087 |
| 0.004748426 | 0.055210087 |
| 0.004761054 | 0.055245587 |
| 0.004790458 | 0.055417814 |
| 0.004801611 | 0.055492918 |
| 0.004806991 | 0.055541617 |
| 0.00483862 | 0.055785275 |
| 0.00488933 | 0.056233785 |
| 0.004892271 | 0.056254034 |
| 0.00490385 | 0.05633277 |
| 0.004906775 | 0.056338094 |
| 0.004912242 | 0.056366502 |
| 0.004915493 | 0.056371348 |
| 0.004946307 | 0.056683772 |
| 0.004951576 | 0.056716865 |
| 0.004954197 | 0.056733243 |
| 0.004966078 | 0.056855621 |
| 0.004969565 | 0.056868207 |
| 0.00497246 | 0.056887668 |
| 0.005016248 | 0.057231365 |
| 0.005018124 | 0.057231365 |
| 0.005027106 | 0.057320075 |
| 0.005030148 | 0.05734104 |
| 0.005044858 | 0.05742626 |
| 0.005049165 | 0.057461548 |
| 0.005051097 | 0.057469807 |
| 0.005060123 | 0.057558747 |
| 0.005079253 | 0.057693675 |
| 0.005093737 | 0.057811084 |
| 0.005108206 | 0.05792584 |
| 0.005143492 | 0.058228926 |
| 0.005150942 | 0.05827171 |
| 0.005171659 | 0.058408969 |
| 0.005177551 | 0.058418527 |
| 0.005178639 | 0.058418527 |
| 0.005193876 | 0.058548811 |
| 0.005210257 | 0.05865537 |

| | |
|---|---|
| 0.00521561 | 0.058691179 |
| 0.005216364 | 0.058691179 |
| 0.005220582 | 0.058723459 |
| 0.005224803 | 0.058723459 |
| 0.005224838 | 0.058723459 |
| 0.005234899 | 0.058767704 |
| 0.005235504 | 0.058767704 |
| 0.005245831 | 0.058869744 |
| 0.005248466 | 0.058883568 |
| 0.005253245 | 0.058883568 |
| 0.005254646 | 0.058885416 |
| 0.005259378 | 0.058924587 |
| 0.005263051 | 0.058942911 |
| 0.005272267 | 0.059012447 |
| 0.005273416 | 0.059012447 |
| 0.00527705 | 0.059039243 |
| 0.005278625 | 0.059043 |
| 0.005291016 | 0.059139925 |
| 0.005294906 | 0.059156546 |
| 0.005312172 | 0.059265101 |
| 0.005313414 | 0.059265101 |
| 0.005317427 | 0.059282096 |
| 0.00532343 | 0.059321246 |
| 0.005335769 | 0.059400567 |
| 0.005345814 | 0.059459382 |
| 0.005358975 | 0.059550142 |
| 0.005366596 | 0.059607017 |
| 0.005378222 | 0.059694381 |
| 0.005390613 | 0.059804038 |
| 0.005420482 | 0.059995657 |
| 0.005422319 | 0.060002037 |
| 0.005425837 | 0.06001308 |
| 0.005436329 | 0.060115168 |
| 0.0054568 | 0.060285551 |
| 0.00546095 | 0.060317413 |
| 0.005491999 | 0.060573718 |
| 0.005500493 | 0.060627633 |
| 0.005505641 | 0.060654367 |
| 0.005521413 | 0.060723095 |
| 0.005522607 | 0.060723095 |
| 0.005524111 | 0.060723095 |
| 0.005531866 | 0.060734478 |
| 0.005537266 | 0.060751755 |
| 0.005548339 | 0.060831205 |
| 0.005576389 | 0.061040374 |

| | |
|---|---|
| 0.005600485 | 0.061158024 |
| 0.005601092 | 0.061158024 |
| 0.005601262 | 0.061158024 |
| 0.005603683 | 0.061170434 |
| 0.005607657 | 0.061199786 |
| 0.005625058 | 0.061327752 |
| 0.005634492 | 0.061354322 |
| 0.005644164 | 0.061429298 |
| 0.005653646 | 0.061490335 |
| 0.005665109 | 0.061572828 |
| 0.00566995 | 0.061603014 |
| 0.005673574 | 0.061608593 |
| 0.00568443 | 0.061642151 |
| 0.005688607 | 0.061659379 |
| 0.005705447 | 0.061782568 |
| 0.005709053 | 0.061782568 |
| 0.005729902 | 0.061963884 |
| 0.005731012 | 0.061963884 |
| 0.005737588 | 0.061995114 |
| 0.005742101 | 0.062013392 |
| 0.005754266 | 0.062130682 |
| 0.005757988 | 0.062156776 |
| 0.005772903 | 0.062275439 |
| 0.005782529 | 0.062351035 |
| 0.005805259 | 0.062553636 |
| 0.005844843 | 0.062880581 |
| 0.00585044 | 0.06289872 |
| 0.005860388 | 0.062976696 |
| 0.005862506 | 0.062985241 |
| 0.005865413 | 0.063002258 |
| 0.005868188 | 0.063017852 |
| 0.005942071 | 0.063549865 |
| 0.005947398 | 0.063581757 |
| 0.005950033 | 0.063595652 |
| 0.005959413 | 0.063632561 |
| 0.005961503 | 0.063632561 |
| 0.005973044 | 0.063727192 |
| 0.005976451 | 0.063735414 |
| 0.005976491 | 0.063735414 |
| 0.00600383 | 0.063955364 |
| 0.006009315 | 0.063985162 |
| 0.006018924 | 0.064073153 |
| 0.006031413 | 0.064163082 |
| 0.006043582 | 0.064235141 |
| 0.006053269 | 0.064283601 |

| | |
|---|---|
| 0.006054232 | 0.064283601 |
| 0.006071217 | 0.064413866 |
| 0.006086045 | 0.064528063 |
| 0.006124184 | 0.064773825 |
| 0.006141326 | 0.064897482 |
| 0.006144384 | 0.064911824 |
| 0.006146757 | 0.064911824 |
| 0.006165688 | 0.065010681 |
| 0.006192014 | 0.06518724 |
| 0.006210897 | 0.06533142 |
| 0.006214654 | 0.06535335 |
| 0.006227383 | 0.065443859 |
| 0.006259261 | 0.0656485 |
| 0.006262041 | 0.065663196 |
| 0.00627202 | 0.065724135 |
| 0.006274172 | 0.065724135 |
| 0.006274753 | 0.065724135 |
| 0.006293438 | 0.065876392 |
| 0.006297167 | 0.065900936 |
| 0.006300858 | 0.065907419 |
| 0.006313753 | 0.066016492 |
| 0.006328723 | 0.066114962 |
| 0.00635258 | 0.06632054 |
| 0.006391695 | 0.06658748 |
| 0.00639408 | 0.066593232 |
| 0.00641011 | 0.066701836 |
| 0.006452749 | 0.067052131 |
| 0.006463205 | 0.067107707 |
| 0.006466173 | 0.06710927 |
| 0.006471517 | 0.067120852 |
| 0.006475668 | 0.067149284 |
| 0.006498445 | 0.067297551 |
| 0.006508583 | 0.067329339 |
| 0.006518123 | 0.067374469 |
| 0.006521868 | 0.067374469 |
| 0.006522182 | 0.067374469 |
| 0.006536692 | 0.06747357 |
| 0.006552048 | 0.067555434 |
| 0.006557389 | 0.067555434 |
| 0.006563379 | 0.0675879 |
| 0.006577643 | 0.067705514 |
| 0.00658629 | 0.067765224 |
| 0.006591621 | 0.067805432 |
| 0.006611544 | 0.067937003 |
| 0.006613322 | 0.067940617 |

| | |
|---|---|
| 0.006654554 | 0.068218153 |
| 0.006663231 | 0.068232574 |
| 0.006674911 | 0.068286253 |
| 0.006687242 | 0.068346194 |
| 0.006731408 | 0.068635573 |
| 0.006762556 | 0.068849991 |
| 0.006780874 | 0.069006987 |
| 0.006792858 | 0.069069906 |
| 0.006803159 | 0.069159884 |
| 0.00682628 | 0.069320949 |
| 0.006833849 | 0.069353444 |
| 0.006838327 | 0.069375097 |
| 0.006856006 | 0.069489467 |
| 0.00685858 | 0.069499315 |
| 0.006864572 | 0.06951998 |
| 0.006864855 | 0.06951998 |
| 0.006866464 | 0.069521488 |
| 0.00688543 | 0.06961867 |
| 0.006886295 | 0.06961867 |
| 0.006892034 | 0.069661907 |
| 0.006895852 | 0.069685709 |
| 0.006914228 | 0.069797362 |
| 0.006925806 | 0.069869804 |
| 0.006972412 | 0.070225379 |
| 0.006977202 | 0.070225379 |
| 0.006978746 | 0.070225379 |
| 0.006996141 | 0.070370693 |
| 0.00700449 | 0.070422171 |
| 0.007017887 | 0.070511798 |
| 0.007021558 | 0.070511798 |
| 0.007022014 | 0.070511798 |
| 0.007030683 | 0.070554221 |
| 0.007045512 | 0.070658367 |
| 0.007057788 | 0.070736955 |
| 0.007065595 | 0.070800148 |
| 0.00710028 | 0.071021264 |
| 0.007126549 | 0.071164783 |
| 0.007136676 | 0.071212749 |
| 0.007149432 | 0.071295228 |
| 0.007164681 | 0.071413113 |
| 0.007166718 | 0.071413113 |
| 0.007167323 | 0.071413113 |
| 0.007168751 | 0.071413113 |
| 0.007179382 | 0.071489115 |
| 0.00718885 | 0.071549942 |

| | |
|---|---|
| 0.007189998 | 0.071549942 |
| 0.007191785 | 0.071552782 |
| 0.007202171 | 0.071626183 |
| 0.007209355 | 0.071682666 |
| 0.007216686 | 0.071695559 |
| 0.007220988 | 0.071695559 |
| 0.00722119 | 0.071695559 |
| 0.007225645 | 0.071724845 |
| 0.007257016 | 0.071875798 |
| 0.007262443 | 0.071880377 |
| 0.007276949 | 0.071988394 |
| 0.007290969 | 0.072018961 |
| 0.007313907 | 0.072179749 |
| 0.007335897 | 0.072320915 |
| 0.007368022 | 0.072533439 |
| 0.007375846 | 0.072555408 |
| 0.007384463 | 0.072579421 |
| 0.0073867 | 0.072579421 |
| 0.007398545 | 0.072653709 |
| 0.007404349 | 0.07267902 |
| 0.007417845 | 0.072738252 |
| 0.007423578 | 0.072764507 |
| 0.007428815 | 0.072779472 |
| 0.007428834 | 0.072779472 |
| 0.007429689 | 0.072779472 |
| 0.007447441 | 0.072893398 |
| 0.007450363 | 0.072907016 |
| 0.007454199 | 0.07291459 |
| 0.007480796 | 0.073084689 |
| 0.007485922 | 0.073119773 |
| 0.007495263 | 0.073177997 |
| 0.007496492 | 0.073177997 |
| 0.007504235 | 0.073223563 |
| 0.007508757 | 0.073252673 |
| 0.007553681 | 0.073550422 |
| 0.007553692 | 0.073550422 |
| 0.007638057 | 0.074026273 |
| 0.007648333 | 0.074068154 |
| 0.007656879 | 0.074092704 |
| 0.007657091 | 0.074092704 |
| 0.007664371 | 0.07413302 |
| 0.007674773 | 0.074203497 |
| 0.007683647 | 0.074261654 |
| 0.007695523 | 0.074298535 |
| 0.007702093 | 0.074346899 |

| | |
|---|---|
| 0.007715324 | 0.074449792 |
| 0.007780962 | 0.074898135 |
| 0.007782436 | 0.074898135 |
| 0.00778725 | 0.074901684 |
| 0.007798576 | 0.074901684 |
| 0.007802029 | 0.074901684 |
| 0.00781431 | 0.074974244 |
| 0.007867748 | 0.075359354 |
| 0.007869086 | 0.075359354 |
| 0.007936946 | 0.075830075 |
| 0.007952705 | 0.075950166 |
| 0.007962928 | 0.075986853 |
| 0.007977618 | 0.076050843 |
| 0.007983771 | 0.076094272 |
| 0.008006023 | 0.076260576 |
| 0.008015199 | 0.076308701 |
| 0.008019465 | 0.076312298 |
| 0.008022505 | 0.076325981 |
| 0.008025403 | 0.076327473 |
| 0.008028523 | 0.076327473 |
| 0.008056811 | 0.076511681 |
| 0.008058089 | 0.076511681 |
| 0.008060828 | 0.076522432 |
| 0.008090788 | 0.076655679 |
| 0.008101733 | 0.076704733 |
| 0.008176141 | 0.077262912 |
| 0.008212693 | 0.077500675 |
| 0.008278416 | 0.077948805 |
| 0.00827982 | 0.077948805 |
| 0.008287967 | 0.077994663 |
| 0.008302212 | 0.078097854 |
| 0.008386167 | 0.078649892 |
| 0.008387864 | 0.078649892 |
| 0.008392076 | 0.078649892 |
| 0.008402555 | 0.078700204 |
| 0.008402598 | 0.078700204 |
| 0.008409808 | 0.078722356 |
| 0.008412356 | 0.078722356 |
| 0.00844353 | 0.07888053 |
| 0.008444409 | 0.07888053 |
| 0.00844922 | 0.078904118 |
| 0.0084648 | 0.079003137 |
| 0.008472049 | 0.079039818 |
| 0.008532602 | 0.079449119 |
| 0.008541097 | 0.079505048 |

| | |
|---|---|
| 0.008544821 | 0.079505048 |
| 0.008576165 | 0.079667842 |
| 0.008580323 | 0.079675567 |
| 0.00858881 | 0.079723107 |
| 0.008598108 | 0.079747246 |
| 0.008601423 | 0.079762459 |
| 0.00860315 | 0.079762939 |
| 0.008605909 | 0.079772992 |
| 0.008637329 | 0.08004866 |
| 0.008646772 | 0.080077207 |
| 0.008647135 | 0.080077207 |
| 0.0086579 | 0.080110366 |
| 0.008672329 | 0.080170224 |
| 0.008685854 | 0.080248528 |
| 0.008698847 | 0.080336632 |
| 0.008703257 | 0.080346972 |
| 0.00870796 | 0.080374811 |
| 0.008711415 | 0.08038613 |
| 0.008717163 | 0.08038613 |
| 0.008736227 | 0.080495318 |
| 0.008784564 | 0.080878102 |
| 0.008822603 | 0.081150297 |
| 0.008832678 | 0.081150297 |
| 0.008834271 | 0.081150297 |
| 0.008836748 | 0.081150297 |
| 0.008842866 | 0.081150297 |
| 0.008846876 | 0.081169364 |
| 0.008858297 | 0.081208501 |
| 0.008860821 | 0.081208501 |
| 0.008868982 | 0.081246974 |
| 0.008891466 | 0.081374681 |
| 0.008898793 | 0.081426096 |
| 0.008904503 | 0.081448484 |
| 0.008911281 | 0.081479798 |
| 0.008912576 | 0.081479798 |
| 0.008938597 | 0.081634365 |
| 0.008962155 | 0.081754564 |
| 0.008972716 | 0.081788237 |
| 0.008991196 | 0.081941007 |
| 0.009078022 | 0.082560746 |
| 0.009102215 | 0.082715358 |
| 0.009140502 | 0.082968302 |
| 0.009143714 | 0.082981642 |
| 0.009157055 | 0.083076533 |
| 0.009172312 | 0.083123454 |

| | |
|---|---|
| 0.009175048 | 0.083123454 |
| 0.009197509 | 0.083258711 |
| 0.009198718 | 0.083258711 |
| 0.009208868 | 0.083334744 |
| 0.009284052 | 0.083728749 |
| 0.00928779 | 0.083744889 |
| 0.009300881 | 0.083775982 |
| 0.009328572 | 0.08392367 |
| 0.009334187 | 0.083931476 |
| 0.00933516 | 0.083931476 |
| 0.009346973 | 0.083962334 |
| 0.009357984 | 0.08404539 |
| 0.009387418 | 0.084198599 |
| 0.009397919 | 0.084276915 |
| 0.009426938 | 0.084425867 |
| 0.009448709 | 0.084541357 |
| 0.009453618 | 0.084569396 |
| 0.009471285 | 0.084607844 |
| 0.009473584 | 0.084607844 |
| 0.009473904 | 0.084607844 |
| 0.009492448 | 0.084741674 |
| 0.00950038 | 0.084783578 |
| 0.009507758 | 0.084814755 |
| 0.009511478 | 0.084832056 |
| 0.009524337 | 0.084883157 |
| 0.009528719 | 0.084906323 |
| 0.009546015 | 0.085028624 |
| 0.009550015 | 0.085032445 |
| 0.009562524 | 0.085112006 |
| 0.009568724 | 0.085150991 |
| 0.009598131 | 0.085283075 |
| 0.009599651 | 0.085283075 |
| 0.009614711 | 0.085384921 |
| 0.009616743 | 0.085384921 |
| 0.009620455 | 0.085388261 |
| 0.009640303 | 0.085516613 |
| 0.009676402 | 0.08570383 |
| 0.009677682 | 0.08570383 |
| 0.009720178 | 0.08598494 |
| 0.00977614 | 0.086351669 |
| 0.009781558 | 0.086383513 |
| 0.009796174 | 0.086496544 |
| 0.009814893 | 0.086632608 |
| 0.009815221 | 0.086632608 |
| 0.009838277 | 0.086691511 |

| | |
|---|---|
| 0.009844788 | 0.086732842 |
| 0.009849818 | 0.086761103 |
| 0.00987828 | 0.08692988 |
| 0.00987993 | 0.08692988 |
| 0.009912132 | 0.087087277 |
| 0.009912447 | 0.087087277 |
| 0.009937739 | 0.087212952 |
| 0.00999896 | 0.087524438 |
| 0.010025086 | 0.087704769 |
| 0.010030829 | 0.08773183 |
| 0.010039231 | 0.08773183 |
| 0.010050325 | 0.087752283 |
| 0.010050784 | 0.087752283 |
| 0.010052912 | 0.087754771 |
| 0.010066158 | 0.087838202 |
| 0.01008165 | 0.087925058 |
| 0.010086448 | 0.087950794 |
| 0.010104675 | 0.088061364 |
| 0.010108322 | 0.088071358 |
| 0.01010952 | 0.088071358 |
| 0.010141265 | 0.088267182 |
| 0.010193697 | 0.088561696 |
| 0.010204609 | 0.088603506 |
| 0.010205951 | 0.088603506 |
| 0.010222712 | 0.088694317 |
| 0.010255589 | 0.08887244 |
| 0.010269252 | 0.088878935 |
| 0.010269401 | 0.088878935 |
| 0.010280027 | 0.088922418 |
| 0.010289681 | 0.088957462 |
| 0.010309676 | 0.089106988 |
| 0.010326646 | 0.089208218 |
| 0.010328052 | 0.089208218 |
| 0.010339173 | 0.089271896 |
| 0.010345782 | 0.089296586 |
| 0.010366202 | 0.08945662 |
| 0.010375767 | 0.089506729 |
| 0.010384247 | 0.089515029 |
| 0.010451901 | 0.089902964 |
| 0.01045687 | 0.089913148 |
| 0.010459083 | 0.089913148 |
| 0.010468853 | 0.08991461 |
| 0.010469121 | 0.08991461 |
| 0.010477413 | 0.089959935 |
| 0.010493702 | 0.090083091 |

| | |
|---|---|
| 0.010498757 | 0.090086346 |
| 0.010528477 | 0.090268184 |
| 0.010564288 | 0.09039622 |
| 0.010573764 | 0.090461048 |
| 0.010613742 | 0.090705308 |
| 0.010640567 | 0.090833485 |
| 0.010669934 | 0.090934053 |
| 0.010680482 | 0.09094928 |
| 0.010694924 | 0.0910397 |
| 0.01070511 | 0.091110125 |
| 0.010710142 | 0.091136668 |
| 0.010723233 | 0.091151611 |
| 0.010761964 | 0.091398014 |
| 0.010784974 | 0.091494183 |
| 0.010789751 | 0.091503469 |
| 0.010797506 | 0.091536639 |
| 0.010863797 | 0.091869684 |
| 0.01089407 | 0.092043976 |
| 0.010934613 | 0.092271935 |
| 0.010948665 | 0.092374146 |
| 0.010977868 | 0.092505824 |
| 0.010990904 | 0.092566536 |
| 0.010996587 | 0.092581657 |
| 0.011001406 | 0.092605862 |
| 0.011016588 | 0.092700888 |
| 0.011033229 | 0.092808127 |
| 0.011042638 | 0.092854466 |
| 0.0110449 | 0.0928571 |
| 0.011067319 | 0.092956586 |
| 0.011070396 | 0.092956586 |
| 0.0110926 | 0.09306099 |
| 0.011177653 | 0.093586129 |
| 0.011180739 | 0.093586129 |
| 0.011190719 | 0.093620298 |
| 0.01122278 | 0.093706604 |
| 0.011226667 | 0.09370706 |
| 0.011232392 | 0.093717877 |
| 0.011233866 | 0.093717877 |
| 0.011241782 | 0.093750965 |
| 0.011243738 | 0.093750965 |
| 0.011259286 | 0.093814905 |
| 0.011338303 | 0.094176725 |
| 0.01139117 | 0.094460355 |
| 0.011398051 | 0.094460355 |
| 0.011420429 | 0.094594917 |

| | |
|---|---|
| 0.011428111 | 0.094642088 |
| 0.011449272 | 0.094800849 |
| 0.011464845 | 0.094847346 |
| 0.011495009 | 0.094981398 |
| 0.011544775 | 0.095296234 |
| 0.011556684 | 0.09535866 |
| 0.011578883 | 0.095517081 |
| 0.011583513 | 0.095518923 |
| 0.011587858 | 0.09553313 |
| 0.01159099 | 0.095542415 |
| 0.011605989 | 0.095632948 |
| 0.011660601 | 0.095950145 |
| 0.011675646 | 0.096024176 |
| 0.011696616 | 0.096113654 |
| 0.011699342 | 0.09611947 |
| 0.011729301 | 0.096204 |
| 0.011777275 | 0.096476827 |
| 0.011783817 | 0.096513815 |
| 0.011798193 | 0.096598341 |
| 0.011864855 | 0.096998841 |
| 0.011879949 | 0.09704253 |
| 0.0119573 | 0.097424558 |
| 0.011975321 | 0.09752883 |
| 0.011978117 | 0.097534926 |
| 0.012007559 | 0.097707847 |
| 0.012043636 | 0.097865471 |
| 0.012053099 | 0.097865471 |
| 0.012157751 | 0.098525983 |
| 0.012165421 | 0.09855061 |
| 0.012166997 | 0.09855061 |
| 0.012204638 | 0.09871541 |
| 0.012205998 | 0.09871541 |
| 0.012218052 | 0.098779349 |
| 0.012235107 | 0.09888189 |
| 0.01223904 | 0.09888189 |
| 0.01226573 | 0.099047122 |
| 0.012297612 | 0.099186849 |
| 0.0123316 | 0.099387934 |
| 0.012332977 | 0.099387934 |
| 0.012359536 | 0.099517771 |
| 0.012389461 | 0.099623985 |
| 0.012403631 | 0.099721092 |
| 0.012413976 | 0.099787416 |
| 0.012419772 | 0.09981716 |
| 0.012442022 | 0.099913236 |

| | |
|---|---|
| 0.012445551 | 0.099923176 |
| 0.012463017 | 0.100029679 |
| 0.012469457 | 0.100047648 |
| 0.012472925 | 0.100058612 |
| 0.012506759 | 0.100262479 |
| 0.012573431 | 0.100593781 |
| 0.012580336 | 0.100632125 |
| 0.012590931 | 0.100699965 |
| 0.012611287 | 0.100824961 |
| 0.012670412 | 0.101070812 |
| 0.012678649 | 0.101112862 |
| 0.012692809 | 0.101191898 |
| 0.012704462 | 0.101250908 |
| 0.012720892 | 0.101328908 |
| 0.012734759 | 0.10139057 |
| 0.012766558 | 0.101609771 |
| 0.012785821 | 0.101661154 |
| 0.012791826 | 0.101665211 |
| 0.012807661 | 0.1017126 |
| 0.0128249 | 0.101784955 |
| 0.012860927 | 0.102019879 |
| 0.01289271 | 0.10210301 |
| 0.012892844 | 0.10210301 |
| 0.012945204 | 0.102380268 |
| 0.012954638 | 0.102407233 |
| 0.012996109 | 0.102715845 |
| 0.013021386 | 0.102796214 |
| 0.013060003 | 0.102964539 |
| 0.01306419 | 0.102980496 |
| 0.013075374 | 0.103034556 |
| 0.013118018 | 0.103285148 |
| 0.01313026 | 0.103326258 |
| 0.013131918 | 0.103326258 |
| 0.013153955 | 0.103394418 |
| 0.013158363 | 0.103394418 |
| 0.013167111 | 0.103394418 |
| 0.013175315 | 0.103394418 |
| 0.013181689 | 0.103410361 |
| 0.013221226 | 0.103579606 |
| 0.013262204 | 0.103768519 |
| 0.013273092 | 0.103819602 |
| 0.013289821 | 0.103911299 |
| 0.013290832 | 0.103911299 |
| 0.013291361 | 0.103911299 |
| 0.013309638 | 0.103951117 |

| | |
|---|---|
| 0.013311321 | 0.103951117 |
| 0.013315701 | 0.103951117 |
| 0.01333477 | 0.103977584 |
| 0.01334125 | 0.10399406 |
| 0.013348593 | 0.104034269 |
| 0.013400318 | 0.104216981 |
| 0.013400483 | 0.104216981 |
| 0.013423117 | 0.104328826 |
| 0.013452179 | 0.104413587 |
| 0.013461312 | 0.104434255 |
| 0.01347951 | 0.104528697 |
| 0.013484954 | 0.104540718 |
| 0.013552089 | 0.10489929 |
| 0.013564504 | 0.104975761 |
| 0.013574474 | 0.104987247 |
| 0.013600001 | 0.105150528 |
| 0.013607758 | 0.105176353 |
| 0.013617104 | 0.105200941 |
| 0.013637148 | 0.105220341 |
| 0.013646588 | 0.105220341 |
| 0.01365354 | 0.105256901 |
| 0.013680502 | 0.105417943 |
| 0.01368107 | 0.105417943 |
| 0.013703712 | 0.105558258 |
| 0.013729851 | 0.10564456 |
| 0.013741993 | 0.105659764 |
| 0.013744712 | 0.105659764 |
| 0.01374573 | 0.105659764 |
| 0.01376407 | 0.105749532 |
| 0.013768881 | 0.105769434 |
| 0.013779585 | 0.105817527 |
| 0.013786236 | 0.105851531 |
| 0.013822008 | 0.106074894 |
| 0.013830922 | 0.106109106 |
| 0.013861174 | 0.106232691 |
| 0.013863868 | 0.106232691 |
| 0.01387729 | 0.106293611 |
| 0.013914173 | 0.106507607 |
| 0.013924173 | 0.106532784 |
| 0.013926675 | 0.106534814 |
| 0.013933907 | 0.106573017 |
| 0.013954855 | 0.106664718 |
| 0.013970543 | 0.106739939 |
| 0.014027067 | 0.106993449 |
| 0.014034295 | 0.10701493 |

| | |
|---|---|
| 0.01403665 | 0.107015129 |
| 0.014148987 | 0.107683344 |
| 0.014150904 | 0.107683344 |
| 0.014152356 | 0.107683344 |
| 0.014157887 | 0.107683344 |
| 0.014177043 | 0.107757743 |
| 0.014185241 | 0.107774648 |
| 0.014251527 | 0.108099613 |
| 0.014294843 | 0.108323475 |
| 0.014296961 | 0.108323475 |
| 0.0143175 | 0.108428731 |
| 0.014335004 | 0.108490901 |
| 0.014365114 | 0.108600691 |
| 0.014367797 | 0.108600691 |
| 0.014384903 | 0.108626888 |
| 0.014404687 | 0.108740305 |
| 0.014409055 | 0.108740305 |
| 0.014422067 | 0.108786791 |
| 0.014440841 | 0.10887667 |
| 0.01448115 | 0.109025583 |
| 0.014481194 | 0.109025583 |
| 0.014502986 | 0.109172392 |
| 0.014529029 | 0.109313954 |
| 0.014533268 | 0.109313954 |
| 0.014558449 | 0.109468779 |
| 0.014561483 | 0.109474309 |
| 0.014565918 | 0.109490372 |
| 0.014574285 | 0.109535975 |
| 0.014587678 | 0.109589089 |
| 0.014635925 | 0.10977656 |
| 0.014636103 | 0.10977656 |
| 0.014646585 | 0.109836706 |
| 0.014700684 | 0.110142132 |
| 0.014703858 | 0.110144799 |
| 0.014727221 | 0.110267783 |
| 0.01473914 | 0.110330256 |
| 0.01477644 | 0.110532039 |
| 0.014791232 | 0.110607944 |
| 0.014810736 | 0.110719027 |
| 0.014857047 | 0.11099554 |
| 0.014882977 | 0.11113696 |
| 0.014886729 | 0.111147556 |
| 0.014901473 | 0.111195626 |
| 0.014902722 | 0.111195626 |
| 0.014971531 | 0.111518474 |

| | |
|---|---|
| 0.015004111 | 0.11167382 |
| 0.015010024 | 0.111700378 |
| 0.01503616 | 0.11182498 |
| 0.015050236 | 0.111868005 |
| 0.01505134 | 0.111868005 |
| 0.015069266 | 0.11194883 |
| 0.015108773 | 0.112119013 |
| 0.015132737 | 0.112210318 |
| 0.015136128 | 0.112217994 |
| 0.015154334 | 0.112300523 |
| 0.015157018 | 0.112302941 |
| 0.015165164 | 0.112322621 |
| 0.015165436 | 0.112322621 |
| 0.015170498 | 0.112332916 |
| 0.015197678 | 0.112481709 |
| 0.015205223 | 0.112494106 |
| 0.015228171 | 0.112602407 |
| 0.015250549 | 0.112697891 |
| 0.015258142 | 0.112719024 |
| 0.015261619 | 0.112727224 |
| 0.015264911 | 0.112734056 |
| 0.015275569 | 0.112795274 |
| 0.015294912 | 0.112885599 |
| 0.015320464 | 0.113039152 |
| 0.015374225 | 0.113312942 |
| 0.015387575 | 0.113393782 |
| 0.015399299 | 0.113445074 |
| 0.015419547 | 0.113576674 |
| 0.015426402 | 0.113609594 |
| 0.015428826 | 0.113609876 |
| 0.015445125 | 0.113631954 |
| 0.015447877 | 0.113631954 |
| 0.01544846 | 0.113631954 |
| 0.015450723 | 0.113631954 |
| 0.015484311 | 0.113789733 |
| 0.015493384 | 0.113821287 |
| 0.015524042 | 0.113993762 |
| 0.015537626 | 0.114013039 |
| 0.015553532 | 0.114087181 |
| 0.015563813 | 0.114110232 |
| 0.015606515 | 0.114229519 |
| 0.015615512 | 0.114243041 |
| 0.01562048 | 0.114243952 |
| 0.015629176 | 0.11427887 |
| 0.015657035 | 0.114458577 |

| | |
|---|---|
| 0.015662301 | 0.114479497 |
| 0.015688837 | 0.114620685 |
| 0.015732792 | 0.11480093 |
| 0.015746694 | 0.114856159 |
| 0.015750007 | 0.114856159 |
| 0.015795608 | 0.114959877 |
| 0.01581932 | 0.11507355 |
| 0.015820813 | 0.11507355 |
| 0.015849267 | 0.115169107 |
| 0.015858304 | 0.115169107 |
| 0.015859363 | 0.115169107 |
| 0.015860629 | 0.115169107 |
| 0.01586445 | 0.115170924 |
| 0.015865715 | 0.115170924 |
| 0.015879403 | 0.115252716 |
| 0.0159496 | 0.115603652 |
| 0.01598433 | 0.115780344 |
| 0.015993945 | 0.115819315 |
| 0.016017392 | 0.115964306 |
| 0.016037104 | 0.116009947 |
| 0.016041803 | 0.116024746 |
| 0.016049501 | 0.116045178 |
| 0.016094287 | 0.116257062 |
| 0.016095177 | 0.116257062 |
| 0.016095892 | 0.116257062 |
| 0.016176388 | 0.116739519 |
| 0.016239938 | 0.117072842 |
| 0.016246586 | 0.117096876 |
| 0.016260425 | 0.117161156 |
| 0.016353979 | 0.117675018 |
| 0.016368951 | 0.117695576 |
| 0.016430332 | 0.117903783 |
| 0.016443696 | 0.117928612 |
| 0.016457281 | 0.117990504 |
| 0.016471641 | 0.118051977 |
| 0.016484283 | 0.118094784 |
| 0.016486703 | 0.118094784 |
| 0.016526962 | 0.11832027 |
| 0.01655808 | 0.118499128 |
| 0.016600733 | 0.118608325 |
| 0.016617016 | 0.118689144 |
| 0.01666144 | 0.118863753 |
| 0.016716069 | 0.119027956 |
| 0.016718591 | 0.119027956 |
| 0.016748051 | 0.119161958 |

| | |
|---|---|
| 0.016748276 | 0.119161958 |
| 0.016807049 | 0.119437378 |
| 0.016828289 | 0.119463237 |
| 0.016889502 | 0.119737471 |
| 0.016909041 | 0.119844711 |
| 0.016915684 | 0.119844711 |
| 0.016918135 | 0.119844711 |
| 0.016932831 | 0.119901867 |
| 0.016946406 | 0.11998015 |
| 0.016955391 | 0.1200217 |
| 0.016972444 | 0.120058981 |
| 0.016972665 | 0.120058981 |
| 0.016983035 | 0.12009666 |
| 0.01699069 | 0.120132955 |
| 0.017022394 | 0.120339256 |
| 0.017041495 | 0.120456408 |
| 0.01705683 | 0.120546913 |
| 0.017076862 | 0.120659515 |
| 0.017100716 | 0.120767469 |
| 0.017124789 | 0.120883702 |
| 0.017163953 | 0.121016669 |
| 0.017168662 | 0.121019429 |
| 0.017169427 | 0.121019429 |
| 0.017184461 | 0.121082468 |
| 0.017288755 | 0.121590599 |
| 0.017350447 | 0.121898474 |
| 0.017382286 | 0.122021619 |
| 0.017383347 | 0.122021619 |
| 0.017448153 | 0.122242306 |
| 0.017480332 | 0.122413732 |
| 0.01751542 | 0.12255133 |
| 0.017530006 | 0.122599352 |
| 0.017613218 | 0.122946609 |
| 0.017629266 | 0.123022573 |
| 0.017644063 | 0.123071729 |
| 0.017699185 | 0.123357271 |
| 0.01770226 | 0.123357271 |
| 0.017728278 | 0.123478315 |
| 0.017760604 | 0.123649218 |
| 0.01776389 | 0.123654017 |
| 0.017826384 | 0.123901392 |
| 0.017827924 | 0.123901392 |
| 0.017828044 | 0.123901392 |
| 0.017867404 | 0.123975942 |
| 0.017875297 | 0.123978227 |

| | |
|---|---|
| 0.017883234 | 0.123997663 |
| 0.017894219 | 0.124035504 |
| 0.017925958 | 0.124106032 |
| 0.017932184 | 0.124107132 |
| 0.017947074 | 0.124143968 |
| 0.017948965 | 0.124143968 |
| 0.017958241 | 0.12419009 |
| 0.017975299 | 0.124290004 |
| 0.01805683 | 0.124673912 |
| 0.018059616 | 0.124673912 |
| 0.018065048 | 0.124693336 |
| 0.018075766 | 0.124749238 |
| 0.018079664 | 0.124758067 |
| 0.018139739 | 0.125009601 |
| 0.018184667 | 0.125210517 |
| 0.0182829 | 0.125723313 |
| 0.018298861 | 0.125805067 |
| 0.018305206 | 0.125805067 |
| 0.018306882 | 0.125805067 |
| 0.018323287 | 0.12587382 |
| 0.018332537 | 0.125901045 |
| 0.018364702 | 0.126049241 |
| 0.018395784 | 0.126208008 |
| 0.018473129 | 0.126592767 |
| 0.018516166 | 0.126812571 |
| 0.018533153 | 0.126876311 |
| 0.01856347 | 0.127011322 |
| 0.018575995 | 0.127047417 |
| 0.01857682 | 0.127047417 |
| 0.018613653 | 0.127166377 |
| 0.018619545 | 0.127175255 |
| 0.018707256 | 0.127627918 |
| 0.018728401 | 0.127735581 |
| 0.018754654 | 0.127856865 |
| 0.018779389 | 0.127941103 |
| 0.01880006 | 0.128022654 |
| 0.018827807 | 0.12812001 |
| 0.018834876 | 0.12814475 |
| 0.018836824 | 0.12814475 |
| 0.01884387 | 0.128156074 |
| 0.018848977 | 0.128172507 |
| 0.018856602 | 0.128204372 |
| 0.018859405 | 0.128204372 |
| 0.018861739 | 0.128204372 |
| 0.018896746 | 0.128369034 |

| | |
|---|---|
| 0.018914826 | 0.128418594 |
| 0.018945045 | 0.128532147 |
| 0.018958797 | 0.1285705 |
| 0.018964805 | 0.128592613 |
| 0.018971416 | 0.128592613 |
| 0.018973072 | 0.128592613 |
| 0.019021533 | 0.128839777 |
| 0.019030683 | 0.128839777 |
| 0.019030966 | 0.128839777 |
| 0.019073385 | 0.129042335 |
| 0.019077988 | 0.129048049 |
| 0.019100648 | 0.1291463 |
| 0.019117261 | 0.129185272 |
| 0.019127864 | 0.129220253 |
| 0.019193836 | 0.129512356 |
| 0.019200474 | 0.129512356 |
| 0.019286069 | 0.129810289 |
| 0.019294572 | 0.129830824 |
| 0.01930025 | 0.129850685 |
| 0.019337284 | 0.129952985 |
| 0.019384848 | 0.130162434 |
| 0.019391489 | 0.130184407 |
| 0.019413697 | 0.130245973 |
| 0.019456463 | 0.130459378 |
| 0.019466046 | 0.130462541 |
| 0.019466064 | 0.130462541 |
| 0.019473966 | 0.130462541 |
| 0.019520707 | 0.130561718 |
| 0.019566191 | 0.130806371 |
| 0.019582927 | 0.130865171 |
| 0.019600855 | 0.130948231 |
| 0.019637579 | 0.131083245 |
| 0.019730207 | 0.131450324 |
| 0.019734986 | 0.13145709 |
| 0.019778472 | 0.13159951 |
| 0.019781559 | 0.131601665 |
| 0.019791439 | 0.131630622 |
| 0.019800744 | 0.131671738 |
| 0.019830814 | 0.131818869 |
| 0.019841125 | 0.131850604 |
| 0.019849989 | 0.131884572 |
| 0.019856885 | 0.131884572 |
| 0.019871567 | 0.13192405 |
| 0.019914126 | 0.132059328 |
| 0.019938848 | 0.132186459 |

| | |
|---|---|
| 0.019961407 | 0.132317594 |
| 0.019975301 | 0.132387615 |
| 0.019985624 | 0.132395872 |
| 0.019998225 | 0.132429516 |
| 0.020043048 | 0.132600397 |
| 0.020099685 | 0.132938158 |
| 0.020116031 | 0.132990864 |
| 0.020141883 | 0.13309295 |
| 0.020143201 | 0.13309295 |
| 0.020193478 | 0.133225837 |
| 0.020194134 | 0.133225837 |
| 0.020196329 | 0.133225837 |
| 0.020210382 | 0.133270311 |
| 0.020211465 | 0.133270311 |
| 0.020248153 | 0.133456793 |
| 0.020304592 | 0.133699265 |
| 0.020331936 | 0.133805324 |
| 0.020356185 | 0.133890902 |
| 0.020366028 | 0.133918654 |
| 0.020422186 | 0.134190406 |
| 0.020427547 | 0.134193488 |
| 0.020430765 | 0.134196116 |
| 0.020540694 | 0.134713803 |
| 0.020559082 | 0.134790045 |
| 0.02056667 | 0.134790045 |
| 0.020569127 | 0.134790045 |
| 0.020613118 | 0.134965762 |
| 0.020621236 | 0.135000357 |
| 0.020638182 | 0.135092731 |
| 0.020751003 | 0.135506235 |
| 0.020752165 | 0.135506235 |
| 0.020809699 | 0.135693253 |
| 0.02082482 | 0.135742339 |
| 0.020825777 | 0.135742339 |
| 0.020850432 | 0.135766657 |
| 0.020890793 | 0.135942861 |
| 0.020907818 | 0.1359573 |
| 0.020914902 | 0.1359573 |
| 0.020915508 | 0.1359573 |
| 0.020994602 | 0.136338955 |
| 0.021007338 | 0.136403065 |
| 0.021058277 | 0.136618417 |
| 0.021060584 | 0.136618417 |
| 0.021135148 | 0.136990157 |
| 0.02114193 | 0.136996831 |

| | |
|---|---|
| 0.021183773 | 0.137174654 |
| 0.021216119 | 0.137328094 |
| 0.021264836 | 0.137587331 |
| 0.021287645 | 0.137697496 |
| 0.021305469 | 0.137794072 |
| 0.021339298 | 0.137956665 |
| 0.021374693 | 0.138021578 |
| 0.021376768 | 0.138021578 |
| 0.021378319 | 0.138021578 |
| 0.021409709 | 0.138149331 |
| 0.02141474 | 0.138163074 |
| 0.021440818 | 0.138275133 |
| 0.02145182 | 0.138289909 |
| 0.021461428 | 0.1383144 |
| 0.021471331 | 0.138333904 |
| 0.021473168 | 0.138333904 |
| 0.021510261 | 0.138535387 |
| 0.02152417 | 0.138562267 |
| 0.021549017 | 0.138627822 |
| 0.021549935 | 0.138627822 |
| 0.021550809 | 0.138627822 |
| 0.021604268 | 0.138840436 |
| 0.021640733 | 0.139018506 |
| 0.021649221 | 0.139033439 |
| 0.021657654 | 0.139033439 |
| 0.021682129 | 0.139134301 |
| 0.02170939 | 0.139235508 |
| 0.021709595 | 0.139235508 |
| 0.021741391 | 0.139369809 |
| 0.02174224 | 0.139369809 |
| 0.021769883 | 0.139490679 |
| 0.021853731 | 0.139914996 |
| 0.021866631 | 0.13995324 |
| 0.021901187 | 0.140064028 |
| 0.021903822 | 0.140064028 |
| 0.021903917 | 0.140064028 |
| 0.021935805 | 0.140124084 |
| 0.021954336 | 0.140128851 |
| 0.021992101 | 0.14028687 |
| 0.02199429 | 0.14028687 |
| 0.02200069 | 0.140290111 |
| 0.022070961 | 0.14062524 |
| 0.022137621 | 0.140974528 |
| 0.02215686 | 0.141078176 |
| 0.022174834 | 0.141154881 |

| | |
|---|---|
| 0.022216842 | 0.141365604 |
| 0.022237074 | 0.141450079 |
| 0.02226557 | 0.141548369 |
| 0.022267746 | 0.141548369 |
| 0.022272314 | 0.141548369 |
| 0.022309133 | 0.141706732 |
| 0.022347608 | 0.141837621 |
| 0.022353132 | 0.141853783 |
| 0.02236316 | 0.141889691 |
| 0.022384202 | 0.141918581 |
| 0.022403562 | 0.141965735 |
| 0.022481725 | 0.142423134 |
| 0.022509293 | 0.142521952 |
| 0.022516523 | 0.142529827 |
| 0.022563883 | 0.14263145 |
| 0.022565263 | 0.14263145 |
| 0.022575676 | 0.142657782 |
| 0.022601964 | 0.142710443 |
| 0.022601988 | 0.142710443 |
| 0.022616963 | 0.142767127 |
| 0.022722774 | 0.14313154 |
| 0.022776868 | 0.143226023 |
| 0.022811765 | 0.143388669 |
| 0.022833875 | 0.143468774 |
| 0.022836559 | 0.143468774 |
| 0.022862795 | 0.143557855 |
| 0.02291029 | 0.143818158 |
| 0.022938178 | 0.143936675 |
| 0.022941258 | 0.143936675 |
| 0.023002702 | 0.1442082 |
| 0.023037417 | 0.144368824 |
| 0.023045292 | 0.144381324 |
| 0.023069529 | 0.144497785 |
| 0.02314541 | 0.144816927 |
| 0.02319021 | 0.145021027 |
| 0.023205804 | 0.145061403 |
| 0.023241899 | 0.145212815 |
| 0.023244537 | 0.145212815 |
| 0.023294577 | 0.145387343 |
| 0.023334609 | 0.145501354 |
| 0.0234005 | 0.145742858 |
| 0.023415791 | 0.145819022 |
| 0.023458523 | 0.146008765 |
| 0.023469443 | 0.146038565 |
| 0.023476733 | 0.14604576 |

| | |
|---|---|
| 0.023487631 | 0.146056355 |
| 0.023522425 | 0.146234503 |
| 0.023568216 | 0.146461827 |
| 0.023608847 | 0.146637933 |
| 0.02362843 | 0.146721149 |
| 0.023659352 | 0.146855721 |
| 0.023686271 | 0.146946204 |
| 0.023777653 | 0.147321227 |
| 0.023844264 | 0.147591128 |
| 0.023856316 | 0.147591128 |
| 0.023856418 | 0.147591128 |
| 0.023861501 | 0.147591128 |
| 0.023913881 | 0.147857513 |
| 0.023939307 | 0.14792565 |
| 0.023956886 | 0.147980274 |
| 0.023978722 | 0.148065394 |
| 0.023989269 | 0.148092925 |
| 0.024016725 | 0.148138189 |
| 0.024122314 | 0.148433264 |
| 0.024154441 | 0.148573365 |
| 0.02417795 | 0.148678491 |
| 0.024187141 | 0.148678491 |
| 0.024207874 | 0.148717122 |
| 0.024209038 | 0.148717122 |
| 0.024221676 | 0.148756385 |
| 0.024251774 | 0.148883633 |
| 0.024257545 | 0.148888791 |
| 0.024261147 | 0.148888791 |
| 0.024264987 | 0.148888791 |
| 0.024281118 | 0.148908914 |
| 0.02428173 | 0.148908914 |
| 0.024284031 | 0.148908914 |
| 0.024296386 | 0.14893189 |
| 0.024297159 | 0.14893189 |
| 0.024303237 | 0.148949976 |
| 0.024330283 | 0.149058193 |
| 0.024364407 | 0.149175982 |
| 0.024405713 | 0.149328225 |
| 0.024578399 | 0.150076333 |
| 0.024633087 | 0.150231782 |
| 0.024635401 | 0.150231782 |
| 0.024645742 | 0.150275603 |
| 0.024690461 | 0.150487814 |
| 0.024700594 | 0.150501671 |
| 0.024728192 | 0.150643336 |

| | |
|---|---|
| 0.02475207 | 0.150711688 |
| 0.024757771 | 0.15071383 |
| 0.024836937 | 0.15101615 |
| 0.024847539 | 0.151027041 |
| 0.024938951 | 0.151327228 |
| 0.024959641 | 0.151433482 |
| 0.02505668 | 0.151813956 |
| 0.0250739 | 0.151875189 |
| 0.025114533 | 0.15192576 |
| 0.02512462 | 0.15192576 |
| 0.025193745 | 0.152133261 |
| 0.025252525 | 0.152395085 |
| 0.025272493 | 0.152492402 |
| 0.02529452 | 0.152552084 |
| 0.025331254 | 0.152657429 |
| 0.02543774 | 0.153105427 |
| 0.025508582 | 0.153415483 |
| 0.025544576 | 0.153535023 |
| 0.025551485 | 0.153557172 |
| 0.025559629 | 0.153574673 |
| 0.025567295 | 0.153574673 |
| 0.025645173 | 0.153925979 |
| 0.025649404 | 0.153931974 |
| 0.025657985 | 0.153961012 |
| 0.025667679 | 0.153983442 |
| 0.025692092 | 0.154052281 |
| 0.025735097 | 0.154182683 |
| 0.02575212 | 0.154218057 |
| 0.025807795 | 0.15445437 |
| 0.025811742 | 0.15445596 |
| 0.025817889 | 0.15445596 |
| 0.025824276 | 0.15445596 |
| 0.025842352 | 0.154502471 |
| 0.025853129 | 0.154512108 |
| 0.02588113 | 0.15462491 |
| 0.025881743 | 0.15462491 |
| 0.026012813 | 0.15521326 |
| 0.026019939 | 0.155236329 |
| 0.026026171 | 0.155254063 |
| 0.026055071 | 0.155387539 |
| 0.026075493 | 0.155450932 |
| 0.026130394 | 0.155700271 |
| 0.026138176 | 0.15572716 |
| 0.026142013 | 0.1557293 |
| 0.026171576 | 0.1557621 |

| | |
|---|---|
| 0.026172283 | 0.1557621 |
| 0.026183176 | 0.1557621 |
| 0.026206126 | 0.155875133 |
| 0.026299181 | 0.156158972 |
| 0.026314717 | 0.156212269 |
| 0.026343931 | 0.156327236 |
| 0.02636586 | 0.156395023 |
| 0.026367405 | 0.156395023 |
| 0.026368489 | 0.156395023 |
| 0.026450558 | 0.156770498 |
| 0.026457319 | 0.156785209 |
| 0.026464123 | 0.156806021 |
| 0.026477683 | 0.156847338 |
| 0.026514797 | 0.156930563 |
| 0.026544494 | 0.157008771 |
| 0.026580044 | 0.157160488 |
| 0.026645724 | 0.157366702 |
| 0.026677794 | 0.157475022 |
| 0.02680631 | 0.157940857 |
| 0.026851431 | 0.158132037 |
| 0.026853333 | 0.158132037 |
| 0.026870616 | 0.158191203 |
| 0.026892119 | 0.158277306 |
| 0.026894123 | 0.158277306 |
| 0.026908795 | 0.158333252 |
| 0.026938122 | 0.158463363 |
| 0.026940136 | 0.158463363 |
| 0.027017574 | 0.15884039 |
| 0.027042675 | 0.15896166 |
| 0.027069961 | 0.159029292 |
| 0.027154438 | 0.159291097 |
| 0.027211213 | 0.159467018 |
| 0.02725042 | 0.159618223 |
| 0.027304381 | 0.159739183 |
| 0.027312959 | 0.159746643 |
| 0.027318113 | 0.159746643 |
| 0.027320556 | 0.159746643 |
| 0.027322656 | 0.159746643 |
| 0.027397808 | 0.160068133 |
| 0.027432877 | 0.160128905 |
| 0.027433226 | 0.160128905 |
| 0.027445186 | 0.160128905 |
| 0.027448556 | 0.160128905 |
| 0.027469741 | 0.160177678 |
| 0.027475426 | 0.160177678 |

| | |
|---|---|
| 0.027489431 | 0.160202609 |
| 0.027491463 | 0.160202609 |
| 0.027501213 | 0.160239815 |
| 0.027539622 | 0.160345912 |
| 0.027609578 | 0.16069429 |
| 0.027667438 | 0.160932715 |
| 0.027682016 | 0.160949798 |
| 0.027689843 | 0.160949798 |
| 0.027698389 | 0.160949798 |
| 0.027700789 | 0.160949798 |
| 0.027717771 | 0.160984695 |
| 0.02773865 | 0.161071527 |
| 0.027752125 | 0.161130131 |
| 0.027837731 | 0.161528708 |
| 0.027935516 | 0.16187916 |
| 0.027994008 | 0.162020974 |
| 0.028016087 | 0.162129059 |
| 0.028039826 | 0.162204571 |
| 0.028065698 | 0.162240836 |
| 0.028069467 | 0.162240836 |
| 0.028076862 | 0.162244199 |
| 0.028087044 | 0.162244937 |
| 0.028088889 | 0.162244937 |
| 0.028102589 | 0.162274735 |
| 0.028115529 | 0.16231226 |
| 0.028115942 | 0.16231226 |
| 0.028126392 | 0.16231226 |
| 0.028128121 | 0.16231226 |
| 0.028161506 | 0.162457367 |
| 0.028275494 | 0.162995645 |
| 0.028289668 | 0.163038724 |
| 0.028306218 | 0.163055182 |
| 0.028312048 | 0.163069042 |
| 0.028321938 | 0.163106282 |
| 0.028335033 | 0.163161964 |
| 0.028384164 | 0.163326395 |
| 0.028432967 | 0.163488124 |
| 0.028468677 | 0.16361502 |
| 0.028578647 | 0.164088637 |
| 0.028585452 | 0.164107919 |
| 0.028602996 | 0.164144835 |
| 0.028614679 | 0.164165451 |
| 0.028625485 | 0.16419481 |
| 0.02863626 | 0.16419481 |
| 0.02867423 | 0.164293471 |

| | |
|---|---|
| 0.02867718 | 0.164293471 |
| 0.028705367 | 0.164400014 |
| 0.028723726 | 0.164485382 |
| 0.0287329 | 0.164515713 |
| 0.028779747 | 0.164645685 |
| 0.028784868 | 0.164645685 |
| 0.028787042 | 0.164645685 |
| 0.028800265 | 0.164666292 |
| 0.028809176 | 0.164677704 |
| 0.028836744 | 0.164756192 |
| 0.028900153 | 0.165019428 |
| 0.028903736 | 0.165019428 |
| 0.029061563 | 0.165682913 |
| 0.029065674 | 0.165686518 |
| 0.029092109 | 0.165779284 |
| 0.029099352 | 0.165779284 |
| 0.029144006 | 0.16592844 |
| 0.02914513 | 0.16592844 |
| 0.029186124 | 0.166055595 |
| 0.029186205 | 0.166055595 |
| 0.029226142 | 0.166203399 |
| 0.029312698 | 0.166536545 |
| 0.0294726 | 0.167265434 |
| 0.029486756 | 0.167325833 |
| 0.029592084 | 0.167758127 |
| 0.029614872 | 0.16777299 |
| 0.02965995 | 0.167928491 |
| 0.029681422 | 0.16801011 |
| 0.029701113 | 0.168021723 |
| 0.029716721 | 0.168070094 |
| 0.029731961 | 0.16810107 |
| 0.029732817 | 0.16810107 |
| 0.029736316 | 0.16810107 |
| 0.0297693 | 0.168248901 |
| 0.029769532 | 0.168248901 |
| 0.029781661 | 0.168297484 |
| 0.029811214 | 0.168404543 |
| 0.029824365 | 0.168450312 |
| 0.029858003 | 0.1685609 |
| 0.029880509 | 0.168615997 |
| 0.029932211 | 0.168727829 |
| 0.029945227 | 0.168776001 |
| 0.029990926 | 0.168934036 |
| 0.030004297 | 0.168961537 |
| 0.030005599 | 0.168961537 |

| | |
|---|---|
| 0.030027355 | 0.169044072 |
| 0.030031598 | 0.169047973 |
| 0.030074607 | 0.169230069 |
| 0.030080108 | 0.169230937 |
| 0.030094586 | 0.169249471 |
| 0.030142386 | 0.169391316 |
| 0.030155911 | 0.169447331 |
| 0.030169335 | 0.169502761 |
| 0.030188799 | 0.169560962 |
| 0.030190374 | 0.169560962 |
| 0.030261261 | 0.169785928 |
| 0.030261476 | 0.169785928 |
| 0.030334692 | 0.170050654 |
| 0.030357047 | 0.170140589 |
| 0.030401673 | 0.170271462 |
| 0.030446567 | 0.170425048 |
| 0.030464014 | 0.170474632 |
| 0.030476933 | 0.170526892 |
| 0.030488148 | 0.1705696 |
| 0.030586468 | 0.171059381 |
| 0.030591891 | 0.171069623 |
| 0.030678024 | 0.171309914 |
| 0.030727718 | 0.17148688 |
| 0.030772472 | 0.171693272 |
| 0.030775515 | 0.171693272 |
| 0.030789497 | 0.171751158 |
| 0.030824647 | 0.171879681 |
| 0.030904512 | 0.172162651 |
| 0.030919512 | 0.172194058 |
| 0.030929408 | 0.172198797 |
| 0.030963737 | 0.172288453 |
| 0.030971035 | 0.172293078 |
| 0.031010166 | 0.172436799 |
| 0.031018767 | 0.172444356 |
| 0.031057098 | 0.172574572 |
| 0.031095892 | 0.17269775 |
| 0.031098536 | 0.17269775 |
| 0.031104233 | 0.17269775 |
| 0.031129345 | 0.172817033 |
| 0.031176837 | 0.172974982 |
| 0.031183219 | 0.172974982 |
| 0.031198644 | 0.17302024 |
| 0.031207192 | 0.173027351 |
| 0.031247017 | 0.17317804 |
| 0.031250952 | 0.17317804 |

| | |
|---|---|
| 0.031263559 | 0.17317804 |
| 0.031307988 | 0.173323902 |
| 0.031321732 | 0.173340426 |
| 0.031329922 | 0.173340426 |
| 0.03135427 | 0.173418889 |
| 0.031365198 | 0.173418889 |
| 0.031404742 | 0.173577075 |
| 0.0314139 | 0.173587403 |
| 0.031447485 | 0.173655059 |
| 0.031453237 | 0.173655059 |
| 0.031496974 | 0.173784223 |
| 0.031561999 | 0.173976797 |
| 0.031563654 | 0.173976797 |
| 0.031563703 | 0.173976797 |
| 0.031605807 | 0.174094363 |
| 0.031676849 | 0.17425194 |
| 0.03169817 | 0.174344602 |
| 0.031720953 | 0.174344602 |
| 0.031722979 | 0.174344602 |
| 0.031740657 | 0.174417409 |
| 0.031763766 | 0.174468101 |
| 0.031801698 | 0.174595902 |
| 0.031821122 | 0.174622028 |
| 0.031827078 | 0.174634592 |
| 0.03183136 | 0.174637967 |
| 0.03184726 | 0.17464601 |
| 0.031878493 | 0.174752035 |
| 0.03193607 | 0.17491021 |
| 0.032013365 | 0.175252935 |
| 0.032041513 | 0.175363149 |
| 0.032060413 | 0.175380913 |
| 0.032073296 | 0.175396992 |
| 0.032117894 | 0.175562831 |
| 0.032122013 | 0.175565196 |
| 0.032137649 | 0.175610351 |
| 0.032193375 | 0.175733398 |
| 0.032247638 | 0.175948935 |
| 0.032292909 | 0.176051731 |
| 0.032294829 | 0.176051731 |
| 0.032391596 | 0.176370702 |
| 0.032408321 | 0.176377996 |
| 0.03241061 | 0.176377996 |
| 0.032502746 | 0.176758612 |
| 0.032633955 | 0.177227045 |
| 0.032648749 | 0.177244679 |

| | |
|---|---|
| 0.03265628 | 0.177264718 |
| 0.032666464 | 0.177299793 |
| 0.032681835 | 0.177354349 |
| 0.032705668 | 0.17745191 |
| 0.032716911 | 0.177456245 |
| 0.032729014 | 0.177456245 |
| 0.032732549 | 0.177456245 |
| 0.03273792 | 0.17746516 |
| 0.032749024 | 0.177505152 |
| 0.032795835 | 0.177637576 |
| 0.032830186 | 0.177709182 |
| 0.032831442 | 0.177709182 |
| 0.03285532 | 0.177798016 |
| 0.03288541 | 0.17789015 |
| 0.032893588 | 0.17789015 |
| 0.032920561 | 0.177908492 |
| 0.032944693 | 0.177978322 |
| 0.032957769 | 0.178028772 |
| 0.03297409 | 0.178096733 |
| 0.033008567 | 0.17820587 |
| 0.033055709 | 0.1783604 |
| 0.033056612 | 0.1783604 |
| 0.033072185 | 0.178424211 |
| 0.033106784 | 0.17852734 |
| 0.033111799 | 0.17852734 |
| 0.033116288 | 0.17852734 |
| 0.033138699 | 0.178580767 |
| 0.033172066 | 0.178703216 |
| 0.033172678 | 0.178703216 |
| 0.033208696 | 0.178836563 |
| 0.033292043 | 0.179088778 |
| 0.033406174 | 0.179450509 |
| 0.033425407 | 0.179475971 |
| 0.033436652 | 0.17951611 |
| 0.033444626 | 0.179518448 |
| 0.03345437 | 0.179550515 |
| 0.03346642 | 0.17959495 |
| 0.033491299 | 0.179694004 |
| 0.033521323 | 0.179740801 |
| 0.033592103 | 0.179960113 |
| 0.033594922 | 0.179960113 |
| 0.033607944 | 0.179963728 |
| 0.033633193 | 0.180023387 |
| 0.033653239 | 0.180090206 |
| 0.033687539 | 0.180207911 |

| | |
|---|---|
| 0.033700262 | 0.180218238 |
| 0.033738722 | 0.180365263 |
| 0.03376182 | 0.180468483 |
| 0.03378492 | 0.180501872 |
| 0.033806051 | 0.180583312 |
| 0.033840651 | 0.180707329 |
| 0.03386784 | 0.180771448 |
| 0.033873738 | 0.180782668 |
| 0.033897062 | 0.180870792 |
| 0.033920353 | 0.180964904 |
| 0.033926905 | 0.180964904 |
| 0.033958247 | 0.181030832 |
| 0.034009023 | 0.181220376 |
| 0.034014026 | 0.18122676 |
| 0.034035904 | 0.181258453 |
| 0.034039003 | 0.181258453 |
| 0.034047888 | 0.181260033 |
| 0.034055758 | 0.181260033 |
| 0.034063489 | 0.181267237 |
| 0.034068906 | 0.181275812 |
| 0.034081689 | 0.181279578 |
| 0.03409735 | 0.181285387 |
| 0.034113154 | 0.181349166 |
| 0.034119969 | 0.181365152 |
| 0.034127017 | 0.181382374 |
| 0.034146451 | 0.181439945 |
| 0.034201843 | 0.181577434 |
| 0.034231267 | 0.181693143 |
| 0.034253381 | 0.181770002 |
| 0.034360722 | 0.182217803 |
| 0.034367607 | 0.182234027 |
| 0.034390919 | 0.182296749 |
| 0.034396355 | 0.182305275 |
| 0.034424869 | 0.182415805 |
| 0.034513313 | 0.182721825 |
| 0.034534481 | 0.182752639 |
| 0.03454015 | 0.18276233 |
| 0.034562876 | 0.182860079 |
| 0.034570389 | 0.182861385 |
| 0.034580182 | 0.182892875 |
| 0.034631569 | 0.183047035 |
| 0.034636233 | 0.183047035 |
| 0.0346404 | 0.183048744 |
| 0.03465497 | 0.183105422 |
| 0.03466685 | 0.18312756 |

| | |
|---|---|
| 0.034700798 | 0.183245915 |
| 0.034720559 | 0.183309618 |
| 0.034736328 | 0.183352219 |
| 0.034763223 | 0.183416963 |
| 0.034767424 | 0.183416963 |
| 0.034810876 | 0.183521635 |
| 0.034825486 | 0.183521635 |
| 0.034826652 | 0.183521635 |
| 0.034830077 | 0.183521635 |
| 0.034860099 | 0.183634252 |
| 0.034902432 | 0.183780889 |
| 0.034991729 | 0.184165739 |
| 0.034994855 | 0.184165739 |
| 0.035038516 | 0.184334408 |
| 0.035047718 | 0.184362455 |
| 0.035051656 | 0.184362806 |
| 0.035069783 | 0.184412087 |
| 0.035109285 | 0.18448256 |
| 0.035181238 | 0.184714743 |
| 0.035184855 | 0.184714743 |
| 0.035188378 | 0.184714743 |
| 0.035223237 | 0.184836611 |
| 0.035294142 | 0.185067339 |
| 0.035294405 | 0.185067339 |
| 0.035337256 | 0.185108613 |
| 0.035400525 | 0.185349667 |
| 0.035423432 | 0.185429398 |
| 0.035428077 | 0.185429398 |
| 0.035429885 | 0.185429398 |
| 0.035484658 | 0.185608744 |
| 0.035518087 | 0.185688248 |
| 0.035532551 | 0.185723094 |
| 0.035543813 | 0.185761569 |
| 0.035596706 | 0.185907009 |
| 0.035598965 | 0.185907009 |
| 0.035612419 | 0.18595688 |
| 0.035640185 | 0.186027066 |
| 0.035641484 | 0.186027066 |
| 0.035729883 | 0.186386324 |
| 0.035734174 | 0.186388296 |
| 0.035741603 | 0.186401216 |
| 0.035759359 | 0.186437982 |
| 0.035783314 | 0.186500326 |
| 0.035855235 | 0.186774281 |
| 0.035934394 | 0.187063872 |

| | |
|---|---|
| 0.035939746 | 0.187071104 |
| 0.035987029 | 0.187203151 |
| 0.036006975 | 0.187216616 |
| 0.036039705 | 0.187305003 |
| 0.036093372 | 0.18744074 |
| 0.036156897 | 0.187647873 |
| 0.036222805 | 0.187867375 |
| 0.036233985 | 0.187904616 |
| 0.036240476 | 0.187917817 |
| 0.036278077 | 0.188051366 |
| 0.036290576 | 0.188095686 |
| 0.036328281 | 0.188240546 |
| 0.036343884 | 0.188247742 |
| 0.036350979 | 0.188247742 |
| 0.036408862 | 0.188421697 |
| 0.036428468 | 0.188461726 |
| 0.036462824 | 0.188578011 |
| 0.036547619 | 0.188929825 |
| 0.036553782 | 0.188929825 |
| 0.036573654 | 0.188966525 |
| 0.036594201 | 0.188970668 |
| 0.036622361 | 0.189024096 |
| 0.036695802 | 0.189299618 |
| 0.036705692 | 0.189299618 |
| 0.03672181 | 0.189355748 |
| 0.036737139 | 0.189359278 |
| 0.036758944 | 0.189375305 |
| 0.036774555 | 0.189378279 |
| 0.036777713 | 0.189378279 |
| 0.036878886 | 0.189745037 |
| 0.036879707 | 0.189745037 |
| 0.036914896 | 0.189864551 |
| 0.036979003 | 0.190021663 |
| 0.03698933 | 0.190021663 |
| 0.037005825 | 0.190085896 |
| 0.037029932 | 0.190168143 |
| 0.03712942 | 0.190503944 |
| 0.037253204 | 0.190862559 |
| 0.037286099 | 0.190915276 |
| 0.037289818 | 0.190915276 |
| 0.037291553 | 0.190915276 |
| 0.037362266 | 0.191133473 |
| 0.03748146 | 0.191584198 |
| 0.0375378 | 0.191743113 |
| 0.037552183 | 0.191796011 |

| | |
|---|---|
| 0.037626151 | 0.191988505 |
| 0.037657457 | 0.192107083 |
| 0.037697213 | 0.192268703 |
| 0.037723996 | 0.192364107 |
| 0.037741247 | 0.192407549 |
| 0.037763065 | 0.192454781 |
| 0.037803333 | 0.192583095 |
| 0.037808546 | 0.192589051 |
| 0.037868445 | 0.19277046 |
| 0.037924722 | 0.192933212 |
| 0.037990532 | 0.193144222 |
| 0.038042816 | 0.193306859 |
| 0.038082319 | 0.19344567 |
| 0.038132123 | 0.193657351 |
| 0.038156168 | 0.193696299 |
| 0.03819525 | 0.193771322 |
| 0.038240574 | 0.193939283 |
| 0.038252735 | 0.193945706 |
| 0.038281319 | 0.193994432 |
| 0.038289345 | 0.193994432 |
| 0.038300236 | 0.193994432 |
| 0.038337187 | 0.194098565 |
| 0.038342955 | 0.194107133 |
| 0.03844856 | 0.194455727 |
| 0.038471273 | 0.194487987 |
| 0.038531304 | 0.194688142 |
| 0.038604032 | 0.194931534 |
| 0.038628888 | 0.195015695 |
| 0.03865925 | 0.195044925 |
| 0.038673071 | 0.195093991 |
| 0.038835769 | 0.195571742 |
| 0.038905981 | 0.195758686 |
| 0.038950603 | 0.195933711 |
| 0.039002955 | 0.195969377 |
| 0.03902828 | 0.196026706 |
| 0.039040037 | 0.196031613 |
| 0.039054177 | 0.196081942 |
| 0.039063865 | 0.196109908 |
| 0.039148102 | 0.196383394 |
| 0.039157696 | 0.196383394 |
| 0.039163566 | 0.196383394 |
| 0.039172868 | 0.196403427 |
| 0.039222428 | 0.196533044 |
| 0.039254233 | 0.196609671 |
| 0.039268862 | 0.19662187 |

| | |
|---|---|
| 0.039269054 | 0.19662187 |
| 0.039275342 | 0.196632681 |
| 0.039281045 | 0.196640567 |
| 0.03937003 | 0.196941121 |
| 0.039462655 | 0.197284534 |
| 0.039467734 | 0.197284821 |
| 0.039573555 | 0.197627105 |
| 0.039645773 | 0.197816601 |
| 0.039666164 | 0.197882099 |
| 0.039680106 | 0.19791019 |
| 0.039711245 | 0.197982568 |
| 0.039743051 | 0.198017692 |
| 0.039785088 | 0.19810991 |
| 0.039788948 | 0.19810991 |
| 0.039790862 | 0.19810991 |
| 0.039806543 | 0.198124203 |
| 0.039844781 | 0.19825828 |
| 0.039903498 | 0.198463231 |
| 0.03991266 | 0.198481901 |
| 0.039958774 | 0.19865517 |
| 0.04000196 | 0.19878687 |
| 0.04003634 | 0.198916209 |
| 0.040056557 | 0.198995892 |
| 0.040159973 | 0.199344592 |
| 0.040285878 | 0.199760074 |
| 0.040303349 | 0.199780954 |
| 0.040404086 | 0.200137848 |
| 0.040560811 | 0.200684576 |
| 0.040621885 | 0.200882415 |
| 0.040628904 | 0.200896266 |
| 0.040653489 | 0.200955239 |
| 0.04081141 | 0.201407452 |
| 0.040816749 | 0.201407452 |
| 0.040859568 | 0.201513786 |
| 0.040885748 | 0.201622028 |
| 0.040916906 | 0.201692515 |
| 0.04101192 | 0.201912914 |
| 0.041012567 | 0.201912914 |
| 0.041087201 | 0.202175858 |
| 0.041121322 | 0.202239281 |
| 0.041129026 | 0.202256286 |
| 0.041153214 | 0.20233345 |
| 0.04117241 | 0.202349 |
| 0.041173371 | 0.202349 |
| 0.041199274 | 0.202425351 |

| | |
|---|---|
| 0.041215343 | 0.202450817 |
| 0.04123255 | 0.202483114 |
| 0.041234672 | 0.202483114 |
| 0.041319375 | 0.202669196 |
| 0.041407068 | 0.203015691 |
| 0.041414556 | 0.203031502 |
| 0.041422209 | 0.203037943 |
| 0.041438391 | 0.203064746 |
| 0.041484638 | 0.203186843 |
| 0.041492729 | 0.203205575 |
| 0.041563274 | 0.203465608 |
| 0.041593347 | 0.203540632 |
| 0.041599453 | 0.203540632 |
| 0.041603881 | 0.203540632 |
| 0.041631489 | 0.203592046 |
| 0.041636904 | 0.20359762 |
| 0.041774769 | 0.204055114 |
| 0.041777593 | 0.204055114 |
| 0.041789036 | 0.204069954 |
| 0.041794874 | 0.204076735 |
| 0.041891413 | 0.204485225 |
| 0.041970408 | 0.20480785 |
| 0.041986595 | 0.204865849 |
| 0.041996857 | 0.204873943 |
| 0.04200685 | 0.204893675 |
| 0.042009506 | 0.204893675 |
| 0.042030976 | 0.204935431 |
| 0.042038393 | 0.204950613 |
| 0.04215483 | 0.205350118 |
| 0.042161267 | 0.205360469 |
| 0.042173976 | 0.205362697 |
| 0.042194684 | 0.205381618 |
| 0.04224035 | 0.205493479 |
| 0.042252562 | 0.205510904 |
| 0.042305334 | 0.205704557 |
| 0.042400342 | 0.206045995 |
| 0.042401512 | 0.206045995 |
| 0.042434366 | 0.206100502 |
| 0.042480475 | 0.206240327 |
| 0.042525394 | 0.206343094 |
| 0.042560286 | 0.206459434 |
| 0.042607644 | 0.206626033 |
| 0.042628461 | 0.206680501 |
| 0.042631895 | 0.206680501 |
| 0.04272433 | 0.206941969 |

| | |
|---|---|
| 0.04272844 | 0.206941969 |
| 0.042737968 | 0.206941969 |
| 0.042768873 | 0.207023795 |
| 0.042769141 | 0.207023795 |
| 0.04277048 | 0.207023795 |
| 0.042772254 | 0.207023795 |
| 0.042777055 | 0.207025994 |
| 0.042781545 | 0.207026688 |
| 0.04279746 | 0.207031821 |
| 0.0428048 | 0.207031821 |
| 0.042857446 | 0.207204504 |
| 0.042891385 | 0.207302925 |
| 0.042895213 | 0.207302925 |
| 0.042965286 | 0.20751226 |
| 0.042994522 | 0.20758279 |
| 0.042998866 | 0.20758279 |
| 0.043036807 | 0.207713216 |
| 0.043062702 | 0.207775031 |
| 0.043077239 | 0.207797669 |
| 0.043085603 | 0.207797669 |
| 0.043116527 | 0.207797669 |
| 0.043117211 | 0.207797669 |
| 0.043119751 | 0.207797669 |
| 0.043183345 | 0.208019942 |
| 0.043188192 | 0.208022249 |
| 0.043197969 | 0.208048305 |
| 0.043221651 | 0.208073486 |
| 0.043299594 | 0.208284999 |
| 0.043306013 | 0.208294835 |
| 0.043538036 | 0.209009781 |
| 0.043570117 | 0.209121635 |
| 0.043588236 | 0.209187515 |
| 0.043595941 | 0.209192842 |
| 0.043619864 | 0.209233885 |
| 0.043763435 | 0.209626987 |
| 0.043798686 | 0.209753646 |
| 0.04393244 | 0.210165153 |
| 0.044017161 | 0.210386666 |
| 0.044024446 | 0.210386666 |
| 0.044025889 | 0.210386666 |
| 0.04412186 | 0.210708093 |
| 0.04416679 | 0.21080982 |
| 0.044172741 | 0.21080982 |
| 0.044213146 | 0.210932536 |
| 0.044238234 | 0.21100995 |

| | |
|---|---|
| 0.044272509 | 0.211088878 |
| 0.044300857 | 0.211181754 |
| 0.044368028 | 0.211369218 |
| 0.044374526 | 0.211369218 |
| 0.044379519 | 0.211369218 |
| 0.044387507 | 0.211383243 |
| 0.044396931 | 0.211401856 |
| 0.044472907 | 0.211662952 |
| 0.04456666 | 0.21200323 |
| 0.044574018 | 0.212017052 |
| 0.04460469 | 0.212099399 |
| 0.044675777 | 0.212301793 |
| 0.044686091 | 0.212302371 |
| 0.044687494 | 0.212302371 |
| 0.044697752 | 0.212329925 |
| 0.044707485 | 0.212354981 |
| 0.044756414 | 0.212478174 |
| 0.044759952 | 0.212478174 |
| 0.044760189 | 0.212478174 |
| 0.044774695 | 0.212525853 |
| 0.044833184 | 0.2126763 |
| 0.044841713 | 0.212695574 |
| 0.044895021 | 0.21289046 |
| 0.045017051 | 0.213314804 |
| 0.04502705 | 0.213340958 |
| 0.045052079 | 0.213438313 |
| 0.045083244 | 0.213530296 |
| 0.045088789 | 0.213530296 |
| 0.045098395 | 0.213530296 |
| 0.045134528 | 0.213680138 |
| 0.045162642 | 0.213770737 |
| 0.045168626 | 0.213777817 |
| 0.0452292 | 0.214018626 |
| 0.04527141 | 0.214123896 |
| 0.045327527 | 0.214320841 |
| 0.045341805 | 0.21436313 |
| 0.045410995 | 0.214605026 |
| 0.045442119 | 0.214724996 |
| 0.045453059 | 0.214739891 |
| 0.045484546 | 0.214842879 |
| 0.045494616 | 0.214850974 |
| 0.045635052 | 0.215258059 |
| 0.045659697 | 0.215352978 |
| 0.045669431 | 0.21537135 |
| 0.045672636 | 0.21537135 |

| | |
|---|---|
| 0.045680143 | 0.215385426 |
| 0.045719672 | 0.21546683 |
| 0.045720028 | 0.21546683 |
| 0.045729495 | 0.215490124 |
| 0.045837475 | 0.215913503 |
| 0.045861678 | 0.215943203 |
| 0.045878467 | 0.215978427 |
| 0.045981305 | 0.216398381 |
| 0.046166336 | 0.216981376 |
| 0.046168964 | 0.216981376 |
| 0.046224104 | 0.217091397 |
| 0.04623298 | 0.217096437 |
| 0.046241317 | 0.217096437 |
| 0.046245276 | 0.217096437 |
| 0.046249115 | 0.217096437 |
| 0.046365963 | 0.217435185 |
| 0.046409427 | 0.21757474 |
| 0.046432034 | 0.217628593 |
| 0.046450516 | 0.217681662 |
| 0.046511906 | 0.217895458 |
| 0.046570829 | 0.218116716 |
| 0.046618098 | 0.218237865 |
| 0.046629422 | 0.218261839 |
| 0.046650901 | 0.218261839 |
| 0.046656808 | 0.218261839 |
| 0.046734114 | 0.218537621 |
| 0.046965325 | 0.21927434 |
| 0.046986753 | 0.219331383 |
| 0.047052146 | 0.219477215 |
| 0.04706211 | 0.219477215 |
| 0.047072517 | 0.219495084 |
| 0.047157421 | 0.219783393 |
| 0.047280933 | 0.220219872 |
| 0.047311057 | 0.220283872 |
| 0.047352988 | 0.220414461 |
| 0.04737429 | 0.220470521 |
| 0.047381774 | 0.220483805 |
| 0.047413756 | 0.220546434 |
| 0.047479652 | 0.220719561 |
| 0.047516518 | 0.220788665 |
| 0.047523392 | 0.220797638 |
| 0.047580256 | 0.220997157 |
| 0.047586277 | 0.221003571 |
| 0.047596417 | 0.221014968 |
| 0.047605531 | 0.221021958 |

| | |
|---|---|
| 0.047609139 | 0.221021958 |
| 0.047655952 | 0.221154633 |
| 0.047692139 | 0.221222024 |
| 0.047701481 | 0.221222024 |
| 0.047834619 | 0.221590655 |
| 0.047862041 | 0.22165747 |
| 0.047906901 | 0.221843652 |
| 0.04792293 | 0.221896302 |
| 0.047963325 | 0.221974589 |
| 0.047965676 | 0.221974589 |
| 0.047974584 | 0.221974589 |
| 0.047995436 | 0.222016185 |
| 0.048013944 | 0.222080228 |
| 0.048079072 | 0.222273534 |
| 0.048108235 | 0.222340883 |
| 0.048210282 | 0.222642403 |
| 0.048252229 | 0.222749724 |
| 0.048278301 | 0.22282225 |
| 0.048291377 | 0.22282225 |
| 0.04829601 | 0.22282225 |
| 0.048384359 | 0.223143396 |
| 0.048432705 | 0.223336747 |
| 0.048442372 | 0.223346064 |
| 0.048448085 | 0.223350782 |
| 0.048479832 | 0.223436287 |
| 0.048480706 | 0.223436287 |
| 0.048520283 | 0.223546006 |
| 0.048523287 | 0.223546006 |
| 0.048558653 | 0.223622412 |
| 0.048563712 | 0.223624086 |
| 0.048590261 | 0.223724708 |
| 0.04864701 | 0.223793445 |
| 0.04864748 | 0.223793445 |
| 0.048742369 | 0.224056828 |
| 0.048749016 | 0.224065757 |
| 0.048819849 | 0.2241966 |
| 0.048827978 | 0.224212313 |
| 0.048854086 | 0.224253628 |
| 0.048856963 | 0.224253628 |
| 0.048860518 | 0.224253628 |
| 0.049036586 | 0.224693618 |
| 0.049189716 | 0.225135363 |
| 0.04921999 | 0.225252276 |
| 0.04922682 | 0.225261889 |
| 0.049236106 | 0.225282737 |

| | |
|---|---|
| 0.04930336 | 0.225512175 |
| 0.04941361 | 0.225769571 |
| 0.049437476 | 0.225829221 |
| 0.049445632 | 0.225829221 |
| 0.049465721 | 0.225844059 |
| 0.049476109 | 0.225844059 |
| 0.049545654 | 0.226004295 |
| 0.049589653 | 0.226112693 |
| 0.049593155 | 0.226112693 |
| 0.049626364 | 0.226239273 |
| 0.049635168 | 0.226239273 |
| 0.049647584 | 0.226249006 |
| 0.049673657 | 0.226328067 |
| 0.049764447 | 0.226568333 |
| 0.049775549 | 0.226592403 |
| 0.049807207 | 0.226654676 |
| 0.049939437 | 0.227029434 |
| 0.049963051 | 0.227078185 |
| 0.049970028 | 0.227078185 |
| 0.049975316 | 0.227078185 |
| 0.049976558 | 0.227078185 |
| 0.049987251 | 0.227099553 |
| 0.049995566 | 0.227099553 |
| 0.050031961 | 0.227137234 |
| 0.050100726 | 0.227399977 |
| 0.050130986 | 0.227454366 |
| 0.05018719 | 0.227622635 |
| 0.050214231 | 0.227723593 |
| 0.050260206 | 0.227893944 |
| 0.050275067 | 0.227912681 |
| 0.050334612 | 0.228079328 |
| 0.050410555 | 0.228255754 |
| 0.050412244 | 0.228255754 |
| 0.050458113 | 0.22830786 |
| 0.050552051 | 0.228580892 |
| 0.050562685 | 0.228585573 |
| 0.05059843 | 0.228702065 |
| 0.050625269 | 0.228779955 |
| 0.050649066 | 0.228802339 |
| 0.050675442 | 0.228812504 |
| 0.050689239 | 0.228812504 |
| 0.050726522 | 0.228935096 |
| 0.050771548 | 0.229116592 |
| 0.050864106 | 0.229382138 |
| 0.050899246 | 0.229460151 |

| | |
|---|---|
| 0.050900676 | 0.229460151 |
| 0.050954724 | 0.229638591 |
| 0.050985769 | 0.229715071 |
| 0.051027514 | 0.229792681 |
| 0.051071112 | 0.229902095 |
| 0.051124186 | 0.230020382 |
| 0.051129226 | 0.230020382 |
| 0.051149997 | 0.23006147 |
| 0.051252747 | 0.230267906 |
| 0.051253859 | 0.230267906 |
| 0.051257913 | 0.230267906 |
| 0.0513807 | 0.230619979 |
| 0.051405245 | 0.230643205 |
| 0.05146482 | 0.230806264 |
| 0.051465818 | 0.230806264 |
| 0.051595212 | 0.231190627 |
| 0.051664224 | 0.231456306 |
| 0.051713717 | 0.231612675 |
| 0.051805579 | 0.232002286 |
| 0.05184179 | 0.232077162 |
| 0.051906273 | 0.232256675 |
| 0.051978013 | 0.232381191 |
| 0.05205136 | 0.232574309 |
| 0.052070041 | 0.232574309 |
| 0.052108482 | 0.2327211 |
| 0.052142936 | 0.232834391 |
| 0.052197396 | 0.232924707 |
| 0.052253509 | 0.233065918 |
| 0.052273846 | 0.233134795 |
| 0.052361846 | 0.23339613 |
| 0.05241752 | 0.233556857 |
| 0.052463913 | 0.23369798 |
| 0.052498818 | 0.233809726 |
| 0.052644845 | 0.234262921 |
| 0.052753009 | 0.234590809 |
| 0.052805007 | 0.23473422 |
| 0.052842425 | 0.234813039 |
| 0.052907887 | 0.234979296 |
| 0.05302594 | 0.235365 |
| 0.053055038 | 0.235420724 |
| 0.053058266 | 0.235420724 |
| 0.05317355 | 0.235756534 |
| 0.053210257 | 0.23585341 |
| 0.053215587 | 0.235855085 |
| 0.05326101 | 0.235998872 |

| | |
|---|---|
| 0.053275639 | 0.236019913 |
| 0.053277554 | 0.236019913 |
| 0.053308369 | 0.236134458 |
| 0.053411077 | 0.236307608 |
| 0.053478248 | 0.236513001 |
| 0.053512198 | 0.236575278 |
| 0.053519701 | 0.236586484 |
| 0.053583214 | 0.236661338 |
| 0.053586324 | 0.236661338 |
| 0.053610967 | 0.236748219 |
| 0.053620803 | 0.236769702 |
| 0.053631589 | 0.236773424 |
| 0.053646467 | 0.236795207 |
| 0.053660933 | 0.236837108 |
| 0.053700384 | 0.236967313 |
| 0.053734914 | 0.237056596 |
| 0.053772485 | 0.237175602 |
| 0.053784383 | 0.237206113 |
| 0.053890867 | 0.237437284 |
| 0.053901912 | 0.237437284 |
| 0.053911188 | 0.237437284 |
| 0.053918255 | 0.237437284 |
| 0.053918587 | 0.237437284 |
| 0.053924198 | 0.237437284 |
| 0.054000597 | 0.237667882 |
| 0.054029833 | 0.237730238 |
| 0.05404159 | 0.237730238 |
| 0.054070642 | 0.237807965 |
| 0.054081582 | 0.237821776 |
| 0.054089412 | 0.237821776 |
| 0.054183075 | 0.237994773 |
| 0.054190649 | 0.238006089 |
| 0.054209852 | 0.238046527 |
| 0.054282029 | 0.238165842 |
| 0.05434725 | 0.23833115 |
| 0.054351367 | 0.23833115 |
| 0.054354735 | 0.23833115 |
| 0.054428118 | 0.23843521 |
| 0.054476833 | 0.238571955 |
| 0.054555117 | 0.238792076 |
| 0.054568034 | 0.238826663 |
| 0.054637956 | 0.239044822 |
| 0.054819296 | 0.239596093 |
| 0.054869733 | 0.239693722 |
| 0.05487545 | 0.239693722 |

| | |
|---|---|
| 0.054982308 | 0.240022215 |
| 0.055098125 | 0.240285541 |
| 0.055178592 | 0.240487168 |
| 0.055179704 | 0.240487168 |
| 0.055207247 | 0.24056318 |
| 0.055343273 | 0.241002073 |
| 0.055343391 | 0.241002073 |
| 0.055352537 | 0.241009791 |
| 0.055522235 | 0.24151589 |
| 0.05557593 | 0.241624962 |
| 0.055577749 | 0.241624962 |
| 0.055630447 | 0.241699624 |
| 0.055651033 | 0.241744966 |
| 0.055669721 | 0.241759989 |
| 0.0556876 | 0.241788397 |
| 0.05568875 | 0.241788397 |
| 0.055729012 | 0.241792799 |
| 0.055744621 | 0.241792799 |
| 0.055775947 | 0.241868434 |
| 0.05583189 | 0.242057174 |
| 0.0558398 | 0.242057174 |
| 0.055886333 | 0.242192757 |
| 0.05589934 | 0.242227084 |
| 0.055948927 | 0.242375802 |
| 0.056030066 | 0.242594903 |
| 0.05604295 | 0.242628634 |
| 0.056084599 | 0.242764811 |
| 0.056309894 | 0.243474473 |
| 0.056318037 | 0.243487577 |
| 0.056335429 | 0.243533267 |
| 0.056352811 | 0.243533267 |
| 0.056371977 | 0.243551423 |
| 0.0563737 | 0.243551423 |
| 0.056422397 | 0.24369543 |
| 0.056477643 | 0.243867696 |
| 0.056494796 | 0.243897538 |
| 0.056537227 | 0.243941141 |
| 0.056543643 | 0.243941141 |
| 0.056558925 | 0.243953262 |
| 0.056623958 | 0.244079036 |
| 0.056642678 | 0.244108275 |
| 0.056688079 | 0.244134472 |
| 0.056703604 | 0.244179256 |
| 0.056781467 | 0.244443627 |
| 0.056800484 | 0.244485917 |

| | |
|---|---|
| 0.056820983 | 0.244552047 |
| 0.056862649 | 0.244671679 |
| 0.056890558 | 0.244718803 |
| 0.056903063 | 0.244750492 |
| 0.056913123 | 0.244771655 |
| 0.056963747 | 0.244809993 |
| 0.056979939 | 0.244837923 |
| 0.057031237 | 0.244869472 |
| 0.057031763 | 0.244869472 |
| 0.057043022 | 0.244869472 |
| 0.057048978 | 0.244869472 |
| 0.057135881 | 0.245071918 |
| 0.057148767 | 0.245084616 |
| 0.057221474 | 0.245145973 |
| 0.057227342 | 0.245145973 |
| 0.057236929 | 0.245145973 |
| 0.057315857 | 0.245351639 |
| 0.057332865 | 0.245402388 |
| 0.057413962 | 0.245639128 |
| 0.057432221 | 0.245695174 |
| 0.057459922 | 0.245769529 |
| 0.057468319 | 0.24578337 |
| 0.057483281 | 0.245803216 |
| 0.057489475 | 0.245807633 |
| 0.05750981 | 0.245850436 |
| 0.057535387 | 0.245893564 |
| 0.057612064 | 0.246154997 |
| 0.057702693 | 0.246409591 |
| 0.057748392 | 0.246507541 |
| 0.057751859 | 0.246507541 |
| 0.057756685 | 0.246507541 |
| 0.057790291 | 0.246563824 |
| 0.057805395 | 0.246582855 |
| 0.057828827 | 0.246616544 |
| 0.057882535 | 0.246779295 |
| 0.057933731 | 0.246842887 |
| 0.05794952 | 0.246888076 |
| 0.05797261 | 0.246964356 |
| 0.057995451 | 0.246995052 |
| 0.058201558 | 0.24757579 |
| 0.058270339 | 0.247767294 |
| 0.058279923 | 0.24778592 |
| 0.05832576 | 0.24793654 |
| 0.058331922 | 0.247940603 |
| 0.058350169 | 0.247973902 |

| | |
|---|---|
| 0.058398496 | 0.24811286 |
| 0.058423219 | 0.248173623 |
| 0.058449862 | 0.248201805 |
| 0.058536891 | 0.24836851 |
| 0.058552596 | 0.248413011 |
| 0.058609871 | 0.248611713 |
| 0.058647257 | 0.248748145 |
| 0.058667365 | 0.248773813 |
| 0.058723491 | 0.248943954 |
| 0.058760755 | 0.248995414 |
| 0.058781178 | 0.249027856 |
| 0.058822262 | 0.249053475 |
| 0.058846359 | 0.249063063 |
| 0.058857316 | 0.249084539 |
| 0.058899561 | 0.24919689 |
| 0.058931575 | 0.24931019 |
| 0.059047013 | 0.249554716 |
| 0.059073608 | 0.249622813 |
| 0.059102166 | 0.249685888 |
| 0.059104262 | 0.249685888 |
| 0.05927228 | 0.250054171 |
| 0.05932237 | 0.250142117 |
| 0.05942801 | 0.250453749 |
| 0.05947485 | 0.250562466 |
| 0.059483941 | 0.250578602 |
| 0.059496969 | 0.25059913 |
| 0.059566143 | 0.250858314 |
| 0.059604609 | 0.250909381 |
| 0.059623369 | 0.250943988 |
| 0.059636423 | 0.250976754 |
| 0.059659498 | 0.250985154 |
| 0.059682792 | 0.251038801 |
| 0.059703172 | 0.251080173 |
| 0.059742406 | 0.251178633 |
| 0.059784713 | 0.251312139 |
| 0.059904747 | 0.251636555 |
| 0.059928901 | 0.251673907 |
| 0.060068341 | 0.252015022 |
| 0.060071922 | 0.252015022 |
| 0.060082002 | 0.252027905 |
| 0.060243827 | 0.252439973 |
| 0.060265324 | 0.252463419 |
| 0.060295148 | 0.252496125 |
| 0.06029603 | 0.252496125 |
| 0.060296326 | 0.252496125 |

| | |
|---|---|
| 0.060326921 | 0.252563186 |
| 0.060331563 | 0.252563186 |
| 0.060338554 | 0.252570252 |
| 0.060400535 | 0.25269644 |
| 0.060452883 | 0.252826328 |
| 0.060476655 | 0.252837221 |
| 0.060507082 | 0.25292003 |
| 0.060530398 | 0.252950889 |
| 0.060548337 | 0.253003653 |
| 0.060582899 | 0.253089212 |
| 0.060585942 | 0.253089212 |
| 0.060612575 | 0.253116164 |
| 0.060648759 | 0.253156744 |
| 0.060655002 | 0.253160615 |
| 0.060682762 | 0.253229691 |
| 0.060687503 | 0.253229691 |
| 0.060706167 | 0.253263194 |
| 0.060790128 | 0.253524644 |
| 0.060811695 | 0.253570181 |
| 0.060811695 | 0.253570181 |
| 0.060845321 | 0.253665976 |
| 0.060879868 | 0.253734686 |
| 0.060883112 | 0.253734686 |
| 0.060917205 | 0.253773537 |
| 0.060924924 | 0.253773537 |
| 0.060939873 | 0.253787265 |
| 0.060949015 | 0.253787265 |
| 0.061021954 | 0.25402435 |
| 0.06104895 | 0.254092312 |
| 0.061207129 | 0.254483801 |
| 0.061220995 | 0.254507193 |
| 0.061228787 | 0.254507193 |
| 0.061467824 | 0.255140203 |
| 0.061471093 | 0.255140203 |
| 0.061523742 | 0.255267971 |
| 0.061524367 | 0.255267971 |
| 0.061545114 | 0.255309569 |
| 0.06158648 | 0.255436669 |
| 0.06165598 | 0.255591378 |
| 0.061747678 | 0.255915188 |
| 0.061923226 | 0.256352999 |
| 0.061925825 | 0.256352999 |
| 0.061955752 | 0.256388497 |
| 0.062072065 | 0.256712941 |
| 0.062139687 | 0.256903375 |

| | |
|---|---|
| 0.062175468 | 0.257006686 |
| 0.062188665 | 0.257038927 |
| 0.062309695 | 0.257338176 |
| 0.062315314 | 0.257339066 |
| 0.062331898 | 0.257385237 |
| 0.062390899 | 0.257571711 |
| 0.062408316 | 0.25760803 |
| 0.06241814 | 0.25760803 |
| 0.062462055 | 0.257766252 |
| 0.062604263 | 0.258174148 |
| 0.062613753 | 0.258190928 |
| 0.062634258 | 0.258218576 |
| 0.062660886 | 0.258227354 |
| 0.062674356 | 0.258227354 |
| 0.062678457 | 0.258227354 |
| 0.062679097 | 0.258227354 |
| 0.062687649 | 0.258227354 |
| 0.06270757 | 0.25826474 |
| 0.062719192 | 0.258290267 |
| 0.062768639 | 0.258426865 |
| 0.062800328 | 0.258465044 |
| 0.062805047 | 0.258465044 |
| 0.062853917 | 0.258633018 |
| 0.062880673 | 0.258686864 |
| 0.062900125 | 0.258722193 |
| 0.062911434 | 0.258727846 |
| 0.062992835 | 0.258924634 |
| 0.063032587 | 0.258979281 |
| 0.063057777 | 0.259057428 |
| 0.063083182 | 0.259097 |
| 0.063101132 | 0.259123719 |
| 0.063106558 | 0.259123719 |
| 0.063140932 | 0.259175488 |
| 0.063179975 | 0.259223781 |
| 0.063199325 | 0.259223781 |
| 0.063270348 | 0.2593884 |
| 0.063290055 | 0.259407527 |
| 0.063348759 | 0.259617178 |
| 0.063411061 | 0.25974702 |
| 0.063446017 | 0.259845509 |
| 0.06346802 | 0.259868587 |
| 0.063517879 | 0.260020172 |
| 0.063559146 | 0.260129885 |
| 0.063598637 | 0.260246781 |
| 0.063604266 | 0.260247456 |

| | |
|---|---|
| 0.063630447 | 0.260332213 |
| 0.063654819 | 0.260374316 |
| 0.063705637 | 0.260476147 |
| 0.063765683 | 0.260505075 |
| 0.063846207 | 0.260744579 |
| 0.063861787 | 0.260785847 |
| 0.06388363 | 0.260830316 |
| 0.063906103 | 0.260883681 |
| 0.063917187 | 0.260883681 |
| 0.063945064 | 0.260904138 |
| 0.063945535 | 0.260904138 |
| 0.063985389 | 0.26095497 |
| 0.064067416 | 0.261155618 |
| 0.064125514 | 0.261281761 |
| 0.064153949 | 0.261329125 |
| 0.064163561 | 0.261334168 |
| 0.064177151 | 0.261356576 |
| 0.064191931 | 0.261394418 |
| 0.064228512 | 0.261521019 |
| 0.064239396 | 0.261536671 |
| 0.064273395 | 0.26159195 |
| 0.064321225 | 0.261697163 |
| 0.064368294 | 0.261866298 |
| 0.064376589 | 0.261877676 |
| 0.064400066 | 0.261950801 |
| 0.064416498 | 0.261995266 |
| 0.064440719 | 0.262047247 |
| 0.064531914 | 0.262330271 |
| 0.064624749 | 0.262581206 |
| 0.064631487 | 0.262581206 |
| 0.064687807 | 0.262629681 |
| 0.064741186 | 0.26279386 |
| 0.064773261 | 0.262823077 |
| 0.064776834 | 0.262823077 |
| 0.064795531 | 0.262833784 |
| 0.064898985 | 0.263133657 |
| 0.064900814 | 0.263133657 |
| 0.064916794 | 0.263166255 |
| 0.064919905 | 0.263166255 |
| 0.06497502 | 0.263322438 |
| 0.065021006 | 0.263419145 |
| 0.065106447 | 0.263651111 |
| 0.065116813 | 0.263651111 |
| 0.065117014 | 0.263651111 |
| 0.065153609 | 0.263754437 |

| | |
|---|---|
| 0.065198403 | 0.263890911 |
| 0.065220282 | 0.263934604 |
| 0.065249686 | 0.264031163 |
| 0.065343981 | 0.264121247 |
| 0.065429592 | 0.264399726 |
| 0.065459479 | 0.264413657 |
| 0.06547292 | 0.264413657 |
| 0.065485178 | 0.264413657 |
| 0.065495355 | 0.264413657 |
| 0.065500273 | 0.264413657 |
| 0.065529809 | 0.264445649 |
| 0.065544922 | 0.264445649 |
| 0.065556423 | 0.264445649 |
| 0.065603357 | 0.264507315 |
| 0.065656592 | 0.26459862 |
| 0.065703832 | 0.2647218 |
| 0.065769813 | 0.264965223 |
| 0.065798525 | 0.265030169 |
| 0.065802628 | 0.265030169 |
| 0.065820569 | 0.265047078 |
| 0.065825565 | 0.265047078 |
| 0.065904031 | 0.265236713 |
| 0.065985214 | 0.26538404 |
| 0.065992059 | 0.265389159 |
| 0.066025369 | 0.265411762 |
| 0.066025543 | 0.265411762 |
| 0.066060145 | 0.265528444 |
| 0.066179569 | 0.26585141 |
| 0.066220988 | 0.265972931 |
| 0.066272266 | 0.266111559 |
| 0.066290644 | 0.26614048 |
| 0.06631021 | 0.266185938 |
| 0.066318734 | 0.266185938 |
| 0.066378462 | 0.266358329 |
| 0.066404066 | 0.266438622 |
| 0.066419781 | 0.266479229 |
| 0.066456346 | 0.266558562 |
| 0.066503868 | 0.266681799 |
| 0.066629838 | 0.267008725 |
| 0.066657875 | 0.267030877 |
| 0.066664379 | 0.267032287 |
| 0.066771294 | 0.267224993 |
| 0.066779597 | 0.267224993 |
| 0.066851203 | 0.267444119 |
| 0.06698399 | 0.267727971 |

| | |
|---|---|
| 0.067007258 | 0.267753561 |
| 0.067092545 | 0.267926365 |
| 0.067118632 | 0.267968343 |
| 0.06715305 | 0.268013645 |
| 0.067178151 | 0.268027657 |
| 0.067179524 | 0.268027657 |
| 0.067184439 | 0.268027657 |
| 0.067294894 | 0.268362548 |
| 0.067318081 | 0.268410068 |
| 0.067429751 | 0.268697854 |
| 0.067449301 | 0.268719543 |
| 0.067477244 | 0.26875219 |
| 0.067551361 | 0.268867514 |
| 0.067572255 | 0.268924738 |
| 0.067577031 | 0.268924738 |
| 0.067643623 | 0.269155716 |
| 0.067646375 | 0.269155716 |
| 0.06789721 | 0.269725617 |
| 0.068041338 | 0.270208023 |
| 0.068112021 | 0.27044362 |
| 0.068156856 | 0.270531425 |
| 0.068203188 | 0.270670215 |
| 0.068238713 | 0.270730153 |
| 0.068246713 | 0.270730153 |
| 0.068320847 | 0.270869523 |
| 0.068480034 | 0.271291041 |
| 0.068511129 | 0.271341577 |
| 0.068530756 | 0.271359688 |
| 0.068554964 | 0.271432976 |
| 0.068652032 | 0.271645424 |
| 0.06865425 | 0.271645424 |
| 0.068676691 | 0.271711643 |
| 0.068713694 | 0.271769389 |
| 0.068722447 | 0.271769389 |
| 0.068846951 | 0.272039159 |
| 0.06886156 | 0.272039159 |
| 0.069049297 | 0.272664858 |
| 0.069101014 | 0.272823842 |
| 0.069171033 | 0.273043131 |
| 0.069173755 | 0.273043131 |
| 0.069184682 | 0.273043532 |
| 0.069218548 | 0.273106767 |
| 0.069237227 | 0.273139844 |
| 0.06926203 | 0.273185714 |
| 0.069355915 | 0.273422245 |

| | |
|---|---|
| 0.06938669 | 0.273520931 |
| 0.069405802 | 0.273550261 |
| 0.069465601 | 0.273717949 |
| 0.069471152 | 0.273717949 |
| 0.069547093 | 0.273852191 |
| 0.069564903 | 0.273852191 |
| 0.069574223 | 0.273852191 |
| 0.069597776 | 0.273914681 |
| 0.069607353 | 0.273914681 |
| 0.069629934 | 0.273958269 |
| 0.069676611 | 0.274066098 |
| 0.069680357 | 0.274066098 |
| 0.069710432 | 0.274116474 |
| 0.069736178 | 0.274138783 |
| 0.069795767 | 0.274203002 |
| 0.069822155 | 0.274284046 |
| 0.069841035 | 0.274312962 |
| 0.069854104 | 0.274341667 |
| 0.069866977 | 0.274348278 |
| 0.069912073 | 0.274491891 |
| 0.069929698 | 0.27451221 |
| 0.070027976 | 0.274670176 |
| 0.07003103 | 0.274670176 |
| 0.070032351 | 0.274670176 |
| 0.07012993 | 0.274926182 |
| 0.070244268 | 0.275238483 |
| 0.070257426 | 0.27524475 |
| 0.07028327 | 0.275323351 |
| 0.070310542 | 0.275384885 |
| 0.07037121 | 0.275554521 |
| 0.070424481 | 0.275659761 |
| 0.070431239 | 0.275659761 |
| 0.070433946 | 0.275659761 |
| 0.070438602 | 0.275659761 |
| 0.070480356 | 0.275777844 |
| 0.070622985 | 0.2761771 |
| 0.070672683 | 0.276303387 |
| 0.070808879 | 0.276544999 |
| 0.070811842 | 0.276544999 |
| 0.070814577 | 0.276544999 |
| 0.070828458 | 0.276549185 |
| 0.070939239 | 0.276882601 |
| 0.070942057 | 0.276882601 |
| 0.07107685 | 0.277321992 |
| 0.071078791 | 0.277321992 |

| | |
|---|---|
| 0.07112326 | 0.277450039 |
| 0.071174431 | 0.277621959 |
| 0.0711826 | 0.277621959 |
| 0.071279899 | 0.277823322 |
| 0.071279938 | 0.277823322 |
| 0.071300383 | 0.277867877 |
| 0.07139383 | 0.278034435 |
| 0.07140013 | 0.278034435 |
| 0.071407173 | 0.278034435 |
| 0.071525644 | 0.278256348 |
| 0.071562836 | 0.278311695 |
| 0.071631548 | 0.278496212 |
| 0.071637036 | 0.278496212 |
| 0.071727593 | 0.278774441 |
| 0.071728012 | 0.278774441 |
| 0.071756781 | 0.278774441 |
| 0.071770083 | 0.278785886 |
| 0.071844587 | 0.279029775 |
| 0.071890641 | 0.279102077 |
| 0.071894044 | 0.279102077 |
| 0.071898364 | 0.279102077 |
| 0.071941981 | 0.279114188 |
| 0.071954228 | 0.279114188 |
| 0.071964044 | 0.279129532 |
| 0.072007785 | 0.279196096 |
| 0.072015253 | 0.279196096 |
| 0.07206867 | 0.279349316 |
| 0.072073496 | 0.279349316 |
| 0.072093664 | 0.279382014 |
| 0.072225774 | 0.279689155 |
| 0.072232949 | 0.279694199 |
| 0.072264284 | 0.279747303 |
| 0.072287568 | 0.279755641 |
| 0.072327739 | 0.279856465 |
| 0.072350413 | 0.279921457 |
| 0.07246666 | 0.280159457 |
| 0.072470752 | 0.280159457 |
| 0.072504279 | 0.280250651 |
| 0.072513166 | 0.280255179 |
| 0.072620032 | 0.280577128 |
| 0.072651266 | 0.280592067 |
| 0.072681193 | 0.280654061 |
| 0.072760652 | 0.280824276 |
| 0.072821993 | 0.280935242 |
| 0.072853735 | 0.280978609 |

| | |
|---|---|
| 0.07291598 | 0.281083554 |
| 0.072916357 | 0.281083554 |
| 0.072929852 | 0.281090074 |
| 0.072953919 | 0.281120024 |
| 0.073066288 | 0.281365467 |
| 0.07307762 | 0.281385639 |
| 0.073157695 | 0.281478521 |
| 0.073159605 | 0.281478521 |
| 0.073177177 | 0.281496601 |
| 0.073222791 | 0.281626573 |
| 0.073305967 | 0.281739112 |
| 0.073311206 | 0.281739112 |
| 0.073355429 | 0.281886316 |
| 0.07338287 | 0.281964758 |
| 0.073405214 | 0.281980546 |
| 0.073447172 | 0.282037009 |
| 0.073547792 | 0.282238391 |
| 0.07368616 | 0.282616412 |
| 0.073707029 | 0.282647743 |
| 0.073875213 | 0.283135608 |
| 0.07390328 | 0.283214636 |
| 0.073941312 | 0.283251204 |
| 0.073950363 | 0.283251204 |
| 0.073962137 | 0.283251204 |
| 0.074027232 | 0.283370437 |
| 0.074071124 | 0.283470102 |
| 0.074089247 | 0.283493899 |
| 0.074112909 | 0.283561659 |
| 0.074125785 | 0.283576113 |
| 0.074148867 | 0.283608103 |
| 0.074191176 | 0.283638042 |
| 0.074198383 | 0.283638042 |
| 0.07421454 | 0.283665655 |
| 0.074335715 | 0.28391277 |
| 0.074340768 | 0.28391277 |
| 0.074400733 | 0.283955742 |
| 0.074407483 | 0.283958751 |
| 0.07441727 | 0.283973346 |
| 0.074479218 | 0.28414986 |
| 0.074487392 | 0.28414986 |
| 0.074528114 | 0.284282433 |
| 0.074552076 | 0.284307688 |
| 0.074566339 | 0.284307688 |
| 0.074596235 | 0.284307688 |
| 0.074602526 | 0.284307688 |

| | |
|---|---|
| 0.074607255 | 0.284307688 |
| 0.074642773 | 0.284355412 |
| 0.074665546 | 0.28439667 |
| 0.074719948 | 0.284503771 |
| 0.074774392 | 0.284583663 |
| 0.074857069 | 0.284753338 |
| 0.074939092 | 0.284891773 |
| 0.074948936 | 0.284904471 |
| 0.074960378 | 0.284904471 |
| 0.074972028 | 0.284926013 |
| 0.075086818 | 0.285248455 |
| 0.075111961 | 0.285275706 |
| 0.075132615 | 0.285331396 |
| 0.075245192 | 0.285553649 |
| 0.075289196 | 0.285584466 |
| 0.075319037 | 0.285652156 |
| 0.075352815 | 0.285712007 |
| 0.075361529 | 0.285722303 |
| 0.075381134 | 0.285773881 |
| 0.075400827 | 0.285803041 |
| 0.075564202 | 0.286285599 |
| 0.07560926 | 0.286365186 |
| 0.075618544 | 0.286377577 |
| 0.075686964 | 0.286591114 |
| 0.075710109 | 0.286633177 |
| 0.075735193 | 0.286705352 |
| 0.07577228 | 0.286720063 |
| 0.075780949 | 0.286720063 |
| 0.076094396 | 0.287516682 |
| 0.07614703 | 0.287594438 |
| 0.076151205 | 0.287594438 |
| 0.076210699 | 0.287773485 |
| 0.0762619 | 0.287829902 |
| 0.076277675 | 0.287843817 |
| 0.076329437 | 0.288010397 |
| 0.076441207 | 0.288293836 |
| 0.076443416 | 0.288293836 |
| 0.076445396 | 0.288293836 |
| 0.076451407 | 0.288293836 |
| 0.076528759 | 0.288560087 |
| 0.076574526 | 0.28866669 |
| 0.076582208 | 0.288672801 |
| 0.076684943 | 0.288880808 |
| 0.076704061 | 0.288903228 |
| 0.076704882 | 0.288903228 |

| | |
|---|---|
| 0.076730604 | 0.288957735 |
| 0.076761946 | 0.288984363 |
| 0.07682537 | 0.289164428 |
| 0.076864959 | 0.28927004 |
| 0.076920778 | 0.289418366 |
| 0.076925842 | 0.289418366 |
| 0.076933887 | 0.289425769 |
| 0.077080822 | 0.289800855 |
| 0.077171054 | 0.290060786 |
| 0.077187644 | 0.290071701 |
| 0.077190853 | 0.290071701 |
| 0.077210546 | 0.290111033 |
| 0.077244769 | 0.290135071 |
| 0.077315802 | 0.29031218 |
| 0.077330567 | 0.290344731 |
| 0.077364632 | 0.290449735 |
| 0.077465247 | 0.290689986 |
| 0.077477566 | 0.290713311 |
| 0.077552719 | 0.290880718 |
| 0.077569649 | 0.290921305 |
| 0.077635859 | 0.291053691 |
| 0.077647725 | 0.291053691 |
| 0.077714599 | 0.291193083 |
| 0.077778519 | 0.291270945 |
| 0.077779072 | 0.291270945 |
| 0.07789924 | 0.291606673 |
| 0.077905459 | 0.291606673 |
| 0.077980588 | 0.291796157 |
| 0.078010127 | 0.291860825 |
| 0.078017061 | 0.291863842 |
| 0.078087176 | 0.291989916 |
| 0.078087546 | 0.291989916 |
| 0.078147052 | 0.292107582 |
| 0.078156078 | 0.29210857 |
| 0.078245168 | 0.292326843 |
| 0.078278262 | 0.292404611 |
| 0.07836729 | 0.292546916 |
| 0.078378203 | 0.292546916 |
| 0.078418826 | 0.29263131 |
| 0.078540638 | 0.292902275 |
| 0.078588157 | 0.292976148 |
| 0.078677957 | 0.293175534 |
| 0.078708966 | 0.293231535 |
| 0.078797351 | 0.293468989 |
| 0.078926382 | 0.293742816 |

| | |
|---|---|
| 0.078952625 | 0.293794567 |
| 0.078963524 | 0.293804597 |
| 0.079000205 | 0.293867409 |
| 0.079003051 | 0.293867409 |
| 0.079052756 | 0.293952367 |
| 0.079087724 | 0.294021625 |
| 0.07921 | 0.294330245 |
| 0.07921019 | 0.294330245 |
| 0.079260355 | 0.294388435 |
| 0.079260564 | 0.294388435 |
| 0.079281927 | 0.29442634 |
| 0.079338918 | 0.294587562 |
| 0.079364751 | 0.294620641 |
| 0.079426298 | 0.294797092 |
| 0.079433335 | 0.294800236 |
| 0.079475408 | 0.294887447 |
| 0.079537062 | 0.295026791 |
| 0.079551176 | 0.295053646 |
| 0.079596558 | 0.295123337 |
| 0.079676178 | 0.295172474 |
| 0.079701577 | 0.295197684 |
| 0.079748649 | 0.295303135 |
| 0.079788986 | 0.295406564 |
| 0.079815014 | 0.295472684 |
| 0.07987098 | 0.295595242 |
| 0.079879173 | 0.29559971 |
| 0.079925596 | 0.295682487 |
| 0.079955393 | 0.295716099 |
| 0.080002599 | 0.2958117 |
| 0.080003296 | 0.2958117 |
| 0.080035672 | 0.2958117 |
| 0.080035955 | 0.2958117 |
| 0.080121997 | 0.29597993 |
| 0.080124073 | 0.29597993 |
| 0.080143918 | 0.296030275 |
| 0.080161403 | 0.2960719 |
| 0.080187318 | 0.296121685 |
| 0.080221822 | 0.296180212 |
| 0.080299315 | 0.296351452 |
| 0.080342488 | 0.296424794 |
| 0.080362755 | 0.296424794 |
| 0.080468894 | 0.2966514 |
| 0.080608375 | 0.296893904 |
| 0.080648468 | 0.296995636 |
| 0.080655577 | 0.296998853 |

| | |
|---|---|
| 0.080692289 | 0.297088098 |
| 0.080895142 | 0.297509164 |
| 0.080899821 | 0.297509164 |
| 0.080900355 | 0.297509164 |
| 0.080939197 | 0.297606046 |
| 0.080973847 | 0.297679762 |
| 0.081123771 | 0.297916714 |
| 0.081193008 | 0.298022757 |
| 0.081196449 | 0.298022757 |
| 0.081255855 | 0.29811615 |
| 0.081281472 | 0.298135335 |
| 0.081316752 | 0.298211515 |
| 0.081386393 | 0.298397626 |
| 0.08139461 | 0.298397626 |
| 0.081409266 | 0.298397626 |
| 0.081416459 | 0.298397626 |
| 0.081418072 | 0.298397626 |
| 0.081423889 | 0.298397626 |
| 0.081542384 | 0.298724223 |
| 0.081600084 | 0.298836387 |
| 0.081852251 | 0.299529556 |
| 0.081865903 | 0.299542984 |
| 0.0818685 | 0.299542984 |
| 0.081973185 | 0.299764809 |
| 0.081994751 | 0.299797638 |
| 0.082114425 | 0.300166071 |
| 0.082153495 | 0.300262798 |
| 0.082216881 | 0.300425301 |
| 0.082257315 | 0.300503885 |
| 0.082334024 | 0.300668805 |
| 0.082370427 | 0.300778681 |
| 0.082554263 | 0.301126741 |
| 0.082581371 | 0.30117949 |
| 0.082605712 | 0.301198489 |
| 0.082611877 | 0.301198489 |
| 0.082620982 | 0.301208628 |
| 0.082700928 | 0.301384737 |
| 0.082718186 | 0.301424563 |
| 0.082772109 | 0.301551843 |
| 0.082859392 | 0.301777492 |
| 0.082877517 | 0.301820422 |
| 0.082898495 | 0.301873736 |
| 0.082909067 | 0.301889155 |
| 0.083004907 | 0.302076466 |
| 0.083090032 | 0.302278544 |

| | |
|---|---|
| 0.083104715 | 0.302294212 |
| 0.08317939 | 0.302434142 |
| 0.08319719 | 0.302475767 |
| 0.083211609 | 0.302482009 |
| 0.083300525 | 0.302708874 |
| 0.083325881 | 0.302745934 |
| 0.083336318 | 0.302750481 |
| 0.083370931 | 0.30285025 |
| 0.08342982 | 0.302974611 |
| 0.08344929 | 0.303022212 |
| 0.08361939 | 0.303465147 |
| 0.083622243 | 0.303465147 |
| 0.083779731 | 0.303763524 |
| 0.083796876 | 0.303797962 |
| 0.083806281 | 0.303808933 |
| 0.083909434 | 0.303992915 |
| 0.083914029 | 0.303992915 |
| 0.083920861 | 0.303992915 |
| 0.084157973 | 0.304550701 |
| 0.084207637 | 0.304675633 |
| 0.084272482 | 0.304826118 |
| 0.084286263 | 0.304852814 |
| 0.084342262 | 0.305032189 |
| 0.084384555 | 0.305138804 |
| 0.084398942 | 0.305167659 |
| 0.084456706 | 0.305330161 |
| 0.084568325 | 0.305548142 |
| 0.084608026 | 0.305587555 |
| 0.084609998 | 0.305587555 |
| 0.084642246 | 0.305587631 |
| 0.084645599 | 0.305587631 |
| 0.084654113 | 0.305587631 |
| 0.084656353 | 0.305587631 |
| 0.084701826 | 0.305625268 |
| 0.084796156 | 0.305884006 |
| 0.084816734 | 0.305888733 |
| 0.084839889 | 0.305902749 |
| 0.084861834 | 0.30595871 |
| 0.084928203 | 0.306160963 |
| 0.084937216 | 0.306160963 |
| 0.085203852 | 0.306565224 |
| 0.085217919 | 0.306592673 |
| 0.085329103 | 0.30687678 |
| 0.085420737 | 0.307104234 |
| 0.085424589 | 0.307104234 |

| | |
|---|---|
| 0.085440531 | 0.307130137 |
| 0.085546948 | 0.307404896 |
| 0.085581967 | 0.307507531 |
| 0.085603817 | 0.307562838 |
| 0.085732049 | 0.307930642 |
| 0.085743881 | 0.307932353 |
| 0.085826411 | 0.308037263 |
| 0.085841796 | 0.308069264 |
| 0.085849878 | 0.308075055 |
| 0.085861257 | 0.308092678 |
| 0.085910653 | 0.30819869 |
| 0.08601368 | 0.30830804 |
| 0.086035155 | 0.308317791 |
| 0.086093132 | 0.308333223 |
| 0.086093157 | 0.308333223 |
| 0.08610227 | 0.308333223 |
| 0.086115118 | 0.308333223 |
| 0.086200764 | 0.308590182 |
| 0.086291881 | 0.308707536 |
| 0.086338788 | 0.308782567 |
| 0.086378065 | 0.308853462 |
| 0.086411145 | 0.308895999 |
| 0.086455716 | 0.308991926 |
| 0.086464729 | 0.30900095 |
| 0.086526359 | 0.309050162 |
| 0.086529264 | 0.309050162 |
| 0.086529666 | 0.309050162 |
| 0.086548068 | 0.309050162 |
| 0.086549879 | 0.309050162 |
| 0.086648034 | 0.309238358 |
| 0.086700352 | 0.30937871 |
| 0.086795139 | 0.309600966 |
| 0.086814705 | 0.309647566 |
| 0.086835173 | 0.309674188 |
| 0.086870068 | 0.309719911 |
| 0.086880509 | 0.309719911 |
| 0.086930189 | 0.309866152 |
| 0.087153179 | 0.310436351 |
| 0.08716926 | 0.310470414 |
| 0.087190099 | 0.310521414 |
| 0.087434801 | 0.310965396 |
| 0.087443933 | 0.310965396 |
| 0.087491802 | 0.311070718 |
| 0.087559984 | 0.311151721 |
| 0.087574088 | 0.311151721 |

| | |
|---|---|
| 0.087583445 | 0.311151721 |
| 0.087587678 | 0.311151721 |
| 0.087600612 | 0.311151721 |
| 0.08767273 | 0.311262118 |
| 0.087681049 | 0.311262118 |
| 0.087689428 | 0.311262118 |
| 0.087692172 | 0.311262118 |
| 0.087775119 | 0.311401157 |
| 0.087777112 | 0.311401157 |
| 0.087800265 | 0.311460093 |
| 0.087823723 | 0.311520104 |
| 0.087831245 | 0.311523587 |
| 0.087874174 | 0.311606231 |
| 0.087925457 | 0.31164886 |
| 0.087936662 | 0.311665381 |
| 0.087950028 | 0.311666366 |
| 0.088130853 | 0.311973689 |
| 0.088131502 | 0.311973689 |
| 0.088139156 | 0.311973689 |
| 0.088212688 | 0.31210948 |
| 0.088226089 | 0.312133707 |
| 0.088251219 | 0.312176232 |
| 0.088314526 | 0.312307389 |
| 0.088355973 | 0.312430762 |
| 0.088444727 | 0.312567996 |
| 0.088454664 | 0.312570876 |
| 0.088523764 | 0.312699053 |
| 0.088621303 | 0.312997168 |
| 0.088643959 | 0.31304999 |
| 0.088649405 | 0.31304999 |
| 0.088749076 | 0.313316847 |
| 0.08876437 | 0.313316847 |
| 0.088764445 | 0.313316847 |
| 0.088915794 | 0.313665099 |
| 0.088927082 | 0.31366899 |
| 0.088930068 | 0.31366899 |
| 0.088953889 | 0.313694298 |
| 0.089000594 | 0.313824437 |
| 0.089073144 | 0.313957102 |
| 0.089077673 | 0.313957102 |
| 0.089094794 | 0.313994211 |
| 0.089131001 | 0.314052097 |
| 0.089220015 | 0.314249495 |
| 0.089251575 | 0.314279591 |
| 0.089254955 | 0.314279591 |

| | |
|---|---|
| 0.089315133 | 0.314421748 |
| 0.089331993 | 0.314426909 |
| 0.089464212 | 0.314737186 |
| 0.089490744 | 0.314783047 |
| 0.089511025 | 0.31480886 |
| 0.089543992 | 0.314855055 |
| 0.089598994 | 0.314978691 |
| 0.089606122 | 0.314980497 |
| 0.089624747 | 0.31499044 |
| 0.089693761 | 0.315113975 |
| 0.089838214 | 0.315433418 |
| 0.089846938 | 0.315433418 |
| 0.089855071 | 0.315433418 |
| 0.089887149 | 0.315502635 |
| 0.089918065 | 0.315567077 |
| 0.089922881 | 0.315567077 |
| 0.089925386 | 0.315567077 |
| 0.090036021 | 0.315769189 |
| 0.090053439 | 0.315807021 |
| 0.090099495 | 0.315922008 |
| 0.090112473 | 0.315925084 |
| 0.090220577 | 0.316230159 |
| 0.090254482 | 0.31632572 |
| 0.090271231 | 0.316335498 |
| 0.090284328 | 0.316335498 |
| 0.090284328 | 0.316335498 |
| 0.090284328 | 0.316335498 |
| 0.090556322 | 0.317032129 |
| 0.09056729 | 0.317032129 |
| 0.090575853 | 0.317032129 |
| 0.090588094 | 0.317042823 |
| 0.090661871 | 0.317182842 |
| 0.090780831 | 0.317447102 |
| 0.090798576 | 0.317447102 |
| 0.090803067 | 0.317447102 |
| 0.090894599 | 0.317649125 |
| 0.091033596 | 0.317968993 |
| 0.091037126 | 0.317968993 |
| 0.091084605 | 0.318025922 |
| 0.091122132 | 0.318032666 |
| 0.091137259 | 0.318043547 |
| 0.091138605 | 0.318043547 |
| 0.091158811 | 0.318067449 |
| 0.091183245 | 0.318080522 |
| 0.091191835 | 0.318080522 |

| | |
|---|---|
| 0.091228142 | 0.318146204 |
| 0.091321454 | 0.318308468 |
| 0.091342892 | 0.318336597 |
| 0.09135077 | 0.318340758 |
| 0.091376806 | 0.318399876 |
| 0.091381105 | 0.318399876 |
| 0.091401371 | 0.31844719 |
| 0.091419993 | 0.318483784 |
| 0.091494111 | 0.318636197 |
| 0.091665871 | 0.318948745 |
| 0.091700001 | 0.319020887 |
| 0.091857721 | 0.319404597 |
| 0.091901279 | 0.319465247 |
| 0.091923995 | 0.319466081 |
| 0.091935005 | 0.319466081 |
| 0.091935291 | 0.319466081 |
| 0.092007842 | 0.319648223 |
| 0.092106318 | 0.319897003 |
| 0.092132758 | 0.319950311 |
| 0.09216919 | 0.320022017 |
| 0.092262025 | 0.320227626 |
| 0.092305596 | 0.320355506 |
| 0.092321596 | 0.320372325 |
| 0.092381127 | 0.32044738 |
| 0.092501755 | 0.32073584 |
| 0.092567503 | 0.32082418 |
| 0.092588523 | 0.320846447 |
| 0.092658296 | 0.321032171 |
| 0.092778094 | 0.321271555 |
| 0.092791368 | 0.321271555 |
| 0.092802128 | 0.321271555 |
| 0.092805637 | 0.321271555 |
| 0.092815428 | 0.321282099 |
| 0.092890053 | 0.321517047 |
| 0.092925364 | 0.321615894 |
| 0.092940131 | 0.321628367 |
| 0.092945713 | 0.321628367 |
| 0.092960727 | 0.321628367 |
| 0.092969486 | 0.321628367 |
| 0.093006222 | 0.321685353 |
| 0.093030485 | 0.321722546 |
| 0.093046331 | 0.321753984 |
| 0.093064434 | 0.321793219 |
| 0.093085842 | 0.321800355 |
| 0.093121282 | 0.321872946 |

| | |
|---|---|
| 0.093390461 | 0.322545867 |
| 0.093601263 | 0.323039665 |
| 0.093705185 | 0.323210954 |
| 0.093758214 | 0.32330636 |
| 0.093762779 | 0.32330636 |
| 0.093779572 | 0.32330636 |
| 0.093886336 | 0.323460974 |
| 0.093949651 | 0.323608884 |
| 0.094022184 | 0.323690201 |
| 0.094031213 | 0.323690201 |
| 0.09405981 | 0.323730784 |
| 0.094165823 | 0.323978593 |
| 0.094165823 | 0.323978593 |
| 0.094240319 | 0.324136627 |
| 0.094245785 | 0.324136627 |
| 0.094332158 | 0.32434 |
| 0.094402061 | 0.324486645 |
| 0.09444602 | 0.324567842 |
| 0.094491013 | 0.324675237 |
| 0.094592681 | 0.324929258 |
| 0.094606519 | 0.324929258 |
| 0.0946102 | 0.324929258 |
| 0.09461086 | 0.324929258 |
| 0.09462461 | 0.324946738 |
| 0.094659216 | 0.325009499 |
| 0.094670182 | 0.325009499 |
| 0.094677765 | 0.325012103 |
| 0.094741836 | 0.32513022 |
| 0.094746306 | 0.32513022 |
| 0.094822674 | 0.325321964 |
| 0.094866679 | 0.325391231 |
| 0.094877024 | 0.325391231 |
| 0.09499229 | 0.325757398 |
| 0.095242551 | 0.326386314 |
| 0.095267424 | 0.326448061 |
| 0.095291324 | 0.326506467 |
| 0.095324249 | 0.32655443 |
| 0.095325892 | 0.32655443 |
| 0.095390669 | 0.326647917 |
| 0.095463872 | 0.326839037 |
| 0.095498586 | 0.326852522 |
| 0.095590535 | 0.326932985 |
| 0.095622179 | 0.326981442 |
| 0.095640272 | 0.327019833 |
| 0.095657048 | 0.327053712 |

| | |
|---|---|
| 0.095693105 | 0.327153507 |
| 0.095739902 | 0.32721954 |
| 0.095910259 | 0.327613706 |
| 0.096013547 | 0.32787246 |
| 0.096057238 | 0.327927609 |
| 0.096105485 | 0.327951273 |
| 0.096205038 | 0.32822044 |
| 0.096244167 | 0.328306902 |
| 0.096298051 | 0.328379773 |
| 0.096300004 | 0.328379773 |
| 0.096351894 | 0.328439141 |
| 0.096422438 | 0.328609049 |
| 0.096474125 | 0.328714633 |
| 0.096577538 | 0.328949323 |
| 0.096587714 | 0.328960458 |
| 0.096608003 | 0.328975669 |
| 0.096613078 | 0.328975669 |
| 0.096626717 | 0.328975669 |
| 0.096713927 | 0.329178461 |
| 0.096780733 | 0.32935877 |
| 0.096845724 | 0.329485947 |
| 0.096947299 | 0.329691821 |
| 0.096959073 | 0.329691821 |
| 0.096963165 | 0.329691821 |
| 0.096994032 | 0.329691821 |
| 0.097005033 | 0.329691821 |
| 0.097008009 | 0.329691821 |
| 0.097017017 | 0.329691821 |
| 0.097040482 | 0.329720134 |
| 0.097040735 | 0.329720134 |
| 0.097046147 | 0.329720134 |
| 0.097065324 | 0.329728233 |
| 0.0970693 | 0.329728233 |
| 0.097086392 | 0.329757907 |
| 0.097091883 | 0.329757907 |
| 0.097178731 | 0.329911713 |
| 0.097366229 | 0.330430481 |
| 0.097387385 | 0.330478731 |
| 0.097402311 | 0.330505831 |
| 0.09741837 | 0.330513229 |
| 0.097622978 | 0.331057732 |
| 0.097642585 | 0.331058448 |
| 0.097646998 | 0.331058448 |
| 0.097659837 | 0.331058448 |
| 0.097693888 | 0.331059713 |

| | |
|---|---|
| 0.097715391 | 0.331061962 |
| 0.097742774 | 0.331080638 |
| 0.097745498 | 0.331080638 |
| 0.097784136 | 0.331164408 |
| 0.097801332 | 0.331188049 |
| 0.097809704 | 0.331188049 |
| 0.09785736 | 0.331294603 |
| 0.097882998 | 0.33130943 |
| 0.09793817 | 0.331426832 |
| 0.097968276 | 0.331458059 |
| 0.098038164 | 0.331600287 |
| 0.098079071 | 0.331667985 |
| 0.098129656 | 0.33180325 |
| 0.098186391 | 0.331883567 |
| 0.098197455 | 0.331883567 |
| 0.098227174 | 0.331956688 |
| 0.098244061 | 0.331990202 |
| 0.098281206 | 0.332048194 |
| 0.098404585 | 0.332344006 |
| 0.098469702 | 0.332516768 |
| 0.09848578 | 0.332547481 |
| 0.098493304 | 0.332549312 |
| 0.098528188 | 0.332596359 |
| 0.098575089 | 0.332687504 |
| 0.098576145 | 0.332687504 |
| 0.09863683 | 0.332772816 |
| 0.098655875 | 0.332772816 |
| 0.09866403 | 0.332772816 |
| 0.098670698 | 0.332772816 |
| 0.098707826 | 0.33282532 |
| 0.098768554 | 0.332912239 |
| 0.098831549 | 0.333077424 |
| 0.09886217 | 0.333135032 |
| 0.098872598 | 0.333135032 |
| 0.098893799 | 0.333145775 |
| 0.09897848 | 0.333360303 |
| 0.099089802 | 0.333564095 |
| 0.099146464 | 0.333625644 |
| 0.099147206 | 0.333625644 |
| 0.099148327 | 0.333625644 |
| 0.099166384 | 0.333662829 |
| 0.099402219 | 0.334273204 |
| 0.099403015 | 0.334273204 |
| 0.099403939 | 0.334273204 |
| 0.099449458 | 0.334363808 |

| | |
|---|---|
| 0.099465984 | 0.334363808 |
| 0.099502418 | 0.334462675 |
| 0.0995341 | 0.334494049 |
| 0.099557535 | 0.33450299 |
| 0.099563575 | 0.33450299 |
| 0.099609076 | 0.334608651 |
| 0.099619655 | 0.334620589 |
| 0.099656334 | 0.334649383 |
| 0.099677165 | 0.334695736 |
| 0.09970658 | 0.33472371 |
| 0.099763601 | 0.33479714 |
| 0.099770926 | 0.334798133 |
| 0.099795949 | 0.334858508 |
| 0.099811101 | 0.334885758 |
| 0.099825599 | 0.334910807 |
| 0.100032228 | 0.33539826 |
| 0.100034272 | 0.33539826 |
| 0.100092543 | 0.335451941 |
| 0.100105048 | 0.335470247 |
| 0.100145887 | 0.335559881 |
| 0.100153527 | 0.335561873 |
| 0.100248117 | 0.33569249 |
| 0.100248898 | 0.33569249 |
| 0.100291228 | 0.335763412 |
| 0.10030429 | 0.335783538 |
| 0.100326229 | 0.335833378 |
| 0.100362885 | 0.335886639 |
| 0.100379532 | 0.335886639 |
| 0.100386238 | 0.335886639 |
| 0.100392443 | 0.335886639 |
| 0.100398559 | 0.335886639 |
| 0.100557574 | 0.336183993 |
| 0.100652841 | 0.336398054 |
| 0.100760173 | 0.336573856 |
| 0.100876999 | 0.336754611 |
| 0.100877192 | 0.336754611 |
| 0.100907246 | 0.33683133 |
| 0.100946942 | 0.336916609 |
| 0.101115545 | 0.337321522 |
| 0.101299042 | 0.337651368 |
| 0.101316869 | 0.337654365 |
| 0.10160629 | 0.33833481 |
| 0.101679601 | 0.338502543 |
| 0.101685092 | 0.338502543 |
| 0.101716057 | 0.338517643 |

| | |
|---|---|
| 0.101718059 | 0.338517643 |
| 0.101802665 | 0.338728205 |
| 0.101964679 | 0.339111826 |
| 0.101982041 | 0.339111826 |
| 0.102114897 | 0.339435107 |
| 0.10214831 | 0.339498782 |
| 0.10217624 | 0.339544223 |
| 0.10234806 | 0.339809179 |
| 0.102361606 | 0.339809179 |
| 0.102374272 | 0.339809179 |
| 0.102377263 | 0.339809179 |
| 0.102396816 | 0.339850393 |
| 0.102430537 | 0.339914938 |
| 0.102507898 | 0.34007609 |
| 0.102514801 | 0.34007609 |
| 0.102534387 | 0.340098652 |
| 0.102581329 | 0.340202004 |
| 0.102672214 | 0.340380714 |
| 0.102755831 | 0.340504804 |
| 0.102792183 | 0.340545584 |
| 0.10288642 | 0.340786671 |
| 0.102920072 | 0.340832538 |
| 0.102955128 | 0.340895708 |
| 0.103056792 | 0.341161175 |
| 0.103082613 | 0.341195743 |
| 0.1031333 | 0.341248415 |
| 0.103175246 | 0.341339777 |
| 0.103234902 | 0.341400426 |
| 0.103236192 | 0.341400426 |
| 0.103262286 | 0.341414405 |
| 0.103352084 | 0.341663753 |
| 0.103436376 | 0.341777742 |
| 0.103443974 | 0.341777742 |
| 0.103459787 | 0.341806277 |
| 0.103483898 | 0.341808853 |
| 0.103489274 | 0.341808853 |
| 0.103502356 | 0.341813945 |
| 0.103524843 | 0.341855212 |
| 0.103598504 | 0.341965078 |
| 0.103608375 | 0.341965078 |
| 0.103689114 | 0.342169246 |
| 0.103705035 | 0.342189256 |
| 0.103730846 | 0.342210634 |
| 0.103740255 | 0.342210634 |
| 0.103759516 | 0.342236511 |

| | |
|---|---|
| 0.103802417 | 0.342320849 |
| 0.103903323 | 0.342465743 |
| 0.103903818 | 0.342465743 |
| 0.104100684 | 0.342931807 |
| 0.104133342 | 0.342937261 |
| 0.104226538 | 0.34307812 |
| 0.104474836 | 0.343444436 |
| 0.104534803 | 0.343564509 |
| 0.104540216 | 0.343564509 |
| 0.104583584 | 0.343659608 |
| 0.104595285 | 0.343674344 |
| 0.104640108 | 0.343721676 |
| 0.104645224 | 0.343721676 |
| 0.104645774 | 0.343721676 |
| 0.10466062 | 0.343723028 |
| 0.104696539 | 0.343794031 |
| 0.104751002 | 0.343868009 |
| 0.10479783 | 0.343944043 |
| 0.104867445 | 0.344050979 |
| 0.104926341 | 0.344152229 |
| 0.105037744 | 0.344391619 |
| 0.105106326 | 0.344484109 |
| 0.105145473 | 0.344555075 |
| 0.105161782 | 0.344584809 |
| 0.105202565 | 0.344694726 |
| 0.105265233 | 0.344876334 |
| 0.105328991 | 0.344962357 |
| 0.105393838 | 0.34509521 |
| 0.105397252 | 0.34509521 |
| 0.10545542 | 0.345200633 |
| 0.10546569 | 0.345200633 |
| 0.105545076 | 0.345365532 |
| 0.105567116 | 0.345413921 |
| 0.105577718 | 0.345424881 |
| 0.105604251 | 0.345487956 |
| 0.105683464 | 0.345627954 |
| 0.105688854 | 0.345627954 |
| 0.105747812 | 0.345720144 |
| 0.105847986 | 0.345839265 |
| 0.105849602 | 0.345839265 |
| 0.105912457 | 0.345903992 |
| 0.105993079 | 0.346107127 |
| 0.105996988 | 0.346107127 |
| 0.106021322 | 0.34616285 |
| 0.106046296 | 0.346196926 |

| | |
|---|---|
| 0.106116351 | 0.346354403 |
| 0.106185093 | 0.346415725 |
| 0.106258775 | 0.346510556 |
| 0.106267342 | 0.346514768 |
| 0.106307626 | 0.346552313 |
| 0.106310642 | 0.346552313 |
| 0.106315238 | 0.346552313 |
| 0.106371901 | 0.346665836 |
| 0.106395255 | 0.346718221 |
| 0.106425271 | 0.346756565 |
| 0.106490017 | 0.34688462 |
| 0.106557853 | 0.347050953 |
| 0.106574466 | 0.347064758 |
| 0.106617585 | 0.34711024 |
| 0.106748089 | 0.347463862 |
| 0.106838376 | 0.347543974 |
| 0.10686322 | 0.347583601 |
| 0.106954612 | 0.347785774 |
| 0.106988862 | 0.34783654 |
| 0.10701204 | 0.34783654 |
| 0.107085277 | 0.347966875 |
| 0.107233007 | 0.348288121 |
| 0.107454802 | 0.348858204 |
| 0.107469343 | 0.348864574 |
| 0.107531911 | 0.348898615 |
| 0.107532037 | 0.348898615 |
| 0.107640426 | 0.349166463 |
| 0.107645058 | 0.349166463 |
| 0.107702923 | 0.349330368 |
| 0.107738158 | 0.349420856 |
| 0.107926776 | 0.349748338 |
| 0.107986184 | 0.349866882 |
| 0.107995044 | 0.349866882 |
| 0.108032131 | 0.349945338 |
| 0.108106033 | 0.350082373 |
| 0.108170734 | 0.350233934 |
| 0.10818141 | 0.350238325 |
| 0.108215524 | 0.350266604 |
| 0.108216425 | 0.350266604 |
| 0.108219562 | 0.350266604 |
| 0.108276756 | 0.350427907 |
| 0.108370407 | 0.350672518 |
| 0.10839575 | 0.350681429 |
| 0.108435837 | 0.350713279 |
| 0.108441859 | 0.350713279 |

| | |
|---|---|
| 0.108458479 | 0.350713279 |
| 0.108497478 | 0.350808511 |
| 0.108552666 | 0.350889248 |
| 0.108566653 | 0.350889248 |
| 0.108710259 | 0.351186564 |
| 0.108747583 | 0.351235666 |
| 0.108904933 | 0.351568333 |
| 0.109046139 | 0.351889726 |
| 0.109060634 | 0.351912658 |
| 0.109095039 | 0.351999826 |
| 0.109166834 | 0.352103443 |
| 0.109171511 | 0.352103443 |
| 0.109222841 | 0.352192291 |
| 0.109228638 | 0.352192291 |
| 0.109238222 | 0.352194805 |
| 0.109244207 | 0.352194805 |
| 0.109256019 | 0.352209045 |
| 0.109284369 | 0.352268284 |
| 0.10930339 | 0.352268284 |
| 0.109303981 | 0.352268284 |
| 0.109382434 | 0.352473423 |
| 0.109427571 | 0.352547313 |
| 0.109558311 | 0.352784362 |
| 0.109560407 | 0.352784362 |
| 0.109589863 | 0.352831501 |
| 0.109604918 | 0.352856121 |
| 0.109762149 | 0.353276889 |
| 0.109765289 | 0.353276889 |
| 0.10981757 | 0.353296849 |
| 0.109835935 | 0.353296849 |
| 0.109848827 | 0.353296849 |
| 0.10986744 | 0.353319146 |
| 0.109887332 | 0.353346546 |
| 0.109890798 | 0.353346546 |
| 0.109919102 | 0.353365985 |
| 0.109943857 | 0.35337401 |
| 0.109966742 | 0.353423715 |
| 0.110067584 | 0.353628495 |
| 0.110120827 | 0.35372797 |
| 0.110129815 | 0.353732984 |
| 0.110233971 | 0.353924318 |
| 0.110279963 | 0.353993177 |
| 0.110329385 | 0.354018586 |
| 0.110337528 | 0.354018586 |
| 0.110442943 | 0.354197098 |

| | |
|---|---|
| 0.110484875 | 0.354197098 |
| 0.11048888 | 0.354197098 |
| 0.110497923 | 0.35419872 |
| 0.110521266 | 0.354249701 |
| 0.110552069 | 0.354324589 |
| 0.110571141 | 0.354338026 |
| 0.110587014 | 0.354365049 |
| 0.1108519 | 0.354998887 |
| 0.110862559 | 0.35500915 |
| 0.11087763 | 0.355013798 |
| 0.110890864 | 0.355013798 |
| 0.11090128 | 0.355013798 |
| 0.111024221 | 0.355296394 |
| 0.111051773 | 0.355328369 |
| 0.111063458 | 0.355341888 |
| 0.111086166 | 0.355388535 |
| 0.111118122 | 0.355445148 |
| 0.111204437 | 0.355581865 |
| 0.111296334 | 0.355800141 |
| 0.111332717 | 0.355866871 |
| 0.111349825 | 0.355866871 |
| 0.111358288 | 0.355866871 |
| 0.111362039 | 0.355866871 |
| 0.111396285 | 0.355922139 |
| 0.11144654 | 0.355942886 |
| 0.111453906 | 0.355942886 |
| 0.111458775 | 0.355942886 |
| 0.11150283 | 0.356006253 |
| 0.111510509 | 0.356006904 |
| 0.111625533 | 0.356326359 |
| 0.111665878 | 0.356405509 |
| 0.111672778 | 0.356405509 |
| 0.111722312 | 0.356471402 |
| 0.111777259 | 0.356619475 |
| 0.111823069 | 0.356737344 |
| 0.111877958 | 0.356821234 |
| 0.111907202 | 0.356866708 |
| 0.111916233 | 0.356868097 |
| 0.111941169 | 0.356868097 |
| 0.112051274 | 0.357182655 |
| 0.112127342 | 0.35734325 |
| 0.112131666 | 0.35734325 |
| 0.112148155 | 0.357347974 |
| 0.112172425 | 0.357378923 |
| 0.112224873 | 0.357520671 |

| | |
|---|---|
| 0.112271309 | 0.357572931 |
| 0.112299532 | 0.357618916 |
| 0.112379131 | 0.357724961 |
| 0.112455633 | 0.357891104 |
| 0.112459277 | 0.357891104 |
| 0.112518889 | 0.35798173 |
| 0.112527442 | 0.35798173 |
| 0.112587333 | 0.358090535 |
| 0.112763794 | 0.358374564 |
| 0.112778933 | 0.358398762 |
| 0.112849498 | 0.358437028 |
| 0.112917077 | 0.358437028 |
| 0.112955768 | 0.358437028 |
| 0.112974544 | 0.358437028 |
| 0.113021665 | 0.358437028 |
| 0.113035117 | 0.358437028 |
| 0.113125029 | 0.358437028 |
| 0.113166697 | 0.358437028 |
| 0.113169187 | 0.358437028 |
| 0.113223825 | 0.358437028 |
| 0.113330422 | 0.358437028 |
| 0.113351364 | 0.358437028 |
| 0.113375686 | 0.358437028 |
| 0.113481059 | 0.358437028 |
| 0.11350215 | 0.358437028 |
| 0.113536364 | 0.358437028 |
| 0.113560353 | 0.358437028 |
| 0.113597412 | 0.358437028 |
| 0.113723359 | 0.358437028 |
| 0.113770065 | 0.358437028 |
| 0.113827148 | 0.358571704 |
| 0.113920872 | 0.358805905 |
| 0.113957177 | 0.358815183 |
| 0.114038707 | 0.35892613 |
| 0.114142346 | 0.359089605 |
| 0.114191421 | 0.35919654 |
| 0.114216079 | 0.359226652 |
| 0.114296814 | 0.359355708 |
| 0.114303563 | 0.359355708 |
| 0.114326385 | 0.35936 |
| 0.114348967 | 0.359383546 |
| 0.114410496 | 0.359505752 |
| 0.114517573 | 0.359763769 |
| 0.11453747 | 0.359763769 |
| 0.114672723 | 0.360044682 |

| | |
|---|---|
| 0.114691138 | 0.360069371 |
| 0.114829176 | 0.360375772 |
| 0.114916938 | 0.360526249 |
| 0.11494828 | 0.360577074 |
| 0.115197466 | 0.361168435 |
| 0.115255175 | 0.361187035 |
| 0.115274619 | 0.361187035 |
| 0.115356874 | 0.361304064 |
| 0.115362104 | 0.361304064 |
| 0.115432007 | 0.361380412 |
| 0.115459721 | 0.361423722 |
| 0.115515505 | 0.361546755 |
| 0.115546132 | 0.361618848 |
| 0.115599555 | 0.361678494 |
| 0.115655155 | 0.361724539 |
| 0.115655852 | 0.361724539 |
| 0.115668199 | 0.361739399 |
| 0.115693193 | 0.361770055 |
| 0.115742818 | 0.361893208 |
| 0.115961318 | 0.362275481 |
| 0.115985982 | 0.362303456 |
| 0.115993094 | 0.362303456 |
| 0.116032336 | 0.362363755 |
| 0.116051549 | 0.362375876 |
| 0.11608427 | 0.362445623 |
| 0.116347425 | 0.363029274 |
| 0.116380988 | 0.363086422 |
| 0.11649238 | 0.363315599 |
| 0.116492589 | 0.363315599 |
| 0.116548077 | 0.363417262 |
| 0.116742265 | 0.363728768 |
| 0.11677699 | 0.363728768 |
| 0.116777308 | 0.363728768 |
| 0.116782542 | 0.363728768 |
| 0.11679983 | 0.363728768 |
| 0.116925251 | 0.363930593 |
| 0.116997428 | 0.364031914 |
| 0.117005771 | 0.364034091 |
| 0.117028875 | 0.364082188 |
| 0.117055844 | 0.364090005 |
| 0.117133004 | 0.364239591 |
| 0.117222172 | 0.364398463 |
| 0.117232671 | 0.364398463 |
| 0.117425695 | 0.364698284 |
| 0.117432661 | 0.364698284 |

| | |
|---|---|
| 0.117433659 | 0.364698284 |
| 0.11752721 | 0.364917427 |
| 0.11755507 | 0.364936083 |
| 0.117555757 | 0.364936083 |
| 0.117604518 | 0.365014686 |
| 0.11769481 | 0.365247021 |
| 0.117729006 | 0.365259994 |
| 0.117865399 | 0.365609962 |
| 0.117937538 | 0.365786085 |
| 0.117973042 | 0.365811379 |
| 0.118118314 | 0.366013074 |
| 0.118219566 | 0.366126957 |
| 0.118224253 | 0.366126957 |
| 0.118284961 | 0.366244546 |
| 0.118292982 | 0.366244546 |
| 0.118337569 | 0.366329162 |
| 0.118345182 | 0.366329162 |
| 0.118358136 | 0.366329162 |
| 0.118379023 | 0.36634559 |
| 0.118379462 | 0.36634559 |
| 0.118432948 | 0.366415877 |
| 0.118553314 | 0.366740627 |
| 0.118588173 | 0.366800812 |
| 0.118603509 | 0.366805651 |
| 0.118664803 | 0.366918689 |
| 0.118755731 | 0.367104512 |
| 0.11876556 | 0.367111068 |
| 0.118805564 | 0.367163233 |
| 0.118842271 | 0.367213169 |
| 0.1188732 | 0.367229285 |
| 0.119069027 | 0.367605128 |
| 0.119069349 | 0.367605128 |
| 0.119096511 | 0.367632998 |
| 0.119145477 | 0.367736481 |
| 0.119251448 | 0.367992013 |
| 0.119267667 | 0.368018217 |
| 0.119396793 | 0.368297347 |
| 0.11945619 | 0.3683546 |
| 0.119469492 | 0.3683546 |
| 0.119580868 | 0.368554844 |
| 0.119598515 | 0.368561926 |
| 0.119700108 | 0.36870761 |
| 0.119720457 | 0.368746442 |
| 0.119754119 | 0.368786716 |
| 0.119837899 | 0.368988705 |

| | |
|---|---|
| 0.119893646 | 0.369051043 |
| 0.119966216 | 0.369203737 |
| 0.119969703 | 0.369203737 |
| 0.12010983 | 0.369539462 |
| 0.120144867 | 0.369599508 |
| 0.120202952 | 0.369682681 |
| 0.120276339 | 0.369836735 |
| 0.12030782 | 0.369863444 |
| 0.120402683 | 0.369994172 |
| 0.120420763 | 0.369994172 |
| 0.120534316 | 0.370301284 |
| 0.12053634 | 0.370301284 |
| 0.120614346 | 0.370445569 |
| 0.120646478 | 0.370496473 |
| 0.120769799 | 0.370755653 |
| 0.120823814 | 0.370897571 |
| 0.120860039 | 0.37096095 |
| 0.12093932 | 0.371113942 |
| 0.120956637 | 0.371113942 |
| 0.121035877 | 0.371309223 |
| 0.121064775 | 0.371373957 |
| 0.121080073 | 0.371396963 |
| 0.121159853 | 0.371569884 |
| 0.121242239 | 0.371798604 |
| 0.121279044 | 0.371851737 |
| 0.121291107 | 0.371852692 |
| 0.12142722 | 0.372126259 |
| 0.121441487 | 0.372144676 |
| 0.121486632 | 0.372210817 |
| 0.121610686 | 0.372426433 |
| 0.121617127 | 0.372426433 |
| 0.121797743 | 0.372630574 |
| 0.121818619 | 0.372630574 |
| 0.121865382 | 0.372672952 |
| 0.121912009 | 0.372735702 |
| 0.121942427 | 0.372769703 |
| 0.121959591 | 0.372798245 |
| 0.122004076 | 0.372891616 |
| 0.122018572 | 0.372891616 |
| 0.122087669 | 0.373046091 |
| 0.122170222 | 0.373187419 |
| 0.122190771 | 0.373193528 |
| 0.122209842 | 0.373219237 |
| 0.122241648 | 0.373248577 |
| 0.122367199 | 0.373397143 |

| | |
|---|---|
| 0.122461564 | 0.373589347 |
| 0.122522773 | 0.373704267 |
| 0.122596469 | 0.373809348 |
| 0.122652656 | 0.373956727 |
| 0.122669588 | 0.373984412 |
| 0.122700393 | 0.374054383 |
| 0.122724059 | 0.374078421 |
| 0.122762257 | 0.374120395 |
| 0.122843653 | 0.374248032 |
| 0.122862979 | 0.374248032 |
| 0.122873926 | 0.374248032 |
| 0.122896511 | 0.37427967 |
| 0.122931714 | 0.37427967 |
| 0.122948057 | 0.374282333 |
| 0.123029051 | 0.37438532 |
| 0.123053525 | 0.37441492 |
| 0.123064896 | 0.374422626 |
| 0.123072908 | 0.374423085 |
| 0.123121272 | 0.374457698 |
| 0.123123597 | 0.374457698 |
| 0.123141509 | 0.374471622 |
| 0.1231439 | 0.374471622 |
| 0.1231596 | 0.374495451 |
| 0.123174773 | 0.374496546 |
| 0.123183549 | 0.374496546 |
| 0.123306559 | 0.374697375 |
| 0.123396295 | 0.374832282 |
| 0.123500712 | 0.375053778 |
| 0.123538722 | 0.37512137 |
| 0.123648288 | 0.375387853 |
| 0.123741453 | 0.375545415 |
| 0.123815194 | 0.375697391 |
| 0.123881984 | 0.375768174 |
| 0.123927546 | 0.375830979 |
| 0.123934521 | 0.375830979 |
| 0.123938131 | 0.375830979 |
| 0.123996571 | 0.375881804 |
| 0.124021987 | 0.375917722 |
| 0.124099864 | 0.376064942 |
| 0.124110839 | 0.376064942 |
| 0.124227528 | 0.376278644 |
| 0.124281725 | 0.376417534 |
| 0.124304506 | 0.376431023 |
| 0.124386348 | 0.37661466 |
| 0.124542558 | 0.376967805 |

| | |
|---|---|
| 0.124561046 | 0.376989766 |
| 0.124589036 | 0.377012646 |
| 0.124650423 | 0.377126373 |
| 0.124655874 | 0.377126373 |
| 0.12468959 | 0.377182359 |
| 0.124745986 | 0.377303636 |
| 0.124803541 | 0.377374024 |
| 0.124954922 | 0.377735834 |
| 0.12503437 | 0.377856089 |
| 0.125049076 | 0.377876551 |
| 0.125094859 | 0.377918989 |
| 0.125128744 | 0.377973406 |
| 0.125137534 | 0.377975986 |
| 0.125294574 | 0.378378333 |
| 0.125330177 | 0.378437859 |
| 0.125349526 | 0.378448297 |
| 0.125451609 | 0.378708487 |
| 0.125542474 | 0.378833162 |
| 0.125551928 | 0.378833162 |
| 0.125603258 | 0.378867699 |
| 0.125656819 | 0.378943666 |
| 0.125832287 | 0.379365828 |
| 0.125839373 | 0.379365828 |
| 0.125850279 | 0.379365828 |
| 0.125857857 | 0.379365828 |
| 0.125908875 | 0.379365828 |
| 0.125914293 | 0.379365828 |
| 0.125924253 | 0.379365828 |
| 0.125965987 | 0.379431474 |
| 0.125969943 | 0.379431474 |
| 0.12599675 | 0.379444658 |
| 0.126006188 | 0.379444658 |
| 0.12606223 | 0.379517436 |
| 0.126103099 | 0.379592485 |
| 0.126132935 | 0.379658299 |
| 0.126374232 | 0.380096341 |
| 0.126397581 | 0.380104943 |
| 0.126416997 | 0.380104943 |
| 0.126478947 | 0.380169976 |
| 0.126486519 | 0.380169976 |
| 0.126496585 | 0.380176238 |
| 0.126527154 | 0.380244116 |
| 0.126551749 | 0.38027267 |
| 0.126610516 | 0.380374633 |
| 0.126731231 | 0.380517428 |

| | |
|---|---|
| 0.126769054 | 0.380517428 |
| 0.126782575 | 0.380517428 |
| 0.126796681 | 0.380517428 |
| 0.126801858 | 0.380517428 |
| 0.126821502 | 0.380536518 |
| 0.1268242 | 0.380536518 |
| 0.126849029 | 0.3805492 |
| 0.126889878 | 0.380613695 |
| 0.126953269 | 0.380659995 |
| 0.127056738 | 0.380910855 |
| 0.127060927 | 0.380910855 |
| 0.127232507 | 0.381173062 |
| 0.127348095 | 0.381435614 |
| 0.127419841 | 0.381530539 |
| 0.127521931 | 0.381620892 |
| 0.12752213 | 0.381620892 |
| 0.127538879 | 0.381632316 |
| 0.127612176 | 0.381722466 |
| 0.127765661 | 0.38207042 |
| 0.12776861 | 0.38207042 |
| 0.127788431 | 0.382105702 |
| 0.127893307 | 0.382347282 |
| 0.127972051 | 0.38255868 |
| 0.128010822 | 0.382650563 |
| 0.1280233 | 0.382663844 |
| 0.128061872 | 0.382755118 |
| 0.128119998 | 0.382847999 |
| 0.128195125 | 0.382872323 |
| 0.128219466 | 0.382872323 |
| 0.128225381 | 0.382872323 |
| 0.128299249 | 0.382995673 |
| 0.12831123 | 0.382995673 |
| 0.128386796 | 0.383197212 |
| 0.128472173 | 0.383379962 |
| 0.128535748 | 0.383506453 |
| 0.128577461 | 0.383598015 |
| 0.128695706 | 0.383758477 |
| 0.128746547 | 0.383763294 |
| 0.128751256 | 0.383763294 |
| 0.128755287 | 0.383763294 |
| 0.128758304 | 0.383763294 |
| 0.128774781 | 0.383778016 |
| 0.128897059 | 0.383998263 |
| 0.128976903 | 0.384128536 |
| 0.128981115 | 0.384128536 |

| | |
|---|---|
| 0.129037483 | 0.384176295 |
| 0.129098797 | 0.384232575 |
| 0.129112859 | 0.384232575 |
| 0.12914076 | 0.384267587 |
| 0.12917231 | 0.384313443 |
| 0.129275819 | 0.384511192 |
| 0.129353945 | 0.38458957 |
| 0.129418331 | 0.384633079 |
| 0.129528158 | 0.38481926 |
| 0.129615217 | 0.38492587 |
| 0.129736989 | 0.385127372 |
| 0.129769256 | 0.385159133 |
| 0.129772976 | 0.385159133 |
| 0.129783144 | 0.385159133 |
| 0.129891914 | 0.385338917 |
| 0.129974685 | 0.385520602 |
| 0.129999418 | 0.385562297 |
| 0.130014992 | 0.385562297 |
| 0.130036447 | 0.385583729 |
| 0.130155423 | 0.385816414 |
| 0.130196282 | 0.385913512 |
| 0.13024284 | 0.386003467 |
| 0.130332903 | 0.386225507 |
| 0.130342087 | 0.386225507 |
| 0.130438156 | 0.386341918 |
| 0.13045836 | 0.386354595 |
| 0.130646866 | 0.386719595 |
| 0.130837947 | 0.387093339 |
| 0.13083815 | 0.387093339 |
| 0.130850908 | 0.387098284 |
| 0.130856077 | 0.387098284 |
| 0.130891547 | 0.387147043 |
| 0.130896946 | 0.387147043 |
| 0.130924194 | 0.387181784 |
| 0.13093308 | 0.387181784 |
| 0.131096914 | 0.387383897 |
| 0.13111008 | 0.387383897 |
| 0.131243544 | 0.387594486 |
| 0.131304195 | 0.387725518 |
| 0.131429714 | 0.388012703 |
| 0.131512331 | 0.388099497 |
| 0.131547247 | 0.388154434 |
| 0.131617866 | 0.38829064 |
| 0.131656997 | 0.388324857 |
| 0.13169229 | 0.388388025 |

| | |
|---|---|
| 0.131815533 | 0.388633055 |
| 0.131857475 | 0.38873265 |
| 0.131884317 | 0.388735701 |
| 0.131887726 | 0.388735701 |
| 0.131919355 | 0.388735701 |
| 0.131922482 | 0.388735701 |
| 0.131974525 | 0.388790761 |
| 0.131975145 | 0.388790761 |
| 0.132000767 | 0.388823125 |
| 0.132031128 | 0.388852676 |
| 0.132045149 | 0.388852676 |
| 0.132080011 | 0.38893129 |
| 0.13212733 | 0.388974333 |
| 0.13214068 | 0.388974333 |
| 0.13217447 | 0.389041067 |
| 0.13224231 | 0.389188868 |
| 0.1322492 | 0.389188868 |
| 0.132272057 | 0.389208036 |
| 0.132272057 | 0.389208036 |
| 0.132305958 | 0.38925124 |
| 0.132311259 | 0.38925124 |
| 0.132425763 | 0.38941972 |
| 0.132551217 | 0.389668342 |
| 0.13259921 | 0.389761313 |
| 0.132642851 | 0.389828813 |
| 0.132734525 | 0.389979541 |
| 0.132909814 | 0.390385182 |
| 0.132926881 | 0.390411233 |
| 0.132947649 | 0.390422574 |
| 0.132950135 | 0.390422574 |
| 0.132985762 | 0.390439706 |
| 0.133043839 | 0.390562064 |
| 0.133079224 | 0.390641863 |
| 0.133213391 | 0.390963408 |
| 0.133299709 | 0.391156741 |
| 0.133304547 | 0.391156741 |
| 0.133312179 | 0.391156741 |
| 0.133330122 | 0.391161373 |
| 0.133521245 | 0.391627721 |
| 0.133521971 | 0.391627721 |
| 0.133540388 | 0.391657618 |
| 0.133551156 | 0.39166508 |
| 0.133645586 | 0.39182138 |
| 0.133701058 | 0.391863401 |
| 0.13372879 | 0.391870232 |

| | |
|---|---|
| 0.133744528 | 0.391870232 |
| 0.133792327 | 0.391962057 |
| 0.133871455 | 0.392169749 |
| 0.1339012 | 0.392178772 |
| 0.133907472 | 0.392178772 |
| 0.133930257 | 0.392197266 |
| 0.133961248 | 0.392215673 |
| 0.134021953 | 0.392345169 |
| 0.134060548 | 0.39237749 |
| 0.134161611 | 0.392550722 |
| 0.134166348 | 0.392550722 |
| 0.134484097 | 0.393243601 |
| 0.13453389 | 0.393243601 |
| 0.134551772 | 0.393243601 |
| 0.134578044 | 0.393272089 |
| 0.134685447 | 0.393513513 |
| 0.134755283 | 0.393645109 |
| 0.13478928 | 0.393720271 |
| 0.134825873 | 0.39376822 |
| 0.134837438 | 0.39376822 |
| 0.134873982 | 0.393822767 |
| 0.134891953 | 0.393849672 |
| 0.134908005 | 0.393849672 |
| 0.135034624 | 0.394044625 |
| 0.135256252 | 0.394401113 |
| 0.135316353 | 0.394401113 |
| 0.135344312 | 0.394401113 |
| 0.135387246 | 0.394401113 |
| 0.135392009 | 0.394401113 |
| 0.135393706 | 0.394401113 |
| 0.135440249 | 0.394484633 |
| 0.135449774 | 0.394488251 |
| 0.135559672 | 0.394643148 |
| 0.135560744 | 0.394643148 |
| 0.135565708 | 0.394643148 |
| 0.135569247 | 0.394643148 |
| 0.135694907 | 0.394898643 |
| 0.135702707 | 0.394898643 |
| 0.135758208 | 0.394975941 |
| 0.135809207 | 0.395076052 |
| 0.135860359 | 0.395200715 |
| 0.135897883 | 0.395212819 |
| 0.135898601 | 0.395212819 |
| 0.135914308 | 0.395212819 |
| 0.135933303 | 0.395219793 |

| | |
|---|---|
| 0.136543253 | 0.396392903 |
| 0.136569827 | 0.396392903 |
| 0.136643064 | 0.396508818 |
| 0.136710208 | 0.396572538 |
| 0.136733025 | 0.396600716 |
| 0.136755697 | 0.396642324 |
| 0.136884872 | 0.396920293 |
| 0.136944565 | 0.396936164 |
| 0.137015313 | 0.39706308 |
| 0.137040896 | 0.397081748 |
| 0.137065613 | 0.397081748 |
| 0.137085894 | 0.397116349 |
| 0.13712084 | 0.397187278 |
| 0.137135398 | 0.397187278 |
| 0.137149472 | 0.397190068 |
| 0.137169286 | 0.397190068 |
| 0.137259442 | 0.39735219 |
| 0.137275767 | 0.39735219 |
| 0.137286189 | 0.397358208 |
| 0.13731344 | 0.397388783 |
| 0.137397063 | 0.397582474 |
| 0.137490519 | 0.397745331 |
| 0.137536856 | 0.397745331 |
| 0.137553165 | 0.397768341 |
| 0.137582186 | 0.397828108 |
| 0.137633008 | 0.39790259 |
| 0.137743451 | 0.398013931 |
| 0.1378418 | 0.398216148 |
| 0.13795159 | 0.398388332 |
| 0.1380149 | 0.398479604 |
| 0.13802235 | 0.398479604 |
| 0.138075425 | 0.398552621 |
| 0.138225017 | 0.398791065 |
| 0.138396519 | 0.399116621 |
| 0.138668097 | 0.399609455 |
| 0.138714983 | 0.39969385 |
| 0.138730952 | 0.39969385 |
| 0.138764458 | 0.399742021 |
| 0.138792775 | 0.399799413 |
| 0.138836936 | 0.399902434 |
| 0.138894093 | 0.39997031 |
| 0.138913995 | 0.399979253 |
| 0.138982462 | 0.400055465 |
| 0.139018993 | 0.400136434 |
| 0.139076612 | 0.400190933 |

| | |
|---|---|
| 0.139124475 | 0.400222916 |
| 0.139170404 | 0.400231561 |
| 0.139192729 | 0.400231561 |
| 0.139194902 | 0.400231561 |
| 0.13929827 | 0.400298092 |
| 0.139325986 | 0.400298092 |
| 0.139472892 | 0.400477077 |
| 0.139473687 | 0.400477077 |
| 0.13954144 | 0.400550877 |
| 0.139553458 | 0.400561232 |
| 0.139617811 | 0.400627592 |
| 0.139636139 | 0.400629541 |
| 0.139647021 | 0.400636629 |
| 0.139687779 | 0.400668598 |
| 0.139699616 | 0.400668598 |
| 0.139700228 | 0.400668598 |
| 0.139748375 | 0.400712429 |
| 0.139778573 | 0.400748501 |
| 0.139826565 | 0.400766733 |
| 0.139827006 | 0.400766733 |
| 0.139910381 | 0.400907007 |
| 0.139945413 | 0.400961329 |
| 0.140069215 | 0.401195364 |
| 0.140099496 | 0.401257964 |
| 0.140145238 | 0.401316574 |
| 0.140275904 | 0.401473499 |
| 0.140306596 | 0.401513886 |
| 0.140306876 | 0.401513886 |
| 0.140318936 | 0.401524271 |
| 0.140367829 | 0.401607771 |
| 0.140421842 | 0.401653084 |
| 0.140500081 | 0.401777086 |
| 0.140540536 | 0.401785688 |
| 0.140592252 | 0.401862558 |
| 0.140617352 | 0.401871144 |
| 0.140793055 | 0.402180221 |
| 0.140823102 | 0.402241926 |
| 0.140846897 | 0.402256593 |
| 0.140901354 | 0.402326196 |
| 0.140939267 | 0.402326196 |
| 0.140939644 | 0.402326196 |
| 0.1409896 | 0.402379656 |
| 0.141051708 | 0.402501591 |
| 0.141124224 | 0.402594994 |
| 0.141177416 | 0.402698496 |

| | |
|---|---|
| 0.141246633 | 0.402871805 |
| 0.141371945 | 0.403010499 |
| 0.141373155 | 0.403010499 |
| 0.141503292 | 0.403193375 |
| 0.141560655 | 0.403332695 |
| 0.141570898 | 0.403337749 |
| 0.141837441 | 0.403822689 |
| 0.141876548 | 0.403894464 |
| 0.141911197 | 0.403968954 |
| 0.142069282 | 0.404265339 |
| 0.142119545 | 0.404320385 |
| 0.142154615 | 0.404371848 |
| 0.14226972 | 0.404602614 |
| 0.142338753 | 0.404750598 |
| 0.142546918 | 0.405081017 |
| 0.142560826 | 0.405081017 |
| 0.14256552 | 0.405081017 |
| 0.142653096 | 0.405254034 |
| 0.142695616 | 0.405280913 |
| 0.14270818 | 0.405280913 |
| 0.14272873 | 0.405280913 |
| 0.142763546 | 0.405353489 |
| 0.142802302 | 0.405415191 |
| 0.142894472 | 0.405562298 |
| 0.14293983 | 0.40556388 |
| 0.142952689 | 0.405576202 |
| 0.143069769 | 0.405787512 |
| 0.143105171 | 0.40585142 |
| 0.14314618 | 0.40588338 |
| 0.143222375 | 0.406026909 |
| 0.143279555 | 0.406068156 |
| 0.143302966 | 0.406086176 |
| 0.143350563 | 0.406148558 |
| 0.14338667 | 0.406226694 |
| 0.143404406 | 0.406226918 |
| 0.143412338 | 0.406226918 |
| 0.143446763 | 0.406276106 |
| 0.143496871 | 0.406393855 |
| 0.143530609 | 0.406465234 |
| 0.143551074 | 0.406493245 |
| 0.14356221 | 0.406493245 |
| 0.143566105 | 0.406493245 |
| 0.14362483 | 0.406611174 |
| 0.143691974 | 0.406777084 |
| 0.143842514 | 0.406962235 |

| | |
|---|---|
| 0.143857515 | 0.406962235 |
| 0.143897939 | 0.407019631 |
| 0.143974651 | 0.407093368 |
| 0.14400117 | 0.40713254 |
| 0.144101003 | 0.407329721 |
| 0.144140663 | 0.40739347 |
| 0.144178529 | 0.407436075 |
| 0.144227074 | 0.407436075 |
| 0.144399283 | 0.407761452 |
| 0.144479619 | 0.407892773 |
| 0.144480044 | 0.407892773 |
| 0.144497621 | 0.407918215 |
| 0.14457778 | 0.40804776 |
| 0.144587977 | 0.408049275 |
| 0.144673883 | 0.408077171 |
| 0.144713061 | 0.408115169 |
| 0.144787474 | 0.408276673 |
| 0.144797972 | 0.408282103 |
| 0.144877617 | 0.408313288 |
| 0.144973954 | 0.408506339 |
| 0.145006251 | 0.408506339 |
| 0.145030402 | 0.408526316 |
| 0.145105416 | 0.408635283 |
| 0.145105416 | 0.408635283 |
| 0.145201358 | 0.408753644 |
| 0.145243661 | 0.408823363 |
| 0.145247446 | 0.408823363 |
| 0.145392734 | 0.409159759 |
| 0.145428733 | 0.409236885 |
| 0.145524209 | 0.409384613 |
| 0.145639721 | 0.409588604 |
| 0.145677282 | 0.409621673 |
| 0.145748375 | 0.409676457 |
| 0.145765495 | 0.409700396 |
| 0.145982127 | 0.41009147 |
| 0.146063833 | 0.410270481 |
| 0.146133984 | 0.410351819 |
| 0.146151463 | 0.410368369 |
| 0.146158243 | 0.410368369 |
| 0.146188127 | 0.41037762 |
| 0.146197154 | 0.41037762 |
| 0.146216414 | 0.41037762 |
| 0.146262562 | 0.410382253 |
| 0.146274799 | 0.410382253 |
| 0.146311071 | 0.410458701 |

| | |
|---|---|
| 0.146395742 | 0.410599493 |
| 0.146449798 | 0.410702734 |
| 0.14652496 | 0.410865131 |
| 0.146575392 | 0.410958156 |
| 0.14660363 | 0.41096475 |
| 0.146707613 | 0.411207832 |
| 0.146750614 | 0.411288903 |
| 0.146761817 | 0.411288903 |
| 0.14690401 | 0.411395147 |
| 0.146979178 | 0.4115324 |
| 0.146998957 | 0.4115324 |
| 0.147004865 | 0.4115324 |
| 0.147017073 | 0.411542386 |
| 0.147085653 | 0.411613397 |
| 0.147169593 | 0.411799907 |
| 0.147187776 | 0.411802398 |
| 0.147211813 | 0.411809229 |
| 0.14726258 | 0.411890705 |
| 0.14732643 | 0.411912687 |
| 0.147330617 | 0.411912687 |
| 0.147333899 | 0.411912687 |
| 0.147376089 | 0.411942073 |
| 0.147423378 | 0.412022157 |
| 0.147459937 | 0.412055528 |
| 0.14751153 | 0.412151335 |
| 0.147691451 | 0.412441589 |
| 0.147745337 | 0.412514196 |
| 0.147944478 | 0.412947193 |
| 0.148105933 | 0.413206097 |
| 0.14817221 | 0.41336679 |
| 0.148223621 | 0.413378791 |
| 0.148232196 | 0.413378791 |
| 0.148249969 | 0.413378791 |
| 0.148348171 | 0.413542777 |
| 0.148412899 | 0.41367479 |
| 0.1484606 | 0.413731006 |
| 0.148464976 | 0.413731006 |
| 0.148507775 | 0.413769733 |
| 0.148529844 | 0.413782807 |
| 0.148668682 | 0.414096926 |
| 0.148743485 | 0.414193001 |
| 0.148746658 | 0.414193001 |
| 0.148837081 | 0.414396332 |
| 0.1488567 | 0.414404906 |
| 0.148880893 | 0.41444563 |

| | |
|---|---|
| 0.148989888 | 0.414556729 |
| 0.149074093 | 0.414668371 |
| 0.14924284 | 0.415010539 |
| 0.149289127 | 0.415072577 |
| 0.149343436 | 0.4151615 |
| 0.149344824 | 0.4151615 |
| 0.14935051 | 0.4151615 |
| 0.149367353 | 0.415168891 |
| 0.149418621 | 0.415287155 |
| 0.149440556 | 0.415312613 |
| 0.149477781 | 0.415354625 |
| 0.149531835 | 0.415384496 |
| 0.149532139 | 0.415384496 |
| 0.149603718 | 0.415413728 |
| 0.149719421 | 0.415613851 |
| 0.149834631 | 0.415860953 |
| 0.149847444 | 0.415872282 |
| 0.149865569 | 0.415884895 |
| 0.149869453 | 0.415884895 |
| 0.149899522 | 0.415898968 |
| 0.149979352 | 0.416028424 |
| 0.150055473 | 0.416110162 |
| 0.150072107 | 0.416132058 |
| 0.150113572 | 0.416187003 |
| 0.150163294 | 0.416187003 |
| 0.150168722 | 0.416187003 |
| 0.150173546 | 0.416187003 |
| 0.15017481 | 0.416187003 |
| 0.150232801 | 0.41626378 |
| 0.150236576 | 0.41626378 |
| 0.150236976 | 0.41626378 |
| 0.150256789 | 0.41626378 |
| 0.150259426 | 0.41626378 |
| 0.150314334 | 0.41626378 |
| 0.150385825 | 0.416310202 |
| 0.15042157 | 0.416325436 |
| 0.150556214 | 0.416583398 |
| 0.150580959 | 0.416621317 |
| 0.150643524 | 0.416721795 |
| 0.150692296 | 0.416784088 |
| 0.150799561 | 0.41700811 |
| 0.150995105 | 0.417306556 |
| 0.151061929 | 0.417467011 |
| 0.151083419 | 0.417502175 |
| 0.151236927 | 0.417853643 |

| | |
|---|---|
| 0.151306089 | 0.418020483 |
| 0.151412136 | 0.418216723 |
| 0.151439612 | 0.41822316 |
| 0.151458003 | 0.41822316 |
| 0.151467262 | 0.41822316 |
| 0.151518865 | 0.418292893 |
| 0.151528068 | 0.418294052 |
| 0.151885669 | 0.41891989 |
| 0.151886717 | 0.41891989 |
| 0.151897976 | 0.418926683 |
| 0.151909437 | 0.418929744 |
| 0.151925474 | 0.418929744 |
| 0.151949632 | 0.418947847 |
| 0.152022448 | 0.419024443 |
| 0.152030201 | 0.419024443 |
| 0.152098214 | 0.41910505 |
| 0.152442867 | 0.419821697 |
| 0.152470322 | 0.419848754 |
| 0.152538031 | 0.419955813 |
| 0.152550849 | 0.419955813 |
| 0.15255329 | 0.419955813 |
| 0.152673873 | 0.420210517 |
| 0.152681105 | 0.420210517 |
| 0.152860621 | 0.420534472 |
| 0.152954294 | 0.420629209 |
| 0.153051461 | 0.420815536 |
| 0.153067975 | 0.420815536 |
| 0.15313456 | 0.420926595 |
| 0.153140188 | 0.420926595 |
| 0.153144562 | 0.420926595 |
| 0.153379488 | 0.42116602 |
| 0.153382002 | 0.42116602 |
| 0.153417783 | 0.421239984 |
| 0.153492153 | 0.421419887 |
| 0.153526044 | 0.421488639 |
| 0.153670341 | 0.421763229 |
| 0.153698423 | 0.42180246 |
| 0.153764101 | 0.421923306 |
| 0.153814424 | 0.422012764 |
| 0.153943336 | 0.422244833 |
| 0.153972769 | 0.422276925 |
| 0.154030214 | 0.422398004 |
| 0.154130334 | 0.422502393 |
| 0.154176115 | 0.422566946 |
| 0.154246233 | 0.422686154 |

| | |
|---|---|
| 0.154287162 | 0.422700238 |
| 0.154319267 | 0.422700238 |
| 0.154321661 | 0.422700238 |
| 0.154340292 | 0.422725004 |
| 0.154502137 | 0.423070965 |
| 0.154547026 | 0.423100746 |
| 0.154548548 | 0.423100746 |
| 0.154665225 | 0.423302416 |
| 0.154666652 | 0.423302416 |
| 0.154696245 | 0.423334755 |
| 0.154726584 | 0.423369127 |
| 0.154770676 | 0.423441118 |
| 0.154812396 | 0.423446461 |
| 0.154860154 | 0.423539941 |
| 0.154877451 | 0.423562925 |
| 0.154896174 | 0.423589807 |
| 0.155005559 | 0.423772676 |
| 0.155024668 | 0.423772676 |
| 0.155099467 | 0.423891389 |
| 0.155117541 | 0.423891389 |
| 0.15522743 | 0.424057405 |
| 0.155318785 | 0.42419299 |
| 0.155321594 | 0.42419299 |
| 0.155391797 | 0.424336054 |
| 0.155433181 | 0.424369515 |
| 0.155448602 | 0.424369515 |
| 0.155511506 | 0.424465887 |
| 0.155524237 | 0.424465887 |
| 0.155528465 | 0.424465887 |
| 0.155573246 | 0.424552505 |
| 0.155591878 | 0.424552505 |
| 0.155649375 | 0.424674198 |
| 0.155676298 | 0.424698994 |
| 0.155768244 | 0.424901154 |
| 0.155911831 | 0.425146729 |
| 0.155937945 | 0.425169252 |
| 0.156133566 | 0.425411447 |
| 0.15613396 | 0.425411447 |
| 0.15616646 | 0.42547566 |
| 0.156200535 | 0.425495476 |
| 0.156237413 | 0.425547254 |
| 0.156321903 | 0.425704345 |
| 0.156408062 | 0.425841581 |
| 0.156432327 | 0.425883298 |
| 0.15665571 | 0.426199102 |

| | |
|---|---|
| 0.156736251 | 0.426320813 |
| 0.156773078 | 0.42637228 |
| 0.156797424 | 0.426389798 |
| 0.156841569 | 0.426412457 |
| 0.156866475 | 0.426431485 |
| 0.156877934 | 0.426438295 |
| 0.156892819 | 0.426454417 |
| 0.156901889 | 0.426454732 |
| 0.156955426 | 0.426527222 |
| 0.15697841 | 0.426541005 |
| 0.157008493 | 0.426596731 |
| 0.157121858 | 0.42675723 |
| 0.157139657 | 0.42675723 |
| 0.157156551 | 0.42675723 |
| 0.157199067 | 0.426810712 |
| 0.157346461 | 0.426950593 |
| 0.157348067 | 0.426950593 |
| 0.157373156 | 0.42698025 |
| 0.157397791 | 0.427022763 |
| 0.157443269 | 0.427091725 |
| 0.15746181 | 0.427091725 |
| 0.157472577 | 0.427091725 |
| 0.157485982 | 0.427091725 |
| 0.157553453 | 0.427201726 |
| 0.157584949 | 0.42722658 |
| 0.157587925 | 0.42722658 |
| 0.157617745 | 0.42722658 |
| 0.157686363 | 0.42722658 |
| 0.157698409 | 0.42722658 |
| 0.157830616 | 0.427434754 |
| 0.157832065 | 0.427434754 |
| 0.157836837 | 0.427434754 |
| 0.157905648 | 0.427475262 |
| 0.157921913 | 0.427494996 |
| 0.157983122 | 0.427603913 |
| 0.158047242 | 0.427639154 |
| 0.158060896 | 0.427639154 |
| 0.158080652 | 0.427639154 |
| 0.158091893 | 0.427639154 |
| 0.158195195 | 0.427772812 |
| 0.158315377 | 0.427854874 |
| 0.158342114 | 0.427877867 |
| 0.158413056 | 0.4279246 |
| 0.158438012 | 0.42796774 |
| 0.158502858 | 0.428049426 |

| | |
|---|---|
| 0.158540464 | 0.428108553 |
| 0.158553064 | 0.428108553 |
| 0.158563887 | 0.428113506 |
| 0.158577619 | 0.428126311 |
| 0.158586991 | 0.428127345 |
| 0.158689062 | 0.428321822 |
| 0.158832551 | 0.428547353 |
| 0.159064641 | 0.428809172 |
| 0.159178961 | 0.428995947 |
| 0.159306258 | 0.429217581 |
| 0.159351936 | 0.42924352 |
| 0.159534934 | 0.429639258 |
| 0.159594982 | 0.429659137 |
| 0.159596443 | 0.429659137 |
| 0.159734274 | 0.429840155 |
| 0.159752425 | 0.429840155 |
| 0.159859535 | 0.430040144 |
| 0.159967336 | 0.430219906 |
| 0.159985931 | 0.430245622 |
| 0.160090386 | 0.430406295 |
| 0.160133196 | 0.43044803 |
| 0.1601335 | 0.43044803 |
| 0.160299394 | 0.430796707 |
| 0.160322788 | 0.430804677 |
| 0.160377037 | 0.430859495 |
| 0.160416352 | 0.430916503 |
| 0.160567164 | 0.431200017 |
| 0.160650056 | 0.431349657 |
| 0.160695619 | 0.43138201 |
| 0.160698335 | 0.43138201 |
| 0.160727992 | 0.431394326 |
| 0.160816641 | 0.431577965 |
| 0.160937608 | 0.431744669 |
| 0.161037976 | 0.43190425 |
| 0.161080553 | 0.431928059 |
| 0.161111842 | 0.431956411 |
| 0.161170307 | 0.432064516 |
| 0.161241685 | 0.432167918 |
| 0.161245174 | 0.432167918 |
| 0.161274546 | 0.432197996 |
| 0.161523468 | 0.432620039 |
| 0.161585154 | 0.432713867 |
| 0.161621956 | 0.432784536 |
| 0.161692727 | 0.432904576 |
| 0.161758429 | 0.433010971 |

| | |
|---|---|
| 0.161759741 | 0.433010971 |
| 0.161828038 | 0.433112902 |
| 0.16184635 | 0.433112902 |
| 0.161877 | 0.433149949 |
| 0.16206713 | 0.433444026 |
| 0.16214606 | 0.433596089 |
| 0.162175387 | 0.433622745 |
| 0.162202994 | 0.433622745 |
| 0.162282055 | 0.433691343 |
| 0.162296193 | 0.433691343 |
| 0.162312187 | 0.433709746 |
| 0.1623396 | 0.433758661 |
| 0.162473623 | 0.434019365 |
| 0.162670184 | 0.434408102 |
| 0.162793773 | 0.43450292 |
| 0.162818855 | 0.43450292 |
| 0.1628958 | 0.434635018 |
| 0.162915408 | 0.434635018 |
| 0.162995732 | 0.434804397 |
| 0.163011476 | 0.434822042 |
| 0.163054294 | 0.434861989 |
| 0.163057005 | 0.434861989 |
| 0.163089769 | 0.434909094 |
| 0.163111753 | 0.434943367 |
| 0.163233051 | 0.435144997 |
| 0.163252195 | 0.435171673 |
| 0.163267561 | 0.435188276 |
| 0.163287379 | 0.435216745 |
| 0.163303225 | 0.435234622 |
| 0.163323058 | 0.435235686 |
| 0.163323444 | 0.435235686 |
| 0.163381957 | 0.435273953 |
| 0.163598231 | 0.435600559 |
| 0.163707121 | 0.435829077 |
| 0.163709287 | 0.435829077 |
| 0.163724379 | 0.435844894 |
| 0.163775486 | 0.43590637 |
| 0.163861253 | 0.436014297 |
| 0.163875828 | 0.436028718 |
| 0.163912398 | 0.436042332 |
| 0.163969758 | 0.436112596 |
| 0.163973615 | 0.436112596 |
| 0.164003826 | 0.436116015 |
| 0.164018479 | 0.436116015 |
| 0.164037627 | 0.436116015 |

| | |
|---|---|
| 0.164052895 | 0.436116015 |
| 0.164099801 | 0.436116015 |
| 0.16410605 | 0.436116015 |
| 0.164305522 | 0.436377287 |
| 0.164380939 | 0.436446146 |
| 0.164510378 | 0.436642945 |
| 0.164625324 | 0.436827138 |
| 0.164706456 | 0.436945045 |
| 0.164732066 | 0.436964305 |
| 0.164761679 | 0.437005833 |
| 0.164829554 | 0.437120007 |
| 0.164851234 | 0.43713433 |
| 0.164907386 | 0.437185864 |
| 0.165060496 | 0.437348331 |
| 0.165076234 | 0.437365699 |
| 0.165102137 | 0.437409994 |
| 0.1652109 | 0.437492292 |
| 0.165210906 | 0.437492292 |
| 0.165269161 | 0.437560416 |
| 0.165449225 | 0.437866763 |
| 0.165514063 | 0.438014019 |
| 0.165535907 | 0.438047485 |
| 0.165645223 | 0.438264241 |
| 0.165654761 | 0.438264241 |
| 0.165709855 | 0.438264241 |
| 0.16571904 | 0.438264241 |
| 0.165739917 | 0.438270781 |
| 0.165815061 | 0.438362416 |
| 0.165827289 | 0.438362416 |
| 0.165829795 | 0.438362416 |
| 0.16586668 | 0.438435584 |
| 0.165885509 | 0.43846102 |
| 0.165904891 | 0.438487915 |
| 0.165978391 | 0.438633493 |
| 0.166162506 | 0.438879582 |
| 0.166189791 | 0.438879582 |
| 0.166233759 | 0.43894302 |
| 0.166246395 | 0.438952049 |
| 0.166272639 | 0.438971543 |
| 0.16630418 | 0.439007271 |
| 0.166428129 | 0.43924549 |
| 0.166431312 | 0.43924549 |
| 0.166550412 | 0.439428898 |
| 0.166565391 | 0.439428898 |
| 0.166656694 | 0.439572375 |

| | |
|---|---|
| 0.166671203 | 0.4395863 |
| 0.166749642 | 0.439700839 |
| 0.166782939 | 0.439759234 |
| 0.166862806 | 0.439848067 |
| 0.166930536 | 0.439891042 |
| 0.166931078 | 0.439891042 |
| 0.166940236 | 0.439891042 |
| 0.167027053 | 0.439971157 |
| 0.167029594 | 0.439971157 |
| 0.16711548 | 0.440124362 |
| 0.167194469 | 0.440259356 |
| 0.16728695 | 0.440451538 |
| 0.167292165 | 0.440451538 |
| 0.16738301 | 0.440512396 |
| 0.167385045 | 0.440512396 |
| 0.167401555 | 0.440512396 |
| 0.167477764 | 0.440615559 |
| 0.167573649 | 0.440794773 |
| 0.167749795 | 0.441018181 |
| 0.167828639 | 0.441112087 |
| 0.16783796 | 0.441112087 |
| 0.167857844 | 0.441128153 |
| 0.16795315 | 0.441305565 |
| 0.167972759 | 0.44133274 |
| 0.168028063 | 0.441453695 |
| 0.168162908 | 0.441650887 |
| 0.168164963 | 0.441650887 |
| 0.16821791 | 0.441692722 |
| 0.168230331 | 0.441692722 |
| 0.16828437 | 0.441697393 |
| 0.168287755 | 0.441697393 |
| 0.168365146 | 0.441763517 |
| 0.168365628 | 0.441763517 |
| 0.16838509 | 0.441782442 |
| 0.168399073 | 0.441787428 |
| 0.168421453 | 0.441787984 |
| 0.168501645 | 0.44186917 |
| 0.168538951 | 0.441912051 |
| 0.168542782 | 0.441912051 |
| 0.168579772 | 0.441976705 |
| 0.168596127 | 0.441995251 |
| 0.168666009 | 0.442087021 |
| 0.16866814 | 0.442087021 |
| 0.168681917 | 0.442087021 |
| 0.168744038 | 0.442115322 |

| | |
|---|---|
| 0.168806496 | 0.442225997 |
| 0.168867316 | 0.44231665 |
| 0.168885102 | 0.44233891 |
| 0.168948297 | 0.442407111 |
| 0.169001501 | 0.442497772 |
| 0.169033106 | 0.442547004 |
| 0.169042985 | 0.442547004 |
| 0.169066429 | 0.442547004 |
| 0.169258762 | 0.442879191 |
| 0.169466533 | 0.443236188 |
| 0.169472118 | 0.443236188 |
| 0.169529125 | 0.44325601 |
| 0.169555212 | 0.44325601 |
| 0.16961798 | 0.443323691 |
| 0.169620966 | 0.443323691 |
| 0.169624113 | 0.443323691 |
| 0.169708344 | 0.443446497 |
| 0.169802154 | 0.443568295 |
| 0.169824128 | 0.443568295 |
| 0.169837263 | 0.443568295 |
| 0.169874739 | 0.44363789 |
| 0.169944527 | 0.443771479 |
| 0.170193435 | 0.444132035 |
| 0.170212316 | 0.444132035 |
| 0.170225096 | 0.444134839 |
| 0.170234812 | 0.444134839 |
| 0.170390641 | 0.44445885 |
| 0.170649728 | 0.444978625 |
| 0.170701868 | 0.44502882 |
| 0.170778893 | 0.44512047 |
| 0.170820257 | 0.445203916 |
| 0.170993079 | 0.445483693 |
| 0.171068795 | 0.445583455 |
| 0.171157176 | 0.445716159 |
| 0.171211296 | 0.445783972 |
| 0.1712635 | 0.445895518 |
| 0.171307229 | 0.445920871 |
| 0.171309973 | 0.445920871 |
| 0.171315544 | 0.445920871 |
| 0.171419181 | 0.446019257 |
| 0.171460864 | 0.446019257 |
| 0.171483056 | 0.446042018 |
| 0.171488345 | 0.446042018 |
| 0.171566617 | 0.446100898 |
| 0.171646575 | 0.446234248 |

| | |
|---|---|
| 0.171700764 | 0.446343952 |
| 0.171914012 | 0.446783185 |
| 0.171939141 | 0.44682411 |
| 0.171970604 | 0.446849219 |
| 0.17197695 | 0.446849219 |
| 0.172035099 | 0.446927162 |
| 0.172046091 | 0.44693134 |
| 0.172202737 | 0.447167534 |
| 0.172213097 | 0.447170056 |
| 0.172334516 | 0.447412155 |
| 0.172358769 | 0.447430284 |
| 0.172381078 | 0.447459864 |
| 0.172405871 | 0.447499833 |
| 0.172418712 | 0.447508775 |
| 0.172455451 | 0.44755667 |
| 0.172593117 | 0.447801852 |
| 0.172613262 | 0.447818488 |
| 0.17269093 | 0.447995584 |
| 0.172753296 | 0.448108558 |
| 0.172877466 | 0.448351413 |
| 0.172882371 | 0.448351413 |
| 0.172899611 | 0.448351413 |
| 0.172903403 | 0.448351413 |
| 0.173006386 | 0.448562715 |
| 0.173010489 | 0.448562715 |
| 0.173019821 | 0.448562715 |
| 0.173022563 | 0.448562715 |
| 0.173032036 | 0.448562856 |
| 0.173069892 | 0.448587741 |
| 0.173268795 | 0.448887812 |
| 0.173307802 | 0.44892254 |
| 0.173321596 | 0.44892254 |
| 0.173399663 | 0.448995867 |
| 0.173449261 | 0.449011198 |
| 0.173450102 | 0.449011198 |
| 0.1734732 | 0.449046583 |
| 0.173557441 | 0.449197514 |
| 0.173579028 | 0.449222865 |
| 0.173632925 | 0.449320649 |
| 0.173785512 | 0.449582105 |
| 0.173801107 | 0.449582105 |
| 0.173857289 | 0.449649802 |
| 0.173899084 | 0.449709056 |
| 0.174005764 | 0.449911646 |
| 0.174018286 | 0.449915665 |

| | |
|---|---|
| 0.174175558 | 0.450204021 |
| 0.174277495 | 0.450371625 |
| 0.174304998 | 0.450416372 |
| 0.174379179 | 0.450487726 |
| 0.17438272 | 0.450487726 |
| 0.174389363 | 0.450487726 |
| 0.174419134 | 0.450496117 |
| 0.174420988 | 0.450496117 |
| 0.17447781 | 0.450520719 |
| 0.174499855 | 0.450534491 |
| 0.174530172 | 0.450534491 |
| 0.174538919 | 0.450534491 |
| 0.174539025 | 0.450534491 |
| 0.174644782 | 0.450683085 |
| 0.174658468 | 0.450683476 |
| 0.174663986 | 0.450683476 |
| 0.174673321 | 0.450683476 |
| 0.174708884 | 0.450726396 |
| 0.174781758 | 0.450821731 |
| 0.174783699 | 0.450821731 |
| 0.174974594 | 0.451069825 |
| 0.175153389 | 0.451310889 |
| 0.175184376 | 0.451365123 |
| 0.175193392 | 0.451365123 |
| 0.175272484 | 0.45146181 |
| 0.175278057 | 0.45146181 |
| 0.175309061 | 0.45146181 |
| 0.17532108 | 0.45146181 |
| 0.17533519 | 0.45146181 |
| 0.175346165 | 0.451465662 |
| 0.175447448 | 0.451509599 |
| 0.175472668 | 0.451509599 |
| 0.175476992 | 0.451509599 |
| 0.17553519 | 0.451601919 |
| 0.175541318 | 0.451601919 |
| 0.175782883 | 0.451801021 |
| 0.175801683 | 0.451801021 |
| 0.175803789 | 0.451801021 |
| 0.17587991 | 0.45193593 |
| 0.175924393 | 0.451977079 |
| 0.17594605 | 0.452008335 |
| 0.175986923 | 0.452014257 |
| 0.176002646 | 0.452014257 |
| 0.176043193 | 0.452053941 |
| 0.176049226 | 0.452053941 |

| | |
|---|---|
| 0.17611821 | 0.452101767 |
| 0.176136678 | 0.452101767 |
| 0.176149079 | 0.452101767 |
| 0.176153285 | 0.452101767 |
| 0.176218162 | 0.452156734 |
| 0.176249673 | 0.45217858 |
| 0.176335625 | 0.452204222 |
| 0.176411424 | 0.452354221 |
| 0.176449871 | 0.452424119 |
| 0.176482094 | 0.452433669 |
| 0.176545254 | 0.452499853 |
| 0.176608642 | 0.452598781 |
| 0.176618867 | 0.452598781 |
| 0.176641444 | 0.45262291 |
| 0.176710407 | 0.452702198 |
| 0.176844084 | 0.452892814 |
| 0.176951414 | 0.453100269 |
| 0.177145886 | 0.453451958 |
| 0.177288279 | 0.453628348 |
| 0.177290992 | 0.453628348 |
| 0.177353184 | 0.453763099 |
| 0.177392953 | 0.453835514 |
| 0.177434955 | 0.453899162 |
| 0.177478302 | 0.453961284 |
| 0.177568919 | 0.4541199 |
| 0.177604796 | 0.454161882 |
| 0.177636622 | 0.454181384 |
| 0.177650178 | 0.454181384 |
| 0.177851508 | 0.454476542 |
| 0.177902348 | 0.454508911 |
| 0.177973977 | 0.45459437 |
| 0.178077336 | 0.454712056 |
| 0.178122915 | 0.454804058 |
| 0.178164224 | 0.454821751 |
| 0.178187143 | 0.454821751 |
| 0.178244149 | 0.454894122 |
| 0.178310415 | 0.45494858 |
| 0.178315818 | 0.45494858 |
| 0.178336953 | 0.45494858 |
| 0.178351607 | 0.45494858 |
| 0.178415303 | 0.454989594 |
| 0.178455106 | 0.455028456 |
| 0.178459204 | 0.455028456 |
| 0.17854045 | 0.455055524 |
| 0.178586378 | 0.455055524 |

| | |
|---|---|
| 0.178622694 | 0.455055524 |
| 0.178663297 | 0.45513347 |
| 0.178672402 | 0.45513347 |
| 0.178727702 | 0.455152619 |
| 0.178727702 | 0.455152619 |
| 0.178844618 | 0.455328627 |
| 0.178894325 | 0.455375726 |
| 0.178904619 | 0.455375726 |
| 0.17902539 | 0.455618373 |
| 0.179088538 | 0.455681686 |
| 0.179124861 | 0.45574976 |
| 0.17914121 | 0.45576701 |
| 0.17929157 | 0.455979048 |
| 0.179308747 | 0.455998382 |
| 0.179340854 | 0.456025004 |
| 0.179398429 | 0.456066601 |
| 0.179402603 | 0.456066601 |
| 0.17942036 | 0.456078058 |
| 0.179439474 | 0.456078058 |
| 0.179465427 | 0.456078058 |
| 0.1796473 | 0.456330638 |
| 0.179651232 | 0.456330638 |
| 0.179659769 | 0.456330638 |
| 0.179786813 | 0.456555921 |
| 0.179812905 | 0.456555921 |
| 0.179830194 | 0.456555921 |
| 0.179834738 | 0.456555921 |
| 0.179874192 | 0.456622175 |
| 0.179889598 | 0.456622175 |
| 0.179920488 | 0.456655648 |
| 0.17997588 | 0.456655648 |
| 0.179979814 | 0.456655648 |
| 0.180014466 | 0.456655648 |
| 0.180019634 | 0.456655648 |
| 0.180027431 | 0.456655648 |
| 0.180067399 | 0.456732705 |
| 0.180092576 | 0.456747916 |
| 0.180152366 | 0.456807972 |
| 0.180155548 | 0.456807972 |
| 0.180173804 | 0.456807972 |
| 0.180185946 | 0.456814439 |
| 0.180273249 | 0.457011447 |
| 0.180295125 | 0.457042575 |
| 0.180360876 | 0.45709838 |
| 0.180365126 | 0.45709838 |

| | |
|---|---|
| 0.180446955 | 0.457208444 |
| 0.180494323 | 0.457219901 |
| 0.180564732 | 0.457336551 |
| 0.180574518 | 0.457337015 |
| 0.180591846 | 0.457353109 |
| 0.180656439 | 0.457386213 |
| 0.180714571 | 0.457386213 |
| 0.180718789 | 0.457386213 |
| 0.18091549 | 0.457665163 |
| 0.180982081 | 0.457784989 |
| 0.180991712 | 0.457785038 |
| 0.181097016 | 0.457919799 |
| 0.1811005 | 0.457919799 |
| 0.18112734 | 0.45793353 |
| 0.181146391 | 0.457939338 |
| 0.181148868 | 0.457939338 |
| 0.1812378 | 0.458049322 |
| 0.181291905 | 0.458104259 |
| 0.18143856 | 0.458266373 |
| 0.181466471 | 0.458266373 |
| 0.181494107 | 0.458301247 |
| 0.18155468 | 0.458360842 |
| 0.181564304 | 0.458360842 |
| 0.181590322 | 0.458398381 |
| 0.181758314 | 0.45865407 |
| 0.181795135 | 0.458661626 |
| 0.181878177 | 0.458711711 |
| 0.181966311 | 0.458853527 |
| 0.181984337 | 0.458853527 |
| 0.1819954 | 0.458853527 |
| 0.181998759 | 0.458853527 |
| 0.182106886 | 0.458972569 |
| 0.182145201 | 0.458997353 |
| 0.182155269 | 0.458997353 |
| 0.182191839 | 0.459065216 |
| 0.182229613 | 0.459087532 |
| 0.182290612 | 0.459144056 |
| 0.182339248 | 0.459193706 |
| 0.18240173 | 0.45926869 |
| 0.182406604 | 0.45926869 |
| 0.182417238 | 0.45926869 |
| 0.182481832 | 0.459325371 |
| 0.182503658 | 0.459325371 |
| 0.1825057 | 0.459325371 |
| 0.182507261 | 0.459325371 |

| | |
|---|---|
| 0.18252044 | 0.459334268 |
| 0.182550207 | 0.459360633 |
| 0.182563512 | 0.459369843 |
| 0.182647897 | 0.459533619 |
| 0.182669783 | 0.45954013 |
| 0.182798579 | 0.459724792 |
| 0.182801102 | 0.459724792 |
| 0.18282169 | 0.459752291 |
| 0.182989842 | 0.460053695 |
| 0.183016554 | 0.460057945 |
| 0.183061241 | 0.460099394 |
| 0.183091281 | 0.460114418 |
| 0.18316987 | 0.460239061 |
| 0.183191863 | 0.460270038 |
| 0.183363012 | 0.460569007 |
| 0.183387361 | 0.460569007 |
| 0.183406711 | 0.460584222 |
| 0.183418365 | 0.460584222 |
| 0.183429157 | 0.460584222 |
| 0.183442629 | 0.460584222 |
| 0.183510119 | 0.460638095 |
| 0.183641741 | 0.460865486 |
| 0.183726582 | 0.460922126 |
| 0.183737644 | 0.460922126 |
| 0.183741732 | 0.460922126 |
| 0.183850517 | 0.461053079 |
| 0.183904459 | 0.461087475 |
| 0.183968669 | 0.461151357 |
| 0.18400766 | 0.461200549 |
| 0.184022541 | 0.461213575 |
| 0.18408015 | 0.461284594 |
| 0.184188424 | 0.461337999 |
| 0.184222551 | 0.46136996 |
| 0.184230245 | 0.46136996 |
| 0.184352843 | 0.461580548 |
| 0.184387595 | 0.461580548 |
| 0.184402308 | 0.461580548 |
| 0.184528833 | 0.461689375 |
| 0.184552175 | 0.461689375 |
| 0.184651539 | 0.461842182 |
| 0.184719009 | 0.461924644 |
| 0.184730108 | 0.461924644 |
| 0.184736635 | 0.461924644 |
| 0.184780521 | 0.461946453 |
| 0.184831822 | 0.46200222 |

| | |
|---|---|
| 0.18509327 | 0.462364271 |
| 0.185296487 | 0.46267777 |
| 0.185316526 | 0.462679292 |
| 0.185341475 | 0.462694985 |
| 0.185346931 | 0.462694985 |
| 0.185351956 | 0.462694985 |
| 0.185377547 | 0.462710362 |
| 0.185409016 | 0.462740407 |
| 0.185435181 | 0.462759423 |
| 0.185526917 | 0.462920621 |
| 0.185652077 | 0.463104356 |
| 0.185665889 | 0.463109578 |
| 0.185702789 | 0.463109578 |
| 0.185717423 | 0.463121824 |
| 0.185853353 | 0.463388004 |
| 0.185926977 | 0.463498778 |
| 0.185988159 | 0.463627031 |
| 0.186084467 | 0.463820012 |
| 0.186262996 | 0.464043734 |
| 0.186313624 | 0.464074 |
| 0.186358802 | 0.464137983 |
| 0.186393914 | 0.46414304 |
| 0.186506294 | 0.464344182 |
| 0.186558121 | 0.464391555 |
| 0.186706569 | 0.464642943 |
| 0.186756259 | 0.464714751 |
| 0.18686199 | 0.464844314 |
| 0.186966525 | 0.465005345 |
| 0.18709438 | 0.465221285 |
| 0.187109693 | 0.465229619 |
| 0.187337663 | 0.465664854 |
| 0.187343403 | 0.465664854 |
| 0.187396671 | 0.465748644 |
| 0.187484289 | 0.465829695 |
| 0.187513329 | 0.465844104 |
| 0.187524484 | 0.465847518 |
| 0.187570081 | 0.465902996 |
| 0.187613937 | 0.465907784 |
| 0.187638685 | 0.465907784 |
| 0.187639131 | 0.465907784 |
| 0.187656351 | 0.465907784 |
| 0.187731852 | 0.465980459 |
| 0.187797691 | 0.465980459 |
| 0.187806432 | 0.465980459 |
| 0.187807621 | 0.465980459 |

| | |
|---|---|
| 0.187917038 | 0.466129435 |
| 0.187966689 | 0.46616796 |
| 0.188034634 | 0.466311968 |
| 0.188133343 | 0.466411259 |
| 0.188230214 | 0.46657732 |
| 0.18826288 | 0.466610959 |
| 0.18839068 | 0.466854828 |
| 0.188412812 | 0.46686109 |
| 0.188437206 | 0.466897242 |
| 0.188456179 | 0.466919962 |
| 0.18851726 | 0.467047 |
| 0.188567356 | 0.467146813 |
| 0.188665919 | 0.467342368 |
| 0.18879085 | 0.467545945 |
| 0.188818969 | 0.467545945 |
| 0.18883637 | 0.467545945 |
| 0.188876042 | 0.467575142 |
| 0.188921512 | 0.467581078 |
| 0.188937529 | 0.467581078 |
| 0.189122533 | 0.467889403 |
| 0.189178969 | 0.467927003 |
| 0.189289122 | 0.467961225 |
| 0.189322383 | 0.46797058 |
| 0.189322383 | 0.46797058 |
| 0.189334512 | 0.467976273 |
| 0.189366718 | 0.468031586 |
| 0.189387164 | 0.468057831 |
| 0.189506076 | 0.468230228 |
| 0.189527006 | 0.468234534 |
| 0.189613158 | 0.468309966 |
| 0.189669139 | 0.468325587 |
| 0.189748345 | 0.468401979 |
| 0.189790952 | 0.468401979 |
| 0.189844237 | 0.468458188 |
| 0.189878051 | 0.468464956 |
| 0.19028183 | 0.469124791 |
| 0.19038376 | 0.469277584 |
| 0.190427965 | 0.469290732 |
| 0.190481623 | 0.469374392 |
| 0.190497097 | 0.469388238 |
| 0.190783825 | 0.469778784 |
| 0.190847202 | 0.469861965 |
| 0.190868818 | 0.469888281 |
| 0.190982517 | 0.470025027 |
| 0.191145659 | 0.470280726 |

| | |
|---|---|
| 0.19118881 | 0.470338297 |
| 0.191304296 | 0.470480997 |
| 0.191306087 | 0.470480997 |
| 0.191330986 | 0.470517937 |
| 0.191353244 | 0.470527094 |
| 0.191354469 | 0.470527094 |
| 0.191393685 | 0.470593024 |
| 0.191401043 | 0.470593024 |
| 0.191467185 | 0.470629838 |
| 0.191472611 | 0.470629838 |
| 0.191508951 | 0.470629838 |
| 0.191512977 | 0.470629838 |
| 0.191513254 | 0.470629838 |
| 0.191515624 | 0.470629838 |
| 0.191649571 | 0.470839477 |
| 0.191714525 | 0.470926179 |
| 0.191763302 | 0.470948837 |
| 0.191867432 | 0.471083111 |
| 0.191934762 | 0.471224131 |
| 0.191997594 | 0.471354092 |
| 0.192103951 | 0.471469385 |
| 0.19222595 | 0.471695879 |
| 0.192261932 | 0.471708265 |
| 0.1922913 | 0.471734715 |
| 0.19233267 | 0.471766532 |
| 0.192400964 | 0.471906516 |
| 0.192592565 | 0.472134996 |
| 0.192626187 | 0.472134996 |
| 0.192646862 | 0.472144914 |
| 0.192680731 | 0.472203621 |
| 0.192736463 | 0.472266301 |
| 0.192778191 | 0.472320947 |
| 0.192793887 | 0.472335107 |
| 0.192975213 | 0.472682086 |
| 0.19299218 | 0.472691181 |
| 0.192998776 | 0.472691181 |
| 0.193033563 | 0.47270262 |
| 0.193047217 | 0.47270262 |
| 0.193103692 | 0.472777953 |
| 0.193127978 | 0.472788806 |
| 0.193200643 | 0.472942383 |
| 0.193266369 | 0.473070271 |
| 0.193412271 | 0.473128026 |
| 0.193428663 | 0.473128026 |
| 0.193441144 | 0.473128026 |

| | |
|---|---|
| 0.193467844 | 0.473128026 |
| 0.193492441 | 0.473145991 |
| 0.193665861 | 0.473491381 |
| 0.193730083 | 0.473539467 |
| 0.193746207 | 0.473544961 |
| 0.19378207 | 0.473562453 |
| 0.19379371 | 0.4735666 |
| 0.19388035 | 0.473636555 |
| 0.193894353 | 0.473636555 |
| 0.193918435 | 0.473636555 |
| 0.194096156 | 0.473787173 |
| 0.194102826 | 0.473787173 |
| 0.194146418 | 0.473804815 |
| 0.194325392 | 0.474111748 |
| 0.194392341 | 0.474226505 |
| 0.19453669 | 0.474428951 |
| 0.194634399 | 0.474556695 |
| 0.194637296 | 0.474556695 |
| 0.194743204 | 0.474735605 |
| 0.1947776 | 0.474758542 |
| 0.19478986 | 0.474758542 |
| 0.194919278 | 0.474864445 |
| 0.194969635 | 0.474921914 |
| 0.194970473 | 0.474921914 |
| 0.195145425 | 0.475222525 |
| 0.195153534 | 0.475222525 |
| 0.195169811 | 0.475222525 |
| 0.195243027 | 0.475352195 |
| 0.195358105 | 0.475488697 |
| 0.195360031 | 0.475488697 |
| 0.195464596 | 0.475648495 |
| 0.195514027 | 0.475695864 |
| 0.195529564 | 0.475709366 |
| 0.195556565 | 0.475750755 |
| 0.195581259 | 0.475786526 |
| 0.195626633 | 0.475859314 |
| 0.195631162 | 0.475859314 |
| 0.195697279 | 0.475905619 |
| 0.195704647 | 0.475905619 |
| 0.195852925 | 0.476179859 |
| 0.195963553 | 0.476300663 |
| 0.196028459 | 0.476387764 |
| 0.196083301 | 0.476470285 |
| 0.196092428 | 0.476470285 |
| 0.196115258 | 0.476501448 |

| | |
|---|---|
| 0.196148867 | 0.476517538 |
| 0.196192158 | 0.476518584 |
| 0.196257606 | 0.476595134 |
| 0.19635271 | 0.476735463 |
| 0.196424093 | 0.476838284 |
| 0.196580964 | 0.477003383 |
| 0.196674868 | 0.477179517 |
| 0.196700467 | 0.477193009 |
| 0.196996631 | 0.477595242 |
| 0.197021141 | 0.477613821 |
| 0.197052163 | 0.477633365 |
| 0.19705247 | 0.477633365 |
| 0.197100399 | 0.477673934 |
| 0.197109119 | 0.477673934 |
| 0.197134012 | 0.477673934 |
| 0.197140724 | 0.477673934 |
| 0.197173039 | 0.477673934 |
| 0.197214869 | 0.47773524 |
| 0.197307603 | 0.477747485 |
| 0.197308103 | 0.477747485 |
| 0.197334384 | 0.477747485 |
| 0.197335557 | 0.477747485 |
| 0.197340296 | 0.477747485 |
| 0.197355754 | 0.477760623 |
| 0.197612445 | 0.478265647 |
| 0.197616351 | 0.478265647 |
| 0.197624623 | 0.478265647 |
| 0.197675646 | 0.478340513 |
| 0.19776084 | 0.478473738 |
| 0.197791174 | 0.478498515 |
| 0.197833638 | 0.478552626 |
| 0.197898012 | 0.4786306 |
| 0.19790607 | 0.4786306 |
| 0.197920679 | 0.478641626 |
| 0.197954399 | 0.478674561 |
| 0.197974486 | 0.4786946 |
| 0.197982788 | 0.4786946 |
| 0.198035381 | 0.478756013 |
| 0.198207715 | 0.479043886 |
| 0.19822278 | 0.479050578 |
| 0.198302507 | 0.479175723 |
| 0.198387873 | 0.479290528 |
| 0.198412026 | 0.47929446 |
| 0.198580368 | 0.479581452 |
| 0.198589777 | 0.479581452 |

| | |
|---|---|
| 0.198641247 | 0.479604921 |
| 0.198677995 | 0.479669329 |
| 0.198776948 | 0.479798196 |
| 0.198889884 | 0.479986244 |
| 0.198932769 | 0.480026221 |
| 0.198944667 | 0.480026221 |
| 0.198955697 | 0.480026221 |
| 0.199119776 | 0.480271379 |
| 0.199138947 | 0.480271379 |
| 0.199139127 | 0.480271379 |
| 0.199168149 | 0.480292847 |
| 0.199239295 | 0.480361067 |
| 0.199246916 | 0.480361067 |
| 0.199308045 | 0.480484118 |
| 0.199497792 | 0.480745401 |
| 0.199617946 | 0.480866114 |
| 0.199647739 | 0.480889235 |
| 0.199666286 | 0.480901267 |
| 0.199771081 | 0.481113333 |
| 0.19980826 | 0.48117854 |
| 0.199822342 | 0.481188123 |
| 0.199860518 | 0.481214305 |
| 0.199867042 | 0.481214305 |
| 0.199921343 | 0.481231896 |
| 0.199924605 | 0.481231896 |
| 0.200006825 | 0.481316006 |
| 0.200146011 | 0.481553629 |
| 0.200180142 | 0.481570538 |
| 0.20024053 | 0.481674464 |
| 0.200246801 | 0.481674464 |
| 0.200362869 | 0.481822263 |
| 0.200368945 | 0.481822263 |
| 0.200471679 | 0.481923213 |
| 0.200580724 | 0.482063791 |
| 0.200594867 | 0.482073455 |
| 0.200664028 | 0.482166674 |
| 0.200710211 | 0.482253316 |
| 0.200750892 | 0.482326729 |
| 0.200787775 | 0.48239101 |
| 0.200854769 | 0.482478948 |
| 0.20091426 | 0.48251147 |
| 0.200980926 | 0.482635939 |
| 0.201033522 | 0.482673222 |
| 0.20107981 | 0.482691805 |
| 0.201185489 | 0.482814149 |

| | |
|---|---|
| 0.201201208 | 0.482814149 |
| 0.201205826 | 0.482814149 |
| 0.201298902 | 0.482839586 |
| 0.201323264 | 0.482859464 |
| 0.20139852 | 0.483005554 |
| 0.201559261 | 0.483220726 |
| 0.201673829 | 0.483420243 |
| 0.201729292 | 0.483482338 |
| 0.201788485 | 0.483567981 |
| 0.201846252 | 0.483567981 |
| 0.20188341 | 0.483597822 |
| 0.20188917 | 0.483597822 |
| 0.201930331 | 0.483647768 |
| 0.201944435 | 0.483657228 |
| 0.202015799 | 0.483706523 |
| 0.202167356 | 0.483826173 |
| 0.202238952 | 0.483837591 |
| 0.202251638 | 0.483837591 |
| 0.202270802 | 0.483837591 |
| 0.202272628 | 0.483837591 |
| 0.202277049 | 0.483837591 |
| 0.202283875 | 0.483837591 |
| 0.202327843 | 0.48389623 |
| 0.202339021 | 0.48389623 |
| 0.202340235 | 0.48389623 |
| 0.202396804 | 0.483932176 |
| 0.202572827 | 0.484153122 |
| 0.202583871 | 0.484153122 |
| 0.202919625 | 0.484700987 |
| 0.202951073 | 0.484751793 |
| 0.203060724 | 0.48495513 |
| 0.203078577 | 0.484959056 |
| 0.203169737 | 0.485110004 |
| 0.203229108 | 0.485196889 |
| 0.203382118 | 0.485437426 |
| 0.203390298 | 0.485437426 |
| 0.203502536 | 0.485630589 |
| 0.203698632 | 0.485952452 |
| 0.203784238 | 0.486075319 |
| 0.203787908 | 0.486075319 |
| 0.203815987 | 0.486075319 |
| 0.203827245 | 0.486075319 |
| 0.203839117 | 0.486075319 |
| 0.203856881 | 0.486075319 |
| 0.203864809 | 0.486075319 |

| | |
|---|---|
| 0.204017257 | 0.486249807 |
| 0.204035564 | 0.486269106 |
| 0.204110829 | 0.486399803 |
| 0.204178751 | 0.486488637 |
| 0.204256512 | 0.486649569 |
| 0.20431269 | 0.486736369 |
| 0.204319669 | 0.486736369 |
| 0.204344042 | 0.486736369 |
| 0.204566742 | 0.487023255 |
| 0.204688347 | 0.487208574 |
| 0.204783282 | 0.487319544 |
| 0.204818429 | 0.48737468 |
| 0.20496965 | 0.487548867 |
| 0.204974141 | 0.487548867 |
| 0.205014514 | 0.487548867 |
| 0.205023756 | 0.487548867 |
| 0.205040651 | 0.487548867 |
| 0.205053675 | 0.487548867 |
| 0.205080951 | 0.487589381 |
| 0.205212439 | 0.487853291 |
| 0.205285976 | 0.487924327 |
| 0.205538459 | 0.488214063 |
| 0.205575692 | 0.488253801 |
| 0.205713823 | 0.488484431 |
| 0.205741988 | 0.488526955 |
| 0.205820556 | 0.488689147 |
| 0.205854015 | 0.488744226 |
| 0.205892496 | 0.488786856 |
| 0.205924565 | 0.488814258 |
| 0.206025251 | 0.488980148 |
| 0.206069977 | 0.48900884 |
| 0.206330863 | 0.489497172 |
| 0.206333852 | 0.489497172 |
| 0.206341941 | 0.489497172 |
| 0.206381467 | 0.489529433 |
| 0.206404373 | 0.489529433 |
| 0.206421139 | 0.489529433 |
| 0.206462394 | 0.489554139 |
| 0.206493803 | 0.48960424 |
| 0.206544063 | 0.489650279 |
| 0.206710342 | 0.489892072 |
| 0.206736556 | 0.489911014 |
| 0.206746624 | 0.489911014 |
| 0.206817185 | 0.489973699 |
| 0.206824517 | 0.489973699 |

| | |
|---|---|
| 0.206860656 | 0.490010567 |
| 0.20689587 | 0.490043154 |
| 0.206896461 | 0.490043154 |
| 0.206923948 | 0.490043154 |
| 0.206930824 | 0.490043154 |
| 0.206982436 | 0.490075163 |
| 0.206990824 | 0.490075163 |
| 0.207150565 | 0.490307124 |
| 0.207252204 | 0.490485341 |
| 0.207287651 | 0.490485341 |
| 0.207464305 | 0.490732677 |
| 0.207535025 | 0.490826827 |
| 0.207570966 | 0.490827293 |
| 0.207653553 | 0.490884004 |
| 0.20768056 | 0.490884004 |
| 0.207699552 | 0.490884004 |
| 0.207703497 | 0.490884004 |
| 0.207764765 | 0.490940003 |
| 0.207768423 | 0.490940003 |
| 0.207812919 | 0.490959011 |
| 0.207813306 | 0.490959011 |
| 0.207945234 | 0.491173801 |
| 0.208034144 | 0.491299756 |
| 0.208098953 | 0.491368229 |
| 0.208104406 | 0.491368229 |
| 0.208159047 | 0.491433495 |
| 0.208194617 | 0.49143505 |
| 0.208370301 | 0.491703529 |
| 0.208449542 | 0.491769164 |
| 0.20849957 | 0.491841596 |
| 0.208590595 | 0.492003975 |
| 0.20871363 | 0.492170735 |
| 0.208742027 | 0.49219053 |
| 0.208812806 | 0.492259937 |
| 0.208971494 | 0.49251214 |
| 0.209012913 | 0.492557373 |
| 0.209072325 | 0.492627894 |
| 0.209118625 | 0.492696748 |
| 0.209122236 | 0.492696748 |
| 0.2091495 | 0.492736606 |
| 0.209296273 | 0.493009228 |
| 0.209353613 | 0.49309552 |
| 0.209470121 | 0.493165099 |
| 0.209508309 | 0.493165099 |
| 0.209531893 | 0.493165099 |

| | |
|---|---|
| 0.209638299 | 0.493351283 |
| 0.209684872 | 0.493394518 |
| 0.209898414 | 0.493670704 |
| 0.209966748 | 0.493739589 |
| 0.209970306 | 0.493739589 |
| 0.209979536 | 0.493739589 |
| 0.210073551 | 0.493911883 |
| 0.210141168 | 0.493975795 |
| 0.210145629 | 0.493975795 |
| 0.210152593 | 0.493975795 |
| 0.210275387 | 0.494191259 |
| 0.210286104 | 0.494192059 |
| 0.210414334 | 0.49439583 |
| 0.210452352 | 0.494436966 |
| 0.210567286 | 0.494560022 |
| 0.210594216 | 0.494583947 |
| 0.210621279 | 0.49461366 |
| 0.210788311 | 0.494908302 |
| 0.21087822 | 0.495070585 |
| 0.210909893 | 0.495105639 |
| 0.210913942 | 0.495105639 |
| 0.210926664 | 0.495111101 |
| 0.211011001 | 0.49516264 |
| 0.211028136 | 0.495178452 |
| 0.211056148 | 0.495219783 |
| 0.211092578 | 0.495256464 |
| 0.211116278 | 0.49528767 |
| 0.211230323 | 0.495506408 |
| 0.211286039 | 0.495588287 |
| 0.211303548 | 0.495604949 |
| 0.211334405 | 0.495619502 |
| 0.211370233 | 0.495619502 |
| 0.21137566 | 0.495619502 |
| 0.211490931 | 0.49575148 |
| 0.211577314 | 0.495880751 |
| 0.211647061 | 0.495883337 |
| 0.21164754 | 0.495883337 |
| 0.21165894 | 0.495883337 |
| 0.211899525 | 0.496342826 |
| 0.211969311 | 0.496457456 |
| 0.212034057 | 0.496514809 |
| 0.212035499 | 0.496514809 |
| 0.212126522 | 0.49662978 |
| 0.21218382 | 0.49663425 |
| 0.212201209 | 0.49663425 |

| | |
|---|---|
| 0.212237663 | 0.496659152 |
| 0.212243133 | 0.496659152 |
| 0.212489629 | 0.497056614 |
| 0.212501766 | 0.497056614 |
| 0.212519934 | 0.497062657 |
| 0.212543577 | 0.497093544 |
| 0.212592339 | 0.497183174 |
| 0.212638592 | 0.497211576 |
| 0.212646242 | 0.497211576 |
| 0.212690361 | 0.497257474 |
| 0.212697194 | 0.497257474 |
| 0.212723066 | 0.497293548 |
| 0.21283487 | 0.497384022 |
| 0.212940556 | 0.497582175 |
| 0.212982744 | 0.497631924 |
| 0.213154074 | 0.497836853 |
| 0.213320179 | 0.49801631 |
| 0.21335639 | 0.49801631 |
| 0.21335639 | 0.49801631 |
| 0.21335639 | 0.49801631 |
| 0.21341406 | 0.498126511 |
| 0.21344743 | 0.498155571 |
| 0.213520539 | 0.498292566 |
| 0.21360739 | 0.498357948 |
| 0.213873839 | 0.498601143 |
| 0.213879131 | 0.498601143 |
| 0.21390251 | 0.49863124 |
| 0.213934757 | 0.498635568 |
| 0.213945842 | 0.498635568 |
| 0.213992589 | 0.498694772 |
| 0.214034542 | 0.498701899 |
| 0.214042002 | 0.498701899 |
| 0.214048003 | 0.498701899 |
| 0.214076885 | 0.498742161 |
| 0.21412175 | 0.498782006 |
| 0.214124276 | 0.498782006 |
| 0.214167237 | 0.498857681 |
| 0.21420111 | 0.498912182 |
| 0.214347158 | 0.499097183 |
| 0.214393835 | 0.499165779 |
| 0.214530186 | 0.499312377 |
| 0.214634772 | 0.499506978 |
| 0.2148593 | 0.499907371 |
| 0.215097518 | 0.500241686 |
| 0.215156952 | 0.500339353 |

| | |
|---|---|
| 0.21517103 | 0.500339353 |
| 0.215211634 | 0.500375407 |
| 0.215253617 | 0.500409235 |
| 0.215264944 | 0.500411141 |
| 0.215345688 | 0.500463617 |
| 0.215407385 | 0.500484629 |
| 0.215440927 | 0.50050266 |
| 0.215598006 | 0.500703166 |
| 0.215611339 | 0.500703166 |
| 0.215644839 | 0.500721203 |
| 0.215748715 | 0.50085124 |
| 0.215808332 | 0.500940804 |
| 0.2158792 | 0.500996194 |
| 0.215885214 | 0.500996194 |
| 0.215899699 | 0.500996194 |
| 0.215905828 | 0.500996194 |
| 0.21598798 | 0.501129384 |
| 0.215994793 | 0.501129384 |
| 0.216034439 | 0.50117598 |
| 0.216072316 | 0.501224023 |
| 0.216077679 | 0.501224023 |
| 0.216144475 | 0.501318979 |
| 0.216227118 | 0.501424138 |
| 0.216270181 | 0.501499582 |
| 0.216496969 | 0.501903287 |
| 0.216577243 | 0.501994259 |
| 0.216580932 | 0.501994259 |
| 0.216617001 | 0.502025198 |
| 0.216654963 | 0.502025198 |
| 0.216702967 | 0.502045372 |
| 0.216705834 | 0.502045372 |
| 0.216786129 | 0.502126137 |
| 0.216848256 | 0.502126137 |
| 0.216848646 | 0.502126137 |
| 0.216855778 | 0.502126137 |
| 0.216856467 | 0.502126137 |
| 0.216890834 | 0.502126137 |
| 0.216942131 | 0.502126137 |
| 0.216952876 | 0.502126137 |
| 0.216971607 | 0.502126137 |
| 0.217018197 | 0.502182766 |
| 0.217089078 | 0.502211563 |
| 0.217115 | 0.502211563 |
| 0.217140851 | 0.502246967 |
| 0.217300159 | 0.502407854 |

| | |
|---|---|
| 0.2173268 | 0.502408673 |
| 0.217402001 | 0.502460576 |
| 0.217662822 | 0.502794997 |
| 0.217720277 | 0.50285369 |
| 0.217751698 | 0.502878346 |
| 0.217980159 | 0.503236683 |
| 0.218100871 | 0.503391818 |
| 0.218129746 | 0.503423452 |
| 0.218173985 | 0.503423452 |
| 0.21823553 | 0.503483027 |
| 0.218247179 | 0.503485512 |
| 0.218260931 | 0.50349285 |
| 0.218285878 | 0.503513372 |
| 0.218371162 | 0.503649557 |
| 0.218548147 | 0.503981035 |
| 0.218557211 | 0.503981035 |
| 0.218627189 | 0.504117991 |
| 0.218647117 | 0.504139536 |
| 0.218749196 | 0.504214329 |
| 0.218786062 | 0.504214329 |
| 0.218798352 | 0.504214329 |
| 0.218802737 | 0.504214329 |
| 0.218803063 | 0.504214329 |
| 0.21881099 | 0.504214329 |
| 0.218814859 | 0.504214329 |
| 0.218828914 | 0.504214329 |
| 0.218865826 | 0.504214329 |
| 0.218998271 | 0.504345422 |
| 0.219021315 | 0.504345422 |
| 0.219037692 | 0.504345422 |
| 0.219097966 | 0.504422045 |
| 0.219146131 | 0.504508545 |
| 0.219218756 | 0.504591537 |
| 0.219280194 | 0.504625948 |
| 0.219281891 | 0.504625948 |
| 0.219305533 | 0.504635409 |
| 0.219317789 | 0.504635409 |
| 0.219317789 | 0.504635409 |
| 0.219361778 | 0.504646607 |
| 0.219385151 | 0.504668497 |
| 0.219428553 | 0.504730396 |
| 0.219439336 | 0.504730396 |
| 0.219484086 | 0.504764095 |
| 0.219489629 | 0.504764095 |
| 0.219511599 | 0.504764095 |

| | |
|---|---|
| 0.219521478 | 0.504764095 |
| 0.219545447 | 0.504769057 |
| 0.219762914 | 0.50497652 |
| 0.219817899 | 0.505078496 |
| 0.21988876 | 0.505192569 |
| 0.219922489 | 0.505221319 |
| 0.220041716 | 0.505446456 |
| 0.220187437 | 0.505683627 |
| 0.220208392 | 0.505707368 |
| 0.220265165 | 0.505806081 |
| 0.220366481 | 0.505910866 |
| 0.220435348 | 0.505990582 |
| 0.220437957 | 0.505990582 |
| 0.220497226 | 0.506012534 |
| 0.220585617 | 0.506134372 |
| 0.220623323 | 0.506172118 |
| 0.220664535 | 0.506242283 |
| 0.220680963 | 0.506255587 |
| 0.220745445 | 0.506354733 |
| 0.220915856 | 0.506623626 |
| 0.221027046 | 0.506805408 |
| 0.221095967 | 0.50691463 |
| 0.2212799 | 0.507142455 |
| 0.221291169 | 0.507142455 |
| 0.22131465 | 0.507171863 |
| 0.221401343 | 0.50734612 |
| 0.221453624 | 0.507395496 |
| 0.221557789 | 0.507455937 |
| 0.221559517 | 0.507455937 |
| 0.221634079 | 0.507488764 |
| 0.221645819 | 0.507491248 |
| 0.221665218 | 0.507511264 |
| 0.221723563 | 0.507563095 |
| 0.221731606 | 0.507563095 |
| 0.221741141 | 0.507563095 |
| 0.221752929 | 0.507565684 |
| 0.221808787 | 0.50762035 |
| 0.22190577 | 0.507744707 |
| 0.222039909 | 0.50797842 |
| 0.22211177 | 0.508045204 |
| 0.222126713 | 0.508054364 |
| 0.222141564 | 0.508054364 |
| 0.22219297 | 0.508117786 |
| 0.222258537 | 0.508139301 |
| 0.222324382 | 0.508189831 |

| | |
|---|---|
| 0.222374218 | 0.508215081 |
| 0.222378111 | 0.508215081 |
| 0.222437131 | 0.508284314 |
| 0.22259942 | 0.50834244 |
| 0.22273352 | 0.508452165 |
| 0.222738068 | 0.508452165 |
| 0.222847024 | 0.508613682 |
| 0.222872899 | 0.508613682 |
| 0.222883967 | 0.508614571 |
| 0.223034405 | 0.50886034 |
| 0.22309683 | 0.50890525 |
| 0.22315558 | 0.508990508 |
| 0.223219868 | 0.509046627 |
| 0.223222937 | 0.509046627 |
| 0.22328427 | 0.509113364 |
| 0.223341553 | 0.509214484 |
| 0.22337462 | 0.509246232 |
| 0.223463173 | 0.509350908 |
| 0.223524796 | 0.509429769 |
| 0.223588405 | 0.509489736 |
| 0.223634176 | 0.509525166 |
| 0.22363584 | 0.509525166 |
| 0.223725603 | 0.50958294 |
| 0.223836777 | 0.509811782 |
| 0.223881111 | 0.509819689 |
| 0.224042662 | 0.510084082 |
| 0.224072835 | 0.510105488 |
| 0.224109753 | 0.510141629 |
| 0.224190614 | 0.510227285 |
| 0.224288176 | 0.510424931 |
| 0.224404127 | 0.510591217 |
| 0.224574573 | 0.510765495 |
| 0.224578121 | 0.510765495 |
| 0.224665649 | 0.51077218 |
| 0.224858725 | 0.51103963 |
| 0.224956454 | 0.511154336 |
| 0.225096515 | 0.511360445 |
| 0.225133553 | 0.511406443 |
| 0.225230433 | 0.511515247 |
| 0.225234089 | 0.511515247 |
| 0.225250578 | 0.511515247 |
| 0.225397631 | 0.511678407 |
| 0.225451295 | 0.511751446 |
| 0.225666319 | 0.512099531 |
| 0.225697772 | 0.512114574 |

| | |
|---|---|
| 0.225731456 | 0.512114574 |
| 0.225741695 | 0.512114574 |
| 0.225773543 | 0.512114574 |
| 0.225800548 | 0.512114574 |
| 0.22592249 | 0.512281186 |
| 0.226069816 | 0.512544191 |
| 0.22608034 | 0.512544191 |
| 0.22616946 | 0.512615763 |
| 0.226398093 | 0.513029462 |
| 0.226402109 | 0.513029462 |
| 0.226421025 | 0.513047916 |
| 0.226477058 | 0.513112117 |
| 0.226522792 | 0.513180849 |
| 0.226534581 | 0.513183147 |
| 0.226673105 | 0.513329158 |
| 0.226674461 | 0.513329158 |
| 0.226701772 | 0.513329158 |
| 0.226709589 | 0.513329158 |
| 0.226776391 | 0.513377785 |
| 0.226890116 | 0.513515196 |
| 0.226897563 | 0.513515196 |
| 0.226913711 | 0.513515196 |
| 0.226968408 | 0.513564126 |
| 0.226972332 | 0.513564126 |
| 0.227002823 | 0.513595138 |
| 0.227007606 | 0.513595138 |
| 0.227040835 | 0.513622124 |
| 0.227068634 | 0.513649328 |
| 0.227119871 | 0.513654641 |
| 0.227145692 | 0.513654641 |
| 0.22715254 | 0.513654641 |
| 0.227174625 | 0.51368019 |
| 0.227326036 | 0.513883811 |
| 0.227353503 | 0.513913873 |
| 0.227458407 | 0.514080886 |
| 0.227488705 | 0.514097487 |
| 0.227750815 | 0.514470122 |
| 0.227828286 | 0.51459631 |
| 0.227981858 | 0.514845861 |
| 0.228057056 | 0.514966501 |
| 0.228091201 | 0.514994769 |
| 0.228158523 | 0.515119436 |
| 0.228168048 | 0.515119436 |
| 0.228288136 | 0.515270066 |
| 0.228299682 | 0.515270066 |

| | |
|---|---|
| 0.228382529 | 0.515408204 |
| 0.228483499 | 0.515550876 |
| 0.228497232 | 0.515550876 |
| 0.228499873 | 0.515550876 |
| 0.228571468 | 0.515621912 |
| 0.228585488 | 0.515621912 |
| 0.228788089 | 0.51590788 |
| 0.228802689 | 0.515916375 |
| 0.228865997 | 0.516010264 |
| 0.229108361 | 0.516336721 |
| 0.229179195 | 0.516367975 |
| 0.229194628 | 0.516367975 |
| 0.229200707 | 0.516367975 |
| 0.229205219 | 0.516367975 |
| 0.229306517 | 0.516465595 |
| 0.229396091 | 0.516600026 |
| 0.229398743 | 0.516600026 |
| 0.229429175 | 0.516644128 |
| 0.229626945 | 0.51698222 |
| 0.229638244 | 0.51698222 |
| 0.229681866 | 0.51698222 |
| 0.229745335 | 0.517063664 |
| 0.229832942 | 0.517186559 |
| 0.22988556 | 0.517238418 |
| 0.229930036 | 0.517238418 |
| 0.230028808 | 0.517398356 |
| 0.230078396 | 0.517450043 |
| 0.230144724 | 0.517545812 |
| 0.230199538 | 0.517595761 |
| 0.230221814 | 0.51759799 |
| 0.230222265 | 0.51759799 |
| 0.230276892 | 0.517623066 |
| 0.230356748 | 0.517729264 |
| 0.230502978 | 0.517904378 |
| 0.230515307 | 0.517904378 |
| 0.230522828 | 0.517904378 |
| 0.230525726 | 0.517904378 |
| 0.230589254 | 0.517958516 |
| 0.230666503 | 0.518083166 |
| 0.230759292 | 0.518180169 |
| 0.230789972 | 0.518189037 |
| 0.230919472 | 0.518309142 |
| 0.231038088 | 0.518504446 |
| 0.231044433 | 0.518504446 |
| 0.23110351 | 0.518526729 |

| | |
|---|---|
| 0.231155312 | 0.518618526 |
| 0.231167058 | 0.518620447 |
| 0.231213769 | 0.518651949 |
| 0.231241085 | 0.518664367 |
| 0.231310748 | 0.518768171 |
| 0.231320042 | 0.518768171 |
| 0.231398573 | 0.518868385 |
| 0.23148018 | 0.518931822 |
| 0.231594677 | 0.519101916 |
| 0.231596673 | 0.519101916 |
| 0.23162145 | 0.519101916 |
| 0.231717229 | 0.51924327 |
| 0.231955878 | 0.519618111 |
| 0.231974887 | 0.519625056 |
| 0.232074846 | 0.519677873 |
| 0.232110118 | 0.519683555 |
| 0.232270844 | 0.519945643 |
| 0.232333742 | 0.520015382 |
| 0.232454239 | 0.52011172 |
| 0.232685297 | 0.520530889 |
| 0.232757428 | 0.520643342 |
| 0.23283951 | 0.520696258 |
| 0.232841285 | 0.520696258 |
| 0.232860098 | 0.520696258 |
| 0.232862166 | 0.520696258 |
| 0.232895155 | 0.520696258 |
| 0.232921228 | 0.520696258 |
| 0.232952285 | 0.520696258 |
| 0.232965582 | 0.520696258 |
| 0.232984415 | 0.520696258 |
| 0.233007125 | 0.52071276 |
| 0.233029238 | 0.520713312 |
| 0.233075354 | 0.520791926 |
| 0.233109411 | 0.520794723 |
| 0.233136541 | 0.520806474 |
| 0.233181528 | 0.520851145 |
| 0.233239097 | 0.520864561 |
| 0.233261693 | 0.520883166 |
| 0.233436143 | 0.521111772 |
| 0.233491845 | 0.521172842 |
| 0.23368771 | 0.521499605 |
| 0.233717157 | 0.521519398 |
| 0.233838943 | 0.521717673 |
| 0.233840199 | 0.521717673 |
| 0.233853762 | 0.521723492 |

| | |
|---|---|
| 0.233888772 | 0.52175272 |
| 0.23397962 | 0.521887846 |
| 0.23398222 | 0.521887846 |
| 0.234013489 | 0.521892179 |
| 0.234038952 | 0.521892179 |
| 0.234163401 | 0.522028631 |
| 0.234539612 | 0.522497175 |
| 0.234563821 | 0.522497175 |
| 0.234589978 | 0.522497175 |
| 0.234609995 | 0.522497175 |
| 0.234616777 | 0.522497175 |
| 0.23467369 | 0.522497175 |
| 0.234676737 | 0.522497175 |
| 0.23471808 | 0.522528285 |
| 0.234731126 | 0.522528285 |
| 0.234741117 | 0.522528285 |
| 0.234757013 | 0.522539246 |
| 0.234791345 | 0.522566819 |
| 0.234875346 | 0.522680503 |
| 0.235120791 | 0.52308006 |
| 0.235157172 | 0.523112126 |
| 0.235182674 | 0.523135725 |
| 0.23525555 | 0.523233215 |
| 0.235284767 | 0.523273762 |
| 0.235315346 | 0.523285343 |
| 0.235322936 | 0.523285343 |
| 0.235439169 | 0.523494925 |
| 0.235532026 | 0.523607345 |
| 0.235723074 | 0.523954957 |
| 0.235967325 | 0.524277655 |
| 0.235978633 | 0.524278321 |
| 0.236140939 | 0.524555631 |
| 0.236155982 | 0.524555631 |
| 0.236221085 | 0.524596738 |
| 0.236324608 | 0.524781178 |
| 0.236451882 | 0.524962417 |
| 0.236594161 | 0.525146925 |
| 0.236601145 | 0.525146925 |
| 0.236620722 | 0.525152518 |
| 0.236681769 | 0.525236547 |
| 0.236789169 | 0.525400046 |
| 0.236808237 | 0.525400046 |
| 0.23685519 | 0.525435171 |
| 0.236863399 | 0.525435171 |
| 0.236903885 | 0.525466528 |

| | |
|---|---|
| 0.236908536 | 0.525466528 |
| 0.236977733 | 0.525509126 |
| 0.236987286 | 0.525509126 |
| 0.237006568 | 0.525509126 |
| 0.237007077 | 0.525509126 |
| 0.237118865 | 0.525708042 |
| 0.23721989 | 0.5258425 |
| 0.237223675 | 0.5258425 |
| 0.237340636 | 0.526019821 |
| 0.237349969 | 0.526019821 |
| 0.237401845 | 0.526041578 |
| 0.237452334 | 0.526093611 |
| 0.237497263 | 0.526130618 |
| 0.237830188 | 0.526549756 |
| 0.238017969 | 0.52691382 |
| 0.23806914 | 0.526956311 |
| 0.238112237 | 0.526978231 |
| 0.23819824 | 0.527095088 |
| 0.238514447 | 0.527672221 |
| 0.238596346 | 0.527776681 |
| 0.238620092 | 0.527776681 |
| 0.238628153 | 0.527776681 |
| 0.238717924 | 0.527912398 |
| 0.238731153 | 0.527912398 |
| 0.238788539 | 0.527939999 |
| 0.238798753 | 0.527939999 |
| 0.238800462 | 0.527939999 |
| 0.238929811 | 0.528125034 |
| 0.238949797 | 0.528136264 |
| 0.239130541 | 0.528341602 |
| 0.239185129 | 0.528378889 |
| 0.239260985 | 0.528383198 |
| 0.239286523 | 0.528383198 |
| 0.239288954 | 0.528383198 |
| 0.239503572 | 0.528680615 |
| 0.23955614 | 0.528724236 |
| 0.239625139 | 0.528788709 |
| 0.239659744 | 0.528830366 |
| 0.239724847 | 0.528900508 |
| 0.239945287 | 0.529190748 |
| 0.239977337 | 0.529236924 |
| 0.240010259 | 0.529260516 |
| 0.240089019 | 0.529383783 |
| 0.240099504 | 0.529383783 |
| 0.240164339 | 0.529479814 |

| | |
|---|---|
| 0.240184643 | 0.529497963 |
| 0.24027527 | 0.529673238 |
| 0.240326784 | 0.529719269 |
| 0.240346437 | 0.529727296 |
| 0.24039088 | 0.529780387 |
| 0.240401741 | 0.529780387 |
| 0.240474985 | 0.529868245 |
| 0.240540056 | 0.529881891 |
| 0.24071449 | 0.530097553 |
| 0.240791658 | 0.530157259 |
| 0.240827712 | 0.530203922 |
| 0.240971241 | 0.530432643 |
| 0.240983046 | 0.530432643 |
| 0.241033231 | 0.53050763 |
| 0.241072197 | 0.530521464 |
| 0.241222402 | 0.530706767 |
| 0.241276329 | 0.530748301 |
| 0.241283561 | 0.530748301 |
| 0.241461246 | 0.531034113 |
| 0.241586172 | 0.531157574 |
| 0.241594084 | 0.531157574 |
| 0.241599238 | 0.531157574 |
| 0.241624525 | 0.531160519 |
| 0.241630248 | 0.531160519 |
| 0.241673395 | 0.531189177 |
| 0.241727137 | 0.531238415 |
| 0.241749608 | 0.531251262 |
| 0.241800594 | 0.531314274 |
| 0.241884176 | 0.531399864 |
| 0.241923547 | 0.531437332 |
| 0.242060638 | 0.531713957 |
| 0.242083889 | 0.531740505 |
| 0.242116946 | 0.531788591 |
| 0.242175011 | 0.531882533 |
| 0.242383571 | 0.532177868 |
| 0.242402402 | 0.532194331 |
| 0.242539672 | 0.532352686 |
| 0.242554375 | 0.532352686 |
| 0.242595148 | 0.532352686 |
| 0.242623794 | 0.532352686 |
| 0.242626278 | 0.532352686 |
| 0.242643402 | 0.532352686 |
| 0.242654779 | 0.532352686 |
| 0.242670296 | 0.532352686 |
| 0.242688374 | 0.532352686 |

| | |
|---|---|
| 0.242694327 | 0.532352686 |
| 0.242700522 | 0.532352686 |
| 0.242702227 | 0.532352686 |
| 0.242709098 | 0.532352686 |
| 0.242872952 | 0.532589441 |
| 0.242941339 | 0.532690353 |
| 0.243067118 | 0.532865874 |
| 0.24310926 | 0.532865874 |
| 0.243110894 | 0.532865874 |
| 0.243129363 | 0.53288183 |
| 0.243141063 | 0.53288295 |
| 0.243265778 | 0.532976066 |
| 0.243292679 | 0.532976066 |
| 0.24338303 | 0.533069835 |
| 0.243794222 | 0.533847697 |
| 0.24380779 | 0.533852862 |
| 0.243852173 | 0.533900954 |
| 0.243958288 | 0.534050771 |
| 0.24402783 | 0.534050771 |
| 0.244055299 | 0.534050771 |
| 0.244057699 | 0.534050771 |
| 0.244078318 | 0.534050771 |
| 0.244088943 | 0.534050771 |
| 0.244264355 | 0.534336691 |
| 0.244276414 | 0.534338527 |
| 0.244287819 | 0.534338934 |
| 0.244397468 | 0.534482607 |
| 0.24483324 | 0.535015953 |
| 0.244853058 | 0.535034713 |
| 0.244989918 | 0.535255501 |
| 0.245001681 | 0.535255501 |
| 0.245009005 | 0.535255501 |
| 0.245079367 | 0.535370764 |
| 0.245085535 | 0.535370764 |
| 0.245285085 | 0.535683819 |
| 0.245368475 | 0.535712358 |
| 0.245383992 | 0.535712358 |
| 0.245410281 | 0.535712358 |
| 0.245489313 | 0.53585955 |
| 0.245541214 | 0.535948277 |
| 0.245803119 | 0.536200482 |
| 0.245816466 | 0.536205038 |
| 0.245845337 | 0.536226233 |
| 0.245983934 | 0.536370005 |
| 0.246002954 | 0.536370005 |

| | |
|---|---|
| 0.246018107 | 0.536374657 |
| 0.246170557 | 0.536619926 |
| 0.246190384 | 0.536619926 |
| 0.246209802 | 0.536619926 |
| 0.24621149 | 0.536619926 |
| 0.246249402 | 0.536657832 |
| 0.246329944 | 0.536710565 |
| 0.246455261 | 0.536812831 |
| 0.246579514 | 0.536921408 |
| 0.246584541 | 0.536921408 |
| 0.246696644 | 0.53703966 |
| 0.246775651 | 0.537074354 |
| 0.246816911 | 0.537075598 |
| 0.246824505 | 0.537075598 |
| 0.246896689 | 0.537116509 |
| 0.246899694 | 0.537116509 |
| 0.246933051 | 0.537147537 |
| 0.246968204 | 0.537167404 |
| 0.246986014 | 0.53718161 |
| 0.247027534 | 0.537222099 |
| 0.247042977 | 0.537222099 |
| 0.24704968 | 0.537222099 |
| 0.247144189 | 0.53732933 |
| 0.247370701 | 0.537576401 |
| 0.247398479 | 0.537602862 |
| 0.247411762 | 0.537602862 |
| 0.247416741 | 0.537602862 |
| 0.24749621 | 0.537608434 |
| 0.247593439 | 0.537765995 |
| 0.247672643 | 0.53781539 |
| 0.247728717 | 0.537878746 |
| 0.247740583 | 0.537878746 |
| 0.247876376 | 0.538012493 |
| 0.247897637 | 0.53801268 |
| 0.247975461 | 0.538072574 |
| 0.247978189 | 0.538072574 |
| 0.248061886 | 0.538169884 |
| 0.24808819 | 0.538202433 |
| 0.248242394 | 0.538438859 |
| 0.248334248 | 0.538586959 |
| 0.248344599 | 0.538586959 |
| 0.248376155 | 0.538630868 |
| 0.248436834 | 0.538737927 |
| 0.248481009 | 0.538749037 |
| 0.248500489 | 0.538749037 |

| | |
|---|---|
| 0.248513206 | 0.538749037 |
| 0.248627406 | 0.538884613 |
| 0.248678034 | 0.538917446 |
| 0.248706578 | 0.538954781 |
| 0.248743728 | 0.539010762 |
| 0.248781537 | 0.539059654 |
| 0.24883031 | 0.539059654 |
| 0.248831492 | 0.539059654 |
| 0.248834201 | 0.539059654 |
| 0.248915441 | 0.539162075 |
| 0.248939902 | 0.539176749 |
| 0.24897333 | 0.539176749 |
| 0.248994683 | 0.539176749 |
| 0.249031479 | 0.539176749 |
| 0.249035424 | 0.539176749 |
| 0.249054183 | 0.539192853 |
| 0.24909874 | 0.539196568 |
| 0.249101184 | 0.539196568 |
| 0.24912641 | 0.539226664 |
| 0.249253552 | 0.539313441 |
| 0.249270073 | 0.539313441 |
| 0.24932945 | 0.53934749 |
| 0.249359598 | 0.539363711 |
| 0.249449484 | 0.539518462 |
| 0.249527438 | 0.539579719 |
| 0.249611261 | 0.539736472 |
| 0.249658576 | 0.539765264 |
| 0.249703568 | 0.539838032 |
| 0.24973075 | 0.539847787 |
| 0.249827948 | 0.53998008 |
| 0.249886414 | 0.540061725 |
| 0.249912083 | 0.540092693 |
| 0.249944041 | 0.540115486 |
| 0.24996967 | 0.540115486 |
| 0.249990106 | 0.540115486 |
| 0.250142142 | 0.540320188 |
| 0.250212347 | 0.540398305 |
| 0.25030119 | 0.540514228 |
| 0.250663479 | 0.541004646 |
| 0.250734188 | 0.541052139 |
| 0.250777347 | 0.541062385 |
| 0.250877205 | 0.541196151 |
| 0.250946249 | 0.541239628 |
| 0.25094863 | 0.541239628 |
| 0.250975146 | 0.541239628 |

| | |
|---|---|
| 0.25098607 | 0.541239628 |
| 0.250988272 | 0.541239628 |
| 0.251009802 | 0.541245056 |
| 0.251013518 | 0.541245056 |
| 0.251037225 | 0.54127167 |
| 0.251116689 | 0.541358062 |
| 0.251134126 | 0.541358062 |
| 0.251180632 | 0.541381502 |
| 0.251190468 | 0.541381502 |
| 0.251281981 | 0.541480721 |
| 0.251373053 | 0.541556169 |
| 0.251373847 | 0.541556169 |
| 0.251442354 | 0.541654757 |
| 0.251466383 | 0.541672592 |
| 0.25147338 | 0.541672592 |
| 0.251515527 | 0.541673502 |
| 0.251519296 | 0.541673502 |
| 0.251671755 | 0.541902974 |
| 0.251682738 | 0.541902974 |
| 0.251726387 | 0.541946845 |
| 0.251741576 | 0.541946845 |
| 0.251795959 | 0.541999734 |
| 0.251874382 | 0.542119539 |
| 0.25201623 | 0.542302308 |
| 0.252059483 | 0.542330263 |
| 0.252267343 | 0.542622019 |
| 0.252284332 | 0.542624789 |
| 0.252291417 | 0.542624789 |
| 0.252393062 | 0.542755698 |
| 0.252526725 | 0.542934738 |
| 0.252575018 | 0.542989545 |
| 0.252628807 | 0.543043344 |
| 0.252634249 | 0.543043344 |
| 0.252729365 | 0.543198766 |
| 0.252892708 | 0.543427226 |
| 0.253003797 | 0.543567839 |
| 0.253025955 | 0.543590923 |
| 0.253080811 | 0.543684249 |
| 0.253154099 | 0.543738869 |
| 0.253283097 | 0.543868511 |
| 0.253286256 | 0.543868511 |
| 0.253292196 | 0.543868511 |
| 0.253349362 | 0.543934863 |
| 0.25335736 | 0.543934863 |
| 0.253542624 | 0.544185424 |

| | |
|---|---|
| 0.253638638 | 0.544317911 |
| 0.253726856 | 0.544433635 |
| 0.253843493 | 0.544585767 |
| 0.25388283 | 0.544629277 |
| 0.253961961 | 0.544741773 |
| 0.253993516 | 0.544784922 |
| 0.254005653 | 0.54478642 |
| 0.254082588 | 0.544804224 |
| 0.254131343 | 0.544878231 |
| 0.254146216 | 0.544878231 |
| 0.254151424 | 0.544878231 |
| 0.25419186 | 0.544915862 |
| 0.254258425 | 0.545034027 |
| 0.254300511 | 0.545069334 |
| 0.254321115 | 0.545069334 |
| 0.254343563 | 0.545069334 |
| 0.254400048 | 0.545092274 |
| 0.254510127 | 0.545205496 |
| 0.254551377 | 0.545222865 |
| 0.254696514 | 0.545457569 |
| 0.254814828 | 0.545661877 |
| 0.254881806 | 0.545738421 |
| 0.25488511 | 0.545738421 |
| 0.25495595 | 0.545841366 |
| 0.25498106 | 0.545858599 |
| 0.254986922 | 0.545858599 |
| 0.255022222 | 0.545860562 |
| 0.255231769 | 0.546174392 |
| 0.255303421 | 0.54626839 |
| 0.255311365 | 0.54626839 |
| 0.255315984 | 0.54626839 |
| 0.255393423 | 0.54640953 |
| 0.255503327 | 0.546521915 |
| 0.255526894 | 0.546547778 |
| 0.255560433 | 0.546562165 |
| 0.255565377 | 0.546562165 |
| 0.255610204 | 0.546627775 |
| 0.255910843 | 0.546974153 |
| 0.25592492 | 0.546974153 |
| 0.255967412 | 0.546974153 |
| 0.255997044 | 0.547012349 |
| 0.256067531 | 0.547094129 |
| 0.256153589 | 0.547175592 |
| 0.256193892 | 0.5472041 |
| 0.256198471 | 0.5472041 |

| | |
|---|---|
| 0.256314257 | 0.547396041 |
| 0.256515919 | 0.54763076 |
| 0.256546191 | 0.547665654 |
| 0.256559979 | 0.547665654 |
| 0.256628601 | 0.547772616 |
| 0.256744936 | 0.547892467 |
| 0.256765344 | 0.547892467 |
| 0.256806434 | 0.547892467 |
| 0.256822966 | 0.547892467 |
| 0.256850394 | 0.547892467 |
| 0.256857308 | 0.547892467 |
| 0.257004564 | 0.548094238 |
| 0.257146228 | 0.548340679 |
| 0.257148027 | 0.548340679 |
| 0.257174176 | 0.548371887 |
| 0.257234016 | 0.548446104 |
| 0.257266559 | 0.548446104 |
| 0.257300128 | 0.548493117 |
| 0.257400771 | 0.548615608 |
| 0.257473431 | 0.548715192 |
| 0.257499108 | 0.548720805 |
| 0.257556735 | 0.548760268 |
| 0.257592818 | 0.548760268 |
| 0.257616424 | 0.548774373 |
| 0.257798798 | 0.549040082 |
| 0.257858702 | 0.549086634 |
| 0.257886056 | 0.549086634 |
| 0.258121942 | 0.549335505 |
| 0.25812354 | 0.549335505 |
| 0.258140983 | 0.549335505 |
| 0.258307924 | 0.549510072 |
| 0.258358085 | 0.549559601 |
| 0.258409214 | 0.549559601 |
| 0.258439201 | 0.549582159 |
| 0.258556817 | 0.549677541 |
| 0.258571307 | 0.549677541 |
| 0.258717052 | 0.549829207 |
| 0.258986251 | 0.550178787 |
| 0.258991925 | 0.550178787 |
| 0.259008371 | 0.550189184 |
| 0.259021223 | 0.550191945 |
| 0.259092336 | 0.550293911 |
| 0.259170977 | 0.550378451 |
| 0.259237221 | 0.550405275 |
| 0.259389075 | 0.550544667 |

| | |
|---|---|
| 0.259400456 | 0.550544667 |
| 0.259418469 | 0.550544667 |
| 0.259445445 | 0.550577383 |
| 0.259494323 | 0.550607507 |
| 0.259565646 | 0.550664204 |
| 0.259578853 | 0.550664204 |
| 0.259649639 | 0.550723765 |
| 0.259653183 | 0.550723765 |
| 0.259681146 | 0.550741374 |
| 0.259684612 | 0.550741374 |
| 0.259704533 | 0.55074367 |
| 0.259821614 | 0.55074367 |
| 0.259822703 | 0.55074367 |
| 0.259824459 | 0.55074367 |
| 0.260026548 | 0.551024889 |
| 0.260082843 | 0.551073359 |
| 0.260093457 | 0.551073359 |
| 0.260115878 | 0.551091588 |
| 0.260198256 | 0.551168045 |
| 0.260297826 | 0.551280886 |
| 0.260333814 | 0.551327782 |
| 0.260343121 | 0.551327782 |
| 0.260401803 | 0.551369842 |
| 0.260579156 | 0.551582375 |
| 0.26070305 | 0.551759888 |
| 0.260709357 | 0.551759888 |
| 0.260728591 | 0.551776077 |
| 0.260757813 | 0.551788882 |
| 0.260783643 | 0.551818582 |
| 0.260847266 | 0.551819459 |
| 0.26093149 | 0.551917648 |
| 0.260938599 | 0.551917648 |
| 0.261028571 | 0.552060433 |
| 0.261037177 | 0.552060433 |
| 0.261068842 | 0.552060433 |
| 0.261077305 | 0.552060433 |
| 0.261244046 | 0.552298953 |
| 0.261272669 | 0.552298953 |
| 0.261287987 | 0.552298953 |
| 0.26133462 | 0.552371365 |
| 0.261415451 | 0.552414699 |
| 0.261499549 | 0.552521112 |
| 0.261509877 | 0.552521112 |
| 0.261546251 | 0.552573451 |
| 0.261602936 | 0.552646167 |

| | |
|---|---|
| 0.261612593 | 0.552646167 |
| 0.261624905 | 0.552646167 |
| 0.261704592 | 0.552711835 |
| 0.261719967 | 0.552718218 |
| 0.261785458 | 0.552776748 |
| 0.26184719 | 0.5528659 |
| 0.26193399 | 0.553000138 |
| 0.262106317 | 0.553192305 |
| 0.262144962 | 0.553249351 |
| 0.262258622 | 0.553342108 |
| 0.262310398 | 0.553402319 |
| 0.262339981 | 0.553440213 |
| 0.262368994 | 0.553469083 |
| 0.262441486 | 0.55353176 |
| 0.262460109 | 0.553546525 |
| 0.262500823 | 0.553580664 |
| 0.262548258 | 0.553609858 |
| 0.262578887 | 0.553649931 |
| 0.262873476 | 0.554050318 |
| 0.262918876 | 0.554121484 |
| 0.262955171 | 0.554152625 |
| 0.262956922 | 0.554152625 |
| 0.263027233 | 0.554276273 |
| 0.263054607 | 0.55427988 |
| 0.263063859 | 0.55427988 |
| 0.263207334 | 0.554508589 |
| 0.263286628 | 0.554591345 |
| 0.263333566 | 0.554602802 |
| 0.26337289 | 0.554661094 |
| 0.263652773 | 0.55498058 |
| 0.263948384 | 0.555455537 |
| 0.264035667 | 0.555556636 |
| 0.264043085 | 0.555556636 |
| 0.264095517 | 0.555576043 |
| 0.264098969 | 0.555576043 |
| 0.264157106 | 0.555649258 |
| 0.264206803 | 0.555704709 |
| 0.264262494 | 0.555772755 |
| 0.264318834 | 0.555817612 |
| 0.264402965 | 0.55588058 |
| 0.264430479 | 0.55588058 |
| 0.264472423 | 0.555886923 |
| 0.264521461 | 0.555886923 |
| 0.264557476 | 0.555886923 |
| 0.264564043 | 0.555886923 |

| | |
|---|---|
| 0.264579575 | 0.555886923 |
| 0.264598772 | 0.555886923 |
| 0.264703508 | 0.555988269 |
| 0.265017865 | 0.556524282 |
| 0.265173698 | 0.556730339 |
| 0.265210518 | 0.556732503 |
| 0.265214309 | 0.556732503 |
| 0.265316634 | 0.556858598 |
| 0.265402696 | 0.557014683 |
| 0.265531607 | 0.557162475 |
| 0.265636671 | 0.557333822 |
| 0.265722254 | 0.557403042 |
| 0.265728181 | 0.557403042 |
| 0.265754216 | 0.55740398 |
| 0.265763739 | 0.55740398 |
| 0.265832583 | 0.557474724 |
| 0.265878634 | 0.557546745 |
| 0.265943731 | 0.557658701 |
| 0.266000525 | 0.557738758 |
| 0.266005331 | 0.557738758 |
| 0.266087579 | 0.557837536 |
| 0.266123291 | 0.557863292 |
| 0.266123291 | 0.557863292 |
| 0.266195939 | 0.557982738 |
| 0.266237242 | 0.558028478 |
| 0.266262032 | 0.558055879 |
| 0.266340545 | 0.558195869 |
| 0.266357478 | 0.558206793 |
| 0.26640638 | 0.558235587 |
| 0.266499269 | 0.558383174 |
| 0.266525859 | 0.558384112 |
| 0.266535467 | 0.558384112 |
| 0.266947724 | 0.559025602 |
| 0.267095931 | 0.559250059 |
| 0.267108444 | 0.559250059 |
| 0.267160589 | 0.559250059 |
| 0.267207189 | 0.559282589 |
| 0.267211357 | 0.559282589 |
| 0.26725387 | 0.559322409 |
| 0.267307611 | 0.559338715 |
| 0.26732654 | 0.559338715 |
| 0.267454099 | 0.559502828 |
| 0.267502132 | 0.559571506 |
| 0.267547217 | 0.559623073 |
| 0.267550284 | 0.559623073 |

| | |
|---|---|
| 0.267611625 | 0.559692531 |
| 0.267671795 | 0.55973024 |
| 0.267683458 | 0.55973024 |
| 0.267815753 | 0.559912634 |
| 0.267909417 | 0.559949316 |
| 0.267917883 | 0.559949316 |
| 0.267941851 | 0.559974836 |
| 0.268021204 | 0.560075937 |
| 0.268034654 | 0.560075937 |
| 0.268065847 | 0.560111085 |
| 0.268277556 | 0.560436474 |
| 0.268339248 | 0.560436474 |
| 0.268409933 | 0.560534942 |
| 0.268433189 | 0.560536003 |
| 0.268445749 | 0.560536003 |
| 0.268694174 | 0.560867833 |
| 0.268769468 | 0.56092897 |
| 0.268804966 | 0.560966354 |
| 0.268818456 | 0.560969926 |
| 0.268861493 | 0.560987797 |
| 0.268872587 | 0.560987797 |
| 0.268966581 | 0.561106935 |
| 0.269088076 | 0.561286637 |
| 0.269361294 | 0.561659749 |
| 0.269419173 | 0.561709004 |
| 0.269420297 | 0.561709004 |
| 0.26953576 | 0.56181255 |
| 0.269572513 | 0.561847824 |
| 0.269581256 | 0.561847824 |
| 0.269633934 | 0.561905591 |
| 0.269644367 | 0.561905591 |
| 0.269702244 | 0.561952435 |
| 0.269874614 | 0.562237795 |
| 0.269898624 | 0.562263221 |
| 0.26992123 | 0.562285721 |
| 0.269937386 | 0.562294781 |
| 0.269963695 | 0.562308126 |
| 0.269967405 | 0.562308126 |
| 0.270033224 | 0.562384051 |
| 0.27005109 | 0.562384051 |
| 0.270154725 | 0.562545254 |
| 0.270175744 | 0.562545254 |
| 0.270272853 | 0.562698251 |
| 0.270391932 | 0.562847751 |
| 0.270439697 | 0.562872689 |

| | |
|---|---|
| 0.270503107 | 0.562872689 |
| 0.2705399 | 0.562872689 |
| 0.270615294 | 0.56288343 |
| 0.270713318 | 0.562953832 |
| 0.270797469 | 0.563052084 |
| 0.270835214 | 0.563081402 |
| 0.270885124 | 0.563111422 |
| 0.270926705 | 0.563118404 |
| 0.2709476 | 0.563118404 |
| 0.271116868 | 0.563415462 |
| 0.271143723 | 0.563415462 |
| 0.271208829 | 0.563415462 |
| 0.271369879 | 0.563532377 |
| 0.271387393 | 0.563532377 |
| 0.271390728 | 0.563532377 |
| 0.271421253 | 0.563532377 |
| 0.271454379 | 0.56355685 |
| 0.271580816 | 0.563711572 |
| 0.271588085 | 0.563711572 |
| 0.271630892 | 0.563721202 |
| 0.271634028 | 0.563721202 |
| 0.271680136 | 0.563735823 |
| 0.271714241 | 0.563737292 |
| 0.271783512 | 0.563797785 |
| 0.271848806 | 0.563800397 |
| 0.271851092 | 0.563800397 |
| 0.27186085 | 0.563800397 |
| 0.271900612 | 0.563800397 |
| 0.272070462 | 0.563906084 |
| 0.272098222 | 0.563906084 |
| 0.272228474 | 0.56404855 |
| 0.272279592 | 0.564114327 |
| 0.272397096 | 0.564308678 |
| 0.272486029 | 0.564429045 |
| 0.272516419 | 0.564457662 |
| 0.272546592 | 0.564478813 |
| 0.272705741 | 0.564633996 |
| 0.272797293 | 0.564646531 |
| 0.272834632 | 0.564666422 |
| 0.272842471 | 0.564666422 |
| 0.272991792 | 0.564926357 |
| 0.273261837 | 0.565187556 |
| 0.273263708 | 0.565187556 |
| 0.273320896 | 0.565263551 |
| 0.273418106 | 0.565389256 |

| | |
|---|---|
| 0.273519443 | 0.565453186 |
| 0.273545555 | 0.565482623 |
| 0.273595993 | 0.56551134 |
| 0.273610699 | 0.56551134 |
| 0.273640596 | 0.56551134 |
| 0.273766856 | 0.565690725 |
| 0.273844811 | 0.565701553 |
| 0.273876212 | 0.565701553 |
| 0.273902821 | 0.565705535 |
| 0.273917079 | 0.565710452 |
| 0.274011922 | 0.565830028 |
| 0.2740225 | 0.565830028 |
| 0.274043325 | 0.565837761 |
| 0.274094774 | 0.565868522 |
| 0.27410774 | 0.565868522 |
| 0.27413802 | 0.565896822 |
| 0.274218664 | 0.565989699 |
| 0.274401575 | 0.566258075 |
| 0.274408138 | 0.566258075 |
| 0.274452578 | 0.566300709 |
| 0.27467259 | 0.566607392 |
| 0.274716466 | 0.566656958 |
| 0.274732312 | 0.566656958 |
| 0.274840178 | 0.566756718 |
| 0.274869838 | 0.566792776 |
| 0.274891279 | 0.566792776 |
| 0.274954636 | 0.566852998 |
| 0.274958279 | 0.566852998 |
| 0.274997699 | 0.566890409 |
| 0.275022298 | 0.566890409 |
| 0.275029168 | 0.566890409 |
| 0.275035939 | 0.566890409 |
| 0.275092048 | 0.566945009 |
| 0.275110045 | 0.566945009 |
| 0.275170922 | 0.567045928 |
| 0.275184182 | 0.567048719 |
| 0.275242739 | 0.567112585 |
| 0.275302078 | 0.567155133 |
| 0.275304539 | 0.567155133 |
| 0.275487987 | 0.567184675 |
| 0.275502109 | 0.567184675 |
| 0.275540031 | 0.567184675 |
| 0.275576706 | 0.567184675 |
| 0.275611377 | 0.567184675 |
| 0.275638361 | 0.567184675 |

| | |
|---|---|
| 0.275680868 | 0.567184675 |
| 0.275803254 | 0.567293526 |
| 0.275849389 | 0.567341073 |
| 0.275869291 | 0.567355848 |
| 0.275982393 | 0.567426748 |
| 0.275997076 | 0.567426748 |
| 0.275998674 | 0.567426748 |
| 0.276023013 | 0.567451462 |
| 0.276058681 | 0.567454462 |
| 0.276090714 | 0.567454462 |
| 0.27626118 | 0.567696044 |
| 0.276411914 | 0.567858786 |
| 0.276452897 | 0.567869495 |
| 0.276570185 | 0.567903364 |
| 0.276575359 | 0.567903364 |
| 0.276594233 | 0.567914287 |
| 0.276606648 | 0.567914287 |
| 0.276770809 | 0.568130477 |
| 0.27683507 | 0.568143264 |
| 0.276836683 | 0.568143264 |
| 0.276889811 | 0.568203327 |
| 0.276942449 | 0.568262377 |
| 0.276967582 | 0.568289465 |
| 0.277016134 | 0.568357468 |
| 0.277024592 | 0.568357468 |
| 0.277062264 | 0.568363895 |
| 0.277063126 | 0.568363895 |
| 0.27714806 | 0.56851282 |
| 0.277284378 | 0.568686502 |
| 0.277292432 | 0.568686502 |
| 0.277341373 | 0.568762382 |
| 0.27735611 | 0.56876261 |
| 0.277451014 | 0.568864758 |
| 0.277500702 | 0.568896318 |
| 0.277614345 | 0.569052653 |
| 0.277664342 | 0.56910615 |
| 0.277732738 | 0.569221839 |
| 0.277806761 | 0.569349052 |
| 0.277961493 | 0.56956813 |
| 0.277980324 | 0.56958221 |
| 0.27802404 | 0.569640197 |
| 0.278032545 | 0.569640197 |
| 0.278061214 | 0.56965298 |
| 0.278071993 | 0.56965298 |
| 0.278130853 | 0.569719072 |

| | |
|---|---|
| 0.278192759 | 0.569772362 |
| 0.278264591 | 0.569870471 |
| 0.278380828 | 0.570034983 |
| 0.278412594 | 0.570051013 |
| 0.278496643 | 0.570174079 |
| 0.278516233 | 0.570179218 |
| 0.278523096 | 0.570179218 |
| 0.278610001 | 0.570260331 |
| 0.278793892 | 0.570479879 |
| 0.278866467 | 0.570531473 |
| 0.278916013 | 0.570531473 |
| 0.278917587 | 0.570531473 |
| 0.278920483 | 0.570531473 |
| 0.278922771 | 0.570531473 |
| 0.27895875 | 0.570580561 |
| 0.278972618 | 0.570584422 |
| 0.279008355 | 0.570609854 |
| 0.279089545 | 0.570652031 |
| 0.279123621 | 0.570697206 |
| 0.279181627 | 0.570781906 |
| 0.279291688 | 0.57079246 |
| 0.279314025 | 0.57079246 |
| 0.279336868 | 0.570814649 |
| 0.279382138 | 0.570828742 |
| 0.279578604 | 0.571112596 |
| 0.279892072 | 0.571517764 |
| 0.279894505 | 0.571517764 |
| 0.279904966 | 0.571517764 |
| 0.27996908 | 0.571542536 |
| 0.280263971 | 0.571850405 |
| 0.280345551 | 0.571914365 |
| 0.280371795 | 0.571914365 |
| 0.280403032 | 0.571914365 |
| 0.280403392 | 0.571914365 |
| 0.280494105 | 0.572050389 |
| 0.280661183 | 0.572268608 |
| 0.280673252 | 0.572268717 |
| 0.280773919 | 0.572332257 |
| 0.280800095 | 0.572332257 |
| 0.280806602 | 0.572332257 |
| 0.280820754 | 0.572332257 |
| 0.280848619 | 0.572332257 |
| 0.280927306 | 0.572443624 |
| 0.281042568 | 0.572556009 |
| 0.281075735 | 0.572557341 |

| | |
|---|---|
| 0.281079287 | 0.572557341 |
| 0.281149059 | 0.572650482 |
| 0.281209827 | 0.572700779 |
| 0.281308925 | 0.572810954 |
| 0.28132406 | 0.572810954 |
| 0.28145458 | 0.573001202 |
| 0.281477652 | 0.573001202 |
| 0.281693917 | 0.573343428 |
| 0.281781294 | 0.573447749 |
| 0.281872233 | 0.5735838 |
| 0.281912363 | 0.57361711 |
| 0.28191915 | 0.57361711 |
| 0.281952337 | 0.57361711 |
| 0.281975879 | 0.57361711 |
| 0.281976532 | 0.57361711 |
| 0.281988402 | 0.57361711 |
| 0.282045293 | 0.57361711 |
| 0.282052849 | 0.57361711 |
| 0.28223225 | 0.573784948 |
| 0.282264754 | 0.573809879 |
| 0.282605724 | 0.574203256 |
| 0.282610673 | 0.574203256 |
| 0.282642332 | 0.574208666 |
| 0.28265862 | 0.574208666 |
| 0.282661561 | 0.574208666 |
| 0.282782561 | 0.574331985 |
| 0.282818465 | 0.574369891 |
| 0.282855326 | 0.574369891 |
| 0.282858267 | 0.574369891 |
| 0.282897366 | 0.574369891 |
| 0.282926029 | 0.574369891 |
| 0.282933128 | 0.574369891 |
| 0.282952377 | 0.574369891 |
| 0.282954217 | 0.574369891 |
| 0.282980239 | 0.574390549 |
| 0.283018784 | 0.574444305 |
| 0.283079918 | 0.574519421 |
| 0.283234301 | 0.574622388 |
| 0.283248237 | 0.574622388 |
| 0.283276762 | 0.574622388 |
| 0.283289239 | 0.574622388 |
| 0.283331368 | 0.574622388 |
| 0.283336743 | 0.574622388 |
| 0.283418978 | 0.57466881 |
| 0.283534102 | 0.574800091 |

| | |
|---|---|
| 0.283534832 | 0.574800091 |
| 0.283546016 | 0.574800091 |
| 0.283764488 | 0.575002181 |
| 0.283782044 | 0.575013292 |
| 0.284017946 | 0.575229436 |
| 0.284273204 | 0.575494342 |
| 0.284467168 | 0.575694855 |
| 0.284474424 | 0.575694855 |
| 0.284561615 | 0.57579855 |
| 0.284580639 | 0.57579855 |
| 0.28461287 | 0.575836739 |
| 0.284623694 | 0.575836739 |
| 0.284692531 | 0.575922663 |
| 0.284848806 | 0.57610697 |
| 0.284854033 | 0.57610697 |
| 0.28491365 | 0.576139124 |
| 0.284967813 | 0.576214743 |
| 0.285097986 | 0.576382597 |
| 0.285101069 | 0.576382597 |
| 0.285122986 | 0.576382597 |
| 0.285167926 | 0.576448054 |
| 0.285234386 | 0.576509001 |
| 0.285258115 | 0.576532494 |
| 0.285305023 | 0.576602833 |
| 0.285336624 | 0.576620524 |
| 0.28542783 | 0.57675313 |
| 0.285446049 | 0.576765473 |
| 0.285592781 | 0.576908043 |
| 0.285600191 | 0.576908043 |
| 0.285601399 | 0.576908043 |
| 0.285650505 | 0.576958296 |
| 0.285734601 | 0.576995332 |
| 0.285741531 | 0.576995332 |
| 0.285758967 | 0.577006076 |
| 0.285874774 | 0.577134286 |
| 0.285883051 | 0.577134286 |
| 0.285907674 | 0.577159531 |
| 0.286034754 | 0.577318185 |
| 0.286187958 | 0.577435246 |
| 0.28618932 | 0.577435246 |
| 0.286213993 | 0.577435246 |
| 0.286259722 | 0.577472107 |
| 0.286416073 | 0.577644156 |
| 0.286426699 | 0.577644156 |
| 0.286508499 | 0.577784658 |

| | |
|---|---|
| 0.286582713 | 0.577909851 |
| 0.286605282 | 0.577930893 |
| 0.286667943 | 0.577983834 |
| 0.286855342 | 0.578239277 |
| 0.286949854 | 0.578331882 |
| 0.28700186 | 0.578371569 |
| 0.287006399 | 0.578371569 |
| 0.287006711 | 0.578371569 |
| 0.287025233 | 0.578371569 |
| 0.287313905 | 0.578566773 |
| 0.287319693 | 0.578566773 |
| 0.287331661 | 0.578566773 |
| 0.28734396 | 0.578566773 |
| 0.287350251 | 0.578566773 |
| 0.287487273 | 0.578655918 |
| 0.287509662 | 0.578676527 |
| 0.287689054 | 0.578933735 |
| 0.28773677 | 0.578986825 |
| 0.287897545 | 0.579187985 |
| 0.288020303 | 0.579361531 |
| 0.2881783 | 0.579630386 |
| 0.288231431 | 0.57966381 |
| 0.288352299 | 0.579686587 |
| 0.288646436 | 0.580078137 |
| 0.288684801 | 0.580078137 |
| 0.288731754 | 0.580078137 |
| 0.288783964 | 0.58013809 |
| 0.288851024 | 0.580219395 |
| 0.288872035 | 0.580219395 |
| 0.28900339 | 0.580432 |
| 0.289090423 | 0.580484404 |
| 0.289136085 | 0.580527142 |
| 0.289164517 | 0.580533647 |
| 0.289188082 | 0.580533647 |
| 0.289241109 | 0.580556212 |
| 0.289346041 | 0.580581558 |
| 0.289378209 | 0.580617529 |
| 0.289401554 | 0.580619564 |
| 0.28953274 | 0.580742013 |
| 0.289543769 | 0.580742013 |
| 0.289561614 | 0.580742013 |
| 0.289621103 | 0.580742013 |
| 0.289634005 | 0.580743433 |
| 0.289700522 | 0.5808279 |
| 0.289729671 | 0.580843941 |

| | |
|---|---|
| 0.28974511 | 0.580843941 |
| 0.289897861 | 0.580945993 |
| 0.289972246 | 0.580945993 |
| 0.28998926 | 0.580945993 |
| 0.290024862 | 0.58096459 |
| 0.290216084 | 0.581274293 |
| 0.290302548 | 0.581325238 |
| 0.290403994 | 0.581476628 |
| 0.290414776 | 0.581476628 |
| 0.290519115 | 0.581587738 |
| 0.290581709 | 0.581615258 |
| 0.290790183 | 0.581880538 |
| 0.290808409 | 0.581880538 |
| 0.290882156 | 0.581947594 |
| 0.290942177 | 0.582018778 |
| 0.291016359 | 0.582067774 |
| 0.291099716 | 0.582187208 |
| 0.291152531 | 0.582243937 |
| 0.291230637 | 0.58232678 |
| 0.291327105 | 0.58238068 |
| 0.291376346 | 0.582382369 |
| 0.291376486 | 0.582382369 |
| 0.291380718 | 0.582382369 |
| 0.291468883 | 0.582433343 |
| 0.291542917 | 0.582510991 |
| 0.291603629 | 0.582547236 |
| 0.29167921 | 0.582547236 |
| 0.291688037 | 0.582547236 |
| 0.291706736 | 0.582547236 |
| 0.291779723 | 0.582602217 |
| 0.291791335 | 0.582602217 |
| 0.291803072 | 0.582602217 |
| 0.291803507 | 0.582602217 |
| 0.291893215 | 0.582697551 |
| 0.291963685 | 0.582813799 |
| 0.292065563 | 0.582968298 |
| 0.292279179 | 0.58332134 |
| 0.292358965 | 0.583382791 |
| 0.292388156 | 0.583402798 |
| 0.292585636 | 0.58363394 |
| 0.292593908 | 0.58363394 |
| 0.292714389 | 0.583676292 |
| 0.292753362 | 0.583698691 |
| 0.292755611 | 0.583698691 |
| 0.292820073 | 0.583764822 |

| | |
|---|---|
| 0.292934202 | 0.583868284 |
| 0.293007963 | 0.583943855 |
| 0.293039526 | 0.583957887 |
| 0.293157296 | 0.584097134 |
| 0.293216769 | 0.584097134 |
| 0.293319875 | 0.584199523 |
| 0.293369352 | 0.584248468 |
| 0.293481047 | 0.584446473 |
| 0.293622871 | 0.584664571 |
| 0.293625886 | 0.584664571 |
| 0.293793273 | 0.584872606 |
| 0.293944251 | 0.585060436 |
| 0.293975694 | 0.585064576 |
| 0.294028724 | 0.585145664 |
| 0.29406271 | 0.585164397 |
| 0.294178608 | 0.585282795 |
| 0.294228578 | 0.585308844 |
| 0.294304756 | 0.585415524 |
| 0.294341355 | 0.585429772 |
| 0.294402647 | 0.585498366 |
| 0.294488925 | 0.585587812 |
| 0.294515931 | 0.585587812 |
| 0.294534629 | 0.585587812 |
| 0.294544208 | 0.585587812 |
| 0.294545985 | 0.585587812 |
| 0.294647454 | 0.585740641 |
| 0.294784499 | 0.585864884 |
| 0.295374383 | 0.586524733 |
| 0.295392798 | 0.586536846 |
| 0.295472524 | 0.586608212 |
| 0.295492246 | 0.586608212 |
| 0.295508295 | 0.586608212 |
| 0.295514957 | 0.586608212 |
| 0.295596139 | 0.586696003 |
| 0.295803197 | 0.586963721 |
| 0.295817293 | 0.586963721 |
| 0.295945882 | 0.587135048 |
| 0.295952949 | 0.587135048 |
| 0.296056777 | 0.587267643 |
| 0.296133385 | 0.587335846 |
| 0.296174023 | 0.58735344 |
| 0.296325701 | 0.587513429 |
| 0.296500405 | 0.587658126 |
| 0.296628458 | 0.587797332 |
| 0.296632349 | 0.587797332 |

| | |
|---|---|
| 0.296668467 | 0.587842546 |
| 0.296762745 | 0.587900372 |
| 0.296805462 | 0.587900372 |
| 0.29682013 | 0.587900372 |
| 0.296838515 | 0.587912336 |
| 0.296891569 | 0.587923411 |
| 0.296918172 | 0.587923411 |
| 0.296949177 | 0.587950166 |
| 0.29701334 | 0.588001395 |
| 0.297027726 | 0.588001395 |
| 0.29707158 | 0.588031591 |
| 0.297307139 | 0.588360789 |
| 0.29735985 | 0.588406563 |
| 0.297426638 | 0.5884666 |
| 0.297446644 | 0.588479384 |
| 0.297748825 | 0.588743103 |
| 0.297755388 | 0.588743103 |
| 0.297775865 | 0.588743103 |
| 0.297778662 | 0.588743103 |
| 0.297837317 | 0.588788723 |
| 0.297866106 | 0.588812246 |
| 0.297938218 | 0.588890418 |
| 0.297958263 | 0.588902723 |
| 0.297969173 | 0.588902723 |
| 0.298031596 | 0.58892834 |
| 0.298083371 | 0.588951432 |
| 0.298210324 | 0.589030684 |
| 0.298237961 | 0.589042853 |
| 0.298283923 | 0.589063858 |
| 0.298364169 | 0.589160847 |
| 0.298391431 | 0.589160847 |
| 0.298398202 | 0.589160847 |
| 0.298401355 | 0.589160847 |
| 0.298518861 | 0.589304392 |
| 0.298568404 | 0.589361162 |
| 0.298590255 | 0.589361162 |
| 0.298751125 | 0.589438153 |
| 0.298755955 | 0.589438153 |
| 0.298767657 | 0.589438153 |
| 0.29877226 | 0.589438153 |
| 0.298819497 | 0.589450697 |
| 0.298978092 | 0.5896 |
| 0.299013582 | 0.589602126 |
| 0.299039621 | 0.589616679 |
| 0.299051164 | 0.589616679 |

| | |
|---|---|
| 0.299102351 | 0.589674132 |
| 0.299168154 | 0.589679376 |
| 0.29927778 | 0.589805081 |
| 0.299280597 | 0.589805081 |
| 0.299292611 | 0.589805081 |
| 0.299373038 | 0.589865947 |
| 0.299550753 | 0.59013777 |
| 0.299560558 | 0.59013777 |
| 0.299620007 | 0.590181648 |
| 0.299801505 | 0.590453303 |
| 0.299872463 | 0.590459144 |
| 0.2998979 | 0.590484818 |
| 0.300063522 | 0.590713238 |
| 0.300108998 | 0.59072951 |
| 0.30012149 | 0.590729687 |
| 0.300164326 | 0.590789584 |
| 0.300231647 | 0.59087325 |
| 0.300406289 | 0.591083377 |
| 0.30041264 | 0.591083377 |
| 0.300538251 | 0.591207934 |
| 0.300639613 | 0.591260802 |
| 0.300817348 | 0.59139801 |
| 0.300926458 | 0.591531817 |
| 0.301004206 | 0.591633734 |
| 0.301015572 | 0.591633734 |
| 0.301107912 | 0.591766384 |
| 0.301122818 | 0.591771259 |
| 0.301346906 | 0.591998938 |
| 0.301350542 | 0.591998938 |
| 0.301386155 | 0.592020062 |
| 0.301426496 | 0.592027911 |
| 0.301452304 | 0.592027911 |
| 0.301481712 | 0.592061252 |
| 0.301631593 | 0.592209081 |
| 0.301664661 | 0.592249591 |
| 0.301714716 | 0.592267878 |
| 0.30177667 | 0.592267878 |
| 0.301817781 | 0.592267878 |
| 0.301830725 | 0.592267878 |
| 0.301861628 | 0.592267878 |
| 0.30202036 | 0.592532694 |
| 0.302133067 | 0.592680571 |
| 0.302201518 | 0.592790432 |
| 0.302233391 | 0.592828537 |
| 0.302343824 | 0.592922758 |

| | |
|---|---|
| 0.302422744 | 0.592974345 |
| 0.30243223 | 0.592974345 |
| 0.302536538 | 0.59309882 |
| 0.302557766 | 0.59309882 |
| 0.302577547 | 0.59309882 |
| 0.302582886 | 0.59309882 |
| 0.302847502 | 0.593488274 |
| 0.302868803 | 0.593488274 |
| 0.302966746 | 0.593509264 |
| 0.303014337 | 0.593578079 |
| 0.303045783 | 0.593615263 |
| 0.303165462 | 0.593727599 |
| 0.303327209 | 0.59394639 |
| 0.303330863 | 0.59394639 |
| 0.30338319 | 0.593952476 |
| 0.303404518 | 0.593952476 |
| 0.303468453 | 0.593993318 |
| 0.303662025 | 0.594235816 |
| 0.303917309 | 0.594539951 |
| 0.304049037 | 0.594675516 |
| 0.304155009 | 0.594809502 |
| 0.304179992 | 0.594814231 |
| 0.304206511 | 0.594831626 |
| 0.304216279 | 0.594831626 |
| 0.304270559 | 0.594888913 |
| 0.304316332 | 0.594905141 |
| 0.304391204 | 0.595027084 |
| 0.304449414 | 0.595116447 |
| 0.30451733 | 0.595144549 |
| 0.304556549 | 0.595144549 |
| 0.304576864 | 0.595144549 |
| 0.304688919 | 0.595218177 |
| 0.304906892 | 0.595380221 |
| 0.304924446 | 0.595385233 |
| 0.304968319 | 0.595422076 |
| 0.305098985 | 0.595522366 |
| 0.305196925 | 0.595551029 |
| 0.305256188 | 0.595610938 |
| 0.305364078 | 0.595662513 |
| 0.305436775 | 0.595726168 |
| 0.3056566 | 0.59598407 |
| 0.305674015 | 0.595984602 |
| 0.305693506 | 0.595984602 |
| 0.305757553 | 0.596012378 |
| 0.305970523 | 0.596254423 |

| | |
|---|---|
| 0.306014418 | 0.596265402 |
| 0.306018211 | 0.596265402 |
| 0.306121844 | 0.596265402 |
| 0.306122655 | 0.596265402 |
| 0.306134407 | 0.596265402 |
| 0.306157532 | 0.596270515 |
| 0.306166575 | 0.596270515 |
| 0.306179704 | 0.596271703 |
| 0.30621799 | 0.596297499 |
| 0.306250093 | 0.59632222 |
| 0.306369688 | 0.596470961 |
| 0.306434569 | 0.596548506 |
| 0.306467366 | 0.596587968 |
| 0.306516459 | 0.596659148 |
| 0.306618592 | 0.596760398 |
| 0.306685985 | 0.596778812 |
| 0.306703235 | 0.596778812 |
| 0.306801741 | 0.596946096 |
| 0.306876751 | 0.596954092 |
| 0.306913724 | 0.596954092 |
| 0.306993671 | 0.597002474 |
| 0.307021492 | 0.597007822 |
| 0.307193131 | 0.597268422 |
| 0.307335088 | 0.597441857 |
| 0.307354472 | 0.597441857 |
| 0.307386591 | 0.597449445 |
| 0.307461359 | 0.597548173 |
| 0.307539967 | 0.597649992 |
| 0.307608325 | 0.597685284 |
| 0.307648552 | 0.597739061 |
| 0.307663585 | 0.597743884 |
| 0.307699256 | 0.597779717 |
| 0.307707131 | 0.597779717 |
| 0.30783299 | 0.597869899 |
| 0.307841425 | 0.597869899 |
| 0.307872541 | 0.597899366 |
| 0.307990541 | 0.598037568 |
| 0.308091046 | 0.59810826 |
| 0.308120762 | 0.598119729 |
| 0.30817909 | 0.59817654 |
| 0.308218748 | 0.59817654 |
| 0.308250152 | 0.59817654 |
| 0.308250246 | 0.59817654 |
| 0.308303566 | 0.598201088 |
| 0.308317207 | 0.598201088 |

| | |
|---|---|
| 0.308320704 | 0.598201088 |
| 0.308383353 | 0.598263705 |
| 0.308484063 | 0.59838596 |
| 0.30849943 | 0.59838939 |
| 0.308514057 | 0.59838939 |
| 0.308523524 | 0.59838939 |
| 0.308634893 | 0.598474 |
| 0.308729043 | 0.598582474 |
| 0.308940986 | 0.598833272 |
| 0.309009831 | 0.598926494 |
| 0.309094055 | 0.59900808 |
| 0.309123727 | 0.599025431 |
| 0.309242099 | 0.599124332 |
| 0.309258956 | 0.599132619 |
| 0.309359027 | 0.59925336 |
| 0.309414597 | 0.599263511 |
| 0.309559041 | 0.59944576 |
| 0.309657643 | 0.599612318 |
| 0.309710958 | 0.599647555 |
| 0.30976814 | 0.599683561 |
| 0.30988787 | 0.599787631 |
| 0.309973762 | 0.599837266 |
| 0.310048943 | 0.599955348 |
| 0.310164729 | 0.60004016 |
| 0.310291127 | 0.600165846 |
| 0.310369068 | 0.60019506 |
| 0.310383928 | 0.60019506 |
| 0.310387056 | 0.60019506 |
| 0.310423035 | 0.600240261 |
| 0.310567498 | 0.600446465 |
| 0.310767837 | 0.600563616 |
| 0.310792827 | 0.600563616 |
| 0.310804628 | 0.600563616 |
| 0.310883712 | 0.600667689 |
| 0.311025826 | 0.600892963 |
| 0.311050925 | 0.600892963 |
| 0.311078949 | 0.600892963 |
| 0.311155454 | 0.600933142 |
| 0.311198946 | 0.600984194 |
| 0.31146108 | 0.601256565 |
| 0.311470218 | 0.601256565 |
| 0.311861369 | 0.601726682 |
| 0.311931806 | 0.601776759 |
| 0.311983143 | 0.601807728 |
| 0.312107728 | 0.601886306 |

| | |
|---|---|
| 0.31211991 | 0.601886306 |
| 0.312121023 | 0.601886306 |
| 0.312167523 | 0.601892558 |
| 0.312177046 | 0.601892558 |
| 0.312181538 | 0.601892558 |
| 0.312200091 | 0.601892558 |
| 0.312320998 | 0.602031433 |
| 0.312434865 | 0.602150183 |
| 0.31252073 | 0.602291297 |
| 0.312731256 | 0.602575109 |
| 0.312788491 | 0.602612251 |
| 0.312890731 | 0.602687328 |
| 0.312923458 | 0.602705057 |
| 0.312925245 | 0.602705057 |
| 0.313474191 | 0.603468636 |
| 0.313479517 | 0.603468636 |
| 0.313486416 | 0.603468636 |
| 0.313560421 | 0.603516803 |
| 0.313632702 | 0.603603859 |
| 0.313726828 | 0.603671642 |
| 0.313730781 | 0.603671642 |
| 0.313757412 | 0.603671642 |
| 0.313760518 | 0.603671642 |
| 0.313769322 | 0.603671642 |
| 0.313824467 | 0.603693255 |
| 0.31385956 | 0.603712508 |
| 0.313934127 | 0.603811237 |
| 0.314067954 | 0.60398026 |
| 0.314069228 | 0.60398026 |
| 0.314173442 | 0.604156277 |
| 0.314226479 | 0.604188378 |
| 0.314261749 | 0.604204115 |
| 0.31439468 | 0.604313319 |
| 0.314555266 | 0.604548794 |
| 0.314622813 | 0.60461206 |
| 0.314627021 | 0.60461206 |
| 0.314631848 | 0.60461206 |
| 0.314845013 | 0.604866777 |
| 0.314846447 | 0.604866777 |
| 0.314945908 | 0.605006606 |
| 0.314974886 | 0.605013467 |
| 0.315020954 | 0.605065788 |
| 0.315027534 | 0.605065788 |
| 0.315069132 | 0.605096882 |
| 0.315085456 | 0.605103831 |

| | |
|---|---|
| 0.315156455 | 0.605206222 |
| 0.315330373 | 0.605354495 |
| 0.315559442 | 0.605672184 |
| 0.315618654 | 0.605723591 |
| 0.31562438 | 0.605723591 |
| 0.315681206 | 0.605735015 |
| 0.315765733 | 0.605821881 |
| 0.315777357 | 0.605821881 |
| 0.315915524 | 0.606062542 |
| 0.315977681 | 0.606157369 |
| 0.316104264 | 0.606305283 |
| 0.316252772 | 0.606489664 |
| 0.316302611 | 0.606557736 |
| 0.316367943 | 0.606557736 |
| 0.316388141 | 0.606557736 |
| 0.316390154 | 0.606557736 |
| 0.316439888 | 0.606628666 |
| 0.316460707 | 0.606644157 |
| 0.31650186 | 0.606698628 |
| 0.316551385 | 0.606758775 |
| 0.316823152 | 0.607192317 |
| 0.316900166 | 0.607279862 |
| 0.316907084 | 0.607279862 |
| 0.317123044 | 0.607586285 |
| 0.317127954 | 0.607586285 |
| 0.317156879 | 0.607603154 |
| 0.317165096 | 0.607603154 |
| 0.317199192 | 0.607644033 |
| 0.31722358 | 0.607649135 |
| 0.317267812 | 0.607702154 |
| 0.31739531 | 0.607897472 |
| 0.317579149 | 0.608176201 |
| 0.317639226 | 0.608217884 |
| 0.317671052 | 0.60825437 |
| 0.31781846 | 0.608487691 |
| 0.317850822 | 0.608525188 |
| 0.317872491 | 0.608542212 |
| 0.317900343 | 0.608571069 |
| 0.31799856 | 0.608661232 |
| 0.318095562 | 0.60875319 |
| 0.318101125 | 0.60875319 |
| 0.31810525 | 0.60875319 |
| 0.318122576 | 0.60875319 |
| 0.318123294 | 0.60875319 |
| 0.318202873 | 0.60877044 |

| | |
|---|---|
| 0.318206685 | 0.60877044 |
| 0.318250791 | 0.608777028 |
| 0.318280241 | 0.60878446 |
| 0.318314713 | 0.608825945 |
| 0.318362142 | 0.608867756 |
| 0.318443017 | 0.608899503 |
| 0.3184525 | 0.608899503 |
| 0.318452975 | 0.608899503 |
| 0.318533059 | 0.608901224 |
| 0.318663923 | 0.609053601 |
| 0.318775863 | 0.609128268 |
| 0.318827712 | 0.609147658 |
| 0.318828245 | 0.609147658 |
| 0.318862095 | 0.609187894 |
| 0.318929234 | 0.609267284 |
| 0.319097731 | 0.609477203 |
| 0.319103104 | 0.609477203 |
| 0.319141062 | 0.609525257 |
| 0.319378215 | 0.609733686 |
| 0.319426297 | 0.609779462 |
| 0.319576772 | 0.609990499 |
| 0.319611608 | 0.610006272 |
| 0.319635547 | 0.610006272 |
| 0.319720264 | 0.610117679 |
| 0.319755932 | 0.610125831 |
| 0.319762968 | 0.610125831 |
| 0.319781348 | 0.610136458 |
| 0.319938526 | 0.610195297 |
| 0.319940306 | 0.610195297 |
| 0.320000726 | 0.610261655 |
| 0.320101159 | 0.61039261 |
| 0.320307488 | 0.610675502 |
| 0.320340775 | 0.610714519 |
| 0.320458181 | 0.610913893 |
| 0.320475982 | 0.610923375 |
| 0.32050021 | 0.610945106 |
| 0.320656055 | 0.611094597 |
| 0.320657762 | 0.611094597 |
| 0.320686452 | 0.611104456 |
| 0.320699666 | 0.611105185 |
| 0.320914025 | 0.611317986 |
| 0.320946635 | 0.611331662 |
| 0.321067901 | 0.611399804 |
| 0.321093271 | 0.611399804 |
| 0.321117797 | 0.611399804 |

| | |
|---|---|
| 0.321133143 | 0.611399804 |
| 0.321136698 | 0.611399804 |
| 0.321221242 | 0.611456495 |
| 0.321272784 | 0.611512224 |
| 0.321317681 | 0.611548801 |
| 0.321365485 | 0.611568573 |
| 0.3213932 | 0.611594771 |
| 0.321415612 | 0.611612983 |
| 0.321487076 | 0.611675655 |
| 0.321617748 | 0.61182651 |
| 0.32165733 | 0.611877367 |
| 0.321780451 | 0.61196934 |
| 0.322028533 | 0.612308656 |
| 0.322038329 | 0.612308656 |
| 0.322290482 | 0.612567994 |
| 0.322306515 | 0.612574022 |
| 0.322332704 | 0.61258243 |
| 0.322336664 | 0.61258243 |
| 0.32236925 | 0.612591976 |
| 0.322380274 | 0.612591976 |
| 0.322475232 | 0.612683225 |
| 0.322477833 | 0.612683225 |
| 0.322491199 | 0.612683225 |
| 0.322503496 | 0.612683225 |
| 0.322518466 | 0.612683225 |
| 0.322531207 | 0.612683225 |
| 0.322565933 | 0.612700315 |
| 0.322604089 | 0.612714337 |
| 0.32261191 | 0.612714337 |
| 0.322673273 | 0.612806443 |
| 0.322749688 | 0.612863079 |
| 0.32276973 | 0.612863079 |
| 0.322818985 | 0.612885538 |
| 0.322923822 | 0.613010176 |
| 0.322930054 | 0.613010176 |
| 0.322935002 | 0.613010176 |
| 0.322964826 | 0.613024861 |
| 0.322990477 | 0.613024861 |
| 0.322996959 | 0.613024861 |
| 0.322998832 | 0.613024861 |
| 0.323075714 | 0.613130659 |
| 0.323210557 | 0.613288813 |
| 0.323235709 | 0.613312103 |
| 0.323293555 | 0.613348555 |
| 0.323324962 | 0.613359274 |

| | |
|---|---|
| 0.323436384 | 0.61343339 |
| 0.32344131 | 0.61343339 |
| 0.323581013 | 0.613644041 |
| 0.323632705 | 0.613649767 |
| 0.32375536 | 0.613809025 |
| 0.323789129 | 0.613814355 |
| 0.3238102 | 0.613814355 |
| 0.323835499 | 0.613814355 |
| 0.323896075 | 0.613880313 |
| 0.323921817 | 0.61389206 |
| 0.324070394 | 0.613990798 |
| 0.324504719 | 0.614593639 |
| 0.324532902 | 0.614604017 |
| 0.324559745 | 0.614604017 |
| 0.324562633 | 0.614604017 |
| 0.324595687 | 0.614604017 |
| 0.324683199 | 0.614687233 |
| 0.324705519 | 0.614705055 |
| 0.324758963 | 0.614757357 |
| 0.324849001 | 0.614903203 |
| 0.32486183 | 0.614903203 |
| 0.324932739 | 0.614975738 |
| 0.32495526 | 0.614975738 |
| 0.324987366 | 0.614975738 |
| 0.325043204 | 0.615050961 |
| 0.325066863 | 0.615062272 |
| 0.32514178 | 0.61513973 |
| 0.325203596 | 0.615187915 |
| 0.325215419 | 0.615187915 |
| 0.325218916 | 0.615187915 |
| 0.325287198 | 0.615292639 |
| 0.325480002 | 0.615535103 |
| 0.325559917 | 0.615637345 |
| 0.325697863 | 0.615751514 |
| 0.325849739 | 0.615988413 |
| 0.325930096 | 0.61606829 |
| 0.325969666 | 0.616085505 |
| 0.326001356 | 0.616105177 |
| 0.326039533 | 0.61612843 |
| 0.326086433 | 0.616132504 |
| 0.326088877 | 0.616132504 |
| 0.326093435 | 0.616132504 |
| 0.326230242 | 0.616266884 |
| 0.326286065 | 0.616319726 |
| 0.326445099 | 0.616444817 |

| | |
|---|---|
| 0.326476074 | 0.61644662 |
| 0.326486509 | 0.61644662 |
| 0.326522679 | 0.616478857 |
| 0.32657269 | 0.616548837 |
| 0.32672376 | 0.616760702 |
| 0.326806805 | 0.616807649 |
| 0.326921347 | 0.616904247 |
| 0.326942283 | 0.616904247 |
| 0.327025732 | 0.616957181 |
| 0.327039636 | 0.616957181 |
| 0.327073968 | 0.616957181 |
| 0.327151994 | 0.617055484 |
| 0.32721008 | 0.617116163 |
| 0.327374108 | 0.617278856 |
| 0.327404084 | 0.617310939 |
| 0.327434549 | 0.617331671 |
| 0.327441003 | 0.617331671 |
| 0.327551129 | 0.617417091 |
| 0.327645355 | 0.617465381 |
| 0.327649962 | 0.617465381 |
| 0.327680315 | 0.617465381 |
| 0.327687799 | 0.617465381 |
| 0.327693429 | 0.617465381 |
| 0.327722795 | 0.617478407 |
| 0.327778132 | 0.617478407 |
| 0.327799716 | 0.617494644 |
| 0.327824572 | 0.617507012 |
| 0.327843989 | 0.617507012 |
| 0.327845179 | 0.617507012 |
| 0.328150518 | 0.617862214 |
| 0.328405494 | 0.618049096 |
| 0.328589176 | 0.61832148 |
| 0.328626627 | 0.618367523 |
| 0.328716149 | 0.618462671 |
| 0.329063093 | 0.618968718 |
| 0.329088983 | 0.618980709 |
| 0.329095461 | 0.618980709 |
| 0.329180541 | 0.619018503 |
| 0.329219011 | 0.619039229 |
| 0.329279964 | 0.619039229 |
| 0.329367831 | 0.619132262 |
| 0.32937152 | 0.619132262 |
| 0.329412303 | 0.619160967 |
| 0.329436491 | 0.619181996 |
| 0.329512509 | 0.619300432 |

| | |
|---|---|
| 0.329638578 | 0.619401927 |
| 0.329677275 | 0.619438993 |
| 0.329793317 | 0.619583698 |
| 0.329858799 | 0.619682274 |
| 0.329970984 | 0.619772591 |
| 0.330068494 | 0.619827228 |
| 0.330078033 | 0.619827228 |
| 0.330079115 | 0.619827228 |
| 0.330111611 | 0.619863811 |
| 0.330287541 | 0.620023046 |
| 0.330427419 | 0.620261181 |
| 0.330505682 | 0.62029284 |
| 0.330588547 | 0.620368035 |
| 0.330617043 | 0.620385303 |
| 0.330664809 | 0.620413365 |
| 0.330707755 | 0.620424102 |
| 0.330709612 | 0.620424102 |
| 0.330933058 | 0.620730781 |
| 0.330972476 | 0.620770532 |
| 0.331023588 | 0.6208175 |
| 0.331067946 | 0.620876243 |
| 0.331128895 | 0.620941645 |
| 0.33116118 | 0.620962467 |
| 0.33120728 | 0.620990833 |
| 0.331340836 | 0.621167881 |
| 0.331527089 | 0.621394752 |
| 0.331669449 | 0.621616027 |
| 0.331697351 | 0.621616027 |
| 0.331845912 | 0.621796572 |
| 0.331868214 | 0.621813896 |
| 0.332033505 | 0.622025714 |
| 0.332050437 | 0.622027894 |
| 0.332158603 | 0.622113247 |
| 0.33231258 | 0.62230375 |
| 0.332427889 | 0.622387315 |
| 0.332500946 | 0.622387315 |
| 0.332508617 | 0.622387315 |
| 0.332530408 | 0.622387315 |
| 0.332568669 | 0.622416221 |
| 0.332659939 | 0.622513651 |
| 0.332766176 | 0.622597449 |
| 0.333268816 | 0.623297457 |
| 0.333275097 | 0.623297457 |
| 0.333545376 | 0.623543607 |
| 0.333562483 | 0.623543607 |

| | |
|---|---|
| 0.333651658 | 0.623658851 |
| 0.333747195 | 0.623709937 |
| 0.333795808 | 0.62373246 |
| 0.333811673 | 0.623734813 |
| 0.333951291 | 0.623862927 |
| 0.334092299 | 0.624017175 |
| 0.334163357 | 0.624100945 |
| 0.334262829 | 0.624237767 |
| 0.334287835 | 0.624259987 |
| 0.334344749 | 0.62434179 |
| 0.334408256 | 0.624410903 |
| 0.334466533 | 0.624410903 |
| 0.33460313 | 0.624477779 |
| 0.33461691 | 0.624477779 |
| 0.334632403 | 0.624477779 |
| 0.334675959 | 0.624477779 |
| 0.334683602 | 0.624477779 |
| 0.334692922 | 0.624477779 |
| 0.334798475 | 0.62457685 |
| 0.334880133 | 0.624680249 |
| 0.334941906 | 0.624771008 |
| 0.334989201 | 0.624834756 |
| 0.335057409 | 0.624893496 |
| 0.335073251 | 0.624893496 |
| 0.335093029 | 0.624893496 |
| 0.335113626 | 0.624893496 |
| 0.335137268 | 0.624893496 |
| 0.335174815 | 0.624893496 |
| 0.335175637 | 0.624893496 |
| 0.335335106 | 0.625040913 |
| 0.335335953 | 0.625040913 |
| 0.335404817 | 0.625120341 |
| 0.335470381 | 0.625218072 |
| 0.335486277 | 0.625223231 |
| 0.335594003 | 0.62529615 |
| 0.335745187 | 0.625363161 |
| 0.335773744 | 0.62536649 |
| 0.335865169 | 0.625464336 |
| 0.33593017 | 0.625525028 |
| 0.335932915 | 0.625525028 |
| 0.335937161 | 0.625525028 |
| 0.336094467 | 0.625744544 |
| 0.336132529 | 0.625790945 |
| 0.336157968 | 0.625813843 |
| 0.336258638 | 0.625878933 |

| | |
|---|---|
| 0.336357152 | 0.625968707 |
| 0.336359442 | 0.625968707 |
| 0.336385637 | 0.625992996 |
| 0.336521722 | 0.626172842 |
| 0.336567888 | 0.626227331 |
| 0.336702566 | 0.626357904 |
| 0.336814174 | 0.626398016 |
| 0.336835809 | 0.626398016 |
| 0.336840019 | 0.626398016 |
| 0.336910961 | 0.626422331 |
| 0.336994323 | 0.626479402 |
| 0.337072665 | 0.626492264 |
| 0.337074858 | 0.626492264 |
| 0.337178594 | 0.626611709 |
| 0.337195101 | 0.626617937 |
| 0.337238575 | 0.626653364 |
| 0.33724048 | 0.626653364 |
| 0.337311145 | 0.626735766 |
| 0.337528994 | 0.627042689 |
| 0.337644318 | 0.627152488 |
| 0.337770661 | 0.627226711 |
| 0.337781121 | 0.627226711 |
| 0.337788357 | 0.627226711 |
| 0.337799256 | 0.627226711 |
| 0.337862458 | 0.627319608 |
| 0.337958853 | 0.627449666 |
| 0.33797213 | 0.627449857 |
| 0.338091574 | 0.627490122 |
| 0.338094855 | 0.627490122 |
| 0.338143428 | 0.627523264 |
| 0.338212515 | 0.627587217 |
| 0.338213544 | 0.627587217 |
| 0.338314725 | 0.627669955 |
| 0.338354706 | 0.627695229 |
| 0.338549417 | 0.627999375 |
| 0.338584451 | 0.628005017 |
| 0.338587625 | 0.628005017 |
| 0.338676598 | 0.628121118 |
| 0.338769337 | 0.628219725 |
| 0.338849104 | 0.628318717 |
| 0.338988746 | 0.628489784 |
| 0.339042307 | 0.628489784 |
| 0.339060124 | 0.628489784 |
| 0.339166568 | 0.628613696 |
| 0.339226776 | 0.628700819 |

| | |
|---|---|
| 0.339275113 | 0.628728864 |
| 0.339306981 | 0.628728864 |
| 0.339317769 | 0.628728864 |
| 0.339330297 | 0.628728864 |
| 0.339350446 | 0.628728864 |
| 0.339361026 | 0.628728864 |
| 0.339413829 | 0.628734332 |
| 0.339461107 | 0.628753195 |
| 0.339465062 | 0.628753195 |
| 0.339536762 | 0.628819006 |
| 0.339572823 | 0.628841981 |
| 0.339588086 | 0.628841981 |
| 0.339659419 | 0.628939743 |
| 0.339705092 | 0.628999861 |
| 0.339789968 | 0.629094798 |
| 0.339878177 | 0.629150907 |
| 0.339892348 | 0.629150907 |
| 0.339986593 | 0.629227538 |
| 0.340125874 | 0.629313835 |
| 0.340191353 | 0.629313835 |
| 0.340207942 | 0.629313835 |
| 0.340237522 | 0.629313835 |
| 0.340275434 | 0.629321921 |
| 0.340374567 | 0.629448838 |
| 0.340398996 | 0.629448838 |
| 0.340449802 | 0.629463538 |
| 0.340470956 | 0.629463538 |
| 0.340556516 | 0.629548408 |
| 0.340778667 | 0.62964137 |
| 0.34079574 | 0.629648487 |
| 0.340845281 | 0.62971559 |
| 0.340987145 | 0.629807169 |
| 0.340990625 | 0.629807169 |
| 0.34100064 | 0.629807169 |
| 0.341088444 | 0.629902757 |
| 0.341092072 | 0.629902757 |
| 0.341121885 | 0.629933388 |
| 0.341149005 | 0.629959043 |
| 0.341181161 | 0.629993996 |
| 0.341239934 | 0.629997988 |
| 0.341262689 | 0.629997988 |
| 0.341326442 | 0.630017999 |
| 0.341414211 | 0.630099627 |
| 0.341436815 | 0.630099627 |
| 0.341732177 | 0.630412438 |

| | |
|---|---|
| 0.341742202 | 0.630412438 |
| 0.34181283 | 0.630450179 |
| 0.341832738 | 0.630450179 |
| 0.341865129 | 0.630474754 |
| 0.341967384 | 0.630590081 |
| 0.342082595 | 0.630729269 |
| 0.342123528 | 0.630731488 |
| 0.342310294 | 0.630867262 |
| 0.342314412 | 0.630867262 |
| 0.342379259 | 0.630958712 |
| 0.342395103 | 0.630963497 |
| 0.342608297 | 0.631169056 |
| 0.342617499 | 0.631169056 |
| 0.342634636 | 0.631169056 |
| 0.342652427 | 0.631169056 |
| 0.342685229 | 0.631205066 |
| 0.342793658 | 0.631282715 |
| 0.342809786 | 0.631288004 |
| 0.342867841 | 0.631321684 |
| 0.342921132 | 0.63138922 |
| 0.342931034 | 0.63138922 |
| 0.342955421 | 0.631409711 |
| 0.342976938 | 0.631424918 |
| 0.343124909 | 0.631624088 |
| 0.343160345 | 0.631641564 |
| 0.343164986 | 0.631641564 |
| 0.343206387 | 0.631676415 |
| 0.343303012 | 0.631749323 |
| 0.343333714 | 0.631764277 |
| 0.343350492 | 0.631770741 |
| 0.343481359 | 0.631828043 |
| 0.343558406 | 0.63185964 |
| 0.343571275 | 0.63185964 |
| 0.343614269 | 0.631865513 |
| 0.343645449 | 0.631866111 |
| 0.343661344 | 0.631866111 |
| 0.343667662 | 0.631866111 |
| 0.343778369 | 0.631923288 |
| 0.343814846 | 0.631965949 |
| 0.343858398 | 0.631997219 |
| 0.343881995 | 0.632016199 |
| 0.343939738 | 0.632021562 |
| 0.343986529 | 0.632061977 |
| 0.344012906 | 0.632067304 |
| 0.344055688 | 0.632086413 |

| | |
|---|---|
| 0.344072374 | 0.632086413 |
| 0.34409273 | 0.632086413 |
| 0.344154495 | 0.632126731 |
| 0.344205971 | 0.63215795 |
| 0.344234225 | 0.63215795 |
| 0.344285608 | 0.632207674 |
| 0.344291483 | 0.632207674 |
| 0.344545587 | 0.6324602 |
| 0.344593992 | 0.632483034 |
| 0.34460264 | 0.632483034 |
| 0.344785965 | 0.632679213 |
| 0.344800673 | 0.632679213 |
| 0.344895097 | 0.632779336 |
| 0.344925603 | 0.632786826 |
| 0.345015182 | 0.632853381 |
| 0.345123166 | 0.632976697 |
| 0.345170366 | 0.632991761 |
| 0.345230352 | 0.633028643 |
| 0.345278014 | 0.633091664 |
| 0.345355625 | 0.633172698 |
| 0.345475524 | 0.633283149 |
| 0.345557056 | 0.633383848 |
| 0.345601477 | 0.633438192 |
| 0.345648836 | 0.633438192 |
| 0.34576617 | 0.6335598 |
| 0.345794558 | 0.633564772 |
| 0.34581899 | 0.633571358 |
| 0.345843253 | 0.633581709 |
| 0.345891155 | 0.633581709 |
| 0.345915374 | 0.633601704 |
| 0.345964194 | 0.633666757 |
| 0.346021568 | 0.633697493 |
| 0.346024264 | 0.633697493 |
| 0.346027663 | 0.633697493 |
| 0.346191211 | 0.633887541 |
| 0.346212547 | 0.633902237 |
| 0.346309863 | 0.633982928 |
| 0.346348228 | 0.634016262 |
| 0.346419388 | 0.63408025 |
| 0.346454361 | 0.63408025 |
| 0.346456219 | 0.63408025 |
| 0.346483154 | 0.634092131 |
| 0.346600789 | 0.63424735 |
| 0.346750926 | 0.634400214 |
| 0.346786103 | 0.63441583 |

| | |
|---|---|
| 0.346957924 | 0.634559543 |
| 0.346986786 | 0.634587959 |
| 0.347095225 | 0.634694967 |
| 0.347098602 | 0.634694967 |
| 0.347112055 | 0.634695199 |
| 0.347226771 | 0.634855445 |
| 0.347239682 | 0.634855445 |
| 0.3472633 | 0.634863422 |
| 0.347270705 | 0.634863422 |
| 0.34734137 | 0.634912787 |
| 0.347364362 | 0.634912787 |
| 0.347432715 | 0.634988983 |
| 0.347506538 | 0.635026431 |
| 0.347664902 | 0.635267073 |
| 0.347729074 | 0.635335581 |
| 0.347775593 | 0.635348993 |
| 0.347776434 | 0.635348993 |
| 0.347886829 | 0.635496469 |
| 0.34789719 | 0.635496469 |
| 0.347917603 | 0.635509384 |
| 0.347987731 | 0.635545951 |
| 0.348161405 | 0.635686534 |
| 0.348290384 | 0.635802766 |
| 0.348291813 | 0.635802766 |
| 0.348348562 | 0.635857617 |
| 0.348450037 | 0.635862053 |
| 0.348455614 | 0.635862053 |
| 0.348458478 | 0.635862053 |
| 0.348471149 | 0.635862053 |
| 0.348555319 | 0.63596588 |
| 0.348578464 | 0.63596588 |
| 0.348597855 | 0.63596588 |
| 0.348605222 | 0.63596588 |
| 0.348608168 | 0.63596588 |
| 0.348737096 | 0.636074857 |
| 0.348748036 | 0.636074857 |
| 0.348871322 | 0.636202262 |
| 0.348923675 | 0.63623952 |
| 0.349057826 | 0.636330269 |
| 0.349061763 | 0.636330269 |
| 0.349246751 | 0.636479893 |
| 0.349338228 | 0.636590596 |
| 0.349395738 | 0.636653089 |
| 0.349491984 | 0.63673102 |
| 0.349502416 | 0.63673102 |

| | |
|---|---|
| 0.349541242 | 0.63673102 |
| 0.34954898 | 0.63673102 |
| 0.349627141 | 0.636821901 |
| 0.349638984 | 0.636821901 |
| 0.349810461 | 0.63701242 |
| 0.349866516 | 0.637017071 |
| 0.349974909 | 0.637092649 |
| 0.350138964 | 0.637220804 |
| 0.35037677 | 0.637526221 |
| 0.350380105 | 0.637526221 |
| 0.350400346 | 0.637526221 |
| 0.350475574 | 0.637553629 |
| 0.350495867 | 0.637553629 |
| 0.350634361 | 0.637732479 |
| 0.350647765 | 0.637732504 |
| 0.350706236 | 0.637746261 |
| 0.350752148 | 0.637751872 |
| 0.350857246 | 0.637845564 |
| 0.351011937 | 0.638078073 |
| 0.3510726 | 0.638098581 |
| 0.351174728 | 0.63822784 |
| 0.351205803 | 0.638229935 |
| 0.351221763 | 0.638229935 |
| 0.351287472 | 0.638229935 |
| 0.351296486 | 0.638229935 |
| 0.351347363 | 0.638244733 |
| 0.351369921 | 0.638244733 |
| 0.351371636 | 0.638244733 |
| 0.351693442 | 0.638588059 |
| 0.351695858 | 0.638588059 |
| 0.351721544 | 0.638588059 |
| 0.351843242 | 0.638735967 |
| 0.351884519 | 0.638762206 |
| 0.35208769 | 0.638910635 |
| 0.35211385 | 0.638910635 |
| 0.352163435 | 0.63897071 |
| 0.352197008 | 0.63897071 |
| 0.352219494 | 0.63897071 |
| 0.352339311 | 0.639027584 |
| 0.352441636 | 0.639149577 |
| 0.352499855 | 0.639149577 |
| 0.352520707 | 0.639149577 |
| 0.352531515 | 0.639149577 |
| 0.352538018 | 0.639149577 |
| 0.35255354 | 0.639149577 |

| | |
|---|---|
| 0.352574995 | 0.639149577 |
| 0.352602511 | 0.639149577 |
| 0.352616859 | 0.639149577 |
| 0.352621292 | 0.639149577 |
| 0.352801266 | 0.639378459 |
| 0.352872632 | 0.639421771 |
| 0.352894723 | 0.639421771 |
| 0.352897272 | 0.639421771 |
| 0.352986604 | 0.639500788 |
| 0.352989611 | 0.639500788 |
| 0.353015476 | 0.639523319 |
| 0.3530816 | 0.639594453 |
| 0.353157646 | 0.639634895 |
| 0.353209269 | 0.639702224 |
| 0.353221683 | 0.639702224 |
| 0.353375563 | 0.639921769 |
| 0.353383217 | 0.639921769 |
| 0.353672055 | 0.640225891 |
| 0.35368021 | 0.640225891 |
| 0.35373222 | 0.640254765 |
| 0.353769697 | 0.640254765 |
| 0.353772276 | 0.640254765 |
| 0.353781409 | 0.640254765 |
| 0.353927465 | 0.640395995 |
| 0.353947044 | 0.640407091 |
| 0.353962373 | 0.640407449 |
| 0.354001569 | 0.64040843 |
| 0.354049077 | 0.640470046 |
| 0.354076453 | 0.640495241 |
| 0.354097456 | 0.640502634 |
| 0.354279284 | 0.640701621 |
| 0.354705474 | 0.641316438 |
| 0.354741561 | 0.641334769 |
| 0.354742544 | 0.641334769 |
| 0.354779678 | 0.641353212 |
| 0.354856657 | 0.641443677 |
| 0.354899457 | 0.641448006 |
| 0.354948183 | 0.641487387 |
| 0.35496308 | 0.641489969 |
| 0.355023807 | 0.641548154 |
| 0.355065607 | 0.641548154 |
| 0.355112218 | 0.641548154 |
| 0.355116509 | 0.641548154 |
| 0.355163852 | 0.641590323 |
| 0.355334812 | 0.641753872 |

| | |
|---|---|
| 0.355492444 | 0.641870566 |
| 0.355848348 | 0.642163886 |
| 0.355870381 | 0.642179314 |
| 0.35591048 | 0.64222734 |
| 0.356097648 | 0.642325836 |
| 0.356113525 | 0.642325836 |
| 0.356244208 | 0.642440094 |
| 0.356273835 | 0.642469197 |
| 0.356344792 | 0.64252417 |
| 0.356487579 | 0.642708632 |
| 0.356643352 | 0.642845737 |
| 0.356654704 | 0.642845737 |
| 0.356672701 | 0.642845737 |
| 0.356685104 | 0.642845737 |
| 0.356851378 | 0.643047948 |
| 0.356857966 | 0.643047948 |
| 0.356899187 | 0.643047948 |
| 0.356905316 | 0.643047948 |
| 0.356919293 | 0.643048805 |
| 0.357002309 | 0.643174041 |
| 0.357052863 | 0.643226049 |
| 0.357071693 | 0.643226049 |
| 0.357191204 | 0.643319675 |
| 0.357221096 | 0.643327885 |
| 0.357257137 | 0.643327885 |
| 0.357263702 | 0.643327885 |
| 0.35728142 | 0.643327885 |
| 0.357331971 | 0.643354242 |
| 0.357405466 | 0.643405531 |
| 0.357418218 | 0.643405531 |
| 0.357428004 | 0.643405531 |
| 0.357560488 | 0.643554167 |
| 0.357564624 | 0.643554167 |
| 0.357604036 | 0.643576459 |
| 0.357686093 | 0.643592462 |
| 0.357750807 | 0.643600836 |
| 0.357752715 | 0.643600836 |
| 0.35779317 | 0.643624992 |
| 0.357838082 | 0.643681471 |
| 0.357851913 | 0.643682041 |
| 0.357917155 | 0.643750767 |
| 0.357961603 | 0.643805663 |
| 0.358071085 | 0.643881732 |
| 0.358109201 | 0.643925961 |
| 0.358153509 | 0.643981319 |

| | |
|---|---|
| 0.35821126 | 0.644057642 |
| 0.358236525 | 0.644057642 |
| 0.358272203 | 0.644097471 |
| 0.358293442 | 0.644098463 |
| 0.358432024 | 0.644223159 |
| 0.358475958 | 0.644244913 |
| 0.35852035 | 0.644276071 |
| 0.358575988 | 0.644283312 |
| 0.358652582 | 0.644283312 |
| 0.358673184 | 0.644283312 |
| 0.35879137 | 0.644416482 |
| 0.358823139 | 0.644416482 |
| 0.35885643 | 0.644418037 |
| 0.358954525 | 0.644432757 |
| 0.358972874 | 0.644432757 |
| 0.359156081 | 0.644664456 |
| 0.359181302 | 0.644685429 |
| 0.359228885 | 0.644746537 |
| 0.359251366 | 0.644747369 |
| 0.359414185 | 0.6449188 |
| 0.359419651 | 0.6449188 |
| 0.359476866 | 0.644980696 |
| 0.359492418 | 0.644980696 |
| 0.359508316 | 0.644980696 |
| 0.359568943 | 0.645040869 |
| 0.359765949 | 0.645199867 |
| 0.359827441 | 0.645285848 |
| 0.359880533 | 0.645330829 |
| 0.360083269 | 0.645598775 |
| 0.360220191 | 0.64574392 |
| 0.360232016 | 0.64574392 |
| 0.360354392 | 0.645890359 |
| 0.360405496 | 0.645935396 |
| 0.360577367 | 0.646193164 |
| 0.360712472 | 0.646386218 |
| 0.360899064 | 0.646568427 |
| 0.361031587 | 0.646666093 |
| 0.361050939 | 0.646676435 |
| 0.361170653 | 0.646710891 |
| 0.361194268 | 0.646714264 |
| 0.361234093 | 0.646761255 |
| 0.361324864 | 0.646850822 |
| 0.361441455 | 0.64700031 |
| 0.361467427 | 0.647008762 |
| 0.361494707 | 0.647033274 |

| | |
|---|---|
| 0.361552925 | 0.647064526 |
| 0.361583696 | 0.64709528 |
| 0.36181928 | 0.647322296 |
| 0.36189577 | 0.647348724 |
| 0.361966892 | 0.647416148 |
| 0.362009743 | 0.647468476 |
| 0.362059912 | 0.647485854 |
| 0.362079109 | 0.647485854 |
| 0.362180752 | 0.647579783 |
| 0.362336577 | 0.647785442 |
| 0.362388335 | 0.647805635 |
| 0.362415122 | 0.647805635 |
| 0.362601795 | 0.647995124 |
| 0.362603524 | 0.647995124 |
| 0.362658553 | 0.648069147 |
| 0.362713136 | 0.648142368 |
| 0.36279514 | 0.648232933 |
| 0.362942228 | 0.648342666 |
| 0.362969141 | 0.648342666 |
| 0.362969929 | 0.648342666 |
| 0.363082394 | 0.648411337 |
| 0.363093526 | 0.648411337 |
| 0.363095631 | 0.648411337 |
| 0.363110272 | 0.648411337 |
| 0.363122329 | 0.648411337 |
| 0.363214321 | 0.648485235 |
| 0.363225883 | 0.648485235 |
| 0.36325598 | 0.648485235 |
| 0.363426409 | 0.648686789 |
| 0.363470765 | 0.648717337 |
| 0.363512797 | 0.648768042 |
| 0.363591313 | 0.648883856 |
| 0.363620956 | 0.64890322 |
| 0.363651566 | 0.64890322 |
| 0.363725694 | 0.64897777 |
| 0.363837196 | 0.649030829 |
| 0.363860872 | 0.649048754 |
| 0.363894552 | 0.649053876 |
| 0.3639029 | 0.649053876 |
| 0.363941795 | 0.649063515 |
| 0.364006282 | 0.649069016 |
| 0.364022141 | 0.649069016 |
| 0.364077187 | 0.649142864 |
| 0.364140056 | 0.649206351 |
| 0.364201485 | 0.649243176 |

| | |
|---|---|
| 0.364201607 | 0.649243176 |
| 0.364326587 | 0.649269897 |
| 0.364376884 | 0.64929624 |
| 0.364419029 | 0.649314818 |
| 0.364594064 | 0.649514316 |
| 0.364661866 | 0.649597386 |
| 0.364819767 | 0.649761455 |
| 0.364909113 | 0.649809165 |
| 0.364935486 | 0.649809165 |
| 0.36496066 | 0.649809165 |
| 0.364963203 | 0.649809165 |
| 0.365094195 | 0.649934261 |
| 0.36518627 | 0.649952409 |
| 0.365383694 | 0.650228675 |
| 0.365541928 | 0.650439566 |
| 0.365603698 | 0.650500871 |
| 0.365681021 | 0.65058984 |
| 0.365889156 | 0.650838575 |
| 0.365999184 | 0.650912738 |
| 0.366085428 | 0.650993192 |
| 0.366175473 | 0.651093818 |
| 0.366236143 | 0.651093818 |
| 0.366237709 | 0.651093818 |
| 0.366259388 | 0.651108053 |
| 0.366273883 | 0.651109519 |
| 0.366306749 | 0.65114364 |
| 0.366368692 | 0.651229443 |
| 0.366390899 | 0.651244611 |
| 0.366509341 | 0.651357901 |
| 0.366601105 | 0.651385916 |
| 0.366658143 | 0.651385916 |
| 0.366697259 | 0.651385916 |
| 0.36670999 | 0.651385916 |
| 0.366777406 | 0.651413238 |
| 0.366900718 | 0.651445738 |
| 0.366994759 | 0.651556576 |
| 0.367081375 | 0.651578313 |
| 0.367084832 | 0.651578313 |
| 0.367217434 | 0.651649348 |
| 0.367269123 | 0.651662564 |
| 0.367376786 | 0.651763711 |
| 0.367422185 | 0.651763711 |
| 0.36774657 | 0.652041712 |
| 0.367756618 | 0.652041712 |
| 0.3678819 | 0.652222233 |

| | |
|---|---|
| 0.367913211 | 0.652222233 |
| 0.367927873 | 0.652223947 |
| 0.367977469 | 0.652279679 |
| 0.368004088 | 0.652279679 |
| 0.368101599 | 0.652316155 |
| 0.368130662 | 0.652316155 |
| 0.368255209 | 0.65238884 |
| 0.36835067 | 0.652408723 |
| 0.368359168 | 0.652408723 |
| 0.36836489 | 0.652408723 |
| 0.368395015 | 0.652408723 |
| 0.368401961 | 0.652408723 |
| 0.368421669 | 0.65241177 |
| 0.368687986 | 0.652648307 |
| 0.368786022 | 0.652694136 |
| 0.368809805 | 0.652694136 |
| 0.368868955 | 0.652727357 |
| 0.368869692 | 0.652727357 |
| 0.368905883 | 0.652742894 |
| 0.369100137 | 0.653038085 |
| 0.369182647 | 0.653084447 |
| 0.369210816 | 0.653085049 |
| 0.369230845 | 0.653085049 |
| 0.369236381 | 0.653085049 |
| 0.369276882 | 0.653128258 |
| 0.369309769 | 0.653128258 |
| 0.369317812 | 0.653128258 |
| 0.369366495 | 0.653128258 |
| 0.36961152 | 0.653311774 |
| 0.369618487 | 0.653311774 |
| 0.369625193 | 0.653311774 |
| 0.369657566 | 0.653320502 |
| 0.369916343 | 0.653656575 |
| 0.369979705 | 0.653744284 |
| 0.37003693 | 0.653748381 |
| 0.370094035 | 0.653776514 |
| 0.37043073 | 0.654152924 |
| 0.370471896 | 0.65417711 |
| 0.370512482 | 0.654184746 |
| 0.370621835 | 0.654281154 |
| 0.370626981 | 0.654281154 |
| 0.37066061 | 0.654313527 |
| 0.370678249 | 0.654313527 |
| 0.370686535 | 0.654313527 |
| 0.370725058 | 0.654338198 |

| | |
|---|---|
| 0.370741728 | 0.654338198 |
| 0.370793436 | 0.65438096 |
| 0.370829085 | 0.654385289 |
| 0.370840187 | 0.654385289 |
| 0.370850848 | 0.654385289 |
| 0.370879933 | 0.654388121 |
| 0.370974008 | 0.654483349 |
| 0.371018166 | 0.654525088 |
| 0.371044567 | 0.654529511 |
| 0.371072688 | 0.654529511 |
| 0.371095583 | 0.654529511 |
| 0.371102 | 0.654529511 |
| 0.371130032 | 0.654529511 |
| 0.371347558 | 0.65480082 |
| 0.371450384 | 0.654918492 |
| 0.37162595 | 0.655145769 |
| 0.371639474 | 0.655145769 |
| 0.371719627 | 0.655259759 |
| 0.371747007 | 0.655259759 |
| 0.371821996 | 0.655259759 |
| 0.371826721 | 0.655259759 |
| 0.372087224 | 0.655515371 |
| 0.37210303 | 0.655515371 |
| 0.372110642 | 0.655515371 |
| 0.372277838 | 0.655712892 |
| 0.372438352 | 0.65586283 |
| 0.37244559 | 0.65586283 |
| 0.372630563 | 0.656104784 |
| 0.372651868 | 0.656104784 |
| 0.372778463 | 0.656296973 |
| 0.372788587 | 0.656296973 |
| 0.372808429 | 0.656307646 |
| 0.372876142 | 0.656341668 |
| 0.37291044 | 0.656341668 |
| 0.372975819 | 0.656429242 |
| 0.373004667 | 0.656434737 |
| 0.373032994 | 0.656460332 |
| 0.373233003 | 0.656715244 |
| 0.373494404 | 0.65693249 |
| 0.373576849 | 0.657032221 |
| 0.373578696 | 0.657032221 |
| 0.373776179 | 0.657307704 |
| 0.373871836 | 0.657426396 |
| 0.373934212 | 0.65748753 |
| 0.374063543 | 0.657520736 |

| | |
|---|---|
| 0.374139297 | 0.657605355 |
| 0.374184196 | 0.657622972 |
| 0.374217794 | 0.657622972 |
| 0.374231988 | 0.657622972 |
| 0.374348065 | 0.657795111 |
| 0.374509563 | 0.6579596 |
| 0.374546144 | 0.6579596 |
| 0.374629344 | 0.658019248 |
| 0.374667063 | 0.658019248 |
| 0.374678732 | 0.658019248 |
| 0.374781068 | 0.658110147 |
| 0.375164833 | 0.658624735 |
| 0.375189067 | 0.658624735 |
| 0.37519088 | 0.658624735 |
| 0.375304188 | 0.658624735 |
| 0.375351844 | 0.658624735 |
| 0.375375368 | 0.658624735 |
| 0.375377134 | 0.658624735 |
| 0.375393756 | 0.658624735 |
| 0.375403769 | 0.658624735 |
| 0.37542806 | 0.658624735 |
| 0.375431464 | 0.658624735 |
| 0.375438363 | 0.658624735 |
| 0.375484754 | 0.658656209 |
| 0.375496655 | 0.658656209 |
| 0.375679364 | 0.65877949 |
| 0.375684074 | 0.65877949 |
| 0.375815483 | 0.658913202 |
| 0.375903966 | 0.659022686 |
| 0.375914278 | 0.659022686 |
| 0.375973112 | 0.659068715 |
| 0.376004142 | 0.659068715 |
| 0.376027596 | 0.659068715 |
| 0.3760374 | 0.659068715 |
| 0.376178129 | 0.659156913 |
| 0.376181779 | 0.659156913 |
| 0.376345276 | 0.659248056 |
| 0.376515296 | 0.659445225 |
| 0.376565727 | 0.659448275 |
| 0.37658202 | 0.659448275 |
| 0.376586268 | 0.659448275 |
| 0.376651645 | 0.659538509 |
| 0.37676375 | 0.659607741 |
| 0.376852759 | 0.659633588 |
| 0.376935774 | 0.659656924 |

| | |
|---|---|
| 0.376936669 | 0.659656924 |
| 0.376940878 | 0.659656924 |
| 0.377049535 | 0.659779501 |
| 0.37711136 | 0.659858288 |
| 0.377143826 | 0.659890852 |
| 0.377211175 | 0.659935959 |
| 0.37725252 | 0.65998036 |
| 0.377291983 | 0.65998036 |
| 0.377322397 | 0.660009322 |
| 0.377402865 | 0.660077348 |
| 0.37765918 | 0.660380136 |
| 0.377753976 | 0.660392193 |
| 0.377798783 | 0.660406027 |
| 0.377873932 | 0.660457967 |
| 0.377875151 | 0.660457967 |
| 0.377885869 | 0.660457967 |
| 0.377887262 | 0.660457967 |
| 0.3779117 | 0.660457967 |
| 0.377927165 | 0.66046076 |
| 0.377951108 | 0.660478368 |
| 0.378023698 | 0.660517369 |
| 0.378044467 | 0.660517369 |
| 0.378068472 | 0.660517369 |
| 0.378072907 | 0.660517369 |
| 0.3781486 | 0.660605339 |
| 0.378252062 | 0.660689146 |
| 0.378286406 | 0.660700673 |
| 0.378335508 | 0.660737972 |
| 0.378372842 | 0.660754715 |
| 0.378500614 | 0.660808238 |
| 0.378564485 | 0.660871296 |
| 0.378646432 | 0.660946988 |
| 0.378662666 | 0.660946988 |
| 0.378714486 | 0.660946988 |
| 0.378742342 | 0.660946988 |
| 0.378768469 | 0.660946988 |
| 0.378775495 | 0.660946988 |
| 0.378776972 | 0.660946988 |
| 0.378821746 | 0.660957003 |
| 0.37896747 | 0.661137917 |
| 0.379019116 | 0.661156042 |
| 0.379179953 | 0.661363936 |
| 0.379245077 | 0.661452692 |
| 0.379320659 | 0.661491661 |
| 0.379322627 | 0.661491661 |

| | |
|---|---|
| 0.379358581 | 0.661530137 |
| 0.379459618 | 0.661657876 |
| 0.379757961 | 0.662065699 |
| 0.379773192 | 0.662065699 |
| 0.37984051 | 0.662065699 |
| 0.379841994 | 0.662065699 |
| 0.379846415 | 0.662065699 |
| 0.379937671 | 0.662200521 |
| 0.380011024 | 0.662237713 |
| 0.380052844 | 0.662237713 |
| 0.380056342 | 0.662237713 |
| 0.380235052 | 0.662423231 |
| 0.38024046 | 0.662423231 |
| 0.380357237 | 0.662541356 |
| 0.380364773 | 0.662541356 |
| 0.380495187 | 0.662687203 |
| 0.380639469 | 0.662879264 |
| 0.380754716 | 0.663035587 |
| 0.380778747 | 0.663035587 |
| 0.380846181 | 0.66308028 |
| 0.380908033 | 0.663139486 |
| 0.380980731 | 0.663217561 |
| 0.381132864 | 0.663358147 |
| 0.38115037 | 0.663358147 |
| 0.381158986 | 0.663358147 |
| 0.381190406 | 0.663365545 |
| 0.381191094 | 0.663365545 |
| 0.38128398 | 0.663455375 |
| 0.381299272 | 0.663455375 |
| 0.381306383 | 0.663455375 |
| 0.381312364 | 0.663455375 |
| 0.381434352 | 0.663505823 |
| 0.381440438 | 0.663505823 |
| 0.381534848 | 0.663551586 |
| 0.38165793 | 0.663717173 |
| 0.381674433 | 0.663721639 |
| 0.382175581 | 0.664200301 |
| 0.382189263 | 0.664200301 |
| 0.382257973 | 0.664232559 |
| 0.382367902 | 0.66433349 |
| 0.38240194 | 0.66433349 |
| 0.382468289 | 0.664374397 |
| 0.38250958 | 0.664395586 |
| 0.382515934 | 0.664395586 |
| 0.382738586 | 0.664640575 |

| | |
|---|---|
| 0.382745704 | 0.664640575 |
| 0.382747119 | 0.664640575 |
| 0.382826602 | 0.664754359 |
| 0.382863682 | 0.66475813 |
| 0.382890396 | 0.66475813 |
| 0.382908136 | 0.66475813 |
| 0.382957968 | 0.664763819 |
| 0.382984697 | 0.664763819 |
| 0.382999542 | 0.664763819 |
| 0.383097722 | 0.664863809 |
| 0.383170569 | 0.664963733 |
| 0.383248973 | 0.665075563 |
| 0.383321721 | 0.665129103 |
| 0.38337599 | 0.665148666 |
| 0.383393882 | 0.665148666 |
| 0.383404511 | 0.665148666 |
| 0.38341679 | 0.665148666 |
| 0.383554334 | 0.665266116 |
| 0.383706474 | 0.665373903 |
| 0.383749438 | 0.665373903 |
| 0.383756183 | 0.665373903 |
| 0.383792758 | 0.665413094 |
| 0.383857754 | 0.665477331 |
| 0.383945635 | 0.66551227 |
| 0.383955624 | 0.66551227 |
| 0.383961748 | 0.66551227 |
| 0.383990255 | 0.665524646 |
| 0.383996835 | 0.665524646 |
| 0.384041909 | 0.665539937 |
| 0.384047282 | 0.665539937 |
| 0.38404758 | 0.665539937 |
| 0.384087483 | 0.665584869 |
| 0.384261649 | 0.665765564 |
| 0.384324553 | 0.665826106 |
| 0.384447967 | 0.66594302 |
| 0.384608734 | 0.666124592 |
| 0.384642646 | 0.666159101 |
| 0.384683913 | 0.666182123 |
| 0.384698815 | 0.666183706 |
| 0.38472736 | 0.666208914 |
| 0.384842889 | 0.666287844 |
| 0.384901748 | 0.666356577 |
| 0.384910572 | 0.666356577 |
| 0.385199121 | 0.666662253 |
| 0.385367152 | 0.666859821 |

| | |
|---|---|
| 0.385473691 | 0.666921924 |
| 0.385475187 | 0.666921924 |
| 0.38568703 | 0.667076946 |
| 0.38568703 | 0.667076946 |
| 0.385762507 | 0.667128211 |
| 0.385824137 | 0.667178187 |
| 0.385857666 | 0.667178187 |
| 0.385907932 | 0.667210117 |
| 0.386003421 | 0.667302538 |
| 0.386089438 | 0.667402143 |
| 0.38611709 | 0.667402143 |
| 0.386219579 | 0.667530842 |
| 0.386323356 | 0.667647093 |
| 0.386328894 | 0.667647093 |
| 0.386589883 | 0.667904246 |
| 0.386675671 | 0.667961584 |
| 0.386710026 | 0.667989935 |
| 0.386737024 | 0.667989935 |
| 0.386741422 | 0.667989935 |
| 0.386759387 | 0.667989935 |
| 0.386827766 | 0.668040605 |
| 0.386917121 | 0.668055144 |
| 0.38692436 | 0.668055144 |
| 0.387071713 | 0.668251875 |
| 0.387232092 | 0.668456065 |
| 0.387346108 | 0.668569086 |
| 0.387426805 | 0.668622547 |
| 0.387551663 | 0.668813793 |
| 0.387584983 | 0.668847058 |
| 0.387644099 | 0.668865873 |
| 0.387680077 | 0.668865873 |
| 0.387680148 | 0.668865873 |
| 0.387886279 | 0.669124554 |
| 0.387978947 | 0.669235932 |
| 0.388000875 | 0.669249517 |
| 0.38806151 | 0.669305625 |
| 0.388083924 | 0.669320045 |
| 0.388179553 | 0.669379543 |
| 0.388200659 | 0.669379543 |
| 0.38820275 | 0.669379543 |
| 0.388228604 | 0.669399889 |
| 0.388279088 | 0.669462699 |
| 0.388343879 | 0.669499653 |
| 0.388429362 | 0.669552125 |
| 0.388755432 | 0.669895978 |

| | |
|---|---|
| 0.388799425 | 0.669943755 |
| 0.388811291 | 0.669943755 |
| 0.388947456 | 0.670105646 |
| 0.389061819 | 0.670181462 |
| 0.389151769 | 0.670287921 |
| 0.389308493 | 0.670479842 |
| 0.389325136 | 0.670479842 |
| 0.389444031 | 0.670509841 |
| 0.389466912 | 0.670515494 |
| 0.389536632 | 0.670562806 |
| 0.389586519 | 0.670576569 |
| 0.38965029 | 0.670646054 |
| 0.389727692 | 0.670673536 |
| 0.389807497 | 0.670762397 |
| 0.389836002 | 0.670787213 |
| 0.389862038 | 0.670807778 |
| 0.389937916 | 0.670889861 |
| 0.390085847 | 0.67107165 |
| 0.390167283 | 0.671116921 |
| 0.390188342 | 0.671126767 |
| 0.39022864 | 0.67117184 |
| 0.3904389 | 0.671398334 |
| 0.390542297 | 0.671425358 |
| 0.390558259 | 0.671425358 |
| 0.390565252 | 0.671425358 |
| 0.390596316 | 0.671425358 |
| 0.390624151 | 0.671425358 |
| 0.390699667 | 0.671496989 |
| 0.390839152 | 0.671639772 |
| 0.390955907 | 0.671779591 |
| 0.390974413 | 0.671779591 |
| 0.390996965 | 0.671779591 |
| 0.391036608 | 0.671785179 |
| 0.39117595 | 0.671840968 |
| 0.39119678 | 0.671840968 |
| 0.391201473 | 0.671840968 |
| 0.391207292 | 0.671840968 |
| 0.391213081 | 0.671840968 |
| 0.39129355 | 0.671867536 |
| 0.391302808 | 0.671867536 |
| 0.391326522 | 0.671867536 |
| 0.391350686 | 0.671867536 |
| 0.391368402 | 0.671867536 |
| 0.391549286 | 0.67203787 |
| 0.391586842 | 0.672051692 |

| | |
|---|---|
| 0.391803405 | 0.672278012 |
| 0.39187366 | 0.672348318 |
| 0.391896617 | 0.672348318 |
| 0.391900847 | 0.672348318 |
| 0.392258158 | 0.672743223 |
| 0.392315315 | 0.672784269 |
| 0.392438375 | 0.672810025 |
| 0.392533314 | 0.672948567 |
| 0.392646742 | 0.673022597 |
| 0.392692076 | 0.673075377 |
| 0.392802665 | 0.673216471 |
| 0.392817797 | 0.673218179 |
| 0.393218274 | 0.673591726 |
| 0.393305547 | 0.673666233 |
| 0.393385491 | 0.673718104 |
| 0.3934591 | 0.673783791 |
| 0.39347321 | 0.673783791 |
| 0.393615979 | 0.67391293 |
| 0.39370803 | 0.674016259 |
| 0.393774501 | 0.674081595 |
| 0.393985466 | 0.67432155 |
| 0.394031599 | 0.67433072 |
| 0.394042967 | 0.67433072 |
| 0.394061616 | 0.67433072 |
| 0.394133179 | 0.67438049 |
| 0.394206109 | 0.674432586 |
| 0.394249335 | 0.674433859 |
| 0.394271476 | 0.674447511 |
| 0.394315458 | 0.674474297 |
| 0.394554833 | 0.674733257 |
| 0.394676753 | 0.674898381 |
| 0.394858653 | 0.675079336 |
| 0.39486762 | 0.675079336 |
| 0.395004805 | 0.67522134 |
| 0.395026109 | 0.67522134 |
| 0.395149554 | 0.6752632 |
| 0.395173559 | 0.6752632 |
| 0.395187839 | 0.6752632 |
| 0.395232226 | 0.675290591 |
| 0.3952586 | 0.675311426 |
| 0.395286735 | 0.675325252 |
| 0.39543785 | 0.675453277 |
| 0.395483875 | 0.675478232 |
| 0.395584597 | 0.675611395 |
| 0.395597068 | 0.675611395 |

| | |
|---|---|
| 0.395755827 | 0.675748854 |
| 0.395836379 | 0.67576526 |
| 0.395851758 | 0.675767292 |
| 0.395946702 | 0.675813453 |
| 0.396009178 | 0.675813453 |
| 0.396015734 | 0.675813453 |
| 0.396018637 | 0.675813453 |
| 0.396080129 | 0.675833111 |
| 0.396146432 | 0.675869843 |
| 0.396153075 | 0.675869843 |
| 0.396236565 | 0.675963854 |
| 0.396339359 | 0.675983417 |
| 0.396358441 | 0.675983417 |
| 0.396404158 | 0.675983417 |
| 0.396459904 | 0.676030069 |
| 0.39650099 | 0.67604293 |
| 0.396530208 | 0.676053131 |
| 0.396562444 | 0.676063886 |
| 0.39661409 | 0.676091106 |
| 0.396645245 | 0.676091106 |
| 0.396651849 | 0.676091106 |
| 0.396722683 | 0.676163442 |
| 0.396759299 | 0.676201651 |
| 0.396837989 | 0.676311562 |
| 0.396876054 | 0.676352231 |
| 0.396901025 | 0.676370585 |
| 0.396974855 | 0.67646774 |
| 0.397027677 | 0.67646774 |
| 0.397126763 | 0.676513812 |
| 0.397245766 | 0.676691731 |
| 0.397321414 | 0.676747979 |
| 0.397579945 | 0.677067259 |
| 0.397804683 | 0.677269361 |
| 0.397815675 | 0.677269361 |
| 0.397838129 | 0.677269361 |
| 0.397844173 | 0.677269361 |
| 0.397892902 | 0.677309598 |
| 0.397926951 | 0.677319142 |
| 0.397953298 | 0.677339781 |
| 0.398011502 | 0.677368956 |
| 0.398167941 | 0.677559831 |
| 0.398251653 | 0.677653858 |
| 0.398337765 | 0.677683626 |
| 0.398372034 | 0.677704248 |
| 0.398380873 | 0.677704248 |

| | |
|---|---|
| 0.39845233 | 0.677798198 |
| 0.398490161 | 0.677798198 |
| 0.398573528 | 0.677910914 |
| 0.398641606 | 0.677976943 |
| 0.398685088 | 0.678003809 |
| 0.398699847 | 0.6780047 |
| 0.398727526 | 0.678027559 |
| 0.398926224 | 0.678195941 |
| 0.399028931 | 0.678322122 |
| 0.399076207 | 0.678342951 |
| 0.399095856 | 0.678342951 |
| 0.399112562 | 0.678342951 |
| 0.399247836 | 0.678465537 |
| 0.39925575 | 0.678465537 |
| 0.39945045 | 0.678602972 |
| 0.399504272 | 0.678645717 |
| 0.399726486 | 0.678904728 |
| 0.399794011 | 0.678927023 |
| 0.399832463 | 0.678927023 |
| 0.399837139 | 0.678927023 |
| 0.399915693 | 0.678977179 |
| 0.399927495 | 0.678977179 |
| 0.400089231 | 0.679058117 |
| 0.400158288 | 0.67912692 |
| 0.40038247 | 0.679289538 |
| 0.400471516 | 0.679370845 |
| 0.400484888 | 0.679370845 |
| 0.400528822 | 0.679392628 |
| 0.40056285 | 0.679401954 |
| 0.400670607 | 0.67948793 |
| 0.400835379 | 0.679624747 |
| 0.400935353 | 0.679651452 |
| 0.400942318 | 0.679651452 |
| 0.40108197 | 0.679798255 |
| 0.401168457 | 0.679848068 |
| 0.401211061 | 0.679882202 |
| 0.401254233 | 0.679882202 |
| 0.40127735 | 0.679887454 |
| 0.40137482 | 0.679966062 |
| 0.401381052 | 0.679966062 |
| 0.401462451 | 0.679976702 |
| 0.401479838 | 0.679976702 |
| 0.401498941 | 0.679976702 |
| 0.401620078 | 0.68002195 |
| 0.401620482 | 0.68002195 |

| | |
|---|---|
| 0.401674358 | 0.680029046 |
| 0.40167504 | 0.680029046 |
| 0.401704733 | 0.68003149 |
| 0.40172061 | 0.68003196 |
| 0.40195815 | 0.68022438 |
| 0.40202949 | 0.68022438 |
| 0.40204238 | 0.68022438 |
| 0.4020475 | 0.68022438 |
| 0.402074493 | 0.680245884 |
| 0.402218415 | 0.680414285 |
| 0.40222332 | 0.680414285 |
| 0.402249985 | 0.680414285 |
| 0.402259748 | 0.680414285 |
| 0.40246312 | 0.680513939 |
| 0.402463797 | 0.680513939 |
| 0.402476917 | 0.680513939 |
| 0.402494206 | 0.680513939 |
| 0.402504413 | 0.680513939 |
| 0.402571037 | 0.68054636 |
| 0.402580745 | 0.68054636 |
| 0.402739004 | 0.680693089 |
| 0.40312061 | 0.681023882 |
| 0.40315596 | 0.681059444 |
| 0.403171945 | 0.681062291 |
| 0.403297292 | 0.681153237 |
| 0.403350191 | 0.681218423 |
| 0.403418176 | 0.681293796 |
| 0.403423429 | 0.681293796 |
| 0.403482711 | 0.681339851 |
| 0.403492772 | 0.681339851 |
| 0.403535998 | 0.681351496 |
| 0.403543431 | 0.681351496 |
| 0.403565855 | 0.681365201 |
| 0.403660749 | 0.681452945 |
| 0.403874169 | 0.68166527 |
| 0.403878483 | 0.68166527 |
| 0.403958447 | 0.681695169 |
| 0.404002193 | 0.681739387 |
| 0.404030169 | 0.681748794 |
| 0.404064757 | 0.681748794 |
| 0.404131859 | 0.681748794 |
| 0.404307639 | 0.681748794 |
| 0.404312964 | 0.681748794 |
| 0.404329412 | 0.681748794 |
| 0.404329616 | 0.681748794 |

| | |
|---|---|
| 0.404375612 | 0.681789764 |
| 0.404436591 | 0.681809998 |
| 0.404442557 | 0.681809998 |
| 0.404444875 | 0.681809998 |
| 0.404672858 | 0.681881053 |
| 0.404673147 | 0.681881053 |
| 0.404757898 | 0.681968078 |
| 0.404782069 | 0.681968078 |
| 0.404831567 | 0.682027345 |
| 0.40486298 | 0.68205614 |
| 0.404915064 | 0.682119757 |
| 0.404962112 | 0.682150757 |
| 0.405001247 | 0.682192552 |
| 0.40506641 | 0.682205803 |
| 0.405221357 | 0.68234898 |
| 0.405223057 | 0.68234898 |
| 0.40524184 | 0.682355484 |
| 0.405257687 | 0.682355484 |
| 0.405269901 | 0.682355484 |
| 0.405477329 | 0.682500273 |
| 0.405532961 | 0.682557103 |
| 0.405558683 | 0.682567944 |
| 0.405568065 | 0.682567944 |
| 0.405587014 | 0.682575714 |
| 0.405651808 | 0.682660636 |
| 0.40574842 | 0.68273032 |
| 0.405922607 | 0.682899187 |
| 0.405937942 | 0.682900865 |
| 0.40601565 | 0.682942806 |
| 0.406064202 | 0.682942806 |
| 0.406076367 | 0.682942806 |
| 0.406092502 | 0.682942806 |
| 0.406130786 | 0.682942806 |
| 0.406139643 | 0.682942806 |
| 0.406159191 | 0.682942806 |
| 0.406179012 | 0.682942806 |
| 0.406192304 | 0.682942806 |
| 0.406350545 | 0.683088289 |
| 0.406522094 | 0.68313555 |
| 0.406687407 | 0.68331691 |
| 0.406807352 | 0.683476361 |
| 0.406811008 | 0.683476361 |
| 0.406837366 | 0.683496533 |
| 0.406872824 | 0.683531993 |
| 0.406906156 | 0.683563878 |

| | |
|---|---|
| 0.40707783 | 0.683779919 |
| 0.407097197 | 0.683788334 |
| 0.407209122 | 0.683879206 |
| 0.407217262 | 0.683879206 |
| 0.407223094 | 0.683879206 |
| 0.407282699 | 0.683919291 |
| 0.407297237 | 0.683919291 |
| 0.407332909 | 0.683919291 |
| 0.407343213 | 0.683919291 |
| 0.407360128 | 0.683919291 |
| 0.407491289 | 0.684036191 |
| 0.407524991 | 0.684036191 |
| 0.407532007 | 0.684036191 |
| 0.407554845 | 0.684050415 |
| 0.407747876 | 0.684302058 |
| 0.407782189 | 0.684335529 |
| 0.408020887 | 0.68463014 |
| 0.408076767 | 0.68466777 |
| 0.408115577 | 0.684677888 |
| 0.408165433 | 0.684692244 |
| 0.408167262 | 0.684692244 |
| 0.408250599 | 0.684783803 |
| 0.408281912 | 0.684794049 |
| 0.408304453 | 0.684801782 |
| 0.408339061 | 0.684801798 |
| 0.408368256 | 0.684801798 |
| 0.408468781 | 0.684908557 |
| 0.408537455 | 0.684999591 |
| 0.408593155 | 0.685039239 |
| 0.408614445 | 0.685039239 |
| 0.408618635 | 0.685039239 |
| 0.408646238 | 0.6850614 |
| 0.408752847 | 0.68516777 |
| 0.408882212 | 0.685336376 |
| 0.408918669 | 0.68534591 |
| 0.408925947 | 0.68534591 |
| 0.40893107 | 0.68534591 |
| 0.409053462 | 0.685478668 |
| 0.409117186 | 0.685537214 |
| 0.409162695 | 0.685565231 |
| 0.409180248 | 0.685570522 |
| 0.409205494 | 0.685588703 |
| 0.409250156 | 0.685616814 |
| 0.409282104 | 0.685620582 |
| 0.40932554 | 0.685623113 |

| | |
|---|---|
| 0.409384548 | 0.685647496 |
| 0.409490326 | 0.685776425 |
| 0.409524366 | 0.685782548 |
| 0.409541943 | 0.685782548 |
| 0.409558502 | 0.685782548 |
| 0.409565977 | 0.685782548 |
| 0.409698459 | 0.685883136 |
| 0.409773604 | 0.685911224 |
| 0.409774924 | 0.685911224 |
| 0.409786843 | 0.685911224 |
| 0.410008213 | 0.686137075 |
| 0.410103329 | 0.686223913 |
| 0.410183278 | 0.686285358 |
| 0.410366574 | 0.686471456 |
| 0.410481681 | 0.686543441 |
| 0.410509926 | 0.686566571 |
| 0.410560797 | 0.686627539 |
| 0.410678394 | 0.686755849 |
| 0.410686257 | 0.686755849 |
| 0.410785942 | 0.686816493 |
| 0.410853347 | 0.686816493 |
| 0.410870705 | 0.686816493 |
| 0.41087567 | 0.686816493 |
| 0.410908291 | 0.686846916 |
| 0.410952577 | 0.686896834 |
| 0.411197032 | 0.687233078 |
| 0.41154848 | 0.687684051 |
| 0.411565383 | 0.687684051 |
| 0.411566568 | 0.687684051 |
| 0.411570286 | 0.687684051 |
| 0.411846301 | 0.687917626 |
| 0.411852062 | 0.687917626 |
| 0.411866612 | 0.687917626 |
| 0.411897015 | 0.687920157 |
| 0.412009615 | 0.688032961 |
| 0.412022342 | 0.688032961 |
| 0.412063443 | 0.688077469 |
| 0.412171701 | 0.688209047 |
| 0.412270659 | 0.688209047 |
| 0.412338847 | 0.688209047 |
| 0.412358989 | 0.688209047 |
| 0.412504974 | 0.688353841 |
| 0.412513156 | 0.688353841 |
| 0.412548109 | 0.68835583 |
| 0.412578304 | 0.688382094 |

| | |
|---|---|
| 0.41265089 | 0.688406737 |
| 0.412798676 | 0.68854026 |
| 0.412803212 | 0.68854026 |
| 0.412859955 | 0.688586674 |
| 0.41300838 | 0.688761865 |
| 0.413216727 | 0.688916339 |
| 0.413337581 | 0.688924144 |
| 0.413655512 | 0.689237722 |
| 0.413858064 | 0.689478733 |
| 0.413873139 | 0.689479729 |
| 0.41395406 | 0.689576605 |
| 0.413960248 | 0.689576605 |
| 0.414201492 | 0.689804456 |
| 0.414205341 | 0.689804456 |
| 0.414247496 | 0.689804456 |
| 0.414317192 | 0.689855265 |
| 0.414323017 | 0.689855265 |
| 0.414588817 | 0.690165033 |
| 0.414642021 | 0.690168672 |
| 0.4147082 | 0.690190108 |
| 0.414733165 | 0.690190108 |
| 0.414738681 | 0.690190108 |
| 0.414748796 | 0.690190108 |
| 0.41482479 | 0.690292453 |
| 0.414868154 | 0.690340492 |
| 0.414990964 | 0.690398262 |
| 0.414998379 | 0.690398262 |
| 0.415033343 | 0.690398275 |
| 0.415096209 | 0.690448716 |
| 0.4151375 | 0.690475073 |
| 0.415185194 | 0.690528294 |
| 0.415354859 | 0.690739434 |
| 0.415412366 | 0.690741708 |
| 0.415565859 | 0.690959607 |
| 0.415619685 | 0.690987494 |
| 0.415669357 | 0.691020632 |
| 0.415736449 | 0.691071253 |
| 0.41583194 | 0.691147367 |
| 0.415897824 | 0.691190972 |
| 0.415940111 | 0.691190972 |
| 0.41605496 | 0.691260751 |
| 0.416073062 | 0.691260751 |
| 0.416227505 | 0.691394649 |
| 0.416335601 | 0.691453445 |
| 0.416563371 | 0.69168719 |

| | |
|---|---|
| 0.416603425 | 0.691729581 |
| 0.4168324 | 0.692037393 |
| 0.41692114 | 0.692112344 |
| 0.417056998 | 0.692193666 |
| 0.417069797 | 0.692193666 |
| 0.417207708 | 0.692282492 |
| 0.417247869 | 0.692282492 |
| 0.417257713 | 0.692282492 |
| 0.417270739 | 0.692282492 |
| 0.417347437 | 0.692361503 |
| 0.417481747 | 0.692439602 |
| 0.41760018 | 0.692587798 |
| 0.417684472 | 0.692679354 |
| 0.417727615 | 0.692726781 |
| 0.417810726 | 0.692782226 |
| 0.41782612 | 0.692782226 |
| 0.417829075 | 0.692782226 |
| 0.41788288 | 0.692816548 |
| 0.417960198 | 0.692919463 |
| 0.418177566 | 0.693135521 |
| 0.418177841 | 0.693135521 |
| 0.418299197 | 0.693218032 |
| 0.418375002 | 0.693317544 |
| 0.418402526 | 0.693339032 |
| 0.418519954 | 0.693365486 |
| 0.418538913 | 0.693365486 |
| 0.418549515 | 0.693365486 |
| 0.418634468 | 0.693433861 |
| 0.418705534 | 0.693527456 |
| 0.418836143 | 0.693668138 |
| 0.418848727 | 0.693668138 |
| 0.41894574 | 0.693780554 |
| 0.419049573 | 0.693859405 |
| 0.419066197 | 0.693859405 |
| 0.419163531 | 0.693972311 |
| 0.419315669 | 0.694127677 |
| 0.419337066 | 0.69413897 |
| 0.419401573 | 0.694163739 |
| 0.419515543 | 0.694269973 |
| 0.4195262 | 0.694269973 |
| 0.419601143 | 0.694310658 |
| 0.419658928 | 0.694320573 |
| 0.419672538 | 0.694320573 |
| 0.419694606 | 0.694320573 |
| 0.419804601 | 0.694406062 |

| | |
|---|---|
| 0.419925806 | 0.694485949 |
| 0.41996023 | 0.694502578 |
| 0.419969467 | 0.694502578 |
| 0.420117867 | 0.694610624 |
| 0.420227215 | 0.694732353 |
| 0.420324409 | 0.694807394 |
| 0.420363254 | 0.694817776 |
| 0.420389424 | 0.694817776 |
| 0.420407663 | 0.694817776 |
| 0.420418222 | 0.694817776 |
| 0.420502561 | 0.694823656 |
| 0.420523993 | 0.694823656 |
| 0.420538491 | 0.694823656 |
| 0.420584673 | 0.694865632 |
| 0.420593076 | 0.694865632 |
| 0.420759785 | 0.694993453 |
| 0.420781033 | 0.694993453 |
| 0.420867004 | 0.695019847 |
| 0.420874995 | 0.695019847 |
| 0.420983015 | 0.695102084 |
| 0.420983255 | 0.695102084 |
| 0.421157946 | 0.695244168 |
| 0.421202425 | 0.695251662 |
| 0.42121728 | 0.695251662 |
| 0.421220877 | 0.695251662 |
| 0.421347053 | 0.695411724 |
| 0.421444224 | 0.695475696 |
| 0.421688912 | 0.695737278 |
| 0.421987979 | 0.695967235 |
| 0.422008581 | 0.695967235 |
| 0.422048957 | 0.695967235 |
| 0.422125799 | 0.696021653 |
| 0.422175075 | 0.696078802 |
| 0.422284936 | 0.696211736 |
| 0.42235594 | 0.696284234 |
| 0.42238256 | 0.696284234 |
| 0.422391015 | 0.696284234 |
| 0.422402007 | 0.696284234 |
| 0.422452944 | 0.696319999 |
| 0.422534287 | 0.696420597 |
| 0.422543221 | 0.696420597 |
| 0.422569166 | 0.696432367 |
| 0.42259423 | 0.696432367 |
| 0.422614008 | 0.696440862 |
| 0.422639236 | 0.696458338 |

| | |
|---|---|
| 0.422671026 | 0.696486627 |
| 0.422736898 | 0.696571072 |
| 0.42276577 | 0.696594545 |
| 0.422821216 | 0.696613604 |
| 0.422864341 | 0.696637681 |
| 0.422970155 | 0.696738477 |
| 0.42319631 | 0.697013133 |
| 0.423217348 | 0.697013133 |
| 0.42322995 | 0.697013133 |
| 0.423267918 | 0.697020796 |
| 0.423329713 | 0.697020796 |
| 0.423351971 | 0.697020796 |
| 0.423356553 | 0.697020796 |
| 0.42335735 | 0.697020796 |
| 0.423370957 | 0.697020796 |
| 0.423375703 | 0.697020796 |
| 0.423450232 | 0.697071208 |
| 0.423522627 | 0.697153581 |
| 0.423529546 | 0.697153581 |
| 0.423672383 | 0.697364597 |
| 0.423782974 | 0.697447501 |
| 0.423829867 | 0.697479184 |
| 0.423985717 | 0.697627271 |
| 0.423993092 | 0.697627271 |
| 0.424278663 | 0.697955834 |
| 0.42433774 | 0.697955834 |
| 0.424357295 | 0.697955834 |
| 0.424368636 | 0.697955834 |
| 0.424432077 | 0.697992515 |
| 0.424449823 | 0.697992946 |
| 0.424496881 | 0.69804623 |
| 0.424532537 | 0.698080761 |
| 0.424567993 | 0.698101993 |
| 0.424574765 | 0.698101993 |
| 0.424700017 | 0.698283831 |
| 0.425038221 | 0.69856826 |
| 0.425057029 | 0.69856826 |
| 0.425063439 | 0.69856826 |
| 0.425078353 | 0.69856826 |
| 0.425156529 | 0.698672624 |
| 0.42520405 | 0.698724072 |
| 0.425358496 | 0.69885875 |
| 0.425502957 | 0.698952519 |
| 0.425540428 | 0.69898158 |
| 0.42555901 | 0.69898158 |

| | |
|---|---|
| 0.425742922 | 0.699015129 |
| 0.425759599 | 0.699015129 |
| 0.425822961 | 0.699044194 |
| 0.425993038 | 0.699246999 |
| 0.426005227 | 0.699246999 |
| 0.426026287 | 0.699254288 |
| 0.426062019 | 0.699254288 |
| 0.426068395 | 0.699254288 |
| 0.426185658 | 0.699398538 |
| 0.426325797 | 0.69953047 |
| 0.426336132 | 0.69953047 |
| 0.426374674 | 0.69953047 |
| 0.426376255 | 0.69953047 |
| 0.426440143 | 0.699535737 |
| 0.426531252 | 0.699604089 |
| 0.4266432 | 0.699691321 |
| 0.426697864 | 0.699756875 |
| 0.426749997 | 0.699818274 |
| 0.426849993 | 0.699943628 |
| 0.427044049 | 0.700107635 |
| 0.427230178 | 0.700316355 |
| 0.427347226 | 0.70040911 |
| 0.427396247 | 0.700419829 |
| 0.427680397 | 0.700716717 |
| 0.427835512 | 0.700886435 |
| 0.427857568 | 0.700886435 |
| 0.428022367 | 0.700965499 |
| 0.428038289 | 0.700965499 |
| 0.428209956 | 0.701101983 |
| 0.42835133 | 0.701237027 |
| 0.428391275 | 0.701278314 |
| 0.428493799 | 0.701422036 |
| 0.428545383 | 0.701453875 |
| 0.428644215 | 0.701475381 |
| 0.428711304 | 0.701531427 |
| 0.428808843 | 0.701545051 |
| 0.428858302 | 0.701545051 |
| 0.428863117 | 0.701545051 |
| 0.428919992 | 0.701608672 |
| 0.428955038 | 0.701608672 |
| 0.428958878 | 0.701608672 |
| 0.428969852 | 0.701608672 |
| 0.429039211 | 0.701663708 |
| 0.429089143 | 0.701697178 |
| 0.429169204 | 0.701755817 |

| | |
|---|---|
| 0.429227719 | 0.701780814 |
| 0.429260381 | 0.701808527 |
| 0.429460122 | 0.702018508 |
| 0.429538139 | 0.702093931 |
| 0.429597232 | 0.70212544 |
| 0.429635416 | 0.702156543 |
| 0.429714183 | 0.702236233 |
| 0.429717966 | 0.702236233 |
| 0.42973421 | 0.702236233 |
| 0.429785903 | 0.702281991 |
| 0.430037126 | 0.702558737 |
| 0.430229316 | 0.702699777 |
| 0.430294221 | 0.702699777 |
| 0.430295115 | 0.702699777 |
| 0.430297971 | 0.702699777 |
| 0.430378488 | 0.702699777 |
| 0.430390414 | 0.702699777 |
| 0.430399448 | 0.702699777 |
| 0.430410438 | 0.702699777 |
| 0.430535044 | 0.702830935 |
| 0.430591268 | 0.702887219 |
| 0.430646249 | 0.702888576 |
| 0.430662548 | 0.702888576 |
| 0.430699853 | 0.702888576 |
| 0.430734138 | 0.702888576 |
| 0.43077055 | 0.702888576 |
| 0.430776968 | 0.702888576 |
| 0.430839939 | 0.702943159 |
| 0.4308904 | 0.70298804 |
| 0.430986998 | 0.703086752 |
| 0.431023114 | 0.703091094 |
| 0.431056573 | 0.703091094 |
| 0.431076941 | 0.703091094 |
| 0.431105305 | 0.703091094 |
| 0.431136004 | 0.703091094 |
| 0.43125765 | 0.703188308 |
| 0.431327809 | 0.703188308 |
| 0.431422556 | 0.703196721 |
| 0.431423524 | 0.703196721 |
| 0.431548131 | 0.703241416 |
| 0.431579376 | 0.703241416 |
| 0.431607404 | 0.703241416 |
| 0.43161268 | 0.703241416 |
| 0.431641122 | 0.703241416 |
| 0.431652098 | 0.703241416 |

| | |
|---|---|
| 0.431676482 | 0.703248017 |
| 0.431709926 | 0.703278446 |
| 0.431784818 | 0.703301069 |
| 0.431829692 | 0.703301069 |
| 0.431837951 | 0.703301069 |
| 0.431881805 | 0.703337133 |
| 0.431991148 | 0.70344641 |
| 0.432046999 | 0.703466631 |
| 0.432117108 | 0.703556731 |
| 0.432229145 | 0.703618877 |
| 0.432309272 | 0.703725262 |
| 0.432412734 | 0.703845566 |
| 0.432492502 | 0.703909945 |
| 0.432662958 | 0.703971752 |
| 0.432717436 | 0.703971752 |
| 0.432726752 | 0.703971752 |
| 0.43276614 | 0.703986644 |
| 0.43278025 | 0.703986644 |
| 0.432905505 | 0.704118247 |
| 0.432931859 | 0.704137063 |
| 0.43303765 | 0.704257647 |
| 0.433077255 | 0.704257647 |
| 0.433086318 | 0.704257647 |
| 0.43309472 | 0.704257647 |
| 0.433214374 | 0.704356024 |
| 0.433285727 | 0.704416902 |
| 0.433296188 | 0.704416902 |
| 0.433394946 | 0.704481267 |
| 0.433448913 | 0.704496857 |
| 0.433477949 | 0.704520007 |
| 0.433495057 | 0.704523769 |
| 0.433605752 | 0.704635618 |
| 0.433608262 | 0.704635618 |
| 0.433679132 | 0.704678653 |
| 0.433747854 | 0.704742233 |
| 0.433765453 | 0.704746784 |
| 0.433933503 | 0.70491865 |
| 0.433945239 | 0.70491865 |
| 0.43399586 | 0.704976836 |
| 0.434147592 | 0.705093807 |
| 0.434208958 | 0.705093807 |
| 0.434217716 | 0.705093807 |
| 0.434250038 | 0.705093807 |
| 0.434269357 | 0.705093807 |
| 0.434307026 | 0.705093807 |

| | |
|---|---|
| 0.434319546 | 0.705093807 |
| 0.434461931 | 0.705228805 |
| 0.434725479 | 0.705416184 |
| 0.434879555 | 0.705570041 |
| 0.434953659 | 0.705605266 |
| 0.435123183 | 0.705704933 |
| 0.435174945 | 0.705704933 |
| 0.435377056 | 0.705872245 |
| 0.435396923 | 0.705880425 |
| 0.43546886 | 0.705946147 |
| 0.435471517 | 0.705946147 |
| 0.43550561 | 0.705953823 |
| 0.435516308 | 0.705953823 |
| 0.435767219 | 0.706143463 |
| 0.435781566 | 0.706143463 |
| 0.435835877 | 0.706183416 |
| 0.436116561 | 0.70642665 |
| 0.436224196 | 0.706548199 |
| 0.43634785 | 0.706682352 |
| 0.436388674 | 0.70668836 |
| 0.436419022 | 0.70668836 |
| 0.436445032 | 0.706689588 |
| 0.436642969 | 0.706889925 |
| 0.436750833 | 0.707016482 |
| 0.436789949 | 0.707027142 |
| 0.436792912 | 0.707027142 |
| 0.436801953 | 0.707027142 |
| 0.43688582 | 0.707066771 |
| 0.437103448 | 0.707322843 |
| 0.437145412 | 0.707366716 |
| 0.437213038 | 0.707405272 |
| 0.437238024 | 0.707405272 |
| 0.437239272 | 0.707405272 |
| 0.437358931 | 0.707440877 |
| 0.437411114 | 0.707440877 |
| 0.437443861 | 0.707440877 |
| 0.437444383 | 0.707440877 |
| 0.437484371 | 0.707440877 |
| 0.43760693 | 0.707584629 |
| 0.437635673 | 0.707607081 |
| 0.43767552 | 0.707647486 |
| 0.437755668 | 0.707688572 |
| 0.437767978 | 0.707688572 |
| 0.437775227 | 0.707688572 |
| 0.43783436 | 0.70773612 |

| | |
|---|---|
| 0.437856791 | 0.707748359 |
| 0.437986051 | 0.707909242 |
| 0.438065731 | 0.707990207 |
| 0.438065874 | 0.707990207 |
| 0.43810672 | 0.708028477 |
| 0.438119286 | 0.708028477 |
| 0.43830944 | 0.708215397 |
| 0.438436974 | 0.708311152 |
| 0.43852287 | 0.708392251 |
| 0.438600056 | 0.708399381 |
| 0.438644234 | 0.708420107 |
| 0.438710715 | 0.708479426 |
| 0.438827736 | 0.708541616 |
| 0.438904998 | 0.708541616 |
| 0.43893117 | 0.708541616 |
| 0.438937632 | 0.708541616 |
| 0.438947346 | 0.708541616 |
| 0.438996101 | 0.708541616 |
| 0.439024761 | 0.708541616 |
| 0.439026593 | 0.708541616 |
| 0.439086258 | 0.708605354 |
| 0.439178873 | 0.708658767 |
| 0.439244325 | 0.708675627 |
| 0.43926372 | 0.708675627 |
| 0.439311564 | 0.708728807 |
| 0.43936621 | 0.708748915 |
| 0.439411458 | 0.708769923 |
| 0.43949229 | 0.708835378 |
| 0.439493025 | 0.708835378 |
| 0.439496687 | 0.708835378 |
| 0.439621346 | 0.708940402 |
| 0.439639437 | 0.708945572 |
| 0.439668297 | 0.708963173 |
| 0.439685674 | 0.708963173 |
| 0.439740849 | 0.708968071 |
| 0.439742704 | 0.708968071 |
| 0.439759096 | 0.708970498 |
| 0.43979613 | 0.709006204 |
| 0.439970675 | 0.709184298 |
| 0.440037966 | 0.709204028 |
| 0.440122144 | 0.70928467 |
| 0.440124534 | 0.70928467 |
| 0.440188152 | 0.70928467 |
| 0.4401988 | 0.70928467 |
| 0.440262804 | 0.709287579 |

| | |
|---|---|
| 0.44028341 | 0.709287579 |
| 0.440344635 | 0.709335318 |
| 0.440345547 | 0.709335318 |
| 0.440469384 | 0.709439924 |
| 0.440482247 | 0.709439924 |
| 0.440755524 | 0.709736056 |
| 0.440827246 | 0.709827548 |
| 0.440892731 | 0.709860996 |
| 0.440966914 | 0.709908438 |
| 0.441029738 | 0.709977086 |
| 0.441161483 | 0.710097602 |
| 0.441183471 | 0.710097602 |
| 0.441203692 | 0.710097602 |
| 0.441426374 | 0.710221622 |
| 0.441429623 | 0.710221622 |
| 0.441730962 | 0.710427482 |
| 0.441743219 | 0.710427482 |
| 0.441924197 | 0.710523532 |
| 0.441930808 | 0.710523532 |
| 0.442271267 | 0.710974909 |
| 0.442364907 | 0.71103427 |
| 0.44236791 | 0.71103427 |
| 0.4424746 | 0.711157756 |
| 0.442511777 | 0.711193507 |
| 0.442716946 | 0.711372983 |
| 0.442927086 | 0.711561132 |
| 0.443448078 | 0.712025542 |
| 0.443627496 | 0.712177596 |
| 0.44368474 | 0.712213451 |
| 0.443731536 | 0.712264564 |
| 0.443851541 | 0.712378996 |
| 0.444018531 | 0.71253094 |
| 0.444106869 | 0.712602852 |
| 0.444132964 | 0.712620716 |
| 0.44428112 | 0.712707654 |
| 0.444306863 | 0.712707654 |
| 0.444396399 | 0.712827269 |
| 0.444467659 | 0.712878587 |
| 0.444479143 | 0.712878587 |
| 0.444582321 | 0.71298142 |
| 0.44461487 | 0.712985603 |
| 0.444761355 | 0.713088404 |
| 0.444808562 | 0.713088404 |
| 0.444819278 | 0.713088404 |
| 0.444827771 | 0.713088404 |

| | |
|---|---|
| 0.444831575 | 0.713088404 |
| 0.444910923 | 0.713100203 |
| 0.444991921 | 0.713182024 |
| 0.445043661 | 0.713240946 |
| 0.445079811 | 0.71327488 |
| 0.445138763 | 0.71329735 |
| 0.445241204 | 0.713393791 |
| 0.445253346 | 0.713393791 |
| 0.445261841 | 0.713393791 |
| 0.445287212 | 0.713393791 |
| 0.445313226 | 0.713407533 |
| 0.445340729 | 0.713407533 |
| 0.445382436 | 0.713407533 |
| 0.445387272 | 0.713407533 |
| 0.445456843 | 0.713494974 |
| 0.445524647 | 0.71357958 |
| 0.445562837 | 0.713586452 |
| 0.445573887 | 0.713586452 |
| 0.445800958 | 0.713806826 |
| 0.445816403 | 0.713806826 |
| 0.44584863 | 0.713834427 |
| 0.445932249 | 0.713934969 |
| 0.44595472 | 0.713934969 |
| 0.446134158 | 0.714128818 |
| 0.44613705 | 0.714128818 |
| 0.446144121 | 0.714128818 |
| 0.446167449 | 0.714128818 |
| 0.446188373 | 0.714138309 |
| 0.446385535 | 0.714357846 |
| 0.446454245 | 0.714386695 |
| 0.446500279 | 0.714397447 |
| 0.446596021 | 0.714502626 |
| 0.446611737 | 0.714503769 |
| 0.446731477 | 0.714538674 |
| 0.44681303 | 0.714574087 |
| 0.446839613 | 0.714574087 |
| 0.446872482 | 0.714574087 |
| 0.446945469 | 0.714597497 |
| 0.447021733 | 0.71467956 |
| 0.447037106 | 0.714680146 |
| 0.447205186 | 0.714908084 |
| 0.447221412 | 0.714908084 |
| 0.447224714 | 0.714908084 |
| 0.44730072 | 0.714957592 |
| 0.447365909 | 0.715013797 |

| | |
|---|---|
| 0.4474422 | 0.71508167 |
| 0.447453418 | 0.71508167 |
| 0.447469355 | 0.715083145 |
| 0.447550101 | 0.715156774 |
| 0.447569068 | 0.715156774 |
| 0.447624765 | 0.715156774 |
| 0.447753592 | 0.715156774 |
| 0.447758574 | 0.715156774 |
| 0.44776834 | 0.715156774 |
| 0.447906877 | 0.715278327 |
| 0.447954291 | 0.715313319 |
| 0.448119938 | 0.715465955 |
| 0.448170334 | 0.71547201 |
| 0.448207758 | 0.71547201 |
| 0.448293922 | 0.715528648 |
| 0.44837483 | 0.715593728 |
| 0.44845234 | 0.715669468 |
| 0.448597272 | 0.715828802 |
| 0.448637708 | 0.715869342 |
| 0.448660197 | 0.715871302 |
| 0.448668998 | 0.715871302 |
| 0.448928839 | 0.716165929 |
| 0.449134299 | 0.716376853 |
| 0.449351599 | 0.716624317 |
| 0.449550217 | 0.716844867 |
| 0.449595251 | 0.716844867 |
| 0.449614989 | 0.716852339 |
| 0.449673375 | 0.716897431 |
| 0.450072261 | 0.717397238 |
| 0.450107382 | 0.717397238 |
| 0.450167975 | 0.717404016 |
| 0.450228979 | 0.717422984 |
| 0.450264382 | 0.717437603 |
| 0.450294999 | 0.717437603 |
| 0.450593077 | 0.717835044 |
| 0.450818751 | 0.717923881 |
| 0.450854831 | 0.717923881 |
| 0.450857423 | 0.717923881 |
| 0.450872555 | 0.717923881 |
| 0.450905097 | 0.717923881 |
| 0.451046221 | 0.718004558 |
| 0.451117459 | 0.718069958 |
| 0.451182556 | 0.718103955 |
| 0.451219255 | 0.718103955 |
| 0.451225406 | 0.718103955 |

| | |
|---|---|
| 0.451339186 | 0.718214241 |
| 0.451602717 | 0.718482183 |
| 0.451774462 | 0.718671724 |
| 0.451848315 | 0.718728866 |
| 0.451895075 | 0.718779239 |
| 0.451972017 | 0.718853607 |
| 0.452020453 | 0.718906636 |
| 0.452066358 | 0.718920719 |
| 0.452083975 | 0.718920719 |
| 0.452166492 | 0.718970839 |
| 0.452333247 | 0.719175289 |
| 0.452357199 | 0.719175289 |
| 0.452411764 | 0.719175289 |
| 0.45241307 | 0.719175289 |
| 0.452471378 | 0.71919864 |
| 0.452475866 | 0.71919864 |
| 0.452547132 | 0.71921037 |
| 0.452591027 | 0.71921037 |
| 0.452639241 | 0.719242289 |
| 0.452761893 | 0.719326116 |
| 0.452833297 | 0.719360956 |
| 0.453043145 | 0.71941778 |
| 0.453051363 | 0.71941778 |
| 0.453066893 | 0.71941778 |
| 0.453201909 | 0.719475464 |
| 0.453206678 | 0.719475464 |
| 0.453259631 | 0.719524267 |
| 0.453284555 | 0.719524267 |
| 0.45328503 | 0.719524267 |
| 0.453389287 | 0.719617809 |
| 0.453438323 | 0.71965007 |
| 0.453500274 | 0.71965007 |
| 0.453641113 | 0.719746735 |
| 0.453704528 | 0.719746735 |
| 0.453710252 | 0.719746735 |
| 0.453737876 | 0.719746735 |
| 0.453740002 | 0.719746735 |
| 0.453740872 | 0.719746735 |
| 0.453757647 | 0.719746735 |
| 0.453835419 | 0.71982215 |
| 0.45400991 | 0.720026971 |
| 0.454046184 | 0.720036547 |
| 0.454122388 | 0.720109438 |
| 0.454185099 | 0.720136951 |
| 0.454202675 | 0.720140844 |

| | |
|---|---|
| 0.454342324 | 0.720338278 |
| 0.45451159 | 0.720446308 |
| 0.454515042 | 0.720446308 |
| 0.454573999 | 0.720489729 |
| 0.454728195 | 0.720601467 |
| 0.454735278 | 0.720601467 |
| 0.454757153 | 0.720612155 |
| 0.454840878 | 0.72069456 |
| 0.454866489 | 0.72069456 |
| 0.454884816 | 0.72069456 |
| 0.45493253 | 0.720698231 |
| 0.454995337 | 0.720725809 |
| 0.455127916 | 0.720846252 |
| 0.455131914 | 0.720846252 |
| 0.455397831 | 0.721197607 |
| 0.45544032 | 0.721197607 |
| 0.455645907 | 0.721341632 |
| 0.4557413 | 0.721341632 |
| 0.455753413 | 0.721341632 |
| 0.455909318 | 0.721453736 |
| 0.456190319 | 0.72175452 |
| 0.456354608 | 0.721894543 |
| 0.456432103 | 0.721993151 |
| 0.456531216 | 0.722077738 |
| 0.456585697 | 0.722077738 |
| 0.456595273 | 0.722077738 |
| 0.45662801 | 0.722077738 |
| 0.45663192 | 0.722077738 |
| 0.456668011 | 0.722077738 |
| 0.456700829 | 0.722077738 |
| 0.456765158 | 0.722088299 |
| 0.456907572 | 0.722141095 |
| 0.456912572 | 0.722141095 |
| 0.456976925 | 0.7221594 |
| 0.457147723 | 0.72223676 |
| 0.457353061 | 0.722394264 |
| 0.457385319 | 0.7223973 |
| 0.457435442 | 0.722452508 |
| 0.457462804 | 0.722461672 |
| 0.457552428 | 0.722517472 |
| 0.457679887 | 0.722608923 |
| 0.457791937 | 0.722660027 |
| 0.457794438 | 0.722660027 |
| 0.457905002 | 0.722714791 |
| 0.457959447 | 0.722752819 |

| | |
|---|---|
| 0.458019381 | 0.722777605 |
| 0.458020679 | 0.722777605 |
| 0.458242325 | 0.722958458 |
| 0.458301295 | 0.722958458 |
| 0.458330983 | 0.722960944 |
| 0.458334196 | 0.722960944 |
| 0.458369622 | 0.722969802 |
| 0.458413898 | 0.722969802 |
| 0.458423148 | 0.722969802 |
| 0.458430891 | 0.722969802 |
| 0.458473614 | 0.723013239 |
| 0.458573054 | 0.723093484 |
| 0.458585229 | 0.723093484 |
| 0.458686908 | 0.723205925 |
| 0.458727871 | 0.723222627 |
| 0.458762652 | 0.72322958 |
| 0.458868585 | 0.72330504 |
| 0.458916524 | 0.72330504 |
| 0.458931104 | 0.72330504 |
| 0.458932013 | 0.72330504 |
| 0.458970511 | 0.723341778 |
| 0.459145307 | 0.723449288 |
| 0.459153813 | 0.723449288 |
| 0.459168139 | 0.723449288 |
| 0.459227466 | 0.723449288 |
| 0.459245632 | 0.723449288 |
| 0.45927405 | 0.723449288 |
| 0.459353043 | 0.723537633 |
| 0.459404258 | 0.723594372 |
| 0.459477787 | 0.723626638 |
| 0.459500712 | 0.723626638 |
| 0.459529109 | 0.723631859 |
| 0.459534415 | 0.723631859 |
| 0.459696987 | 0.723733834 |
| 0.459705544 | 0.723733834 |
| 0.459785222 | 0.723811423 |
| 0.459920595 | 0.723928819 |
| 0.460058189 | 0.724049678 |
| 0.460299184 | 0.724320796 |
| 0.460302911 | 0.724320796 |
| 0.460404028 | 0.724380967 |
| 0.460406351 | 0.724380967 |
| 0.460442783 | 0.724380967 |
| 0.460446836 | 0.724380967 |
| 0.460559541 | 0.724470508 |

| | |
|---|---|
| 0.460614972 | 0.724495022 |
| 0.46073431 | 0.724610936 |
| 0.460807501 | 0.724675687 |
| 0.460858277 | 0.724675687 |
| 0.46087887 | 0.724675687 |
| 0.460906223 | 0.724677932 |
| 0.461116393 | 0.724828887 |
| 0.461193718 | 0.72485739 |
| 0.461227039 | 0.72485739 |
| 0.461275983 | 0.724864434 |
| 0.461350163 | 0.724885334 |
| 0.461387047 | 0.724895457 |
| 0.461561722 | 0.725002478 |
| 0.46160328 | 0.725043844 |
| 0.461667777 | 0.725073412 |
| 0.461731281 | 0.725101416 |
| 0.461762937 | 0.72510331 |
| 0.461804605 | 0.725132648 |
| 0.461848541 | 0.725148673 |
| 0.461859302 | 0.725148673 |
| 0.461867953 | 0.725148673 |
| 0.461936142 | 0.725207918 |
| 0.46200246 | 0.725288126 |
| 0.46220988 | 0.725398556 |
| 0.462231173 | 0.725408068 |
| 0.462253675 | 0.725419479 |
| 0.462305367 | 0.725476696 |
| 0.462375752 | 0.725500096 |
| 0.462379654 | 0.725500096 |
| 0.462381211 | 0.725500096 |
| 0.462570144 | 0.725698073 |
| 0.462712058 | 0.725851822 |
| 0.46273821 | 0.725868939 |
| 0.462759662 | 0.725878682 |
| 0.462873404 | 0.725948243 |
| 0.462901933 | 0.725948243 |
| 0.463005644 | 0.726023554 |
| 0.463019704 | 0.726023554 |
| 0.463077641 | 0.726090496 |
| 0.463098587 | 0.726099434 |
| 0.463188822 | 0.72614355 |
| 0.463261585 | 0.726153626 |
| 0.463318239 | 0.726153626 |
| 0.4633499 | 0.726153626 |
| 0.463362535 | 0.726153626 |

| | |
|---|---|
| 0.463414835 | 0.726153626 |
| 0.463425411 | 0.726153626 |
| 0.463426011 | 0.726153626 |
| 0.463483824 | 0.726153626 |
| 0.46357883 | 0.726254693 |
| 0.463761539 | 0.726456424 |
| 0.464010932 | 0.726542475 |
| 0.464029351 | 0.726542475 |
| 0.46403656 | 0.726542475 |
| 0.464055724 | 0.726542475 |
| 0.464065635 | 0.726542475 |
| 0.464067622 | 0.726542475 |
| 0.464180882 | 0.726599486 |
| 0.464186395 | 0.726599486 |
| 0.464224684 | 0.726599486 |
| 0.464234128 | 0.726599486 |
| 0.464378033 | 0.726660419 |
| 0.464501349 | 0.7267676 |
| 0.4647832 | 0.727083412 |
| 0.464824776 | 0.727083412 |
| 0.464850775 | 0.727083412 |
| 0.464898007 | 0.727083412 |
| 0.465032715 | 0.727192866 |
| 0.465091277 | 0.727260564 |
| 0.465108561 | 0.727263714 |
| 0.465139352 | 0.727267161 |
| 0.465210236 | 0.727351057 |
| 0.465235842 | 0.727367214 |
| 0.465354995 | 0.727457982 |
| 0.465517607 | 0.727621258 |
| 0.465589932 | 0.727658059 |
| 0.465618241 | 0.727678424 |
| 0.466073442 | 0.728150889 |
| 0.46619402 | 0.728261648 |
| 0.466296969 | 0.728313017 |
| 0.466350796 | 0.728341933 |
| 0.466418586 | 0.728341933 |
| 0.466455109 | 0.728341933 |
| 0.466455701 | 0.728341933 |
| 0.466538381 | 0.72840593 |
| 0.466568532 | 0.72840593 |
| 0.466626097 | 0.72841695 |
| 0.466716329 | 0.728475984 |
| 0.466763066 | 0.728487499 |
| 0.466844423 | 0.7285906 |

| | |
|---|---|
| 0.466862492 | 0.728594924 |
| 0.467037402 | 0.728700748 |
| 0.46712151 | 0.728784228 |
| 0.46720522 | 0.728843203 |
| 0.467225216 | 0.728847601 |
| 0.467277427 | 0.728847601 |
| 0.467284555 | 0.728847601 |
| 0.467308544 | 0.728859203 |
| 0.4673226 | 0.728859203 |
| 0.467460198 | 0.729029703 |
| 0.467462534 | 0.729029703 |
| 0.46758418 | 0.729052306 |
| 0.467627781 | 0.729075092 |
| 0.46773046 | 0.729145819 |
| 0.467844316 | 0.729243045 |
| 0.467925748 | 0.729323309 |
| 0.468022433 | 0.729401323 |
| 0.468094845 | 0.729442567 |
| 0.468172078 | 0.729487224 |
| 0.468184768 | 0.729487224 |
| 0.468387539 | 0.729662415 |
| 0.468404449 | 0.729662415 |
| 0.468585978 | 0.729825841 |
| 0.468660405 | 0.729917891 |
| 0.468728292 | 0.729952011 |
| 0.468879838 | 0.730106693 |
| 0.468888937 | 0.730106693 |
| 0.468935501 | 0.730131455 |
| 0.469109258 | 0.730330368 |
| 0.469170194 | 0.730401361 |
| 0.469263705 | 0.73049828 |
| 0.469278464 | 0.73049828 |
| 0.469531001 | 0.73079002 |
| 0.4695426 | 0.73079002 |
| 0.469573229 | 0.730813808 |
| 0.469620303 | 0.730833598 |
| 0.4696281 | 0.730833598 |
| 0.469714017 | 0.730891187 |
| 0.469715392 | 0.730891187 |
| 0.46973037 | 0.730891187 |
| 0.469829401 | 0.730981048 |
| 0.469834166 | 0.730981048 |
| 0.469916059 | 0.730983153 |
| 0.469927608 | 0.730983153 |
| 0.470076256 | 0.731158305 |

| | |
|---|---|
| 0.470086263 | 0.731158305 |
| 0.47019731 | 0.731235504 |
| 0.470257353 | 0.731265164 |
| 0.470443971 | 0.731356411 |
| 0.470486143 | 0.731374226 |
| 0.47053745 | 0.731417106 |
| 0.470544442 | 0.731417106 |
| 0.470656087 | 0.731542898 |
| 0.470789756 | 0.731666947 |
| 0.470815542 | 0.731666947 |
| 0.471048252 | 0.731818082 |
| 0.471138022 | 0.731838184 |
| 0.471185768 | 0.73187106 |
| 0.471209743 | 0.731877982 |
| 0.471320438 | 0.732002168 |
| 0.471471143 | 0.732212349 |
| 0.471569155 | 0.732284548 |
| 0.471609862 | 0.732284548 |
| 0.471679854 | 0.732325481 |
| 0.471739444 | 0.732325481 |
| 0.471757412 | 0.732325481 |
| 0.471757585 | 0.732325481 |
| 0.471804693 | 0.732340598 |
| 0.471815124 | 0.732340598 |
| 0.47193156 | 0.732425848 |
| 0.471992893 | 0.732448928 |
| 0.472038705 | 0.732448928 |
| 0.472136972 | 0.732489412 |
| 0.472144577 | 0.732489412 |
| 0.472378329 | 0.732665604 |
| 0.472472165 | 0.732715698 |
| 0.472578304 | 0.732735443 |
| 0.472587397 | 0.732735443 |
| 0.472666529 | 0.732780113 |
| 0.472752192 | 0.732815903 |
| 0.472785576 | 0.732843789 |
| 0.47285971 | 0.732887155 |
| 0.472890833 | 0.732891579 |
| 0.472946508 | 0.732926279 |
| 0.47298785 | 0.732966498 |
| 0.473108586 | 0.733129742 |
| 0.473176663 | 0.733205847 |
| 0.473188488 | 0.733205847 |
| 0.473314764 | 0.733268156 |
| 0.473318502 | 0.733268156 |

| | |
|---|---|
| 0.473370141 | 0.733272611 |
| 0.473395765 | 0.733288454 |
| 0.473499716 | 0.733401768 |
| 0.473580859 | 0.733479742 |
| 0.473601165 | 0.73348734 |
| 0.473775581 | 0.733614332 |
| 0.47386718 | 0.733689874 |
| 0.474045035 | 0.733912261 |
| 0.474074982 | 0.733934768 |
| 0.474102408 | 0.733944808 |
| 0.474161742 | 0.733973651 |
| 0.474279592 | 0.734059464 |
| 0.474355055 | 0.734081494 |
| 0.474380027 | 0.734081494 |
| 0.474395113 | 0.734081494 |
| 0.474400955 | 0.734081494 |
| 0.474729077 | 0.734421425 |
| 0.474772049 | 0.734440963 |
| 0.47483494 | 0.734490538 |
| 0.474861779 | 0.734508198 |
| 0.474989449 | 0.734644956 |
| 0.475029289 | 0.734648004 |
| 0.475047857 | 0.734652865 |
| 0.475173822 | 0.734752237 |
| 0.475278121 | 0.734841939 |
| 0.475359338 | 0.734882288 |
| 0.475435027 | 0.73492816 |
| 0.475764244 | 0.735283242 |
| 0.475802632 | 0.735317152 |
| 0.475827742 | 0.735317152 |
| 0.475832503 | 0.735317152 |
| 0.475872989 | 0.735355856 |
| 0.47598839 | 0.735486457 |
| 0.476062877 | 0.735577687 |
| 0.476085673 | 0.735584287 |
| 0.47609895 | 0.735584287 |
| 0.476154433 | 0.735623692 |
| 0.476187015 | 0.735646456 |
| 0.476332986 | 0.735756345 |
| 0.476527847 | 0.735952058 |
| 0.476536954 | 0.735952058 |
| 0.47657992 | 0.735970684 |
| 0.476696017 | 0.736054504 |
| 0.476720192 | 0.736064481 |
| 0.476781637 | 0.736079469 |

| | |
|---|---|
| 0.47681142 | 0.736079469 |
| 0.476846632 | 0.736095176 |
| 0.476902128 | 0.736095176 |
| 0.476907822 | 0.736095176 |
| 0.476939362 | 0.736096147 |
| 0.476964991 | 0.736104205 |
| 0.476986899 | 0.736104205 |
| 0.47699095 | 0.736104205 |
| 0.477125705 | 0.736243091 |
| 0.477228148 | 0.736327102 |
| 0.477267989 | 0.736358986 |
| 0.477375331 | 0.736433532 |
| 0.477389902 | 0.736433532 |
| 0.477517731 | 0.736559154 |
| 0.477570919 | 0.736593483 |
| 0.477670139 | 0.736689557 |
| 0.477714576 | 0.736689557 |
| 0.477760047 | 0.736718181 |
| 0.477959052 | 0.73697733 |
| 0.478195411 | 0.737187781 |
| 0.478283638 | 0.73728686 |
| 0.478387891 | 0.737328244 |
| 0.478420109 | 0.73735404 |
| 0.478436462 | 0.737355382 |
| 0.478452399 | 0.737356083 |
| 0.478717925 | 0.737622084 |
| 0.478740045 | 0.737632303 |
| 0.478878051 | 0.737786588 |
| 0.478902143 | 0.737786588 |
| 0.479042119 | 0.737869132 |
| 0.479083938 | 0.737869132 |
| 0.479100375 | 0.737869132 |
| 0.479101983 | 0.737869132 |
| 0.479151899 | 0.737884939 |
| 0.479215667 | 0.737935419 |
| 0.479440587 | 0.73815966 |
| 0.479454282 | 0.73815966 |
| 0.479600998 | 0.738298327 |
| 0.479660025 | 0.738306057 |
| 0.479682017 | 0.738306057 |
| 0.479688887 | 0.738306057 |
| 0.479738111 | 0.738329598 |
| 0.479773403 | 0.738340696 |
| 0.479857504 | 0.738389845 |
| 0.479898081 | 0.738389845 |

| | |
|---|---|
| 0.479904112 | 0.738389845 |
| 0.47994733 | 0.738393626 |
| 0.480037453 | 0.738435104 |
| 0.480056706 | 0.738435104 |
| 0.480209587 | 0.738537909 |
| 0.480211686 | 0.738537909 |
| 0.480232787 | 0.738546512 |
| 0.480272784 | 0.738572491 |
| 0.480489594 | 0.738798318 |
| 0.480561386 | 0.738837147 |
| 0.48061372 | 0.738869871 |
| 0.4806626 | 0.738869871 |
| 0.480695796 | 0.738876834 |
| 0.480763571 | 0.738930124 |
| 0.480789101 | 0.738930124 |
| 0.480792525 | 0.738930124 |
| 0.480838706 | 0.738960203 |
| 0.480913013 | 0.739002098 |
| 0.480916938 | 0.739002098 |
| 0.480960212 | 0.739033363 |
| 0.481149865 | 0.739243614 |
| 0.481167238 | 0.739243614 |
| 0.481186683 | 0.739249642 |
| 0.481267215 | 0.73933422 |
| 0.481467257 | 0.739537557 |
| 0.481526801 | 0.739585403 |
| 0.481636542 | 0.739702157 |
| 0.48171138 | 0.73974553 |
| 0.481823715 | 0.739894182 |
| 0.481856211 | 0.739901098 |
| 0.481904919 | 0.73992112 |
| 0.481912637 | 0.73992112 |
| 0.481971454 | 0.73992112 |
| 0.482113276 | 0.739945163 |
| 0.482167051 | 0.739945163 |
| 0.482167578 | 0.739945163 |
| 0.482172304 | 0.739945163 |
| 0.482373328 | 0.740102809 |
| 0.482448043 | 0.740184677 |
| 0.482536949 | 0.740257054 |
| 0.482550283 | 0.740257054 |
| 0.482604731 | 0.740259234 |
| 0.482643676 | 0.740270204 |
| 0.482721373 | 0.740328631 |
| 0.482743947 | 0.740328631 |

| | |
|---|---|
| 0.482899637 | 0.740379948 |
| 0.483077607 | 0.740578015 |
| 0.483175223 | 0.740606237 |
| 0.483189317 | 0.740606237 |
| 0.483228892 | 0.74064306 |
| 0.483354057 | 0.740735064 |
| 0.483365262 | 0.740735064 |
| 0.483372724 | 0.740735064 |
| 0.48339019 | 0.740735064 |
| 0.483395172 | 0.740735064 |
| 0.483397789 | 0.740735064 |
| 0.483502244 | 0.740832396 |
| 0.483571943 | 0.740832396 |
| 0.483574196 | 0.740832396 |
| 0.48360179 | 0.740832396 |
| 0.483717077 | 0.740832396 |
| 0.483728264 | 0.740832396 |
| 0.483732451 | 0.740832396 |
| 0.483914854 | 0.74102671 |
| 0.483932682 | 0.741030184 |
| 0.483955209 | 0.741039534 |
| 0.48407212 | 0.741124561 |
| 0.484100992 | 0.74114494 |
| 0.484141607 | 0.741183296 |
| 0.484217927 | 0.741234572 |
| 0.484224716 | 0.741234572 |
| 0.484336057 | 0.741309476 |
| 0.484348548 | 0.741309476 |
| 0.484484233 | 0.741442601 |
| 0.484497177 | 0.741442601 |
| 0.484524814 | 0.741442601 |
| 0.484585818 | 0.741442601 |
| 0.484591204 | 0.741442601 |
| 0.484628109 | 0.741475247 |
| 0.484846228 | 0.741718287 |
| 0.484849254 | 0.741718287 |
| 0.484894528 | 0.741741154 |
| 0.48489535 | 0.741741154 |
| 0.484989624 | 0.741813888 |
| 0.485056252 | 0.741863564 |
| 0.485147795 | 0.741935845 |
| 0.485240717 | 0.741935845 |
| 0.485343163 | 0.742000793 |
| 0.485375022 | 0.742000793 |
| 0.485390014 | 0.742000793 |

| | |
|---|---|
| 0.485393267 | 0.742000793 |
| 0.48540783 | 0.742000793 |
| 0.485445765 | 0.742025186 |
| 0.485454947 | 0.742025186 |
| 0.485500791 | 0.742071443 |
| 0.485680535 | 0.742281994 |
| 0.485696313 | 0.742281994 |
| 0.485752287 | 0.742330322 |
| 0.48580664 | 0.742372151 |
| 0.48582281 | 0.742373043 |
| 0.485947862 | 0.742492664 |
| 0.485975998 | 0.742509692 |
| 0.486208566 | 0.742628932 |
| 0.486266033 | 0.742645257 |
| 0.486317465 | 0.742676176 |
| 0.486372871 | 0.742736974 |
| 0.486664982 | 0.743034729 |
| 0.486666009 | 0.743034729 |
| 0.486681389 | 0.743034729 |
| 0.486698864 | 0.743034729 |
| 0.48671666 | 0.743034729 |
| 0.486736391 | 0.743034729 |
| 0.486770668 | 0.743034729 |
| 0.486819353 | 0.743085229 |
| 0.486846389 | 0.743092274 |
| 0.486868138 | 0.743092274 |
| 0.486874134 | 0.743092274 |
| 0.486911168 | 0.743092274 |
| 0.486917582 | 0.743092274 |
| 0.486940577 | 0.743103556 |
| 0.486987936 | 0.743120024 |
| 0.487011991 | 0.743120024 |
| 0.487071746 | 0.743184662 |
| 0.487882261 | 0.743978462 |
| 0.488042587 | 0.744189162 |
| 0.488108964 | 0.744218895 |
| 0.488309648 | 0.74443522 |
| 0.48835928 | 0.74443522 |
| 0.488359915 | 0.74443522 |
| 0.488403133 | 0.744464624 |
| 0.488415739 | 0.744464624 |
| 0.488510791 | 0.744536751 |
| 0.488555715 | 0.744536751 |
| 0.488644109 | 0.744536751 |
| 0.488708161 | 0.744597059 |

| | |
|---|---|
| 0.488911874 | 0.744846966 |
| 0.489176999 | 0.745028574 |
| 0.489276526 | 0.745068198 |
| 0.489285793 | 0.745068198 |
| 0.489291747 | 0.745068198 |
| 0.489359949 | 0.745100585 |
| 0.489519212 | 0.745211632 |
| 0.489607103 | 0.745262472 |
| 0.489631137 | 0.745275237 |
| 0.489782902 | 0.745452002 |
| 0.489841634 | 0.745452002 |
| 0.489856832 | 0.745452002 |
| 0.489913628 | 0.745505625 |
| 0.489923374 | 0.745505625 |
| 0.489986229 | 0.745529811 |
| 0.490078631 | 0.745598945 |
| 0.490118573 | 0.745612077 |
| 0.490167989 | 0.745645626 |
| 0.49018822 | 0.745645626 |
| 0.49025546 | 0.745701226 |
| 0.490271919 | 0.745702445 |
| 0.490319136 | 0.745738636 |
| 0.490343948 | 0.745740555 |
| 0.490509504 | 0.745788055 |
| 0.490707673 | 0.745974542 |
| 0.490745854 | 0.745974542 |
| 0.490758593 | 0.745974542 |
| 0.490921561 | 0.746166306 |
| 0.491154577 | 0.746282311 |
| 0.491262818 | 0.746304825 |
| 0.491310051 | 0.746304825 |
| 0.491312874 | 0.746304825 |
| 0.491339284 | 0.746304825 |
| 0.491444962 | 0.746366366 |
| 0.491576222 | 0.746470499 |
| 0.491733824 | 0.746552086 |
| 0.491792432 | 0.746560788 |
| 0.491868957 | 0.74662936 |
| 0.491996805 | 0.746704433 |
| 0.492110236 | 0.746805194 |
| 0.492245463 | 0.746939009 |
| 0.492462959 | 0.747116592 |
| 0.49261736 | 0.74726808 |
| 0.492656766 | 0.747277423 |
| 0.492806914 | 0.747457562 |

| | |
|---|---|
| 0.492875848 | 0.747492234 |
| 0.492879192 | 0.747492234 |
| 0.493089703 | 0.747640618 |
| 0.49332725 | 0.747818102 |
| 0.493756379 | 0.748063518 |
| 0.49378649 | 0.748063518 |
| 0.493787576 | 0.748063518 |
| 0.49385784 | 0.748117641 |
| 0.49394937 | 0.748170369 |
| 0.494056946 | 0.748209805 |
| 0.494147685 | 0.748299635 |
| 0.494344728 | 0.748506516 |
| 0.494411652 | 0.748560014 |
| 0.494464836 | 0.748613285 |
| 0.494501322 | 0.748639896 |
| 0.49459081 | 0.748690488 |
| 0.494722361 | 0.748741362 |
| 0.494842864 | 0.748854924 |
| 0.494859093 | 0.748854924 |
| 0.494876013 | 0.748854924 |
| 0.495086019 | 0.749146424 |
| 0.495193798 | 0.749240581 |
| 0.495318153 | 0.749364311 |
| 0.495356319 | 0.749364311 |
| 0.495480284 | 0.749392489 |
| 0.495559614 | 0.749392489 |
| 0.495561686 | 0.749392489 |
| 0.495649099 | 0.749490153 |
| 0.495682431 | 0.749490153 |
| 0.495693814 | 0.749490153 |
| 0.495713432 | 0.749490153 |
| 0.495782385 | 0.74953621 |
| 0.49578617 | 0.74953621 |
| 0.495798366 | 0.74953621 |
| 0.495823882 | 0.749550992 |
| 0.495942025 | 0.749682001 |
| 0.496174703 | 0.749879339 |
| 0.496296191 | 0.749994217 |
| 0.496306039 | 0.749994217 |
| 0.496362329 | 0.750031684 |
| 0.496568943 | 0.750193191 |
| 0.496602477 | 0.750193191 |
| 0.496649095 | 0.75022698 |
| 0.496741889 | 0.750281374 |
| 0.496764304 | 0.750282029 |

| | |
|---|---|
| 0.496786859 | 0.750283912 |
| 0.496814223 | 0.750283912 |
| 0.496987511 | 0.750428767 |
| 0.497085537 | 0.750529193 |
| 0.497215796 | 0.750607255 |
| 0.497276357 | 0.750632568 |
| 0.497402996 | 0.750757578 |
| 0.497410195 | 0.750757578 |
| 0.497573825 | 0.750943496 |
| 0.497580676 | 0.750943496 |
| 0.497608 | 0.750960936 |
| 0.497681711 | 0.75098959 |
| 0.497705827 | 0.75098959 |
| 0.497759518 | 0.751046811 |
| 0.497853625 | 0.751108965 |
| 0.497863794 | 0.751108965 |
| 0.497987667 | 0.751224459 |
| 0.498083361 | 0.751321221 |
| 0.498131423 | 0.751369921 |
| 0.498282196 | 0.751502142 |
| 0.498463473 | 0.751667301 |
| 0.498510199 | 0.75167939 |
| 0.498544364 | 0.75168331 |
| 0.498597477 | 0.751739594 |
| 0.498620124 | 0.751749942 |
| 0.49866961 | 0.751753533 |
| 0.49871621 | 0.751799612 |
| 0.498773776 | 0.751834984 |
| 0.498871143 | 0.751914172 |
| 0.498919444 | 0.751924901 |
| 0.498925625 | 0.751924901 |
| 0.499030101 | 0.751928718 |
| 0.499036303 | 0.751928718 |
| 0.499071063 | 0.751928718 |
| 0.499111855 | 0.751928718 |
| 0.499128538 | 0.751928718 |
| 0.499194944 | 0.751973862 |
| 0.499321808 | 0.75211739 |
| 0.499381411 | 0.75218338 |
| 0.499531999 | 0.75233882 |
| 0.499588658 | 0.752343773 |
| 0.499623406 | 0.752343773 |
| 0.499657725 | 0.75235838 |
| 0.499682171 | 0.75235838 |
| 0.499688269 | 0.75235838 |

| | |
|---|---|
| 0.499835449 | 0.75235838 |
| 0.499883158 | 0.75235838 |
| 0.499895984 | 0.75235838 |
| 0.499909151 | 0.75235838 |
| 0.499984041 | 0.752359224 |
| 0.500057749 | 0.752369307 |
| 0.500076401 | 0.752373603 |
| 0.500152119 | 0.752452698 |
| 0.50017637 | 0.752452698 |
| 0.50021357 | 0.752484893 |
| 0.500294247 | 0.752541861 |
| 0.500336964 | 0.752561447 |
| 0.500457787 | 0.752634734 |
| 0.50050653 | 0.752634734 |
| 0.500588603 | 0.752692451 |
| 0.500617673 | 0.752712397 |
| 0.500675925 | 0.752776218 |
| 0.500772182 | 0.752849643 |
| 0.500830398 | 0.752884898 |
| 0.500846542 | 0.752884898 |
| 0.500858864 | 0.752884898 |
| 0.500928908 | 0.752942659 |
| 0.501009891 | 0.753006152 |
| 0.501018581 | 0.753006152 |
| 0.501088448 | 0.75306363 |
| 0.501159745 | 0.753123251 |
| 0.50121056 | 0.753128323 |
| 0.501435582 | 0.753323843 |
| 0.501654387 | 0.753573133 |
| 0.50173089 | 0.753573133 |
| 0.501733073 | 0.753573133 |
| 0.501896989 | 0.753734592 |
| 0.501925597 | 0.753734592 |
| 0.502215024 | 0.753968728 |
| 0.502228943 | 0.753968728 |
| 0.502295066 | 0.753996198 |
| 0.502353702 | 0.753998157 |
| 0.502360296 | 0.753998157 |
| 0.502382425 | 0.753998157 |
| 0.50245345 | 0.754068078 |
| 0.502479117 | 0.754077652 |
| 0.502491496 | 0.754077652 |
| 0.502531219 | 0.754096748 |
| 0.502578771 | 0.754101614 |
| 0.502602066 | 0.754101614 |

| | |
|---|---|
| 0.502710041 | 0.754109182 |
| 0.502738595 | 0.754109182 |
| 0.502768526 | 0.754109182 |
| 0.502940928 | 0.754299329 |
| 0.502960429 | 0.754299329 |
| 0.503228559 | 0.754541826 |
| 0.503347148 | 0.754648314 |
| 0.503362959 | 0.754648314 |
| 0.503600416 | 0.754821465 |
| 0.503614407 | 0.754821465 |
| 0.503622195 | 0.754821465 |
| 0.503672614 | 0.754827425 |
| 0.503758711 | 0.754885194 |
| 0.504088528 | 0.755209196 |
| 0.504101781 | 0.755209196 |
| 0.504177112 | 0.755225552 |
| 0.504263566 | 0.755225552 |
| 0.504268829 | 0.755225552 |
| 0.50427127 | 0.755225552 |
| 0.504406953 | 0.755326475 |
| 0.504426487 | 0.755326475 |
| 0.504433812 | 0.755326475 |
| 0.504485032 | 0.755361548 |
| 0.504500562 | 0.755361548 |
| 0.504540151 | 0.755366962 |
| 0.504602313 | 0.75543628 |
| 0.504652118 | 0.755447724 |
| 0.504673405 | 0.755447724 |
| 0.504750307 | 0.755521196 |
| 0.504754214 | 0.755521196 |
| 0.504828772 | 0.7555268 |
| 0.504837275 | 0.7555268 |
| 0.50504935 | 0.755725452 |
| 0.505127334 | 0.755794651 |
| 0.505211587 | 0.755873221 |
| 0.505462123 | 0.756084546 |
| 0.505583206 | 0.756167923 |
| 0.505645115 | 0.756221245 |
| 0.505763305 | 0.75636603 |
| 0.505826092 | 0.756412423 |
| 0.506000795 | 0.756576228 |
| 0.506043134 | 0.756576228 |
| 0.506073755 | 0.756576228 |
| 0.506153002 | 0.756616208 |
| 0.506172683 | 0.756621882 |

| | |
|---|---|
| 0.506235827 | 0.756692518 |
| 0.506297101 | 0.756760357 |
| 0.506454007 | 0.756908114 |
| 0.506457157 | 0.756908114 |
| 0.506459524 | 0.756908114 |
| 0.506654047 | 0.757127555 |
| 0.506740211 | 0.757185042 |
| 0.506839299 | 0.757285585 |
| 0.506886435 | 0.757332253 |
| 0.507073911 | 0.757446028 |
| 0.507203692 | 0.757592368 |
| 0.507219897 | 0.757592813 |
| 0.507292097 | 0.757629373 |
| 0.507416417 | 0.757743758 |
| 0.507441349 | 0.757757231 |
| 0.507561622 | 0.757841789 |
| 0.507661689 | 0.757923266 |
| 0.507663932 | 0.757923266 |
| 0.507834897 | 0.758017282 |
| 0.507838315 | 0.758017282 |
| 0.508032027 | 0.758140102 |
| 0.508127997 | 0.758244574 |
| 0.508190992 | 0.758282289 |
| 0.508314013 | 0.758411531 |
| 0.50832538 | 0.758411531 |
| 0.508399475 | 0.758462546 |
| 0.508421869 | 0.758462546 |
| 0.508565647 | 0.758603653 |
| 0.50870574 | 0.758717992 |
| 0.50877958 | 0.758740054 |
| 0.508781435 | 0.758740054 |
| 0.508784536 | 0.758740054 |
| 0.508894099 | 0.758784649 |
| 0.509033092 | 0.758872743 |
| 0.509087635 | 0.758872743 |
| 0.509194142 | 0.758994411 |
| 0.509325377 | 0.759108509 |
| 0.509388938 | 0.759165972 |
| 0.509441734 | 0.759220899 |
| 0.509680904 | 0.759407706 |
| 0.509694643 | 0.759407706 |
| 0.509839634 | 0.75951758 |
| 0.509855083 | 0.75951758 |
| 0.509911912 | 0.75951758 |
| 0.51011089 | 0.759742679 |

| | |
|---|---|
| 0.510174399 | 0.759768912 |
| 0.51023386 | 0.759822013 |
| 0.51024658 | 0.759822013 |
| 0.510285843 | 0.759822013 |
| 0.510597794 | 0.760182592 |
| 0.510664146 | 0.760223069 |
| 0.510672867 | 0.760223069 |
| 0.510741035 | 0.76027702 |
| 0.510889991 | 0.760388221 |
| 0.510912745 | 0.760390029 |
| 0.510939343 | 0.760391918 |
| 0.510948824 | 0.760391918 |
| 0.511106477 | 0.760490012 |
| 0.511128038 | 0.760496566 |
| 0.511350783 | 0.760726367 |
| 0.511517652 | 0.760857027 |
| 0.511581628 | 0.760857027 |
| 0.511603557 | 0.760857027 |
| 0.511629049 | 0.760857027 |
| 0.511647066 | 0.760857027 |
| 0.511674023 | 0.760857307 |
| 0.511690551 | 0.760858129 |
| 0.511840281 | 0.760989124 |
| 0.511868753 | 0.760989124 |
| 0.511983353 | 0.761079657 |
| 0.512113171 | 0.761218006 |
| 0.512265136 | 0.761306279 |
| 0.512267133 | 0.761306279 |
| 0.512293164 | 0.761306279 |
| 0.51229565 | 0.761306279 |
| 0.512466512 | 0.761465153 |
| 0.512524868 | 0.761514849 |
| 0.512703606 | 0.76168691 |
| 0.512718378 | 0.76168691 |
| 0.512769912 | 0.761725854 |
| 0.512951314 | 0.761784763 |
| 0.512956572 | 0.761784763 |
| 0.513035615 | 0.761784763 |
| 0.513038723 | 0.761784763 |
| 0.513101235 | 0.761784763 |
| 0.513135728 | 0.761784763 |
| 0.513145401 | 0.761784763 |
| 0.513166935 | 0.761784763 |
| 0.513172795 | 0.761784763 |
| 0.51322828 | 0.761784763 |

| | |
|---|---|
| 0.513235921 | 0.761784763 |
| 0.513369379 | 0.761927187 |
| 0.513417694 | 0.761928841 |
| 0.513418488 | 0.761928841 |
| 0.513550148 | 0.762035422 |
| 0.513554306 | 0.762035422 |
| 0.513612166 | 0.762093914 |
| 0.513807981 | 0.762153849 |
| 0.513832518 | 0.762153849 |
| 0.513874219 | 0.762153849 |
| 0.513898886 | 0.762153849 |
| 0.513903316 | 0.762153849 |
| 0.51391177 | 0.762153849 |
| 0.513922163 | 0.762153849 |
| 0.513948176 | 0.762168694 |
| 0.514000892 | 0.762199405 |
| 0.514188673 | 0.762382913 |
| 0.514220384 | 0.762406196 |
| 0.514255465 | 0.76241074 |
| 0.514329268 | 0.762420616 |
| 0.514361117 | 0.762424977 |
| 0.514460719 | 0.762548882 |
| 0.514808518 | 0.762945678 |
| 0.514982891 | 0.762999031 |
| 0.514988701 | 0.762999031 |
| 0.515022056 | 0.763024713 |
| 0.515150959 | 0.763120734 |
| 0.515333744 | 0.76329654 |
| 0.5154141 | 0.76336808 |
| 0.515643784 | 0.763637018 |
| 0.515693614 | 0.763687067 |
| 0.515886676 | 0.763827481 |
| 0.515942636 | 0.763827481 |
| 0.516065344 | 0.763912202 |
| 0.516085189 | 0.763912202 |
| 0.516173282 | 0.763912202 |
| 0.516174349 | 0.763912202 |
| 0.51622026 | 0.763920654 |
| 0.516259438 | 0.763954893 |
| 0.516350536 | 0.763994749 |
| 0.516383078 | 0.764017025 |
| 0.516501755 | 0.764123539 |
| 0.516535043 | 0.764149049 |
| 0.5165739 | 0.764162004 |
| 0.516591934 | 0.764162004 |

| | |
|---|---|
| 0.516628668 | 0.764177766 |
| 0.516638386 | 0.764177766 |
| 0.516650725 | 0.764177766 |
| 0.516768027 | 0.764272784 |
| 0.516795201 | 0.764272784 |
| 0.516874692 | 0.764316894 |
| 0.516885394 | 0.764316894 |
| 0.516886332 | 0.764316894 |
| 0.51703851 | 0.764490178 |
| 0.517087 | 0.764514406 |
| 0.517151337 | 0.764585792 |
| 0.517266924 | 0.764709206 |
| 0.517440786 | 0.76489501 |
| 0.517508521 | 0.764923913 |
| 0.517554599 | 0.764960308 |
| 0.51765352 | 0.765067003 |
| 0.517829365 | 0.765161343 |
| 0.517835407 | 0.765161343 |
| 0.517845571 | 0.765161343 |
| 0.517845876 | 0.765161343 |
| 0.518127975 | 0.765364465 |
| 0.51821033 | 0.765402333 |
| 0.518228908 | 0.765402333 |
| 0.518367761 | 0.765457519 |
| 0.518391475 | 0.765468804 |
| 0.518443206 | 0.765493611 |
| 0.518524878 | 0.765570847 |
| 0.518546429 | 0.765578932 |
| 0.518604631 | 0.7655874 |
| 0.518624677 | 0.7655874 |
| 0.518657555 | 0.7655874 |
| 0.518700417 | 0.765616412 |
| 0.518717915 | 0.765618512 |
| 0.51874526 | 0.765635145 |
| 0.518793007 | 0.765658161 |
| 0.518838109 | 0.765700998 |
| 0.519114079 | 0.765858426 |
| 0.519129483 | 0.765858426 |
| 0.519157031 | 0.765858426 |
| 0.519174808 | 0.765858426 |
| 0.519262845 | 0.765858426 |
| 0.519362502 | 0.765858426 |
| 0.51936382 | 0.765858426 |
| 0.519427939 | 0.765890876 |
| 0.519433119 | 0.765890876 |

| | |
|---|---|
| 0.519496539 | 0.765960673 |
| 0.51954038 | 0.7660016 |
| 0.519654028 | 0.766079836 |
| 0.519772891 | 0.766175844 |
| 0.519841676 | 0.766175844 |
| 0.519846775 | 0.766175844 |
| 0.519884333 | 0.766175844 |
| 0.519996529 | 0.766238826 |
| 0.520023103 | 0.766238826 |
| 0.520063124 | 0.766238826 |
| 0.520087399 | 0.766238826 |
| 0.520146078 | 0.766252185 |
| 0.520193058 | 0.766252185 |
| 0.520197593 | 0.766252185 |
| 0.520209086 | 0.766252185 |
| 0.520559606 | 0.766544108 |
| 0.520575277 | 0.766544108 |
| 0.520587849 | 0.766544108 |
| 0.520591054 | 0.766544108 |
| 0.520644002 | 0.766544108 |
| 0.52066411 | 0.766544108 |
| 0.520667772 | 0.766544108 |
| 0.520795518 | 0.766618084 |
| 0.520866729 | 0.766675641 |
| 0.521002946 | 0.766849821 |
| 0.521029654 | 0.766849821 |
| 0.521055092 | 0.766857974 |
| 0.521178714 | 0.766976469 |
| 0.521193674 | 0.766976469 |
| 0.521293523 | 0.767042963 |
| 0.521359882 | 0.76706951 |
| 0.521554371 | 0.767293949 |
| 0.521746801 | 0.767471881 |
| 0.521753512 | 0.767471881 |
| 0.521831405 | 0.76750245 |
| 0.522092303 | 0.767815088 |
| 0.522124578 | 0.767815088 |
| 0.52224037 | 0.767866823 |
| 0.522261776 | 0.76787459 |
| 0.522300321 | 0.767907556 |
| 0.522401116 | 0.767986924 |
| 0.522437399 | 0.767986924 |
| 0.522460065 | 0.767986924 |
| 0.522561113 | 0.768039186 |
| 0.52259949 | 0.768039186 |

| | |
|---|---|
| 0.522838051 | 0.768210138 |
| 0.522876228 | 0.768210138 |
| 0.522877265 | 0.768210138 |
| 0.522994895 | 0.768312088 |
| 0.5230234 | 0.768322642 |
| 0.523051898 | 0.768322642 |
| 0.523057357 | 0.768322642 |
| 0.523066608 | 0.768322642 |
| 0.52311504 | 0.768346389 |
| 0.52328014 | 0.768524565 |
| 0.523284757 | 0.768524565 |
| 0.523322921 | 0.768556915 |
| 0.523407987 | 0.768563348 |
| 0.523530394 | 0.768673945 |
| 0.523547864 | 0.768673945 |
| 0.523656154 | 0.768772871 |
| 0.523706695 | 0.768772871 |
| 0.523712092 | 0.768772871 |
| 0.523760651 | 0.768820456 |
| 0.523925813 | 0.768971691 |
| 0.523928263 | 0.768971691 |
| 0.523997516 | 0.769002239 |
| 0.524153205 | 0.769183322 |
| 0.524199337 | 0.769227317 |
| 0.524396436 | 0.769374316 |
| 0.524460651 | 0.769380144 |
| 0.524465025 | 0.769380144 |
| 0.524506678 | 0.769387787 |
| 0.524547548 | 0.769387787 |
| 0.524580469 | 0.769387787 |
| 0.524656298 | 0.769447437 |
| 0.524697125 | 0.769459924 |
| 0.524760943 | 0.76951882 |
| 0.524784272 | 0.76951882 |
| 0.524785758 | 0.76951882 |
| 0.524826522 | 0.769540435 |
| 0.524832814 | 0.769540435 |
| 0.524994218 | 0.76964684 |
| 0.524996697 | 0.76964684 |
| 0.525002342 | 0.76964684 |
| 0.525115779 | 0.769765749 |
| 0.525181555 | 0.769791091 |
| 0.525226089 | 0.769832676 |
| 0.52540457 | 0.770002102 |
| 0.525432013 | 0.770016016 |

| | |
|---|---|
| 0.525593156 | 0.77010999 |
| 0.525684845 | 0.770173253 |
| 0.525718255 | 0.77019851 |
| 0.525805065 | 0.7702783 |
| 0.526011163 | 0.770508366 |
| 0.526026823 | 0.770508366 |
| 0.52610014 | 0.770566449 |
| 0.526115015 | 0.770566449 |
| 0.526150423 | 0.770577415 |
| 0.52615486 | 0.770577415 |
| 0.526252866 | 0.770627325 |
| 0.526253661 | 0.770627325 |
| 0.526350572 | 0.77066729 |
| 0.526429423 | 0.77066729 |
| 0.526457165 | 0.77066729 |
| 0.526480944 | 0.77066729 |
| 0.526500947 | 0.77066729 |
| 0.526507491 | 0.77066729 |
| 0.52654112 | 0.770692827 |
| 0.526600704 | 0.770720101 |
| 0.526624483 | 0.770720101 |
| 0.526674422 | 0.770769503 |
| 0.526733558 | 0.770813493 |
| 0.527032274 | 0.771174724 |
| 0.527100404 | 0.771250721 |
| 0.527178589 | 0.771282267 |
| 0.527203413 | 0.771282267 |
| 0.527216421 | 0.771282267 |
| 0.527441653 | 0.77151301 |
| 0.527479494 | 0.771520971 |
| 0.527549752 | 0.771552649 |
| 0.527656441 | 0.771630949 |
| 0.527670778 | 0.771630949 |
| 0.527700499 | 0.771630949 |
| 0.527764213 | 0.771653042 |
| 0.527943298 | 0.771820415 |
| 0.527980877 | 0.771820415 |
| 0.52800727 | 0.771820415 |
| 0.528010347 | 0.771820415 |
| 0.528039733 | 0.771820415 |
| 0.528041408 | 0.771820415 |
| 0.528056947 | 0.771820415 |
| 0.528182607 | 0.771885328 |
| 0.528235461 | 0.771901441 |
| 0.528300178 | 0.771962711 |

| | |
|---|---|
| 0.528352896 | 0.772016056 |
| 0.528459891 | 0.772077647 |
| 0.528498506 | 0.772094399 |
| 0.528609148 | 0.772167774 |
| 0.528640383 | 0.772175536 |
| 0.528739818 | 0.772265421 |
| 0.528766784 | 0.772265421 |
| 0.528800492 | 0.772267295 |
| 0.528910685 | 0.772367039 |
| 0.528917442 | 0.772367039 |
| 0.528959587 | 0.77238122 |
| 0.528991903 | 0.772385173 |
| 0.529040293 | 0.772428021 |
| 0.529121053 | 0.772455071 |
| 0.529192242 | 0.772531461 |
| 0.529235202 | 0.772543394 |
| 0.529252783 | 0.772543394 |
| 0.529265299 | 0.772543394 |
| 0.529425316 | 0.772729598 |
| 0.529449204 | 0.772740783 |
| 0.529553216 | 0.772828274 |
| 0.529704529 | 0.772863328 |
| 0.529731406 | 0.772863328 |
| 0.529756415 | 0.772863328 |
| 0.52976035 | 0.772863328 |
| 0.529872391 | 0.772882925 |
| 0.529876642 | 0.772882925 |
| 0.529895888 | 0.772882925 |
| 0.529913842 | 0.772882925 |
| 0.529919833 | 0.772882925 |
| 0.529971916 | 0.772911548 |
| 0.5300444 | 0.77296992 |
| 0.530215704 | 0.773099174 |
| 0.530221737 | 0.773099174 |
| 0.530346858 | 0.773195469 |
| 0.530361408 | 0.773195469 |
| 0.530532027 | 0.773280376 |
| 0.530659456 | 0.773351388 |
| 0.53066321 | 0.773351388 |
| 0.53066321 | 0.773351388 |
| 0.53069744 | 0.773377607 |
| 0.530752589 | 0.773424117 |
| 0.530761833 | 0.773424117 |
| 0.530998482 | 0.773589586 |
| 0.531005328 | 0.773589586 |

| | |
|---|---|
| 0.531071364 | 0.773631498 |
| 0.53111225 | 0.773650695 |
| 0.531150711 | 0.773673827 |
| 0.531276631 | 0.773795477 |
| 0.531319992 | 0.773811304 |
| 0.531342579 | 0.773820536 |
| 0.531421054 | 0.773887496 |
| 0.531786785 | 0.774183379 |
| 0.5318275 | 0.774192011 |
| 0.53183286 | 0.774192011 |
| 0.531941043 | 0.774286144 |
| 0.532070039 | 0.774406374 |
| 0.532109667 | 0.774433236 |
| 0.532121016 | 0.774433236 |
| 0.532306189 | 0.774601312 |
| 0.532403388 | 0.774702153 |
| 0.532441768 | 0.774710661 |
| 0.532585615 | 0.774868128 |
| 0.532592072 | 0.774868128 |
| 0.5325988 | 0.774868128 |
| 0.532660474 | 0.774930961 |
| 0.532697256 | 0.774940353 |
| 0.532777273 | 0.774999732 |
| 0.53278689 | 0.774999732 |
| 0.532917129 | 0.775095316 |
| 0.532951105 | 0.775120233 |
| 0.533162539 | 0.775339465 |
| 0.533179506 | 0.775339465 |
| 0.533212526 | 0.775339465 |
| 0.533259191 | 0.775339465 |
| 0.533297815 | 0.775339465 |
| 0.533314508 | 0.775339465 |
| 0.533329754 | 0.775339465 |
| 0.533411405 | 0.775434497 |
| 0.533495468 | 0.775509359 |
| 0.533619514 | 0.775547653 |
| 0.53366786 | 0.775570584 |
| 0.533735044 | 0.775610443 |
| 0.533744142 | 0.775610443 |
| 0.533776245 | 0.775633427 |
| 0.533818443 | 0.775671079 |
| 0.534018731 | 0.775843759 |
| 0.534135972 | 0.775857829 |
| 0.534165424 | 0.775857829 |
| 0.534165858 | 0.775857829 |

| | |
|---|---|
| 0.534170665 | 0.775857829 |
| 0.53417185 | 0.775857829 |
| 0.534524821 | 0.776084187 |
| 0.534526416 | 0.776084187 |
| 0.534633164 | 0.776144879 |
| 0.534642509 | 0.776144879 |
| 0.534804276 | 0.776227319 |
| 0.534878189 | 0.776287283 |
| 0.53494602 | 0.776338415 |
| 0.535021431 | 0.776400539 |
| 0.535126584 | 0.776482157 |
| 0.535163413 | 0.776511939 |
| 0.535293975 | 0.77664979 |
| 0.535299776 | 0.77664979 |
| 0.535314046 | 0.77664979 |
| 0.535354806 | 0.77664979 |
| 0.535372567 | 0.77664979 |
| 0.535417749 | 0.776691677 |
| 0.535491896 | 0.776775577 |
| 0.535546255 | 0.776807111 |
| 0.535609763 | 0.776830763 |
| 0.535641843 | 0.776830763 |
| 0.535644115 | 0.776830763 |
| 0.535704436 | 0.776850268 |
| 0.535760608 | 0.776857775 |
| 0.535853087 | 0.77692091 |
| 0.535929903 | 0.776963872 |
| 0.535958007 | 0.776963872 |
| 0.536173205 | 0.777122502 |
| 0.536350971 | 0.777193249 |
| 0.536434149 | 0.77726648 |
| 0.536506418 | 0.777347546 |
| 0.536528868 | 0.777356425 |
| 0.536796317 | 0.777460109 |
| 0.536933429 | 0.77758287 |
| 0.537000093 | 0.777605649 |
| 0.537027421 | 0.777605649 |
| 0.537134258 | 0.777689415 |
| 0.537186814 | 0.777694583 |
| 0.537246266 | 0.777705529 |
| 0.537265314 | 0.777705529 |
| 0.537290442 | 0.777705529 |
| 0.537353726 | 0.777715359 |
| 0.537567354 | 0.777920164 |
| 0.537574251 | 0.777920164 |

| | |
|---|---|
| 0.537620304 | 0.777920164 |
| 0.537759469 | 0.778026982 |
| 0.537780385 | 0.778033608 |
| 0.537812121 | 0.77803756 |
| 0.537822576 | 0.77803756 |
| 0.537832124 | 0.77803756 |
| 0.537880088 | 0.77805968 |
| 0.537979911 | 0.77810603 |
| 0.538001768 | 0.77810603 |
| 0.538020157 | 0.77810603 |
| 0.538234726 | 0.778289015 |
| 0.538464834 | 0.778482255 |
| 0.538537769 | 0.778482255 |
| 0.538547819 | 0.778482255 |
| 0.538575086 | 0.778482255 |
| 0.538583896 | 0.778482255 |
| 0.538666161 | 0.778482255 |
| 0.538666171 | 0.778482255 |
| 0.538669402 | 0.778482255 |
| 0.538675798 | 0.778482255 |
| 0.538682391 | 0.778482255 |
| 0.538695741 | 0.778482255 |
| 0.53870837 | 0.778482255 |
| 0.538752029 | 0.778517033 |
| 0.538860694 | 0.778579566 |
| 0.539045378 | 0.77870249 |
| 0.539048127 | 0.77870249 |
| 0.53906022 | 0.77870249 |
| 0.539134281 | 0.778712752 |
| 0.539171593 | 0.778712752 |
| 0.539196086 | 0.778712752 |
| 0.539205904 | 0.778712752 |
| 0.539398206 | 0.77881278 |
| 0.539595614 | 0.778986431 |
| 0.539648581 | 0.778986431 |
| 0.539649055 | 0.778986431 |
| 0.539695099 | 0.779028898 |
| 0.539826051 | 0.779090504 |
| 0.539828556 | 0.779090504 |
| 0.539881652 | 0.779104285 |
| 0.53991091 | 0.779104285 |
| 0.540024338 | 0.779165536 |
| 0.540031148 | 0.779165536 |
| 0.54004204 | 0.779165536 |
| 0.540051515 | 0.779165536 |

| | |
|---|---|
| 0.540098329 | 0.779185869 |
| 0.540134419 | 0.779214332 |
| 0.540337952 | 0.779374248 |
| 0.540421758 | 0.779416379 |
| 0.540529429 | 0.779471207 |
| 0.540584343 | 0.779471207 |
| 0.540590704 | 0.779471207 |
| 0.540637198 | 0.77949105 |
| 0.540713301 | 0.779577175 |
| 0.540793025 | 0.779611229 |
| 0.540885244 | 0.77968346 |
| 0.540936889 | 0.779720803 |
| 0.540943892 | 0.779720803 |
| 0.541000887 | 0.779779355 |
| 0.541046918 | 0.779822103 |
| 0.54124623 | 0.779992238 |
| 0.54132148 | 0.780005389 |
| 0.541413192 | 0.780090337 |
| 0.541609597 | 0.780312313 |
| 0.541794483 | 0.780437851 |
| 0.541801859 | 0.780437851 |
| 0.541896529 | 0.780527006 |
| 0.541914957 | 0.780529944 |
| 0.542086893 | 0.780649568 |
| 0.542123449 | 0.780649568 |
| 0.542133626 | 0.780649568 |
| 0.542329954 | 0.780797096 |
| 0.542410977 | 0.780866536 |
| 0.542433326 | 0.780875106 |
| 0.542511451 | 0.780940364 |
| 0.542530361 | 0.780943981 |
| 0.542568381 | 0.780975106 |
| 0.542678103 | 0.780991419 |
| 0.542910259 | 0.781168861 |
| 0.542934889 | 0.781168861 |
| 0.542947239 | 0.781168861 |
| 0.542949016 | 0.781168861 |
| 0.54301885 | 0.781201753 |
| 0.543069077 | 0.781216965 |
| 0.543121249 | 0.781251456 |
| 0.543158042 | 0.781257189 |
| 0.543280505 | 0.781315525 |
| 0.54335412 | 0.781374035 |
| 0.543393985 | 0.781389875 |
| 0.543428405 | 0.781389875 |

| | |
|---|---|
| 0.543447167 | 0.781389875 |
| 0.543475143 | 0.78140651 |
| 0.543577662 | 0.781466559 |
| 0.543585815 | 0.781466559 |
| 0.543609557 | 0.781466559 |
| 0.543615356 | 0.781466559 |
| 0.543667054 | 0.7814937 |
| 0.543738623 | 0.781541059 |
| 0.543868004 | 0.781604132 |
| 0.543891577 | 0.781604132 |
| 0.543960512 | 0.78167961 |
| 0.544253184 | 0.781935035 |
| 0.544356452 | 0.78205981 |
| 0.544551629 | 0.782245828 |
| 0.544593947 | 0.782267065 |
| 0.544629965 | 0.782287573 |
| 0.544670485 | 0.782300274 |
| 0.544671659 | 0.782300274 |
| 0.544778835 | 0.782407019 |
| 0.544837127 | 0.782443548 |
| 0.544896104 | 0.782477159 |
| 0.544944427 | 0.782477159 |
| 0.544959106 | 0.782477159 |
| 0.544998245 | 0.782509765 |
| 0.545034623 | 0.782538405 |
| 0.545148759 | 0.782641684 |
| 0.545425831 | 0.78288811 |
| 0.545428736 | 0.78288811 |
| 0.545442747 | 0.78288811 |
| 0.545621627 | 0.783074208 |
| 0.545667243 | 0.783074208 |
| 0.545698749 | 0.783074208 |
| 0.545706866 | 0.783074208 |
| 0.54578326 | 0.783117505 |
| 0.54585228 | 0.783169349 |
| 0.545893427 | 0.783170087 |
| 0.545972408 | 0.783170087 |
| 0.545990832 | 0.783170087 |
| 0.54601742 | 0.783170087 |
| 0.546024406 | 0.783170087 |
| 0.546042095 | 0.783170087 |
| 0.546129696 | 0.783170087 |
| 0.546179412 | 0.783170087 |
| 0.546229939 | 0.783170087 |
| 0.54623536 | 0.783170087 |

| | |
|---|---|
| 0.546250518 | 0.783170087 |
| 0.54625389 | 0.783170087 |
| 0.546256466 | 0.783170087 |
| 0.546435994 | 0.78327536 |
| 0.546558558 | 0.783306106 |
| 0.546568351 | 0.783306106 |
| 0.546577249 | 0.783306106 |
| 0.546582765 | 0.783306106 |
| 0.546598034 | 0.783306106 |
| 0.546605464 | 0.783306106 |
| 0.546656158 | 0.783331613 |
| 0.54673877 | 0.783355717 |
| 0.546772832 | 0.783380954 |
| 0.546915269 | 0.783502581 |
| 0.546953865 | 0.783522475 |
| 0.546975014 | 0.783529205 |
| 0.547020805 | 0.783547666 |
| 0.547069758 | 0.783570653 |
| 0.547162224 | 0.783627403 |
| 0.547177101 | 0.783627403 |
| 0.547240982 | 0.783627403 |
| 0.547241107 | 0.783627403 |
| 0.547251353 | 0.783627403 |
| 0.54725746 | 0.783627403 |
| 0.547316745 | 0.783660755 |
| 0.54732938 | 0.783660755 |
| 0.547346568 | 0.783660755 |
| 0.54743169 | 0.783711948 |
| 0.547484411 | 0.783763866 |
| 0.547677113 | 0.783969039 |
| 0.547705442 | 0.783986028 |
| 0.547780522 | 0.784022807 |
| 0.547821718 | 0.784034646 |
| 0.547914891 | 0.784120868 |
| 0.547982417 | 0.78417038 |
| 0.548086174 | 0.784224605 |
| 0.548145131 | 0.78426184 |
| 0.548297996 | 0.78440986 |
| 0.54833896 | 0.784414487 |
| 0.54846593 | 0.784414487 |
| 0.548484011 | 0.784416791 |
| 0.548598107 | 0.784487425 |
| 0.548613938 | 0.784487425 |
| 0.548615757 | 0.784487425 |
| 0.548658144 | 0.784500928 |

| | |
|---|---|
| 0.548789466 | 0.784641586 |
| 0.548848848 | 0.78467747 |
| 0.548887652 | 0.78467747 |
| 0.54889694 | 0.78467747 |
| 0.549168634 | 0.784924511 |
| 0.549286056 | 0.784981746 |
| 0.5493078 | 0.784981746 |
| 0.549345982 | 0.784981746 |
| 0.549357015 | 0.784981746 |
| 0.549445522 | 0.784995142 |
| 0.549491433 | 0.785020143 |
| 0.549501336 | 0.785020143 |
| 0.549603899 | 0.785080823 |
| 0.549676971 | 0.785152269 |
| 0.549690664 | 0.785152269 |
| 0.549970346 | 0.785410425 |
| 0.549991475 | 0.785417048 |
| 0.550015807 | 0.785428245 |
| 0.550125604 | 0.785487629 |
| 0.550139854 | 0.785487629 |
| 0.55016574 | 0.785501041 |
| 0.550187618 | 0.785508729 |
| 0.550380875 | 0.785562703 |
| 0.550381979 | 0.785562703 |
| 0.550441048 | 0.785564991 |
| 0.550441449 | 0.785564991 |
| 0.55069709 | 0.785648689 |
| 0.550698046 | 0.785648689 |
| 0.551079704 | 0.786010075 |
| 0.55110783 | 0.786021401 |
| 0.551297805 | 0.786151099 |
| 0.551316999 | 0.786154931 |
| 0.551392187 | 0.786192471 |
| 0.551545509 | 0.78631598 |
| 0.551593696 | 0.786326331 |
| 0.551610124 | 0.786326331 |
| 0.551834724 | 0.78654243 |
| 0.551883193 | 0.78654243 |
| 0.551886009 | 0.78654243 |
| 0.551886009 | 0.78654243 |
| 0.551886009 | 0.78654243 |
| 0.551906789 | 0.786542941 |
| 0.551956435 | 0.786568429 |
| 0.551972329 | 0.786568429 |
| 0.551997271 | 0.786568429 |

| | |
|---|---|
| 0.552003342 | 0.786568429 |
| 0.552048294 | 0.786603962 |
| 0.552075572 | 0.786603962 |
| 0.552125357 | 0.786624618 |
| 0.552173269 | 0.786669347 |
| 0.552305432 | 0.786810565 |
| 0.552347187 | 0.78682298 |
| 0.552558612 | 0.787074516 |
| 0.552613453 | 0.787074516 |
| 0.552618622 | 0.787074516 |
| 0.552648125 | 0.787076986 |
| 0.552690521 | 0.787076986 |
| 0.552875804 | 0.787222815 |
| 0.552907554 | 0.787244487 |
| 0.552990449 | 0.787338976 |
| 0.553081952 | 0.787398641 |
| 0.553109694 | 0.787404133 |
| 0.553179244 | 0.787442999 |
| 0.553255523 | 0.787504506 |
| 0.553461349 | 0.787674987 |
| 0.553528425 | 0.787699113 |
| 0.553557631 | 0.787699113 |
| 0.553790387 | 0.787861978 |
| 0.553800101 | 0.787861978 |
| 0.55381209 | 0.787861978 |
| 0.553919785 | 0.787881888 |
| 0.553930708 | 0.787881888 |
| 0.553987314 | 0.787910347 |
| 0.554101559 | 0.788011489 |
| 0.554163028 | 0.788042594 |
| 0.554452489 | 0.788336529 |
| 0.554505669 | 0.788353063 |
| 0.554530329 | 0.788353063 |
| 0.5545614 | 0.788373704 |
| 0.55462525 | 0.788440938 |
| 0.554682 | 0.788498079 |
| 0.554758758 | 0.78856012 |
| 0.554789616 | 0.788560363 |
| 0.554792044 | 0.788560363 |
| 0.554880046 | 0.788607812 |
| 0.554891658 | 0.788607812 |
| 0.55497029 | 0.78862941 |
| 0.554991207 | 0.78862941 |
| 0.555034507 | 0.78862941 |
| 0.555060662 | 0.788631783 |

| | |
|---|---|
| 0.55507411 | 0.788631783 |
| 0.555100229 | 0.788645366 |
| 0.555149304 | 0.788652676 |
| 0.555163408 | 0.788652676 |
| 0.55517346 | 0.788652676 |
| 0.555188169 | 0.788652676 |
| 0.555217623 | 0.788670994 |
| 0.555239795 | 0.788678966 |
| 0.555316554 | 0.788705979 |
| 0.555344169 | 0.788709608 |
| 0.555416028 | 0.788788143 |
| 0.555582816 | 0.788977959 |
| 0.555695704 | 0.789052487 |
| 0.555739461 | 0.789071852 |
| 0.55574834 | 0.789071852 |
| 0.555773381 | 0.789083882 |
| 0.555893205 | 0.789193774 |
| 0.555900492 | 0.789193774 |
| 0.555946856 | 0.789224295 |
| 0.556000651 | 0.789265387 |
| 0.556053357 | 0.789316679 |
| 0.556076815 | 0.789326452 |
| 0.556106124 | 0.78934453 |
| 0.556168812 | 0.789386458 |
| 0.556225666 | 0.789420102 |
| 0.556344445 | 0.789476989 |
| 0.55645618 | 0.78955533 |
| 0.556577756 | 0.789660961 |
| 0.556710689 | 0.789802508 |
| 0.556765705 | 0.789833504 |
| 0.556984292 | 0.790096526 |
| 0.557064253 | 0.790186418 |
| 0.557217235 | 0.790309269 |
| 0.55732821 | 0.790359095 |
| 0.557355833 | 0.790359095 |
| 0.557751736 | 0.790705349 |
| 0.557762128 | 0.790705349 |
| 0.557788743 | 0.790719542 |
| 0.558058039 | 0.791010741 |
| 0.558256969 | 0.791128383 |
| 0.558312889 | 0.791128383 |
| 0.55847706 | 0.791271365 |
| 0.558496752 | 0.791275725 |
| 0.55872536 | 0.791411275 |
| 0.559055045 | 0.791607432 |

| | |
|---|---|
| 0.559079916 | 0.791607432 |
| 0.559175349 | 0.791695483 |
| 0.559415058 | 0.791900166 |
| 0.559538556 | 0.791979572 |
| 0.559636473 | 0.792068592 |
| 0.559696445 | 0.792068592 |
| 0.559713246 | 0.792068592 |
| 0.559739279 | 0.792068592 |
| 0.559771707 | 0.792068592 |
| 0.559808659 | 0.792068592 |
| 0.559846025 | 0.792068592 |
| 0.559984364 | 0.792181523 |
| 0.560016984 | 0.792185342 |
| 0.560072374 | 0.792185342 |
| 0.56019316 | 0.792227418 |
| 0.560271543 | 0.792305376 |
| 0.560314398 | 0.792318478 |
| 0.560320587 | 0.792318478 |
| 0.560391294 | 0.792345889 |
| 0.56044901 | 0.792365858 |
| 0.560464044 | 0.792365858 |
| 0.560503054 | 0.792397488 |
| 0.560546739 | 0.792435724 |
| 0.560822129 | 0.792613298 |
| 0.560872598 | 0.792614954 |
| 0.560935487 | 0.792653903 |
| 0.560950717 | 0.792653903 |
| 0.560995478 | 0.792676402 |
| 0.561025115 | 0.792676402 |
| 0.561113092 | 0.792690928 |
| 0.561313662 | 0.792808094 |
| 0.561342821 | 0.792808094 |
| 0.561427508 | 0.792904188 |
| 0.561560074 | 0.792942132 |
| 0.561570386 | 0.792942132 |
| 0.561663556 | 0.792942132 |
| 0.561692762 | 0.792942132 |
| 0.561695684 | 0.792942132 |
| 0.561745947 | 0.792954186 |
| 0.561803596 | 0.792990449 |
| 0.561838237 | 0.792990449 |
| 0.561977756 | 0.793116858 |
| 0.562013431 | 0.793143703 |
| 0.562102646 | 0.793152097 |
| 0.562178392 | 0.793188703 |

| | |
|---|---|
| 0.562319675 | 0.7932395 |
| 0.562338338 | 0.7932395 |
| 0.562344929 | 0.7932395 |
| 0.562366017 | 0.7932395 |
| 0.562480719 | 0.793307036 |
| 0.562493653 | 0.793307036 |
| 0.56256224 | 0.793307036 |
| 0.562631556 | 0.793381291 |
| 0.562749426 | 0.793424178 |
| 0.56281576 | 0.793424178 |
| 0.562878538 | 0.793424178 |
| 0.563151666 | 0.79371521 |
| 0.563238861 | 0.793802887 |
| 0.563247207 | 0.793802887 |
| 0.563458511 | 0.793936169 |
| 0.563667894 | 0.794019361 |
| 0.563706275 | 0.794019361 |
| 0.563719949 | 0.794019361 |
| 0.563720318 | 0.794019361 |
| 0.563865757 | 0.794128746 |
| 0.564093875 | 0.794235824 |
| 0.564103502 | 0.794235824 |
| 0.564140798 | 0.794235824 |
| 0.564141254 | 0.794235824 |
| 0.564163625 | 0.794235824 |
| 0.564166101 | 0.794235824 |
| 0.564188095 | 0.794235824 |
| 0.564248337 | 0.794297152 |
| 0.564282256 | 0.794321422 |
| 0.564373053 | 0.794378798 |
| 0.564782375 | 0.794640095 |
| 0.564808964 | 0.794640095 |
| 0.564880204 | 0.794693374 |
| 0.565041901 | 0.794850418 |
| 0.565134718 | 0.794887569 |
| 0.56527708 | 0.795063832 |
| 0.565331764 | 0.795074496 |
| 0.565358303 | 0.795074496 |
| 0.565404141 | 0.795080692 |
| 0.5654961 | 0.795137067 |
| 0.565519644 | 0.795146698 |
| 0.565562757 | 0.795156031 |
| 0.565656713 | 0.795187376 |
| 0.565745587 | 0.795187376 |
| 0.565761232 | 0.795187376 |

| | |
|---|---|
| 0.565915825 | 0.79532807 |
| 0.566193848 | 0.795565685 |
| 0.566216853 | 0.795565685 |
| 0.566222968 | 0.795565685 |
| 0.566276159 | 0.795589824 |
| 0.566323053 | 0.795589824 |
| 0.56633594 | 0.795589824 |
| 0.566486032 | 0.795730262 |
| 0.566589521 | 0.79585182 |
| 0.566613538 | 0.79585182 |
| 0.566643519 | 0.79585759 |
| 0.566849142 | 0.795981995 |
| 0.56689357 | 0.795981995 |
| 0.56689622 | 0.795981995 |
| 0.566901997 | 0.795981995 |
| 0.566918326 | 0.795981995 |
| 0.566932648 | 0.795981995 |
| 0.567003261 | 0.796034203 |
| 0.567090622 | 0.796086451 |
| 0.567186226 | 0.796126799 |
| 0.567407154 | 0.796334875 |
| 0.56746193 | 0.7963495 |
| 0.567611836 | 0.796410533 |
| 0.567616558 | 0.796410533 |
| 0.56776139 | 0.796490205 |
| 0.567762875 | 0.796490205 |
| 0.567901753 | 0.796638101 |
| 0.568113717 | 0.796724243 |
| 0.568163141 | 0.796744398 |
| 0.568334054 | 0.796843845 |
| 0.568349579 | 0.796843845 |
| 0.56841832 | 0.796913409 |
| 0.568600948 | 0.797059067 |
| 0.56863044 | 0.797059067 |
| 0.568653706 | 0.797059067 |
| 0.568753961 | 0.797176131 |
| 0.568824149 | 0.797232936 |
| 0.568827968 | 0.797232936 |
| 0.568942098 | 0.797322504 |
| 0.569044239 | 0.797388922 |
| 0.569061901 | 0.797388922 |
| 0.569210457 | 0.797481655 |
| 0.56935715 | 0.797481655 |
| 0.569404039 | 0.797481655 |
| 0.569430252 | 0.797481655 |

| | |
|---|---|
| 0.569448512 | 0.797481655 |
| 0.569460195 | 0.797481655 |
| 0.569490303 | 0.797481655 |
| 0.569511351 | 0.797481655 |
| 0.569602253 | 0.797484373 |
| 0.569610167 | 0.797484373 |
| 0.5696705 | 0.797545398 |
| 0.569739859 | 0.797619055 |
| 0.569844602 | 0.797652862 |
| 0.569952425 | 0.79768986 |
| 0.569954298 | 0.79768986 |
| 0.57014124 | 0.797852643 |
| 0.570319308 | 0.798013852 |
| 0.570361603 | 0.798013852 |
| 0.570376073 | 0.798013852 |
| 0.570388688 | 0.798013852 |
| 0.570396079 | 0.798013852 |
| 0.570407239 | 0.798013852 |
| 0.570570503 | 0.798143631 |
| 0.570583793 | 0.798143631 |
| 0.570656597 | 0.79822082 |
| 0.570682354 | 0.79822082 |
| 0.570708734 | 0.79822082 |
| 0.570732751 | 0.79822082 |
| 0.570756294 | 0.79822082 |
| 0.570797254 | 0.798231225 |
| 0.570855328 | 0.798286849 |
| 0.570897053 | 0.798286849 |
| 0.570904075 | 0.798286849 |
| 0.571036338 | 0.798401469 |
| 0.571129807 | 0.79842781 |
| 0.571131225 | 0.79842781 |
| 0.571155763 | 0.79842781 |
| 0.57130893 | 0.798548171 |
| 0.571332577 | 0.798557789 |
| 0.571411605 | 0.798621373 |
| 0.571467959 | 0.798653261 |
| 0.571720385 | 0.798823275 |
| 0.571756689 | 0.798823275 |
| 0.571764569 | 0.798823275 |
| 0.571848811 | 0.798904207 |
| 0.571939826 | 0.798984486 |
| 0.572058618 | 0.799056685 |
| 0.57208113 | 0.799064695 |
| 0.572118493 | 0.799091264 |

| | |
|---|---|
| 0.572143105 | 0.799091264 |
| 0.572150486 | 0.799091264 |
| 0.572184571 | 0.79911198 |
| 0.572198876 | 0.79911198 |
| 0.572296835 | 0.799225351 |
| 0.572340001 | 0.799253718 |
| 0.572386826 | 0.799257282 |
| 0.572474126 | 0.799349493 |
| 0.57248643 | 0.799349493 |
| 0.5725888 | 0.79942212 |
| 0.572917153 | 0.799786771 |
| 0.572954083 | 0.799814881 |
| 0.573079363 | 0.799872545 |
| 0.573126336 | 0.799914664 |
| 0.573257894 | 0.800051389 |
| 0.573285528 | 0.800066267 |
| 0.573374157 | 0.800079078 |
| 0.573379744 | 0.800079078 |
| 0.573419926 | 0.800079078 |
| 0.573420518 | 0.800079078 |
| 0.573521261 | 0.800114154 |
| 0.573576713 | 0.800144639 |
| 0.573887882 | 0.800463535 |
| 0.573891796 | 0.800463535 |
| 0.573917831 | 0.800463535 |
| 0.573937111 | 0.800463535 |
| 0.574013367 | 0.800532253 |
| 0.574036548 | 0.800532253 |
| 0.574056272 | 0.800532253 |
| 0.574120538 | 0.800598433 |
| 0.574203679 | 0.800667485 |
| 0.574255849 | 0.80069958 |
| 0.574313301 | 0.80075001 |
| 0.574353881 | 0.800759705 |
| 0.574379716 | 0.800772283 |
| 0.574470707 | 0.800794688 |
| 0.574538306 | 0.800801782 |
| 0.57454494 | 0.800801782 |
| 0.574578411 | 0.800801782 |
| 0.574635151 | 0.800801782 |
| 0.574777346 | 0.800896371 |
| 0.574788226 | 0.800896371 |
| 0.575013193 | 0.800994784 |
| 0.575018265 | 0.800994784 |
| 0.575318377 | 0.801283435 |

| | |
|---|---|
| 0.575355317 | 0.801311451 |
| 0.575408708 | 0.801329454 |
| 0.575504437 | 0.801398108 |
| 0.575815639 | 0.801645683 |
| 0.575830946 | 0.801645683 |
| 0.575851904 | 0.801651427 |
| 0.575930756 | 0.801723709 |
| 0.575960406 | 0.801723709 |
| 0.575994287 | 0.801723709 |
| 0.576035754 | 0.801723709 |
| 0.57603923 | 0.801723709 |
| 0.576090709 | 0.801723709 |
| 0.576139748 | 0.801723709 |
| 0.576169956 | 0.801723709 |
| 0.57617316 | 0.801723709 |
| 0.576193007 | 0.801726985 |
| 0.576248489 | 0.801726985 |
| 0.576271481 | 0.801726985 |
| 0.576290174 | 0.801726985 |
| 0.576421363 | 0.801806563 |
| 0.576434726 | 0.801806563 |
| 0.576656086 | 0.802008932 |
| 0.57666441 | 0.802008932 |
| 0.576756256 | 0.802066404 |
| 0.576785726 | 0.802083966 |
| 0.576900793 | 0.802197135 |
| 0.577021657 | 0.80231835 |
| 0.577045786 | 0.802323192 |
| 0.577103921 | 0.802323192 |
| 0.577106797 | 0.802323192 |
| 0.577209522 | 0.802323192 |
| 0.577243337 | 0.802323192 |
| 0.577323687 | 0.802373809 |
| 0.577342127 | 0.802373809 |
| 0.577347942 | 0.802373809 |
| 0.577459651 | 0.802484084 |
| 0.577644424 | 0.802597249 |
| 0.577662554 | 0.802597249 |
| 0.577677236 | 0.802597249 |
| 0.577721156 | 0.802619665 |
| 0.577732155 | 0.802619665 |
| 0.577833595 | 0.802697408 |
| 0.577870847 | 0.802725743 |
| 0.577910793 | 0.802756461 |
| 0.57792667 | 0.802756461 |

| | |
|---|---|
| 0.577957028 | 0.802775216 |
| 0.578140566 | 0.802984631 |
| 0.578187354 | 0.802984631 |
| 0.578192095 | 0.802984631 |
| 0.578378413 | 0.803046436 |
| 0.57838767 | 0.803046436 |
| 0.578429678 | 0.803080407 |
| 0.578480729 | 0.803081053 |
| 0.5785095 | 0.803097586 |
| 0.578559434 | 0.803135764 |
| 0.578806231 | 0.803283943 |
| 0.578944589 | 0.803373753 |
| 0.579081185 | 0.803469661 |
| 0.579122466 | 0.803503531 |
| 0.579286903 | 0.803661443 |
| 0.579371737 | 0.80372397 |
| 0.579394925 | 0.80372397 |
| 0.579598646 | 0.803930009 |
| 0.57967963 | 0.804004959 |
| 0.57969884 | 0.804004959 |
| 0.579703326 | 0.804004959 |
| 0.579924915 | 0.804195194 |
| 0.579943375 | 0.804197377 |
| 0.579988047 | 0.804223507 |
| 0.580051533 | 0.804277107 |
| 0.580128152 | 0.804359927 |
| 0.580162812 | 0.80436115 |
| 0.580208048 | 0.804374352 |
| 0.580256779 | 0.804374352 |
| 0.580293435 | 0.804401754 |
| 0.580352917 | 0.80443738 |
| 0.580398931 | 0.804452801 |
| 0.580448495 | 0.804452801 |
| 0.580634427 | 0.804663656 |
| 0.58072278 | 0.804739266 |
| 0.580760524 | 0.804768154 |
| 0.580795755 | 0.80479356 |
| 0.580982941 | 0.805006096 |
| 0.581038391 | 0.805059506 |
| 0.581117504 | 0.805108547 |
| 0.581121475 | 0.805108547 |
| 0.581165276 | 0.805118201 |
| 0.581270379 | 0.805134092 |
| 0.581325367 | 0.805134092 |
| 0.58133317 | 0.805134092 |

| | |
|---|---|
| 0.581395644 | 0.805156301 |
| 0.581465216 | 0.805229235 |
| 0.581535204 | 0.805255914 |
| 0.581582559 | 0.805289258 |
| 0.581593101 | 0.805289258 |
| 0.581810118 | 0.805402427 |
| 0.581887506 | 0.805467175 |
| 0.581913822 | 0.80547575 |
| 0.581953496 | 0.805507255 |
| 0.582056528 | 0.805597884 |
| 0.582151418 | 0.805687547 |
| 0.582274592 | 0.805811186 |
| 0.582340796 | 0.805840056 |
| 0.582399806 | 0.805867389 |
| 0.58246669 | 0.805913109 |
| 0.582499475 | 0.805935058 |
| 0.582555408 | 0.805989031 |
| 0.58264633 | 0.806046266 |
| 0.582681397 | 0.806046266 |
| 0.582797488 | 0.806072675 |
| 0.582802036 | 0.806072675 |
| 0.582841074 | 0.806098454 |
| 0.58287968 | 0.806098454 |
| 0.582919186 | 0.806106173 |
| 0.58307459 | 0.806135916 |
| 0.583098306 | 0.806135916 |
| 0.583113219 | 0.806135916 |
| 0.583134588 | 0.806135916 |
| 0.583135502 | 0.806135916 |
| 0.583302821 | 0.806216028 |
| 0.583329176 | 0.806216028 |
| 0.583330074 | 0.806216028 |
| 0.583342507 | 0.806216028 |
| 0.583378674 | 0.806222081 |
| 0.583384038 | 0.806222081 |
| 0.583454665 | 0.806270011 |
| 0.583503912 | 0.806270011 |
| 0.583554151 | 0.806270011 |
| 0.583695738 | 0.806403827 |
| 0.583701798 | 0.806403827 |
| 0.583752205 | 0.80642278 |
| 0.583766313 | 0.80642278 |
| 0.583858249 | 0.806502997 |
| 0.583892059 | 0.806526308 |
| 0.584001288 | 0.806630397 |

| | |
|---|---|
| 0.584077716 | 0.806665778 |
| 0.584117956 | 0.806697961 |
| 0.58415314 | 0.806723159 |
| 0.584342725 | 0.806914786 |
| 0.584431525 | 0.80695954 |
| 0.584437905 | 0.80695954 |
| 0.584442908 | 0.80695954 |
| 0.584523997 | 0.807003907 |
| 0.584525874 | 0.807003907 |
| 0.58467473 | 0.807115833 |
| 0.584857663 | 0.80722798 |
| 0.585016728 | 0.807344358 |
| 0.585022428 | 0.807344358 |
| 0.58504369 | 0.807344358 |
| 0.585200502 | 0.807443777 |
| 0.585320266 | 0.807562234 |
| 0.585360154 | 0.807570479 |
| 0.585567327 | 0.807692521 |
| 0.585623202 | 0.807695691 |
| 0.585647262 | 0.807695691 |
| 0.585705803 | 0.807695691 |
| 0.585709057 | 0.807695691 |
| 0.585733852 | 0.807695691 |
| 0.585756841 | 0.807695691 |
| 0.585776304 | 0.807695691 |
| 0.585811995 | 0.807702513 |
| 0.585869408 | 0.807734905 |
| 0.585904099 | 0.80775765 |
| 0.585995726 | 0.807792141 |
| 0.586069662 | 0.807866068 |
| 0.586155057 | 0.807912986 |
| 0.586168208 | 0.807912986 |
| 0.586235813 | 0.807956722 |
| 0.586275487 | 0.807956722 |
| 0.586362561 | 0.8079938 |
| 0.586497929 | 0.808063436 |
| 0.58660372 | 0.808162435 |
| 0.586771447 | 0.808321183 |
| 0.586818751 | 0.808321183 |
| 0.586849724 | 0.808337676 |
| 0.586904337 | 0.808366144 |
| 0.587000144 | 0.808474721 |
| 0.587035853 | 0.808487492 |
| 0.587095815 | 0.808489587 |
| 0.587393503 | 0.808841486 |

| | |
|---|---|
| 0.587419083 | 0.808841486 |
| 0.587419283 | 0.808841486 |
| 0.58753673 | 0.808979817 |
| 0.587556353 | 0.808983447 |
| 0.587617398 | 0.80900806 |
| 0.58763621 | 0.80900806 |
| 0.58765904 | 0.80900806 |
| 0.587697539 | 0.809037508 |
| 0.587899213 | 0.809236314 |
| 0.587905628 | 0.809236314 |
| 0.587945671 | 0.809236314 |
| 0.587972497 | 0.809236314 |
| 0.587977883 | 0.809236314 |
| 0.588111397 | 0.809373292 |
| 0.588190582 | 0.809449421 |
| 0.588227592 | 0.809449421 |
| 0.588251692 | 0.809449421 |
| 0.588336531 | 0.809519384 |
| 0.588406986 | 0.80956955 |
| 0.588496694 | 0.809600308 |
| 0.588525198 | 0.809615251 |
| 0.588569732 | 0.809639868 |
| 0.588602767 | 0.809639868 |
| 0.588621592 | 0.809639868 |
| 0.588627623 | 0.809639868 |
| 0.588749466 | 0.809751374 |
| 0.588760164 | 0.809751374 |
| 0.588883963 | 0.809804715 |
| 0.588982088 | 0.80988834 |
| 0.589012794 | 0.80988834 |
| 0.589093155 | 0.809951524 |
| 0.589108908 | 0.809951524 |
| 0.589290957 | 0.810154699 |
| 0.589391081 | 0.810244739 |
| 0.589452305 | 0.810275801 |
| 0.58955725 | 0.810319458 |
| 0.589596362 | 0.810319458 |
| 0.589598557 | 0.810319458 |
| 0.589677472 | 0.810381145 |
| 0.589809959 | 0.810516444 |
| 0.58990433 | 0.810622739 |
| 0.589923176 | 0.810625248 |
| 0.589962607 | 0.810630072 |
| 0.590113155 | 0.810745954 |
| 0.590321033 | 0.81095232 |

| | |
|---|---|
| 0.590415158 | 0.810997108 |
| 0.590443048 | 0.811012028 |
| 0.590531076 | 0.81108379 |
| 0.590546383 | 0.81108379 |
| 0.590616889 | 0.811153007 |
| 0.590707514 | 0.811208621 |
| 0.590739818 | 0.811208621 |
| 0.590772458 | 0.811208621 |
| 0.590825404 | 0.811208621 |
| 0.590859362 | 0.811208621 |
| 0.590917488 | 0.811219178 |
| 0.590951548 | 0.811219178 |
| 0.591038609 | 0.811315306 |
| 0.591112142 | 0.81134609 |
| 0.591216259 | 0.811395463 |
| 0.59124066 | 0.811396269 |
| 0.59126712 | 0.811396269 |
| 0.591453662 | 0.811616443 |
| 0.591481557 | 0.811616443 |
| 0.591563135 | 0.811616443 |
| 0.591583541 | 0.811616443 |
| 0.591665709 | 0.811616443 |
| 0.591671656 | 0.811616443 |
| 0.591679625 | 0.811616443 |
| 0.591686845 | 0.811616443 |
| 0.591733126 | 0.811616443 |
| 0.591780074 | 0.811616443 |
| 0.591882615 | 0.811678472 |
| 0.592004538 | 0.811715693 |
| 0.592040886 | 0.811715693 |
| 0.592090403 | 0.811715693 |
| 0.59209474 | 0.811715693 |
| 0.592096123 | 0.811715693 |
| 0.592276212 | 0.811884126 |
| 0.592288281 | 0.811884126 |
| 0.592388239 | 0.811916303 |
| 0.592395383 | 0.811916303 |
| 0.592396992 | 0.811916303 |
| 0.592513094 | 0.812005327 |
| 0.592590039 | 0.812048361 |
| 0.592731962 | 0.812126705 |
| 0.592755129 | 0.812126705 |
| 0.592860611 | 0.812154409 |
| 0.59288887 | 0.812156088 |
| 0.592895942 | 0.812156088 |

| | |
|---|---|
| 0.593014204 | 0.812195522 |
| 0.59302705 | 0.812195522 |
| 0.593104764 | 0.812255243 |
| 0.593136433 | 0.812275257 |
| 0.593162678 | 0.812287842 |
| 0.593187525 | 0.812298512 |
| 0.593247718 | 0.81234629 |
| 0.593274083 | 0.81234629 |
| 0.593279076 | 0.81234629 |
| 0.593365624 | 0.812385308 |
| 0.593387367 | 0.812385308 |
| 0.59342496 | 0.812411074 |
| 0.593477727 | 0.81242046 |
| 0.593481274 | 0.81242046 |
| 0.593562537 | 0.812446561 |
| 0.593591642 | 0.812446561 |
| 0.593620828 | 0.812446561 |
| 0.593621888 | 0.812446561 |
| 0.593631817 | 0.812446561 |
| 0.59367329 | 0.812446561 |
| 0.593787833 | 0.812525677 |
| 0.593859927 | 0.812540254 |
| 0.593885998 | 0.812540254 |
| 0.593892934 | 0.812540254 |
| 0.593953127 | 0.812599263 |
| 0.594165865 | 0.81274134 |
| 0.594177279 | 0.81274134 |
| 0.594208288 | 0.81274134 |
| 0.594210557 | 0.81274134 |
| 0.594348216 | 0.812882936 |
| 0.594371742 | 0.812890473 |
| 0.594453043 | 0.812944939 |
| 0.594529026 | 0.812944939 |
| 0.594538196 | 0.812944939 |
| 0.594651329 | 0.813013371 |
| 0.594683163 | 0.813013371 |
| 0.594685624 | 0.813013371 |
| 0.594822633 | 0.813013371 |
| 0.594849795 | 0.813013371 |
| 0.594851429 | 0.813013371 |
| 0.59493969 | 0.813013371 |
| 0.59504402 | 0.81306408 |
| 0.595098776 | 0.813115569 |
| 0.595192698 | 0.81322057 |
| 0.595218423 | 0.813224004 |

| | |
|---|---|
| 0.595273718 | 0.813261277 |
| 0.595400062 | 0.813370819 |
| 0.59542221 | 0.813370819 |
| 0.595601127 | 0.813421285 |
| 0.595603157 | 0.813421285 |
| 0.595752263 | 0.813506089 |
| 0.595774348 | 0.813506089 |
| 0.595790216 | 0.813506089 |
| 0.595795224 | 0.813506089 |
| 0.595811607 | 0.813506089 |
| 0.595846346 | 0.81352852 |
| 0.595920244 | 0.813536341 |
| 0.596109712 | 0.813656529 |
| 0.596138372 | 0.81367233 |
| 0.596177956 | 0.813703037 |
| 0.596242527 | 0.813744529 |
| 0.596301563 | 0.813801781 |
| 0.596384467 | 0.813891602 |
| 0.59654745 | 0.814032438 |
| 0.596686505 | 0.814118089 |
| 0.596687428 | 0.814118089 |
| 0.596713134 | 0.814118089 |
| 0.596771437 | 0.814144817 |
| 0.596792183 | 0.814144817 |
| 0.596823519 | 0.814164172 |
| 0.596906021 | 0.814230075 |
| 0.596954516 | 0.814238702 |
| 0.59696075 | 0.814238702 |
| 0.596963634 | 0.814238702 |
| 0.597010244 | 0.8142417 |
| 0.597034217 | 0.8142417 |
| 0.597071758 | 0.814269582 |
| 0.597135069 | 0.81428679 |
| 0.597152765 | 0.81428679 |
| 0.597375363 | 0.814520367 |
| 0.597402391 | 0.8145339 |
| 0.597432313 | 0.814551379 |
| 0.597538945 | 0.814650121 |
| 0.597603845 | 0.814668641 |
| 0.597662686 | 0.814725534 |
| 0.597687144 | 0.814735557 |
| 0.597799268 | 0.814818435 |
| 0.598011861 | 0.815014906 |
| 0.598056684 | 0.815029349 |
| 0.598145641 | 0.815103932 |

| | |
|---|---|
| 0.59820662 | 0.815122361 |
| 0.598256787 | 0.815138778 |
| 0.598317712 | 0.815193098 |
| 0.598367558 | 0.815213477 |
| 0.598380077 | 0.815213477 |
| 0.598501589 | 0.815285736 |
| 0.598580781 | 0.815328317 |
| 0.598729526 | 0.815432975 |
| 0.598867282 | 0.815597267 |
| 0.599118779 | 0.815836452 |
| 0.599136005 | 0.815836452 |
| 0.599179945 | 0.815836452 |
| 0.599225211 | 0.815851437 |
| 0.599367044 | 0.815978112 |
| 0.599369649 | 0.815978112 |
| 0.599441124 | 0.816028766 |
| 0.599579981 | 0.816112225 |
| 0.599610728 | 0.816119689 |
| 0.599766654 | 0.816242343 |
| 0.599769396 | 0.816242343 |
| 0.599830836 | 0.816279309 |
| 0.599930355 | 0.816344762 |
| 0.599950144 | 0.816348366 |
| 0.599997934 | 0.81639007 |
| 0.60003176 | 0.816412605 |
| 0.600100205 | 0.816412605 |
| 0.600235373 | 0.816503202 |
| 0.600273364 | 0.81650824 |
| 0.60046874 | 0.816610738 |
| 0.600520762 | 0.816658168 |
| 0.600622149 | 0.816713106 |
| 0.600697796 | 0.816713106 |
| 0.600737714 | 0.816743316 |
| 0.600988665 | 0.817014455 |
| 0.601106039 | 0.817017445 |
| 0.601269496 | 0.817093111 |
| 0.601310804 | 0.817125932 |
| 0.601343153 | 0.817134769 |
| 0.601386982 | 0.817136192 |
| 0.601475379 | 0.817212919 |
| 0.601494927 | 0.817212919 |
| 0.601640092 | 0.817333829 |
| 0.601652565 | 0.817333829 |
| 0.601724682 | 0.817342665 |
| 0.601788331 | 0.817357617 |

| | |
|---|---|
| 0.601902946 | 0.817405197 |
| 0.601911052 | 0.817405197 |
| 0.601985823 | 0.817471187 |
| 0.602028712 | 0.817476943 |
| 0.602159306 | 0.817578444 |
| 0.602343413 | 0.817682105 |
| 0.602346686 | 0.817682105 |
| 0.602373085 | 0.817682105 |
| 0.602389653 | 0.817682105 |
| 0.60250954 | 0.817803445 |
| 0.602538825 | 0.817803445 |
| 0.602571511 | 0.817803445 |
| 0.60260933 | 0.817803445 |
| 0.602624461 | 0.817803445 |
| 0.602868431 | 0.817918387 |
| 0.602873479 | 0.817918387 |
| 0.602875542 | 0.817918387 |
| 0.602890018 | 0.817918387 |
| 0.602955628 | 0.817960798 |
| 0.603174893 | 0.818162717 |
| 0.603190745 | 0.818162717 |
| 0.603205884 | 0.818162717 |
| 0.603216903 | 0.818162717 |
| 0.603287277 | 0.818200961 |
| 0.603337189 | 0.818223917 |
| 0.603623862 | 0.81841848 |
| 0.603712226 | 0.8184976 |
| 0.603729747 | 0.818498055 |
| 0.603813749 | 0.818575909 |
| 0.603991271 | 0.818738338 |
| 0.603999155 | 0.818738338 |
| 0.604060782 | 0.8187837 |
| 0.60425256 | 0.818872253 |
| 0.604290166 | 0.818884872 |
| 0.604336407 | 0.818907468 |
| 0.604392183 | 0.818953213 |
| 0.60442442 | 0.818954019 |
| 0.604693205 | 0.819174653 |
| 0.604746647 | 0.819177997 |
| 0.604835187 | 0.819247418 |
| 0.604850126 | 0.819247418 |
| 0.604897114 | 0.819287762 |
| 0.6049639 | 0.819308319 |
| 0.604987997 | 0.819314164 |
| 0.605002621 | 0.819314164 |

| | |
|---|---|
| 0.605107308 | 0.819432635 |
| 0.605142025 | 0.819456349 |
| 0.605214861 | 0.819484951 |
| 0.605249179 | 0.819484951 |
| 0.605499715 | 0.819800862 |
| 0.605586 | 0.819824462 |
| 0.605631842 | 0.819863218 |
| 0.605667446 | 0.819882049 |
| 0.605680181 | 0.819882049 |
| 0.60582975 | 0.819998259 |
| 0.605832053 | 0.819998259 |
| 0.60590834 | 0.820036102 |
| 0.605920823 | 0.820036102 |
| 0.605974286 | 0.820047091 |
| 0.606139853 | 0.820224537 |
| 0.606227688 | 0.820273476 |
| 0.606294491 | 0.820317256 |
| 0.606325016 | 0.820335252 |
| 0.606443327 | 0.8204021 |
| 0.606518937 | 0.820411173 |
| 0.606617516 | 0.820497911 |
| 0.606668909 | 0.82050007 |
| 0.606696393 | 0.82050007 |
| 0.606704241 | 0.82050007 |
| 0.606722478 | 0.82050007 |
| 0.60675631 | 0.82050834 |
| 0.606776122 | 0.82050834 |
| 0.606780277 | 0.82050834 |
| 0.606866362 | 0.820554855 |
| 0.606971861 | 0.820604316 |
| 0.607128026 | 0.82074555 |
| 0.607243001 | 0.820863136 |
| 0.607283722 | 0.820863136 |
| 0.607360251 | 0.8209293 |
| 0.607612904 | 0.821035701 |
| 0.607627154 | 0.821035701 |
| 0.607633492 | 0.821035701 |
| 0.607635719 | 0.821035701 |
| 0.607690793 | 0.821063529 |
| 0.607765728 | 0.821084168 |
| 0.607775028 | 0.821084168 |
| 0.607844371 | 0.821131265 |
| 0.607946102 | 0.821201291 |
| 0.608072829 | 0.821266679 |
| 0.608085528 | 0.821266679 |

| | |
|---|---|
| 0.608140115 | 0.821266679 |
| 0.608168005 | 0.821266679 |
| 0.608483973 | 0.821502119 |
| 0.608576711 | 0.821584598 |
| 0.60861558 | 0.821613782 |
| 0.608653508 | 0.821629238 |
| 0.608857196 | 0.821699518 |
| 0.608864967 | 0.821699518 |
| 0.608871342 | 0.821699518 |
| 0.60897033 | 0.821766587 |
| 0.609025435 | 0.821817663 |
| 0.609077973 | 0.821841987 |
| 0.609114394 | 0.821844564 |
| 0.609198918 | 0.821897281 |
| 0.609205236 | 0.821897281 |
| 0.609250679 | 0.821928511 |
| 0.609498002 | 0.82209003 |
| 0.609498002 | 0.82209003 |
| 0.609503455 | 0.82209003 |
| 0.609608697 | 0.822154534 |
| 0.609608995 | 0.822154534 |
| 0.609620328 | 0.822154534 |
| 0.609653004 | 0.822175321 |
| 0.609729563 | 0.822255286 |
| 0.609794264 | 0.822265896 |
| 0.609852626 | 0.822304824 |
| 0.610103522 | 0.82250341 |
| 0.61023737 | 0.822660568 |
| 0.61033147 | 0.822717563 |
| 0.610398506 | 0.822761353 |
| 0.610541669 | 0.822842327 |
| 0.61055275 | 0.822842327 |
| 0.610587989 | 0.822842327 |
| 0.610596794 | 0.822842327 |
| 0.610618998 | 0.822848966 |
| 0.610713725 | 0.822887305 |
| 0.61076238 | 0.822890804 |
| 0.610843695 | 0.8229422 |
| 0.610922283 | 0.823012324 |
| 0.610930306 | 0.823012324 |
| 0.611014135 | 0.823078691 |
| 0.611071194 | 0.823103372 |
| 0.611084304 | 0.823103372 |
| 0.611340231 | 0.823354952 |
| 0.61146478 | 0.823441034 |

| | |
|---|---|
| 0.611693662 | 0.823557132 |
| 0.611753127 | 0.823586396 |
| 0.611754172 | 0.823586396 |
| 0.611918093 | 0.823690841 |
| 0.612030162 | 0.823771669 |
| 0.612047812 | 0.823772146 |
| 0.612179271 | 0.823855952 |
| 0.612231228 | 0.823879314 |
| 0.612280897 | 0.823903367 |
| 0.612291064 | 0.823903367 |
| 0.61251813 | 0.824079123 |
| 0.612561772 | 0.824097362 |
| 0.612566293 | 0.824097362 |
| 0.612684759 | 0.82414034 |
| 0.61274328 | 0.824193713 |
| 0.612781989 | 0.824193713 |
| 0.612882216 | 0.824266266 |
| 0.612988971 | 0.824363284 |
| 0.613052233 | 0.824376492 |
| 0.613090125 | 0.824376492 |
| 0.613105442 | 0.824376492 |
| 0.613129724 | 0.824376492 |
| 0.613140314 | 0.824376492 |
| 0.613217022 | 0.824390637 |
| 0.613481847 | 0.824621003 |
| 0.613485078 | 0.824621003 |
| 0.613509577 | 0.824621003 |
| 0.613596064 | 0.824706235 |
| 0.613818172 | 0.824837125 |
| 0.613862801 | 0.824839712 |
| 0.61393214 | 0.824863068 |
| 0.613967829 | 0.824887748 |
| 0.614131353 | 0.825060899 |
| 0.614321312 | 0.825148572 |
| 0.614331131 | 0.825148572 |
| 0.614369864 | 0.825148572 |
| 0.614462262 | 0.825218747 |
| 0.614523501 | 0.825237616 |
| 0.614688071 | 0.825334572 |
| 0.614690128 | 0.825334572 |
| 0.614701687 | 0.825334572 |
| 0.614722438 | 0.825334572 |
| 0.614902232 | 0.825434526 |
| 0.614905922 | 0.825434526 |
| 0.614929397 | 0.825434526 |

| | |
|---|---|
| 0.614995903 | 0.825482559 |
| 0.61504314 | 0.825494143 |
| 0.615121363 | 0.825573003 |
| 0.615340423 | 0.825730259 |
| 0.61539758 | 0.82578369 |
| 0.615453672 | 0.82581143 |
| 0.615461226 | 0.82581143 |
| 0.61547027 | 0.82581143 |
| 0.61563598 | 0.82589417 |
| 0.615799844 | 0.825991261 |
| 0.615826038 | 0.825991261 |
| 0.615829754 | 0.825991261 |
| 0.615976694 | 0.826124988 |
| 0.616086225 | 0.82619507 |
| 0.616153302 | 0.82619507 |
| 0.616173088 | 0.82619507 |
| 0.616229235 | 0.82619507 |
| 0.616236509 | 0.82619507 |
| 0.616240859 | 0.82619507 |
| 0.616241914 | 0.82619507 |
| 0.61633796 | 0.826285446 |
| 0.616361371 | 0.826285446 |
| 0.616426788 | 0.826349883 |
| 0.616498354 | 0.826403497 |
| 0.616501486 | 0.826403497 |
| 0.616538662 | 0.826406812 |
| 0.616680919 | 0.826550969 |
| 0.616783142 | 0.826618192 |
| 0.61697569 | 0.826744226 |
| 0.616981925 | 0.826744226 |
| 0.617039498 | 0.826775641 |
| 0.617117349 | 0.826826728 |
| 0.617160085 | 0.826826728 |
| 0.617249039 | 0.826889783 |
| 0.617340873 | 0.826889783 |
| 0.617375068 | 0.826889783 |
| 0.617377396 | 0.826889783 |
| 0.617385111 | 0.826889783 |
| 0.61743654 | 0.826911212 |
| 0.617517562 | 0.826911212 |
| 0.617517742 | 0.826911212 |
| 0.617602774 | 0.826948758 |
| 0.617711074 | 0.827026468 |
| 0.617712906 | 0.827026468 |
| 0.617777165 | 0.827033731 |

| | |
|---|---|
| 0.617876253 | 0.827033731 |
| 0.617883826 | 0.827033731 |
| 0.618011641 | 0.827124162 |
| 0.618165834 | 0.827260788 |
| 0.618198882 | 0.827281769 |
| 0.618285816 | 0.827305123 |
| 0.618323682 | 0.827332547 |
| 0.618366656 | 0.827349415 |
| 0.618420693 | 0.827354494 |
| 0.61846148 | 0.827354494 |
| 0.618476703 | 0.827354494 |
| 0.61847998 | 0.827354494 |
| 0.618510779 | 0.827354494 |
| 0.618526564 | 0.827354494 |
| 0.618531172 | 0.827354494 |
| 0.618632985 | 0.827360649 |
| 0.618657375 | 0.827360649 |
| 0.618743336 | 0.827372438 |
| 0.618789288 | 0.827372438 |
| 0.618802251 | 0.827372438 |
| 0.618804947 | 0.827372438 |
| 0.618808093 | 0.827372438 |
| 0.618928754 | 0.827454291 |
| 0.618935879 | 0.827454291 |
| 0.619113427 | 0.827563075 |
| 0.619122029 | 0.827563075 |
| 0.619236489 | 0.827563075 |
| 0.619265101 | 0.827563075 |
| 0.619273334 | 0.827563075 |
| 0.619293989 | 0.827563075 |
| 0.619295264 | 0.827563075 |
| 0.619520691 | 0.827697968 |
| 0.619523681 | 0.827697968 |
| 0.619532486 | 0.827697968 |
| 0.619829757 | 0.827975313 |
| 0.619901908 | 0.827991467 |
| 0.61991269 | 0.827991467 |
| 0.619935766 | 0.827991467 |
| 0.62014179 | 0.828134003 |
| 0.620191318 | 0.828156332 |
| 0.620262743 | 0.828228487 |
| 0.620324159 | 0.828271952 |
| 0.620330076 | 0.828271952 |
| 0.620419719 | 0.828335255 |
| 0.620437651 | 0.828335255 |

| | |
|---|---|
| 0.620599367 | 0.828335255 |
| 0.620617691 | 0.828335255 |
| 0.6206244 | 0.828335255 |
| 0.620630305 | 0.828335255 |
| 0.620772658 | 0.828444838 |
| 0.620794736 | 0.828451088 |
| 0.620873412 | 0.828490213 |
| 0.620932471 | 0.828542034 |
| 0.621069195 | 0.828631609 |
| 0.621377775 | 0.828904216 |
| 0.621461882 | 0.828969754 |
| 0.621523518 | 0.829007655 |
| 0.621525108 | 0.829007655 |
| 0.621571507 | 0.829046325 |
| 0.621702655 | 0.82914663 |
| 0.62200768 | 0.829480001 |
| 0.622018566 | 0.829480001 |
| 0.62206314 | 0.82948938 |
| 0.622067195 | 0.82948938 |
| 0.622224334 | 0.829584976 |
| 0.622310709 | 0.829584976 |
| 0.622324748 | 0.829584976 |
| 0.622383323 | 0.829584976 |
| 0.622434075 | 0.829584976 |
| 0.622476272 | 0.829584976 |
| 0.622482462 | 0.829584976 |
| 0.622489471 | 0.829584976 |
| 0.622552938 | 0.829635099 |
| 0.622663799 | 0.829681329 |
| 0.622678529 | 0.829681329 |
| 0.622698374 | 0.829681329 |
| 0.622709572 | 0.829681329 |
| 0.622735837 | 0.829693113 |
| 0.6228668 | 0.829795487 |
| 0.62288503 | 0.829795487 |
| 0.622899789 | 0.829795487 |
| 0.623097685 | 0.829893762 |
| 0.623117422 | 0.829893762 |
| 0.623182659 | 0.829893762 |
| 0.623205062 | 0.829900391 |
| 0.623392798 | 0.829997239 |
| 0.623396661 | 0.829997239 |
| 0.623417205 | 0.829997239 |
| 0.623444162 | 0.830009926 |
| 0.623598812 | 0.830120352 |

| | |
|---|---|
| 0.623610695 | 0.830120352 |
| 0.623631683 | 0.830120352 |
| 0.623739748 | 0.830217791 |
| 0.623947667 | 0.830358306 |
| 0.624036462 | 0.830403861 |
| 0.624100385 | 0.830442518 |
| 0.62432079 | 0.830666171 |
| 0.624493855 | 0.830764631 |
| 0.62449945 | 0.830764631 |
| 0.624550623 | 0.830786296 |
| 0.624610629 | 0.830836546 |
| 0.624623289 | 0.830836546 |
| 0.624700147 | 0.830892518 |
| 0.624752598 | 0.830892518 |
| 0.625077408 | 0.831152487 |
| 0.625111795 | 0.831152487 |
| 0.625211157 | 0.831183792 |
| 0.625215935 | 0.831183792 |
| 0.625251502 | 0.831207874 |
| 0.625281073 | 0.831223983 |
| 0.625450605 | 0.831379736 |
| 0.625526114 | 0.831442861 |
| 0.625562135 | 0.831458371 |
| 0.625612139 | 0.831501628 |
| 0.625635529 | 0.831509511 |
| 0.625671114 | 0.831511912 |
| 0.625687586 | 0.831511912 |
| 0.625689712 | 0.831511912 |
| 0.625715918 | 0.831523536 |
| 0.625893411 | 0.831666587 |
| 0.625955935 | 0.831703259 |
| 0.625990272 | 0.831725679 |
| 0.626066882 | 0.831781058 |
| 0.62609884 | 0.831800314 |
| 0.626200867 | 0.831843045 |
| 0.626226084 | 0.831853341 |
| 0.626266902 | 0.831868856 |
| 0.62637985 | 0.831964781 |
| 0.626435801 | 0.832015893 |
| 0.626511072 | 0.832092661 |
| 0.626537642 | 0.832095679 |
| 0.626566651 | 0.832095679 |
| 0.626573109 | 0.832095679 |
| 0.626624819 | 0.832100844 |
| 0.626627142 | 0.832100844 |

| | |
|---|---|
| 0.626639531 | 0.832100844 |
| 0.626764724 | 0.83224388 |
| 0.626815321 | 0.832287861 |
| 0.626887843 | 0.83232214 |
| 0.62693573 | 0.832350645 |
| 0.627004686 | 0.832400061 |
| 0.627025077 | 0.832400126 |
| 0.627050978 | 0.832400126 |
| 0.627057167 | 0.832400126 |
| 0.627173548 | 0.832482527 |
| 0.627196012 | 0.832482527 |
| 0.627238461 | 0.832482527 |
| 0.627283943 | 0.832515532 |
| 0.627402084 | 0.832625921 |
| 0.627470766 | 0.832693867 |
| 0.627613132 | 0.83281318 |
| 0.627669573 | 0.832864869 |
| 0.627714301 | 0.832901015 |
| 0.627824777 | 0.832972703 |
| 0.6278375 | 0.832972703 |
| 0.627855777 | 0.832972703 |
| 0.627937908 | 0.833035256 |
| 0.628002249 | 0.833068406 |
| 0.628015371 | 0.833068406 |
| 0.628080595 | 0.83310852 |
| 0.628208832 | 0.83313784 |
| 0.628215773 | 0.83313784 |
| 0.628305741 | 0.833198316 |
| 0.628418923 | 0.833203747 |
| 0.628420861 | 0.833203747 |
| 0.628432297 | 0.833203747 |
| 0.628734932 | 0.83348898 |
| 0.628777154 | 0.833500537 |
| 0.62877865 | 0.833500537 |
| 0.628878746 | 0.833586821 |
| 0.629038987 | 0.833741467 |
| 0.629246011 | 0.833861178 |
| 0.629278318 | 0.833861178 |
| 0.629278318 | 0.833861178 |
| 0.629397935 | 0.833950075 |
| 0.629489133 | 0.834024506 |
| 0.629576527 | 0.834117093 |
| 0.629605995 | 0.834118446 |
| 0.629773086 | 0.834181688 |
| 0.629946268 | 0.834328445 |

| | |
|---|---|
| 0.630272562 | 0.83459814 |
| 0.630461401 | 0.834784865 |
| 0.630483683 | 0.834784865 |
| 0.630513819 | 0.834801559 |
| 0.630640494 | 0.83483958 |
| 0.630657357 | 0.83483958 |
| 0.630684006 | 0.83483958 |
| 0.630747746 | 0.83483958 |
| 0.630757282 | 0.83483958 |
| 0.630845264 | 0.83483958 |
| 0.630879814 | 0.83483958 |
| 0.630916303 | 0.83483958 |
| 0.630951763 | 0.83483958 |
| 0.630963225 | 0.83483958 |
| 0.631020032 | 0.834891548 |
| 0.631051471 | 0.834909951 |
| 0.631183513 | 0.834991866 |
| 0.63124265 | 0.835000519 |
| 0.63126546 | 0.835005317 |
| 0.631316406 | 0.835005317 |
| 0.631342137 | 0.835016161 |
| 0.631379246 | 0.835018864 |
| 0.631523703 | 0.835100592 |
| 0.631705405 | 0.835218292 |
| 0.631765745 | 0.835251697 |
| 0.631802092 | 0.835253379 |
| 0.631851281 | 0.83528787 |
| 0.632040819 | 0.83528787 |
| 0.632079766 | 0.83528787 |
| 0.632089722 | 0.83528787 |
| 0.632126329 | 0.83528787 |
| 0.632126329 | 0.83528787 |
| 0.632246451 | 0.835357403 |
| 0.632282627 | 0.835357403 |
| 0.632297014 | 0.835357403 |
| 0.632301728 | 0.835357403 |
| 0.632356423 | 0.835360146 |
| 0.632438312 | 0.835444301 |
| 0.632487107 | 0.835463265 |
| 0.632550711 | 0.835500935 |
| 0.632593832 | 0.835534719 |
| 0.632651148 | 0.835587249 |
| 0.632691563 | 0.835588361 |
| 0.632873484 | 0.835647557 |
| 0.632888293 | 0.835647557 |

| | |
|---|---|
| 0.632894126 | 0.835647557 |
| 0.632939232 | 0.835666477 |
| 0.633417087 | 0.836065615 |
| 0.633568426 | 0.836186338 |
| 0.633702457 | 0.836202969 |
| 0.633755615 | 0.83621588 |
| 0.633759178 | 0.83621588 |
| 0.634010674 | 0.836385518 |
| 0.634198646 | 0.836523194 |
| 0.634202857 | 0.836523194 |
| 0.634249313 | 0.836538134 |
| 0.634315683 | 0.836556172 |
| 0.63437238 | 0.836572879 |
| 0.634487088 | 0.836623281 |
| 0.634545198 | 0.836650356 |
| 0.634650053 | 0.836714305 |
| 0.634768608 | 0.836714305 |
| 0.634782867 | 0.836714305 |
| 0.634821592 | 0.836714305 |
| 0.634834372 | 0.836714305 |
| 0.634991346 | 0.836857744 |
| 0.635001197 | 0.836857744 |
| 0.635025773 | 0.836862078 |
| 0.63509428 | 0.836909056 |
| 0.635197752 | 0.836954666 |
| 0.635307692 | 0.83703544 |
| 0.635400833 | 0.837106455 |
| 0.635468688 | 0.837162918 |
| 0.635538829 | 0.837169452 |
| 0.635553307 | 0.837169452 |
| 0.635555129 | 0.837169452 |
| 0.635571694 | 0.837169452 |
| 0.635738141 | 0.837310693 |
| 0.635743791 | 0.837310693 |
| 0.635851388 | 0.837413368 |
| 0.635864691 | 0.837413368 |
| 0.636059586 | 0.837580455 |
| 0.636102409 | 0.837613687 |
| 0.636215496 | 0.837669959 |
| 0.636891484 | 0.838267975 |
| 0.636968746 | 0.838274763 |
| 0.636974062 | 0.838274763 |
| 0.637068734 | 0.838320742 |
| 0.637077267 | 0.838320742 |
| 0.637107476 | 0.838334514 |

| | |
|---|---|
| 0.637142911 | 0.838339061 |
| 0.63714866 | 0.838339061 |
| 0.637199503 | 0.838362958 |
| 0.63729543 | 0.838466007 |
| 0.637341234 | 0.838503107 |
| 0.637511874 | 0.83868127 |
| 0.637570377 | 0.838735066 |
| 0.637834128 | 0.838888795 |
| 0.637838692 | 0.838888795 |
| 0.637838692 | 0.838888795 |
| 0.637838692 | 0.838888795 |
| 0.637845759 | 0.838888795 |
| 0.638088051 | 0.839068434 |
| 0.638168024 | 0.83912726 |
| 0.638237013 | 0.839194805 |
| 0.63827949 | 0.83920749 |
| 0.638293438 | 0.83920749 |
| 0.638405394 | 0.839304193 |
| 0.639071383 | 0.839967236 |
| 0.639324075 | 0.840198571 |
| 0.639335596 | 0.840198571 |
| 0.639466723 | 0.840205168 |
| 0.639471017 | 0.840205168 |
| 0.639478829 | 0.840205168 |
| 0.639554617 | 0.840214512 |
| 0.639582707 | 0.840214512 |
| 0.639682077 | 0.840214512 |
| 0.639710902 | 0.840214512 |
| 0.639727197 | 0.840214512 |
| 0.639753481 | 0.840214512 |
| 0.639804692 | 0.840249403 |
| 0.639815333 | 0.840249403 |
| 0.639844925 | 0.84025433 |
| 0.639927302 | 0.84025433 |
| 0.63993891 | 0.84025433 |
| 0.639942581 | 0.84025433 |
| 0.639990887 | 0.840294591 |
| 0.640120589 | 0.840372217 |
| 0.640253765 | 0.840454387 |
| 0.640313337 | 0.84046064 |
| 0.640343686 | 0.84046064 |
| 0.640344848 | 0.84046064 |
| 0.640364409 | 0.84046064 |
| 0.640395814 | 0.840474877 |
| 0.640468337 | 0.840488666 |

| | |
|---|---|
| 0.640473999 | 0.840488666 |
| 0.640546319 | 0.84056041 |
| 0.640652402 | 0.840676456 |
| 0.640757536 | 0.840718431 |
| 0.640772651 | 0.840718431 |
| 0.640835993 | 0.840778376 |
| 0.640865789 | 0.840794307 |
| 0.640904502 | 0.840800527 |
| 0.640940713 | 0.840823119 |
| 0.641266808 | 0.841053531 |
| 0.641292943 | 0.841053531 |
| 0.641420546 | 0.841105075 |
| 0.641468651 | 0.841143508 |
| 0.641711376 | 0.841301138 |
| 0.6418081 | 0.841381623 |
| 0.641998584 | 0.841515524 |
| 0.642126098 | 0.841659503 |
| 0.642291148 | 0.841672833 |
| 0.642295318 | 0.841672833 |
| 0.642354633 | 0.841704242 |
| 0.642377666 | 0.841711266 |
| 0.642724541 | 0.841981921 |
| 0.642761914 | 0.841981921 |
| 0.642783582 | 0.841981921 |
| 0.642796368 | 0.841981921 |
| 0.64286235 | 0.842022032 |
| 0.642887195 | 0.842024674 |
| 0.642966087 | 0.842088432 |
| 0.643020718 | 0.842113668 |
| 0.643039285 | 0.842114827 |
| 0.64307495 | 0.842132774 |
| 0.64313978 | 0.842173932 |
| 0.643260204 | 0.842260753 |
| 0.643276854 | 0.842260753 |
| 0.64329219 | 0.842260753 |
| 0.64338876 | 0.842273836 |
| 0.64340333 | 0.842273836 |
| 0.643408292 | 0.842273836 |
| 0.643428057 | 0.84227656 |
| 0.643575378 | 0.842353648 |
| 0.643725937 | 0.842425367 |
| 0.643743989 | 0.842425367 |
| 0.643749548 | 0.842425367 |
| 0.64387845 | 0.842484318 |
| 0.644130318 | 0.842644919 |

| | |
|---|---|
| 0.644225639 | 0.842644919 |
| 0.644233245 | 0.842644919 |
| 0.644287006 | 0.842644919 |
| 0.644385814 | 0.842650207 |
| 0.644459381 | 0.842677127 |
| 0.644475974 | 0.842677127 |
| 0.644486415 | 0.842677127 |
| 0.644635694 | 0.842722447 |
| 0.6446682 | 0.842741811 |
| 0.644710457 | 0.842773918 |
| 0.64480009 | 0.842844821 |
| 0.644833428 | 0.842865266 |
| 0.644855936 | 0.842871554 |
| 0.644883175 | 0.842884024 |
| 0.644928742 | 0.842897318 |
| 0.645030439 | 0.84297546 |
| 0.645178192 | 0.843069383 |
| 0.645257648 | 0.843069383 |
| 0.645283249 | 0.843069383 |
| 0.64536955 | 0.843126386 |
| 0.6454289 | 0.843164125 |
| 0.645483523 | 0.843205895 |
| 0.645505845 | 0.843207677 |
| 0.645648285 | 0.843216796 |
| 0.645676059 | 0.843216796 |
| 0.645725983 | 0.843216796 |
| 0.645731236 | 0.843216796 |
| 0.645731632 | 0.843216796 |
| 0.64575299 | 0.843216796 |
| 0.645759084 | 0.843216796 |
| 0.645775141 | 0.843216796 |
| 0.645850011 | 0.843271917 |
| 0.645950799 | 0.843356509 |
| 0.645988385 | 0.843356509 |
| 0.646232952 | 0.843461825 |
| 0.646240654 | 0.843461825 |
| 0.646246152 | 0.843461825 |
| 0.64637324 | 0.843558347 |
| 0.646415086 | 0.843585983 |
| 0.646661752 | 0.843737294 |
| 0.64666825 | 0.843737294 |
| 0.64674612 | 0.843737294 |
| 0.646772615 | 0.843737294 |
| 0.64679794 | 0.843737294 |
| 0.646811522 | 0.843737294 |

| | |
|---|---|
| 0.64684077 | 0.843746867 |
| 0.646859972 | 0.843746867 |
| 0.646897936 | 0.843746867 |
| 0.646904401 | 0.843746867 |
| 0.647050476 | 0.843826137 |
| 0.647255438 | 0.843945556 |
| 0.647318686 | 0.843945556 |
| 0.647348704 | 0.843945556 |
| 0.647409243 | 0.843945556 |
| 0.647435064 | 0.843945556 |
| 0.647533299 | 0.844008441 |
| 0.647605038 | 0.844022372 |
| 0.647675934 | 0.844055757 |
| 0.64775854 | 0.84409412 |
| 0.647809973 | 0.844138046 |
| 0.647873579 | 0.844174736 |
| 0.647931592 | 0.844221657 |
| 0.648113873 | 0.844320675 |
| 0.64816563 | 0.844320675 |
| 0.648210326 | 0.844320675 |
| 0.648333032 | 0.844354175 |
| 0.648349529 | 0.844354175 |
| 0.648370632 | 0.844354175 |
| 0.648389614 | 0.844354175 |
| 0.648496163 | 0.84440842 |
| 0.648595112 | 0.844467999 |
| 0.648645799 | 0.844510907 |
| 0.648827327 | 0.844654888 |
| 0.648845322 | 0.844655228 |
| 0.648897601 | 0.844677109 |
| 0.649313209 | 0.845033342 |
| 0.649771767 | 0.845581256 |
| 0.649787482 | 0.845581256 |
| 0.649869439 | 0.845610837 |
| 0.649881233 | 0.845610837 |
| 0.649940204 | 0.845618262 |
| 0.650127046 | 0.845722737 |
| 0.650304851 | 0.845824198 |
| 0.650400395 | 0.845824198 |
| 0.650439959 | 0.845829459 |
| 0.650554818 | 0.845916876 |
| 0.650631519 | 0.845916876 |
| 0.65064079 | 0.845916876 |
| 0.650824925 | 0.846088496 |
| 0.650850108 | 0.846088496 |

| | |
|---|---|
| 0.650870299 | 0.846088754 |
| 0.650971672 | 0.846197435 |
| 0.651052163 | 0.846255871 |
| 0.651168882 | 0.846292427 |
| 0.651186902 | 0.846292427 |
| 0.651320839 | 0.846397206 |
| 0.651347913 | 0.846409294 |
| 0.651399867 | 0.846453713 |
| 0.651522859 | 0.846507992 |
| 0.651530507 | 0.846507992 |
| 0.651563657 | 0.84652797 |
| 0.651720661 | 0.846600968 |
| 0.651782395 | 0.846600968 |
| 0.651784539 | 0.846600968 |
| 0.651815852 | 0.846600968 |
| 0.651860331 | 0.846600968 |
| 0.651868437 | 0.846600968 |
| 0.651904254 | 0.846600968 |
| 0.651971337 | 0.846641914 |
| 0.652012671 | 0.846650278 |
| 0.65213896 | 0.846755566 |
| 0.652139041 | 0.846755566 |
| 0.652428109 | 0.846922802 |
| 0.652444196 | 0.846922802 |
| 0.652523641 | 0.846966545 |
| 0.652554203 | 0.846983132 |
| 0.652599388 | 0.846995583 |
| 0.652634932 | 0.846995583 |
| 0.652698924 | 0.84705555 |
| 0.652897746 | 0.847151991 |
| 0.652967192 | 0.847195936 |
| 0.6532134 | 0.847414266 |
| 0.653259883 | 0.847414266 |
| 0.653312002 | 0.847414266 |
| 0.653316254 | 0.847414266 |
| 0.653365204 | 0.847431403 |
| 0.653480635 | 0.847515786 |
| 0.653513273 | 0.847525169 |
| 0.653570724 | 0.847534958 |
| 0.653584394 | 0.847534958 |
| 0.653702978 | 0.847596421 |
| 0.653766983 | 0.847656333 |
| 0.653803257 | 0.84766641 |
| 0.653810351 | 0.84766641 |
| 0.654069405 | 0.847840715 |

| | |
|---|---|
| 0.654092843 | 0.847841324 |
| 0.654201983 | 0.847841324 |
| 0.654217416 | 0.847841324 |
| 0.654220981 | 0.847841324 |
| 0.654221885 | 0.847841324 |
| 0.654253313 | 0.847841324 |
| 0.654294225 | 0.847855228 |
| 0.65454278 | 0.84811164 |
| 0.654581006 | 0.848134544 |
| 0.654703498 | 0.848157544 |
| 0.654705607 | 0.848157544 |
| 0.654786091 | 0.84819826 |
| 0.654807739 | 0.84819826 |
| 0.65494538 | 0.848283575 |
| 0.655002164 | 0.84829758 |
| 0.655157615 | 0.848445607 |
| 0.65528769 | 0.848542328 |
| 0.655451415 | 0.848657887 |
| 0.655561695 | 0.848710214 |
| 0.65559554 | 0.848710214 |
| 0.655668457 | 0.848758469 |
| 0.655712494 | 0.848792405 |
| 0.65589547 | 0.84898194 |
| 0.655932106 | 0.84898194 |
| 0.655961438 | 0.84898194 |
| 0.655980871 | 0.84898194 |
| 0.655993935 | 0.84898194 |
| 0.656002231 | 0.84898194 |
| 0.656189606 | 0.849065531 |
| 0.656190902 | 0.849065531 |
| 0.656293658 | 0.849100837 |
| 0.656318484 | 0.849100837 |
| 0.656421055 | 0.849100837 |
| 0.656447961 | 0.849100837 |
| 0.656515417 | 0.849100837 |
| 0.656518542 | 0.849100837 |
| 0.656528402 | 0.849100837 |
| 0.656535639 | 0.849100837 |
| 0.656539809 | 0.849100837 |
| 0.656557465 | 0.849100837 |
| 0.656584212 | 0.849100837 |
| 0.65667057 | 0.849100837 |
| 0.6566796 | 0.849100837 |
| 0.656736133 | 0.849100837 |
| 0.656736142 | 0.849100837 |

| | |
|---|---|
| 0.656740452 | 0.849100837 |
| 0.656753032 | 0.849100837 |
| 0.656831999 | 0.849133781 |
| 0.656870024 | 0.849150467 |
| 0.656880565 | 0.849150467 |
| 0.657023711 | 0.849243311 |
| 0.657082098 | 0.849272683 |
| 0.657196221 | 0.849351297 |
| 0.657214256 | 0.849351297 |
| 0.657274029 | 0.84938245 |
| 0.657383681 | 0.849407314 |
| 0.657532756 | 0.849467636 |
| 0.657536143 | 0.849467636 |
| 0.65763387 | 0.849570844 |
| 0.657726943 | 0.849573047 |
| 0.657742872 | 0.849573047 |
| 0.657773213 | 0.849573047 |
| 0.657809827 | 0.849573047 |
| 0.65783006 | 0.849573047 |
| 0.657861413 | 0.849573047 |
| 0.657889124 | 0.849573047 |
| 0.657958071 | 0.84961474 |
| 0.65797111 | 0.84961474 |
| 0.658106342 | 0.849663453 |
| 0.658218676 | 0.849684352 |
| 0.658221265 | 0.849684352 |
| 0.658251935 | 0.849700913 |
| 0.658331016 | 0.849743948 |
| 0.658393602 | 0.849743948 |
| 0.658397493 | 0.849743948 |
| 0.658463843 | 0.849743948 |
| 0.658494625 | 0.849743948 |
| 0.658511319 | 0.849743948 |
| 0.658517215 | 0.849743948 |
| 0.658689226 | 0.849919855 |
| 0.658942046 | 0.850092552 |
| 0.658973365 | 0.850092552 |
| 0.65900852 | 0.850092552 |
| 0.659021263 | 0.850092552 |
| 0.659036646 | 0.850092552 |
| 0.659171905 | 0.850137125 |
| 0.659275318 | 0.850186712 |
| 0.659369663 | 0.850244997 |
| 0.65939986 | 0.850260915 |
| 0.659540232 | 0.850395869 |

| | |
|---|---|
| 0.659695264 | 0.850480642 |
| 0.659771905 | 0.85051038 |
| 0.659977943 | 0.85063784 |
| 0.660032925 | 0.850662665 |
| 0.660083091 | 0.85068128 |
| 0.660171062 | 0.850748611 |
| 0.660249253 | 0.850757295 |
| 0.660305986 | 0.850784363 |
| 0.660350271 | 0.850818406 |
| 0.660413506 | 0.850830827 |
| 0.660456195 | 0.850851209 |
| 0.660536578 | 0.850861097 |
| 0.660544191 | 0.850861097 |
| 0.660600325 | 0.850887377 |
| 0.660759092 | 0.850918111 |
| 0.660762178 | 0.850918111 |
| 0.660796231 | 0.850918111 |
| 0.660824728 | 0.850918111 |
| 0.660825756 | 0.850918111 |
| 0.660825764 | 0.850918111 |
| 0.660900749 | 0.850952148 |
| 0.660969597 | 0.851017788 |
| 0.661043257 | 0.85102486 |
| 0.661046564 | 0.85102486 |
| 0.661169901 | 0.851152293 |
| 0.661258185 | 0.851228271 |
| 0.661306656 | 0.851266269 |
| 0.661361915 | 0.851292774 |
| 0.661385556 | 0.851300196 |
| 0.661420744 | 0.851322481 |
| 0.661449021 | 0.851330044 |
| 0.661462943 | 0.851330044 |
| 0.6614881 | 0.851330044 |
| 0.661608235 | 0.851353347 |
| 0.661765592 | 0.851490203 |
| 0.661871304 | 0.851495064 |
| 0.661943224 | 0.851495064 |
| 0.661979267 | 0.851495064 |
| 0.662199806 | 0.851620051 |
| 0.662206248 | 0.851620051 |
| 0.662273382 | 0.851664926 |
| 0.662276906 | 0.851664926 |
| 0.662459698 | 0.851712084 |
| 0.662501841 | 0.851712084 |
| 0.662544716 | 0.851712084 |

| | |
|---|---|
| 0.66258149 | 0.851712084 |
| 0.662649576 | 0.851712084 |
| 0.662652855 | 0.851712084 |
| 0.662720503 | 0.851712084 |
| 0.662745674 | 0.851712084 |
| 0.662755374 | 0.851712084 |
| 0.662803322 | 0.851712084 |
| 0.662813625 | 0.851712084 |
| 0.662832184 | 0.851712084 |
| 0.66286296 | 0.851712888 |
| 0.662882578 | 0.851712888 |
| 0.662886458 | 0.851712888 |
| 0.662937661 | 0.851755698 |
| 0.663002237 | 0.851768708 |
| 0.663508057 | 0.85219058 |
| 0.663516737 | 0.85219058 |
| 0.66380682 | 0.852246369 |
| 0.663868046 | 0.852246369 |
| 0.663953559 | 0.852303135 |
| 0.664010404 | 0.852303135 |
| 0.66402587 | 0.852303135 |
| 0.664096024 | 0.852370208 |
| 0.664153965 | 0.852378717 |
| 0.664184759 | 0.852392216 |
| 0.664264869 | 0.852472055 |
| 0.664308642 | 0.85250526 |
| 0.664585391 | 0.85269958 |
| 0.664637619 | 0.85274362 |
| 0.664686174 | 0.852755378 |
| 0.664700499 | 0.852755378 |
| 0.664824213 | 0.852825352 |
| 0.664827387 | 0.852825352 |
| 0.664960744 | 0.852951397 |
| 0.665150498 | 0.853125865 |
| 0.665209627 | 0.853152214 |
| 0.665242169 | 0.853152214 |
| 0.665253107 | 0.853152214 |
| 0.665289394 | 0.853152214 |
| 0.665354542 | 0.853152214 |
| 0.665379087 | 0.853152214 |
| 0.665403913 | 0.853152214 |
| 0.665445917 | 0.853155801 |
| 0.665453254 | 0.853155801 |
| 0.665623606 | 0.85322717 |
| 0.66572044 | 0.853305363 |

| | |
|---|---|
| 0.665741082 | 0.853308857 |
| 0.665890945 | 0.853435362 |
| 0.666065279 | 0.853521849 |
| 0.666130844 | 0.853566126 |
| 0.666207565 | 0.853566126 |
| 0.666210629 | 0.853566126 |
| 0.666258421 | 0.853586468 |
| 0.666417491 | 0.853719994 |
| 0.666426706 | 0.853719994 |
| 0.666444653 | 0.853719994 |
| 0.666456199 | 0.853719994 |
| 0.666605089 | 0.853864788 |
| 0.66665187 | 0.853878781 |
| 0.666781863 | 0.854010963 |
| 0.666803328 | 0.854010963 |
| 0.666906581 | 0.854109921 |
| 0.666956118 | 0.854126579 |
| 0.666972711 | 0.854126579 |
| 0.667266917 | 0.854367864 |
| 0.667285095 | 0.854368171 |
| 0.66734108 | 0.854408972 |
| 0.667352841 | 0.854408972 |
| 0.667438436 | 0.854469662 |
| 0.667454067 | 0.854469662 |
| 0.667605869 | 0.854549148 |
| 0.667747771 | 0.854638909 |
| 0.667904306 | 0.854747377 |
| 0.667979405 | 0.854820515 |
| 0.668232336 | 0.854999941 |
| 0.668292722 | 0.855014707 |
| 0.668359515 | 0.855021168 |
| 0.668481576 | 0.855021168 |
| 0.668588692 | 0.855021168 |
| 0.668709819 | 0.855021168 |
| 0.668734542 | 0.855021168 |
| 0.668751954 | 0.855021168 |
| 0.6687833 | 0.855021168 |
| 0.668814406 | 0.855021168 |
| 0.668834992 | 0.855021168 |
| 0.668938491 | 0.855067668 |
| 0.668944534 | 0.855067668 |
| 0.668971216 | 0.855078825 |
| 0.669099971 | 0.855198708 |
| 0.669100919 | 0.855198708 |
| 0.669198236 | 0.855208754 |

| | |
|---|---|
| 0.669212736 | 0.855208754 |
| 0.669216517 | 0.855208754 |
| 0.669458384 | 0.855372905 |
| 0.669470687 | 0.855372905 |
| 0.669544319 | 0.855409652 |
| 0.6695889 | 0.855409652 |
| 0.669631054 | 0.855409652 |
| 0.669638805 | 0.855409652 |
| 0.669881301 | 0.855622096 |
| 0.669947309 | 0.855683459 |
| 0.670150761 | 0.855851508 |
| 0.670242483 | 0.855890339 |
| 0.670253049 | 0.855890339 |
| 0.670404707 | 0.855988989 |
| 0.670430715 | 0.855988989 |
| 0.670438139 | 0.855988989 |
| 0.670707297 | 0.856143971 |
| 0.670765755 | 0.856154811 |
| 0.670806927 | 0.856161473 |
| 0.670921262 | 0.856284453 |
| 0.670986779 | 0.856322177 |
| 0.671047246 | 0.856353451 |
| 0.671099629 | 0.856374408 |
| 0.671276668 | 0.856490157 |
| 0.671312857 | 0.856508812 |
| 0.671430881 | 0.856544348 |
| 0.671458682 | 0.856544348 |
| 0.671597941 | 0.856627423 |
| 0.67160367 | 0.856627423 |
| 0.671658537 | 0.856668929 |
| 0.671797271 | 0.856713756 |
| 0.671895276 | 0.856761696 |
| 0.67191191 | 0.856761696 |
| 0.671913153 | 0.856761696 |
| 0.671916271 | 0.856761696 |
| 0.671970603 | 0.856796779 |
| 0.672068145 | 0.856796779 |
| 0.672078251 | 0.856796779 |
| 0.672148952 | 0.856841041 |
| 0.672276402 | 0.856886481 |
| 0.672285383 | 0.856886481 |
| 0.672384582 | 0.85695792 |
| 0.672455124 | 0.857001959 |
| 0.672540329 | 0.857072818 |
| 0.672641184 | 0.857140307 |

| | |
|---|---|
| 0.672653724 | 0.857140307 |
| 0.672671663 | 0.857140307 |
| 0.672695851 | 0.857148196 |
| 0.672793978 | 0.857180285 |
| 0.672811023 | 0.857180285 |
| 0.672905651 | 0.857212479 |
| 0.672915772 | 0.857212479 |
| 0.672986435 | 0.857252017 |
| 0.673108576 | 0.857331592 |
| 0.673112424 | 0.857331592 |
| 0.67335223 | 0.85745693 |
| 0.673452129 | 0.857537664 |
| 0.673455723 | 0.857537664 |
| 0.673513561 | 0.857570655 |
| 0.673577004 | 0.857598747 |
| 0.673601822 | 0.857598747 |
| 0.673747026 | 0.857730337 |
| 0.673886391 | 0.857844344 |
| 0.673940789 | 0.857844344 |
| 0.674113278 | 0.858004919 |
| 0.674146914 | 0.858004919 |
| 0.674160876 | 0.858004919 |
| 0.674218458 | 0.858032326 |
| 0.674231088 | 0.858032326 |
| 0.674250219 | 0.858032326 |
| 0.674254474 | 0.858032326 |
| 0.674351173 | 0.858119776 |
| 0.674405265 | 0.858132502 |
| 0.674583286 | 0.858292864 |
| 0.674658405 | 0.858339945 |
| 0.674898243 | 0.858550971 |
| 0.67490822 | 0.858550971 |
| 0.674914404 | 0.858550971 |
| 0.67493446 | 0.858553552 |
| 0.67496381 | 0.858567955 |
| 0.675054295 | 0.858607315 |
| 0.675076831 | 0.858607315 |
| 0.675181022 | 0.8586551 |
| 0.675196691 | 0.8586551 |
| 0.67520039 | 0.8586551 |
| 0.675212607 | 0.8586551 |
| 0.675273429 | 0.858709519 |
| 0.675429084 | 0.858859185 |
| 0.675471867 | 0.858859185 |
| 0.675479018 | 0.858859185 |

| | |
|---|---|
| 0.675556762 | 0.858917512 |
| 0.675831569 | 0.859166937 |
| 0.676125482 | 0.859357076 |
| 0.676208601 | 0.859385137 |
| 0.676219737 | 0.859385137 |
| 0.676301408 | 0.859443064 |
| 0.676463355 | 0.85949812 |
| 0.676471057 | 0.85949812 |
| 0.676550865 | 0.859554521 |
| 0.676566989 | 0.859554521 |
| 0.676603596 | 0.859574794 |
| 0.676667948 | 0.859628408 |
| 0.676762217 | 0.859684583 |
| 0.677032291 | 0.85991704 |
| 0.677035488 | 0.85991704 |
| 0.677089168 | 0.859934241 |
| 0.677116604 | 0.859934241 |
| 0.677157364 | 0.859934241 |
| 0.677302027 | 0.860072086 |
| 0.677397591 | 0.860147571 |
| 0.677462607 | 0.860173619 |
| 0.677612937 | 0.860235596 |
| 0.677685123 | 0.860235596 |
| 0.677714903 | 0.860235596 |
| 0.677929173 | 0.860380234 |
| 0.677937848 | 0.860380234 |
| 0.678003259 | 0.860380234 |
| 0.678003301 | 0.860380234 |
| 0.678134142 | 0.860486353 |
| 0.678180254 | 0.860521939 |
| 0.678212127 | 0.860525776 |
| 0.678219414 | 0.860525776 |
| 0.678422975 | 0.860682553 |
| 0.678498549 | 0.860719433 |
| 0.678537884 | 0.86072675 |
| 0.678558534 | 0.86072675 |
| 0.678672672 | 0.860779829 |
| 0.678728118 | 0.860827228 |
| 0.678840511 | 0.860874219 |
| 0.678859935 | 0.860874219 |
| 0.678889584 | 0.860874219 |
| 0.678913785 | 0.860874219 |
| 0.678930097 | 0.860874219 |
| 0.678985068 | 0.860874219 |
| 0.679066918 | 0.860874219 |

| | |
|---|---|
| 0.679095477 | 0.860874219 |
| 0.679115819 | 0.860874219 |
| 0.679133889 | 0.860874219 |
| 0.679174904 | 0.860874219 |
| 0.679224237 | 0.860874219 |
| 0.679230942 | 0.860874219 |
| 0.679314342 | 0.860897009 |
| 0.679363263 | 0.860897009 |
| 0.679429391 | 0.860897009 |
| 0.679434288 | 0.860897009 |
| 0.679436218 | 0.860897009 |
| 0.679476346 | 0.860897009 |
| 0.679524246 | 0.860897009 |
| 0.679568501 | 0.860930174 |
| 0.679636387 | 0.860947827 |
| 0.679636665 | 0.860947827 |
| 0.679822032 | 0.861136837 |
| 0.679840762 | 0.861137659 |
| 0.679923608 | 0.86119883 |
| 0.679983644 | 0.861227024 |
| 0.680119102 | 0.861284067 |
| 0.680154309 | 0.86130575 |
| 0.680397513 | 0.86148518 |
| 0.680431421 | 0.86148518 |
| 0.680703484 | 0.861689339 |
| 0.680710515 | 0.861689339 |
| 0.680831889 | 0.861751365 |
| 0.680975328 | 0.861853598 |
| 0.680987336 | 0.861853598 |
| 0.681083159 | 0.861853598 |
| 0.681193997 | 0.861929598 |
| 0.681306423 | 0.861992017 |
| 0.681363824 | 0.861992017 |
| 0.681367029 | 0.861992017 |
| 0.681375935 | 0.861992017 |
| 0.681422306 | 0.861992017 |
| 0.681434301 | 0.861992017 |
| 0.68143829 | 0.861992017 |
| 0.681554899 | 0.862041767 |
| 0.681568119 | 0.862041767 |
| 0.681608981 | 0.862057797 |
| 0.681672762 | 0.862105436 |
| 0.681872201 | 0.862266078 |
| 0.681917271 | 0.862300177 |
| 0.681946882 | 0.862314725 |

| | |
|---|---|
| 0.68205061 | 0.862422991 |
| 0.682148882 | 0.862497054 |
| 0.682156614 | 0.862497054 |
| 0.682207663 | 0.862515125 |
| 0.682244451 | 0.862515125 |
| 0.682260522 | 0.862515125 |
| 0.68233383 | 0.86253783 |
| 0.682340644 | 0.86253783 |
| 0.682409177 | 0.862601568 |
| 0.682546653 | 0.862693454 |
| 0.682702869 | 0.862743832 |
| 0.682741752 | 0.862770077 |
| 0.682781949 | 0.862775089 |
| 0.682867595 | 0.862796107 |
| 0.682934162 | 0.862796107 |
| 0.682943508 | 0.862796107 |
| 0.682973175 | 0.862810701 |
| 0.683046954 | 0.862816063 |
| 0.683081078 | 0.862816063 |
| 0.683174771 | 0.862816063 |
| 0.683176697 | 0.862816063 |
| 0.683200849 | 0.86281967 |
| 0.683215785 | 0.86281967 |
| 0.683331565 | 0.862897244 |
| 0.683560831 | 0.86309522 |
| 0.683636519 | 0.863111088 |
| 0.683677811 | 0.863111088 |
| 0.683682132 | 0.863111088 |
| 0.683841322 | 0.863205301 |
| 0.683847381 | 0.863205301 |
| 0.683941458 | 0.863232538 |
| 0.683994375 | 0.863276449 |
| 0.684076211 | 0.863313402 |
| 0.684091341 | 0.863313402 |
| 0.684113908 | 0.863313402 |
| 0.684266989 | 0.863313402 |
| 0.684269715 | 0.863313402 |
| 0.684373808 | 0.863313402 |
| 0.684374514 | 0.863313402 |
| 0.684513099 | 0.863450628 |
| 0.684576274 | 0.863461711 |
| 0.684626054 | 0.86350163 |
| 0.684697931 | 0.863534311 |
| 0.684706358 | 0.863534311 |
| 0.68475612 | 0.863574202 |

| | |
|---|---|
| 0.68494314 | 0.86368249 |
| 0.684950791 | 0.86368249 |
| 0.684990501 | 0.863709694 |
| 0.6850252 | 0.86373058 |
| 0.68507398 | 0.863746352 |
| 0.685174334 | 0.863804277 |
| 0.685263086 | 0.863864436 |
| 0.685280274 | 0.863864436 |
| 0.685294604 | 0.863864436 |
| 0.685493209 | 0.864069052 |
| 0.685580197 | 0.86415583 |
| 0.685658366 | 0.864224513 |
| 0.685681882 | 0.864224513 |
| 0.685743277 | 0.864269894 |
| 0.685864555 | 0.864331625 |
| 0.685892599 | 0.864331625 |
| 0.685906945 | 0.864331625 |
| 0.686056822 | 0.864394358 |
| 0.686105279 | 0.864395377 |
| 0.686156832 | 0.864402249 |
| 0.68627283 | 0.864473445 |
| 0.686286507 | 0.864473445 |
| 0.686322252 | 0.864473445 |
| 0.686381609 | 0.864478916 |
| 0.686421928 | 0.864478916 |
| 0.686439178 | 0.864478916 |
| 0.686444994 | 0.864478916 |
| 0.686567237 | 0.864530518 |
| 0.686608613 | 0.864559312 |
| 0.686695817 | 0.864578135 |
| 0.686828034 | 0.864629832 |
| 0.686884249 | 0.864629832 |
| 0.686936683 | 0.864629832 |
| 0.686936756 | 0.864629832 |
| 0.686978912 | 0.864629832 |
| 0.687000682 | 0.864629832 |
| 0.687023325 | 0.864629832 |
| 0.687037767 | 0.864629832 |
| 0.687045996 | 0.864629832 |
| 0.687073416 | 0.864629832 |
| 0.687081808 | 0.864629832 |
| 0.687264876 | 0.864700549 |
| 0.687288842 | 0.864700549 |
| 0.68735787 | 0.864701755 |
| 0.687372051 | 0.864701755 |

| | |
|---|---|
| 0.687403767 | 0.864715122 |
| 0.687563256 | 0.864870063 |
| 0.687660234 | 0.864969205 |
| 0.687712501 | 0.865007763 |
| 0.687825868 | 0.865063313 |
| 0.688004601 | 0.865147481 |
| 0.688031015 | 0.865147481 |
| 0.688210485 | 0.865210838 |
| 0.688223153 | 0.865210838 |
| 0.688233829 | 0.865210838 |
| 0.688295903 | 0.865266035 |
| 0.688372237 | 0.865297096 |
| 0.688459729 | 0.865334939 |
| 0.688498567 | 0.865360918 |
| 0.688668402 | 0.865505181 |
| 0.688675312 | 0.865505181 |
| 0.688704209 | 0.865505181 |
| 0.688950177 | 0.865722919 |
| 0.689003314 | 0.865731685 |
| 0.689113006 | 0.86579047 |
| 0.68913958 | 0.865801018 |
| 0.689163285 | 0.865807961 |
| 0.689216843 | 0.865829568 |
| 0.689273282 | 0.86587763 |
| 0.689326198 | 0.865921265 |
| 0.68935855 | 0.865939066 |
| 0.68950756 | 0.866034878 |
| 0.689544561 | 0.866035674 |
| 0.689646961 | 0.866045552 |
| 0.689647872 | 0.866045552 |
| 0.689667669 | 0.866045552 |
| 0.689852336 | 0.86617097 |
| 0.689881063 | 0.866184203 |
| 0.690060909 | 0.866250338 |
| 0.690367607 | 0.866475949 |
| 0.690368128 | 0.866475949 |
| 0.690399627 | 0.866492649 |
| 0.690417933 | 0.866492791 |
| 0.69044109 | 0.86649902 |
| 0.690675182 | 0.866747125 |
| 0.690710017 | 0.866767182 |
| 0.690727563 | 0.866767182 |
| 0.690769605 | 0.866797102 |
| 0.690821672 | 0.866836455 |
| 0.690871171 | 0.866836455 |

| | |
|---|---|
| 0.690932524 | 0.866864501 |
| 0.691050258 | 0.866917331 |
| 0.69120973 | 0.866917331 |
| 0.69121678 | 0.866917331 |
| 0.691226113 | 0.866917331 |
| 0.691229463 | 0.866917331 |
| 0.691450144 | 0.867103812 |
| 0.691450977 | 0.867103812 |
| 0.691523835 | 0.867172346 |
| 0.691554694 | 0.86718821 |
| 0.691617035 | 0.867233047 |
| 0.691626868 | 0.867233047 |
| 0.691646333 | 0.867234623 |
| 0.691740886 | 0.86726185 |
| 0.691827167 | 0.867324362 |
| 0.692093355 | 0.86755773 |
| 0.69225723 | 0.867703637 |
| 0.692353191 | 0.86780108 |
| 0.692599763 | 0.867950241 |
| 0.692669537 | 0.867966676 |
| 0.692681056 | 0.867966676 |
| 0.692733313 | 0.867966676 |
| 0.692738382 | 0.867966676 |
| 0.692787892 | 0.867966676 |
| 0.692938352 | 0.868100453 |
| 0.693033065 | 0.868163288 |
| 0.693133233 | 0.868207571 |
| 0.693235185 | 0.868312438 |
| 0.693299253 | 0.868355152 |
| 0.693353046 | 0.868391552 |
| 0.693427684 | 0.868439356 |
| 0.693647223 | 0.868645779 |
| 0.693919651 | 0.868911748 |
| 0.693935404 | 0.868911748 |
| 0.694021129 | 0.868931239 |
| 0.694072468 | 0.868940852 |
| 0.694083541 | 0.868940852 |
| 0.694106983 | 0.868947358 |
| 0.694309353 | 0.868995122 |
| 0.694344742 | 0.868995122 |
| 0.694346722 | 0.868995122 |
| 0.694349583 | 0.868995122 |
| 0.69435196 | 0.868995122 |
| 0.694354993 | 0.868995122 |
| 0.694364105 | 0.868995122 |

| | |
|---|---|
| 0.694373635 | 0.868995122 |
| 0.694382664 | 0.868995122 |
| 0.694400578 | 0.868995122 |
| 0.69444094 | 0.869022797 |
| 0.69448637 | 0.869036883 |
| 0.69448869 | 0.869036883 |
| 0.694609377 | 0.869135065 |
| 0.69461964 | 0.869135065 |
| 0.694689192 | 0.869176796 |
| 0.694705415 | 0.869176796 |
| 0.694718194 | 0.869176796 |
| 0.694740632 | 0.869176796 |
| 0.694851762 | 0.869176796 |
| 0.694889649 | 0.869176796 |
| 0.694890169 | 0.869176796 |
| 0.695130091 | 0.869294319 |
| 0.695132209 | 0.869294319 |
| 0.695173175 | 0.869294319 |
| 0.695291557 | 0.869335263 |
| 0.695348288 | 0.869335263 |
| 0.695356046 | 0.869335263 |
| 0.695365993 | 0.869335263 |
| 0.695481073 | 0.869434031 |
| 0.695499618 | 0.869434031 |
| 0.695511117 | 0.869434031 |
| 0.695520612 | 0.869434031 |
| 0.695536272 | 0.869434031 |
| 0.69562519 | 0.869485222 |
| 0.695697933 | 0.869485222 |
| 0.695717076 | 0.869485222 |
| 0.695770888 | 0.869499097 |
| 0.695855267 | 0.869558912 |
| 0.695992476 | 0.869596645 |
| 0.696003249 | 0.869596645 |
| 0.696013272 | 0.869596645 |
| 0.696109771 | 0.869694396 |
| 0.696186752 | 0.869721007 |
| 0.696197653 | 0.869721007 |
| 0.696371294 | 0.869884219 |
| 0.696715793 | 0.870138808 |
| 0.696751337 | 0.870138808 |
| 0.696757799 | 0.870138808 |
| 0.696805462 | 0.870152698 |
| 0.696858189 | 0.87018745 |
| 0.696893991 | 0.870187609 |

| | |
|---|---|
| 0.696926643 | 0.870187609 |
| 0.696941589 | 0.870187609 |
| 0.696978931 | 0.870187609 |
| 0.696997857 | 0.870187609 |
| 0.697052113 | 0.870209724 |
| 0.697087741 | 0.870217219 |
| 0.697158627 | 0.870251453 |
| 0.697279736 | 0.870349473 |
| 0.697308705 | 0.870349473 |
| 0.697516945 | 0.870493398 |
| 0.697543607 | 0.870503862 |
| 0.69760241 | 0.870554434 |
| 0.697770145 | 0.870617734 |
| 0.69784433 | 0.870618392 |
| 0.697877219 | 0.87062363 |
| 0.697932762 | 0.870670116 |
| 0.698152044 | 0.870807333 |
| 0.698241435 | 0.870807333 |
| 0.698251675 | 0.870807333 |
| 0.698253957 | 0.870807333 |
| 0.698353118 | 0.870852353 |
| 0.698420629 | 0.870913737 |
| 0.698536252 | 0.870997323 |
| 0.698544236 | 0.870997323 |
| 0.698720213 | 0.871150461 |
| 0.698845437 | 0.871196254 |
| 0.6988459 | 0.871196254 |
| 0.698891336 | 0.871204169 |
| 0.698931467 | 0.871231391 |
| 0.699063068 | 0.871281415 |
| 0.699063068 | 0.871281415 |
| 0.699238633 | 0.871477426 |
| 0.699482994 | 0.871667928 |
| 0.699788343 | 0.871979996 |
| 0.699895958 | 0.872022832 |
| 0.700039678 | 0.872133453 |
| 0.70021827 | 0.872287496 |
| 0.700326042 | 0.872358088 |
| 0.70033192 | 0.872358088 |
| 0.700362684 | 0.872358088 |
| 0.700403871 | 0.872358088 |
| 0.700412234 | 0.872358088 |
| 0.700496443 | 0.872360216 |
| 0.700563294 | 0.872420657 |
| 0.700653818 | 0.872441076 |

| | |
|---|---|
| 0.700666022 | 0.872441076 |
| 0.700899554 | 0.872656865 |
| 0.70099827 | 0.872716298 |
| 0.701202871 | 0.872769825 |
| 0.701227069 | 0.872769825 |
| 0.701274843 | 0.872781879 |
| 0.701311255 | 0.872783969 |
| 0.701349824 | 0.872783969 |
| 0.701388614 | 0.872789611 |
| 0.701391009 | 0.872789611 |
| 0.701521422 | 0.872865385 |
| 0.701613763 | 0.872907152 |
| 0.701726582 | 0.872983645 |
| 0.701730234 | 0.872983645 |
| 0.701754668 | 0.872991239 |
| 0.7018335 | 0.872998096 |
| 0.702206258 | 0.873267649 |
| 0.702209093 | 0.873267649 |
| 0.702233397 | 0.873267649 |
| 0.702233559 | 0.873267649 |
| 0.702320193 | 0.873306977 |
| 0.702380124 | 0.873318683 |
| 0.702384617 | 0.873318683 |
| 0.702464298 | 0.873344028 |
| 0.702488605 | 0.873344028 |
| 0.702570036 | 0.873344028 |
| 0.702678151 | 0.873387241 |
| 0.702766384 | 0.873430107 |
| 0.702934737 | 0.87359217 |
| 0.703007227 | 0.873659461 |
| 0.703085107 | 0.87371065 |
| 0.703119632 | 0.873712562 |
| 0.703218187 | 0.873739508 |
| 0.703218402 | 0.873739508 |
| 0.703326616 | 0.873797349 |
| 0.703382327 | 0.873829202 |
| 0.703446545 | 0.873857212 |
| 0.703522903 | 0.873889868 |
| 0.703586395 | 0.873907894 |
| 0.703592462 | 0.873907894 |
| 0.703661201 | 0.87397048 |
| 0.703810552 | 0.87401922 |
| 0.703872172 | 0.874061286 |
| 0.70388113 | 0.874061286 |
| 0.703932177 | 0.874075183 |

| | |
|---|---|
| 0.703947378 | 0.874075183 |
| 0.704089888 | 0.874195473 |
| 0.704123319 | 0.874195473 |
| 0.704138137 | 0.874195473 |
| 0.704175845 | 0.874195473 |
| 0.704337229 | 0.874238013 |
| 0.704353854 | 0.874238013 |
| 0.704383197 | 0.874238248 |
| 0.704390756 | 0.874238248 |
| 0.704431914 | 0.874251866 |
| 0.704571218 | 0.874276924 |
| 0.704572219 | 0.874276924 |
| 0.704585643 | 0.874276924 |
| 0.704619212 | 0.874276924 |
| 0.704623623 | 0.874276924 |
| 0.704633531 | 0.874276924 |
| 0.704675802 | 0.874276924 |
| 0.704704649 | 0.874276924 |
| 0.7047228 | 0.874276924 |
| 0.704744309 | 0.874276924 |
| 0.70475234 | 0.874276924 |
| 0.704776802 | 0.874284498 |
| 0.704841333 | 0.874302321 |
| 0.704846717 | 0.874302321 |
| 0.704858302 | 0.874302321 |
| 0.70486567 | 0.874302321 |
| 0.704882955 | 0.874302321 |
| 0.704970706 | 0.874342851 |
| 0.705039936 | 0.874360406 |
| 0.70515422 | 0.874360704 |
| 0.705168685 | 0.874360704 |
| 0.705220415 | 0.874402081 |
| 0.705311821 | 0.874492648 |
| 0.705410888 | 0.874579809 |
| 0.705542753 | 0.87469128 |
| 0.705563852 | 0.87469128 |
| 0.705694349 | 0.874764548 |
| 0.705696422 | 0.874764548 |
| 0.705784322 | 0.874850738 |
| 0.705914456 | 0.874965729 |
| 0.705940024 | 0.874965729 |
| 0.705950575 | 0.874965729 |
| 0.705971045 | 0.874968329 |
| 0.706085317 | 0.875041644 |
| 0.706290261 | 0.875176101 |

| | |
|---|---|
| 0.706319798 | 0.875176101 |
| 0.706359194 | 0.875176101 |
| 0.706429925 | 0.875211548 |
| 0.706462374 | 0.875211548 |
| 0.706476211 | 0.875211548 |
| 0.70655317 | 0.875279817 |
| 0.706612683 | 0.875330774 |
| 0.706782581 | 0.875495695 |
| 0.707107416 | 0.875668978 |
| 0.707149755 | 0.875668978 |
| 0.707150261 | 0.875668978 |
| 0.707179016 | 0.875668978 |
| 0.707179875 | 0.875668978 |
| 0.707222061 | 0.875698448 |
| 0.70734835 | 0.875752478 |
| 0.707356597 | 0.875752478 |
| 0.707413342 | 0.875764293 |
| 0.70744995 | 0.875764293 |
| 0.707499039 | 0.875764293 |
| 0.707500257 | 0.875764293 |
| 0.707545103 | 0.875764293 |
| 0.707593904 | 0.875764293 |
| 0.707632184 | 0.875764293 |
| 0.707661386 | 0.875764293 |
| 0.707707812 | 0.875793601 |
| 0.707815457 | 0.87580716 |
| 0.707843135 | 0.87580716 |
| 0.707921889 | 0.875843708 |
| 0.707965032 | 0.875843708 |
| 0.707968443 | 0.875843708 |
| 0.708081012 | 0.875942191 |
| 0.708216624 | 0.876057035 |
| 0.70822903 | 0.876057035 |
| 0.708332355 | 0.876119752 |
| 0.708394177 | 0.876170293 |
| 0.708483721 | 0.876230919 |
| 0.708509398 | 0.876230919 |
| 0.708753635 | 0.876354303 |
| 0.708821817 | 0.876354303 |
| 0.708835928 | 0.876354303 |
| 0.708837357 | 0.876354303 |
| 0.708881939 | 0.876386671 |
| 0.708910869 | 0.876396098 |
| 0.708926367 | 0.876396098 |
| 0.708994244 | 0.87645726 |

| | |
|---|---|
| 0.709022126 | 0.876468978 |
| 0.709057561 | 0.876490032 |
| 0.709177673 | 0.876546114 |
| 0.709179608 | 0.876546114 |
| 0.70928212 | 0.876546499 |
| 0.709347081 | 0.876546499 |
| 0.709360903 | 0.876546499 |
| 0.70938322 | 0.876551335 |
| 0.709432299 | 0.876580084 |
| 0.709576377 | 0.876638204 |
| 0.709612156 | 0.876638204 |
| 0.709625448 | 0.876638204 |
| 0.709664006 | 0.876638204 |
| 0.709673296 | 0.876638204 |
| 0.709701858 | 0.87664257 |
| 0.709750399 | 0.876663873 |
| 0.709792862 | 0.876675865 |
| 0.709811371 | 0.876675865 |
| 0.70981533 | 0.876675865 |
| 0.709906544 | 0.876723241 |
| 0.709983415 | 0.87676978 |
| 0.710031795 | 0.87680679 |
| 0.710077084 | 0.876839982 |
| 0.710119035 | 0.876869051 |
| 0.710290962 | 0.87705861 |
| 0.710356343 | 0.87709085 |
| 0.710401955 | 0.87709085 |
| 0.710457944 | 0.87709085 |
| 0.710504688 | 0.87709085 |
| 0.710544529 | 0.87709085 |
| 0.710722828 | 0.87709085 |
| 0.710784539 | 0.87709085 |
| 0.710800771 | 0.87709085 |
| 0.710857994 | 0.87709085 |
| 0.710898722 | 0.87709085 |
| 0.710920563 | 0.87709085 |
| 0.710953505 | 0.87709085 |
| 0.710981877 | 0.87709085 |
| 0.711050917 | 0.87709085 |
| 0.71105128 | 0.87709085 |
| 0.711059663 | 0.87709085 |
| 0.711062599 | 0.87709085 |
| 0.711241185 | 0.877179511 |
| 0.711248407 | 0.877179511 |
| 0.711254506 | 0.877179511 |

| | |
|---|---|
| 0.711288762 | 0.877199044 |
| 0.711364891 | 0.87723041 |
| 0.711403143 | 0.87723041 |
| 0.711427489 | 0.877233838 |
| 0.711705204 | 0.877451536 |
| 0.71171458 | 0.877451536 |
| 0.711778948 | 0.877475217 |
| 0.711800547 | 0.877475217 |
| 0.711816212 | 0.877475217 |
| 0.71185876 | 0.877493003 |
| 0.71193202 | 0.877515174 |
| 0.711971143 | 0.877520991 |
| 0.712144694 | 0.877586029 |
| 0.71215734 | 0.877586029 |
| 0.712165498 | 0.877586029 |
| 0.712205496 | 0.877586029 |
| 0.712315614 | 0.877615926 |
| 0.712345444 | 0.877615926 |
| 0.712400001 | 0.877637735 |
| 0.712476841 | 0.877709694 |
| 0.712621404 | 0.877815485 |
| 0.712695016 | 0.877815485 |
| 0.712702692 | 0.877815485 |
| 0.712758178 | 0.87782922 |
| 0.712833659 | 0.877876779 |
| 0.712863956 | 0.87789139 |
| 0.712976535 | 0.877947836 |
| 0.713038445 | 0.877975504 |
| 0.713119349 | 0.878001583 |
| 0.713270384 | 0.878142138 |
| 0.713343798 | 0.878180306 |
| 0.713410791 | 0.878180306 |
| 0.713421535 | 0.878180306 |
| 0.713424031 | 0.878180306 |
| 0.713430458 | 0.878180306 |
| 0.71350392 | 0.878183887 |
| 0.713516372 | 0.878183887 |
| 0.713543999 | 0.878183887 |
| 0.713686854 | 0.878337006 |
| 0.713832035 | 0.878368236 |
| 0.713844917 | 0.878368236 |
| 0.713868496 | 0.878368236 |
| 0.713917878 | 0.878368236 |
| 0.713970036 | 0.878368236 |
| 0.714002349 | 0.878368236 |

| | |
|---|---|
| 0.714031653 | 0.878368236 |
| 0.714063878 | 0.878368236 |
| 0.714136411 | 0.878368236 |
| 0.714193698 | 0.878378967 |
| 0.714254951 | 0.878404663 |
| 0.714267226 | 0.878404663 |
| 0.714310418 | 0.878404663 |
| 0.714332018 | 0.878404663 |
| 0.714376452 | 0.878413943 |
| 0.714544239 | 0.878506853 |
| 0.714658407 | 0.878571883 |
| 0.71467092 | 0.878571883 |
| 0.714742251 | 0.878614215 |
| 0.714788914 | 0.878621681 |
| 0.714804166 | 0.878621681 |
| 0.714840564 | 0.878621681 |
| 0.714916529 | 0.878679858 |
| 0.714924794 | 0.878679858 |
| 0.715021041 | 0.878775473 |
| 0.715049403 | 0.878787653 |
| 0.715109136 | 0.878838386 |
| 0.71514067 | 0.878840086 |
| 0.715147425 | 0.878840086 |
| 0.715218383 | 0.878876264 |
| 0.715232223 | 0.878876264 |
| 0.715319239 | 0.878915159 |
| 0.715481056 | 0.879057321 |
| 0.715521929 | 0.879057321 |
| 0.715539156 | 0.879057321 |
| 0.715563754 | 0.879057321 |
| 0.715613298 | 0.879057321 |
| 0.715637968 | 0.879057321 |
| 0.715816795 | 0.879164906 |
| 0.71581742 | 0.879164906 |
| 0.71584412 | 0.879164906 |
| 0.715882864 | 0.879164906 |
| 0.715910035 | 0.879164906 |
| 0.715951497 | 0.879164906 |
| 0.715956266 | 0.879164906 |
| 0.715965524 | 0.879164906 |
| 0.716038666 | 0.879189219 |
| 0.716104009 | 0.879221614 |
| 0.716263288 | 0.879352217 |
| 0.716344722 | 0.879403806 |
| 0.716566713 | 0.879500439 |

| | |
|---|---|
| 0.716574895 | 0.879500439 |
| 0.716593306 | 0.879500439 |
| 0.71661152 | 0.879500439 |
| 0.716662567 | 0.879500439 |
| 0.716749304 | 0.879500439 |
| 0.716940672 | 0.879618403 |
| 0.716944313 | 0.879618403 |
| 0.716982655 | 0.879633279 |
| 0.717013911 | 0.879635074 |
| 0.717045935 | 0.879635074 |
| 0.717124465 | 0.879635074 |
| 0.717152462 | 0.879635074 |
| 0.717196287 | 0.879635074 |
| 0.717197817 | 0.879635074 |
| 0.717199393 | 0.879635074 |
| 0.71723494 | 0.879635074 |
| 0.717276129 | 0.879662938 |
| 0.717494473 | 0.879885399 |
| 0.71768795 | 0.880036153 |
| 0.717709792 | 0.880036153 |
| 0.717872565 | 0.88011268 |
| 0.717883079 | 0.88011268 |
| 0.718021884 | 0.880237536 |
| 0.718090074 | 0.88025411 |
| 0.718146298 | 0.88025411 |
| 0.718241951 | 0.880348699 |
| 0.718321981 | 0.880385785 |
| 0.718334762 | 0.880385785 |
| 0.718353766 | 0.880385785 |
| 0.718364633 | 0.880385785 |
| 0.718510248 | 0.880541584 |
| 0.718602483 | 0.880563986 |
| 0.718656636 | 0.880566846 |
| 0.718694406 | 0.880566846 |
| 0.718735412 | 0.880566846 |
| 0.718798867 | 0.880566846 |
| 0.718808194 | 0.880566846 |
| 0.718879416 | 0.88060366 |
| 0.718893714 | 0.88060366 |
| 0.719031673 | 0.88073018 |
| 0.719166513 | 0.88077945 |
| 0.719169018 | 0.88077945 |
| 0.719221405 | 0.88077945 |
| 0.71925844 | 0.88077945 |
| 0.719272503 | 0.88077945 |

| | |
|---|---|
| 0.719295574 | 0.88077945 |
| 0.719296023 | 0.88077945 |
| 0.719312986 | 0.88077945 |
| 0.719317179 | 0.88077945 |
| 0.719353863 | 0.880782211 |
| 0.719455726 | 0.880795304 |
| 0.719457024 | 0.880795304 |
| 0.719621771 | 0.880864055 |
| 0.719628537 | 0.880864055 |
| 0.719678043 | 0.880864055 |
| 0.719680525 | 0.880864055 |
| 0.719719446 | 0.880867504 |
| 0.7197792 | 0.880895903 |
| 0.719779641 | 0.880895903 |
| 0.719814476 | 0.8809159 |
| 0.719857704 | 0.88092353 |
| 0.719980999 | 0.880953905 |
| 0.720165735 | 0.881057069 |
| 0.720168017 | 0.881057069 |
| 0.720189649 | 0.881057069 |
| 0.720245735 | 0.881060227 |
| 0.720246895 | 0.881060227 |
| 0.720296982 | 0.881069332 |
| 0.72030861 | 0.881069332 |
| 0.720456973 | 0.881160791 |
| 0.720458613 | 0.881160791 |
| 0.720511342 | 0.881202652 |
| 0.720697945 | 0.881317714 |
| 0.720754092 | 0.881344961 |
| 0.720757236 | 0.881344961 |
| 0.720891514 | 0.881430232 |
| 0.720936609 | 0.881446259 |
| 0.721192285 | 0.881632828 |
| 0.721196275 | 0.881632828 |
| 0.721316695 | 0.881757404 |
| 0.721368266 | 0.881761545 |
| 0.721548605 | 0.881900414 |
| 0.72156251 | 0.881900414 |
| 0.721564873 | 0.881900414 |
| 0.721581818 | 0.881900414 |
| 0.721647459 | 0.881958007 |
| 0.721690642 | 0.88198815 |
| 0.721767261 | 0.882020926 |
| 0.721784677 | 0.882020926 |
| 0.721791538 | 0.882020926 |

| | |
|---|---|
| 0.721962979 | 0.88211726 |
| 0.722115357 | 0.882245908 |
| 0.722145355 | 0.882245908 |
| 0.722159673 | 0.882245908 |
| 0.722194853 | 0.882245908 |
| 0.722197939 | 0.882245908 |
| 0.722254171 | 0.882269343 |
| 0.722404313 | 0.882407484 |
| 0.722452505 | 0.88242706 |
| 0.722610968 | 0.882507917 |
| 0.722641617 | 0.882516277 |
| 0.722696592 | 0.882532494 |
| 0.722755834 | 0.882542614 |
| 0.722857583 | 0.88256174 |
| 0.722864151 | 0.88256174 |
| 0.722933956 | 0.882609601 |
| 0.72303568 | 0.882663616 |
| 0.72304422 | 0.882663616 |
| 0.723062996 | 0.882663616 |
| 0.723067922 | 0.882663616 |
| 0.723095856 | 0.882663616 |
| 0.723372925 | 0.882888685 |
| 0.723456114 | 0.882909742 |
| 0.723538481 | 0.882909742 |
| 0.723660688 | 0.883033799 |
| 0.723686887 | 0.883033799 |
| 0.723698537 | 0.883033799 |
| 0.723806477 | 0.883078389 |
| 0.723898542 | 0.883082916 |
| 0.723906744 | 0.883082916 |
| 0.723911453 | 0.883082916 |
| 0.724049619 | 0.883082916 |
| 0.724081332 | 0.883082916 |
| 0.724099263 | 0.883082916 |
| 0.724101215 | 0.883082916 |
| 0.724266528 | 0.883218985 |
| 0.724279731 | 0.883218985 |
| 0.72430722 | 0.883218985 |
| 0.724329089 | 0.883218985 |
| 0.724483072 | 0.883292937 |
| 0.724513973 | 0.883308 |
| 0.724559526 | 0.883340924 |
| 0.724695043 | 0.883438399 |
| 0.724738816 | 0.883446433 |
| 0.724967823 | 0.883635125 |

| | |
|---|---|
| 0.724995601 | 0.883640118 |
| 0.725015909 | 0.883640118 |
| 0.72502758 | 0.883640118 |
| 0.725153016 | 0.883747508 |
| 0.725207859 | 0.883769374 |
| 0.725261268 | 0.883805582 |
| 0.725286582 | 0.883805582 |
| 0.725324491 | 0.883805582 |
| 0.725358588 | 0.883805582 |
| 0.725370972 | 0.883805582 |
| 0.725414914 | 0.883805582 |
| 0.725446411 | 0.883805582 |
| 0.725463391 | 0.883805582 |
| 0.7255921 | 0.883861716 |
| 0.725629508 | 0.883861716 |
| 0.725654793 | 0.883861716 |
| 0.725814625 | 0.883965968 |
| 0.725891661 | 0.884024472 |
| 0.725904364 | 0.884024472 |
| 0.725942314 | 0.884024472 |
| 0.726119129 | 0.88404817 |
| 0.726141562 | 0.88404817 |
| 0.726175832 | 0.88404817 |
| 0.726179111 | 0.88404817 |
| 0.726209165 | 0.884056156 |
| 0.726222795 | 0.884056156 |
| 0.726302761 | 0.884097326 |
| 0.726455407 | 0.884141331 |
| 0.726471276 | 0.884141331 |
| 0.72649414 | 0.884141331 |
| 0.726535951 | 0.884141331 |
| 0.726545874 | 0.884141331 |
| 0.726555974 | 0.884141331 |
| 0.726578163 | 0.884141331 |
| 0.726580222 | 0.884141331 |
| 0.726589785 | 0.884141331 |
| 0.726667778 | 0.88415207 |
| 0.726732632 | 0.8841596 |
| 0.726733322 | 0.8841596 |
| 0.726734748 | 0.8841596 |
| 0.726901646 | 0.884249213 |
| 0.726919803 | 0.884249213 |
| 0.727112255 | 0.884429012 |
| 0.727206165 | 0.884484601 |
| 0.727301399 | 0.884577842 |

| | |
|---|---|
| 0.727372446 | 0.884641663 |
| 0.727406252 | 0.884660187 |
| 0.727613862 | 0.884756539 |
| 0.727684071 | 0.884769703 |
| 0.727685191 | 0.884769703 |
| 0.72774349 | 0.884774504 |
| 0.727763793 | 0.884774504 |
| 0.727889716 | 0.884841459 |
| 0.727928473 | 0.884850383 |
| 0.727934215 | 0.884850383 |
| 0.728092141 | 0.884997177 |
| 0.728120734 | 0.885009346 |
| 0.728168722 | 0.885011072 |
| 0.728177901 | 0.885011072 |
| 0.728355705 | 0.885104339 |
| 0.728396732 | 0.885104339 |
| 0.728440481 | 0.885104339 |
| 0.728470704 | 0.885118481 |
| 0.728504032 | 0.885136394 |
| 0.72858188 | 0.885153358 |
| 0.728592334 | 0.885153358 |
| 0.728645176 | 0.885173372 |
| 0.728645979 | 0.885173372 |
| 0.728720679 | 0.885173807 |
| 0.72877925 | 0.885222376 |
| 0.728918437 | 0.885269021 |
| 0.728949218 | 0.885269021 |
| 0.728980839 | 0.885269021 |
| 0.72900259 | 0.885269021 |
| 0.729049695 | 0.885302513 |
| 0.729349622 | 0.88544722 |
| 0.729352082 | 0.88544722 |
| 0.729394108 | 0.88544722 |
| 0.72941055 | 0.88544722 |
| 0.729454124 | 0.885475289 |
| 0.72950516 | 0.88548112 |
| 0.729626763 | 0.885551031 |
| 0.72970711 | 0.885558938 |
| 0.730034599 | 0.885866082 |
| 0.730078471 | 0.885896747 |
| 0.730263242 | 0.885962948 |
| 0.730319087 | 0.885997518 |
| 0.730328943 | 0.885997518 |
| 0.730503618 | 0.886119038 |
| 0.730522139 | 0.886119038 |

| | |
|---|---|
| 0.730668256 | 0.886204816 |
| 0.730684358 | 0.886204816 |
| 0.730696636 | 0.886204816 |
| 0.730876787 | 0.886354693 |
| 0.730935943 | 0.886368837 |
| 0.730956505 | 0.886368837 |
| 0.730970243 | 0.886368837 |
| 0.731078602 | 0.886437479 |
| 0.731090415 | 0.886437479 |
| 0.731238026 | 0.886535828 |
| 0.731319819 | 0.886612423 |
| 0.731445782 | 0.886742561 |
| 0.731607206 | 0.886809793 |
| 0.731721049 | 0.886846905 |
| 0.731809847 | 0.886912161 |
| 0.731941663 | 0.886940528 |
| 0.731960742 | 0.886940528 |
| 0.731961764 | 0.886940528 |
| 0.732027089 | 0.88699573 |
| 0.732187255 | 0.887054395 |
| 0.732510175 | 0.88733276 |
| 0.7325675 | 0.887365446 |
| 0.732574422 | 0.887365446 |
| 0.732702866 | 0.887453319 |
| 0.732881383 | 0.887577068 |
| 0.732892792 | 0.887577068 |
| 0.732898216 | 0.887577068 |
| 0.733000613 | 0.887655933 |
| 0.73306929 | 0.887655933 |
| 0.733157559 | 0.887720964 |
| 0.733166148 | 0.887720964 |
| 0.733274057 | 0.887760129 |
| 0.733287495 | 0.887760129 |
| 0.733320267 | 0.887760129 |
| 0.733373023 | 0.887778497 |
| 0.733381427 | 0.887778497 |
| 0.733473359 | 0.887867217 |
| 0.733867354 | 0.88818902 |
| 0.733869744 | 0.88818902 |
| 0.733963027 | 0.888249143 |
| 0.734003224 | 0.888249143 |
| 0.734012672 | 0.888249143 |
| 0.734112615 | 0.888337733 |
| 0.734123183 | 0.888337733 |
| 0.734152388 | 0.888344848 |

| | |
|---|---|
| 0.734192626 | 0.888344848 |
| 0.734248432 | 0.888344848 |
| 0.734251525 | 0.888344848 |
| 0.734646391 | 0.888627722 |
| 0.734757623 | 0.88872159 |
| 0.734791329 | 0.888739788 |
| 0.734898672 | 0.888830889 |
| 0.735035369 | 0.888830889 |
| 0.735120464 | 0.888860738 |
| 0.73515261 | 0.888860738 |
| 0.735265341 | 0.888974472 |
| 0.735306022 | 0.88900109 |
| 0.735361053 | 0.88903066 |
| 0.73547015 | 0.889109243 |
| 0.735524627 | 0.889152531 |
| 0.735630073 | 0.88925743 |
| 0.735673655 | 0.889279777 |
| 0.735739738 | 0.889290459 |
| 0.735759001 | 0.889290459 |
| 0.735769427 | 0.889290459 |
| 0.735931349 | 0.889378091 |
| 0.735939086 | 0.889378091 |
| 0.735953974 | 0.889378091 |
| 0.736084709 | 0.889468372 |
| 0.736160645 | 0.889514995 |
| 0.736421707 | 0.889717027 |
| 0.73646265 | 0.889717027 |
| 0.736488314 | 0.889717027 |
| 0.736495974 | 0.889717027 |
| 0.736599446 | 0.889748851 |
| 0.736606185 | 0.889748851 |
| 0.736640894 | 0.889756685 |
| 0.736759791 | 0.889832595 |
| 0.736849571 | 0.889895895 |
| 0.736910617 | 0.889913773 |
| 0.736978169 | 0.889913773 |
| 0.737008374 | 0.889913773 |
| 0.737013855 | 0.889913773 |
| 0.737097102 | 0.88992728 |
| 0.737147279 | 0.88992728 |
| 0.737167125 | 0.88992728 |
| 0.737194558 | 0.88992728 |
| 0.737206211 | 0.88992728 |
| 0.737259892 | 0.88992728 |
| 0.737299415 | 0.88992728 |

| | |
|---|---|
| 0.737305205 | 0.88992728 |
| 0.737325805 | 0.88992728 |
| 0.737347162 | 0.88992728 |
| 0.737367923 | 0.88992728 |
| 0.73739327 | 0.88992728 |
| 0.737467796 | 0.889965507 |
| 0.737549401 | 0.890018885 |
| 0.737626225 | 0.890031375 |
| 0.737680626 | 0.890031375 |
| 0.737703006 | 0.890031375 |
| 0.737707613 | 0.890031375 |
| 0.737762401 | 0.890059206 |
| 0.737869938 | 0.890112498 |
| 0.73804308 | 0.890186081 |
| 0.738219778 | 0.890354108 |
| 0.738245242 | 0.890362273 |
| 0.738345289 | 0.890460386 |
| 0.738635803 | 0.890652879 |
| 0.738696411 | 0.89070341 |
| 0.738800393 | 0.890752158 |
| 0.738830354 | 0.890752158 |
| 0.738853452 | 0.890756739 |
| 0.738946416 | 0.890777768 |
| 0.73898129 | 0.890777768 |
| 0.739017261 | 0.890777768 |
| 0.739029695 | 0.890777768 |
| 0.739265501 | 0.890934181 |
| 0.739278527 | 0.890934181 |
| 0.739280636 | 0.890934181 |
| 0.739345022 | 0.89095988 |
| 0.739353582 | 0.89095988 |
| 0.739517832 | 0.891062206 |
| 0.739662991 | 0.891109914 |
| 0.739669394 | 0.891109914 |
| 0.739679725 | 0.891109914 |
| 0.739688398 | 0.891109914 |
| 0.739752423 | 0.891163777 |
| 0.739770531 | 0.891163777 |
| 0.739824653 | 0.891206433 |
| 0.739845002 | 0.891208405 |
| 0.739959827 | 0.891245889 |
| 0.739962178 | 0.891245889 |
| 0.74007956 | 0.891267258 |
| 0.740104882 | 0.891267258 |
| 0.740116613 | 0.891267258 |

| | |
|---|---|
| 0.740118422 | 0.891267258 |
| 0.740170856 | 0.89128934 |
| 0.740174187 | 0.89128934 |
| 0.740284069 | 0.891376583 |
| 0.740361206 | 0.891401855 |
| 0.740388466 | 0.891412141 |
| 0.740454379 | 0.891446428 |
| 0.740740793 | 0.8916327 |
| 0.740788161 | 0.891645423 |
| 0.740855822 | 0.891659259 |
| 0.741024663 | 0.891772327 |
| 0.741024663 | 0.891772327 |
| 0.741081922 | 0.891796167 |
| 0.741105183 | 0.891801626 |
| 0.741154955 | 0.891837614 |
| 0.741173745 | 0.891837614 |
| 0.741323264 | 0.891951372 |
| 0.741546157 | 0.892089478 |
| 0.741550432 | 0.892089478 |
| 0.741596484 | 0.892122344 |
| 0.741653048 | 0.892145322 |
| 0.741673665 | 0.892147589 |
| 0.741750038 | 0.892185247 |
| 0.741789109 | 0.892185247 |
| 0.741849917 | 0.892185247 |
| 0.741963951 | 0.892185247 |
| 0.741972512 | 0.892185247 |
| 0.742126604 | 0.892185247 |
| 0.742140365 | 0.892185247 |
| 0.742161769 | 0.892185247 |
| 0.742167369 | 0.892185247 |
| 0.742193355 | 0.892185247 |
| 0.742194405 | 0.892185247 |
| 0.742273092 | 0.892201877 |
| 0.74228079 | 0.892201877 |
| 0.742377885 | 0.892273546 |
| 0.742497098 | 0.892352828 |
| 0.742637697 | 0.89240235 |
| 0.742653687 | 0.89240235 |
| 0.742672786 | 0.892402785 |
| 0.742759866 | 0.89243498 |
| 0.742789255 | 0.89243498 |
| 0.742845224 | 0.892452374 |
| 0.742942148 | 0.892546304 |
| 0.742964591 | 0.892549829 |

| | |
|---|---|
| 0.743001303 | 0.892549829 |
| 0.743059745 | 0.892576666 |
| 0.743061126 | 0.892576666 |
| 0.743119879 | 0.892624729 |
| 0.743147735 | 0.892635675 |
| 0.743211574 | 0.892687811 |
| 0.743266113 | 0.892687811 |
| 0.743445673 | 0.892858437 |
| 0.74360707 | 0.892984252 |
| 0.743612213 | 0.892984252 |
| 0.743706121 | 0.89304134 |
| 0.743756147 | 0.893051154 |
| 0.744138932 | 0.893341542 |
| 0.744320961 | 0.893459042 |
| 0.744353784 | 0.893475923 |
| 0.744401975 | 0.893495364 |
| 0.744407501 | 0.893495364 |
| 0.744441526 | 0.893508313 |
| 0.744460116 | 0.893508313 |
| 0.744493332 | 0.893508313 |
| 0.744681005 | 0.893581756 |
| 0.744691803 | 0.893581756 |
| 0.744754595 | 0.893581756 |
| 0.744799129 | 0.893581756 |
| 0.744812927 | 0.893581756 |
| 0.744834088 | 0.893581756 |
| 0.744843888 | 0.893581756 |
| 0.744963206 | 0.893625866 |
| 0.745043269 | 0.893664267 |
| 0.745129899 | 0.893664267 |
| 0.745216902 | 0.893742972 |
| 0.745263203 | 0.893742972 |
| 0.745283665 | 0.893742972 |
| 0.745448707 | 0.893827046 |
| 0.745491664 | 0.893827046 |
| 0.74550193 | 0.893827046 |
| 0.745680113 | 0.893827046 |
| 0.745707813 | 0.893827046 |
| 0.745711176 | 0.893827046 |
| 0.745737363 | 0.893827046 |
| 0.745864724 | 0.893898475 |
| 0.745869473 | 0.893898475 |
| 0.745993114 | 0.893979167 |
| 0.746263311 | 0.894116286 |
| 0.746276494 | 0.894116286 |

| | |
|---|---|
| 0.746325984 | 0.894153087 |
| 0.7463523 | 0.894162122 |
| 0.746408108 | 0.894206489 |
| 0.746775553 | 0.894511689 |
| 0.746897235 | 0.894592796 |
| 0.747004989 | 0.894592796 |
| 0.747140712 | 0.894611592 |
| 0.747190042 | 0.894648167 |
| 0.747367998 | 0.894793757 |
| 0.747411673 | 0.894801059 |
| 0.747519493 | 0.894832487 |
| 0.747533851 | 0.894832487 |
| 0.747569427 | 0.894832487 |
| 0.747569443 | 0.894832487 |
| 0.74767617 | 0.894892767 |
| 0.747721706 | 0.894924778 |
| 0.74785637 | 0.89498328 |
| 0.747879366 | 0.89498328 |
| 0.747890381 | 0.89498328 |
| 0.747902126 | 0.89498328 |
| 0.747948627 | 0.895016439 |
| 0.747986623 | 0.895034442 |
| 0.748048624 | 0.895034442 |
| 0.748092501 | 0.895034442 |
| 0.748114013 | 0.895034442 |
| 0.748143066 | 0.895046717 |
| 0.748162134 | 0.895047046 |
| 0.748280418 | 0.8951211 |
| 0.748329934 | 0.895132876 |
| 0.748334928 | 0.895132876 |
| 0.74835034 | 0.895132876 |
| 0.748365441 | 0.895132876 |
| 0.748423087 | 0.895134386 |
| 0.748609104 | 0.895266937 |
| 0.748716258 | 0.89536611 |
| 0.748755182 | 0.89536611 |
| 0.748776854 | 0.89536611 |
| 0.748798638 | 0.89536611 |
| 0.748840037 | 0.89536611 |
| 0.748842426 | 0.89536611 |
| 0.74887996 | 0.895384422 |
| 0.749007743 | 0.895410569 |
| 0.749011214 | 0.895410569 |
| 0.749041658 | 0.895424488 |
| 0.749124464 | 0.895495505 |

| | |
|---|---|
| 0.749145166 | 0.895495505 |
| 0.749248612 | 0.895495505 |
| 0.749435055 | 0.895557591 |
| 0.749558011 | 0.895561167 |
| 0.749559017 | 0.895561167 |
| 0.749569672 | 0.895561167 |
| 0.749633411 | 0.895587586 |
| 0.74963834 | 0.895587586 |
| 0.749726635 | 0.895587586 |
| 0.749728899 | 0.895587586 |
| 0.749767965 | 0.895587586 |
| 0.749795713 | 0.895587586 |
| 0.749798631 | 0.895587586 |
| 0.749893778 | 0.895627917 |
| 0.749971065 | 0.895681234 |
| 0.750006113 | 0.895690172 |
| 0.750034969 | 0.895690172 |
| 0.750083805 | 0.895721789 |
| 0.750186293 | 0.895758582 |
| 0.750327596 | 0.895867134 |
| 0.750448581 | 0.895895855 |
| 0.750541314 | 0.89595787 |
| 0.750568408 | 0.895967757 |
| 0.750592243 | 0.895973753 |
| 0.75062149 | 0.895986208 |
| 0.750732626 | 0.896052881 |
| 0.750752133 | 0.896052881 |
| 0.750752602 | 0.896052881 |
| 0.751001796 | 0.896238005 |
| 0.751132187 | 0.896331713 |
| 0.751212059 | 0.896354171 |
| 0.751342219 | 0.896380617 |
| 0.75134944 | 0.896380617 |
| 0.751353856 | 0.896380617 |
| 0.751375826 | 0.896380617 |
| 0.751639604 | 0.896476877 |
| 0.751712358 | 0.896476877 |
| 0.751740301 | 0.896476877 |
| 0.751795797 | 0.896489194 |
| 0.751884093 | 0.896544265 |
| 0.751895156 | 0.896544265 |
| 0.751954638 | 0.896544265 |
| 0.752016613 | 0.896544265 |
| 0.7520419 | 0.896544265 |
| 0.752068016 | 0.896544265 |

| | |
|---|---|
| 0.752119695 | 0.896544265 |
| 0.752133101 | 0.896544265 |
| 0.752189553 | 0.896544265 |
| 0.752288565 | 0.896605378 |
| 0.752354116 | 0.896661065 |
| 0.752435628 | 0.896724847 |
| 0.752468364 | 0.896729908 |
| 0.752606307 | 0.896782101 |
| 0.752787509 | 0.896928069 |
| 0.752809509 | 0.896928069 |
| 0.75285466 | 0.896943388 |
| 0.75294287 | 0.897003604 |
| 0.753156164 | 0.897048936 |
| 0.753162805 | 0.897048936 |
| 0.753162805 | 0.897048936 |
| 0.753169271 | 0.897048936 |
| 0.753375214 | 0.897198297 |
| 0.753390577 | 0.897198297 |
| 0.753412189 | 0.897198297 |
| 0.753445379 | 0.897198297 |
| 0.753582934 | 0.897294792 |
| 0.753737707 | 0.897380552 |
| 0.753826502 | 0.897427757 |
| 0.754182813 | 0.897739742 |
| 0.754280222 | 0.897788378 |
| 0.75431425 | 0.897806443 |
| 0.754375661 | 0.897850464 |
| 0.754388939 | 0.897850464 |
| 0.754441223 | 0.897867816 |
| 0.754462275 | 0.897870434 |
| 0.754557822 | 0.897939269 |
| 0.75460569 | 0.897973795 |
| 0.754641827 | 0.89799436 |
| 0.75494503 | 0.898242947 |
| 0.754997131 | 0.898269867 |
| 0.755022039 | 0.898269867 |
| 0.755043097 | 0.898269867 |
| 0.755120572 | 0.898326079 |
| 0.755128069 | 0.898326079 |
| 0.755188255 | 0.898375238 |
| 0.755298992 | 0.898462089 |
| 0.755395675 | 0.89855282 |
| 0.755466178 | 0.898561819 |
| 0.755472545 | 0.898561819 |
| 0.755499244 | 0.898565618 |

| | |
|---|---|
| 0.755550632 | 0.898565618 |
| 0.755574692 | 0.898565618 |
| 0.755681124 | 0.898595613 |
| 0.755767363 | 0.898618699 |
| 0.755770269 | 0.898618699 |
| 0.755790247 | 0.898620018 |
| 0.756049545 | 0.898838563 |
| 0.75617012 | 0.898958529 |
| 0.756261714 | 0.898984437 |
| 0.756266623 | 0.898984437 |
| 0.756309233 | 0.898986075 |
| 0.756324628 | 0.898986075 |
| 0.756537744 | 0.899194509 |
| 0.756751121 | 0.899315435 |
| 0.756870943 | 0.899410973 |
| 0.757068226 | 0.899484261 |
| 0.757096639 | 0.899484261 |
| 0.757104991 | 0.899484261 |
| 0.757114116 | 0.899484261 |
| 0.757145773 | 0.899484261 |
| 0.757240935 | 0.899518273 |
| 0.757253816 | 0.899518273 |
| 0.757273459 | 0.899518273 |
| 0.757292377 | 0.899518273 |
| 0.757379404 | 0.899518273 |
| 0.757392121 | 0.899518273 |
| 0.757551599 | 0.899636205 |
| 0.757615193 | 0.899666862 |
| 0.757653713 | 0.899676272 |
| 0.757757225 | 0.899738295 |
| 0.757781458 | 0.899738295 |
| 0.757795136 | 0.899738295 |
| 0.757807588 | 0.899738295 |
| 0.757852495 | 0.899740896 |
| 0.757867695 | 0.899740896 |
| 0.757960491 | 0.89975129 |
| 0.758005956 | 0.89975129 |
| 0.758087338 | 0.899819261 |
| 0.758112587 | 0.899819261 |
| 0.758121391 | 0.899819261 |
| 0.758154809 | 0.899820394 |
| 0.758166301 | 0.899820394 |
| 0.758339086 | 0.899930805 |
| 0.758357851 | 0.899930805 |
| 0.758390937 | 0.899930805 |

| | |
|---|---|
| 0.758551571 | 0.899991301 |
| 0.758626822 | 0.899991301 |
| 0.75863419 | 0.899991301 |
| 0.758677835 | 0.899991301 |
| 0.758697331 | 0.899991301 |
| 0.75875058 | 0.899991301 |
| 0.758843703 | 0.899991301 |
| 0.758866556 | 0.899991301 |
| 0.75889815 | 0.899991301 |
| 0.758905841 | 0.899991301 |
| 0.758968022 | 0.899991301 |
| 0.758968051 | 0.899991301 |
| 0.759016562 | 0.899991301 |
| 0.759023697 | 0.899991301 |
| 0.759041102 | 0.899991301 |
| 0.759068627 | 0.900001531 |
| 0.759195398 | 0.900060509 |
| 0.75921286 | 0.900060509 |
| 0.759286757 | 0.90012571 |
| 0.75945027 | 0.900274735 |
| 0.759494939 | 0.900305278 |
| 0.75957834 | 0.900346045 |
| 0.759670238 | 0.900401026 |
| 0.759819291 | 0.900446238 |
| 0.759836468 | 0.900446238 |
| 0.759865689 | 0.900446238 |
| 0.759871341 | 0.900446238 |
| 0.759999602 | 0.900544884 |
| 0.760210577 | 0.900638751 |
| 0.760227635 | 0.900638751 |
| 0.76027712 | 0.900672049 |
| 0.760404473 | 0.900725758 |
| 0.760417017 | 0.900725758 |
| 0.7604612 | 0.900741272 |
| 0.760467939 | 0.900741272 |
| 0.760524517 | 0.900763483 |
| 0.760553476 | 0.900775381 |
| 0.760641226 | 0.900818722 |
| 0.760739979 | 0.900884255 |
| 0.760976305 | 0.901009755 |
| 0.761013259 | 0.901009755 |
| 0.761018445 | 0.901009755 |
| 0.761028232 | 0.901009755 |
| 0.761035136 | 0.901009755 |
| 0.761119432 | 0.901048102 |

| | |
|---|---|
| 0.761143201 | 0.901048102 |
| 0.761277456 | 0.901139838 |
| 0.761331382 | 0.901181273 |
| 0.761407867 | 0.901217199 |
| 0.761628527 | 0.901309588 |
| 0.761673708 | 0.901309588 |
| 0.761723649 | 0.901309588 |
| 0.762085479 | 0.90159635 |
| 0.762090221 | 0.90159635 |
| 0.762098512 | 0.90159635 |
| 0.76223096 | 0.901598867 |
| 0.762285717 | 0.901598867 |
| 0.762306738 | 0.901598867 |
| 0.762310647 | 0.901598867 |
| 0.76232972 | 0.901598867 |
| 0.762370986 | 0.901598867 |
| 0.762395032 | 0.901598867 |
| 0.762632354 | 0.901739271 |
| 0.762635854 | 0.901739271 |
| 0.762707947 | 0.901776318 |
| 0.762786789 | 0.901776318 |
| 0.762789684 | 0.901776318 |
| 0.762791982 | 0.901776318 |
| 0.76283471 | 0.901776318 |
| 0.762873227 | 0.901796058 |
| 0.762945373 | 0.901821144 |
| 0.763107817 | 0.901891101 |
| 0.763124057 | 0.901891101 |
| 0.763283922 | 0.90194582 |
| 0.763320463 | 0.90194582 |
| 0.763387029 | 0.902000448 |
| 0.763472402 | 0.902056564 |
| 0.763540714 | 0.902114896 |
| 0.763675035 | 0.902207424 |
| 0.763675858 | 0.902207424 |
| 0.763723354 | 0.902222865 |
| 0.763864502 | 0.902299486 |
| 0.763867455 | 0.902299486 |
| 0.763990375 | 0.902377046 |
| 0.763996999 | 0.902377046 |
| 0.764008903 | 0.902377046 |
| 0.764203179 | 0.90251698 |
| 0.764241142 | 0.902539435 |
| 0.76441952 | 0.902663482 |
| 0.764421992 | 0.902663482 |

| | |
|---|---|
| 0.764447041 | 0.902670681 |
| 0.764603424 | 0.902728612 |
| 0.764647735 | 0.902728612 |
| 0.764770891 | 0.902739744 |
| 0.764795432 | 0.902740002 |
| 0.764810236 | 0.902740002 |
| 0.76487524 | 0.902740002 |
| 0.764997751 | 0.902776941 |
| 0.765052455 | 0.902776941 |
| 0.765072083 | 0.902776941 |
| 0.765142355 | 0.902776941 |
| 0.765157102 | 0.902776941 |
| 0.765200461 | 0.902776941 |
| 0.765324385 | 0.902823439 |
| 0.765362162 | 0.902823439 |
| 0.765362162 | 0.902823439 |
| 0.765531559 | 0.902930202 |
| 0.765546623 | 0.902930202 |
| 0.76558286 | 0.90293732 |
| 0.765666678 | 0.902957044 |
| 0.765675419 | 0.902957044 |
| 0.765703346 | 0.90296762 |
| 0.765899852 | 0.903154627 |
| 0.766062441 | 0.903205903 |
| 0.766086182 | 0.903205903 |
| 0.766212549 | 0.903211736 |
| 0.766647589 | 0.903489862 |
| 0.766665147 | 0.903489862 |
| 0.766702296 | 0.903489862 |
| 0.766748896 | 0.903489862 |
| 0.766749384 | 0.903489862 |
| 0.766770771 | 0.903489862 |
| 0.766836233 | 0.90352059 |
| 0.766887363 | 0.90355848 |
| 0.767007115 | 0.90363251 |
| 0.767031729 | 0.903639156 |
| 0.767178929 | 0.903767127 |
| 0.7672177 | 0.903767127 |
| 0.767235234 | 0.903767127 |
| 0.767301227 | 0.903776715 |
| 0.767320161 | 0.903776715 |
| 0.767338254 | 0.903776715 |
| 0.767396945 | 0.903780362 |
| 0.767617612 | 0.903945675 |
| 0.767666817 | 0.903945675 |

| | |
|---|---|
| 0.767696661 | 0.903945675 |
| 0.767733016 | 0.903945675 |
| 0.767972315 | 0.904143408 |
| 0.768037063 | 0.904153432 |
| 0.768142954 | 0.904237454 |
| 0.768147126 | 0.904237454 |
| 0.768297753 | 0.904325361 |
| 0.768421235 | 0.904353742 |
| 0.768439395 | 0.904353742 |
| 0.768474656 | 0.904353742 |
| 0.768637828 | 0.904353742 |
| 0.768638293 | 0.904353742 |
| 0.768738817 | 0.904353742 |
| 0.768739606 | 0.904353742 |
| 0.768812022 | 0.904381718 |
| 0.768820354 | 0.904381718 |
| 0.768918875 | 0.9043896 |
| 0.769066253 | 0.904451511 |
| 0.769078899 | 0.904451511 |
| 0.769098989 | 0.904451511 |
| 0.769310189 | 0.904600444 |
| 0.769346953 | 0.904605102 |
| 0.769373512 | 0.904605102 |
| 0.769377592 | 0.904605102 |
| 0.769409118 | 0.904605102 |
| 0.769458375 | 0.904640682 |
| 0.769600278 | 0.904673523 |
| 0.769638228 | 0.904689602 |
| 0.769651947 | 0.904689602 |
| 0.769792815 | 0.904788194 |
| 0.76990764 | 0.904900824 |
| 0.770135633 | 0.905079451 |
| 0.770272917 | 0.905196116 |
| 0.770394431 | 0.905298166 |
| 0.770397771 | 0.905298166 |
| 0.770642626 | 0.905474192 |
| 0.770740318 | 0.90548667 |
| 0.770870401 | 0.905584795 |
| 0.7709166 | 0.905584795 |
| 0.770926901 | 0.905584795 |
| 0.771171403 | 0.905766996 |
| 0.771173095 | 0.905766996 |
| 0.771174696 | 0.905766996 |
| 0.771295293 | 0.905851188 |
| 0.771305839 | 0.905851188 |

| | |
|---|---|
| 0.771365191 | 0.90589095 |
| 0.771503871 | 0.905972061 |
| 0.771544417 | 0.905997336 |
| 0.771616602 | 0.906037422 |
| 0.771754246 | 0.906142807 |
| 0.771840371 | 0.906186886 |
| 0.772029587 | 0.906308285 |
| 0.772033958 | 0.906308285 |
| 0.772052133 | 0.906308285 |
| 0.772056601 | 0.906308285 |
| 0.772187728 | 0.906411603 |
| 0.772216533 | 0.906411603 |
| 0.772232118 | 0.906411603 |
| 0.772295123 | 0.906411603 |
| 0.772315898 | 0.906411603 |
| 0.772377109 | 0.906448342 |
| 0.772385266 | 0.906448342 |
| 0.772477565 | 0.906511987 |
| 0.772603573 | 0.906592844 |
| 0.772752452 | 0.906684961 |
| 0.772752485 | 0.906684961 |
| 0.772774898 | 0.906684961 |
| 0.77294978 | 0.906792378 |
| 0.7732249 | 0.906996116 |
| 0.773232899 | 0.906996116 |
| 0.773301091 | 0.907053765 |
| 0.773395642 | 0.907104928 |
| 0.773420894 | 0.907104928 |
| 0.77346086 | 0.907107125 |
| 0.773592554 | 0.907172221 |
| 0.773629347 | 0.907193029 |
| 0.773709204 | 0.907264335 |
| 0.773778814 | 0.907276391 |
| 0.773795683 | 0.907276391 |
| 0.773832203 | 0.907287231 |
| 0.773892516 | 0.907300582 |
| 0.77398726 | 0.907311049 |
| 0.774019311 | 0.907311049 |
| 0.774113751 | 0.907311049 |
| 0.774128807 | 0.907311049 |
| 0.774159272 | 0.907311049 |
| 0.774163576 | 0.907311049 |
| 0.774278208 | 0.907338176 |
| 0.77434755 | 0.907341501 |
| 0.774351476 | 0.907341501 |

| | |
|---|---|
| 0.77435367 | 0.907341501 |
| 0.774384642 | 0.907341501 |
| 0.774550149 | 0.907468449 |
| 0.774550149 | 0.907468449 |
| 0.774687753 | 0.907607341 |
| 0.774761377 | 0.907671269 |
| 0.774810307 | 0.907706265 |
| 0.7749755 | 0.90779254 |
| 0.775002589 | 0.90779254 |
| 0.775003057 | 0.90779254 |
| 0.775017374 | 0.90779254 |
| 0.775132999 | 0.907879066 |
| 0.775155566 | 0.907879066 |
| 0.775167494 | 0.907879066 |
| 0.775206865 | 0.907902852 |
| 0.775269049 | 0.907931202 |
| 0.775343689 | 0.907994737 |
| 0.775428286 | 0.908002128 |
| 0.775457878 | 0.908002128 |
| 0.775485242 | 0.908002128 |
| 0.775604268 | 0.908060481 |
| 0.775609605 | 0.908060481 |
| 0.775646514 | 0.908060481 |
| 0.77570683 | 0.908108772 |
| 0.775737374 | 0.908122208 |
| 0.775771267 | 0.908139563 |
| 0.775938602 | 0.908290801 |
| 0.776028613 | 0.908373839 |
| 0.776269071 | 0.908553165 |
| 0.776332062 | 0.908570172 |
| 0.77636627 | 0.908570172 |
| 0.776386645 | 0.908570172 |
| 0.776408178 | 0.908570172 |
| 0.776424328 | 0.908570172 |
| 0.776497339 | 0.908616907 |
| 0.776503356 | 0.908616907 |
| 0.776659295 | 0.908746485 |
| 0.776671335 | 0.908746485 |
| 0.77685067 | 0.908903926 |
| 0.776871122 | 0.908903926 |
| 0.776985821 | 0.908980464 |
| 0.77708336 | 0.909005261 |
| 0.777109345 | 0.909013331 |
| 0.777261285 | 0.909131926 |
| 0.777359713 | 0.909131926 |

| | |
|---|---|
| 0.77736344 | 0.909131926 |
| 0.777544914 | 0.909234347 |
| 0.77761973 | 0.909234347 |
| 0.777651085 | 0.909234347 |
| 0.777708474 | 0.909234347 |
| 0.777740419 | 0.909234347 |
| 0.777765921 | 0.909234347 |
| 0.777861418 | 0.909312398 |
| 0.777909028 | 0.90931342 |
| 0.777919569 | 0.90931342 |
| 0.77798315 | 0.909327252 |
| 0.77798868 | 0.909327252 |
| 0.7780997 | 0.909390062 |
| 0.778220184 | 0.909440316 |
| 0.778261884 | 0.909440316 |
| 0.778267756 | 0.909440316 |
| 0.778272536 | 0.909440316 |
| 0.778302272 | 0.909440316 |
| 0.778351693 | 0.909440316 |
| 0.778353769 | 0.909440316 |
| 0.778489072 | 0.909440316 |
| 0.77849812 | 0.909440316 |
| 0.77851831 | 0.909440316 |
| 0.778539753 | 0.909440316 |
| 0.778845104 | 0.909606798 |
| 0.778858102 | 0.909606798 |
| 0.778987514 | 0.909668453 |
| 0.779093914 | 0.909748347 |
| 0.779293798 | 0.909854756 |
| 0.779299664 | 0.909854756 |
| 0.779396448 | 0.909912818 |
| 0.77957361 | 0.909951531 |
| 0.779723357 | 0.910037109 |
| 0.779797867 | 0.910085441 |
| 0.77996505 | 0.91019811 |
| 0.78003863 | 0.91024893 |
| 0.780067433 | 0.910260239 |
| 0.780103548 | 0.910280078 |
| 0.780234079 | 0.91036922 |
| 0.780259697 | 0.91036922 |
| 0.780306444 | 0.91036922 |
| 0.780351906 | 0.91036922 |
| 0.78037405 | 0.91036922 |
| 0.780398721 | 0.91036922 |
| 0.780433299 | 0.91036922 |

| | |
|---|---|
| 0.780441413 | 0.91036922 |
| 0.780493643 | 0.91036922 |
| 0.780498589 | 0.91036922 |
| 0.780513056 | 0.91036922 |
| 0.780551784 | 0.91036922 |
| 0.780639551 | 0.91036922 |
| 0.78064213 | 0.91036922 |
| 0.780643637 | 0.91036922 |
| 0.780696034 | 0.91036922 |
| 0.780750256 | 0.910387867 |
| 0.780770467 | 0.910389146 |
| 0.78080209 | 0.91040373 |
| 0.780974196 | 0.910537528 |
| 0.781018957 | 0.910557211 |
| 0.781120282 | 0.910564037 |
| 0.781209634 | 0.910609175 |
| 0.781226843 | 0.910609175 |
| 0.781287198 | 0.910657238 |
| 0.781347944 | 0.910661182 |
| 0.781566767 | 0.91073133 |
| 0.781641562 | 0.91073133 |
| 0.781652411 | 0.91073133 |
| 0.781663196 | 0.91073133 |
| 0.781736672 | 0.91073133 |
| 0.781755289 | 0.91073133 |
| 0.781757691 | 0.91073133 |
| 0.781789485 | 0.91073133 |
| 0.781805421 | 0.91073133 |
| 0.782112822 | 0.910830041 |
| 0.782127458 | 0.910830041 |
| 0.782170719 | 0.910830041 |
| 0.782174895 | 0.910830041 |
| 0.782230893 | 0.910865523 |
| 0.782484953 | 0.911049989 |
| 0.782625796 | 0.911147151 |
| 0.782715985 | 0.911208824 |
| 0.782732516 | 0.911208824 |
| 0.782838544 | 0.911208824 |
| 0.782850931 | 0.911208824 |
| 0.782873924 | 0.911208824 |
| 0.782890499 | 0.911208824 |
| 0.782908337 | 0.911208824 |
| 0.782992862 | 0.911208824 |
| 0.783032299 | 0.911208824 |
| 0.783034856 | 0.911208824 |

| | |
|---|---|
| 0.78303526 | 0.911208824 |
| 0.783042429 | 0.911208824 |
| 0.78307558 | 0.911208824 |
| 0.783161881 | 0.911222801 |
| 0.783170385 | 0.911222801 |
| 0.783182385 | 0.911222801 |
| 0.78321858 | 0.911222801 |
| 0.783226485 | 0.911222801 |
| 0.783279841 | 0.911247154 |
| 0.783338159 | 0.911263684 |
| 0.783402602 | 0.91127811 |
| 0.783589066 | 0.911355225 |
| 0.783621811 | 0.911366784 |
| 0.783656412 | 0.911366784 |
| 0.783706054 | 0.911380008 |
| 0.783763808 | 0.911385009 |
| 0.783799752 | 0.911385009 |
| 0.783806033 | 0.911385009 |
| 0.783931093 | 0.911463666 |
| 0.783988589 | 0.911508264 |
| 0.78404988 | 0.91154563 |
| 0.784080016 | 0.91154563 |
| 0.7841043 | 0.91154563 |
| 0.784118432 | 0.91154563 |
| 0.784192981 | 0.91154563 |
| 0.78429195 | 0.911616959 |
| 0.784452908 | 0.911658447 |
| 0.784461261 | 0.911658447 |
| 0.78448036 | 0.911658447 |
| 0.784500226 | 0.911658447 |
| 0.784838021 | 0.911939312 |
| 0.784903731 | 0.911973988 |
| 0.784906161 | 0.911973988 |
| 0.785181575 | 0.912186773 |
| 0.785185062 | 0.912186773 |
| 0.7852265 | 0.912212663 |
| 0.785403165 | 0.912282959 |
| 0.785421744 | 0.912282959 |
| 0.78548053 | 0.912282959 |
| 0.785509204 | 0.912282959 |
| 0.785514552 | 0.912282959 |
| 0.785585594 | 0.912282959 |
| 0.785585694 | 0.912282959 |
| 0.785589535 | 0.912282959 |
| 0.785686039 | 0.912323702 |

| | |
|---|---|
| 0.785711033 | 0.912330482 |
| 0.785779775 | 0.912348637 |
| 0.785784137 | 0.912348637 |
| 0.785954646 | 0.912413151 |
| 0.786032561 | 0.912452792 |
| 0.786046268 | 0.912452792 |
| 0.786239133 | 0.912587702 |
| 0.786395822 | 0.912725084 |
| 0.786504019 | 0.912806173 |
| 0.786537754 | 0.912823081 |
| 0.786576144 | 0.912845391 |
| 0.786609146 | 0.912855221 |
| 0.786632129 | 0.912855221 |
| 0.786674566 | 0.912855221 |
| 0.786677443 | 0.912855221 |
| 0.786680448 | 0.912855221 |
| 0.787152657 | 0.913158437 |
| 0.787224905 | 0.913197763 |
| 0.787261058 | 0.913217459 |
| 0.787396843 | 0.913307903 |
| 0.787409222 | 0.913307903 |
| 0.787415733 | 0.913307903 |
| 0.787489085 | 0.913370739 |
| 0.787512329 | 0.913375455 |
| 0.787562754 | 0.913411695 |
| 0.787747603 | 0.913578485 |
| 0.787830544 | 0.913578485 |
| 0.787840836 | 0.913578485 |
| 0.787926973 | 0.913589394 |
| 0.787974897 | 0.913611676 |
| 0.788076717 | 0.913696288 |
| 0.788202851 | 0.913714404 |
| 0.788209372 | 0.913714404 |
| 0.788241895 | 0.913714404 |
| 0.788244842 | 0.913714404 |
| 0.78832256 | 0.913714404 |
| 0.788344452 | 0.913717542 |
| 0.788441186 | 0.913735589 |
| 0.788467906 | 0.913735589 |
| 0.788482322 | 0.913735589 |
| 0.788513007 | 0.913735589 |
| 0.788551092 | 0.913735589 |
| 0.788628848 | 0.913735589 |
| 0.788629215 | 0.913735589 |
| 0.788639115 | 0.913735589 |

| | |
|---|---|
| 0.78869406 | 0.913735589 |
| 0.788705142 | 0.913735589 |
| 0.788712048 | 0.913735589 |
| 0.788723663 | 0.913735589 |
| 0.788858386 | 0.913735589 |
| 0.788863385 | 0.913735589 |
| 0.788876415 | 0.913735589 |
| 0.7888868 | 0.913735589 |
| 0.78889721 | 0.913735589 |
| 0.788929469 | 0.913750731 |
| 0.788970162 | 0.91377564 |
| 0.789223936 | 0.913936209 |
| 0.789295445 | 0.913974572 |
| 0.789475813 | 0.914009323 |
| 0.789619625 | 0.914127707 |
| 0.789944424 | 0.914343953 |
| 0.789980471 | 0.914343953 |
| 0.790038555 | 0.914343953 |
| 0.790065778 | 0.914343953 |
| 0.790167063 | 0.914343953 |
| 0.790183504 | 0.914343953 |
| 0.790201207 | 0.914343953 |
| 0.790228773 | 0.914343953 |
| 0.790526083 | 0.914576864 |
| 0.790669791 | 0.914629768 |
| 0.790745737 | 0.914652456 |
| 0.790749639 | 0.914652456 |
| 0.790900207 | 0.914743053 |
| 0.791088071 | 0.914759202 |
| 0.791163496 | 0.914759202 |
| 0.791168738 | 0.914759202 |
| 0.791176522 | 0.914759202 |
| 0.79117969 | 0.914759202 |
| 0.791198064 | 0.914759202 |
| 0.791281654 | 0.914782002 |
| 0.791344191 | 0.914782002 |
| 0.791387385 | 0.914782002 |
| 0.791434655 | 0.914802913 |
| 0.791457377 | 0.914802913 |
| 0.791600866 | 0.914892424 |
| 0.791619127 | 0.914892424 |
| 0.791703966 | 0.914949369 |
| 0.791758669 | 0.914949369 |
| 0.791782498 | 0.914949369 |
| 0.792146467 | 0.915206554 |

| | |
|---|---|
| 0.792222805 | 0.915272548 |
| 0.792338188 | 0.915338436 |
| 0.792366328 | 0.915338436 |
| 0.792429076 | 0.915372145 |
| 0.792446622 | 0.915372145 |
| 0.792462768 | 0.915372145 |
| 0.792567827 | 0.915471295 |
| 0.792630382 | 0.915521347 |
| 0.792808806 | 0.91563861 |
| 0.792901842 | 0.915686849 |
| 0.793042649 | 0.915686849 |
| 0.793121145 | 0.915686849 |
| 0.793130028 | 0.915686849 |
| 0.793130226 | 0.915686849 |
| 0.793144712 | 0.915686849 |
| 0.793173811 | 0.915686849 |
| 0.793177419 | 0.915686849 |
| 0.793432611 | 0.915853611 |
| 0.79369938 | 0.916022982 |
| 0.793884163 | 0.916214041 |
| 0.793932172 | 0.916224637 |
| 0.793980651 | 0.91623358 |
| 0.794152513 | 0.916279506 |
| 0.794330023 | 0.916373317 |
| 0.794483122 | 0.916505539 |
| 0.794662084 | 0.916650499 |
| 0.794735351 | 0.916707691 |
| 0.79480344 | 0.916719624 |
| 0.794931661 | 0.916800906 |
| 0.794955564 | 0.916801494 |
| 0.795089669 | 0.916894326 |
| 0.795252013 | 0.917005429 |
| 0.795340497 | 0.917037531 |
| 0.795447248 | 0.917129033 |
| 0.795557967 | 0.917198934 |
| 0.795584907 | 0.917198934 |
| 0.795687795 | 0.917273226 |
| 0.795771769 | 0.917303339 |
| 0.795812701 | 0.91732832 |
| 0.795873492 | 0.917335358 |
| 0.795876589 | 0.917335358 |
| 0.795927965 | 0.917350173 |
| 0.796100997 | 0.917460793 |
| 0.79628619 | 0.917628168 |
| 0.796463943 | 0.91770142 |

| | |
|---|---|
| 0.796497969 | 0.917718424 |
| 0.796636158 | 0.917811034 |
| 0.796709295 | 0.917873091 |
| 0.796734754 | 0.917880219 |
| 0.796817293 | 0.917908698 |
| 0.796993189 | 0.917996297 |
| 0.797017722 | 0.917996297 |
| 0.797084735 | 0.917996297 |
| 0.797143907 | 0.917996297 |
| 0.797180082 | 0.918015759 |
| 0.797227459 | 0.91804812 |
| 0.797296886 | 0.918083673 |
| 0.797414583 | 0.918135632 |
| 0.79747178 | 0.918135632 |
| 0.797476953 | 0.918135632 |
| 0.797515772 | 0.91814186 |
| 0.797520917 | 0.91814186 |
| 0.797550728 | 0.918153985 |
| 0.797600196 | 0.918188739 |
| 0.797621036 | 0.918190537 |
| 0.797769956 | 0.918287296 |
| 0.797818537 | 0.918287296 |
| 0.797848149 | 0.918287296 |
| 0.797871554 | 0.918287296 |
| 0.797881705 | 0.918287296 |
| 0.797883748 | 0.918287296 |
| 0.797929712 | 0.918287296 |
| 0.798110259 | 0.918420733 |
| 0.798195284 | 0.918452001 |
| 0.79830438 | 0.918510314 |
| 0.798321071 | 0.918510314 |
| 0.798321313 | 0.918510314 |
| 0.798343057 | 0.918510314 |
| 0.79838096 | 0.918510314 |
| 0.798439607 | 0.918524395 |
| 0.798478377 | 0.918524395 |
| 0.798485407 | 0.918524395 |
| 0.798561807 | 0.918524395 |
| 0.798589316 | 0.918524395 |
| 0.798610631 | 0.918524395 |
| 0.798651389 | 0.918524395 |
| 0.79869197 | 0.918535302 |
| 0.798739859 | 0.918568196 |
| 0.79876605 | 0.918576136 |
| 0.798880047 | 0.918611242 |

| | |
|---|---|
| 0.798889284 | 0.918611242 |
| 0.798893015 | 0.918611242 |
| 0.798918335 | 0.918618178 |
| 0.799126721 | 0.918746882 |
| 0.799202738 | 0.918767743 |
| 0.799422523 | 0.918931507 |
| 0.799460957 | 0.918931507 |
| 0.799488319 | 0.91894078 |
| 0.799572844 | 0.919015755 |
| 0.799609595 | 0.91902171 |
| 0.799670221 | 0.91902171 |
| 0.799678141 | 0.91902171 |
| 0.799693802 | 0.91902171 |
| 0.799896295 | 0.919099176 |
| 0.799961672 | 0.919138222 |
| 0.799968874 | 0.919138222 |
| 0.800034282 | 0.919191198 |
| 0.800128808 | 0.919229055 |
| 0.800185357 | 0.919229055 |
| 0.800229328 | 0.919229055 |
| 0.800240959 | 0.919229055 |
| 0.800252405 | 0.919229055 |
| 0.80037133 | 0.919296125 |
| 0.800376533 | 0.919296125 |
| 0.800401855 | 0.91930304 |
| 0.800477373 | 0.919347666 |
| 0.80071629 | 0.919398926 |
| 0.800736293 | 0.919398926 |
| 0.800767057 | 0.919409314 |
| 0.800904689 | 0.919525772 |
| 0.800944623 | 0.919549454 |
| 0.801062527 | 0.919618314 |
| 0.801111384 | 0.919620754 |
| 0.801161197 | 0.919620754 |
| 0.801207633 | 0.919651891 |
| 0.801323304 | 0.919690742 |
| 0.801334314 | 0.919690742 |
| 0.801336813 | 0.919690742 |
| 0.801424625 | 0.919745798 |
| 0.801600247 | 0.919788948 |
| 0.801604473 | 0.919788948 |
| 0.801611659 | 0.919788948 |
| 0.801662536 | 0.919819355 |
| 0.801832899 | 0.919904359 |
| 0.801917045 | 0.919960829 |

| | |
|---|---|
| 0.801962951 | 0.919960829 |
| 0.80196745 | 0.919960829 |
| 0.801971396 | 0.919960829 |
| 0.801978996 | 0.919960829 |
| 0.802119754 | 0.920022124 |
| 0.802150244 | 0.920022124 |
| 0.802171758 | 0.920022124 |
| 0.802187346 | 0.920022124 |
| 0.80221753 | 0.920022124 |
| 0.802225603 | 0.920022124 |
| 0.802291787 | 0.920033602 |
| 0.802293563 | 0.920033602 |
| 0.802338273 | 0.920040568 |
| 0.802391136 | 0.920056881 |
| 0.80242913 | 0.920075665 |
| 0.802465472 | 0.920075665 |
| 0.802548725 | 0.920122286 |
| 0.802602308 | 0.920128109 |
| 0.802637362 | 0.920128109 |
| 0.802639249 | 0.920128109 |
| 0.802685087 | 0.920128109 |
| 0.802724388 | 0.920128867 |
| 0.802760689 | 0.920148332 |
| 0.802904846 | 0.920289003 |
| 0.802960473 | 0.920289003 |
| 0.803076642 | 0.920289003 |
| 0.803100287 | 0.920293955 |
| 0.803234025 | 0.920370794 |
| 0.803308128 | 0.920370794 |
| 0.803370442 | 0.920370794 |
| 0.803640453 | 0.920562714 |
| 0.803775337 | 0.920646641 |
| 0.803965201 | 0.92080731 |
| 0.803980287 | 0.92080731 |
| 0.804017247 | 0.920813 |
| 0.804126018 | 0.920915427 |
| 0.804166229 | 0.920922276 |
| 0.80417067 | 0.920922276 |
| 0.804219563 | 0.920956123 |
| 0.804239388 | 0.920956681 |
| 0.804482408 | 0.921105525 |
| 0.804530545 | 0.92112202 |
| 0.804588934 | 0.921157624 |
| 0.804625552 | 0.921176776 |
| 0.804676459 | 0.921176776 |

| | |
|---|---|
| 0.804757705 | 0.921217979 |
| 0.804815033 | 0.921240719 |
| 0.804835599 | 0.921240719 |
| 0.804932891 | 0.921261573 |
| 0.804973975 | 0.921261573 |
| 0.804984478 | 0.921261573 |
| 0.805009769 | 0.921261573 |
| 0.805018112 | 0.921261573 |
| 0.805052965 | 0.9212655 |
| 0.805070212 | 0.9212655 |
| 0.805143515 | 0.921268961 |
| 0.805213041 | 0.921268961 |
| 0.805260191 | 0.921277148 |
| 0.805287265 | 0.921277148 |
| 0.805292984 | 0.921277148 |
| 0.805447045 | 0.921391631 |
| 0.805511477 | 0.921413402 |
| 0.805532614 | 0.921413402 |
| 0.805566856 | 0.921413402 |
| 0.805601262 | 0.921430627 |
| 0.805622076 | 0.921432305 |
| 0.805704587 | 0.921482418 |
| 0.805775966 | 0.921525326 |
| 0.805784551 | 0.921525326 |
| 0.805858196 | 0.921525326 |
| 0.805932125 | 0.921549772 |
| 0.80597574 | 0.921552007 |
| 0.805996347 | 0.921552007 |
| 0.805997624 | 0.921552007 |
| 0.806124501 | 0.921652822 |
| 0.806170816 | 0.921683649 |
| 0.806289403 | 0.9217971 |
| 0.806367661 | 0.92186444 |
| 0.806430984 | 0.921892575 |
| 0.806551512 | 0.921954101 |
| 0.806562235 | 0.921954101 |
| 0.80661325 | 0.92196816 |
| 0.806686896 | 0.921982683 |
| 0.8067422 | 0.921982683 |
| 0.806745255 | 0.921982683 |
| 0.806745255 | 0.921982683 |
| 0.806792343 | 0.921982683 |
| 0.806824338 | 0.921982683 |
| 0.806829902 | 0.921982683 |
| 0.806854384 | 0.921982683 |

| | |
|---|---|
| 0.806858257 | 0.921982683 |
| 0.806880586 | 0.921986076 |
| 0.806981886 | 0.922043909 |
| 0.806989277 | 0.922043909 |
| 0.807133491 | 0.922164438 |
| 0.807232895 | 0.922220498 |
| 0.807260493 | 0.922220498 |
| 0.807481465 | 0.922362867 |
| 0.80756217 | 0.922366586 |
| 0.807889446 | 0.922629739 |
| 0.807946514 | 0.92265474 |
| 0.807950083 | 0.92265474 |
| 0.808040012 | 0.922703924 |
| 0.808105682 | 0.92274393 |
| 0.808151957 | 0.922755589 |
| 0.808174017 | 0.922755589 |
| 0.808215363 | 0.922780676 |
| 0.808371516 | 0.92289259 |
| 0.8084587 | 0.922947878 |
| 0.808551213 | 0.922986188 |
| 0.808574395 | 0.922986188 |
| 0.808576934 | 0.922986188 |
| 0.808656327 | 0.923024923 |
| 0.808711496 | 0.923024923 |
| 0.808893449 | 0.923167167 |
| 0.808902769 | 0.923167167 |
| 0.808950721 | 0.923189739 |
| 0.809016741 | 0.923189739 |
| 0.809085167 | 0.923220471 |
| 0.809375925 | 0.923447217 |
| 0.809499296 | 0.923447217 |
| 0.809510217 | 0.923447217 |
| 0.80952999 | 0.923447217 |
| 0.809538879 | 0.923447217 |
| 0.809560594 | 0.923447217 |
| 0.80957813 | 0.923447217 |
| 0.809658173 | 0.923469228 |
| 0.809697653 | 0.923469228 |
| 0.809698457 | 0.923469228 |
| 0.809729742 | 0.923469228 |
| 0.809775437 | 0.923499228 |
| 0.809872679 | 0.923499559 |
| 0.810028434 | 0.923593292 |
| 0.810146327 | 0.923634333 |
| 0.810177661 | 0.923634333 |

| | |
|---|---|
| 0.810211098 | 0.923634333 |
| 0.810319405 | 0.923699345 |
| 0.81046597 | 0.923844305 |
| 0.810500875 | 0.923857758 |
| 0.810516568 | 0.923857758 |
| 0.810561197 | 0.923886518 |
| 0.810628768 | 0.923919313 |
| 0.810659889 | 0.923932673 |
| 0.810724985 | 0.923975199 |
| 0.810736002 | 0.923975199 |
| 0.81079969 | 0.923981273 |
| 0.810817878 | 0.923981273 |
| 0.810916912 | 0.923990351 |
| 0.810924697 | 0.923990351 |
| 0.811097837 | 0.923990351 |
| 0.811150966 | 0.923990351 |
| 0.811156064 | 0.923990351 |
| 0.811270107 | 0.923990351 |
| 0.811315476 | 0.923990351 |
| 0.811327948 | 0.923990351 |
| 0.811373297 | 0.923990351 |
| 0.811425485 | 0.92400922 |
| 0.811462736 | 0.924029546 |
| 0.811492007 | 0.9240381 |
| 0.811584922 | 0.9240381 |
| 0.811589625 | 0.9240381 |
| 0.811679291 | 0.924069077 |
| 0.811716133 | 0.924075061 |
| 0.811883111 | 0.924176811 |
| 0.811883129 | 0.924176811 |
| 0.811944785 | 0.924224906 |
| 0.812202738 | 0.924244323 |
| 0.812237008 | 0.924244323 |
| 0.812319049 | 0.924244323 |
| 0.812366754 | 0.924244323 |
| 0.812368547 | 0.924244323 |
| 0.812369367 | 0.924244323 |
| 0.812409903 | 0.924268363 |
| 0.812450479 | 0.924292446 |
| 0.812529026 | 0.924359725 |
| 0.81260132 | 0.924409526 |
| 0.812670843 | 0.924410658 |
| 0.812823493 | 0.924465772 |
| 0.812847627 | 0.924465772 |
| 0.812856462 | 0.924465772 |

| | |
|---|---|
| 0.812891206 | 0.924465772 |
| 0.813153441 | 0.924540069 |
| 0.813189347 | 0.924558821 |
| 0.813469862 | 0.924767374 |
| 0.813523712 | 0.924784443 |
| 0.813558856 | 0.924800556 |
| 0.813576721 | 0.924800556 |
| 0.813597174 | 0.924801733 |
| 0.813689563 | 0.924829031 |
| 0.813698862 | 0.924829031 |
| 0.813785868 | 0.924839103 |
| 0.813804816 | 0.924839103 |
| 0.814023556 | 0.924994799 |
| 0.814051938 | 0.924994799 |
| 0.814134865 | 0.925026782 |
| 0.814144764 | 0.925026782 |
| 0.814170233 | 0.925033653 |
| 0.814263414 | 0.925073317 |
| 0.814398105 | 0.925164545 |
| 0.81440199 | 0.925164545 |
| 0.814610787 | 0.925288881 |
| 0.814628007 | 0.925288881 |
| 0.81467521 | 0.925320428 |
| 0.814742843 | 0.925322074 |
| 0.814836088 | 0.925370747 |
| 0.814859076 | 0.925374788 |
| 0.814895184 | 0.925393728 |
| 0.814933684 | 0.925409585 |
| 0.815039242 | 0.92546047 |
| 0.815093088 | 0.925486066 |
| 0.815193547 | 0.925578064 |
| 0.81523474 | 0.925580926 |
| 0.815313993 | 0.925597422 |
| 0.815457663 | 0.925701397 |
| 0.81551639 | 0.925745998 |
| 0.815594862 | 0.92581301 |
| 0.815754999 | 0.925899214 |
| 0.815768569 | 0.925899214 |
| 0.815786593 | 0.925899214 |
| 0.815787446 | 0.925899214 |
| 0.815842629 | 0.925923183 |
| 0.815847448 | 0.925923183 |
| 0.815933511 | 0.925954663 |
| 0.816037442 | 0.925991551 |
| 0.816165005 | 0.925991551 |

| | |
|---|---|
| 0.81617079 | 0.925991551 |
| 0.816372339 | 0.926121654 |
| 0.816430012 | 0.926157115 |
| 0.816464122 | 0.926159594 |
| 0.816573413 | 0.926165307 |
| 0.816623423 | 0.926165307 |
| 0.816644174 | 0.926165307 |
| 0.816801663 | 0.926299803 |
| 0.816905821 | 0.926312872 |
| 0.816913033 | 0.926312872 |
| 0.816921664 | 0.926312872 |
| 0.816926402 | 0.926312872 |
| 0.816978344 | 0.926345767 |
| 0.817018309 | 0.926369027 |
| 0.817092401 | 0.926430981 |
| 0.817137325 | 0.926459861 |
| 0.817342557 | 0.926645205 |
| 0.817417878 | 0.926653073 |
| 0.817443933 | 0.926653073 |
| 0.817598063 | 0.926805735 |
| 0.817643162 | 0.926834798 |
| 0.8177639 | 0.926887264 |
| 0.817788191 | 0.926887264 |
| 0.817812401 | 0.926887264 |
| 0.81781532 | 0.926887264 |
| 0.817956887 | 0.926954105 |
| 0.817977569 | 0.926954105 |
| 0.817988461 | 0.926954105 |
| 0.817996118 | 0.926954105 |
| 0.818001429 | 0.926954105 |
| 0.818062765 | 0.927001554 |
| 0.81815558 | 0.927084671 |
| 0.818196914 | 0.927087394 |
| 0.818220497 | 0.927092059 |
| 0.818379332 | 0.927184295 |
| 0.818435701 | 0.927225603 |
| 0.818468584 | 0.927240801 |
| 0.81868916 | 0.927387303 |
| 0.818705019 | 0.927387303 |
| 0.81871972 | 0.927387303 |
| 0.818753676 | 0.927387303 |
| 0.818806733 | 0.927403288 |
| 0.818990609 | 0.92752976 |
| 0.819107366 | 0.927567324 |
| 0.819311449 | 0.927732255 |

| | |
|---|---|
| 0.819434561 | 0.927815664 |
| 0.819464949 | 0.927817833 |
| 0.819563198 | 0.927864167 |
| 0.819580709 | 0.927864167 |
| 0.8195838 | 0.927864167 |
| 0.81968547 | 0.927898221 |
| 0.819857781 | 0.928041986 |
| 0.819958052 | 0.928111371 |
| 0.820121852 | 0.928174638 |
| 0.820155497 | 0.928174638 |
| 0.820177746 | 0.928174638 |
| 0.820186565 | 0.928174638 |
| 0.82026038 | 0.928174638 |
| 0.820337241 | 0.928187626 |
| 0.820487993 | 0.92830244 |
| 0.820497187 | 0.92830244 |
| 0.820612826 | 0.928318144 |
| 0.820615395 | 0.928318144 |
| 0.82065574 | 0.928318144 |
| 0.820695101 | 0.928318144 |
| 0.820717518 | 0.928318144 |
| 0.820720073 | 0.928318144 |
| 0.820782672 | 0.928318144 |
| 0.820892029 | 0.928318144 |
| 0.820900084 | 0.928318144 |
| 0.820939809 | 0.928318144 |
| 0.820944539 | 0.928318144 |
| 0.820956006 | 0.928318144 |
| 0.820957705 | 0.928318144 |
| 0.821024812 | 0.928348062 |
| 0.821139415 | 0.928433562 |
| 0.821248706 | 0.92853509 |
| 0.821284722 | 0.928553767 |
| 0.821337394 | 0.928569233 |
| 0.821439375 | 0.928598806 |
| 0.821461039 | 0.928598806 |
| 0.821537039 | 0.928662677 |
| 0.821757036 | 0.928704007 |
| 0.821875089 | 0.928758242 |
| 0.822015212 | 0.928894547 |
| 0.822160473 | 0.9289944 |
| 0.822162092 | 0.9289944 |
| 0.822253077 | 0.929039806 |
| 0.822260797 | 0.929039806 |
| 0.822336189 | 0.929075107 |

| | |
|---|---|
| 0.8223532 | 0.929075107 |
| 0.822383334 | 0.929075107 |
| 0.822395692 | 0.929075107 |
| 0.822433934 | 0.929075107 |
| 0.822525616 | 0.929102593 |
| 0.822530953 | 0.929102593 |
| 0.822579254 | 0.92911308 |
| 0.822612464 | 0.929128556 |
| 0.822662502 | 0.929163037 |
| 0.822771954 | 0.929207926 |
| 0.822799796 | 0.929207926 |
| 0.822937958 | 0.929274579 |
| 0.823059472 | 0.929279072 |
| 0.823082175 | 0.929279072 |
| 0.823087593 | 0.929279072 |
| 0.823096933 | 0.929279072 |
| 0.823313061 | 0.929409198 |
| 0.823382943 | 0.929409198 |
| 0.823492222 | 0.92943895 |
| 0.823561694 | 0.92943895 |
| 0.823613056 | 0.929443157 |
| 0.823675195 | 0.929469234 |
| 0.823792049 | 0.929579073 |
| 0.823896135 | 0.929608425 |
| 0.823948423 | 0.929614838 |
| 0.82394989 | 0.929614838 |
| 0.824043411 | 0.929630035 |
| 0.824051921 | 0.929630035 |
| 0.824230682 | 0.929769896 |
| 0.824476805 | 0.929911118 |
| 0.824497624 | 0.929912578 |
| 0.82457611 | 0.92992323 |
| 0.824637623 | 0.92992323 |
| 0.824647522 | 0.92992323 |
| 0.824704509 | 0.929926472 |
| 0.824705195 | 0.929926472 |
| 0.824821798 | 0.929981549 |
| 0.824950157 | 0.930029348 |
| 0.82497168 | 0.930029348 |
| 0.824972177 | 0.930029348 |
| 0.825213451 | 0.930147273 |
| 0.82521349 | 0.930147273 |
| 0.825241465 | 0.930150741 |
| 0.82529123 | 0.930150741 |
| 0.825300073 | 0.930150741 |

| | |
|---|---|
| 0.825376559 | 0.930198991 |
| 0.825415911 | 0.930221329 |
| 0.825570711 | 0.930351756 |
| 0.825609838 | 0.930371222 |
| 0.825712427 | 0.930371222 |
| 0.825724293 | 0.930371222 |
| 0.825742778 | 0.930371222 |
| 0.825754311 | 0.930371222 |
| 0.825763796 | 0.930371222 |
| 0.82590388 | 0.930462009 |
| 0.825931357 | 0.930462009 |
| 0.825942057 | 0.930462009 |
| 0.826051167 | 0.930484733 |
| 0.826051312 | 0.930484733 |
| 0.826055967 | 0.930484733 |
| 0.82605963 | 0.930484733 |
| 0.826142765 | 0.930519767 |
| 0.826148509 | 0.930519767 |
| 0.826384951 | 0.930674845 |
| 0.826415063 | 0.930674845 |
| 0.826432443 | 0.930674845 |
| 0.826460944 | 0.930674845 |
| 0.82647796 | 0.930674845 |
| 0.826525402 | 0.930674845 |
| 0.826541344 | 0.930674845 |
| 0.826567133 | 0.93068188 |
| 0.826691192 | 0.930799561 |
| 0.82676292 | 0.930818211 |
| 0.826766388 | 0.930818211 |
| 0.826809407 | 0.930844639 |
| 0.826835421 | 0.930851923 |
| 0.826859353 | 0.930856862 |
| 0.827006393 | 0.930962544 |
| 0.827011869 | 0.930962544 |
| 0.827094657 | 0.931025783 |
| 0.827126692 | 0.931025783 |
| 0.82718789 | 0.931050661 |
| 0.827218093 | 0.931062652 |
| 0.827440755 | 0.931200903 |
| 0.827455223 | 0.931200903 |
| 0.827458236 | 0.931200903 |
| 0.82750519 | 0.93122757 |
| 0.827536164 | 0.93122757 |
| 0.827542052 | 0.93122757 |
| 0.827634032 | 0.93122757 |

| | |
|---|---|
| 0.827689711 | 0.93122757 |
| 0.827694757 | 0.93122757 |
| 0.827735645 | 0.93122757 |
| 0.827757386 | 0.93122757 |
| 0.82792374 | 0.93132956 |
| 0.82792757 | 0.93132956 |
| 0.827944098 | 0.93132956 |
| 0.828032238 | 0.931376621 |
| 0.828092646 | 0.931408663 |
| 0.828239711 | 0.931530079 |
| 0.828285473 | 0.931559549 |
| 0.828441785 | 0.931608056 |
| 0.828462528 | 0.931608056 |
| 0.828463535 | 0.931608056 |
| 0.828719635 | 0.93180151 |
| 0.828808315 | 0.931851384 |
| 0.828854334 | 0.931858947 |
| 0.828885532 | 0.931858947 |
| 0.828940268 | 0.931858947 |
| 0.829076593 | 0.931955076 |
| 0.829107505 | 0.931955893 |
| 0.829209738 | 0.932004819 |
| 0.829329923 | 0.932014001 |
| 0.829348914 | 0.932014001 |
| 0.829354891 | 0.932014001 |
| 0.829390484 | 0.932032008 |
| 0.829534787 | 0.932084207 |
| 0.829620082 | 0.932123641 |
| 0.829687305 | 0.932123641 |
| 0.829746665 | 0.932123641 |
| 0.829763553 | 0.932123641 |
| 0.829793043 | 0.932123641 |
| 0.829813746 | 0.932123641 |
| 0.82985221 | 0.932123641 |
| 0.829862712 | 0.932123641 |
| 0.829881169 | 0.932123641 |
| 0.829918863 | 0.932123641 |
| 0.82993893 | 0.932123641 |
| 0.829950853 | 0.932123641 |
| 0.829969844 | 0.932123641 |
| 0.830042033 | 0.932128478 |
| 0.830070426 | 0.932128478 |
| 0.830096055 | 0.932128478 |
| 0.830102616 | 0.932128478 |
| 0.830236485 | 0.932220502 |

| | |
|---|---|
| 0.830300452 | 0.932247274 |
| 0.830334523 | 0.932247274 |
| 0.830384755 | 0.932247478 |
| 0.830457479 | 0.932275182 |
| 0.830468156 | 0.932275182 |
| 0.830594726 | 0.932342733 |
| 0.830608351 | 0.932342733 |
| 0.830676868 | 0.932356044 |
| 0.830778866 | 0.932436753 |
| 0.830788284 | 0.932436753 |
| 0.830818664 | 0.932448876 |
| 0.830843874 | 0.932455197 |
| 0.830875044 | 0.932466064 |
| 0.830892714 | 0.932466064 |
| 0.830942555 | 0.932500025 |
| 0.831048596 | 0.932597052 |
| 0.831231077 | 0.932703487 |
| 0.831280526 | 0.932703487 |
| 0.831329016 | 0.932707457 |
| 0.831362399 | 0.932707457 |
| 0.831465363 | 0.932801 |
| 0.831549249 | 0.932873135 |
| 0.831694134 | 0.932973946 |
| 0.831746485 | 0.932973946 |
| 0.831784776 | 0.932983527 |
| 0.831906684 | 0.93301822 |
| 0.831941605 | 0.933027593 |
| 0.832090444 | 0.933103702 |
| 0.83210265 | 0.933103702 |
| 0.832123329 | 0.933103702 |
| 0.832216947 | 0.933125342 |
| 0.83222344 | 0.933125342 |
| 0.832224687 | 0.933125342 |
| 0.83231824 | 0.933163153 |
| 0.832365109 | 0.933163153 |
| 0.832375968 | 0.933163153 |
| 0.832466945 | 0.933196865 |
| 0.8324688 | 0.933196865 |
| 0.832576201 | 0.933223146 |
| 0.832586236 | 0.933223146 |
| 0.832678157 | 0.933260286 |
| 0.83271923 | 0.933269294 |
| 0.832725385 | 0.933269294 |
| 0.832750664 | 0.933275664 |
| 0.832851304 | 0.933322566 |

| | |
|---|---|
| 0.83292822 | 0.933369157 |
| 0.832962067 | 0.933369157 |
| 0.832990867 | 0.933369157 |
| 0.833082409 | 0.933422448 |
| 0.833144927 | 0.933422448 |
| 0.833165952 | 0.933422448 |
| 0.833172843 | 0.933422448 |
| 0.833195216 | 0.933422448 |
| 0.833283383 | 0.933433395 |
| 0.833673996 | 0.933698929 |
| 0.833684141 | 0.933698929 |
| 0.833716471 | 0.933698929 |
| 0.833803047 | 0.933745254 |
| 0.8338478 | 0.933745254 |
| 0.83389145 | 0.933745254 |
| 0.833903066 | 0.933745254 |
| 0.833914678 | 0.933745254 |
| 0.833943183 | 0.933748204 |
| 0.833956605 | 0.933748204 |
| 0.833976129 | 0.933748204 |
| 0.834319224 | 0.933982606 |
| 0.834370619 | 0.933990581 |
| 0.834427934 | 0.933990581 |
| 0.834586403 | 0.934106526 |
| 0.83469841 | 0.934106526 |
| 0.834725638 | 0.934106526 |
| 0.834745712 | 0.934106526 |
| 0.834746413 | 0.934106526 |
| 0.834771991 | 0.934106526 |
| 0.834784622 | 0.934106526 |
| 0.834844198 | 0.934106526 |
| 0.834856029 | 0.934106526 |
| 0.834866218 | 0.934106526 |
| 0.834892274 | 0.934106526 |
| 0.834904165 | 0.934106526 |
| 0.834966077 | 0.934144275 |
| 0.83505892 | 0.934144275 |
| 0.835077333 | 0.934144275 |
| 0.835172365 | 0.934194192 |
| 0.835304554 | 0.93428635 |
| 0.835405081 | 0.934293848 |
| 0.835426059 | 0.93429537 |
| 0.835524167 | 0.934361208 |
| 0.835583443 | 0.934405557 |
| 0.835634231 | 0.934418471 |

| | |
|---|---|
| 0.835724811 | 0.93442483 |
| 0.835778337 | 0.93442483 |
| 0.835800941 | 0.93442483 |
| 0.835940786 | 0.93442483 |
| 0.83598287 | 0.93442483 |
| 0.836006212 | 0.93442483 |
| 0.836046173 | 0.93442483 |
| 0.8360466 | 0.93442483 |
| 0.836082853 | 0.93442483 |
| 0.836102046 | 0.93442483 |
| 0.8361174 | 0.93442483 |
| 0.836130433 | 0.93442483 |
| 0.836146537 | 0.93442483 |
| 0.836150029 | 0.93442483 |
| 0.836174425 | 0.934429637 |
| 0.836195244 | 0.934429637 |
| 0.836213189 | 0.934429637 |
| 0.836237252 | 0.934434602 |
| 0.836415336 | 0.934569478 |
| 0.836427244 | 0.934569478 |
| 0.836436445 | 0.934569478 |
| 0.836682618 | 0.934734889 |
| 0.836753031 | 0.934791626 |
| 0.836885408 | 0.934798895 |
| 0.836921818 | 0.934798895 |
| 0.836975642 | 0.93479912 |
| 0.837082752 | 0.934874903 |
| 0.837182389 | 0.934964256 |
| 0.837231741 | 0.934997447 |
| 0.837277266 | 0.935024613 |
| 0.837360183 | 0.935024613 |
| 0.837416268 | 0.935024613 |
| 0.837426064 | 0.935024613 |
| 0.837549048 | 0.935103524 |
| 0.837605249 | 0.935129517 |
| 0.837676501 | 0.935165222 |
| 0.837743193 | 0.935173915 |
| 0.837809726 | 0.935204346 |
| 0.838107375 | 0.935454854 |
| 0.838369017 | 0.935631296 |
| 0.838433218 | 0.935659096 |
| 0.838502356 | 0.935696725 |
| 0.838512948 | 0.935696725 |
| 0.838536925 | 0.935696725 |
| 0.838895867 | 0.935822156 |

| | |
|---|---|
| 0.838908776 | 0.935822156 |
| 0.838913367 | 0.935822156 |
| 0.838980855 | 0.935875519 |
| 0.839008124 | 0.935879947 |
| 0.839024124 | 0.935879947 |
| 0.839353975 | 0.936104809 |
| 0.839383213 | 0.936104809 |
| 0.839481705 | 0.936149176 |
| 0.83962699 | 0.936223507 |
| 0.839711901 | 0.936236356 |
| 0.839736802 | 0.936236356 |
| 0.839824661 | 0.936272872 |
| 0.839848187 | 0.936272872 |
| 0.839926529 | 0.936319768 |
| 0.840017723 | 0.936368034 |
| 0.840031852 | 0.936368034 |
| 0.840067974 | 0.936386383 |
| 0.840101312 | 0.936401628 |
| 0.840143248 | 0.936426456 |
| 0.840215242 | 0.936440955 |
| 0.840266215 | 0.936453937 |
| 0.840337517 | 0.936489574 |
| 0.840413455 | 0.936530373 |
| 0.840565129 | 0.9366358 |
| 0.84073378 | 0.9366358 |
| 0.840750007 | 0.9366358 |
| 0.840772975 | 0.9366358 |
| 0.840800789 | 0.9366358 |
| 0.840801714 | 0.9366358 |
| 0.841001308 | 0.93680726 |
| 0.841169699 | 0.936934663 |
| 0.841525946 | 0.937221875 |
| 0.841576391 | 0.937256139 |
| 0.841608855 | 0.93725978 |
| 0.841624898 | 0.93725978 |
| 0.841638698 | 0.93725978 |
| 0.84167805 | 0.937270422 |
| 0.841703497 | 0.937270422 |
| 0.841713496 | 0.937270422 |
| 0.841798929 | 0.937270422 |
| 0.841825501 | 0.937270422 |
| 0.841904087 | 0.937270422 |
| 0.842042835 | 0.937312146 |
| 0.842050208 | 0.937312146 |
| 0.842100076 | 0.937312146 |

| | |
|---|---|
| 0.842159409 | 0.937312146 |
| 0.842196192 | 0.937312146 |
| 0.842197409 | 0.937312146 |
| 0.842244327 | 0.937315531 |
| 0.842423955 | 0.937412655 |
| 0.842473896 | 0.937444611 |
| 0.842523724 | 0.937453079 |
| 0.842585649 | 0.937455921 |
| 0.842619128 | 0.937455921 |
| 0.842639683 | 0.937455921 |
| 0.842656991 | 0.937455921 |
| 0.84266121 | 0.937455921 |
| 0.842702102 | 0.937479516 |
| 0.842759611 | 0.937492061 |
| 0.842795633 | 0.937492061 |
| 0.842862905 | 0.937505116 |
| 0.84289812 | 0.937522391 |
| 0.843014583 | 0.937541696 |
| 0.843115717 | 0.937611139 |
| 0.84341738 | 0.937883208 |
| 0.843537752 | 0.937961294 |
| 0.843568439 | 0.937961294 |
| 0.843729336 | 0.937982679 |
| 0.84373436 | 0.937982679 |
| 0.843962245 | 0.938179123 |
| 0.844208581 | 0.938388268 |
| 0.844259182 | 0.938421458 |
| 0.844314929 | 0.938421458 |
| 0.844317255 | 0.938421458 |
| 0.844531451 | 0.93849757 |
| 0.844531749 | 0.93849757 |
| 0.844561691 | 0.93849757 |
| 0.844660539 | 0.938531054 |
| 0.844719993 | 0.938540548 |
| 0.845090763 | 0.938815744 |
| 0.845106248 | 0.938815744 |
| 0.845123908 | 0.938815744 |
| 0.845138885 | 0.938815744 |
| 0.845205097 | 0.938815744 |
| 0.845218394 | 0.938815744 |
| 0.8452479 | 0.938815744 |
| 0.845267184 | 0.938815744 |
| 0.845380581 | 0.938828799 |
| 0.845392928 | 0.938828799 |
| 0.845448025 | 0.938828799 |

| | |
|---|---|
| 0.84545252 | 0.938828799 |
| 0.845616245 | 0.938966821 |
| 0.845652307 | 0.938984973 |
| 0.845685465 | 0.938999898 |
| 0.845733401 | 0.93900347 |
| 0.845742986 | 0.93900347 |
| 0.845762659 | 0.93900347 |
| 0.845791923 | 0.939008646 |
| 0.845929413 | 0.939118906 |
| 0.845948237 | 0.939118906 |
| 0.846004697 | 0.93913712 |
| 0.846006236 | 0.93913712 |
| 0.846132326 | 0.939211418 |
| 0.846191579 | 0.939255298 |
| 0.846341628 | 0.939356173 |
| 0.846366424 | 0.939361802 |
| 0.846425983 | 0.939400907 |
| 0.846441105 | 0.939400907 |
| 0.846476412 | 0.939418201 |
| 0.84659522 | 0.939509057 |
| 0.846607457 | 0.939509057 |
| 0.846617459 | 0.939509057 |
| 0.846650802 | 0.939524168 |
| 0.846691872 | 0.939547851 |
| 0.846741852 | 0.939581421 |
| 0.846766455 | 0.939586831 |
| 0.846916696 | 0.939665971 |
| 0.847101157 | 0.939804952 |
| 0.847268732 | 0.939968971 |
| 0.84732907 | 0.939981333 |
| 0.847469444 | 0.940019997 |
| 0.847472623 | 0.940019997 |
| 0.847499387 | 0.94002779 |
| 0.847581119 | 0.940034765 |
| 0.847638464 | 0.940034765 |
| 0.847678618 | 0.940034765 |
| 0.84768357 | 0.940034765 |
| 0.847715538 | 0.940034765 |
| 0.847725359 | 0.940034765 |
| 0.847742524 | 0.940034765 |
| 0.847807976 | 0.940073423 |
| 0.847859997 | 0.940087339 |
| 0.847918177 | 0.940087339 |
| 0.848108978 | 0.940100741 |
| 0.848144915 | 0.940100741 |

| | |
|---|---|
| 0.848151557 | 0.940100741 |
| 0.848153396 | 0.940100741 |
| 0.848157321 | 0.940100741 |
| 0.848243378 | 0.940174246 |
| 0.848294654 | 0.940183941 |
| 0.848371061 | 0.940206364 |
| 0.848435689 | 0.940234231 |
| 0.848600791 | 0.940329677 |
| 0.848666381 | 0.940380478 |
| 0.848761048 | 0.940421363 |
| 0.848762515 | 0.940421363 |
| 0.848907787 | 0.940538562 |
| 0.849015656 | 0.940582157 |
| 0.84902613 | 0.940582157 |
| 0.849154692 | 0.94068082 |
| 0.849247071 | 0.940719396 |
| 0.849268521 | 0.940719396 |
| 0.849545345 | 0.940916612 |
| 0.849584844 | 0.940938478 |
| 0.84979905 | 0.941131946 |
| 0.849882732 | 0.94119517 |
| 0.849895661 | 0.94119517 |
| 0.850080823 | 0.941310196 |
| 0.850150168 | 0.941310196 |
| 0.850156227 | 0.941310196 |
| 0.850179266 | 0.941310196 |
| 0.850266588 | 0.941315161 |
| 0.850376941 | 0.941377082 |
| 0.85051539 | 0.941474523 |
| 0.850523502 | 0.941474523 |
| 0.850813119 | 0.941641913 |
| 0.850899277 | 0.941650625 |
| 0.851014867 | 0.941722887 |
| 0.851027181 | 0.941722887 |
| 0.851045692 | 0.941722887 |
| 0.851370677 | 0.941816029 |
| 0.851443433 | 0.941858004 |
| 0.851539073 | 0.941876296 |
| 0.851667456 | 0.941974546 |
| 0.851748791 | 0.941997774 |
| 0.851882046 | 0.941998224 |
| 0.851891853 | 0.941998224 |
| 0.85197053 | 0.941998224 |
| 0.851980917 | 0.941998224 |
| 0.851994082 | 0.941998224 |

| | |
|---|---|
| 0.852035852 | 0.941998224 |
| 0.852045097 | 0.941998224 |
| 0.852059847 | 0.941998224 |
| 0.852084437 | 0.941998224 |
| 0.85215852 | 0.942045351 |
| 0.852166625 | 0.942045351 |
| 0.852292559 | 0.942140831 |
| 0.852493845 | 0.942182078 |
| 0.852709568 | 0.942182078 |
| 0.852716654 | 0.942182078 |
| 0.852727618 | 0.942182078 |
| 0.852770506 | 0.942182078 |
| 0.852826616 | 0.942182078 |
| 0.852848459 | 0.942182078 |
| 0.852860358 | 0.942182078 |
| 0.852864056 | 0.942182078 |
| 0.85290587 | 0.942182078 |
| 0.852929409 | 0.942182078 |
| 0.853013463 | 0.942182078 |
| 0.853013571 | 0.942182078 |
| 0.853027421 | 0.942182078 |
| 0.853055883 | 0.942182078 |
| 0.853070094 | 0.942182078 |
| 0.853080177 | 0.942182078 |
| 0.853101391 | 0.942182078 |
| 0.853189401 | 0.94223558 |
| 0.853235662 | 0.94226482 |
| 0.853291079 | 0.94229111 |
| 0.853362987 | 0.942339879 |
| 0.853426094 | 0.942385464 |
| 0.853551758 | 0.942397531 |
| 0.853593007 | 0.942401212 |
| 0.853722058 | 0.942474166 |
| 0.853808453 | 0.942525848 |
| 0.853899215 | 0.942563515 |
| 0.854012356 | 0.942575601 |
| 0.854020188 | 0.942575601 |
| 0.854047452 | 0.942575601 |
| 0.854091012 | 0.942575601 |
| 0.854118307 | 0.942583882 |
| 0.85430665 | 0.942748043 |
| 0.854481461 | 0.942860071 |
| 0.854565441 | 0.942860071 |
| 0.854569797 | 0.942860071 |
| 0.854578662 | 0.942860071 |

| | |
|---|---|
| 0.854587122 | 0.942860071 |
| 0.854591624 | 0.942860071 |
| 0.854624944 | 0.942860071 |
| 0.854632539 | 0.942860071 |
| 0.854645729 | 0.942860071 |
| 0.854728953 | 0.942908203 |
| 0.854793396 | 0.942940778 |
| 0.854798079 | 0.942940778 |
| 0.854857926 | 0.942984955 |
| 0.854909786 | 0.943003387 |
| 0.854914234 | 0.943003387 |
| 0.855239914 | 0.943190438 |
| 0.855285427 | 0.943190438 |
| 0.855347467 | 0.943190438 |
| 0.855366279 | 0.943190438 |
| 0.855370893 | 0.943190438 |
| 0.855378764 | 0.943190438 |
| 0.855420762 | 0.943212591 |
| 0.85545416 | 0.943227579 |
| 0.855581333 | 0.943240954 |
| 0.855610657 | 0.943240954 |
| 0.855627798 | 0.943240954 |
| 0.855822778 | 0.943371025 |
| 0.855887814 | 0.943371025 |
| 0.855911993 | 0.943371025 |
| 0.855977779 | 0.943371025 |
| 0.855987766 | 0.943371025 |
| 0.856070937 | 0.943405306 |
| 0.856253964 | 0.943585172 |
| 0.856305321 | 0.943619933 |
| 0.856329845 | 0.943625125 |
| 0.856359462 | 0.943631989 |
| 0.856496154 | 0.943663434 |
| 0.856521169 | 0.943663434 |
| 0.856626329 | 0.943734582 |
| 0.856656158 | 0.94374261 |
| 0.856723025 | 0.943765322 |
| 0.856734493 | 0.943765322 |
| 0.856900035 | 0.943907511 |
| 0.856903835 | 0.943907511 |
| 0.856969152 | 0.943907511 |
| 0.856974929 | 0.943907511 |
| 0.857107475 | 0.943909397 |
| 0.857128448 | 0.943909397 |
| 0.857142745 | 0.943909397 |

| | |
|---|---|
| 0.857204796 | 0.943931533 |
| 0.85743257 | 0.944019137 |
| 0.857440608 | 0.944019137 |
| 0.85753708 | 0.944052192 |
| 0.857550166 | 0.944052192 |
| 0.857574485 | 0.944052192 |
| 0.857592695 | 0.944052192 |
| 0.857679005 | 0.944052192 |
| 0.85770665 | 0.944052192 |
| 0.857707487 | 0.944052192 |
| 0.857728315 | 0.944052192 |
| 0.857728315 | 0.944052192 |
| 0.857830212 | 0.944110578 |
| 0.858095585 | 0.944276738 |
| 0.858138567 | 0.944276738 |
| 0.858211317 | 0.944276738 |
| 0.858240652 | 0.944276738 |
| 0.858264099 | 0.944276738 |
| 0.858320228 | 0.944289056 |
| 0.858425579 | 0.944294358 |
| 0.858427723 | 0.944294358 |
| 0.858762472 | 0.944513781 |
| 0.858835584 | 0.944547243 |
| 0.8589308 | 0.944589849 |
| 0.858953103 | 0.944592566 |
| 0.859029591 | 0.944647331 |
| 0.859042571 | 0.944647331 |
| 0.859120379 | 0.944655055 |
| 0.859126591 | 0.944655055 |
| 0.859169056 | 0.944655055 |
| 0.859211963 | 0.944655055 |
| 0.859277368 | 0.944655055 |
| 0.859285947 | 0.944655055 |
| 0.8593507 | 0.944655055 |
| 0.859366946 | 0.944655055 |
| 0.859656361 | 0.944864154 |
| 0.859820008 | 0.94490128 |
| 0.859829017 | 0.94490128 |
| 0.859933866 | 0.944971484 |
| 0.859992006 | 0.944971484 |
| 0.859994179 | 0.944971484 |
| 0.860017039 | 0.944971484 |
| 0.860076387 | 0.944971484 |
| 0.860110873 | 0.944971484 |
| 0.860128034 | 0.944971484 |

| | |
|---|---|
| 0.860170478 | 0.944971484 |
| 0.860181119 | 0.944971484 |
| 0.860268325 | 0.944971484 |
| 0.860275753 | 0.944971484 |
| 0.86028377 | 0.944971484 |
| 0.860319439 | 0.944971484 |
| 0.860331085 | 0.944971484 |
| 0.860385882 | 0.944971484 |
| 0.860428609 | 0.944971484 |
| 0.860606458 | 0.945047184 |
| 0.860616592 | 0.945047184 |
| 0.860906695 | 0.945244136 |
| 0.860958264 | 0.945244136 |
| 0.861043656 | 0.945276403 |
| 0.861163411 | 0.945294061 |
| 0.86124577 | 0.945345755 |
| 0.861312054 | 0.945353403 |
| 0.861312286 | 0.945353403 |
| 0.861348672 | 0.94536541 |
| 0.861362924 | 0.94536541 |
| 0.8614252 | 0.945385113 |
| 0.861440425 | 0.945385113 |
| 0.861475359 | 0.945401301 |
| 0.861675484 | 0.945512355 |
| 0.861745347 | 0.945539069 |
| 0.861842748 | 0.945565183 |
| 0.861995943 | 0.945667906 |
| 0.862273668 | 0.945873806 |
| 0.862346507 | 0.945916576 |
| 0.862484109 | 0.945963768 |
| 0.862545469 | 0.946009281 |
| 0.86269173 | 0.946065086 |
| 0.86269922 | 0.946065086 |
| 0.862702462 | 0.946065086 |
| 0.86272296 | 0.946065086 |
| 0.86274842 | 0.946065086 |
| 0.862866151 | 0.946121332 |
| 0.862906797 | 0.946122336 |
| 0.862965081 | 0.946143099 |
| 0.862965466 | 0.946143099 |
| 0.863094281 | 0.946256811 |
| 0.863206448 | 0.946256811 |
| 0.863210203 | 0.946256811 |
| 0.863225337 | 0.946256811 |
| 0.863232324 | 0.946256811 |

| | |
|---|---|
| 0.86324506 | 0.946256811 |
| 0.863251096 | 0.946256811 |
| 0.863329211 | 0.946256811 |
| 0.863351668 | 0.946256811 |
| 0.863375958 | 0.946256811 |
| 0.863586534 | 0.946256811 |
| 0.863615539 | 0.946256811 |
| 0.863637444 | 0.946256811 |
| 0.863665223 | 0.946256811 |
| 0.863747653 | 0.946303585 |
| 0.863843906 | 0.946365499 |
| 0.863942255 | 0.94641176 |
| 0.863954539 | 0.94641176 |
| 0.863956517 | 0.94641176 |
| 0.86398228 | 0.94641176 |
| 0.864010319 | 0.94641176 |
| 0.864199847 | 0.946494184 |
| 0.864239919 | 0.946516306 |
| 0.864320751 | 0.946546076 |
| 0.864387174 | 0.946546076 |
| 0.864419912 | 0.946546076 |
| 0.864420859 | 0.946546076 |
| 0.864422851 | 0.946546076 |
| 0.864433586 | 0.946546076 |
| 0.86463554 | 0.946605948 |
| 0.864643581 | 0.946605948 |
| 0.864662047 | 0.946605948 |
| 0.864753446 | 0.946665112 |
| 0.864865834 | 0.946688129 |
| 0.864873857 | 0.946688129 |
| 0.864896878 | 0.94669157 |
| 0.865026587 | 0.946746513 |
| 0.865063606 | 0.946765272 |
| 0.865103112 | 0.946772614 |
| 0.865158843 | 0.946772614 |
| 0.865382349 | 0.94695956 |
| 0.86540019 | 0.94695956 |
| 0.865468934 | 0.947012404 |
| 0.865488251 | 0.947012404 |
| 0.865799775 | 0.947266221 |
| 0.865841942 | 0.947268835 |
| 0.865878834 | 0.947287436 |
| 0.865914618 | 0.947304824 |
| 0.865967457 | 0.947312801 |
| 0.865999145 | 0.947312801 |

| | |
|---|---|
| 0.866063361 | 0.947312801 |
| 0.866151333 | 0.947346183 |
| 0.866199298 | 0.947376888 |
| 0.866344051 | 0.947489447 |
| 0.866361894 | 0.947489447 |
| 0.866514573 | 0.947634663 |
| 0.866611723 | 0.947654015 |
| 0.86663507 | 0.947654015 |
| 0.86675608 | 0.947657059 |
| 0.866791127 | 0.947657059 |
| 0.866841834 | 0.947657059 |
| 0.866921414 | 0.947709647 |
| 0.867063793 | 0.947725389 |
| 0.867342577 | 0.947856863 |
| 0.867425283 | 0.947856863 |
| 0.86752735 | 0.947856863 |
| 0.867531293 | 0.947856863 |
| 0.867539799 | 0.947856863 |
| 0.86760032 | 0.947886138 |
| 0.867751081 | 0.948007357 |
| 0.867955196 | 0.948117282 |
| 0.867963413 | 0.948117282 |
| 0.868087298 | 0.948170485 |
| 0.868155564 | 0.948170485 |
| 0.868157946 | 0.948170485 |
| 0.868181846 | 0.948170485 |
| 0.868217751 | 0.948170485 |
| 0.86830906 | 0.948170485 |
| 0.868755259 | 0.948450626 |
| 0.86876686 | 0.948450626 |
| 0.868780661 | 0.948450626 |
| 0.868794076 | 0.948450626 |
| 0.868873828 | 0.948515949 |
| 0.86889431 | 0.948516567 |
| 0.86892273 | 0.94852585 |
| 0.868954243 | 0.94853851 |
| 0.868990315 | 0.948556145 |
| 0.869018612 | 0.948565293 |
| 0.869097777 | 0.948623248 |
| 0.869111542 | 0.948623248 |
| 0.869147081 | 0.948633761 |
| 0.869352373 | 0.948741497 |
| 0.869547058 | 0.948810315 |
| 0.869548715 | 0.948810315 |
| 0.869649952 | 0.948867917 |

| | |
|---|---|
| 0.869734084 | 0.948933044 |
| 0.869789081 | 0.948949571 |
| 0.869853635 | 0.948998261 |
| 0.86990821 | 0.949014324 |
| 0.869977279 | 0.949067934 |
| 0.870120388 | 0.949137092 |
| 0.870417543 | 0.94938074 |
| 0.870469894 | 0.949403487 |
| 0.87062701 | 0.949499671 |
| 0.87067109 | 0.949499671 |
| 0.870756009 | 0.949525974 |
| 0.870869373 | 0.949584368 |
| 0.870921649 | 0.949589608 |
| 0.870940428 | 0.949589608 |
| 0.871116904 | 0.949659117 |
| 0.871126918 | 0.949659117 |
| 0.87113732 | 0.949659117 |
| 0.87118861 | 0.949680371 |
| 0.871196697 | 0.949680371 |
| 0.871260766 | 0.949699001 |
| 0.87135611 | 0.949699001 |
| 0.87137331 | 0.949699001 |
| 0.871424422 | 0.949732973 |
| 0.871515882 | 0.94975655 |
| 0.871726681 | 0.949850768 |
| 0.871743039 | 0.949850768 |
| 0.871770627 | 0.949850768 |
| 0.871953282 | 0.949945907 |
| 0.871958873 | 0.949945907 |
| 0.872170061 | 0.950097778 |
| 0.872198019 | 0.950097778 |
| 0.872239742 | 0.950121496 |
| 0.87233093 | 0.950155633 |
| 0.872377107 | 0.950184199 |
| 0.872478041 | 0.950250672 |
| 0.872499979 | 0.950252835 |
| 0.872650966 | 0.950259164 |
| 0.872660859 | 0.950259164 |
| 0.872680433 | 0.950259164 |
| 0.87271364 | 0.950259164 |
| 0.872734419 | 0.950259164 |
| 0.87273616 | 0.950259164 |
| 0.872747322 | 0.950259164 |
| 0.872765392 | 0.950259164 |
| 0.872803683 | 0.950259164 |

| | |
|---|---|
| 0.872812018 | 0.950259164 |
| 0.872828168 | 0.950259164 |
| 0.872937036 | 0.950280415 |
| 0.872948651 | 0.950280415 |
| 0.872964258 | 0.950280415 |
| 0.873081532 | 0.95036923 |
| 0.873186362 | 0.950372137 |
| 0.873188198 | 0.950372137 |
| 0.873223642 | 0.950388995 |
| 0.873270507 | 0.950418281 |
| 0.873326028 | 0.950456989 |
| 0.873377574 | 0.950469649 |
| 0.873456581 | 0.950512192 |
| 0.873714027 | 0.950599185 |
| 0.873716775 | 0.950599185 |
| 0.873763215 | 0.950599185 |
| 0.873788893 | 0.950599185 |
| 0.873802661 | 0.950599185 |
| 0.873815952 | 0.950599185 |
| 0.873923076 | 0.950672293 |
| 0.874262875 | 0.950808509 |
| 0.874265231 | 0.950808509 |
| 0.874272443 | 0.950808509 |
| 0.874272443 | 0.950808509 |
| 0.874505431 | 0.950936554 |
| 0.874533742 | 0.950940391 |
| 0.874557782 | 0.950940391 |
| 0.874588825 | 0.950940391 |
| 0.874654599 | 0.950990197 |
| 0.874803161 | 0.951086588 |
| 0.874837159 | 0.951101839 |
| 0.874877158 | 0.951123614 |
| 0.874964511 | 0.951193263 |
| 0.875018564 | 0.951193263 |
| 0.875021111 | 0.951193263 |
| 0.87513674 | 0.951297245 |
| 0.875361037 | 0.951430198 |
| 0.875373828 | 0.951430198 |
| 0.875636626 | 0.95161726 |
| 0.875653016 | 0.95161726 |
| 0.875683266 | 0.95161726 |
| 0.875710863 | 0.95161726 |
| 0.875773975 | 0.951642416 |
| 0.875876779 | 0.951709623 |
| 0.87593581 | 0.951709702 |

| | |
|---|---|
| 0.876117214 | 0.951885084 |
| 0.876175485 | 0.951911669 |
| 0.876207749 | 0.951911669 |
| 0.876243281 | 0.951911669 |
| 0.87628295 | 0.951911669 |
| 0.876330145 | 0.951911669 |
| 0.876423982 | 0.951979497 |
| 0.876453819 | 0.951990195 |
| 0.876517964 | 0.952038155 |
| 0.876597161 | 0.952059039 |
| 0.876666919 | 0.95209432 |
| 0.876702026 | 0.95210624 |
| 0.876734563 | 0.95210624 |
| 0.876740575 | 0.95210624 |
| 0.876913736 | 0.952227521 |
| 0.87693223 | 0.952227521 |
| 0.876980568 | 0.952253175 |
| 0.876995843 | 0.952253175 |
| 0.877078938 | 0.95230097 |
| 0.877116826 | 0.95230097 |
| 0.87711984 | 0.95230097 |
| 0.877210295 | 0.952364369 |
| 0.877224615 | 0.952364369 |
| 0.877238224 | 0.952364369 |
| 0.877289475 | 0.952389394 |
| 0.877324357 | 0.952389394 |
| 0.877341412 | 0.952389394 |
| 0.877358445 | 0.952389394 |
| 0.877361258 | 0.952389394 |
| 0.877393268 | 0.952402434 |
| 0.87747998 | 0.952462305 |
| 0.877635993 | 0.952579066 |
| 0.877764476 | 0.95269681 |
| 0.877937305 | 0.952819252 |
| 0.878028659 | 0.952853259 |
| 0.87809308 | 0.952878564 |
| 0.878116135 | 0.952878564 |
| 0.878167912 | 0.952878564 |
| 0.878198651 | 0.952878564 |
| 0.878203224 | 0.952878564 |
| 0.878266099 | 0.952915516 |
| 0.8783035 | 0.952929494 |
| 0.87868416 | 0.953169063 |
| 0.878749739 | 0.953201399 |
| 0.878830279 | 0.953267052 |

| | |
|---|---|
| 0.878896607 | 0.953313109 |
| 0.878994376 | 0.953319292 |
| 0.879023234 | 0.953319292 |
| 0.879030395 | 0.953319292 |
| 0.879101678 | 0.953330788 |
| 0.879109221 | 0.953330788 |
| 0.879277229 | 0.953416137 |
| 0.879284787 | 0.953416137 |
| 0.879288017 | 0.953416137 |
| 0.879322587 | 0.953431915 |
| 0.879394903 | 0.953466912 |
| 0.879457756 | 0.953491647 |
| 0.879518066 | 0.953523209 |
| 0.879543129 | 0.953523209 |
| 0.879546929 | 0.953523209 |
| 0.879716295 | 0.953590158 |
| 0.879739427 | 0.953590158 |
| 0.879746118 | 0.953590158 |
| 0.879748839 | 0.953590158 |
| 0.879790548 | 0.953600679 |
| 0.879991099 | 0.953666403 |
| 0.879999393 | 0.953666403 |
| 0.8801325 | 0.953778848 |
| 0.880335945 | 0.953900897 |
| 0.880589762 | 0.954057116 |
| 0.880729394 | 0.954142936 |
| 0.880733268 | 0.954142936 |
| 0.88073962 | 0.954142936 |
| 0.880830752 | 0.954161828 |
| 0.880905992 | 0.954161828 |
| 0.880915295 | 0.954161828 |
| 0.880923015 | 0.954161828 |
| 0.881021891 | 0.954209978 |
| 0.881149709 | 0.954263891 |
| 0.881151813 | 0.954263891 |
| 0.881223772 | 0.954319451 |
| 0.881243191 | 0.954319451 |
| 0.88128292 | 0.954333567 |
| 0.881325894 | 0.954333567 |
| 0.881410782 | 0.954333567 |
| 0.881425882 | 0.954333567 |
| 0.881520296 | 0.954402527 |
| 0.881589398 | 0.95442716 |
| 0.881704651 | 0.95442716 |
| 0.881767651 | 0.95442716 |

| | |
|---|---|
| 0.881784492 | 0.95442716 |
| 0.881862149 | 0.95442716 |
| 0.881863682 | 0.95442716 |
| 0.882010973 | 0.954521499 |
| 0.88207482 | 0.954528359 |
| 0.882123489 | 0.954537682 |
| 0.882126137 | 0.954537682 |
| 0.882254566 | 0.9545899 |
| 0.882276461 | 0.954591903 |
| 0.882317125 | 0.954614214 |
| 0.88234935 | 0.954619472 |
| 0.882436642 | 0.954620377 |
| 0.882436959 | 0.954620377 |
| 0.882438257 | 0.954620377 |
| 0.882443083 | 0.954620377 |
| 0.882488591 | 0.954633408 |
| 0.882586859 | 0.954667492 |
| 0.882610564 | 0.954671451 |
| 0.882676436 | 0.954686563 |
| 0.882709818 | 0.954686563 |
| 0.882835351 | 0.954686563 |
| 0.882859067 | 0.954686563 |
| 0.882863428 | 0.954686563 |
| 0.882996758 | 0.954749932 |
| 0.883042448 | 0.954749932 |
| 0.883043956 | 0.954749932 |
| 0.883243814 | 0.954900986 |
| 0.883334268 | 0.954955423 |
| 0.883454444 | 0.954958496 |
| 0.883471277 | 0.954958496 |
| 0.883574001 | 0.955041167 |
| 0.883617404 | 0.955066405 |
| 0.883843379 | 0.95510974 |
| 0.883846397 | 0.95510974 |
| 0.883857605 | 0.95510974 |
| 0.883872074 | 0.95510974 |
| 0.883881071 | 0.95510974 |
| 0.883883903 | 0.95510974 |
| 0.883898144 | 0.95510974 |
| 0.884008494 | 0.955207309 |
| 0.884068406 | 0.955234773 |
| 0.88429813 | 0.955404028 |
| 0.884412684 | 0.955448962 |
| 0.88448104 | 0.955480803 |
| 0.884482282 | 0.955480803 |

| | |
|---|---|
| 0.884529739 | 0.955510397 |
| 0.884672885 | 0.955572963 |
| 0.884687976 | 0.955572963 |
| 0.884733477 | 0.955589609 |
| 0.884860469 | 0.955640394 |
| 0.884866804 | 0.955640394 |
| 0.884891087 | 0.955640639 |
| 0.884935667 | 0.955667113 |
| 0.884969924 | 0.955667829 |
| 0.885059658 | 0.955692664 |
| 0.8851929 | 0.955755917 |
| 0.885198506 | 0.955755917 |
| 0.88524637 | 0.955785929 |
| 0.885368628 | 0.95580956 |
| 0.885373244 | 0.95580956 |
| 0.885408738 | 0.95580956 |
| 0.885484654 | 0.955869846 |
| 0.885520198 | 0.95588655 |
| 0.885547229 | 0.955894064 |
| 0.885607084 | 0.955937008 |
| 0.885642925 | 0.955947327 |
| 0.885657051 | 0.955947327 |
| 0.885825839 | 0.956035143 |
| 0.885851847 | 0.956035143 |
| 0.885853305 | 0.956035143 |
| 0.885867786 | 0.956035143 |
| 0.88587866 | 0.956035143 |
| 0.885907236 | 0.956044319 |
| 0.886070066 | 0.956138815 |
| 0.886143392 | 0.956138815 |
| 0.886377826 | 0.956292145 |
| 0.886516099 | 0.956397994 |
| 0.886574665 | 0.956417848 |
| 0.886712857 | 0.956540032 |
| 0.886728095 | 0.956540032 |
| 0.88680665 | 0.95658144 |
| 0.886871426 | 0.956629647 |
| 0.886917835 | 0.956636374 |
| 0.887017221 | 0.956721906 |
| 0.88720819 | 0.956834923 |
| 0.887278625 | 0.956834923 |
| 0.88730055 | 0.956834923 |
| 0.887357866 | 0.956850958 |
| 0.887398982 | 0.956873629 |
| 0.887449067 | 0.95690597 |

| | |
|---|---|
| 0.887595861 | 0.956982063 |
| 0.887636027 | 0.956999232 |
| 0.888094043 | 0.957303037 |
| 0.88810563 | 0.957303037 |
| 0.888118813 | 0.957303037 |
| 0.88830838 | 0.957442365 |
| 0.888347149 | 0.957449281 |
| 0.888449948 | 0.957486608 |
| 0.888473663 | 0.9574905 |
| 0.888563007 | 0.957515371 |
| 0.888565561 | 0.957515371 |
| 0.888729901 | 0.957593299 |
| 0.8889125 | 0.957700871 |
| 0.888949925 | 0.957700871 |
| 0.888967918 | 0.957700871 |
| 0.889012151 | 0.957700871 |
| 0.88907036 | 0.957700871 |
| 0.889070408 | 0.957700871 |
| 0.889071038 | 0.957700871 |
| 0.889396843 | 0.957900171 |
| 0.889480355 | 0.957913693 |
| 0.889544102 | 0.957943799 |
| 0.889607242 | 0.957975132 |
| 0.889717886 | 0.958072617 |
| 0.889781578 | 0.95810842 |
| 0.889791369 | 0.95810842 |
| 0.889882636 | 0.958156176 |
| 0.88991407 | 0.958156176 |
| 0.889985712 | 0.958179958 |
| 0.890049384 | 0.958179958 |
| 0.890079108 | 0.958179958 |
| 0.890079108 | 0.958179958 |
| 0.890079108 | 0.958179958 |
| 0.890217906 | 0.958255577 |
| 0.890260635 | 0.958255577 |
| 0.890467719 | 0.958372658 |
| 0.890651016 | 0.958429373 |
| 0.890652899 | 0.958429373 |
| 0.890712538 | 0.958471894 |
| 0.890786368 | 0.958480971 |
| 0.890811648 | 0.958480971 |
| 0.890882173 | 0.958481188 |
| 0.890923179 | 0.958501971 |
| 0.891113916 | 0.958608774 |
| 0.891121524 | 0.958608774 |

| | |
|---|---|
| 0.891165617 | 0.958618077 |
| 0.891211312 | 0.958640404 |
| 0.891244336 | 0.958654275 |
| 0.891383202 | 0.958760338 |
| 0.891485413 | 0.958816879 |
| 0.891515865 | 0.958816879 |
| 0.891516297 | 0.958816879 |
| 0.891850938 | 0.959029158 |
| 0.891858848 | 0.959029158 |
| 0.891860601 | 0.959029158 |
| 0.891874765 | 0.959029158 |
| 0.891993055 | 0.959069741 |
| 0.892027877 | 0.959069869 |
| 0.892053586 | 0.959069869 |
| 0.892146308 | 0.959126255 |
| 0.892232521 | 0.959138518 |
| 0.892297928 | 0.959138518 |
| 0.892359325 | 0.959138759 |
| 0.892410289 | 0.959158722 |
| 0.892418176 | 0.959158722 |
| 0.892455279 | 0.959162332 |
| 0.892586969 | 0.959162332 |
| 0.892587139 | 0.959162332 |
| 0.892624473 | 0.959163996 |
| 0.892656123 | 0.959176364 |
| 0.892754959 | 0.959260923 |
| 0.892798919 | 0.959264876 |
| 0.892952919 | 0.959293348 |
| 0.892984884 | 0.959293348 |
| 0.892993405 | 0.959293348 |
| 0.893071457 | 0.959319516 |
| 0.893237715 | 0.959388514 |
| 0.893747956 | 0.959760802 |
| 0.894074681 | 0.959943047 |
| 0.894176686 | 0.960008859 |
| 0.894252442 | 0.960068551 |
| 0.894370092 | 0.96010829 |
| 0.894417129 | 0.960115502 |
| 0.894484662 | 0.960122613 |
| 0.894497043 | 0.960122613 |
| 0.894504864 | 0.960122613 |
| 0.894505749 | 0.960122613 |
| 0.894601839 | 0.960183932 |
| 0.894644442 | 0.960208019 |
| 0.894703065 | 0.960237236 |

| | |
|---|---|
| 0.894803301 | 0.960237236 |
| 0.894840847 | 0.960237236 |
| 0.894856258 | 0.960237236 |
| 0.894907371 | 0.960237236 |
| 0.894993557 | 0.960253336 |
| 0.895027346 | 0.960253336 |
| 0.895059017 | 0.960253336 |
| 0.895066386 | 0.960253336 |
| 0.895069742 | 0.960253336 |
| 0.895101316 | 0.960265579 |
| 0.895217945 | 0.960304175 |
| 0.895245144 | 0.960311722 |
| 0.895523239 | 0.960498373 |
| 0.895563443 | 0.960501725 |
| 0.895611925 | 0.960532092 |
| 0.895705961 | 0.960589683 |
| 0.895773909 | 0.960619292 |
| 0.895799381 | 0.960624977 |
| 0.895924249 | 0.960671255 |
| 0.895949348 | 0.960671255 |
| 0.896037006 | 0.960682558 |
| 0.89620731 | 0.960737948 |
| 0.896236115 | 0.960747202 |
| 0.896274623 | 0.960766858 |
| 0.896352618 | 0.960802252 |
| 0.896368162 | 0.960802252 |
| 0.89641437 | 0.960805531 |
| 0.896442327 | 0.960805531 |
| 0.896472089 | 0.960805531 |
| 0.896512682 | 0.960827415 |
| 0.89656669 | 0.960830565 |
| 0.89659426 | 0.960830565 |
| 0.896608477 | 0.960830565 |
| 0.896812537 | 0.960994592 |
| 0.896875606 | 0.961021744 |
| 0.897107621 | 0.961248727 |
| 0.897137409 | 0.96124963 |
| 0.897163393 | 0.96124963 |
| 0.897375596 | 0.96134121 |
| 0.897439733 | 0.961388294 |
| 0.897558237 | 0.961410943 |
| 0.897561807 | 0.961410943 |
| 0.897733751 | 0.961475928 |
| 0.897767527 | 0.961475928 |
| 0.89777011 | 0.961475928 |

| | |
|---|---|
| 0.897814534 | 0.961475928 |
| 0.897846156 | 0.961475928 |
| 0.897873698 | 0.961475928 |
| 0.897925174 | 0.961475928 |
| 0.897925291 | 0.961475928 |
| 0.898026106 | 0.961527897 |
| 0.898060474 | 0.961527897 |
| 0.898072817 | 0.961527897 |
| 0.898153855 | 0.96156934 |
| 0.898202445 | 0.96157082 |
| 0.898237603 | 0.96157082 |
| 0.898256186 | 0.96157082 |
| 0.898369919 | 0.961588669 |
| 0.898389825 | 0.961588669 |
| 0.898393884 | 0.961588669 |
| 0.898395011 | 0.961588669 |
| 0.89841419 | 0.961588669 |
| 0.898570051 | 0.961677833 |
| 0.898578263 | 0.961677833 |
| 0.89863484 | 0.961716772 |
| 0.89872862 | 0.961784993 |
| 0.899026472 | 0.961962971 |
| 0.899110788 | 0.962031576 |
| 0.899136693 | 0.96203768 |
| 0.899166821 | 0.962048303 |
| 0.899202194 | 0.962064537 |
| 0.899297921 | 0.962068491 |
| 0.899313914 | 0.962068491 |
| 0.899331424 | 0.962068491 |
| 0.899348188 | 0.962068491 |
| 0.899359667 | 0.962068491 |
| 0.89938769 | 0.962068491 |
| 0.899429482 | 0.962091588 |
| 0.899555787 | 0.962147261 |
| 0.899575918 | 0.962147261 |
| 0.899576738 | 0.962147261 |
| 0.899733646 | 0.962177571 |
| 0.899771533 | 0.962177571 |
| 0.899790707 | 0.962177571 |
| 0.89993682 | 0.96226687 |
| 0.900046615 | 0.962341059 |
| 0.90017587 | 0.962400969 |
| 0.900183475 | 0.962400969 |
| 0.90049494 | 0.962669132 |
| 0.900608055 | 0.962702262 |

| | |
|---|---|
| 0.900624209 | 0.962702262 |
| 0.900689584 | 0.962720981 |
| 0.90070515 | 0.962720981 |
| 0.900769912 | 0.962744668 |
| 0.90091475 | 0.962836972 |
| 0.900953296 | 0.962856561 |
| 0.900980891 | 0.962864446 |
| 0.901009204 | 0.962873098 |
| 0.901266812 | 0.963105172 |
| 0.901290976 | 0.963108713 |
| 0.901329891 | 0.963108713 |
| 0.90136115 | 0.963119545 |
| 0.901469545 | 0.963202515 |
| 0.901510615 | 0.963202515 |
| 0.901635624 | 0.963239942 |
| 0.901767731 | 0.963316251 |
| 0.901888374 | 0.963380304 |
| 0.902156693 | 0.963491595 |
| 0.902201854 | 0.963491595 |
| 0.902222769 | 0.963491595 |
| 0.902269781 | 0.963506791 |
| 0.902424738 | 0.96355917 |
| 0.90244022 | 0.96355917 |
| 0.902461031 | 0.96355979 |
| 0.902586482 | 0.963585725 |
| 0.902648717 | 0.963608967 |
| 0.902772063 | 0.963675842 |
| 0.902851257 | 0.963738778 |
| 0.903093965 | 0.963933042 |
| 0.903259908 | 0.963963906 |
| 0.903267508 | 0.963963906 |
| 0.903295532 | 0.963963906 |
| 0.903315658 | 0.963963906 |
| 0.903453999 | 0.964037025 |
| 0.903454523 | 0.964037025 |
| 0.903673566 | 0.964163463 |
| 0.903719681 | 0.964168752 |
| 0.903771388 | 0.964202319 |
| 0.903797531 | 0.964208517 |
| 0.903817688 | 0.964208517 |
| 0.903910111 | 0.964263917 |
| 0.903944362 | 0.964278857 |
| 0.903997734 | 0.964297659 |
| 0.904002481 | 0.964297659 |
| 0.9042761 | 0.964531018 |

| | |
|---|---|
| 0.904282004 | 0.964531018 |
| 0.904541769 | 0.964663728 |
| 0.90456098 | 0.964663728 |
| 0.904689081 | 0.964679328 |
| 0.904694181 | 0.964679328 |
| 0.904714148 | 0.964679328 |
| 0.904764789 | 0.964679328 |
| 0.904869475 | 0.964679328 |
| 0.904886913 | 0.964679328 |
| 0.9051905 | 0.964863673 |
| 0.905203479 | 0.964863673 |
| 0.905238494 | 0.964863673 |
| 0.905368991 | 0.964863673 |
| 0.905377099 | 0.964863673 |
| 0.905601966 | 0.964964302 |
| 0.905683151 | 0.965004682 |
| 0.905891387 | 0.965120577 |
| 0.905942995 | 0.965120577 |
| 0.905959357 | 0.965120577 |
| 0.905967807 | 0.965120577 |
| 0.90597859 | 0.965120577 |
| 0.906061906 | 0.96514654 |
| 0.906072121 | 0.96514654 |
| 0.90617917 | 0.965192784 |
| 0.906208137 | 0.965202052 |
| 0.906357882 | 0.965296783 |
| 0.906401703 | 0.965321867 |
| 0.90646849 | 0.965336654 |
| 0.906517853 | 0.965336654 |
| 0.906575002 | 0.965336654 |
| 0.906585282 | 0.965336654 |
| 0.906587337 | 0.965336654 |
| 0.906676372 | 0.965398515 |
| 0.906779799 | 0.965411857 |
| 0.906799276 | 0.965411857 |
| 0.906810524 | 0.965411857 |
| 0.906837354 | 0.96541884 |
| 0.906937634 | 0.965470871 |
| 0.907011969 | 0.965518407 |
| 0.907094502 | 0.965538736 |
| 0.907112158 | 0.965538736 |
| 0.907155868 | 0.965563682 |
| 0.907224413 | 0.96561506 |
| 0.907274448 | 0.965629394 |
| 0.907340231 | 0.965652012 |

| | |
|---|---|
| 0.90758335 | 0.965781268 |
| 0.907687668 | 0.965869484 |
| 0.90779683 | 0.965904195 |
| 0.907913746 | 0.965960731 |
| 0.907914253 | 0.965960731 |
| 0.908010031 | 0.966007143 |
| 0.90804333 | 0.966007143 |
| 0.908121515 | 0.966051761 |
| 0.908307573 | 0.966163367 |
| 0.908396496 | 0.966214796 |
| 0.908812374 | 0.96650605 |
| 0.908860554 | 0.966522698 |
| 0.908878511 | 0.966522698 |
| 0.909086559 | 0.966581148 |
| 0.909104293 | 0.966581148 |
| 0.909104293 | 0.966581148 |
| 0.90954389 | 0.967024887 |
| 0.909581667 | 0.967043464 |
| 0.909689282 | 0.9671147 |
| 0.90973015 | 0.96713656 |
| 0.909911939 | 0.967199626 |
| 0.909931628 | 0.967199626 |
| 0.910011029 | 0.967224749 |
| 0.910031811 | 0.967224749 |
| 0.910036497 | 0.967224749 |
| 0.910253053 | 0.967357799 |
| 0.910262578 | 0.967357799 |
| 0.910341812 | 0.967370147 |
| 0.910495718 | 0.967438018 |
| 0.910533636 | 0.967438018 |
| 0.910536736 | 0.967438018 |
| 0.910541848 | 0.967438018 |
| 0.910628878 | 0.96748732 |
| 0.910702241 | 0.96754368 |
| 0.910888107 | 0.967633221 |
| 0.911008187 | 0.96769603 |
| 0.911180803 | 0.967793049 |
| 0.911233307 | 0.967827232 |
| 0.911449852 | 0.967856992 |
| 0.911474123 | 0.967856992 |
| 0.911536915 | 0.967856992 |
| 0.91155319 | 0.967856992 |
| 0.911656399 | 0.967856992 |
| 0.911662532 | 0.967856992 |
| 0.911708402 | 0.967856992 |

| | |
|---|---|
| 0.911741315 | 0.967858933 |
| 0.911768597 | 0.967858933 |
| 0.911846812 | 0.967896056 |
| 0.91194404 | 0.967977687 |
| 0.912267587 | 0.968105751 |
| 0.912280524 | 0.968105751 |
| 0.912488299 | 0.968202375 |
| 0.912531044 | 0.968202375 |
| 0.912535864 | 0.968202375 |
| 0.912557975 | 0.968202375 |
| 0.912562299 | 0.968202375 |
| 0.91260372 | 0.968217443 |
| 0.912637489 | 0.968217443 |
| 0.912734992 | 0.968217864 |
| 0.912758511 | 0.968217864 |
| 0.912767388 | 0.968217864 |
| 0.912768117 | 0.968217864 |
| 0.912780189 | 0.968217864 |
| 0.912802802 | 0.968220286 |
| 0.912931318 | 0.968291909 |
| 0.913122988 | 0.968408937 |
| 0.913345221 | 0.968601488 |
| 0.913601238 | 0.968756099 |
| 0.913741981 | 0.968801185 |
| 0.913832256 | 0.968801185 |
| 0.913966704 | 0.968801185 |
| 0.914011481 | 0.968801185 |
| 0.91402172 | 0.968801185 |
| 0.914236827 | 0.968935144 |
| 0.914331516 | 0.968936589 |
| 0.914358738 | 0.968936589 |
| 0.914373254 | 0.968936589 |
| 0.914504312 | 0.96898923 |
| 0.914603954 | 0.969073249 |
| 0.914650153 | 0.96907908 |
| 0.914719286 | 0.969085435 |
| 0.914728748 | 0.969085435 |
| 0.914823814 | 0.969085435 |
| 0.914861632 | 0.969085435 |
| 0.91489312 | 0.969085435 |
| 0.914909637 | 0.969085435 |
| 0.914957578 | 0.969102979 |
| 0.915010766 | 0.969137277 |
| 0.915030656 | 0.969137277 |
| 0.915105909 | 0.969195427 |

| | |
|---|---|
| 0.915173771 | 0.969219488 |
| 0.915208536 | 0.969219488 |
| 0.915223362 | 0.969219488 |
| 0.915226506 | 0.969219488 |
| 0.91562759 | 0.969510788 |
| 0.915696498 | 0.969521598 |
| 0.915892917 | 0.969632361 |
| 0.915996932 | 0.969702214 |
| 0.916022094 | 0.969702214 |
| 0.916063357 | 0.969713493 |
| 0.916091224 | 0.96971984 |
| 0.916126209 | 0.96971984 |
| 0.916130434 | 0.96971984 |
| 0.916151602 | 0.969720694 |
| 0.916382196 | 0.969835448 |
| 0.916628167 | 0.969964226 |
| 0.916702368 | 0.969980309 |
| 0.916799901 | 0.970041753 |
| 0.916803969 | 0.970041753 |
| 0.91694028 | 0.970057805 |
| 0.917018958 | 0.970057805 |
| 0.91702456 | 0.970057805 |
| 0.917040073 | 0.970057805 |
| 0.917042127 | 0.970057805 |
| 0.917075182 | 0.970057805 |
| 0.917101492 | 0.970057805 |
| 0.917153502 | 0.970069729 |
| 0.917263873 | 0.970104402 |
| 0.917345917 | 0.970104402 |
| 0.917374499 | 0.97010958 |
| 0.917412579 | 0.970128309 |
| 0.917512999 | 0.970144855 |
| 0.917517987 | 0.970144855 |
| 0.917530074 | 0.970144855 |
| 0.917607253 | 0.970174267 |
| 0.917651137 | 0.970186705 |
| 0.917697778 | 0.970192943 |
| 0.917864446 | 0.970299294 |
| 0.91793588 | 0.970312324 |
| 0.918241977 | 0.970502536 |
| 0.91829604 | 0.970502536 |
| 0.918296277 | 0.970502536 |
| 0.918592759 | 0.970602753 |
| 0.918594383 | 0.970602753 |
| 0.918634816 | 0.970602753 |

| | |
|---|---|
| 0.918640759 | 0.970602753 |
| 0.918643807 | 0.970602753 |
| 0.918697506 | 0.970602753 |
| 0.918839507 | 0.970615014 |
| 0.91889979 | 0.970615014 |
| 0.918902793 | 0.970615014 |
| 0.919007694 | 0.970615114 |
| 0.919023942 | 0.970615114 |
| 0.919046721 | 0.970615114 |
| 0.91905268 | 0.970615114 |
| 0.919054943 | 0.970615114 |
| 0.919105095 | 0.970625391 |
| 0.919208563 | 0.970669732 |
| 0.919451588 | 0.970788549 |
| 0.919468928 | 0.970788549 |
| 0.919543091 | 0.970813414 |
| 0.919556497 | 0.970813414 |
| 0.919770618 | 0.970921865 |
| 0.919776416 | 0.970921865 |
| 0.919793891 | 0.970921865 |
| 0.919926202 | 0.970967692 |
| 0.920013428 | 0.970967692 |
| 0.920035619 | 0.970967692 |
| 0.920038573 | 0.970967692 |
| 0.920251441 | 0.971125042 |
| 0.920439601 | 0.971242254 |
| 0.920444084 | 0.971242254 |
| 0.920643299 | 0.971430941 |
| 0.920687281 | 0.971455828 |
| 0.920799896 | 0.971500958 |
| 0.920861463 | 0.971500958 |
| 0.920956329 | 0.97150823 |
| 0.920987203 | 0.97150823 |
| 0.921071949 | 0.971517334 |
| 0.921110021 | 0.971520184 |
| 0.921198377 | 0.971520184 |
| 0.921199724 | 0.971520184 |
| 0.921217441 | 0.971520184 |
| 0.921352829 | 0.971641451 |
| 0.921381701 | 0.971650384 |
| 0.921497591 | 0.971720352 |
| 0.92158139 | 0.971774899 |
| 0.921638256 | 0.971784212 |
| 0.92165695 | 0.971784212 |
| 0.921695577 | 0.971784212 |

| | |
|---|---|
| 0.921817412 | 0.971794049 |
| 0.921852274 | 0.971794049 |
| 0.921865697 | 0.971794049 |
| 0.921877271 | 0.971794049 |
| 0.921885211 | 0.971794049 |
| 0.922107278 | 0.971884627 |
| 0.922165156 | 0.971884627 |
| 0.922226922 | 0.971884627 |
| 0.922231291 | 0.971884627 |
| 0.922242197 | 0.971884627 |
| 0.922259012 | 0.971884627 |
| 0.922266622 | 0.971884627 |
| 0.922320591 | 0.971884627 |
| 0.922341407 | 0.971884627 |
| 0.922358633 | 0.971884627 |
| 0.922407599 | 0.971884627 |
| 0.922464183 | 0.971884627 |
| 0.922484075 | 0.971884627 |
| 0.92255477 | 0.971884627 |
| 0.922572984 | 0.971884627 |
| 0.922635226 | 0.971884627 |
| 0.92264454 | 0.971884627 |
| 0.922742551 | 0.971942246 |
| 0.922947813 | 0.972096586 |
| 0.922999055 | 0.972129058 |
| 0.923031387 | 0.972131393 |
| 0.923162758 | 0.972131393 |
| 0.923206385 | 0.972131393 |
| 0.92321893 | 0.972131393 |
| 0.923225797 | 0.972131393 |
| 0.923344813 | 0.972170662 |
| 0.923380264 | 0.972170662 |
| 0.923408508 | 0.972170662 |
| 0.923445455 | 0.972170662 |
| 0.923454251 | 0.972170662 |
| 0.923456344 | 0.972170662 |
| 0.923581413 | 0.972235167 |
| 0.923589726 | 0.972235167 |
| 0.92372497 | 0.972291575 |
| 0.923814386 | 0.97231305 |
| 0.923827034 | 0.97231305 |
| 0.923888414 | 0.972356164 |
| 0.924058215 | 0.972491893 |
| 0.924115522 | 0.972509224 |
| 0.92427879 | 0.97259508 |

| | |
|---|---|
| 0.924302805 | 0.972598862 |
| 0.924471815 | 0.972737093 |
| 0.924579264 | 0.972782305 |
| 0.924624996 | 0.972808524 |
| 0.924645035 | 0.972808524 |
| 0.924714024 | 0.972838126 |
| 0.924745238 | 0.972849476 |
| 0.924909602 | 0.972937854 |
| 0.924912982 | 0.972937854 |
| 0.924998938 | 0.972937854 |
| 0.9250831 | 0.972968505 |
| 0.92522389 | 0.973023002 |
| 0.925249168 | 0.973023002 |
| 0.92531599 | 0.973063031 |
| 0.925389044 | 0.973106443 |
| 0.925400208 | 0.973106443 |
| 0.925418567 | 0.973106443 |
| 0.925519204 | 0.973147809 |
| 0.925655658 | 0.973263565 |
| 0.925801427 | 0.9733586 |
| 0.925974353 | 0.973450759 |
| 0.926075644 | 0.973496473 |
| 0.92610859 | 0.973509619 |
| 0.926150414 | 0.973510611 |
| 0.926187552 | 0.973527903 |
| 0.926246557 | 0.973527903 |
| 0.926251254 | 0.973527903 |
| 0.926289509 | 0.973527903 |
| 0.926373679 | 0.973534838 |
| 0.926420217 | 0.973534838 |
| 0.926420463 | 0.973534838 |
| 0.926422873 | 0.973534838 |
| 0.926439194 | 0.973534838 |
| 0.9265251 | 0.973603631 |
| 0.926657867 | 0.97369117 |
| 0.926706462 | 0.97369117 |
| 0.926712305 | 0.97369117 |
| 0.926876215 | 0.97369117 |
| 0.926924668 | 0.97369117 |
| 0.926935243 | 0.97369117 |
| 0.926983761 | 0.973698901 |
| 0.927019144 | 0.973714592 |
| 0.92709957 | 0.97375612 |
| 0.927362969 | 0.973902502 |
| 0.927461242 | 0.973964161 |

| | |
|---|---|
| 0.927505135 | 0.973988779 |
| 0.927664528 | 0.974024565 |
| 0.927679719 | 0.974024565 |
| 0.927680587 | 0.974024565 |
| 0.927740568 | 0.974024565 |
| 0.927779079 | 0.974040212 |
| 0.927810666 | 0.974051902 |
| 0.928047252 | 0.974239913 |
| 0.928181663 | 0.974281531 |
| 0.928193045 | 0.974281531 |
| 0.928256869 | 0.974296821 |
| 0.928288641 | 0.974296821 |
| 0.928315283 | 0.974296821 |
| 0.928383035 | 0.974330641 |
| 0.928410493 | 0.974337987 |
| 0.928439518 | 0.974346978 |
| 0.928704854 | 0.974555293 |
| 0.928761246 | 0.974555293 |
| 0.928779943 | 0.974555293 |
| 0.928801715 | 0.974555293 |
| 0.928860646 | 0.974595655 |
| 0.928993742 | 0.974614812 |
| 0.928999311 | 0.974614812 |
| 0.929052583 | 0.974614812 |
| 0.929071785 | 0.974614812 |
| 0.92908968 | 0.974614812 |
| 0.929158178 | 0.974620344 |
| 0.929249504 | 0.974638643 |
| 0.929307118 | 0.97464138 |
| 0.92933096 | 0.97464138 |
| 0.92933397 | 0.97464138 |
| 0.929500128 | 0.974790047 |
| 0.929656252 | 0.974893505 |
| 0.929734193 | 0.9749537 |
| 0.9297769 | 0.974955623 |
| 0.929861912 | 0.975001834 |
| 0.92999243 | 0.975098471 |
| 0.929997885 | 0.975098471 |
| 0.93003597 | 0.975098471 |
| 0.930105056 | 0.975123813 |
| 0.930133224 | 0.975123813 |
| 0.930141328 | 0.975123813 |
| 0.930142038 | 0.975123813 |
| 0.930338506 | 0.97530601 |
| 0.930366159 | 0.97530601 |

| | |
|---|---|
| 0.930439053 | 0.975349315 |
| 0.930489444 | 0.97538067 |
| 0.930533286 | 0.975405159 |
| 0.930690927 | 0.975505992 |
| 0.930690927 | 0.975505992 |
| 0.930765657 | 0.975541382 |
| 0.930892024 | 0.975609419 |
| 0.930951507 | 0.975634104 |
| 0.931030305 | 0.975634104 |
| 0.931032729 | 0.975634104 |
| 0.931075891 | 0.975634104 |
| 0.931094123 | 0.975634104 |
| 0.931127607 | 0.975641627 |
| 0.93115654 | 0.975650479 |
| 0.931247142 | 0.975723945 |
| 0.93127471 | 0.975731364 |
| 0.931313448 | 0.97573892 |
| 0.931322896 | 0.97573892 |
| 0.931366401 | 0.975763035 |
| 0.931448609 | 0.975827697 |
| 0.931545123 | 0.975864411 |
| 0.931628258 | 0.975930035 |
| 0.931662169 | 0.975944093 |
| 0.931876921 | 0.97596949 |
| 0.931889167 | 0.97596949 |
| 0.931899467 | 0.97596949 |
| 0.931900934 | 0.97596949 |
| 0.931941941 | 0.97596949 |
| 0.931988632 | 0.975985545 |
| 0.932019004 | 0.975995891 |
| 0.932129491 | 0.97604721 |
| 0.932205629 | 0.976062557 |
| 0.932338639 | 0.976158903 |
| 0.932419493 | 0.976188417 |
| 0.932448814 | 0.976188417 |
| 0.932534712 | 0.976239274 |
| 0.932557148 | 0.976239274 |
| 0.932590145 | 0.976250444 |
| 0.932610551 | 0.976250444 |
| 0.932721927 | 0.976278599 |
| 0.932739939 | 0.976278599 |
| 0.932778277 | 0.97629727 |
| 0.932899909 | 0.976358448 |
| 0.932926631 | 0.976358448 |
| 0.93293792 | 0.976358448 |

| | |
|---|---|
| 0.932967712 | 0.976358448 |
| 0.933025935 | 0.976363374 |
| 0.933147931 | 0.976448127 |
| 0.933184492 | 0.976464931 |
| 0.933250711 | 0.976498855 |
| 0.933304621 | 0.976526268 |
| 0.933472011 | 0.976594136 |
| 0.933601375 | 0.976666279 |
| 0.933644363 | 0.976672167 |
| 0.933716042 | 0.976690688 |
| 0.933930949 | 0.976829758 |
| 0.933933121 | 0.976829758 |
| 0.934040481 | 0.976829758 |
| 0.934045073 | 0.976829758 |
| 0.934068601 | 0.976829758 |
| 0.934086389 | 0.976829758 |
| 0.934092484 | 0.976829758 |
| 0.934105945 | 0.976829758 |
| 0.934211851 | 0.976860858 |
| 0.934343041 | 0.976906849 |
| 0.934406775 | 0.976906849 |
| 0.934427476 | 0.976907046 |
| 0.934448498 | 0.97690758 |
| 0.934576235 | 0.976966054 |
| 0.934578386 | 0.976966054 |
| 0.934586482 | 0.976966054 |
| 0.934666554 | 0.976981325 |
| 0.934796038 | 0.976981325 |
| 0.934801544 | 0.976981325 |
| 0.934831148 | 0.976985936 |
| 0.934939806 | 0.977078053 |
| 0.935033708 | 0.977111866 |
| 0.935165607 | 0.977206818 |
| 0.935305745 | 0.97728893 |
| 0.935394787 | 0.977344592 |
| 0.935400058 | 0.977344592 |
| 0.935453139 | 0.977378612 |
| 0.935473887 | 0.977378849 |
| 0.935573889 | 0.977419005 |
| 0.935744277 | 0.977554128 |
| 0.935808302 | 0.977563015 |
| 0.935855722 | 0.977563015 |
| 0.935863026 | 0.977563015 |
| 0.93586954 | 0.977563015 |
| 0.935890638 | 0.977563015 |

| | |
|---|---|
| 0.936027206 | 0.977678138 |
| 0.936303865 | 0.977881325 |
| 0.936372801 | 0.977931877 |
| 0.936489709 | 0.97800553 |
| 0.936504928 | 0.97800553 |
| 0.936567336 | 0.978041347 |
| 0.936587118 | 0.978041347 |
| 0.936627268 | 0.978041347 |
| 0.936791922 | 0.978041347 |
| 0.936829536 | 0.978041347 |
| 0.93685359 | 0.978041347 |
| 0.93690328 | 0.978041347 |
| 0.936973196 | 0.978041347 |
| 0.936991005 | 0.978041347 |
| 0.93704741 | 0.978055445 |
| 0.93709395 | 0.978055445 |
| 0.937136223 | 0.978055445 |
| 0.937165157 | 0.978055445 |
| 0.937219049 | 0.978055445 |
| 0.937230404 | 0.978055445 |
| 0.937408003 | 0.978173255 |
| 0.937425449 | 0.978173255 |
| 0.937473573 | 0.978202039 |
| 0.937674584 | 0.978283512 |
| 0.937736519 | 0.978283512 |
| 0.93785284 | 0.978362002 |
| 0.938100962 | 0.978533907 |
| 0.938107382 | 0.978533907 |
| 0.938120356 | 0.978533907 |
| 0.93816939 | 0.978563622 |
| 0.938305764 | 0.978617918 |
| 0.938331948 | 0.978617918 |
| 0.938369971 | 0.978617918 |
| 0.938385825 | 0.978617918 |
| 0.938540563 | 0.978757858 |
| 0.93870275 | 0.978905561 |
| 0.938738883 | 0.978921807 |
| 0.938779438 | 0.978942664 |
| 0.938879177 | 0.979025235 |
| 0.938940617 | 0.979045834 |
| 0.938952347 | 0.979045834 |
| 0.938960601 | 0.979045834 |
| 0.939024884 | 0.979091426 |
| 0.939189663 | 0.97913313 |
| 0.93920879 | 0.97913313 |

| | |
|---|---|
| 0.939295566 | 0.97920216 |
| 0.939328362 | 0.979214916 |
| 0.93942156 | 0.979269204 |
| 0.939485659 | 0.979293975 |
| 0.939500853 | 0.979293975 |
| 0.939555959 | 0.979302133 |
| 0.939661041 | 0.979316214 |
| 0.93978087 | 0.979316214 |
| 0.939850761 | 0.979316214 |
| 0.939921643 | 0.979324246 |
| 0.939988423 | 0.979324246 |
| 0.940021543 | 0.97933733 |
| 0.940149317 | 0.979441363 |
| 0.940200287 | 0.979441363 |
| 0.940207152 | 0.979441363 |
| 0.940221019 | 0.979441363 |
| 0.940224224 | 0.979441363 |
| 0.940311296 | 0.979446376 |
| 0.94037265 | 0.979488862 |
| 0.940396976 | 0.979492778 |
| 0.940498516 | 0.979512854 |
| 0.940694883 | 0.979649058 |
| 0.940711572 | 0.979649058 |
| 0.940788315 | 0.979682527 |
| 0.940792203 | 0.979682527 |
| 0.940825514 | 0.979682527 |
| 0.940867129 | 0.979682527 |
| 0.94089885 | 0.979694137 |
| 0.941056986 | 0.97969621 |
| 0.941057888 | 0.97969621 |
| 0.941076624 | 0.97969621 |
| 0.941123068 | 0.97969621 |
| 0.94113098 | 0.97969621 |
| 0.941198287 | 0.97969621 |
| 0.941206238 | 0.97969621 |
| 0.94122299 | 0.97969621 |
| 0.941240393 | 0.97969621 |
| 0.941258384 | 0.97969621 |
| 0.941311852 | 0.979698192 |
| 0.941387703 | 0.979722452 |
| 0.941416804 | 0.979722452 |
| 0.94149924 | 0.979722452 |
| 0.941602434 | 0.979722452 |
| 0.941620499 | 0.979722452 |
| 0.941633183 | 0.979722452 |

| | |
|---|---|
| 0.941740552 | 0.979722452 |
| 0.94178245 | 0.979722452 |
| 0.941782665 | 0.979722452 |
| 0.942026168 | 0.97978068 |
| 0.942027375 | 0.97978068 |
| 0.942068555 | 0.97978068 |
| 0.942077855 | 0.97978068 |
| 0.942128445 | 0.97978068 |
| 0.942139795 | 0.97978068 |
| 0.942173801 | 0.97978068 |
| 0.942195991 | 0.97978068 |
| 0.942213374 | 0.97978068 |
| 0.942299209 | 0.979845518 |
| 0.942325546 | 0.979851512 |
| 0.94248014 | 0.979869569 |
| 0.942549743 | 0.979876631 |
| 0.942613688 | 0.979876631 |
| 0.942672288 | 0.979876631 |
| 0.942676766 | 0.979876631 |
| 0.942678887 | 0.979876631 |
| 0.94272648 | 0.979884211 |
| 0.942759219 | 0.979884211 |
| 0.942791992 | 0.979884211 |
| 0.942823095 | 0.979884211 |
| 0.942829885 | 0.979884211 |
| 0.942870912 | 0.979884211 |
| 0.942871347 | 0.979884211 |
| 0.943031361 | 0.979991227 |
| 0.943082567 | 0.979991227 |
| 0.943097779 | 0.979991227 |
| 0.943139067 | 0.979991288 |
| 0.943142246 | 0.979991288 |
| 0.94320339 | 0.979998579 |
| 0.94323618 | 0.980003687 |
| 0.943303596 | 0.980003687 |
| 0.943315536 | 0.980003687 |
| 0.943658465 | 0.980167529 |
| 0.943716095 | 0.980184636 |
| 0.943853598 | 0.980272278 |
| 0.943862223 | 0.980272278 |
| 0.944021372 | 0.980337044 |
| 0.94403688 | 0.980337044 |
| 0.944068669 | 0.980337044 |
| 0.944108612 | 0.980350964 |
| 0.944246808 | 0.980350964 |

| | |
|---|---|
| 0.944247751 | 0.980350964 |
| 0.944258848 | 0.980350964 |
| 0.94439462 | 0.980350964 |
| 0.944419312 | 0.980350964 |
| 0.944608562 | 0.980448189 |
| 0.944628594 | 0.980448189 |
| 0.944852887 | 0.980616877 |
| 0.944945475 | 0.980645389 |
| 0.944959751 | 0.980645389 |
| 0.94496272 | 0.980645389 |
| 0.944991344 | 0.980653726 |
| 0.945024278 | 0.980666535 |
| 0.945145501 | 0.980686675 |
| 0.945167232 | 0.980686675 |
| 0.945278812 | 0.980759716 |
| 0.945330518 | 0.980768944 |
| 0.945349485 | 0.980768944 |
| 0.945447112 | 0.980789436 |
| 0.945451609 | 0.980789436 |
| 0.945495668 | 0.980795659 |
| 0.945498795 | 0.980795659 |
| 0.945553615 | 0.980831163 |
| 0.945611543 | 0.980856554 |
| 0.945703478 | 0.980901162 |
| 0.945852446 | 0.980976548 |
| 0.945936599 | 0.980976548 |
| 0.945939642 | 0.980976548 |
| 0.945961813 | 0.980976548 |
| 0.945966399 | 0.980976548 |
| 0.945997211 | 0.980976548 |
| 0.946083576 | 0.981004675 |
| 0.946106142 | 0.981004675 |
| 0.946170532 | 0.981024737 |
| 0.946273597 | 0.981086545 |
| 0.946313072 | 0.981106114 |
| 0.946478931 | 0.981192636 |
| 0.946574648 | 0.981244396 |
| 0.946590668 | 0.981244396 |
| 0.946815367 | 0.981405672 |
| 0.946817905 | 0.981405672 |
| 0.946873397 | 0.981411383 |
| 0.947046496 | 0.981495275 |
| 0.947173912 | 0.98157965 |
| 0.947257114 | 0.981593406 |
| 0.947289233 | 0.981605333 |

| | |
|---|---|
| 0.947331024 | 0.981605924 |
| 0.947390139 | 0.981633195 |
| 0.947460397 | 0.981633195 |
| 0.947550397 | 0.981654613 |
| 0.947605155 | 0.981654613 |
| 0.947605741 | 0.981654613 |
| 0.947625384 | 0.981654613 |
| 0.947681398 | 0.981654613 |
| 0.947687183 | 0.981654613 |
| 0.947839861 | 0.981722196 |
| 0.947877791 | 0.981722196 |
| 0.947891515 | 0.981722196 |
| 0.947896716 | 0.981722196 |
| 0.947965945 | 0.981772546 |
| 0.948013012 | 0.981799942 |
| 0.948054505 | 0.981821564 |
| 0.94811519 | 0.98184534 |
| 0.948207368 | 0.981873121 |
| 0.948284763 | 0.981916273 |
| 0.948300939 | 0.981916273 |
| 0.948310891 | 0.981916273 |
| 0.948367681 | 0.981933898 |
| 0.948379521 | 0.981933898 |
| 0.948410382 | 0.981933898 |
| 0.948462506 | 0.981945172 |
| 0.948498087 | 0.981960664 |
| 0.948634501 | 0.982059196 |
| 0.948676531 | 0.982060014 |
| 0.948724729 | 0.982088562 |
| 0.948774474 | 0.982097366 |
| 0.948817413 | 0.982120467 |
| 0.948859591 | 0.982142781 |
| 0.948896292 | 0.982159423 |
| 0.949048359 | 0.98225278 |
| 0.949075268 | 0.982254168 |
| 0.949175728 | 0.982283881 |
| 0.949257974 | 0.982302411 |
| 0.949262998 | 0.982302411 |
| 0.949324865 | 0.982302411 |
| 0.949327055 | 0.982302411 |
| 0.949455613 | 0.982321494 |
| 0.94949865 | 0.982321494 |
| 0.949506631 | 0.982321494 |
| 0.949529243 | 0.98232355 |
| 0.949590772 | 0.982331185 |

| | |
|---|---|
| 0.949610488 | 0.982331185 |
| 0.949696662 | 0.982331185 |
| 0.949751888 | 0.982331185 |
| 0.949785939 | 0.982331185 |
| 0.949815191 | 0.982331185 |
| 0.949868431 | 0.982331185 |
| 0.94989274 | 0.982331185 |
| 0.949921367 | 0.982331185 |
| 0.949944967 | 0.982331185 |
| 0.950015789 | 0.982331185 |
| 0.950022694 | 0.982331185 |
| 0.950058776 | 0.982331185 |
| 0.950084007 | 0.982331185 |
| 0.950114565 | 0.982331185 |
| 0.950128916 | 0.982331185 |
| 0.950145686 | 0.982331185 |
| 0.950145686 | 0.982331185 |
| 0.950170034 | 0.982331185 |
| 0.950176014 | 0.982331185 |
| 0.950220547 | 0.982334577 |
| 0.95033785 | 0.982421242 |
| 0.950381765 | 0.982421242 |
| 0.950632729 | 0.98249462 |
| 0.950643534 | 0.98249462 |
| 0.950673386 | 0.982504152 |
| 0.950743403 | 0.982533873 |
| 0.950818122 | 0.982589769 |
| 0.95088083 | 0.982594459 |
| 0.950886497 | 0.982594459 |
| 0.950905184 | 0.982594459 |
| 0.950968128 | 0.98263818 |
| 0.951016953 | 0.982645776 |
| 0.951037374 | 0.982645776 |
| 0.951102629 | 0.982653807 |
| 0.951169203 | 0.982653807 |
| 0.951185335 | 0.982653807 |
| 0.951238211 | 0.982653807 |
| 0.951277896 | 0.982653807 |
| 0.951303975 | 0.982653807 |
| 0.951308775 | 0.982653807 |
| 0.951317812 | 0.982653807 |
| 0.951334 | 0.982653807 |
| 0.951462263 | 0.982713758 |
| 0.951594514 | 0.98273108 |
| 0.951695439 | 0.982750069 |

| | |
|---|---|
| 0.951970726 | 0.982885164 |
| 0.95202294 | 0.982917767 |
| 0.952064997 | 0.982939881 |
| 0.95224326 | 0.983059993 |
| 0.952309977 | 0.98310756 |
| 0.952372616 | 0.983150914 |
| 0.952493068 | 0.983232636 |
| 0.952557952 | 0.983278302 |
| 0.952792496 | 0.983406262 |
| 0.952823309 | 0.983406262 |
| 0.952826451 | 0.983406262 |
| 0.95287027 | 0.983430176 |
| 0.952936762 | 0.983434868 |
| 0.953010282 | 0.983472751 |
| 0.953054233 | 0.983472751 |
| 0.953123494 | 0.983475101 |
| 0.953126826 | 0.983475101 |
| 0.953140944 | 0.983475101 |
| 0.953183001 | 0.98349719 |
| 0.953209906 | 0.983503644 |
| 0.953289057 | 0.983564002 |
| 0.953700344 | 0.983732621 |
| 0.953770811 | 0.983763433 |
| 0.953771527 | 0.983763433 |
| 0.953861712 | 0.983771233 |
| 0.953951257 | 0.983777604 |
| 0.953959539 | 0.983777604 |
| 0.954010331 | 0.983777604 |
| 0.954068226 | 0.983777604 |
| 0.954095104 | 0.983777604 |
| 0.954303894 | 0.983907684 |
| 0.95448574 | 0.983988665 |
| 0.954567259 | 0.984051404 |
| 0.95458841 | 0.984051908 |
| 0.954655203 | 0.984099462 |
| 0.954839384 | 0.984206054 |
| 0.955023043 | 0.984306794 |
| 0.955031357 | 0.984306794 |
| 0.95514567 | 0.984306794 |
| 0.955411124 | 0.984516446 |
| 0.955461024 | 0.984525263 |
| 0.955502402 | 0.984546599 |
| 0.955546984 | 0.984549936 |
| 0.955685224 | 0.984621725 |
| 0.95571427 | 0.984621725 |

| | |
|---|---|
| 0.955877415 | 0.984733711 |
| 0.956140191 | 0.98484156 |
| 0.956219217 | 0.984880358 |
| 0.956269683 | 0.984894774 |
| 0.956295941 | 0.984894774 |
| 0.95635729 | 0.984894774 |
| 0.956389029 | 0.984906164 |
| 0.956463188 | 0.984938774 |
| 0.956526968 | 0.984943163 |
| 0.956627035 | 0.984943163 |
| 0.956637403 | 0.984943163 |
| 0.956651907 | 0.984943163 |
| 0.956675329 | 0.984943163 |
| 0.956734043 | 0.984943163 |
| 0.956734065 | 0.984943163 |
| 0.956779617 | 0.984943163 |
| 0.956871734 | 0.984943163 |
| 0.956987192 | 0.984943163 |
| 0.956991635 | 0.984943163 |
| 0.957017791 | 0.984943163 |
| 0.957030608 | 0.984943163 |
| 0.957141774 | 0.984943163 |
| 0.957147458 | 0.984943163 |
| 0.957237898 | 0.984943163 |
| 0.957312734 | 0.984943163 |
| 0.957318391 | 0.984943163 |
| 0.957319905 | 0.984943163 |
| 0.95737061 | 0.984943163 |
| 0.957381545 | 0.984943163 |
| 0.9573973 | 0.984943163 |
| 0.957410632 | 0.984943163 |
| 0.957417725 | 0.984943163 |
| 0.957431305 | 0.984943163 |
| 0.957519198 | 0.985000826 |
| 0.957547894 | 0.985000826 |
| 0.957682433 | 0.985053899 |
| 0.957691275 | 0.985053899 |
| 0.957722329 | 0.985053899 |
| 0.957729706 | 0.985053899 |
| 0.95775361 | 0.985053899 |
| 0.95777286 | 0.985053899 |
| 0.957915137 | 0.985135804 |
| 0.958046183 | 0.985195362 |
| 0.958080603 | 0.985195362 |
| 0.95808723 | 0.985195362 |

| | |
|---|---|
| 0.95812173 | 0.985195362 |
| 0.958171764 | 0.985195362 |
| 0.958182521 | 0.985195362 |
| 0.958224728 | 0.985195362 |
| 0.958235633 | 0.985195362 |
| 0.958241962 | 0.985195362 |
| 0.958345779 | 0.985233502 |
| 0.95836213 | 0.985233502 |
| 0.958378328 | 0.985233502 |
| 0.958382492 | 0.985233502 |
| 0.958567298 | 0.985376236 |
| 0.95861591 | 0.985376236 |
| 0.958635491 | 0.985376236 |
| 0.958678205 | 0.985376236 |
| 0.958868868 | 0.985499516 |
| 0.958990566 | 0.985593921 |
| 0.959069761 | 0.985609775 |
| 0.959070633 | 0.985609775 |
| 0.959147095 | 0.985638643 |
| 0.959168917 | 0.985638643 |
| 0.959224984 | 0.985664095 |
| 0.959235955 | 0.985664095 |
| 0.959294062 | 0.985695111 |
| 0.95933216 | 0.985695111 |
| 0.959344191 | 0.985695111 |
| 0.959465732 | 0.98575133 |
| 0.95951036 | 0.985775916 |
| 0.959653284 | 0.985850437 |
| 0.959665693 | 0.985850437 |
| 0.959802135 | 0.985919036 |
| 0.959912273 | 0.985944521 |
| 0.960036693 | 0.986040145 |
| 0.960063649 | 0.986046567 |
| 0.960108847 | 0.986051896 |
| 0.960132525 | 0.986051896 |
| 0.960148445 | 0.986051896 |
| 0.960230493 | 0.986111601 |
| 0.960319276 | 0.986160249 |
| 0.960409229 | 0.986197737 |
| 0.960470566 | 0.986230551 |
| 0.960625893 | 0.986287017 |
| 0.960717321 | 0.986287017 |
| 0.960735019 | 0.986287017 |
| 0.960738923 | 0.986287017 |
| 0.960760072 | 0.986287017 |

| | |
|---|---|
| 0.96077406 | 0.986287017 |
| 0.960803656 | 0.98629614 |
| 0.960851865 | 0.986303112 |
| 0.96105969 | 0.986430615 |
| 0.961061128 | 0.986430615 |
| 0.96109993 | 0.986430615 |
| 0.961105392 | 0.986430615 |
| 0.961146806 | 0.986430615 |
| 0.961199325 | 0.986447169 |
| 0.961222972 | 0.986450181 |
| 0.961328088 | 0.9865204 |
| 0.961340095 | 0.9865204 |
| 0.961357348 | 0.9865204 |
| 0.961374249 | 0.9865204 |
| 0.961422336 | 0.986527234 |
| 0.961470842 | 0.986549343 |
| 0.961588235 | 0.986631751 |
| 0.961627279 | 0.986631751 |
| 0.961627771 | 0.986631751 |
| 0.961708287 | 0.986648485 |
| 0.961737165 | 0.986648485 |
| 0.961737882 | 0.986648485 |
| 0.961745716 | 0.986648485 |
| 0.961942437 | 0.986784536 |
| 0.962179631 | 0.986953847 |
| 0.962310808 | 0.987007532 |
| 0.962325609 | 0.987007532 |
| 0.962368964 | 0.987025155 |
| 0.962448841 | 0.987025155 |
| 0.962459623 | 0.987025155 |
| 0.962461055 | 0.987025155 |
| 0.962479079 | 0.987025155 |
| 0.962613387 | 0.987111374 |
| 0.962687012 | 0.987144365 |
| 0.96273402 | 0.987171314 |
| 0.962898655 | 0.987318871 |
| 0.963050237 | 0.987345382 |
| 0.963101999 | 0.987345382 |
| 0.963110842 | 0.987345382 |
| 0.963118753 | 0.987345382 |
| 0.963154497 | 0.987345382 |
| 0.963175682 | 0.987345382 |
| 0.963223346 | 0.987345382 |
| 0.963227903 | 0.987345382 |
| 0.963250217 | 0.987345382 |

| | |
|---|---|
| 0.963256203 | 0.987345382 |
| 0.963610632 | 0.987559898 |
| 0.963725643 | 0.987612958 |
| 0.963745351 | 0.987612958 |
| 0.963798351 | 0.987646019 |
| 0.963890709 | 0.987719411 |
| 0.963957256 | 0.98776015 |
| 0.963992683 | 0.98776015 |
| 0.964058193 | 0.9877673 |
| 0.964058908 | 0.9877673 |
| 0.96406188 | 0.9877673 |
| 0.964217414 | 0.987862903 |
| 0.964413111 | 0.988038572 |
| 0.964473641 | 0.988061649 |
| 0.964516992 | 0.988084807 |
| 0.964593603 | 0.988132636 |
| 0.964636903 | 0.988132636 |
| 0.964665427 | 0.988132636 |
| 0.96479308 | 0.988159142 |
| 0.964902664 | 0.988224828 |
| 0.964945059 | 0.988225045 |
| 0.965006265 | 0.988238802 |
| 0.965060704 | 0.988253951 |
| 0.965119192 | 0.988276574 |
| 0.965253984 | 0.9883721 |
| 0.965402395 | 0.988460312 |
| 0.965562066 | 0.988577237 |
| 0.965690012 | 0.988577237 |
| 0.965697844 | 0.988577237 |
| 0.965732806 | 0.988577237 |
| 0.965745045 | 0.988577237 |
| 0.965924085 | 0.988596878 |
| 0.965966132 | 0.988596878 |
| 0.966052076 | 0.988596878 |
| 0.966079526 | 0.988596878 |
| 0.966393191 | 0.988596878 |
| 0.966449346 | 0.988596878 |
| 0.966449346 | 0.988596878 |
| 0.966449346 | 0.988596878 |
| 0.966449346 | 0.988596878 |
| 0.966449346 | 0.988596878 |
| 0.966449346 | 0.988596878 |
| 0.966449346 | 0.988596878 |
| 0.966449346 | 0.988596878 |
| 0.966449346 | 0.988596878 |

| | |
|---|---|
| 0.966449346 | 0.988596878 |
| 0.966449346 | 0.988596878 |
| 0.966449346 | 0.988596878 |
| 0.966449346 | 0.988596878 |
| 0.966449346 | 0.988596878 |
| 0.966449346 | 0.988596878 |
| 0.966449346 | 0.988596878 |
| 0.966449346 | 0.988596878 |
| 0.966449346 | 0.988596878 |
| 0.966449346 | 0.988596878 |
| 0.966449346 | 0.988596878 |
| 0.966644377 | 0.988711703 |
| 0.966704213 | 0.988725423 |
| 0.966722799 | 0.988725423 |
| 0.96677847 | 0.988725423 |
| 0.966782349 | 0.988725423 |
| 0.966895764 | 0.988756485 |
| 0.966935165 | 0.988775547 |
| 0.967004286 | 0.988806706 |
| 0.967013989 | 0.988806706 |
| 0.96702792 | 0.988806706 |
| 0.967281899 | 0.988917785 |
| 0.967343089 | 0.988937886 |
| 0.967390534 | 0.988965161 |
| 0.967471981 | 0.989027196 |
| 0.967544316 | 0.989033073 |
| 0.967552105 | 0.989033073 |
| 0.967560795 | 0.989033073 |
| 0.967613539 | 0.98906576 |
| 0.967716152 | 0.98910985 |
| 0.967739744 | 0.98910985 |
| 0.967764105 | 0.989113523 |
| 0.967833295 | 0.989157397 |
| 0.967848571 | 0.989157397 |
| 0.967933468 | 0.989189602 |
| 0.96796316 | 0.989189602 |
| 0.967987018 | 0.989192759 |
| 0.968049276 | 0.989213931 |
| 0.968079049 | 0.98922313 |
| 0.968124972 | 0.989227608 |
| 0.968181847 | 0.989264499 |
| 0.968208849 | 0.989270865 |
| 0.968319085 | 0.989341051 |
| 0.968389932 | 0.989356156 |
| 0.968393728 | 0.989356156 |

| | |
|---|---|
| 0.96846985 | 0.989362866 |
| 0.968527395 | 0.989362866 |
| 0.9686575 | 0.9894672 |
| 0.968671083 | 0.9894672 |
| 0.968736449 | 0.9894862 |
| 0.968760188 | 0.9894862 |
| 0.968772787 | 0.9894862 |
| 0.968906901 | 0.989587208 |
| 0.968913236 | 0.989587208 |
| 0.969011907 | 0.989609147 |
| 0.969017113 | 0.989609147 |
| 0.969029209 | 0.989609147 |
| 0.969043052 | 0.989609147 |
| 0.96908157 | 0.989609147 |
| 0.969097928 | 0.989609147 |
| 0.96910421 | 0.989609147 |
| 0.969148185 | 0.989609147 |
| 0.969201463 | 0.989609147 |
| 0.969226519 | 0.989609147 |
| 0.969255728 | 0.989609147 |
| 0.969287948 | 0.989609147 |
| 0.969344587 | 0.989627051 |
| 0.969347333 | 0.989627051 |
| 0.96936782 | 0.989627051 |
| 0.969415085 | 0.989631666 |
| 0.969419794 | 0.989631666 |
| 0.969465125 | 0.98963361 |
| 0.969534385 | 0.989669813 |
| 0.969593631 | 0.989678124 |
| 0.969656306 | 0.989678124 |
| 0.969675756 | 0.989678124 |
| 0.969727267 | 0.989678124 |
| 0.9697991 | 0.989685623 |
| 0.969820012 | 0.989685623 |
| 0.969884111 | 0.989723797 |
| 0.969905387 | 0.989723797 |
| 0.969933654 | 0.989723797 |
| 0.969974548 | 0.989723797 |
| 0.969986131 | 0.989723797 |
| 0.970084756 | 0.989786128 |
| 0.970167242 | 0.989827879 |
| 0.970218415 | 0.989831055 |
| 0.970293922 | 0.989831055 |
| 0.970315836 | 0.989831055 |
| 0.970491859 | 0.989919707 |

| | |
|---|---|
| 0.970493448 | 0.989919707 |
| 0.970539634 | 0.989924367 |
| 0.970795472 | 0.990021622 |
| 0.970821801 | 0.990021622 |
| 0.970830097 | 0.990021622 |
| 0.970854554 | 0.990021622 |
| 0.970857872 | 0.990021622 |
| 0.970906954 | 0.990021622 |
| 0.970914418 | 0.990021622 |
| 0.971307654 | 0.990273613 |
| 0.97134867 | 0.990290759 |
| 0.97138287 | 0.990304428 |
| 0.971413669 | 0.990314628 |
| 0.971454218 | 0.990334769 |
| 0.97158826 | 0.990407817 |
| 0.971644262 | 0.990443706 |
| 0.971739329 | 0.990478386 |
| 0.971742866 | 0.990478386 |
| 0.971800781 | 0.990478386 |
| 0.97182386 | 0.990478386 |
| 0.971873501 | 0.990481713 |
| 0.97190334 | 0.990481713 |
| 0.971910311 | 0.990481713 |
| 0.972167763 | 0.990659527 |
| 0.972167992 | 0.990659527 |
| 0.972254796 | 0.990726785 |
| 0.972380114 | 0.990796499 |
| 0.972429143 | 0.990819648 |
| 0.972485249 | 0.990829537 |
| 0.972507526 | 0.990829537 |
| 0.972666379 | 0.990911737 |
| 0.972697434 | 0.990911737 |
| 0.972706774 | 0.990911737 |
| 0.972778385 | 0.990949495 |
| 0.972797113 | 0.990949495 |
| 0.972835867 | 0.990956406 |
| 0.972884576 | 0.990956406 |
| 0.972894655 | 0.990956406 |
| 0.972919509 | 0.990956406 |
| 0.972967565 | 0.990956406 |
| 0.972972605 | 0.990956406 |
| 0.973090074 | 0.991026643 |
| 0.973134926 | 0.991043355 |
| 0.9731481 | 0.991043355 |
| 0.973169952 | 0.991044418 |

| | |
|---|---|
| 0.973583176 | 0.991251936 |
| 0.973611326 | 0.991251936 |
| 0.973638745 | 0.991251936 |
| 0.973723508 | 0.991251936 |
| 0.973758894 | 0.991251936 |
| 0.973777317 | 0.991251936 |
| 0.97385573 | 0.991281263 |
| 0.973860421 | 0.991281263 |
| 0.973904531 | 0.99128379 |
| 0.974070691 | 0.99141054 |
| 0.974145639 | 0.991423261 |
| 0.974266711 | 0.991504106 |
| 0.974404351 | 0.991613428 |
| 0.974415774 | 0.991613428 |
| 0.974465832 | 0.991643181 |
| 0.974492531 | 0.991649163 |
| 0.974622056 | 0.991756754 |
| 0.974719196 | 0.991759558 |
| 0.97471957 | 0.991759558 |
| 0.974741587 | 0.991759558 |
| 0.974798825 | 0.991771024 |
| 0.974820521 | 0.991771024 |
| 0.974936129 | 0.991827523 |
| 0.975033902 | 0.991869222 |
| 0.975213549 | 0.991953639 |
| 0.975279697 | 0.991953639 |
| 0.975282073 | 0.991953639 |
| 0.975405976 | 0.991998715 |
| 0.975440062 | 0.991998715 |
| 0.975511839 | 0.992008164 |
| 0.975549109 | 0.992010601 |
| 0.975570317 | 0.992010601 |
| 0.975576722 | 0.992010601 |
| 0.975662224 | 0.992068369 |
| 0.975675192 | 0.992068369 |
| 0.975699209 | 0.992071609 |
| 0.975885619 | 0.99217642 |
| 0.976042794 | 0.992228653 |
| 0.976061994 | 0.992228653 |
| 0.97611774 | 0.992264142 |
| 0.97631244 | 0.992419699 |
| 0.976448191 | 0.992476813 |
| 0.976464482 | 0.992476813 |
| 0.976517596 | 0.992501142 |
| 0.976766422 | 0.992673152 |

| | |
|---|---|
| 0.976895964 | 0.992747366 |
| 0.976932542 | 0.992753396 |
| 0.977011142 | 0.992782055 |
| 0.977098303 | 0.992790755 |
| 0.977166999 | 0.992790755 |
| 0.97732944 | 0.992790755 |
| 0.977357921 | 0.992790755 |
| 0.977413384 | 0.992790755 |
| 0.977440316 | 0.992790755 |
| 0.977460087 | 0.992790755 |
| 0.977583689 | 0.992790755 |
| 0.977589068 | 0.992790755 |
| 0.977612632 | 0.992790755 |
| 0.97762387 | 0.992790755 |
| 0.977647826 | 0.992790755 |
| 0.977660944 | 0.992790755 |
| 0.977669448 | 0.992790755 |
| 0.977719093 | 0.992790755 |
| 0.977724047 | 0.992790755 |
| 0.977746696 | 0.992790755 |
| 0.978031055 | 0.992996875 |
| 0.978080219 | 0.993008603 |
| 0.978208841 | 0.993075681 |
| 0.978324624 | 0.993140935 |
| 0.978359523 | 0.993140935 |
| 0.978377381 | 0.993140935 |
| 0.978453605 | 0.993175973 |
| 0.978476254 | 0.993177796 |
| 0.978567228 | 0.993227801 |
| 0.978588725 | 0.993228454 |
| 0.978688315 | 0.993308366 |
| 0.978798543 | 0.993313272 |
| 0.978899908 | 0.993313272 |
| 0.978901709 | 0.993313272 |
| 0.979002917 | 0.993352477 |
| 0.979028117 | 0.993356884 |
| 0.979221685 | 0.993384211 |
| 0.979292074 | 0.993384211 |
| 0.979302769 | 0.993384211 |
| 0.979348698 | 0.993384211 |
| 0.979376387 | 0.993384211 |
| 0.979384628 | 0.993384211 |
| 0.979388771 | 0.993384211 |
| 0.97943721 | 0.993412185 |
| 0.979467221 | 0.993421468 |

| | |
|---|---|
| 0.979849996 | 0.99370389 |
| 0.979891207 | 0.993724523 |
| 0.97993318 | 0.993745929 |
| 0.9801796 | 0.993901764 |
| 0.980198953 | 0.993901764 |
| 0.980232928 | 0.993901764 |
| 0.980366781 | 0.99399516 |
| 0.980590178 | 0.994112798 |
| 0.980601437 | 0.994112798 |
| 0.980688992 | 0.994170409 |
| 0.980765733 | 0.994170409 |
| 0.980813121 | 0.994193728 |
| 0.980867241 | 0.994227426 |
| 0.980917044 | 0.994251274 |
| 0.980948171 | 0.994251274 |
| 0.981016023 | 0.994251274 |
| 0.981089242 | 0.994291655 |
| 0.981185929 | 0.994320848 |
| 0.981243627 | 0.994354991 |
| 0.981441866 | 0.994437652 |
| 0.981453915 | 0.994437652 |
| 0.981476228 | 0.994437652 |
| 0.981492236 | 0.994437652 |
| 0.981664787 | 0.99451085 |
| 0.98176202 | 0.994561997 |
| 0.981857214 | 0.994561997 |
| 0.981937871 | 0.994592948 |
| 0.98206599 | 0.994638109 |
| 0.982076206 | 0.994638109 |
| 0.982128646 | 0.994638109 |
| 0.982254982 | 0.99471978 |
| 0.982282732 | 0.99471978 |
| 0.982351281 | 0.99471978 |
| 0.982403966 | 0.99471978 |
| 0.982420765 | 0.99471978 |
| 0.982435985 | 0.99471978 |
| 0.982443172 | 0.99471978 |
| 0.982474301 | 0.99471978 |
| 0.982489347 | 0.99471978 |
| 0.982599122 | 0.99471978 |
| 0.982617854 | 0.99471978 |
| 0.982627002 | 0.99471978 |
| 0.982980289 | 0.994869108 |
| 0.983013397 | 0.994869108 |
| 0.983046068 | 0.994869108 |

| | |
|---|---|
| 0.983188068 | 0.99494939 |
| 0.983267897 | 0.994967213 |
| 0.983303873 | 0.994967213 |
| 0.983349263 | 0.994967213 |
| 0.98335754 | 0.994967213 |
| 0.983393697 | 0.994967213 |
| 0.983452942 | 0.994990728 |
| 0.983479613 | 0.994990728 |
| 0.983553291 | 0.995021973 |
| 0.983636777 | 0.995021973 |
| 0.983645079 | 0.995021973 |
| 0.983709716 | 0.995021973 |
| 0.9837533 | 0.995021973 |
| 0.983756219 | 0.995021973 |
| 0.983849667 | 0.995046493 |
| 0.984032766 | 0.995190975 |
| 0.984063613 | 0.995201038 |
| 0.984175129 | 0.995271549 |
| 0.984261039 | 0.995273589 |
| 0.98434554 | 0.995273589 |
| 0.984366551 | 0.995273589 |
| 0.984431705 | 0.995273589 |
| 0.98446568 | 0.995273589 |
| 0.984478702 | 0.995273589 |
| 0.98451763 | 0.995273589 |
| 0.984607494 | 0.995273589 |
| 0.98462645 | 0.995273589 |
| 0.984697276 | 0.995273589 |
| 0.984704395 | 0.995273589 |
| 0.984761548 | 0.995273589 |
| 0.98477444 | 0.995273589 |
| 0.984835924 | 0.995273589 |
| 0.984846647 | 0.995273589 |
| 0.984940363 | 0.995273589 |
| 0.98497655 | 0.995273589 |
| 0.984982895 | 0.995273589 |
| 0.984985509 | 0.995273589 |
| 0.98499076 | 0.995273589 |
| 0.98508774 | 0.995273589 |
| 0.985088209 | 0.995273589 |
| 0.98513842 | 0.995273589 |
| 0.985270805 | 0.995343681 |
| 0.985298735 | 0.995343681 |
| 0.985313077 | 0.995343681 |
| 0.985355372 | 0.995343681 |

| | |
|---|---|
| 0.985378897 | 0.995343681 |
| 0.985378956 | 0.995343681 |
| 0.985505235 | 0.995369692 |
| 0.98555882 | 0.995370818 |
| 0.985617693 | 0.995370818 |
| 0.985617693 | 0.995370818 |
| 0.985622121 | 0.995370818 |
| 0.985656709 | 0.995374922 |
| 0.98574005 | 0.995416873 |
| 0.985766382 | 0.995422358 |
| 0.985950035 | 0.995517802 |
| 0.986072889 | 0.995563002 |
| 0.986154532 | 0.995624325 |
| 0.986207719 | 0.995656916 |
| 0.986248783 | 0.995677268 |
| 0.98631464 | 0.995701543 |
| 0.986459588 | 0.995805655 |
| 0.986641858 | 0.995888404 |
| 0.986704475 | 0.995888404 |
| 0.986741641 | 0.995888404 |
| 0.986774797 | 0.995888404 |
| 0.986845577 | 0.995888404 |
| 0.986889909 | 0.995888404 |
| 0.986904997 | 0.995888404 |
| 0.986906516 | 0.995888404 |
| 0.986917944 | 0.995888404 |
| 0.987094895 | 0.995963012 |
| 0.987097185 | 0.995963012 |
| 0.987126984 | 0.995963012 |
| 0.987138261 | 0.995963012 |
| 0.987207515 | 0.995966948 |
| 0.987252803 | 0.995966948 |
| 0.987267633 | 0.995966948 |
| 0.987306679 | 0.995985241 |
| 0.987378704 | 0.996028965 |
| 0.987425201 | 0.996028965 |
| 0.987437815 | 0.996028965 |
| 0.987502281 | 0.996028965 |
| 0.987542277 | 0.996028965 |
| 0.987559153 | 0.996028965 |
| 0.987655213 | 0.996071355 |
| 0.987673511 | 0.996071355 |
| 0.987673661 | 0.996071355 |
| 0.987684838 | 0.996071355 |
| 0.987820597 | 0.99616103 |

| | |
|---|---|
| 0.987828843 | 0.99616103 |
| 0.987836506 | 0.99616103 |
| 0.987909348 | 0.9961923 |
| 0.98798968 | 0.996217625 |
| 0.988192689 | 0.996260347 |
| 0.988215924 | 0.996260347 |
| 0.988218065 | 0.996260347 |
| 0.988281691 | 0.996260347 |
| 0.988307848 | 0.996260347 |
| 0.988344881 | 0.996260347 |
| 0.988446393 | 0.996260347 |
| 0.988450993 | 0.996260347 |
| 0.988470791 | 0.996260347 |
| 0.988570561 | 0.996260347 |
| 0.988581626 | 0.996260347 |
| 0.988633734 | 0.996260347 |
| 0.988712102 | 0.996260347 |
| 0.988778379 | 0.996260347 |
| 0.988813523 | 0.996260347 |
| 0.988868461 | 0.996260347 |
| 0.988876001 | 0.996260347 |
| 0.988884249 | 0.996260347 |
| 0.988922153 | 0.996260347 |
| 0.989023651 | 0.996260347 |
| 0.98902758 | 0.996260347 |
| 0.989059827 | 0.996260347 |
| 0.989064564 | 0.996260347 |
| 0.989195099 | 0.996267392 |
| 0.989197067 | 0.996267392 |
| 0.989233228 | 0.996282744 |
| 0.98925975 | 0.996288387 |
| 0.989326875 | 0.996300513 |
| 0.989496729 | 0.996391685 |
| 0.989508764 | 0.996391685 |
| 0.989542769 | 0.996403758 |
| 0.989675882 | 0.996468615 |
| 0.98972074 | 0.996468615 |
| 0.98973162 | 0.996468615 |
| 0.98987499 | 0.996526037 |
| 0.989914194 | 0.996526037 |
| 0.990133947 | 0.996578742 |
| 0.990224432 | 0.996591176 |
| 0.990239085 | 0.996591176 |
| 0.990373852 | 0.996668369 |
| 0.990374643 | 0.996668369 |

| | |
|---|---|
| 0.990390407 | 0.996668369 |
| 0.990439642 | 0.996696856 |
| 0.990480856 | 0.996717272 |
| 0.990546438 | 0.996741146 |
| 0.990723347 | 0.996888804 |
| 0.990865311 | 0.996935636 |
| 0.990891049 | 0.99694047 |
| 0.991185667 | 0.99709982 |
| 0.991216481 | 0.99709982 |
| 0.991216916 | 0.99709982 |
| 0.991263652 | 0.997104713 |
| 0.991356339 | 0.997132848 |
| 0.991425625 | 0.997132848 |
| 0.991438185 | 0.997132848 |
| 0.991450885 | 0.997132848 |
| 0.99149489 | 0.997132848 |
| 0.991500985 | 0.997132848 |
| 0.991751515 | 0.997300566 |
| 0.991888789 | 0.997388951 |
| 0.991898669 | 0.997388951 |
| 0.991979754 | 0.997424784 |
| 0.992009531 | 0.997430495 |
| 0.992078289 | 0.997430495 |
| 0.992096332 | 0.997430495 |
| 0.992125637 | 0.997430495 |
| 0.992132032 | 0.997430495 |
| 0.992162587 | 0.997440159 |
| 0.992428819 | 0.997612803 |
| 0.992438631 | 0.997612803 |
| 0.992439049 | 0.997612803 |
| 0.992661046 | 0.99777278 |
| 0.992776026 | 0.997790669 |
| 0.992853838 | 0.997790669 |
| 0.992858413 | 0.997790669 |
| 0.992882489 | 0.997790669 |
| 0.992917637 | 0.997790669 |
| 0.992931303 | 0.997790669 |
| 0.992974092 | 0.997790669 |
| 0.993009384 | 0.997790669 |
| 0.993027625 | 0.997790669 |
| 0.993147451 | 0.997861559 |
| 0.993326141 | 0.997941742 |
| 0.993371019 | 0.99795987 |
| 0.99357605 | 0.997973589 |
| 0.993600929 | 0.997973589 |

| | |
|---|---|
| 0.993616689 | 0.997973589 |
| 0.993773176 | 0.997973589 |
| 0.993830249 | 0.997973589 |
| 0.993831058 | 0.997973589 |
| 0.99384594 | 0.997973589 |
| 0.993853796 | 0.997973589 |
| 0.993873551 | 0.997973589 |
| 0.99388757 | 0.997973589 |
| 0.993961214 | 0.997992207 |
| 0.993972594 | 0.997992207 |
| 0.993989928 | 0.997992207 |
| 0.99403215 | 0.998006786 |
| 0.994187119 | 0.998021877 |
| 0.994294324 | 0.998061054 |
| 0.994309969 | 0.998061054 |
| 0.994347781 | 0.998062283 |
| 0.994353105 | 0.998062283 |
| 0.994414539 | 0.998090119 |
| 0.994425084 | 0.998090119 |
| 0.99450732 | 0.998090119 |
| 0.994527532 | 0.998090119 |
| 0.994555787 | 0.998090809 |
| 0.994570132 | 0.998090809 |
| 0.994700056 | 0.99820016 |
| 0.994788373 | 0.998267753 |
| 0.994934702 | 0.998351488 |
| 0.99501082 | 0.998385799 |
| 0.995186621 | 0.998498882 |
| 0.995205667 | 0.998498882 |
| 0.995244135 | 0.998498882 |
| 0.995244821 | 0.998498882 |
| 0.995272921 | 0.998498882 |
| 0.995275699 | 0.998498882 |
| 0.995302826 | 0.998498882 |
| 0.995384742 | 0.998498892 |
| 0.995508003 | 0.998534633 |
| 0.995560463 | 0.998550582 |
| 0.995573402 | 0.998550582 |
| 0.995635827 | 0.998552308 |
| 0.995681798 | 0.998554158 |
| 0.995776142 | 0.998610382 |
| 0.995802656 | 0.998612282 |
| 0.995867219 | 0.998615972 |
| 0.996069279 | 0.99875348 |
| 0.996301615 | 0.998839412 |

| | |
|---|---|
| 0.996440992 | 0.99891368 |
| 0.996448015 | 0.99891368 |
| 0.996472415 | 0.99891368 |
| 0.996501234 | 0.99891368 |
| 0.996501706 | 0.99891368 |
| 0.996589694 | 0.99895983 |
| 0.996759547 | 0.999045982 |
| 0.996856203 | 0.999049484 |
| 0.996873425 | 0.999049484 |
| 0.996920916 | 0.999049484 |
| 0.996968615 | 0.999049484 |
| 0.997013312 | 0.999049484 |
| 0.997021354 | 0.999049484 |
| 0.997046972 | 0.999049484 |
| 0.997053227 | 0.999049484 |
| 0.997141321 | 0.999049484 |
| 0.997260656 | 0.999065119 |
| 0.997278493 | 0.999065119 |
| 0.997318246 | 0.999065119 |
| 0.99732697 | 0.999065119 |
| 0.997488945 | 0.999107496 |
| 0.997586909 | 0.999168552 |
| 0.997612023 | 0.999168552 |
| 0.997666707 | 0.999168552 |
| 0.997679191 | 0.999168552 |
| 0.997700019 | 0.999168552 |
| 0.997864535 | 0.999261241 |
| 0.997948193 | 0.999269059 |
| 0.998282067 | 0.999409117 |
| 0.998300669 | 0.999409117 |
| 0.998346441 | 0.999409117 |
| 0.998346961 | 0.999409117 |
| 0.998353807 | 0.999409117 |
| 0.998374215 | 0.999409117 |
| 0.99838486 | 0.999409117 |
| 0.998516712 | 0.99948205 |
| 0.998704385 | 0.999606879 |
| 0.998785222 | 0.999660234 |
| 0.998900455 | 0.999698081 |
| 0.998958626 | 0.999735291 |
| 0.999016578 | 0.99977228 |
| 0.999095875 | 0.99980962 |
| 0.999156009 | 0.999827781 |
| 0.999208899 | 0.999832579 |
| 0.99927254 | 0.999832579 |

| | |
|---|---|
| 0.999321475 | 0.999832579 |
| 0.999339967 | 0.999832579 |
| 0.999373238 | 0.999832579 |
| 0.999382343 | 0.999832579 |
| 0.999391726 | 0.999832579 |
| 0.999433304 | 0.999846197 |
| 0.999476699 | 0.999854581 |
| 0.999538346 | 0.999895249 |
| 0.999676666 | 0.99996483 |
| 0.999749598 | 0.99996483 |
| 0.999771682 | 0.99996483 |
| 0.999778311 | 0.99996483 |
| 0.99981786 | 0.99996483 |
| 0.999860555 | 0.999983411 |
| 0.99987843 | 0.999983411 |
| 0.99990178 | 0.999985765 |
| 0.999993464 | 0.999993464 |
| 0.99998868 | 0.999993464 |
| 0.999963819 | 0.999993464 |
| 0.999935375 | 0.999993464 |
| 0.999806373 | 0.99996483 |
| 0.9997653 | 0.99996483 |
| 0.999709374 | 0.99996483 |
| 0.999672831 | 0.99996483 |
| 0.999609514 | 0.99994544 |
| 0.999447325 | 0.999846197 |
| 0.999375619 | 0.999832579 |
| 0.999364497 | 0.999832579 |
| 0.999361366 | 0.999832579 |
| 0.999189988 | 0.999832579 |
| 0.999148602 | 0.999827781 |
| 0.999055865 | 0.999790588 |
| 0.998896811 | 0.999698081 |
| 0.99882066 | 0.999660234 |
| 0.998815555 | 0.999660234 |
| 0.998700735 | 0.999606879 |
| 0.998661426 | 0.999605896 |
| 0.99850704 | 0.99948205 |
| 0.998401881 | 0.999409117 |
| 0.998352105 | 0.999409117 |
| 0.998179124 | 0.999375177 |
| 0.998119193 | 0.999336181 |
| 0.998061746 | 0.999299672 |
| 0.998048694 | 0.999299672 |
| 0.998041001 | 0.999299672 |

| | |
|---|---|
| 0.997968228 | 0.999269059 |
| 0.997938171 | 0.999269059 |
| 0.997897477 | 0.999261241 |
| 0.997886756 | 0.999261241 |
| 0.997869506 | 0.999261241 |
| 0.997729131 | 0.999176696 |
| 0.997688892 | 0.999168552 |
| 0.997654041 | 0.999168552 |
| 0.997492209 | 0.999107496 |
| 0.997482956 | 0.999107496 |
| 0.997424499 | 0.999102711 |
| 0.997414436 | 0.999102711 |
| 0.997345016 | 0.999065119 |
| 0.997327336 | 0.999065119 |
| 0.997203549 | 0.999049484 |
| 0.997187495 | 0.999049484 |
| 0.997178049 | 0.999049484 |
| 0.997176535 | 0.999049484 |
| 0.997174792 | 0.999049484 |
| 0.997123086 | 0.999049484 |
| 0.997112918 | 0.999049484 |
| 0.997071217 | 0.999049484 |
| 0.996929323 | 0.999049484 |
| 0.996840637 | 0.999049484 |
| 0.996825346 | 0.999049484 |
| 0.996800267 | 0.999049484 |
| 0.996755701 | 0.999045982 |
| 0.996730793 | 0.999045982 |
| 0.996619252 | 0.998968433 |
| 0.996574373 | 0.99895983 |
| 0.996410935 | 0.99891368 |
| 0.996301785 | 0.998839412 |
| 0.996285754 | 0.998839412 |
| 0.996228596 | 0.998829113 |
| 0.996220831 | 0.998829113 |
| 0.996205704 | 0.998829113 |
| 0.99618995 | 0.998829113 |
| 0.995998213 | 0.998703249 |
| 0.995963935 | 0.998689903 |
| 0.995869238 | 0.998615972 |
| 0.995865145 | 0.998615972 |
| 0.995779794 | 0.998610382 |
| 0.995663366 | 0.998554158 |
| 0.99563802 | 0.998552308 |
| 0.995626244 | 0.998552308 |

| | |
|---|---|
| 0.995515569 | 0.998534633 |
| 0.995439758 | 0.99850065 |
| 0.995417041 | 0.998498892 |
| 0.995403723 | 0.998498892 |
| 0.995401199 | 0.998498892 |
| 0.995379943 | 0.998498892 |
| 0.995312206 | 0.998498882 |
| 0.995280931 | 0.998498882 |
| 0.994965715 | 0.998361574 |
| 0.994844636 | 0.998282145 |
| 0.99482578 | 0.998282145 |
| 0.994519157 | 0.998090119 |
| 0.994515478 | 0.998090119 |
| 0.994514214 | 0.998090119 |
| 0.994259068 | 0.998052031 |
| 0.994210417 | 0.998024229 |
| 0.994186741 | 0.998021877 |
| 0.994185664 | 0.998021877 |
| 0.99415323 | 0.998021877 |
| 0.994106246 | 0.998021877 |
| 0.994105511 | 0.998021877 |
| 0.994046359 | 0.998006786 |
| 0.993966323 | 0.997992207 |
| 0.993837661 | 0.997973589 |
| 0.993831561 | 0.997973589 |
| 0.993795385 | 0.997973589 |
| 0.993761371 | 0.997973589 |
| 0.993714209 | 0.997973589 |
| 0.993704363 | 0.997973589 |
| 0.993629373 | 0.997973589 |
| 0.993622857 | 0.997973589 |
| 0.993614825 | 0.997973589 |
| 0.993563117 | 0.997973589 |
| 0.993499958 | 0.997973589 |
| 0.993498681 | 0.997973589 |
| 0.993425265 | 0.997972266 |
| 0.993406998 | 0.997972266 |
| 0.993332021 | 0.997941742 |
| 0.993291285 | 0.997941742 |
| 0.993264949 | 0.99793751 |
| 0.993200764 | 0.997894074 |
| 0.993055945 | 0.997790669 |
| 0.993042933 | 0.997790669 |
| 0.992990649 | 0.997790669 |
| 0.992988444 | 0.997790669 |

| | |
|---|---|
| 0.992939487 | 0.997790669 |
| 0.992858342 | 0.997790669 |
| 0.99280888 | 0.997790669 |
| 0.992785938 | 0.997790669 |
| 0.992693766 | 0.997784611 |
| 0.992589144 | 0.997721565 |
| 0.992470855 | 0.997623719 |
| 0.992319667 | 0.997555961 |
| 0.992191802 | 0.997448475 |
| 0.99212747 | 0.997430495 |
| 0.99209524 | 0.997430495 |
| 0.991923175 | 0.997388951 |
| 0.991921293 | 0.997388951 |
| 0.991603378 | 0.997172655 |
| 0.991565473 | 0.99715559 |
| 0.991545205 | 0.99715559 |
| 0.991462213 | 0.997132848 |
| 0.99142107 | 0.997132848 |
| 0.991374948 | 0.997132848 |
| 0.991362155 | 0.997132848 |
| 0.99124862 | 0.997104713 |
| 0.991187967 | 0.99709982 |
| 0.991093043 | 0.997059448 |
| 0.990966684 | 0.996953386 |
| 0.990946207 | 0.996953386 |
| 0.990941122 | 0.996953386 |
| 0.990838299 | 0.996929518 |
| 0.990795055 | 0.996907068 |
| 0.990774706 | 0.996907068 |
| 0.99073504 | 0.996888804 |
| 0.990538147 | 0.996741146 |
| 0.990250925 | 0.996591176 |
| 0.990231216 | 0.996591176 |
| 0.990203198 | 0.996591176 |
| 0.990118975 | 0.996578742 |
| 0.990057666 | 0.996544085 |
| 0.990043165 | 0.996544085 |
| 0.990025528 | 0.996544085 |
| 0.990012545 | 0.996544085 |
| 0.990011646 | 0.996544085 |
| 0.989998404 | 0.996544085 |
| 0.989908843 | 0.996526037 |
| 0.989872812 | 0.996526037 |
| 0.989822272 | 0.996517753 |
| 0.989793517 | 0.996509868 |

| | |
|---|---|
| 0.989720318 | 0.996468615 |
| 0.989583517 | 0.996403758 |
| 0.989568269 | 0.996403758 |
| 0.989444475 | 0.99636908 |
| 0.989433206 | 0.99636908 |
| 0.989334546 | 0.996300513 |
| 0.989333006 | 0.996300513 |
| 0.989196107 | 0.996267392 |
| 0.989164722 | 0.996267392 |
| 0.989143042 | 0.996267392 |
| 0.989121694 | 0.996267392 |
| 0.989052984 | 0.996260347 |
| 0.988994911 | 0.996260347 |
| 0.988981854 | 0.996260347 |
| 0.988979662 | 0.996260347 |
| 0.988932874 | 0.996260347 |
| 0.988900165 | 0.996260347 |
| 0.988854145 | 0.996260347 |
| 0.988788835 | 0.996260347 |
| 0.98878106 | 0.996260347 |
| 0.988766616 | 0.996260347 |
| 0.988683581 | 0.996260347 |
| 0.988673381 | 0.996260347 |
| 0.988665085 | 0.996260347 |
| 0.988648983 | 0.996260347 |
| 0.988599076 | 0.996260347 |
| 0.988558604 | 0.996260347 |
| 0.98843158 | 0.996260347 |
| 0.988398033 | 0.996260347 |
| 0.988382202 | 0.996260347 |
| 0.988341717 | 0.996260347 |
| 0.988309778 | 0.996260347 |
| 0.988297714 | 0.996260347 |
| 0.988281708 | 0.996260347 |
| 0.98821472 | 0.996260347 |
| 0.988208945 | 0.996260347 |
| 0.988150571 | 0.996260347 |
| 0.987997214 | 0.996217625 |
| 0.987981964 | 0.996217625 |
| 0.987862207 | 0.996165855 |
| 0.987538018 | 0.996028965 |
| 0.987492176 | 0.996028965 |
| 0.987480573 | 0.996028965 |
| 0.987441668 | 0.996028965 |
| 0.987264838 | 0.995966948 |

| | |
|---|---|
| 0.987249662 | 0.995966948 |
| 0.987192308 | 0.995966948 |
| 0.987084672 | 0.995963012 |
| 0.987008249 | 0.995937328 |
| 0.986946849 | 0.995896472 |
| 0.986908465 | 0.995888404 |
| 0.986884733 | 0.995888404 |
| 0.986854918 | 0.995888404 |
| 0.986791289 | 0.995888404 |
| 0.986757436 | 0.995888404 |
| 0.986698968 | 0.995888404 |
| 0.986628788 | 0.995888404 |
| 0.986592234 | 0.995888404 |
| 0.986544705 | 0.995870471 |
| 0.98634089 | 0.995706937 |
| 0.986294975 | 0.995701543 |
| 0.986013545 | 0.99552419 |
| 0.985986316 | 0.995517802 |
| 0.985974452 | 0.995517802 |
| 0.985875793 | 0.995469522 |
| 0.985866148 | 0.995469522 |
| 0.985802024 | 0.995437243 |
| 0.985704425 | 0.995402003 |
| 0.985631745 | 0.995370818 |
| 0.985623318 | 0.995370818 |
| 0.985497806 | 0.995369692 |
| 0.985475897 | 0.995369692 |
| 0.985442294 | 0.995369446 |
| 0.985395887 | 0.995343681 |
| 0.985370433 | 0.995343681 |
| 0.985287846 | 0.995343681 |
| 0.985134783 | 0.995273589 |
| 0.985102274 | 0.995273589 |
| 0.985058155 | 0.995273589 |
| 0.985043011 | 0.995273589 |
| 0.985010982 | 0.995273589 |
| 0.985008227 | 0.995273589 |
| 0.984944463 | 0.995273589 |
| 0.984922776 | 0.995273589 |
| 0.984903332 | 0.995273589 |
| 0.984889033 | 0.995273589 |
| 0.98487918 | 0.995273589 |
| 0.984870326 | 0.995273589 |
| 0.984814758 | 0.995273589 |
| 0.984587395 | 0.995273589 |

| | |
|---|---|
| 0.984545612 | 0.995273589 |
| 0.984516919 | 0.995273589 |
| 0.984487571 | 0.995273589 |
| 0.984442913 | 0.995273589 |
| 0.984420247 | 0.995273589 |
| 0.984412799 | 0.995273589 |
| 0.98433347 | 0.995273589 |
| 0.984274488 | 0.995273589 |
| 0.984256971 | 0.995273589 |
| 0.984097676 | 0.995214354 |
| 0.983869013 | 0.995046493 |
| 0.983814148 | 0.995033265 |
| 0.983782091 | 0.995021973 |
| 0.983762262 | 0.995021973 |
| 0.983739803 | 0.995021973 |
| 0.983723064 | 0.995021973 |
| 0.983637072 | 0.995021973 |
| 0.983616071 | 0.995021973 |
| 0.983584446 | 0.995021973 |
| 0.983478751 | 0.994990728 |
| 0.983388034 | 0.994967213 |
| 0.983380777 | 0.994967213 |
| 0.983372959 | 0.994967213 |
| 0.983274199 | 0.994967213 |
| 0.98314053 | 0.994922424 |
| 0.9831345 | 0.994922424 |
| 0.98304389 | 0.994869108 |
| 0.983041626 | 0.994869108 |
| 0.983023027 | 0.994869108 |
| 0.983000858 | 0.994869108 |
| 0.982969166 | 0.994869108 |
| 0.9828997 | 0.994869108 |
| 0.982860761 | 0.994869108 |
| 0.982734267 | 0.994764934 |
| 0.982695464 | 0.994746798 |
| 0.98265009 | 0.994722009 |
| 0.982606092 | 0.99471978 |
| 0.982600198 | 0.99471978 |
| 0.982565705 | 0.99471978 |
| 0.982544423 | 0.99471978 |
| 0.982505755 | 0.99471978 |
| 0.982484705 | 0.99471978 |
| 0.982403911 | 0.99471978 |
| 0.982149568 | 0.994638148 |
| 0.982124974 | 0.994638109 |

| | |
|---|---|
| 0.982124974 | 0.994638109 |
| 0.98208049 | 0.994638109 |
| 0.981959642 | 0.994593846 |
| 0.981900039 | 0.99457578 |
| 0.98186555 | 0.994561997 |
| 0.98186163 | 0.994561997 |
| 0.981815242 | 0.994561997 |
| 0.981814543 | 0.994561997 |
| 0.981689768 | 0.99451085 |
| 0.981678674 | 0.99451085 |
| 0.981591967 | 0.994475231 |
| 0.98157696 | 0.994475231 |
| 0.981564871 | 0.994475231 |
| 0.981481781 | 0.994437652 |
| 0.981377715 | 0.994427395 |
| 0.981322394 | 0.994392495 |
| 0.98126696 | 0.994357479 |
| 0.981189057 | 0.994320848 |
| 0.981160803 | 0.994320848 |
| 0.98109762 | 0.994291655 |
| 0.981008514 | 0.994251274 |
| 0.980992733 | 0.994251274 |
| 0.980971518 | 0.994251274 |
| 0.980769242 | 0.994170409 |
| 0.9807613 | 0.994170409 |
| 0.980751605 | 0.994170409 |
| 0.980608042 | 0.994112798 |
| 0.980558601 | 0.994112798 |
| 0.980483392 | 0.994071069 |
| 0.980432592 | 0.994040726 |
| 0.980316561 | 0.993965403 |
| 0.980232533 | 0.993901764 |
| 0.980115437 | 0.99386727 |
| 0.980013393 | 0.993784953 |
| 0.979995534 | 0.993784953 |
| 0.979735389 | 0.993608819 |
| 0.979632695 | 0.993525827 |
| 0.979618406 | 0.993525827 |
| 0.979508555 | 0.993442236 |
| 0.979350691 | 0.993384211 |
| 0.979323035 | 0.993384211 |
| 0.979301806 | 0.993384211 |
| 0.979280158 | 0.993384211 |
| 0.979265465 | 0.993384211 |
| 0.979246774 | 0.993384211 |

| | |
|---|---|
| 0.979226777 | 0.993384211 |
| 0.979226302 | 0.993384211 |
| 0.979149689 | 0.993384211 |
| 0.978957178 | 0.99332723 |
| 0.978934612 | 0.993325495 |
| 0.978897446 | 0.993313272 |
| 0.978876974 | 0.993313272 |
| 0.978821045 | 0.993313272 |
| 0.978801109 | 0.993313272 |
| 0.978761984 | 0.993313272 |
| 0.978731194 | 0.99330955 |
| 0.978724267 | 0.99330955 |
| 0.978501952 | 0.993182714 |
| 0.978443151 | 0.993175973 |
| 0.978352554 | 0.993140935 |
| 0.978306208 | 0.993140935 |
| 0.978201659 | 0.993075681 |
| 0.97811631 | 0.993024077 |
| 0.978047817 | 0.992996875 |
| 0.97804201 | 0.992996875 |
| 0.977831143 | 0.992840383 |
| 0.97776142 | 0.992790755 |
| 0.977713288 | 0.992790755 |
| 0.977656864 | 0.992790755 |
| 0.977623243 | 0.992790755 |
| 0.977617337 | 0.992790755 |
| 0.977608191 | 0.992790755 |
| 0.97754803 | 0.992790755 |
| 0.977518088 | 0.992790755 |
| 0.977502788 | 0.992790755 |
| 0.977411382 | 0.992790755 |
| 0.977334333 | 0.992790755 |
| 0.977322471 | 0.992790755 |
| 0.977264557 | 0.992790755 |
| 0.977248261 | 0.992790755 |
| 0.977149263 | 0.992790755 |
| 0.977119944 | 0.992790755 |
| 0.977096214 | 0.992790755 |
| 0.977059403 | 0.992790755 |
| 0.977023279 | 0.992782055 |
| 0.976975135 | 0.992775497 |
| 0.976905763 | 0.992747366 |
| 0.976791742 | 0.992673856 |
| 0.976770206 | 0.992673152 |
| 0.976752885 | 0.992673152 |

| | |
|---|---|
| 0.97668018 | 0.992645205 |
| 0.97647282 | 0.992476813 |
| 0.97644792 | 0.992476813 |
| 0.976410019 | 0.992476813 |
| 0.976266764 | 0.99239445 |
| 0.976060495 | 0.992228653 |
| 0.976040167 | 0.992228653 |
| 0.976017609 | 0.992228653 |
| 0.975984413 | 0.992228653 |
| 0.975836397 | 0.992147555 |
| 0.975833383 | 0.992147555 |
| 0.97573931 | 0.992091203 |
| 0.975485992 | 0.992003061 |
| 0.975464499 | 0.992002385 |
| 0.975438959 | 0.991998715 |
| 0.975435556 | 0.991998715 |
| 0.975326877 | 0.991968334 |
| 0.9752916 | 0.991953639 |
| 0.975268877 | 0.991953639 |
| 0.975259564 | 0.991953639 |
| 0.975219659 | 0.991953639 |
| 0.975075772 | 0.991882397 |
| 0.975041995 | 0.991869222 |
| 0.974974549 | 0.991842981 |
| 0.974938529 | 0.991827523 |
| 0.974868298 | 0.991798445 |
| 0.974807283 | 0.991771024 |
| 0.97474678 | 0.991759558 |
| 0.97473314 | 0.991759558 |
| 0.974639908 | 0.991756754 |
| 0.974184746 | 0.991441875 |
| 0.974142463 | 0.991423261 |
| 0.974123147 | 0.991423261 |
| 0.973976835 | 0.991336198 |
| 0.973883628 | 0.991283698 |
| 0.973789984 | 0.991251936 |
| 0.973739902 | 0.991251936 |
| 0.97371553 | 0.991251936 |
| 0.973686788 | 0.991251936 |
| 0.973680261 | 0.991251936 |
| 0.973561658 | 0.991251936 |
| 0.973506947 | 0.991218047 |
| 0.973453263 | 0.991184576 |
| 0.97341305 | 0.991164821 |
| 0.973383693 | 0.991156119 |

| | |
|---|---|
| 0.97330313 | 0.991095274 |
| 0.97328997 | 0.991095274 |
| 0.973263472 | 0.991095274 |
| 0.973191224 | 0.991044889 |
| 0.973086854 | 0.991026643 |
| 0.972979495 | 0.990956406 |
| 0.972971958 | 0.990956406 |
| 0.972806258 | 0.990949495 |
| 0.972630755 | 0.99089788 |
| 0.972544061 | 0.990830752 |
| 0.972522065 | 0.990829537 |
| 0.972460851 | 0.990829537 |
| 0.97238562 | 0.990796499 |
| 0.972294899 | 0.990746451 |
| 0.972071611 | 0.990603702 |
| 0.971969761 | 0.990521104 |
| 0.97184572 | 0.99047947 |
| 0.971823859 | 0.990478386 |
| 0.971796953 | 0.990478386 |
| 0.97167936 | 0.990458285 |
| 0.971558602 | 0.990398783 |
| 0.971522966 | 0.990383654 |
| 0.971311059 | 0.990273613 |
| 0.971301762 | 0.990273613 |
| 0.971282197 | 0.990273613 |
| 0.971281437 | 0.990273613 |
| 0.971026566 | 0.990089536 |
| 0.970990509 | 0.990073968 |
| 0.970918385 | 0.990021622 |
| 0.970888266 | 0.990021622 |
| 0.970871866 | 0.990021622 |
| 0.970855178 | 0.990021622 |
| 0.970658346 | 0.989989615 |
| 0.970641686 | 0.989989615 |
| 0.970552802 | 0.989924367 |
| 0.970506664 | 0.989919707 |
| 0.970461705 | 0.989919707 |
| 0.970362466 | 0.989857422 |
| 0.970303065 | 0.989831055 |
| 0.970286427 | 0.989831055 |
| 0.970243419 | 0.989831055 |
| 0.9701894 | 0.989829282 |
| 0.97010869 | 0.989789344 |
| 0.970002884 | 0.989723797 |
| 0.969960344 | 0.989723797 |

| | |
|---|---|
| 0.969810584 | 0.989685623 |
| 0.969750324 | 0.989678124 |
| 0.969749196 | 0.989678124 |
| 0.969701581 | 0.989678124 |
| 0.969659774 | 0.989678124 |
| 0.969654557 | 0.989678124 |
| 0.969608494 | 0.989678124 |
| 0.96947814 | 0.98963361 |
| 0.969434677 | 0.989631666 |
| 0.969280546 | 0.989609147 |
| 0.969242231 | 0.989609147 |
| 0.969169522 | 0.989609147 |
| 0.969149676 | 0.989609147 |
| 0.969006831 | 0.989609147 |
| 0.968718592 | 0.9894862 |
| 0.968523342 | 0.989362866 |
| 0.968514096 | 0.989362866 |
| 0.968485477 | 0.989362866 |
| 0.968454355 | 0.989362866 |
| 0.968396189 | 0.989356156 |
| 0.968275693 | 0.989317939 |
| 0.968108776 | 0.989227608 |
| 0.968011834 | 0.989196894 |
| 0.967957375 | 0.989189602 |
| 0.967947399 | 0.989189602 |
| 0.967732328 | 0.98910985 |
| 0.967650749 | 0.989082567 |
| 0.967492811 | 0.989027262 |
| 0.967321783 | 0.988937332 |
| 0.967255213 | 0.98891173 |
| 0.96724959 | 0.98891173 |
| 0.967208463 | 0.98890639 |
| 0.967113044 | 0.988830058 |
| 0.96707552 | 0.988812919 |
| 0.967069431 | 0.988812919 |
| 0.966872936 | 0.988754371 |
| 0.966860745 | 0.988754371 |
| 0.966806596 | 0.988728989 |
| 0.96676791 | 0.988725423 |
| 0.966697928 | 0.988725423 |
| 0.966511358 | 0.988596878 |
| 0.966507365 | 0.988596878 |
| 0.966500284 | 0.988596878 |
| 0.966285266 | 0.988596878 |
| 0.966283683 | 0.988596878 |

| | |
|---|---|
| 0.966163952 | 0.988596878 |
| 0.966163771 | 0.988596878 |
| 0.965937965 | 0.988596878 |
| 0.965863936 | 0.988596878 |
| 0.965786429 | 0.988577237 |
| 0.96578603 | 0.988577237 |
| 0.965771543 | 0.988577237 |
| 0.965704951 | 0.988577237 |
| 0.965635032 | 0.988577237 |
| 0.96562397 | 0.988577237 |
| 0.965608482 | 0.988577237 |
| 0.96554134 | 0.988577237 |
| 0.965378888 | 0.988457493 |
| 0.965307286 | 0.988405429 |
| 0.96515333 | 0.988290283 |
| 0.965076349 | 0.988253951 |
| 0.965020056 | 0.988238802 |
| 0.964965123 | 0.988225045 |
| 0.964942711 | 0.988225045 |
| 0.96481778 | 0.988159142 |
| 0.964814298 | 0.988159142 |
| 0.964796534 | 0.988159142 |
| 0.964755377 | 0.988159142 |
| 0.964688164 | 0.988132636 |
| 0.96467852 | 0.988132636 |
| 0.964633077 | 0.988132636 |
| 0.964430369 | 0.988038572 |
| 0.964205013 | 0.987862903 |
| 0.964105647 | 0.987790893 |
| 0.963986293 | 0.98776015 |
| 0.963703566 | 0.987612638 |
| 0.963650641 | 0.98757965 |
| 0.963534593 | 0.987503219 |
| 0.963521418 | 0.987503219 |
| 0.963457091 | 0.987466286 |
| 0.963454479 | 0.987466286 |
| 0.963320553 | 0.987368842 |
| 0.963287597 | 0.987356312 |
| 0.963138967 | 0.987345382 |
| 0.963133963 | 0.987345382 |
| 0.963089753 | 0.987345382 |
| 0.963070626 | 0.987345382 |
| 0.962984313 | 0.987345382 |
| 0.962975256 | 0.987345382 |
| 0.962669318 | 0.987144365 |

| | |
|---|---|
| 0.962591691 | 0.98711038 |
| 0.962487859 | 0.987025155 |
| 0.962391161 | 0.987025155 |
| 0.962309762 | 0.987007532 |
| 0.962297816 | 0.987007532 |
| 0.962190376 | 0.986953847 |
| 0.962121548 | 0.986925758 |
| 0.96206295 | 0.986886905 |
| 0.961809072 | 0.986668979 |
| 0.961799878 | 0.986668979 |
| 0.961747663 | 0.986648485 |
| 0.96148531 | 0.986549343 |
| 0.961419048 | 0.986527234 |
| 0.961162483 | 0.986430615 |
| 0.961159441 | 0.986430615 |
| 0.96104495 | 0.986430615 |
| 0.961015823 | 0.986430615 |
| 0.960835264 | 0.986303112 |
| 0.960750627 | 0.986287017 |
| 0.960707781 | 0.986287017 |
| 0.96070562 | 0.986287017 |
| 0.960704265 | 0.986287017 |
| 0.96054901 | 0.986268571 |
| 0.960498397 | 0.986237865 |
| 0.960417903 | 0.986197737 |
| 0.960347411 | 0.986167877 |
| 0.960305194 | 0.986160249 |
| 0.960151652 | 0.986051896 |
| 0.959922889 | 0.985944521 |
| 0.959905183 | 0.985944521 |
| 0.959836532 | 0.985919609 |
| 0.959815273 | 0.985919036 |
| 0.95971972 | 0.985863409 |
| 0.959704649 | 0.985863409 |
| 0.959632227 | 0.985850437 |
| 0.959560586 | 0.985806251 |
| 0.959448347 | 0.98575133 |
| 0.959448343 | 0.98575133 |
| 0.959348923 | 0.985695111 |
| 0.959169795 | 0.985638643 |
| 0.95907962 | 0.985609775 |
| 0.95900211 | 0.985593921 |
| 0.958859588 | 0.985499516 |
| 0.958772995 | 0.985443511 |
| 0.958686851 | 0.985376236 |

| | |
|---|---|
| 0.958669432 | 0.985376236 |
| 0.958643528 | 0.985376236 |
| 0.958628517 | 0.985376236 |
| 0.958289126 | 0.985222584 |
| 0.958223904 | 0.985195362 |
| 0.958177558 | 0.985195362 |
| 0.95810855 | 0.985195362 |
| 0.958101113 | 0.985195362 |
| 0.957892568 | 0.985133866 |
| 0.957799384 | 0.985059303 |
| 0.957773447 | 0.985053899 |
| 0.957610918 | 0.985035641 |
| 0.957556391 | 0.985000826 |
| 0.957438294 | 0.984943163 |
| 0.957417158 | 0.984943163 |
| 0.957412372 | 0.984943163 |
| 0.957373064 | 0.984943163 |
| 0.957351859 | 0.984943163 |
| 0.957274324 | 0.984943163 |
| 0.957183726 | 0.984943163 |
| 0.957147786 | 0.984943163 |
| 0.957111229 | 0.984943163 |
| 0.957111035 | 0.984943163 |
| 0.957089214 | 0.984943163 |
| 0.957053249 | 0.984943163 |
| 0.957035329 | 0.984943163 |
| 0.957027313 | 0.984943163 |
| 0.956989025 | 0.984943163 |
| 0.956927557 | 0.984943163 |
| 0.956877961 | 0.984943163 |
| 0.956700999 | 0.984943163 |
| 0.956696589 | 0.984943163 |
| 0.956631002 | 0.984943163 |
| 0.956581595 | 0.984943163 |
| 0.956482736 | 0.984938774 |
| 0.95644996 | 0.984938774 |
| 0.956345803 | 0.984894774 |
| 0.956317696 | 0.984894774 |
| 0.956285263 | 0.984894774 |
| 0.956186898 | 0.98486837 |
| 0.956134714 | 0.98484156 |
| 0.956130779 | 0.98484156 |
| 0.956117609 | 0.98484156 |
| 0.956081893 | 0.98484156 |
| 0.955979219 | 0.984782244 |

| | |
|---|---|
| 0.95591143 | 0.984733711 |
| 0.955900529 | 0.984733711 |
| 0.955803021 | 0.984685929 |
| 0.955720027 | 0.984621725 |
| 0.955612307 | 0.984574641 |
| 0.955583321 | 0.984566075 |
| 0.955535929 | 0.984549936 |
| 0.955459392 | 0.984525263 |
| 0.955370945 | 0.984496343 |
| 0.955222878 | 0.984365061 |
| 0.955138777 | 0.984306794 |
| 0.955120567 | 0.984306794 |
| 0.955112628 | 0.984306794 |
| 0.955091937 | 0.984306794 |
| 0.955066379 | 0.984306794 |
| 0.95506606 | 0.984306794 |
| 0.954940551 | 0.984287091 |
| 0.954841265 | 0.984206054 |
| 0.954762909 | 0.984167887 |
| 0.954684381 | 0.98410824 |
| 0.954485739 | 0.983988665 |
| 0.954445756 | 0.983988665 |
| 0.954409848 | 0.983974323 |
| 0.954340447 | 0.983924071 |
| 0.954267478 | 0.983891438 |
| 0.95423082 | 0.983874941 |
| 0.95421207 | 0.983874941 |
| 0.95409369 | 0.983777604 |
| 0.954037753 | 0.983777604 |
| 0.954013324 | 0.983777604 |
| 0.953999839 | 0.983777604 |
| 0.95396377 | 0.983777604 |
| 0.953934155 | 0.983777604 |
| 0.95384812 | 0.983771233 |
| 0.953844981 | 0.983771233 |
| 0.953808877 | 0.983771233 |
| 0.95367664 | 0.983729475 |
| 0.953645667 | 0.983718832 |
| 0.953630837 | 0.983718832 |
| 0.953622432 | 0.983718832 |
| 0.953505691 | 0.983638354 |
| 0.953472632 | 0.983625556 |
| 0.953456852 | 0.983625556 |
| 0.95345484 | 0.983625556 |
| 0.953445663 | 0.983625556 |

| | |
|---|---|
| 0.953353046 | 0.983587408 |
| 0.953345844 | 0.983587408 |
| 0.953076718 | 0.983472751 |
| 0.953065054 | 0.983472751 |
| 0.953044004 | 0.983472751 |
| 0.95291916 | 0.983434868 |
| 0.952913399 | 0.983434868 |
| 0.952762962 | 0.983404667 |
| 0.952710972 | 0.983372317 |
| 0.952685814 | 0.983367662 |
| 0.95267091 | 0.983367662 |
| 0.952438231 | 0.983197339 |
| 0.95222257 | 0.983059943 |
| 0.952129042 | 0.982984694 |
| 0.951894612 | 0.982827885 |
| 0.95187939 | 0.982827885 |
| 0.951833303 | 0.982807196 |
| 0.951805667 | 0.982799967 |
| 0.951756502 | 0.982770508 |
| 0.951729437 | 0.982763868 |
| 0.95167628 | 0.982750069 |
| 0.951647081 | 0.982742748 |
| 0.951622734 | 0.982738914 |
| 0.951580421 | 0.98273108 |
| 0.951537975 | 0.982715308 |
| 0.951522318 | 0.982715308 |
| 0.951514988 | 0.982715308 |
| 0.951474573 | 0.982713758 |
| 0.951427071 | 0.982707317 |
| 0.951372272 | 0.982672028 |
| 0.951333048 | 0.982653807 |
| 0.951309559 | 0.982653807 |
| 0.951216314 | 0.982653807 |
| 0.951162266 | 0.982653807 |
| 0.951109202 | 0.982653807 |
| 0.951008556 | 0.982645776 |
| 0.950849025 | 0.982594459 |
| 0.950694241 | 0.982504385 |
| 0.950636976 | 0.98249462 |
| 0.950600325 | 0.98249462 |
| 0.950490025 | 0.982421242 |
| 0.950490025 | 0.982421242 |
| 0.950490025 | 0.982421242 |
| 0.950490025 | 0.982421242 |
| 0.950490025 | 0.982421242 |

| | |
|---|---|
| 0.950481166 | 0.982421242 |
| 0.950400449 | 0.982421242 |
| 0.950218432 | 0.982334577 |
| 0.950021068 | 0.982331185 |
| 0.949969485 | 0.982331185 |
| 0.949954969 | 0.982331185 |
| 0.94990322 | 0.982331185 |
| 0.949788896 | 0.982331185 |
| 0.949784503 | 0.982331185 |
| 0.949769121 | 0.982331185 |
| 0.949735155 | 0.982331185 |
| 0.949704999 | 0.982331185 |
| 0.949643638 | 0.982331185 |
| 0.949603771 | 0.982331185 |
| 0.949475322 | 0.982321494 |
| 0.949442989 | 0.982321494 |
| 0.949438499 | 0.982321494 |
| 0.949364436 | 0.982302411 |
| 0.94934418 | 0.982302411 |
| 0.949298551 | 0.982302411 |
| 0.949274423 | 0.982302411 |
| 0.949181531 | 0.982283881 |
| 0.949177082 | 0.982283881 |
| 0.949090948 | 0.982254168 |
| 0.94901233 | 0.982236836 |
| 0.948989171 | 0.982234211 |
| 0.948756843 | 0.982097366 |
| 0.948663982 | 0.982060014 |
| 0.948523033 | 0.981965145 |
| 0.948455195 | 0.981945172 |
| 0.948392596 | 0.981933898 |
| 0.948190803 | 0.981873121 |
| 0.948189229 | 0.981873121 |
| 0.948118693 | 0.98184534 |
| 0.947875431 | 0.981722196 |
| 0.947826983 | 0.981722196 |
| 0.947750021 | 0.981698353 |
| 0.947675188 | 0.981654613 |
| 0.947653373 | 0.981654613 |
| 0.947614024 | 0.981654613 |
| 0.947509297 | 0.981654613 |
| 0.947459528 | 0.981633195 |
| 0.947459093 | 0.981633195 |
| 0.947402177 | 0.981633195 |
| 0.947311823 | 0.981605924 |

| | |
|---|---|
| 0.947241854 | 0.981593406 |
| 0.947229632 | 0.981593406 |
| 0.947182009 | 0.98157965 |
| 0.947059375 | 0.981495275 |
| 0.946953457 | 0.981428218 |
| 0.946926068 | 0.981421187 |
| 0.946902846 | 0.981418477 |
| 0.946875395 | 0.981411383 |
| 0.946828673 | 0.981405672 |
| 0.94661691 | 0.981250241 |
| 0.94650138 | 0.98119455 |
| 0.946407117 | 0.981139543 |
| 0.946399796 | 0.981139543 |
| 0.946354062 | 0.981127254 |
| 0.946199153 | 0.981030718 |
| 0.946172786 | 0.981024737 |
| 0.946112242 | 0.981004675 |
| 0.946023324 | 0.980976548 |
| 0.946021579 | 0.980976548 |
| 0.945959688 | 0.980976548 |
| 0.945921017 | 0.980976548 |
| 0.945882694 | 0.980976548 |
| 0.945859244 | 0.980976548 |
| 0.945654783 | 0.980872016 |
| 0.945619281 | 0.980856554 |
| 0.945435877 | 0.980789436 |
| 0.945377633 | 0.980776783 |
| 0.945331262 | 0.980768944 |
| 0.945267581 | 0.980759716 |
| 0.945158297 | 0.980686675 |
| 0.945154126 | 0.980686675 |
| 0.945125168 | 0.980686675 |
| 0.945076316 | 0.980686675 |
| 0.944942867 | 0.980645389 |
| 0.944773776 | 0.980556138 |
| 0.944663802 | 0.980463365 |
| 0.944597111 | 0.980448189 |
| 0.944541141 | 0.980421518 |
| 0.944452585 | 0.980350964 |
| 0.944439299 | 0.980350964 |
| 0.944429081 | 0.980350964 |
| 0.944414737 | 0.980350964 |
| 0.944363446 | 0.980350964 |
| 0.944323603 | 0.980350964 |
| 0.944304609 | 0.980350964 |

| | |
|---|---|
| 0.944280683 | 0.980350964 |
| 0.944249565 | 0.980350964 |
| 0.944248696 | 0.980350964 |
| 0.94419544 | 0.980350964 |
| 0.944132208 | 0.980350964 |
| 0.944067825 | 0.980337044 |
| 0.943949416 | 0.980298703 |
| 0.943933843 | 0.980298703 |
| 0.943919709 | 0.980298703 |
| 0.943744285 | 0.980192539 |
| 0.943697802 | 0.980184636 |
| 0.943624808 | 0.980153945 |
| 0.943619868 | 0.980153945 |
| 0.943567172 | 0.980136831 |
| 0.943536748 | 0.980126604 |
| 0.943517401 | 0.980126604 |
| 0.943474897 | 0.980105109 |
| 0.943472409 | 0.980105109 |
| 0.943407187 | 0.980077524 |
| 0.943313074 | 0.980003687 |
| 0.94328842 | 0.980003687 |
| 0.943207737 | 0.979998579 |
| 0.943159567 | 0.979991288 |
| 0.94308051 | 0.979991227 |
| 0.943069629 | 0.979991227 |
| 0.942996949 | 0.979991227 |
| 0.942834205 | 0.979884211 |
| 0.942760799 | 0.979884211 |
| 0.942664538 | 0.979876631 |
| 0.942621103 | 0.979876631 |
| 0.942600156 | 0.979876631 |
| 0.942571908 | 0.979876631 |
| 0.942486928 | 0.979869569 |
| 0.942479071 | 0.979869569 |
| 0.942436886 | 0.979869569 |
| 0.942432939 | 0.979869569 |
| 0.942396544 | 0.979869569 |
| 0.942387337 | 0.979869569 |
| 0.942216283 | 0.97978068 |
| 0.942202293 | 0.97978068 |
| 0.942169369 | 0.97978068 |
| 0.942019244 | 0.97978068 |
| 0.942013367 | 0.97978068 |
| 0.941955542 | 0.97978068 |
| 0.941912213 | 0.97978068 |

| | |
|---|---|
| 0.941853319 | 0.979745507 |
| 0.941844317 | 0.979745507 |
| 0.941790014 | 0.979722452 |
| 0.941783131 | 0.979722452 |
| 0.941774143 | 0.979722452 |
| 0.941771321 | 0.979722452 |
| 0.941737019 | 0.979722452 |
| 0.941711001 | 0.979722452 |
| 0.941603693 | 0.979722452 |
| 0.94158499 | 0.979722452 |
| 0.941552455 | 0.979722452 |
| 0.941517897 | 0.979722452 |
| 0.941499802 | 0.979722452 |
| 0.941432897 | 0.979722452 |
| 0.941410431 | 0.979722452 |
| 0.941314149 | 0.979698192 |
| 0.941271104 | 0.97969621 |
| 0.941269918 | 0.97969621 |
| 0.941237414 | 0.97969621 |
| 0.941126668 | 0.97969621 |
| 0.941094193 | 0.97969621 |
| 0.941042828 | 0.97969621 |
| 0.940968374 | 0.97969621 |
| 0.940923529 | 0.97969621 |
| 0.940858291 | 0.979682527 |
| 0.940839232 | 0.979682527 |
| 0.940671406 | 0.979649058 |
| 0.940523406 | 0.979517357 |
| 0.940467682 | 0.97950216 |
| 0.940439142 | 0.979493855 |
| 0.940432973 | 0.979493855 |
| 0.940288812 | 0.979444377 |
| 0.940269791 | 0.979444377 |
| 0.940260983 | 0.979444377 |
| 0.939983979 | 0.979324246 |
| 0.939964657 | 0.979324246 |
| 0.939932105 | 0.979324246 |
| 0.939894126 | 0.979324246 |
| 0.939857339 | 0.979316214 |
| 0.939834552 | 0.979316214 |
| 0.939826494 | 0.979316214 |
| 0.939815194 | 0.979316214 |
| 0.939780856 | 0.979316214 |
| 0.939736293 | 0.979316214 |
| 0.939694022 | 0.979316214 |

| | |
|---|---|
| 0.93969264 | 0.979316214 |
| 0.939674934 | 0.979316214 |
| 0.939658067 | 0.979316214 |
| 0.939585756 | 0.979311758 |
| 0.93952757 | 0.979293975 |
| 0.939509447 | 0.979293975 |
| 0.939353855 | 0.979220059 |
| 0.939194473 | 0.97913313 |
| 0.939183789 | 0.97913313 |
| 0.939151974 | 0.97913313 |
| 0.939101112 | 0.979128035 |
| 0.939049385 | 0.979095537 |
| 0.938373969 | 0.978617918 |
| 0.938328055 | 0.978617918 |
| 0.938322936 | 0.978617918 |
| 0.938190021 | 0.97856371 |
| 0.938073195 | 0.978533907 |
| 0.938057608 | 0.978533907 |
| 0.937786287 | 0.978314003 |
| 0.937736519 | 0.978283512 |
| 0.937713565 | 0.978283512 |
| 0.937699477 | 0.978283512 |
| 0.937696002 | 0.978283512 |
| 0.93762367 | 0.978283512 |
| 0.937542228 | 0.978230813 |
| 0.937502223 | 0.978210502 |
| 0.937334367 | 0.978121073 |
| 0.937257214 | 0.978061992 |
| 0.937224294 | 0.978055445 |
| 0.937210796 | 0.978055445 |
| 0.937165355 | 0.978055445 |
| 0.937107707 | 0.978055445 |
| 0.937013423 | 0.978043312 |
| 0.93696148 | 0.978041347 |
| 0.936954459 | 0.978041347 |
| 0.936938881 | 0.978041347 |
| 0.936905713 | 0.978041347 |
| 0.936897825 | 0.978041347 |
| 0.936874605 | 0.978041347 |
| 0.936824964 | 0.978041347 |
| 0.936774172 | 0.978041347 |
| 0.936732561 | 0.978041347 |
| 0.936730622 | 0.978041347 |
| 0.9367204 | 0.978041347 |
| 0.936675619 | 0.978041347 |

| | |
|---|---|
| 0.936670077 | 0.978041347 |
| 0.936449715 | 0.977990759 |
| 0.936245903 | 0.977842232 |
| 0.936207373 | 0.977823434 |
| 0.936205706 | 0.977823434 |
| 0.935896462 | 0.977563015 |
| 0.935792378 | 0.977563015 |
| 0.935717108 | 0.977547187 |
| 0.935531142 | 0.977395786 |
| 0.935506729 | 0.97739172 |
| 0.935259776 | 0.977262337 |
| 0.935222983 | 0.977245332 |
| 0.935158008 | 0.977206818 |
| 0.935032838 | 0.977111866 |
| 0.934984377 | 0.977103193 |
| 0.934806223 | 0.976981325 |
| 0.934797175 | 0.976981325 |
| 0.934781245 | 0.976981325 |
| 0.934777733 | 0.976981325 |
| 0.934777619 | 0.976981325 |
| 0.934736477 | 0.976981325 |
| 0.934734616 | 0.976981325 |
| 0.934528543 | 0.976966054 |
| 0.934393954 | 0.976906849 |
| 0.934386434 | 0.976906849 |
| 0.934359359 | 0.976906849 |
| 0.934260233 | 0.976860858 |
| 0.934255647 | 0.976860858 |
| 0.934251416 | 0.976860858 |
| 0.934246642 | 0.976860858 |
| 0.93420674 | 0.976860858 |
| 0.934107429 | 0.976829758 |
| 0.93404344 | 0.976829758 |
| 0.933949987 | 0.976829758 |
| 0.933906344 | 0.976829758 |
| 0.933728356 | 0.976690688 |
| 0.933712101 | 0.976690688 |
| 0.933649129 | 0.976672167 |
| 0.933602487 | 0.976666279 |
| 0.933501507 | 0.976603542 |
| 0.933407336 | 0.976547925 |
| 0.933401664 | 0.976547925 |
| 0.933400336 | 0.976547925 |
| 0.933396303 | 0.976547925 |
| 0.933257918 | 0.976498855 |

| | |
|---|---|
| 0.933047335 | 0.976364314 |
| 0.933025533 | 0.976363374 |
| 0.932980228 | 0.976358448 |
| 0.932954798 | 0.976358448 |
| 0.932875459 | 0.976358448 |
| 0.932713449 | 0.976278599 |
| 0.93271041 | 0.976278599 |
| 0.932684794 | 0.976278599 |
| 0.932558885 | 0.976239274 |
| 0.932430807 | 0.976188417 |
| 0.932408059 | 0.976188417 |
| 0.932232361 | 0.976069088 |
| 0.932178563 | 0.976055675 |
| 0.9321586 | 0.976055675 |
| 0.932124604 | 0.97604721 |
| 0.932105192 | 0.97604721 |
| 0.931952809 | 0.97596949 |
| 0.931942599 | 0.97596949 |
| 0.931860338 | 0.97596949 |
| 0.931855996 | 0.97596949 |
| 0.931831758 | 0.97596949 |
| 0.931827044 | 0.97596949 |
| 0.931727022 | 0.975969096 |
| 0.931705996 | 0.975968537 |
| 0.931544294 | 0.975864411 |
| 0.931515642 | 0.975864411 |
| 0.931099942 | 0.975634104 |
| 0.931084573 | 0.975634104 |
| 0.9310435 | 0.975634104 |
| 0.931022445 | 0.975634104 |
| 0.930888897 | 0.975609419 |
| 0.930857086 | 0.975609419 |
| 0.93072681 | 0.975522134 |
| 0.930570616 | 0.975422822 |
| 0.930377264 | 0.97530601 |
| 0.930034545 | 0.975098471 |
| 0.929832261 | 0.974992209 |
| 0.929754596 | 0.9749537 |
| 0.929622803 | 0.974879893 |
| 0.92951666 | 0.974790047 |
| 0.929313628 | 0.97464138 |
| 0.929223354 | 0.974632678 |
| 0.92919113 | 0.974620344 |
| 0.929172243 | 0.974620344 |
| 0.929160314 | 0.974620344 |

| | |
|---|---|
| 0.929104002 | 0.974614812 |
| 0.929075367 | 0.974614812 |
| 0.929053369 | 0.974614812 |
| 0.929037245 | 0.974614812 |
| 0.928963386 | 0.974614812 |
| 0.928901357 | 0.974614812 |
| 0.928800534 | 0.974555293 |
| 0.928765632 | 0.974555293 |
| 0.928662484 | 0.974538022 |
| 0.928491707 | 0.974380278 |
| 0.928330353 | 0.974296821 |
| 0.928298945 | 0.974296821 |
| 0.928224005 | 0.974292555 |
| 0.9281857 | 0.974281531 |
| 0.928168547 | 0.974281531 |
| 0.928166172 | 0.974281531 |
| 0.928051118 | 0.974239913 |
| 0.927900086 | 0.974124306 |
| 0.927743724 | 0.974024565 |
| 0.927717438 | 0.974024565 |
| 0.9277161 | 0.974024565 |
| 0.92767874 | 0.974024565 |
| 0.927662041 | 0.974024565 |
| 0.927635458 | 0.974024565 |
| 0.927382079 | 0.973902502 |
| 0.927381446 | 0.973902502 |
| 0.927375104 | 0.973902502 |
| 0.927243194 | 0.973842542 |
| 0.927221569 | 0.973841306 |
| 0.92716746 | 0.97380595 |
| 0.927043824 | 0.973719041 |
| 0.926970764 | 0.973698901 |
| 0.926935513 | 0.97369117 |
| 0.926901279 | 0.97369117 |
| 0.926850781 | 0.97369117 |
| 0.926822306 | 0.97369117 |
| 0.926817537 | 0.97369117 |
| 0.926795378 | 0.97369117 |
| 0.926781091 | 0.97369117 |
| 0.926759865 | 0.97369117 |
| 0.926720937 | 0.97369117 |
| 0.926646341 | 0.97369117 |
| 0.926381923 | 0.973534838 |
| 0.92635025 | 0.973534838 |
| 0.926283869 | 0.973527903 |

| | |
|---|---|
| 0.926245523 | 0.973527903 |
| 0.926143731 | 0.973510611 |
| 0.926065954 | 0.973496473 |
| 0.925991279 | 0.973450759 |
| 0.925948939 | 0.973449222 |
| 0.925929711 | 0.973449222 |
| 0.925923336 | 0.973449222 |
| 0.925702048 | 0.973275601 |
| 0.925670165 | 0.973263565 |
| 0.925499914 | 0.973147809 |
| 0.925486105 | 0.973147809 |
| 0.925257495 | 0.973023002 |
| 0.925253232 | 0.973023002 |
| 0.925240365 | 0.973023002 |
| 0.925230014 | 0.973023002 |
| 0.925081136 | 0.972968505 |
| 0.925013101 | 0.972937854 |
| 0.924998938 | 0.972937854 |
| 0.924965027 | 0.972937854 |
| 0.92496199 | 0.972937854 |
| 0.924960689 | 0.972937854 |
| 0.924789211 | 0.972874246 |
| 0.924667014 | 0.972810159 |
| 0.924553357 | 0.972776538 |
| 0.92454594 | 0.972776538 |
| 0.92447502 | 0.972737093 |
| 0.92424539 | 0.972581423 |
| 0.924232811 | 0.972581423 |
| 0.924201857 | 0.97257859 |
| 0.924084339 | 0.972497897 |
| 0.924053237 | 0.972491893 |
| 0.923786052 | 0.972312891 |
| 0.92375092 | 0.9722974 |
| 0.923720417 | 0.972291575 |
| 0.923640043 | 0.972245156 |
| 0.923619066 | 0.972244563 |
| 0.923512644 | 0.972197 |
| 0.923467213 | 0.972170662 |
| 0.92342138 | 0.972170662 |
| 0.923372224 | 0.972170662 |
| 0.92325084 | 0.97213627 |
| 0.923209588 | 0.972131393 |
| 0.92317143 | 0.972131393 |
| 0.92314892 | 0.972131393 |
| 0.923141981 | 0.972131393 |

| | |
|---|---|
| 0.923091419 | 0.972131393 |
| 0.923050286 | 0.972131393 |
| 0.922908263 | 0.972076426 |
| 0.922760462 | 0.971942246 |
| 0.922741449 | 0.971942246 |
| 0.92263153 | 0.971884627 |
| 0.922614971 | 0.971884627 |
| 0.922587729 | 0.971884627 |
| 0.922580357 | 0.971884627 |
| 0.922562583 | 0.971884627 |
| 0.922549207 | 0.971884627 |
| 0.922489415 | 0.971884627 |
| 0.922443352 | 0.971884627 |
| 0.922438327 | 0.971884627 |
| 0.922227394 | 0.971884627 |
| 0.922223018 | 0.971884627 |
| 0.922182503 | 0.971884627 |
| 0.92209191 | 0.971884627 |
| 0.922073793 | 0.971884627 |
| 0.922023147 | 0.971884627 |
| 0.921963604 | 0.971855176 |
| 0.921817412 | 0.971794049 |
| 0.921810945 | 0.971794049 |
| 0.92178008 | 0.971794049 |
| 0.921712646 | 0.971784212 |
| 0.921700852 | 0.971784212 |
| 0.921626725 | 0.971784212 |
| 0.921509258 | 0.971720352 |
| 0.921501453 | 0.971720352 |
| 0.921213538 | 0.971520184 |
| 0.921195351 | 0.971520184 |
| 0.921187724 | 0.971520184 |
| 0.921042919 | 0.97150823 |
| 0.921042269 | 0.97150823 |
| 0.921011484 | 0.97150823 |
| 0.920997395 | 0.97150823 |
| 0.920926068 | 0.97150823 |
| 0.920918263 | 0.97150823 |
| 0.92087284 | 0.971500958 |
| 0.920870683 | 0.971500958 |
| 0.920853963 | 0.971500958 |
| 0.920769683 | 0.971499727 |
| 0.920712356 | 0.971460763 |
| 0.920431856 | 0.971242254 |
| 0.920413589 | 0.971242254 |

| | |
|---|---|
| 0.92020802 | 0.971100737 |
| 0.920101484 | 0.971009824 |
| 0.920041174 | 0.970967692 |
| 0.920036838 | 0.970967692 |
| 0.919984449 | 0.970967692 |
| 0.919921216 | 0.970967692 |
| 0.919898173 | 0.970967692 |
| 0.919832779 | 0.970941396 |
| 0.919664531 | 0.970849867 |
| 0.919609615 | 0.970813414 |
| 0.919598113 | 0.970813414 |
| 0.919570367 | 0.970813414 |
| 0.919484147 | 0.970788549 |
| 0.919402816 | 0.97076724 |
| 0.919352037 | 0.970735146 |
| 0.919312347 | 0.970714757 |
| 0.919259601 | 0.970680583 |
| 0.919243437 | 0.970680583 |
| 0.919195566 | 0.970669732 |
| 0.919125814 | 0.970625391 |
| 0.919118232 | 0.970625391 |
| 0.918971716 | 0.970615114 |
| 0.918932572 | 0.970615014 |
| 0.918926201 | 0.970615014 |
| 0.918873751 | 0.970615014 |
| 0.918798688 | 0.970602753 |
| 0.918787839 | 0.970602753 |
| 0.918783372 | 0.970602753 |
| 0.918759315 | 0.970602753 |
| 0.918746621 | 0.970602753 |
| 0.918730707 | 0.970602753 |
| 0.918696673 | 0.970602753 |
| 0.918672189 | 0.970602753 |
| 0.918576225 | 0.970602753 |
| 0.918462118 | 0.97057012 |
| 0.918451619 | 0.97057012 |
| 0.91835855 | 0.970503743 |
| 0.91834636 | 0.970503743 |
| 0.91833264 | 0.970503743 |
| 0.918285149 | 0.970502536 |
| 0.918230836 | 0.970502536 |
| 0.918185792 | 0.970493447 |
| 0.918160287 | 0.970488027 |
| 0.917984106 | 0.970323339 |
| 0.917953312 | 0.970312324 |

| | |
|---|---|
| 0.91790036 | 0.970299422 |
| 0.917879866 | 0.970299294 |
| 0.917797084 | 0.970254855 |
| 0.917781907 | 0.970254855 |
| 0.917692402 | 0.970192943 |
| 0.917619002 | 0.970174267 |
| 0.91761363 | 0.970174267 |
| 0.917528735 | 0.970144855 |
| 0.91743795 | 0.970133598 |
| 0.917349234 | 0.970104402 |
| 0.917311783 | 0.970104402 |
| 0.917297379 | 0.970104402 |
| 0.917292611 | 0.970104402 |
| 0.917284482 | 0.970104402 |
| 0.917195678 | 0.970092795 |
| 0.917153215 | 0.970069729 |
| 0.917100853 | 0.970057805 |
| 0.917002583 | 0.970057805 |
| 0.916986858 | 0.970057805 |
| 0.916941147 | 0.970057805 |
| 0.9168952 | 0.970057805 |
| 0.916890084 | 0.970057805 |
| 0.91682154 | 0.970041753 |
| 0.916666803 | 0.969964226 |
| 0.916661337 | 0.969964226 |
| 0.916655175 | 0.969964226 |
| 0.91661531 | 0.969964226 |
| 0.916569765 | 0.969964226 |
| 0.916550522 | 0.969964226 |
| 0.916512098 | 0.969951374 |
| 0.916341376 | 0.969813798 |
| 0.916325255 | 0.969813798 |
| 0.916300269 | 0.969813393 |
| 0.916218415 | 0.96974831 |
| 0.916211843 | 0.96974831 |
| 0.916032341 | 0.969702214 |
| 0.915905278 | 0.969632361 |
| 0.915838754 | 0.96960504 |
| 0.915752513 | 0.969535288 |
| 0.915737796 | 0.969535288 |
| 0.915698869 | 0.969521598 |
| 0.915649251 | 0.969512169 |
| 0.915550616 | 0.969450837 |
| 0.915497758 | 0.96941642 |
| 0.915458154 | 0.969396036 |

| | |
|---|---|
| 0.91533831 | 0.969290682 |
| 0.915314573 | 0.969287097 |
| 0.915230379 | 0.969219488 |
| 0.914920666 | 0.969085435 |
| 0.914907675 | 0.969085435 |
| 0.914904187 | 0.969085435 |
| 0.914867937 | 0.969085435 |
| 0.914845514 | 0.969085435 |
| 0.914754285 | 0.969085435 |
| 0.914692284 | 0.969085435 |
| 0.91463528 | 0.96907908 |
| 0.914449933 | 0.968953167 |
| 0.914444596 | 0.968953167 |
| 0.91443923 | 0.968953167 |
| 0.914362539 | 0.968936589 |
| 0.914324455 | 0.968936589 |
| 0.914321062 | 0.968936589 |
| 0.914249825 | 0.968935144 |
| 0.914239101 | 0.968935144 |
| 0.914159043 | 0.968903613 |
| 0.914103592 | 0.968866402 |
| 0.914018011 | 0.968801185 |
| 0.913990522 | 0.968801185 |
| 0.913926418 | 0.968801185 |
| 0.913924298 | 0.968801185 |
| 0.913921868 | 0.968801185 |
| 0.913909523 | 0.968801185 |
| 0.913849648 | 0.968801185 |
| 0.913800794 | 0.968801185 |
| 0.913796806 | 0.968801185 |
| 0.913778357 | 0.968801185 |
| 0.91376744 | 0.968801185 |
| 0.913762343 | 0.968801185 |
| 0.913741817 | 0.968801185 |
| 0.913613054 | 0.968756099 |
| 0.91358715 | 0.968756099 |
| 0.913565973 | 0.968756099 |
| 0.913519643 | 0.968743322 |
| 0.913474056 | 0.968716548 |
| 0.913295507 | 0.968570333 |
| 0.913093635 | 0.968399371 |
| 0.913012599 | 0.96833499 |
| 0.913002171 | 0.96833499 |
| 0.912857715 | 0.968235405 |
| 0.912846673 | 0.968235405 |

| | |
|---|---|
| 0.912707097 | 0.968217864 |
| 0.912694486 | 0.968217864 |
| 0.912631615 | 0.968217443 |
| 0.912468917 | 0.968202375 |
| 0.912453632 | 0.968202375 |
| 0.912406763 | 0.968188332 |
| 0.912317624 | 0.968115313 |
| 0.912288287 | 0.968105751 |
| 0.912244778 | 0.968105751 |
| 0.912224566 | 0.968105751 |
| 0.912190625 | 0.968105751 |
| 0.912178242 | 0.968105751 |
| 0.912042729 | 0.967996146 |
| 0.912041043 | 0.967996146 |
| 0.912015446 | 0.967996146 |
| 0.911989894 | 0.967996146 |
| 0.911831864 | 0.967896056 |
| 0.911771196 | 0.967858933 |
| 0.911695999 | 0.967856992 |
| 0.911691495 | 0.967856992 |
| 0.911661202 | 0.967856992 |
| 0.911654402 | 0.967856992 |
| 0.911627855 | 0.967856992 |
| 0.911610848 | 0.967856992 |
| 0.911591784 | 0.967856992 |
| 0.911575023 | 0.967856992 |
| 0.911573791 | 0.967856992 |
| 0.911555069 | 0.967856992 |
| 0.911431941 | 0.967856992 |
| 0.911422696 | 0.967856992 |
| 0.911417447 | 0.967856992 |
| 0.911359392 | 0.967856992 |
| 0.911283155 | 0.967856992 |
| 0.911083383 | 0.967711158 |
| 0.911082529 | 0.967711158 |
| 0.911058446 | 0.967711158 |
| 0.910975278 | 0.967682656 |
| 0.910924844 | 0.967650665 |
| 0.910869252 | 0.967633221 |
| 0.910845234 | 0.967630843 |
| 0.910839845 | 0.967630843 |
| 0.910794715 | 0.967620341 |
| 0.910586445 | 0.967463819 |
| 0.910457529 | 0.967434758 |
| 0.91043785 | 0.967434758 |

| | |
|---|---|
| 0.9103561 | 0.967370147 |
| 0.910303858 | 0.967357799 |
| 0.910284019 | 0.967357799 |
| 0.910282592 | 0.967357799 |
| 0.91019472 | 0.967349741 |
| 0.910133555 | 0.967306321 |
| 0.910025698 | 0.967224749 |
| 0.909885282 | 0.967193535 |
| 0.909878172 | 0.967193535 |
| 0.909875949 | 0.967193535 |
| 0.909866254 | 0.967193535 |
| 0.909821432 | 0.967193535 |
| 0.909686274 | 0.9671147 |
| 0.909106233 | 0.966581148 |
| 0.909104293 | 0.966581148 |
| 0.909026134 | 0.966581148 |
| 0.908942166 | 0.96653617 |
| 0.90894066 | 0.96653617 |
| 0.908888909 | 0.966522698 |
| 0.908660393 | 0.966365999 |
| 0.908570487 | 0.966291961 |
| 0.908567285 | 0.966291961 |
| 0.908511879 | 0.966272784 |
| 0.908481358 | 0.966261901 |
| 0.908452041 | 0.966252297 |
| 0.9083938 | 0.966214796 |
| 0.908187972 | 0.966057724 |
| 0.908169619 | 0.966057724 |
| 0.908142652 | 0.966052669 |
| 0.90805929 | 0.966007143 |
| 0.908043975 | 0.966007143 |
| 0.907985576 | 0.966007143 |
| 0.907885129 | 0.965960731 |
| 0.907800273 | 0.965904195 |
| 0.907767183 | 0.965904195 |
| 0.90770681 | 0.965869484 |
| 0.907542897 | 0.965759798 |
| 0.907532001 | 0.965759798 |
| 0.907509685 | 0.965759798 |
| 0.90747889 | 0.965756421 |
| 0.907443054 | 0.965739863 |
| 0.907327148 | 0.965652012 |
| 0.907278429 | 0.965629394 |
| 0.907069663 | 0.965536661 |
| 0.907066497 | 0.965536661 |

| | |
|---|---|
| 0.906947042 | 0.965470871 |
| 0.906885823 | 0.96544886 |
| 0.906763327 | 0.965411857 |
| 0.906729889 | 0.965411857 |
| 0.906719317 | 0.965411857 |
| 0.906598006 | 0.965336654 |
| 0.906569903 | 0.965336654 |
| 0.906533333 | 0.965336654 |
| 0.906501933 | 0.965336654 |
| 0.906347929 | 0.965296783 |
| 0.906245717 | 0.965220492 |
| 0.906162054 | 0.965192784 |
| 0.906095225 | 0.96514654 |
| 0.906079647 | 0.96514654 |
| 0.905989793 | 0.965120577 |
| 0.905947987 | 0.965120577 |
| 0.905888074 | 0.965120577 |
| 0.905862337 | 0.965120577 |
| 0.905698644 | 0.965004682 |
| 0.905620223 | 0.964964302 |
| 0.905587248 | 0.964964302 |
| 0.905579189 | 0.964964302 |
| 0.905445332 | 0.964864295 |
| 0.905434517 | 0.964864295 |
| 0.90540423 | 0.964863673 |
| 0.905389415 | 0.964863673 |
| 0.905382155 | 0.964863673 |
| 0.90536717 | 0.964863673 |
| 0.905353058 | 0.964863673 |
| 0.905265377 | 0.964863673 |
| 0.905254046 | 0.964863673 |
| 0.905225233 | 0.964863673 |
| 0.905117383 | 0.96483864 |
| 0.905067811 | 0.964807391 |
| 0.905015713 | 0.964773448 |
| 0.90498049 | 0.964757494 |
| 0.904870686 | 0.964679328 |
| 0.904866023 | 0.964679328 |
| 0.904819281 | 0.964679328 |
| 0.904808916 | 0.964679328 |
| 0.904796506 | 0.964679328 |
| 0.904743362 | 0.964679328 |
| 0.904712628 | 0.964679328 |
| 0.904692196 | 0.964679328 |
| 0.904588716 | 0.964663728 |

| | |
|---|---|
| 0.90458392 | 0.964663728 |
| 0.904426132 | 0.964576738 |
| 0.904391445 | 0.964561344 |
| 0.90438012 | 0.964561344 |
| 0.90436073 | 0.964561344 |
| 0.904328072 | 0.964558553 |
| 0.904066252 | 0.964344085 |
| 0.903874952 | 0.964248009 |
| 0.90369448 | 0.964163463 |
| 0.903678451 | 0.964163463 |
| 0.903566559 | 0.964091774 |
| 0.903516802 | 0.964060282 |
| 0.903490838 | 0.964054175 |
| 0.90334552 | 0.963963906 |
| 0.903342058 | 0.963963906 |
| 0.903325105 | 0.963963906 |
| 0.903303852 | 0.963963906 |
| 0.903240417 | 0.963963906 |
| 0.903237763 | 0.963963906 |
| 0.903171348 | 0.963963906 |
| 0.903039495 | 0.963896505 |
| 0.902875909 | 0.963743493 |
| 0.902725703 | 0.963647953 |
| 0.902669611 | 0.963609673 |
| 0.902623662 | 0.963603819 |
| 0.90257688 | 0.963585725 |
| 0.902570898 | 0.963585725 |
| 0.902549638 | 0.963585725 |
| 0.902520857 | 0.963585725 |
| 0.902419548 | 0.96355917 |
| 0.902395725 | 0.96355917 |
| 0.902365465 | 0.96355917 |
| 0.902322856 | 0.963541864 |
| 0.902235321 | 0.963491595 |
| 0.902224063 | 0.963491595 |
| 0.902192361 | 0.963491595 |
| 0.902186974 | 0.963491595 |
| 0.902071938 | 0.963468339 |
| 0.902069274 | 0.963468339 |
| 0.902016937 | 0.963452807 |
| 0.901999446 | 0.963452807 |
| 0.901976065 | 0.963452365 |
| 0.901872895 | 0.963380304 |
| 0.901868181 | 0.963380304 |
| 0.90175913 | 0.963316251 |

| | |
|---|---|
| 0.901683165 | 0.963269124 |
| 0.901618796 | 0.963239942 |
| 0.901585204 | 0.96322929 |
| 0.901551383 | 0.963214764 |
| 0.901519695 | 0.963202515 |
| 0.901517732 | 0.963202515 |
| 0.90133079 | 0.963108713 |
| 0.901233517 | 0.963091203 |
| 0.90089097 | 0.962833163 |
| 0.900787954 | 0.962744668 |
| 0.900765862 | 0.962744668 |
| 0.900677387 | 0.962720981 |
| 0.900626997 | 0.962702262 |
| 0.900586766 | 0.962702262 |
| 0.900535382 | 0.962690758 |
| 0.900493587 | 0.962669132 |
| 0.900334546 | 0.962540875 |
| 0.900140497 | 0.962398228 |
| 0.900108134 | 0.96238523 |
| 0.900020558 | 0.962334803 |
| 0.899883718 | 0.962231693 |
| 0.8998129 | 0.962177571 |
| 0.89979309 | 0.962177571 |
| 0.899683933 | 0.962147675 |
| 0.899680774 | 0.962147675 |
| 0.899673559 | 0.962147675 |
| 0.899659958 | 0.962147675 |
| 0.899654772 | 0.962147675 |
| 0.899582538 | 0.962147261 |
| 0.899507243 | 0.962147261 |
| 0.899373762 | 0.962068491 |
| 0.899354978 | 0.962068491 |
| 0.899349314 | 0.962068491 |
| 0.898988004 | 0.961943421 |
| 0.898987189 | 0.961943421 |
| 0.898844642 | 0.961833238 |
| 0.898779362 | 0.961784993 |
| 0.898759625 | 0.961784993 |
| 0.898742763 | 0.961784993 |
| 0.898551263 | 0.961677833 |
| 0.898546139 | 0.961677833 |
| 0.898410427 | 0.961588669 |
| 0.898315057 | 0.961588669 |
| 0.898234157 | 0.96157082 |
| 0.898193043 | 0.96157082 |

| | |
|---|---|
| 0.898112597 | 0.961546783 |
| 0.898074768 | 0.961527897 |
| 0.897987471 | 0.961520892 |
| 0.897903924 | 0.961475928 |
| 0.897842777 | 0.961475928 |
| 0.897828057 | 0.961475928 |
| 0.897809707 | 0.961475928 |
| 0.897708716 | 0.961475928 |
| 0.897648229 | 0.961460266 |
| 0.897620577 | 0.961452271 |
| 0.897549275 | 0.961410943 |
| 0.897531373 | 0.961410943 |
| 0.897490073 | 0.961410943 |
| 0.897357271 | 0.96134121 |
| 0.897344417 | 0.96134121 |
| 0.897304581 | 0.961330003 |
| 0.897274425 | 0.961319319 |
| 0.897218713 | 0.961281255 |
| 0.897169012 | 0.96124963 |
| 0.896830089 | 0.960994592 |
| 0.896716625 | 0.96091625 |
| 0.896616491 | 0.960830565 |
| 0.896534285 | 0.960828947 |
| 0.896457202 | 0.960805531 |
| 0.896423679 | 0.960805531 |
| 0.896363624 | 0.960802252 |
| 0.896199384 | 0.960737948 |
| 0.896115299 | 0.960682558 |
| 0.896100514 | 0.960682558 |
| 0.896093586 | 0.960682558 |
| 0.89607318 | 0.960682558 |
| 0.896027821 | 0.960682558 |
| 0.895988187 | 0.960676033 |
| 0.89596356 | 0.960671255 |
| 0.895960106 | 0.960671255 |
| 0.895929403 | 0.960671255 |
| 0.895831366 | 0.960637648 |
| 0.895726792 | 0.960590392 |
| 0.895667488 | 0.960570052 |
| 0.895540151 | 0.960498373 |
| 0.895520212 | 0.960498373 |
| 0.895395134 | 0.960407721 |
| 0.895383822 | 0.960407721 |
| 0.895336641 | 0.960388239 |
| 0.895208205 | 0.960304175 |

| | |
|---|---|
| 0.895199536 | 0.960304175 |
| 0.89516611 | 0.960304175 |
| 0.895033814 | 0.960253336 |
| 0.894933765 | 0.960237236 |
| 0.894914065 | 0.960237236 |
| 0.894874975 | 0.960237236 |
| 0.89487383 | 0.960237236 |
| 0.894861721 | 0.960237236 |
| 0.894819255 | 0.960237236 |
| 0.894765332 | 0.960237236 |
| 0.894735801 | 0.960237236 |
| 0.894524549 | 0.960122613 |
| 0.894410873 | 0.960115502 |
| 0.894364087 | 0.96010829 |
| 0.894351055 | 0.96010829 |
| 0.894339843 | 0.96010829 |
| 0.894142333 | 0.959993618 |
| 0.894075076 | 0.959943047 |
| 0.894040409 | 0.959943047 |
| 0.894016765 | 0.959943047 |
| 0.893980415 | 0.959927972 |
| 0.893955773 | 0.959923154 |
| 0.893784426 | 0.959760802 |
| 0.893767022 | 0.959760802 |
| 0.893695032 | 0.959729724 |
| 0.893635281 | 0.959687198 |
| 0.893613535 | 0.959685483 |
| 0.893522527 | 0.959609385 |
| 0.893309633 | 0.959402379 |
| 0.893276579 | 0.959388514 |
| 0.893259483 | 0.959388514 |
| 0.893233829 | 0.959388514 |
| 0.893221306 | 0.959388514 |
| 0.893111625 | 0.959319516 |
| 0.893109084 | 0.959319516 |
| 0.893065721 | 0.959319516 |
| 0.893006695 | 0.959293348 |
| 0.892991342 | 0.959293348 |
| 0.892990005 | 0.959293348 |
| 0.892983389 | 0.959293348 |
| 0.892927615 | 0.959293348 |
| 0.892875372 | 0.959293348 |
| 0.892780763 | 0.959264876 |
| 0.892602786 | 0.959162332 |
| 0.892590451 | 0.959162332 |

| | |
|---|---|
| 0.89255876 | 0.959162332 |
| 0.892555273 | 0.959162332 |
| 0.892492568 | 0.959162332 |
| 0.892488121 | 0.959162332 |
| 0.892359025 | 0.959138759 |
| 0.892358267 | 0.959138759 |
| 0.892298685 | 0.959138518 |
| 0.892295557 | 0.959138518 |
| 0.892280153 | 0.959138518 |
| 0.892247201 | 0.959138518 |
| 0.892235882 | 0.959138518 |
| 0.892144416 | 0.959126255 |
| 0.892052666 | 0.959069869 |
| 0.891964272 | 0.959060444 |
| 0.891929138 | 0.959044318 |
| 0.891916004 | 0.959044318 |
| 0.891830147 | 0.959029158 |
| 0.891791509 | 0.959029158 |
| 0.891776095 | 0.959029158 |
| 0.891753585 | 0.959029158 |
| 0.891405784 | 0.958762975 |
| 0.891360475 | 0.958757545 |
| 0.891170428 | 0.958618077 |
| 0.891066921 | 0.958593337 |
| 0.891034546 | 0.958580161 |
| 0.89094174 | 0.958501971 |
| 0.89087247 | 0.958481188 |
| 0.890841721 | 0.958480971 |
| 0.890834977 | 0.958480971 |
| 0.890808362 | 0.958480971 |
| 0.890781588 | 0.958480971 |
| 0.890646919 | 0.958429373 |
| 0.890592705 | 0.958429373 |
| 0.890533413 | 0.95838741 |
| 0.890527632 | 0.95838741 |
| 0.89047946 | 0.958372658 |
| 0.890468119 | 0.958372658 |
| 0.890310313 | 0.958255577 |
| 0.890304022 | 0.958255577 |
| 0.89029925 | 0.958255577 |
| 0.890294731 | 0.958255577 |
| 0.890236034 | 0.958255577 |
| 0.890157052 | 0.958242206 |
| 0.890040523 | 0.958179958 |
| 0.890006457 | 0.958179958 |

| | |
|---|---|
| 0.889916191 | 0.958156176 |
| 0.889837488 | 0.958136418 |
| 0.889558033 | 0.957943799 |
| 0.88948985 | 0.957913693 |
| 0.889466871 | 0.957913693 |
| 0.889430296 | 0.957913693 |
| 0.889289885 | 0.957806634 |
| 0.889289141 | 0.957806634 |
| 0.88925066 | 0.957806634 |
| 0.889200482 | 0.95777532 |
| 0.889184508 | 0.95777532 |
| 0.889175624 | 0.95777532 |
| 0.889012583 | 0.957700871 |
| 0.8889772 | 0.957700871 |
| 0.88887887 | 0.957688812 |
| 0.888841693 | 0.957670422 |
| 0.888784777 | 0.957630762 |
| 0.888705116 | 0.957588258 |
| 0.888669443 | 0.957571483 |
| 0.888597264 | 0.957515371 |
| 0.88858886 | 0.957515371 |
| 0.888587546 | 0.957515371 |
| 0.888446813 | 0.957486608 |
| 0.888386071 | 0.957461099 |
| 0.888355003 | 0.957449281 |
| 0.888257528 | 0.957409222 |
| 0.88815352 | 0.957318782 |
| 0.888067631 | 0.957303037 |
| 0.888043072 | 0.957303037 |
| 0.88799231 | 0.957275006 |
| 0.887960471 | 0.95726235 |
| 0.887870522 | 0.957187047 |
| 0.887788949 | 0.957120771 |
| 0.887683389 | 0.95702863 |
| 0.887600009 | 0.956982063 |
| 0.887538367 | 0.956958929 |
| 0.887498242 | 0.956937329 |
| 0.887322944 | 0.956834964 |
| 0.887302816 | 0.956834923 |
| 0.887274002 | 0.956834923 |
| 0.887257059 | 0.956834923 |
| 0.887251203 | 0.956834923 |
| 0.887207149 | 0.956834923 |
| 0.887062734 | 0.956749328 |
| 0.886910657 | 0.956636374 |

| | |
|---|---|
| 0.886791224 | 0.95658144 |
| 0.886571652 | 0.956417848 |
| 0.886436248 | 0.956333511 |
| 0.886328289 | 0.956260362 |
| 0.886195555 | 0.956138815 |
| 0.886188237 | 0.956138815 |
| 0.886162809 | 0.956138815 |
| 0.88613345 | 0.956138815 |
| 0.88612504 | 0.956138815 |
| 0.886099447 | 0.956138815 |
| 0.886097203 | 0.956138815 |
| 0.88605181 | 0.956138815 |
| 0.885742721 | 0.955996749 |
| 0.885676859 | 0.955947327 |
| 0.885392421 | 0.95580956 |
| 0.885341726 | 0.95580956 |
| 0.885330842 | 0.95580956 |
| 0.885297498 | 0.95580956 |
| 0.885147224 | 0.95574388 |
| 0.885137439 | 0.95574388 |
| 0.885016593 | 0.955667829 |
| 0.885014382 | 0.955667829 |
| 0.884982905 | 0.955667829 |
| 0.884870796 | 0.955640394 |
| 0.884763579 | 0.955589609 |
| 0.884763432 | 0.955589609 |
| 0.884637249 | 0.955561513 |
| 0.884609237 | 0.955552927 |
| 0.884576417 | 0.955539147 |
| 0.884370644 | 0.955425216 |
| 0.884330972 | 0.955404028 |
| 0.884316926 | 0.955404028 |
| 0.884282082 | 0.955404028 |
| 0.884242136 | 0.955394742 |
| 0.884074024 | 0.955234773 |
| 0.883797251 | 0.95510974 |
| 0.883750846 | 0.95510974 |
| 0.883719187 | 0.95510974 |
| 0.88370629 | 0.95510974 |
| 0.883658007 | 0.955088617 |
| 0.883477465 | 0.954958496 |
| 0.883453676 | 0.954958496 |
| 0.883402745 | 0.954958496 |
| 0.883398745 | 0.954958496 |
| 0.883370479 | 0.954958496 |

| | |
|---|---|
| 0.883317326 | 0.954955423 |
| 0.883224289 | 0.954900986 |
| 0.883155775 | 0.954849155 |
| 0.883006672 | 0.954749932 |
| 0.882967448 | 0.954749932 |
| 0.882885121 | 0.954686563 |
| 0.882866972 | 0.954686563 |
| 0.882823757 | 0.954686563 |
| 0.882817881 | 0.954686563 |
| 0.882808517 | 0.954686563 |
| 0.882732774 | 0.954686563 |
| 0.882692954 | 0.954686563 |
| 0.882640882 | 0.954682563 |
| 0.882541206 | 0.954639791 |
| 0.882515261 | 0.954633408 |
| 0.882508374 | 0.954633408 |
| 0.882362071 | 0.954619472 |
| 0.882192937 | 0.954544903 |
| 0.882178992 | 0.954544903 |
| 0.882152783 | 0.954544828 |
| 0.882077438 | 0.954528359 |
| 0.88206925 | 0.954528359 |
| 0.881958925 | 0.95448686 |
| 0.88188648 | 0.954430145 |
| 0.881855928 | 0.95442716 |
| 0.88185387 | 0.95442716 |
| 0.881853191 | 0.95442716 |
| 0.881846448 | 0.95442716 |
| 0.881814989 | 0.95442716 |
| 0.881797667 | 0.95442716 |
| 0.881793368 | 0.95442716 |
| 0.881717836 | 0.95442716 |
| 0.881671703 | 0.95442716 |
| 0.881662169 | 0.95442716 |
| 0.881436565 | 0.954333567 |
| 0.881409489 | 0.954333567 |
| 0.881381903 | 0.954333567 |
| 0.881381364 | 0.954333567 |
| 0.881326664 | 0.954333567 |
| 0.881115766 | 0.954263891 |
| 0.881099976 | 0.954263891 |
| 0.880995783 | 0.954203401 |
| 0.880937365 | 0.954161828 |
| 0.880923318 | 0.954161828 |
| 0.880910474 | 0.954161828 |

| | |
|---|---|
| 0.880866516 | 0.954161828 |
| 0.880840582 | 0.954161828 |
| 0.880600307 | 0.954057116 |
| 0.880531064 | 0.954025502 |
| 0.88046858 | 0.953979505 |
| 0.880441875 | 0.953972271 |
| 0.880420649 | 0.953970976 |
| 0.880311986 | 0.953896639 |
| 0.880234902 | 0.953834812 |
| 0.880194296 | 0.953812512 |
| 0.880143204 | 0.953778848 |
| 0.879962903 | 0.953666403 |
| 0.879923499 | 0.953649254 |
| 0.879861466 | 0.953603724 |
| 0.879838635 | 0.953600679 |
| 0.879828296 | 0.953600679 |
| 0.879818698 | 0.953600679 |
| 0.879625541 | 0.953543317 |
| 0.879607313 | 0.953543317 |
| 0.879593016 | 0.953543317 |
| 0.879422895 | 0.953475556 |
| 0.879379251 | 0.953466912 |
| 0.879183473 | 0.953367894 |
| 0.879176569 | 0.953367894 |
| 0.879070612 | 0.953330788 |
| 0.879038571 | 0.953319292 |
| 0.879022289 | 0.953319292 |
| 0.87896595 | 0.953319292 |
| 0.878912773 | 0.953313109 |
| 0.878699916 | 0.953169063 |
| 0.87862398 | 0.953130109 |
| 0.878604558 | 0.953130109 |
| 0.878492898 | 0.953031325 |
| 0.878447051 | 0.953003296 |
| 0.878438361 | 0.953003296 |
| 0.878431244 | 0.953003296 |
| 0.878318999 | 0.952929494 |
| 0.878212034 | 0.952878564 |
| 0.878132013 | 0.952878564 |
| 0.878052836 | 0.952857784 |
| 0.878012883 | 0.952853259 |
| 0.877990348 | 0.952853259 |
| 0.877904797 | 0.952805683 |
| 0.877844496 | 0.952761948 |
| 0.877508419 | 0.952462305 |

| | |
|---|---|
| 0.877494796 | 0.952462305 |
| 0.877111829 | 0.95230097 |
| 0.876819545 | 0.952148577 |
| 0.876792631 | 0.952141061 |
| 0.876669627 | 0.95209432 |
| 0.876568333 | 0.95204944 |
| 0.876541666 | 0.952042187 |
| 0.876341551 | 0.951911669 |
| 0.876330586 | 0.951911669 |
| 0.876317782 | 0.951911669 |
| 0.87627836 | 0.951911669 |
| 0.876264644 | 0.951911669 |
| 0.875930643 | 0.951709702 |
| 0.875895772 | 0.951709623 |
| 0.875835427 | 0.951687479 |
| 0.875736233 | 0.951623117 |
| 0.875704812 | 0.95161726 |
| 0.875638908 | 0.95161726 |
| 0.875476171 | 0.951492483 |
| 0.875402354 | 0.951433968 |
| 0.875378909 | 0.951430198 |
| 0.875349005 | 0.951430198 |
| 0.875279578 | 0.951409085 |
| 0.875164648 | 0.95130587 |
| 0.875020497 | 0.951193263 |
| 0.874754827 | 0.95105575 |
| 0.874717906 | 0.951037318 |
| 0.874585366 | 0.950940391 |
| 0.874492778 | 0.950936554 |
| 0.874417999 | 0.950884902 |
| 0.8743845 | 0.950870185 |
| 0.874307823 | 0.950808509 |
| 0.874301012 | 0.950808509 |
| 0.874291272 | 0.950808509 |
| 0.874265652 | 0.950808509 |
| 0.874216496 | 0.950808509 |
| 0.874172199 | 0.950808509 |
| 0.874154488 | 0.950808509 |
| 0.874089545 | 0.950808509 |
| 0.873945442 | 0.95067491 |
| 0.873842252 | 0.950606084 |
| 0.873788388 | 0.950599185 |
| 0.873711514 | 0.950599185 |
| 0.873703866 | 0.950599185 |
| 0.873634062 | 0.95059673 |

| | |
|---|---|
| 0.873597085 | 0.950578213 |
| 0.873566449 | 0.950566595 |
| 0.873556899 | 0.950566595 |
| 0.873501818 | 0.9505397 |
| 0.87341473 | 0.950488367 |
| 0.87336292 | 0.950469649 |
| 0.873175799 | 0.950372137 |
| 0.873125665 | 0.95036923 |
| 0.873119461 | 0.95036923 |
| 0.873103094 | 0.95036923 |
| 0.872944164 | 0.950280415 |
| 0.872898623 | 0.950280415 |
| 0.872844975 | 0.950259164 |
| 0.872820788 | 0.950259164 |
| 0.872703547 | 0.950259164 |
| 0.872634033 | 0.950259164 |
| 0.872557426 | 0.950259164 |
| 0.872535459 | 0.950259164 |
| 0.872419213 | 0.95020833 |
| 0.872311574 | 0.950155633 |
| 0.872264689 | 0.950126941 |
| 0.872183535 | 0.950097778 |
| 0.872173847 | 0.950097778 |
| 0.871988149 | 0.949956072 |
| 0.871950026 | 0.949945907 |
| 0.871883619 | 0.94992911 |
| 0.871847906 | 0.949911931 |
| 0.871771826 | 0.949850768 |
| 0.871755871 | 0.949850768 |
| 0.871585645 | 0.94975655 |
| 0.871571588 | 0.94975655 |
| 0.87155106 | 0.94975655 |
| 0.871534335 | 0.94975655 |
| 0.871527631 | 0.94975655 |
| 0.871476746 | 0.94975655 |
| 0.871359086 | 0.949699001 |
| 0.871341975 | 0.949699001 |
| 0.871337811 | 0.949699001 |
| 0.871295322 | 0.949699001 |
| 0.871259137 | 0.949699001 |
| 0.87106261 | 0.949642882 |
| 0.871033798 | 0.949633208 |
| 0.871004806 | 0.949623338 |
| 0.87095393 | 0.949589608 |
| 0.87090312 | 0.949589608 |

| | |
|---|---|
| 0.870836573 | 0.949570343 |
| 0.87079372 | 0.949545355 |
| 0.870733505 | 0.949523175 |
| 0.870692016 | 0.949499671 |
| 0.87067369 | 0.949499671 |
| 0.870656431 | 0.949499671 |
| 0.870582291 | 0.949488717 |
| 0.87048421 | 0.949403487 |
| 0.870423487 | 0.94938074 |
| 0.870399744 | 0.94938074 |
| 0.870265161 | 0.94927327 |
| 0.870100816 | 0.949137092 |
| 0.870090412 | 0.949137092 |
| 0.870083246 | 0.949137092 |
| 0.869901083 | 0.949014324 |
| 0.869767414 | 0.948947669 |
| 0.86965447 | 0.948867917 |
| 0.869561833 | 0.948810315 |
| 0.869552149 | 0.948810315 |
| 0.869528236 | 0.948810315 |
| 0.869450725 | 0.948797765 |
| 0.869379242 | 0.948741497 |
| 0.869362165 | 0.948741497 |
| 0.869252237 | 0.948668109 |
| 0.869237911 | 0.948668109 |
| 0.869195978 | 0.948650187 |
| 0.86916101 | 0.948633761 |
| 0.868705155 | 0.948440511 |
| 0.868682554 | 0.948437577 |
| 0.868659793 | 0.948434469 |
| 0.868494134 | 0.948275335 |
| 0.868429031 | 0.94822599 |
| 0.868420155 | 0.94822599 |
| 0.86833838 | 0.948170485 |
| 0.868336461 | 0.948170485 |
| 0.868315077 | 0.948170485 |
| 0.868261798 | 0.948170485 |
| 0.868256211 | 0.948170485 |
| 0.868251739 | 0.948170485 |
| 0.86824748 | 0.948170485 |
| 0.868217756 | 0.948170485 |
| 0.86806049 | 0.948170485 |
| 0.86799105 | 0.948117282 |
| 0.867988629 | 0.948117282 |
| 0.867963413 | 0.948117282 |

| | |
|---|---|
| 0.867935066 | 0.948117282 |
| 0.867786807 | 0.948024641 |
| 0.867728821 | 0.948004785 |
| 0.867553622 | 0.947856863 |
| 0.867539261 | 0.947856863 |
| 0.867538446 | 0.947856863 |
| 0.867528477 | 0.947856863 |
| 0.867478359 | 0.947856863 |
| 0.867455159 | 0.947856863 |
| 0.867454299 | 0.947856863 |
| 0.867268106 | 0.94780583 |
| 0.867263694 | 0.94780583 |
| 0.867209322 | 0.947785084 |
| 0.867170143 | 0.947764013 |
| 0.867132246 | 0.947744343 |
| 0.867113354 | 0.947744343 |
| 0.867075106 | 0.947725389 |
| 0.867056839 | 0.947725389 |
| 0.867052542 | 0.947725389 |
| 0.867018481 | 0.947725389 |
| 0.866965026 | 0.947713817 |
| 0.866945396 | 0.947713817 |
| 0.866853412 | 0.947657059 |
| 0.866845692 | 0.947657059 |
| 0.86682646 | 0.947657059 |
| 0.866820527 | 0.947657059 |
| 0.866773013 | 0.947657059 |
| 0.8667302 | 0.947657059 |
| 0.866704524 | 0.947657059 |
| 0.866651653 | 0.947654015 |
| 0.866623123 | 0.947654015 |
| 0.866615396 | 0.947654015 |
| 0.866588883 | 0.947654015 |
| 0.866351302 | 0.947489447 |
| 0.866100922 | 0.947312801 |
| 0.866087677 | 0.947312801 |
| 0.866074133 | 0.947312801 |
| 0.866040856 | 0.947312801 |
| 0.866002496 | 0.947312801 |
| 0.865978923 | 0.947312801 |
| 0.865836445 | 0.947268835 |
| 0.865739937 | 0.947222513 |
| 0.865715661 | 0.947217712 |
| 0.865529185 | 0.947035437 |
| 0.865189587 | 0.946772614 |

| | |
|---|---|
| 0.865187556 | 0.946772614 |
| 0.8651641 | 0.946772614 |
| 0.865152503 | 0.946772614 |
| 0.865025829 | 0.946746513 |
| 0.865010453 | 0.946746513 |
| 0.864955516 | 0.946733996 |
| 0.864848002 | 0.946688129 |
| 0.864810198 | 0.946683721 |
| 0.864791313 | 0.946683721 |
| 0.864679526 | 0.946605948 |
| 0.864622016 | 0.946605948 |
| 0.864609225 | 0.946605948 |
| 0.864530092 | 0.946572903 |
| 0.864485717 | 0.946546076 |
| 0.864469535 | 0.946546076 |
| 0.864459352 | 0.946546076 |
| 0.864365812 | 0.946546076 |
| 0.864260962 | 0.946517587 |
| 0.864132026 | 0.946441669 |
| 0.864111052 | 0.946440463 |
| 0.864064975 | 0.94641176 |
| 0.864056565 | 0.94641176 |
| 0.864025848 | 0.94641176 |
| 0.863997166 | 0.94641176 |
| 0.863824699 | 0.946365499 |
| 0.863691274 | 0.946263585 |
| 0.863659136 | 0.946256811 |
| 0.863649028 | 0.946256811 |
| 0.86364072 | 0.946256811 |
| 0.863602145 | 0.946256811 |
| 0.863578619 | 0.946256811 |
| 0.863563107 | 0.946256811 |
| 0.863534883 | 0.946256811 |
| 0.863532883 | 0.946256811 |
| 0.863518309 | 0.946256811 |
| 0.863481977 | 0.946256811 |
| 0.863472629 | 0.946256811 |
| 0.863411955 | 0.946256811 |
| 0.863364719 | 0.946256811 |
| 0.863335727 | 0.946256811 |
| 0.86329517 | 0.946256811 |
| 0.86315833 | 0.946256811 |
| 0.862898542 | 0.946122336 |
| 0.862864011 | 0.946121332 |
| 0.862796588 | 0.94608862 |

| | |
|---|---|
| 0.862755262 | 0.946065086 |
| 0.86270874 | 0.946065086 |
| 0.862610143 | 0.946058428 |
| 0.862465707 | 0.945963768 |
| 0.862443193 | 0.945962461 |
| 0.862386534 | 0.945922099 |
| 0.862361638 | 0.945916576 |
| 0.862282926 | 0.945873806 |
| 0.862281373 | 0.945873806 |
| 0.862175128 | 0.94582091 |
| 0.862092556 | 0.945752111 |
| 0.861950452 | 0.945639781 |
| 0.861867798 | 0.945570884 |
| 0.861799094 | 0.945539069 |
| 0.861798091 | 0.945539069 |
| 0.861780065 | 0.945539069 |
| 0.861740249 | 0.945539069 |
| 0.86163588 | 0.945490682 |
| 0.86156909 | 0.945439174 |
| 0.861547028 | 0.945436748 |
| 0.861494876 | 0.945401301 |
| 0.861422778 | 0.945385113 |
| 0.861272805 | 0.945353403 |
| 0.861178827 | 0.945294061 |
| 0.861155701 | 0.945294061 |
| 0.861142352 | 0.945294061 |
| 0.861133843 | 0.945294061 |
| 0.861106829 | 0.945294061 |
| 0.861004607 | 0.945255323 |
| 0.86097457 | 0.945244136 |
| 0.860963514 | 0.945244136 |
| 0.860947393 | 0.945244136 |
| 0.86089556 | 0.945244136 |
| 0.860855711 | 0.945244136 |
| 0.860821549 | 0.94522866 |
| 0.860681252 | 0.945096396 |
| 0.860554278 | 0.945022333 |
| 0.860551067 | 0.945022333 |
| 0.86048302 | 0.944987659 |
| 0.860478199 | 0.944987659 |
| 0.86040948 | 0.944971484 |
| 0.860392747 | 0.944971484 |
| 0.860370008 | 0.944971484 |
| 0.860359071 | 0.944971484 |
| 0.860357136 | 0.944971484 |

| | |
|---|---|
| 0.860336844 | 0.944971484 |
| 0.860288061 | 0.944971484 |
| 0.860278879 | 0.944971484 |
| 0.860268755 | 0.944971484 |
| 0.860217687 | 0.944971484 |
| 0.860093792 | 0.944971484 |
| 0.859812679 | 0.94490128 |
| 0.859775759 | 0.94490128 |
| 0.859740045 | 0.944890715 |
| 0.859715161 | 0.94488517 |
| 0.859677628 | 0.944865723 |
| 0.859650769 | 0.944864154 |
| 0.859523394 | 0.944761613 |
| 0.859483637 | 0.944739716 |
| 0.859392325 | 0.944661149 |
| 0.859360583 | 0.944655055 |
| 0.859352652 | 0.944655055 |
| 0.859336134 | 0.944655055 |
| 0.859285518 | 0.944655055 |
| 0.859240864 | 0.944655055 |
| 0.859230618 | 0.944655055 |
| 0.859186005 | 0.944655055 |
| 0.859130625 | 0.944655055 |
| 0.858872225 | 0.944547243 |
| 0.858855756 | 0.944547243 |
| 0.858807508 | 0.944541502 |
| 0.85867425 | 0.94443856 |
| 0.858637324 | 0.944419756 |
| 0.858536654 | 0.944330837 |
| 0.858483663 | 0.944294358 |
| 0.858474816 | 0.944294358 |
| 0.858464958 | 0.944294358 |
| 0.858447286 | 0.944294358 |
| 0.858386264 | 0.944294358 |
| 0.858371664 | 0.944294358 |
| 0.858289205 | 0.944276738 |
| 0.858284059 | 0.944276738 |
| 0.858277723 | 0.944276738 |
| 0.858230957 | 0.944276738 |
| 0.858224504 | 0.944276738 |
| 0.858204295 | 0.944276738 |
| 0.858120252 | 0.944276738 |
| 0.858053531 | 0.944276738 |
| 0.857993638 | 0.944235113 |
| 0.857971212 | 0.944232252 |

| | |
|---|---|
| 0.857840831 | 0.944110578 |
| 0.857759179 | 0.944064345 |
| 0.857707401 | 0.944052192 |
| 0.857620425 | 0.944052192 |
| 0.857532793 | 0.944052192 |
| 0.857508889 | 0.944052192 |
| 0.857435463 | 0.944019137 |
| 0.857396018 | 0.944019137 |
| 0.857390308 | 0.944019137 |
| 0.857275762 | 0.943946749 |
| 0.857272483 | 0.943946749 |
| 0.857222304 | 0.943931533 |
| 0.857165514 | 0.943912646 |
| 0.8571213 | 0.943909397 |
| 0.8571187 | 0.943909397 |
| 0.857102894 | 0.943909397 |
| 0.857070176 | 0.943909397 |
| 0.857002301 | 0.943907511 |
| 0.856998749 | 0.943907511 |
| 0.85694842 | 0.943907511 |
| 0.856674244 | 0.94374261 |
| 0.856627327 | 0.943734582 |
| 0.856523119 | 0.943663434 |
| 0.856492724 | 0.943663434 |
| 0.856418475 | 0.943635458 |
| 0.856414595 | 0.943635458 |
| 0.8563757 | 0.943631989 |
| 0.856055078 | 0.943405306 |
| 0.856000215 | 0.943371025 |
| 0.855985967 | 0.943371025 |
| 0.855933327 | 0.943371025 |
| 0.855887818 | 0.943371025 |
| 0.855878816 | 0.943371025 |
| 0.855839915 | 0.943371025 |
| 0.855816631 | 0.943371025 |
| 0.855644533 | 0.943240954 |
| 0.855601471 | 0.943240954 |
| 0.855565433 | 0.943240954 |
| 0.855548988 | 0.943240954 |
| 0.855535381 | 0.943240954 |
| 0.855489619 | 0.943240954 |
| 0.855380867 | 0.943190438 |
| 0.855372764 | 0.943190438 |
| 0.855371994 | 0.943190438 |
| 0.855331384 | 0.943190438 |

| | |
|---|---|
| 0.855288967 | 0.943190438 |
| 0.855272055 | 0.943190438 |
| 0.855261987 | 0.943190438 |
| 0.855238831 | 0.943190438 |
| 0.854979734 | 0.943053794 |
| 0.854683702 | 0.942880123 |
| 0.854621786 | 0.942860071 |
| 0.854571986 | 0.942860071 |
| 0.854472463 | 0.942860071 |
| 0.854296798 | 0.942748043 |
| 0.854081627 | 0.942575601 |
| 0.854074559 | 0.942575601 |
| 0.85407101 | 0.942575601 |
| 0.854044131 | 0.942575601 |
| 0.854003796 | 0.942575601 |
| 0.853901946 | 0.942563515 |
| 0.853861852 | 0.942562948 |
| 0.853760413 | 0.942494661 |
| 0.853691996 | 0.942462824 |
| 0.853686729 | 0.942462824 |
| 0.853596613 | 0.942401212 |
| 0.853553704 | 0.942397531 |
| 0.853527463 | 0.942397531 |
| 0.853500385 | 0.942397531 |
| 0.853491262 | 0.942397531 |
| 0.853443841 | 0.942385464 |
| 0.853299038 | 0.94229111 |
| 0.853124316 | 0.942185549 |
| 0.853092363 | 0.942182078 |
| 0.852954434 | 0.942182078 |
| 0.852941789 | 0.942182078 |
| 0.852910479 | 0.942182078 |
| 0.852902076 | 0.942182078 |
| 0.852871863 | 0.942182078 |
| 0.852782073 | 0.942182078 |
| 0.852695558 | 0.942182078 |
| 0.852694973 | 0.942182078 |
| 0.852674962 | 0.942182078 |
| 0.852631283 | 0.942182078 |
| 0.852624331 | 0.942182078 |
| 0.852598075 | 0.942182078 |
| 0.852571343 | 0.942182078 |
| 0.852523351 | 0.942182078 |
| 0.852509438 | 0.942182078 |
| 0.852489553 | 0.942182078 |

| | |
|---|---|
| 0.852460539 | 0.942182078 |
| 0.85245956 | 0.942182078 |
| 0.852455296 | 0.942182078 |
| 0.852424014 | 0.942182078 |
| 0.852253639 | 0.942119675 |
| 0.852065906 | 0.941998224 |
| 0.852030397 | 0.941998224 |
| 0.852017634 | 0.941998224 |
| 0.851983458 | 0.941998224 |
| 0.851967198 | 0.941998224 |
| 0.851931512 | 0.941998224 |
| 0.851805901 | 0.941998224 |
| 0.851767572 | 0.941997774 |
| 0.851766491 | 0.941997774 |
| 0.851711953 | 0.941997774 |
| 0.851650166 | 0.941974546 |
| 0.851512367 | 0.941868631 |
| 0.851484122 | 0.941859263 |
| 0.851478114 | 0.941859263 |
| 0.851385712 | 0.941816029 |
| 0.851345137 | 0.941814895 |
| 0.851340357 | 0.941814895 |
| 0.851293497 | 0.941801519 |
| 0.851263182 | 0.941789857 |
| 0.851236209 | 0.941781893 |
| 0.851231817 | 0.941781893 |
| 0.851201352 | 0.941781893 |
| 0.85114111 | 0.941742308 |
| 0.851115345 | 0.941735678 |
| 0.851084012 | 0.941722887 |
| 0.85107498 | 0.941722887 |
| 0.851054254 | 0.941722887 |
| 0.850900077 | 0.941650625 |
| 0.850896844 | 0.941650625 |
| 0.85083715 | 0.941646627 |
| 0.85075049 | 0.941594478 |
| 0.850724187 | 0.941587247 |
| 0.850671085 | 0.941550355 |
| 0.850653479 | 0.941550355 |
| 0.850610317 | 0.941526856 |
| 0.850585702 | 0.941521492 |
| 0.850395943 | 0.941377082 |
| 0.850336549 | 0.941355093 |
| 0.850280713 | 0.941315161 |
| 0.850274734 | 0.941315161 |

| | |
|---|---|
| 0.850216936 | 0.941310196 |
| 0.850210648 | 0.941310196 |
| 0.850166078 | 0.941310196 |
| 0.85014221 | 0.941310196 |
| 0.850109712 | 0.941310196 |
| 0.850051649 | 0.941310196 |
| 0.849991999 | 0.94127997 |
| 0.849745156 | 0.941094143 |
| 0.849461097 | 0.940845182 |
| 0.849356038 | 0.940750698 |
| 0.84932928 | 0.940742939 |
| 0.849295871 | 0.940727811 |
| 0.849261246 | 0.940719396 |
| 0.849184671 | 0.940692151 |
| 0.849115246 | 0.940659002 |
| 0.848951245 | 0.940542952 |
| 0.848944507 | 0.940542952 |
| 0.848808355 | 0.940450275 |
| 0.848687425 | 0.940381918 |
| 0.848590932 | 0.940329677 |
| 0.848527831 | 0.940292584 |
| 0.848466469 | 0.940246464 |
| 0.848416505 | 0.940234231 |
| 0.848331087 | 0.940183941 |
| 0.848328307 | 0.940183941 |
| 0.848315336 | 0.940183941 |
| 0.848153971 | 0.940100741 |
| 0.848076892 | 0.940100741 |
| 0.848029346 | 0.940100741 |
| 0.847988515 | 0.940088661 |
| 0.847967583 | 0.940087339 |
| 0.84795336 | 0.940087339 |
| 0.847951973 | 0.940087339 |
| 0.847922883 | 0.940087339 |
| 0.847916226 | 0.940087339 |
| 0.847816863 | 0.940073423 |
| 0.847673913 | 0.940034765 |
| 0.84766878 | 0.940034765 |
| 0.847647467 | 0.940034765 |
| 0.847591875 | 0.940034765 |
| 0.847535211 | 0.940034765 |
| 0.847469566 | 0.940019997 |
| 0.847453091 | 0.940019997 |
| 0.847358821 | 0.939981333 |
| 0.847345827 | 0.939981333 |

| | |
|---|---|
| 0.847336954 | 0.939981333 |
| 0.847040007 | 0.939759002 |
| 0.846976597 | 0.939710541 |
| 0.846861597 | 0.939626728 |
| 0.846832149 | 0.939615943 |
| 0.846804144 | 0.939606761 |
| 0.84630679 | 0.939339396 |
| 0.846272353 | 0.939323064 |
| 0.846060685 | 0.939153784 |
| 0.846035099 | 0.939147271 |
| 0.845950392 | 0.939118906 |
| 0.845767545 | 0.93900347 |
| 0.845575124 | 0.938943052 |
| 0.845449656 | 0.938828799 |
| 0.845440713 | 0.938828799 |
| 0.845370842 | 0.938828799 |
| 0.845359242 | 0.938828799 |
| 0.84528307 | 0.938815744 |
| 0.845235339 | 0.938815744 |
| 0.845225631 | 0.938815744 |
| 0.845175498 | 0.938815744 |
| 0.845108105 | 0.938815744 |
| 0.844857566 | 0.93862771 |
| 0.844840072 | 0.93862771 |
| 0.844823996 | 0.93862771 |
| 0.844691743 | 0.938531054 |
| 0.844678952 | 0.938531054 |
| 0.844654554 | 0.938531054 |
| 0.844637419 | 0.938531054 |
| 0.844563081 | 0.93849757 |
| 0.844542564 | 0.93849757 |
| 0.844523329 | 0.93849757 |
| 0.844480867 | 0.93849757 |
| 0.84442502 | 0.938497432 |
| 0.844399318 | 0.938490766 |
| 0.844284779 | 0.938421458 |
| 0.84408865 | 0.938276855 |
| 0.84398103 | 0.938179123 |
| 0.843964239 | 0.938179123 |
| 0.843745227 | 0.937982679 |
| 0.843742907 | 0.937982679 |
| 0.843721865 | 0.937982679 |
| 0.843721314 | 0.937982679 |
| 0.84371838 | 0.937982679 |
| 0.843676426 | 0.937982679 |

| | |
|---|---|
| 0.843562816 | 0.937961294 |
| 0.843486503 | 0.93793588 |
| 0.843419442 | 0.937883208 |
| 0.843316864 | 0.937812932 |
| 0.843033588 | 0.937541696 |
| 0.84300098 | 0.937541696 |
| 0.842991503 | 0.937541696 |
| 0.842988778 | 0.937541696 |
| 0.842969738 | 0.937541696 |
| 0.842831483 | 0.937492061 |
| 0.842828056 | 0.937492061 |
| 0.842815034 | 0.937492061 |
| 0.842808435 | 0.937492061 |
| 0.842634766 | 0.937455921 |
| 0.842540555 | 0.937453079 |
| 0.842516838 | 0.937453079 |
| 0.842425495 | 0.937412655 |
| 0.842344795 | 0.937366657 |
| 0.842279171 | 0.937315531 |
| 0.84226326 | 0.937315531 |
| 0.842229055 | 0.937315531 |
| 0.842179104 | 0.937312146 |
| 0.842145604 | 0.937312146 |
| 0.842110684 | 0.937312146 |
| 0.842081596 | 0.937312146 |
| 0.842046917 | 0.937312146 |
| 0.841981727 | 0.937312146 |
| 0.841954839 | 0.937305014 |
| 0.841896471 | 0.937270422 |
| 0.841881044 | 0.937270422 |
| 0.841876505 | 0.937270422 |
| 0.841856167 | 0.937270422 |
| 0.841836601 | 0.937270422 |
| 0.841757924 | 0.937270422 |
| 0.84174721 | 0.937270422 |
| 0.841497468 | 0.937212074 |
| 0.841452775 | 0.937184213 |
| 0.84130642 | 0.93704312 |
| 0.841298349 | 0.93704312 |
| 0.841088288 | 0.936865894 |
| 0.841015979 | 0.93680726 |
| 0.840948664 | 0.936776097 |
| 0.840803053 | 0.9366358 |
| 0.840789867 | 0.9366358 |
| 0.840759813 | 0.9366358 |

| | |
|---|---|
| 0.840715891 | 0.9366358 |
| 0.840689797 | 0.9366358 |
| 0.84068818 | 0.9366358 |
| 0.840657062 | 0.9366358 |
| 0.840609518 | 0.9366358 |
| 0.840589021 | 0.9366358 |
| 0.840397724 | 0.936530373 |
| 0.840333453 | 0.936489574 |
| 0.84023718 | 0.936443492 |
| 0.840211728 | 0.936440955 |
| 0.840211135 | 0.936440955 |
| 0.840017352 | 0.936368034 |
| 0.839929573 | 0.936319768 |
| 0.839844813 | 0.936272872 |
| 0.839824258 | 0.936272872 |
| 0.839722493 | 0.936236356 |
| 0.839682162 | 0.936236356 |
| 0.839680905 | 0.936236356 |
| 0.839605718 | 0.936221707 |
| 0.839597016 | 0.936221707 |
| 0.839516385 | 0.93616593 |
| 0.839480295 | 0.936149176 |
| 0.839402609 | 0.936104809 |
| 0.83939308 | 0.936104809 |
| 0.839260392 | 0.936033878 |
| 0.839250574 | 0.936033878 |
| 0.839209169 | 0.936020587 |
| 0.839193728 | 0.936020587 |
| 0.839051445 | 0.935888503 |
| 0.838896957 | 0.935822156 |
| 0.838893892 | 0.935822156 |
| 0.838881115 | 0.935822156 |
| 0.838826723 | 0.935822156 |
| 0.838793586 | 0.935822156 |
| 0.838753 | 0.935818613 |
| 0.838703639 | 0.935785461 |
| 0.838692705 | 0.935785461 |
| 0.838652181 | 0.935771892 |
| 0.838635825 | 0.935771892 |
| 0.838545523 | 0.935696725 |
| 0.83841905 | 0.935659096 |
| 0.838349999 | 0.935631296 |
| 0.838336241 | 0.935631296 |
| 0.83831364 | 0.935631296 |
| 0.838179158 | 0.935507095 |

| | |
|---|---|
| 0.838112711 | 0.935454854 |
| 0.838036624 | 0.935413774 |
| 0.838013711 | 0.935410121 |
| 0.837768379 | 0.935180111 |
| 0.837724693 | 0.935173915 |
| 0.837717435 | 0.935173915 |
| 0.837643121 | 0.935149877 |
| 0.837562333 | 0.935103524 |
| 0.837488566 | 0.935065005 |
| 0.837432756 | 0.935024613 |
| 0.837381369 | 0.935024613 |
| 0.837364658 | 0.935024613 |
| 0.837337424 | 0.935024613 |
| 0.837331925 | 0.935024613 |
| 0.837082348 | 0.934874903 |
| 0.836955812 | 0.934798895 |
| 0.836942912 | 0.934798895 |
| 0.836935631 | 0.934798895 |
| 0.836912076 | 0.934798895 |
| 0.836870763 | 0.934798895 |
| 0.836842542 | 0.934798895 |
| 0.836823398 | 0.934798895 |
| 0.836784952 | 0.934798895 |
| 0.836672864 | 0.934734889 |
| 0.836650152 | 0.934734889 |
| 0.836630704 | 0.934734889 |
| 0.836512584 | 0.934632624 |
| 0.83638046 | 0.934569478 |
| 0.836122822 | 0.93442483 |
| 0.836085535 | 0.93442483 |
| 0.836015038 | 0.93442483 |
| 0.836006508 | 0.93442483 |
| 0.835978786 | 0.93442483 |
| 0.835974521 | 0.93442483 |
| 0.835963005 | 0.93442483 |
| 0.835892484 | 0.93442483 |
| 0.835789859 | 0.93442483 |
| 0.83575098 | 0.93442483 |
| 0.835689605 | 0.93442483 |
| 0.835685099 | 0.93442483 |
| 0.835603183 | 0.934405691 |
| 0.835519539 | 0.934361208 |
| 0.835387445 | 0.934293848 |
| 0.835359143 | 0.93428635 |
| 0.835345064 | 0.93428635 |

| | |
|---|---|
| 0.835331418 | 0.93428635 |
| 0.835198284 | 0.934194192 |
| 0.835190701 | 0.934194192 |
| 0.835094816 | 0.934144275 |
| 0.835064022 | 0.934144275 |
| 0.835054419 | 0.934144275 |
| 0.835005703 | 0.934144275 |
| 0.834983472 | 0.934144275 |
| 0.834886258 | 0.934106526 |
| 0.83488621 | 0.934106526 |
| 0.834769213 | 0.934106526 |
| 0.834725671 | 0.934106526 |
| 0.834711029 | 0.934106526 |
| 0.834631283 | 0.934106526 |
| 0.834559596 | 0.934106526 |
| 0.834414364 | 0.933990581 |
| 0.834389836 | 0.933990581 |
| 0.834362814 | 0.933990581 |
| 0.834322758 | 0.933982606 |
| 0.834311965 | 0.933982606 |
| 0.8342297 | 0.933944288 |
| 0.834192647 | 0.933924759 |
| 0.834136171 | 0.933883484 |
| 0.834065152 | 0.933825924 |
| 0.833882882 | 0.933745254 |
| 0.833843493 | 0.933745254 |
| 0.833758272 | 0.933723787 |
| 0.833701137 | 0.933698929 |
| 0.833620142 | 0.933678868 |
| 0.833619273 | 0.933678868 |
| 0.833496262 | 0.933584029 |
| 0.833463471 | 0.933569255 |
| 0.83345008 | 0.933569255 |
| 0.833393821 | 0.93353515 |
| 0.833275339 | 0.933433395 |
| 0.833275322 | 0.933433395 |
| 0.833224893 | 0.933433395 |
| 0.833189447 | 0.933422448 |
| 0.833189272 | 0.933422448 |
| 0.833083814 | 0.933422448 |
| 0.832983904 | 0.933369157 |
| 0.832969258 | 0.933369157 |
| 0.832811981 | 0.93330046 |
| 0.832799294 | 0.93330046 |
| 0.832662367 | 0.933260286 |

| | |
|---|---|
| 0.832659119 | 0.933260286 |
| 0.832567505 | 0.933223146 |
| 0.832565706 | 0.933223146 |
| 0.832484415 | 0.933196865 |
| 0.832460979 | 0.933196865 |
| 0.832370721 | 0.933163153 |
| 0.832360422 | 0.933163153 |
| 0.832285757 | 0.933163153 |
| 0.832192586 | 0.933125342 |
| 0.832127019 | 0.933103702 |
| 0.832091175 | 0.933103702 |
| 0.832043333 | 0.933103702 |
| 0.831913657 | 0.93301822 |
| 0.831905788 | 0.93301822 |
| 0.831883023 | 0.93301822 |
| 0.831840239 | 0.93301822 |
| 0.831756646 | 0.932973946 |
| 0.831748292 | 0.932973946 |
| 0.831724376 | 0.932973946 |
| 0.831585456 | 0.93289178 |
| 0.831349374 | 0.932707457 |
| 0.831342257 | 0.932707457 |
| 0.831276345 | 0.932703487 |
| 0.831273706 | 0.932703487 |
| 0.831255952 | 0.932703487 |
| 0.831192281 | 0.932703487 |
| 0.831075576 | 0.932605354 |
| 0.830677221 | 0.932356044 |
| 0.83062621 | 0.932342733 |
| 0.83055298 | 0.932326454 |
| 0.83053902 | 0.932326454 |
| 0.83043961 | 0.932275182 |
| 0.830365 | 0.932247274 |
| 0.830363032 | 0.932247274 |
| 0.830310219 | 0.932247274 |
| 0.830243287 | 0.932220502 |
| 0.830127812 | 0.932134794 |
| 0.830091371 | 0.932128478 |
| 0.830047512 | 0.932128478 |
| 0.82998088 | 0.932123641 |
| 0.829950887 | 0.932123641 |
| 0.829879276 | 0.932123641 |
| 0.829779778 | 0.932123641 |
| 0.82977841 | 0.932123641 |
| 0.829762226 | 0.932123641 |

| | |
|---|---|
| 0.829738102 | 0.932123641 |
| 0.829564243 | 0.932095315 |
| 0.829520738 | 0.932084207 |
| 0.829501927 | 0.932084207 |
| 0.829471523 | 0.932079092 |
| 0.829463987 | 0.932079092 |
| 0.829348639 | 0.932014001 |
| 0.829295934 | 0.932014001 |
| 0.829284521 | 0.932014001 |
| 0.829257972 | 0.932014001 |
| 0.829201356 | 0.932004819 |
| 0.829132549 | 0.931962048 |
| 0.82908721 | 0.931955076 |
| 0.828962561 | 0.931858947 |
| 0.828950776 | 0.931858947 |
| 0.82894492 | 0.931858947 |
| 0.828937208 | 0.931858947 |
| 0.828818873 | 0.931851384 |
| 0.828735385 | 0.93180151 |
| 0.828646764 | 0.93174586 |
| 0.828642412 | 0.93174586 |
| 0.828552849 | 0.931684253 |
| 0.828465526 | 0.931608056 |
| 0.828422395 | 0.931608056 |
| 0.82837827 | 0.931608056 |
| 0.828336666 | 0.931595126 |
| 0.828233886 | 0.931530079 |
| 0.828044602 | 0.931376621 |
| 0.828041198 | 0.931376621 |
| 0.827862033 | 0.931303235 |
| 0.82777522 | 0.93122757 |
| 0.827753137 | 0.93122757 |
| 0.827743322 | 0.93122757 |
| 0.827736496 | 0.93122757 |
| 0.827692048 | 0.93122757 |
| 0.827614816 | 0.93122757 |
| 0.827570608 | 0.93122757 |
| 0.827416076 | 0.931200903 |
| 0.827377608 | 0.931198179 |
| 0.827288093 | 0.931119435 |
| 0.827165232 | 0.93104716 |
| 0.827125766 | 0.931025783 |
| 0.826991447 | 0.930962544 |
| 0.826761552 | 0.930818211 |
| 0.826512783 | 0.930674845 |

| | |
|---|---|
| 0.826496842 | 0.930674845 |
| 0.826385649 | 0.930674845 |
| 0.826342476 | 0.930670949 |
| 0.826188704 | 0.930519767 |
| 0.82618491 | 0.930519767 |
| 0.826171072 | 0.930519767 |
| 0.826059913 | 0.930484733 |
| 0.825890236 | 0.930462009 |
| 0.825856828 | 0.930454029 |
| 0.82574574 | 0.930371222 |
| 0.825679143 | 0.930371222 |
| 0.825664468 | 0.930371222 |
| 0.825519957 | 0.930316572 |
| 0.825314216 | 0.930150741 |
| 0.825304067 | 0.930150741 |
| 0.825185743 | 0.930147273 |
| 0.825156917 | 0.930147273 |
| 0.825100671 | 0.930108157 |
| 0.825067032 | 0.930092251 |
| 0.825004249 | 0.93004349 |
| 0.824871198 | 0.929981549 |
| 0.82486821 | 0.929981549 |
| 0.824850455 | 0.929981549 |
| 0.824766387 | 0.929951437 |
| 0.824742769 | 0.929946823 |
| 0.82466327 | 0.92992323 |
| 0.824642312 | 0.92992323 |
| 0.824631382 | 0.92992323 |
| 0.824618494 | 0.92992323 |
| 0.824593661 | 0.92992323 |
| 0.824412134 | 0.929860198 |
| 0.824352247 | 0.929814671 |
| 0.824315701 | 0.929795469 |
| 0.824273507 | 0.929769896 |
| 0.824271853 | 0.929769896 |
| 0.824260538 | 0.929769896 |
| 0.824159718 | 0.929729622 |
| 0.82402854 | 0.929630035 |
| 0.823979893 | 0.929614838 |
| 0.823968177 | 0.929614838 |
| 0.823881899 | 0.929608425 |
| 0.823852554 | 0.929603297 |
| 0.823833273 | 0.929603297 |
| 0.82363456 | 0.929445402 |
| 0.823608545 | 0.929443157 |

| | |
|---|---|
| 0.823570298 | 0.92943895 |
| 0.823557562 | 0.92943895 |
| 0.823524179 | 0.92943895 |
| 0.823513256 | 0.92943895 |
| 0.823429114 | 0.929411755 |
| 0.823408337 | 0.92941033 |
| 0.82338782 | 0.929409198 |
| 0.823382256 | 0.929409198 |
| 0.823361003 | 0.929409198 |
| 0.823267995 | 0.929384077 |
| 0.823255986 | 0.929384077 |
| 0.823240358 | 0.929384077 |
| 0.82318891 | 0.929360884 |
| 0.823091935 | 0.929279072 |
| 0.82308616 | 0.929279072 |
| 0.822997403 | 0.929279072 |
| 0.822956373 | 0.929274579 |
| 0.82293281 | 0.929274579 |
| 0.822928558 | 0.929274579 |
| 0.82291468 | 0.929274579 |
| 0.822790219 | 0.929207926 |
| 0.822779783 | 0.929207926 |
| 0.822746309 | 0.929207926 |
| 0.822573231 | 0.92911308 |
| 0.822467606 | 0.929075107 |
| 0.822463464 | 0.929075107 |
| 0.822447884 | 0.929075107 |
| 0.822414673 | 0.929075107 |
| 0.822224022 | 0.929039806 |
| 0.822145441 | 0.9289944 |
| 0.821847727 | 0.928749359 |
| 0.821804045 | 0.92872203 |
| 0.821768597 | 0.928704007 |
| 0.821749898 | 0.928704007 |
| 0.821720072 | 0.928704007 |
| 0.821677261 | 0.928688932 |
| 0.821661027 | 0.928688932 |
| 0.821640083 | 0.928688932 |
| 0.821623821 | 0.928688932 |
| 0.821606672 | 0.928688932 |
| 0.821601317 | 0.928688932 |
| 0.821456901 | 0.928598806 |
| 0.821446171 | 0.928598806 |
| 0.821421959 | 0.928598806 |
| 0.8213168 | 0.928567992 |

| | |
|---|---|
| 0.821058366 | 0.928363962 |
| 0.820978861 | 0.928318144 |
| 0.82096001 | 0.928318144 |
| 0.820932182 | 0.928318144 |
| 0.820913041 | 0.928318144 |
| 0.820871222 | 0.928318144 |
| 0.82080117 | 0.928318144 |
| 0.820790393 | 0.928318144 |
| 0.820790118 | 0.928318144 |
| 0.820771394 | 0.928318144 |
| 0.820626684 | 0.928318144 |
| 0.82056474 | 0.928318144 |
| 0.820437459 | 0.928278967 |
| 0.820323652 | 0.928187626 |
| 0.820286784 | 0.928174638 |
| 0.820276076 | 0.928174638 |
| 0.820250552 | 0.928174638 |
| 0.820238978 | 0.928174638 |
| 0.820196842 | 0.928174638 |
| 0.82017771 | 0.928174638 |
| 0.820047611 | 0.928146568 |
| 0.820034567 | 0.928146568 |
| 0.820002761 | 0.928139918 |
| 0.819945723 | 0.928111371 |
| 0.819716888 | 0.927904555 |
| 0.81969181 | 0.927898221 |
| 0.819654053 | 0.927898221 |
| 0.81960896 | 0.927870596 |
| 0.819532406 | 0.927864167 |
| 0.819443553 | 0.927815664 |
| 0.819333948 | 0.927735675 |
| 0.819308385 | 0.927732255 |
| 0.819148013 | 0.927591298 |
| 0.819073605 | 0.927551147 |
| 0.819035244 | 0.92752976 |
| 0.819023601 | 0.92752976 |
| 0.819016042 | 0.92752976 |
| 0.818872187 | 0.927433312 |
| 0.818829229 | 0.927406713 |
| 0.818797819 | 0.927403288 |
| 0.818741911 | 0.927387303 |
| 0.818645443 | 0.927374986 |
| 0.818635246 | 0.927374986 |
| 0.81856332 | 0.927326068 |
| 0.818379771 | 0.927184295 |

| | |
|---|---|
| 0.818285486 | 0.927121582 |
| 0.818257133 | 0.927111514 |
| 0.818179352 | 0.927087394 |
| 0.817968701 | 0.926954105 |
| 0.817825676 | 0.926887264 |
| 0.817825676 | 0.926887264 |
| 0.817716352 | 0.926887264 |
| 0.81743762 | 0.926653073 |
| 0.817424822 | 0.926653073 |
| 0.817359167 | 0.926645205 |
| 0.817357833 | 0.926645205 |
| 0.816929884 | 0.926312872 |
| 0.816839664 | 0.926312872 |
| 0.816664284 | 0.92616606 |
| 0.816640153 | 0.926165307 |
| 0.816638881 | 0.926165307 |
| 0.816616314 | 0.926165307 |
| 0.816590213 | 0.926165307 |
| 0.816578647 | 0.926165307 |
| 0.816577558 | 0.926165307 |
| 0.81644249 | 0.926157115 |
| 0.816273254 | 0.926031306 |
| 0.816271114 | 0.926031306 |
| 0.816261379 | 0.926031306 |
| 0.816179884 | 0.925991551 |
| 0.816136685 | 0.925991551 |
| 0.81610897 | 0.925991551 |
| 0.816107555 | 0.925991551 |
| 0.816080365 | 0.925991551 |
| 0.816070446 | 0.925991551 |
| 0.816031929 | 0.925991551 |
| 0.815973934 | 0.925978473 |
| 0.815924751 | 0.925954663 |
| 0.815921777 | 0.925954663 |
| 0.81577919 | 0.925899214 |
| 0.815647381 | 0.925850561 |
| 0.815437153 | 0.925700179 |
| 0.815327202 | 0.925597422 |
| 0.815273803 | 0.925580926 |
| 0.815256328 | 0.925580926 |
| 0.815243607 | 0.925580926 |
| 0.815051114 | 0.92546047 |
| 0.814978674 | 0.925422342 |
| 0.814948009 | 0.925409585 |
| 0.814773801 | 0.925322074 |

| | |
|---|---|
| 0.814764214 | 0.925322074 |
| 0.814761547 | 0.925322074 |
| 0.814752083 | 0.925322074 |
| 0.814577983 | 0.925276196 |
| 0.814574091 | 0.925276196 |
| 0.814548309 | 0.925276196 |
| 0.814432489 | 0.925177125 |
| 0.814318794 | 0.925114166 |
| 0.814253699 | 0.925073317 |
| 0.814214204 | 0.925061543 |
| 0.814092387 | 0.925011407 |
| 0.814058349 | 0.924994799 |
| 0.81392309 | 0.924907306 |
| 0.813862434 | 0.924860446 |
| 0.813830688 | 0.924846437 |
| 0.813767676 | 0.924839103 |
| 0.813762035 | 0.924839103 |
| 0.813745218 | 0.924839103 |
| 0.813646971 | 0.924814192 |
| 0.813642059 | 0.924814192 |
| 0.813489708 | 0.924767861 |
| 0.81342427 | 0.924737616 |
| 0.813314938 | 0.924635394 |
| 0.81327341 | 0.924610252 |
| 0.813214932 | 0.924565839 |
| 0.813068685 | 0.924465772 |
| 0.813057309 | 0.924465772 |
| 0.813055328 | 0.924465772 |
| 0.813050782 | 0.924465772 |
| 0.813021351 | 0.924465772 |
| 0.813018365 | 0.924465772 |
| 0.812992473 | 0.924465772 |
| 0.812990769 | 0.924465772 |
| 0.812983998 | 0.924465772 |
| 0.812899491 | 0.924465772 |
| 0.812878915 | 0.924465772 |
| 0.812850665 | 0.924465772 |
| 0.81283909 | 0.924465772 |
| 0.812753928 | 0.924465772 |
| 0.812654103 | 0.924410658 |
| 0.812631029 | 0.924409526 |
| 0.812622164 | 0.924409526 |
| 0.812363232 | 0.924244323 |
| 0.812334081 | 0.924244323 |
| 0.81233364 | 0.924244323 |

| | |
|---|---|
| 0.812270039 | 0.924244323 |
| 0.812266085 | 0.924244323 |
| 0.812168724 | 0.924244323 |
| 0.812155231 | 0.924244323 |
| 0.812126229 | 0.924244323 |
| 0.812125236 | 0.924244323 |
| 0.812092727 | 0.924244323 |
| 0.812048564 | 0.92423259 |
| 0.812043774 | 0.92423259 |
| 0.812024424 | 0.92423259 |
| 0.812005782 | 0.92423259 |
| 0.811971471 | 0.92423259 |
| 0.811815089 | 0.924143535 |
| 0.811814881 | 0.924143535 |
| 0.811691474 | 0.924069077 |
| 0.811688962 | 0.924069077 |
| 0.81160606 | 0.9240381 |
| 0.81160606 | 0.9240381 |
| 0.811581586 | 0.9240381 |
| 0.811575442 | 0.9240381 |
| 0.811389515 | 0.923990351 |
| 0.811376644 | 0.923990351 |
| 0.811367562 | 0.923990351 |
| 0.811361178 | 0.923990351 |
| 0.811351731 | 0.923990351 |
| 0.811307355 | 0.923990351 |
| 0.811302823 | 0.923990351 |
| 0.811298606 | 0.923990351 |
| 0.811289992 | 0.923990351 |
| 0.811285757 | 0.923990351 |
| 0.811261907 | 0.923990351 |
| 0.811137767 | 0.923990351 |
| 0.811087143 | 0.923990351 |
| 0.811047337 | 0.923990351 |
| 0.810996896 | 0.923990351 |
| 0.81098219 | 0.923990351 |
| 0.810962564 | 0.923990351 |
| 0.810919785 | 0.923990351 |
| 0.810867092 | 0.923990351 |
| 0.810862336 | 0.923990351 |
| 0.810818933 | 0.923981273 |
| 0.810784375 | 0.923981273 |
| 0.810589122 | 0.923896236 |
| 0.81030611 | 0.923699345 |
| 0.810223588 | 0.923634333 |

| | |
|---|---|
| 0.810203624 | 0.923634333 |
| 0.810153514 | 0.923634333 |
| 0.810107802 | 0.923634333 |
| 0.810100088 | 0.923634333 |
| 0.810032448 | 0.923593292 |
| 0.809966147 | 0.923561916 |
| 0.809946917 | 0.923561916 |
| 0.809870372 | 0.923499559 |
| 0.809859963 | 0.923499559 |
| 0.809839633 | 0.923499559 |
| 0.809816132 | 0.923499559 |
| 0.809718881 | 0.923469228 |
| 0.809626074 | 0.923461563 |
| 0.809594107 | 0.923447217 |
| 0.809520299 | 0.923447217 |
| 0.809496553 | 0.923447217 |
| 0.809476109 | 0.923447217 |
| 0.80946492 | 0.923447217 |
| 0.809398322 | 0.923447217 |
| 0.809304696 | 0.923404599 |
| 0.80925277 | 0.923367472 |
| 0.809179121 | 0.923305558 |
| 0.80903885 | 0.923189739 |
| 0.809038837 | 0.923189739 |
| 0.809017075 | 0.923189739 |
| 0.808976341 | 0.923189739 |
| 0.808739371 | 0.923024923 |
| 0.808721211 | 0.923024923 |
| 0.808702967 | 0.923024923 |
| 0.808684912 | 0.923024923 |
| 0.808589156 | 0.922986188 |
| 0.808483421 | 0.922953976 |
| 0.808439156 | 0.922947689 |
| 0.80833778 | 0.922876197 |
| 0.808267631 | 0.92281823 |
| 0.808158655 | 0.922755589 |
| 0.808051273 | 0.922703924 |
| 0.808045867 | 0.922703924 |
| 0.807845317 | 0.922601465 |
| 0.807812575 | 0.922586195 |
| 0.807802448 | 0.922586195 |
| 0.807664665 | 0.92246151 |
| 0.807562187 | 0.922366586 |
| 0.80755817 | 0.922366586 |
| 0.807511506 | 0.922366586 |

| | |
|---|---|
| 0.807409705 | 0.922303017 |
| 0.807392779 | 0.922303017 |
| 0.807320997 | 0.922245926 |
| 0.807279374 | 0.922220498 |
| 0.807242506 | 0.922220498 |
| 0.807207402 | 0.922220498 |
| 0.807112352 | 0.922162409 |
| 0.806981079 | 0.922043909 |
| 0.806825284 | 0.921982683 |
| 0.806801212 | 0.921982683 |
| 0.806644236 | 0.92198145 |
| 0.806601617 | 0.92196816 |
| 0.806477748 | 0.921901779 |
| 0.806457405 | 0.921900651 |
| 0.806425924 | 0.921892575 |
| 0.806096865 | 0.92164335 |
| 0.805937622 | 0.921549772 |
| 0.805887733 | 0.921536976 |
| 0.805846236 | 0.921525326 |
| 0.805838226 | 0.921525326 |
| 0.805819823 | 0.921525326 |
| 0.805671244 | 0.921466413 |
| 0.805560443 | 0.921413402 |
| 0.805472586 | 0.921394088 |
| 0.805451092 | 0.921391631 |
| 0.805374515 | 0.92134829 |
| 0.805227797 | 0.921268961 |
| 0.805217602 | 0.921268961 |
| 0.805175102 | 0.921268961 |
| 0.805174258 | 0.921268961 |
| 0.805173701 | 0.921268961 |
| 0.805089369 | 0.9212655 |
| 0.805027907 | 0.921261573 |
| 0.804979567 | 0.921261573 |
| 0.804926891 | 0.921261573 |
| 0.804923141 | 0.921261573 |
| 0.804820471 | 0.921240719 |
| 0.804735558 | 0.921214768 |
| 0.804683028 | 0.921176776 |
| 0.804663588 | 0.921176776 |
| 0.804538495 | 0.92112202 |
| 0.804485408 | 0.921105525 |
| 0.804422533 | 0.921077822 |
| 0.804368911 | 0.921038567 |
| 0.804341991 | 0.921029885 |

| | |
|---|---|
| 0.804293754 | 0.920996793 |
| 0.803992945 | 0.92080731 |
| 0.803954092 | 0.92080731 |
| 0.803703121 | 0.920586064 |
| 0.803663407 | 0.920562714 |
| 0.80366231 | 0.920562714 |
| 0.803578489 | 0.920531862 |
| 0.803554123 | 0.920526091 |
| 0.803492728 | 0.920477899 |
| 0.803379911 | 0.920370794 |
| 0.803369273 | 0.920370794 |
| 0.803347044 | 0.920370794 |
| 0.803342751 | 0.920370794 |
| 0.803289531 | 0.920370794 |
| 0.803266885 | 0.920370794 |
| 0.803244256 | 0.920370794 |
| 0.803232054 | 0.920370794 |
| 0.803075212 | 0.920289003 |
| 0.80305615 | 0.920289003 |
| 0.803043348 | 0.920289003 |
| 0.803030334 | 0.920289003 |
| 0.803021553 | 0.920289003 |
| 0.802962496 | 0.920289003 |
| 0.802925684 | 0.920289003 |
| 0.802706667 | 0.920128867 |
| 0.802672417 | 0.920128109 |
| 0.802649188 | 0.920128109 |
| 0.802564092 | 0.920122286 |
| 0.802525355 | 0.920122173 |
| 0.802457871 | 0.920075665 |
| 0.802370267 | 0.920055103 |
| 0.802333491 | 0.920040568 |
| 0.802263099 | 0.920033602 |
| 0.802221802 | 0.920022124 |
| 0.802162775 | 0.920022124 |
| 0.80211276 | 0.920022124 |
| 0.80204718 | 0.920016885 |
| 0.801833195 | 0.919904359 |
| 0.801810993 | 0.919904359 |
| 0.801795765 | 0.919904359 |
| 0.801698754 | 0.919838751 |
| 0.801616723 | 0.919788948 |
| 0.801612053 | 0.919788948 |
| 0.801547904 | 0.919788948 |
| 0.801526436 | 0.919788948 |

| | |
|---|---|
| 0.801519888 | 0.919788948 |
| 0.80137924 | 0.919715875 |
| 0.801338031 | 0.919690742 |
| 0.801298512 | 0.919690742 |
| 0.801159628 | 0.919620754 |
| 0.801156975 | 0.919620754 |
| 0.801107369 | 0.919620754 |
| 0.801031198 | 0.919604514 |
| 0.801024466 | 0.919604514 |
| 0.80078395 | 0.919409314 |
| 0.800728634 | 0.919398926 |
| 0.800720511 | 0.919398926 |
| 0.800711018 | 0.919398926 |
| 0.80065177 | 0.919398926 |
| 0.800603836 | 0.919379826 |
| 0.80058841 | 0.919379826 |
| 0.800577974 | 0.919379826 |
| 0.800555712 | 0.919379826 |
| 0.800535153 | 0.919379826 |
| 0.800479316 | 0.919347666 |
| 0.800351633 | 0.919296125 |
| 0.800260237 | 0.919229055 |
| 0.800224697 | 0.919229055 |
| 0.800163324 | 0.919229055 |
| 0.800116765 | 0.919229055 |
| 0.800083459 | 0.919225525 |
| 0.799876138 | 0.919098189 |
| 0.799857474 | 0.919098189 |
| 0.799839988 | 0.919098189 |
| 0.799800931 | 0.919078294 |
| 0.799748681 | 0.919040425 |
| 0.799736191 | 0.919040425 |
| 0.799684474 | 0.91902171 |
| 0.799672238 | 0.91902171 |
| 0.799443339 | 0.918931507 |
| 0.799380894 | 0.918906011 |
| 0.799357233 | 0.918900991 |
| 0.799249329 | 0.918799126 |
| 0.799195063 | 0.918767743 |
| 0.799187688 | 0.918767743 |
| 0.799078934 | 0.918714119 |
| 0.799066062 | 0.918714119 |
| 0.799004383 | 0.918672761 |
| 0.798957559 | 0.9186411 |
| 0.798884457 | 0.918611242 |

| | |
|---|---|
| 0.798856841 | 0.918611242 |
| 0.7986632 | 0.918524395 |
| 0.798660126 | 0.918524395 |
| 0.798639087 | 0.918524395 |
| 0.798638446 | 0.918524395 |
| 0.79862559 | 0.918524395 |
| 0.798606339 | 0.918524395 |
| 0.798447782 | 0.918524395 |
| 0.798375495 | 0.918510314 |
| 0.79833288 | 0.918510314 |
| 0.798171498 | 0.918446821 |
| 0.798153126 | 0.918446821 |
| 0.798076887 | 0.918404521 |
| 0.797958843 | 0.918290866 |
| 0.797936459 | 0.918287296 |
| 0.797920856 | 0.918287296 |
| 0.797892599 | 0.918287296 |
| 0.797877442 | 0.918287296 |
| 0.797781814 | 0.918287296 |
| 0.797472679 | 0.918135632 |
| 0.797470579 | 0.918135632 |
| 0.797343612 | 0.918093083 |
| 0.797320636 | 0.918088824 |
| 0.797273267 | 0.918078672 |
| 0.797129861 | 0.917996297 |
| 0.797128593 | 0.917996297 |
| 0.797123534 | 0.917996297 |
| 0.797110623 | 0.917996297 |
| 0.797063674 | 0.917996297 |
| 0.797005137 | 0.917996297 |
| 0.797001837 | 0.917996297 |
| 0.796868392 | 0.917923158 |
| 0.796849588 | 0.917923158 |
| 0.796793985 | 0.91790405 |
| 0.79676142 | 0.917888736 |
| 0.796602928 | 0.91779495 |
| 0.79653698 | 0.91774117 |
| 0.796426589 | 0.917680582 |
| 0.796361834 | 0.917628168 |
| 0.796356883 | 0.917628168 |
| 0.796355933 | 0.917628168 |
| 0.796308773 | 0.917628168 |
| 0.79624728 | 0.917607172 |
| 0.796069309 | 0.917446474 |
| 0.796001844 | 0.917390922 |

| | |
|---|---|
| 0.795990847 | 0.917390922 |
| 0.795900775 | 0.917341034 |
| 0.795851138 | 0.917335358 |
| 0.795732637 | 0.917280431 |
| 0.795719262 | 0.917280431 |
| 0.795687867 | 0.917273226 |
| 0.795582205 | 0.917198934 |
| 0.795561424 | 0.917198934 |
| 0.795348632 | 0.917037531 |
| 0.795333776 | 0.917037531 |
| 0.795263028 | 0.917005429 |
| 0.795227857 | 0.917005429 |
| 0.795136012 | 0.916925567 |
| 0.795010589 | 0.91682533 |
| 0.79497067 | 0.916801494 |
| 0.794927053 | 0.916800906 |
| 0.794856308 | 0.9167584 |
| 0.794798931 | 0.916719624 |
| 0.79476273 | 0.91671707 |
| 0.794666522 | 0.916650499 |
| 0.794648799 | 0.916650499 |
| 0.794452463 | 0.916492369 |
| 0.794316143 | 0.916373317 |
| 0.794315795 | 0.916373317 |
| 0.794232239 | 0.916327096 |
| 0.794186227 | 0.916296208 |
| 0.79409347 | 0.91623358 |
| 0.794088038 | 0.91623358 |
| 0.794068618 | 0.91623358 |
| 0.794064854 | 0.91623358 |
| 0.794027895 | 0.91623358 |
| 0.793997866 | 0.91623358 |
| 0.793951057 | 0.916224637 |
| 0.793949346 | 0.916224637 |
| 0.793623242 | 0.915957305 |
| 0.793580548 | 0.915930225 |
| 0.79355925 | 0.915927841 |
| 0.793546861 | 0.915927841 |
| 0.793516123 | 0.915922456 |
| 0.793437249 | 0.915853611 |
| 0.793379277 | 0.915831088 |
| 0.793300025 | 0.915761799 |
| 0.793245743 | 0.915721332 |
| 0.793239821 | 0.915721332 |
| 0.793158621 | 0.915686849 |

| | |
|---|---|
| 0.793093396 | 0.915686849 |
| 0.79303235 | 0.915686849 |
| 0.793011189 | 0.915686849 |
| 0.793007932 | 0.915686849 |
| 0.792967911 | 0.915686849 |
| 0.792957518 | 0.915686849 |
| 0.792918001 | 0.915686849 |
| 0.792854728 | 0.915669442 |
| 0.792767021 | 0.915612555 |
| 0.792718461 | 0.915578672 |
| 0.792690515 | 0.915568598 |
| 0.792375929 | 0.915338436 |
| 0.792304525 | 0.915322554 |
| 0.792286617 | 0.915322554 |
| 0.792115603 | 0.915193097 |
| 0.792089857 | 0.915185552 |
| 0.792027089 | 0.91513523 |
| 0.79195238 | 0.915071108 |
| 0.791879686 | 0.915009312 |
| 0.791823844 | 0.914966985 |
| 0.791789388 | 0.914949369 |
| 0.791743764 | 0.914949369 |
| 0.791723593 | 0.914949369 |
| 0.791624851 | 0.914892424 |
| 0.791489777 | 0.914802913 |
| 0.791483595 | 0.914802913 |
| 0.791394857 | 0.914782002 |
| 0.791375327 | 0.914782002 |
| 0.791373003 | 0.914782002 |
| 0.791325256 | 0.914782002 |
| 0.791240685 | 0.914759202 |
| 0.791238972 | 0.914759202 |
| 0.791216787 | 0.914759202 |
| 0.791175239 | 0.914759202 |
| 0.791156067 | 0.914759202 |
| 0.791082489 | 0.914759202 |
| 0.791074605 | 0.914759202 |
| 0.791054917 | 0.914759202 |
| 0.791045668 | 0.914759202 |
| 0.790980729 | 0.914759202 |
| 0.790935121 | 0.914759202 |
| 0.790891974 | 0.914743053 |
| 0.790891875 | 0.914743053 |
| 0.790764263 | 0.914652456 |
| 0.790687035 | 0.914629768 |

| | |
|---|---|
| 0.790625762 | 0.914603317 |
| 0.790602062 | 0.914598115 |
| 0.790572042 | 0.914585602 |
| 0.790552425 | 0.914585124 |
| 0.790351363 | 0.914396939 |
| 0.790320696 | 0.91438367 |
| 0.790311198 | 0.91438367 |
| 0.790261421 | 0.914359515 |
| 0.790217838 | 0.914343953 |
| 0.790180441 | 0.914343953 |
| 0.790166357 | 0.914343953 |
| 0.79014133 | 0.914343953 |
| 0.790098776 | 0.914343953 |
| 0.790097005 | 0.914343953 |
| 0.790077186 | 0.914343953 |
| 0.790046011 | 0.914343953 |
| 0.790032574 | 0.914343953 |
| 0.789990075 | 0.914343953 |
| 0.789866852 | 0.914325019 |
| 0.789841562 | 0.914317966 |
| 0.789684341 | 0.914158186 |
| 0.789648882 | 0.914139357 |
| 0.789569509 | 0.914091909 |
| 0.789478983 | 0.914009323 |
| 0.789475769 | 0.914009323 |
| 0.789464626 | 0.914009323 |
| 0.789458109 | 0.914009323 |
| 0.789419425 | 0.914009323 |
| 0.789398401 | 0.914009323 |
| 0.789386657 | 0.914009323 |
| 0.789271764 | 0.913969372 |
| 0.789153024 | 0.913876312 |
| 0.789140308 | 0.913876312 |
| 0.789088647 | 0.913846201 |
| 0.789042277 | 0.913814719 |
| 0.789002336 | 0.913790683 |
| 0.788840986 | 0.913735589 |
| 0.788836695 | 0.913735589 |
| 0.78878932 | 0.913735589 |
| 0.788788421 | 0.913735589 |
| 0.788767477 | 0.913735589 |
| 0.788733488 | 0.913735589 |
| 0.788611795 | 0.913735589 |
| 0.788547684 | 0.913735589 |
| 0.788539405 | 0.913735589 |

| | |
|---|---|
| 0.788402863 | 0.913735589 |
| 0.788380938 | 0.913735589 |
| 0.78832013 | 0.913714404 |
| 0.788287043 | 0.913714404 |
| 0.788285133 | 0.913714404 |
| 0.788278041 | 0.913714404 |
| 0.788218303 | 0.913714404 |
| 0.788143727 | 0.913714404 |
| 0.788115804 | 0.913714404 |
| 0.787984556 | 0.913611676 |
| 0.787921517 | 0.913589394 |
| 0.787904513 | 0.913589394 |
| 0.787896201 | 0.913589394 |
| 0.787819934 | 0.913578485 |
| 0.787804357 | 0.913578485 |
| 0.787779895 | 0.913578485 |
| 0.787591748 | 0.91342308 |
| 0.787297001 | 0.91323691 |
| 0.787220553 | 0.913197763 |
| 0.787129193 | 0.913153459 |
| 0.787064108 | 0.913100194 |
| 0.786969111 | 0.913012225 |
| 0.786967734 | 0.913012225 |
| 0.786881289 | 0.912954814 |
| 0.786799167 | 0.912881773 |
| 0.786785254 | 0.912881773 |
| 0.786780008 | 0.912881773 |
| 0.786729177 | 0.912867284 |
| 0.786714626 | 0.912867284 |
| 0.78643854 | 0.912752421 |
| 0.78639117 | 0.912725084 |
| 0.786237275 | 0.912587702 |
| 0.786104869 | 0.912476338 |
| 0.786093235 | 0.912476338 |
| 0.786011338 | 0.912452792 |
| 0.785938262 | 0.912413151 |
| 0.785916399 | 0.912413151 |
| 0.785889755 | 0.912404537 |
| 0.785850647 | 0.912381375 |
| 0.785841423 | 0.912381375 |
| 0.785762747 | 0.912348637 |
| 0.785678732 | 0.912323702 |
| 0.785632523 | 0.912306046 |
| 0.785593487 | 0.912282959 |
| 0.785524796 | 0.912282959 |

| | |
|---|---|
| 0.785500266 | 0.912282959 |
| 0.785476324 | 0.912282959 |
| 0.785460919 | 0.912282959 |
| 0.78545369 | 0.912282959 |
| 0.7854407 | 0.912282959 |
| 0.785351229 | 0.912282959 |
| 0.785071154 | 0.912098938 |
| 0.785021394 | 0.912063375 |
| 0.784934439 | 0.911984595 |
| 0.784602649 | 0.911688064 |
| 0.784575095 | 0.911678289 |
| 0.784549334 | 0.911670598 |
| 0.78451975 | 0.911658464 |
| 0.784500594 | 0.911658447 |
| 0.784419513 | 0.911658447 |
| 0.784371725 | 0.911642154 |
| 0.784356631 | 0.911642154 |
| 0.784352071 | 0.911642154 |
| 0.784292626 | 0.911616959 |
| 0.784192031 | 0.91154563 |
| 0.784170994 | 0.91154563 |
| 0.784147925 | 0.91154563 |
| 0.784063506 | 0.91154563 |
| 0.783923129 | 0.911463666 |
| 0.783863233 | 0.911429267 |
| 0.783780854 | 0.911385009 |
| 0.783752738 | 0.911385009 |
| 0.783684328 | 0.911376996 |
| 0.783646989 | 0.911366784 |
| 0.783558637 | 0.911342089 |
| 0.783536629 | 0.911338747 |
| 0.783515546 | 0.911336482 |
| 0.783509268 | 0.911336482 |
| 0.783497605 | 0.911336482 |
| 0.783407961 | 0.91127811 |
| 0.783385572 | 0.91127811 |
| 0.783307529 | 0.91125031 |
| 0.783285683 | 0.911247154 |
| 0.783216871 | 0.911222801 |
| 0.783112738 | 0.911222801 |
| 0.783080546 | 0.911208824 |
| 0.783005353 | 0.911208824 |
| 0.782986918 | 0.911208824 |
| 0.782916178 | 0.911208824 |
| 0.782820924 | 0.911208824 |

| | |
|---|---|
| 0.782789816 | 0.911208824 |
| 0.782767163 | 0.911208824 |
| 0.782752783 | 0.911208824 |
| 0.782556624 | 0.911088891 |
| 0.782553212 | 0.911088891 |
| 0.782484258 | 0.911049989 |
| 0.782352311 | 0.910940092 |
| 0.782325467 | 0.910931106 |
| 0.782276523 | 0.910896386 |
| 0.782181298 | 0.910830041 |
| 0.782178231 | 0.910830041 |
| 0.782113242 | 0.910830041 |
| 0.782022224 | 0.910800686 |
| 0.782013004 | 0.910800686 |
| 0.782012283 | 0.910800686 |
| 0.781981599 | 0.910800686 |
| 0.781952198 | 0.910800686 |
| 0.781947166 | 0.910800686 |
| 0.78187126 | 0.910758493 |
| 0.78183855 | 0.910742665 |
| 0.781809697 | 0.91073133 |
| 0.781788312 | 0.91073133 |
| 0.781755937 | 0.91073133 |
| 0.781636846 | 0.91073133 |
| 0.781616313 | 0.91073133 |
| 0.781567706 | 0.91073133 |
| 0.781539983 | 0.91073133 |
| 0.781526565 | 0.91073133 |
| 0.781466364 | 0.910710054 |
| 0.781449832 | 0.910710054 |
| 0.781416048 | 0.910695985 |
| 0.78139155 | 0.910689719 |
| 0.781331878 | 0.910661182 |
| 0.781323632 | 0.910661182 |
| 0.781174767 | 0.910593045 |
| 0.781130764 | 0.910564037 |
| 0.781114377 | 0.910564037 |
| 0.781071706 | 0.910562053 |
| 0.781048433 | 0.910557211 |
| 0.781036882 | 0.910557211 |
| 0.780869579 | 0.910437842 |
| 0.780843043 | 0.910429191 |
| 0.78071815 | 0.910372719 |
| 0.78069324 | 0.91036922 |
| 0.780693119 | 0.91036922 |

| | |
|---|---|
| 0.780626276 | 0.91036922 |
| 0.78061136 | 0.91036922 |
| 0.780561197 | 0.91036922 |
| 0.780559162 | 0.91036922 |
| 0.780489384 | 0.91036922 |
| 0.780359112 | 0.91036922 |
| 0.780326578 | 0.91036922 |
| 0.780314191 | 0.91036922 |
| 0.780140251 | 0.910300604 |
| 0.779975968 | 0.91019811 |
| 0.779973225 | 0.91019811 |
| 0.779830333 | 0.910095057 |
| 0.779802985 | 0.910085441 |
| 0.779721106 | 0.910037109 |
| 0.779710466 | 0.910037109 |
| 0.779676418 | 0.910037109 |
| 0.779556114 | 0.909951531 |
| 0.779526756 | 0.909941443 |
| 0.779483129 | 0.909912818 |
| 0.779467987 | 0.909912818 |
| 0.779455885 | 0.909912818 |
| 0.779441176 | 0.909912818 |
| 0.779426457 | 0.909912818 |
| 0.779386184 | 0.909912818 |
| 0.779249806 | 0.909841154 |
| 0.779235562 | 0.909841154 |
| 0.779209586 | 0.909838803 |
| 0.779150252 | 0.909791827 |
| 0.779006394 | 0.909668453 |
| 0.778984667 | 0.909668453 |
| 0.778948831 | 0.909668146 |
| 0.778906633 | 0.909641171 |
| 0.778852592 | 0.909606798 |
| 0.778798031 | 0.909603557 |
| 0.77879523 | 0.909603557 |
| 0.778759591 | 0.909603272 |
| 0.77872139 | 0.90958096 |
| 0.778670885 | 0.909544274 |
| 0.77856279 | 0.909440316 |
| 0.778544558 | 0.909440316 |
| 0.778523361 | 0.909440316 |
| 0.778493438 | 0.909440316 |
| 0.778479767 | 0.909440316 |
| 0.778466873 | 0.909440316 |
| 0.778425408 | 0.909440316 |

| | |
|---|---|
| 0.778409558 | 0.909440316 |
| 0.778406979 | 0.909440316 |
| 0.778343316 | 0.909440316 |
| 0.778239709 | 0.909440316 |
| 0.778089144 | 0.909390062 |
| 0.778037551 | 0.909362057 |
| 0.777973664 | 0.909327252 |
| 0.777918241 | 0.90931342 |
| 0.777775559 | 0.909234347 |
| 0.777762467 | 0.909234347 |
| 0.777729177 | 0.909234347 |
| 0.777719655 | 0.909234347 |
| 0.777704294 | 0.909234347 |
| 0.77769371 | 0.909234347 |
| 0.777665956 | 0.909234347 |
| 0.777653898 | 0.909234347 |
| 0.777561062 | 0.909234347 |
| 0.777493239 | 0.909234347 |
| 0.77744554 | 0.909205617 |
| 0.777337932 | 0.909131926 |
| 0.777333381 | 0.909131926 |
| 0.777292363 | 0.909131926 |
| 0.777274999 | 0.909131926 |
| 0.777270914 | 0.909131926 |
| 0.777067034 | 0.909005261 |
| 0.777065565 | 0.909005261 |
| 0.777046163 | 0.909005261 |
| 0.776901313 | 0.908903926 |
| 0.776891634 | 0.908903926 |
| 0.776740559 | 0.908805154 |
| 0.776552106 | 0.908651626 |
| 0.776425263 | 0.908570172 |
| 0.776300111 | 0.908557659 |
| 0.776277195 | 0.908553165 |
| 0.776188075 | 0.908493511 |
| 0.776171028 | 0.908493511 |
| 0.776142439 | 0.908484749 |
| 0.775877968 | 0.908242147 |
| 0.775643534 | 0.908060481 |
| 0.775548178 | 0.908034639 |
| 0.775501345 | 0.908002128 |
| 0.775495826 | 0.908002128 |
| 0.775452879 | 0.908002128 |
| 0.775380645 | 0.907994737 |
| 0.775366235 | 0.907994737 |

| | |
|---|---|
| 0.775269199 | 0.907931202 |
| 0.775064033 | 0.907824866 |
| 0.775011209 | 0.90779254 |
| 0.774979459 | 0.90779254 |
| 0.774891454 | 0.907779001 |
| 0.774476821 | 0.907427183 |
| 0.774380729 | 0.907341501 |
| 0.774286549 | 0.907338176 |
| 0.774281443 | 0.907338176 |
| 0.774260286 | 0.907338176 |
| 0.774188398 | 0.907312454 |
| 0.774168148 | 0.907311049 |
| 0.774146663 | 0.907311049 |
| 0.77412818 | 0.907311049 |
| 0.774105436 | 0.907311049 |
| 0.774051656 | 0.907311049 |
| 0.774034286 | 0.907311049 |
| 0.773960175 | 0.907311049 |
| 0.773916066 | 0.907305858 |
| 0.773872875 | 0.90729989 |
| 0.773843028 | 0.907287231 |
| 0.773793508 | 0.907276391 |
| 0.773730527 | 0.907267002 |
| 0.773567469 | 0.907165141 |
| 0.773548763 | 0.907165141 |
| 0.773526079 | 0.907161276 |
| 0.773458932 | 0.907107125 |
| 0.773409842 | 0.907104928 |
| 0.773366726 | 0.907104928 |
| 0.773229499 | 0.906996116 |
| 0.77303249 | 0.906828039 |
| 0.77298305 | 0.906792378 |
| 0.772972864 | 0.906792378 |
| 0.772948108 | 0.906792378 |
| 0.772947466 | 0.906792378 |
| 0.772796299 | 0.906684961 |
| 0.772786183 | 0.906684961 |
| 0.772717638 | 0.906684961 |
| 0.772598249 | 0.906592844 |
| 0.772564254 | 0.90659138 |
| 0.772410859 | 0.906456041 |
| 0.772309331 | 0.906411603 |
| 0.772272731 | 0.906411603 |
| 0.772259111 | 0.906411603 |
| 0.772139576 | 0.906383349 |

| | |
|---|---|
| 0.771966686 | 0.906292086 |
| 0.771858051 | 0.906186886 |
| 0.771851984 | 0.906186886 |
| 0.771763429 | 0.906142807 |
| 0.771760284 | 0.906142807 |
| 0.771579606 | 0.906016319 |
| 0.771483065 | 0.905969966 |
| 0.771452446 | 0.905956347 |
| 0.771377737 | 0.90589095 |
| 0.771252288 | 0.905832968 |
| 0.771177099 | 0.905766996 |
| 0.771099436 | 0.905765127 |
| 0.770898624 | 0.905584795 |
| 0.770819053 | 0.905547447 |
| 0.770748307 | 0.90548667 |
| 0.770743752 | 0.90548667 |
| 0.770731775 | 0.90548667 |
| 0.770711736 | 0.90548667 |
| 0.770574256 | 0.905416196 |
| 0.770556786 | 0.905416196 |
| 0.770534502 | 0.90541416 |
| 0.770520544 | 0.90541416 |
| 0.770253165 | 0.905195239 |
| 0.770085298 | 0.905042628 |
| 0.770067826 | 0.905042628 |
| 0.769959728 | 0.904939713 |
| 0.769790935 | 0.904788194 |
| 0.769687139 | 0.90470864 |
| 0.769598219 | 0.904673523 |
| 0.769596716 | 0.904673523 |
| 0.769589019 | 0.904673523 |
| 0.769535602 | 0.904673523 |
| 0.769527809 | 0.904673523 |
| 0.769407449 | 0.904605102 |
| 0.769263019 | 0.904567311 |
| 0.769161954 | 0.904470801 |
| 0.769126559 | 0.904451511 |
| 0.769110491 | 0.904451511 |
| 0.769053829 | 0.904451511 |
| 0.768959977 | 0.9043896 |
| 0.768951446 | 0.9043896 |
| 0.768951386 | 0.9043896 |
| 0.768905799 | 0.9043896 |
| 0.768878896 | 0.9043896 |
| 0.76885649 | 0.9043896 |

| | |
|---|---|
| 0.768762118 | 0.904357886 |
| 0.768724899 | 0.904353742 |
| 0.768719851 | 0.904353742 |
| 0.76866644 | 0.904353742 |
| 0.768664118 | 0.904353742 |
| 0.768637382 | 0.904353742 |
| 0.768621136 | 0.904353742 |
| 0.768613625 | 0.904353742 |
| 0.768583471 | 0.904353742 |
| 0.768569979 | 0.904353742 |
| 0.768500623 | 0.904353742 |
| 0.768475745 | 0.904353742 |
| 0.768465411 | 0.904353742 |
| 0.768439017 | 0.904353742 |
| 0.768397057 | 0.904353742 |
| 0.768331546 | 0.904342787 |
| 0.768263908 | 0.904307872 |
| 0.768241922 | 0.904304343 |
| 0.768168822 | 0.904240644 |
| 0.768037781 | 0.904153432 |
| 0.768031443 | 0.904153432 |
| 0.767952993 | 0.90414301 |
| 0.767841539 | 0.904034139 |
| 0.767747423 | 0.903945675 |
| 0.767724888 | 0.903945675 |
| 0.767720085 | 0.903945675 |
| 0.767701457 | 0.903945675 |
| 0.76769578 | 0.903945675 |
| 0.767538788 | 0.903901139 |
| 0.767522186 | 0.903901139 |
| 0.767398279 | 0.903780362 |
| 0.767357482 | 0.903777011 |
| 0.767309475 | 0.903776715 |
| 0.767304782 | 0.903776715 |
| 0.767210728 | 0.903767127 |
| 0.767194911 | 0.903767127 |
| 0.766980601 | 0.903623626 |
| 0.766976659 | 0.903623626 |
| 0.766791184 | 0.903489862 |
| 0.766784807 | 0.903489862 |
| 0.766763531 | 0.903489862 |
| 0.766762118 | 0.903489862 |
| 0.766708352 | 0.903489862 |
| 0.766666848 | 0.903489862 |
| 0.76666526 | 0.903489862 |

| | |
|---|---|
| 0.766653741 | 0.903489862 |
| 0.766572406 | 0.903489862 |
| 0.766500551 | 0.903482694 |
| 0.766454949 | 0.903451302 |
| 0.766295729 | 0.903285979 |
| 0.766213782 | 0.903211736 |
| 0.766212852 | 0.903211736 |
| 0.766209475 | 0.903211736 |
| 0.766152275 | 0.903211736 |
| 0.766145914 | 0.903211736 |
| 0.766095048 | 0.903205903 |
| 0.766091515 | 0.903205903 |
| 0.766077269 | 0.903205903 |
| 0.766008872 | 0.903205903 |
| 0.765995372 | 0.903205903 |
| 0.765954353 | 0.903196533 |
| 0.765788395 | 0.903045556 |
| 0.765649159 | 0.902957044 |
| 0.765647075 | 0.902957044 |
| 0.765557866 | 0.902930202 |
| 0.765508277 | 0.902930202 |
| 0.765502859 | 0.902930202 |
| 0.765372565 | 0.902823439 |
| 0.765359147 | 0.902823439 |
| 0.7652772 | 0.902822749 |
| 0.765264171 | 0.902822749 |
| 0.765198521 | 0.902776941 |
| 0.765184924 | 0.902776941 |
| 0.765169676 | 0.902776941 |
| 0.765137966 | 0.902776941 |
| 0.765072562 | 0.902776941 |
| 0.765057759 | 0.902776941 |
| 0.765038255 | 0.902776941 |
| 0.764922744 | 0.902740002 |
| 0.764911076 | 0.902740002 |
| 0.764908062 | 0.902740002 |
| 0.764871051 | 0.902740002 |
| 0.764860156 | 0.902740002 |
| 0.764768238 | 0.902739744 |
| 0.764767782 | 0.902739744 |
| 0.764766481 | 0.902739744 |
| 0.764707329 | 0.902739744 |
| 0.764681484 | 0.902739744 |
| 0.764640235 | 0.902728612 |
| 0.764622319 | 0.902728612 |

| | |
|---|---|
| 0.764617242 | 0.902728612 |
| 0.764611167 | 0.902728612 |
| 0.764563571 | 0.902728612 |
| 0.764528382 | 0.902728612 |
| 0.764283602 | 0.902544819 |
| 0.764282016 | 0.902544819 |
| 0.764161756 | 0.902490439 |
| 0.764158848 | 0.902490439 |
| 0.764141358 | 0.902490439 |
| 0.763917799 | 0.902336574 |
| 0.763800687 | 0.902265373 |
| 0.763798691 | 0.902265373 |
| 0.763726815 | 0.902222865 |
| 0.763620906 | 0.902187261 |
| 0.763460241 | 0.902056564 |
| 0.763321859 | 0.90194582 |
| 0.763295207 | 0.90194582 |
| 0.763261664 | 0.90194582 |
| 0.763260226 | 0.90194582 |
| 0.763186021 | 0.901919571 |
| 0.763159235 | 0.901910295 |
| 0.76309585 | 0.901891101 |
| 0.763089496 | 0.901891101 |
| 0.763000587 | 0.901834693 |
| 0.762979775 | 0.901832475 |
| 0.762951254 | 0.901821144 |
| 0.762903451 | 0.901809403 |
| 0.762837594 | 0.901776318 |
| 0.762837168 | 0.901776318 |
| 0.762760559 | 0.901776318 |
| 0.762759726 | 0.901776318 |
| 0.762529913 | 0.901658775 |
| 0.762473711 | 0.901614705 |
| 0.762447756 | 0.901606398 |
| 0.762432498 | 0.901606398 |
| 0.762403527 | 0.901598867 |
| 0.762376255 | 0.901598867 |
| 0.762371261 | 0.901598867 |
| 0.76226547 | 0.901598867 |
| 0.762251592 | 0.901598867 |
| 0.762246963 | 0.901598867 |
| 0.762244756 | 0.901598867 |
| 0.762186896 | 0.901598867 |
| 0.762133336 | 0.901598867 |
| 0.761986144 | 0.901530594 |

| | |
|---|---|
| 0.76197098 | 0.901530594 |
| 0.761915821 | 0.901492182 |
| 0.761837122 | 0.901421461 |
| 0.761709499 | 0.901309588 |
| 0.761696233 | 0.901309588 |
| 0.761688252 | 0.901309588 |
| 0.761668625 | 0.901309588 |
| 0.761649806 | 0.901309588 |
| 0.761534627 | 0.901265057 |
| 0.761529191 | 0.901265057 |
| 0.761457754 | 0.901218868 |
| 0.761444544 | 0.901218868 |
| 0.761418499 | 0.901217199 |
| 0.761378036 | 0.9012141 |
| 0.761275357 | 0.901139838 |
| 0.761261605 | 0.901139838 |
| 0.761129418 | 0.901048102 |
| 0.76107526 | 0.901034861 |
| 0.760976227 | 0.901009755 |
| 0.760961453 | 0.901009755 |
| 0.760848069 | 0.900945053 |
| 0.760790137 | 0.900898853 |
| 0.760778688 | 0.900898853 |
| 0.76067985 | 0.900835449 |
| 0.760646813 | 0.900818722 |
| 0.760574548 | 0.900777938 |
| 0.76051025 | 0.900763483 |
| 0.760382251 | 0.900725758 |
| 0.760372976 | 0.900725758 |
| 0.76033816 | 0.900721957 |
| 0.760230102 | 0.900638751 |
| 0.760210056 | 0.900638751 |
| 0.760177145 | 0.900638751 |
| 0.76014096 | 0.900638751 |
| 0.760103993 | 0.900623765 |
| 0.760020865 | 0.900547673 |
| 0.759947616 | 0.900505688 |
| 0.759878539 | 0.900446238 |
| 0.759809696 | 0.900446238 |
| 0.759801759 | 0.900446238 |
| 0.759765819 | 0.900446238 |
| 0.759697847 | 0.900411342 |
| 0.759665664 | 0.900401026 |
| 0.759586042 | 0.900346045 |
| 0.759547569 | 0.900345257 |

| | |
|---|---|
| 0.759366696 | 0.900198069 |
| 0.759190496 | 0.900060509 |
| 0.759173912 | 0.900060509 |
| 0.759098328 | 0.90001434 |
| 0.759019122 | 0.899991301 |
| 0.758994596 | 0.899991301 |
| 0.758992183 | 0.899991301 |
| 0.758989191 | 0.899991301 |
| 0.758897734 | 0.899991301 |
| 0.758829999 | 0.899991301 |
| 0.758787752 | 0.899991301 |
| 0.758764767 | 0.899991301 |
| 0.758719818 | 0.899991301 |
| 0.758704499 | 0.899991301 |
| 0.758697521 | 0.899991301 |
| 0.758636684 | 0.899991301 |
| 0.758591268 | 0.899991301 |
| 0.758510741 | 0.899989812 |
| 0.758454455 | 0.899945448 |
| 0.758453372 | 0.899945448 |
| 0.758404323 | 0.899930805 |
| 0.758370349 | 0.899930805 |
| 0.758285814 | 0.899902284 |
| 0.758265605 | 0.899900724 |
| 0.758179025 | 0.899820394 |
| 0.758007449 | 0.89975129 |
| 0.757970615 | 0.89975129 |
| 0.757960062 | 0.89975129 |
| 0.757946669 | 0.89975129 |
| 0.757885344 | 0.899740896 |
| 0.757855359 | 0.899740896 |
| 0.757758127 | 0.899738295 |
| 0.757660898 | 0.899676272 |
| 0.757608023 | 0.899666862 |
| 0.757486557 | 0.899581395 |
| 0.757414519 | 0.899518273 |
| 0.757407041 | 0.899518273 |
| 0.757352341 | 0.899518273 |
| 0.7573468 | 0.899518273 |
| 0.757341815 | 0.899518273 |
| 0.757178134 | 0.899484261 |
| 0.757177246 | 0.899484261 |
| 0.757174757 | 0.899484261 |
| 0.757174396 | 0.899484261 |
| 0.75717297 | 0.899484261 |

| | |
|---|---|
| 0.757154171 | 0.899484261 |
| 0.757016372 | 0.899484261 |
| 0.756954234 | 0.899484261 |
| 0.756821905 | 0.89937514 |
| 0.756752781 | 0.899315435 |
| 0.756656495 | 0.899245887 |
| 0.756619994 | 0.899224945 |
| 0.756615505 | 0.899224945 |
| 0.756612858 | 0.899224945 |
| 0.756394847 | 0.899047102 |
| 0.75630624 | 0.898986075 |
| 0.756207078 | 0.898958529 |
| 0.756207015 | 0.898958529 |
| 0.755989951 | 0.89879015 |
| 0.755956293 | 0.89877257 |
| 0.755908219 | 0.898737849 |
| 0.755763295 | 0.898618699 |
| 0.755694251 | 0.898595613 |
| 0.755693407 | 0.898595613 |
| 0.755627495 | 0.898583539 |
| 0.755600478 | 0.898573846 |
| 0.755573672 | 0.898565618 |
| 0.755538735 | 0.898565618 |
| 0.755477165 | 0.898561819 |
| 0.755412999 | 0.89855282 |
| 0.755214406 | 0.898383907 |
| 0.755032157 | 0.898269867 |
| 0.754908637 | 0.898222086 |
| 0.754895641 | 0.898222086 |
| 0.75489365 | 0.898222086 |
| 0.754745322 | 0.898095077 |
| 0.75449673 | 0.897889002 |
| 0.754438318 | 0.897867816 |
| 0.754263286 | 0.897788378 |
| 0.754232698 | 0.897776685 |
| 0.754085218 | 0.897646006 |
| 0.754078315 | 0.897646006 |
| 0.75405163 | 0.897646006 |
| 0.753993834 | 0.897604528 |
| 0.753776228 | 0.897390336 |
| 0.753749168 | 0.897380552 |
| 0.753696672 | 0.897362917 |
| 0.753682804 | 0.897362917 |
| 0.753651496 | 0.897353995 |
| 0.753534677 | 0.897259764 |

| | |
|---|---|
| 0.753492784 | 0.897232313 |
| 0.753434594 | 0.897198297 |
| 0.753324637 | 0.897166675 |
| 0.753268382 | 0.897122112 |
| 0.753231309 | 0.897100391 |
| 0.753162805 | 0.897048936 |
| 0.75315562 | 0.897048936 |
| 0.75309341 | 0.897048936 |
| 0.753081755 | 0.897048936 |
| 0.753032044 | 0.897048936 |
| 0.752993699 | 0.89704172 |
| 0.752940826 | 0.897003604 |
| 0.752822969 | 0.896928069 |
| 0.752798973 | 0.896928069 |
| 0.752787586 | 0.896928069 |
| 0.752573107 | 0.896764978 |
| 0.752572691 | 0.896764978 |
| 0.752561508 | 0.896764978 |
| 0.752540137 | 0.896764978 |
| 0.752445289 | 0.896724847 |
| 0.752265522 | 0.896600351 |
| 0.752199641 | 0.896544265 |
| 0.752196115 | 0.896544265 |
| 0.752180421 | 0.896544265 |
| 0.752113097 | 0.896544265 |
| 0.752075262 | 0.896544265 |
| 0.752051215 | 0.896544265 |
| 0.752024563 | 0.896544265 |
| 0.752008273 | 0.896544265 |
| 0.751990082 | 0.896544265 |
| 0.75192496 | 0.896544265 |
| 0.751766645 | 0.896476877 |
| 0.751757988 | 0.896476877 |
| 0.751732175 | 0.896476877 |
| 0.751695367 | 0.896476877 |
| 0.751692579 | 0.896476877 |
| 0.75168984 | 0.896476877 |
| 0.751659883 | 0.896476877 |
| 0.751607479 | 0.896476877 |
| 0.751558176 | 0.896475187 |
| 0.75150646 | 0.89643595 |
| 0.751505864 | 0.89643595 |
| 0.751423481 | 0.896381869 |
| 0.751403611 | 0.896380617 |
| 0.75138529 | 0.896380617 |

| | |
|---|---|
| 0.751369979 | 0.896380617 |
| 0.751303052 | 0.896380617 |
| 0.751287021 | 0.896380617 |
| 0.751174417 | 0.896331713 |
| 0.751161981 | 0.896331713 |
| 0.751155474 | 0.896331713 |
| 0.751084062 | 0.896313721 |
| 0.750929017 | 0.896173606 |
| 0.750910878 | 0.896173606 |
| 0.750899749 | 0.896173606 |
| 0.750881101 | 0.896173606 |
| 0.750684296 | 0.89603872 |
| 0.750470553 | 0.895895855 |
| 0.750466025 | 0.895895855 |
| 0.750448952 | 0.895895855 |
| 0.750405138 | 0.895895855 |
| 0.750391342 | 0.895895855 |
| 0.750333634 | 0.895867134 |
| 0.750236825 | 0.895796462 |
| 0.750158039 | 0.895747303 |
| 0.750117865 | 0.895721789 |
| 0.750113806 | 0.895721789 |
| 0.750033735 | 0.895690172 |
| 0.749907617 | 0.895627917 |
| 0.749895551 | 0.895627917 |
| 0.749861936 | 0.895627917 |
| 0.749774217 | 0.895587586 |
| 0.749748054 | 0.895587586 |
| 0.749747048 | 0.895587586 |
| 0.749693708 | 0.895587586 |
| 0.749540979 | 0.895561167 |
| 0.749510657 | 0.895561167 |
| 0.749488909 | 0.895561167 |
| 0.749470183 | 0.895561167 |
| 0.749434598 | 0.895557591 |
| 0.749367506 | 0.89552181 |
| 0.749334975 | 0.895505403 |
| 0.749307889 | 0.895495505 |
| 0.749289339 | 0.895495505 |
| 0.749279965 | 0.895495505 |
| 0.749268089 | 0.895495505 |
| 0.74923811 | 0.895495505 |
| 0.74921514 | 0.895495505 |
| 0.749207208 | 0.895495505 |
| 0.749184874 | 0.895495505 |

| | |
|---|---|
| 0.74900895 | 0.895410569 |
| 0.748950895 | 0.895405886 |
| 0.748914142 | 0.895384422 |
| 0.748909959 | 0.895384422 |
| 0.748840883 | 0.89536611 |
| 0.748753988 | 0.89536611 |
| 0.748561842 | 0.895232897 |
| 0.748507475 | 0.895190355 |
| 0.748487403 | 0.895188829 |
| 0.748406217 | 0.895134386 |
| 0.748393628 | 0.895134386 |
| 0.748203294 | 0.895051322 |
| 0.748184827 | 0.895051322 |
| 0.748104281 | 0.895034442 |
| 0.748074454 | 0.895034442 |
| 0.748046487 | 0.895034442 |
| 0.748023787 | 0.895034442 |
| 0.747824255 | 0.894980043 |
| 0.747809017 | 0.894980043 |
| 0.747785822 | 0.894979025 |
| 0.747650069 | 0.894884015 |
| 0.747613898 | 0.89486321 |
| 0.747547693 | 0.894832487 |
| 0.747506117 | 0.894832487 |
| 0.747453288 | 0.894828388 |
| 0.747407244 | 0.894801059 |
| 0.747319437 | 0.894758109 |
| 0.747211449 | 0.894651307 |
| 0.747137503 | 0.894611592 |
| 0.747087447 | 0.894592796 |
| 0.747085907 | 0.894592796 |
| 0.747085907 | 0.894592796 |
| 0.747074692 | 0.894592796 |
| 0.747070073 | 0.894592796 |
| 0.74703021 | 0.894592796 |
| 0.747006609 | 0.894592796 |
| 0.746998921 | 0.894592796 |
| 0.746994441 | 0.894592796 |
| 0.746943632 | 0.894592796 |
| 0.746881899 | 0.894592796 |
| 0.746623435 | 0.89435197 |
| 0.746507137 | 0.894235152 |
| 0.746500509 | 0.894235152 |
| 0.746496372 | 0.894235152 |
| 0.746463699 | 0.894235152 |

| | |
|---|---|
| 0.74625294 | 0.894116286 |
| 0.746251693 | 0.894116286 |
| 0.746218628 | 0.894116286 |
| 0.746187103 | 0.894116286 |
| 0.746072907 | 0.894007305 |
| 0.746042308 | 0.893993132 |
| 0.746014579 | 0.893982396 |
| 0.745935552 | 0.893932679 |
| 0.745932321 | 0.893932679 |
| 0.745773455 | 0.893828384 |
| 0.745753571 | 0.893827046 |
| 0.745738829 | 0.893827046 |
| 0.745727677 | 0.893827046 |
| 0.745681009 | 0.893827046 |
| 0.745676878 | 0.893827046 |
| 0.745646625 | 0.893827046 |
| 0.745635982 | 0.893827046 |
| 0.745605283 | 0.893827046 |
| 0.745580064 | 0.893827046 |
| 0.745539377 | 0.893827046 |
| 0.7454763 | 0.893827046 |
| 0.745470116 | 0.893827046 |
| 0.745432067 | 0.893827046 |
| 0.745308116 | 0.893742972 |
| 0.74530217 | 0.893742972 |
| 0.745286069 | 0.893742972 |
| 0.745126426 | 0.893664267 |
| 0.745116424 | 0.893664267 |
| 0.745115304 | 0.893664267 |
| 0.745100663 | 0.893664267 |
| 0.745093526 | 0.893664267 |
| 0.74496654 | 0.893625866 |
| 0.744892243 | 0.893581756 |
| 0.744880258 | 0.893581756 |
| 0.744877542 | 0.893581756 |
| 0.744861484 | 0.893581756 |
| 0.744815046 | 0.893581756 |
| 0.744770397 | 0.893581756 |
| 0.744731553 | 0.893581756 |
| 0.744706758 | 0.893581756 |
| 0.74465996 | 0.893581756 |
| 0.744536845 | 0.893515505 |
| 0.744519633 | 0.893515505 |
| 0.744475173 | 0.893508313 |
| 0.744309573 | 0.893459042 |

| | |
|---|---|
| 0.744224188 | 0.893387912 |
| 0.744217595 | 0.893387912 |
| 0.744148046 | 0.893341542 |
| 0.744142873 | 0.893341542 |
| 0.744142237 | 0.893341542 |
| 0.744137892 | 0.893341542 |
| 0.744128603 | 0.893341542 |
| 0.743862544 | 0.893133874 |
| 0.743858826 | 0.893133874 |
| 0.743729223 | 0.89304134 |
| 0.743720176 | 0.89304134 |
| 0.743625431 | 0.892984252 |
| 0.743504607 | 0.8929067 |
| 0.743442849 | 0.892858437 |
| 0.743263199 | 0.892687811 |
| 0.743256924 | 0.892687811 |
| 0.742982905 | 0.892549829 |
| 0.742812008 | 0.89243498 |
| 0.742795727 | 0.89243498 |
| 0.742790318 | 0.89243498 |
| 0.742762342 | 0.89243498 |
| 0.742631856 | 0.89240235 |
| 0.742603409 | 0.89240235 |
| 0.742562059 | 0.892382307 |
| 0.742518793 | 0.892352828 |
| 0.74250954 | 0.892352828 |
| 0.742478054 | 0.892352828 |
| 0.742313134 | 0.892218236 |
| 0.742242909 | 0.892201379 |
| 0.742210756 | 0.892185247 |
| 0.742159511 | 0.892185247 |
| 0.742149598 | 0.892185247 |
| 0.742128992 | 0.892185247 |
| 0.742088195 | 0.892185247 |
| 0.742086223 | 0.892185247 |
| 0.742086152 | 0.892185247 |
| 0.742069619 | 0.892185247 |
| 0.74206407 | 0.892185247 |
| 0.742027824 | 0.892185247 |
| 0.741941361 | 0.892185247 |
| 0.741901165 | 0.892185247 |
| 0.741837454 | 0.892185247 |
| 0.741829356 | 0.892185247 |
| 0.741774285 | 0.892185247 |
| 0.741766777 | 0.892185247 |

| | |
|---|---|
| 0.741639609 | 0.892145322 |
| 0.741546443 | 0.892089478 |
| 0.741472495 | 0.892063317 |
| 0.741380422 | 0.891975076 |
| 0.741348364 | 0.891959039 |
| 0.741225928 | 0.891856789 |
| 0.741191267 | 0.891837614 |
| 0.74108044 | 0.891796167 |
| 0.740979184 | 0.891762662 |
| 0.740894604 | 0.891683403 |
| 0.740855809 | 0.891659259 |
| 0.740829445 | 0.891659259 |
| 0.740758869 | 0.8916327 |
| 0.7406524 | 0.89154961 |
| 0.74064707 | 0.89154961 |
| 0.740605817 | 0.891538602 |
| 0.740530795 | 0.891470822 |
| 0.740507638 | 0.891465479 |
| 0.7404827 | 0.891457991 |
| 0.740416996 | 0.891423956 |
| 0.740333816 | 0.891391411 |
| 0.740306052 | 0.891380516 |
| 0.740217379 | 0.891318815 |
| 0.740025824 | 0.891245889 |
| 0.740025503 | 0.891245889 |
| 0.74000269 | 0.891245889 |
| 0.739979961 | 0.891245889 |
| 0.739952057 | 0.891245889 |
| 0.739894818 | 0.891245871 |
| 0.739682289 | 0.891109914 |
| 0.739656465 | 0.891109914 |
| 0.739637007 | 0.891109914 |
| 0.739494941 | 0.891057166 |
| 0.739440241 | 0.891013798 |
| 0.739376788 | 0.89095988 |
| 0.739367643 | 0.89095988 |
| 0.739236229 | 0.890934181 |
| 0.739168616 | 0.890889353 |
| 0.739121167 | 0.890854709 |
| 0.739038627 | 0.890777768 |
| 0.738991362 | 0.890777768 |
| 0.738964554 | 0.890777768 |
| 0.73892869 | 0.890777768 |
| 0.738871559 | 0.890756739 |
| 0.7388128 | 0.890752158 |

| | |
|---|---|
| 0.738806134 | 0.890752158 |
| 0.738730007 | 0.890721369 |
| 0.738585747 | 0.890615068 |
| 0.738551394 | 0.890596193 |
| 0.738484527 | 0.890538108 |
| 0.738455998 | 0.890526253 |
| 0.738423338 | 0.890509415 |
| 0.738397669 | 0.890501007 |
| 0.738192441 | 0.890343683 |
| 0.738022756 | 0.890184111 |
| 0.738013741 | 0.890184111 |
| 0.73797351 | 0.8901698 |
| 0.737939274 | 0.890151047 |
| 0.737922678 | 0.890151047 |
| 0.737856341 | 0.890112498 |
| 0.737826868 | 0.890105632 |
| 0.737769696 | 0.890059206 |
| 0.737709252 | 0.890031375 |
| 0.737651921 | 0.890031375 |
| 0.737627677 | 0.890031375 |
| 0.737588567 | 0.890031375 |
| 0.737500377 | 0.889982276 |
| 0.737463525 | 0.889965507 |
| 0.737398748 | 0.88992728 |
| 0.737373173 | 0.88992728 |
| 0.737289463 | 0.88992728 |
| 0.737263866 | 0.88992728 |
| 0.73720014 | 0.88992728 |
| 0.73719885 | 0.88992728 |
| 0.737165067 | 0.88992728 |
| 0.737151121 | 0.88992728 |
| 0.736981783 | 0.889913773 |
| 0.736976834 | 0.889913773 |
| 0.736929734 | 0.889913773 |
| 0.736901362 | 0.889913773 |
| 0.736812506 | 0.889873696 |
| 0.736715689 | 0.889801895 |
| 0.73666195 | 0.889759554 |
| 0.736615726 | 0.889748851 |
| 0.736573454 | 0.889748851 |
| 0.736560038 | 0.889748851 |
| 0.736450119 | 0.889717027 |
| 0.73642953 | 0.889717027 |
| 0.73636221 | 0.889690834 |
| 0.736303064 | 0.889641941 |

| | |
|---|---|
| 0.73618183 | 0.889518026 |
| 0.736107604 | 0.889473471 |
| 0.7360826 | 0.889468372 |
| 0.736043208 | 0.889463358 |
| 0.735908295 | 0.889378091 |
| 0.735854896 | 0.889348622 |
| 0.735846957 | 0.889348622 |
| 0.735754548 | 0.889290459 |
| 0.735685903 | 0.889279777 |
| 0.735400219 | 0.889047269 |
| 0.735367813 | 0.88903066 |
| 0.735148723 | 0.888860738 |
| 0.735123392 | 0.888860738 |
| 0.735053273 | 0.888830889 |
| 0.735025001 | 0.888830889 |
| 0.735017618 | 0.888830889 |
| 0.735000334 | 0.888830889 |
| 0.734993926 | 0.888830889 |
| 0.734979421 | 0.888830889 |
| 0.734964757 | 0.888830889 |
| 0.734927843 | 0.888830889 |
| 0.73466136 | 0.888627722 |
| 0.734571703 | 0.888564409 |
| 0.734563772 | 0.888564409 |
| 0.734426092 | 0.888433402 |
| 0.734405268 | 0.888430777 |
| 0.734367531 | 0.888407692 |
| 0.734364112 | 0.888407692 |
| 0.734329012 | 0.888406225 |
| 0.734259628 | 0.888344848 |
| 0.734221084 | 0.888344848 |
| 0.734207567 | 0.888344848 |
| 0.733983216 | 0.888249143 |
| 0.733945339 | 0.888249143 |
| 0.733698069 | 0.888026377 |
| 0.7336553 | 0.887997178 |
| 0.733655126 | 0.887997178 |
| 0.733532987 | 0.887894263 |
| 0.73350057 | 0.88787759 |
| 0.733328973 | 0.887760129 |
| 0.73332014 | 0.887760129 |
| 0.733241543 | 0.887760129 |
| 0.733205259 | 0.887745752 |
| 0.733075164 | 0.887655933 |
| 0.733043019 | 0.887655933 |

| | |
|---|---|
| 0.733042848 | 0.887655933 |
| 0.733030447 | 0.887655933 |
| 0.732792991 | 0.887517337 |
| 0.732736163 | 0.887471079 |
| 0.732618492 | 0.887373692 |
| 0.732604621 | 0.887373692 |
| 0.73243779 | 0.887267643 |
| 0.732430783 | 0.887267643 |
| 0.732427791 | 0.887267643 |
| 0.732335001 | 0.887210823 |
| 0.732174721 | 0.887054395 |
| 0.73217312 | 0.887054395 |
| 0.732138529 | 0.887054395 |
| 0.732137644 | 0.887054395 |
| 0.732074006 | 0.887030012 |
| 0.731962908 | 0.886940528 |
| 0.731956459 | 0.886940528 |
| 0.731902311 | 0.886940528 |
| 0.731827766 | 0.886912161 |
| 0.73173668 | 0.886846905 |
| 0.73173654 | 0.886846905 |
| 0.731687022 | 0.886846905 |
| 0.731685999 | 0.886846905 |
| 0.731612958 | 0.886809793 |
| 0.731570701 | 0.88680371 |
| 0.73154032 | 0.886789453 |
| 0.731524352 | 0.886789453 |
| 0.731522157 | 0.886789453 |
| 0.731209518 | 0.886523833 |
| 0.731182887 | 0.886514112 |
| 0.731101069 | 0.886437479 |
| 0.730988623 | 0.886368837 |
| 0.730974104 | 0.886368837 |
| 0.730883907 | 0.886354693 |
| 0.730777882 | 0.886271248 |
| 0.730704498 | 0.886204816 |
| 0.730623788 | 0.886197198 |
| 0.730621333 | 0.886197198 |
| 0.730404738 | 0.886021762 |
| 0.730397519 | 0.886021762 |
| 0.730393576 | 0.886021762 |
| 0.730239837 | 0.88595712 |
| 0.73023532 | 0.88595712 |
| 0.730232035 | 0.88595712 |
| 0.730229161 | 0.88595712 |

| | |
|---|---|
| 0.730210462 | 0.88595712 |
| 0.730178914 | 0.88595712 |
| 0.730020402 | 0.885866082 |
| 0.729963168 | 0.885824543 |
| 0.729815761 | 0.885668228 |
| 0.729685165 | 0.885554871 |
| 0.729663408 | 0.885551031 |
| 0.729660518 | 0.885551031 |
| 0.729603474 | 0.885545991 |
| 0.72955011 | 0.885503787 |
| 0.729512843 | 0.88548112 |
| 0.729470855 | 0.885475289 |
| 0.729393825 | 0.88544722 |
| 0.72937917 | 0.88544722 |
| 0.729295123 | 0.885442511 |
| 0.729192751 | 0.885340789 |
| 0.729174958 | 0.885340789 |
| 0.729174259 | 0.885340789 |
| 0.729162161 | 0.885340789 |
| 0.729142946 | 0.885340789 |
| 0.729130568 | 0.885340789 |
| 0.729003528 | 0.885269021 |
| 0.728977136 | 0.885269021 |
| 0.728950525 | 0.885269021 |
| 0.728950525 | 0.885269021 |
| 0.728873902 | 0.885269021 |
| 0.728830325 | 0.885261837 |
| 0.728716757 | 0.885173807 |
| 0.728709159 | 0.885173807 |
| 0.728695223 | 0.885173807 |
| 0.728564974 | 0.885153358 |
| 0.728562294 | 0.885153358 |
| 0.728433874 | 0.885104339 |
| 0.728412002 | 0.885104339 |
| 0.728404767 | 0.885104339 |
| 0.728354281 | 0.885104339 |
| 0.728311303 | 0.885104339 |
| 0.728285126 | 0.885096218 |
| 0.728225582 | 0.885046437 |
| 0.728143651 | 0.885011072 |
| 0.728049705 | 0.884968182 |
| 0.727841106 | 0.884804951 |
| 0.727797483 | 0.884774504 |
| 0.727785212 | 0.884774504 |
| 0.727719226 | 0.884769703 |

| | |
|---|---|
| 0.727703217 | 0.884769703 |
| 0.727698546 | 0.884769703 |
| 0.727615515 | 0.884756539 |
| 0.727607224 | 0.884756539 |
| 0.727509084 | 0.884694884 |
| 0.727474225 | 0.884675082 |
| 0.727466778 | 0.884675082 |
| 0.727435488 | 0.884673152 |
| 0.727141891 | 0.884429012 |
| 0.72713451 | 0.884429012 |
| 0.727076623 | 0.884417386 |
| 0.726897526 | 0.884249213 |
| 0.726890155 | 0.884249213 |
| 0.726868966 | 0.884249213 |
| 0.726792163 | 0.884206865 |
| 0.726672867 | 0.88415207 |
| 0.726668767 | 0.88415207 |
| 0.726615119 | 0.884149568 |
| 0.726557597 | 0.884141331 |
| 0.726423438 | 0.884141331 |
| 0.726419243 | 0.884141331 |
| 0.726402491 | 0.884141331 |
| 0.726312303 | 0.884097326 |
| 0.726248598 | 0.884064971 |
| 0.726143118 | 0.88404817 |
| 0.726141059 | 0.88404817 |
| 0.72611554 | 0.88404817 |
| 0.726103944 | 0.88404817 |
| 0.726085501 | 0.88404817 |
| 0.726003788 | 0.884038106 |
| 0.725974031 | 0.884024472 |
| 0.725973211 | 0.884024472 |
| 0.725921576 | 0.884024472 |
| 0.725768103 | 0.883931912 |
| 0.725718368 | 0.883893942 |
| 0.725705214 | 0.883893942 |
| 0.7256547 | 0.883861716 |
| 0.725648668 | 0.883861716 |
| 0.725515923 | 0.883805582 |
| 0.725501325 | 0.883805582 |
| 0.725493996 | 0.883805582 |
| 0.725432743 | 0.883805582 |
| 0.725415583 | 0.883805582 |
| 0.725392161 | 0.883805582 |
| 0.725337356 | 0.883805582 |

| | |
|---|---|
| 0.72517136 | 0.883747508 |
| 0.725154155 | 0.883747508 |
| 0.72491453 | 0.88359278 |
| 0.724836885 | 0.883520751 |
| 0.724806902 | 0.883506816 |
| 0.724730625 | 0.883446433 |
| 0.724695127 | 0.883438399 |
| 0.724616865 | 0.883388216 |
| 0.724482477 | 0.883292937 |
| 0.724415065 | 0.883255243 |
| 0.724398542 | 0.883255243 |
| 0.724367244 | 0.88324216 |
| 0.724329694 | 0.883218985 |
| 0.724147998 | 0.883110481 |
| 0.724106854 | 0.883082916 |
| 0.724100363 | 0.883082916 |
| 0.724044578 | 0.883082916 |
| 0.724036089 | 0.883082916 |
| 0.724004939 | 0.883082916 |
| 0.723967048 | 0.883082916 |
| 0.723930165 | 0.883082916 |
| 0.72389708 | 0.883082916 |
| 0.723840938 | 0.883082916 |
| 0.723763196 | 0.883048206 |
| 0.72374802 | 0.883048206 |
| 0.723714308 | 0.883033799 |
| 0.723535179 | 0.882909742 |
| 0.723520473 | 0.882909742 |
| 0.723513917 | 0.882909742 |
| 0.723492673 | 0.882909742 |
| 0.723484585 | 0.882909742 |
| 0.723393422 | 0.882891077 |
| 0.723257503 | 0.882770433 |
| 0.723251363 | 0.882770433 |
| 0.723207513 | 0.882754664 |
| 0.723115894 | 0.882665454 |
| 0.723087322 | 0.882663616 |
| 0.722940415 | 0.882609601 |
| 0.722833325 | 0.88256174 |
| 0.722792989 | 0.882542729 |
| 0.722778692 | 0.882542729 |
| 0.722729016 | 0.882532494 |
| 0.722721815 | 0.882532494 |
| 0.722688437 | 0.882532494 |
| 0.722616242 | 0.882507917 |

| | |
|---|---|
| 0.722600364 | 0.882507917 |
| 0.722589923 | 0.882507917 |
| 0.722572964 | 0.882507917 |
| 0.722457395 | 0.88242706 |
| 0.722321914 | 0.882329464 |
| 0.72222225 | 0.882252978 |
| 0.722177419 | 0.882245908 |
| 0.722109463 | 0.882245908 |
| 0.721873749 | 0.882030864 |
| 0.721864751 | 0.882030864 |
| 0.721849666 | 0.882030864 |
| 0.721814852 | 0.882026784 |
| 0.721758475 | 0.882020926 |
| 0.721505589 | 0.881897773 |
| 0.721375624 | 0.881761545 |
| 0.721336368 | 0.881758821 |
| 0.721121486 | 0.881586658 |
| 0.721087121 | 0.881567275 |
| 0.721017439 | 0.881504713 |
| 0.720951121 | 0.881446259 |
| 0.720900998 | 0.881430232 |
| 0.720868303 | 0.881430232 |
| 0.720825836 | 0.881406215 |
| 0.720663814 | 0.881298605 |
| 0.720661768 | 0.881298605 |
| 0.720653192 | 0.881298605 |
| 0.720634519 | 0.881298605 |
| 0.7204458 | 0.881160791 |
| 0.720391266 | 0.881146305 |
| 0.720309836 | 0.881069332 |
| 0.720207316 | 0.881057069 |
| 0.720196055 | 0.881057069 |
| 0.720172929 | 0.881057069 |
| 0.720063537 | 0.881016958 |
| 0.719993507 | 0.880953905 |
| 0.719938575 | 0.880931956 |
| 0.719935286 | 0.880931956 |
| 0.719931702 | 0.880931956 |
| 0.719887677 | 0.880931956 |
| 0.719837008 | 0.880920838 |
| 0.719698133 | 0.880864055 |
| 0.719697037 | 0.880864055 |
| 0.719690887 | 0.880864055 |
| 0.719541154 | 0.880830374 |
| 0.719526887 | 0.880830374 |

| | |
|---|---|
| 0.719513866 | 0.880830374 |
| 0.719455456 | 0.880795304 |
| 0.719442337 | 0.880795304 |
| 0.719374499 | 0.880784835 |
| 0.719333115 | 0.88077945 |
| 0.719307421 | 0.88077945 |
| 0.719301884 | 0.88077945 |
| 0.719188626 | 0.88077945 |
| 0.719083866 | 0.880768627 |
| 0.719033984 | 0.88073018 |
| 0.718830914 | 0.88057203 |
| 0.718798096 | 0.880566846 |
| 0.71878058 | 0.880566846 |
| 0.718718709 | 0.880566846 |
| 0.718700967 | 0.880566846 |
| 0.718674425 | 0.880566846 |
| 0.718673678 | 0.880566846 |
| 0.718593097 | 0.880563986 |
| 0.718589979 | 0.880563986 |
| 0.718577475 | 0.880563986 |
| 0.718361346 | 0.880385785 |
| 0.718144368 | 0.88025411 |
| 0.718124787 | 0.88025411 |
| 0.718115747 | 0.88025411 |
| 0.718110495 | 0.88025411 |
| 0.717992489 | 0.880224157 |
| 0.717849368 | 0.88011268 |
| 0.717841813 | 0.88011268 |
| 0.717779538 | 0.880076358 |
| 0.717745163 | 0.880056867 |
| 0.717700196 | 0.880036153 |
| 0.717545085 | 0.879902154 |
| 0.717525283 | 0.879900527 |
| 0.717300159 | 0.879669757 |
| 0.717221997 | 0.879635074 |
| 0.717207346 | 0.879635074 |
| 0.717197137 | 0.879635074 |
| 0.717090024 | 0.879635074 |
| 0.717058602 | 0.879635074 |
| 0.716993376 | 0.879633279 |
| 0.716910119 | 0.879618403 |
| 0.716896586 | 0.879618403 |
| 0.716894424 | 0.879618403 |
| 0.716755832 | 0.879500439 |
| 0.71673123 | 0.879500439 |

| | |
|---|---|
| 0.716722999 | 0.879500439 |
| 0.716710543 | 0.879500439 |
| 0.716707415 | 0.879500439 |
| 0.716693206 | 0.879500439 |
| 0.71666984 | 0.879500439 |
| 0.716586513 | 0.879500439 |
| 0.716559234 | 0.879500439 |
| 0.716464122 | 0.879482377 |
| 0.716462074 | 0.879482377 |
| 0.716450939 | 0.879482377 |
| 0.716284235 | 0.879352217 |
| 0.716282589 | 0.879352217 |
| 0.716191211 | 0.879306012 |
| 0.71609476 | 0.879221614 |
| 0.716040704 | 0.879189219 |
| 0.715993579 | 0.879176688 |
| 0.715946896 | 0.879164906 |
| 0.715904072 | 0.879164906 |
| 0.71589306 | 0.879164906 |
| 0.715885799 | 0.879164906 |
| 0.715733049 | 0.87915144 |
| 0.7156239 | 0.879057321 |
| 0.715616968 | 0.879057321 |
| 0.715592361 | 0.879057321 |
| 0.715576242 | 0.879057321 |
| 0.715450541 | 0.879053813 |
| 0.715292349 | 0.878904794 |
| 0.715269535 | 0.878899436 |
| 0.715172118 | 0.878847755 |
| 0.714826463 | 0.878621681 |
| 0.714782177 | 0.878621681 |
| 0.71470413 | 0.878590032 |
| 0.714649511 | 0.878571883 |
| 0.714647473 | 0.878571883 |
| 0.714514817 | 0.878493358 |
| 0.714480307 | 0.878473605 |
| 0.714455097 | 0.878465287 |
| 0.714439522 | 0.878465287 |
| 0.714370752 | 0.878413943 |
| 0.714321426 | 0.878404663 |
| 0.714288004 | 0.878404663 |
| 0.714200464 | 0.878378967 |
| 0.714177825 | 0.878378967 |
| 0.714135596 | 0.878368236 |
| 0.714125001 | 0.878368236 |

| | |
|---|---|
| 0.714098767 | 0.878368236 |
| 0.714076102 | 0.878368236 |
| 0.714068557 | 0.878368236 |
| 0.714056007 | 0.878368236 |
| 0.714034132 | 0.878368236 |
| 0.71403291 | 0.878368236 |
| 0.714028835 | 0.878368236 |
| 0.713952527 | 0.878368236 |
| 0.713872614 | 0.878368236 |
| 0.713771979 | 0.878368236 |
| 0.713760328 | 0.878368236 |
| 0.713725541 | 0.878361922 |
| 0.713531374 | 0.878183887 |
| 0.713494962 | 0.878183887 |
| 0.713476376 | 0.878183887 |
| 0.713390435 | 0.878180306 |
| 0.713379886 | 0.878180306 |
| 0.713201656 | 0.878080221 |
| 0.71311329 | 0.878001583 |
| 0.713068223 | 0.87798403 |
| 0.713042864 | 0.877975504 |
| 0.712983526 | 0.877947836 |
| 0.712973752 | 0.877947836 |
| 0.712954611 | 0.877947836 |
| 0.712830804 | 0.877876779 |
| 0.712728595 | 0.877815485 |
| 0.712716253 | 0.877815485 |
| 0.712702297 | 0.877815485 |
| 0.7126724 | 0.877815485 |
| 0.712517063 | 0.877713839 |
| 0.712500161 | 0.877713839 |
| 0.712369872 | 0.877623319 |
| 0.712339971 | 0.877615926 |
| 0.712310965 | 0.877615926 |
| 0.712247472 | 0.877586029 |
| 0.712242293 | 0.877586029 |
| 0.712229413 | 0.877586029 |
| 0.712226942 | 0.877586029 |
| 0.71221015 | 0.877586029 |
| 0.712188506 | 0.877586029 |
| 0.712071867 | 0.877586029 |
| 0.712022658 | 0.877558759 |
| 0.711973589 | 0.877520991 |
| 0.711931188 | 0.877515174 |
| 0.711905756 | 0.877515174 |

| | |
|---|---|
| 0.711825907 | 0.877475217 |
| 0.711817534 | 0.877475217 |
| 0.711705686 | 0.877451536 |
| 0.711622354 | 0.877405972 |
| 0.711506212 | 0.877285483 |
| 0.711465407 | 0.877257882 |
| 0.711406289 | 0.87723041 |
| 0.711382258 | 0.87723041 |
| 0.71131199 | 0.877204975 |
| 0.711252986 | 0.877179511 |
| 0.711216615 | 0.877179511 |
| 0.711115714 | 0.877121905 |
| 0.711072121 | 0.87709085 |
| 0.710999493 | 0.87709085 |
| 0.710994611 | 0.87709085 |
| 0.710983923 | 0.87709085 |
| 0.710969743 | 0.87709085 |
| 0.71094991 | 0.87709085 |
| 0.710944866 | 0.87709085 |
| 0.710939278 | 0.87709085 |
| 0.710909171 | 0.87709085 |
| 0.710903844 | 0.87709085 |
| 0.710901774 | 0.87709085 |
| 0.710871742 | 0.87709085 |
| 0.710869301 | 0.87709085 |
| 0.710851791 | 0.87709085 |
| 0.710802173 | 0.87709085 |
| 0.710676122 | 0.87709085 |
| 0.710660354 | 0.87709085 |
| 0.710649103 | 0.87709085 |
| 0.710569221 | 0.87709085 |
| 0.710566353 | 0.87709085 |
| 0.710461582 | 0.87709085 |
| 0.710435255 | 0.87709085 |
| 0.71041654 | 0.87709085 |
| 0.710353495 | 0.87709085 |
| 0.709949393 | 0.876750498 |
| 0.709908913 | 0.876723241 |
| 0.709903297 | 0.876723241 |
| 0.709714746 | 0.87664257 |
| 0.709674398 | 0.876638204 |
| 0.709632616 | 0.876638204 |
| 0.709580489 | 0.876638204 |
| 0.709547917 | 0.876638204 |
| 0.709476438 | 0.876598291 |

| | |
|---|---|
| 0.709443297 | 0.876580084 |
| 0.7093578 | 0.876546499 |
| 0.709334479 | 0.876546499 |
| 0.709306227 | 0.876546499 |
| 0.709267809 | 0.876546499 |
| 0.709262115 | 0.876546499 |
| 0.709256427 | 0.876546499 |
| 0.709194952 | 0.876546114 |
| 0.709177505 | 0.876546114 |
| 0.70911334 | 0.876536232 |
| 0.708806063 | 0.876354303 |
| 0.708798388 | 0.876354303 |
| 0.708740122 | 0.876354303 |
| 0.708719795 | 0.876354303 |
| 0.708600418 | 0.876243337 |
| 0.708571979 | 0.876230919 |
| 0.708557131 | 0.876230919 |
| 0.708536511 | 0.876230919 |
| 0.708528091 | 0.876230919 |
| 0.708452509 | 0.876219685 |
| 0.708334918 | 0.876119752 |
| 0.708275041 | 0.876091196 |
| 0.70813477 | 0.87598594 |
| 0.707983012 | 0.875843708 |
| 0.707983012 | 0.875843708 |
| 0.707983012 | 0.875843708 |
| 0.707840237 | 0.87580716 |
| 0.707833737 | 0.87580716 |
| 0.707798257 | 0.87580716 |
| 0.707740234 | 0.875793601 |
| 0.707730278 | 0.875793601 |
| 0.707658343 | 0.875764293 |
| 0.707656104 | 0.875764293 |
| 0.707616689 | 0.875764293 |
| 0.707602204 | 0.875764293 |
| 0.707567748 | 0.875764293 |
| 0.707562076 | 0.875764293 |
| 0.707511284 | 0.875764293 |
| 0.707492286 | 0.875764293 |
| 0.707357635 | 0.875752478 |
| 0.707353935 | 0.875752478 |
| 0.707241334 | 0.875699546 |
| 0.707158101 | 0.875668978 |
| 0.707145717 | 0.875668978 |
| 0.707128022 | 0.875668978 |

| | |
|---|---|
| 0.707057319 | 0.875668978 |
| 0.707028671 | 0.875668978 |
| 0.706948066 | 0.875609589 |
| 0.706907768 | 0.875582447 |
| 0.706857907 | 0.875543459 |
| 0.7068493 | 0.875543459 |
| 0.706736263 | 0.87546109 |
| 0.706479685 | 0.875211548 |
| 0.706448258 | 0.875211548 |
| 0.706358409 | 0.875176101 |
| 0.706352179 | 0.875176101 |
| 0.706335495 | 0.875176101 |
| 0.706202217 | 0.875118203 |
| 0.706166797 | 0.875097079 |
| 0.706116569 | 0.875057604 |
| 0.706054066 | 0.875025683 |
| 0.70600137 | 0.874983144 |
| 0.705828988 | 0.874883333 |
| 0.705664716 | 0.874764548 |
| 0.705659411 | 0.874764548 |
| 0.705557828 | 0.87469128 |
| 0.705418846 | 0.874579809 |
| 0.705158175 | 0.874360704 |
| 0.705127946 | 0.874360704 |
| 0.705105401 | 0.874360704 |
| 0.705088543 | 0.874360704 |
| 0.705068076 | 0.874360704 |
| 0.705026624 | 0.874360406 |
| 0.705006897 | 0.874360406 |
| 0.704925119 | 0.874309079 |
| 0.704913909 | 0.874309079 |
| 0.704746018 | 0.874276924 |
| 0.704725739 | 0.874276924 |
| 0.704721809 | 0.874276924 |
| 0.704714727 | 0.874276924 |
| 0.704569059 | 0.874276924 |
| 0.704438441 | 0.874251866 |
| 0.704332555 | 0.874238013 |
| 0.704332 | 0.874238013 |
| 0.704331769 | 0.874238013 |
| 0.704296203 | 0.874238013 |
| 0.704229774 | 0.874220701 |
| 0.704191096 | 0.874195473 |
| 0.704188564 | 0.874195473 |
| 0.704119045 | 0.874195473 |

| | |
|---|---|
| 0.70406785 | 0.874195473 |
| 0.703929746 | 0.874075183 |
| 0.703794861 | 0.87401922 |
| 0.703792739 | 0.87401922 |
| 0.703731096 | 0.873988915 |
| 0.703705557 | 0.873979988 |
| 0.703701585 | 0.873979988 |
| 0.703556292 | 0.873907894 |
| 0.703511724 | 0.873889868 |
| 0.703459918 | 0.873857212 |
| 0.703417286 | 0.873849838 |
| 0.703338341 | 0.873797349 |
| 0.703276618 | 0.873766251 |
| 0.703265174 | 0.873766251 |
| 0.70319953 | 0.873739508 |
| 0.703143388 | 0.873714684 |
| 0.703123335 | 0.873712562 |
| 0.703058691 | 0.87370062 |
| 0.702785994 | 0.873430107 |
| 0.702775451 | 0.873430107 |
| 0.702758 | 0.873430107 |
| 0.702661194 | 0.873387241 |
| 0.702648408 | 0.873387241 |
| 0.702591275 | 0.873347636 |
| 0.702562196 | 0.873344028 |
| 0.702535386 | 0.873344028 |
| 0.702523827 | 0.873344028 |
| 0.702499244 | 0.873344028 |
| 0.70247634 | 0.873344028 |
| 0.702421357 | 0.873341564 |
| 0.702372585 | 0.873318683 |
| 0.702295172 | 0.873298664 |
| 0.702273431 | 0.873294431 |
| 0.702205116 | 0.873267649 |
| 0.702166967 | 0.873267649 |
| 0.701990373 | 0.873102015 |
| 0.701949872 | 0.873074442 |
| 0.701942346 | 0.873074442 |
| 0.701899159 | 0.873056966 |
| 0.701831024 | 0.872998096 |
| 0.701818386 | 0.872998096 |
| 0.701793627 | 0.872998096 |
| 0.701676088 | 0.872961889 |
| 0.701576293 | 0.872883337 |
| 0.701543538 | 0.872865385 |

| | |
|---|---|
| 0.701532572 | 0.872865385 |
| 0.701500406 | 0.872865385 |
| 0.701444908 | 0.872833877 |
| 0.701335504 | 0.872783969 |
| 0.701314778 | 0.872783969 |
| 0.701253655 | 0.872778317 |
| 0.701228507 | 0.872769825 |
| 0.701186855 | 0.872769825 |
| 0.701156221 | 0.872769825 |
| 0.701112626 | 0.872739631 |
| 0.701107876 | 0.872739631 |
| 0.701101163 | 0.872739631 |
| 0.701038909 | 0.872716298 |
| 0.701032688 | 0.872716298 |
| 0.701029812 | 0.872716298 |
| 0.701015019 | 0.872716298 |
| 0.700787594 | 0.872540279 |
| 0.700709437 | 0.872465775 |
| 0.700671282 | 0.872441076 |
| 0.700621152 | 0.872441076 |
| 0.700599057 | 0.872441076 |
| 0.700476812 | 0.872358579 |
| 0.700468604 | 0.872358579 |
| 0.700439785 | 0.872358088 |
| 0.700422442 | 0.872358088 |
| 0.700392493 | 0.872358088 |
| 0.700254698 | 0.872310059 |
| 0.700187679 | 0.872272203 |
| 0.700172501 | 0.872272203 |
| 0.699999316 | 0.872105981 |
| 0.699977192 | 0.87210123 |
| 0.699868772 | 0.872011772 |
| 0.699826776 | 0.871982259 |
| 0.699826776 | 0.871982259 |
| 0.699732105 | 0.871932732 |
| 0.699522385 | 0.871694208 |
| 0.699436414 | 0.871632688 |
| 0.699431382 | 0.871632688 |
| 0.699396405 | 0.871628443 |
| 0.699289072 | 0.871517483 |
| 0.699027429 | 0.871281415 |
| 0.698985471 | 0.871253102 |
| 0.698961998 | 0.871246645 |
| 0.698866695 | 0.871196254 |
| 0.698857526 | 0.871196254 |

| | |
|---|---|
| 0.698767247 | 0.871163491 |
| 0.698739376 | 0.871151548 |
| 0.698666802 | 0.871106672 |
| 0.698560811 | 0.870997323 |
| 0.698541516 | 0.870997323 |
| 0.698341282 | 0.870852353 |
| 0.698284108 | 0.870811899 |
| 0.698262163 | 0.870807333 |
| 0.698255374 | 0.870807333 |
| 0.698216409 | 0.870807333 |
| 0.698215525 | 0.870807333 |
| 0.698168271 | 0.870807333 |
| 0.698127154 | 0.870807333 |
| 0.698122526 | 0.870807333 |
| 0.698106136 | 0.870807333 |
| 0.69785474 | 0.870618392 |
| 0.697831806 | 0.870618392 |
| 0.697799373 | 0.870617734 |
| 0.697794809 | 0.870617734 |
| 0.697775964 | 0.870617734 |
| 0.697749893 | 0.870617734 |
| 0.697732173 | 0.870617734 |
| 0.697708041 | 0.870617734 |
| 0.697697793 | 0.870617734 |
| 0.697506698 | 0.870493398 |
| 0.697470258 | 0.870480752 |
| 0.697346796 | 0.870349473 |
| 0.697346392 | 0.870349473 |
| 0.697339301 | 0.870349473 |
| 0.697268086 | 0.870349473 |
| 0.697112931 | 0.870217219 |
| 0.697098481 | 0.870217219 |
| 0.69703527 | 0.870209724 |
| 0.696993643 | 0.870187609 |
| 0.696951284 | 0.870187609 |
| 0.696869833 | 0.87018745 |
| 0.696782331 | 0.870146628 |
| 0.696738769 | 0.870138808 |
| 0.696718337 | 0.870138808 |
| 0.696707193 | 0.870138808 |
| 0.696670808 | 0.870138808 |
| 0.696640477 | 0.870138808 |
| 0.696486845 | 0.869982925 |
| 0.696399863 | 0.869897091 |
| 0.696298202 | 0.869815728 |

| | |
|---|---|
| 0.696204115 | 0.869721007 |
| 0.6961494 | 0.869721007 |
| 0.695969863 | 0.869596645 |
| 0.695959834 | 0.869596645 |
| 0.695951759 | 0.869596645 |
| 0.695947721 | 0.869596645 |
| 0.695841439 | 0.869558912 |
| 0.695743246 | 0.869487368 |
| 0.695723273 | 0.869485222 |
| 0.695689412 | 0.869485222 |
| 0.695688629 | 0.869485222 |
| 0.695640887 | 0.869485222 |
| 0.695595195 | 0.869484865 |
| 0.695340189 | 0.869335263 |
| 0.695314527 | 0.869335263 |
| 0.695280968 | 0.869335263 |
| 0.695210362 | 0.869300425 |
| 0.695187226 | 0.869294319 |
| 0.695180676 | 0.869294319 |
| 0.695176733 | 0.869294319 |
| 0.695065729 | 0.869279331 |
| 0.695056824 | 0.869279331 |
| 0.695000833 | 0.86924382 |
| 0.694938594 | 0.869188802 |
| 0.694931714 | 0.869188802 |
| 0.694892495 | 0.869176796 |
| 0.694856027 | 0.869176796 |
| 0.694853496 | 0.869176796 |
| 0.694827114 | 0.869176796 |
| 0.694819698 | 0.869176796 |
| 0.694766715 | 0.869176796 |
| 0.694621897 | 0.869135065 |
| 0.694287312 | 0.868995122 |
| 0.694231348 | 0.868995122 |
| 0.694226956 | 0.868995122 |
| 0.694173709 | 0.868995122 |
| 0.694041387 | 0.868933759 |
| 0.693987318 | 0.868911748 |
| 0.693978157 | 0.868911748 |
| 0.693955694 | 0.868911748 |
| 0.693867202 | 0.868875565 |
| 0.69367978 | 0.86866371 |
| 0.693599124 | 0.868608385 |
| 0.693567087 | 0.868591104 |
| 0.693378254 | 0.868400287 |

| | |
|---|---|
| 0.693305751 | 0.868355152 |
| 0.69311115 | 0.868202745 |
| 0.693082528 | 0.868189726 |
| 0.693043195 | 0.868163288 |
| 0.693037777 | 0.868163288 |
| 0.692873737 | 0.868042336 |
| 0.69279512 | 0.867966676 |
| 0.692788537 | 0.867966676 |
| 0.69277386 | 0.867966676 |
| 0.692768097 | 0.867966676 |
| 0.69274748 | 0.867966676 |
| 0.692538791 | 0.867896669 |
| 0.692524947 | 0.867896669 |
| 0.692491233 | 0.867882742 |
| 0.692452546 | 0.867857093 |
| 0.692439715 | 0.867857093 |
| 0.692424827 | 0.867857093 |
| 0.692215581 | 0.867674267 |
| 0.692104393 | 0.86755773 |
| 0.692065311 | 0.867554406 |
| 0.692011107 | 0.867509292 |
| 0.691910855 | 0.867406445 |
| 0.691796993 | 0.867309363 |
| 0.69172484 | 0.86726185 |
| 0.691712193 | 0.86726185 |
| 0.691698988 | 0.86726185 |
| 0.691302444 | 0.866963201 |
| 0.691276047 | 0.866952926 |
| 0.691205104 | 0.866917331 |
| 0.691151159 | 0.866917331 |
| 0.691144268 | 0.866917331 |
| 0.691119261 | 0.866917331 |
| 0.691118214 | 0.866917331 |
| 0.691097911 | 0.866917331 |
| 0.691050661 | 0.866917331 |
| 0.690994688 | 0.866896821 |
| 0.690968106 | 0.866886307 |
| 0.690925932 | 0.866864501 |
| 0.690873769 | 0.866836455 |
| 0.69086181 | 0.866836455 |
| 0.690568705 | 0.866636339 |
| 0.690219973 | 0.866335661 |
| 0.690184402 | 0.866313844 |
| 0.690154894 | 0.866299638 |
| 0.69011198 | 0.866268601 |

| | |
|---|---|
| 0.690079242 | 0.866250338 |
| 0.6900748 | 0.866250338 |
| 0.690034588 | 0.866250338 |
| 0.690022184 | 0.866250338 |
| 0.689965732 | 0.866222003 |
| 0.689937159 | 0.866208965 |
| 0.689909792 | 0.866197439 |
| 0.689839452 | 0.86617097 |
| 0.689819478 | 0.86617097 |
| 0.689741519 | 0.866100328 |
| 0.689679713 | 0.866045552 |
| 0.689669803 | 0.866045552 |
| 0.689635569 | 0.866045552 |
| 0.689631014 | 0.866045552 |
| 0.689530475 | 0.866035674 |
| 0.689501761 | 0.866034878 |
| 0.689457247 | 0.866017362 |
| 0.689414453 | 0.865986448 |
| 0.689186746 | 0.865814596 |
| 0.689099247 | 0.86579047 |
| 0.689029862 | 0.865731685 |
| 0.689017334 | 0.865731685 |
| 0.688998876 | 0.865731685 |
| 0.688858543 | 0.865630613 |
| 0.688821118 | 0.865606422 |
| 0.688803878 | 0.865606422 |
| 0.688702863 | 0.865505181 |
| 0.68864382 | 0.865505181 |
| 0.688438636 | 0.865331264 |
| 0.688404958 | 0.86531177 |
| 0.688375117 | 0.865297096 |
| 0.688318499 | 0.865271601 |
| 0.688196696 | 0.865210838 |
| 0.688159608 | 0.865208882 |
| 0.688092606 | 0.865147481 |
| 0.688084819 | 0.865147481 |
| 0.688072308 | 0.865147481 |
| 0.688058868 | 0.865147481 |
| 0.687993567 | 0.865147481 |
| 0.687957166 | 0.865137062 |
| 0.687949335 | 0.865137062 |
| 0.687875547 | 0.865080106 |
| 0.687861932 | 0.865080106 |
| 0.687808195 | 0.865063313 |
| 0.687773664 | 0.865043344 |

| | |
|---|---|
| 0.687727212 | 0.865007763 |
| 0.687529826 | 0.864850854 |
| 0.687374986 | 0.864701755 |
| 0.687374143 | 0.864701755 |
| 0.687301404 | 0.864700549 |
| 0.687276488 | 0.864700549 |
| 0.687234207 | 0.864700549 |
| 0.687217971 | 0.864700549 |
| 0.687150709 | 0.864648 |
| 0.687139856 | 0.864648 |
| 0.68710952 | 0.864641859 |
| 0.687022527 | 0.864629832 |
| 0.68690033 | 0.864629832 |
| 0.68689797 | 0.864629832 |
| 0.686889397 | 0.864629832 |
| 0.686839311 | 0.864629832 |
| 0.686833686 | 0.864629832 |
| 0.686818818 | 0.864629832 |
| 0.686744759 | 0.864616898 |
| 0.686662715 | 0.864559312 |
| 0.686658523 | 0.864559312 |
| 0.686635129 | 0.864559312 |
| 0.686552464 | 0.864530518 |
| 0.68652435 | 0.864522229 |
| 0.686515559 | 0.864522229 |
| 0.686453653 | 0.864478916 |
| 0.686444815 | 0.864478916 |
| 0.686356791 | 0.864478916 |
| 0.686314273 | 0.864473445 |
| 0.686204576 | 0.864416667 |
| 0.686196315 | 0.864416667 |
| 0.686133228 | 0.864395377 |
| 0.686132065 | 0.864395377 |
| 0.686107916 | 0.864395377 |
| 0.686059821 | 0.864394358 |
| 0.685998443 | 0.864362757 |
| 0.685955587 | 0.864331625 |
| 0.685945398 | 0.864331625 |
| 0.685932742 | 0.864331625 |
| 0.68590117 | 0.864331625 |
| 0.685877121 | 0.864331625 |
| 0.685872212 | 0.864331625 |
| 0.685689124 | 0.864224513 |
| 0.685414299 | 0.863992452 |
| 0.685206673 | 0.863822182 |

| | |
|---|---|
| 0.685156596 | 0.863804277 |
| 0.68513512 | 0.86380057 |
| 0.685066389 | 0.863746352 |
| 0.684932913 | 0.86368249 |
| 0.684916924 | 0.86368249 |
| 0.684876116 | 0.863679794 |
| 0.684801492 | 0.863608555 |
| 0.684686294 | 0.863534311 |
| 0.684559489 | 0.863461711 |
| 0.684556416 | 0.863461711 |
| 0.684386184 | 0.863313402 |
| 0.684381335 | 0.863313402 |
| 0.684368485 | 0.863313402 |
| 0.684362408 | 0.863313402 |
| 0.684337437 | 0.863313402 |
| 0.684329979 | 0.863313402 |
| 0.684319267 | 0.863313402 |
| 0.684315134 | 0.863313402 |
| 0.68421684 | 0.863313402 |
| 0.684205345 | 0.863313402 |
| 0.684155972 | 0.863313402 |
| 0.684154656 | 0.863313402 |
| 0.684082446 | 0.863313402 |
| 0.683938671 | 0.863232538 |
| 0.683921186 | 0.863232538 |
| 0.683901531 | 0.863232538 |
| 0.683777433 | 0.86316276 |
| 0.683765036 | 0.86316276 |
| 0.683727404 | 0.863145361 |
| 0.683676151 | 0.863111088 |
| 0.683623216 | 0.863111088 |
| 0.683612406 | 0.863111088 |
| 0.683545772 | 0.86309522 |
| 0.683526259 | 0.86309522 |
| 0.683473158 | 0.863053162 |
| 0.683299922 | 0.862880164 |
| 0.68327307 | 0.862869134 |
| 0.683157442 | 0.862816063 |
| 0.683133156 | 0.862816063 |
| 0.683130589 | 0.862816063 |
| 0.683127902 | 0.862816063 |
| 0.683104981 | 0.862816063 |
| 0.683031309 | 0.862816063 |
| 0.683002578 | 0.862816063 |
| 0.682923062 | 0.862796107 |

| | |
|---|---|
| 0.682897226 | 0.862796107 |
| 0.682872278 | 0.862796107 |
| 0.682854108 | 0.862796107 |
| 0.68280961 | 0.862787151 |
| 0.682773505 | 0.862775089 |
| 0.682683289 | 0.86274198 |
| 0.682682937 | 0.86274198 |
| 0.682673511 | 0.86274198 |
| 0.682652472 | 0.86274198 |
| 0.682572437 | 0.862693454 |
| 0.682565138 | 0.862693454 |
| 0.682476427 | 0.862640786 |
| 0.682433625 | 0.862609578 |
| 0.682316597 | 0.86253783 |
| 0.682268352 | 0.862515125 |
| 0.682241404 | 0.862515125 |
| 0.68216351 | 0.862497054 |
| 0.681842935 | 0.862251963 |
| 0.681742986 | 0.862148461 |
| 0.681696772 | 0.862112909 |
| 0.681616993 | 0.862057797 |
| 0.681539276 | 0.862041767 |
| 0.681498032 | 0.862021798 |
| 0.681497311 | 0.862021798 |
| 0.681421704 | 0.861992017 |
| 0.681343396 | 0.861992017 |
| 0.681333382 | 0.861992017 |
| 0.68120797 | 0.861929598 |
| 0.68113697 | 0.861885557 |
| 0.681093618 | 0.861853598 |
| 0.681064257 | 0.861853598 |
| 0.681061951 | 0.861853598 |
| 0.681049434 | 0.861853598 |
| 0.681035399 | 0.861853598 |
| 0.680996279 | 0.861853598 |
| 0.680957307 | 0.861853598 |
| 0.68081875 | 0.861751365 |
| 0.680764215 | 0.86171151 |
| 0.680738718 | 0.861702138 |
| 0.680685483 | 0.861689339 |
| 0.680631509 | 0.861658034 |
| 0.680476882 | 0.86148518 |
| 0.680472405 | 0.86148518 |
| 0.680468407 | 0.86148518 |
| 0.680429974 | 0.86148518 |

| | |
|---|---|
| 0.680371548 | 0.86148518 |
| 0.680312731 | 0.861437653 |
| 0.680312485 | 0.861437653 |
| 0.680293137 | 0.861437653 |
| 0.680073223 | 0.861248867 |
| 0.680043673 | 0.861234345 |
| 0.680035604 | 0.861234345 |
| 0.680017279 | 0.861234345 |
| 0.679943301 | 0.86119883 |
| 0.67992681 | 0.86119883 |
| 0.679664277 | 0.860959905 |
| 0.679607912 | 0.860947827 |
| 0.679523039 | 0.860897009 |
| 0.679500925 | 0.860897009 |
| 0.679498613 | 0.860897009 |
| 0.67948861 | 0.860897009 |
| 0.679472759 | 0.860897009 |
| 0.679434592 | 0.860897009 |
| 0.679423618 | 0.860897009 |
| 0.679253202 | 0.860874219 |
| 0.679239332 | 0.860874219 |
| 0.67922717 | 0.860874219 |
| 0.679220093 | 0.860874219 |
| 0.679200303 | 0.860874219 |
| 0.679173089 | 0.860874219 |
| 0.679155186 | 0.860874219 |
| 0.679144928 | 0.860874219 |
| 0.679132926 | 0.860874219 |
| 0.679130526 | 0.860874219 |
| 0.679094419 | 0.860874219 |
| 0.679053511 | 0.860874219 |
| 0.679022915 | 0.860874219 |
| 0.678950921 | 0.860874219 |
| 0.678662535 | 0.860779829 |
| 0.678638052 | 0.860779829 |
| 0.678579128 | 0.86072995 |
| 0.678551997 | 0.86072675 |
| 0.678451406 | 0.860682553 |
| 0.678448998 | 0.860682553 |
| 0.678412261 | 0.860682553 |
| 0.678391781 | 0.860682553 |
| 0.678256487 | 0.860549888 |
| 0.678064931 | 0.860421455 |
| 0.678014382 | 0.860380234 |
| 0.678009304 | 0.860380234 |

| | |
|---|---|
| 0.677970276 | 0.860380234 |
| 0.677897191 | 0.860380234 |
| 0.677797655 | 0.860288586 |
| 0.677754402 | 0.860256613 |
| 0.677719782 | 0.860235596 |
| 0.677682085 | 0.860235596 |
| 0.677676449 | 0.860235596 |
| 0.677637175 | 0.860235596 |
| 0.677620969 | 0.860235596 |
| 0.677529837 | 0.860177894 |
| 0.677508409 | 0.860173619 |
| 0.677497316 | 0.860173619 |
| 0.677495355 | 0.860173619 |
| 0.677476827 | 0.860173619 |
| 0.677393296 | 0.860147571 |
| 0.677253162 | 0.860032965 |
| 0.677154854 | 0.859934241 |
| 0.677124209 | 0.859934241 |
| 0.677119999 | 0.859934241 |
| 0.676978868 | 0.859890989 |
| 0.676961832 | 0.859890989 |
| 0.6768441 | 0.859765668 |
| 0.676711363 | 0.859642913 |
| 0.676681896 | 0.859628408 |
| 0.67656959 | 0.859554521 |
| 0.676456007 | 0.85949812 |
| 0.676450694 | 0.85949812 |
| 0.676449285 | 0.85949812 |
| 0.676378088 | 0.859494642 |
| 0.676346198 | 0.859477051 |
| 0.676283468 | 0.859443064 |
| 0.676210176 | 0.859385137 |
| 0.67619138 | 0.859385137 |
| 0.676093133 | 0.859338893 |
| 0.676058788 | 0.859318172 |
| 0.676035395 | 0.85931137 |
| 0.676034286 | 0.85931137 |
| 0.67594726 | 0.859245207 |
| 0.675929134 | 0.859245098 |
| 0.675876105 | 0.859200621 |
| 0.675809902 | 0.859162326 |
| 0.675781136 | 0.859148688 |
| 0.675625939 | 0.858974309 |
| 0.675563231 | 0.858917512 |
| 0.675481289 | 0.858859185 |

| | |
|---|---|
| 0.675400844 | 0.858848614 |
| 0.67517205 | 0.8586551 |
| 0.675084891 | 0.858607315 |
| 0.675072356 | 0.858607315 |
| 0.675054606 | 0.858607315 |
| 0.67473928 | 0.858396979 |
| 0.674703896 | 0.858374891 |
| 0.67460479 | 0.85829466 |
| 0.674585357 | 0.858292864 |
| 0.674475256 | 0.858198633 |
| 0.674377246 | 0.858119776 |
| 0.674362763 | 0.858119776 |
| 0.674133676 | 0.858004919 |
| 0.674042351 | 0.857945776 |
| 0.67394465 | 0.857844344 |
| 0.673937494 | 0.857844344 |
| 0.673935526 | 0.857844344 |
| 0.67386171 | 0.857844344 |
| 0.673669333 | 0.857654352 |
| 0.67360765 | 0.857598747 |
| 0.673580607 | 0.857598747 |
| 0.673469645 | 0.857537664 |
| 0.673323352 | 0.857443083 |
| 0.673319002 | 0.857443083 |
| 0.673316925 | 0.857443083 |
| 0.673268353 | 0.857441823 |
| 0.67324519 | 0.857435251 |
| 0.673244517 | 0.857435251 |
| 0.673127796 | 0.857331592 |
| 0.673070566 | 0.857327481 |
| 0.672993321 | 0.857252017 |
| 0.672926285 | 0.857212479 |
| 0.672898344 | 0.857212479 |
| 0.672890587 | 0.857212479 |
| 0.672808584 | 0.857180285 |
| 0.672794573 | 0.857180285 |
| 0.672752807 | 0.857180285 |
| 0.672607575 | 0.857127441 |
| 0.672546715 | 0.857072818 |
| 0.672443398 | 0.857001959 |
| 0.672310539 | 0.856886481 |
| 0.67230336 | 0.856886481 |
| 0.672273199 | 0.856886481 |
| 0.672264067 | 0.856886481 |
| 0.672231476 | 0.856886481 |

| | |
|---|---|
| 0.67211444 | 0.856819979 |
| 0.672072175 | 0.856796779 |
| 0.672066995 | 0.856796779 |
| 0.672059417 | 0.856796779 |
| 0.672032596 | 0.856796779 |
| 0.671924809 | 0.856761696 |
| 0.671796142 | 0.856713756 |
| 0.67178293 | 0.856713756 |
| 0.6717753 | 0.856713756 |
| 0.671728273 | 0.856713756 |
| 0.671672185 | 0.856668929 |
| 0.671550679 | 0.856605715 |
| 0.671484585 | 0.856544348 |
| 0.671470623 | 0.856544348 |
| 0.671465055 | 0.856544348 |
| 0.671408434 | 0.856544348 |
| 0.671370284 | 0.85653619 |
| 0.671364055 | 0.85653619 |
| 0.671280253 | 0.856490157 |
| 0.671253568 | 0.856490157 |
| 0.67124441 | 0.856490157 |
| 0.671155083 | 0.856422225 |
| 0.671075067 | 0.85636601 |
| 0.671014715 | 0.856334883 |
| 0.670965656 | 0.856318166 |
| 0.670794842 | 0.856161473 |
| 0.670756557 | 0.856154811 |
| 0.67072131 | 0.856143971 |
| 0.670696503 | 0.856143971 |
| 0.670693986 | 0.856143971 |
| 0.670674226 | 0.856143971 |
| 0.670665646 | 0.856143971 |
| 0.670644363 | 0.856143971 |
| 0.670640357 | 0.856143971 |
| 0.670564602 | 0.856127501 |
| 0.670424318 | 0.855988989 |
| 0.670340673 | 0.855956333 |
| 0.670339112 | 0.855956333 |
| 0.670232412 | 0.855890339 |
| 0.670194143 | 0.855883961 |
| 0.670129755 | 0.855847631 |
| 0.670038012 | 0.855753409 |
| 0.669984461 | 0.855707962 |
| 0.669816186 | 0.855561871 |
| 0.669713815 | 0.855454054 |

| | |
|---|---|
| 0.669661093 | 0.855409652 |
| 0.669650511 | 0.855409652 |
| 0.669634094 | 0.855409652 |
| 0.669619912 | 0.855409652 |
| 0.669561527 | 0.855409652 |
| 0.669468728 | 0.855372905 |
| 0.669404318 | 0.855356946 |
| 0.669378613 | 0.855347049 |
| 0.669359827 | 0.855345992 |
| 0.669280561 | 0.855267649 |
| 0.669213334 | 0.855208754 |
| 0.669169165 | 0.855208754 |
| 0.669127018 | 0.855208754 |
| 0.668899114 | 0.855055508 |
| 0.668875211 | 0.855047902 |
| 0.668836346 | 0.855021168 |
| 0.668830315 | 0.855021168 |
| 0.66882403 | 0.855021168 |
| 0.66882209 | 0.855021168 |
| 0.668804466 | 0.855021168 |
| 0.668769603 | 0.855021168 |
| 0.668735927 | 0.855021168 |
| 0.668711425 | 0.855021168 |
| 0.668682575 | 0.855021168 |
| 0.668622883 | 0.855021168 |
| 0.668617182 | 0.855021168 |
| 0.668605096 | 0.855021168 |
| 0.66859196 | 0.855021168 |
| 0.668525168 | 0.855021168 |
| 0.668520822 | 0.855021168 |
| 0.668493914 | 0.855021168 |
| 0.668440347 | 0.855021168 |
| 0.668430286 | 0.855021168 |
| 0.66842949 | 0.855021168 |
| 0.668382768 | 0.855021168 |
| 0.668322047 | 0.855021168 |
| 0.668267523 | 0.855005435 |
| 0.668245277 | 0.854999941 |
| 0.668214778 | 0.854999941 |
| 0.668156176 | 0.854954843 |
| 0.668102829 | 0.854909549 |
| 0.668097886 | 0.854909549 |
| 0.668032765 | 0.854865831 |
| 0.667890434 | 0.854747377 |
| 0.667849372 | 0.854723009 |

| | |
|---|---|
| 0.667795019 | 0.854676413 |
| 0.667708687 | 0.854611852 |
| 0.667700498 | 0.854611852 |
| 0.667632892 | 0.854560772 |
| 0.667580038 | 0.85453905 |
| 0.667543477 | 0.854515218 |
| 0.667534208 | 0.854515218 |
| 0.667528732 | 0.854515218 |
| 0.667435086 | 0.854469662 |
| 0.667114351 | 0.854195483 |
| 0.66709952 | 0.854195483 |
| 0.667057142 | 0.854168159 |
| 0.667007464 | 0.85412751 |
| 0.666988803 | 0.854126579 |
| 0.666921995 | 0.854109921 |
| 0.666808863 | 0.854010963 |
| 0.666628568 | 0.853871899 |
| 0.666529036 | 0.853790334 |
| 0.666294793 | 0.85360507 |
| 0.66626235 | 0.853586468 |
| 0.666204305 | 0.853566126 |
| 0.666193804 | 0.853566126 |
| 0.66615487 | 0.853566126 |
| 0.666068545 | 0.853521849 |
| 0.666052529 | 0.853521849 |
| 0.666031497 | 0.853521849 |
| 0.666025716 | 0.853521849 |
| 0.66591397 | 0.853438584 |
| 0.665893537 | 0.853435362 |
| 0.665761327 | 0.853311842 |
| 0.665671039 | 0.853265007 |
| 0.665589734 | 0.853206714 |
| 0.665581746 | 0.853206714 |
| 0.665571586 | 0.853206714 |
| 0.665507952 | 0.853170768 |
| 0.665488998 | 0.853169435 |
| 0.66546045 | 0.853155801 |
| 0.665398976 | 0.853152214 |
| 0.665390331 | 0.853152214 |
| 0.665315716 | 0.853152214 |
| 0.665313613 | 0.853152214 |
| 0.665285328 | 0.853152214 |
| 0.66524522 | 0.853152214 |
| 0.665004473 | 0.852961544 |
| 0.664983005 | 0.852956979 |

| | |
|---|---|
| 0.664844573 | 0.852825352 |
| 0.664762754 | 0.852789304 |
| 0.664750345 | 0.852789304 |
| 0.664664059 | 0.85275457 |
| 0.664550488 | 0.852677769 |
| 0.664514645 | 0.85265475 |
| 0.664504192 | 0.85265475 |
| 0.664499859 | 0.85265475 |
| 0.664438817 | 0.852626365 |
| 0.664388177 | 0.852584354 |
| 0.664156344 | 0.852378717 |
| 0.664136515 | 0.852378717 |
| 0.664019129 | 0.852303135 |
| 0.66400637 | 0.852303135 |
| 0.663966508 | 0.852303135 |
| 0.663874279 | 0.852246369 |
| 0.663848001 | 0.852246369 |
| 0.663833095 | 0.852246369 |
| 0.66380682 | 0.852246369 |
| 0.663771436 | 0.852246369 |
| 0.663754069 | 0.852246369 |
| 0.663737198 | 0.852246369 |
| 0.663734849 | 0.852246369 |
| 0.663720762 | 0.852246369 |
| 0.66370276 | 0.852246369 |
| 0.66363253 | 0.852211672 |
| 0.66359961 | 0.852192375 |
| 0.663580319 | 0.85219058 |
| 0.663562918 | 0.85219058 |
| 0.663558258 | 0.85219058 |
| 0.66352737 | 0.85219058 |
| 0.663477976 | 0.85219058 |
| 0.663439913 | 0.852171114 |
| 0.663439648 | 0.852171114 |
| 0.663261842 | 0.851988342 |
| 0.663148038 | 0.851865131 |
| 0.663099265 | 0.851825454 |
| 0.663083218 | 0.851825454 |
| 0.663019323 | 0.851768708 |
| 0.662995006 | 0.851768708 |
| 0.66297081 | 0.851768708 |
| 0.662821993 | 0.851712084 |
| 0.662816795 | 0.851712084 |
| 0.662800152 | 0.851712084 |
| 0.66277855 | 0.851712084 |

| | |
|---|---|
| 0.662754257 | 0.851712084 |
| 0.662722601 | 0.851712084 |
| 0.662699898 | 0.851712084 |
| 0.662625274 | 0.851712084 |
| 0.662597094 | 0.851712084 |
| 0.662593433 | 0.851712084 |
| 0.662567065 | 0.851712084 |
| 0.662445235 | 0.851712084 |
| 0.662428764 | 0.851712084 |
| 0.662428052 | 0.851712084 |
| 0.662345015 | 0.851683523 |
| 0.66232826 | 0.851683523 |
| 0.662304491 | 0.851677403 |
| 0.662158473 | 0.851604601 |
| 0.662080315 | 0.851527076 |
| 0.662037547 | 0.851495064 |
| 0.662019733 | 0.851495064 |
| 0.662018097 | 0.851495064 |
| 0.661991438 | 0.851495064 |
| 0.661976285 | 0.851495064 |
| 0.661968983 | 0.851495064 |
| 0.661926125 | 0.851495064 |
| 0.66191133 | 0.851495064 |
| 0.66188816 | 0.851495064 |
| 0.66182808 | 0.851495064 |
| 0.6618038 | 0.851493356 |
| 0.661796305 | 0.851493356 |
| 0.661647202 | 0.851360872 |
| 0.661623479 | 0.851353347 |
| 0.661602162 | 0.851353347 |
| 0.661579083 | 0.851353347 |
| 0.661571929 | 0.851353347 |
| 0.661528609 | 0.851346278 |
| 0.66149812 | 0.851330044 |
| 0.66132345 | 0.851266269 |
| 0.661181292 | 0.851152293 |
| 0.661034636 | 0.85102486 |
| 0.661010764 | 0.85102486 |
| 0.660866287 | 0.850930781 |
| 0.660838581 | 0.850918111 |
| 0.660728524 | 0.850918111 |
| 0.660725896 | 0.850918111 |
| 0.660689819 | 0.850918111 |
| 0.660652455 | 0.850908498 |
| 0.660626444 | 0.850898009 |

| | |
|---|---|
| 0.660569805 | 0.850871078 |
| 0.660531466 | 0.850861097 |
| 0.660482921 | 0.850851209 |
| 0.660469502 | 0.850851209 |
| 0.660387427 | 0.850820245 |
| 0.660369759 | 0.850820245 |
| 0.660282783 | 0.850777483 |
| 0.660244987 | 0.850757295 |
| 0.660219711 | 0.850757295 |
| 0.660210349 | 0.850757295 |
| 0.660116898 | 0.850701829 |
| 0.660052189 | 0.850664473 |
| 0.660026962 | 0.850662665 |
| 0.659961319 | 0.85063784 |
| 0.659948691 | 0.85063784 |
| 0.659937188 | 0.85063784 |
| 0.659899617 | 0.850628967 |
| 0.659856892 | 0.850596915 |
| 0.659737429 | 0.850488957 |
| 0.659732779 | 0.850488957 |
| 0.659675458 | 0.850478128 |
| 0.659635476 | 0.850449604 |
| 0.659618361 | 0.850449604 |
| 0.65960589 | 0.850449604 |
| 0.65942043 | 0.850264419 |
| 0.659324537 | 0.850209826 |
| 0.659288762 | 0.850186712 |
| 0.659277945 | 0.850186712 |
| 0.659227316 | 0.850176534 |
| 0.659178909 | 0.850137125 |
| 0.659129307 | 0.850119195 |
| 0.659103796 | 0.850109312 |
| 0.659076371 | 0.850096961 |
| 0.659055103 | 0.850092552 |
| 0.659041439 | 0.850092552 |
| 0.659037677 | 0.850092552 |
| 0.65898437 | 0.850092552 |
| 0.658882496 | 0.850077113 |
| 0.658862992 | 0.850074978 |
| 0.658743582 | 0.849943939 |
| 0.65871093 | 0.849924834 |
| 0.658687199 | 0.849919855 |
| 0.658512162 | 0.849743948 |
| 0.658492868 | 0.849743948 |
| 0.658465091 | 0.849743948 |

| | |
|---|---|
| 0.658395122 | 0.849743948 |
| 0.658390003 | 0.849743948 |
| 0.658350542 | 0.849743948 |
| 0.65821998 | 0.849684352 |
| 0.658151555 | 0.849663453 |
| 0.658149213 | 0.849663453 |
| 0.658148409 | 0.849663453 |
| 0.658131139 | 0.849663453 |
| 0.658115867 | 0.849663453 |
| 0.658081479 | 0.849663453 |
| 0.657997166 | 0.849625349 |
| 0.657903145 | 0.849573047 |
| 0.657900781 | 0.849573047 |
| 0.657881817 | 0.849573047 |
| 0.657854316 | 0.849573047 |
| 0.657840728 | 0.849573047 |
| 0.657840333 | 0.849573047 |
| 0.657700874 | 0.849573047 |
| 0.657691463 | 0.849573047 |
| 0.657453781 | 0.849407314 |
| 0.65745015 | 0.849407314 |
| 0.657446299 | 0.849407314 |
| 0.657433054 | 0.849407314 |
| 0.657413354 | 0.849407314 |
| 0.657384599 | 0.849407314 |
| 0.657344975 | 0.849407314 |
| 0.657299781 | 0.849392681 |
| 0.657238002 | 0.849358939 |
| 0.657205576 | 0.849351297 |
| 0.657140834 | 0.84932555 |
| 0.65706655 | 0.849272683 |
| 0.657014278 | 0.849243311 |
| 0.65694618 | 0.84918919 |
| 0.656907998 | 0.849162882 |
| 0.656815811 | 0.849133781 |
| 0.656812679 | 0.849133781 |
| 0.656745394 | 0.849100837 |
| 0.656706403 | 0.849100837 |
| 0.65666578 | 0.849100837 |
| 0.656639159 | 0.849100837 |
| 0.656614967 | 0.849100837 |
| 0.656572216 | 0.849100837 |
| 0.656555418 | 0.849100837 |
| 0.65638774 | 0.849100837 |
| 0.656354585 | 0.849100837 |

| | |
|---|---|
| 0.656343605 | 0.849100837 |
| 0.656322331 | 0.849100837 |
| 0.656309938 | 0.849100837 |
| 0.656286819 | 0.849100837 |
| 0.656169231 | 0.849065531 |
| 0.656105264 | 0.849023921 |
| 0.656099016 | 0.849023921 |
| 0.656095792 | 0.849023921 |
| 0.656019345 | 0.84898194 |
| 0.655928589 | 0.84898194 |
| 0.655924519 | 0.84898194 |
| 0.655643969 | 0.848749839 |
| 0.655580054 | 0.848710214 |
| 0.655540491 | 0.848708156 |
| 0.655500366 | 0.848679279 |
| 0.655466025 | 0.848657887 |
| 0.655443156 | 0.848657887 |
| 0.655385372 | 0.848622671 |
| 0.655337623 | 0.848583915 |
| 0.655250256 | 0.848516924 |
| 0.655233031 | 0.848516924 |
| 0.655159555 | 0.848445607 |
| 0.655009627 | 0.84829758 |
| 0.654984421 | 0.84829758 |
| 0.654913428 | 0.848265258 |
| 0.654884244 | 0.848250528 |
| 0.654831825 | 0.848205699 |
| 0.654808273 | 0.84819826 |
| 0.654771854 | 0.84819826 |
| 0.654699086 | 0.848157544 |
| 0.654687658 | 0.848157544 |
| 0.654676822 | 0.848157544 |
| 0.654614826 | 0.84815529 |
| 0.654545522 | 0.84811164 |
| 0.654387421 | 0.847952923 |
| 0.654265694 | 0.847841324 |
| 0.654258508 | 0.847841324 |
| 0.654241703 | 0.847841324 |
| 0.654163131 | 0.847841324 |
| 0.654156476 | 0.847841324 |
| 0.654056194 | 0.847840715 |
| 0.654051194 | 0.847840715 |
| 0.654038189 | 0.847840715 |
| 0.653967073 | 0.847800364 |
| 0.653908382 | 0.847747353 |

| | |
|---|---|
| 0.653875643 | 0.847727985 |
| 0.653697007 | 0.847596421 |
| 0.653683455 | 0.847596421 |
| 0.653627892 | 0.847568286 |
| 0.653577847 | 0.847534958 |
| 0.65352346 | 0.847525169 |
| 0.653397778 | 0.847431403 |
| 0.653383771 | 0.847431403 |
| 0.653331187 | 0.847414266 |
| 0.653329583 | 0.847414266 |
| 0.653328041 | 0.847414266 |
| 0.653227302 | 0.847414266 |
| 0.653201183 | 0.847414266 |
| 0.653034467 | 0.847237062 |
| 0.653016846 | 0.847237062 |
| 0.652933365 | 0.847175127 |
| 0.652843725 | 0.847104974 |
| 0.652842455 | 0.847104974 |
| 0.65282808 | 0.847104974 |
| 0.652798892 | 0.847104974 |
| 0.652787262 | 0.847104974 |
| 0.652747253 | 0.847095189 |
| 0.652621005 | 0.846995583 |
| 0.652585945 | 0.846995583 |
| 0.652509789 | 0.846966545 |
| 0.652454375 | 0.846922802 |
| 0.652450876 | 0.846922802 |
| 0.652379999 | 0.846918588 |
| 0.652351649 | 0.846904869 |
| 0.652282725 | 0.846838472 |
| 0.652254085 | 0.846824374 |
| 0.652213251 | 0.846794441 |
| 0.652170336 | 0.846761807 |
| 0.65214775 | 0.846755566 |
| 0.652013331 | 0.846650278 |
| 0.651927642 | 0.846608256 |
| 0.651893408 | 0.846600968 |
| 0.651881818 | 0.846600968 |
| 0.651865777 | 0.846600968 |
| 0.651818207 | 0.846600968 |
| 0.651780041 | 0.846600968 |
| 0.651677454 | 0.84658344 |
| 0.651676096 | 0.84658344 |
| 0.651638196 | 0.846578624 |
| 0.651637473 | 0.846578624 |

| | |
|---|---|
| 0.651500648 | 0.846507992 |
| 0.651492721 | 0.846507992 |
| 0.651439793 | 0.846482498 |
| 0.651310503 | 0.846397206 |
| 0.651254311 | 0.846356938 |
| 0.651173115 | 0.846292427 |
| 0.65116416 | 0.846292427 |
| 0.651149798 | 0.846292427 |
| 0.651076997 | 0.846265054 |
| 0.650993675 | 0.846202941 |
| 0.650852336 | 0.846088496 |
| 0.650710167 | 0.845972952 |
| 0.650649273 | 0.845916876 |
| 0.650621963 | 0.845916876 |
| 0.650610274 | 0.845916876 |
| 0.650561497 | 0.845916876 |
| 0.65047985 | 0.845858238 |
| 0.650426626 | 0.845829459 |
| 0.650389045 | 0.845824198 |
| 0.650375112 | 0.845824198 |
| 0.650352659 | 0.845824198 |
| 0.650339023 | 0.845824198 |
| 0.650317877 | 0.845824198 |
| 0.650220082 | 0.845751361 |
| 0.650214087 | 0.845751361 |
| 0.650208356 | 0.845751361 |
| 0.650158956 | 0.845741147 |
| 0.650057758 | 0.8456557 |
| 0.650040662 | 0.8456557 |
| 0.650017258 | 0.84564921 |
| 0.650011729 | 0.84564921 |
| 0.649965742 | 0.845628387 |
| 0.649923299 | 0.845618262 |
| 0.649900294 | 0.845612538 |
| 0.649878805 | 0.845610837 |
| 0.649823278 | 0.845604734 |
| 0.649576165 | 0.845352461 |
| 0.64930104 | 0.845033342 |
| 0.649173725 | 0.844897989 |
| 0.649165819 | 0.844897989 |
| 0.649165614 | 0.844897989 |
| 0.649050258 | 0.844806554 |
| 0.649037937 | 0.844806554 |
| 0.649027178 | 0.844806554 |
| 0.648870481 | 0.844664891 |

| | |
|---|---|
| 0.648780834 | 0.84461745 |
| 0.64877746 | 0.84461745 |
| 0.648687581 | 0.844542218 |
| 0.648588524 | 0.844467999 |
| 0.648571453 | 0.844467999 |
| 0.648467359 | 0.844393999 |
| 0.648419047 | 0.844354175 |
| 0.648415641 | 0.844354175 |
| 0.648378568 | 0.844354175 |
| 0.6483367 | 0.844354175 |
| 0.648294802 | 0.844354175 |
| 0.648288939 | 0.844354175 |
| 0.648216043 | 0.844320675 |
| 0.64821364 | 0.844320675 |
| 0.648164149 | 0.844320675 |
| 0.648156685 | 0.844320675 |
| 0.648122531 | 0.844320675 |
| 0.648119379 | 0.844320675 |
| 0.648081691 | 0.844320675 |
| 0.648068701 | 0.844320675 |
| 0.647945041 | 0.844221657 |
| 0.647847477 | 0.844163821 |
| 0.647730273 | 0.844080379 |
| 0.647705983 | 0.844071821 |
| 0.647663924 | 0.844055757 |
| 0.647614873 | 0.844022372 |
| 0.647605071 | 0.844022372 |
| 0.647560066 | 0.844020232 |
| 0.647499011 | 0.843986847 |
| 0.647449613 | 0.843945556 |
| 0.647446447 | 0.843945556 |
| 0.647416797 | 0.843945556 |
| 0.64736891 | 0.843945556 |
| 0.647335241 | 0.843945556 |
| 0.647319178 | 0.843945556 |
| 0.647312288 | 0.843945556 |
| 0.64727806 | 0.843945556 |
| 0.647226856 | 0.843945556 |
| 0.647126372 | 0.843847545 |
| 0.647092238 | 0.843826137 |
| 0.647076465 | 0.843826137 |
| 0.647059851 | 0.843826137 |
| 0.647056084 | 0.843826137 |
| 0.647052971 | 0.843826137 |
| 0.646967989 | 0.843802733 |

| | |
|---|---|
| 0.646907439 | 0.843746867 |
| 0.646788073 | 0.843737294 |
| 0.646743973 | 0.843737294 |
| 0.646740451 | 0.843737294 |
| 0.646697675 | 0.843737294 |
| 0.646678058 | 0.843737294 |
| 0.646657752 | 0.843737294 |
| 0.646596929 | 0.843734676 |
| 0.646535411 | 0.843677517 |
| 0.646447553 | 0.843585983 |
| 0.646447553 | 0.843585983 |
| 0.646341859 | 0.843540507 |
| 0.64627013 | 0.843470007 |
| 0.646204968 | 0.843461825 |
| 0.646198611 | 0.843461825 |
| 0.646191633 | 0.843461825 |
| 0.646191074 | 0.843461825 |
| 0.646179642 | 0.843461825 |
| 0.646084883 | 0.843436251 |
| 0.646027572 | 0.843384551 |
| 0.645978372 | 0.843356509 |
| 0.645969355 | 0.843356509 |
| 0.645852767 | 0.843271917 |
| 0.645765648 | 0.843216796 |
| 0.64571891 | 0.843216796 |
| 0.645668979 | 0.843216796 |
| 0.645652258 | 0.843216796 |
| 0.645595383 | 0.843216796 |
| 0.645564602 | 0.843216796 |
| 0.645520295 | 0.843207677 |
| 0.645433844 | 0.843164125 |
| 0.645308215 | 0.843069383 |
| 0.645301261 | 0.843069383 |
| 0.645272341 | 0.843069383 |
| 0.645269753 | 0.843069383 |
| 0.645250686 | 0.843069383 |
| 0.645247291 | 0.843069383 |
| 0.645108685 | 0.843016829 |
| 0.645064134 | 0.842981739 |
| 0.645041629 | 0.84297546 |
| 0.645026003 | 0.84297546 |
| 0.644903951 | 0.842888048 |
| 0.644750146 | 0.842802668 |
| 0.64461788 | 0.84272229 |
| 0.644607142 | 0.84272229 |

| | |
|---|---|
| 0.644601404 | 0.84272229 |
| 0.644591458 | 0.84272229 |
| 0.64454205 | 0.842715684 |
| 0.644494867 | 0.842677127 |
| 0.644440987 | 0.842677127 |
| 0.644381027 | 0.842650207 |
| 0.644372153 | 0.842650207 |
| 0.644353161 | 0.842650207 |
| 0.644311001 | 0.842644919 |
| 0.6442956 | 0.842644919 |
| 0.644260327 | 0.842644919 |
| 0.644244606 | 0.842644919 |
| 0.644238765 | 0.842644919 |
| 0.644237894 | 0.842644919 |
| 0.644176113 | 0.842644919 |
| 0.644171447 | 0.842644919 |
| 0.644167544 | 0.842644919 |
| 0.644142069 | 0.842644919 |
| 0.644034504 | 0.84260724 |
| 0.643977171 | 0.842555374 |
| 0.643923514 | 0.842508316 |
| 0.643887483 | 0.842484318 |
| 0.643876126 | 0.842484318 |
| 0.643808409 | 0.842450287 |
| 0.643781547 | 0.842438282 |
| 0.643753989 | 0.842425367 |
| 0.643729754 | 0.842425367 |
| 0.643632938 | 0.842382686 |
| 0.643601613 | 0.842364836 |
| 0.643572605 | 0.842353648 |
| 0.643569519 | 0.842353648 |
| 0.643478333 | 0.842296072 |
| 0.64346695 | 0.842296072 |
| 0.643393175 | 0.842273836 |
| 0.643385131 | 0.842273836 |
| 0.643332152 | 0.842273836 |
| 0.643220969 | 0.842236964 |
| 0.643155149 | 0.842173932 |
| 0.643088353 | 0.842132774 |
| 0.642989247 | 0.842095608 |
| 0.642899726 | 0.842024674 |
| 0.642835505 | 0.842010028 |
| 0.642778531 | 0.841981921 |
| 0.642744318 | 0.841981921 |
| 0.642727994 | 0.841981921 |

| | |
|---|---|
| 0.642653136 | 0.841956397 |
| 0.642527916 | 0.8418155 |
| 0.642524641 | 0.8418155 |
| 0.642506004 | 0.8418155 |
| 0.642447703 | 0.841779877 |
| 0.642339386 | 0.841704242 |
| 0.642292607 | 0.841672833 |
| 0.642258436 | 0.841672833 |
| 0.642215779 | 0.841661234 |
| 0.642207306 | 0.841661234 |
| 0.642202343 | 0.841661234 |
| 0.642167184 | 0.841661234 |
| 0.64216288 | 0.841661234 |
| 0.641993215 | 0.841515524 |
| 0.641917558 | 0.841455633 |
| 0.64191026 | 0.841455633 |
| 0.641892949 | 0.841455633 |
| 0.641738372 | 0.841313371 |
| 0.641686836 | 0.841292123 |
| 0.641659827 | 0.841279871 |
| 0.641606424 | 0.841233012 |
| 0.641573841 | 0.841213449 |
| 0.641507267 | 0.841149316 |
| 0.641485177 | 0.841143508 |
| 0.641392007 | 0.84109081 |
| 0.64136001 | 0.841072008 |
| 0.64133297 | 0.841059707 |
| 0.641313319 | 0.841057094 |
| 0.641288128 | 0.841053531 |
| 0.641221527 | 0.841029347 |
| 0.641203654 | 0.841029067 |
| 0.641132672 | 0.840959123 |
| 0.641060313 | 0.84088737 |
| 0.641047572 | 0.84088737 |
| 0.641015908 | 0.840875441 |
| 0.640970234 | 0.840838687 |
| 0.640905837 | 0.840800527 |
| 0.640765603 | 0.840718431 |
| 0.640729787 | 0.840718431 |
| 0.640676242 | 0.840684577 |
| 0.64046482 | 0.840488666 |
| 0.64041055 | 0.840474877 |
| 0.640352229 | 0.84046064 |
| 0.640333212 | 0.84046064 |
| 0.640252632 | 0.840454387 |

| | |
|---|---|
| 0.640201746 | 0.84043243 |
| 0.640152991 | 0.84039159 |
| 0.640095344 | 0.84036224 |
| 0.640093614 | 0.84036224 |
| 0.64005685 | 0.840358032 |
| 0.639938541 | 0.84025433 |
| 0.639902475 | 0.84025433 |
| 0.639860024 | 0.84025433 |
| 0.639742421 | 0.840214512 |
| 0.639735813 | 0.840214512 |
| 0.639725267 | 0.840214512 |
| 0.639698341 | 0.840214512 |
| 0.639677733 | 0.840214512 |
| 0.639660623 | 0.840214512 |
| 0.639640258 | 0.840214512 |
| 0.639538067 | 0.840214512 |
| 0.639499388 | 0.840205168 |
| 0.639497724 | 0.840205168 |
| 0.639463599 | 0.840205168 |
| 0.639453139 | 0.840205168 |
| 0.639408023 | 0.840205168 |
| 0.639367535 | 0.840205168 |
| 0.639283242 | 0.840176136 |
| 0.639249643 | 0.840155162 |
| 0.639102515 | 0.839984973 |
| 0.638938227 | 0.839815398 |
| 0.638894316 | 0.839780857 |
| 0.638845799 | 0.839740261 |
| 0.638617522 | 0.839463367 |
| 0.638614461 | 0.839463367 |
| 0.638610752 | 0.839463367 |
| 0.638588368 | 0.839463367 |
| 0.638566547 | 0.839463367 |
| 0.638408319 | 0.839304193 |
| 0.638299522 | 0.83920749 |
| 0.63814499 | 0.839120139 |
| 0.638025223 | 0.839008981 |
| 0.638022135 | 0.839008981 |
| 0.638021425 | 0.839008981 |
| 0.63801781 | 0.839008981 |
| 0.637864681 | 0.838890516 |
| 0.63778722 | 0.838888795 |
| 0.637747095 | 0.838888795 |
| 0.637659501 | 0.838805973 |
| 0.637628371 | 0.83878819 |

| | |
|---|---|
| 0.637406963 | 0.838566417 |
| 0.637163738 | 0.838339061 |
| 0.637079407 | 0.838320742 |
| 0.63701142 | 0.838300763 |
| 0.636943316 | 0.838274763 |
| 0.636931112 | 0.838274763 |
| 0.636898502 | 0.838267975 |
| 0.636854448 | 0.838256324 |
| 0.636734842 | 0.838122055 |
| 0.63673399 | 0.838122055 |
| 0.636656029 | 0.838064639 |
| 0.636626015 | 0.838048292 |
| 0.636613152 | 0.838048292 |
| 0.636567626 | 0.838017754 |
| 0.636404241 | 0.837825823 |
| 0.636359908 | 0.837790616 |
| 0.636302314 | 0.837737949 |
| 0.636261217 | 0.837706998 |
| 0.636207454 | 0.837669959 |
| 0.636187566 | 0.837669959 |
| 0.636161325 | 0.837668108 |
| 0.636009792 | 0.837538042 |
| 0.635897644 | 0.837413513 |
| 0.635879952 | 0.837413368 |
| 0.635749244 | 0.837310693 |
| 0.635702025 | 0.837310693 |
| 0.635535793 | 0.837169452 |
| 0.635478845 | 0.837162918 |
| 0.635329354 | 0.83703544 |
| 0.635327052 | 0.83703544 |
| 0.635288251 | 0.83703544 |
| 0.635145565 | 0.836909056 |
| 0.635141556 | 0.836909056 |
| 0.635139477 | 0.836909056 |
| 0.635039628 | 0.836862078 |
| 0.6349845 | 0.836857744 |
| 0.634839652 | 0.836714305 |
| 0.634830546 | 0.836714305 |
| 0.634794403 | 0.836714305 |
| 0.634786293 | 0.836714305 |
| 0.634742584 | 0.836714305 |
| 0.634733997 | 0.836714305 |
| 0.63469574 | 0.836714305 |
| 0.634684167 | 0.836714305 |
| 0.634586928 | 0.836682213 |

| | |
|---|---|
| 0.634507097 | 0.836623281 |
| 0.634494036 | 0.836623281 |
| 0.634437441 | 0.836600919 |
| 0.63443445 | 0.836600919 |
| 0.634381046 | 0.836572879 |
| 0.63433852 | 0.836563125 |
| 0.634288314 | 0.836543242 |
| 0.634286341 | 0.836543242 |
| 0.634227867 | 0.836533015 |
| 0.634195392 | 0.836523194 |
| 0.634133418 | 0.836501103 |
| 0.634112127 | 0.836496186 |
| 0.634007503 | 0.836385518 |
| 0.633876391 | 0.836254699 |
| 0.633874538 | 0.836254699 |
| 0.633860415 | 0.836254699 |
| 0.633832486 | 0.836254699 |
| 0.633802428 | 0.836249778 |
| 0.633714279 | 0.836202969 |
| 0.633704262 | 0.836202969 |
| 0.633678707 | 0.836202969 |
| 0.63366445 | 0.836202969 |
| 0.633639684 | 0.836202969 |
| 0.633630584 | 0.836202969 |
| 0.633578776 | 0.836186338 |
| 0.633545547 | 0.836186338 |
| 0.633520261 | 0.836178624 |
| 0.63341213 | 0.836065615 |
| 0.633408947 | 0.836065615 |
| 0.63329447 | 0.835973273 |
| 0.633135215 | 0.835786216 |
| 0.633094705 | 0.835755904 |
| 0.633062884 | 0.835737062 |
| 0.63305526 | 0.835737062 |
| 0.633032164 | 0.835737062 |
| 0.632965866 | 0.835678475 |
| 0.632907357 | 0.835647557 |
| 0.632845693 | 0.835647557 |
| 0.632813815 | 0.835639881 |
| 0.63276375 | 0.835596937 |
| 0.632761663 | 0.835596937 |
| 0.632756108 | 0.835596937 |
| 0.632704623 | 0.835588361 |
| 0.632682697 | 0.835588361 |
| 0.632516968 | 0.835479536 |

| | |
|---|---|
| 0.63245521 | 0.835444301 |
| 0.632339132 | 0.835360146 |
| 0.632328048 | 0.835360146 |
| 0.632284899 | 0.835357403 |
| 0.632276828 | 0.835357403 |
| 0.632190424 | 0.835349388 |
| 0.632119404 | 0.83528787 |
| 0.632072569 | 0.83528787 |
| 0.632068106 | 0.83528787 |
| 0.632036805 | 0.83528787 |
| 0.632011782 | 0.83528787 |
| 0.631986114 | 0.83528787 |
| 0.631965907 | 0.83528787 |
| 0.631962736 | 0.83528787 |
| 0.631941072 | 0.83528787 |
| 0.631912978 | 0.83528787 |
| 0.631880811 | 0.83528787 |
| 0.631795921 | 0.835253379 |
| 0.631727095 | 0.835223783 |
| 0.631669856 | 0.835194476 |
| 0.631581635 | 0.835101015 |
| 0.631563781 | 0.835100592 |
| 0.631558371 | 0.835100592 |
| 0.631546873 | 0.835100592 |
| 0.631508302 | 0.835100592 |
| 0.631460533 | 0.835079992 |
| 0.631424714 | 0.835055809 |
| 0.631367614 | 0.835018864 |
| 0.631302739 | 0.835005317 |
| 0.631292611 | 0.835005317 |
| 0.631225319 | 0.835000519 |
| 0.631213994 | 0.835000519 |
| 0.631176018 | 0.834991866 |
| 0.631096131 | 0.834922651 |
| 0.631090563 | 0.834922651 |
| 0.630962544 | 0.83483958 |
| 0.630922532 | 0.83483958 |
| 0.630893425 | 0.83483958 |
| 0.630879638 | 0.83483958 |
| 0.630843723 | 0.83483958 |
| 0.63082025 | 0.83483958 |
| 0.630804895 | 0.83483958 |
| 0.630804534 | 0.83483958 |
| 0.630727455 | 0.83483958 |
| 0.630684676 | 0.83483958 |

| | |
|---|---|
| 0.630648428 | 0.83483958 |
| 0.630607405 | 0.83483958 |
| 0.630585147 | 0.83483958 |
| 0.630553467 | 0.834830844 |
| 0.630475085 | 0.834784865 |
| 0.630329721 | 0.834650623 |
| 0.630264434 | 0.83459814 |
| 0.630117292 | 0.834438934 |
| 0.630087997 | 0.83442334 |
| 0.630087146 | 0.83442334 |
| 0.630016128 | 0.834374565 |
| 0.629982422 | 0.834353127 |
| 0.629901455 | 0.834292295 |
| 0.629841844 | 0.83423654 |
| 0.629782916 | 0.834181688 |
| 0.629775212 | 0.834181688 |
| 0.629742952 | 0.834181688 |
| 0.629722847 | 0.834181688 |
| 0.629682486 | 0.834164657 |
| 0.629635507 | 0.834125624 |
| 0.629612575 | 0.834118446 |
| 0.629417024 | 0.833952167 |
| 0.629315052 | 0.833863454 |
| 0.629302251 | 0.833863454 |
| 0.629247701 | 0.833861178 |
| 0.629203532 | 0.833854878 |
| 0.629202262 | 0.833854878 |
| 0.629154089 | 0.833835758 |
| 0.629065438 | 0.833741467 |
| 0.629059397 | 0.833741467 |
| 0.629025184 | 0.833741467 |
| 0.628801949 | 0.833508222 |
| 0.628695342 | 0.833459697 |
| 0.62862788 | 0.83339346 |
| 0.62860662 | 0.833388472 |
| 0.62859925 | 0.833388472 |
| 0.628416984 | 0.833203747 |
| 0.628376282 | 0.833203747 |
| 0.628371001 | 0.833203747 |
| 0.628355624 | 0.833203747 |
| 0.628301308 | 0.833198316 |
| 0.62822515 | 0.83313784 |
| 0.628222205 | 0.83313784 |
| 0.628196821 | 0.83313784 |
| 0.628143582 | 0.83313784 |

| | |
|---|---|
| 0.628127305 | 0.83313784 |
| 0.628041264 | 0.833079551 |
| 0.62795559 | 0.83303551 |
| 0.627880172 | 0.832981865 |
| 0.627852677 | 0.832972703 |
| 0.627845691 | 0.832972703 |
| 0.627557232 | 0.832762205 |
| 0.627533049 | 0.832753317 |
| 0.627363648 | 0.832598113 |
| 0.627241596 | 0.832482527 |
| 0.62723821 | 0.832482527 |
| 0.627231259 | 0.832482527 |
| 0.627142006 | 0.832482527 |
| 0.626949987 | 0.832350645 |
| 0.626893565 | 0.83232214 |
| 0.62685432 | 0.83231644 |
| 0.626583229 | 0.832095679 |
| 0.626356346 | 0.831956764 |
| 0.626272696 | 0.831868856 |
| 0.626193183 | 0.831843045 |
| 0.62616288 | 0.831838986 |
| 0.626153826 | 0.831838986 |
| 0.626010583 | 0.831729462 |
| 0.625935527 | 0.831699346 |
| 0.625870789 | 0.831659731 |
| 0.625866175 | 0.831659731 |
| 0.625736149 | 0.831527221 |
| 0.625533009 | 0.831442861 |
| 0.625402317 | 0.831338753 |
| 0.625314049 | 0.831244619 |
| 0.625184051 | 0.831183792 |
| 0.625153525 | 0.831170424 |
| 0.625122583 | 0.831152487 |
| 0.625093575 | 0.831152487 |
| 0.625055459 | 0.831152487 |
| 0.624946224 | 0.831034002 |
| 0.624942221 | 0.831034002 |
| 0.624876613 | 0.830987838 |
| 0.624875362 | 0.830987838 |
| 0.624836784 | 0.830981278 |
| 0.624749225 | 0.830892518 |
| 0.624740937 | 0.830892518 |
| 0.624729599 | 0.830892518 |
| 0.624547858 | 0.830786296 |
| 0.62441289 | 0.830695886 |

| | |
|---|---|
| 0.624382855 | 0.830679131 |
| 0.624382484 | 0.830679131 |
| 0.624345373 | 0.830675674 |
| 0.624202344 | 0.830531779 |
| 0.62418808 | 0.830531779 |
| 0.624079053 | 0.830437333 |
| 0.624028605 | 0.830403861 |
| 0.623967359 | 0.830358306 |
| 0.623964654 | 0.830358306 |
| 0.623958785 | 0.830358306 |
| 0.623932191 | 0.830358306 |
| 0.623901578 | 0.830358306 |
| 0.623773807 | 0.830239922 |
| 0.623718871 | 0.830213204 |
| 0.623631559 | 0.830120352 |
| 0.623551734 | 0.830106732 |
| 0.623531421 | 0.830102893 |
| 0.623407864 | 0.829997239 |
| 0.623385999 | 0.829997239 |
| 0.623362399 | 0.829997239 |
| 0.62335293 | 0.829997239 |
| 0.623287596 | 0.829987092 |
| 0.623181854 | 0.829893762 |
| 0.623163247 | 0.829893762 |
| 0.62314133 | 0.829893762 |
| 0.623129413 | 0.829893762 |
| 0.623111184 | 0.829893762 |
| 0.623026053 | 0.829870842 |
| 0.622965994 | 0.829814051 |
| 0.622955944 | 0.829814051 |
| 0.622933742 | 0.829814051 |
| 0.622878678 | 0.829795487 |
| 0.622859398 | 0.829795487 |
| 0.622656614 | 0.829681329 |
| 0.622629337 | 0.829681329 |
| 0.622613305 | 0.829681329 |
| 0.622497908 | 0.829584976 |
| 0.622489981 | 0.829584976 |
| 0.622451053 | 0.829584976 |
| 0.622426975 | 0.829584976 |
| 0.622396841 | 0.829584976 |
| 0.622328554 | 0.829584976 |
| 0.622296054 | 0.829584976 |
| 0.622281145 | 0.829584976 |
| 0.622237413 | 0.829584976 |

| | |
|---|---|
| 0.622136184 | 0.829497495 |
| 0.622112681 | 0.82948938 |
| 0.622095878 | 0.82948938 |
| 0.622085748 | 0.82948938 |
| 0.62199493 | 0.829480001 |
| 0.621716347 | 0.82914663 |
| 0.621690297 | 0.82914663 |
| 0.621627475 | 0.829097755 |
| 0.621461754 | 0.828969754 |
| 0.621377941 | 0.828904216 |
| 0.621329864 | 0.828886514 |
| 0.621234749 | 0.82878284 |
| 0.621219112 | 0.82878284 |
| 0.621205539 | 0.82878284 |
| 0.621069071 | 0.828631609 |
| 0.621049115 | 0.828631609 |
| 0.62100042 | 0.828609486 |
| 0.62087624 | 0.828490213 |
| 0.620829224 | 0.828473898 |
| 0.620689009 | 0.828356416 |
| 0.62065576 | 0.828335255 |
| 0.620638707 | 0.828335255 |
| 0.620594337 | 0.828335255 |
| 0.620588581 | 0.828335255 |
| 0.620556295 | 0.828335255 |
| 0.620553187 | 0.828335255 |
| 0.620531714 | 0.828335255 |
| 0.620485704 | 0.828335255 |
| 0.620473722 | 0.828335255 |
| 0.620358262 | 0.828286365 |
| 0.620157208 | 0.828134003 |
| 0.620122772 | 0.828134003 |
| 0.620084834 | 0.828107013 |
| 0.619980929 | 0.827991467 |
| 0.619970738 | 0.827991467 |
| 0.619968354 | 0.827991467 |
| 0.61995502 | 0.827991467 |
| 0.619904345 | 0.827991467 |
| 0.619795323 | 0.827952539 |
| 0.619682521 | 0.827825071 |
| 0.619621239 | 0.827766423 |
| 0.619610114 | 0.827766423 |
| 0.619535239 | 0.827697968 |
| 0.619532543 | 0.827697968 |
| 0.619498236 | 0.827697968 |

| | |
|---|---|
| 0.61941149 | 0.827671942 |
| 0.619401941 | 0.827671942 |
| 0.619231493 | 0.827563075 |
| 0.619222636 | 0.827563075 |
| 0.619176622 | 0.827563075 |
| 0.619160802 | 0.827563075 |
| 0.619151943 | 0.827563075 |
| 0.619143515 | 0.827563075 |
| 0.619022865 | 0.827500897 |
| 0.618992064 | 0.827482949 |
| 0.618983223 | 0.827482949 |
| 0.618874647 | 0.827418882 |
| 0.618822537 | 0.827372438 |
| 0.618794561 | 0.827372438 |
| 0.618750652 | 0.827372438 |
| 0.61869867 | 0.827372438 |
| 0.618646688 | 0.827360649 |
| 0.618611077 | 0.827360649 |
| 0.618609499 | 0.827360649 |
| 0.618595384 | 0.827360649 |
| 0.618557931 | 0.827360649 |
| 0.618467053 | 0.827354494 |
| 0.618412748 | 0.827354494 |
| 0.618371032 | 0.827349415 |
| 0.61828057 | 0.827305123 |
| 0.618252491 | 0.827305123 |
| 0.618252255 | 0.827305123 |
| 0.618056432 | 0.827137622 |
| 0.618030385 | 0.827126005 |
| 0.617960341 | 0.827078745 |
| 0.617909343 | 0.827033731 |
| 0.617898732 | 0.827033731 |
| 0.617855025 | 0.827033731 |
| 0.617846046 | 0.827033731 |
| 0.617823146 | 0.827033731 |
| 0.617802551 | 0.827033731 |
| 0.617772518 | 0.827033731 |
| 0.617745949 | 0.827033731 |
| 0.617680803 | 0.827026468 |
| 0.617578515 | 0.826939524 |
| 0.617547822 | 0.826921674 |
| 0.617522646 | 0.826911212 |
| 0.617485684 | 0.826911212 |
| 0.617466012 | 0.826911212 |
| 0.617419696 | 0.826911212 |

| | |
|---|---|
| 0.617371429 | 0.826889783 |
| 0.617370108 | 0.826889783 |
| 0.617345569 | 0.826889783 |
| 0.617277447 | 0.826889783 |
| 0.617181788 | 0.826826728 |
| 0.617178932 | 0.826826728 |
| 0.617138931 | 0.826826728 |
| 0.617073589 | 0.826798059 |
| 0.616998694 | 0.826744226 |
| 0.616939248 | 0.826734351 |
| 0.616922747 | 0.826734351 |
| 0.616851321 | 0.826663042 |
| 0.616830881 | 0.826658911 |
| 0.616752514 | 0.826600404 |
| 0.61673821 | 0.826600404 |
| 0.616647102 | 0.826528904 |
| 0.616527644 | 0.826406812 |
| 0.616345149 | 0.826285446 |
| 0.616234456 | 0.82619507 |
| 0.616193423 | 0.82619507 |
| 0.616177626 | 0.82619507 |
| 0.616021212 | 0.82613173 |
| 0.616004551 | 0.82613173 |
| 0.615981493 | 0.826124988 |
| 0.615851159 | 0.82599671 |
| 0.615712937 | 0.825904355 |
| 0.615704724 | 0.825904355 |
| 0.615674048 | 0.825898712 |
| 0.615657157 | 0.825898712 |
| 0.615628347 | 0.82589417 |
| 0.615596282 | 0.82588744 |
| 0.615563599 | 0.825866857 |
| 0.615529623 | 0.825844537 |
| 0.615491912 | 0.825817204 |
| 0.615302726 | 0.825702937 |
| 0.615288464 | 0.825702937 |
| 0.61522208 | 0.825641241 |
| 0.615156518 | 0.825576518 |
| 0.615136564 | 0.825573003 |
| 0.615017178 | 0.825482559 |
| 0.615006272 | 0.825482559 |
| 0.614912953 | 0.825434526 |
| 0.614905009 | 0.825434526 |
| 0.614886113 | 0.825434526 |
| 0.614800442 | 0.825401004 |

| | |
|---|---|
| 0.61473363 | 0.825334572 |
| 0.614706264 | 0.825334572 |
| 0.614577097 | 0.825263996 |
| 0.614540125 | 0.825237616 |
| 0.614534427 | 0.825237616 |
| 0.614474093 | 0.825218747 |
| 0.614453947 | 0.825218747 |
| 0.614353458 | 0.825148572 |
| 0.614345075 | 0.825148572 |
| 0.614341564 | 0.825148572 |
| 0.614333807 | 0.825148572 |
| 0.61431105 | 0.825148572 |
| 0.614232515 | 0.825148572 |
| 0.614215584 | 0.825148572 |
| 0.614038058 | 0.824958831 |
| 0.613900673 | 0.824844059 |
| 0.613884173 | 0.824844059 |
| 0.613860424 | 0.824839712 |
| 0.613826238 | 0.824837125 |
| 0.613801289 | 0.824837125 |
| 0.613785976 | 0.824837125 |
| 0.613753806 | 0.824832882 |
| 0.613711731 | 0.824799611 |
| 0.613624937 | 0.824706235 |
| 0.613611252 | 0.824706235 |
| 0.613507201 | 0.824621003 |
| 0.613387685 | 0.824550248 |
| 0.613362992 | 0.824540326 |
| 0.613300161 | 0.824479134 |
| 0.613204734 | 0.824390637 |
| 0.613196554 | 0.824390637 |
| 0.613154573 | 0.824376492 |
| 0.613107964 | 0.824376492 |
| 0.613035689 | 0.824376492 |
| 0.613010832 | 0.824369406 |
| 0.612975525 | 0.824363284 |
| 0.61282086 | 0.824207022 |
| 0.612793658 | 0.824193713 |
| 0.612785214 | 0.824193713 |
| 0.612670581 | 0.82414034 |
| 0.612667878 | 0.82414034 |
| 0.612656004 | 0.82414034 |
| 0.612610381 | 0.824133396 |
| 0.612486977 | 0.824060489 |
| 0.612467239 | 0.824057212 |

| | |
|---|---|
| 0.612454219 | 0.824057212 |
| 0.612318298 | 0.823903367 |
| 0.612311893 | 0.823903367 |
| 0.612219229 | 0.823879314 |
| 0.612140838 | 0.823827508 |
| 0.612089084 | 0.823781135 |
| 0.61206746 | 0.823775311 |
| 0.612001483 | 0.823756349 |
| 0.611981115 | 0.823752214 |
| 0.611918226 | 0.823690841 |
| 0.611912419 | 0.823690841 |
| 0.611906642 | 0.823690841 |
| 0.611879849 | 0.823690841 |
| 0.611697851 | 0.823557132 |
| 0.611666071 | 0.823557132 |
| 0.611654958 | 0.823557132 |
| 0.611644214 | 0.823557132 |
| 0.611552653 | 0.823478037 |
| 0.611548096 | 0.823478037 |
| 0.611490594 | 0.823441034 |
| 0.611479274 | 0.823441034 |
| 0.611464545 | 0.823441034 |
| 0.611360602 | 0.823359105 |
| 0.611235585 | 0.823237295 |
| 0.611225534 | 0.823237295 |
| 0.611180669 | 0.82320989 |
| 0.611042473 | 0.823093584 |
| 0.610977215 | 0.823052237 |
| 0.610792302 | 0.82289624 |
| 0.61077099 | 0.822890804 |
| 0.610733837 | 0.822887305 |
| 0.610718616 | 0.822887305 |
| 0.610709582 | 0.822887305 |
| 0.610655174 | 0.822874434 |
| 0.610590107 | 0.822842327 |
| 0.610527846 | 0.822842327 |
| 0.610519197 | 0.822842327 |
| 0.610489956 | 0.822842327 |
| 0.610377317 | 0.822756077 |
| 0.610300555 | 0.822699175 |
| 0.610281352 | 0.822696573 |
| 0.610075104 | 0.82248838 |
| 0.609988422 | 0.822394796 |
| 0.6099317 | 0.822341602 |
| 0.609907501 | 0.822332255 |

| | |
|---|---|
| 0.609896648 | 0.822332255 |
| 0.60980649 | 0.822265896 |
| 0.609801516 | 0.822265896 |
| 0.609752162 | 0.822262478 |
| 0.60955782 | 0.822140074 |
| 0.609425852 | 0.822055202 |
| 0.609383814 | 0.822021778 |
| 0.609346304 | 0.82199446 |
| 0.609293143 | 0.821946028 |
| 0.6092629 | 0.821928511 |
| 0.609168042 | 0.821893664 |
| 0.609113813 | 0.821844564 |
| 0.609074448 | 0.821841987 |
| 0.608917524 | 0.82171861 |
| 0.608886123 | 0.821699518 |
| 0.60888353 | 0.821699518 |
| 0.60886281 | 0.821699518 |
| 0.608785628 | 0.821699518 |
| 0.608757157 | 0.821688454 |
| 0.608696034 | 0.821629238 |
| 0.608689741 | 0.821629238 |
| 0.608679549 | 0.821629238 |
| 0.608498367 | 0.821502119 |
| 0.608457292 | 0.821493237 |
| 0.608402321 | 0.821442306 |
| 0.608358503 | 0.82140643 |
| 0.608336639 | 0.821400196 |
| 0.608302106 | 0.821376854 |
| 0.608203551 | 0.821267062 |
| 0.608194453 | 0.821267062 |
| 0.608168781 | 0.821266679 |
| 0.608150599 | 0.821266679 |
| 0.608108398 | 0.821266679 |
| 0.608035965 | 0.821250324 |
| 0.608035541 | 0.821250324 |
| 0.607965178 | 0.821201291 |
| 0.6079567 | 0.821201291 |
| 0.607906505 | 0.821191909 |
| 0.607832145 | 0.821131265 |
| 0.60774122 | 0.821084168 |
| 0.607726703 | 0.821084168 |
| 0.607672075 | 0.821061531 |
| 0.60762579 | 0.821035701 |
| 0.607568501 | 0.821035701 |
| 0.607542342 | 0.821035701 |

| | |
|---|---|
| 0.607467713 | 0.820971731 |
| 0.607462687 | 0.820971731 |
| 0.607442503 | 0.820971731 |
| 0.607384607 | 0.8209293 |
| 0.60736907 | 0.8209293 |
| 0.607283948 | 0.820863136 |
| 0.607260697 | 0.820863136 |
| 0.607112149 | 0.82074555 |
| 0.607080067 | 0.820727308 |
| 0.606971563 | 0.820604316 |
| 0.606915234 | 0.820574345 |
| 0.606908266 | 0.820574345 |
| 0.606862852 | 0.820554855 |
| 0.606823418 | 0.82054338 |
| 0.606717887 | 0.82050007 |
| 0.606717136 | 0.82050007 |
| 0.606573912 | 0.820462234 |
| 0.6065147 | 0.820411173 |
| 0.606501091 | 0.820411173 |
| 0.606475127 | 0.820411173 |
| 0.60643771 | 0.8204021 |
| 0.606416041 | 0.8204021 |
| 0.606389815 | 0.820399617 |
| 0.606273179 | 0.820311725 |
| 0.606212821 | 0.820273476 |
| 0.606161174 | 0.820230083 |
| 0.606031755 | 0.82010156 |
| 0.605948948 | 0.820036102 |
| 0.605938623 | 0.820036102 |
| 0.605897859 | 0.820036102 |
| 0.605834899 | 0.819998259 |
| 0.60577264 | 0.8199839 |
| 0.605579897 | 0.819824462 |
| 0.605561214 | 0.819824462 |
| 0.605522693 | 0.819808667 |
| 0.605234455 | 0.819484951 |
| 0.605232347 | 0.819484951 |
| 0.605176697 | 0.819479999 |
| 0.604941308 | 0.819301021 |
| 0.604915046 | 0.81928875 |
| 0.604826419 | 0.819247418 |
| 0.604747273 | 0.819177997 |
| 0.604713489 | 0.819177997 |
| 0.604601755 | 0.819074064 |
| 0.604589618 | 0.819074064 |

| | |
|---|---|
| 0.60455341 | 0.819055164 |
| 0.604551977 | 0.819055164 |
| 0.604471118 | 0.818990268 |
| 0.604427168 | 0.818954019 |
| 0.604341228 | 0.818907468 |
| 0.60426366 | 0.818872253 |
| 0.604235423 | 0.818872253 |
| 0.604198334 | 0.818853624 |
| 0.604183266 | 0.818853624 |
| 0.604146104 | 0.818829439 |
| 0.604110621 | 0.818804649 |
| 0.604102094 | 0.818804649 |
| 0.604010126 | 0.818738338 |
| 0.604000212 | 0.818738338 |
| 0.603821547 | 0.818575909 |
| 0.603636685 | 0.81841848 |
| 0.603617184 | 0.81841848 |
| 0.603612067 | 0.81841848 |
| 0.603473273 | 0.818290102 |
| 0.603471904 | 0.818290102 |
| 0.603463122 | 0.818290102 |
| 0.603379437 | 0.818232751 |
| 0.603355743 | 0.818223917 |
| 0.603345693 | 0.818223917 |
| 0.6032419 | 0.818162717 |
| 0.603237929 | 0.818162717 |
| 0.603237639 | 0.818162717 |
| 0.603045136 | 0.818058922 |
| 0.602938596 | 0.817960798 |
| 0.602834115 | 0.817918387 |
| 0.602797251 | 0.817909024 |
| 0.602723885 | 0.817832777 |
| 0.602703064 | 0.817827825 |
| 0.60269037 | 0.817827825 |
| 0.602660733 | 0.817816985 |
| 0.602633584 | 0.817803445 |
| 0.602586786 | 0.817803445 |
| 0.602558995 | 0.817803445 |
| 0.602540858 | 0.817803445 |
| 0.602371722 | 0.817682105 |
| 0.602337394 | 0.817682105 |
| 0.602261408 | 0.817647846 |
| 0.602195525 | 0.817581706 |
| 0.602175956 | 0.817578444 |
| 0.602093178 | 0.817512666 |

| | |
|---|---|
| 0.602082991 | 0.817512666 |
| 0.60203254 | 0.817476943 |
| 0.601993973 | 0.817471187 |
| 0.601882518 | 0.817405197 |
| 0.601859424 | 0.817405007 |
| 0.601835264 | 0.817395504 |
| 0.601790207 | 0.817357617 |
| 0.601776018 | 0.817357617 |
| 0.601727715 | 0.817342665 |
| 0.601706465 | 0.817342665 |
| 0.60168394 | 0.817342665 |
| 0.60156363 | 0.817259635 |
| 0.601527351 | 0.817233659 |
| 0.601487461 | 0.817212919 |
| 0.601372622 | 0.817136192 |
| 0.601351621 | 0.817134769 |
| 0.601251246 | 0.817091625 |
| 0.601228815 | 0.817084456 |
| 0.601198353 | 0.817066373 |
| 0.601196875 | 0.817066373 |
| 0.601128043 | 0.817017445 |
| 0.601122956 | 0.817017445 |
| 0.601106039 | 0.817017445 |
| 0.601106039 | 0.817017445 |
| 0.601054767 | 0.817017445 |
| 0.601022916 | 0.817014455 |
| 0.601006656 | 0.817014455 |
| 0.600910255 | 0.816931257 |
| 0.600831069 | 0.816846921 |
| 0.600698346 | 0.816713106 |
| 0.600673483 | 0.816713106 |
| 0.600659414 | 0.816713106 |
| 0.600655986 | 0.816713106 |
| 0.600613704 | 0.816713106 |
| 0.600558175 | 0.816685727 |
| 0.600442611 | 0.816598521 |
| 0.600438229 | 0.816598521 |
| 0.600435667 | 0.816598521 |
| 0.600374595 | 0.816575972 |
| 0.600343387 | 0.816556845 |
| 0.6002935 | 0.81651231 |
| 0.600253131 | 0.816504038 |
| 0.600222457 | 0.816503202 |
| 0.600218903 | 0.816503202 |
| 0.600180288 | 0.816498232 |

| | |
|---|---|
| 0.60009791 | 0.816412605 |
| 0.600068787 | 0.816412605 |
| 0.600051512 | 0.816412605 |
| 0.599928267 | 0.816344762 |
| 0.599904231 | 0.816344762 |
| 0.599811847 | 0.816276791 |
| 0.59968575 | 0.81617515 |
| 0.599668408 | 0.816174871 |
| 0.599588109 | 0.816112225 |
| 0.599538417 | 0.816091234 |
| 0.599497044 | 0.81605824 |
| 0.599476634 | 0.816053781 |
| 0.59941047 | 0.81601036 |
| 0.599288988 | 0.815914946 |
| 0.599223693 | 0.815851437 |
| 0.599173574 | 0.815836452 |
| 0.599148817 | 0.815836452 |
| 0.599142197 | 0.815836452 |
| 0.599126135 | 0.815836452 |
| 0.59888474 | 0.815597721 |
| 0.598707422 | 0.815426189 |
| 0.598638634 | 0.815355817 |
| 0.598626313 | 0.815355817 |
| 0.598584205 | 0.815328317 |
| 0.598534424 | 0.815307145 |
| 0.598480869 | 0.815280829 |
| 0.598447228 | 0.81525832 |
| 0.598419581 | 0.815243976 |
| 0.598330886 | 0.815193098 |
| 0.598249235 | 0.815138778 |
| 0.598210508 | 0.815122361 |
| 0.598188798 | 0.815122361 |
| 0.598096716 | 0.815060582 |
| 0.598031841 | 0.815018815 |
| 0.597984535 | 0.815000986 |
| 0.597976634 | 0.815000986 |
| 0.597880537 | 0.814905885 |
| 0.597772838 | 0.814805728 |
| 0.597714445 | 0.814749453 |
| 0.597596049 | 0.814668641 |
| 0.597574534 | 0.814668641 |
| 0.597492286 | 0.814609828 |
| 0.597361543 | 0.814520367 |
| 0.597219639 | 0.814354665 |
| 0.597140108 | 0.81428679 |

| | |
|---|---|
| 0.597139356 | 0.81428679 |
| 0.597030907 | 0.8142417 |
| 0.597016056 | 0.8142417 |
| 0.596881987 | 0.81422061 |
| 0.596792237 | 0.814144817 |
| 0.596721363 | 0.814118089 |
| 0.596670464 | 0.814118089 |
| 0.596581388 | 0.814043713 |
| 0.596556033 | 0.814032438 |
| 0.596548494 | 0.814032438 |
| 0.596480279 | 0.813999033 |
| 0.596242313 | 0.813744529 |
| 0.596088933 | 0.813651485 |
| 0.596000902 | 0.81355464 |
| 0.595970415 | 0.813536341 |
| 0.595966535 | 0.813536341 |
| 0.595965745 | 0.813536341 |
| 0.59589636 | 0.81352852 |
| 0.595886394 | 0.81352852 |
| 0.595868885 | 0.81352852 |
| 0.595780524 | 0.813506089 |
| 0.595746513 | 0.813506089 |
| 0.595670664 | 0.813476894 |
| 0.595612864 | 0.813421285 |
| 0.595610912 | 0.813421285 |
| 0.595571162 | 0.813421285 |
| 0.595540499 | 0.813421285 |
| 0.595500624 | 0.813407945 |
| 0.595470282 | 0.813389827 |
| 0.595444728 | 0.81337825 |
| 0.595414714 | 0.813370819 |
| 0.595383615 | 0.813370819 |
| 0.595229361 | 0.813224004 |
| 0.595015794 | 0.813048837 |
| 0.594992394 | 0.813040189 |
| 0.594955698 | 0.813013371 |
| 0.59495375 | 0.813013371 |
| 0.594944033 | 0.813013371 |
| 0.594923799 | 0.813013371 |
| 0.594908814 | 0.813013371 |
| 0.594877061 | 0.813013371 |
| 0.594840382 | 0.813013371 |
| 0.594797229 | 0.813013371 |
| 0.594795655 | 0.813013371 |
| 0.594795648 | 0.813013371 |

| | |
|---|---|
| 0.594776108 | 0.813013371 |
| 0.594757126 | 0.813013371 |
| 0.594607775 | 0.812981125 |
| 0.594574406 | 0.812958839 |
| 0.594547171 | 0.812944939 |
| 0.59454438 | 0.812944939 |
| 0.594525988 | 0.812944939 |
| 0.594486478 | 0.812944939 |
| 0.594387863 | 0.812890473 |
| 0.594299071 | 0.812839063 |
| 0.594171228 | 0.81274134 |
| 0.59413603 | 0.81274134 |
| 0.594126228 | 0.81274134 |
| 0.594047898 | 0.812682232 |
| 0.594038502 | 0.812682232 |
| 0.593889117 | 0.812540254 |
| 0.593814039 | 0.812525677 |
| 0.593807652 | 0.812525677 |
| 0.593705044 | 0.812446561 |
| 0.593689209 | 0.812446561 |
| 0.593654232 | 0.812446561 |
| 0.593639427 | 0.812446561 |
| 0.593532694 | 0.812444146 |
| 0.593515669 | 0.812444146 |
| 0.593440302 | 0.812411074 |
| 0.593383 | 0.812385308 |
| 0.593339688 | 0.812385308 |
| 0.593290641 | 0.81234629 |
| 0.593051209 | 0.812205255 |
| 0.593013692 | 0.812195522 |
| 0.593008636 | 0.812195522 |
| 0.592975454 | 0.812195522 |
| 0.592919061 | 0.812164398 |
| 0.592852439 | 0.812154409 |
| 0.592829085 | 0.812154409 |
| 0.59280453 | 0.812147663 |
| 0.592785086 | 0.812144386 |
| 0.592748107 | 0.812126705 |
| 0.592716868 | 0.812126705 |
| 0.592700113 | 0.812126705 |
| 0.592625634 | 0.81206609 |
| 0.592595646 | 0.812048361 |
| 0.592531419 | 0.812007075 |
| 0.592459237 | 0.811954882 |
| 0.592431175 | 0.811939788 |

| | |
|---|---|
| 0.592392714 | 0.811916303 |
| 0.592316179 | 0.811899 |
| 0.592197445 | 0.811806342 |
| 0.592167734 | 0.811788978 |
| 0.592097233 | 0.811715693 |
| 0.592072261 | 0.811715693 |
| 0.59203063 | 0.811715693 |
| 0.592003928 | 0.811715693 |
| 0.591973636 | 0.811715693 |
| 0.591908862 | 0.811691094 |
| 0.591878294 | 0.811678472 |
| 0.591831449 | 0.811655046 |
| 0.591786261 | 0.811616443 |
| 0.591754283 | 0.811616443 |
| 0.591748141 | 0.811616443 |
| 0.591713368 | 0.811616443 |
| 0.591706529 | 0.811616443 |
| 0.591689857 | 0.811616443 |
| 0.591638337 | 0.811616443 |
| 0.591561363 | 0.811616443 |
| 0.591514736 | 0.811616443 |
| 0.591502293 | 0.811616443 |
| 0.591486122 | 0.811616443 |
| 0.591267955 | 0.811396269 |
| 0.591201538 | 0.811395463 |
| 0.591163316 | 0.811369565 |
| 0.591150353 | 0.811369565 |
| 0.591109872 | 0.81134609 |
| 0.591069871 | 0.811334835 |
| 0.590936488 | 0.811219178 |
| 0.59089276 | 0.811208621 |
| 0.590879792 | 0.811208621 |
| 0.590873657 | 0.811208621 |
| 0.590863014 | 0.811208621 |
| 0.59085126 | 0.811208621 |
| 0.590813291 | 0.811208621 |
| 0.590754545 | 0.811208621 |
| 0.59073569 | 0.811208621 |
| 0.590630842 | 0.811153007 |
| 0.590502626 | 0.811070471 |
| 0.590365525 | 0.81095232 |
| 0.590362652 | 0.81095232 |
| 0.590337513 | 0.81095232 |
| 0.590290312 | 0.810942561 |
| 0.590225266 | 0.81087659 |

| | |
|---|---|
| 0.590071758 | 0.810712466 |
| 0.589998543 | 0.810635258 |
| 0.589977748 | 0.810630072 |
| 0.589956722 | 0.810630072 |
| 0.589741342 | 0.810445535 |
| 0.5896463 | 0.810361689 |
| 0.589577854 | 0.810319458 |
| 0.589498741 | 0.810275801 |
| 0.589495838 | 0.810275801 |
| 0.589477738 | 0.810275801 |
| 0.589420866 | 0.810262298 |
| 0.589308573 | 0.810154699 |
| 0.589308189 | 0.810154699 |
| 0.589109764 | 0.809951524 |
| 0.58899617 | 0.80988834 |
| 0.588986964 | 0.80988834 |
| 0.588866692 | 0.809804347 |
| 0.58885657 | 0.809804347 |
| 0.588849038 | 0.809804347 |
| 0.588785394 | 0.809762691 |
| 0.588653221 | 0.809651052 |
| 0.588628091 | 0.809639868 |
| 0.588497337 | 0.809600308 |
| 0.588485619 | 0.809600308 |
| 0.588438704 | 0.809589801 |
| 0.588399773 | 0.80956955 |
| 0.588324166 | 0.809519384 |
| 0.588235328 | 0.809449421 |
| 0.588224756 | 0.809449421 |
| 0.58805586 | 0.809320246 |
| 0.587955674 | 0.809236314 |
| 0.58794163 | 0.809236314 |
| 0.587833359 | 0.809201092 |
| 0.587659161 | 0.80900806 |
| 0.587619655 | 0.80900806 |
| 0.587235951 | 0.808659187 |
| 0.587077318 | 0.808487492 |
| 0.587071694 | 0.808487492 |
| 0.58706157 | 0.808487492 |
| 0.586874983 | 0.80834909 |
| 0.586820779 | 0.808321183 |
| 0.586812883 | 0.808321183 |
| 0.586673019 | 0.80821115 |
| 0.586632886 | 0.808179239 |
| 0.586603401 | 0.808162435 |

| | |
|---|---|
| 0.586447465 | 0.808017283 |
| 0.586424536 | 0.808009066 |
| 0.586408955 | 0.808009066 |
| 0.586407998 | 0.808009066 |
| 0.586318689 | 0.807956722 |
| 0.586307481 | 0.807956722 |
| 0.586305204 | 0.807956722 |
| 0.586287893 | 0.807956722 |
| 0.58622117 | 0.807956722 |
| 0.586130254 | 0.807907435 |
| 0.58608328 | 0.807866068 |
| 0.585978759 | 0.807792131 |
| 0.585964553 | 0.807792131 |
| 0.585919826 | 0.80775765 |
| 0.585852756 | 0.807734905 |
| 0.585790089 | 0.807695691 |
| 0.585748131 | 0.807695691 |
| 0.585743838 | 0.807695691 |
| 0.585707845 | 0.807695691 |
| 0.585615519 | 0.807695691 |
| 0.585597711 | 0.807695691 |
| 0.585531753 | 0.807666844 |
| 0.585491938 | 0.807635316 |
| 0.585486304 | 0.807635316 |
| 0.585466548 | 0.807635316 |
| 0.585461104 | 0.807635316 |
| 0.585445667 | 0.807635316 |
| 0.585355662 | 0.807570479 |
| 0.585306534 | 0.807562234 |
| 0.585198624 | 0.807443777 |
| 0.585167081 | 0.807443777 |
| 0.585089426 | 0.807360686 |
| 0.585071185 | 0.807358908 |
| 0.585040514 | 0.807344358 |
| 0.584964828 | 0.807329099 |
| 0.584945104 | 0.807325271 |
| 0.584829562 | 0.807212587 |
| 0.584817905 | 0.807212587 |
| 0.584788999 | 0.807203387 |
| 0.584732684 | 0.807149046 |
| 0.584709643 | 0.807140634 |
| 0.584617648 | 0.807060427 |
| 0.58461589 | 0.807060427 |
| 0.584567159 | 0.807037512 |
| 0.584480425 | 0.806987946 |

| | |
|---|---|
| 0.584366624 | 0.806924392 |
| 0.584302579 | 0.806882743 |
| 0.584279759 | 0.806874625 |
| 0.584062816 | 0.806665778 |
| 0.584050023 | 0.806665778 |
| 0.583928961 | 0.806553889 |
| 0.583787457 | 0.806428598 |
| 0.583736077 | 0.80642278 |
| 0.583672476 | 0.806403827 |
| 0.583542452 | 0.806270011 |
| 0.583521923 | 0.806270011 |
| 0.583521846 | 0.806270011 |
| 0.583517797 | 0.806270011 |
| 0.583504162 | 0.806270011 |
| 0.583345802 | 0.806216028 |
| 0.58334314 | 0.806216028 |
| 0.583319054 | 0.806216028 |
| 0.583212299 | 0.80619528 |
| 0.583157012 | 0.806142252 |
| 0.583094468 | 0.806135916 |
| 0.583057736 | 0.806135916 |
| 0.583015375 | 0.806133641 |
| 0.582998201 | 0.806133299 |
| 0.582963609 | 0.80610887 |
| 0.582959521 | 0.80610887 |
| 0.582927808 | 0.806106173 |
| 0.582888375 | 0.806098454 |
| 0.582868611 | 0.806098454 |
| 0.582801062 | 0.806072675 |
| 0.582778106 | 0.806072675 |
| 0.582776982 | 0.806072675 |
| 0.582704616 | 0.806054974 |
| 0.58266898 | 0.806046266 |
| 0.582651213 | 0.806046266 |
| 0.582651057 | 0.806046266 |
| 0.582458446 | 0.805913109 |
| 0.582377917 | 0.805860515 |
| 0.582346212 | 0.805840056 |
| 0.582301351 | 0.805824803 |
| 0.582246759 | 0.805796082 |
| 0.582069716 | 0.805597884 |
| 0.582027241 | 0.805585917 |
| 0.581890715 | 0.805467175 |
| 0.581730383 | 0.805315456 |
| 0.581730383 | 0.805315456 |

| | |
|---|---|
| 0.581730383 | 0.805315456 |
| 0.581724508 | 0.805315456 |
| 0.5816906 | 0.805315456 |
| 0.58167034 | 0.805315456 |
| 0.581640532 | 0.805315456 |
| 0.581527121 | 0.805255914 |
| 0.581491068 | 0.805241622 |
| 0.581395125 | 0.805156301 |
| 0.581345797 | 0.805134092 |
| 0.581325046 | 0.805134092 |
| 0.581312261 | 0.805134092 |
| 0.581310401 | 0.805134092 |
| 0.581304872 | 0.805134092 |
| 0.581250168 | 0.805134092 |
| 0.581195513 | 0.805134092 |
| 0.581151053 | 0.805118201 |
| 0.581124499 | 0.805108547 |
| 0.580941458 | 0.804972036 |
| 0.580672456 | 0.804692944 |
| 0.580624513 | 0.804663656 |
| 0.580436932 | 0.804452801 |
| 0.580424495 | 0.804452801 |
| 0.58040954 | 0.804452801 |
| 0.580330093 | 0.804429155 |
| 0.580256618 | 0.804374352 |
| 0.580223032 | 0.804374352 |
| 0.580189774 | 0.804374352 |
| 0.580152547 | 0.80436115 |
| 0.57999599 | 0.804223507 |
| 0.579871206 | 0.804144127 |
| 0.57977082 | 0.804028327 |
| 0.579763114 | 0.804028327 |
| 0.579735853 | 0.804026659 |
| 0.579518274 | 0.803841939 |
| 0.579477528 | 0.803808832 |
| 0.579399475 | 0.80372397 |
| 0.579385576 | 0.80372397 |
| 0.579205015 | 0.803571244 |
| 0.579158978 | 0.803530781 |
| 0.579050588 | 0.803450614 |
| 0.578989001 | 0.803388567 |
| 0.578980769 | 0.803388567 |
| 0.57894113 | 0.803373753 |
| 0.57891017 | 0.803372804 |
| 0.578848951 | 0.803311256 |

| | |
|---|---|
| 0.578812403 | 0.803283943 |
| 0.578805401 | 0.803283943 |
| 0.578801969 | 0.803283943 |
| 0.578705013 | 0.803228527 |
| 0.57864465 | 0.803168151 |
| 0.578644602 | 0.803168151 |
| 0.578587591 | 0.803135764 |
| 0.578586611 | 0.803135764 |
| 0.578471051 | 0.803081053 |
| 0.578449493 | 0.803081053 |
| 0.578388349 | 0.803046436 |
| 0.578386797 | 0.803046436 |
| 0.578380755 | 0.803046436 |
| 0.578338568 | 0.803046436 |
| 0.578313307 | 0.803046436 |
| 0.57825439 | 0.803024313 |
| 0.578253176 | 0.803024313 |
| 0.578191254 | 0.802984631 |
| 0.578171959 | 0.802984631 |
| 0.577771176 | 0.802634109 |
| 0.577743926 | 0.802619665 |
| 0.577671548 | 0.802597249 |
| 0.577578103 | 0.802553165 |
| 0.577557523 | 0.802547984 |
| 0.577511719 | 0.80250775 |
| 0.577477839 | 0.802484084 |
| 0.57747493 | 0.802484084 |
| 0.577347569 | 0.802373809 |
| 0.577244137 | 0.802323192 |
| 0.577230786 | 0.802323192 |
| 0.577193091 | 0.802323192 |
| 0.577181972 | 0.802323192 |
| 0.577148166 | 0.802323192 |
| 0.577147946 | 0.802323192 |
| 0.577140934 | 0.802323192 |
| 0.577129415 | 0.802323192 |
| 0.576995996 | 0.802306093 |
| 0.576818555 | 0.802106199 |
| 0.576706292 | 0.80202034 |
| 0.576695835 | 0.80202034 |
| 0.576653427 | 0.802008932 |
| 0.57660594 | 0.801997872 |
| 0.576548477 | 0.801941366 |
| 0.576363485 | 0.801754299 |
| 0.576357073 | 0.801754299 |

| | |
|---|---|
| 0.576310183 | 0.801726985 |
| 0.576306501 | 0.801726985 |
| 0.576255822 | 0.801726985 |
| 0.576221863 | 0.801726985 |
| 0.576143607 | 0.801723709 |
| 0.576128633 | 0.801723709 |
| 0.576127577 | 0.801723709 |
| 0.576123589 | 0.801723709 |
| 0.576035866 | 0.801723709 |
| 0.575989147 | 0.801723709 |
| 0.575955362 | 0.801723709 |
| 0.57581128 | 0.801645683 |
| 0.575778606 | 0.801643115 |
| 0.575770127 | 0.801643115 |
| 0.575650131 | 0.801511103 |
| 0.57562591 | 0.80150081 |
| 0.575599928 | 0.801488065 |
| 0.575518498 | 0.801398108 |
| 0.575435544 | 0.801329454 |
| 0.575425618 | 0.801329454 |
| 0.575397533 | 0.801329454 |
| 0.57531089 | 0.801283435 |
| 0.57530586 | 0.801283435 |
| 0.575303672 | 0.801283435 |
| 0.575255941 | 0.801283435 |
| 0.575027036 | 0.800994784 |
| 0.575011369 | 0.800994784 |
| 0.57500905 | 0.800994784 |
| 0.574986319 | 0.800994784 |
| 0.574961677 | 0.800994784 |
| 0.574956123 | 0.800994784 |
| 0.574904705 | 0.800994784 |
| 0.574860505 | 0.800973648 |
| 0.574692379 | 0.800809676 |
| 0.5746699 | 0.800801782 |
| 0.574655214 | 0.800801782 |
| 0.57464407 | 0.800801782 |
| 0.574635198 | 0.800801782 |
| 0.574634735 | 0.800801782 |
| 0.574616314 | 0.800801782 |
| 0.57451879 | 0.800801782 |
| 0.574479856 | 0.800794688 |
| 0.574477393 | 0.800794688 |
| 0.574417863 | 0.800778585 |
| 0.574398713 | 0.800775327 |

| | |
|---|---|
| 0.57434063 | 0.800759705 |
| 0.57426032 | 0.80069958 |
| 0.574146999 | 0.800611891 |
| 0.574041758 | 0.800532253 |
| 0.573939767 | 0.800463535 |
| 0.573897963 | 0.800463535 |
| 0.573846767 | 0.800463535 |
| 0.573677122 | 0.80026127 |
| 0.573555085 | 0.800137903 |
| 0.573520266 | 0.800114154 |
| 0.573506266 | 0.800114154 |
| 0.573445722 | 0.800079078 |
| 0.573445254 | 0.800079078 |
| 0.573358967 | 0.800079078 |
| 0.573326043 | 0.800076165 |
| 0.573302152 | 0.800066267 |
| 0.573157854 | 0.799935213 |
| 0.5730722 | 0.799872545 |
| 0.573030165 | 0.799850756 |
| 0.573027459 | 0.799850756 |
| 0.572987918 | 0.79983867 |
| 0.572818378 | 0.79967232 |
| 0.572769083 | 0.799626942 |
| 0.572664845 | 0.799504854 |
| 0.572541252 | 0.799379169 |
| 0.572513562 | 0.799363942 |
| 0.57236953 | 0.799256564 |
| 0.572350711 | 0.799253718 |
| 0.572011899 | 0.799014862 |
| 0.571995264 | 0.799014862 |
| 0.571983868 | 0.799014862 |
| 0.571901668 | 0.798954616 |
| 0.571774108 | 0.798823275 |
| 0.571769318 | 0.798823275 |
| 0.571736625 | 0.798823275 |
| 0.571708042 | 0.798823275 |
| 0.571697665 | 0.798823275 |
| 0.571696516 | 0.798823275 |
| 0.571682155 | 0.798823275 |
| 0.57151713 | 0.798698543 |
| 0.57142938 | 0.798622779 |
| 0.571383719 | 0.798605833 |
| 0.571305614 | 0.798548171 |
| 0.571253113 | 0.798517023 |
| 0.571203455 | 0.798471044 |

| | |
|---|---|
| 0.571143842 | 0.79842781 |
| 0.571090955 | 0.79842781 |
| 0.571068768 | 0.798423373 |
| 0.570998538 | 0.798372056 |
| 0.570997198 | 0.798372056 |
| 0.570879645 | 0.798286849 |
| 0.570779416 | 0.798229717 |
| 0.57074382 | 0.79822082 |
| 0.57072414 | 0.79822082 |
| 0.570529965 | 0.798115217 |
| 0.570514178 | 0.798115217 |
| 0.570456426 | 0.798059223 |
| 0.570383167 | 0.798013852 |
| 0.570264082 | 0.797977654 |
| 0.570246597 | 0.797976632 |
| 0.570138502 | 0.797852643 |
| 0.57005501 | 0.797778854 |
| 0.569979827 | 0.797697076 |
| 0.569957922 | 0.79768986 |
| 0.569946138 | 0.79768986 |
| 0.569931589 | 0.79768986 |
| 0.569847747 | 0.797652862 |
| 0.569825684 | 0.797652862 |
| 0.569812719 | 0.797652862 |
| 0.569811115 | 0.797652862 |
| 0.569600893 | 0.797484373 |
| 0.569585142 | 0.797484373 |
| 0.569541248 | 0.797481655 |
| 0.569533723 | 0.797481655 |
| 0.569492969 | 0.797481655 |
| 0.569479211 | 0.797481655 |
| 0.5694778 | 0.797481655 |
| 0.569467686 | 0.797481655 |
| 0.569421432 | 0.797481655 |
| 0.56936532 | 0.797481655 |
| 0.569351696 | 0.797481655 |
| 0.569335665 | 0.797481655 |
| 0.5693217 | 0.797481655 |
| 0.569321111 | 0.797481655 |
| 0.569321111 | 0.797481655 |
| 0.569264473 | 0.797481655 |
| 0.569197101 | 0.797481655 |
| 0.56916436 | 0.797481655 |
| 0.569073203 | 0.797388922 |
| 0.569072149 | 0.797388922 |

| | |
|---|---|
| 0.569007646 | 0.797388922 |
| 0.568933676 | 0.797322504 |
| 0.568874198 | 0.797274268 |
| 0.568653556 | 0.797059067 |
| 0.568613914 | 0.797059067 |
| 0.568584516 | 0.797059067 |
| 0.568485811 | 0.796964464 |
| 0.56843266 | 0.796913409 |
| 0.568305787 | 0.79682936 |
| 0.568294609 | 0.79682936 |
| 0.568270092 | 0.796826229 |
| 0.568202089 | 0.796754332 |
| 0.568178275 | 0.796744398 |
| 0.568173394 | 0.796744398 |
| 0.568113 | 0.796724243 |
| 0.568083885 | 0.796724243 |
| 0.568027204 | 0.796673292 |
| 0.567992092 | 0.796647506 |
| 0.567975927 | 0.796647506 |
| 0.567953951 | 0.796640931 |
| 0.56794428 | 0.796640931 |
| 0.56792935 | 0.796640931 |
| 0.567810259 | 0.796533215 |
| 0.567704372 | 0.796455053 |
| 0.567667665 | 0.796427015 |
| 0.567639196 | 0.796410533 |
| 0.567629384 | 0.796410533 |
| 0.567624864 | 0.796410533 |
| 0.5676047 | 0.796410533 |
| 0.56754936 | 0.796410533 |
| 0.567480806 | 0.796352525 |
| 0.567434788 | 0.796334875 |
| 0.567430388 | 0.796334875 |
| 0.567404764 | 0.796334875 |
| 0.567319212 | 0.796266528 |
| 0.567262085 | 0.796209811 |
| 0.567152077 | 0.796102327 |
| 0.567148103 | 0.796102327 |
| 0.567146615 | 0.796102327 |
| 0.567074893 | 0.796086451 |
| 0.567051288 | 0.796078164 |
| 0.566992188 | 0.796034203 |
| 0.566881906 | 0.795981995 |
| 0.566833025 | 0.795981995 |
| 0.566833022 | 0.795981995 |

| | |
|---|---|
| 0.566767945 | 0.79596193 |
| 0.566735584 | 0.795939953 |
| 0.566730069 | 0.795939953 |
| 0.566622701 | 0.79585182 |
| 0.56648012 | 0.795730262 |
| 0.566463306 | 0.795730262 |
| 0.56633029 | 0.795589824 |
| 0.56631234 | 0.795589824 |
| 0.566235236 | 0.795565685 |
| 0.566184243 | 0.795565685 |
| 0.566175891 | 0.795565685 |
| 0.56607777 | 0.795485245 |
| 0.566006423 | 0.795408454 |
| 0.565965566 | 0.795374506 |
| 0.565802383 | 0.795192106 |
| 0.565782321 | 0.795187376 |
| 0.565775996 | 0.795187376 |
| 0.565762451 | 0.795187376 |
| 0.565726624 | 0.795187376 |
| 0.565716645 | 0.795187376 |
| 0.565671571 | 0.795187376 |
| 0.565609876 | 0.795156194 |
| 0.565593064 | 0.795156031 |
| 0.565576957 | 0.795156031 |
| 0.56554077 | 0.795152927 |
| 0.56548347 | 0.795137067 |
| 0.565422618 | 0.795080692 |
| 0.565406618 | 0.795080692 |
| 0.565368131 | 0.795074496 |
| 0.565358303 | 0.795074496 |
| 0.56530619 | 0.795074496 |
| 0.56513507 | 0.794887569 |
| 0.565105536 | 0.794887569 |
| 0.565062253 | 0.794855571 |
| 0.565037453 | 0.794850418 |
| 0.564936345 | 0.794748879 |
| 0.564858436 | 0.794686224 |
| 0.564807657 | 0.794640095 |
| 0.564799985 | 0.794640095 |
| 0.5647917 | 0.794640095 |
| 0.564782212 | 0.794640095 |
| 0.564720914 | 0.794640095 |
| 0.564670697 | 0.794609877 |
| 0.56466763 | 0.794609877 |
| 0.564638674 | 0.794609877 |

| | |
|---|---|
| 0.564524579 | 0.79447468 |
| 0.564452708 | 0.794397007 |
| 0.56444459 | 0.794397007 |
| 0.564424918 | 0.794397007 |
| 0.564413245 | 0.794397007 |
| 0.564368641 | 0.794378798 |
| 0.564366644 | 0.794378798 |
| 0.564175442 | 0.794235824 |
| 0.564102743 | 0.794235824 |
| 0.564091882 | 0.794235824 |
| 0.564091411 | 0.794235824 |
| 0.564064722 | 0.794235824 |
| 0.563911945 | 0.794128746 |
| 0.563908503 | 0.794128746 |
| 0.563881318 | 0.794128746 |
| 0.563851964 | 0.794128746 |
| 0.56379912 | 0.79408726 |
| 0.56373424 | 0.794019361 |
| 0.563669904 | 0.794019361 |
| 0.563667511 | 0.794019361 |
| 0.563635774 | 0.794019361 |
| 0.563592275 | 0.794007255 |
| 0.563567635 | 0.793996029 |
| 0.563525439 | 0.793960067 |
| 0.563507587 | 0.793958402 |
| 0.563475137 | 0.793936169 |
| 0.563424192 | 0.79391136 |
| 0.563379025 | 0.793871204 |
| 0.56335333 | 0.793858484 |
| 0.563320148 | 0.793835213 |
| 0.563286652 | 0.793811498 |
| 0.56328182 | 0.793811498 |
| 0.563117459 | 0.793690486 |
| 0.563115562 | 0.793690486 |
| 0.56309747 | 0.793690486 |
| 0.562868501 | 0.793424178 |
| 0.562848968 | 0.793424178 |
| 0.562839877 | 0.793424178 |
| 0.562827996 | 0.793424178 |
| 0.562797078 | 0.793424178 |
| 0.562789168 | 0.793424178 |
| 0.562739766 | 0.793424178 |
| 0.562713132 | 0.793424178 |
| 0.562672533 | 0.793392093 |
| 0.562654693 | 0.793390428 |

| | |
|---|---|
| 0.562552222 | 0.793307036 |
| 0.562543745 | 0.793307036 |
| 0.562538977 | 0.793307036 |
| 0.562515995 | 0.793307036 |
| 0.562449026 | 0.793307036 |
| 0.562381107 | 0.7932395 |
| 0.56234957 | 0.7932395 |
| 0.562323398 | 0.7932395 |
| 0.562235047 | 0.79319792 |
| 0.562234186 | 0.79319792 |
| 0.562209908 | 0.79319792 |
| 0.562178551 | 0.793188703 |
| 0.5621756 | 0.793188703 |
| 0.562087768 | 0.793152097 |
| 0.562087543 | 0.793152097 |
| 0.562067068 | 0.793152097 |
| 0.562031709 | 0.793145995 |
| 0.561976097 | 0.793116858 |
| 0.561958836 | 0.793116858 |
| 0.561835456 | 0.792990449 |
| 0.561814202 | 0.792990449 |
| 0.561720759 | 0.792942132 |
| 0.561718592 | 0.792942132 |
| 0.5617164 | 0.792942132 |
| 0.561686952 | 0.792942132 |
| 0.561665033 | 0.792942132 |
| 0.561607205 | 0.792942132 |
| 0.561582944 | 0.792942132 |
| 0.561553446 | 0.792942132 |
| 0.561523261 | 0.792942132 |
| 0.561514961 | 0.792942132 |
| 0.561466342 | 0.79293552 |
| 0.561327152 | 0.792808094 |
| 0.561297487 | 0.792808094 |
| 0.561253279 | 0.792775666 |
| 0.561248232 | 0.792775666 |
| 0.561198223 | 0.792744922 |
| 0.561153693 | 0.79270553 |
| 0.561126713 | 0.792690928 |
| 0.561113821 | 0.792690928 |
| 0.561082539 | 0.792690928 |
| 0.561054534 | 0.79268301 |
| 0.561033213 | 0.792676402 |
| 0.561019028 | 0.792676402 |
| 0.560915137 | 0.792650661 |

| | |
|---|---|
| 0.560873227 | 0.792614954 |
| 0.560846962 | 0.792614954 |
| 0.5608123 | 0.792613298 |
| 0.560808196 | 0.792613298 |
| 0.560747367 | 0.792578194 |
| 0.560726718 | 0.792572529 |
| 0.560669226 | 0.792514787 |
| 0.56066438 | 0.792514787 |
| 0.560631793 | 0.792508916 |
| 0.56058517 | 0.792466531 |
| 0.560438096 | 0.792365858 |
| 0.56040001 | 0.792345889 |
| 0.560385917 | 0.792345889 |
| 0.560330716 | 0.792318478 |
| 0.560199781 | 0.792227418 |
| 0.560188746 | 0.792227418 |
| 0.560136785 | 0.792208898 |
| 0.560108931 | 0.792193029 |
| 0.560086863 | 0.792185342 |
| 0.560083105 | 0.792185342 |
| 0.560053075 | 0.792185342 |
| 0.560025277 | 0.792185342 |
| 0.559887904 | 0.792068592 |
| 0.559878399 | 0.792068592 |
| 0.559873563 | 0.792068592 |
| 0.559856162 | 0.792068592 |
| 0.559819319 | 0.792068592 |
| 0.559805903 | 0.792068592 |
| 0.559755747 | 0.792068592 |
| 0.559746704 | 0.792068592 |
| 0.559682266 | 0.792068592 |
| 0.559663759 | 0.792068592 |
| 0.55954229 | 0.791979572 |
| 0.559466339 | 0.791919141 |
| 0.559465725 | 0.791919141 |
| 0.55941968 | 0.791900166 |
| 0.559383651 | 0.791896238 |
| 0.559367715 | 0.791896238 |
| 0.559331663 | 0.791869715 |
| 0.559270171 | 0.791806196 |
| 0.559130255 | 0.791655171 |
| 0.559068525 | 0.791607432 |
| 0.559011293 | 0.791580866 |
| 0.559006629 | 0.791580866 |
| 0.558950206 | 0.791541432 |

| | |
|---|---|
| 0.558908989 | 0.791506598 |
| 0.55889482 | 0.791506598 |
| 0.55885614 | 0.791478824 |
| 0.558825376 | 0.79145879 |
| 0.558808935 | 0.79145879 |
| 0.558801455 | 0.79145879 |
| 0.558800708 | 0.79145879 |
| 0.558676253 | 0.791365253 |
| 0.558663716 | 0.791365253 |
| 0.558663278 | 0.791365253 |
| 0.558632413 | 0.791365253 |
| 0.558600097 | 0.791351522 |
| 0.558556639 | 0.791313494 |
| 0.558531777 | 0.791301811 |
| 0.558359532 | 0.791128383 |
| 0.558356222 | 0.791128383 |
| 0.558338124 | 0.791128383 |
| 0.558319132 | 0.791128383 |
| 0.558299301 | 0.791128383 |
| 0.558295629 | 0.791128383 |
| 0.558289046 | 0.791128383 |
| 0.558280573 | 0.791128383 |
| 0.558185635 | 0.791117344 |
| 0.558111946 | 0.791036445 |
| 0.558108072 | 0.791036445 |
| 0.558060594 | 0.791010741 |
| 0.558034893 | 0.791010741 |
| 0.557817972 | 0.790737441 |
| 0.557706031 | 0.790672891 |
| 0.557647916 | 0.790614035 |
| 0.557545931 | 0.790492977 |
| 0.557533331 | 0.790492977 |
| 0.557522151 | 0.790492977 |
| 0.557510592 | 0.790492977 |
| 0.557385124 | 0.790359095 |
| 0.557377572 | 0.790359095 |
| 0.557346032 | 0.790359095 |
| 0.557343636 | 0.790359095 |
| 0.557318774 | 0.790359095 |
| 0.557316397 | 0.790359095 |
| 0.55720123 | 0.790309269 |
| 0.55719701 | 0.790309269 |
| 0.557151216 | 0.790286237 |
| 0.556953568 | 0.790076475 |
| 0.556756529 | 0.789833504 |

| | |
|---|---|
| 0.556612047 | 0.789686087 |
| 0.556551612 | 0.789647391 |
| 0.556470149 | 0.78955533 |
| 0.556412298 | 0.78952029 |
| 0.556372384 | 0.789487175 |
| 0.556348629 | 0.789476989 |
| 0.556330098 | 0.789476989 |
| 0.55628974 | 0.789463988 |
| 0.556280612 | 0.789463988 |
| 0.556206566 | 0.789416519 |
| 0.55613607 | 0.789363511 |
| 0.555955133 | 0.789224295 |
| 0.555826678 | 0.789136028 |
| 0.555701566 | 0.789052487 |
| 0.55563837 | 0.789009799 |
| 0.555617893 | 0.789004246 |
| 0.555440175 | 0.788798915 |
| 0.555325053 | 0.788705979 |
| 0.555323146 | 0.788705979 |
| 0.555310995 | 0.788705979 |
| 0.555158551 | 0.788652676 |
| 0.555039322 | 0.78862941 |
| 0.555036393 | 0.78862941 |
| 0.555029169 | 0.78862941 |
| 0.554949612 | 0.78862941 |
| 0.554922324 | 0.788627862 |
| 0.554838436 | 0.788579236 |
| 0.554821467 | 0.788578651 |
| 0.554700995 | 0.788501545 |
| 0.554522409 | 0.788353063 |
| 0.554472985 | 0.788342135 |
| 0.55439254 | 0.788274823 |
| 0.554384786 | 0.788274823 |
| 0.554318743 | 0.788216957 |
| 0.554316743 | 0.788216957 |
| 0.55412461 | 0.788011489 |
| 0.554118024 | 0.788011489 |
| 0.554068178 | 0.788001825 |
| 0.553964809 | 0.787901868 |
| 0.553934218 | 0.787881888 |
| 0.553933597 | 0.787881888 |
| 0.553933209 | 0.787881888 |
| 0.553911468 | 0.787881888 |
| 0.553820967 | 0.787861978 |
| 0.553801025 | 0.787861978 |

| | |
|---|---|
| 0.553733225 | 0.787854821 |
| 0.553677428 | 0.787798968 |
| 0.553657535 | 0.787794198 |
| 0.553611025 | 0.787751554 |
| 0.553547757 | 0.787699113 |
| 0.553545197 | 0.787699113 |
| 0.553474523 | 0.787674987 |
| 0.55340953 | 0.787629562 |
| 0.553390056 | 0.787625383 |
| 0.553337386 | 0.787573955 |
| 0.55330293 | 0.787548448 |
| 0.553238419 | 0.787503697 |
| 0.553163901 | 0.787442999 |
| 0.553118875 | 0.787404133 |
| 0.553070988 | 0.787398641 |
| 0.553034266 | 0.787377824 |
| 0.552823389 | 0.787171715 |
| 0.55274883 | 0.787089081 |
| 0.552733502 | 0.787089081 |
| 0.552732179 | 0.787089081 |
| 0.552690758 | 0.787076986 |
| 0.552658161 | 0.787076986 |
| 0.55262292 | 0.787074516 |
| 0.552617318 | 0.787074516 |
| 0.552507095 | 0.787027229 |
| 0.552340532 | 0.78682298 |
| 0.552219025 | 0.786711003 |
| 0.552101265 | 0.786613826 |
| 0.552077826 | 0.786603962 |
| 0.551919397 | 0.786542941 |
| 0.551886009 | 0.78654243 |
| 0.551737906 | 0.786472563 |
| 0.55161881 | 0.786326331 |
| 0.551575443 | 0.786326331 |
| 0.551538938 | 0.78631598 |
| 0.551494768 | 0.786290715 |
| 0.551409354 | 0.786192471 |
| 0.551406031 | 0.786192471 |
| 0.551358538 | 0.786190625 |
| 0.551296573 | 0.786151099 |
| 0.551288 | 0.786151099 |
| 0.551229421 | 0.786124195 |
| 0.551186723 | 0.78608684 |
| 0.551138609 | 0.78604176 |
| 0.551083386 | 0.786010075 |

| | |
|---|---|
| 0.551011933 | 0.785955237 |
| 0.550899922 | 0.785819 |
| 0.550827427 | 0.785739124 |
| 0.550801055 | 0.785725038 |
| 0.550750348 | 0.785676236 |
| 0.550716216 | 0.785651078 |
| 0.550678122 | 0.785648689 |
| 0.550638325 | 0.785634089 |
| 0.550636174 | 0.785634089 |
| 0.550619993 | 0.785634089 |
| 0.550611848 | 0.785634089 |
| 0.550604619 | 0.785634089 |
| 0.550586187 | 0.785634089 |
| 0.550538823 | 0.785633333 |
| 0.550530914 | 0.785633333 |
| 0.550495588 | 0.785618714 |
| 0.550406857 | 0.785562703 |
| 0.550401802 | 0.785562703 |
| 0.550400501 | 0.785562703 |
| 0.550358914 | 0.785562703 |
| 0.550355349 | 0.785562703 |
| 0.550332215 | 0.785562703 |
| 0.550323259 | 0.785562703 |
| 0.550278466 | 0.785562703 |
| 0.550224861 | 0.785538353 |
| 0.550086244 | 0.785458178 |
| 0.550075553 | 0.785458178 |
| 0.550039037 | 0.785437869 |
| 0.549937159 | 0.785386581 |
| 0.549806347 | 0.785223309 |
| 0.549795563 | 0.785223309 |
| 0.549737781 | 0.785172474 |
| 0.549728107 | 0.785172474 |
| 0.549607676 | 0.785080823 |
| 0.549547278 | 0.785041638 |
| 0.549515748 | 0.785020143 |
| 0.549448801 | 0.784995142 |
| 0.54943204 | 0.784995142 |
| 0.549425488 | 0.784995142 |
| 0.549420498 | 0.784995142 |
| 0.549342775 | 0.784981746 |
| 0.549312172 | 0.784981746 |
| 0.549308082 | 0.784981746 |
| 0.54926809 | 0.784981746 |
| 0.549186483 | 0.784926467 |

| | |
|---|---|
| 0.549165373 | 0.784924511 |
| 0.549100885 | 0.784874785 |
| 0.549089781 | 0.784874785 |
| 0.549040032 | 0.784834912 |
| 0.548948822 | 0.784728084 |
| 0.548865966 | 0.78467747 |
| 0.54883137 | 0.78467747 |
| 0.548705481 | 0.78454506 |
| 0.54863434 | 0.784490444 |
| 0.548573207 | 0.784487425 |
| 0.548544445 | 0.784479665 |
| 0.548454894 | 0.784414487 |
| 0.548439586 | 0.784414487 |
| 0.548436685 | 0.784414487 |
| 0.54842176 | 0.784414487 |
| 0.548391817 | 0.784414487 |
| 0.548391491 | 0.784414487 |
| 0.548391255 | 0.784414487 |
| 0.548382303 | 0.784414487 |
| 0.5482632 | 0.784383642 |
| 0.548200719 | 0.784317812 |
| 0.548130581 | 0.78426184 |
| 0.54803741 | 0.78417839 |
| 0.548026959 | 0.78417839 |
| 0.548018558 | 0.78417839 |
| 0.547936595 | 0.784128368 |
| 0.547899459 | 0.784120868 |
| 0.547801945 | 0.784029908 |
| 0.547763989 | 0.784022704 |
| 0.547746301 | 0.78402095 |
| 0.547547537 | 0.783807116 |
| 0.54754073 | 0.783807116 |
| 0.54740029 | 0.783690552 |
| 0.547373759 | 0.783676127 |
| 0.547337477 | 0.783660755 |
| 0.547232016 | 0.783627403 |
| 0.547163127 | 0.783627403 |
| 0.547093477 | 0.783581061 |
| 0.54706749 | 0.783570653 |
| 0.546995792 | 0.783535403 |
| 0.546923526 | 0.783502581 |
| 0.546843508 | 0.783435079 |
| 0.546799731 | 0.783395926 |
| 0.546731031 | 0.783355717 |
| 0.546718821 | 0.783355717 |

| | |
|---|---|
| 0.546706292 | 0.783355717 |
| 0.546632262 | 0.78332094 |
| 0.546499464 | 0.783295614 |
| 0.546495861 | 0.783295614 |
| 0.546479898 | 0.783295614 |
| 0.546435564 | 0.78327536 |
| 0.546388326 | 0.783254178 |
| 0.546387456 | 0.783254178 |
| 0.546341026 | 0.783233522 |
| 0.546292039 | 0.783186868 |
| 0.54626389 | 0.783170087 |
| 0.54623754 | 0.783170087 |
| 0.546202476 | 0.783170087 |
| 0.546194332 | 0.783170087 |
| 0.546181061 | 0.783170087 |
| 0.546112325 | 0.783170087 |
| 0.546083499 | 0.783170087 |
| 0.546067058 | 0.783170087 |
| 0.546043548 | 0.783170087 |
| 0.546000449 | 0.783170087 |
| 0.54594385 | 0.783170087 |
| 0.545923643 | 0.783170087 |
| 0.54581102 | 0.783133743 |
| 0.545736644 | 0.783074208 |
| 0.545727325 | 0.783074208 |
| 0.545675944 | 0.783074208 |
| 0.545664019 | 0.783074208 |
| 0.545659295 | 0.783074208 |
| 0.545459008 | 0.78288811 |
| 0.545351473 | 0.782828132 |
| 0.545337252 | 0.782828132 |
| 0.545248679 | 0.782727758 |
| 0.545208039 | 0.782693009 |
| 0.545155854 | 0.782641684 |
| 0.545085165 | 0.782587379 |
| 0.54492946 | 0.782477159 |
| 0.544926946 | 0.782477159 |
| 0.544882438 | 0.782477159 |
| 0.54482085 | 0.782443548 |
| 0.544731031 | 0.782361955 |
| 0.544599262 | 0.782267065 |
| 0.544504416 | 0.782201599 |
| 0.544501902 | 0.782201599 |
| 0.544415591 | 0.78212118 |
| 0.544177607 | 0.781850037 |

| | |
|---|---|
| 0.544116203 | 0.781785398 |
| 0.544100848 | 0.781785398 |
| 0.544095344 | 0.781785398 |
| 0.544051369 | 0.781762998 |
| 0.544026903 | 0.781751427 |
| 0.543877238 | 0.781604132 |
| 0.543873088 | 0.781604132 |
| 0.543851184 | 0.781604132 |
| 0.543788961 | 0.781574579 |
| 0.543749229 | 0.781541059 |
| 0.54371136 | 0.7815338 |
| 0.543663127 | 0.7814937 |
| 0.543596989 | 0.781466559 |
| 0.543546388 | 0.781466559 |
| 0.54343306 | 0.781389875 |
| 0.543403344 | 0.781389875 |
| 0.543323865 | 0.78135412 |
| 0.543280623 | 0.781315525 |
| 0.543277969 | 0.781315525 |
| 0.543192179 | 0.781259102 |
| 0.543188627 | 0.781259102 |
| 0.543148031 | 0.781257189 |
| 0.543080868 | 0.781216965 |
| 0.543051978 | 0.781216965 |
| 0.543021085 | 0.781201753 |
| 0.543013897 | 0.781201753 |
| 0.542934165 | 0.781168861 |
| 0.542894316 | 0.781168861 |
| 0.542891424 | 0.781168861 |
| 0.542867796 | 0.781168861 |
| 0.542767648 | 0.781096685 |
| 0.542671742 | 0.780991419 |
| 0.542665429 | 0.780991419 |
| 0.542656632 | 0.780991419 |
| 0.542610664 | 0.780988761 |
| 0.542605316 | 0.780988761 |
| 0.542505015 | 0.780940364 |
| 0.542357487 | 0.780813133 |
| 0.542237736 | 0.780687928 |
| 0.54223589 | 0.780687928 |
| 0.542196018 | 0.780675064 |
| 0.542181001 | 0.780675064 |
| 0.542145529 | 0.780649568 |
| 0.542141909 | 0.780649568 |
| 0.542132491 | 0.780649568 |

| | |
|---|---|
| 0.542127793 | 0.780649568 |
| 0.54197523 | 0.780569546 |
| 0.541932053 | 0.780530963 |
| 0.541866904 | 0.78050794 |
| 0.541798901 | 0.780437851 |
| 0.541724915 | 0.780397826 |
| 0.541692712 | 0.780375039 |
| 0.541651247 | 0.780338907 |
| 0.541616403 | 0.780312313 |
| 0.541523381 | 0.780225499 |
| 0.541361039 | 0.780038791 |
| 0.541295976 | 0.779992238 |
| 0.541295203 | 0.779992238 |
| 0.541283453 | 0.779992238 |
| 0.541271859 | 0.779992238 |
| 0.541232323 | 0.779992238 |
| 0.541219003 | 0.779992238 |
| 0.541144096 | 0.779938564 |
| 0.540871087 | 0.77968346 |
| 0.540802398 | 0.779611229 |
| 0.540801213 | 0.779611229 |
| 0.540742749 | 0.779596032 |
| 0.540607198 | 0.779471392 |
| 0.540562259 | 0.779471207 |
| 0.540551599 | 0.779471207 |
| 0.540485334 | 0.779437261 |
| 0.540464016 | 0.779430119 |
| 0.54045074 | 0.779430119 |
| 0.540418338 | 0.779416379 |
| 0.54040965 | 0.779416379 |
| 0.540343454 | 0.779374248 |
| 0.540334887 | 0.779374248 |
| 0.540267697 | 0.779335784 |
| 0.540234312 | 0.779311231 |
| 0.54019928 | 0.779284299 |
| 0.540071663 | 0.779171002 |
| 0.540010972 | 0.779165536 |
| 0.539998367 | 0.779165536 |
| 0.539904548 | 0.779104285 |
| 0.539884542 | 0.779104285 |
| 0.539835928 | 0.779090504 |
| 0.539810858 | 0.779090504 |
| 0.539746581 | 0.779055987 |
| 0.539746305 | 0.779055987 |
| 0.539649323 | 0.778986431 |

| | |
|---|---|
| 0.539633648 | 0.778986431 |
| 0.539562119 | 0.7789786 |
| 0.539554601 | 0.7789786 |
| 0.53943167 | 0.778837486 |
| 0.539391416 | 0.77881278 |
| 0.539390201 | 0.77881278 |
| 0.539382454 | 0.77881278 |
| 0.539337002 | 0.77881278 |
| 0.539302902 | 0.778793224 |
| 0.539293407 | 0.778793224 |
| 0.539214476 | 0.778712752 |
| 0.539172604 | 0.778712752 |
| 0.539137443 | 0.778712752 |
| 0.539094272 | 0.778704443 |
| 0.539081874 | 0.778704443 |
| 0.538992046 | 0.778674862 |
| 0.53898442 | 0.778674862 |
| 0.538952616 | 0.778665136 |
| 0.538940507 | 0.778665136 |
| 0.538858164 | 0.778579566 |
| 0.538848562 | 0.778579566 |
| 0.538814635 | 0.778579566 |
| 0.538711617 | 0.778482255 |
| 0.538597245 | 0.778482255 |
| 0.538573068 | 0.778482255 |
| 0.538551066 | 0.778482255 |
| 0.538548865 | 0.778482255 |
| 0.538507054 | 0.778482255 |
| 0.538409297 | 0.778470539 |
| 0.538405008 | 0.778470539 |
| 0.53840311 | 0.778470539 |
| 0.538224749 | 0.778289015 |
| 0.538114274 | 0.778162092 |
| 0.538086827 | 0.778146029 |
| 0.53805261 | 0.778120174 |
| 0.538026493 | 0.77810603 |
| 0.538013945 | 0.77810603 |
| 0.537921111 | 0.778071757 |
| 0.537913569 | 0.778071757 |
| 0.537873771 | 0.77805968 |
| 0.537712826 | 0.777983133 |
| 0.537677659 | 0.777955884 |
| 0.537662644 | 0.777955884 |
| 0.537613549 | 0.777920164 |
| 0.537588272 | 0.777920164 |

| | |
|---|---|
| 0.537547133 | 0.777920164 |
| 0.537427523 | 0.777782997 |
| 0.537364458 | 0.777715359 |
| 0.537346077 | 0.777715359 |
| 0.537308679 | 0.777705529 |
| 0.537297072 | 0.777705529 |
| 0.537270393 | 0.777705529 |
| 0.537241415 | 0.777705529 |
| 0.537182763 | 0.777694583 |
| 0.537157611 | 0.777694583 |
| 0.537081251 | 0.777636308 |
| 0.537068768 | 0.777636308 |
| 0.5370203 | 0.777605649 |
| 0.536962711 | 0.77758287 |
| 0.536949708 | 0.77758287 |
| 0.536935875 | 0.77758287 |
| 0.536864049 | 0.777534561 |
| 0.536738706 | 0.777400309 |
| 0.53669456 | 0.777360009 |
| 0.536690371 | 0.777360009 |
| 0.536687533 | 0.777360009 |
| 0.536683138 | 0.777360009 |
| 0.536671038 | 0.777360009 |
| 0.536662828 | 0.777360009 |
| 0.536636885 | 0.777360009 |
| 0.536633247 | 0.777360009 |
| 0.536584558 | 0.777360009 |
| 0.536567723 | 0.777360009 |
| 0.536417086 | 0.777265403 |
| 0.536350969 | 0.777193249 |
| 0.536335603 | 0.777193249 |
| 0.536305571 | 0.777193249 |
| 0.536280152 | 0.777185211 |
| 0.536247558 | 0.777161623 |
| 0.536244318 | 0.777161623 |
| 0.53618793 | 0.777122502 |
| 0.536139567 | 0.777099704 |
| 0.536132359 | 0.777099704 |
| 0.53612265 | 0.777099704 |
| 0.536098894 | 0.777099704 |
| 0.535995526 | 0.776985509 |
| 0.535964286 | 0.776963872 |
| 0.535942548 | 0.776963872 |
| 0.535929903 | 0.776963872 |
| 0.535835935 | 0.776919693 |

| | |
|---|---|
| 0.535788281 | 0.776874249 |
| 0.535748014 | 0.776857775 |
| 0.535738197 | 0.776857775 |
| 0.535706497 | 0.776850268 |
| 0.535660942 | 0.776831512 |
| 0.535632135 | 0.776830763 |
| 0.535604138 | 0.776830763 |
| 0.535518115 | 0.776789952 |
| 0.535365597 | 0.77664979 |
| 0.535330348 | 0.77664979 |
| 0.535086702 | 0.776447942 |
| 0.535062957 | 0.776437143 |
| 0.534970921 | 0.776350896 |
| 0.534902287 | 0.776298603 |
| 0.534858917 | 0.77628297 |
| 0.53480323 | 0.776227319 |
| 0.534753482 | 0.776200905 |
| 0.534716806 | 0.776171324 |
| 0.534686249 | 0.776150625 |
| 0.534676298 | 0.776150625 |
| 0.534649698 | 0.776144879 |
| 0.534628851 | 0.776144879 |
| 0.534622581 | 0.776144879 |
| 0.534520269 | 0.776084187 |
| 0.534496892 | 0.776084187 |
| 0.534492761 | 0.776084187 |
| 0.534474418 | 0.776084187 |
| 0.534464816 | 0.776084187 |
| 0.534387444 | 0.776048018 |
| 0.534377595 | 0.776048018 |
| 0.534313604 | 0.775988109 |
| 0.534300015 | 0.775988109 |
| 0.534275553 | 0.775980171 |
| 0.534175028 | 0.775857829 |
| 0.534105996 | 0.775857829 |
| 0.534097795 | 0.775857829 |
| 0.534041026 | 0.775852482 |
| 0.533969865 | 0.775796429 |
| 0.533947165 | 0.775787114 |
| 0.533911653 | 0.775759184 |
| 0.533894771 | 0.775758321 |
| 0.533698671 | 0.775591695 |
| 0.533644086 | 0.775559698 |
| 0.533599512 | 0.775542249 |
| 0.533576251 | 0.775532107 |

| | |
|---|---|
| 0.533574024 | 0.775532107 |
| 0.53355362 | 0.775532107 |
| 0.533533442 | 0.775532107 |
| 0.533441929 | 0.775455201 |
| 0.53331679 | 0.775339465 |
| 0.533313105 | 0.775339465 |
| 0.533236394 | 0.775339465 |
| 0.533226734 | 0.775339465 |
| 0.533220786 | 0.775339465 |
| 0.53309506 | 0.77528225 |
| 0.533014391 | 0.775188604 |
| 0.532917698 | 0.775095316 |
| 0.532820786 | 0.775001697 |
| 0.532820307 | 0.775001697 |
| 0.532749767 | 0.774993072 |
| 0.532674529 | 0.774930961 |
| 0.532433993 | 0.774710661 |
| 0.532317823 | 0.774601312 |
| 0.532179481 | 0.774447328 |
| 0.532172063 | 0.774447328 |
| 0.532145168 | 0.77444472 |
| 0.532014836 | 0.774349691 |
| 0.531954919 | 0.774286144 |
| 0.531876972 | 0.774220012 |
| 0.531841481 | 0.774192011 |
| 0.531757956 | 0.774165072 |
| 0.531673003 | 0.774065055 |
| 0.531650012 | 0.774055244 |
| 0.531624616 | 0.77404193 |
| 0.531578023 | 0.773997752 |
| 0.531560225 | 0.7739955 |
| 0.53155616 | 0.7739955 |
| 0.531493406 | 0.77394553 |
| 0.531460832 | 0.773921758 |
| 0.531365153 | 0.77382975 |
| 0.531301731 | 0.773808371 |
| 0.53124813 | 0.773777628 |
| 0.531160619 | 0.773673827 |
| 0.531082828 | 0.773631498 |
| 0.531049446 | 0.773630196 |
| 0.530968454 | 0.773583193 |
| 0.530936321 | 0.77356004 |
| 0.530872697 | 0.773491003 |
| 0.530852821 | 0.773485707 |
| 0.530838809 | 0.773485707 |

| | |
|---|---|
| 0.530789017 | 0.773440064 |
| 0.530637287 | 0.773351388 |
| 0.530592879 | 0.773343546 |
| 0.530533302 | 0.773280376 |
| 0.530523253 | 0.773280376 |
| 0.530508088 | 0.773280376 |
| 0.530488313 | 0.773280376 |
| 0.530452362 | 0.773280376 |
| 0.530426518 | 0.773266721 |
| 0.53027428 | 0.773115778 |
| 0.530246658 | 0.773099174 |
| 0.530245971 | 0.773099174 |
| 0.530162028 | 0.773070446 |
| 0.530148168 | 0.773070446 |
| 0.530074171 | 0.772989667 |
| 0.530000814 | 0.772930025 |
| 0.529949042 | 0.772901856 |
| 0.529886687 | 0.772882925 |
| 0.529825483 | 0.772882925 |
| 0.529816349 | 0.772882925 |
| 0.529806607 | 0.772882925 |
| 0.529750824 | 0.772863328 |
| 0.529733801 | 0.772863328 |
| 0.529721353 | 0.772863328 |
| 0.529682479 | 0.772863328 |
| 0.529646549 | 0.772863328 |
| 0.529642388 | 0.772863328 |
| 0.52960598 | 0.772863328 |
| 0.529557829 | 0.772828274 |
| 0.529540269 | 0.772828274 |
| 0.529404419 | 0.772722779 |
| 0.529257438 | 0.772543394 |
| 0.529123695 | 0.772455071 |
| 0.529072815 | 0.772428146 |
| 0.529068735 | 0.772428146 |
| 0.528994729 | 0.772385173 |
| 0.52895695 | 0.77238122 |
| 0.528845867 | 0.77230988 |
| 0.528782994 | 0.772265421 |
| 0.528781523 | 0.772265421 |
| 0.528686977 | 0.772219912 |
| 0.528618857 | 0.772167774 |
| 0.528595674 | 0.772167774 |
| 0.528541655 | 0.772126048 |
| 0.528503779 | 0.772094399 |

| | |
|---|---|
| 0.528455108 | 0.772077647 |
| 0.528407664 | 0.77204871 |
| 0.528385948 | 0.772040666 |
| 0.52824204 | 0.771901441 |
| 0.5281986 | 0.771885328 |
| 0.528140506 | 0.771847797 |
| 0.528135421 | 0.771847797 |
| 0.528111602 | 0.771847797 |
| 0.528107441 | 0.771847797 |
| 0.52803989 | 0.771820415 |
| 0.528035092 | 0.771820415 |
| 0.528008341 | 0.771820415 |
| 0.527822324 | 0.771714316 |
| 0.527738594 | 0.771639273 |
| 0.527731921 | 0.771639273 |
| 0.527686138 | 0.771630949 |
| 0.527665123 | 0.771630949 |
| 0.527653366 | 0.771630949 |
| 0.52752011 | 0.77153299 |
| 0.527509482 | 0.77153299 |
| 0.527461287 | 0.771518035 |
| 0.527388792 | 0.771459381 |
| 0.5273467 | 0.771421502 |
| 0.527282134 | 0.771350745 |
| 0.527219129 | 0.771282267 |
| 0.52721549 | 0.771282267 |
| 0.527166455 | 0.771282267 |
| 0.526998374 | 0.771148813 |
| 0.526901898 | 0.77103133 |
| 0.52673685 | 0.770813493 |
| 0.526623932 | 0.770720101 |
| 0.526603764 | 0.770720101 |
| 0.526491296 | 0.77066729 |
| 0.52648377 | 0.77066729 |
| 0.526465562 | 0.77066729 |
| 0.526433431 | 0.77066729 |
| 0.526432595 | 0.77066729 |
| 0.526432391 | 0.77066729 |
| 0.526412087 | 0.77066729 |
| 0.526272541 | 0.770631278 |
| 0.526230082 | 0.770627325 |
| 0.526202591 | 0.770623621 |
| 0.526078514 | 0.770560382 |
| 0.52599286 | 0.770506012 |
| 0.525824704 | 0.770283377 |

| | |
|---|---|
| 0.525787659 | 0.770276495 |
| 0.525674666 | 0.770173253 |
| 0.525652928 | 0.770173253 |
| 0.525587305 | 0.77010999 |
| 0.525586161 | 0.77010999 |
| 0.525499956 | 0.770044502 |
| 0.525490286 | 0.770044502 |
| 0.525478347 | 0.770044502 |
| 0.525406352 | 0.770002102 |
| 0.525361461 | 0.7699837 |
| 0.525259931 | 0.769858585 |
| 0.525167671 | 0.769791091 |
| 0.525132408 | 0.769766434 |
| 0.525042277 | 0.769681693 |
| 0.524993129 | 0.76964684 |
| 0.524967659 | 0.76964684 |
| 0.524909611 | 0.769629346 |
| 0.524676863 | 0.769453903 |
| 0.524647613 | 0.769447437 |
| 0.524583317 | 0.769387787 |
| 0.524579401 | 0.769387787 |
| 0.524575996 | 0.769387787 |
| 0.5245139 | 0.769387787 |
| 0.524448663 | 0.769380144 |
| 0.524412762 | 0.769374569 |
| 0.524383943 | 0.769374316 |
| 0.52438152 | 0.769374316 |
| 0.524346607 | 0.769372311 |
| 0.524332371 | 0.769372311 |
| 0.524256565 | 0.769287593 |
| 0.524138419 | 0.769183322 |
| 0.523986758 | 0.769002239 |
| 0.52398008 | 0.769002239 |
| 0.523927465 | 0.768971691 |
| 0.523785366 | 0.768833039 |
| 0.52370764 | 0.768772871 |
| 0.523705096 | 0.768772871 |
| 0.523677709 | 0.768772871 |
| 0.523533974 | 0.768673945 |
| 0.523523347 | 0.768673945 |
| 0.523406222 | 0.768563348 |
| 0.523386092 | 0.768563348 |
| 0.523359779 | 0.768563348 |
| 0.523339868 | 0.768558104 |
| 0.523208527 | 0.768460004 |

| | |
|---|---|
| 0.523102734 | 0.768346389 |
| 0.522925649 | 0.768234057 |
| 0.522893238 | 0.768210138 |
| 0.522885914 | 0.768210138 |
| 0.522818344 | 0.768210138 |
| 0.522812646 | 0.768210138 |
| 0.522759203 | 0.768179092 |
| 0.522756153 | 0.768179092 |
| 0.522731544 | 0.768179092 |
| 0.522678396 | 0.768131448 |
| 0.522594997 | 0.768039186 |
| 0.522580297 | 0.768039186 |
| 0.522541229 | 0.768039186 |
| 0.522483304 | 0.767986924 |
| 0.522474003 | 0.767986924 |
| 0.522472932 | 0.767986924 |
| 0.522464104 | 0.767986924 |
| 0.522324821 | 0.767919872 |
| 0.522223315 | 0.767865452 |
| 0.522216906 | 0.767865452 |
| 0.522210633 | 0.767865452 |
| 0.522177838 | 0.767865452 |
| 0.522121434 | 0.767815088 |
| 0.522115841 | 0.767815088 |
| 0.521959795 | 0.767667577 |
| 0.521794507 | 0.767471881 |
| 0.521778905 | 0.767471881 |
| 0.521771263 | 0.767471881 |
| 0.521735687 | 0.767471881 |
| 0.521664526 | 0.7674229 |
| 0.52156076 | 0.767293949 |
| 0.521456688 | 0.767188239 |
| 0.521327161 | 0.767045063 |
| 0.521322621 | 0.767045063 |
| 0.521275426 | 0.767040032 |
| 0.521216126 | 0.766976469 |
| 0.521212441 | 0.766976469 |
| 0.521172172 | 0.766976469 |
| 0.521033452 | 0.766849821 |
| 0.520866813 | 0.766675641 |
| 0.52078513 | 0.766618084 |
| 0.520782314 | 0.766618084 |
| 0.520723086 | 0.766582542 |
| 0.520680884 | 0.766544108 |
| 0.520646823 | 0.766544108 |

| | |
|---|---|
| 0.520644002 | 0.766544108 |
| 0.520568387 | 0.766544108 |
| 0.520522782 | 0.766544108 |
| 0.52043106 | 0.766460624 |
| 0.520415223 | 0.766460624 |
| 0.520365415 | 0.766411343 |
| 0.520365218 | 0.766411343 |
| 0.520267837 | 0.766315024 |
| 0.520208583 | 0.766252185 |
| 0.520189899 | 0.766252185 |
| 0.520136777 | 0.766252185 |
| 0.520086305 | 0.766238826 |
| 0.520077993 | 0.766238826 |
| 0.520054009 | 0.766238826 |
| 0.519977887 | 0.766238826 |
| 0.519915953 | 0.766175844 |
| 0.519914281 | 0.766175844 |
| 0.519886813 | 0.766175844 |
| 0.519864327 | 0.766175844 |
| 0.51986402 | 0.766175844 |
| 0.519825396 | 0.766175844 |
| 0.519791141 | 0.766175844 |
| 0.519673869 | 0.766079836 |
| 0.519657925 | 0.766079836 |
| 0.519573402 | 0.766002861 |
| 0.519571099 | 0.766002861 |
| 0.519378951 | 0.765858426 |
| 0.519368375 | 0.765858426 |
| 0.519366796 | 0.765858426 |
| 0.519312211 | 0.765858426 |
| 0.519305974 | 0.765858426 |
| 0.519277254 | 0.765858426 |
| 0.5192729 | 0.765858426 |
| 0.519235323 | 0.765858426 |
| 0.519227222 | 0.765858426 |
| 0.51919335 | 0.765858426 |
| 0.519182084 | 0.765858426 |
| 0.519165826 | 0.765858426 |
| 0.519147852 | 0.765858426 |
| 0.519128066 | 0.765858426 |
| 0.51908157 | 0.765858426 |
| 0.51905796 | 0.765858426 |
| 0.518915154 | 0.765719792 |
| 0.518904228 | 0.765719792 |
| 0.518895457 | 0.765719792 |

| | |
|---|---|
| 0.518866074 | 0.765718543 |
| 0.51876342 | 0.765638222 |
| 0.518664687 | 0.7655874 |
| 0.51863913 | 0.7655874 |
| 0.518610666 | 0.7655874 |
| 0.518601321 | 0.7655874 |
| 0.518456493 | 0.765493611 |
| 0.518426859 | 0.765493611 |
| 0.518345416 | 0.765448255 |
| 0.518345364 | 0.765448255 |
| 0.518316162 | 0.765448255 |
| 0.518282828 | 0.765427029 |
| 0.518274644 | 0.765427029 |
| 0.518233965 | 0.765402333 |
| 0.518223774 | 0.765402333 |
| 0.518195225 | 0.765402333 |
| 0.518125699 | 0.765364465 |
| 0.518121968 | 0.765364465 |
| 0.518097293 | 0.765364465 |
| 0.51806321 | 0.765363748 |
| 0.518062934 | 0.765363748 |
| 0.518061037 | 0.765363748 |
| 0.517973819 | 0.765302904 |
| 0.517935845 | 0.765270537 |
| 0.517833993 | 0.765161343 |
| 0.517817019 | 0.765161343 |
| 0.517745101 | 0.765154867 |
| 0.517725114 | 0.765149072 |
| 0.517565267 | 0.764960308 |
| 0.517505351 | 0.764923913 |
| 0.517458504 | 0.76489746 |
| 0.517383893 | 0.764834648 |
| 0.517320436 | 0.764764577 |
| 0.517207156 | 0.764644583 |
| 0.517059248 | 0.764497107 |
| 0.516973951 | 0.764418453 |
| 0.516889219 | 0.764316894 |
| 0.51678689 | 0.764272784 |
| 0.516771407 | 0.764272784 |
| 0.516675125 | 0.764190124 |
| 0.516587061 | 0.764162004 |
| 0.516445174 | 0.764063565 |
| 0.516397674 | 0.764017025 |
| 0.516345462 | 0.763994749 |
| 0.516339806 | 0.763994749 |

| | |
|---|---|
| 0.516326072 | 0.763994749 |
| 0.516205149 | 0.763920654 |
| 0.516182469 | 0.763912202 |
| 0.516168976 | 0.763912202 |
| 0.516159696 | 0.763912202 |
| 0.516146335 | 0.763912202 |
| 0.516134459 | 0.763912202 |
| 0.516081758 | 0.763912202 |
| 0.515975433 | 0.763843154 |
| 0.515948808 | 0.763827481 |
| 0.515935171 | 0.763827481 |
| 0.515912392 | 0.763827481 |
| 0.515906936 | 0.763827481 |
| 0.515852725 | 0.763827481 |
| 0.515828423 | 0.763815455 |
| 0.515760709 | 0.763738933 |
| 0.51574443 | 0.763738572 |
| 0.515558824 | 0.763534939 |
| 0.515499496 | 0.763470816 |
| 0.515389324 | 0.763355123 |
| 0.515317518 | 0.763296244 |
| 0.515235443 | 0.763198409 |
| 0.515208418 | 0.763182114 |
| 0.515149273 | 0.763120734 |
| 0.515097343 | 0.763088779 |
| 0.515044531 | 0.763034276 |
| 0.514964236 | 0.762999031 |
| 0.514960447 | 0.762999031 |
| 0.514952638 | 0.762999031 |
| 0.514939095 | 0.762999031 |
| 0.514926499 | 0.762999031 |
| 0.514901651 | 0.762999031 |
| 0.514857716 | 0.762994849 |
| 0.514773735 | 0.76291787 |
| 0.514706314 | 0.762841687 |
| 0.514544956 | 0.762626272 |
| 0.514498958 | 0.762581827 |
| 0.514342167 | 0.762420616 |
| 0.514323982 | 0.762420616 |
| 0.51431341 | 0.762420616 |
| 0.514284275 | 0.762420616 |
| 0.514254625 | 0.76241074 |
| 0.514125182 | 0.762312508 |
| 0.514121015 | 0.762312508 |
| 0.514033951 | 0.762224696 |

| | |
|---|---|
| 0.513989914 | 0.762199405 |
| 0.513878307 | 0.762153849 |
| 0.513859761 | 0.762153849 |
| 0.51385639 | 0.762153849 |
| 0.513779295 | 0.762153849 |
| 0.513745877 | 0.762153466 |
| 0.513734255 | 0.762153466 |
| 0.51372052 | 0.762153466 |
| 0.513677585 | 0.762123371 |
| 0.513641729 | 0.762093914 |
| 0.513629095 | 0.762093914 |
| 0.513483885 | 0.761978408 |
| 0.513473107 | 0.761978408 |
| 0.513415332 | 0.761928841 |
| 0.513360944 | 0.761927187 |
| 0.513241428 | 0.761784763 |
| 0.513196609 | 0.761784763 |
| 0.513189819 | 0.761784763 |
| 0.513118481 | 0.761784763 |
| 0.513112849 | 0.761784763 |
| 0.513099398 | 0.761784763 |
| 0.513090279 | 0.761784763 |
| 0.513081624 | 0.761784763 |
| 0.513026416 | 0.761784763 |
| 0.51300956 | 0.761784763 |
| 0.512976685 | 0.761784763 |
| 0.512944904 | 0.761784763 |
| 0.512935878 | 0.761784763 |
| 0.512908031 | 0.761784763 |
| 0.512832182 | 0.761770837 |
| 0.512820767 | 0.761770837 |
| 0.512727703 | 0.76168691 |
| 0.512713808 | 0.76168691 |
| 0.512602935 | 0.76159658 |
| 0.512531935 | 0.761514849 |
| 0.512434581 | 0.761441464 |
| 0.512413817 | 0.761434366 |
| 0.51233262 | 0.761337463 |
| 0.512286446 | 0.761306279 |
| 0.512156335 | 0.761218006 |
| 0.512147958 | 0.761218006 |
| 0.512140216 | 0.761218006 |
| 0.512000975 | 0.761082099 |
| 0.51196524 | 0.761076486 |
| 0.511930912 | 0.76104921 |

| | |
|---|---|
| 0.511874516 | 0.760989124 |
| 0.511874241 | 0.760989124 |
| 0.511724021 | 0.760860388 |
| 0.511714685 | 0.760860388 |
| 0.511657859 | 0.760857027 |
| 0.511651541 | 0.760857027 |
| 0.511613488 | 0.760857027 |
| 0.511604005 | 0.760857027 |
| 0.51154815 | 0.760857027 |
| 0.511519586 | 0.760857027 |
| 0.511482312 | 0.760857027 |
| 0.511453879 | 0.760838764 |
| 0.51136235 | 0.760726367 |
| 0.511300643 | 0.76068209 |
| 0.511245604 | 0.760623966 |
| 0.511181524 | 0.760552387 |
| 0.511107666 | 0.760490012 |
| 0.511058349 | 0.760464148 |
| 0.511043391 | 0.760464148 |
| 0.510995823 | 0.760418624 |
| 0.510961911 | 0.760391918 |
| 0.510895565 | 0.760388221 |
| 0.510889919 | 0.760388221 |
| 0.510859017 | 0.760388221 |
| 0.510790874 | 0.760327446 |
| 0.510727695 | 0.76027702 |
| 0.510657476 | 0.760223069 |
| 0.510513043 | 0.760080174 |
| 0.510390339 | 0.759921242 |
| 0.510378932 | 0.759921242 |
| 0.510291787 | 0.759822013 |
| 0.510268182 | 0.759822013 |
| 0.510176361 | 0.759768912 |
| 0.510147487 | 0.759768912 |
| 0.510064537 | 0.759697399 |
| 0.510007489 | 0.759636185 |
| 0.509904075 | 0.75951758 |
| 0.509901538 | 0.75951758 |
| 0.509878034 | 0.75951758 |
| 0.509873474 | 0.75951758 |
| 0.509856758 | 0.75951758 |
| 0.509735085 | 0.759444205 |
| 0.509684825 | 0.759407706 |
| 0.509673889 | 0.759407706 |
| 0.509604911 | 0.759369035 |

| | |
|---|---|
| 0.509593225 | 0.759369035 |
| 0.50956541 | 0.75935769 |
| 0.509473643 | 0.759244695 |
| 0.509334443 | 0.759108509 |
| 0.509282982 | 0.75907932 |
| 0.509276244 | 0.75907932 |
| 0.509096584 | 0.758872743 |
| 0.509081012 | 0.758872743 |
| 0.509071482 | 0.758872743 |
| 0.509066483 | 0.758872743 |
| 0.508999027 | 0.758869828 |
| 0.508973633 | 0.758855723 |
| 0.508932628 | 0.758818341 |
| 0.508854471 | 0.758749316 |
| 0.508840214 | 0.758749316 |
| 0.508836129 | 0.758749316 |
| 0.508817787 | 0.758749316 |
| 0.508760843 | 0.758740054 |
| 0.50870602 | 0.758717992 |
| 0.508688694 | 0.758717992 |
| 0.508664051 | 0.758717992 |
| 0.508515771 | 0.758553014 |
| 0.508447286 | 0.75847461 |
| 0.508423274 | 0.758462546 |
| 0.508383363 | 0.758462546 |
| 0.508220381 | 0.758302384 |
| 0.508149795 | 0.758244574 |
| 0.508143097 | 0.758244574 |
| 0.507944253 | 0.758032867 |
| 0.507937015 | 0.758032867 |
| 0.507927368 | 0.758032867 |
| 0.507922875 | 0.758032867 |
| 0.507905944 | 0.758032867 |
| 0.507896616 | 0.758032867 |
| 0.507834872 | 0.758017282 |
| 0.507826555 | 0.758017282 |
| 0.50778112 | 0.758017282 |
| 0.507764444 | 0.758017282 |
| 0.507708898 | 0.757966637 |
| 0.50759139 | 0.757862476 |
| 0.507541842 | 0.757836013 |
| 0.507521397 | 0.757829244 |
| 0.507481503 | 0.757793433 |
| 0.507357633 | 0.757679731 |
| 0.507342937 | 0.757679731 |

| | |
|---|---|
| 0.507279272 | 0.757629373 |
| 0.50726081 | 0.757629373 |
| 0.507174206 | 0.757572085 |
| 0.50704243 | 0.757422759 |
| 0.507014971 | 0.757405496 |
| 0.507001678 | 0.757405496 |
| 0.506938506 | 0.757338778 |
| 0.506920015 | 0.757334909 |
| 0.506911622 | 0.757334909 |
| 0.506774908 | 0.75721313 |
| 0.506706701 | 0.757158725 |
| 0.506705187 | 0.757158725 |
| 0.50660545 | 0.757078688 |
| 0.506505778 | 0.756953488 |
| 0.506438201 | 0.756908114 |
| 0.50613832 | 0.756616208 |
| 0.506094485 | 0.756576228 |
| 0.506086094 | 0.756576228 |
| 0.506085315 | 0.756576228 |
| 0.506060749 | 0.756576228 |
| 0.506056706 | 0.756576228 |
| 0.505988543 | 0.756576228 |
| 0.505860202 | 0.756439682 |
| 0.505822472 | 0.756412423 |
| 0.505650613 | 0.756221245 |
| 0.505543392 | 0.756132121 |
| 0.505518047 | 0.756117958 |
| 0.505479833 | 0.756084546 |
| 0.505466579 | 0.756084546 |
| 0.505459978 | 0.756084546 |
| 0.505444857 | 0.756084546 |
| 0.505431549 | 0.756084546 |
| 0.505388344 | 0.756084546 |
| 0.505350982 | 0.756058027 |
| 0.505154073 | 0.755810915 |
| 0.505094901 | 0.755769867 |
| 0.504984068 | 0.755651508 |
| 0.504926131 | 0.755588552 |
| 0.50491602 | 0.755588552 |
| 0.504889828 | 0.755581708 |
| 0.504819318 | 0.7555268 |
| 0.504813068 | 0.7555268 |
| 0.504784512 | 0.7555268 |
| 0.504670699 | 0.755447724 |
| 0.504670349 | 0.755447724 |

| | |
|---|---|
| 0.504520676 | 0.755361548 |
| 0.504520061 | 0.755361548 |
| 0.504422435 | 0.755326475 |
| 0.504341027 | 0.755282524 |
| 0.504337191 | 0.755282524 |
| 0.504245812 | 0.755225552 |
| 0.504244564 | 0.755225552 |
| 0.504217909 | 0.755225552 |
| 0.504166063 | 0.755225552 |
| 0.504139793 | 0.75521863 |
| 0.504132817 | 0.75521863 |
| 0.504079852 | 0.755209196 |
| 0.504073014 | 0.755209196 |
| 0.50402427 | 0.755188093 |
| 0.503901293 | 0.755027587 |
| 0.503891531 | 0.755027587 |
| 0.503821257 | 0.754955166 |
| 0.503725482 | 0.754859151 |
| 0.503724898 | 0.754859151 |
| 0.503652788 | 0.754821465 |
| 0.503637317 | 0.754821465 |
| 0.503569222 | 0.754814985 |
| 0.503556642 | 0.754814985 |
| 0.503501449 | 0.754760906 |
| 0.503494856 | 0.754760906 |
| 0.503449565 | 0.754730639 |
| 0.503419269 | 0.754708978 |
| 0.503288376 | 0.754584004 |
| 0.503266495 | 0.754574952 |
| 0.503195561 | 0.754516103 |
| 0.503154453 | 0.754478217 |
| 0.503129528 | 0.754464595 |
| 0.503047987 | 0.754366073 |
| 0.502987641 | 0.754299329 |
| 0.502982724 | 0.754299329 |
| 0.502911252 | 0.754279773 |
| 0.502781678 | 0.754109182 |
| 0.502779074 | 0.754109182 |
| 0.502763255 | 0.754109182 |
| 0.502746952 | 0.754109182 |
| 0.502707353 | 0.754109182 |
| 0.502698672 | 0.754109182 |
| 0.502674087 | 0.754109182 |
| 0.502662322 | 0.754109182 |
| 0.502602464 | 0.754101614 |

| | |
|---|---|
| 0.502582956 | 0.754101614 |
| 0.502535887 | 0.754096748 |
| 0.502391028 | 0.753998157 |
| 0.502326398 | 0.753996198 |
| 0.50232553 | 0.753996198 |
| 0.502308477 | 0.753996198 |
| 0.502288513 | 0.753996198 |
| 0.502174307 | 0.753934234 |
| 0.50217279 | 0.753934234 |
| 0.502161323 | 0.753934234 |
| 0.502142987 | 0.753934234 |
| 0.502124143 | 0.753934234 |
| 0.502103832 | 0.753934234 |
| 0.502067126 | 0.753915913 |
| 0.50193055 | 0.753734592 |
| 0.501924422 | 0.753734592 |
| 0.501815231 | 0.75365647 |
| 0.501743919 | 0.753573133 |
| 0.501721213 | 0.753573133 |
| 0.501709482 | 0.753573133 |
| 0.501699593 | 0.753573133 |
| 0.501645155 | 0.753573133 |
| 0.501602354 | 0.75355062 |
| 0.501385982 | 0.753273089 |
| 0.501383789 | 0.753273089 |
| 0.50133288 | 0.75324083 |
| 0.501300778 | 0.753216359 |
| 0.501276871 | 0.7532042 |
| 0.501191876 | 0.753124009 |
| 0.501181959 | 0.753124009 |
| 0.501145747 | 0.753123251 |
| 0.501043094 | 0.753019231 |
| 0.500951847 | 0.752953375 |
| 0.500908515 | 0.752935769 |
| 0.500805908 | 0.752876581 |
| 0.500745079 | 0.752832662 |
| 0.500725407 | 0.75282685 |
| 0.500554669 | 0.75266519 |
| 0.500518612 | 0.752634734 |
| 0.500513982 | 0.752634734 |
| 0.500489998 | 0.752634734 |
| 0.500468643 | 0.752634734 |
| 0.500457139 | 0.752634734 |
| 0.500380978 | 0.752594103 |
| 0.500343465 | 0.752561447 |

| | |
|---|---|
| 0.500298842 | 0.752541861 |
| 0.500264832 | 0.752538239 |
| 0.500163678 | 0.752452698 |
| 0.500050965 | 0.752369307 |
| 0.500026654 | 0.752369307 |
| 0.500008071 | 0.752365867 |
| 0.49998786 | 0.752359224 |
| 0.499987199 | 0.752359224 |
| 0.499958001 | 0.752359224 |
| 0.499924111 | 0.75235838 |
| 0.499883936 | 0.75235838 |
| 0.499875159 | 0.75235838 |
| 0.4998719 | 0.75235838 |
| 0.499865753 | 0.75235838 |
| 0.499830589 | 0.75235838 |
| 0.49982267 | 0.75235838 |
| 0.499796359 | 0.75235838 |
| 0.499788409 | 0.75235838 |
| 0.499786237 | 0.75235838 |
| 0.499784049 | 0.75235838 |
| 0.499630067 | 0.752343773 |
| 0.499623406 | 0.752343773 |
| 0.499583053 | 0.752343773 |
| 0.499565058 | 0.752343773 |
| 0.499520258 | 0.75233882 |
| 0.499452519 | 0.752266694 |
| 0.49929648 | 0.752103025 |
| 0.499179927 | 0.751973862 |
| 0.499133399 | 0.751928718 |
| 0.499120681 | 0.751928718 |
| 0.499104046 | 0.751928718 |
| 0.499047887 | 0.751928718 |
| 0.499035609 | 0.751928718 |
| 0.499016185 | 0.751928718 |
| 0.499008195 | 0.751928718 |
| 0.499006821 | 0.751928718 |
| 0.498914504 | 0.751924901 |
| 0.498849148 | 0.751904815 |
| 0.498787032 | 0.751834984 |
| 0.498773185 | 0.751834984 |
| 0.498669859 | 0.751753533 |
| 0.498639238 | 0.751753533 |
| 0.498536033 | 0.75168331 |
| 0.498503224 | 0.75167939 |
| 0.498470616 | 0.751667301 |

| | |
|---|---|
| 0.498463348 | 0.751667301 |
| 0.498355541 | 0.751565162 |
| 0.498314733 | 0.751527416 |
| 0.498262327 | 0.751495973 |
| 0.498213277 | 0.751445791 |
| 0.498202486 | 0.751445791 |
| 0.4980085 | 0.751232092 |
| 0.497963096 | 0.751211186 |
| 0.497934834 | 0.751192345 |
| 0.497837435 | 0.751108965 |
| 0.49780612 | 0.751093331 |
| 0.497695278 | 0.75098959 |
| 0.497646443 | 0.750971361 |
| 0.497644407 | 0.750971361 |
| 0.497439536 | 0.750778071 |
| 0.497295849 | 0.750632568 |
| 0.497293587 | 0.750632568 |
| 0.497256543 | 0.750632568 |
| 0.497216039 | 0.750607255 |
| 0.497214514 | 0.750607255 |
| 0.497212028 | 0.750607255 |
| 0.497191106 | 0.750607255 |
| 0.497076172 | 0.750529193 |
| 0.496880403 | 0.750290827 |
| 0.496877666 | 0.750290827 |
| 0.496852587 | 0.750290827 |
| 0.496828564 | 0.750283912 |
| 0.496816281 | 0.750283912 |
| 0.496748117 | 0.750281374 |
| 0.496701061 | 0.750257887 |
| 0.496687694 | 0.750257887 |
| 0.496610975 | 0.750193191 |
| 0.496581594 | 0.750193191 |
| 0.496543302 | 0.750186139 |
| 0.496477231 | 0.750110113 |
| 0.496469907 | 0.750110113 |
| 0.496453817 | 0.750110113 |
| 0.496412203 | 0.750083248 |
| 0.496339224 | 0.750020566 |
| 0.496222533 | 0.749915615 |
| 0.496182785 | 0.749879339 |
| 0.496151846 | 0.749879339 |
| 0.496102904 | 0.749829996 |
| 0.496056538 | 0.749783712 |
| 0.4959969 | 0.749717361 |

| | |
|---|---|
| 0.495984986 | 0.749717361 |
| 0.49591261 | 0.749661331 |
| 0.49572069 | 0.749490153 |
| 0.495713319 | 0.749490153 |
| 0.495560156 | 0.749392489 |
| 0.495534378 | 0.749392489 |
| 0.495520216 | 0.749392489 |
| 0.495503661 | 0.749392489 |
| 0.49549633 | 0.749392489 |
| 0.495496058 | 0.749392489 |
| 0.495437506 | 0.749392489 |
| 0.495385712 | 0.749364311 |
| 0.495370273 | 0.749364311 |
| 0.495361406 | 0.749364311 |
| 0.495337133 | 0.749364311 |
| 0.495256515 | 0.749311669 |
| 0.495141252 | 0.749184879 |
| 0.495100108 | 0.749146424 |
| 0.494870471 | 0.748854924 |
| 0.494778425 | 0.748802416 |
| 0.494688527 | 0.748713947 |
| 0.494685774 | 0.748713947 |
| 0.494670229 | 0.748713947 |
| 0.494627235 | 0.748692557 |
| 0.494619572 | 0.748692557 |
| 0.494594428 | 0.748690488 |
| 0.494572877 | 0.748690488 |
| 0.49451385 | 0.748639896 |
| 0.494413932 | 0.748560014 |
| 0.494347166 | 0.748506516 |
| 0.494257634 | 0.74841854 |
| 0.494172272 | 0.748313074 |
| 0.494140441 | 0.748299635 |
| 0.494025585 | 0.748186101 |
| 0.493999488 | 0.748170369 |
| 0.493995161 | 0.748170369 |
| 0.493992026 | 0.748170369 |
| 0.493955515 | 0.748170369 |
| 0.493945921 | 0.748170369 |
| 0.493870426 | 0.748117641 |
| 0.493823059 | 0.748093478 |
| 0.493768297 | 0.748063518 |
| 0.493734764 | 0.748063518 |
| 0.49371683 | 0.748063518 |
| 0.493708022 | 0.748063518 |

| | |
|---|---|
| 0.493699986 | 0.748063518 |
| 0.493680626 | 0.748063518 |
| 0.493662624 | 0.748063518 |
| 0.493647166 | 0.748063518 |
| 0.493621767 | 0.748063518 |
| 0.493599232 | 0.748063518 |
| 0.493561991 | 0.748031088 |
| 0.493548005 | 0.748031088 |
| 0.493409843 | 0.747848092 |
| 0.493400516 | 0.747848092 |
| 0.493375115 | 0.747843054 |
| 0.493364083 | 0.747843054 |
| 0.493310985 | 0.747817247 |
| 0.493308866 | 0.747817247 |
| 0.493273314 | 0.747807745 |
| 0.493186773 | 0.747700349 |
| 0.493131677 | 0.747640618 |
| 0.493130115 | 0.747640618 |
| 0.493112432 | 0.747640618 |
| 0.493063946 | 0.747633129 |
| 0.493019379 | 0.747589353 |
| 0.492978531 | 0.747551212 |
| 0.492923942 | 0.747492234 |
| 0.492917253 | 0.747492234 |
| 0.492907549 | 0.747492234 |
| 0.492847103 | 0.747492234 |
| 0.492711357 | 0.747336427 |
| 0.492634917 | 0.74726808 |
| 0.492634119 | 0.74726808 |
| 0.492536036 | 0.747189483 |
| 0.492472301 | 0.747116592 |
| 0.492463691 | 0.747116592 |
| 0.492454392 | 0.747116592 |
| 0.492419951 | 0.747116592 |
| 0.492376398 | 0.747090086 |
| 0.492318082 | 0.7470254 |
| 0.49222952 | 0.746938614 |
| 0.492189365 | 0.746901478 |
| 0.492044809 | 0.746729698 |
| 0.492019505 | 0.74671509 |
| 0.491960308 | 0.746672836 |
| 0.491952735 | 0.746672836 |
| 0.491925599 | 0.746667745 |
| 0.491901001 | 0.746654204 |
| 0.491858635 | 0.74662936 |

| | |
|---|---|
| 0.491782349 | 0.746560788 |
| 0.491780043 | 0.746560788 |
| 0.491739675 | 0.746552086 |
| 0.491676168 | 0.746503264 |
| 0.491675913 | 0.746503264 |
| 0.4916299 | 0.746480609 |
| 0.491618857 | 0.746480609 |
| 0.491611736 | 0.746480609 |
| 0.491556837 | 0.746464863 |
| 0.491531235 | 0.746449784 |
| 0.491517688 | 0.746449784 |
| 0.491437206 | 0.746366366 |
| 0.491374488 | 0.746306931 |
| 0.491357431 | 0.746304825 |
| 0.491345576 | 0.746304825 |
| 0.491334021 | 0.746304825 |
| 0.491324112 | 0.746304825 |
| 0.491322325 | 0.746304825 |
| 0.491310286 | 0.746304825 |
| 0.491288271 | 0.746304825 |
| 0.491231591 | 0.746304825 |
| 0.491096699 | 0.746218175 |
| 0.49108168 | 0.746218175 |
| 0.491080094 | 0.746218175 |
| 0.491065576 | 0.746218175 |
| 0.491048478 | 0.746218175 |
| 0.49103989 | 0.746218175 |
| 0.49101489 | 0.746218175 |
| 0.491012337 | 0.746218175 |
| 0.490998005 | 0.746218175 |
| 0.490893132 | 0.746146907 |
| 0.490764069 | 0.745974542 |
| 0.490725557 | 0.745974542 |
| 0.490712986 | 0.745974542 |
| 0.490594138 | 0.745859068 |
| 0.49053177 | 0.745788055 |
| 0.490519111 | 0.745788055 |
| 0.490513433 | 0.745788055 |
| 0.490485103 | 0.745788055 |
| 0.490477046 | 0.745788055 |
| 0.490471112 | 0.745788055 |
| 0.490422048 | 0.745787888 |
| 0.490419465 | 0.745787888 |
| 0.490399086 | 0.745787888 |
| 0.490327028 | 0.745738636 |

| | |
|---|---|
| 0.49020325 | 0.745645626 |
| 0.49018797 | 0.745645626 |
| 0.490101304 | 0.745609621 |
| 0.490035692 | 0.745557434 |
| 0.490015564 | 0.745550627 |
| 0.489980168 | 0.745529811 |
| 0.489965277 | 0.745529811 |
| 0.489850806 | 0.745452002 |
| 0.489826653 | 0.745452002 |
| 0.489810452 | 0.745452002 |
| 0.489729519 | 0.745401163 |
| 0.489574618 | 0.745236842 |
| 0.48954393 | 0.745213947 |
| 0.489526763 | 0.745211632 |
| 0.489520638 | 0.745211632 |
| 0.489462235 | 0.745184856 |
| 0.489418733 | 0.745142446 |
| 0.489387377 | 0.745118526 |
| 0.489327952 | 0.745075685 |
| 0.489315889 | 0.745075685 |
| 0.489263657 | 0.745068198 |
| 0.489203153 | 0.745028574 |
| 0.489199385 | 0.745028574 |
| 0.489174828 | 0.745028574 |
| 0.489158124 | 0.745028574 |
| 0.489133666 | 0.745028574 |
| 0.48911634 | 0.745028574 |
| 0.489101593 | 0.745028574 |
| 0.489063414 | 0.74500634 |
| 0.489015264 | 0.744956818 |
| 0.488948299 | 0.744878631 |
| 0.488888775 | 0.744835599 |
| 0.48871657 | 0.744597059 |
| 0.488645721 | 0.744536751 |
| 0.488634655 | 0.744536751 |
| 0.488627393 | 0.744536751 |
| 0.48862705 | 0.744536751 |
| 0.488583151 | 0.744536751 |
| 0.48858199 | 0.744536751 |
| 0.488557087 | 0.744536751 |
| 0.488511646 | 0.744536751 |
| 0.488426441 | 0.744464624 |
| 0.488360258 | 0.74443522 |
| 0.488336765 | 0.74443522 |
| 0.48831361 | 0.74443522 |

| | |
|---|---|
| 0.488286036 | 0.74443522 |
| 0.488104007 | 0.744218895 |
| 0.488098761 | 0.744218895 |
| 0.487888788 | 0.743978462 |
| 0.487858093 | 0.743978462 |
| 0.487799839 | 0.743914279 |
| 0.487726856 | 0.743826794 |
| 0.487726516 | 0.743826794 |
| 0.487711312 | 0.743826794 |
| 0.487683572 | 0.743826794 |
| 0.487673215 | 0.743826794 |
| 0.487595464 | 0.74374549 |
| 0.487548056 | 0.743696995 |
| 0.487520354 | 0.743678557 |
| 0.487515551 | 0.743678557 |
| 0.487487046 | 0.743675386 |
| 0.487422867 | 0.743601297 |
| 0.487401085 | 0.743591885 |
| 0.487321389 | 0.743494115 |
| 0.487138066 | 0.743238232 |
| 0.487089719 | 0.743188276 |
| 0.48701378 | 0.743120024 |
| 0.486959765 | 0.743109027 |
| 0.486911547 | 0.743092274 |
| 0.486769638 | 0.743034729 |
| 0.486749859 | 0.743034729 |
| 0.486719515 | 0.743034729 |
| 0.486632538 | 0.743034729 |
| 0.486424818 | 0.742768669 |
| 0.486419409 | 0.742768669 |
| 0.486310772 | 0.742676176 |
| 0.48625235 | 0.742645257 |
| 0.486229773 | 0.742637508 |
| 0.486196936 | 0.742628932 |
| 0.486195568 | 0.742628932 |
| 0.486178414 | 0.742628932 |
| 0.486175225 | 0.742628932 |
| 0.48617094 | 0.742628932 |
| 0.48607977 | 0.742575096 |
| 0.48607045 | 0.742575096 |
| 0.486066966 | 0.742575096 |
| 0.485990187 | 0.742509692 |
| 0.485917524 | 0.742470129 |
| 0.485840917 | 0.742376893 |
| 0.485763681 | 0.742330322 |

| | |
|---|---|
| 0.485700885 | 0.742281994 |
| 0.485595214 | 0.742191947 |
| 0.485363403 | 0.742000793 |
| 0.48531955 | 0.742000793 |
| 0.48527085 | 0.741958098 |
| 0.485231792 | 0.741935845 |
| 0.485222362 | 0.741935845 |
| 0.485220968 | 0.741935845 |
| 0.48522057 | 0.741935845 |
| 0.485203813 | 0.741935845 |
| 0.485181278 | 0.741935845 |
| 0.485068832 | 0.741863564 |
| 0.485008071 | 0.741818279 |
| 0.484955474 | 0.741785476 |
| 0.484916527 | 0.741749725 |
| 0.484801237 | 0.741692477 |
| 0.484711095 | 0.741578391 |
| 0.484560688 | 0.741442601 |
| 0.484552603 | 0.741442601 |
| 0.484531084 | 0.741442601 |
| 0.484507365 | 0.741442601 |
| 0.484478572 | 0.741442601 |
| 0.484308067 | 0.741295164 |
| 0.484270863 | 0.741262041 |
| 0.484237354 | 0.741234572 |
| 0.48423053 | 0.741234572 |
| 0.484032029 | 0.741087004 |
| 0.483996486 | 0.741056408 |
| 0.483969906 | 0.741039534 |
| 0.483892081 | 0.741015663 |
| 0.483796899 | 0.740893727 |
| 0.483741295 | 0.740832396 |
| 0.483735647 | 0.740832396 |
| 0.483728333 | 0.740832396 |
| 0.483701675 | 0.740832396 |
| 0.483693589 | 0.740832396 |
| 0.483680713 | 0.740832396 |
| 0.483647494 | 0.740832396 |
| 0.483592156 | 0.740832396 |
| 0.483588867 | 0.740832396 |
| 0.483580379 | 0.740832396 |
| 0.483521096 | 0.740832396 |
| 0.483276136 | 0.740691633 |
| 0.483179996 | 0.740606237 |
| 0.483172493 | 0.740606237 |

| | |
|---|---|
| 0.483163474 | 0.740606237 |
| 0.483163339 | 0.740606237 |
| 0.483007176 | 0.740493877 |
| 0.482982742 | 0.740480252 |
| 0.482901772 | 0.740379948 |
| 0.482897767 | 0.740379948 |
| 0.482852778 | 0.740376333 |
| 0.482834861 | 0.740372696 |
| 0.482828562 | 0.740372696 |
| 0.482808936 | 0.740372696 |
| 0.482767618 | 0.740341093 |
| 0.482743781 | 0.740328631 |
| 0.482711835 | 0.740328631 |
| 0.482620981 | 0.740259234 |
| 0.482616549 | 0.740259234 |
| 0.482577837 | 0.740259234 |
| 0.482557388 | 0.740257054 |
| 0.482535466 | 0.740257054 |
| 0.482379142 | 0.740102809 |
| 0.482376778 | 0.740102809 |
| 0.482374949 | 0.740102809 |
| 0.482365602 | 0.740102809 |
| 0.482198711 | 0.739945163 |
| 0.482189164 | 0.739945163 |
| 0.482179674 | 0.739945163 |
| 0.482079831 | 0.73992962 |
| 0.482074898 | 0.73992962 |
| 0.482056604 | 0.73992962 |
| 0.482049024 | 0.73992962 |
| 0.482031363 | 0.73992962 |
| 0.482020577 | 0.73992962 |
| 0.481981078 | 0.73992112 |
| 0.481971454 | 0.73992112 |
| 0.481971454 | 0.73992112 |
| 0.481936293 | 0.73992112 |
| 0.481859289 | 0.739901098 |
| 0.481674619 | 0.739712929 |
| 0.481670406 | 0.739712929 |
| 0.481616627 | 0.739695424 |
| 0.481529464 | 0.739585403 |
| 0.481370823 | 0.739413281 |
| 0.481353622 | 0.739410707 |
| 0.481288306 | 0.73933422 |
| 0.481282667 | 0.73933422 |
| 0.481159522 | 0.739243614 |

| | |
|---|---|
| 0.481044094 | 0.739125949 |
| 0.480968319 | 0.739033363 |
| 0.480893597 | 0.739002098 |
| 0.480843128 | 0.738960203 |
| 0.480743925 | 0.738926965 |
| 0.480675753 | 0.738869871 |
| 0.480648085 | 0.738869871 |
| 0.480631426 | 0.738869871 |
| 0.480620997 | 0.738869871 |
| 0.480535817 | 0.738821686 |
| 0.480505415 | 0.738798793 |
| 0.480472267 | 0.738795528 |
| 0.480433517 | 0.738759795 |
| 0.480296202 | 0.738572491 |
| 0.480294099 | 0.738572491 |
| 0.480174252 | 0.738528034 |
| 0.480137515 | 0.73849538 |
| 0.480101549 | 0.738463909 |
| 0.480067318 | 0.738435104 |
| 0.480046764 | 0.738435104 |
| 0.480027365 | 0.738435104 |
| 0.480008939 | 0.738435104 |
| 0.479929369 | 0.738389845 |
| 0.47991812 | 0.738389845 |
| 0.479879061 | 0.738389845 |
| 0.479870665 | 0.738389845 |
| 0.479838735 | 0.738389845 |
| 0.479750689 | 0.738329598 |
| 0.479707959 | 0.738311552 |
| 0.479647775 | 0.738306057 |
| 0.479630132 | 0.738306057 |
| 0.479606356 | 0.738298327 |
| 0.479531177 | 0.738230319 |
| 0.479513119 | 0.738226381 |
| 0.479443758 | 0.73815966 |
| 0.479428594 | 0.73815966 |
| 0.479362796 | 0.73811425 |
| 0.479357514 | 0.73811425 |
| 0.479173925 | 0.737894998 |
| 0.47914983 | 0.737884939 |
| 0.47911065 | 0.737869132 |
| 0.47908245 | 0.737869132 |
| 0.479055447 | 0.737869132 |
| 0.479053788 | 0.737869132 |
| 0.47894825 | 0.737809888 |

| | |
|---|---|
| 0.478918949 | 0.737788614 |
| 0.478888907 | 0.737786588 |
| 0.478855345 | 0.737786087 |
| 0.478716584 | 0.737622084 |
| 0.478664889 | 0.73758809 |
| 0.478604828 | 0.737519401 |
| 0.478588941 | 0.737518783 |
| 0.478545651 | 0.737475934 |
| 0.478370005 | 0.737324539 |
| 0.478355393 | 0.737324539 |
| 0.478353834 | 0.737324539 |
| 0.478343064 | 0.737324539 |
| 0.478203886 | 0.737187781 |
| 0.478125914 | 0.737115299 |
| 0.478116864 | 0.737115299 |
| 0.478093898 | 0.73711366 |
| 0.47802512 | 0.737031479 |
| 0.47801128 | 0.737031479 |
| 0.477939878 | 0.736971625 |
| 0.477726016 | 0.736689557 |
| 0.477703994 | 0.736689557 |
| 0.477696231 | 0.736689557 |
| 0.47764251 | 0.736680046 |
| 0.477537203 | 0.736565334 |
| 0.477497722 | 0.736552145 |
| 0.477468501 | 0.736530925 |
| 0.477362709 | 0.736433532 |
| 0.477295195 | 0.736358986 |
| 0.477285935 | 0.736358986 |
| 0.477224423 | 0.736327102 |
| 0.477142782 | 0.736243091 |
| 0.47713402 | 0.736243091 |
| 0.477026603 | 0.736135373 |
| 0.476931279 | 0.736096147 |
| 0.476897389 | 0.736095176 |
| 0.476878586 | 0.736095176 |
| 0.47682037 | 0.736079469 |
| 0.476807683 | 0.736079469 |
| 0.476770468 | 0.736079469 |
| 0.476733387 | 0.736064481 |
| 0.476677405 | 0.736049629 |
| 0.476660823 | 0.736047889 |
| 0.476638639 | 0.736037496 |
| 0.476558298 | 0.735961156 |
| 0.476495561 | 0.735935859 |

| | |
|---|---|
| 0.476433794 | 0.735864324 |
| 0.47636637 | 0.735784047 |
| 0.476315577 | 0.735753315 |
| 0.476281934 | 0.735725209 |
| 0.476281234 | 0.735725209 |
| 0.476200059 | 0.735646456 |
| 0.476113482 | 0.735584287 |
| 0.475959083 | 0.735465033 |
| 0.47562056 | 0.735085035 |
| 0.475584835 | 0.735053672 |
| 0.475511024 | 0.734963444 |
| 0.475472765 | 0.73492816 |
| 0.475469679 | 0.73492816 |
| 0.475444488 | 0.73492816 |
| 0.475381368 | 0.734882288 |
| 0.475368598 | 0.734882288 |
| 0.475334348 | 0.734881159 |
| 0.475332098 | 0.734881159 |
| 0.4752296 | 0.734790773 |
| 0.475214312 | 0.734790773 |
| 0.475170286 | 0.734752237 |
| 0.475161646 | 0.734752237 |
| 0.475100859 | 0.734710976 |
| 0.475019831 | 0.734648004 |
| 0.474996468 | 0.734644956 |
| 0.474957783 | 0.734632838 |
| 0.474822305 | 0.734490538 |
| 0.474743998 | 0.734421425 |
| 0.474722662 | 0.734421425 |
| 0.474685185 | 0.734402005 |
| 0.474632518 | 0.734344376 |
| 0.474629835 | 0.734344376 |
| 0.474526214 | 0.734227607 |
| 0.474460875 | 0.734150361 |
| 0.474331083 | 0.734068773 |
| 0.474320359 | 0.734068773 |
| 0.474294242 | 0.734059464 |
| 0.474259562 | 0.734053498 |
| 0.474245457 | 0.734053498 |
| 0.474181685 | 0.733980666 |
| 0.474149983 | 0.733973651 |
| 0.474112289 | 0.733944808 |
| 0.473901682 | 0.733714173 |
| 0.473870582 | 0.733689874 |
| 0.473809841 | 0.733643528 |

| | |
|---|---|
| 0.47375831 | 0.733611439 |
| 0.473705628 | 0.733553711 |
| 0.473701929 | 0.733553711 |
| 0.473700451 | 0.733553711 |
| 0.473658613 | 0.733552459 |
| 0.47354917 | 0.733454514 |
| 0.473471148 | 0.73338137 |
| 0.47336172 | 0.733272611 |
| 0.473336473 | 0.733268156 |
| 0.473335298 | 0.733268156 |
| 0.473307498 | 0.733268156 |
| 0.473305569 | 0.733268156 |
| 0.473241039 | 0.733263418 |
| 0.472923117 | 0.732913879 |
| 0.47289334 | 0.732891579 |
| 0.472856039 | 0.732887155 |
| 0.472800955 | 0.732843789 |
| 0.472743102 | 0.732815903 |
| 0.472706857 | 0.732793329 |
| 0.472689395 | 0.732790112 |
| 0.47266756 | 0.732780113 |
| 0.472610341 | 0.73273911 |
| 0.472592591 | 0.732735443 |
| 0.472577967 | 0.732735443 |
| 0.472563737 | 0.732735443 |
| 0.472532699 | 0.732735443 |
| 0.472503899 | 0.732735443 |
| 0.472455963 | 0.732714431 |
| 0.472409799 | 0.732666693 |
| 0.472397463 | 0.732666693 |
| 0.472328989 | 0.732612935 |
| 0.472273313 | 0.732550435 |
| 0.472218776 | 0.732489697 |
| 0.47220959 | 0.732489697 |
| 0.472195921 | 0.732489697 |
| 0.472183835 | 0.732489697 |
| 0.472157074 | 0.732489412 |
| 0.472148608 | 0.732489412 |
| 0.472109451 | 0.732489412 |
| 0.47208012 | 0.732489327 |
| 0.472028154 | 0.732448928 |
| 0.472013396 | 0.732448928 |
| 0.472006253 | 0.732448928 |
| 0.47200211 | 0.732448928 |
| 0.471918137 | 0.732425848 |

| | |
|---|---|
| 0.471852596 | 0.732351026 |
| 0.471844925 | 0.732351026 |
| 0.471774681 | 0.732325557 |
| 0.471759256 | 0.732325481 |
| 0.471749173 | 0.732325481 |
| 0.471739735 | 0.732325481 |
| 0.47170417 | 0.732325481 |
| 0.471609884 | 0.732284548 |
| 0.471609741 | 0.732284548 |
| 0.471555643 | 0.732284548 |
| 0.471516826 | 0.73225942 |
| 0.471273782 | 0.731953577 |
| 0.47118992 | 0.73187106 |
| 0.471127789 | 0.731838184 |
| 0.471121639 | 0.731838184 |
| 0.471121042 | 0.731838184 |
| 0.471101709 | 0.731838184 |
| 0.471031954 | 0.731816633 |
| 0.470998833 | 0.731789046 |
| 0.470926456 | 0.731700463 |
| 0.470914413 | 0.731700463 |
| 0.470907137 | 0.731700463 |
| 0.47089375 | 0.731700463 |
| 0.470863932 | 0.7316988 |
| 0.470828072 | 0.731666947 |
| 0.470810912 | 0.731666947 |
| 0.470772616 | 0.731666947 |
| 0.470682035 | 0.731559354 |
| 0.470598595 | 0.731477408 |
| 0.470465995 | 0.731366777 |
| 0.470429538 | 0.731356411 |
| 0.470424681 | 0.731356411 |
| 0.470384106 | 0.73133496 |
| 0.470353952 | 0.731311951 |
| 0.470350593 | 0.731311951 |
| 0.470293152 | 0.731265164 |
| 0.470286502 | 0.731265164 |
| 0.47028027 | 0.731265164 |
| 0.470248145 | 0.731265164 |
| 0.470173245 | 0.731221954 |
| 0.470143502 | 0.731199575 |
| 0.470123663 | 0.731192597 |
| 0.469952629 | 0.730998199 |
| 0.469925908 | 0.730983153 |
| 0.469924414 | 0.730983153 |

| | |
|---|---|
| 0.469915459 | 0.730983153 |
| 0.469896547 | 0.730983153 |
| 0.469821018 | 0.730981048 |
| 0.469710479 | 0.730891187 |
| 0.469631979 | 0.730833598 |
| 0.469450797 | 0.730694895 |
| 0.469435025 | 0.730694227 |
| 0.469298023 | 0.730504851 |
| 0.469188807 | 0.730406462 |
| 0.469039568 | 0.730245742 |
| 0.468967614 | 0.730157586 |
| 0.468916406 | 0.730125594 |
| 0.468793339 | 0.73000557 |
| 0.468772885 | 0.729997587 |
| 0.468702275 | 0.729935362 |
| 0.468693098 | 0.729935362 |
| 0.468547933 | 0.729790451 |
| 0.468539409 | 0.729790451 |
| 0.46852961 | 0.729790451 |
| 0.468515601 | 0.729790451 |
| 0.468398525 | 0.729662415 |
| 0.46837084 | 0.729662415 |
| 0.46831566 | 0.729619559 |
| 0.468247298 | 0.729536918 |
| 0.468206275 | 0.729496868 |
| 0.468150515 | 0.729481583 |
| 0.468133121 | 0.729478346 |
| 0.468080984 | 0.729442567 |
| 0.468064684 | 0.729442567 |
| 0.468017807 | 0.729401323 |
| 0.467993096 | 0.729401323 |
| 0.467926436 | 0.729323309 |
| 0.467791815 | 0.729185075 |
| 0.467751322 | 0.729145819 |
| 0.467748755 | 0.729145819 |
| 0.467743461 | 0.729145819 |
| 0.467735091 | 0.729145819 |
| 0.46762941 | 0.729075092 |
| 0.467555051 | 0.729030755 |
| 0.467539388 | 0.7290302 |
| 0.46753854 | 0.7290302 |
| 0.467516577 | 0.7290302 |
| 0.467496171 | 0.7290302 |
| 0.467485935 | 0.7290302 |
| 0.467284014 | 0.728847601 |

| | |
|---|---|
| 0.467260007 | 0.728847601 |
| 0.46719865 | 0.728843203 |
| 0.46719039 | 0.728843203 |
| 0.467083988 | 0.728749561 |
| 0.467033789 | 0.728700748 |
| 0.467010009 | 0.728700748 |
| 0.46697301 | 0.728671895 |
| 0.466928633 | 0.728626519 |
| 0.466900008 | 0.728605724 |
| 0.466884572 | 0.728605508 |
| 0.466740392 | 0.728475984 |
| 0.466727347 | 0.728475984 |
| 0.466719319 | 0.728475984 |
| 0.466718985 | 0.728475984 |
| 0.466616367 | 0.72841695 |
| 0.466606605 | 0.72841695 |
| 0.466573156 | 0.72840593 |
| 0.466546725 | 0.72840593 |
| 0.466480154 | 0.728356236 |
| 0.466451102 | 0.728341933 |
| 0.466450427 | 0.728341933 |
| 0.466438063 | 0.728341933 |
| 0.466390436 | 0.728341933 |
| 0.466341091 | 0.728341933 |
| 0.466299553 | 0.728313017 |
| 0.466239761 | 0.728267393 |
| 0.466222174 | 0.728263808 |
| 0.466205501 | 0.728261648 |
| 0.466188656 | 0.728261648 |
| 0.466105591 | 0.728177228 |
| 0.466064609 | 0.728150889 |
| 0.466056417 | 0.728150889 |
| 0.465945308 | 0.728022347 |
| 0.465788957 | 0.727801931 |
| 0.465742909 | 0.727753856 |
| 0.465742174 | 0.727753856 |
| 0.465705413 | 0.727743018 |
| 0.465652649 | 0.727684441 |
| 0.465647304 | 0.727684441 |
| 0.465563988 | 0.727641389 |
| 0.46553583 | 0.727621258 |
| 0.465525715 | 0.727621258 |
| 0.465400587 | 0.727481498 |
| 0.465400129 | 0.727481498 |
| 0.465302023 | 0.727399049 |

| | |
|---|---|
| 0.46528845 | 0.727399049 |
| 0.465251708 | 0.727368143 |
| 0.465141306 | 0.727267161 |
| 0.464996893 | 0.727160724 |
| 0.464964996 | 0.727134718 |
| 0.464916922 | 0.727083412 |
| 0.46490468 | 0.727083412 |
| 0.464885991 | 0.727083412 |
| 0.464883992 | 0.727083412 |
| 0.464869346 | 0.727083412 |
| 0.464818252 | 0.727083412 |
| 0.464734368 | 0.727036649 |
| 0.464607428 | 0.726861938 |
| 0.464575321 | 0.726835583 |
| 0.464563172 | 0.726835583 |
| 0.464461003 | 0.726728348 |
| 0.464457557 | 0.726728348 |
| 0.464387074 | 0.726660419 |
| 0.464385364 | 0.726660419 |
| 0.464379098 | 0.726660419 |
| 0.464354183 | 0.726660419 |
| 0.464306181 | 0.72665321 |
| 0.464280775 | 0.726637325 |
| 0.464241341 | 0.726599486 |
| 0.464231714 | 0.726599486 |
| 0.464173153 | 0.726599486 |
| 0.464135604 | 0.726599486 |
| 0.464088918 | 0.726551932 |
| 0.464028645 | 0.726542475 |
| 0.464008349 | 0.726542475 |
| 0.464005881 | 0.726542475 |
| 0.463968995 | 0.726542475 |
| 0.463967818 | 0.726542475 |
| 0.463931656 | 0.726542475 |
| 0.463915437 | 0.726542475 |
| 0.463863984 | 0.726534134 |
| 0.463861984 | 0.726534134 |
| 0.463785317 | 0.726458702 |
| 0.46376861 | 0.726456424 |
| 0.463698992 | 0.726395155 |
| 0.46366009 | 0.726358105 |
| 0.463528359 | 0.726199512 |
| 0.463482 | 0.726153626 |
| 0.463474622 | 0.726153626 |
| 0.463473723 | 0.726153626 |

| | |
|---|---|
| 0.463421645 | 0.726153626 |
| 0.463420801 | 0.726153626 |
| 0.463371105 | 0.726153626 |
| 0.463310836 | 0.726153626 |
| 0.463286321 | 0.726153626 |
| 0.463236847 | 0.726148877 |
| 0.463218202 | 0.72614355 |
| 0.463207976 | 0.72614355 |
| 0.463165576 | 0.726132752 |
| 0.463131991 | 0.726104 |
| 0.463129002 | 0.726104 |
| 0.462956318 | 0.725971967 |
| 0.462926379 | 0.72594892 |
| 0.462910705 | 0.725948243 |
| 0.462906942 | 0.725948243 |
| 0.462900494 | 0.725948243 |
| 0.462885195 | 0.725948243 |
| 0.462866919 | 0.725948243 |
| 0.462632043 | 0.725750208 |
| 0.462583573 | 0.725698073 |
| 0.462548362 | 0.725690643 |
| 0.46254776 | 0.725690643 |
| 0.46243514 | 0.725560811 |
| 0.462342191 | 0.725500096 |
| 0.462200709 | 0.725398556 |
| 0.462146755 | 0.725347289 |
| 0.46214135 | 0.725347289 |
| 0.462130869 | 0.725347289 |
| 0.462126433 | 0.725347289 |
| 0.462121566 | 0.725347289 |
| 0.462089184 | 0.725347289 |
| 0.462039698 | 0.725322677 |
| 0.461895548 | 0.725168093 |
| 0.46181207 | 0.725132648 |
| 0.461762536 | 0.72510331 |
| 0.461722707 | 0.725101416 |
| 0.461720974 | 0.725101416 |
| 0.461656592 | 0.725073412 |
| 0.461620526 | 0.72504702 |
| 0.461494246 | 0.724920398 |
| 0.461488348 | 0.724920398 |
| 0.461480923 | 0.724920398 |
| 0.461474302 | 0.724920398 |
| 0.461443096 | 0.724920398 |
| 0.461426439 | 0.724920398 |

| | |
|---|---|
| 0.461384454 | 0.724895457 |
| 0.461344492 | 0.724885334 |
| 0.461336589 | 0.724885334 |
| 0.461292899 | 0.724867099 |
| 0.461256282 | 0.72485739 |
| 0.461249368 | 0.72485739 |
| 0.461234872 | 0.72485739 |
| 0.461211174 | 0.72485739 |
| 0.461175129 | 0.72485739 |
| 0.46114449 | 0.724849129 |
| 0.461087245 | 0.724806992 |
| 0.461053898 | 0.724778493 |
| 0.461013961 | 0.724739632 |
| 0.461009748 | 0.724739632 |
| 0.460961912 | 0.724705651 |
| 0.460955601 | 0.724705651 |
| 0.46091385 | 0.724677932 |
| 0.460881991 | 0.724675687 |
| 0.460879624 | 0.724675687 |
| 0.460852014 | 0.724675687 |
| 0.460765762 | 0.724636474 |
| 0.460708065 | 0.724593587 |
| 0.460693217 | 0.724593587 |
| 0.460610518 | 0.724495022 |
| 0.460568964 | 0.724470508 |
| 0.460495127 | 0.724402214 |
| 0.460488634 | 0.724402214 |
| 0.460451202 | 0.724380967 |
| 0.460370214 | 0.724373193 |
| 0.460321705 | 0.724320796 |
| 0.460313279 | 0.724320796 |
| 0.460298575 | 0.724320796 |
| 0.460178303 | 0.724214786 |
| 0.460057653 | 0.724049678 |
| 0.460056918 | 0.724049678 |
| 0.459964174 | 0.723973486 |
| 0.459908822 | 0.723928819 |
| 0.459839529 | 0.723849063 |
| 0.459812755 | 0.723830841 |
| 0.459731522 | 0.723750809 |
| 0.459697708 | 0.723733834 |
| 0.45966209 | 0.723733834 |
| 0.459643046 | 0.723731134 |
| 0.459623581 | 0.723724413 |
| 0.459600408 | 0.723711851 |

| | |
|---|---|
| 0.459492633 | 0.723626638 |
| 0.459473069 | 0.723626638 |
| 0.459461768 | 0.723626638 |
| 0.459281766 | 0.723449288 |
| 0.459267842 | 0.723449288 |
| 0.459263547 | 0.723449288 |
| 0.459219443 | 0.723449288 |
| 0.459199224 | 0.723449288 |
| 0.459177068 | 0.723449288 |
| 0.45914918 | 0.723449288 |
| 0.459092642 | 0.723449288 |
| 0.459077913 | 0.723449288 |
| 0.459022091 | 0.723399131 |
| 0.458899195 | 0.72330504 |
| 0.458898962 | 0.72330504 |
| 0.458856411 | 0.72330504 |
| 0.458817152 | 0.723291556 |
| 0.458750423 | 0.72322958 |
| 0.458703149 | 0.72320759 |
| 0.458667627 | 0.723199466 |
| 0.458568998 | 0.723093484 |
| 0.458534585 | 0.723085448 |
| 0.458398247 | 0.722969802 |
| 0.458388503 | 0.722969802 |
| 0.458302261 | 0.722958458 |
| 0.458296535 | 0.722958458 |
| 0.458289826 | 0.722958458 |
| 0.458264662 | 0.722958458 |
| 0.458254354 | 0.722958458 |
| 0.458229567 | 0.722958458 |
| 0.458192486 | 0.722958458 |
| 0.458056512 | 0.722786254 |
| 0.458038422 | 0.722781656 |
| 0.457995193 | 0.722777605 |
| 0.457944553 | 0.722752819 |
| 0.457889835 | 0.722714791 |
| 0.457879024 | 0.722714791 |
| 0.457861979 | 0.722714791 |
| 0.457835574 | 0.72270101 |
| 0.457777497 | 0.722660027 |
| 0.457716547 | 0.722608923 |
| 0.457714324 | 0.722608923 |
| 0.457686912 | 0.722608923 |
| 0.457649051 | 0.722598175 |
| 0.457593475 | 0.722534377 |

| | |
|---|---|
| 0.457568325 | 0.72251862 |
| 0.457537742 | 0.722517472 |
| 0.457526358 | 0.722517472 |
| 0.457471584 | 0.722461672 |
| 0.457384179 | 0.7223973 |
| 0.457327191 | 0.722377358 |
| 0.457281711 | 0.722329477 |
| 0.457249813 | 0.722303046 |
| 0.457192687 | 0.72223676 |
| 0.457182448 | 0.72223676 |
| 0.457179357 | 0.72223676 |
| 0.457165823 | 0.72223676 |
| 0.457156565 | 0.72223676 |
| 0.45714685 | 0.72223676 |
| 0.457125355 | 0.72223676 |
| 0.457079129 | 0.72223676 |
| 0.457055184 | 0.722235144 |
| 0.457051372 | 0.722235144 |
| 0.456964863 | 0.7221594 |
| 0.456962104 | 0.7221594 |
| 0.456919854 | 0.722141095 |
| 0.456883542 | 0.722141095 |
| 0.456867033 | 0.722141095 |
| 0.456819851 | 0.722102856 |
| 0.456817691 | 0.722102856 |
| 0.456803471 | 0.722102856 |
| 0.456765106 | 0.722088299 |
| 0.456728156 | 0.722077738 |
| 0.456714697 | 0.722077738 |
| 0.456705353 | 0.722077738 |
| 0.456698539 | 0.722077738 |
| 0.456654463 | 0.722077738 |
| 0.456636619 | 0.722077738 |
| 0.456611437 | 0.722077738 |
| 0.456542798 | 0.722077738 |
| 0.456491239 | 0.722062711 |
| 0.456339609 | 0.721894543 |
| 0.456336545 | 0.721894543 |
| 0.456275678 | 0.721841612 |
| 0.456241678 | 0.721811798 |
| 0.456141772 | 0.721701687 |
| 0.456030009 | 0.721548827 |
| 0.456021863 | 0.721548827 |
| 0.456020584 | 0.721548827 |
| 0.455935599 | 0.721471353 |

| | |
|---|---|
| 0.455880999 | 0.721432892 |
| 0.455854429 | 0.721414816 |
| 0.455831408 | 0.721402355 |
| 0.455800305 | 0.721377102 |
| 0.455762748 | 0.721341632 |
| 0.455723139 | 0.721341632 |
| 0.455723063 | 0.721341632 |
| 0.455722348 | 0.721341632 |
| 0.455687044 | 0.721341632 |
| 0.455676076 | 0.721341632 |
| 0.455669583 | 0.721341632 |
| 0.45566082 | 0.721341632 |
| 0.455626765 | 0.721341632 |
| 0.455609926 | 0.721341632 |
| 0.455579242 | 0.721341632 |
| 0.455469783 | 0.721213491 |
| 0.45544461 | 0.721197607 |
| 0.455418041 | 0.721197607 |
| 0.455412052 | 0.721197607 |
| 0.4551948 | 0.720921878 |
| 0.455128905 | 0.720846252 |
| 0.455058446 | 0.720801803 |
| 0.454979178 | 0.720724184 |
| 0.454967032 | 0.720724184 |
| 0.454925347 | 0.720698231 |
| 0.454912381 | 0.720698231 |
| 0.454882458 | 0.72069456 |
| 0.454804092 | 0.720662558 |
| 0.454732385 | 0.720601467 |
| 0.454722907 | 0.720601467 |
| 0.454658709 | 0.720576031 |
| 0.454657121 | 0.720576031 |
| 0.45456608 | 0.720489729 |
| 0.45451635 | 0.720446308 |
| 0.454507273 | 0.720446308 |
| 0.454457989 | 0.720446308 |
| 0.454412182 | 0.720401072 |
| 0.454367653 | 0.720354456 |
| 0.454178577 | 0.720136951 |
| 0.45414872 | 0.720127217 |
| 0.454074434 | 0.720057371 |
| 0.454042724 | 0.720036547 |
| 0.453986456 | 0.720013751 |
| 0.453911447 | 0.719918761 |
| 0.453790921 | 0.719775543 |

| | |
|---|---|
| 0.453752397 | 0.719746735 |
| 0.453744824 | 0.719746735 |
| 0.453722459 | 0.719746735 |
| 0.453703329 | 0.719746735 |
| 0.453573437 | 0.719718211 |
| 0.453532551 | 0.719677311 |
| 0.453497867 | 0.71965007 |
| 0.453489029 | 0.71965007 |
| 0.453485589 | 0.71965007 |
| 0.453480782 | 0.71965007 |
| 0.453357982 | 0.719592101 |
| 0.453318873 | 0.719554006 |
| 0.453208965 | 0.719475464 |
| 0.453131702 | 0.719424748 |
| 0.453112209 | 0.719417782 |
| 0.453109571 | 0.719417782 |
| 0.453086186 | 0.719417782 |
| 0.45306156 | 0.71941778 |
| 0.453016386 | 0.71941778 |
| 0.452975377 | 0.719392386 |
| 0.45295334 | 0.719381377 |
| 0.452940508 | 0.719381377 |
| 0.452917816 | 0.719372935 |
| 0.452907558 | 0.719372935 |
| 0.452898699 | 0.719372935 |
| 0.452870575 | 0.719369873 |
| 0.452866739 | 0.719369873 |
| 0.452834754 | 0.719360956 |
| 0.452813505 | 0.719360956 |
| 0.452767512 | 0.719326116 |
| 0.452750996 | 0.719326116 |
| 0.45268623 | 0.71926896 |
| 0.452668385 | 0.719264601 |
| 0.452604053 | 0.71921037 |
| 0.452583666 | 0.71921037 |
| 0.452562329 | 0.71921037 |
| 0.452557458 | 0.71921037 |
| 0.452536879 | 0.71921037 |
| 0.452515366 | 0.71921037 |
| 0.452430974 | 0.719175289 |
| 0.452425526 | 0.719175289 |
| 0.452416029 | 0.719175289 |
| 0.452376575 | 0.719175289 |
| 0.452199449 | 0.718999237 |
| 0.452148177 | 0.71896572 |

| | |
|---|---|
| 0.452130104 | 0.718960985 |
| 0.452089687 | 0.718920719 |
| 0.452049164 | 0.718920719 |
| 0.451912535 | 0.718783005 |
| 0.451797302 | 0.718671724 |
| 0.451796089 | 0.718671724 |
| 0.451690323 | 0.718573553 |
| 0.451649802 | 0.718533089 |
| 0.451559481 | 0.718437395 |
| 0.451484994 | 0.718342881 |
| 0.45147293 | 0.718342881 |
| 0.451426224 | 0.718297366 |
| 0.451420314 | 0.718297366 |
| 0.451343822 | 0.718214241 |
| 0.45124436 | 0.718103955 |
| 0.451234513 | 0.718103955 |
| 0.451227415 | 0.718103955 |
| 0.451172245 | 0.718103955 |
| 0.451065779 | 0.718011693 |
| 0.451033393 | 0.718004558 |
| 0.451013021 | 0.717999705 |
| 0.450960243 | 0.717939681 |
| 0.45095265 | 0.717939681 |
| 0.450946325 | 0.717939681 |
| 0.450894865 | 0.717923881 |
| 0.450875508 | 0.717923881 |
| 0.450832318 | 0.717923881 |
| 0.450823879 | 0.717923881 |
| 0.450822032 | 0.717923881 |
| 0.450797485 | 0.717923881 |
| 0.450788476 | 0.717923881 |
| 0.45071176 | 0.71790405 |
| 0.450682618 | 0.71788164 |
| 0.450666898 | 0.71788061 |
| 0.45065002 | 0.717877734 |
| 0.450639695 | 0.717877734 |
| 0.450483991 | 0.717685267 |
| 0.450343982 | 0.717486212 |
| 0.450298407 | 0.717437603 |
| 0.450268782 | 0.717437603 |
| 0.450202012 | 0.717404016 |
| 0.450194399 | 0.717404016 |
| 0.450177322 | 0.717404016 |
| 0.450165521 | 0.717404016 |
| 0.450136348 | 0.717404016 |

| | |
|---|---|
| 0.450107382 | 0.717397238 |
| 0.450107382 | 0.717397238 |
| 0.45010226 | 0.717397238 |
| 0.449941342 | 0.717252612 |
| 0.449804833 | 0.717059003 |
| 0.449702626 | 0.716920065 |
| 0.449665321 | 0.716897431 |
| 0.449589143 | 0.716844867 |
| 0.449579714 | 0.716844867 |
| 0.449542304 | 0.716844867 |
| 0.449448889 | 0.716731475 |
| 0.44943687 | 0.716731475 |
| 0.449278027 | 0.716530977 |
| 0.449218765 | 0.716460455 |
| 0.449170638 | 0.716407688 |
| 0.449136264 | 0.716376853 |
| 0.449105515 | 0.716375791 |
| 0.44909708 | 0.716375791 |
| 0.449067079 | 0.716362465 |
| 0.448903844 | 0.716150043 |
| 0.44889466 | 0.716150043 |
| 0.448772498 | 0.715988468 |
| 0.448691327 | 0.715882947 |
| 0.448589278 | 0.715828802 |
| 0.448582349 | 0.715828802 |
| 0.448394644 | 0.715601371 |
| 0.448319029 | 0.715528648 |
| 0.448308183 | 0.715528648 |
| 0.448238474 | 0.71547201 |
| 0.448232454 | 0.71547201 |
| 0.448229634 | 0.71547201 |
| 0.448208765 | 0.71547201 |
| 0.448144547 | 0.715465955 |
| 0.448131033 | 0.715465955 |
| 0.448128615 | 0.715465955 |
| 0.448128447 | 0.715465955 |
| 0.448026202 | 0.71539692 |
| 0.447958827 | 0.715313319 |
| 0.447895856 | 0.715278327 |
| 0.447863171 | 0.715256497 |
| 0.447785713 | 0.715156774 |
| 0.447779655 | 0.715156774 |
| 0.447760262 | 0.715156774 |
| 0.447726078 | 0.715156774 |
| 0.447724288 | 0.715156774 |

| | |
|---|---|
| 0.447703105 | 0.715156774 |
| 0.447681625 | 0.715156774 |
| 0.447676893 | 0.715156774 |
| 0.447660555 | 0.715156774 |
| 0.44763688 | 0.715156774 |
| 0.447588 | 0.715156774 |
| 0.447503118 | 0.715113106 |
| 0.447430224 | 0.71508167 |
| 0.447318328 | 0.714961743 |
| 0.447300223 | 0.714957592 |
| 0.447274922 | 0.714957592 |
| 0.4469554 | 0.714597497 |
| 0.446937183 | 0.714597497 |
| 0.446895748 | 0.714574087 |
| 0.44688554 | 0.714574087 |
| 0.446879034 | 0.714574087 |
| 0.446848327 | 0.714574087 |
| 0.44676858 | 0.714538674 |
| 0.446761503 | 0.714538674 |
| 0.446740615 | 0.714538674 |
| 0.446740114 | 0.714538674 |
| 0.446689683 | 0.714532462 |
| 0.446681543 | 0.714532462 |
| 0.446670121 | 0.714532462 |
| 0.446650683 | 0.714532462 |
| 0.446537828 | 0.714433523 |
| 0.44647856 | 0.714386695 |
| 0.446469865 | 0.714386695 |
| 0.446440572 | 0.714386695 |
| 0.446410864 | 0.714374377 |
| 0.446343189 | 0.714314081 |
| 0.446323982 | 0.714307345 |
| 0.446215792 | 0.714158194 |
| 0.446160981 | 0.714128818 |
| 0.446011828 | 0.713999709 |
| 0.445956397 | 0.713934969 |
| 0.445808423 | 0.713806826 |
| 0.445788173 | 0.713806826 |
| 0.445770283 | 0.713806826 |
| 0.445656814 | 0.713671264 |
| 0.445620143 | 0.713636535 |
| 0.44554875 | 0.713586452 |
| 0.44537168 | 0.713407533 |
| 0.445358986 | 0.713407533 |
| 0.44528882 | 0.713393791 |

| | |
|---|---|
| 0.445257034 | 0.713393791 |
| 0.445126711 | 0.71329735 |
| 0.44510852 | 0.713296886 |
| 0.44497883 | 0.713182024 |
| 0.444888589 | 0.713088404 |
| 0.444887482 | 0.713088404 |
| 0.444887029 | 0.713088404 |
| 0.444869121 | 0.713088404 |
| 0.444834391 | 0.713088404 |
| 0.444747485 | 0.713088404 |
| 0.444742971 | 0.713088404 |
| 0.444736701 | 0.713088404 |
| 0.444722615 | 0.713088404 |
| 0.444602499 | 0.712985603 |
| 0.444524805 | 0.712913186 |
| 0.444488264 | 0.712878587 |
| 0.444437017 | 0.712868413 |
| 0.444302608 | 0.712707654 |
| 0.444294326 | 0.712707654 |
| 0.44427383 | 0.712707654 |
| 0.444263383 | 0.712707654 |
| 0.444170319 | 0.712632636 |
| 0.444154647 | 0.712631497 |
| 0.444092086 | 0.712602852 |
| 0.444032131 | 0.71253094 |
| 0.443980731 | 0.712496471 |
| 0.443942044 | 0.712458393 |
| 0.443914062 | 0.712437492 |
| 0.443862656 | 0.712378996 |
| 0.443860908 | 0.712378996 |
| 0.443802864 | 0.712355048 |
| 0.443672813 | 0.712213451 |
| 0.443632497 | 0.712177596 |
| 0.443623328 | 0.712177596 |
| 0.443588292 | 0.712177596 |
| 0.44353358 | 0.712114813 |
| 0.44352413 | 0.712114813 |
| 0.443411109 | 0.711990185 |
| 0.443402166 | 0.711990185 |
| 0.443367335 | 0.711967904 |
| 0.443364589 | 0.711967904 |
| 0.443310087 | 0.711923982 |
| 0.443273836 | 0.71188977 |
| 0.443226255 | 0.711837358 |
| 0.443200273 | 0.711819634 |

| | |
|---|---|
| 0.443157566 | 0.711775045 |
| 0.443061985 | 0.711645526 |
| 0.443045625 | 0.711643249 |
| 0.443026785 | 0.711636987 |
| 0.443009569 | 0.711633335 |
| 0.443005765 | 0.711633335 |
| 0.442934741 | 0.711561132 |
| 0.442851807 | 0.711475899 |
| 0.442800932 | 0.711418162 |
| 0.442757875 | 0.711372983 |
| 0.442744604 | 0.711372983 |
| 0.442738405 | 0.711372983 |
| 0.442692914 | 0.711364601 |
| 0.442628646 | 0.711285328 |
| 0.442602048 | 0.711266584 |
| 0.442559987 | 0.711222989 |
| 0.44255045 | 0.711222989 |
| 0.442393104 | 0.711050768 |
| 0.442344932 | 0.71103427 |
| 0.442291491 | 0.710983423 |
| 0.442170874 | 0.710837515 |
| 0.442083405 | 0.71072089 |
| 0.442045155 | 0.710683385 |
| 0.441897324 | 0.710517668 |
| 0.4418765 | 0.710508172 |
| 0.441854658 | 0.710497039 |
| 0.441807743 | 0.710445587 |
| 0.441801214 | 0.710445587 |
| 0.441766651 | 0.710427482 |
| 0.441756365 | 0.710427482 |
| 0.441752187 | 0.710427482 |
| 0.441722719 | 0.710427482 |
| 0.441658189 | 0.71039698 |
| 0.441621229 | 0.710361522 |
| 0.441599152 | 0.710350001 |
| 0.441571603 | 0.710329677 |
| 0.441548768 | 0.710316936 |
| 0.441514558 | 0.710285892 |
| 0.441505345 | 0.710285892 |
| 0.441448706 | 0.710227933 |
| 0.441429871 | 0.710221622 |
| 0.441409245 | 0.710221622 |
| 0.441395587 | 0.710221622 |
| 0.441353598 | 0.710218867 |
| 0.441353598 | 0.710218867 |

| | |
|---|---|
| 0.441353598 | 0.710218867 |
| 0.441257514 | 0.71013623 |
| 0.441243613 | 0.71013623 |
| 0.441200446 | 0.710097602 |
| 0.441191381 | 0.710097602 |
| 0.44111908 | 0.710081402 |
| 0.44103937 | 0.709977086 |
| 0.440942218 | 0.709892676 |
| 0.440924916 | 0.709888816 |
| 0.440885484 | 0.709860996 |
| 0.440845656 | 0.709833194 |
| 0.440692917 | 0.709659236 |
| 0.440687007 | 0.709659236 |
| 0.440655592 | 0.709647119 |
| 0.440651394 | 0.709647119 |
| 0.44054826 | 0.70952225 |
| 0.440456491 | 0.709439924 |
| 0.440450482 | 0.709439924 |
| 0.440357724 | 0.709335318 |
| 0.440272187 | 0.709287579 |
| 0.440260355 | 0.709287579 |
| 0.440222035 | 0.70928467 |
| 0.440216365 | 0.70928467 |
| 0.440206842 | 0.70928467 |
| 0.440166327 | 0.70928467 |
| 0.440140546 | 0.70928467 |
| 0.440022111 | 0.709202474 |
| 0.439995944 | 0.709184298 |
| 0.439992667 | 0.709184298 |
| 0.439948919 | 0.709180503 |
| 0.439876564 | 0.70908787 |
| 0.439838025 | 0.709049744 |
| 0.439709895 | 0.708963173 |
| 0.439706876 | 0.708963173 |
| 0.439542113 | 0.70883663 |
| 0.439532755 | 0.70883663 |
| 0.439513376 | 0.70883663 |
| 0.439383553 | 0.708748915 |
| 0.439381525 | 0.708748915 |
| 0.439374878 | 0.708748915 |
| 0.439257623 | 0.708675627 |
| 0.439230072 | 0.708675627 |
| 0.439218797 | 0.708675627 |
| 0.439161371 | 0.708654536 |
| 0.439127527 | 0.708623931 |

| | |
|---|---|
| 0.439116359 | 0.708623931 |
| 0.439031886 | 0.708541616 |
| 0.439016808 | 0.708541616 |
| 0.439013825 | 0.708541616 |
| 0.439004586 | 0.708541616 |
| 0.438979894 | 0.708541616 |
| 0.438963481 | 0.708541616 |
| 0.438948941 | 0.708541616 |
| 0.438893362 | 0.708541616 |
| 0.438875592 | 0.708541616 |
| 0.438864227 | 0.708541616 |
| 0.438764729 | 0.708541616 |
| 0.438698434 | 0.708479426 |
| 0.438616527 | 0.708399381 |
| 0.438615283 | 0.708399381 |
| 0.438604605 | 0.708399381 |
| 0.438555768 | 0.70839734 |
| 0.438553815 | 0.70839734 |
| 0.438491635 | 0.708365821 |
| 0.438442923 | 0.708311152 |
| 0.438408403 | 0.708303436 |
| 0.438364003 | 0.708255728 |
| 0.43832417 | 0.708215397 |
| 0.438323148 | 0.708215397 |
| 0.438238876 | 0.708149656 |
| 0.438183211 | 0.708083731 |
| 0.438176764 | 0.708083731 |
| 0.437962411 | 0.707895057 |
| 0.437830661 | 0.70773612 |
| 0.437766821 | 0.707688572 |
| 0.437733056 | 0.707688572 |
| 0.437503172 | 0.707440877 |
| 0.43750164 | 0.707440877 |
| 0.437468217 | 0.707440877 |
| 0.437460829 | 0.707440877 |
| 0.437414393 | 0.707440877 |
| 0.437378922 | 0.707440877 |
| 0.437370806 | 0.707440877 |
| 0.437359488 | 0.707440877 |
| 0.437349492 | 0.707440877 |
| 0.437315401 | 0.707440877 |
| 0.437286773 | 0.707440877 |
| 0.437243505 | 0.707405272 |
| 0.437161673 | 0.707368995 |
| 0.436957948 | 0.70711142 |

| | |
|---|---|
| 0.436928447 | 0.707087705 |
| 0.436917634 | 0.707087705 |
| 0.436852219 | 0.707036417 |
| 0.436851575 | 0.707036417 |
| 0.43682809 | 0.707036417 |
| 0.436692313 | 0.706945777 |
| 0.436630028 | 0.706889925 |
| 0.436568737 | 0.706817801 |
| 0.436554217 | 0.706817801 |
| 0.436506865 | 0.70676568 |
| 0.436429436 | 0.70668836 |
| 0.436426053 | 0.70668836 |
| 0.436413149 | 0.70668836 |
| 0.436351536 | 0.706682352 |
| 0.43626122 | 0.706584136 |
| 0.436224134 | 0.706548199 |
| 0.436119486 | 0.70642665 |
| 0.436100943 | 0.70642665 |
| 0.436097315 | 0.70642665 |
| 0.436050617 | 0.706411195 |
| 0.436014037 | 0.706375962 |
| 0.435978169 | 0.706341881 |
| 0.43593893 | 0.706302334 |
| 0.435892203 | 0.706250653 |
| 0.435820614 | 0.70618271 |
| 0.435759469 | 0.706143463 |
| 0.435712112 | 0.706102989 |
| 0.435703075 | 0.706102989 |
| 0.435693394 | 0.706102989 |
| 0.435685973 | 0.706102989 |
| 0.435599038 | 0.706015837 |
| 0.435593865 | 0.706015837 |
| 0.435586451 | 0.706015837 |
| 0.435481928 | 0.705946147 |
| 0.435306364 | 0.705781657 |
| 0.435258564 | 0.705728183 |
| 0.435256772 | 0.705728183 |
| 0.43521459 | 0.705704933 |
| 0.435210299 | 0.705704933 |
| 0.435206189 | 0.705704933 |
| 0.435199088 | 0.705704933 |
| 0.435187919 | 0.705704933 |
| 0.435179775 | 0.705704933 |
| 0.435167085 | 0.705704933 |
| 0.435083423 | 0.705704933 |

| | |
|---|---|
| 0.435075648 | 0.705704933 |
| 0.434990157 | 0.705605266 |
| 0.434989593 | 0.705605266 |
| 0.434987078 | 0.705605266 |
| 0.434937363 | 0.705605266 |
| 0.434920832 | 0.705605266 |
| 0.434873582 | 0.705570041 |
| 0.434819432 | 0.705520564 |
| 0.434784451 | 0.70548784 |
| 0.434673399 | 0.705355707 |
| 0.434661811 | 0.705355707 |
| 0.434661811 | 0.705355707 |
| 0.434661811 | 0.705355707 |
| 0.434661331 | 0.705355707 |
| 0.434645583 | 0.705355707 |
| 0.434626527 | 0.705355707 |
| 0.43457353 | 0.705355707 |
| 0.434533151 | 0.705320373 |
| 0.434400169 | 0.705152585 |
| 0.434354412 | 0.705102341 |
| 0.434336762 | 0.705097722 |
| 0.434303811 | 0.705093807 |
| 0.434302538 | 0.705093807 |
| 0.434281198 | 0.705093807 |
| 0.434278128 | 0.705093807 |
| 0.434230431 | 0.705093807 |
| 0.43421551 | 0.705093807 |
| 0.434212457 | 0.705093807 |
| 0.434160924 | 0.705093807 |
| 0.434143757 | 0.705093807 |
| 0.434019904 | 0.704991849 |
| 0.433932668 | 0.70491865 |
| 0.433828034 | 0.704800374 |
| 0.433792249 | 0.704766278 |
| 0.433708974 | 0.704703101 |
| 0.433678515 | 0.704678653 |
| 0.433668864 | 0.704678653 |
| 0.433588516 | 0.704635618 |
| 0.433434663 | 0.704496857 |
| 0.433430561 | 0.704496857 |
| 0.433380734 | 0.704481267 |
| 0.433365844 | 0.704481267 |
| 0.433330739 | 0.704449026 |
| 0.433276207 | 0.704416902 |
| 0.433187072 | 0.704335678 |

| | |
|---|---|
| 0.433162478 | 0.704319734 |
| 0.43314506 | 0.704315459 |
| 0.433073645 | 0.704257647 |
| 0.432974439 | 0.704182271 |
| 0.432890951 | 0.704118247 |
| 0.432831862 | 0.704046553 |
| 0.432776724 | 0.703986644 |
| 0.432710385 | 0.703971752 |
| 0.432706928 | 0.703971752 |
| 0.432681826 | 0.703971752 |
| 0.43266914 | 0.703971752 |
| 0.432653025 | 0.703971752 |
| 0.432618939 | 0.703971752 |
| 0.43261201 | 0.703971752 |
| 0.432607203 | 0.703971752 |
| 0.432589203 | 0.703971752 |
| 0.432511404 | 0.703909945 |
| 0.432507625 | 0.703909945 |
| 0.432452744 | 0.703886635 |
| 0.432370451 | 0.703800795 |
| 0.432202472 | 0.703599506 |
| 0.432193142 | 0.703599506 |
| 0.432165273 | 0.703587046 |
| 0.432157544 | 0.703587046 |
| 0.43201981 | 0.70344641 |
| 0.432012857 | 0.70344641 |
| 0.432009219 | 0.70344641 |
| 0.431893627 | 0.703337133 |
| 0.431841948 | 0.703301069 |
| 0.431837637 | 0.703301069 |
| 0.431804395 | 0.703301069 |
| 0.431801857 | 0.703301069 |
| 0.431751624 | 0.703301069 |
| 0.431657664 | 0.703241416 |
| 0.43163286 | 0.703241416 |
| 0.431599819 | 0.703241416 |
| 0.431587363 | 0.703241416 |
| 0.431559022 | 0.703241416 |
| 0.431496493 | 0.703241416 |
| 0.431489455 | 0.703241416 |
| 0.431440825 | 0.703200855 |
| 0.431404355 | 0.703196721 |
| 0.43137407 | 0.703188308 |
| 0.431359932 | 0.703188308 |
| 0.431352044 | 0.703188308 |

| | |
|---|---|
| 0.431345538 | 0.703188308 |
| 0.431336108 | 0.703188308 |
| 0.431304232 | 0.703188308 |
| 0.431293677 | 0.703188308 |
| 0.431293037 | 0.703188308 |
| 0.431273605 | 0.703188308 |
| 0.431193456 | 0.703158622 |
| 0.431137283 | 0.703091094 |
| 0.431132806 | 0.703091094 |
| 0.431130534 | 0.703091094 |
| 0.431087955 | 0.703091094 |
| 0.431012576 | 0.703091094 |
| 0.430936316 | 0.703028154 |
| 0.430896967 | 0.70298804 |
| 0.430828431 | 0.702943159 |
| 0.430756872 | 0.702888576 |
| 0.430729344 | 0.702888576 |
| 0.430709342 | 0.702888576 |
| 0.430704071 | 0.702888576 |
| 0.430680138 | 0.702888576 |
| 0.430663506 | 0.702888576 |
| 0.430599038 | 0.702887219 |
| 0.430510055 | 0.702814232 |
| 0.430482142 | 0.702792752 |
| 0.430369703 | 0.702699777 |
| 0.430359884 | 0.702699777 |
| 0.430326333 | 0.702699777 |
| 0.430310573 | 0.702699777 |
| 0.430307011 | 0.702699777 |
| 0.430301056 | 0.702699777 |
| 0.430292808 | 0.702699777 |
| 0.430266179 | 0.702699777 |
| 0.430240691 | 0.702699777 |
| 0.43017571 | 0.702699777 |
| 0.430127953 | 0.702672156 |
| 0.430043775 | 0.702558737 |
| 0.429940913 | 0.702438883 |
| 0.42992613 | 0.702438827 |
| 0.429911263 | 0.702438633 |
| 0.429865929 | 0.702388658 |
| 0.429743155 | 0.702236233 |
| 0.429601643 | 0.70212544 |
| 0.429575966 | 0.70212544 |
| 0.429503104 | 0.702060759 |
| 0.429462516 | 0.702018508 |

| | |
|---|---|
| 0.429379538 | 0.701931052 |
| 0.429337707 | 0.70188676 |
| 0.429284387 | 0.701823682 |
| 0.429228696 | 0.701780814 |
| 0.42921957 | 0.701780814 |
| 0.429142286 | 0.701735894 |
| 0.429131397 | 0.701735894 |
| 0.429078138 | 0.701697178 |
| 0.428999054 | 0.701622127 |
| 0.428976096 | 0.701608672 |
| 0.428863547 | 0.701545051 |
| 0.428862517 | 0.701545051 |
| 0.428827287 | 0.701545051 |
| 0.428818029 | 0.701545051 |
| 0.428796241 | 0.701545051 |
| 0.428737381 | 0.701531427 |
| 0.428726176 | 0.701531427 |
| 0.428686218 | 0.701520015 |
| 0.428631171 | 0.701475381 |
| 0.428611238 | 0.701469621 |
| 0.428606392 | 0.701469621 |
| 0.42859965 | 0.701469621 |
| 0.428557433 | 0.701453875 |
| 0.428538028 | 0.701453875 |
| 0.428337378 | 0.701237027 |
| 0.428254551 | 0.701126793 |
| 0.428250729 | 0.701126793 |
| 0.428186406 | 0.701087525 |
| 0.428176348 | 0.701087525 |
| 0.428147272 | 0.701071655 |
| 0.428137972 | 0.701071655 |
| 0.428135106 | 0.701071655 |
| 0.428028891 | 0.700965499 |
| 0.428011687 | 0.700965499 |
| 0.42798625 | 0.700965499 |
| 0.427979378 | 0.700965499 |
| 0.427931933 | 0.700935935 |
| 0.427905506 | 0.700916754 |
| 0.427872281 | 0.700886435 |
| 0.427869574 | 0.700886435 |
| 0.427817794 | 0.700886435 |
| 0.427705557 | 0.700733834 |
| 0.427531227 | 0.700496413 |
| 0.427502844 | 0.700474007 |
| 0.427496798 | 0.700474007 |

| | |
|---|---|
| 0.42748757 | 0.700474007 |
| 0.427438423 | 0.700440744 |
| 0.427426005 | 0.700440744 |
| 0.427385429 | 0.700419829 |
| 0.427360294 | 0.70040911 |
| 0.427344893 | 0.70040911 |
| 0.427314987 | 0.700407162 |
| 0.427289666 | 0.700389762 |
| 0.42717126 | 0.700243877 |
| 0.427137615 | 0.700212825 |
| 0.427085337 | 0.700151223 |
| 0.426929313 | 0.699943628 |
| 0.426926304 | 0.699943628 |
| 0.426917866 | 0.699943628 |
| 0.426894213 | 0.699943628 |
| 0.426876307 | 0.699943628 |
| 0.426864767 | 0.699943628 |
| 0.426640016 | 0.699691321 |
| 0.426630274 | 0.699691321 |
| 0.426612679 | 0.699691321 |
| 0.426523017 | 0.699604089 |
| 0.426492672 | 0.699588996 |
| 0.426445516 | 0.699535737 |
| 0.426443899 | 0.699535737 |
| 0.426429749 | 0.699535737 |
| 0.426383554 | 0.69953047 |
| 0.426341881 | 0.69953047 |
| 0.426313338 | 0.69953047 |
| 0.426267221 | 0.699508285 |
| 0.426140843 | 0.69934909 |
| 0.426043931 | 0.699254288 |
| 0.425911445 | 0.699141255 |
| 0.425892491 | 0.699134239 |
| 0.42580883 | 0.699044194 |
| 0.425775902 | 0.699015129 |
| 0.425773192 | 0.699015129 |
| 0.425732043 | 0.699015129 |
| 0.425719003 | 0.699015129 |
| 0.425692986 | 0.699015129 |
| 0.425692747 | 0.699015129 |
| 0.425673413 | 0.699015129 |
| 0.425667226 | 0.699015129 |
| 0.425656283 | 0.699015129 |
| 0.425642648 | 0.699015129 |
| 0.425618244 | 0.699015129 |

| | |
|---|---|
| 0.425564677 | 0.69898158 |
| 0.425457871 | 0.698902563 |
| 0.425446224 | 0.698902563 |
| 0.425437477 | 0.698902563 |
| 0.425433146 | 0.698902563 |
| 0.425372505 | 0.69885875 |
| 0.425361679 | 0.69885875 |
| 0.42533953 | 0.69885875 |
| 0.42527289 | 0.698791511 |
| 0.425217178 | 0.698724072 |
| 0.425070409 | 0.69856826 |
| 0.425061548 | 0.69856826 |
| 0.425040784 | 0.69856826 |
| 0.425022363 | 0.69856826 |
| 0.42501697 | 0.69856826 |
| 0.424827288 | 0.698372541 |
| 0.424808889 | 0.698366399 |
| 0.424794143 | 0.698366264 |
| 0.424760073 | 0.698334357 |
| 0.424745816 | 0.698334357 |
| 0.424434905 | 0.697992515 |
| 0.424409005 | 0.697992515 |
| 0.424367434 | 0.697955834 |
| 0.42435555 | 0.697955834 |
| 0.424344611 | 0.697955834 |
| 0.424296461 | 0.697955834 |
| 0.424235632 | 0.697929894 |
| 0.424231828 | 0.697929894 |
| 0.424108605 | 0.697769121 |
| 0.42407992 | 0.697746031 |
| 0.423967644 | 0.697627271 |
| 0.423952781 | 0.697627271 |
| 0.423950881 | 0.697627271 |
| 0.42379597 | 0.697447501 |
| 0.423771301 | 0.697447501 |
| 0.423708637 | 0.697376067 |
| 0.42370518 | 0.697376067 |
| 0.423410594 | 0.697030049 |
| 0.423410502 | 0.697030049 |
| 0.423364062 | 0.697020796 |
| 0.423278882 | 0.697020796 |
| 0.423239335 | 0.697013133 |
| 0.423223966 | 0.697013133 |
| 0.42319283 | 0.697013133 |
| 0.423016587 | 0.696790861 |

| | |
|---|---|
| 0.422928853 | 0.696694538 |
| 0.422879711 | 0.696637681 |
| 0.422872362 | 0.696637681 |
| 0.422815074 | 0.696613604 |
| 0.422787082 | 0.696605561 |
| 0.422583646 | 0.696432367 |
| 0.422425517 | 0.696298888 |
| 0.422379007 | 0.696284234 |
| 0.42219807 | 0.696092619 |
| 0.422110779 | 0.696020984 |
| 0.422068067 | 0.69597465 |
| 0.422035921 | 0.695967235 |
| 0.422020308 | 0.695967235 |
| 0.422000157 | 0.695967235 |
| 0.421980019 | 0.695967235 |
| 0.421975069 | 0.695967235 |
| 0.42196179 | 0.695967235 |
| 0.42194633 | 0.695967235 |
| 0.421845177 | 0.695872134 |
| 0.421805433 | 0.695830673 |
| 0.42180088 | 0.695830673 |
| 0.421781713 | 0.695830673 |
| 0.421704994 | 0.695737278 |
| 0.421697856 | 0.695737278 |
| 0.42169309 | 0.695737278 |
| 0.421688891 | 0.695737278 |
| 0.421656285 | 0.695737278 |
| 0.421559472 | 0.695641777 |
| 0.421410406 | 0.695443985 |
| 0.421403404 | 0.695443985 |
| 0.42137658 | 0.695436357 |
| 0.421284069 | 0.695331867 |
| 0.421204108 | 0.695251662 |
| 0.421127815 | 0.695218524 |
| 0.421042686 | 0.695102084 |
| 0.421040561 | 0.695102084 |
| 0.421039222 | 0.695102084 |
| 0.421032541 | 0.695102084 |
| 0.421015253 | 0.695102084 |
| 0.420890722 | 0.695019847 |
| 0.420878026 | 0.695019847 |
| 0.420853825 | 0.695019847 |
| 0.42080298 | 0.694995442 |
| 0.420787184 | 0.694993453 |
| 0.420758479 | 0.694993453 |

| | |
|---|---|
| 0.420756794 | 0.694993453 |
| 0.420728888 | 0.694993453 |
| 0.420680023 | 0.694961064 |
| 0.420661049 | 0.694953824 |
| 0.420529351 | 0.694823656 |
| 0.420517779 | 0.694823656 |
| 0.420489194 | 0.694823656 |
| 0.420482652 | 0.694823656 |
| 0.420449929 | 0.694823656 |
| 0.420381755 | 0.694817776 |
| 0.420359585 | 0.694817776 |
| 0.42029678 | 0.694785837 |
| 0.420249839 | 0.694732353 |
| 0.420248073 | 0.694732353 |
| 0.420142746 | 0.694627645 |
| 0.420054977 | 0.694530756 |
| 0.420024292 | 0.694504131 |
| 0.420020513 | 0.694504131 |
| 0.420006882 | 0.694504131 |
| 0.419979607 | 0.694502578 |
| 0.419917536 | 0.694485949 |
| 0.419856124 | 0.694418933 |
| 0.41985499 | 0.694418933 |
| 0.419852571 | 0.694418933 |
| 0.41980354 | 0.694406062 |
| 0.419750767 | 0.694365245 |
| 0.419734255 | 0.694362048 |
| 0.41968102 | 0.694320573 |
| 0.419676983 | 0.694320573 |
| 0.419654575 | 0.694320573 |
| 0.419575646 | 0.694292588 |
| 0.419563785 | 0.694292588 |
| 0.419532824 | 0.694269973 |
| 0.419424904 | 0.694163739 |
| 0.41942345 | 0.694163739 |
| 0.419386444 | 0.694163739 |
| 0.419357085 | 0.694147982 |
| 0.419261238 | 0.694061695 |
| 0.419248067 | 0.694061695 |
| 0.419212787 | 0.694029735 |
| 0.419108271 | 0.693904943 |
| 0.419065883 | 0.693859405 |
| 0.419028917 | 0.693859405 |
| 0.418960521 | 0.693780909 |
| 0.418874038 | 0.693685936 |

| | |
|---|---|
| 0.418801649 | 0.693638411 |
| 0.418737581 | 0.693556417 |
| 0.418596899 | 0.693395747 |
| 0.418595804 | 0.693395747 |
| 0.418522569 | 0.693365486 |
| 0.418503083 | 0.693365486 |
| 0.418498683 | 0.693365486 |
| 0.418461204 | 0.693363893 |
| 0.418456402 | 0.693363893 |
| 0.418451995 | 0.693363893 |
| 0.418300397 | 0.693218032 |
| 0.418295054 | 0.693218032 |
| 0.418255332 | 0.693215711 |
| 0.418227076 | 0.693193003 |
| 0.418142056 | 0.69312445 |
| 0.41809295 | 0.693067173 |
| 0.418077842 | 0.69306625 |
| 0.41801296 | 0.692982814 |
| 0.417883574 | 0.692816548 |
| 0.41783378 | 0.692782226 |
| 0.417753383 | 0.69274539 |
| 0.417647159 | 0.692641592 |
| 0.417597083 | 0.692587798 |
| 0.417466683 | 0.692438731 |
| 0.417460415 | 0.692438731 |
| 0.417413874 | 0.692399366 |
| 0.417391681 | 0.692386667 |
| 0.417367989 | 0.69237148 |
| 0.417290007 | 0.692290343 |
| 0.417265214 | 0.692282492 |
| 0.417257702 | 0.692282492 |
| 0.417208516 | 0.692282492 |
| 0.417180233 | 0.692282492 |
| 0.417105869 | 0.692201861 |
| 0.417086397 | 0.692193666 |
| 0.417081381 | 0.692193666 |
| 0.4170304 | 0.692193666 |
| 0.417006086 | 0.69218098 |
| 0.416981735 | 0.692164684 |
| 0.416976643 | 0.692164684 |
| 0.416882344 | 0.692072063 |
| 0.416851049 | 0.692044232 |
| 0.416813598 | 0.692030302 |
| 0.41668724 | 0.69184463 |
| 0.416433424 | 0.691495526 |

| | |
|---|---|
| 0.416427026 | 0.691495526 |
| 0.416395888 | 0.691481415 |
| 0.416371785 | 0.691465498 |
| 0.416350009 | 0.691453445 |
| 0.416349739 | 0.691453445 |
| 0.416330351 | 0.691453445 |
| 0.416283019 | 0.691438632 |
| 0.416274812 | 0.691438632 |
| 0.416221866 | 0.691394649 |
| 0.416196447 | 0.691391287 |
| 0.416189107 | 0.691391287 |
| 0.416160853 | 0.691380388 |
| 0.416074327 | 0.691260751 |
| 0.416038062 | 0.691260751 |
| 0.415996189 | 0.691227384 |
| 0.415969593 | 0.691207305 |
| 0.41594525 | 0.691190972 |
| 0.415942429 | 0.691190972 |
| 0.41591805 | 0.691190972 |
| 0.415904898 | 0.691190972 |
| 0.415772608 | 0.69107287 |
| 0.415771576 | 0.69107287 |
| 0.415742615 | 0.691071253 |
| 0.415683144 | 0.691020632 |
| 0.415666162 | 0.691020632 |
| 0.415618132 | 0.690987494 |
| 0.415573896 | 0.690959607 |
| 0.415413835 | 0.690741708 |
| 0.415395618 | 0.690741708 |
| 0.415368959 | 0.690739434 |
| 0.415284528 | 0.690647259 |
| 0.415198495 | 0.690528294 |
| 0.415107156 | 0.690448716 |
| 0.415082421 | 0.690448716 |
| 0.415018839 | 0.690398262 |
| 0.41501783 | 0.690398262 |
| 0.414974226 | 0.690398262 |
| 0.414972906 | 0.690398262 |
| 0.414938898 | 0.690398262 |
| 0.414890697 | 0.690353884 |
| 0.414719652 | 0.690190108 |
| 0.414674034 | 0.690186273 |
| 0.414648968 | 0.690168672 |
| 0.41464001 | 0.690168672 |
| 0.414636553 | 0.690168672 |

| | |
|---|---|
| 0.414583329 | 0.690165033 |
| 0.414544993 | 0.690140324 |
| 0.41449989 | 0.690089358 |
| 0.414453844 | 0.690036819 |
| 0.414330314 | 0.689855265 |
| 0.414256347 | 0.689804456 |
| 0.414253552 | 0.689804456 |
| 0.414215346 | 0.689804456 |
| 0.414182867 | 0.689804456 |
| 0.414178793 | 0.689804456 |
| 0.414156619 | 0.689804456 |
| 0.414148975 | 0.689804456 |
| 0.414058232 | 0.689715704 |
| 0.413785626 | 0.689382166 |
| 0.413757468 | 0.689359369 |
| 0.41369947 | 0.689286851 |
| 0.413648611 | 0.689237722 |
| 0.413635168 | 0.689237722 |
| 0.413537047 | 0.68911266 |
| 0.413470075 | 0.689025167 |
| 0.413464142 | 0.689025167 |
| 0.413441395 | 0.689025167 |
| 0.413418621 | 0.689011746 |
| 0.413351593 | 0.688924144 |
| 0.413344564 | 0.688924144 |
| 0.413321709 | 0.688924144 |
| 0.413312025 | 0.688924144 |
| 0.413281623 | 0.688924144 |
| 0.413266639 | 0.688924144 |
| 0.413257904 | 0.688924144 |
| 0.413243449 | 0.688924144 |
| 0.413194709 | 0.688903745 |
| 0.413164661 | 0.688877763 |
| 0.413164634 | 0.688877763 |
| 0.413149974 | 0.688877763 |
| 0.413124161 | 0.688877763 |
| 0.413106439 | 0.688877153 |
| 0.413104256 | 0.688877153 |
| 0.412986162 | 0.688748931 |
| 0.412940723 | 0.688697266 |
| 0.412838262 | 0.688574607 |
| 0.412779768 | 0.68854026 |
| 0.41271125 | 0.688459202 |
| 0.412675715 | 0.688424037 |
| 0.412641428 | 0.688406737 |

| | |
|---|---|
| 0.412614396 | 0.688394081 |
| 0.412609543 | 0.688394081 |
| 0.412543879 | 0.68835583 |
| 0.412518011 | 0.688353841 |
| 0.412489654 | 0.688353841 |
| 0.41242636 | 0.688297366 |
| 0.41235644 | 0.688209047 |
| 0.412353907 | 0.688209047 |
| 0.41233903 | 0.688209047 |
| 0.412314547 | 0.688209047 |
| 0.412300482 | 0.688209047 |
| 0.412277617 | 0.688209047 |
| 0.412235736 | 0.688209047 |
| 0.412234796 | 0.688209047 |
| 0.41222732 | 0.688209047 |
| 0.412189592 | 0.688209047 |
| 0.412155937 | 0.688207791 |
| 0.41202176 | 0.688032961 |
| 0.411979419 | 0.688032961 |
| 0.411892983 | 0.687920157 |
| 0.411866496 | 0.687917626 |
| 0.411770785 | 0.687854062 |
| 0.41172282 | 0.687798062 |
| 0.411704506 | 0.687791592 |
| 0.411659996 | 0.687741358 |
| 0.41161541 | 0.687690992 |
| 0.411596816 | 0.687684051 |
| 0.411585923 | 0.687684051 |
| 0.411499411 | 0.687666051 |
| 0.411446615 | 0.687601949 |
| 0.411335515 | 0.687440402 |
| 0.411185474 | 0.687233078 |
| 0.41110086 | 0.687120571 |
| 0.410863295 | 0.686816493 |
| 0.410828878 | 0.686816493 |
| 0.410822623 | 0.686816493 |
| 0.410814564 | 0.686816493 |
| 0.410751511 | 0.68680179 |
| 0.410709615 | 0.686755849 |
| 0.410706983 | 0.686755849 |
| 0.410705435 | 0.686755849 |
| 0.410452163 | 0.68651818 |
| 0.410437959 | 0.68651818 |
| 0.410414741 | 0.686503806 |
| 0.410383676 | 0.686475953 |

| | |
|---|---|
| 0.410299463 | 0.686383298 |
| 0.410299002 | 0.686383298 |
| 0.410222832 | 0.686303317 |
| 0.410210787 | 0.686303317 |
| 0.410162025 | 0.686273907 |
| 0.410125378 | 0.686236698 |
| 0.410050787 | 0.686160102 |
| 0.410024228 | 0.686139768 |
| 0.409968835 | 0.686095285 |
| 0.409955385 | 0.686095285 |
| 0.409921681 | 0.686064586 |
| 0.409850277 | 0.685969186 |
| 0.409828082 | 0.685956144 |
| 0.409756563 | 0.685911224 |
| 0.409712458 | 0.685883136 |
| 0.409707802 | 0.685883136 |
| 0.409692586 | 0.685883136 |
| 0.40959825 | 0.685788366 |
| 0.409589251 | 0.685788366 |
| 0.409550946 | 0.685782548 |
| 0.409421395 | 0.685685096 |
| 0.409355593 | 0.685623113 |
| 0.409351081 | 0.685623113 |
| 0.409348176 | 0.685623113 |
| 0.409328608 | 0.685623113 |
| 0.409280207 | 0.685620582 |
| 0.409251063 | 0.685616814 |
| 0.40916256 | 0.685565231 |
| 0.409106726 | 0.685537214 |
| 0.409032786 | 0.685468139 |
| 0.408955329 | 0.685362449 |
| 0.408801742 | 0.685225613 |
| 0.408737954 | 0.68516692 |
| 0.408690225 | 0.685111026 |
| 0.408615986 | 0.685039239 |
| 0.408409811 | 0.684833788 |
| 0.408376355 | 0.684801798 |
| 0.408363785 | 0.684801798 |
| 0.408362147 | 0.684801798 |
| 0.408285465 | 0.684794049 |
| 0.408244095 | 0.684783803 |
| 0.40814004 | 0.684692244 |
| 0.408105825 | 0.684677888 |
| 0.408080794 | 0.68466777 |
| 0.408029616 | 0.68463014 |

| | |
|---|---|
| 0.407958085 | 0.684558353 |
| 0.407929723 | 0.684534879 |
| 0.407893857 | 0.684498809 |
| 0.407745021 | 0.684302058 |
| 0.40770349 | 0.68427579 |
| 0.407509424 | 0.684036191 |
| 0.407468345 | 0.684025761 |
| 0.407390562 | 0.683919291 |
| 0.407390166 | 0.683919291 |
| 0.407378919 | 0.683919291 |
| 0.407344247 | 0.683919291 |
| 0.407264692 | 0.683919291 |
| 0.407187911 | 0.683879206 |
| 0.407157924 | 0.683866217 |
| 0.407055522 | 0.683766563 |
| 0.406928314 | 0.68357699 |
| 0.406682064 | 0.68331691 |
| 0.406594695 | 0.68320934 |
| 0.406556454 | 0.683169186 |
| 0.40651468 | 0.68313555 |
| 0.406508553 | 0.68313555 |
| 0.406499199 | 0.68313555 |
| 0.406473284 | 0.68313555 |
| 0.406471787 | 0.68313555 |
| 0.406464303 | 0.68313555 |
| 0.406417682 | 0.683128814 |
| 0.406389786 | 0.683106033 |
| 0.406383371 | 0.683106033 |
| 0.406336074 | 0.683088074 |
| 0.406294216 | 0.683041815 |
| 0.406254412 | 0.682999008 |
| 0.406245636 | 0.682999008 |
| 0.406191924 | 0.682942806 |
| 0.406191415 | 0.682942806 |
| 0.40612085 | 0.682942806 |
| 0.406112943 | 0.682942806 |
| 0.406074465 | 0.682942806 |
| 0.406054066 | 0.682942806 |
| 0.405960136 | 0.682914079 |
| 0.40589679 | 0.682879876 |
| 0.405844321 | 0.682815723 |
| 0.405834946 | 0.682815723 |
| 0.405778587 | 0.682753365 |
| 0.405750556 | 0.68273032 |
| 0.405739746 | 0.68273032 |

| | |
|---|---|
| 0.405703194 | 0.682722988 |
| 0.405484866 | 0.682500273 |
| 0.405472836 | 0.682500273 |
| 0.405467775 | 0.682500273 |
| 0.405451553 | 0.682500273 |
| 0.405400024 | 0.682478066 |
| 0.405381704 | 0.682471348 |
| 0.405368746 | 0.682471348 |
| 0.405330227 | 0.682432929 |
| 0.405220465 | 0.68234898 |
| 0.405181344 | 0.68234898 |
| 0.405162535 | 0.682343566 |
| 0.405045781 | 0.682195184 |
| 0.405038666 | 0.682195184 |
| 0.405038626 | 0.682195184 |
| 0.404961668 | 0.682150757 |
| 0.404779159 | 0.681968078 |
| 0.40474831 | 0.681968078 |
| 0.404663924 | 0.681881053 |
| 0.404630713 | 0.681881053 |
| 0.404623387 | 0.681881053 |
| 0.404614133 | 0.681881053 |
| 0.404593588 | 0.681881053 |
| 0.40455797 | 0.681855844 |
| 0.404535085 | 0.681841402 |
| 0.40452016 | 0.681840376 |
| 0.404513098 | 0.681840376 |
| 0.404484537 | 0.681828592 |
| 0.404477246 | 0.681828592 |
| 0.404422423 | 0.681809998 |
| 0.404336997 | 0.681748794 |
| 0.404329612 | 0.681748794 |
| 0.404328188 | 0.681748794 |
| 0.404326956 | 0.681748794 |
| 0.404315627 | 0.681748794 |
| 0.404306017 | 0.681748794 |
| 0.404297188 | 0.681748794 |
| 0.404290349 | 0.681748794 |
| 0.404263531 | 0.681748794 |
| 0.404178678 | 0.681748794 |
| 0.404145005 | 0.681748794 |
| 0.404133996 | 0.681748794 |
| 0.404129423 | 0.681748794 |
| 0.404112721 | 0.681748794 |
| 0.404109638 | 0.681748794 |

| | |
|---|---|
| 0.404101073 | 0.681748794 |
| 0.403961676 | 0.681695169 |
| 0.403951385 | 0.681695169 |
| 0.403944298 | 0.681695169 |
| 0.403886708 | 0.68166527 |
| 0.403881257 | 0.68166527 |
| 0.403881209 | 0.68166527 |
| 0.403845057 | 0.68166527 |
| 0.403698906 | 0.681493205 |
| 0.403648983 | 0.681452945 |
| 0.403590922 | 0.681383368 |
| 0.403532787 | 0.681351496 |
| 0.403493617 | 0.681339851 |
| 0.403284392 | 0.681153237 |
| 0.403254094 | 0.681128586 |
| 0.403226052 | 0.681105376 |
| 0.403188277 | 0.681065724 |
| 0.403115384 | 0.681023882 |
| 0.40306786 | 0.680983077 |
| 0.403035468 | 0.680952508 |
| 0.403004154 | 0.680923756 |
| 0.402959223 | 0.680871995 |
| 0.402941733 | 0.680866598 |
| 0.402928957 | 0.680866598 |
| 0.40292053 | 0.680866598 |
| 0.402893342 | 0.680857297 |
| 0.402883655 | 0.680857297 |
| 0.402816364 | 0.680775521 |
| 0.402806895 | 0.680775521 |
| 0.402737298 | 0.680693089 |
| 0.402686717 | 0.680653025 |
| 0.40267768 | 0.680653025 |
| 0.402634893 | 0.680613737 |
| 0.402579606 | 0.68054636 |
| 0.402543227 | 0.68054636 |
| 0.402431183 | 0.680510917 |
| 0.402392379 | 0.680469459 |
| 0.402385015 | 0.680469459 |
| 0.402382386 | 0.680469459 |
| 0.402376206 | 0.680469459 |
| 0.402353397 | 0.680469459 |
| 0.402328936 | 0.680469459 |
| 0.402307751 | 0.680469459 |
| 0.402251817 | 0.680414285 |
| 0.402092119 | 0.68025154 |

| | |
|---|---|
| 0.402040842 | 0.68022438 |
| 0.40201448 | 0.68022438 |
| 0.401999682 | 0.68022438 |
| 0.401971652 | 0.68022438 |
| 0.401921808 | 0.68020503 |
| 0.401890729 | 0.680176601 |
| 0.401865257 | 0.680157661 |
| 0.401850483 | 0.680156828 |
| 0.401844505 | 0.680156828 |
| 0.401769331 | 0.680067808 |
| 0.401733874 | 0.68003196 |
| 0.40170504 | 0.68003149 |
| 0.401642292 | 0.68002195 |
| 0.401639094 | 0.68002195 |
| 0.401628677 | 0.68002195 |
| 0.40157124 | 0.68002195 |
| 0.401569031 | 0.68002195 |
| 0.401535919 | 0.680011066 |
| 0.40150135 | 0.679976702 |
| 0.401479142 | 0.679976702 |
| 0.401458465 | 0.679976702 |
| 0.401409406 | 0.679966062 |
| 0.401397903 | 0.679966062 |
| 0.401326628 | 0.67992257 |
| 0.401315549 | 0.67992257 |
| 0.401259975 | 0.679882202 |
| 0.401241288 | 0.679882202 |
| 0.401241288 | 0.679882202 |
| 0.401142845 | 0.679828855 |
| 0.401140884 | 0.679828855 |
| 0.401127327 | 0.679828855 |
| 0.40100281 | 0.679688274 |
| 0.400966815 | 0.679651452 |
| 0.400964362 | 0.679651452 |
| 0.400948381 | 0.679651452 |
| 0.400928514 | 0.679651452 |
| 0.400916271 | 0.679651452 |
| 0.400905196 | 0.679651452 |
| 0.400836902 | 0.679624747 |
| 0.400819282 | 0.679624747 |
| 0.400797678 | 0.679624747 |
| 0.400739841 | 0.679556947 |
| 0.400733048 | 0.679556947 |
| 0.40065221 | 0.679480925 |
| 0.400598333 | 0.679413746 |

| | |
|---|---|
| 0.400586062 | 0.679413746 |
| 0.400547879 | 0.679400757 |
| 0.400516522 | 0.679392628 |
| 0.400487451 | 0.679370845 |
| 0.400437224 | 0.679358232 |
| 0.400335688 | 0.679234364 |
| 0.400320959 | 0.679233569 |
| 0.400304696 | 0.679230175 |
| 0.400275723 | 0.67920521 |
| 0.400271026 | 0.67920521 |
| 0.40023862 | 0.679190649 |
| 0.400229761 | 0.679190649 |
| 0.400229747 | 0.679190649 |
| 0.400148739 | 0.67912692 |
| 0.400043752 | 0.679005124 |
| 0.400034079 | 0.679005124 |
| 0.400032761 | 0.679005124 |
| 0.400016251 | 0.679005124 |
| 0.400013873 | 0.679005124 |
| 0.399988996 | 0.679005124 |
| 0.399946241 | 0.678984801 |
| 0.399896123 | 0.678972323 |
| 0.399855187 | 0.678927023 |
| 0.39985489 | 0.678927023 |
| 0.399832593 | 0.678927023 |
| 0.399830742 | 0.678927023 |
| 0.399742274 | 0.678904728 |
| 0.399739108 | 0.678904728 |
| 0.399728508 | 0.678904728 |
| 0.39959401 | 0.678749739 |
| 0.399536599 | 0.678676425 |
| 0.399495617 | 0.678645717 |
| 0.399450612 | 0.678602972 |
| 0.399378652 | 0.678529128 |
| 0.399358295 | 0.678518747 |
| 0.399316462 | 0.678471876 |
| 0.399300367 | 0.678468733 |
| 0.399291614 | 0.678468733 |
| 0.399282835 | 0.678468733 |
| 0.399236883 | 0.678465537 |
| 0.399180823 | 0.678410828 |
| 0.399126641 | 0.678342951 |
| 0.399118028 | 0.678342951 |
| 0.399074392 | 0.678342951 |
| 0.398949017 | 0.678210482 |

| | |
|---|---|
| 0.398886034 | 0.678151822 |
| 0.398878268 | 0.678151822 |
| 0.39887342 | 0.678151822 |
| 0.398849862 | 0.678151822 |
| 0.398772071 | 0.678054889 |
| 0.398750396 | 0.678042241 |
| 0.398655055 | 0.677976943 |
| 0.398604644 | 0.677939626 |
| 0.398493026 | 0.677798198 |
| 0.398469166 | 0.677798198 |
| 0.398340292 | 0.677683626 |
| 0.398332052 | 0.677683626 |
| 0.398314949 | 0.677683626 |
| 0.39831163 | 0.677683626 |
| 0.398186946 | 0.677567961 |
| 0.398152022 | 0.67755695 |
| 0.398075903 | 0.67745162 |
| 0.398013107 | 0.677368956 |
| 0.398001919 | 0.677368956 |
| 0.397913459 | 0.677319142 |
| 0.397855044 | 0.677269361 |
| 0.397853682 | 0.677269361 |
| 0.397817391 | 0.677269361 |
| 0.397804901 | 0.677269361 |
| 0.397725911 | 0.677243188 |
| 0.397716228 | 0.677243188 |
| 0.397630012 | 0.677128311 |
| 0.397542104 | 0.677027026 |
| 0.397520503 | 0.677014447 |
| 0.397452155 | 0.67692225 |
| 0.397415277 | 0.676883646 |
| 0.397314609 | 0.676747979 |
| 0.397308918 | 0.676747979 |
| 0.397127116 | 0.676513812 |
| 0.397110528 | 0.676513812 |
| 0.397092351 | 0.676513812 |
| 0.397043257 | 0.67646774 |
| 0.397039327 | 0.67646774 |
| 0.39702086 | 0.67646774 |
| 0.39701493 | 0.67646774 |
| 0.396666046 | 0.676091106 |
| 0.39665653 | 0.676091106 |
| 0.396617706 | 0.676091106 |
| 0.396564906 | 0.676063886 |
| 0.39651003 | 0.67604293 |

| | |
|---|---|
| 0.396497694 | 0.67604293 |
| 0.396458618 | 0.676030069 |
| 0.396395433 | 0.675983417 |
| 0.39638139 | 0.675983417 |
| 0.396378003 | 0.675983417 |
| 0.396359267 | 0.675983417 |
| 0.396353872 | 0.675983417 |
| 0.396349051 | 0.675983417 |
| 0.396346835 | 0.675983417 |
| 0.396312003 | 0.675983417 |
| 0.396171126 | 0.675876427 |
| 0.396140391 | 0.675869843 |
| 0.396088975 | 0.675833111 |
| 0.39608234 | 0.675833111 |
| 0.396034885 | 0.675813453 |
| 0.396027728 | 0.675813453 |
| 0.396009412 | 0.675813453 |
| 0.395976658 | 0.675813453 |
| 0.395939464 | 0.675813453 |
| 0.395891259 | 0.67578628 |
| 0.395890121 | 0.67578628 |
| 0.395831461 | 0.67576526 |
| 0.395829048 | 0.67576526 |
| 0.395815961 | 0.67576526 |
| 0.395785318 | 0.67576526 |
| 0.395729159 | 0.675727543 |
| 0.395672182 | 0.675654477 |
| 0.395671755 | 0.675654477 |
| 0.395626122 | 0.675624272 |
| 0.395604397 | 0.675611395 |
| 0.395460165 | 0.675461959 |
| 0.395440899 | 0.675453277 |
| 0.395427305 | 0.675453277 |
| 0.39541544 | 0.675453277 |
| 0.395407465 | 0.675453277 |
| 0.39529505 | 0.675325252 |
| 0.395215566 | 0.675286352 |
| 0.395183925 | 0.6752632 |
| 0.395181333 | 0.6752632 |
| 0.395165986 | 0.6752632 |
| 0.3951248 | 0.6752632 |
| 0.395117057 | 0.6752632 |
| 0.395049922 | 0.67522134 |
| 0.39503626 | 0.67522134 |
| 0.395028684 | 0.67522134 |

| | |
|---|---|
| 0.394990423 | 0.67522134 |
| 0.394903244 | 0.675116006 |
| 0.394791461 | 0.674997592 |
| 0.394738422 | 0.674931137 |
| 0.394734693 | 0.674931137 |
| 0.3947284 | 0.674931137 |
| 0.394566022 | 0.674733257 |
| 0.394506025 | 0.674679108 |
| 0.394468804 | 0.674639677 |
| 0.394463452 | 0.674639677 |
| 0.394439323 | 0.674637709 |
| 0.394393815 | 0.674584099 |
| 0.39430814 | 0.674474297 |
| 0.394235474 | 0.674433859 |
| 0.394220521 | 0.674433017 |
| 0.394195652 | 0.674432586 |
| 0.394175333 | 0.674428386 |
| 0.394121695 | 0.67438049 |
| 0.394079674 | 0.674337392 |
| 0.394051173 | 0.67433072 |
| 0.394003671 | 0.674328477 |
| 0.393928123 | 0.674247635 |
| 0.393914876 | 0.674247635 |
| 0.393898387 | 0.6742452 |
| 0.393856648 | 0.674197986 |
| 0.393734929 | 0.674038082 |
| 0.393671717 | 0.67397832 |
| 0.393619373 | 0.67391293 |
| 0.39361604 | 0.67391293 |
| 0.393558362 | 0.673881146 |
| 0.393487981 | 0.67378486 |
| 0.393463691 | 0.673783791 |
| 0.393392413 | 0.673718104 |
| 0.393366134 | 0.673718104 |
| 0.393355068 | 0.673718104 |
| 0.393247904 | 0.673591726 |
| 0.39324278 | 0.673591726 |
| 0.393237691 | 0.673591726 |
| 0.393221035 | 0.673591726 |
| 0.393184946 | 0.673591726 |
| 0.393175613 | 0.673591726 |
| 0.393126049 | 0.67355257 |
| 0.393105259 | 0.67354118 |
| 0.393101851 | 0.67354118 |
| 0.393081956 | 0.67354118 |

| | |
|---|---|
| 0.39307962 | 0.67354118 |
| 0.392938863 | 0.673352975 |
| 0.392905524 | 0.673320069 |
| 0.392846217 | 0.67324266 |
| 0.392766407 | 0.673178555 |
| 0.392647152 | 0.673022597 |
| 0.392626255 | 0.673022597 |
| 0.392574945 | 0.67297149 |
| 0.392571496 | 0.67297149 |
| 0.392424184 | 0.672809915 |
| 0.392416309 | 0.672809915 |
| 0.392390147 | 0.672799998 |
| 0.392384475 | 0.672799998 |
| 0.392382098 | 0.672799998 |
| 0.392338595 | 0.672784269 |
| 0.392335994 | 0.672784269 |
| 0.392281744 | 0.672759448 |
| 0.392257553 | 0.672743223 |
| 0.392176607 | 0.672651803 |
| 0.392128149 | 0.672592911 |
| 0.392041995 | 0.672469354 |
| 0.392029904 | 0.672469354 |
| 0.392019133 | 0.672469354 |
| 0.392015388 | 0.672469354 |
| 0.39198236 | 0.672463938 |
| 0.391819331 | 0.672281118 |
| 0.39178209 | 0.672265658 |
| 0.391739179 | 0.672216244 |
| 0.39173133 | 0.672216244 |
| 0.391722023 | 0.672216244 |
| 0.391654291 | 0.67214323 |
| 0.391564678 | 0.67203787 |
| 0.391553699 | 0.67203787 |
| 0.391517829 | 0.672030116 |
| 0.391476153 | 0.671982799 |
| 0.391394897 | 0.671867536 |
| 0.391392575 | 0.671867536 |
| 0.391382055 | 0.671867536 |
| 0.391366818 | 0.671867536 |
| 0.391289233 | 0.671867536 |
| 0.391259884 | 0.671867536 |
| 0.391224251 | 0.671840968 |
| 0.391221186 | 0.671840968 |
| 0.391219166 | 0.671840968 |
| 0.391200424 | 0.671840968 |

| | |
|---|---|
| 0.391150202 | 0.671840968 |
| 0.391057627 | 0.671797058 |
| 0.39101925 | 0.671779591 |
| 0.391016625 | 0.671779591 |
| 0.391003303 | 0.671779591 |
| 0.390986529 | 0.671779591 |
| 0.390835228 | 0.671639772 |
| 0.390773104 | 0.671574734 |
| 0.390719682 | 0.671507156 |
| 0.390663626 | 0.671459276 |
| 0.390629795 | 0.671425358 |
| 0.390618933 | 0.671425358 |
| 0.39059519 | 0.671425358 |
| 0.390577461 | 0.671425358 |
| 0.390536742 | 0.671425358 |
| 0.390536415 | 0.671425358 |
| 0.390459755 | 0.671399622 |
| 0.390444908 | 0.671398334 |
| 0.390412717 | 0.671391459 |
| 0.390329038 | 0.671271795 |
| 0.390326403 | 0.671271795 |
| 0.390245651 | 0.671176861 |
| 0.390168527 | 0.671116921 |
| 0.390144217 | 0.671116921 |
| 0.390137417 | 0.671116921 |
| 0.390063678 | 0.671057751 |
| 0.390021149 | 0.671008823 |
| 0.389931727 | 0.670889861 |
| 0.389765338 | 0.670714086 |
| 0.389713235 | 0.670672891 |
| 0.389684624 | 0.670647886 |
| 0.389672576 | 0.670647886 |
| 0.389655397 | 0.670646054 |
| 0.389586866 | 0.670576569 |
| 0.389584928 | 0.670576569 |
| 0.389513719 | 0.670547599 |
| 0.389499531 | 0.670547413 |
| 0.389449551 | 0.670509841 |
| 0.38942356 | 0.670509841 |
| 0.38939648 | 0.670491185 |
| 0.389386084 | 0.670491185 |
| 0.389364303 | 0.67048426 |
| 0.389363636 | 0.67048426 |
| 0.389333582 | 0.670479842 |
| 0.389252272 | 0.670412539 |

| | |
|---|---|
| 0.389187386 | 0.670325026 |
| 0.38909007 | 0.670205887 |
| 0.389036593 | 0.670162247 |
| 0.389024822 | 0.670162247 |
| 0.389023864 | 0.670162247 |
| 0.388986283 | 0.670148299 |
| 0.388922207 | 0.670086385 |
| 0.388887614 | 0.670051023 |
| 0.388723147 | 0.669864582 |
| 0.388700244 | 0.669849351 |
| 0.388691236 | 0.669849351 |
| 0.388644116 | 0.669801096 |
| 0.388603124 | 0.669754685 |
| 0.388598544 | 0.669754685 |
| 0.388591379 | 0.669754685 |
| 0.388534686 | 0.669709438 |
| 0.388414302 | 0.669550399 |
| 0.388379527 | 0.669514685 |
| 0.38835675 | 0.669499653 |
| 0.388346969 | 0.669499653 |
| 0.388342982 | 0.669499653 |
| 0.388197238 | 0.669379543 |
| 0.388188012 | 0.669379543 |
| 0.388177224 | 0.669379543 |
| 0.388016795 | 0.669252739 |
| 0.387909618 | 0.669140578 |
| 0.387869321 | 0.669119537 |
| 0.387752806 | 0.668942764 |
| 0.387723386 | 0.66891624 |
| 0.387658723 | 0.668865873 |
| 0.387644273 | 0.668865873 |
| 0.387635896 | 0.668865873 |
| 0.38738179 | 0.668569086 |
| 0.387376387 | 0.668569086 |
| 0.387354482 | 0.668569086 |
| 0.387347707 | 0.668569086 |
| 0.387301406 | 0.668551486 |
| 0.387161376 | 0.668358218 |
| 0.387134864 | 0.668336674 |
| 0.387020986 | 0.668188519 |
| 0.386929707 | 0.668055144 |
| 0.386919408 | 0.668055144 |
| 0.386869583 | 0.668048209 |
| 0.386851153 | 0.668040605 |
| 0.386841854 | 0.668040605 |

| | |
|---|---|
| 0.386779735 | 0.667989935 |
| 0.38677134 | 0.667989935 |
| 0.38667917 | 0.667961584 |
| 0.386658182 | 0.667961584 |
| 0.386641388 | 0.667961584 |
| 0.386568768 | 0.667891994 |
| 0.386449602 | 0.667710328 |
| 0.386448346 | 0.667710328 |
| 0.386442604 | 0.667710328 |
| 0.386424857 | 0.667710328 |
| 0.38642331 | 0.667710328 |
| 0.386401329 | 0.667710328 |
| 0.386323072 | 0.667647093 |
| 0.386168188 | 0.667466241 |
| 0.386110698 | 0.667402143 |
| 0.386105343 | 0.667402143 |
| 0.385993201 | 0.667302538 |
| 0.38593639 | 0.667235097 |
| 0.385875456 | 0.667178187 |
| 0.385867839 | 0.667178187 |
| 0.385864814 | 0.667178187 |
| 0.385809917 | 0.667178187 |
| 0.385762279 | 0.667128211 |
| 0.385717875 | 0.667099472 |
| 0.385690845 | 0.667076946 |
| 0.385673208 | 0.667076946 |
| 0.385605213 | 0.667025732 |
| 0.385578755 | 0.667004191 |
| 0.385549091 | 0.6669771 |
| 0.385542483 | 0.6669771 |
| 0.385533531 | 0.6669771 |
| 0.385419584 | 0.666874174 |
| 0.385418619 | 0.666874174 |
| 0.38539813 | 0.666874174 |
| 0.385369283 | 0.666859821 |
| 0.38533249 | 0.66684461 |
| 0.385289495 | 0.666794432 |
| 0.385184742 | 0.666661592 |
| 0.385172182 | 0.666661592 |
| 0.385154008 | 0.666656852 |
| 0.385118162 | 0.666619034 |
| 0.385098188 | 0.666608687 |
| 0.385087933 | 0.666608687 |
| 0.384992469 | 0.666474133 |
| 0.384838113 | 0.666287844 |

| | |
|---|---|
| 0.38479203 | 0.666248228 |
| 0.384781152 | 0.666248228 |
| 0.384780853 | 0.666248228 |
| 0.384680743 | 0.666182123 |
| 0.384574713 | 0.666089893 |
| 0.384529498 | 0.6660358 |
| 0.384464739 | 0.665947852 |
| 0.384402013 | 0.665887637 |
| 0.384390365 | 0.665887637 |
| 0.384344483 | 0.665836414 |
| 0.384296363 | 0.665801487 |
| 0.384223303 | 0.665723343 |
| 0.384187299 | 0.665685178 |
| 0.384158701 | 0.665659843 |
| 0.384103058 | 0.665587642 |
| 0.383914656 | 0.665503306 |
| 0.38390516 | 0.665503306 |
| 0.383891612 | 0.665503306 |
| 0.383854762 | 0.665477331 |
| 0.383740574 | 0.665373903 |
| 0.383714431 | 0.665373903 |
| 0.383705281 | 0.665373903 |
| 0.383649194 | 0.665333727 |
| 0.383637857 | 0.665333727 |
| 0.38359613 | 0.665290153 |
| 0.38357556 | 0.665278704 |
| 0.383513472 | 0.665219466 |
| 0.383505407 | 0.665219466 |
| 0.383495402 | 0.665219466 |
| 0.383494942 | 0.665219466 |
| 0.383392129 | 0.665148666 |
| 0.383367263 | 0.665148666 |
| 0.383277594 | 0.665076765 |
| 0.383271211 | 0.665076765 |
| 0.38309903 | 0.664863809 |
| 0.383085916 | 0.664863809 |
| 0.382994792 | 0.664763819 |
| 0.382989331 | 0.664763819 |
| 0.382926477 | 0.66475813 |
| 0.382916048 | 0.66475813 |
| 0.382895488 | 0.66475813 |
| 0.382875226 | 0.66475813 |
| 0.38267365 | 0.664585688 |
| 0.382668769 | 0.664585688 |
| 0.38265329 | 0.664585688 |

| | |
|---|---|
| 0.382522338 | 0.664395586 |
| 0.382436791 | 0.664343913 |
| 0.382416843 | 0.66433349 |
| 0.382411737 | 0.66433349 |
| 0.38239298 | 0.66433349 |
| 0.382375288 | 0.66433349 |
| 0.382315153 | 0.664302217 |
| 0.382261117 | 0.664232559 |
| 0.382222039 | 0.664213122 |
| 0.382200715 | 0.664200301 |
| 0.382200426 | 0.664200301 |
| 0.382199829 | 0.664200301 |
| 0.382134251 | 0.664200301 |
| 0.382116866 | 0.6642 |
| 0.382078086 | 0.66415683 |
| 0.382067399 | 0.66415683 |
| 0.381998692 | 0.664067296 |
| 0.381973553 | 0.664047832 |
| 0.381946092 | 0.664024329 |
| 0.381943622 | 0.664024329 |
| 0.381928328 | 0.664024329 |
| 0.381918891 | 0.664024329 |
| 0.381912087 | 0.664024329 |
| 0.381885055 | 0.664024329 |
| 0.381714233 | 0.663766613 |
| 0.381607931 | 0.663654456 |
| 0.381518543 | 0.663547459 |
| 0.381509706 | 0.663547459 |
| 0.38149964 | 0.663547459 |
| 0.381452809 | 0.663505823 |
| 0.381404949 | 0.66349527 |
| 0.381392587 | 0.66349527 |
| 0.381367314 | 0.663478267 |
| 0.381366127 | 0.663478267 |
| 0.381355721 | 0.663478267 |
| 0.381245299 | 0.663435634 |
| 0.38109797 | 0.663324672 |
| 0.381071768 | 0.663303308 |
| 0.381058146 | 0.663303308 |
| 0.381025317 | 0.663270934 |
| 0.380948351 | 0.663185434 |
| 0.380885952 | 0.663125283 |
| 0.380831318 | 0.663078642 |
| 0.380802651 | 0.663052968 |
| 0.380767706 | 0.663035587 |

| | |
|---|---|
| 0.380647217 | 0.662879264 |
| 0.3805273 | 0.662718898 |
| 0.380432617 | 0.662602459 |
| 0.380411851 | 0.662590522 |
| 0.380369713 | 0.662541356 |
| 0.380304048 | 0.662499667 |
| 0.380246262 | 0.662423231 |
| 0.38020682 | 0.662423231 |
| 0.380199385 | 0.662423231 |
| 0.380102271 | 0.662293511 |
| 0.380043202 | 0.662237713 |
| 0.380018528 | 0.662237713 |
| 0.380007585 | 0.662237713 |
| 0.379974545 | 0.662237713 |
| 0.379826973 | 0.662065699 |
| 0.379817814 | 0.662065699 |
| 0.379793769 | 0.662065699 |
| 0.379714544 | 0.662029673 |
| 0.379532814 | 0.661737053 |
| 0.379525766 | 0.661737053 |
| 0.379389986 | 0.661560681 |
| 0.379294836 | 0.661491638 |
| 0.379258616 | 0.661452692 |
| 0.379174508 | 0.661363936 |
| 0.379069719 | 0.661220096 |
| 0.378994844 | 0.661137917 |
| 0.378981309 | 0.661137917 |
| 0.378873474 | 0.660998826 |
| 0.37885124 | 0.660984248 |
| 0.378815258 | 0.660957003 |
| 0.378788251 | 0.660946988 |
| 0.378788194 | 0.660946988 |
| 0.378748161 | 0.660946988 |
| 0.378747938 | 0.660946988 |
| 0.378730207 | 0.660946988 |
| 0.378606033 | 0.660919604 |
| 0.378560683 | 0.660871296 |
| 0.378488926 | 0.660808238 |
| 0.378477498 | 0.660808238 |
| 0.378469494 | 0.660808238 |
| 0.378442647 | 0.660803928 |
| 0.378423734 | 0.660795131 |
| 0.378413256 | 0.660795131 |
| 0.378367938 | 0.660754715 |
| 0.378306842 | 0.660712136 |

| | |
|---|---|
| 0.378285191 | 0.660700673 |
| 0.378220512 | 0.660658268 |
| 0.378210403 | 0.660658268 |
| 0.378171966 | 0.660621927 |
| 0.378084374 | 0.660517369 |
| 0.378037725 | 0.660517369 |
| 0.378035372 | 0.660517369 |
| 0.378015448 | 0.660517369 |
| 0.377907309 | 0.660457967 |
| 0.377798599 | 0.660406027 |
| 0.377763137 | 0.660392193 |
| 0.377761683 | 0.660392193 |
| 0.377748657 | 0.660392193 |
| 0.3777182 | 0.660392193 |
| 0.37770744 | 0.660392193 |
| 0.377696873 | 0.660392193 |
| 0.37761096 | 0.660320062 |
| 0.377610803 | 0.660320062 |
| 0.377608296 | 0.660320062 |
| 0.377453414 | 0.660117275 |
| 0.377440518 | 0.660117275 |
| 0.377400262 | 0.660077348 |
| 0.377343364 | 0.660021756 |
| 0.377291726 | 0.65998036 |
| 0.377268314 | 0.65998036 |
| 0.377193728 | 0.659929678 |
| 0.377188938 | 0.659929678 |
| 0.37705248 | 0.659779501 |
| 0.376975244 | 0.659692826 |
| 0.376897024 | 0.659652895 |
| 0.376891937 | 0.659652895 |
| 0.376858293 | 0.659633588 |
| 0.376840211 | 0.659633588 |
| 0.376825699 | 0.659633588 |
| 0.376808504 | 0.659633588 |
| 0.376774279 | 0.659607741 |
| 0.376720147 | 0.659561461 |
| 0.376717341 | 0.659561461 |
| 0.376712086 | 0.659561461 |
| 0.376706333 | 0.659561461 |
| 0.376551457 | 0.659448275 |
| 0.376541543 | 0.659448275 |
| 0.376509917 | 0.659445225 |
| 0.376451924 | 0.659382729 |
| 0.376396755 | 0.659310344 |

| | |
|---|---|
| 0.376347353 | 0.659248056 |
| 0.376347209 | 0.659248056 |
| 0.37634379 | 0.659248056 |
| 0.376330059 | 0.659248056 |
| 0.376293628 | 0.659248056 |
| 0.376255539 | 0.659232702 |
| 0.376237494 | 0.659225338 |
| 0.376184602 | 0.659156913 |
| 0.376177132 | 0.659156913 |
| 0.376108755 | 0.65912101 |
| 0.3761085 | 0.65912101 |
| 0.376098441 | 0.65912101 |
| 0.376017317 | 0.659068715 |
| 0.376011623 | 0.659068715 |
| 0.375945802 | 0.659053693 |
| 0.375824157 | 0.658913202 |
| 0.375794959 | 0.658910521 |
| 0.375748867 | 0.658853961 |
| 0.375692564 | 0.65877949 |
| 0.37567247 | 0.65877949 |
| 0.375639462 | 0.65877949 |
| 0.375593629 | 0.658727271 |
| 0.375582335 | 0.658727271 |
| 0.375578896 | 0.658727271 |
| 0.375568202 | 0.658727271 |
| 0.375497793 | 0.658656209 |
| 0.375411575 | 0.658624735 |
| 0.375400946 | 0.658624735 |
| 0.375324481 | 0.658624735 |
| 0.375324466 | 0.658624735 |
| 0.375314949 | 0.658624735 |
| 0.375269998 | 0.658624735 |
| 0.375267643 | 0.658624735 |
| 0.375217264 | 0.658624735 |
| 0.375208482 | 0.658624735 |
| 0.375093281 | 0.658528741 |
| 0.375040483 | 0.65846032 |
| 0.374930498 | 0.658291485 |
| 0.374902076 | 0.65826585 |
| 0.374884123 | 0.658258595 |
| 0.374785762 | 0.658110147 |
| 0.374732004 | 0.658064273 |
| 0.374697756 | 0.658028392 |
| 0.37467785 | 0.658019248 |
| 0.374668442 | 0.658019248 |

| | |
|---|---|
| 0.374637073 | 0.658019248 |
| 0.374617184 | 0.658019248 |
| 0.374548065 | 0.6579596 |
| 0.37452351 | 0.6579596 |
| 0.374456495 | 0.657895805 |
| 0.374422897 | 0.657861043 |
| 0.374408082 | 0.657859283 |
| 0.374357749 | 0.657795111 |
| 0.374232167 | 0.657622972 |
| 0.374212769 | 0.657622972 |
| 0.374189551 | 0.657622972 |
| 0.37412476 | 0.657604072 |
| 0.3740559 | 0.657520736 |
| 0.374049883 | 0.657520736 |
| 0.374043503 | 0.657520736 |
| 0.374034282 | 0.657520736 |
| 0.373996452 | 0.657520736 |
| 0.373976891 | 0.657514023 |
| 0.373975847 | 0.657514023 |
| 0.373908011 | 0.657465733 |
| 0.373855084 | 0.657421212 |
| 0.373776735 | 0.657307704 |
| 0.373631754 | 0.657101272 |
| 0.373493986 | 0.65693249 |
| 0.373463595 | 0.656926823 |
| 0.373456093 | 0.656926823 |
| 0.373440072 | 0.656926823 |
| 0.373428913 | 0.656926823 |
| 0.373417372 | 0.656926823 |
| 0.373339813 | 0.656830382 |
| 0.373328339 | 0.656830382 |
| 0.373266223 | 0.656749434 |
| 0.373178553 | 0.656643697 |
| 0.373134603 | 0.656590621 |
| 0.37308255 | 0.656523282 |
| 0.372987762 | 0.656429242 |
| 0.372904702 | 0.656341668 |
| 0.372901064 | 0.656341668 |
| 0.372839968 | 0.656314649 |
| 0.372828104 | 0.656314649 |
| 0.372642258 | 0.656104784 |
| 0.372581894 | 0.656054344 |
| 0.372564725 | 0.656048366 |
| 0.372382802 | 0.655800758 |
| 0.372352657 | 0.655771918 |

| | |
|---|---|
| 0.372351321 | 0.655771918 |
| 0.372307417 | 0.65574074 |
| 0.372171542 | 0.655549911 |
| 0.372163263 | 0.655549911 |
| 0.372159983 | 0.655549911 |
| 0.372105958 | 0.655515371 |
| 0.372064094 | 0.655515371 |
| 0.372053286 | 0.655515371 |
| 0.372018316 | 0.6554982 |
| 0.371936176 | 0.655377714 |
| 0.371897703 | 0.655334169 |
| 0.371884795 | 0.655334169 |
| 0.37182796 | 0.655259759 |
| 0.371814195 | 0.655259759 |
| 0.371785754 | 0.655259759 |
| 0.371783102 | 0.655259759 |
| 0.371759719 | 0.655259759 |
| 0.371619572 | 0.655145769 |
| 0.371578351 | 0.655110762 |
| 0.371455545 | 0.654918492 |
| 0.371373018 | 0.65482147 |
| 0.371281895 | 0.654709275 |
| 0.371234422 | 0.6546498 |
| 0.371181635 | 0.65458095 |
| 0.371138723 | 0.654529511 |
| 0.371132405 | 0.654529511 |
| 0.371113989 | 0.654529511 |
| 0.371026274 | 0.654525088 |
| 0.37097513 | 0.654483349 |
| 0.37095828 | 0.654483349 |
| 0.370874954 | 0.654388121 |
| 0.370841356 | 0.654385289 |
| 0.370784052 | 0.65438096 |
| 0.3707358 | 0.654338198 |
| 0.370584707 | 0.654255026 |
| 0.370552563 | 0.654222528 |
| 0.370517427 | 0.654184746 |
| 0.370512743 | 0.654184746 |
| 0.370460802 | 0.65417711 |
| 0.370413449 | 0.654146662 |
| 0.370360373 | 0.654077184 |
| 0.370290864 | 0.653978676 |
| 0.370264505 | 0.653956374 |
| 0.370240693 | 0.653938568 |
| 0.370227433 | 0.653938568 |

| | |
|---|---|
| 0.370212109 | 0.653936584 |
| 0.370125598 | 0.653808021 |
| 0.370073416 | 0.65376434 |
| 0.370056721 | 0.653759096 |
| 0.37003541 | 0.653748381 |
| 0.370025932 | 0.653748381 |
| 0.370011622 | 0.653748381 |
| 0.369844453 | 0.653553791 |
| 0.369759389 | 0.653427718 |
| 0.369730223 | 0.65340042 |
| 0.369674948 | 0.653326979 |
| 0.369641812 | 0.653316903 |
| 0.369603545 | 0.653311774 |
| 0.369602452 | 0.653311774 |
| 0.369591285 | 0.653311774 |
| 0.369579712 | 0.653311774 |
| 0.369563927 | 0.653311774 |
| 0.369473389 | 0.653237401 |
| 0.369465083 | 0.653237401 |
| 0.369401606 | 0.65315898 |
| 0.369370517 | 0.653128258 |
| 0.369368029 | 0.653128258 |
| 0.369330209 | 0.653128258 |
| 0.369293251 | 0.653128258 |
| 0.369194903 | 0.653084447 |
| 0.369188052 | 0.653084447 |
| 0.369157076 | 0.653084447 |
| 0.369127711 | 0.65306261 |
| 0.36894663 | 0.65279074 |
| 0.368890471 | 0.652739874 |
| 0.368855392 | 0.652727357 |
| 0.368801918 | 0.652694136 |
| 0.368800072 | 0.652694136 |
| 0.368791046 | 0.652694136 |
| 0.368751498 | 0.652694136 |
| 0.36873311 | 0.652694136 |
| 0.368657714 | 0.652618977 |
| 0.368617781 | 0.65257254 |
| 0.368591571 | 0.652550396 |
| 0.368576527 | 0.652548018 |
| 0.368559264 | 0.652541712 |
| 0.368521473 | 0.65249906 |
| 0.368470409 | 0.652432901 |
| 0.368444777 | 0.65241177 |
| 0.368439878 | 0.65241177 |

| | |
|---|---|
| 0.368377663 | 0.652408723 |
| 0.368373094 | 0.652408723 |
| 0.368302116 | 0.652401713 |
| 0.368281151 | 0.65238884 |
| 0.368280007 | 0.65238884 |
| 0.368268805 | 0.65238884 |
| 0.368265074 | 0.65238884 |
| 0.368210938 | 0.652385787 |
| 0.368157941 | 0.652316155 |
| 0.368152557 | 0.652316155 |
| 0.368137193 | 0.652316155 |
| 0.368120034 | 0.652316155 |
| 0.36809011 | 0.652316155 |
| 0.36808853 | 0.652316155 |
| 0.368052277 | 0.652316155 |
| 0.368014094 | 0.652279679 |
| 0.368001205 | 0.652279679 |
| 0.36791068 | 0.652222233 |
| 0.367896726 | 0.652222233 |
| 0.367726628 | 0.652037086 |
| 0.367705009 | 0.652023026 |
| 0.367621506 | 0.651899228 |
| 0.367609176 | 0.651899228 |
| 0.367591816 | 0.651895123 |
| 0.367577459 | 0.651893935 |
| 0.36757151 | 0.651893935 |
| 0.367502069 | 0.651808775 |
| 0.367462976 | 0.651763711 |
| 0.367450105 | 0.651763711 |
| 0.367446458 | 0.651763711 |
| 0.367408339 | 0.651763711 |
| 0.367400981 | 0.651763711 |
| 0.367397073 | 0.651763711 |
| 0.367342736 | 0.651744616 |
| 0.367288685 | 0.651672994 |
| 0.367254313 | 0.651660563 |
| 0.367248905 | 0.651660563 |
| 0.367220628 | 0.651649348 |
| 0.367218478 | 0.651649348 |
| 0.367195946 | 0.651649348 |
| 0.367195858 | 0.651649348 |
| 0.367166315 | 0.651649348 |
| 0.367143153 | 0.651649348 |
| 0.367047735 | 0.651561032 |
| 0.367031545 | 0.651556576 |

| | |
|---|---|
| 0.367030461 | 0.651556576 |
| 0.367009959 | 0.651556576 |
| 0.36694962 | 0.651508277 |
| 0.366898577 | 0.651445738 |
| 0.366885264 | 0.651445738 |
| 0.36687721 | 0.651445738 |
| 0.366872503 | 0.651445738 |
| 0.366841765 | 0.651445738 |
| 0.366811389 | 0.651432845 |
| 0.366786672 | 0.651413238 |
| 0.366758176 | 0.651411212 |
| 0.366747501 | 0.651411212 |
| 0.36671658 | 0.651385916 |
| 0.366686283 | 0.651385916 |
| 0.366670275 | 0.651385916 |
| 0.36666896 | 0.651385916 |
| 0.366641317 | 0.651385916 |
| 0.3666264 | 0.651385916 |
| 0.366588977 | 0.651385916 |
| 0.366577757 | 0.651385916 |
| 0.366572069 | 0.651385916 |
| 0.366529186 | 0.651368864 |
| 0.366442429 | 0.651263287 |
| 0.366438866 | 0.651263287 |
| 0.366429888 | 0.651263287 |
| 0.366237709 | 0.651093818 |
| 0.366237709 | 0.651093818 |
| 0.366237709 | 0.651093818 |
| 0.366234456 | 0.651093818 |
| 0.366045187 | 0.650945938 |
| 0.366040197 | 0.650945938 |
| 0.365946798 | 0.650843875 |
| 0.365938489 | 0.650843875 |
| 0.365930156 | 0.650843875 |
| 0.365929905 | 0.650843875 |
| 0.365845043 | 0.650784413 |
| 0.365819583 | 0.650763428 |
| 0.365724796 | 0.650619112 |
| 0.365712094 | 0.650619112 |
| 0.365632658 | 0.650528095 |
| 0.365568926 | 0.650463303 |
| 0.36541672 | 0.650241066 |
| 0.365396104 | 0.650228675 |
| 0.365371363 | 0.650228675 |
| 0.365234471 | 0.650013907 |

| | |
|---|---|
| 0.365168432 | 0.64994495 |
| 0.365155953 | 0.64994495 |
| 0.365144022 | 0.64994495 |
| 0.365130465 | 0.64994495 |
| 0.365116245 | 0.64994495 |
| 0.36501028 | 0.649809165 |
| 0.365000122 | 0.649809165 |
| 0.365000122 | 0.649809165 |
| 0.365000122 | 0.649809165 |
| 0.365000122 | 0.649809165 |
| 0.36496797 | 0.649809165 |
| 0.364966624 | 0.649809165 |
| 0.364927251 | 0.649809165 |
| 0.364753551 | 0.649667815 |
| 0.364739601 | 0.649667264 |
| 0.364686731 | 0.649597386 |
| 0.3646769 | 0.649597386 |
| 0.364599182 | 0.649514316 |
| 0.364529466 | 0.649438704 |
| 0.364493909 | 0.649399649 |
| 0.364440156 | 0.649328169 |
| 0.364416747 | 0.649314818 |
| 0.364381336 | 0.64929624 |
| 0.364339288 | 0.649269897 |
| 0.364333233 | 0.649269897 |
| 0.364314453 | 0.649269897 |
| 0.364289025 | 0.649269897 |
| 0.364260999 | 0.649251847 |
| 0.364242797 | 0.649243702 |
| 0.36423673 | 0.649243702 |
| 0.364226908 | 0.649243702 |
| 0.364170687 | 0.649236658 |
| 0.364112225 | 0.649181033 |
| 0.364009694 | 0.649069016 |
| 0.363981146 | 0.64906882 |
| 0.363979712 | 0.64906882 |
| 0.363950915 | 0.649063515 |
| 0.363918255 | 0.649053876 |
| 0.363909439 | 0.649053876 |
| 0.36382172 | 0.649027532 |
| 0.363817936 | 0.649027532 |
| 0.363797067 | 0.649027532 |
| 0.363794294 | 0.649027532 |
| 0.363792739 | 0.649027532 |
| 0.363670287 | 0.64890322 |

| | |
|---|---|
| 0.36366649 | 0.64890322 |
| 0.36366339 | 0.64890322 |
| 0.363448692 | 0.648702251 |
| 0.363326021 | 0.648531909 |
| 0.363296822 | 0.648504094 |
| 0.363291406 | 0.648504094 |
| 0.363259025 | 0.648485235 |
| 0.363238548 | 0.648485235 |
| 0.363181925 | 0.648469128 |
| 0.363174257 | 0.648469128 |
| 0.363116731 | 0.648411337 |
| 0.363106373 | 0.648411337 |
| 0.362999973 | 0.648363012 |
| 0.362974969 | 0.648342666 |
| 0.362938019 | 0.648342666 |
| 0.362929663 | 0.648342666 |
| 0.362924796 | 0.648342666 |
| 0.362864781 | 0.648316048 |
| 0.36280465 | 0.648232933 |
| 0.362777059 | 0.648232272 |
| 0.362589407 | 0.647995124 |
| 0.362534673 | 0.647945019 |
| 0.362531216 | 0.647945019 |
| 0.362429482 | 0.647805635 |
| 0.362417968 | 0.647805635 |
| 0.362408723 | 0.647805635 |
| 0.362361048 | 0.647804874 |
| 0.362291458 | 0.647729092 |
| 0.362253592 | 0.647685706 |
| 0.362137283 | 0.64752637 |
| 0.362101029 | 0.647485854 |
| 0.362097353 | 0.647485854 |
| 0.362064229 | 0.647485854 |
| 0.362042297 | 0.647485854 |
| 0.361915604 | 0.647348724 |
| 0.361911868 | 0.647348724 |
| 0.361904836 | 0.647348724 |
| 0.361867163 | 0.647348724 |
| 0.361837201 | 0.647330041 |
| 0.361817554 | 0.647322296 |
| 0.36179288 | 0.647322296 |
| 0.361732445 | 0.647239881 |
| 0.361676838 | 0.647164698 |
| 0.361645695 | 0.647133285 |
| 0.361629364 | 0.647128376 |

| | |
|---|---|
| 0.361609029 | 0.647116301 |
| 0.361537394 | 0.647061045 |
| 0.361522124 | 0.64705803 |
| 0.36144912 | 0.64700031 |
| 0.36138869 | 0.646940769 |
| 0.36129387 | 0.646819649 |
| 0.361251051 | 0.646767303 |
| 0.361178806 | 0.646710891 |
| 0.361175113 | 0.646710891 |
| 0.361163021 | 0.646710891 |
| 0.361110033 | 0.646684998 |
| 0.361094426 | 0.646681366 |
| 0.361090926 | 0.646681366 |
| 0.361076677 | 0.646681366 |
| 0.361018859 | 0.646666093 |
| 0.360949909 | 0.646568427 |
| 0.360948386 | 0.646568427 |
| 0.360941004 | 0.646568427 |
| 0.360915725 | 0.646568427 |
| 0.360911715 | 0.646568427 |
| 0.360852819 | 0.646540411 |
| 0.360816746 | 0.6465001 |
| 0.360762799 | 0.646427759 |
| 0.360735102 | 0.646402449 |
| 0.360682807 | 0.646357378 |
| 0.360577603 | 0.646193164 |
| 0.360406644 | 0.645935396 |
| 0.36034611 | 0.645890359 |
| 0.360257419 | 0.645765153 |
| 0.360215591 | 0.64574392 |
| 0.360168133 | 0.645702314 |
| 0.36012631 | 0.645651639 |
| 0.359972572 | 0.645424602 |
| 0.35993068 | 0.645373786 |
| 0.359893173 | 0.645330829 |
| 0.359869912 | 0.645330829 |
| 0.359742043 | 0.645181287 |
| 0.359736245 | 0.645181287 |
| 0.359732296 | 0.645181287 |
| 0.359679529 | 0.645142053 |
| 0.359658676 | 0.645128945 |
| 0.359634446 | 0.64510978 |
| 0.359594747 | 0.645062862 |
| 0.3595521 | 0.64503495 |
| 0.359495558 | 0.644980696 |

| | |
|---|---|
| 0.359313147 | 0.644776279 |
| 0.359283498 | 0.644747369 |
| 0.359280971 | 0.644747369 |
| 0.359271953 | 0.644747369 |
| 0.359079076 | 0.644550528 |
| 0.359004366 | 0.644440711 |
| 0.359000129 | 0.644440711 |
| 0.35897148 | 0.644432757 |
| 0.358948285 | 0.644432757 |
| 0.358931874 | 0.644432757 |
| 0.358905754 | 0.644432757 |
| 0.358893436 | 0.644432757 |
| 0.358888157 | 0.644432757 |
| 0.358849507 | 0.644418037 |
| 0.358828503 | 0.644416482 |
| 0.358819418 | 0.644416482 |
| 0.358815712 | 0.644416482 |
| 0.358785142 | 0.644416482 |
| 0.358669464 | 0.644283312 |
| 0.358649721 | 0.644283312 |
| 0.358647718 | 0.644283312 |
| 0.358625465 | 0.644283312 |
| 0.358604462 | 0.644283312 |
| 0.358603127 | 0.644283312 |
| 0.358585919 | 0.644283312 |
| 0.358573624 | 0.644283312 |
| 0.358499269 | 0.644262497 |
| 0.358450327 | 0.644223159 |
| 0.358443245 | 0.644223159 |
| 0.358401705 | 0.644208703 |
| 0.358385899 | 0.644204605 |
| 0.358336127 | 0.644139449 |
| 0.358299802 | 0.644098463 |
| 0.358230702 | 0.644057642 |
| 0.358054397 | 0.643876035 |
| 0.358037947 | 0.643870764 |
| 0.358025298 | 0.643870764 |
| 0.357974711 | 0.643805663 |
| 0.357874564 | 0.643698472 |
| 0.357789176 | 0.643624992 |
| 0.357750137 | 0.643600836 |
| 0.357718281 | 0.643600836 |
| 0.357694007 | 0.643592462 |
| 0.357692681 | 0.643592462 |
| 0.357690193 | 0.643592462 |

| | |
|---|---|
| 0.357637452 | 0.643587959 |
| 0.357620262 | 0.643581342 |
| 0.357591303 | 0.643576459 |
| 0.357455829 | 0.643406982 |
| 0.357446683 | 0.643406982 |
| 0.357361389 | 0.643358567 |
| 0.357355047 | 0.643358567 |
| 0.357303824 | 0.643327885 |
| 0.357295386 | 0.643327885 |
| 0.357282577 | 0.643327885 |
| 0.357264399 | 0.643327885 |
| 0.357166821 | 0.643300087 |
| 0.357142298 | 0.643280244 |
| 0.357120163 | 0.643264703 |
| 0.357104348 | 0.643260544 |
| 0.357068855 | 0.643226049 |
| 0.356892283 | 0.643047948 |
| 0.356868634 | 0.643047948 |
| 0.356830365 | 0.643047948 |
| 0.356793688 | 0.643017102 |
| 0.356667825 | 0.642845737 |
| 0.35665729 | 0.642845737 |
| 0.356631354 | 0.642845737 |
| 0.356631123 | 0.642845737 |
| 0.356501665 | 0.642709698 |
| 0.356449955 | 0.642665127 |
| 0.356434619 | 0.642661806 |
| 0.356311549 | 0.642488553 |
| 0.356301796 | 0.642488553 |
| 0.356182954 | 0.642353953 |
| 0.356153876 | 0.642325836 |
| 0.356142497 | 0.642325836 |
| 0.35612776 | 0.642325836 |
| 0.356126851 | 0.642325836 |
| 0.356116864 | 0.642325836 |
| 0.356102337 | 0.642325836 |
| 0.356094361 | 0.642325836 |
| 0.356078695 | 0.642325836 |
| 0.356068434 | 0.642325836 |
| 0.356051589 | 0.642325836 |
| 0.356012826 | 0.642325836 |
| 0.35600704 | 0.642325836 |
| 0.355815161 | 0.642128327 |
| 0.355809 | 0.642128327 |
| 0.355786265 | 0.642124842 |

| | |
|---|---|
| 0.355726181 | 0.642040731 |
| 0.355684338 | 0.641989539 |
| 0.355683484 | 0.641989539 |
| 0.355659268 | 0.641989539 |
| 0.355659006 | 0.641989539 |
| 0.355600946 | 0.641936332 |
| 0.355598579 | 0.641936332 |
| 0.355578841 | 0.641936332 |
| 0.355570224 | 0.641936332 |
| 0.355510606 | 0.641870566 |
| 0.355499965 | 0.641870566 |
| 0.355489603 | 0.641870566 |
| 0.3554589 | 0.641870566 |
| 0.355440971 | 0.641866502 |
| 0.355435124 | 0.641866502 |
| 0.355351652 | 0.641753872 |
| 0.355341285 | 0.641753872 |
| 0.355222849 | 0.641594243 |
| 0.355207207 | 0.641590323 |
| 0.355195019 | 0.641590323 |
| 0.355185074 | 0.641590323 |
| 0.3551721 | 0.641590323 |
| 0.355114358 | 0.641548154 |
| 0.355098958 | 0.641548154 |
| 0.355065916 | 0.641548154 |
| 0.355060578 | 0.641548154 |
| 0.354985721 | 0.641506542 |
| 0.354946477 | 0.641487387 |
| 0.354895873 | 0.641448006 |
| 0.354873085 | 0.641448006 |
| 0.354804514 | 0.641373766 |
| 0.354757001 | 0.64133656 |
| 0.354522232 | 0.641009467 |
| 0.354460253 | 0.640921734 |
| 0.354370855 | 0.640784415 |
| 0.354360064 | 0.640784415 |
| 0.354298163 | 0.640701621 |
| 0.354285102 | 0.640701621 |
| 0.354269486 | 0.640701621 |
| 0.354223492 | 0.640663892 |
| 0.354183748 | 0.640616337 |
| 0.354107436 | 0.640502634 |
| 0.353987581 | 0.640407449 |
| 0.353976701 | 0.640407449 |
| 0.353903749 | 0.640377413 |

| | |
|---|---|
| 0.353871148 | 0.640342749 |
| 0.353859755 | 0.640342749 |
| 0.353823404 | 0.64030501 |
| 0.353782196 | 0.640254765 |
| 0.353699029 | 0.640225891 |
| 0.353699029 | 0.640225891 |
| 0.353617151 | 0.640175006 |
| 0.353522703 | 0.640028348 |
| 0.353501168 | 0.640013691 |
| 0.353484471 | 0.64000779 |
| 0.353479889 | 0.64000779 |
| 0.353464296 | 0.64000779 |
| 0.353413963 | 0.639953115 |
| 0.353364508 | 0.639921769 |
| 0.353133293 | 0.63961511 |
| 0.353112849 | 0.639602406 |
| 0.353095685 | 0.63959564 |
| 0.353062984 | 0.639585057 |
| 0.35298494 | 0.639500788 |
| 0.352905719 | 0.639421771 |
| 0.352879485 | 0.639421771 |
| 0.352822222 | 0.639392107 |
| 0.352739127 | 0.639290171 |
| 0.352738687 | 0.639290171 |
| 0.352653945 | 0.639184436 |
| 0.352602331 | 0.639149577 |
| 0.352599397 | 0.639149577 |
| 0.352527094 | 0.639149577 |
| 0.35251386 | 0.639149577 |
| 0.352483346 | 0.639149577 |
| 0.352372708 | 0.639063819 |
| 0.352323278 | 0.639022841 |
| 0.352290503 | 0.638987729 |
| 0.352270398 | 0.638975597 |
| 0.35226377 | 0.638975597 |
| 0.352240872 | 0.63897071 |
| 0.352234217 | 0.63897071 |
| 0.352214547 | 0.63897071 |
| 0.352189654 | 0.63897071 |
| 0.352110028 | 0.638910635 |
| 0.352106605 | 0.638910635 |
| 0.352095712 | 0.638910635 |
| 0.352089882 | 0.638910635 |
| 0.352063509 | 0.638910635 |
| 0.352049724 | 0.638910635 |

| | |
|---|---|
| 0.352043109 | 0.638910635 |
| 0.352024847 | 0.638910635 |
| 0.351989189 | 0.638910635 |
| 0.351876911 | 0.638762206 |
| 0.351769188 | 0.638625872 |
| 0.351744189 | 0.63860483 |
| 0.351716615 | 0.638588059 |
| 0.351700836 | 0.638588059 |
| 0.351585918 | 0.638463511 |
| 0.351540723 | 0.638405781 |
| 0.351540723 | 0.638405781 |
| 0.351496767 | 0.638374639 |
| 0.351452824 | 0.638319173 |
| 0.351446584 | 0.638319173 |
| 0.351430389 | 0.638319173 |
| 0.3513683 | 0.638244733 |
| 0.351316276 | 0.638241543 |
| 0.351278812 | 0.638229935 |
| 0.351274905 | 0.638229935 |
| 0.351258384 | 0.638229935 |
| 0.351254014 | 0.638229935 |
| 0.351219133 | 0.638229935 |
| 0.351147697 | 0.638203066 |
| 0.351076809 | 0.638098581 |
| 0.3510574 | 0.638098581 |
| 0.35104566 | 0.638098581 |
| 0.350894702 | 0.637889307 |
| 0.35080026 | 0.637766309 |
| 0.350791385 | 0.637766309 |
| 0.350777665 | 0.637766309 |
| 0.350736725 | 0.637748176 |
| 0.350722281 | 0.637746261 |
| 0.350712553 | 0.637746261 |
| 0.350690524 | 0.637746261 |
| 0.350680226 | 0.637746261 |
| 0.350579581 | 0.637657198 |
| 0.350547686 | 0.637623535 |
| 0.350487211 | 0.637553629 |
| 0.350475083 | 0.637553629 |
| 0.350465002 | 0.637553629 |
| 0.350463448 | 0.637553629 |
| 0.350400485 | 0.637526221 |
| 0.350377448 | 0.637526221 |
| 0.350350339 | 0.637526221 |
| 0.350271248 | 0.637437192 |

| | |
|---|---|
| 0.350128847 | 0.637220804 |
| 0.350119645 | 0.637220804 |
| 0.35006443 | 0.637158199 |
| 0.350053696 | 0.637158199 |
| 0.350014505 | 0.637116027 |
| 0.350003716 | 0.637116027 |
| 0.349938733 | 0.637051144 |
| 0.349926602 | 0.637051144 |
| 0.349920675 | 0.637051144 |
| 0.349880246 | 0.637017718 |
| 0.349864369 | 0.637017071 |
| 0.34985215 | 0.637017071 |
| 0.349846765 | 0.637017071 |
| 0.3497792 | 0.636979848 |
| 0.349732551 | 0.63691925 |
| 0.349726496 | 0.63691925 |
| 0.349724495 | 0.63691925 |
| 0.349587853 | 0.636777477 |
| 0.349544866 | 0.63673102 |
| 0.34951678 | 0.63673102 |
| 0.349482111 | 0.63673102 |
| 0.349412625 | 0.636653089 |
| 0.34940896 | 0.636653089 |
| 0.349264115 | 0.636479893 |
| 0.349252606 | 0.636479893 |
| 0.349229954 | 0.636479893 |
| 0.349210915 | 0.636479893 |
| 0.349206639 | 0.636479893 |
| 0.349193665 | 0.636479893 |
| 0.34916168 | 0.636463692 |
| 0.349140353 | 0.636449176 |
| 0.349051356 | 0.636330269 |
| 0.349044474 | 0.636330269 |
| 0.349009003 | 0.636330269 |
| 0.349004947 | 0.636330269 |
| 0.34893183 | 0.63623952 |
| 0.348927502 | 0.63623952 |
| 0.348867156 | 0.636202262 |
| 0.348856788 | 0.636202262 |
| 0.348851891 | 0.636202262 |
| 0.34871759 | 0.636068043 |
| 0.348678533 | 0.636021159 |
| 0.348671211 | 0.636021159 |
| 0.348631168 | 0.635983478 |
| 0.348534627 | 0.635953516 |

| | |
|---|---|
| 0.348462862 | 0.635862053 |
| 0.348458628 | 0.635862053 |
| 0.348445117 | 0.635862053 |
| 0.348440778 | 0.635862053 |
| 0.348371944 | 0.635862053 |
| 0.348308652 | 0.635809136 |
| 0.348290795 | 0.635802766 |
| 0.348289346 | 0.635802766 |
| 0.348273352 | 0.635802766 |
| 0.348159611 | 0.635686534 |
| 0.348138831 | 0.635686534 |
| 0.348110076 | 0.635665922 |
| 0.34806735 | 0.635612272 |
| 0.348018535 | 0.635547499 |
| 0.348004343 | 0.635545951 |
| 0.347998451 | 0.635545951 |
| 0.347997289 | 0.635545951 |
| 0.347996894 | 0.635545951 |
| 0.347802268 | 0.635371816 |
| 0.347771103 | 0.635348993 |
| 0.347698421 | 0.635303947 |
| 0.347536337 | 0.63505652 |
| 0.347483124 | 0.63500801 |
| 0.34747261 | 0.63500801 |
| 0.347455109 | 0.635005544 |
| 0.347428338 | 0.634988983 |
| 0.347363001 | 0.634912787 |
| 0.34732511 | 0.634912787 |
| 0.347312982 | 0.634912787 |
| 0.347234269 | 0.634855445 |
| 0.347070661 | 0.634692611 |
| 0.34700291 | 0.63459308 |
| 0.346957155 | 0.634559543 |
| 0.346956243 | 0.634559543 |
| 0.346944023 | 0.634559543 |
| 0.346903776 | 0.634557981 |
| 0.34688006 | 0.63453897 |
| 0.346824006 | 0.634460801 |
| 0.346770406 | 0.634411482 |
| 0.346738273 | 0.634400214 |
| 0.34669042 | 0.634338249 |
| 0.346664952 | 0.63431602 |
| 0.346650451 | 0.634313856 |
| 0.346571441 | 0.634218014 |
| 0.346489338 | 0.634092131 |

| | |
|---|---|
| 0.346443756 | 0.63408025 |
| 0.346441482 | 0.63408025 |
| 0.346354696 | 0.634016262 |
| 0.346295757 | 0.633981474 |
| 0.346269912 | 0.633958527 |
| 0.346252178 | 0.633950427 |
| 0.346159381 | 0.633853627 |
| 0.346131714 | 0.633827334 |
| 0.346061748 | 0.63372358 |
| 0.346034197 | 0.633697493 |
| 0.345881321 | 0.633581709 |
| 0.345865967 | 0.633581709 |
| 0.34585408 | 0.633581709 |
| 0.345802092 | 0.633564772 |
| 0.345772774 | 0.6335598 |
| 0.345767932 | 0.6335598 |
| 0.34568705 | 0.633475842 |
| 0.345653205 | 0.633438192 |
| 0.345639508 | 0.633438192 |
| 0.345615174 | 0.633438192 |
| 0.345546497 | 0.633383848 |
| 0.345456702 | 0.63327302 |
| 0.345428931 | 0.633246486 |
| 0.34542101 | 0.633246486 |
| 0.345362092 | 0.633172698 |
| 0.345336954 | 0.633172698 |
| 0.345220296 | 0.633028643 |
| 0.345217504 | 0.633028643 |
| 0.345156818 | 0.632991288 |
| 0.345135571 | 0.632976697 |
| 0.345052206 | 0.632872546 |
| 0.345032689 | 0.632861121 |
| 0.345012845 | 0.632853381 |
| 0.344959958 | 0.632800828 |
| 0.344941231 | 0.632790848 |
| 0.344925752 | 0.632786826 |
| 0.344828063 | 0.632680721 |
| 0.344819371 | 0.632680721 |
| 0.344788933 | 0.632679213 |
| 0.344765532 | 0.632679213 |
| 0.344748022 | 0.632679213 |
| 0.344691086 | 0.632599991 |
| 0.344614079 | 0.632483034 |
| 0.344583447 | 0.632483034 |
| 0.344548519 | 0.6324602 |

| | |
|---|---|
| 0.344478227 | 0.632379914 |
| 0.344461491 | 0.632373567 |
| 0.344428679 | 0.632337703 |
| 0.344370852 | 0.632255911 |
| 0.344347669 | 0.632237719 |
| 0.34434714 | 0.632237719 |
| 0.344325214 | 0.632237719 |
| 0.344237857 | 0.63215795 |
| 0.344234739 | 0.63215795 |
| 0.34421382 | 0.63215795 |
| 0.344143902 | 0.632126731 |
| 0.344128541 | 0.632126731 |
| 0.344079775 | 0.632086413 |
| 0.344039747 | 0.632086413 |
| 0.344015971 | 0.632067304 |
| 0.343951264 | 0.632021562 |
| 0.343942404 | 0.632021562 |
| 0.343930775 | 0.632021562 |
| 0.343903395 | 0.632021562 |
| 0.343834004 | 0.631976771 |
| 0.34377457 | 0.631923288 |
| 0.343771375 | 0.631923288 |
| 0.343761593 | 0.631923288 |
| 0.343730398 | 0.631923288 |
| 0.343689044 | 0.631881031 |
| 0.343666894 | 0.631866111 |
| 0.343608357 | 0.631865513 |
| 0.343591703 | 0.631865513 |
| 0.343546594 | 0.63185964 |
| 0.343514295 | 0.631828043 |
| 0.343510523 | 0.631828043 |
| 0.343500852 | 0.631828043 |
| 0.343478634 | 0.631828043 |
| 0.343467771 | 0.631828043 |
| 0.343457748 | 0.631828043 |
| 0.343418526 | 0.631822696 |
| 0.34339266 | 0.631799514 |
| 0.343383915 | 0.631799514 |
| 0.343312322 | 0.631749323 |
| 0.343290884 | 0.631749323 |
| 0.343255568 | 0.631718108 |
| 0.343233169 | 0.631701296 |
| 0.343174189 | 0.631641564 |
| 0.343112603 | 0.631624088 |
| 0.343014477 | 0.631469618 |

| | |
|---|---|
| 0.342864058 | 0.631321684 |
| 0.342853443 | 0.631321684 |
| 0.342788571 | 0.631282715 |
| 0.34278223 | 0.631282715 |
| 0.342755571 | 0.631282715 |
| 0.342736898 | 0.631275824 |
| 0.342647097 | 0.631169056 |
| 0.342640388 | 0.631169056 |
| 0.342625022 | 0.631169056 |
| 0.342618182 | 0.631169056 |
| 0.342601477 | 0.631169056 |
| 0.342599335 | 0.631169056 |
| 0.342419806 | 0.630984605 |
| 0.342316389 | 0.630867262 |
| 0.342315817 | 0.630867262 |
| 0.342308399 | 0.630867262 |
| 0.342285452 | 0.630867262 |
| 0.342205623 | 0.630809583 |
| 0.342164084 | 0.630757426 |
| 0.342154861 | 0.630757426 |
| 0.342113974 | 0.630731488 |
| 0.342096832 | 0.630731102 |
| 0.342010402 | 0.630620574 |
| 0.342004018 | 0.630620574 |
| 0.341910181 | 0.63050901 |
| 0.341905462 | 0.63050901 |
| 0.341838567 | 0.630450179 |
| 0.341778394 | 0.630412438 |
| 0.34177012 | 0.630412438 |
| 0.341759469 | 0.630412438 |
| 0.341695851 | 0.630382254 |
| 0.341655714 | 0.630332623 |
| 0.341593223 | 0.630241746 |
| 0.341575507 | 0.630233475 |
| 0.341561359 | 0.630231785 |
| 0.341543953 | 0.630224085 |
| 0.341483497 | 0.630136943 |
| 0.341481235 | 0.630136943 |
| 0.341434057 | 0.630099627 |
| 0.341429604 | 0.630099627 |
| 0.341402425 | 0.630099627 |
| 0.341303537 | 0.630000139 |
| 0.341301209 | 0.630000139 |
| 0.34129135 | 0.630000139 |
| 0.341250424 | 0.629997988 |

| | |
|---|---|
| 0.341243799 | 0.629997988 |
| 0.34122227 | 0.629997988 |
| 0.341199855 | 0.629997988 |
| 0.341019291 | 0.629817193 |
| 0.340963591 | 0.629807169 |
| 0.34094641 | 0.629804704 |
| 0.340942531 | 0.629804704 |
| 0.340912145 | 0.629790261 |
| 0.340867877 | 0.629732908 |
| 0.340763894 | 0.6296385 |
| 0.340760236 | 0.6296385 |
| 0.340754838 | 0.6296385 |
| 0.340746699 | 0.6296385 |
| 0.340743194 | 0.6296385 |
| 0.340703177 | 0.6296385 |
| 0.340690715 | 0.6296385 |
| 0.340655834 | 0.629609818 |
| 0.34065326 | 0.629609818 |
| 0.340639479 | 0.629609818 |
| 0.34061287 | 0.629603709 |
| 0.340609876 | 0.629603709 |
| 0.340502691 | 0.62947334 |
| 0.340493001 | 0.62947334 |
| 0.340468941 | 0.629463538 |
| 0.340423356 | 0.629448838 |
| 0.340410458 | 0.629448838 |
| 0.34038588 | 0.629448838 |
| 0.340341279 | 0.629419256 |
| 0.340260478 | 0.629318699 |
| 0.340244635 | 0.629313835 |
| 0.340235062 | 0.629313835 |
| 0.34020464 | 0.629313835 |
| 0.340191498 | 0.629313835 |
| 0.340189091 | 0.629313835 |
| 0.340177553 | 0.629313835 |
| 0.340125338 | 0.629313835 |
| 0.340107754 | 0.629313835 |
| 0.340099074 | 0.629313835 |
| 0.340059773 | 0.629289615 |
| 0.340034444 | 0.629267192 |
| 0.340004293 | 0.629235845 |
| 0.339982555 | 0.629227538 |
| 0.33995257 | 0.629213471 |
| 0.339921249 | 0.629179951 |
| 0.339882325 | 0.629150907 |

| | |
|---|---|
| 0.339842454 | 0.629131907 |
| 0.339838908 | 0.629131907 |
| 0.339795991 | 0.629094798 |
| 0.339747691 | 0.629054282 |
| 0.339593419 | 0.628841981 |
| 0.339541402 | 0.628819006 |
| 0.339495839 | 0.628783525 |
| 0.339466262 | 0.628753195 |
| 0.339416474 | 0.628734332 |
| 0.339390447 | 0.628734332 |
| 0.339373919 | 0.628728864 |
| 0.3393641 | 0.628728864 |
| 0.339334064 | 0.628728864 |
| 0.339302613 | 0.628728864 |
| 0.339125697 | 0.628562405 |
| 0.33909758 | 0.628534751 |
| 0.339053904 | 0.628489784 |
| 0.339052412 | 0.628489784 |
| 0.33903596 | 0.628489784 |
| 0.339025634 | 0.628489784 |
| 0.338931432 | 0.628422444 |
| 0.33892332 | 0.628422444 |
| 0.338795601 | 0.628243968 |
| 0.338759102 | 0.628219725 |
| 0.338709943 | 0.628158501 |
| 0.338658664 | 0.628112316 |
| 0.338558212 | 0.627999375 |
| 0.338419996 | 0.627791896 |
| 0.338348536 | 0.627695229 |
| 0.338292116 | 0.627652459 |
| 0.338257995 | 0.627613602 |
| 0.338233675 | 0.627592926 |
| 0.338217421 | 0.627587217 |
| 0.338114624 | 0.62749426 |
| 0.338099219 | 0.627490122 |
| 0.338081079 | 0.627490122 |
| 0.338078759 | 0.627490122 |
| 0.338052702 | 0.627490122 |
| 0.338016151 | 0.627482663 |
| 0.338005771 | 0.627482663 |
| 0.337880756 | 0.627329128 |
| 0.337751999 | 0.627226711 |
| 0.337720326 | 0.627202416 |
| 0.337717296 | 0.627202416 |
| 0.337703034 | 0.627202416 |

| | |
|---|---|
| 0.337653938 | 0.627152488 |
| 0.337650797 | 0.627152488 |
| 0.337586723 | 0.627101012 |
| 0.337558936 | 0.627073854 |
| 0.337453688 | 0.626927243 |
| 0.337374531 | 0.626804634 |
| 0.337334615 | 0.626754923 |
| 0.337257837 | 0.626661165 |
| 0.337169789 | 0.626611709 |
| 0.337104039 | 0.62652205 |
| 0.337066676 | 0.626492264 |
| 0.337040218 | 0.626492264 |
| 0.337033071 | 0.626492264 |
| 0.337002168 | 0.626479402 |
| 0.336976963 | 0.626479402 |
| 0.336971222 | 0.626479402 |
| 0.336918857 | 0.626422331 |
| 0.336904637 | 0.626422331 |
| 0.336880771 | 0.626422331 |
| 0.336871735 | 0.626422331 |
| 0.336823154 | 0.626398016 |
| 0.33681106 | 0.626398016 |
| 0.336754823 | 0.626361867 |
| 0.336750716 | 0.626361867 |
| 0.33672639 | 0.626357904 |
| 0.336722214 | 0.626357904 |
| 0.336718753 | 0.626357904 |
| 0.336691225 | 0.626357904 |
| 0.33666161 | 0.626357904 |
| 0.336577303 | 0.626227331 |
| 0.33645303 | 0.626069486 |
| 0.336413468 | 0.626020327 |
| 0.336333254 | 0.625968707 |
| 0.336304138 | 0.625939161 |
| 0.336257216 | 0.625878933 |
| 0.336234241 | 0.625878933 |
| 0.336210213 | 0.625862178 |
| 0.33620966 | 0.625862178 |
| 0.336066876 | 0.625717636 |
| 0.336018833 | 0.625652643 |
| 0.335828899 | 0.625421246 |
| 0.335786366 | 0.62536649 |
| 0.335761671 | 0.62536649 |
| 0.335708515 | 0.62531931 |
| 0.335697194 | 0.62531931 |

| | |
|---|---|
| 0.335669823 | 0.62529615 |
| 0.335669759 | 0.62529615 |
| 0.335660981 | 0.62529615 |
| 0.335660651 | 0.62529615 |
| 0.335654125 | 0.62529615 |
| 0.335620065 | 0.62529615 |
| 0.335612574 | 0.62529615 |
| 0.335609419 | 0.62529615 |
| 0.335603648 | 0.62529615 |
| 0.33551676 | 0.625255575 |
| 0.335390284 | 0.625117719 |
| 0.335286797 | 0.624998211 |
| 0.335235938 | 0.624927865 |
| 0.335222357 | 0.624927008 |
| 0.33519126 | 0.624893496 |
| 0.335190712 | 0.624893496 |
| 0.335147956 | 0.624893496 |
| 0.335146992 | 0.624893496 |
| 0.335122473 | 0.624893496 |
| 0.335112312 | 0.624893496 |
| 0.33485289 | 0.624653894 |
| 0.334793768 | 0.62457685 |
| 0.334787962 | 0.62457685 |
| 0.334724427 | 0.624512096 |
| 0.334683626 | 0.624477779 |
| 0.334677788 | 0.624477779 |
| 0.334665124 | 0.624477779 |
| 0.334602034 | 0.624477779 |
| 0.334601343 | 0.624477779 |
| 0.334551622 | 0.624477779 |
| 0.33451172 | 0.62443324 |
| 0.334486644 | 0.624410903 |
| 0.334485923 | 0.624410903 |
| 0.334482408 | 0.624410903 |
| 0.334480892 | 0.624410903 |
| 0.334448054 | 0.624410903 |
| 0.334440691 | 0.624410903 |
| 0.33425693 | 0.624237767 |
| 0.33415433 | 0.624100945 |
| 0.334057016 | 0.623975742 |
| 0.334022329 | 0.623935422 |
| 0.333992398 | 0.623903982 |
| 0.333957321 | 0.623862927 |
| 0.333927014 | 0.62385525 |
| 0.333906288 | 0.623841001 |

| | |
|---|---|
| 0.333845844 | 0.62375254 |
| 0.33382326 | 0.623734813 |
| 0.333786549 | 0.62373246 |
| 0.333757563 | 0.623709937 |
| 0.333745822 | 0.623709937 |
| 0.333715484 | 0.623704718 |
| 0.333693205 | 0.623687555 |
| 0.333674219 | 0.623676544 |
| 0.333612333 | 0.623609819 |
| 0.333563819 | 0.623543607 |
| 0.33345372 | 0.6234112 |
| 0.333453402 | 0.6234112 |
| 0.333450377 | 0.6234112 |
| 0.333393754 | 0.6233725 |
| 0.333371408 | 0.623355191 |
| 0.333360645 | 0.623355191 |
| 0.333336483 | 0.62333883 |
| 0.333316983 | 0.62332684 |
| 0.333295779 | 0.623311662 |
| 0.333184751 | 0.623177432 |
| 0.333183637 | 0.623177432 |
| 0.333159689 | 0.623177432 |
| 0.333034528 | 0.622969854 |
| 0.333026736 | 0.622969854 |
| 0.332991175 | 0.622937692 |
| 0.332984898 | 0.622937692 |
| 0.332783154 | 0.622597449 |
| 0.33277203 | 0.622597449 |
| 0.332762375 | 0.622597449 |
| 0.332745676 | 0.622597449 |
| 0.332676634 | 0.622520432 |
| 0.332652562 | 0.622513651 |
| 0.33262328 | 0.622493966 |
| 0.332540156 | 0.622387315 |
| 0.332537114 | 0.622387315 |
| 0.332514531 | 0.622387315 |
| 0.332474213 | 0.622387315 |
| 0.332465786 | 0.622387315 |
| 0.332441599 | 0.622387315 |
| 0.332402536 | 0.622374319 |
| 0.332399809 | 0.622374319 |
| 0.33236328 | 0.622349753 |
| 0.332332206 | 0.622316032 |
| 0.332275785 | 0.622259312 |
| 0.332272565 | 0.622259312 |

| | |
|---|---|
| 0.332225366 | 0.622213823 |
| 0.332133735 | 0.622091136 |
| 0.332122762 | 0.622091136 |
| 0.332090679 | 0.622059417 |
| 0.332060789 | 0.622027894 |
| 0.331987195 | 0.621963422 |
| 0.331952471 | 0.621922832 |
| 0.331926224 | 0.621898122 |
| 0.331757916 | 0.621656145 |
| 0.331747599 | 0.621656145 |
| 0.331710817 | 0.621616805 |
| 0.331694326 | 0.621616027 |
| 0.331677716 | 0.621616027 |
| 0.331480444 | 0.621331776 |
| 0.331386911 | 0.621180903 |
| 0.331381165 | 0.621180903 |
| 0.331370903 | 0.621180903 |
| 0.331296877 | 0.621109917 |
| 0.33126127 | 0.621067611 |
| 0.33119643 | 0.620990833 |
| 0.331166075 | 0.620962467 |
| 0.33109315 | 0.620899061 |
| 0.331001233 | 0.620800021 |
| 0.330938249 | 0.620730781 |
| 0.330829671 | 0.620576003 |
| 0.330825203 | 0.620576003 |
| 0.330784113 | 0.620539426 |
| 0.330700414 | 0.620424102 |
| 0.3306482 | 0.620406643 |
| 0.330623801 | 0.620385303 |
| 0.330583467 | 0.620368035 |
| 0.330573272 | 0.620368035 |
| 0.330509404 | 0.62029284 |
| 0.330485264 | 0.62029284 |
| 0.330472501 | 0.62029284 |
| 0.3304449 | 0.620269549 |
| 0.33025473 | 0.61998589 |
| 0.330233638 | 0.619970731 |
| 0.330200438 | 0.619932838 |
| 0.330195463 | 0.619932838 |
| 0.330175736 | 0.619932838 |
| 0.330153479 | 0.619917988 |
| 0.330074955 | 0.619827228 |
| 0.330070275 | 0.619827228 |
| 0.329984953 | 0.619772591 |

| | |
|---|---|
| 0.329975457 | 0.619772591 |
| 0.329945112 | 0.619771085 |
| 0.329942049 | 0.619771085 |
| 0.329927173 | 0.619771085 |
| 0.329773736 | 0.619571349 |
| 0.329766327 | 0.619571349 |
| 0.329644543 | 0.619401927 |
| 0.329640317 | 0.619401927 |
| 0.329578386 | 0.619350923 |
| 0.3295651 | 0.619350392 |
| 0.32953468 | 0.619317662 |
| 0.329384031 | 0.619132262 |
| 0.329322538 | 0.619089975 |
| 0.329282546 | 0.619039229 |
| 0.329275545 | 0.619039229 |
| 0.329266463 | 0.619039229 |
| 0.329236011 | 0.619039229 |
| 0.329227346 | 0.619039229 |
| 0.329174581 | 0.619018503 |
| 0.32916552 | 0.619018503 |
| 0.329125942 | 0.61898915 |
| 0.329118824 | 0.61898915 |
| 0.329017555 | 0.618907504 |
| 0.328939707 | 0.618785505 |
| 0.328784567 | 0.618518093 |
| 0.328748126 | 0.618473969 |
| 0.328737672 | 0.618473969 |
| 0.328697204 | 0.618451456 |
| 0.328657085 | 0.618400402 |
| 0.328551306 | 0.618274647 |
| 0.328449209 | 0.618106941 |
| 0.328388239 | 0.618041043 |
| 0.328335205 | 0.617965652 |
| 0.328323014 | 0.617965652 |
| 0.328320871 | 0.617965652 |
| 0.328303749 | 0.617965652 |
| 0.32826938 | 0.61793944 |
| 0.328249478 | 0.617926399 |
| 0.328246964 | 0.617926399 |
| 0.328234335 | 0.617926399 |
| 0.32820324 | 0.617912627 |
| 0.328180573 | 0.617894376 |
| 0.328135198 | 0.617857796 |
| 0.328116679 | 0.617847352 |
| 0.328086678 | 0.617815286 |

| | |
|---|---|
| 0.328061004 | 0.617791367 |
| 0.328055361 | 0.617791367 |
| 0.328011935 | 0.617747811 |
| 0.32794698 | 0.617649905 |
| 0.327889299 | 0.617565691 |
| 0.327776942 | 0.617478407 |
| 0.327766253 | 0.617478407 |
| 0.327748305 | 0.617478407 |
| 0.327738182 | 0.617478407 |
| 0.327675945 | 0.617465381 |
| 0.327660055 | 0.617465381 |
| 0.327654386 | 0.617465381 |
| 0.327564993 | 0.617418787 |
| 0.327524488 | 0.617391308 |
| 0.327486012 | 0.617343213 |
| 0.32747618 | 0.617343213 |
| 0.327475366 | 0.617343213 |
| 0.327372734 | 0.617278856 |
| 0.327359361 | 0.617278856 |
| 0.327313028 | 0.617236996 |
| 0.327297726 | 0.617232579 |
| 0.327227349 | 0.617124295 |
| 0.32720947 | 0.617116163 |
| 0.327136614 | 0.617050912 |
| 0.32706317 | 0.616957181 |
| 0.327053666 | 0.616957181 |
| 0.327041967 | 0.616957181 |
| 0.327025061 | 0.616957181 |
| 0.327002751 | 0.616957181 |
| 0.326942257 | 0.616904247 |
| 0.3268786 | 0.616857402 |
| 0.326868176 | 0.616857402 |
| 0.326830708 | 0.616815904 |
| 0.326813384 | 0.616807649 |
| 0.326813179 | 0.616807649 |
| 0.326767559 | 0.616794489 |
| 0.32676018 | 0.616794489 |
| 0.326682008 | 0.61670633 |
| 0.32666518 | 0.616699007 |
| 0.326492661 | 0.61644662 |
| 0.326452877 | 0.616444817 |
| 0.32642419 | 0.616439528 |
| 0.326409447 | 0.616436128 |
| 0.326353811 | 0.616355499 |
| 0.32634425 | 0.616355499 |

| | |
|---|---|
| 0.326320807 | 0.616342048 |
| 0.326296048 | 0.616319726 |
| 0.326242193 | 0.616266884 |
| 0.326210919 | 0.616256691 |
| 0.326208448 | 0.616256691 |
| 0.326156124 | 0.616202059 |
| 0.326145478 | 0.616202059 |
| 0.326083378 | 0.616132504 |
| 0.326021084 | 0.616118013 |
| 0.325978011 | 0.616085505 |
| 0.325961068 | 0.616085505 |
| 0.325886762 | 0.616010828 |
| 0.32586197 | 0.615988413 |
| 0.325856193 | 0.615988413 |
| 0.325681551 | 0.615745117 |
| 0.325647638 | 0.61570544 |
| 0.325637624 | 0.61570544 |
| 0.325610432 | 0.615683978 |
| 0.325597133 | 0.615683276 |
| 0.325504173 | 0.615556373 |
| 0.325473523 | 0.615535103 |
| 0.325402228 | 0.615436897 |
| 0.325379509 | 0.615418365 |
| 0.325312932 | 0.615316879 |
| 0.325190118 | 0.615187915 |
| 0.325119597 | 0.615122196 |
| 0.32507501 | 0.615062272 |
| 0.324990538 | 0.614975738 |
| 0.324964802 | 0.614975738 |
| 0.324939917 | 0.614975738 |
| 0.324930449 | 0.614975738 |
| 0.324756229 | 0.614757357 |
| 0.32466176 | 0.614671078 |
| 0.324613434 | 0.614604017 |
| 0.324606803 | 0.614604017 |
| 0.324599577 | 0.614604017 |
| 0.32458122 | 0.614604017 |
| 0.324419105 | 0.614455926 |
| 0.324407566 | 0.614455926 |
| 0.324407374 | 0.614455926 |
| 0.32434151 | 0.614382257 |
| 0.32433606 | 0.614382257 |
| 0.32431231 | 0.614375816 |
| 0.324306049 | 0.614375816 |
| 0.324181647 | 0.614177147 |

| | |
|---|---|
| 0.32404713 | 0.613971144 |
| 0.324021288 | 0.613946606 |
| 0.324013067 | 0.613946606 |
| 0.323969868 | 0.613898023 |
| 0.323953832 | 0.61389206 |
| 0.323949644 | 0.61389206 |
| 0.323931487 | 0.61389206 |
| 0.323881873 | 0.613877823 |
| 0.323831269 | 0.613814355 |
| 0.323828364 | 0.613814355 |
| 0.323799046 | 0.613814355 |
| 0.323730272 | 0.613785894 |
| 0.323717614 | 0.613785894 |
| 0.323616801 | 0.613644041 |
| 0.323609321 | 0.613644041 |
| 0.323596728 | 0.613644041 |
| 0.32354115 | 0.61359831 |
| 0.32343157 | 0.61343339 |
| 0.323385655 | 0.613401116 |
| 0.323369855 | 0.613395575 |
| 0.323360215 | 0.613395575 |
| 0.323321304 | 0.613359274 |
| 0.323292976 | 0.613348555 |
| 0.323263909 | 0.613341175 |
| 0.323204386 | 0.613288813 |
| 0.323184041 | 0.613287367 |
| 0.323177949 | 0.613287367 |
| 0.323007094 | 0.613024861 |
| 0.322830737 | 0.612885538 |
| 0.32279875 | 0.612873673 |
| 0.322780303 | 0.612863079 |
| 0.322764475 | 0.612863079 |
| 0.322753777 | 0.612863079 |
| 0.322691277 | 0.612816198 |
| 0.322595982 | 0.612714337 |
| 0.322562926 | 0.612700315 |
| 0.322523668 | 0.612683225 |
| 0.322408004 | 0.612620228 |
| 0.322377304 | 0.612591976 |
| 0.322236826 | 0.612490454 |
| 0.322204911 | 0.612454233 |
| 0.322188665 | 0.612447796 |
| 0.322164078 | 0.612425503 |
| 0.322104538 | 0.61233676 |
| 0.322096222 | 0.61233676 |

| | |
|---|---|
| 0.322081399 | 0.61233676 |
| 0.322073058 | 0.61233676 |
| 0.321964592 | 0.612217336 |
| 0.321941627 | 0.612198111 |
| 0.32187206 | 0.612090261 |
| 0.321864281 | 0.612090261 |
| 0.321782774 | 0.61196934 |
| 0.32173597 | 0.611929199 |
| 0.321710564 | 0.611905314 |
| 0.321703912 | 0.611905314 |
| 0.321692382 | 0.611905314 |
| 0.321612431 | 0.61182651 |
| 0.321598118 | 0.61182651 |
| 0.321514075 | 0.611702586 |
| 0.321456794 | 0.611642472 |
| 0.321437797 | 0.611630761 |
| 0.321366592 | 0.611568573 |
| 0.321362914 | 0.611568573 |
| 0.321314283 | 0.611548801 |
| 0.321230667 | 0.611456495 |
| 0.32122902 | 0.611456495 |
| 0.321228657 | 0.611456495 |
| 0.321162211 | 0.611423936 |
| 0.321131077 | 0.611399804 |
| 0.321073952 | 0.611399804 |
| 0.321063636 | 0.611399804 |
| 0.321032902 | 0.611397714 |
| 0.320994338 | 0.611348717 |
| 0.320972548 | 0.611331662 |
| 0.320971419 | 0.611331662 |
| 0.320966622 | 0.611331662 |
| 0.320884282 | 0.611285776 |
| 0.32085541 | 0.611255225 |
| 0.320852042 | 0.611255225 |
| 0.320849599 | 0.611255225 |
| 0.320755521 | 0.611138264 |
| 0.320749587 | 0.611138264 |
| 0.320721312 | 0.611121982 |
| 0.320668448 | 0.611094597 |
| 0.320656403 | 0.611094597 |
| 0.320626979 | 0.611094597 |
| 0.320580182 | 0.611048638 |
| 0.320520027 | 0.610958431 |
| 0.320295093 | 0.610675502 |
| 0.320179762 | 0.610480864 |

| | |
|---|---|
| 0.320174651 | 0.610480864 |
| 0.320107842 | 0.61039261 |
| 0.320029203 | 0.610291525 |
| 0.319976323 | 0.610239553 |
| 0.319930614 | 0.610195297 |
| 0.319925728 | 0.610195297 |
| 0.319896984 | 0.610195297 |
| 0.319854733 | 0.610154259 |
| 0.319843089 | 0.610154259 |
| 0.319832586 | 0.610154259 |
| 0.319826484 | 0.610154259 |
| 0.319819547 | 0.610154259 |
| 0.319742801 | 0.610125831 |
| 0.319649076 | 0.610006272 |
| 0.319640466 | 0.610006272 |
| 0.319607245 | 0.610006272 |
| 0.319552151 | 0.609967949 |
| 0.319499283 | 0.609891477 |
| 0.319427799 | 0.609779462 |
| 0.319360017 | 0.609723383 |
| 0.319309217 | 0.609650835 |
| 0.319232935 | 0.609529626 |
| 0.319230096 | 0.609529626 |
| 0.319230053 | 0.609529626 |
| 0.319230002 | 0.609529626 |
| 0.319217799 | 0.609529626 |
| 0.319210815 | 0.609529626 |
| 0.319157371 | 0.609529626 |
| 0.318985959 | 0.60930233 |
| 0.318982143 | 0.60930233 |
| 0.318976623 | 0.60930233 |
| 0.318883855 | 0.609205028 |
| 0.318792517 | 0.609128268 |
| 0.31878568 | 0.609128268 |
| 0.318732954 | 0.609087766 |
| 0.318726253 | 0.609087766 |
| 0.318721396 | 0.609087766 |
| 0.318719146 | 0.609087766 |
| 0.31864542 | 0.609042677 |
| 0.318614729 | 0.609008456 |
| 0.31859683 | 0.608998685 |
| 0.318519374 | 0.608899503 |
| 0.318517515 | 0.608899503 |
| 0.318506121 | 0.608899503 |
| 0.318490657 | 0.608899503 |

| | |
|---|---|
| 0.318471609 | 0.608899503 |
| 0.318467993 | 0.608899503 |
| 0.318467409 | 0.608899503 |
| 0.318455625 | 0.608899503 |
| 0.318353068 | 0.608867756 |
| 0.318264885 | 0.608779537 |
| 0.318247617 | 0.608777028 |
| 0.318229638 | 0.608777028 |
| 0.318209001 | 0.60877044 |
| 0.318207787 | 0.60877044 |
| 0.318160543 | 0.60877044 |
| 0.318139209 | 0.608759185 |
| 0.318081785 | 0.60875319 |
| 0.31797998 | 0.608650129 |
| 0.317950108 | 0.608617411 |
| 0.317928332 | 0.608600187 |
| 0.317706984 | 0.608298715 |
| 0.317623583 | 0.608212383 |
| 0.317600338 | 0.608192324 |
| 0.317481972 | 0.608014552 |
| 0.317455103 | 0.607987542 |
| 0.317358726 | 0.607851847 |
| 0.317227373 | 0.607649135 |
| 0.317130776 | 0.607586285 |
| 0.317044876 | 0.607495024 |
| 0.316928596 | 0.60729665 |
| 0.316863651 | 0.607245498 |
| 0.31676238 | 0.607100277 |
| 0.31661083 | 0.60683424 |
| 0.316558718 | 0.606758775 |
| 0.316387605 | 0.606557736 |
| 0.316353821 | 0.606557736 |
| 0.316350517 | 0.606557736 |
| 0.316346512 | 0.606557736 |
| 0.316182667 | 0.606379635 |
| 0.316131167 | 0.606305283 |
| 0.316125189 | 0.606305283 |
| 0.316122845 | 0.606305283 |
| 0.31612053 | 0.606305283 |
| 0.316094042 | 0.606305283 |
| 0.315759038 | 0.605821881 |
| 0.31569589 | 0.605738786 |
| 0.315679914 | 0.605735015 |
| 0.315656407 | 0.605735015 |
| 0.315656031 | 0.605735015 |

| | |
|---|---|
| 0.315588064 | 0.605702709 |
| 0.315492289 | 0.605567696 |
| 0.315487143 | 0.605567696 |
| 0.315439897 | 0.60551594 |
| 0.315389189 | 0.605443003 |
| 0.315313507 | 0.605346518 |
| 0.315284656 | 0.605315528 |
| 0.315239201 | 0.605252658 |
| 0.315215069 | 0.605230722 |
| 0.315199212 | 0.605224675 |
| 0.315193154 | 0.605224675 |
| 0.315164187 | 0.605206222 |
| 0.315062024 | 0.605096882 |
| 0.314974598 | 0.605013467 |
| 0.314916411 | 0.604974342 |
| 0.314847718 | 0.604866777 |
| 0.31476928 | 0.604789273 |
| 0.31472345 | 0.604725612 |
| 0.314659235 | 0.60462662 |
| 0.314638963 | 0.60461206 |
| 0.314513539 | 0.604492992 |
| 0.314459769 | 0.604414037 |
| 0.314380927 | 0.604311274 |
| 0.314337239 | 0.604251682 |
| 0.314323261 | 0.604249201 |
| 0.314302615 | 0.604233901 |
| 0.314294476 | 0.604233901 |
| 0.314240878 | 0.604188378 |
| 0.314237368 | 0.604188378 |
| 0.314192413 | 0.604168366 |
| 0.31403292 | 0.603959209 |
| 0.313943303 | 0.603811237 |
| 0.313866618 | 0.603712508 |
| 0.313831258 | 0.603693255 |
| 0.313805098 | 0.603691699 |
| 0.313796482 | 0.603691699 |
| 0.313752686 | 0.603671642 |
| 0.313702075 | 0.603671642 |
| 0.313677396 | 0.603665481 |
| 0.313574796 | 0.603516803 |
| 0.313565395 | 0.603516803 |
| 0.313542135 | 0.603516803 |
| 0.313508757 | 0.603487252 |
| 0.313402035 | 0.603379364 |
| 0.313399516 | 0.603379364 |

| | |
|---|---|
| 0.31336355 | 0.603354051 |
| 0.313326253 | 0.603306629 |
| 0.313278483 | 0.603239036 |
| 0.313224708 | 0.603159874 |
| 0.313203989 | 0.603144363 |
| 0.313103817 | 0.602975839 |
| 0.313099918 | 0.602975839 |
| 0.312968657 | 0.602764294 |
| 0.312875386 | 0.602682144 |
| 0.312874856 | 0.602682144 |
| 0.312852745 | 0.602682144 |
| 0.312823604 | 0.602655521 |
| 0.312777613 | 0.602612251 |
| 0.31274695 | 0.602580969 |
| 0.312713557 | 0.602565382 |
| 0.312647985 | 0.602463407 |
| 0.312630782 | 0.602454632 |
| 0.312534129 | 0.602292747 |
| 0.312429908 | 0.602150183 |
| 0.312387231 | 0.602107109 |
| 0.31233968 | 0.602039821 |
| 0.312322688 | 0.602031433 |
| 0.312241772 | 0.601924183 |
| 0.312226911 | 0.601919899 |
| 0.31218856 | 0.601892558 |
| 0.312162703 | 0.601892558 |
| 0.312062003 | 0.601845597 |
| 0.31203223 | 0.601812546 |
| 0.312019169 | 0.601811727 |
| 0.312017302 | 0.601811727 |
| 0.311991824 | 0.601807728 |
| 0.311970354 | 0.601807728 |
| 0.311937864 | 0.601776759 |
| 0.311908171 | 0.601768227 |
| 0.311876462 | 0.601731426 |
| 0.311855082 | 0.601726682 |
| 0.311754601 | 0.601569418 |
| 0.311754109 | 0.601569418 |
| 0.311722442 | 0.601556109 |
| 0.311639065 | 0.601419578 |
| 0.311633822 | 0.601419578 |
| 0.311612388 | 0.601416834 |
| 0.311569969 | 0.601359333 |
| 0.311512927 | 0.601273604 |
| 0.311499365 | 0.601271794 |

| | |
|---|---|
| 0.311478851 | 0.601256565 |
| 0.311373046 | 0.601125418 |
| 0.311345924 | 0.601097422 |
| 0.311340472 | 0.601097422 |
| 0.311322867 | 0.601097422 |
| 0.311289407 | 0.601061404 |
| 0.311261217 | 0.60103134 |
| 0.31125626 | 0.60103134 |
| 0.311211629 | 0.600984319 |
| 0.311159893 | 0.600933142 |
| 0.311140744 | 0.600933142 |
| 0.311129054 | 0.600933142 |
| 0.311088622 | 0.600892963 |
| 0.311065373 | 0.600892963 |
| 0.311040832 | 0.600892963 |
| 0.310982638 | 0.600834453 |
| 0.310847347 | 0.600621794 |
| 0.310783496 | 0.600563616 |
| 0.310751028 | 0.600557508 |
| 0.310728167 | 0.600537694 |
| 0.310725049 | 0.600537694 |
| 0.310692939 | 0.600518348 |
| 0.31068985 | 0.600518348 |
| 0.310686447 | 0.600518348 |
| 0.310670265 | 0.600518348 |
| 0.310611883 | 0.600459152 |
| 0.310600621 | 0.600459152 |
| 0.310582625 | 0.600451336 |
| 0.31048857 | 0.600318237 |
| 0.310450458 | 0.600268917 |
| 0.310340914 | 0.600178938 |
| 0.310324482 | 0.600171531 |
| 0.310315253 | 0.600171531 |
| 0.31029634 | 0.600165846 |
| 0.310254898 | 0.600134434 |
| 0.310254893 | 0.600134434 |
| 0.310220496 | 0.600116636 |
| 0.310168365 | 0.60004016 |
| 0.310153935 | 0.60004016 |
| 0.310152515 | 0.60004016 |
| 0.310125182 | 0.60004016 |
| 0.310106377 | 0.60004016 |
| 0.309975325 | 0.599837266 |
| 0.30995147 | 0.599837266 |
| 0.309917643 | 0.599798756 |

| | |
|---|---|
| 0.309899301 | 0.599787631 |
| 0.309890938 | 0.599787631 |
| 0.309878754 | 0.599787631 |
| 0.309877317 | 0.599787631 |
| 0.309835593 | 0.599786196 |
| 0.309769983 | 0.599683561 |
| 0.309761523 | 0.599683561 |
| 0.309713612 | 0.599647555 |
| 0.309673015 | 0.599617705 |
| 0.309491099 | 0.599338561 |
| 0.309477039 | 0.599335703 |
| 0.309476649 | 0.599335703 |
| 0.309407401 | 0.599263511 |
| 0.309402312 | 0.599263511 |
| 0.309385198 | 0.599263511 |
| 0.309330796 | 0.599223046 |
| 0.309273512 | 0.599136448 |
| 0.309230175 | 0.599124332 |
| 0.309201428 | 0.599094277 |
| 0.3091636 | 0.599045352 |
| 0.309140741 | 0.599025431 |
| 0.309132142 | 0.599025431 |
| 0.309081377 | 0.599007883 |
| 0.309026805 | 0.598926494 |
| 0.309023767 | 0.598926494 |
| 0.30888338 | 0.598745981 |
| 0.30886415 | 0.598733073 |
| 0.308851037 | 0.598732024 |
| 0.308779703 | 0.598618103 |
| 0.308748757 | 0.598582474 |
| 0.308743064 | 0.598582474 |
| 0.308664778 | 0.598492749 |
| 0.308642543 | 0.598474 |
| 0.308636309 | 0.598474 |
| 0.308599428 | 0.598463495 |
| 0.308594588 | 0.598463495 |
| 0.308588831 | 0.598463495 |
| 0.308437393 | 0.598319799 |
| 0.308397437 | 0.598266659 |
| 0.308338516 | 0.598201088 |
| 0.308329079 | 0.598201088 |
| 0.308278682 | 0.598201088 |
| 0.308250506 | 0.59817654 |
| 0.30824257 | 0.59817654 |
| 0.308229497 | 0.59817654 |

| | |
|---|---|
| 0.308178654 | 0.59817654 |
| 0.308102296 | 0.59810826 |
| 0.308082368 | 0.59810826 |
| 0.308034881 | 0.598050519 |
| 0.30803127 | 0.598050519 |
| 0.308005325 | 0.598041894 |
| 0.307976258 | 0.598034216 |
| 0.30792501 | 0.597959083 |
| 0.307881705 | 0.597899366 |
| 0.307827245 | 0.597869899 |
| 0.307819647 | 0.597869899 |
| 0.307819647 | 0.597869899 |
| 0.307792331 | 0.597869899 |
| 0.307785162 | 0.597869899 |
| 0.307602938 | 0.597685284 |
| 0.307573654 | 0.597666684 |
| 0.307564056 | 0.597666684 |
| 0.307490488 | 0.597578221 |
| 0.30746248 | 0.597548173 |
| 0.307370143 | 0.597441857 |
| 0.307360334 | 0.597441857 |
| 0.307351657 | 0.597441857 |
| 0.307296283 | 0.597420201 |
| 0.307220319 | 0.597296899 |
| 0.307187432 | 0.597268422 |
| 0.307124044 | 0.597182855 |
| 0.307010242 | 0.597007822 |
| 0.306982367 | 0.597002474 |
| 0.306974131 | 0.597002474 |
| 0.30695915 | 0.597002474 |
| 0.306918656 | 0.596954092 |
| 0.306914079 | 0.596954092 |
| 0.306898575 | 0.596954092 |
| 0.306893128 | 0.596954092 |
| 0.306861343 | 0.596954092 |
| 0.306861301 | 0.596954092 |
| 0.306835712 | 0.596954092 |
| 0.306698508 | 0.596778812 |
| 0.306693923 | 0.596778812 |
| 0.306672367 | 0.596778812 |
| 0.306643876 | 0.596778812 |
| 0.306604383 | 0.596757129 |
| 0.30659306 | 0.596757129 |
| 0.306539793 | 0.596680183 |
| 0.306390366 | 0.596486836 |

| | |
|---|---|
| 0.306350753 | 0.59645848 |
| 0.306288545 | 0.596361741 |
| 0.306255727 | 0.59632222 |
| 0.306203265 | 0.596293204 |
| 0.30613891 | 0.596265402 |
| 0.306120894 | 0.596265402 |
| 0.306098937 | 0.596265402 |
| 0.306091681 | 0.596265402 |
| 0.306069442 | 0.596265402 |
| 0.306047745 | 0.596265402 |
| 0.306032888 | 0.596265402 |
| 0.306016302 | 0.596265402 |
| 0.305939792 | 0.596218933 |
| 0.305875486 | 0.596118003 |
| 0.305850672 | 0.596094036 |
| 0.305840551 | 0.596094036 |
| 0.305783746 | 0.596012378 |
| 0.305781347 | 0.596012378 |
| 0.305766806 | 0.596012378 |
| 0.305730415 | 0.596005995 |
| 0.305706927 | 0.595984602 |
| 0.305701003 | 0.595984602 |
| 0.305654349 | 0.59598407 |
| 0.305642392 | 0.59598407 |
| 0.305583056 | 0.595913859 |
| 0.305491464 | 0.59575964 |
| 0.305480887 | 0.59575964 |
| 0.305473118 | 0.59575964 |
| 0.305420445 | 0.595718713 |
| 0.30541432 | 0.595718713 |
| 0.305366618 | 0.595662513 |
| 0.305360529 | 0.595662513 |
| 0.305302661 | 0.595610938 |
| 0.305299977 | 0.595610938 |
| 0.305291315 | 0.595610938 |
| 0.305271656 | 0.595610938 |
| 0.305222501 | 0.595576534 |
| 0.305169733 | 0.595522366 |
| 0.305169479 | 0.595522366 |
| 0.305167554 | 0.595522366 |
| 0.305156512 | 0.595522366 |
| 0.305150843 | 0.595522366 |
| 0.305147189 | 0.595522366 |
| 0.305090803 | 0.595522366 |
| 0.305087128 | 0.595522366 |

| | |
|---|---|
| 0.305041381 | 0.595491489 |
| 0.305007071 | 0.595448917 |
| 0.304983006 | 0.595426342 |
| 0.304954153 | 0.595418827 |
| 0.304909379 | 0.595380221 |
| 0.304891256 | 0.595380221 |
| 0.304885094 | 0.595380221 |
| 0.304817218 | 0.59529789 |
| 0.304811414 | 0.59529789 |
| 0.304779731 | 0.595273497 |
| 0.304772261 | 0.595273497 |
| 0.304730681 | 0.595226515 |
| 0.304721785 | 0.595226515 |
| 0.304717079 | 0.595226515 |
| 0.3046448 | 0.595156401 |
| 0.304626237 | 0.595144549 |
| 0.304623205 | 0.595144549 |
| 0.304609275 | 0.595144549 |
| 0.304598944 | 0.595144549 |
| 0.304593476 | 0.595144549 |
| 0.304590856 | 0.595144549 |
| 0.304560687 | 0.595144549 |
| 0.304543274 | 0.595144549 |
| 0.304537541 | 0.595144549 |
| 0.304502228 | 0.595144549 |
| 0.304305616 | 0.594905141 |
| 0.304298047 | 0.594905141 |
| 0.304242284 | 0.594858051 |
| 0.304182405 | 0.594814231 |
| 0.304137389 | 0.594799466 |
| 0.304112047 | 0.594774329 |
| 0.304035781 | 0.59467401 |
| 0.304033796 | 0.59467401 |
| 0.303994608 | 0.594642319 |
| 0.303950832 | 0.594581109 |
| 0.303886265 | 0.59450364 |
| 0.30384126 | 0.594440013 |
| 0.30384123 | 0.594440013 |
| 0.303840501 | 0.594440013 |
| 0.303758524 | 0.594351393 |
| 0.303745931 | 0.59435117 |
| 0.303689557 | 0.594265275 |
| 0.303615496 | 0.594169176 |
| 0.303563753 | 0.594092326 |
| 0.303517256 | 0.594025739 |

| | |
|---|---|
| 0.303488744 | 0.593994346 |
| 0.303475747 | 0.593993318 |
| 0.303461659 | 0.593993318 |
| 0.303417468 | 0.593952476 |
| 0.303411912 | 0.593952476 |
| 0.303387276 | 0.593952476 |
| 0.303369744 | 0.593952476 |
| 0.303339534 | 0.59394639 |
| 0.303338507 | 0.59394639 |
| 0.303295217 | 0.59394639 |
| 0.303163431 | 0.593727599 |
| 0.303147477 | 0.593727599 |
| 0.303140321 | 0.593727599 |
| 0.30308259 | 0.593662946 |
| 0.302953252 | 0.593507242 |
| 0.302948419 | 0.593507242 |
| 0.302936831 | 0.593507242 |
| 0.302926074 | 0.593507242 |
| 0.302902941 | 0.59350633 |
| 0.302893371 | 0.59350633 |
| 0.302864492 | 0.593488274 |
| 0.302792052 | 0.593411121 |
| 0.302789015 | 0.593411121 |
| 0.302690488 | 0.5932609 |
| 0.302636883 | 0.593180248 |
| 0.30257585 | 0.59309882 |
| 0.302513087 | 0.593084048 |
| 0.302459426 | 0.593003255 |
| 0.302415866 | 0.592974345 |
| 0.302376876 | 0.592939044 |
| 0.302356122 | 0.592922758 |
| 0.302317129 | 0.592895117 |
| 0.302312161 | 0.592895117 |
| 0.302303763 | 0.592895117 |
| 0.302278751 | 0.592893093 |
| 0.30210057 | 0.592641233 |
| 0.302092652 | 0.592641233 |
| 0.301872945 | 0.592267878 |
| 0.301871088 | 0.592267878 |
| 0.301870457 | 0.592267878 |
| 0.301869818 | 0.592267878 |
| 0.301866491 | 0.592267878 |
| 0.301860088 | 0.592267878 |
| 0.301860082 | 0.592267878 |
| 0.301854726 | 0.592267878 |

| | |
|---|---|
| 0.301841409 | 0.592267878 |
| 0.301752538 | 0.592267878 |
| 0.301683526 | 0.592262214 |
| 0.301605757 | 0.592182767 |
| 0.30159159 | 0.592179365 |
| 0.301547694 | 0.592117586 |
| 0.301539692 | 0.592117586 |
| 0.301497823 | 0.592068478 |
| 0.301443443 | 0.592027911 |
| 0.301427956 | 0.592027911 |
| 0.301409654 | 0.592027911 |
| 0.3013663 | 0.592005477 |
| 0.30133435 | 0.591998938 |
| 0.301313072 | 0.591998584 |
| 0.301291234 | 0.591980099 |
| 0.301278708 | 0.59197991 |
| 0.301253686 | 0.591955165 |
| 0.301242454 | 0.591955165 |
| 0.301174882 | 0.591849154 |
| 0.301087821 | 0.591751316 |
| 0.300965434 | 0.591584016 |
| 0.300868476 | 0.591442253 |
| 0.300863781 | 0.591442253 |
| 0.300821135 | 0.59139801 |
| 0.300812197 | 0.59139801 |
| 0.300807165 | 0.59139801 |
| 0.300801853 | 0.59139801 |
| 0.300752519 | 0.59138517 |
| 0.300737752 | 0.591380549 |
| 0.300697559 | 0.591325928 |
| 0.30069733 | 0.591325928 |
| 0.300613202 | 0.591233275 |
| 0.30060934 | 0.591233275 |
| 0.300606037 | 0.591233275 |
| 0.300600525 | 0.591233275 |
| 0.300556474 | 0.591219362 |
| 0.300517667 | 0.591191859 |
| 0.300499501 | 0.591180541 |
| 0.300437701 | 0.591083377 |
| 0.300437016 | 0.591083377 |
| 0.300416185 | 0.591083377 |
| 0.300404591 | 0.591083377 |
| 0.300366224 | 0.591083377 |
| 0.300268279 | 0.590920925 |
| 0.300229607 | 0.59087325 |

| | |
|---|---|
| 0.300107418 | 0.59072951 |
| 0.30008112 | 0.590723465 |
| 0.300015823 | 0.59064375 |
| 0.300012114 | 0.59064375 |
| 0.299979954 | 0.590621963 |
| 0.299860967 | 0.590459144 |
| 0.299845198 | 0.590454281 |
| 0.299844156 | 0.590454281 |
| 0.299819906 | 0.590453303 |
| 0.299816655 | 0.590453303 |
| 0.299805083 | 0.590453303 |
| 0.299649391 | 0.590215118 |
| 0.299612437 | 0.590181648 |
| 0.299588283 | 0.590167975 |
| 0.299521874 | 0.59011038 |
| 0.29939833 | 0.589891377 |
| 0.299352166 | 0.589849226 |
| 0.29934506 | 0.589849226 |
| 0.299326534 | 0.589847525 |
| 0.299233 | 0.589760814 |
| 0.299179298 | 0.589679376 |
| 0.29915596 | 0.589679376 |
| 0.299139942 | 0.589675015 |
| 0.29913052 | 0.589675015 |
| 0.299118974 | 0.589675015 |
| 0.299060829 | 0.589616679 |
| 0.299016309 | 0.589602126 |
| 0.298994028 | 0.589602126 |
| 0.298969523 | 0.5896 |
| 0.298920428 | 0.589535104 |
| 0.29889113 | 0.589501733 |
| 0.298852877 | 0.589450697 |
| 0.298852302 | 0.589450697 |
| 0.298839082 | 0.589450697 |
| 0.298827641 | 0.589450697 |
| 0.298795237 | 0.589450697 |
| 0.298765046 | 0.589438153 |
| 0.298761547 | 0.589438153 |
| 0.298749093 | 0.589438153 |
| 0.29873297 | 0.589438153 |
| 0.298693547 | 0.589438153 |
| 0.298637773 | 0.589392559 |
| 0.298611248 | 0.589364631 |
| 0.298597116 | 0.589361162 |
| 0.298590937 | 0.589361162 |

| | |
|---|---|
| 0.29849955 | 0.589290697 |
| 0.298487359 | 0.589290697 |
| 0.298409035 | 0.589160847 |
| 0.298322791 | 0.589112678 |
| 0.298285701 | 0.589063858 |
| 0.298259587 | 0.589061139 |
| 0.298219432 | 0.589030684 |
| 0.298218163 | 0.589030684 |
| 0.298175369 | 0.589016934 |
| 0.298132388 | 0.588956459 |
| 0.29812594 | 0.588956459 |
| 0.298121378 | 0.588956459 |
| 0.298092746 | 0.588951432 |
| 0.298070681 | 0.588951432 |
| 0.298064592 | 0.588951432 |
| 0.298026523 | 0.58892834 |
| 0.298024134 | 0.58892834 |
| 0.298002657 | 0.58892834 |
| 0.297886305 | 0.588812246 |
| 0.297884006 | 0.588812246 |
| 0.297815899 | 0.58877082 |
| 0.297789518 | 0.588743103 |
| 0.297770029 | 0.588743103 |
| 0.297749936 | 0.588743103 |
| 0.297739098 | 0.588743103 |
| 0.297717742 | 0.588743103 |
| 0.297665004 | 0.588716878 |
| 0.297658449 | 0.588716878 |
| 0.297622697 | 0.588682097 |
| 0.297610033 | 0.588681495 |
| 0.297562164 | 0.588611254 |
| 0.297513548 | 0.588539531 |
| 0.297479211 | 0.588496049 |
| 0.297458431 | 0.588479384 |
| 0.297427258 | 0.5884666 |
| 0.29737982 | 0.588421632 |
| 0.297331984 | 0.588375869 |
| 0.29731201 | 0.588360789 |
| 0.29727357 | 0.58833361 |
| 0.297159441 | 0.588132177 |
| 0.297144263 | 0.588126576 |
| 0.297097649 | 0.588058754 |
| 0.297043978 | 0.588001395 |
| 0.297042631 | 0.588001395 |
| 0.297027038 | 0.588001395 |

| | |
|---|---|
| 0.296956374 | 0.587950166 |
| 0.296916156 | 0.587923411 |
| 0.296911957 | 0.587923411 |
| 0.296900682 | 0.587923411 |
| 0.296870053 | 0.587923411 |
| 0.296813858 | 0.587900372 |
| 0.296795137 | 0.587900372 |
| 0.296784508 | 0.587900372 |
| 0.296772098 | 0.587900372 |
| 0.29675517 | 0.587900372 |
| 0.296736175 | 0.587900372 |
| 0.296679851 | 0.587842546 |
| 0.296578262 | 0.587739063 |
| 0.296565453 | 0.587738133 |
| 0.296547666 | 0.587727338 |
| 0.296437688 | 0.587558275 |
| 0.296425529 | 0.587558275 |
| 0.296424389 | 0.587558275 |
| 0.296413531 | 0.587558275 |
| 0.296407851 | 0.587558275 |
| 0.296401653 | 0.587558275 |
| 0.296395461 | 0.587558275 |
| 0.296328713 | 0.587513429 |
| 0.296292475 | 0.587490496 |
| 0.296218392 | 0.587368056 |
| 0.296199296 | 0.587354645 |
| 0.296191432 | 0.587354645 |
| 0.296156787 | 0.587343714 |
| 0.296140488 | 0.587335846 |
| 0.296100123 | 0.587304704 |
| 0.296070191 | 0.587269793 |
| 0.29605159 | 0.587267643 |
| 0.296029491 | 0.587262437 |
| 0.295950731 | 0.587135048 |
| 0.295950404 | 0.587135048 |
| 0.295806329 | 0.586963721 |
| 0.295745388 | 0.5868944 |
| 0.295671666 | 0.58677255 |
| 0.295654231 | 0.5867624 |
| 0.29561499 | 0.586708969 |
| 0.295587099 | 0.586696003 |
| 0.295560252 | 0.586673674 |
| 0.295485562 | 0.586608212 |
| 0.295455899 | 0.586608212 |
| 0.295420267 | 0.586566934 |

| | |
|---|---|
| 0.295300988 | 0.586403442 |
| 0.295287169 | 0.58640045 |
| 0.295281769 | 0.58640045 |
| 0.295275318 | 0.58640045 |
| 0.295262157 | 0.58640045 |
| 0.29522532 | 0.586375425 |
| 0.295169635 | 0.586289273 |
| 0.295130192 | 0.586235378 |
| 0.295080654 | 0.586161424 |
| 0.295022363 | 0.586070076 |
| 0.295018842 | 0.586070076 |
| 0.295014817 | 0.586070076 |
| 0.294988084 | 0.586070076 |
| 0.294976178 | 0.586070076 |
| 0.294962249 | 0.586070076 |
| 0.294957561 | 0.586070076 |
| 0.294906223 | 0.586010459 |
| 0.294826729 | 0.58587694 |
| 0.294812977 | 0.585874057 |
| 0.29479606 | 0.585864884 |
| 0.294742318 | 0.585806969 |
| 0.294741084 | 0.585806969 |
| 0.294696148 | 0.585764093 |
| 0.294696148 | 0.585764093 |
| 0.294660055 | 0.585741244 |
| 0.294628442 | 0.585727296 |
| 0.294502917 | 0.585587812 |
| 0.294490328 | 0.585587812 |
| 0.294420408 | 0.585509239 |
| 0.294373744 | 0.585465332 |
| 0.294343572 | 0.585429772 |
| 0.294311825 | 0.585415524 |
| 0.294233614 | 0.585308844 |
| 0.294195942 | 0.585282795 |
| 0.294191897 | 0.585282795 |
| 0.294182715 | 0.585282795 |
| 0.294107626 | 0.585204876 |
| 0.294093224 | 0.585200668 |
| 0.29404867 | 0.585160907 |
| 0.293965326 | 0.585064576 |
| 0.293949045 | 0.585060436 |
| 0.293936306 | 0.585060436 |
| 0.293879987 | 0.584996327 |
| 0.293874131 | 0.584996327 |
| 0.293792254 | 0.584872606 |

| | |
|---|---|
| 0.293772396 | 0.584872606 |
| 0.29372011 | 0.584800287 |
| 0.293645736 | 0.58467665 |
| 0.293627394 | 0.584664571 |
| 0.293332509 | 0.584199523 |
| 0.293320456 | 0.584199523 |
| 0.293267186 | 0.584142707 |
| 0.293242418 | 0.584117801 |
| 0.293219778 | 0.584097134 |
| 0.293219069 | 0.584097134 |
| 0.293208234 | 0.584097134 |
| 0.2931874 | 0.584097134 |
| 0.293175224 | 0.584097134 |
| 0.293140418 | 0.584097134 |
| 0.293081948 | 0.584017987 |
| 0.293025108 | 0.583953588 |
| 0.292957784 | 0.583868284 |
| 0.292947761 | 0.583868284 |
| 0.292927434 | 0.583868284 |
| 0.292908614 | 0.583868019 |
| 0.292901525 | 0.583868019 |
| 0.29285238 | 0.583804793 |
| 0.292812002 | 0.583764822 |
| 0.29276239 | 0.583698691 |
| 0.29270823 | 0.583676292 |
| 0.292707864 | 0.583676292 |
| 0.292701445 | 0.583676292 |
| 0.292673382 | 0.583676292 |
| 0.292655247 | 0.583676292 |
| 0.292625494 | 0.583645647 |
| 0.29260737 | 0.58363394 |
| 0.292606657 | 0.58363394 |
| 0.292529365 | 0.58357611 |
| 0.292477263 | 0.583496612 |
| 0.292473886 | 0.583496612 |
| 0.292427224 | 0.583445662 |
| 0.292393491 | 0.583402798 |
| 0.292347454 | 0.583382791 |
| 0.29231098 | 0.583335918 |
| 0.292300972 | 0.583335918 |
| 0.29226165 | 0.583310799 |
| 0.29222182 | 0.583255747 |
| 0.292062572 | 0.582968298 |
| 0.291872739 | 0.582681097 |
| 0.291850073 | 0.582660271 |

| | |
|---|---|
| 0.291808761 | 0.582602217 |
| 0.291730975 | 0.582569021 |
| 0.291707834 | 0.582547236 |
| 0.291702168 | 0.582547236 |
| 0.291696536 | 0.582547236 |
| 0.291682992 | 0.582547236 |
| 0.291676165 | 0.582547236 |
| 0.291645768 | 0.582547236 |
| 0.291598781 | 0.582547236 |
| 0.2915558 | 0.582512294 |
| 0.291542609 | 0.582510991 |
| 0.291479596 | 0.582433343 |
| 0.291463888 | 0.582433343 |
| 0.29146012 | 0.582433343 |
| 0.291458733 | 0.582433343 |
| 0.291394844 | 0.582386162 |
| 0.291348868 | 0.582382369 |
| 0.291330961 | 0.58238068 |
| 0.291324605 | 0.58238068 |
| 0.291318935 | 0.58238068 |
| 0.291286671 | 0.58238068 |
| 0.291245381 | 0.582331812 |
| 0.291221847 | 0.58232678 |
| 0.291180663 | 0.582275745 |
| 0.291116909 | 0.582197145 |
| 0.291027776 | 0.582067774 |
| 0.291016359 | 0.582067774 |
| 0.291016359 | 0.582067774 |
| 0.290957121 | 0.582024227 |
| 0.290929549 | 0.582017963 |
| 0.290847979 | 0.581903663 |
| 0.290824203 | 0.581880538 |
| 0.290820378 | 0.581880538 |
| 0.290813358 | 0.581880538 |
| 0.290773419 | 0.581880538 |
| 0.290752367 | 0.581880538 |
| 0.290692571 | 0.581788254 |
| 0.290621091 | 0.581669638 |
| 0.290563951 | 0.581604157 |
| 0.290550478 | 0.581601632 |
| 0.290537505 | 0.581600107 |
| 0.290506819 | 0.581587569 |
| 0.29048869 | 0.581575721 |
| 0.290439697 | 0.581502079 |
| 0.290353169 | 0.581402161 |

| | |
|---|---|
| 0.290293716 | 0.581325238 |
| 0.290273652 | 0.581316256 |
| 0.290259927 | 0.581313215 |
| 0.290258404 | 0.581313215 |
| 0.290210301 | 0.581274293 |
| 0.290128477 | 0.581147704 |
| 0.290019629 | 0.58096459 |
| 0.289991182 | 0.580945993 |
| 0.289983777 | 0.580945993 |
| 0.289978028 | 0.580945993 |
| 0.289945128 | 0.580945993 |
| 0.289942577 | 0.580945993 |
| 0.289932671 | 0.580945993 |
| 0.289890165 | 0.580945993 |
| 0.289869459 | 0.580945993 |
| 0.289865907 | 0.580945993 |
| 0.289832529 | 0.58092138 |
| 0.289802215 | 0.580885068 |
| 0.289790436 | 0.580885068 |
| 0.289774876 | 0.580879163 |
| 0.289735398 | 0.580843941 |
| 0.289696457 | 0.5808279 |
| 0.28961414 | 0.580742013 |
| 0.289611651 | 0.580742013 |
| 0.289608095 | 0.580742013 |
| 0.289605438 | 0.580742013 |
| 0.289581258 | 0.580742013 |
| 0.289546679 | 0.580742013 |
| 0.28954169 | 0.580742013 |
| 0.289515918 | 0.580742013 |
| 0.289419198 | 0.580630502 |
| 0.289388349 | 0.580617529 |
| 0.289338494 | 0.580581558 |
| 0.28932184 | 0.580581558 |
| 0.28929684 | 0.580556212 |
| 0.289288427 | 0.580556212 |
| 0.289279798 | 0.580556212 |
| 0.289265493 | 0.580556212 |
| 0.289262013 | 0.580556212 |
| 0.28925407 | 0.580556212 |
| 0.289243357 | 0.580556212 |
| 0.289176949 | 0.580533647 |
| 0.289155561 | 0.580533647 |
| 0.289126465 | 0.580527142 |
| 0.289076753 | 0.580481428 |

| | |
|---|---|
| 0.289061318 | 0.580474908 |
| 0.289056859 | 0.580474908 |
| 0.289046591 | 0.580474908 |
| 0.288951695 | 0.580352649 |
| 0.288873166 | 0.580219395 |
| 0.288860716 | 0.580219395 |
| 0.28874194 | 0.580078137 |
| 0.288741692 | 0.580078137 |
| 0.288738194 | 0.580078137 |
| 0.288704913 | 0.580078137 |
| 0.288702123 | 0.580078137 |
| 0.288697222 | 0.580078137 |
| 0.288676851 | 0.580078137 |
| 0.288538011 | 0.579913063 |
| 0.288529947 | 0.579913063 |
| 0.288481521 | 0.579848467 |
| 0.288478558 | 0.579848467 |
| 0.288424181 | 0.57978215 |
| 0.288413611 | 0.57978215 |
| 0.288351803 | 0.579686587 |
| 0.288351461 | 0.579686587 |
| 0.28834527 | 0.579686587 |
| 0.28832668 | 0.579686587 |
| 0.28832319 | 0.579686587 |
| 0.288286773 | 0.579686587 |
| 0.288266668 | 0.57968572 |
| 0.288253676 | 0.579684069 |
| 0.288219068 | 0.579663425 |
| 0.288193136 | 0.579635746 |
| 0.288037446 | 0.579371545 |
| 0.2879882 | 0.579321422 |
| 0.287914902 | 0.57919844 |
| 0.287842884 | 0.579102482 |
| 0.287842772 | 0.579102482 |
| 0.287776805 | 0.579018457 |
| 0.287761436 | 0.579011994 |
| 0.287698231 | 0.578933735 |
| 0.287691539 | 0.578933735 |
| 0.287682403 | 0.578933735 |
| 0.28765616 | 0.578933735 |
| 0.287485311 | 0.578655918 |
| 0.287464824 | 0.578655918 |
| 0.287416358 | 0.578586536 |
| 0.287394392 | 0.578566773 |
| 0.28739343 | 0.578566773 |

| | |
|---|---|
| 0.287378461 | 0.578566773 |
| 0.287366009 | 0.578566773 |
| 0.287361542 | 0.578566773 |
| 0.287287637 | 0.578566773 |
| 0.287274197 | 0.578566773 |
| 0.287264373 | 0.578566773 |
| 0.287253255 | 0.578566773 |
| 0.287250856 | 0.578566773 |
| 0.287228327 | 0.578566773 |
| 0.28722288 | 0.578566773 |
| 0.287222633 | 0.578566773 |
| 0.28715379 | 0.57852257 |
| 0.287138739 | 0.578516719 |
| 0.287094528 | 0.578452115 |
| 0.287046895 | 0.578380609 |
| 0.287030265 | 0.578371569 |
| 0.286935116 | 0.578326651 |
| 0.28689655 | 0.578273393 |
| 0.286892223 | 0.578273393 |
| 0.286846246 | 0.578239277 |
| 0.286798757 | 0.578174155 |
| 0.286777058 | 0.578154883 |
| 0.286692796 | 0.578009473 |
| 0.286634566 | 0.577941005 |
| 0.286617755 | 0.577931576 |
| 0.286392685 | 0.577624477 |
| 0.286381481 | 0.577624477 |
| 0.286295752 | 0.577477884 |
| 0.286283943 | 0.577477884 |
| 0.286268639 | 0.577472107 |
| 0.286236469 | 0.57745613 |
| 0.28620457 | 0.577435246 |
| 0.286200656 | 0.577435246 |
| 0.286155146 | 0.577435246 |
| 0.28613536 | 0.577423363 |
| 0.286102854 | 0.57738223 |
| 0.286098421 | 0.57738223 |
| 0.286051756 | 0.577328036 |
| 0.286020156 | 0.577313187 |
| 0.285998011 | 0.577292956 |
| 0.285991128 | 0.577292956 |
| 0.285827932 | 0.577071933 |
| 0.285823939 | 0.577071933 |
| 0.285812342 | 0.577071933 |
| 0.285729173 | 0.576995332 |

| | |
|---|---|
| 0.285725656 | 0.576995332 |
| 0.285713407 | 0.576995332 |
| 0.285681227 | 0.576995332 |
| 0.28564126 | 0.576958296 |
| 0.28556939 | 0.576908043 |
| 0.285555388 | 0.576908043 |
| 0.285544258 | 0.576908043 |
| 0.285475417 | 0.576800343 |
| 0.28536347 | 0.576647545 |
| 0.285337991 | 0.576620524 |
| 0.285231825 | 0.576509001 |
| 0.285180025 | 0.576448054 |
| 0.285123438 | 0.576382597 |
| 0.285064697 | 0.576361706 |
| 0.284993639 | 0.576242499 |
| 0.284918319 | 0.576139124 |
| 0.284887332 | 0.576125386 |
| 0.284881395 | 0.576125386 |
| 0.284827266 | 0.576101762 |
| 0.284806602 | 0.576084431 |
| 0.284714993 | 0.575923592 |
| 0.284702441 | 0.575922663 |
| 0.284682708 | 0.575922663 |
| 0.284577081 | 0.57579855 |
| 0.284493126 | 0.575694855 |
| 0.284481324 | 0.575694855 |
| 0.284441994 | 0.575689225 |
| 0.284434275 | 0.575689225 |
| 0.284428455 | 0.575689225 |
| 0.28440283 | 0.575683352 |
| 0.284386802 | 0.575675374 |
| 0.284329919 | 0.575584693 |
| 0.284272052 | 0.575494342 |
| 0.28422782 | 0.575451385 |
| 0.284222661 | 0.575451385 |
| 0.28421169 | 0.575451385 |
| 0.284200965 | 0.575451385 |
| 0.284174965 | 0.575442222 |
| 0.28413777 | 0.575391368 |
| 0.284089832 | 0.575318752 |
| 0.284040777 | 0.575243871 |
| 0.284021572 | 0.575229436 |
| 0.284020671 | 0.575229436 |
| 0.284006674 | 0.575229436 |
| 0.283997212 | 0.575229436 |

| | |
|---|---|
| 0.283990087 | 0.575229436 |
| 0.28398591 | 0.575229436 |
| 0.283930635 | 0.575216486 |
| 0.2838995 | 0.575177874 |
| 0.28384114 | 0.575084102 |
| 0.283829938 | 0.575084102 |
| 0.283758372 | 0.575002181 |
| 0.283684815 | 0.574889656 |
| 0.283655105 | 0.574853908 |
| 0.283625775 | 0.574818927 |
| 0.283604412 | 0.574800091 |
| 0.283593258 | 0.574800091 |
| 0.283584454 | 0.574800091 |
| 0.283573991 | 0.574800091 |
| 0.283557166 | 0.574800091 |
| 0.283480844 | 0.574745312 |
| 0.283462701 | 0.574732996 |
| 0.283413943 | 0.57466881 |
| 0.283396929 | 0.57466881 |
| 0.283368278 | 0.574639402 |
| 0.283347823 | 0.574622388 |
| 0.283345993 | 0.574622388 |
| 0.283343204 | 0.574622388 |
| 0.283343201 | 0.574622388 |
| 0.283342703 | 0.574622388 |
| 0.283299296 | 0.574622388 |
| 0.283277832 | 0.574622388 |
| 0.283276976 | 0.574622388 |
| 0.283276168 | 0.574622388 |
| 0.283269548 | 0.574622388 |
| 0.283233914 | 0.574622388 |
| 0.28315534 | 0.574622388 |
| 0.283068059 | 0.574519421 |
| 0.282958002 | 0.574369891 |
| 0.282941672 | 0.574369891 |
| 0.282918822 | 0.574369891 |
| 0.282900656 | 0.574369891 |
| 0.28288473 | 0.574369891 |
| 0.282778416 | 0.574331985 |
| 0.282771487 | 0.574331985 |
| 0.282752107 | 0.574331985 |
| 0.282714218 | 0.574291139 |
| 0.282658865 | 0.574208666 |
| 0.282592256 | 0.574203256 |
| 0.282442028 | 0.573934059 |

| | |
|---|---|
| 0.282422842 | 0.573919559 |
| 0.2824009 | 0.573899457 |
| 0.28236029 | 0.573841413 |
| 0.282359852 | 0.573841413 |
| 0.282320677 | 0.573809879 |
| 0.282320052 | 0.573809879 |
| 0.282310553 | 0.573809879 |
| 0.282309752 | 0.573809879 |
| 0.282274383 | 0.573809879 |
| 0.282236126 | 0.573784948 |
| 0.282204489 | 0.573769613 |
| 0.282182657 | 0.573749719 |
| 0.282178159 | 0.573749719 |
| 0.282163894 | 0.573749719 |
| 0.282128694 | 0.573713475 |
| 0.282069262 | 0.57361711 |
| 0.282068569 | 0.57361711 |
| 0.282056036 | 0.57361711 |
| 0.28203505 | 0.57361711 |
| 0.28202294 | 0.57361711 |
| 0.282016725 | 0.57361711 |
| 0.281931148 | 0.57361711 |
| 0.281849071 | 0.573561173 |
| 0.281769299 | 0.573447749 |
| 0.28175736 | 0.573447749 |
| 0.281657422 | 0.573293646 |
| 0.281558945 | 0.573117696 |
| 0.281530951 | 0.573085207 |
| 0.281471781 | 0.573001202 |
| 0.281379236 | 0.572874315 |
| 0.281338464 | 0.572815792 |
| 0.281320643 | 0.572810954 |
| 0.281301357 | 0.572810954 |
| 0.281243189 | 0.572744232 |
| 0.281206642 | 0.572700779 |
| 0.281205306 | 0.572700779 |
| 0.281142609 | 0.572650482 |
| 0.281059587 | 0.572557341 |
| 0.281016092 | 0.57252656 |
| 0.281009634 | 0.57252656 |
| 0.281001079 | 0.57252656 |
| 0.280973934 | 0.572514142 |
| 0.280865468 | 0.572342103 |
| 0.280843877 | 0.572332257 |
| 0.280839976 | 0.572332257 |

| | |
|---|---|
| 0.280835883 | 0.572332257 |
| 0.280835188 | 0.572332257 |
| 0.280785192 | 0.572332257 |
| 0.280778819 | 0.572332257 |
| 0.280778561 | 0.572332257 |
| 0.280645186 | 0.572260489 |
| 0.280629229 | 0.572252454 |
| 0.280624263 | 0.572252454 |
| 0.280530932 | 0.572100998 |
| 0.280421249 | 0.571926295 |
| 0.280402311 | 0.571914365 |
| 0.280392207 | 0.571914365 |
| 0.280338001 | 0.571914365 |
| 0.280337758 | 0.571914365 |
| 0.280327507 | 0.571914365 |
| 0.280247093 | 0.571840464 |
| 0.280242787 | 0.571840464 |
| 0.280242214 | 0.571840464 |
| 0.280191658 | 0.571800844 |
| 0.280110595 | 0.57165991 |
| 0.280078467 | 0.571618835 |
| 0.280066338 | 0.571618576 |
| 0.280055755 | 0.571618576 |
| 0.280051533 | 0.571618576 |
| 0.280041731 | 0.571618576 |
| 0.28002424 | 0.571618576 |
| 0.279958641 | 0.571542536 |
| 0.279932945 | 0.571517764 |
| 0.279928498 | 0.571517764 |
| 0.279912746 | 0.571517764 |
| 0.279900908 | 0.571517764 |
| 0.279866298 | 0.571517764 |
| 0.279828602 | 0.571500721 |
| 0.279815187 | 0.571497831 |
| 0.279711598 | 0.57131076 |
| 0.279698224 | 0.571307943 |
| 0.279633561 | 0.57120036 |
| 0.279544823 | 0.571068084 |
| 0.279533148 | 0.571068084 |
| 0.279476599 | 0.570977692 |
| 0.279391707 | 0.570828742 |
| 0.279386523 | 0.570828742 |
| 0.279357924 | 0.570828742 |
| 0.279305479 | 0.57079246 |
| 0.279295739 | 0.57079246 |

| | |
|---|---|
| 0.279294524 | 0.57079246 |
| 0.279281164 | 0.57079246 |
| 0.279275819 | 0.57079246 |
| 0.279260516 | 0.57079246 |
| 0.27925743 | 0.57079246 |
| 0.279204426 | 0.570788909 |
| 0.279189017 | 0.570781906 |
| 0.279072792 | 0.570642274 |
| 0.27907231 | 0.570642274 |
| 0.279032975 | 0.570609854 |
| 0.279026906 | 0.570609854 |
| 0.279009337 | 0.570609854 |
| 0.278876786 | 0.570531473 |
| 0.278873524 | 0.570531473 |
| 0.27883395 | 0.57052131 |
| 0.278801723 | 0.570479879 |
| 0.27870318 | 0.570327246 |
| 0.278701054 | 0.570327246 |
| 0.278681618 | 0.570327246 |
| 0.278662429 | 0.570317369 |
| 0.278622586 | 0.570260331 |
| 0.278621519 | 0.570260331 |
| 0.278594511 | 0.570260331 |
| 0.27853729 | 0.570183766 |
| 0.278464119 | 0.570131999 |
| 0.278410009 | 0.570051013 |
| 0.278299677 | 0.569893314 |
| 0.278292528 | 0.569893314 |
| 0.278210972 | 0.569785163 |
| 0.278166706 | 0.569743505 |
| 0.278144195 | 0.569721898 |
| 0.278098277 | 0.569676846 |
| 0.278074666 | 0.56965298 |
| 0.277928349 | 0.569524718 |
| 0.277915075 | 0.56952202 |
| 0.277885296 | 0.5694855 |
| 0.277636805 | 0.5690742 |
| 0.277589299 | 0.569025804 |
| 0.277547473 | 0.568964555 |
| 0.277502241 | 0.568896318 |
| 0.277465953 | 0.568870898 |
| 0.277415904 | 0.568817258 |
| 0.277400428 | 0.568810014 |
| 0.277365368 | 0.56876261 |
| 0.277286433 | 0.568686502 |

| | |
|---|---|
| 0.277250365 | 0.568673692 |
| 0.27717334 | 0.568540189 |
| 0.277063526 | 0.568363895 |
| 0.276940976 | 0.568262377 |
| 0.276848758 | 0.568143562 |
| 0.276820505 | 0.568143264 |
| 0.27681302 | 0.568143264 |
| 0.276783805 | 0.568132666 |
| 0.276733948 | 0.568079295 |
| 0.276713852 | 0.568062527 |
| 0.276665533 | 0.567987816 |
| 0.276617793 | 0.567914287 |
| 0.276576701 | 0.567903364 |
| 0.276572357 | 0.567903364 |
| 0.276570181 | 0.567903364 |
| 0.276555543 | 0.567903364 |
| 0.276506001 | 0.567903364 |
| 0.276491505 | 0.567899814 |
| 0.27647536 | 0.567891145 |
| 0.276449546 | 0.567869495 |
| 0.276434849 | 0.567869495 |
| 0.2764118 | 0.567858786 |
| 0.276394827 | 0.567858786 |
| 0.276392104 | 0.567858786 |
| 0.276360779 | 0.567851711 |
| 0.276313267 | 0.567778582 |
| 0.276186227 | 0.567566509 |
| 0.276179462 | 0.567566509 |
| 0.276148674 | 0.567538315 |
| 0.276095959 | 0.567454462 |
| 0.276087601 | 0.567454462 |
| 0.276085434 | 0.567454462 |
| 0.276051163 | 0.567454462 |
| 0.275999077 | 0.567426748 |
| 0.275951246 | 0.567426388 |
| 0.275914823 | 0.567375989 |
| 0.275908428 | 0.567375989 |
| 0.275887586 | 0.567368974 |
| 0.275850195 | 0.567341073 |
| 0.275786207 | 0.567282962 |
| 0.275774972 | 0.567282962 |
| 0.275736096 | 0.567228882 |
| 0.275712497 | 0.567204833 |
| 0.275690789 | 0.567184675 |
| 0.275690789 | 0.567184675 |

| | |
|---|---|
| 0.275690789 | 0.567184675 |
| 0.275690789 | 0.567184675 |
| 0.275690789 | 0.567184675 |
| 0.275690789 | 0.567184675 |
| 0.275690789 | 0.567184675 |
| 0.275690789 | 0.567184675 |
| 0.275690789 | 0.567184675 |
| 0.275690789 | 0.567184675 |
| 0.275690789 | 0.567184675 |
| 0.275690789 | 0.567184675 |
| 0.275690789 | 0.567184675 |
| 0.275690789 | 0.567184675 |
| 0.275690789 | 0.567184675 |
| 0.275641947 | 0.567184675 |
| 0.275612674 | 0.567184675 |
| 0.275599575 | 0.567184675 |
| 0.275585132 | 0.567184675 |
| 0.275513401 | 0.567184675 |
| 0.275431816 | 0.567184675 |
| 0.275380564 | 0.567184675 |
| 0.275372213 | 0.567184675 |
| 0.275350579 | 0.567184675 |
| 0.275307273 | 0.567155133 |
| 0.275250898 | 0.567112585 |
| 0.275214579 | 0.567086821 |
| 0.275101299 | 0.566945009 |
| 0.275060443 | 0.566916381 |
| 0.275024623 | 0.566890409 |
| 0.274936501 | 0.566852998 |
| 0.274893366 | 0.566792776 |
| 0.274840178 | 0.566756718 |
| 0.274824974 | 0.566756718 |
| 0.274784603 | 0.566715728 |
| 0.274782377 | 0.566715728 |
| 0.274727124 | 0.566656958 |
| 0.274657611 | 0.566601032 |
| 0.274653908 | 0.566601032 |
| 0.274647357 | 0.566601032 |
| 0.274508538 | 0.5663671 |
| 0.274484654 | 0.566342356 |
| 0.274430084 | 0.566278827 |
| 0.274355856 | 0.566199252 |
| 0.274338399 | 0.56618776 |
| 0.274312268 | 0.566158362 |
| 0.274193474 | 0.565962233 |

| | |
|---|---|
| 0.274161153 | 0.565920047 |
| 0.274112429 | 0.565868522 |
| 0.274093928 | 0.565868522 |
| 0.274050006 | 0.565837761 |
| 0.273989158 | 0.565810243 |
| 0.273971628 | 0.565798575 |
| 0.273889015 | 0.565701553 |
| 0.273884123 | 0.565701553 |
| 0.273881201 | 0.565701553 |
| 0.273875493 | 0.565701553 |
| 0.273822532 | 0.565701553 |
| 0.273819368 | 0.565701553 |
| 0.273788752 | 0.565690725 |
| 0.273786041 | 0.565690725 |
| 0.273729413 | 0.565641737 |
| 0.27369853 | 0.565602459 |
| 0.273642563 | 0.56551134 |
| 0.273626357 | 0.56551134 |
| 0.273623877 | 0.56551134 |
| 0.273583115 | 0.56551134 |
| 0.273499804 | 0.565437131 |
| 0.27349919 | 0.565437131 |
| 0.273496145 | 0.565437131 |
| 0.273446642 | 0.565400852 |
| 0.273429163 | 0.565389256 |
| 0.273351875 | 0.565278528 |
| 0.273333082 | 0.565264207 |
| 0.273272283 | 0.565187556 |
| 0.273261511 | 0.565187556 |
| 0.27325907 | 0.565187556 |
| 0.273239057 | 0.565187556 |
| 0.273234115 | 0.565187556 |
| 0.273224082 | 0.565187556 |
| 0.27315188 | 0.565134864 |
| 0.273081413 | 0.565013618 |
| 0.273065135 | 0.565004484 |
| 0.273037898 | 0.564972671 |
| 0.273030928 | 0.564972671 |
| 0.272880884 | 0.564721381 |
| 0.272838175 | 0.564666422 |
| 0.27279669 | 0.564646531 |
| 0.272796397 | 0.564646531 |
| 0.27276765 | 0.564646531 |
| 0.272763747 | 0.564646531 |
| 0.272749653 | 0.564646531 |

| | |
|---|---|
| 0.272746136 | 0.564646531 |
| 0.27270825 | 0.564633996 |
| 0.272695225 | 0.564633996 |
| 0.272662722 | 0.564613368 |
| 0.272614689 | 0.564538449 |
| 0.272614328 | 0.564538449 |
| 0.272564157 | 0.564482893 |
| 0.272550335 | 0.564478813 |
| 0.272490752 | 0.564429045 |
| 0.272473588 | 0.564429045 |
| 0.272343938 | 0.564223096 |
| 0.272236001 | 0.56404855 |
| 0.27214998 | 0.563919385 |
| 0.272136105 | 0.56391517 |
| 0.27211988 | 0.563906084 |
| 0.272119474 | 0.563906084 |
| 0.272114601 | 0.563906084 |
| 0.272078106 | 0.563906084 |
| 0.272052106 | 0.563906084 |
| 0.271986014 | 0.563800397 |
| 0.271980011 | 0.563800397 |
| 0.271978219 | 0.563800397 |
| 0.271972177 | 0.563800397 |
| 0.271963317 | 0.563800397 |
| 0.271958683 | 0.563800397 |
| 0.271927577 | 0.563800397 |
| 0.271923412 | 0.563800397 |
| 0.271916715 | 0.563800397 |
| 0.27186977 | 0.563800397 |
| 0.271844693 | 0.563800397 |
| 0.271830098 | 0.563800397 |
| 0.271826562 | 0.563800397 |
| 0.271764561 | 0.563783027 |
| 0.271730678 | 0.563737292 |
| 0.271721094 | 0.563737292 |
| 0.271694461 | 0.563735823 |
| 0.271692196 | 0.563735823 |
| 0.271651906 | 0.563721202 |
| 0.271643808 | 0.563721202 |
| 0.27162129 | 0.563721202 |
| 0.271530589 | 0.563641361 |
| 0.271497704 | 0.563597662 |
| 0.271470779 | 0.563566332 |
| 0.271430759 | 0.563532377 |
| 0.271415504 | 0.563532377 |

| | |
|---|---|
| 0.271415266 | 0.563532377 |
| 0.271385605 | 0.563532377 |
| 0.271359511 | 0.563532377 |
| 0.271320963 | 0.563525512 |
| 0.271307812 | 0.563522771 |
| 0.271282385 | 0.563494534 |
| 0.27128082 | 0.563494534 |
| 0.271247106 | 0.563470403 |
| 0.271205911 | 0.563415462 |
| 0.271189958 | 0.563415462 |
| 0.271187379 | 0.563415462 |
| 0.271171629 | 0.563415462 |
| 0.271157816 | 0.563415462 |
| 0.271133838 | 0.563415462 |
| 0.271097946 | 0.563406287 |
| 0.270940744 | 0.563118404 |
| 0.270929319 | 0.563118404 |
| 0.270928755 | 0.563118404 |
| 0.270879595 | 0.563111422 |
| 0.270864213 | 0.563111422 |
| 0.270810844 | 0.563055313 |
| 0.270797032 | 0.563052084 |
| 0.270732926 | 0.562967039 |
| 0.270714754 | 0.562953832 |
| 0.270709678 | 0.562953832 |
| 0.270705965 | 0.562953832 |
| 0.270694586 | 0.562953832 |
| 0.270621807 | 0.56288343 |
| 0.270593006 | 0.562872689 |
| 0.270589732 | 0.562872689 |
| 0.27058936 | 0.562872689 |
| 0.270586498 | 0.562872689 |
| 0.270583026 | 0.562872689 |
| 0.270582807 | 0.562872689 |
| 0.270580174 | 0.562872689 |
| 0.270571893 | 0.562872689 |
| 0.270564185 | 0.562872689 |
| 0.270525481 | 0.562872689 |
| 0.27045871 | 0.562872689 |
| 0.27043664 | 0.562872689 |
| 0.27041472 | 0.562870586 |
| 0.270322902 | 0.562728653 |
| 0.270298558 | 0.562702571 |
| 0.27029642 | 0.562702571 |
| 0.270233364 | 0.562640632 |

| | |
|---|---|
| 0.270170371 | 0.562545254 |
| 0.270088368 | 0.562437091 |
| 0.270046048 | 0.562384051 |
| 0.269987126 | 0.56232461 |
| 0.269871646 | 0.562237795 |
| 0.269829383 | 0.562192747 |
| 0.269696551 | 0.561952435 |
| 0.26967892 | 0.561952435 |
| 0.269614487 | 0.561892494 |
| 0.269540739 | 0.56181255 |
| 0.269519767 | 0.56181255 |
| 0.269459656 | 0.561717295 |
| 0.269455203 | 0.561717295 |
| 0.269450648 | 0.561717295 |
| 0.269402046 | 0.561709004 |
| 0.269345646 | 0.561651712 |
| 0.269251436 | 0.561479843 |
| 0.269218801 | 0.56143637 |
| 0.269178839 | 0.561377615 |
| 0.269143036 | 0.561327525 |
| 0.269125508 | 0.56131555 |
| 0.269117489 | 0.56131555 |
| 0.269019207 | 0.561167562 |
| 0.269018518 | 0.561167562 |
| 0.268912042 | 0.561017733 |
| 0.268885914 | 0.560987797 |
| 0.268876456 | 0.560987797 |
| 0.268857106 | 0.560987797 |
| 0.268775275 | 0.56092897 |
| 0.268761947 | 0.56092897 |
| 0.268731506 | 0.56091136 |
| 0.268698876 | 0.560867833 |
| 0.268669407 | 0.560855484 |
| 0.268645178 | 0.560829486 |
| 0.268629568 | 0.56082148 |
| 0.268615916 | 0.560817562 |
| 0.268594736 | 0.560797927 |
| 0.268557802 | 0.560745394 |
| 0.26844398 | 0.560536003 |
| 0.268395056 | 0.56052845 |
| 0.268337898 | 0.560436474 |
| 0.268329793 | 0.560436474 |
| 0.268306999 | 0.560436474 |
| 0.268301566 | 0.560436474 |
| 0.268287167 | 0.560436474 |

| | |
|---|---|
| 0.268149172 | 0.560211459 |
| 0.268139953 | 0.560211459 |
| 0.268110258 | 0.560179306 |
| 0.268037265 | 0.560075937 |
| 0.268033405 | 0.560075937 |
| 0.267891319 | 0.559942942 |
| 0.267872816 | 0.55992884 |
| 0.267853307 | 0.559912634 |
| 0.267851219 | 0.559912634 |
| 0.267836545 | 0.559912634 |
| 0.267823871 | 0.559912634 |
| 0.267805246 | 0.559912634 |
| 0.26774628 | 0.559836336 |
| 0.267683786 | 0.55973024 |
| 0.267630497 | 0.559692531 |
| 0.267620804 | 0.559692531 |
| 0.267569248 | 0.559638161 |
| 0.267457553 | 0.559502828 |
| 0.26744744 | 0.559502828 |
| 0.26738247 | 0.559419474 |
| 0.267332126 | 0.559338715 |
| 0.267328465 | 0.559338715 |
| 0.26732554 | 0.559338715 |
| 0.267273898 | 0.559338715 |
| 0.267247503 | 0.559322409 |
| 0.267207138 | 0.559282589 |
| 0.267156021 | 0.559250059 |
| 0.267142629 | 0.559250059 |
| 0.267136224 | 0.559250059 |
| 0.267134601 | 0.559250059 |
| 0.267126815 | 0.559250059 |
| 0.267126588 | 0.559250059 |
| 0.266930156 | 0.55901339 |
| 0.266903482 | 0.558982109 |
| 0.266822214 | 0.558836482 |
| 0.266804834 | 0.558824655 |
| 0.266789679 | 0.558817488 |
| 0.266680756 | 0.558613906 |
| 0.266559789 | 0.558385076 |
| 0.266548684 | 0.558385076 |
| 0.266535881 | 0.558384112 |
| 0.266500261 | 0.558383174 |
| 0.266404753 | 0.558235587 |
| 0.266378355 | 0.558225984 |
| 0.266203703 | 0.557982738 |

| | |
|---|---|
| 0.266070893 | 0.557827109 |
| 0.266028194 | 0.557762141 |
| 0.265828218 | 0.557474724 |
| 0.265809991 | 0.557474724 |
| 0.265747653 | 0.55740398 |
| 0.265721048 | 0.557403042 |
| 0.265690919 | 0.557398529 |
| 0.2656819 | 0.557398529 |
| 0.265583015 | 0.557245793 |
| 0.265521188 | 0.557162475 |
| 0.265487521 | 0.557119059 |
| 0.265478568 | 0.557119059 |
| 0.265476881 | 0.557119059 |
| 0.265305224 | 0.556858598 |
| 0.265245007 | 0.556757335 |
| 0.265221487 | 0.556732503 |
| 0.265196935 | 0.556732503 |
| 0.265135273 | 0.556674203 |
| 0.265098551 | 0.556621641 |
| 0.265041272 | 0.556525908 |
| 0.265028812 | 0.556524282 |
| 0.26501385 | 0.556524282 |
| 0.264968385 | 0.556470997 |
| 0.26495784 | 0.556470997 |
| 0.2649467 | 0.556470997 |
| 0.264680761 | 0.555965008 |
| 0.264636619 | 0.555896805 |
| 0.264620779 | 0.55588805 |
| 0.264608571 | 0.555886923 |
| 0.26460789 | 0.555886923 |
| 0.264593959 | 0.555886923 |
| 0.264585318 | 0.555886923 |
| 0.26453608 | 0.555886923 |
| 0.264527702 | 0.555886923 |
| 0.264526239 | 0.555886923 |
| 0.264465479 | 0.555886923 |
| 0.264460668 | 0.555886923 |
| 0.264421353 | 0.55588058 |
| 0.264416205 | 0.55588058 |
| 0.264410846 | 0.55588058 |
| 0.264402965 | 0.55588058 |
| 0.264344921 | 0.555847926 |
| 0.264295945 | 0.55579402 |
| 0.264285317 | 0.55579402 |
| 0.264244249 | 0.555758925 |

| | |
|---|---|
| 0.264198909 | 0.555704709 |
| 0.264126232 | 0.555608854 |
| 0.264094865 | 0.555576043 |
| 0.264072987 | 0.555576043 |
| 0.264017667 | 0.555552241 |
| 0.263993981 | 0.555526945 |
| 0.263943994 | 0.555455537 |
| 0.263910619 | 0.555425145 |
| 0.263858508 | 0.555340013 |
| 0.263844722 | 0.555335538 |
| 0.263732349 | 0.555123552 |
| 0.263627483 | 0.554951872 |
| 0.263587117 | 0.554891425 |
| 0.263573482 | 0.554887249 |
| 0.263567238 | 0.554887249 |
| 0.263499961 | 0.554781514 |
| 0.263488794 | 0.554781514 |
| 0.263476828 | 0.554781514 |
| 0.263451913 | 0.554753923 |
| 0.26341713 | 0.554705207 |
| 0.26340962 | 0.554705207 |
| 0.263316482 | 0.554591345 |
| 0.263313974 | 0.554591345 |
| 0.263299024 | 0.554591345 |
| 0.263238396 | 0.554524971 |
| 0.263231542 | 0.554524971 |
| 0.263186764 | 0.554489783 |
| 0.263148441 | 0.554433569 |
| 0.263041196 | 0.55427988 |
| 0.262828784 | 0.553980637 |
| 0.262778331 | 0.553898808 |
| 0.262706872 | 0.553772691 |
| 0.262695758 | 0.553772691 |
| 0.26266875 | 0.553741348 |
| 0.262646216 | 0.553718355 |
| 0.262645038 | 0.553718355 |
| 0.262642002 | 0.553718355 |
| 0.262538151 | 0.553609858 |
| 0.262511166 | 0.553580664 |
| 0.262508905 | 0.553580664 |
| 0.262415009 | 0.553500427 |
| 0.262392046 | 0.553476503 |
| 0.262376907 | 0.553469083 |
| 0.26229581 | 0.553396055 |
| 0.262249586 | 0.553342108 |

| | |
|---|---|
| 0.262212507 | 0.553293832 |
| 0.262202228 | 0.553293832 |
| 0.262197298 | 0.553293832 |
| 0.262191199 | 0.553293832 |
| 0.262074557 | 0.553149786 |
| 0.262073249 | 0.553149786 |
| 0.262067229 | 0.553149786 |
| 0.261983513 | 0.553031149 |
| 0.261975956 | 0.553031149 |
| 0.261953496 | 0.553016806 |
| 0.261930636 | 0.553000138 |
| 0.26179336 | 0.552776748 |
| 0.261790421 | 0.552776748 |
| 0.261730825 | 0.552718218 |
| 0.261700368 | 0.552711835 |
| 0.261696765 | 0.552711835 |
| 0.261670533 | 0.552711835 |
| 0.261627084 | 0.552646167 |
| 0.261493678 | 0.552521112 |
| 0.261424715 | 0.552414699 |
| 0.26142378 | 0.552414699 |
| 0.261419607 | 0.552414699 |
| 0.261379889 | 0.552414699 |
| 0.261346314 | 0.552371567 |
| 0.261288765 | 0.552298953 |
| 0.261285434 | 0.552298953 |
| 0.261236278 | 0.552298953 |
| 0.261221382 | 0.552298953 |
| 0.261155483 | 0.552188776 |
| 0.261083192 | 0.552060433 |
| 0.261056738 | 0.552060433 |
| 0.260946136 | 0.551917648 |
| 0.260889949 | 0.551872334 |
| 0.260853367 | 0.551819459 |
| 0.260836139 | 0.551819459 |
| 0.260818194 | 0.551818582 |
| 0.260813807 | 0.551818582 |
| 0.260799023 | 0.551818582 |
| 0.260743432 | 0.551782967 |
| 0.260701246 | 0.551759888 |
| 0.260650912 | 0.551709746 |
| 0.260552184 | 0.551549796 |
| 0.260551026 | 0.551549796 |
| 0.260520074 | 0.55153085 |
| 0.26050098 | 0.551514944 |

| | |
|---|---|
| 0.260457501 | 0.551447405 |
| 0.26040929 | 0.551369842 |
| 0.260374971 | 0.5513462 |
| 0.26036444 | 0.5513462 |
| 0.260275097 | 0.551257261 |
| 0.260241873 | 0.551211407 |
| 0.260239797 | 0.551211407 |
| 0.260152294 | 0.551095196 |
| 0.260141265 | 0.551095196 |
| 0.260135622 | 0.551095196 |
| 0.260095703 | 0.551073359 |
| 0.26005318 | 0.551056808 |
| 0.260015542 | 0.551024889 |
| 0.259931845 | 0.550873225 |
| 0.259921166 | 0.550873225 |
| 0.259893603 | 0.550841202 |
| 0.2598919 | 0.550841202 |
| 0.25979531 | 0.55074367 |
| 0.259780535 | 0.55074367 |
| 0.259776374 | 0.55074367 |
| 0.259774062 | 0.55074367 |
| 0.259754144 | 0.55074367 |
| 0.259742655 | 0.55074367 |
| 0.259736384 | 0.55074367 |
| 0.259725752 | 0.55074367 |
| 0.259652489 | 0.550723765 |
| 0.259592836 | 0.55066934 |
| 0.259573927 | 0.550664204 |
| 0.2595616 | 0.550664204 |
| 0.259524488 | 0.550646981 |
| 0.25948222 | 0.550606358 |
| 0.259473505 | 0.550606358 |
| 0.259415236 | 0.550544667 |
| 0.259401812 | 0.550544667 |
| 0.259371821 | 0.550544667 |
| 0.259355921 | 0.550544667 |
| 0.259353766 | 0.550544667 |
| 0.259329797 | 0.550544667 |
| 0.25929181 | 0.550496637 |
| 0.259229142 | 0.550405275 |
| 0.259216729 | 0.550405275 |
| 0.259203394 | 0.550405275 |
| 0.259178363 | 0.550378451 |
| 0.259148329 | 0.55036375 |
| 0.259133245 | 0.550356258 |

| | |
|---|---|
| 0.259058826 | 0.550247278 |
| 0.258944217 | 0.550126515 |
| 0.25889318 | 0.550042624 |
| 0.258854994 | 0.549986027 |
| 0.258830713 | 0.549958972 |
| 0.258786134 | 0.549888784 |
| 0.258784791 | 0.549888784 |
| 0.258737108 | 0.549833671 |
| 0.258723463 | 0.549829207 |
| 0.258675886 | 0.549777162 |
| 0.258662274 | 0.549772764 |
| 0.258658046 | 0.549772764 |
| 0.258651625 | 0.549772764 |
| 0.25859444 | 0.549702181 |
| 0.258570753 | 0.549677541 |
| 0.258570474 | 0.549677541 |
| 0.258547961 | 0.549677541 |
| 0.258490162 | 0.549627703 |
| 0.258476549 | 0.549623296 |
| 0.258445664 | 0.549582159 |
| 0.258411978 | 0.549559601 |
| 0.258393597 | 0.549559601 |
| 0.258387893 | 0.549559601 |
| 0.258386296 | 0.549559601 |
| 0.258380194 | 0.549559601 |
| 0.258289917 | 0.549496309 |
| 0.258258881 | 0.549454825 |
| 0.258250434 | 0.549454825 |
| 0.258232572 | 0.549447942 |
| 0.258173075 | 0.549345889 |
| 0.258157494 | 0.549337279 |
| 0.258145126 | 0.549335505 |
| 0.25813113 | 0.549335505 |
| 0.258114675 | 0.549335505 |
| 0.258107513 | 0.549335505 |
| 0.25801879 | 0.549238442 |
| 0.257968865 | 0.549156713 |
| 0.257968685 | 0.549156713 |
| 0.257952302 | 0.549156713 |
| 0.257901358 | 0.549086634 |
| 0.25789744 | 0.549086634 |
| 0.257885643 | 0.549086634 |
| 0.257847878 | 0.549086634 |
| 0.257823233 | 0.549067569 |
| 0.257755438 | 0.548972286 |

| | |
|---|---|
| 0.257714669 | 0.548910001 |
| 0.25769749 | 0.548897959 |
| 0.25764755 | 0.548816131 |
| 0.257598281 | 0.548760268 |
| 0.257598281 | 0.548760268 |
| 0.257598281 | 0.548760268 |
| 0.257549717 | 0.548760268 |
| 0.257536849 | 0.548760268 |
| 0.257493085 | 0.548720805 |
| 0.257468815 | 0.548715192 |
| 0.257403665 | 0.548615608 |
| 0.257366388 | 0.548585255 |
| 0.257333948 | 0.548540659 |
| 0.25726574 | 0.548446104 |
| 0.257260724 | 0.548446104 |
| 0.257256007 | 0.548446104 |
| 0.25700944 | 0.548094238 |
| 0.256974073 | 0.548067895 |
| 0.256953787 | 0.548049172 |
| 0.256890404 | 0.547938523 |
| 0.256845602 | 0.547892467 |
| 0.256837933 | 0.547892467 |
| 0.256826115 | 0.547892467 |
| 0.25680609 | 0.547892467 |
| 0.25680244 | 0.547892467 |
| 0.25680029 | 0.547892467 |
| 0.256707661 | 0.547867708 |
| 0.256696782 | 0.547867708 |
| 0.256660107 | 0.547815313 |
| 0.256566991 | 0.547665654 |
| 0.256516149 | 0.54763076 |
| 0.256515712 | 0.54763076 |
| 0.256461494 | 0.547587714 |
| 0.256446296 | 0.547579814 |
| 0.256377108 | 0.547456624 |
| 0.256374162 | 0.547456624 |
| 0.256339764 | 0.547425968 |
| 0.256251104 | 0.54728571 |
| 0.256201403 | 0.5472041 |
| 0.256133492 | 0.547157202 |
| 0.256092479 | 0.547094129 |
| 0.256083078 | 0.547094129 |
| 0.256069898 | 0.547094129 |
| 0.256008257 | 0.547012349 |
| 0.255962836 | 0.546974153 |

| | |
|---|---|
| 0.255952443 | 0.546974153 |
| 0.255945561 | 0.546974153 |
| 0.255939384 | 0.546974153 |
| 0.255929827 | 0.546974153 |
| 0.255905294 | 0.546974153 |
| 0.255757807 | 0.54674703 |
| 0.255754562 | 0.54674703 |
| 0.255742442 | 0.54674703 |
| 0.25572393 | 0.54674703 |
| 0.255707914 | 0.546738533 |
| 0.255690793 | 0.54672647 |
| 0.255677409 | 0.546722397 |
| 0.255652949 | 0.546694639 |
| 0.255568048 | 0.546562165 |
| 0.255500479 | 0.546521915 |
| 0.255479033 | 0.546519039 |
| 0.255466131 | 0.546515988 |
| 0.255414213 | 0.546429463 |
| 0.255237648 | 0.546174392 |
| 0.255235092 | 0.546174392 |
| 0.255197458 | 0.546162009 |
| 0.255176039 | 0.546140709 |
| 0.255146304 | 0.54610161 |
| 0.2550177 | 0.545860562 |
| 0.25501678 | 0.545860562 |
| 0.254896407 | 0.545738421 |
| 0.254857704 | 0.545729156 |
| 0.254760269 | 0.545569574 |
| 0.254668138 | 0.545421324 |
| 0.254610782 | 0.545323007 |
| 0.254552578 | 0.545222865 |
| 0.254548638 | 0.545222865 |
| 0.254508072 | 0.545205496 |
| 0.254506324 | 0.545205496 |
| 0.254486262 | 0.545205496 |
| 0.254473663 | 0.545205496 |
| 0.254395899 | 0.545092274 |
| 0.254378516 | 0.545092274 |
| 0.254368851 | 0.545092274 |
| 0.254342333 | 0.545069334 |
| 0.254319836 | 0.545069334 |
| 0.254292128 | 0.545069334 |
| 0.254166064 | 0.544885088 |
| 0.254068976 | 0.544799563 |
| 0.254065623 | 0.544799563 |

| | |
|---|---|
| 0.254049813 | 0.544799563 |
| 0.254048237 | 0.544799563 |
| 0.254046556 | 0.544799563 |
| 0.253935344 | 0.544709211 |
| 0.253886644 | 0.544629277 |
| 0.253780635 | 0.54447544 |
| 0.253774039 | 0.54447544 |
| 0.253764071 | 0.54447544 |
| 0.253718849 | 0.544433635 |
| 0.253660983 | 0.544341338 |
| 0.25362687 | 0.544317184 |
| 0.253568807 | 0.544217095 |
| 0.253526429 | 0.544175188 |
| 0.253506363 | 0.544156642 |
| 0.253469231 | 0.544101459 |
| 0.253433611 | 0.544049517 |
| 0.253409054 | 0.544021322 |
| 0.253313969 | 0.543890741 |
| 0.25325741 | 0.543867376 |
| 0.253217866 | 0.543806975 |
| 0.253174736 | 0.543738869 |
| 0.253170884 | 0.543738869 |
| 0.253138187 | 0.54373393 |
| 0.253127093 | 0.54373393 |
| 0.253100472 | 0.543701961 |
| 0.252977575 | 0.54353602 |
| 0.25296113 | 0.543525208 |
| 0.252946414 | 0.543518108 |
| 0.252838147 | 0.543334495 |
| 0.252826574 | 0.543334141 |
| 0.252769909 | 0.543236879 |
| 0.252745102 | 0.543208077 |
| 0.252664725 | 0.543084342 |
| 0.252611465 | 0.543043344 |
| 0.252557886 | 0.542977224 |
| 0.252513563 | 0.542930949 |
| 0.252496045 | 0.542917794 |
| 0.252451871 | 0.542847318 |
| 0.252397867 | 0.542755698 |
| 0.252359389 | 0.542721964 |
| 0.252316615 | 0.542654478 |
| 0.252202031 | 0.542506036 |
| 0.252184676 | 0.542493206 |
| 0.252136217 | 0.542413461 |
| 0.252119812 | 0.542402668 |

| | |
|---|---|
| 0.252099263 | 0.542382962 |
| 0.252063382 | 0.542330263 |
| 0.252041356 | 0.542330263 |
| 0.251972957 | 0.54223369 |
| 0.251970628 | 0.54223369 |
| 0.251923929 | 0.54217718 |
| 0.251919364 | 0.54217718 |
| 0.251841413 | 0.542073077 |
| 0.251760009 | 0.541946845 |
| 0.251756504 | 0.541946845 |
| 0.251737236 | 0.541946845 |
| 0.251663937 | 0.541902974 |
| 0.251624094 | 0.541850194 |
| 0.251601788 | 0.541826657 |
| 0.251510683 | 0.541673502 |
| 0.251502404 | 0.541673502 |
| 0.251423669 | 0.541639004 |
| 0.251328061 | 0.541506517 |
| 0.251313632 | 0.541499926 |
| 0.251305048 | 0.541499926 |
| 0.251225744 | 0.541384034 |
| 0.251223342 | 0.541384034 |
| 0.251206899 | 0.541384034 |
| 0.251188202 | 0.541381502 |
| 0.251166771 | 0.541381502 |
| 0.25112843 | 0.541358062 |
| 0.251110695 | 0.541358062 |
| 0.251087789 | 0.541356185 |
| 0.250972596 | 0.541239628 |
| 0.250941391 | 0.541239628 |
| 0.250931323 | 0.541239628 |
| 0.250836602 | 0.541133073 |
| 0.250799593 | 0.541077741 |
| 0.250781115 | 0.541062385 |
| 0.25075741 | 0.541060261 |
| 0.250742285 | 0.541052139 |
| 0.250737735 | 0.541052139 |
| 0.25071681 | 0.541052139 |
| 0.250712538 | 0.541052139 |
| 0.250590981 | 0.540872684 |
| 0.250570013 | 0.540851939 |
| 0.250541292 | 0.540814455 |
| 0.250473084 | 0.540691727 |
| 0.250471163 | 0.540691727 |
| 0.250457305 | 0.540691727 |

| | |
|---|---|
| 0.250445127 | 0.540691727 |
| 0.250384842 | 0.54059925 |
| 0.250381138 | 0.54059925 |
| 0.250322765 | 0.540514228 |
| 0.250314613 | 0.540514228 |
| 0.250282332 | 0.540500438 |
| 0.250265552 | 0.540488707 |
| 0.250209252 | 0.540398305 |
| 0.250182506 | 0.540382866 |
| 0.250103073 | 0.540260299 |
| 0.250033253 | 0.540133975 |
| 0.250013354 | 0.540115486 |
| 0.250013065 | 0.540115486 |
| 0.250000307 | 0.540115486 |
| 0.249974648 | 0.540115486 |
| 0.249962872 | 0.540115486 |
| 0.249837299 | 0.53998008 |
| 0.249821405 | 0.53998008 |
| 0.249781645 | 0.539933299 |
| 0.249720874 | 0.539847787 |
| 0.249648885 | 0.539765264 |
| 0.249646972 | 0.539765264 |
| 0.249516689 | 0.539579719 |
| 0.249476455 | 0.539518462 |
| 0.249474656 | 0.539518462 |
| 0.249456676 | 0.539518462 |
| 0.249342972 | 0.539352245 |
| 0.249302385 | 0.539313441 |
| 0.249300538 | 0.539313441 |
| 0.249291074 | 0.539313441 |
| 0.249264275 | 0.539313441 |
| 0.249264275 | 0.539313441 |
| 0.249264275 | 0.539313441 |
| 0.249264275 | 0.539313441 |
| 0.249264275 | 0.539313441 |
| 0.249188428 | 0.539311881 |
| 0.249158734 | 0.539272122 |
| 0.249097483 | 0.539196568 |
| 0.249082272 | 0.539196568 |
| 0.249034856 | 0.539176749 |
| 0.2490184 | 0.539176749 |
| 0.248988345 | 0.539176749 |
| 0.248974834 | 0.539176749 |
| 0.248960046 | 0.539176749 |
| 0.248893001 | 0.539137988 |

| | |
|---|---|
| 0.248868765 | 0.539110009 |
| 0.248800965 | 0.539059654 |
| 0.248789121 | 0.539059654 |
| 0.248651569 | 0.538884613 |
| 0.248646028 | 0.538884613 |
| 0.24864158 | 0.538884613 |
| 0.248607179 | 0.538884613 |
| 0.248561317 | 0.538811609 |
| 0.24852114 | 0.538749037 |
| 0.248519123 | 0.538749037 |
| 0.248483251 | 0.538749037 |
| 0.248478269 | 0.538749037 |
| 0.248292571 | 0.538523169 |
| 0.248239042 | 0.538438859 |
| 0.248217093 | 0.538433024 |
| 0.248161038 | 0.538335949 |
| 0.24804521 | 0.53815822 |
| 0.248017369 | 0.538122328 |
| 0.24798314 | 0.538072574 |
| 0.247921647 | 0.53801268 |
| 0.24791423 | 0.53801268 |
| 0.247907479 | 0.53801268 |
| 0.247864964 | 0.538012244 |
| 0.247801947 | 0.537899973 |
| 0.247780874 | 0.537878746 |
| 0.247772166 | 0.537878746 |
| 0.247770309 | 0.537878746 |
| 0.247748163 | 0.537878746 |
| 0.247687577 | 0.537823299 |
| 0.247653396 | 0.537798116 |
| 0.24764548 | 0.537798116 |
| 0.247637427 | 0.537798116 |
| 0.247620309 | 0.537798116 |
| 0.247509609 | 0.537608434 |
| 0.247499672 | 0.537608434 |
| 0.247498284 | 0.537608434 |
| 0.247490947 | 0.537608434 |
| 0.24748649 | 0.537608434 |
| 0.247463714 | 0.537608434 |
| 0.247455465 | 0.537608434 |
| 0.24733753 | 0.537528841 |
| 0.247320329 | 0.537515985 |
| 0.247299901 | 0.537496115 |
| 0.247287668 | 0.537494057 |
| 0.24727183 | 0.537484161 |

| | |
|---|---|
| 0.247269612 | 0.537484161 |
| 0.247248854 | 0.537483278 |
| 0.247224855 | 0.53745564 |
| 0.247201728 | 0.537429895 |
| 0.247090975 | 0.537238158 |
| 0.247072309 | 0.537222099 |
| 0.247069218 | 0.537222099 |
| 0.247051152 | 0.537222099 |
| 0.246967376 | 0.537167404 |
| 0.246936514 | 0.537147537 |
| 0.246871896 | 0.537105102 |
| 0.246856315 | 0.537095738 |
| 0.246848644 | 0.537095738 |
| 0.246819597 | 0.537075598 |
| 0.246790103 | 0.537074354 |
| 0.246783012 | 0.537074354 |
| 0.246756915 | 0.537074354 |
| 0.246744526 | 0.537073331 |
| 0.246717781 | 0.53703966 |
| 0.246712977 | 0.53703966 |
| 0.246704597 | 0.53703966 |
| 0.246682682 | 0.53703966 |
| 0.246651429 | 0.537017948 |
| 0.246598037 | 0.536926246 |
| 0.246549765 | 0.536894776 |
| 0.246545513 | 0.536894776 |
| 0.246537059 | 0.536894776 |
| 0.246529926 | 0.536894776 |
| 0.246482961 | 0.536847486 |
| 0.246455779 | 0.536812831 |
| 0.246430748 | 0.53680741 |
| 0.246390834 | 0.536745014 |
| 0.246388187 | 0.536745014 |
| 0.246353216 | 0.536712163 |
| 0.246343388 | 0.536712163 |
| 0.24632781 | 0.536710565 |
| 0.246326423 | 0.536710565 |
| 0.24631989 | 0.536710565 |
| 0.246299191 | 0.536710565 |
| 0.246220741 | 0.536619926 |
| 0.246190698 | 0.536619926 |
| 0.246144736 | 0.536601608 |
| 0.2461341 | 0.536601608 |
| 0.246004711 | 0.536370005 |
| 0.245983934 | 0.536370005 |

| | |
|---|---|
| 0.245978893 | 0.536370005 |
| 0.245972217 | 0.536370005 |
| 0.245919254 | 0.536330996 |
| 0.245882428 | 0.536275238 |
| 0.2458487 | 0.536226233 |
| 0.245794351 | 0.536200482 |
| 0.245782455 | 0.536200482 |
| 0.245781605 | 0.536200482 |
| 0.245773981 | 0.536200482 |
| 0.245739283 | 0.536184022 |
| 0.24573334 | 0.536184022 |
| 0.245681449 | 0.536106954 |
| 0.245639216 | 0.536039356 |
| 0.245624763 | 0.536032376 |
| 0.245595712 | 0.535993539 |
| 0.245593392 | 0.535993539 |
| 0.245562962 | 0.535971182 |
| 0.245410633 | 0.535712358 |
| 0.245404655 | 0.535712358 |
| 0.245394061 | 0.535712358 |
| 0.24538775 | 0.535712358 |
| 0.245326938 | 0.53570153 |
| 0.245312127 | 0.535693749 |
| 0.245303421 | 0.535693749 |
| 0.245247338 | 0.535625945 |
| 0.245196768 | 0.535540055 |
| 0.245176388 | 0.535520101 |
| 0.245120042 | 0.535421586 |
| 0.245010292 | 0.535255501 |
| 0.244944595 | 0.535210176 |
| 0.244799855 | 0.534967545 |
| 0.244758745 | 0.534902252 |
| 0.24469821 | 0.534794495 |
| 0.244682762 | 0.534785273 |
| 0.244585807 | 0.534597899 |
| 0.244585799 | 0.534597899 |
| 0.244571834 | 0.534597899 |
| 0.244530594 | 0.534550814 |
| 0.244530594 | 0.534550814 |
| 0.244530594 | 0.534550814 |
| 0.244530594 | 0.534550814 |
| 0.244506182 | 0.534550814 |
| 0.244486282 | 0.534550814 |
| 0.244461906 | 0.534547873 |
| 0.244443557 | 0.534532291 |

| | |
|---|---|
| 0.244409614 | 0.534482607 |
| 0.244403172 | 0.534482607 |
| 0.244341602 | 0.534407489 |
| 0.244331191 | 0.534407489 |
| 0.244217722 | 0.534259219 |
| 0.244113929 | 0.534056689 |
| 0.244100011 | 0.534050771 |
| 0.244098324 | 0.534050771 |
| 0.244097348 | 0.534050771 |
| 0.244081749 | 0.534050771 |
| 0.244011869 | 0.534050771 |
| 0.243998199 | 0.534050771 |
| 0.243998163 | 0.534050771 |
| 0.243974186 | 0.534050771 |
| 0.243927834 | 0.53401751 |
| 0.243910492 | 0.53400409 |
| 0.24383868 | 0.533895955 |
| 0.243713505 | 0.533695485 |
| 0.243700258 | 0.533691014 |
| 0.243549559 | 0.533385516 |
| 0.243414064 | 0.533113291 |
| 0.243379116 | 0.533069835 |
| 0.243368071 | 0.533069835 |
| 0.243306646 | 0.532976066 |
| 0.243300262 | 0.532976066 |
| 0.243295431 | 0.532976066 |
| 0.243219016 | 0.53290666 |
| 0.243213698 | 0.53290666 |
| 0.243191701 | 0.532895842 |
| 0.243178593 | 0.532891637 |
| 0.243172397 | 0.532891637 |
| 0.243156195 | 0.532891593 |
| 0.243099659 | 0.532865874 |
| 0.243085367 | 0.532865874 |
| 0.243023242 | 0.532796352 |
| 0.243018346 | 0.532796352 |
| 0.24297224 | 0.532733583 |
| 0.242924446 | 0.532677836 |
| 0.242819355 | 0.532496427 |
| 0.242815877 | 0.532496427 |
| 0.242805637 | 0.532496427 |
| 0.242785335 | 0.532495379 |
| 0.242692715 | 0.532352686 |
| 0.242677824 | 0.532352686 |
| 0.242642434 | 0.532352686 |

| | |
|---|---|
| 0.242635741 | 0.532352686 |
| 0.242553288 | 0.532352686 |
| 0.242533773 | 0.532352686 |
| 0.24250838 | 0.532352686 |
| 0.242413417 | 0.532194331 |
| 0.242324139 | 0.532071908 |
| 0.242318787 | 0.532071908 |
| 0.242230091 | 0.531914451 |
| 0.242218847 | 0.531914286 |
| 0.242194497 | 0.531885338 |
| 0.242182052 | 0.531882533 |
| 0.24191499 | 0.531437332 |
| 0.241879107 | 0.531399864 |
| 0.241856982 | 0.531389143 |
| 0.241814665 | 0.531320679 |
| 0.241764281 | 0.531258993 |
| 0.241732608 | 0.531238415 |
| 0.241716614 | 0.531238415 |
| 0.241676744 | 0.531189177 |
| 0.241644073 | 0.531166395 |
| 0.241606603 | 0.531157574 |
| 0.241590205 | 0.531157574 |
| 0.241560267 | 0.531157574 |
| 0.241546717 | 0.531157574 |
| 0.241492456 | 0.531078229 |
| 0.241394694 | 0.530912264 |
| 0.241392612 | 0.530912264 |
| 0.241361969 | 0.530889323 |
| 0.241286712 | 0.530748301 |
| 0.241273475 | 0.530748301 |
| 0.241223258 | 0.530706767 |
| 0.241210746 | 0.530706767 |
| 0.241207065 | 0.530706767 |
| 0.241120818 | 0.530579432 |
| 0.241099846 | 0.530557796 |
| 0.241054772 | 0.53050763 |
| 0.241050657 | 0.53050763 |
| 0.240987287 | 0.530432643 |
| 0.240983338 | 0.530432643 |
| 0.24090066 | 0.530340008 |
| 0.240795386 | 0.530157259 |
| 0.240772803 | 0.530156555 |
| 0.240734877 | 0.530097553 |
| 0.240729491 | 0.530097553 |
| 0.240685654 | 0.530062686 |

| | |
|---|---|
| 0.240677785 | 0.530062686 |
| 0.240664077 | 0.530062686 |
| 0.240607098 | 0.529963198 |
| 0.240582847 | 0.529934292 |
| 0.240547932 | 0.529881891 |
| 0.24053983 | 0.529881891 |
| 0.240535836 | 0.529881891 |
| 0.240535022 | 0.529881891 |
| 0.240534471 | 0.529881891 |
| 0.240445853 | 0.529828566 |
| 0.240415207 | 0.529785548 |
| 0.240355405 | 0.529727296 |
| 0.240329518 | 0.529719269 |
| 0.240297427 | 0.529697563 |
| 0.240165293 | 0.529479814 |
| 0.240076237 | 0.529381497 |
| 0.240003505 | 0.529260516 |
| 0.239890104 | 0.529093544 |
| 0.239841354 | 0.529010521 |
| 0.239830128 | 0.52901026 |
| 0.239794328 | 0.52895579 |
| 0.239759049 | 0.528902465 |
| 0.23975153 | 0.528902465 |
| 0.239737317 | 0.528902465 |
| 0.239694747 | 0.528858598 |
| 0.239671243 | 0.528831239 |
| 0.239629763 | 0.528788709 |
| 0.239623857 | 0.528788709 |
| 0.239591185 | 0.528777079 |
| 0.239550912 | 0.528724236 |
| 0.239514175 | 0.528680615 |
| 0.239440686 | 0.528567395 |
| 0.239438485 | 0.528567395 |
| 0.239434336 | 0.528567395 |
| 0.239430509 | 0.528567395 |
| 0.239428054 | 0.528567395 |
| 0.239348086 | 0.528485456 |
| 0.239290679 | 0.528383198 |
| 0.239270589 | 0.528383198 |
| 0.239261532 | 0.528383198 |
| 0.23924265 | 0.528383198 |
| 0.239233689 | 0.528383198 |
| 0.23922223 | 0.528383198 |
| 0.239188881 | 0.528378889 |
| 0.239149815 | 0.528341602 |

| | |
|---|---|
| 0.23914412 | 0.528341602 |
| 0.239138947 | 0.528341602 |
| 0.239101126 | 0.528332066 |
| 0.239054563 | 0.528253685 |
| 0.239036114 | 0.528237426 |
| 0.238985019 | 0.528149018 |
| 0.238982402 | 0.528149018 |
| 0.238957069 | 0.528136264 |
| 0.238898997 | 0.528081431 |
| 0.238889909 | 0.528081431 |
| 0.238832998 | 0.527984553 |
| 0.23880176 | 0.527939999 |
| 0.238797364 | 0.527939999 |
| 0.238733853 | 0.527912398 |
| 0.238724658 | 0.527912398 |
| 0.238612597 | 0.527776681 |
| 0.238584245 | 0.527776681 |
| 0.238536663 | 0.527696857 |
| 0.23850138 | 0.527667823 |
| 0.23849158 | 0.527667823 |
| 0.238484429 | 0.527667823 |
| 0.23825431 | 0.527194667 |
| 0.238170754 | 0.527058753 |
| 0.238159732 | 0.527058753 |
| 0.238094659 | 0.526963814 |
| 0.23808569 | 0.526963814 |
| 0.238064573 | 0.526956311 |
| 0.238027817 | 0.52691382 |
| 0.237902348 | 0.526685034 |
| 0.237784962 | 0.5264741 |
| 0.237761794 | 0.526447278 |
| 0.237742585 | 0.52642922 |
| 0.237740434 | 0.52642922 |
| 0.237715664 | 0.526418557 |
| 0.237697929 | 0.526403761 |
| 0.237604871 | 0.526222143 |
| 0.237552963 | 0.526131648 |
| 0.237549705 | 0.526131648 |
| 0.237538847 | 0.526131648 |
| 0.237538806 | 0.526131648 |
| 0.237519737 | 0.526131648 |
| 0.237469512 | 0.526093611 |
| 0.237462525 | 0.526093611 |
| 0.237457616 | 0.526093611 |
| 0.237397359 | 0.526041578 |

| | |
|---|---|
| 0.237369937 | 0.526019821 |
| 0.237365268 | 0.526019821 |
| 0.237344515 | 0.526019821 |
| 0.237324519 | 0.526019821 |
| 0.237183973 | 0.52580344 |
| 0.237154385 | 0.525762318 |
| 0.237044501 | 0.525567639 |
| 0.23698261 | 0.525509126 |
| 0.236954161 | 0.525509126 |
| 0.236923103 | 0.525469714 |
| 0.236910634 | 0.525466528 |
| 0.236852879 | 0.525435171 |
| 0.236814471 | 0.525400046 |
| 0.236805742 | 0.525400046 |
| 0.2367062 | 0.525257708 |
| 0.236685636 | 0.525236547 |
| 0.236625717 | 0.525152518 |
| 0.236580511 | 0.525146925 |
| 0.236555638 | 0.525119336 |
| 0.236507605 | 0.525037182 |
| 0.236502672 | 0.525037182 |
| 0.236431623 | 0.52494191 |
| 0.23636614 | 0.524820986 |
| 0.236337193 | 0.524781178 |
| 0.236330733 | 0.524781178 |
| 0.236214812 | 0.524596738 |
| 0.236207964 | 0.524596738 |
| 0.236169534 | 0.524555631 |
| 0.236168043 | 0.524555631 |
| 0.236076574 | 0.524446988 |
| 0.236050877 | 0.524414364 |
| 0.235953689 | 0.524271816 |
| 0.235935165 | 0.524255113 |
| 0.235911696 | 0.524227422 |
| 0.235871262 | 0.524162029 |
| 0.235831574 | 0.524098287 |
| 0.235808434 | 0.524071315 |
| 0.235759362 | 0.523986707 |
| 0.235750506 | 0.523986707 |
| 0.23565506 | 0.523828225 |
| 0.235638935 | 0.523816829 |
| 0.235533705 | 0.523607345 |
| 0.235463544 | 0.523500244 |
| 0.235461905 | 0.523500244 |
| 0.235415862 | 0.523467542 |

| | |
|---|---|
| 0.235310573 | 0.523285343 |
| 0.235236391 | 0.523215036 |
| 0.235189748 | 0.523135725 |
| 0.235146071 | 0.523111865 |
| 0.235093799 | 0.523044442 |
| 0.235063029 | 0.523000414 |
| 0.235040061 | 0.522973745 |
| 0.234967104 | 0.522835838 |
| 0.234901771 | 0.522714883 |
| 0.234836106 | 0.522617598 |
| 0.234807888 | 0.522579219 |
| 0.234774708 | 0.522554211 |
| 0.234741117 | 0.522528285 |
| 0.234683259 | 0.522497175 |
| 0.234681474 | 0.522497175 |
| 0.234678792 | 0.522497175 |
| 0.234641127 | 0.522497175 |
| 0.234638021 | 0.522497175 |
| 0.234625354 | 0.522497175 |
| 0.234624207 | 0.522497175 |
| 0.234615102 | 0.522497175 |
| 0.234575987 | 0.522497175 |
| 0.234495279 | 0.522469436 |
| 0.234469672 | 0.522436822 |
| 0.234438407 | 0.522391598 |
| 0.234418679 | 0.522372078 |
| 0.234417603 | 0.522372078 |
| 0.23440503 | 0.522372078 |
| 0.234373309 | 0.522344295 |
| 0.23429736 | 0.522199464 |
| 0.23428817 | 0.522199464 |
| 0.234270864 | 0.522189285 |
| 0.234256285 | 0.522181226 |
| 0.234221235 | 0.522127534 |
| 0.234165907 | 0.522028631 |
| 0.234152127 | 0.522028631 |
| 0.234146633 | 0.522028631 |
| 0.234140989 | 0.522028631 |
| 0.234097395 | 0.521998064 |
| 0.234030849 | 0.521892179 |
| 0.234025628 | 0.521892179 |
| 0.23400584 | 0.521892179 |
| 0.233956012 | 0.521878271 |
| 0.233867508 | 0.52172972 |
| 0.233826486 | 0.521717673 |

| | |
|---|---|
| 0.23371848 | 0.521519398 |
| 0.233615022 | 0.521361823 |
| 0.233582571 | 0.52131383 |
| 0.233568812 | 0.52130755 |
| 0.233497514 | 0.521172842 |
| 0.23344827 | 0.521111772 |
| 0.233440086 | 0.521111772 |
| 0.233352227 | 0.520970633 |
| 0.233314538 | 0.520910908 |
| 0.233294311 | 0.520890167 |
| 0.233280239 | 0.520883166 |
| 0.233277802 | 0.520883166 |
| 0.233233954 | 0.520864561 |
| 0.233211218 | 0.520851145 |
| 0.233204055 | 0.520851145 |
| 0.233203412 | 0.520851145 |
| 0.233186484 | 0.520851145 |
| 0.233131678 | 0.520806474 |
| 0.233102083 | 0.520794723 |
| 0.233093253 | 0.520794723 |
| 0.233027463 | 0.520713312 |
| 0.232988808 | 0.520696258 |
| 0.232982701 | 0.520696258 |
| 0.232976489 | 0.520696258 |
| 0.232969541 | 0.520696258 |
| 0.232956997 | 0.520696258 |
| 0.232900723 | 0.520696258 |
| 0.23289785 | 0.520696258 |
| 0.232855699 | 0.520696258 |
| 0.232853557 | 0.520696258 |
| 0.23283749 | 0.520696258 |
| 0.232709696 | 0.52056102 |
| 0.232678234 | 0.520530889 |
| 0.232639476 | 0.520477279 |
| 0.232630941 | 0.520477279 |
| 0.232421762 | 0.520063485 |
| 0.232420986 | 0.520063485 |
| 0.232417884 | 0.520063485 |
| 0.232396827 | 0.520063485 |
| 0.232356591 | 0.520015382 |
| 0.232352241 | 0.520015382 |
| 0.232349926 | 0.520015382 |
| 0.232293318 | 0.519971512 |
| 0.232267166 | 0.519945643 |
| 0.232223988 | 0.519889625 |

| | |
|---|---|
| 0.232214369 | 0.519889625 |
| 0.232104905 | 0.519683555 |
| 0.232086977 | 0.519680604 |
| 0.232058955 | 0.519666722 |
| 0.232042482 | 0.519654266 |
| 0.232039814 | 0.519654266 |
| 0.232031029 | 0.519654266 |
| 0.232014348 | 0.519654266 |
| 0.23201098 | 0.519654266 |
| 0.231960876 | 0.519618111 |
| 0.231951992 | 0.519618111 |
| 0.231874579 | 0.519498097 |
| 0.231813267 | 0.519385165 |
| 0.231775085 | 0.519324049 |
| 0.231732088 | 0.519252136 |
| 0.231716928 | 0.51924327 |
| 0.231655487 | 0.519153767 |
| 0.231612499 | 0.519101916 |
| 0.231608813 | 0.519101916 |
| 0.231548777 | 0.519061169 |
| 0.231443577 | 0.518874188 |
| 0.23144293 | 0.518874188 |
| 0.2314192 | 0.518868385 |
| 0.231414148 | 0.518868385 |
| 0.231373539 | 0.518839283 |
| 0.231369504 | 0.518839283 |
| 0.231284221 | 0.518736689 |
| 0.231235301 | 0.518664367 |
| 0.231211378 | 0.518651949 |
| 0.231197408 | 0.518651949 |
| 0.231097792 | 0.518526729 |
| 0.23109005 | 0.518526729 |
| 0.231060918 | 0.518504446 |
| 0.231059556 | 0.518504446 |
| 0.231030695 | 0.518504446 |
| 0.230911417 | 0.518309142 |
| 0.230849689 | 0.518201354 |
| 0.230846283 | 0.518201354 |
| 0.230839368 | 0.518201354 |
| 0.23081611 | 0.51819925 |
| 0.230800681 | 0.518189037 |
| 0.230787325 | 0.518189037 |
| 0.230764091 | 0.518180169 |
| 0.230745424 | 0.518180169 |
| 0.230744786 | 0.518180169 |

| | |
|---|---|
| 0.230737552 | 0.518180169 |
| 0.230660584 | 0.518083166 |
| 0.230576902 | 0.517955199 |
| 0.230571653 | 0.517955199 |
| 0.23053253 | 0.517904378 |
| 0.230493412 | 0.517904378 |
| 0.23046684 | 0.517903377 |
| 0.230405473 | 0.517789906 |
| 0.230384385 | 0.517766946 |
| 0.230320738 | 0.51767276 |
| 0.230291806 | 0.517632161 |
| 0.230257289 | 0.517603432 |
| 0.230256252 | 0.517603432 |
| 0.230253798 | 0.517603432 |
| 0.230174695 | 0.517564335 |
| 0.230163451 | 0.517563489 |
| 0.230092722 | 0.517453304 |
| 0.230080407 | 0.517450043 |
| 0.230035698 | 0.517398356 |
| 0.230005044 | 0.517378276 |
| 0.229932009 | 0.517238418 |
| 0.229930269 | 0.517238418 |
| 0.22992817 | 0.517238418 |
| 0.229913531 | 0.517238418 |
| 0.229912506 | 0.517238418 |
| 0.229805052 | 0.517148233 |
| 0.22977856 | 0.517113049 |
| 0.229745759 | 0.517063664 |
| 0.229687862 | 0.51698222 |
| 0.229677047 | 0.51698222 |
| 0.229676411 | 0.51698222 |
| 0.229664748 | 0.51698222 |
| 0.229663005 | 0.51698222 |
| 0.229627586 | 0.51698222 |
| 0.229443549 | 0.516652068 |
| 0.229341907 | 0.516520878 |
| 0.229275027 | 0.516419093 |
| 0.229259576 | 0.516408712 |
| 0.229243984 | 0.516398016 |
| 0.229232904 | 0.51639748 |
| 0.229208965 | 0.516367975 |
| 0.22916009 | 0.516367975 |
| 0.229155186 | 0.516367975 |
| 0.229125729 | 0.516351429 |
| 0.229104592 | 0.516336721 |

| | |
|---|---|
| 0.229099427 | 0.516336721 |
| 0.229073265 | 0.516330923 |
| 0.229027077 | 0.516251247 |
| 0.229017211 | 0.516251247 |
| 0.228948898 | 0.516123879 |
| 0.228922814 | 0.516089503 |
| 0.228912659 | 0.516089503 |
| 0.228862714 | 0.516010264 |
| 0.228739662 | 0.515823099 |
| 0.228656363 | 0.515659671 |
| 0.228640932 | 0.515649289 |
| 0.228637026 | 0.515649289 |
| 0.228618199 | 0.515646854 |
| 0.228610674 | 0.515646854 |
| 0.228583233 | 0.515621912 |
| 0.228558054 | 0.515621912 |
| 0.228513969 | 0.515558256 |
| 0.228482207 | 0.515550876 |
| 0.228459523 | 0.515550876 |
| 0.228320319 | 0.515292225 |
| 0.228290248 | 0.515270066 |
| 0.228285391 | 0.515270066 |
| 0.228208969 | 0.51516298 |
| 0.228184983 | 0.51513325 |
| 0.228073313 | 0.514978792 |
| 0.227998795 | 0.514859352 |
| 0.22798201 | 0.514845861 |
| 0.227965956 | 0.514845861 |
| 0.227861197 | 0.514646239 |
| 0.227780326 | 0.514512383 |
| 0.227739044 | 0.514467933 |
| 0.227624081 | 0.514232621 |
| 0.2275974 | 0.514196734 |
| 0.227593292 | 0.514196734 |
| 0.22758556 | 0.514196734 |
| 0.227567273 | 0.514196734 |
| 0.227548946 | 0.514184829 |
| 0.227512926 | 0.51412783 |
| 0.227473541 | 0.514087612 |
| 0.227459771 | 0.514080886 |
| 0.22741174 | 0.514021117 |
| 0.227329414 | 0.513883811 |
| 0.227247113 | 0.513746534 |
| 0.227225336 | 0.513721689 |
| 0.227221562 | 0.513721689 |

| | |
|---|---|
| 0.227210185 | 0.513721689 |
| 0.227145169 | 0.513654641 |
| 0.227140176 | 0.513654641 |
| 0.22710057 | 0.513654641 |
| 0.227092197 | 0.513654641 |
| 0.227074697 | 0.513649328 |
| 0.227041102 | 0.513622124 |
| 0.226964196 | 0.513564126 |
| 0.226918362 | 0.513515196 |
| 0.226837619 | 0.513430059 |
| 0.226808139 | 0.513387731 |
| 0.226792966 | 0.513377785 |
| 0.226790444 | 0.513377785 |
| 0.2267543 | 0.513363457 |
| 0.226749035 | 0.513363457 |
| 0.226717594 | 0.513329158 |
| 0.226713958 | 0.513329158 |
| 0.226694399 | 0.513329158 |
| 0.226656185 | 0.513329158 |
| 0.226632809 | 0.513329158 |
| 0.226629087 | 0.513329158 |
| 0.226563941 | 0.513225248 |
| 0.226481679 | 0.513112117 |
| 0.226469493 | 0.513112117 |
| 0.22636719 | 0.512999152 |
| 0.22630993 | 0.512893791 |
| 0.226176489 | 0.512615763 |
| 0.226149441 | 0.512603247 |
| 0.226123532 | 0.512568913 |
| 0.226109232 | 0.512560895 |
| 0.226099297 | 0.512560895 |
| 0.225942818 | 0.512281186 |
| 0.225942811 | 0.512281186 |
| 0.225880641 | 0.512213363 |
| 0.225831074 | 0.512125349 |
| 0.22581557 | 0.512114574 |
| 0.225811394 | 0.512114574 |
| 0.225795249 | 0.512114574 |
| 0.225788907 | 0.512114574 |
| 0.225781422 | 0.512114574 |
| 0.225720257 | 0.512114574 |
| 0.225718582 | 0.512114574 |
| 0.225688523 | 0.512114574 |
| 0.225669157 | 0.512099531 |
| 0.22554207 | 0.511859914 |

| | |
|---|---|
| 0.22554207 | 0.511859914 |
| 0.225520053 | 0.511858729 |
| 0.22549932 | 0.511836064 |
| 0.225428909 | 0.511725021 |
| 0.22537681 | 0.511655529 |
| 0.225365486 | 0.511654208 |
| 0.225362516 | 0.511654208 |
| 0.225293486 | 0.511539514 |
| 0.225275578 | 0.51152324 |
| 0.225273054 | 0.51152324 |
| 0.2252458 | 0.511515247 |
| 0.22516839 | 0.511426148 |
| 0.225162367 | 0.511426148 |
| 0.225138239 | 0.511406443 |
| 0.225037155 | 0.511249981 |
| 0.22500936 | 0.51121122 |
| 0.224988119 | 0.511187347 |
| 0.224962858 | 0.511154336 |
| 0.224946256 | 0.511154336 |
| 0.224904677 | 0.511095289 |
| 0.224895635 | 0.511095289 |
| 0.224843271 | 0.511028894 |
| 0.224802725 | 0.510961124 |
| 0.224799534 | 0.510961124 |
| 0.22478153 | 0.510961124 |
| 0.22469181 | 0.510782151 |
| 0.224676699 | 0.51077218 |
| 0.224674119 | 0.51077218 |
| 0.224669022 | 0.51077218 |
| 0.224638101 | 0.51077218 |
| 0.224624467 | 0.51077218 |
| 0.224617566 | 0.51077218 |
| 0.224598689 | 0.510765495 |
| 0.224592537 | 0.510765495 |
| 0.224580066 | 0.510765495 |
| 0.224565925 | 0.510765495 |
| 0.224557949 | 0.510765495 |
| 0.224519257 | 0.510755572 |
| 0.224488827 | 0.510710742 |
| 0.22443836 | 0.51062032 |
| 0.22441659 | 0.510595182 |
| 0.224376083 | 0.510551799 |
| 0.224359482 | 0.510538416 |
| 0.22432967 | 0.510494968 |
| 0.224168047 | 0.510200306 |

| | |
|---|---|
| 0.224147106 | 0.510177023 |
| 0.224129711 | 0.510161812 |
| 0.224110133 | 0.510141629 |
| 0.224052722 | 0.510084082 |
| 0.224030798 | 0.510082934 |
| 0.2239508 | 0.509925168 |
| 0.223910779 | 0.509858418 |
| 0.223883066 | 0.509819689 |
| 0.223872345 | 0.509819689 |
| 0.223856769 | 0.509819689 |
| 0.223719436 | 0.50958294 |
| 0.223680894 | 0.509529842 |
| 0.223678335 | 0.509529842 |
| 0.223675375 | 0.509529842 |
| 0.223656083 | 0.509529842 |
| 0.223636048 | 0.509525166 |
| 0.223551393 | 0.509429769 |
| 0.223544305 | 0.509429769 |
| 0.223534945 | 0.509429769 |
| 0.223495856 | 0.509400704 |
| 0.223463313 | 0.509350908 |
| 0.223442392 | 0.509350908 |
| 0.223439489 | 0.509350908 |
| 0.223350003 | 0.509214484 |
| 0.223273688 | 0.509113364 |
| 0.223264002 | 0.509113364 |
| 0.223215102 | 0.509046627 |
| 0.223214785 | 0.509046627 |
| 0.223138313 | 0.5089755 |
| 0.223090465 | 0.50890525 |
| 0.223085356 | 0.50890525 |
| 0.223059834 | 0.50889398 |
| 0.22299143 | 0.508786665 |
| 0.222966589 | 0.508754359 |
| 0.222936943 | 0.508711085 |
| 0.222869231 | 0.508613682 |
| 0.222868083 | 0.508613682 |
| 0.222863419 | 0.508613682 |
| 0.222763185 | 0.508485129 |
| 0.222715364 | 0.508449078 |
| 0.222713396 | 0.508449078 |
| 0.222658856 | 0.508368808 |
| 0.222638746 | 0.508347264 |
| 0.222630955 | 0.508347264 |
| 0.222615287 | 0.50834244 |

| | |
|---|---|
| 0.222611768 | 0.50834244 |
| 0.222601185 | 0.50834244 |
| 0.222591355 | 0.50834244 |
| 0.222591344 | 0.50834244 |
| 0.22256938 | 0.50834244 |
| 0.22253928 | 0.508339486 |
| 0.222505179 | 0.50828597 |
| 0.222493783 | 0.508284314 |
| 0.222493524 | 0.508284314 |
| 0.22249092 | 0.508284314 |
| 0.222482588 | 0.508284314 |
| 0.222455932 | 0.508284314 |
| 0.222444523 | 0.508284314 |
| 0.222428597 | 0.508284314 |
| 0.222355442 | 0.508212047 |
| 0.222346132 | 0.508212047 |
| 0.222322801 | 0.508189831 |
| 0.222315131 | 0.508189831 |
| 0.222297527 | 0.508189831 |
| 0.222259599 | 0.508139301 |
| 0.222255059 | 0.508139301 |
| 0.222248779 | 0.508139301 |
| 0.222241225 | 0.508139301 |
| 0.222196845 | 0.508117786 |
| 0.222147776 | 0.508054364 |
| 0.222096426 | 0.508034507 |
| 0.222086258 | 0.508034507 |
| 0.222057675 | 0.507994663 |
| 0.221987911 | 0.507883857 |
| 0.221928429 | 0.50777216 |
| 0.221857593 | 0.507658864 |
| 0.221853869 | 0.507658864 |
| 0.221833191 | 0.507651805 |
| 0.221789384 | 0.507600336 |
| 0.221784361 | 0.507600336 |
| 0.221726401 | 0.507563095 |
| 0.221721082 | 0.507563095 |
| 0.221609088 | 0.507455937 |
| 0.221608827 | 0.507455937 |
| 0.221593865 | 0.507455937 |
| 0.22158898 | 0.507455937 |
| 0.221579953 | 0.507455937 |
| 0.221579499 | 0.507455937 |
| 0.221571435 | 0.507455937 |
| 0.221530273 | 0.507455937 |

| | |
|---|---|
| 0.221491616 | 0.50743091 |
| 0.221480885 | 0.507430735 |
| 0.221454851 | 0.507395496 |
| 0.221451949 | 0.507395496 |
| 0.22128403 | 0.507142455 |
| 0.221271846 | 0.507142455 |
| 0.221232571 | 0.507105769 |
| 0.22121745 | 0.507095513 |
| 0.221200016 | 0.507079956 |
| 0.221193914 | 0.507079956 |
| 0.221113866 | 0.506931265 |
| 0.221093697 | 0.50691463 |
| 0.221023296 | 0.506805408 |
| 0.221001875 | 0.506796491 |
| 0.220831923 | 0.50645553 |
| 0.220826615 | 0.50645553 |
| 0.220813844 | 0.50645553 |
| 0.220806562 | 0.50645553 |
| 0.220718384 | 0.506317044 |
| 0.220597361 | 0.506136935 |
| 0.220547704 | 0.506071762 |
| 0.220540042 | 0.506071762 |
| 0.220500643 | 0.506012534 |
| 0.22048726 | 0.506012534 |
| 0.220486389 | 0.506012534 |
| 0.220479323 | 0.506012534 |
| 0.220400038 | 0.505952312 |
| 0.220371361 | 0.505910866 |
| 0.220323027 | 0.505848669 |
| 0.220302125 | 0.505825063 |
| 0.220283237 | 0.505806081 |
| 0.220278144 | 0.505806081 |
| 0.220172183 | 0.505672978 |
| 0.220153304 | 0.505654004 |
| 0.220120337 | 0.505602666 |
| 0.220015232 | 0.505409996 |
| 0.219921059 | 0.505221319 |
| 0.21984089 | 0.505106955 |
| 0.219757603 | 0.50497652 |
| 0.219716239 | 0.504917989 |
| 0.219695101 | 0.504893774 |
| 0.219694585 | 0.504893774 |
| 0.219690951 | 0.504893774 |
| 0.219688628 | 0.504893774 |
| 0.219688045 | 0.504893774 |

| | |
|---|---|
| 0.219670207 | 0.504893774 |
| 0.219667186 | 0.504893774 |
| 0.219633918 | 0.504893774 |
| 0.219559644 | 0.50477733 |
| 0.219532691 | 0.504764095 |
| 0.219522776 | 0.504764095 |
| 0.21950524 | 0.504764095 |
| 0.219443851 | 0.504730396 |
| 0.219365039 | 0.504646607 |
| 0.219352743 | 0.504646607 |
| 0.219348251 | 0.504646607 |
| 0.219245749 | 0.504591537 |
| 0.219244041 | 0.504591537 |
| 0.219238396 | 0.504591537 |
| 0.21918116 | 0.504540412 |
| 0.21916338 | 0.504523867 |
| 0.219054095 | 0.504345422 |
| 0.219047561 | 0.504345422 |
| 0.219036745 | 0.504345422 |
| 0.219036462 | 0.504345422 |
| 0.219015348 | 0.504345422 |
| 0.219013625 | 0.504345422 |
| 0.21900551 | 0.504345422 |
| 0.218987069 | 0.504345422 |
| 0.218882814 | 0.50421919 |
| 0.218870116 | 0.504214329 |
| 0.218859612 | 0.504214329 |
| 0.218851723 | 0.504214329 |
| 0.218821294 | 0.504214329 |
| 0.218799626 | 0.504214329 |
| 0.218778279 | 0.504214329 |
| 0.21873581 | 0.504214329 |
| 0.218699167 | 0.504210728 |
| 0.218670357 | 0.504168713 |
| 0.21850558 | 0.503910771 |
| 0.218460563 | 0.50383135 |
| 0.218352669 | 0.503631292 |
| 0.218290972 | 0.503513372 |
| 0.218204568 | 0.50343598 |
| 0.218198369 | 0.50343598 |
| 0.218177997 | 0.503423452 |
| 0.218160315 | 0.503423452 |
| 0.218146549 | 0.503423452 |
| 0.218137835 | 0.503423452 |
| 0.218054145 | 0.503308363 |

| | |
|---|---|
| 0.218043277 | 0.503307671 |
| 0.218030476 | 0.503302515 |
| 0.218027251 | 0.503302515 |
| 0.217980825 | 0.503236683 |
| 0.217898436 | 0.50309525 |
| 0.217855323 | 0.503020092 |
| 0.217819794 | 0.502962441 |
| 0.217805293 | 0.502953343 |
| 0.217781166 | 0.502922014 |
| 0.217730464 | 0.50285369 |
| 0.217703178 | 0.502839441 |
| 0.217680577 | 0.502811623 |
| 0.217646701 | 0.502782143 |
| 0.217632417 | 0.502773532 |
| 0.217597579 | 0.502717433 |
| 0.217596784 | 0.502717433 |
| 0.217551688 | 0.502660179 |
| 0.217502463 | 0.502570823 |
| 0.2174888 | 0.502563634 |
| 0.217483905 | 0.502563634 |
| 0.217473708 | 0.502563634 |
| 0.217414162 | 0.5024643 |
| 0.217380022 | 0.50243416 |
| 0.217375555 | 0.50243416 |
| 0.217370772 | 0.50243416 |
| 0.217358352 | 0.50243416 |
| 0.217315896 | 0.502407854 |
| 0.217306621 | 0.502407854 |
| 0.21727293 | 0.50238168 |
| 0.217261381 | 0.502379365 |
| 0.217223368 | 0.502315855 |
| 0.217194124 | 0.502272617 |
| 0.217193969 | 0.502272617 |
| 0.21716346 | 0.502250479 |
| 0.217158215 | 0.502250479 |
| 0.217113976 | 0.502211563 |
| 0.217113757 | 0.502211563 |
| 0.217084061 | 0.502211563 |
| 0.217052653 | 0.502189298 |
| 0.217049172 | 0.502189298 |
| 0.21703666 | 0.502189298 |
| 0.216983182 | 0.502126137 |
| 0.21697914 | 0.502126137 |
| 0.216968257 | 0.502126137 |
| 0.216967032 | 0.502126137 |

| | |
|---|---|
| 0.216932846 | 0.502126137 |
| 0.216915907 | 0.502126137 |
| 0.216911753 | 0.502126137 |
| 0.216896743 | 0.502126137 |
| 0.216873821 | 0.502126137 |
| 0.216847279 | 0.502126137 |
| 0.216838974 | 0.502126137 |
| 0.21679788 | 0.502126137 |
| 0.21675549 | 0.502111563 |
| 0.216737633 | 0.502094617 |
| 0.216692201 | 0.502045372 |
| 0.216688728 | 0.502045372 |
| 0.216647729 | 0.502025198 |
| 0.216645204 | 0.502025198 |
| 0.216635479 | 0.502025198 |
| 0.21658891 | 0.501994259 |
| 0.216586161 | 0.501994259 |
| 0.216550844 | 0.501994259 |
| 0.216408756 | 0.501723205 |
| 0.216374006 | 0.501667062 |
| 0.216292605 | 0.501502745 |
| 0.216286438 | 0.501502745 |
| 0.21621402 | 0.501418179 |
| 0.216160719 | 0.501318979 |
| 0.216158292 | 0.501318979 |
| 0.216111583 | 0.501278252 |
| 0.216035922 | 0.50117598 |
| 0.215955903 | 0.501087975 |
| 0.215871515 | 0.500996194 |
| 0.215870056 | 0.500996194 |
| 0.215833212 | 0.500974141 |
| 0.215769457 | 0.500874978 |
| 0.215737234 | 0.500849001 |
| 0.215716189 | 0.500824557 |
| 0.215683105 | 0.500772155 |
| 0.215650646 | 0.500721203 |
| 0.215635394 | 0.500721203 |
| 0.215601205 | 0.500703166 |
| 0.215599786 | 0.500703166 |
| 0.215546945 | 0.500651273 |
| 0.215505642 | 0.500579751 |
| 0.215497631 | 0.500579751 |
| 0.215464341 | 0.50053264 |
| 0.21544062 | 0.50050266 |
| 0.215412149 | 0.500484629 |

| | |
|---|---|
| 0.21540351 | 0.500484629 |
| 0.215400924 | 0.500484629 |
| 0.215382187 | 0.500484629 |
| 0.215350565 | 0.500463617 |
| 0.21531845 | 0.500437821 |
| 0.215302795 | 0.500425855 |
| 0.215288024 | 0.500415946 |
| 0.215287677 | 0.500415946 |
| 0.215231995 | 0.500383395 |
| 0.215218053 | 0.500375407 |
| 0.215211976 | 0.500375407 |
| 0.215168191 | 0.500339353 |
| 0.215078062 | 0.500220863 |
| 0.21507509 | 0.500220863 |
| 0.215072471 | 0.500220863 |
| 0.215008729 | 0.50013288 |
| 0.214930458 | 0.499975234 |
| 0.214894077 | 0.49991502 |
| 0.214891853 | 0.49991502 |
| 0.214878642 | 0.49991502 |
| 0.214811729 | 0.499821105 |
| 0.214807165 | 0.499821105 |
| 0.214714777 | 0.49964434 |
| 0.214691377 | 0.499614297 |
| 0.21455496 | 0.499345636 |
| 0.214514631 | 0.499300574 |
| 0.214471975 | 0.499225686 |
| 0.214457841 | 0.499217185 |
| 0.21445663 | 0.499217185 |
| 0.214438453 | 0.499217185 |
| 0.214435336 | 0.499217185 |
| 0.214353893 | 0.499097183 |
| 0.214352555 | 0.499097183 |
| 0.214325627 | 0.499097183 |
| 0.214325218 | 0.499097183 |
| 0.214253178 | 0.498984651 |
| 0.21422473 | 0.498942795 |
| 0.214086227 | 0.498742161 |
| 0.214039179 | 0.498701899 |
| 0.214008513 | 0.498701899 |
| 0.213989254 | 0.498694772 |
| 0.213946246 | 0.498635568 |
| 0.213916575 | 0.498635568 |
| 0.213804468 | 0.498475885 |
| 0.213803503 | 0.498475885 |

| | |
|---|---|
| 0.213766922 | 0.498437149 |
| 0.21373851 | 0.498395301 |
| 0.213737863 | 0.498395301 |
| 0.213721793 | 0.498395301 |
| 0.213721282 | 0.498395301 |
| 0.213697641 | 0.498395301 |
| 0.213686931 | 0.498395301 |
| 0.213680507 | 0.498395301 |
| 0.213656342 | 0.498374504 |
| 0.213643643 | 0.49836929 |
| 0.213642998 | 0.49836929 |
| 0.213637845 | 0.49836929 |
| 0.213603389 | 0.498357948 |
| 0.213594732 | 0.498357948 |
| 0.213591823 | 0.498357948 |
| 0.213590272 | 0.498357948 |
| 0.213527054 | 0.498292566 |
| 0.213428078 | 0.498134817 |
| 0.21335639 | 0.49801631 |
| 0.21335639 | 0.49801631 |
| 0.213342382 | 0.49801631 |
| 0.213255242 | 0.497951043 |
| 0.213241221 | 0.497942716 |
| 0.213239205 | 0.497942716 |
| 0.213217787 | 0.497936822 |
| 0.213212673 | 0.497936822 |
| 0.213129266 | 0.497803323 |
| 0.213113551 | 0.497791031 |
| 0.213096076 | 0.497774624 |
| 0.21305006 | 0.497691545 |
| 0.213043624 | 0.497691545 |
| 0.213022325 | 0.497675574 |
| 0.213016096 | 0.497675574 |
| 0.212974626 | 0.497631924 |
| 0.212908742 | 0.497532243 |
| 0.212823067 | 0.497380843 |
| 0.212810477 | 0.497375826 |
| 0.212786405 | 0.497343973 |
| 0.212774593 | 0.497340771 |
| 0.212770864 | 0.497340771 |
| 0.212738343 | 0.497304852 |
| 0.212693428 | 0.497257474 |
| 0.212637435 | 0.497211576 |
| 0.212605684 | 0.497189969 |
| 0.212506914 | 0.497056614 |

| | |
|---|---|
| 0.212472639 | 0.497049678 |
| 0.212466466 | 0.497049678 |
| 0.21241894 | 0.496972877 |
| 0.212356583 | 0.496851394 |
| 0.212272795 | 0.496679754 |
| 0.212264631 | 0.496679754 |
| 0.212228502 | 0.496659152 |
| 0.21219386 | 0.49663425 |
| 0.212181827 | 0.49663425 |
| 0.212171951 | 0.49663425 |
| 0.212162382 | 0.49663425 |
| 0.212136734 | 0.49662978 |
| 0.212086289 | 0.496560499 |
| 0.21205802 | 0.496518721 |
| 0.212047218 | 0.496517839 |
| 0.211995871 | 0.496470834 |
| 0.211983527 | 0.496466336 |
| 0.211932402 | 0.496395421 |
| 0.211884454 | 0.496331934 |
| 0.211768859 | 0.496085557 |
| 0.211718113 | 0.495991076 |
| 0.211661712 | 0.495883337 |
| 0.211649139 | 0.495883337 |
| 0.211616942 | 0.495883337 |
| 0.211611743 | 0.495883337 |
| 0.211611196 | 0.495883337 |
| 0.211537032 | 0.495810738 |
| 0.211516972 | 0.49578812 |
| 0.211476993 | 0.495743207 |
| 0.211456806 | 0.495720283 |
| 0.211421209 | 0.495661231 |
| 0.211393003 | 0.495619502 |
| 0.211384063 | 0.495619502 |
| 0.21136628 | 0.495619502 |
| 0.211342062 | 0.495619502 |
| 0.211315521 | 0.495608623 |
| 0.211245075 | 0.495516606 |
| 0.211148753 | 0.495339458 |
| 0.211081294 | 0.495254387 |
| 0.21100928 | 0.49516264 |
| 0.210973121 | 0.49512254 |
| 0.210971533 | 0.49512254 |
| 0.210956368 | 0.49512254 |
| 0.210948729 | 0.49512254 |
| 0.210803173 | 0.494918798 |

| | |
|---|---|
| 0.210770377 | 0.494890591 |
| 0.210749032 | 0.494864869 |
| 0.21068718 | 0.494744027 |
| 0.210598241 | 0.494583947 |
| 0.210529026 | 0.494494546 |
| 0.210502149 | 0.494455802 |
| 0.210492642 | 0.494455802 |
| 0.210485506 | 0.494455802 |
| 0.210476186 | 0.494455802 |
| 0.210452604 | 0.494436966 |
| 0.210404406 | 0.49439583 |
| 0.210368732 | 0.494337459 |
| 0.210321772 | 0.494251493 |
| 0.210249909 | 0.494155764 |
| 0.210201825 | 0.494067134 |
| 0.210138084 | 0.493975795 |
| 0.210128521 | 0.493975795 |
| 0.210038603 | 0.493854094 |
| 0.209941947 | 0.49372433 |
| 0.20991329 | 0.493681312 |
| 0.209821525 | 0.493514236 |
| 0.209815796 | 0.493514236 |
| 0.209757427 | 0.493412209 |
| 0.209752754 | 0.493412209 |
| 0.209747373 | 0.493412209 |
| 0.209718951 | 0.493394808 |
| 0.209708469 | 0.493394518 |
| 0.209698585 | 0.493394518 |
| 0.20969388 | 0.493394518 |
| 0.209680212 | 0.493394518 |
| 0.209617105 | 0.493325784 |
| 0.209538475 | 0.493165099 |
| 0.209537419 | 0.493165099 |
| 0.209531459 | 0.493165099 |
| 0.209489005 | 0.493165099 |
| 0.209480626 | 0.493165099 |
| 0.209477667 | 0.493165099 |
| 0.209473978 | 0.493165099 |
| 0.209460986 | 0.493165099 |
| 0.20943418 | 0.493165099 |
| 0.209424416 | 0.493165099 |
| 0.20940124 | 0.49315892 |
| 0.209382188 | 0.493138437 |
| 0.209325586 | 0.493053892 |
| 0.209203441 | 0.492814933 |

| | |
|---|---|
| 0.20919786 | 0.492814933 |
| 0.209055684 | 0.492613055 |
| 0.209021712 | 0.492557373 |
| 0.209020048 | 0.492557373 |
| 0.208965945 | 0.49251214 |
| 0.208927306 | 0.492456735 |
| 0.208893427 | 0.492401249 |
| 0.208882967 | 0.492400964 |
| 0.208810475 | 0.492259937 |
| 0.20877064 | 0.492209262 |
| 0.208755375 | 0.492197636 |
| 0.208723298 | 0.492170735 |
| 0.208692873 | 0.492147725 |
| 0.208642511 | 0.492053323 |
| 0.208626649 | 0.492040281 |
| 0.208624119 | 0.492040281 |
| 0.208538539 | 0.491905551 |
| 0.208501098 | 0.491841596 |
| 0.208449743 | 0.491769164 |
| 0.208419336 | 0.491746146 |
| 0.208413996 | 0.491746146 |
| 0.208404127 | 0.491746146 |
| 0.208359699 | 0.491702874 |
| 0.208260934 | 0.491494152 |
| 0.208251011 | 0.491494152 |
| 0.208249402 | 0.491494152 |
| 0.208230561 | 0.491494152 |
| 0.208183639 | 0.491433495 |
| 0.208176564 | 0.491433495 |
| 0.208172207 | 0.491433495 |
| 0.208152644 | 0.491433495 |
| 0.208090405 | 0.491368229 |
| 0.208050007 | 0.491312856 |
| 0.207970497 | 0.491173801 |
| 0.207969721 | 0.491173801 |
| 0.207950956 | 0.491173801 |
| 0.207852568 | 0.490992673 |
| 0.207850814 | 0.490992673 |
| 0.207817701 | 0.490959011 |
| 0.20779324 | 0.490959011 |
| 0.207755493 | 0.490940003 |
| 0.207749133 | 0.490940003 |
| 0.207692961 | 0.490884004 |
| 0.207679968 | 0.490884004 |
| 0.207675036 | 0.490884004 |

| | |
|---|---|
| 0.207671246 | 0.490884004 |
| 0.207644012 | 0.490884004 |
| 0.207616744 | 0.490884004 |
| 0.207576445 | 0.490827293 |
| 0.207570584 | 0.490827293 |
| 0.207550175 | 0.490827293 |
| 0.207520381 | 0.490816565 |
| 0.20750602 | 0.490806974 |
| 0.207372926 | 0.490540894 |
| 0.20736065 | 0.490536218 |
| 0.207354168 | 0.490536218 |
| 0.207352816 | 0.490536218 |
| 0.207344225 | 0.490536218 |
| 0.207300773 | 0.490492022 |
| 0.207284696 | 0.490485341 |
| 0.207280952 | 0.490485341 |
| 0.207265563 | 0.490485341 |
| 0.207203065 | 0.490407015 |
| 0.207139634 | 0.490305618 |
| 0.207135173 | 0.490305618 |
| 0.207041253 | 0.490121466 |
| 0.207034015 | 0.490121466 |
| 0.207015034 | 0.490108118 |
| 0.206988857 | 0.490075163 |
| 0.206946437 | 0.490043154 |
| 0.206943295 | 0.490043154 |
| 0.206930359 | 0.490043154 |
| 0.206852177 | 0.490010567 |
| 0.206809121 | 0.489973699 |
| 0.206789965 | 0.48996496 |
| 0.206776278 | 0.489956904 |
| 0.206718058 | 0.489892072 |
| 0.206717876 | 0.489892072 |
| 0.206702291 | 0.489892072 |
| 0.206647354 | 0.489822003 |
| 0.206594428 | 0.489720924 |
| 0.206587737 | 0.489720924 |
| 0.206523492 | 0.489625881 |
| 0.206505218 | 0.48960693 |
| 0.206460406 | 0.489554139 |
| 0.206441632 | 0.489553656 |
| 0.206420306 | 0.489529433 |
| 0.206406605 | 0.489529433 |
| 0.206368113 | 0.489525556 |
| 0.206345869 | 0.489497172 |

| | |
|---|---|
| 0.206276202 | 0.489429417 |
| 0.2060942 | 0.489021945 |
| 0.20607841 | 0.48900884 |
| 0.206075268 | 0.48900884 |
| 0.206042512 | 0.488996747 |
| 0.206017138 | 0.488980148 |
| 0.205943277 | 0.488834311 |
| 0.205919377 | 0.488814258 |
| 0.205889359 | 0.488786856 |
| 0.205691422 | 0.488455593 |
| 0.205644282 | 0.488367999 |
| 0.205595052 | 0.488275433 |
| 0.205551158 | 0.488219878 |
| 0.20548747 | 0.488117293 |
| 0.205486168 | 0.488117293 |
| 0.205470039 | 0.488117293 |
| 0.205465235 | 0.488117293 |
| 0.205454863 | 0.488117293 |
| 0.205453295 | 0.488117293 |
| 0.205445646 | 0.488117293 |
| 0.205435653 | 0.488117293 |
| 0.205351541 | 0.487989115 |
| 0.20531702 | 0.487931425 |
| 0.205303788 | 0.487924327 |
| 0.205303202 | 0.487924327 |
| 0.205301005 | 0.487924327 |
| 0.205266032 | 0.487924327 |
| 0.205243929 | 0.487903798 |
| 0.205201475 | 0.487851577 |
| 0.205052618 | 0.487548867 |
| 0.205050547 | 0.487548867 |
| 0.205007724 | 0.487548867 |
| 0.20499719 | 0.487548867 |
| 0.204991132 | 0.487548867 |
| 0.204956035 | 0.487548867 |
| 0.204948142 | 0.487548867 |
| 0.20492646 | 0.487548867 |
| 0.204888423 | 0.487496674 |
| 0.204826918 | 0.48737468 |
| 0.204751838 | 0.487269065 |
| 0.204750073 | 0.487269065 |
| 0.204721895 | 0.487246499 |
| 0.204695731 | 0.487208574 |
| 0.204693071 | 0.487208574 |
| 0.204659032 | 0.487194268 |

| | |
|---|---|
| 0.2045889 | 0.487051663 |
| 0.204563476 | 0.487023255 |
| 0.204556164 | 0.487023255 |
| 0.204508004 | 0.486956441 |
| 0.204471781 | 0.486894531 |
| 0.204458108 | 0.486886315 |
| 0.204397471 | 0.486766256 |
| 0.204393757 | 0.486766256 |
| 0.204388738 | 0.486766256 |
| 0.20438226 | 0.486766256 |
| 0.204335342 | 0.486736369 |
| 0.204325535 | 0.486736369 |
| 0.204174438 | 0.486488637 |
| 0.204139441 | 0.486443646 |
| 0.204062248 | 0.486308365 |
| 0.20394737 | 0.486107567 |
| 0.20394246 | 0.486107567 |
| 0.203913429 | 0.486075319 |
| 0.203909628 | 0.486075319 |
| 0.203908918 | 0.486075319 |
| 0.203905807 | 0.486075319 |
| 0.203893895 | 0.486075319 |
| 0.203867358 | 0.486075319 |
| 0.203853761 | 0.486075319 |
| 0.203843831 | 0.486075319 |
| 0.203760497 | 0.486075319 |
| 0.203693574 | 0.485952452 |
| 0.203688496 | 0.485952452 |
| 0.20366982 | 0.485952452 |
| 0.203669448 | 0.485952452 |
| 0.20352973 | 0.485671148 |
| 0.203448968 | 0.485527084 |
| 0.203436045 | 0.485520572 |
| 0.203391014 | 0.485437426 |
| 0.203356201 | 0.485427315 |
| 0.203282953 | 0.485276787 |
| 0.203248752 | 0.485219465 |
| 0.203218672 | 0.485196296 |
| 0.203172344 | 0.485110004 |
| 0.203100236 | 0.484986461 |
| 0.203066751 | 0.48495513 |
| 0.203013343 | 0.484876209 |
| 0.202846144 | 0.48454977 |
| 0.202824038 | 0.484521264 |
| 0.202724428 | 0.4843076 |

| | |
|---|---|
| 0.202714145 | 0.484307328 |
| 0.202710173 | 0.484307328 |
| 0.202650407 | 0.48420363 |
| 0.202630182 | 0.484179596 |
| 0.202627082 | 0.484179596 |
| 0.202598772 | 0.484153122 |
| 0.202596466 | 0.484153122 |
| 0.202535049 | 0.484098003 |
| 0.202501336 | 0.484041712 |
| 0.202498877 | 0.484041712 |
| 0.202479292 | 0.484037607 |
| 0.202431987 | 0.483948812 |
| 0.202422714 | 0.483948812 |
| 0.202404706 | 0.483932176 |
| 0.202404706 | 0.483932176 |
| 0.202359192 | 0.48389623 |
| 0.202351303 | 0.48389623 |
| 0.20228232 | 0.483837591 |
| 0.202278119 | 0.483837591 |
| 0.202272874 | 0.483837591 |
| 0.202237498 | 0.483837591 |
| 0.202215238 | 0.483837591 |
| 0.202148063 | 0.483804312 |
| 0.202147048 | 0.483804312 |
| 0.202117889 | 0.483780716 |
| 0.202111898 | 0.483780716 |
| 0.202079891 | 0.483738389 |
| 0.202079211 | 0.483738389 |
| 0.202059678 | 0.483738389 |
| 0.202051943 | 0.483738389 |
| 0.202028616 | 0.483712894 |
| 0.202005148 | 0.483705338 |
| 0.201991082 | 0.483695977 |
| 0.201974546 | 0.483680697 |
| 0.201974476 | 0.483680697 |
| 0.201911002 | 0.483625793 |
| 0.201857304 | 0.483570134 |
| 0.201837101 | 0.483567981 |
| 0.201835416 | 0.483567981 |
| 0.20183526 | 0.483567981 |
| 0.201815339 | 0.483567981 |
| 0.201800678 | 0.483567981 |
| 0.201764088 | 0.483541398 |
| 0.201693234 | 0.483420243 |
| 0.201690936 | 0.483420243 |

| | |
|---|---|
| 0.201626207 | 0.483332563 |
| 0.201608481 | 0.483314397 |
| 0.201550564 | 0.483220726 |
| 0.201530661 | 0.483200805 |
| 0.201524254 | 0.483200805 |
| 0.201497871 | 0.483170833 |
| 0.201487689 | 0.483170744 |
| 0.201445187 | 0.483093149 |
| 0.201327466 | 0.482859464 |
| 0.201283752 | 0.482827563 |
| 0.201271247 | 0.482821885 |
| 0.201257885 | 0.482814149 |
| 0.201257546 | 0.482814149 |
| 0.201253257 | 0.482814149 |
| 0.201230691 | 0.482814149 |
| 0.201217394 | 0.482814149 |
| 0.201213923 | 0.482814149 |
| 0.201195586 | 0.482814149 |
| 0.20112313 | 0.482734027 |
| 0.201105013 | 0.48271487 |
| 0.201085269 | 0.482691805 |
| 0.20107572 | 0.482691805 |
| 0.201065893 | 0.482691805 |
| 0.20103699 | 0.482673222 |
| 0.201024675 | 0.482673222 |
| 0.20100886 | 0.482673222 |
| 0.200918963 | 0.48251147 |
| 0.200905695 | 0.48251147 |
| 0.200897912 | 0.48251147 |
| 0.200867675 | 0.482485614 |
| 0.200850542 | 0.482478948 |
| 0.200846656 | 0.482478948 |
| 0.200649982 | 0.482157249 |
| 0.200634557 | 0.482144511 |
| 0.200580532 | 0.482063791 |
| 0.200514113 | 0.481952342 |
| 0.200511099 | 0.481952342 |
| 0.200481756 | 0.481923213 |
| 0.200475524 | 0.481923213 |
| 0.200451298 | 0.481922966 |
| 0.200440877 | 0.481922241 |
| 0.2004163 | 0.481887476 |
| 0.200415482 | 0.481887476 |
| 0.200359986 | 0.481822263 |
| 0.20032382 | 0.481786728 |

| | |
|---|---|
| 0.200312413 | 0.481783621 |
| 0.200303234 | 0.481783621 |
| 0.200183373 | 0.481570538 |
| 0.200164704 | 0.481570538 |
| 0.200122525 | 0.481521445 |
| 0.200065636 | 0.481408885 |
| 0.200057924 | 0.481408885 |
| 0.200006457 | 0.481316006 |
| 0.199974696 | 0.481287322 |
| 0.199941562 | 0.481231896 |
| 0.199938614 | 0.481231896 |
| 0.199914989 | 0.481231896 |
| 0.199883734 | 0.481214305 |
| 0.199873757 | 0.481214305 |
| 0.19984711 | 0.481214305 |
| 0.199672928 | 0.480901267 |
| 0.199631296 | 0.48087395 |
| 0.199616387 | 0.480866114 |
| 0.199602404 | 0.480866114 |
| 0.199596069 | 0.480866114 |
| 0.199564856 | 0.480835502 |
| 0.199562347 | 0.480835502 |
| 0.199507272 | 0.480745401 |
| 0.199497579 | 0.480745401 |
| 0.199454866 | 0.480692081 |
| 0.199448534 | 0.480692081 |
| 0.199446296 | 0.480692081 |
| 0.199430921 | 0.480692081 |
| 0.19941204 | 0.480686164 |
| 0.199347001 | 0.480553707 |
| 0.199226472 | 0.48036041 |
| 0.199190425 | 0.480297811 |
| 0.199178282 | 0.480292847 |
| 0.199169864 | 0.480292847 |
| 0.19910627 | 0.480265097 |
| 0.199096537 | 0.480265097 |
| 0.199082902 | 0.480257376 |
| 0.198990846 | 0.480059621 |
| 0.198971951 | 0.480038352 |
| 0.198956844 | 0.480026221 |
| 0.198901878 | 0.479990866 |
| 0.198838368 | 0.479886235 |
| 0.198830278 | 0.479886235 |
| 0.198781741 | 0.479798196 |
| 0.198778572 | 0.479798196 |

| | |
|---|---|
| 0.198746933 | 0.479787124 |
| 0.198728688 | 0.479767399 |
| 0.198621457 | 0.479581452 |
| 0.198617429 | 0.479581452 |
| 0.19860163 | 0.479581452 |
| 0.198596684 | 0.479581452 |
| 0.198525217 | 0.47949493 |
| 0.198446379 | 0.479328822 |
| 0.198427866 | 0.479308414 |
| 0.198406007 | 0.47929446 |
| 0.198390271 | 0.479290528 |
| 0.198385983 | 0.479290528 |
| 0.198324932 | 0.479205597 |
| 0.198290108 | 0.479170072 |
| 0.1982306 | 0.479050578 |
| 0.198178508 | 0.478997604 |
| 0.19808905 | 0.478805683 |
| 0.198058449 | 0.478756013 |
| 0.198049674 | 0.478756013 |
| 0.198046434 | 0.478756013 |
| 0.197999744 | 0.478711294 |
| 0.197933652 | 0.478648696 |
| 0.197867528 | 0.478585992 |
| 0.197867171 | 0.478585992 |
| 0.197807508 | 0.478513724 |
| 0.197781694 | 0.478498515 |
| 0.19775462 | 0.478473738 |
| 0.197719643 | 0.478422671 |
| 0.197655663 | 0.478316461 |
| 0.197591135 | 0.478257508 |
| 0.197415467 | 0.4778566 |
| 0.197389343 | 0.477817648 |
| 0.197329549 | 0.477747485 |
| 0.19731489 | 0.477747485 |
| 0.197311244 | 0.477747485 |
| 0.197284445 | 0.477747485 |
| 0.19725545 | 0.477747485 |
| 0.197238953 | 0.477744984 |
| 0.197238049 | 0.477744984 |
| 0.197179532 | 0.477673934 |
| 0.197164424 | 0.477673934 |
| 0.197163936 | 0.477673934 |
| 0.197152748 | 0.477673934 |
| 0.197125775 | 0.477673934 |
| 0.197095242 | 0.477673934 |

| | |
|---|---|
| 0.197024351 | 0.477613821 |
| 0.196971417 | 0.477558424 |
| 0.19695858 | 0.477551611 |
| 0.196941906 | 0.477535494 |
| 0.196858961 | 0.477358675 |
| 0.196858739 | 0.477358675 |
| 0.196826794 | 0.477329279 |
| 0.196810857 | 0.477314936 |
| 0.196806882 | 0.477314936 |
| 0.196748365 | 0.477211977 |
| 0.196731446 | 0.477195241 |
| 0.196730621 | 0.477195241 |
| 0.196714818 | 0.477195241 |
| 0.196689848 | 0.477191553 |
| 0.196619316 | 0.477069038 |
| 0.196582241 | 0.477003383 |
| 0.196568893 | 0.477003383 |
| 0.196561937 | 0.477003383 |
| 0.196514251 | 0.4769356 |
| 0.19650982 | 0.4769356 |
| 0.196465746 | 0.47686648 |
| 0.196459928 | 0.47686648 |
| 0.196451142 | 0.47686648 |
| 0.196377794 | 0.476750189 |
| 0.196361719 | 0.476735463 |
| 0.196283905 | 0.476595134 |
| 0.196273943 | 0.476595134 |
| 0.196266381 | 0.476595134 |
| 0.196257909 | 0.476595134 |
| 0.196237925 | 0.476595134 |
| 0.196192347 | 0.476518584 |
| 0.196171906 | 0.476517538 |
| 0.196167454 | 0.476517538 |
| 0.196155542 | 0.476517538 |
| 0.196129994 | 0.476512943 |
| 0.196046517 | 0.476407341 |
| 0.195982617 | 0.476300663 |
| 0.195980441 | 0.476300663 |
| 0.195960967 | 0.476300663 |
| 0.195959916 | 0.476300663 |
| 0.19594301 | 0.476300663 |
| 0.195932627 | 0.476300663 |
| 0.195901133 | 0.476272754 |
| 0.195824348 | 0.476134685 |
| 0.195823194 | 0.476134685 |

| | |
|---|---|
| 0.195710153 | 0.475905619 |
| 0.19570336 | 0.475905619 |
| 0.195691422 | 0.475905619 |
| 0.195642176 | 0.4758618 |
| 0.195511061 | 0.475695864 |
| 0.195478024 | 0.475656867 |
| 0.19541958 | 0.475563248 |
| 0.195378961 | 0.475488697 |
| 0.195377463 | 0.475488697 |
| 0.195357426 | 0.475488697 |
| 0.195351395 | 0.475488697 |
| 0.195330916 | 0.475488697 |
| 0.1953239 | 0.475488697 |
| 0.195215383 | 0.475309188 |
| 0.195165517 | 0.475222525 |
| 0.195052862 | 0.475034909 |
| 0.195020548 | 0.474980498 |
| 0.194990682 | 0.474932046 |
| 0.194976551 | 0.474921914 |
| 0.194923046 | 0.474864445 |
| 0.194905405 | 0.474864445 |
| 0.194903832 | 0.474864445 |
| 0.194897992 | 0.474864445 |
| 0.194872188 | 0.474861995 |
| 0.194865525 | 0.474861995 |
| 0.194818327 | 0.474779331 |
| 0.194804578 | 0.474770117 |
| 0.19478986 | 0.474758542 |
| 0.194750546 | 0.474735605 |
| 0.194739773 | 0.474735605 |
| 0.194677527 | 0.474630488 |
| 0.194631981 | 0.474556695 |
| 0.194555018 | 0.474428951 |
| 0.194548998 | 0.474428951 |
| 0.194546891 | 0.474428951 |
| 0.194491239 | 0.474370575 |
| 0.194441261 | 0.474272966 |
| 0.194429458 | 0.474268466 |
| 0.194416936 | 0.474262211 |
| 0.194373653 | 0.474205202 |
| 0.194322793 | 0.474111748 |
| 0.19432125 | 0.474111748 |
| 0.194282228 | 0.474079295 |
| 0.194192592 | 0.473884848 |
| 0.194149847 | 0.473804815 |

| | |
|---|---|
| 0.194149319 | 0.473804815 |
| 0.194146362 | 0.473804815 |
| 0.194096216 | 0.473787173 |
| 0.19409127 | 0.473787173 |
| 0.194088517 | 0.473787173 |
| 0.194083039 | 0.473787173 |
| 0.194076989 | 0.473787173 |
| 0.194032914 | 0.473786495 |
| 0.194000654 | 0.473732011 |
| 0.193993304 | 0.473732011 |
| 0.193945982 | 0.473647077 |
| 0.193931729 | 0.473636555 |
| 0.193931679 | 0.473636555 |
| 0.193920875 | 0.473636555 |
| 0.193900967 | 0.473636555 |
| 0.193899106 | 0.473636555 |
| 0.193874652 | 0.473636555 |
| 0.193873023 | 0.473636555 |
| 0.193870021 | 0.473636555 |
| 0.193776504 | 0.473562453 |
| 0.193755027 | 0.473544961 |
| 0.193732894 | 0.473539467 |
| 0.193730875 | 0.473539467 |
| 0.193717495 | 0.473539467 |
| 0.193673454 | 0.473491381 |
| 0.193659126 | 0.473491381 |
| 0.193520304 | 0.47318983 |
| 0.19347516 | 0.473128026 |
| 0.193456129 | 0.473128026 |
| 0.193453713 | 0.473128026 |
| 0.193430573 | 0.473128026 |
| 0.193430162 | 0.473128026 |
| 0.193429338 | 0.473128026 |
| 0.193418553 | 0.473128026 |
| 0.193412271 | 0.473128026 |
| 0.19337523 | 0.473128026 |
| 0.193371386 | 0.473128026 |
| 0.193336555 | 0.473128026 |
| 0.19333075 | 0.473128026 |
| 0.193328711 | 0.473128026 |
| 0.193303974 | 0.47312238 |
| 0.193272752 | 0.473070271 |
| 0.19312373 | 0.472788806 |
| 0.193068209 | 0.472715381 |
| 0.193053072 | 0.47270262 |

| | |
|---|---|
| 0.193035908 | 0.47270262 |
| 0.193024231 | 0.47270262 |
| 0.192951755 | 0.472648935 |
| 0.192928094 | 0.472615282 |
| 0.192836395 | 0.472414947 |
| 0.192745971 | 0.472266301 |
| 0.192730489 | 0.472266301 |
| 0.19271414 | 0.472261195 |
| 0.192632902 | 0.472134996 |
| 0.192632619 | 0.472134996 |
| 0.192620903 | 0.472134996 |
| 0.19260319 | 0.472134996 |
| 0.192584278 | 0.472134996 |
| 0.192555814 | 0.472116127 |
| 0.192537192 | 0.472094771 |
| 0.192516476 | 0.472068279 |
| 0.192511927 | 0.472068279 |
| 0.19248997 | 0.472051893 |
| 0.192465118 | 0.472015251 |
| 0.192459304 | 0.472015251 |
| 0.192333986 | 0.471766532 |
| 0.192320635 | 0.471766532 |
| 0.192270614 | 0.471708265 |
| 0.192265605 | 0.471708265 |
| 0.192255508 | 0.471708265 |
| 0.192164146 | 0.471568517 |
| 0.19216363 | 0.471568517 |
| 0.192100149 | 0.471469385 |
| 0.192099466 | 0.471469385 |
| 0.192076698 | 0.471469385 |
| 0.192060732 | 0.471460493 |
| 0.192046541 | 0.471449957 |
| 0.191822097 | 0.470996083 |
| 0.191814768 | 0.470996083 |
| 0.1918123 | 0.470996083 |
| 0.191795749 | 0.470996083 |
| 0.191757339 | 0.470948837 |
| 0.191739504 | 0.470938958 |
| 0.191724677 | 0.470926828 |
| 0.191696041 | 0.470905063 |
| 0.191661292 | 0.470843986 |
| 0.191641217 | 0.470839477 |
| 0.191608289 | 0.470786627 |
| 0.191598038 | 0.470785729 |
| 0.191524713 | 0.470629838 |

| | |
|---|---|
| 0.191514018 | 0.470629838 |
| 0.191484406 | 0.470629838 |
| 0.191443336 | 0.470629838 |
| 0.191425099 | 0.470627875 |
| 0.191296882 | 0.470480997 |
| 0.191256176 | 0.470431125 |
| 0.191248002 | 0.470431125 |
| 0.191240385 | 0.470431125 |
| 0.19117575 | 0.470330463 |
| 0.191126909 | 0.470258888 |
| 0.191101449 | 0.470220537 |
| 0.1911011 | 0.470220537 |
| 0.191085505 | 0.470220537 |
| 0.191016635 | 0.470084704 |
| 0.190977563 | 0.470025027 |
| 0.190940913 | 0.469971208 |
| 0.190940815 | 0.469971208 |
| 0.19088749 | 0.469888281 |
| 0.190878771 | 0.469888281 |
| 0.190840374 | 0.469861965 |
| 0.190829613 | 0.469861965 |
| 0.190751731 | 0.469724042 |
| 0.190705938 | 0.469635558 |
| 0.190695927 | 0.46963519 |
| 0.190693147 | 0.46963519 |
| 0.190687662 | 0.46963519 |
| 0.190678548 | 0.46963519 |
| 0.190678548 | 0.46963519 |
| 0.190672961 | 0.46963519 |
| 0.190647886 | 0.46963519 |
| 0.190600868 | 0.469571051 |
| 0.19058457 | 0.469555187 |
| 0.190570274 | 0.469544255 |
| 0.190438097 | 0.469291419 |
| 0.190412777 | 0.469277584 |
| 0.19040672 | 0.469277584 |
| 0.190401141 | 0.469277584 |
| 0.190391189 | 0.469277584 |
| 0.190384683 | 0.469277584 |
| 0.190292897 | 0.469127792 |
| 0.190234886 | 0.469033333 |
| 0.190232721 | 0.469033333 |
| 0.190148216 | 0.468868189 |
| 0.190145187 | 0.468868189 |
| 0.190141973 | 0.468868189 |

| | |
|---|---|
| 0.190091911 | 0.468802166 |
| 0.190055988 | 0.468737846 |
| 0.190037389 | 0.468716247 |
| 0.190013239 | 0.468680953 |
| 0.189955775 | 0.46856348 |
| 0.189935212 | 0.468537025 |
| 0.189911567 | 0.468502964 |
| 0.189886324 | 0.468464956 |
| 0.189883783 | 0.468464956 |
| 0.189860202 | 0.468464956 |
| 0.189843439 | 0.468458188 |
| 0.189836667 | 0.468458188 |
| 0.189791954 | 0.468401979 |
| 0.189789986 | 0.468401979 |
| 0.189772102 | 0.468401979 |
| 0.189767202 | 0.468401979 |
| 0.189729681 | 0.46839392 |
| 0.189692168 | 0.468325587 |
| 0.189691096 | 0.468325587 |
| 0.189690155 | 0.468325587 |
| 0.189680403 | 0.468325587 |
| 0.189649014 | 0.468325587 |
| 0.18964004 | 0.468325587 |
| 0.189617011 | 0.468309966 |
| 0.189601385 | 0.468309966 |
| 0.189580081 | 0.46829161 |
| 0.189578216 | 0.46829161 |
| 0.189553027 | 0.468273356 |
| 0.189527481 | 0.468234534 |
| 0.189473212 | 0.468173316 |
| 0.189428113 | 0.468086164 |
| 0.18942719 | 0.468086164 |
| 0.18939886 | 0.468062448 |
| 0.189307131 | 0.46797058 |
| 0.189270254 | 0.46793887 |
| 0.189270163 | 0.46793887 |
| 0.189246292 | 0.467928209 |
| 0.189245794 | 0.467928209 |
| 0.189226155 | 0.467927003 |
| 0.189224173 | 0.467927003 |
| 0.189209427 | 0.467927003 |
| 0.189203607 | 0.467927003 |
| 0.189202792 | 0.467927003 |
| 0.189193385 | 0.467927003 |
| 0.189190076 | 0.467927003 |

| | |
|---|---|
| 0.189177655 | 0.467927003 |
| 0.189046365 | 0.467725257 |
| 0.189034248 | 0.467719576 |
| 0.189027762 | 0.467719576 |
| 0.188999425 | 0.467682009 |
| 0.18896225 | 0.467614315 |
| 0.188939002 | 0.467581078 |
| 0.188920793 | 0.467581078 |
| 0.18890752 | 0.467581078 |
| 0.188889877 | 0.467580986 |
| 0.188877699 | 0.467575142 |
| 0.188851285 | 0.467558358 |
| 0.188836454 | 0.467545945 |
| 0.188829914 | 0.467545945 |
| 0.18882109 | 0.467545945 |
| 0.188799296 | 0.467545945 |
| 0.18874801 | 0.467497086 |
| 0.188726998 | 0.467469354 |
| 0.188614204 | 0.467238568 |
| 0.188407089 | 0.46686109 |
| 0.1883571 | 0.466795899 |
| 0.188349677 | 0.466795899 |
| 0.188239511 | 0.46657732 |
| 0.188163573 | 0.466437646 |
| 0.188161921 | 0.466437646 |
| 0.188132617 | 0.466411259 |
| 0.188127598 | 0.466411259 |
| 0.188063919 | 0.466311968 |
| 0.188058937 | 0.466311968 |
| 0.188054168 | 0.466311968 |
| 0.187944474 | 0.466137138 |
| 0.187931581 | 0.466129435 |
| 0.187930202 | 0.466129435 |
| 0.187920307 | 0.466129435 |
| 0.187916071 | 0.466129435 |
| 0.187822598 | 0.465980459 |
| 0.187821646 | 0.465980459 |
| 0.187790211 | 0.465980459 |
| 0.187789464 | 0.465980459 |
| 0.187780954 | 0.465980459 |
| 0.187777997 | 0.465980459 |
| 0.187744116 | 0.465980459 |
| 0.187729819 | 0.465980459 |
| 0.187721361 | 0.465980459 |
| 0.187670182 | 0.465917836 |

| | |
|---|---|
| 0.187652516 | 0.465907784 |
| 0.187650339 | 0.465907784 |
| 0.187629877 | 0.465907784 |
| 0.18760069 | 0.465907784 |
| 0.187576164 | 0.465902996 |
| 0.187564403 | 0.465902996 |
| 0.187497748 | 0.465829695 |
| 0.187493903 | 0.465829695 |
| 0.187462088 | 0.465813996 |
| 0.187438142 | 0.465778796 |
| 0.187421862 | 0.465762641 |
| 0.187420198 | 0.465762641 |
| 0.187379067 | 0.465729195 |
| 0.187332668 | 0.465664854 |
| 0.187247264 | 0.465498769 |
| 0.187190454 | 0.465381831 |
| 0.187190165 | 0.465381831 |
| 0.187096573 | 0.465221285 |
| 0.187071628 | 0.465207833 |
| 0.186980439 | 0.465005345 |
| 0.186976567 | 0.465005345 |
| 0.186917726 | 0.464922213 |
| 0.186876647 | 0.464844314 |
| 0.186868152 | 0.464844314 |
| 0.186859972 | 0.464844314 |
| 0.18678985 | 0.464725498 |
| 0.186782548 | 0.464725498 |
| 0.186767748 | 0.46471906 |
| 0.186732712 | 0.464680437 |
| 0.186707889 | 0.464642943 |
| 0.186675771 | 0.464611567 |
| 0.186666741 | 0.464611567 |
| 0.186594757 | 0.464458476 |
| 0.186557658 | 0.464391555 |
| 0.186519591 | 0.464344182 |
| 0.186517744 | 0.464344182 |
| 0.186500561 | 0.464344182 |
| 0.186399814 | 0.46414304 |
| 0.186390891 | 0.46414304 |
| 0.186384628 | 0.46414304 |
| 0.18634023 | 0.464116 |
| 0.186305294 | 0.464074 |
| 0.186293444 | 0.464072277 |
| 0.186272243 | 0.464043734 |
| 0.186261076 | 0.464043734 |

| | |
|---|---|
| 0.186255299 | 0.464043734 |
| 0.186237372 | 0.464043734 |
| 0.186177518 | 0.463929087 |
| 0.186164587 | 0.463921138 |
| 0.186164544 | 0.463921138 |
| 0.186127983 | 0.463878465 |
| 0.186119607 | 0.463878465 |
| 0.186085052 | 0.463820012 |
| 0.185886742 | 0.463422731 |
| 0.18588355 | 0.463422731 |
| 0.185786607 | 0.463245836 |
| 0.185762326 | 0.463209543 |
| 0.185699345 | 0.463109578 |
| 0.185693154 | 0.463109578 |
| 0.185690278 | 0.463109578 |
| 0.185603553 | 0.463007564 |
| 0.185595796 | 0.463007564 |
| 0.185580001 | 0.462997314 |
| 0.185565711 | 0.462985915 |
| 0.185529821 | 0.462920621 |
| 0.185477656 | 0.462838956 |
| 0.185436067 | 0.462759423 |
| 0.185396301 | 0.462732924 |
| 0.185374102 | 0.462710362 |
| 0.185311343 | 0.462679292 |
| 0.185279639 | 0.462659957 |
| 0.185246746 | 0.462602075 |
| 0.185223043 | 0.462567137 |
| 0.185217886 | 0.462567137 |
| 0.185212987 | 0.462567137 |
| 0.185212038 | 0.462567137 |
| 0.185106872 | 0.462373996 |
| 0.185081235 | 0.462358458 |
| 0.185078401 | 0.462358458 |
| 0.185028182 | 0.462274419 |
| 0.185023556 | 0.462274419 |
| 0.185010772 | 0.462274419 |
| 0.184998442 | 0.462272865 |
| 0.184931809 | 0.462130608 |
| 0.184928934 | 0.462130608 |
| 0.184922319 | 0.462130608 |
| 0.184919528 | 0.462130608 |
| 0.18486102 | 0.462050686 |
| 0.18483193 | 0.46200222 |
| 0.184799292 | 0.46196913 |

| | |
|---|---|
| 0.184764887 | 0.461931615 |
| 0.184761644 | 0.461931615 |
| 0.184742701 | 0.461924644 |
| 0.184710922 | 0.461924644 |
| 0.184707663 | 0.461924644 |
| 0.184621217 | 0.461790595 |
| 0.184571056 | 0.461689375 |
| 0.184568938 | 0.461689375 |
| 0.184556661 | 0.461689375 |
| 0.184554307 | 0.461689375 |
| 0.184544325 | 0.461689375 |
| 0.184507585 | 0.461689375 |
| 0.184504103 | 0.461689375 |
| 0.184441645 | 0.461583849 |
| 0.184436572 | 0.461583849 |
| 0.184426532 | 0.461583849 |
| 0.184411251 | 0.461580548 |
| 0.184393624 | 0.461580548 |
| 0.184388122 | 0.461580548 |
| 0.184384084 | 0.461580548 |
| 0.184376902 | 0.461580548 |
| 0.184334959 | 0.461580548 |
| 0.184330416 | 0.461580548 |
| 0.184218129 | 0.46136996 |
| 0.184185469 | 0.461337999 |
| 0.184167258 | 0.461333509 |
| 0.184166554 | 0.461333509 |
| 0.184145167 | 0.4613267 |
| 0.184132847 | 0.461320101 |
| 0.184113927 | 0.461296964 |
| 0.184099304 | 0.461284594 |
| 0.184097961 | 0.461284594 |
| 0.184071119 | 0.461284594 |
| 0.184057099 | 0.461275913 |
| 0.183995091 | 0.461193316 |
| 0.183965567 | 0.461151357 |
| 0.183924511 | 0.461089203 |
| 0.183914763 | 0.461089035 |
| 0.183881751 | 0.461054813 |
| 0.183876471 | 0.461054813 |
| 0.183870848 | 0.461054813 |
| 0.183852018 | 0.461053079 |
| 0.183848532 | 0.461053079 |
| 0.183821287 | 0.46104884 |
| 0.183811311 | 0.4610481 |

| | |
|---|---|
| 0.183800934 | 0.461046351 |
| 0.183736916 | 0.460922126 |
| 0.183728036 | 0.460922126 |
| 0.183727484 | 0.460922126 |
| 0.1836987 | 0.460922126 |
| 0.183679728 | 0.460922126 |
| 0.183641225 | 0.460865486 |
| 0.183600766 | 0.460811218 |
| 0.183577112 | 0.460776132 |
| 0.183512445 | 0.460638095 |
| 0.183494392 | 0.460638095 |
| 0.183481368 | 0.460632918 |
| 0.183453073 | 0.460586163 |
| 0.183436608 | 0.460584222 |
| 0.183388218 | 0.460569007 |
| 0.183385001 | 0.460569007 |
| 0.18337704 | 0.460569007 |
| 0.183297644 | 0.460462942 |
| 0.183259069 | 0.460390318 |
| 0.183236319 | 0.460357448 |
| 0.183162555 | 0.460239061 |
| 0.183126183 | 0.460177847 |
| 0.183075642 | 0.460099394 |
| 0.183071688 | 0.460099394 |
| 0.183054153 | 0.460099394 |
| 0.183020511 | 0.460057945 |
| 0.183008501 | 0.460057945 |
| 0.182980476 | 0.460053695 |
| 0.182915076 | 0.459914282 |
| 0.182909505 | 0.459914282 |
| 0.18287548 | 0.459863279 |
| 0.182786728 | 0.459724792 |
| 0.182724301 | 0.459604452 |
| 0.18271121 | 0.459595797 |
| 0.1826885 | 0.459562942 |
| 0.182667406 | 0.45954013 |
| 0.182593996 | 0.459422276 |
| 0.182544587 | 0.459360633 |
| 0.182497724 | 0.459325371 |
| 0.182480165 | 0.459325371 |
| 0.182440433 | 0.459302807 |
| 0.182413471 | 0.45926869 |
| 0.182385845 | 0.45926869 |
| 0.18233096 | 0.459193706 |
| 0.182304648 | 0.459155129 |

| | |
|---|---|
| 0.182289536 | 0.459144056 |
| 0.182267392 | 0.459134135 |
| 0.182253855 | 0.459124319 |
| 0.182228578 | 0.459087532 |
| 0.182220388 | 0.459087532 |
| 0.182152212 | 0.458997353 |
| 0.18213931 | 0.458997353 |
| 0.182050019 | 0.458853527 |
| 0.182044312 | 0.458853527 |
| 0.1820409 | 0.458853527 |
| 0.182039476 | 0.458853527 |
| 0.182031473 | 0.458853527 |
| 0.182018567 | 0.458853527 |
| 0.182014268 | 0.458853527 |
| 0.181988874 | 0.458853527 |
| 0.181871455 | 0.458711711 |
| 0.181868862 | 0.458711711 |
| 0.181858312 | 0.458711711 |
| 0.181855485 | 0.458711711 |
| 0.181810167 | 0.458661626 |
| 0.18180623 | 0.458661626 |
| 0.18180029 | 0.458661626 |
| 0.181768652 | 0.45865407 |
| 0.181767936 | 0.45865407 |
| 0.181721768 | 0.45860866 |
| 0.181708538 | 0.458599573 |
| 0.181682272 | 0.458557582 |
| 0.181631029 | 0.458452541 |
| 0.181620861 | 0.458451175 |
| 0.181565827 | 0.458360842 |
| 0.181564162 | 0.458360842 |
| 0.181541938 | 0.458360842 |
| 0.181470674 | 0.458266373 |
| 0.18146483 | 0.458266373 |
| 0.181447501 | 0.458266373 |
| 0.181414436 | 0.458245842 |
| 0.181382813 | 0.458190264 |
| 0.181359059 | 0.458154559 |
| 0.181349343 | 0.458154316 |
| 0.181334157 | 0.458140252 |
| 0.181310292 | 0.458104259 |
| 0.181307621 | 0.458104259 |
| 0.181302799 | 0.458104259 |
| 0.181250079 | 0.458049322 |
| 0.18124405 | 0.458049322 |

| | |
|---|---|
| 0.181230208 | 0.458049322 |
| 0.181195668 | 0.458009026 |
| 0.181159962 | 0.457943077 |
| 0.181118205 | 0.45793353 |
| 0.18110268 | 0.457919799 |
| 0.181091081 | 0.457919799 |
| 0.18104041 | 0.457859581 |
| 0.181012479 | 0.457813251 |
| 0.180928579 | 0.457673967 |
| 0.180910264 | 0.457665163 |
| 0.180898264 | 0.457665163 |
| 0.180897554 | 0.457665163 |
| 0.180841751 | 0.457575841 |
| 0.180815462 | 0.457533631 |
| 0.180768858 | 0.457440009 |
| 0.180756919 | 0.457434101 |
| 0.180732818 | 0.457397412 |
| 0.180708634 | 0.457386213 |
| 0.180708625 | 0.457386213 |
| 0.180698728 | 0.457386213 |
| 0.180690203 | 0.457386213 |
| 0.18067798 | 0.457386213 |
| 0.180660442 | 0.457386213 |
| 0.180660001 | 0.457386213 |
| 0.180633728 | 0.457386213 |
| 0.180600078 | 0.457353109 |
| 0.180509077 | 0.457219901 |
| 0.180504262 | 0.457219901 |
| 0.180499724 | 0.457219901 |
| 0.180467549 | 0.457211978 |
| 0.180463256 | 0.457211978 |
| 0.180430866 | 0.457192002 |
| 0.180409422 | 0.457161987 |
| 0.180401505 | 0.457161987 |
| 0.180363253 | 0.45709838 |
| 0.180363075 | 0.45709838 |
| 0.180307042 | 0.457048458 |
| 0.180170836 | 0.456807972 |
| 0.180161257 | 0.456807972 |
| 0.180132975 | 0.456807972 |
| 0.18007966 | 0.45673948 |
| 0.180019325 | 0.456655648 |
| 0.18000104 | 0.456655648 |
| 0.179996723 | 0.456655648 |
| 0.179985143 | 0.456655648 |

| | |
|---|---|
| 0.17997342 | 0.456655648 |
| 0.179943554 | 0.456655648 |
| 0.179927673 | 0.456655648 |
| 0.179880439 | 0.456622175 |
| 0.179827655 | 0.456555921 |
| 0.179815035 | 0.456555921 |
| 0.179802493 | 0.456555921 |
| 0.179723339 | 0.456443422 |
| 0.179678769 | 0.456354562 |
| 0.179642462 | 0.456330638 |
| 0.179617099 | 0.456319602 |
| 0.179604784 | 0.456312655 |
| 0.179594143 | 0.456309963 |
| 0.179593704 | 0.456309963 |
| 0.179577691 | 0.456309963 |
| 0.179474142 | 0.456078058 |
| 0.179469136 | 0.456078058 |
| 0.179468497 | 0.456078058 |
| 0.179448589 | 0.456078058 |
| 0.179384325 | 0.456066601 |
| 0.179373276 | 0.456065076 |
| 0.17934794 | 0.456025004 |
| 0.179341558 | 0.456025004 |
| 0.179285818 | 0.455979048 |
| 0.179232104 | 0.455876499 |
| 0.179224753 | 0.455876499 |
| 0.179211952 | 0.455873933 |
| 0.179207514 | 0.455873933 |
| 0.179158799 | 0.455787412 |
| 0.179086237 | 0.455681686 |
| 0.179086196 | 0.455681686 |
| 0.179042972 | 0.455638772 |
| 0.179011934 | 0.455608476 |
| 0.178910924 | 0.455375726 |
| 0.178896802 | 0.455375726 |
| 0.178878388 | 0.455375726 |
| 0.17883391 | 0.455325704 |
| 0.178797423 | 0.455257144 |
| 0.178786038 | 0.455252494 |
| 0.178784057 | 0.455252494 |
| 0.178727702 | 0.455152619 |
| 0.178727702 | 0.455152619 |
| 0.178703256 | 0.455152619 |
| 0.178618783 | 0.455055524 |
| 0.178610933 | 0.455055524 |

| | |
|---|---|
| 0.178606773 | 0.455055524 |
| 0.17860377 | 0.455055524 |
| 0.178596457 | 0.455055524 |
| 0.178588734 | 0.455055524 |
| 0.178578869 | 0.455055524 |
| 0.178562647 | 0.455055524 |
| 0.178544142 | 0.455055524 |
| 0.178520457 | 0.455055524 |
| 0.178496589 | 0.455050692 |
| 0.178482722 | 0.455039698 |
| 0.178475758 | 0.455039698 |
| 0.178456871 | 0.455028456 |
| 0.178410318 | 0.454989594 |
| 0.178385043 | 0.454961147 |
| 0.178370564 | 0.45494858 |
| 0.178368104 | 0.45494858 |
| 0.178352107 | 0.45494858 |
| 0.178347991 | 0.45494858 |
| 0.178337582 | 0.45494858 |
| 0.178321528 | 0.45494858 |
| 0.178282595 | 0.45494858 |
| 0.178234967 | 0.454894122 |
| 0.178226816 | 0.454894122 |
| 0.178179226 | 0.454821751 |
| 0.178174625 | 0.454821751 |
| 0.17817392 | 0.454821751 |
| 0.178163695 | 0.454821751 |
| 0.178069788 | 0.454712056 |
| 0.178032228 | 0.454645627 |
| 0.178028869 | 0.454645627 |
| 0.178010898 | 0.45463991 |
| 0.177994902 | 0.454623434 |
| 0.177966207 | 0.45459437 |
| 0.177928301 | 0.454526453 |
| 0.177924313 | 0.454526453 |
| 0.17789302 | 0.454508911 |
| 0.177885667 | 0.454508911 |
| 0.17788314 | 0.454508911 |
| 0.177822284 | 0.454426244 |
| 0.177816817 | 0.454426244 |
| 0.177815303 | 0.454426244 |
| 0.177806244 | 0.454426244 |
| 0.177717003 | 0.454254692 |
| 0.17768916 | 0.454207902 |
| 0.177687466 | 0.454207902 |

| | |
|---|---|
| 0.177667004 | 0.454200021 |
| 0.177643657 | 0.454181384 |
| 0.177613942 | 0.454161882 |
| 0.177603651 | 0.454161882 |
| 0.177504452 | 0.453979409 |
| 0.177488693 | 0.453963481 |
| 0.177447873 | 0.453907827 |
| 0.177400545 | 0.453835514 |
| 0.177283587 | 0.453628348 |
| 0.177269815 | 0.453628348 |
| 0.177268785 | 0.453628348 |
| 0.177259216 | 0.453628348 |
| 0.17723026 | 0.453619173 |
| 0.177205177 | 0.45357935 |
| 0.177057789 | 0.453250811 |
| 0.177033164 | 0.453212133 |
| 0.177031427 | 0.453212133 |
| 0.177013186 | 0.45320971 |
| 0.176963517 | 0.453106898 |
| 0.176865116 | 0.452903645 |
| 0.176851377 | 0.452892814 |
| 0.176808331 | 0.452831274 |
| 0.176798928 | 0.452831274 |
| 0.176777091 | 0.452799963 |
| 0.176746403 | 0.452745708 |
| 0.176740519 | 0.452745708 |
| 0.176673218 | 0.452631272 |
| 0.176662909 | 0.452629208 |
| 0.176655295 | 0.452629208 |
| 0.176622524 | 0.452598781 |
| 0.176612571 | 0.452598781 |
| 0.176545915 | 0.452499853 |
| 0.176537481 | 0.452499853 |
| 0.176501239 | 0.452458394 |
| 0.176469904 | 0.45242677 |
| 0.176459841 | 0.452425327 |
| 0.176413112 | 0.452354221 |
| 0.176330784 | 0.452204222 |
| 0.176326403 | 0.452204222 |
| 0.176305419 | 0.452199806 |
| 0.176304719 | 0.452199806 |
| 0.176290308 | 0.452199806 |
| 0.176287403 | 0.452199806 |
| 0.176269213 | 0.452199806 |
| 0.176234761 | 0.452164679 |

| | |
|---|---|
| 0.17622217 | 0.452156734 |
| 0.176213985 | 0.452156734 |
| 0.176181266 | 0.452124852 |
| 0.176178634 | 0.452124852 |
| 0.176152874 | 0.452101767 |
| 0.176149172 | 0.452101767 |
| 0.176143251 | 0.452101767 |
| 0.176099697 | 0.452101767 |
| 0.176090235 | 0.452101767 |
| 0.176047024 | 0.452053941 |
| 0.1760053 | 0.452014257 |
| 0.176001456 | 0.452014257 |
| 0.175989093 | 0.452014257 |
| 0.175979761 | 0.452014257 |
| 0.175917583 | 0.451977079 |
| 0.17591147 | 0.451977079 |
| 0.175827138 | 0.451824706 |
| 0.175808435 | 0.451801021 |
| 0.175782398 | 0.451801021 |
| 0.175747024 | 0.451765086 |
| 0.175739592 | 0.451765086 |
| 0.175720295 | 0.451745142 |
| 0.175696259 | 0.451707733 |
| 0.175693281 | 0.451707733 |
| 0.17566832 | 0.451684665 |
| 0.175667348 | 0.451684665 |
| 0.175647051 | 0.451678748 |
| 0.175622386 | 0.451639707 |
| 0.175621759 | 0.451639707 |
| 0.175611294 | 0.451639707 |
| 0.175610215 | 0.451639707 |
| 0.175585331 | 0.451639707 |
| 0.175580643 | 0.451639707 |
| 0.175580077 | 0.451639707 |
| 0.175535414 | 0.451601919 |
| 0.175473504 | 0.451509599 |
| 0.175467027 | 0.451509599 |
| 0.1754549 | 0.451509599 |
| 0.175411715 | 0.451487989 |
| 0.175407148 | 0.451487989 |
| 0.175398758 | 0.451487989 |
| 0.175390182 | 0.451487989 |
| 0.175383569 | 0.451487989 |
| 0.175372176 | 0.451487989 |
| 0.175332898 | 0.45146181 |

| | |
|---|---|
| 0.175320209 | 0.45146181 |
| 0.175319529 | 0.45146181 |
| 0.175295927 | 0.45146181 |
| 0.175293558 | 0.45146181 |
| 0.175262214 | 0.45146181 |
| 0.175143618 | 0.451310127 |
| 0.175106319 | 0.451238431 |
| 0.17509282 | 0.451228058 |
| 0.175083423 | 0.451228058 |
| 0.175048384 | 0.451162366 |
| 0.175035952 | 0.451154741 |
| 0.175012434 | 0.451118536 |
| 0.174991136 | 0.45108805 |
| 0.174944596 | 0.451016904 |
| 0.174940596 | 0.451016904 |
| 0.174937965 | 0.451016904 |
| 0.174909658 | 0.451000068 |
| 0.174909464 | 0.451000068 |
| 0.174904599 | 0.451000068 |
| 0.174870337 | 0.450971926 |
| 0.174835964 | 0.450907697 |
| 0.174794373 | 0.450824845 |
| 0.174762885 | 0.450816874 |
| 0.174735697 | 0.450771155 |
| 0.174699438 | 0.450726396 |
| 0.174626892 | 0.450661337 |
| 0.174586034 | 0.45058031 |
| 0.174576615 | 0.45058031 |
| 0.174575835 | 0.45058031 |
| 0.174539901 | 0.450534491 |
| 0.174531095 | 0.450534491 |
| 0.174465771 | 0.450514057 |
| 0.174464903 | 0.450514057 |
| 0.174448762 | 0.450514057 |
| 0.174447374 | 0.450514057 |
| 0.174420207 | 0.450496117 |
| 0.174377486 | 0.450487726 |
| 0.174364893 | 0.450487726 |
| 0.17433791 | 0.450476976 |
| 0.174278226 | 0.450371625 |
| 0.174248746 | 0.450344313 |
| 0.174202105 | 0.450248205 |
| 0.174122914 | 0.450092377 |
| 0.174096651 | 0.450048915 |
| 0.174060335 | 0.449979458 |

| | |
|---|---|
| 0.174026212 | 0.449915665 |
| 0.173979182 | 0.449867339 |
| 0.17395757 | 0.449835877 |
| 0.173888614 | 0.449706398 |
| 0.173837627 | 0.449623366 |
| 0.17383105 | 0.449623366 |
| 0.173802795 | 0.449582105 |
| 0.173769779 | 0.449569952 |
| 0.173736907 | 0.449509326 |
| 0.173721855 | 0.4494948 |
| 0.173680882 | 0.4494132 |
| 0.17363568 | 0.449320649 |
| 0.173559801 | 0.449197514 |
| 0.173544242 | 0.449197514 |
| 0.173425325 | 0.448995867 |
| 0.173417038 | 0.448995867 |
| 0.173414792 | 0.448995867 |
| 0.17340909 | 0.448995867 |
| 0.173357781 | 0.448943018 |
| 0.17335489 | 0.448943018 |
| 0.173333076 | 0.448927861 |
| 0.173313576 | 0.44892254 |
| 0.173286778 | 0.448905594 |
| 0.173270488 | 0.448887812 |
| 0.173267111 | 0.448887812 |
| 0.173203914 | 0.448788576 |
| 0.17319654 | 0.448788576 |
| 0.173176821 | 0.448767204 |
| 0.173156627 | 0.44873929 |
| 0.17309252 | 0.448597564 |
| 0.173087429 | 0.448597564 |
| 0.173066613 | 0.448587741 |
| 0.173054813 | 0.448587741 |
| 0.172895128 | 0.448351413 |
| 0.172869003 | 0.448351413 |
| 0.17274477 | 0.448108558 |
| 0.172597447 | 0.447801852 |
| 0.172585351 | 0.447801852 |
| 0.172487217 | 0.44758901 |
| 0.172477791 | 0.447588938 |
| 0.17245596 | 0.44755667 |
| 0.172360288 | 0.447430284 |
| 0.172289026 | 0.447318438 |
| 0.172255497 | 0.447255768 |
| 0.172173059 | 0.447114846 |

| | |
|---|---|
| 0.172163607 | 0.44711468 |
| 0.172135559 | 0.447066215 |
| 0.172120197 | 0.447050698 |
| 0.172092986 | 0.447004399 |
| 0.172061621 | 0.446947304 |
| 0.172025137 | 0.446925661 |
| 0.17201591 | 0.446925661 |
| 0.171972417 | 0.446849219 |
| 0.171756338 | 0.44639777 |
| 0.171753251 | 0.44639777 |
| 0.171738841 | 0.44639777 |
| 0.171707516 | 0.446343952 |
| 0.171641502 | 0.446234248 |
| 0.171633791 | 0.446234248 |
| 0.171567181 | 0.446100898 |
| 0.171563772 | 0.446100898 |
| 0.171548509 | 0.446100898 |
| 0.171532024 | 0.446100898 |
| 0.171517504 | 0.446093499 |
| 0.171458133 | 0.446019257 |
| 0.171456157 | 0.446019257 |
| 0.171438215 | 0.446019257 |
| 0.171424684 | 0.446019257 |
| 0.171421272 | 0.446019257 |
| 0.171357503 | 0.445920871 |
| 0.171350953 | 0.445920871 |
| 0.171340029 | 0.445920871 |
| 0.171336691 | 0.445920871 |
| 0.1713362 | 0.445920871 |
| 0.171301847 | 0.445920871 |
| 0.17120646 | 0.445783972 |
| 0.171205306 | 0.445783972 |
| 0.171143278 | 0.445704335 |
| 0.171117189 | 0.445660762 |
| 0.171102366 | 0.445646527 |
| 0.171026196 | 0.445496863 |
| 0.171025172 | 0.445496863 |
| 0.171005124 | 0.445490705 |
| 0.17097781 | 0.44546828 |
| 0.170903995 | 0.445300322 |
| 0.1708797 | 0.445261379 |
| 0.170861583 | 0.445238531 |
| 0.170844062 | 0.445217234 |
| 0.170836936 | 0.445217234 |
| 0.170743153 | 0.445051673 |

| | |
|---|---|
| 0.170737129 | 0.445051673 |
| 0.170723323 | 0.445048697 |
| 0.170706354 | 0.44502882 |
| 0.170671987 | 0.444987942 |
| 0.170664915 | 0.444987942 |
| 0.170550747 | 0.444744876 |
| 0.170529853 | 0.444714741 |
| 0.170432141 | 0.444484262 |
| 0.170413065 | 0.44445885 |
| 0.170410921 | 0.44445885 |
| 0.17040384 | 0.44445885 |
| 0.170317433 | 0.444306753 |
| 0.170242208 | 0.444134839 |
| 0.170213157 | 0.444132035 |
| 0.170201752 | 0.444132035 |
| 0.170167978 | 0.444111473 |
| 0.17010823 | 0.443979871 |
| 0.170068072 | 0.443899384 |
| 0.170054417 | 0.443888071 |
| 0.170043027 | 0.443882667 |
| 0.170025064 | 0.443860103 |
| 0.170004008 | 0.443829463 |
| 0.170001169 | 0.443829463 |
| 0.169967594 | 0.44378305 |
| 0.169954803 | 0.44377398 |
| 0.169921132 | 0.443734716 |
| 0.169838777 | 0.443568295 |
| 0.169824128 | 0.443568295 |
| 0.169815478 | 0.443568295 |
| 0.169809695 | 0.443568295 |
| 0.169741499 | 0.443484469 |
| 0.169717693 | 0.443446596 |
| 0.169689161 | 0.443420699 |
| 0.169655306 | 0.443356556 |
| 0.169638816 | 0.443337791 |
| 0.169590078 | 0.44330772 |
| 0.169560989 | 0.44325601 |
| 0.169554039 | 0.44325601 |
| 0.169549189 | 0.44325601 |
| 0.169537713 | 0.44325601 |
| 0.169512784 | 0.44325601 |
| 0.169491568 | 0.443244832 |
| 0.169478956 | 0.443236188 |
| 0.169470159 | 0.443236188 |
| 0.169436567 | 0.443222682 |

| | |
|---|---|
| 0.169434785 | 0.443222682 |
| 0.169434093 | 0.443222682 |
| 0.169353782 | 0.443079135 |
| 0.169289215 | 0.44293454 |
| 0.16923921 | 0.442852364 |
| 0.169153789 | 0.442653159 |
| 0.169153416 | 0.442653159 |
| 0.169137128 | 0.442653159 |
| 0.169130494 | 0.442653159 |
| 0.169091802 | 0.442588221 |
| 0.169066763 | 0.442547004 |
| 0.169065635 | 0.442547004 |
| 0.168964553 | 0.442425354 |
| 0.168931691 | 0.442387949 |
| 0.168927774 | 0.442387949 |
| 0.168903396 | 0.442362499 |
| 0.168865453 | 0.44231665 |
| 0.168814136 | 0.442225997 |
| 0.16879009 | 0.442211652 |
| 0.168741372 | 0.442115322 |
| 0.16872237 | 0.442107192 |
| 0.168721254 | 0.442107192 |
| 0.168718658 | 0.442107192 |
| 0.168686825 | 0.442087021 |
| 0.168669795 | 0.442087021 |
| 0.168640317 | 0.442086767 |
| 0.168545833 | 0.441912051 |
| 0.168482628 | 0.441843628 |
| 0.168478578 | 0.441843628 |
| 0.168457611 | 0.441826679 |
| 0.168449735 | 0.441826679 |
| 0.168424306 | 0.441787984 |
| 0.168405542 | 0.441787428 |
| 0.168368601 | 0.441763517 |
| 0.168348246 | 0.441763517 |
| 0.168321716 | 0.441737842 |
| 0.168310902 | 0.441733803 |
| 0.168273137 | 0.441697393 |
| 0.168270523 | 0.441697393 |
| 0.16825082 | 0.441697393 |
| 0.168243538 | 0.441697393 |
| 0.168223878 | 0.441692722 |
| 0.168186578 | 0.441650887 |
| 0.168182931 | 0.441650887 |
| 0.168180125 | 0.441650887 |

| | |
|---|---|
| 0.168160723 | 0.441650887 |
| 0.168116381 | 0.441612645 |
| 0.168099383 | 0.441592351 |
| 0.168091186 | 0.441592351 |
| 0.167920048 | 0.441242932 |
| 0.167891082 | 0.44119116 |
| 0.167842468 | 0.441112087 |
| 0.167825146 | 0.441112087 |
| 0.167813007 | 0.441112087 |
| 0.167781463 | 0.441073454 |
| 0.167751178 | 0.441018181 |
| 0.167750975 | 0.441018181 |
| 0.167746593 | 0.441018181 |
| 0.167686592 | 0.440945745 |
| 0.167680998 | 0.440945745 |
| 0.167668625 | 0.440945745 |
| 0.167612414 | 0.440823703 |
| 0.16760525 | 0.440823703 |
| 0.167593105 | 0.440821606 |
| 0.167509245 | 0.440649701 |
| 0.16748785 | 0.440617754 |
| 0.167469596 | 0.440615559 |
| 0.167466813 | 0.440615559 |
| 0.167457832 | 0.440615559 |
| 0.167401386 | 0.440512396 |
| 0.167398699 | 0.440512396 |
| 0.16739507 | 0.440512396 |
| 0.16734226 | 0.440502394 |
| 0.167337246 | 0.440502394 |
| 0.167312092 | 0.440471671 |
| 0.167295196 | 0.440451538 |
| 0.167153444 | 0.440175666 |
| 0.167146553 | 0.440175666 |
| 0.167114935 | 0.440124362 |
| 0.16705869 | 0.440023463 |
| 0.166996535 | 0.439932739 |
| 0.166995829 | 0.439932739 |
| 0.166980712 | 0.439932739 |
| 0.166962775 | 0.439916801 |
| 0.166943763 | 0.439891042 |
| 0.166921693 | 0.439891042 |
| 0.166915178 | 0.439891042 |
| 0.166902623 | 0.439891042 |
| 0.166835639 | 0.439800796 |
| 0.166834488 | 0.439800796 |

| | |
|---|---|
| 0.166809705 | 0.439781114 |
| 0.16680442 | 0.439781114 |
| 0.16675156 | 0.439700839 |
| 0.166705255 | 0.439627419 |
| 0.166685765 | 0.439600361 |
| 0.166641646 | 0.439557025 |
| 0.16660358 | 0.439480958 |
| 0.166596409 | 0.439480958 |
| 0.166561703 | 0.439428898 |
| 0.166519588 | 0.439381075 |
| 0.16651946 | 0.439381075 |
| 0.166480912 | 0.439327704 |
| 0.166458388 | 0.439292604 |
| 0.16639546 | 0.439199544 |
| 0.166326512 | 0.439041888 |
| 0.166281429 | 0.438971543 |
| 0.166266339 | 0.438971543 |
| 0.16621662 | 0.438922098 |
| 0.166191304 | 0.438879582 |
| 0.166187032 | 0.438879582 |
| 0.166175614 | 0.438879582 |
| 0.166165038 | 0.438879582 |
| 0.166147949 | 0.438879582 |
| 0.166140373 | 0.438879582 |
| 0.166125377 | 0.438879582 |
| 0.166112028 | 0.438879582 |
| 0.166095285 | 0.438869356 |
| 0.166004378 | 0.438653491 |
| 0.166002971 | 0.438653491 |
| 0.165954662 | 0.438595123 |
| 0.165787993 | 0.4383249 |
| 0.165779611 | 0.4383249 |
| 0.165771159 | 0.4383249 |
| 0.165730542 | 0.438270322 |
| 0.165705012 | 0.438264241 |
| 0.16570074 | 0.438264241 |
| 0.165695137 | 0.438264241 |
| 0.165682126 | 0.438264241 |
| 0.165674453 | 0.438264241 |
| 0.16566193 | 0.438264241 |
| 0.165569973 | 0.43811329 |
| 0.165445229 | 0.437866763 |
| 0.165419646 | 0.43783714 |
| 0.165410416 | 0.43783704 |
| 0.165409955 | 0.43783704 |

| | |
|---|---|
| 0.165378264 | 0.437800599 |
| 0.165337253 | 0.437716362 |
| 0.165234244 | 0.437492292 |
| 0.165227745 | 0.437492292 |
| 0.16522621 | 0.437492292 |
| 0.165209582 | 0.437492292 |
| 0.165203198 | 0.437492292 |
| 0.165150327 | 0.437440332 |
| 0.165145575 | 0.437440332 |
| 0.165132449 | 0.437440332 |
| 0.165129351 | 0.437440332 |
| 0.164999737 | 0.437211665 |
| 0.164997906 | 0.437211665 |
| 0.164996968 | 0.437211665 |
| 0.164974448 | 0.437211665 |
| 0.16497377 | 0.437211665 |
| 0.16496318 | 0.437211665 |
| 0.16494711 | 0.437211665 |
| 0.164941149 | 0.437211665 |
| 0.164939836 | 0.437211665 |
| 0.164893158 | 0.437172479 |
| 0.164889508 | 0.437172479 |
| 0.164868071 | 0.437154638 |
| 0.164836654 | 0.437120007 |
| 0.164814339 | 0.437109507 |
| 0.164766073 | 0.437005833 |
| 0.164723513 | 0.436964305 |
| 0.164671714 | 0.436877214 |
| 0.164671335 | 0.436877214 |
| 0.164663391 | 0.436877214 |
| 0.164606069 | 0.436800381 |
| 0.164587349 | 0.436775042 |
| 0.164546922 | 0.43669209 |
| 0.164519236 | 0.436642945 |
| 0.164468859 | 0.436557897 |
| 0.164466866 | 0.436557897 |
| 0.164457401 | 0.436557897 |
| 0.16444146 | 0.436557897 |
| 0.164391165 | 0.436448966 |
| 0.164345842 | 0.436377287 |
| 0.16434468 | 0.436377287 |
| 0.164336084 | 0.436377287 |
| 0.164333549 | 0.436377287 |
| 0.164319906 | 0.436377287 |
| 0.164287923 | 0.436369466 |

| | |
|---|---|
| 0.164245185 | 0.436280282 |
| 0.164223836 | 0.436247903 |
| 0.164223368 | 0.436247903 |
| 0.164210526 | 0.436247903 |
| 0.164198904 | 0.436247903 |
| 0.16413756 | 0.436116015 |
| 0.164135254 | 0.436116015 |
| 0.164134279 | 0.436116015 |
| 0.164130853 | 0.436116015 |
| 0.164121719 | 0.436116015 |
| 0.164114743 | 0.436116015 |
| 0.164106256 | 0.436116015 |
| 0.164068519 | 0.436116015 |
| 0.164039513 | 0.436116015 |
| 0.164028409 | 0.436116015 |
| 0.163996462 | 0.436116015 |
| 0.163980607 | 0.436112596 |
| 0.163931486 | 0.436055007 |
| 0.163917566 | 0.436042332 |
| 0.163897406 | 0.436037412 |
| 0.163890229 | 0.436037412 |
| 0.163859084 | 0.436014297 |
| 0.163817029 | 0.435945333 |
| 0.163793235 | 0.43590637 |
| 0.163785924 | 0.43590637 |
| 0.163782441 | 0.43590637 |
| 0.163766911 | 0.43590637 |
| 0.163605157 | 0.435600559 |
| 0.163591493 | 0.435600559 |
| 0.163589674 | 0.435600559 |
| 0.1635353 | 0.435511952 |
| 0.163528582 | 0.435511952 |
| 0.163498578 | 0.435462854 |
| 0.163462004 | 0.435389792 |
| 0.163449689 | 0.43538134 |
| 0.163430975 | 0.435355839 |
| 0.163411122 | 0.435327303 |
| 0.163376565 | 0.435273953 |
| 0.163365185 | 0.435273953 |
| 0.163344221 | 0.43524646 |
| 0.163331039 | 0.435235686 |
| 0.163217653 | 0.435128305 |
| 0.163215215 | 0.435128305 |
| 0.163192159 | 0.435109051 |
| 0.163190516 | 0.435109051 |

| | |
|---|---|
| 0.163062974 | 0.434861989 |
| 0.163050251 | 0.434861989 |
| 0.16292311 | 0.434635018 |
| 0.162913763 | 0.434635018 |
| 0.162898831 | 0.434635018 |
| 0.162897587 | 0.434635018 |
| 0.162811735 | 0.43450292 |
| 0.162809922 | 0.43450292 |
| 0.162809428 | 0.43450292 |
| 0.162797222 | 0.43450292 |
| 0.162792289 | 0.43450292 |
| 0.162781743 | 0.43450292 |
| 0.162767546 | 0.43450292 |
| 0.162756863 | 0.43450292 |
| 0.162755107 | 0.43450292 |
| 0.162718972 | 0.43450292 |
| 0.162673872 | 0.434408102 |
| 0.162651974 | 0.434398339 |
| 0.162640245 | 0.434391372 |
| 0.162626477 | 0.43437896 |
| 0.162619004 | 0.43437896 |
| 0.162464891 | 0.434019365 |
| 0.162460286 | 0.434019365 |
| 0.16243434 | 0.433987453 |
| 0.162289057 | 0.433691343 |
| 0.162281936 | 0.433691343 |
| 0.162266855 | 0.433691343 |
| 0.162246976 | 0.433681487 |
| 0.162215895 | 0.433622745 |
| 0.162211264 | 0.433622745 |
| 0.162196585 | 0.433622745 |
| 0.162184753 | 0.433622745 |
| 0.162151299 | 0.433596089 |
| 0.162148169 | 0.433596089 |
| 0.162067047 | 0.433444026 |
| 0.162055087 | 0.433444026 |
| 0.162005958 | 0.433353428 |
| 0.16199488 | 0.433348132 |
| 0.161989055 | 0.433348132 |
| 0.161962539 | 0.433310294 |
| 0.161908261 | 0.433189412 |
| 0.161884416 | 0.433149949 |
| 0.161852383 | 0.433112902 |
| 0.161840698 | 0.433112902 |
| 0.161789263 | 0.433041319 |

| | |
|---|---|
| 0.161777358 | 0.43303379 |
| 0.16171798 | 0.432947849 |
| 0.161689291 | 0.432904576 |
| 0.161629717 | 0.432784536 |
| 0.161583757 | 0.432713867 |
| 0.161531949 | 0.432620039 |
| 0.161477902 | 0.432523933 |
| 0.161450691 | 0.432475369 |
| 0.161449636 | 0.432475369 |
| 0.161447344 | 0.432475369 |
| 0.161411542 | 0.432443467 |
| 0.161395809 | 0.432425642 |
| 0.161392967 | 0.432425642 |
| 0.161375509 | 0.432419904 |
| 0.161308834 | 0.432265561 |
| 0.161266653 | 0.432197996 |
| 0.161205391 | 0.432109925 |
| 0.161188193 | 0.432088146 |
| 0.161147917 | 0.432028811 |
| 0.161109963 | 0.431956411 |
| 0.161083129 | 0.431928059 |
| 0.16108229 | 0.431928059 |
| 0.161050113 | 0.431912481 |
| 0.160998829 | 0.431823575 |
| 0.160968262 | 0.431765905 |
| 0.16095128 | 0.431744669 |
| 0.160949156 | 0.431744669 |
| 0.160896305 | 0.43167014 |
| 0.160892308 | 0.43167014 |
| 0.160878783 | 0.43167014 |
| 0.160849518 | 0.431617556 |
| 0.160837613 | 0.431609928 |
| 0.160739154 | 0.431394326 |
| 0.160735223 | 0.431394326 |
| 0.160729035 | 0.431394326 |
| 0.160677765 | 0.431375417 |
| 0.160666791 | 0.431370271 |
| 0.160597692 | 0.431233368 |
| 0.160589699 | 0.431233368 |
| 0.160542078 | 0.431156962 |
| 0.160532331 | 0.431155098 |
| 0.160516642 | 0.431137271 |
| 0.160489807 | 0.431089507 |
| 0.160399607 | 0.430895826 |
| 0.160366365 | 0.430855129 |

| | |
|---|---|
| 0.160355938 | 0.430851418 |
| 0.160329496 | 0.430804677 |
| 0.160327754 | 0.430804677 |
| 0.160286517 | 0.430786409 |
| 0.160164588 | 0.430483004 |
| 0.160145381 | 0.430455673 |
| 0.160118767 | 0.43044803 |
| 0.160090863 | 0.430406295 |
| 0.160078984 | 0.430406295 |
| 0.160046282 | 0.430359319 |
| 0.160044708 | 0.430359319 |
| 0.159965551 | 0.430219906 |
| 0.159908703 | 0.430110793 |
| 0.159896983 | 0.430103557 |
| 0.159864378 | 0.430040144 |
| 0.159820593 | 0.42997093 |
| 0.159771898 | 0.429864207 |
| 0.159753933 | 0.429840155 |
| 0.159741301 | 0.429840155 |
| 0.159727503 | 0.429840155 |
| 0.1596543 | 0.429693452 |
| 0.159645824 | 0.429693452 |
| 0.15964369 | 0.429693452 |
| 0.159623423 | 0.429683195 |
| 0.159619979 | 0.429683195 |
| 0.159581868 | 0.429659137 |
| 0.159574562 | 0.429659137 |
| 0.15956155 | 0.429659137 |
| 0.159549864 | 0.429655173 |
| 0.159530198 | 0.429639258 |
| 0.159519077 | 0.429639258 |
| 0.159506406 | 0.429635313 |
| 0.159339937 | 0.429235475 |
| 0.1593396 | 0.429235475 |
| 0.159335405 | 0.429235475 |
| 0.15930362 | 0.429217581 |
| 0.159249194 | 0.429112383 |
| 0.159218509 | 0.429053975 |
| 0.159209938 | 0.429053975 |
| 0.159168457 | 0.428991914 |
| 0.159158954 | 0.428990576 |
| 0.159144743 | 0.428976552 |
| 0.159102148 | 0.42888601 |
| 0.159057462 | 0.428809172 |
| 0.159034758 | 0.428777152 |

| | |
|---|---|
| 0.159023183 | 0.428770219 |
| 0.159020864 | 0.428770219 |
| 0.15899032 | 0.428730153 |
| 0.158971049 | 0.428702461 |
| 0.158962981 | 0.428702461 |
| 0.158947659 | 0.428687929 |
| 0.158935731 | 0.428680035 |
| 0.158912287 | 0.428641076 |
| 0.158875529 | 0.4285662 |
| 0.158869778 | 0.4285662 |
| 0.158869778 | 0.4285662 |
| 0.158860212 | 0.4285662 |
| 0.158817346 | 0.428530607 |
| 0.158783536 | 0.428463653 |
| 0.158779825 | 0.428463653 |
| 0.158772387 | 0.428463653 |
| 0.158703995 | 0.428321822 |
| 0.158702281 | 0.428321822 |
| 0.158680979 | 0.428321822 |
| 0.158678294 | 0.428321822 |
| 0.158550446 | 0.428108553 |
| 0.158504203 | 0.428049426 |
| 0.158499924 | 0.428049426 |
| 0.158492172 | 0.428049426 |
| 0.158398186 | 0.427908701 |
| 0.158393362 | 0.427908701 |
| 0.158392555 | 0.427908701 |
| 0.15835982 | 0.427877867 |
| 0.158353335 | 0.427877867 |
| 0.158325256 | 0.427857295 |
| 0.158314283 | 0.427854874 |
| 0.158306353 | 0.427854874 |
| 0.15830532 | 0.427854874 |
| 0.158286668 | 0.427854874 |
| 0.158260466 | 0.427827859 |
| 0.158251511 | 0.427827859 |
| 0.158244238 | 0.427827859 |
| 0.158230299 | 0.427819157 |
| 0.158226032 | 0.427819157 |
| 0.158179939 | 0.427755845 |
| 0.158172823 | 0.427755845 |
| 0.158130729 | 0.427671337 |
| 0.158114812 | 0.427652573 |
| 0.158109279 | 0.427652573 |
| 0.15808706 | 0.427639154 |

| | |
|---|---|
| 0.158085446 | 0.427639154 |
| 0.158070346 | 0.427639154 |
| 0.158066722 | 0.427639154 |
| 0.158017744 | 0.427632882 |
| 0.158012165 | 0.427632882 |
| 0.157989083 | 0.427603913 |
| 0.157931013 | 0.427495331 |
| 0.157898723 | 0.427475262 |
| 0.157894053 | 0.427475262 |
| 0.157890685 | 0.427475262 |
| 0.157888578 | 0.427475262 |
| 0.15785705 | 0.427465189 |
| 0.157753627 | 0.427282302 |
| 0.157730853 | 0.427244914 |
| 0.157725386 | 0.427244914 |
| 0.157706144 | 0.42722658 |
| 0.157698908 | 0.42722658 |
| 0.157684233 | 0.42722658 |
| 0.157680348 | 0.42722658 |
| 0.157669535 | 0.42722658 |
| 0.15766716 | 0.42722658 |
| 0.157642835 | 0.42722658 |
| 0.157641832 | 0.42722658 |
| 0.157614424 | 0.42722658 |
| 0.157609933 | 0.42722658 |
| 0.157597564 | 0.42722658 |
| 0.157532369 | 0.427168875 |
| 0.157509061 | 0.427129991 |
| 0.157480372 | 0.427091725 |
| 0.157480131 | 0.427091725 |
| 0.157479809 | 0.427091725 |
| 0.157353261 | 0.426950593 |
| 0.157304547 | 0.426891376 |
| 0.157295061 | 0.426889958 |
| 0.157292022 | 0.426889958 |
| 0.157279966 | 0.426889958 |
| 0.157238977 | 0.426810712 |
| 0.157230213 | 0.426810712 |
| 0.157228396 | 0.426810712 |
| 0.157222594 | 0.426810712 |
| 0.157203419 | 0.426810712 |
| 0.157167432 | 0.426762444 |
| 0.157155264 | 0.42675723 |
| 0.157135249 | 0.42675723 |
| 0.15712381 | 0.42675723 |

| | |
|---|---|
| 0.157111617 | 0.42675723 |
| 0.157108514 | 0.42675723 |
| 0.157083751 | 0.426754196 |
| 0.157016833 | 0.426596731 |
| 0.156970496 | 0.426541005 |
| 0.156948052 | 0.426527222 |
| 0.156928901 | 0.42650381 |
| 0.156851228 | 0.426414376 |
| 0.156826222 | 0.426395071 |
| 0.156821453 | 0.426395071 |
| 0.156814244 | 0.426395071 |
| 0.15678717 | 0.426386258 |
| 0.156749054 | 0.426331289 |
| 0.156733135 | 0.426320813 |
| 0.156694291 | 0.42625537 |
| 0.1566932 | 0.42625537 |
| 0.15665414 | 0.426199102 |
| 0.156601876 | 0.426101323 |
| 0.156592346 | 0.426099734 |
| 0.156544385 | 0.425993569 |
| 0.156540488 | 0.425993569 |
| 0.156537604 | 0.425993569 |
| 0.156528809 | 0.425993569 |
| 0.156509626 | 0.425993569 |
| 0.156509626 | 0.425993569 |
| 0.156494543 | 0.425993569 |
| 0.156447642 | 0.425900647 |
| 0.15639907 | 0.425841443 |
| 0.156359145 | 0.425757075 |
| 0.156357386 | 0.425757075 |
| 0.156312314 | 0.425702573 |
| 0.156285468 | 0.4256538 |
| 0.156223158 | 0.425532762 |
| 0.156195847 | 0.425495476 |
| 0.156187382 | 0.425495476 |
| 0.156113821 | 0.425405249 |
| 0.156101474 | 0.425395944 |
| 0.156061858 | 0.425312322 |
| 0.156040894 | 0.425279523 |
| 0.156040238 | 0.425279523 |
| 0.156030191 | 0.425279523 |
| 0.156007437 | 0.425261344 |
| 0.155998712 | 0.425261344 |
| 0.15596806 | 0.425202678 |
| 0.155949513 | 0.425176452 |

| | |
|---|---|
| 0.155936307 | 0.425169252 |
| 0.155904611 | 0.425146729 |
| 0.155846637 | 0.425017624 |
| 0.155809051 | 0.424939454 |
| 0.155804747 | 0.424939454 |
| 0.155779401 | 0.424907253 |
| 0.15574283 | 0.424856165 |
| 0.155669129 | 0.424698994 |
| 0.155595859 | 0.424552505 |
| 0.155586701 | 0.424552505 |
| 0.155512575 | 0.424465887 |
| 0.15546197 | 0.424381684 |
| 0.155444551 | 0.424369515 |
| 0.155431147 | 0.424369515 |
| 0.155429934 | 0.424369515 |
| 0.155355458 | 0.424261147 |
| 0.155291096 | 0.424158343 |
| 0.155281366 | 0.424156092 |
| 0.155238243 | 0.424062621 |
| 0.155198314 | 0.424002187 |
| 0.15515806 | 0.423916527 |
| 0.155142592 | 0.423898585 |
| 0.155131058 | 0.423891389 |
| 0.155128754 | 0.423891389 |
| 0.155117469 | 0.423891389 |
| 0.155103718 | 0.423891389 |
| 0.155097955 | 0.423891389 |
| 0.155025329 | 0.423772676 |
| 0.155021939 | 0.423772676 |
| 0.155021331 | 0.423772676 |
| 0.155015531 | 0.423772676 |
| 0.154915333 | 0.423617878 |
| 0.154817083 | 0.423446461 |
| 0.15481541 | 0.423446461 |
| 0.154791595 | 0.423446461 |
| 0.154790622 | 0.423446461 |
| 0.154752436 | 0.423415539 |
| 0.154707859 | 0.423342211 |
| 0.154685136 | 0.423328679 |
| 0.15464113 | 0.423281213 |
| 0.154583598 | 0.423148056 |
| 0.154572126 | 0.423140972 |
| 0.154548363 | 0.423100746 |
| 0.154545569 | 0.423100746 |
| 0.154461283 | 0.422983413 |

| | |
|---|---|
| 0.154425243 | 0.422909036 |
| 0.154387088 | 0.422828859 |
| 0.154322375 | 0.422700238 |
| 0.154297854 | 0.422700238 |
| 0.154282614 | 0.422700238 |
| 0.154265415 | 0.422690077 |
| 0.154263352 | 0.422690077 |
| 0.1542347 | 0.422678869 |
| 0.15420442 | 0.422620206 |
| 0.154143691 | 0.422502393 |
| 0.154141 | 0.422502393 |
| 0.154117883 | 0.422502393 |
| 0.15409054 | 0.422453957 |
| 0.154083955 | 0.422453957 |
| 0.154059257 | 0.422416823 |
| 0.154043524 | 0.422398004 |
| 0.154039378 | 0.422398004 |
| 0.153967299 | 0.422276925 |
| 0.153904903 | 0.422163728 |
| 0.153885658 | 0.422135252 |
| 0.153860163 | 0.422089625 |
| 0.15385799 | 0.422089625 |
| 0.153790745 | 0.421972106 |
| 0.153711204 | 0.42180246 |
| 0.15370468 | 0.42180246 |
| 0.153635877 | 0.421692942 |
| 0.153615448 | 0.42166117 |
| 0.153611547 | 0.42166117 |
| 0.153570505 | 0.421586399 |
| 0.153375549 | 0.42116602 |
| 0.153368783 | 0.42116602 |
| 0.153356647 | 0.42116602 |
| 0.153350662 | 0.42116602 |
| 0.153347017 | 0.42116602 |
| 0.153342786 | 0.42116602 |
| 0.153333183 | 0.42116602 |
| 0.153330344 | 0.42116602 |
| 0.153302112 | 0.42116602 |
| 0.153294001 | 0.42116602 |
| 0.153249409 | 0.421093268 |
| 0.153192657 | 0.420961613 |
| 0.153187456 | 0.420961613 |
| 0.153183988 | 0.420961613 |
| 0.153165339 | 0.420959409 |
| 0.153116898 | 0.420923433 |

| | |
|---|---|
| 0.153068813 | 0.420815536 |
| 0.153061333 | 0.420815536 |
| 0.153059382 | 0.420815536 |
| 0.152956879 | 0.420629209 |
| 0.152952117 | 0.420629209 |
| 0.152938883 | 0.420629209 |
| 0.152923987 | 0.420629209 |
| 0.152899665 | 0.420593297 |
| 0.152898508 | 0.420593297 |
| 0.152786056 | 0.420353619 |
| 0.152764541 | 0.420318706 |
| 0.152730719 | 0.420249924 |
| 0.152722306 | 0.420249924 |
| 0.152721458 | 0.420249924 |
| 0.152691309 | 0.420214318 |
| 0.152651456 | 0.420177478 |
| 0.152648654 | 0.420177478 |
| 0.152540322 | 0.419955813 |
| 0.152503005 | 0.419914473 |
| 0.152458012 | 0.419839128 |
| 0.15239809 | 0.419722654 |
| 0.152380546 | 0.419698604 |
| 0.152368947 | 0.419690927 |
| 0.152348728 | 0.419659506 |
| 0.15232954 | 0.419630922 |
| 0.152323639 | 0.419630922 |
| 0.152243749 | 0.41944311 |
| 0.152164661 | 0.419249469 |
| 0.152103445 | 0.41910505 |
| 0.152083131 | 0.419097568 |
| 0.152069404 | 0.41908399 |
| 0.152046162 | 0.419044185 |
| 0.152025037 | 0.419024443 |
| 0.152016119 | 0.419024443 |
| 0.152004227 | 0.419024443 |
| 0.151962302 | 0.418958526 |
| 0.151949162 | 0.418947847 |
| 0.151920581 | 0.418929744 |
| 0.151869092 | 0.418919799 |
| 0.15182673 | 0.418827203 |
| 0.151826152 | 0.418827203 |
| 0.151816365 | 0.418827203 |
| 0.151720522 | 0.418606957 |
| 0.15171718 | 0.418606957 |
| 0.151666098 | 0.418505288 |

| | |
|---|---|
| 0.151617529 | 0.418395506 |
| 0.151613772 | 0.418395506 |
| 0.151607847 | 0.418395506 |
| 0.151599583 | 0.418395506 |
| 0.151591921 | 0.418395506 |
| 0.151554514 | 0.418342811 |
| 0.151509163 | 0.418290353 |
| 0.151503569 | 0.418290353 |
| 0.151464245 | 0.41822316 |
| 0.151447002 | 0.41822316 |
| 0.151421025 | 0.418216723 |
| 0.151413228 | 0.418216723 |
| 0.151361941 | 0.418126278 |
| 0.151325999 | 0.418051238 |
| 0.151203789 | 0.417786325 |
| 0.151094078 | 0.417507407 |
| 0.150988132 | 0.417306556 |
| 0.150974712 | 0.417298624 |
| 0.150971944 | 0.417298624 |
| 0.150950154 | 0.41727918 |
| 0.150946169 | 0.41727918 |
| 0.150931718 | 0.417276655 |
| 0.150922454 | 0.417275265 |
| 0.150863642 | 0.417136876 |
| 0.150856889 | 0.417136876 |
| 0.150794043 | 0.41700811 |
| 0.150756178 | 0.416936558 |
| 0.150681562 | 0.416778602 |
| 0.150660892 | 0.416745633 |
| 0.150641468 | 0.416721795 |
| 0.150593805 | 0.416632654 |
| 0.150558507 | 0.416583398 |
| 0.150551354 | 0.416583398 |
| 0.150455555 | 0.416371104 |
| 0.150448162 | 0.416371104 |
| 0.150413138 | 0.416325436 |
| 0.150398583 | 0.416310202 |
| 0.150398279 | 0.416310202 |
| 0.150389094 | 0.416310202 |
| 0.150346853 | 0.41626378 |
| 0.150343501 | 0.41626378 |
| 0.150334037 | 0.41626378 |
| 0.150325504 | 0.41626378 |
| 0.150318452 | 0.41626378 |
| 0.150308086 | 0.41626378 |

| | |
|---|---|
| 0.1503061 | 0.41626378 |
| 0.1502813 | 0.41626378 |
| 0.150269662 | 0.41626378 |
| 0.150266706 | 0.41626378 |
| 0.150179306 | 0.416187003 |
| 0.150149154 | 0.416187003 |
| 0.150148244 | 0.416187003 |
| 0.150131674 | 0.416187003 |
| 0.15009152 | 0.41616166 |
| 0.150049087 | 0.416110162 |
| 0.150041646 | 0.416110162 |
| 0.150026858 | 0.416103493 |
| 0.14999781 | 0.416047154 |
| 0.149982322 | 0.416028424 |
| 0.149966601 | 0.416028424 |
| 0.149959789 | 0.416028424 |
| 0.149900722 | 0.415898968 |
| 0.149891124 | 0.415898968 |
| 0.149802754 | 0.41579671 |
| 0.149793956 | 0.415796523 |
| 0.14969362 | 0.415566453 |
| 0.149654733 | 0.415482715 |
| 0.149639273 | 0.41546401 |
| 0.149616393 | 0.415424704 |
| 0.14960371 | 0.415413728 |
| 0.149595216 | 0.415413728 |
| 0.149588405 | 0.415413728 |
| 0.14956611 | 0.415406176 |
| 0.149559051 | 0.415406176 |
| 0.149551666 | 0.415406176 |
| 0.149518793 | 0.415384496 |
| 0.149498131 | 0.415362705 |
| 0.149488619 | 0.415360507 |
| 0.149467618 | 0.415350618 |
| 0.149445221 | 0.415312613 |
| 0.149355978 | 0.4151615 |
| 0.149265186 | 0.415030241 |
| 0.149249386 | 0.415010539 |
| 0.149167073 | 0.414830093 |
| 0.149149799 | 0.414806275 |
| 0.149138792 | 0.414799886 |
| 0.149089429 | 0.41468681 |
| 0.149058833 | 0.414650138 |
| 0.149019838 | 0.414565878 |
| 0.149007845 | 0.414556729 |

| | |
|---|---|
| 0.149001155 | 0.414556729 |
| 0.148997955 | 0.414556729 |
| 0.148976649 | 0.414556729 |
| 0.148966521 | 0.414556729 |
| 0.14896437 | 0.414556729 |
| 0.148963419 | 0.414556729 |
| 0.148955905 | 0.414556729 |
| 0.148919228 | 0.414528117 |
| 0.148857562 | 0.414404906 |
| 0.148825925 | 0.414389497 |
| 0.148720933 | 0.414169798 |
| 0.148716625 | 0.414169798 |
| 0.148707868 | 0.414169798 |
| 0.148595166 | 0.413916363 |
| 0.148592166 | 0.413916363 |
| 0.148522478 | 0.413782807 |
| 0.148494237 | 0.413756217 |
| 0.148492876 | 0.413756217 |
| 0.148467815 | 0.413731006 |
| 0.148465203 | 0.413731006 |
| 0.148378753 | 0.41360382 |
| 0.148334161 | 0.413527927 |
| 0.148320357 | 0.413513648 |
| 0.148263652 | 0.413379753 |
| 0.148254627 | 0.413378791 |
| 0.148248794 | 0.413378791 |
| 0.148235237 | 0.413378791 |
| 0.148232325 | 0.413378791 |
| 0.14820106 | 0.413378791 |
| 0.148182107 | 0.413370188 |
| 0.148102194 | 0.413206097 |
| 0.148068333 | 0.413149597 |
| 0.147997669 | 0.412976621 |
| 0.147983105 | 0.412960177 |
| 0.147978353 | 0.412960177 |
| 0.147971691 | 0.412960177 |
| 0.147952441 | 0.412947193 |
| 0.147928355 | 0.412928366 |
| 0.147905881 | 0.412889831 |
| 0.147896684 | 0.412888357 |
| 0.147766 | 0.412547704 |
| 0.14773208 | 0.412501364 |
| 0.147702012 | 0.412441589 |
| 0.147700996 | 0.412441589 |
| 0.147668645 | 0.412420957 |

| | |
|---|---|
| 0.147663251 | 0.412420957 |
| 0.147653953 | 0.412420957 |
| 0.14764677 | 0.412420957 |
| 0.147612026 | 0.412359551 |
| 0.147540309 | 0.412183381 |
| 0.147539211 | 0.412183381 |
| 0.147499297 | 0.412141333 |
| 0.147439344 | 0.412022157 |
| 0.147437224 | 0.412022157 |
| 0.147403825 | 0.41199542 |
| 0.147375856 | 0.411942073 |
| 0.147348278 | 0.411912687 |
| 0.14734763 | 0.411912687 |
| 0.147336972 | 0.411912687 |
| 0.147333899 | 0.411912687 |
| 0.147330412 | 0.411912687 |
| 0.14731415 | 0.411912687 |
| 0.147228096 | 0.411818437 |
| 0.147216158 | 0.411809229 |
| 0.147207555 | 0.411809229 |
| 0.147179821 | 0.411802398 |
| 0.147138121 | 0.411736035 |
| 0.147076193 | 0.411611108 |
| 0.147059794 | 0.411589401 |
| 0.147051619 | 0.411589401 |
| 0.147048443 | 0.411589401 |
| 0.146988176 | 0.4115324 |
| 0.146978263 | 0.4115324 |
| 0.146939684 | 0.411470857 |
| 0.146889681 | 0.411379209 |
| 0.146873803 | 0.411358932 |
| 0.146865044 | 0.411358588 |
| 0.146864757 | 0.411358588 |
| 0.146847005 | 0.411356447 |
| 0.146840985 | 0.411356447 |
| 0.146840939 | 0.411356447 |
| 0.146836338 | 0.411356447 |
| 0.146800203 | 0.411322113 |
| 0.146783373 | 0.411299154 |
| 0.146780572 | 0.411299154 |
| 0.146762444 | 0.411288903 |
| 0.146653011 | 0.41107898 |
| 0.146600434 | 0.41096475 |
| 0.146592051 | 0.41096475 |
| 0.146544584 | 0.410895968 |

| | |
|---|---|
| 0.146465222 | 0.410721804 |
| 0.146404846 | 0.410600848 |
| 0.146353723 | 0.410505816 |
| 0.146345163 | 0.410505816 |
| 0.146323427 | 0.410469187 |
| 0.146275204 | 0.410382253 |
| 0.146256381 | 0.410382253 |
| 0.146254337 | 0.410382253 |
| 0.14623047 | 0.41037762 |
| 0.146223499 | 0.41037762 |
| 0.146218548 | 0.41037762 |
| 0.146217093 | 0.41037762 |
| 0.146205423 | 0.41037762 |
| 0.146135117 | 0.410351819 |
| 0.146129089 | 0.410351819 |
| 0.146088864 | 0.410294513 |
| 0.146071693 | 0.410270481 |
| 0.1460697 | 0.410270481 |
| 0.145980364 | 0.41009147 |
| 0.145920846 | 0.409967682 |
| 0.145911651 | 0.409966031 |
| 0.145909621 | 0.409966031 |
| 0.145904725 | 0.409966031 |
| 0.145845438 | 0.409852529 |
| 0.145822997 | 0.409813648 |
| 0.14580865 | 0.409797507 |
| 0.145745081 | 0.409676457 |
| 0.145717728 | 0.409638663 |
| 0.145710456 | 0.409638663 |
| 0.145696037 | 0.409626044 |
| 0.145686073 | 0.409622209 |
| 0.145664598 | 0.409610192 |
| 0.145658492 | 0.409610192 |
| 0.145628192 | 0.409580364 |
| 0.145621766 | 0.409580364 |
| 0.145605366 | 0.40956454 |
| 0.145572927 | 0.409497479 |
| 0.145521333 | 0.409384613 |
| 0.145517458 | 0.409384613 |
| 0.145496515 | 0.409379246 |
| 0.1454635 | 0.409310535 |
| 0.145349462 | 0.409062153 |
| 0.145283083 | 0.408899503 |
| 0.145205512 | 0.408753644 |
| 0.145204985 | 0.408753644 |

| | |
|---|---|
| 0.145183448 | 0.408753644 |
| 0.145172106 | 0.408753644 |
| 0.145170005 | 0.408753644 |
| 0.145111986 | 0.408635283 |
| 0.145105416 | 0.408635283 |
| 0.145055196 | 0.408571991 |
| 0.145014733 | 0.408506339 |
| 0.145011661 | 0.408506339 |
| 0.144992334 | 0.408506339 |
| 0.144985114 | 0.408506339 |
| 0.144974069 | 0.408506339 |
| 0.144889378 | 0.408322272 |
| 0.144868723 | 0.408312385 |
| 0.144855136 | 0.408298253 |
| 0.144845272 | 0.408294613 |
| 0.144844624 | 0.408294613 |
| 0.144843822 | 0.408294613 |
| 0.144819348 | 0.408294031 |
| 0.144808477 | 0.408287553 |
| 0.144759701 | 0.408222528 |
| 0.144704472 | 0.408115111 |
| 0.144693789 | 0.408109148 |
| 0.144666923 | 0.408077171 |
| 0.144664977 | 0.408077171 |
| 0.144644204 | 0.408065957 |
| 0.144643752 | 0.408065957 |
| 0.144641492 | 0.408065957 |
| 0.144638781 | 0.408065957 |
| 0.144615895 | 0.408065957 |
| 0.144595452 | 0.408049275 |
| 0.144545817 | 0.407981724 |
| 0.144531375 | 0.407965138 |
| 0.144518435 | 0.407952792 |
| 0.144442295 | 0.40783455 |
| 0.144415515 | 0.407783111 |
| 0.144368722 | 0.407699326 |
| 0.144329226 | 0.407611958 |
| 0.144321645 | 0.407611958 |
| 0.144311719 | 0.407610856 |
| 0.144250236 | 0.40746136 |
| 0.14423273 | 0.407436075 |
| 0.144223917 | 0.407436075 |
| 0.144223515 | 0.407436075 |
| 0.144205789 | 0.407436075 |
| 0.144202712 | 0.407436075 |

| | |
|---|---|
| 0.144197915 | 0.407436075 |
| 0.144191145 | 0.407436075 |
| 0.144129189 | 0.407385216 |
| 0.144091239 | 0.407326297 |
| 0.144059083 | 0.407259569 |
| 0.144005601 | 0.40713254 |
| 0.143957278 | 0.407068412 |
| 0.143931639 | 0.407020078 |
| 0.143922934 | 0.407019631 |
| 0.143918132 | 0.407019631 |
| 0.143914419 | 0.407019631 |
| 0.143899225 | 0.407019631 |
| 0.143859915 | 0.406962235 |
| 0.143843714 | 0.406962235 |
| 0.143835528 | 0.406962235 |
| 0.143830213 | 0.406962235 |
| 0.143806905 | 0.406957306 |
| 0.143777241 | 0.406897536 |
| 0.143765563 | 0.406888665 |
| 0.143727644 | 0.40680552 |
| 0.143722915 | 0.40680552 |
| 0.143705158 | 0.406790226 |
| 0.143607133 | 0.406585241 |
| 0.14344454 | 0.406276106 |
| 0.143403424 | 0.406226918 |
| 0.143321991 | 0.406091765 |
| 0.143320866 | 0.406091765 |
| 0.143297282 | 0.406086176 |
| 0.143277988 | 0.406068156 |
| 0.143255204 | 0.406047467 |
| 0.143251717 | 0.406047467 |
| 0.143246734 | 0.406047467 |
| 0.143210457 | 0.40601729 |
| 0.14318155 | 0.405959499 |
| 0.14313447 | 0.405874339 |
| 0.143117866 | 0.40585142 |
| 0.143117136 | 0.40585142 |
| 0.143048935 | 0.405752583 |
| 0.143001515 | 0.405642236 |
| 0.142984697 | 0.405618688 |
| 0.142971864 | 0.405606442 |
| 0.142937049 | 0.40556388 |
| 0.142927907 | 0.40556388 |
| 0.142913727 | 0.405562298 |
| 0.142905954 | 0.405562298 |

| | |
|---|---|
| 0.142902799 | 0.405562298 |
| 0.142902799 | 0.405562298 |
| 0.142889471 | 0.405562298 |
| 0.142830851 | 0.405472071 |
| 0.14279013 | 0.405404801 |
| 0.142729476 | 0.405280913 |
| 0.142722151 | 0.405280913 |
| 0.142701233 | 0.405280913 |
| 0.142675056 | 0.405271361 |
| 0.142660448 | 0.405254034 |
| 0.142590253 | 0.405102957 |
| 0.14257759 | 0.405091146 |
| 0.142557282 | 0.405081017 |
| 0.142538462 | 0.405081017 |
| 0.142494999 | 0.40500145 |
| 0.142484361 | 0.404995385 |
| 0.14244121 | 0.404896898 |
| 0.14243108 | 0.404892267 |
| 0.142422102 | 0.404890911 |
| 0.142409571 | 0.404879455 |
| 0.142389274 | 0.404845918 |
| 0.142380125 | 0.404844072 |
| 0.142296853 | 0.404655613 |
| 0.142243669 | 0.404552682 |
| 0.142239132 | 0.404552682 |
| 0.142163475 | 0.4043729 |
| 0.142137256 | 0.40434662 |
| 0.142115319 | 0.404320385 |
| 0.14208322 | 0.404265339 |
| 0.142076901 | 0.404265339 |
| 0.142037922 | 0.404208898 |
| 0.142027715 | 0.404204004 |
| 0.141982115 | 0.404098373 |
| 0.141977919 | 0.404098373 |
| 0.141973501 | 0.404098373 |
| 0.141842857 | 0.403822689 |
| 0.141818668 | 0.4038021 |
| 0.14181114 | 0.4038021 |
| 0.141748108 | 0.403649454 |
| 0.141740191 | 0.403649454 |
| 0.141712594 | 0.403596587 |
| 0.141640986 | 0.403416771 |
| 0.141633161 | 0.403416771 |
| 0.14161912 | 0.40340274 |
| 0.141609825 | 0.40340039 |

| | |
|---|---|
| 0.141609265 | 0.40340039 |
| 0.141490847 | 0.403182037 |
| 0.141486869 | 0.403182037 |
| 0.141459991 | 0.403142353 |
| 0.141411163 | 0.403027316 |
| 0.141396801 | 0.403010499 |
| 0.141393787 | 0.403010499 |
| 0.141392076 | 0.403010499 |
| 0.141359389 | 0.403010499 |
| 0.1413178 | 0.402930009 |
| 0.141311585 | 0.402930009 |
| 0.141303734 | 0.402930009 |
| 0.141293081 | 0.402930009 |
| 0.141290613 | 0.402930009 |
| 0.141267131 | 0.40290614 |
| 0.141148731 | 0.402640789 |
| 0.141120937 | 0.402594994 |
| 0.141118949 | 0.402594994 |
| 0.141094331 | 0.402582056 |
| 0.141057679 | 0.402501591 |
| 0.141001532 | 0.402389597 |
| 0.140978783 | 0.402372895 |
| 0.140953974 | 0.402326196 |
| 0.140946594 | 0.402326196 |
| 0.140938019 | 0.402326196 |
| 0.140934634 | 0.402326196 |
| 0.140904056 | 0.402326196 |
| 0.140884864 | 0.402321826 |
| 0.140853575 | 0.402256593 |
| 0.140834525 | 0.40225043 |
| 0.140775948 | 0.402155476 |
| 0.140735528 | 0.402064123 |
| 0.140721853 | 0.402049172 |
| 0.140717852 | 0.402049172 |
| 0.140714716 | 0.402049172 |
| 0.140668182 | 0.40196817 |
| 0.140659263 | 0.401966802 |
| 0.140615475 | 0.401871144 |
| 0.140597472 | 0.401862558 |
| 0.140553706 | 0.401785688 |
| 0.140552537 | 0.401785688 |
| 0.140551133 | 0.401785688 |
| 0.140547077 | 0.401785688 |
| 0.140538538 | 0.401785688 |
| 0.140466713 | 0.401705785 |

| | |
|---|---|
| 0.140442927 | 0.401661881 |
| 0.140431418 | 0.401653084 |
| 0.140427684 | 0.401653084 |
| 0.140403755 | 0.401646322 |
| 0.140381846 | 0.401607771 |
| 0.140376965 | 0.401607771 |
| 0.140345405 | 0.401575884 |
| 0.140255952 | 0.401440518 |
| 0.140243029 | 0.401427653 |
| 0.140238948 | 0.401427653 |
| 0.140231985 | 0.401427653 |
| 0.140227998 | 0.401427653 |
| 0.140208351 | 0.401424895 |
| 0.140201409 | 0.401424895 |
| 0.140174772 | 0.401377013 |
| 0.140124194 | 0.40128044 |
| 0.140110811 | 0.40126624 |
| 0.140046776 | 0.401155218 |
| 0.140002438 | 0.401052336 |
| 0.140000808 | 0.401052336 |
| 0.139958426 | 0.400974491 |
| 0.139918035 | 0.400907007 |
| 0.139890722 | 0.400876979 |
| 0.139862409 | 0.400819962 |
| 0.139860182 | 0.400819962 |
| 0.139812915 | 0.400766733 |
| 0.139797401 | 0.400754232 |
| 0.139791372 | 0.400754232 |
| 0.139771785 | 0.400748501 |
| 0.139749164 | 0.400712429 |
| 0.139728366 | 0.40070104 |
| 0.13971387 | 0.400683596 |
| 0.1396838 | 0.400668598 |
| 0.139680867 | 0.400668598 |
| 0.139627048 | 0.400627592 |
| 0.139626623 | 0.400627592 |
| 0.139589076 | 0.400591042 |
| 0.139577789 | 0.400582787 |
| 0.139569078 | 0.400581925 |
| 0.139511397 | 0.400488777 |
| 0.139505636 | 0.400488777 |
| 0.139500613 | 0.400488777 |
| 0.139499383 | 0.400488777 |
| 0.139473305 | 0.400477077 |
| 0.1394545 | 0.400477077 |

| | |
|---|---|
| 0.139429433 | 0.400446577 |
| 0.13942891 | 0.400446577 |
| 0.139409685 | 0.400438156 |
| 0.13939468 | 0.400419207 |
| 0.139373566 | 0.400382702 |
| 0.139345373 | 0.400325859 |
| 0.139327303 | 0.400298092 |
| 0.139324059 | 0.400298092 |
| 0.139305933 | 0.400298092 |
| 0.139304615 | 0.400298092 |
| 0.139292907 | 0.400298092 |
| 0.139286321 | 0.400298092 |
| 0.139286077 | 0.400298092 |
| 0.139284126 | 0.400298092 |
| 0.139279443 | 0.400298092 |
| 0.139275784 | 0.400298092 |
| 0.139248567 | 0.400298092 |
| 0.139186774 | 0.400231561 |
| 0.139183779 | 0.400231561 |
| 0.139173953 | 0.400231561 |
| 0.139147464 | 0.400231561 |
| 0.139139413 | 0.400231561 |
| 0.139124669 | 0.400222916 |
| 0.139124187 | 0.400222916 |
| 0.139118737 | 0.400222916 |
| 0.139079941 | 0.400190933 |
| 0.139052656 | 0.400160776 |
| 0.139041532 | 0.400152943 |
| 0.139027508 | 0.400136761 |
| 0.138956136 | 0.400003861 |
| 0.138940309 | 0.399982478 |
| 0.138938054 | 0.399982478 |
| 0.138927783 | 0.399982478 |
| 0.138909286 | 0.399979253 |
| 0.138873347 | 0.399934751 |
| 0.138861642 | 0.399925225 |
| 0.138850938 | 0.399918578 |
| 0.138742117 | 0.39970184 |
| 0.138728714 | 0.39969385 |
| 0.138699829 | 0.399676715 |
| 0.138571678 | 0.399355762 |
| 0.13856622 | 0.399355762 |
| 0.138562108 | 0.399355762 |
| 0.138543382 | 0.399346708 |
| 0.138531859 | 0.399337663 |

| | |
|---|---|
| 0.138523589 | 0.399337663 |
| 0.138514147 | 0.399334943 |
| 0.138504458 | 0.399331184 |
| 0.13848044 | 0.399286107 |
| 0.138443959 | 0.399205089 |
| 0.138437808 | 0.399205089 |
| 0.138334604 | 0.398962224 |
| 0.138305864 | 0.398903494 |
| 0.138297227 | 0.398902739 |
| 0.13828171 | 0.398882139 |
| 0.138280926 | 0.398882139 |
| 0.138247198 | 0.398830898 |
| 0.138224493 | 0.398791065 |
| 0.138204422 | 0.39877996 |
| 0.138160103 | 0.398676234 |
| 0.138127557 | 0.398606468 |
| 0.138123671 | 0.398606468 |
| 0.138115368 | 0.398606468 |
| 0.138090195 | 0.3985711 |
| 0.138053888 | 0.39851461 |
| 0.138033395 | 0.398479604 |
| 0.138027622 | 0.398479604 |
| 0.137988425 | 0.398446388 |
| 0.137981765 | 0.398446388 |
| 0.137937678 | 0.39837231 |
| 0.137936684 | 0.39837231 |
| 0.137932706 | 0.39837231 |
| 0.13792159 | 0.39837231 |
| 0.13791101 | 0.39837231 |
| 0.137806917 | 0.398139523 |
| 0.137775514 | 0.398072942 |
| 0.137746732 | 0.398013931 |
| 0.137736746 | 0.398013931 |
| 0.137728913 | 0.398013931 |
| 0.137724526 | 0.398013931 |
| 0.137717192 | 0.398013931 |
| 0.137680652 | 0.39796786 |
| 0.137651983 | 0.39790914 |
| 0.137648525 | 0.39790914 |
| 0.137623909 | 0.397900437 |
| 0.13761365 | 0.397894932 |
| 0.13752979 | 0.397745331 |
| 0.137523703 | 0.397745331 |
| 0.137514863 | 0.397745331 |
| 0.137507473 | 0.397745331 |

| | |
|---|---|
| 0.137497282 | 0.397745331 |
| 0.137478198 | 0.397744755 |
| 0.137471441 | 0.397744755 |
| 0.137455237 | 0.397726644 |
| 0.13734836 | 0.39746569 |
| 0.137312242 | 0.397388783 |
| 0.137267995 | 0.39735219 |
| 0.137209788 | 0.397233638 |
| 0.137203967 | 0.397233638 |
| 0.137193152 | 0.397233638 |
| 0.13716972 | 0.397190068 |
| 0.137167984 | 0.397190068 |
| 0.13713489 | 0.397187278 |
| 0.137061644 | 0.397081748 |
| 0.137057222 | 0.397081748 |
| 0.137047256 | 0.397081748 |
| 0.137017484 | 0.39706308 |
| 0.137017152 | 0.39706308 |
| 0.136948686 | 0.396936164 |
| 0.136944452 | 0.396936164 |
| 0.13692474 | 0.396936164 |
| 0.136922425 | 0.396936164 |
| 0.13692221 | 0.396936164 |
| 0.136897869 | 0.396933813 |
| 0.136858729 | 0.396868648 |
| 0.136831588 | 0.396814105 |
| 0.136826929 | 0.396814105 |
| 0.136714983 | 0.396572538 |
| 0.136707339 | 0.396572538 |
| 0.136684265 | 0.396555889 |
| 0.136667389 | 0.396531084 |
| 0.136657219 | 0.396525733 |
| 0.136631977 | 0.396500803 |
| 0.136624271 | 0.396500803 |
| 0.13658938 | 0.396425496 |
| 0.136568979 | 0.396392903 |
| 0.136562733 | 0.396392903 |
| 0.136561702 | 0.396392903 |
| 0.136539506 | 0.396392903 |
| 0.136535183 | 0.396392903 |
| 0.136534948 | 0.396392903 |
| 0.136505915 | 0.396392903 |
| 0.136479278 | 0.396347473 |
| 0.136392706 | 0.396120215 |
| 0.136362961 | 0.39605798 |

| | |
|---|---|
| 0.136342456 | 0.396022574 |
| 0.136250344 | 0.395779161 |
| 0.136212674 | 0.395693873 |
| 0.136202599 | 0.395688738 |
| 0.136163525 | 0.395599353 |
| 0.136160293 | 0.395599353 |
| 0.136157799 | 0.395599353 |
| 0.136088695 | 0.395454319 |
| 0.13608645 | 0.395454319 |
| 0.136041267 | 0.395364744 |
| 0.135999007 | 0.395266047 |
| 0.135990674 | 0.395265947 |
| 0.135986784 | 0.395265947 |
| 0.135983645 | 0.395265947 |
| 0.135959179 | 0.395246771 |
| 0.135950736 | 0.39524635 |
| 0.135932573 | 0.395219793 |
| 0.135909228 | 0.395212819 |
| 0.135881476 | 0.395212819 |
| 0.135877663 | 0.395212819 |
| 0.13579122 | 0.395047855 |
| 0.135752347 | 0.394975941 |
| 0.135732541 | 0.39494952 |
| 0.135706765 | 0.394898643 |
| 0.135687253 | 0.394898643 |
| 0.135683721 | 0.394898643 |
| 0.13566438 | 0.394895943 |
| 0.135532704 | 0.394633252 |
| 0.13552701 | 0.394633252 |
| 0.1354908 | 0.394559483 |
| 0.135471778 | 0.394528211 |
| 0.13543366 | 0.394484633 |
| 0.135395011 | 0.394401113 |
| 0.135395011 | 0.394401113 |
| 0.13538581 | 0.394401113 |
| 0.135385794 | 0.394401113 |
| 0.135372277 | 0.394401113 |
| 0.135330311 | 0.394401113 |
| 0.135328853 | 0.394401113 |
| 0.135321102 | 0.394401113 |
| 0.135310675 | 0.394401113 |
| 0.135297221 | 0.394401113 |
| 0.135285631 | 0.394401113 |
| 0.135271321 | 0.394401113 |
| 0.135233015 | 0.394363386 |

| | |
|---|---|
| 0.135231652 | 0.394363386 |
| 0.135224764 | 0.394363386 |
| 0.135205512 | 0.394355621 |
| 0.135201409 | 0.394355621 |
| 0.135189922 | 0.394355621 |
| 0.135155743 | 0.394282896 |
| 0.135148307 | 0.394282896 |
| 0.135082397 | 0.394117208 |
| 0.135077399 | 0.394117208 |
| 0.135040972 | 0.394044625 |
| 0.135036629 | 0.394044625 |
| 0.13502424 | 0.394044625 |
| 0.135013355 | 0.394044625 |
| 0.134978927 | 0.393984272 |
| 0.134973232 | 0.393984272 |
| 0.13496442 | 0.393984272 |
| 0.134906393 | 0.393849672 |
| 0.134847033 | 0.39376822 |
| 0.134842934 | 0.39376822 |
| 0.134799049 | 0.393724659 |
| 0.134748111 | 0.393645109 |
| 0.134718437 | 0.393585756 |
| 0.1346718 | 0.393497782 |
| 0.134626375 | 0.393389187 |
| 0.134560039 | 0.393243601 |
| 0.134558755 | 0.393243601 |
| 0.134548147 | 0.393243601 |
| 0.134545027 | 0.393243601 |
| 0.134530468 | 0.393243601 |
| 0.134527966 | 0.393243601 |
| 0.134527436 | 0.393243601 |
| 0.134491413 | 0.393243601 |
| 0.134479254 | 0.393243601 |
| 0.134458768 | 0.393237196 |
| 0.134456215 | 0.393237196 |
| 0.134419913 | 0.393171847 |
| 0.134336394 | 0.392951692 |
| 0.134294374 | 0.392852907 |
| 0.134275639 | 0.392822227 |
| 0.13417922 | 0.392564269 |
| 0.13414272 | 0.392529819 |
| 0.134139681 | 0.392529819 |
| 0.134083854 | 0.392405783 |
| 0.134065948 | 0.39237749 |
| 0.134062702 | 0.39237749 |

| | |
|---|---|
| 0.134055664 | 0.39237749 |
| 0.134007212 | 0.392326129 |
| 0.133955425 | 0.392215673 |
| 0.133947331 | 0.392215673 |
| 0.133926457 | 0.392197266 |
| 0.133903354 | 0.392178772 |
| 0.133886278 | 0.392178772 |
| 0.133783637 | 0.391960708 |
| 0.133739556 | 0.391870232 |
| 0.133725251 | 0.391870232 |
| 0.133710907 | 0.391868153 |
| 0.133689352 | 0.391853208 |
| 0.133686768 | 0.391853208 |
| 0.133665107 | 0.391830371 |
| 0.133654048 | 0.391822071 |
| 0.13363677 | 0.391819652 |
| 0.133623859 | 0.39180592 |
| 0.133596359 | 0.391749401 |
| 0.133594437 | 0.391749401 |
| 0.133493359 | 0.391592029 |
| 0.133348877 | 0.391192298 |
| 0.13332033 | 0.391156741 |
| 0.133306792 | 0.391156741 |
| 0.133120672 | 0.390715368 |
| 0.133103108 | 0.390687892 |
| 0.133039728 | 0.390562064 |
| 0.132985753 | 0.390439706 |
| 0.132983957 | 0.390439706 |
| 0.132955335 | 0.390422574 |
| 0.132841708 | 0.390209202 |
| 0.132832794 | 0.390207085 |
| 0.132781014 | 0.390079037 |
| 0.132738958 | 0.389979541 |
| 0.13269206 | 0.38988986 |
| 0.132672723 | 0.389857093 |
| 0.132654914 | 0.389828813 |
| 0.132648026 | 0.389828813 |
| 0.132639006 | 0.389828813 |
| 0.132593771 | 0.389761313 |
| 0.132530588 | 0.389631748 |
| 0.132514183 | 0.389607567 |
| 0.13247236 | 0.38950865 |
| 0.132458844 | 0.389492954 |
| 0.132408299 | 0.389392407 |
| 0.132407267 | 0.389392407 |

| | |
|---|---|
| 0.132352652 | 0.389276833 |
| 0.132349206 | 0.389276833 |
| 0.132344845 | 0.389276833 |
| 0.132323039 | 0.389261851 |
| 0.132292789 | 0.389244989 |
| 0.132196248 | 0.389081122 |
| 0.132143631 | 0.388974333 |
| 0.132134497 | 0.388974333 |
| 0.132115387 | 0.388974333 |
| 0.132096982 | 0.388957216 |
| 0.132038754 | 0.388852676 |
| 0.132034809 | 0.388852676 |
| 0.132002459 | 0.388823125 |
| 0.131950751 | 0.388766991 |
| 0.131931968 | 0.388735701 |
| 0.131930064 | 0.388735701 |
| 0.131915485 | 0.388735701 |
| 0.131909952 | 0.388735701 |
| 0.131892692 | 0.388735701 |
| 0.13179417 | 0.388594128 |
| 0.131759688 | 0.388516509 |
| 0.131720822 | 0.388425955 |
| 0.131699805 | 0.388388025 |
| 0.131662078 | 0.388324857 |
| 0.131642496 | 0.388315197 |
| 0.131638056 | 0.388315197 |
| 0.131589895 | 0.38823217 |
| 0.131562005 | 0.388173933 |
| 0.131539591 | 0.388154434 |
| 0.131507571 | 0.388099497 |
| 0.131500664 | 0.388099497 |
| 0.131475209 | 0.388062083 |
| 0.131459693 | 0.388040333 |
| 0.131457094 | 0.388040333 |
| 0.131434038 | 0.388012703 |
| 0.131393338 | 0.387940643 |
| 0.131376531 | 0.387915067 |
| 0.131269193 | 0.387646195 |
| 0.131243544 | 0.387594486 |
| 0.131232259 | 0.387594486 |
| 0.131225091 | 0.387594486 |
| 0.131182047 | 0.387508983 |
| 0.131170121 | 0.387497789 |
| 0.131139293 | 0.387430748 |
| 0.1311153 | 0.387383897 |

| | |
|---|---|
| 0.131105466 | 0.387383897 |
| 0.131080282 | 0.387376557 |
| 0.131065136 | 0.387355832 |
| 0.131058146 | 0.387355832 |
| 0.131050973 | 0.387355832 |
| 0.131040972 | 0.387355832 |
| 0.131012948 | 0.387297726 |
| 0.130987876 | 0.387247645 |
| 0.13097276 | 0.387226993 |
| 0.130966774 | 0.387226993 |
| 0.130955329 | 0.387223534 |
| 0.130929753 | 0.387181784 |
| 0.130882299 | 0.387147043 |
| 0.130820408 | 0.387088939 |
| 0.13082034 | 0.387088939 |
| 0.13081192 | 0.387088939 |
| 0.130759584 | 0.386981093 |
| 0.130685102 | 0.386784699 |
| 0.130658414 | 0.386729743 |
| 0.130589208 | 0.386572951 |
| 0.130566535 | 0.386529858 |
| 0.130525148 | 0.386431356 |
| 0.130499401 | 0.386379147 |
| 0.130482996 | 0.386354595 |
| 0.130478821 | 0.386354595 |
| 0.130477594 | 0.386354595 |
| 0.130467644 | 0.386354595 |
| 0.130401402 | 0.386257079 |
| 0.130396322 | 0.386257079 |
| 0.130389229 | 0.386257079 |
| 0.13037852 | 0.386257079 |
| 0.130364057 | 0.386242547 |
| 0.130359607 | 0.386242547 |
| 0.130334265 | 0.386225507 |
| 0.130221232 | 0.385963445 |
| 0.13014687 | 0.385815075 |
| 0.130138096 | 0.385813077 |
| 0.130135423 | 0.385813077 |
| 0.130055089 | 0.385614997 |
| 0.130021124 | 0.385562297 |
| 0.130020853 | 0.385562297 |
| 0.129916737 | 0.385372724 |
| 0.12989725 | 0.385338917 |
| 0.12986565 | 0.385293174 |
| 0.129839373 | 0.38523921 |

| | |
|---|---|
| 0.129804297 | 0.385159133 |
| 0.12980342 | 0.385159133 |
| 0.129792516 | 0.385159133 |
| 0.129787159 | 0.385159133 |
| 0.129733502 | 0.385127372 |
| 0.129699688 | 0.385064645 |
| 0.12969153 | 0.38506443 |
| 0.129689336 | 0.38506443 |
| 0.129676383 | 0.38506443 |
| 0.129620616 | 0.38492587 |
| 0.129618263 | 0.38492587 |
| 0.129609862 | 0.38492587 |
| 0.129588227 | 0.384925687 |
| 0.129586051 | 0.384925687 |
| 0.129561298 | 0.384893707 |
| 0.129492404 | 0.38473704 |
| 0.129484808 | 0.38473704 |
| 0.129441262 | 0.384633079 |
| 0.129438244 | 0.384633079 |
| 0.129438202 | 0.384633079 |
| 0.129412711 | 0.384633079 |
| 0.129408374 | 0.384633079 |
| 0.129387912 | 0.384618531 |
| 0.129383956 | 0.384618531 |
| 0.129368658 | 0.384609306 |
| 0.12933262 | 0.384550175 |
| 0.129330683 | 0.384550175 |
| 0.129308152 | 0.384525433 |
| 0.12930277 | 0.384525433 |
| 0.129300863 | 0.384525433 |
| 0.129279142 | 0.384511192 |
| 0.129256485 | 0.384491826 |
| 0.129210075 | 0.384377779 |
| 0.129197646 | 0.384364813 |
| 0.129153512 | 0.384281523 |
| 0.129127373 | 0.384251757 |
| 0.129107753 | 0.384232575 |
| 0.129090656 | 0.384232575 |
| 0.129089552 | 0.384232575 |
| 0.129059237 | 0.384193029 |
| 0.129053398 | 0.384193029 |
| 0.129022841 | 0.384156717 |
| 0.12901477 | 0.384156701 |
| 0.129014206 | 0.384156701 |
| 0.129006548 | 0.384156701 |

| | |
|---|---|
| 0.128980384 | 0.384128536 |
| 0.128931643 | 0.384053247 |
| 0.128924891 | 0.384053247 |
| 0.128888577 | 0.383997014 |
| 0.128866959 | 0.383956626 |
| 0.128847564 | 0.383922857 |
| 0.128834489 | 0.383907916 |
| 0.128806329 | 0.383848018 |
| 0.128761783 | 0.383763294 |
| 0.128760853 | 0.383763294 |
| 0.128753384 | 0.383763294 |
| 0.128707989 | 0.383763294 |
| 0.128684951 | 0.383750431 |
| 0.12867715 | 0.383750431 |
| 0.128669854 | 0.383750431 |
| 0.128666691 | 0.383750431 |
| 0.128657978 | 0.383750431 |
| 0.1286148 | 0.383661347 |
| 0.128608842 | 0.383661347 |
| 0.128538718 | 0.383506453 |
| 0.128527283 | 0.383506453 |
| 0.12844961 | 0.383336648 |
| 0.128412737 | 0.383250622 |
| 0.128307145 | 0.382995673 |
| 0.128269228 | 0.382942299 |
| 0.12823775 | 0.382872323 |
| 0.128235263 | 0.382872323 |
| 0.128229658 | 0.382872323 |
| 0.128225814 | 0.382872323 |
| 0.128213447 | 0.382872323 |
| 0.128211125 | 0.382872323 |
| 0.128189917 | 0.382872323 |
| 0.12817098 | 0.382872323 |
| 0.128154633 | 0.382872323 |
| 0.128133141 | 0.382847999 |
| 0.128130058 | 0.382847999 |
| 0.128101663 | 0.382825996 |
| 0.128083018 | 0.382794296 |
| 0.127873047 | 0.382310709 |
| 0.127832292 | 0.382212854 |
| 0.127740388 | 0.382034 |
| 0.127685506 | 0.381893845 |
| 0.127622708 | 0.381729996 |
| 0.127605064 | 0.381722466 |
| 0.127589199 | 0.381701682 |

| | |
|---|---|
| 0.127588245 | 0.381701682 |
| 0.127572727 | 0.381700356 |
| 0.127541973 | 0.381632316 |
| 0.127516732 | 0.381620892 |
| 0.127514703 | 0.381620892 |
| 0.127468894 | 0.381557478 |
| 0.127466463 | 0.381557478 |
| 0.127454967 | 0.381557478 |
| 0.127453316 | 0.381557478 |
| 0.127444297 | 0.381557478 |
| 0.127404844 | 0.38150962 |
| 0.127388148 | 0.38148361 |
| 0.12736798 | 0.381447196 |
| 0.127359214 | 0.381444929 |
| 0.127319819 | 0.381374906 |
| 0.127314541 | 0.381374906 |
| 0.127236428 | 0.381173062 |
| 0.127228277 | 0.381173062 |
| 0.12721199 | 0.381171781 |
| 0.127195129 | 0.381145238 |
| 0.127186906 | 0.381144578 |
| 0.127149444 | 0.38105629 |
| 0.127143694 | 0.38105629 |
| 0.12712801 | 0.381040007 |
| 0.127118191 | 0.381034557 |
| 0.127109234 | 0.381031689 |
| 0.126973569 | 0.380696894 |
| 0.126939375 | 0.380642296 |
| 0.126930073 | 0.380638369 |
| 0.126924819 | 0.380638369 |
| 0.126909262 | 0.380623894 |
| 0.12690832 | 0.380623894 |
| 0.126860387 | 0.3805492 |
| 0.126858753 | 0.3805492 |
| 0.126842074 | 0.3805492 |
| 0.12679425 | 0.380517428 |
| 0.126787301 | 0.380517428 |
| 0.12678216 | 0.380517428 |
| 0.12677013 | 0.380517428 |
| 0.126754903 | 0.380517428 |
| 0.126747011 | 0.380517428 |
| 0.126736968 | 0.380517428 |
| 0.126726292 | 0.380517428 |
| 0.126709301 | 0.380517428 |
| 0.126705747 | 0.380517428 |

| | |
|---|---|
| 0.126636705 | 0.38038133 |
| 0.126630712 | 0.38038133 |
| 0.126624741 | 0.38038133 |
| 0.126596114 | 0.380355356 |
| 0.12657594 | 0.380318736 |
| 0.126552624 | 0.38027267 |
| 0.126485853 | 0.380169976 |
| 0.126476007 | 0.380169976 |
| 0.126457508 | 0.380169976 |
| 0.126431235 | 0.380123758 |
| 0.126412779 | 0.380104943 |
| 0.126411022 | 0.380104943 |
| 0.126387937 | 0.380104943 |
| 0.126371108 | 0.380096341 |
| 0.126338669 | 0.380037376 |
| 0.126326455 | 0.380024637 |
| 0.126263906 | 0.379860466 |
| 0.126255559 | 0.379859349 |
| 0.126243977 | 0.379848499 |
| 0.126243966 | 0.379848499 |
| 0.126230682 | 0.379848499 |
| 0.126185787 | 0.379745387 |
| 0.126165543 | 0.379708456 |
| 0.126149589 | 0.379684434 |
| 0.126079722 | 0.379546103 |
| 0.12604039 | 0.379475671 |
| 0.126022901 | 0.379447003 |
| 0.126017608 | 0.379447003 |
| 0.126004172 | 0.379444658 |
| 0.125987356 | 0.379444658 |
| 0.125953553 | 0.3794301 |
| 0.125919843 | 0.379365828 |
| 0.125908245 | 0.379365828 |
| 0.125889791 | 0.379365828 |
| 0.125889059 | 0.379365828 |
| 0.125884736 | 0.379365828 |
| 0.125863466 | 0.379365828 |
| 0.125758223 | 0.379177443 |
| 0.125737741 | 0.37913969 |
| 0.125692043 | 0.379025893 |
| 0.125623674 | 0.378867699 |
| 0.125617715 | 0.378867699 |
| 0.125615858 | 0.378867699 |
| 0.125606547 | 0.378867699 |
| 0.125582994 | 0.378864969 |

| | |
|---|---|
| 0.125564497 | 0.378833162 |
| 0.125556729 | 0.378833162 |
| 0.125540041 | 0.378833162 |
| 0.125528834 | 0.378833162 |
| 0.125518879 | 0.378833162 |
| 0.125483197 | 0.378755828 |
| 0.125467685 | 0.378733009 |
| 0.125427964 | 0.37866111 |
| 0.125338405 | 0.378438709 |
| 0.125322002 | 0.378437166 |
| 0.125160638 | 0.377997826 |
| 0.125159652 | 0.377997826 |
| 0.125114227 | 0.377953525 |
| 0.125090736 | 0.377918989 |
| 0.125074039 | 0.377904032 |
| 0.125073117 | 0.377904032 |
| 0.124992315 | 0.377752965 |
| 0.124984058 | 0.377751983 |
| 0.124977201 | 0.377751983 |
| 0.124974809 | 0.377751983 |
| 0.124922056 | 0.377660451 |
| 0.124848485 | 0.377461994 |
| 0.124821277 | 0.377403691 |
| 0.124797095 | 0.377374024 |
| 0.124792922 | 0.377374024 |
| 0.124775757 | 0.377361886 |
| 0.124748575 | 0.377303636 |
| 0.124692637 | 0.377182359 |
| 0.124658292 | 0.377126373 |
| 0.124605843 | 0.377039548 |
| 0.124583967 | 0.377012646 |
| 0.124565644 | 0.376989766 |
| 0.124532755 | 0.376962091 |
| 0.124511469 | 0.376921614 |
| 0.124510216 | 0.376921614 |
| 0.124488077 | 0.376898711 |
| 0.124364421 | 0.37657221 |
| 0.12430989 | 0.376431023 |
| 0.124305547 | 0.376431023 |
| 0.124227967 | 0.376278644 |
| 0.124184627 | 0.37619522 |
| 0.12414689 | 0.376104822 |
| 0.124127304 | 0.376069408 |
| 0.124117933 | 0.376064942 |
| 0.124108028 | 0.376064942 |

| | |
|---|---|
| 0.124087223 | 0.376064942 |
| 0.124050967 | 0.375981635 |
| 0.124002245 | 0.375881804 |
| 0.123991638 | 0.375881804 |
| 0.123968326 | 0.375850753 |
| 0.123957166 | 0.375840841 |
| 0.123949329 | 0.375840841 |
| 0.123904505 | 0.375800795 |
| 0.12388586 | 0.375768174 |
| 0.123877573 | 0.375768174 |
| 0.12386207 | 0.375767808 |
| 0.123839825 | 0.375724255 |
| 0.123831736 | 0.375723648 |
| 0.123810058 | 0.375697391 |
| 0.123804793 | 0.375697391 |
| 0.123727131 | 0.375525876 |
| 0.123703924 | 0.375479369 |
| 0.123668009 | 0.375394281 |
| 0.12365801 | 0.375387853 |
| 0.12365546 | 0.375387853 |
| 0.123549209 | 0.375129298 |
| 0.123525712 | 0.375105783 |
| 0.123474873 | 0.374999219 |
| 0.123445004 | 0.374932417 |
| 0.123413921 | 0.374861916 |
| 0.123391678 | 0.374832282 |
| 0.123381067 | 0.374832282 |
| 0.123361567 | 0.374798506 |
| 0.123346122 | 0.374775485 |
| 0.123312546 | 0.374697375 |
| 0.123297634 | 0.374697375 |
| 0.123269938 | 0.37463961 |
| 0.123258683 | 0.374629311 |
| 0.123236898 | 0.374587001 |
| 0.123224011 | 0.374571737 |
| 0.123217879 | 0.374571737 |
| 0.123179836 | 0.374496546 |
| 0.123117647 | 0.374457698 |
| 0.123099472 | 0.374456061 |
| 0.123084255 | 0.374433688 |
| 0.123054501 | 0.37441492 |
| 0.123027244 | 0.37438532 |
| 0.123012646 | 0.374383239 |
| 0.123002969 | 0.374377711 |
| 0.122967314 | 0.374293108 |

| | |
|---|---|
| 0.122963085 | 0.374293108 |
| 0.122939324 | 0.37427967 |
| 0.122937959 | 0.37427967 |
| 0.122930006 | 0.37427967 |
| 0.122913214 | 0.37427967 |
| 0.122909173 | 0.37427967 |
| 0.122870243 | 0.374248032 |
| 0.12286803 | 0.374248032 |
| 0.122865955 | 0.374248032 |
| 0.122799342 | 0.374164435 |
| 0.122796123 | 0.374164435 |
| 0.122769178 | 0.374120395 |
| 0.122739696 | 0.374078421 |
| 0.122738523 | 0.374078421 |
| 0.122723962 | 0.374078421 |
| 0.122587056 | 0.373804579 |
| 0.122553555 | 0.373726351 |
| 0.122548748 | 0.373726351 |
| 0.122545003 | 0.373726351 |
| 0.122500674 | 0.373660792 |
| 0.12249956 | 0.373660792 |
| 0.12245675 | 0.373589347 |
| 0.122441431 | 0.373575785 |
| 0.122380251 | 0.373413044 |
| 0.122360155 | 0.373397143 |
| 0.122352849 | 0.373397143 |
| 0.122341201 | 0.373389579 |
| 0.122318569 | 0.373344429 |
| 0.122313937 | 0.373344429 |
| 0.1222935 | 0.373315763 |
| 0.122281916 | 0.373304329 |
| 0.1222564 | 0.373250358 |
| 0.12224798 | 0.373248577 |
| 0.122223179 | 0.373220706 |
| 0.122214862 | 0.373219237 |
| 0.122187773 | 0.373193528 |
| 0.1221731 | 0.373187419 |
| 0.122146451 | 0.373153883 |
| 0.122104982 | 0.373051124 |
| 0.122095599 | 0.373046387 |
| 0.12207193 | 0.37302193 |
| 0.122021455 | 0.372891616 |
| 0.122013024 | 0.372891616 |
| 0.121939735 | 0.372769703 |
| 0.121915652 | 0.372735702 |

| | |
|---|---|
| 0.121900892 | 0.372735702 |
| 0.121871653 | 0.372672952 |
| 0.121865187 | 0.372672952 |
| 0.121834323 | 0.372630574 |
| 0.121834152 | 0.372630574 |
| 0.121833889 | 0.372630574 |
| 0.121824126 | 0.372630574 |
| 0.121817909 | 0.372630574 |
| 0.121807558 | 0.372630574 |
| 0.121777603 | 0.372630574 |
| 0.121759733 | 0.372617799 |
| 0.121756143 | 0.372617799 |
| 0.121754113 | 0.372617799 |
| 0.121743572 | 0.372617799 |
| 0.12172455 | 0.372605898 |
| 0.121717172 | 0.372605898 |
| 0.121681035 | 0.372520579 |
| 0.121675649 | 0.372520579 |
| 0.121652413 | 0.372480841 |
| 0.121626824 | 0.372426433 |
| 0.121624979 | 0.372426433 |
| 0.121621879 | 0.372426433 |
| 0.121612399 | 0.372426433 |
| 0.12153337 | 0.372283882 |
| 0.121531528 | 0.372283882 |
| 0.121493887 | 0.372210817 |
| 0.121485411 | 0.372210817 |
| 0.121448857 | 0.372144676 |
| 0.121415954 | 0.372115678 |
| 0.12140384 | 0.372102496 |
| 0.12138961 | 0.372082826 |
| 0.121337425 | 0.371946808 |
| 0.121328155 | 0.37194233 |
| 0.121282988 | 0.371851737 |
| 0.12125985 | 0.371828667 |
| 0.121107054 | 0.371431879 |
| 0.121097251 | 0.371425732 |
| 0.120980777 | 0.3711641 |
| 0.120956505 | 0.371113942 |
| 0.120946438 | 0.371113942 |
| 0.120915298 | 0.37108273 |
| 0.120894596 | 0.371043106 |
| 0.120840183 | 0.370923911 |
| 0.120762288 | 0.370755653 |
| 0.120757458 | 0.370755653 |

| | |
|---|---|
| 0.120743373 | 0.370746221 |
| 0.120736917 | 0.370746221 |
| 0.120637255 | 0.370492039 |
| 0.120597068 | 0.370416391 |
| 0.120580446 | 0.370389224 |
| 0.120544042 | 0.370301284 |
| 0.120419432 | 0.369994172 |
| 0.120409753 | 0.369994172 |
| 0.120391322 | 0.369994172 |
| 0.120339386 | 0.369863444 |
| 0.120334505 | 0.369863444 |
| 0.120334505 | 0.369863444 |
| 0.12033293 | 0.369863444 |
| 0.120315136 | 0.369863444 |
| 0.12028757 | 0.369847391 |
| 0.120254639 | 0.369793885 |
| 0.120218279 | 0.369705944 |
| 0.120186717 | 0.369656622 |
| 0.12017841 | 0.369654942 |
| 0.120175528 | 0.369654942 |
| 0.120129991 | 0.369577615 |
| 0.120076277 | 0.369460097 |
| 0.120031645 | 0.36934663 |
| 0.120013262 | 0.369313927 |
| 0.11994598 | 0.369178441 |
| 0.11989684 | 0.369051043 |
| 0.119885006 | 0.369051043 |
| 0.119853345 | 0.368988705 |
| 0.119850701 | 0.368988705 |
| 0.119785302 | 0.368850753 |
| 0.119756763 | 0.368786716 |
| 0.119730835 | 0.368754558 |
| 0.119684234 | 0.368682557 |
| 0.119670728 | 0.368664799 |
| 0.119642057 | 0.368600314 |
| 0.119628054 | 0.368581016 |
| 0.119622898 | 0.368581016 |
| 0.119607488 | 0.368565336 |
| 0.119598643 | 0.368561926 |
| 0.1195729 | 0.368554135 |
| 0.119515313 | 0.368400481 |
| 0.119514014 | 0.368400481 |
| 0.119511543 | 0.368400481 |
| 0.119508029 | 0.368400481 |
| 0.119468532 | 0.3683546 |

| | |
|---|---|
| 0.11945606 | 0.3683546 |
| 0.119445289 | 0.3683546 |
| 0.11943025 | 0.368352836 |
| 0.11942707 | 0.368352836 |
| 0.119379003 | 0.368266324 |
| 0.119369534 | 0.368260967 |
| 0.119297596 | 0.368062876 |
| 0.119294784 | 0.368062876 |
| 0.11922502 | 0.367934298 |
| 0.119161098 | 0.36776086 |
| 0.11911571 | 0.367668432 |
| 0.119080867 | 0.367608534 |
| 0.119072046 | 0.367605128 |
| 0.119058708 | 0.367605128 |
| 0.119042469 | 0.367605128 |
| 0.119028843 | 0.36759089 |
| 0.118999651 | 0.367524569 |
| 0.1189524 | 0.367402462 |
| 0.118938731 | 0.367384067 |
| 0.118890328 | 0.367258376 |
| 0.118872971 | 0.367229285 |
| 0.118855729 | 0.367222952 |
| 0.118844852 | 0.367213169 |
| 0.118839301 | 0.367213169 |
| 0.118787168 | 0.367130205 |
| 0.118774225 | 0.367114026 |
| 0.118749601 | 0.367104512 |
| 0.118715518 | 0.367027847 |
| 0.118709433 | 0.367027847 |
| 0.118630001 | 0.366834895 |
| 0.118612842 | 0.366805651 |
| 0.118607139 | 0.366805651 |
| 0.118580603 | 0.366800812 |
| 0.118546635 | 0.366740627 |
| 0.118422134 | 0.366406221 |
| 0.118404418 | 0.366375208 |
| 0.11840026 | 0.366375208 |
| 0.11835877 | 0.366329162 |
| 0.118316717 | 0.366294221 |
| 0.118287153 | 0.366244546 |
| 0.118253637 | 0.366194147 |
| 0.118214974 | 0.366126957 |
| 0.118195842 | 0.366110391 |
| 0.118191266 | 0.366110391 |
| 0.118180109 | 0.366109278 |

| | |
|---|---|
| 0.118177468 | 0.366109278 |
| 0.118149914 | 0.36606336 |
| 0.118140254 | 0.366057244 |
| 0.118108327 | 0.366005941 |
| 0.118083338 | 0.365952311 |
| 0.118070584 | 0.365936599 |
| 0.118068817 | 0.365936599 |
| 0.118064188 | 0.365936599 |
| 0.118036211 | 0.365901499 |
| 0.118020464 | 0.365876499 |
| 0.118003504 | 0.365847733 |
| 0.117984097 | 0.365811379 |
| 0.1179823 | 0.365811379 |
| 0.117971799 | 0.365811379 |
| 0.117969141 | 0.365811379 |
| 0.117929478 | 0.365784908 |
| 0.117860661 | 0.365609962 |
| 0.117756943 | 0.365321125 |
| 0.117729569 | 0.365259994 |
| 0.117716752 | 0.365259994 |
| 0.117702381 | 0.365247021 |
| 0.117613568 | 0.365018989 |
| 0.117590759 | 0.36499577 |
| 0.117568716 | 0.364951132 |
| 0.117556205 | 0.364936083 |
| 0.11748581 | 0.364812667 |
| 0.117480979 | 0.364812667 |
| 0.117420557 | 0.364698284 |
| 0.117404957 | 0.364698284 |
| 0.117402526 | 0.364698284 |
| 0.11739929 | 0.364698284 |
| 0.117355749 | 0.364622757 |
| 0.117328461 | 0.364561756 |
| 0.117309656 | 0.364527109 |
| 0.117292013 | 0.364496064 |
| 0.117257309 | 0.364411996 |
| 0.117245303 | 0.364398463 |
| 0.117240225 | 0.364398463 |
| 0.117235488 | 0.364398463 |
| 0.117205004 | 0.364392109 |
| 0.117202187 | 0.364392109 |
| 0.117179424 | 0.364360152 |
| 0.117077255 | 0.364090005 |
| 0.117075974 | 0.364090005 |
| 0.11707529 | 0.364090005 |

| | |
|---|---|
| 0.117070632 | 0.364090005 |
| 0.117042028 | 0.364090005 |
| 0.116975327 | 0.363986925 |
| 0.116966421 | 0.363982994 |
| 0.116943387 | 0.363935096 |
| 0.116934299 | 0.363930593 |
| 0.116933794 | 0.363930593 |
| 0.116923404 | 0.363930593 |
| 0.116915278 | 0.363930593 |
| 0.116898459 | 0.363930593 |
| 0.116890144 | 0.363930593 |
| 0.116860007 | 0.363865818 |
| 0.116821472 | 0.363769616 |
| 0.116800718 | 0.363728768 |
| 0.116785911 | 0.363728768 |
| 0.116768298 | 0.363728768 |
| 0.116754935 | 0.363728768 |
| 0.116734193 | 0.363728768 |
| 0.116706031 | 0.363671693 |
| 0.116698513 | 0.363671693 |
| 0.116682799 | 0.363646884 |
| 0.116660879 | 0.363602361 |
| 0.116649702 | 0.363591319 |
| 0.116633753 | 0.3635654 |
| 0.116628094 | 0.3635654 |
| 0.11660299 | 0.363517091 |
| 0.116574359 | 0.363451623 |
| 0.116556958 | 0.36342116 |
| 0.116545594 | 0.363417262 |
| 0.116509605 | 0.363344876 |
| 0.116457975 | 0.363255222 |
| 0.116410621 | 0.363131297 |
| 0.11639267 | 0.363099084 |
| 0.116368363 | 0.363070816 |
| 0.116343312 | 0.363029274 |
| 0.116321654 | 0.362996423 |
| 0.116311093 | 0.362987252 |
| 0.116279918 | 0.36291374 |
| 0.116269175 | 0.362903992 |
| 0.116241769 | 0.36284223 |
| 0.11624037 | 0.36284223 |
| 0.116160111 | 0.362634871 |
| 0.116158675 | 0.362634871 |
| 0.116054323 | 0.362375876 |
| 0.116035224 | 0.362363755 |

| | |
|---|---|
| 0.116013292 | 0.36234278 |
| 0.115990439 | 0.362303456 |
| 0.115956358 | 0.362275481 |
| 0.115947027 | 0.362275481 |
| 0.115928296 | 0.3622436 |
| 0.115903017 | 0.362188374 |
| 0.115895381 | 0.362188275 |
| 0.11586496 | 0.362116967 |
| 0.115838426 | 0.362057796 |
| 0.11581918 | 0.3620214 |
| 0.115800875 | 0.361987942 |
| 0.115775915 | 0.361933674 |
| 0.115762858 | 0.361916609 |
| 0.115747774 | 0.361893208 |
| 0.115690116 | 0.361770055 |
| 0.115625942 | 0.361678494 |
| 0.115618676 | 0.361678494 |
| 0.115616223 | 0.361678494 |
| 0.115616223 | 0.361678494 |
| 0.115616223 | 0.361678494 |
| 0.115604995 | 0.361678494 |
| 0.115559346 | 0.361636439 |
| 0.115509177 | 0.361546755 |
| 0.115461018 | 0.361423722 |
| 0.115430591 | 0.361380412 |
| 0.115428087 | 0.361380412 |
| 0.115422143 | 0.361380412 |
| 0.115410129 | 0.361380412 |
| 0.115387287 | 0.361359174 |
| 0.115351993 | 0.361304064 |
| 0.115333769 | 0.361286597 |
| 0.115300888 | 0.361207352 |
| 0.115298135 | 0.361207352 |
| 0.115279235 | 0.361187035 |
| 0.115271472 | 0.361187035 |
| 0.115267748 | 0.361187035 |
| 0.115262502 | 0.361187035 |
| 0.115259974 | 0.361187035 |
| 0.115255933 | 0.361187035 |
| 0.115213908 | 0.361172436 |
| 0.115211992 | 0.361172436 |
| 0.115174905 | 0.361121473 |
| 0.115171769 | 0.361121473 |
| 0.115064669 | 0.360823345 |
| 0.115041088 | 0.360773155 |

| | |
|---|---|
| 0.114999938 | 0.360667854 |
| 0.114981966 | 0.360635238 |
| 0.114977805 | 0.360635238 |
| 0.114945595 | 0.360577074 |
| 0.114868157 | 0.360396949 |
| 0.114855687 | 0.360381567 |
| 0.114846274 | 0.360375772 |
| 0.114843905 | 0.360375772 |
| 0.114833288 | 0.360375772 |
| 0.114755617 | 0.360186225 |
| 0.114743718 | 0.360172613 |
| 0.114719662 | 0.360120836 |
| 0.114695707 | 0.360069371 |
| 0.114661274 | 0.360032469 |
| 0.114659435 | 0.360032469 |
| 0.114659009 | 0.360032469 |
| 0.114650203 | 0.360032469 |
| 0.114560321 | 0.359810366 |
| 0.114537931 | 0.359763769 |
| 0.114524237 | 0.359763769 |
| 0.114517279 | 0.359763769 |
| 0.11446715 | 0.359660045 |
| 0.114398805 | 0.359492733 |
| 0.114364735 | 0.359409385 |
| 0.114336699 | 0.359368703 |
| 0.114322985 | 0.35936 |
| 0.114309929 | 0.359355708 |
| 0.114291457 | 0.359355708 |
| 0.114290087 | 0.359355708 |
| 0.114276924 | 0.359355708 |
| 0.114226615 | 0.359236067 |
| 0.114203658 | 0.359211309 |
| 0.114187108 | 0.35919654 |
| 0.114124061 | 0.359055798 |
| 0.114121332 | 0.359055798 |
| 0.11409181 | 0.35900176 |
| 0.114081551 | 0.358993196 |
| 0.114068275 | 0.358975138 |
| 0.114045166 | 0.35892613 |
| 0.114034198 | 0.35892613 |
| 0.114014531 | 0.358900865 |
| 0.114004096 | 0.358891736 |
| 0.113972819 | 0.358816992 |
| 0.11397164 | 0.358816992 |
| 0.113954153 | 0.358815183 |

| | |
|---|---|
| 0.113950791 | 0.358815183 |
| 0.113933891 | 0.358813027 |
| 0.113924095 | 0.358805905 |
| 0.11386007 | 0.35865169 |
| 0.11377687 | 0.358437028 |
| 0.113770065 | 0.358437028 |
| 0.113770065 | 0.358437028 |
| 0.113770065 | 0.358437028 |
| 0.113770065 | 0.358437028 |
| 0.113770065 | 0.358437028 |
| 0.113770065 | 0.358437028 |
| 0.113770065 | 0.358437028 |
| 0.113770065 | 0.358437028 |
| 0.113770065 | 0.358437028 |
| 0.113770065 | 0.358437028 |
| 0.113770065 | 0.358437028 |
| 0.113770065 | 0.358437028 |
| 0.113770065 | 0.358437028 |
| 0.113770065 | 0.358437028 |
| 0.113770065 | 0.358437028 |
| 0.113770065 | 0.358437028 |
| 0.113770065 | 0.358437028 |
| 0.113770065 | 0.358437028 |
| 0.113770065 | 0.358437028 |
| 0.113770065 | 0.358437028 |
| 0.113770065 | 0.358437028 |
| 0.113770065 | 0.358437028 |
| 0.113770065 | 0.358437028 |
| 0.113770065 | 0.358437028 |
| 0.113770065 | 0.358437028 |
| 0.113770065 | 0.358437028 |
| 0.113770065 | 0.358437028 |
| 0.113770065 | 0.358437028 |
| 0.113770065 | 0.358437028 |
| 0.113770065 | 0.358437028 |
| 0.113770065 | 0.358437028 |
| 0.113770065 | 0.358437028 |
| 0.113770065 | 0.358437028 |
| 0.113770065 | 0.358437028 |
| 0.113770065 | 0.358437028 |
| 0.113770065 | 0.358437028 |
| 0.113770065 | 0.358437028 |
| 0.113770065 | 0.358437028 |
| 0.113770065 | 0.358437028 |
| 0.113770065 | 0.358437028 |

| | |
|---|---|
| 0.113770065 | 0.358437028 |
| 0.113770065 | 0.358437028 |
| 0.113770065 | 0.358437028 |
| 0.113770065 | 0.358437028 |
| 0.113770065 | 0.358437028 |
| 0.113770065 | 0.358437028 |
| 0.113770065 | 0.358437028 |
| 0.113770065 | 0.358437028 |
| 0.113770065 | 0.358437028 |
| 0.113770065 | 0.358437028 |
| 0.113770065 | 0.358437028 |
| 0.113770065 | 0.358437028 |
| 0.113770065 | 0.358437028 |
| 0.113770065 | 0.358437028 |
| 0.113770065 | 0.358437028 |
| 0.113770065 | 0.358437028 |
| 0.113770065 | 0.358437028 |
| 0.113770065 | 0.358437028 |
| 0.113770065 | 0.358437028 |
| 0.113770065 | 0.358437028 |
| 0.113770065 | 0.358437028 |
| 0.113770065 | 0.358437028 |
| 0.113770065 | 0.358437028 |
| 0.113770065 | 0.358437028 |
| 0.113770065 | 0.358437028 |
| 0.113770065 | 0.358437028 |
| 0.113770065 | 0.358437028 |
| 0.113770065 | 0.358437028 |
| 0.113770065 | 0.358437028 |
| 0.113770065 | 0.358437028 |
| 0.113770065 | 0.358437028 |
| 0.113770065 | 0.358437028 |
| 0.113770065 | 0.358437028 |
| 0.113770065 | 0.358437028 |
| 0.113770065 | 0.358437028 |
| 0.113770065 | 0.358437028 |
| 0.113770065 | 0.358437028 |
| 0.113770065 | 0.358437028 |
| 0.113770065 | 0.358437028 |
| 0.113770065 | 0.358437028 |
| 0.113770065 | 0.358437028 |
| 0.113764581 | 0.358437028 |
| 0.113715656 | 0.358437028 |
| 0.113663317 | 0.358437028 |
| 0.113639706 | 0.358437028 |
| 0.113563506 | 0.358437028 |

| | |
|---|---|
| 0.113524368 | 0.358437028 |
| 0.113520342 | 0.358437028 |
| 0.113516818 | 0.358437028 |
| 0.113483649 | 0.358437028 |
| 0.1134447 | 0.358437028 |
| 0.113428886 | 0.358437028 |
| 0.11341633 | 0.358437028 |
| 0.113403002 | 0.358437028 |
| 0.113389868 | 0.358437028 |
| 0.113338263 | 0.358437028 |
| 0.113322784 | 0.358437028 |
| 0.113316088 | 0.358437028 |
| 0.113262761 | 0.358437028 |
| 0.113234929 | 0.358437028 |
| 0.113166426 | 0.358437028 |
| 0.113142181 | 0.358437028 |
| 0.113131656 | 0.358437028 |
| 0.113108583 | 0.358437028 |
| 0.113064378 | 0.358437028 |
| 0.113043302 | 0.358437028 |
| 0.113017136 | 0.358437028 |
| 0.112989914 | 0.358437028 |
| 0.112929274 | 0.358437028 |
| 0.112840899 | 0.358437028 |
| 0.112820245 | 0.358437028 |
| 0.112741856 | 0.358328754 |
| 0.112739645 | 0.358328754 |
| 0.112691931 | 0.358217886 |
| 0.112686337 | 0.358217886 |
| 0.112682666 | 0.358217886 |
| 0.112682602 | 0.358217886 |
| 0.112680375 | 0.358217886 |
| 0.112652579 | 0.358212338 |
| 0.112617712 | 0.358125376 |
| 0.112599237 | 0.358090535 |
| 0.112597873 | 0.358090535 |
| 0.112553726 | 0.358017517 |
| 0.112536368 | 0.357986214 |
| 0.112522979 | 0.35798173 |
| 0.11250887 | 0.35798173 |
| 0.112488598 | 0.357953795 |
| 0.112461383 | 0.357891104 |
| 0.11242917 | 0.357860328 |
| 0.112366748 | 0.35770945 |
| 0.112363111 | 0.35770945 |

| | |
|---|---|
| 0.112361547 | 0.35770945 |
| 0.112357675 | 0.35770945 |
| 0.112324345 | 0.357670092 |
| 0.112300764 | 0.357618916 |
| 0.112271253 | 0.357572931 |
| 0.112253738 | 0.357564792 |
| 0.112253119 | 0.357564792 |
| 0.112172875 | 0.357378923 |
| 0.112140121 | 0.357346283 |
| 0.112115333 | 0.35733902 |
| 0.112092158 | 0.357289065 |
| 0.111945102 | 0.356868097 |
| 0.111942774 | 0.356868097 |
| 0.111926277 | 0.356868097 |
| 0.111891627 | 0.356840936 |
| 0.111844164 | 0.356737344 |
| 0.111844142 | 0.356737344 |
| 0.111831631 | 0.356737344 |
| 0.111723363 | 0.356471402 |
| 0.111707532 | 0.356468652 |
| 0.111699122 | 0.356465698 |
| 0.111645055 | 0.356364792 |
| 0.111555262 | 0.35612591 |
| 0.111475509 | 0.355942886 |
| 0.111474959 | 0.355942886 |
| 0.111471477 | 0.355942886 |
| 0.111458169 | 0.355942886 |
| 0.111445242 | 0.355942886 |
| 0.111427285 | 0.355942886 |
| 0.111401754 | 0.355922139 |
| 0.111372613 | 0.355876782 |
| 0.111344097 | 0.355866871 |
| 0.111306409 | 0.355808467 |
| 0.111294736 | 0.355800141 |
| 0.111269422 | 0.355761867 |
| 0.111205658 | 0.355581865 |
| 0.111178279 | 0.355542059 |
| 0.111177432 | 0.355542059 |
| 0.111171085 | 0.355542059 |
| 0.111167462 | 0.355542059 |
| 0.111092962 | 0.355388535 |
| 0.111049403 | 0.355328369 |
| 0.111026859 | 0.355296394 |
| 0.111022531 | 0.355296394 |
| 0.110976793 | 0.355207777 |

| | |
|---|---|
| 0.110963979 | 0.355190631 |
| 0.110897726 | 0.355013798 |
| 0.110888781 | 0.355013798 |
| 0.1108506 | 0.354998887 |
| 0.11072932 | 0.354654024 |
| 0.110712185 | 0.354622989 |
| 0.11068049 | 0.354545311 |
| 0.110625092 | 0.354391689 |
| 0.110616632 | 0.354388424 |
| 0.110607826 | 0.354384051 |
| 0.110607824 | 0.354384051 |
| 0.110569953 | 0.354338026 |
| 0.11048998 | 0.354197098 |
| 0.110487985 | 0.354197098 |
| 0.110475223 | 0.354197098 |
| 0.110461034 | 0.354197098 |
| 0.110450538 | 0.354197098 |
| 0.110429751 | 0.354194743 |
| 0.110420426 | 0.354188688 |
| 0.110406179 | 0.354166842 |
| 0.110377369 | 0.354098271 |
| 0.110360879 | 0.354069222 |
| 0.110337663 | 0.354018586 |
| 0.110330124 | 0.354018586 |
| 0.110300013 | 0.353993177 |
| 0.110299011 | 0.353993177 |
| 0.110297533 | 0.353993177 |
| 0.110291763 | 0.353993177 |
| 0.110270981 | 0.353993177 |
| 0.110221408 | 0.353907841 |
| 0.110218434 | 0.353907841 |
| 0.110201711 | 0.353892313 |
| 0.110198907 | 0.353892313 |
| 0.110147334 | 0.353765397 |
| 0.110100896 | 0.353687804 |
| 0.110084879 | 0.353660204 |
| 0.110022781 | 0.353508398 |
| 0.110019928 | 0.353508398 |
| 0.110005005 | 0.353498978 |
| 0.109975142 | 0.353426863 |
| 0.109934402 | 0.353367468 |
| 0.109927916 | 0.353367468 |
| 0.109910839 | 0.353363274 |
| 0.109901426 | 0.353356862 |
| 0.109853088 | 0.353296849 |

| | |
|---|---|
| 0.109851747 | 0.353296849 |
| 0.109844143 | 0.353296849 |
| 0.109828035 | 0.353296849 |
| 0.109817692 | 0.353296849 |
| 0.109808993 | 0.353296849 |
| 0.109795561 | 0.353296849 |
| 0.109777081 | 0.353290967 |
| 0.10967281 | 0.353026962 |
| 0.109651396 | 0.352981888 |
| 0.109569366 | 0.35278936 |
| 0.109542699 | 0.352775044 |
| 0.10953172 | 0.35276354 |
| 0.109519949 | 0.352749482 |
| 0.109489218 | 0.352674353 |
| 0.109478729 | 0.352664418 |
| 0.109468606 | 0.35265566 |
| 0.109398249 | 0.352476687 |
| 0.109390282 | 0.352474863 |
| 0.109350165 | 0.352393282 |
| 0.109300119 | 0.352268284 |
| 0.10920916 | 0.352177174 |
| 0.109184421 | 0.352121237 |
| 0.109155578 | 0.352099743 |
| 0.109140164 | 0.352073868 |
| 0.109113203 | 0.352010737 |
| 0.109104106 | 0.352005233 |
| 0.108998043 | 0.351758356 |
| 0.108991584 | 0.351758356 |
| 0.108961891 | 0.351689346 |
| 0.10895175 | 0.351680446 |
| 0.108909635 | 0.351568333 |
| 0.10888869 | 0.351548375 |
| 0.108881553 | 0.351548375 |
| 0.108879017 | 0.351548375 |
| 0.108878791 | 0.351548375 |
| 0.108828266 | 0.351448593 |
| 0.108803787 | 0.351393364 |
| 0.108743143 | 0.351235666 |
| 0.108739081 | 0.351235666 |
| 0.108665321 | 0.351065203 |
| 0.108651473 | 0.351044277 |
| 0.108637335 | 0.351022413 |
| 0.108619207 | 0.350987648 |
| 0.108597869 | 0.35094251 |
| 0.108585127 | 0.350925143 |

| | |
|---|---|
| 0.108564219 | 0.350889248 |
| 0.108540974 | 0.350877688 |
| 0.108529972 | 0.350865936 |
| 0.108518584 | 0.350852937 |
| 0.108460661 | 0.350713279 |
| 0.108459106 | 0.350713279 |
| 0.108455574 | 0.350713279 |
| 0.108414047 | 0.350705413 |
| 0.108399269 | 0.350681429 |
| 0.10839417 | 0.350681429 |
| 0.108374426 | 0.350672518 |
| 0.108318409 | 0.35053889 |
| 0.108211925 | 0.350266604 |
| 0.1081727 | 0.350233934 |
| 0.108111188 | 0.350082373 |
| 0.108108934 | 0.350082373 |
| 0.108101114 | 0.350082373 |
| 0.108042498 | 0.349955118 |
| 0.108000565 | 0.349866882 |
| 0.107970638 | 0.349841322 |
| 0.107956391 | 0.349818956 |
| 0.107927254 | 0.349748338 |
| 0.107902894 | 0.349716989 |
| 0.107891642 | 0.349704317 |
| 0.10788463 | 0.349704317 |
| 0.10787706 | 0.349704317 |
| 0.107873065 | 0.349704317 |
| 0.107860566 | 0.349698787 |
| 0.107843073 | 0.349665876 |
| 0.107792347 | 0.349525197 |
| 0.107790184 | 0.349525197 |
| 0.107784372 | 0.349525197 |
| 0.10757256 | 0.348978839 |
| 0.107566972 | 0.348978839 |
| 0.10753318 | 0.348898615 |
| 0.107525609 | 0.348898615 |
| 0.107504079 | 0.348898615 |
| 0.107493214 | 0.348887782 |
| 0.107493017 | 0.348887782 |
| 0.107471414 | 0.348864574 |
| 0.107453178 | 0.348858204 |
| 0.107360654 | 0.348600073 |
| 0.107294718 | 0.348409731 |
| 0.107286059 | 0.34840537 |
| 0.107257096 | 0.348335064 |

| | |
|---|---|
| 0.107235329 | 0.348288121 |
| 0.107198805 | 0.348216989 |
| 0.107190478 | 0.348213689 |
| 0.107141082 | 0.348076965 |
| 0.107134249 | 0.348076965 |
| 0.107115918 | 0.348042695 |
| 0.107055477 | 0.347893778 |
| 0.1070376 | 0.34785942 |
| 0.107026518 | 0.347847139 |
| 0.107015954 | 0.34783654 |
| 0.107005713 | 0.34783654 |
| 0.107005184 | 0.34783654 |
| 0.106994229 | 0.34783654 |
| 0.106956521 | 0.347785774 |
| 0.106911934 | 0.347688269 |
| 0.106896579 | 0.347662071 |
| 0.106865154 | 0.347583601 |
| 0.106835282 | 0.347543974 |
| 0.106827422 | 0.347543974 |
| 0.106806251 | 0.347510679 |
| 0.106795544 | 0.347499583 |
| 0.106780081 | 0.347473006 |
| 0.106777604 | 0.347473006 |
| 0.10677211 | 0.347473006 |
| 0.106762309 | 0.347473006 |
| 0.106740976 | 0.347463862 |
| 0.10673864 | 0.347463862 |
| 0.106604677 | 0.347091942 |
| 0.106604516 | 0.347091942 |
| 0.106581825 | 0.347064991 |
| 0.10656294 | 0.347050953 |
| 0.106538513 | 0.34701886 |
| 0.106474282 | 0.346857087 |
| 0.106438787 | 0.346765176 |
| 0.106428863 | 0.346756565 |
| 0.106408895 | 0.346738946 |
| 0.106365963 | 0.346665836 |
| 0.106341448 | 0.346614027 |
| 0.106312011 | 0.346552313 |
| 0.106300382 | 0.346552313 |
| 0.106222421 | 0.346415725 |
| 0.106219975 | 0.346415725 |
| 0.106214888 | 0.346415725 |
| 0.10620503 | 0.346415725 |
| 0.106203964 | 0.346415725 |

| | |
|---|---|
| 0.106200411 | 0.346415725 |
| 0.106193187 | 0.346415725 |
| 0.106182488 | 0.346415725 |
| 0.106169538 | 0.346415725 |
| 0.106140937 | 0.346387172 |
| 0.106133294 | 0.346385964 |
| 0.106083806 | 0.346271911 |
| 0.106070717 | 0.346252914 |
| 0.106037189 | 0.346190925 |
| 0.105957845 | 0.346026762 |
| 0.105912989 | 0.345903992 |
| 0.105907036 | 0.345903992 |
| 0.105900888 | 0.345903992 |
| 0.105888233 | 0.345903992 |
| 0.105864013 | 0.345862626 |
| 0.10583732 | 0.345839265 |
| 0.105821136 | 0.345817431 |
| 0.1058098 | 0.345804114 |
| 0.10580673 | 0.345804114 |
| 0.105800015 | 0.345804114 |
| 0.105789312 | 0.345804114 |
| 0.105778662 | 0.345797264 |
| 0.105734108 | 0.34569907 |
| 0.105723271 | 0.345687368 |
| 0.105702476 | 0.345643104 |
| 0.105690586 | 0.345627954 |
| 0.105654756 | 0.34558197 |
| 0.105635743 | 0.345543511 |
| 0.105628037 | 0.345542035 |
| 0.105519483 | 0.345305511 |
| 0.105511539 | 0.345303242 |
| 0.105498885 | 0.345285556 |
| 0.105465057 | 0.345200633 |
| 0.105460263 | 0.345200633 |
| 0.105452946 | 0.345200633 |
| 0.105384303 | 0.34509521 |
| 0.105334948 | 0.344962357 |
| 0.105332238 | 0.344962357 |
| 0.105310041 | 0.344951947 |
| 0.105303814 | 0.344951947 |
| 0.105292285 | 0.344941235 |
| 0.1051196 | 0.344493993 |
| 0.105109351 | 0.344484109 |
| 0.105098064 | 0.344484109 |
| 0.105070387 | 0.344427511 |

| | |
|---|---|
| 0.105057117 | 0.344407718 |
| 0.105045963 | 0.344394856 |
| 0.105025993 | 0.344376797 |
| 0.105017485 | 0.344372608 |
| 0.105007965 | 0.344365101 |
| 0.104950539 | 0.344200476 |
| 0.104928602 | 0.344152229 |
| 0.104928369 | 0.344152229 |
| 0.104910774 | 0.344152229 |
| 0.104868836 | 0.344050979 |
| 0.104860072 | 0.344050979 |
| 0.104814577 | 0.343944043 |
| 0.104810411 | 0.343944043 |
| 0.104805254 | 0.343944043 |
| 0.104776513 | 0.343913921 |
| 0.104755306 | 0.343868009 |
| 0.104746392 | 0.343868009 |
| 0.104722655 | 0.343831924 |
| 0.104710424 | 0.343815469 |
| 0.104696677 | 0.343794031 |
| 0.104655911 | 0.343723028 |
| 0.104623964 | 0.343721151 |
| 0.104605524 | 0.343684277 |
| 0.104554805 | 0.343588745 |
| 0.104528283 | 0.343564509 |
| 0.104522523 | 0.343564509 |
| 0.104473216 | 0.343444436 |
| 0.104470258 | 0.343444436 |
| 0.104465479 | 0.343444436 |
| 0.104458239 | 0.343444436 |
| 0.104449101 | 0.343444436 |
| 0.104422124 | 0.343413376 |
| 0.104421753 | 0.343413376 |
| 0.10441965 | 0.343413376 |
| 0.104419066 | 0.343413376 |
| 0.104407972 | 0.343413376 |
| 0.10440037 | 0.343413376 |
| 0.104386462 | 0.343413376 |
| 0.104377706 | 0.343413376 |
| 0.104368264 | 0.343413376 |
| 0.104358081 | 0.343413376 |
| 0.104303896 | 0.343261586 |
| 0.104277483 | 0.343198376 |
| 0.104245066 | 0.343115394 |
| 0.104201443 | 0.343019223 |

| | |
|---|---|
| 0.104200384 | 0.343019223 |
| 0.104190986 | 0.343019223 |
| 0.104187315 | 0.343019223 |
| 0.104184702 | 0.343019223 |
| 0.104150339 | 0.342969521 |
| 0.104131184 | 0.342937261 |
| 0.104128763 | 0.342937261 |
| 0.104128183 | 0.342937261 |
| 0.104102884 | 0.342931807 |
| 0.104081902 | 0.342910124 |
| 0.104039951 | 0.342795623 |
| 0.104001951 | 0.342694128 |
| 0.103995795 | 0.342694128 |
| 0.103967551 | 0.342628187 |
| 0.103911068 | 0.342465743 |
| 0.103906137 | 0.342465743 |
| 0.103898048 | 0.342465743 |
| 0.103859114 | 0.342412989 |
| 0.103856676 | 0.342412989 |
| 0.10384786 | 0.342412989 |
| 0.103819328 | 0.342352915 |
| 0.103769657 | 0.342236511 |
| 0.103768821 | 0.342236511 |
| 0.103737039 | 0.342210634 |
| 0.103724446 | 0.342210634 |
| 0.103691787 | 0.342169246 |
| 0.103640962 | 0.342048929 |
| 0.103607562 | 0.341965078 |
| 0.103567675 | 0.341901826 |
| 0.103560536 | 0.341901826 |
| 0.103552758 | 0.341899979 |
| 0.103550204 | 0.341899979 |
| 0.10350517 | 0.341813945 |
| 0.103483009 | 0.341808853 |
| 0.103468507 | 0.341808853 |
| 0.103442407 | 0.341777742 |
| 0.103424314 | 0.341777742 |
| 0.103419752 | 0.341777742 |
| 0.103377111 | 0.341675343 |
| 0.103374108 | 0.341675343 |
| 0.103364995 | 0.341675343 |
| 0.103318224 | 0.341575529 |
| 0.10326232 | 0.341414405 |
| 0.103243755 | 0.341400426 |
| 0.10324099 | 0.341400426 |

| | |
|---|---|
| 0.10322952 | 0.341400426 |
| 0.103218816 | 0.341400426 |
| 0.103212648 | 0.341400426 |
| 0.103154812 | 0.341295883 |
| 0.103129584 | 0.341248415 |
| 0.103115037 | 0.3412354 |
| 0.10309589 | 0.341195743 |
| 0.103089231 | 0.341195743 |
| 0.103069261 | 0.341178739 |
| 0.103030121 | 0.341096593 |
| 0.102994916 | 0.341003743 |
| 0.10295039 | 0.340895708 |
| 0.102921737 | 0.340832538 |
| 0.102920764 | 0.340832538 |
| 0.102863526 | 0.340734534 |
| 0.102848735 | 0.340709238 |
| 0.102782143 | 0.340536009 |
| 0.10276645 | 0.340507702 |
| 0.102758426 | 0.340504804 |
| 0.102749231 | 0.340504804 |
| 0.102705946 | 0.340401954 |
| 0.102692864 | 0.340382282 |
| 0.102685244 | 0.340380714 |
| 0.102683399 | 0.340380714 |
| 0.102676711 | 0.340380714 |
| 0.102670586 | 0.340380714 |
| 0.102661038 | 0.340380714 |
| 0.10264984 | 0.340380714 |
| 0.102543025 | 0.340098652 |
| 0.102539938 | 0.340098652 |
| 0.102500644 | 0.34007609 |
| 0.102469574 | 0.339997102 |
| 0.10243992 | 0.339922389 |
| 0.102430077 | 0.339914938 |
| 0.102370016 | 0.339809179 |
| 0.10236576 | 0.339809179 |
| 0.1023484 | 0.339809179 |
| 0.102346321 | 0.339809179 |
| 0.102339848 | 0.339809179 |
| 0.102332284 | 0.339809179 |
| 0.102312405 | 0.339809179 |
| 0.102265034 | 0.339697084 |
| 0.102256704 | 0.339693106 |
| 0.102256299 | 0.339693106 |
| 0.10223975 | 0.339684171 |

| | |
|---|---|
| 0.102212424 | 0.339617072 |
| 0.102210654 | 0.339617072 |
| 0.102172239 | 0.339544223 |
| 0.102123937 | 0.339441466 |
| 0.102112686 | 0.339435107 |
| 0.10206257 | 0.339308532 |
| 0.102023135 | 0.339201108 |
| 0.102020586 | 0.339201108 |
| 0.101980789 | 0.339111826 |
| 0.10197142 | 0.339111826 |
| 0.101959661 | 0.339111826 |
| 0.101912427 | 0.338998686 |
| 0.101887399 | 0.338939103 |
| 0.101840376 | 0.338806341 |
| 0.101823074 | 0.338772446 |
| 0.101785487 | 0.338694712 |
| 0.101745143 | 0.338584121 |
| 0.101711769 | 0.338517643 |
| 0.101710277 | 0.338517643 |
| 0.101650208 | 0.338433727 |
| 0.101628328 | 0.338384533 |
| 0.101569782 | 0.338236891 |
| 0.101569601 | 0.338236891 |
| 0.10150987 | 0.338084655 |
| 0.101477826 | 0.338001567 |
| 0.101442075 | 0.337906121 |
| 0.101424261 | 0.337870412 |
| 0.101377576 | 0.337738515 |
| 0.101377339 | 0.337738515 |
| 0.101368737 | 0.337738515 |
| 0.101352812 | 0.337726886 |
| 0.101352242 | 0.337726886 |
| 0.101316172 | 0.337654365 |
| 0.101301791 | 0.337651368 |
| 0.101299742 | 0.337651368 |
| 0.101298469 | 0.337651368 |
| 0.101281607 | 0.337651368 |
| 0.101227983 | 0.337523465 |
| 0.101206291 | 0.337474762 |
| 0.101203044 | 0.337474762 |
| 0.101197223 | 0.337474762 |
| 0.101174646 | 0.337440116 |
| 0.101157002 | 0.337404896 |
| 0.101124923 | 0.337321522 |
| 0.101122083 | 0.337321522 |

| | |
|---|---|
| 0.101113823 | 0.337321522 |
| 0.101107504 | 0.337321522 |
| 0.101105929 | 0.337321522 |
| 0.101070992 | 0.337283363 |
| 0.101058817 | 0.337266363 |
| 0.100922372 | 0.336858212 |
| 0.100840167 | 0.336678208 |
| 0.100832107 | 0.336674899 |
| 0.100829269 | 0.336674899 |
| 0.100803017 | 0.336624968 |
| 0.100794143 | 0.336618935 |
| 0.100773578 | 0.336573856 |
| 0.10077051 | 0.336573856 |
| 0.100736539 | 0.336520946 |
| 0.10072258 | 0.336497917 |
| 0.100716795 | 0.336497917 |
| 0.100698281 | 0.336463941 |
| 0.100675575 | 0.336411677 |
| 0.100664435 | 0.336398054 |
| 0.100662788 | 0.336398054 |
| 0.100658927 | 0.336398054 |
| 0.100626669 | 0.336366254 |
| 0.100571108 | 0.336204125 |
| 0.100558027 | 0.336183993 |
| 0.100549522 | 0.336183993 |
| 0.100515759 | 0.336113462 |
| 0.100502468 | 0.336092618 |
| 0.1004688 | 0.336003618 |
| 0.100467943 | 0.336003618 |
| 0.100452373 | 0.335995868 |
| 0.100424516 | 0.335926284 |
| 0.100412572 | 0.335909922 |
| 0.100394369 | 0.335886639 |
| 0.100385791 | 0.335886639 |
| 0.100376233 | 0.335886639 |
| 0.100270251 | 0.335716782 |
| 0.100267501 | 0.335716782 |
| 0.100236764 | 0.33569249 |
| 0.100229441 | 0.33569249 |
| 0.10022335 | 0.33569249 |
| 0.10019439 | 0.335627953 |
| 0.100177542 | 0.335595118 |
| 0.100161018 | 0.335563365 |
| 0.100122756 | 0.335505979 |
| 0.100091826 | 0.335451941 |

| | |
|---|---|
| 0.100086667 | 0.335451941 |
| 0.100084639 | 0.335451941 |
| 0.100076386 | 0.335451941 |
| 0.100043627 | 0.335406012 |
| 0.100016635 | 0.335386345 |
| 0.099997552 | 0.335345966 |
| 0.099993487 | 0.335345966 |
| 0.099956686 | 0.335256134 |
| 0.099931771 | 0.335196173 |
| 0.099911772 | 0.335152695 |
| 0.099846352 | 0.334956837 |
| 0.099755542 | 0.334793688 |
| 0.099745747 | 0.334784403 |
| 0.099729572 | 0.334753704 |
| 0.099722307 | 0.334752912 |
| 0.099706132 | 0.33472371 |
| 0.099692695 | 0.33472371 |
| 0.099655796 | 0.334649383 |
| 0.099649329 | 0.334649383 |
| 0.099630013 | 0.334631781 |
| 0.099592368 | 0.334576123 |
| 0.099546867 | 0.334494049 |
| 0.099544947 | 0.334494049 |
| 0.099533897 | 0.334494049 |
| 0.099518894 | 0.334494049 |
| 0.099463061 | 0.334363808 |
| 0.099458246 | 0.334363808 |
| 0.099422819 | 0.334313089 |
| 0.099324992 | 0.33407849 |
| 0.099292935 | 0.333994252 |
| 0.099282154 | 0.333981575 |
| 0.09928069 | 0.333981575 |
| 0.099209922 | 0.33378574 |
| 0.099123491 | 0.333612784 |
| 0.099102021 | 0.333564095 |
| 0.099099106 | 0.333564095 |
| 0.099089718 | 0.333564095 |
| 0.099056045 | 0.333503622 |
| 0.099031523 | 0.333444633 |
| 0.099025983 | 0.333444633 |
| 0.099009908 | 0.333418997 |
| 0.09899844 | 0.333403952 |
| 0.098978345 | 0.333360303 |
| 0.098913686 | 0.333189201 |
| 0.098883615 | 0.333135032 |

| | |
|---|---|
| 0.098879286 | 0.333135032 |
| 0.098868751 | 0.333135032 |
| 0.09880184 | 0.333000867 |
| 0.098765919 | 0.332912239 |
| 0.098748509 | 0.33289179 |
| 0.098747055 | 0.33289179 |
| 0.09873041 | 0.3328779 |
| 0.098707715 | 0.33282532 |
| 0.098702296 | 0.33282532 |
| 0.098671294 | 0.332772816 |
| 0.098666032 | 0.332772816 |
| 0.098651802 | 0.332772816 |
| 0.09863438 | 0.332772816 |
| 0.098599707 | 0.332719871 |
| 0.098588539 | 0.332705758 |
| 0.098536463 | 0.332600718 |
| 0.098515384 | 0.332576706 |
| 0.09850464 | 0.332564011 |
| 0.098426058 | 0.332392958 |
| 0.098393235 | 0.33232924 |
| 0.098373042 | 0.332284602 |
| 0.098351954 | 0.332236933 |
| 0.098334043 | 0.332199992 |
| 0.098282138 | 0.332048194 |
| 0.098280913 | 0.332048194 |
| 0.098198569 | 0.331883567 |
| 0.098194078 | 0.331883567 |
| 0.098179743 | 0.331883567 |
| 0.098168475 | 0.331883567 |
| 0.098133004 | 0.33180325 |
| 0.098078876 | 0.331667985 |
| 0.098056786 | 0.331639722 |
| 0.098037695 | 0.331600287 |
| 0.098011559 | 0.331557392 |
| 0.097996938 | 0.331531479 |
| 0.097964744 | 0.331458059 |
| 0.09795562 | 0.331458059 |
| 0.09791716 | 0.331379278 |
| 0.097889565 | 0.33130943 |
| 0.097882779 | 0.33130943 |
| 0.097876116 | 0.33130943 |
| 0.097822623 | 0.331200546 |
| 0.097811978 | 0.331188049 |
| 0.097753861 | 0.331085419 |
| 0.097726082 | 0.331061962 |

| | |
|---|---|
| 0.097722264 | 0.331061962 |
| 0.097704565 | 0.331059713 |
| 0.09770342 | 0.331059713 |
| 0.097683339 | 0.331058448 |
| 0.097683161 | 0.331058448 |
| 0.097680359 | 0.331058448 |
| 0.097678562 | 0.331058448 |
| 0.097668924 | 0.331058448 |
| 0.097627519 | 0.331057732 |
| 0.097574811 | 0.330926122 |
| 0.097572922 | 0.330926122 |
| 0.097566722 | 0.330926122 |
| 0.097557311 | 0.330926122 |
| 0.097461588 | 0.330636303 |
| 0.097410024 | 0.330508457 |
| 0.097307145 | 0.330253501 |
| 0.097298028 | 0.330246093 |
| 0.097287598 | 0.330234225 |
| 0.097233594 | 0.330074438 |
| 0.097162638 | 0.329880592 |
| 0.09715895 | 0.329880592 |
| 0.097140049 | 0.329850929 |
| 0.097118488 | 0.329801227 |
| 0.09709884 | 0.329758017 |
| 0.097061913 | 0.329728233 |
| 0.097017046 | 0.329691821 |
| 0.097015648 | 0.329691821 |
| 0.097000405 | 0.329691821 |
| 0.096987953 | 0.329691821 |
| 0.096960749 | 0.329691821 |
| 0.096951187 | 0.329691821 |
| 0.09694458 | 0.329691821 |
| 0.096890694 | 0.329591677 |
| 0.096852694 | 0.329485947 |
| 0.096847211 | 0.329485947 |
| 0.096835421 | 0.329485947 |
| 0.096806605 | 0.329423277 |
| 0.096758746 | 0.329307473 |
| 0.096698742 | 0.329150299 |
| 0.096691378 | 0.329148757 |
| 0.096662448 | 0.329073795 |
| 0.096621574 | 0.328975669 |
| 0.096614323 | 0.328975669 |
| 0.09656257 | 0.328921867 |
| 0.096558102 | 0.328921867 |

| | |
|---|---|
| 0.096533383 | 0.328869489 |
| 0.096516595 | 0.328835817 |
| 0.096463974 | 0.328703563 |
| 0.096437549 | 0.328637031 |
| 0.096419385 | 0.328609049 |
| 0.096418662 | 0.328609049 |
| 0.09634646 | 0.328439141 |
| 0.096339509 | 0.328439141 |
| 0.096334747 | 0.328439141 |
| 0.09630998 | 0.328390279 |
| 0.096271745 | 0.328330425 |
| 0.096258851 | 0.32830996 |
| 0.096253312 | 0.32830996 |
| 0.096212498 | 0.32822238 |
| 0.096138152 | 0.328015742 |
| 0.096136677 | 0.328015742 |
| 0.09610455 | 0.327951273 |
| 0.096100812 | 0.327951273 |
| 0.096099343 | 0.327951273 |
| 0.096076922 | 0.327947792 |
| 0.09606533 | 0.327931727 |
| 0.096044892 | 0.327908967 |
| 0.096029795 | 0.327880928 |
| 0.096025423 | 0.327880928 |
| 0.095968735 | 0.327742934 |
| 0.095949604 | 0.327701096 |
| 0.095918474 | 0.327618272 |
| 0.095905011 | 0.327613706 |
| 0.095821495 | 0.327357458 |
| 0.095812446 | 0.327350026 |
| 0.095807535 | 0.327350026 |
| 0.095800552 | 0.327350026 |
| 0.095788303 | 0.327337986 |
| 0.095752477 | 0.327239038 |
| 0.095732486 | 0.327217677 |
| 0.095713888 | 0.327177588 |
| 0.095703048 | 0.327164014 |
| 0.095607505 | 0.32695474 |
| 0.09559428 | 0.326932985 |
| 0.095583745 | 0.326932985 |
| 0.095550164 | 0.326852522 |
| 0.095547295 | 0.326852522 |
| 0.095546937 | 0.326852522 |
| 0.095541257 | 0.326852522 |
| 0.095537459 | 0.326852522 |

| | |
|---|---|
| 0.095532983 | 0.326852522 |
| 0.095527431 | 0.326852522 |
| 0.095502722 | 0.326852522 |
| 0.095490841 | 0.326852522 |
| 0.095489982 | 0.326852522 |
| 0.095479258 | 0.326852522 |
| 0.09543439 | 0.326761588 |
| 0.095394333 | 0.326647917 |
| 0.095394149 | 0.326647917 |
| 0.095389582 | 0.326647917 |
| 0.095359683 | 0.326623202 |
| 0.095338052 | 0.326572598 |
| 0.095299133 | 0.326509732 |
| 0.095210735 | 0.326300761 |
| 0.095205493 | 0.326300761 |
| 0.095205203 | 0.326300761 |
| 0.09516156 | 0.326202649 |
| 0.095137723 | 0.326144413 |
| 0.095102861 | 0.326048372 |
| 0.095059727 | 0.325923952 |
| 0.095043726 | 0.325892552 |
| 0.09499747 | 0.325757398 |
| 0.094876595 | 0.325391231 |
| 0.094862262 | 0.325391231 |
| 0.094843346 | 0.325369448 |
| 0.09478205 | 0.325206015 |
| 0.09476306 | 0.325164285 |
| 0.09473682 | 0.32513022 |
| 0.094716482 | 0.325098148 |
| 0.094703908 | 0.325078418 |
| 0.094664635 | 0.325009499 |
| 0.094635818 | 0.324961795 |
| 0.094612698 | 0.324929258 |
| 0.094610813 | 0.324929258 |
| 0.094598952 | 0.324929258 |
| 0.094465849 | 0.324612194 |
| 0.094446128 | 0.324567842 |
| 0.094432752 | 0.324567842 |
| 0.094371803 | 0.324406054 |
| 0.094364494 | 0.324404341 |
| 0.094340593 | 0.324345587 |
| 0.094331511 | 0.32434 |
| 0.094315131 | 0.324328284 |
| 0.094283519 | 0.32424299 |
| 0.094229065 | 0.324125935 |

| | |
|---|---|
| 0.094227133 | 0.324125935 |
| 0.094218836 | 0.324125935 |
| 0.094153199 | 0.323978593 |
| 0.094144676 | 0.323976048 |
| 0.09407821 | 0.323770714 |
| 0.094050779 | 0.323723094 |
| 0.094034426 | 0.323690201 |
| 0.094020762 | 0.323690201 |
| 0.094009967 | 0.323690201 |
| 0.094007459 | 0.323690201 |
| 0.093999037 | 0.323690201 |
| 0.093986288 | 0.323688259 |
| 0.093966303 | 0.323642834 |
| 0.093936224 | 0.323586038 |
| 0.09389735 | 0.323475523 |
| 0.093882284 | 0.323460974 |
| 0.093875291 | 0.323460974 |
| 0.093856319 | 0.323427747 |
| 0.093850328 | 0.323427747 |
| 0.093840415 | 0.323419745 |
| 0.093825671 | 0.323392332 |
| 0.093822506 | 0.323392332 |
| 0.093797735 | 0.323342844 |
| 0.093780363 | 0.32330636 |
| 0.093768463 | 0.32330636 |
| 0.093723822 | 0.323228421 |
| 0.093718473 | 0.323228421 |
| 0.093688201 | 0.323175777 |
| 0.09368733 | 0.323175777 |
| 0.093676078 | 0.323175777 |
| 0.093672723 | 0.323175777 |
| 0.093649666 | 0.323136467 |
| 0.09363597 | 0.323112617 |
| 0.093630736 | 0.323112617 |
| 0.093582824 | 0.322999433 |
| 0.093566163 | 0.322965331 |
| 0.093562864 | 0.322965331 |
| 0.093552935 | 0.322965331 |
| 0.093525893 | 0.322896537 |
| 0.093497093 | 0.322820504 |
| 0.093475592 | 0.322769669 |
| 0.093440294 | 0.322671178 |
| 0.093425933 | 0.32264498 |
| 0.09337372 | 0.322511433 |
| 0.093347379 | 0.322443835 |

| | |
|---|---|
| 0.09332425 | 0.322387324 |
| 0.093314424 | 0.322376763 |
| 0.093209515 | 0.322037692 |
| 0.093197303 | 0.322018859 |
| 0.093192516 | 0.322018859 |
| 0.093191938 | 0.322018859 |
| 0.093183966 | 0.322018859 |
| 0.093129155 | 0.321876797 |
| 0.093093525 | 0.321800355 |
| 0.093086811 | 0.321800355 |
| 0.093084736 | 0.321800355 |
| 0.093017197 | 0.321699954 |
| 0.0929981 | 0.321680624 |
| 0.092990831 | 0.321678843 |
| 0.092968613 | 0.321628367 |
| 0.092957455 | 0.321628367 |
| 0.092800669 | 0.321271555 |
| 0.092799016 | 0.321271555 |
| 0.092792984 | 0.321271555 |
| 0.092792879 | 0.321271555 |
| 0.092747045 | 0.321255528 |
| 0.092728758 | 0.321215547 |
| 0.092669352 | 0.321033111 |
| 0.09266234 | 0.321032171 |
| 0.092652043 | 0.321032171 |
| 0.092575361 | 0.32082418 |
| 0.092568779 | 0.32082418 |
| 0.092537277 | 0.320762219 |
| 0.092522932 | 0.32073584 |
| 0.092519003 | 0.32073584 |
| 0.092514145 | 0.32073584 |
| 0.092511469 | 0.32073584 |
| 0.092501558 | 0.32073584 |
| 0.092450264 | 0.320623953 |
| 0.092417075 | 0.320532191 |
| 0.092385894 | 0.32044738 |
| 0.09236175 | 0.320410303 |
| 0.092356028 | 0.320410303 |
| 0.09235493 | 0.320410303 |
| 0.092347233 | 0.320410303 |
| 0.092323895 | 0.320372325 |
| 0.092254372 | 0.320224399 |
| 0.09224902 | 0.320224399 |
| 0.092231794 | 0.320192698 |
| 0.092211907 | 0.320146998 |

| | |
|---|---|
| 0.092150121 | 0.319979133 |
| 0.092135102 | 0.319950311 |
| 0.092104742 | 0.319897003 |
| 0.092033727 | 0.31969151 |
| 0.092027113 | 0.31969151 |
| 0.091996181 | 0.319631028 |
| 0.09198112 | 0.319602013 |
| 0.091926897 | 0.319466081 |
| 0.091916572 | 0.319466081 |
| 0.091901513 | 0.319465247 |
| 0.091894653 | 0.319465247 |
| 0.091863946 | 0.319404597 |
| 0.091862891 | 0.319404597 |
| 0.091814247 | 0.319301726 |
| 0.091803296 | 0.319286956 |
| 0.091790959 | 0.319267363 |
| 0.091771243 | 0.319222101 |
| 0.091714962 | 0.319049632 |
| 0.091681172 | 0.318978681 |
| 0.09164723 | 0.318907183 |
| 0.091631751 | 0.318876618 |
| 0.091628376 | 0.318876618 |
| 0.091619555 | 0.318876618 |
| 0.091614379 | 0.318876618 |
| 0.091589115 | 0.318821429 |
| 0.09157048 | 0.318779858 |
| 0.091565269 | 0.318779858 |
| 0.091563862 | 0.318779858 |
| 0.091512308 | 0.318647222 |
| 0.091502452 | 0.318636197 |
| 0.091500316 | 0.318636197 |
| 0.091450625 | 0.318525592 |
| 0.091441208 | 0.318516085 |
| 0.091425248 | 0.318483784 |
| 0.091337417 | 0.318336597 |
| 0.091316624 | 0.318308468 |
| 0.091284498 | 0.318226233 |
| 0.091280313 | 0.318226233 |
| 0.091272742 | 0.318226233 |
| 0.091262919 | 0.318220885 |
| 0.091239894 | 0.31816389 |
| 0.091221427 | 0.318146083 |
| 0.09119595 | 0.318080522 |
| 0.091181635 | 0.318080522 |
| 0.091177419 | 0.318080522 |

| | |
|---|---|
| 0.091156084 | 0.318067449 |
| 0.091113522 | 0.318025922 |
| 0.091112369 | 0.318025922 |
| 0.091102438 | 0.318025922 |
| 0.091095443 | 0.318025922 |
| 0.091094412 | 0.318025922 |
| 0.091075271 | 0.318025922 |
| 0.091054731 | 0.317983844 |
| 0.09105017 | 0.317983844 |
| 0.091033508 | 0.317968993 |
| 0.091030743 | 0.317968993 |
| 0.091012002 | 0.317968993 |
| 0.090962794 | 0.31782591 |
| 0.090912142 | 0.317672237 |
| 0.090898858 | 0.317649125 |
| 0.090839147 | 0.317487052 |
| 0.090827843 | 0.317470841 |
| 0.090821779 | 0.317470841 |
| 0.090807721 | 0.317447102 |
| 0.090803905 | 0.317447102 |
| 0.090783418 | 0.317447102 |
| 0.090779969 | 0.317447102 |
| 0.090773175 | 0.317447102 |
| 0.090727076 | 0.317351533 |
| 0.090723441 | 0.317351533 |
| 0.09066553 | 0.317182842 |
| 0.090652362 | 0.317182842 |
| 0.09064633 | 0.317182842 |
| 0.090638243 | 0.317180571 |
| 0.090592222 | 0.317042823 |
| 0.090573943 | 0.317032129 |
| 0.090568496 | 0.317032129 |
| 0.090498944 | 0.316879371 |
| 0.090446251 | 0.316718153 |
| 0.090444375 | 0.316718153 |
| 0.090368402 | 0.316492086 |
| 0.090354178 | 0.316465541 |
| 0.09031431 | 0.316349167 |
| 0.090303765 | 0.316335498 |
| 0.090300781 | 0.316335498 |
| 0.090294225 | 0.316335498 |
| 0.090201863 | 0.316187836 |
| 0.090120272 | 0.315925084 |
| 0.090116324 | 0.315925084 |
| 0.090094613 | 0.315922008 |

| | |
|---|---|
| 0.090007784 | 0.315693404 |
| 0.09000605 | 0.315693404 |
| 0.089974133 | 0.315621863 |
| 0.089973928 | 0.315621863 |
| 0.089955933 | 0.315604509 |
| 0.089939147 | 0.315568864 |
| 0.089937357 | 0.315568864 |
| 0.089860806 | 0.315433418 |
| 0.089857451 | 0.315433418 |
| 0.089847358 | 0.315433418 |
| 0.089843384 | 0.315433418 |
| 0.08979617 | 0.315369235 |
| 0.089757056 | 0.31525511 |
| 0.089737244 | 0.315208771 |
| 0.08972096 | 0.315174818 |
| 0.089697024 | 0.315113975 |
| 0.089675893 | 0.315086225 |
| 0.089667803 | 0.315081041 |
| 0.089640669 | 0.315008939 |
| 0.089628791 | 0.31499044 |
| 0.08962454 | 0.31499044 |
| 0.089595472 | 0.314978691 |
| 0.089578565 | 0.314953368 |
| 0.089541175 | 0.314855055 |
| 0.089521565 | 0.314822682 |
| 0.089497076 | 0.314783047 |
| 0.089492427 | 0.314783047 |
| 0.089461212 | 0.314737186 |
| 0.089429423 | 0.314661283 |
| 0.089370568 | 0.314477434 |
| 0.089355441 | 0.314447435 |
| 0.089343006 | 0.314426909 |
| 0.089338079 | 0.314426909 |
| 0.089329104 | 0.314426909 |
| 0.089305304 | 0.314410387 |
| 0.089296941 | 0.314404185 |
| 0.089237121 | 0.314263263 |
| 0.089227123 | 0.31425129 |
| 0.089210528 | 0.314239317 |
| 0.08916839 | 0.314114121 |
| 0.089163491 | 0.314114121 |
| 0.089152454 | 0.314104447 |
| 0.089122868 | 0.314046672 |
| 0.089110918 | 0.314027797 |
| 0.089073547 | 0.313957102 |

| | |
|---|---|
| 0.089070585 | 0.313957102 |
| 0.089024369 | 0.313862136 |
| 0.089007086 | 0.313824437 |
| 0.088957003 | 0.313694298 |
| 0.088942094 | 0.313688176 |
| 0.088904844 | 0.313649704 |
| 0.088870927 | 0.313553275 |
| 0.088848482 | 0.313497307 |
| 0.088841856 | 0.313497153 |
| 0.088840566 | 0.313497153 |
| 0.088805076 | 0.313413808 |
| 0.088779889 | 0.313348139 |
| 0.088763979 | 0.313316847 |
| 0.088688362 | 0.313141122 |
| 0.088668089 | 0.313092754 |
| 0.088561559 | 0.312809359 |
| 0.088502423 | 0.312646854 |
| 0.088494786 | 0.312643067 |
| 0.088478937 | 0.312610262 |
| 0.088462168 | 0.312574201 |
| 0.088447286 | 0.312567996 |
| 0.088442885 | 0.312567996 |
| 0.088402634 | 0.312479756 |
| 0.088401344 | 0.312479756 |
| 0.088399195 | 0.312479756 |
| 0.088370249 | 0.312434849 |
| 0.088366663 | 0.312434849 |
| 0.088312788 | 0.312307389 |
| 0.088304406 | 0.312307389 |
| 0.088294326 | 0.312305522 |
| 0.088246469 | 0.312176232 |
| 0.088208065 | 0.31210948 |
| 0.088184274 | 0.31205531 |
| 0.088168869 | 0.312023981 |
| 0.088148108 | 0.311973689 |
| 0.088146387 | 0.311973689 |
| 0.088133863 | 0.311973689 |
| 0.088121757 | 0.311973689 |
| 0.088112217 | 0.311973689 |
| 0.088106692 | 0.311973689 |
| 0.08810181 | 0.311973689 |
| 0.088062776 | 0.31190346 |
| 0.08805378 | 0.311894792 |
| 0.088029013 | 0.311830258 |
| 0.087999029 | 0.311747231 |

| | |
|---|---|
| 0.087985671 | 0.311723092 |
| 0.087974296 | 0.31170598 |
| 0.087971682 | 0.31170598 |
| 0.087944608 | 0.311666366 |
| 0.087920975 | 0.31164886 |
| 0.087916573 | 0.31164886 |
| 0.087914769 | 0.31164886 |
| 0.087905189 | 0.31164886 |
| 0.087896737 | 0.31164886 |
| 0.087873533 | 0.311606231 |
| 0.087843247 | 0.311542952 |
| 0.087752134 | 0.311358926 |
| 0.087741094 | 0.311342949 |
| 0.087740846 | 0.311342949 |
| 0.087707397 | 0.311269762 |
| 0.087702853 | 0.311269762 |
| 0.087680632 | 0.311262118 |
| 0.087675381 | 0.311262118 |
| 0.087630473 | 0.311182266 |
| 0.087615339 | 0.311151721 |
| 0.087609143 | 0.311151721 |
| 0.087605883 | 0.311151721 |
| 0.087603151 | 0.311151721 |
| 0.087600203 | 0.311151721 |
| 0.087590661 | 0.311151721 |
| 0.087587519 | 0.311151721 |
| 0.087574092 | 0.311151721 |
| 0.087535634 | 0.311151721 |
| 0.087530259 | 0.311151721 |
| 0.087494559 | 0.311070718 |
| 0.087494332 | 0.311070718 |
| 0.087445347 | 0.310965396 |
| 0.087404659 | 0.310890343 |
| 0.08739066 | 0.310863767 |
| 0.087355196 | 0.310760825 |
| 0.087334757 | 0.310711321 |
| 0.087331392 | 0.310711321 |
| 0.087330411 | 0.310711321 |
| 0.087325184 | 0.310711321 |
| 0.08732403 | 0.310711321 |
| 0.087287359 | 0.310658725 |
| 0.087272563 | 0.310629277 |
| 0.087260941 | 0.310611125 |
| 0.087259332 | 0.310611125 |
| 0.087244062 | 0.310597469 |

| | |
|---|---|
| 0.087227684 | 0.310562376 |
| 0.087222038 | 0.310562376 |
| 0.087214735 | 0.310562376 |
| 0.087203633 | 0.310546392 |
| 0.087074095 | 0.310177853 |
| 0.087069889 | 0.310177853 |
| 0.087057749 | 0.310166026 |
| 0.087045322 | 0.310144949 |
| 0.087023678 | 0.310091032 |
| 0.087013493 | 0.310077938 |
| 0.087006 | 0.310074437 |
| 0.086979444 | 0.310002991 |
| 0.086934544 | 0.309866152 |
| 0.086879753 | 0.309719911 |
| 0.086866104 | 0.309719911 |
| 0.086847832 | 0.309696142 |
| 0.086824586 | 0.309659618 |
| 0.086786681 | 0.309593982 |
| 0.086749898 | 0.309485948 |
| 0.086727029 | 0.309427537 |
| 0.086710811 | 0.309392853 |
| 0.086671839 | 0.309300141 |
| 0.08664318 | 0.309238358 |
| 0.086615343 | 0.309168022 |
| 0.086609071 | 0.309168022 |
| 0.086608963 | 0.309168022 |
| 0.0866042 | 0.309168022 |
| 0.086580967 | 0.309137994 |
| 0.086544798 | 0.309050162 |
| 0.086538778 | 0.309050162 |
| 0.086532408 | 0.309050162 |
| 0.086529666 | 0.309050162 |
| 0.086507827 | 0.309050162 |
| 0.086489438 | 0.309050162 |
| 0.08644313 | 0.308970128 |
| 0.086415905 | 0.308895999 |
| 0.086402108 | 0.30889305 |
| 0.086400937 | 0.30889305 |
| 0.086372318 | 0.308853462 |
| 0.086346918 | 0.308788457 |
| 0.086333477 | 0.308782567 |
| 0.086321324 | 0.308766483 |
| 0.086314099 | 0.308763827 |
| 0.08628204 | 0.308695515 |
| 0.086271492 | 0.308680967 |

| | |
|---|---|
| 0.086266444 | 0.308680967 |
| 0.086260626 | 0.308680967 |
| 0.086260193 | 0.308680967 |
| 0.086253791 | 0.308680967 |
| 0.086221821 | 0.308619168 |
| 0.086208846 | 0.308595919 |
| 0.086196353 | 0.308590182 |
| 0.086116038 | 0.308333223 |
| 0.086110298 | 0.308333223 |
| 0.086103643 | 0.308333223 |
| 0.086084362 | 0.308333223 |
| 0.086061416 | 0.308323084 |
| 0.086057112 | 0.308323084 |
| 0.086054386 | 0.308323084 |
| 0.086040518 | 0.308317791 |
| 0.08602488 | 0.308308146 |
| 0.086018377 | 0.30830804 |
| 0.086005168 | 0.308307101 |
| 0.085999665 | 0.308307101 |
| 0.085999165 | 0.308307101 |
| 0.085998294 | 0.308307101 |
| 0.085994241 | 0.308307101 |
| 0.085980556 | 0.308307101 |
| 0.085972053 | 0.308307101 |
| 0.085963951 | 0.308307101 |
| 0.085937616 | 0.308250552 |
| 0.085916686 | 0.30819869 |
| 0.085902256 | 0.308193353 |
| 0.085885764 | 0.308157395 |
| 0.085825191 | 0.308037263 |
| 0.085816301 | 0.308037263 |
| 0.085814141 | 0.308037263 |
| 0.085807813 | 0.308037263 |
| 0.085797321 | 0.308037263 |
| 0.085762268 | 0.307946288 |
| 0.085753715 | 0.307938789 |
| 0.085745457 | 0.307932353 |
| 0.085703109 | 0.307849911 |
| 0.085695833 | 0.307846992 |
| 0.085645361 | 0.30768889 |
| 0.085539339 | 0.307400747 |
| 0.085470394 | 0.307176158 |
| 0.085464719 | 0.307176158 |
| 0.085444691 | 0.307130137 |
| 0.085405502 | 0.307081976 |

| | |
|---|---|
| 0.085369888 | 0.306977099 |
| 0.085338947 | 0.306889009 |
| 0.085310029 | 0.306831349 |
| 0.085296843 | 0.306807093 |
| 0.085287303 | 0.30679595 |
| 0.085235897 | 0.306634192 |
| 0.085186996 | 0.306527733 |
| 0.085184968 | 0.306527733 |
| 0.085142949 | 0.30641554 |
| 0.085136645 | 0.30641554 |
| 0.08513095 | 0.30641554 |
| 0.085121094 | 0.306406344 |
| 0.085117487 | 0.306406344 |
| 0.085107265 | 0.306402882 |
| 0.085095329 | 0.30638307 |
| 0.08506728 | 0.306305234 |
| 0.08506728 | 0.306305234 |
| 0.085066266 | 0.306305234 |
| 0.085057638 | 0.306305234 |
| 0.085043917 | 0.306305234 |
| 0.085036418 | 0.306305234 |
| 0.085036043 | 0.306305234 |
| 0.085010294 | 0.306262125 |
| 0.085006678 | 0.306262125 |
| 0.084993276 | 0.306247144 |
| 0.084986223 | 0.306244901 |
| 0.084986223 | 0.306244901 |
| 0.084986223 | 0.306244901 |
| 0.084972684 | 0.306244901 |
| 0.084937087 | 0.306160963 |
| 0.084839159 | 0.305902749 |
| 0.084834838 | 0.305902749 |
| 0.084810699 | 0.305888733 |
| 0.084808051 | 0.305888733 |
| 0.084757287 | 0.305766951 |
| 0.084756426 | 0.305766951 |
| 0.084705179 | 0.305625268 |
| 0.084701172 | 0.305625268 |
| 0.084690654 | 0.305625268 |
| 0.084670643 | 0.305593263 |
| 0.084662666 | 0.305587631 |
| 0.08465323 | 0.305587631 |
| 0.08463832 | 0.305587631 |
| 0.084618088 | 0.305587631 |
| 0.084611315 | 0.305587555 |

| | |
|---|---|
| 0.084609259 | 0.305587555 |
| 0.084600506 | 0.305587555 |
| 0.084554121 | 0.305520001 |
| 0.084530087 | 0.305456331 |
| 0.08449909 | 0.305367492 |
| 0.084491123 | 0.305361869 |
| 0.084490681 | 0.305361869 |
| 0.084485899 | 0.305361869 |
| 0.084473731 | 0.305361869 |
| 0.084419668 | 0.305219427 |
| 0.08436879 | 0.305104964 |
| 0.084228592 | 0.304690501 |
| 0.084226649 | 0.304690501 |
| 0.084211687 | 0.304675633 |
| 0.084175068 | 0.304589421 |
| 0.08414393 | 0.304523019 |
| 0.084139598 | 0.304523019 |
| 0.084128481 | 0.304513388 |
| 0.08412151 | 0.304511299 |
| 0.08409713 | 0.304446185 |
| 0.084062056 | 0.304342343 |
| 0.084051938 | 0.304328848 |
| 0.084028308 | 0.30426642 |
| 0.083993709 | 0.304164264 |
| 0.083978051 | 0.304130684 |
| 0.083973784 | 0.304130684 |
| 0.083963737 | 0.304125097 |
| 0.08391741 | 0.303992915 |
| 0.083910823 | 0.303992915 |
| 0.083909806 | 0.303992915 |
| 0.083883013 | 0.303992915 |
| 0.083880884 | 0.303992915 |
| 0.083872666 | 0.303992915 |
| 0.083841565 | 0.303913709 |
| 0.083780999 | 0.303763524 |
| 0.083773577 | 0.303763524 |
| 0.083757126 | 0.303746336 |
| 0.083747158 | 0.303733316 |
| 0.083720969 | 0.303661458 |
| 0.083712612 | 0.303654269 |
| 0.083700089 | 0.303631972 |
| 0.083698941 | 0.303631972 |
| 0.083691477 | 0.303631972 |
| 0.083659584 | 0.303554395 |
| 0.083635075 | 0.303488589 |

| | |
|---|---|
| 0.083589757 | 0.303393489 |
| 0.08357178 | 0.303351358 |
| 0.083501995 | 0.303121152 |
| 0.083492963 | 0.303111468 |
| 0.08348418 | 0.303102688 |
| 0.083456234 | 0.303024325 |
| 0.083405598 | 0.302909745 |
| 0.083382857 | 0.30285025 |
| 0.08338167 | 0.30285025 |
| 0.083328709 | 0.302745934 |
| 0.083305798 | 0.302708874 |
| 0.083289702 | 0.302696576 |
| 0.083265477 | 0.302631629 |
| 0.083235311 | 0.302545077 |
| 0.083206275 | 0.302482009 |
| 0.083168677 | 0.302418277 |
| 0.083167898 | 0.302418277 |
| 0.083123726 | 0.302300986 |
| 0.083115517 | 0.302294212 |
| 0.083112889 | 0.302294212 |
| 0.083092168 | 0.302278544 |
| 0.083047567 | 0.302162451 |
| 0.08302777 | 0.3021135 |
| 0.0830263 | 0.3021135 |
| 0.082999828 | 0.302076466 |
| 0.082982352 | 0.302040541 |
| 0.082978722 | 0.302040541 |
| 0.082953582 | 0.301981981 |
| 0.082932069 | 0.301926743 |
| 0.082926958 | 0.301926743 |
| 0.082817089 | 0.301646491 |
| 0.082802672 | 0.301617045 |
| 0.082781443 | 0.301562778 |
| 0.082763767 | 0.301544516 |
| 0.082736918 | 0.301469756 |
| 0.082694891 | 0.301384737 |
| 0.08267466 | 0.301335123 |
| 0.082670433 | 0.301335123 |
| 0.082663574 | 0.301335123 |
| 0.082609671 | 0.301198489 |
| 0.082600351 | 0.301198489 |
| 0.082567115 | 0.301150559 |
| 0.082549035 | 0.301126741 |
| 0.082539816 | 0.301120171 |
| 0.082537703 | 0.301120171 |

| | |
|---|---|
| 0.08253442 | 0.301120171 |
| 0.082526542 | 0.301120171 |
| 0.082521843 | 0.301120171 |
| 0.08250384 | 0.301104246 |
| 0.08248362 | 0.30105352 |
| 0.082465496 | 0.301010438 |
| 0.082464922 | 0.301010438 |
| 0.082461004 | 0.301010438 |
| 0.0824606 | 0.301010438 |
| 0.082420306 | 0.300937743 |
| 0.082318693 | 0.30063587 |
| 0.082298345 | 0.300584607 |
| 0.082295137 | 0.300584607 |
| 0.082272953 | 0.30053796 |
| 0.082236469 | 0.300450774 |
| 0.082228023 | 0.300442965 |
| 0.082180204 | 0.300314322 |
| 0.08217164 | 0.300306069 |
| 0.082135958 | 0.300221738 |
| 0.082105118 | 0.300155085 |
| 0.082061083 | 0.300017134 |
| 0.081981758 | 0.299773144 |
| 0.081968674 | 0.299764809 |
| 0.081966526 | 0.299764809 |
| 0.081948055 | 0.299741956 |
| 0.081933844 | 0.299712995 |
| 0.0819216 | 0.299691225 |
| 0.081899316 | 0.299632717 |
| 0.081834685 | 0.299488284 |
| 0.081818801 | 0.299453165 |
| 0.081762884 | 0.299271508 |
| 0.081697369 | 0.299054692 |
| 0.081694518 | 0.299054692 |
| 0.081694468 | 0.299054692 |
| 0.081680746 | 0.299054692 |
| 0.081665816 | 0.299031124 |
| 0.081621211 | 0.298890774 |
| 0.08158707 | 0.298811703 |
| 0.081585391 | 0.298811703 |
| 0.08155064 | 0.298724223 |
| 0.081549561 | 0.298724223 |
| 0.081547119 | 0.298724223 |
| 0.081540733 | 0.298724223 |
| 0.08151318 | 0.298701872 |
| 0.081402869 | 0.298397626 |

| | |
|---|---|
| 0.081395003 | 0.298397626 |
| 0.081341905 | 0.298280791 |
| 0.081305713 | 0.298193993 |
| 0.081283459 | 0.298135335 |
| 0.081272245 | 0.298135335 |
| 0.081259451 | 0.29811615 |
| 0.081256379 | 0.29811615 |
| 0.081252876 | 0.29811615 |
| 0.081216353 | 0.298049872 |
| 0.081209769 | 0.298048676 |
| 0.081191941 | 0.298022757 |
| 0.081170072 | 0.297994837 |
| 0.081165428 | 0.297994837 |
| 0.081156944 | 0.297992582 |
| 0.081156397 | 0.297992582 |
| 0.081109732 | 0.297888127 |
| 0.081088437 | 0.297832884 |
| 0.081085562 | 0.297832884 |
| 0.0810734 | 0.29782359 |
| 0.081072532 | 0.29782359 |
| 0.081062097 | 0.29782359 |
| 0.0810355 | 0.29775326 |
| 0.081033237 | 0.29775326 |
| 0.08102129 | 0.297746988 |
| 0.081019507 | 0.297746988 |
| 0.081012261 | 0.297746988 |
| 0.08098566 | 0.29768496 |
| 0.080977996 | 0.297679762 |
| 0.080951398 | 0.297627932 |
| 0.080922038 | 0.297565929 |
| 0.080891005 | 0.297509164 |
| 0.080868054 | 0.29748226 |
| 0.080858555 | 0.297470292 |
| 0.080852662 | 0.297470292 |
| 0.080798364 | 0.297294783 |
| 0.080787993 | 0.297279592 |
| 0.080786688 | 0.297279592 |
| 0.080768401 | 0.297253431 |
| 0.080760888 | 0.297248751 |
| 0.080722857 | 0.297131734 |
| 0.080716963 | 0.297131734 |
| 0.08069957 | 0.297091939 |
| 0.080675271 | 0.297048406 |
| 0.080616802 | 0.296901982 |
| 0.080598735 | 0.296881359 |

| | |
|---|---|
| 0.080588502 | 0.296866626 |
| 0.080583501 | 0.296866626 |
| 0.080571806 | 0.29685104 |
| 0.080566985 | 0.29685104 |
| 0.080543123 | 0.296791283 |
| 0.080539654 | 0.296791283 |
| 0.080537282 | 0.296791283 |
| 0.080525521 | 0.296791283 |
| 0.080480247 | 0.2966514 |
| 0.080478451 | 0.2966514 |
| 0.080463078 | 0.2966514 |
| 0.080462428 | 0.2966514 |
| 0.080425477 | 0.296564278 |
| 0.080417232 | 0.296556834 |
| 0.080392482 | 0.29648852 |
| 0.080369742 | 0.296427609 |
| 0.080361429 | 0.296424794 |
| 0.080353 | 0.296424794 |
| 0.080351674 | 0.296424794 |
| 0.080333688 | 0.296424794 |
| 0.080316541 | 0.296392059 |
| 0.080295813 | 0.296351452 |
| 0.080295007 | 0.296351452 |
| 0.080277378 | 0.29633938 |
| 0.0802721 | 0.29633938 |
| 0.080206218 | 0.296145556 |
| 0.08019847 | 0.296139909 |
| 0.080173068 | 0.29609202 |
| 0.080101007 | 0.29594063 |
| 0.080086852 | 0.295911287 |
| 0.080053688 | 0.2958117 |
| 0.080053541 | 0.2958117 |
| 0.080051635 | 0.2958117 |
| 0.080041614 | 0.2958117 |
| 0.080035672 | 0.2958117 |
| 0.080029819 | 0.2958117 |
| 0.080027714 | 0.2958117 |
| 0.079959517 | 0.295716099 |
| 0.079952461 | 0.295716099 |
| 0.079940696 | 0.295715381 |
| 0.079914304 | 0.295663682 |
| 0.079894653 | 0.29561394 |
| 0.0798846 | 0.29559971 |
| 0.079868155 | 0.295595242 |
| 0.079851274 | 0.295568246 |

| | |
|---|---|
| 0.079819253 | 0.295472684 |
| 0.079755947 | 0.295307197 |
| 0.079733022 | 0.295268226 |
| 0.079730277 | 0.295268226 |
| 0.079695854 | 0.295197684 |
| 0.079690664 | 0.295197684 |
| 0.079665738 | 0.295156756 |
| 0.079655775 | 0.295142803 |
| 0.079652082 | 0.295142803 |
| 0.079648548 | 0.295142803 |
| 0.079638491 | 0.295142803 |
| 0.079632115 | 0.295142803 |
| 0.079619538 | 0.295123337 |
| 0.079614972 | 0.295123337 |
| 0.079614691 | 0.295123337 |
| 0.079610833 | 0.295123337 |
| 0.079594192 | 0.295123337 |
| 0.079591279 | 0.295123337 |
| 0.079576525 | 0.295123337 |
| 0.079537741 | 0.295026791 |
| 0.079531151 | 0.295026791 |
| 0.079530391 | 0.295026791 |
| 0.079453631 | 0.294829615 |
| 0.079447352 | 0.294829286 |
| 0.07938755 | 0.294676241 |
| 0.079366385 | 0.294620641 |
| 0.079360423 | 0.294620641 |
| 0.07930422 | 0.294481686 |
| 0.079283134 | 0.29442634 |
| 0.079233913 | 0.294335352 |
| 0.07923262 | 0.294335352 |
| 0.0792237 | 0.294335352 |
| 0.079213998 | 0.294330245 |
| 0.07920429 | 0.294330245 |
| 0.079159454 | 0.294219394 |
| 0.079122145 | 0.294103677 |
| 0.079107754 | 0.294073135 |
| 0.079084893 | 0.294021625 |
| 0.079056751 | 0.293952367 |
| 0.0790544 | 0.293952367 |
| 0.079051348 | 0.293952367 |
| 0.079027874 | 0.293936788 |
| 0.078974195 | 0.293805966 |
| 0.078967658 | 0.293804597 |
| 0.078946183 | 0.29379355 |

| | |
|---|---|
| 0.078868072 | 0.293548741 |
| 0.078860269 | 0.293542636 |
| 0.078854759 | 0.293542636 |
| 0.078840651 | 0.29351549 |
| 0.078828493 | 0.293493171 |
| 0.078824251 | 0.293493171 |
| 0.078824145 | 0.293493171 |
| 0.078821682 | 0.293493171 |
| 0.078797296 | 0.293468989 |
| 0.078782382 | 0.293459135 |
| 0.078747434 | 0.2933519 |
| 0.078705336 | 0.293231535 |
| 0.078681623 | 0.293175534 |
| 0.078663392 | 0.293153471 |
| 0.078632833 | 0.293062519 |
| 0.078621705 | 0.293043977 |
| 0.078598468 | 0.292980295 |
| 0.078591204 | 0.292976148 |
| 0.078562767 | 0.292915994 |
| 0.078561651 | 0.292915994 |
| 0.078549669 | 0.292913019 |
| 0.078530659 | 0.292887995 |
| 0.078516438 | 0.29285789 |
| 0.078495558 | 0.29280294 |
| 0.078486783 | 0.292793141 |
| 0.078453522 | 0.292691988 |
| 0.07843536 | 0.292647153 |
| 0.078431569 | 0.292647153 |
| 0.078402628 | 0.292593793 |
| 0.078383925 | 0.292546916 |
| 0.078382701 | 0.292546916 |
| 0.078372602 | 0.292546916 |
| 0.078361428 | 0.292546916 |
| 0.078331589 | 0.29248911 |
| 0.078320088 | 0.292469096 |
| 0.078315574 | 0.292469096 |
| 0.078300892 | 0.292443269 |
| 0.078294674 | 0.292442981 |
| 0.078271123 | 0.292400877 |
| 0.078208808 | 0.292213926 |
| 0.07819652 | 0.292190935 |
| 0.078175378 | 0.292134855 |
| 0.078172096 | 0.292134855 |
| 0.07814968 | 0.292107582 |
| 0.078140593 | 0.292107582 |

| | |
|---|---|
| 0.078127803 | 0.292094581 |
| 0.078125504 | 0.292094581 |
| 0.078072557 | 0.291979714 |
| 0.078066304 | 0.291979258 |
| 0.0780646 | 0.291979258 |
| 0.078032412 | 0.291898341 |
| 0.077994241 | 0.291824316 |
| 0.077951987 | 0.291712053 |
| 0.077943949 | 0.291704891 |
| 0.07791609 | 0.291623547 |
| 0.077905245 | 0.291606673 |
| 0.077887754 | 0.291606673 |
| 0.077863571 | 0.291541531 |
| 0.07779909 | 0.291323001 |
| 0.077776582 | 0.291270945 |
| 0.077760958 | 0.291270945 |
| 0.077741101 | 0.291220342 |
| 0.077727711 | 0.291193083 |
| 0.077721768 | 0.291193083 |
| 0.077720196 | 0.291193083 |
| 0.077692317 | 0.291152097 |
| 0.077672967 | 0.291102487 |
| 0.077663841 | 0.29109119 |
| 0.077644318 | 0.291053691 |
| 0.077638699 | 0.291053691 |
| 0.077619636 | 0.291040027 |
| 0.077618676 | 0.291040027 |
| 0.077617178 | 0.291040027 |
| 0.077551611 | 0.290880718 |
| 0.077545438 | 0.290880718 |
| 0.077535326 | 0.290880718 |
| 0.077518828 | 0.290845221 |
| 0.077420927 | 0.290546566 |
| 0.077420577 | 0.290546566 |
| 0.077389431 | 0.290474147 |
| 0.077380492 | 0.290463486 |
| 0.077380391 | 0.290463486 |
| 0.077280501 | 0.290202508 |
| 0.077278952 | 0.290202508 |
| 0.077250359 | 0.290135071 |
| 0.077244795 | 0.290135071 |
| 0.077225685 | 0.290111033 |
| 0.077223584 | 0.290111033 |
| 0.07722084 | 0.290111033 |
| 0.077175768 | 0.290060786 |

| | |
|---|---|
| 0.077152543 | 0.29001927 |
| 0.077120202 | 0.289920584 |
| 0.077082269 | 0.289800855 |
| 0.077081709 | 0.289800855 |
| 0.07706073 | 0.289788503 |
| 0.07700449 | 0.289599877 |
| 0.077002706 | 0.289599877 |
| 0.076985649 | 0.28957475 |
| 0.076941658 | 0.289432142 |
| 0.076922353 | 0.289418366 |
| 0.076868197 | 0.28927004 |
| 0.07682799 | 0.289164428 |
| 0.076818676 | 0.289164428 |
| 0.076758948 | 0.288984363 |
| 0.076751951 | 0.288984363 |
| 0.076743467 | 0.288983327 |
| 0.076710064 | 0.288903228 |
| 0.076685915 | 0.288880808 |
| 0.07666884 | 0.288862183 |
| 0.076666989 | 0.288862183 |
| 0.076654918 | 0.288855432 |
| 0.076654007 | 0.288855432 |
| 0.076633227 | 0.288819396 |
| 0.076602732 | 0.288727312 |
| 0.076541832 | 0.288566282 |
| 0.07653413 | 0.288560087 |
| 0.076419134 | 0.288263432 |
| 0.076407315 | 0.288241678 |
| 0.076395497 | 0.288219926 |
| 0.076333912 | 0.288010397 |
| 0.076275587 | 0.287843817 |
| 0.076257595 | 0.287829902 |
| 0.076256838 | 0.287829902 |
| 0.076247263 | 0.287829902 |
| 0.076239474 | 0.287829902 |
| 0.076217209 | 0.287775252 |
| 0.076175553 | 0.287663579 |
| 0.076147918 | 0.287594438 |
| 0.076137125 | 0.287594438 |
| 0.076132893 | 0.287594438 |
| 0.076130246 | 0.287594438 |
| 0.076067831 | 0.287439113 |
| 0.076052435 | 0.28740374 |
| 0.076046186 | 0.287402929 |
| 0.076041477 | 0.287402929 |

| | |
|---|---|
| 0.076024002 | 0.287364692 |
| 0.07601187 | 0.287341641 |
| 0.075978109 | 0.287236816 |
| 0.075962365 | 0.28720009 |
| 0.07594029 | 0.287139425 |
| 0.075923353 | 0.287098177 |
| 0.075862597 | 0.286891212 |
| 0.07585322 | 0.286878527 |
| 0.07584102 | 0.286855167 |
| 0.075838486 | 0.286855167 |
| 0.075831261 | 0.286855167 |
| 0.075810527 | 0.286808163 |
| 0.07578122 | 0.286720063 |
| 0.075780317 | 0.286720063 |
| 0.07577526 | 0.286720063 |
| 0.075772912 | 0.286720063 |
| 0.075750798 | 0.286720063 |
| 0.07570052 | 0.286619658 |
| 0.075634682 | 0.286415917 |
| 0.075584849 | 0.286295501 |
| 0.075581341 | 0.286295501 |
| 0.075576546 | 0.286295501 |
| 0.075554401 | 0.286271236 |
| 0.075453594 | 0.285912033 |
| 0.075444895 | 0.285901816 |
| 0.075419477 | 0.285828237 |
| 0.075410915 | 0.285818532 |
| 0.075398129 | 0.285803041 |
| 0.075346433 | 0.285710555 |
| 0.075334026 | 0.285686255 |
| 0.075297281 | 0.285592387 |
| 0.075275723 | 0.285556104 |
| 0.075269674 | 0.2855559 |
| 0.075263085 | 0.285553649 |
| 0.07526292 | 0.285553649 |
| 0.075260187 | 0.285553649 |
| 0.075230895 | 0.2855225 |
| 0.075219892 | 0.285503491 |
| 0.07520379 | 0.285465123 |
| 0.075201989 | 0.285465123 |
| 0.075201989 | 0.285465123 |
| 0.075196184 | 0.285465123 |
| 0.075163545 | 0.285403345 |
| 0.075153703 | 0.285388726 |
| 0.07510759 | 0.285275706 |

| | |
|---|---|
| 0.075100396 | 0.285275706 |
| 0.075072074 | 0.285215192 |
| 0.07505946 | 0.285190019 |
| 0.075044088 | 0.285154363 |
| 0.075031555 | 0.285129489 |
| 0.074957864 | 0.284904471 |
| 0.074934702 | 0.284891773 |
| 0.074923773 | 0.284879016 |
| 0.074915292 | 0.28486951 |
| 0.074914647 | 0.28486951 |
| 0.074911356 | 0.28486951 |
| 0.074876831 | 0.284791475 |
| 0.074860826 | 0.284753338 |
| 0.074852932 | 0.284753338 |
| 0.074838063 | 0.284734976 |
| 0.074813334 | 0.284663629 |
| 0.074811575 | 0.284663629 |
| 0.074804331 | 0.284663629 |
| 0.074752979 | 0.284524905 |
| 0.074741453 | 0.284503771 |
| 0.074738423 | 0.284503771 |
| 0.074737944 | 0.284503771 |
| 0.074725292 | 0.284503771 |
| 0.074711115 | 0.284501979 |
| 0.074708242 | 0.284501979 |
| 0.074698246 | 0.284498468 |
| 0.074652165 | 0.284368443 |
| 0.074630335 | 0.284330772 |
| 0.074624287 | 0.284330472 |
| 0.074612337 | 0.284307688 |
| 0.074606172 | 0.284307688 |
| 0.074604134 | 0.284307688 |
| 0.074599487 | 0.284307688 |
| 0.074585393 | 0.284307688 |
| 0.074571374 | 0.284307688 |
| 0.074566007 | 0.284307688 |
| 0.074549162 | 0.284307688 |
| 0.074486057 | 0.28414986 |
| 0.07445447 | 0.284092537 |
| 0.074384812 | 0.283917729 |
| 0.074377551 | 0.28391277 |
| 0.074377506 | 0.28391277 |
| 0.074376884 | 0.28391277 |
| 0.074372483 | 0.28391277 |
| 0.074371637 | 0.28391277 |

| | |
|---|---|
| 0.074369975 | 0.28391277 |
| 0.074356763 | 0.28391277 |
| 0.074349448 | 0.28391277 |
| 0.074340427 | 0.28391277 |
| 0.07433597 | 0.28391277 |
| 0.074335746 | 0.28391277 |
| 0.074239865 | 0.283682769 |
| 0.07422943 | 0.283665655 |
| 0.074224572 | 0.283665655 |
| 0.074222797 | 0.283665655 |
| 0.07419505 | 0.283638042 |
| 0.074193099 | 0.283638042 |
| 0.074186368 | 0.283638042 |
| 0.074174052 | 0.283638042 |
| 0.074173267 | 0.283638042 |
| 0.074139996 | 0.283596948 |
| 0.074128595 | 0.283576113 |
| 0.074083153 | 0.283493359 |
| 0.074068724 | 0.283470102 |
| 0.074049528 | 0.283433003 |
| 0.073991836 | 0.283257709 |
| 0.073984189 | 0.283251204 |
| 0.073982413 | 0.283251204 |
| 0.073973381 | 0.283251204 |
| 0.073971434 | 0.283251204 |
| 0.073966075 | 0.283251204 |
| 0.073946073 | 0.283251204 |
| 0.073923869 | 0.28322517 |
| 0.073917882 | 0.283225016 |
| 0.073915237 | 0.283225016 |
| 0.073876713 | 0.283135608 |
| 0.073861173 | 0.283121617 |
| 0.073858938 | 0.283121617 |
| 0.073810431 | 0.282972663 |
| 0.073777558 | 0.282869406 |
| 0.073715644 | 0.282654778 |
| 0.073707875 | 0.282647743 |
| 0.073687837 | 0.282616412 |
| 0.073687749 | 0.282616412 |
| 0.073687219 | 0.282616412 |
| 0.073682453 | 0.282616412 |
| 0.073655979 | 0.282608017 |
| 0.073567515 | 0.282291334 |
| 0.073542935 | 0.282238391 |
| 0.073532713 | 0.282226007 |

| | |
|---|---|
| 0.073510247 | 0.282162519 |
| 0.073502511 | 0.282155563 |
| 0.073471178 | 0.282058018 |
| 0.073463793 | 0.282052403 |
| 0.073460402 | 0.282052403 |
| 0.07344794 | 0.282037009 |
| 0.073430296 | 0.282014734 |
| 0.073425662 | 0.282014734 |
| 0.073409553 | 0.281980546 |
| 0.073393602 | 0.281964758 |
| 0.073388588 | 0.281964758 |
| 0.073301274 | 0.281739112 |
| 0.073296399 | 0.281739112 |
| 0.07327131 | 0.281676705 |
| 0.073260944 | 0.281659587 |
| 0.073251596 | 0.281646383 |
| 0.07324566 | 0.281646297 |
| 0.073245491 | 0.281646297 |
| 0.073239576 | 0.281646297 |
| 0.073220556 | 0.281626573 |
| 0.073171288 | 0.281496601 |
| 0.073160657 | 0.281478521 |
| 0.073146903 | 0.281478521 |
| 0.073141903 | 0.281478521 |
| 0.073136928 | 0.281478521 |
| 0.073117201 | 0.281447743 |
| 0.073110882 | 0.281446167 |
| 0.073089251 | 0.281385639 |
| 0.073086783 | 0.281385639 |
| 0.073063419 | 0.281365467 |
| 0.073061235 | 0.281365467 |
| 0.07304244 | 0.281341874 |
| 0.073036776 | 0.281341874 |
| 0.073014872 | 0.281281186 |
| 0.073011443 | 0.281281186 |
| 0.072966306 | 0.281139574 |
| 0.07295533 | 0.281120024 |
| 0.072937744 | 0.281097744 |
| 0.072925821 | 0.281090074 |
| 0.072913898 | 0.281083554 |
| 0.072905057 | 0.281083554 |
| 0.072891249 | 0.281077768 |
| 0.072885658 | 0.281077768 |
| 0.072843087 | 0.280960292 |
| 0.072842543 | 0.280960292 |

| | |
|---|---|
| 0.072824795 | 0.280935242 |
| 0.072796122 | 0.280870133 |
| 0.072789874 | 0.28086878 |
| 0.072782462 | 0.280862934 |
| 0.072778999 | 0.280862934 |
| 0.072760128 | 0.280824276 |
| 0.072748308 | 0.280822149 |
| 0.072727995 | 0.280766492 |
| 0.072724319 | 0.280766492 |
| 0.072719866 | 0.280766492 |
| 0.072667623 | 0.280624414 |
| 0.072653356 | 0.280592067 |
| 0.072651645 | 0.280592067 |
| 0.07265004 | 0.280592067 |
| 0.072638262 | 0.280592067 |
| 0.072587331 | 0.280473537 |
| 0.072570479 | 0.280431172 |
| 0.072551215 | 0.280379482 |
| 0.07250611 | 0.280250651 |
| 0.07246605 | 0.280159457 |
| 0.072459695 | 0.280159457 |
| 0.072441327 | 0.280136657 |
| 0.072436463 | 0.280136657 |
| 0.07240321 | 0.280034732 |
| 0.072394729 | 0.280024668 |
| 0.072386938 | 0.280017276 |
| 0.072370937 | 0.279978118 |
| 0.072295808 | 0.279755641 |
| 0.072293517 | 0.279755641 |
| 0.072287829 | 0.279755641 |
| 0.072285222 | 0.279755641 |
| 0.072248567 | 0.279709193 |
| 0.072244315 | 0.279709193 |
| 0.072225168 | 0.279689155 |
| 0.072189869 | 0.279595583 |
| 0.072183467 | 0.279593526 |
| 0.072170568 | 0.2795663 |
| 0.072163747 | 0.279562614 |
| 0.072156593 | 0.279557637 |
| 0.072115044 | 0.279419393 |
| 0.07210641 | 0.279408674 |
| 0.072081614 | 0.279358045 |
| 0.072049804 | 0.279302949 |
| 0.072016378 | 0.279196096 |
| 0.071986053 | 0.279146696 |

| | |
|---|---|
| 0.071979115 | 0.279142523 |
| 0.071979033 | 0.279142523 |
| 0.071953104 | 0.279114188 |
| 0.071950723 | 0.279114188 |
| 0.071948819 | 0.279114188 |
| 0.071930816 | 0.279114188 |
| 0.071921393 | 0.279114188 |
| 0.071914202 | 0.279114188 |
| 0.07191325 | 0.279114188 |
| 0.071893386 | 0.279102077 |
| 0.071874589 | 0.279100781 |
| 0.071852102 | 0.279036209 |
| 0.071832993 | 0.279007498 |
| 0.071761284 | 0.278774441 |
| 0.071756792 | 0.278774441 |
| 0.071748699 | 0.278774441 |
| 0.071741927 | 0.278774441 |
| 0.071741267 | 0.278774441 |
| 0.071681844 | 0.278647665 |
| 0.071591695 | 0.27836539 |
| 0.071579962 | 0.278342493 |
| 0.071566198 | 0.278311695 |
| 0.071546418 | 0.278280215 |
| 0.071541335 | 0.278280215 |
| 0.071528597 | 0.278256348 |
| 0.071521997 | 0.278256348 |
| 0.0715086 | 0.27824674 |
| 0.071499388 | 0.278233628 |
| 0.071480647 | 0.278183429 |
| 0.071477545 | 0.278183429 |
| 0.071451615 | 0.278115894 |
| 0.071442789 | 0.278104268 |
| 0.071442203 | 0.278104268 |
| 0.071428885 | 0.278095603 |
| 0.071407336 | 0.278034435 |
| 0.071400216 | 0.278034435 |
| 0.071391412 | 0.278034435 |
| 0.071378779 | 0.278034435 |
| 0.071377493 | 0.278034435 |
| 0.071362228 | 0.278034435 |
| 0.071361944 | 0.278034435 |
| 0.071350663 | 0.278034435 |
| 0.071283117 | 0.277823322 |
| 0.071276427 | 0.277823322 |
| 0.071264821 | 0.277823322 |

| | |
|---|---|
| 0.071255009 | 0.277823322 |
| 0.071205164 | 0.277655832 |
| 0.071190648 | 0.277621959 |
| 0.071187428 | 0.277621959 |
| 0.071108307 | 0.277414429 |
| 0.071051224 | 0.277259866 |
| 0.070963753 | 0.276941222 |
| 0.070942918 | 0.276882601 |
| 0.070918783 | 0.276856465 |
| 0.070857991 | 0.276641819 |
| 0.070825655 | 0.276549185 |
| 0.070815773 | 0.276544999 |
| 0.070795851 | 0.276544999 |
| 0.07078704 | 0.276544999 |
| 0.070780578 | 0.276543607 |
| 0.070745008 | 0.276427312 |
| 0.070736689 | 0.276417485 |
| 0.070725751 | 0.27639742 |
| 0.070723699 | 0.27639742 |
| 0.070701527 | 0.276348101 |
| 0.070701372 | 0.276348101 |
| 0.070694386 | 0.276348101 |
| 0.070667658 | 0.276303387 |
| 0.07066254 | 0.276303387 |
| 0.07061896 | 0.2761771 |
| 0.070616653 | 0.2761771 |
| 0.070586068 | 0.276100745 |
| 0.070580449 | 0.276100745 |
| 0.070560311 | 0.276045336 |
| 0.070526353 | 0.275935153 |
| 0.070459702 | 0.275719678 |
| 0.070413185 | 0.275650894 |
| 0.070411262 | 0.275650894 |
| 0.070387581 | 0.275595964 |
| 0.070342623 | 0.27546523 |
| 0.070329818 | 0.275437732 |
| 0.070298458 | 0.275360198 |
| 0.070254019 | 0.27524475 |
| 0.070217156 | 0.275154887 |
| 0.070207299 | 0.275138899 |
| 0.070167574 | 0.275005845 |
| 0.070163758 | 0.275005845 |
| 0.070140865 | 0.274946419 |
| 0.070074734 | 0.274732414 |
| 0.070053423 | 0.274671471 |

| | |
|---|---|
| 0.070047325 | 0.274670176 |
| 0.070045293 | 0.274670176 |
| 0.070038978 | 0.274670176 |
| 0.070007087 | 0.274648069 |
| 0.069998795 | 0.274638157 |
| 0.069992223 | 0.274634997 |
| 0.069957525 | 0.274521466 |
| 0.069954525 | 0.274521466 |
| 0.069943639 | 0.27451221 |
| 0.069941031 | 0.27451221 |
| 0.069936885 | 0.27451221 |
| 0.069915408 | 0.274491891 |
| 0.069867308 | 0.274348278 |
| 0.069835375 | 0.274312962 |
| 0.069783713 | 0.274178262 |
| 0.069780531 | 0.274178262 |
| 0.069766474 | 0.274155763 |
| 0.069766005 | 0.274155763 |
| 0.069750641 | 0.274138783 |
| 0.069749549 | 0.274138783 |
| 0.06974951 | 0.274138783 |
| 0.069729344 | 0.274138783 |
| 0.06972479 | 0.274138783 |
| 0.069694545 | 0.274076634 |
| 0.069688014 | 0.274073581 |
| 0.069672025 | 0.274066098 |
| 0.069641226 | 0.273980067 |
| 0.069622006 | 0.273949709 |
| 0.069606406 | 0.273914681 |
| 0.069573702 | 0.273852191 |
| 0.069568177 | 0.273852191 |
| 0.06956423 | 0.273852191 |
| 0.069550139 | 0.273852191 |
| 0.069549138 | 0.273852191 |
| 0.069534941 | 0.273852191 |
| 0.069531818 | 0.273852191 |
| 0.069501017 | 0.273790316 |
| 0.069490328 | 0.273770854 |
| 0.069424988 | 0.273581325 |
| 0.069411361 | 0.273550261 |
| 0.069405073 | 0.273550261 |
| 0.069341725 | 0.273388937 |
| 0.069310737 | 0.273289382 |
| 0.069299991 | 0.273269638 |
| 0.069272973 | 0.273185714 |

| | |
|---|---|
| 0.069271205 | 0.273185714 |
| 0.069252004 | 0.273170877 |
| 0.069238401 | 0.273139844 |
| 0.069217328 | 0.273106767 |
| 0.069193191 | 0.273051959 |
| 0.069185322 | 0.273043532 |
| 0.069138221 | 0.272948112 |
| 0.069082642 | 0.272773916 |
| 0.069030818 | 0.272614493 |
| 0.068974014 | 0.272412752 |
| 0.068953785 | 0.272355443 |
| 0.068923651 | 0.272258996 |
| 0.068862286 | 0.272039159 |
| 0.068861564 | 0.272039159 |
| 0.068844205 | 0.272039159 |
| 0.068842704 | 0.272039159 |
| 0.068840828 | 0.272039159 |
| 0.068826637 | 0.272039159 |
| 0.06880533 | 0.271994643 |
| 0.068771532 | 0.271883601 |
| 0.068750036 | 0.271821181 |
| 0.068747794 | 0.271821181 |
| 0.068725524 | 0.271769389 |
| 0.068717695 | 0.271769389 |
| 0.068714884 | 0.271769389 |
| 0.068714626 | 0.271769389 |
| 0.068651196 | 0.271645424 |
| 0.068647894 | 0.271645424 |
| 0.068627712 | 0.271630687 |
| 0.068615843 | 0.27160628 |
| 0.068605315 | 0.271587179 |
| 0.068576094 | 0.271494069 |
| 0.068529176 | 0.271359688 |
| 0.068514788 | 0.271341577 |
| 0.06849543 | 0.271310033 |
| 0.068484939 | 0.271291041 |
| 0.068418295 | 0.271072135 |
| 0.068384495 | 0.27096076 |
| 0.068382879 | 0.27096076 |
| 0.068357265 | 0.270897943 |
| 0.068346375 | 0.270877325 |
| 0.06833909 | 0.270870991 |
| 0.068333032 | 0.270869523 |
| 0.068332197 | 0.270869523 |
| 0.068330305 | 0.270869523 |

| | |
|---|---|
| 0.068324764 | 0.270869523 |
| 0.068300928 | 0.270854981 |
| 0.068287157 | 0.270822919 |
| 0.068278544 | 0.270811309 |
| 0.068273806 | 0.270811309 |
| 0.068244624 | 0.270730153 |
| 0.06824396 | 0.270730153 |
| 0.068242349 | 0.270730153 |
| 0.068184412 | 0.27061825 |
| 0.068145628 | 0.270509403 |
| 0.068141198 | 0.270509403 |
| 0.068118075 | 0.270445111 |
| 0.068092114 | 0.270387119 |
| 0.068034114 | 0.270201862 |
| 0.067990786 | 0.270052301 |
| 0.067920134 | 0.269794182 |
| 0.06786305 | 0.269612402 |
| 0.067860846 | 0.269612402 |
| 0.067851493 | 0.269611468 |
| 0.06783424 | 0.269565402 |
| 0.067830685 | 0.269565402 |
| 0.067817879 | 0.269545365 |
| 0.067794341 | 0.269474302 |
| 0.06776714 | 0.269388664 |
| 0.067763596 | 0.269388664 |
| 0.067759146 | 0.269388664 |
| 0.067730032 | 0.269308585 |
| 0.06771194 | 0.269259129 |
| 0.067711447 | 0.269259129 |
| 0.067694843 | 0.2692361 |
| 0.067691518 | 0.2692361 |
| 0.067683441 | 0.269235718 |
| 0.067669207 | 0.269201582 |
| 0.067658883 | 0.269182994 |
| 0.067543725 | 0.268859591 |
| 0.067543221 | 0.268859591 |
| 0.067542455 | 0.268859591 |
| 0.067527962 | 0.268859591 |
| 0.067519609 | 0.268853477 |
| 0.067508644 | 0.26883229 |
| 0.067497099 | 0.26880879 |
| 0.067467408 | 0.268735487 |
| 0.067464523 | 0.268735487 |
| 0.06745212 | 0.268719543 |
| 0.06744643 | 0.268719543 |

| | |
|---|---|
| 0.067428044 | 0.268697854 |
| 0.067415841 | 0.268687388 |
| 0.067405559 | 0.268668888 |
| 0.067380618 | 0.268591957 |
| 0.067357807 | 0.268523501 |
| 0.067336663 | 0.268461681 |
| 0.067315514 | 0.268410068 |
| 0.067291666 | 0.268362548 |
| 0.06727994 | 0.268347852 |
| 0.067198525 | 0.268045574 |
| 0.067188406 | 0.268027657 |
| 0.067174083 | 0.268027657 |
| 0.067156757 | 0.268013645 |
| 0.067145631 | 0.268013645 |
| 0.067131386 | 0.26797976 |
| 0.067122899 | 0.267968343 |
| 0.067101133 | 0.267926365 |
| 0.067100888 | 0.267926365 |
| 0.06709438 | 0.267926365 |
| 0.067090589 | 0.267926365 |
| 0.06708266 | 0.267926365 |
| 0.067045391 | 0.267838522 |
| 0.06704446 | 0.267838522 |
| 0.067028263 | 0.267815022 |
| 0.066996481 | 0.267732958 |
| 0.066990276 | 0.267730627 |
| 0.066972889 | 0.267706067 |
| 0.066969041 | 0.267706067 |
| 0.066961276 | 0.267704584 |
| 0.066943271 | 0.267655069 |
| 0.066941471 | 0.267655069 |
| 0.066923255 | 0.267619971 |
| 0.066920968 | 0.267619971 |
| 0.066917896 | 0.267619971 |
| 0.066880012 | 0.267514432 |
| 0.066865482 | 0.267478777 |
| 0.066809945 | 0.267301515 |
| 0.066806344 | 0.267301515 |
| 0.066774409 | 0.267224993 |
| 0.066755859 | 0.267197352 |
| 0.066751592 | 0.267197352 |
| 0.066728071 | 0.267131027 |
| 0.066716391 | 0.26710672 |
| 0.066692193 | 0.267032287 |
| 0.066690566 | 0.267032287 |

| | |
|---|---|
| 0.066689671 | 0.267032287 |
| 0.066687378 | 0.267032287 |
| 0.066679749 | 0.267032287 |
| 0.0666582 | 0.267030877 |
| 0.066643404 | 0.267016522 |
| 0.066635852 | 0.267008725 |
| 0.066631211 | 0.267008725 |
| 0.066620651 | 0.267008725 |
| 0.066620158 | 0.267008725 |
| 0.066605153 | 0.266998034 |
| 0.066579346 | 0.266917045 |
| 0.066549321 | 0.266819137 |
| 0.066525132 | 0.266744609 |
| 0.066503442 | 0.266681799 |
| 0.066491198 | 0.266675899 |
| 0.066448389 | 0.266549097 |
| 0.066427623 | 0.266488242 |
| 0.06634097 | 0.266230316 |
| 0.066331122 | 0.266213227 |
| 0.066318572 | 0.266185938 |
| 0.066288895 | 0.26614048 |
| 0.066236759 | 0.265991409 |
| 0.066228668 | 0.265981346 |
| 0.066185295 | 0.26585199 |
| 0.066168629 | 0.265829886 |
| 0.066145495 | 0.265759363 |
| 0.066127885 | 0.265711024 |
| 0.066121643 | 0.26570836 |
| 0.066103818 | 0.265659146 |
| 0.06608181 | 0.265593111 |
| 0.066023478 | 0.265411762 |
| 0.066009559 | 0.265411762 |
| 0.066008108 | 0.265411762 |
| 0.065983038 | 0.26538404 |
| 0.065968356 | 0.265361055 |
| 0.065965074 | 0.265361055 |
| 0.065963842 | 0.265361055 |
| 0.065945868 | 0.265337828 |
| 0.065940813 | 0.265337828 |
| 0.065939568 | 0.265337828 |
| 0.065879595 | 0.265160776 |
| 0.0658749 | 0.265160776 |
| 0.065865145 | 0.265147431 |
| 0.06586311 | 0.265147431 |
| 0.065829086 | 0.265047078 |

| | |
|---|---|
| 0.065821776 | 0.265047078 |
| 0.065787941 | 0.265015839 |
| 0.065673938 | 0.264623744 |
| 0.065673839 | 0.264623744 |
| 0.065649082 | 0.264590743 |
| 0.065636169 | 0.264561087 |
| 0.065617274 | 0.264507315 |
| 0.065613403 | 0.264507315 |
| 0.065606168 | 0.264507315 |
| 0.065593502 | 0.264501035 |
| 0.065588208 | 0.264501035 |
| 0.065583942 | 0.264501035 |
| 0.06556311 | 0.264445649 |
| 0.06555973 | 0.264445649 |
| 0.065556047 | 0.264445649 |
| 0.065552994 | 0.264445649 |
| 0.065535107 | 0.264445649 |
| 0.065525302 | 0.264445649 |
| 0.065505212 | 0.264413657 |
| 0.065497929 | 0.264413657 |
| 0.065494291 | 0.264413657 |
| 0.065490066 | 0.264413657 |
| 0.065485739 | 0.264413657 |
| 0.065477794 | 0.264413657 |
| 0.06544154 | 0.264403141 |
| 0.065439565 | 0.264403141 |
| 0.065425536 | 0.264399726 |
| 0.065356931 | 0.264150928 |
| 0.065344041 | 0.264121247 |
| 0.065337963 | 0.264121247 |
| 0.065332899 | 0.264121247 |
| 0.065328361 | 0.264121247 |
| 0.065322499 | 0.264121247 |
| 0.065320224 | 0.264121247 |
| 0.06529071 | 0.264062555 |
| 0.065289631 | 0.264062555 |
| 0.065287381 | 0.264062555 |
| 0.065278172 | 0.264062555 |
| 0.065276324 | 0.264062555 |
| 0.06526868 | 0.264062555 |
| 0.065207824 | 0.263906614 |
| 0.065187109 | 0.26386762 |
| 0.065137712 | 0.263712496 |
| 0.065089918 | 0.263608632 |
| 0.065059005 | 0.263505843 |

| | |
|---|---|
| 0.065052118 | 0.263500358 |
| 0.06505057 | 0.263500358 |
| 0.065020257 | 0.263419145 |
| 0.064999699 | 0.263377631 |
| 0.064997033 | 0.263377631 |
| 0.064973067 | 0.263322438 |
| 0.064961427 | 0.26331216 |
| 0.064898381 | 0.263133657 |
| 0.064824442 | 0.262891152 |
| 0.064804777 | 0.262833784 |
| 0.064804468 | 0.262833784 |
| 0.064801079 | 0.262833784 |
| 0.064780064 | 0.262823077 |
| 0.064758661 | 0.262803406 |
| 0.064750792 | 0.26279386 |
| 0.064748115 | 0.26279386 |
| 0.064693796 | 0.262629681 |
| 0.064689273 | 0.262629681 |
| 0.064689054 | 0.262629681 |
| 0.064679853 | 0.262629681 |
| 0.064674222 | 0.262629681 |
| 0.064668726 | 0.262629681 |
| 0.064667578 | 0.262629681 |
| 0.064637749 | 0.262581206 |
| 0.064637498 | 0.262581206 |
| 0.064621403 | 0.262581206 |
| 0.06461551 | 0.262581206 |
| 0.064605441 | 0.262581206 |
| 0.064556795 | 0.262409018 |
| 0.064503578 | 0.262237466 |
| 0.064451287 | 0.262047247 |
| 0.064449285 | 0.262047247 |
| 0.064423929 | 0.262003114 |
| 0.064300367 | 0.261634649 |
| 0.064294465 | 0.261632988 |
| 0.064285331 | 0.261618172 |
| 0.064268897 | 0.26159195 |
| 0.064259932 | 0.261581862 |
| 0.064243339 | 0.261536671 |
| 0.064166161 | 0.261334168 |
| 0.064144731 | 0.261313925 |
| 0.06413135 | 0.261281761 |
| 0.064130869 | 0.261281761 |
| 0.064129688 | 0.261281761 |
| 0.064122758 | 0.261281761 |

| | |
|---|---|
| 0.064101309 | 0.26127112 |
| 0.064067488 | 0.261155618 |
| 0.064067178 | 0.261155618 |
| 0.064040486 | 0.261112595 |
| 0.064029939 | 0.261091944 |
| 0.063991906 | 0.2609592 |
| 0.063982622 | 0.26095497 |
| 0.063962759 | 0.260907359 |
| 0.063962248 | 0.260907359 |
| 0.063962015 | 0.260907359 |
| 0.063945228 | 0.260904138 |
| 0.063924088 | 0.260883681 |
| 0.063919525 | 0.260883681 |
| 0.06391537 | 0.260883681 |
| 0.06387399 | 0.260813317 |
| 0.063817492 | 0.26064966 |
| 0.063789101 | 0.260556049 |
| 0.063781496 | 0.260547328 |
| 0.063763036 | 0.260505075 |
| 0.063760775 | 0.260505075 |
| 0.063760468 | 0.260505075 |
| 0.063756846 | 0.260505075 |
| 0.063750176 | 0.260505075 |
| 0.063743858 | 0.260505075 |
| 0.063720874 | 0.26047842 |
| 0.063714849 | 0.260476147 |
| 0.063709825 | 0.260476147 |
| 0.063702331 | 0.260476147 |
| 0.063673921 | 0.260398233 |
| 0.063668231 | 0.260397322 |
| 0.063657138 | 0.260374316 |
| 0.063642752 | 0.260360191 |
| 0.063568882 | 0.260147375 |
| 0.063527549 | 0.260022909 |
| 0.063521421 | 0.260020172 |
| 0.063495079 | 0.259957029 |
| 0.063464529 | 0.259868587 |
| 0.063460001 | 0.259868587 |
| 0.063423547 | 0.259775822 |
| 0.063410071 | 0.25974702 |
| 0.063408921 | 0.25974702 |
| 0.063398811 | 0.25974702 |
| 0.063382196 | 0.259718134 |
| 0.063352108 | 0.259617178 |
| 0.063279942 | 0.2593884 |

| | |
|---|---|
| 0.063279772 | 0.2593884 |
| 0.063251388 | 0.259338312 |
| 0.063250521 | 0.259338312 |
| 0.063222385 | 0.259264036 |
| 0.063215068 | 0.259256356 |
| 0.063201683 | 0.259223781 |
| 0.063198282 | 0.259223781 |
| 0.063198175 | 0.259223781 |
| 0.063195847 | 0.259223781 |
| 0.063191151 | 0.259223781 |
| 0.06318298 | 0.259223781 |
| 0.063181453 | 0.259223781 |
| 0.063130467 | 0.259154864 |
| 0.063130203 | 0.259154864 |
| 0.063115354 | 0.259137498 |
| 0.063093992 | 0.259116802 |
| 0.06308373 | 0.259097 |
| 0.063065314 | 0.259066045 |
| 0.063033318 | 0.258979281 |
| 0.063013682 | 0.258943283 |
| 0.063012279 | 0.258943283 |
| 0.063009592 | 0.258943283 |
| 0.062992359 | 0.258924634 |
| 0.062992099 | 0.258924634 |
| 0.062988918 | 0.258924634 |
| 0.062954149 | 0.258854983 |
| 0.062928568 | 0.25877214 |
| 0.062912365 | 0.258727846 |
| 0.062886632 | 0.258689031 |
| 0.062876721 | 0.258686864 |
| 0.062856724 | 0.258633018 |
| 0.0628024 | 0.258465044 |
| 0.062787633 | 0.25846038 |
| 0.062783252 | 0.25846038 |
| 0.062741619 | 0.258337953 |
| 0.062724818 | 0.258291103 |
| 0.062706202 | 0.25826474 |
| 0.062684662 | 0.258227354 |
| 0.062664406 | 0.258227354 |
| 0.062660278 | 0.258227354 |
| 0.062642145 | 0.258218576 |
| 0.062641354 | 0.258218576 |
| 0.062634412 | 0.258218576 |
| 0.062599663 | 0.258174148 |
| 0.062567707 | 0.258068084 |

| | |
|---|---|
| 0.062507056 | 0.257840253 |
| 0.062504149 | 0.257840253 |
| 0.062486643 | 0.257800705 |
| 0.062485311 | 0.257800705 |
| 0.062482727 | 0.257800705 |
| 0.062418306 | 0.25760803 |
| 0.062393282 | 0.257571711 |
| 0.062358035 | 0.257470839 |
| 0.06229385 | 0.257295048 |
| 0.062285316 | 0.257282109 |
| 0.062268551 | 0.257235171 |
| 0.062259341 | 0.257219433 |
| 0.062241666 | 0.257168718 |
| 0.062240744 | 0.257168718 |
| 0.062226607 | 0.257151115 |
| 0.062197045 | 0.257051256 |
| 0.062161565 | 0.256971517 |
| 0.06211644 | 0.256829558 |
| 0.062095879 | 0.256766836 |
| 0.062085766 | 0.256747312 |
| 0.062067442 | 0.256712941 |
| 0.062047283 | 0.256655022 |
| 0.062036 | 0.256630641 |
| 0.062005755 | 0.256527803 |
| 0.061966699 | 0.256388497 |
| 0.061966571 | 0.256388497 |
| 0.061963254 | 0.256388497 |
| 0.061944882 | 0.25638732 |
| 0.061942569 | 0.25638732 |
| 0.06190437 | 0.256308739 |
| 0.061900658 | 0.256308739 |
| 0.061887557 | 0.256283688 |
| 0.061862376 | 0.256201687 |
| 0.061845452 | 0.256153867 |
| 0.061828049 | 0.256104062 |
| 0.061820461 | 0.256094904 |
| 0.061808436 | 0.256067363 |
| 0.061789773 | 0.256012311 |
| 0.061769519 | 0.255950661 |
| 0.061768247 | 0.255950661 |
| 0.061750212 | 0.255915188 |
| 0.061726346 | 0.255860808 |
| 0.061637455 | 0.255536828 |
| 0.061634758 | 0.255536828 |
| 0.061630837 | 0.255536828 |

| | |
|---|---|
| 0.061621053 | 0.255535559 |
| 0.061609258 | 0.255508894 |
| 0.06157216 | 0.255399517 |
| 0.06154338 | 0.255309569 |
| 0.0615105 | 0.255254909 |
| 0.061502974 | 0.255245918 |
| 0.061472144 | 0.255140203 |
| 0.061463109 | 0.255140203 |
| 0.061460147 | 0.255140203 |
| 0.061459068 | 0.255140203 |
| 0.061441085 | 0.255140203 |
| 0.061432014 | 0.255129274 |
| 0.061425041 | 0.255122564 |
| 0.061414761 | 0.255102112 |
| 0.061379052 | 0.254976023 |
| 0.06135809 | 0.25491118 |
| 0.061357013 | 0.25491118 |
| 0.061322029 | 0.254805816 |
| 0.06131277 | 0.254789571 |
| 0.06130963 | 0.254789571 |
| 0.061239314 | 0.254528737 |
| 0.061223472 | 0.254507193 |
| 0.061204248 | 0.254483801 |
| 0.061196233 | 0.254482929 |
| 0.061186883 | 0.254466267 |
| 0.061186073 | 0.254466267 |
| 0.061149113 | 0.254353606 |
| 0.061129013 | 0.25429221 |
| 0.061119434 | 0.25427457 |
| 0.061100742 | 0.254219013 |
| 0.061066121 | 0.254097164 |
| 0.061064946 | 0.254097164 |
| 0.061057182 | 0.254097164 |
| 0.061032358 | 0.254045457 |
| 0.060998392 | 0.253948464 |
| 0.060964539 | 0.253829712 |
| 0.060948104 | 0.253787265 |
| 0.060929733 | 0.253773537 |
| 0.060920436 | 0.253773537 |
| 0.060919756 | 0.253773537 |
| 0.060912997 | 0.253773537 |
| 0.060892544 | 0.25375179 |
| 0.060867575 | 0.253714337 |
| 0.06085867 | 0.253699422 |
| 0.060841341 | 0.253665976 |

| | |
|---|---|
| 0.060784271 | 0.253522418 |
| 0.060777496 | 0.253516359 |
| 0.060763319 | 0.253479424 |
| 0.06070031 | 0.253260944 |
| 0.060674223 | 0.253218649 |
| 0.060646954 | 0.253156744 |
| 0.060629428 | 0.253120421 |
| 0.060627403 | 0.253120421 |
| 0.060617779 | 0.253116164 |
| 0.060617297 | 0.253116164 |
| 0.060602294 | 0.253116164 |
| 0.060590069 | 0.253089212 |
| 0.060585008 | 0.253089212 |
| 0.060522515 | 0.252940137 |
| 0.060516947 | 0.252939066 |
| 0.060497964 | 0.25290411 |
| 0.060470877 | 0.252835262 |
| 0.060467876 | 0.252835262 |
| 0.060458124 | 0.252826328 |
| 0.060436916 | 0.252782031 |
| 0.060433823 | 0.252782031 |
| 0.060416047 | 0.252739136 |
| 0.060391494 | 0.252680813 |
| 0.06038015 | 0.252655543 |
| 0.060363999 | 0.252610157 |
| 0.060362392 | 0.252610157 |
| 0.060359871 | 0.252610157 |
| 0.060328538 | 0.252563186 |
| 0.060299639 | 0.252496125 |
| 0.060292679 | 0.252496125 |
| 0.060261669 | 0.252463419 |
| 0.060252596 | 0.252454509 |
| 0.060239443 | 0.252439973 |
| 0.060233638 | 0.252439973 |
| 0.060202771 | 0.252334538 |
| 0.060202472 | 0.252334538 |
| 0.060200005 | 0.252334538 |
| 0.060186103 | 0.252331295 |
| 0.060165091 | 0.25226541 |
| 0.060157435 | 0.252255515 |
| 0.06015248 | 0.252255515 |
| 0.060115885 | 0.252125685 |
| 0.060087279 | 0.252027905 |
| 0.060073625 | 0.252015022 |
| 0.060058865 | 0.252015022 |

| | |
|---|---|
| 0.060020081 | 0.251879144 |
| 0.06001091 | 0.25186285 |
| 0.059999112 | 0.251835526 |
| 0.059989591 | 0.251817754 |
| 0.059970364 | 0.251759234 |
| 0.059967905 | 0.251759234 |
| 0.059967759 | 0.251759234 |
| 0.059937671 | 0.251688542 |
| 0.059914723 | 0.251636555 |
| 0.059910425 | 0.251636555 |
| 0.059907126 | 0.251636555 |
| 0.059867087 | 0.251525208 |
| 0.05983944 | 0.251431231 |
| 0.059838223 | 0.251431231 |
| 0.059806774 | 0.251338325 |
| 0.059803348 | 0.251338325 |
| 0.059795394 | 0.251334854 |
| 0.059753284 | 0.251202196 |
| 0.059738403 | 0.251178633 |
| 0.059717083 | 0.251116504 |
| 0.059702074 | 0.251080173 |
| 0.059677166 | 0.251037307 |
| 0.059658767 | 0.250985154 |
| 0.059656821 | 0.250985154 |
| 0.059655021 | 0.250985154 |
| 0.059619844 | 0.250943988 |
| 0.059602237 | 0.250909381 |
| 0.05959213 | 0.2509012 |
| 0.059585938 | 0.250897308 |
| 0.059581112 | 0.250897308 |
| 0.059499338 | 0.25059913 |
| 0.059471804 | 0.250562466 |
| 0.059444748 | 0.250479961 |
| 0.059433938 | 0.250456569 |
| 0.059422487 | 0.250452635 |
| 0.059364806 | 0.250231667 |
| 0.059341373 | 0.250155032 |
| 0.059334305 | 0.250147379 |
| 0.059331785 | 0.250147379 |
| 0.059322553 | 0.250142117 |
| 0.059306943 | 0.250120586 |
| 0.059295115 | 0.25009285 |
| 0.059280695 | 0.250054171 |
| 0.05927944 | 0.250054171 |
| 0.05927013 | 0.250054171 |

| | |
|---|---|
| 0.05926752 | 0.250054171 |
| 0.059258631 | 0.250054171 |
| 0.059253728 | 0.250054171 |
| 0.059246163 | 0.250054171 |
| 0.059210837 | 0.24993659 |
| 0.059192259 | 0.249880317 |
| 0.059187689 | 0.249880317 |
| 0.059182677 | 0.249880317 |
| 0.059155313 | 0.249790771 |
| 0.05915215 | 0.249790771 |
| 0.059149667 | 0.249790771 |
| 0.059133903 | 0.249766798 |
| 0.059130074 | 0.249766798 |
| 0.059101021 | 0.249685888 |
| 0.059057028 | 0.249574896 |
| 0.059039176 | 0.24954374 |
| 0.059038882 | 0.24954374 |
| 0.059006673 | 0.249450632 |
| 0.059004668 | 0.249450632 |
| 0.05899616 | 0.249450473 |
| 0.058987854 | 0.249437498 |
| 0.05896206 | 0.249350563 |
| 0.058959121 | 0.249350563 |
| 0.058947808 | 0.249334568 |
| 0.058937151 | 0.249311635 |
| 0.058889431 | 0.249176167 |
| 0.058880514 | 0.249160571 |
| 0.058847012 | 0.249063063 |
| 0.058837117 | 0.249063063 |
| 0.058829059 | 0.249053475 |
| 0.058824829 | 0.249053475 |
| 0.05882231 | 0.249053475 |
| 0.058813137 | 0.249053475 |
| 0.05881301 | 0.249053475 |
| 0.058811224 | 0.249053475 |
| 0.0587866 | 0.249028674 |
| 0.058781153 | 0.249027856 |
| 0.058763064 | 0.248995414 |
| 0.058751072 | 0.248988905 |
| 0.058742125 | 0.24897314 |
| 0.058736851 | 0.248972945 |
| 0.058724785 | 0.248943954 |
| 0.058706148 | 0.248909259 |
| 0.058668979 | 0.248773813 |
| 0.058667452 | 0.248773813 |

| | |
|---|---|
| 0.058600168 | 0.248592696 |
| 0.058536818 | 0.24836851 |
| 0.058534919 | 0.24836851 |
| 0.058522689 | 0.24836851 |
| 0.05850177 | 0.248307977 |
| 0.058500997 | 0.248307977 |
| 0.058495741 | 0.248307977 |
| 0.058461122 | 0.248201805 |
| 0.058457547 | 0.248201805 |
| 0.058456672 | 0.248201805 |
| 0.058454868 | 0.248201805 |
| 0.05845069 | 0.248201805 |
| 0.058404068 | 0.2481144 |
| 0.05839556 | 0.24811286 |
| 0.058366632 | 0.248021734 |
| 0.058338369 | 0.247945877 |
| 0.058322927 | 0.24793654 |
| 0.058261056 | 0.247749938 |
| 0.058214905 | 0.24757579 |
| 0.058214413 | 0.24757579 |
| 0.058206319 | 0.24757579 |
| 0.058197441 | 0.24757579 |
| 0.058115647 | 0.247264062 |
| 0.05808879 | 0.247171877 |
| 0.058078191 | 0.247148857 |
| 0.058068604 | 0.247130142 |
| 0.058055456 | 0.247096265 |
| 0.058040334 | 0.247053982 |
| 0.058040185 | 0.247053982 |
| 0.058032135 | 0.247053982 |
| 0.058031145 | 0.247053982 |
| 0.058005746 | 0.246995052 |
| 0.058001614 | 0.246995052 |
| 0.05798596 | 0.246977047 |
| 0.057980816 | 0.246977047 |
| 0.057928592 | 0.246842887 |
| 0.05792309 | 0.246841713 |
| 0.057921714 | 0.246841713 |
| 0.057912409 | 0.246840369 |
| 0.057908394 | 0.246840369 |
| 0.057907356 | 0.246840369 |
| 0.057865122 | 0.246727141 |
| 0.057852306 | 0.246694582 |
| 0.057819992 | 0.246600946 |
| 0.057811146 | 0.246585298 |

| | |
|---|---|
| 0.0578008 | 0.246582855 |
| 0.05779058 | 0.246563824 |
| 0.057779062 | 0.246558857 |
| 0.057771722 | 0.246549626 |
| 0.057749837 | 0.246507541 |
| 0.057726178 | 0.246465682 |
| 0.057722206 | 0.246465682 |
| 0.057702116 | 0.246409591 |
| 0.05767715 | 0.246344687 |
| 0.057664526 | 0.246312857 |
| 0.057662145 | 0.246312857 |
| 0.057629023 | 0.24620537 |
| 0.057573269 | 0.246011313 |
| 0.05755951 | 0.245974589 |
| 0.057535027 | 0.245893564 |
| 0.057522519 | 0.245882699 |
| 0.057502498 | 0.245841244 |
| 0.057482826 | 0.245803216 |
| 0.057450051 | 0.24574938 |
| 0.057386776 | 0.245544872 |
| 0.057364603 | 0.245472053 |
| 0.057364534 | 0.245472053 |
| 0.05734191 | 0.245419045 |
| 0.057301808 | 0.245313547 |
| 0.05730129 | 0.245313547 |
| 0.057300105 | 0.245313547 |
| 0.057281226 | 0.24529158 |
| 0.057259961 | 0.245222566 |
| 0.057236527 | 0.245145973 |
| 0.057234466 | 0.245145973 |
| 0.057231459 | 0.245145973 |
| 0.057209917 | 0.245145973 |
| 0.057208916 | 0.245145973 |
| 0.057187413 | 0.245110234 |
| 0.057177411 | 0.24508942 |
| 0.057176583 | 0.24508942 |
| 0.057167519 | 0.24508942 |
| 0.057167311 | 0.24508942 |
| 0.057155706 | 0.245084616 |
| 0.057154377 | 0.245084616 |
| 0.057137308 | 0.245071918 |
| 0.057136597 | 0.245071918 |
| 0.057124626 | 0.245071918 |
| 0.057117539 | 0.245071918 |
| 0.057109524 | 0.245063094 |

| | |
|---|---|
| 0.057084004 | 0.244975659 |
| 0.057076852 | 0.244967037 |
| 0.057043889 | 0.244869472 |
| 0.057037467 | 0.244869472 |
| 0.057030368 | 0.244869472 |
| 0.057029736 | 0.244869472 |
| 0.057024364 | 0.244869472 |
| 0.057018047 | 0.244869472 |
| 0.057013014 | 0.244869472 |
| 0.057012871 | 0.244869472 |
| 0.056968298 | 0.244809993 |
| 0.056964514 | 0.244809993 |
| 0.056960158 | 0.244809993 |
| 0.056946598 | 0.244805093 |
| 0.056941526 | 0.244805093 |
| 0.056940312 | 0.244805093 |
| 0.056938314 | 0.244805093 |
| 0.056918965 | 0.244774677 |
| 0.056882806 | 0.24470756 |
| 0.056869328 | 0.244671679 |
| 0.056867617 | 0.244671679 |
| 0.056841026 | 0.244616206 |
| 0.056785527 | 0.244443627 |
| 0.056754696 | 0.244355081 |
| 0.056729634 | 0.244269258 |
| 0.056680894 | 0.244125606 |
| 0.056671745 | 0.244108275 |
| 0.056668093 | 0.244108275 |
| 0.05666569 | 0.244108275 |
| 0.056663081 | 0.244108275 |
| 0.056655289 | 0.244108275 |
| 0.056652478 | 0.244108275 |
| 0.056648358 | 0.244108275 |
| 0.05660352 | 0.24401302 |
| 0.056598652 | 0.24401302 |
| 0.056593937 | 0.24401302 |
| 0.056576907 | 0.243964542 |
| 0.056576208 | 0.243964542 |
| 0.05656625 | 0.243962765 |
| 0.056550993 | 0.243941141 |
| 0.056550275 | 0.243941141 |
| 0.056546372 | 0.243941141 |
| 0.05653665 | 0.243941141 |
| 0.056523241 | 0.243941141 |
| 0.056520463 | 0.243941141 |

| | |
|---|---|
| 0.056503734 | 0.243914015 |
| 0.056486466 | 0.243883684 |
| 0.056476193 | 0.243867696 |
| 0.056430725 | 0.243709297 |
| 0.056410663 | 0.243666848 |
| 0.056394568 | 0.24361942 |
| 0.056373714 | 0.243551423 |
| 0.056354172 | 0.243533267 |
| 0.056346767 | 0.243533267 |
| 0.056339745 | 0.243533267 |
| 0.056300995 | 0.243458098 |
| 0.056280822 | 0.243392962 |
| 0.056259735 | 0.243323865 |
| 0.056229676 | 0.243215945 |
| 0.056177861 | 0.243013897 |
| 0.056175491 | 0.243013897 |
| 0.056160555 | 0.242983172 |
| 0.056158394 | 0.242983172 |
| 0.0561537 | 0.242983172 |
| 0.056144186 | 0.242978568 |
| 0.056121274 | 0.242901484 |
| 0.056083613 | 0.242764811 |
| 0.056020676 | 0.242576301 |
| 0.056017941 | 0.242576301 |
| 0.056004609 | 0.242550835 |
| 0.055999349 | 0.242550108 |
| 0.055967835 | 0.242435662 |
| 0.055918053 | 0.242264085 |
| 0.055907163 | 0.242238943 |
| 0.05586441 | 0.24211978 |
| 0.055861108 | 0.24211978 |
| 0.055834909 | 0.242057174 |
| 0.055786766 | 0.241893329 |
| 0.055753428 | 0.241792799 |
| 0.055752507 | 0.241792799 |
| 0.055741837 | 0.241792799 |
| 0.055739893 | 0.241792799 |
| 0.055738139 | 0.241792799 |
| 0.055721796 | 0.241792799 |
| 0.055712813 | 0.241792798 |
| 0.05571272 | 0.241792798 |
| 0.055708694 | 0.241792798 |
| 0.055696569 | 0.241788397 |
| 0.055693358 | 0.241788397 |
| 0.055668193 | 0.241759989 |

| | |
|---|---|
| 0.05565611 | 0.241744966 |
| 0.05565492 | 0.241744966 |
| 0.055626751 | 0.241699624 |
| 0.055623376 | 0.241699624 |
| 0.055604548 | 0.241653234 |
| 0.055599859 | 0.241653234 |
| 0.055593911 | 0.241651108 |
| 0.055592302 | 0.241651108 |
| 0.055566606 | 0.241620629 |
| 0.055561242 | 0.241619363 |
| 0.055546486 | 0.241577254 |
| 0.055533886 | 0.241544508 |
| 0.055499558 | 0.241439299 |
| 0.055482258 | 0.241386089 |
| 0.055465286 | 0.241334292 |
| 0.055443894 | 0.241263256 |
| 0.055438702 | 0.241262705 |
| 0.055393601 | 0.241088461 |
| 0.05537761 | 0.241040891 |
| 0.055375956 | 0.241040891 |
| 0.055370862 | 0.241040891 |
| 0.055355284 | 0.241009791 |
| 0.055300289 | 0.240858424 |
| 0.055282701 | 0.240803841 |
| 0.055277232 | 0.240802044 |
| 0.055267211 | 0.240780413 |
| 0.055232096 | 0.24064944 |
| 0.055185963 | 0.240492438 |
| 0.055157734 | 0.240435418 |
| 0.055154119 | 0.240435418 |
| 0.055142442 | 0.240412769 |
| 0.055116172 | 0.240320233 |
| 0.055104661 | 0.240292042 |
| 0.0550827 | 0.240240268 |
| 0.055082075 | 0.240240268 |
| 0.055048738 | 0.240136126 |
| 0.055044738 | 0.240136126 |
| 0.055044382 | 0.240136126 |
| 0.055040003 | 0.240136126 |
| 0.055021346 | 0.240104604 |
| 0.055009623 | 0.240075442 |
| 0.055004806 | 0.240075442 |
| 0.055001849 | 0.240075442 |
| 0.054981417 | 0.240022215 |
| 0.054941339 | 0.239887343 |

| | |
|---|---|
| 0.054941269 | 0.239887343 |
| 0.054917923 | 0.239829079 |
| 0.05491165 | 0.239823672 |
| 0.054876863 | 0.239693722 |
| 0.054849111 | 0.239638438 |
| 0.054835343 | 0.239600266 |
| 0.054834543 | 0.239600266 |
| 0.05483383 | 0.239600266 |
| 0.054757141 | 0.239346397 |
| 0.054750587 | 0.239339716 |
| 0.054745556 | 0.239339691 |
| 0.054744024 | 0.239339691 |
| 0.054729615 | 0.239313934 |
| 0.054726237 | 0.239313934 |
| 0.054705601 | 0.239252862 |
| 0.0546932 | 0.239220592 |
| 0.054676083 | 0.239167691 |
| 0.054674703 | 0.239167691 |
| 0.054635103 | 0.239044822 |
| 0.054634208 | 0.239044822 |
| 0.054623703 | 0.239044822 |
| 0.054555014 | 0.238792076 |
| 0.054554738 | 0.238792076 |
| 0.05451886 | 0.238699188 |
| 0.054484791 | 0.238571955 |
| 0.054480525 | 0.238571955 |
| 0.054467232 | 0.238560864 |
| 0.054444336 | 0.238482517 |
| 0.05442853 | 0.23843521 |
| 0.054423783 | 0.23843521 |
| 0.054413682 | 0.23843521 |
| 0.054397875 | 0.238388626 |
| 0.054396973 | 0.238388626 |
| 0.054393683 | 0.238388626 |
| 0.054388049 | 0.238388626 |
| 0.054377452 | 0.238386864 |
| 0.054367578 | 0.238365518 |
| 0.054343158 | 0.23833115 |
| 0.054338745 | 0.23833115 |
| 0.054326621 | 0.238317582 |
| 0.054315809 | 0.238292102 |
| 0.054276976 | 0.23816561 |
| 0.054271283 | 0.238162573 |
| 0.054259419 | 0.238132451 |
| 0.054252288 | 0.238123097 |

| | |
|---|---|
| 0.054236109 | 0.238074025 |
| 0.054235061 | 0.238074025 |
| 0.054230429 | 0.238074025 |
| 0.054227149 | 0.238074025 |
| 0.054203762 | 0.23804173 |
| 0.054166387 | 0.237943415 |
| 0.054161749 | 0.237943415 |
| 0.054159829 | 0.237943415 |
| 0.054154631 | 0.237943415 |
| 0.054135223 | 0.237894286 |
| 0.054134259 | 0.237894286 |
| 0.054129606 | 0.237894286 |
| 0.054128876 | 0.237894286 |
| 0.054107675 | 0.237861015 |
| 0.054093756 | 0.237821776 |
| 0.0540887 | 0.237821776 |
| 0.054050019 | 0.237739218 |
| 0.054042986 | 0.237730238 |
| 0.054033817 | 0.237730238 |
| 0.054029732 | 0.237730238 |
| 0.05400386 | 0.237667882 |
| 0.053980181 | 0.237607588 |
| 0.053978063 | 0.237607588 |
| 0.053966571 | 0.237591597 |
| 0.053926535 | 0.237437284 |
| 0.053919168 | 0.237437284 |
| 0.053912347 | 0.237437284 |
| 0.053885819 | 0.237437284 |
| 0.053882935 | 0.237437284 |
| 0.053871185 | 0.237435031 |
| 0.053859185 | 0.237404108 |
| 0.053846093 | 0.237368371 |
| 0.053823047 | 0.237288742 |
| 0.053817554 | 0.237286486 |
| 0.05380897 | 0.237270606 |
| 0.053790943 | 0.237213079 |
| 0.053745661 | 0.237079244 |
| 0.053735549 | 0.237056596 |
| 0.05371798 | 0.237022997 |
| 0.053667784 | 0.2368454 |
| 0.053642265 | 0.236795207 |
| 0.053627634 | 0.236773424 |
| 0.053576829 | 0.236661338 |
| 0.053575301 | 0.236661338 |
| 0.053565227 | 0.236655944 |

| | |
|---|---|
| 0.053557539 | 0.236643931 |
| 0.053556341 | 0.236643931 |
| 0.053554376 | 0.236643931 |
| 0.053547727 | 0.236643931 |
| 0.053541981 | 0.236643931 |
| 0.053512139 | 0.236575278 |
| 0.053504946 | 0.236575278 |
| 0.053492341 | 0.236553363 |
| 0.053448539 | 0.236403561 |
| 0.053424436 | 0.236318903 |
| 0.05342353 | 0.236318903 |
| 0.05341196 | 0.236307608 |
| 0.053410874 | 0.236307608 |
| 0.053409926 | 0.236307608 |
| 0.053403024 | 0.236307608 |
| 0.053400047 | 0.236307608 |
| 0.053398654 | 0.236307608 |
| 0.053393494 | 0.236307608 |
| 0.053392912 | 0.236307608 |
| 0.053378625 | 0.236307608 |
| 0.053351143 | 0.236258011 |
| 0.05334898 | 0.236258011 |
| 0.053336763 | 0.236238258 |
| 0.053262895 | 0.235998872 |
| 0.053256487 | 0.235998872 |
| 0.053192246 | 0.235795524 |
| 0.053182246 | 0.23577314 |
| 0.053159745 | 0.235717267 |
| 0.053148961 | 0.235691392 |
| 0.053148085 | 0.235691392 |
| 0.05311738 | 0.23559522 |
| 0.053104806 | 0.235561385 |
| 0.053071623 | 0.235436121 |
| 0.05307002 | 0.235436121 |
| 0.05305787 | 0.235420724 |
| 0.053035668 | 0.235386244 |
| 0.053022308 | 0.235365 |
| 0.052993193 | 0.235263499 |
| 0.052946376 | 0.235077564 |
| 0.052922853 | 0.234995034 |
| 0.05291802 | 0.234995034 |
| 0.052909442 | 0.234979296 |
| 0.052880262 | 0.234893512 |
| 0.052873488 | 0.23488533 |
| 0.052860202 | 0.234848213 |

| | |
|---|---|
| 0.052848998 | 0.234820342 |
| 0.052838757 | 0.234813039 |
| 0.052817343 | 0.234745391 |
| 0.052809901 | 0.23473422 |
| 0.052785101 | 0.234667787 |
| 0.052781662 | 0.234667787 |
| 0.052773911 | 0.234661846 |
| 0.052749679 | 0.234590809 |
| 0.052724243 | 0.23450668 |
| 0.052717988 | 0.234500757 |
| 0.052703998 | 0.234460426 |
| 0.052697305 | 0.234452549 |
| 0.052670208 | 0.234353887 |
| 0.052644049 | 0.234262921 |
| 0.052622321 | 0.234206457 |
| 0.05262066 | 0.234206457 |
| 0.052597289 | 0.234138804 |
| 0.052595533 | 0.234138804 |
| 0.05258244 | 0.234116469 |
| 0.052562372 | 0.234049 |
| 0.052549362 | 0.234012945 |
| 0.052491555 | 0.233799243 |
| 0.052443848 | 0.233630449 |
| 0.05242579 | 0.233571854 |
| 0.052415593 | 0.233556857 |
| 0.052390348 | 0.233479471 |
| 0.052373023 | 0.233424102 |
| 0.052360467 | 0.23339613 |
| 0.052335896 | 0.233324133 |
| 0.052309038 | 0.233226228 |
| 0.052299776 | 0.233206762 |
| 0.05229754 | 0.233206762 |
| 0.052248319 | 0.233064595 |
| 0.052241748 | 0.233057113 |
| 0.052225364 | 0.233005846 |
| 0.05221951 | 0.233001556 |
| 0.052182911 | 0.232881889 |
| 0.052171562 | 0.232853058 |
| 0.052167903 | 0.232853058 |
| 0.052160016 | 0.232845165 |
| 0.052159684 | 0.232845165 |
| 0.052155281 | 0.232845165 |
| 0.052112678 | 0.2327211 |
| 0.052069828 | 0.232574309 |
| 0.052067259 | 0.232574309 |

| | |
|---|---|
| 0.052066198 | 0.232574309 |
| 0.052063813 | 0.232574309 |
| 0.052040608 | 0.232573712 |
| 0.052016058 | 0.23248581 |
| 0.051988489 | 0.232384399 |
| 0.051986416 | 0.232384399 |
| 0.051973911 | 0.232381191 |
| 0.051959161 | 0.232340527 |
| 0.051943819 | 0.232293732 |
| 0.051943763 | 0.232293732 |
| 0.051940998 | 0.232293732 |
| 0.051940482 | 0.232293732 |
| 0.051927038 | 0.232293732 |
| 0.051913395 | 0.232266721 |
| 0.051880822 | 0.232164608 |
| 0.051879383 | 0.232164608 |
| 0.051868553 | 0.232153331 |
| 0.051865255 | 0.232153331 |
| 0.051830447 | 0.232048194 |
| 0.051827334 | 0.232048194 |
| 0.051811965 | 0.232009068 |
| 0.051710318 | 0.231612675 |
| 0.051671531 | 0.231467268 |
| 0.051644567 | 0.231390009 |
| 0.051586354 | 0.231172683 |
| 0.051563663 | 0.231092745 |
| 0.051562477 | 0.231092745 |
| 0.051562063 | 0.231092745 |
| 0.05153854 | 0.231045373 |
| 0.051527268 | 0.231016584 |
| 0.051519896 | 0.231005282 |
| 0.051492868 | 0.230905832 |
| 0.051446617 | 0.230763613 |
| 0.051443645 | 0.230763613 |
| 0.051421846 | 0.230695953 |
| 0.051405156 | 0.230643205 |
| 0.051394687 | 0.230639287 |
| 0.051386674 | 0.230625057 |
| 0.051380396 | 0.230619979 |
| 0.051372737 | 0.230619979 |
| 0.051355382 | 0.230571528 |
| 0.051316431 | 0.230418371 |
| 0.051315913 | 0.230418371 |
| 0.051285789 | 0.230324212 |
| 0.051284075 | 0.230324212 |

| | |
|---|---|
| 0.051273411 | 0.230312059 |
| 0.051258747 | 0.230267906 |
| 0.05123167 | 0.230233134 |
| 0.051223455 | 0.230217937 |
| 0.051210391 | 0.230180947 |
| 0.051208884 | 0.230180947 |
| 0.051201137 | 0.230180947 |
| 0.051196802 | 0.230180947 |
| 0.051192844 | 0.230180947 |
| 0.051189129 | 0.230180947 |
| 0.051187899 | 0.230180947 |
| 0.051136032 | 0.230020382 |
| 0.051135463 | 0.230020382 |
| 0.051133079 | 0.230020382 |
| 0.05110039 | 0.229968657 |
| 0.05108826 | 0.229935793 |
| 0.051085562 | 0.229935793 |
| 0.051071118 | 0.229902095 |
| 0.051068055 | 0.229902095 |
| 0.051041457 | 0.229833738 |
| 0.051013813 | 0.229752706 |
| 0.051013053 | 0.229752706 |
| 0.051007712 | 0.229752706 |
| 0.050998661 | 0.229749642 |
| 0.050986164 | 0.229715071 |
| 0.050971319 | 0.229691644 |
| 0.05092987 | 0.229548304 |
| 0.050909648 | 0.229478877 |
| 0.050885075 | 0.229433256 |
| 0.05087629 | 0.229415362 |
| 0.050854081 | 0.229358648 |
| 0.050849658 | 0.229358648 |
| 0.050840221 | 0.229339571 |
| 0.050839272 | 0.229339571 |
| 0.050830057 | 0.229337164 |
| 0.050786645 | 0.229163007 |
| 0.050700837 | 0.22884086 |
| 0.05068975 | 0.228812504 |
| 0.050683539 | 0.228812504 |
| 0.050677977 | 0.228812504 |
| 0.05067311 | 0.228812504 |
| 0.050669742 | 0.228812504 |
| 0.050667864 | 0.228812504 |
| 0.050639307 | 0.228779955 |
| 0.050636532 | 0.228779955 |

| | |
|---|---|
| 0.050630249 | 0.228779955 |
| 0.050602859 | 0.228702065 |
| 0.050592006 | 0.22869642 |
| 0.050558307 | 0.228585573 |
| 0.050532792 | 0.228515504 |
| 0.050499506 | 0.228386664 |
| 0.050490484 | 0.228367549 |
| 0.050472921 | 0.228309794 |
| 0.050470009 | 0.228309794 |
| 0.050462981 | 0.228308197 |
| 0.050457043 | 0.22830786 |
| 0.05045685 | 0.22830786 |
| 0.050456269 | 0.22830786 |
| 0.050434589 | 0.228288173 |
| 0.050433279 | 0.228288173 |
| 0.050429569 | 0.228288173 |
| 0.050413049 | 0.228255754 |
| 0.05040428 | 0.228255754 |
| 0.050365646 | 0.228127875 |
| 0.050361135 | 0.228127875 |
| 0.050360574 | 0.228127875 |
| 0.050342928 | 0.228090053 |
| 0.050335772 | 0.228079328 |
| 0.050323978 | 0.228069285 |
| 0.050307309 | 0.228015435 |
| 0.050281462 | 0.227919976 |
| 0.050273136 | 0.227912681 |
| 0.050261364 | 0.227893944 |
| 0.050168034 | 0.227557423 |
| 0.050164119 | 0.227557423 |
| 0.050136827 | 0.2274592 |
| 0.050129455 | 0.227454366 |
| 0.050110404 | 0.227404307 |
| 0.050104675 | 0.227399977 |
| 0.050087098 | 0.227363536 |
| 0.050032476 | 0.227137234 |
| 0.05003079 | 0.227137234 |
| 0.050030161 | 0.227137234 |
| 0.050026527 | 0.227137234 |
| 0.050010414 | 0.227137234 |
| 0.049990965 | 0.227099553 |
| 0.049975342 | 0.227078185 |
| 0.049941995 | 0.227029434 |
| 0.049905408 | 0.226906432 |
| 0.049898363 | 0.22689606 |

| | |
|---|---|
| 0.04988757 | 0.226868643 |
| 0.049886036 | 0.226868643 |
| 0.049863459 | 0.22680231 |
| 0.049859257 | 0.22680231 |
| 0.049848652 | 0.226778272 |
| 0.049832881 | 0.226728183 |
| 0.049820868 | 0.226695184 |
| 0.04980663 | 0.226654676 |
| 0.049784752 | 0.226595795 |
| 0.049779249 | 0.226592403 |
| 0.049752838 | 0.226537133 |
| 0.049745248 | 0.226524229 |
| 0.049732176 | 0.22648636 |
| 0.049713961 | 0.226425059 |
| 0.049709776 | 0.226425059 |
| 0.049695611 | 0.226384778 |
| 0.049691575 | 0.226384778 |
| 0.049651555 | 0.226249006 |
| 0.049645653 | 0.226249006 |
| 0.049634404 | 0.226239273 |
| 0.049562941 | 0.226018213 |
| 0.049561863 | 0.226018213 |
| 0.04955126 | 0.226008222 |
| 0.049541205 | 0.226004295 |
| 0.049537264 | 0.226004295 |
| 0.049530084 | 0.225998208 |
| 0.049519815 | 0.225973001 |
| 0.049514278 | 0.225969387 |
| 0.04950626 | 0.225954445 |
| 0.049477332 | 0.225844059 |
| 0.04947444 | 0.225844059 |
| 0.04946441 | 0.225844059 |
| 0.049460375 | 0.225844059 |
| 0.049445591 | 0.225829221 |
| 0.049437511 | 0.225829221 |
| 0.04940541 | 0.225753763 |
| 0.04940169 | 0.225753763 |
| 0.049391275 | 0.225732486 |
| 0.049378577 | 0.225696111 |
| 0.049369142 | 0.225674647 |
| 0.049363323 | 0.225669705 |
| 0.049353698 | 0.225647364 |
| 0.049322756 | 0.225527545 |
| 0.049316501 | 0.225520597 |
| 0.049314279 | 0.225520597 |

| | |
|---|---|
| 0.04930519 | 0.225512175 |
| 0.049302877 | 0.225512175 |
| 0.049285916 | 0.225488982 |
| 0.049173893 | 0.225084575 |
| 0.049168745 | 0.225082642 |
| 0.049128023 | 0.224917847 |
| 0.049121015 | 0.224907383 |
| 0.049110672 | 0.224881642 |
| 0.049109327 | 0.224881642 |
| 0.049098875 | 0.224870863 |
| 0.0490769 | 0.224791836 |
| 0.049072487 | 0.224791836 |
| 0.049062358 | 0.22476846 |
| 0.049059791 | 0.22476846 |
| 0.049036449 | 0.224693618 |
| 0.049027121 | 0.224693482 |
| 0.049020457 | 0.224684563 |
| 0.049020363 | 0.224684563 |
| 0.049018504 | 0.224684563 |
| 0.04900681 | 0.224684563 |
| 0.049003199 | 0.224684563 |
| 0.048979332 | 0.224604144 |
| 0.048979084 | 0.224604144 |
| 0.048972097 | 0.224604144 |
| 0.048961077 | 0.224585303 |
| 0.048956804 | 0.224585303 |
| 0.048926754 | 0.224471101 |
| 0.048898373 | 0.224362506 |
| 0.048891732 | 0.224353649 |
| 0.048875404 | 0.224300332 |
| 0.04884528 | 0.224248518 |
| 0.048838497 | 0.224238995 |
| 0.048815202 | 0.2241966 |
| 0.048810972 | 0.2241966 |
| 0.048810426 | 0.2241966 |
| 0.048809226 | 0.2241966 |
| 0.048795872 | 0.224194579 |
| 0.048788405 | 0.224181897 |
| 0.048771687 | 0.2241267 |
| 0.048763894 | 0.224112513 |
| 0.048738521 | 0.224056828 |
| 0.048733529 | 0.224056828 |
| 0.048728013 | 0.224055715 |
| 0.048719924 | 0.224040148 |
| 0.048717258 | 0.224040148 |

| | |
|---|---|
| 0.048712028 | 0.224040148 |
| 0.048658664 | 0.223823276 |
| 0.048644486 | 0.223793445 |
| 0.048644321 | 0.223793445 |
| 0.048636386 | 0.223793445 |
| 0.04860992 | 0.223728707 |
| 0.0486047 | 0.223726303 |
| 0.048604353 | 0.223726303 |
| 0.04859822 | 0.223726303 |
| 0.048539337 | 0.223555071 |
| 0.048538528 | 0.223555071 |
| 0.048534469 | 0.223555071 |
| 0.048513387 | 0.223543644 |
| 0.048504262 | 0.223523219 |
| 0.048473529 | 0.223436287 |
| 0.048435662 | 0.223336747 |
| 0.04834848 | 0.222999522 |
| 0.048332603 | 0.222947884 |
| 0.048323427 | 0.222927144 |
| 0.048293018 | 0.22282225 |
| 0.048289698 | 0.22282225 |
| 0.048267835 | 0.22280017 |
| 0.048248583 | 0.222749724 |
| 0.048243604 | 0.222749724 |
| 0.048225713 | 0.222692073 |
| 0.048202083 | 0.222626124 |
| 0.048185437 | 0.222570827 |
| 0.048166044 | 0.222502829 |
| 0.048137228 | 0.222391287 |
| 0.048129085 | 0.222375238 |
| 0.048119647 | 0.222353199 |
| 0.048112313 | 0.222340883 |
| 0.048104543 | 0.222340883 |
| 0.04808977 | 0.222301415 |
| 0.048076973 | 0.222273534 |
| 0.048066394 | 0.222258074 |
| 0.048028727 | 0.222105466 |
| 0.048018959 | 0.222081855 |
| 0.047981783 | 0.221974589 |
| 0.047978487 | 0.221974589 |
| 0.047974273 | 0.221974589 |
| 0.047966204 | 0.221974589 |
| 0.04796147 | 0.221974589 |
| 0.047945759 | 0.221974589 |
| 0.047842961 | 0.221590655 |

| | |
|---|---|
| 0.047840049 | 0.221590655 |
| 0.047834752 | 0.221590655 |
| 0.047829798 | 0.221590655 |
| 0.0478242 | 0.221590655 |
| 0.04779496 | 0.221497575 |
| 0.04777242 | 0.22141466 |
| 0.047771973 | 0.22141466 |
| 0.047771895 | 0.22141466 |
| 0.047740407 | 0.221330904 |
| 0.047721198 | 0.221263387 |
| 0.047712693 | 0.221245491 |
| 0.047702987 | 0.221222024 |
| 0.047696882 | 0.221222024 |
| 0.047693289 | 0.221222024 |
| 0.047692317 | 0.221222024 |
| 0.047683075 | 0.221222024 |
| 0.047654487 | 0.221154633 |
| 0.047639383 | 0.221120834 |
| 0.04761344 | 0.221021958 |
| 0.047598012 | 0.221014968 |
| 0.047570688 | 0.220974269 |
| 0.047569259 | 0.220974269 |
| 0.047516825 | 0.220788665 |
| 0.047507295 | 0.220787464 |
| 0.04748805 | 0.220719561 |
| 0.047485719 | 0.220719561 |
| 0.047484248 | 0.220719561 |
| 0.047475533 | 0.220719561 |
| 0.047454785 | 0.220672629 |
| 0.047447458 | 0.220660098 |
| 0.047424354 | 0.220574191 |
| 0.047409137 | 0.220546434 |
| 0.04740502 | 0.220546434 |
| 0.047401373 | 0.220546434 |
| 0.047359751 | 0.220424399 |
| 0.047334604 | 0.220350423 |
| 0.047322959 | 0.220317747 |
| 0.047305074 | 0.22027755 |
| 0.047304148 | 0.22027755 |
| 0.04728344 | 0.220219872 |
| 0.047280981 | 0.220219872 |
| 0.047279484 | 0.220219872 |
| 0.047264533 | 0.220217952 |
| 0.047215853 | 0.220012662 |
| 0.047167484 | 0.219808782 |

| | |
|---|---|
| 0.04715072 | 0.219773667 |
| 0.047130535 | 0.219701087 |
| 0.047110669 | 0.219629975 |
| 0.047103175 | 0.219616536 |
| 0.047064077 | 0.219477215 |
| 0.047054995 | 0.219477215 |
| 0.047054581 | 0.219477215 |
| 0.047049428 | 0.219477215 |
| 0.047045634 | 0.219477215 |
| 0.047027744 | 0.219458198 |
| 0.047013197 | 0.219411811 |
| 0.046998715 | 0.219365718 |
| 0.046985537 | 0.219331383 |
| 0.046959038 | 0.219266479 |
| 0.046958459 | 0.219266479 |
| 0.046951729 | 0.219266479 |
| 0.046942026 | 0.219251533 |
| 0.046930541 | 0.219219386 |
| 0.046902904 | 0.219111783 |
| 0.046890724 | 0.219076369 |
| 0.04686663 | 0.218985283 |
| 0.046857006 | 0.218961798 |
| 0.046788078 | 0.218661149 |
| 0.046787765 | 0.218661149 |
| 0.046785278 | 0.218661149 |
| 0.046782085 | 0.218661149 |
| 0.046758583 | 0.218609112 |
| 0.046758334 | 0.218609112 |
| 0.046713844 | 0.218464283 |
| 0.046682666 | 0.218339913 |
| 0.046678258 | 0.218339913 |
| 0.046655475 | 0.218261839 |
| 0.046654396 | 0.218261839 |
| 0.046648668 | 0.218261839 |
| 0.046636098 | 0.218261839 |
| 0.046619607 | 0.218237865 |
| 0.046618631 | 0.218237865 |
| 0.046601663 | 0.218218209 |
| 0.046592255 | 0.218195607 |
| 0.046554689 | 0.218062566 |
| 0.046514438 | 0.217895458 |
| 0.046502925 | 0.217884389 |
| 0.046481835 | 0.217807001 |
| 0.046434622 | 0.217628593 |
| 0.046432872 | 0.217628593 |

| | |
|---|---|
| 0.046403543 | 0.217568573 |
| 0.046382339 | 0.217490563 |
| 0.046347888 | 0.217371823 |
| 0.046347888 | 0.217371823 |
| 0.04634717 | 0.217371823 |
| 0.046330104 | 0.217352627 |
| 0.04632698 | 0.217352627 |
| 0.046285034 | 0.217183973 |
| 0.046278639 | 0.217175363 |
| 0.046259735 | 0.217108041 |
| 0.046252704 | 0.217096437 |
| 0.046235015 | 0.217096437 |
| 0.046224281 | 0.217091397 |
| 0.046212539 | 0.217079059 |
| 0.046205475 | 0.217067288 |
| 0.046201985 | 0.217067288 |
| 0.046195389 | 0.217062725 |
| 0.046193411 | 0.217062725 |
| 0.046123433 | 0.216810182 |
| 0.046092041 | 0.216684003 |
| 0.046091395 | 0.216684003 |
| 0.046084803 | 0.216684003 |
| 0.046076191 | 0.216673651 |
| 0.046054666 | 0.216593815 |
| 0.046035599 | 0.216525527 |
| 0.046030251 | 0.216521753 |
| 0.046026126 | 0.216521753 |
| 0.046020817 | 0.216520146 |
| 0.046004561 | 0.216465048 |
| 0.046003446 | 0.216465048 |
| 0.045969343 | 0.216363467 |
| 0.045938318 | 0.216238808 |
| 0.045867814 | 0.215949623 |
| 0.045861916 | 0.215943203 |
| 0.045859117 | 0.215943203 |
| 0.045850897 | 0.215943203 |
| 0.045833642 | 0.215913503 |
| 0.045807914 | 0.215816946 |
| 0.045738942 | 0.215513316 |
| 0.045718543 | 0.21546683 |
| 0.045693028 | 0.215403526 |
| 0.045686025 | 0.215391833 |
| 0.045630505 | 0.21525793 |
| 0.045620244 | 0.215230844 |
| 0.045600728 | 0.215160082 |

| | |
|---|---|
| 0.045577502 | 0.2150718 |
| 0.045565802 | 0.215037895 |
| 0.045558898 | 0.21502662 |
| 0.045551375 | 0.215012422 |
| 0.045542625 | 0.214992428 |
| 0.045535937 | 0.214982164 |
| 0.045521646 | 0.214936 |
| 0.045503198 | 0.214870199 |
| 0.045488391 | 0.214842879 |
| 0.045462596 | 0.214763646 |
| 0.045445398 | 0.214724996 |
| 0.045407933 | 0.214605026 |
| 0.045405439 | 0.214605026 |
| 0.045347948 | 0.214370888 |
| 0.045328361 | 0.214320841 |
| 0.045277716 | 0.214123896 |
| 0.045277493 | 0.214123896 |
| 0.045270114 | 0.214123896 |
| 0.045237481 | 0.214018626 |
| 0.045236757 | 0.214018626 |
| 0.045219305 | 0.213996405 |
| 0.045158611 | 0.213770737 |
| 0.045096163 | 0.213530296 |
| 0.045093203 | 0.213530296 |
| 0.045060517 | 0.213457056 |
| 0.045013709 | 0.213314804 |
| 0.045000985 | 0.213281113 |
| 0.044959962 | 0.213107891 |
| 0.044937733 | 0.213023727 |
| 0.044929318 | 0.213005038 |
| 0.044907656 | 0.212923538 |
| 0.04489621 | 0.21289046 |
| 0.044877483 | 0.212844043 |
| 0.044825977 | 0.212663292 |
| 0.044811952 | 0.212617937 |
| 0.044810904 | 0.212617937 |
| 0.04479818 | 0.21259495 |
| 0.04478509 | 0.21255401 |
| 0.044738804 | 0.212440182 |
| 0.044734254 | 0.212439756 |
| 0.044729273 | 0.212437284 |
| 0.044678457 | 0.212301793 |
| 0.044677927 | 0.212301793 |
| 0.044632816 | 0.212148414 |
| 0.044619575 | 0.212106649 |

| | |
|---|---|
| 0.044618176 | 0.212106649 |
| 0.044617853 | 0.212106649 |
| 0.044595072 | 0.212074839 |
| 0.044579485 | 0.212021882 |
| 0.044536402 | 0.211880456 |
| 0.044502191 | 0.211738845 |
| 0.044492774 | 0.211715193 |
| 0.04448106 | 0.211680602 |
| 0.044431257 | 0.211485856 |
| 0.044421456 | 0.211460336 |
| 0.044405043 | 0.211403336 |
| 0.044400293 | 0.211401856 |
| 0.044380124 | 0.211369218 |
| 0.044364423 | 0.211369218 |
| 0.044348868 | 0.211326016 |
| 0.044340852 | 0.211308961 |
| 0.044333141 | 0.211293358 |
| 0.044308786 | 0.211198416 |
| 0.04428403 | 0.211122673 |
| 0.044270418 | 0.211088878 |
| 0.044266473 | 0.211088878 |
| 0.04426465 | 0.211088878 |
| 0.044217818 | 0.210933697 |
| 0.044191849 | 0.210852054 |
| 0.044178571 | 0.21080982 |
| 0.044177648 | 0.21080982 |
| 0.044161551 | 0.21080982 |
| 0.044151102 | 0.210784336 |
| 0.044150064 | 0.210784336 |
| 0.044147175 | 0.210784336 |
| 0.044111712 | 0.210680758 |
| 0.04410231 | 0.210656977 |
| 0.044076722 | 0.210555873 |
| 0.044056902 | 0.210482304 |
| 0.044046631 | 0.210454346 |
| 0.044028049 | 0.210386666 |
| 0.044009032 | 0.210380224 |
| 0.043985886 | 0.210290682 |
| 0.043981298 | 0.210289857 |
| 0.043979515 | 0.210289857 |
| 0.0439606 | 0.210233103 |
| 0.043941978 | 0.210165153 |
| 0.043940762 | 0.210165153 |
| 0.043933684 | 0.210165153 |
| 0.043930085 | 0.210165153 |

| | |
|---|---|
| 0.043927729 | 0.210165153 |
| 0.043921388 | 0.210165153 |
| 0.043915762 | 0.210165153 |
| 0.043891117 | 0.210090677 |
| 0.043870715 | 0.210014128 |
| 0.043855371 | 0.209961776 |
| 0.043853861 | 0.209961776 |
| 0.043827982 | 0.209872841 |
| 0.043773908 | 0.209656064 |
| 0.043742116 | 0.209545942 |
| 0.043736552 | 0.209540363 |
| 0.043734434 | 0.209540363 |
| 0.043727734 | 0.209540274 |
| 0.043712632 | 0.209488986 |
| 0.043709772 | 0.209488986 |
| 0.043697074 | 0.209456574 |
| 0.043690136 | 0.209444401 |
| 0.043687717 | 0.209444401 |
| 0.043643612 | 0.209263496 |
| 0.043641132 | 0.209263496 |
| 0.043637291 | 0.209263496 |
| 0.043635561 | 0.209263496 |
| 0.043615181 | 0.209232496 |
| 0.043602523 | 0.209192842 |
| 0.043601706 | 0.209192842 |
| 0.043562559 | 0.209106431 |
| 0.043504749 | 0.208871035 |
| 0.043504294 | 0.208871035 |
| 0.043497854 | 0.208871035 |
| 0.043483442 | 0.208831895 |
| 0.043478145 | 0.208827514 |
| 0.043474842 | 0.208827514 |
| 0.043469717 | 0.208827514 |
| 0.043447615 | 0.208744031 |
| 0.043415423 | 0.208610408 |
| 0.043413546 | 0.208610408 |
| 0.043410704 | 0.208610408 |
| 0.043399653 | 0.208597766 |
| 0.043395348 | 0.208597766 |
| 0.043394903 | 0.208597766 |
| 0.043392903 | 0.208597766 |
| 0.043389589 | 0.208597766 |
| 0.043336073 | 0.208397329 |
| 0.04332301 | 0.20835555 |
| 0.043290605 | 0.208262794 |

| | |
|---|---|
| 0.043276304 | 0.208215025 |
| 0.043247305 | 0.208096525 |
| 0.043240241 | 0.208083559 |
| 0.043233779 | 0.208073486 |
| 0.043229675 | 0.208073486 |
| 0.043226525 | 0.208073486 |
| 0.043216778 | 0.208073486 |
| 0.043211305 | 0.208070448 |
| 0.043208223 | 0.208070448 |
| 0.043172631 | 0.207989367 |
| 0.043163629 | 0.207967038 |
| 0.043145886 | 0.20790258 |
| 0.043116075 | 0.207797669 |
| 0.043111174 | 0.207797669 |
| 0.043107536 | 0.207797669 |
| 0.043100861 | 0.207797669 |
| 0.043100695 | 0.207797669 |
| 0.043096025 | 0.207797669 |
| 0.043094025 | 0.207797669 |
| 0.043052777 | 0.207748187 |
| 0.043045774 | 0.207735441 |
| 0.043015852 | 0.207633118 |
| 0.043001067 | 0.20758279 |
| 0.042998421 | 0.20758279 |
| 0.042969028 | 0.20751226 |
| 0.042952457 | 0.207474307 |
| 0.04293105 | 0.20739194 |
| 0.042915375 | 0.207337245 |
| 0.042911417 | 0.207337245 |
| 0.04290687 | 0.207337245 |
| 0.042882659 | 0.207284317 |
| 0.042878887 | 0.207284317 |
| 0.042840478 | 0.207143495 |
| 0.042824569 | 0.207087596 |
| 0.042808688 | 0.207031821 |
| 0.042808017 | 0.207031821 |
| 0.042801072 | 0.207031821 |
| 0.042799976 | 0.207031821 |
| 0.042737351 | 0.206941969 |
| 0.042736284 | 0.206941969 |
| 0.042704487 | 0.206885019 |
| 0.042689127 | 0.206831643 |
| 0.042679957 | 0.206808252 |
| 0.042667571 | 0.206769273 |
| 0.042660733 | 0.206757173 |

| | |
|---|---|
| 0.042651823 | 0.206735024 |
| 0.042649224 | 0.206735024 |
| 0.042627112 | 0.206680501 |
| 0.04259688 | 0.20659487 |
| 0.042587781 | 0.206571772 |
| 0.042533957 | 0.206352731 |
| 0.042527638 | 0.206343094 |
| 0.042520587 | 0.206343094 |
| 0.042508037 | 0.206311044 |
| 0.042494683 | 0.20626725 |
| 0.042492301 | 0.20626725 |
| 0.042476221 | 0.206240327 |
| 0.042472432 | 0.206240327 |
| 0.042445005 | 0.206131156 |
| 0.042429553 | 0.206098148 |
| 0.042422377 | 0.206084308 |
| 0.042415437 | 0.206071612 |
| 0.042412253 | 0.206071612 |
| 0.042399283 | 0.206045995 |
| 0.042389342 | 0.206045995 |
| 0.042371356 | 0.205983524 |
| 0.042329246 | 0.205799814 |
| 0.042288427 | 0.205643343 |
| 0.042285168 | 0.205643343 |
| 0.042244797 | 0.205494121 |
| 0.042215573 | 0.205393919 |
| 0.042213012 | 0.205393919 |
| 0.042205589 | 0.205387308 |
| 0.042204691 | 0.205387308 |
| 0.042195795 | 0.205381618 |
| 0.042191858 | 0.205381618 |
| 0.042178972 | 0.205362697 |
| 0.042175977 | 0.205362697 |
| 0.042173643 | 0.205362697 |
| 0.042149249 | 0.205343931 |
| 0.042136435 | 0.205302504 |
| 0.042131508 | 0.2052995 |
| 0.042130821 | 0.2052995 |
| 0.042112181 | 0.205247323 |
| 0.042109926 | 0.205247323 |
| 0.042063254 | 0.205050827 |
| 0.04203081 | 0.204935431 |
| 0.042027157 | 0.204935431 |
| 0.041996285 | 0.204873943 |
| 0.041943825 | 0.204699106 |

| | |
|---|---|
| 0.041898651 | 0.204499597 |
| 0.041886926 | 0.204484279 |
| 0.04188066 | 0.204474646 |
| 0.041789201 | 0.204069954 |
| 0.041744433 | 0.203934979 |
| 0.041737014 | 0.203919654 |
| 0.041732063 | 0.20391638 |
| 0.041727057 | 0.20391284 |
| 0.041694008 | 0.203772243 |
| 0.041677464 | 0.203712292 |
| 0.0416692 | 0.203692803 |
| 0.041667389 | 0.203692803 |
| 0.04166038 | 0.2036915 |
| 0.04162825 | 0.203592046 |
| 0.041625422 | 0.203592046 |
| 0.041613192 | 0.203565272 |
| 0.041602729 | 0.203540632 |
| 0.041589013 | 0.203540632 |
| 0.041567186 | 0.203465608 |
| 0.041544654 | 0.203397128 |
| 0.041530901 | 0.203350699 |
| 0.041523797 | 0.203336815 |
| 0.041483202 | 0.203186843 |
| 0.041471219 | 0.203162902 |
| 0.041466401 | 0.203160198 |
| 0.041452067 | 0.203110864 |
| 0.04143714 | 0.203064746 |
| 0.041424396 | 0.203037943 |
| 0.04139848 | 0.202994482 |
| 0.041382628 | 0.202937649 |
| 0.04134868 | 0.202792047 |
| 0.041316837 | 0.202669196 |
| 0.041304993 | 0.20264039 |
| 0.041301768 | 0.20264039 |
| 0.041301649 | 0.20264039 |
| 0.041275261 | 0.202557121 |
| 0.041274547 | 0.202557121 |
| 0.041261876 | 0.202533171 |
| 0.041256515 | 0.202527731 |
| 0.041244661 | 0.202490408 |
| 0.041240676 | 0.202490408 |
| 0.041225775 | 0.202481179 |
| 0.041208192 | 0.202436569 |
| 0.041201658 | 0.202425351 |
| 0.04119467 | 0.202425351 |

| | |
|---|---|
| 0.041167914 | 0.202349 |
| 0.041167896 | 0.202349 |
| 0.041148885 | 0.202333051 |
| 0.041118063 | 0.202239281 |
| 0.041112781 | 0.202239042 |
| 0.041112292 | 0.202239042 |
| 0.041106442 | 0.202239042 |
| 0.041081392 | 0.20216816 |
| 0.041080694 | 0.20216816 |
| 0.041071252 | 0.202160038 |
| 0.041053225 | 0.202092189 |
| 0.041010375 | 0.201912914 |
| 0.041009839 | 0.201912914 |
| 0.0410016 | 0.201912914 |
| 0.040989786 | 0.201905111 |
| 0.04097149 | 0.201835868 |
| 0.040953982 | 0.201770487 |
| 0.040948327 | 0.2017635 |
| 0.040940073 | 0.201743701 |
| 0.040934169 | 0.201735477 |
| 0.040926983 | 0.201720937 |
| 0.040916982 | 0.201692515 |
| 0.040898074 | 0.201641053 |
| 0.04089416 | 0.201641053 |
| 0.040853159 | 0.201503047 |
| 0.040835663 | 0.20143761 |
| 0.040825502 | 0.201408349 |
| 0.040823202 | 0.201408349 |
| 0.040816863 | 0.201407452 |
| 0.040815081 | 0.201407452 |
| 0.040778651 | 0.201302324 |
| 0.040775058 | 0.201302324 |
| 0.040741204 | 0.201159166 |
| 0.040725873 | 0.201104321 |
| 0.040712761 | 0.201060419 |
| 0.040711962 | 0.201060419 |
| 0.040710399 | 0.201060419 |
| 0.040697956 | 0.201049846 |
| 0.040697493 | 0.201049846 |
| 0.040688693 | 0.201045793 |
| 0.040674372 | 0.200995881 |
| 0.040664091 | 0.200965928 |
| 0.040660124 | 0.200965928 |
| 0.040648754 | 0.200952689 |
| 0.040633553 | 0.200898396 |

| | |
|---|---|
| 0.040618472 | 0.200882415 |
| 0.04060873 | 0.20085907 |
| 0.040602881 | 0.200850998 |
| 0.040565153 | 0.200685211 |
| 0.040541642 | 0.200610574 |
| 0.04054113 | 0.200610574 |
| 0.040534341 | 0.200610574 |
| 0.040528867 | 0.200609887 |
| 0.040495657 | 0.200466338 |
| 0.040488613 | 0.200452306 |
| 0.040478551 | 0.200423323 |
| 0.040458987 | 0.200347285 |
| 0.040457742 | 0.200347285 |
| 0.040412812 | 0.200160255 |
| 0.040399376 | 0.20013533 |
| 0.040392356 | 0.20012137 |
| 0.040388618 | 0.20012137 |
| 0.040368074 | 0.200042687 |
| 0.040328868 | 0.1998692 |
| 0.040306868 | 0.199780954 |
| 0.040294802 | 0.199762729 |
| 0.040293578 | 0.199762729 |
| 0.040285402 | 0.199760074 |
| 0.040275733 | 0.199751365 |
| 0.040273118 | 0.199751365 |
| 0.040272157 | 0.199751365 |
| 0.040267192 | 0.199751365 |
| 0.040265751 | 0.199751365 |
| 0.040261055 | 0.199751365 |
| 0.040220564 | 0.199602461 |
| 0.040198067 | 0.199511602 |
| 0.040160232 | 0.199344592 |
| 0.040148259 | 0.199326708 |
| 0.040142582 | 0.1993193 |
| 0.040116995 | 0.199213026 |
| 0.040113304 | 0.199213026 |
| 0.04009696 | 0.199155065 |
| 0.040071198 | 0.199047864 |
| 0.040029251 | 0.198901733 |
| 0.039984741 | 0.198722034 |
| 0.039979036 | 0.198714417 |
| 0.03996657 | 0.19867319 |
| 0.039919754 | 0.198481901 |
| 0.039918663 | 0.198481901 |
| 0.039894984 | 0.198441607 |

| | |
|---|---|
| 0.039876374 | 0.198369758 |
| 0.03985878 | 0.198302949 |
| 0.039845639 | 0.19825828 |
| 0.039810373 | 0.198124203 |
| 0.039804709 | 0.198124203 |
| 0.039799362 | 0.198124203 |
| 0.039785473 | 0.19810991 |
| 0.03976033 | 0.198040706 |
| 0.039755415 | 0.198036938 |
| 0.039747394 | 0.198017692 |
| 0.039747048 | 0.198017692 |
| 0.039736323 | 0.198017692 |
| 0.039726207 | 0.197994983 |
| 0.039724171 | 0.197994983 |
| 0.039719897 | 0.197994983 |
| 0.039698006 | 0.197937285 |
| 0.039696972 | 0.197937285 |
| 0.039694425 | 0.197937285 |
| 0.039677075 | 0.19791019 |
| 0.039648881 | 0.197816601 |
| 0.039639768 | 0.197812579 |
| 0.039638136 | 0.197812579 |
| 0.039631026 | 0.197810403 |
| 0.039611387 | 0.197733104 |
| 0.039603276 | 0.197713337 |
| 0.039591013 | 0.197672833 |
| 0.039577815 | 0.19762766 |
| 0.039564721 | 0.197603711 |
| 0.039553708 | 0.197569423 |
| 0.039530656 | 0.197474989 |
| 0.039515471 | 0.197419838 |
| 0.039511433 | 0.197419838 |
| 0.039501711 | 0.197392506 |
| 0.03949963 | 0.197392506 |
| 0.039484933 | 0.197350079 |
| 0.039463535 | 0.197284534 |
| 0.039460661 | 0.197284534 |
| 0.03944057 | 0.197231838 |
| 0.039437892 | 0.197231838 |
| 0.039396784 | 0.197054256 |
| 0.039369069 | 0.196941121 |
| 0.039365998 | 0.196941121 |
| 0.039349167 | 0.196898802 |
| 0.039317378 | 0.196760405 |
| 0.039314292 | 0.196760405 |

| | |
|---|---|
| 0.039308121 | 0.196755424 |
| 0.039259457 | 0.196615159 |
| 0.03925353 | 0.196609671 |
| 0.039246503 | 0.196609671 |
| 0.039230748 | 0.196554054 |
| 0.039209371 | 0.196488289 |
| 0.039196706 | 0.196445493 |
| 0.039193782 | 0.196445493 |
| 0.039180065 | 0.196403427 |
| 0.039178917 | 0.196403427 |
| 0.039163699 | 0.196383394 |
| 0.039163065 | 0.196383394 |
| 0.03913868 | 0.196361311 |
| 0.039134192 | 0.196359479 |
| 0.039127576 | 0.196346966 |
| 0.03909503 | 0.196204317 |
| 0.039091631 | 0.196204317 |
| 0.039068218 | 0.196111089 |
| 0.039035774 | 0.196030872 |
| 0.039030828 | 0.196026706 |
| 0.039025845 | 0.196026706 |
| 0.039012028 | 0.195994288 |
| 0.038996202 | 0.19595612 |
| 0.038995403 | 0.19595612 |
| 0.038989053 | 0.19595612 |
| 0.03898293 | 0.195951452 |
| 0.038982307 | 0.195951452 |
| 0.038973792 | 0.195946879 |
| 0.038972327 | 0.195946879 |
| 0.038971787 | 0.195946879 |
| 0.038966938 | 0.195946879 |
| 0.038964055 | 0.195946879 |
| 0.038931236 | 0.195856976 |
| 0.038907589 | 0.195758686 |
| 0.038887529 | 0.195699106 |
| 0.038880585 | 0.195684839 |
| 0.038873347 | 0.195669087 |
| 0.038856125 | 0.19560307 |
| 0.03885598 | 0.19560307 |
| 0.038841689 | 0.195571742 |
| 0.038841292 | 0.195571742 |
| 0.038836319 | 0.195571742 |
| 0.038834958 | 0.195571742 |
| 0.038820884 | 0.195570365 |
| 0.038818297 | 0.195570365 |

| | |
|---|---|
| 0.038812706 | 0.195570365 |
| 0.038806993 | 0.195562445 |
| 0.038793403 | 0.195514644 |
| 0.038786399 | 0.195500035 |
| 0.038781881 | 0.195497949 |
| 0.038777744 | 0.195497791 |
| 0.038758325 | 0.195420572 |
| 0.038736631 | 0.19533187 |
| 0.038735061 | 0.19533187 |
| 0.038686794 | 0.195121881 |
| 0.038683522 | 0.195121881 |
| 0.038656826 | 0.195044925 |
| 0.038652341 | 0.195044925 |
| 0.038646209 | 0.195041113 |
| 0.03864502 | 0.195041113 |
| 0.038642649 | 0.195041113 |
| 0.038623375 | 0.195008532 |
| 0.038597208 | 0.194917744 |
| 0.038582741 | 0.194865344 |
| 0.038579942 | 0.194865344 |
| 0.038540548 | 0.19469354 |
| 0.038538543 | 0.19469354 |
| 0.038526003 | 0.194682008 |
| 0.038522089 | 0.194682008 |
| 0.038498974 | 0.194586716 |
| 0.038493191 | 0.194578139 |
| 0.038467787 | 0.194487987 |
| 0.038465927 | 0.194487987 |
| 0.038465814 | 0.194487987 |
| 0.038444816 | 0.194455727 |
| 0.038429749 | 0.194401873 |
| 0.038400987 | 0.194277014 |
| 0.038396664 | 0.194275779 |
| 0.038389185 | 0.194258574 |
| 0.038382624 | 0.194246014 |
| 0.038356941 | 0.194136666 |
| 0.038348759 | 0.194115883 |
| 0.038323682 | 0.194050817 |
| 0.038308473 | 0.193994432 |
| 0.038308322 | 0.193994432 |
| 0.0383047 | 0.193994432 |
| 0.038303203 | 0.193994432 |
| 0.038289911 | 0.193994432 |
| 0.038285249 | 0.193994432 |
| 0.038262518 | 0.19394731 |

| | |
|---|---|
| 0.03825897 | 0.19394731 |
| 0.038254057 | 0.193945706 |
| 0.038252122 | 0.193945706 |
| 0.038233412 | 0.193923611 |
| 0.038208103 | 0.193815882 |
| 0.038191809 | 0.193771322 |
| 0.038185051 | 0.193760861 |
| 0.038178459 | 0.193748051 |
| 0.03816696 | 0.193710339 |
| 0.038160127 | 0.193696299 |
| 0.038158496 | 0.193696299 |
| 0.038140395 | 0.193658058 |
| 0.038140303 | 0.193658058 |
| 0.038122601 | 0.193629641 |
| 0.03805974 | 0.193351597 |
| 0.038059624 | 0.193351597 |
| 0.038035404 | 0.193289817 |
| 0.038025276 | 0.19325897 |
| 0.038014907 | 0.193226893 |
| 0.03800505 | 0.193197407 |
| 0.037987267 | 0.193144222 |
| 0.0379811 | 0.193137504 |
| 0.037976854 | 0.193136532 |
| 0.037976295 | 0.193136532 |
| 0.037972072 | 0.193136532 |
| 0.037915489 | 0.192906844 |
| 0.03789964 | 0.192846814 |
| 0.037899007 | 0.192846814 |
| 0.037898592 | 0.192846814 |
| 0.037889224 | 0.192846814 |
| 0.037867989 | 0.19277046 |
| 0.037866216 | 0.19277046 |
| 0.037845954 | 0.192717764 |
| 0.037836904 | 0.192692285 |
| 0.037818767 | 0.192620514 |
| 0.037774105 | 0.192454781 |
| 0.037771716 | 0.192454781 |
| 0.037766688 | 0.192454781 |
| 0.037748675 | 0.192407549 |
| 0.037744784 | 0.192407549 |
| 0.037739479 | 0.192407549 |
| 0.037713216 | 0.192329727 |
| 0.037682729 | 0.192215416 |
| 0.037634216 | 0.192009085 |
| 0.037621114 | 0.191983368 |

| | |
|---|---|
| 0.037618417 | 0.191983368 |
| 0.037616631 | 0.191983368 |
| 0.037613415 | 0.191983368 |
| 0.037612396 | 0.191983368 |
| 0.03759492 | 0.191952537 |
| 0.03758719 | 0.191933643 |
| 0.037572517 | 0.191879291 |
| 0.037531795 | 0.191733004 |
| 0.037504412 | 0.19161367 |
| 0.037496525 | 0.191593929 |
| 0.037491959 | 0.191591153 |
| 0.037489857 | 0.191591153 |
| 0.037482553 | 0.191584198 |
| 0.037458755 | 0.191503661 |
| 0.037456255 | 0.191503661 |
| 0.037427799 | 0.191386494 |
| 0.03739454 | 0.191236952 |
| 0.037384827 | 0.191207815 |
| 0.037383131 | 0.191207815 |
| 0.037358123 | 0.191132808 |
| 0.037354786 | 0.191132808 |
| 0.037353447 | 0.191132808 |
| 0.037340049 | 0.191101926 |
| 0.037333613 | 0.191089522 |
| 0.037308514 | 0.190981578 |
| 0.037284306 | 0.190915276 |
| 0.037282931 | 0.190915276 |
| 0.037276988 | 0.190915276 |
| 0.037269757 | 0.190915276 |
| 0.037236329 | 0.190796625 |
| 0.037232982 | 0.190796625 |
| 0.037230968 | 0.190796625 |
| 0.037223444 | 0.190792183 |
| 0.037219346 | 0.190791712 |
| 0.03720647 | 0.190746236 |
| 0.037205272 | 0.190746236 |
| 0.037192596 | 0.190716166 |
| 0.037179218 | 0.190668098 |
| 0.037173813 | 0.190660909 |
| 0.037158692 | 0.190603878 |
| 0.037137527 | 0.190515836 |
| 0.037131209 | 0.190503944 |
| 0.037123287 | 0.190503944 |
| 0.037121648 | 0.190503944 |
| 0.037082854 | 0.190337869 |

| | |
|---|---|
| 0.037077448 | 0.190330631 |
| 0.037063751 | 0.190280832 |
| 0.037053745 | 0.19024997 |
| 0.037033815 | 0.190168143 |
| 0.037021678 | 0.190146818 |
| 0.036989326 | 0.190021663 |
| 0.036984574 | 0.190021663 |
| 0.036981734 | 0.190021663 |
| 0.036972598 | 0.190021663 |
| 0.036972052 | 0.190021663 |
| 0.036963016 | 0.190021663 |
| 0.036961329 | 0.190021663 |
| 0.036953848 | 0.190021663 |
| 0.036925061 | 0.189896329 |
| 0.036909516 | 0.189857382 |
| 0.03689523 | 0.189804397 |
| 0.036852017 | 0.189643544 |
| 0.036849349 | 0.189643544 |
| 0.036835957 | 0.18960187 |
| 0.036815518 | 0.189517151 |
| 0.036798535 | 0.189450201 |
| 0.036780588 | 0.189378279 |
| 0.036778372 | 0.189378279 |
| 0.036775843 | 0.189378279 |
| 0.03676013 | 0.189375305 |
| 0.036758258 | 0.189375305 |
| 0.036757535 | 0.189375305 |
| 0.036753806 | 0.189375305 |
| 0.036736171 | 0.189359278 |
| 0.036732373 | 0.189359278 |
| 0.036724527 | 0.189355748 |
| 0.036704535 | 0.189299618 |
| 0.03670119 | 0.189299618 |
| 0.036653448 | 0.189112095 |
| 0.036643498 | 0.189081246 |
| 0.036628454 | 0.189024096 |
| 0.036626744 | 0.189024096 |
| 0.036616994 | 0.189024096 |
| 0.036609165 | 0.189006471 |
| 0.036598262 | 0.188970668 |
| 0.036594337 | 0.188970668 |
| 0.036592479 | 0.188970668 |
| 0.036588872 | 0.188970668 |
| 0.036585016 | 0.188970668 |
| 0.036566124 | 0.188948115 |

| | |
|---|---|
| 0.036563682 | 0.188948115 |
| 0.03655465 | 0.188929825 |
| 0.036552831 | 0.188929825 |
| 0.036527418 | 0.188871062 |
| 0.036514541 | 0.188824977 |
| 0.036460886 | 0.188578011 |
| 0.036433416 | 0.188466854 |
| 0.036428236 | 0.188461726 |
| 0.036421847 | 0.188461726 |
| 0.036392536 | 0.188357675 |
| 0.036383475 | 0.18833124 |
| 0.036379132 | 0.18832923 |
| 0.036376132 | 0.18832923 |
| 0.036356754 | 0.18825431 |
| 0.036351533 | 0.188247742 |
| 0.036349316 | 0.188247742 |
| 0.03634271 | 0.188247742 |
| 0.036330382 | 0.188240546 |
| 0.03632446 | 0.188240546 |
| 0.036271238 | 0.188036381 |
| 0.036252175 | 0.187958019 |
| 0.036222859 | 0.187867375 |
| 0.036211038 | 0.18784698 |
| 0.036192298 | 0.187770214 |
| 0.036181603 | 0.187735176 |
| 0.036176888 | 0.187731169 |
| 0.036148385 | 0.187624143 |
| 0.036130682 | 0.187552694 |
| 0.036118284 | 0.187508775 |
| 0.036118089 | 0.187508775 |
| 0.036112322 | 0.187508775 |
| 0.036083153 | 0.187408104 |
| 0.03607832 | 0.187403441 |
| 0.036060503 | 0.187331323 |
| 0.03605716 | 0.187331323 |
| 0.036053067 | 0.187331323 |
| 0.036048062 | 0.187327992 |
| 0.036037275 | 0.187305003 |
| 0.036030719 | 0.187299177 |
| 0.036025182 | 0.187290838 |
| 0.036005786 | 0.187216616 |
| 0.03600513 | 0.187216616 |
| 0.036004451 | 0.187216616 |
| 0.035988663 | 0.187203151 |
| 0.03597527 | 0.18717437 |

| | |
|---|---|
| 0.03597493 | 0.18717437 |
| 0.035958557 | 0.1871283 |
| 0.035943638 | 0.187071104 |
| 0.035918138 | 0.186999691 |
| 0.035903898 | 0.186945989 |
| 0.035890582 | 0.186897086 |
| 0.035885665 | 0.186891913 |
| 0.035862657 | 0.186792514 |
| 0.035849747 | 0.186766123 |
| 0.035835054 | 0.18671 |
| 0.035819821 | 0.186651054 |
| 0.03578698 | 0.186500326 |
| 0.035781684 | 0.186500326 |
| 0.035773066 | 0.186489035 |
| 0.035758908 | 0.186437982 |
| 0.035744479 | 0.186401216 |
| 0.035715181 | 0.186330041 |
| 0.035697989 | 0.186260748 |
| 0.035674789 | 0.186160092 |
| 0.035663401 | 0.186121062 |
| 0.035635253 | 0.186027066 |
| 0.035617708 | 0.185964107 |
| 0.035595889 | 0.185907009 |
| 0.035593711 | 0.185907009 |
| 0.035590291 | 0.185907009 |
| 0.03557908 | 0.185905086 |
| 0.035554232 | 0.185795636 |
| 0.035526808 | 0.185713457 |
| 0.035505182 | 0.185641159 |
| 0.03549803 | 0.185624141 |
| 0.035491455 | 0.185610136 |
| 0.035487291 | 0.185608744 |
| 0.035483651 | 0.185608744 |
| 0.035476173 | 0.185608744 |
| 0.03546809 | 0.185589839 |
| 0.035433535 | 0.185429398 |
| 0.035402732 | 0.185349667 |
| 0.035389935 | 0.185323415 |
| 0.035376877 | 0.185275402 |
| 0.035368462 | 0.185251703 |
| 0.035332097 | 0.185101947 |
| 0.035323845 | 0.18507907 |
| 0.035320921 | 0.18507907 |
| 0.035319391 | 0.18507907 |
| 0.035313425 | 0.18507907 |

| | |
|---|---|
| 0.035312995 | 0.18507907 |
| 0.035311614 | 0.18507907 |
| 0.035309557 | 0.18507907 |
| 0.035289757 | 0.185067339 |
| 0.035275532 | 0.18502949 |
| 0.035273099 | 0.18502949 |
| 0.035263974 | 0.185009616 |
| 0.03525788 | 0.184998021 |
| 0.035213271 | 0.18480468 |
| 0.035209899 | 0.18480468 |
| 0.035182276 | 0.184714743 |
| 0.035177526 | 0.184714743 |
| 0.035168938 | 0.184714491 |
| 0.035152151 | 0.184646686 |
| 0.035125639 | 0.184527777 |
| 0.035119068 | 0.18451361 |
| 0.035103694 | 0.184473534 |
| 0.035100979 | 0.184473534 |
| 0.035094827 | 0.184467647 |
| 0.035091955 | 0.184467647 |
| 0.035083557 | 0.184449117 |
| 0.035072641 | 0.184412087 |
| 0.035056568 | 0.184368284 |
| 0.035022217 | 0.184269017 |
| 0.035019502 | 0.184269017 |
| 0.034994691 | 0.184165739 |
| 0.034937996 | 0.183927482 |
| 0.03493634 | 0.183927482 |
| 0.034886032 | 0.183714846 |
| 0.034868411 | 0.183642355 |
| 0.034863017 | 0.183634252 |
| 0.034858021 | 0.183634252 |
| 0.034828575 | 0.183521635 |
| 0.034825924 | 0.183521635 |
| 0.034821796 | 0.183521635 |
| 0.034807385 | 0.183521635 |
| 0.034801585 | 0.183521635 |
| 0.034776125 | 0.18341999 |
| 0.0347717 | 0.183416963 |
| 0.034770684 | 0.183416963 |
| 0.034747157 | 0.183368738 |
| 0.034744491 | 0.183368738 |
| 0.034730734 | 0.183343013 |
| 0.034719481 | 0.183309618 |
| 0.034693179 | 0.183225994 |

| | |
|---|---|
| 0.034674951 | 0.18315004 |
| 0.034661871 | 0.183121565 |
| 0.034633434 | 0.183047035 |
| 0.034626646 | 0.183047035 |
| 0.034623212 | 0.183047035 |
| 0.034613773 | 0.183029883 |
| 0.034593892 | 0.182945067 |
| 0.034566303 | 0.182860079 |
| 0.034525189 | 0.182723767 |
| 0.034521601 | 0.182723767 |
| 0.034519025 | 0.182723767 |
| 0.034513125 | 0.182721825 |
| 0.034503902 | 0.182712626 |
| 0.034501609 | 0.182712626 |
| 0.034458989 | 0.182515377 |
| 0.034456651 | 0.182515377 |
| 0.034452724 | 0.182515377 |
| 0.034443001 | 0.182491578 |
| 0.034407157 | 0.182342238 |
| 0.034386764 | 0.182295017 |
| 0.034385173 | 0.182295017 |
| 0.034344033 | 0.182149581 |
| 0.03434122 | 0.182149581 |
| 0.034289138 | 0.181898951 |
| 0.034266746 | 0.181800412 |
| 0.034263777 | 0.181800412 |
| 0.034243214 | 0.181736301 |
| 0.034214383 | 0.181623765 |
| 0.034194781 | 0.181560183 |
| 0.034193472 | 0.181560183 |
| 0.034169844 | 0.181468237 |
| 0.034165733 | 0.181466643 |
| 0.034162083 | 0.181466643 |
| 0.034160746 | 0.181466643 |
| 0.034149278 | 0.181439945 |
| 0.034146406 | 0.181439945 |
| 0.034097088 | 0.181285387 |
| 0.034095973 | 0.181285387 |
| 0.034084839 | 0.181279578 |
| 0.034084765 | 0.181279578 |
| 0.034080474 | 0.181279578 |
| 0.034058329 | 0.181260033 |
| 0.034056174 | 0.181260033 |
| 0.034048505 | 0.181260033 |
| 0.034028278 | 0.181241867 |

| | |
|---|---|
| 0.034026974 | 0.181241867 |
| 0.034025864 | 0.181241867 |
| 0.034000321 | 0.18119428 |
| 0.033976246 | 0.181086244 |
| 0.033962173 | 0.181031494 |
| 0.033957786 | 0.181030832 |
| 0.033944027 | 0.180995541 |
| 0.033935488 | 0.180970272 |
| 0.033930682 | 0.180964904 |
| 0.033926413 | 0.180964904 |
| 0.033897846 | 0.180870792 |
| 0.03386397 | 0.180771051 |
| 0.033852826 | 0.180731816 |
| 0.033845233 | 0.180711538 |
| 0.033839788 | 0.180707329 |
| 0.033811209 | 0.180590611 |
| 0.033787016 | 0.180501872 |
| 0.033786687 | 0.180501872 |
| 0.033782454 | 0.180501872 |
| 0.033777225 | 0.180501872 |
| 0.033727794 | 0.180327082 |
| 0.033703652 | 0.180218238 |
| 0.033697688 | 0.180218238 |
| 0.03369037 | 0.180207911 |
| 0.033679353 | 0.180189458 |
| 0.03367771 | 0.180189458 |
| 0.033642767 | 0.180054395 |
| 0.033622552 | 0.179986655 |
| 0.03361593 | 0.179971437 |
| 0.033610712 | 0.179963728 |
| 0.033603114 | 0.17996351 |
| 0.033602015 | 0.17996351 |
| 0.033594678 | 0.179960113 |
| 0.033585558 | 0.179960113 |
| 0.033574869 | 0.179933644 |
| 0.033550975 | 0.179825826 |
| 0.033548194 | 0.179825826 |
| 0.033533103 | 0.179770495 |
| 0.03352379 | 0.179740801 |
| 0.033511732 | 0.179716614 |
| 0.033509886 | 0.179716614 |
| 0.033500686 | 0.179697844 |
| 0.033496494 | 0.179695598 |
| 0.033492424 | 0.179694004 |
| 0.033441378 | 0.179518448 |

| | |
|---|---|
| 0.033421432 | 0.179474859 |
| 0.033413679 | 0.179453459 |
| 0.033409362 | 0.179450509 |
| 0.033400983 | 0.17944598 |
| 0.033399614 | 0.17944598 |
| 0.033388111 | 0.179417305 |
| 0.033382264 | 0.179406128 |
| 0.033380815 | 0.179406128 |
| 0.033371669 | 0.179389673 |
| 0.033350067 | 0.179293786 |
| 0.03334643 | 0.179293786 |
| 0.033329747 | 0.179225003 |
| 0.033315053 | 0.17916622 |
| 0.03330367 | 0.17912523 |
| 0.033293133 | 0.179088778 |
| 0.033283463 | 0.179077213 |
| 0.033281118 | 0.179077213 |
| 0.033275856 | 0.179076747 |
| 0.033254603 | 0.178982594 |
| 0.033250436 | 0.178980395 |
| 0.033249814 | 0.178980395 |
| 0.033235469 | 0.178940274 |
| 0.033222461 | 0.178890465 |
| 0.033208182 | 0.178836563 |
| 0.033198934 | 0.178824432 |
| 0.033148174 | 0.178611617 |
| 0.033137149 | 0.178580767 |
| 0.033128387 | 0.178565604 |
| 0.033117539 | 0.17852734 |
| 0.033109359 | 0.17852734 |
| 0.033088531 | 0.178471969 |
| 0.033086273 | 0.178471969 |
| 0.033053543 | 0.1783604 |
| 0.033049662 | 0.1783604 |
| 0.033041213 | 0.17835814 |
| 0.033009263 | 0.17820587 |
| 0.032994563 | 0.178166902 |
| 0.03298755 | 0.178149228 |
| 0.032940578 | 0.177976279 |
| 0.03292606 | 0.177918023 |
| 0.032916163 | 0.177904911 |
| 0.032915524 | 0.177904911 |
| 0.032910219 | 0.177904911 |
| 0.032908421 | 0.177904911 |
| 0.032906667 | 0.177904911 |

| | |
|---|---|
| 0.032894756 | 0.17789015 |
| 0.032892599 | 0.17789015 |
| 0.032886263 | 0.17789015 |
| 0.032873011 | 0.17787354 |
| 0.032838249 | 0.177725826 |
| 0.032827955 | 0.177709182 |
| 0.03281947 | 0.17770497 |
| 0.032804728 | 0.17764534 |
| 0.032803814 | 0.17764534 |
| 0.032779925 | 0.177571597 |
| 0.032777977 | 0.177571597 |
| 0.032775713 | 0.177571597 |
| 0.032771816 | 0.177571597 |
| 0.032770024 | 0.177571597 |
| 0.032730133 | 0.177456245 |
| 0.032724456 | 0.177456245 |
| 0.032718723 | 0.177456245 |
| 0.032715681 | 0.177456245 |
| 0.032683963 | 0.177354349 |
| 0.032648867 | 0.177244679 |
| 0.032648107 | 0.177244679 |
| 0.032634455 | 0.177227045 |
| 0.032633643 | 0.177227045 |
| 0.032626546 | 0.177227045 |
| 0.032609592 | 0.177172831 |
| 0.032577857 | 0.177020606 |
| 0.032574573 | 0.177020606 |
| 0.032559608 | 0.176961822 |
| 0.032557214 | 0.176961822 |
| 0.032531742 | 0.176850736 |
| 0.032517366 | 0.176792764 |
| 0.032507884 | 0.176761387 |
| 0.032503662 | 0.176758612 |
| 0.032501851 | 0.176758612 |
| 0.032447094 | 0.176511449 |
| 0.032442551 | 0.176506896 |
| 0.032439105 | 0.176506896 |
| 0.032411452 | 0.176377996 |
| 0.032399043 | 0.176370927 |
| 0.03239896 | 0.176370927 |
| 0.03237535 | 0.176302402 |
| 0.032367932 | 0.176282158 |
| 0.032338057 | 0.1761396 |
| 0.032331725 | 0.176125248 |
| 0.032328138 | 0.176125248 |

| | |
|---|---|
| 0.032322551 | 0.176115564 |
| 0.032314713 | 0.176093003 |
| 0.032312378 | 0.176093003 |
| 0.03230257 | 0.17606712 |
| 0.03229605 | 0.176051731 |
| 0.032293263 | 0.176051731 |
| 0.032292069 | 0.176051731 |
| 0.032286097 | 0.176051731 |
| 0.032272433 | 0.176043886 |
| 0.03225867 | 0.175988968 |
| 0.032237742 | 0.175915095 |
| 0.032204776 | 0.17575534 |
| 0.032203685 | 0.17575534 |
| 0.032188144 | 0.175724982 |
| 0.032184292 | 0.175724094 |
| 0.032184023 | 0.175724094 |
| 0.032165918 | 0.175664049 |
| 0.03216325 | 0.175664049 |
| 0.032159785 | 0.175664049 |
| 0.032159343 | 0.175664049 |
| 0.032152763 | 0.175664049 |
| 0.032126109 | 0.175567433 |
| 0.032113167 | 0.175557138 |
| 0.032099556 | 0.175502876 |
| 0.032076507 | 0.175396992 |
| 0.032074349 | 0.175396992 |
| 0.032062519 | 0.175380913 |
| 0.032057168 | 0.175380913 |
| 0.032052263 | 0.175380913 |
| 0.032044544 | 0.175363149 |
| 0.032036065 | 0.175357045 |
| 0.032009715 | 0.175252935 |
| 0.031997634 | 0.175207094 |
| 0.031951091 | 0.174972357 |
| 0.031935542 | 0.17491021 |
| 0.031917622 | 0.17484939 |
| 0.031903483 | 0.174792039 |
| 0.031901601 | 0.174792039 |
| 0.03189521 | 0.174786924 |
| 0.031885174 | 0.174752035 |
| 0.031884816 | 0.174752035 |
| 0.031864685 | 0.174700052 |
| 0.031863053 | 0.174700052 |
| 0.031847494 | 0.17464601 |
| 0.03184712 | 0.17464601 |

| | |
|---|---|
| 0.031840275 | 0.17464601 |
| 0.031817696 | 0.174622028 |
| 0.031815287 | 0.174622028 |
| 0.031813399 | 0.174622028 |
| 0.031778616 | 0.174489294 |
| 0.03177364 | 0.174482089 |
| 0.031767664 | 0.174469386 |
| 0.031760772 | 0.174468101 |
| 0.031748916 | 0.174426763 |
| 0.031743551 | 0.174417409 |
| 0.03172117 | 0.174344602 |
| 0.031719407 | 0.174344602 |
| 0.031719145 | 0.174344602 |
| 0.031718205 | 0.174344602 |
| 0.031716968 | 0.174344602 |
| 0.031674353 | 0.17425194 |
| 0.031672411 | 0.17425194 |
| 0.031668256 | 0.17425194 |
| 0.03166682 | 0.17425194 |
| 0.031659947 | 0.17425194 |
| 0.031649641 | 0.174223006 |
| 0.031624256 | 0.174103392 |
| 0.031620433 | 0.174102469 |
| 0.031615305 | 0.174094363 |
| 0.031611918 | 0.174094363 |
| 0.031608704 | 0.174094363 |
| 0.03160525 | 0.174094363 |
| 0.031600095 | 0.174094363 |
| 0.03159685 | 0.174094363 |
| 0.031577383 | 0.174026383 |
| 0.031564732 | 0.173976797 |
| 0.031554105 | 0.173976797 |
| 0.03154786 | 0.173976797 |
| 0.03153052 | 0.173908984 |
| 0.0315196 | 0.173868887 |
| 0.031500603 | 0.173784223 |
| 0.031488646 | 0.173758515 |
| 0.03148175 | 0.173740591 |
| 0.031471606 | 0.173704737 |
| 0.031460547 | 0.173663824 |
| 0.031455313 | 0.173655059 |
| 0.031452293 | 0.173655059 |
| 0.031452112 | 0.173655059 |
| 0.031448393 | 0.173655059 |
| 0.031444875 | 0.173655059 |

| | |
|---|---|
| 0.03142574 | 0.173632682 |
| 0.031412863 | 0.173587403 |
| 0.031386778 | 0.17349792 |
| 0.031385872 | 0.17349792 |
| 0.031363818 | 0.173418889 |
| 0.031358813 | 0.173418889 |
| 0.031355319 | 0.173418889 |
| 0.03133281 | 0.173340426 |
| 0.031331389 | 0.173340426 |
| 0.031323011 | 0.173340426 |
| 0.031318598 | 0.173340426 |
| 0.031302521 | 0.173313781 |
| 0.031283931 | 0.173230995 |
| 0.031272806 | 0.173189526 |
| 0.031267096 | 0.17317804 |
| 0.031263772 | 0.17317804 |
| 0.03126218 | 0.17317804 |
| 0.031258996 | 0.17317804 |
| 0.031248329 | 0.17317804 |
| 0.031225977 | 0.173111351 |
| 0.031202753 | 0.173022878 |
| 0.031193705 | 0.173012996 |
| 0.031180953 | 0.172974982 |
| 0.031178506 | 0.172974982 |
| 0.031175489 | 0.172974982 |
| 0.031150357 | 0.172893375 |
| 0.031138758 | 0.172849143 |
| 0.031101642 | 0.17269775 |
| 0.031090441 | 0.17269775 |
| 0.031075414 | 0.172638368 |
| 0.031060307 | 0.172574572 |
| 0.031053283 | 0.172574572 |
| 0.031050549 | 0.172574572 |
| 0.031030609 | 0.172490055 |
| 0.03101759 | 0.172444356 |
| 0.031003836 | 0.172421729 |
| 0.030988026 | 0.172353931 |
| 0.030973468 | 0.172293078 |
| 0.030965401 | 0.172288453 |
| 0.030951467 | 0.17225117 |
| 0.03093844 | 0.172198797 |
| 0.030935954 | 0.172198797 |
| 0.030933418 | 0.172198797 |
| 0.030926161 | 0.172198797 |
| 0.030915918 | 0.172194058 |

| | |
|---|---|
| 0.030906643 | 0.172162651 |
| 0.030902808 | 0.172162651 |
| 0.030876959 | 0.17205767 |
| 0.030846151 | 0.17190611 |
| 0.0308407 | 0.17189585 |
| 0.030836703 | 0.171893689 |
| 0.030836667 | 0.171893689 |
| 0.030826973 | 0.171879681 |
| 0.030824264 | 0.171879681 |
| 0.030804151 | 0.171812779 |
| 0.030752527 | 0.171605228 |
| 0.03072232 | 0.171476848 |
| 0.030714366 | 0.171452547 |
| 0.030685547 | 0.171311751 |
| 0.030685519 | 0.171311751 |
| 0.030676168 | 0.171309914 |
| 0.030675484 | 0.171309914 |
| 0.030670865 | 0.171309914 |
| 0.030657988 | 0.171278358 |
| 0.030653563 | 0.171273725 |
| 0.030648027 | 0.171262888 |
| 0.030646244 | 0.171262888 |
| 0.030619647 | 0.171144457 |
| 0.030610063 | 0.171110973 |
| 0.030607098 | 0.171110973 |
| 0.030602609 | 0.171109467 |
| 0.030566349 | 0.170966939 |
| 0.030543271 | 0.170857924 |
| 0.030453314 | 0.170434783 |
| 0.030447997 | 0.170425048 |
| 0.030440424 | 0.17042272 |
| 0.030436749 | 0.170422179 |
| 0.030415709 | 0.170324392 |
| 0.030402682 | 0.170271462 |
| 0.030379207 | 0.170180014 |
| 0.030373482 | 0.170167958 |
| 0.030371606 | 0.170167958 |
| 0.030370585 | 0.170167958 |
| 0.030357879 | 0.170140589 |
| 0.030331673 | 0.170050654 |
| 0.030321822 | 0.170018531 |
| 0.030304013 | 0.169938682 |
| 0.030287227 | 0.16986455 |
| 0.030274826 | 0.169814998 |
| 0.030266079 | 0.169785928 |

| | |
|---|---|
| 0.030263991 | 0.169785928 |
| 0.030246709 | 0.169757247 |
| 0.030244785 | 0.169757247 |
| 0.03022842 | 0.169694596 |
| 0.030225683 | 0.169694596 |
| 0.03021556 | 0.1696624 |
| 0.030210209 | 0.169652357 |
| 0.030177766 | 0.169530131 |
| 0.030139549 | 0.169391316 |
| 0.030135165 | 0.169390709 |
| 0.03012923 | 0.16937734 |
| 0.030110332 | 0.169291088 |
| 0.030099377 | 0.169249471 |
| 0.030097679 | 0.169249471 |
| 0.030091034 | 0.169249471 |
| 0.030081868 | 0.169230937 |
| 0.030058043 | 0.16915685 |
| 0.030050265 | 0.16913306 |
| 0.030012441 | 0.168980083 |
| 0.029993621 | 0.168934036 |
| 0.029970044 | 0.168841173 |
| 0.029955786 | 0.168780813 |
| 0.029951388 | 0.168776001 |
| 0.029949758 | 0.168776001 |
| 0.02993115 | 0.168727829 |
| 0.029913447 | 0.168661982 |
| 0.02991333 | 0.168661982 |
| 0.029900894 | 0.168631127 |
| 0.029900398 | 0.168631127 |
| 0.029894423 | 0.168631127 |
| 0.029892646 | 0.168631127 |
| 0.029890669 | 0.168631127 |
| 0.029879774 | 0.168615997 |
| 0.029869906 | 0.168596114 |
| 0.029860128 | 0.1685609 |
| 0.029849307 | 0.168539768 |
| 0.029846203 | 0.168539768 |
| 0.02982639 | 0.168450312 |
| 0.029800282 | 0.168362758 |
| 0.02978993 | 0.168324239 |
| 0.029726527 | 0.16810107 |
| 0.029708707 | 0.168044727 |
| 0.029699837 | 0.168021723 |
| 0.029699225 | 0.168021723 |
| 0.029695035 | 0.168021723 |

| | |
|---|---|
| 0.029690531 | 0.168021723 |
| 0.029667301 | 0.16795014 |
| 0.029656127 | 0.167926808 |
| 0.029652306 | 0.167925131 |
| 0.029628594 | 0.167810802 |
| 0.029622264 | 0.167794903 |
| 0.029613599 | 0.16777299 |
| 0.029609903 | 0.16777299 |
| 0.029608301 | 0.16777299 |
| 0.029598159 | 0.167758127 |
| 0.029595354 | 0.167758127 |
| 0.029580906 | 0.167720216 |
| 0.029553592 | 0.167585297 |
| 0.029546556 | 0.167565352 |
| 0.029516093 | 0.167412523 |
| 0.029505191 | 0.167370622 |
| 0.02949666 | 0.16734216 |
| 0.029496128 | 0.16734216 |
| 0.029459506 | 0.167211047 |
| 0.029448333 | 0.167167549 |
| 0.029398464 | 0.166904355 |
| 0.029392241 | 0.166888921 |
| 0.029345257 | 0.166642011 |
| 0.0293323 | 0.166588296 |
| 0.029331056 | 0.166588296 |
| 0.029330118 | 0.166588296 |
| 0.029311172 | 0.166536545 |
| 0.029307391 | 0.166536545 |
| 0.029283757 | 0.166431684 |
| 0.029268769 | 0.166366351 |
| 0.02925767 | 0.166323115 |
| 0.029256898 | 0.166323115 |
| 0.02924406 | 0.16628544 |
| 0.029223947 | 0.166203399 |
| 0.029202514 | 0.166108701 |
| 0.029190077 | 0.16605779 |
| 0.029162121 | 0.165958219 |
| 0.029154477 | 0.165934548 |
| 0.02915446 | 0.165934548 |
| 0.029146436 | 0.16592844 |
| 0.029123918 | 0.165859719 |
| 0.029110681 | 0.165804159 |
| 0.0291079 | 0.165804159 |
| 0.029098585 | 0.165779284 |
| 0.02909439 | 0.165779284 |

| | |
|---|---|
| 0.029081602 | 0.165757473 |
| 0.029057501 | 0.165679589 |
| 0.029056855 | 0.165679589 |
| 0.029048653 | 0.165668804 |
| 0.029045471 | 0.165668804 |
| 0.029027935 | 0.165590307 |
| 0.029017458 | 0.165550367 |
| 0.028984843 | 0.165384106 |
| 0.028980449 | 0.165378843 |
| 0.028965938 | 0.165315843 |
| 0.028934146 | 0.16515419 |
| 0.028907071 | 0.165019428 |
| 0.028893316 | 0.165000235 |
| 0.028885107 | 0.164973139 |
| 0.028843893 | 0.164757507 |
| 0.028840571 | 0.164757507 |
| 0.028831545 | 0.164746251 |
| 0.02882762 | 0.164743587 |
| 0.028823058 | 0.164737281 |
| 0.0288048 | 0.164672453 |
| 0.028798302 | 0.164666292 |
| 0.028789747 | 0.164645685 |
| 0.028788753 | 0.164645685 |
| 0.028781413 | 0.164645685 |
| 0.028767966 | 0.164639742 |
| 0.028751385 | 0.164564619 |
| 0.028739614 | 0.164517019 |
| 0.028735932 | 0.164515713 |
| 0.028699951 | 0.164388762 |
| 0.028679865 | 0.164293471 |
| 0.028677546 | 0.164293471 |
| 0.028666022 | 0.164293218 |
| 0.028651427 | 0.164229331 |
| 0.028642406 | 0.164197385 |
| 0.028638509 | 0.16419481 |
| 0.028632988 | 0.16419481 |
| 0.028628901 | 0.16419481 |
| 0.028616154 | 0.164165451 |
| 0.028605668 | 0.164144835 |
| 0.028602797 | 0.164144835 |
| 0.028600126 | 0.164144835 |
| 0.028553514 | 0.163964097 |
| 0.028524009 | 0.163814418 |
| 0.028517933 | 0.163799279 |
| 0.028502171 | 0.163728489 |

| | |
|---|---|
| 0.028501084 | 0.163728489 |
| 0.028492276 | 0.163711141 |
| 0.028477777 | 0.163647574 |
| 0.028462919 | 0.163601671 |
| 0.028451399 | 0.163555195 |
| 0.028436299 | 0.163488124 |
| 0.028435904 | 0.163488124 |
| 0.028415694 | 0.163428842 |
| 0.028408309 | 0.163406105 |
| 0.028404279 | 0.163402655 |
| 0.028395867 | 0.163374001 |
| 0.028380869 | 0.163326395 |
| 0.028365812 | 0.163260245 |
| 0.028361687 | 0.163256232 |
| 0.028356914 | 0.163248488 |
| 0.028356116 | 0.163248488 |
| 0.028305665 | 0.163055182 |
| 0.028301683 | 0.163055182 |
| 0.028294021 | 0.163044084 |
| 0.028278771 | 0.162995645 |
| 0.028208342 | 0.162629064 |
| 0.028208331 | 0.162629064 |
| 0.028207278 | 0.162629064 |
| 0.028202709 | 0.162629064 |
| 0.028185621 | 0.162576791 |
| 0.028158283 | 0.162457367 |
| 0.028129536 | 0.16231226 |
| 0.028123386 | 0.16231226 |
| 0.028095562 | 0.162253827 |
| 0.028090616 | 0.162244937 |
| 0.028082204 | 0.162244937 |
| 0.028075783 | 0.162244199 |
| 0.028069042 | 0.162240836 |
| 0.028057545 | 0.162230991 |
| 0.028049685 | 0.162205238 |
| 0.028046164 | 0.162204571 |
| 0.028045509 | 0.162204571 |
| 0.028027012 | 0.162152881 |
| 0.028023242 | 0.162150766 |
| 0.027983717 | 0.1619811 |
| 0.027983264 | 0.1619811 |
| 0.027982895 | 0.1619811 |
| 0.027970457 | 0.161963402 |
| 0.027968527 | 0.161963402 |
| 0.02795951 | 0.161939386 |

| | |
|---|---|
| 0.027956852 | 0.161939386 |
| 0.027953832 | 0.161939386 |
| 0.027943955 | 0.161908362 |
| 0.027921046 | 0.161815 |
| 0.027904459 | 0.161738551 |
| 0.027904202 | 0.161738551 |
| 0.027880891 | 0.161641288 |
| 0.027880873 | 0.161641288 |
| 0.027880288 | 0.161641288 |
| 0.027869407 | 0.161633736 |
| 0.027862036 | 0.161610665 |
| 0.027860094 | 0.161610665 |
| 0.027859649 | 0.161610665 |
| 0.027827957 | 0.161491669 |
| 0.027823266 | 0.161484121 |
| 0.027811785 | 0.16143716 |
| 0.027798948 | 0.161382311 |
| 0.027720316 | 0.160984695 |
| 0.02771835 | 0.160984695 |
| 0.027718023 | 0.160984695 |
| 0.027696642 | 0.160949798 |
| 0.027694534 | 0.160949798 |
| 0.027686274 | 0.160949798 |
| 0.027674921 | 0.16093693 |
| 0.027671668 | 0.16093693 |
| 0.027667197 | 0.160932715 |
| 0.027638014 | 0.16080084 |
| 0.027636432 | 0.16080084 |
| 0.027619054 | 0.160729799 |
| 0.027600453 | 0.160660814 |
| 0.027574533 | 0.160529555 |
| 0.027535879 | 0.160343722 |
| 0.027528211 | 0.16031867 |
| 0.027523994 | 0.160313719 |
| 0.027519697 | 0.160308293 |
| 0.027507531 | 0.160257024 |
| 0.027483258 | 0.160193994 |
| 0.027477096 | 0.160177678 |
| 0.027472228 | 0.160177678 |
| 0.027471512 | 0.160177678 |
| 0.027463182 | 0.160177678 |
| 0.027445686 | 0.160128905 |
| 0.027445366 | 0.160128905 |
| 0.027436754 | 0.160128905 |
| 0.02743007 | 0.160128905 |

| | |
|---|---|
| 0.027420647 | 0.160122996 |
| 0.027416251 | 0.160116957 |
| 0.027409047 | 0.160094518 |
| 0.027402676 | 0.160076935 |
| 0.027395376 | 0.160068133 |
| 0.027383007 | 0.160020918 |
| 0.027381347 | 0.160020918 |
| 0.02737317 | 0.160002696 |
| 0.027362891 | 0.159962245 |
| 0.027319743 | 0.159746643 |
| 0.027304611 | 0.159739183 |
| 0.027302173 | 0.159739183 |
| 0.027298445 | 0.159739183 |
| 0.027296241 | 0.159739183 |
| 0.027294347 | 0.159739183 |
| 0.027286744 | 0.159739183 |
| 0.027281711 | 0.159739183 |
| 0.027266738 | 0.159674529 |
| 0.027254053 | 0.159619871 |
| 0.027248776 | 0.159618223 |
| 0.027242593 | 0.159611633 |
| 0.027215232 | 0.159470949 |
| 0.027202736 | 0.159436958 |
| 0.027194151 | 0.159406258 |
| 0.027184656 | 0.159370215 |
| 0.027175183 | 0.159334288 |
| 0.027170457 | 0.159326194 |
| 0.027168813 | 0.159326194 |
| 0.027168241 | 0.159326194 |
| 0.027129052 | 0.159161781 |
| 0.027125403 | 0.159159981 |
| 0.027122797 | 0.159159981 |
| 0.027120163 | 0.159159981 |
| 0.027116215 | 0.159159981 |
| 0.027114839 | 0.159159981 |
| 0.027114343 | 0.159159981 |
| 0.027096744 | 0.159109436 |
| 0.027084119 | 0.159054911 |
| 0.02708183 | 0.159054911 |
| 0.027073078 | 0.159029292 |
| 0.027071005 | 0.159029292 |
| 0.027070098 | 0.159029292 |
| 0.027065404 | 0.159029292 |
| 0.027044877 | 0.15896166 |
| 0.027001013 | 0.15876262 |

| | |
|---|---|
| 0.02697296 | 0.158617257 |
| 0.026959254 | 0.158556235 |
| 0.026911367 | 0.158333252 |
| 0.026895212 | 0.158277306 |
| 0.026857246 | 0.158132037 |
| 0.026856421 | 0.158132037 |
| 0.026840447 | 0.158111311 |
| 0.026808195 | 0.157940857 |
| 0.026793265 | 0.157891959 |
| 0.026779027 | 0.157827586 |
| 0.026778496 | 0.157827586 |
| 0.0267727 | 0.157827586 |
| 0.026747396 | 0.157699708 |
| 0.026742738 | 0.157691763 |
| 0.02672467 | 0.157604735 |
| 0.026717987 | 0.157584837 |
| 0.026708858 | 0.157550507 |
| 0.026702959 | 0.157535217 |
| 0.026697304 | 0.157521367 |
| 0.026686142 | 0.157475022 |
| 0.026684327 | 0.157475022 |
| 0.026683255 | 0.157475022 |
| 0.02668152 | 0.157475022 |
| 0.026655464 | 0.157391491 |
| 0.026647961 | 0.157366702 |
| 0.026645973 | 0.157366702 |
| 0.026639598 | 0.157366702 |
| 0.026616546 | 0.157259177 |
| 0.026609015 | 0.157234191 |
| 0.026605918 | 0.157234191 |
| 0.026601824 | 0.157230711 |
| 0.026595392 | 0.157212205 |
| 0.026584272 | 0.157165978 |
| 0.026579542 | 0.157160488 |
| 0.026549586 | 0.157019387 |
| 0.026541249 | 0.157008771 |
| 0.026539949 | 0.157008771 |
| 0.02653284 | 0.156998334 |
| 0.026526716 | 0.156981596 |
| 0.026513488 | 0.156930563 |
| 0.02651304 | 0.156930563 |
| 0.026505473 | 0.156930563 |
| 0.026493592 | 0.156883042 |
| 0.026486448 | 0.156860241 |
| 0.026485383 | 0.156860241 |

| | |
|---|---|
| 0.026477109 | 0.156847338 |
| 0.026451545 | 0.156770498 |
| 0.026447553 | 0.156770498 |
| 0.026443792 | 0.156770498 |
| 0.026438391 | 0.156770498 |
| 0.026375601 | 0.156417723 |
| 0.026354557 | 0.15637081 |
| 0.026335249 | 0.156295193 |
| 0.026321693 | 0.156234208 |
| 0.026307983 | 0.156191763 |
| 0.026292077 | 0.156136254 |
| 0.026287476 | 0.156128397 |
| 0.026275451 | 0.156076439 |
| 0.026273376 | 0.156076439 |
| 0.026270691 | 0.156076439 |
| 0.026258818 | 0.15603602 |
| 0.026255753 | 0.15603602 |
| 0.026251793 | 0.156033216 |
| 0.026231272 | 0.1559307 |
| 0.026230906 | 0.1559307 |
| 0.026227428 | 0.1559307 |
| 0.026223137 | 0.1559307 |
| 0.026208833 | 0.155875133 |
| 0.026183286 | 0.1557621 |
| 0.026179925 | 0.1557621 |
| 0.026175825 | 0.1557621 |
| 0.026169584 | 0.1557621 |
| 0.026159763 | 0.15575835 |
| 0.026154683 | 0.155747572 |
| 0.026151601 | 0.155747572 |
| 0.026145075 | 0.1557293 |
| 0.02611572 | 0.155632305 |
| 0.026099911 | 0.155557558 |
| 0.02609897 | 0.155557558 |
| 0.026073585 | 0.155450932 |
| 0.026068391 | 0.15544751 |
| 0.026038441 | 0.155307806 |
| 0.025991938 | 0.155108136 |
| 0.025961154 | 0.154943847 |
| 0.025957781 | 0.154943129 |
| 0.0259433 | 0.154876105 |
| 0.025934788 | 0.154844696 |
| 0.025926311 | 0.154813491 |
| 0.025913167 | 0.154754405 |
| 0.025905185 | 0.154726139 |

| | |
|---|---|
| 0.025893438 | 0.154675376 |
| 0.025871186 | 0.154600623 |
| 0.025848968 | 0.154506628 |
| 0.025845029 | 0.154502471 |
| 0.025831661 | 0.154461334 |
| 0.025829205 | 0.154461334 |
| 0.025823802 | 0.15445596 |
| 0.025816155 | 0.15445596 |
| 0.025800631 | 0.1544309 |
| 0.025782981 | 0.15434465 |
| 0.025781414 | 0.15434465 |
| 0.025780152 | 0.15434465 |
| 0.025745092 | 0.15419536 |
| 0.025742488 | 0.15419536 |
| 0.025736501 | 0.154182683 |
| 0.025733768 | 0.154182683 |
| 0.025732767 | 0.154182683 |
| 0.025731204 | 0.154182683 |
| 0.025707179 | 0.154103938 |
| 0.025702222 | 0.15409362 |
| 0.025685176 | 0.154030202 |
| 0.025680973 | 0.154024392 |
| 0.025677394 | 0.154022326 |
| 0.025660708 | 0.153961012 |
| 0.025628918 | 0.153847801 |
| 0.025623034 | 0.153831875 |
| 0.025591662 | 0.153662897 |
| 0.02558297 | 0.153630076 |
| 0.025574897 | 0.153600958 |
| 0.025565993 | 0.153574673 |
| 0.025563677 | 0.153574673 |
| 0.025541776 | 0.153535023 |
| 0.025541365 | 0.153535023 |
| 0.025523789 | 0.153468187 |
| 0.025519959 | 0.153464531 |
| 0.025500598 | 0.153386836 |
| 0.025496879 | 0.153383837 |
| 0.025469848 | 0.153240584 |
| 0.025459031 | 0.153194852 |
| 0.025450289 | 0.1531616 |
| 0.025435849 | 0.153105427 |
| 0.025423399 | 0.153057796 |
| 0.025409373 | 0.152992693 |
| 0.025402506 | 0.152970688 |
| 0.025400199 | 0.152970688 |

| | |
|---|---|
| 0.025384616 | 0.152901619 |
| 0.025378515 | 0.152884204 |
| 0.025348755 | 0.152724246 |
| 0.025336112 | 0.152667383 |
| 0.02532663 | 0.152648875 |
| 0.025317966 | 0.15261597 |
| 0.025308088 | 0.152575736 |
| 0.025306032 | 0.152575736 |
| 0.02530388 | 0.152575736 |
| 0.025294556 | 0.152552084 |
| 0.02529437 | 0.152552084 |
| 0.025275055 | 0.152492402 |
| 0.025248853 | 0.152392235 |
| 0.025247967 | 0.152392235 |
| 0.025214936 | 0.152226108 |
| 0.025196363 | 0.152133261 |
| 0.025194582 | 0.152133261 |
| 0.025188216 | 0.152133261 |
| 0.025186966 | 0.152133261 |
| 0.025182679 | 0.152133261 |
| 0.025181499 | 0.152133261 |
| 0.025178822 | 0.152133261 |
| 0.025165289 | 0.152099898 |
| 0.025164074 | 0.152099898 |
| 0.025161611 | 0.152099898 |
| 0.025126907 | 0.15192576 |
| 0.025122303 | 0.15192576 |
| 0.025119879 | 0.15192576 |
| 0.025118928 | 0.15192576 |
| 0.02511663 | 0.15192576 |
| 0.02510858 | 0.15192576 |
| 0.025101493 | 0.15192576 |
| 0.025101449 | 0.15192576 |
| 0.025090559 | 0.151898825 |
| 0.025090559 | 0.151898825 |
| 0.02509002 | 0.151898825 |
| 0.025082906 | 0.151898825 |
| 0.025067389 | 0.151855068 |
| 0.025057415 | 0.151813956 |
| 0.025053533 | 0.151813956 |
| 0.025044475 | 0.151793487 |
| 0.025040645 | 0.151789587 |
| 0.025038867 | 0.151789587 |
| 0.025009197 | 0.151637559 |
| 0.024988581 | 0.151531847 |

| | |
|---|---|
| 0.024984429 | 0.151525967 |
| 0.0249811 | 0.151525067 |
| 0.024969391 | 0.151473338 |
| 0.024934685 | 0.151320623 |
| 0.024930837 | 0.151316553 |
| 0.024929647 | 0.151316553 |
| 0.024921915 | 0.151300962 |
| 0.024919189 | 0.151300962 |
| 0.024909704 | 0.151265395 |
| 0.024903371 | 0.151246219 |
| 0.024901015 | 0.151246219 |
| 0.024896278 | 0.151241713 |
| 0.02489501 | 0.151241713 |
| 0.02488986 | 0.151241306 |
| 0.024866792 | 0.151120418 |
| 0.024848256 | 0.151027041 |
| 0.024844793 | 0.151027041 |
| 0.024836952 | 0.15101615 |
| 0.02482568 | 0.150986166 |
| 0.024814555 | 0.15093778 |
| 0.024810088 | 0.150929883 |
| 0.024789836 | 0.150825948 |
| 0.02478908 | 0.150825948 |
| 0.024775887 | 0.150779601 |
| 0.024771921 | 0.150774733 |
| 0.02475875 | 0.15071383 |
| 0.024751885 | 0.150711688 |
| 0.024746417 | 0.150711688 |
| 0.024732108 | 0.150647925 |
| 0.024701778 | 0.150501671 |
| 0.024693184 | 0.150487814 |
| 0.024682966 | 0.150464049 |
| 0.024652155 | 0.150295466 |
| 0.024630004 | 0.150231782 |
| 0.024614204 | 0.150160202 |
| 0.024613094 | 0.150160202 |
| 0.024610114 | 0.150160202 |
| 0.024600587 | 0.150134826 |
| 0.024593858 | 0.150112994 |
| 0.024591436 | 0.150112994 |
| 0.024583977 | 0.150091154 |
| 0.024575041 | 0.150075071 |
| 0.024571789 | 0.150074452 |
| 0.024537608 | 0.149884914 |
| 0.024536682 | 0.149884914 |

| | |
|---|---|
| 0.024511511 | 0.149763917 |
| 0.024498319 | 0.149702517 |
| 0.024486846 | 0.149651611 |
| 0.024481256 | 0.149636647 |
| 0.024481046 | 0.149636647 |
| 0.024474668 | 0.14963479 |
| 0.024460731 | 0.149568782 |
| 0.024455298 | 0.149554759 |
| 0.02445488 | 0.149554759 |
| 0.024431486 | 0.149447521 |
| 0.024412933 | 0.149353212 |
| 0.024401871 | 0.1493239 |
| 0.024401576 | 0.1493239 |
| 0.024390522 | 0.149292817 |
| 0.024370003 | 0.149186391 |
| 0.02436517 | 0.149175982 |
| 0.024354307 | 0.149147818 |
| 0.024337985 | 0.149067031 |
| 0.024334106 | 0.149062442 |
| 0.024312601 | 0.148969027 |
| 0.024310853 | 0.148969027 |
| 0.024296736 | 0.14893189 |
| 0.024280587 | 0.148908914 |
| 0.024268245 | 0.148888791 |
| 0.024267935 | 0.148888791 |
| 0.024235672 | 0.148803965 |
| 0.024234631 | 0.148803965 |
| 0.024217739 | 0.148751387 |
| 0.024206234 | 0.148717122 |
| 0.024206068 | 0.148717122 |
| 0.024204171 | 0.148717122 |
| 0.024187059 | 0.148678491 |
| 0.024186794 | 0.148678491 |
| 0.024177502 | 0.148678491 |
| 0.024140047 | 0.148504009 |
| 0.024138601 | 0.148504009 |
| 0.024118252 | 0.148427442 |
| 0.024117335 | 0.148427442 |
| 0.024110166 | 0.148416039 |
| 0.024109857 | 0.148416039 |
| 0.024108508 | 0.148416039 |
| 0.024084611 | 0.148316234 |
| 0.02408345 | 0.148316234 |
| 0.024082351 | 0.148316234 |
| 0.024082247 | 0.148316234 |

| | |
|---|---|
| 0.024080047 | 0.148316234 |
| 0.024079558 | 0.148316234 |
| 0.024074137 | 0.148316234 |
| 0.024064319 | 0.148316234 |
| 0.024061707 | 0.148316234 |
| 0.02404694 | 0.14825678 |
| 0.024025328 | 0.148142711 |
| 0.024021484 | 0.148138189 |
| 0.024020147 | 0.148138189 |
| 0.024018189 | 0.148138189 |
| 0.024017832 | 0.148138189 |
| 0.024009931 | 0.148138189 |
| 0.023997217 | 0.148103594 |
| 0.02399314 | 0.148097624 |
| 0.023981701 | 0.148065394 |
| 0.023965178 | 0.148001753 |
| 0.023958593 | 0.147980274 |
| 0.023943957 | 0.147928245 |
| 0.023940431 | 0.14792565 |
| 0.023937466 | 0.14792565 |
| 0.023934042 | 0.14792565 |
| 0.023922204 | 0.147889773 |
| 0.023911738 | 0.147857513 |
| 0.023880337 | 0.147688455 |
| 0.023858794 | 0.147591128 |
| 0.023854748 | 0.147591128 |
| 0.023852092 | 0.147591128 |
| 0.023836791 | 0.147572448 |
| 0.023832499 | 0.14756506 |
| 0.023828653 | 0.147560427 |
| 0.023816209 | 0.147502549 |
| 0.0238134 | 0.147502549 |
| 0.02381214 | 0.147502549 |
| 0.023774594 | 0.147321227 |
| 0.023774577 | 0.147321227 |
| 0.023761139 | 0.147276387 |
| 0.023750319 | 0.147228484 |
| 0.02374124 | 0.147191361 |
| 0.023740643 | 0.147191361 |
| 0.023740515 | 0.147191361 |
| 0.023703391 | 0.147014119 |
| 0.023689929 | 0.146949762 |
| 0.023672782 | 0.146881659 |
| 0.023671938 | 0.146881659 |
| 0.023669642 | 0.146881659 |

| | |
|---|---|
| 0.023656141 | 0.146854931 |
| 0.023653396 | 0.146854931 |
| 0.02361806 | 0.146675884 |
| 0.02360887 | 0.146637933 |
| 0.023600173 | 0.146622152 |
| 0.023575279 | 0.146486604 |
| 0.023545865 | 0.146342019 |
| 0.023542568 | 0.146340626 |
| 0.023496982 | 0.146095401 |
| 0.023487636 | 0.146056355 |
| 0.023482275 | 0.146056355 |
| 0.023474211 | 0.14604576 |
| 0.023468099 | 0.146038565 |
| 0.02344435 | 0.145939622 |
| 0.023440086 | 0.145932153 |
| 0.023437568 | 0.145932153 |
| 0.023395406 | 0.145730188 |
| 0.023385085 | 0.145684947 |
| 0.023380863 | 0.145677706 |
| 0.023369974 | 0.145628912 |
| 0.023369248 | 0.145628912 |
| 0.023347286 | 0.14552561 |
| 0.023341057 | 0.145505829 |
| 0.023337284 | 0.145501354 |
| 0.023335154 | 0.145501354 |
| 0.023335154 | 0.145501354 |
| 0.023327052 | 0.145501354 |
| 0.023324721 | 0.145501354 |
| 0.023322232 | 0.145501354 |
| 0.023302491 | 0.145417681 |
| 0.023283861 | 0.145339506 |
| 0.023270079 | 0.145272521 |
| 0.023266577 | 0.145269696 |
| 0.023265118 | 0.145269696 |
| 0.023260277 | 0.14526845 |
| 0.023255819 | 0.145259658 |
| 0.023245271 | 0.145212815 |
| 0.023235607 | 0.145209584 |
| 0.023210174 | 0.145069679 |
| 0.023203258 | 0.145061403 |
| 0.023202576 | 0.145061403 |
| 0.02318845 | 0.145021027 |
| 0.023173663 | 0.144955614 |
| 0.023154401 | 0.144854153 |
| 0.023140272 | 0.1448038 |

| | |
|---|---|
| 0.023134878 | 0.144789069 |
| 0.023133583 | 0.144789069 |
| 0.023132152 | 0.144789069 |
| 0.023106852 | 0.144670703 |
| 0.023105121 | 0.144670703 |
| 0.023099186 | 0.144660744 |
| 0.023070132 | 0.144497785 |
| 0.023045475 | 0.144381324 |
| 0.023017338 | 0.144261974 |
| 0.023012117 | 0.144248235 |
| 0.022998718 | 0.144202205 |
| 0.022977547 | 0.14408843 |
| 0.022975615 | 0.14408843 |
| 0.022969596 | 0.144076511 |
| 0.022946252 | 0.143949042 |
| 0.022940602 | 0.143936675 |
| 0.022932883 | 0.143936675 |
| 0.022886008 | 0.14368467 |
| 0.022853212 | 0.143516602 |
| 0.022842934 | 0.143470978 |
| 0.022840628 | 0.143470978 |
| 0.022822654 | 0.143419255 |
| 0.022820783 | 0.143419255 |
| 0.022785406 | 0.143241888 |
| 0.022780568 | 0.143230376 |
| 0.022772283 | 0.143216099 |
| 0.022771825 | 0.143216099 |
| 0.022765016 | 0.143208215 |
| 0.022758503 | 0.143186157 |
| 0.022752903 | 0.143169838 |
| 0.022749661 | 0.143168356 |
| 0.022743264 | 0.14314701 |
| 0.022741778 | 0.14314701 |
| 0.022740184 | 0.14314701 |
| 0.022736058 | 0.14314701 |
| 0.022732762 | 0.14314701 |
| 0.022732093 | 0.14314701 |
| 0.022720303 | 0.14313154 |
| 0.022718153 | 0.14313154 |
| 0.02271513 | 0.14313154 |
| 0.022694095 | 0.143026551 |
| 0.022689291 | 0.1430152 |
| 0.022684847 | 0.143006118 |
| 0.022665285 | 0.142901709 |
| 0.022662879 | 0.142901709 |

| | |
|---|---|
| 0.022661253 | 0.142901709 |
| 0.022650441 | 0.142864859 |
| 0.022647048 | 0.14286238 |
| 0.022635965 | 0.142811382 |
| 0.022630637 | 0.142796681 |
| 0.022629957 | 0.142796681 |
| 0.022619958 | 0.142767127 |
| 0.02260564 | 0.142714582 |
| 0.022600207 | 0.142710443 |
| 0.022590907 | 0.14269723 |
| 0.022587056 | 0.142691832 |
| 0.022581414 | 0.142675114 |
| 0.022568514 | 0.14263145 |
| 0.022568036 | 0.14263145 |
| 0.022555992 | 0.142628016 |
| 0.022551074 | 0.142615858 |
| 0.022548315 | 0.142615858 |
| 0.022543652 | 0.142606786 |
| 0.022534847 | 0.142570027 |
| 0.022534685 | 0.142570027 |
| 0.022526276 | 0.142553673 |
| 0.022519642 | 0.142530628 |
| 0.022514861 | 0.142529827 |
| 0.022508836 | 0.142521952 |
| 0.022507837 | 0.142521952 |
| 0.022507381 | 0.142521952 |
| 0.022471172 | 0.142375218 |
| 0.02240252 | 0.141965735 |
| 0.022393932 | 0.141942484 |
| 0.022392322 | 0.141942484 |
| 0.022380251 | 0.141912424 |
| 0.022379999 | 0.141912424 |
| 0.022375411 | 0.141912424 |
| 0.022369488 | 0.141900851 |
| 0.022364749 | 0.141889691 |
| 0.022339242 | 0.141803424 |
| 0.02233541 | 0.141797995 |
| 0.022332623 | 0.141797995 |
| 0.022327452 | 0.141785273 |
| 0.022327006 | 0.141785273 |
| 0.022292292 | 0.141618649 |
| 0.022291073 | 0.141618649 |
| 0.02227922 | 0.14157337 |
| 0.022271189 | 0.141548369 |
| 0.02226493 | 0.141548369 |

| | |
|---|---|
| 0.022252331 | 0.141515793 |
| 0.022239028 | 0.141450079 |
| 0.022229431 | 0.141426813 |
| 0.022201245 | 0.141285234 |
| 0.022189524 | 0.141229512 |
| 0.022165593 | 0.141114919 |
| 0.022116246 | 0.140857244 |
| 0.022110735 | 0.140840973 |
| 0.022090701 | 0.140732182 |
| 0.022058742 | 0.140566188 |
| 0.022051157 | 0.140536659 |
| 0.022049006 | 0.140536659 |
| 0.022030273 | 0.140441146 |
| 0.022013271 | 0.140351545 |
| 0.022000015 | 0.140290111 |
| 0.021987543 | 0.140281406 |
| 0.021984225 | 0.140279028 |
| 0.021961054 | 0.140149956 |
| 0.021954805 | 0.140128851 |
| 0.021949138 | 0.140128851 |
| 0.021947666 | 0.140128851 |
| 0.021945404 | 0.140128851 |
| 0.021939347 | 0.140124084 |
| 0.021937235 | 0.140124084 |
| 0.021927868 | 0.140107132 |
| 0.021924643 | 0.140105324 |
| 0.021924523 | 0.140105324 |
| 0.021922821 | 0.140105324 |
| 0.021913826 | 0.140092592 |
| 0.021906417 | 0.140064028 |
| 0.021902967 | 0.140064028 |
| 0.021891711 | 0.140064013 |
| 0.021880676 | 0.140012221 |
| 0.02186852 | 0.13995324 |
| 0.021867888 | 0.13995324 |
| 0.021851172 | 0.139914996 |
| 0.021849688 | 0.139914996 |
| 0.02184814 | 0.139914996 |
| 0.021831132 | 0.139845504 |
| 0.021774971 | 0.139504513 |
| 0.021751426 | 0.139391168 |
| 0.021748217 | 0.139389358 |
| 0.021730543 | 0.139332335 |
| 0.021730196 | 0.139332335 |
| 0.021694197 | 0.139174238 |

| | |
|---|---|
| 0.021692468 | 0.139174238 |
| 0.021677315 | 0.13912215 |
| 0.02167521 | 0.13912215 |
| 0.021657309 | 0.139033439 |
| 0.021652953 | 0.139033439 |
| 0.021644555 | 0.139024306 |
| 0.021619382 | 0.138900083 |
| 0.021614185 | 0.138885428 |
| 0.021602047 | 0.138840436 |
| 0.021594199 | 0.138813186 |
| 0.02158291 | 0.138759346 |
| 0.021572443 | 0.13871078 |
| 0.021568724 | 0.13870559 |
| 0.021559581 | 0.138665518 |
| 0.021539479 | 0.138611105 |
| 0.021537455 | 0.138611105 |
| 0.021526072 | 0.138562267 |
| 0.021519662 | 0.138558463 |
| 0.021515771 | 0.138552138 |
| 0.021497621 | 0.138472708 |
| 0.02146939 | 0.138333904 |
| 0.021461149 | 0.1383144 |
| 0.021448176 | 0.138285133 |
| 0.021444245 | 0.138278506 |
| 0.021433814 | 0.13824868 |
| 0.021427589 | 0.138227248 |
| 0.021399248 | 0.13810054 |
| 0.021390502 | 0.138062805 |
| 0.021386781 | 0.138057499 |
| 0.021370293 | 0.138021578 |
| 0.021364159 | 0.13800499 |
| 0.021363875 | 0.13800499 |
| 0.021360233 | 0.13800499 |
| 0.02135846 | 0.13800499 |
| 0.021357786 | 0.13800499 |
| 0.02134739 | 0.137990246 |
| 0.021333553 | 0.13793825 |
| 0.021322965 | 0.137888507 |
| 0.021273815 | 0.13762673 |
| 0.021254935 | 0.137541955 |
| 0.021225382 | 0.137369379 |
| 0.021209122 | 0.137301464 |
| 0.021194445 | 0.137225098 |
| 0.021180861 | 0.137174446 |
| 0.021172399 | 0.137138289 |

| | |
|---|---|
| 0.021151448 | 0.137021218 |
| 0.021146934 | 0.137010614 |
| 0.021140656 | 0.136996831 |
| 0.021133145 | 0.136990157 |
| 0.02112257 | 0.136945907 |
| 0.021091397 | 0.13676242 |
| 0.021077587 | 0.136691478 |
| 0.021076756 | 0.136691478 |
| 0.021058552 | 0.136618417 |
| 0.02105093 | 0.136611613 |
| 0.021041306 | 0.136567766 |
| 0.021035265 | 0.136547166 |
| 0.021024805 | 0.136497869 |
| 0.020994346 | 0.136338955 |
| 0.020970797 | 0.136221513 |
| 0.020970433 | 0.136221513 |
| 0.020950597 | 0.136127433 |
| 0.020937293 | 0.136059553 |
| 0.020918704 | 0.1359573 |
| 0.020918544 | 0.1359573 |
| 0.020913213 | 0.1359573 |
| 0.020911102 | 0.1359573 |
| 0.020908882 | 0.1359573 |
| 0.020894841 | 0.135950629 |
| 0.020869697 | 0.135824143 |
| 0.020867057 | 0.135824143 |
| 0.020859469 | 0.135794687 |
| 0.020852313 | 0.135766657 |
| 0.020850189 | 0.135766657 |
| 0.020844736 | 0.135766657 |
| 0.020842973 | 0.135766657 |
| 0.020836122 | 0.135754036 |
| 0.020833668 | 0.135754036 |
| 0.020829053 | 0.135745115 |
| 0.02082286 | 0.135742339 |
| 0.020809094 | 0.135693253 |
| 0.020808449 | 0.135693253 |
| 0.020796891 | 0.135665458 |
| 0.020791338 | 0.135647816 |
| 0.020790963 | 0.135647816 |
| 0.020790179 | 0.135647816 |
| 0.020783816 | 0.135647816 |
| 0.02077537 | 0.135617951 |
| 0.020757933 | 0.135522698 |
| 0.020752566 | 0.135506235 |

| | |
|---|---|
| 0.02075119 | 0.135506235 |
| 0.020738091 | 0.135486015 |
| 0.020738051 | 0.135486015 |
| 0.020735658 | 0.135486015 |
| 0.020730428 | 0.135486015 |
| 0.02072192 | 0.135454695 |
| 0.020711469 | 0.135404962 |
| 0.020704794 | 0.135379901 |
| 0.020702925 | 0.135379901 |
| 0.020687887 | 0.135306511 |
| 0.020676718 | 0.13525203 |
| 0.020664317 | 0.135189481 |
| 0.020653365 | 0.135136392 |
| 0.020648794 | 0.135125052 |
| 0.020644315 | 0.135114303 |
| 0.020611478 | 0.134965762 |
| 0.020602338 | 0.134932282 |
| 0.02058552 | 0.134840674 |
| 0.02057501 | 0.134790373 |
| 0.020573946 | 0.134790373 |
| 0.0205693 | 0.134790045 |
| 0.020567271 | 0.134790045 |
| 0.020550756 | 0.134761233 |
| 0.020536024 | 0.134701726 |
| 0.020526064 | 0.134654939 |
| 0.020524827 | 0.134654939 |
| 0.020519383 | 0.1346482 |
| 0.02051452 | 0.134634842 |
| 0.020483403 | 0.134449149 |
| 0.020480963 | 0.134449149 |
| 0.020449931 | 0.134266452 |
| 0.020444912 | 0.134252012 |
| 0.020439195 | 0.134232975 |
| 0.02042426 | 0.134190406 |
| 0.020423746 | 0.134190406 |
| 0.020413371 | 0.13417439 |
| 0.020391766 | 0.134050884 |
| 0.020374223 | 0.133954049 |
| 0.020361929 | 0.133910192 |
| 0.020354911 | 0.133890902 |
| 0.020351609 | 0.133890902 |
| 0.020347613 | 0.133889993 |
| 0.020327248 | 0.133792955 |
| 0.020325232 | 0.133792955 |
| 0.020308065 | 0.13370365 |

| | |
|---|---|
| 0.02029509 | 0.133655176 |
| 0.020288689 | 0.133631503 |
| 0.020271549 | 0.133537076 |
| 0.020260721 | 0.133484208 |
| 0.020259626 | 0.133484208 |
| 0.020253423 | 0.133473054 |
| 0.020233421 | 0.133378152 |
| 0.020230127 | 0.133374897 |
| 0.020205442 | 0.133267495 |
| 0.020179698 | 0.13317147 |
| 0.020178188 | 0.13317147 |
| 0.020170982 | 0.133150849 |
| 0.020169885 | 0.133150849 |
| 0.020156622 | 0.13309295 |
| 0.020155806 | 0.13309295 |
| 0.020154952 | 0.13309295 |
| 0.020153951 | 0.13309295 |
| 0.020146091 | 0.13309295 |
| 0.020137822 | 0.13309295 |
| 0.020131997 | 0.133077951 |
| 0.020112746 | 0.13298761 |
| 0.020111172 | 0.13298761 |
| 0.020073644 | 0.132784366 |
| 0.020028455 | 0.132522257 |
| 0.020021047 | 0.132491651 |
| 0.020012692 | 0.132454767 |
| 0.020011652 | 0.132454767 |
| 0.020004379 | 0.132436552 |
| 0.020000536 | 0.132429516 |
| 0.019993166 | 0.132417541 |
| 0.019987115 | 0.132395872 |
| 0.019985571 | 0.132395872 |
| 0.019977529 | 0.132387615 |
| 0.019921616 | 0.132090602 |
| 0.0199083 | 0.132039073 |
| 0.019904018 | 0.132029062 |
| 0.019897786 | 0.132006106 |
| 0.019889722 | 0.131970996 |
| 0.019885884 | 0.131963909 |
| 0.019884685 | 0.131963909 |
| 0.019876105 | 0.131935783 |
| 0.019863569 | 0.131889334 |
| 0.019860082 | 0.131884572 |
| 0.019859546 | 0.131884572 |
| 0.019858796 | 0.131884572 |

| | |
|---|---|
| 0.019836549 | 0.131838589 |
| 0.019822818 | 0.13178411 |
| 0.019803151 | 0.131671738 |
| 0.01978998 | 0.131630622 |
| 0.019770804 | 0.13156687 |
| 0.019770228 | 0.13156687 |
| 0.019765545 | 0.13156687 |
| 0.019760511 | 0.131553518 |
| 0.019754329 | 0.131530746 |
| 0.019752578 | 0.131530746 |
| 0.019740635 | 0.131476329 |
| 0.019731211 | 0.131450324 |
| 0.019729332 | 0.131450324 |
| 0.019726725 | 0.131450324 |
| 0.019724448 | 0.131450324 |
| 0.019719363 | 0.131450324 |
| 0.019690868 | 0.131291749 |
| 0.019690141 | 0.131291749 |
| 0.019683949 | 0.131282372 |
| 0.019681695 | 0.131282372 |
| 0.01966676 | 0.131204481 |
| 0.019660305 | 0.131179792 |
| 0.019653305 | 0.131151455 |
| 0.019645017 | 0.131114521 |
| 0.019632079 | 0.131064907 |
| 0.019630561 | 0.131064907 |
| 0.019624979 | 0.131054244 |
| 0.019617638 | 0.131023591 |
| 0.019612371 | 0.131006788 |
| 0.01959146 | 0.130903827 |
| 0.019575232 | 0.130832105 |
| 0.019568635 | 0.130806371 |
| 0.019560663 | 0.130789795 |
| 0.019542709 | 0.130688093 |
| 0.01952107 | 0.130561718 |
| 0.019518182 | 0.130561718 |
| 0.019517786 | 0.130561718 |
| 0.019515181 | 0.130561718 |
| 0.019505221 | 0.130547378 |
| 0.019501382 | 0.130540034 |
| 0.019499527 | 0.130540034 |
| 0.019490038 | 0.130500778 |
| 0.019485106 | 0.130486097 |
| 0.019482283 | 0.130485543 |
| 0.019476109 | 0.130462541 |

| | |
|---|---|
| 0.019474738 | 0.130462541 |
| 0.019472023 | 0.130462541 |
| 0.019470988 | 0.130462541 |
| 0.019438247 | 0.130355592 |
| 0.019434844 | 0.130351125 |
| 0.019425182 | 0.130304666 |
| 0.019406963 | 0.130219134 |
| 0.019401048 | 0.130197789 |
| 0.01939632 | 0.130184407 |
| 0.019395381 | 0.130184407 |
| 0.019381856 | 0.130160695 |
| 0.019378644 | 0.130157479 |
| 0.019374099 | 0.130145305 |
| 0.019369427 | 0.130132277 |
| 0.019341239 | 0.129961231 |
| 0.019332229 | 0.129937351 |
| 0.019330043 | 0.129937351 |
| 0.019327788 | 0.129937351 |
| 0.019326494 | 0.129937351 |
| 0.019322742 | 0.129937351 |
| 0.019315629 | 0.129917449 |
| 0.019310312 | 0.129900032 |
| 0.019291522 | 0.129828642 |
| 0.019285712 | 0.129810289 |
| 0.019281718 | 0.129810289 |
| 0.01927959 | 0.129810289 |
| 0.019276336 | 0.129810289 |
| 0.01927625 | 0.129810289 |
| 0.019265576 | 0.129782404 |
| 0.019257729 | 0.129747897 |
| 0.019248572 | 0.129704547 |
| 0.019240829 | 0.129670717 |
| 0.019230389 | 0.1296187 |
| 0.019225426 | 0.12960359 |
| 0.019222618 | 0.129603006 |
| 0.019219649 | 0.129601332 |
| 0.019215367 | 0.129590805 |
| 0.019201015 | 0.129512356 |
| 0.019200969 | 0.129512356 |
| 0.019190667 | 0.129512356 |
| 0.019177796 | 0.129447403 |
| 0.019169932 | 0.129412668 |
| 0.019165267 | 0.129399516 |
| 0.019164156 | 0.129399516 |
| 0.019156774 | 0.129378854 |

| | |
|---|---|
| 0.019149705 | 0.129349454 |
| 0.019122988 | 0.129205638 |
| 0.019116964 | 0.129185272 |
| 0.019115546 | 0.129185272 |
| 0.019103544 | 0.129147549 |
| 0.019094433 | 0.129122611 |
| 0.019087447 | 0.129093703 |
| 0.019074433 | 0.129042335 |
| 0.019070561 | 0.129042335 |
| 0.019046476 | 0.12890813 |
| 0.01904338 | 0.128905491 |
| 0.019027575 | 0.128839777 |
| 0.019023893 | 0.128839777 |
| 0.019020859 | 0.128839777 |
| 0.019019181 | 0.128839777 |
| 0.018975558 | 0.128592613 |
| 0.018970251 | 0.128592613 |
| 0.018953583 | 0.128553448 |
| 0.018952915 | 0.128553448 |
| 0.018941893 | 0.12852907 |
| 0.018938648 | 0.128525364 |
| 0.018938642 | 0.128525364 |
| 0.018927597 | 0.128486984 |
| 0.018914381 | 0.128418594 |
| 0.018909144 | 0.128416627 |
| 0.018903352 | 0.1283956 |
| 0.018884733 | 0.128305732 |
| 0.018873858 | 0.128250144 |
| 0.018865693 | 0.12821295 |
| 0.01884074 | 0.128153089 |
| 0.018818902 | 0.128077715 |
| 0.01881502 | 0.128069593 |
| 0.018809241 | 0.128048563 |
| 0.018805152 | 0.128039027 |
| 0.018789492 | 0.127968989 |
| 0.018782711 | 0.127941103 |
| 0.018782214 | 0.127941103 |
| 0.01877878 | 0.127941103 |
| 0.018765903 | 0.127899779 |
| 0.018756922 | 0.127856865 |
| 0.018756599 | 0.127856865 |
| 0.018741371 | 0.127805741 |
| 0.018713603 | 0.127652932 |
| 0.01870335 | 0.127619547 |
| 0.018698477 | 0.127604583 |

| | |
|---|---|
| 0.018688182 | 0.127552597 |
| 0.018681086 | 0.127522442 |
| 0.018670806 | 0.127470541 |
| 0.018655449 | 0.127383949 |
| 0.018625001 | 0.127194284 |
| 0.018615575 | 0.127166377 |
| 0.018604205 | 0.127125179 |
| 0.018602624 | 0.127125179 |
| 0.018598989 | 0.127125179 |
| 0.018597728 | 0.127125179 |
| 0.018595059 | 0.127125179 |
| 0.018591878 | 0.127125179 |
| 0.018569961 | 0.127036991 |
| 0.018563542 | 0.127011322 |
| 0.018559268 | 0.127011322 |
| 0.018553695 | 0.126998681 |
| 0.018525848 | 0.126844531 |
| 0.018518518 | 0.126812571 |
| 0.018509962 | 0.126790445 |
| 0.018489033 | 0.126665299 |
| 0.018487837 | 0.126665299 |
| 0.018468518 | 0.126579378 |
| 0.018457156 | 0.126519711 |
| 0.018436934 | 0.126399285 |
| 0.018425883 | 0.126341714 |
| 0.018419016 | 0.126312811 |
| 0.018413192 | 0.126291055 |
| 0.018398476 | 0.126208295 |
| 0.018384722 | 0.126150289 |
| 0.018367879 | 0.126052879 |
| 0.018361604 | 0.126046142 |
| 0.018344052 | 0.125943806 |
| 0.018343429 | 0.125943806 |
| 0.01833016 | 0.125901045 |
| 0.018310637 | 0.125805067 |
| 0.018308525 | 0.125805067 |
| 0.018308092 | 0.125805067 |
| 0.018238515 | 0.125436211 |
| 0.018235259 | 0.125431926 |
| 0.018226402 | 0.125389116 |
| 0.018216722 | 0.125340627 |
| 0.01820508 | 0.12527862 |
| 0.018194471 | 0.125223707 |
| 0.018192797 | 0.125223707 |
| 0.018187854 | 0.125214357 |

| | |
|---|---|
| 0.018175869 | 0.125168032 |
| 0.018166993 | 0.125125 |
| 0.018161075 | 0.125102333 |
| 0.018154975 | 0.125078401 |
| 0.018146826 | 0.125040347 |
| 0.01813672 | 0.125006882 |
| 0.018136489 | 0.125006882 |
| 0.018131334 | 0.125005943 |
| 0.018130981 | 0.125005943 |
| 0.018114642 | 0.124927028 |
| 0.01810941 | 0.12490903 |
| 0.018108385 | 0.12490903 |
| 0.01808904 | 0.124804677 |
| 0.018058184 | 0.124673912 |
| 0.018051564 | 0.124672543 |
| 0.018047984 | 0.124665893 |
| 0.018046896 | 0.124665893 |
| 0.018039613 | 0.124644228 |
| 0.018025853 | 0.124567222 |
| 0.018008098 | 0.124462587 |
| 0.018005243 | 0.124460913 |
| 0.018004522 | 0.124460913 |
| 0.017939786 | 0.124116532 |
| 0.017937268 | 0.124116532 |
| 0.017933216 | 0.124107132 |
| 0.017927846 | 0.124106032 |
| 0.017927358 | 0.124106032 |
| 0.017910608 | 0.124040793 |
| 0.017908555 | 0.124040793 |
| 0.017905182 | 0.12403929 |
| 0.017903616 | 0.12403929 |
| 0.017901294 | 0.12403929 |
| 0.017898253 | 0.12403929 |
| 0.017892988 | 0.124035504 |
| 0.017883552 | 0.123997663 |
| 0.017875543 | 0.123978227 |
| 0.017870144 | 0.12397689 |
| 0.017865231 | 0.123975942 |
| 0.017862237 | 0.123975942 |
| 0.01785922 | 0.123973336 |
| 0.017851863 | 0.123940334 |
| 0.017851464 | 0.123940334 |
| 0.017847321 | 0.123940334 |
| 0.01784727 | 0.123940334 |
| 0.017845179 | 0.123940334 |

| | |
|---|---|
| 0.017843979 | 0.123940334 |
| 0.017834646 | 0.123929194 |
| 0.017827467 | 0.123901392 |
| 0.017826451 | 0.123901392 |
| 0.017825649 | 0.123901392 |
| 0.017820676 | 0.123901392 |
| 0.017807889 | 0.123887866 |
| 0.017792269 | 0.123797279 |
| 0.017776258 | 0.123703951 |
| 0.017767593 | 0.123661717 |
| 0.017757546 | 0.123646 |
| 0.017745923 | 0.12358314 |
| 0.017725205 | 0.123474965 |
| 0.01770572 | 0.123357271 |
| 0.017705519 | 0.123357271 |
| 0.017697397 | 0.123357271 |
| 0.017688926 | 0.123330478 |
| 0.01767428 | 0.123246404 |
| 0.017669949 | 0.123234243 |
| 0.01763981 | 0.123060092 |
| 0.017638213 | 0.123060092 |
| 0.017617363 | 0.122957521 |
| 0.01761197 | 0.122946609 |
| 0.017605959 | 0.122931973 |
| 0.017593814 | 0.122865188 |
| 0.017580348 | 0.122789153 |
| 0.017579437 | 0.122789153 |
| 0.017571832 | 0.12276568 |
| 0.017566663 | 0.122747573 |
| 0.017562067 | 0.122733467 |
| 0.017553166 | 0.122689262 |
| 0.017549826 | 0.122683924 |
| 0.017548362 | 0.122683924 |
| 0.017543515 | 0.122675817 |
| 0.017522495 | 0.12256482 |
| 0.017518748 | 0.122556613 |
| 0.017507575 | 0.12251444 |
| 0.017495836 | 0.122450281 |
| 0.017492871 | 0.122447528 |
| 0.017490231 | 0.122447043 |
| 0.017484753 | 0.122426689 |
| 0.017464374 | 0.122319959 |
| 0.017454451 | 0.122268444 |
| 0.01744212 | 0.122218016 |
| 0.017440952 | 0.122218016 |

| | |
|---|---|
| 0.017430796 | 0.122174614 |
| 0.017424895 | 0.122151234 |
| 0.017421184 | 0.122143194 |
| 0.017414604 | 0.122115038 |
| 0.01741348 | 0.122115038 |
| 0.017398216 | 0.122036056 |
| 0.017396028 | 0.122036056 |
| 0.017395878 | 0.122036056 |
| 0.017394463 | 0.122036056 |
| 0.017389421 | 0.122036056 |
| 0.017382545 | 0.122021619 |
| 0.017371788 | 0.121994417 |
| 0.017355886 | 0.121900723 |
| 0.017353561 | 0.121900723 |
| 0.01732818 | 0.121759989 |
| 0.017326065 | 0.121759989 |
| 0.01730247 | 0.121615221 |
| 0.017300169 | 0.121615221 |
| 0.017295719 | 0.121603664 |
| 0.01729559 | 0.121603664 |
| 0.017284362 | 0.121577654 |
| 0.017282328 | 0.121577654 |
| 0.017279676 | 0.121577654 |
| 0.017278827 | 0.121577654 |
| 0.017272059 | 0.121562929 |
| 0.017253773 | 0.121452178 |
| 0.017251714 | 0.121452178 |
| 0.017251177 | 0.121452178 |
| 0.01723066 | 0.121343286 |
| 0.017217624 | 0.121269411 |
| 0.017209761 | 0.121231959 |
| 0.017185997 | 0.121082468 |
| 0.017179524 | 0.121072683 |
| 0.017161727 | 0.121016669 |
| 0.017159731 | 0.121016669 |
| 0.017154001 | 0.121000241 |
| 0.017147659 | 0.120973423 |
| 0.017143211 | 0.120959958 |
| 0.017137818 | 0.120939823 |
| 0.01713397 | 0.120930584 |
| 0.017123895 | 0.120883702 |
| 0.017114117 | 0.12084419 |
| 0.01709886 | 0.120767469 |
| 0.017082583 | 0.120675196 |
| 0.01707783 | 0.120659515 |

| | |
|---|---|
| 0.016977443 | 0.120074942 |
| 0.016962355 | 0.1200217 |
| 0.016962069 | 0.1200217 |
| 0.016960307 | 0.1200217 |
| 0.016923897 | 0.119856424 |
| 0.016919727 | 0.119844711 |
| 0.016914295 | 0.119844711 |
| 0.016913133 | 0.119844711 |
| 0.016885231 | 0.119725012 |
| 0.016884886 | 0.119725012 |
| 0.016872404 | 0.119669689 |
| 0.016866153 | 0.119643173 |
| 0.016862085 | 0.119632135 |
| 0.016850711 | 0.119569254 |
| 0.016848837 | 0.119569254 |
| 0.016830754 | 0.119463237 |
| 0.016823838 | 0.119449751 |
| 0.016822737 | 0.119449751 |
| 0.016819488 | 0.119449751 |
| 0.016818634 | 0.119449751 |
| 0.016817151 | 0.119449751 |
| 0.016810204 | 0.119441977 |
| 0.016795797 | 0.119375231 |
| 0.016794165 | 0.119375231 |
| 0.016790445 | 0.119372822 |
| 0.016781466 | 0.119326794 |
| 0.016770906 | 0.119269515 |
| 0.016758022 | 0.119195683 |
| 0.016757771 | 0.119195683 |
| 0.01674241 | 0.119155825 |
| 0.016737282 | 0.119137127 |
| 0.016719445 | 0.119027956 |
| 0.016718845 | 0.119027956 |
| 0.016715795 | 0.119027956 |
| 0.01671203 | 0.119027956 |
| 0.0166989 | 0.118988406 |
| 0.016690698 | 0.118947757 |
| 0.016686858 | 0.118938188 |
| 0.016686663 | 0.118938188 |
| 0.016686062 | 0.118938188 |
| 0.016678719 | 0.118933574 |
| 0.016670867 | 0.118895384 |
| 0.016668871 | 0.118895384 |
| 0.016658224 | 0.118858614 |
| 0.0166467 | 0.118794185 |

| | |
|---|---|
| 0.016644419 | 0.118794185 |
| 0.016630849 | 0.11871664 |
| 0.016625156 | 0.118693797 |
| 0.016622013 | 0.118689144 |
| 0.016621985 | 0.118689144 |
| 0.01661864 | 0.118689144 |
| 0.016598174 | 0.118607836 |
| 0.016594729 | 0.118601014 |
| 0.016590939 | 0.118591725 |
| 0.016583897 | 0.118559179 |
| 0.016573889 | 0.118505424 |
| 0.016571078 | 0.118503114 |
| 0.016570046 | 0.118503114 |
| 0.016569937 | 0.118503114 |
| 0.016569689 | 0.118503114 |
| 0.016567952 | 0.118503114 |
| 0.016552161 | 0.118474566 |
| 0.01652812 | 0.11832027 |
| 0.016520329 | 0.11830006 |
| 0.016507042 | 0.118222691 |
| 0.016485493 | 0.118094784 |
| 0.016473291 | 0.118051977 |
| 0.016462436 | 0.118009702 |
| 0.016449105 | 0.11794964 |
| 0.016440073 | 0.117920389 |
| 0.016439418 | 0.117920389 |
| 0.016435515 | 0.117920389 |
| 0.016423276 | 0.117870909 |
| 0.016416498 | 0.117840019 |
| 0.016411733 | 0.117823576 |
| 0.016397588 | 0.117739775 |
| 0.016395723 | 0.117739775 |
| 0.016394733 | 0.117739775 |
| 0.016393854 | 0.117739775 |
| 0.016389847 | 0.117739775 |
| 0.016386334 | 0.117739775 |
| 0.016381985 | 0.117734238 |
| 0.016380708 | 0.117734238 |
| 0.016376067 | 0.117727233 |
| 0.016369192 | 0.117695576 |
| 0.016365072 | 0.117695576 |
| 0.016360039 | 0.117683062 |
| 0.016357462 | 0.117682299 |
| 0.01635339 | 0.117675018 |
| 0.016332896 | 0.117558839 |

| | |
|---|---|
| 0.016332388 | 0.117558839 |
| 0.016318226 | 0.117488761 |
| 0.016315631 | 0.11748784 |
| 0.016311928 | 0.117478936 |
| 0.016292209 | 0.117354661 |
| 0.016271771 | 0.117225176 |
| 0.016249781 | 0.117102183 |
| 0.016240793 | 0.117072842 |
| 0.016239922 | 0.117072842 |
| 0.016235833 | 0.117072842 |
| 0.016228939 | 0.117058255 |
| 0.016228768 | 0.117058255 |
| 0.016214258 | 0.116987797 |
| 0.016177396 | 0.116739519 |
| 0.016174015 | 0.116739519 |
| 0.016170802 | 0.116739519 |
| 0.016126439 | 0.116442378 |
| 0.016123596 | 0.116439503 |
| 0.016083788 | 0.116222512 |
| 0.016065335 | 0.116106786 |
| 0.016059593 | 0.116082904 |
| 0.016055365 | 0.116069955 |
| 0.016046021 | 0.11603763 |
| 0.016037321 | 0.116009947 |
| 0.016033632 | 0.116009947 |
| 0.016030171 | 0.116009947 |
| 0.01602696 | 0.116005473 |
| 0.016018837 | 0.115964306 |
| 0.015986133 | 0.115780344 |
| 0.015964306 | 0.115657439 |
| 0.015962055 | 0.115657439 |
| 0.015960857 | 0.115657439 |
| 0.015936527 | 0.115526482 |
| 0.015934849 | 0.115526482 |
| 0.015919178 | 0.115435852 |
| 0.015909605 | 0.115384007 |
| 0.015903974 | 0.115360733 |
| 0.015903157 | 0.115360733 |
| 0.015897267 | 0.115347214 |
| 0.015894187 | 0.11534244 |
| 0.015860176 | 0.115169107 |
| 0.015853968 | 0.115169107 |
| 0.015848097 | 0.115169107 |
| 0.015842586 | 0.115160999 |
| 0.015836151 | 0.115131788 |

| | |
|---|---|
| 0.015832022 | 0.115119348 |
| 0.015826516 | 0.115096883 |
| 0.015820892 | 0.11507355 |
| 0.015802795 | 0.114994607 |
| 0.015788401 | 0.114924983 |
| 0.015786614 | 0.114924983 |
| 0.015783897 | 0.114924983 |
| 0.015782718 | 0.114924983 |
| 0.015780055 | 0.114924983 |
| 0.015774789 | 0.114913715 |
| 0.015771571 | 0.114907849 |
| 0.015765466 | 0.114880942 |
| 0.015762782 | 0.114878962 |
| 0.015761681 | 0.114878962 |
| 0.015757802 | 0.114877826 |
| 0.015754686 | 0.114872691 |
| 0.015749639 | 0.114856159 |
| 0.01573945 | 0.114831919 |
| 0.01573176 | 0.11480093 |
| 0.015724494 | 0.114775551 |
| 0.0157229 | 0.114775551 |
| 0.015714401 | 0.11473705 |
| 0.015701502 | 0.11466045 |
| 0.015700054 | 0.11466045 |
| 0.015694257 | 0.114642698 |
| 0.015687374 | 0.114620685 |
| 0.015671277 | 0.114527529 |
| 0.015630053 | 0.11427887 |
| 0.015617957 | 0.114243041 |
| 0.015616405 | 0.114243041 |
| 0.015614874 | 0.114243041 |
| 0.015597226 | 0.114179077 |
| 0.015593266 | 0.114167637 |
| 0.015591125 | 0.114167637 |
| 0.015590608 | 0.114167637 |
| 0.015586304 | 0.114167637 |
| 0.015585787 | 0.114167637 |
| 0.015576241 | 0.114130713 |
| 0.015576196 | 0.114130713 |
| 0.015570159 | 0.114121262 |
| 0.01556715 | 0.114116768 |
| 0.015563862 | 0.114110232 |
| 0.015560014 | 0.114110232 |
| 0.015550152 | 0.114079961 |
| 0.015538636 | 0.114013039 |

| | |
|---|---|
| 0.015536406 | 0.114013039 |
| 0.01553024 | 0.114004125 |
| 0.015528639 | 0.114004125 |
| 0.015523304 | 0.113993762 |
| 0.015517116 | 0.113978048 |
| 0.015492425 | 0.113821287 |
| 0.015477688 | 0.113758622 |
| 0.015475522 | 0.113758622 |
| 0.015473559 | 0.113758622 |
| 0.015467072 | 0.113733248 |
| 0.015450911 | 0.113631954 |
| 0.015450703 | 0.113631954 |
| 0.015446571 | 0.113631954 |
| 0.015442488 | 0.113631954 |
| 0.015396461 | 0.113441712 |
| 0.015367801 | 0.113283124 |
| 0.015361741 | 0.113255978 |
| 0.015357539 | 0.11324253 |
| 0.015355218 | 0.11324253 |
| 0.015352526 | 0.113240631 |
| 0.015333908 | 0.113120824 |
| 0.015307382 | 0.112960133 |
| 0.015294756 | 0.112885599 |
| 0.015294287 | 0.112885599 |
| 0.015254675 | 0.112710895 |
| 0.015248324 | 0.112697891 |
| 0.01524503 | 0.112692075 |
| 0.015232375 | 0.11261601 |
| 0.015226692 | 0.112602407 |
| 0.015220853 | 0.112583251 |
| 0.015206439 | 0.112494106 |
| 0.015204165 | 0.112494106 |
| 0.015181691 | 0.112380854 |
| 0.015174702 | 0.112346579 |
| 0.015166749 | 0.112322621 |
| 0.015154261 | 0.112300523 |
| 0.015140644 | 0.112233998 |
| 0.015120598 | 0.112137764 |
| 0.015116435 | 0.112124346 |
| 0.015113948 | 0.112123366 |
| 0.015111007 | 0.112119013 |
| 0.015110367 | 0.112119013 |
| 0.015102337 | 0.112107075 |
| 0.015098811 | 0.112098376 |
| 0.015089109 | 0.112043812 |

| | |
|---|---|
| 0.015082758 | 0.112014112 |
| 0.015078841 | 0.112002487 |
| 0.015062677 | 0.111917338 |
| 0.01505391 | 0.111869648 |
| 0.015046133 | 0.111864216 |
| 0.015039518 | 0.111832495 |
| 0.015025774 | 0.111765191 |
| 0.015019354 | 0.111734889 |
| 0.015017724 | 0.111734889 |
| 0.01499127 | 0.111595687 |
| 0.014988817 | 0.111594867 |
| 0.014985246 | 0.111585728 |
| 0.014985078 | 0.111585728 |
| 0.014962501 | 0.111468645 |
| 0.014956572 | 0.111441905 |
| 0.014955684 | 0.111441905 |
| 0.014946651 | 0.111402836 |
| 0.01494418 | 0.111401855 |
| 0.014942435 | 0.111401855 |
| 0.014930929 | 0.111337928 |
| 0.014914899 | 0.111235812 |
| 0.014911868 | 0.111230626 |
| 0.014904841 | 0.111195626 |
| 0.014903598 | 0.111195626 |
| 0.014902897 | 0.111195626 |
| 0.014880442 | 0.111135455 |
| 0.014860633 | 0.111004919 |
| 0.014855325 | 0.11099554 |
| 0.014816852 | 0.110729991 |
| 0.014815702 | 0.110729991 |
| 0.014807892 | 0.110715143 |
| 0.014786113 | 0.110587028 |
| 0.014773727 | 0.110529105 |
| 0.014760067 | 0.110444258 |
| 0.014744408 | 0.110344427 |
| 0.014740198 | 0.110330256 |
| 0.01471807 | 0.110216595 |
| 0.014711466 | 0.110184464 |
| 0.014701189 | 0.110142132 |
| 0.014689829 | 0.11009166 |
| 0.014686552 | 0.110084418 |
| 0.01467388 | 0.110006753 |
| 0.014651371 | 0.109855299 |
| 0.01463626 | 0.10977656 |
| 0.014631066 | 0.10977656 |

| | |
|---|---|
| 0.014627561 | 0.10977656 |
| 0.014624519 | 0.109774934 |
| 0.014619357 | 0.109753488 |
| 0.014598475 | 0.109613992 |
| 0.014598144 | 0.109613992 |
| 0.014596923 | 0.109613992 |
| 0.014588255 | 0.109589089 |
| 0.014576719 | 0.10953698 |
| 0.014552723 | 0.109443004 |
| 0.014531991 | 0.109313954 |
| 0.014527301 | 0.109313954 |
| 0.014510216 | 0.109209555 |
| 0.014477448 | 0.109025583 |
| 0.014472388 | 0.109010981 |
| 0.014452884 | 0.108881286 |
| 0.014451795 | 0.108881286 |
| 0.014450968 | 0.108881286 |
| 0.014447319 | 0.108881286 |
| 0.014445085 | 0.108881286 |
| 0.014432817 | 0.108833404 |
| 0.014425918 | 0.108798604 |
| 0.01441445 | 0.108746553 |
| 0.014413467 | 0.108746553 |
| 0.014407826 | 0.108740305 |
| 0.014404551 | 0.108740305 |
| 0.01438051 | 0.108610937 |
| 0.014379302 | 0.108610937 |
| 0.014374593 | 0.108600691 |
| 0.014373609 | 0.108600691 |
| 0.014373377 | 0.108600691 |
| 0.014373323 | 0.108600691 |
| 0.014371738 | 0.108600691 |
| 0.014370068 | 0.108600691 |
| 0.014358199 | 0.108600691 |
| 0.014353912 | 0.108599485 |
| 0.014340922 | 0.108518444 |
| 0.014333813 | 0.108490901 |
| 0.014333394 | 0.108490901 |
| 0.014328751 | 0.108490901 |
| 0.014317683 | 0.108428731 |
| 0.014313325 | 0.108428731 |
| 0.014283006 | 0.108252188 |
| 0.01426755 | 0.10815225 |
| 0.014264485 | 0.108146235 |
| 0.014262267 | 0.108146235 |

| | |
|---|---|
| 0.014256777 | 0.108122219 |
| 0.014249094 | 0.108098373 |
| 0.014240099 | 0.108047345 |
| 0.01423489 | 0.10802503 |
| 0.014225836 | 0.107973529 |
| 0.014219219 | 0.107940508 |
| 0.014202349 | 0.107829631 |
| 0.014197301 | 0.107808489 |
| 0.014195182 | 0.107808489 |
| 0.014194987 | 0.107808489 |
| 0.014186056 | 0.107774648 |
| 0.014183905 | 0.107774648 |
| 0.014165868 | 0.107689991 |
| 0.014165397 | 0.107689991 |
| 0.014160472 | 0.107683344 |
| 0.014159298 | 0.107683344 |
| 0.014157171 | 0.107683344 |
| 0.014157066 | 0.107683344 |
| 0.014140242 | 0.107667039 |
| 0.014128139 | 0.107592084 |
| 0.014123696 | 0.107575445 |
| 0.014105142 | 0.107451308 |
| 0.014097885 | 0.10741321 |
| 0.014086975 | 0.107347254 |
| 0.014076845 | 0.107287231 |
| 0.014062339 | 0.107193823 |
| 0.014034377 | 0.10701493 |
| 0.014025297 | 0.106993449 |
| 0.01401327 | 0.106922464 |
| 0.014012442 | 0.106922464 |
| 0.013998433 | 0.106843487 |
| 0.013998068 | 0.106843487 |
| 0.013996305 | 0.106843487 |
| 0.013995207 | 0.106843487 |
| 0.013990588 | 0.106843487 |
| 0.013980627 | 0.106793153 |
| 0.013971419 | 0.106739939 |
| 0.013963007 | 0.106709903 |
| 0.013953873 | 0.106664718 |
| 0.013941969 | 0.106600446 |
| 0.013937174 | 0.10658089 |
| 0.01392305 | 0.106532784 |
| 0.013919138 | 0.106528487 |
| 0.01390769 | 0.10647509 |
| 0.013890809 | 0.106362954 |

| | |
|---|---|
| 0.013885073 | 0.106336128 |
| 0.013876968 | 0.106293611 |
| 0.013864875 | 0.106232691 |
| 0.013858887 | 0.106232691 |
| 0.013850965 | 0.106194447 |
| 0.013839726 | 0.106125361 |
| 0.013835824 | 0.106112527 |
| 0.013833258 | 0.106109931 |
| 0.013826144 | 0.106089541 |
| 0.01382119 | 0.106074894 |
| 0.013811763 | 0.106030436 |
| 0.013778488 | 0.105817527 |
| 0.013758522 | 0.105723963 |
| 0.013749801 | 0.105674002 |
| 0.013737098 | 0.10564456 |
| 0.013736738 | 0.10564456 |
| 0.013734373 | 0.10564456 |
| 0.013734121 | 0.10564456 |
| 0.013730398 | 0.10564456 |
| 0.013725645 | 0.10564456 |
| 0.013725498 | 0.10564456 |
| 0.013712804 | 0.105594132 |
| 0.01371272 | 0.105594132 |
| 0.013699748 | 0.105544795 |
| 0.013657535 | 0.105270661 |
| 0.013646391 | 0.105220341 |
| 0.013645143 | 0.105220341 |
| 0.013642703 | 0.105220341 |
| 0.013640954 | 0.105220341 |
| 0.013637346 | 0.105220341 |
| 0.013634883 | 0.105220341 |
| 0.013627981 | 0.105213141 |
| 0.013625529 | 0.105211262 |
| 0.01362491 | 0.105211262 |
| 0.013619775 | 0.105200941 |
| 0.0136192 | 0.105200941 |
| 0.013618612 | 0.105200941 |
| 0.013606937 | 0.105176353 |
| 0.01359875 | 0.105150528 |
| 0.013572683 | 0.104987247 |
| 0.013571384 | 0.104987247 |
| 0.013566376 | 0.104975761 |
| 0.01354233 | 0.10484079 |
| 0.013538992 | 0.104831992 |
| 0.01353023 | 0.104781179 |

| | |
|---|---|
| 0.013528316 | 0.104781179 |
| 0.01350131 | 0.104591233 |
| 0.013497648 | 0.104579874 |
| 0.013493025 | 0.104561066 |
| 0.013492842 | 0.104561066 |
| 0.013486009 | 0.104540718 |
| 0.013480069 | 0.104528697 |
| 0.013478103 | 0.104528697 |
| 0.013456455 | 0.104413587 |
| 0.013454823 | 0.104413587 |
| 0.013451967 | 0.104413587 |
| 0.013448358 | 0.104413587 |
| 0.013448193 | 0.104413587 |
| 0.013445839 | 0.104413587 |
| 0.013440751 | 0.104410806 |
| 0.013431826 | 0.104358495 |
| 0.013430361 | 0.104358495 |
| 0.013423626 | 0.104328826 |
| 0.013411315 | 0.104267171 |
| 0.013405921 | 0.104242251 |
| 0.013395399 | 0.104211478 |
| 0.013388363 | 0.104173754 |
| 0.013386121 | 0.104173329 |
| 0.013385607 | 0.104173329 |
| 0.013383674 | 0.104173329 |
| 0.013380902 | 0.104173329 |
| 0.013376026 | 0.10416285 |
| 0.013373973 | 0.10416285 |
| 0.013366801 | 0.10412506 |
| 0.013361356 | 0.104099672 |
| 0.013355549 | 0.104071454 |
| 0.013340716 | 0.10399406 |
| 0.013334022 | 0.103977584 |
| 0.013332353 | 0.103977584 |
| 0.013332195 | 0.103977584 |
| 0.01332889 | 0.103977584 |
| 0.013324556 | 0.103977584 |
| 0.01331828 | 0.103951117 |
| 0.013318266 | 0.103951117 |
| 0.013314489 | 0.103951117 |
| 0.013310441 | 0.103951117 |
| 0.013306185 | 0.103951117 |
| 0.013302264 | 0.103951117 |
| 0.013294651 | 0.103919968 |
| 0.013269961 | 0.103812163 |

| | |
|---|---|
| 0.013257615 | 0.103749659 |
| 0.01325662 | 0.103749659 |
| 0.013241858 | 0.103660419 |
| 0.013236427 | 0.103634933 |
| 0.013227185 | 0.103579606 |
| 0.013226036 | 0.103579606 |
| 0.013224331 | 0.103579606 |
| 0.013223456 | 0.103579606 |
| 0.013218167 | 0.103579606 |
| 0.013217365 | 0.103579606 |
| 0.01320185 | 0.103500329 |
| 0.013197601 | 0.103484051 |
| 0.013192949 | 0.103464609 |
| 0.013187853 | 0.103441681 |
| 0.013178743 | 0.103404279 |
| 0.013175213 | 0.103394418 |
| 0.013175071 | 0.103394418 |
| 0.013174686 | 0.103394418 |
| 0.013174605 | 0.103394418 |
| 0.013167286 | 0.103394418 |
| 0.013165396 | 0.103394418 |
| 0.013159788 | 0.103394418 |
| 0.013154218 | 0.103394418 |
| 0.013151707 | 0.103394418 |
| 0.013143805 | 0.103368563 |
| 0.013142618 | 0.103368563 |
| 0.013139766 | 0.103368563 |
| 0.013128345 | 0.103326258 |
| 0.013121396 | 0.103294669 |
| 0.013103541 | 0.103188218 |
| 0.013089463 | 0.103094401 |
| 0.013082582 | 0.10305725 |
| 0.013081531 | 0.10305725 |
| 0.013067874 | 0.102992491 |
| 0.013055898 | 0.102949211 |
| 0.013054638 | 0.102949211 |
| 0.013048859 | 0.102927793 |
| 0.013047592 | 0.102927793 |
| 0.01304323 | 0.10291748 |
| 0.013041542 | 0.10291748 |
| 0.013026737 | 0.102821409 |
| 0.01301321 | 0.102748699 |
| 0.013011866 | 0.102748699 |
| 0.013011209 | 0.102748699 |
| 0.013010623 | 0.102748699 |

| | |
|---|---|
| 0.013006109 | 0.102748699 |
| 0.013004727 | 0.102748699 |
| 0.012954912 | 0.102407233 |
| 0.012947201 | 0.102380268 |
| 0.012942104 | 0.102373958 |
| 0.012935963 | 0.102342373 |
| 0.012928625 | 0.102301319 |
| 0.012923594 | 0.102278502 |
| 0.012920386 | 0.102270108 |
| 0.012902298 | 0.102143907 |
| 0.012901298 | 0.102143907 |
| 0.012890557 | 0.10210301 |
| 0.012882155 | 0.102069269 |
| 0.012879779 | 0.102067424 |
| 0.012876459 | 0.102058102 |
| 0.012872664 | 0.102045002 |
| 0.012871075 | 0.102045002 |
| 0.012870741 | 0.102045002 |
| 0.012869375 | 0.102045002 |
| 0.012859239 | 0.102019879 |
| 0.012840077 | 0.10188843 |
| 0.012818966 | 0.101754817 |
| 0.012817043 | 0.101754817 |
| 0.012809377 | 0.1017126 |
| 0.012806336 | 0.1017126 |
| 0.012801651 | 0.101702127 |
| 0.012795996 | 0.101674158 |
| 0.012792735 | 0.101665211 |
| 0.012788211 | 0.101663184 |
| 0.012777441 | 0.101611486 |
| 0.012775476 | 0.101611486 |
| 0.01277445 | 0.101611486 |
| 0.012772 | 0.101611486 |
| 0.012771727 | 0.101611486 |
| 0.012754788 | 0.101533059 |
| 0.01272506 | 0.101330288 |
| 0.01272276 | 0.101328908 |
| 0.01271356 | 0.101289514 |
| 0.012712813 | 0.101289514 |
| 0.012694962 | 0.101192128 |
| 0.012683811 | 0.101137099 |
| 0.012671254 | 0.101070812 |
| 0.012670852 | 0.101070812 |
| 0.012669177 | 0.101070812 |
| 0.012665724 | 0.101070812 |

| | |
|---|---|
| 0.012663673 | 0.101070812 |
| 0.012643595 | 0.100951637 |
| 0.012642622 | 0.100951637 |
| 0.012637183 | 0.100934277 |
| 0.012627046 | 0.100870232 |
| 0.012623516 | 0.100858949 |
| 0.01262005 | 0.100848176 |
| 0.012615028 | 0.100824961 |
| 0.012613882 | 0.100824961 |
| 0.012596661 | 0.100728872 |
| 0.012569146 | 0.100576396 |
| 0.012564886 | 0.100559206 |
| 0.012562255 | 0.100555046 |
| 0.012558676 | 0.100543294 |
| 0.012556118 | 0.100539715 |
| 0.012554991 | 0.100539715 |
| 0.012531986 | 0.100380239 |
| 0.012528945 | 0.100372758 |
| 0.012521932 | 0.100333456 |
| 0.012516465 | 0.100306529 |
| 0.012515813 | 0.100306529 |
| 0.012504565 | 0.100261769 |
| 0.01250184 | 0.100256798 |
| 0.012490034 | 0.100178992 |
| 0.012465622 | 0.100033727 |
| 0.012447908 | 0.099925251 |
| 0.012442216 | 0.099913236 |
| 0.012437832 | 0.099911726 |
| 0.012427778 | 0.099847803 |
| 0.012426049 | 0.099847803 |
| 0.012383724 | 0.09959467 |
| 0.012382071 | 0.09959467 |
| 0.012377084 | 0.099574899 |
| 0.012376772 | 0.099574899 |
| 0.012369142 | 0.099544629 |
| 0.012364386 | 0.099523172 |
| 0.012363715 | 0.099523172 |
| 0.012358408 | 0.099517771 |
| 0.012349799 | 0.099473004 |
| 0.012349684 | 0.099473004 |
| 0.012340382 | 0.099430788 |
| 0.012318101 | 0.099301658 |
| 0.012315338 | 0.099296192 |
| 0.012313172 | 0.099295537 |
| 0.012295646 | 0.099186849 |

| | |
|---|---|
| 0.012288239 | 0.099144837 |
| 0.012280059 | 0.099095621 |
| 0.012276637 | 0.099084801 |
| 0.012276123 | 0.099084801 |
| 0.012275971 | 0.099084801 |
| 0.012264905 | 0.099047122 |
| 0.012263431 | 0.099047122 |
| 0.012238498 | 0.09888189 |
| 0.012230701 | 0.098864829 |
| 0.012215997 | 0.098779349 |
| 0.012201638 | 0.09871368 |
| 0.012201032 | 0.09871368 |
| 0.012195417 | 0.098696879 |
| 0.012193587 | 0.098696879 |
| 0.01219211 | 0.098696879 |
| 0.012191883 | 0.098696879 |
| 0.012186127 | 0.098688774 |
| 0.012166076 | 0.09855061 |
| 0.012157046 | 0.098525983 |
| 0.012154156 | 0.098525983 |
| 0.012148625 | 0.098502313 |
| 0.012144679 | 0.09848708 |
| 0.012129967 | 0.098384525 |
| 0.012117304 | 0.098298562 |
| 0.012104372 | 0.098210375 |
| 0.012103111 | 0.098210375 |
| 0.012095474 | 0.098171635 |
| 0.012077789 | 0.098044806 |
| 0.012053643 | 0.097865471 |
| 0.012052213 | 0.097865471 |
| 0.01204995 | 0.097865471 |
| 0.012049013 | 0.097865471 |
| 0.012039272 | 0.097848874 |
| 0.012035864 | 0.097837873 |
| 0.012029013 | 0.097798878 |
| 0.012025654 | 0.097788254 |
| 0.012018656 | 0.097748042 |
| 0.012018389 | 0.097748042 |
| 0.012017717 | 0.097748042 |
| 0.012006117 | 0.097707847 |
| 0.011993338 | 0.097625484 |
| 0.011984811 | 0.097572754 |
| 0.011973774 | 0.09752883 |
| 0.011958426 | 0.097424558 |
| 0.011956312 | 0.097424558 |

| | |
|---|---|
| 0.011951311 | 0.097416579 |
| 0.011949438 | 0.097416579 |
| 0.011946518 | 0.097410853 |
| 0.011945017 | 0.097410853 |
| 0.011935946 | 0.09735799 |
| 0.011918103 | 0.097229097 |
| 0.011915862 | 0.097227471 |
| 0.011905087 | 0.097156198 |
| 0.011898339 | 0.097117773 |
| 0.011894633 | 0.097104165 |
| 0.011893966 | 0.097104165 |
| 0.011891188 | 0.097104165 |
| 0.011880971 | 0.09704253 |
| 0.011875175 | 0.097028471 |
| 0.011872104 | 0.097020025 |
| 0.011871623 | 0.097020025 |
| 0.011865439 | 0.096998841 |
| 0.011857574 | 0.096967834 |
| 0.011856946 | 0.096967834 |
| 0.011845568 | 0.096902929 |
| 0.011830109 | 0.096793095 |
| 0.011824339 | 0.096762508 |
| 0.011818946 | 0.096734994 |
| 0.011812925 | 0.096702331 |
| 0.011786746 | 0.096521205 |
| 0.01177702 | 0.096476827 |
| 0.011776367 | 0.096476827 |
| 0.011768134 | 0.096451714 |
| 0.01173589 | 0.096204 |
| 0.011735176 | 0.096204 |
| 0.011732564 | 0.096204 |
| 0.01173233 | 0.096204 |
| 0.011728687 | 0.096204 |
| 0.011724196 | 0.096204 |
| 0.011721439 | 0.096201442 |
| 0.011717569 | 0.096186259 |
| 0.011716278 | 0.096186259 |
| 0.011712094 | 0.096174469 |
| 0.011706596 | 0.096145892 |
| 0.0117014 | 0.096119794 |
| 0.011696567 | 0.096113654 |
| 0.011696458 | 0.096113654 |
| 0.011694683 | 0.096113654 |
| 0.011692763 | 0.096113654 |
| 0.011674655 | 0.096024176 |

| | |
|---|---|
| 0.011671054 | 0.09601957 |
| 0.011651393 | 0.095890941 |
| 0.011634536 | 0.095768763 |
| 0.011634267 | 0.095768763 |
| 0.011633545 | 0.095768763 |
| 0.011631869 | 0.095768763 |
| 0.011626622 | 0.095768763 |
| 0.011612793 | 0.095672459 |
| 0.011600384 | 0.0956033 |
| 0.011584129 | 0.095518923 |
| 0.011579894 | 0.095517081 |
| 0.011552306 | 0.095339055 |
| 0.011545117 | 0.095296234 |
| 0.011542478 | 0.095296234 |
| 0.011526988 | 0.095196094 |
| 0.01152499 | 0.095196094 |
| 0.011517515 | 0.095150857 |
| 0.011493404 | 0.094981398 |
| 0.011485018 | 0.094931788 |
| 0.011478536 | 0.094894675 |
| 0.011474188 | 0.094875204 |
| 0.011469437 | 0.094852385 |
| 0.011469097 | 0.094852385 |
| 0.011459915 | 0.094823029 |
| 0.011455629 | 0.094804035 |
| 0.011454502 | 0.094804035 |
| 0.011453919 | 0.094804035 |
| 0.011406961 | 0.094499794 |
| 0.011404265 | 0.0944939 |
| 0.011398233 | 0.094460355 |
| 0.011397024 | 0.094460355 |
| 0.011392958 | 0.094460355 |
| 0.011387551 | 0.094454005 |
| 0.011380904 | 0.09441531 |
| 0.01136632 | 0.094310754 |
| 0.011366035 | 0.094310754 |
| 0.011359209 | 0.094284599 |
| 0.011349353 | 0.094219213 |
| 0.011347771 | 0.094219213 |
| 0.011341552 | 0.094187288 |
| 0.011332799 | 0.094147429 |
| 0.011315831 | 0.09402287 |
| 0.011314969 | 0.09402287 |
| 0.011312473 | 0.09402287 |
| 0.011309372 | 0.09401841 |

| | |
|---|---|
| 0.011301898 | 0.093972674 |
| 0.011300623 | 0.093972674 |
| 0.01129857 | 0.093972674 |
| 0.011291861 | 0.093938425 |
| 0.011284021 | 0.093889599 |
| 0.011283025 | 0.093889599 |
| 0.01127913 | 0.093881707 |
| 0.011273385 | 0.093850292 |
| 0.011272732 | 0.093850292 |
| 0.011268157 | 0.093839574 |
| 0.011266903 | 0.093839574 |
| 0.011262877 | 0.093828416 |
| 0.01125731 | 0.093814857 |
| 0.011256206 | 0.093814857 |
| 0.011254618 | 0.093814857 |
| 0.011243632 | 0.093750965 |
| 0.011230964 | 0.093717877 |
| 0.011224645 | 0.093706604 |
| 0.011224434 | 0.093706604 |
| 0.011222592 | 0.093706604 |
| 0.011214635 | 0.093688723 |
| 0.01121358 | 0.093688723 |
| 0.011212577 | 0.093688723 |
| 0.011212127 | 0.093688723 |
| 0.011208206 | 0.093688723 |
| 0.01120342 | 0.09367719 |
| 0.011201066 | 0.093673951 |
| 0.011194638 | 0.093636636 |
| 0.011185562 | 0.0935936 |
| 0.011185475 | 0.0935936 |
| 0.011179126 | 0.093586129 |
| 0.011167053 | 0.093520875 |
| 0.011146433 | 0.093364607 |
| 0.011142492 | 0.093348016 |
| 0.011137719 | 0.093324447 |
| 0.01113211 | 0.093293858 |
| 0.011129489 | 0.093288312 |
| 0.011118241 | 0.093210428 |
| 0.011112931 | 0.093182318 |
| 0.011111633 | 0.093182318 |
| 0.011107333 | 0.09316818 |
| 0.011087532 | 0.093034867 |
| 0.011082572 | 0.093009629 |
| 0.011081127 | 0.093009629 |
| 0.0110729 | 0.09296122 |

| | |
|---|---|
| 0.011069892 | 0.092956586 |
| 0.011059457 | 0.092913872 |
| 0.011058138 | 0.092913872 |
| 0.011056579 | 0.092913872 |
| 0.011054742 | 0.092913872 |
| 0.011040095 | 0.092849483 |
| 0.011023109 | 0.092739378 |
| 0.011014466 | 0.09269941 |
| 0.010995727 | 0.092581657 |
| 0.010983633 | 0.092521664 |
| 0.010980763 | 0.092513851 |
| 0.010973435 | 0.092484831 |
| 0.010969474 | 0.092467811 |
| 0.010962226 | 0.092423075 |
| 0.010960216 | 0.09242249 |
| 0.010958814 | 0.09242249 |
| 0.010954232 | 0.092404748 |
| 0.010929032 | 0.092241184 |
| 0.010927672 | 0.092241184 |
| 0.010919816 | 0.092196076 |
| 0.010917635 | 0.092194005 |
| 0.010916759 | 0.092194005 |
| 0.010914338 | 0.092194005 |
| 0.010889513 | 0.092021795 |
| 0.010881882 | 0.09197363 |
| 0.010877551 | 0.09195334 |
| 0.010873815 | 0.09193808 |
| 0.010859797 | 0.091852166 |
| 0.010858819 | 0.091852166 |
| 0.010855832 | 0.091851256 |
| 0.010852877 | 0.091842565 |
| 0.010832754 | 0.091688565 |
| 0.010829548 | 0.091677724 |
| 0.01082413 | 0.091648151 |
| 0.010820513 | 0.091633812 |
| 0.01081643 | 0.091615524 |
| 0.010810971 | 0.091585579 |
| 0.010809701 | 0.091585579 |
| 0.010807642 | 0.091585579 |
| 0.01080069 | 0.091547334 |
| 0.010794423 | 0.091526794 |
| 0.010786735 | 0.091494183 |
| 0.010784223 | 0.091494183 |
| 0.010771683 | 0.091415352 |
| 0.010771127 | 0.091415352 |

| | |
|---|---|
| 0.010770224 | 0.091415352 |
| 0.010766848 | 0.091415352 |
| 0.010758184 | 0.091382204 |
| 0.010746951 | 0.091303077 |
| 0.010733146 | 0.091202058 |
| 0.010725295 | 0.091151611 |
| 0.010725058 | 0.091151611 |
| 0.01072396 | 0.091151611 |
| 0.010722202 | 0.091151611 |
| 0.010716414 | 0.091151611 |
| 0.010713725 | 0.091150876 |
| 0.010692024 | 0.091031286 |
| 0.010679269 | 0.09094928 |
| 0.010674875 | 0.090934053 |
| 0.010674623 | 0.090934053 |
| 0.010674244 | 0.090934053 |
| 0.010674131 | 0.090934053 |
| 0.010667622 | 0.090934053 |
| 0.010659063 | 0.090896898 |
| 0.010649719 | 0.090833485 |
| 0.010648247 | 0.090833485 |
| 0.010647469 | 0.090833485 |
| 0.010646028 | 0.090833485 |
| 0.010644863 | 0.090833485 |
| 0.010643892 | 0.090833485 |
| 0.010636893 | 0.090833485 |
| 0.01063424 | 0.090831595 |
| 0.010630413 | 0.09081519 |
| 0.01062386 | 0.090775491 |
| 0.010607989 | 0.090672418 |
| 0.010604884 | 0.090662146 |
| 0.01058818 | 0.090535595 |
| 0.010582565 | 0.090503829 |
| 0.010582248 | 0.090503829 |
| 0.010557741 | 0.09035643 |
| 0.010556452 | 0.09035643 |
| 0.010555238 | 0.09035643 |
| 0.010554846 | 0.09035643 |
| 0.010544899 | 0.090311433 |
| 0.010544325 | 0.090311433 |
| 0.010537367 | 0.090279395 |
| 0.01053557 | 0.090279395 |
| 0.010535424 | 0.090279395 |
| 0.010531295 | 0.090276091 |
| 0.010526422 | 0.09026682 |

| | |
|---|---|
| 0.010521385 | 0.090239867 |
| 0.010514475 | 0.090196847 |
| 0.010499702 | 0.090086346 |
| 0.01049554 | 0.090083091 |
| 0.010470246 | 0.08991461 |
| 0.010468327 | 0.08991461 |
| 0.010466399 | 0.08991461 |
| 0.010460637 | 0.089913148 |
| 0.010455414 | 0.089913148 |
| 0.010450969 | 0.089902964 |
| 0.010436194 | 0.089800288 |
| 0.010435171 | 0.089800288 |
| 0.01043456 | 0.089800288 |
| 0.01043258 | 0.089800288 |
| 0.01042757 | 0.089790976 |
| 0.010404526 | 0.089608747 |
| 0.010404263 | 0.089608747 |
| 0.010400052 | 0.089602615 |
| 0.010397071 | 0.089593145 |
| 0.010393606 | 0.08957949 |
| 0.010384239 | 0.089515029 |
| 0.010384193 | 0.089515029 |
| 0.010383165 | 0.089515029 |
| 0.010371276 | 0.089484198 |
| 0.010342364 | 0.089283262 |
| 0.01033278 | 0.089232876 |
| 0.01031259 | 0.089106988 |
| 0.01031171 | 0.089106988 |
| 0.01028812 | 0.088957462 |
| 0.010286203 | 0.088957462 |
| 0.010275254 | 0.088897277 |
| 0.010272996 | 0.088893895 |
| 0.010266034 | 0.088878935 |
| 0.010264888 | 0.088878935 |
| 0.010263876 | 0.088878935 |
| 0.010260061 | 0.088878852 |
| 0.010259013 | 0.088878852 |
| 0.01024818 | 0.088824398 |
| 0.010248059 | 0.088824398 |
| 0.010242789 | 0.08881 |
| 0.010234843 | 0.088757261 |
| 0.010230109 | 0.088732364 |
| 0.01022386 | 0.088694317 |
| 0.010210179 | 0.088607905 |
| 0.010208328 | 0.088607905 |

| | |
|---|---|
| 0.010200938 | 0.088592282 |
| 0.010195986 | 0.088565429 |
| 0.010192322 | 0.088561696 |
| 0.010179294 | 0.088468836 |
| 0.01017647 | 0.088460438 |
| 0.010169535 | 0.08841629 |
| 0.01016681 | 0.088408741 |
| 0.010166054 | 0.088408741 |
| 0.01016513 | 0.088408741 |
| 0.010155462 | 0.088358458 |
| 0.010150799 | 0.088334019 |
| 0.010136677 | 0.088243376 |
| 0.010133482 | 0.088231686 |
| 0.010129169 | 0.08821026 |
| 0.010120058 | 0.088147039 |
| 0.010091747 | 0.087964793 |
| 0.010091635 | 0.087964793 |
| 0.010081439 | 0.087925058 |
| 0.010071238 | 0.087866434 |
| 0.01005886 | 0.087790606 |
| 0.010045689 | 0.087739963 |
| 0.010045278 | 0.087739963 |
| 0.010043153 | 0.087739963 |
| 0.010037525 | 0.08773183 |
| 0.01003563 | 0.08773183 |
| 0.010034785 | 0.08773183 |
| 0.010034135 | 0.08773183 |
| 0.010024143 | 0.087704769 |
| 0.010021457 | 0.087704769 |
| 0.009997515 | 0.087524438 |
| 0.009996771 | 0.087524438 |
| 0.00999299 | 0.087520434 |
| 0.009988804 | 0.087499864 |
| 0.009968165 | 0.087335137 |
| 0.009965323 | 0.087326299 |
| 0.00996516 | 0.087326299 |
| 0.009962631 | 0.087326299 |
| 0.009954887 | 0.087283025 |
| 0.009951698 | 0.087271131 |
| 0.00995021 | 0.087271131 |
| 0.009948756 | 0.087271131 |
| 0.009947443 | 0.087271131 |
| 0.009933981 | 0.087196042 |
| 0.009927174 | 0.087152356 |
| 0.009924273 | 0.087142952 |

| | |
|---|---|
| 0.009923185 | 0.087142952 |
| 0.009915138 | 0.08709485 |
| 0.00990197 | 0.087027331 |
| 0.0098972 | 0.087001468 |
| 0.009893094 | 0.08698143 |
| 0.009887102 | 0.086944791 |
| 0.009884507 | 0.08693803 |
| 0.009882448 | 0.086935972 |
| 0.009872991 | 0.086900932 |
| 0.009862331 | 0.086823153 |
| 0.009859476 | 0.086814064 |
| 0.009852875 | 0.086771978 |
| 0.009833321 | 0.086663876 |
| 0.00983289 | 0.086663876 |
| 0.009832639 | 0.086663876 |
| 0.009831834 | 0.086663876 |
| 0.009829182 | 0.086663876 |
| 0.009827262 | 0.086663876 |
| 0.009822567 | 0.086663876 |
| 0.009817855 | 0.0866398 |
| 0.009773478 | 0.086344172 |
| 0.009772434 | 0.086344172 |
| 0.00977035 | 0.086344172 |
| 0.009764865 | 0.086316112 |
| 0.009764663 | 0.086316112 |
| 0.009738251 | 0.086112827 |
| 0.009737556 | 0.086112827 |
| 0.009704746 | 0.085864375 |
| 0.009703461 | 0.085864375 |
| 0.00970154 | 0.085864375 |
| 0.009694417 | 0.08582082 |
| 0.009679402 | 0.08570383 |
| 0.009676947 | 0.08570383 |
| 0.009675884 | 0.08570383 |
| 0.009674376 | 0.08570383 |
| 0.009672752 | 0.08570383 |
| 0.009670557 | 0.08570383 |
| 0.009662758 | 0.085683891 |
| 0.009646101 | 0.085552111 |
| 0.00963998 | 0.085516613 |
| 0.00962962 | 0.085453684 |
| 0.009618286 | 0.085384921 |
| 0.009613921 | 0.085384921 |
| 0.009597706 | 0.085283075 |
| 0.0095802 | 0.085157929 |

| | |
|---|---|
| 0.009580093 | 0.085157929 |
| 0.009579313 | 0.085157929 |
| 0.009578564 | 0.085157929 |
| 0.009575797 | 0.085157929 |
| 0.00957048 | 0.085150991 |
| 0.009559457 | 0.085100605 |
| 0.009547967 | 0.085030113 |
| 0.00954372 | 0.08502408 |
| 0.009523503 | 0.084883157 |
| 0.009519716 | 0.084873742 |
| 0.00951622 | 0.084858456 |
| 0.009506558 | 0.084814755 |
| 0.009500702 | 0.084783578 |
| 0.009485892 | 0.084699022 |
| 0.009470303 | 0.084607844 |
| 0.009469344 | 0.084607844 |
| 0.00946585 | 0.084607844 |
| 0.009463332 | 0.084607844 |
| 0.009462884 | 0.084607844 |
| 0.009458605 | 0.084598122 |
| 0.009435825 | 0.08444195 |
| 0.009432636 | 0.084429274 |
| 0.009431264 | 0.084429274 |
| 0.00943042 | 0.084429274 |
| 0.009425461 | 0.084425867 |
| 0.009417674 | 0.084374638 |
| 0.009410519 | 0.084326395 |
| 0.009408822 | 0.084326395 |
| 0.009405192 | 0.084310388 |
| 0.0094033 | 0.084309297 |
| 0.009379962 | 0.084147567 |
| 0.009377153 | 0.084138215 |
| 0.009376919 | 0.084138215 |
| 0.009375263 | 0.084138215 |
| 0.009370105 | 0.08412252 |
| 0.009363331 | 0.084077556 |
| 0.009346416 | 0.083962334 |
| 0.009342692 | 0.083955547 |
| 0.009341182 | 0.083955547 |
| 0.009336489 | 0.083931476 |
| 0.009331469 | 0.083931476 |
| 0.009315948 | 0.083825929 |
| 0.009311072 | 0.08379789 |
| 0.009308702 | 0.083792395 |
| 0.009308523 | 0.083792395 |

| | |
|---|---|
| 0.009303361 | 0.083775982 |
| 0.009302117 | 0.083775982 |
| 0.009296575 | 0.083762391 |
| 0.009294106 | 0.083755985 |
| 0.009291116 | 0.083744889 |
| 0.009290372 | 0.083744889 |
| 0.009274482 | 0.083658287 |
| 0.009264305 | 0.083582315 |
| 0.009262967 | 0.083582315 |
| 0.009261492 | 0.083582315 |
| 0.009259574 | 0.083582315 |
| 0.009258703 | 0.083582315 |
| 0.009257816 | 0.083582315 |
| 0.009257228 | 0.083582315 |
| 0.009253909 | 0.083582315 |
| 0.0092485 | 0.083566361 |
| 0.009243777 | 0.083539542 |
| 0.009234347 | 0.083470158 |
| 0.009231688 | 0.083461957 |
| 0.009227879 | 0.083443364 |
| 0.00922751 | 0.083443364 |
| 0.009216975 | 0.083376425 |
| 0.009213224 | 0.083358322 |
| 0.009198585 | 0.083258711 |
| 0.009194251 | 0.083258711 |
| 0.009191082 | 0.083252885 |
| 0.009173959 | 0.083123454 |
| 0.009172856 | 0.083123454 |
| 0.009172591 | 0.083123454 |
| 0.00917215 | 0.083123454 |
| 0.009165636 | 0.083123454 |
| 0.009157658 | 0.083076533 |
| 0.009138699 | 0.082967747 |
| 0.009135404 | 0.082953644 |
| 0.00912515 | 0.082876337 |
| 0.00912137 | 0.082857807 |
| 0.00911623 | 0.082826917 |
| 0.009090258 | 0.082622462 |
| 0.009084528 | 0.082586147 |
| 0.009080784 | 0.082567869 |
| 0.009078267 | 0.082560746 |
| 0.009072171 | 0.082536827 |
| 0.009068693 | 0.082520951 |
| 0.009067284 | 0.082520951 |
| 0.009066748 | 0.082520951 |

| | |
|---|---|
| 0.009025736 | 0.082177159 |
| 0.009019358 | 0.082134794 |
| 0.009016272 | 0.082122395 |
| 0.009015635 | 0.082122395 |
| 0.009006109 | 0.082061209 |
| 0.008971193 | 0.081788237 |
| 0.008968222 | 0.081778575 |
| 0.00896563 | 0.081770597 |
| 0.008954398 | 0.081699447 |
| 0.008953622 | 0.081699447 |
| 0.008949951 | 0.08169017 |
| 0.008947614 | 0.081684496 |
| 0.008940409 | 0.081634365 |
| 0.008939172 | 0.081634365 |
| 0.008937735 | 0.081634365 |
| 0.008929479 | 0.081597145 |
| 0.008924707 | 0.081569187 |
| 0.008913216 | 0.081479798 |
| 0.00890466 | 0.081448484 |
| 0.00888451 | 0.081326654 |
| 0.008883124 | 0.081326654 |
| 0.008881947 | 0.081326654 |
| 0.008871895 | 0.081258024 |
| 0.008864872 | 0.081224945 |
| 0.008861372 | 0.081208501 |
| 0.008858568 | 0.081208501 |
| 0.008856814 | 0.081208501 |
| 0.008850486 | 0.081186843 |
| 0.008843094 | 0.081150297 |
| 0.00884271 | 0.081150297 |
| 0.008842647 | 0.081150297 |
| 0.008841158 | 0.081150297 |
| 0.008840859 | 0.081150297 |
| 0.008839706 | 0.081150297 |
| 0.00883566 | 0.081150297 |
| 0.008831683 | 0.081150297 |
| 0.008831156 | 0.081150297 |
| 0.008815197 | 0.081113099 |
| 0.008814074 | 0.081113099 |
| 0.008811831 | 0.081113099 |
| 0.008771934 | 0.080777437 |
| 0.00877111 | 0.080777437 |
| 0.008754639 | 0.08064936 |
| 0.008730447 | 0.08045762 |
| 0.008723154 | 0.080405974 |

| | |
|---|---|
| 0.008719314 | 0.08038613 |
| 0.008718201 | 0.08038613 |
| 0.008717171 | 0.08038613 |
| 0.008716868 | 0.08038613 |
| 0.00870045 | 0.080336632 |
| 0.008696605 | 0.080332271 |
| 0.008684355 | 0.080248528 |
| 0.008679609 | 0.080221953 |
| 0.008669837 | 0.080162748 |
| 0.008662489 | 0.080110366 |
| 0.008662483 | 0.080110366 |
| 0.008661911 | 0.080110366 |
| 0.00865892 | 0.080110366 |
| 0.008658124 | 0.080110366 |
| 0.008650909 | 0.080096583 |
| 0.008645868 | 0.080077207 |
| 0.008644986 | 0.080077207 |
| 0.008597222 | 0.079747246 |
| 0.008595849 | 0.079747246 |
| 0.008591307 | 0.079730747 |
| 0.008582016 | 0.079675567 |
| 0.008580402 | 0.079675567 |
| 0.00857532 | 0.079667842 |
| 0.00856321 | 0.079578536 |
| 0.008555123 | 0.079518901 |
| 0.008554196 | 0.079518901 |
| 0.008553448 | 0.079518901 |
| 0.0085498 | 0.079515981 |
| 0.008546955 | 0.079505048 |
| 0.008546395 | 0.079505048 |
| 0.008546012 | 0.079505048 |
| 0.008530106 | 0.079441408 |
| 0.0085177 | 0.079341386 |
| 0.008510316 | 0.07928811 |
| 0.008505942 | 0.07926286 |
| 0.008502167 | 0.079243187 |
| 0.008490917 | 0.079153829 |
| 0.008488465 | 0.079146464 |
| 0.008488218 | 0.079146464 |
| 0.008486791 | 0.079146464 |
| 0.008466542 | 0.079003917 |
| 0.008460641 | 0.078979804 |
| 0.00845097 | 0.078904989 |
| 0.008445038 | 0.07888053 |
| 0.008440339 | 0.07888053 |

| | |
|---|---|
| 0.008434168 | 0.078840846 |
| 0.00842946 | 0.078812311 |
| 0.008427319 | 0.07880776 |
| 0.008424346 | 0.078795426 |
| 0.008417579 | 0.078747598 |
| 0.008413228 | 0.078722356 |
| 0.008411131 | 0.078722356 |
| 0.008410641 | 0.078722356 |
| 0.008393924 | 0.078649892 |
| 0.008393299 | 0.078649892 |
| 0.008393258 | 0.078649892 |
| 0.008391002 | 0.078649892 |
| 0.008383274 | 0.078649892 |
| 0.008368626 | 0.078536463 |
| 0.008362339 | 0.078492932 |
| 0.008355607 | 0.078445202 |
| 0.008355044 | 0.078445202 |
| 0.008351626 | 0.078438748 |
| 0.00834417 | 0.07838418 |
| 0.008326922 | 0.078237587 |
| 0.008321498 | 0.078202048 |
| 0.008321187 | 0.078202048 |
| 0.008318289 | 0.078202048 |
| 0.008311638 | 0.078155654 |
| 0.008309275 | 0.078148861 |
| 0.008293905 | 0.078035131 |
| 0.008283492 | 0.077967961 |
| 0.008269174 | 0.077879369 |
| 0.008269161 | 0.077879369 |
| 0.008264377 | 0.07786498 |
| 0.008260074 | 0.077839837 |
| 0.008236954 | 0.077637327 |
| 0.00822898 | 0.077577519 |
| 0.008223521 | 0.077541404 |
| 0.008220743 | 0.07753056 |
| 0.008218852 | 0.077528077 |
| 0.008214385 | 0.077501293 |
| 0.008206412 | 0.077456757 |
| 0.008201745 | 0.077428052 |
| 0.008201677 | 0.077428052 |
| 0.008200479 | 0.077428052 |
| 0.008194288 | 0.077403677 |
| 0.008192537 | 0.077402488 |
| 0.008165952 | 0.077181939 |
| 0.008150385 | 0.077050096 |

| | |
|---|---|
| 0.008134922 | 0.076919186 |
| 0.008130192 | 0.076889723 |
| 0.008125722 | 0.076862712 |
| 0.008112196 | 0.076750014 |
| 0.008108327 | 0.076728648 |
| 0.008104419 | 0.076706907 |
| 0.008102579 | 0.076704733 |
| 0.008098962 | 0.076700985 |
| 0.008092985 | 0.076659627 |
| 0.008090959 | 0.076655679 |
| 0.008085676 | 0.076636117 |
| 0.008085017 | 0.076636117 |
| 0.008083966 | 0.076636117 |
| 0.008080562 | 0.0766334 |
| 0.008080405 | 0.0766334 |
| 0.00807929 | 0.0766334 |
| 0.008078772 | 0.0766334 |
| 0.008077479 | 0.0766334 |
| 0.008056752 | 0.076511681 |
| 0.008051075 | 0.076490831 |
| 0.008046945 | 0.076466848 |
| 0.00804659 | 0.076466848 |
| 0.008029073 | 0.076327473 |
| 0.008026396 | 0.076327473 |
| 0.008018747 | 0.076312298 |
| 0.008015882 | 0.076308701 |
| 0.008009487 | 0.076278315 |
| 0.008003873 | 0.076255347 |
| 0.007987574 | 0.076115279 |
| 0.007975201 | 0.076043027 |
| 0.007973073 | 0.076037951 |
| 0.007972982 | 0.076037951 |
| 0.007965602 | 0.075997136 |
| 0.007958223 | 0.075957175 |
| 0.007957372 | 0.075957175 |
| 0.00795632 | 0.075957175 |
| 0.007939976 | 0.075843806 |
| 0.007932651 | 0.075804244 |
| 0.007932125 | 0.075804244 |
| 0.007928094 | 0.075791112 |
| 0.007924 | 0.075767181 |
| 0.007922971 | 0.075767181 |
| 0.007904494 | 0.075611034 |
| 0.007894511 | 0.075530711 |
| 0.007888795 | 0.075491185 |

| | |
|---|---|
| 0.007885673 | 0.075476477 |
| 0.007885041 | 0.075476477 |
| 0.007870272 | 0.075359354 |
| 0.007868705 | 0.075359354 |
| 0.007868132 | 0.075359354 |
| 0.007862368 | 0.075359354 |
| 0.007855046 | 0.075304404 |
| 0.007853535 | 0.075304404 |
| 0.007838345 | 0.07517456 |
| 0.007837891 | 0.07517456 |
| 0.007807709 | 0.074926003 |
| 0.007805789 | 0.074922678 |
| 0.007801191 | 0.074901684 |
| 0.007800438 | 0.074901684 |
| 0.007799955 | 0.074901684 |
| 0.007798135 | 0.074901684 |
| 0.007798071 | 0.074901684 |
| 0.007794382 | 0.074901684 |
| 0.007792112 | 0.074901684 |
| 0.007789458 | 0.074901684 |
| 0.007785788 | 0.074901684 |
| 0.007782789 | 0.074898135 |
| 0.007778942 | 0.074898135 |
| 0.007778934 | 0.074898135 |
| 0.007768664 | 0.074837812 |
| 0.00775094 | 0.074682182 |
| 0.007744728 | 0.074637431 |
| 0.007734251 | 0.07455154 |
| 0.007733135 | 0.07455154 |
| 0.007731409 | 0.07455154 |
| 0.007728775 | 0.074544024 |
| 0.007722121 | 0.074494923 |
| 0.007715879 | 0.074449792 |
| 0.007693974 | 0.074298535 |
| 0.007691018 | 0.074285158 |
| 0.007687025 | 0.074261654 |
| 0.007686934 | 0.074261654 |
| 0.007685959 | 0.074261654 |
| 0.007668742 | 0.074160234 |
| 0.007663854 | 0.07413302 |
| 0.007656426 | 0.074092704 |
| 0.007651866 | 0.074087297 |
| 0.00764467 | 0.074047727 |
| 0.007642295 | 0.074039783 |
| 0.007639346 | 0.074026273 |

| | |
|---|---|
| 0.00763515 | 0.074015727 |
| 0.007632047 | 0.074000716 |
| 0.007630169 | 0.073997572 |
| 0.007629008 | 0.073997572 |
| 0.007623916 | 0.073967049 |
| 0.007623645 | 0.073967049 |
| 0.007619955 | 0.073958748 |
| 0.007619381 | 0.073958748 |
| 0.007603884 | 0.073832864 |
| 0.007602025 | 0.073829863 |
| 0.007599426 | 0.073819674 |
| 0.007599401 | 0.073819674 |
| 0.007588292 | 0.073741603 |
| 0.007584622 | 0.073720983 |
| 0.00757856 | 0.073677096 |
| 0.00757783 | 0.073677096 |
| 0.007576913 | 0.073677096 |
| 0.007572395 | 0.073662269 |
| 0.007565313 | 0.073608411 |
| 0.007562969 | 0.07360064 |
| 0.007557469 | 0.073562142 |
| 0.00755472 | 0.073550422 |
| 0.007551182 | 0.073550422 |
| 0.007545668 | 0.073522403 |
| 0.007534743 | 0.073430977 |
| 0.007534483 | 0.073430977 |
| 0.007533052 | 0.073430977 |
| 0.00752299 | 0.073361478 |
| 0.007519837 | 0.073345751 |
| 0.00749804 | 0.073178104 |
| 0.007495017 | 0.073177997 |
| 0.007479533 | 0.073084689 |
| 0.007479325 | 0.073084689 |
| 0.007464224 | 0.072967693 |
| 0.007463453 | 0.072967693 |
| 0.007457265 | 0.072929599 |
| 0.007453976 | 0.07291459 |
| 0.007442965 | 0.072864564 |
| 0.007442879 | 0.072864564 |
| 0.007434288 | 0.072809545 |
| 0.007421174 | 0.072755924 |
| 0.007414767 | 0.072723044 |
| 0.007410167 | 0.072692896 |
| 0.007408979 | 0.072692896 |
| 0.0074057 | 0.07267902 |

| | |
|---|---|
| 0.00740191 | 0.072671775 |
| 0.007397214 | 0.072653709 |
| 0.007387932 | 0.072579421 |
| 0.007387384 | 0.072579421 |
| 0.007386711 | 0.072579421 |
| 0.007381458 | 0.072579421 |
| 0.007376347 | 0.072555408 |
| 0.007375209 | 0.072555408 |
| 0.007371072 | 0.072548475 |
| 0.007361614 | 0.072485342 |
| 0.007354246 | 0.072427761 |
| 0.007352676 | 0.07242728 |
| 0.00734698 | 0.072386143 |
| 0.007342433 | 0.072356314 |
| 0.007337323 | 0.072320915 |
| 0.007329323 | 0.072271979 |
| 0.007325342 | 0.072247685 |
| 0.00732221 | 0.072231751 |
| 0.007321312 | 0.072231751 |
| 0.007305116 | 0.07210793 |
| 0.00730003 | 0.072072665 |
| 0.007294922 | 0.072037167 |
| 0.007291566 | 0.072018961 |
| 0.00728807 | 0.072014298 |
| 0.007286103 | 0.072009804 |
| 0.007282668 | 0.071990795 |
| 0.007282007 | 0.071990795 |
| 0.007281295 | 0.071990795 |
| 0.007277891 | 0.071988394 |
| 0.007273063 | 0.071970532 |
| 0.007261714 | 0.071880377 |
| 0.007261267 | 0.071880377 |
| 0.007257453 | 0.071875798 |
| 0.00725624 | 0.071875798 |
| 0.007253145 | 0.071875798 |
| 0.007250655 | 0.071868247 |
| 0.007248068 | 0.071857563 |
| 0.007239872 | 0.071791256 |
| 0.007239366 | 0.071791256 |
| 0.00723936 | 0.071791256 |
| 0.007232994 | 0.071767872 |
| 0.007227627 | 0.071729567 |
| 0.007220928 | 0.071695559 |
| 0.007215514 | 0.071695559 |
| 0.007214653 | 0.071695559 |

| | |
|---|---|
| 0.007212454 | 0.071695559 |
| 0.007193868 | 0.071558552 |
| 0.007187527 | 0.071549942 |
| 0.007170616 | 0.071416754 |
| 0.007167078 | 0.071413113 |
| 0.007143713 | 0.071253111 |
| 0.007141751 | 0.07124847 |
| 0.007130374 | 0.071164783 |
| 0.007129153 | 0.071164783 |
| 0.007128791 | 0.071164783 |
| 0.007121123 | 0.071132075 |
| 0.007120619 | 0.071132075 |
| 0.007118788 | 0.071132075 |
| 0.007118026 | 0.071132075 |
| 0.007105754 | 0.071038149 |
| 0.007102574 | 0.071021264 |
| 0.007101181 | 0.071021264 |
| 0.007100378 | 0.071021264 |
| 0.007096802 | 0.071021264 |
| 0.007095103 | 0.071021112 |
| 0.007087572 | 0.070960649 |
| 0.007082721 | 0.070926988 |
| 0.007078141 | 0.070896026 |
| 0.007072937 | 0.070858812 |
| 0.007057804 | 0.070736955 |
| 0.007051478 | 0.070703312 |
| 0.00703624 | 0.070580248 |
| 0.007032591 | 0.070558499 |
| 0.007026763 | 0.070529749 |
| 0.007025524 | 0.070529749 |
| 0.007020724 | 0.070511798 |
| 0.007015178 | 0.070502609 |
| 0.007005695 | 0.070422171 |
| 0.006999924 | 0.07039387 |
| 0.006987077 | 0.070294363 |
| 0.00697772 | 0.070225379 |
| 0.006976741 | 0.070225379 |
| 0.006973736 | 0.070225379 |
| 0.006962208 | 0.070147894 |
| 0.006946681 | 0.070006261 |
| 0.00694488 | 0.070002923 |
| 0.006939914 | 0.069967672 |
| 0.00693359 | 0.069918717 |
| 0.00693035 | 0.069900838 |
| 0.006918189 | 0.069807752 |

| | |
|---|---|
| 0.006917012 | 0.069807752 |
| 0.006910787 | 0.069777408 |
| 0.006907322 | 0.069757217 |
| 0.006904824 | 0.069746773 |
| 0.006903571 | 0.069746773 |
| 0.006880287 | 0.069587472 |
| 0.006878592 | 0.06958511 |
| 0.006878092 | 0.06958511 |
| 0.006875886 | 0.06958511 |
| 0.006872947 | 0.06957234 |
| 0.006859896 | 0.069499315 |
| 0.006850985 | 0.069453353 |
| 0.006845595 | 0.069413489 |
| 0.006840352 | 0.069375097 |
| 0.006839651 | 0.069375097 |
| 0.00683378 | 0.069353444 |
| 0.006831283 | 0.069353444 |
| 0.0068257 | 0.069320949 |
| 0.006825641 | 0.069320949 |
| 0.00681815 | 0.069282702 |
| 0.006810301 | 0.069217718 |
| 0.006790875 | 0.069064494 |
| 0.006789314 | 0.069063361 |
| 0.006782918 | 0.069013038 |
| 0.006779187 | 0.06900456 |
| 0.006758944 | 0.068827926 |
| 0.006753295 | 0.06878511 |
| 0.006751684 | 0.068783415 |
| 0.006747357 | 0.068754042 |
| 0.006746046 | 0.068754042 |
| 0.006741532 | 0.068724092 |
| 0.00672322 | 0.068566764 |
| 0.006723076 | 0.068566764 |
| 0.006722663 | 0.068566764 |
| 0.006718297 | 0.068560601 |
| 0.006716703 | 0.068559031 |
| 0.006706269 | 0.068467192 |
| 0.006699826 | 0.068416084 |
| 0.006699091 | 0.068416084 |
| 0.006692411 | 0.068369677 |
| 0.00669201 | 0.068369677 |
| 0.006684914 | 0.068337065 |
| 0.00668293 | 0.068331451 |
| 0.006681076 | 0.068327164 |
| 0.006675642 | 0.068286253 |

| | |
|---|---|
| 0.006675476 | 0.068286253 |
| 0.006674858 | 0.068286253 |
| 0.006666263 | 0.06824895 |
| 0.006663109 | 0.068232574 |
| 0.006662893 | 0.068232574 |
| 0.006657526 | 0.068218153 |
| 0.00665637 | 0.068218153 |
| 0.006656151 | 0.068218153 |
| 0.006655671 | 0.068218153 |
| 0.006644967 | 0.068162796 |
| 0.006643785 | 0.068162796 |
| 0.006630622 | 0.068044951 |
| 0.006630096 | 0.068044951 |
| 0.006627211 | 0.068039274 |
| 0.00662248 | 0.068005357 |
| 0.006618077 | 0.067974801 |
| 0.006603321 | 0.067867145 |
| 0.00660323 | 0.067867145 |
| 0.006600002 | 0.067862328 |
| 0.006597703 | 0.067853337 |
| 0.006582643 | 0.067742342 |
| 0.006567702 | 0.067617802 |
| 0.00656185 | 0.06758677 |
| 0.006556466 | 0.067555434 |
| 0.006555741 | 0.067555434 |
| 0.006553583 | 0.067555434 |
| 0.006551906 | 0.067555434 |
| 0.006546879 | 0.067534805 |
| 0.006544861 | 0.067528617 |
| 0.006542779 | 0.067521759 |
| 0.006527627 | 0.067394601 |
| 0.006525034 | 0.067382439 |
| 0.006522848 | 0.067374469 |
| 0.006518997 | 0.067374469 |
| 0.006515826 | 0.067374469 |
| 0.006514662 | 0.067374469 |
| 0.006507922 | 0.067329339 |
| 0.006505938 | 0.067329339 |
| 0.006504916 | 0.067329339 |
| 0.006502538 | 0.0673253 |
| 0.006498369 | 0.067297551 |
| 0.006487326 | 0.067211631 |
| 0.006484276 | 0.067194652 |
| 0.006480065 | 0.067165626 |
| 0.00647821 | 0.06716102 |

| | |
|---|---|
| 0.006470177 | 0.067120852 |
| 0.006468971 | 0.067120852 |
| 0.006466023 | 0.06710927 |
| 0.006457153 | 0.06705949 |
| 0.006455036 | 0.067052131 |
| 0.006453662 | 0.067052131 |
| 0.006450072 | 0.06704443 |
| 0.00643651 | 0.066918063 |
| 0.006433553 | 0.06690193 |
| 0.006428188 | 0.066860739 |
| 0.006427751 | 0.066860739 |
| 0.006403471 | 0.066647318 |
| 0.006402746 | 0.066647318 |
| 0.006397287 | 0.066612061 |
| 0.00639213 | 0.06658748 |
| 0.006388363 | 0.066577367 |
| 0.006386086 | 0.066568203 |
| 0.006372995 | 0.066446282 |
| 0.006371951 | 0.066446282 |
| 0.006370857 | 0.066446282 |
| 0.00636987 | 0.066446282 |
| 0.006369584 | 0.066446282 |
| 0.006364912 | 0.066434733 |
| 0.00635006 | 0.066308771 |
| 0.006347647 | 0.066298111 |
| 0.006327224 | 0.066113796 |
| 0.006321964 | 0.066073334 |
| 0.006315628 | 0.066021599 |
| 0.006301938 | 0.065907419 |
| 0.006301559 | 0.065907419 |
| 0.006292297 | 0.065876392 |
| 0.006286554 | 0.065833267 |
| 0.006267175 | 0.065688107 |
| 0.006266422 | 0.065688107 |
| 0.006256247 | 0.065631333 |
| 0.006253768 | 0.065619781 |
| 0.006248496 | 0.065578917 |
| 0.006240016 | 0.065504357 |
| 0.006237276 | 0.065490027 |
| 0.006236344 | 0.065490027 |
| 0.006235607 | 0.065490027 |
| 0.006233616 | 0.065490027 |
| 0.006223266 | 0.065415023 |
| 0.006219639 | 0.065391333 |
| 0.006211197 | 0.06533142 |

| | |
|---|---|
| 0.00620959 | 0.06533142 |
| 0.006204031 | 0.065299316 |
| 0.006185998 | 0.0651383 |
| 0.006185328 | 0.0651383 |
| 0.006183435 | 0.0651383 |
| 0.00618328 | 0.0651383 |
| 0.006176135 | 0.065092007 |
| 0.006168775 | 0.065028829 |
| 0.006160283 | 0.064968066 |
| 0.006158841 | 0.064967246 |
| 0.00615848 | 0.064967246 |
| 0.006158346 | 0.064967246 |
| 0.006156998 | 0.064967246 |
| 0.006152953 | 0.064962691 |
| 0.006146772 | 0.064911824 |
| 0.006135102 | 0.064846091 |
| 0.006133976 | 0.064846091 |
| 0.006126675 | 0.064785775 |
| 0.006116668 | 0.064708699 |
| 0.006114274 | 0.064697739 |
| 0.006114084 | 0.064697739 |
| 0.006112219 | 0.064697739 |
| 0.006111482 | 0.064697739 |
| 0.00610589 | 0.064666492 |
| 0.006095565 | 0.064571506 |
| 0.006094078 | 0.064570111 |
| 0.00609295 | 0.064570111 |
| 0.00609263 | 0.064570111 |
| 0.006085138 | 0.064528063 |
| 0.006074805 | 0.064437579 |
| 0.006067382 | 0.06438752 |
| 0.006065982 | 0.064387015 |
| 0.00605489 | 0.064283601 |
| 0.006050669 | 0.064281777 |
| 0.006046184 | 0.064248464 |
| 0.006042649 | 0.064235141 |
| 0.006041243 | 0.064235141 |
| 0.006041233 | 0.064235141 |
| 0.006024695 | 0.064105934 |
| 0.006024171 | 0.064105934 |
| 0.006007324 | 0.063978266 |
| 0.005997768 | 0.063905082 |
| 0.005992538 | 0.063863637 |
| 0.005986773 | 0.063816482 |
| 0.005981926 | 0.063779084 |

| | |
|---|---|
| 0.005968968 | 0.063697973 |
| 0.005960754 | 0.063632561 |
| 0.005959114 | 0.063632561 |
| 0.005954953 | 0.063619681 |
| 0.005953288 | 0.063616169 |
| 0.005943081 | 0.063549865 |
| 0.005933063 | 0.063471252 |
| 0.005926385 | 0.063414054 |
| 0.005925769 | 0.063414054 |
| 0.005924857 | 0.063414054 |
| 0.005922865 | 0.063414054 |
| 0.005921851 | 0.063414054 |
| 0.005920678 | 0.063414054 |
| 0.005919992 | 0.063414054 |
| 0.005916313 | 0.06340599 |
| 0.005915124 | 0.06340599 |
| 0.005902921 | 0.063290957 |
| 0.005889089 | 0.063156863 |
| 0.005887526 | 0.063154326 |
| 0.005886566 | 0.063154326 |
| 0.005883832 | 0.063143141 |
| 0.005881862 | 0.063136231 |
| 0.005870885 | 0.063032603 |
| 0.005858366 | 0.062969168 |
| 0.005850491 | 0.06289872 |
| 0.005847288 | 0.062892677 |
| 0.005841575 | 0.062859619 |
| 0.005840286 | 0.062859619 |
| 0.005835354 | 0.062821072 |
| 0.005831853 | 0.062797573 |
| 0.005830255 | 0.062794565 |
| 0.005811057 | 0.062601951 |
| 0.005799725 | 0.062508147 |
| 0.005785848 | 0.062372704 |
| 0.005778737 | 0.06232425 |
| 0.005765948 | 0.062214496 |
| 0.005761646 | 0.062182166 |
| 0.005741635 | 0.062013392 |
| 0.005737805 | 0.061995114 |
| 0.005737405 | 0.061995114 |
| 0.005730964 | 0.061963884 |
| 0.005711755 | 0.06179776 |
| 0.005707978 | 0.061782568 |
| 0.005707322 | 0.061782568 |
| 0.005703093 | 0.061774208 |

| | |
|---|---|
| 0.005699442 | 0.061748709 |
| 0.005693353 | 0.061696771 |
| 0.00568696 | 0.061655548 |
| 0.005683357 | 0.061642151 |
| 0.005680611 | 0.061628805 |
| 0.005679977 | 0.061628805 |
| 0.005679526 | 0.061628805 |
| 0.005677468 | 0.061628805 |
| 0.005671766 | 0.061603014 |
| 0.005671725 | 0.061603014 |
| 0.005660956 | 0.061541733 |
| 0.005656137 | 0.061503391 |
| 0.005649583 | 0.06146019 |
| 0.005647173 | 0.061448006 |
| 0.005637827 | 0.061374351 |
| 0.005634699 | 0.061354322 |
| 0.005634683 | 0.061354322 |
| 0.005630784 | 0.061353772 |
| 0.005628158 | 0.061339196 |
| 0.00562582 | 0.061327752 |
| 0.005616967 | 0.061259287 |
| 0.005613044 | 0.061230523 |
| 0.005610287 | 0.061214467 |
| 0.005597353 | 0.061157406 |
| 0.005597198 | 0.061157406 |
| 0.005595189 | 0.061157406 |
| 0.005594789 | 0.061157406 |
| 0.005592543 | 0.061157406 |
| 0.005591715 | 0.061157406 |
| 0.005589029 | 0.061150624 |
| 0.005587086 | 0.061143419 |
| 0.005573696 | 0.061024924 |
| 0.005568201 | 0.060978782 |
| 0.005564845 | 0.06095604 |
| 0.005560053 | 0.060917561 |
| 0.005558963 | 0.060917561 |
| 0.005555738 | 0.060898309 |
| 0.005543679 | 0.060794111 |
| 0.005541811 | 0.060787618 |
| 0.005534654 | 0.060737089 |
| 0.005534275 | 0.060737089 |
| 0.005531202 | 0.060734478 |
| 0.005529435 | 0.060734478 |
| 0.005529279 | 0.060734478 |
| 0.005526576 | 0.060732406 |

| | |
|---|---|
| 0.005524454 | 0.060723095 |
| 0.00552171 | 0.060723095 |
| 0.005521133 | 0.060723095 |
| 0.005516449 | 0.06071919 |
| 0.005513298 | 0.060698531 |
| 0.005509291 | 0.060668443 |
| 0.00550674 | 0.060654367 |
| 0.005503798 | 0.06065002 |
| 0.00549566 | 0.060588381 |
| 0.005493058 | 0.060573718 |
| 0.005492413 | 0.060573718 |
| 0.005489622 | 0.060573718 |
| 0.005489485 | 0.060573718 |
| 0.005483302 | 0.060536215 |
| 0.005473496 | 0.060441969 |
| 0.005443812 | 0.060156018 |
| 0.005439661 | 0.060124101 |
| 0.005439643 | 0.060124101 |
| 0.005425616 | 0.06001308 |
| 0.005412966 | 0.059926391 |
| 0.005411823 | 0.059926391 |
| 0.005410636 | 0.059926391 |
| 0.005410403 | 0.059926391 |
| 0.005406232 | 0.059907538 |
| 0.005402217 | 0.059876981 |
| 0.005396992 | 0.059832994 |
| 0.005396558 | 0.059832994 |
| 0.005392582 | 0.059811945 |
| 0.005381152 | 0.059712987 |
| 0.005374996 | 0.059672483 |
| 0.005373594 | 0.059670825 |
| 0.005364724 | 0.059600123 |
| 0.005357712 | 0.059550004 |
| 0.00535566 | 0.059541086 |
| 0.005348318 | 0.059473349 |
| 0.005344302 | 0.059456451 |
| 0.00534316 | 0.059456451 |
| 0.005336784 | 0.059400567 |
| 0.005333237 | 0.059388843 |
| 0.005332706 | 0.059388843 |
| 0.005325598 | 0.059331522 |
| 0.005323213 | 0.059321246 |
| 0.005316899 | 0.059282096 |
| 0.005313288 | 0.059265101 |
| 0.005309731 | 0.059265101 |

| | |
|---|---|
| 0.005305796 | 0.059235614 |
| 0.005300085 | 0.059185738 |
| 0.005298556 | 0.059182539 |
| 0.005294987 | 0.059156546 |
| 0.005290315 | 0.059139925 |
| 0.005287748 | 0.059131147 |
| 0.005270744 | 0.059010274 |
| 0.005263489 | 0.058942911 |
| 0.005252977 | 0.058883568 |
| 0.005252897 | 0.058883568 |
| 0.005249829 | 0.058883568 |
| 0.005230318 | 0.058737176 |
| 0.005230142 | 0.058737176 |
| 0.005229511 | 0.058737176 |
| 0.005225398 | 0.058723459 |
| 0.005224323 | 0.058723459 |
| 0.005210718 | 0.05865537 |
| 0.005208477 | 0.05865537 |
| 0.005198648 | 0.058561019 |
| 0.005197587 | 0.058561019 |
| 0.005197473 | 0.058561019 |
| 0.005192834 | 0.058548811 |
| 0.005182828 | 0.05845194 |
| 0.005177083 | 0.058418527 |
| 0.005175095 | 0.058418527 |
| 0.005173994 | 0.058418527 |
| 0.005170772 | 0.058408969 |
| 0.005165701 | 0.058369353 |
| 0.005160334 | 0.058322555 |
| 0.005159404 | 0.058322555 |
| 0.005156859 | 0.058310947 |
| 0.005155581 | 0.058310346 |
| 0.005148295 | 0.058255602 |
| 0.005147532 | 0.058255602 |
| 0.005134571 | 0.058141754 |
| 0.005126202 | 0.058060785 |
| 0.005121289 | 0.058018946 |
| 0.005120959 | 0.058018946 |
| 0.005111757 | 0.05793852 |
| 0.00511147 | 0.05793852 |
| 0.005100902 | 0.057856786 |
| 0.005100285 | 0.057856786 |
| 0.005094445 | 0.057811084 |
| 0.005087327 | 0.057757834 |
| 0.005087253 | 0.057757834 |

| | |
|---|---|
| 0.005076985 | 0.057681675 |
| 0.005073672 | 0.057657786 |
| 0.005072237 | 0.057655231 |
| 0.005065646 | 0.057594062 |
| 0.005064582 | 0.057594062 |
| 0.005043605 | 0.057425721 |
| 0.005043485 | 0.057425721 |
| 0.00504327 | 0.057425721 |
| 0.005035594 | 0.057375656 |
| 0.005034709 | 0.057375656 |
| 0.005017628 | 0.057231365 |
| 0.005016016 | 0.057231365 |
| 0.005010748 | 0.057202033 |
| 0.005009603 | 0.057202033 |
| 0.005006304 | 0.05717871 |
| 0.004999105 | 0.057110192 |
| 0.004995864 | 0.057086851 |
| 0.004992007 | 0.057056466 |
| 0.00498962 | 0.057042881 |
| 0.004975406 | 0.056894033 |
| 0.004974572 | 0.056894033 |
| 0.004968436 | 0.056868207 |
| 0.004948183 | 0.056691637 |
| 0.004944025 | 0.056671261 |
| 0.004927028 | 0.056490026 |
| 0.004914577 | 0.056371348 |
| 0.004912704 | 0.056366502 |
| 0.004907862 | 0.056338094 |
| 0.004907289 | 0.056338094 |
| 0.004902289 | 0.056328429 |
| 0.004898388 | 0.056297184 |
| 0.004896784 | 0.056292331 |
| 0.004870719 | 0.056033271 |
| 0.004868982 | 0.056026816 |
| 0.004867715 | 0.05602578 |
| 0.004855442 | 0.055898023 |
| 0.004855175 | 0.055898023 |
| 0.004851445 | 0.055879029 |
| 0.004850902 | 0.055879029 |
| 0.004846074 | 0.055844177 |
| 0.004842856 | 0.055820596 |
| 0.004835771 | 0.055765922 |
| 0.004832567 | 0.055742472 |
| 0.004832352 | 0.055742472 |
| 0.004830862 | 0.055742472 |

| | |
|---|---|
| 0.00482614 | 0.055708817 |
| 0.004824203 | 0.055699959 |
| 0.004817608 | 0.055637297 |
| 0.004810818 | 0.055572356 |
| 0.004801213 | 0.055492918 |
| 0.004795552 | 0.055449805 |
| 0.004792187 | 0.055424359 |
| 0.004788172 | 0.055404822 |
| 0.00478464 | 0.055377413 |
| 0.004781257 | 0.055351707 |
| 0.004779354 | 0.055343136 |
| 0.004775101 | 0.055307334 |
| 0.004773409 | 0.055301182 |
| 0.004772048 | 0.055298869 |
| 0.004771868 | 0.055298869 |
| 0.004765131 | 0.055245587 |
| 0.004764085 | 0.055245587 |
| 0.004763456 | 0.055245587 |
| 0.00476339 | 0.055245587 |
| 0.004763251 | 0.055245587 |
| 0.004763158 | 0.055245587 |
| 0.004761207 | 0.055245587 |
| 0.004758287 | 0.055245587 |
| 0.004757438 | 0.055245587 |
| 0.004756977 | 0.055245587 |
| 0.004749317 | 0.055210087 |
| 0.004735676 | 0.055118779 |
| 0.004732654 | 0.055097066 |
| 0.004732374 | 0.055097066 |
| 0.004731453 | 0.055097066 |
| 0.004727982 | 0.05509655 |
| 0.004726643 | 0.055094428 |
| 0.004724089 | 0.055078137 |
| 0.004717298 | 0.055012431 |
| 0.004713473 | 0.054981286 |
| 0.004708039 | 0.054931355 |
| 0.00469436 | 0.054812028 |
| 0.004693207 | 0.054812006 |
| 0.004685084 | 0.054730559 |
| 0.004681251 | 0.054699201 |
| 0.004680141 | 0.054699201 |
| 0.0046726 | 0.054665171 |
| 0.004662339 | 0.054571942 |
| 0.004658615 | 0.054541752 |
| 0.004655269 | 0.054529391 |

| | |
|---|---|
| 0.004653181 | 0.054518342 |
| 0.00464725 | 0.054462248 |
| 0.004639447 | 0.054384182 |
| 0.004636727 | 0.054379067 |
| 0.004629696 | 0.054309987 |
| 0.004626377 | 0.054284424 |
| 0.004626183 | 0.054284424 |
| 0.004618753 | 0.054221676 |
| 0.004606992 | 0.054096949 |
| 0.004592634 | 0.053974398 |
| 0.004590382 | 0.053974398 |
| 0.004580138 | 0.053874589 |
| 0.004578754 | 0.053871622 |
| 0.004576341 | 0.053856535 |
| 0.004568875 | 0.053795258 |
| 0.004568812 | 0.053795258 |
| 0.004567279 | 0.053795258 |
| 0.004564668 | 0.053785608 |
| 0.004561359 | 0.053762987 |
| 0.004560653 | 0.053762987 |
| 0.004556393 | 0.053741293 |
| 0.00455497 | 0.05373781 |
| 0.004550514 | 0.053711849 |
| 0.004549992 | 0.053711849 |
| 0.004549723 | 0.053711849 |
| 0.004543247 | 0.053679295 |
| 0.004538693 | 0.053652108 |
| 0.004514471 | 0.053418816 |
| 0.004511592 | 0.053398011 |
| 0.004510091 | 0.053393515 |
| 0.00450992 | 0.053393515 |
| 0.004500512 | 0.053306608 |
| 0.004494588 | 0.053249684 |
| 0.004489629 | 0.053204175 |
| 0.004489361 | 0.053204175 |
| 0.004486949 | 0.053204175 |
| 0.004483264 | 0.053181672 |
| 0.004481975 | 0.053179627 |
| 0.004479836 | 0.053167489 |
| 0.004478314 | 0.053167489 |
| 0.004455313 | 0.05295563 |
| 0.004451462 | 0.052923057 |
| 0.004449734 | 0.052923057 |
| 0.004449011 | 0.052923057 |
| 0.004445151 | 0.052900846 |

| | |
|---|---|
| 0.004439652 | 0.052848613 |
| 0.004429948 | 0.052779627 |
| 0.004424092 | 0.052729285 |
| 0.004418982 | 0.052681563 |
| 0.004407827 | 0.052574899 |
| 0.004406289 | 0.052574171 |
| 0.004405254 | 0.052574171 |
| 0.004401205 | 0.052548553 |
| 0.004399381 | 0.052539948 |
| 0.004398446 | 0.052539948 |
| 0.004397156 | 0.052539732 |
| 0.004386302 | 0.052449516 |
| 0.004383062 | 0.052423931 |
| 0.004382804 | 0.052423931 |
| 0.004370486 | 0.052312926 |
| 0.004370312 | 0.052312926 |
| 0.004369322 | 0.052312926 |
| 0.004368892 | 0.052312926 |
| 0.004368227 | 0.052312926 |
| 0.004359299 | 0.052257822 |
| 0.004358047 | 0.052257822 |
| 0.004357065 | 0.052257822 |
| 0.004356841 | 0.052257822 |
| 0.004354491 | 0.052257822 |
| 0.004352682 | 0.052257419 |
| 0.004343063 | 0.052168234 |
| 0.004342131 | 0.052168234 |
| 0.004339164 | 0.052147711 |
| 0.004338402 | 0.052147711 |
| 0.004326392 | 0.0520336 |
| 0.004323335 | 0.052009965 |
| 0.004320423 | 0.051988066 |
| 0.00431877 | 0.051981314 |
| 0.004308419 | 0.05188295 |
| 0.004296587 | 0.051779742 |
| 0.004295988 | 0.051779742 |
| 0.004295763 | 0.051779742 |
| 0.004294144 | 0.051779742 |
| 0.004290615 | 0.051760161 |
| 0.004284992 | 0.051705426 |
| 0.004283923 | 0.051705426 |
| 0.004281081 | 0.051684428 |
| 0.004280948 | 0.051684428 |
| 0.00427962 | 0.051684428 |
| 0.00427796 | 0.051684428 |

| | |
|---|---|
| 0.004273966 | 0.051650897 |
| 0.004271804 | 0.051637869 |
| 0.004259029 | 0.05153577 |
| 0.004257452 | 0.051529779 |
| 0.004256328 | 0.051529779 |
| 0.004252289 | 0.051506553 |
| 0.004249103 | 0.051481051 |
| 0.004241362 | 0.051413421 |
| 0.004235716 | 0.051358054 |
| 0.004224954 | 0.051240613 |
| 0.004219858 | 0.051204887 |
| 0.004217823 | 0.051204887 |
| 0.004214523 | 0.051192323 |
| 0.004207341 | 0.051118119 |
| 0.004206844 | 0.051118119 |
| 0.004205954 | 0.051118119 |
| 0.004202593 | 0.051112585 |
| 0.004199395 | 0.051086741 |
| 0.00419909 | 0.051086741 |
| 0.004198463 | 0.051086741 |
| 0.004195156 | 0.051074309 |
| 0.004185568 | 0.05098364 |
| 0.004180491 | 0.050947858 |
| 0.004178052 | 0.050931172 |
| 0.004174897 | 0.05090574 |
| 0.004167769 | 0.050844854 |
| 0.004154939 | 0.05071432 |
| 0.00414911 | 0.050656159 |
| 0.004147694 | 0.050656159 |
| 0.004144431 | 0.050640597 |
| 0.004131239 | 0.050515669 |
| 0.00412582 | 0.050475328 |
| 0.00411842 | 0.050423642 |
| 0.004118294 | 0.050423642 |
| 0.00411702 | 0.050423642 |
| 0.004112883 | 0.050394724 |
| 0.004097829 | 0.050236112 |
| 0.00409421 | 0.050204669 |
| 0.004084753 | 0.0501016 |
| 0.004084459 | 0.0501016 |
| 0.004069697 | 0.049955529 |
| 0.004069348 | 0.049955529 |
| 0.004061482 | 0.049906146 |
| 0.004059291 | 0.049892087 |
| 0.004045345 | 0.04974636 |

| | |
|---|---|
| 0.004042253 | 0.049721171 |
| 0.004034458 | 0.049650933 |
| 0.004025494 | 0.049591882 |
| 0.004019198 | 0.04952713 |
| 0.004018864 | 0.04952713 |
| 0.003991047 | 0.049256698 |
| 0.003989789 | 0.04925393 |
| 0.003987943 | 0.049243904 |
| 0.003985059 | 0.049221064 |
| 0.003978738 | 0.04915573 |
| 0.003974013 | 0.049122838 |
| 0.003959108 | 0.048951305 |
| 0.003955231 | 0.048916076 |
| 0.003954824 | 0.048916076 |
| 0.003954796 | 0.048916076 |
| 0.003951073 | 0.048902749 |
| 0.003942143 | 0.048817588 |
| 0.003939901 | 0.04880251 |
| 0.003934531 | 0.048761361 |
| 0.003932185 | 0.048744973 |
| 0.003929366 | 0.048722705 |
| 0.003923304 | 0.048710943 |
| 0.003921392 | 0.048708752 |
| 0.00391972 | 0.048704537 |
| 0.003918392 | 0.04870075 |
| 0.0039132 | 0.048648905 |
| 0.003910859 | 0.048632503 |
| 0.003907644 | 0.048605216 |
| 0.003906449 | 0.048603048 |
| 0.003898468 | 0.048516422 |
| 0.003872469 | 0.048205462 |
| 0.003870609 | 0.048194897 |
| 0.003867211 | 0.048177786 |
| 0.003865328 | 0.048166927 |
| 0.003862094 | 0.048139213 |
| 0.003861516 | 0.048139213 |
| 0.003855963 | 0.04808797 |
| 0.003855366 | 0.04808797 |
| 0.003855192 | 0.04808797 |
| 0.003845204 | 0.04801666 |
| 0.003836966 | 0.047938924 |
| 0.003834019 | 0.047914672 |
| 0.003832941 | 0.047913779 |
| 0.003825008 | 0.047827157 |
| 0.003824205 | 0.047827157 |

| | |
|---|---|
| 0.003823414 | 0.047827157 |
| 0.00381976 | 0.047811751 |
| 0.00381639 | 0.047782126 |
| 0.003813079 | 0.047765788 |
| 0.003809411 | 0.047732392 |
| 0.003804963 | 0.047689205 |
| 0.003802596 | 0.047672081 |
| 0.003787634 | 0.047534561 |
| 0.003783607 | 0.047496532 |
| 0.003781253 | 0.047479501 |
| 0.003780597 | 0.047479501 |
| 0.003776437 | 0.047456566 |
| 0.003776345 | 0.047456566 |
| 0.003773493 | 0.047444599 |
| 0.003773274 | 0.047444599 |
| 0.003771529 | 0.047444599 |
| 0.003768983 | 0.047428865 |
| 0.003768487 | 0.047428865 |
| 0.003761898 | 0.047376589 |
| 0.003757599 | 0.047357279 |
| 0.003757421 | 0.047357279 |
| 0.003756999 | 0.047357279 |
| 0.003751362 | 0.047341318 |
| 0.003750435 | 0.047341318 |
| 0.003749618 | 0.047341318 |
| 0.003739553 | 0.047242354 |
| 0.003734675 | 0.047205779 |
| 0.003734436 | 0.047205779 |
| 0.003728286 | 0.047162564 |
| 0.00372517 | 0.047135669 |
| 0.003721562 | 0.047102532 |
| 0.003712351 | 0.047010936 |
| 0.003711559 | 0.047010936 |
| 0.003706492 | 0.046961715 |
| 0.003706043 | 0.046961715 |
| 0.003694533 | 0.046860059 |
| 0.003693811 | 0.046860059 |
| 0.003693702 | 0.046860059 |
| 0.003692774 | 0.046860059 |
| 0.003692692 | 0.046860059 |
| 0.003690082 | 0.046860059 |
| 0.003687946 | 0.046851381 |
| 0.003687257 | 0.046851381 |
| 0.003676844 | 0.046735264 |
| 0.003664124 | 0.046598465 |

| | |
|---|---|
| 0.00365854 | 0.046539867 |
| 0.003655142 | 0.046509064 |
| 0.003654871 | 0.046509064 |
| 0.003653226 | 0.046509064 |
| 0.003650691 | 0.04648969 |
| 0.003644891 | 0.046428249 |
| 0.003644247 | 0.046428249 |
| 0.00364218 | 0.046428249 |
| 0.003640573 | 0.046428249 |
| 0.003629777 | 0.046310046 |
| 0.003619284 | 0.046188543 |
| 0.003616843 | 0.046169762 |
| 0.003615656 | 0.046166982 |
| 0.003613982 | 0.046157989 |
| 0.003611083 | 0.046133324 |
| 0.003606873 | 0.046093254 |
| 0.003603151 | 0.046081438 |
| 0.00359972 | 0.0460623 |
| 0.003599135 | 0.0460623 |
| 0.003597111 | 0.046053661 |
| 0.003594392 | 0.046031216 |
| 0.00359137 | 0.046004887 |
| 0.003589275 | 0.045990428 |
| 0.003586451 | 0.045966598 |
| 0.003583651 | 0.045943078 |
| 0.003581851 | 0.045932365 |
| 0.003581356 | 0.045932365 |
| 0.003578293 | 0.045911461 |
| 0.003575044 | 0.045894501 |
| 0.003573719 | 0.04588986 |
| 0.003568657 | 0.045837221 |
| 0.003567204 | 0.045830925 |
| 0.003563346 | 0.04579371 |
| 0.003563315 | 0.04579371 |
| 0.003554266 | 0.045714028 |
| 0.003553711 | 0.045714028 |
| 0.003551363 | 0.045701364 |
| 0.003548069 | 0.045671319 |
| 0.00354039 | 0.045584789 |
| 0.003539531 | 0.045584789 |
| 0.00352936 | 0.045491972 |
| 0.00352913 | 0.045491972 |
| 0.003526843 | 0.045484143 |
| 0.00352636 | 0.045484143 |
| 0.003523121 | 0.045473079 |

| | |
|---|---|
| 0.003520796 | 0.0454554 |
| 0.00351931 | 0.045448526 |
| 0.003513682 | 0.045388154 |
| 0.003510395 | 0.045358003 |
| 0.003508225 | 0.045342256 |
| 0.003506491 | 0.045332154 |
| 0.003495758 | 0.04520566 |
| 0.00348609 | 0.04510513 |
| 0.003482127 | 0.04507835 |
| 0.003479317 | 0.04506647 |
| 0.003473168 | 0.045023569 |
| 0.003468448 | 0.044986871 |
| 0.003464865 | 0.044952636 |
| 0.003463256 | 0.044944009 |
| 0.003460633 | 0.044922201 |
| 0.003459793 | 0.044922201 |
| 0.003447029 | 0.044782229 |
| 0.003446938 | 0.044782229 |
| 0.003443021 | 0.044766796 |
| 0.0034383 | 0.044717622 |
| 0.003421763 | 0.044526857 |
| 0.003417026 | 0.044489524 |
| 0.003406714 | 0.044392784 |
| 0.003406419 | 0.044392784 |
| 0.003398223 | 0.044317403 |
| 0.003397495 | 0.044317403 |
| 0.003395937 | 0.04431186 |
| 0.003395216 | 0.04431186 |
| 0.003394802 | 0.04431186 |
| 0.003392621 | 0.044305236 |
| 0.003391732 | 0.044305236 |
| 0.003387453 | 0.044273932 |
| 0.003387099 | 0.044273932 |
| 0.003375703 | 0.044156715 |
| 0.003371729 | 0.044116848 |
| 0.003366333 | 0.044058355 |
| 0.003354246 | 0.043924297 |
| 0.003346615 | 0.043848479 |
| 0.003345208 | 0.04384211 |
| 0.003345135 | 0.04384211 |
| 0.003338733 | 0.043786694 |
| 0.00333641 | 0.043774992 |
| 0.003335165 | 0.043770707 |
| 0.003334893 | 0.043770707 |
| 0.003333581 | 0.043770707 |

| | |
|---|---|
| 0.003332654 | 0.043770707 |
| 0.00332673 | 0.043708187 |
| 0.003323517 | 0.043678238 |
| 0.003320727 | 0.043665453 |
| 0.003317776 | 0.043638692 |
| 0.003313164 | 0.043590072 |
| 0.003309894 | 0.043559085 |
| 0.003305029 | 0.043507074 |
| 0.003304982 | 0.043507074 |
| 0.003302437 | 0.043497004 |
| 0.0032989 | 0.043462435 |
| 0.003290423 | 0.043374747 |
| 0.003289428 | 0.043373637 |
| 0.003289219 | 0.043373637 |
| 0.003282809 | 0.04332234 |
| 0.003279618 | 0.043292226 |
| 0.003278912 | 0.043292226 |
| 0.003278358 | 0.043292226 |
| 0.0032776 | 0.043292226 |
| 0.003277314 | 0.043292226 |
| 0.003277118 | 0.043292226 |
| 0.003277071 | 0.043292226 |
| 0.003276297 | 0.043292226 |
| 0.003270418 | 0.04327875 |
| 0.003267353 | 0.043262221 |
| 0.003262562 | 0.043222821 |
| 0.003258734 | 0.043184122 |
| 0.003257225 | 0.043176137 |
| 0.003253932 | 0.043144499 |
| 0.003253325 | 0.043144499 |
| 0.003249162 | 0.043105242 |
| 0.003244935 | 0.043061159 |
| 0.003242671 | 0.043043107 |
| 0.003241853 | 0.043043107 |
| 0.003224392 | 0.042848248 |
| 0.0032214 | 0.042820429 |
| 0.003218387 | 0.042804275 |
| 0.00321564 | 0.042779685 |
| 0.003207777 | 0.042687001 |
| 0.003206972 | 0.042687001 |
| 0.003198229 | 0.042585152 |
| 0.003195741 | 0.042574422 |
| 0.003187505 | 0.042488468 |
| 0.003186259 | 0.042483746 |
| 0.00318335 | 0.042480641 |

| | |
|---|---|
| 0.003180433 | 0.042453605 |
| 0.003177335 | 0.042436042 |
| 0.00317569 | 0.042425968 |
| 0.003170358 | 0.042366624 |
| 0.00316697 | 0.042333218 |
| 0.003166567 | 0.042333218 |
| 0.003166522 | 0.042333218 |
| 0.003156062 | 0.042246698 |
| 0.003144422 | 0.042150124 |
| 0.003138793 | 0.042098368 |
| 0.003136674 | 0.042081793 |
| 0.003130125 | 0.042005765 |
| 0.003127816 | 0.041998441 |
| 0.003118727 | 0.041923686 |
| 0.003116425 | 0.041904563 |
| 0.003112553 | 0.041864323 |
| 0.003107136 | 0.041803261 |
| 0.003106287 | 0.041803261 |
| 0.003103785 | 0.041793596 |
| 0.003101412 | 0.041785281 |
| 0.003100317 | 0.041782336 |
| 0.00308676 | 0.041623191 |
| 0.003080183 | 0.041558042 |
| 0.003079589 | 0.041558042 |
| 0.003077054 | 0.041551131 |
| 0.003076131 | 0.041550456 |
| 0.003075115 | 0.041548511 |
| 0.003063931 | 0.041444436 |
| 0.003060451 | 0.041409122 |
| 0.003058392 | 0.041393023 |
| 0.003056149 | 0.041374421 |
| 0.00305064 | 0.04132333 |
| 0.003044965 | 0.04125819 |
| 0.003042503 | 0.041236564 |
| 0.003038004 | 0.041187303 |
| 0.003035835 | 0.041169621 |
| 0.003031394 | 0.041128195 |
| 0.00302834 | 0.041103098 |
| 0.003025693 | 0.041078868 |
| 0.003024659 | 0.041076544 |
| 0.003020323 | 0.041041056 |
| 0.003019593 | 0.041041056 |
| 0.003017417 | 0.041036693 |
| 0.003017041 | 0.041036693 |
| 0.003016515 | 0.041036693 |

| | |
|---|---|
| 0.003013746 | 0.041033648 |
| 0.003008843 | 0.040978605 |
| 0.003004235 | 0.040939255 |
| 0.003003871 | 0.040939255 |
| 0.00300089 | 0.040917093 |
| 0.003000302 | 0.040917093 |
| 0.002991351 | 0.040833782 |
| 0.002989434 | 0.04082793 |
| 0.002982861 | 0.040752933 |
| 0.002981635 | 0.040747865 |
| 0.002970537 | 0.040642858 |
| 0.002968922 | 0.040632433 |
| 0.002967649 | 0.040626675 |
| 0.002967399 | 0.040626675 |
| 0.002966038 | 0.040626675 |
| 0.0029601 | 0.040569981 |
| 0.002958322 | 0.040557279 |
| 0.002957513 | 0.040557279 |
| 0.002950166 | 0.040480431 |
| 0.002948064 | 0.040463237 |
| 0.002947554 | 0.040463237 |
| 0.002947199 | 0.040463237 |
| 0.002942861 | 0.040438447 |
| 0.002936693 | 0.040376982 |
| 0.002932261 | 0.040327688 |
| 0.002924907 | 0.040261436 |
| 0.002924632 | 0.040261436 |
| 0.002923848 | 0.040261436 |
| 0.002921278 | 0.04025896 |
| 0.002921264 | 0.04025896 |
| 0.0029106 | 0.040145707 |
| 0.002906308 | 0.040109743 |
| 0.002901381 | 0.04005393 |
| 0.002894028 | 0.039986624 |
| 0.002893654 | 0.039986624 |
| 0.002890683 | 0.039975192 |
| 0.002886107 | 0.039935097 |
| 0.002883688 | 0.039913229 |
| 0.002879069 | 0.039872474 |
| 0.002875825 | 0.039839129 |
| 0.002874358 | 0.039830389 |
| 0.002871371 | 0.039813751 |
| 0.0028681 | 0.039801567 |
| 0.002865354 | 0.039786652 |
| 0.002857309 | 0.039686514 |

| | |
|---|---|
| 0.002855576 | 0.039678253 |
| 0.002854101 | 0.039676665 |
| 0.002849651 | 0.039626377 |
| 0.002839487 | 0.039508106 |
| 0.002839124 | 0.039508106 |
| 0.002836692 | 0.039492287 |
| 0.002835061 | 0.039481131 |
| 0.00283422 | 0.039480955 |
| 0.002830857 | 0.039457195 |
| 0.002830673 | 0.039457195 |
| 0.002821421 | 0.039360227 |
| 0.00282031 | 0.039356259 |
| 0.002817665 | 0.039330872 |
| 0.002814747 | 0.039324722 |
| 0.002812678 | 0.039315266 |
| 0.002810761 | 0.039303609 |
| 0.002804217 | 0.03923515 |
| 0.002798193 | 0.03918376 |
| 0.002796394 | 0.039171725 |
| 0.002795017 | 0.03916395 |
| 0.002792472 | 0.039139814 |
| 0.002788154 | 0.039090788 |
| 0.002786977 | 0.039085797 |
| 0.002785966 | 0.03908312 |
| 0.002784921 | 0.039079975 |
| 0.002778623 | 0.039013893 |
| 0.002773399 | 0.038964214 |
| 0.002773183 | 0.038964214 |
| 0.002769302 | 0.038929622 |
| 0.002764743 | 0.038877008 |
| 0.002761125 | 0.038837595 |
| 0.00275993 | 0.038837595 |
| 0.002759452 | 0.038837595 |
| 0.002756433 | 0.038817456 |
| 0.002753883 | 0.03880191 |
| 0.002750866 | 0.038773456 |
| 0.002749854 | 0.038770662 |
| 0.002749654 | 0.038770662 |
| 0.002747041 | 0.038753945 |
| 0.002744966 | 0.038736152 |
| 0.002742425 | 0.038711753 |
| 0.002742335 | 0.038711753 |
| 0.002740879 | 0.038711753 |
| 0.002738163 | 0.038708968 |
| 0.00273393 | 0.0386606 |

| | |
|---|---|
| 0.002725394 | 0.038551337 |
| 0.002716473 | 0.038445237 |
| 0.002715551 | 0.038445237 |
| 0.002711971 | 0.038417197 |
| 0.002708878 | 0.03838612 |
| 0.002704746 | 0.038361808 |
| 0.002702152 | 0.038336423 |
| 0.002700863 | 0.038329555 |
| 0.00269901 | 0.038314679 |
| 0.002697338 | 0.038306296 |
| 0.002694843 | 0.038289762 |
| 0.002691285 | 0.038250618 |
| 0.002690685 | 0.038250618 |
| 0.002688118 | 0.038239837 |
| 0.002684929 | 0.038217307 |
| 0.002683395 | 0.038206899 |
| 0.002674782 | 0.038107045 |
| 0.002670313 | 0.038054763 |
| 0.002670305 | 0.038054763 |
| 0.00266645 | 0.038033844 |
| 0.002666141 | 0.038033844 |
| 0.002662998 | 0.038007376 |
| 0.00266276 | 0.038007376 |
| 0.002662247 | 0.038007376 |
| 0.002651799 | 0.037881594 |
| 0.002650467 | 0.037873931 |
| 0.002649755 | 0.037873931 |
| 0.002640127 | 0.037782845 |
| 0.002629468 | 0.037652933 |
| 0.002614012 | 0.03747669 |
| 0.00260918 | 0.037429945 |
| 0.002608721 | 0.037429945 |
| 0.002605593 | 0.037401017 |
| 0.002604932 | 0.037401017 |
| 0.002595773 | 0.037282531 |
| 0.002585721 | 0.037149365 |
| 0.002583669 | 0.037131082 |
| 0.002580641 | 0.037109963 |
| 0.002571566 | 0.037001802 |
| 0.002558892 | 0.036852837 |
| 0.002552577 | 0.036773008 |
| 0.002552048 | 0.036773008 |
| 0.002550083 | 0.036768699 |
| 0.002549866 | 0.036768699 |
| 0.002548578 | 0.036768699 |

| | |
|---|---|
| 0.002545156 | 0.036743903 |
| 0.002544099 | 0.036739781 |
| 0.002543229 | 0.036738355 |
| 0.002538701 | 0.036695202 |
| 0.002534765 | 0.036660562 |
| 0.002530462 | 0.036609446 |
| 0.002529289 | 0.036603604 |
| 0.002525665 | 0.036562264 |
| 0.002525247 | 0.036562264 |
| 0.002519122 | 0.036500834 |
| 0.002517146 | 0.036483299 |
| 0.00251212 | 0.036432618 |
| 0.002510783 | 0.036424332 |
| 0.002507183 | 0.036383182 |
| 0.002506251 | 0.036380747 |
| 0.002500791 | 0.036323632 |
| 0.002499281 | 0.036312775 |
| 0.002498299 | 0.036309583 |
| 0.002497404 | 0.036307654 |
| 0.002492361 | 0.036245398 |
| 0.002492171 | 0.036245398 |
| 0.002491898 | 0.036245398 |
| 0.002484169 | 0.036170445 |
| 0.002483129 | 0.036166361 |
| 0.002482551 | 0.036166361 |
| 0.002480712 | 0.036166361 |
| 0.002477585 | 0.036151938 |
| 0.002474537 | 0.036118536 |
| 0.00247446 | 0.036118536 |
| 0.00246883 | 0.036057336 |
| 0.002464639 | 0.036018208 |
| 0.002463889 | 0.036018208 |
| 0.002463331 | 0.036018208 |
| 0.00246327 | 0.036018208 |
| 0.002462318 | 0.036018208 |
| 0.002451218 | 0.035899894 |
| 0.002449695 | 0.035898967 |
| 0.002446072 | 0.035856905 |
| 0.002443714 | 0.035833366 |
| 0.002440693 | 0.035811114 |
| 0.002438976 | 0.035796951 |
| 0.00243493 | 0.035748585 |
| 0.002432918 | 0.03573006 |
| 0.002431765 | 0.03572414 |
| 0.002427316 | 0.035669792 |

| | |
|---|---|
| 0.002424737 | 0.035642881 |
| 0.002422778 | 0.035625087 |
| 0.002422327 | 0.035625087 |
| 0.002419686 | 0.035612609 |
| 0.002417308 | 0.035588605 |
| 0.002417061 | 0.035588605 |
| 0.002415483 | 0.035583732 |
| 0.002413821 | 0.035576821 |
| 0.00241155 | 0.035558793 |
| 0.002411106 | 0.035558793 |
| 0.002405647 | 0.035515727 |
| 0.002402329 | 0.035477748 |
| 0.002396897 | 0.035430484 |
| 0.002396678 | 0.035430484 |
| 0.002395732 | 0.035430484 |
| 0.002392672 | 0.035400988 |
| 0.002390862 | 0.035385197 |
| 0.002389627 | 0.035377925 |
| 0.002387648 | 0.035359614 |
| 0.002386228 | 0.03534957 |
| 0.002385243 | 0.035345975 |
| 0.002384076 | 0.035342927 |
| 0.002383327 | 0.035342927 |
| 0.002373823 | 0.035220577 |
| 0.002373797 | 0.035220577 |
| 0.002367753 | 0.035152417 |
| 0.002364896 | 0.035120956 |
| 0.002363827 | 0.035116033 |
| 0.002359069 | 0.035067218 |
| 0.002357728 | 0.035058226 |
| 0.002355043 | 0.035040192 |
| 0.002352985 | 0.035020501 |
| 0.002352044 | 0.035017438 |
| 0.002348738 | 0.034979158 |
| 0.002348369 | 0.034979158 |
| 0.002345462 | 0.03497412 |
| 0.002344755 | 0.03497412 |
| 0.002341919 | 0.034943168 |
| 0.002336981 | 0.034880409 |
| 0.002336817 | 0.034880409 |
| 0.002336437 | 0.034880409 |
| 0.002332192 | 0.034841685 |
| 0.002330708 | 0.03483044 |
| 0.002330695 | 0.03483044 |
| 0.00232844 | 0.034818409 |

| | |
|---|---|
| 0.002327873 | 0.034818409 |
| 0.002322901 | 0.034779249 |
| 0.002319947 | 0.034745956 |
| 0.00231776 | 0.034724115 |
| 0.002317578 | 0.034724115 |
| 0.002315991 | 0.034719455 |
| 0.002315007 | 0.034715635 |
| 0.002314257 | 0.034715315 |
| 0.00230987 | 0.034660425 |
| 0.002307845 | 0.034640957 |
| 0.002307712 | 0.034640957 |
| 0.002307142 | 0.034640957 |
| 0.002304422 | 0.034622308 |
| 0.002302886 | 0.034610152 |
| 0.002301376 | 0.034598376 |
| 0.002292967 | 0.034482831 |
| 0.002291607 | 0.034473273 |
| 0.002289987 | 0.034459789 |
| 0.002281485 | 0.0343427 |
| 0.002272151 | 0.034213001 |
| 0.002266257 | 0.03414585 |
| 0.002265715 | 0.03414585 |
| 0.002262932 | 0.034117328 |
| 0.002262126 | 0.034115986 |
| 0.002262031 | 0.034115986 |
| 0.002260422 | 0.034111888 |
| 0.0022571 | 0.034072556 |
| 0.002253039 | 0.034022028 |
| 0.002252714 | 0.034022028 |
| 0.002251482 | 0.034020089 |
| 0.002239475 | 0.033870903 |
| 0.002236492 | 0.033836535 |
| 0.002235461 | 0.033831686 |
| 0.002227165 | 0.033719481 |
| 0.002219321 | 0.033630162 |
| 0.002217161 | 0.033608128 |
| 0.002216606 | 0.033608128 |
| 0.002215844 | 0.033608128 |
| 0.002215408 | 0.033608128 |
| 0.00221245 | 0.033590163 |
| 0.002211341 | 0.033584029 |
| 0.002201096 | 0.033449779 |
| 0.002200165 | 0.033449779 |
| 0.002197011 | 0.033419687 |
| 0.00218891 | 0.033338755 |

| | |
|---|---|
| 0.002184415 | 0.033291883 |
| 0.00218249 | 0.03328385 |
| 0.002181466 | 0.033278887 |
| 0.002180155 | 0.033269535 |
| 0.002175868 | 0.033214757 |
| 0.002174994 | 0.033212074 |
| 0.002174546 | 0.033212074 |
| 0.002173293 | 0.033211659 |
| 0.002170598 | 0.0331875 |
| 0.002161315 | 0.033056187 |
| 0.002157187 | 0.033003638 |
| 0.002153971 | 0.032975621 |
| 0.00215077 | 0.032937207 |
| 0.002145049 | 0.03286016 |
| 0.002143511 | 0.032847174 |
| 0.002140163 | 0.032806415 |
| 0.002139981 | 0.032806415 |
| 0.002138814 | 0.032806415 |
| 0.002127966 | 0.032672034 |
| 0.00211737 | 0.03253032 |
| 0.002114622 | 0.032509077 |
| 0.002113908 | 0.032508594 |
| 0.002112309 | 0.032494497 |
| 0.002111861 | 0.032494497 |
| 0.002108337 | 0.032454345 |
| 0.002106609 | 0.032438239 |
| 0.002102275 | 0.032381964 |
| 0.002101811 | 0.032381964 |
| 0.002097457 | 0.032328675 |
| 0.002096946 | 0.032328675 |
| 0.002096024 | 0.032328675 |
| 0.002095564 | 0.032328675 |
| 0.002094111 | 0.032328675 |
| 0.002090284 | 0.032280784 |
| 0.002089755 | 0.032280784 |
| 0.002088421 | 0.032272948 |
| 0.002088306 | 0.032272948 |
| 0.002087926 | 0.032272948 |
| 0.002083529 | 0.032228718 |
| 0.002082443 | 0.032222394 |
| 0.002081216 | 0.032213871 |
| 0.002074592 | 0.032121777 |
| 0.002068755 | 0.032052244 |
| 0.002067699 | 0.032046311 |
| 0.002062541 | 0.031976766 |

| | |
|---|---|
| 0.002051128 | 0.03184129 |
| 0.002048677 | 0.031823989 |
| 0.002047092 | 0.031809745 |
| 0.002043622 | 0.031766177 |
| 0.002043024 | 0.031766177 |
| 0.002042873 | 0.031766177 |
| 0.002038344 | 0.031735932 |
| 0.002035737 | 0.0317057 |
| 0.002035361 | 0.0317057 |
| 0.002026817 | 0.031608127 |
| 0.00202316 | 0.031571764 |
| 0.002021062 | 0.031549367 |
| 0.00202007 | 0.031544228 |
| 0.002019125 | 0.031539804 |
| 0.002017433 | 0.031523719 |
| 0.002013273 | 0.031469036 |
| 0.002010928 | 0.0314427 |
| 0.002004679 | 0.03135528 |
| 0.002003098 | 0.031343636 |
| 0.001999872 | 0.031310955 |
| 0.001999482 | 0.031310955 |
| 0.00199796 | 0.031310955 |
| 0.00198933 | 0.031225497 |
| 0.001985623 | 0.031180107 |
| 0.001978634 | 0.031090858 |
| 0.001977649 | 0.031085646 |
| 0.001972946 | 0.031032193 |
| 0.001972122 | 0.031029483 |
| 0.001966389 | 0.030949511 |
| 0.001965627 | 0.030947734 |
| 0.001963827 | 0.030929622 |
| 0.001960682 | 0.030890305 |
| 0.001956772 | 0.030843662 |
| 0.001953881 | 0.030813735 |
| 0.001941846 | 0.030674724 |
| 0.001934231 | 0.030584871 |
| 0.00193311 | 0.030584871 |
| 0.001919507 | 0.03040251 |
| 0.001916027 | 0.030367589 |
| 0.00191602 | 0.030367589 |
| 0.001911082 | 0.030309394 |
| 0.001910705 | 0.030309394 |
| 0.001910286 | 0.030309394 |
| 0.001910053 | 0.030309394 |
| 0.001906211 | 0.030282565 |

| | |
|---|---|
| 0.001903823 | 0.030254718 |
| 0.001902149 | 0.030238198 |
| 0.001900825 | 0.030227237 |
| 0.00190068 | 0.030227237 |
| 0.001900553 | 0.030227237 |
| 0.00189908 | 0.030227237 |
| 0.001896123 | 0.030202888 |
| 0.001890963 | 0.03013077 |
| 0.001888549 | 0.030102386 |
| 0.001887435 | 0.030094699 |
| 0.001885243 | 0.030069818 |
| 0.00188004 | 0.029996879 |
| 0.001877893 | 0.029972663 |
| 0.001869069 | 0.029861841 |
| 0.00186284 | 0.029772314 |
| 0.001859749 | 0.029732889 |
| 0.001853486 | 0.029662632 |
| 0.001847837 | 0.029602057 |
| 0.001846264 | 0.02958682 |
| 0.0018454 | 0.029582921 |
| 0.001843733 | 0.029575983 |
| 0.001842645 | 0.029575983 |
| 0.001840106 | 0.029557757 |
| 0.001839566 | 0.029557757 |
| 0.001837958 | 0.029543176 |
| 0.001833092 | 0.029474911 |
| 0.001824747 | 0.029350629 |
| 0.001824515 | 0.029350629 |
| 0.001821859 | 0.029323987 |
| 0.001821782 | 0.029323987 |
| 0.001821166 | 0.029323987 |
| 0.001813965 | 0.029236448 |
| 0.001812275 | 0.029219098 |
| 0.001810932 | 0.029217233 |
| 0.001801433 | 0.02907382 |
| 0.00179498 | 0.029009003 |
| 0.001793582 | 0.028996243 |
| 0.001792501 | 0.028988604 |
| 0.001787085 | 0.028910828 |
| 0.001786466 | 0.028910828 |
| 0.001784665 | 0.028910828 |
| 0.001781107 | 0.028892637 |
| 0.001779362 | 0.028874161 |
| 0.001779334 | 0.028874161 |
| 0.001762795 | 0.028654145 |

| | |
|---|---|
| 0.001757149 | 0.028572118 |
| 0.001752863 | 0.028512157 |
| 0.00175062 | 0.028485403 |
| 0.001749675 | 0.028479757 |
| 0.00174719 | 0.028458764 |
| 0.001743139 | 0.028402487 |
| 0.001741338 | 0.028382862 |
| 0.00173669 | 0.028316789 |
| 0.001730371 | 0.028233079 |
| 0.001727048 | 0.028198195 |
| 0.001724819 | 0.028174955 |
| 0.001724729 | 0.028174955 |
| 0.00171248 | 0.02803723 |
| 0.001711418 | 0.028032863 |
| 0.001710379 | 0.02803174 |
| 0.001709973 | 0.02803174 |
| 0.001706373 | 0.027985342 |
| 0.001703814 | 0.027952989 |
| 0.001701539 | 0.02792529 |
| 0.001694331 | 0.027826173 |
| 0.001687166 | 0.027720059 |
| 0.001686778 | 0.027720059 |
| 0.001684062 | 0.027695725 |
| 0.001674592 | 0.027568541 |
| 0.001673744 | 0.027564109 |
| 0.001673599 | 0.027564109 |
| 0.001670813 | 0.027534876 |
| 0.001670124 | 0.027533054 |
| 0.001668288 | 0.02752039 |
| 0.001665246 | 0.027481175 |
| 0.00166226 | 0.027441406 |
| 0.001661931 | 0.027441406 |
| 0.001658049 | 0.027390885 |
| 0.001657132 | 0.027390885 |
| 0.00165464 | 0.027363036 |
| 0.001648825 | 0.027276344 |
| 0.001648021 | 0.02727251 |
| 0.001642252 | 0.027195951 |
| 0.00163896 | 0.027156071 |
| 0.001636721 | 0.027132647 |
| 0.001629052 | 0.027042895 |
| 0.001627476 | 0.027035864 |
| 0.001625663 | 0.027015168 |
| 0.001625199 | 0.027015168 |
| 0.001620887 | 0.026954612 |

| | |
|---|---|
| 0.00162086 | 0.026954612 |
| 0.001611186 | 0.026821382 |
| 0.001607504 | 0.026797555 |
| 0.001605355 | 0.026780467 |
| 0.001605051 | 0.026780467 |
| 0.001592773 | 0.026617191 |
| 0.001592489 | 0.026617191 |
| 0.00158778 | 0.026561706 |
| 0.001586298 | 0.02654625 |
| 0.001581011 | 0.026488541 |
| 0.001580706 | 0.026488541 |
| 0.001577849 | 0.026469962 |
| 0.001577336 | 0.026469962 |
| 0.001576958 | 0.026469962 |
| 0.001576159 | 0.026469571 |
| 0.001574339 | 0.026457665 |
| 0.001572622 | 0.02643814 |
| 0.00157047 | 0.026411294 |
| 0.001567443 | 0.02636969 |
| 0.001564625 | 0.026331592 |
| 0.001554645 | 0.02620283 |
| 0.001553297 | 0.02620283 |
| 0.001550499 | 0.026167828 |
| 0.001548111 | 0.02614164 |
| 0.001540022 | 0.026046389 |
| 0.001539917 | 0.026046389 |
| 0.001531226 | 0.025916028 |
| 0.001528892 | 0.025904133 |
| 0.001527927 | 0.025897 |
| 0.001527458 | 0.025897 |
| 0.001522219 | 0.025827827 |
| 0.001517322 | 0.025753923 |
| 0.001515692 | 0.025735419 |
| 0.001508967 | 0.025639511 |
| 0.001505831 | 0.025595368 |
| 0.001505709 | 0.025595368 |
| 0.001503393 | 0.025572174 |
| 0.001503018 | 0.025572174 |
| 0.001500181 | 0.025535788 |
| 0.001495814 | 0.025490197 |
| 0.001494619 | 0.025486652 |
| 0.001492361 | 0.025460557 |
| 0.001491817 | 0.025460557 |
| 0.001490393 | 0.025460172 |
| 0.001490384 | 0.025460172 |

| | |
|---|---|
| 0.001489054 | 0.02545556 |
| 0.001487991 | 0.025446519 |
| 0.001485158 | 0.025425458 |
| 0.00147764 | 0.025305843 |
| 0.001477542 | 0.025305843 |
| 0.001471604 | 0.025229688 |
| 0.001470812 | 0.025225202 |
| 0.001469789 | 0.025216729 |
| 0.001468211 | 0.02520089 |
| 0.001467064 | 0.025197206 |
| 0.00145979 | 0.025108497 |
| 0.001459735 | 0.025108497 |
| 0.001458111 | 0.025097739 |
| 0.001457413 | 0.025096839 |
| 0.001457287 | 0.025096839 |
| 0.001456824 | 0.025096839 |
| 0.001454683 | 0.025084059 |
| 0.001454027 | 0.025081827 |
| 0.001452307 | 0.025070331 |
| 0.001452235 | 0.025070331 |
| 0.001451499 | 0.025070331 |
| 0.00145144 | 0.025070331 |
| 0.001449392 | 0.025056335 |
| 0.001447454 | 0.02503192 |
| 0.001440491 | 0.024947727 |
| 0.001439353 | 0.02493709 |
| 0.001439214 | 0.02493709 |
| 0.001436991 | 0.024914296 |
| 0.001435648 | 0.024900077 |
| 0.001434652 | 0.024891857 |
| 0.00143385 | 0.024887014 |
| 0.001433834 | 0.024887014 |
| 0.001428184 | 0.024815796 |
| 0.001426813 | 0.024801034 |
| 0.001426354 | 0.024801034 |
| 0.001423559 | 0.024771584 |
| 0.001418883 | 0.024708286 |
| 0.00141837 | 0.024708286 |
| 0.001415268 | 0.024663366 |
| 0.001412014 | 0.024615662 |
| 0.001409987 | 0.024607348 |
| 0.001408304 | 0.024595997 |
| 0.001406579 | 0.024588941 |
| 0.001402527 | 0.024540103 |
| 0.001399773 | 0.02450091 |

| | |
|---|---|
| 0.001397041 | 0.024462087 |
| 0.001392274 | 0.024396553 |
| 0.001391514 | 0.024396553 |
| 0.001390962 | 0.024396553 |
| 0.001390111 | 0.024396553 |
| 0.001389101 | 0.024394797 |
| 0.001387998 | 0.02438442 |
| 0.001387414 | 0.024383161 |
| 0.001387315 | 0.024383161 |
| 0.00138603 | 0.024376836 |
| 0.001385483 | 0.024376212 |
| 0.001383174 | 0.024346425 |
| 0.001382471 | 0.024346425 |
| 0.001378912 | 0.024314491 |
| 0.001377332 | 0.024295621 |
| 0.001371457 | 0.024200952 |
| 0.001369423 | 0.024174025 |
| 0.001365649 | 0.024152165 |
| 0.001362172 | 0.024117526 |
| 0.001356982 | 0.02404116 |
| 0.001356609 | 0.02404116 |
| 0.001354328 | 0.024014361 |
| 0.001352242 | 0.023986306 |
| 0.001351254 | 0.023986306 |
| 0.001350277 | 0.023978239 |
| 0.001348174 | 0.023958767 |
| 0.001345352 | 0.02391753 |
| 0.001343921 | 0.023901025 |
| 0.001343849 | 0.023901025 |
| 0.001343767 | 0.023901025 |
| 0.001342663 | 0.023901025 |
| 0.001340839 | 0.023881873 |
| 0.001339707 | 0.023870635 |
| 0.001335345 | 0.023810742 |
| 0.001333941 | 0.023799176 |
| 0.001332514 | 0.023786976 |
| 0.001332337 | 0.023786976 |
| 0.001327229 | 0.02371041 |
| 0.001325569 | 0.023689631 |
| 0.001325529 | 0.023689631 |
| 0.001325338 | 0.023689631 |
| 0.001324002 | 0.02368829 |
| 0.001322659 | 0.023673166 |
| 0.001319793 | 0.023630743 |
| 0.001318375 | 0.023623109 |

| | |
|---|---|
| 0.00131679 | 0.023605654 |
| 0.001314347 | 0.023577555 |
| 0.001311349 | 0.023532636 |
| 0.001309353 | 0.023505677 |
| 0.00130858 | 0.023505066 |
| 0.001305476 | 0.023480333 |
| 0.001305068 | 0.023480333 |
| 0.001304029 | 0.023472024 |
| 0.001298393 | 0.023379423 |
| 0.00129694 | 0.023370931 |
| 0.001294708 | 0.023344816 |
| 0.001293175 | 0.023329565 |
| 0.001292392 | 0.023324273 |
| 0.001286176 | 0.023229701 |
| 0.001284897 | 0.023215391 |
| 0.001282865 | 0.023192799 |
| 0.001279543 | 0.023153795 |
| 0.001278093 | 0.023136345 |
| 0.00127461 | 0.02308207 |
| 0.001273164 | 0.02306773 |
| 0.001270462 | 0.023033222 |
| 0.001269762 | 0.023029311 |
| 0.001263235 | 0.022937124 |
| 0.001261456 | 0.02291357 |
| 0.001260847 | 0.022911241 |
| 0.001258108 | 0.022870186 |
| 0.001255018 | 0.022822739 |
| 0.0012529 | 0.02280218 |
| 0.001252585 | 0.02280218 |
| 0.00125003 | 0.022775512 |
| 0.001249015 | 0.022765716 |
| 0.001248106 | 0.022758733 |
| 0.001247392 | 0.022758733 |
| 0.001241383 | 0.022678697 |
| 0.001239624 | 0.022655243 |
| 0.001234147 | 0.022589827 |
| 0.001223776 | 0.02241722 |
| 0.001223208 | 0.022415433 |
| 0.001221299 | 0.022389068 |
| 0.001217776 | 0.022333079 |
| 0.00121706 | 0.022328559 |
| 0.001216251 | 0.022322313 |
| 0.001212767 | 0.022266953 |
| 0.001212416 | 0.022266953 |
| 0.001210145 | 0.022235956 |

| | |
|---|---|
| 0.001207606 | 0.022197857 |
| 0.001205915 | 0.022175335 |
| 0.001204491 | 0.022166269 |
| 0.001199878 | 0.022098455 |
| 0.001196428 | 0.022051957 |
| 0.001194121 | 0.022017965 |
| 0.001191336 | 0.021975117 |
| 0.001191161 | 0.021975117 |
| 0.001189037 | 0.021949717 |
| 0.001185478 | 0.021901001 |
| 0.001184688 | 0.0218949 |
| 0.001182337 | 0.021876903 |
| 0.001182259 | 0.021876903 |
| 0.001181966 | 0.021876903 |
| 0.00118083 | 0.021876903 |
| 0.001174145 | 0.021784581 |
| 0.00117218 | 0.0217566 |
| 0.001169099 | 0.02170787 |
| 0.001167125 | 0.021696582 |
| 0.001164373 | 0.021653886 |
| 0.001163243 | 0.021641326 |
| 0.001161544 | 0.021626598 |
| 0.001159612 | 0.021599083 |
| 0.001159605 | 0.021599083 |
| 0.001156382 | 0.02155577 |
| 0.001150248 | 0.021485378 |
| 0.001148368 | 0.021465207 |
| 0.001146522 | 0.02143911 |
| 0.00114414 | 0.021419785 |
| 0.001143515 | 0.02141651 |
| 0.001143146 | 0.02141651 |
| 0.001138152 | 0.021349659 |
| 0.001135889 | 0.021315595 |
| 0.001133787 | 0.02128454 |
| 0.001133217 | 0.021282233 |
| 0.001132425 | 0.02127574 |
| 0.001131509 | 0.021266931 |
| 0.001131329 | 0.021266931 |
| 0.001130093 | 0.021257078 |
| 0.001129746 | 0.021257078 |
| 0.001128725 | 0.021248127 |
| 0.001128569 | 0.021248127 |
| 0.001128357 | 0.021248127 |
| 0.001123129 | 0.021176271 |
| 0.001120753 | 0.021139834 |

| | |
|---|---|
| 0.001120322 | 0.021139834 |
| 0.001115812 | 0.021063336 |
| 0.001114846 | 0.021061781 |
| 0.001114306 | 0.021059933 |
| 0.001113828 | 0.02105926 |
| 0.001112221 | 0.021037223 |
| 0.001108998 | 0.020984595 |
| 0.001105398 | 0.020933124 |
| 0.001103877 | 0.020926429 |
| 0.001102865 | 0.020918413 |
| 0.001096743 | 0.020818892 |
| 0.001095096 | 0.0207959 |
| 0.001091432 | 0.020742866 |
| 0.001089623 | 0.02071676 |
| 0.001088895 | 0.020711184 |
| 0.001087385 | 0.020699004 |
| 0.001087053 | 0.020699004 |
| 0.001086884 | 0.020699004 |
| 0.001085337 | 0.020684808 |
| 0.001082602 | 0.020657489 |
| 0.001082315 | 0.020657489 |
| 0.00108177 | 0.020657489 |
| 0.001079784 | 0.020636795 |
| 0.00107976 | 0.020636795 |
| 0.001078788 | 0.020634315 |
| 0.001075892 | 0.020616188 |
| 0.001073919 | 0.020590792 |
| 0.001073069 | 0.020582779 |
| 0.001066936 | 0.020479869 |
| 0.001066107 | 0.020473977 |
| 0.001059581 | 0.020365082 |
| 0.001058866 | 0.020359556 |
| 0.001055258 | 0.020298379 |
| 0.001055131 | 0.020298379 |
| 0.001054234 | 0.020295067 |
| 0.001054061 | 0.020295067 |
| 0.001053709 | 0.020295067 |
| 0.001052724 | 0.020295067 |
| 0.001052498 | 0.020295067 |
| 0.001052217 | 0.020295067 |
| 0.001051889 | 0.020295067 |
| 0.001050523 | 0.020289242 |
| 0.001046458 | 0.020218922 |
| 0.001046432 | 0.020218922 |
| 0.001040358 | 0.020125564 |

| | |
|---|---|
| 0.001039082 | 0.020117228 |
| 0.001034902 | 0.020052586 |
| 0.001032574 | 0.02002378 |
| 0.001026425 | 0.01992076 |
| 0.001024397 | 0.019897618 |
| 0.001023809 | 0.019894317 |
| 0.001021454 | 0.019856652 |
| 0.001018001 | 0.019797613 |
| 0.001017488 | 0.019795707 |
| 0.00101709 | 0.019795707 |
| 0.001015794 | 0.019786998 |
| 0.001015468 | 0.019786998 |
| 0.001013407 | 0.019756663 |
| 0.001013091 | 0.019756663 |
| 0.001013052 | 0.019756663 |
| 0.001007648 | 0.019692748 |
| 0.001005312 | 0.019655164 |
| 0.00100442 | 0.019653868 |
| 0.000997229 | 0.019524171 |
| 0.000993235 | 0.019458994 |
| 0.000989184 | 0.019395586 |
| 0.000986811 | 0.019357025 |
| 0.000983699 | 0.019304265 |
| 0.000983058 | 0.019304265 |
| 0.000980294 | 0.019276823 |
| 0.000980084 | 0.019276823 |
| 0.000980052 | 0.019276823 |
| 0.000977631 | 0.0192483 |
| 0.000973384 | 0.019180547 |
| 0.000971684 | 0.019154966 |
| 0.000970711 | 0.019143715 |
| 0.000967096 | 0.019080308 |
| 0.000965443 | 0.019063389 |
| 0.000964864 | 0.019059964 |
| 0.000960597 | 0.01899143 |
| 0.000955971 | 0.018915683 |
| 0.000955641 | 0.018915683 |
| 0.000954259 | 0.018905357 |
| 0.000954122 | 0.018905357 |
| 0.000952663 | 0.01888946 |
| 0.000951629 | 0.018887995 |
| 0.000949481 | 0.018857772 |
| 0.000949297 | 0.018857772 |
| 0.000949027 | 0.018857772 |
| 0.000946702 | 0.018841861 |

| | |
|---|---|
| 0.000942449 | 0.018780761 |
| 0.000939846 | 0.018736732 |
| 0.000939308 | 0.01873612 |
| 0.000930899 | 0.018589494 |
| 0.00093006 | 0.018580518 |
| 0.000924995 | 0.018494852 |
| 0.000919568 | 0.018409522 |
| 0.000918977 | 0.018405428 |
| 0.000917192 | 0.018385136 |
| 0.000913815 | 0.018332883 |
| 0.000913556 | 0.018332883 |
| 0.000911856 | 0.018309017 |
| 0.000909341 | 0.018266214 |
| 0.000908505 | 0.018257126 |
| 0.000907481 | 0.018244242 |
| 0.000907192 | 0.018244242 |
| 0.000905376 | 0.018225006 |
| 0.000900779 | 0.018140121 |
| 0.000898556 | 0.018116943 |
| 0.000898414 | 0.018116943 |
| 0.000895786 | 0.018082407 |
| 0.000895432 | 0.018082407 |
| 0.000891425 | 0.018012675 |
| 0.000890965 | 0.018012675 |
| 0.000890738 | 0.018012675 |
| 0.000888633 | 0.01799443 |
| 0.000885261 | 0.017941424 |
| 0.000882135 | 0.017885679 |
| 0.000880385 | 0.017857794 |
| 0.000874143 | 0.017738728 |
| 0.000874115 | 0.017738728 |
| 0.000872067 | 0.017719256 |
| 0.000870287 | 0.01769819 |
| 0.000869112 | 0.017696956 |
| 0.000867338 | 0.017668398 |
| 0.000866341 | 0.01765563 |
| 0.000865229 | 0.017640532 |
| 0.000862454 | 0.017599006 |
| 0.000862201 | 0.017599006 |
| 0.000861527 | 0.017595165 |
| 0.000860648 | 0.017584753 |
| 0.000860605 | 0.017584753 |
| 0.000859206 | 0.017570371 |
| 0.000857817 | 0.0175495 |
| 0.000855465 | 0.017508918 |

| | |
|---|---|
| 0.000853453 | 0.017490272 |
| 0.000852098 | 0.017470032 |
| 0.000851505 | 0.017465386 |
| 0.000849581 | 0.017433422 |
| 0.000848873 | 0.017426415 |
| 0.000847234 | 0.017400274 |
| 0.000845581 | 0.017373802 |
| 0.000844089 | 0.017350647 |
| 0.000843616 | 0.017348402 |
| 0.000843099 | 0.017345263 |
| 0.000841916 | 0.017328406 |
| 0.000838812 | 0.017279445 |
| 0.000835062 | 0.017209659 |
| 0.000834272 | 0.01720081 |
| 0.000833391 | 0.017190091 |
| 0.000831037 | 0.017156385 |
| 0.00083068 | 0.017156385 |
| 0.000830413 | 0.017156385 |
| 0.000830029 | 0.017156385 |
| 0.000826254 | 0.017094698 |
| 0.00082416 | 0.017058786 |
| 0.000822162 | 0.017024823 |
| 0.000822129 | 0.017024823 |
| 0.000821264 | 0.017021027 |
| 0.000816978 | 0.016946951 |
| 0.000816428 | 0.016942918 |
| 0.000816184 | 0.016942918 |
| 0.000813617 | 0.0169067 |
| 0.000811651 | 0.016895322 |
| 0.000809956 | 0.016867407 |
| 0.000809805 | 0.016867407 |
| 0.000809567 | 0.016867407 |
| 0.000809322 | 0.016867407 |
| 0.000807721 | 0.016857734 |
| 0.000807037 | 0.016850833 |
| 0.000806235 | 0.016841467 |
| 0.000803549 | 0.016792722 |
| 0.000801007 | 0.016746947 |
| 0.000799622 | 0.016725337 |
| 0.00079899 | 0.01671946 |
| 0.000796221 | 0.016690863 |
| 0.000794238 | 0.016656617 |
| 0.000792268 | 0.016622619 |
| 0.000789713 | 0.016583627 |
| 0.000788946 | 0.016574823 |

| | |
|---|---|
| 0.000788051 | 0.01656333 |
| 0.000782883 | 0.016469243 |
| 0.000781846 | 0.016454702 |
| 0.000780579 | 0.016435286 |
| 0.000778603 | 0.016415464 |
| 0.000776337 | 0.016374925 |
| 0.000776296 | 0.016374925 |
| 0.000774173 | 0.016343771 |
| 0.000773207 | 0.016330621 |
| 0.000772747 | 0.016328144 |
| 0.000771316 | 0.016305138 |
| 0.000770001 | 0.01628457 |
| 0.000768216 | 0.01626125 |
| 0.000766409 | 0.016230208 |
| 0.00076567 | 0.016221774 |
| 0.000764176 | 0.01619733 |
| 0.000763542 | 0.016191104 |
| 0.000757967 | 0.01609437 |
| 0.000756814 | 0.016077057 |
| 0.000755751 | 0.016061657 |
| 0.000750609 | 0.015966611 |
| 0.000745213 | 0.015858926 |
| 0.000742967 | 0.015818192 |
| 0.000742179 | 0.015808477 |
| 0.000741715 | 0.015805667 |
| 0.000736536 | 0.015728607 |
| 0.000736098 | 0.015728607 |
| 0.000734464 | 0.015707372 |
| 0.000733271 | 0.015688896 |
| 0.000732732 | 0.015684419 |
| 0.000732715 | 0.015684419 |
| 0.00072892 | 0.015623894 |
| 0.000728261 | 0.015616793 |
| 0.000727221 | 0.015601503 |
| 0.000725991 | 0.015582148 |
| 0.000724873 | 0.015565167 |
| 0.000724827 | 0.015565167 |
| 0.000724594 | 0.015565167 |
| 0.00072344 | 0.015555423 |
| 0.000720964 | 0.015523221 |
| 0.000720618 | 0.015522792 |
| 0.000718055 | 0.015481571 |
| 0.000715889 | 0.015441877 |
| 0.000715428 | 0.015438923 |
| 0.000715369 | 0.015438923 |

| | |
|---|---|
| 0.0007151 | 0.015438923 |
| 0.000714253 | 0.015434543 |
| 0.000711505 | 0.015382146 |
| 0.000709009 | 0.015349773 |
| 0.000704773 | 0.015271251 |
| 0.000704727 | 0.015271251 |
| 0.000699501 | 0.015170818 |
| 0.00069752 | 0.015141658 |
| 0.000697397 | 0.015141658 |
| 0.00069508 | 0.015102444 |
| 0.000691115 | 0.015030008 |
| 0.00068862 | 0.014982592 |
| 0.000688551 | 0.014982592 |
| 0.000688213 | 0.014982592 |
| 0.000687736 | 0.014982592 |
| 0.000686005 | 0.014966197 |
| 0.000685891 | 0.014966197 |
| 0.000683534 | 0.014926481 |
| 0.000680773 | 0.014878174 |
| 0.000678411 | 0.014838699 |
| 0.000676582 | 0.014808617 |
| 0.000676013 | 0.014802966 |
| 0.000674655 | 0.014780023 |
| 0.000674204 | 0.014777601 |
| 0.000667449 | 0.014662635 |
| 0.000666953 | 0.014658507 |
| 0.00066627 | 0.014650241 |
| 0.000666029 | 0.014650241 |
| 0.00066462 | 0.014634228 |
| 0.000663946 | 0.014626162 |
| 0.000663102 | 0.014614324 |
| 0.000659471 | 0.014561259 |
| 0.000659313 | 0.014561259 |
| 0.00065557 | 0.014515487 |
| 0.000654189 | 0.014505144 |
| 0.000652955 | 0.014484522 |
| 0.000651935 | 0.014475384 |
| 0.000651055 | 0.014462598 |
| 0.000650842 | 0.014462598 |
| 0.000649493 | 0.01444681 |
| 0.000649394 | 0.01444681 |
| 0.000644704 | 0.014354994 |
| 0.000644523 | 0.014354994 |
| 0.000644301 | 0.014354994 |
| 0.000643095 | 0.014339268 |

| | |
|---|---|
| 0.00064152 | 0.014310858 |
| 0.000638577 | 0.014271946 |
| 0.000637615 | 0.014257121 |
| 0.00063648 | 0.014238436 |
| 0.000633823 | 0.014192325 |
| 0.000626173 | 0.014045882 |
| 0.000619289 | 0.01389956 |
| 0.000618223 | 0.013888718 |
| 0.000618192 | 0.013888718 |
| 0.000615144 | 0.013832604 |
| 0.000614271 | 0.013819507 |
| 0.000613531 | 0.013809373 |
| 0.000612773 | 0.013798844 |
| 0.000612287 | 0.013794416 |
| 0.000611723 | 0.013788223 |
| 0.000611382 | 0.013787075 |
| 0.000610685 | 0.013777866 |
| 0.000610468 | 0.013777866 |
| 0.000608113 | 0.013745888 |
| 0.000606999 | 0.013733748 |
| 0.000605512 | 0.013706612 |
| 0.000605499 | 0.013706612 |
| 0.00060539 | 0.013706612 |
| 0.000601386 | 0.013632666 |
| 0.0006006 | 0.013621317 |
| 0.000600034 | 0.013618674 |
| 0.000598523 | 0.01359363 |
| 0.000596047 | 0.013543869 |
| 0.000595845 | 0.013543869 |
| 0.00059083 | 0.013457422 |
| 0.000590455 | 0.013455304 |
| 0.000590051 | 0.01345253 |
| 0.000588956 | 0.013434013 |
| 0.000588382 | 0.013427353 |
| 0.000587582 | 0.013419599 |
| 0.000586728 | 0.013408878 |
| 0.000585422 | 0.013385444 |
| 0.000585318 | 0.013385444 |
| 0.000584204 | 0.01337045 |
| 0.000583552 | 0.013361929 |
| 0.000580334 | 0.013294651 |
| 0.000579764 | 0.013294382 |
| 0.000579221 | 0.013288326 |
| 0.000579208 | 0.013288326 |
| 0.000578457 | 0.013288326 |

| | |
|---|---|
| 0.00057816 | 0.013288326 |
| 0.000577465 | 0.013280023 |
| 0.000575378 | 0.01323842 |
| 0.000574412 | 0.013222575 |
| 0.000572002 | 0.013199007 |
| 0.000570699 | 0.013175325 |
| 0.000567548 | 0.013108924 |
| 0.000566348 | 0.013087563 |
| 0.000565067 | 0.013070628 |
| 0.000563289 | 0.013054883 |
| 0.000562366 | 0.013046177 |
| 0.000560273 | 0.013003968 |
| 0.000558473 | 0.012968512 |
| 0.000557627 | 0.012955168 |
| 0.000556932 | 0.012945334 |
| 0.000551833 | 0.012839356 |
| 0.000550618 | 0.012817339 |
| 0.000550088 | 0.012811277 |
| 0.000544846 | 0.012707817 |
| 0.000544329 | 0.012701995 |
| 0.000543577 | 0.012690661 |
| 0.00054223 | 0.012665411 |
| 0.00054113 | 0.012652124 |
| 0.000540888 | 0.012652124 |
| 0.000539833 | 0.012639267 |
| 0.000538149 | 0.012613392 |
| 0.000535585 | 0.012559485 |
| 0.000535289 | 0.012558721 |
| 0.000533439 | 0.01252148 |
| 0.000532524 | 0.012506173 |
| 0.000531053 | 0.012492124 |
| 0.000528198 | 0.012441387 |
| 0.00052469 | 0.012377113 |
| 0.000523889 | 0.012364353 |
| 0.000522749 | 0.012343554 |
| 0.00052256 | 0.012343554 |
| 0.00052247 | 0.012343554 |
| 0.000521002 | 0.012320631 |
| 0.000518665 | 0.012281333 |
| 0.000518193 | 0.012278601 |
| 0.000515254 | 0.012233303 |
| 0.000515252 | 0.012233303 |
| 0.000514397 | 0.012231241 |
| 0.000512927 | 0.01220239 |
| 0.000511689 | 0.012179026 |

| | |
|---|---|
| 0.000511516 | 0.012179026 |
| 0.000511241 | 0.012179026 |
| 0.000510553 | 0.012170231 |
| 0.000509918 | 0.012161184 |
| 0.0005092 | 0.012150128 |
| 0.000508349 | 0.012135905 |
| 0.000506992 | 0.012109578 |
| 0.000504458 | 0.012061156 |
| 0.000502791 | 0.012033392 |
| 0.000502654 | 0.012033392 |
| 0.000501518 | 0.012014982 |
| 0.000500595 | 0.011998922 |
| 0.000498188 | 0.011965302 |
| 0.000496973 | 0.011942142 |
| 0.000496739 | 0.011942142 |
| 0.000496445 | 0.011941497 |
| 0.000492605 | 0.011878956 |
| 0.000488713 | 0.011797229 |
| 0.000485963 | 0.01174276 |
| 0.000485231 | 0.011736964 |
| 0.000483013 | 0.011701143 |
| 0.000482655 | 0.01169843 |
| 0.00048225 | 0.01169456 |
| 0.00048166 | 0.011686215 |
| 0.000479731 | 0.011645331 |
| 0.000477333 | 0.011593026 |
| 0.000477216 | 0.011593026 |
| 0.000471605 | 0.011477329 |
| 0.000470774 | 0.011462951 |
| 0.000470372 | 0.011459038 |
| 0.00047005 | 0.011457045 |
| 0.000467532 | 0.011419048 |
| 0.00046556 | 0.011376709 |
| 0.000464202 | 0.011355193 |
| 0.000463665 | 0.011347883 |
| 0.000462524 | 0.011325766 |
| 0.000459925 | 0.011273713 |
| 0.000459029 | 0.011257557 |
| 0.000458949 | 0.011257557 |
| 0.000458517 | 0.011256592 |
| 0.000458335 | 0.011256592 |
| 0.00045744 | 0.011247557 |
| 0.000456653 | 0.011236163 |
| 0.000452504 | 0.01116075 |
| 0.000452132 | 0.011157365 |

| | |
|---|---|
| 0.000451574 | 0.011149361 |
| 0.000449125 | 0.011100407 |
| 0.00044906 | 0.011100407 |
| 0.000448141 | 0.011087597 |
| 0.000446693 | 0.011057502 |
| 0.000445212 | 0.011026577 |
| 0.00044442 | 0.011012695 |
| 0.000443593 | 0.010997916 |
| 0.000441708 | 0.010956898 |
| 0.000438515 | 0.010889027 |
| 0.000437922 | 0.010879972 |
| 0.00043491 | 0.010827748 |
| 0.000434344 | 0.010819312 |
| 0.000429086 | 0.010721972 |
| 0.000428583 | 0.010715032 |
| 0.000426439 | 0.010667024 |
| 0.00042541 | 0.01064688 |
| 0.00042206 | 0.010585278 |
| 0.000417349 | 0.010478158 |
| 0.000416065 | 0.010451434 |
| 0.000415862 | 0.010451434 |
| 0.000413907 | 0.010408216 |
| 0.000411758 | 0.010359648 |
| 0.000409713 | 0.010323782 |
| 0.000409489 | 0.010323782 |
| 0.000406498 | 0.01026527 |
| 0.000405757 | 0.010251975 |
| 0.000404122 | 0.010216103 |
| 0.00040412 | 0.010216103 |
| 0.000403593 | 0.010213557 |
| 0.000401725 | 0.010171701 |
| 0.000401259 | 0.010165309 |
| 0.000400884 | 0.010161211 |
| 0.000399694 | 0.010136424 |
| 0.000398823 | 0.010119738 |
| 0.000395839 | 0.010072966 |
| 0.000394581 | 0.01004958 |
| 0.000392602 | 0.010004518 |
| 0.000390876 | 0.009971211 |
| 0.000390047 | 0.009955399 |
| 0.000389822 | 0.009955399 |
| 0.000387775 | 0.009923986 |
| 0.00038764 | 0.009923986 |
| 0.000387623 | 0.009923986 |
| 0.000387229 | 0.009923986 |

| | |
|---|---|
| 0.000385361 | 0.009888774 |
| 0.000384498 | 0.009871956 |
| 0.000384389 | 0.009871956 |
| 0.000384388 | 0.009871956 |
| 0.000383565 | 0.009871956 |
| 0.000382881 | 0.009862343 |
| 0.000382753 | 0.009862343 |
| 0.000382677 | 0.009862343 |
| 0.000381364 | 0.00983924 |
| 0.00038112 | 0.00983924 |
| 0.000380763 | 0.00983924 |
| 0.000380591 | 0.00983924 |
| 0.000380366 | 0.00983924 |
| 0.000379325 | 0.009818534 |
| 0.000376345 | 0.009751996 |
| 0.000372098 | 0.0096472 |
| 0.000371282 | 0.009631287 |
| 0.000370452 | 0.009615006 |
| 0.000369352 | 0.009591684 |
| 0.000367579 | 0.009556053 |
| 0.0003671 | 0.009548806 |
| 0.00036635 | 0.009534505 |
| 0.000366041 | 0.009531664 |
| 0.000365322 | 0.009518157 |
| 0.000364737 | 0.009510747 |
| 0.000363807 | 0.009494261 |
| 0.000363591 | 0.009493839 |
| 0.000362129 | 0.009466048 |
| 0.000359502 | 0.009418044 |
| 0.000359345 | 0.009418044 |
| 0.000359318 | 0.009418044 |
| 0.000357937 | 0.009397729 |
| 0.000357895 | 0.009397729 |
| 0.000355263 | 0.009337798 |
| 0.00035476 | 0.009329744 |
| 0.000352214 | 0.009267897 |
| 0.000352072 | 0.009267897 |
| 0.000351593 | 0.009266918 |
| 0.000351226 | 0.009262374 |
| 0.000350986 | 0.009261171 |
| 0.000349754 | 0.00924167 |
| 0.000348935 | 0.009232639 |
| 0.000347755 | 0.00921163 |
| 0.000347751 | 0.00921163 |
| 0.000346367 | 0.009185091 |

| | |
|---|---|
| 0.000346026 | 0.009181167 |
| 0.000344338 | 0.009141459 |
| 0.000343619 | 0.009127461 |
| 0.000342925 | 0.009114122 |
| 0.000342888 | 0.009114122 |
| 0.000342379 | 0.009109765 |
| 0.00034157 | 0.009093312 |
| 0.000341034 | 0.009084135 |
| 0.000338995 | 0.009039927 |
| 0.000338687 | 0.009039927 |
| 0.000338672 | 0.009039927 |
| 0.00033863 | 0.009039927 |
| 0.000338401 | 0.009039927 |
| 0.000338355 | 0.009039927 |
| 0.000337754 | 0.009039927 |
| 0.000337679 | 0.009039927 |
| 0.000336322 | 0.009024702 |
| 0.000335493 | 0.009012151 |
| 0.000334925 | 0.00900197 |
| 0.000334713 | 0.009001345 |
| 0.000333434 | 0.008977089 |
| 0.000333096 | 0.008973063 |
| 0.000333002 | 0.008973063 |
| 0.000331064 | 0.008933463 |
| 0.000330367 | 0.00891973 |
| 0.000329794 | 0.008914364 |
| 0.000326711 | 0.008841064 |
| 0.000325147 | 0.008808759 |
| 0.000324355 | 0.008792281 |
| 0.000324336 | 0.008792281 |
| 0.000323829 | 0.008788032 |
| 0.000322049 | 0.008744719 |
| 0.000321083 | 0.008733432 |
| 0.000320354 | 0.008718576 |
| 0.000320015 | 0.008714324 |
| 0.000318871 | 0.00868813 |
| 0.00031847 | 0.00868218 |
| 0.00031807 | 0.008676255 |
| 0.000317461 | 0.008664591 |
| 0.000317153 | 0.008661148 |
| 0.000314365 | 0.008589933 |
| 0.000313628 | 0.008574718 |
| 0.000312287 | 0.008542959 |
| 0.000311318 | 0.008531154 |
| 0.000310903 | 0.008529058 |

| | |
|---|---|
| 0.00031007 | 0.008511632 |
| 0.000308124 | 0.008472884 |
| 0.000307264 | 0.008454123 |
| 0.000306881 | 0.008448463 |
| 0.000306689 | 0.008448059 |
| 0.000306639 | 0.008448059 |
| 0.000305971 | 0.00843802 |
| 0.000304136 | 0.00839228 |
| 0.000302588 | 0.008359254 |
| 0.000302558 | 0.008359254 |
| 0.00030235 | 0.008359254 |
| 0.000301896 | 0.008359254 |
| 0.000301246 | 0.008346388 |
| 0.000300017 | 0.008317169 |
| 0.000299994 | 0.008317169 |
| 0.000299659 | 0.008316948 |
| 0.000297108 | 0.008270243 |
| 0.000296918 | 0.008269768 |
| 0.000295777 | 0.00824281 |
| 0.00029544 | 0.00824281 |
| 0.000295408 | 0.00824281 |
| 0.000294682 | 0.008236389 |
| 0.000291128 | 0.008146626 |
| 0.000290723 | 0.008140073 |
| 0.000290269 | 0.008136939 |
| 0.000287377 | 0.00807963 |
| 0.000286975 | 0.008073094 |
| 0.000284775 | 0.008015936 |
| 0.000281855 | 0.007947834 |
| 0.0002794 | 0.007897331 |
| 0.000279176 | 0.007895675 |
| 0.000278403 | 0.007883188 |
| 0.000276455 | 0.007832668 |
| 0.000274179 | 0.007782082 |
| 0.000272426 | 0.007736923 |
| 0.000270605 | 0.007694386 |
| 0.000268965 | 0.007652317 |
| 0.000268106 | 0.007632447 |
| 0.000268024 | 0.007632447 |
| 0.000266788 | 0.007608583 |
| 0.000264452 | 0.007555509 |
| 0.000264023 | 0.007553842 |
| 0.000263747 | 0.007553497 |
| 0.000262588 | 0.007524826 |
| 0.000261843 | 0.007517055 |

| | |
|---|---|
| 0.000259804 | 0.007463024 |
| 0.000259151 | 0.007448747 |
| 0.000258275 | 0.007428066 |
| 0.00025774 | 0.007417144 |
| 0.000257 | 0.007409298 |
| 0.000256868 | 0.007409298 |
| 0.000256561 | 0.007409298 |
| 0.000255288 | 0.007386752 |
| 0.000255256 | 0.007386752 |
| 0.000255108 | 0.007386752 |
| 0.000254782 | 0.007385584 |
| 0.000252314 | 0.007327421 |
| 0.000252247 | 0.007327421 |
| 0.000251486 | 0.007312297 |
| 0.00025141 | 0.007312297 |
| 0.000250997 | 0.007306977 |
| 0.000249715 | 0.007274101 |
| 0.0002497 | 0.007274101 |
| 0.000249458 | 0.007274101 |
| 0.000249365 | 0.007274101 |
| 0.000248831 | 0.007270589 |
| 0.000247031 | 0.007229798 |
| 0.000246635 | 0.007224169 |
| 0.000246301 | 0.007218825 |
| 0.000241983 | 0.007105375 |
| 0.000241319 | 0.007101417 |
| 0.000239977 | 0.007068278 |
| 0.000239315 | 0.007053126 |
| 0.000239233 | 0.007053126 |
| 0.000239164 | 0.007053126 |
| 0.000238261 | 0.007035138 |
| 0.000238088 | 0.007034369 |
| 0.000237377 | 0.00701773 |
| 0.000235876 | 0.006982027 |
| 0.000235486 | 0.006974796 |
| 0.000234926 | 0.0069669 |
| 0.000234348 | 0.006958413 |
| 0.000233603 | 0.006949298 |
| 0.000233534 | 0.006949298 |
| 0.000232399 | 0.006922112 |
| 0.000231478 | 0.006899007 |
| 0.000230197 | 0.006873714 |
| 0.000229208 | 0.00684848 |
| 0.000229099 | 0.00684848 |
| 0.000228326 | 0.006835006 |

| | |
|---|---|
| 0.000227553 | 0.006820439 |
| 0.000225448 | 0.006765858 |
| 0.000225368 | 0.006765858 |
| 0.000224885 | 0.006761727 |
| 0.000224774 | 0.006761727 |
| 0.000224339 | 0.006753864 |
| 0.000224268 | 0.006753864 |
| 0.000223151 | 0.006726584 |
| 0.000222965 | 0.006725244 |
| 0.000222342 | 0.006710705 |
| 0.000221418 | 0.00669128 |
| 0.000220608 | 0.006671031 |
| 0.000219568 | 0.006643827 |
| 0.000219213 | 0.006637294 |
| 0.000216565 | 0.006582229 |
| 0.00021592 | 0.00656681 |
| 0.000214752 | 0.006535458 |
| 0.000212908 | 0.006483485 |
| 0.000211958 | 0.00645869 |
| 0.00021164 | 0.006457249 |
| 0.000211151 | 0.006446481 |
| 0.000209954 | 0.006418147 |
| 0.000209914 | 0.006418147 |
| 0.000209212 | 0.006403695 |
| 0.000207748 | 0.006367058 |
| 0.000207576 | 0.006365874 |
| 0.000207432 | 0.006365581 |
| 0.000206836 | 0.006355613 |
| 0.000206781 | 0.006355613 |
| 0.000206173 | 0.00634737 |
| 0.000205097 | 0.006318326 |
| 0.000203653 | 0.006281991 |
| 0.000203633 | 0.006281991 |
| 0.000203139 | 0.006277246 |
| 0.00020272 | 0.006269453 |
| 0.000202696 | 0.006269453 |
| 0.000202485 | 0.006269453 |
| 0.000202174 | 0.006268848 |
| 0.000200804 | 0.006234482 |
| 0.000200548 | 0.006234482 |
| 0.000200318 | 0.00623159 |
| 0.000200276 | 0.00623159 |
| 0.000199938 | 0.006227893 |
| 0.000199313 | 0.006216556 |
| 0.000198876 | 0.006206998 |

| | |
|---|---|
| 0.00019869 | 0.00620619 |
| 0.000198641 | 0.00620619 |
| 0.000197493 | 0.00618408 |
| 0.000197443 | 0.00618408 |
| 0.00019742 | 0.00618408 |
| 0.000197372 | 0.00618408 |
| 0.000196291 | 0.006162656 |
| 0.000195046 | 0.006127611 |
| 0.00019464 | 0.006118905 |
| 0.000194516 | 0.006118905 |
| 0.000194119 | 0.006110564 |
| 0.000194114 | 0.006110564 |
| 0.000192717 | 0.006078498 |
| 0.000191566 | 0.006050208 |
| 0.000190669 | 0.006033873 |
| 0.000190158 | 0.006021716 |
| 0.000189699 | 0.006011189 |
| 0.000189446 | 0.006007155 |
| 0.000188868 | 0.005993183 |
| 0.000186375 | 0.005929527 |
| 0.000181435 | 0.005783927 |
| 0.000180707 | 0.005764577 |
| 0.000180431 | 0.005759647 |
| 0.000180015 | 0.005750211 |
| 0.000179628 | 0.005745575 |
| 0.000177678 | 0.005687015 |
| 0.000177648 | 0.005687015 |
| 0.000176314 | 0.005650963 |
| 0.000175852 | 0.005643729 |
| 0.000175768 | 0.005643729 |
| 0.000175757 | 0.005643729 |
| 0.000174131 | 0.005599819 |
| 0.000172808 | 0.005561016 |
| 0.000172368 | 0.005550633 |
| 0.000172354 | 0.005550633 |
| 0.000171561 | 0.005539604 |
| 0.000171198 | 0.00553164 |
| 0.000170105 | 0.005503807 |
| 0.000169984 | 0.005503606 |
| 0.000169953 | 0.005503606 |
| 0.000169917 | 0.005503606 |
| 0.000169637 | 0.005503606 |
| 0.00016936 | 0.00550212 |
| 0.00016827 | 0.005470431 |
| 0.00016796 | 0.005467822 |

| | |
|---|---|
| 0.000166206 | 0.00541442 |
| 0.000166092 | 0.005414402 |
| 0.000165995 | 0.005414402 |
| 0.000165381 | 0.005402329 |
| 0.000165332 | 0.005402329 |
| 0.000164989 | 0.005400639 |
| 0.000164724 | 0.005395658 |
| 0.000164491 | 0.005391749 |
| 0.000163439 | 0.005360943 |
| 0.000163116 | 0.005354035 |
| 0.000162691 | 0.005343798 |
| 0.000162613 | 0.005343798 |
| 0.000162367 | 0.005340512 |
| 0.000162363 | 0.005340512 |
| 0.000162294 | 0.005340512 |
| 0.000162126 | 0.005340512 |
| 0.000161365 | 0.005322236 |
| 0.000161052 | 0.005315605 |
| 0.000160733 | 0.005308755 |
| 0.000159579 | 0.005275921 |
| 0.000159354 | 0.00527419 |
| 0.000158179 | 0.005242599 |
| 0.000156864 | 0.005206243 |
| 0.000155815 | 0.005182253 |
| 0.000155805 | 0.005182253 |
| 0.000155153 | 0.005167467 |
| 0.000155045 | 0.005167466 |
| 0.000153655 | 0.005128343 |
| 0.000153374 | 0.005122545 |
| 0.000153337 | 0.005122545 |
| 0.000152998 | 0.00511716 |
| 0.00015231 | 0.005097732 |
| 0.00015118 | 0.005067489 |
| 0.00015065 | 0.00505637 |
| 0.000150196 | 0.005044713 |
| 0.000149704 | 0.005031731 |
| 0.000148713 | 0.005001938 |
| 0.000148701 | 0.005001938 |
| 0.000148271 | 0.004994147 |
| 0.000148057 | 0.004990446 |
| 0.000147305 | 0.004968619 |
| 0.000146164 | 0.004933622 |
| 0.00014609 | 0.004933622 |
| 0.000146011 | 0.004933622 |
| 0.000145502 | 0.004921752 |

| | |
|---|---|
| 0.000145344 | 0.004921752 |
| 0.00014525 | 0.004921752 |
| 0.000144923 | 0.004921752 |
| 0.000144258 | 0.00490755 |
| 0.000144058 | 0.004904257 |
| 0.000144043 | 0.004904257 |
| 0.000143214 | 0.004882493 |
| 0.000142394 | 0.004858018 |
| 0.000142175 | 0.004854012 |
| 0.000141845 | 0.00484622 |
| 0.000140511 | 0.004810999 |
| 0.000140166 | 0.004802653 |
| 0.000139506 | 0.004783479 |
| 0.000139232 | 0.004780979 |
| 0.000138874 | 0.004772116 |
| 0.00013865 | 0.004767872 |
| 0.000137213 | 0.004730159 |
| 0.000135601 | 0.004683288 |
| 0.000135548 | 0.004683288 |
| 0.000135181 | 0.00467558 |
| 0.000134584 | 0.004658316 |
| 0.000134495 | 0.004658316 |
| 0.000133425 | 0.004625757 |
| 0.000132841 | 0.004611375 |
| 0.00013144 | 0.004566085 |
| 0.000130706 | 0.004547217 |
| 0.000129561 | 0.004510687 |
| 0.000129129 | 0.004498928 |
| 0.000129082 | 0.004498928 |
| 0.000128157 | 0.004474892 |
| 0.000127763 | 0.004466337 |
| 0.000127561 | 0.004463879 |
| 0.000126831 | 0.004448155 |
| 0.000126491 | 0.004442753 |
| 0.000126425 | 0.004442753 |
| 0.000125837 | 0.004429583 |
| 0.000125764 | 0.004429583 |
| 0.000125724 | 0.004429583 |
| 0.000125609 | 0.004429583 |
| 0.000125321 | 0.004424518 |
| 0.000125245 | 0.004424518 |
| 0.000125237 | 0.004424518 |
| 0.000124908 | 0.004423568 |
| 0.000124686 | 0.004421745 |
| 0.000124431 | 0.004417541 |

| | |
|---|---|
| 0.000123475 | 0.004391889 |
| 0.000121663 | 0.004340397 |
| 0.000121501 | 0.004337878 |
| 0.000121284 | 0.004333396 |
| 0.000120874 | 0.004321984 |
| 0.000120724 | 0.004319844 |
| 0.000120539 | 0.004316491 |
| 0.00011845 | 0.004251251 |
| 0.000118406 | 0.004251251 |
| 0.000117124 | 0.004213181 |
| 0.000115396 | 0.004154155 |
| 0.000115268 | 0.004152685 |
| 0.000115127 | 0.004150754 |
| 0.000114175 | 0.004122687 |
| 0.000113074 | 0.00409533 |
| 0.000112825 | 0.00408944 |
| 0.000112593 | 0.004084135 |
| 0.000112325 | 0.004077504 |
| 0.000111901 | 0.004065224 |
| 0.00011164 | 0.004058843 |
| 0.000111445 | 0.00405486 |
| 0.000111061 | 0.004043951 |
| 0.000110925 | 0.004042085 |
| 0.000110834 | 0.004041874 |
| 0.00011051 | 0.004033145 |
| 0.000109867 | 0.004012765 |
| 0.000109133 | 0.00398909 |
| 0.000108043 | 0.003955249 |
| 0.000107873 | 0.003952068 |
| 0.000107416 | 0.003938333 |
| 0.000107034 | 0.003927355 |
| 0.000106666 | 0.003916888 |
| 0.000106361 | 0.003908703 |
| 0.000106236 | 0.003907101 |
| 0.000105923 | 0.00389862 |
| 0.000105634 | 0.003894 |
| 0.00010544 | 0.003889847 |
| 0.000104974 | 0.003875671 |
| 0.000104842 | 0.003873774 |
| 0.000104663 | 0.00387019 |
| 0.000104395 | 0.003863273 |
| 0.000104317 | 0.003863273 |
| 0.000102979 | 0.003825743 |
| 0.000102925 | 0.003825743 |
| 0.000102622 | 0.003819573 |

| | |
|---|---|
| 0.000102434 | 0.003817396 |
| 0.000100803 | 0.003762492 |
| 0.000100153 | 0.003744094 |
| 0.000100131 | 0.003744094 |
| 9.98E-05 | 0.00373674 |
| 9.97E-05 | 0.00373581 |
| 9.95E-05 | 0.003734268 |
| 9.84E-05 | 0.003699312 |
| 9.80E-05 | 0.00368733 |
| 9.77E-05 | 0.003678518 |
| 9.75E-05 | 0.003675437 |
| 9.71E-05 | 0.003662159 |
| 9.67E-05 | 0.003648261 |
| 9.66E-05 | 0.003648261 |
| 9.55E-05 | 0.003612787 |
| 9.46E-05 | 0.003588164 |
| 9.46E-05 | 0.003588164 |
| 9.45E-05 | 0.003588164 |
| 9.43E-05 | 0.003587787 |
| 9.40E-05 | 0.003578654 |
| 9.33E-05 | 0.003556724 |
| 9.32E-05 | 0.003556724 |
| 9.26E-05 | 0.003540479 |
| 9.26E-05 | 0.003540479 |
| 9.24E-05 | 0.003537092 |
| 9.21E-05 | 0.003530524 |
| 9.18E-05 | 0.003520341 |
| 9.18E-05 | 0.003520341 |
| 9.08E-05 | 0.00348693 |
| 8.92E-05 | 0.00342838 |
| 8.88E-05 | 0.003417541 |
| 8.78E-05 | 0.003387052 |
| 8.74E-05 | 0.00337671 |
| 8.72E-05 | 0.003370019 |
| 8.64E-05 | 0.003342628 |
| 8.63E-05 | 0.00334032 |
| 8.55E-05 | 0.003313388 |
| 8.49E-05 | 0.003296241 |
| 8.49E-05 | 0.003296241 |
| 8.48E-05 | 0.003295458 |
| 8.46E-05 | 0.003292429 |
| 8.40E-05 | 0.003272482 |
| 8.39E-05 | 0.0032708 |
| 8.30E-05 | 0.003238534 |
| 8.18E-05 | 0.003196328 |

| | |
|---|---|
| 8.08E-05 | 0.00316204 |
| 8.08E-05 | 0.00316204 |
| 8.06E-05 | 0.003159139 |
| 7.98E-05 | 0.003136933 |
| 7.93E-05 | 0.003122895 |
| 7.85E-05 | 0.003092281 |
| 7.82E-05 | 0.003082111 |
| 7.80E-05 | 0.003076078 |
| 7.73E-05 | 0.00305201 |
| 7.72E-05 | 0.003050512 |
| 7.64E-05 | 0.003023879 |
| 7.64E-05 | 0.003023879 |
| 7.64E-05 | 0.003023879 |
| 7.61E-05 | 0.003020979 |
| 7.55E-05 | 0.002999316 |
| 7.50E-05 | 0.002983012 |
| 7.48E-05 | 0.002980584 |
| 7.47E-05 | 0.002977286 |
| 7.43E-05 | 0.002964298 |
| 7.43E-05 | 0.002964298 |
| 7.41E-05 | 0.002962665 |
| 7.38E-05 | 0.002957528 |
| 7.38E-05 | 0.002957528 |
| 7.35E-05 | 0.002948226 |
| 7.23E-05 | 0.002904863 |
| 7.17E-05 | 0.002887542 |
| 7.16E-05 | 0.002886965 |
| 7.12E-05 | 0.00287359 |
| 7.09E-05 | 0.002865209 |
| 7.02E-05 | 0.002838211 |
| 6.98E-05 | 0.002830676 |
| 6.97E-05 | 0.002830676 |
| 6.95E-05 | 0.002824933 |
| 6.94E-05 | 0.002821578 |
| 6.94E-05 | 0.002821578 |
| 6.93E-05 | 0.002821578 |
| 6.92E-05 | 0.002821578 |
| 6.78E-05 | 0.002774827 |
| 6.78E-05 | 0.002774827 |
| 6.66E-05 | 0.002736256 |
| 6.51E-05 | 0.002680938 |
| 6.45E-05 | 0.002662358 |
| 6.43E-05 | 0.002662358 |
| 6.40E-05 | 0.002654421 |
| 6.35E-05 | 0.002633347 |

| | |
|---|---|
| 6.33E-05 | 0.002629361 |
| 6.32E-05 | 0.002629361 |
| 6.26E-05 | 0.002612407 |
| 6.23E-05 | 0.002600976 |
| 6.12E-05 | 0.002564708 |
| 6.11E-05 | 0.00256012 |
| 6.09E-05 | 0.00255698 |
| 6.06E-05 | 0.002546698 |
| 5.98E-05 | 0.002517468 |
| 5.98E-05 | 0.002516428 |
| 5.86E-05 | 0.002470231 |
| 5.84E-05 | 0.00246378 |
| 5.75E-05 | 0.002431103 |
| 5.72E-05 | 0.002422525 |
| 5.65E-05 | 0.002397774 |
| 5.64E-05 | 0.002395878 |
| 5.64E-05 | 0.002395878 |
| 5.60E-05 | 0.002384551 |
| 5.57E-05 | 0.002376639 |
| 5.55E-05 | 0.002366785 |
| 5.55E-05 | 0.002366785 |
| 5.47E-05 | 0.002340777 |
| 5.45E-05 | 0.00233436 |
| 5.43E-05 | 0.002327707 |
| 5.37E-05 | 0.002307105 |
| 5.31E-05 | 0.00228287 |
| 5.21E-05 | 0.002245737 |
| 5.20E-05 | 0.002243172 |
| 5.18E-05 | 0.002235411 |
| 5.16E-05 | 0.002232685 |
| 5.16E-05 | 0.002232473 |
| 5.16E-05 | 0.002232473 |
| 5.15E-05 | 0.002232473 |
| 5.14E-05 | 0.002232473 |
| 5.11E-05 | 0.002226304 |
| 5.08E-05 | 0.002214004 |
| 5.07E-05 | 0.002209768 |
| 5.03E-05 | 0.002195173 |
| 5.01E-05 | 0.002191109 |
| 4.98E-05 | 0.002179111 |
| 4.94E-05 | 0.002167701 |
| 4.93E-05 | 0.002163786 |
| 4.92E-05 | 0.002160304 |
| 4.92E-05 | 0.002160304 |
| 4.87E-05 | 0.002146749 |

| | |
|---|---|
| 4.81E-05 | 0.002123916 |
| 4.81E-05 | 0.002123695 |
| 4.80E-05 | 0.002123695 |
| 4.77E-05 | 0.002113341 |
| 4.75E-05 | 0.00210553 |
| 4.67E-05 | 0.002072788 |
| 4.66E-05 | 0.002072788 |
| 4.66E-05 | 0.002072788 |
| 4.64E-05 | 0.002068459 |
| 4.62E-05 | 0.002058216 |
| 4.58E-05 | 0.002047986 |
| 4.54E-05 | 0.00203215 |
| 4.47E-05 | 0.0020016 |
| 4.43E-05 | 0.001987543 |
| 4.38E-05 | 0.001969992 |
| 4.37E-05 | 0.00196382 |
| 4.36E-05 | 0.001961583 |
| 4.31E-05 | 0.001942092 |
| 4.31E-05 | 0.001942092 |
| 4.30E-05 | 0.001942092 |
| 4.29E-05 | 0.00193923 |
| 4.29E-05 | 0.00193923 |
| 4.25E-05 | 0.001923548 |
| 4.16E-05 | 0.001888948 |
| 4.15E-05 | 0.001886732 |
| 4.13E-05 | 0.001885617 |
| 4.07E-05 | 0.001864761 |
| 4.06E-05 | 0.001861249 |
| 4.05E-05 | 0.001859544 |
| 4.05E-05 | 0.001859544 |
| 4.04E-05 | 0.001859527 |
| 4.00E-05 | 0.00184757 |
| 3.97E-05 | 0.001833275 |
| 3.95E-05 | 0.001824392 |
| 3.93E-05 | 0.001820775 |
| 3.88E-05 | 0.001801202 |
| 3.87E-05 | 0.001798143 |
| 3.84E-05 | 0.001782765 |
| 3.83E-05 | 0.001782765 |
| 3.82E-05 | 0.001777891 |
| 3.80E-05 | 0.001773466 |
| 3.77E-05 | 0.001759778 |
| 3.76E-05 | 0.001757164 |
| 3.74E-05 | 0.001749488 |
| 3.70E-05 | 0.001735049 |

| | |
|---|---|
| 3.69E-05 | 0.001733373 |
| 3.67E-05 | 0.001727214 |
| 3.67E-05 | 0.00172672 |
| 3.65E-05 | 0.001721442 |
| 3.62E-05 | 0.001710273 |
| 3.58E-05 | 0.001694599 |
| 3.55E-05 | 0.001684339 |
| 3.54E-05 | 0.001678197 |
| 3.50E-05 | 0.001667265 |
| 3.50E-05 | 0.001666067 |
| 3.48E-05 | 0.001660037 |
| 3.44E-05 | 0.00164194 |
| 3.34E-05 | 0.001605723 |
| 3.29E-05 | 0.001582369 |
| 3.26E-05 | 0.001574475 |
| 3.25E-05 | 0.001573332 |
| 3.25E-05 | 0.001573332 |
| 3.24E-05 | 0.001571514 |
| 3.20E-05 | 0.001554123 |
| 3.20E-05 | 0.001552555 |
| 3.19E-05 | 0.001551387 |
| 3.19E-05 | 0.001551387 |
| 3.16E-05 | 0.001543504 |
| 3.14E-05 | 0.001537819 |
| 3.14E-05 | 0.001537819 |
| 3.13E-05 | 0.001535965 |
| 3.13E-05 | 0.001535919 |
| 3.11E-05 | 0.001529845 |
| 3.10E-05 | 0.001526693 |
| 3.08E-05 | 0.001522141 |
| 3.07E-05 | 0.001515513 |
| 3.04E-05 | 0.001503864 |
| 3.04E-05 | 0.001503075 |
| 3.02E-05 | 0.001494369 |
| 3.00E-05 | 0.001491192 |
| 2.99E-05 | 0.001486532 |
| 2.99E-05 | 0.001486532 |
| 2.98E-05 | 0.001485226 |
| 2.96E-05 | 0.001479581 |
| 2.91E-05 | 0.001461342 |
| 2.90E-05 | 0.001453225 |
| 2.87E-05 | 0.001441909 |
| 2.82E-05 | 0.001420502 |
| 2.81E-05 | 0.001415657 |
| 2.79E-05 | 0.001405708 |

| | |
|---|---|
| 2.77E-05 | 0.001401449 |
| 2.72E-05 | 0.001384553 |
| 2.70E-05 | 0.001376294 |
| 2.69E-05 | 0.001371289 |
| 2.65E-05 | 0.001363706 |
| 2.64E-05 | 0.001358898 |
| 2.62E-05 | 0.001351549 |
| 2.57E-05 | 0.00132812 |
| 2.56E-05 | 0.001325226 |
| 2.53E-05 | 0.001313798 |
| 2.51E-05 | 0.001305654 |
| 2.51E-05 | 0.001303661 |
| 2.50E-05 | 0.001303597 |
| 2.50E-05 | 0.001303597 |
| 2.50E-05 | 0.001303597 |
| 2.50E-05 | 0.001303369 |
| 2.49E-05 | 0.001300909 |
| 2.48E-05 | 0.001297956 |
| 2.46E-05 | 0.001289276 |
| 2.45E-05 | 0.00128755 |
| 2.41E-05 | 0.001268558 |
| 2.37E-05 | 0.001250429 |
| 2.37E-05 | 0.001250141 |
| 2.37E-05 | 0.001250141 |
| 2.36E-05 | 0.001248793 |
| 2.33E-05 | 0.001234517 |
| 2.31E-05 | 0.001228917 |
| 2.29E-05 | 0.001218255 |
| 2.27E-05 | 0.001211605 |
| 2.26E-05 | 0.001208688 |
| 2.19E-05 | 0.001172303 |
| 2.17E-05 | 0.001166433 |
| 2.16E-05 | 0.001163962 |
| 2.15E-05 | 0.001159158 |
| 2.14E-05 | 0.001156475 |
| 2.14E-05 | 0.001156475 |
| 2.14E-05 | 0.001155043 |
| 2.13E-05 | 0.001154242 |
| 2.12E-05 | 0.001149402 |
| 2.12E-05 | 0.001149402 |
| 2.11E-05 | 0.001147673 |
| 2.10E-05 | 0.001142647 |
| 2.09E-05 | 0.00114254 |
| 2.09E-05 | 0.001141251 |
| 2.08E-05 | 0.00113735 |

| | |
|---|---|
| 2.08E-05 | 0.00113735 |
| 2.07E-05 | 0.00113735 |
| 2.07E-05 | 0.001136173 |
| 2.06E-05 | 0.001136173 |
| 2.05E-05 | 0.001128659 |
| 2.01E-05 | 0.001112641 |
| 2.00E-05 | 0.00110831 |
| 1.99E-05 | 0.00110215 |
| 1.96E-05 | 0.001093026 |
| 1.96E-05 | 0.001091613 |
| 1.94E-05 | 0.001082437 |
| 1.89E-05 | 0.00105724 |
| 1.87E-05 | 0.001051138 |
| 1.86E-05 | 0.001046175 |
| 1.85E-05 | 0.001042972 |
| 1.84E-05 | 0.001038506 |
| 1.83E-05 | 0.001031117 |
| 1.81E-05 | 0.001022015 |
| 1.81E-05 | 0.001022015 |
| 1.78E-05 | 0.001012109 |
| 1.78E-05 | 0.001012109 |
| 1.78E-05 | 0.001012109 |
| 1.78E-05 | 0.001012109 |
| 1.76E-05 | 0.001005611 |
| 1.74E-05 | 0.000996505 |
| 1.71E-05 | 0.000985009 |
| 1.71E-05 | 0.000985009 |
| 1.71E-05 | 0.000985009 |
| 1.70E-05 | 0.000982542 |
| 1.69E-05 | 0.000981694 |
| 1.67E-05 | 0.000970012 |
| 1.67E-05 | 0.000968363 |
| 1.66E-05 | 0.000967295 |
| 1.65E-05 | 0.00096146 |
| 1.64E-05 | 0.000960991 |
| 1.64E-05 | 0.000960991 |
| 1.64E-05 | 0.000960991 |
| 1.64E-05 | 0.000960902 |
| 1.63E-05 | 0.000956928 |
| 1.62E-05 | 0.000953635 |
| 1.56E-05 | 0.000925196 |
| 1.56E-05 | 0.000925196 |
| 1.56E-05 | 0.000925196 |
| 1.55E-05 | 0.000924922 |
| 1.54E-05 | 0.000917512 |

| | |
|---|---|
| 1.54E-05 | 0.000917512 |
| 1.53E-05 | 0.000914032 |
| 1.53E-05 | 0.000912481 |
| 1.52E-05 | 0.000911596 |
| 1.51E-05 | 0.000905293 |
| 1.51E-05 | 0.000904271 |
| 1.50E-05 | 0.000904271 |
| 1.50E-05 | 0.000904271 |
| 1.50E-05 | 0.000904271 |
| 1.49E-05 | 0.000898139 |
| 1.48E-05 | 0.000895361 |
| 1.47E-05 | 0.000892211 |
| 1.46E-05 | 0.000889532 |
| 1.44E-05 | 0.000879179 |
| 1.43E-05 | 0.000875046 |
| 1.43E-05 | 0.000874888 |
| 1.42E-05 | 0.000871396 |
| 1.42E-05 | 0.000871396 |
| 1.41E-05 | 0.000868516 |
| 1.41E-05 | 0.000868516 |
| 1.41E-05 | 0.000868516 |
| 1.37E-05 | 0.000852709 |
| 1.35E-05 | 0.000843787 |
| 1.35E-05 | 0.000843787 |
| 1.35E-05 | 0.00084356 |
| 1.34E-05 | 0.000842 |
| 1.33E-05 | 0.000834171 |
| 1.26E-05 | 0.000797236 |
| 1.24E-05 | 0.000789162 |
| 1.24E-05 | 0.000788404 |
| 1.22E-05 | 0.000778644 |
| 1.22E-05 | 0.000778644 |
| 1.22E-05 | 0.000778644 |
| 1.21E-05 | 0.000776344 |
| 1.21E-05 | 0.000774628 |
| 1.20E-05 | 0.000773439 |
| 1.19E-05 | 0.000769017 |
| 1.19E-05 | 0.000769017 |
| 1.17E-05 | 0.00075514 |
| 1.16E-05 | 0.00075205 |
| 1.16E-05 | 0.00075205 |
| 1.15E-05 | 0.000750224 |
| 1.14E-05 | 0.000744884 |
| 1.14E-05 | 0.000744884 |
| 1.12E-05 | 0.000736367 |

| | |
|---|---|
| 1.11E-05 | 0.000730761 |
| 1.09E-05 | 0.000720913 |
| 1.09E-05 | 0.000719446 |
| 1.08E-05 | 0.00071419 |
| 1.08E-05 | 0.000714069 |
| 1.07E-05 | 0.000712248 |
| 1.06E-05 | 0.000707565 |
| 1.06E-05 | 0.000707354 |
| 1.05E-05 | 0.000700671 |
| 1.05E-05 | 0.000700498 |
| 1.05E-05 | 0.000700498 |
| 1.03E-05 | 0.000695207 |
| 1.03E-05 | 0.000695207 |
| 1.03E-05 | 0.000694541 |
| 1.00E-05 | 0.000678497 |
| 9.80E-06 | 0.000662931 |
| 9.80E-06 | 0.000662931 |
| 9.69E-06 | 0.000659107 |
| 9.33E-06 | 0.000638702 |
| 9.18E-06 | 0.00062984 |
| 9.13E-06 | 0.000627266 |
| 9.10E-06 | 0.000626292 |
| 9.09E-06 | 0.000626292 |
| 8.93E-06 | 0.000617572 |
| 8.93E-06 | 0.000617572 |
| 8.64E-06 | 0.000600728 |
| 8.60E-06 | 0.000599885 |
| 8.13E-06 | 0.000571798 |
| 8.13E-06 | 0.000571798 |
| 8.10E-06 | 0.000571175 |
| 7.85E-06 | 0.000554414 |
| 7.82E-06 | 0.000554136 |
| 7.75E-06 | 0.000549947 |
| 7.65E-06 | 0.000544321 |
| 7.55E-06 | 0.000538791 |
| 7.52E-06 | 0.000537568 |
| 7.51E-06 | 0.000537568 |
| 7.47E-06 | 0.000535645 |
| 7.40E-06 | 0.000531318 |
| 7.33E-06 | 0.0005292 |
| 7.22E-06 | 0.000523233 |
| 7.11E-06 | 0.000516211 |
| 7.08E-06 | 0.000514639 |
| 7.07E-06 | 0.000514581 |
| 6.99E-06 | 0.000509784 |

| | |
|---|---|
| 6.89E-06 | 0.00050425 |
| 6.81E-06 | 0.00049925 |
| 6.73E-06 | 0.000494072 |
| 6.71E-06 | 0.000493419 |
| 6.67E-06 | 0.000492064 |
| 6.52E-06 | 0.000481776 |
| 6.49E-06 | 0.000479711 |
| 6.48E-06 | 0.000479711 |
| 6.46E-06 | 0.000479711 |
| 6.33E-06 | 0.000472101 |
| 6.20E-06 | 0.000463181 |
| 6.09E-06 | 0.00045641 |
| 5.99E-06 | 0.000449538 |
| 5.97E-06 | 0.00044944 |
| 5.91E-06 | 0.000446827 |
| 5.88E-06 | 0.000445184 |
| 5.76E-06 | 0.000438442 |
| 5.66E-06 | 0.000432196 |
| 5.64E-06 | 0.000431255 |
| 5.60E-06 | 0.00042852 |
| 5.56E-06 | 0.000426469 |
| 5.47E-06 | 0.000420115 |
| 5.46E-06 | 0.000420115 |
| 5.39E-06 | 0.000415867 |
| 5.35E-06 | 0.00041363 |
| 5.26E-06 | 0.000407095 |
| 5.23E-06 | 0.000405828 |
| 5.22E-06 | 0.000405183 |
| 5.17E-06 | 0.000403438 |
| 5.14E-06 | 0.000402357 |
| 5.14E-06 | 0.000402265 |
| 5.12E-06 | 0.000401544 |
| 5.11E-06 | 0.000401544 |
| 5.08E-06 | 0.000401044 |
| 4.99E-06 | 0.000395006 |
| 4.97E-06 | 0.000394048 |
| 4.97E-06 | 0.000394048 |
| 4.89E-06 | 0.000389252 |
| 4.79E-06 | 0.000383488 |
| 4.74E-06 | 0.000379691 |
| 4.73E-06 | 0.000379502 |
| 4.65E-06 | 0.000375074 |
| 4.62E-06 | 0.000372606 |
| 4.58E-06 | 0.00037067 |
| 4.54E-06 | 0.000367889 |

| | |
|---|---|
| 4.53E-06 | 0.000367509 |
| 4.51E-06 | 0.00036653 |
| 4.51E-06 | 0.00036653 |
| 4.49E-06 | 0.000366167 |
| 4.43E-06 | 0.00036244 |
| 4.38E-06 | 0.000359897 |
| 4.32E-06 | 0.00035549 |
| 4.32E-06 | 0.00035549 |
| 4.31E-06 | 0.00035549 |
| 4.27E-06 | 0.00035389 |
| 4.23E-06 | 0.000351168 |
| 4.22E-06 | 0.000351168 |
| 4.21E-06 | 0.000351168 |
| 4.17E-06 | 0.000349891 |
| 4.12E-06 | 0.00034607 |
| 4.09E-06 | 0.000343885 |
| 4.05E-06 | 0.000341927 |
| 4.00E-06 | 0.000339205 |
| 3.96E-06 | 0.000336678 |
| 3.94E-06 | 0.000336067 |
| 3.94E-06 | 0.000336067 |
| 3.89E-06 | 0.000333412 |
| 3.88E-06 | 0.0003326 |
| 3.85E-06 | 0.000331723 |
| 3.80E-06 | 0.000327472 |
| 3.76E-06 | 0.000326018 |
| 3.73E-06 | 0.000323271 |
| 3.72E-06 | 0.000323271 |
| 3.69E-06 | 0.000321572 |
| 3.67E-06 | 0.000320336 |
| 3.66E-06 | 0.000320205 |
| 3.65E-06 | 0.000319598 |
| 3.61E-06 | 0.000316557 |
| 3.59E-06 | 0.000315719 |
| 3.49E-06 | 0.0003071 |
| 3.47E-06 | 0.00030626 |
| 3.47E-06 | 0.00030626 |
| 3.45E-06 | 0.000305642 |
| 3.43E-06 | 0.000305117 |
| 3.40E-06 | 0.000304173 |
| 3.38E-06 | 0.00030354 |
| 3.35E-06 | 0.000300776 |
| 3.34E-06 | 0.000300322 |
| 3.29E-06 | 0.000296709 |
| 3.28E-06 | 0.000295985 |

| | |
|---|---|
| 3.24E-06 | 0.000293174 |
| 3.23E-06 | 0.000292925 |
| 3.21E-06 | 0.000292572 |
| 3.18E-06 | 0.00029005 |
| 3.12E-06 | 0.000285351 |
| 3.06E-06 | 0.000280544 |
| 3.05E-06 | 0.000280458 |
| 3.04E-06 | 0.000280458 |
| 3.03E-06 | 0.000279554 |
| 2.99E-06 | 0.000276442 |
| 2.96E-06 | 0.000274685 |
| 2.94E-06 | 0.00027314 |
| 2.92E-06 | 0.000271837 |
| 2.91E-06 | 0.000271679 |
| 2.88E-06 | 0.000270152 |
| 2.86E-06 | 0.000268476 |
| 2.85E-06 | 0.00026785 |
| 2.81E-06 | 0.000264544 |
| 2.76E-06 | 0.000261332 |
| 2.72E-06 | 0.000257803 |
| 2.71E-06 | 0.000257803 |
| 2.71E-06 | 0.000257803 |
| 2.67E-06 | 0.000255413 |
| 2.67E-06 | 0.000255413 |
| 2.62E-06 | 0.000252981 |
| 2.54E-06 | 0.000245724 |
| 2.50E-06 | 0.0002416 |
| 2.43E-06 | 0.000236514 |
| 2.41E-06 | 0.000235519 |
| 2.35E-06 | 0.000230218 |
| 2.31E-06 | 0.000226516 |
| 2.29E-06 | 0.000225207 |
| 2.26E-06 | 0.000222756 |
| 2.23E-06 | 0.00022029 |
| 2.23E-06 | 0.00022029 |
| 2.22E-06 | 0.00022029 |
| 2.17E-06 | 0.00021708 |
| 2.15E-06 | 0.000215905 |
| 2.14E-06 | 0.000215215 |
| 2.14E-06 | 0.000215215 |
| 2.12E-06 | 0.000214574 |
| 2.11E-06 | 0.000214221 |
| 2.10E-06 | 0.000213998 |
| 2.06E-06 | 0.000210783 |
| 2.05E-06 | 0.000210783 |

| | |
|---|---|
| 1.95E-06 | 0.000203044 |
| 1.95E-06 | 0.000203044 |
| 1.94E-06 | 0.000202416 |
| 1.88E-06 | 0.000197315 |
| 1.85E-06 | 0.000195256 |
| 1.84E-06 | 0.000194429 |
| 1.84E-06 | 0.000194429 |
| 1.84E-06 | 0.000194429 |
| 1.84E-06 | 0.000194429 |
| 1.84E-06 | 0.000194429 |
| 1.82E-06 | 0.000194429 |
| 1.82E-06 | 0.000194429 |
| 1.82E-06 | 0.000194429 |
| 1.82E-06 | 0.000194429 |
| 1.81E-06 | 0.000194429 |
| 1.76E-06 | 0.000190667 |
| 1.75E-06 | 0.000190319 |
| 1.75E-06 | 0.000190245 |
| 1.73E-06 | 0.000189236 |
| 1.71E-06 | 0.000186923 |
| 1.65E-06 | 0.000181183 |
| 1.65E-06 | 0.000181183 |
| 1.64E-06 | 0.000181123 |
| 1.62E-06 | 0.000179717 |
| 1.61E-06 | 0.000178843 |
| 1.61E-06 | 0.000178843 |
| 1.58E-06 | 0.000176681 |
| 1.54E-06 | 0.000173117 |
| 1.54E-06 | 0.000173117 |
| 1.51E-06 | 0.000171626 |
| 1.49E-06 | 0.000169791 |
| 1.49E-06 | 0.000169791 |
| 1.48E-06 | 0.000169176 |
| 1.45E-06 | 0.000166964 |
| 1.42E-06 | 0.000164186 |
| 1.41E-06 | 0.000164099 |
| 1.37E-06 | 0.000159405 |
| 1.35E-06 | 0.000158649 |
| 1.34E-06 | 0.000157501 |
| 1.29E-06 | 0.000151987 |
| 1.27E-06 | 0.00015166 |
| 1.25E-06 | 0.000150966 |
| 1.21E-06 | 0.000147662 |
| 1.21E-06 | 0.000147662 |
| 1.21E-06 | 0.000147662 |

| | |
|---|---|
| 1.20E-06 | 0.000147662 |
| 1.19E-06 | 0.000147245 |
| 1.19E-06 | 0.000147201 |
| 1.13E-06 | 0.000140335 |
| 1.10E-06 | 0.000137757 |
| 1.09E-06 | 0.000136906 |
| 1.08E-06 | 0.000136906 |
| 1.07E-06 | 0.000136617 |
| 1.04E-06 | 0.000133968 |
| 1.04E-06 | 0.000133968 |
| 1.02E-06 | 0.000132314 |
| 1.01E-06 | 0.000131648 |
| 9.69E-07 | 0.000127202 |
| 9.63E-07 | 0.00012702 |
| 9.39E-07 | 0.000125251 |
| 9.14E-07 | 0.000122599 |
| 9.02E-07 | 0.00012166 |
| 9.00E-07 | 0.00012166 |
| 8.81E-07 | 0.00011953 |
| 8.36E-07 | 0.000114479 |
| 8.28E-07 | 0.00011429 |
| 8.19E-07 | 0.000113751 |
| 8.10E-07 | 0.000113118 |
| 8.08E-07 | 0.000113118 |
| 7.94E-07 | 0.00011195 |
| 7.87E-07 | 0.00011149 |
| 7.83E-07 | 0.00011149 |
| 7.80E-07 | 0.00011149 |
| 7.59E-07 | 0.000109236 |
| 7.57E-07 | 0.000109211 |
| 7.44E-07 | 0.000107991 |
| 7.38E-07 | 0.000107616 |
| 7.30E-07 | 0.000106987 |
| 7.25E-07 | 0.000106624 |
| 7.20E-07 | 0.00010654 |
| 6.87E-07 | 0.000102413 |
| 6.82E-07 | 0.000102205 |
| 6.45E-07 | 9.70E-05 |
| 6.45E-07 | 9.70E-05 |
| 6.40E-07 | 9.67E-05 |
| 6.33E-07 | 9.60E-05 |
| 6.18E-07 | 9.47E-05 |
| 6.12E-07 | 9.40E-05 |
| 6.08E-07 | 9.37E-05 |
| 5.99E-07 | 9.29E-05 |

| | |
|---|---|
| 5.84E-07 | 9.15E-05 |
| 5.68E-07 | 8.93E-05 |
| 5.24E-07 | 8.35E-05 |
| 5.11E-07 | 8.17E-05 |
| 5.07E-07 | 8.12E-05 |
| 5.05E-07 | 8.12E-05 |
| 4.98E-07 | 8.04E-05 |
| 4.98E-07 | 8.04E-05 |
| 4.96E-07 | 8.04E-05 |
| 4.92E-07 | 8.02E-05 |
| 4.87E-07 | 7.98E-05 |
| 4.70E-07 | 7.79E-05 |
| 4.63E-07 | 7.71E-05 |
| 4.61E-07 | 7.71E-05 |
| 4.43E-07 | 7.52E-05 |
| 4.23E-07 | 7.25E-05 |
| 4.17E-07 | 7.17E-05 |
| 4.11E-07 | 7.12E-05 |
| 4.08E-07 | 7.10E-05 |
| 4.02E-07 | 7.03E-05 |
| 3.82E-07 | 6.76E-05 |
| 3.78E-07 | 6.73E-05 |
| 3.77E-07 | 6.73E-05 |
| 3.75E-07 | 6.73E-05 |
| 3.67E-07 | 6.69E-05 |
| 3.65E-07 | 6.68E-05 |
| 3.45E-07 | 6.40E-05 |
| 3.39E-07 | 6.36E-05 |
| 3.39E-07 | 6.36E-05 |
| 3.36E-07 | 6.36E-05 |
| 3.31E-07 | 6.30E-05 |
| 3.29E-07 | 6.29E-05 |
| 3.19E-07 | 6.12E-05 |
| 3.18E-07 | 6.12E-05 |
| 3.06E-07 | 5.99E-05 |
| 3.04E-07 | 5.99E-05 |
| 2.99E-07 | 5.94E-05 |
| 2.94E-07 | 5.88E-05 |
| 2.90E-07 | 5.83E-05 |
| 2.90E-07 | 5.83E-05 |
| 2.87E-07 | 5.82E-05 |
| 2.74E-07 | 5.60E-05 |
| 2.67E-07 | 5.49E-05 |
| 2.62E-07 | 5.43E-05 |
| 2.51E-07 | 5.26E-05 |

| | |
|---|---|
| 2.45E-07 | 5.15E-05 |
| 2.41E-07 | 5.13E-05 |
| 2.36E-07 | 5.05E-05 |
| 2.36E-07 | 5.05E-05 |
| 2.33E-07 | 5.02E-05 |
| 2.30E-07 | 4.99E-05 |
| 2.29E-07 | 4.99E-05 |
| 2.28E-07 | 4.99E-05 |
| 2.21E-07 | 4.94E-05 |
| 2.21E-07 | 4.94E-05 |
| 2.13E-07 | 4.83E-05 |
| 2.11E-07 | 4.81E-05 |
| 2.10E-07 | 4.81E-05 |
| 2.10E-07 | 4.81E-05 |
| 2.01E-07 | 4.66E-05 |
| 1.99E-07 | 4.65E-05 |
| 1.96E-07 | 4.60E-05 |
| 1.94E-07 | 4.60E-05 |
| 1.90E-07 | 4.57E-05 |
| 1.89E-07 | 4.56E-05 |
| 1.83E-07 | 4.45E-05 |
| 1.79E-07 | 4.38E-05 |
| 1.75E-07 | 4.31E-05 |
| 1.62E-07 | 4.04E-05 |
| 1.57E-07 | 3.96E-05 |
| 1.57E-07 | 3.96E-05 |
| 1.56E-07 | 3.96E-05 |
| 1.53E-07 | 3.91E-05 |
| 1.53E-07 | 3.91E-05 |
| 1.51E-07 | 3.91E-05 |
| 1.48E-07 | 3.89E-05 |
| 1.44E-07 | 3.80E-05 |
| 1.38E-07 | 3.66E-05 |
| 1.33E-07 | 3.60E-05 |
| 1.29E-07 | 3.50E-05 |
| 1.28E-07 | 3.50E-05 |
| 1.26E-07 | 3.48E-05 |
| 1.25E-07 | 3.48E-05 |
| 1.22E-07 | 3.44E-05 |
| 1.16E-07 | 3.32E-05 |
| 1.13E-07 | 3.28E-05 |
| 1.06E-07 | 3.13E-05 |
| 1.04E-07 | 3.07E-05 |
| 1.01E-07 | 3.02E-05 |
| 1.00E-07 | 3.00E-05 |

| | |
|---|---|
| 9.91E-08 | 2.99E-05 |
| 9.90E-08 | 2.99E-05 |
| 9.76E-08 | 2.98E-05 |
| 8.94E-08 | 2.79E-05 |
| 8.58E-08 | 2.72E-05 |
| 8.50E-08 | 2.72E-05 |
| 8.13E-08 | 2.62E-05 |
| 8.04E-08 | 2.61E-05 |
| 6.66E-08 | 2.24E-05 |
| 6.14E-08 | 2.09E-05 |
| 5.74E-08 | 2.00E-05 |
| 5.52E-08 | 1.93E-05 |
| 4.89E-08 | 1.73E-05 |
| 4.74E-08 | 1.70E-05 |
| 4.70E-08 | 1.70E-05 |
| 4.62E-08 | 1.68E-05 |
| 4.56E-08 | 1.67E-05 |
| 4.43E-08 | 1.63E-05 |
| 4.39E-08 | 1.63E-05 |
| 4.30E-08 | 1.63E-05 |
| 4.20E-08 | 1.60E-05 |
| 4.04E-08 | 1.56E-05 |
| 3.55E-08 | 1.41E-05 |
| 3.39E-08 | 1.36E-05 |
| 3.21E-08 | 1.30E-05 |
| 3.19E-08 | 1.30E-05 |
| 3.15E-08 | 1.30E-05 |
| 2.83E-08 | 1.20E-05 |
| 2.66E-08 | 1.15E-05 |
| 2.55E-08 | 1.12E-05 |
| 2.50E-08 | 1.11E-05 |
| 2.39E-08 | 1.08E-05 |
| 2.36E-08 | 1.08E-05 |
| 2.36E-08 | 1.08E-05 |
| 2.07E-08 | 9.96E-06 |
| 1.94E-08 | 9.45E-06 |
| 1.50E-08 | 7.58E-06 |
| 1.49E-08 | 7.58E-06 |
| 1.43E-08 | 7.39E-06 |
| 1.35E-08 | 7.06E-06 |
| 1.28E-08 | 6.92E-06 |
| 1.10E-08 | 6.15E-06 |
| 1.10E-08 | 6.15E-06 |
| 9.34E-09 | 5.43E-06 |
| 8.66E-09 | 5.09E-06 |

| | |
|---|---|
| 7.92E-09 | 4.84E-06 |
| 7.17E-09 | 4.44E-06 |
| 6.30E-09 | 3.95E-06 |
| 6.29E-09 | 3.95E-06 |
| 6.00E-09 | 3.92E-06 |
| 5.88E-09 | 3.92E-06 |
| 5.86E-09 | 3.92E-06 |
| 4.78E-09 | 3.56E-06 |
| 4.12E-09 | 3.16E-06 |
| 3.48E-09 | 2.76E-06 |
| 3.01E-09 | 2.43E-06 |
| 1.40E-09 | 1.28E-06 |
| 1.04E-09 | 1.03E-06 |
| 1.00E-09 | 1.02E-06 |
| 9.81E-10 | 1.02E-06 |
| 8.15E-10 | 9.46E-07 |
| 6.29E-10 | 7.80E-07 |
| 1.29E-10 | 2.28E-07 |
| 2.48E-11 | 7.81E-08 |
| 7.29E-12 | 3.16E-08 |
