## Supplementary Table S6 for "Deep-learning quantified cell-type-specific nuclear morphology predicts genomic instability and prognosis in multiple cancer types"

**Table S6.** Association between OS and CAF nuHIFs.

| | N | coef | coef upper 95% | coef lower 95% | HR | HR upper 95% | HR lower 95% | HR p | corrected HR p |
|---|---|---|---|---|---|---|---|---|---|
| MEAN[FIBROBLAST_NUCLEUS_AREA]_H & E | 869 | 0.22883764114300303 | 0.38886355426388514 | 0.068811728 | 1.2571379149646529 | 1.4753032387628406 | 1.0712345067221318 | 0.005066804 | 0.038001029 |
| MEAN[FIBROBLAST_NUCLEUS_CIRCULARITY]_H & E | 869 | 0.21907000287810188 | 0.3980153129908672 | 0.040124693 | 1.2449184215098092 | 1.488866828736869 | 1.0409405638577625 | 0.016419974883189647 | 0.061574906 |
| MEAN[FIBROBLAST_NUCLEUS_ECCENTRICITY]_H & E | 869 | -0.201185114 | -0.023726668 | -0.378643559 | 0.8177610387204644 | 0.9765525960950849 | 0.6847896561057932 | 0.026282196919975798 | 0.078846591 |
| MEAN[FIBROBLAST_NUCLEUS_MAJOR_AXIS_LENGTH]_H & E | 869 | 0.20331182 | 0.364797601 | 0.041826039 | 1.225454530418393 | 1.4402224797281344 | 1.042713071945969 | 0.013601861706664872 | 0.061574906 |
| MEAN[FIBROBLAST_NUCLEUS_MEAN_GRAYSCALE_CHANNEL_GRAY]_H & E | 869 | -0.025002789 | 0.17335573651486583 | -0.223361314 | 0.975307192 | 1.1892891034024098 | 0.7998258085287959 | 0.8048691693576534 | 0.8623598243117715 |
| MEAN[FIBROBLAST_NUCLEUS_MEAN_HSV_CHANNEL_SATURATION]_H & E | 869 | 0.19417616938038595 | 0.3973391154018358 | -0.008986777 | 1.2143101883873708 | 1.4878604008872796 | 0.991053484 | 0.061031622 | 0.1307820473065336 |
| MEAN[FIBROBLAST_NUCLEUS_MEAN_LAB_CHANNEL_A]_H & E | 869 | 0.27105931360613184 | 0.4390716636178024 | 0.10304696359446125 | 1.3113528491442898 | 1.5512664527705908 | 1.1085434690394589 | 0.001566529 | 0.03748984 |
| MEAN[FIBROBLAST_NUCLEUS_MEAN_LAB_CHANNEL_B]_H & E | 869 | -0.207261655 | 0.005935926 | -0.420459235 | 0.8128069474833756 | 1.0059535782370452 | 0.6567451502434684 | 0.056728882 | 0.1307820473065336 |
| MEAN[FIBROBLAST_NUCLEUS_MINOR_AXIS_LENGTH]_H & E | 869 | 0.25794498530620336 | 0.4323645183140181 | 0.083525452 | 1.2942676129291597 | 1.5408966978677008 | 1.0871128844622713 | 0.003748984 | 0.03748984 |
| MEAN[FIBROBLAST_NUCLEUS_PERIMETER]_H & E | 869 | 0.21367787153220524 | 0.37241064443956773 | 0.054945099 | 1.2382237234325115 | 1.451228797986814 | 1.0564826107351017 | 0.008329767 | 0.049978601223705045 |
| MEAN[FIBROBLAST_NUCLEUS_SOLIDITY]_H & E | 869 | 0.28353611501255366 | 0.4696642095882806 | 0.09740802 | 1.3278168335695544 | 1.5994570207022087 | 1.1023100469036837 | 0.002829404 | 0.03748984 |
| MEAN[FIBROBLAST_NUCLEUS_STD_GRAYSCALE_CHANNEL_GRAY]_H & E | 869 | -0.086441717 | 0.093634616 | -0.26651805 | 0.9171890040219934 | 1.0981584235961206 | 0.7660421766324718 | 0.3467879001470321 | 0.5201818502205482 |
| MEAN[FIBROBLAST_NUCLEUS_STD_HSV_CHANNEL_SATURATION]_H & E | 869 | -0.007210141 | 0.19303030336580507 | -0.207450584 | 0.9928157901559341 | 1.2129195485069875 | 0.8126533984848819 | 0.9437374130810634 | 0.9437374130810634 |
| MEAN[FIBROBLAST_NUCLEUS_STD_LAB_CHANNEL_A]_H & E | 869 | 0.036191887 | 0.19725711954995448 | -0.124873347 | 1.0368547858539063 | 1.2180571867618757 | 0.8826086809652564 | 0.6596399370471133 | 0.791360028 |
| MEAN[FIBROBLAST_NUCLEUS_STD_LAB_CHANNEL_B]_H & E | 869 | 0.10983618811670914 | 0.2799909414001165 | -0.060318565 | 1.1160952258223442 | 1.3231178266881598 | 0.9414645679904485 | 0.2058097965823532 | 0.34301632763725537 |
| STD[FIBROBLAST_NUCLEUS_AREA]_H & E | 869 | 0.20493439396244956 | 0.3704244400976033 | 0.039444347827295834 | 1.2274445345493412 | 1.4483492217071858 | 1.040232606069397 | 0.015219263918093577 | 0.061574906 |
| STD[FIBROBLAST_NUCLEUS_CIRCULARITY]_H & E | 869 | -0.045248494 | 0.11071997374886619 | -0.201216962 | 0.9557599517838258 | 1.1170820507536607 | 0.8177349952203835 | 0.5696200185994056 | 0.712025023 |
| STD[FIBROBLAST_NUCLEUS_ECCENTRICITY]_H & E | 869 | 0.17699824163198102 | 0.36214657145449025 | -0.008150088 | 1.1936289942862435 | 1.4364094631560342 | 0.9918830337349436 | 0.060973459 | 0.1307820473065336 |
| STD[FIBROBLAST_NUCLEUS_MAJOR_AXIS_LENGTH]_H & E | 869 | 0.13235315554075103 | 0.2887277468683976 | -0.024021436 | 1.1415113791600096 | 1.3347282950996295 | 0.9762647825297824 | 0.097139027 | 0.18213567539329337 |
| STD[FIBROBLAST_NUCLEUS_MEAN_GRAYSCALE_CHANNEL_GRAY]_H & E | 869 | -0.039188571 | 0.15438780909782746 | -0.232764951 | 0.9615693678234865 | 1.1669433502859266 | 0.7923397900248614 | 0.6915260739282402 | 0.791360028 |
| STD[FIBROBLAST_NUCLEUS_MEAN_HSV_CHANNEL_SATURATION]_H & E | 869 | -0.040197584 | 0.17338889593133883 | -0.253784064 | 0.9605996213955943 | 1.1893285401889433 | 0.7758593201494735 | 0.7122240256390263 | 0.791360028 |
| STD[FIBROBLAST_NUCLEUS_MEAN_LAB_CHANNEL_A]_H & E | 869 | -0.054944079 | 0.10363661062249398 | -0.213524769 | 0.9465380776420471 | 1.1091973111505076 | 0.8077321531702957 | 0.4970894052285969 | 0.6778491889480868 |
| STD[FIBROBLAST_NUCLEUS_MEAN_LAB_CHANNEL_B]_H & E | 869 | 0.015400586005668591 | 0.18688443856223602 | -0.156083267 | 1.0155197861616163 | 1.205487969077389 | 0.8554879538739965 | 0.8602782016967794 | 0.8899429672725304 |
| STD[FIBROBLAST_NUCLEUS_MINOR_AXIS_LENGTH]_H & E | 869 | 0.1955175423269703 | 0.37360701922568895 | 0.017428065 | 1.2159401241543137 | 1.4529660505249724 | 1.0175808202773946 | 0.031415585448690705 | 0.085678869 |
| STD[FIBROBLAST_NUCLEUS_PERIMETER]_H & E | 869 | 0.13057095025932353 | 0.28363539272045407 | -0.022493492 | 1.1394787833407667 | 1.3279486627250425 | 0.9777576002217757 | 0.094536417 | 0.18213567539329337 |
| STD[FIBROBLAST_NUCLEUS_SOLIDITY]_H & E | 869 | -0.10954533 | 0.056931802 | -0.276022462 | 0.8962415370053036 | 1.058583614750257 | 0.7587958867501771 | 0.19715620815246307 | 0.34301632763725537 |
| STD[FIBROBLAST_NUCLEUS_STD_GRAYSCALE_CHANNEL_GRAY]_H & E | 869 | 0.051534771 | 0.22411834302407632 | -0.1210488 | 1.0528857960571798 | 1.2512190837171504 | 0.8859907221408423 | 0.5583727701402865 | 0.712025023 |
| STD[FIBROBLAST_NUCLEUS_STD_HSV_CHANNEL_SATURATION]_H & E | 869 | 0.08323761 | 0.28854760923606965 | -0.122072389 | 1.086800012434261 | 1.3344878819591264 | 0.8850842956274116 | 0.4268366037891962 | 0.6097665768417089 |
| STD[FIBROBLAST_NUCLEUS_STD_LAB_CHANNEL_A]_H & E | 869 | 0.1908178298474283 | 0.35025301745986576 | 0.031382642 | 1.2102389625821446 | 1.4194266428864273 | 1.031880269327239 | 0.018988525287551253 | 0.063295084 |
| STD[FIBROBLAST_NUCLEUS_STD_LAB_CHANNEL_B]_H & E | 869 | 0.11507774784973267 | 0.301285996 | -0.0711305 | 1.1219606642101796 | 1.3515958370087215 | 0.9313403441822118 | 0.22579240190880523 | 0.3565143188033767 |
