## Supplementary Table S5 for "Deep-learning quantified cell-type-specific nuclear morphology predicts genomic instability and prognosis in multiple cancer types"

**Table S5**. Association between PFS and CAF nuHIFs.

| | N | coef | coef upper 95% | coef lower 95% | HR | HR upper 95% | HR lower 95% | HR p | corrected HR p |
|---|---|---|---|---|---|---|---|---|---|
| MEAN[FIBROBLAST_NUCLEUS_AREA]_H & E | 869 | 0.2227848577696467 | 0.36764703963953405 | 0.077922676 | 1.2495517134728442 | 1.4443321569963432 | 1.0810390650651935 | 0.002576186 | 0.014201608156398616 |
| MEAN[FIBROBLAST_NUCLEUS_CIRCULARITY]_H & E | 869 | 0.2379840056235913 | 0.3917310382564573 | 0.084236973 | 1.2686889007748074 | 1.479539718299954 | 1.0878866630215556 | 0.002414823 | 0.014201608156398616 |
| MEAN[FIBROBLAST_NUCLEUS_ECCENTRICITY]_H & E | 869 | -0.232814576 | -0.079922456 | -0.385706695 | 0.7923004717427704 | 0.9231879310552198 | 0.679969935 | 0.002840322 | 0.014201608156398616 |
| MEAN[FIBROBLAST_NUCLEUS_MAJOR_AXIS_LENGTH]_H & E | 869 | 0.1596697811624682 | 0.30582347879333094 | 0.013516084 | 1.1731234195715599 | 1.357742615028446 | 1.013607838712816 | 0.032256411 | 0.087972031 |
| MEAN[FIBROBLAST_NUCLEUS_MEAN_GRAYSCALE_CHANNEL_GRAY]_H & E | 869 | 0.065036912 | 0.23587606144379297 | -0.105802238 | 1.0671984156037477 | 1.266017392075273 | 0.8996025373713299 | 0.45558256193526336 | 0.706641603 |
| MEAN[FIBROBLAST_NUCLEUS_MEAN_HSV_CHANNEL_SATURATION]_H & E | 869 | 0.093603 | 0.2708139525931644 | -0.083607953 | 1.0981237043088956 | 1.3110311337506875 | 0.9197917874880968 | 0.3005498807499173 | 0.5635310264060949 |
| MEAN[FIBROBLAST_NUCLEUS_MEAN_LAB_CHANNEL_A]_H & E | 869 | 0.22598340100368636 | 0.3739866939367227 | 0.077980108 | 1.253554857353036 | 1.4535178097282275 | 1.081101153268431 | 0.002765834 | 0.014201608156398616 |
| MEAN[FIBROBLAST_NUCLEUS_MEAN_LAB_CHANNEL_B]_H & E | 869 | -0.087323609 | 0.09259106 | -0.267238278 | 0.9163804985842817 | 1.097013030994594 | 0.7654906500283085 | 0.3414568905805737 | 0.5690948176342896 |
| MEAN[FIBROBLAST_NUCLEUS_MINOR_AXIS_LENGTH]_H & E | 869 | 0.25856325882051084 | 0.41417469008707103 | 0.10295182755395066 | 1.2950680717404022 | 1.5131214311126082 | 1.1084380116195607 | 0.001127284 | 0.014201608156398616 |
| MEAN[FIBROBLAST_NUCLEUS_PERIMETER]_H & E | 869 | 0.1916653687161907 | 0.3363776101145092 | 0.046953127 | 1.211265121936967 | 1.399867529 | 1.0480728819412624 | 0.009434596 | 0.029272096 |
| MEAN[FIBROBLAST_NUCLEUS_SOLIDITY]_H & E | 869 | 0.23388500818645439 | 0.3924566430942096 | 0.075313373 | 1.2634991917902814 | 1.4806136690620133 | 1.0782219839062084 | 0.003842062 | 0.014407733128836036 |
| MEAN[FIBROBLAST_NUCLEUS_STD_GRAYSCALE_CHANNEL_GRAY]_H & E | 869 | -0.100422646 | 0.061730903 | -0.262576194 | 0.904455073 | 1.0636760738864273 | 0.769067764 | 0.2248175305373894 | 0.4496350610747788 |
| MEAN[FIBROBLAST_NUCLEUS_STD_HSV_CHANNEL_SATURATION]_H & E | 869 | -0.063958176 | 0.10997800088400203 | -0.237894352 | 0.9380442318908352 | 1.1162535135982288 | 0.788285968 | 0.4710944016778187 | 0.706641603 |
| MEAN[FIBROBLAST_NUCLEUS_STD_LAB_CHANNEL_A]_H & E | 869 | -0.04800152 | 0.095205591 | -0.19120863 | 0.9531323384853171 | 1.0998849580170822 | 0.8259602497922148 | 0.5112070555191986 | 0.7302957935988552 |
| MEAN[FIBROBLAST_NUCLEUS_STD_LAB_CHANNEL_B]_H & E | 869 | -0.004129406 | 0.14309559784559614 | -0.151354409 | 0.995879109 | 1.153840101021556 | 0.8595430149040563 | 0.9561595083241762 | 0.9841195145055311 |
| STD[FIBROBLAST_NUCLEUS_AREA]_H & E | 869 | 0.221291306 | 0.36962627760696515 | 0.072956335 | 1.2476868365896514 | 1.447193664907468 | 1.0756835660268225 | 0.003456281 | 0.014407733128836036 |
| STD[FIBROBLAST_NUCLEUS_CIRCULARITY]_H & E | 869 | -0.038712564 | 0.098087109 | -0.175512236 | 0.9620271911117604 | 1.1030588669466053 | 0.8390271309819253 | 0.5791376225077487 | 0.7882303635937439 |
| STD[FIBROBLAST_NUCLEUS_ECCENTRICITY]_H & E | 869 | 0.21153111646859174 | 0.37195813372737674 | 0.051104099 | 1.2355684115562173 | 1.4505722499685936 | 1.0524324449668792 | 0.009757365 | 0.029272096 |
| STD[FIBROBLAST_NUCLEUS_MAJOR_AXIS_LENGTH]_H & E | 869 | 0.10443350641121861 | 0.24426013927258378 | -0.035393126 | 1.1100815781041382 | 1.2766764009655678 | 0.9652258858346427 | 0.14323357938941214 | 0.30692909869159746 |
| STD[FIBROBLAST_NUCLEUS_MEAN_GRAYSCALE_CHANNEL_GRAY]_H & E | 869 | 0.035925979 | 0.2024523615004806 | -0.130600404 | 1.0365791145573062 | 1.2244017550701274 | 0.8775683767905637 | 0.6724137176737344 | 0.8093532720532981 |
| STD[FIBROBLAST_NUCLEUS_MEAN_HSV_CHANNEL_SATURATION]_H & E | 869 | 0.033666146 | 0.2150916134249105 | -0.147759321 | 1.0342392643464278 | 1.2399754901641105 | 0.8626387089105061 | 0.716082078 | 0.8093532720532981 |
| STD[FIBROBLAST_NUCLEUS_MEAN_LAB_CHANNEL_A]_H & E | 869 | 0.001391296 | 0.13838964765197345 | -0.135607055 | 1.0013922646339017 | 1.1484229434090478 | 0.8731856790424986 | 0.9841195145055311 | 0.9841195145055311 |
| STD[FIBROBLAST_NUCLEUS_MEAN_LAB_CHANNEL_B]_H & E | 869 | 0.072875222 | 0.21780131163288005 | -0.072050868 | 1.0755963178503476 | 1.2433400058801847 | 0.9304835632263185 | 0.3243509251454352 | 0.5690948176342896 |
| STD[FIBROBLAST_NUCLEUS_MINOR_AXIS_LENGTH]_H & E | 869 | 0.24653509947949367 | 0.4043060535781936 | 0.088764145 | 1.2795840951651138 | 1.4982624253714385 | 1.0928228786045986 | 0.002193739 | 0.014201608156398616 |
| STD[FIBROBLAST_NUCLEUS_PERIMETER]_H & E | 869 | 0.14010603208743908 | 0.2799843364818857 | 0.000227728 | 1.1503957712556263 | 1.323109087631965 | 1.000227753624912 | 0.049628179 | 0.12407044865062226 |
| STD[FIBROBLAST_NUCLEUS_SOLIDITY]_H & E | 869 | -0.039343112 | 0.10945890951897358 | -0.188145133 | 0.9614207777286736 | 1.1156742264026356 | 0.8284944565124571 | 0.6043099454218703 | 0.7882303635937439 |
| STD[FIBROBLAST_NUCLEUS_STD_GRAYSCALE_CHANNEL_GRAY]_H & E | 869 | 0.013555342438795838 | 0.16222127221449137 | -0.135110587 | 1.0136476326300115 | 1.1761204553085303 | 0.8736192951145373 | 0.858165973 | 0.9194635425950196 |
| STD[FIBROBLAST_NUCLEUS_STD_HSV_CHANNEL_SATURATION]_H & E | 869 | 0.036932767 | 0.2163253989581214 | -0.142459865 | 1.0376232561382357 | 1.241506298135589 | 0.8672223598831302 | 0.6865723719041734 | 0.8093532720532981 |
| STD[FIBROBLAST_NUCLEUS_STD_LAB_CHANNEL_A]_H & E | 869 | 0.12493456522363297 | 0.268161737 | -0.018292606 | 1.133074308177063 | 1.3075586032597502 | 0.9818736878410397 | 0.087332016 | 0.20153542221466528 |
| STD[FIBROBLAST_NUCLEUS_STD_LAB_CHANNEL_B]_H & E | 869 | -0.02847243 | 0.13224182575055837 | -0.189186686 | 0.97192909 | 1.1413843020115324 | 0.8276319853440098 | 0.7284179448479683 | 0.8093532720532981 |
