## Supplementary Table S4 for "Deep-learning quantified cell-type-specific nuclear morphology predicts genomic instability and prognosis in multiple cancer types"

**Table S4.** Top nuHIF Mean Importance for WGD Prediction

| BRCA | Mean Importance Score |
|---|---|
| Standard deviation of cancer cell nucleus major axis length | 0.0685 |
| Mean of cancer cell nucleus major axis length | 0.0655 |
| Standard deviation of cancer cell nucleus minor axis length | 0.0572 |
| Standard deciation of cancer cell nucleus area | 0.0521 |
| Mean of cancer cell nucleus perimeter | 0.0519 |
| **LUAD** | **Mean Importance Score** |
| Mean cancer cell nucleus perimeter | 0.0479 |
| Standard deviation of cancer cell nucleus minor axis length | 0.0441 |
| Mean cancer cell nucleus major axis length | 0.0425 |
| Mean of the cancer cell nucleus saturation (in HSV colorspace) standard deviation | 0.0367 |
| Mean cancer cell nucleus eccentricity | 0.0355 |
| **PRAD** | **Mean Importance Score** |
| Standard deviation of cancer cell nucleus major axis length | 0.0618 |
| Mean of the cancer cell nucleus saturation (in HSV colorspace) mean value | 0.0544 |
| Mean of the cancer cell nucleus grayscale value mean value | 0.0481 |
| Slide-level standard deviation of the cancer cell nucleus grayscale value standard deviation | 0.046 |
| Mean cancer nucleus circularity | 0.0436 |
