## Supplementary Table S3 for "Deep-learning quantified cell-type-specific nuclear morphology predicts genomic instability and prognosis in multiple cancer types"

**Table S3.** nuHIFs extracted from the nuclear segmentation model.

| nuHIF Name | Descriptor |
|---|---|
| MEAN[CELL TYPE X_NUCLEUS_AREA]_H & E | Slide-level mean of the area of each nucleus of cell type X, in um^2. |
| MEAN[CELL TYPE X_NUCLEUS_CIRCULARITY]_H & E | Slide-level mean of the circularity of each nucleus of cell type X. |
| MEAN[CELL TYPE X_NUCLEUS_ECCENTRICITY]_H & E | Slide-level mean of the eccentricity of each nucleus of cell type X. |
| MEAN[CELL TYPE X_NUCLEUS_MAJOR_AXIS_LENGTH]_H & E | Slide-level mean of the major axis length of each nucleus of cell type X, in um. |
| MEAN[CELL TYPE X_NUCLEUS_MEAN_GRAYSCALE_CHANNEL_GRAY]_H & E | Slide-level mean of the (mean grayscale intensity value of each nucleus) of cell type X. |
| MEAN[CELL TYPE X_NUCLEUS_MEAN_HSV_CHANNEL_SATURATION]_H & E | Slide-level mean of the (mean saturation value in HSV colorspace of each nucleus) of cell type X. |
| MEAN[CELL TYPE X_NUCLEUS_MEAN_LAB_CHANNEL_A]_H & E | Slide-level mean of the (mean A value in LAB colorspace of each nucleus) of cell type X. |
| MEAN[CELL TYPE X_NUCLEUS_MEAN_LAB_CHANNEL_B]_H & E | Slide-level mean of the (mean B value in LAB colorspace of each nucleus) of cell type X. |
| MEAN[CELL TYPE X_NUCLEUS_MINOR_AXIS_LENGTH]_H & E | Slide-level mean of the minor axis length of each nucleus of cell type X, in um. |
| MEAN[CELL TYPE X_NUCLEUS_PERIMETER]_H & E | Slide-level mean of the perimeter length of each nucleus of cell type X, in um. |
| MEAN[CELL TYPE X_NUCLEUS_SOLIDITY]_H & E | Slide-level mean of the solidity of each nucleus of cell type X. |
| MEAN[CELL TYPE X_NUCLEUS_STD_GRAYSCALE_CHANNEL_GRAY]_H & E | Slide-level mean of the (standard deviation of the grayscale intensity value of each nucleus) of cell type X. |
| MEAN[CELL TYPE X_NUCLEUS_STD_HSV_CHANNEL_SATURATION]_H & E | Slide-level mean of the (standard deviation of the saturation value in HSV colorspace of each nucleus) of cell type X. |
| MEAN[CELL TYPE X_NUCLEUS_STD_LAB_CHANNEL_A]_H & E | Slide-level mean of the (standard deviation of the A value in LAB colorspace of each nucleus) of cell type X. |
| MEAN[CELL TYPE X_NUCLEUS_STD_LAB_CHANNEL_B]_H & E | Slide-level mean of the (standard deviation of the B value in LAB colorspace of each nucleus) of cell type X. |
| STD[CELL TYPE X_NUCLEUS_AREA]_H & E | Slide-level standard deviation of the area of each nucleus of cell type X, in um^2. |
| STD[CELL TYPE X_NUCLEUS_CIRCULARITY]_H & E | Slide-level standard deviation of the circularity of each nucleus of cell type X. |
| STD[CELL TYPE X_NUCLEUS_ECCENTRICITY]_H & E | Slide-level standard deviation of the eccentricity of each nucleus of cell type X. |
| STD[CELL TYPE X_NUCLEUS_MAJOR_AXIS_LENGTH]_H & E | Slide-level standard deviation of the major axis length of each nucleus of cell type X, in um. |
| STD[CELL TYPE X_NUCLEUS_MEAN_GRAYSCALE_CHANNEL_GRAY]_H & E | Slide-level standard deviation of the (mean grayscale intensity value of each nucleus) of cell type X. |
| STD[CELL TYPE X_NUCLEUS_MEAN_HSV_CHANNEL_SATURATION]_H & E | Slide-level standard deviation of the (mean saturation value in HSV colorspace of each nucleus) of cell type X. |
| STD[CELL TYPE X_NUCLEUS_MEAN_LAB_CHANNEL_A]_H & E | Slide-level standard deviation of the (mean A value in LAB colorspace of each nucleus) of cell type X. |
| STD[CELL TYPE X_NUCLEUS_MEAN_LAB_CHANNEL_B]_H & E | Slide-level standard deviation of the (mean B value in LAB colorspace of each nucleus) of cell type X. |
| STD[CELL TYPE X_NUCLEUS_MINOR_AXIS_LENGTH]_H & E | Slide-level standard deviation of the minor axis length of each nucleus of cell type X, in um. |
| STD[CELL TYPE X_NUCLEUS_PERIMETER]_H & E | Slide-level standard deviation of the perimeter length of each nucleus of cell type X, in um. |
| STD[CELL TYPE X_NUCLEUS_SOLIDITY]_H & E | Slide-level standard deviation of the solidity of each nucleus of cell type X. |
| STD[CELL TYPE X_NUCLEUS_STD_GRAYSCALE_CHANNEL_GRAY]_H & E | Slide-level standard deviation of the (standard deviation of the grayscale intensity value of each nucleus) of cell type X. |
| STD[CELL TYPE X_NUCLEUS_STD_HSV_CHANNEL_SATURATION]_H & E | Slide-level standard deviation of the (standard deviation of the saturation value in HSV colorspace of each nucleus) of cell type X. |
| STD[CELL TYPE X_NUCLEUS_STD_LAB_CHANNEL_A]_H & E | Slide-level standard deviation of the (standard deviation of the A value in LAB colorspace of each nucleus) of cell type X. |
| STD[CELL TYPE X_NUCLEUS_STD_LAB_CHANNEL_B]_H & E | Slide-level standard deviation of the (standard deviation of the B value in LAB colorspace of each nucleus) of cell type X. |
