## Supplementary Table S2 for "Deep-learning quantified cell-type-specific nuclear morphology predicts genomic instability and prognosis in multiple cancer types"

**Table S2. Summary table of cell model performance characteristics.**

| | Model vs. Avg Pathologist [95% CI] | Δrecall [95% CI] | Model vs. Avg Pathologist [95% CI] | ΔPrecision [95% CI] | Model vs. Avg Pathologist [95% CI] | ΔF1 [95% CI] |
|---|---|---|---|---|---|---|
| | | | BRCA | | | |
| | Model vs. Avg Pathologist [95% CI] | Δrecall [95% CI] | Model vs. Avg Pathologist [95% CI] | ΔPrecision [95% CI] | Model vs. Avg Pathologist [95% CI] | ΔF1 [95% CI] |
| **Cancer Cells (N = 486 cells)** | 0.36 [0.404, 0.248] | -0.204 [-0.141, -0.271] | -0.057 [0, -0.137] | -0.057 [0, -0.137] | 0.446 [0.508, 0.364] | -0.101 [-0.064, -0.189] |
| **Fibroblasts (N = 476 cells)** | 0.233 [0.278, 0.176] | -0.159 [-0.112, -0.228] | -0.042 [0.002, -0.087] | -0.002 [0.032, -0.043] | 0.278 [0.328, 0.257] | -0.077 [-0.041, -0.129] |
| **Lymphocytes (N = 1,624 cells)** | 0.589 [0.603, 0.483] | 0.003 [0.033, -0.064] | -0.052 [-0.016, -0.077] | -0.042 [0.002, -0.087] | 0.566 [0.592, 0.513] | 0.013 [0.033, -0.021] |
| **Macrophages (N = 299 cells)** | 0.144 [0.18, 0.099] | 0.013 [0.057, -0.026] | -0.002 [0.032, -0.043] | -0.275 [-0.2, -0.35] | 0.213 [0.3, 0.205] | -0.009 [0.061, -0.078] |
| **Plasma Cells (N = 194 cells)** | 0.354 [0.423, 0.194] | -0.108 [-0.001, -0.216] | -0.275 [-0.2, -0.35] | -0.052 [-0.016, -0.077] | 0.387 [0.495, 0.342] | -0.179 [-0.127, -0.282] |
| | | | LUAD | | | |
| | Model vs. Avg Pathologist [95% CI] | Δrecall [95% CI] | Model vs. Avg Pathologist [95% CI] | ΔPrecision [95% CI] | Model vs. Avg Pathologist [95% CI] | ΔF1 [95% CI] |
| **Cancer Cells (N = 5,492 cells)** | 0.547 [0.506, 0.586] | -0.132 [-0.161, -0.0991] | 0.717 [0.679, 0.745] | 0.0377 [0.00369, 0.0640] | 0.608 [0.570, 0.638] | -0.051 [-0.0743, -0.0273] |
| **Fibroblasts (N = 5,960 cells)** | 0.451 [0.412, 0.486] | 0.0284 [-0.00804, 0.0623] | 0.337 [0.319, 0.356] | -0.0863 [-0.103, -0.0675] | 0.306 [0.285, 0.326] | 0.0308 [0.0171, 0.0485] |
| **Lymphocytes (N = 8,940 cells)** | 0.485 [0.420, 0.495] | -0.113 [-0.156, -0.0691] | 0.549 [0.521, 0.575] | -0.0218 [-0.0414, -0.00144] | 0.477 [0.445, 0.500] | -0.0509 [-0.0836, -0.0226] |
| **Macrophages (N = 2,100 cells)** | 0.164 [0.136, 0.195] | -0.108 [-0.140, -0.0748] | 0.307 [0.281, 0.333] | 0.0349 [0.00816, 0.0595] | 0.185 [0.161, 0.210] | -0.0121 [-0.0393, 0.0125] |
| **Plasma Cells (N = 4,697 cells)** | 0.563 [0.513, 0.597] | 0.0449 [0.00601, 0.0754] | 0.409 [0.380, 0.435] | -0.11 [-0.131, -0.0868] | 0.425 [0.389, 0.454] | -0.00294 [-0.0208, 0.0159] |
| | | | PRAD | | | |
| | Model vs. Avg Pathologist [95% CI] | Δrecall [95% CI] | Model vs. Avg Pathologist [95% CI] | ΔPrecision [95% CI] | Model vs. Avg Pathologist [95% CI] | ΔF1 [95% CI] |
| **Cancer Cells (N = 2,539 cells)** | 0.615 [0.553, 0.641] | -0.02 [-0.050, 0.008] | 0.567 [0.514, 0.609] | -0.082 [-0.122, -0.037] | 0.588 [0.533, 0.611] | -0.042 [-0.075, -0.013] |
| **Fibroblasts (N = 2,844 cells)** | 0.451 [0.422, 0.484] | 0.02 [-0.004, 0.062] | 0.380 [0.355, 0.399] | -0.04 [-0.067, -0.027] | 0.359 [0.339, 0.382] | 0.014 [-0.009, 0.033] |
| **Lymphocytes (N = 975 cells)** | 0.491 [0.423, 0.534] | 0.14 [0.106, 0.211] | 0.226 [0.183, 0.251] | -0.128 [-0.165, -0.073] | 0.325 [0.296, 0.373] | -0.006 [-0.045, 0.029] |
| **Macrophages (N = 336 cells)** | 0.115 [0.069, 0.150] | 0.01 [-0.048, 0.083] | 0.079 [0.060, 0.096] | -0.025 [-0.08, 0.028] | 0.099 [0.089, 0.137] | -0.042 [-0.142, 0.032] |
| **Plasma Cells (N = 169 cells)** | 0.365 [0.217, 0.452] | 0.18 [0.071, 0.259] | 0.104 [0.051, 0.137] | -0.082 [-0.147, -0.025] | 0.315 [0.239, 0.424] | 0.016 [-0.174, 0.075] |
