## Supplementary Table S1 for "Deep-learning quantified cell-type-specific nuclear morphology predicts genomic instability and prognosis in multiple cancer types"

**Table S1. Summary of agreement (ICC) between model-derived cell counts and pathologists' cell counts.**

| | BRCA (n=177 frames) | | | LUAD (n=834 frames) | | | PRAD (n=409 frames) | | |
|---|---|---|---|---|---|---|---|---|---|
| | Number of Cells | Model vs. Consensus [95% CI] | ΔICC [95% CI] | Number of Cells | Model vs. Consensus [95% CI] | ΔICC [95% CI] | Number of Cells | Model vs. Consensus [95% CI] | ΔICC [95% CI] |
| Cancer Cells | 486 | 0.7 [0.615, 0.768] | -0.018 [-0.209, -0.011] | 5492 | 0.745 [0.704, 0.780] | -0.0199 [-0.0634, 0.0322] | 2539 | 0.839 [0.805, 0.867] | 0.002 [-0.030, 0.068] |
| Fibroblasts | 476 | 0.401 [0.266, 0.52] | -0.058 [-0.093, -0.027] | 5960 | 0.517 [0.442, 0.581] | 0.00818 [-0.0435, 0.0505 | 2844 | 0.37 [0.282, 0.451] | 0.038 [-0.024, 0.068] |
| Lymphocytes | 1624 | 0.898 0.865, 0.923] | 0.043 [-0.025, 0.075] | 8940 | 0.631 [0.579, 0.676] | -0.120 [-0.188, -0.0137] | 975 | 0.59 [0.432, 0.698] | 0.022 [-0.026, 0.063] |
| Macrophages | 299 | 0.18 [0.039, 0.315] | 0.049 [0.04, 0.087] | 2100 | 0.398 [0.336, 0.457] | 0.00524 [-0.0570, 0.0833 | 336 | 0.214 [0.117, 0.307] | -0.003 [-0.076, 0.076] |
| Plasma Cells | 194 | 0.531 [0.403, 0.636] | -0.102 [-0.169, -0.033] | 4697 | 0.688 [0.593, 0.758] | 0.0441 [0.0190, 0.0783] | 169 | 0.228 [0.135, 0.318] | 0.076 [-0.006, 0.097] |
