## Supplementary figure S1 through S6 for "Deep-learning quantified cell-type-specific nuclear morphology predicts genomic instability and prognosis in multiple cancer types"

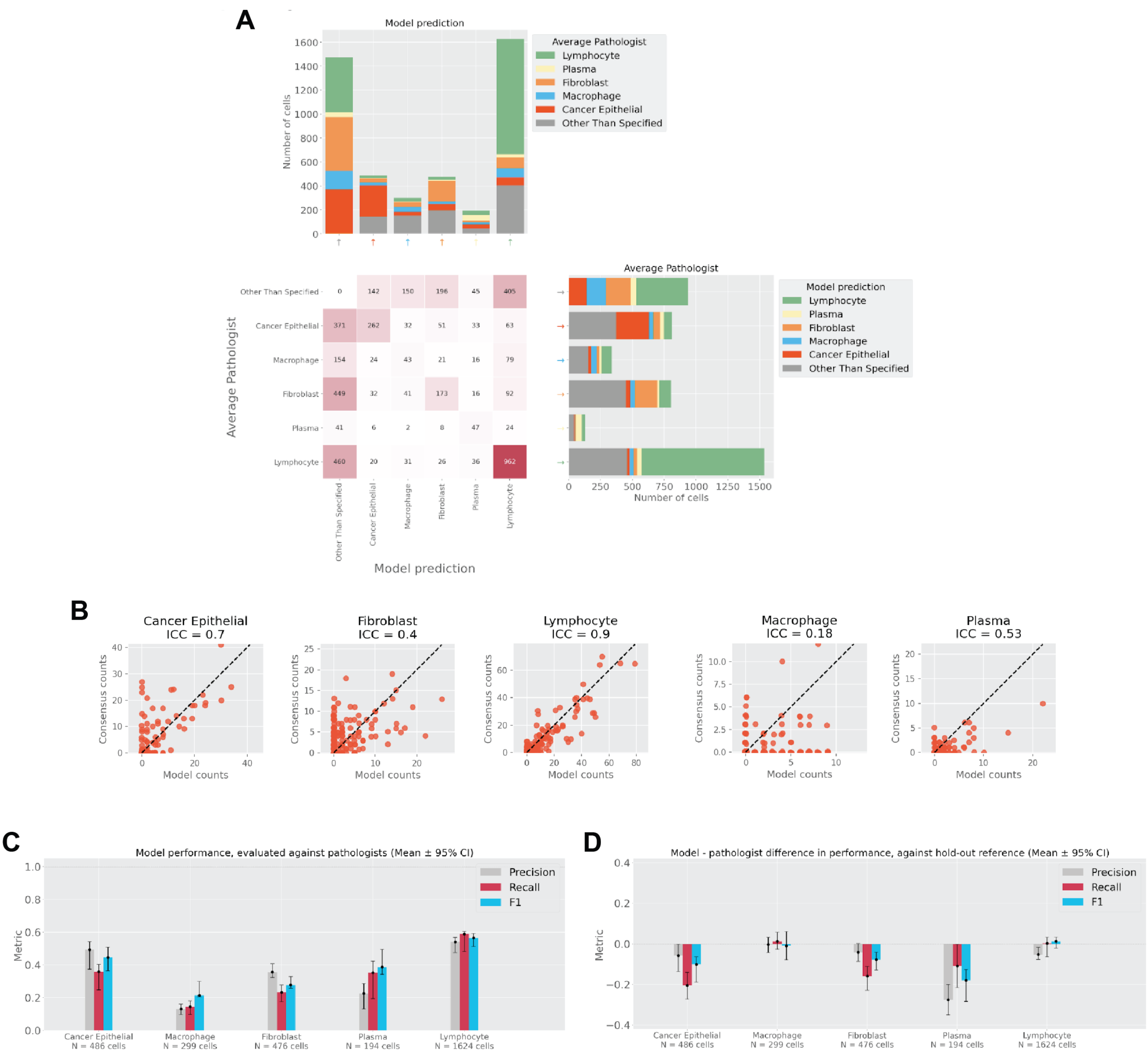


**Supplementary Figure S1. Performance of cell classification model in breast cancer.** A) Comparison of model-predicted cell types to average pathologist annotations. The bar graph at the top left depicts the breakdown of average pathologist annotations for each class of model prediction (precision). The bar graph at the bottom right shows the breakdown of model predictions for each class of average pathologist annotation (recall). "Other Than Specified" refers to predictions of classes other than those listed, or background class. B) Agreement between model-derived cell counts and pathologist consensus counts . C) Precision, Recall, and F1 scores of model predictions compared to pathologists' annotations in nested pairwise fashion32. D) Difference in the nested pairwise metric comparing mean difference between model and individual pathologist performance. Positive values indicate that the model out-performed pathologists when evaluated against held-out pathologists, while negative values indicate the model under-performed pathologists. Confidence intervals were obtained by bootstrapping.

### Supplementary Figure S2

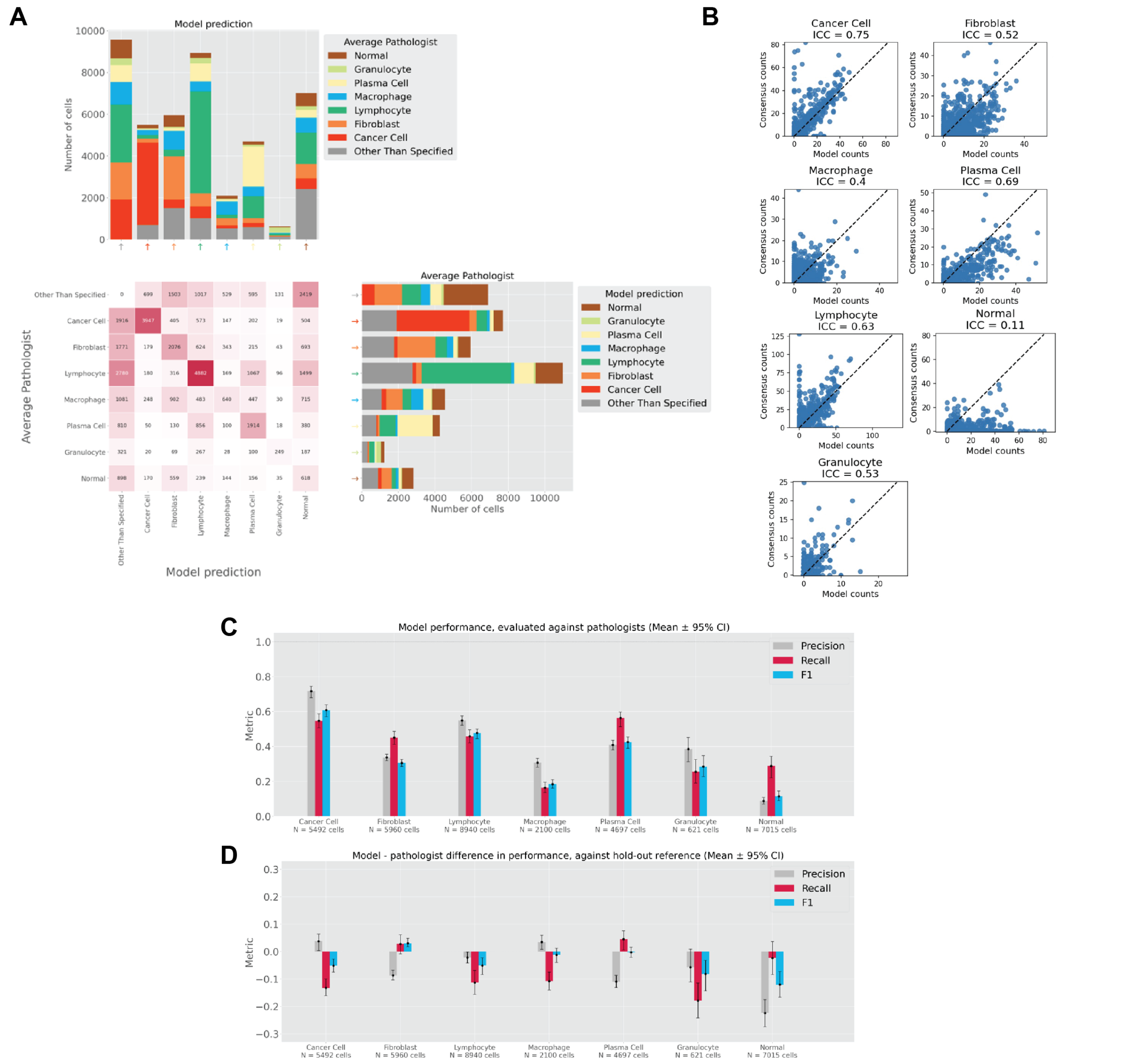


**Supplementary Figure S2. Performance of cell classification model in NSCLC.** A) Comparison of model-predicted cell types to average pathologist annotations. The bar graph at the top left depicts the breakdown of average pathologist annotations for each class of model prediction (precision). The bar graph at the bottom right shows the breakdown of model predictions for each class of average pathologist annotation (recall). “Other Than Specified” refers to predictions of classes other than those listed, or background class. B) Agreement between model-derived cell counts and pathologist consensus counts . C) Precision, Recall, and F1 scores of model predictions compared to pathologists’ annotations in nested pairwise fashion32. D) Difference in the nested pairwise metric comparing mean difference between model and individual pathologist performance. Positive values indicate that the model out-performed pathologists when evaluated against held-out pathologists, while negative values indicate the model under-performed pathologists. Confidence intervals were obtained by bootstrapping.

### Supplementary Figure S3

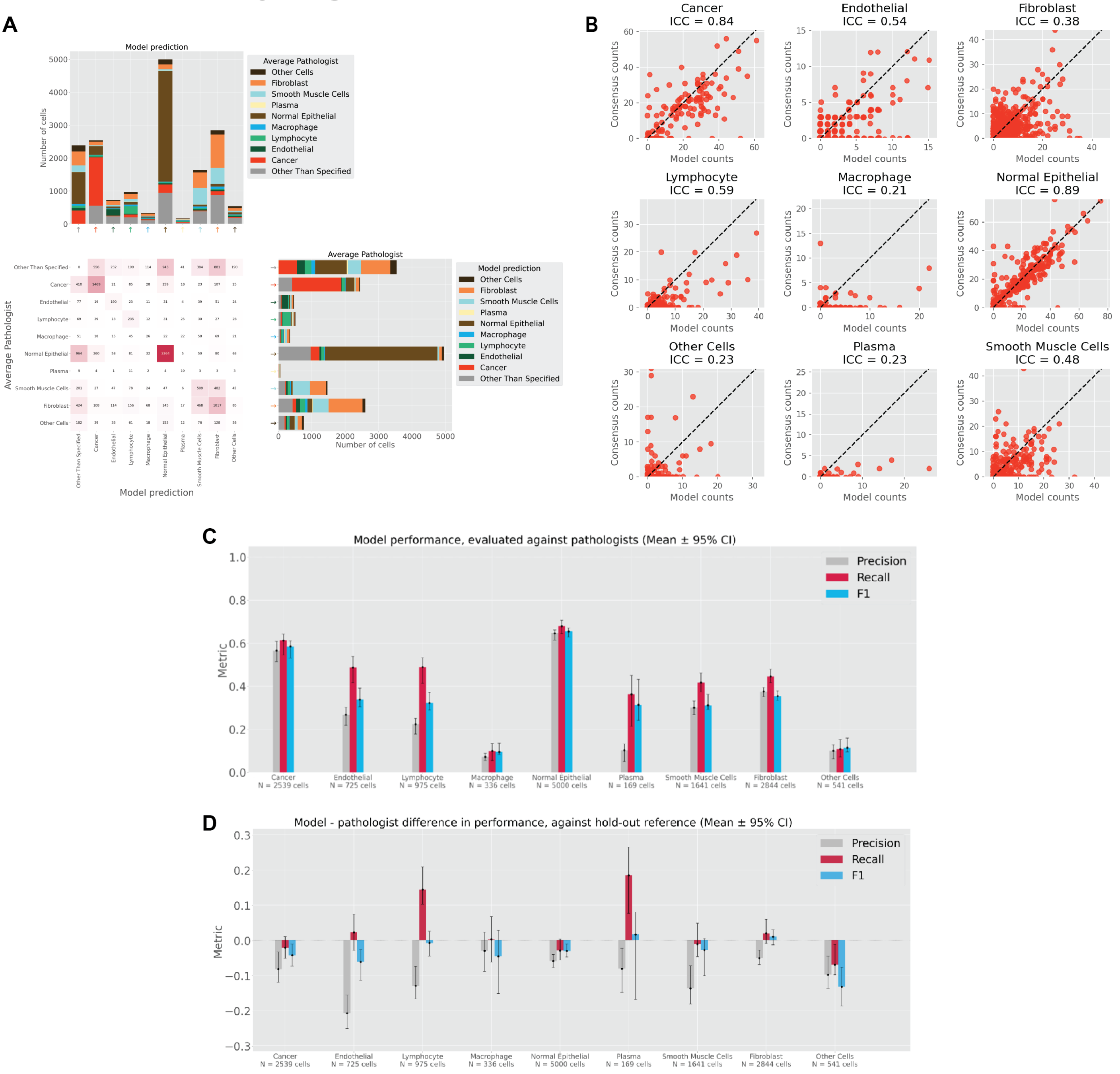


**Supplementary Figure S3. Performance of cell classification model in prostate cancer.** A) Comparison of model-predicted cell types to average pathologist annotations. The bar graph at the top left depicts the breakdown of average pathologist annotations for each class of model prediction (precision). The bar graph at the bottom right shows the breakdown of model predictions for each class of average pathologist annotation (recall). "Other Than Specified" refers to predictions of classes other than those listed, or background class. B) Agreement between model-derived cell counts and pathologist consensus counts . C) Precision, Recall, and F1 scores of model predictions compared to pathologists' annotations in nested pairwise fashion32. D) Difference in the nested pairwise metric comparing mean difference between model and individual pathologist performance. Positive values indicate that the model out-performed pathologists when evaluated against held-out pathologists, while negative values indicate the model under-performed pathologists. Confidence intervals were obtained by bootstrapping.

### Supplementary Figure S4

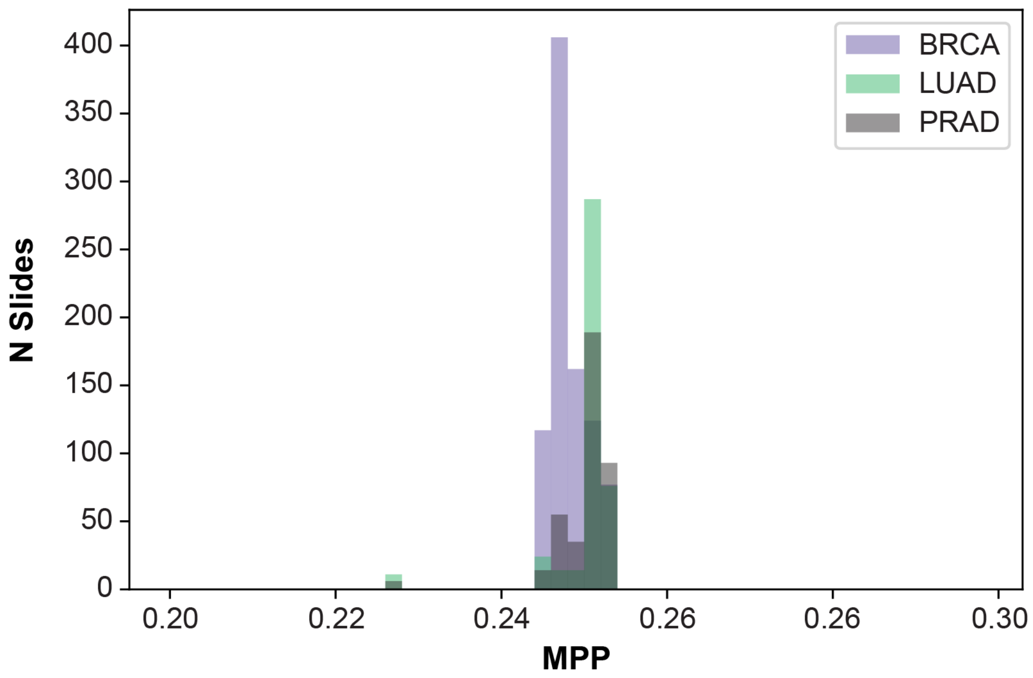


**Supplementary Figure S4. Distribution of pixel sizes (in MPP) across the three TCGA datasets (BRCA, LUAD, PRAD) used in this study.**

### Supplementary Figure S5

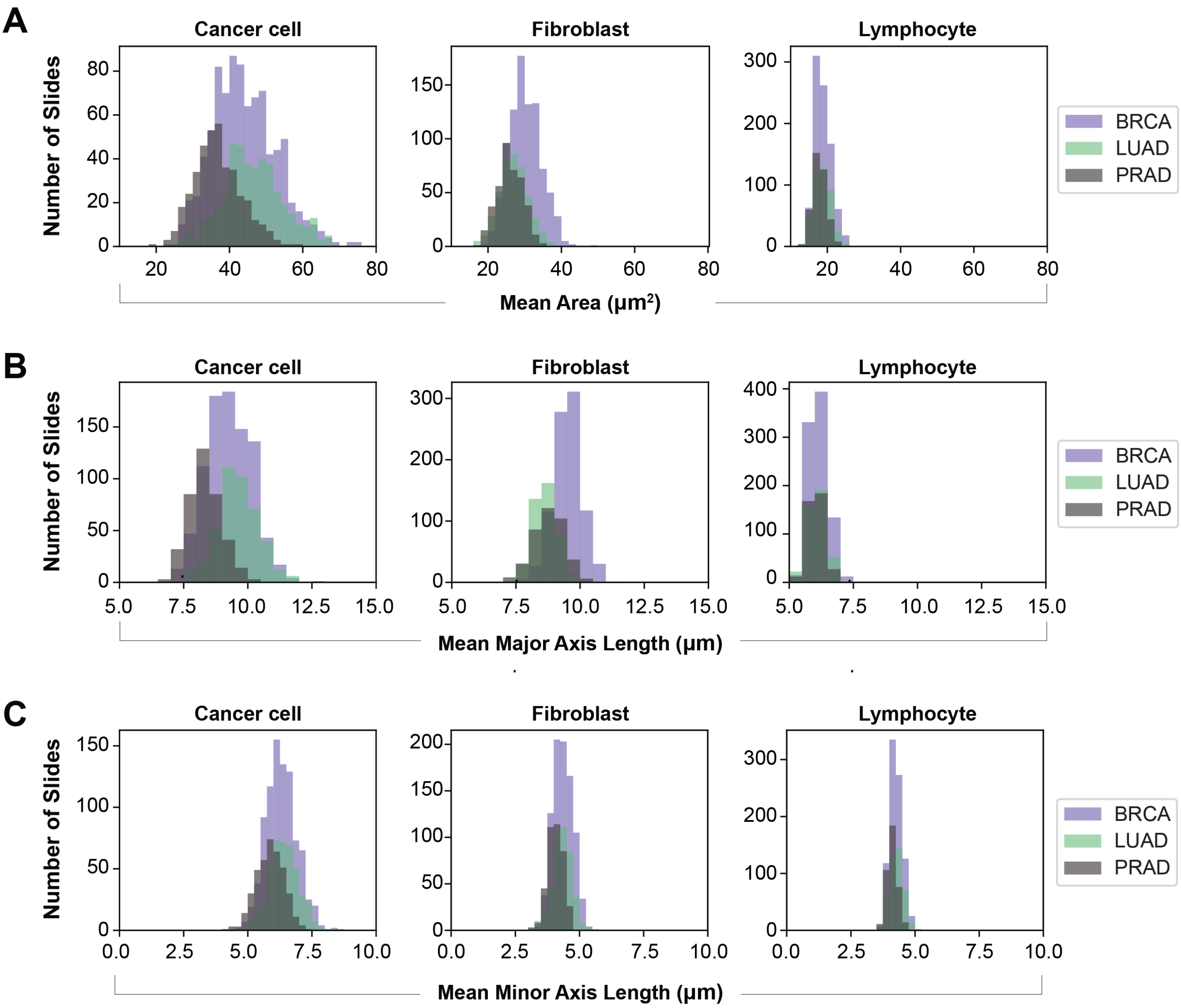


**Supplementary Figure S5. Distributions of nuclear size features in BRCA, LUAD, and PRAD datasets.** A) Distribution of area. B) Distribution of major axis length. C) Distribution of minor axis length.

### Supplementary Figure S6

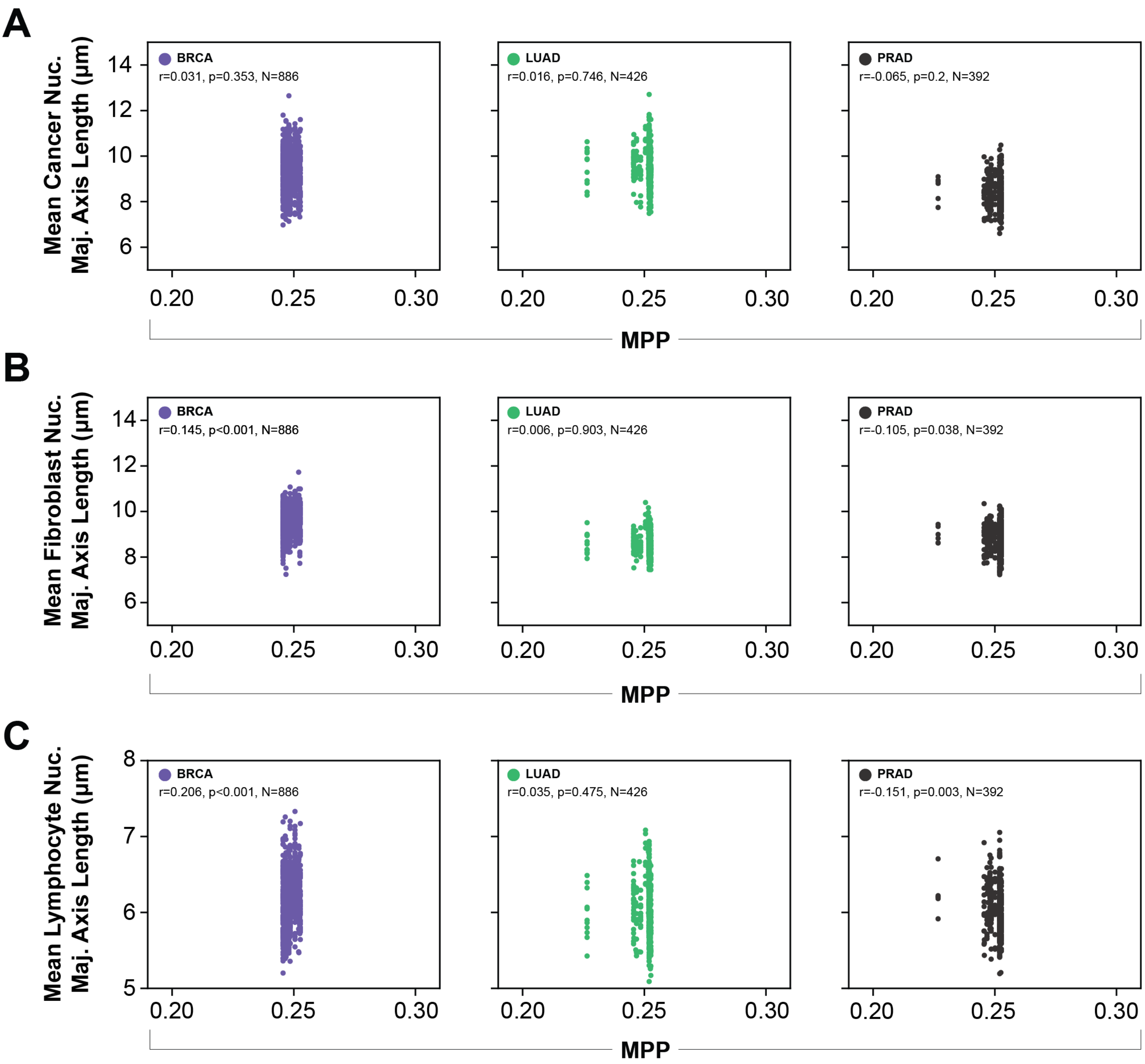


**Supplementary Figure S6. Pearson correlation between nuclear size (using major axis length as a representative feature) and MPP for each cell type within BRCA, LUAD, and PRAD datasets.** A) Correlation between major axis length and MPP in cancer epithelial cells. B) Correlation between major axis length and MPP in fibroblasts. C) Correlation between major axis length and MPP in lymphocytes.
